# HIF-1α coordinates adrenal steroidogenesis through direct transcriptional control and regulation of miRNA biogenesis

**DOI:** 10.64898/2026.02.24.707817

**Authors:** Barbara Krystyna Stepien, Anupam Sinha, Stephen Ariyeloye, Anja Krüger, Peter Mirtschink, Rafał Bartoszewski, Ben Wielockx

**Affiliations:** Institute of Clinical Chemistry and Laboratory Medicine, University Carl Gustav Carus and Medical Faculty, Technische Universität Dresden, 01307 Dresden, Germany; Department of Biophysics, Faculty of Biotechnology, University of Wroclaw, 50-383 Wroclaw, Poland; Experimental Centre, Faculty of Medicine, Technische Universität Dresden, 01307 Dresden, Germany

**Keywords:** CUT&Tag, HIF, MicroRNA, Adrenal steroidogenesis, RISC, Microprocessor complex, Argonaute, Drosha

## Abstract

**Background:** Adrenal steroid hormone production is essential for systemic stress adaptation and metabolic homeostasis, and it is tightly regulated by oxygen availability. Previously, we demonstrated that acute hypoxia suppresses adrenal steroidogenesis through HIF-1α-dependent induction of microRNAs (miRNAs) that target key steroidogenic enzymes. However, the mechanisms by which HIF-1α controls miRNA expression and activity in this context remain unclear.

**Methods:** To address this issue, we mapped the genome-wide HIF-1α binding landscape in murine adrenocortical cells using Cleavage Under Targets & Tagmentation (CUT&Tag). We integrated this data with gene expression analyses following pharmacological HIF-1α stabilization, physiological hypoxia, and genetic HIF-1α depletion to distinguish HIF-1α-dependent effects from broader hypoxia-driven responses.

**Results:** We detected HIF-1α binding at loci encoding steroidogenic enzymes and steroidogenesis-associated miRNAs. Unexpectedly, we also detected binding at genes involved in miRNA biogenesis and function, including components of the nuclear microprocessor complex and the cytoplasmic RNA-induced silencing complex (RISC). Functional analyses revealed that hypoxia broadly represses the expression of miRNA-processing genes through both HIF-1α-dependent and -independent mechanisms. Notably, HIF-1α selectively modulated or counteracted this repression in a gene-specific manner, indicating a regulatory role beyond direct transcriptional activation.

**Conclusions:** These findings reveal an unrecognized layer of hypoxia-driven cell communication, wherein HIF-1α coordinates the transcriptional and post-transcriptional regulation of adrenal steroidogenesis by shaping the miRNA-processing landscape. This work extends our understanding of how oxygen-sensitive signaling pathways integrate gene expression and RNA-based regulatory mechanisms to control endocrine function.

## Background

Steroid hormones are a diverse class of cholesterol-derived molecules that include sex hormones such as estradiol and testosterone, mineralocorticoids such as aldosterone, and glucocorticoids, which are represented primarily by cortisol in humans and corticosterone in rodents (1,2). These hormones regulate a broad range of physiological functions, including metabolism, electrolyte homeostasis, stress responses, immune regulation, and fertility. Dysregulation of steroidogenesis has been linked to numerous human disorders, including common conditions such as hypertension and polycystic ovary syndrome (1), while overproduction manifests as Cushing’s disease and related syndromes (2). The adrenal cortex is the primary site of non-sex steroid hormone production in mammals, with distinct cortical zones synthesizing mineralocorticoids and glucocorticoids that regulate vital physiological processes (1–3). Steroid hormone production is a tightly regulated process that responds to a variety of internal and external stimuli to maintain homeostasis. One such stimulus that can modulate steroidogenesis in the adrenal gland is hypoxia, a state of reduced tissue oxygenation that activates the hypoxia-inducible factor (HIF) signaling pathway (reviewed in (4,5)). Mammals express three HIF-α family members, HIF-1α, HIF-2α, and HIF-3α, whose protein levels are tightly regulated by oxygen availability. Under normoxic conditions, HIF-α proteins undergo oxygen-dependent hydroxylation of conserved prolyl residues by prolyl hydroxylase domain (PHD) enzymes, which targets them for ubiquitination by the von Hippel–Lindau (VHL) ubiquitin ligase complex and subsequent proteasomal degradation. In contrast, reduced oxygen availability leads to PHD inactivation and stabilization of HIF-α proteins. Stabilized HIF-α subunits accumulate and translocate to the nucleus, where they heterodimerize with the constitutively expressed HIF-1β subunit, bind conserved hypoxia-response elements in target genes, and regulate transcription (4–7).

Previously, we showed that HIF-1α, but not HIF-2α, modulates adrenal steroidogenesis by repressing the expression of steroidogenic enzymes and reducing overall steroid hormone production (8). More recently, we demonstrated that stabilization of HIF-1α increases the expression of specific miRNAs that target key enzymes in the steroidogenic pathway, both in adrenocortical cells *in vitro* and in the adrenal gland *in vivo*. These changes were associated with altered expression of steroidogenic enzymes and corresponding changes in steroid hormone output (9). However, it remains largely unclear whether HIF regulates these miRNAs through direct binding to regulatory DNA elements or via indirect mechanisms, and how such regulation contributes to HIF-dependent transcriptional control of steroidogenesis.

In the present study, we employed genome-wide mapping of HIF-1α binding using the CUT&Tag method (10) in a murine adrenocortical cell line to further elucidate how HIF-1α shapes the transcriptional landscape underlying steroidogenesis. This analysis revealed extensive HIF-1α occupancy across the genome, including binding at multiple miRNA gene loci, such as *miR-6924*, which we previously identified as a hypoxia-responsive regulator of steroidogenic enzymes (9). Notably, HIF-1α binding was also detected at genomic regions associated with genes involved in miRNA biogenesis and function, pointing to a broader role for HIF-1α in coordinating miRNA homeostasis beyond the regulation of individual miRNA genes. Consistent with this, mRNA expression analyses confirmed HIF-1α-dependent regulation of several miRNA-processing components. Together, these findings indicate that the influence of HIF-1α on the adrenal miRNA landscape extends beyond specific miRNA targets and includes modulation of global miRNA biogenesis, thereby adding an additional regulatory layer to HIF-mediated control of steroidogenesis.

## Methods

### Cell culture

The murine Y-1 adrenocortical cell line (11,12) was cultured in Dulbeccós Modified Eagle’s Medium (DMEM/F12; Thermo Fisher Scientific, #31330-038), which was supplemented with 10% Fetal Calf Serum (FCS; Thermo Fisher Scientific, #10500-064), 2.5% horse serum (Biowest, #S0910-500), and 1% penicillin-streptomycin (Thermo Fisher Scientific, #15140122). During hypoxia experiments the medium was additionally supplemented with 2.5% UltroSerG (Pall Life Sciences, #15950-017) and 1% Insulin-Transferrin-Selenium-Ethanolamine (ITS-X) (Thermo Fisher Scientific, # 51500056). Unless otherwise indicated, cells were grown at 37°C in a humidified incubator under normoxia (atmospheric O_2_ concentration) with 5% CO_2_. For HIF hydroxylase (PHD) inhibition, dimethyloxalylglycine (DMOG; Biomol, Cay71210-100) in 1x PBS was added to the growth medium at 1 mM final concentration for 24 or 48 hours (solvent only was used as control). DMOG is known to stabilise HIF-α proteins via the inhibition of prolyl hydroxylase domain enzymes, which under normoxic conditions target HIF for rapid degradation. For the hypoxia treatment cells were grown in an atmosphere containing 5% O_2_ and 5% CO_2_ (Whitley H35 Hypoxystation; Don Whitley Scientific Limited, United Kingdom) for 24 or 48 hours. For the shRNA treatment, the Y-1 cells were transduced with the lentivirus carrying either shHIF-1α or scramble (shScr) as described previously (9). At the end of each experiment, the cells were immediately collected for RNA isolation by removing the medium and resuspending the cells in an appropriate lysis buffer.

### CUT&Tag assay

The CUT&Tag assay was performed as previously described (13). Nuclear preparation was performed based on the protocol described by Kaya-Okur and colleagues (14). Briefly, Y-1 cells were cultured for 24 hours in the presence of 1 mM DMOG or under control conditions. The cells were washed with 1x PBS and treated with trypsin-EDTA (Thermo Fisher Scientific, #15400054) for 4 minutes at 37°C in a cell culture incubator. Trypsin was quenched with cell culture medium and detached cells were transferred to a 15 ml tube and pelleted for 5 minutes at 491 g. The cells were then resuspended in fresh cell culture medium and washed with 1x PBS. Cell pellet was resuspended in ice-cold NE1 buffer (20 mM HEPES-KOH pH 7.9, 10 mM KCl, 0.1% Triton X-100, 20% glycerol, 0.5 mM spermidine) supplemented with Roche Complete Protease Inhibitor EDTA-Free tablets (Merck, #11836170001) at ½ volume of the starting culture and incubated for 10 minutes on ice. The cell nuclei were pelleted by centrifugation (4 minutes, 1300 g, 4°C) and resuspended in ice-cold 1x PBS. Nuclei were crosslinked with 0.1% formaldehyde (16% (w/v) formaldehyde; Thermo Fisher Scientific, #28906) for 2 minutes at room temperature and the reaction was quenched by glycine solution (70 mM final concentration). Fixed nuclei were then washed with 0.5 ml of nuclei wash buffer (20 mM HEPES-NaOH pH 7.5, 150 mM NaCl, 0.5 mM spermidine) supplemented with Roche Complete Protease Inhibitor EDTA-Free tablets and counted. The nuclear pellet was then resuspended in nuclei wash buffer at a concentration of 1 million nuclei per ml, aliquoted and stored at -80°C until use.

The frozen nuclei were thawed at room temperature for 30 minutes and bound to ConA-coated magnetic beads (BioMag Plus Concanavalin A; Bangs Laboratories, #BP531), which were pre-equilibrated with binding buffer (20 mM HEPES-KOH pH 7.9, 10 mM KCl, 1 mM CaCl_2_, 1 mM MnCl_2_) in 0.5 ml tubes (3.5 µl beads per 50,000 nuclei) for 10 minutes rotating at room temperature. The supernatant was removed and anti-HIF-1α primary antibody solution (Cayman Chemicals, #10006421) in antibody buffer (CUT&Tag wash buffer with 0.1% BSA) was added at 1:50 dilution. The beads were rotated with the primary Ab solution for 2 hours at RT followed by overnight incubation at 4°C. Next, the supernatant was removed and guinea pig α-rabbit secondary antibody (Antibodies online, #ABIN101961) solution, 1:100 in CUT&Tag wash buffer (20 mM HEPES-NaOH pH 7.5, 150 mM NaCl, 0.5 mM spermidine, Roche Complete Protease Inhibitor EDTA-Free tablets; Merck, #11836170001) was added to the beads. The beads were rotated for 1 hour at RT and the supernatant was removed. The beads were then washed with 0.5 ml of CUT&Tag wash buffer and the pA-Tn5 adapter complex (Epicypher, #15-1117) in 300 wash buffer (20 mM HEPES-NaOH pH 7.5, 300 mM NaCl, 0.5 mM spermidine, Roche Complete Protease Inhibitor EDTA-Free tablets; Merck, #11836170001) was added at 1:20. The complex was rotated with the beads for 2 hours at RT and the supernatant was removed. The beads were washed with 0.5 ml of 300 wash buffer and resuspended in CUT&Tag tagmentation buffer (300 wash buffer with 10 mM MgCl_2_). The samples were incubated for 1 hour at 37°C and cooled down to 8°C in a PCR cycler. The supernatant was removed and beads were washed with TAPS wash buffer (10 mM TAPS pH 8.5, 0.2 mM EDTA). The Tn5 complex was released with 5 µl SDS release solution (10 mM TAPS pH 8.5, 0.1% SDS) at 58°C for 1 h in a PCR cycler. Subsequently, 15 µl Triton neutralization solution (0.67% Triton X-100) was added per sample. PCR reactions (gap filling: 5 minutes, 58°C; 5 minutes, 72°C; initial denaturation: 30 s, 98°C; 11 cycles of denaturation 10 s, 98°C; and annealing and elongation 10s, 60°C; final elongation: 1 minute, 72°C; hold 8°C) were run with barcoded primers (15) using the NEBNext High Fidelity 2x PCR mix (New England Biolabs, #M0541L). DNA was cleaned up with SPRI beads (AMPure XP Bead-Based Reagent; Beckman Coulter, #A63880) for 10 minutes at RT, washed twice with 80% ethanol and eluted with 10 mM Tris-HCl pH 8 for 1 hour at RT. The libraries were subjected to size selection to enrich for fractions between 170 and 700 bp (Fragment Analyzer, Advanced Analytical) and sequenced (Illumina next generation sequencing) at the DRESDEN-concept Genome Center c/o Center for Molecular and Cellular Bioengineering (CMCB), Technische Universität Dresden.

### Bioinformatic analysis

An in-house pipeline was created to perform the CUT&TAG analyses. Briefly, bowtie2 v2.4.2 (16) was run using the parameters “--end-to-end --very-sensitive --no-mixed --no-discordant --phred33 -I 10 -X 700 -p 2” using the Mus_musculus.GRCm39.111.gtf and Mus_musculus.GRCm39.dna_sm.primary_assembly.fa files from ENSEMBL (17) database. samtools v1.15.1 (18) was used to filter, sort and index the bam files. bamCoverage command from deepTools v3.5.6 (19) command line software package was used to create normalized bigwig files from the filtered and sorted bam files. macs2 v2.2.7.1 (20) was run using the following parameters: “callpeak -f BAMPE -g 2654621783 --keep-dup all --nomodel --pvalue 0.1“. The resultant .narrowPeak files were sorted and merged using bedtools v2.17.0 (21) to create a reference bed file containing all the accessible regions across all the samples. This was further used to generate per-region counts table using featureCounts of the Rsubread v2.10.2 (22) package of R. The counts table was used as input for DESeq2 v1.46.0 (23), another R package, to perform differential accessibility analyses. Homer2 (24) was used to predict the motifs in the differentially accessible regions. To extract a list of HIF-1α target regions used for downstream analysis the results were filtered according to the following criteria: p-value < 0.05 and log2FoldChange > 2. Accordingly, 9903 HIF-1α-bound gene regions, represent 6989 unique genes, were found to be enriched in DMOG-treated cells (Additional file 1). In contrast, 3455 gene regions representing 2792 genes, were depleted compared to control cells. miRNet platform (25) was used to identify the network of miRNA-regulated genes for the putative HIF-1α miRNA targets. Over-Representation analysis (KEGG pathways) was done using the WebGestalt platform (26,27) using a list of 6989 unique CUT&Tag target gene IDs. Pathways with FDR < 0.05 were considered significant (Additional file 3). ggplot2 v3.5.2 (28) was used for plotting. The data has been submitted to GEO under the accession number GSE311437.

### RNA isolation and reverse transcription

RNA was isolated from Y-1 cells using either the RNA Easy Plus micro kit (Qiagen, #74034) or the RNeasy Plus Mini kit (Qiagen, #74134) following the manufacturer’s instructions. Reverse transcription was performed using the iScript cDNA Synthesis Kit (Bio-Rad, 1708890) for protein coding genes or Mir-X miRNA First-Strand Synthesis Kit (Takara, #638313) for miRNA genes.

### Real-time quantitative PCR

For gene expression analysis, quantitative real-time PCR was performed using the Real-Time PCR Detection System-CFX384 (Bio-Rad, 1855484). Reactions were set up using the SsoFast EvaGreen Supermix (Bio-Rad, 1725201) for protein coding genes or the Mir-X miRNA qRT-PCR TB Green Kit (Takara, #638316) for miRNA genes according to the manufacturers’ protocols. mRNA expression levels were calculated relative to beta-2 microglobulin (*B2M*) and eukaryotic translation elongation factor 2 (*EF2*) housekeeping genes using the 2(-ddCt) method, where ddCT was calculated by subtracting the average control dCT from dCT of each sample. miRNA expression levels were calculated with the 2(-ddCt) method using the product of U6 control primers as a reference (Mir-X miRNA qRT-PCR TB Green Kit; Takara, #638316). Primer sequences and references (8,29,30) are listed in Additional file 6. Melting curves for the products of primers used in this study can be seen in Additional file 11. The additional analysis of the product of the Mtdh primers was done *in silico* using the uMelt Quartz v4.0 software (Blake & Delcourt (1998) method) (31) and the presence of a single product was confirmed with agarose gel electrophoresis (Additional file 11).

### Western blotting

Protein lysates were prepared in modified RIPA buffer (150mM NaCl, 50 mM Tris-HCl pH 8.0, 1 mM EDTA, 0.1% SDS, 1% NP-40) with Roche Complete Protease Inhibitor EDTA-Free tablets (Merck, #11836170001). Protein concentration was measured with the BCA assay (Pierce BCA protein assay kit, Thermo Scientific, #23227) and samples were adjusted to the same amount of protein (25 μg) per lane. Protein samples were prepared in NuPAGE LDS Sample Buffer (4x) (Thermo Fisher Scientific, NP0007) with NuPAGE Sample Reducing Agent (10x) (Thermo Fisher Scientific, NP0004) with 500 mM DTT and loaded onto a 4-12% Bis-Tris gradient gels (Thermo Fisher Scientific, NP0321BOX). The gels were run using the XCell SureLock Mini-Cell Electrophoresis (ThermoFisher Scientific, EI0001) system in 1x NuPAGE MOPS SDS Running Buffer (Thermo Fisher Scientific, NP0001) at 150 V, room temperature. After electrophoresis the proteins were transferred onto Amersham Protran Supported Nitrocellulose Western Blotting Membranes, Cytiva, pore size 0.2 µm (Avantor, #10600015) using the XCell II Blot Module (Thermo Fisher Scientific, EI9051) system at 30 V for 2 hours on ice in 1x NuPAGE Transfer Buffer (Thermo Fisher Scientific, NP00061) with 20% methanol (concentration for transferring 2 gels). The membranes were cut at the height of 70 kDa band (PageRuler Plus Prestained Protein Ladder, 10 bis 250 kDa, Thermo Fisher Scientific, #26619) and each half was blocked in 5% skimmed milk in PBST (1x PBS, 0.05% TritonX-100) for 1 hour at room temperature and then incubated with primary antibodies either against AGO2 (Abcam, ab186733) at 1:1000, AGO4 (Abcam, ab25921) at 1:1000 or tubulin α (Sigma Aldrich, #T5168) at 1:5000 in 5% skimmed milk in PBST overnight at 4°C. The membranes were then washed 3 times 10 minutes in PBST and then incubated with HRP-conjugated secondary antibodies (goat anti-rabbit, R&D Systems, HAF008 or horse anti-mouse, Cell Signalling, #7076), for 1 h at room temperature, washed 3 times 10 minutes in PBST and developed with SuperSignal West Femto Maximum Sensitivity Substrate (ThermoFisher Scientific, #34095) or SuperSignal West Pico PLUS Chemiluminescent Substrate (ThermoFisher Scientific, #34577). Imaging was done with Fusion FX Vilber Lourmat (Peqlab) imager. For quantification AGO2 and AGO4 signals were normalized to respective tubulin α signal intensity using Fiji software (32).

### Statistical analysis and data visualization

To assess statistical significance between two experimental groups, a Mann–Whitney U-test was used for non-normally distributed data, while an unpaired t-test with Welch’s correction, accounting for unequal variances, was applied for normally distributed data. Normality was tested using the Shapiro-Wilk test. Statistical differences presented in the figures were considered significant at p-values below 0.05. All statistical analyses were performed using GraphPad Prism v10.02 for Mac or higher (GraphPad Software, La Jolla, California, USA, www.graphpad.com). Graphical elements were generated using Affinity Designer 2 (Serif, Nottingham, UK) and BioRender.com (BioRender, Toronto, Canada).

## Results

### CUT&Tag reveals a HIF-1α DNA binding profile in Y-1 cells

To gain insight into the direct regulation of steroidogenesis by HIF-1α, we performed a genome-wide CUT&Tag analysis in the murine adrenocortical cell line Y-1. Cells were incubated for 24 hours in the presence of DMOG, a hypoxia-mimicking agent previously shown to stabilize and promote accumulation of HIF-1α protein in Y-1 cells (9). Nuclei were subsequently isolated, lightly fixed, and processed for CUT&Tag library preparation, followed by sequencing to generate a genome-wide HIF-1α binding profile (Figure 1A). In DMOG-treated cells, we identified 9,903 HIF-1α–associated regions corresponding to 6,989 unique genes, defined as the putative HIF-1α target list (Additional file 1). Notably, 28.8% of these binding sites were located within promoter regions (Additional file 2 and Figure 1B).

**Figure 1.**
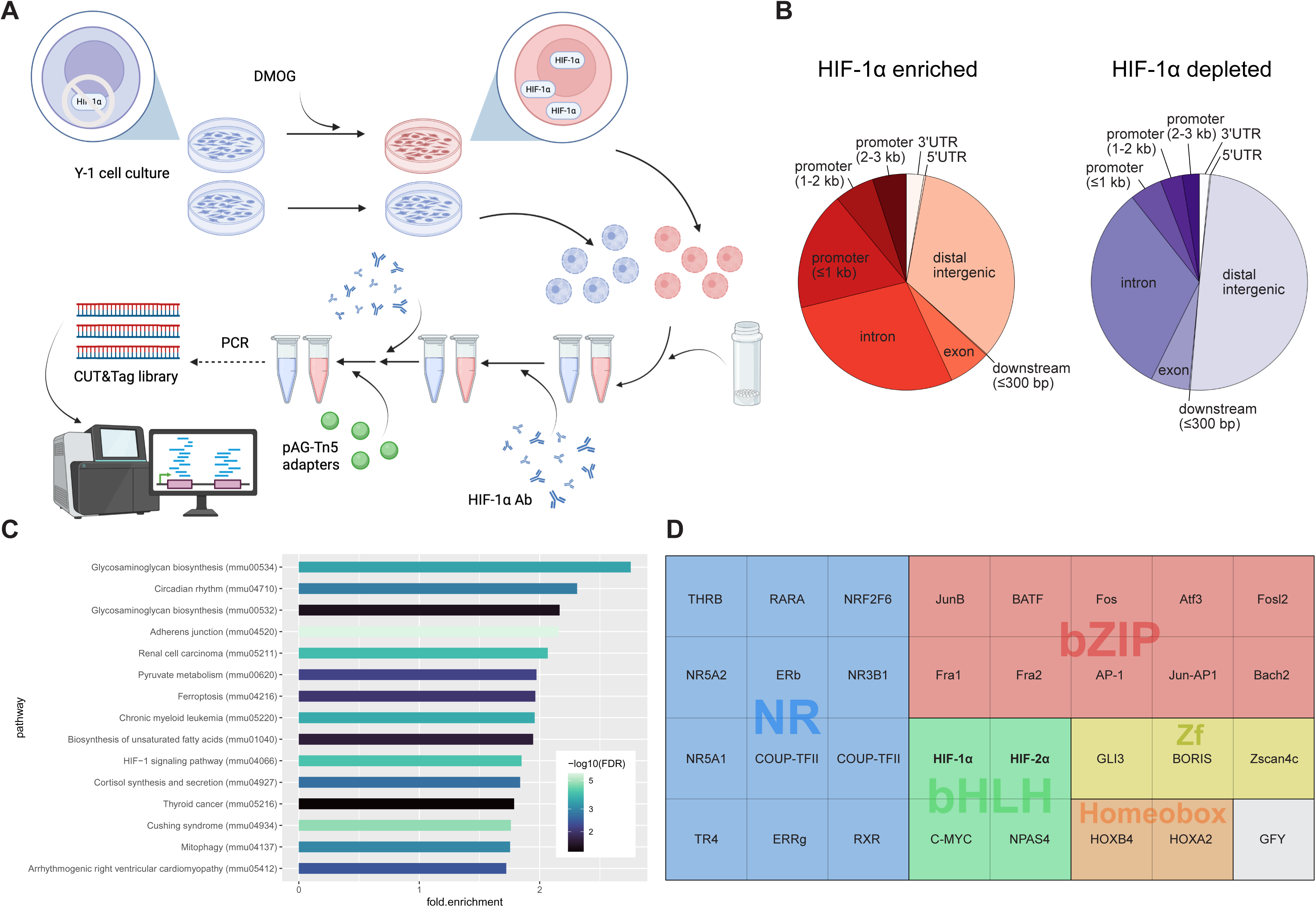
Genome-wide HIF-1α binding profile in Y-1 adrenocortical cells. A) Schematic representation of the CUT&Tag protocol. Under standard culture conditions (21% O□), Y-1 cells express undetectable levels of HIF-1α protein. For the CUT&Tag procedure, the cells were either left untreated as a control or treated with DMOG for 24 hours to stabilize HIF-1α. This treatment leads to nuclear accumulation of HIF-1α and its binding to target genes. The nuclei were then isolated, lightly fixed with formaldehyde, and bound to concanavalin A-coated beads. The HIF-1α-bound chromatin regions were labeled on the beads using a primary anti-HIF-1α antibody, a secondary antibody, and protein A-Tn5 adapters. The tagged DNA fragments were then released, PCR-amplified, and subjected to Illumina next-generation sequencing. B) The genomic distribution of enriched (left) and depleted (right) HIF-1α-bound sequences in DMOG-treated versus control cells (see also Additional file 2). C) Top 15 KEGG pathways enriched among HIF-1α-bound genes. Bar length indicates the pathway enrichment ratio, and colors represent FDR (see also Additional file 3). D) Motif enrichment analysis of HIF-1α-bound sequences revealed conserved binding motifs for HIF-1/2α and 30 additional transcription factors (p < 0.01, Benjamini < 0.1) that are grouped into six major families (see Additional file 4).

To assess the functional relevance of the identified HIF-1α target genes, we performed over-representation analysis using the WebGestalt platform (26,27). KEGG pathway analysis revealed a strong enrichment for the HIF-1 signaling pathway, with 51 of 114 annotated genes present in the target list (enrichment ratio 1.85), confirming the robustness of the dataset (Figure 1C; Additional file 3). Several pathways related to HIF activity were also enriched, including carbon metabolism, glycolysis, mito- and autophagy, and cancer-related pathways. Genes within these categories included canonical HIF targets such as Erythropoietin (*Epo*) and *Vegfa*, as well as multiple glycolytic and glucose metabolism–related enzymes (*Hk1, Pfkp, Pfkl, Pgk1, Eno1, Pdk1, Pdha1/2, Pdhb, Ldha,* and *Ldhc*). Importantly, beyond hypoxia-related pathways, we also observed significant enrichment in categories directly linked to steroid hormone biosynthesis, including Cushing syndrome and cortisol synthesis and secretion (Figure 1C).

Motif enrichment analysis of HIF-1α–bound regions using HOMER identified 31 significantly enriched transcription factor motifs, including the expected canonical HIF-1α binding motif (Figure 1D; Additional file 4). Notably, motifs for NR5A1 (SF-1) and NR5A2 (LRH-1), key regulators of adrenal development and steroidogenesis (1), were also significantly enriched, supporting a role for HIF-1α within transcriptional networks governing steroid hormone production. In addition, HIF-1α–bound regions were enriched for motifs recognized by members of the AP-1 transcription complex, some of which have been described as HIF-1α cofactors in cancer cells (33), as well as motifs associated with nuclear receptor superfamily members.

### HIF-1**α** directly binds predicted miRNA gene regions

We previously demonstrated that HIF-1α upregulates the expression of multiple miRNAs that modulate steroidogenic enzyme expression under DMOG treatment or hypoxic conditions, both in Y-1 cells and *in vivo* (9). To assess whether HIF-1α directly regulates miRNA gene expression, we analyzed our CUT&Tag dataset for HIF-1α binding in proximity to annotated miRNA loci. This analysis identified 154 enriched HIF-1α–bound regions associated with 122 miRNA-containing transcripts (Additional file 1).

To explore the potential regulatory impact of these miRNAs, we used the miRNet platform (25) to identify predicted target genes of the HIF-1α-associated miRNAs. This analysis yielded a large interaction network comprising more than 13,000 genes, of which 5,257 were predicted to be regulated by at least five distinct miRNAs (Additional file 5). Notably, 12 of the miRNAs identified correspond to previously described hypoxamiRs (hypoxia-regulated microRNAs) regulated by hypoxia and HIF signaling (9,34) and 10 have been implicated in the regulation of steroidogenesis (Table 1). Among these candidates, *miR-29b* emerged as a prominent potential direct HIF-1α target, with four distinct HIF-1α–bound regions detected in its vicinity. To determine whether pharmacological HIF-1α stabilization affects *miR-29b* expression in adrenocortical cells, we measured *miR-29b* levels by qPCR following 24 hours of DMOG treatment under normoxic conditions. Under these conditions, DMOG induced an approximately two-fold increase in *miR-29b* expression in Y-1 cells (Figure 2).

**Figure 2.**
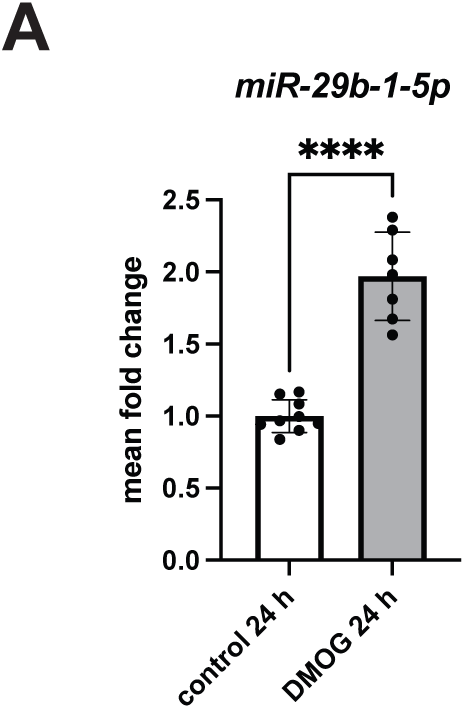
Pharmacological HIF-1α stabilization increases mir-29b expression. A) qPCR analysis of *miR-29b* RNA expression in Y-1 cells treated with DMOG for 24 h under normoxia. Bar heights represent mean fold change relative to control; error bars denote standard deviation of the mean. Statistical significance was determined using an unpaired *t*-test with Welch’s correction (**** p < 0.0001).

**Table 1.**
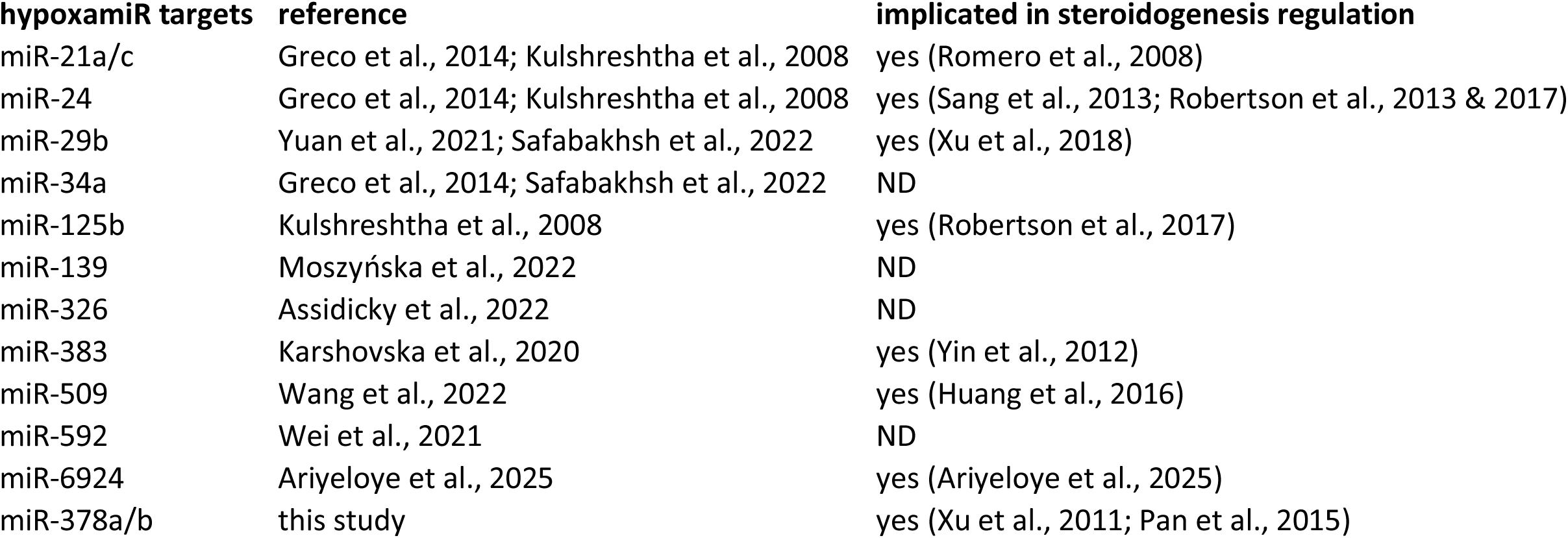
Hypoxia-regulated genes enriched in the HIF-1α CUT&Tag dataset and their reported involvement in steroidogenesis. A list of miR genes enriched in HIF-1α CUT&Tag dataset reported to be regulated by hypoxia (hypoxamiRs) and/or implicated in steroidogenesis regulation. ND – no data available. Citations can be found in the reference list (9,34,36,38–43,52–61).

Consistent with our previous findings, *miR-6924*, one of the hypoxia induced miRNAs reported by our group to target *Cyp11a1* mRNA expression (9) was also identified among the HIF-1α–associated miRNA loci. In the present study, CUT&Tag mapping provides complementary genomic evidence by revealing HIF-1α binding 325 nucleotides upstream of the *miR-6924* transcriptional start site within its 1 kb promoter region, including two canonical HIF-1α binding motifs. Together, these findings support direct HIF-1α–dependent transcriptional regulation of steroidogenesis-relevant miRNAs.

### HIF-1**α** binds within loci of genes crucial for miRNA biogenesis and function

To investigate mechanisms by which HIF-1α could influence miRNA levels beyond direct transcriptional regulation of individual miRNA genes, we examined our CUT&Tag dataset for HIF-1α binding at loci encoding components of the miRNA-processing machinery. This analysis revealed HIF-1α association with multiple genes encoding both the nuclear microprocessor complex (MC) and RISC (Figure 3 and Additional file 1). As these complexes are essential for miRNA maturation and function (Figure 3A), their association with HIF-1α points to a potential role in modulating miRNA biogenesis and activity. Notably, HIF-1α binding was detected at loci encoding the core MC components *Drosha* and *Dgcr8*, all four Argonaute orthologues (*Ago1–4*), and several additional RISC-associated factors. Canonical HIF-1α binding motifs (A/GCGTG) were identified within the majority of these HIF-1α–bound regions (Figure 3B). However, no canonical motif was detected in the regions associated with *Drosha* or *Tnrc6a/b*. Nevertheless, the absence of a consensus motif does not exclude functional HIF-1α binding, as non-canonical or indirect binding mechanisms have been described previously (35).

**Figure 3.**
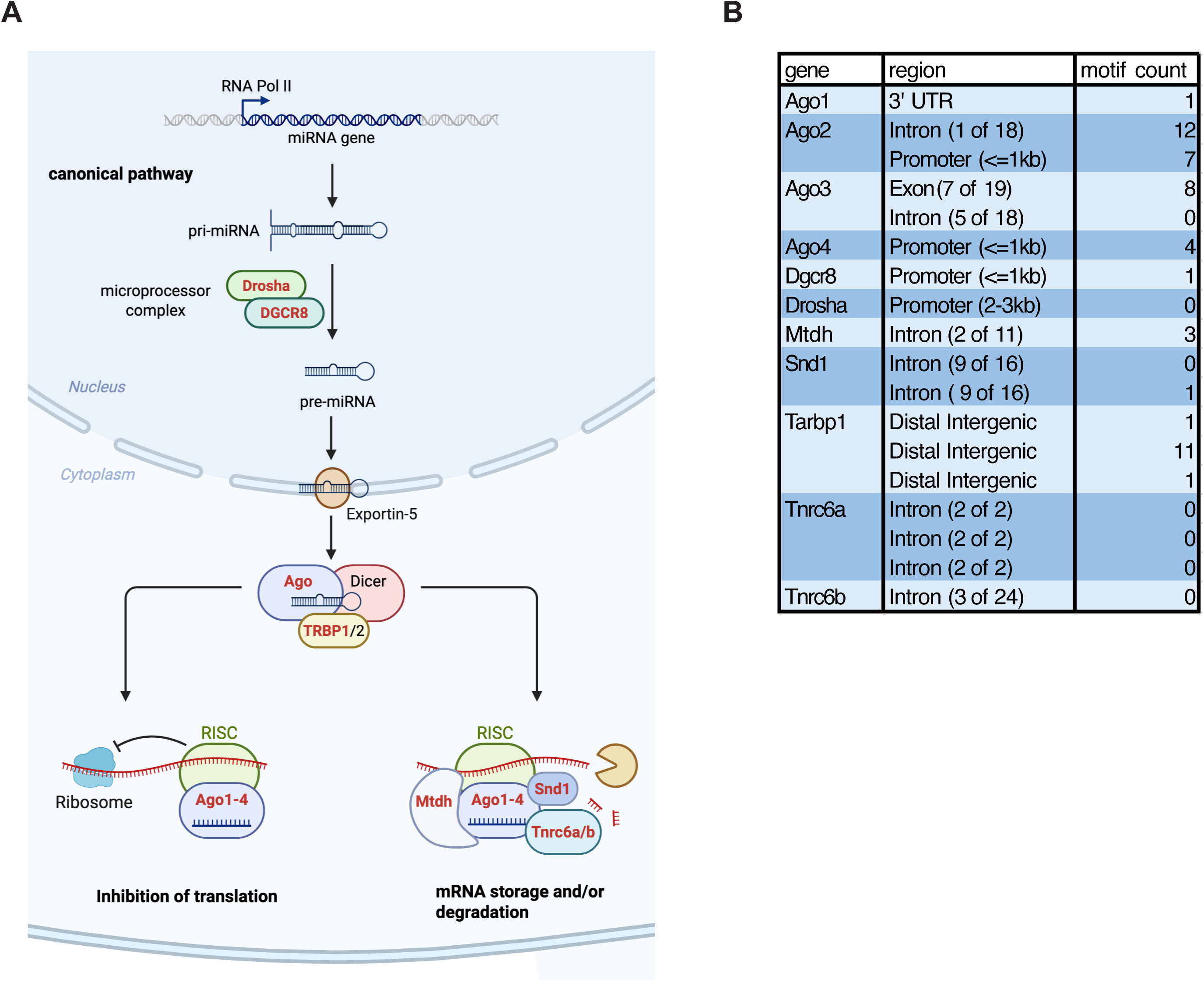
HIF-1α binding within the miRNA processing gene regions revealed with CUT&Tag analysis. A) Schematic depiction of the miRNA processing pathway. The pathway components identified in the HIF-1α CUT&Tag assay are depicted with protein names highlighted in red. Among the genes bound by HIF-1α are both members of the nuclear microprocessor complex (*Drosha* and *Dgcr8*) as well as a number of RISC components (*Ago1-4*, *Snd1*, *Mtdh*, *Tarbp1* and *Tnrc6a/b*). B) Identified miRNA biogenesis and function genes with the assigned location of the putative HIF-1α binding region and a number of canonical HIF-1α binding sites detected within each region.

### HIF-1**α** modulates hypoxic repression of miRNA biogenesis and function

To determine whether HIF-1α binding to genes encoding components of the miRNA-processing machinery is associated with functional changes in their expression, we next examined the effects of pharmacological HIF-1α stabilization, physiological hypoxia, and genetic HIF-1α depletion on MC and RISC gene expression.

To assess whether pharmacological HIF-1α stabilization affects the expression of miRNA-processing genes identified as potential HIF-1α targets in the CUT&Tag dataset, we performed qPCR analysis on Y-1 cells treated with DMOG for 24 hours under normoxic conditions (21% O□) (Figure 4A and Table 2). DMOG treatment did not alter *Drosha* mRNA levels but led to a significant reduction in *Dgcr8* expression (Figure 4B). In contrast, DMOG exerted gene-specific effects on components of the RISC complex: *Ago2* and *Ago4* were upregulated, whereas *Mtdh* and *Tnrc6a* were repressed, while the remaining RISC genes were unaffected (Figure 4C). As expected, DMOG did not affect *Hif1a* mRNA expression (Additional file 7A), consistent with HIF-1α regulation occurring at the level of protein stabilization. To determine whether DMOG-induced changes in mRNA abundance were reflected at the protein level, we performed quantitative Western blot analysis of AGO2 and AGO4. Consistent with the qPCR results, protein levels of both AGO2 and AGO4 increased by approximately 30% in DMOG-treated Y-1 cells (Figure 4D and Additional file 8).

**Figure 4.**
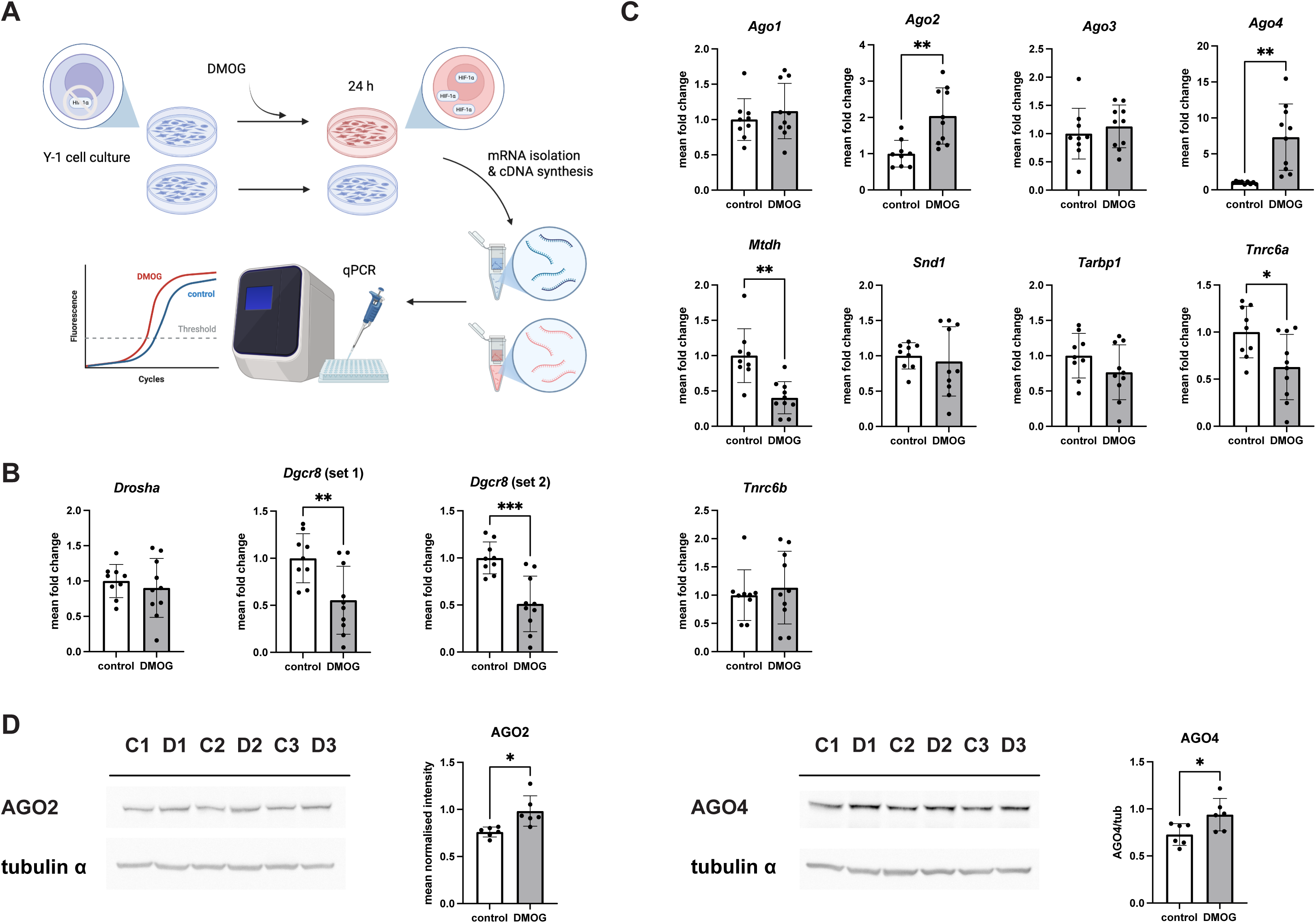
Effects of DMOG-induced HIF-1α stabilization on MC and RISC gene expression. A) Schematic depiction of the experimental setup. Y-1 cells cultures were treated with DMOG or vehicle for 24 h, collected and used for RNA and protein expression analysis. B) qPCR analysis of *Drosha* and *Dgcr8* mRNA expression in Y-1 cells treated with DMOG for 24 h under normoxia. ‘Set 1’ and ‘set 2’ refer to two independent primer pairs used for *Dgcr8* quantification. n = 9 (control) and n = 10 (DMOG). C) qPCR analysis of RISC component genes (*Ago1–4, Mtdh, Snd1, Tarbp1, Tnrc6a,* and *Tnrc6b*) in extracts from control and DMOG-treated Y-1 cells (24 h treatment). n = 9 (control) and n = 10 (DMOG). Bar heights represent mean fold change relative to control; error bars denote standard deviation of the mean. Statistical significance was determined using a Mann–Whitney U-test or unpaired *t*-test with Welch’s correction (* p < 0.05, ** p < 0.01, *** p < 0.001, **** p < 0.0001). D) Western blot analysis of AGO2 and AGO4 protein expression in Y-1 cells treated with DMOG for 24 h under normoxia. Representative image of AGO2, AGO4 and respective loading controls (tubulin α) blots are shown (C1-3 control Y-1 cell extracts, D1-3 DMOG-treated Y-1 cell extracts). Quantification is shown on the right of each gel set. Bar heights represent mean normalized AGO2 or AGO4 signal intensity; error bars denote standard deviation of the mean. N = 6 samples were analyzed per condition. Statistical significance was determined using an unpaired *t*-test with Welch’s correction (* p < 0.05).

**Table 2.**
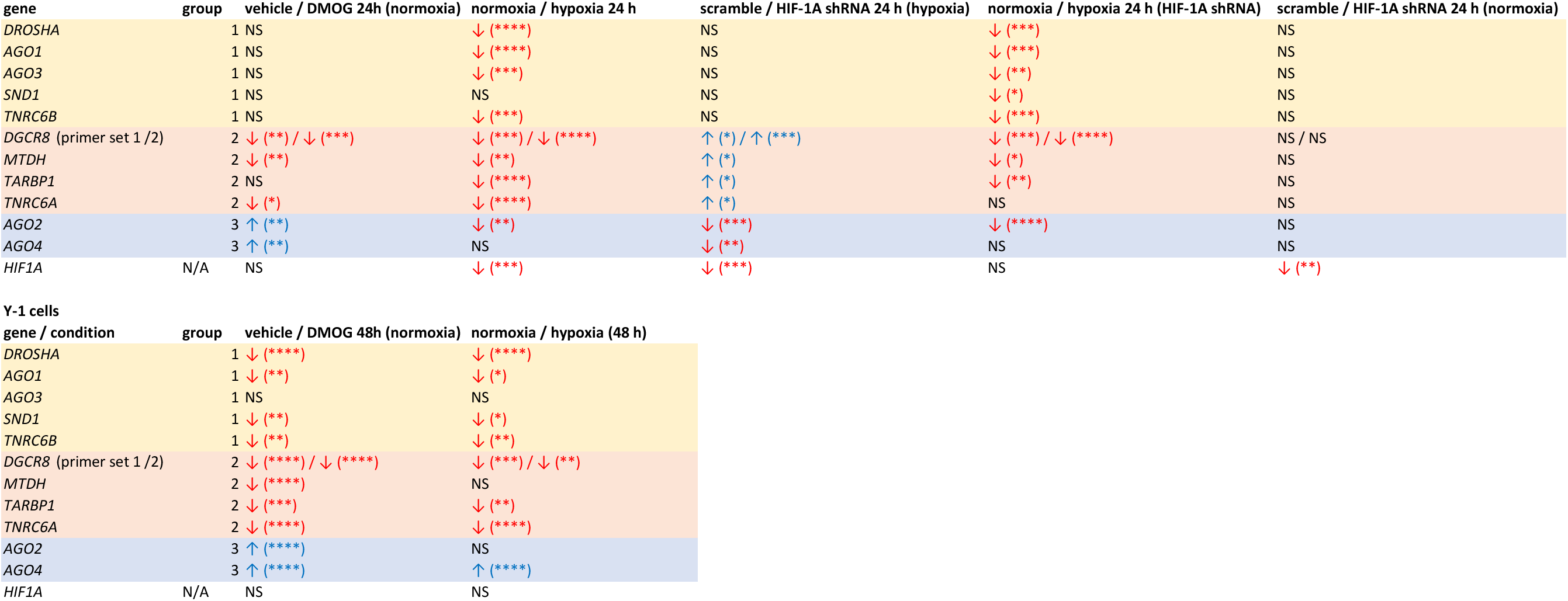
Summary of qPCR analysis of miRNA-processing gene expression in response to hypoxia and HIF-1α manipulation. Summary of qPCR analysis of miRNA-processing gene expression upon hypoxia or HIF-1α manipulation. Arrows indicate a decrease (red arrow down) or increase (blue arrow up) in gene expression upon treatment. Statistical significance was determined using a Mann–Whitney U-test or unpaired *t*-test with Welch’s correction (* p < 0.05, ** p < 0.01, *** p < 0.001, **** p < 0.0001, NS – not significant).

We next asked whether physiological stabilization of HIF-1α during hypoxia induces similar transcriptional effects. Y-1 cells were exposed to acute hypoxia (5% O□for 24 hours), a condition previously shown to stabilize HIF-1α but not HIF-2α protein in this cell line (Figure 5A) (9). In contrast to DMOG treatment, acute hypoxia resulted in a broad repression of genes involved in miRNA biogenesis and function. Both microprocessor components, *Drosha* and *Dgcr8*, were reduced by approximately 50% (Figure 5B), and most Argonaute orthologues were repressed, with the exception of *Ago4* (Figure 5C). Virtually all of the remaining RISC genes were also significantly downregulated, as was *Hif1a* mRNA itself (Additional file 7B). A summary of these results is provided in Table 2.

**Figure 5.**
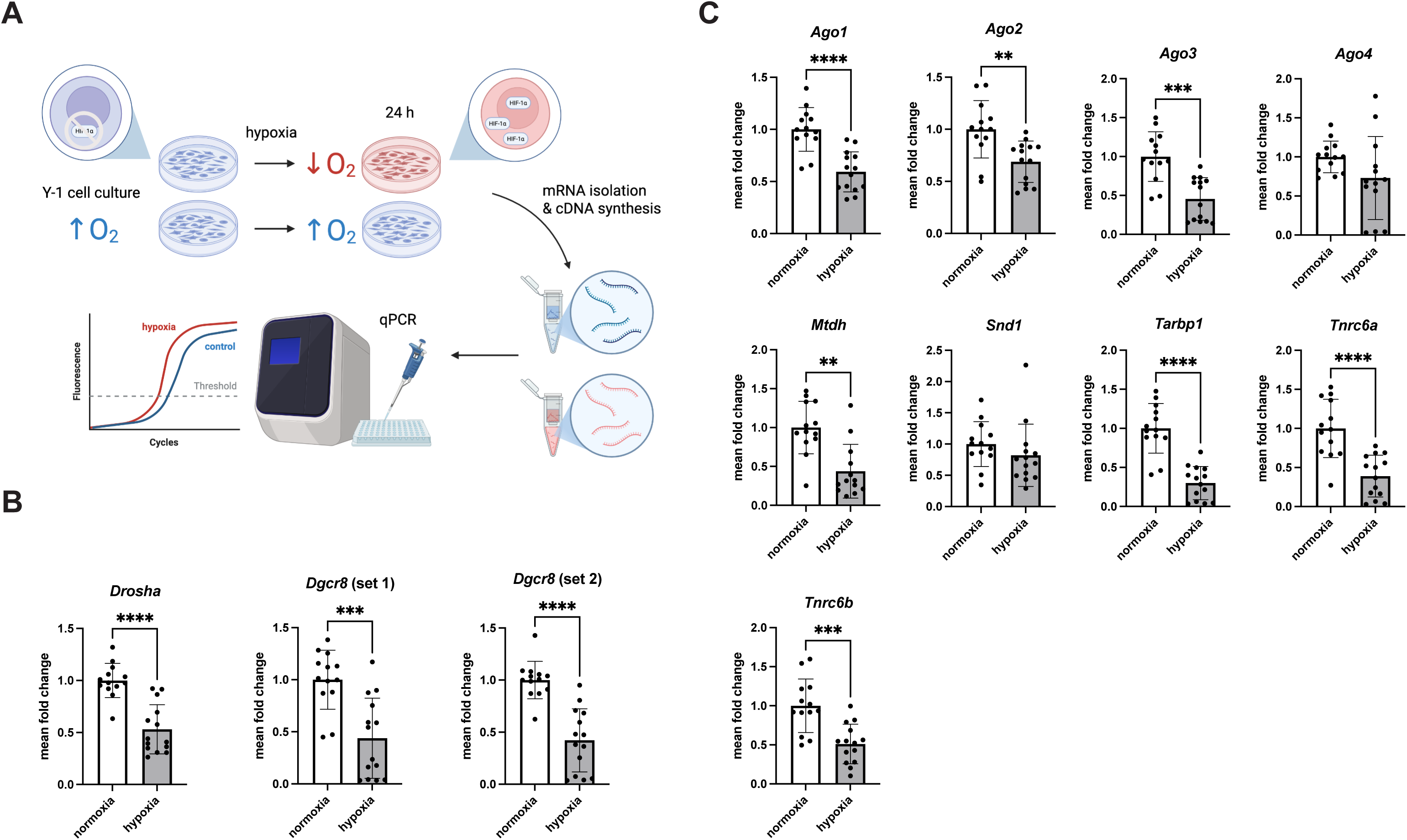
Effects of acute hypoxia on MC and RISC gene expression in Y-1 cells. A) Schematic depiction of the experimental setup. Y-1 cells were cultured in hypoxic (5% O_2_) or control (atmospheric oxygen concentration) conditions for 24 h, collected and used for RNA expression analysis. B) qPCR analysis of mRNA expression of the microprocessor complex components *Drosha* and *Dgcr8* in extracts from Y-1 cell grown under normoxia (21% O_2_) and hypoxia (5% O_2_) for 24 h. ‘Set 1’ and ‘set 2’ refer to two independent primer pairs used for *Dgcr8* qPCR. Normoxia: n = 13; hypoxia n = 14. C) qPCR analysis of mRNA expression of the RISC components *Ago1-4*, *Mtdh*, *Snd1*, *Tarbp1*, *Tnrc6a* and *Tnrc6b* in extracts from Y-1 cell grown under normoxia (21% O_2_) and hypoxia (5% O_2_) for 24 h. Normoxia: n = 13; hypoxia n = 14 (n = 13 for *Ago4* and *Mthd*). Bar heights represent mean fold change relative to control; error bars denote standard deviation of the mean. Statistical significance was determined using a Mann–Whitney U-test or unpaired *t*-test with Welch’s correction (* p < 0.05, ** p < 0.01, *** p < 0.001, **** p < 0.0001).

To determine whether these hypoxia-induced expression changes depend on HIF-1α, we performed shRNA-mediated *Hif1a* knockdown (KD) in Y-1 cells cultured under acute hypoxia (Figure 6A). As expected, *Hif1a* mRNA levels were significantly reduced compared to scrambled controls (Additional file 7C), consistent with previously confirmed protein depletion (9). Under hypoxia, HIF-1α depletion had no effect on *Drosha* expression but resulted in a significant increase in *Dgcr8* mRNA levels (Figure 6B). Similarly, knockdown of *Hif1a* led to upregulation of the RISC genes *Mtdh*, *Tarbp1*, and *Tnrc6a* (Figure 6C), indicating that their repression by hypoxia is at least partially HIF-1α dependent. In contrast, *Ago1*, *Ago3*, *Tnrc6b*, and *Snd1* were unaffected by *Hif1a* knockdown. Notably, *Ago2* and *Ago4* expression decreased upon HIF-1α depletion, suggesting that HIF-1α counteracts, rather than mediates, their hypoxia-induced repression. Under normoxia, *Hif1a* knockdown had no effect on the expression of any analyzed genes as expected due to HIF-1α protein destabilization already in the control condition (Additional file 9).

**Figure 6.**
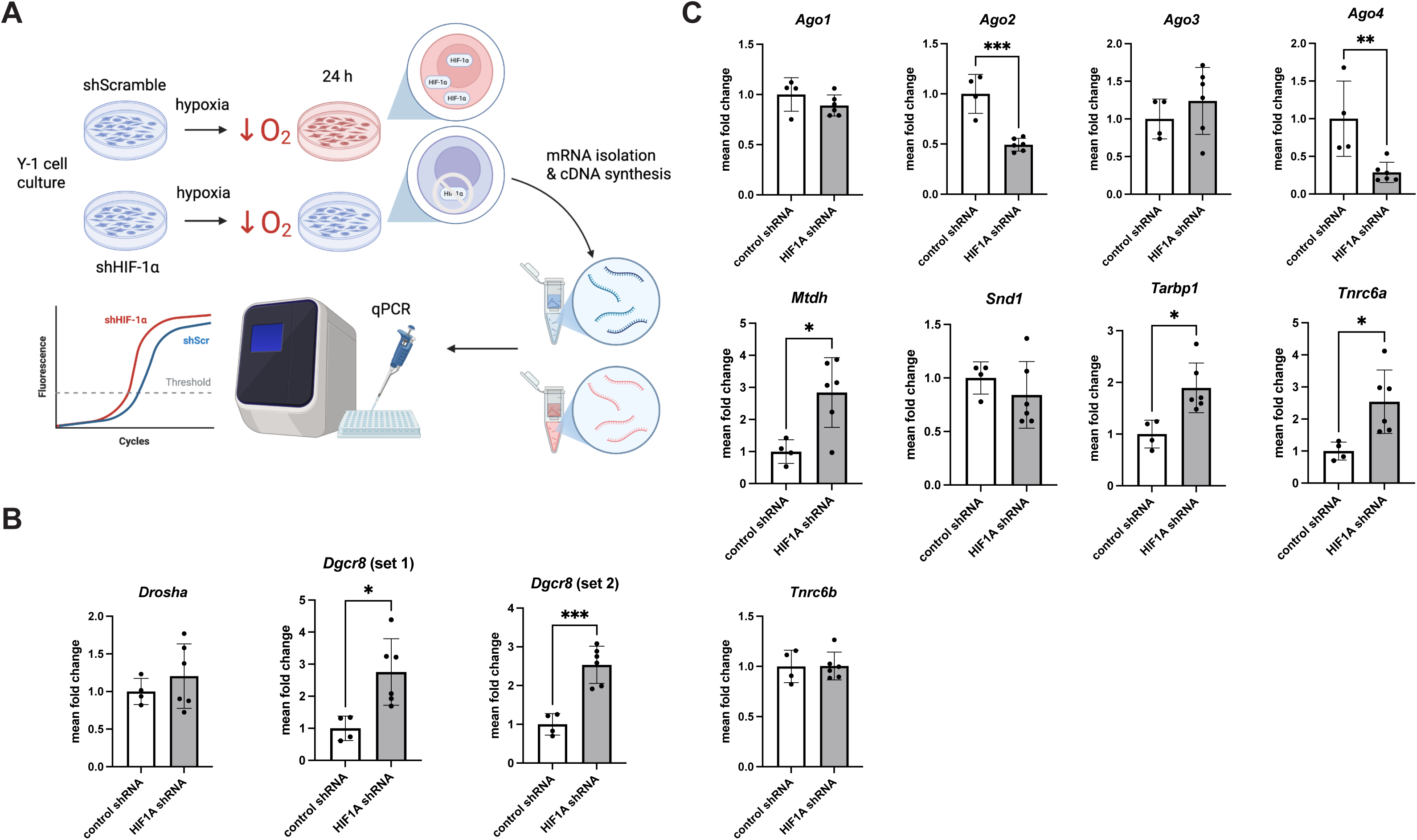
Effects of HIF-1α knockdown on MC and RISC gene expression under hypoxia. A) Schematic depiction of the experimental setup. Y-1 cells were transfected with shRNA against *HifI1a* or control shRNA (scramble) and then cultured in hypoxic (5% O_2_) conditions for 24 h, collected and used for RNA expression analysis. B) qPCR analysis of mRNA expression of the microprocessor complex components *Drosha* and *Dgcr8* in extracts from Y-1 cells treated with scramble (control) or *Hif1a* shRNA (*HifI1a* KD) and grown under hypoxia (5% O_2_) for 24 h. ‘Set 1’ and ‘set 2’ refer to two independent primer pairs used for *Dgcr8* qPCR. Control: n = 4; *HifI1a* KD n = 6. C) qPCR analysis of mRNA expression of the RISC components *Ago1-4*, *Mtdh*, *Snd1*, *Tarbp1*, *Tnrc6a* and *Tnrc6b* in extracts from Y-1 cells treated with scramble (control) or *Hif1a* shRNA (*HifI1a* KD) and grown under hypoxia (5% O_2_) for 24 h. Control: n = 4; *HifI1a* KD n = 6. Bar heights represent mean fold change relative to control; error bars denote standard deviation of the mean. Statistical significance was determined using a Mann–Whitney U-test or unpaired *t*-test with Welch’s correction (* p < 0.05, ** p < 0.01, *** p < 0.001, **** p < 0.0001).

To further delineate the contribution of HIF-1α to hypoxia-driven repression, we compared MC and RISC gene expression in *Hif1a*-depleted Y-1 cells cultured under normoxia or hypoxia (Figure 7A). With the exception of Ago4 and Tnrc6a, all analyzed genes remained significantly repressed by hypoxia despite HIF-1α depletion (Figure 7B-C and Table 2). These findings indicate that HIF-1α contributes to, but is not strictly required for, hypoxia-induced repression of a subset of miRNA biogenesis and function genes, while hypoxia itself exerts a broader suppressive effect on the miRNA biogenesis.

**Figure 7.**
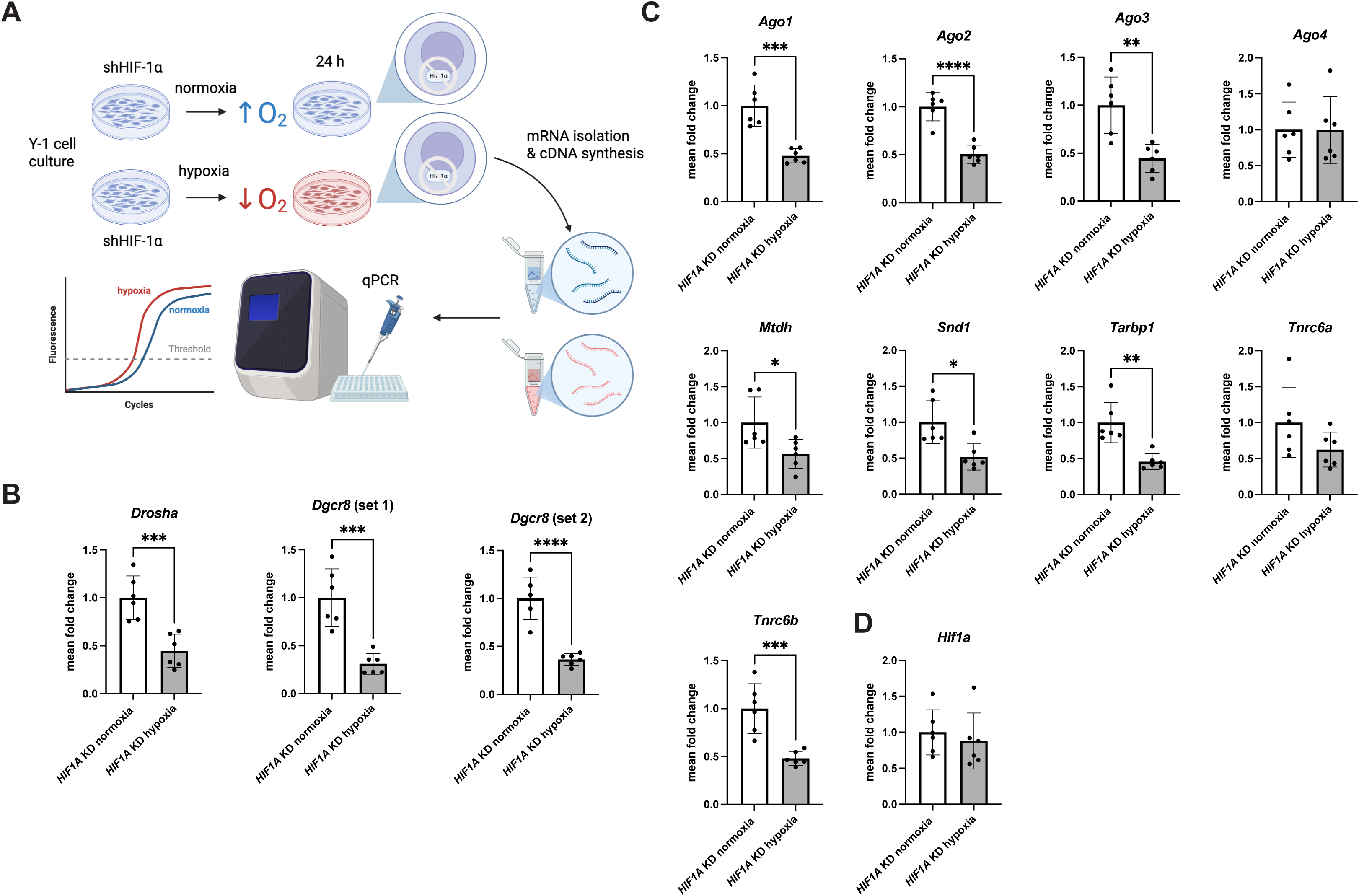
Comparison of HIF-1α knockdown in Y-1 cells under normoxia and hypoxia. A) Schematic depiction of the experimental setup. Y-1 cells were transfected with shRNA against *HifI1a* and then cultured in either hypoxic (5% O_2_) or control (atmospheric oxygen concentration) conditions for 24 h, collected and used for RNA expression analysis. B) qPCR analysis of mRNA expression of the microprocessor complex components *Drosha* and *Dgcr8* in extracts from Y-1 cells treated with *Hif1a* shRNA (*Hif1a* KD) and grown under either normoxia (21% O_2_) or hypoxia (5% O_2_) for 24 h. ‘Set 1’ and ‘set 2’ refer to two independent primer pairs used for *Dgcr8* qPCR. Normoxia: n = 6; hypoxia n = 6. C) qPCR analysis of mRNA expression of the RISC components *Ago1-4*, *Mtdh*, *Snd1*, *Tarbp1*, *Tnrc6a* and *Tnrc6b* in extracts from Y-1 cells treated with *Hif1a* shRNA (*Hif1a* KD) under either normoxia (21% O_2_) or hypoxia (5% O_2_) for 24 h. Normoxia: n = 6; hypoxia n = 6. D) qPCR analysis of the *Hif1a* mRNA expression in extracts from Y-1 cells treated with *Hif1a* shRNA (*Hif1a* KD) and grown under either normoxia (21% O_2_) or hypoxia (5% O_2_) for 24 h. Normoxia: n = 6; hypoxia n = 6. Bar heights represent mean fold change relative to control; error bars denote standard deviation of the mean. Statistical significance was determined using a Mann–Whitney U-test or unpaired *t*-test with Welch’s correction (* p < 0.05, ** p < 0.01, *** p < 0.001, **** p < 0.0001).

In summary, combined pharmacological and genetic manipulation of HIF-1α reveals a modulatory role for HIF-1α in shaping the transcriptional response of miRNA-processing genes to hypoxia. Whereas a subset of genes is repressed independently of HIF-1α (Table 2; group 1), others display HIF-1α-dependent repression (Table 2; group 2) or counter-regulation (Table 2; group 3). Notably, hypoxia-induced repression of *Dgcr8* and the RISC components *Mtdh*, *Tarbp1*, and *Tnrc6a* was partially mimicked by DMOG and alleviated by *Hif1a* knockdown, while repression of *Ago2* and *Ago4* was antagonized by HIF-1α stabilization and enhanced by its depletion. Together, these data define distinct classes of miRNA-processing genes whose expression is differentially shaped by hypoxia and HIF-1α activity (Table 2 and Figure 8).

**Figure 8.**
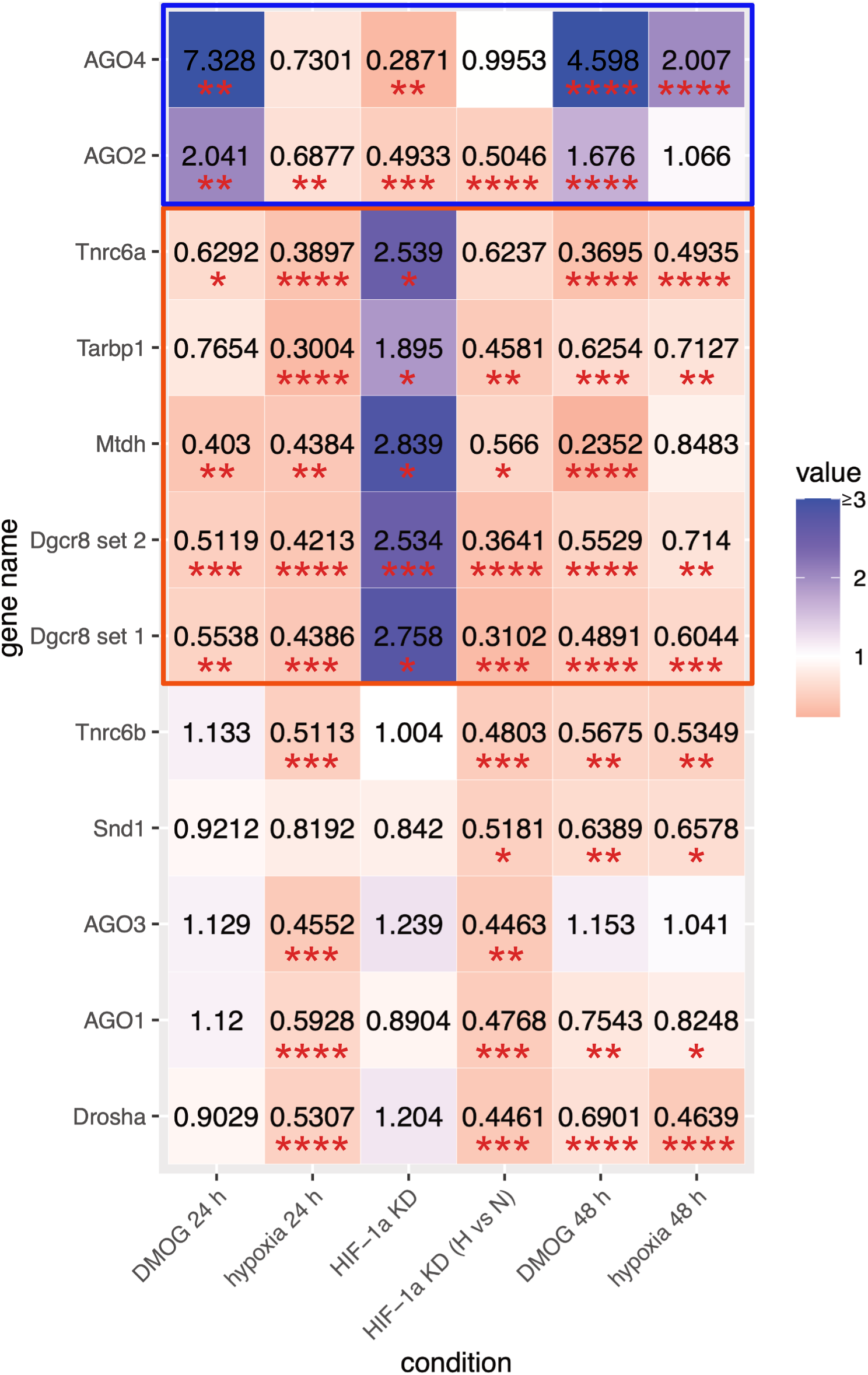
Effects of hypoxia and HIF-1α manipulation on the expression of miRNA biogenesis and function genes. Summary of the MC and RISC gene expression analysis. Fold-change in expression is indicated by the color code with no change (normalized to 1) depicted in white. Maximum value of 3 was set for the color scale to avoid the disproportionate effect of the most upregulated gene (*Ago4*). Three classes of genes can be distinguished – upregulated in the presence of HIF-1α stabilization (group 3; blue outline), downregulated in the presence of HIF-1α stabilization (group 2; red outline) and unaffected by HIF-1α (group 1; no outline). Red stars indicate significantly changed expression determined using a Mann–Whitney U-test or unpaired *t*-test with Welch’s correction (* p < 0.05, ** p < 0.01, *** p < 0.001, **** p < 0.0001). Hypoxia exerts a generalized suppressive effect on the expression of most of the analyzed genes.

### Prolonged hypoxia induces partial adaptation of gene expression of RISC components

To determine whether the hypoxia-induced changes in miRNA biogenesis gene expression persist or adapt over time, we extended hypoxic exposure to 48 hours (Figure 9A). qPCR analysis of Y-1 cells cultured under normoxic (21% O□) or hypoxic (5% O□) conditions revealed that prolonged hypoxia sustained repression of the microprocessor components *Drosha* and *Dgcr8* (Figure 9B). In contrast, several RISC genes exhibited differential responses over time. While *Ago1, Tarbp1, Tnrc6a,* and *Tnrc6b* remained repressed after 48 hours of hypoxia, *Ago2, Ago3,* and *Mtdh*, which were suppressed after 24 hours, returned to baseline expression levels (Figure 9C). Notably, *Ago4* expression, which was unaffected by acute hypoxia, was significantly increased following prolonged hypoxic exposure. Prolonged hypoxia had little effect on *Hif1a* mRNA levels (Additional file 7D). Together, these data indicate that sustained low oxygen conditions trigger compensatory transcriptional adaptation in a subset of RISC genes, whereas repression of core microprocessor genes persists (Table 2 and Figure 8).

**Figure 9.**
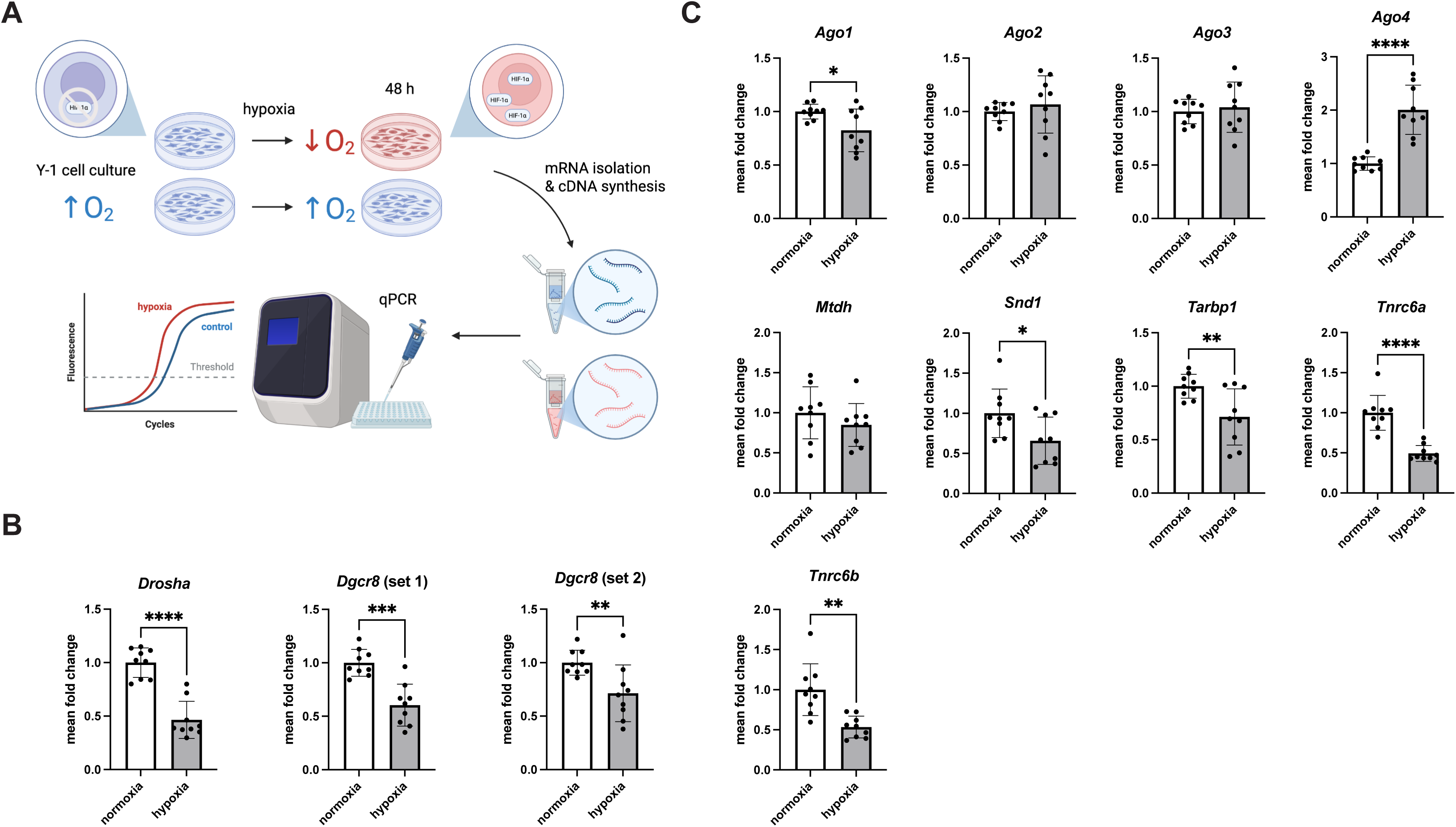
Effects of prolonged hypoxia on MC and RISC gene expression in Y-1 cells. A) Schematic depiction of the experimental setup. Y-1 cells were cultured in hypoxic (5% O_2_) or control (atmospheric oxygen concentration) conditions for 48 h, collected and used for RNA expression analysis. B) qPCR analysis of mRNA expression of the microprocessor complex components *Drosha* and *Dgcr8* in extracts from Y-1 cell grown under normoxia (21% O_2_) and hypoxia (5% O_2_) for 48 h. ‘Set 1’ and ‘set 2’ refer to two independent primer pairs used for *Dgcr8* qPCR. Normoxia: n = 9; hypoxia n = 9. C) qPCR analysis of mRNA expression of the RISC components *Ago1-4*, *Mtdh*, *Snd1*, *Tarbp1*, *Tnrc6a* and *Tnrc6b* in extracts from Y-1 cell grown under normoxia (21% O_2_) and hypoxia (5% O_2_) for 48 h. Normoxia: n = 9; hypoxia n = 9. Bar heights represent mean fold change relative to control; error bars denote standard deviation of the mean. Statistical significance was determined using a Mann–Whitney U-test or unpaired *t*-test with Welch’s correction (* p < 0.05, ** p < 0.01, *** p < 0.001, **** p < 0.0001).

To assess whether prolonged pharmacological stabilization of HIF-1α elicits a similar transcriptional response, we treated Y-1 cells with DMOG for 48 hours under normoxic conditions. Overall, extended DMOG treatment largely recapitulated the effects observed under prolonged hypoxia, maintaining repression of *Drosha, Dgcr8, Ago1, Tarbp1, Tnrc6a,* and *Tnrc6b*. However, distinct differences were observed: *Mtdh* remained repressed, and *Ago2* expression was induced following 48-hour DMOG treatment, consistent with the effects observed after 24 hours of DMOG exposure. This contrasts with their normalization under prolonged hypoxia (Additional file 10).

Taken together, these findings indicate that acute hypoxia induces widespread repression of miRNA biogenesis and RISC genes through a combination of HIF-1α-dependent and - independent mechanisms, whereas prolonged hypoxia is accompanied by partial transcriptional adaptation that selectively restores expression of specific RISC components but not microprocessor genes. The overlapping response to extended pharmacological HIF-1α stabilization underscores a sustained yet gene-specific role for HIF signaling in regulating the miRNA-processing machinery over time.

## Discussion

In this study, we examined how HIF-1α regulates adrenal steroidogenesis and miRNA biogenesis under altered hypoxic signaling. Using complementary pharmacological, physiological, and genetic approaches in murine adrenocortical cells, we combined genome-wide CUT&Tag profiling with transcriptional analyses to distinguish direct HIF-1α–dependent effects from broader, HIF-independent hypoxia-driven responses. These data reveal that HIF-1α directly targets steroidogenic genes, steroidogenesis-associated miRNAs, and components of the miRNA-processing machinery, while hypoxia additionally imposes a more general repression of miRNA biogenesis that is only partially modulated by HIF-1α. Together, our findings link oxygen-sensitive transcriptional control to post-transcriptional regulation in the adrenal cortex.

One established mechanism by which HIF-1α represses gene expression is through the induction of miRNAs that inhibit translation or promote degradation of specific target mRNAs. Numerous hypoxamiRs, miRNAs induced by hypoxia and/or HIF-1α stabilization, have been identified across diverse cell types, and this list continues to expand (3,4,34,36). In our CUT&Tag dataset, we detected HIF-1α binding at 12 hypoxamiRs loci, eight of which have also been implicated in the regulation of steroid hormone biosynthesis in the adrenal gland or gonads (9,37–43). Among these, *miR-29b* emerged as a strong candidate for direct HIF-1α regulation, with multiple HIF-1α binding sites detected in its genomic vicinity. Consistent with this, we observed a significant induction of *miR-29b* expression in Y-1 cells following DMOG treatment.

Interestingly, several miRNA genes previously shown to be regulated by HIF-1α and to target steroidogenic enzymes (9) were not detected among direct HIF-1α binding sites in our dataset. This observation suggests that their regulation may occur indirectly, either through downstream transcription factors or through modulation of the miRNA-processing machinery itself. Indeed, hypoxia and HIF signaling have been reported to affect the expression of key miRNA-processing components such as *Drosha* and *Dicer* in various cancer and immune cell types (44–48). However, whether such mechanisms operate in the context of adrenal steroidogenesis has remained unclear. Our data demonstrate that both hypoxia and HIF-1α regulate the expression of multiple components of the nuclear microprocessor complex and the cytoplasmic RISC in adrenocortical cells.

Based on their transcriptional responses to hypoxia and HIF-1α manipulation, we identified three distinct gene groups: (1) genes repressed independently of HIF-1α: *Drosha*, *Ago1*, *Ago3*, *Snd1* and *Tnrc6b*, (2) genes repressed by both HIF-1α-dependent and -independent mechanisms: *Dgcr8*, *Mtdh*, *Tarbp1* and *Tnrc6a*, and (3) genes whose repression by hypoxia is counteracted by HIF-1α: *Ago2* and *Ago4* (overview in Table 2 and Figure 8). The widespread hypoxia-induced repression of miRNA biogenesis genes, together with the modulatory role of HIF-1α, suggests that HIF signaling may fine-tune the miRNA system to permit the activity of specific miRNAs while globally constraining miRNA production. Notably, maintenance of *Ago2* and *Ago4* expression by HIF-1α despite repression of microprocessor components may help preserve a functional pool of existing miRNAs under conditions of reduced miRNA biogenesis.

Although our study focused on HIF-1α–mediated regulation of miRNA processing in adrenocortical cells in the context of steroidogenesis, the implications of these findings likely extend beyond the adrenal gland. miRNAs are critical regulators of physiological brain development and responses to perinatal ischemia (49) and hypoxia-induced miRNAs play central roles in the pathogenesis of glioblastoma multiforme (50). Given the importance of HIF-1α signaling in both brain development (6) and tumor progression (50,51), it is plausible that HIF-dependent control of miRNA metabolism represents a broader adaptive mechanism across tissues. The ability to generate genome-wide HIF-1α binding profiles using CUT&Tag provides a powerful framework for exploring such regulatory principles in diverse physiological and pathological contexts.

## Conclusions

Our study identifies HIF-1α as a key integrator of oxygen-sensitive transcriptional and post-transcriptional regulation in adrenal steroidogenesis. By linking direct gene regulation to modulation of miRNA biogenesis and function, these findings reveal an additional layer through which hypoxic signaling reshapes endocrine cell communication and hormone output.

## List of abbreviations

CUT&Tag: Cleavage Under Targets & Tagmentation
DMOG: dimethyloxalylglycine
HIF: hypoxia-inducible factor
KD: knockdown
MC: microprocessor complex
miRNA: microRNA
PHD: prolyl hydroxylase domain
RISC: RNA-induced silencing complex
VHL: von Hippel–Lindau

## Additional files

Tables: Additional files 1-6

Figures: Additional files 7-11

## Statements & Declarations

## Ethics approval and consent to participate

Not applicable

## Consent for publication

Not applicable

## Availability of data and materials

Source data for this study are openly available at GEO under the accession number GSE311437. Materials are available upon reasonable request.

## Authors’ contributions

BKS designed and performed the majority of experiments, analyzed data, and wrote the manuscript. SA performed experiments and contributed to the discussion. AK performed experiments. AS performed bioinformatic analysis of the CUT&Tag data and contributed to the discussion. PM and RB provided tools and contributed to the discussion. BW supervised the overall study and co-wrote the manuscript. All authors read and approved the final manuscript

## Acknowledgements

The authors would like to thank Prof. Mareike Albert and her group (particularly Janine Epperlein and Nora Ditzer) for fruitful discussions and help in establishing the CUT&Tag protocol for HIF-1α.

## Additional files

**Additional file 1.xlsx List of HIF-1**α **CUT&Tag target genes in DMOG-treated Y-1 cells compared to controls.**

List of all regions significantly (p < 0.05; log2-fold change > 2 or <-2) enriched (positive value of log2FoldChange) or depleted (negative value of log2FoldChange) in the HIF-1α CUT&Tag dataset from DMOG-treated compared to control Y-1 cells. The HIF-1α CUT&Tag-enriched target regions map to 6989 unique genes and include 122 miRNA-associated genes.

**Additional file 2.xlsx Genomic distribution of HIF-1α–bound sequences enriched or depleted in DMOG-treated versus control Y-1 cells.**

Genomic distribution of HIF-1α-bound sequences enriched (DMOG UP) or depleted (DMOG DOWN) in the dataset obtained from DMOG-treated compared to control cells. Note that in the DMOG UP condition more binding can be detected in promoter regions (28.8% compared to 10.7%) while in DMOG down bound sequences are predominantly located to distal intergenic regions (49.6% *vs* 33.7%).

**Additional file 3.xlsx KEGG pathway analysis of HIF-1α target genes identified by CUT&Tag.**

Over-Representation analysis (KEGG pathways) of the HIF-1α target gene dataset. The analysis was performed using the WebGestalt platform (26,27).

**Additional file 4.xlsx Transcription factor motif enrichment analysis of HIF-1α–bound sequences.**

Transcription factor motif analysis. As expected, the analysis identified enrichment of the HIF-1α targets sequences in canonical HIF-1/2α motifs. Moreover, HIF-1α-bound sequences were enriched in motifs for several transcription factors, primarily members of the bZIP and NR superfamilies. The analysis was performed using HOMER.

**Additional file 5.xlsx**

miRNA-regulated genes of the putative HIF-1α target miRNAs. The analysis was performed using miRNet.

**Additional file 6.xlsx**

qPCR primer sequences and references.

**Additional file 7.pdf**

**qPCR analysis of *Hif1a* mRNA expression under different experimental conditions.**

A) qPCR analysis of the *Hif1a* mRNA expression in extracts from control Y-1 cells and cells treated with DMOG inhibitor for 24 h. Control: n = 3; DMOG n = 4. B) qPCR analysis of the *Hif1a* mRNA expression in extracts from Y-1 cell grown under normoxia (21% O_2_) or hypoxia (5% O_2_) for 24 h. Normoxia: n = 11; hypoxia n = 14. C) qPCR analysis of the *Hif1a* mRNA expression in extracts from Y-1 cells treated with scramble (control) or *Hif1a* shRNA (*Hif1a* KD) and grown under hypoxia (5% O_2_) for 24 h. Control: n = 4; *Hif1a* KD n = 6. D) qPCR analysis of the *Hif1a* mRNA expression in extracts from Y-1 cell grown under normoxia (21% O_2_) or hypoxia (5% O_2_) for 48 h. Normoxia: n = 9; hypoxia n = 9. Bar heights represent mean fold change relative to control; error bars denote standard deviation of the mean. Statistical significance was determined using a Mann–Whitney U-test or unpaired *t*-test with Welch’s correction (* p < 0.05, ** p < 0.01, *** p < 0.001, **** p < 0.0001).

**Additional file 8.pdf**

**Western blot analysis of AGO2 and AGO4 protein expression in Y-1 cells treated with DMOG for 24 h.**

Uncropped Western blots for AGO2, AGO4 and tubulin α (visualized merged with the marker image) and Ponceau staining of the respective membranes. Lane description: M, marker; C1-C6, control Y-1 cell extracts; D1-D6, DMOG-treated Y-1 cell extracts; E, empty lane; N/A, unrelated data (not analyzed).

**Additional file 9.pdf**

**qPCR analysis of miRNA-processing gene expression upon HIF-1**α **depletion under normoxia.**

A) qPCR analysis of mRNA expression of the microprocessor complex components *Drosha* and *Dgcr8* in extracts from Y-1 cells treated with scramble (control) or *Hif1a* shRNA (*Hif1a* KD) and grown under normoxia (21% O_2_) for 24 h. Control: n = 4; *Hif1a* KD n = 6. B) qPCR analysis of mRNA expression of the RISC components *Ago1-4*, *Mtdh*, *Snd1*, *Tarbp1*, *Tnrc6a* and *Tnrc6b* in extracts from Y-1 cells treated with scramble (control) or *Hif1a* shRNA (*Hif1a* KD) and grown under normoxia (21% O_2_) for 24 h. Control: n = 4; *HIF1A* KD n = 6. C) qPCR analysis of the *Hif1a* mRNA expression in extracts from Y-1 cells treated with scramble (control) or *Hif1a* shRNA (*Hif1a* KD) and grown under normoxia (21% O_2_) for 24 h. Control: n = 4; *Hif1a* KD n = 6. Bar heights represent mean fold change relative to control; error bars denote standard deviation of the mean. Statistical significance was determined using a Mann–Whitney U-test or unpaired *t*-test with Welch’s correction (* p < 0.05, ** p < 0.01, *** p < 0.001, **** p < 0.0001).

**Additional file 10.pdf**

**Effects of prolonged DMOG-induced HIF-1**α **stabilization on miRNA-processing gene expression.**

A) qPCR analysis of mRNA expression of the microprocessor complex components *Drosha* and *Dgcr8* in extracts from control Y-1 cells and cells treated with DMOG inhibitor for 48 h. ‘Set 1’ and ‘set 2’ refer to two independent primer pairs used for *Dgcr8* qPCR. Control: n = 9; DMOG n = 9. B) qPCR analysis of mRNA expression of the RISC components *Ago1-4*, *Mtdh*, *Snd1*, *Tarbp1*, *Tnrc6a* and *Tnrc6b* in extracts from control Y-1 cells and cells treated with DMOG inhibitor for 48 h. Control: n = 9; DMOG n = 9. C) qPCR analysis of the *Hif1a* mRNA expression in extracts from control Y-1 cells and cells treated with DMOG inhibitor for 48 h. Control: n = 9; DMOG n = 9. Bar heights represent mean fold change relative to control; error bars denote standard deviation of the mean. Statistical significance was determined using a Mann–Whitney U-test or unpaired *t*-test with Welch’s correction (* p < 0.05, ** p < 0.01, *** p < 0.001, **** p < 0.0001).

**Additional file 11.pdf**

**Melting curves for the products of qPCR primers used in this study.**

Melting curves and derivative plots for the qPCR primers used in this study. Note that for Mtdh product two distinct peaks can be seen. This is in agreement with the uMelt Quartz (31) analysis of the product sequence which also predicts two distinct peaks for this product. Additionally, agarose gel electrophoresis shows the presence of a single band at ∼200 bp, in agreement with the predicted product size (199 bp). The DNA size marker in the flanking wells is GeneRuler DNA Ladder Mix (Thermo Fisher Scientific, SM0331).

