## Supplementary Figures + Table for "HIF-1α coordinates adrenal steroidogenesis through direct transcriptional control and regulation of miRNA biogenesis"

Add. Figure 1

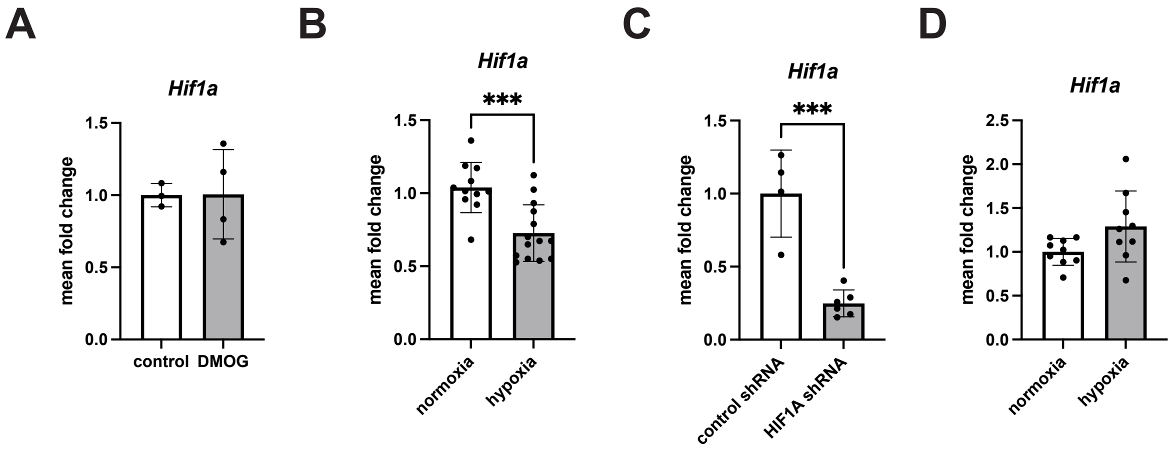

A
Hif1a
mean fold change
1.5
1.0
0.5
0.0
control
DMOG
B
Hif1a
***
mean fold change
1.5
1.0
0.5
0.0
normoxia
hypoxia
C
Hif1a
***
mean fold change
1.5
1.0
0.5
0.0
control shRNA
HIF1A shRNA
D
Hif1a
mean fold change
2.5
2.0
1.5
1.0
0.5
0.0
normoxia
hypoxia

Add. Figure 2

**uncropped blot AGO2 - gel 1**

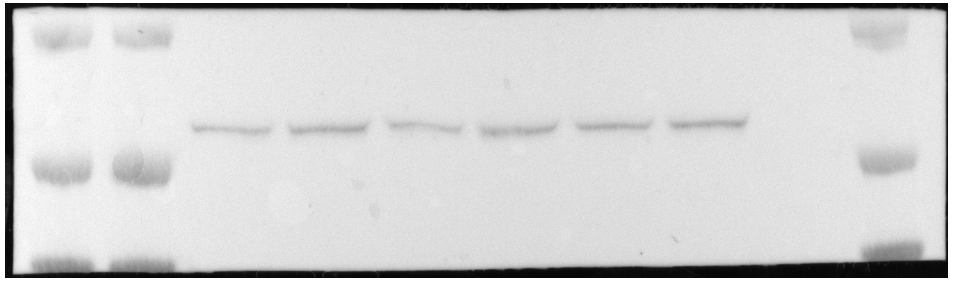

**uncropped blot AGO2 - gel 2**

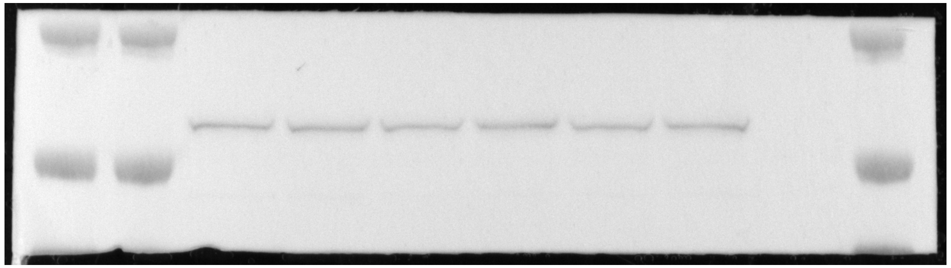

**uncropped blot tubulin α - gel 1**

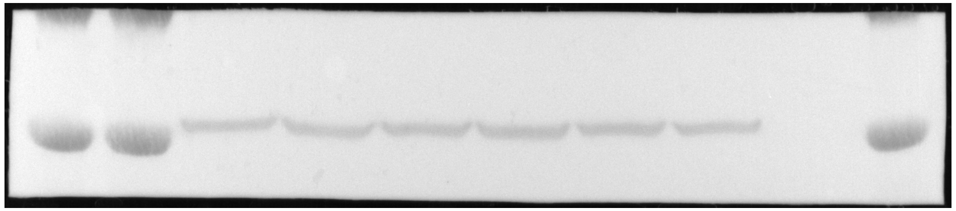

**uncropped blot tubulin α - gel 2**

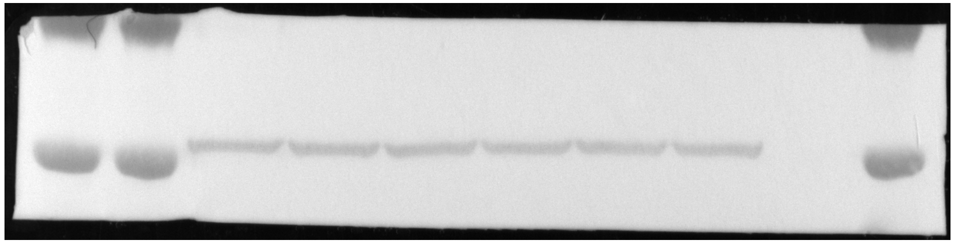

**Ponceau staining - gel 1**

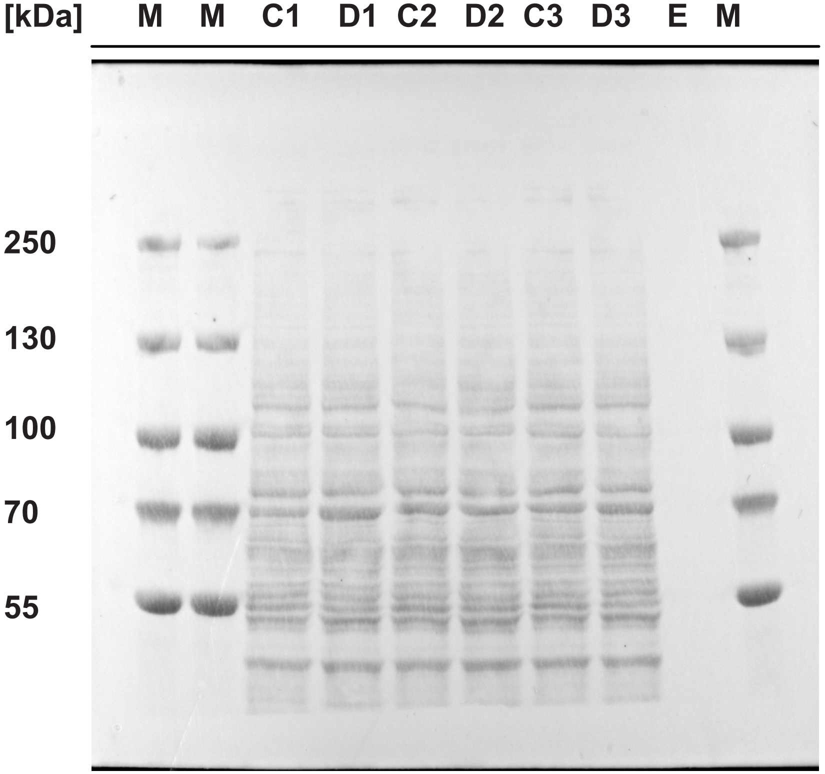


**Ponceau staining - gel 2**

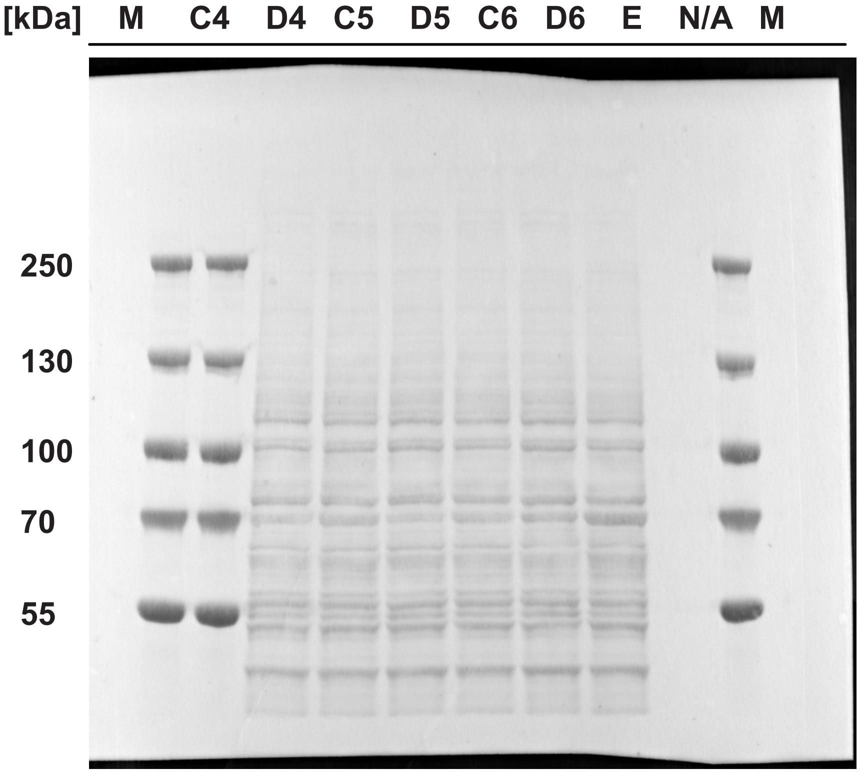

uncropped blot AGO4 - gel 1

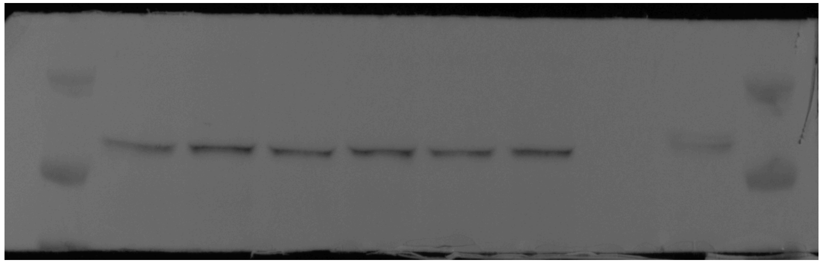

uncropped blot tubulin α - gel 1

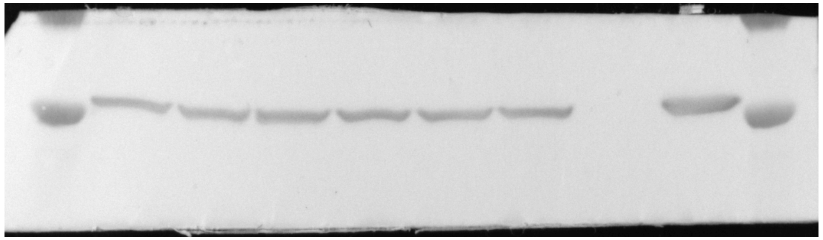

Ponceau staining - gel 1

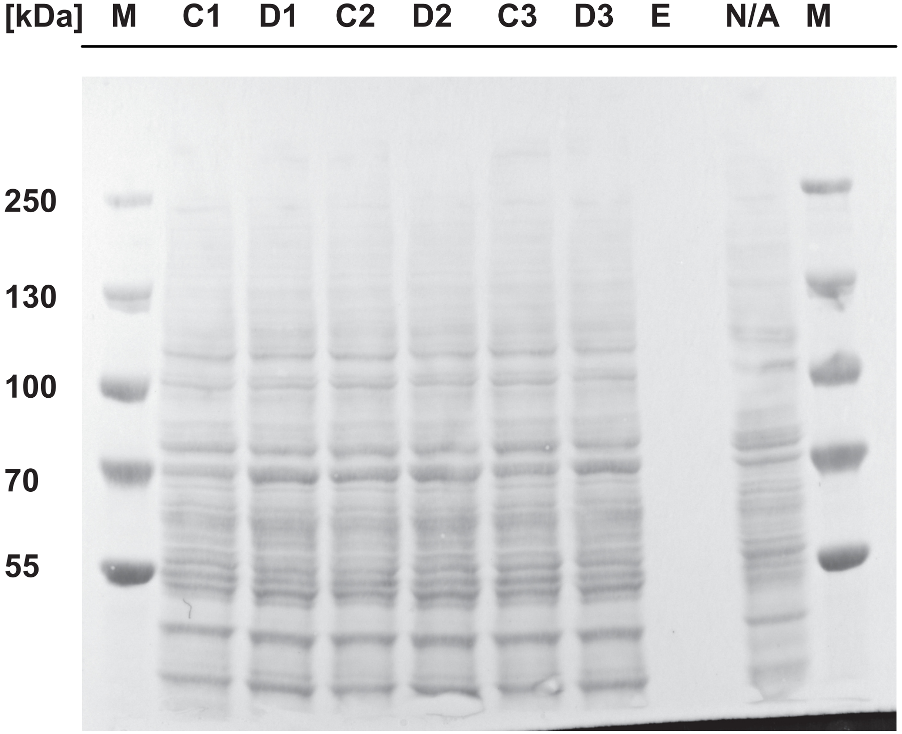


uncropped blot AGO4 - gel 2

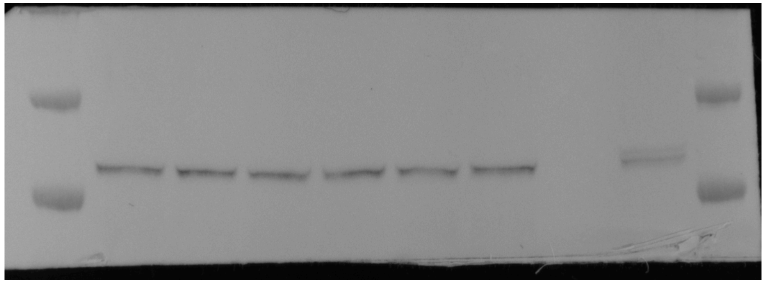

uncropped blot tubulin α - gel 2

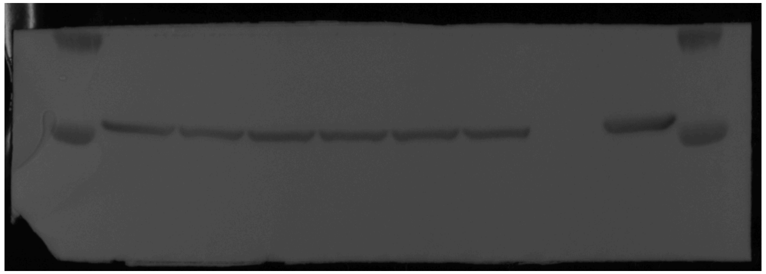

Ponceau staining - gel 2

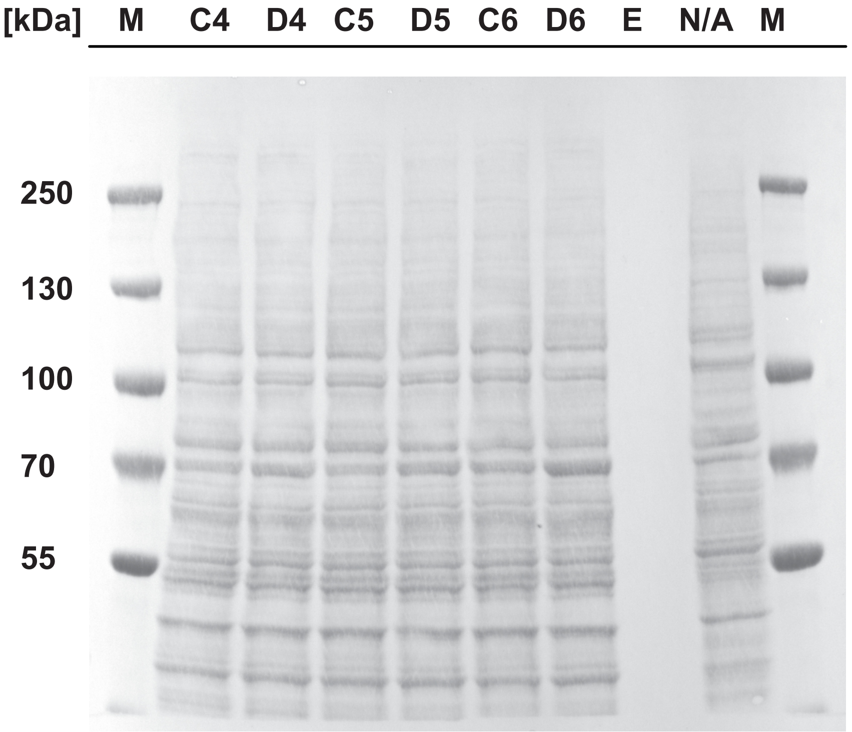

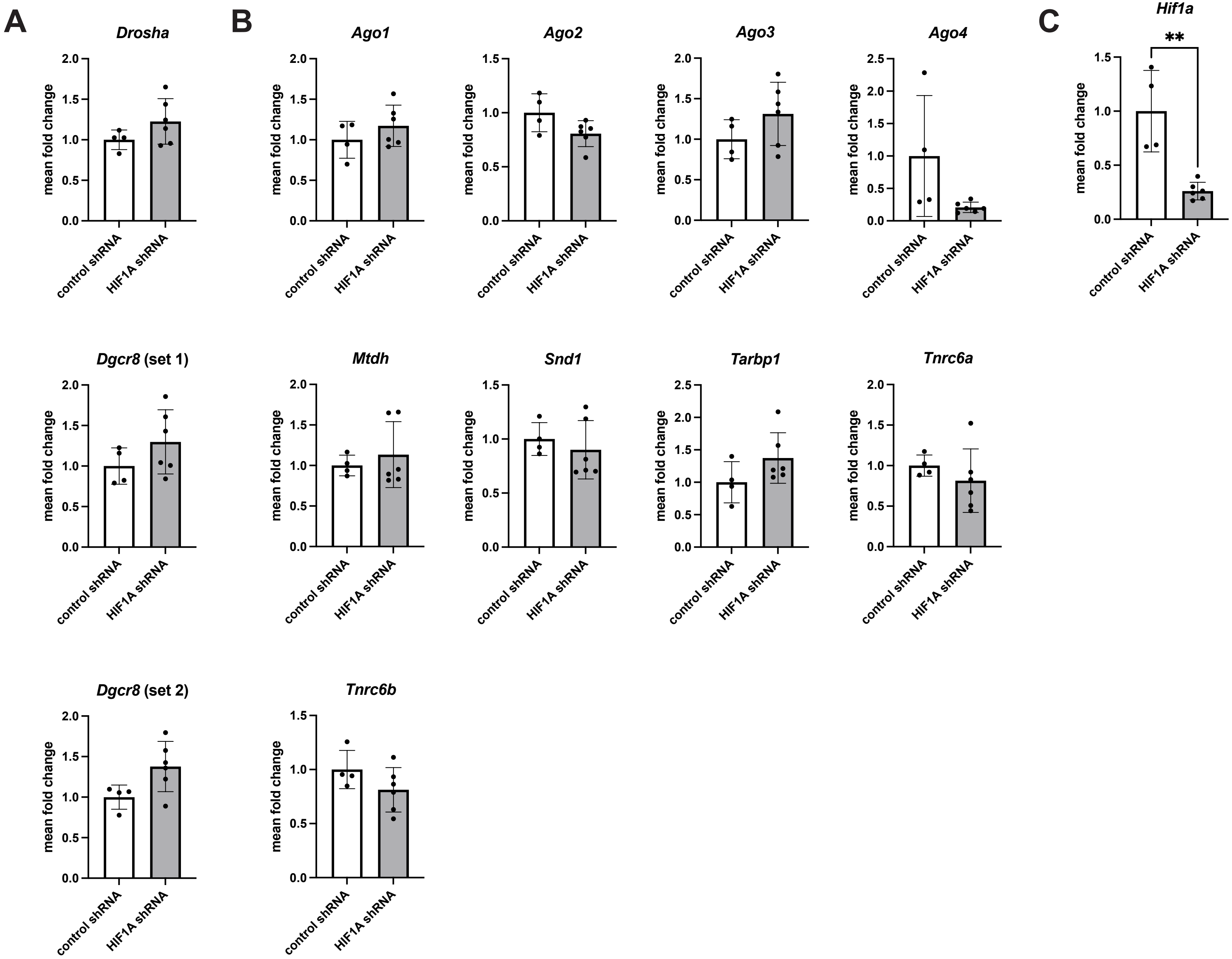

Add. Figure 3
A
B
C
Drosha
Ago1
Ago2
Ago3
Ago4
Hif1a
**
Dgcr8 (set 1)
Mtdh
Snd1
Tarbp1
Tnrc6a
Dgcr8 (set 2)
Tnrc6b
mean fold change
control shRNA
HIF1A shRNA

Add. Figure 4

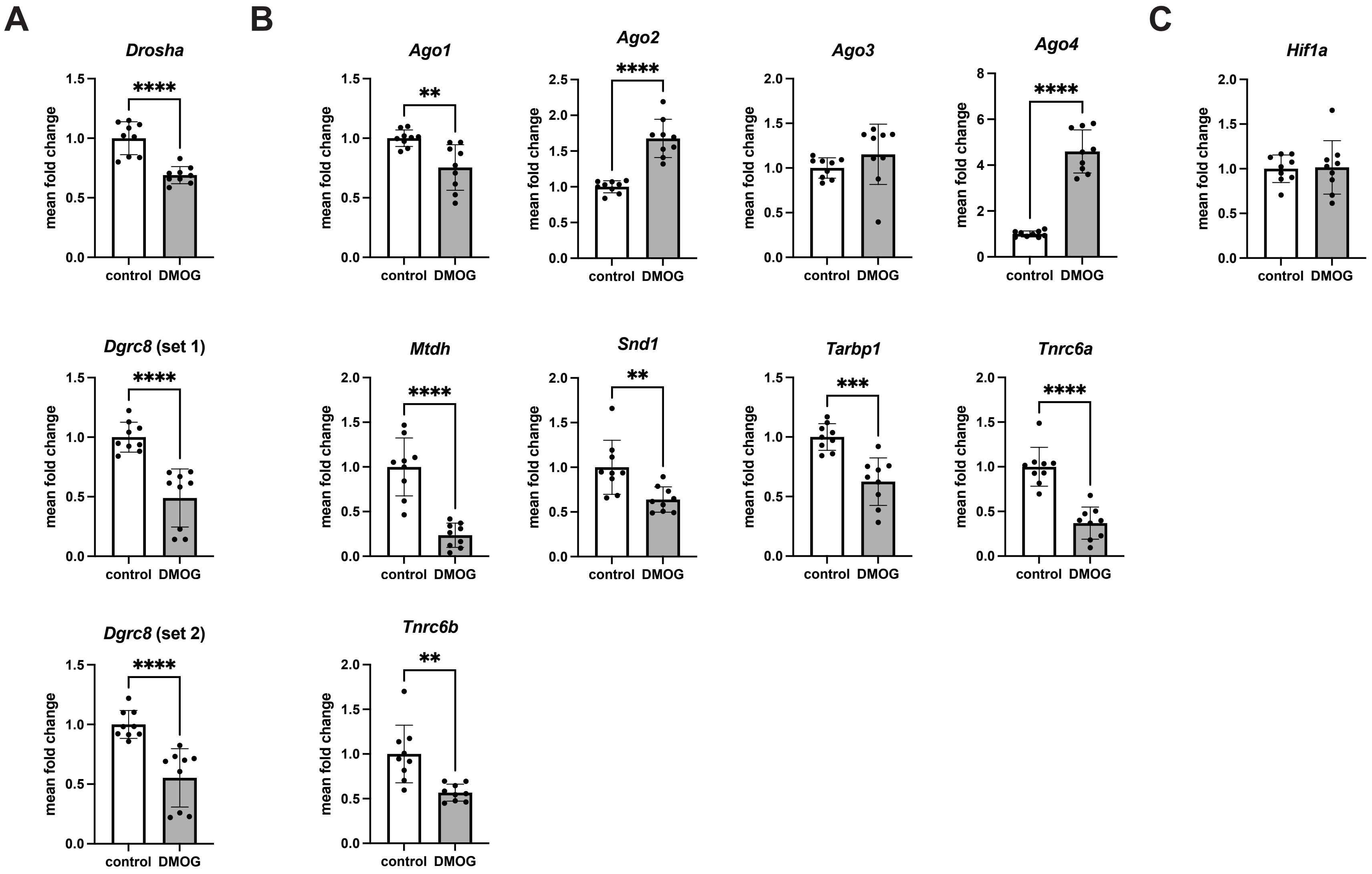

A
B
C
Drosha
Ago1
Ago2
Ago3
Ago4
Hif1a
Dgrc8 (set 1)
Mtdh
Snd1
Tarbp1
Tnrc6a
Dgrc8 (set 2)
Tnrc6b
mean fold change
control
DMOG
****
**
***
0.0
0.5
1.0
1.5
2.0
2.5
0
2
4
6
8

Add. Figure 5

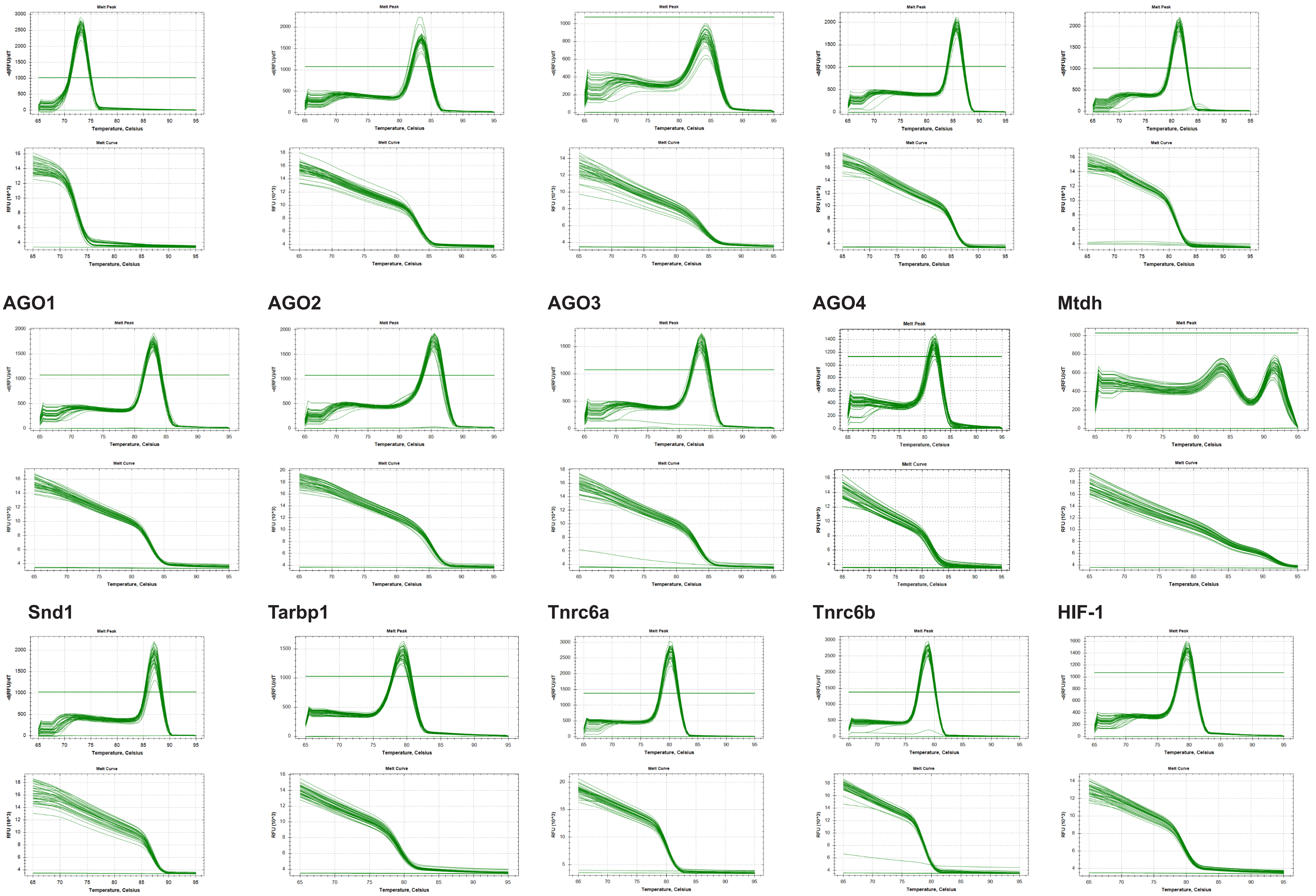
B2M
EF2
Drosha
Dgcr8 (set 1)
Dgcr8 (set 2)
AGO1
AGO2
AGO3
AGO4
Mtdh
Snd1
Tarbp1
Tnrc6a
Tnrc6b
HIF-1
Melt Peak
Melt Curve
-d(RFU)/dT
RFU (10^3)
Temperature, Celsius

### Mtdh

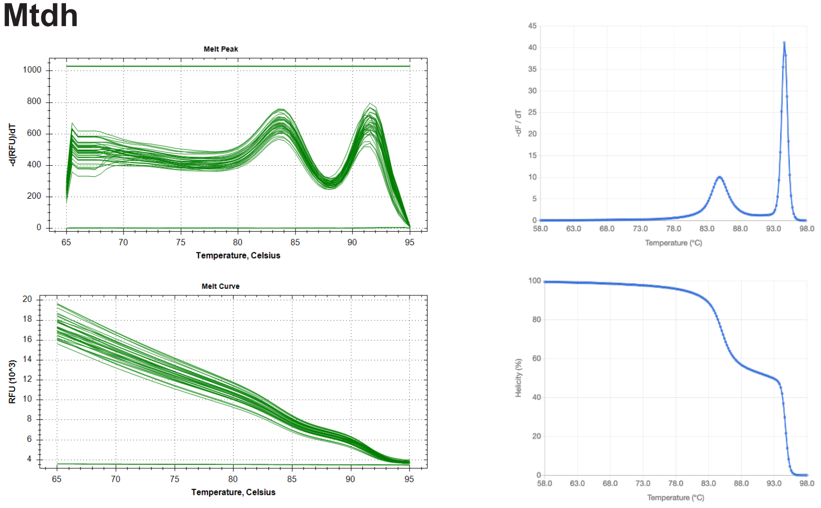
Melt Peak
-d(RFU)/dT
Temperature, Celsius
-dF / dT
Temperature (°C)
Melt Curve
RFU (10^3)
Temperature, Celsius
Helicity (%)
Temperature (°C)

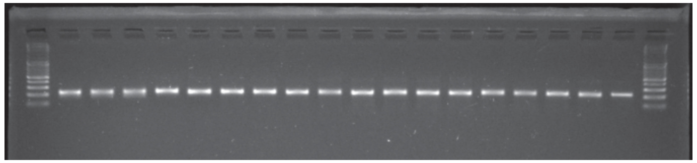

### Add. Table 1

| seqnames | start | end | width | strand | baseMean | log2FoldChange | lfcSE | stat | pvalue | padj | weight | annotation | geneChr | geneStart | geneEnd | geneLength | geneStran | geneId | transcriptId | distanceToTSS | ENSEMBL | SYMBOL | GENENAME |
|---|---|---|---|---|---|---|---|---|---|---|---|---|---|---|---|---|---|---|---|---|---|---|---|
| chr1 | 98461716 | 98463466 | 1751 | * | 10,2933 | -7,234096799 | 1,718724 | -4,20899 | 2,57E-05 | 0,070291925 | 1,425929 | Distal Intergenic | 1 | 98427614 | 98437108 | 9495 | 1 | 70866 | ENSMUST00000162782.2 | 34102 | ENSMUSG00000026336 | Slco6d1 | solute carrier organic anion transporter family, member 6d1 |
| chr3 | 1,41E+08 | 1,41E+08 | 1143 | * | 9,526937 | -6,883236778 | 1,747678 | -3,9385 | 8,2E-05 | 0,143127763 | 1,449706 | Distal Intergenic | 3 | 1,41E+08 | 1,41E+08 | 1372 | 1 | 75852 | ENSMUST00000196030.2 | 39752 | | 4930590L14Rik | RIKEN cDNA 4930590L14 gene |
| chr14 | 1,19E+08 | 1,19E+08 | 1113 | * | 7,803541 | -6,85080636 | 1,898795 | -3,60797 | 0,000309 | 0,301441602 | 1,360622 | Promoter (2-3kb) | 14 | 1,19E+08 | 1,19E+08 | 2208 | 2 | 239273 | ENSMUST00000228691.2 | 2534 | ENSMUSG00000032849 | Abcc4 | ATP-binding cassette, sub-family C (CFTR/MRP), member 4 |
| chr1 | 99825683 | 99826653 | 971 | * | 7,930322 | -6,809364533 | 1,791107 | -3,80176 | 0,000144 | 0,205744841 | 1,360622 | Intron (ENSMUST0000 | 1 | 99700490 | 1E+08 | 712085 | 1 | 241175 | ENSMUST00000086738.10 | 125193 | ENSMUSG00000067028 | Cntnap5b | contactin associated protein-like 5B |
| chr12 | 1,18E+08 | 1,18E+08 | 1043 | * | 9,943865 | -6,740770883 | 1,824 | -3,6956 | 0,000219 | 0,284660899 | 1,086118 | Intron (ENSMUST0000 | 12 | 1,18E+08 | 1,18E+08 | 39649 | 2 | 20688 | ENSMUST00000221844.2 | -3082 | ENSMUSG00000025323 | Sp4 | trans-acting transcription factor 4 |
| chr14 | 10532375 | 10533094 | 720 | * | 7,227145 | -6,698579707 | 1,817941 | -3,68471 | 0,000229 | 1 | 0 | Intron (ENSMUST0000 | 14 | 10347631 | 10378603 | 30973 | 2 | 19270 | ENSMUST00000248717.1 | -153772 | | Ptprg | protein tyrosine phosphatase, receptor type, G |
| chr2 | 57858072 | 57858826 | 755 | * | 9,214919 | -6,697431427 | 1,736811 | -3,85617 | 0,000115 | 0,133396034 | 2,237403 | Distal Intergenic | 2 | 57872204 | 57876029 | 3826 | 2 | 77767 | ENSMUST00000209502.2 | 17203 | ENSMUSG00000026830 | Ermn | ermin, ERM-like protein |
| chr14 | 5129371 | 5129998 | 628 | * | 5,356618 | -6,69679438 | 1,996713 | -3,35391 | 0,000797 | 1 | 0 | Distal Intergenic | 14 | 5363820 | 5364135 | 316 | 2 | 1E+08 | ENSMUST00000225472.2 | 234137 | | Pou5f1-rs6 | POU domain, class 5, transcription factor 1, related sequence 6 |
| chr7 | 1,15E+08 | 1,15E+08 | 1036 | * | 10,59508 | -6,651699274 | 1,694081 | -3,92644 | 8,62E-05 | 0,216838386 | 0,738441 | Distal Intergenic | 7 | 1,15E+08 | 1,15E+08 | 1803 | 1 | 1,03E+08 | ENSMUST00000206921.2 | 46060 | | Gm34225 | predicted gene, 34225 |
| chr13 | 90390600 | 90391633 | 1034 | * | 10,72658 | -6,648738517 | 1,700319 | -3,91029 | 9,22E-05 | 0,225004052 | 0,738441 | Distal Intergenic | 13 | 90247100 | 90247233 | 134 | 1 | 1,15E+08 | ENSMUST00000157683.3 | 143500 | ENSMUSG00000088308 | Gm24507 | predicted gene, 24507 |
| chrX | 1,09E+08 | 1,09E+08 | 532 | * | 3,479436 | -6,62492065 | 2,510257 | -2,63914 | 0,008312 | 1 | 0 | Distal Intergenic | 20 | 1,09E+08 | 1,09E+08 | 2406 | 1 | 634340 | ENSMUST00000122296.2 | -361528 | ENSMUSG00000082853 | Gm7134 | kinesin family member 18B pseudogene |
| chrX | 41580792 | 41582387 | 1596 | * | 8,813108 | -6,575805102 | 1,867409 | -3,52135 | 0,000429 | 0,337054778 | 1,425929 | Distal Intergenic | 20 | 41591330 | 41612574 | 21245 | 1 | 20400 | ENSMUST00000129855.8 | -8943 | ENSMUSG00000005696 | Sh2d1a | SH2 domain containing 1A |
| chr19 | 30403930 | 30404851 | 922 | * | 7,706113 | -6,574539997 | 1,828663 | -3,59527 | 0,000324 | 0,306095127 | 1,360622 | Distal Intergenic | 19 | 30516328 | 30516945 | 618 | 2 | 546723 | ENSMUST00000078012.6 | 112094 | | Ppp1r2-ps3 | protein phosphatase 1, regulatory (inhibitor) subunit 2, pseudogene 3 |
| chr4 | 1,31E+08 | 1,31E+08 | 475 | * | 5,231839 | -6,56603295 | 2,034558 | -3,22725 | 0,00125 | 1 | 0 | Distal Intergenic | 4 | 1,31E+08 | 1,31E+08 | 9643 | 2 | 77835 | ENSMUST00000153810.2 | -37843 | ENSMUSG00000086262 | A930031H19Rik | RIKEN cDNA A930031H19 gene |
| chr9 | 68128157 | 68128884 | 728 | * | 6,702324 | -6,551672879 | 1,880195 | -3,48457 | 0,000493 | 1 | 0 | Distal Intergenic | 9 | 68187856 | 68187985 | 130 | 1 | 1,15E+08 | ENSMUST00000158185.3 | -58972 | ENSMUSG00000088810 | Gm22382 | predicted gene, 22382 |
| chr9 | 1,17E+08 | 1,17E+08 | 927 | * | 7,091102 | -6,538204443 | 1,808078 | -3,61611 | 0,000299 | 1 | 0 | Intron (ENSMUST0000 | 9 | 1,16E+08 | 1,17E+08 | 239998 | 2 | 207181 | ENSMUST00000172469.3 | -118085 | ENSMUSG00000039607 | Rbms3 | RNA binding motif, single stranded interacting protein |
| chr14 | 1,17E+08 | 1,17E+08 | 1211 | * | 5,206273 | -6,52578439 | 2,118265 | -3,08072 | 0,002065 | 1 | 0 | Distal Intergenic | 14 | 1,17E+08 | 1,18E+08 | 701884 | 1 | 23888 | ENSMUST00000123239.2 | -302222 | ENSMUSG00000058571 | Gpc6 | glypican 6 |
| chr5 | 4246685 | 4247775 | 1091 | * | 6,939471 | -6,514835705 | 1,826492 | -3,56686 | 0,000361 | 1 | 0 | 3' UTR | 5 | 4242367 | 4247651 | 5285 | 1 | 208595 | ENSMUST00000177258.2 | 4318 | ENSMUSG00000053178 | Mterf1b | mitochondrial transcription termination factor 1b |
| chr3 | 47136200 | 47137231 | 1032 | * | 9,704386 | -6,514723459 | 1,784184 | -3,65137 | 0,000261 | 0,274904537 | 1,425929 | Distal Intergenic | 3 | 47155062 | 47155199 | 138 | 1 | 1,15E+08 | ENSMUST00000158114.3 | -17831 | ENSMUSG00000088739 | Gm23891 | predicted gene, 23891 |
| chrX | 93140887 | 93142118 | 1232 | * | 5,838147 | -6,506992819 | 2,050652 | -3,17313 | 0,001508 | 1 | 0 | 3' UTR | 20 | 93118238 | 93166506 | 48269 | 2 | 22764 | ENSMUST00000113926.8 | 24388 | ENSMUSG00000079509 | Zfx | zinc finger protein X-linked |
| chr19 | 53431876 | 53432296 | 421 | * | 4,786822 | -6,50694904 | 2,075939 | -3,13446 | 0,001722 | 1 | 0 | Exon (ENSMUST00000 | 19 | 53432445 | 53439922 | 7478 | 2 | 381232 | ENSMUST00000180667.3 | 7626 | | Mirt1 | myocardial infarction associated transcript 1 |
| chr7 | 68168143 | 68168896 | 754 | * | 8,699484 | -6,493376672 | 1,74135 | -3,72893 | 0,000192 | 1 | 0 | Distal Intergenic | 7 | 68082833 | 68140636 | 57804 | 1 | 1,03E+08 | ENSMUST00000207356.2 | 85310 | | Gm34549 | predicted gene, 34549 |
| chr4 | 1,36E+08 | 1,36E+08 | 1309 | * | 10,51736 | -6,486527004 | 2,315121 | -2,80181 | 0,005082 | 0,802490409 | 1,173325 | Intron (ENSMUST0000 | 4 | 1,36E+08 | 1,36E+08 | 32578 | 2 | 13844 | ENSMUST00000156558.2 | -36867 | ENSMUSG00000028664 | Ephb2 | Eph receptor B2 |
| chrX | 1,01E+08 | 1,01E+08 | 831 | * | 6,70022 | -6,485231633 | 1,90274 | -3,40837 | 0,000654 | 1 | 0 | Intron (ENSMUST0000 | 20 | 1,01E+08 | 1,01E+08 | 116085 | 1 | 1E+08 | ENSMUST00000155477.2 | 19710 | ENSMUSG00000079481 | Nhsl2 | NHS-like 2 |
| chr7 | 1,41E+08 | 1,41E+08 | 544 | * | 5,932268 | -6,477119612 | 1,912049 | -3,38753 | 0,000705 | 1 | 0 | Distal Intergenic | 7 | 1,41E+08 | 1,41E+08 | 30260 | 1 | 17833 | ENSMUST00000155534.9 | -5870 | ENSMUSG00000037974 | Muc5ac | mucin 5, subtypes A and C, tracheobronchial/gastric |
| chr10 | 82297364 | 82298662 | 1299 | * | 9,45936 | -6,473505911 | 1,765103 | -3,66749 | 0,000245 | 0,263251859 | 1,449706 | Distal Intergenic | 10 | 82322371 | 82328452 | 6082 | 2 | 432480 | ENSMUST00000177934.3 | 29790 | ENSMUSG00000094186 | Gm1553 | predicted gene 1553 |
| chrX | 43839005 | 43840350 | 1346 | * | 6,640367 | -6,472226795 | 1,856264 | -3,4867 | 0,000489 | 1 | 0 | Distal Intergenic | 20 | 43875447 | 43879038 | 3592 | 2 | 245404 | ENSMUST00000115056.2 | 38688 | ENSMUSG00000045284 | Dcaf12l1 | DDB1 and CUL4 associated factor 12-like 1 |
| chr15 | 11097532 | 11098801 | 1270 | * | 9,580245 | -6,469021048 | 1,840933 | -3,51399 | 0,000441 | 0,339611164 | 1,425929 | Intron (ENSMUST0000 | 15 | 11064904 | 11217207 | 152304 | 1 | 239337 | ENSMUST00000228940.2 | 32628 | ENSMUSG00000047497 | Adamts12 | a disintegrin-like and metallopeptidase (reprolysin type) with thrombospondin type 1 motif, 12 |
| chrX | 48313949 | 48314853 | 905 | * | 5,61214 | -6,463255245 | 2,038032 | -3,17132 | 0,001517 | 1 | 0 | Intron (ENSMUST0000 | 20 | 48098584 | 48377089 | 278506 | 2 | 209224 | ENSMUST00000114914.8 | 62236 | ENSMUSG00000031109 | Enox2 | ecto-NOX disulfide-thiol exchanger 2 |
| chr2 | 1,33E+08 | 1,33E+08 | 1224 | * | 10,99348 | -6,453384787 | 1,740715 | -3,70732 | 0,000209 | 0,259991115 | 1,2763 | Promoter (<=1kb) | 2 | 1,33E+08 | 1,33E+08 | 103 | 1 | 1,15E+08 | ENSMUST00000082479.4 | -340 | ENSMUSG00000064413 | Gm22245 | predicted gene, 22245 |
| chr14 | 82937947 | 82938754 | 808 | * | 6,135141 | -6,450351368 | 1,966904 | -3,27944 | 0,00104 | 1 | 0 | Distal Intergenic | 14 | 83643071 | 83643204 | 134 | 1 | 1,15E+08 | ENSMUST00000157384.3 | -704317 | ENSMUSG00000088009 | Gm24774 | predicted gene, 24774 |
| chr4 | 1,48E+08 | 1,48E+08 | 803 | * | 8,194788 | -6,442177176 | 1,772872 | -3,63375 | 0,000279 | 0,282588188 | 1,425929 | Intron (ENSMUST0000 | 4 | 1,48E+08 | 1,48E+08 | 47702 | 2 | 242748 | ENSMUST00000047720.9 | 7849 | ENSMUSG00000041544 | Disp3 | dispatched RND transporter family member 3 |
| chr9 | 64407613 | 64408274 | 662 | * | 7,589091 | -6,441756449 | 1,7995 | -3,57975 | 0,000344 | 1 | 0 | Promoter (<=1kb) | 9 | 64408618 | 64416354 | 7737 | 1 | 214058 | ENSMUST00000148963.8 | -344 | ENSMUSG00000036466 | Megf11 | multiple EGF-like-domains 11 |
| chr13 | 52479459 | 52480089 | 631 | * | 6,473222 | -6,44141447 | 1,968991 | -3,27143 | 0,00107 | 1 | 0 | Distal Intergenic | 13 | 52334447 | 52335673 | 1227 | 1 | 1,19E+08 | ENSMUST00000222876.2 | 145012 | | Gm48199 | predicted gene, 48199 |
| chr2 | 67831502 | 67832096 | 595 | * | 5,880205 | -6,431429252 | 2,099533 | -3,06327 | 0,002189 | 1 | 0 | Intron (ENSMUST0000 | 2 | 67935015 | 67950858 | 15844 | 1 | 26877 | ENSMUST00000042456.4 | -102919 | ENSMUSG00000034780 | B3galt1 | UDP-Gal:betaGlcNAc beta 1,3-galactosyltransferase, polypeptide 1 |
| chr3 | 1,12E+08 | 1,12E+08 | 504 | * | 6,042921 | -6,42019766 | 2,082296 | -3,08323 | 0,002048 | 1 | 0 | Distal Intergenic | 3 | 1,12E+08 | 1,12E+08 | 127 | 2 | 1,15E+08 | ENSMUST00000158392.3 | -6820 | ENSMUSG00000089017 | Gm25519 | predicted gene, 25519 |
| chr1 | 1,74E+08 | 1,74E+08 | 1129 | * | 14,14651 | -6,415208191 | 1,69981 | -3,77407 | 0,000161 | 0,247743734 | 1,075735 | Intron (ENSMUST0000 | 1 | 1,74E+08 | 1,74E+08 | 45508 | 1 | 54418 | ENSMUST00000191821.2 | 21838 | ENSMUSG00000028354 | Fmn2 | formin 2 |
| chr2 | 1,8E+08 | 1,8E+08 | 529 | * | 5,598886 | -6,41414454 | 1,941369 | -3,30393 | 0,000953 | 1 | 0 | Distal Intergenic | 2 | 1,8E+08 | 1,8E+08 | 3143 | 1 | 228993 | ENSMUST00000140955.2 | 16261 | ENSMUSG00000023393 | Slc17a9 | solute carrier family 17, member 9 |
| chr14 | 67168730 | 67169565 | 836 | * | 5,884227 | -6,411807035 | 1,898485 | -3,37733 | 0,000732 | 1 | 0 | Distal Intergenic | 14 | 67151765 | 67153484 | 1720 | 1 | 239157 | ENSMUST00000165831.2 | 16965 | ENSMUSG00000046204 | Pnma2 | paraneoplastic antigen MA2 |
| chr16 | 16007763 | 16008318 | 556 | * | 4,809733 | -6,406536082 | 2,06827 | -3,09753 | 0,001951 | 1 | 0 | Distal Intergenic | 16 | 16031182 | 16045151 | 13970 | 1 | 67451 | ENSMUST00000162150.8 | -22864 | ENSMUSG00000041957 | Pkp2 | plakophilin 2 |
| chrX | 59044137 | 59045111 | 975 | * | 5,60207 | -6,40229647 | 1,972918 | -3,24509 | 0,001174 | 1 | 0 | Promoter (<=1kb) | 20 | 59044824 | 59076119 | 31296 | 1 | 14071 | ENSMUST00000033477.5 | 0 | ENSMUSG00000031138 | F9 | coagulation factor IX |
| chr3 | 71576577 | 71576995 | 419 | * | 7,102001 | -6,396984422 | 2,100959 | -3,04479 | 0,002328 | 0,646404087 | 1,360622 | Distal Intergenic | 3 | 70709254 | 70714652 | 5399 | 2 | 625901 | ENSMUST00000242834.1 | -861925 | | Gm6634 | predicted gene 6634 |
| chr18 | 6994512 | 6995339 | 828 | * | 6,312617 | -6,395866389 | 1,940155 | -3,29657 | 0,000979 | 1 | 0 | Promoter (1-2kb) | 18 | 6936952 | 6992988 | 56037 | 2 | 210719 | ENSMUST00000188926.2 | -1524 | ENSMUSG00000061013 | Mkx | mohawk homeobox |
| chr10 | 28116779 | 28117480 | 702 | * | 7,958274 | -6,391752369 | 1,960971 | -3,25948 | 0,001116 | 1 | 0 | Intron (ENSMUST0000 | 10 | 27951006 | 28140669 | 189664 | 1 | 19272 | ENSMUST00000219761.2 | 165773 | ENSMUSG00000019889 | Ptprk | protein tyrosine phosphatase, receptor type, K |
| chr3 | 1,27E+08 | 1,27E+08 | 1135 | * | 8,220117 | -6,389532847 | 1,781468 | -3,58667 | 0,000335 | 0,304413931 | 1,425929 | Distal Intergenic | 3 | 1,27E+08 | 1,27E+08 | 6025 | 2 | 109676 | ENSMUST00000182118.9 | 12458 | ENSMUSG00000032826 | Ank2 | ankyrin 2, brain |
| chrX | 1,08E+08 | 1,08E+08 | 528 | * | 4,01402 | -6,385271283 | 2,216329 | -2,88101 | 0,003964 | 1 | 0 | Intron (ENSMUST0000 | 20 | 1,08E+08 | 1,08E+08 | 92730 | 1 | 69964 | ENSMUST00000207966.2 | -278811 | ENSMUSG00000086629 | 2810403D21Rik | RIKEN cDNA 2810403D21 gene |
| chr3 | 46833216 | 46834125 | 910 | * | 6,540405 | -6,383355223 | 1,991503 | -3,20529 | 0,001349 | 1 | 0 | Distal Intergenic | 3 | 46855103 | 46855217 | 115 | 2 | 1,02E+08 | ENSMUST00000183392.3 | 21092 | ENSMUSG00000098417 | Mir6379 | microRNA 6379 |
| chr7 | 58925728 | 58926595 | 868 | * | 6,908589 | -6,38062128 | 1,837894 | -3,4717 | 0,000517 | 1 | 0 | Exon (ENSMUST00000 | 7 | 58935926 | 58954680 | 18755 | 1 | 22215 | ENSMUST00000202247.4 | -9331 | ENSMUSG00000025326 | Ube3a | ubiquitin protein ligase E3A |
| chrX | 1,13E+08 | 1,13E+08 | 1406 | * | 5,97576 | -6,379278984 | 1,954028 | -3,26468 | 0,001096 | 1 | 0 | Downstream (<=300bp | 20 | 1,13E+08 | 1,13E+08 | 85905 | 1 | 237010 | ENSMUST00000133447.2 | 86511 | ENSMUSG00000025597 | Klhl4 | kelch-like 4 |
| chr18 | 78572496 | 78572849 | 354 | * | 5,199085 | -6,378342731 | 2,109968 | -3,02296 | 0,002503 | 1 | 0 | Intron (ENSMUST0000 | 18 | 78576673 | 78576779 | 107 | 1 | 1,15E+08 | ENSMUST00000240445.1 | -3824 | ENSMUSG00000119659 | Gm26430 | predicted gene, 26430 |
| chrX | 1,52E+08 | 1,52E+08 | 629 | * | 5,758526 | -6,376205171 | 2,051459 | -3,10813 | 0,001883 | 1 | 0 | Distal Intergenic | 20 | 1,52E+08 | 1,52E+08 | 11127 | 2 | 207474 | ENSMUST00000039424.14 | 29520 | ENSMUSG00000041633 | Kctd12b | potassium channel tetramerisation domain containing 12b |
| chr1 | 1,92E+08 | 1,92E+08 | 878 | * | 6,375694 | -6,371979232 | 1,959276 | -3,25221 | 0,001145 | 1 | 0 | Exon (ENSMUST00000 | 1 | 1,92E+08 | 1,92E+08 | 11607 | 1 | 18005 | ENSMUST00000027931.8 | -3822 | ENSMUSG00000026622 | Nek2 | NIMA (never in mitosis gene a)-related expressed kinase 2 |
| chr3 | 12166500 | 12167414 | 915 | * | 5,944704 | -6,367462015 | 1,93153 | -3,29659 | 0,000979 | 1 | 0 | Distal Intergenic | 3 | 11888289 | 11909419 | 21131 | 2 | 1E+08 | ENSMUST00000192022.2 | -257081 | ENSMUSG00000104015 | Gm10745 | predicted gene 10745 |
| chr1 | 71671021 | 71671866 | 846 | * | 9,129457 | -6,357808593 | 1,725086 | -3,6855 | 0,000228 | 0,202462686 | 2,237403 | Exon (ENSMUST00000 | 1 | 71643256 | 71663503 | 20248 | 2 | 14268 | ENSMUST00000189160.2 | -7518 | ENSMUSG00000026193 | Fn1 | fibronectin 1 |
| chr5 | 53915482 | 53916158 | 677 | * | 8,126684 | -6,356051057 | 1,82091 | -3,49059 | 0,000482 | 0,352527836 | 1,449706 | Distal Intergenic | 5 | 53855827 | 53865047 | 9221 | 2 | 12425 | ENSMUST00000031093.5 | -50435 | ENSMUSG00000029193 | Cckar | cholecystokinin A receptor |
| chrX | 41269183 | 41270634 | 1452 | * | 5,374153 | -6,354755969 | 2,035692 | -3,12167 | 0,001798 | 1 | 0 | Intron (ENSMUST0000 | 20 | 41241057 | 41309906 | 68850 | 1 | 20843 | ENSMUST00000128799.3 | 28126 | ENSMUSG00000025862 | Stag2 | stromal antigen 2 |
| chr11 | 42862200 | 42862767 | 568 | * | 5,317643 | -6,354607582 | 1,927203 | -3,29732 | 0,000976 | 1 | 0 | Distal Intergenic | 11 | 43040704 | 43050302 | 9599 | 1 | 319767 | ENSMUST00000148911.2 | -177937 | ENSMUSG00000055415 | Atp10b | ATPase, class V, type 10B |
| chr2 | 1,1E+08 | 1,1E+08 | 642 | * | 3,851698 | -6,349419672 | 2,307203 | -2,752 | 0,005923 | 1 | 0 | Distal Intergenic | 2 | 1,1E+08 | 1,1E+08 | 2267 | 2 | 67606 | ENSMUST00000099626.5 | -9446 | ENSMUSG00000074971 | Fibin | fin bud initiation factor homolog (zebrafish) |
| chr3 | 69719884 | 69720285 | 402 | * | 5,894619 | -6,345100235 | 1,899639 | -3,34016 | 0,000837 | 1 | 0 | Distal Intergenic | 3 | 69727911 | 69767255 | 39345 | 2 | 66183 | ENSMUST00000171529.4 | 46970 | ENSMUSG00000043461 | Sptssb | serine palmitoyltransferase, small subunit B |
| chr10 | 97340299 | 97341557 | 1259 | * | 9,50256 | -6,344935498 | 1,74454 | -3,63703 | 0,000276 | 0,302119236 | 1,211315 | Intron (ENSMUST0000 | 10 | 97319199 | 97353578 | 34380 | 1 | 13179 | ENSMUST00000219539.2 | 21100 | ENSMUSG00000019929 | Dcn | decorin |
| chr3 | 1,38E+08 | 1,38E+08 | 656 | * | 5,727334 | -6,343220305 | 1,899983 | -3,33857 | 0,000842 | 1 | 0 | Promoter (2-3kb) | 3 | 1,38E+08 | 1,38E+08 | 3366 | 1 | 56692 | ENSMUST00000200107.2 | -2140 | ENSMUSG00000091512 | Lamtor3 | late endosomal/lysosomal adaptor, MAPK and MTOR activator 3 |
| chr1 | 1,49E+08 | 1,49E+08 | 627 | * | 4,338167 | -6,333255123 | 2,251655 | -2,81271 | 0,004913 | 1 | 0 | Distal Intergenic | 1 | 1,48E+08 | 1,48E+08 | 129 | 2 | 1,15E+08 | ENSMUST00000157957.3 | -544502 | ENSMUSG00000088582 | Gm22966 | predicted gene, 22966 |
| chr15 | 94627145 | 94627399 | 255 | * | 4,427249 | -6,33323951 | 2,224356 | -2,84722 | 0,00441 | 1 | 0 | Intron (ENSMUST0000 | 15 | 94579136 | 94579267 | 132 | 2 | 1,15E+08 | ENSMUST00000178490.3 | -47878 | ENSMUSG00000119001 | Gm25546 | predicted gene, 25546 |
| chr10 | 85082596 | 85083545 | 950 | * | 6,435514 | -6,328922897 | 1,900795 | -3,32962 | 0,00087 | 1 | 0 | Intron (ENSMUST0000 | 10 | 84967564 | 85020928 | 53365 | 2 | 12952 | ENSMUST00000020227.11 | -61668 | ENSMUSG00000020038 | Cry1 | cryptochrome 1 (photolyase-like) |
| chr13 | 82448730 | 82449819 | 1090 | * | 6,302433 | -6,327919411 | 1,909668 | -3,31362 | 0,000921 | 1 | 0 | Distal Intergenic | 13 | 82198870 | 82199000 | 131 | 1 | 1,15E+08 | ENSMUST00000158298.3 | 249860 | ENSMUSG00000088923 | Gm24295 | predicted gene, 24295 |
| chr7 | 1,29E+08 | 1,29E+08 | 611 | * | 5,982604 | -6,325335385 | 1,970038 | -3,21077 | 0,001324 | 1 | 0 | Distal Intergenic | 7 | 1,29E+08 | 1,29E+08 | 25976 | 1 | 1,03E+08 | ENSMUST00000205836.2 | 65656 | | Gm33248 | predicted gene, 33248 |
| chr14 | 22642984 | 22643824 | 841 | * | 5,822331 | -6,322327614 | 1,932236 | -3,27203 | 0,001068 | 1 | 0 | Promoter (2-3kb) | 14 | 22646581 | 22649336 | 2756 | 1 | 76633 | ENSMUST00000226003.2 | -2757 | ENSMUSG00000063458 | Lrmda | leucine rich melanocyte differentiation associated |
| chrX | 93023373 | 93024223 | 851 | * | 6,260152 | -6,321125337 | 2,039834 | -3,09884 | 0,001943 | 1 | 0 | Distal Intergenic | 20 | 93013402 | 93018677 | 5276 | 1 | 385493 | ENSMUST00000120624.2 | 9971 | ENSMUSG00000081044 | AU015836 | expressed sequence AU015836 |
| chr2 | 58092900 | 58093756 | 857 | * | 6,206428 | -6,321080346 | 1,95256 | -3,23733 | 0,001207 | 1 | 0 | Distal Intergenic | 2 | 58049988 | 58085544 | 35557 | 2 | 227929 | ENSMUST00000153052.2 | -7356 | ENSMUSG00000026832 | Cytip | cytohesin 1 interacting protein |
| chr2 | 1,32E+08 | 1,32E+08 | 628 | * | 4,155346 | -6,319003924 | 2,261272 | -2,79445 | 0,005199 | 1 | 0 | Distal Intergenic | 2 | 1,32E+08 | 1,32E+08 | 14283 | 2 | 246313 | ENSMUST00000110156.2 | -5662 | ENSMUSG00000050558 | Prokr2 | prokineticin receptor 2 |
| chr9 | 22879610 | 22880468 | 859 | * | 6,167548 | -6,31605217 | 2,027462 | -3,11525 | 0,001838 | 1 | 0 | Distal Intergenic | 9 | 23134372 | 23396496 | 262125 | 1 | 73230 | ENSMUST00000071982.7 | -253904 | ENSMUSG00000031963 | Bmper | BMP-binding endothelial regulator |
| chr18 | 13427423 | 13428097 | 675 | * | 6,274508 | -6,315084645 | 1,85534 | -3,40373 | 0,000665 | 1 | 0 | Intron (ENSMUST0000 | 18 | 13342759 | 13342865 | 107 | 2 | 1,15E+08 | ENSMUST00000083216.3 | -84558 | ENSMUSG00000119747 | Gm22251 | predicted gene, 22251 |
| chr8 | 1,04E+08 | 1,04E+08 | 268 | * | 4,09089 | -6,313482261 | 2,311896 | -2,73087 | 0,006317 | 1 | 0 | Intron (ENSMUST0000 | 8 | 1,04E+08 | 1,04E+08 | 68740 | 1 | 1,03E+08 | ENSMUST00000209324.2 | 23647 | | Gm32531 | predicted gene, 32531 |
| chrX | 14898383 | 14899052 | 670 | * | 4,803463 | -6,313479426 | 2,090934 | -3,01945 | 0,002532 | 1 | 0 | Distal Intergenic | 20 | 14977071 | 14977619 | 549 | 2 | 67395 | ENSMUST00000173100.2 | 78567 | ENSMUSG00000045217 | Ppp1r2-ps9 | protein phosphatase 1, regulatory (inhibitor) subunit 2, pseudogene 9 |
| chr10 | 54604812 | 54605735 | 924 | * | 8,234681 | -6,312121966 | 1,790258 | -3,52582 | 0,000422 | 0,332694435 | 1,449706 | Distal Intergenic | 10 | 54589072 | 54589133 | 62 | 2 | 1,15E+08 | ENSMUST00000158362.3 | -15679 | ENSMUSG00000088987 | Gm25602 | predicted gene, 25602 |
| chr5 | 50099442 | 50100456 | 1015 | * | 9,469459 | -6,308656046 | 1,738615 | -3,62855 | 0,000285 | 0,226551964 | 2,237403 | Distal Intergenic | 5 | 50123452 | 50129417 | 5966 | 2 | 70693 | ENSMUST00000196177.2 | 28961 | ENSMUSG00000029090 | Adgra3 | adhesion G protein-coupled receptor A3 |
| chr8 | 52727095 | 52728185 | 1091 | * | 5,91151 | -6,308590286 | 1,869265 | -3,3749 | 0,000738 | 1 | 0 | Distal Intergenic | 8 | 52789950 | 52790140 | 191 | 2 | 1,15E+08 | ENSMUST00000082748.3 | 61955 | ENSMUSG00000064682 | Gm25813 | predicted gene, 25813 |
| chr1 | 1,89E+08 | 1,89E+08 | 1165 | * | 6,143486 | -6,305644511 | 1,912153 | -3,29767 | 0,000975 | 1 | 0 | Intron (ENSMUST0000 | 1 | 1,89E+08 | 1,89E+08 | 192461 | 2 | 16526 | ENSMUST00000194402.6 | 25361 | ENSMUSG00000037624 | Kcnk2 | potassium channel, subfamily K, member 2 |
| chrX | 23286116 | 23286826 | 711 | * | 5,321868 | -6,304146708 | 1,987226 | -3,17234 | 0,001512 | 1 | 0 | Distal Intergenic | 20 | 23085510 | 23231321 | 145812 | 2 | 67455 | ENSMUST00000115313.8 | -54795 | ENSMUSG00000036782 | Klhl13 | kelch-like 13 |
| chr1 | 49752185 | 49752912 | 728 | * | 5,311009 | -6,303787789 | 2,049668 | -3,07552 | 0,002101 | 1 | 0 | Distal Intergenic | 1 | 49283775 | 49379602 | 95828 | 1 | 442837 | ENSMUST00000188673.7 | 468410 | ENSMUSG00000051616 | C230029F24Rik | RIKEN cDNA C230029F24 gene |
| chr18 | 40476984 | 40477779 | 796 | * | 7,989251 | -6,30161161 | 1,792703 | -3,51515 | 0,00044 | 1 | 0 | Intron (ENSMUST0000 | 18 | 40541138 | 40664683 | 123546 | 1 | 383348 | ENSMUST00000236301.2 | -63359 | ENSMUSG00000051401 | Kctd16 | potassium channel tetramerisation domain containing 16 |
| chr9 | 42161533 | 42162606 | 1074 | * | 8,304677 | -6,299101724 | 1,783271 | -3,53233 | 0,000412 | 0,337696055 | 1,360622 | Distal Intergenic | 9 | 42166029 | 42175496 | 9468 | 2 | 235293 | ENSMUST00000169609.2 | 12890 | ENSMUSG00000032018 | Sc5d | sterol-C5-desaturase |
| chr14 | 1,23E+08 | 1,23E+08 | 1033 | * | 5,48728 | -6,298677116 | 2,014975 | -3,12593 | 0,001772 | 1 | 0 | Intron (ENSMUST0000 | 14 | 1,22E+08 | 1,23E+08 | 144187 | 1 | 69634 | ENSMUST00000226136.2 | 32685 | ENSMUSG00000025545 | Clybl | citrate lyase beta like |
| chr7 | 1,4E+08 | 1,4E+08 | 568 | * | 4,903498 | -6,290892375 | 2,127226 | -2,95732 | 0,003103 | 1 | 0 | Promoter (1-2kb) | 7 | 1,4E+08 | 1,4E+08 | 3826 | 2 | 74237 | ENSMUST00000211261.2 | 1239 | ENSMUSG00000025474 | Tubgcp2 | tubulin, gamma complex associated protein 2 |
| chr15 | 65185831 | 65186402 | 572 | * | 4,805138 | -6,288378824 | 2,138629 | -2,94038 | 0,003278 | 1 | 0 | Distal Intergenic | 15 | 65566398 | 65566586 | 189 | 1 | 1,15E+08 | ENSMUST00000104595.4 | -379996 | ENSMUSG00000077783 | Gm23907 | predicted gene, 23907 |
| chr6 | 1,04E+08 | 1,04E+08 | 369 | * | 6,486872 | -6,287646612 | 1,858814 | -3,38261 | 0,000718 | 1 | 0 | Distal Intergenic | 6 | 1,04E+08 | 1,05E+08 | 150160 | 1 | 53870 | ENSMUST00000161446.2 | -82806 | ENSMUSG00000030092 | Cntn6 | contactin 6 |
| chr4 | 27939503 | 27940138 | 636 | * | 4,651243 | -6,286077123 | 2,082664 | -3,01829 | 0,002542 | 1 | 0 | Distal Intergenic | 4 | 27700671 | 27700777 | 107 | 1 | 1,15E+08 | ENSMUST00000179794.2 | 238832 | ENSMUSG00000095003 | Gm25173 | predicted gene, 25173 |
| chrX | 40339063 | 40340025 | 963 | * | 5,440333 | -6,286014432 | 1,929066 | -3,25858 | 0,00112 | 1 | 0 | Distal Intergenic | 20 | 40461136 | 40461391 | 256 | 1 | 1,08E+08 | ENSMUST00000121134.2 | -121111 | ENSMUSG00000083515 | Gm14577 | predicted gene 14577 |
| chr14 | 80431322 | 80431825 | 504 | * | 3,856375 | -6,282676247 | 2,291845 | -2,74132 | 0,006119 | 1 | 0 | Distal Intergenic | 14 | 80237742 | 80259370 | 21629 | 1 | 380924 | ENSMUST00000088735.4 | 193580 | ENSMUSG00000022026 | Olfm4 | olfactomedin 4 |
| chr14 | 82257863 | 82259303 | 1441 | * | 8,257814 | -6,278881022 | 1,97627 | -3,17714 | 0,001487 | 0,474484649 | 2,237403 | Distal Intergenic | 14 | 81923250 | 81958568 | 35319 | 1 | 1,05E+08 | ENSMUST00000227374.2 | 334613 | | Gm41219 | predicted gene, 41219 |
| chr8 | 68916504 | 68916975 | 472 | * | 5,736937 | -6,276620273 | 1,98672 | -3,15929 | 0,001582 | 1 | 0 | Intron (ENSMUST0000 | 8 | 68914477 | 68982539 | 68063 | 2 | 234356 | ENSMUST00000211871.2 | 65564 | ENSMUSG00000036356 | Csgalnact1 | chondroitin sulfate N-acetylgalactosaminyltransferase 1 |
| chr14 | 27811451 | 27812542 | 1092 | * | 11,52623 | -6,271364973 | 1,699181 | -3,69082 | 0,000224 | 0,269133964 | 1,2763 | Intron (ENSMUST0000 | 14 | 27768385 | 28200281 | 431897 | 1 | 238988 | ENSMUST00000209800.2 | 43066 | ENSMUSG00000040640 | Erc2 | ELKS/RAB6-interacting/CAST family member 2 |
| chr14 | 1,08E+08 | 1,08E+08 | 834 | * | 3,555222 | -6,270793128 | 2,322331 | -2,70022 | 0,006929 | 1 | 0 | Distal Intergenic | 14 | 1,07E+08 | 1,07E+08 | 127 | 1 | 1,15E+08 | ENSMUST00000104036.3 | 739478 | ENSMUSG00000077224 | Gm23437 | predicted gene, 23437 |
| chr14 | 1,23E+08 | 1,23E+08 | 1083 | * | 7,076252 | -6,266318544 | 1,834453 | -3,41591 | 0,000636 | 1 | 0 | Intron (ENSMUST0000 | 14 | 1,22E+08 | 1,23E+08 | 144187 | 1 | 69634 | ENSMUST00000226136.2 | 13957 | ENSMUSG00000025545 | Clybl | citrate lyase beta like |
| chr3 | 91238443 | 91239217 | 775 | * | 5,834024 | -6,263720639 | 2,053558 | -3,05018 | 0,002287 | 1 | 0 | Distal Intergenic | 3 | 90995444 | 90998110 | 2667 | 2 | 229550 | ENSMUST00000166173.3 | -240333 | ENSMUSG00000091175 | S100a7l2 | S100 calcium binding protein A7 like 2 |
| chr9 | 67592329 | 67593750 | 1422 | * | 10,73815 | -6,263285937 | 1,734918 | -3,61013 | 0,000306 | 0,396006019 | 0,744375 | Distal Intergenic | 9 | 67634999 | 67635062 | 64 | 2 | 1,02E+08 | ENSMUST00000184532.3 | 41312 | ENSMUSG00000098937 | Mir7241 | microRNA 7241 |
| chr14 | 1,17E+08 | 1,17E+08 | 1199 | * | 6,390691 | -6,259819423 | 1,996568 | -3,13529 | 0,001717 | 1 | 0 | Intron (ENSMUST0000 | 14 | 1,17E+08 | 1,18E+08 | 701884 | 1 | 23888 | ENSMUST00000123239.2 | -554609 | ENSMUSG00000058571 | Gpc6 | glypican 6 |
| chr6 | 1,21E+08 | 1,21E+08 | 1274 | * | 12,2458 | -6,25753789 | 1,688055 | -3,70695 | 0,00021 | 0,302171532 | 0,918255 | Promoter (<=1kb) | 6 | 1,21E+08 | 1,21E+08 | 916 | 1 | 24110 | ENSMUST00000204710.2 | 398 | ENSMUSG00000030107 | Usp18 | ubiquitin specific peptidase 18 |
| chr3 | 38995307 | 38996314 | 1008 | * | 5,625851 | -6,256108363 | 1,907734 | -3,27934 | 0,00104 | 1 | 0 | Intron (ENSMUST0000 | 3 | 38945091 | 39006578 | 61488 | 1 | 329628 | ENSMUST00000129182.2 | 50216 | ENSMUSG00000046743 | Fat4 | FAT atypical cadherin 4 |
| chrX | 86618941 | 86619851 | 911 | * | 5,295186 | -6,253795613 | 1,962429 | -3,18676 | 0,001439 | 1 | 0 | Intron (ENSMUST0000 | 20 | 85790848 | 86933841 | 1142994 | 2 | 331461 | ENSMUST00000078875.8 | 313990 | ENSMUSG00000052372 | Il1rapl1 | interleukin 1 receptor accessory protein-like 1 |

| | | | | | | | | | | | | | | | | | | | | | | | |
|---|---|---|---|---|---|---|---|---|---|---|---|---|---|---|---|---|---|---|---|---|---|---|---|
| chr1 | 41574622 | 41575631 | 1010 | * | 9,189244 | -6,248840641 | 1,746472 | -3,57798 | 0,000346 | 0,308275655 | 1,425929 | Distal Intergenic | 1 | 41782848 | 41978749 | 195902 | 1 | 1,03E+08 | ENSMUST00000247631.1 | -207217 | | Gm29260 | predicted gene 29260 |
| chr18 | 70098982 | 70099436 | 455 | * | 5,945158 | -6,245769756 | 1,965361 | -3,17792 | 0,001483 | 1 | 0 | Intron (ENSMUST0000 | 18 | 70058647 | 70102555 | 43909 | 1 | 381175 | ENSMUST00000080050.6 | 40335 | ENSMUSG00000038903 | Ccdc68 | coiled-coil domain containing 68 |
| chr3 | 46998452 | 46999502 | 1051 | * | 9,442625 | -6,240035854 | 1,759665 | -3,54615 | 0,000391 | 0,323988916 | 1,425929 | Distal Intergenic | 3 | 46855103 | 46855217 | 115 | 2 | 1,02E+08 | ENSMUST00000183392.3 | -143235 | ENSMUSG00000098417 | Mir6379 | microRNA 6379 |
| chr18 | 45558241 | 45559102 | 862 | * | 8,97813 | -6,239556735 | 1,855454 | -3,36282 | 0,000772 | 0,466212802 | 1,211315 | Intron (ENSMUST0000 | 18 | 45692536 | 45804316 | 111781 | 1 | 140492 | ENSMUST00000184101.8 | -133434 | ENSMUSG00000054477 | Kcnn2 | potassium intermediate/small conductance calcium-activated channel, subfamily N, member 2 |
| chr1 | 49006744 | 49007580 | 837 | * | 5,769625 | -6,238638103 | 1,949324 | -3,20041 | 0,001372 | 1 | 0 | Distal Intergenic | 1 | 49283775 | 49377405 | 93631 | 1 | 442837 | ENSMUST00000056879.9 | -276195 | ENSMUSG00000051616 | C230029F24Rik | RIKEN cDNA C230029F24 gene |
| chrX | 90385692 | 90386540 | 849 | * | 5,960449 | -6,236832605 | 1,954067 | -3,19172 | 0,001414 | 1 | 0 | Distal Intergenic | 20 | 90375361 | 90376503 | 1143 | 1 | 17146 | ENSMUST00000101397.2 | 10331 | ENSMUSG00000073069 | Mageb2 | MAGE family member B2 |
| chr16 | 36239874 | 36240443 | 570 | * | 5,438633 | -6,235403524 | 1,994017 | -3,12706 | 0,001766 | 1 | 0 | Promoter (<=1kb) | 16 | 36239449 | 36239552 | 104 | 1 | 1,15E+08 | ENSMUST00000239658.1 | 425 | ENSMUSG00000119903 | Gm22263 | predicted gene, 22263 |
| chr11 | 43808077 | 43808924 | 848 | * | 7,53712 | -6,232891764 | 1,842329 | -3,38316 | 0,000717 | 1 | 0 | Distal Intergenic | 11 | 43666850 | 43792037 | 125188 | 2 | 11548 | ENSMUST00000139906.2 | -16040 | ENSMUSG00000050541 | Adra1b | adrenergic receptor, alpha 1b |
| chr18 | 67860173 | 67861160 | 988 | * | 4,137521 | -6,232357732 | 2,085753 | -2,98806 | 0,002808 | 1 | 0 | Promoter (2-3kb) | 18 | 67804282 | 67857665 | 53384 | 2 | 19255 | ENSMUST00000122412.2 | -2508 | ENSMUSG00000024539 | Ptpn2 | protein tyrosine phosphatase, non-receptor type 2 |
| chr13 | 17982081 | 17982876 | 796 | * | 3,526191 | -6,229786188 | 2,333358 | -2,66988 | 0,007588 | 1 | 0 | Promoter (1-2kb) | 13 | 17984176 | 17984282 | 107 | 2 | 1,15E+08 | ENSMUST00000083232.3 | 1406 | ENSMUSG00000118773 | Gm24018 | predicted gene, 24018 |
| chr14 | 99617455 | 99618846 | 1392 | * | 8,081858 | -6,227518899 | 1,846608 | -3,37241 | 0,000745 | 1 | 0 | Distal Intergenic | 14 | 99536590 | 99542762 | 6173 | 1 | 12224 | ENSMUST00000226784.2 | 80865 | ENSMUSG00000005148 | Klf5 | Kruppel-like factor 5 |
| chr16 | 47005940 | 47006762 | 823 | * | 7,647192 | -6,226618424 | 1,812862 | -3,43469 | 0,000593 | 1 | 0 | Distal Intergenic | 16 | 47326682 | 47326788 | 107 | 2 | 1,15E+08 | ENSMUST00000157107.4 | 320026 | ENSMUSG00000119721 | Gm24651 | predicted gene, 24651 |
| chr17 | 13732689 | 13733222 | 534 | * | 8,237347 | -6,223310448 | 1,761703 | -3,53255 | 0,000412 | 0,275349208 | 2,237403 | Intron (ENSMUST0000 | 17 | 13741718 | 13748695 | 6978 | 1 | 272667 | ENSMUST00000144015.2 | -8496 | ENSMUSG00000079711 | Smok4a | sperm motility kinase 4A |
| chr18 | 65810846 | 65811540 | 695 | * | 8,993821 | -6,220505717 | 1,911417 | -3,25439 | 0,001136 | 0,511299448 | 1,360622 | Intron (ENSMUST0000 | 18 | 65831339 | 65884608 | 53270 | 1 | 319888 | ENSMUST00000115097.8 | -19799 | ENSMUSG00000046610 | Oacyl | O-acyltransferase like |
| chr5 | 58801175 | 58802104 | 930 | * | 3,691237 | -6,218621095 | 2,300051 | -2,70369 | 0,006857 | 1 | 0 | Distal Intergenic | 5 | 58600582 | 58610342 | 9761 | 2 | 78163 | ENSMUST00000199751.2 | -190833 | ENSMUSG00000105234 | 4930459L07Rik | RIKEN cDNA 4930459L07 gene |
| chr6 | 1,01E+08 | 1,01E+08 | 632 | * | 6,922868 | -6,216060488 | 1,846777 | -3,3659 | 0,000763 | 1 | 0 | Distal Intergenic | 6 | 1,01E+08 | 1,01E+08 | 6027 | 1 | 73395 | ENSMUST00000187986.2 | 34816 | ENSMUSG00000101553 | 1700049E22Rik | RIKEN cDNA 1700049E22 gene |
| chr14 | 78793278 | 78793987 | 710 | * | 4,313553 | -6,213944841 | 2,129799 | -2,91762 | 0,003527 | 1 | 0 | Distal Intergenic | 14 | 78729686 | 78774248 | 44563 | 2 | 219181 | ENSMUST00000123853.9 | -19030 | ENSMUSG00000022016 | Akap11 | A kinase (PRKA) anchor protein 11 |
| chr3 | 1,05E+08 | 1,05E+08 | 1407 | * | 9,280007 | -6,213688364 | 1,842469 | -3,37248 | 0,000745 | 0,352774915 | 2,237403 | Distal Intergenic | 3 | 1,05E+08 | 1,05E+08 | 10133 | 2 | 78635 | ENSMUST00000143893.2 | 174721 | ENSMUSG00000074346 | Kcnd3os | potassium voltage-gated channel, Shal-related family, member 3, opposite strand |
| chr10 | 1,25E+08 | 1,25E+08 | 741 | * | 4,902119 | -6,21120427 | 2,078242 | -2,98868 | 0,002802 | 1 | 0 | Exon (ENSMUST00000 | 10 | 1,25E+08 | 1,25E+08 | 23819 | 1 | 1,03E+08 | ENSMUST00000219518.2 | -140355 | | Gm36719 | predicted gene, 36719 |
| chr18 | 43736009 | 43736437 | 429 | * | 4,516262 | -6,21040437 | 2,290713 | -2,71112 | 0,006706 | 1 | 0 | Intron (ENSMUST0000 | 18 | 43702690 | 43707310 | 4621 | 2 | 76217 | ENSMUST00000235168.2 | -28699 | ENSMUSG00000024502 | Jakmip2 | janus kinase and microtubule interacting protein 2 |
| chr16 | 78536601 | 78537324 | 724 | * | 5,666692 | -6,208817901 | 1,952125 | -3,18054 | 0,00147 | 1 | 0 | Distal Intergenic | 16 | 78491922 | 78505971 | 14050 | 1 | 78812 | ENSMUST00000231496.2 | 44679 | ENSMUSG00000116786 | 4930570E03Rik | RIKEN cDNA 4930570E03 gene |
| chr15 | 11540563 | 11541238 | 676 | * | 5,425393 | -6,208745208 | 2,017648 | -3,07722 | 0,002089 | 1 | 0 | Distal Intergenic | 15 | 11382387 | 11399751 | 17365 | 2 | 110960 | ENSMUST00000022849.7 | -140812 | ENSMUSG00000022241 | Tars | threonyl-tRNA synthetase |
| chr16 | 27546228 | 27546909 | 682 | * | 4,190141 | -6,208243407 | 2,222318 | -2,79359 | 0,005213 | 1 | 0 | Distal Intergenic | 16 | 27668682 | 27744880 | 76199 | 1 | 1E+08 | ENSMUST00000231532.2 | -121773 | ENSMUSG00000116654 | Gm10823 | predicted gene 10823 |
| chrX | 84624052 | 84624774 | 723 | * | 4,988418 | -6,203095623 | 2,17844 | -2,84749 | 0,004406 | 1 | 0 | Intron (ENSMUST0000 | 20 | 84617756 | 84657589 | 39834 | 1 | 66724 | ENSMUST00000146063.2 | 6296 | ENSMUSG00000035476 | Tab3 | TGF-beta activated kinase 1/MAP3K7 binding protein 3 |
| chr5 | 59254472 | 59255371 | 900 | * | 5,229661 | -6,197883472 | 2,15208 | -2,87995 | 0,003977 | 1 | 0 | Distal Intergenic | 5 | 59372289 | 59372395 | 107 | 2 | 1,15E+08 | ENSMUST00000239835.1 | 117024 | ENSMUSG00000119859 | Gm22308 | predicted gene, 22308 |
| chr1 | 1,22E+08 | 1,22E+08 | 544 | * | 6,305314 | -6,196330001 | 1,938009 | -3,19727 | 0,001387 | 1 | 0 | Distal Intergenic | 1 | 1,21E+08 | 1,21E+08 | 14155 | 2 | 66942 | ENSMUST00000001724.12 | -316363 | ENSMUSG00000001674 | Ddx18 | DEAD box helicase 18 |
| chr15 | 3681963 | 3682523 | 561 | * | 5,5668 | -6,195209932 | 1,953057 | -3,17206 | 0,001514 | 1 | 0 | Distal Intergenic | 15 | 3632754 | 3632879 | 126 | 2 | 1,15E+08 | ENSMUST00000116945.4 | -49084 | ENSMUSG00000080595 | Gm22031 | predicted gene, 22031 |
| chr7 | 26613214 | 26613680 | 467 | * | 5,503914 | -6,193163338 | 1,929216 | -3,2102 | 0,001326 | 1 | 0 | Distal Intergenic | 7 | 26631480 | 26638805 | 7326 | 2 | 233005 | ENSMUST00000170227.3 | 25125 | ENSMUSG00000091867 | Cyp2a22 | cytochrome P450, family 2, subfamily a, polypeptide 22 |
| chr3 | 1,32E+08 | 1,32E+08 | 688 | * | 4,432683 | -6,192574792 | 2,175665 | -2,84629 | 0,004423 | 1 | 0 | Exon (ENSMUST00000 | 3 | 1,32E+08 | 1,32E+08 | 10365 | 2 | 13722 | ENSMUST00000197793.5 | -12615 | ENSMUSG00000028029 | Aimp1 | aminoacyl tRNA synthetase complex-interacting multifunctional protein 1 |
| chr1 | 39197517 | 39198147 | 631 | * | 7,080168 | -6,191426545 | 1,839565 | -3,3657 | 0,000763 | 1 | 0 | Distal Intergenic | 1 | 39225977 | 39226073 | 97 | 2 | 1,02E+08 | ENSMUST00000184059.3 | 27926 | ENSMUSG00000098833 | Mir6349 | microRNA 6349 |
| chr1 | 1,03E+08 | 1,03E+08 | 802 | * | 6,668517 | -6,189894933 | 1,83019 | -3,38211 | 0,000719 | 1 | 0 | Intron (ENSMUST0000 | 1 | 1,03E+08 | 1,03E+08 | 134 | 2 | 1,15E+08 | ENSMUST00000157283.3 | -423922 | ENSMUSG00000087908 | Gm22034 | predicted gene, 22034 |
| chr2 | 1,26E+08 | 1,26E+08 | 1431 | * | 7,774793 | -6,189327327 | 1,914353 | -3,23312 | 0,001224 | 0,525184597 | 1,360622 | Intron (ENSMUST0000 | 2 | 1,26E+08 | 1,26E+08 | 55549 | 1 | 14178 | ENSMUST00000110442.2 | 9494 | ENSMUSG00000027208 | Fgf7 | fibroblast growth factor 7 |
| chr4 | 48253143 | 48254066 | 924 | * | 7,223783 | -6,188407249 | 1,833939 | -3,37438 | 0,00074 | 1 | 0 | Intron (ENSMUST0000 | 4 | 48238249 | 48244474 | 6226 | 2 | 76299 | ENSMUST00000132927.8 | -8669 | ENSMUSG00000028343 | Erp44 | endoplasmic reticulum protein 44 |
| chr1 | 1,66E+08 | 1,66E+08 | 776 | * | 6,015461 | -6,186406008 | 1,872092 | -3,30454 | 0,000951 | 1 | 0 | Promoter (<=1kb) | 1 | 1,66E+08 | 1,66E+08 | 18723 | 2 | 226604 | ENSMUST00000046662.8 | 0 | ENSMUSG00000051081 | Gm4847 | predicted gene 4847 |
| chr19 | 54693766 | 54694669 | 904 | * | 8,268658 | -6,186083055 | 1,818564 | -3,40163 | 0,00067 | 0,416350185 | 1,425929 | Distal Intergenic | 19 | 55058166 | 55087883 | 29718 | 2 | 14732 | ENSMUST00000061856.6 | 393214 | ENSMUSG00000024978 | Gpam | glycerol-3-phosphate acyltransferase, mitochondrial |
| chr4 | 34688360 | 34688934 | 575 | * | 5,808416 | -6,184663396 | 1,896835 | -3,26052 | 0,001112 | 1 | 0 | Promoter (<=1kb) | 4 | 34663257 | 34687438 | 24182 | 2 | 24060 | ENSMUST00000029970.14 | -922 | ENSMUSG00000028293 | Slc35a1 | solute carrier family 35 (CMP-sialic acid transporter), member 1 |
| chr14 | 93364872 | 93365359 | 488 | * | 4,792863 | -6,184624673 | 1,976809 | -3,12859 | 0,001756 | 1 | 0 | Intron (ENSMUST0000 | 14 | 93375466 | 93375625 | 160 | 2 | 1,15E+08 | ENSMUST00000083080.3 | 10266 | ENSMUSG00000065014 | Gm23509 | predicted gene, 23509 |
| chr14 | 71485618 | 71486983 | 1366 | * | 5,613702 | -6,176361661 | 2,124236 | -2,90757 | 0,003643 | 1 | 0 | Promoter (1-2kb) | 14 | 71488713 | 71508487 | 19775 | 1 | 1E+08 | ENSMUST00000228402.2 | -1730 | ENSMUSG00000115139 | Gm4251 | predicted gene 4251 |
| chr2 | 8876056 | 8876768 | 713 | * | 5,221721 | -6,175669075 | 2,021878 | -3,05442 | 0,002255 | 1 | 0 | Distal Intergenic | 2 | 9194326 | 9202288 | 7963 | 1 | 73438 | ENSMUST00000132414.2 | -317558 | ENSMUSG00000085580 | 1700061F12Rik | RIKEN cDNA 1700061F12 gene |
| chr13 | 1,02E+08 | 1,02E+08 | 305 | * | 4,649589 | -6,173870286 | 2,03359 | -3,03595 | 0,002398 | 1 | 0 | Distal Intergenic | 13 | 1,02E+08 | 1,02E+08 | 4376 | 2 | 1,18E+08 | ENSMUST00000225534.2 | -39081 | ENSMUSG00000114652 | Gm41043 | predicted gene, 41043 |
| chrX | 56314716 | 56315511 | 796 | * | 4,138462 | -6,173770334 | 2,152571 | -2,86809 | 0,00413 | 1 | 0 | Intron (ENSMUST0000 | 20 | 56278800 | 56326954 | 48155 | 2 | 73341 | ENSMUST00000114735.9 | 11443 | ENSMUSG00000031133 | Arhgef6 | Rac/Cdc42 guanine nucleotide exchange factor (GEF) 6 |
| chr3 | 17875025 | 17875548 | 524 | * | 4,701407 | -6,173601284 | 2,326573 | -2,65352 | 0,007966 | 1 | 0 | Distal Intergenic | 3 | 17857785 | 17857911 | 127 | 2 | 1,15E+08 | ENSMUST00000157268.3 | -17114 | ENSMUSG00000087893 | Gm23441 | predicted gene, 23441 |
| chr14 | 74543975 | 74545050 | 1076 | * | 5,501817 | -6,172019549 | 1,92838 | -3,20062 | 0,001371 | 1 | 0 | Intron (ENSMUST0000 | 14 | 74592967 | 74607181 | 14215 | 1 | 74425 | ENSMUST00000226927.2 | -47917 | ENSMUSG00000115845 | 4933402J15Rik | RIKEN cDNA 4933402J15 gene |
| chr5 | 78830109 | 78830851 | 743 | * | 5,179851 | -6,171677543 | 1,986544 | -3,10674 | 0,001892 | 1 | 0 | Distal Intergenic | 5 | 78927706 | 78927841 | 136 | 1 | 1,15E+08 | ENSMUST00000103990.3 | -96855 | ENSMUSG00000077178 | Gm25756 | predicted gene, 25756 |
| chr4 | 24986966 | 24987556 | 591 | * | 3,733679 | -6,171102963 | 2,253597 | -2,73833 | 0,006175 | 1 | 0 | Intron (ENSMUST0000 | 4 | 24973419 | 25009233 | 35815 | 1 | 81006 | ENSMUST00000038920.2 | 13547 | ENSMUSG00000040372 | Gpr63 | G protein-coupled receptor 63 |
| chr13 | 10588006 | 10588782 | 777 | * | 8,04413 | -6,170479668 | 1,83111 | -3,3698 | 0,000752 | 1 | 0 | Exon (ENSMUST00000 | 13 | 10604695 | 10726784 | 122090 | 1 | 1,05E+08 | ENSMUST00000222066.2 | -15913 | | Gm40666 | predicted gene, 40666 |
| chr18 | 83297022 | 83297521 | 500 | * | 7,30295 | -6,167414914 | 1,845932 | -3,34108 | 0,000835 | 1 | 0 | Distal Intergenic | 18 | 83310896 | 83324586 | 13691 | 2 | 70960 | ENSMUST00000235143.2 | 27065 | ENSMUSG00000118333 | 4921531P14Rik | RIKEN cDNA 4921531P14 gene |
| chr5 | 62636614 | 62637365 | 752 | * | 7,480434 | -6,16733116 | 1,853898 | -3,32668 | 0,000879 | 1 | 0 | Distal Intergenic | 5 | 62680751 | 62680882 | 132 | 2 | 1,15E+08 | ENSMUST00000104352.4 | 43517 | ENSMUSG00000119858 | Gm24079 | predicted gene, 24079 |
| chr7 | 15847672 | 15848348 | 677 | * | 6,295249 | -6,166521916 | 1,95256 | -3,15817 | 0,001588 | 1 | 0 | Promoter (<=1kb) | 7 | 15847569 | 15850484 | 2916 | 1 | 108124 | ENSMUST00000150208.8 | 103 | ENSMUSG00000006024 | Napa | N-ethylmaleimide sensitive fusion protein attachment protein alpha |
| chr4 | 68505215 | 68506193 | 979 | * | 4,353914 | -6,165571778 | 2,147158 | -2,8715 | 0,004085 | 1 | 0 | Distal Intergenic | 4 | 68744818 | 68747595 | 2778 | 2 | 56710 | ENSMUST00000140572.2 | 241402 | ENSMUSG00000028351 | Brinp1 | bone morphogenic protein/retinoic acid inducible neural specific 1 |
| chr14 | 1,23E+08 | 1,23E+08 | 655 | * | 4,972099 | -6,165413994 | 1,958617 | -3,14784 | 0,001645 | 1 | 0 | Intron (ENSMUST0000 | 14 | 1,23E+08 | 1,23E+08 | 6306 | 1 | 380934 | ENSMUST00000226639.2 | -15425 | ENSMUSG00000115016 | AA536875 | expressed sequence AA536875 |
| chr7 | 83197247 | 83198290 | 1044 | * | 7,885535 | -6,165143297 | 1,798923 | -3,42713 | 0,00061 | 1 | 0 | Promoter (1-2kb) | 7 | 83192078 | 83199464 | 7387 | 2 | 1E+08 | ENSMUST00000208877.2 | 1174 | | Gm10610 | predicted gene 10610 |
| chrX | 1,22E+08 | 1,22E+08 | 452 | * | 3,771721 | -6,162631232 | 2,296223 | -2,68381 | 0,007279 | 1 | 0 | Distal Intergenic | 20 | 1,22E+08 | 1,22E+08 | 42157 | 1 | 73061 | ENSMUST00000113343.8 | -145811 | ENSMUSG00000079450 | Cldn34c1 | claudin 34C1 |
| chr4 | 70821169 | 70821713 | 545 | * | 4,595527 | -6,160154224 | 2,134543 | -2,88594 | 0,003903 | 1 | 0 | Distal Intergenic | 4 | 70993346 | 71049370 | 56025 | 2 | 1,03E+08 | ENSMUST00000143796.2 | 227657 | ENSMUSG00000084937 | Gm11228 | predicted gene 11228 |
| chr1 | 83554282 | 83555414 | 1133 | * | 8,367214 | -6,159881029 | 1,779701 | -3,46119 | 0,000538 | 0,376805967 | 1,425929 | Distal Intergenic | 1 | 83231860 | 83385921 | 154062 | 2 | 77629 | ENSMUST00000160953.8 | -168361 | ENSMUSG00000026163 | Sphkap | SPHK1 interactor, AKAP domain containing |
| chr1 | 1,55E+08 | 1,55E+08 | 985 | * | 4,99488 | -6,15736842 | 2,143446 | -2,87265 | 0,00407 | 1 | 0 | Intron (ENSMUST0000 | 1 | 1,54E+08 | 1,55E+08 | 465675 | 2 | 12290 | ENSMUST00000211821.2 | -5874 | ENSMUSG00000004110 | Cacna1e | calcium channel, voltage-dependent, R type, alpha 1E subunit |
| chr11 | 45410716 | 45411297 | 582 | * | 6,347289 | -6,154638533 | 1,866492 | -3,29744 | 0,000976 | 1 | 0 | Distal Intergenic | 11 | 45530210 | 45530341 | 132 | 1 | 1,15E+08 | ENSMUST00000104132.3 | -118913 | ENSMUSG00000077320 | Gm22284 | predicted gene, 22284 |
| chr3 | 9073562 | 9074325 | 764 | * | 8,111553 | -6,153807123 | 1,764586 | -3,4874 | 0,000488 | 1 | 0 | Distal Intergenic | 3 | 8990653 | 9069783 | 79131 | 2 | 21985 | ENSMUST00000120143.8 | -3779 | ENSMUSG00000027506 | Tpd52 | tumor protein D52 |
| chr5 | 1,29E+08 | 1,29E+08 | 466 | * | 5,66913 | -6,153509036 | 1,956909 | -3,1445 | 0,001664 | 1 | 0 | Distal Intergenic | 5 | 1,3E+08 | 1,3E+08 | 70164 | 1 | 231769 | ENSMUST00000053737.9 | -102148 | ENSMUSG00000029439 | Sfswap | splicing factor SWAP |
| chr8 | 46808956 | 46809451 | 496 | * | 6,62456 | -6,151403685 | 1,971246 | -3,12057 | 0,001805 | 1 | 0 | Distal Intergenic | 8 | 46816288 | 46820241 | 3954 | 1 | 1,03E+08 | ENSMUST00000211698.2 | -6837 | ENSMUSG00000110035 | Gm30931 | predicted gene, 30931 |
| chr14 | 1,14E+08 | 1,14E+08 | 751 | * | 6,638644 | -6,151102993 | 1,880935 | -3,27024 | 0,001075 | 1 | 0 | Distal Intergenic | 14 | 1,15E+08 | 1,15E+08 | 29363 | 1 | 75822 | ENSMUST00000226121.2 | -633074 | ENSMUSG00000115320 | 4930524C18Rik | RIKEN cDNA 4930524C18 gene |
| chr18 | 7809969 | 7811190 | 1222 | * | 8,475192 | -6,14600011 | 1,771573 | -3,46923 | 0,000522 | 0,367692073 | 1,449706 | Distal Intergenic | 18 | 7835177 | 7835283 | 107 | 1 | 1,15E+08 | ENSMUST00000082888.2 | -23987 | ENSMUSG00000064822 | Gm24292 | predicted gene, 24292 |
| chr8 | 1,03E+08 | 1,03E+08 | 392 | * | 5,260125 | -6,144091743 | 2,059917 | -2,98269 | 0,002857 | 1 | 0 | Distal Intergenic | 8 | 1,03E+08 | 1,03E+08 | 2716 | 2 | 12552 | ENSMUST00000210578.2 | 623403 | ENSMUSG00000031673 | Cdh11 | cadherin 11 |
| chr3 | 1,2E+08 | 1,2E+08 | 1008 | * | 7,513064 | -6,143993945 | 1,833494 | -3,35098 | 0,000805 | 1 | 0 | Distal Intergenic | 3 | 1,2E+08 | 1,2E+08 | 23918 | 2 | 56195 | ENSMUST00000199026.5 | -241608 | ENSMUSG00000028134 | Ptbp2 | polypyrimidine tract binding protein 2 |
| chr2 | 1,39E+08 | 1,39E+08 | 720 | * | 5,153897 | -6,1436538 | 2,038549 | -3,01374 | 0,00258 | 1 | 0 | Intron (ENSMUST0000 | 2 | 1,39E+08 | 1,39E+08 | 143144 | 1 | 228677 | ENSMUST00000110083.8 | -98323 | ENSMUSG00000039092 | Sptlc3 | serine palmitoyltransferase, long chain base subunit 3 |
| chr14 | 1,17E+08 | 1,17E+08 | 691 | * | 3,657486 | -6,140938025 | 2,372588 | -2,58829 | 0,009645 | 1 | 0 | Intron (ENSMUST0000 | 14 | 1,17E+08 | 1,18E+08 | 701884 | 1 | 23888 | ENSMUST00000123239.2 | -401060 | ENSMUSG00000058571 | Gpc6 | glypican 6 |
| chrX | 1,18E+08 | 1,18E+08 | 817 | * | 4,067905 | -6,140910296 | 2,331555 | -2,63383 | 0,008443 | 1 | 0 | Distal Intergenic | 20 | 1,17E+08 | 1,17E+08 | 1030 | 2 | 245583 | ENSMUST00000073857.6 | -601305 | ENSMUSG00000100133 | Tgif2lx1 | TGFB-induced factor homeobox 2-like, X-linked 1 |
| chr3 | 33423655 | 33424278 | 624 | * | 5,662336 | -6,140285263 | 1,889487 | -3,24971 | 0,001155 | 1 | 0 | Distal Intergenic | 3 | 33447206 | 33447348 | 143 | 2 | 1,15E+08 | ENSMUST00000157635.3 | 23070 | ENSMUSG00000088260 | Gm24780 | predicted gene, 24780 |
| chr8 | 37912974 | 37913635 | 662 | * | 7,232677 | -6,138154817 | 2,040016 | -3,00888 | 0,002622 | 1 | 0 | Distal Intergenic | 8 | 37784631 | 37784732 | 102 | 2 | 1,15E+08 | ENSMUST00000083343.3 | -128242 | ENSMUSG00000065277 | Gm25126 | predicted gene, 25126 |
| chrX | 25608491 | 25609351 | 861 | * | 4,534948 | -6,137217869 | 2,336612 | -2,62655 | 0,008626 | 1 | 0 | Distal Intergenic | 20 | 25577812 | 25598961 | 21150 | 2 | 1E+08 | ENSMUST00000118537.2 | -9530 | ENSMUSG00000080959 | Gm2003 | predicted pseudogene 2003 |
| chr1 | 3655675 | 3656455 | 781 | * | 6,372359 | -6,136514557 | 1,870185 | -3,28123 | 0,001034 | 1 | 0 | Intron (ENSMUST0000 | 1 | 3284705 | 3741721 | 457017 | 2 | 497097 | ENSMUST00000070533.5 | 85266 | ENSMUSG00000051951 | Xkr4 | X-linked Kx blood group related 4 |
| chrX | 1,3E+08 | 1,3E+08 | 883 | * | 4,465373 | -6,135240025 | 2,218964 | -2,76491 | 0,005694 | 1 | 0 | Distal Intergenic | 20 | 1,3E+08 | 1,3E+08 | 129 | 2 | 1,15E+08 | ENSMUST00000104191.3 | -491997 | ENSMUSG00000077379 | Gm25170 | predicted gene, 25170 |
| chr12 | 1,11E+08 | 1,11E+08 | 600 | * | 3,988725 | -6,135143854 | 2,173966 | -2,8221 | 0,004771 | 1 | 0 | Intron (ENSMUST0000 | 12 | 1,11E+08 | 1,11E+08 | 24642 | 2 | 78062 | ENSMUST00000121608.2 | -23030 | ENSMUSG00000083193 | 4930595D18Rik | RIKEN cDNA 4930595D18 gene |
| chr14 | 12947127 | 12947539 | 413 | * | 4,635543 | -6,132532348 | 2,085278 | -2,94087 | 0,003273 | 1 | 0 | Distal Intergenic | 14 | 12583005 | 12585578 | 2574 | 2 | 1E+08 | ENSMUST00000224435.2 | -361549 | ENSMUSG00000089686 | Mnd1-ps | Mnd1 retrotransposed pseudogene |
| chr14 | 68462301 | 68463258 | 958 | * | 5,337809 | -6,131442865 | 1,922293 | -3,18965 | 0,001424 | 1 | 0 | Distal Intergenic | 14 | 68356732 | 68362295 | 5564 | 2 | 18040 | ENSMUST00000022638.6 | -100006 | ENSMUSG00000022054 | Nefm | neurofilament, medium polypeptide |
| chr7 | 41058110 | 41058591 | 482 | * | 3,6233 | -6,129647736 | 2,304596 | -2,65975 | 0,00782 | 1 | 0 | Intron (ENSMUST0000 | 7 | 41043122 | 41046871 | 3750 | 1 | 1,03E+08 | ENSMUST00000170929.2 | 14988 | ENSMUSG00000091864 | Gm17102 | predicted gene 17102 |
| chr10 | 46713609 | 46714197 | 589 | * | 3,939489 | -6,129166135 | 2,296948 | -2,6684 | 0,007621 | 1 | 0 | Promoter (<=1kb) | 10 | 46714442 | 46714603 | 162 | 2 | 1,15E+08 | ENSMUST00000083784.3 | 406 | ENSMUSG00000065718 | Gm25650 | predicted gene, 25650 |
| chr2 | 69759914 | 69761055 | 1142 | * | 8,093495 | -6,127151733 | 1,78897 | -3,42496 | 0,000615 | 0,410018041 | 1,360622 | Intron (ENSMUST0000 | 2 | 69766313 | 69775645 | 9333 | 1 | 68795 | ENSMUST00000142563.2 | -5258 | ENSMUSG00000044308 | Ubr3 | ubiquitin protein ligase E3 component n-recognin 3 |
| chr14 | 3725711 | 3726658 | 948 | * | 4,796779 | -6,126252417 | 1,985599 | -3,08534 | 0,002033 | 1 | 0 | Intron (ENSMUST0000 | 14 | 3655393 | 3655527 | 135 | 2 | 1,15E+08 | ENSMUST00000104501.3 | -70184 | ENSMUSG00000077689 | Gm22499 | predicted gene, 22499 |
| chr1 | 13687757 | 13688978 | 1222 | * | 6,605215 | -6,125390234 | 1,835378 | -3,3374 | 0,000846 | 1 | 0 | Distal Intergenic | 1 | 13700219 | 13708635 | 8417 | 2 | 212442 | ENSMUST00000156222.2 | 19657 | ENSMUSG00000025937 | Lactb2 | lactamase, beta 2 |
| chr14 | 61722507 | 61723415 | 909 | * | 7,499386 | -6,121738103 | 1,859125 | -3,29281 | 0,000992 | 1 | 0 | Distal Intergenic | 14 | 61730137 | 61737831 | 7695 | 1 | 75349 | ENSMUST00000228251.2 | -6722 | ENSMUSG00000115656 | 4930556J02Rik | RIKEN cDNA 4930556J02 gene |
| chr1 | 1,77E+08 | 1,77E+08 | 837 | * | 8,212976 | -6,11533391 | 1,855277 | -3,29618 | 0,00098 | 0,476080922 | 1,449706 | Distal Intergenic | 1 | 1,77E+08 | 1,77E+08 | 4887 | 1 | 30928 | ENSMUST00000195002.2 | -77720 | ENSMUSG00000063659 | Zbtb18 | zinc finger and BTB domain containing 18 |
| chrX | 50010730 | 50011401 | 672 | * | 5,665024 | -6,1149536 | 1,930967 | -3,16678 | 0,001541 | 1 | 0 | Intron (ENSMUST0000 | 20 | 49984057 | 50031587 | 47531 | 2 | 385354 | ENSMUST00000060650.7 | 20186 | ENSMUSG00000036131 | Frmd7 | FERM domain containing 7 |
| chrX | 1,04E+08 | 1,04E+08 | 563 | * | 3,7848 | -6,113153073 | 2,332302 | -2,62108 | 0,008765 | 1 | 0 | Distal Intergenic | 20 | 1,04E+08 | 1,04E+08 | 2316 | 2 | 272790 | ENSMUST00000033575.7 | -23754 | ENSMUSG00000031224 | Magee2 | MAGE family member E2 |
| chrX | 85229938 | 85230972 | 1035 | * | 5,859924 | -6,110739472 | 1,956329 | -3,12358 | 0,001787 | 1 | 0 | Distal Intergenic | 20 | 85235370 | 85239553 | 4184 | 1 | 11614 | ENSMUST00000026036.5 | -4398 | ENSMUSG00000025056 | Nr0b1 | nuclear receptor subfamily 0, group B, member 1 |
| chrX | 92666407 | 92667397 | 991 | * | 6,115674 | -6,109710864 | 2,06604 | -2,95721 | 0,003104 | 1 | 0 | Exon (ENSMUST00000 | 20 | 92666904 | 92675721 | 8818 | 2 | 18968 | ENSMUST00000124236.2 | 8324 | ENSMUSG00000006678 | Pola1 | polymerase (DNA directed), alpha 1 |
| chrX | 1,38E+08 | 1,38E+08 | 896 | * | 6,10477 | -6,109117852 | 1,850059 | -3,30212 | 0,00096 | 1 | 0 | Intron (ENSMUST0000 | 20 | 1,38E+08 | 1,38E+08 | 74212 | 1 | 102871 | ENSMUST00000113030.3 | 20325 | ENSMUSG00000042498 | Radx | RPA1 related single stranded DNA binding protein, X-linked |
| chr19 | 26032720 | 26033570 | 851 | * | 7,024892 | -6,103535348 | 1,823915 | -3,34639 | 0,000819 | 1 | 0 | Distal Intergenic | 19 | 26015816 | 26023831 | 8016 | 2 | 73433 | ENSMUST00000237184.2 | -8889 | ENSMUSG00000118016 | 1700048O14Rik | RIKEN cDNA 1700048O14 gene |
| chr19 | 37140193 | 37140761 | 569 | * | 6,4433 | -6,103516562 | 1,915267 | -3,18677 | 0,001439 | 1 | 0 | Intron (ENSMUST0000 | 19 | 37151243 | 37173940 | 22698 | 1 | 319931 | ENSMUST00000180609.2 | -10482 | ENSMUSG00000097591 | A330032B11Rik | RIKEN cDNA A330032B11 gene |
| chr3 | 6063023 | 6063683 | 661 | * | 6,112086 | -6,10287245 | 1,970049 | -3,09783 | 0,001949 | 1 | 0 | Distal Intergenic | 3 | 5625560 | 5641299 | 15740 | 2 | 19302 | ENSMUST00000059021.10 | -421724 | ENSMUSG00000040374 | Pex2 | peroxisomal biogenesis factor 2 |
| chr16 | 7880399 | 7881482 | 1084 | * | 6,56549 | -6,101822454 | 1,923982 | -3,17145 | 0,001517 | 1 | 0 | Distal Intergenic | 16 | 8154857 | 8154960 | 104 | 1 | 1,15E+08 | ENSMUST00000158968.3 | -273375 | ENSMUSG00000089593 | Gm26159 | predicted gene, 26159 |
| chrX | 1,18E+08 | 1,18E+08 | 914 | * | 4,217601 | -6,101296155 | 2,188424 | -2,78799 | 0,005304 | 1 | 0 | Distal Intergenic | 20 | 1,17E+08 | 1,17E+08 | 1030 | 2 | 245583 | ENSMUST00000073857.6 | -655335 | ENSMUSG00000100133 | Tgif2lx1 | TGFB-induced factor homeobox 2-like, X-linked 1 |
| chr1 | 86409893 | 86410956 | 1064 | * | 4,197739 | -6,101118008 | 2,211997 | -2,75819 | 0,005812 | 1 | 0 | Distal Intergenic | 1 | 86454448 | 86458434 | 3987 | 1 | 19231 | ENSMUST00000045897.15 | -43492 | ENSMUSG00000026238 | Ptma | prothymosin alpha |
| chr13 | 93458496 | 93460595 | 2100 | * | 12,65637 | -6,096766659 | 1,816877 | -3,35563 | 0,000792 | 0,554723541 | 0,738441 | Intron (ENSMUST0000 | 13 | 93444542 | 93541637 | 97096 | 1 | 26556 | ENSMUST00000079086.8 | 13954 | ENSMUSG00000007617 | Homer1 | homer scaffolding protein 1 |
| chr1 | 1,28E+08 | 1,28E+08 | 777 | * | 6,746576 | -6,094136086 | 1,853156 | -3,28852 | 0,001007 | 1 | 0 | Promoter (1-2kb) | 1 | 1,28E+08 | 1,28E+08 | 7828 | 2 | 226409 | ENSMUST00000143389.2 | -1125 | ENSMUSG00000036086 | Zranb3 | zinc finger, RAN-binding domain containing 3 |
| chr17 | 55024339 | 55024995 | 657 | * | 5,844808 | -6,094101816 | 1,940251 | -3,14088 | 0,001684 | 1 | 0 | Distal Intergenic | 17 | 54958964 | 54959100 | 137 | 1 | 1,15E+08 | ENSMUST00000158889.3 | 65375 | ENSMUSG00000089514 | Gm26291 | predicted gene, 26291 |
| chr19 | 3107511 | 3108913 | 1403 | * | 9,526363 | -6,092753788 | 1,731724 | -3,51832 | 0,000434 | 0,337020859 | 1,449706 | Intron (ENSMUST0000 | 19 | 3186984 | 3187090 | 107 | 2 | 1,15E+08 | ENSMUST00000101978.3 | 78177 | ENSMUSG00000118966 | Gm25431 | predicted gene, 25431 |
| chr14 | 3388549 | 3389553 | 1005 | * | 5,763105 | -6,092676821 | 2,019137 | -3,01747 | 0,002549 | 1 | 0 | Distal Intergenic | 14 | 3575429 | 3839361 | 263933 | 1 | 218793 | ENSMUST00000124353.2 | -185876 | ENSMUSG00000058317 | Ube2e2 | ubiquitin-conjugating enzyme E2E 2 |
| chr8 | 87564361 | 87565030 | 670 | * | 5,800534 | -6,088343499 | 2,000346 | -3,04365 | 0,002337 | 1 | 0 | Intron (ENSMUST0000 | 8 | 87534788 | 87570673 | 35886 | 2 | 1,05E+08 | ENSMUST00000209389.2 | 5643 | | Gm39215 | predicted gene, 39215 |
| chr18 | 59623884 | 59624660 | 777 | * | 5,494793 | -6,084421207 | 2,059578 | -2,95421 | 0,003135 | 1 | 0 | Intron (ENSMUST0000 | 18 | 59712702 | 59712807 | 106 | 1 | 1,02E+08 | ENSMUST00000183687.3 | -88042 | ENSMUSG00000098287 | Mir6355 | microRNA 6355 |
| chr14 | 1,16E+08 | 1,16E+08 | 905 | * | 4,291805 | -6,084020935 | 2,203964 | -2,76049 | 0,005771 | 1 | 0 | Intron (ENSMUST0000 | 14 | 1,16E+08 | 1,16E+08 | 13002 | 2 | 75088 | ENSMUST00000176582.2 | -22486 | ENSMUSG00000093551 | 4930505G20Rik | RIKEN cDNA 4930505G20 gene |

| | | | | | | | | | | | | | | | | | | | | | | | |
|---|---|---|---|---|---|---|---|---|---|---|---|---|---|---|---|---|---|---|---|---|---|---|---|
| chr6 | 67329354 | 67329604 | 251 | * | 4,404225 | -6,083895971 | 2,208241 | -2,75509 | 0,005868 | 1 | 0 | Intron (ENSMUST0000( | 6 | 67268302 | 67316678 | 48377 | 2 | 16162 | ENSMUST00000117441.8 | -12676 | ENSMUSG00000018341 | Il12rb2 | interleukin 12 receptor, beta 2 |
| chrX | 11541231 | 11541971 | 741 | * | 6,266883 | -6,083246484 | 1,904508 | -3,19413 | 0,001403 | 1 | 0 | Distal Intergenic | 20 | 11716723 | 11716829 | 107 | 1 | 1,15E+08 | ENSMUST00000083891.3 | -174752 | ENSMUSG00000065825 | Gm26314 | predicted gene, 26314 |
| chr1 | 1,87E+08 | 1,87E+08 | 667 | * | 5,679551 | -6,081378251 | 1,906342 | -3,19008 | 0,001422 | 1 | 0 | Distal Intergenic | 1 | 1,87E+08 | 1,87E+08 | 3212 | 1 | 110957 | ENSMUST00000045108.2 | 75791 | ENSMUSG00000039224 | D1Pas1 | DNA segment, Chr 1, Pasteur Institute 1 |
| chr2 | 90760647 | 90761394 | 748 | * | 5,375172 | -6,081281018 | 2,020007 | -3,01053 | 0,002608 | 1 | 0 | Distal Intergenic | 2 | 90770727 | 90818099 | 47373 | 1 | 13046 | ENSMUST00000150546.2 | -9333 | ENSMUSG00000005506 | Celf1 | CUGBP, Elav-like family member 1 |
| chr14 | 98799165 | 98799754 | 590 | * | 4,187555 | -6,081182422 | 2,157318 | -2,81886 | 0,004819 | 1 | 0 | Distal Intergenic | 14 | 98861626 | 98861732 | 107 | 1 | 1,15E+08 | ENSMUST00000180098.3 | -61872 | ENSMUSG00000119462 | Gm26139 | predicted gene, 26139 |
| chr4 | 7690748 | 7691197 | 450 | * | 3,790555 | -6,080767683 | 2,279662 | -2,6674 | 0,007644 | 1 | 0 | Distal Intergenic | 4 | 7820916 | 7821047 | 132 | 1 | 1,15E+08 | ENSMUST00000179446.2 | -129719 | ENSMUSG00000094771 | Gm23860 | predicted gene, 23860 |
| chr3 | 5053611 | 5054354 | 744 | * | 7,994704 | -6,080623357 | 1,784305 | -3,40784 | 0,000655 | 1 | 0 | Distal Intergenic | 3 | 5155336 | 5159840 | 4505 | 1 | 1,01E+08 | ENSMUST00000181749.3 | -100982 | ENSMUSG00000096922 | Gm17308 | predicted gene, 17308 |
| chrX | 94123645 | 94124448 | 804 | * | 5,04505 | -6,078490276 | 2,154083 | -2,82185 | 0,004775 | 1 | 0 | Promoter (<=1kb) | 20 | 94095210 | 94125235 | 30026 | 2 | 236915 | ENSMUST00000196354.5 | 787 | ENSMUSG00000025656 | Arhgef9 | CDC42 guanine nucleotide exchange factor (GEF) 9 |
| chr5 | 21596084 | 21596596 | 513 | * | 4,473863 | -6,078399217 | 2,141542 | -2,83833 | 0,004535 | 1 | 0 | Intron (ENSMUST0000( | 5 | 21577640 | 21583372 | 5733 | 1 | 14190 | ENSMUST00000035799.6 | 18444 | ENSMUSG00000039899 | Fgl2 | fibrinogen-like protein 2 |
| chr19 | 42471138 | 42472181 | 1044 | * | 10,24512 | -6,076449939 | 1,719 | -3,53488 | 0,000408 | 0,360585824 | 1,173325 | Distal Intergenic | 19 | 42507198 | 42580782 | 73585 | 1 | 52013 | ENSMUST00000026188.10 | -35017 | ENSMUSG00000025184 | R3hcc1l | R3H domain and coiled-coil containing 1 like |
| chr10 | 50318207 | 50319209 | 1003 | * | 4,992831 | -6,075560459 | 2,009397 | -3,02357 | 0,002498 | 1 | 0 | Distal Intergenic | 10 | 50468765 | 50727581 | 258817 | 1 | 77987 | ENSMUST00000035606.10 | -149556 | ENSMUSG00000038774 | Ascc3 | activating signal cointegrator 1 complex subunit 3 |
| chrX | 1,06E+08 | 1,06E+08 | 642 | * | 3,55791 | -6,075267365 | 2,312779 | -2,62683 | 0,008619 | 1 | 0 | Distal Intergenic | 20 | 1,06E+08 | 1,06E+08 | 107 | 2 | 1,15E+08 | ENSMUST00000083799.3 | -78062 | ENSMUSG00000065733 | Gm24491 | predicted gene, 24491 |
| chr13 | 73619683 | 73620759 | 1077 | * | 8,083944 | -6,073938572 | 1,895581 | -3,20426 | 0,001354 | 0,54122637 | 1,360622 | Promoter (<=1kb) | 13 | 73619870 | 73663244 | 43375 | 1 | 210992 | ENSMUST00000223060.2 | 0 | ENSMUSG00000021608 | Lpcat1 | lysophosphatidylcholine acyltransferase 1 |
| chr14 | 92177433 | 92178173 | 741 | * | 4,757449 | -6,072784503 | 2,095091 | -2,89858 | 0,003749 | 1 | 0 | Distal Intergenic | 14 | 92605936 | 92606065 | 130 | 2 | 1,15E+08 | ENSMUST00000104110.3 | 427892 | ENSMUSG00000077298 | Gm24127 | predicted gene, 24127 |
| chr3 | 1,52E+08 | 1,52E+08 | 466 | * | 4,794042 | -6,070380894 | 2,324662 | -2,6113 | 0,00902 | 1 | 0 | Promoter (1-2kb) | 3 | 1,52E+08 | 1,52E+08 | 3144 | 1 | 329777 | ENSMUST00000160651.2 | 1498 | ENSMUSG00000039047 | Pigk | phosphatidylinositol glycan anchor biosynthesis, class K |
| chr4 | 82150955 | 82151622 | 668 | * | 7,499255 | -6,069293229 | 1,807029 | -3,35871 | 0,000783 | 1 | 0 | Distal Intergenic | 4 | 81952907 | 81984130 | 31224 | 2 | 1,03E+08 | ENSMUST00000243987.1 | -166825 | ENSMUSG00000087110 | Gm11264 | predicted gene 11264 |
| chr15 | 4988658 | 4989313 | 656 | * | 7,980348 | -6,064833373 | 1,813951 | -3,34344 | 0,000827 | 1 | 0 | Intron (ENSMUST0000( | 15 | 4977964 | 4991554 | 13591 | 1 | 223825 | ENSMUST00000226849.2 | 10694 | ENSMUSG00000022155 | Mroh2b | maestro heat-like repeat family member 2B |
| chr18 | 87882311 | 87883320 | 1010 | * | 5,75935 | -6,064440507 | 1,893355 | -3,20301 | 0,00136 | 1 | 0 | Distal Intergenic | 18 | 87943555 | 87943688 | 134 | 2 | 1,15E+08 | ENSMUST00000104453.3 | 60368 | ENSMUSG00000077641 | Gm24987 | predicted gene, 24987 |
| chr16 | 75555943 | 75556412 | 470 | * | 5,128309 | -6,064095487 | 1,9937 | -3,04163 | 0,002353 | 1 | 0 | Exon (ENSMUST000002 | 16 | 75556076 | 75563670 | 7595 | 2 | 110920 | ENSMUST00000137806.2 | 7258 | ENSMUSG00000032932 | Hspa13 | heat shock protein 70 family, member 13 |
| chr7 | 1,25E+08 | 1,25E+08 | 542 | * | 4,877101 | -6,063248105 | 2,055516 | -2,94974 | 0,00318 | 1 | 0 | Distal Intergenic | 7 | 1,25E+08 | 1,25E+08 | 54248 | 2 | 71208 | ENSMUST00000205794.2 | 166353 | ENSMUSG00000045928 | 4933440M02Rik | RIKEN cDNA 4933440M02 gene |
| chr7 | 83775727 | 83776401 | 675 | * | 5,201907 | -6,06314646 | 2,030775 | -2,98563 | 0,00283 | 1 | 0 | Intron (ENSMUST0000( | 7 | 83763459 | 83797023 | 33565 | 2 | 70178 | ENSMUST00000131505.2 | 20622 | ENSMUSG00000038459 | Abhd17c | abhydrolase domain containing 17C |
| chr14 | 82982153 | 82982760 | 608 | * | 4,072708 | -6,060747003 | 2,192955 | -2,76374 | 0,005714 | 1 | 0 | Distal Intergenic | 14 | 83643071 | 83643204 | 134 | 1 | 1,15E+08 | ENSMUST00000157384.3 | -660311 | ENSMUSG00000088009 | Gm24774 | predicted gene, 24774 |
| chr14 | 70133031 | 70133737 | 707 | * | 5,064338 | -6,060638279 | 1,980301 | -3,06046 | 0,00221 | 1 | 0 | Intron (ENSMUST0000( | 14 | 70077869 | 70297367 | 219499 | 1 | 73523 | ENSMUST00000022678.5 | 55162 | ENSMUSG00000022085 | Pebp4 | phosphatidylethanolamine binding protein 4 |
| chr1 | 1,54E+08 | 1,54E+08 | 1035 | * | 7,518115 | -6,059296745 | 1,806324 | -3,35449 | 0,000795 | 1 | 0 | Promoter (<=1kb) | 1 | 1,54E+08 | 1,54E+08 | 1074 | 1 | 14645 | ENSMUST00000154576.2 | 758 | ENSMUSG00000026473 | Glul | glutamate-ammonia ligase (glutamine synthetase) |
| chr19 | 36436578 | 36436844 | 267 | * | 4,39524 | -6,055303009 | 2,10104 | -2,88205 | 0,003951 | 1 | 0 | 3' UTR | 19 | 36417054 | 36423690 | 6637 | 1 | 76073 | ENSMUST00000225185.2 | 19524 | ENSMUSG00000024805 | Pcgf5 | polycomb group ring finger 5 |
| chr2 | 68583006 | 68583467 | 462 | * | 4,607394 | -6,054905496 | 2,184856 | -2,77131 | 0,005583 | 1 | 0 | Distal Intergenic | 2 | 68546635 | 68547654 | 1020 | 1 | 75721 | ENSMUST00000135023.2 | 36371 | ENSMUSG00000079324 | 4932414N04Rik | RIKEN cDNA 4932414N04 gene |
| chr14 | 1,15E+08 | 1,15E+08 | 1112 | * | 6,721197 | -6,054829158 | 1,847665 | -3,27702 | 0,001049 | 1 | 0 | Distal Intergenic | 14 | 1,15E+08 | 1,15E+08 | 29363 | 1 | 75822 | ENSMUST00000226121.2 | -542897 | ENSMUSG00000115320 | 4930524C18Rik | RIKEN cDNA 4930524C18 gene |
| chr3 | 80789716 | 80790415 | 700 | * | 6,01045 | -6,052275593 | 2,064072 | -2,9322 | 0,003366 | 1 | 0 | Intron (ENSMUST0000( | 3 | 80752268 | 80819396 | 67129 | 2 | 14658 | ENSMUST00000194085.6 | 28981 | ENSMUSG00000028020 | Glrb | glycine receptor, beta subunit |
| chr19 | 41196120 | 41196816 | 697 | * | 6,897614 | -6,051008688 | 1,912564 | -3,16382 | 0,001557 | 1 | 0 | Promoter (<=1kb) | 19 | 41071192 | 41195274 | 124083 | 2 | 24087 | ENSMUST00000025986.15 | -846 | ENSMUSG00000025013 | Tll2 | tolloid-like 2 |
| chr9 | 49848536 | 49849447 | 912 | * | 5,341583 | -6,049554249 | 2,084306 | -2,90243 | 0,003703 | 1 | 0 | Distal Intergenic | 9 | 49737169 | 49737275 | 107 | 2 | 1,15E+08 | ENSMUST00000178565.3 | -111261 | ENSMUSG00000119677 | Gm25633 | predicted gene, 25633 |
| chr7 | 65426228 | 65427259 | 1032 | * | 8,086771 | -6,049217409 | 1,78391 | -3,39099 | 0,000696 | 1 | 0 | Distal Intergenic | 7 | 65452758 | 65455998 | 3241 | 1 | 1,01E+08 | ENSMUST00000243649.1 | -25499 | ENSMUSG00000100094 | 1810008I18Rik | RIKEN cDNA 1810008I18 gene |
| chr7 | 79230385 | 79231401 | 1017 | * | 10,21579 | -6,049211731 | 1,788874 | -3,38157 | 0,000721 | 0,454149353 | 1,211315 | Distal Intergenic | 7 | 79243117 | 79244677 | 1561 | 2 | 56315 | ENSMUST00000172788.3 | 13276 | ENSMUSG00000030549 | Rhcg | Rhesus blood group-associated C glycoprotein |
| chr17 | 20876717 | 20877456 | 740 | * | 6,369847 | -6,04913767 | 1,856529 | -3,2583 | 0,001121 | 1 | 0 | Distal Intergenic | 17 | 20907770 | 20908666 | 897 | 1 | 171225 | ENSMUST00000061756.4 | -30314 | ENSMUSG00000042848 | Vmn1r226 | vomeronasal 1 receptor 226 |
| chr2 | 1,13E+08 | 1,13E+08 | 694 | * | 5,293836 | -6,048301138 | 2,060298 | -2,93564 | 0,003329 | 1 | 0 | Intron (ENSMUST0000( | 2 | 1,13E+08 | 1,13E+08 | 9792 | 1 | 14260 | ENSMUST00000154834.2 | -10009 | ENSMUSG00000044042 | Fmn1 | formin 1 |
| chr14 | 83857460 | 83859064 | 1605 | * | 8,423021 | -6,045942697 | 1,816997 | -3,32744 | 0,000876 | 0,467714818 | 1,360622 | Distal Intergenic | 14 | 83643071 | 83643204 | 134 | 1 | 1,15E+08 | ENSMUST00000157384.3 | 214389 | ENSMUSG00000088009 | Gm24774 | predicted gene, 24774 |
| chr14 | 1,16E+08 | 1,16E+08 | 861 | * | 5,2579 | -6,045673229 | 2,042028 | -2,96062 | 0,00307 | 1 | 0 | Intron (ENSMUST0000( | 14 | 1,16E+08 | 1,16E+08 | 13002 | 2 | 75088 | ENSMUST00000176582.2 | -508382 | ENSMUSG00000093551 | 4930505G20Rik | RIKEN cDNA 4930505G20 gene |
| chr3 | 50279821 | 50280449 | 629 | * | 6,255455 | -6,044183601 | 1,979917 | -3,05275 | 0,002268 | 1 | 0 | Intron (ENSMUST0000( | 3 | 50319387 | 50398060 | 78674 | 2 | 26570 | ENSMUST00000029297.6 | 117611 | ENSMUSG00000027737 | Slc7a11 | solute carrier family 7 (cationic amino acid transporter, y+ system), member 11 |
| chr5 | 44474809 | 44475408 | 600 | * | 4,695979 | -6,043980203 | 2,042645 | -2,9589 | 0,003087 | 1 | 0 | Distal Intergenic | 5 | 44375171 | 44383968 | 8798 | 2 | 231225 | ENSMUST00000196151.2 | -90841 | ENSMUSG00000046985 | Tapt1 | transmembrane anterior posterior transformation 1 |
| chr18 | 34488243 | 34488662 | 420 | * | 4,237677 | -6,040051148 | 2,196671 | -2,74964 | 0,005966 | 1 | 0 | Intron (ENSMUST0000( | 18 | 34480045 | 34505550 | 25506 | 2 | 13476 | ENSMUST00000236887.2 | 16888 | ENSMUSG00000005873 | Reep5 | receptor accessory protein 5 |
| chr9 | 55645674 | 55646741 | 1068 | * | 7,689722 | -6,037977292 | 1,784525 | -3,38352 | 0,000716 | 0,434788971 | 1,360622 | Intron (ENSMUST0000( | 9 | 55650913 | 55699594 | 48682 | 2 | 244891 | ENSMUST00000215323.2 | 52853 | ENSMUSG00000034007 | Scaper | S phase cyclin A-associated protein in the ER |
| chrX | 97718833 | 97720124 | 1292 | * | 5,353064 | -6,0375638 | 2,116505 | -2,85261 | 0,004336 | 1 | 0 | Intron (ENSMUST0000( | 20 | 97601863 | 97622095 | 20233 | 2 | 94190 | ENSMUST00000147529.2 | -96738 | ENSMUSG00000031214 | Ophn1 | oligophrenin 1 |
| chrX | 59580814 | 59581559 | 746 | * | 4,438667 | -6,037242469 | 2,054396 | -2,93869 | 0,003296 | 1 | 0 | Promoter (1-2kb) | 20 | 59481241 | 59582855 | 101615 | 2 | 631784 | ENSMUST00000166562.3 | 1296 | ENSMUSG00000079583 | Gm7073 | predicted gene 7073 |
| chr8 | 1,05E+08 | 1,05E+08 | 355 | * | 6,378418 | -6,036186989 | 2,003302 | -3,01312 | 0,002586 | 1 | 0 | Distal Intergenic | 8 | 1,05E+08 | 1,05E+08 | 42887 | 1 | 12562 | ENSMUST00000034339.10 | -218202 | ENSMUSG00000031871 | Cdh5 | cadherin 5 |
| chr12 | 66232200 | 66233220 | 1021 | * | 4,539902 | -6,03396005 | 2,100322 | -2,87287 | 0,004068 | 1 | 0 | Distal Intergenic | 12 | 66330153 | 66331175 | 1023 | 2 | 238217 | ENSMUST00000081908.8 | 97955 | ENSMUSG00000060499 | Rpl10l | ribosomal protein L10-like |
| chr3 | 40435102 | 40436370 | 1269 | * | 7,145851 | -6,03339188 | 1,83274 | -3,29201 | 0,000995 | 1 | 0 | Distal Intergenic | 3 | 40562579 | 40563923 | 1345 | 2 | 1E+08 | ENSMUST00000202778.2 | 127553 | ENSMUSG00000037884 | 1700017G19Rik | RIKEN cDNA 1700017G19 gene |
| chr9 | 13015671 | 13016502 | 832 | * | 6,875478 | -6,033328505 | 1,816476 | -3,32145 | 0,000896 | 1 | 0 | Distal Intergenic | 9 | 13091272 | 13105860 | 14589 | 2 | 78219 | ENSMUST00000216694.2 | 89358 | ENSMUSG00000111830 | 4930568E12Rik | RIKEN cDNA 4930568E12 gene |
| chr19 | 27185774 | 27186780 | 1007 | * | 5,140188 | -6,031949281 | 2,074357 | -2,90787 | 0,003639 | 1 | 0 | Distal Intergenic | 19 | 27193884 | 27227155 | 33272 | 1 | 22359 | ENSMUST00000164509.8 | -7104 | ENSMUSG00000024924 | Vldlr | very low density lipoprotein receptor |
| chr12 | 36635529 | 36636501 | 973 | * | 8,901755 | -6,03191228 | 1,761044 | -3,42519 | 0,000614 | 0,400053238 | 1,449706 | Intron (ENSMUST0000( | 12 | 36432179 | 36738591 | 306413 | 1 | 75847 | ENSMUST00000221177.2 | 203350 | ENSMUSG00000043153 | Crppa | CDP-L-ribitol pyrophosphorylase A |
| chr3 | 1,42E+08 | 1,42E+08 | 577 | * | 4,72536 | -6,031462915 | 1,9969 | -3,02041 | 0,002524 | 1 | 0 | Exon (ENSMUST00000: | 3 | 1,42E+08 | 1,42E+08 | 43477 | 1 | 229905 | ENSMUST00000106218.8 | -37399 | ENSMUSG00000040213 | Kyat3 | kynurenine aminotransferase 3 |
| chr4 | 20620351 | 20620892 | 542 | * | 3,940814 | -6,03127005 | 2,284341 | -2,64027 | 0,008284 | 1 | 0 | Intron (ENSMUST0000( | 4 | 20118874 | 20778567 | 659694 | 2 | 269513 | ENSMUST00000119374.8 | 157675 | ENSMUSG00000055761 | Nkain3 | Na+/K+ transporting ATPase interacting 3 |
| chr3 | 1,33E+08 | 1,33E+08 | 857 | * | 7,206215 | -6,031264307 | 1,887402 | -3,19554 | 0,001396 | 1 | 0 | Intron (ENSMUST0000( | 3 | 1,32E+08 | 1,33E+08 | 51597 | 1 | 271981 | ENSMUST00000196491.2 | 26499 | ENSMUSG00000028030 | Tbck | TBC1 domain containing kinase |
| chr11 | 45607856 | 45608349 | 494 | * | 4,174766 | -6,031123476 | 2,166721 | -2,78353 | 0,005377 | 1 | 0 | Intron (ENSMUST0000( | 11 | 45511198 | 45644439 | 133242 | 2 | 1,05E+08 | ENSMUST00000245312.1 | 36090 | | Gm39816 | predicted gene, 39816 |
| chrX | 85674129 | 85674852 | 724 | * | 3,744209 | -6,030970712 | 2,273424 | -2,65281 | 0,007982 | 1 | 0 | Distal Intergenic | 20 | 85293862 | 85348699 | 54838 | 2 | 434903 | ENSMUST00000096378.3 | -325430 | ENSMUSG00000035427 | Mageb4 | MAGE family member B4 |
| chr2 | 97344197 | 97345099 | 903 | * | 7,877956 | -6,030664182 | 1,83702 | -3,28285 | 0,001028 | 1 | 0 | Intron (ENSMUST0000( | 2 | 97298434 | 97459408 | 160975 | 1 | 241568 | ENSMUST00000170144.2 | 45763 | ENSMUSG00000050587 | Lrrc4c | leucine rich repeat containing 4C |
| chr7 | 1,09E+08 | 1,09E+08 | 419 | * | 5,873944 | -6,029666703 | 1,927865 | -3,12764 | 0,001762 | 1 | 0 | Distal Intergenic | 7 | 1,09E+08 | 1,09E+08 | 36533 | 2 | 109594 | ENSMUST00000207178.2 | -15835 | ENSMUSG00000036111 | Lmo1 | LIM domain only 1 |
| chr2 | 20143342 | 20143814 | 473 | * | 7,121796 | -6,026449635 | 1,812814 | -3,32436 | 0,000886 | 0,470097239 | 1,360622 | Intron (ENSMUST0000( | 2 | 20285330 | 20285436 | 107 | 1 | 1,15E+08 | ENSMUST00000083319.3 | -141516 | ENSMUSG00000118880 | Gm23970 | predicted gene, 23970 |
| chr16 | 38716599 | 38717582 | 984 | * | 6,901595 | -6,025898598 | 1,923619 | -3,13258 | 0,001733 | 1 | 0 | Intron (ENSMUST0000( | 16 | 38713033 | 38744830 | 31798 | 1 | 207683 | ENSMUST00000210186.2 | 3566 | ENSMUSG00000022790 | Igsf11 | immunoglobulin superfamily, member 11 |
| chr16 | 93912331 | 93912820 | 490 | * | 5,078177 | -6,025489618 | 2,02034 | -2,98241 | 0,00286 | 1 | 0 | Intron (ENSMUST0000( | 16 | 93926040 | 93926574 | 535 | 1 | 20465 | ENSMUST00000231295.2 | -13220 | ENSMUSG00000062713 | Sim2 | single-minded family bHLH transcription factor 2 |
| chr18 | 87923112 | 87923808 | 697 | * | 5,160425 | -6,024783636 | 2,25473 | -2,67206 | 0,007539 | 1 | 0 | Distal Intergenic | 18 | 87943555 | 87943688 | 134 | 2 | 1,15E+08 | ENSMUST00000104453.3 | 19880 | ENSMUSG00000077641 | Gm24987 | predicted gene, 24987 |
| chrX | 86027259 | 86028323 | 1065 | * | 6,231481 | -6,024382126 | 2,008717 | -2,99912 | 0,002708 | 1 | 0 | Intron (ENSMUST0000( | 20 | 85293862 | 85348699 | 54838 | 2 | 434903 | ENSMUST00000096378.3 | -678560 | ENSMUSG00000035427 | Mageb4 | MAGE family member B4 |
| chr9 | 99951224 | 99952072 | 849 | * | 6,63695 | -6,024053671 | 1,944558 | -3,0979 | 0,001949 | 1 | 0 | Distal Intergenic | 9 | 99905701 | 99916703 | 11003 | 1 | 108970 | ENSMUST00000215896.2 | 45523 | ENSMUSG00000101584 | 4930519F24Rik | RIKEN cDNA 4930519F24 gene |
| chr14 | 3536453 | 3537384 | 932 | * | 3,631204 | -6,022044147 | 2,30552 | -2,61201 | 0,009001 | 1 | 0 | Distal Intergenic | 14 | 3575429 | 3839361 | 263933 | 1 | 218793 | ENSMUST00000124353.2 | -38045 | ENSMUSG00000058317 | Ube2e2 | ubiquitin-conjugating enzyme E2E 2 |
| chrX | 89192327 | 89193768 | 1442 | * | 12,45609 | -6,020702462 | 1,760192 | -3,42048 | 0,000625 | 0,477253929 | 0,918255 | Distal Intergenic | 20 | 88973704 | 88976458 | 2755 | 2 | 245511 | ENSMUST00000088146.3 | -215869 | ENSMUSG00000035387 | 4930415L06Rik | RIKEN cDNA 4930415L06 gene |
| chr4 | 1,02E+08 | 1,02E+08 | 491 | * | 6,166797 | -6,020150151 | 1,892052 | -3,18181 | 0,001464 | 1 | 0 | Promoter (1-2kb) | 4 | 1,02E+08 | 1,02E+08 | 3428 | 1 | 381536 | ENSMUST00000106914.8 | -1310 | ENSMUSG00000078625 | Gm12789 | predicted gene 12789 |
| chrX | 20262120 | 20263053 | 934 | * | 5,886886 | -6,019893272 | 1,938768 | -3,10501 | 0,001903 | 1 | 0 | Intron (ENSMUST0000( | 20 | 20291927 | 20386178 | 94252 | 1 | 382207 | ENSMUST00000115384.9 | -28874 | ENSMUSG00000037315 | Jade3 | jade family PHD finger 3 |
| chr1 | 73500136 | 73500682 | 547 | * | 5,319421 | -6,01943903 | 2,052526 | -2,9327 | 0,00336 | 1 | 0 | Distal Intergenic | 1 | 73434493 | 73446728 | 12236 | 2 | 723792 | ENSMUST00000190487.2 | -53408 | | Pinc | pregnancy induced noncoding RNA |
| chr1 | 1,23E+08 | 1,23E+08 | 556 | * | 4,363172 | -6,019426451 | 2,055472 | -2,92849 | 0,003406 | 1 | 0 | Intron (ENSMUST0000( | 1 | 1,23E+08 | 1,23E+08 | 131 | 2 | 1,15E+08 | ENSMUST00000104284.3 | -129865 | ENSMUSG00000077472 | Gm24791 | predicted gene, 24791 |
| chr2 | 1,3E+08 | 1,3E+08 | 666 | * | 4,677322 | -6,018551416 | 2,040417 | -2,94967 | 0,003181 | 1 | 0 | Promoter (<=1kb) | 2 | 1,3E+08 | 1,3E+08 | 812 | 1 | 75656 | ENSMUST00000110281.2 | -941 | ENSMUSG00000027409 | 1700020A23Rik | RIKEN cDNA 1700020A23 gene |
| chr12 | 81826180 | 81827220 | 1041 | * | 7,374771 | -6,017245957 | 2,006356 | -2,99909 | 0,002708 | 0,671214802 | 1,360622 | Promoter (<=1kb) | 12 | 81768813 | 81827648 | 58836 | 2 | 338372 | ENSMUST00000222322.2 | 428 | ENSMUSG00000042724 | Map3k9 | mitogen-activated protein kinase kinase kinase 9 |
| chr16 | 76862935 | 76863242 | 308 | * | 4,022465 | -6,016005682 | 2,146545 | -2,80265 | 0,005069 | 1 | 0 | Intron (ENSMUST0000( | 16 | 76878390 | 76890816 | 12427 | 1 | 30940 | ENSMUST00000232538.2 | -15148 | ENSMUSG00000022867 | Usp25 | ubiquitin specific peptidase 25 |
| chr12 | 1,02E+08 | 1,02E+08 | 543 | * | 4,15292 | -6,015975121 | 2,30251 | -2,61279 | 0,008981 | 1 | 0 | Exon (ENSMUST00000: | 12 | 1,02E+08 | 1,02E+08 | 60504 | 2 | 74413 | ENSMUST00000160715.8 | 58042 | ENSMUSG00000021187 | Tc2n | tandem C2 domains, nuclear |
| chrX | 1,3E+08 | 1,3E+08 | 1106 | * | 5,156023 | -6,01399288 | 1,939596 | -3,10064 | 0,001931 | 1 | 0 | Distal Intergenic | 20 | 1,3E+08 | 1,3E+08 | 129 | 2 | 1,15E+08 | ENSMUST00000104191.3 | -182865 | ENSMUSG00000077379 | Gm25170 | predicted gene, 25170 |
| chr7 | 69174371 | 69174989 | 619 | * | 4,815045 | -6,013417417 | 2,06232 | -2,91585 | 0,003547 | 1 | 0 | Intron (ENSMUST0000( | 7 | 69148327 | 69148433 | 107 | 2 | 1,15E+08 | ENSMUST00000240350.1 | -25938 | ENSMUSG00000118803 | Gm23815 | predicted gene, 23815 |
| chr16 | 42946360 | 42946858 | 499 | * | 4,602963 | -6,013085215 | 2,163573 | -2,77924 | 0,005449 | 1 | 0 | Promoter (<=1kb) | 16 | 42946596 | 42946702 | 107 | 1 | 1,15E+08 | ENSMUST00000239901.1 | 0 | ENSMUSG00000118809 | Gm25873 | predicted gene, 25873 |
| chr10 | 39859569 | 39859981 | 413 | * | 3,65641 | -6,012754157 | 2,301132 | -2,61295 | 0,008976 | 1 | 0 | Promoter (2-3kb) | 10 | 39844151 | 39862635 | 18485 | 2 | 215928 | ENSMUST00000170579.9 | 2654 | ENSMUSG00000038528 | Mfsd4b5 | major facilitator superfamily domain containing 4B5 |
| chrX | 1,32E+08 | 1,32E+08 | 547 | * | 5,147377 | -6,011293328 | 2,003171 | -3,00089 | 0,002692 | 1 | 0 | Distal Intergenic | 20 | 1,31E+08 | 1,31E+08 | 131 | 2 | 1,15E+08 | ENSMUST00000104440.3 | -133724 | ENSMUSG00000077628 | Gm23768 | predicted gene, 23768 |
| chr8 | 23308974 | 23309550 | 577 | * | 5,193187 | -6,011287758 | 2,003972 | -2,99969 | 0,002703 | 1 | 0 | Distal Intergenic | 8 | 23295782 | 23299553 | 3772 | 1 | 73299 | ENSMUST00000186344.2 | 13192 | | 1700041G16Rik | RIKEN cDNA 1700041G16 gene |
| chr7 | 1,12E+08 | 1,12E+08 | 984 | * | 7,023663 | -6,010225404 | 1,869309 | -3,21521 | 0,001303 | 0,537150047 | 1,360622 | Intron (ENSMUST0000( | 7 | 1,12E+08 | 1,12E+08 | 67832 | 1 | 21676 | ENSMUST00000171373.3 | 8560 | ENSMUSG00000055320 | Tead1 | TEA domain family member 1 |
| chr4 | 51376554 | 51377483 | 930 | * | 5,142026 | -6,009099371 | 2,043588 | -2,94046 | 0,003277 | 1 | 0 | Distal Intergenic | 4 | 51216678 | 51250622 | 33945 | 1 | 74914 | ENSMUST00000156384.4 | 159876 | ENSMUSG00000039555 | Cylc2 | cylicin, basic protein of sperm head cytoskeleton 2 |
| chr14 | 1,11E+08 | 1,11E+08 | 684 | * | 4,547814 | -6,007149456 | 2,124554 | -2,82749 | 0,004691 | 1 | 0 | Distal Intergenic | 14 | 1,11E+08 | 1,11E+08 | 103 | 1 | 1,15E+08 | ENSMUST00000122492.3 | -27965 | ENSMUSG00000084441 | Gm26255 | predicted gene, 26255 |
| chr14 | 89481810 | 89482885 | 1076 | * | 4,48635 | -6,007001144 | 2,330918 | -2,5771 | 0,009963 | 1 | 0 | Distal Intergenic | 14 | 89944241 | 89944372 | 132 | 1 | 1,15E+08 | ENSMUST00000104220.3 | -461356 | ENSMUSG00000077408 | Gm25415 | predicted gene, 25415 |
| chr1 | 1,93E+08 | 1,93E+08 | 805 | * | 5,242096 | -6,006309989 | 1,957303 | -3,06867 | 0,00215 | 1 | 0 | Intron (ENSMUST0000( | 1 | 1,93E+08 | 1,93E+08 | 11269 | 2 | 214791 | ENSMUST00000155579.8 | -80454 | ENSMUSG00000016262 | Sertad4 | SERTA domain containing 4 |
| chr17 | 61228481 | 61229166 | 686 | * | 5,984362 | -6,002510934 | 1,894326 | -3,16868 | 0,001531 | 1 | 0 | Distal Intergenic | 17 | 61250872 | 61265873 | 15002 | 2 | 1,03E+08 | ENSMUST00000233618.2 | 36707 | | Gm33890 | predicted gene, 33890 |
| chr14 | 87052055 | 87053120 | 1066 | * | 4,047957 | -6,000749619 | 2,219936 | -2,70312 | 0,006869 | 1 | 0 | Intron (ENSMUST0000( | 14 | 86893608 | 87027325 | 133718 | 2 | 56419 | ENSMUST00000227638.2 | -24730 | ENSMUSG00000022021 | Diaph3 | diaphanous related formin 3 |
| chr14 | 3819793 | 3820591 | 799 | * | 5,210966 | -6,000350549 | 1,965772 | -3,05241 | 0,00227 | 1 | 0 | Intron (ENSMUST0000( | 14 | 3804546 | 3882808 | 78263 | 1 | 218793 | ENSMUST00000135381.8 | 15247 | ENSMUSG00000058317 | Ube2e2 | ubiquitin-conjugating enzyme E2E 2 |
| chr8 | 1,07E+08 | 1,07E+08 | 600 | * | 6,442547 | -6,000101449 | 1,937977 | -3,09606 | 0,001961 | 1 | 0 | Intron (ENSMUST0000( | 8 | 1,07E+08 | 1,07E+08 | 85441 | 2 | 58994 | ENSMUST00000067512.8 | 27468 | ENSMUSG00000031906 | Smpd3 | sphingomyelin phosphodiesterase 3, neutral |
| chr3 | 32667447 | 32668182 | 736 | * | 5,408777 | -5,995794708 | 1,987547 | -3,01668 | 0,002556 | 1 | 0 | Promoter (2-3kb) | 3 | 32645283 | 32670628 | 25346 | 2 | 14696 | ENSMUST00000193050.2 | 2446 | ENSMUSG00000027669 | Gnb4 | guanine nucleotide binding protein (G protein), beta 4 |
| chr14 | 62218644 | 62219516 | 873 | * | 7,600518 | -5,995049512 | 1,810548 | -3,31118 | 0,000929 | 0,477253929 | 1,360622 | Distal Intergenic | 14 | 62350569 | 62408575 | 58007 | 2 | 1,03E+08 | ENSMUST00000226702.2 | 189059 | | Gm35419 | predicted gene, 35419 |
| chr10 | 1,14E+08 | 1,14E+08 | 572 | * | 4,336289 | -5,994576307 | 2,162551 | -2,77199 | 0,005571 | 1 | 0 | Intron (ENSMUST0000( | 10 | 1,14E+08 | 1,15E+08 | 402550 | 2 | 237553 | ENSMUST00000061632.9 | 295819 | ENSMUSG00000050663 | Trhde | TRH-degrading enzyme |
| chr15 | 1E+08 | 1E+08 | 451 | * | 5,4056 | -5,994430901 | 2,024469 | -2,96099 | 0,003067 | 1 | 0 | Intron (ENSMUST0000( | 15 | 1E+08 | 1E+08 | 5650 | 2 | 18174 | ENSMUST00000124324.2 | 13331 | ENSMUSG00000023030 | Slc11a2 | solute carrier family 11 (proton-coupled divalent metal ion transporters), member 2 |
| chr10 | 1,18E+08 | 1,18E+08 | 1034 | * | 7,732836 | -5,993132709 | 1,778603 | -3,36957 | 0,000753 | 1 | 0 | 3' UTR | 10 | 1,18E+08 | 1,18E+08 | 1572 | 1 | 15978 | ENSMUST00000220309.2 | 3543 | ENSMUSG00000055170 | Ifng | interferon gamma |
| chr9 | 1,14E+08 | 1,14E+08 | 1300 | * | 6,270725 | -5,992562305 | 1,866664 | -3,21031 | 0,001326 | 1 | 0 | Promoter (1-2kb) | 9 | 1,14E+08 | 1,14E+08 | 22657 | 2 | 78893 | ENSMUST00000216785.2 | -1244 | ENSMUSG00000056167 | Cnot10 | CCR4-NOT transcription complex, subunit 10 |
| chr11 | 53911329 | 53912174 | 846 | * | 8,420468 | -5,992251171 | 1,787058 | -3,35314 | 0,000799 | 0,452016789 | 1,360622 | Intron (ENSMUST0000( | 11 | 53905086 | 53918778 | 13693 | 2 | 30805 | ENSMUST00000154369.2 | 6604 | ENSMUSG00000020334 | Slc22a4 | solute carrier family 22 (organic cation transporter), member 4 |
| chr5 | 73123775 | 73124471 | 697 | * | 6,212607 | -5,992229616 | 1,942669 | -3,08453 | 0,002039 | 1 | 0 | Intron (ENSMUST0000( | 5 | 73105842 | 73132045 | 26204 | 1 | 75991 | ENSMUST00000145939.2 | 17933 | ENSMUSG00000036087 | Slain2 | SLAIN motif family, member 2 |
| chr16 | 71628863 | 71629351 | 489 | * | 4,868095 | -5,992047257 | 1,993606 | -3,00563 | 0,00265 | 1 | 0 | Distal Intergenic | 16 | 71460676 | 71460806 | 131 | 1 | 1,15E+08 | ENSMUST00000083990.3 | 168187 | ENSMUSG00000065924 | Gm22797 | predicted gene, 22797 |
| chr5 | 1,21E+08 | 1,21E+08 | 593 | * | 4,301221 | -5,991697469 | 2,130843 | -2,81189 | 0,004925 | 1 | 0 | Distal Intergenic | 5 | 1,21E+08 | 1,21E+08 | 7117 | 1 | 100535 | ENSMUST00000044224.5 | 25312 | ENSMUSG00000032623 | Oas1d | 2'-5' oligoadenylate synthetase 1D |
| chr14 | 90776637 | 90777516 | 880 | * | 3,442794 | -5,988158293 | 2,301927 | -2,60137 | 0,009285 | 1 | 0 | Distal Intergenic | 14 | 90905695 | 90928881 | 23187 | 2 | 1,03E+08 | ENSMUST00000226296.2 | 151365 | | Gm33246 | predicted gene, 33246 |
| chr15 | 54393404 | 54393770 | 367 | * | 8,778064 | -5,988048042 | 1,779356 | -3,36529 | 0,000765 | 1 | 0 | Distal Intergenic | 15 | 54434588 | 54466243 | 31656 | 1 | 105853 | ENSMUST00000025356.4 | -40818 | ENSMUSG00000024479 | Mal2 | mal, T cell differentiation protein 2 |
| chrX | 37366409 | 37367197 | 789 | * | 4,420076 | -5,987950725 | 2,128345 | -2,81343 | 0,004902 | 1 | 0 | Distal Intergenic | 20 | 37286183 | 37341316 | 55134 | 2 | 245386 | ENSMUST00000155848.8 | -25093 | ENSMUSG00000036502 | Tmem255a | transmembrane protein 255A |
| chr10 | 44546422 | 44547344 | 923 | * | 9,57518 | -5,986977819 | 1,76384 | -3,39429 | 0,000688 | 0,451739959 | 1,173325 | Intron (ENSMUST0000( | 10 | 44608449 | 44608563 | 115 | 1 | 1,15E+08 | ENSMUST00000158983.4 | -61105 | ENSMUSG00000089608 | Gm22503 | predicted gene, 22503 |
| chr10 | 57421719 | 57422410 | 692 | * | 6,54409 | -5,986852317 | 2,041145 | -2,93308 | 0,003356 | 1 | 0 | Distal Intergenic | 10 | 57391870 | 57408626 | 16757 | 2 | 56442 | ENSMUST00000020027.11 | -13093 | ENSMUSG00000019877 | Serinc1 | serine incorporator 1 |

| | | | | | | | | | | | | | | | | | | | | | | | |
|---|---|---|---|---|---|---|---|---|---|---|---|---|---|---|---|---|---|---|---|---|---|---|---|
| chr14 | 9200784 | 9201811 | 1028 | * | 6,492549 | -5,986325146 | 1,920608 | -3,11689 | 0,001828 | 1 | 0 | Intron (ENSMUST0000( | 14 | 9185133 | 9206630 | 21498 | 1 | 328354 | ENSMUST00000140967.2 | 15651 | ENSMUSG00000051729 | Gm5087 | predicted gene 5087 |
| chr4 | 1,49E+08 | 1,49E+08 | 612 | * | 5,721531 | -5,986096841 | 1,969962 | -3,03869 | 0,002376 | 1 | 0 | Promoter (2-3kb) | 4 | 1,49E+08 | 1,49E+08 | 104 | 1 | 1,15E+08 | ENSMUST00000158456.3 | 2102 | ENSMUSG00000089081 | Gm24002 | predicted gene, 24002 |
| chr5 | 21613852 | 21614569 | 718 | * | 4,800448 | -5,985866873 | 2,028853 | -2,95037 | 0,003174 | 1 | 0 | Intron (ENSMUST0000( | 5 | 21497959 | 21629675 | 131717 | 2 | 75172 | ENSMUST00000115245.8 | 15106 | ENSMUSG00000064280 | Ccdc146 | coiled-coil domain containing 146 |
| chr1 | 88572585 | 88573741 | 1157 | * | 12,97923 | -5,983378225 | 1,671211 | -3,58027 | 0,000343 | 0,414202366 | 0,738441 | Distal Intergenic | 1 | 88625947 | 88629943 | 3997 | 2 | 320982 | ENSMUST00000159814.2 | 56202 | ENSMUSG00000049866 | Arl4c | ADP-ribosylation factor-like 4C |
| chrX | 26802517 | 26803192 | 676 | * | 4,324982 | -5,982914845 | 2,097755 | -2,85206 | 0,004344 | 1 | 0 | Distal Intergenic | 20 | 26868043 | 26890969 | 22927 | 2 | 619991 | ENSMUST00000167438.2 | 87777 | ENSMUSG00000080725 | Gm6121 | predicted gene 6121 |
| chr14 | 55582305 | 55582586 | 282 | * | 4,910001 | -5,982267986 | 2,199847 | -2,7194 | 0,00654 | 1 | 0 | Distal Intergenic | 14 | 55472111 | 55478705 | 6595 | 1 | 71412 | ENSMUST00000165432.2 | 110194 | ENSMUSG00000022209 | Dhrs2 | dehydrogenase/reductase member 2 |
| chrX | 99304471 | 99305017 | 547 | * | 3,773695 | -5,981998472 | 2,289277 | -2,61305 | 0,008974 | 1 | 0 | Intron (ENSMUST0000( | 20 | 99424703 | 99424791 | 89 | 1 | 751534 | ENSMUST00000102443.3 | -119686 | ENSMUSG00000076398 | Mir676 | microRNA 676 |
| chr11 | 1,09E+08 | 1,09E+08 | 779 | * | 8,91928 | -5,981942231 | 1,747267 | -3,4236 | 0,000618 | 0,410415129 | 1,360622 | Promoter (1-2kb) | 11 | 1,09E+08 | 1,09E+08 | 29577 | 1 | 74008 | ENSMUST00000106696.2 | -1335 | ENSMUSG00000020604 | Arsg | arylsulfatase G |
| chr2 | 1,34E+08 | 1,34E+08 | 676 | * | 4,896985 | -5,979999508 | 1,977032 | -3,02474 | 0,002489 | 1 | 0 | Distal Intergenic | 2 | 1,33E+08 | 1,33E+08 | 107 | 2 | 1,15E+08 | ENSMUST00000239857.1 | -145597 | ENSMUSG00000119228 | Gm25258 | predicted gene, 25258 |
| chr11 | 11368491 | 11369158 | 668 | * | 5,346933 | -5,979935211 | 1,990041 | -3,00493 | 0,002656 | 1 | 0 | Intron (ENSMUST0000( | 11 | 11377385 | 11377512 | 128 | 2 | 1,15E+08 | ENSMUST00000082832.3 | 8354 | ENSMUSG00000064766 | Gm26458 | predicted gene, 26458 |
| chr12 | 1,07E+08 | 1,07E+08 | 662 | * | 4,889612 | -5,978645871 | 2,03548 | -2,93722 | 0,003312 | 1 | 0 | Distal Intergenic | 12 | 1,07E+08 | 1,07E+08 | 20609 | 2 | 71033 | ENSMUST00000124363.8 | 42393 | ENSMUSG00000085820 | 4933406K04Rik | RIKEN cDNA 4933406K04 gene |
| chr4 | 80777067 | 80778060 | 994 | * | 6,97134 | -5,978005048 | 1,89446 | -3,15552 | 0,001602 | 0,575442234 | 1,360622 | Distal Intergenic | 4 | 80764825 | 80769237 | 4413 | 1 | 22178 | ENSMUST00000133932.2 | 12242 | ENSMUSG00000005994 | Tyrp1 | tyrosinase-related protein 1 |
| chr13 | 13877377 | 13877999 | 623 | * | 6,405085 | -5,977779639 | 1,860135 | -3,21363 | 0,001311 | 1 | 0 | Intron (ENSMUST0000( | 13 | 13883936 | 13922114 | 38179 | 1 | 17101 | ENSMUST00000221717.2 | -5937 | ENSMUSG00000019726 | Lyst | lysosomal trafficking regulator |
| chr16 | 70555263 | 70555725 | 463 | * | 3,988248 | -5,976094556 | 2,103956 | -2,84041 | 0,004506 | 1 | 0 | Distal Intergenic | 16 | 70414927 | 70421712 | 6786 | 1 | 52645 | ENSMUST00000231858.2 | 140336 | ENSMUSG00000116792 | D16Ertd519e | DNA segment, Chr 16, ERATO Doi 519, expressed |
| chr1 | 55663111 | 55664184 | 1074 | * | 8,107715 | -5,972468469 | 1,809106 | -3,30134 | 0,000962 | 0,474289497 | 1,449706 | Intron (ENSMUST0000( | 1 | 55741184 | 55774386 | 33203 | 1 | 227120 | ENSMUST00000187059.2 | -77000 | ENSMUSG00000038349 | Plcl1 | phospholipase C-like 1 |
| chr17 | 5326628 | 5327126 | 499 | * | 5,260118 | -5,972233864 | 1,955561 | -3,05398 | 0,002258 | 1 | 0 | Intron (ENSMUST0000( | 17 | 5263228 | 5267593 | 4366 | 2 | 1,03E+08 | ENSMUST00000247135.1 | -59035 | | Gm34854 | predicted gene, 34854 |
| chr19 | 39437027 | 39437680 | 654 | * | 5,690304 | -5,972118544 | 1,926525 | -3,09994 | 0,001936 | 1 | 0 | Intron (ENSMUST0000( | 19 | 39378000 | 39451519 | 73520 | 2 | 13097 | ENSMUST00000035488.3 | 13839 | ENSMUSG00000032808 | Cyp2c38 | cytochrome P450, family 2, subfamily c, polypeptide 38 |
| chr17 | 3880416 | 3880874 | 459 | * | 4,619798 | -5,969922711 | 2,089991 | -2,85643 | 0,004284 | 1 | 0 | Distal Intergenic | 17 | 3685515 | 3746536 | 61022 | 2 | 224480 | ENSMUST00000115800.2 | -133880 | ENSMUSG00000023802 | Nox3 | NADPH oxidase 3 |
| chrX | 46061941 | 46062534 | 594 | * | 4,066958 | -5,969797996 | 2,231049 | -2,67578 | 0,007456 | 1 | 0 | Distal Intergenic | 20 | 45417884 | 45419222 | 1339 | 1 | 73360 | ENSMUST00000059466.3 | 644057 | ENSMUSG00000046615 | Actrt1 | actin-related protein T1 |
| chr1 | 62646093 | 62647013 | 921 | * | 6,611389 | -5,967692781 | 1,896617 | -3,14649 | 0,001652 | 1 | 0 | Intron (ENSMUST0000( | 1 | 62742444 | 62779091 | 36648 | 1 | 18187 | ENSMUST00000126344.2 | -95431 | ENSMUSG00000025969 | Nrp2 | neuropilin 2 |
| chr2 | 7699773 | 7700781 | 1009 | * | 4,709738 | -5,966991894 | 2,138185 | -2,79068 | 0,00526 | 1 | 0 | Distal Intergenic | 2 | 7398887 | 7514374 | 115488 | 2 | 14007 | ENSMUST00000182037.2 | -185399 | ENSMUSG00000002107 | Celf2 | CUGBP, Elav-like family member 2 |
| chr16 | 73207530 | 73208091 | 562 | * | 3,842488 | -5,966975663 | 2,141705 | -2,78609 | 0,005335 | 1 | 0 | Intron (ENSMUST0000( | 16 | 73219681 | 73219847 | 167 | 2 | 1,15E+08 | ENSMUST00000082896.3 | 11756 | ENSMUSG00000064830 | Gm25699 | predicted gene, 25699 |
| chr9 | 86292784 | 86293385 | 602 | * | 4,902654 | -5,966647887 | 2,222511 | -2,68464 | 0,007261 | 1 | 0 | Intron (ENSMUST0000( | 9 | 86189359 | 86347003 | 157645 | 2 | 70348 | ENSMUST00000034986.14 | 53618 | ENSMUSG00000032415 | Ube2cbp | ubiquitin-conjugating enzyme E2C binding protein |
| chr1 | 1,14E+08 | 1,14E+08 | 740 | * | 5,076084 | -5,966231478 | 1,997353 | -2,98707 | 0,002817 | 1 | 0 | Distal Intergenic | 1 | 1,16E+08 | 1,16E+08 | 15632 | 2 | 1,08E+08 | ENSMUST00000186137.7 | 1433081 | | Gm50605 | predicted gene, 50605 |
| chr14 | 37255991 | 37256887 | 897 | * | 6,31559 | -5,963983675 | 1,934699 | -3,08264 | 0,002052 | 1 | 0 | Distal Intergenic | 14 | 37323064 | 37328745 | 5682 | 2 | 1,03E+08 | ENSMUST00000224767.2 | 71858 | ENSMUSG00000114334 | Gm31393 | predicted gene, 31393 |
| chrX | 1,17E+08 | 1,17E+08 | 1456 | * | 6,02229 | -5,963718969 | 2,088326 | -2,85574 | 0,004294 | 1 | 0 | Distal Intergenic | 20 | 1,17E+08 | 1,17E+08 | 1030 | 1 | 1E+08 | ENSMUST00000072518.8 | -43307 | ENSMUSG00000100194 | Tgif2lx2 | TGFB-induced factor homeobox 2-like, X-linked 2 |
| chr1 | 1,86E+08 | 1,86E+08 | 745 | * | 4,223689 | -5,963600981 | 2,167048 | -2,75195 | 0,005924 | 1 | 0 | Distal Intergenic | 1 | 1,86E+08 | 1,86E+08 | 46635 | 2 | 320449 | ENSMUST00000193084.2 | 161233 | ENSMUSG00000103690 | A730004F24Rik | RIKEN cDNA A730004F24 gene |
| chr15 | 4228541 | 4229147 | 607 | * | 4,399099 | -5,963594615 | 2,171141 | -2,74676 | 0,006019 | 1 | 0 | Distal Intergenic | 15 | 4172294 | 4183441 | 11148 | 1 | 67041 | ENSMUST00000133130.2 | 56247 | ENSMUSG00000022186 | Oxct1 | 3-oxoacid CoA transferase 1 |
| chr14 | 1,17E+08 | 1,17E+08 | 656 | * | 4,223528 | -5,963546869 | 2,217234 | -2,68963 | 0,007153 | 1 | 0 | Intron (ENSMUST0000( | 14 | 1,17E+08 | 1,18E+08 | 701884 | 1 | 23888 | ENSMUST00000123239.2 | -646209 | ENSMUSG00000058571 | Gpc6 | glypican 6 |
| chrX | 76794685 | 76795706 | 1022 | * | 6,521043 | -5,962398012 | 1,992446 | -2,9925 | 0,002767 | 1 | 0 | Distal Intergenic | 20 | 76815061 | 76839578 | 24518 | 2 | 19108 | ENSMUST00000114044.2 | 43872 | ENSMUSG00000035725 | Prkx | protein kinase, X-linked |
| chr3 | 1,03E+08 | 1,03E+08 | 823 | * | 6,177816 | -5,961754034 | 1,967494 | -3,03013 | 0,002445 | 1 | 0 | Exon (ENSMUST00000( | 3 | 1,03E+08 | 1,03E+08 | 15179 | 2 | 20957 | ENSMUST00000198651.2 | -18061 | ENSMUSG00000027855 | Sycp1 | synaptonemal complex protein 1 |
| chr18 | 20926239 | 20926699 | 461 | * | 5,204876 | -5,961540917 | 1,996615 | -2,98582 | 0,002828 | 1 | 0 | Distal Intergenic | 18 | 20817656 | 20879461 | 61806 | 2 | 56386 | ENSMUST00000070080.6 | -46778 | ENSMUSG00000056124 | B4galt6 | UDP-Gal:betaGlcNAc beta 1,4-galactosyltransferase, polypeptide 6 |
| chr12 | 68074131 | 68074559 | 429 | * | 5,294573 | -5,958028386 | 2,118305 | -2,81264 | 0,004914 | 1 | 0 | Distal Intergenic | 12 | 67659070 | 67659255 | 186 | 1 | 1,15E+08 | ENSMUST00000122499.3 | 415061 | ENSMUSG00000084448 | Gm26256 | predicted gene, 26256 |
| chr12 | 88658500 | 88658836 | 337 | * | 5,116162 | -5,957786398 | 1,973549 | -3,01882 | 0,002538 | 1 | 0 | Intron (ENSMUST0000( | 12 | 88689646 | 89228341 | 538696 | 1 | 18191 | ENSMUST00000167734.8 | -30810 | ENSMUSG00000066392 | Nrxn3 | neurexin III |
| chr8 | 83234793 | 83235300 | 508 | * | 3,558047 | -5,957367166 | 2,290183 | -2,60126 | 0,009288 | 1 | 0 | Distal Intergenic | 8 | 83135720 | 83135855 | 136 | 1 | 1,15E+08 | ENSMUST00000083248.3 | 99073 | ENSMUSG00000065182 | Gm25138 | predicted gene, 25138 |
| chr1 | 1,02E+08 | 1,02E+08 | 448 | * | 4,807583 | -5,955177246 | 2,02303 | -2,94369 | 0,003243 | 1 | 0 | Distal Intergenic | 1 | 1,02E+08 | 1,02E+08 | 132 | 2 | 1,15E+08 | ENSMUST00000104107.3 | 29845 | ENSMUSG00000077295 | Gm24122 | predicted gene, 24122 |
| chr19 | 42579729 | 42580687 | 959 | * | 7,488306 | -5,95363304 | 1,845806 | -3,22549 | 0,001258 | 1 | 0 | 3' UTR | 19 | 42569600 | 42573210 | 3611 | 1 | 52013 | ENSMUST00000162238.2 | 10129 | ENSMUSG00000025184 | R3hcc1l | R3H domain and coiled-coil containing 1 like |
| chr5 | 28759271 | 28759855 | 585 | * | 4,363142 | -5,948214731 | 2,137989 | -2,78215 | 0,0054 | 1 | 0 | Intron (ENSMUST0000( | 5 | 28661813 | 28672254 | 10442 | 2 | 20423 | ENSMUST00000002708.5 | -87017 | ENSMUSG00000002633 | Shh | sonic hedgehog |
| chr4 | 17455955 | 17456665 | 711 | * | 4,603353 | -5,94688519 | 2,091001 | -2,84404 | 0,004455 | 1 | 0 | Distal Intergenic | 4 | 17546447 | 17636191 | 89745 | 2 | 1,03E+08 | ENSMUST00000245354.1 | 179526 | | Gm30092 | predicted gene, 30092 |
| chr4 | 70267137 | 70267776 | 640 | * | 6,089135 | -5,946114945 | 2,056383 | -2,89154 | 0,003834 | 1 | 0 | 3' UTR | 4 | 70317123 | 70321778 | 4656 | 2 | 214444 | ENSMUST00000126416.2 | 54002 | ENSMUSG00000039298 | Cdk5rap2 | CDK5 regulatory subunit associated protein 2 |
| chr10 | 63132615 | 63133260 | 646 | * | 5,089731 | -5,945208209 | 2,019415 | -2,94402 | 0,00324 | 1 | 0 | Intron (ENSMUST0000( | 10 | 63139486 | 63143688 | 4203 | 1 | 67345 | ENSMUST00000219279.2 | -6226 | ENSMUSG00000020064 | Herc4 | hect domain and RLD 4 |
| chr8 | 4901203 | 4902232 | 1030 | * | 8,716927 | -5,942654786 | 1,826825 | -3,253 | 0,001142 | 0,504427013 | 1,425929 | Distal Intergenic | 8 | 4813468 | 4829567 | 16100 | 2 | 20419 | ENSMUST00000208381.2 | -71636 | ENSMUSG00000022322 | Shcbp1 | Shc SH2-domain binding protein 1 |
| chr4 | 43804961 | 43805537 | 577 | * | 4,836383 | -5,942109254 | 2,105865 | -2,8217 | 0,004777 | 1 | 0 | Distal Intergenic | 4 | 43820335 | 43821433 | 1099 | 2 | 29846 | ENSMUST00000079465.5 | 15896 | ENSMUSG00000110970 | Olfr156 | olfactory receptor 156 |
| chr13 | 15711050 | 15711546 | 497 | * | 4,464723 | -5,942009122 | 2,12088 | -2,80167 | 0,005084 | 1 | 0 | Intron (ENSMUST0000( | 13 | 15638565 | 15904611 | 266047 | 1 | 14634 | ENSMUST00000110510.4 | 72485 | ENSMUSG00000021318 | Gli3 | GLI-Kruppel family member GLI3 |
| chr9 | 94021120 | 94021881 | 762 | * | 7,279106 | -5,941518266 | 1,789782 | -3,31969 | 0,000901 | 1 | 0 | Exon (ENSMUST000002 | 9 | 94166549 | 94166705 | 157 | 1 | 1,15E+08 | ENSMUST00000082955.3 | -144668 | ENSMUSG00000064889 | Gm24200 | predicted gene, 24200 |
| chr11 | 99241779 | 99242434 | 656 | * | 4,920102 | -5,94138091 | 2,113153 | -2,81162 | 0,004929 | 1 | 0 | Promoter (<=1kb) | 11 | 99236391 | 99241930 | 5540 | 2 | 16675 | ENSMUST00000017732.3 | 0 | ENSMUSG00000017588 | Krt27 | keratin 27 |
| chrX | 71275069 | 71276058 | 990 | * | 5,846848 | -5,935137043 | 2,014693 | -2,94593 | 0,00322 | 1 | 0 | Exon (ENSMUST000002 | 20 | 71265623 | 71266570 | 948 | 1 | 17140 | ENSMUST00000048675.2 | 9446 | ENSMUSG00000033343 | Magea4 | MAGE family member A4 |
| chr9 | 24924910 | 24925675 | 766 | * | 4,041133 | -5,932225334 | 2,111031 | -2,81011 | 0,004952 | 1 | 0 | Distal Intergenic | 9 | 24885504 | 24909716 | 24213 | 1 | 1,03E+08 | ENSMUST00000214469.2 | 39406 | ENSMUSG00000111866 | E130101E03Rik | RIKEN cDNA E130101E03 gene |
| chr11 | 54351020 | 54351636 | 617 | * | 4,526952 | -5,932045411 | 2,154885 | -2,75284 | 0,005908 | 1 | 0 | Promoter (2-3kb) | 11 | 54348988 | 54349095 | 108 | 1 | 1,15E+08 | ENSMUST00000157812.4 | 2032 | ENSMUSG00000088437 | Gm24198 | predicted gene, 24198 |
| chrX | 1,65E+08 | 1,65E+08 | 649 | * | 5,399535 | -5,931342897 | 2,039782 | -2,90783 | 0,003639 | 1 | 0 | Intron (ENSMUST0000( | 20 | 1,65E+08 | 1,65E+08 | 62714 | 2 | 54156 | ENSMUST00000000412.3 | 27941 | ENSMUSG00000000402 | Egfl6 | EGF-like-domain, multiple 6 |
| chr16 | 36622249 | 36622926 | 678 | * | 6,263778 | -5,930689579 | 1,867706 | -3,17539 | 0,001496 | 1 | 0 | Intron (ENSMUST0000( | 16 | 36613246 | 36629067 | 15822 | 2 | 106389 | ENSMUST00000114825.3 | 6141 | ENSMUSG00000022838 | Eaf2 | ELL associated factor 2 |
| chrX | 57536214 | 57537054 | 841 | * | 5,133156 | -5,929927007 | 2,049031 | -2,89402 | 0,003803 | 1 | 0 | Distal Intergenic | 20 | 57403312 | 57403418 | 107 | 2 | 1,15E+08 | ENSMUST00000240146.1 | -132796 | ENSMUSG00000119633 | Gm25851 | predicted gene, 25851 |
| chr2 | 35653016 | 35653457 | 442 | * | 8,579819 | -5,929486786 | 1,786607 | -3,31885 | 0,000904 | 0,46544192 | 1,425929 | Exon (ENSMUST000002 | 2 | 35600339 | 35621006 | 20668 | 1 | 69601 | ENSMUST00000156669.2 | 52677 | ENSMUSG00000026883 | Dab2ip | disabled 2 interacting protein |
| chr13 | 45012961 | 45013618 | 658 | * | 4,449875 | -5,929063627 | 2,159665 | -2,74536 | 0,006044 | 1 | 0 | Intron (ENSMUST0000( | 13 | 44994259 | 45049957 | 55699 | 1 | 16468 | ENSMUST00000173003.2 | 18702 | ENSMUSG00000038518 | Jarid2 | jumonji, AT rich interactive domain 2 |
| chr14 | 1,18E+08 | 1,18E+08 | 539 | * | 4,60523 | -5,926182852 | 2,049548 | -2,89146 | 0,003835 | 1 | 0 | Intron (ENSMUST0000( | 14 | 1,17E+08 | 1,18E+08 | 1049151 | 1 | 23888 | ENSMUST00000078849.12 | 441313 | ENSMUSG00000058571 | Gpc6 | glypican 6 |
| chrX | 42007554 | 42008397 | 844 | * | 4,398257 | -5,92584102 | 2,114693 | -2,80222 | 0,005075 | 1 | 0 | Intron (ENSMUST0000( | 20 | 41770634 | 41770763 | 130 | 2 | 1,15E+08 | ENSMUST00000104370.3 | -236791 | ENSMUSG00000077558 | Gm23705 | predicted gene, 23705 |
| chr13 | 22298254 | 22298902 | 649 | * | 3,979278 | -5,925292233 | 2,244142 | -2,64034 | 0,008282 | 1 | 0 | Distal Intergenic | 13 | 22285798 | 22286934 | 1137 | 2 | 252906 | ENSMUST00000186062.3 | -11320 | ENSMUSG00000099611 | Vmn1r189 | vomeronasal 1 receptor 189 |
| chr8 | 19726568 | 19727072 | 505 | * | 6,026498 | -5,924670177 | 1,913356 | -3,09648 | 0,001958 | 1 | 0 | Distal Intergenic | 8 | 19644916 | 19715338 | 70423 | 2 | 1,03E+08 | ENSMUST00000212862.2 | -11230 | | Gm35998 | predicted gene, 35998 |
| chr1 | 34230779 | 34231585 | 807 | * | 7,841247 | -5,924161887 | 1,795669 | -3,29914 | 0,00097 | 1 | 0 | Exon (ENSMUST000001 | 1 | 34204937 | 34217352 | 12416 | 2 | 1,08E+08 | ENSMUST00000181692.3 | -13427 | ENSMUSG00000096992 | Gm26788 | predicted gene, 26788 |
| chr12 | 4418905 | 4419968 | 1064 | * | 7,6915 | -5,923260274 | 1,815891 | -3,2619 | 0,001107 | 1 | 0 | Intron (ENSMUST0000( | 12 | 4324813 | 4404436 | 79624 | 2 | 17977 | ENSMUST00000217674.2 | -14469 | ENSMUSG00000020647 | Ncoa1 | nuclear receptor coactivator 1 |
| chr8 | 99642266 | 99642807 | 542 | * | 6,552225 | -5,922643793 | 1,96389 | -3,01577 | 0,002563 | 1 | 0 | Distal Intergenic | 8 | 99738518 | 99759372 | 20855 | 1 | 1,01E+08 | ENSMUST00000161244.2 | -95711 | ENSMUSG00000089819 | Gm15679 | predicted gene 15679 |
| chr11 | 74625217 | 74626107 | 891 | * | 5,457166 | -5,92251014 | 2,013661 | -2,94117 | 0,00327 | 1 | 0 | Distal Intergenic | 11 | 74619229 | 74619335 | 107 | 1 | 1,15E+08 | ENSMUST00000158264.3 | 5988 | ENSMUSG00000088889 | Gm24211 | predicted gene, 24211 |
| chr14 | 1,2E+08 | 1,2E+08 | 441 | * | 4,303495 | -5,922305383 | 2,102158 | -2,81725 | 0,004844 | 1 | 0 | Intron (ENSMUST0000( | 14 | 1,19E+08 | 1,2E+08 | 731475 | 1 | 50787 | ENSMUST00000065904.5 | 126243 | ENSMUSG00000053465 | Hs6st3 | heparan sulfate 6-O-sulfotransferase 3 |
| chr10 | 64413397 | 64414341 | 945 | * | 4,905411 | -5,922301453 | 2,103008 | -2,81611 | 0,004861 | 1 | 0 | Intron (ENSMUST0000( | 10 | 63765669 | 63926056 | 160388 | 2 | 216028 | ENSMUST00000131945.2 | -487341 | ENSMUSG00000042846 | Lrrtm3 | leucine rich repeat transmembrane neuronal 3 |
| chr10 | 18560031 | 18560915 | 885 | * | 6,288558 | -5,921060278 | 1,904157 | -3,10954 | 0,001874 | 1 | 0 | Intron (ENSMUST0000( | 10 | 18619217 | 18713727 | 94511 | 1 | 1,03E+08 | ENSMUST00000215162.2 | -58302 | ENSMUSG00000111474 | Gm32926 | predicted gene, 32926 |
| chr4 | 1,15E+08 | 1,15E+08 | 507 | * | 4,227858 | -5,919089981 | 2,187373 | -2,70603 | 0,006809 | 1 | 0 | Distal Intergenic | 4 | 1,15E+08 | 1,15E+08 | 27959 | 2 | 81906 | ENSMUST00000106545.8 | -34573 | ENSMUSG00000047155 | Cyp4x1 | cytochrome P450, family 4, subfamily x, polypeptide 1 |
| chr19 | 28305298 | 28306654 | 1357 | * | 8,345003 | -5,918991035 | 1,805695 | -3,27796 | 0,001046 | 0,415338694 | 2,237403 | Promoter (1-2kb) | 19 | 28239747 | 28303406 | 63660 | 2 | 226075 | ENSMUST00000159520.8 | -1892 | ENSMUSG00000052942 | Glis3 | GLIS family zinc finger 3 |
| chr15 | 25364703 | 25365274 | 572 | * | 4,734777 | -5,918879927 | 2,262983 | -2,61552 | 0,008909 | 1 | 0 | 5' UTR | 15 | 25285888 | 25332745 | 46858 | 2 | 78115 | ENSMUST00000226199.2 | -31958 | | 4930445E18Rik | RIKEN cDNA 4930445E18 gene |
| chr14 | 78811468 | 78812233 | 766 | * | 4,517113 | -5,918756282 | 2,154629 | -2,747 | 0,006014 | 1 | 0 | Intron (ENSMUST0000( | 14 | 78841150 | 78841271 | 122 | 1 | 1,15E+08 | ENSMUST00000122498.3 | -28917 | ENSMUSG00000084447 | Gm26251 | predicted gene, 26251 |
| chr19 | 30088465 | 30089412 | 948 | * | 7,874859 | -5,917992887 | 1,778764 | -3,32702 | 0,000878 | 1 | 0 | Intron (ENSMUST0000( | 19 | 30069361 | 30070339 | 979 | 1 | 109113 | ENSMUST00000143769.2 | 19104 | ENSMUSG00000024817 | Uhrf2 | ubiquitin-like, containing PHD and RING finger domains 2 |
| chrX | 1,3E+08 | 1,3E+08 | 702 | * | 4,96707 | -5,914586217 | 2,00306 | -2,95278 | 0,003149 | 1 | 0 | Distal Intergenic | 20 | 1,3E+08 | 1,3E+08 | 129 | 2 | 1,15E+08 | ENSMUST00000104191.3 | -514655 | ENSMUSG00000077379 | Gm25170 | predicted gene, 25170 |
| chr9 | 55472631 | 55473261 | 631 | * | 5,235535 | -5,914517284 | 1,990788 | -2,97094 | 0,002969 | 1 | 0 | Intron (ENSMUST0000( | 9 | 55449620 | 55451564 | 1945 | 1 | 104360 | ENSMUST00000165302.2 | 23011 | ENSMUSG00000032318 | Isl2 | insulin related protein 2 (islet 2) |
| chr17 | 36697305 | 36698001 | 697 | * | 6,075504 | -5,914194205 | 1,961321 | -3,01541 | 0,002566 | 1 | 0 | Distal Intergenic | 17 | 36675895 | 36679309 | 3415 | 2 | 110696 | ENSMUST00000073546.5 | -17996 | ENSMUSG00000058124 | H2-M10.3 | histocompatibility 2, M region locus 10.3 |
| chrX | 1,64E+08 | 1,64E+08 | 843 | * | 5,183424 | -5,913950768 | 1,987321 | -2,97584 | 0,002922 | 1 | 0 | Distal Intergenic | 20 | 1,64E+08 | 1,64E+08 | 16561 | 1 | 237211 | ENSMUST00000057150.8 | 49388 | ENSMUSG00000047757 | Fancb | Fanconi anemia, complementation group B |
| chr2 | 1,13E+08 | 1,13E+08 | 652 | * | 6,604601 | -5,913686368 | 1,880895 | -3,14408 | 0,001666 | 1 | 0 | Intron (ENSMUST0000( | 2 | 1,12E+08 | 1,13E+08 | 398952 | 2 | 20192 | ENSMUST00000091818.6 | -27480 | ENSMUSG00000057378 | Ryr3 | ryanodine receptor 3 |
| chr12 | 14695657 | 14696013 | 357 | * | 4,637115 | -5,913153622 | 2,093176 | -2,82497 | 0,004729 | 1 | 0 | Distal Intergenic | 12 | 14556588 | 14556694 | 107 | 1 | 1,15E+08 | ENSMUST00000240467.1 | 139069 | ENSMUSG00000119624 | Gm25414 | predicted gene, 25414 |
| chr11 | 31211731 | 31212129 | 399 | * | 4,506126 | -5,912439866 | 2,231692 | -2,64931 | 0,008066 | 1 | 0 | Distal Intergenic | 11 | 31315441 | 31318166 | 2726 | 2 | 20856 | ENSMUST00000149467.2 | 106037 | ENSMUSG00000020303 | Stc2 | stanniocalcin 2 |
| chr5 | 24077027 | 24077384 | 358 | * | 4,122951 | -5,912407262 | 2,27257 | -2,60164 | 0,009278 | 1 | 0 | Exon (ENSMUST000002 | 5 | 24057208 | 24057278 | 71 | 1 | 1E+08 | ENSMUST00000175580.3 | 19819 | | Snord93 | small nucleolar RNA, C/D box 93 |
| chr6 | 1,15E+08 | 1,15E+08 | 686 | * | 9,590989 | -5,91141987 | 1,774478 | -3,33136 | 0,000864 | 0,454309106 | 1,449706 | Exon (ENSMUST000002 | 6 | 1,15E+08 | 1,15E+08 | 129449 | 1 | 19016 | ENSMUST00000171644.8 | -43907 | ENSMUSG00000000440 | Pparg | peroxisome proliferator activated receptor gamma |
| chr7 | 1,15E+08 | 1,15E+08 | 642 | * | 5,269796 | -5,909878108 | 1,990915 | -2,96842 | 0,002993 | 1 | 0 | Distal Intergenic | 7 | 1,15E+08 | 1,15E+08 | 1803 | 1 | 1,03E+08 | ENSMUST00000206921.2 | 159161 | | Gm34225 | predicted gene, 34225 |
| chrX | 62134450 | 62135518 | 1069 | * | 6,011342 | -5,909416179 | 2,121392 | -2,78563 | 0,005342 | 1 | 0 | Distal Intergenic | 20 | 61554145 | 61570617 | 16473 | 2 | 75545 | ENSMUST00000129479.2 | -563833 | ENSMUSG00000085757 | Tslrn1 | testis specific long noncoding RNA 1 |
| chr15 | 5425090 | 5425562 | 473 | * | 4,628419 | -5,909306595 | 2,160095 | -2,73567 | 0,006225 | 1 | 0 | Distal Intergenic | 15 | 5525747 | 5624465 | 98719 | 1 | 71432 | ENSMUST00000211750.2 | -100185 | ENSMUSG00000110018 | 5430437J10Rik | RIKEN cDNA 5430437J10 gene |
| chrX | 29329583 | 29330394 | 812 | * | 3,617206 | -5,909083575 | 2,254476 | -2,62104 | 0,008766 | 1 | 0 | Distal Intergenic | 20 | 29185123 | 29206497 | 21375 | 2 | 1,03E+08 | ENSMUST00000119298.2 | -123086 | | Gm7421 | predicted gene 7421 |
| chr16 | 6071013 | 6071816 | 804 | * | 4,443544 | -5,90856472 | 2,086628 | -2,83163 | 0,004631 | 1 | 0 | Intron (ENSMUST0000( | 16 | 6167002 | 7229390 | 1062389 | 1 | 268859 | ENSMUST00000056416.9 | -95186 | ENSMUSG00000008658 | Rbfox1 | RNA binding protein, fox-1 homolog (C. elegans) 1 |
| chr9 | 49015171 | 49015849 | 679 | * | 5,76525 | -5,907058195 | 2,04877 | -2,88322 | 0,003936 | 1 | 0 | Promoter (1-2kb) | 9 | 49014121 | 49028291 | 14171 | 1 | 80893 | ENSMUST00000167095.8 | 1050 | ENSMUSG00000032268 | Tmprss5 | transmembrane protease, serine 5 (spinesin) |
| chr16 | 54050737 | 54051701 | 965 | * | 7,396619 | -5,906800723 | 1,817407 | -3,25012 | 0,001154 | 0,514210241 | 1,360622 | Distal Intergenic | 16 | 54174825 | 54174956 | 132 | 1 | 1,15E+08 | ENSMUST00000082793.3 | -123124 | ENSMUSG00000064727 | Gm25853 | predicted gene, 25853 |
| chr2 | 17643492 | 17643899 | 408 | * | 3,975566 | -5,905805453 | 2,26672 | -2,60544 | 0,009176 | 1 | 0 | Intron (ENSMUST0000( | 2 | 17568325 | 17568437 | 113 | 1 | 1,02E+08 | ENSMUST00000184784.3 | 75167 | ENSMUSG00000099161 | Mir6419 | microRNA 6419 |
| chr15 | 3523700 | 3524582 | 883 | * | 10,85085 | -5,905744876 | 1,747143 | -3,38023 | 0,000724 | 0,446344042 | 1,2763 | Intron (ENSMUST0000( | 15 | 3357021 | 3501126 | 144106 | 2 | 14600 | ENSMUST00000110697.9 | -22574 | ENSMUSG00000055737 | Ghr | growth hormone receptor |
| chr11 | 52056856 | 52057606 | 751 | * | 5,502309 | -5,904439963 | 2,033383 | -2,90375 | 0,003687 | 1 | 0 | Promoter (1-2kb) | 11 | 52045549 | 52058986 | 13438 | 2 | 257871 | ENSMUST00000203403.3 | 1380 | | Olfr1372-ps1 | olfactory receptor 1372, pseudogene 1 |
| chr4 | 1,23E+08 | 1,23E+08 | 690 | * | 4,761166 | -5,903120485 | 2,072095 | -2,84887 | 0,004388 | 1 | 0 | Exon (ENSMUST000001 | 4 | 1,23E+08 | 1,23E+08 | 8191 | 2 | 11426 | ENSMUST00000106213.9 | 21980 | ENSMUSG00000028649 | Macf1 | microtubule-actin crosslinking factor 1 |
| chr6 | 1,18E+08 | 1,18E+08 | 810 | * | 9,49339 | -5,902024851 | 1,840123 | -3,20741 | 0,001339 | 0,563858922 | 1,211315 | Exon (ENSMUST00000( | 6 | 1,18E+08 | 1,18E+08 | 22642 | 2 | 213895 | ENSMUST00000205207.3 | 6923 | ENSMUSG00000030138 | Bms1 | BMS1, ribosome biogenesis factor |
| chr9 | 24208915 | 24209519 | 605 | * | 4,720342 | -5,901897091 | 2,032937 | -2,90314 | 0,003694 | 1 | 0 | Promoter (2-3kb) | 9 | 24211630 | 24226167 | 14538 | 1 | 319239 | ENSMUST00000154337.2 | -2111 | ENSMUSG00000043659 | Npsr1 | neuropeptide S receptor 1 |
| chr18 | 83199389 | 83199863 | 475 | * | 4,931546 | -5,899998187 | 2,131933 | -2,76744 | 0,00565 | 1 | 0 | Distal Intergenic | 18 | 83093433 | 83126015 | 32583 | 2 | 76995 | ENSMUST00000235874.2 | -73374 | | 1700095A13Rik | RIKEN cDNA 1700095A13 gene |
| chr14 | 1,1E+08 | 1,1E+08 | 1285 | * | 7,261582 | -5,899647935 | 1,790588 | -3,29481 | 0,000985 | 0,487259032 | 1,360622 | Distal Intergenic | 14 | 1,09E+08 | 1,09E+08 | 5359 | 2 | 76965 | ENSMUST00000100322.4 | -349471 | ENSMUSG00000075478 | Slitrk1 | SLIT and NTRK-like family, member 1 |
| chr10 | 1,16E+08 | 1,16E+08 | 610 | * | 4,614515 | -5,89934837 | 2,073867 | -2,84461 | 0,004447 | 1 | 0 | Intron (ENSMUST0000( | 10 | 1,16E+08 | 1,16E+08 | 16481 | 1 | 216350 | ENSMUST00000217900.2 | -12294 | ENSMUSG00000034127 | Tspan8 | tetraspanin 8 |
| chrX | 1,51E+08 | 1,51E+08 | 696 | * | 5,034932 | -5,899311582 | 2,137461 | -2,75996 | 0,005781 | 1 | 0 | Distal Intergenic | 20 | 1,51E+08 | 1,51E+08 | 119 | 1 | 1,15E+08 | ENSMUST00000093707.3 | 4376 | ENSMUSG00000118781 | n-R5s13 | nuclear encoded rRNA 5S 13 |
| chr5 | 32474504 | 32474937 | 434 | * | 5,284779 | -5,897141425 | 1,982145 | -2,97513 | 0,002929 | 1 | 0 | Exon (ENSMUST000002 | 5 | 32510913 | 32523864 | 12952 | 1 | 665270 | ENSMUST00000202453.4 | -35976 | ENSMUSG00000029134 | Plb1 | phospholipase B1 |
| chr9 | 85049490 | 85050028 | 539 | * | 5,663186 | -5,894927722 | 1,908719 | -3,08842 | 0,002012 | 1 | 0 | Distal Intergenic | 9 | 85202492 | 85209177 | 6686 | 2 | 212943 | ENSMUST00000034802.15 | 159149 | ENSMUSG00000032265 | Tent5a | terminal nucleotidyltransferase 5A |
| chr10 | 26265515 | 26265948 | 434 | * | 5,315834 | -5,893724689 | 2,100183 | -2,80629 | 0,005012 | 1 | 0 | Distal Intergenic | 10 | 26209756 | 26251967 | 42212 | 2 | 237339 | ENSMUST00000217699.2 | -13548 | ENSMUSG00000039089 | L3mbtl3 | L3MBTL3 histone methyl-lysine binding protein |
| chr2 | 1,15E+08 | 1,15E+08 | 594 | * | 4,884078 | -5,893162509 | 2,178484 | -2,70517 | 0,006827 | 1 | 0 | Intron (ENSMUST0000( | 2 | 1,15E+08 | 1,15E+08 | 189880 | 1 | 1,03E+08 | ENSMUST00000144368.2 | 131144 | | Gm13974 | predicted gene 13974 |
| chr18 | 12424390 | 12424970 | 581 | * | 4,580427 | -5,893124999 | 2,184048 | -2,69826 | 0,00697 | 1 | 0 | Intron (ENSMUST0000( | 18 | 12395169 | 12429048 | 33880 | 2 | 225187 | ENSMUST00000234212.2 | 4078 | ENSMUSG00000057766 | Ankrd29 | ankyrin repeat domain 29 |

| | | | | | | | | | | | | | | | | | | | | | | | |
|---|---|---|---|---|---|---|---|---|---|---|---|---|---|---|---|---|---|---|---|---|---|---|---|
| chrX | 1,45E+08 | 1,45E+08 | 791 | * | 5,48889 | -5,89309145 | 2,190409 | -2,69041 | 0,007136 | 1 | 0 | Distal Intergenic | 20 | 1,46E+08 | 1,46E+08 | 26816 | 1 | 15560 | ENSMUST00000149260.8 | -670446 | ENSMUSG00000041380 | Htr2c | 5-hydroxytryptamine (serotonin) receptor 2C |
| chr3 | 73399427 | 73399931 | 505 | * | 4,295496 | -5,892785961 | 2,125683 | -2,77218 | 0,005568 | 1 | 0 | Intron (ENSMUST0000( | 3 | 73258539 | 73388984 | 130446 | 2 | 74693 | ENSMUST00000194514.2 | -10443 | ENSMUSG00000102362 | 4930509J09Rik | RIKEN cDNA 4930509J09 gene |
| chr3 | 56547811 | 56548558 | 748 | * | 4,027443 | -5,892595283 | 2,278725 | -2,58592 | 0,009712 | 1 | 0 | Distal Intergenic | 3 | 56352703 | 56352833 | 131 | 2 | 1,15E+08 | ENSMUST00000104522.3 | -194978 | ENSMUSG00000119391 | Gm25727 | predicted gene, 25727 |
| chr14 | 73791771 | 73792391 | 621 | * | 4,858365 | -5,890584083 | 2,001397 | -2,94324 | 0,003248 | 1 | 0 | Promoter (1-2kb) | 14 | 73790206 | 73833223 | 43018 | 1 | 20916 | ENSMUST00000022706.7 | 1565 | ENSMUSG00000022110 | Sucla2 | succinate-Coenzyme A ligase, ADP-forming, beta subunit |
| chr10 | 68823014 | 68823432 | 419 | * | 4,370296 | -5,889022813 | 2,068878 | -2,84648 | 0,004421 | 1 | 0 | Distal Intergenic | 10 | 68856570 | 68869718 | 13149 | 2 | 1,03E+08 | ENSMUST00000219854.2 | 46286 | ENSMUSG00000112331 | Gm33035 | predicted gene, 33035 |
| chr2 | 1,27E+08 | 1,27E+08 | 668 | * | 6,342582 | -5,888726066 | 1,957971 | -3,00757 | 0,002633 | 1 | 0 | Exon (ENSMUST00000: | 2 | 1,27E+08 | 1,27E+08 | 26082 | 2 | 66552 | ENSMUST00000143700.2 | 4735 | ENSMUSG00000027366 | Sppl2a | signal peptide peptidase like 2A |
| chr4 | 1,1E+08 | 1,1E+08 | 1095 | * | 9,124204 | -5,887580888 | 1,74609 | -3,37187 | 0,000747 | 0,45991801 | 1,211315 | Intron (ENSMUST0000( | 4 | 1,1E+08 | 1,1E+08 | 6349 | 1 | 14084 | ENSMUST00000143217.2 | 29691 | ENSMUSG00000010517 | Faf1 | Fas-associated factor 1 |
| chr14 | 72452261 | 72453349 | 1089 | * | 6,490842 | -5,886973363 | 1,851156 | -3,18016 | 0,001472 | 1 | 0 | Distal Intergenic | 14 | 72663453 | 72729278 | 65826 | 2 | 675947 | ENSMUST00000208955.2 | 275929 | | Gm9195 | predicted gene 9195 |
| chr9 | 36173902 | 36174584 | 683 | * | 5,826587 | -5,886802519 | 1,894624 | -3,10711 | 0,001889 | 1 | 0 | Exon (ENSMUST00000: | 9 | 36167227 | 36169619 | 2393 | 2 | 1,05E+08 | ENSMUST00000181639.3 | -4283 | | Gm3867 | predicted gene 3867 |
| chr11 | 96039619 | 96040096 | 478 | * | 4,47799 | -5,886596798 | 2,180495 | -2,69966 | 0,006941 | 1 | 0 | Intron (ENSMUST0000( | 11 | 96025045 | 96056277 | 31233 | 1 | 237930 | ENSMUST00000107680.2 | 14574 | ENSMUSG00000038756 | Ttll6 | tubulin tyrosine ligase-like family, member 6 |
| chr2 | 50239158 | 50239640 | 483 | * | 4,682257 | -5,883528238 | 2,091635 | -2,81289 | 0,00491 | 1 | 0 | Intron (ENSMUST0000( | 2 | 50270412 | 50302622 | 32211 | 1 | 1,03E+08 | ENSMUST00000227437.2 | -30772 | | Gm48908 | predicted gene 48908 |
| chr9 | 70913064 | 70913537 | 474 | * | 4,093372 | -5,882647409 | 2,192566 | -2,683 | 0,007297 | 1 | 0 | Distal Intergenic | 9 | 70705454 | 70859508 | 154055 | 2 | 15450 | ENSMUST00000216798.2 | -53556 | ENSMUSG00000032207 | Lipc | lipase, hepatic |
| chr1 | 31547586 | 31548752 | 1167 | * | 8,506806 | -5,882408943 | 1,796095 | -3,27511 | 0,001056 | 1 | 0 | Distal Intergenic | 1 | 31695713 | 31695819 | 107 | 2 | 1,15E+08 | ENSMUST00000157995.3 | 147067 | ENSMUSG00000088620 | Gm23032 | predicted gene, 23032 |
| chr9 | 62399548 | 62400073 | 526 | * | 5,345108 | -5,882357234 | 1,99527 | -2,94815 | 0,003197 | 1 | 0 | Intron (ENSMUST0000( | 9 | 62335218 | 62417780 | 82563 | 2 | 235431 | ENSMUST00000164246.9 | 17707 | ENSMUSG00000041729 | Coro2b | coronin, actin binding protein, 2B |
| chr5 | 49024207 | 49025092 | 886 | * | 8,055775 | -5,8798135 | 1,82348 | -3,2245 | 0,001262 | 1 | 0 | Intron (ENSMUST0000( | 5 | 48546844 | 49046899 | 500056 | 2 | 80334 | ENSMUST00000176978.8 | 21807 | ENSMUSG00000029088 | Kcnip4 | Kv channel interacting protein 4 |
| chr14 | 70015040 | 70015411 | 372 | * | 4,067629 | -5,879503816 | 2,207819 | -2,66304 | 0,007744 | 1 | 0 | Promoter (<=1kb) | 14 | 70014918 | 70020123 | 5206 | 1 | 21933 | ENSMUST00000224235.2 | 122 | ENSMUSG00000022074 | Tnfrsf10b | tumor necrosis factor receptor superfamily, member 10b |
| chr6 | 1,37E+08 | 1,37E+08 | 439 | * | 4,759705 | -5,876628894 | 2,149245 | -2,73428 | 0,006252 | 1 | 0 | Intron (ENSMUST0000( | 6 | 1,37E+08 | 1,37E+08 | 84458 | 2 | 14917 | ENSMUST00000078095.11 | 8449 | ENSMUSG00000042638 | Gucy2c | guanylate cyclase 2c |
| chrX | 36383240 | 36384375 | 1136 | * | 4,008881 | -5,876178282 | 2,244079 | -2,61853 | 0,008831 | 1 | 0 | Distal Intergenic | 20 | 36390448 | 36414399 | 23952 | 1 | 67050 | ENSMUST00000016553.5 | -6073 | ENSMUSG00000016409 | Nkap | NFKB activating protein |
| chr16 | 9066167 | 9066733 | 567 | * | 4,463033 | -5,875586033 | 2,187506 | -2,68597 | 0,007232 | 1 | 0 | Distal Intergenic | 16 | 8867836 | 8873262 | 5427 | 1 | 1,03E+08 | ENSMUST00000229201.2 | 198331 | | Gm30567 | predicted gene, 30567 |
| chr4 | 1,2E+08 | 1,2E+08 | 932 | * | 9,663525 | -5,874961361 | 1,73509 | -3,38597 | 0,000709 | 0,422040194 | 1,449706 | Distal Intergenic | 4 | 1,2E+08 | 1,2E+08 | 2175 | 2 | 14916 | ENSMUST00000106309.9 | 4658 | ENSMUSG00000032978 | Guca2b | guanylate cyclase activator 2b (retina) |
| chr14 | 1,08E+08 | 1,08E+08 | 975 | * | 7,412184 | -5,873002316 | 1,919805 | -3,05917 | 0,00222 | 1 | 0 | Distal Intergenic | 14 | 1,09E+08 | 1,09E+08 | 5359 | 2 | 76965 | ENSMUST00000100322.4 | 923385 | ENSMUSG00000075478 | Slitrk1 | SLIT and NTRK-like family, member 1 |
| chr17 | 13066755 | 13067364 | 610 | * | 4,581695 | -5,872339757 | 2,149682 | -2,73172 | 0,0063 | 1 | 0 | Promoter (<=1kb) | 17 | 13060002 | 13067007 | 7006 | 2 | 17171 | ENSMUST00000165020.8 | 0 | ENSMUSG00000068037 | Mas1 | MAS1 oncogene |
| chr1 | 1,29E+08 | 1,29E+08 | 530 | * | 5,186326 | -5,87178825 | 2,022638 | -2,90304 | 0,003696 | 1 | 0 | Intron (ENSMUST0000( | 1 | 1,29E+08 | 1,3E+08 | 355528 | 1 | 210417 | ENSMUST00000140629.2 | 27426 | ENSMUSG00000042581 | Thsd7b | thrombospondin, type I, domain containing 7B |
| chrX | 66778934 | 66779680 | 747 | * | 6,031433 | -5,870994398 | 2,014437 | -2,91446 | 0,003563 | 1 | 0 | Distal Intergenic | 20 | 66739304 | 66749956 | 10653 | 1 | 666842 | ENSMUST00000114672.2 | 39630 | ENSMUSG00000079577 | Gm14692 | predicted gene 14692 |
| chr12 | 30633416 | 30633585 | 170 | * | 4,829555 | -5,870361777 | 2,093114 | -2,80461 | 0,005038 | 1 | 0 | Promoter (<=1kb) | 12 | 30634425 | 30641214 | 6790 | 1 | 211986 | ENSMUST00000057151.10 | -840 | ENSMUSG00000043061 | Tmem18 | transmembrane protein 18 |
| chrX | 1,27E+08 | 1,27E+08 | 760 | * | 4,060295 | -5,869593498 | 2,226151 | -2,63666 | 0,008373 | 1 | 0 | Distal Intergenic | 20 | 1,27E+08 | 1,27E+08 | 107 | 2 | 1,15E+08 | ENSMUST00000240076.1 | -372728 | ENSMUSG00000119321 | Gm25061 | predicted gene, 25061 |
| chr2 | 1,18E+08 | 1,18E+08 | 504 | * | 4,174034 | -5,869355142 | 2,219783 | -2,64411 | 0,008191 | 1 | 0 | Distal Intergenic | 2 | 1,18E+08 | 1,18E+08 | 43381 | 1 | 12236 | ENSMUST00000038341.8 | -13479 | ENSMUSG00000040084 | Bub1b | BUB1B, mitotic checkpoint serine/threonine kinase |
| chr1 | 1,86E+08 | 1,86E+08 | 1085 | * | 4,372769 | -5,869354068 | 2,208346 | -2,65781 | 0,007865 | 1 | 0 | Promoter (<=1kb) | 1 | 1,86E+08 | 1,86E+08 | 46635 | 2 | 320449 | ENSMUST00000193084.2 | 816 | ENSMUSG00000103690 | A730004F24Rik | RIKEN cDNA A730004F24 gene |
| chrX | 55913793 | 55914455 | 663 | * | 4,075616 | -5,869334931 | 2,12952 | -2,75618 | 0,005848 | 1 | 0 | Promoter (1-2kb) | 20 | 55915523 | 56025697 | 110175 | 1 | 236798 | ENSMUST00000154818.8 | -1068 | ENSMUSG00000053852 | Adgrg4 | adhesion G protein-coupled receptor G4 |
| chrX | 1,28E+08 | 1,28E+08 | 568 | * | 5,053297 | -5,869272566 | 1,942557 | -3,02142 | 0,002516 | 1 | 0 | Distal Intergenic | 20 | 1,28E+08 | 1,28E+08 | 132 | 2 | 1,15E+08 | ENSMUST00000158659.3 | -115138 | ENSMUSG00000089284 | Gm26029 | predicted gene, 26029 |
| chr6 | 16129142 | 16130045 | 904 | * | 5,748142 | -5,866889234 | 2,010365 | -2,91832 | 0,003519 | 1 | 0 | Distal Intergenic | 6 | 16468235 | 16469903 | 1669 | 2 | 1,03E+08 | ENSMUST00000203052.2 | 339858 | | Gm36406 | predicted gene, 36406 |
| chr10 | 83877646 | 83878296 | 651 | * | 4,140612 | -5,865583932 | 2,121955 | -2,76424 | 0,005706 | 1 | 0 | Intron (ENSMUST0000( | 10 | 83928130 | 83932113 | 3984 | 1 | 1,03E+08 | ENSMUST00000213796.2 | -49834 | ENSMUSG00000111524 | Gm34113 | predicted gene, 34113 |
| chr16 | 54817140 | 54817743 | 604 | * | 4,787288 | -5,865210568 | 2,024669 | -2,89687 | 0,003769 | 1 | 0 | Distal Intergenic | 16 | 54792794 | 54792900 | 107 | 1 | 1,15E+08 | ENSMUST00000082615.3 | 24346 | ENSMUSG00000064549 | Gm22977 | predicted gene, 22977 |
| chr13 | 73720173 | 73720754 | 582 | * | 4,626098 | -5,863032633 | 2,204358 | -2,65975 | 0,00782 | 1 | 0 | Intron (ENSMUST0000( | 13 | 73752125 | 73768724 | 16600 | 1 | 218335 | ENSMUST00000022102.9 | -31371 | ENSMUSG00000021610 | Clptm1l | CLPTM1-like |
| chr7 | 47731375 | 47731794 | 420 | * | 3,692673 | -5,862639613 | 2,26049 | -2,59353 | 0,0095 | 1 | 0 | Distal Intergenic | 7 | 47670719 | 47677345 | 6627 | 2 | 404242 | ENSMUST00000094390.3 | -54030 | ENSMUSG00000070552 | Mrgprx1 | MAS-related GPR, member X1 |
| chr6 | 6004026 | 6004640 | 615 | * | 5,783272 | -5,860966602 | 1,912569 | -3,06445 | 0,002181 | 1 | 0 | Promoter (1-2kb) | 6 | 6006066 | 6027787 | 21722 | 1 | 13426 | ENSMUST00000203520.2 | -1426 | ENSMUSG00000029757 | Dync1i1 | dynein cytoplasmic 1 intermediate chain 1 |
| chr1 | 1,4E+08 | 1,4E+08 | 646 | * | 5,018481 | -5,859965616 | 2,077697 | -2,82041 | 0,004796 | 1 | 0 | Intron (ENSMUST0000( | 1 | 1,4E+08 | 1,41E+08 | 61834 | 1 | 240776 | ENSMUST00000193606.2 | -18441 | ENSMUSG00000052726 | Kcnt2 | potassium channel, subfamily T, member 2 |
| chr3 | 1,08E+08 | 1,08E+08 | 489 | * | 3,976087 | -5,859283268 | 2,272839 | -2,57796 | 0,009939 | 1 | 0 | Intron (ENSMUST0000( | 3 | 1,08E+08 | 1,08E+08 | 5972 | 1 | 20661 | ENSMUST00000129755.3 | -18927 | ENSMUSG00000068747 | Sort1 | sortilin 1 |
| chr1 | 1,69E+08 | 1,69E+08 | 1889 | * | 8,738543 | -5,857095704 | 1,789478 | -3,27308 | 0,001064 | 0,490699003 | 1,449706 | Distal Intergenic | 1 | 1,68E+08 | 1,68E+08 | 120 | 1 | 1,02E+08 | ENSMUST00000183804.3 | 218144 | ENSMUSG00000098407 | Mir6348 | microRNA 6348 |
| chr1 | 1,64E+08 | 1,64E+08 | 1043 | * | 7,61227 | -5,856513941 | 1,80231 | -3,24945 | 0,001156 | 1 | 0 | Distal Intergenic | 1 | 1,64E+08 | 1,64E+08 | 22977 | 2 | 240879 | ENSMUST00000159679.8 | 51571 | ENSMUSG00000040113 | Mettl11b | methyltransferase like 11B |
| chr19 | 21885217 | 21886343 | 1127 | * | 6,934358 | -5,85636043 | 1,863243 | -3,1431 | 0,001672 | 1 | 0 | Distal Intergenic | 19 | 21755752 | 21785266 | 29515 | 1 | 83921 | ENSMUST00000238079.2 | 129465 | ENSMUSG00000024754 | Cemip2 | cell migration inducing hyaluronidase 2 |
| chr4 | 3980539 | 3981484 | 946 | * | 7,034206 | -5,856346432 | 1,870171 | -3,13145 | 0,001739 | 1 | 0 | Distal Intergenic | 4 | 3946808 | 3949872 | 3065 | 1 | 66433 | ENSMUST00000151519.2 | 33731 | ENSMUSG00000042198 | Chchd7 | coiled-coil-helix-coiled-coil-helix domain containing 7 |
| chr9 | 99164885 | 99165593 | 709 | * | 6,837582 | -5,856320497 | 1,882834 | -3,11038 | 0,001868 | 1 | 0 | Intron (ENSMUST0000( | 9 | 99173541 | 99178404 | 4864 | 1 | 68121 | ENSMUST00000187883.2 | -7948 | ENSMUSG00000056267 | Cep70 | centrosomal protein 70 |
| chr18 | 32795175 | 32795972 | 798 | * | 6,436917 | -5,856317272 | 1,884079 | -3,10832 | 0,001882 | 1 | 0 | Distal Intergenic | 18 | 32754973 | 32755109 | 137 | 2 | 1,15E+08 | ENSMUST00000157090.3 | -40066 | ENSMUSG00000087715 | Gm25826 | predicted gene, 25826 |
| chr18 | 45166336 | 45166642 | 307 | * | 3,943941 | -5,85545863 | 2,185315 | -2,67946 | 0,007374 | 1 | 0 | Distal Intergenic | 18 | 45177929 | 45262810 | 84882 | 1 | 1,03E+08 | ENSMUST00000238022.2 | -11287 | | Gm31706 | predicted gene, 31706 |
| chrX | 1,3E+08 | 1,3E+08 | 1041 | * | 6,309243 | -5,85410841 | 1,953852 | -2,99619 | 0,002734 | 1 | 0 | Distal Intergenic | 20 | 1,3E+08 | 1,3E+08 | 129 | 2 | 1,15E+08 | ENSMUST00000104191.3 | -321878 | ENSMUSG00000077379 | Gm25170 | predicted gene, 25170 |
| chr1 | 1,54E+08 | 1,54E+08 | 573 | * | 5,306722 | -5,853987861 | 1,96947 | -2,97237 | 0,002955 | 1 | 0 | Distal Intergenic | 1 | 1,54E+08 | 1,54E+08 | 4503 | 1 | 1,01E+08 | ENSMUST00000086195.7 | -15868 | ENSMUSG00000066797 | Zfp648 | zinc finger protein 648 |
| chr5 | 1,27E+08 | 1,27E+08 | 626 | * | 5,794974 | -5,851752117 | 1,968666 | -2,97245 | 0,002954 | 1 | 0 | Promoter (2-3kb) | 5 | 1,27E+08 | 1,27E+08 | 1126 | 2 | 1,05E+08 | ENSMUST00000196498.2 | 2112 | | Gm40330 | predicted gene, 40330 |
| chr1 | 1,72E+08 | 1,72E+08 | 792 | * | 6,833218 | -5,851444767 | 1,896254 | -3,08579 | 0,00203 | 1 | 0 | Promoter (<=1kb) | 1 | 1,72E+08 | 1,72E+08 | 4267 | 2 | 93840 | ENSMUST00000148363.2 | -211 | ENSMUSG00000026556 | Vangl2 | VANGL planar cell polarity 2 |
| chr1 | 49114903 | 49115506 | 604 | * | 5,665355 | -5,850684886 | 2,067596 | -2,8297 | 0,004659 | 1 | 0 | Distal Intergenic | 1 | 49283775 | 49377405 | 93631 | 1 | 442837 | ENSMUST00000056879.9 | -168269 | ENSMUSG00000051616 | C230029F24Rik | RIKEN cDNA C230029F24 gene |
| chr7 | 51502853 | 51503799 | 947 | * | 6,363466 | -5,849766318 | 1,900186 | -3,07852 | 0,00208 | 1 | 0 | Distal Intergenic | 7 | 51511763 | 51644723 | 132961 | 1 | 14453 | ENSMUST00000129604.8 | -7964 | ENSMUSG00000030498 | Gas2 | growth arrest specific 2 |
| chr3 | 20100574 | 20101321 | 748 | * | 5,674674 | -5,849677706 | 1,929273 | -3,03206 | 0,002429 | 1 | 0 | Distal Intergenic | 3 | 20111975 | 20171445 | 59471 | 1 | 20585 | ENSMUST00000143005.8 | -10654 | ENSMUSG00000002428 | Hltf | helicase-like transcription factor |
| chr7 | 1,25E+08 | 1,25E+08 | 418 | * | 4,685518 | -5,849265257 | 2,141074 | -2,73193 | 0,006296 | 1 | 0 | Distal Intergenic | 7 | 1,25E+08 | 1,25E+08 | 5795 | 2 | 233863 | ENSMUST00000206042.2 | 13575 | ENSMUSG00000032777 | Gtf3c1 | general transcription factor III C 1 |
| chr12 | 76405547 | 76405986 | 440 | * | 3,974761 | -5,848688739 | 2,185046 | -2,67669 | 0,007435 | 1 | 0 | Promoter (1-2kb) | 12 | 76394556 | 76403914 | 9359 | 2 | 109929 | ENSMUST00000176102.8 | -1633 | ENSMUSG00000056459 | Zbtb25 | zinc finger and BTB domain containing 25 |
| chr17 | 85291089 | 85291550 | 462 | * | 4,512611 | -5,848624771 | 2,170461 | -2,69465 | 0,007046 | 1 | 0 | Intron (ENSMUST0000( | 17 | 85301101 | 85331419 | 30319 | 1 | 19043 | ENSMUST00000112307.4 | -9551 | ENSMUSG00000061130 | Ppm1b | protein phosphatase 1B, magnesium dependent, beta isoform |
| chr7 | 57780498 | 57781414 | 917 | * | 5,159554 | -5,848432772 | 2,132136 | -2,74299 | 0,006088 | 1 | 0 | Distal Intergenic | 7 | 57797844 | 57797975 | 132 | 2 | 1,15E+08 | ENSMUST00000104456.3 | 16561 | ENSMUSG00000119548 | Gm26288 | predicted gene, 26288 |
| chr7 | 1,37E+08 | 1,37E+08 | 471 | * | 4,806886 | -5,848015017 | 2,06129 | -2,83707 | 0,004553 | 1 | 0 | Distal Intergenic | 7 | 1,37E+08 | 1,37E+08 | 4481 | 1 | 30926 | ENSMUST00000211598.2 | 203264 | ENSMUSG00000031068 | Glrx3 | glutaredoxin 3 |
| chr2 | 44408028 | 44408355 | 328 | * | 3,548184 | -5,845620263 | 2,235435 | -2,61498 | 0,008923 | 1 | 0 | Distal Intergenic | 2 | 44455145 | 44461359 | 6215 | 2 | 227835 | ENSMUST00000134813.2 | 53004 | ENSMUSG00000036890 | Gtdc1 | glycosyltransferase-like domain containing 1 |
| chr13 | 20923489 | 20924566 | 1078 | * | 6,658688 | -5,845525379 | 1,949858 | -2,99792 | 0,002718 | 1 | 0 | Distal Intergenic | 13 | 20978283 | 21208422 | 230140 | 1 | 27052 | ENSMUST00000021757.5 | -53717 | ENSMUSG00000021322 | Aoah | acyloxyacyl hydrolase |
| chr16 | 37321306 | 37322092 | 787 | * | 4,304122 | -5,845512663 | 2,149503 | -2,71947 | 0,006539 | 1 | 0 | Distal Intergenic | 16 | 37352882 | 37360138 | 7257 | 2 | 74197 | ENSMUST00000130350.2 | 38046 | ENSMUSG00000022828 | Gtf2e1 | general transcription factor II E, polypeptide 1 (alpha subunit) |
| chr12 | 50107625 | 50108595 | 971 | * | 6,240369 | -5,843436055 | 1,85588 | -3,14861 | 0,001641 | 1 | 0 | Distal Intergenic | 12 | 50167092 | 50192019 | 24928 | 1 | 1,05E+08 | ENSMUST00000218806.2 | -58497 | ENSMUSG00000112353 | Gm40418 | predicted gene, 40418 |
| chrX | 50086791 | 50087665 | 875 | * | 6,0386 | -5,843322904 | 1,87832 | -3,11093 | 0,001865 | 1 | 0 | Distal Intergenic | 20 | 50092789 | 50106895 | 14107 | 2 | 72065 | ENSMUST00000053593.8 | 19230 | ENSMUSG00000050029 | Rap2c | RAP2C, member of RAS oncogene family |
| chr2 | 33024060 | 33024648 | 589 | * | 5,080654 | -5,842953249 | 1,968509 | -2,96821 | 0,002995 | 1 | 0 | Promoter (2-3kb) | 2 | 32944183 | 33021666 | 77484 | 2 | 99326 | ENSMUST00000127509.2 | -2394 | ENSMUSG00000038860 | Garnl3 | GTPase activating RANGAP domain-like 3 |
| chr1 | 1,71E+08 | 1,71E+08 | 603 | * | 5,831957 | -5,842862165 | 1,999439 | -2,92225 | 0,003475 | 1 | 0 | Intron (ENSMUST0000( | 1 | 1,71E+08 | 1,71E+08 | 163098 | 2 | 226641 | ENSMUST00000027974.7 | -3687 | ENSMUSG00000026663 | Atf6 | activating transcription factor 6 |
| chr7 | 72588445 | 72589052 | 608 | * | 4,498689 | -5,842599934 | 2,097921 | -2,78495 | 0,005354 | 1 | 0 | Distal Intergenic | 7 | 72747497 | 72751816 | 4320 | 2 | 625243 | ENSMUST00000205529.2 | 162764 | ENSMUSG00000108383 | Gm6567 | predicted gene 6567 |
| chr11 | 6338454 | 6339070 | 617 | * | 4,936676 | -5,839929061 | 2,076407 | -2,81252 | 0,004916 | 1 | 0 | Promoter (<=1kb) | 11 | 6339074 | 6356158 | 17085 | 1 | 52915 | ENSMUST00000109787.8 | -4 | ENSMUSG00000041164 | Zmiz2 | zinc finger, MIZ-type containing 2 |
| chr18 | 33232145 | 33232911 | 767 | * | 6,114318 | -5,839866429 | 1,886576 | -3,09548 | 0,001965 | 1 | 0 | Intron (ENSMUST0000( | 18 | 33336458 | 33346819 | 10362 | 2 | 170459 | ENSMUST00000119991.8 | 113908 | ENSMUSG00000024378 | Stard4 | StAR-related lipid transfer (START) domain containing 4 |
| chr14 | 45819782 | 45820571 | 790 | * | 6,193845 | -5,838614492 | 2,047993 | -2,8509 | 0,00436 | 1 | 0 | Intron (ENSMUST0000( | 14 | 45802988 | 45823577 | 20590 | 1 | 1,15E+08 | ENSMUST00000228948.2 | 16794 | | Gm15601 | predicted gene 15601 |
| chr11 | 18606070 | 18607162 | 1093 | * | 11,73883 | -5,837219944 | 1,672773 | -3,48955 | 0,000484 | 0,470097239 | 0,744375 | Intron (ENSMUST0000( | 11 | 18688695 | 18688796 | 102 | 1 | 1,15E+08 | ENSMUST00000083450.4 | -81533 | ENSMUSG00000065384 | Gm23093 | predicted gene, 23093 |
| chr4 | 10918226 | 10918708 | 483 | * | 4,589247 | -5,835645165 | 2,179879 | -2,67705 | 0,007427 | 1 | 0 | Distal Intergenic | 4 | 10879539 | 10891778 | 12240 | 1 | 67157 | ENSMUST00000142026.2 | 38687 | ENSMUSG00000059482 | Cfap418 | cilia and flagella associated protein 418 |
| chr2 | 1,3E+08 | 1,3E+08 | 337 | * | 3,721973 | -5,835535056 | 2,227236 | -2,62008 | 0,008791 | 1 | 0 | Distal Intergenic | 2 | 1,3E+08 | 1,3E+08 | 9962 | 2 | 241638 | ENSMUST00000045761.7 | -4843 | ENSMUSG00000037703 | Lzts3 | leucine zipper, putative tumor suppressor family member 3 |
| chr8 | 1,29E+08 | 1,29E+08 | 572 | * | 4,967359 | -5,834381011 | 2,031221 | -2,87235 | 0,004074 | 1 | 0 | Distal Intergenic | 8 | 1,28E+08 | 1,28E+08 | 31125 | 1 | 93742 | ENSMUST00000159940.2 | 198596 | ENSMUSG00000025812 | Pard3 | par-3 family cell polarity regulator |
| chr8 | 9128299 | 9128747 | 449 | * | 4,509513 | -5,832795355 | 2,132707 | -2,73493 | 0,006239 | 1 | 0 | Distal Intergenic | 8 | 9175412 | 9182288 | 6877 | 2 | 71249 | ENSMUST00000208031.2 | 53541 | ENSMUSG00000108848 | 4933430N04Rik | RIKEN cDNA 4933430N04 gene |
| chr13 | 1,15E+08 | 1,15E+08 | 802 | * | 7,487459 | -5,831740807 | 1,907876 | -3,05667 | 0,002238 | 0,641187423 | 1,360622 | Distal Intergenic | 13 | 1,15E+08 | 1,15E+08 | 10488 | 1 | 17434 | ENSMUST00000166104.9 | -90465 | ENSMUSG00000015536 | Mocs2 | molybdenum cofactor synthesis 2 |
| chrX | 69676040 | 69677114 | 1075 | * | 5,016904 | -5,830124383 | 2,028783 | -2,87371 | 0,004057 | 1 | 0 | Distal Intergenic | 20 | 69737082 | 69737213 | 132 | 1 | 1,15E+08 | ENSMUST00000104087.3 | -59968 | ENSMUSG00000077275 | Gm22785 | predicted gene, 22785 |
| chr1 | 29357831 | 29358865 | 1035 | * | 6,547165 | -5,829562523 | 1,982558 | -2,94042 | 0,003278 | 1 | 0 | Distal Intergenic | 1 | 29460569 | 29460693 | 125 | 1 | 1,02E+08 | ENSMUST00000183870.3 | -101704 | ENSMUSG00000099257 | Mir6342 | microRNA 6342 |
| chrX | 57108237 | 57109047 | 811 | * | 4,183956 | -5,828712925 | 2,217196 | -2,62887 | 0,008567 | 1 | 0 | Distal Intergenic | 20 | 57076370 | 57087096 | 10727 | 1 | 22773 | ENSMUST00000088629.4 | 31867 | ENSMUSG00000067860 | Zic3 | zinc finger protein of the cerebellum 3 |
| chr14 | 78966210 | 78967326 | 1117 | * | 6,911183 | -5,825102373 | 1,829498 | -3,18399 | 0,001453 | 1 | 0 | Promoter (2-3kb) | 14 | 78796190 | 78970216 | 174027 | 2 | 380921 | ENSMUST00000226342.3 | 2890 | ENSMUSG00000034731 | Dgkh | diacylglycerol kinase, eta |
| chr13 | 47261957 | 47262492 | 536 | * | 6,100613 | -5,825099066 | 1,996455 | -2,91772 | 0,003526 | 1 | 0 | Promoter (2-3kb) | 13 | 47242071 | 47259677 | 17607 | 2 | 110052 | ENSMUST00000135278.8 | -2280 | ENSMUSG00000021377 | Dek | DEK proto-oncogene (DNA binding) |
| chr2 | 90441137 | 90441869 | 733 | * | 5,530275 | -5,825093345 | 1,965726 | -2,96333 | 0,003043 | 1 | 0 | Distal Intergenic | 2 | 90263233 | 90410939 | 147707 | 2 | 19271 | ENSMUST00000168621.4 | -30198 | ENSMUSG00000025314 | Ptprj | protein tyrosine phosphatase, receptor type, J |
| chr4 | 75615574 | 75615969 | 396 | * | 5,188652 | -5,825092552 | 2,012982 | -2,89376 | 0,003807 | 1 | 0 | Distal Intergenic | 4 | 75957574 | 76002694 | 45121 | 2 | 19266 | ENSMUST00000212365.2 | 386725 | ENSMUSG00000028399 | Ptprd | protein tyrosine phosphatase, receptor type, D |
| chr7 | 1,17E+08 | 1,17E+08 | 330 | * | 3,866253 | -5,825084761 | 2,158521 | -2,69865 | 0,006962 | 1 | 0 | Distal Intergenic | 7 | 1,18E+08 | 1,18E+08 | 5566 | 2 | 267019 | ENSMUST00000134270.2 | 279824 | ENSMUSG00000008683 | Rps15a | ribosomal protein S15A |
| chr3 | 20339578 | 20341022 | 1445 | * | 8,943207 | -5,824757173 | 1,781937 | -3,26878 | 0,00108 | 0,524368688 | 1,211315 | Distal Intergenic | 3 | 20302428 | 20329897 | 27470 | 2 | 76703 | ENSMUST00000011607.6 | -9681 | ENSMUSG00000011463 | Cpb1 | carboxypeptidase B1 (tissue) |
| chr18 | 42945144 | 42945535 | 392 | * | 4,249603 | -5,824262009 | 2,146677 | -2,71315 | 0,006665 | 1 | 0 | Intron (ENSMUST0000( | 18 | 42891533 | 43031880 | 140348 | 2 | 72930 | ENSMUST00000155262.2 | 86345 | ENSMUSG00000024500 | Ppp2r2b | protein phosphatase 2, regulatory subunit B, beta |
| chr11 | 23317874 | 23318658 | 785 | * | 5,176988 | -5,823730065 | 2,073713 | -2,80836 | 0,004979 | 1 | 0 | Intron (ENSMUST0000( | 11 | 23364976 | 23370652 | 5677 | 1 | 17847 | ENSMUST00000155984.2 | -46318 | ENSMUSG00000056342 | Usp34 | ubiquitin specific peptidase 34 |
| chr9 | 95040197 | 95040690 | 494 | * | 5,297921 | -5,823019766 | 1,955401 | -2,97792 | 0,002902 | 1 | 0 | Intron (ENSMUST0000( | 9 | 95003739 | 95006604 | 2866 | 1 | 331004 | ENSMUST00000162329.2 | 36458 | ENSMUSG00000031129 | Slc9a9 | solute carrier family 9 (sodium/hydrogen exchanger), member 9 |
| chr1 | 1,37E+08 | 1,37E+08 | 869 | * | 6,107421 | -5,822277762 | 1,934957 | -3,009 | 0,002621 | 1 | 0 | Distal Intergenic | 1 | 1,37E+08 | 1,37E+08 | 104 | 2 | 1,15E+08 | ENSMUST00000104019.3 | -227173 | ENSMUSG00000077207 | Gm25609 | predicted gene, 25609 |
| chr14 | 1,01E+08 | 1,01E+08 | 611 | * | 4,281354 | -5,822009696 | 2,164263 | -2,69007 | 0,007144 | 1 | 0 | Distal Intergenic | 14 | 1,01E+08 | 1,01E+08 | 101 | 2 | 1,15E+08 | ENSMUST00000157535.4 | -99044 | ENSMUSG00000088160 | Gm23538 | predicted gene, 23538 |
| chr3 | 1,16E+08 | 1,16E+08 | 449 | * | 4,580491 | -5,821434055 | 2,219815 | -2,62249 | 0,008729 | 1 | 0 | Distal Intergenic | 3 | 1,16E+08 | 1,16E+08 | 41252 | 1 | 69740 | ENSMUST00000189799.7 | -60394 | ENSMUSG00000033554 | Dph5 | diphthamide biosynthesis 5 |
| chr1 | 1,91E+08 | 1,91E+08 | 941 | * | 3,864346 | -5,821401523 | 2,205135 | -2,63993 | 0,008292 | 1 | 0 | Intron (ENSMUST0000( | 1 | 1,91E+08 | 1,91E+08 | 11406 | 2 | 66208 | ENSMUST00000046770.10 | 9517 | ENSMUSG00000037499 | Nenf | neuron derived neurotrophic factor |
| chrX | 58985488 | 58986230 | 743 | * | 5,541287 | -5,820712001 | 1,973702 | -2,94913 | 0,003187 | 1 | 0 | Distal Intergenic | 20 | 59044824 | 59076119 | 31296 | 1 | 14071 | ENSMUST00000033477.5 | -58594 | ENSMUSG00000031138 | F9 | coagulation factor IX |
| chr7 | 24300342 | 24301501 | 1160 | * | 7,15036 | -5,819737403 | 1,85317 | -3,14042 | 0,001687 | 0,586565597 | 1,360622 | Intron (ENSMUST0000( | 7 | 24310188 | 24328722 | 18535 | 1 | 232970 | ENSMUST00000206422.2 | -8687 | ENSMUSG00000074277 | Phldb3 | pleckstrin homology like domain, family B, member 3 |
| chr4 | 47876665 | 47877698 | 1034 | * | 7,803791 | -5,818630124 | 1,792695 | -3,24575 | 0,001171 | 1 | 0 | Intron (ENSMUST0000( | 4 | 48045153 | 48086445 | 41293 | 1 | 18124 | ENSMUST00000030025.10 | -167455 | ENSMUSG00000028341 | Nr4a3 | nuclear receptor subfamily 4, group A, member 3 |
| chr19 | 45268585 | 45269439 | 855 | * | 7,734256 | -5,818599018 | 1,807909 | -3,21841 | 0,001289 | 1 | 0 | Distal Intergenic | 19 | 45288909 | 45291776 | 2868 | 2 | 1,03E+08 | ENSMUST00000187954.3 | 22337 | ENSMUSG00000100417 | Gm29595 | predicted gene 29595 |
| chr7 | 58295253 | 58295846 | 594 | * | 5,603805 | -5,818434227 | 1,905559 | -3,0534 | 0,002263 | 1 | 0 | Distal Intergenic | 7 | 58305914 | 58308626 | 2713 | 1 | 11982 | ENSMUST00000207668.2 | -10068 | ENSMUSG00000025324 | Atp10a | ATPase, class V, type 10A |
| chr14 | 71778894 | 71780005 | 1112 | * | 4,751135 | -5,816925192 | 2,079059 | -2,79786 | 0,005144 | 1 | 0 | Distal Intergenic | 14 | 71658452 | 71659051 | 600 | 1 | 73857 | ENSMUST00000226329.2 | 120442 | ENSMUSG00000115286 | 4930429C20Rik | RIKEN cDNA 4930429C20 gene |
| chr7 | 72606983 | 72607725 | 743 | * | 6,639174 | -5,813558959 | 1,92528 | -3,01959 | 0,002531 | 1 | 0 | Distal Intergenic | 7 | 72747497 | 72751816 | 4320 | 2 | 625243 | ENSMUST00000205529.2 | 144091 | ENSMUSG00000108383 | Gm6567 | predicted gene 6567 |
| chr1 | 1,21E+08 | 1,21E+08 | 760 | * | 6,571244 | -5,813362845 | 1,890687 | -3,07474 | 0,002107 | 1 | 0 | Promoter (2-3kb) | 1 | 1,21E+08 | 1,21E+08 | 5123 | 1 | 69904 | ENSMUST00000188147.2 | -2137 | | 2610027F03Rik | RIKEN cDNA 2610027F03 gene |
| chr14 | 30520470 | 30521390 | 921 | * | 5,624642 | -5,813192266 | 2,038642 | -2,8515 | 0,004351 | 1 | 0 | Promoter (2-3kb) | 14 | 30518308 | 30520063 | 1756 | 1 | 54650 | ENSMUST00000227445.2 | 2162 | ENSMUSG00000006527 | Sfmbt1 | Scm-like with four mbt domains 1 |
| chr1 | 91406242 | 91407217 | 976 | * | 8,07404 | -5,81279064 | 1,812701 | -3,2067 | 0,001343 | 1 | 0 | Distal Intergenic | 1 | 91410746 | 91412496 | 1751 | 1 | 1,03E+08 | ENSMUST00000187902.2 | -3529 | | Gm28382 | predicted gene 28382 |
| chrX | 1,4E+08 | 1,4E+08 | 597 | * | 6,125544 | -5,811600078 | 1,970162 | -2,94981 | 0,00318 | 1 | 0 | Distal Intergenic | 20 | 1,4E+08 | 1,4E+08 | 14110 | 1 | 23947 | ENSMUST00000128809.2 | -8368 | ENSMUSG00000000266 | Mid2 | midline 2 |

| | | | | | | | | | | | | | | | | | | | | | | | |
|---|---|---|---|---|---|---|---|---|---|---|---|---|---|---|---|---|---|---|---|---|---|---|---|
| chr14 | 22279820 | 22280756 | 937 | * | 7,468545 | -5,811529014 | 1,98219 | -2,93187 | 0,003369 | 0,706976265 | 1,360622 | Intron (ENSMUST0000 | 14 | 22360233 | 22634507 | 274275 | 1 | 76633 | ENSMUST00000159107.2 | -79477 | ENSMUSG00000063458 | Lrmda | leucine rich melanocyte differentiation associated |
| chr12 | 1,15E+08 | 1,15E+08 | 675 | * | 4,40407 | -5,810456886 | 2,146212 | -2,70731 | 0,006783 | 1 | 0 | Distal Intergenic | 12 | 1,15E+08 | 1,15E+08 | 25225 | 1 | 1,03E+08 | ENSMUST00000195683.2 | -182201 | ENSMUSG00000103168 | Gm30948 | predicted gene, 30948 |
| chr10 | 1,11E+08 | 1,11E+08 | 455 | * | 5,1144 | -5,808938673 | 2,036209 | -2,85282 | 0,004333 | 1 | 0 | Distal Intergenic | 10 | 1,11E+08 | 1,11E+08 | 3049 | 1 | 71769 | ENSMUST00000040454.5 | 3869 | ENSMUSG00000035759 | Bbs10 | Bardet-Biedl syndrome 10 (human) |
| chr7 | 61650931 | 61651282 | 352 | * | 4,402465 | -5,808422838 | 2,122389 | -2,73674 | 0,006205 | 1 | 0 | Distal Intergenic | 7 | 61632037 | 61632107 | 71 | 2 | 1,01E+08 | ENSMUST00000175109.3 | -18824 | ENSMUSG00000092850 | Mir344g | microRNA 344g |
| chr4 | 94975989 | 94976569 | 581 | * | 4,190401 | -5,807587652 | 2,240211 | -2,59243 | 0,00953 | 1 | 0 | Intron (ENSMUST0000 | 4 | 94981747 | 95049471 | 67725 | 1 | 230451 | ENSMUST00000130684.8 | -5178 | | Junos | jun proto-oncogene, opposite strand |
| chr17 | 12374086 | 12374940 | 855 | * | 4,203944 | -5,807571431 | 2,229384 | -2,60501 | 0,009187 | 1 | 0 | Intron (ENSMUST0000 | 17 | 12370659 | 12437966 | 67308 | 1 | 68262 | ENSMUST00000170858.8 | 3427 | ENSMUSG00000023827 | Agpat4 | 1-acylglycerol-3-phosphate O-acyltransferase 4 (lysophosphatidic acid acyltransferase, delta) |
| chrX | 65315454 | 65316076 | 623 | * | 4,22512 | -5,807456629 | 2,162553 | -2,68546 | 0,007243 | 1 | 0 | Distal Intergenic | 20 | 65692924 | 65704999 | 12076 | 1 | 245450 | ENSMUST00000166241.2 | -376848 | ENSMUSG00000036790 | Slitrk2 | SLIT and NTRK-like family, member 2 |
| chr2 | 27442798 | 27443223 | 426 | * | 5,440529 | -5,807070028 | 1,99779 | -2,90675 | 0,003652 | 1 | 0 | Distal Intergenic | 2 | 27430438 | 27433374 | 2937 | 1 | 1,05E+08 | ENSMUST00000142511.2 | 12360 | | Gm13421 | predicted gene 13421 |
| chr12 | 6094914 | 6096074 | 1161 | * | 6,532434 | -5,806416746 | 1,845699 | -3,14592 | 0,001656 | 1 | 0 | Distal Intergenic | 12 | 5858234 | 5906142 | 47909 | 1 | 1,03E+08 | ENSMUST00000219472.2 | 236680 | | Gm32442 | predicted gene, 32442 |
| chr14 | 86288742 | 86289763 | 1022 | * | 4,796978 | -5,806296556 | 2,118557 | -2,74068 | 0,006131 | 1 | 0 | Distal Intergenic | 14 | 86410363 | 86417021 | 6659 | 1 | 1,03E+08 | ENSMUST00000227116.2 | -120600 | | Gm32729 | predicted gene, 32729 |
| chr14 | 22252720 | 22253348 | 629 | * | 5,671658 | -5,805927493 | 2,081735 | -2,78899 | 0,005287 | 1 | 0 | Intron (ENSMUST0000 | 14 | 22360233 | 22634507 | 274275 | 1 | 76633 | ENSMUST00000159107.2 | -106885 | ENSMUSG00000063458 | Lrmda | leucine rich melanocyte differentiation associated |
| chr5 | 53140650 | 53141351 | 702 | * | 5,168201 | -5,804800601 | 1,973498 | -2,94138 | 0,003268 | 1 | 0 | Promoter (1-2kb) | 5 | 53139379 | 53153095 | 13717 | 1 | 1,03E+08 | ENSMUST00000199474.2 | 1271 | | Gm30301 | predicted gene, 30301 |
| chr2 | 53042687 | 53043597 | 911 | * | 8,508078 | -5,803536997 | 1,914556 | -3,03127 | 0,002435 | 0,654756507 | 1,360622 | 3' UTR | 2 | 53029184 | 53035321 | 6138 | 2 | 56194 | ENSMUST00000133384.2 | -7366 | ENSMUSG00000061136 | Prpf40a | pre-mRNA processing factor 40A |
| chr2 | 3815911 | 3816467 | 557 | * | 5,399199 | -5,802901402 | 2,043068 | -2,84029 | 0,004507 | 1 | 0 | Distal Intergenic | 2 | 3856067 | 3878612 | 22546 | 1 | 1,03E+08 | ENSMUST00000141222.2 | -39600 | | Gm36031 | predicted gene, 36031 |
| chr13 | 75610069 | 75610620 | 552 | * | 5,218521 | -5,80276252 | 2,024207 | -2,86668 | 0,004148 | 1 | 0 | Distal Intergenic | 13 | 75793015 | 75793290 | 276 | 1 | 751556 | ENSMUST00000099371.5 | -182395 | ENSMUSG00000076236 | Mir682 | microRNA 682 |
| chr19 | 26209312 | 26210535 | 1224 | * | 6,216929 | -5,802240132 | 2,017138 | -2,87647 | 0,004021 | 1 | 0 | Distal Intergenic | 19 | 26015816 | 26023831 | 8016 | 2 | 73433 | ENSMUST00000237184.2 | -185481 | ENSMUSG00000118016 | 1700048O14Rik | RIKEN cDNA 1700048O14 gene |
| chr14 | 1,12E+08 | 1,12E+08 | 490 | * | 4,853269 | -5,802065114 | 2,039198 | -2,84527 | 0,004437 | 1 | 0 | Distal Intergenic | 14 | 1,12E+08 | 1,12E+08 | 90973 | 2 | 1,03E+08 | ENSMUST00000242291.1 | 175815 | | Gm30439 | predicted gene, 30439 |
| chr3 | 72750631 | 72751030 | 400 | * | 4,295166 | -5,801543774 | 2,116686 | -2,74086 | 0,006128 | 1 | 0 | Distal Intergenic | 3 | 72795893 | 72874199 | 78307 | 2 | 69983 | ENSMUST00000094190.5 | 123169 | ENSMUSG00000027790 | Sis | sucrase isomaltase (alpha-glucosidase) |
| chr3 | 57321754 | 57322304 | 551 | * | 4,637967 | -5,800204629 | 1,991118 | -2,91304 | 0,003579 | 1 | 0 | Distal Intergenic | 3 | 57332735 | 57349098 | 16364 | 1 | 229302 | ENSMUST00000029377.8 | -10431 | ENSMUSG00000027801 | Tm4sf4 | transmembrane 4 superfamily member 4 |
| chr3 | 92410577 | 92411205 | 629 | * | 5,642101 | -5,799960628 | 1,900534 | -3,05175 | 0,002275 | 1 | 0 | Promoter (2-3kb) | 3 | 92407570 | 92407800 | 231 | 2 | 229562 | ENSMUST00000062129.2 | -2777 | ENSMUSG00000045566 | Sprr4 | small proline-rich protein 4 |
| chr16 | 19925326 | 19925966 | 641 | * | 6,831493 | -5,799882694 | 1,877639 | -3,08892 | 0,002009 | 1 | 0 | 5' UTR | 16 | 19916342 | 19925882 | 9541 | 1 | 75785 | ENSMUST00000231842.2 | 8984 | ENSMUSG00000062901 | Klhl24 | kelch-like 24 |
| chr1 | 1,82E+08 | 1,82E+08 | 1987 | * | 13,6985 | -5,798628324 | 1,681962 | -3,44754 | 0,000566 | 0,456474583 | 0,938682 | Distal Intergenic | 1 | 1,82E+08 | 1,82E+08 | 921 | 2 | 625823 | ENSMUST00000169123.4 | 13552 | ENSMUSG00000091013 | Vmn1r1 | vomeronasal 1 receptor 1 |
| chr14 | 1,19E+08 | 1,19E+08 | 895 | * | 6,375423 | -5,798289352 | 1,906207 | -3,04179 | 0,002352 | 1 | 0 | Exon (ENSMUST00000 | 14 | 1,19E+08 | 1,19E+08 | 3679 | 2 | 66435 | ENSMUST00000173470.2 | -5969 | ENSMUSG00000042104 | Uggt2 | UDP-glucose glycoprotein glucosyltransferase 2 |
| chr17 | 5748864 | 5749640 | 777 | * | 6,080422 | -5,798279635 | 1,908353 | -3,03837 | 0,002379 | 1 | 0 | Intron (ENSMUST0000 | 17 | 5848932 | 5853168 | 4237 | 1 | 78512 | ENSMUST00000180924.8 | -99292 | ENSMUSG00000096965 | 3300005D01Rik | RIKEN cDNA 3300005D01 gene |
| chr14 | 67464199 | 67464922 | 724 | * | 4,651265 | -5,79666075 | 2,175721 | -2,66425 | 0,007716 | 1 | 0 | Distal Intergenic | 14 | 67469886 | 67470578 | 693 | 2 | 1,05E+08 | ENSMUST00000175861.2 | 5656 | | Gm20675 | predicted gene 20675 |
| chr3 | 83988169 | 83988743 | 575 | * | 5,242952 | -5,796001973 | 2,045244 | -2,83389 | 0,004598 | 1 | 0 | Distal Intergenic | 3 | 83805516 | 83947482 | 141967 | 2 | 229473 | ENSMUST00000052342.9 | -40687 | ENSMUSG00000033767 | Tmem131l | transmembrane 131 like |
| chr1 | 23207756 | 23208332 | 577 | * | 4,33533 | -5,793448952 | 2,092315 | -2,76892 | 0,005624 | 1 | 0 | Intron (ENSMUST0000 | 1 | 23139554 | 23141334 | 1781 | 2 | 66755 | ENSMUST00000073179.6 | -66422 | ENSMUSG00000073730 | Ppp1r14bl | protein phosphatase 1, regulatory inhibitor subunit 14B like |
| chr12 | 1,09E+08 | 1,09E+08 | 1271 | * | 9,580303 | -5,793045922 | 1,834381 | -3,15804 | 0,001588 | 0,596263346 | 1,211315 | Promoter (<=1kb) | 12 | 1,09E+08 | 1,09E+08 | 5568 | 1 | 13386 | ENSMUST00000124293.3 | 477 | ENSMUSG00000040856 | Dlk1 | delta like non-canonical Notch ligand 1 |
| chr12 | 17390169 | 17391354 | 1186 | * | 7,736525 | -5,792850569 | 1,772536 | -3,26811 | 0,001083 | 1 | 0 | Distal Intergenic | 12 | 17394402 | 17398374 | 3973 | 2 | 1,08E+08 | ENSMUST00000221894.2 | 7020 | | Gm46336 | predicted gene, 46336 |
| chr16 | 77012174 | 77013193 | 1020 | * | 9,884156 | -5,792559791 | 1,708477 | -3,39048 | 0,000698 | 0,422040194 | 1,425929 | Distal Intergenic | 16 | 77033184 | 77256431 | 223248 | 1 | 77994 | ENSMUST00000244279.1 | -19991 | ENSMUSG00000090386 | Mir99ahg | Mir99a and Mirlet7c-1 host gene (non-protein coding) |
| chr4 | 1,24E+08 | 1,24E+08 | 1059 | * | 10,18723 | -5,792556067 | 1,788781 | -3,23827 | 0,001203 | 0,54670137 | 1,173325 | Exon (ENSMUST00000 | 4 | 1,24E+08 | 1,24E+08 | 5456 | 1 | 108809 | ENSMUST00000133499.2 | 4726 | | 4933407E24Rik | RIKEN cDNA 4933407E24 gene |
| chr1 | 85558483 | 85558770 | 288 | * | 7,453717 | -5,791994666 | 1,844475 | -3,14019 | 0,001688 | 0,586565597 | 1,360622 | Promoter (1-2kb) | 1 | 85560199 | 85572663 | 12465 | 1 | 434484 | ENSMUST00000161497.2 | -1429 | ENSMUSG00000070031 | Sp140 | Sp140 nuclear body protein |
| chr2 | 1,79E+08 | 1,79E+08 | 678 | * | 4,175456 | -5,790373117 | 2,178492 | -2,65797 | 0,007861 | 1 | 0 | Distal Intergenic | 2 | 1,78E+08 | 1,78E+08 | 1713 | 1 | 1,19E+08 | ENSMUST00000246055.1 | 413378 | | Gm54095 | predicted gene, 54095 |
| chr18 | 8003628 | 8004360 | 733 | * | 4,575804 | -5,789534232 | 2,148827 | -2,69428 | 0,007054 | 1 | 0 | Distal Intergenic | 18 | 7910619 | 7918503 | 7885 | 1 | 225131 | ENSMUST00000167542.2 | 93009 | ENSMUSG00000024283 | Wac | WW domain containing adaptor with coiled-coil |
| chr1 | 1,76E+08 | 1,76E+08 | 735 | * | 6,642889 | -5,787575022 | 1,857757 | -3,11536 | 0,001837 | 1 | 0 | Distal Intergenic | 1 | 1,76E+08 | 1,76E+08 | 97163 | 2 | 1,03E+08 | ENSMUST00000194241.2 | -61560 | ENSMUSG00000104021 | Gm36536 | predicted gene, 36536 |
| chr19 | 30920051 | 30920781 | 731 | * | 4,403802 | -5,786483534 | 2,137164 | -2,70755 | 0,006778 | 1 | 0 | Intron (ENSMUST0000 | 19 | 31060237 | 31064469 | 4233 | 1 | 83410 | ENSMUST00000066039.8 | -139456 | ENSMUSG00000053536 | Cstf2t | cleavage stimulation factor, 3' pre-RNA subunit 2, tau |
| chr7 | 24340646 | 24341362 | 717 | * | 5,159136 | -5,786423646 | 1,953731 | -2,96173 | 0,003059 | 1 | 0 | Downstream (<=300bp | 7 | 24330570 | 24337341 | 6772 | 2 | 1,03E+08 | ENSMUST00000180699.2 | -3305 | | Gm26550 | predicted gene, 26550 |
| chr5 | 60014739 | 60015154 | 416 | * | 4,612872 | -5,783916126 | 2,035251 | -2,84187 | 0,004485 | 1 | 0 | Distal Intergenic | 5 | 60120467 | 60143341 | 22875 | 2 | 71057 | ENSMUST00000198882.2 | 128187 | ENSMUSG00000106013 | 4933402J10Rik | RIKEN cDNA 4933402J10 gene |
| chr3 | 24672861 | 24673517 | 657 | * | 4,666977 | -5,783343646 | 2,174704 | -2,65937 | 0,007829 | 1 | 0 | Intron (ENSMUST0000 | 3 | 24616238 | 24616344 | 107 | 1 | 1,15E+08 | ENSMUST00000093715.3 | 56623 | ENSMUSG00000070227 | Gm24704 | predicted gene, 24704 |
| chr14 | 7061540 | 7062058 | 519 | * | 3,586675 | -5,783286522 | 2,192967 | -2,6372 | 0,008359 | 1 | 0 | Intron (ENSMUST0000 | 14 | 7033258 | 7112248 | 78991 | 1 | 218763 | ENSMUST00000163937.3 | 28282 | ENSMUSG00000045201 | Lrrc3b | leucine rich repeat containing 3B |
| chr9 | 72908134 | 72908963 | 830 | * | 7,722679 | -5,782287862 | 1,871569 | -3,08954 | 0,002005 | 0,621084104 | 1,360622 | 3' UTR | 9 | 72917170 | 72919259 | 2090 | 1 | 72278 | ENSMUST00000131924.2 | -8207 | ENSMUSG00000034563 | Ccpg1 | cell cycle progression 1 |
| chr17 | 24884839 | 24885377 | 539 | * | 4,688447 | -5,781896316 | 2,008741 | -2,87837 | 0,003997 | 1 | 0 | Distal Intergenic | 17 | 24888661 | 24900990 | 12330 | 1 | 213753 | ENSMUST00000234956.2 | -3284 | ENSMUSG00000041130 | Zfp598 | zinc finger protein 598 |
| chr14 | 9228270 | 9229282 | 1013 | * | 4,572265 | -5,780335669 | 2,157005 | -2,6798 | 0,007367 | 1 | 0 | Intron (ENSMUST0000 | 14 | 9185133 | 9206630 | 21498 | 1 | 328354 | ENSMUST00000140967.2 | 43137 | ENSMUSG00000051729 | Gm5087 | predicted gene 5087 |
| chr11 | 80550651 | 80551713 | 1063 | * | 8,325365 | -5,779698524 | 1,768299 | -3,26851 | 0,001081 | 0,503036965 | 1,360622 | Intron (ENSMUST0000 | 11 | 80564493 | 80670829 | 106337 | 2 | 338367 | ENSMUST00000125944.2 | 119116 | ENSMUSG00000035441 | Myo1d | myosin ID |
| chr14 | 89542507 | 89543874 | 1368 | * | 6,984531 | -5,779658718 | 1,809332 | -3,19436 | 0,001401 | 1 | 0 | Distal Intergenic | 14 | 89944241 | 89944372 | 132 | 1 | 1,15E+08 | ENSMUST00000104220.3 | -400367 | ENSMUSG00000077408 | Gm25415 | predicted gene, 25415 |
| chr19 | 23251551 | 23252136 | 586 | * | 4,383261 | -5,77942812 | 2,171414 | -2,6616 | 0,007777 | 1 | 0 | Promoter (<=1kb) | 19 | 23186087 | 23251261 | 65175 | 2 | 226026 | ENSMUST00000223934.2 | -290 | ENSMUSG00000024943 | Smc5 | structural maintenance of chromosomes 5 |
| chr17 | 40430723 | 40431281 | 559 | * | 3,467891 | -5,779412167 | 2,215944 | -2,6081 | 0,009105 | 1 | 0 | Distal Intergenic | 17 | 40517906 | 40519531 | 1626 | 2 | 18663 | ENSMUST00000033585.7 | 88250 | ENSMUSG00000031233 | Pgk2 | phosphoglycerate kinase 2 |
| chr6 | 1,05E+08 | 1,05E+08 | 288 | * | 4,031761 | -5,779411129 | 2,194515 | -2,63357 | 0,008449 | 1 | 0 | Distal Intergenic | 6 | 1,05E+08 | 1,05E+08 | 139 | 1 | 1,15E+08 | ENSMUST00000158962.4 | 12984 | ENSMUSG00000089587 | Gm23339 | predicted gene, 23339 |
| chr6 | 17398241 | 17398651 | 411 | * | 5,641967 | -5,77740354 | 2,016748 | -2,86471 | 0,004174 | 1 | 0 | Distal Intergenic | 6 | 17436305 | 17436411 | 107 | 1 | 1,15E+08 | ENSMUST00000240512.1 | -37654 | ENSMUSG00000119734 | Gm25021 | predicted gene, 25021 |
| chr11 | 38926197 | 38926928 | 732 | * | 7,541089 | -5,77685259 | 1,875499 | -3,08017 | 0,002069 | 1 | 0 | Distal Intergenic | 11 | 38688998 | 38689104 | 107 | 2 | 1,15E+08 | ENSMUST00000240209.1 | -237093 | ENSMUSG00000119681 | Gm23520 | predicted gene, 23520 |
| chr14 | 63482918 | 63483549 | 632 | * | 3,843325 | -5,776139965 | 2,214141 | -2,60875 | 0,009087 | 1 | 0 | Promoter (<=1kb) | 14 | 63436371 | 63482720 | 46350 | 2 | 14463 | ENSMUST00000067417.10 | -198 | ENSMUSG00000021944 | Gata4 | GATA binding protein 4 |
| chr8 | 76219260 | 76219713 | 454 | * | 4,007842 | -5,776118153 | 2,218519 | -2,60359 | 0,009225 | 1 | 0 | Intron (ENSMUST0000 | 8 | 76175611 | 76711131 | 535521 | 1 | 71831 | ENSMUST00000121983.2 | 43649 | ENSMUSG00000031620 | Iqcm | IQ motif containing M |
| chr14 | 1,07E+08 | 1,07E+08 | 728 | * | 3,4302 | -5,775616745 | 2,228477 | -2,59173 | 0,009549 | 1 | 0 | Distal Intergenic | 14 | 1,07E+08 | 1,07E+08 | 5957 | 1 | 1,03E+08 | ENSMUST00000227226.2 | 31885 | | Gm36107 | predicted gene, 36107 |
| chr14 | 68676982 | 68677726 | 745 | * | 6,434836 | -5,774700738 | 1,963368 | -2,94122 | 0,003269 | 1 | 0 | Distal Intergenic | 14 | 68699080 | 68733866 | 34787 | 1 | 1,03E+08 | ENSMUST00000225482.2 | -21354 | | Gm31227 | predicted gene, 31227 |
| chr5 | 82569060 | 82569740 | 681 | * | 4,821212 | -5,774085435 | 2,066568 | -2,79405 | 0,005205 | 1 | 0 | Distal Intergenic | 5 | 82270274 | 82272560 | 2287 | 2 | 67327 | ENSMUST00000185410.2 | -296500 | ENSMUSG00000099863 | 1700031L13Rik | RIKEN cDNA 1700031L13 gene |
| chr19 | 8207675 | 8208868 | 1194 | * | 9,655353 | -5,773552293 | 1,750685 | -3,29788 | 0,000974 | 0,477253929 | 1,425929 | Distal Intergenic | 19 | 8137668 | 8196264 | 58597 | 2 | 236293 | ENSMUST00000140910.8 | -11411 | ENSMUSG00000075044 | Slc22a29 | solute carrier family 22. member 29 |
| chr10 | 72925483 | 72926105 | 623 | * | 4,107252 | -5,771270368 | 2,13152 | -2,70758 | 0,006777 | 1 | 0 | Promoter (2-3kb) | 10 | 72853596 | 72922541 | 68946 | 2 | 327768 | ENSMUST00000219536.3 | -2942 | ENSMUSG00000111994 | A330049N07Rik | RIKEN cDNA A330049N07 gene |
| chr14 | 1,18E+08 | 1,18E+08 | 1052 | * | 4,525681 | -5,771180416 | 2,118444 | -2,72425 | 0,006445 | 1 | 0 | Intron (ENSMUST0000 | 14 | 1,18E+08 | 1,18E+08 | 105 | 2 | 1,02E+08 | ENSMUST00000183564.3 | 297953 | ENSMUSG00000098648 | Mir6239 | microRNA 6239 |
| chr2 | 12429079 | 12429483 | 405 | * | 4,373537 | -5,768370637 | 2,17438 | -2,65288 | 0,007981 | 1 | 0 | Distal Intergenic | 2 | 12352074 | 12424281 | 72208 | 2 | 66960 | ENSMUST00000028105.13 | -4798 | ENSMUSG00000026767 | Mindy3 | MINDY lysine 48 deubiquitinase 3 |
| chr4 | 11665739 | 11666358 | 620 | * | 6,271875 | -5,767361881 | 1,886415 | -3,05731 | 0,002233 | 1 | 0 | Distal Intergenic | 4 | 11704457 | 11714752 | 10296 | 1 | 14579 | ENSMUST00000108304.9 | -38099 | ENSMUSG00000028214 | Gem | GTP binding protein (gene overexpressed in skeletal muscle) |
| chr14 | 86089941 | 86091403 | 1463 | * | 7,81362 | -5,767173789 | 1,85586 | -3,10755 | 0,001886 | 1 | 0 | Distal Intergenic | 14 | 86410363 | 86417021 | 6659 | 1 | 1,03E+08 | ENSMUST00000227116.2 | -318960 | | Gm32729 | predicted gene, 32729 |
| chr6 | 43788873 | 43789715 | 843 | * | 6,211715 | -5,766958583 | 2,076106 | -2,77778 | 0,005473 | 1 | 0 | Distal Intergenic | 6 | 43864128 | 43864258 | 131 | 1 | 1,15E+08 | ENSMUST00000103958.3 | -74413 | ENSMUSG00000077146 | Gm22939 | predicted gene, 22939 |
| chr1 | 1,88E+08 | 1,88E+08 | 1056 | * | 5,292901 | -5,765742639 | 1,931088 | -2,98575 | 0,002829 | 1 | 0 | Intron (ENSMUST0000 | 1 | 1,88E+08 | 1,88E+08 | 131 | 2 | 1,15E+08 | ENSMUST00000104047.3 | -37468 | ENSMUSG00000077235 | Gm25095 | predicted gene, 25095 |
| chrX | 1,58E+08 | 1,58E+08 | 283 | * | 4,527793 | -5,765666406 | 2,225588 | -2,59063 | 0,00958 | 1 | 0 | Distal Intergenic | 20 | 1,58E+08 | 1,58E+08 | 25152 | 1 | 382252 | ENSMUST00000144598.8 | -19486 | ENSMUSG00000044150 | Bclaf3 | Bclaf1 and Thrap3 family member 3 |
| chr2 | 1,02E+08 | 1,02E+08 | 1089 | * | 8,31717 | -5,765600077 | 1,813529 | -3,17922 | 0,001477 | 0,547949061 | 1,425929 | Intron (ENSMUST0000 | 2 | 1,02E+08 | 1,02E+08 | 3844 | 1 | 1,03E+08 | ENSMUST00000146134.2 | 10012 | | Gm13919 | predicted gene 13919 |
| chr15 | 33670599 | 33671183 | 585 | * | 5,110917 | -5,76552323 | 2,010903 | -2,86713 | 0,004142 | 1 | 0 | Distal Intergenic | 15 | 33684021 | 33688030 | 4010 | 2 | 239364 | ENSMUST00000042021.5 | 16847 | ENSMUSG00000038984 | Tspyl5 | testis-specific protein, Y-encoded-like 5 |
| chr2 | 1,5E+08 | 1,5E+08 | 589 | * | 4,370824 | -5,765413557 | 2,06037 | -2,79824 | 0,005138 | 1 | 0 | Intron (ENSMUST0000 | 2 | 1,5E+08 | 1,5E+08 | 22653 | 1 | 245174 | ENSMUST00000073782.4 | 14733 | ENSMUSG00000060336 | Zfp937 | zinc finger protein 937 |
| chr14 | 1,18E+08 | 1,18E+08 | 886 | * | 4,165116 | -5,765187101 | 2,18853 | -2,63427 | 0,008432 | 1 | 0 | Intron (ENSMUST0000 | 14 | 1,18E+08 | 1,18E+08 | 105 | 2 | 1,02E+08 | ENSMUST00000183564.3 | 510402 | ENSMUSG00000098648 | Mir6239 | microRNA 6239 |
| chr2 | 1,67E+08 | 1,67E+08 | 736 | * | 6,065295 | -5,763040207 | 2,021955 | -2,85023 | 0,004369 | 1 | 0 | Intron (ENSMUST0000 | 2 | 1,67E+08 | 1,67E+08 | 1442 | 2 | 407243 | ENSMUST00000142148.2 | -7467 | ENSMUSG00000090213 | Peds1 | plasmanylethanolamine desaturase 1 |
| chr18 | 44244920 | 44245772 | 853 | * | 4,318783 | -5,762101188 | 2,166917 | -2,65912 | 0,007834 | 1 | 0 | Distal Intergenic | 18 | 44249330 | 44252634 | 3305 | 1 | 1,15E+08 | ENSMUST00000212114.2 | -3558 | | Gm37797 | predicted gene, 37797 |
| chr14 | 1,07E+08 | 1,07E+08 | 1037 | * | 6,35102 | -5,760334282 | 1,858533 | -3,0994 | 0,001939 | 1 | 0 | Distal Intergenic | 14 | 1,07E+08 | 1,07E+08 | 127 | 1 | 1,15E+08 | ENSMUST00000104036.3 | -60818 | ENSMUSG00000077224 | Gm23437 | predicted gene, 23437 |
| chr11 | 1,13E+08 | 1,13E+08 | 963 | * | 9,135255 | -5,759456345 | 1,732461 | -3,32444 | 0,000886 | 0,461973573 | 1,425929 | Intron (ENSMUST0000 | 11 | 1,13E+08 | 1,13E+08 | 320904 | 2 | 69806 | ENSMUST00000071539.10 | 38641 | ENSMUSG00000041654 | Slc39a11 | solute carrier family 39 (metal ion transporter), member 11 |
| chr13 | 76966096 | 76966854 | 759 | * | 6,853843 | -5,759313276 | 1,836392 | -3,13621 | 0,001711 | 1 | 0 | Intron (ENSMUST0000 | 13 | 76973326 | 76976195 | 2870 | 1 | 78771 | ENSMUST00000155275.2 | -6472 | ENSMUSG00000021596 | Mctp1 | multiple C2 domains, transmembrane 1 |
| chr18 | 31974197 | 31974975 | 779 | * | 5,212905 | -5,759238487 | 2,139575 | -2,69177 | 0,007107 | 1 | 0 | 3' UTR | 18 | 31937229 | 31968475 | 31247 | 1 | 74320 | ENSMUST00000234344.2 | 36968 | ENSMUSG00000024400 | Wdr33 | WD repeat domain 33 |
| chr18 | 23375188 | 23375915 | 728 | * | 6,141585 | -5,759092969 | 2,046625 | -2,81395 | 0,004894 | 1 | 0 | Distal Intergenic | 18 | 23349726 | 23350668 | 943 | 1 | 665797 | ENSMUST00000234562.2 | 25462 | | Gm7788 | glyceraldehyde-3-phosphate dehydrogenase pseudogene |
| chr4 | 30237243 | 30237994 | 752 | * | 7,88502 | -5,758979218 | 1,893235 | -3,04187 | 0,002351 | 0,649217039 | 1,360622 | Distal Intergenic | 4 | 31174909 | 31175012 | 104 | 2 | 1,15E+08 | ENSMUST00000083119.2 | 937018 | ENSMUSG00000065053 | Gm25705 | predicted gene, 25705 |
| chr16 | 10123647 | 10124794 | 1148 | * | 8,766619 | -5,758874352 | 1,813237 | -3,17602 | 0,001493 | 0,562910619 | 1,360622 | Intron (ENSMUST0000 | 16 | 10099613 | 10131832 | 32220 | 2 | 13731 | ENSMUST00000078357.5 | 7038 | ENSMUSG00000022505 | Emp2 | epithelial membrane protein 2 |
| chr11 | 28385423 | 28386102 | 680 | * | 4,771619 | -5,758298704 | 2,075289 | -2,7747 | 0,005525 | 1 | 0 | Promoter (1-2kb) | 11 | 28338817 | 28384110 | 45294 | 2 | 216613 | ENSMUST00000160016.8 | -1313 | ENSMUSG00000032878 | Ccdc85a | coiled-coil domain containing 85A |
| chr2 | 42323632 | 42323987 | 356 | * | 3,998782 | -5,758101791 | 2,15254 | -2,67503 | 0,007472 | 1 | 0 | Intron (ENSMUST0000 | 2 | 42337451 | 42337580 | 130 | 2 | 1,02E+08 | ENSMUST00000183581.3 | 13593 | ENSMUSG00000098660 | Mir6336 | microRNA 6336 |
| chr10 | 1,06E+08 | 1,06E+08 | 685 | * | 4,658724 | -5,757956891 | 2,22543 | -2,58735 | 0,009672 | 1 | 0 | Distal Intergenic | 10 | 1,06E+08 | 1,06E+08 | 1479 | 1 | 52713 | ENSMUST00000155406.2 | 143985 | ENSMUSG00000019897 | Ccdc59 | coiled-coil domain containing 59 |
| chrX | 1,53E+08 | 1,53E+08 | 418 | * | 4,020167 | -5,757942248 | 2,231846 | -2,5799 | 0,009883 | 1 | 0 | Distal Intergenic | 20 | 1,53E+08 | 1,53E+08 | 9675 | 2 | 78092 | ENSMUST00000178698.2 | 17092 | ENSMUSG00000094802 | Samt1 | spermatogenesis associated multipass transmembrane protein 1 |
| chr13 | 5713275 | 5714121 | 847 | * | 4,217143 | -5,757212562 | 2,177131 | -2,6444 | 0,008183 | 1 | 0 | Intron (ENSMUST0000 | 13 | 5590073 | 5774444 | 184372 | 2 | 1,03E+08 | ENSMUST00000223426.3 | 60323 | | Gm35330 | predicted gene, 35330 |
| chr3 | 43523243 | 43523726 | 484 | * | 4,370772 | -5,755510092 | 2,081467 | -2,76512 | 0,00569 | 1 | 0 | Distal Intergenic | 3 | 45281480 | 45332640 | 51161 | 2 | 72511 | ENSMUST00000191843.2 | 1808914 | ENSMUSG00000097040 | 2610316D01Rik | RIKEN cDNA 2610316D01 gene |
| chr16 | 56554680 | 56555029 | 350 | * | 3,824211 | -5,754005175 | 2,14464 | -2,68297 | 0,007297 | 1 | 0 | Intron (ENSMUST0000 | 16 | 56510695 | 56537813 | 27119 | 2 | 21787 | ENSMUST00000065515.14 | -16867 | ENSMUSG00000022757 | Tfg | Trk-fused gene |
| chr7 | 76180810 | 76181656 | 847 | * | 5,820811 | -5,753946147 | 2,097921 | -2,74269 | 0,006094 | 1 | 0 | Intron (ENSMUST0000 | 7 | 76060810 | 76416356 | 355547 | 1 | 244071 | ENSMUST00000107442.3 | 120000 | ENSMUSG00000025754 | Agbl1 | ATP/GTP binding protein-like 1 |
| chr3 | 42869711 | 42870096 | 386 | * | 4,608607 | -5,753928439 | 2,081696 | -2,76406 | 0,005709 | 1 | 0 | Distal Intergenic | 3 | 41705631 | 41710927 | 5297 | 1 | 73852 | ENSMUST00000195030.2 | 1164080 | ENSMUSG00000025766 | D3Ertd751e | DNA segment, Chr 3, ERATO Doi 751, expressed |
| chr10 | 18003878 | 18005050 | 1173 | * | 8,757595 | -5,752981623 | 1,749755 | -3,28788 | 0,001009 | 0,492617108 | 1,360622 | Exon (ENSMUST00000 | 10 | 17997932 | 17998857 | 926 | 1 | 19707 | ENSMUST00000150533.2 | 5946 | ENSMUSG00000019854 | Reps1 | RalBP1 associated Eps domain containing protein |
| chrX | 55792353 | 55793403 | 1051 | * | 5,721732 | -5,751913883 | 1,915168 | -3,00335 | 0,00267 | 1 | 0 | Promoter (1-2kb) | 20 | 55795370 | 55795475 | 106 | 1 | 1,15E+08 | ENSMUST00000158319.3 | -1967 | ENSMUSG00000088944 | Gm23258 | predicted gene, 23258 |
| chr2 | 18768634 | 18769292 | 659 | * | 5,003887 | -5,751509022 | 1,991767 | -2,88764 | 0,003881 | 1 | 0 | Promoter (2-3kb) | 2 | 18747526 | 18765673 | 18148 | 2 | 1E+08 | ENSMUST00000174811.2 | -2961 | ENSMUSG00000092353 | Gm3363 | predicted gene 3363 |
| chr4 | 56796384 | 56796924 | 541 | * | 4,677182 | -5,751056481 | 2,11221 | -2,72277 | 0,006474 | 1 | 0 | Exon (ENSMUST00000 | 4 | 56749680 | 56802331 | 52652 | 2 | 230233 | ENSMUST00000030140.3 | 5407 | ENSMUSG00000028431 | Elp1 | elongator complex protein 1 |
| chr14 | 72725821 | 72726557 | 737 | * | 4,26948 | -5,750883427 | 2,17196 | -2,64779 | 0,008102 | 1 | 0 | Promoter (2-3kb) | 14 | 72663453 | 72729278 | 65826 | 2 | 675947 | ENSMUST00000208955.2 | 2721 | | Gm9195 | predicted gene 9195 |
| chrX | 44757775 | 44758346 | 572 | * | 4,836121 | -5,749332971 | 2,044754 | -2,81175 | 0,004927 | 1 | 0 | Distal Intergenic | 20 | 44884238 | 44884344 | 107 | 2 | 1,15E+08 | ENSMUST00000082530.3 | 125998 | ENSMUSG00000119236 | Gm25078 | predicted gene, 25078 |
| chr9 | 24502525 | 24503473 | 949 | * | 7,6924 | -5,748969882 | 1,989291 | -2,88996 | 0,003853 | 1 | 0 | Intron (ENSMUST0000 | 9 | 24536492 | 24537120 | 629 | 1 | 235067 | ENSMUST00000053956.8 | -33019 | ENSMUSG00000047995 | Cypt4 | cysteine-rich perinuclear theca 4 |
| chr13 | 6262113 | 6263184 | 1072 | * | 6,805159 | -5,748024717 | 1,886583 | -3,04679 | 0,002313 | 1 | 0 | Distal Intergenic | 13 | 6330414 | 6344750 | 14337 | 2 | 1,03E+08 | ENSMUST00000221255.2 | 81566 | ENSMUSG00000114097 | Gm35615 | predicted gene, 35615 |
| chrX | 1,2E+08 | 1,2E+08 | 894 | * | 6,383256 | -5,745255008 | 1,875865 | -3,06272 | 0,002193 | 1 | 0 | Intron (ENSMUST0000 | 20 | 1,2E+08 | 1,2E+08 | 128 | 1 | 1,15E+08 | ENSMUST00000104247.3 | -115632 | ENSMUSG00000077435 | Gm22590 | predicted gene, 22590 |
| chr1 | 1,74E+08 | 1,74E+08 | 490 | * | 4,612151 | -5,744033173 | 2,072388 | -2,7717 | 0,005576 | 1 | 0 | Intron (ENSMUST0000 | 1 | 1,74E+08 | 1,74E+08 | 32072 | 1 | 20739 | ENSMUST00000156092.2 | 12308 | ENSMUSG00000026532 | Spta1 | spectrin alpha, erythrocytic 1 |
| chrX | 99508147 | 99508751 | 605 | * | 4,365332 | -5,743840549 | 2,12196 | -2,70686 | 0,006792 | 1 | 0 | Distal Intergenic | 20 | 99445832 | 99486323 | 40492 | 2 | 245532 | ENSMUST00000147103.2 | -21824 | ENSMUSG00000031220 | Awat2 | acyl-CoA wax alcohol acyltransferase 2 |
| chr4 | 69827555 | 69828003 | 449 | * | 4,933496 | -5,741594146 | 2,043587 | -2,80957 | 0,004961 | 1 | 0 | Distal Intergenic | 4 | 70160708 | 70162089 | 1382 | 2 | 214444 | ENSMUST00000124251.2 | 334086 | ENSMUSG00000039298 | Cdk5rap2 | CDK5 regulatory subunit associated protein 2 |
| chr2 | 78486304 | 78486892 | 589 | * | 4,523696 | -5,741085468 | 2,0981 | -2,73633 | 0,006213 | 1 | 0 | Distal Intergenic | 2 | 78698468 | 78700951 | 2484 | 1 | 22193 | ENSMUST00000124120.2 | -211576 | ENSMUSG00000027011 | Ube2e3 | ubiquitin-conjugating enzyme E2E 3 |
| chr6 | 15660588 | 15661084 | 497 | * | 4,895803 | -5,739586021 | 2,036609 | -2,81821 | 0,004829 | 1 | 0 | Distal Intergenic | 6 | 15720660 | 15802164 | 81505 | 1 | 16543 | ENSMUST00000101663.10 | -59576 | ENSMUSG00000041390 | Mdfic | MyoD family inhibitor domain containing |
| chr4 | 51996510 | 51997044 | 535 | * | 4,115146 | -5,739575197 | 2,120873 | -2,70623 | 0,006805 | 1 | 0 | Intron (ENSMUST0000 | 4 | 51968094 | 52051110 | 83017 | 2 | 329844 | ENSMUST00000141452.2 | 54066 | ENSMUSG00000085515 | C630028M04Rik | RIKEN cDNA C630028M04 gene |

| | | | | | | | | | | | | | | | | | | | | | | | |
|---|---|---|---|---|---|---|---|---|---|---|---|---|---|---|---|---|---|---|---|---|---|---|---|
| chrX | 7150963 | 7151989 | 1027 | * | 4,011593 | -5,738884341 | 2,211395 | -2,59514 | 0,009455 | 1 | 0 | Intron (ENSMUST00000 | 20 | 7079230 | 7185409 | 106180 | 2 | 12728 | ENSMUST00000132126.2 | 33420 | ENSMUSG00000004317 | Clcn5 | chloride channel, voltage-sensitive 5 |
| chr7 | 96371511 | 96372045 | 535 | * | 5,919634 | -5,738102029 | 1,896079 | -3,0263 | 0,002476 | 1 | 0 | Intron (ENSMUST00000 | 7 | 96427141 | 96446940 | 19800 | 1 | 23966 | ENSMUST00000140140.2 | -55096 | ENSMUSG00000048078 | Tenm4 | teneurin transmembrane protein 4 |
| chr13 | 32717255 | 32718519 | 1265 | * | 11,46059 | -5,737946093 | 1,69967 | -3,37592 | 0,000736 | 0,504427013 | 0,918255 | Exon (ENSMUST000002 | 13 | 32003562 | 32522723 | 519162 | 2 | 218138 | ENSMUST00000041859.9 | -194532 | ENSMUSG00000038372 | Gmds | GDP-mannose 4, 6-dehydratase |
| chr9 | 47874205 | 47874705 | 501 | * | 5,84374 | -5,737606589 | 1,953424 | -2,93721 | 0,003312 | 1 | 0 | Distal Intergenic | 9 | 48043802 | 48062728 | 18927 | 2 | 1,05E+08 | ENSMUST00000213821.2 | 188023 | ENSMUSG00000110924 | Gm40518 | predicted gene, 40518 |
| chr13 | 49185797 | 49186451 | 655 | * | 4,987712 | -5,73695364 | 2,054573 | -2,79229 | 0,005234 | 1 | 0 | Distal Intergenic | 13 | 49191085 | 49214359 | 23275 | 2 | 75607 | ENSMUST00000159196.8 | 27908 | ENSMUSG00000037989 | Wnk2 | WNK lysine deficient protein kinase 2 |
| chr7 | 40106798 | 40107321 | 524 | * | 4,368827 | -5,736810722 | 2,080852 | -2,75695 | 0,005834 | 1 | 0 | Intron (ENSMUST00000 | 7 | 40324161 | 40332685 | 8525 | 1 | 71029 | ENSMUST00000206289.2 | -216840 | ENSMUSG00000102004 | 4933402C06Rik | RIKEN cDNA 4933402C06 gene |
| chr7 | 1,37E+08 | 1,37E+08 | 781 | * | 5,081384 | -5,735493629 | 2,180781 | -2,63002 | 0,008538 | 1 | 0 | Intron (ENSMUST00000 | 7 | 1,37E+08 | 1,37E+08 | 15206 | 2 | 13593 | ENSMUST00000209864.2 | 10212 | ENSMUSG00000010476 | Ebf3 | early B cell factor 3 |
| chr14 | 1,23E+08 | 1,23E+08 | 725 | * | 4,600927 | -5,735474929 | 2,17761 | -2,63384 | 0,008443 | 1 | 0 | Intron (ENSMUST00000 | 14 | 1,23E+08 | 1,23E+08 | 61409 | 1 | 110821 | ENSMUST00000143240.2 | 4497 | ENSMUSG00000041650 | Pcca | propionyl-Coenzyme A carboxylase, alpha polypeptide |
| chr12 | 86646456 | 86647510 | 1055 | * | 8,77067 | -5,734477179 | 1,839775 | -3,11695 | 0,001827 | 0,594430593 | 1,425929 | Distal Intergenic | 12 | 86701899 | 86702005 | 107 | 1 | 1,15E+08 | ENSMUST00000101884.3 | -54389 | ENSMUSG00000075832 | Gm22004 | predicted gene, 22004 |
| chr3 | 1,02E+08 | 1,02E+08 | 813 | * | 5,311286 | -5,73438613 | 1,936272 | -2,96156 | 0,003061 | 1 | 0 | Distal Intergenic | 3 | 1,02E+08 | 1,02E+08 | 26376 | 1 | 319951 | ENSMUST00000198712.6 | -18251 | ENSMUSG00000105891 | A230001M10Rik | RIKEN cDNA A230001M10 gene |
| chr10 | 12095677 | 12096120 | 444 | * | 5,05389 | -5,733317361 | 2,164191 | -2,64917 | 0,008069 | 1 | 0 | Distal Intergenic | 10 | 12239154 | 12239317 | 164 | 1 | 1,15E+08 | ENSMUST00000083162.3 | -143034 | ENSMUSG00000065096 | Gm23680 | predicted gene, 23680 |
| chr1 | 1,63E+08 | 1,63E+08 | 318 | * | 5,745803 | -5,732576517 | 1,903541 | -3,01153 | 0,002599 | 1 | 0 | Intron (ENSMUST00000 | 1 | 1,63E+08 | 1,63E+08 | 134 | 2 | 1,15E+08 | ENSMUST00000104420.3 | 27609 | ENSMUSG00000077608 | Gm22671 | predicted gene, 22671 |
| chr11 | 16283666 | 16284492 | 827 | * | 5,395565 | -5,732429503 | 2,01259 | -2,84829 | 0,004396 | 1 | 0 | Intron (ENSMUST00000 | 11 | 16213218 | 16234551 | 21334 | 1 | 211739 | ENSMUST00000207256.2 | 70448 | ENSMUSG00000048834 | Vstm2a | V-set and transmembrane domain containing 2A |
| chr10 | 11812863 | 11813434 | 572 | * | 5,434568 | -5,730612881 | 1,942886 | -2,94954 | 0,003183 | 1 | 0 | Distal Intergenic | 10 | 11523749 | 11523878 | 130 | 1 | 1,15E+08 | ENSMUST00000104351.3 | 289114 | ENSMUSG00000077539 | Gm22546 | predicted gene, 22546 |
| chr16 | 40711075 | 40712355 | 1281 | * | 8,09228 | -5,73038043 | 1,969586 | -2,90943 | 0,003621 | 1 | 0 | Intron (ENSMUST00000 | 16 | 41353220 | 41966576 | 613357 | 1 | 268890 | ENSMUST00000078873.11 | -640865 | ENSMUSG00000061080 | Lsamp | limbic system-associated membrane protein |
| chr15 | 93353872 | 93354269 | 398 | * | 5,647382 | -5,730087343 | 2,017153 | -2,84068 | 0,004502 | 1 | 0 | Intron (ENSMUST00000 | 15 | 93330826 | 93342896 | 12071 | 1 | 223828 | ENSMUST00000229721.2 | 23046 | ENSMUSG00000036167 | Pphln1 | periphilin 1 |
| chrX | 1,46E+08 | 1,46E+08 | 786 | * | 4,356656 | -5,729555734 | 2,0878 | -2,7443 | 0,006064 | 1 | 0 | Intron (ENSMUST00000 | 20 | 1,46E+08 | 1,46E+08 | 128 | 1 | 1E+08 | ENSMUST00000104579.3 | -8194 | ENSMUSG00000077767 | Snora35 | small nucleolar RNA, H/ACA box 35 |
| chr5 | 77188464 | 77189133 | 670 | * | 6,601457 | -5,728592842 | 1,900145 | -3,01482 | 0,002571 | 1 | 0 | Intron (ENSMUST00000 | 5 | 77184901 | 77195342 | 10442 | 1 | 71868 | ENSMUST00000118941.2 | 3563 | ENSMUSG00000029248 | Thegl | theg spermatid protein like |
| chr4 | 29587741 | 29588698 | 958 | * | 6,344321 | -5,72817541 | 1,975829 | -2,89913 | 0,003742 | 1 | 0 | Distal Intergenic | 4 | 29068079 | 29088811 | 20733 | 1 | 1,03E+08 | ENSMUST00000154813.2 | 519662 | | Gm11917 | predicted gene 11917 |
| chr3 | 58126490 | 58127142 | 653 | * | 4,896861 | -5,727887136 | 2,080924 | -2,75257 | 0,005913 | 1 | 0 | Distal Intergenic | 3 | 58071353 | 58075037 | 3685 | 1 | 1,05E+08 | ENSMUST00000200072.2 | 55137 | | Gm40056 | predicted gene, 40056 |
| chr2 | 1,78E+08 | 1,78E+08 | 595 | * | 6,380013 | -5,727837665 | 1,998946 | -2,86543 | 0,004164 | 1 | 0 | Distal Intergenic | 2 | 1,78E+08 | 1,78E+08 | 1713 | 1 | 1,19E+08 | ENSMUST00000246055.1 | 253875 | | Gm54095 | predicted gene, 54095 |
| chr16 | 52110438 | 52111082 | 645 | * | 4,929208 | -5,726806848 | 2,083194 | -2,74905 | 0,005977 | 1 | 0 | Intron (ENSMUST00000 | 16 | 52071450 | 52117287 | 45838 | 2 | 11658 | ENSMUST00000167115.8 | 6205 | ENSMUSG00000022636 | Alcam | activated leukocyte cell adhesion molecule |
| chr2 | 1,5E+08 | 1,5E+08 | 976 | * | 12,99333 | -5,726595649 | 1,655104 | -3,45996 | 0,00054 | 0,452373067 | 0,918255 | Distal Intergenic | 2 | 1,5E+08 | 1,5E+08 | 22303 | 2 | 104348 | ENSMUST00000089207.13 | -12872 | ENSMUSG00000068134 | Zfp120 | zinc finger protein 120 |
| chr9 | 72768481 | 72769038 | 558 | * | 5,348651 | -5,72542919 | 1,933701 | -2,96087 | 0,003068 | 1 | 0 | Intron (ENSMUST00000 | 9 | 72817909 | 72819587 | 1679 | 1 | 235472 | ENSMUST00000184274.2 | -48871 | ENSMUSG00000036030 | Prtg | protogenin |
| chr13 | 7716415 | 7717129 | 715 | * | 4,354496 | -5,725016144 | 2,148127 | -2,66512 | 0,007696 | 1 | 0 | Distal Intergenic | 13 | 7496545 | 7515379 | 18835 | 1 | 1,03E+08 | ENSMUST00000221383.2 | 219870 | | Gm36074 | predicted gene, 36074 |
| chr16 | 23109643 | 23110340 | 698 | * | 6,309207 | -5,724111931 | 1,891751 | -3,02583 | 0,00248 | 1 | 0 | Promoter (<=1kb) | 16 | 23109220 | 23177345 | 68126 | 1 | 20440 | ENSMUST00000115335.2 | 423 | ENSMUSG00000022885 | St6gal1 | beta galactoside alpha 2,6 sialyltransferase 1 |
| chr7 | 1,38E+08 | 1,38E+08 | 500 | * | 4,894134 | -5,723080334 | 2,042409 | -2,80212 | 0,005077 | 1 | 0 | Intron (ENSMUST00000 | 7 | 1,38E+08 | 1,38E+08 | 188755 | 2 | 70571 | ENSMUST00000160436.2 | 43776 | ENSMUSG00000091002 | Tcerg1l | transcription elongation regulator 1-like |
| chrX | 24516497 | 24517324 | 828 | * | 5,343552 | -5,723020828 | 2,035695 | -2,81134 | 0,004934 | 1 | 0 | Distal Intergenic | 20 | 24418489 | 24441134 | 22646 | 2 | 1E+08 | ENSMUST00000119285.2 | -75363 | ENSMUSG00000081218 | Gm4297 | predicted gene 4297 |
| chrX | 51941198 | 51942330 | 1133 | * | 4,840282 | -5,722916951 | 2,004746 | -2,85468 | 0,004308 | 1 | 0 | Distal Intergenic | 20 | 51967120 | 51967610 | 491 | 1 | 624936 | ENSMUST00000119119.2 | -24790 | | Pou5f1-rs8 | POU domain, class 5, transcription factor 1, related sequence 8 |
| chr4 | 1,1E+08 | 1,1E+08 | 861 | * | 5,121683 | -5,722835998 | 2,01467 | -2,84058 | 0,004503 | 1 | 0 | Intron (ENSMUST00000 | 4 | 1,1E+08 | 1,1E+08 | 84594 | 1 | 14084 | ENSMUST00000155204.2 | -11875 | ENSMUSG00000010517 | Faf1 | Fas-associated factor 1 |
| chr10 | 55011665 | 55012317 | 653 | * | 3,697589 | -5,720874466 | 2,210487 | -2,58806 | 0,009652 | 1 | 0 | Distal Intergenic | 10 | 54589072 | 54589133 | 62 | 2 | 1,15E+08 | ENSMUST00000158362.3 | -422532 | ENSMUSG00000088987 | Gm25602 | predicted gene, 25602 |
| chr1 | 1,65E+08 | 1,65E+08 | 526 | * | 3,98597 | -5,720839163 | 2,18823 | -2,61437 | 0,008939 | 1 | 0 | Exon (ENSMUST00000 | 1 | 1,65E+08 | 1,65E+08 | 1524 | 2 | 68481 | ENSMUST00000192716.2 | 20415 | ENSMUSG00000026566 | Mpzl1 | myelin protein zero-like 1 |
| chr18 | 27563696 | 27564401 | 706 | * | 5,42446 | -5,719308522 | 2,10508 | -2,71691 | 0,006589 | 1 | 0 | Distal Intergenic | 18 | 27942785 | 27960738 | 17954 | 1 | 1,03E+08 | ENSMUST00000234286.2 | -378384 | | Gm33674 | predicted gene, 33674 |
| chr14 | 33457222 | 33458723 | 1502 | * | 14,97029 | -5,718974077 | 1,571978 | -3,63807 | 0,000275 | 0,227352887 | 2,141612 | Distal Intergenic | 14 | 33530032 | 33533240 | 3209 | 2 | 218902 | ENSMUST00000228030.2 | 74517 | ENSMUSG00000115751 | 4930503F20Rik | RIKEN cDNA 4930503F20 gene |
| chr5 | 21448679 | 21449164 | 486 | * | 5,210321 | -5,718089363 | 1,98955 | -2,87406 | 0,004052 | 1 | 0 | Intron (ENSMUST00000 | 5 | 21456157 | 21463281 | 7125 | 1 | 212167 | ENSMUST00000196035.2 | -6993 | ENSMUSG00000039934 | Gsap | gamma-secretase activating protein |
| chr1 | 1,5E+08 | 1,5E+08 | 611 | * | 5,483715 | -5,717970396 | 2,027555 | -2,82013 | 0,0048 | 1 | 0 | Promoter (1-2kb) | 1 | 1,5E+08 | 1,5E+08 | 1616 | 1 | 108989 | ENSMUST00000147769.2 | 1614 | ENSMUSG00000006005 | Tpr | translocated promoter region, nuclear basket protein |
| chr17 | 19965531 | 19966059 | 529 | * | 5,046231 | -5,717751625 | 2,114757 | -2,70374 | 0,006856 | 1 | 0 | Distal Intergenic | 17 | 19993625 | 20032798 | 39174 | 1 | 627636 | ENSMUST00000172203.2 | -27566 | ENSMUSG00000091771 | Vmn2r103 | vomeronasal 2, receptor 103 |
| chr17 | 64293786 | 64294369 | 584 | * | 3,846644 | -5,717606031 | 2,186277 | -2,61522 | 0,008917 | 1 | 0 | Intron (ENSMUST00000 | 17 | 64244967 | 64446192 | 201226 | 1 | 14158 | ENSMUST00000038080.7 | 48819 | ENSMUSG00000000127 | Fer | fer (fms/fps related) protein kinase |
| chr2 | 1,54E+08 | 1,54E+08 | 703 | * | 5,809733 | -5,715237938 | 2,058874 | -2,77591 | 0,005505 | 1 | 0 | Distal Intergenic | 2 | 1,54E+08 | 1,54E+08 | 1481 | 1 | 19194 | ENSMUST00000132933.2 | 33532 | ENSMUSG00000042459 | Bpifa2 | BPI fold containing family A, member 2 |
| chr9 | 64694723 | 64695575 | 853 | * | 5,283347 | -5,71293938 | 1,931806 | -2,9573 | 0,003103 | 1 | 0 | Distal Intergenic | 9 | 64701734 | 64703685 | 1952 | 2 | 1,03E+08 | ENSMUST00000241365.1 | 8110 | | Gm36459 | predicted gene, 36459 |
| chr7 | 1,09E+08 | 1,09E+08 | 750 | * | 5,445605 | -5,712890154 | 1,916447 | -2,98098 | 0,002873 | 1 | 0 | Intron (ENSMUST00000 | 7 | 1,09E+08 | 1,09E+08 | 102080 | 2 | 76954 | ENSMUST00000208734.2 | 5453 | ENSMUSG00000031024 | Denn2b | DENN domain containing 2B |
| chr14 | 84847272 | 84848458 | 1187 | * | 6,20421 | -5,711617994 | 2,139999 | -2,66898 | 0,007608 | 1 | 0 | Distal Intergenic | 14 | 84696164 | 84713864 | 17701 | 2 | 319903 | ENSMUST00000228125.2 | -133408 | ENSMUSG00000115529 | 9630013A20Rik | RIKEN cDNA 9630013A20 gene |
| chr13 | 92013282 | 92014054 | 773 | * | 6,411977 | -5,711224267 | 1,917829 | -2,97796 | 0,002902 | 1 | 0 | 5' UTR | 13 | 92001506 | 92025004 | 23499 | 2 | 76722 | ENSMUST00000022122.4 | 10950 | ENSMUSG00000021622 | Ckmt2 | creatine kinase, mitochondrial 2 |
| chr4 | 1,19E+08 | 1,19E+08 | 638 | * | 5,284293 | -5,711209914 | 1,919956 | -2,97466 | 0,002933 | 1 | 0 | Intron (ENSMUST00000 | 4 | 1,19E+08 | 1,19E+08 | 16251 | 2 | 73332 | ENSMUST00000238485.2 | -14660 | ENSMUSG00000028637 | Ccdc30 | coiled-coil domain containing 30 |
| chr12 | 15385258 | 15385692 | 435 | * | 4,256674 | -5,710586316 | 2,060329 | -2,77169 | 0,005577 | 1 | 0 | Distal Intergenic | 12 | 15118118 | 15187519 | 69402 | 1 | 1,03E+08 | ENSMUST00000222816.2 | 267140 | | Gm36128 | predicted gene, 36128 |
| chr2 | 32653705 | 32654109 | 405 | * | 4,260705 | -5,710479677 | 2,091402 | -2,73045 | 0,006325 | 1 | 0 | Intron (ENSMUST00000 | 2 | 32650307 | 32651801 | 1495 | 1 | 30933 | ENSMUST00000149987.2 | 3398 | ENSMUSG00000009563 | Tor2a | torsin family 2, member A |
| chr3 | 56061136 | 56061970 | 835 | * | 7,450983 | -5,710373575 | 1,868117 | -3,05675 | 0,002237 | 1 | 0 | Intron (ENSMUST00000 | 3 | 55532616 | 56091122 | 558507 | 2 | 26422 | ENSMUST00000029374.8 | 29152 | ENSMUSG00000027799 | Nbea | neurobeachin |
| chr9 | 9087726 | 9088537 | 812 | * | 6,96516 | -5,708992181 | 1,959484 | -2,91352 | 0,003574 | 1 | 0 | Intron (ENSMUST00000 | 9 | 9236293 | 9260888 | 24596 | 1 | 1,01E+08 | ENSMUST00000189204.2 | -147756 | ENSMUSG00000085253 | Gm16833 | predicted gene, 16833 |
| chr3 | 74925232 | 74925781 | 550 | * | 4,671516 | -5,708299143 | 2,019003 | -2,82729 | 0,004694 | 1 | 0 | Distal Intergenic | 3 | 74968668 | 75051010 | 82343 | 2 | 213234 | ENSMUST00000107776.8 | 125229 | ENSMUSG00000034151 | Zbbx | zinc finger, B-box domain containing |
| chr14 | 90089818 | 90090607 | 790 | * | 4,762345 | -5,7078905 | 2,063517 | -2,7661 | 0,005673 | 1 | 0 | Distal Intergenic | 14 | 89944241 | 89944372 | 132 | 1 | 1,15E+08 | ENSMUST00000104220.3 | 145577 | ENSMUSG00000077408 | Gm25415 | predicted gene, 25415 |
| chr9 | 1,05E+08 | 1,05E+08 | 1084 | * | 6,470447 | -5,701747902 | 1,961137 | -2,90737 | 0,003645 | 1 | 0 | Intron (ENSMUST00000 | 9 | 1,05E+08 | 1,05E+08 | 149743 | 1 | 74020 | ENSMUST00000213452.2 | -11207 | ENSMUSG00000032564 | Cpne4 | copine IV |
| chr12 | 72271410 | 72271894 | 485 | * | 4,684516 | -5,701601003 | 2,133428 | -2,67251 | 0,007529 | 1 | 0 | Intron (ENSMUST00000 | 12 | 72258526 | 72282513 | 23988 | 2 | 104001 | ENSMUST00000150156.2 | 10619 | ENSMUSG00000021087 | Rtn1 | reticulon 1 |
| chr4 | 82541717 | 82542214 | 498 | * | 5,341379 | -5,701200341 | 2,004018 | -2,84488 | 0,004443 | 1 | 0 | Intron (ENSMUST00000 | 4 | 82503285 | 82505556 | 2272 | 2 | 1,05E+08 | ENSMUST00000137477.2 | -36161 | | Gm11267 | predicted gene 11267 |
| chr1 | 54003765 | 54004468 | 704 | * | 6,828215 | -5,699455456 | 1,854632 | -3,07309 | 0,002119 | 1 | 0 | Intron (ENSMUST00000 | 1 | 53878176 | 53905671 | 27496 | 2 | 329152 | ENSMUST00000152870.2 | -98094 | ENSMUSG00000042807 | Hecw2 | HECT, C2 and WW domain containing E3 ubiquitin protein ligase 2 |
| chrX | 69262193 | 69263123 | 931 | * | 4,13147 | -5,698575206 | 2,145459 | -2,65611 | 0,007905 | 1 | 0 | Distal Intergenic | 20 | 68972865 | 69187194 | 214330 | 2 | 74305 | ENSMUST00000188663.2 | -74999 | ENSMUSG00000100511 | 1700111N16Rik | RIKEN cDNA 1700111N16 gene |
| chr9 | 36596356 | 36596947 | 592 | * | 4,484343 | -5,698551086 | 2,045389 | -2,78605 | 0,005335 | 1 | 0 | Intron (ENSMUST00000 | 9 | 36547049 | 36604265 | 57217 | 2 | 235973 | ENSMUST00000216487.2 | 7318 | ENSMUSG00000070313 | Pate14 | prostate and testis expressed 14 |
| chr11 | 52392755 | 52393458 | 704 | * | 6,826908 | -5,6985366 | 1,987525 | -2,86715 | 0,004142 | 1 | 0 | Distal Intergenic | 11 | 52287473 | 52294101 | 6629 | 1 | 213673 | ENSMUST00000136303.2 | 105282 | ENSMUSG00000036275 | 9530068E07Rik | RIKEN cDNA 9530068E07 gene |
| chr16 | 57548117 | 57548593 | 477 | * | 5,322022 | -5,698524894 | 1,981033 | -2,87654 | 0,004021 | 1 | 0 | Intron (ENSMUST00000 | 16 | 57444621 | 57575100 | 130480 | 2 | 12837 | ENSMUST00000089332.5 | 26507 | ENSMUSG00000068196 | Col8a1 | collagen, type VIII, alpha 1 |
| chr17 | 29278373 | 29279037 | 665 | * | 5,326964 | -5,698513734 | 1,940091 | -2,93724 | 0,003311 | 1 | 0 | Promoter (1-2kb) | 17 | 29277203 | 29278729 | 1527 | 1 | 1,05E+08 | ENSMUST00000233408.2 | 1170 | | Gm41556 | predicted gene, 41556 |
| chr7 | 1,23E+08 | 1,23E+08 | 479 | * | 7,054874 | -5,698456525 | 1,838056 | -3,10026 | 0,001933 | 1 | 0 | Distal Intergenic | 7 | 1,23E+08 | 1,23E+08 | 6236 | 2 | 210162 | ENSMUST00000128217.2 | -145213 | ENSMUSG00000030757 | Zkscan2 | zinc finger with KRAB and SCAN domains 2 |
| chr13 | 35413968 | 35415250 | 1283 | * | 8,000996 | -5,698384808 | 1,761052 | -3,23578 | 0,001213 | 1 | 0 | Distal Intergenic | 13 | 35394643 | 35396031 | 1389 | 2 | 75543 | ENSMUST00000220954.2 | -17937 | ENSMUSG00000113770 | 1700019C18Rik | RIKEN cDNA 1700019C18 gene |
| chr17 | 86060267 | 86060775 | 509 | * | 8,299495 | -5,698325306 | 1,78156 | -3,1985 | 0,001381 | 1 | 0 | Distal Intergenic | 17 | 86001057 | 86020456 | 19400 | 1 | 1,03E+08 | ENSMUST00000235073.2 | 59210 | | CJ186046Rik | Riken cDNA CJ186046 gene |
| chr4 | 1,01E+08 | 1,01E+08 | 570 | * | 5,418778 | -5,696459829 | 1,967281 | -2,8956 | 0,003784 | 1 | 0 | Promoter (<=1kb) | 4 | 1,01E+08 | 1,01E+08 | 164 | 1 | 1,15E+08 | ENSMUST00000083345.3 | -387 | ENSMUSG00000065279 | Gm25124 | predicted gene, 25124 |
| chr14 | 79764021 | 79765004 | 984 | * | 4,636186 | -5,695746469 | 2,088859 | -2,72673 | 0,006397 | 1 | 0 | Intron (ENSMUST00000 | 14 | 79753138 | 79776267 | 23130 | 1 | 13709 | ENSMUST00000227104.2 | 10883 | ENSMUSG00000036461 | Elf1 | E74 like ETS transcription factor 1 |
| chr5 | 92856662 | 92857385 | 724 | * | 6,114875 | -5,693334641 | 1,974848 | -2,88292 | 0,00394 | 1 | 0 | Intron (ENSMUST00000 | 5 | 92831484 | 93112621 | 281138 | 1 | 27428 | ENSMUST00000168878.8 | 25178 | ENSMUSG00000029381 | Shroom3 | shroom family member 3 |
| chr1 | 56494618 | 56495376 | 759 | * | 4,364944 | -5,692683689 | 2,115935 | -2,69039 | 0,007137 | 1 | 0 | Distal Intergenic | 1 | 56675203 | 56676594 | 1392 | 2 | 71066 | ENSMUST00000062085.6 | 181218 | ENSMUSG00000045336 | Hsfy2 | heat shock transcription factor, Y-linked 2 |
| chr14 | 1,19E+08 | 1,19E+08 | 558 | * | 6,830108 | -5,692303015 | 1,855152 | -3,06838 | 0,002152 | 1 | 0 | Promoter (<=1kb) | 14 | 1,19E+08 | 1,19E+08 | 2923 | 1 | 1,03E+08 | ENSMUST00000227592.2 | 0 | | Gm32093 | predicted gene, 32093 |
| chr3 | 74780308 | 74781009 | 702 | * | 5,443757 | -5,691266962 | 1,913355 | -2,9745 | 0,002935 | 1 | 0 | Distal Intergenic | 3 | 74968668 | 75051010 | 82343 | 2 | 213234 | ENSMUST00000107776.8 | 270001 | ENSMUSG00000034151 | Zbbx | zinc finger, B-box domain containing |
| chrX | 81865405 | 81866397 | 993 | * | 4,883253 | -5,691083305 | 2,19876 | -2,58832 | 0,009645 | 1 | 0 | Distal Intergenic | 20 | 81992476 | 82536345 | 543870 | 1 | 13405 | ENSMUST00000127295.8 | -126079 | ENSMUSG00000045103 | Dmd | dystrophin, muscular dystrophy |
| chr8 | 26263668 | 26264272 | 605 | * | 4,745679 | -5,689824275 | 2,150807 | -2,64544 | 0,008159 | 1 | 0 | Intron (ENSMUST00000 | 8 | 26259539 | 26267610 | 8072 | 2 | 67384 | ENSMUST00000210103.2 | 3338 | ENSMUSG00000037316 | Bag4 | BCL2-associated athanogene 4 |
| chr4 | 5214355 | 5214972 | 618 | * | 4,90314 | -5,689598913 | 2,121213 | -2,68224 | 0,007313 | 1 | 0 | Distal Intergenic | 4 | 5632328 | 5642284 | 9957 | 1 | 73872 | ENSMUST00000156903.8 | -417356 | ENSMUSG00000086066 | 4930423M02Rik | RIKEN cDNA 4930423M02 gene |
| chr1 | 87843611 | 87844343 | 733 | * | 6,441884 | -5,689452065 | 1,935551 | -2,93945 | 0,003288 | 1 | 0 | Promoter (1-2kb) | 1 | 87845620 | 87849693 | 4074 | 1 | 227333 | ENSMUST00000191589.2 | -1277 | ENSMUSG00000070738 | Dgkd | diacylglycerol kinase, delta |
| chr11 | 43771825 | 43772415 | 591 | * | 4,895587 | -5,689358288 | 2,092637 | -2,71875 | 0,006553 | 1 | 0 | Intron (ENSMUST00000 | 11 | 43665433 | 43792037 | 126605 | 2 | 11548 | ENSMUST00000067258.9 | 19622 | ENSMUSG00000050541 | Adra1b | adrenergic receptor, alpha 1b |
| chr3 | 68122994 | 68123606 | 613 | * | 4,99511 | -5,688857593 | 2,040156 | -2,78844 | 0,005296 | 1 | 0 | Intron (ENSMUST00000 | 3 | 67972135 | 68533797 | 561663 | 1 | 30953 | ENSMUST00000169909.10 | 150859 | ENSMUSG00000027777 | Schip1 | schwannomin interacting protein 1 |
| chr2 | 1,55E+08 | 1,55E+08 | 325 | * | 4,677243 | -5,68811477 | 2,128667 | -2,67215 | 0,007537 | 1 | 0 | Intron (ENSMUST00000 | 2 | 1,55E+08 | 1,55E+08 | 3427 | 2 | 228812 | ENSMUST00000125334.2 | -20919 | ENSMUSG00000038383 | Pigu | phosphatidylinositol glycan anchor biosynthesis, class U |
| chr3 | 1,4E+08 | 1,4E+08 | 948 | * | 5,17495 | -5,687946174 | 2,164554 | -2,62777 | 0,008595 | 1 | 0 | Distal Intergenic | 3 | 1,41E+08 | 1,41E+08 | 3470 | 2 | 1,03E+08 | ENSMUST00000199504.2 | 322772 | | Gm32754 | predicted gene, 32754 |
| chr2 | 72996180 | 72996720 | 541 | * | 4,02904 | -5,686871309 | 2,2058 | -2,57814 | 0,009933 | 1 | 0 | Intron (ENSMUST00000 | 2 | 72923546 | 73043304 | 119759 | 2 | 67059 | ENSMUST00000112055.8 | 46584 | ENSMUSG00000027108 | Ola1 | Obg-like ATPase 1 |
| chr7 | 47278262 | 47278785 | 524 | * | 4,95556 | -5,68671092 | 2,151627 | -2,64298 | 0,008218 | 1 | 0 | Distal Intergenic | 7 | 47238698 | 47251120 | 12423 | 2 | 233222 | ENSMUST00000176369.2 | -27142 | ENSMUSG00000078698 | Mrgpra3 | MAS-related GPR, member A3 |
| chr15 | 4390389 | 4391008 | 620 | * | 5,215713 | -5,685976208 | 2,048834 | -2,77522 | 0,005516 | 1 | 0 | Distal Intergenic | 15 | 4404973 | 4549896 | 144924 | 1 | 239318 | ENSMUST00000130332.2 | -13965 | ENSMUSG00000049148 | Plcxd3 | phosphatidylinositol-specific phospholipase C, X domain containing 3 |
| chr4 | 86160192 | 86160516 | 325 | * | 4,254167 | -5,684497396 | 2,200226 | -2,5836 | 0,009778 | 1 | 0 | Intron (ENSMUST00000 | 4 | 86194726 | 86195339 | 614 | 1 | 77739 | ENSMUST00000136418.2 | -34210 | ENSMUSG00000066113 | Adamtsl1 | ADAMTS-like 1 |
| chr14 | 79155493 | 79156433 | 941 | * | 6,244331 | -5,684155151 | 1,857532 | -3,06006 | 0,002213 | 1 | 0 | Intron (ENSMUST00000 | 14 | 79086694 | 79302708 | 216015 | 1 | 219189 | ENSMUST00000227255.2 | 68799 | ENSMUSG00000058997 | Vwa8 | von Willebrand factor A domain containing 8 |
| chr1 | 1,85E+08 | 1,85E+08 | 793 | * | 4,577716 | -5,683573574 | 2,178695 | -2,60871 | 0,009089 | 1 | 0 | Intron (ENSMUST00000 | 1 | 1,85E+08 | 1,85E+08 | 21565 | 2 | 66112 | ENSMUST00000189492.2 | 11300 | ENSMUSG00000026621 | Mtarc1 | mitochondrial amidoxime reducing component 1 |
| chr11 | 9065393 | 9066032 | 640 | * | 5,772705 | -5,682154194 | 1,936576 | -2,93412 | 0,003345 | 1 | 0 | Promoter (2-3kb) | 11 | 9068103 | 9086164 | 18062 | 1 | 22271 | ENSMUST00000020677.8 | -2071 | ENSMUSG00000020407 | Upp1 | uridine phosphorylase 1 |
| chr10 | 84864748 | 84865239 | 492 | * | 4,499891 | -5,682053062 | 2,118128 | -2,68258 | 0,007306 | 1 | 0 | Distal Intergenic | 10 | 84844580 | 84854201 | 9622 | 1 | 237422 | ENSMUST00000214820.2 | 20168 | ENSMUSG00000035620 | Ric8b | RIC8 guanine nucleotide exchange factor B |
| chr5 | 1,2E+08 | 1,2E+08 | 1258 | * | 9,683207 | -5,681662225 | 1,878065 | -3,02527 | 0,002484 | 0,649854434 | 1,425929 | Distal Intergenic | 5 | 1,2E+08 | 1,2E+08 | 9160 | 2 | 1,03E+08 | ENSMUST00000201435.2 | 124767 | ENSMUSG00000106938 | Gm31314 | predicted gene, 31314 |
| chr14 | 1,11E+08 | 1,11E+08 | 947 | * | 3,91784 | -5,68055489 | 2,14122 | -2,65295 | 0,007979 | 1 | 0 | Distal Intergenic | 14 | 1,11E+08 | 1,11E+08 | 6570 | 2 | 239250 | ENSMUST00000078386.4 | -37567 | ENSMUSG00000045871 | Slitrk6 | SLIT and NTRK-like family, member 6 |
| chr13 | 42019245 | 42020107 | 863 | * | 7,872241 | -5,679614491 | 1,897257 | -2,99359 | 0,002757 | 1 | 0 | Promoter (<=1kb) | 13 | 42019380 | 42066583 | 47204 | 1 | 1E+08 | ENSMUST00000221285.3 | 0 | ENSMUSG00000113856 | Gm10790 | predicted gene 10790 |
| chr14 | 97528710 | 97529602 | 893 | * | 5,096381 | -5,678343145 | 2,2018 | -2,57896 | 0,00991 | 1 | 0 | Distal Intergenic | 14 | 97573931 | 97574068 | 138 | 2 | 1,15E+08 | ENSMUST00000157913.3 | 44466 | ENSMUSG00000088538 | Gm23276 | predicted gene, 23276 |
| chr13 | 1,15E+08 | 1,15E+08 | 1010 | * | 6,804086 | -5,676884742 | 1,871925 | -3,03265 | 0,002424 | 1 | 0 | Promoter (<=1kb) | 13 | 1,15E+08 | 1,15E+08 | 148869 | 2 | 109700 | ENSMUST00000061673.9 | -544 | ENSMUSG00000042284 | Itga1 | integrin alpha 1 |
| chrX | 1,61E+08 | 1,61E+08 | 759 | * | 6,264807 | -5,676865495 | 1,87613 | -3,02584 | 0,002479 | 1 | 0 | Distal Intergenic | 20 | 1,61E+08 | 1,61E+08 | 6360 | 2 | 194590 | ENSMUST00000155043.2 | 166142 | ENSMUSG00000040855 | Reps2 | RALBP1 associated Eps domain containing protein 2 |
| chrX | 67717576 | 67718447 | 872 | * | 5,868513 | -5,676789886 | 1,892389 | -2,9998 | 0,002702 | 1 | 0 | Distal Intergenic | 20 | 67722147 | 67761569 | 39423 | 1 | 14265 | ENSMUST00000088546.12 | -3700 | ENSMUSG00000000838 | Fmr1 | FMRP translational regulator 1 |
| chr11 | 1,03E+08 | 1,03E+08 | 644 | * | 5,428782 | -5,676129732 | 2,117578 | -2,68048 | 0,007352 | 1 | 0 | Promoter (<=1kb) | 11 | 1,03E+08 | 1,03E+08 | 5323 | 2 | 14615 | ENSMUST00000107075.9 | 0 | ENSMUSG00000034520 | Gjc1 | gap junction protein, gamma 1 |
| chr7 | 25643997 | 25644447 | 451 | * | 3,942082 | -5,676028868 | 2,171537 | -2,61383 | 0,008953 | 1 | 0 | Distal Intergenic | 7 | 25616191 | 25616855 | 665 | 1 | 13088 | ENSMUST00000144140.2 | 27806 | ENSMUSG00000030483 | Cyp2b10 | cytochrome P450, family 2, subfamily b, polypeptide 10 |
| chr9 | 82340157 | 82340477 | 321 | * | 4,463681 | -5,674115287 | 2,06427 | -2,74873 | 0,005983 | 1 | 0 | Distal Intergenic | 9 | 82321467 | 82321527 | 61 | 2 | 1,15E+08 | ENSMUST00000158441.3 | -18630 | ENSMUSG00000089066 | Gm22836 | predicted gene, 22836 |
| chr3 | 1,16E+08 | 1,16E+08 | 708 | * | 4,119143 | -5,672892648 | 2,162171 | -2,6237 | 0,008698 | 1 | 0 | Intron (ENSMUST00000 | 3 | 1,16E+08 | 1,16E+08 | 3850 | 2 | 64378 | ENSMUST00000090473.7 | -82316 | ENSMUSG00000068696 | Gpr88 | G-protein coupled receptor 88 |
| chr1 | 22481502 | 22482281 | 780 | * | 4,277893 | -5,672819783 | 2,175096 | -2,60808 | 0,009105 | 1 | 0 | Intron (ENSMUST00000 | 1 | 22507272 | 22548503 | 41232 | 2 | 116837 | ENSMUST00000239339.2 | 66222 | ENSMUSG00000041670 | Rims1 | regulating synaptic membrane exocytosis 1 |
| chr12 | 55192497 | 55192947 | 451 | * | 3,799903 | -5,671652757 | 2,177784 | -2,60432 | 0,009206 | 1 | 0 | Intron (ENSMUST00000 | 12 | 55188894 | 55201681 | 12788 | 2 | 1E+08 | ENSMUST00000219562.3 | 8734 | | Gm2436 | predicted gene 2436 |
| chr10 | 27620987 | 27621616 | 630 | * | 5,254374 | -5,671259393 | 2,001737 | -2,83317 | 0,004609 | 1 | 0 | Exon (ENSMUST000002 | 10 | 27522309 | 27574323 | 52015 | 2 | 1,03E+08 | ENSMUST00000218769.2 | -46664 | | Gm30165 | predicted gene, 30165 |
| chrX | 1,12E+08 | 1,12E+08 | 1177 | * | 5,640064 | -5,67119895 | 1,982548 | -2,86056 | 0,004229 | 1 | 0 | Exon (ENSMUST00000 | 20 | 1,12E+08 | 1,12E+08 | 70 | 2 | 723850 | ENSMUST00000083576.3 | 5283 | ENSMUSG00000065510 | Mir361 | microRNA 361 |
| chr3 | 47584240 | 47585415 | 1176 | * | 6,955717 | -5,669928959 | 1,83019 | -3,098 | 0,001948 | 1 | 0 | Distal Intergenic | 3 | 47980448 | 47981195 | 748 | 2 | 666202 | ENSMUST00000193787.2 | 395780 | ENSMUSG00000103257 | Gm7977 | tyrosine 3-monooxygenase/tryptophan 5-monooxygenase activation protein, theta polypeptide pseudogene |
| chr18 | 21542383 | 21543316 | 934 | * | 8,357183 | -5,669728194 | 1,857401 | -3,05251 | 0,002269 | 1 | 0 | Distal Intergenic | 18 | 21513882 | 21513988 | 107 | 2 | 1,15E+08 | ENSMUST00000240423.1 | -28395 | ENSMUSG00000119907 | Gm22886 | predicted gene, 22886 |

| | | | | | | | | | | | | | | | | | | | | | | | |
|---|---|---|---|---|---|---|---|---|---|---|---|---|---|---|---|---|---|---|---|---|---|---|---|
| chr14 | 48401123 | 48401664 | 542 | * | 6,530659 | -5,669726365 | 1,85746 | -3,05241 | 0,00227 | 1 | 0 | Intron (ENSMUST0000 | 14 | 48405730 | 48519032 | 113303 | 1 | 93834 | ENSMUST00000227362.2 | -4066 | ENSMUSG00000021846 | Peli2 | pellino 2 |
| chr5 | 1,46E+08 | 1,46E+08 | 762 | * | 6,59865 | -5,66908531 | 1,972748 | -2,8737 | 0,004057 | 1 | 0 | Distal Intergenic | 5 | 1,46E+08 | 1,46E+08 | 26088 | 2 | 53973 | ENSMUST00000094111.6 | -10897 | ENSMUSG00000075551 | Cyp3a41a | cytochrome P450, family 3, subfamily a, polypeptide 41A |
| chr14 | 92755162 | 92755969 | 808 | * | 3,97819 | -5,66729746 | 2,185065 | -2,59365 | 0,009496 | 1 | 0 | Distal Intergenic | 14 | 92605936 | 92606065 | 130 | 2 | 1,15E+08 | ENSMUST00000104110.3 | -149097 | ENSMUSG00000077298 | Gm24127 | predicted gene, 24127 |
| chrX | 1,57E+08 | 1,57E+08 | 527 | * | 4,40921 | -5,665663234 | 2,112438 | -2,68205 | 0,007317 | 1 | 0 | Intron (ENSMUST0000 | 20 | 1,57E+08 | 1,57E+08 | 31858 | 1 | 66106 | ENSMUST00000126418.2 | 3571 | ENSMUSG00000041476 | Smpx | small muscle protein, X-linked |
| chr4 | 81493398 | 81494449 | 1052 | * | 5,733896 | -5,665356929 | 1,93234 | -2,93186 | 0,003369 | 1 | 0 | Distal Intergenic | 4 | 81334777 | 81361052 | 26276 | 2 | 17475 | ENSMUST00000142273.2 | -132346 | ENSMUSG00000028402 | Mpdz | multiple PDZ domain crumbs cell polarity complex component |
| chr16 | 61264730 | 61265245 | 516 | * | 3,963859 | -5,663986424 | 2,140652 | -2,64592 | 0,008147 | 1 | 0 | Distal Intergenic | 16 | 60890381 | 60890457 | 77 | 1 | 1,15E+08 | ENSMUST00000158376.3 | 374349 | ENSMUSG00000089001 | Gm23852 | predicted gene, 23852 |
| chrX | 1,25E+08 | 1,25E+08 | 893 | * | 4,042292 | -5,663965768 | 2,12371 | -2,66701 | 0,007653 | 1 | 0 | Distal Intergenic | 20 | 1,25E+08 | 1,25E+08 | 399 | 2 | 626048 | ENSMUST00000118529.2 | -128144 | ENSMUSG00000096437 | Rps12-ps16 | ribosomal protein S12, pseudogene 16 |
| chr1 | 49512281 | 49513158 | 878 | * | 5,70322 | -5,663695264 | 1,964376 | -2,8832 | 0,003937 | 1 | 0 | Distal Intergenic | 1 | 49283775 | 49379602 | 95828 | 1 | 442837 | ENSMUST00000188673.7 | 228506 | ENSMUSG00000051616 | C230029F24Rik | RIKEN cDNA C230029F24 gene |
| chr3 | 1,45E+08 | 1,45E+08 | 782 | * | 5,691005 | -5,663592781 | 1,922206 | -2,9464 | 0,003215 | 1 | 0 | 3' UTR | 3 | 1,45E+08 | 1,45E+08 | 7034 | 1 | 71355 | ENSMUST00000142441.2 | 5538 | ENSMUSG00000028197 | Col24a1 | collagen, type XXIV, alpha 1 |
| chr7 | 62845434 | 62846270 | 837 | * | 7,035313 | -5,662794461 | 1,82398 | -3,10464 | 0,001905 | 1 | 0 | Intron (ENSMUST0000 | 7 | 62748440 | 62862261 | 113822 | 2 | 11441 | ENSMUST00000032738.7 | 15991 | ENSMUSG00000030525 | Chrna7 | cholinergic receptor, nicotinic, alpha polypeptide 7 |
| chr12 | 3533298 | 3534088 | 791 | * | 5,569909 | -5,661856789 | 2,07849 | -2,72402 | 0,006449 | 1 | 0 | Intron (ENSMUST0000 | 12 | 3524659 | 3547061 | 22403 | 1 | 75302 | ENSMUST00000219208.2 | 8639 | ENSMUSG00000037486 | Asxl2 | ASXL transcriptional regulator 2 |
| chr1 | 1,94E+08 | 1,94E+08 | 790 | * | 4,918395 | -5,661479176 | 2,004108 | -2,82494 | 0,004729 | 1 | 0 | Distal Intergenic | 1 | 1,94E+08 | 1,94E+08 | 197045 | 1 | 18845 | ENSMUST00000027952.12 | -76150 | ENSMUSG00000026640 | Plxna2 | plexin A2 |
| chr7 | 82107731 | 82108662 | 932 | * | 7,313936 | -5,661187056 | 1,874731 | -3,01973 | 0,00253 | 1 | 0 | Intron (ENSMUST0000 | 7 | 82057682 | 82115052 | 57371 | 1 | 269959 | ENSMUST00000174495.2 | 50049 | ENSMUSG00000070469 | Adamtsl3 | ADAMTS-like 3 |
| chr1 | 43368694 | 43369168 | 475 | * | 5,017713 | -5,659318985 | 2,119441 | -2,67019 | 0,007581 | 1 | 0 | Distal Intergenic | 1 | 43289316 | 43342085 | 52770 | 2 | 1,03E+08 | ENSMUST00000189899.2 | -26609 | | Gm29040 | predicted gene 29040 |
| chr13 | 1,16E+08 | 1,16E+08 | 796 | * | 4,991024 | -5,658724214 | 2,045525 | -2,76639 | 0,005668 | 1 | 0 | Distal Intergenic | 13 | 1,16E+08 | 1,16E+08 | 3957 | 2 | 16392 | ENSMUST00000176444.2 | 152553 | ENSMUSG00000042258 | Isl1 | ISL1 transcription factor, LIM/homeodomain |
| chr3 | 93395036 | 93395475 | 440 | * | 4,061414 | -5,656380123 | 2,194275 | -2,57779 | 0,009943 | 1 | 0 | Distal Intergenic | 3 | 93376061 | 93379287 | 3227 | 1 | 71325 | ENSMUST00000029516.3 | 18975 | ENSMUSG00000027908 | Tchhl1 | trichohyalin-like 1 |
| chr10 | 74881169 | 74881535 | 367 | * | 5,600864 | -5,656221005 | 1,95848 | -2,88807 | 0,003876 | 1 | 0 | Intron (ENSMUST0000 | 10 | 74874055 | 74886522 | 12468 | 1 | 76877 | ENSMUST00000151432.2 | 7114 | ENSMUSG00000020175 | Rab36 | RAB36, member RAS oncogene family |
| chrX | 63085678 | 63086484 | 807 | * | 4,628515 | -5,655150546 | 2,176868 | -2,59784 | 0,009381 | 1 | 0 | Distal Intergenic | 20 | 63195011 | 63195804 | 794 | 2 | 627470 | ENSMUST00000114682.4 | 109320 | ENSMUSG00000079579 | Gm6760 | predicted gene 6760 |
| chr11 | 98128891 | 98130141 | 1251 | * | 6,297944 | -5,654706939 | 1,905152 | -2,96811 | 0,002996 | 1 | 0 | Promoter (2-3kb) | 11 | 98132563 | 98169330 | 36768 | 1 | 69131 | ENSMUST00000147441.8 | -2422 | ENSMUSG00000003119 | Cdk12 | cyclin-dependent kinase 12 |
| chr14 | 1,02E+08 | 1,02E+08 | 931 | * | 4,578866 | -5,65332545 | 2,13146 | -2,65233 | 0,007994 | 1 | 0 | Promoter (1-2kb) | 14 | 1,02E+08 | 1,02E+08 | 631 | 2 | 66200 | ENSMUST00000131022.2 | -1718 | ENSMUSG00000075486 | Commd6 | COMM domain containing 6 |
| chr15 | 65536093 | 65536525 | 433 | * | 3,704764 | -5,653180605 | 2,16685 | -2,60894 | 0,009082 | 1 | 0 | Distal Intergenic | 15 | 65566398 | 65566586 | 189 | 1 | 1,15E+08 | ENSMUST00000104595.4 | -29873 | ENSMUSG00000077783 | Gm23907 | predicted gene, 23907 |
| chr19 | 10553081 | 10553771 | 691 | * | 4,906775 | -5,651299819 | 2,05 | -2,75673 | 0,005838 | 1 | 0 | Promoter (<=1kb) | 19 | 10551041 | 10554445 | 3405 | 2 | 72982 | ENSMUST00000236958.2 | 674 | ENSMUSG00000024666 | Tmem138 | transmembrane protein 138 |
| chr12 | 45663158 | 45663651 | 494 | * | 4,707479 | -5,651120011 | 2,020809 | -2,79646 | 0,005167 | 1 | 0 | Distal Intergenic | 12 | 45513079 | 45542564 | 29486 | 1 | 1,03E+08 | ENSMUST00000219610.2 | 150079 | ENSMUSG00000112054 | Gm33680 | predicted gene, 33680 |
| chr14 | 1,01E+08 | 1,01E+08 | 737 | * | 4,757176 | -5,650762079 | 2,083481 | -2,71217 | 0,006684 | 1 | 0 | Distal Intergenic | 14 | 1,01E+08 | 1,01E+08 | 2633 | 2 | 76627 | ENSMUST00000187304.2 | -3738 | ENSMUSG00000042888 | Prr30 | proline rich 30 |
| chr18 | 40019001 | 40019633 | 633 | * | 5,346126 | -5,65071166 | 1,964428 | -2,87652 | 0,004021 | 1 | 0 | Distal Intergenic | 18 | 40039233 | 40062427 | 23195 | 2 | 1,05E+08 | ENSMUST00000237624.2 | 42794 | | Gm41708 | predicted gene, 41708 |
| chr4 | 98068455 | 98069221 | 767 | * | 6,852111 | -5,650644647 | 1,955231 | -2,89001 | 0,003852 | 1 | 0 | Distal Intergenic | 4 | 98209635 | 98211087 | 1453 | 1 | 78687 | ENSMUST00000131324.2 | -140414 | | 0610025J13Rik | RIKEN cDNA 0610025J13 gene |
| chr16 | 74265185 | 74265817 | 633 | * | 4,422125 | -5,649976408 | 2,169752 | -2,60397 | 0,009215 | 1 | 0 | Distal Intergenic | 16 | 74146683 | 74208713 | 62031 | 2 | 268902 | ENSMUST00000156010.2 | -56472 | ENSMUSG00000052516 | Robo2 | roundabout guidance receptor 2 |
| chr17 | 45713068 | 45714101 | 1034 | * | 6,446477 | -5,64956392 | 1,850533 | -3,05294 | 0,002266 | 1 | 0 | Intron (ENSMUST0000 | 17 | 45719298 | 45726610 | 7313 | 2 | 71702 | ENSMUST00000232900.2 | 12509 | ENSMUSG00000023932 | Cdc5l | cell division cycle 5-like (S. pombe) |
| chrX | 1,59E+08 | 1,59E+08 | 548 | * | 5,412484 | -5,648707559 | 2,024736 | -2,78985 | 0,005273 | 1 | 0 | Intron (ENSMUST0000 | 20 | 1,59E+08 | 1,59E+08 | 134919 | 1 | 270672 | ENSMUST00000033665.9 | 68349 | ENSMUSG00000031303 | Map3k15 | mitogen-activated protein kinase kinase kinase 15 |
| chr2 | 1,02E+08 | 1,02E+08 | 630 | * | 4,263003 | -5,648697817 | 2,067374 | -2,73231 | 0,006289 | 1 | 0 | Distal Intergenic | 2 | 1,02E+08 | 1,02E+08 | 9137 | 1 | 76501 | ENSMUST00000156799.2 | 18367 | ENSMUSG00000027163 | Commd9 | COMM domain containing 9 |
| chr4 | 27879572 | 27880333 | 762 | * | 4,981954 | -5,646493818 | 1,994055 | -2,83166 | 0,004631 | 1 | 0 | Distal Intergenic | 4 | 27700671 | 27700777 | 107 | 1 | 1,15E+08 | ENSMUST00000179794.2 | 178901 | ENSMUSG00000095003 | Gm25173 | predicted gene, 25173 |
| chr13 | 85577948 | 85578406 | 459 | * | 4,874284 | -5,646280312 | 2,023745 | -2,79002 | 0,005271 | 1 | 0 | Distal Intergenic | 13 | 85363614 | 85437249 | 73636 | 2 | 218397 | ENSMUST00000223598.2 | -140699 | ENSMUSG00000021549 | Rasa1 | RAS p21 protein activator 1 |
| chr15 | 5581906 | 5582297 | 392 | * | 4,073032 | -5,645859324 | 2,09296 | -2,69755 | 0,006985 | 1 | 0 | Intron (ENSMUST0000 | 15 | 5525799 | 5624464 | 98666 | 1 | 71432 | ENSMUST00000228069.2 | 56107 | ENSMUSG00000110018 | 5430437J10Rik | RIKEN cDNA 5430437J10 gene |
| chrX | 36856800 | 36857345 | 546 | * | 4,398677 | -5,645710517 | 2,12122 | -2,66154 | 0,007778 | 1 | 0 | Promoter (1-2kb) | 20 | 36850650 | 36854930 | 4281 | 2 | 622301 | ENSMUST00000115162.2 | -1870 | ENSMUSG00000079627 | Rhox2h | reproductive homeobox 2H |
| chr13 | 71651676 | 71652130 | 455 | * | 4,415582 | -5,645640427 | 2,135859 | -2,64326 | 0,008211 | 1 | 0 | Intron (ENSMUST0000 | 13 | 71635726 | 71760828 | 125103 | 1 | 1,03E+08 | ENSMUST00000221719.2 | 15950 | | 1700112M02Rik | RIKEN cDNA 1700112M02 gene |
| chr5 | 5066666 | 5067202 | 537 | * | 5,213884 | -5,645554004 | 2,154427 | -2,62044 | 0,008782 | 1 | 0 | Intron (ENSMUST0000 | 5 | 4981690 | 5086516 | 104827 | 2 | 18647 | ENSMUST00000156660.2 | 19314 | ENSMUSG00000028926 | Cdk14 | cyclin-dependent kinase 14 |
| chr9 | 54647554 | 54648327 | 774 | * | 6,498259 | -5,645524482 | 1,989695 | -2,83738 | 0,004549 | 1 | 0 | Distal Intergenic | 9 | 54624448 | 54641803 | 17356 | 2 | 66317 | ENSMUST00000121204.8 | -5751 | ENSMUSG00000061559 | Wdr61 | WD repeat domain 61 |
| chr10 | 98734570 | 98734901 | 332 | * | 4,912188 | -5,645480701 | 2,166275 | -2,60608 | 0,009159 | 1 | 0 | Distal Intergenic | 10 | 98750268 | 98822984 | 72717 | 1 | 67972 | ENSMUST00000219557.2 | -15367 | ENSMUSG00000019943 | Atp2b1 | ATPase, Ca++ transporting, plasma membrane 1 |
| chr2 | 72289118 | 72289893 | 776 | * | 5,753378 | -5,643753277 | 2,032466 | -2,7768 | 0,00549 | 1 | 0 | Distal Intergenic | 2 | 72306503 | 72317237 | 10735 | 1 | 66953 | ENSMUST00000102691.11 | -16610 | ENSMUSG00000055612 | Cdca7 | cell division cycle associated 7 |
| chr1 | 67307194 | 67308108 | 915 | * | 5,77752 | -5,642627438 | 1,894786 | -2,97798 | 0,002902 | 1 | 0 | Intron (ENSMUST0000 | 1 | 67296383 | 67296511 | 129 | 1 | 1,15E+08 | ENSMUST00000104557.3 | 10811 | ENSMUSG00000077745 | Gm26342 | predicted gene, 26342 |
| chr7 | 58931696 | 58932867 | 1172 | * | 5,959322 | -5,639254706 | 1,960073 | -2,87706 | 0,004014 | 1 | 0 | Exon (ENSMUST00000 | 7 | 58935926 | 58954680 | 18755 | 1 | 22215 | ENSMUST00000202247.4 | -3059 | ENSMUSG00000025326 | Ube3a | ubiquitin protein ligase E3A |
| chr14 | 1,17E+08 | 1,17E+08 | 487 | * | 4,508493 | -5,638283383 | 2,078053 | -2,71325 | 0,006663 | 1 | 0 | Distal Intergenic | 14 | 1,17E+08 | 1,18E+08 | 701884 | 1 | 23888 | ENSMUST00000123239.2 | -84404 | ENSMUSG00000058571 | Gpc6 | glypican 6 |
| chrX | 1,65E+08 | 1,65E+08 | 784 | * | 7,389011 | -5,637675019 | 1,941664 | -2,90353 | 0,00369 | 1 | 0 | Distal Intergenic | 20 | 1,65E+08 | 1,65E+08 | 150078 | 1 | 14758 | ENSMUST00000112235.8 | -19894 | ENSMUSG00000031342 | Gpm6b | glycoprotein m6b |
| chr9 | 6370019 | 6370557 | 539 | * | 5,036749 | -5,635977708 | 2,04614 | -2,75444 | 0,005879 | 1 | 0 | Intron (ENSMUST0000 | 9 | 6262733 | 6269846 | 7114 | 2 | 71829 | ENSMUST00000051706.6 | -100173 | ENSMUSG00000047619 | Ddi1 | DNA-damage inducible 1 |
| chr12 | 27329694 | 27330335 | 642 | * | 5,565796 | -5,634996255 | 2,126143 | -2,65034 | 0,008041 | 1 | 0 | Distal Intergenic | 12 | 27384263 | 27392573 | 8311 | 2 | 20666 | ENSMUST00000079063.7 | 62238 | ENSMUSG00000063632 | Sox11 | SRY (sex determining region Y)-box 11 |
| chr16 | 12087102 | 12088226 | 1125 | * | 8,167092 | -5,634030091 | 1,779693 | -3,16573 | 0,001547 | 1 | 0 | 3' UTR | 16 | 11802584 | 11813260 | 10677 | 1 | 72555 | ENSMUST00000229804.2 | 284518 | ENSMUSG00000022494 | Shisa9 | shisa family member 9 |
| chr9 | 22398964 | 22399371 | 408 | * | 4,648671 | -5,633216929 | 2,152801 | -2,61669 | 0,008879 | 1 | 0 | Promoter (2-3kb) | 9 | 22402125 | 22799576 | 397452 | 1 | 319845 | ENSMUST00000147405.8 | -2754 | ENSMUSG00000035919 | Bbs9 | Bardet-Biedl syndrome 9 (human) |
| chr8 | 41710058 | 41710448 | 391 | * | 4,946108 | -5,631752525 | 2,161519 | -2,60546 | 0,009175 | 1 | 0 | Intron (ENSMUST0000 | 8 | 41692815 | 41785129 | 92315 | 1 | 18536 | ENSMUST00000211247.2 | 17243 | ENSMUSG00000031592 | Pcm1 | pericentriolar material 1 |
| chr19 | 29246388 | 29247074 | 687 | * | 5,176092 | -5,631213293 | 2,088017 | -2,69692 | 0,006998 | 1 | 0 | Intron (ENSMUST0000 | 19 | 29229841 | 29289740 | 59900 | 1 | 16452 | ENSMUST00000238009.2 | 16547 | ENSMUSG00000024789 | Jak2 | Janus kinase 2 |
| chr12 | 97838873 | 97839480 | 608 | * | 4,786621 | -5,63116614 | 2,011038 | -2,80013 | 0,005108 | 1 | 0 | Promoter (<=1kb) | 12 | 97836038 | 97838779 | 2742 | 2 | 1,03E+08 | ENSMUST00000218234.2 | -94 | | Gm35326 | predicted gene, 35326 |
| chr5 | 48907813 | 48908297 | 485 | * | 4,002492 | -5,630135229 | 2,144446 | -2,62545 | 0,008653 | 1 | 0 | Intron (ENSMUST0000 | 5 | 48553376 | 48911906 | 358531 | 2 | 80334 | ENSMUST00000101214.4 | 3609 | ENSMUSG00000029088 | Kcnip4 | Kv channel interacting protein 4 |
| chr1 | 50038466 | 50039609 | 1144 | * | 7,705373 | -5,62963499 | 1,941144 | -2,90016 | 0,00373 | 1 | 0 | Distal Intergenic | 1 | 49283775 | 49379602 | 95828 | 1 | 442837 | ENSMUST00000188673.7 | 754691 | ENSMUSG00000051616 | C230029F24Rik | RIKEN cDNA C230029F24 gene |
| chr4 | 63965927 | 63966470 | 544 | * | 4,460351 | -5,628498469 | 2,127086 | -2,64611 | 0,008142 | 1 | 0 | Promoter (<=1kb) | 4 | 63878022 | 63965252 | 87231 | 2 | 21923 | ENSMUST00000107377.10 | -675 | ENSMUSG00000028364 | Tnc | tenascin C |
| chrX | 1,58E+08 | 1,58E+08 | 773 | * | 4,411422 | -5,628428328 | 2,099893 | -2,68034 | 0,007355 | 1 | 0 | Intron (ENSMUST0000 | 20 | 1,58E+08 | 1,58E+08 | 50597 | 1 | 110651 | ENSMUST00000156648.8 | 3937 | ENSMUSG00000031309 | Rps6ka3 | ribosomal protein S6 kinase polypeptide 3 |
| chr19 | 17508161 | 17509105 | 945 | * | 4,811054 | -5,628169355 | 2,016657 | -2,79084 | 0,005257 | 1 | 0 | Intron (ENSMUST0000 | 19 | 17374801 | 17377325 | 2525 | 1 | 54391 | ENSMUST00000146171.2 | 133360 | ENSMUSG00000024712 | Rfk | riboflavin kinase |
| chr1 | 1,5E+08 | 1,5E+08 | 789 | * | 8,53266 | -5,627251768 | 1,772106 | -3,17546 | 0,001496 | 0,547497234 | 1,449706 | Distal Intergenic | 1 | 1,5E+08 | 1,5E+08 | 465 | 2 | 1E+08 | ENSMUST00000121422.2 | 4716 | ENSMUSG00000031167 | Rbm3-ps | RNA binding motif (RNP1, RRM) protein 3 |
| chr11 | 1,06E+08 | 1,06E+08 | 1258 | * | 5,854319 | -5,626922523 | 2,143607 | -2,62498 | 0,008665 | 1 | 0 | Intron (ENSMUST0000 | 11 | 1,06E+08 | 1,06E+08 | 27637 | 1 | 77097 | ENSMUST00000146162.8 | -30063 | ENSMUSG00000053580 | Tanc2 | tetratricopeptide repeat, ankyrin repeat and coiled-coil containing 2 |
| chr4 | 1,37E+08 | 1,37E+08 | 775 | * | 6,073186 | -5,626683723 | 2,122642 | -2,65079 | 0,00803 | 1 | 0 | Intron (ENSMUST0000 | 4 | 1,36E+08 | 1,37E+08 | 181198 | 2 | 13844 | ENSMUST00000105846.9 | 61716 | ENSMUSG00000028664 | Ephb2 | Eph receptor B2 |
| chr1 | 21195629 | 21196236 | 608 | * | 5,657807 | -5,626284004 | 1,905543 | -2,95259 | 0,003151 | 1 | 0 | Distal Intergenic | 1 | 21066523 | 21149453 | 82931 | 2 | 170829 | ENSMUST00000037998.6 | -46176 | ENSMUSG00000041779 | Tram2 | translocating chain-associating membrane protein 2 |
| chr1 | 93817055 | 93817817 | 763 | * | 5,0021 | -5,625926247 | 1,984008 | -2,83564 | 0,004573 | 1 | 0 | Distal Intergenic | 1 | 93789548 | 93801546 | 11999 | 1 | 381334 | ENSMUST00000187896.2 | 27507 | ENSMUSG00000094651 | Gal3st2 | galactose-3-O-sulfotransferase 2 |
| chrX | 1,09E+08 | 1,09E+08 | 1169 | * | 5,194842 | -5,62580242 | 2,018671 | -2,78688 | 0,005322 | 1 | 0 | Distal Intergenic | 20 | 1,08E+08 | 1,08E+08 | 3682 | 1 | 622976 | ENSMUST00000060013.4 | 385872 | ENSMUSG00000048621 | Gm6377 | predicted gene 6377 |
| chr15 | 40960941 | 40961552 | 612 | * | 4,596343 | -5,625510377 | 2,113844 | -2,66127 | 0,007785 | 1 | 0 | Intron (ENSMUST0000 | 15 | 41036883 | 41037264 | 382 | 1 | 1E+08 | ENSMUST00000060066.6 | -75331 | | 4930555K19Rik | leucine rich repeat and sterile alpha motif containing 1 pseudogene |
| chr7 | 1,21E+08 | 1,21E+08 | 328 | * | 3,669695 | -5,625354657 | 2,182364 | -2,57764 | 0,009948 | 1 | 0 | Intron (ENSMUST0000 | 7 | 1,21E+08 | 1,21E+08 | 492 | 2 | 668548 | ENSMUST00000077747.6 | 4810 | | Gm9234 | predicted pseudogene 9234 |
| chr9 | 1,21E+08 | 1,21E+08 | 496 | * | 4,613519 | -5,623215483 | 2,060924 | -2,72849 | 0,006362 | 1 | 0 | Distal Intergenic | 9 | 1,21E+08 | 1,21E+08 | 20487 | 2 | 1,19E+08 | ENSMUST00000215630.2 | -7668 | ENSMUSG00000111598 | Gm47095 | predicted gene, 47095 |
| chr2 | 1,01E+08 | 1,01E+08 | 770 | * | 4,698262 | -5,622902892 | 2,024971 | -2,77678 | 0,00549 | 1 | 0 | Intron (ENSMUST0000 | 2 | 1,01E+08 | 1,01E+08 | 48757 | 2 | 68170 | ENSMUST00000136601.9 | 10142 | ENSMUSG00000027165 | Iftap | intraflagellar transport associated protein |
| chr2 | 7394487 | 7395215 | 729 | * | 4,7017 | -5,622812525 | 2,084054 | -2,69802 | 0,006975 | 1 | 0 | Intron (ENSMUST0000 | 2 | 6558700 | 7400704 | 842005 | 2 | 14007 | ENSMUST00000182404.8 | 5489 | ENSMUSG00000002107 | Celf2 | CUGBP, Elav-like family member 2 |
| chr13 | 1,18E+08 | 1,18E+08 | 599 | * | 5,184596 | -5,620743662 | 2,163331 | -2,59819 | 0,009372 | 1 | 0 | Intron (ENSMUST0000 | 13 | 1,18E+08 | 1,18E+08 | 130 | 1 | 1,15E+08 | ENSMUST00000157530.3 | -151221 | ENSMUSG00000088155 | Gm26349 | predicted gene, 26349 |
| chr2 | 1,11E+08 | 1,11E+08 | 567 | * | 4,931694 | -5,620535777 | 2,050292 | -2,74133 | 0,006119 | 1 | 0 | Promoter (1-2kb) | 2 | 1,11E+08 | 1,11E+08 | 1043 | 2 | 258398 | ENSMUST00000207786.3 | 1908 | ENSMUSG00000108919 | Olfr1295 | olfactory receptor 1295 |
| chr12 | 1,14E+08 | 1,14E+08 | 492 | * | 5,655514 | -5,620124988 | 1,906683 | -2,94759 | 0,003203 | 1 | 0 | Distal Intergenic | 12 | 1,14E+08 | 1,14E+08 | 50281 | 1 | 1,19E+08 | ENSMUST00000247233.1 | -375490 | | Gm53771 | predicted gene, 53771 |
| chr10 | 24935033 | 24935822 | 790 | * | 7,856332 | -5,619724137 | 1,819033 | -3,0894 | 0,002006 | 0,621084104 | 1,360622 | Distal Intergenic | 10 | 24803468 | 24804659 | 1192 | 1 | 1,05E+08 | ENSMUST00000244341.1 | 131565 | | Gm40616 | predicted gene, 40616 |
| chr1 | 1,47E+08 | 1,47E+08 | 900 | * | 4,927644 | -5,61825532 | 1,984524 | -2,83103 | 0,00464 | 1 | 0 | Exon (ENSMUST00000 | 1 | 1,48E+08 | 1,48E+08 | 129 | 2 | 1,15E+08 | ENSMUST00000157957.3 | 806106 | ENSMUSG00000088582 | Gm22966 | predicted gene, 22966 |
| chr7 | 1,25E+08 | 1,25E+08 | 525 | * | 4,855974 | -5,617947241 | 2,053031 | -2,73642 | 0,006211 | 1 | 0 | Intron (ENSMUST0000 | 7 | 1,25E+08 | 1,25E+08 | 53779 | 1 | 233865 | ENSMUST00000148701.4 | 29343 | ENSMUSG00000032743 | Katnip | katanin interacting protein |
| chr8 | 41730226 | 41731098 | 873 | * | 4,249277 | -5,617678568 | 2,126417 | -2,64185 | 0,008245 | 1 | 0 | Intron (ENSMUST0000 | 8 | 41692815 | 41785129 | 92315 | 1 | 18536 | ENSMUST00000211247.2 | 37411 | ENSMUSG00000031592 | Pcm1 | pericentriolar material 1 |
| chr10 | 24362904 | 24363863 | 960 | * | 5,901329 | -5,616148998 | 1,984763 | -2,82963 | 0,00466 | 1 | 0 | Distal Intergenic | 10 | 24455946 | 24469656 | 13711 | 2 | 1,03E+08 | ENSMUST00000141076.2 | 105793 | | Gm15270 | predicted gene 15270 |
| chr2 | 1,3E+08 | 1,3E+08 | 644 | * | 4,545175 | -5,614735599 | 2,119383 | -2,64923 | 0,008068 | 1 | 0 | Distal Intergenic | 2 | 1,3E+08 | 1,3E+08 | 1859 | 2 | 18610 | ENSMUST00000152113.2 | -29318 | ENSMUSG00000027400 | Pdyn | prodynorphin |
| chr14 | 1,06E+08 | 1,06E+08 | 826 | * | 5,286061 | -5,614587032 | 1,964679 | -2,85776 | 0,004266 | 1 | 0 | Exon (ENSMUST00000 | 14 | 1,06E+08 | 1,06E+08 | 4873 | 2 | 24064 | ENSMUST00000022709.6 | 43216 | ENSMUSG00000022114 | Spry2 | sprouty RTK signaling antagonist 2 |
| chr4 | 16503076 | 16503390 | 315 | * | 4,651739 | -5,614377926 | 2,164187 | -2,59422 | 0,009481 | 1 | 0 | Intron (ENSMUST0000 | 4 | 16675923 | 16676029 | 107 | 1 | 1,15E+08 | ENSMUST00000179782.2 | -172533 | ENSMUSG00000096423 | Gm24978 | predicted gene, 24978 |
| chr10 | 91662577 | 91663742 | 1166 | * | 7,483351 | -5,612953525 | 1,842215 | -3,04685 | 0,002313 | 1 | 0 | Distal Intergenic | 10 | 91724691 | 91789442 | 64752 | 1 | 1,03E+08 | ENSMUST00000217529.2 | -60949 | ENSMUSG00000111657 | Gm31592 | predicted gene, 31592 |
| chrX | 99091313 | 99092222 | 910 | * | 4,852176 | -5,612857602 | 2,139272 | -2,62372 | 0,008697 | 1 | 0 | Intron (ENSMUST0000 | 20 | 99019387 | 99440762 | 421376 | 1 | 13607 | ENSMUST00000071453.3 | 71926 | ENSMUSG00000059327 | Eda | ectodysplasin-A |
| chr9 | 60137666 | 60137907 | 242 | * | 4,201624 | -5,61285115 | 2,135414 | -2,62846 | 0,008577 | 1 | 0 | Intron (ENSMUST0000 | 9 | 59877002 | 60048503 | 171502 | 2 | 207596 | ENSMUST00000034829.6 | -89163 | ENSMUSG00000032289 | Thsd4 | thrombospondin, type I, domain containing 4 |
| chr1 | 59092138 | 59092598 | 461 | * | 4,441459 | -5,612850553 | 2,135275 | -2,62863 | 0,008573 | 1 | 0 | Intron (ENSMUST0000 | 1 | 59075214 | 59077065 | 1852 | 1 | 1,05E+08 | ENSMUST00000191356.2 | 16924 | | Gm29018 | predicted gene 29018 |
| chr3 | 1,34E+08 | 1,34E+08 | 599 | * | 4,764008 | -5,612742588 | 2,036593 | -2,75595 | 0,005852 | 1 | 0 | Intron (ENSMUST0000 | 3 | 1,34E+08 | 1,34E+08 | 19493 | 2 | 75208 | ENSMUST00000198231.2 | -158673 | ENSMUSG00000105765 | 4930539C22Rik | RIKEN cDNA 4930539C22 gene |
| chr8 | 1,15E+08 | 1,15E+08 | 281 | * | 5,330561 | -5,612741203 | 2,033903 | -2,75959 | 0,005787 | 1 | 0 | Intron (ENSMUST0000 | 8 | 1,15E+08 | 1,15E+08 | 5955 | 1 | 270097 | ENSMUST00000124143.2 | -61306 | ENSMUSG00000046844 | Vat1l | vesicle amine transport protein 1 like |
| chrX | 50411148 | 50411765 | 618 | * | 5,000252 | -5,612662358 | 1,982003 | -2,83181 | 0,004628 | 1 | 0 | Distal Intergenic | 20 | 50253294 | 50295409 | 42116 | 2 | 171170 | ENSMUST00000156028.2 | -115739 | ENSMUSG00000036109 | Mbnl3 | muscleblind like splicing factor 3 |
| chr1 | 1,71E+08 | 1,71E+08 | 643 | * | 5,241137 | -5,612594905 | 1,940721 | -2,89202 | 0,003828 | 1 | 0 | Promoter (<=1kb) | 1 | 1,71E+08 | 1,71E+08 | 13187 | 1 | 21945 | ENSMUST00000111300.8 | -324 | ENSMUSG00000013973 | Dedd | death effector domain-containing |
| chr7 | 86113123 | 86114232 | 1110 | * | 7,342729 | -5,612544228 | 1,915031 | -2,93078 | 0,003381 | 1 | 0 | Promoter (<=1kb) | 7 | 86114621 | 86115613 | 993 | 1 | 257929 | ENSMUST00000057734.4 | -389 | ENSMUSG00000020168 | Olfr299 | olfactory receptor 299 |
| chr9 | 1,04E+08 | 1,04E+08 | 901 | * | 6,673667 | -5,611539643 | 1,864897 | -3,00904 | 0,002621 | 1 | 0 | Distal Intergenic | 9 | 1,04E+08 | 1,04E+08 | 624 | 2 | 1,03E+08 | ENSMUST00000216588.3 | 10658 | ENSMUSG00000111035 | Gm33054 | predicted gene, 33054 |
| chr3 | 78338922 | 78339489 | 568 | * | 6,060544 | -5,61117184 | 1,903625 | -2,94762 | 0,003202 | 1 | 0 | Distal Intergenic | 3 | 78887229 | 78887358 | 130 | 1 | 1,15E+08 | ENSMUST00000240248.1 | -547740 | ENSMUSG00000119445 | Gm23440 | predicted gene, 23440 |
| chr18 | 65458416 | 65458922 | 507 | * | 5,161797 | -5,610401153 | 2,010156 | -2,79103 | 0,005254 | 1 | 0 | Intron (ENSMUST0000 | 18 | 65398600 | 65437527 | 38928 | 2 | 225638 | ENSMUST00000235298.2 | -20889 | ENSMUSG00000032845 | Alpk2 | alpha-kinase 2 |
| chr9 | 34336100 | 34336900 | 801 | * | 5,273838 | -5,61019507 | 2,057043 | -2,72731 | 0,006385 | 1 | 0 | Distal Intergenic | 9 | 34397190 | 34400454 | 3265 | 1 | 67703 | ENSMUST00000184203.8 | -60290 | ENSMUSG00000032036 | Kirrel3 | kirre like nephrin family adhesion molecule 3 |
| chr10 | 15592264 | 15593264 | 1001 | * | 6,281287 | -5,609991619 | 2,090068 | -2,68412 | 0,007272 | 1 | 0 | Intron (ENSMUST0000 | 10 | 15610660 | 15614570 | 3911 | 2 | 1,03E+08 | ENSMUST00000218861.2 | 21306 | | Gm32283 | predicted gene, 32283 |
| chr5 | 1,38E+08 | 1,38E+08 | 1082 | * | 6,290302 | -5,609056145 | 1,946132 | -2,88216 | 0,00395 | 1 | 0 | Promoter (1-2kb) | 5 | 1,38E+08 | 1,38E+08 | 19373 | 1 | 74570 | ENSMUST00000110963.8 | 1602 | ENSMUSG00000029729 | Zkscan1 | zinc finger with KRAB and SCAN domains 1 |
| chr5 | 1,15E+08 | 1,15E+08 | 517 | * | 5,306266 | -5,607835223 | 2,100391 | -2,6699 | 0,007587 | 1 | 0 | Promoter (1-2kb) | 5 | 1,15E+08 | 1,15E+08 | 8153 | 1 | 17855 | ENSMUST00000139420.3 | 1277 | ENSMUSG00000041939 | Mvk | mevalonate kinase |
| chr4 | 1,03E+08 | 1,03E+08 | 434 | * | 4,634293 | -5,607522968 | 2,050269 | -2,73502 | 0,006238 | 1 | 0 | Promoter (1-2kb) | 4 | 1,03E+08 | 1,03E+08 | 1723 | 1 | 73094 | ENSMUST00000141624.2 | -1743 | ENSMUSG00000028524 | Sgip1 | SH3-domain GRB2-like (endophilin) interacting protein 1 |
| chr7 | 1,09E+08 | 1,09E+08 | 474 | * | 4,449497 | -5,606665786 | 2,144805 | -2,61407 | 0,008947 | 1 | 0 | Intron (ENSMUST0000 | 7 | 1,09E+08 | 1,09E+08 | 41834 | 2 | 56788 | ENSMUST00000132113.8 | -9728 | ENSMUSG00000007279 | Scube2 | signal peptide, CUB domain, EGF-like 2 |
| chr1 | 1,57E+08 | 1,57E+08 | 740 | * | 5,370379 | -5,606613403 | 1,938984 | -2,89152 | 0,003834 | 1 | 0 | Intron (ENSMUST0000 | 1 | 1,57E+08 | 1,57E+08 | 38184 | 2 | 78255 | ENSMUST00000189648.2 | 36051 | ENSMUSG00000026594 | Ralgps2 | Ral GEF with PH domain and SH3 binding motif 2 |
| chr1 | 30691241 | 30691990 | 750 | * | 6,100305 | -5,606547015 | 1,932143 | -2,90172 | 0,003711 | 1 | 0 | Distal Intergenic | 1 | 30845286 | 30848186 | 2901 | 2 | 213109 | ENSMUST00000190190.2 | 156196 | ENSMUSG00000048874 | Phf3 | PHD finger protein 3 |
| chr15 | 37868802 | 37869739 | 938 | * | 6,173517 | -5,606040939 | 1,887551 | -2,97001 | 0,002978 | 1 | 0 | Distal Intergenic | 15 | 37893239 | 37899960 | 6722 | 2 | 1,03E+08 | ENSMUST00000226812.2 | 30221 | ENSMUSG00000115326 | Gm35019 | predicted gene, 35019 |
| chr2 | 15954904 | 15955465 | 562 | * | 4,490205 | -5,604958722 | 2,133878 | -2,62665 | 0,008623 | 1 | 0 | Intron (ENSMUST0000 | 2 | 16361115 | 16760650 | 399536 | 1 | 67448 | ENSMUST00000028081.13 | -405650 | ENSMUSG00000026748 | Plxdc2 | plexin domain containing 2 |
| chr1 | 1,11E+08 | 1,11E+08 | 649 | * | 4,470533 | -5,604939701 | 2,059717 | -2,72122 | 0,006504 | 1 | 0 | Distal Intergenic | 1 | 1,11E+08 | 1,11E+08 | 27232 | 2 | 227485 | ENSMUST00000187555.2 | -389148 | ENSMUSG00000047216 | Cdh19 | cadherin 19, type 2 |
| chr16 | 35206864 | 35207348 | 485 | * | 4,828703 | -5,602635018 | 2,010527 | -2,78665 | 0,005326 | 1 | 0 | Distal Intergenic | 16 | 35202415 | 35202617 | 203 | 1 | 1,15E+08 | ENSMUST00000082529.3 | 4449 | ENSMUSG00000064463 | Gm25079 | predicted gene, 25079 |

| | | | | | | | | | | | | | | | | | | | | | | | |
|---|---|---|---|---|---|---|---|---|---|---|---|---|---|---|---|---|---|---|---|---|---|---|---|
| chr19 | 58410341 | 58410904 | 564 | * | 4,830454 | -5,601806085 | 2,157995 | -2,59584 | 0,009436 | 1 | 0 | Intron (ENSMUST00000 | 19 | 58225124 | 58442898 | 217775 | 2 | 14585 | ENSMUST00000169850.8 | 31994 | ENSMUSG00000025089 | Gfra1 | glial cell line derived neurotrophic factor family receptor alpha 1 |
| chrX | 1,22E+08 | 1,22E+08 | 1036 | * | 5,934171 | -5,600111928 | 2,026679 | -2,7632 | 0,005724 | 1 | 0 | Distal Intergenic | 20 | 1,22E+08 | 1,22E+08 | 42157 | 1 | 73061 | ENSMUST00000113343.8 | -144655 | ENSMUSG00000079450 | Cldn34c1 | claudin 34C1 |
| chr2 | 8780097 | 8780697 | 601 | * | 4,827019 | -5,598967936 | 2,124612 | -2,63529 | 0,008407 | 1 | 0 | Distal Intergenic | 2 | 9194326 | 9202288 | 7963 | 1 | 73438 | ENSMUST00000132414.2 | -413629 | ENSMUSG00000085580 | 1700061F12Rik | RIKEN cDNA 1700061F12 gene |
| chr1 | 1,57E+08 | 1,57E+08 | 823 | * | 8,650056 | -5,597646309 | 1,826545 | -3,06461 | 0,00218 | 0,539620856 | 2,237403 | Intron (ENSMUST00000 | 1 | 1,57E+08 | 1,57E+08 | 80511 | 2 | 226525 | ENSMUST00000134543.8 | -34157 | ENSMUSG00000070565 | Rasal2 | RAS protein activator like 2 |
| chrX | 26049521 | 26050738 | 1218 | * | 5,507556 | -5,597240649 | 1,896311 | -2,95165 | 0,003161 | 1 | 0 | Distal Intergenic | 20 | 25983514 | 26006341 | 22828 | 2 | 382275 | ENSMUST00000089165.9 | -43180 | ENSMUSG00000079655 | Gm5168 | predicted gene 5168 |
| chr4 | 1,21E+08 | 1,21E+08 | 762 | * | 5,871491 | -5,597227245 | 1,908433 | -2,93289 | 0,003358 | 1 | 0 | Exon (ENSMUST00000 | 4 | 1,21E+08 | 1,21E+08 | 132 | 1 | 1,15E+08 | ENSMUST00000157793.3 | -29341 | ENSMUSG00000088418 | Gm26162 | predicted gene, 26162 |
| chr8 | 63095072 | 63096003 | 932 | * | 5,801881 | -5,597205108 | 1,925369 | -2,90708 | 0,003648 | 1 | 0 | Distal Intergenic | 8 | 63404043 | 63809886 | 405844 | 1 | 72902 | ENSMUST00000117377.8 | -308040 | ENSMUSG00000054162 | Spock3 | sparc/osteonectin, cwcv and kazal-like domains proteoglycan 3 |
| chr5 | 1,47E+08 | 1,47E+08 | 700 | * | 6,331323 | -5,597109672 | 2,029542 | -2,75782 | 0,005819 | 1 | 0 | Distal Intergenic | 5 | 1,47E+08 | 1,47E+08 | 875 | 1 | 75949 | ENSMUST00000197707.2 | -18166 | | 4930573C15Rik | RIKEN cDNA 4930573C15 gene |
| chrX | 69431866 | 69432447 | 582 | * | 3,725722 | -5,597011478 | 2,166119 | -2,58389 | 0,009769 | 1 | 0 | Promoter (2-3kb) | 20 | 69429483 | 69433023 | 3541 | 1 | 68618 | ENSMUST00000053981.6 | 2383 | ENSMUSG00000045237 | Eola1 | endothelium and lymphocyte associated ASCH domain 1 |
| chr1 | 69290881 | 69291398 | 518 | * | 5,259633 | -5,59699812 | 2,168688 | -2,58082 | 0,009857 | 1 | 0 | Distal Intergenic | 1 | 68071345 | 69147218 | 1075874 | 2 | 13869 | ENSMUST00000121473.8 | -143663 | ENSMUSG00000062209 | Erbb4 | erb-b2 receptor tyrosine kinase 4 |
| chr18 | 69963610 | 69964305 | 696 | * | 6,853329 | -5,596059587 | 1,892191 | -2,95745 | 0,003102 | 1 | 0 | Distal Intergenic | 18 | 69891395 | 69912832 | 21438 | 2 | 1,03E+08 | ENSMUST00000244028.1 | -50778 | | Gm31502 | predicted gene, 31502 |
| chr7 | 66428136 | 66428828 | 693 | * | 4,505014 | -5,595184061 | 2,135024 | -2,62067 | 0,008776 | 1 | 0 | Intron (ENSMUST00000 | 7 | 66413917 | 66471590 | 57674 | 1 | 545975 | ENSMUST00000208521.2 | 14219 | ENSMUSG00000030510 | Cers3 | ceramide synthase 3 |
| chr14 | 96096710 | 96097750 | 1041 | * | 4,33924 | -5,594187662 | 2,099776 | -2,66418 | 0,007718 | 1 | 0 | Distal Intergenic | 14 | 96118690 | 96120216 | 1527 | 1 | 66732 | ENSMUST00000228913.2 | -20940 | ENSMUSG00000034689 | Spertl | spermatid associated like |
| chr14 | 1,04E+08 | 1,04E+08 | 969 | * | 5,350125 | -5,593994586 | 2,021228 | -2,76762 | 0,005647 | 1 | 0 | Promoter (2-3kb) | 14 | 1,04E+08 | 1,04E+08 | 49515 | 2 | 73877 | ENSMUST00000228770.2 | -2065 | ENSMUSG00000115727 | 4930432J09Rik | RIKEN cDNA 4930432J09 gene |
| chrX | 1,15E+08 | 1,15E+08 | 547 | * | 4,353462 | -5,593822445 | 2,162851 | -2,58632 | 0,009701 | 1 | 0 | Distal Intergenic | 20 | 1,15E+08 | 1,15E+08 | 164 | 1 | 1,15E+08 | ENSMUST00000082617.3 | -232468 | ENSMUSG00000064551 | Gm24624 | predicted gene, 24624 |
| chr16 | 88505302 | 88505917 | 616 | * | 4,061204 | -5,593806023 | 2,166449 | -2,58202 | 0,009823 | 1 | 0 | Promoter (<=1kb) | 16 | 88504057 | 88504850 | 794 | 2 | 69661 | ENSMUST00000050882.4 | -452 | ENSMUSG00000050704 | 2310061N02Rik | RIKEN cDNA 2310061N02 gene |
| chr19 | 11253208 | 11254012 | 805 | * | 7,325566 | -5,592667666 | 1,93111 | -2,89609 | 0,003778 | 1 | 0 | Promoter (2-3kb) | 19 | 11250602 | 11256807 | 6206 | 2 | 269063 | ENSMUST00000191110.7 | 2795 | ENSMUSG00000054523 | Ms4a5 | membrane-spanning 4-domains, subfamily A, member 5 |
| chr7 | 92009936 | 92010599 | 664 | * | 3,731853 | -5,592195821 | 2,156612 | -2,59305 | 0,009513 | 1 | 0 | Intron (ENSMUST00000 | 7 | 92076877 | 92087272 | 10396 | 1 | 23859 | ENSMUST00000207891.2 | -66278 | ENSMUSG00000052572 | Dlg2 | discs large MAGUK scaffold protein 2 |
| chr6 | 42036817 | 42037409 | 593 | * | 5,77318 | -5,591553578 | 2,081437 | -2,68639 | 0,007223 | 1 | 0 | Distal Intergenic | 6 | 42015328 | 42019785 | 4458 | 1 | 20939 | ENSMUST00000024059.5 | 21489 | ENSMUSG00000023289 | Sva | seminal vesicle antigen |
| chr14 | 51679759 | 51681165 | 1407 | * | 6,703647 | -5,591455303 | 1,991087 | -2,80824 | 0,004981 | 1 | 0 | Intron (ENSMUST00000 | 14 | 51689314 | 51698311 | 8998 | 1 | 22301 | ENSMUST00000239412.3 | -8149 | ENSMUSG00000070448 | Vmn2r89 | vomeronasal 2, receptor 89 |
| chr2 | 10341589 | 10342279 | 691 | * | 4,930586 | -5,591424228 | 1,986593 | -2,81458 | 0,004884 | 1 | 0 | Distal Intergenic | 2 | 10375321 | 10443948 | 68628 | 1 | 353282 | ENSMUST00000114862.8 | -33042 | ENSMUSG00000061186 | Sfmbt2 | Scm-like with four mbt domains 2 |
| chr16 | 77952672 | 77953114 | 443 | * | 5,300062 | -5,589118538 | 2,090672 | -2,67336 | 0,00751 | 1 | 0 | Distal Intergenic | 16 | 78037146 | 78037276 | 131 | 1 | 1,15E+08 | ENSMUST00000104263.3 | -84032 | ENSMUSG00000077451 | Gm25916 | predicted gene, 25916 |
| chr17 | 41866750 | 41867255 | 506 | * | 4,615729 | -5,589113636 | 2,094176 | -2,66888 | 0,00761 | 1 | 0 | Distal Intergenic | 17 | 41266079 | 41269872 | 3794 | 1 | 17850 | ENSMUST00000233222.2 | 600671 | ENSMUSG00000023921 | Mmut | methylmalonyl-Coenzyme A mutase |
| chr7 | 1,39E+08 | 1,39E+08 | 773 | * | 6,832402 | -5,588878285 | 1,862716 | -3,00039 | 0,002696 | 1 | 0 | Intron (ENSMUST00000 | 7 | 1,39E+08 | 1,39E+08 | 84880 | 2 | 57740 | ENSMUST00000016125.12 | 42644 | ENSMUSG00000015981 | Stk32c | serine/threonine kinase 32C |
| chr16 | 18946953 | 18947755 | 803 | * | 4,272532 | -5,587427537 | 2,16783 | -2,57743 | 0,009954 | 1 | 0 | Distal Intergenic | 16 | 18882081 | 18925646 | 43566 | 1 | 1,05E+08 | ENSMUST00000242960.1 | 64872 | | LOC105246083 | uncharacterized LOC105246083 |
| chr14 | 63319134 | 63320034 | 901 | * | 6,340312 | -5,585832819 | 1,999845 | -2,79313 | 0,00522 | 1 | 0 | Distal Intergenic | 14 | 63336985 | 63337091 | 107 | 1 | 1,15E+08 | ENSMUST00000082465.3 | -16951 | ENSMUSG00000064399 | Gm23629 | predicted gene, 23629 |
| chr11 | 79597115 | 79598150 | 1036 | * | 6,153642 | -5,585535739 | 1,976974 | -2,8253 | 0,004724 | 1 | 0 | Promoter (1-2kb) | 11 | 79595456 | 79596246 | 791 | 1 | 1,03E+08 | ENSMUST00000137738.2 | 1659 | | Gm11205 | predicted gene 11205 |
| chr10 | 1,14E+08 | 1,14E+08 | 941 | * | 4,945852 | -5,583890255 | 2,067508 | -2,70078 | 0,006918 | 1 | 0 | Distal Intergenic | 10 | 1,13E+08 | 1,13E+08 | 11133 | 2 | 1,05E+08 | ENSMUST00000218612.2 | -292160 | | Gm40765 | predicted gene, 40765 |
| chrX | 23791036 | 23792082 | 1047 | * | 6,338184 | -5,582904689 | 1,876341 | -2,97542 | 0,002926 | 1 | 0 | Distal Intergenic | 20 | 23759023 | 23773736 | 14714 | 1 | 236749 | ENSMUST00000163718.3 | 32013 | ENSMUSG00000068113 | Tesl1 | testin LIM domain protein like 1 |
| chr18 | 55972574 | 55973727 | 1154 | * | 8,742114 | -5,582259246 | 1,808745 | -3,08626 | 0,002027 | 0,623120685 | 1,360622 | Distal Intergenic | 18 | 56326071 | 56326205 | 135 | 1 | 1,15E+08 | ENSMUST00000157815.3 | -352344 | ENSMUSG00000088440 | Gm23555 | predicted gene, 23555 |
| chr3 | 92482233 | 92482834 | 602 | * | 5,375264 | -5,581671557 | 1,933481 | -2,88685 | 0,003891 | 1 | 0 | Promoter (1-2kb) | 3 | 92478209 | 92481042 | 2834 | 2 | 16447 | ENSMUST00000053107.6 | -1191 | ENSMUSG00000049128 | Ivl | involucrin |
| chr1 | 75245412 | 75246020 | 609 | * | 4,414159 | -5,5810914 | 2,120271 | -2,63225 | 0,008482 | 1 | 0 | Distal Intergenic | 1 | 75235203 | 75241146 | 5944 | 2 | 19275 | ENSMUST00000185849.7 | -4266 | ENSMUSG00000026204 | Ptprn | protein tyrosine phosphatase, receptor type, N |
| chrX | 67054341 | 67055090 | 750 | * | 3,935323 | -5,581070332 | 2,126672 | -2,62432 | 0,008682 | 1 | 0 | Promoter (<=1kb) | 20 | 67053711 | 67053785 | 75 | 2 | 1E+08 | ENSMUST00000196089.3 | -556 | ENSMUSG00000106233 | Mir509 | microRNA 509 |
| chr11 | 85378515 | 85379593 | 1079 | * | 6,032809 | -5,579494345 | 1,977597 | -2,82135 | 0,004782 | 1 | 0 | Intron (ENSMUST00000 | 11 | 85383394 | 85386740 | 3347 | 1 | 192197 | ENSMUST00000140215.2 | -3801 | ENSMUSG00000059439 | Bcas3 | BCAS3 microtubule associated cell migration factor |
| chr5 | 1,38E+08 | 1,38E+08 | 735 | * | 4,724512 | -5,579337239 | 2,165872 | -2,57602 | 0,009994 | 1 | 0 | Distal Intergenic | 5 | 1,38E+08 | 1,38E+08 | 14788 | 1 | 72154 | ENSMUST00000100524.3 | 24081 | ENSMUSG00000036898 | Zfp157 | zinc finger protein 157 |
| chr12 | 73967637 | 73968466 | 830 | * | 5,014789 | -5,578966104 | 2,024682 | -2,75548 | 0,005861 | 1 | 0 | Intron (ENSMUST00000 | 12 | 73973376 | 73975306 | 1931 | 1 | 15251 | ENSMUST00000149740.2 | -4910 | ENSMUSG00000021109 | Hif1a | hypoxia inducible factor 1, alpha subunit |
| chr9 | 11228729 | 11229491 | 763 | * | 5,73912 | -5,577510848 | 1,967277 | -2,83514 | 0,004581 | 1 | 0 | Distal Intergenic | 9 | 9660974 | 10904780 | 1243807 | 2 | 244682 | ENSMUST00000160216.8 | -323949 | ENSMUSG00000039488 | Cntn5 | contactin 5 |
| chr12 | 34463025 | 34463732 | 708 | * | 6,252382 | -5,577234513 | 1,946019 | -2,86597 | 0,004157 | 1 | 0 | Intron (ENSMUST00000 | 12 | 34423826 | 34487306 | 63481 | 2 | 79221 | ENSMUST00000211107.2 | 23574 | ENSMUSG00000004698 | Hdac9 | histone deacetylase 9 |
| chr6 | 58034172 | 58034475 | 304 | * | 5,869955 | -5,576921631 | 1,924408 | -2,89799 | 0,003756 | 1 | 0 | Distal Intergenic | 6 | 57985168 | 57986187 | 1020 | 2 | 171190 | ENSMUST00000049694.4 | -47985 | ENSMUSG00000048697 | Vmn1r26 | vomeronasal 1 receptor 26 |
| chr7 | 1,07E+08 | 1,07E+08 | 594 | * | 7,154898 | -5,57670281 | 1,91021 | -2,91942 | 0,003507 | 1 | 0 | Promoter (<=1kb) | 7 | 1,07E+08 | 1,07E+08 | 9578 | 2 | 320635 | ENSMUST00000052438.8 | 0 | ENSMUSG00000048065 | Cyb5r2 | cytochrome b5 reductase 2 |
| chrX | 60337168 | 60337834 | 667 | * | 4,481579 | -5,576077663 | 2,139892 | -2,60577 | 0,009167 | 1 | 0 | Distal Intergenic | 20 | 60331794 | 60336090 | 4297 | 1 | 245440 | ENSMUST00000232598.2 | 5374 | ENSMUSG00000118608 | Gm4988 | predicted gene 4988 |
| chr11 | 26759806 | 26760332 | 527 | * | 5,168842 | -5,5759711 | 2,096474 | -2,65969 | 0,007821 | 1 | 0 | Distal Intergenic | 11 | 26497848 | 26543999 | 46152 | 2 | 69922 | ENSMUST00000156264.2 | -215807 | ENSMUSG00000064090 | Vrk2 | vaccinia related kinase 2 |
| chr4 | 53133638 | 53134393 | 756 | * | 7,483488 | -5,575816229 | 1,860255 | -2,99734 | 0,002723 | 1 | 0 | Intron (ENSMUST00000 | 4 | 53030787 | 53159895 | 129109 | 2 | 11303 | ENSMUST00000030010.4 | 25502 | ENSMUSG00000015243 | Abca1 | ATP-binding cassette, sub-family A (ABC1), member 1 |
| chr5 | 78488523 | 78489351 | 829 | * | 6,242001 | -5,575247333 | 1,890195 | -2,94956 | 0,003182 | 1 | 0 | Distal Intergenic | 5 | 78927706 | 78927841 | 136 | 1 | 1,15E+08 | ENSMUST00000103990.3 | -438355 | ENSMUSG00000077178 | Gm25756 | predicted gene, 25756 |
| chr9 | 48640005 | 48640443 | 439 | * | 6,864608 | -5,575182254 | 1,876924 | -2,97038 | 0,002974 | 1 | 0 | Intron (ENSMUST00000 | 9 | 48565597 | 48747245 | 181649 | 2 | 235320 | ENSMUST00000093852.5 | 106802 | ENSMUSG00000066687 | Zbtb16 | zinc finger and BTB domain containing 16 |
| chr13 | 48136982 | 48138422 | 1441 | * | 8,908151 | -5,575126959 | 1,828244 | -3,04944 | 0,002293 | 0,644341614 | 1,360622 | Intron (ENSMUST00000 | 13 | 48113806 | 48172084 | 58279 | 2 | 70971 | ENSMUST00000061899.6 | 33662 | ENSMUSG00000047324 | 4931429P17Rik | RIKEN cDNA 4931429P17 gene |
| chr16 | 27532640 | 27533417 | 778 | * | 7,912403 | -5,574918808 | 1,833826 | -3,04005 | 0,002365 | 1 | 0 | Distal Intergenic | 16 | 27668682 | 27744880 | 76199 | 1 | 1E+08 | ENSMUST00000231532.2 | -135265 | ENSMUSG00000116654 | Gm10823 | predicted gene 10823 |
| chr15 | 32579248 | 32580059 | 812 | * | 6,44177 | -5,574412108 | 1,854023 | -3,00666 | 0,002641 | 1 | 0 | Intron (ENSMUST00000 | 15 | 32601821 | 32689782 | 87962 | 1 | 20356 | ENSMUST00000227429.2 | -21762 | ENSMUSG00000022231 | Sema5a | sema domain, seven thrombospondin repeats (type 1 and type 1-like), transmembrane domain (TM) and short cytoplasmic domain, (semaphorin) 5A |
| chr8 | 1,28E+08 | 1,28E+08 | 690 | * | 5,282919 | -5,573849546 | 1,93565 | -2,87958 | 0,003982 | 1 | 0 | Intron (ENSMUST00000 | 8 | 1,28E+08 | 1,28E+08 | 26768 | 1 | 93742 | ENSMUST00000159511.2 | 28117 | ENSMUSG00000025812 | Pard3 | par-3 family cell polarity regulator |
| chr10 | 63295019 | 63295677 | 659 | * | 5,07519 | -5,573569028 | 1,988525 | -2,80287 | 0,005065 | 1 | 0 | Promoter (1-2kb) | 10 | 63293289 | 64839446 | 1546158 | 1 | 216033 | ENSMUST00000105440.8 | 1730 | ENSMUSG00000060843 | Ctnna3 | catenin (cadherin associated protein), alpha 3 |
| chr19 | 29543152 | 29543657 | 506 | * | 5,123832 | -5,573281972 | 2,055241 | -2,71174 | 0,006693 | 1 | 0 | Intron (ENSMUST00000 | 19 | 29572928 | 29582585 | 9658 | 1 | 226089 | ENSMUST00000160452.2 | -29271 | ENSMUSG00000038658 | Ric1 | RAB6A GEF complex partner 1 |
| chr14 | 1,04E+08 | 1,04E+08 | 642 | * | 6,20302 | -5,571418613 | 1,993607 | -2,79464 | 0,005196 | 1 | 0 | Distal Intergenic | 14 | 1,04E+08 | 1,04E+08 | 6460 | 2 | 70898 | ENSMUST00000160117.2 | -29157 | ENSMUSG00000089707 | Slain1os | SLAIN motif family, member 1, opposite strand |
| chr12 | 1,12E+08 | 1,12E+08 | 772 | * | 4,603435 | -5,570264259 | 2,100056 | -2,65244 | 0,007991 | 1 | 0 | Intron (ENSMUST00000 | 12 | 1,12E+08 | 1,12E+08 | 22724 | 1 | 17169 | ENSMUST00000222870.2 | 5291 | ENSMUSG00000007411 | Mark3 | MAP/microtubule affinity regulating kinase 3 |
| chr5 | 1,49E+08 | 1,49E+08 | 1051 | * | 7,315258 | -5,568674981 | 1,897781 | -2,93431 | 0,003343 | 1 | 0 | Distal Intergenic | 5 | 1,49E+08 | 1,49E+08 | 5118 | 1 | 78015 | ENSMUST00000155504.2 | 15328 | ENSMUSG00000085740 | 4930505K14Rik | RIKEN cDNA 4930505K14 gene |
| chr3 | 1,35E+08 | 1,35E+08 | 606 | * | 4,720743 | -5,567870382 | 2,062806 | -2,69917 | 0,006951 | 1 | 0 | Promoter (1-2kb) | 3 | 1,35E+08 | 1,35E+08 | 3821 | 1 | 74446 | ENSMUST00000159276.3 | -1511 | ENSMUSG00000050150 | Slc9b1 | solute carrier family 9, subfamily B (NHA1, cation proton antiporter 1), member 1 |
| chr9 | 96365474 | 96366183 | 710 | * | 5,459042 | -5,567606358 | 1,941163 | -2,86818 | 0,004128 | 1 | 0 | Distal Intergenic | 9 | 96352990 | 96360728 | 7739 | 2 | 19823 | ENSMUST00000057500.6 | -4746 | ENSMUSG00000051234 | Rnf7 | ring finger protein 7 |
| chr14 | 80022277 | 80023308 | 1032 | * | 6,5149 | -5,566863384 | 1,84105 | -3,02374 | 0,002497 | 1 | 0 | Distal Intergenic | 14 | 80004220 | 80008752 | 4533 | 2 | 18530 | ENSMUST00000195355.2 | -13525 | ENSMUSG00000036422 | Pcdh8 | protocadherin 8 |
| chr3 | 35729462 | 35730262 | 801 | * | 6,115293 | -5,565631562 | 1,9894 | -2,79764 | 0,005148 | 1 | 0 | Distal Intergenic | 3 | 35651301 | 35720312 | 69012 | 2 | 625963 | ENSMUST00000198751.5 | -9150 | ENSMUSG00000105662 | Gm6639 | predicted gene 6639 |
| chr7 | 1,17E+08 | 1,17E+08 | 679 | * | 4,864606 | -5,565296113 | 2,050469 | -2,71416 | 0,006644 | 1 | 0 | Distal Intergenic | 7 | 1,17E+08 | 1,17E+08 | 49233 | 1 | 1,05E+08 | ENSMUST00000206031.2 | -47297 | | Gm39075 | predicted gene, 39075 |
| chr11 | 61422325 | 61423367 | 1043 | * | 6,425243 | -5,564170116 | 2,002546 | -2,77855 | 0,00546 | 1 | 0 | Exon (ENSMUST00000 | 11 | 61423490 | 61435139 | 11650 | 2 | 13855 | ENSMUST00000153984.2 | 11772 | ENSMUSG00000001036 | Epn2 | epsin 2 |
| chr12 | 21996456 | 21997219 | 764 | * | 4,279049 | -5,562870199 | 2,127851 | -2,61431 | 0,008941 | 1 | 0 | Distal Intergenic | 12 | 21826160 | 21826708 | 549 | 2 | 434693 | ENSMUST00000164169.2 | -169748 | | Mrto4-ps2 | mRNA turnover 4, pseudogene 2 |
| chr1 | 1,74E+08 | 1,74E+08 | 648 | * | 5,596313 | -5,562739955 | 2,105809 | -2,64162 | 0,008251 | 1 | 0 | Intron (ENSMUST00000 | 1 | 1,74E+08 | 1,74E+08 | 18321 | 2 | 226691 | ENSMUST00000042610.15 | 4477 | ENSMUSG00000073490 | Ifi207 | interferon activated gene 207 |
| chr17 | 17008617 | 17009264 | 648 | * | 5,0381 | -5,562366129 | 2,0523 | -2,71031 | 0,006722 | 1 | 0 | Distal Intergenic | 17 | 17171801 | 17194425 | 22625 | 1 | 213811 | ENSMUST00000190415.3 | -162537 | ENSMUSG00000060149 | BC002059 | cDNA sequence BC002059 |
| chr14 | 1,03E+08 | 1,03E+08 | 740 | * | 5,105983 | -5,562128399 | 2,023141 | -2,74925 | 0,005973 | 1 | 0 | Distal Intergenic | 14 | 1,03E+08 | 1,03E+08 | 16595 | 2 | 1,03E+08 | ENSMUST00000227348.2 | -55008 | ENSMUSG00000115816 | Gm34589 | predicted gene, 34589 |
| chr3 | 87779994 | 87780461 | 468 | * | 4,496306 | -5,562124995 | 2,108801 | -2,63758 | 0,00835 | 1 | 0 | Intron (ENSMUST00000 | 3 | 87766210 | 87792915 | 26706 | 2 | 94315 | ENSMUST00000005015.10 | 12454 | ENSMUSG00000004895 | Prcc | papillary renal cell carcinoma (translocation-associated) |
| chr2 | 1,79E+08 | 1,79E+08 | 411 | * | 4,768604 | -5,56197747 | 2,124253 | -2,61832 | 0,008836 | 1 | 0 | Intron (ENSMUST00000 | 2 | 1,79E+08 | 1,79E+08 | 20349 | 1 | 12561 | ENSMUST00000136411.2 | -85481 | ENSMUSG00000000305 | Cdh4 | cadherin 4 |
| chr15 | 79724182 | 79724928 | 747 | * | 7,335633 | -5,560276039 | 1,86077 | -2,98816 | 0,002807 | 1 | 0 | Distal Intergenic | 15 | 79708097 | 79718889 | 10793 | 2 | 494448 | ENSMUST00000109627.8 | -5293 | ENSMUSG00000089715 | Cbx6 | chromobox 6 |
| chrX | 1,03E+08 | 1,03E+08 | 947 | * | 5,10413 | -5,55984972 | 2,128241 | -2,61242 | 0,00899 | 1 | 0 | Intron (ENSMUST00000 | 20 | 1,03E+08 | 1,03E+08 | 133190 | 2 | 11306 | ENSMUST00000033695.6 | 94758 | ENSMUSG00000031333 | Abcb7 | ATP-binding cassette, sub-family B (MDR/TAP), member 7 |
| chr5 | 47195779 | 47196511 | 733 | * | 4,210123 | -5,559808878 | 2,068956 | -2,68725 | 0,007204 | 1 | 0 | Distal Intergenic | 5 | 48140480 | 48465075 | 324596 | 1 | 20563 | ENSMUST00000173107.8 | -943969 | ENSMUSG00000031558 | Slit2 | slit guidance ligand 2 |
| chr12 | 12496408 | 12497150 | 743 | * | 6,18207 | -5,559607763 | 1,871132 | -2,97125 | 0,002966 | 1 | 0 | Distal Intergenic | 12 | 12401523 | 12442616 | 41094 | 2 | 70858 | ENSMUST00000220545.2 | -53792 | ENSMUSG00000114136 | 4921511I17Rik | RIKEN cDNA 4921511I17 gene |
| chr10 | 74297617 | 74298369 | 753 | * | 6,29107 | -5,559593417 | 1,862179 | -2,98553 | 0,002831 | 1 | 0 | Intron (ENSMUST00000 | 10 | 74221705 | 74232556 | 10852 | 1 | 11994 | ENSMUST00000139106.3 | 75912 | ENSMUSG00000052613 | Pcdh15 | protocadherin 15 |
| chrX | 1,03E+08 | 1,03E+08 | 903 | * | 4,701253 | -5,557077336 | 2,080709 | -2,67076 | 0,007568 | 1 | 0 | Promoter (<=1kb) | 20 | 1,03E+08 | 1,03E+08 | 27806 | 2 | 20502 | ENSMUST00000148684.3 | -268 | ENSMUSG00000033965 | Slc16a2 | solute carrier family 16 (monocarboxylic acid transporters), member 2 |
| chr4 | 62796682 | 62797068 | 387 | * | 4,232239 | -5,556758053 | 2,143525 | -2,59235 | 0,009532 | 1 | 0 | Distal Intergenic | 4 | 62857578 | 62857711 | 134 | 1 | 1,15E+08 | ENSMUST00000157342.3 | -60510 | ENSMUSG00000087967 | Gm25391 | predicted gene, 25391 |
| chr14 | 72692677 | 72693470 | 794 | * | 5,933863 | -5,554837149 | 2,041462 | -2,72101 | 0,006508 | 1 | 0 | Exon (ENSMUST00000 | 14 | 72663453 | 72729278 | 65826 | 2 | 675947 | ENSMUST00000208955.2 | 35808 | | Gm9195 | predicted gene 9195 |
| chr3 | 1,46E+08 | 1,46E+08 | 864 | * | 7,647742 | -5,554703968 | 1,925673 | -2,88455 | 0,00392 | 1 | 0 | Distal Intergenic | 3 | 1,46E+08 | 1,46E+08 | 10095 | 1 | 12042 | ENSMUST00000029842.9 | -18926 | ENSMUSG00000028191 | Bcl10 | B cell leukemia/lymphoma 10 |
| chr10 | 19456833 | 19457775 | 943 | * | 5,130609 | -5,554607889 | 2,109291 | -2,6334 | 0,008453 | 1 | 0 | Distal Intergenic | 10 | 19467697 | 19485977 | 18281 | 1 | 15979 | ENSMUST00000020188.13 | -9922 | ENSMUSG00000020009 | Ifngr1 | interferon gamma receptor 1 |
| chr5 | 1,24E+08 | 1,24E+08 | 1119 | * | 9,2782 | -5,552154349 | 1,840007 | -3,01746 | 0,002549 | 0,681421811 | 1,211315 | Promoter (<=1kb) | 5 | 1,24E+08 | 1,24E+08 | 34269 | 2 | 56325 | ENSMUST00000031354.11 | 0 | ENSMUSG00000029408 | Abcb9 | ATP-binding cassette, sub-family B (MDR/TAP), member 9 |
| chr19 | 30431554 | 30432885 | 1332 | * | 6,268922 | -5,551781257 | 1,87174 | -2,96611 | 0,003016 | 1 | 0 | Distal Intergenic | 19 | 30516328 | 30516945 | 618 | 2 | 546723 | ENSMUST00000078012.6 | 84060 | | Ppp1r2-ps3 | protein phosphatase 1, regulatory (inhibitor) subunit 2, pseudogene 3 |
| chrX | 1,54E+08 | 1,54E+08 | 635 | * | 5,41413 | -5,551708714 | 2,054733 | -2,70191 | 0,006894 | 1 | 0 | Promoter (<=1kb) | 20 | 1,54E+08 | 1,54E+08 | 110 | 1 | 1,15E+08 | ENSMUST00000082990.4 | 606 | ENSMUSG00002075311 | Gm24268 | predicted gene, 24268 |
| chr1 | 1,28E+08 | 1,28E+08 | 601 | * | 4,17352 | -5,551696564 | 2,092194 | -2,65353 | 0,007965 | 1 | 0 | Intron (ENSMUST00000 | 1 | 1,28E+08 | 1,28E+08 | 27369 | 1 | 226412 | ENSMUST00000189317.3 | 7721 | ENSMUSG00000056211 | R3hdm1 | R3H domain containing 1 |
| chr2 | 1,47E+08 | 1,47E+08 | 431 | * | 5,013072 | -5,551694173 | 2,116573 | -2,62296 | 0,008717 | 1 | 0 | Promoter (<=1kb) | 2 | 1,47E+08 | 1,47E+08 | 7939 | 2 | 18088 | ENSMUST00000109969.3 | -309 | ENSMUSG00000027434 | Nkx2-2 | NK2 homeobox 2 |
| chr6 | 1,41E+08 | 1,41E+08 | 495 | * | 6,441468 | -5,551119489 | 1,862068 | -2,98116 | 0,002872 | 1 | 0 | Intron (ENSMUST00000 | 6 | 1,41E+08 | 1,41E+08 | 86800 | 1 | 1,03E+08 | ENSMUST00000204355.2 | 27934 | ENSMUSG00000107638 | Gm30524 | predicted gene, 30524 |
| chr19 | 19193113 | 19194185 | 1073 | * | 7,330771 | -5,547680067 | 1,984838 | -2,79503 | 0,00519 | 1 | 0 | Distal Intergenic | 19 | 19253263 | 19253394 | 132 | 1 | 1,15E+08 | ENSMUST00000158282.3 | -59078 | ENSMUSG00000088907 | Gm25437 | predicted gene, 25437 |
| chr9 | 47005744 | 47006336 | 593 | * | 5,369619 | -5,546527506 | 2,058519 | -2,69443 | 0,007051 | 1 | 0 | Intron (ENSMUST00000 | 9 | 46997245 | 47077192 | 79948 | 2 | 1,03E+08 | ENSMUST00000214227.2 | 70856 | | Gm31698 | predicted gene, 31698 |
| chr14 | 68696982 | 68697767 | 786 | * | 4,478887 | -5,546170745 | 2,060747 | -2,69134 | 0,007117 | 1 | 0 | Promoter (1-2kb) | 14 | 68699080 | 68733866 | 34787 | 1 | 1,03E+08 | ENSMUST00000225482.2 | -1313 | | Gm31227 | predicted gene, 31227 |
| chr12 | 33894103 | 33894618 | 516 | * | 4,927754 | -5,543693187 | 1,979797 | -2,80013 | 0,005108 | 1 | 0 | Distal Intergenic | 12 | 33978424 | 33979309 | 886 | 1 | 114712 | ENSMUST00000061035.4 | -83806 | ENSMUSG00000046518 | Ferd3l | Fer3 like bHLH transcription factor |
| chr2 | 1,71E+08 | 1,71E+08 | 708 | * | 5,484235 | -5,543629463 | 2,019767 | -2,74469 | 0,006057 | 1 | 0 | Distal Intergenic | 2 | 1,71E+08 | 1,71E+08 | 264 | 2 | 1,15E+08 | ENSMUST00000157880.3 | -222706 | ENSMUSG00000088505 | Gm26489 | predicted gene, 26489 |
| chr10 | 1,26E+08 | 1,26E+08 | 688 | * | 5,153029 | -5,54360746 | 2,031594 | -2,7287 | 0,006358 | 1 | 0 | Distal Intergenic | 10 | 1,26E+08 | 1,26E+08 | 3088 | 1 | 66334 | ENSMUST00000218075.2 | 84797 | | 1700025K04Rik | RIKEN cDNA 1700025K04 gene |
| chr10 | 54719003 | 54719382 | 380 | * | 4,629152 | -5,54353186 | 2,089591 | -2,65293 | 0,00798 | 1 | 0 | Distal Intergenic | 10 | 54589072 | 54589133 | 62 | 2 | 1,15E+08 | ENSMUST00000158362.3 | -129870 | ENSMUSG00000088987 | Gm25602 | predicted gene, 25602 |
| chr8 | 36326920 | 36327228 | 309 | * | 5,032502 | -5,543530347 | 2,086926 | -2,65631 | 0,0079 | 1 | 0 | Distal Intergenic | 8 | 36331239 | 36350111 | 18873 | 2 | 330737 | ENSMUST00000210717.2 | 22883 | ENSMUSG00000109761 | 5430403N17Rik | RIKEN cDNA 5430403N17 gene |
| chr10 | 72943327 | 72944501 | 1175 | * | 8,337955 | -5,54322516 | 1,768301 | -3,13477 | 0,00172 | 0,577046464 | 1,449706 | Intron (ENSMUST00000 | 10 | 72935285 | 73319952 | 384668 | 1 | 11994 | ENSMUST00000156999.2 | 8042 | ENSMUSG00000052613 | Pcdh15 | protocadherin 15 |
| chr12 | 1,14E+08 | 1,14E+08 | 1123 | * | 6,3865 | -5,542920739 | 1,885805 | -2,93929 | 0,00329 | 1 | 0 | Distal Intergenic | 12 | 1,14E+08 | 1,14E+08 | 2558 | 1 | 238406 | ENSMUST00000053086.3 | 373919 | ENSMUSG00000043945 | Adam6a | a disintegrin and metallopeptidase domain 6A |
| chr17 | 9113448 | 9113791 | 344 | * | 4,676014 | -5,540847255 | 2,075568 | -2,66956 | 0,007595 | 1 | 0 | Promoter (1-2kb) | 17 | 9112123 | 9149362 | 37240 | 1 | 23984 | ENSMUST00000136160.2 | 1325 | ENSMUSG00000023868 | Pde10a | phosphodiesterase 10A |
| chr4 | 1,19E+08 | 1,19E+08 | 405 | * | 4,908325 | -5,540759778 | 2,087413 | -2,65437 | 0,007946 | 1 | 0 | Intron (ENSMUST00000 | 4 | 1,19E+08 | 1,19E+08 | 17961 | 2 | 194237 | ENSMUST00000126728.2 | 10399 | ENSMUSG00000048899 | Rimkla | ribosomal modification protein rimK-like family member A |
| chr4 | 1,19E+08 | 1,19E+08 | 600 | * | 6,662341 | -5,53859859 | 2,045937 | -2,70712 | 0,006787 | 1 | 0 | Distal Intergenic | 4 | 1,19E+08 | 1,19E+08 | 948 | 1 | 258331 | ENSMUST00000094830.2 | -10852 | ENSMUSG00000073768 | Olfr1330 | olfactory receptor 1330 |
| chr11 | 68259779 | 68260511 | 733 | * | 5,890728 | -5,538233616 | 2,121361 | -2,6107 | 0,009036 | 1 | 0 | Intron (ENSMUST00000 | 11 | 68100190 | 68277652 | 177463 | 2 | 18208 | ENSMUST00000021284.4 | 17141 | ENSMUSG00000020902 | Ntn1 | netrin 1 |
| chr1 | 1,15E+08 | 1,15E+08 | 652 | * | 6,201627 | -5,537545121 | 1,973317 | -2,80621 | 0,005013 | 1 | 0 | Distal Intergenic | 1 | 1,16E+08 | 1,16E+08 | 15632 | 2 | 1,08E+08 | ENSMUST00000186137.7 | 427170 | | Gm50605 | predicted gene, 50605 |
| chrX | 76057556 | 76058562 | 1007 | * | 5,497731 | -5,537489022 | 1,964889 | -2,81822 | 0,004829 | 1 | 0 | Distal Intergenic | 20 | 75873640 | 75873778 | 139 | 1 | 1,15E+08 | ENSMUST00000157905.4 | 183916 | ENSMUSG00000088530 | Gm23280 | predicted gene, 23280 |
| chr9 | 30986952 | 30987933 | 982 | * | 7,63046 | -5,536419891 | 1,804927 | -3,06739 | 0,002159 | 1 | 0 | 3' UTR | 9 | 30991913 | 30993771 | 1859 | 2 | 1,05E+08 | ENSMUST00000181215.2 | 5838 | | Gm17508 | predicted gene, 17508 |
| chrX | 11560646 | 11562100 | 1455 | * | 8,262936 | -5,536236074 | 1,835012 | -3,017 | 0,002553 | 0,651089638 | 1,449706 | Distal Intergenic | 20 | 11716723 | 11716829 | 107 | 1 | 1,15E+08 | ENSMUST00000083891.3 | -154623 | ENSMUSG00000065825 | Gm26314 | predicted gene, 26314 |
| chr15 | 68926355 | 68927399 | 1045 | * | 7,425634 | -5,536197885 | 1,841489 | -3,00637 | 0,002644 | 0,669377201 | 1,360622 | Intron (ENSMUST00000 | 15 | 68801917 | 68896614 | 94698 | 1 | 13992 | ENSMUST00000230073.2 | 124438 | ENSMUSG00000022332 | Khdrbs3 | KH domain containing, RNA binding, signal transduction associated 3 |

| | | | | | | | | | | | | | | | | | | | | | | | |
|---|---|---|---|---|---|---|---|---|---|---|---|---|---|---|---|---|---|---|---|---|---|---|---|
| chr19 | 8138368 | 8139746 | 1379 | * | 9,455857 | -5,53591705 | 1,791408 | -3,09026 | 0,002 | 0,620861959 | 1,360622 | Exon (ENSMUST000001 | 19 | 8137658 | 8147409 | 9752 | 2 | 236293 | ENSMUST00000142517.3 | 7663 | ENSMUSG00000075044 | Slc22a29 | solute carrier family 22. member 29 |
| chr17 | 31614167 | 31615041 | 875 | * | 7,794711 | -5,535876929 | 1,788452 | -3,09535 | 0,001966 | 1 | 0 | Intron (ENSMUST00000 | 17 | 31605268 | 31674096 | 68829 | 1 | 18585 | ENSMUST00000141314.8 | 8899 | ENSMUSG00000041119 | Pde9a | phosphodiesterase 9A |
| chr3 | 83264563 | 83265539 | 977 | * | 4,977432 | -5,535476245 | 2,025207 | -2,73329 | 0,006271 | 1 | 0 | Downstream (<=300bp | 3 | 83462670 | 83481429 | 18760 | 2 | 69460 | ENSMUST00000185938.2 | 215890 | ENSMUSG00000101334 | 1700028M03Rik | RIKEN cDNA 1700028M03 gene |
| chr13 | 5417664 | 5418168 | 505 | * | 5,024469 | -5,535463229 | 2,031712 | -2,72453 | 0,006439 | 1 | 0 | Distal Intergenic | 13 | 5504908 | 5549701 | 44794 | 1 | 1,05E+08 | ENSMUST00000220659.2 | -86740 | | Gm40660 | predicted gene, 40660 |
| chr3 | 86089752 | 86090542 | 791 | * | 6,87766 | -5,535347966 | 1,868178 | -2,96297 | 0,003047 | 1 | 0 | Distal Intergenic | 3 | 86098723 | 86098833 | 111 | 1 | 1,15E+08 | ENSMUST00000158769.3 | -8181 | ENSMUSG00000089394 | Gm23025 | predicted gene, 23025 |
| chr14 | 94390324 | 94391277 | 954 | * | 6,429019 | -5,533706237 | 1,975372 | -2,80135 | 0,005089 | 1 | 0 | Distal Intergenic | 14 | 94123549 | 94128115 | 4567 | 2 | 211712 | ENSMUST00000194056.2 | -262209 | ENSMUSG00000055421 | Pcdh9 | protocadherin 9 |
| chr18 | 71411946 | 71412945 | 1000 | * | 5,76099 | -5,533281183 | 2,012247 | -2,7498 | 0,005963 | 1 | 0 | Intron (ENSMUST00000 | 18 | 70750844 | 70758848 | 8005 | 1 | 17191 | ENSMUST00000128555.2 | 661102 | ENSMUSG00000024513 | Mbd2 | methyl-CpG binding domain protein 2 |
| chr8 | 37395784 | 37396409 | 626 | * | 5,515479 | -5,532221489 | 2,128524 | -2,59909 | 0,009347 | 1 | 0 | Intron (ENSMUST00000 | 8 | 37316280 | 37419511 | 103232 | 2 | 50768 | ENSMUST00000036104.11 | 23102 | ENSMUSG00000031523 | Dlc1 | deleted in liver cancer 1 |
| chr2 | 82058355 | 82059676 | 1322 | * | 13,00822 | -5,53199177 | 1,626577 | -3,401 | 0,000671 | 0,527676616 | 0,738441 | Intron (ENSMUST00000 | 2 | 82065517 | 82087224 | 21708 | 1 | 241514 | ENSMUST00000127187.2 | -5841 | ENSMUSG00000070866 | Zfp804a | zinc finger protein 804A |
| chr11 | 1,16E+08 | 1,16E+08 | 1044 | * | 9,404691 | -5,529639963 | 1,785028 | -3,09779 | 0,00195 | 0,637143933 | 1,211315 | Promoter (2-3kb) | 11 | 1,16E+08 | 1,16E+08 | 2362 | 2 | 53413 | ENSMUST00000155288.2 | -2243 | ENSMUSG00000020792 | Exoc7 | exocyst complex component 7 |
| chrX | 61217197 | 61217955 | 759 | * | 4,986367 | -5,529538142 | 1,999672 | -2,76522 | 0,005688 | 1 | 0 | Distal Intergenic | 20 | 61327105 | 61335684 | 8580 | 1 | 74437 | ENSMUST00000178085.8 | -109150 | ENSMUSG00000045330 | Magec2 | MAGE family member C2 |
| chrX | 10124076 | 10125201 | 1126 | * | 8,631616 | -5,528422322 | 1,817399 | -3,04194 | 0,002351 | 0,550608817 | 2,237403 | Intron (ENSMUST00000 | 20 | 10118606 | 10183103 | 64498 | 1 | 18416 | ENSMUST00000115528.3 | 5470 | ENSMUSG00000031173 | Otc | ornithine transcarbamylase |
| chr19 | 21447810 | 21448838 | 1029 | * | 5,943809 | -5,528121786 | 1,870459 | -2,95549 | 0,003122 | 1 | 0 | Promoter (1-2kb) | 19 | 21396089 | 21449952 | 53864 | 2 | 14544 | ENSMUST00000136258.2 | 1114 | ENSMUSG00000058624 | Gda | guanine deaminase |
| chr10 | 1,09E+08 | 1,09E+08 | 439 | * | 5,162042 | -5,527466262 | 1,985131 | -2,78443 | 0,005362 | 1 | 0 | Distal Intergenic | 10 | 1,09E+08 | 1,09E+08 | 106 | 2 | 1,15E+08 | ENSMUST00000083868.3 | 109311 | ENSMUSG00000065802 | Gm23041 | predicted gene, 23041 |
| chr9 | 3657634 | 3658162 | 529 | * | 4,568163 | -5,524609953 | 2,057745 | -2,68479 | 0,007258 | 1 | 0 | Intron (ENSMUST00000 | 9 | 3532888 | 3761177 | 228290 | 1 | 234889 | ENSMUST00000213060.2 | 124746 | ENSMUSG00000041624 | Gucy1a2 | guanylate cyclase 1, soluble, alpha 2 |
| chr1 | 13045316 | 13046002 | 687 | * | 4,815959 | -5,524025029 | 2,053203 | -2,69044 | 0,007136 | 1 | 0 | Intron (ENSMUST00000 | 1 | 12949907 | 13061138 | 111232 | 2 | 240726 | ENSMUST00000146763.8 | 15136 | ENSMUSG00000025938 | Slco5a1 | solute carrier organic anion transporter family, member 5A1 |
| chrX | 70244727 | 70245526 | 800 | * | 4,514956 | -5,521651505 | 2,136707 | -2,58419 | 0,009761 | 1 | 0 | Distal Intergenic | 20 | 70254373 | 70358901 | 104529 | 1 | 17772 | ENSMUST00000171933.8 | -8847 | ENSMUSG00000031337 | Mtm1 | X-linked myotubular myopathy gene 1 |
| chr5 | 1,48E+08 | 1,48E+08 | 554 | * | 4,641881 | -5,521361978 | 2,006665 | -2,75151 | 0,005932 | 1 | 0 | Distal Intergenic | 5 | 1,47E+08 | 1,48E+08 | 164408 | 2 | 14254 | ENSMUST00000031653.12 | -54497 | ENSMUSG00000029648 | Flt1 | FMS-like tyrosine kinase 1 |
| chr2 | 1,26E+08 | 1,26E+08 | 489 | * | 5,051147 | -5,521293313 | 1,982787 | -2,78461 | 0,005359 | 1 | 0 | Distal Intergenic | 2 | 1,26E+08 | 1,26E+08 | 45885 | 2 | 70354 | ENSMUST00000053699.13 | -13137 | ENSMUSG00000035093 | Secisbp2l | SECIS binding protein 2-like |
| chr7 | 66466097 | 66467194 | 1098 | * | 5,758665 | -5,52110493 | 1,927914 | -2,86377 | 0,004186 | 1 | 0 | Intron (ENSMUST00000 | 7 | 66489483 | 66619303 | 129821 | 1 | 233332 | ENSMUST00000156396.8 | -22289 | ENSMUSG00000058145 | Adamts17 | a disintegrin-like and metallopeptidase (reprolysin type) with thrombospondin type 1 motif, 17 |
| chr8 | 17966260 | 17967302 | 1043 | * | 7,959792 | -5,520920449 | 1,802476 | -3,06296 | 0,002192 | 1 | 0 | Distal Intergenic | 8 | 16394877 | 17585602 | 1190726 | 2 | 94109 | ENSMUST00000137947.2 | -380658 | ENSMUSG00000060924 | Csmd1 | CUB and Sushi multiple domains 1 |
| chr17 | 14571540 | 14572381 | 842 | * | 8,303418 | -5,520801071 | 1,811825 | -3,04709 | 0,002311 | 0,547497234 | 2,237403 | Intron (ENSMUST00000 | 17 | 14609375 | 14619204 | 9830 | 2 | 74917 | ENSMUST00000233310.2 | 46823 | ENSMUSG00000117328 | 4930474M22Rik | RIKEN cDNA 4930474M22 gene |
| chr7 | 34458159 | 34458773 | 615 | * | 5,199305 | -5,519501985 | 1,954423 | -2,82411 | 0,004741 | 1 | 0 | Intron (ENSMUST00000 | 7 | 34373893 | 34453237 | 79345 | 2 | 68947 | ENSMUST00000238947.2 | -4922 | ENSMUSG00000060402 | Chst8 | carbohydrate sulfotransferase 8 |
| chr14 | 59394962 | 59395666 | 705 | * | 5,500061 | -5,519442286 | 1,963471 | -2,81106 | 0,004938 | 1 | 0 | Distal Intergenic | 14 | 59378775 | 59380342 | 1568 | 2 | 67932 | ENSMUST00000162271.2 | -14620 | ENSMUSG00000021977 | 1700129C05Rik | RIKEN cDNA 1700129C05 gene |
| chr11 | 66914290 | 66915007 | 718 | * | 4,722673 | -5,519194516 | 2,004658 | -2,75319 | 0,005902 | 1 | 0 | Promoter (<=1kb) | 11 | 66915980 | 66925616 | 9637 | 1 | 338369 | ENSMUST00000079077.12 | -973 | ENSMUSG00000050270 | Tmem220 | transmembrane protein 220 |
| chr2 | 1,41E+08 | 1,41E+08 | 688 | * | 4,822391 | -5,519160298 | 2,010198 | -2,74558 | 0,00604 | 1 | 0 | Intron (ENSMUST00000 | 2 | 1,41E+08 | 1,41E+08 | 56561 | 2 | 1,03E+08 | ENSMUST00000151538.2 | 12596 | | Macrod2os1 | mono-ADP ribosylhydrolase 2, opposite strand 1 |
| chr16 | 23071708 | 23072688 | 981 | * | 7,056092 | -5,518323472 | 1,925668 | -2,86567 | 0,004161 | 1 | 0 | Intron (ENSMUST00000 | 16 | 23088379 | 23139929 | 51551 | 1 | 20440 | ENSMUST00000128050.8 | -15691 | ENSMUSG00000022885 | St6gal1 | beta galactoside alpha 2,6 sialyltransferase 1 |
| chr2 | 38189201 | 38189550 | 350 | * | 5,419003 | -5,516940542 | 2,012121 | -2,74185 | 0,006109 | 1 | 0 | Distal Intergenic | 2 | 37706928 | 38177402 | 470475 | 2 | 227801 | ENSMUST00000140552.8 | -11799 | ENSMUSG00000035392 | Dennd1a | DENN/MADD domain containing 1A |
| chr13 | 82858220 | 82859004 | 785 | * | 4,411683 | -5,515920672 | 2,097659 | -2,62956 | 0,00855 | 1 | 0 | Distal Intergenic | 13 | 82846394 | 82846452 | 59 | 2 | 1,01E+08 | ENSMUST00000198795.3 | -11768 | ENSMUSG00000106655 | Mir3961 | microRNA 3961 |
| chr3 | 19455748 | 19456319 | 572 | * | 5,468674 | -5,515337563 | 2,011493 | -2,74191 | 0,006108 | 1 | 0 | Distal Intergenic | 3 | 19277272 | 19365486 | 88215 | 2 | 18583 | ENSMUST00000099195.10 | -90262 | ENSMUSG00000069094 | Pde7a | phosphodiesterase 7A |
| chr1 | 1,74E+08 | 1,74E+08 | 713 | * | 5,659416 | -5,514885067 | 1,968343 | -2,80179 | 0,005082 | 1 | 0 | Distal Intergenic | 1 | 1,74E+08 | 1,74E+08 | 20177 | 2 | 26388 | ENSMUST00000000266.9 | -21066 | ENSMUSG00000026535 | Ifi202b | interferon activated gene 202B |
| chr5 | 39790080 | 39790944 | 865 | * | 7,468673 | -5,512629395 | 1,802483 | -3,05835 | 0,002226 | 1 | 0 | Intron (ENSMUST00000 | 5 | 39772417 | 39801940 | 29524 | 2 | 15476 | ENSMUST00000152057.2 | 10996 | ENSMUSG00000051022 | Hs3st1 | heparan sulfate (glucosamine) 3-O-sulfotransferase 1 |
| chr13 | 92375886 | 92376774 | 889 | * | 6,478658 | -5,51225465 | 1,863855 | -2,95745 | 0,003102 | 1 | 0 | Promoter (<=1kb) | 13 | 92348388 | 92377638 | 29251 | 2 | 17686 | ENSMUST00000191242.7 | 864 | ENSMUSG00000014850 | Msh3 | mutS homolog 3 |
| chr16 | 65414281 | 65415054 | 774 | * | 6,911343 | -5,511865314 | 1,898656 | -2,90303 | 0,003696 | 1 | 0 | Distal Intergenic | 16 | 65461988 | 65551514 | 89527 | 1 | 1,05E+08 | ENSMUST00000241896.1 | -46934 | | Gm41469 | predicted gene, 41469 |
| chr3 | 28682473 | 28683320 | 848 | * | 6,859822 | -5,511742872 | 1,910848 | -2,88445 | 0,003921 | 1 | 0 | Promoter (<=1kb) | 3 | 28684021 | 28692618 | 8598 | 1 | 665113 | ENSMUST00000162225.2 | -701 | ENSMUSG00000027692 | Tnik | TRAF2 and NCK interacting kinase |
| chr1 | 1,56E+08 | 1,56E+08 | 562 | * | 4,542433 | -5,510742216 | 2,037687 | -2,70441 | 0,006843 | 1 | 0 | Intron (ENSMUST00000 | 1 | 1,56E+08 | 1,56E+08 | 1630 | 1 | 240832 | ENSMUST00000125371.2 | 35413 | ENSMUSG00000050565 | Tor1aip2 | torsin A interacting protein 2 |
| chr19 | 57906219 | 57906535 | 317 | * | 4,221375 | -5,510525212 | 2,075436 | -2,65512 | 0,007928 | 1 | 0 | Intron (ENSMUST00000 | 19 | 58039599 | 58039668 | 70 | 1 | 1,01E+08 | ENSMUST00000176631.3 | -133064 | ENSMUSG00000093757 | Mir5623 | microRNA 5623 |
| chrX | 95309406 | 95310341 | 936 | * | 6,311773 | -5,509271138 | 1,977057 | -2,7866 | 0,005326 | 1 | 0 | Intron (ENSMUST00000 | 20 | 95286423 | 95286532 | 110 | 1 | 723814 | ENSMUST00000102112.3 | 22983 | ENSMUSG00000076066 | Mir223 | microRNA 223 |
| chr1 | 1,87E+08 | 1,87E+08 | 834 | * | 4,630921 | -5,507900282 | 2,133825 | -2,58123 | 0,009845 | 1 | 0 | Intron (ENSMUST00000 | 1 | 1,87E+08 | 1,87E+08 | 3497 | 2 | 1,05E+08 | ENSMUST00000162615.2 | 20305 | ENSMUSG00000089947 | Gm15509 | predicted gene 15509 |
| chr7 | 1,37E+08 | 1,37E+08 | 742 | * | 6,2264 | -5,506360648 | 1,942039 | -2,83535 | 0,004578 | 1 | 0 | Distal Intergenic | 7 | 1,37E+08 | 1,37E+08 | 4481 | 1 | 30926 | ENSMUST00000211598.2 | 65477 | ENSMUSG00000031068 | Glrx3 | glutaredoxin 3 |
| chr2 | 95167373 | 95167882 | 510 | * | 5,766682 | -5,506318764 | 1,938511 | -2,84049 | 0,004504 | 1 | 0 | Distal Intergenic | 2 | 95224587 | 95227996 | 3410 | 1 | 1,03E+08 | ENSMUST00000148637.2 | -56705 | | Gm13794 | predicted gene 13794 |
| chr14 | 87455946 | 87456549 | 604 | * | 5,23678 | -5,504831475 | 2,010416 | -2,73816 | 0,006178 | 1 | 0 | Distal Intergenic | 14 | 87202887 | 87378671 | 175785 | 2 | 56419 | ENSMUST00000228000.2 | -77275 | ENSMUSG00000022021 | Diaph3 | diaphanous related formin 3 |
| chr10 | 35900609 | 35901109 | 501 | * | 5,847732 | -5,50481787 | 1,947557 | -2,82652 | 0,004706 | 1 | 0 | Distal Intergenic | 10 | 35714486 | 35730471 | 15986 | 1 | 1,05E+08 | ENSMUST00000213846.2 | 186123 | | Gm40623 | predicted gene, 40623 |
| chr11 | 81126093 | 81126900 | 808 | * | 6,148775 | -5,504816999 | 1,945691 | -2,82924 | 0,004666 | 1 | 0 | Intron (ENSMUST00000 | 11 | 80770995 | 81044147 | 273153 | 2 | 11418 | ENSMUST00000021045.13 | -81946 | ENSMUSG00000020704 | Asic2 | acid-sensing (proton-gated) ion channel 2 |
| chrX | 1,33E+08 | 1,33E+08 | 583 | * | 5,49647 | -5,504800994 | 1,90016 | -2,89702 | 0,003767 | 1 | 0 | Distal Intergenic | 20 | 1,33E+08 | 1,33E+08 | 14598 | 1 | 64103 | ENSMUST00000033602.9 | -27030 | ENSMUSG00000031250 | Tnmd | tenomodulin |
| chr18 | 6159602 | 6160816 | 1215 | * | 9,455797 | -5,504700128 | 1,731683 | -3,17882 | 0,001479 | 0,559413722 | 1,360622 | Distal Intergenic | 18 | 6025217 | 6136098 | 110882 | 2 | 75415 | ENSMUST00000182559.8 | -23504 | ENSMUSG00000041225 | Arhgap12 | Rho GTPase activating protein 12 |
| chr7 | 13498999 | 13500019 | 1021 | * | 7,397948 | -5,504423492 | 1,812215 | -3,0374 | 0,002386 | 1 | 0 | Intron (ENSMUST00000 | 7 | 13467430 | 13513561 | 46132 | 1 | 1E+08 | ENSMUST00000086148.8 | 31569 | ENSMUSG00000070811 | Sult2a2 | sulfotransferase family 2A, dehydroepiandrosterone (DHEA)-preferring, member 2 |
| chr5 | 1,51E+08 | 1,51E+08 | 567 | * | 4,469809 | -5,501996562 | 2,101801 | -2,61775 | 0,008851 | 1 | 0 | Intron (ENSMUST00000 | 5 | 1,51E+08 | 1,51E+08 | 364618 | 2 | 243362 | ENSMUST00000202365.2 | 9209 | ENSMUSG00000016128 | Stard13 | StAR-related lipid transfer (START) domain containing 13 |
| chr1 | 1,05E+08 | 1,05E+08 | 451 | * | 4,365221 | -5,501904684 | 2,117066 | -2,59884 | 0,009354 | 1 | 0 | Distal Intergenic | 1 | 1,05E+08 | 1,05E+08 | 226953 | 1 | 23836 | ENSMUST00000062528.9 | -146389 | ENSMUSG00000050840 | Cdh20 | cadherin 20 |
| chr16 | 53858450 | 53858847 | 398 | * | 4,130351 | -5,499394769 | 2,127067 | -2,58544 | 0,009726 | 1 | 0 | Distal Intergenic | 16 | 54174825 | 54174956 | 132 | 1 | 1,15E+08 | ENSMUST00000082793.3 | -315978 | ENSMUSG00000064727 | Gm25853 | predicted gene, 25853 |
| chr10 | 19905104 | 19905821 | 718 | * | 4,10171 | -5,499273732 | 2,102719 | -2,61532 | 0,008914 | 1 | 0 | Intron (ENSMUST00000 | 10 | 19975271 | 19981183 | 5913 | 1 | 26408 | ENSMUST00000138994.2 | -69450 | ENSMUSG00000071369 | Map3k5 | mitogen-activated protein kinase kinase kinase 5 |
| chr10 | 1,15E+08 | 1,15E+08 | 610 | * | 4,960772 | -5,498855604 | 2,12699 | -2,58528 | 0,00973 | 1 | 0 | Intron (ENSMUST00000 | 10 | 1,14E+08 | 1,15E+08 | 402550 | 2 | 237553 | ENSMUST00000061632.9 | 39793 | ENSMUSG00000050663 | Trhde | TRH-degrading enzyme |
| chr9 | 60956352 | 60957220 | 869 | * | 6,382637 | -5,498340728 | 1,967163 | -2,79506 | 0,005189 | 1 | 0 | Exon (ENSMUST000002 | 9 | 60956291 | 60962951 | 6661 | 2 | 1,05E+08 | ENSMUST00000217239.2 | 5731 | | Gm39348 | predicted gene, 39348 |
| chr19 | 42374147 | 42374852 | 706 | * | 6,168398 | -5,497346612 | 1,8728 | -2,93536 | 0,003332 | 1 | 0 | Intron (ENSMUST00000 | 19 | 42398833 | 42402685 | 3853 | 1 | 1,03E+08 | ENSMUST00000235253.2 | -23981 | | Gm34299 | predicted gene, 34299 |
| chr19 | 21012655 | 21013998 | 1344 | * | 7,085793 | -5,497051875 | 1,850971 | -2,96982 | 0,00298 | 1 | 0 | Distal Intergenic | 19 | 21081866 | 21084521 | 2656 | 1 | 75354 | ENSMUST00000236278.2 | -67868 | ENSMUSG00000118047 | 4930554I06Rik | RIKEN cDNA 4930554I06 gene |
| chr16 | 35932363 | 35933730 | 1368 | * | 8,826742 | -5,496894064 | 1,826467 | -3,00958 | 0,002616 | 0,683710007 | 1,211315 | Distal Intergenic | 16 | 35942539 | 35951575 | 9037 | 2 | 209294 | ENSMUST00000161638.2 | 17845 | ENSMUSG00000034362 | Csta1 | cystatin A1 |
| chr13 | 78441031 | 78441838 | 808 | * | 6,965299 | -5,496756235 | 1,842862 | -2,98273 | 0,002857 | 0,681316955 | 1,360622 | Distal Intergenic | 13 | 78350672 | 78383779 | 33108 | 1 | 320174 | ENSMUST00000134705.3 | 90359 | ENSMUSG00000087143 | A830082K12Rik | RIKEN cDNA A830082K12 gene |
| chr7 | 38689985 | 38690917 | 933 | * | 5,888869 | -5,496620066 | 1,918337 | -2,8653 | 0,004166 | 1 | 0 | Distal Intergenic | 7 | 38492062 | 38492168 | 107 | 1 | 1,15E+08 | ENSMUST00000239705.1 | 197923 | ENSMUSG00000119867 | Gm24797 | predicted gene, 24797 |
| chr10 | 1,11E+08 | 1,11E+08 | 502 | * | 5,508717 | -5,494814416 | 1,963647 | -2,79827 | 0,005138 | 1 | 0 | Promoter (2-3kb) | 10 | 1,11E+08 | 1,11E+08 | 132498 | 1 | 237542 | ENSMUST00000105275.9 | -2190 | ENSMUSG00000020189 | Osbpl8 | oxysterol binding protein-like 8 |
| chr17 | 30613514 | 30614448 | 935 | * | 6,384316 | -5,494518395 | 1,991609 | -2,75883 | 0,005801 | 1 | 0 | Promoter (<=1kb) | 17 | 30614594 | 30615007 | 414 | 2 | 623169 | ENSMUST00000235315.2 | 559 | | Gm6402 | ribosomal protein S17 pseudogene |
| chr13 | 62445044 | 62445654 | 611 | * | 4,244238 | -5,493689405 | 2,085453 | -2,63429 | 0,008431 | 1 | 0 | Distal Intergenic | 13 | 62484277 | 62488541 | 4265 | 2 | 1,01E+08 | ENSMUST00000222522.2 | 42887 | | Gm19792 | predicted gene, 19792 |
| chr3 | 35082132 | 35082900 | 769 | * | 5,354134 | -5,490861886 | 2,117596 | -2,59297 | 0,009515 | 1 | 0 | Distal Intergenic | 3 | 34976694 | 34976800 | 107 | 2 | 1,02E+08 | ENSMUST00000183357.3 | -105332 | ENSMUSG00000098862 | Mir6378 | microRNA 6378 |
| chr4 | 1,21E+08 | 1,21E+08 | 432 | * | 4,587274 | -5,490717009 | 2,087226 | -2,63063 | 0,008523 | 1 | 0 | Intron (ENSMUST00000 | 4 | 1,21E+08 | 1,21E+08 | 51111 | 2 | 60613 | ENSMUST00000030376.8 | 39697 | ENSMUSG00000028631 | Kcnq4 | potassium voltage-gated channel, subfamily Q, member 4 |
| chr16 | 45482187 | 45482736 | 550 | * | 5,488462 | -5,490583668 | 2,060611 | -2,66454 | 0,007709 | 1 | 0 | Exon (ENSMUST000001 | 16 | 45447211 | 45474431 | 27221 | 2 | 212998 | ENSMUST00000036732.9 | -7756 | ENSMUSG00000033187 | BC016579 | cDNA sequence, BC016579 |
| chr1 | 1,2E+08 | 1,2E+08 | 895 | * | 5,687884 | -5,49028434 | 2,007224 | -2,73526 | 0,006233 | 1 | 0 | Intron (ENSMUST00000 | 1 | 1,2E+08 | 1,2E+08 | 16346 | 2 | 226352 | ENSMUST00000189042.2 | 8332 | ENSMUSG00000026383 | Epb41l5 | erythrocyte membrane protein band 4.1 like 5 |
| chr10 | 19838404 | 19839276 | 873 | * | 6,957432 | -5,489996439 | 1,964564 | -2,79451 | 0,005198 | 1 | 0 | Intron (ENSMUST00000 | 10 | 19826801 | 19899445 | 72645 | 1 | 26408 | ENSMUST00000152533.2 | 11603 | ENSMUSG00000071369 | Map3k5 | mitogen-activated protein kinase kinase kinase 5 |
| chr11 | 55284658 | 55285676 | 1019 | * | 6,446002 | -5,48834458 | 1,934772 | -2,83669 | 0,004558 | 1 | 0 | 3' UTR | 11 | 55286212 | 55290400 | 4189 | 2 | 20692 | ENSMUST00000123775.8 | 4724 | ENSMUSG00000018593 | Sparc | secreted acidic cysteine rich glycoprotein |
| chr1 | 1,75E+08 | 1,75E+08 | 291 | * | 4,964108 | -5,488252834 | 1,964386 | -2,79388 | 0,005208 | 1 | 0 | Distal Intergenic | 1 | 1,75E+08 | 1,75E+08 | 88035 | 2 | 23893 | ENSMUST00000055294.4 | -54523 | ENSMUSG00000050069 | Grem2 | gremlin 2, DAN family BMP antagonist |
| chr11 | 6024902 | 6025248 | 347 | * | 5,061874 | -5,488206632 | 1,980577 | -2,77101 | 0,005588 | 1 | 0 | Distal Intergenic | 11 | 5942820 | 6016362 | 73543 | 2 | 12323 | ENSMUST00000123391.8 | -8540 | ENSMUSG00000057897 | Camk2b | calcium/calmodulin-dependent protein kinase II, beta |
| chr4 | 55753050 | 55754048 | 999 | * | 5,753942 | -5,488175113 | 1,993675 | -2,75279 | 0,005909 | 1 | 0 | Distal Intergenic | 4 | 55605149 | 55636901 | 31753 | 1 | 1,03E+08 | ENSMUST00000123620.2 | 147901 | | Gm12506 | predicted gene 12506 |
| chr1 | 1,8E+08 | 1,8E+08 | 410 | * | 4,312713 | -5,48804323 | 2,047833 | -2,67993 | 0,007364 | 1 | 0 | Promoter (1-2kb) | 1 | 1,8E+08 | 1,8E+08 | 3062 | 2 | 19165 | ENSMUST00000130711.8 | -1488 | ENSMUSG00000010609 | Psen2 | presenilin 2 |
| chr16 | 54168475 | 54169615 | 1141 | * | 6,666956 | -5,487636344 | 1,890037 | -2,90345 | 0,003691 | 1 | 0 | Distal Intergenic | 16 | 54174825 | 54174956 | 132 | 1 | 1,15E+08 | ENSMUST00000082793.3 | -5210 | ENSMUSG00000064727 | Gm25853 | predicted gene, 25853 |
| chr12 | 1,04E+08 | 1,04E+08 | 601 | * | 6,07422 | -5,48583654 | 2,023109 | -2,71159 | 0,006696 | 1 | 0 | Promoter (<=1kb) | 12 | 1,04E+08 | 1,04E+08 | 9715 | 2 | 380780 | ENSMUST00000120251.9 | -64 | ENSMUSG00000063232 | Serpina11 | serine (or cysteine) peptidase inhibitor, clade A (alpha-1 antiproteinase, antitrypsin), member 11 |
| chr1 | 1,39E+08 | 1,39E+08 | 744 | * | 4,641341 | -5,484995283 | 2,113016 | -2,59581 | 0,009437 | 1 | 0 | Distal Intergenic | 1 | 1,39E+08 | 1,39E+08 | 680 | 2 | 75546 | ENSMUST00000186955.3 | -10980 | ENSMUSG00000099808 | 1700019P21Rik | RIKEN cDNA 1700019P21 gene |
| chr14 | 22823005 | 22823918 | 914 | * | 5,499878 | -5,481714272 | 1,978996 | -2,76995 | 0,005607 | 1 | 0 | Intron (ENSMUST00000 | 14 | 22646581 | 22649336 | 2756 | 1 | 76633 | ENSMUST00000226003.2 | 176424 | ENSMUSG00000063458 | Lrmda | leucine rich melanocyte differentiation associated |
| chrX | 1,09E+08 | 1,09E+08 | 829 | * | 5,457026 | -5,481485049 | 1,940123 | -2,82533 | 0,004723 | 1 | 0 | Distal Intergenic | 20 | 1,09E+08 | 1,09E+08 | 2406 | 1 | 634340 | ENSMUST00000122296.2 | -306910 | ENSMUSG00000082853 | Gm7134 | kinesin family member 18B pseudogene |
| chr14 | 1,07E+08 | 1,07E+08 | 1440 | * | 9,036369 | -5,48134149 | 1,762598 | -3,10981 | 0,001872 | 0,62786102 | 1,211315 | Distal Intergenic | 14 | 1,07E+08 | 1,07E+08 | 127 | 1 | 1,15E+08 | ENSMUST00000104036.3 | 141871 | ENSMUSG00000077224 | Gm23437 | predicted gene, 23437 |
| chr19 | 52684942 | 52685631 | 690 | * | 7,453033 | -5,48122448 | 1,902368 | -2,88126 | 0,003961 | 0,731734453 | 1,360622 | Distal Intergenic | 19 | 52911612 | 52915362 | 3751 | 1 | 1E+08 | ENSMUST00000236393.2 | -225981 | ENSMUSG00000117896 | AA387883 | expressed sequence AA387883 |
| chr3 | 66574204 | 66575061 | 858 | * | 8,787757 | -5,481069225 | 1,820491 | -3,01076 | 0,002606 | 0,654833721 | 1,449706 | Distal Intergenic | 3 | 66880113 | 66885780 | 5668 | 2 | 20429 | ENSMUST00000162060.9 | 310719 | ENSMUSG00000027833 | Shox2 | short stature homeobox 2 |
| chr10 | 41850572 | 41851415 | 844 | * | 6,869674 | -5,480622909 | 1,840849 | -2,97723 | 0,002909 | 1 | 0 | Promoter (<=1kb) | 10 | 41792047 | 41852118 | 60072 | 2 | 213402 | ENSMUST00000160367.8 | 703 | ENSMUSG00000071324 | Armc2 | armadillo repeat containing 2 |
| chr16 | 34501344 | 34502243 | 900 | * | 5,590074 | -5,479858649 | 1,975922 | -2,77332 | 0,005549 | 1 | 0 | Distal Intergenic | 16 | 34510986 | 34544937 | 33952 | 1 | 239839 | ENSMUST00000231364.2 | -8743 | ENSMUSG00000022833 | Ccdc14 | coiled-coil domain containing 14 |
| chr14 | 1,14E+08 | 1,14E+08 | 737 | * | 5,150535 | -5,479834942 | 1,961666 | -2,79346 | 0,005215 | 1 | 0 | Distal Intergenic | 14 | 1,15E+08 | 1,15E+08 | 29363 | 1 | 75822 | ENSMUST00000226121.2 | -1121906 | ENSMUSG00000115320 | 4930524C18Rik | RIKEN cDNA 4930524C18 gene |
| chr9 | 12762763 | 12763394 | 632 | * | 5,296851 | -5,479675669 | 2,019974 | -2,71275 | 0,006673 | 1 | 0 | Distal Intergenic | 9 | 13091272 | 13105860 | 14589 | 2 | 78219 | ENSMUST00000216694.2 | 342466 | ENSMUSG00000111830 | 4930568E12Rik | RIKEN cDNA 4930568E12 gene |
| chr2 | 71603948 | 71604577 | 630 | * | 4,474099 | -5,479381743 | 2,114903 | -2,59084 | 0,009574 | 1 | 0 | Intron (ENSMUST00000 | 2 | 71617273 | 71651287 | 34015 | 1 | 16403 | ENSMUST00000152009.2 | -12696 | ENSMUSG00000027111 | Itga6 | integrin alpha 6 |
| chr17 | 50076965 | 50077466 | 502 | * | 4,413866 | -5,479364728 | 2,120139 | -2,58444 | 0,009754 | 1 | 0 | Intron (ENSMUST00000 | 17 | 50060939 | 50066061 | 5123 | 1 | 319991 | ENSMUST00000162419.2 | 16026 | ENSMUSG00000023999 | Kif6 | kinesin family member 6 |
| chr1 | 1,64E+08 | 1,64E+08 | 548 | * | 6,348566 | -5,478602867 | 1,937955 | -2,827 | 0,004699 | 1 | 0 | Intron (ENSMUST00000 | 1 | 1,64E+08 | 1,64E+08 | 27482 | 1 | 1,19E+08 | ENSMUST00000194406.2 | 31438 | | Gm37754 | predicted gene, 37754 |
| chr8 | 1,09E+08 | 1,09E+08 | 656 | * | 5,816482 | -5,478500998 | 1,944157 | -2,81793 | 0,004833 | 1 | 0 | Distal Intergenic | 8 | 1,09E+08 | 1,1E+08 | 1018987 | 1 | 11906 | ENSMUST00000220518.2 | -122678 | ENSMUSG00000038872 | Zfhx3 | zinc finger homeobox 3 |
| chrX | 1,07E+08 | 1,07E+08 | 593 | * | 4,490737 | -5,473645814 | 2,085486 | -2,62464 | 0,008674 | 1 | 0 | Distal Intergenic | 20 | 1,07E+08 | 1,07E+08 | 80320 | 1 | 213449 | ENSMUST00000101292.9 | -32015 | ENSMUSG00000073007 | Tent5d | terminal nucleotidyltransferase 5D |
| chr11 | 38840117 | 38840728 | 612 | * | 4,578744 | -5,47349865 | 2,037235 | -2,68673 | 0,007216 | 1 | 0 | Distal Intergenic | 11 | 38688998 | 38689104 | 107 | 2 | 1,15E+08 | ENSMUST00000240209.1 | -151013 | ENSMUSG00000119681 | Gm23520 | predicted gene, 23520 |
| chr9 | 69594678 | 69595455 | 778 | * | 6,281756 | -5,472848294 | 1,887792 | -2,89907 | 0,003743 | 1 | 0 | Distal Intergenic | 9 | 69666022 | 69738127 | 72106 | 1 | 319970 | ENSMUST00000180488.8 | -70567 | ENSMUSG00000097623 | B230323A14Rik | RIKEN cDNA B230323A14 gene |
| chr6 | 94015596 | 94016558 | 963 | * | 6,333477 | -5,472810979 | 1,880986 | -2,90954 | 0,00362 | 1 | 0 | Intron (ENSMUST00000 | 6 | 93655017 | 93890671 | 235655 | 2 | 14924 | ENSMUST00000093769.8 | -124925 | ENSMUSG00000045095 | Magi1 | membrane associated guanylate kinase, WW and PDZ domain containing 1 |
| chr2 | 76711249 | 76711802 | 554 | * | 6,01273 | -5,464734545 | 1,944998 | -2,80963 | 0,00496 | 1 | 0 | Exon (ENSMUST00000 | 2 | 76719095 | 76719851 | 757 | 2 | 22138 | ENSMUST00000139832.2 | 8049 | ENSMUSG00000051747 | Ttn | titin |
| chrX | 49581169 | 49582057 | 889 | * | 5,50914 | -5,464675222 | 1,925767 | -2,83766 | 0,004545 | 1 | 0 | Distal Intergenic | 20 | 49626209 | 49651481 | 25273 | 1 | 1,03E+08 | ENSMUST00000241980.1 | -44152 | | Gm35612 | predicted gene, 35612 |
| chr13 | 23257481 | 23258011 | 531 | * | 6,110037 | -5,464579649 | 1,897455 | -2,87995 | 0,003977 | 1 | 0 | Promoter (1-2kb) | 13 | 23259913 | 23260960 | 1048 | 1 | 171253 | ENSMUST00000072972.5 | -1902 | ENSMUSG00000099917 | Vmn1r215 | vomeronasal 1 receptor 215 |
| chrX | 49562538 | 49563296 | 759 | * | 5,461244 | -5,463268143 | 1,943363 | -2,81124 | 0,004935 | 1 | 0 | Distal Intergenic | 20 | 49514308 | 49515459 | 1152 | 2 | 258289 | ENSMUST00000219631.2 | -47079 | ENSMUSG00000048173 | Olfr1324 | olfactory receptor 1324 |
| chr1 | 19048984 | 19049967 | 984 | * | 5,771691 | -5,463053496 | 1,969742 | -2,77349 | 0,005546 | 1 | 0 | Distal Intergenic | 1 | 19107490 | 19107601 | 112 | 1 | 1,08E+08 | ENSMUST00000185386.2 | -57523 | | Gm28343 | predicted gene 28343 |
| chr7 | 1,14E+08 | 1,14E+08 | 830 | * | 7,303983 | -5,462706044 | 1,902117 | -2,87191 | 0,00408 | 1 | 0 | Intron (ENSMUST00000 | 7 | 1,14E+08 | 1,14E+08 | 36905 | 1 | 233752 | ENSMUST00000206274.3 | 3512 | ENSMUSG00000048782 | Insc | INSC spindle orientation adaptor protein |
| chr15 | 98128616 | 98129330 | 715 | * | 6,207788 | -5,462291832 | 2,116158 | -2,58123 | 0,009845 | 1 | 0 | Promoter (2-3kb) | 15 | 98124873 | 98132131 | 7259 | 2 | 629524 | ENSMUST00000206621.2 | 2801 | ENSMUSG00000059460 | Olfr286 | olfactory receptor 286 |
| chr5 | 6692364 | 6692981 | 618 | * | 4,422498 | -5,462238735 | 2,102539 | -2,59793 | 0,009379 | 1 | 0 | Distal Intergenic | 5 | 6819030 | 6926523 | 107494 | 2 | 207618 | ENSMUST00000164784.2 | 233542 | ENSMUSG00000092094 | Zfp804b | zinc finger protein 804B |
| chr4 | 65857047 | 65857942 | 896 | * | 7,605092 | -5,460899558 | 2,025879 | -2,69557 | 0,007027 | 1 | 0 | Intron (ENSMUST00000 | 4 | 65523486 | 65531993 | 8508 | 1 | 69807 | ENSMUST00000156922.2 | 333561 | ENSMUSG00000051675 | Trim32 | tripartite motif-containing 32 |
| chr3 | 1,54E+08 | 1,54E+08 | 424 | * | 5,909222 | -5,458262653 | 1,98966 | -2,74331 | 0,006082 | 1 | 0 | Intron (ENSMUST00000 | 3 | 1,54E+08 | 1,54E+08 | 3044 | 1 | 242259 | ENSMUST00000128362.2 | -3409 | ENSMUSG00000028360 | Slc44a5 | solute carrier family 44, member 5 |

| | | | | | | | | | | | | | | | | | | | | | | | |
|---|---|---|---|---|---|---|---|---|---|---|---|---|---|---|---|---|---|---|---|---|---|---|---|
| chr18 | 85371748 | 85372338 | 591 | * | 5,435487 | -5,457695373 | 1,941029 | -2,81175 | 0,004927 | 1 | 0 | Distal Intergenic | 18 | 84983635 | 84999597 | 15963 | 1 | 50764 | ENSMUST00000225015.2 | 388113 | ENSMUSG00000034391 | Fbxo15 | F-box protein 15 |
| chr3 | 1,07E+08 | 1,07E+08 | 386 | * | 6,637391 | -5,456252767 | 1,919654 | -2,84231 | 0,004479 | 1 | 0 | Distal Intergenic | 3 | 1,07E+08 | 1,07E+08 | 21250 | 2 | 99738 | ENSMUST00000009617.10 | 60187 | ENSMUSG00000027895 | Kcnc4 | potassium voltage gated channel, Shaw-related subfamily, member 4 |
| chr18 | 3675563 | 3676028 | 466 | * | 5,993068 | -5,455428899 | 1,891922 | -2,88354 | 0,003932 | 1 | 0 | Distal Intergenic | 18 | 3507923 | 3516404 | 8482 | 1 | 68010 | ENSMUST00000025075.3 | 167640 | ENSMUSG00000024232 | Bambi | BMP and activin membrane-bound inhibitor |
| chr10 | 1,29E+08 | 1,29E+08 | 508 | * | 6,250238 | -5,451916704 | 2,02293 | -2,69506 | 0,007038 | 1 | 0 | Distal Intergenic | 10 | 1,29E+08 | 1,29E+08 | 954 | 2 | 258540 | ENSMUST00000078914.3 | 7210 | ENSMUSG00000061367 | Olfr771 | olfactory receptor 771 |
| chrX | 50852240 | 50853562 | 1323 | * | 5,839786 | -5,451417111 | 2,054701 | -2,65314 | 0,007975 | 1 | 0 | Intron (ENSMUST0000 | 20 | 50842836 | 50890021 | 47186 | 2 | 83563 | ENSMUST00000069509.4 | 36459 | ENSMUSG00000055780 | Usp26 | ubiquitin specific peptidase 26 |
| chr11 | 1,13E+08 | 1,13E+08 | 340 | * | 5,351645 | -5,450962714 | 2,087565 | -2,61116 | 0,009024 | 1 | 0 | Intron (ENSMUST0000 | 11 | 1,13E+08 | 1,13E+08 | 3690 | 2 | 319871 | ENSMUST00000242930.1 | -158262 | ENSMUSG00000085421 | 4732490B19Rik | RIKEN cDNA 4732490B19 gene |
| chr3 | 15919375 | 15920269 | 895 | * | 8,98531 | -5,450788366 | 1,702727 | -3,20121 | 0,001369 | 0,533987851 | 1,449706 | Distal Intergenic | 3 | 15849309 | 15902692 | 53384 | 2 | 1E+08 | ENSMUST00000050623.11 | -16683 | ENSMUSG00000074677 | Sirpb1c | signal-regulatory protein beta 1C |
| chr1 | 1,56E+08 | 1,56E+08 | 461 | * | 5,057164 | -5,450203574 | 2,052953 | -2,65481 | 0,007935 | 1 | 0 | Intron (ENSMUST0000 | 1 | 1,56E+08 | 1,56E+08 | 43997 | 2 | 16872 | ENSMUST00000027740.14 | 58238 | ENSMUSG00000026468 | Lhx4 | LIM homeobox protein 4 |
| chr11 | 72531176 | 72531918 | 743 | * | 5,222453 | -5,447765389 | 1,984517 | -2,74513 | 0,006049 | 1 | 0 | Intron (ENSMUST0000 | 11 | 72506832 | 72506976 | 145 | 2 | 1,15E+08 | ENSMUST00000082633.3 | -24200 | ENSMUSG00000064567 | Gm24143 | predicted gene, 24143 |
| chr6 | 1,32E+08 | 1,32E+08 | 617 | * | 3,988252 | -5,447703986 | 2,108066 | -2,58422 | 0,00976 | 1 | 0 | Distal Intergenic | 6 | 1,32E+08 | 1,32E+08 | 3087 | 2 | 1,01E+08 | ENSMUST00000100870.6 | 36517 | ENSMUSG00000079276 | Gm4736 | predicted gene 4736 |
| chr3 | 62374691 | 62375436 | 746 | * | 5,641317 | -5,446426424 | 1,938738 | -2,80926 | 0,004966 | 1 | 0 | 3' UTR | 3 | 62378066 | 62380008 | 1943 | 2 | 72162 | ENSMUST00000160858.2 | 4572 | ENSMUSG00000027770 | Dhx36 | DEAH (Asp-Glu-Ala-His) box polypeptide 36 |
| chr4 | 1,5E+08 | 1,5E+08 | 1030 | * | 6,351538 | -5,446337826 | 1,946178 | -2,79848 | 0,005134 | 1 | 0 | Distal Intergenic | 4 | 1,5E+08 | 1,51E+08 | 340321 | 1 | 68703 | ENSMUST00000105682.9 | -4172 | ENSMUSG00000039852 | Rere | arginine glutamic acid dipeptide (RE) repeats |
| chr6 | 1,19E+08 | 1,19E+08 | 638 | * | 4,272039 | -5,444939509 | 2,094337 | -2,59984 | 0,009327 | 1 | 0 | Intron (ENSMUST0000 | 6 | 1,19E+08 | 1,19E+08 | 188006 | 2 | 12288 | ENSMUST00000220022.2 | -76084 | ENSMUSG00000051331 | Cacna1c | calcium channel, voltage-dependent, L type, alpha 1C subunit |
| chr12 | 74840079 | 74840675 | 597 | * | 4,168574 | -5,441787921 | 2,103843 | -2,58659 | 0,009693 | 1 | 0 | Distal Intergenic | 12 | 74872119 | 74874496 | 2378 | 1 | 1,05E+08 | ENSMUST00000223498.2 | -31444 | | Gm39473 | predicted gene, 39473 |
| chrX | 72245484 | 72246671 | 1188 | * | 6,866242 | -5,441702492 | 1,926435 | -2,82475 | 0,004732 | 1 | 0 | 3' UTR | 20 | 72235828 | 72246526 | 10699 | 1 | 574437 | ENSMUST00000101486.5 | 9656 | ENSMUSG00000073125 | Xlr3b | X-linked lymphocyte-regulated 3B |
| chr2 | 1,74E+08 | 1,74E+08 | 515 | * | 4,9019 | -5,44110143 | 1,996039 | -2,72595 | 0,006412 | 1 | 0 | Distal Intergenic | 2 | 1,74E+08 | 1,74E+08 | 9926 | 1 | 56491 | ENSMUST00000154306.2 | 84577 | ENSMUSG00000054455 | Vapb | vesicle-associated membrane protein, associated protein B and C |
| chr13 | 69126364 | 69127076 | 713 | * | 7,379673 | -5,439514719 | 1,848145 | -2,94323 | 0,003248 | 1 | 0 | Intron (ENSMUST0000 | 13 | 69146912 | 69179593 | 32682 | 1 | 1,03E+08 | ENSMUST00000221765.2 | -19836 | | Gm35161 | predicted gene, 35161 |
| chr17 | 3600877 | 3601574 | 698 | * | 5,907445 | -5,439341269 | 1,912311 | -2,84438 | 0,00445 | 1 | 0 | Exon (ENSMUST00000 | 17 | 3607803 | 3610141 | 2339 | 1 | 68230 | ENSMUST00000241894.1 | -6229 | ENSMUSG00000116852 | 1700102H20Rik | RIKEN cDNA 1700102H20 gene |
| chr14 | 76197502 | 76198774 | 1273 | * | 5,233653 | -5,439234621 | 1,953853 | -2,78385 | 0,005372 | 1 | 0 | Intron (ENSMUST0000 | 14 | 76192449 | 76203646 | 11198 | 1 | 67516 | ENSMUST00000050120.4 | 5053 | ENSMUSG00000046523 | Kctd4 | potassium channel tetramerisation domain containing 4 |
| chrX | 20090467 | 20091254 | 788 | * | 4,64746 | -5,439098925 | 2,017262 | -2,69628 | 0,007012 | 1 | 0 | Intron (ENSMUST0000 | 20 | 19971993 | 20028439 | 56447 | 2 | 236727 | ENSMUST00000224372.2 | -62028 | ENSMUSG00000037341 | Slc9a7 | solute carrier family 9 (sodium/hydrogen exchanger), member 7 |
| chr9 | 29738844 | 29739300 | 457 | * | 4,787097 | -5,439019718 | 2,063417 | -2,63593 | 0,008391 | 1 | 0 | Intron (ENSMUST0000 | 9 | 29322884 | 29874415 | 551532 | 2 | 235106 | ENSMUST00000146651.2 | 135115 | ENSMUSG00000059974 | Ntm | neurotrimin |
| chr15 | 58637891 | 58638226 | 336 | * | 5,729128 | -5,437721442 | 1,953869 | -2,78305 | 0,005385 | 1 | 0 | Distal Intergenic | 15 | 58691172 | 58691327 | 156 | 2 | 1,15E+08 | ENSMUST00000083115.4 | 53101 | ENSMUSG00000065049 | Gm24041 | predicted gene, 24041 |
| chrX | 1,17E+08 | 1,17E+08 | 824 | * | 5,169581 | -5,436330837 | 2,080342 | -2,61319 | 0,00897 | 1 | 0 | Distal Intergenic | 20 | 1,17E+08 | 1,17E+08 | 1030 | 1 | 1E+08 | ENSMUST00000072518.8 | -264278 | ENSMUSG00000100194 | Tgif2lx2 | TGFB-induced factor homeobox 2-like, X-linked 2 |
| chr10 | 63704690 | 63705460 | 771 | * | 4,513773 | -5,436304591 | 2,083373 | -2,60938 | 0,009071 | 1 | 0 | Intron (ENSMUST0000 | 10 | 63656834 | 63913266 | 256433 | 1 | 216033 | ENSMUST00000135474.2 | 47856 | ENSMUSG00000060843 | Ctnna3 | catenin (cadherin associated protein), alpha 3 |
| chr18 | 40048607 | 40049119 | 513 | * | 5,15879 | -5,435443806 | 2,072202 | -2,62303 | 0,008715 | 1 | 0 | Intron (ENSMUST0000 | 18 | 40039233 | 40062427 | 23195 | 2 | 1,05E+08 | ENSMUST00000237624.2 | 13308 | | Gm41708 | predicted gene, 41708 |
| chr4 | 1,38E+08 | 1,38E+08 | 1008 | * | 6,729096 | -5,432819221 | 1,912637 | -2,84049 | 0,004504 | 1 | 0 | Intron (ENSMUST0000 | 4 | 1,38E+08 | 1,38E+08 | 924 | 2 | 73563 | ENSMUST00000124620.2 | 62779 | ENSMUSG00000084804 | 1700095J12Rik | RIKEN cDNA 1700095J12 gene |
| chr19 | 8089771 | 8090674 | 904 | * | 5,50204 | -5,432707899 | 2,045279 | -2,65622 | 0,007902 | 1 | 0 | Intron (ENSMUST0000 | 19 | 8039574 | 8109346 | 69773 | 2 | 434674 | ENSMUST00000065651.5 | 18672 | ENSMUSG00000063590 | Slc22a28 | solute carrier family 22, member 28 |
| chr16 | 20076034 | 20076735 | 702 | * | 5,227472 | -5,432656143 | 2,035733 | -2,66865 | 0,007616 | 1 | 0 | Distal Intergenic | 16 | 20052053 | 20060108 | 8056 | 2 | 208158 | ENSMUST00000040880.9 | -15926 | ENSMUSG00000041205 | Map6d1 | MAP6 domain containing 1 |
| chr13 | 43802798 | 43803483 | 686 | * | 5,795805 | -5,430783309 | 1,922392 | -2,82501 | 0,004728 | 1 | 0 | Intron (ENSMUST0000 | 13 | 43769459 | 43821542 | 52084 | 1 | 328234 | ENSMUST00000161817.2 | 33339 | ENSMUSG00000044164 | Rnf182 | ring finger protein 182 |
| chr11 | 8333394 | 8334110 | 717 | * | 5,911629 | -5,430749984 | 1,929616 | -2,81442 | 0,004887 | 1 | 0 | Exon (ENSMUST00000 | 11 | 8401070 | 8418675 | 17606 | 2 | 319939 | ENSMUST00000129074.2 | 84565 | ENSMUSG00000020422 | Tns3 | tensin 3 |
| chr2 | 73099288 | 73099914 | 627 | * | 4,642878 | -5,430298995 | 2,050059 | -2,64885 | 0,008077 | 1 | 0 | Promoter (2-3kb) | 2 | 73102269 | 73107815 | 5547 | 1 | 381373 | ENSMUST00000090813.6 | -2355 | ENSMUSG00000068859 | Sp9 | trans-acting transcription factor 9 |
| chr2 | 3098648 | 3099549 | 902 | * | 5,677221 | -5,429001257 | 1,96361 | -2,76481 | 0,005696 | 1 | 0 | Distal Intergenic | 2 | 3115261 | 3228287 | 113027 | 1 | 269233 | ENSMUST00000072955.12 | -15712 | ENSMUSG00000050530 | Fam171a1 | family with sequence similarity 171, member A1 |
| chr14 | 73841403 | 73842133 | 731 | * | 4,336807 | -5,427362273 | 2,101333 | -2,58282 | 0,0098 | 1 | 0 | Distal Intergenic | 14 | 73827948 | 73828473 | 526 | 1 | 20916 | ENSMUST00000161082.2 | 13455 | ENSMUSG00000022110 | Sucla2 | succinate-Coenzyme A ligase, ADP-forming, beta subunit |
| chrX | 1,05E+08 | 1,05E+08 | 801 | * | 4,972303 | -5,426253436 | 2,041692 | -2,65772 | 0,007867 | 1 | 0 | Distal Intergenic | 20 | 1,05E+08 | 1,05E+08 | 16600 | 1 | 18655 | ENSMUST00000081593.13 | -19627 | ENSMUSG00000062070 | Pgk1 | phosphoglycerate kinase 1 |
| chr7 | 83004905 | 83005463 | 559 | * | 4,32949 | -5,423984421 | 2,063392 | -2,62867 | 0,008572 | 1 | 0 | Promoter (<=1kb) | 7 | 82976946 | 83005785 | 28840 | 2 | 1,03E+08 | ENSMUST00000208251.2 | 322 | | Gm29976 | predicted gene, 29976 |
| chr1 | 1,48E+08 | 1,48E+08 | 553 | * | 6,097834 | -5,423679562 | 1,992074 | -2,72263 | 0,006476 | 1 | 0 | Distal Intergenic | 1 | 1,48E+08 | 1,48E+08 | 129 | 2 | 1,15E+08 | ENSMUST00000157957.3 | 188564 | ENSMUSG00000088582 | Gm22966 | predicted gene, 22966 |
| chr17 | 28281753 | 28282433 | 681 | * | 7,058768 | -5,423380758 | 1,934627 | -2,80332 | 0,005058 | 1 | 0 | Downstream (<=300bp | 17 | 28285771 | 28287450 | 1680 | 2 | 21463 | ENSMUST00000123628.2 | 5017 | ENSMUSG00000062859 | Tcp11 | t-complex protein 11 |
| chr19 | 28222228 | 28223012 | 785 | * | 6,643509 | -5,42293451 | 1,869595 | -2,90059 | 0,003725 | 1 | 0 | Distal Intergenic | 19 | 28241351 | 28261306 | 19956 | 2 | 226075 | ENSMUST00000159639.2 | 38294 | ENSMUSG00000052942 | Glis3 | GLIS family zinc finger 3 |
| chr4 | 1,36E+08 | 1,36E+08 | 999 | * | 6,352349 | -5,42292195 | 1,868058 | -2,90297 | 0,003696 | 1 | 0 | Distal Intergenic | 4 | 1,36E+08 | 1,36E+08 | 2933 | 2 | 52830 | ENSMUST00000030436.12 | -6142 | ENSMUSG00000028675 | Pnrc2 | proline-rich nuclear receptor coactivator 2 |
| chr10 | 72411935 | 72412791 | 857 | * | 6,627741 | -5,422910675 | 1,83615 | -2,95341 | 0,003143 | 1 | 0 | Distal Intergenic | 10 | 72363382 | 72363514 | 133 | 1 | 1,15E+08 | ENSMUST00000104454.3 | 48553 | ENSMUSG00000077642 | Gm24984 | predicted gene, 24984 |
| chr9 | 14895722 | 14896511 | 790 | * | 6,420733 | -5,422649774 | 1,864745 | -2,90798 | 0,003638 | 1 | 0 | Distal Intergenic | 9 | 14919214 | 14921804 | 2591 | 2 | 55991 | ENSMUST00000166933.2 | 25293 | ENSMUSG00000031934 | Panx1 | pannexin 1 |
| chr19 | 30876196 | 30877223 | 1028 | * | 7,426182 | -5,422245651 | 1,871346 | -2,89751 | 0,003761 | 1 | 0 | Intron (ENSMUST0000 | 19 | 30686285 | 30758852 | 72568 | 2 | 19091 | ENSMUST00000182401.2 | -117344 | ENSMUSG00000052920 | Prkg1 | protein kinase, cGMP-dependent, type I |
| chr3 | 86819857 | 86820864 | 1008 | * | 6,792197 | -5,421788132 | 1,994282 | -2,71867 | 0,006555 | 1 | 0 | Intron (ENSMUST0000 | 3 | 86694519 | 86827477 | 132959 | 2 | 70762 | ENSMUST00000193632.6 | 6613 | ENSMUSG00000028078 | Dclk2 | doublecortin-like kinase 2 |
| chr12 | 1,16E+08 | 1,16E+08 | 556 | * | 4,781379 | -5,421669979 | 2,017324 | -2,68755 | 0,007198 | 1 | 0 | Intron (ENSMUST0000 | 12 | 1,16E+08 | 1,16E+08 | 268 | 2 | 627375 | ENSMUST00000189273.2 | -12195 | | Gm6750 | predicted gene 6750 |
| chr12 | 39301368 | 39302010 | 643 | * | 4,585632 | -5,421426068 | 2,072357 | -2,61607 | 0,008895 | 1 | 0 | Distal Intergenic | 12 | 39111906 | 39204760 | 92855 | 2 | 1,03E+08 | ENSMUST00000222945.2 | -96608 | ENSMUSG00000113272 | Gm34809 | predicted gene, 34809 |
| chrX | 1,63E+08 | 1,63E+08 | 602 | * | 5,458061 | -5,419221538 | 2,041764 | -2,65419 | 0,00795 | 1 | 0 | Promoter (1-2kb) | 20 | 1,63E+08 | 1,63E+08 | 57014 | 2 | 12169 | ENSMUST00000112263.2 | -1635 | ENSMUSG00000031377 | Bmx | BMX non-receptor tyrosine kinase |
| chr9 | 94036347 | 94036704 | 358 | * | 4,613882 | -5,418750115 | 2,081048 | -2,60386 | 0,009218 | 1 | 0 | Distal Intergenic | 9 | 94166549 | 94166705 | 157 | 1 | 1,15E+08 | ENSMUST00000082955.3 | -129845 | ENSMUSG00000064889 | Gm24200 | predicted gene, 24200 |
| chrX | 1,46E+08 | 1,46E+08 | 797 | * | 5,210425 | -5,416847482 | 1,995464 | -2,71458 | 0,006636 | 1 | 0 | Distal Intergenic | 20 | 1,46E+08 | 1,46E+08 | 83702 | 2 | 210297 | ENSMUST00000136789.8 | -37525 | ENSMUSG00000031290 | Lrch2 | leucine-rich repeats and calponin homology (CH) domain containing 2 |
| chr16 | 36971454 | 36972578 | 1125 | * | 8,609613 | -5,416028458 | 1,741929 | -3,10921 | 0,001876 | 0,512062732 | 2,237403 | Intron (ENSMUST0000 | 16 | 36977114 | 37024938 | 47825 | 2 | 207227 | ENSMUST00000149984.2 | 52360 | ENSMUSG00000022829 | Stxbp5l | syntaxin binding protein 5-like |
| chr3 | 1,35E+08 | 1,35E+08 | 1113 | * | 12,03999 | -5,415679777 | 1,622125 | -3,33863 | 0,000842 | 0,472504624 | 1,2763 | Distal Intergenic | 3 | 1,35E+08 | 1,35E+08 | 80958 | 2 | 18033 | ENSMUST00000138602.3 | -62527 | ENSMUSG00000028163 | Nfkb1 | nuclear factor of kappa light polypeptide gene enhancer in B cells 1, p105 |
| chr12 | 3339383 | 3340180 | 798 | * | 4,994399 | -5,415113399 | 2,05768 | -2,63166 | 0,008497 | 1 | 0 | Intron (ENSMUST0000 | 12 | 3297428 | 3359969 | 62542 | 2 | 19325 | ENSMUST00000021001.10 | 19789 | ENSMUSG00000020671 | Rab10 | RAB10, member RAS oncogene family |
| chr14 | 4545661 | 4546383 | 723 | * | 4,747892 | -5,414907593 | 1,987583 | -2,72437 | 0,006442 | 1 | 0 | Intron (ENSMUST0000 | 14 | 4458435 | 4506875 | 48441 | 2 | 21834 | ENSMUST00000224934.2 | -38786 | ENSMUSG00000021779 | Thrb | thyroid hormone receptor beta |
| chr14 | 66994963 | 66995634 | 672 | * | 5,041548 | -5,414857516 | 1,973265 | -2,74411 | 0,006068 | 1 | 0 | Intron (ENSMUST0000 | 14 | 67040313 | 67106137 | 65825 | 2 | 12934 | ENSMUST00000022629.9 | 110503 | ENSMUSG00000022048 | Dpysl2 | dihydropyrimidinase-like 2 |
| chr3 | 61822342 | 61822917 | 576 | * | 6,232273 | -5,414477677 | 1,882643 | -2,876 | 0,004028 | 1 | 0 | Distal Intergenic | 3 | 62245765 | 62369642 | 123878 | 1 | 622434 | ENSMUST00000079300.13 | -422848 | ENSMUSG00000036885 | Arhgef26 | Rho guanine nucleotide exchange factor (GEF) 26 |
| chr8 | 57255350 | 57255857 | 508 | * | 6,327043 | -5,41363214 | 1,917854 | -2,82276 | 0,004761 | 1 | 0 | Distal Intergenic | 8 | 57325115 | 57341925 | 16811 | 2 | 1,03E+08 | ENSMUST00000209983.2 | 86068 | ENSMUSG00000110000 | Gm33501 | predicted gene, 33501 |
| chr5 | 89752093 | 89753132 | 1040 | * | 8,682068 | -5,413581755 | 1,755206 | -3,0843 | 0,00204 | 0,528429778 | 2,237403 | Distal Intergenic | 5 | 89675288 | 89731599 | 56312 | 1 | 104443 | ENSMUST00000048557.3 | 76805 | ENSMUSG00000035528 | Npffr2 | neuropeptide FF receptor 2 |
| chr4 | 1,44E+08 | 1,44E+08 | 250 | * | 4,642527 | -5,412602294 | 2,066264 | -2,61951 | 0,008806 | 1 | 0 | Distal Intergenic | 4 | 1,44E+08 | 1,44E+08 | 6512 | 2 | 626943 | ENSMUST00000105762.3 | -4895 | ENSMUSG00000078509 | Pramel14 | PRAME like 14 |
| chr12 | 1,03E+08 | 1,03E+08 | 1707 | * | 14,24638 | -5,410075857 | 1,627564 | -3,32403 | 0,000887 | 0,579872171 | 0,738441 | Intron (ENSMUST0000 | 12 | 1,03E+08 | 1,03E+08 | 11766 | 1 | 217843 | ENSMUST00000178467.2 | 8819 | ENSMUSG00000021198 | Unc79 | unc-79 homolog |
| chr9 | 46689875 | 46690374 | 500 | * | 5,813635 | -5,408473351 | 2,036196 | -2,65617 | 0,007903 | 1 | 0 | Distal Intergenic | 9 | 46741637 | 46748914 | 7278 | 1 | 1,03E+08 | ENSMUST00000215504.2 | -51263 | | Gm31614 | predicted gene, 31614 |
| chr3 | 1,09E+08 | 1,09E+08 | 1135 | * | 7,508264 | -5,407962722 | 1,739113 | -3,10961 | 0,001873 | 1 | 0 | Distal Intergenic | 3 | 1,09E+08 | 1,1E+08 | 345046 | 1 | 57257 | ENSMUST00000046864.14 | -17074 | ENSMUSG00000033721 | Vav3 | vav 3 oncogene |
| chr4 | 1,29E+08 | 1,29E+08 | 589 | * | 7,583602 | -5,406326096 | 1,881145 | -2,87396 | 0,004054 | 1 | 0 | Promoter (<=1kb) | 4 | 1,29E+08 | 1,29E+08 | 5917 | 1 | 66260 | ENSMUST00000030577.11 | 30 | ENSMUSG00000028786 | Tmem54 | transmembrane protein 54 |
| chr8 | 32998482 | 32999375 | 894 | * | 4,313758 | -5,40623166 | 2,065638 | -2,61722 | 0,008865 | 1 | 0 | Intron (ENSMUST0000 | 8 | 32339339 | 32782941 | 443603 | 2 | 211323 | ENSMUST00000238824.2 | -215541 | ENSMUSG00000062991 | Nrg1 | neuregulin 1 |
| chr12 | 1,1E+08 | 1,1E+08 | 831 | * | 5,913189 | -5,406163673 | 2,016665 | -2,68074 | 0,007346 | 1 | 0 | Distal Intergenic | 12 | 1,1E+08 | 1,1E+08 | 375 | 1 | 629595 | ENSMUST00000222676.2 | 58340 | | Gm6988 | predicted gene 6988 |
| chr1 | 58238215 | 58239385 | 1171 | * | 6,119347 | -5,405817426 | 1,859463 | -2,90719 | 0,003647 | 1 | 0 | 3' UTR | 1 | 58249556 | 58307756 | 58201 | 1 | 71872 | ENSMUST00000040442.6 | -10171 | ENSMUSG00000038242 | Aox4 | aldehyde oxidase 4 |
| chr1 | 1,71E+08 | 1,71E+08 | 549 | * | 5,816659 | -5,40509469 | 1,909101 | -2,83122 | 0,004637 | 1 | 0 | Promoter (<=1kb) | 1 | 1,71E+08 | 1,71E+08 | 4523 | 1 | 1,03E+08 | ENSMUST00000194503.2 | 987 | | Gm10522 | predicted gene 10522 |
| chr10 | 24483567 | 24484272 | 706 | * | 5,809007 | -5,404859478 | 1,929766 | -2,80079 | 0,005098 | 1 | 0 | Distal Intergenic | 10 | 24425265 | 24471747 | 46483 | 2 | 1,03E+08 | ENSMUST00000136908.9 | -11820 | | Gm15270 | predicted gene 15270 |
| chrX | 88517989 | 88518955 | 967 | * | 5,332268 | -5,404656764 | 1,949963 | -2,77167 | 0,005577 | 1 | 0 | Distal Intergenic | 20 | 88447454 | 88453295 | 5842 | 2 | 18630 | ENSMUST00000113960.3 | -64694 | ENSMUSG00000035395 | Dcaf8l | DDB1 and CUL4 associated factor 8 like |
| chr10 | 1,16E+08 | 1,16E+08 | 442 | * | 5,408342 | -5,404380741 | 1,979227 | -2,73055 | 0,006323 | 1 | 0 | Distal Intergenic | 10 | 1,15E+08 | 1,15E+08 | 137470 | 2 | 14160 | ENSMUST00000020350.15 | -86725 | ENSMUSG00000020140 | Lgr5 | leucine rich repeat containing G protein coupled receptor 5 |
| chr13 | 48887759 | 48888232 | 474 | * | 5,782965 | -5,404324965 | 1,98603 | -2,72117 | 0,006505 | 1 | 0 | Distal Intergenic | 13 | 48816474 | 48819983 | 3510 | 1 | 12022 | ENSMUST00000021813.5 | 71285 | ENSMUSG00000021381 | Barx1 | BarH-like homeobox 1 |
| chr3 | 30702978 | 30703719 | 742 | * | 6,030452 | -5,399133553 | 1,994079 | -2,70758 | 0,006778 | 1 | 0 | Promoter (<=1kb) | 3 | 30702035 | 30726448 | 24414 | 1 | 68307 | ENSMUST00000108265.3 | 943 | ENSMUSG00000027703 | Lrriq4 | leucine-rich repeats and IQ motif containing 4 |
| chr14 | 81851899 | 81853034 | 1136 | * | 6,087749 | -5,397686747 | 1,883007 | -2,86653 | 0,00415 | 1 | 0 | Distal Intergenic | 14 | 81923250 | 81958568 | 35319 | 1 | 1,05E+08 | ENSMUST00000227374.2 | -70216 | | Gm41219 | predicted gene, 41219 |
| chr7 | 1,43E+08 | 1,43E+08 | 1007 | * | 6,937058 | -5,397290582 | 1,859695 | -2,90224 | 0,003705 | 1 | 0 | Distal Intergenic | 7 | 1,43E+08 | 1,43E+08 | 94 | 1 | 1,02E+08 | ENSMUST00000183699.3 | 9460 | ENSMUSG00000099321 | Mir6404 | microRNA 6404 |
| chrX | 82171363 | 82172185 | 823 | * | 4,55431 | -5,397278339 | 2,093881 | -2,57764 | 0,009948 | 1 | 0 | Intron (ENSMUST0000 | 20 | 82087008 | 82773937 | 686930 | 1 | 13405 | ENSMUST00000147740.2 | 84355 | ENSMUSG00000045103 | Dmd | dystrophin, muscular dystrophy |
| chrX | 1,51E+08 | 1,51E+08 | 580 | * | 5,116083 | -5,397261936 | 1,944025 | -2,77633 | 0,005498 | 1 | 0 | Distal Intergenic | 20 | 1,51E+08 | 1,51E+08 | 28844 | 2 | 72997 | ENSMUST00000149098.8 | -4700 | ENSMUSG00000087403 | Kantr | Kdm5c adjacent non-coding transcript |
| chr2 | 73084555 | 73085517 | 963 | * | 7,088011 | -5,396813176 | 1,834676 | -2,94156 | 0,003266 | 0,698702076 | 1,360622 | Distal Intergenic | 2 | 73102269 | 73107815 | 5547 | 1 | 381373 | ENSMUST00000090813.6 | -16752 | ENSMUSG00000068859 | Sp9 | trans-acting transcription factor 9 |
| chr9 | 29588152 | 29588763 | 612 | * | 5,159825 | -5,395701684 | 1,967507 | -2,7424 | 0,006099 | 1 | 0 | Intron (ENSMUST0000 | 9 | 29501780 | 29503806 | 2027 | 2 | 1,05E+08 | ENSMUST00000141752.2 | -84346 | | Gm15521 | predicted gene 15521 |
| chr5 | 67052694 | 67053202 | 509 | * | 5,060525 | -5,390558604 | 2,036442 | -2,64705 | 0,00812 | 1 | 0 | Promoter (2-3kb) | 5 | 67055774 | 67126496 | 70723 | 1 | 77569 | ENSMUST00000122812.5 | -2572 | ENSMUSG00000037736 | Limch1 | LIM and calponin homology domains 1 |
| chr7 | 1,06E+08 | 1,06E+08 | 820 | * | 6,885324 | -5,390526075 | 2,0309 | -2,65426 | 0,007948 | 0,85617852 | 1,360622 | Distal Intergenic | 7 | 1,06E+08 | 1,06E+08 | 136 | 1 | 1,15E+08 | ENSMUST00000104589.3 | 94465 | ENSMUSG00000077777 | Gm22372 | predicted gene, 22372 |
| chr2 | 1,2E+08 | 1,2E+08 | 919 | * | 5,346561 | -5,390041536 | 1,975515 | -2,72842 | 0,006364 | 1 | 0 | Promoter (<=1kb) | 2 | 1,2E+08 | 1,2E+08 | 1819 | 1 | 29808 | ENSMUST00000131396.2 | -421 | ENSMUSG00000033943 | Mga | MAX gene associated |
| chr10 | 32311481 | 32311779 | 299 | * | 5,478854 | -5,38977457 | 1,948935 | -2,7655 | 0,005684 | 1 | 0 | Intron (ENSMUST0000 | 10 | 32205761 | 32286331 | 80571 | 2 | 432450 | ENSMUST00000092603.5 | -25150 | ENSMUSG00000069670 | Nkain2 | Na+/K+ transporting ATPase interacting 2 |
| chr6 | 1,46E+08 | 1,46E+08 | 662 | * | 6,538931 | -5,389376965 | 1,913956 | -2,81583 | 0,004865 | 1 | 0 | Intron (ENSMUST0000 | 6 | 1,46E+08 | 1,46E+08 | 3799 | 2 | 16439 | ENSMUST00000144557.2 | -43607 | ENSMUSG00000030287 | Itpr2 | inositol 1,4,5-triphosphate receptor 2 |
| chr12 | 1,05E+08 | 1,05E+08 | 751 | * | 6,372811 | -5,388408422 | 1,97314 | -2,73088 | 0,006317 | 1 | 0 | Distal Intergenic | 12 | 1,05E+08 | 1,05E+08 | 8376 | 2 | 1E+08 | ENSMUST00000222652.2 | -22616 | ENSMUSG00000113314 | Gm2721 | predicted gene 2721 |
| chr15 | 5609130 | 5609635 | 506 | * | 4,480483 | -5,388300486 | 2,04502 | -2,63484 | 0,008418 | 1 | 0 | Intron (ENSMUST0000 | 15 | 5525799 | 5624464 | 98666 | 1 | 71432 | ENSMUST00000228069.2 | 83331 | ENSMUSG00000110018 | 5430437J10Rik | RIKEN cDNA 5430437J10 gene |
| chrX | 95233086 | 95233861 | 776 | * | 4,662216 | -5,388287422 | 2,059693 | -2,61606 | 0,008895 | 1 | 0 | Distal Intergenic | 20 | 95237657 | 95238094 | 438 | 1 | 1,01E+08 | ENSMUST00000120611.2 | -3796 | ENSMUSG00000081848 | Gm14925 | predicted gene 14925 |
| chr1 | 89499621 | 89499921 | 301 | * | 4,326774 | -5,388261462 | 2,079611 | -2,59099 | 0,00957 | 1 | 0 | Intron (ENSMUST0000 | 1 | 89507958 | 89822999 | 315042 | 1 | 347722 | ENSMUST00000074945.8 | -8037 | ENSMUSG00000055013 | Agap1 | ArfGAP with GTPase domain, ankyrin repeat and PH domain 1 |
| chr10 | 1,03E+08 | 1,03E+08 | 695 | * | 4,350143 | -5,388254574 | 2,08275 | -2,58709 | 0,009679 | 1 | 0 | Distal Intergenic | 10 | 1,03E+08 | 1,03E+08 | 14424 | 2 | 216285 | ENSMUST00000164226.3 | 129593 | ENSMUSG00000036602 | Alx1 | ALX homeobox 1 |
| chr16 | 5614035 | 5614949 | 915 | * | 8,604814 | -5,386917617 | 1,77084 | -3,04201 | 0,00235 | 1 | 0 | Distal Intergenic | 16 | 5703219 | 7170909 | 1467691 | 1 | 268859 | ENSMUST00000230658.2 | -88270 | ENSMUSG00000008658 | Rbfox1 | RNA binding protein, fox-1 homolog (C. elegans) 1 |
| chr5 | 79933079 | 79933834 | 756 | * | 5,426373 | -5,385845233 | 1,996541 | -2,69759 | 0,006984 | 1 | 0 | Distal Intergenic | 5 | 80246755 | 80246861 | 107 | 1 | 1,15E+08 | ENSMUST00000239632.1 | -312921 | ENSMUSG00000119601 | Gm24304 | predicted gene, 24304 |
| chr4 | 1,25E+08 | 1,25E+08 | 838 | * | 4,984329 | -5,385787885 | 2,065917 | -2,60697 | 0,009135 | 1 | 0 | Intron (ENSMUST0000 | 4 | 1,25E+08 | 1,25E+08 | 20995 | 1 | 230737 | ENSMUST00000106176.8 | 4607 | ENSMUSG00000028869 | Gnl2 | guanine nucleotide binding protein-like 2 (nucleolar) |
| chr5 | 73003430 | 73003878 | 449 | * | 4,594424 | -5,381744882 | 2,070108 | -2,59974 | 0,009329 | 1 | 0 | Promoter (1-2kb) | 5 | 72980805 | 73001792 | 20988 | 2 | 21682 | ENSMUST00000202547.2 | -1638 | ENSMUSG00000029217 | Tec | tec protein tyrosine kinase |
| chr8 | 1,11E+08 | 1,11E+08 | 834 | * | 9,60924 | -5,38174425 | 1,795419 | -2,99749 | 0,002722 | 0,662625192 | 1,449706 | Intron (ENSMUST0000 | 8 | 1,11E+08 | 1,11E+08 | 6513 | 1 | 11765 | ENSMUST00000172892.2 | 5063 | ENSMUSG00000031731 | Ap1g1 | adaptor protein complex AP-1, gamma 1 subunit |
| chr2 | 1,35E+08 | 1,35E+08 | 527 | * | 4,6615 | -5,381516608 | 2,032654 | -2,64753 | 0,008108 | 1 | 0 | Promoter (1-2kb) | 2 | 1,35E+08 | 1,35E+08 | 5541 | 1 | 1,03E+08 | ENSMUST00000247656.1 | -1358 | ENSMUSG00000084762 | Platr3 | pluripotency associated transcript 3 |
| chr1 | 1,82E+08 | 1,82E+08 | 1156 | * | 8,883372 | -5,381134753 | 1,815673 | -2,96371 | 0,00304 | 0,604345495 | 2,237403 | Distal Intergenic | 1 | 1,82E+08 | 1,82E+08 | 7183 | 1 | 27058 | ENSMUST00000111018.2 | -5757 | ENSMUSG00000026511 | Srp9 | signal recognition particle 9 |
| chr11 | 12278490 | 12278944 | 455 | * | 6,249042 | -5,380447129 | 1,903368 | -2,8268 | 0,004702 | 1 | 0 | Intron (ENSMUST0000 | 11 | 12272275 | 12301028 | 28754 | 1 | 1,01E+08 | ENSMUST00000174550.2 | 6215 | | Gm12002 | predicted gene 12002 |
| chr19 | 23738732 | 23739488 | 757 | * | 7,184626 | -5,379594038 | 1,834005 | -2,93325 | 0,003354 | 1 | 0 | Promoter (2-3kb) | 19 | 23736251 | 23926960 | 190710 | 1 | 319924 | ENSMUST00000025830.9 | 2481 | ENSMUSG00000024897 | Apba1 | amyloid beta (A4) precursor protein binding, family A, member 1 |
| chr13 | 6877990 | 6878753 | 764 | * | 5,241382 | -5,379582721 | 1,947425 | -2,76241 | 0,005738 | 1 | 0 | Distal Intergenic | 13 | 6712601 | 6712930 | 330 | 2 | 1E+08 | ENSMUST00000089635.6 | -165060 | | Gm10029 | predicted gene 10029 |
| chr16 | 74097642 | 74098233 | 592 | * | 5,39281 | -5,379424434 | 1,994529 | -2,69709 | 0,006995 | 1 | 0 | Intron (ENSMUST0000 | 16 | 74138878 | 74138955 | 78 | 2 | 751561 | ENSMUST00000199719.3 | 40722 | ENSMUSG00000104758 | Mir691 | microRNA 691 |
| chrX | 1,38E+08 | 1,38E+08 | 550 | * | 5,925353 | -5,37685445 | 2,0566 | -2,61444 | 0,008937 | 1 | 0 | Intron (ENSMUST0000 | 20 | 1,38E+08 | 1,38E+08 | 21170 | 1 | 245631 | ENSMUST00000113042.3 | 37760 | ENSMUSG00000042515 | Pwwp3b | PWWP domain containing 3B |
| chr8 | 88860877 | 88861599 | 723 | * | 4,558004 | -5,376614494 | 2,078935 | -2,58624 | 0,009703 | 1 | 0 | Promoter (2-3kb) | 8 | 88864483 | 88898655 | 34173 | 1 | 234549 | ENSMUST00000034079.14 | -2884 | ENSMUSG00000031657 | Heatr3 | HEAT repeat containing 3 |
| chr14 | 32588625 | 32589013 | 389 | * | 4,748608 | -5,376598618 | 2,080256 | -2,58458 | 0,00975 | 1 | 0 | Intron (ENSMUST0000 | 14 | 32578713 | 32661448 | 82736 | 1 | 320736 | ENSMUST00000053175.13 | 9912 | ENSMUSG00000050666 | Vstm4 | V-set and transmembrane domain containing 4 |
| chr7 | 61616498 | 61617169 | 672 | * | 5,467964 | -5,37486133 | 2,047993 | -2,62445 | 0,008679 | 1 | 0 | Intron (ENSMUST0000 | 7 | 61593649 | 61632051 | 38403 | 2 | 320026 | ENSMUST00000207694.2 | 14882 | ENSMUSG00000109321 | A330076H08Rik | RIKEN cDNA A330076H08 gene |

| | | | | | | | | | | | | | | | | | | | | | | | |
|---|---|---|---|---|---|---|---|---|---|---|---|---|---|---|---|---|---|---|---|---|---|---|---|
| chr11 | 98411293 | 98411952 | 660 | * | 4,949718 | -5,374859418 | 2,047706 | -2,62482 | 0,008669 | 1 | 0 | Intron (ENSMUST00000 | 11 | 98355728 | 98436626 | 80899 | 2 | 22780 | ENSMUST00000103141.4 | 24674 | ENSMUSG00000018168 | Ikzf3 | IKAROS family zinc finger 3 |
| chrX | 51672848 | 51673722 | 875 | * | 5,321318 | -5,374504938 | 2,023185 | -2,65646 | 0,007897 | 1 | 0 | Exon (ENSMUST00000 | 20 | 51664720 | 51707924 | 43205 | 1 | 1,01E+08 | ENSMUST00000114854.8 | 8128 | ENSMUSG00000055110 | A630012P03Rik | RIKEN cDNA A630012P03 gene |
| chr2 | 13847066 | 13847954 | 889 | * | 9,310574 | -5,372477616 | 1,820525 | -2,95106 | 0,003167 | 0,693003605 | 1,360622 | Distal Intergenic | 2 | 13655832 | 13798875 | 143044 | 2 | 241230 | ENSMUST00000003509.10 | -48191 | ENSMUSG00000003418 | St8sia6 | ST8 alpha-N-acetyl-neuraminide alpha-2,8-sialyltransferase 6 |
| chrX | 1,04E+08 | 1,04E+08 | 863 | * | 5,050829 | -5,372447942 | 2,032641 | -2,64309 | 0,008215 | 1 | 0 | Distal Intergenic | 20 | 1,04E+08 | 1,04E+08 | 111 | 2 | 1,15E+08 | ENSMUST00000158335.3 | 27605 | ENSMUSG00000088960 | Gm22090 | predicted gene, 22090 |
| chr1 | 1,7E+08 | 1,7E+08 | 712 | * | 4,753702 | -5,372317356 | 2,00813 | -2,67528 | 0,007467 | 1 | 0 | Intron (ENSMUST00000 | 1 | 1,7E+08 | 1,7E+08 | 2988 | 1 | 67303 | ENSMUST00000242643.1 | 10156 | ENSMUSG00000097503 | 3110045C21Rik | RIKEN cDNA 3110045C21 gene |
| chr16 | 13824729 | 13825387 | 659 | * | 5,897957 | -5,37157145 | 1,899081 | -2,82851 | 0,004677 | 1 | 0 | Intron (ENSMUST00000 | 16 | 13799563 | 13804752 | 5190 | 2 | 74482 | ENSMUST00000230206.2 | -19977 | ENSMUSG00000065968 | Ifitm7 | interferon induced transmembrane protein 7 |
| chr11 | 45874822 | 45875658 | 837 | * | 6,180104 | -5,371244735 | 1,863517 | -2,88232 | 0,003948 | 1 | 0 | Exon (ENSMUST00000 | 11 | 45871137 | 45908821 | 37685 | 1 | 214105 | ENSMUST00000049038.4 | 3685 | ENSMUSG00000040489 | Sox30 | SRY (sex determining region Y)-box 30 |
| chr2 | 1,64E+08 | 1,64E+08 | 532 | * | 5,391923 | -5,370737658 | 1,938026 | -2,77124 | 0,005584 | 1 | 0 | Distal Intergenic | 2 | 1,64E+08 | 1,64E+08 | 1737 | 2 | 53878 | ENSMUST00000109374.2 | 5535 | ENSMUSG00000040132 | Semg1 | semenogelin 1 |
| chr19 | 56181047 | 56181592 | 546 | * | 5,658826 | -5,370606256 | 1,961498 | -2,73801 | 0,006181 | 1 | 0 | Distal Intergenic | 19 | 56265950 | 56266077 | 128 | 2 | 1,15E+08 | ENSMUST00000104035.3 | 84485 | ENSMUSG00000077223 | Gm22271 | predicted gene, 22271 |
| chr3 | 1,37E+08 | 1,37E+08 | 771 | * | 4,64564 | -5,37009902 | 2,07529 | -2,58764 | 0,009664 | 1 | 0 | Intron (ENSMUST00000 | 3 | 1,37E+08 | 1,37E+08 | 86151 | 1 | 59308 | ENSMUST00000197511.2 | 76581 | ENSMUSG00000054690 | Emcn | endomucin |
| chr14 | 1,07E+08 | 1,07E+08 | 652 | * | 4,538028 | -5,370067458 | 2,083936 | -2,57689 | 0,009969 | 1 | 0 | Distal Intergenic | 14 | 1,07E+08 | 1,07E+08 | 127 | 1 | 1,15E+08 | ENSMUST00000104036.3 | 148014 | ENSMUSG00000077224 | Gm23437 | predicted gene, 23437 |
| chr9 | 1,23E+08 | 1,23E+08 | 638 | * | 5,030216 | -5,368182639 | 2,025999 | -2,64965 | 0,008058 | 1 | 0 | Exon (ENSMUST00000 | 9 | 1,23E+08 | 1,23E+08 | 6580 | 1 | 22646 | ENSMUST00000148851.2 | 12662 | ENSMUSG00000057895 | Zfp105 | zinc finger protein 105 |
| chr5 | 80434090 | 80434831 | 742 | * | 5,217802 | -5,363420537 | 2,057648 | -2,60658 | 0,009145 | 1 | 0 | Distal Intergenic | 5 | 80246755 | 80246861 | 107 | 1 | 1,15E+08 | ENSMUST00000239632.1 | 187335 | ENSMUSG00000119601 | Gm24304 | predicted gene, 24304 |
| chr11 | 12636029 | 12636532 | 504 | * | 4,845919 | -5,363387189 | 2,046739 | -2,62045 | 0,008781 | 1 | 0 | Distal Intergenic | 11 | 12642813 | 12642942 | 130 | 1 | 1,15E+08 | ENSMUST00000157884.3 | -6281 | ENSMUSG00000088509 | Gm22789 | predicted gene, 22789 |
| chrX | 7880847 | 7881998 | 1152 | * | 6,609117 | -5,362828839 | 1,921264 | -2,7913 | 0,00525 | 1 | 0 | Promoter (2-3kb) | 20 | 7870439 | 7884731 | 14293 | 2 | 69824 | ENSMUST00000033503.3 | 2733 | ENSMUSG00000031163 | Glod5 | glyoxalase domain containing 5 |
| chr17 | 31267738 | 31268338 | 601 | * | 5,429282 | -5,362814088 | 1,919202 | -2,79429 | 0,005201 | 1 | 0 | Distal Intergenic | 17 | 31276649 | 31324579 | 47931 | 1 | 11307 | ENSMUST00000236391.2 | -8311 | ENSMUSG00000024030 | Abcg1 | ATP binding cassette subfamily G member 1 |
| chr10 | 78293486 | 78294459 | 974 | * | 8,146723 | -5,362518802 | 1,871546 | -2,86529 | 0,004166 | 0,730607562 | 1,449706 | Intron (ENSMUST00000 | 10 | 78272578 | 78300809 | 28232 | 2 | 216134 | ENSMUST00000041616.15 | 6350 | ENSMUSG00000032788 | Pdxk | pyridoxal (pyridoxine, vitamin B6) kinase |
| chr8 | 10479314 | 10480413 | 1100 | * | 7,636696 | -5,362040361 | 1,812288 | -2,95871 | 0,003089 | 1 | 0 | Intron (ENSMUST00000 | 8 | 10322572 | 10684741 | 362170 | 1 | 244281 | ENSMUST00000042103.9 | 156742 | ENSMUSG00000039057 | Myo16 | myosin XVI |
| chr2 | 1,54E+08 | 1,54E+08 | 1154 | * | 8,417522 | -5,36177058 | 1,784952 | -3,00387 | 0,002666 | 0,669586355 | 1,360622 | Distal Intergenic | 2 | 1,54E+08 | 1,54E+08 | 3564 | 1 | 71425 | ENSMUST00000126084.2 | 13338 | ENSMUSG00000067998 | Bpifb9a | BPI fold containing family B, member 9A |
| chr1 | 60416858 | 60417689 | 832 | * | 5,485087 | -5,361745822 | 1,945923 | -2,75537 | 0,005862 | 1 | 0 | Intron (ENSMUST00000 | 1 | 60448778 | 60520317 | 71540 | 1 | 329165 | ENSMUST00000052332.15 | -31089 | ENSMUSG00000026782 | Abi2 | abl interactor 2 |
| chrX | 38830340 | 38830935 | 596 | * | 5,924563 | -5,354070819 | 1,992678 | -2,68687 | 0,007212 | 1 | 0 | Distal Intergenic | 20 | 38951930 | 38952460 | 531 | 2 | 1,01E+08 | ENSMUST00000179096.3 | 121525 | ENSMUSG00000095240 | Cypt14 | cysteine-rich perinuclear theca 14 |
| chr12 | 4705319 | 4705956 | 638 | * | 5,926617 | -5,353975964 | 1,959361 | -2,73251 | 0,006285 | 1 | 0 | Exon (ENSMUST00000 | 12 | 4714535 | 4724167 | 9633 | 1 | 20403 | ENSMUST00000218985.2 | -8579 | ENSMUSG00000020640 | Itsn2 | intersectin 2 |
| chr7 | 11122482 | 11123222 | 741 | * | 5,327987 | -5,353930005 | 1,942785 | -2,7558 | 0,005855 | 1 | 0 | Distal Intergenic | 7 | 11131842 | 11136245 | 4404 | 1 | 665902 | ENSMUST00000145237.8 | -8620 | ENSMUSG00000070828 | Zscan4f | zinc finger and SCAN domain containing 4F |
| chr2 | 1,13E+08 | 1,13E+08 | 980 | * | 7,070546 | -5,353830006 | 1,855571 | -2,88527 | 0,003911 | 1 | 0 | Intron (ENSMUST00000 | 2 | 1,13E+08 | 1,14E+08 | 268792 | 1 | 14260 | ENSMUST00000081349.9 | -9710 | ENSMUSG00000044042 | Fmn1 | formin 1 |
| chr5 | 22641687 | 22642677 | 991 | * | 8,801575 | -5,35361959 | 1,870829 | -2,86163 | 0,004215 | 0,731734453 | 1,449706 | Distal Intergenic | 5 | 22733319 | 22733717 | 399 | 2 | 1E+08 | ENSMUST00000118224.2 | 91040 | | Gm15421 | ribosomal protein L22 like 1 pseudogene |
| chr7 | 16073575 | 16074315 | 741 | * | 7,04827 | -5,353456413 | 1,827261 | -2,92977 | 0,003392 | 0,708539289 | 1,360622 | Intron (ENSMUST00000 | 7 | 16047442 | 16051622 | 4181 | 1 | 170770 | ENSMUST00000147267.2 | 26133 | ENSMUSG00000002083 | Bbc3 | BCL2 binding component 3 |
| chr18 | 83010308 | 83011070 | 763 | * | 5,545412 | -5,352738868 | 1,947694 | -2,74824 | 0,005992 | 1 | 0 | Intron (ENSMUST00000 | 18 | 83005933 | 83019825 | 13893 | 1 | 329003 | ENSMUST00000236468.2 | 4375 | ENSMUSG00000058881 | Zfp516 | zinc finger protein 516 |
| chr2 | 1,32E+08 | 1,32E+08 | 1048 | * | 7,666931 | -5,352446473 | 1,910541 | -2,80153 | 0,005086 | 1 | 0 | Distal Intergenic | 2 | 1,32E+08 | 1,32E+08 | 28502 | 1 | 19122 | ENSMUST00000091288.13 | -12226 | ENSMUSG00000079037 | Prnp | prion protein |
| chr5 | 1,48E+08 | 1,48E+08 | 981 | * | 5,569663 | -5,352191767 | 2,010384 | -2,66227 | 0,007761 | 1 | 0 | Distal Intergenic | 5 | 1,48E+08 | 1,48E+08 | 15996 | 1 | 66537 | ENSMUST00000031654.10 | -7156 | ENSMUSG00000029649 | Pomp | proteasome maturation protein |
| chr11 | 32280341 | 32281124 | 784 | * | 6,438178 | -5,3519602 | 2,041904 | -2,62106 | 0,008766 | 1 | 0 | Distal Intergenic | 11 | 32297820 | 32378173 | 80354 | 1 | 268396 | ENSMUST00000038753.6 | -16696 | ENSMUSG00000040711 | Sh3pxd2b | SH3 and PX domains 2B |
| chr1 | 1,86E+08 | 1,86E+08 | 850 | * | 5,071948 | -5,351724572 | 2,076981 | -2,57668 | 0,009975 | 1 | 0 | Distal Intergenic | 1 | 1,86E+08 | 1,86E+08 | 28756 | 2 | 226791 | ENSMUST00000195481.2 | 50267 | ENSMUSG00000039246 | Lyplal1 | lysophospholipase-like 1 |
| chr7 | 6207243 | 6208189 | 947 | * | 10,71447 | -5,349748824 | 1,743099 | -3,0691 | 0,002147 | 0,731734453 | 0,738441 | Promoter (1-2kb) | 7 | 6205800 | 6214365 | 8566 | 1 | 232836 | ENSMUST00000130076.2 | 1443 | ENSMUSG00000034660 | Galp | galanin-like peptide |
| chr14 | 58185911 | 58186265 | 355 | * | 4,578089 | -5,347353107 | 2,05776 | -2,59863 | 0,00936 | 1 | 0 | Intron (ENSMUST00000 | 14 | 58161403 | 58191651 | 30249 | 2 | 68514 | ENSMUST00000224607.2 | 5386 | ENSMUSG00000021973 | Micu2 | mitochondrial calcium uptake 2 |
| chr6 | 1,22E+08 | 1,22E+08 | 356 | * | 5,579377 | -5,346691605 | 1,996134 | -2,67852 | 0,007395 | 1 | 0 | Distal Intergenic | 6 | 1,22E+08 | 1,22E+08 | 12287 | 2 | 50928 | ENSMUST00000032207.9 | 15019 | ENSMUSG00000030114 | Klrg1 | killer cell lectin-like receptor subfamily G, member 1 |
| chr9 | 84089027 | 84089501 | 475 | * | 4,336252 | -5,344960895 | 2,037809 | -2,6229 | 0,008719 | 1 | 0 | Distal Intergenic | 9 | 84070408 | 84070592 | 185 | 1 | 1,15E+08 | ENSMUST00000104478.4 | 18619 | ENSMUSG00000077666 | Gm26146 | predicted gene, 26146 |
| chr12 | 5642969 | 5643419 | 451 | * | 5,516059 | -5,344924976 | 1,994656 | -2,67962 | 0,007371 | 1 | 0 | Distal Intergenic | 12 | 5461641 | 5466631 | 4991 | 1 | 72669 | ENSMUST00000163627.3 | 181328 | ENSMUSG00000037735 | 2810032G03Rik | RIKEN cDNA 2810032G03 gene |
| chr7 | 30027350 | 30028067 | 718 | * | 5,827226 | -5,344167584 | 1,911166 | -2,79629 | 0,005169 | 1 | 0 | Distal Intergenic | 7 | 30020831 | 30023320 | 2490 | 2 | 68332 | ENSMUST00000098586.5 | -4030 | ENSMUSG00000074211 | Sdhaf1 | succinate dehydrogenase complex assembly factor 1 |
| chr1 | 1,82E+08 | 1,82E+08 | 850 | * | 5,708662 | -5,343811399 | 1,93851 | -2,75666 | 0,00584 | 1 | 0 | Intron (ENSMUST00000 | 1 | 1,82E+08 | 1,82E+08 | 4475 | 1 | 240960 | ENSMUST00000160793.2 | 25229 | ENSMUSG00000047369 | Dnah14 | dynein, axonemal, heavy chain 14 |
| chr7 | 71731654 | 71732077 | 424 | * | 4,775813 | -5,343105514 | 2,002176 | -2,66865 | 0,007616 | 1 | 0 | Intron (ENSMUST00000 | 7 | 71730161 | 71760331 | 30171 | 2 | 244049 | ENSMUST00000206466.2 | 28254 | ENSMUSG00000032776 | Mctp2 | multiple C2 domains, transmembrane 2 |
| chrX | 1,34E+08 | 1,34E+08 | 667 | * | 5,264932 | -5,342988534 | 2,013751 | -2,65325 | 0,007972 | 1 | 0 | Distal Intergenic | 20 | 1,34E+08 | 1,34E+08 | 5075 | 2 | 67416 | ENSMUST00000168264.2 | -45821 | ENSMUSG00000033436 | Armcx2 | armadillo repeat containing, X-linked 2 |
| chrX | 1,14E+08 | 1,14E+08 | 656 | * | 4,840636 | -5,342518434 | 2,069933 | -2,58101 | 0,009851 | 1 | 0 | Distal Intergenic | 20 | 1,14E+08 | 1,14E+08 | 381 | 1 | 1,15E+08 | ENSMUST00000185033.3 | -189275 | ENSMUSG00000098405 | Gm27414 | predicted gene, 27414 |
| chr2 | 7135468 | 7136395 | 928 | * | 10,44969 | -5,34212206 | 1,652587 | -3,23258 | 0,001227 | 0,600272691 | 0,918255 | Intron (ENSMUST00000 | 2 | 6548241 | 7086113 | 537873 | 2 | 14007 | ENSMUST00000183209.8 | -49355 | ENSMUSG00000002107 | Celf2 | CUGBP, Elav-like family member 2 |
| chr17 | 27975554 | 27976289 | 736 | * | 6,759356 | -5,34045297 | 2,047062 | -2,60884 | 0,009085 | 1 | 0 | Intron (ENSMUST00000 | 17 | 27933633 | 27947929 | 14297 | 2 | 30051 | ENSMUST00000114870.9 | -27625 | ENSMUSG00000024215 | Spdef | SAM pointed domain containing ets transcription factor |
| chr9 | 58218088 | 58218833 | 746 | * | 5,395051 | -5,33760448 | 2,009835 | -2,65574 | 0,007913 | 1 | 0 | Promoter (1-2kb) | 9 | 58195021 | 58220469 | 25449 | 2 | 16949 | ENSMUST00000061799.10 | 1636 | ENSMUSG00000032334 | Loxl1 | lysyl oxidase-like 1 |
| chrX | 96219868 | 96220997 | 1130 | * | 7,370442 | -5,337583955 | 1,766955 | -3,02078 | 0,002521 | 1 | 0 | Distal Intergenic | 20 | 96116202 | 96125710 | 9509 | 1 | 245526 | ENSMUST00000033556.4 | 103666 | ENSMUSG00000031212 | Pgr15l | G protein-coupled receptor 15-like |
| chr9 | 68564281 | 68564950 | 670 | * | 5,939557 | -5,337518997 | 2,001072 | -2,66733 | 0,007646 | 1 | 0 | Promoter (1-2kb) | 9 | 68562585 | 68571462 | 8878 | 1 | 19883 | ENSMUST00000143507.2 | 1696 | ENSMUSG00000032238 | Rora | RAR-related orphan receptor alpha |
| chr18 | 85894972 | 85896232 | 1261 | * | 8,972344 | -5,33732779 | 1,732704 | -3,08035 | 0,002068 | 0,646266933 | 1,211315 | Distal Intergenic | 18 | 86367711 | 86367817 | 107 | 1 | 1,15E+08 | ENSMUST00000082733.3 | -471479 | ENSMUSG00000119275 | Gm25453 | predicted gene, 25453 |
| chr1 | 1,93E+08 | 1,93E+08 | 1001 | * | 6,29844 | -5,336609096 | 1,923691 | -2,77415 | 0,005535 | 1 | 0 | Intron (ENSMUST00000 | 1 | 1,93E+08 | 1,93E+08 | 129274 | 1 | 1,03E+08 | ENSMUST00000246715.1 | 88263 | | Gm31870 | predicted gene, 31870 |
| chr16 | 87330201 | 87330786 | 586 | * | 6,995277 | -5,33630315 | 1,849952 | -2,88456 | 0,00392 | 1 | 0 | Distal Intergenic | 16 | 87324918 | 87325053 | 136 | 2 | 1,15E+08 | ENSMUST00000104391.3 | -5148 | ENSMUSG00000077579 | Gm24891 | predicted gene, 24891 |
| chr12 | 10381346 | 10381862 | 517 | * | 5,328316 | -5,335795076 | 1,911846 | -2,79091 | 0,005256 | 1 | 0 | Distal Intergenic | 12 | 10419973 | 10425935 | 5963 | 1 | 70881 | ENSMUST00000143739.9 | -38111 | ENSMUSG00000020622 | Nt5c1b | 5'-nucleotidase, cytosolic IB |
| chr10 | 24349935 | 24350500 | 566 | * | 5,102853 | -5,335750274 | 1,961953 | -2,71961 | 0,006536 | 1 | 0 | Distal Intergenic | 10 | 24455946 | 24469656 | 13711 | 2 | 1,03E+08 | ENSMUST00000141076.2 | 119156 | | Gm15270 | predicted gene 15270 |
| chr10 | 90510480 | 90510994 | 515 | * | 4,286714 | -5,335679461 | 2,053494 | -2,59834 | 0,009368 | 1 | 0 | Intron (ENSMUST00000 | 10 | 90412854 | 90805682 | 392829 | 1 | 77531 | ENSMUST00000099364.12 | 97626 | ENSMUSG00000058589 | Anks1b | ankyrin repeat and sterile alpha motif domain containing 1B |
| chr1 | 79353993 | 79354549 | 557 | * | 4,351237 | -5,335674188 | 2,066719 | -2,58171 | 0,009831 | 1 | 0 | Distal Intergenic | 1 | 79412386 | 79417759 | 5374 | 2 | 20254 | ENSMUST00000185234.2 | 63210 | ENSMUSG00000050711 | Scg2 | secretogranin II |
| chr3 | 70576129 | 70576691 | 563 | * | 6,586397 | -5,334550262 | 1,952502 | -2,73216 | 0,006292 | 1 | 0 | Distal Intergenic | 3 | 70697557 | 70714624 | 17068 | 2 | 625901 | ENSMUST00000246772.1 | 137933 | | Gm6634 | predicted gene 6634 |
| chr4 | 94471321 | 94472377 | 1057 | * | 5,760342 | -5,333577412 | 2,033162 | -2,62329 | 0,008708 | 1 | 0 | Promoter (<=1kb) | 4 | 94459924 | 94471735 | 11812 | 2 | 18786 | ENSMUST00000127656.3 | 0 | ENSMUSG00000028577 | Plaa | phospholipase A2, activating protein |
| chr13 | 20930801 | 20931572 | 772 | * | 6,982573 | -5,333406549 | 2,049865 | -2,60183 | 0,009273 | 1 | 0 | Distal Intergenic | 13 | 20978283 | 21208422 | 230140 | 1 | 27052 | ENSMUST00000021757.5 | -46711 | ENSMUSG00000021322 | Aoah | acyloxyacyl hydrolase |
| chr13 | 95070295 | 95071202 | 908 | * | 7,045948 | -5,32900419 | 1,953548 | -2,72786 | 0,006375 | 1 | 0 | Distal Intergenic | 13 | 95112852 | 95151039 | 38188 | 1 | 218460 | ENSMUST00000160115.8 | -41650 | ENSMUSG00000042015 | Wdr41 | WD repeat domain 41 |
| chr9 | 1,02E+08 | 1,02E+08 | 715 | * | 5,158523 | -5,328069059 | 1,98264 | -2,68736 | 0,007202 | 1 | 0 | Intron (ENSMUST00000 | 9 | 1,02E+08 | 1,02E+08 | 17941 | 2 | 270190 | ENSMUST00000217184.2 | -20053 | ENSMUSG00000032537 | Ephb1 | Eph receptor B1 |
| chr5 | 1,3E+08 | 1,3E+08 | 1416 | * | 8,77101 | -5,327260876 | 1,791699 | -2,9733 | 0,002946 | 1 | 0 | Promoter (<=1kb) | 5 | 1,3E+08 | 1,3E+08 | 2933 | 1 | 231769 | ENSMUST00000200500.2 | 0 | ENSMUSG00000029439 | Sfswap | splicing factor SWAP |
| chr3 | 1,02E+08 | 1,02E+08 | 493 | * | 7,231748 | -5,327103776 | 1,819929 | -2,92709 | 0,003421 | 1 | 0 | Promoter (2-3kb) | 3 | 1,02E+08 | 1,02E+08 | 1535 | 1 | 12373 | ENSMUST00000159833.2 | 2114 | ENSMUSG00000027861 | Casq2 | calsequestrin 2 |
| chr10 | 47820240 | 47820702 | 463 | * | 6,053042 | -5,326688402 | 1,920115 | -2,77415 | 0,005535 | 1 | 0 | Exon (ENSMUST00000 | 10 | 48495408 | 48495491 | 84 | 2 | 1,15E+08 | ENSMUST00000083696.4 | 674789 | | Gm24256 | predicted gene, 24256 |
| chr9 | 78230257 | 78230860 | 604 | * | 5,114244 | -5,326604283 | 1,946325 | -2,73675 | 0,006205 | 1 | 0 | Distal Intergenic | 9 | 78234871 | 78236878 | 2008 | 1 | 382088 | ENSMUST00000187667.7 | -4011 | ENSMUSG00000038750 | Omt2b | oocyte maturation, beta |
| chr7 | 1,22E+08 | 1,22E+08 | 865 | * | 6,641955 | -5,326573831 | 1,856353 | -2,86938 | 0,004113 | 1 | 0 | Intron (ENSMUST00000 | 7 | 1,22E+08 | 1,22E+08 | 41259 | 1 | 18751 | ENSMUST00000149583.2 | 10097 | ENSMUSG00000052889 | Prkcb | protein kinase C, beta |
| chr17 | 43291110 | 43291485 | 376 | * | 5,553028 | -5,326516286 | 1,978262 | -2,69252 | 0,007091 | 1 | 0 | Distal Intergenic | 17 | 43327446 | 43400080 | 72635 | 1 | 94185 | ENSMUST00000024708.6 | -35961 | ENSMUSG00000023915 | Tnfrsf21 | tumor necrosis factor receptor superfamily, member 21 |
| chr5 | 1,24E+08 | 1,24E+08 | 786 | * | 5,576623 | -5,326449597 | 2,004637 | -2,65706 | 0,007882 | 1 | 0 | Exon (ENSMUST00000 | 5 | 1,24E+08 | 1,24E+08 | 63133 | 1 | 208104 | ENSMUST00000068237.12 | 3733 | ENSMUSG00000038342 | Mlxip | MLX interacting protein |
| chr7 | 1,22E+08 | 1,22E+08 | 612 | * | 4,814236 | -5,326423764 | 2,014458 | -2,6441 | 0,008191 | 1 | 0 | Intron (ENSMUST00000 | 7 | 1,22E+08 | 1,22E+08 | 28097 | 1 | 18751 | ENSMUST00000127910.2 | -26547 | ENSMUSG00000052889 | Prkcb | protein kinase C, beta |
| chrX | 89876073 | 89876805 | 733 | * | 4,460592 | -5,326339507 | 2,051003 | -2,59694 | 0,009406 | 1 | 0 | Distal Intergenic | 20 | 89935748 | 89936740 | 993 | 1 | 212952 | ENSMUST00000172155.2 | -58943 | ENSMUSG00000091497 | Mageb11 | MAGE family member B11 |
| chr5 | 1,46E+08 | 1,46E+08 | 474 | * | 5,111418 | -5,326326634 | 2,059697 | -2,58598 | 0,00971 | 1 | 0 | Distal Intergenic | 5 | 1,46E+08 | 1,46E+08 | 2738 | 2 | 433961 | ENSMUST00000199463.2 | 14036 | ENSMUSG00000104752 | Gm5565 | predicted gene 5565 |
| chr7 | 1,25E+08 | 1,25E+08 | 635 | * | 4,895998 | -5,32435814 | 2,027219 | -2,62644 | 0,008628 | 1 | 0 | Distal Intergenic | 7 | 1,24E+08 | 1,24E+08 | 83266 | 2 | 73076 | ENSMUST00000130240.2 | -270430 | ENSMUSG00000086254 | 3100003L05Rik | RIKEN cDNA 3100003L05 gene |
| chr4 | 72942370 | 72942988 | 619 | * | 4,981329 | -5,324293862 | 2,031746 | -2,62055 | 0,008779 | 1 | 0 | Distal Intergenic | 4 | 73032894 | 73033020 | 127 | 1 | 1,15E+08 | ENSMUST00000158569.3 | -89906 | ENSMUSG00000089194 | Gm25769 | predicted gene, 25769 |
| chr4 | 65359375 | 65360110 | 736 | * | 5,114133 | -5,323979537 | 2,059672 | -2,58487 | 0,009742 | 1 | 0 | Intron (ENSMUST00000 | 4 | 65523223 | 65534475 | 11253 | 1 | 69807 | ENSMUST00000050850.14 | -163113 | ENSMUSG00000051675 | Trim32 | tripartite motif-containing 32 |
| chrX | 1E+08 | 1E+08 | 1023 | * | 5,14541 | -5,323941296 | 2,063176 | -2,58046 | 0,009867 | 1 | 0 | Distal Intergenic | 20 | 1E+08 | 1E+08 | 664 | 2 | 11989 | ENSMUST00000144410.2 | 8086 | ENSMUSG00000031297 | Slc7a3 | solute carrier family 7 (cationic amino acid transporter, y+ system), member 3 |
| chr1 | 1,89E+08 | 1,89E+08 | 1050 | * | 5,88982 | -5,321480451 | 2,050426 | -2,5953 | 0,009451 | 1 | 0 | Distal Intergenic | 1 | 1,89E+08 | 1,89E+08 | 5193 | 2 | 16526 | ENSMUST00000193686.2 | 37556 | ENSMUSG00000037624 | Kcnk2 | potassium channel, subfamily K, member 2 |
| chr5 | 1,46E+08 | 1,46E+08 | 480 | * | 4,703592 | -5,319287186 | 2,064137 | -2,577 | 0,009966 | 1 | 0 | Distal Intergenic | 5 | 1,46E+08 | 1,46E+08 | 25539 | 2 | 13112 | ENSMUST00000035918.8 | 36342 | ENSMUSG00000056035 | Cyp3a11 | cytochrome P450, family 3, subfamily a, polypeptide 11 |
| chrX | 99599691 | 99601267 | 1577 | * | 9,221656 | -5,319025093 | 1,748239 | -3,0425 | 0,002346 | 0,641187423 | 1,425929 | Distal Intergenic | 20 | 99615857 | 99621811 | 5955 | 1 | 245533 | ENSMUST00000096361.5 | -14590 | ENSMUSG00000015665 | Awat1 | acyl-CoA wax alcohol acyltransferase 1 |
| chr17 | 41256061 | 41256682 | 622 | * | 5,493437 | -5,318620633 | 1,963758 | -2,70839 | 0,006761 | 1 | 0 | Intron (ENSMUST00000 | 17 | 41266079 | 41269872 | 3794 | 1 | 17850 | ENSMUST00000233222.2 | -9397 | ENSMUSG00000023921 | Mmut | methylmalonyl-Coenzyme A mutase |
| chrX | 16866945 | 16867872 | 928 | * | 6,760816 | -5,317749025 | 1,872552 | -2,83984 | 0,004514 | 1 | 0 | Distal Intergenic | 20 | 16751760 | 16778013 | 26254 | 2 | 17986 | ENSMUST00000040134.8 | -88932 | ENSMUSG00000040138 | Ndp | Norrie disease (pseudoglioma) (human) |
| chr9 | 48415329 | 48415962 | 634 | * | 5,122625 | -5,317284611 | 1,969866 | -2,69931 | 0,006948 | 1 | 0 | Distal Intergenic | 9 | 48406742 | 48407272 | 531 | 1 | 434402 | ENSMUST00000048824.9 | 8587 | ENSMUSG00000042293 | Gm5617 | predicted gene 5617 |
| chr9 | 14898544 | 14899083 | 540 | * | 6,341581 | -5,315257968 | 2,011318 | -2,64267 | 0,008225 | 1 | 0 | Distal Intergenic | 9 | 14919214 | 14921804 | 2591 | 2 | 55991 | ENSMUST00000166933.2 | 22721 | ENSMUSG00000031934 | Panx1 | pannexin 1 |
| chr4 | 21338733 | 21339368 | 636 | * | 4,499932 | -5,309767156 | 2,05569 | -2,58296 | 0,009796 | 1 | 0 | Distal Intergenic | 4 | 21679747 | 21685781 | 6035 | 2 | 230025 | ENSMUST00000149516.8 | 346413 | ENSMUSG00000040478 | Prdm13 | PR domain containing 13 |
| chr13 | 95702502 | 95703310 | 809 | * | 5,377548 | -5,309631418 | 2,024781 | -2,62232 | 0,008733 | 1 | 0 | Distal Intergenic | 13 | 95648240 | 95661735 | 13496 | 2 | 14063 | ENSMUST00000022185.10 | -40767 | ENSMUSG00000021678 | F2rl1 | coagulation factor II (thrombin) receptor-like 1 |
| chr13 | 33490876 | 33491556 | 681 | * | 6,14719 | -5,309535252 | 2,005557 | -2,64741 | 0,008111 | 1 | 0 | Distal Intergenic | 13 | 33508060 | 33519353 | 11294 | 1 | 20709 | ENSMUST00000075515.7 | -16504 | ENSMUSG00000038327 | Serpinb9f | serine (or cysteine) peptidase inhibitor, clade B, member 9f |
| chr18 | 69511177 | 69511675 | 499 | * | 5,473187 | -5,309183244 | 1,945166 | -2,72942 | 0,006345 | 1 | 0 | Intron (ENSMUST00000 | 18 | 69482015 | 69776003 | 293989 | 1 | 21413 | ENSMUST00000201124.4 | 29162 | ENSMUSG00000053477 | Tcf4 | transcription factor 4 |
| chr13 | 51742247 | 51743088 | 842 | * | 7,658512 | -5,308025971 | 1,811985 | -2,9294 | 0,003396 | 1 | 0 | Distal Intergenic | 13 | 51585077 | 51723523 | 138447 | 2 | 20418 | ENSMUST00000239056.2 | -18724 | ENSMUSG00000021448 | Shc3 | src homology 2 domain-containing transforming protein C3 |
| chr10 | 12736042 | 12736638 | 597 | * | 4,700274 | -5,307635456 | 2,027455 | -2,61788 | 0,008848 | 1 | 0 | Promoter (<=1kb) | 10 | 12257932 | 12737479 | 479548 | 2 | 22288 | ENSMUST00000076817.5 | 841 | ENSMUSG00000019820 | Utrn | utrophin |
| chr19 | 23410232 | 23410879 | 648 | * | 4,483074 | -5,307543911 | 2,044357 | -2,59619 | 0,009426 | 1 | 0 | Intron (ENSMUST00000 | 19 | 23279973 | 23425762 | 145790 | 2 | 71738 | ENSMUST00000036069.8 | 14883 | ENSMUSG00000033207 | Mamdc2 | MAM domain containing 2 |
| chr16 | 69636309 | 69637012 | 704 | * | 5,978394 | -5,306199481 | 1,993969 | -2,66112 | 0,007788 | 1 | 0 | Distal Intergenic | 16 | 69654022 | 69657374 | 3353 | 2 | 224318 | ENSMUST00000166256.2 | 20362 | ENSMUSG00000063163 | Speer2 | spermatogenesis associated glutamate (E)-rich protein 2 |
| chr12 | 31863738 | 31864436 | 699 | * | 4,723814 | -5,300187769 | 2,034564 | -2,60507 | 0,009185 | 1 | 0 | Intron (ENSMUST00000 | 12 | 31908457 | 31908591 | 135 | 1 | 1,15E+08 | ENSMUST00000157617.3 | -44021 | ENSMUSG00000088242 | Gm25913 | predicted gene, 25913 |
| chr5 | 76265511 | 76266272 | 762 | * | 4,959547 | -5,300007382 | 1,982747 | -2,67306 | 0,007516 | 1 | 0 | Distal Intergenic | 5 | 76288118 | 76299362 | 11245 | 1 | 57357 | ENSMUST00000138699.8 | -21846 | ENSMUSG00000029233 | Srd5a3 | steroid 5 alpha-reductase 3 |
| chr12 | 86430139 | 86430448 | 310 | * | 4,819396 | -5,299996225 | 1,979492 | -2,67745 | 0,007418 | 1 | 0 | Intron (ENSMUST00000 | 12 | 86407891 | 86568402 | 160512 | 1 | 26380 | ENSMUST00000116402.10 | 22248 | ENSMUSG00000021255 | Esrrb | estrogen related receptor, beta |
| chr9 | 71572720 | 71573517 | 798 | * | 5,945705 | -5,29977839 | 1,927828 | -2,74909 | 0,005976 | 1 | 0 | Intron (ENSMUST00000 | 9 | 71539915 | 71555632 | 15718 | 2 | 68178 | ENSMUST00000146567.2 | -17088 | ENSMUSG00000032232 | Cgnl1 | cingulin-like 1 |
| chrX | 80335882 | 80336813 | 932 | * | 6,727192 | -5,299166516 | 1,915413 | -2,76659 | 0,005665 | 1 | 0 | Distal Intergenic | 20 | 80463181 | 80501727 | 38547 | 2 | 245492 | ENSMUST00000080083.3 | 164914 | ENSMUSG00000060673 | 4930595M18Rik | RIKEN cDNA 4930595M18 gene |
| chr4 | 1,31E+08 | 1,31E+08 | 1099 | * | 16,69197 | -5,297670068 | 1,646017 | -3,21848 | 0,001289 | 0,542967057 | 1,2763 | Promoter (<=1kb) | 4 | 1,31E+08 | 1,31E+08 | 1919 | 1 | 16792 | ENSMUST00000125332.2 | 620 | ENSMUSG00000028581 | Laptm5 | lysosomal-associated protein transmembrane 5 |
| chr10 | 4391205 | 4391872 | 668 | * | 5,37734 | -5,290630564 | 2,023082 | -2,61513 | 0,008919 | 1 | 0 | Intron (ENSMUST00000 | 10 | 4384581 | 4390469 | 5889 | 1 | 73419 | ENSMUST00000131636.2 | 6624 | ENSMUSG00000061759 | Armt1 | acidic residue methyltransferase 1 |
| chr17 | 43582852 | 43583531 | 680 | * | 5,15756 | -5,290514675 | 1,96199 | -2,6965 | 0,007007 | 1 | 0 | Promoter (1-2kb) | 17 | 43581220 | 43635628 | 54409 | 1 | 77596 | ENSMUST00000047399.6 | 1632 | ENSMUSG00000041293 | Adgrf1 | adhesion G protein-coupled receptor F1 |
| chr11 | 69178248 | 69179119 | 872 | * | 5,679955 | -5,290413386 | 1,917619 | -2,75884 | 0,005801 | 1 | 0 | Distal Intergenic | 11 | 69190313 | 69191316 | 1004 | 2 | 216846 | ENSMUST00000156671.8 | 12197 | ENSMUSG00000032782 | Cntrob | centrobin, centrosomal BRCA2 interacting protein |
| chr2 | 70182822 | 70183809 | 988 | * | 6,621286 | -5,290363605 | 1,897798 | -2,78763 | 0,005309 | 1 | 0 | Intron (ENSMUST00000 | 2 | 70305267 | 70308073 | 2807 | 1 | 64406 | ENSMUST00000100043.3 | -121458 | ENSMUSG00000075304 | Sp5 | trans-acting transcription factor 5 |
| chr12 | 74413943 | 74414853 | 911 | * | 6,274569 | -5,290344137 | 1,883862 | -2,80824 | 0,004981 | 1 | 0 | Distal Intergenic | 12 | 74360975 | 74363936 | 2962 | 2 | 1,03E+08 | ENSMUST00000223381.2 | -50007 | | Gm34552 | predicted gene, 34552 |
| chr18 | 48167841 | 48168578 | 738 | * | 7,538871 | -5,290212914 | 1,892829 | -2,79487 | 0,005192 | 0,782360218 | 1,360622 | Intron (ENSMUST00000 | 18 | 48178402 | 48181479 | 3078 | 1 | 433182 | ENSMUST00000076155.6 | -9824 | ENSMUSG00000059040 | Eno1b | enolase 1B, retrotransposed |
| chr12 | 1,06E+08 | 1,06E+08 | 803 | * | 7,525976 | -5,290048557 | 1,804991 | -2,93079 | 0,003381 | 1 | 0 | Distal Intergenic | 12 | 1,06E+08 | 1,06E+08 | 64993 | 1 | 22367 | ENSMUST00000072040.7 | -31051 | ENSMUSG00000021115 | Vrk1 | vaccinia related kinase 1 |
| chr8 | 67203988 | 67204468 | 481 | * | 6,594239 | -5,288966874 | 1,995717 | -2,65016 | 0,008045 | 1 | 0 | Distal Intergenic | 8 | 67150074 | 67159444 | 9371 | 1 | 18166 | ENSMUST00000039303.7 | 53914 | ENSMUSG00000036437 | Npy1r | neuropeptide Y receptor Y1 |

| | | | | | | | | | | | | | | | | | | | | | | | |
|---|---|---|---|---|---|---|---|---|---|---|---|---|---|---|---|---|---|---|---|---|---|---|---|
| chr3 | 46326655 | 46327396 | 742 | * | 6,713824 | -5,286243746 | 1,785491 | -2,96067 | 0,00307 | 1 | 0 | Distal Intergenic | 3 | 46401491 | 46402355 | 865 | 2 | 241989 | ENSMUST00000195537.2 | 74959 | ENSMUSG00000090919 | Pabpc4l | poly(A) binding protein, cytoplasmic 4-like |
| chr3 | 81507434 | 81507982 | 549 | * | 4,850416 | -5,282760813 | 2,00535 | -2,63433 | 0,00843 | 1 | 0 | Distal Intergenic | 3 | 81709332 | 81712468 | 3137 | 1 | 320566 | ENSMUST00000193672.2 | -201350 | ENSMUSG00000103695 | A830029E22Rik | RIKEN cDNA A830029E22 gene |
| chr5 | 1,5E+08 | 1,5E+08 | 690 | * | 5,370232 | -5,282710558 | 2,001072 | -2,63994 | 0,008292 | 1 | 0 | Distal Intergenic | 5 | 1,5E+08 | 1,5E+08 | 863 | 1 | 1,03E+08 | ENSMUST00000201713.2 | 10694 | | Gm36241 | predicted gene, 36241 |
| chr14 | 80374506 | 80375397 | 892 | * | 7,127041 | -5,281051874 | 1,949554 | -2,70885 | 0,006752 | 1 | 0 | Distal Intergenic | 14 | 80237742 | 80259370 | 21629 | 1 | 380924 | ENSMUST00000088735.4 | 136764 | ENSMUSG00000022026 | Olfm4 | olfactomedin 4 |
| chr18 | 6009083 | 6009617 | 535 | * | 4,264267 | -5,281051856 | 2,049384 | -2,5769 | 0,009969 | 1 | 0 | Distal Intergenic | 18 | 6032960 | 6034707 | 1748 | 2 | 75415 | ENSMUST00000182343.2 | 25090 | ENSMUSG00000041225 | Arhgap12 | Rho GTPase activating protein 12 |
| chr8 | 98995392 | 98995934 | 543 | * | 5,352955 | -5,281051801 | 1,964325 | -2,68848 | 0,007178 | 1 | 0 | Distal Intergenic | 8 | 99401709 | 99401834 | 126 | 2 | 1,15E+08 | ENSMUST00000116830.3 | 405900 | ENSMUSG00000080480 | Gm23494 | predicted gene, 23494 |
| chr7 | 1,01E+08 | 1,01E+08 | 751 | * | 5,238486 | -5,281047668 | 1,932355 | -2,73296 | 0,006277 | 1 | 0 | Exon (ENSMUST00000: | 7 | 1,01E+08 | 1,01E+08 | 32248 | 1 | 207278 | ENSMUST00000130426.2 | 32114 | ENSMUSG00000030691 | Fchsd2 | FCH and double SH3 domains 2 |
| chr11 | 25257058 | 25258060 | 1003 | * | 4,716408 | -5,281046247 | 2,00984 | -2,6276 | 0,008599 | 1 | 0 | Intron (ENSMUST0000( | 11 | 25251263 | 25272786 | 21524 | 1 | 1,03E+08 | ENSMUST00000145127.2 | 5795 | | Gm12069 | predicted gene 12069 |
| chr19 | 55781332 | 55782438 | 1107 | * | 8,289824 | -5,280173881 | 1,84413 | -2,86323 | 0,004193 | 0,744054256 | 1,360622 | Intron (ENSMUST0000( | 19 | 55731277 | 55899113 | 167837 | 1 | 21416 | ENSMUST00000133008.8 | 50055 | ENSMUSG00000024985 | Tcf7l2 | transcription factor 7 like 2, T cell specific, HMG box |
| chr9 | 1,03E+08 | 1,03E+08 | 632 | * | 4,7952 | -5,279160272 | 2,02287 | -2,60974 | 0,009061 | 1 | 0 | 3' UTR | 9 | 1,03E+08 | 1,03E+08 | 3674 | 1 | 235559 | ENSMUST00000185972.2 | 3156 | ENSMUSG00000032555 | Topbp1 | topoisomerase (DNA) II binding protein 1 |
| chrX | 12258971 | 12259799 | 829 | * | 4,583894 | -5,273427889 | 2,03641 | -2,58957 | 0,00961 | 1 | 0 | Distal Intergenic | 20 | 12206021 | 12221235 | 15215 | 2 | 1E+08 | ENSMUST00000146535.8 | -37736 | ENSMUSG00000087591 | Gm14635 | predicted gene 14635 |
| chr4 | 1,29E+08 | 1,29E+08 | 704 | * | 5,818357 | -5,272980244 | 1,990045 | -2,64968 | 0,008057 | 1 | 0 | Distal Intergenic | 4 | 1,29E+08 | 1,29E+08 | 4191 | 1 | 1,05E+08 | ENSMUST00000154515.2 | -13247 | | Gm15904 | predicted gene 15904 |
| chr9 | 99110655 | 99111516 | 862 | * | 7,777501 | -5,272354588 | 1,860941 | -2,83317 | 0,004609 | 1 | 0 | 3' UTR | 9 | 99111521 | 99118423 | 6903 | 2 | 623459 | ENSMUST00000188555.2 | 6907 | ENSMUSG00000096316 | Faiml | Fas apoptotic inhibitory molecule like |
| chr4 | 56797501 | 56798394 | 894 | * | 6,879953 | -5,271781282 | 1,886837 | -2,79398 | 0,005206 | 0,782859869 | 1,360622 | Intron (ENSMUST0000( | 4 | 56749680 | 56802331 | 52652 | 2 | 230233 | ENSMUST00000030140.3 | 3937 | ENSMUSG00000028431 | Elp1 | elongator complex protein 1 |
| chr17 | 31195410 | 31196842 | 1433 | * | 12,18294 | -5,271758474 | 1,634347 | -3,22561 | 0,001257 | 0,606153628 | 0,918255 | Intron (ENSMUST0000( | 17 | 31173653 | 31229684 | 56032 | 1 | 52020 | ENSMUST00000237889.2 | 21757 | ENSMUSG00000054134 | Umodl1 | uromodulin-like 1 |
| chr12 | 28868014 | 28868493 | 480 | * | 5,022008 | -5,269432601 | 2,032862 | -2,59213 | 0,009539 | 1 | 0 | Intron (ENSMUST0000( | 12 | 28860137 | 28861190 | 1054 | 1 | 380752 | ENSMUST00000222829.2 | 7877 | ENSMUSG00000036613 | Eipr1 | EARP complex and GARP complex interacting protein 1 |
| chr14 | 1,21E+08 | 1,21E+08 | 1048 | * | 6,865374 | -5,262629913 | 1,919131 | -2,74219 | 0,006103 | 1 | 0 | Intron (ENSMUST0000( | 14 | 1,21E+08 | 1,21E+08 | 35211 | 1 | 105559 | ENSMUST00000227153.2 | 8611 | ENSMUSG00000022139 | Mbnl2 | muscleblind like splicing factor 2 |
| chr13 | 32815795 | 32816838 | 1044 | * | 7,036191 | -5,262521379 | 1,857906 | -2,8325 | 0,004619 | 1 | 0 | Intron (ENSMUST0000( | 13 | 32901558 | 32960396 | 58839 | 2 | 238564 | ENSMUST00000230119.2 | 143558 | ENSMUSG00000044951 | Mylk4 | myosin light chain kinase family, member 4 |
| chr6 | 82655305 | 82655713 | 409 | * | 5,973113 | -5,262354142 | 1,896081 | -2,77539 | 0,005514 | 1 | 0 | Promoter (1-2kb) | 6 | 82657113 | 82671803 | 14691 | 1 | 1,03E+08 | ENSMUST00000203711.2 | -1400 | | Gm32425 | predicted gene, 32425 |
| chr11 | 54060367 | 54061292 | 926 | * | 6,356825 | -5,262331588 | 1,888669 | -2,78626 | 0,005332 | 1 | 0 | Distal Intergenic | 11 | 54018519 | 54022491 | 3973 | 1 | 18452 | ENSMUST00000139315.2 | 41848 | ENSMUSG00000018906 | P4ha2 | procollagen-proline, 2-oxoglutarate 4-dioxygenase (proline 4-hydroxylase), alpha II polypeptide |
| chr1 | 1,86E+08 | 1,86E+08 | 995 | * | 6,047019 | -5,261866314 | 1,989748 | -2,64449 | 0,008181 | 1 | 0 | Distal Intergenic | 1 | 1,86E+08 | 1,86E+08 | 46635 | 2 | 320449 | ENSMUST00000193084.2 | 120678 | ENSMUSG00000103690 | A730004F24Rik | RIKEN cDNA A730004F24 gene |
| chr1 | 27227490 | 27227994 | 505 | * | 4,677363 | -5,261736069 | 2,024576 | -2,59893 | 0,009351 | 1 | 0 | Distal Intergenic | 1 | 26720895 | 26726541 | 5647 | 2 | 210940 | ENSMUST00000097801.4 | -500949 | ENSMUSG00000073722 | 4931408C20Rik | RIKEN cDNA 4931408C20 gene |
| chr8 | 5042996 | 5043321 | 326 | * | 5,156843 | -5,261687632 | 2,03872 | -2,58088 | 0,009855 | 1 | 0 | Distal Intergenic | 8 | 5133219 | 5155351 | 22133 | 2 | 20494 | ENSMUST00000023835.3 | 112030 | ENSMUSG00000023073 | Slc10a2 | solute carrier family 10, member 2 |
| chr12 | 97261761 | 97262204 | 444 | * | 5,075302 | -5,261679631 | 2,040523 | -2,57859 | 0,00992 | 1 | 0 | Distal Intergenic | 12 | 97279623 | 97282627 | 3005 | 1 | 1,05E+08 | ENSMUST00000218233.2 | -17419 | | Gm40552 | predicted gene, 40552 |
| chr10 | 99234047 | 99235864 | 1818 | * | 12,65133 | -5,259250214 | 1,668794 | -3,15153 | 0,001624 | 0,684274806 | 0,738441 | Promoter (2-3kb) | 10 | 99238652 | 99249134 | 10483 | 1 | 1,19E+08 | ENSMUST00000218725.2 | -2788 | | LOC118567817 | uncharacterized LOC118567817 |
| chrX | 1,38E+08 | 1,38E+08 | 1086 | * | 8,497159 | -5,258122759 | 1,690555 | -3,11029 | 0,001869 | 0,594676756 | 1,449706 | Intron (ENSMUST0000( | 20 | 1,38E+08 | 1,38E+08 | 94663 | 1 | 27206 | ENSMUST00000064937.14 | -68996 | ENSMUSG00000052854 | Nrk | Nik related kinase |
| chrX | 97793189 | 97793925 | 737 | * | 6,366392 | -5,255962895 | 2,039708 | -2,57682 | 0,009971 | 1 | 0 | Intron (ENSMUST0000( | 20 | 97858381 | 97858487 | 107 | 1 | 1,15E+08 | ENSMUST00000157991.3 | -64456 | ENSMUSG00000088616 | Gm26000 | predicted gene, 26000 |
| chr6 | 76202654 | 76203222 | 569 | * | 7,08013 | -5,253028342 | 1,914099 | -2,74439 | 0,006062 | 1 | 0 | Distal Intergenic | 6 | 76177101 | 76177261 | 161 | 2 | 1,15E+08 | ENSMUST00000082556.3 | -25393 | ENSMUSG00000064490 | Gm22283 | predicted gene, 22283 |
| chr13 | 37511821 | 37512412 | 592 | * | 5,609372 | -5,252781393 | 1,903941 | -2,7589 | 0,0058 | 1 | 0 | Distal Intergenic | 13 | 37529184 | 37603012 | 73829 | 1 | 17084 | ENSMUST00000021860.7 | -16772 | ENSMUSG00000021423 | Ly86 | lymphocyte antigen 86 |
| chr9 | 1,02E+08 | 1,02E+08 | 598 | * | 6,663002 | -5,25261359 | 1,925643 | -2,72772 | 0,006377 | 1 | 0 | Intron (ENSMUST0000( | 9 | 1,02E+08 | 1,02E+08 | 17607 | 2 | 270190 | ENSMUST00000215514.2 | 8082 | ENSMUSG00000032537 | Ephb1 | Eph receptor B1 |
| chr8 | 31629903 | 31630381 | 479 | * | 7,403282 | -5,252116063 | 1,830763 | -2,86881 | 0,00412 | 1 | 0 | Distal Intergenic | 8 | 31639892 | 31639995 | 104 | 2 | 1E+08 | ENSMUST00000157627.3 | 9614 | ENSMUSG00000088252 | Snord13 | small nucleolar RNA, C/D box 13 |
| chr12 | 1,14E+08 | 1,14E+08 | 444 | * | 4,4906 | -5,244041028 | 2,027869 | -2,58599 | 0,00971 | 1 | 0 | Distal Intergenic | 12 | 1,14E+08 | 1,14E+08 | 2558 | 1 | 238406 | ENSMUST00000053086.3 | 39639 | ENSMUSG00000043945 | Adam6a | a disintegrin and metallopeptidase domain 6A |
| chr9 | 5183377 | 5183977 | 601 | * | 5,455569 | -5,244028241 | 2,025099 | -2,58952 | 0,009611 | 1 | 0 | Distal Intergenic | 9 | 5111399 | 5111533 | 135 | 1 | 1,15E+08 | ENSMUST00000122741.2 | 71978 | ENSMUSG00000084690 | Gm24896 | predicted gene, 24896 |
| chr9 | 38496574 | 38497224 | 651 | * | 6,601725 | -5,243851983 | 1,919538 | -2,73183 | 0,006298 | 1 | 0 | 3' UTR | 9 | 38491474 | 38498172 | 6699 | 1 | 258806 | ENSMUST00000217160.3 | 5100 | ENSMUSG00000111448 | Olfr912 | olfactory receptor 912 |
| chr2 | 1,56E+08 | 1,56E+08 | 1036 | * | 6,236187 | -5,243031157 | 1,925821 | -2,72249 | 0,006479 | 1 | 0 | Intron (ENSMUST0000( | 2 | 1,56E+08 | 1,56E+08 | 3195 | 1 | 68295 | ENSMUST00000126250.2 | 24898 | ENSMUSG00000027628 | Aar2 | AAR2 splicing factor homolog |
| chrX | 25500145 | 25501227 | 1083 | * | 4,826513 | -5,242488818 | 1,986571 | -2,63896 | 0,008316 | 1 | 0 | Distal Intergenic | 20 | 25503596 | 25525752 | 22157 | 2 | 1E+08 | ENSMUST00000120268.2 | 24525 | ENSMUSG00000073267 | Gm1993 | predicted gene 1993 |
| chr7 | 1,44E+08 | 1,44E+08 | 1180 | * | 9,364821 | -5,241762453 | 1,745825 | -3,00246 | 0,002678 | 0,579150837 | 2,237403 | Promoter (1-2kb) | 7 | 1,44E+08 | 1,44E+08 | 5809 | 1 | 320688 | ENSMUST00000209134.2 | -1048 | | D930030F02Rik | RIKEN cDNA D930030F02 gene |
| chr4 | 56482845 | 56483522 | 678 | * | 4,999432 | -5,236031694 | 2,029924 | -2,57942 | 0,009897 | 1 | 0 | Distal Intergenic | 4 | 56678904 | 56679035 | 132 | 2 | 1,15E+08 | ENSMUST00000104475.2 | 195513 | ENSMUSG00000077663 | Gm26144 | predicted gene, 26144 |
| chr13 | 64018546 | 64019257 | 712 | * | 4,95505 | -5,233959229 | 1,960135 | -2,6702 | 0,007581 | 1 | 0 | Promoter (<=1kb) | 13 | 64018323 | 64020311 | 1989 | 1 | 76251 | ENSMUST00000109776.3 | 223 | ENSMUSG00000021470 | Ercc6l2 | excision repair cross-complementing rodent repair deficiency, complementation group 6 like 2 |
| chr14 | 61777285 | 61778079 | 795 | * | 6,307757 | -5,233865504 | 1,892111 | -2,76615 | 0,005672 | 1 | 0 | Exon (ENSMUST00000( | 14 | 61771754 | 61794207 | 22454 | 2 | 66674 | ENSMUST00000100496.5 | 16128 | ENSMUSG00000021930 | Spryd7 | SPRY domain containing 7 |
| chr7 | 39778677 | 39779256 | 580 | * | 5,477899 | -5,23379182 | 1,912804 | -2,73619 | 0,006216 | 1 | 0 | Distal Intergenic | 7 | 39279816 | 39279926 | 111 | 1 | 1,15E+08 | ENSMUST00000093724.4 | 498861 | ENSMUSG00000070236 | Gm26327 | predicted gene, 26327 |
| chrX | 88592363 | 88593179 | 817 | * | 5,804933 | -5,23369784 | 1,890109 | -2,76899 | 0,005623 | 1 | 0 | Distal Intergenic | 20 | 88447454 | 88453295 | 5842 | 2 | 18630 | ENSMUST00000113960.3 | -139068 | ENSMUSG00000035395 | Dcaf8l | DDB1 and CUL4 associated factor 8 like |
| chr15 | 41378522 | 41379074 | 553 | * | 7,072902 | -5,233320796 | 1,81736 | -2,87963 | 0,003981 | 1 | 0 | Intron (ENSMUST0000( | 15 | 41312423 | 41724443 | 412021 | 1 | 170719 | ENSMUST00000230203.2 | 66099 | ENSMUSG00000022307 | Oxr1 | oxidation resistance 1 |
| chr8 | 1,17E+08 | 1,17E+08 | 456 | * | 6,487448 | -5,232599245 | 1,83501 | -2,85154 | 0,004351 | 1 | 0 | Intron (ENSMUST0000( | 8 | 1,17E+08 | 1,17E+08 | 10901 | 1 | 75465 | ENSMUST00000078170.7 | 90990 | ENSMUSG00000034467 | Dynlrb2 | dynein light chain roadblock-type 2 |
| chr17 | 27451691 | 27452124 | 434 | * | 5,290254 | -5,232487225 | 2,010864 | -2,60211 | 0,009265 | 1 | 0 | Distal Intergenic | 17 | 27454043 | 27461613 | 7571 | 2 | 319720 | ENSMUST00000244703.1 | 9489 | ENSMUSG00000117916 | 9630028I04Rik | RIKEN cDNA 9630028I04 gene |
| chr1 | 20812865 | 20813615 | 751 | * | 4,765865 | -5,225946915 | 2,020785 | -2,5861 | 0,009707 | 1 | 0 | Distal Intergenic | 1 | 20801129 | 20804720 | 3592 | 1 | 16171 | ENSMUST00000027061.5 | 11736 | ENSMUSG00000025929 | Il17a | interleukin 17A |
| chr2 | 1,25E+08 | 1,25E+08 | 380 | * | 5,273431 | -5,225812969 | 2,010221 | -2,59962 | 0,009333 | 1 | 0 | Intron (ENSMUST0000( | 2 | 1,25E+08 | 1,25E+08 | 77403 | 1 | 20495 | ENSMUST00000028630.9 | 4866 | ENSMUSG00000027202 | Slc12a1 | solute carrier family 12, member 1 |
| chr19 | 52627483 | 52628161 | 679 | * | 5,391448 | -5,225227519 | 1,968635 | -2,65424 | 0,007949 | 1 | 0 | Distal Intergenic | 19 | 52911612 | 52915362 | 3751 | 1 | 1E+08 | ENSMUST00000236393.2 | -283451 | ENSMUSG00000117896 | AA387883 | expressed sequence AA387883 |
| chr3 | 37716638 | 37717607 | 970 | * | 5,151209 | -5,224163472 | 1,96294 | -2,6614 | 0,007782 | 1 | 0 | Distal Intergenic | 3 | 37692728 | 37694805 | 2078 | 2 | 1,19E+08 | ENSMUST00000195887.2 | -21833 | | Gm42922 | predicted gene 42922 |
| chr4 | 71617785 | 71618284 | 500 | * | 5,578552 | -5,224117099 | 1,921482 | -2,7188 | 0,006552 | 1 | 0 | Distal Intergenic | 4 | 71673880 | 71676202 | 2323 | 2 | 1,03E+08 | ENSMUST00000084489.3 | 57918 | ENSMUSG00000066141 | Gm11232 | predicted gene 11232 |
| chr13 | 94862544 | 94863010 | 467 | * | 4,889047 | -5,222643518 | 2,007915 | -2,60103 | 0,009294 | 1 | 0 | Distal Intergenic | 13 | 94837526 | 94839226 | 1701 | 2 | 1,03E+08 | ENSMUST00000220503.2 | -23318 | | Gm32351 | predicted gene, 32351 |
| chr14 | 1,21E+08 | 1,21E+08 | 664 | * | 5,008623 | -5,218266432 | 1,987318 | -2,62578 | 0,008645 | 1 | 0 | Distal Intergenic | 14 | 1,21E+08 | 1,21E+08 | 36776 | 1 | 70572 | ENSMUST00000032898.9 | -37721 | ENSMUSG00000030662 | Ipo5 | importin 5 |
| chr10 | 21071970 | 21073268 | 1299 | * | 9,031365 | -5,214582923 | 1,79791 | -2,90036 | 0,003727 | 0,723174878 | 1,360622 | Distal Intergenic | 10 | 21053526 | 21065081 | 11556 | 2 | 1,05E+08 | ENSMUST00000213692.2 | -6889 | | Gm40608 | predicted gene, 40608 |
| chr18 | 36594125 | 36594817 | 693 | * | 4,767643 | -5,214266797 | 1,993212 | -2,61601 | 0,008896 | 1 | 0 | Distal Intergenic | 18 | 36536729 | 36587577 | 50849 | 2 | 67199 | ENSMUST00000237792.2 | -6548 | ENSMUSG00000024346 | Pfdn1 | prefoldin 1 |
| chr1 | 98586474 | 98586847 | 374 | * | 4,598656 | -5,214246156 | 2,012049 | -2,59151 | 0,009556 | 1 | 0 | Distal Intergenic | 1 | 98517243 | 98534869 | 17627 | 1 | 666182 | ENSMUST00000190991.2 | 69231 | | Gm7967 | predicted gene 7967 |
| chr15 | 80806670 | 80807274 | 605 | * | 5,256938 | -5,212680157 | 2,01096 | -2,59214 | 0,009538 | 1 | 0 | Promoter (<=1kb) | 15 | 80807265 | 80814233 | 6969 | 1 | 213988 | ENSMUST00000228525.2 | 0 | ENSMUSG00000047888 | Tnrc6b | trinucleotide repeat containing 6b |
| chr19 | 5218714 | 5220660 | 1947 | * | 12,15201 | -5,208616526 | 1,705859 | -3,05337 | 0,002263 | 0,652564605 | 1,2763 | Intron (ENSMUST0000( | 19 | 5191577 | 5193527 | 1951 | 2 | 107975 | ENSMUST00000237967.2 | -25187 | ENSMUSG00000024855 | Pacs1 | phosphofurin acidic cluster sorting protein 1 |
| chr9 | 1,24E+08 | 1,24E+08 | 1382 | * | 9,829615 | -5,207762164 | 1,769478 | -2,94311 | 0,003249 | 0,73764377 | 1,086118 | Promoter (<=1kb) | 9 | 1,24E+08 | 1,24E+08 | 103527 | 1 | 12769 | ENSMUST00000163559.8 | 0 | ENSMUSG00000029530 | Ccr9 | chemokine (C-C motif) receptor 9 |
| chr3 | 1,33E+08 | 1,33E+08 | 633 | * | 8,138181 | -5,207371589 | 1,730061 | -3,00994 | 0,002613 | 0,658947078 | 1,425929 | Intron (ENSMUST0000( | 3 | 1,33E+08 | 1,33E+08 | 29992 | 1 | 74776 | ENSMUST00000106315.4 | 26471 | ENSMUSG00000028013 | Ppa2 | pyrophosphatase (inorganic) 2 |
| chr17 | 47421694 | 47422208 | 515 | * | 5,06903 | -5,205662303 | 1,998944 | -2,60421 | 0,009209 | 1 | 0 | Distal Intergenic | 17 | 47451801 | 47640462 | 188662 | 1 | 224829 | ENSMUST00000188947.7 | -29593 | ENSMUSG00000064043 | Trerf1 | transcriptional regulating factor 1 |
| chr14 | 39136714 | 39137115 | 402 | * | 5,43616 | -5,205660902 | 1,999002 | -2,60413 | 0,009211 | 1 | 0 | Intron (ENSMUST0000( | 14 | 38731144 | 39194623 | 463480 | 2 | 18183 | ENSMUST00000176122.2 | 57508 | ENSMUSG00000041014 | Nrg3 | neuregulin 3 |
| chr15 | 36293310 | 36294254 | 945 | * | 7,062145 | -5,204998023 | 1,830532 | -2,84343 | 0,004463 | 1 | 0 | Distal Intergenic | 15 | 36240079 | 36283293 | 43215 | 2 | 30945 | ENSMUST00000022890.10 | -10017 | ENSMUSG00000022280 | Rnf19a | ring finger protein 19A |
| chr7 | 4223790 | 4224748 | 959 | * | 6,87442 | -5,204750976 | 1,875548 | -2,77506 | 0,005519 | 1 | 0 | Distal Intergenic | 7 | 4240753 | 4246462 | 5710 | 1 | 232801 | ENSMUST00000117550.2 | -16005 | ENSMUSG00000070873 | Lilra5 | leukocyte immunoglobulin-like receptor, subfamily A (with TM domain), member 5 |
| chr3 | 1,32E+08 | 1,32E+08 | 786 | * | 6,466479 | -5,204680226 | 1,889932 | -2,7539 | 0,005889 | 1 | 0 | Distal Intergenic | 3 | 1,32E+08 | 1,32E+08 | 95013 | 1 | 56811 | ENSMUST00000029665.7 | 115670 | ENSMUSG00000028031 | Dkk2 | dickkopf WNT signaling pathway inhibitor 2 |
| chrX | 1,42E+08 | 1,42E+08 | 749 | * | 5,980756 | -5,204623042 | 1,902677 | -2,73542 | 0,00623 | 1 | 0 | Distal Intergenic | 20 | 1,42E+08 | 1,42E+08 | 104426 | 2 | 83453 | ENSMUST00000063029.13 | 111807 | ENSMUSG00000031283 | Chrdl1 | chordin-like 1 |
| chr9 | 78196628 | 78197412 | 785 | * | 5,248318 | -5,204434322 | 1,949101 | -2,67017 | 0,007581 | 1 | 0 | Promoter (<=1kb) | 9 | 78197205 | 78212807 | 15603 | 1 | 1E+08 | ENSMUST00000119823.8 | 0 | ENSMUSG00000074179 | Gsta5 | glutathione S-transferase alpha 5 |
| chr14 | 1,22E+08 | 1,22E+08 | 831 | * | 6,2459 | -5,204397939 | 1,960055 | -2,65523 | 0,007925 | 1 | 0 | Distal Intergenic | 14 | 1,22E+08 | 1,22E+08 | 3747 | 1 | 223262 | ENSMUST00000045976.7 | 11632 | ENSMUSG00000071229 | Timm8a2 | translocase of inner mitochondrial membrane 8A2 |
| chr18 | 86189389 | 86190078 | 690 | * | 5,50836 | -5,204307811 | 1,98594 | -2,62058 | 0,008778 | 1 | 0 | Distal Intergenic | 18 | 86367711 | 86367817 | 107 | 1 | 1,15E+08 | ENSMUST00000082733.3 | -177633 | ENSMUSG00000119275 | Gm25453 | predicted gene, 25453 |
| chr14 | 21893634 | 21894569 | 936 | * | 5,976932 | -5,203955134 | 1,939173 | -2,68359 | 0,007284 | 1 | 0 | Promoter (1-2kb) | 14 | 21892343 | 21895587 | 3245 | 1 | 22334 | ENSMUST00000142084.3 | 1291 | ENSMUSG00000021771 | Vdac2 | voltage-dependent anion channel 2 |
| chr6 | 10644899 | 10645490 | 592 | * | 5,913341 | -5,203246674 | 1,977189 | -2,63164 | 0,008497 | 1 | 0 | Distal Intergenic | 6 | 10840587 | 10840693 | 107 | 2 | 1,15E+08 | ENSMUST00000240049.1 | 195203 | ENSMUSG00000118743 | Gm26058 | predicted gene, 26058 |
| chr1 | 14575400 | 14576160 | 761 | * | 5,56398 | -5,202828962 | 2,00198 | -2,59884 | 0,009354 | 1 | 0 | Distal Intergenic | 1 | 14239182 | 14380459 | 141278 | 2 | 14048 | ENSMUST00000168081.9 | -194941 | ENSMUSG00000025932 | Eya1 | EYA transcriptional coactivator and phosphatase 1 |
| chr5 | 42598462 | 42599666 | 1205 | * | 9,304414 | -5,196214036 | 1,729773 | -3,00399 | 0,002665 | 0,661775851 | 1,425929 | Distal Intergenic | 5 | 42225303 | 42374141 | 148839 | 1 | 433882 | ENSMUST00000087332.5 | 373159 | ENSMUSG00000067285 | Gm16223 | predicted gene 16223 |
| chr12 | 86053733 | 86054596 | 864 | * | 7,200585 | -5,195469848 | 1,827774 | -2,84251 | 0,004476 | 1 | 0 | Intron (ENSMUST0000( | 12 | 86008775 | 86100296 | 91522 | 1 | 320244 | ENSMUST00000220906.2 | 44958 | ENSMUSG00000012609 | Ttll5 | tubulin tyrosine ligase-like family, member 5 |
| chrX | 1,39E+08 | 1,39E+08 | 744 | * | 5,836843 | -5,195382916 | 1,975404 | -2,63004 | 0,008538 | 1 | 0 | Promoter (<=1kb) | 20 | 1,39E+08 | 1,39E+08 | 93205 | 1 | 245643 | ENSMUST00000208130.2 | 612 | ENSMUSG00000042425 | Frmpd3 | FERM and PDZ domain containing 3 |
| chr18 | 66389728 | 66390341 | 614 | * | 5,11607 | -5,195244895 | 1,960473 | -2,65 | 0,008049 | 1 | 0 | Intron (ENSMUST0000( | 18 | 66178373 | 66424517 | 246145 | 2 | 320924 | ENSMUST00000146610.2 | 34176 | ENSMUSG00000046318 | Ccbe1 | collagen and calcium binding EGF domains 1 |
| chr14 | 4439700 | 4440575 | 876 | * | 5,188387 | -5,195142569 | 1,949547 | -2,66479 | 0,007704 | 1 | 0 | Intron (ENSMUST0000( | 14 | 4431611 | 4488167 | 56557 | 2 | 21834 | ENSMUST00000022304.12 | 47592 | ENSMUSG00000021779 | Thrb | thyroid hormone receptor beta |
| chr3 | 56180359 | 56181144 | 786 | * | 6,371493 | -5,194517057 | 1,891854 | -2,74573 | 0,006038 | 1 | 0 | Distal Intergenic | 3 | 55532616 | 56091122 | 558507 | 2 | 26422 | ENSMUST00000029374.8 | -89237 | ENSMUSG00000027799 | Nbea | neurobeachin |
| chr9 | 71288989 | 71290072 | 1084 | * | 7,317721 | -5,194482292 | 1,832191 | -2,83512 | 0,004581 | 1 | 0 | Distal Intergenic | 9 | 71385753 | 71392240 | 6488 | 2 | 28015 | ENSMUST00000163972.2 | 102168 | ENSMUSG00000032199 | Polr2m | polymerase (RNA) II (DNA directed) polypeptide M |
| chr3 | 99226942 | 99227648 | 707 | * | 5,596867 | -5,194393062 | 1,966968 | -2,64081 | 0,008271 | 1 | 0 | Intron (ENSMUST0000( | 3 | 99203826 | 99223556 | 19731 | 1 | 21384 | ENSMUST00000150756.3 | 23116 | ENSMUSG00000027868 | Tbx15 | T-box 15 |
| chr14 | 84785416 | 84786028 | 613 | * | 4,822645 | -5,19430596 | 1,98236 | -2,62026 | 0,008786 | 1 | 0 | Distal Intergenic | 14 | 84696164 | 84713864 | 17701 | 2 | 319903 | ENSMUST00000228125.2 | -71552 | ENSMUSG00000115529 | 9630013A20Rik | RIKEN cDNA 9630013A20 gene |
| chr1 | 1,42E+08 | 1,42E+08 | 782 | * | 5,179448 | -5,194263699 | 1,990113 | -2,61003 | 0,009053 | 1 | 0 | Distal Intergenic | 1 | 1,41E+08 | 1,41E+08 | 107 | 1 | 1,15E+08 | ENSMUST00000239951.1 | 900439 | ENSMUSG00000118759 | Gm25457 | predicted gene, 25457 |
| chr1 | 80830716 | 80831222 | 507 | * | 4,675529 | -5,194193129 | 2,003426 | -2,59265 | 0,009524 | 1 | 0 | Distal Intergenic | 1 | 80560934 | 80736244 | 175311 | 2 | 210293 | ENSMUST00000191505.2 | -94472 | ENSMUSG00000038608 | Dock10 | dedicator of cytokinesis 10 |
| chrX | 12319328 | 12320428 | 1101 | * | 9,791581 | -5,193697589 | 1,696861 | -3,06077 | 0,002208 | 0,627862701 | 1,425929 | Distal Intergenic | 20 | 12206021 | 12221235 | 15215 | 2 | 1E+08 | ENSMUST00000146535.8 | -98093 | ENSMUSG00000087591 | Gm14635 | predicted gene 14635 |
| chr17 | 64838204 | 64839285 | 1082 | * | 9,388594 | -5,188709475 | 1,698118 | -3,05557 | 0,002246 | 0,631820259 | 1,425929 | Intron (ENSMUST0000( | 17 | 64821134 | 64862498 | 41365 | 2 | 1E+08 | ENSMUST00000168379.3 | 23213 | ENSMUSG00000073377 | AU016765 | expressed sequence AU016765 |
| chr12 | 21209896 | 21210592 | 697 | * | 9,869622 | -5,187599108 | 1,736061 | -2,98814 | 0,002807 | 0,6980047 | 1,173325 | Intron (ENSMUST0000( | 12 | 21161955 | 21318180 | 156226 | 1 | 211914 | ENSMUST00000050990.10 | 47941 | ENSMUSG00000052632 | Asap2 | ArfGAP with SH3 domain, ankyrin repeat and PH domain 2 |
| chr12 | 52243881 | 52244478 | 598 | * | 7,269156 | -5,186085162 | 1,814214 | -2,85859 | 0,004255 | 1 | 0 | Intron (ENSMUST0000( | 12 | 52222870 | 52317862 | 94993 | 1 | 76826 | ENSMUST00000162111.2 | 21011 | ENSMUSG00000035142 | Nubpl | nucleotide binding protein-like |
| chr2 | 1,24E+08 | 1,24E+08 | 633 | * | 6,391098 | -5,185701768 | 1,948904 | -2,66083 | 0,007795 | 1 | 0 | Intron (ENSMUST0000( | 2 | 1,24E+08 | 1,24E+08 | 364697 | 1 | 214968 | ENSMUST00000151199.2 | 31963 | ENSMUSG00000027200 | Sema6d | sema domain, transmembrane domain (TM), and cytoplasmic domain, (semaphorin) 6D |
| chr19 | 3393045 | 3393621 | 577 | * | 5,821232 | -5,184935112 | 1,904578 | -2,72235 | 0,006482 | 1 | 0 | Promoter (1-2kb) | 19 | 3391627 | 3414538 | 22912 | 1 | 12894 | ENSMUST00000238015.2 | 1418 | ENSMUSG00000024900 | Cpt1a | carnitine palmitoyltransferase 1a, liver |
| chr16 | 26809855 | 26810911 | 1057 | * | 7,346807 | -5,184073627 | 1,93986 | -2,6724 | 0,007531 | 1 | 0 | Promoter (<=1kb) | 16 | 26777855 | 26810402 | 32548 | 2 | 239789 | ENSMUST00000231299.2 | 0 | ENSMUSG00000068428 | Gmnc | geminin coiled-coil domain containing |
| chr11 | 63911990 | 63912919 | 930 | * | 7,613877 | -5,183871376 | 1,821369 | -2,84614 | 0,004425 | 1 | 0 | Intron (ENSMUST0000( | 11 | 63855052 | 63858866 | 3815 | 1 | 78883 | ENSMUST00000146915.2 | 56938 | ENSMUSG00000086359 | 9630013K17Rik | RIKEN cDNA 9630013K17 gene |
| chr9 | 59024005 | 59024889 | 885 | * | 8,253083 | -5,183505081 | 1,79456 | -2,88845 | 0,003871 | 0,722439117 | 1,425929 | Distal Intergenic | 9 | 58781970 | 58943724 | 161755 | 2 | 18007 | ENSMUST00000068664.7 | -80281 | ENSMUSG00000032340 | Neo1 | neogenin |
| chr7 | 1,2E+08 | 1,2E+08 | 1538 | * | 9,196621 | -5,183460077 | 1,791428 | -2,89348 | 0,00381 | 0,727652793 | 1,360622 | Exon (ENSMUST00000( | 7 | 1,2E+08 | 1,2E+08 | 123113 | 1 | 233810 | ENSMUST00000120490.8 | 4069 | ENSMUSG00000051900 | Abca16 | ATP-binding cassette, sub-family A (ABC1), member 16 |
| chr5 | 1,2E+08 | 1,2E+08 | 736 | * | 5,084083 | -5,175245529 | 2,001186 | -2,58609 | 0,009707 | 1 | 0 | Distal Intergenic | 5 | 1,2E+08 | 1,2E+08 | 538 | 2 | 100993 | ENSMUST00000200851.2 | 14807 | | | |
| chr14 | 83984665 | 83985575 | 911 | * | 6,193635 | -5,17500669 | 1,947771 | -2,65689 | 0,007887 | 1 | 0 | Distal Intergenic | 14 | 84283871 | 84312730 | 28860 | 2 | 1,03E+08 | ENSMUST00000228897.2 | 327155 | ENSMUSG00000115043 | Gm32455 | predicted gene, 32455 |
| chr1 | 84838243 | 84838807 | 565 | * | 5,352179 | -5,174918979 | 1,930436 | -2,6807 | 0,007347 | 1 | 0 | Intron (ENSMUST0000( | 1 | 84844157 | 84878208 | 34052 | 1 | 66153 | ENSMUST00000139562.8 | -5350 | ENSMUSG00000073633 | Fbxo36 | F-box protein 36 |
| chr15 | 80842761 | 80843324 | 564 | * | 5,567573 | -5,174904457 | 1,928603 | -2,68324 | 0,007291 | 1 | 0 | Promoter (2-3kb) | 15 | 80845414 | 80848045 | 2632 | 1 | 11564 | ENSMUST00000163575.2 | -2090 | ENSMUSG00000022407 | Adsl | adenylosuccinate lyase |
| chr15 | 25900236 | 25900833 | 598 | * | 5,791189 | -5,174862676 | 1,919733 | -2,69562 | 0,007026 | 1 | 0 | Intron (ENSMUST0000( | 15 | 25910510 | 25966778 | 56269 | 1 | 66270 | ENSMUST00000228600.2 | -9677 | ENSMUSG00000022270 | Retreg1 | reticulophagy regulator 1 |
| chr14 | 75497458 | 75498373 | 916 | * | 6,170804 | -5,174853401 | 1,91812 | -2,69788 | 0,006978 | 1 | 0 | Exon (ENSMUST00000( | 14 | 75509789 | 75520986 | 11198 | 1 | 56373 | ENSMUST00000227817.2 | -11416 | ENSMUSG00000021999 | Cpb2 | carboxypeptidase B2 (plasma) |
| chr4 | 22295241 | 22295883 | 643 | * | 6,024731 | -5,174847878 | 1,917398 | -2,69889 | 0,006957 | 1 | 0 | Distal Intergenic | 4 | 22357543 | 22376725 | 19183 | 1 | 269514 | ENSMUST00000131972.8 | -61660 | ENSMUSG00000040410 | Fbxl4 | F-box and leucine-rich repeat protein 4 |
| chr5 | 1,32E+08 | 1,32E+08 | 634 | * | 5,755467 | -5,174055358 | 1,988078 | -2,60254 | 0,009254 | 1 | 0 | Intron (ENSMUST0000( | 5 | 1,32E+08 | 1,32E+08 | 418371 | 2 | 319974 | ENSMUST00000160135.4 | 187927 | ENSMUSG00000029673 | Auts2 | autism susceptibility candidate 2 |
| chr7 | 79806051 | 79806575 | 525 | * | 5,003543 | -5,17387765 | 2,006409 | -2,57868 | 0,009918 | 1 | 0 | Promoter (1-2kb) | 7 | 79804917 | 79806519 | 1603 | 1 | 1,15E+08 | ENSMUST00000205955.2 | 1134 | | Gm45202 | predicted gene 45202 |

| | | | | | | | | | | | | | | | | | | | | | | | |
|---|---|---|---|---|---|---|---|---|---|---|---|---|---|---|---|---|---|---|---|---|---|---|---|
| chr14 | 72441407 | 72442166 | 760 | * | 4,764732 | -5,173873504 | 2,006975 | -2,57795 | 0,009939 | 1 | 0 | Distal Intergenic | 14 | 72663453 | 72729278 | 65826 | 2 | 675947 | ENSMUST00000208955.2 | 287112 | | Gm9195 | predicted gene 9195 |
| chr18 | 73029840 | 73030728 | 889 | * | 9,469746 | -5,170964255 | 1,674441 | -3,08817 | 0,002014 | 0,61020361 | 1,449706 | Distal Intergenic | 18 | 73041961 | 73054161 | 12201 | 2 | 1,03E+08 | ENSMUST00000236243.2 | 23433 | ENSMUSG00000118316 | Gm31908 | predicted gene, 31908 |
| chr6 | 74141270 | 74141924 | 655 | * | 5,75115 | -5,164966786 | 1,99575 | -2,58798 | 0,009654 | 1 | 0 | Distal Intergenic | 6 | 74317933 | 74318064 | 132 | 1 | 1,15E+08 | ENSMUST00000240164.1 | -176009 | ENSMUSG00000118786 | Gm26179 | predicted gene, 26179 |
| chr11 | 71039081 | 71039543 | 463 | * | 5,47636 | -5,164840626 | 1,937611 | -2,66557 | 0,007686 | 1 | 0 | Distal Intergenic | 11 | 70983062 | 71035530 | 52469 | 2 | 195046 | ENSMUST00000108518.3 | -3551 | ENSMUSG00000069830 | Nlrp1a | NLR family, pyrin domain containing 1A |
| chr16 | 36262468 | 36263049 | 582 | * | 5,357466 | -5,16482919 | 1,93394 | -2,67063 | 0,007571 | 1 | 0 | Distal Intergenic | 16 | 36259006 | 36259108 | 103 | 1 | 1,15E+08 | ENSMUST00000158492.3 | 3462 | ENSMUSG00000089117 | Gm25903 | predicted gene, 25903 |
| chr1 | 1,86E+08 | 1,86E+08 | 830 | * | 5,829798 | -5,164765834 | 1,910913 | -2,70277 | 0,006876 | 1 | 0 | Distal Intergenic | 1 | 1,85E+08 | 1,85E+08 | 630 | 1 | 114668 | ENSMUST00000192616.2 | 307782 | ENSMUSG00000102185 | 5033404E19Rik | NSA2 ribosome biogenesis homolog pseudogene |
| chr14 | 64296864 | 64297922 | 1059 | * | 6,59441 | -5,164662132 | 1,874839 | -2,75472 | 0,005874 | 1 | 0 | Distal Intergenic | 14 | 64312894 | 64322857 | 9964 | 2 | 71037 | ENSMUST00000089338.6 | 24935 | ENSMUSG00000034623 | Prss55 | protease, serine 55 |
| chr5 | 24283614 | 24284821 | 1208 | * | 7,76813 | -5,164296238 | 1,784599 | -2,89381 | 0,003806 | 1 | 0 | Distal Intergenic | 5 | 24305603 | 24365783 | 60181 | 1 | 52323 | ENSMUST00000030841.10 | -20782 | ENSMUSG00000028986 | Klhl7 | kelch-like 7 |
| chr7 | 29669713 | 29670595 | 883 | * | 5,84463 | -5,164015342 | 1,975618 | -2,61387 | 0,008952 | 1 | 0 | Exon (ENSMUST000002 | 7 | 29683380 | 29724536 | 41157 | 1 | 243905 | ENSMUST00000148442.8 | -12785 | ENSMUSG00000074221 | Zfp568 | zinc finger protein 568 |
| chr8 | 1,25E+08 | 1,25E+08 | 1066 | * | 10,13214 | -5,163098798 | 1,851318 | -2,78888 | 0,005289 | 0,824134241 | 1,086118 | Intron (ENSMUST0000 | 8 | 1,25E+08 | 1,25E+08 | 23194 | 2 | 209966 | ENSMUST00000136892.3 | -19739 | ENSMUSG00000050751 | Pgbd5 | piggyBac transposable element derived 5 |
| chr9 | 52559799 | 52560590 | 792 | * | 7,445211 | -5,155529548 | 1,833294 | -2,81217 | 0,004921 | 1 | 0 | Distal Intergenic | 9 | 52584344 | 52590729 | 6386 | 2 | 330941 | ENSMUST00000098768.3 | 30139 | ENSMUSG00000078307 | AI593442 | expressed sequence AI593442 |
| chr17 | 66328261 | 66329174 | 914 | * | 8,172985 | -5,154683503 | 1,876298 | -2,74726 | 0,006009 | 1 | 0 | Intron (ENSMUST0000 | 17 | 66282843 | 66289949 | 7107 | 2 | 106585 | ENSMUST00000146090.2 | -38312 | ENSMUSG00000034647 | Ankrd12 | ankyrin repeat domain 12 |
| chr11 | 86769718 | 86770476 | 759 | * | 5,879125 | -5,154405371 | 1,93798 | -2,65968 | 0,007822 | 1 | 0 | Distal Intergenic | 11 | 86659672 | 86698572 | 38901 | 2 | 67487 | ENSMUST00000018569.14 | -71146 | ENSMUSG00000018425 | Dhx40 | DEAH (Asp-Glu-Ala-His) box polypeptide 40 |
| chr17 | 52902972 | 52903503 | 532 | * | 5,369899 | -5,153474292 | 1,997221 | -2,58032 | 0,009871 | 1 | 0 | Intron (ENSMUST0000 | 17 | 52900009 | 52909688 | 9680 | 2 | 1,08E+08 | ENSMUST00000183438.2 | 6185 | ENSMUSG00000098552 | Gm27217 | predicted gene 27217 |
| chr10 | 80138454 | 80139309 | 856 | * | 7,648013 | -5,152417592 | 1,883546 | -2,73549 | 0,006229 | 0,814343635 | 1,360622 | Promoter (<=1kb) | 10 | 80137654 | 80154090 | 16437 | 1 | 23805 | ENSMUST00000020349.7 | 800 | ENSMUSG00000020135 | Apc2 | APC regulator of WNT signaling pathway 2 |
| chr19 | 20960023 | 20960670 | 648 | * | 6,624388 | -5,149313737 | 1,789069 | -2,87821 | 0,003999 | 1 | 0 | Distal Intergenic | 19 | 20760822 | 20931566 | 170745 | 2 | 13409 | ENSMUST00000039500.4 | -28457 | ENSMUSG00000024749 | Tmc1 | transmembrane channel-like gene family 1 |
| chr4 | 1,21E+08 | 1,21E+08 | 505 | * | 5,706145 | -5,146941727 | 1,925947 | -2,67242 | 0,007531 | 1 | 0 | Promoter (2-3kb) | 4 | 1,21E+08 | 1,21E+08 | 3576 | 2 | 69469 | ENSMUST00000106268.4 | 2060 | ENSMUSG00000078577 | Tmco2 | transmembrane and coiled-coil domains 2 |
| chrX | 50113462 | 50114291 | 830 | * | 5,581204 | -5,145719153 | 1,916743 | -2,68462 | 0,007261 | 1 | 0 | Distal Intergenic | 20 | 50092789 | 50106895 | 14107 | 2 | 72065 | ENSMUST00000053593.8 | -6567 | ENSMUSG00000050029 | Rap2c | RAP2C, member of RAS oncogene family |
| chr5 | 33652898 | 33653703 | 806 | * | 7,023096 | -5,144874869 | 1,821662 | -2,82428 | 0,004739 | 0,764849314 | 1,360622 | Distal Intergenic | 5 | 33588078 | 33591320 | 3243 | 2 | 672284 | ENSMUST00000173348.2 | -61578 | ENSMUSG00000029112 | Nkx1-1 | NK1 homeobox 1 |
| chr16 | 67079015 | 67079872 | 858 | * | 7,220192 | -5,144870268 | 1,822241 | -2,82338 | 0,004752 | 1 | 0 | Intron (ENSMUST0000 | 16 | 66459979 | 66750222 | 290244 | 2 | 239857 | ENSMUST00000128168.3 | -328793 | ENSMUSG00000064115 | Cadm2 | cell adhesion molecule 2 |
| chr11 | 51407667 | 51408722 | 1056 | * | 8,119814 | -5,14485488 | 1,826037 | -2,8175 | 0,00484 | 0,683757236 | 2,237403 | Exon (ENSMUST00000 | 11 | 51465259 | 51471126 | 5868 | 1 | 237759 | ENSMUST00000151098.2 | -56537 | ENSMUSG00000063564 | Col23a1 | collagen, type XXIII, alpha 1 |
| chr2 | 68117424 | 68117993 | 570 | * | 5,916937 | -5,14459142 | 1,889742 | -2,72238 | 0,006481 | 1 | 0 | Intron (ENSMUST0000 | 2 | 68042084 | 68081700 | 39617 | 2 | 53416 | ENSMUST00000144457.2 | -35724 | ENSMUSG00000027030 | Stk39 | serine/threonine kinase 39 |
| chr13 | 87883056 | 87883738 | 683 | * | 5,932197 | -5,144492562 | 1,919544 | -2,68006 | 0,007361 | 1 | 0 | Distal Intergenic | 13 | 88669309 | 88726287 | 56979 | 1 | 1,03E+08 | ENSMUST00000224910.2 | -785571 | ENSMUSG00000114273 | Gm34585 | predicted gene, 34585 |
| chr9 | 53994443 | 53995395 | 953 | * | 6,067549 | -5,144395449 | 1,954693 | -2,63182 | 0,008493 | 1 | 0 | Distal Intergenic | 9 | 53930689 | 53975672 | 44984 | 2 | 244882 | ENSMUST00000098760.5 | -18771 | ENSMUSG00000074345 | Tnfaip8l3 | tumor necrosis factor, alpha-induced protein 8-like 3 |
| chr12 | 53082173 | 53082788 | 616 | * | 5,749595 | -5,144275579 | 1,913663 | -2,68818 | 0,007184 | 1 | 0 | Intron (ENSMUST0000 | 12 | 53093491 | 53093609 | 119 | 2 | 1,15E+08 | ENSMUST00000083299.3 | 10821 | ENSMUSG00000119139 | n-R5s58 | nuclear encoded rRNA 5S 58 |
| chr1 | 1,55E+08 | 1,55E+08 | 773 | * | 9,994709 | -5,142766286 | 1,679193 | -3,06264 | 0,002194 | 0,627785088 | 1,425929 | Intron (ENSMUST0000 | 1 | 1,54E+08 | 1,55E+08 | 335626 | 2 | 12290 | ENSMUST00000187541.7 | -33457 | ENSMUSG00000004110 | Cacna1e | calcium channel, voltage-dependent, R type, alpha 1E subunit |
| chr3 | 1,4E+08 | 1,4E+08 | 1092 | * | 8,637896 | -5,140578882 | 1,789521 | -2,8726 | 0,004071 | 0,73764377 | 1,360622 | Distal Intergenic | 3 | 1,4E+08 | 1,4E+08 | 122 | 1 | 1,02E+08 | ENSMUST00000184427.3 | 4612 | ENSMUSG00000099169 | Mir7j | microRNA 7j |
| chr8 | 68562923 | 68564476 | 1554 | * | 13,66128 | -5,13894826 | 1,607198 | -3,19746 | 0,001386 | 0,662625192 | 0,738441 | Intron (ENSMUST0000 | 8 | 68552106 | 68573965 | 21860 | 2 | 234353 | ENSMUST00000185416.3 | 9489 | ENSMUSG00000030465 | Psd3 | pleckstrin and Sec7 domain containing 3 |
| chr6 | 67536325 | 67536790 | 466 | * | 7,394708 | -5,1345429 | 1,877992 | -2,73406 | 0,006256 | 1 | 0 | Distal Intergenic | 6 | 67542175 | 67543430 | 1256 | 2 | 75070 | ENSMUST00000199440.2 | 6640 | | 4930515G16Rik | myc induced nuclear antigen pseudogene |
| chr2 | 1,71E+08 | 1,71E+08 | 493 | * | 6,032799 | -5,134463367 | 1,88905 | -2,71801 | 0,006568 | 1 | 0 | Distal Intergenic | 2 | 1,71E+08 | 1,71E+08 | 264 | 2 | 1,15E+08 | ENSMUST00000157880.3 | 182638 | ENSMUSG00000088505 | Gm26489 | predicted gene, 26489 |
| chr10 | 1,08E+08 | 1,08E+08 | 487 | * | 5,559244 | -5,13416045 | 1,937235 | -2,65025 | 0,008043 | 1 | 0 | Intron (ENSMUST0000 | 10 | 1,08E+08 | 1,08E+08 | 6695 | 1 | 1,03E+08 | ENSMUST00000220211.2 | 72395 | ENSMUSG00000112532 | Gm36283 | predicted gene, 36283 |
| chr6 | 1,29E+08 | 1,29E+08 | 476 | * | 7,179035 | -5,13270337 | 1,832928 | -2,80027 | 0,005106 | 1 | 0 | Promoter (<=1kb) | 6 | 1,29E+08 | 1,29E+08 | 1678 | 1 | 330428 | ENSMUST00000204692.2 | -172 | ENSMUSG00000030160 | Tmem52b | transmembrane protein 52B |
| chr13 | 20547443 | 20548273 | 831 | * | 8,811857 | -5,129756634 | 1,717821 | -2,9862 | 0,002825 | 0,693376834 | 1,211315 | Intron (ENSMUST0000 | 13 | 20467087 | 20557301 | 90215 | 1 | 140580 | ENSMUST00000221595.3 | 80356 | ENSMUSG00000041112 | Elmo1 | engulfment and cell motility 1 |
| chr4 | 79095146 | 79096077 | 932 | * | 10,90311 | -5,129166674 | 1,637438 | -3,13243 | 0,001734 | 0,604345495 | 1,2763 | Distal Intergenic | 4 | 78130313 | 78133846 | 3534 | 1 | 1,03E+08 | ENSMUST00000246656.1 | 964833 | | B230208B08Rik | RIKEN cDNA B230208B08 gene |
| chr11 | 78649065 | 78650582 | 1518 | * | 10,37853 | -5,126422877 | 1,66402 | -3,08075 | 0,002065 | 0,725842761 | 0,744375 | Distal Intergenic | 11 | 78641332 | 78642555 | 1224 | 2 | 69109 | ENSMUST00000059468.6 | -6510 | ENSMUSG00000049489 | Ccnq | cyclin Q |
| chr2 | 1,26E+08 | 1,26E+08 | 1109 | * | 6,744697 | -5,124340164 | 1,883375 | -2,72083 | 0,006512 | 1 | 0 | Intron (ENSMUST0000 | 2 | 1,26E+08 | 1,26E+08 | 1447 | 1 | 14178 | ENSMUST00000138576.2 | -60348 | ENSMUSG00000027208 | Fgf7 | fibroblast growth factor 7 |
| chr7 | 1,43E+08 | 1,43E+08 | 835 | * | 6,600919 | -5,124169654 | 1,899713 | -2,69734 | 0,00699 | 1 | 0 | Intron (ENSMUST0000 | 7 | 1,43E+08 | 1,43E+08 | 590 | 2 | 12577 | ENSMUST00000208118.2 | 46991 | ENSMUSG00000037664 | Cdkn1c | cyclin-dependent kinase inhibitor 1C (P57) |
| chr10 | 1,23E+08 | 1,23E+08 | 475 | * | 5,231166 | -5,12408157 | 1,955646 | -2,62015 | 0,008789 | 1 | 0 | Intron (ENSMUST0000 | 10 | 1,23E+08 | 1,23E+08 | 46082 | 1 | 319468 | ENSMUST00000162969.2 | 14218 | ENSMUSG00000034613 | Ppm1h | protein phosphatase 1H (PP2C domain containing) |
| chr1 | 1,61E+08 | 1,61E+08 | 714 | * | 5,641714 | -5,12389466 | 1,928525 | -2,6569 | 0,007886 | 1 | 0 | Distal Intergenic | 1 | 1,61E+08 | 1,61E+08 | 10636 | 1 | 240873 | ENSMUST00000086084.2 | 50643 | ENSMUSG00000066755 | Tnfsf18 | tumor necrosis factor (ligand) superfamily, member 18 |
| chr12 | 64338521 | 64339204 | 684 | * | 5,86259 | -5,123736106 | 1,919638 | -2,66912 | 0,007605 | 1 | 0 | Distal Intergenic | 12 | 64518104 | 64521464 | 3361 | 2 | 623046 | ENSMUST00000059833.8 | 182260 | ENSMUSG00000043060 | Fscb | fibrous sheath CABYR binding protein |
| chr19 | 60278903 | 60279495 | 593 | * | 6,450401 | -5,123321014 | 1,882024 | -2,72224 | 0,006484 | 1 | 0 | Distal Intergenic | 19 | 60187867 | 60215133 | 27267 | 2 | 76539 | ENSMUST00000237339.2 | -63770 | ENSMUSG00000057858 | Fam204a | family with sequence similarity 204, member A |
| chr1 | 1,78E+08 | 1,78E+08 | 1596 | * | 10,44924 | -5,118164039 | 1,707082 | -2,9982 | 0,002716 | 0,775744126 | 0,738441 | Intron (ENSMUST0000 | 1 | 1,78E+08 | 1,78E+08 | 107 | 2 | 1,15E+08 | ENSMUST00000240272.1 | 5840 | ENSMUSG00000119789 | Gm25312 | predicted gene, 25312 |
| chrX | 1,51E+08 | 1,51E+08 | 796 | * | 6,604787 | -5,114997291 | 1,83479 | -2,78778 | 0,005307 | 1 | 0 | Distal Intergenic | 20 | 1,51E+08 | 1,51E+08 | 2598 | 1 | 15108 | ENSMUST00000026289.10 | -10386 | ENSMUSG00000025260 | Hsd17b10 | hydroxysteroid (17-beta) dehydrogenase 10 |
| chr2 | 1,18E+08 | 1,18E+08 | 978 | * | 5,367392 | -5,113684385 | 1,97226 | -2,5928 | 0,00952 | 1 | 0 | Intron (ENSMUST0000 | 2 | 1,18E+08 | 1,18E+08 | 11635 | 2 | 71313 | ENSMUST00000231041.2 | -13943 | ENSMUSG00000027344 | Fsip1 | fibrous sheath-interacting protein 1 |
| chr15 | 63370305 | 63371050 | 746 | * | 7,142328 | -5,113196535 | 1,901287 | -2,68933 | 0,007159 | 1 | 0 | Intron (ENSMUST0000 | 15 | 63353063 | 63353125 | 63 | 2 | 1,15E+08 | ENSMUST00000239577.1 | -17180 | ENSMUSG00000119866 | Gm25702 | predicted gene, 25702 |
| chr11 | 89574175 | 89574760 | 586 | * | 5,444629 | -5,113038756 | 1,973163 | -2,59129 | 0,009562 | 1 | 0 | Intron (ENSMUST0000 | 11 | 89311813 | 89587671 | 275859 | 2 | 382543 | ENSMUST00000128717.9 | 12911 | ENSMUSG00000047773 | Ankfn1 | ankyrin-repeat and fibronectin type III domain containing 1 |
| chr6 | 6841549 | 6842639 | 1091 | * | 10,68498 | -5,108680381 | 1,798948 | -2,83982 | 0,004514 | 0,767135014 | 1,2763 | Intron (ENSMUST0000 | 6 | 6824188 | 6830791 | 6604 | 2 | 320038 | ENSMUST00000210702.2 | -10758 | ENSMUSG00000090063 | Dlx6os1 | distal-less homeobox 6, opposite strand 1 |
| chr6 | 1,33E+08 | 1,33E+08 | 736 | * | 8,318784 | -5,108678723 | 1,701471 | -3,00251 | 0,002678 | 1 | 0 | Distal Intergenic | 6 | 1,34E+08 | 1,34E+08 | 3574 | 1 | 545884 | ENSMUST00000178578.4 | -76680 | ENSMUSG00000095412 | Gm5885 | predicted gene 5885 |
| chr8 | 11648575 | 11649449 | 875 | * | 9,315839 | -5,107241752 | 1,747047 | -2,92336 | 0,003463 | 0,730469267 | 1,211315 | Distal Intergenic | 8 | 11661585 | 11674626 | 13042 | 2 | 102334 | ENSMUST00000211690.2 | 25177 | ENSMUSG00000031508 | Ankrd10 | ankyrin repeat domain 10 |
| chr14 | 85682448 | 85683283 | 836 | * | 6,193815 | -5,103798944 | 1,886299 | -2,70572 | 0,006816 | 1 | 0 | Distal Intergenic | 14 | 86410363 | 86417021 | 6659 | 1 | 1,03E+08 | ENSMUST00000227116.2 | -727080 | | Gm32729 | predicted gene, 32729 |
| chr13 | 92327071 | 92327823 | 753 | * | 5,699579 | -5,103423889 | 1,959894 | -2,60393 | 0,009216 | 1 | 0 | Distal Intergenic | 13 | 92348765 | 92372107 | 23343 | 2 | 17686 | ENSMUST00000190761.2 | 44284 | ENSMUSG00000014850 | Msh3 | mutS homolog 3 |
| chr9 | 62663577 | 62664203 | 627 | * | 5,357333 | -5,10305571 | 1,964741 | -2,59732 | 0,009396 | 1 | 0 | Intron (ENSMUST0000 | 9 | 62699103 | 62719216 | 20114 | 2 | 14155 | ENSMUST00000034775.10 | 55013 | ENSMUSG00000032244 | Fem1b | fem 1 homolog b |
| chr7 | 10601700 | 10602288 | 589 | * | 5,5185 | -5,10302564 | 1,956228 | -2,6086 | 0,009091 | 1 | 0 | Distal Intergenic | 7 | 10634667 | 10638954 | 4288 | 2 | 665780 | ENSMUST00000210847.2 | 36666 | ENSMUSG00000095339 | Zscan4b | zinc finger and SCAN domain containing 4B |
| chr16 | 71708132 | 71708764 | 633 | * | 5,549227 | -5,103000367 | 1,935443 | -2,63661 | 0,008374 | 1 | 0 | Distal Intergenic | 16 | 71460676 | 71460806 | 131 | 1 | 1,15E+08 | ENSMUST00000083990.3 | 247456 | ENSMUSG00000065924 | Gm22797 | predicted gene, 22797 |
| chr9 | 83311658 | 83312490 | 833 | * | 5,387836 | -5,102877753 | 1,943966 | -2,62498 | 0,008665 | 1 | 0 | Promoter (1-2kb) | 9 | 83313600 | 83336626 | 23027 | 1 | 1E+08 | ENSMUST00000188217.2 | -1110 | ENSMUSG00000099718 | Gm2087 | predicted gene 2087 |
| chr8 | 1,17E+08 | 1,17E+08 | 409 | * | 5,783219 | -5,10273057 | 1,888145 | -2,70251 | 0,006882 | 1 | 0 | Distal Intergenic | 8 | 1,17E+08 | 1,17E+08 | 11426 | 1 | 1,03E+08 | ENSMUST00000241934.1 | 329569 | | Gm30925 | predicted gene, 30925 |
| chr7 | 75200645 | 75201226 | 582 | * | 6,036576 | -5,102599044 | 1,964466 | -2,59745 | 0,009392 | 1 | 0 | Intron (ENSMUST0000 | 7 | 75259787 | 75347091 | 87305 | 1 | 75547 | ENSMUST00000207923.2 | -58561 | ENSMUSG00000066406 | Akap13 | A kinase (PRKA) anchor protein 13 |
| chr12 | 61999494 | 62000389 | 896 | * | 6,494491 | -5,099016442 | 1,855896 | -2,74747 | 0,006006 | 1 | 0 | Distal Intergenic | 12 | 61570734 | 61904714 | 333981 | 1 | 238205 | ENSMUST00000119481.2 | 428760 | ENSMUSG00000035653 | Lrfn5 | leucine rich repeat and fibronectin type III domain containing 5 |
| chrX | 36654571 | 36655496 | 926 | * | 6,760501 | -5,093910667 | 1,866533 | -2,72908 | 0,006351 | 1 | 0 | Promoter (2-3kb) | 20 | 36651650 | 36655617 | 3968 | 1 | 1E+08 | ENSMUST00000151387.2 | 2921 | ENSMUSG00000079636 | Rhox3c | reproductive homeobox 3C |
| chr2 | 1,32E+08 | 1,32E+08 | 639 | * | 5,829673 | -5,092328963 | 1,945094 | -2,61804 | 0,008844 | 1 | 0 | Distal Intergenic | 2 | 1,31E+08 | 1,32E+08 | 68942 | 2 | 1,19E+08 | ENSMUST00000138080.2 | -56102 | | Gm14280 | predicted gene 14280 |
| chr3 | 1,25E+08 | 1,25E+08 | 762 | * | 6,191455 | -5,092310575 | 1,886781 | -2,69894 | 0,006956 | 1 | 0 | Distal Intergenic | 3 | 1,24E+08 | 1,24E+08 | 7885 | 2 | 384775 | ENSMUST00000180033.8 | -220032 | ENSMUSG00000039174 | 1700003H04Rik | RIKEN cDNA 1700003H04 gene |
| chr1 | 49715043 | 49715730 | 688 | * | 6,410783 | -5,092205478 | 1,952172 | -2,60848 | 0,009094 | 1 | 0 | Distal Intergenic | 1 | 49283775 | 49379602 | 95828 | 1 | 442837 | ENSMUST00000188673.7 | 431268 | ENSMUSG00000051616 | C230029F24Rik | RIKEN cDNA C230029F24 gene |
| chr10 | 53870656 | 53871448 | 793 | * | 6,276035 | -5,091154657 | 1,900799 | -2,67843 | 0,007397 | 1 | 0 | Intron (ENSMUST0000 | 10 | 53794815 | 53853211 | 58397 | 2 | 17155 | ENSMUST00000219234.2 | -17445 | ENSMUSG00000003746 | Man1a | mannosidase 1, alpha |
| chr4 | 27563723 | 27564601 | 879 | * | 7,142069 | -5,091066792 | 1,738086 | -2,92912 | 0,003399 | 1 | 0 | Distal Intergenic | 4 | 27700671 | 27700777 | 107 | 1 | 1,15E+08 | ENSMUST00000179794.2 | -136070 | ENSMUSG00000095003 | Gm25173 | predicted gene, 25173 |
| chr11 | 1,07E+08 | 1,07E+08 | 994 | * | 5,693897 | -5,088739215 | 1,82692 | -2,78542 | 0,005346 | 1 | 0 | Intron (ENSMUST0000 | 11 | 1,07E+08 | 1,07E+08 | 98251 | 2 | 66313 | ENSMUST00000139297.8 | 24133 | ENSMUSG00000018363 | Smurf2 | SMAD specific E3 ubiquitin protein ligase 2 |
| chr11 | 1,11E+08 | 1,11E+08 | 1060 | * | 8,027048 | -5,086148856 | 1,796117 | -2,83175 | 0,004629 | 1 | 0 | Distal Intergenic | 11 | 1,11E+08 | 1,11E+08 | 2269 | 2 | 1,03E+08 | ENSMUST00000247083.1 | 186808 | | Gm30158 | predicted gene, 30158 |
| chr14 | 69328437 | 69329478 | 1042 | * | 7,114333 | -5,085738077 | 1,789694 | -2,84168 | 0,004488 | 1 | 0 | Distal Intergenic | 14 | 69256883 | 69267101 | 10219 | 2 | 1,05E+08 | ENSMUST00000224911.2 | -61336 | | Gm41192 | predicted gene, 41192 |
| chr9 | 84351306 | 84352402 | 1097 | * | 8,750688 | -5,083845232 | 1,858054 | -2,73611 | 0,006217 | 0,813839948 | 1,360622 | Distal Intergenic | 9 | 84460013 | 84495737 | 35725 | 1 | 1,05E+08 | ENSMUST00000214502.2 | -107611 | | Gm39388 | predicted gene, 39388 |
| chr15 | 45132889 | 45133522 | 634 | * | 6,37277 | -5,082491245 | 1,910261 | -2,66063 | 0,0078 | 1 | 0 | Distal Intergenic | 15 | 44976181 | 44978316 | 2136 | 2 | 67498 | ENSMUST00000228536.2 | -154573 | ENSMUSG00000022342 | Kcnv1 | potassium channel, subfamily V, member 1 |
| chr10 | 67804189 | 67805051 | 863 | * | 7,451463 | -5,082102393 | 1,794781 | -2,8316 | 0,004632 | 1 | 0 | Intron (ENSMUST0000 | 10 | 67798309 | 67809538 | 11230 | 2 | 75067 | ENSMUST00000219339.2 | 4487 | ENSMUSG00000112899 | 4930521O17Rik | RIKEN cDNA 4930521O17 gene |
| chr19 | 28043831 | 28044717 | 887 | * | 8,109262 | -5,082055472 | 1,808947 | -2,8094 | 0,004963 | 0,764729825 | 1,425929 | Distal Intergenic | 19 | 27782543 | 27988566 | 206024 | 2 | 19726 | ENSMUST00000173161.2 | -55265 | ENSMUSG00000040929 | Rfx3 | regulatory factor X, 3 (influences HLA class II expression) |
| chr13 | 1,06E+08 | 1,06E+08 | 894 | * | 8,067644 | -5,081642533 | 1,843772 | -2,75611 | 0,005849 | 1 | 0 | Distal Intergenic | 13 | 1,07E+08 | 1,07E+08 | 14510 | 2 | 1,03E+08 | ENSMUST00000225713.2 | 401647 | | Gm30411 | predicted gene, 30411 |
| chr1 | 1,14E+08 | 1,14E+08 | 683 | * | 6,022551 | -5,080995512 | 1,904528 | -2,66785 | 0,007634 | 1 | 0 | Distal Intergenic | 1 | 1,16E+08 | 1,16E+08 | 15632 | 2 | 1,08E+08 | ENSMUST00000186137.7 | 1310267 | | Gm50605 | predicted gene, 50605 |
| chr5 | 16677257 | 16678728 | 1472 | * | 10,8774 | -5,076997892 | 1,668253 | -3,0433 | 0,00234 | 0,748382289 | 0,744375 | Distal Intergenic | 5 | 16758493 | 16778432 | 19940 | 1 | 15234 | ENSMUST00000195957.5 | -79765 | ENSMUSG00000028864 | Hgf | hepatocyte growth factor |
| chr10 | 69575793 | 69576911 | 1119 | * | 10,16974 | -5,076102047 | 1,712817 | -2,9636 | 0,003041 | 0,714462637 | 1,173325 | Intron (ENSMUST0000 | 10 | 69542308 | 69658465 | 116158 | 1 | 11735 | ENSMUST00000182162.8 | 33485 | ENSMUSG00000069601 | Ank3 | ankyrin 3, epithelial |
| chr4 | 1,37E+08 | 1,37E+08 | 1214 | * | 9,598378 | -5,073924534 | 1,846407 | -2,748 | 0,005996 | 0,826891088 | 1,211315 | Distal Intergenic | 4 | 1,37E+08 | 1,37E+08 | 22238 | 1 | 22417 | ENSMUST00000045747.5 | -17274 | ENSMUSG00000036856 | Wnt4 | wingless-type MMTV integration site family, member 4 |
| chr4 | 1,44E+08 | 1,44E+08 | 524 | * | 6,602497 | -5,073285075 | 1,851968 | -2,7394 | 0,006155 | 1 | 0 | Distal Intergenic | 4 | 1,44E+08 | 1,44E+08 | 5107 | 2 | 100012 | ENSMUST00000050933.9 | 22971 | ENSMUSG00000050810 | Oog3 | oogenesin 3 |
| chr5 | 1,09E+08 | 1,09E+08 | 1438 | * | 9,353148 | -5,072453181 | 1,751505 | -2,89605 | 0,003779 | 0,746743246 | 1,211315 | Distal Intergenic | 5 | 1,09E+08 | 1,09E+08 | 10071 | 1 | 69587 | ENSMUST00000151897.2 | 25921 | ENSMUSG00000033623 | Pcgf3 | polycomb group ring finger 3 |
| chr1 | 1,7E+08 | 1,7E+08 | 625 | * | 5,293918 | -5,071147498 | 1,95796 | -2,59002 | 0,009597 | 1 | 0 | Exon (ENSMUST00000 | 1 | 1,7E+08 | 1,7E+08 | 2988 | 1 | 67303 | ENSMUST00000242643.1 | 18656 | ENSMUSG00000097503 | 3110045C21Rik | RIKEN cDNA 3110045C21 gene |
| chr14 | 1,01E+08 | 1,01E+08 | 1445 | * | 9,001416 | -5,070700361 | 1,721806 | -2,94499 | 0,00323 | 0,685026746 | 1,449706 | Distal Intergenic | 14 | 1,01E+08 | 1,01E+08 | 101 | 2 | 1,15E+08 | ENSMUST00000157535.4 | 4679 | ENSMUSG00000088160 | Gm23538 | predicted gene, 23538 |
| chr3 | 42900043 | 42900810 | 768 | * | 6,403041 | -5,070225575 | 1,820545 | -2,785 | 0,005353 | 1 | 0 | Distal Intergenic | 3 | 41705631 | 41710927 | 5297 | 1 | 73852 | ENSMUST00000195030.2 | 1194412 | ENSMUSG00000025766 | D3Ertd751e | DNA segment, Chr 3, ERATO Doi 751, expressed |
| chr11 | 1,03E+08 | 1,03E+08 | 1089 | * | 6,986981 | -5,069341732 | 1,85158 | -2,73785 | 0,006184 | 1 | 0 | Promoter (1-2kb) | 11 | 1,03E+08 | 1,03E+08 | 5058 | 2 | 14615 | ENSMUST00000155089.2 | -1824 | ENSMUSG00000034520 | Gjc1 | gap junction protein, gamma 1 |
| chr15 | 11282547 | 11283573 | 1027 | * | 8,045347 | -5,065831064 | 1,746988 | -2,89975 | 0,003735 | 1 | 0 | Intron (ENSMUST0000 | 15 | 11361697 | 11361823 | 127 | 2 | 1,15E+08 | ENSMUST00000158679.3 | 78250 | ENSMUSG00000089304 | Gm24844 | predicted gene, 24844 |
| chr14 | 84545973 | 84547009 | 1037 | * | 7,000351 | -5,065797166 | 1,816081 | -2,78941 | 0,00528 | 1 | 0 | Distal Intergenic | 14 | 84681003 | 84776442 | 95440 | 1 | 219228 | ENSMUST00000071370.7 | -133994 | ENSMUSG00000035566 | Pcdh17 | protocadherin 17 |
| chr15 | 86750652 | 86751045 | 394 | * | 5,644851 | -5,064266698 | 1,909359 | -2,65234 | 0,007994 | 1 | 0 | Distal Intergenic | 15 | 86622027 | 86650341 | 28315 | 1 | 75160 | ENSMUST00000230110.2 | 128625 | | 4930543I03Rik | RIKEN cDNA 4930543I03 gene |
| chr14 | 23321687 | 23322910 | 1224 | * | 12,03247 | -5,063172739 | 1,654913 | -3,05948 | 0,002217 | 0,73688097 | 0,744375 | Distal Intergenic | 14 | 23088262 | 23144639 | 56378 | 2 | 791307 | ENSMUST00000090409.2 | -177048 | ENSMUSG00000068680 | Gm10248 | predicted gene 10248 |
| chr1 | 84121980 | 84122620 | 641 | * | 4,782106 | -5,062529846 | 1,961683 | -2,58071 | 0,00986 | 1 | 0 | Intron (ENSMUST0000 | 1 | 84196570 | 84196682 | 113 | 2 | 1,02E+08 | ENSMUST00000183596.3 | 74062 | ENSMUSG00000098851 | Mir6353 | microRNA 6353 |
| chr15 | 99532399 | 99533137 | 739 | * | 7,81789 | -5,061523206 | 1,819391 | -2,78199 | 0,005403 | 1 | 0 | Promoter (<=1kb) | 15 | 99530040 | 99532655 | 2616 | 2 | 26934 | ENSMUST00000168976.2 | 0 | ENSMUSG00000023015 | Racgap1 | Rac GTPase-activating protein 1 |
| chr10 | 52924908 | 52925562 | 655 | * | 5,909357 | -5,061290293 | 1,922517 | -2,63264 | 0,008472 | 1 | 0 | Intron (ENSMUST0000 | 10 | 53057276 | 53096559 | 39284 | 1 | 1,05E+08 | ENSMUST00000219795.2 | -131714 | | Gm40649 | predicted gene, 40649 |
| chr5 | 1,21E+08 | 1,21E+08 | 713 | * | 6,846943 | -5,060364813 | 1,931555 | -2,61984 | 0,008797 | 1 | 0 | 3' UTR | 5 | 1,21E+08 | 1,21E+08 | 2660 | 2 | 23960 | ENSMUST00000161809.2 | 6327 | ENSMUSG00000066861 | Oas1g | 2'-5' oligoadenylate synthetase 1G |
| chr5 | 74108803 | 74109303 | 501 | * | 5,36344 | -5,060250412 | 1,945357 | -2,60119 | 0,00929 | 1 | 0 | Distal Intergenic | 5 | 74070239 | 74077764 | 7526 | 1 | 69402 | ENSMUST00000202487.2 | 38564 | ENSMUSG00000107027 | 1700019F05Rik | RIKEN cDNA 1700019F05 gene |
| chr14 | 27073169 | 27073896 | 728 | * | 6,302279 | -5,059510421 | 1,896154 | -2,6683 | 0,007624 | 1 | 0 | Intron (ENSMUST0000 | 14 | 27083346 | 27101604 | 18259 | 1 | 71704 | ENSMUST00000225494.2 | -9450 | ENSMUSG00000021895 | Arhgef3 | Rho guanine nucleotide exchange factor (GEF) 3 |
| chr2 | 71656139 | 71657171 | 1033 | * | 6,38346 | -5,057393657 | 1,8022 | -2,80623 | 0,005012 | 1 | 0 | Promoter (<=1kb) | 2 | 71657106 | 71685295 | 28190 | 1 | 16403 | ENSMUST00000155249.2 | 0 | ENSMUSG00000027111 | Itga6 | integrin alpha 6 |
| chr3 | 37988978 | 37990161 | 1184 | * | 8,573581 | -5,057147877 | 1,704702 | -2,96659 | 0,003011 | 0,602555268 | 2,237403 | Exon (ENSMUST00000 | 3 | 37952962 | 38046877 | 93916 | 1 | 626410 | ENSMUST00000196893.2 | 36016 | ENSMUSG00000106461 | Gm20755 | predicted gene, 20755 |
| chr6 | 91306450 | 91307404 | 955 | * | 9,382682 | -5,056642979 | 1,752461 | -2,88545 | 0,003909 | 0,730469267 | 1,360622 | Distal Intergenic | 6 | 91242861 | 91249413 | 6553 | 1 | 14115 | ENSMUST00000137029.2 | 63589 | ENSMUSG00000064080 | Fbln2 | fibulin 2 |
| chr17 | 60180580 | 60181510 | 931 | * | 8,784166 | -5,055213469 | 1,929927 | -2,61938 | 0,008809 | 0,861293497 | 1,449706 | Intron (ENSMUST0000 | 17 | 59876554 | 59876660 | 107 | 2 | 1,15E+08 | ENSMUST00000104374.3 | -303920 | ENSMUSG00000077562 | Gm23203 | predicted gene, 23203 |
| chr11 | 1,04E+08 | 1,04E+08 | 1267 | * | 9,056421 | -5,0515334 | 1,802293 | -2,80284 | 0,005066 | 0,796850747 | 1,211315 | Promoter (2-3kb) | 11 | 1,04E+08 | 1,04E+08 | 96636 | 1 | 17762 | ENSMUST00000100347.11 | 2994 | ENSMUSG00000018411 | Mapt | microtubule-associated protein tau |
| chr5 | 1,22E+08 | 1,22E+08 | 903 | * | 6,574701 | -5,049586328 | 1,930498 | -2,61569 | 0,008905 | 1 | 0 | Intron (ENSMUST0000 | 5 | 1,22E+08 | 1,22E+08 | 6108 | 2 | 654470 | ENSMUST00000176677.8 | 16368 | ENSMUSG00000038593 | Tctn1 | tectonic family member 1 |
| chr12 | 1,08E+08 | 1,08E+08 | 555 | * | 5,16727 | -5,049356742 | 1,951117 | -2,58793 | 0,009655 | 1 | 0 | Distal Intergenic | 12 | 1,08E+08 | 1,08E+08 | 93200 | 2 | 58208 | ENSMUST00000066060.11 | -25382 | ENSMUSG00000048251 | Bcl11b | B cell leukemia/lymphoma 11B |
| chr14 | 4026050 | 4026773 | 724 | * | 6,347861 | -5,048878438 | 1,906561 | -2,64816 | 0,008093 | 1 | 0 | Distal Intergenic | 14 | 4137837 | 4186974 | 49138 | 1 | 22194 | ENSMUST00000022296.7 | -111064 | ENSMUSG00000021774 | Ube2e1 | ubiquitin-conjugating enzyme E2E 1 |
| chr1 | 53853924 | 53855026 | 1103 | * | 9,217506 | -5,04778386 | 1,827919 | -2,76149 | 0,005754 | 0,713797407 | 2,237403 | Intron (ENSMUST0000 | 1 | 53871851 | 53880330 | 8480 | 2 | 329152 | ENSMUST00000150677.8 | 25304 | ENSMUSG00000042807 | Hecw2 | HECT, C2 and WW domain containing E3 ubiquitin protein ligase 2 |

| | | | | | | | | | | | | | | | | | | | | | | | |
|---|---|---|---|---|---|---|---|---|---|---|---|---|---|---|---|---|---|---|---|---|---|---|---|
| chr14 | 5844774 | 5845622 | 849 | * | 5,745092 | -5,038544317 | 1,943752 | -2,59217 | 0,009537 | 1 | 0 | Intron (ENSMUST0000( | 14 | 5893936 | 6038426 | 144491 | 1 | 218772 | ENSMUST00000225921.2 | -48314 | ENSMUSG00000017491 | Rarb | retinoic acid receptor, beta |
| chr10 | 1,21E+08 | 1,21E+08 | 581 | * | 5,547542 | -5,038158294 | 1,925371 | -2,61672 | 0,008878 | 1 | 0 | Promoter (1-2kb) | 10 | 1,21E+08 | 1,21E+08 | 626 | 1 | 1,08E+08 | ENSMUST00000219800.2 | -1376 | | Gm46204 | predicted gene, 46204 |
| chr10 | 73747001 | 73748034 | 1034 | * | 8,962729 | -5,034621715 | 1,75486 | -2,86896 | 0,004118 | 0,7591629 | 1,211315 | Intron (ENSMUST0000( | 10 | 73782888 | 74480050 | 697163 | 1 | 11994 | ENSMUST00000191709.6 | -34854 | ENSMUSG00000052613 | Pcdh15 | protocadherin 15 |
| chr10 | 85235983 | 85237431 | 1449 | * | 11,14422 | -5,031866633 | 1,780392 | -2,82627 | 0,004709 | 0,791242424 | 1,173325 | Intron (ENSMUST0000( | 10 | 85222691 | 85476458 | 253768 | 1 | 74007 | ENSMUST00000020231.10 | 13292 | ENSMUSG00000020042 | Btbd11 | BTB (POZ) domain containing 11 |
| chr1 | 41919767 | 41920769 | 1003 | * | 10,18386 | -5,031561809 | 1,668028 | -3,01647 | 0,002557 | 0,651501713 | 1,449706 | Intron (ENSMUST0000( | 1 | 41957512 | 41959245 | 1734 | 1 | 1,03E+08 | ENSMUST00000187227.2 | -36743 | | Gm29260 | predicted gene 29260 |
| chr10 | 85351617 | 85352302 | 686 | * | 8,07465 | -5,030976575 | 1,730137 | -2,90785 | 0,003639 | 0,718390388 | 1,360622 | Intron (ENSMUST0000( | 10 | 85434265 | 85496156 | 61892 | 1 | 74007 | ENSMUST00000105306.3 | -81963 | ENSMUSG00000020042 | Btbd11 | BTB (POZ) domain containing 11 |
| chr10 | 1,09E+08 | 1,09E+08 | 1114 | * | 8,151088 | -5,030744077 | 1,726455 | -2,91392 | 0,003569 | 0,634611828 | 2,237403 | Intron (ENSMUST0000( | 10 | 1,09E+08 | 1,09E+08 | 1397 | 2 | 368203 | ENSMUST00000218023.2 | -37260 | ENSMUSG00000112039 | Gm5136 | predicted gene 5136 |
| chr11 | 57046764 | 57047262 | 499 | * | 6,397478 | -5,028256521 | 1,908085 | -2,63524 | 0,008408 | 1 | 0 | Intron (ENSMUST0000( | 11 | 57067344 | 57080619 | 13276 | 1 | 14799 | ENSMUST00000173531.2 | -20082 | ENSMUSG00000020524 | Gria1 | glutamate receptor, ionotropic, AMPA1 (alpha 1) |
| chrX | 1,56E+08 | 1,56E+08 | 1061 | * | 5,34486 | -5,027259816 | 1,949401 | -2,57887 | 0,009912 | 1 | 0 | Exon (ENSMUST00000: | 20 | 1,56E+08 | 1,56E+08 | 50892 | 2 | 270669 | ENSMUST00000058098.15 | -13316 | ENSMUSG00000046873 | Mbtps2 | membrane-bound transcription factor peptidase, site 2 |
| chr13 | 19851947 | 19852539 | 593 | * | 6,580215 | -5,026890781 | 1,837587 | -2,73559 | 0,006227 | 1 | 0 | Intron (ENSMUST0000( | 13 | 19829248 | 19859901 | 30654 | 2 | 73412 | ENSMUST00000223466.2 | 7362 | ENSMUSG00000041138 | Nme8 | NME/NM23 family member 8 |
| chr9 | 91414846 | 91415491 | 646 | * | 6,14905 | -5,026515 | 1,927801 | -2,60738 | 0,009124 | 1 | 0 | Intron (ENSMUST0000( | 9 | 91286527 | 91444871 | 158345 | 1 | 1,03E+08 | ENSMUST00000191272.2 | 128319 | | Gm29478 | predicted gene 29478 |
| chr1 | 87601485 | 87602330 | 846 | * | 6,302948 | -5,0264644 | 1,925624 | -2,6103 | 0,009046 | 1 | 0 | Promoter (1-2kb) | 1 | 87603961 | 87648224 | 44264 | 1 | 16331 | ENSMUST00000167032.2 | -1631 | ENSMUSG00000026288 | Inpp5d | inositol polyphosphate-5-phosphatase D |
| chr2 | 92230326 | 92231327 | 1002 | * | 9,255214 | -5,026394313 | 1,744949 | -2,88054 | 0,00397 | 0,723027638 | 1,449706 | Promoter (<=1kb) | 2 | 92214025 | 92231608 | 17584 | 2 | 19099 | ENSMUST00000050312.3 | 281 | ENSMUSG00000027223 | Mapk8ip1 | mitogen-activated protein kinase 8 interacting protein 1 |
| chr17 | 63739635 | 63740785 | 1151 | * | 10,21703 | -5,024620388 | 1,705658 | -2,94586 | 0,003221 | 0,685026746 | 1,449706 | Intron (ENSMUST0000( | 17 | 63365933 | 63758706 | 392774 | 2 | 50758 | ENSMUST00000169134.2 | 17921 | ENSMUSG00000023965 | Fbxl17 | F-box and leucine-rich repeat protein 17 |
| chr14 | 1,22E+08 | 1,22E+08 | 657 | * | 7,506222 | -5,01884639 | 1,853017 | -2,70847 | 0,006759 | 1 | 0 | Intron (ENSMUST0000( | 14 | 1,22E+08 | 1,22E+08 | 195666 | 2 | 105445 | ENSMUST00000239077.2 | -25158 | ENSMUSG00000025558 | Dock9 | dedicator of cytokinesis 9 |
| chr9 | 47487325 | 47488498 | 1174 | * | 8,305687 | -5,016817825 | 1,71124 | -2,93169 | 0,003371 | 0,707074539 | 1,360622 | Intron (ENSMUST0000( | 9 | 47441671 | 47761688 | 320018 | 1 | 54725 | ENSMUST00000034581.4 | 45654 | ENSMUSG00000032076 | Cadm1 | cell adhesion molecule 1 |
| chr11 | 30612011 | 30612529 | 519 | * | 5,73146 | -5,016171234 | 1,920573 | -2,61181 | 0,009006 | 1 | 0 | Distal Intergenic | 11 | 30455991 | 30599587 | 143597 | 2 | 75572 | ENSMUST00000074613.4 | -12424 | ENSMUSG00000060923 | Acyp2 | acylphosphatase 2, muscle type |
| chrX | 52386414 | 52387706 | 1293 | * | 5,744811 | -5,016021935 | 1,920134 | -2,61233 | 0,008993 | 1 | 0 | Exon (ENSMUST00000: | 20 | 52362301 | 52420062 | 57762 | 1 | 73866 | ENSMUST00000124137.8 | 24113 | ENSMUSG00000036013 | Fam122c | family with sequence similarity 122, member C |
| chr14 | 33753292 | 33754057 | 766 | * | 6,043592 | -5,015095777 | 1,913422 | -2,62101 | 0,008767 | 1 | 0 | Intron (ENSMUST0000( | 14 | 33762995 | 33768938 | 5944 | 1 | 432838 | ENSMUST00000227522.2 | -8938 | ENSMUSG00000072624 | Gm5460 | predicted gene 5460 |
| chr7 | 1,19E+08 | 1,19E+08 | 994 | * | 9,870962 | -5,013400506 | 1,669168 | -3,00353 | 0,002669 | 0,705098751 | 1,086118 | Promoter (<=1kb) | 7 | 1,19E+08 | 1,19E+08 | 9041 | 2 | 1,05E+08 | ENSMUST00000207060.2 | 450 | ENSMUSG00000109093 | Gm39079 | predicted gene, 39079 |
| chrX | 1,5E+08 | 1,5E+08 | 1056 | * | 9,387507 | -5,011628911 | 1,732156 | -2,89329 | 0,003812 | 0,748382289 | 1,211315 | Promoter (2-3kb) | 20 | 1,5E+08 | 1,5E+08 | 1072 | 1 | 14163 | ENSMUST00000125620.2 | -2500 | ENSMUSG00000025265 | Fgd1 | FYVE, RhoGEF and PH domain containing 1 |
| chr14 | 6309143 | 6310262 | 1120 | * | 6,492389 | -5,006755523 | 1,784301 | -2,806 | 0,005016 | 1 | 0 | Distal Intergenic | 14 | 6226476 | 6227493 | 1018 | 1 | 71147 | ENSMUST00000148121.3 | 82667 | ENSMUSG00000021786 | Oxsm | 3-oxoacyl-ACP synthase, mitochondrial |
| chr5 | 1,18E+08 | 1,18E+08 | 836 | * | 6,550288 | -5,005306336 | 1,874239 | -2,67058 | 0,007572 | 1 | 0 | Distal Intergenic | 5 | 1,18E+08 | 1,18E+08 | 132 | 1 | 1,15E+08 | ENSMUST00000104098.4 | 19557 | ENSMUSG00000119733 | Gm25076 | predicted gene, 25076 |
| chr3 | 55152196 | 55153090 | 895 | * | 6,070475 | -5,003986688 | 1,914563 | -2,61364 | 0,008958 | 1 | 0 | Promoter (1-2kb) | 3 | 55154700 | 55163812 | 9113 | 1 | 13175 | ENSMUST00000198154.2 | -1610 | ENSMUSG00000027797 | Dclk1 | doublecortin-like kinase 1 |
| chr5 | 49602978 | 49604108 | 1131 | * | 11,53803 | -5,001957698 | 1,677708 | -2,98142 | 0,002869 | 0,686261859 | 1,2763 | Intron (ENSMUST0000( | 5 | 48546849 | 49682106 | 1135258 | 2 | 80334 | ENSMUST00000176191.8 | 77998 | ENSMUSG00000029088 | Kcnip4 | Kv channel interacting protein 4 |
| chr6 | 1,39E+08 | 1,39E+08 | 736 | * | 9,205488 | -5,00081261 | 1,790722 | -2,79262 | 0,005228 | 0,802490409 | 1,211315 | Distal Intergenic | 6 | 1,39E+08 | 1,39E+08 | 167 | 2 | 1,15E+08 | ENSMUST00000158278.3 | -52302 | ENSMUSG00000088903 | Gm25434 | predicted gene, 25434 |
| chr13 | 1,07E+08 | 1,07E+08 | 757 | * | 7,306457 | -4,997633965 | 1,904947 | -2,6235 | 0,008703 | 0,87277522 | 1,360622 | Distal Intergenic | 13 | 1,07E+08 | 1,07E+08 | 22618 | 1 | 1E+08 | ENSMUST00000224838.2 | 104442 | ENSMUSG00000071181 | 3830408C21Rik | RIKEN cDNA 3830408C21 gene |
| chr10 | 1,14E+08 | 1,14E+08 | 920 | * | 6,9919 | -4,997553307 | 1,757146 | -2,84413 | 0,004453 | 0,75147965 | 1,360622 | Distal Intergenic | 10 | 1,13E+08 | 1,13E+08 | 11133 | 2 | 1,05E+08 | ENSMUST00000218612.2 | -332951 | | Gm40765 | predicted gene, 40765 |
| chr5 | 50642649 | 50643241 | 593 | * | 6,077758 | -4,997048335 | 1,917401 | -2,60616 | 0,009156 | 1 | 0 | Intron (ENSMUST0000( | 5 | 50684283 | 50700877 | 16595 | 2 | 1,03E+08 | ENSMUST00000199991.2 | 57636 | | Gm36814 | predicted gene, 36814 |
| chr10 | 1,08E+08 | 1,08E+08 | 664 | * | 7,165894 | -4,99634816 | 1,769679 | -2,82331 | 0,004753 | 1 | 0 | Distal Intergenic | 10 | 1,08E+08 | 1,08E+08 | 31190 | 1 | 628877 | ENSMUST00000219670.2 | 107117 | ENSMUSG00000112910 | Gm29685 | predicted gene, 29685 |
| chr1 | 29832455 | 29833413 | 959 | * | 6,930393 | -4,994912275 | 1,789626 | -2,79104 | 0,005254 | 1 | 0 | Distal Intergenic | 1 | 29460569 | 29460693 | 125 | 1 | 1,02E+08 | ENSMUST00000183870.3 | 371886 | ENSMUSG00000099257 | Mir6342 | microRNA 6342 |
| chr19 | 30067964 | 30068742 | 779 | * | 6,34375 | -4,99444855 | 1,86423 | -2,67909 | 0,007382 | 1 | 0 | Promoter (<=1kb) | 19 | 30069361 | 30070339 | 979 | 1 | 109113 | ENSMUST00000143769.2 | -619 | ENSMUSG00000024817 | Uhrf2 | ubiquitin-like, containing PHD and RING finger domains 2 |
| chr5 | 1,16E+08 | 1,16E+08 | 637 | * | 6,218201 | -4,994068923 | 1,907147 | -2,61861 | 0,008829 | 1 | 0 | Distal Intergenic | 5 | 1,16E+08 | 1,16E+08 | 44490 | 2 | 545798 | ENSMUST00000111997.3 | -19239 | ENSMUSG00000079278 | Tmem233 | transmembrane protein 233 |
| chr18 | 88869044 | 88870055 | 1012 | * | 5,676129 | -4,99386078 | 1,933993 | -2,58215 | 0,009819 | 1 | 0 | Exon (ENSMUST00000: | 18 | 88888719 | 88905461 | 16743 | 2 | 54607 | ENSMUST00000236108.2 | 35406 | ENSMUSG00000056153 | Socs6 | suppressor of cytokine signaling 6 |
| chr10 | 95281043 | 95281583 | 541 | * | 6,284169 | -4,992605026 | 1,903078 | -2,62344 | 0,008705 | 1 | 0 | Exon (ENSMUST00000: | 10 | 95253242 | 95279547 | 26306 | 1 | 70523 | ENSMUST00000244592.1 | 27801 | ENSMUSG00000097766 | 5730420D15Rik | RIKEN cDNA 5730420D15 gene |
| chr4 | 1,16E+08 | 1,16E+08 | 1195 | * | 9,991117 | -4,992414735 | 1,710263 | -2,91909 | 0,003511 | 0,749931768 | 1,086118 | Exon (ENSMUST00000: | 4 | 1,16E+08 | 1,16E+08 | 1198 | 2 | 17776 | ENSMUST00000128713.2 | 16107 | ENSMUSG00000003810 | Mast2 | microtubule associated serine/threonine kinase 2 |
| chr6 | 1,43E+08 | 1,43E+08 | 1540 | * | 11,56522 | -4,988085466 | 1,728348 | -2,88604 | 0,003901 | 0,742014238 | 1,2763 | Distal Intergenic | 6 | 1,43E+08 | 1,43E+08 | 9988 | 1 | 52285 | ENSMUST00000160951.2 | 77103 | | D6Ertd474e | DNA segment, Chr 6, ERATO Doi 474, expressed |
| chr7 | 1,4E+08 | 1,4E+08 | 896 | * | 7,155182 | -4,985063097 | 1,824236 | -2,73269 | 0,006282 | 1 | 0 | Distal Intergenic | 7 | 1,4E+08 | 1,4E+08 | 930 | 2 | 259028 | ENSMUST00000073226.2 | -13643 | ENSMUSG00000063823 | Olfr532 | olfactory receptor 532 |
| chr7 | 1,15E+08 | 1,15E+08 | 1338 | * | 7,944319 | -4,984298025 | 1,813707 | -2,74813 | 0,005994 | 1 | 0 | Intron (ENSMUST0000( | 7 | 1,15E+08 | 1,15E+08 | 65136 | 2 | 20679 | ENSMUST00000206427.2 | -48681 | ENSMUSG00000051910 | Sox6 | SRY (sex determining region Y)-box 6 |
| chr5 | 1,51E+08 | 1,51E+08 | 909 | * | 8,329847 | -4,98260373 | 1,71206 | -2,9103 | 0,003611 | 0,708532921 | 1,449706 | Intron (ENSMUST0000( | 5 | 1,51E+08 | 1,52E+08 | 44832 | 2 | 632671 | ENSMUST00000232853.2 | 60647 | ENSMUSG00000091794 | Vmn2r18 | vomeronasal 2, receptor 18 |
| chr12 | 84083168 | 84084005 | 838 | * | 6,533687 | -4,981514803 | 1,913168 | -2,6038 | 0,00922 | 1 | 0 | Promoter (1-2kb) | 12 | 84085153 | 84095375 | 10223 | 1 | 171282 | ENSMUST00000021652.5 | -1148 | ENSMUSG00000052392 | Acot4 | acyl-CoA thioesterase 4 |
| chr5 | 1,16E+08 | 1,16E+08 | 1279 | * | 9,857617 | -4,980407311 | 1,770779 | -2,81255 | 0,004915 | 0,792545355 | 1,211315 | Distal Intergenic | 5 | 1,16E+08 | 1,16E+08 | 80425 | 1 | 12704 | ENSMUST00000147330.8 | -48012 | ENSMUSG00000029516 | Cit | citron |
| chr12 | 28584496 | 28585316 | 821 | * | 7,437298 | -4,980153318 | 1,823206 | -2,73154 | 0,006304 | 0,816770951 | 1,360622 | Intron (ENSMUST0000( | 12 | 28566055 | 28574748 | 8694 | 2 | 68511 | ENSMUST00000189747.2 | -9748 | ENSMUSG00000020633 | Dcdc2c | doublecortin domain containing 2C |
| chr11 | 50319398 | 50320544 | 1147 | * | 8,661387 | -4,979701411 | 1,80018 | -2,76622 | 0,005671 | 1 | 0 | Promoter (1-2kb) | 11 | 50280113 | 50321952 | 41840 | 2 | 216724 | ENSMUST00000020643.4 | 1408 | ENSMUSG00000020375 | Rufy1 | RUN and FYVE domain containing 1 |
| chr17 | 30969384 | 30969885 | 502 | * | 7,990666 | -4,979517236 | 1,791496 | -2,77953 | 0,005444 | 1 | 0 | Exon (ENSMUST00000: | 17 | 30998707 | 31011740 | 13034 | 1 | 13417 | ENSMUST00000237094.2 | -28822 | ENSMUSG00000033826 | Dnah8 | dynein, axonemal, heavy chain 8 |
| chr16 | 13997585 | 13999092 | 1508 | * | 7,747633 | -4,978293858 | 1,760662 | -2,82751 | 0,004691 | 0,762875156 | 1,360622 | Promoter (2-3kb) | 16 | 14001437 | 14008384 | 6948 | 1 | 67203 | ENSMUST00000149232.2 | -2345 | ENSMUSG00000022678 | Nde1 | nudE neurodevelopment protein 1 |
| chr7 | 1,02E+08 | 1,02E+08 | 1302 | * | 8,617747 | -4,974450214 | 1,731015 | -2,87372 | 0,004057 | 1 | 0 | Intron (ENSMUST0000( | 7 | 1,02E+08 | 1,02E+08 | 31190 | 2 | 269966 | ENSMUST00000211764.2 | 11437 | ENSMUSG00000063550 | Nup98 | nucleoporin 98 |
| chr1 | 1,67E+08 | 1,67E+08 | 1418 | * | 11,30445 | -4,974028253 | 1,640015 | -3,03292 | 0,002422 | 0,751210692 | 0,744375 | Distal Intergenic | 1 | 1,67E+08 | 1,67E+08 | 25706 | 2 | 1,01E+08 | ENSMUST00000180995.3 | 148007 | ENSMUSG00000102548 | Gm16701 | predicted gene, 16701 |
| chr11 | 99685843 | 99686663 | 821 | * | 5,595316 | -4,970268774 | 1,921668 | -2,58643 | 0,009697 | 1 | 0 | Promoter (1-2kb) | 11 | 99683215 | 99684293 | 1079 | 2 | 670464 | ENSMUST00000105058.2 | -1550 | ENSMUSG00000078261 | Gm11596 | predicted gene 11596 |
| chr14 | 62432400 | 62433595 | 1196 | * | 6,226865 | -4,970142247 | 1,907189 | -2,606 | 0,009161 | 1 | 0 | Distal Intergenic | 14 | 62350569 | 62408575 | 58007 | 2 | 1,03E+08 | ENSMUST00000226702.2 | -23825 | | Gm35419 | predicted gene, 35419 |
| chr18 | 3282499 | 3283671 | 1173 | * | 6,826323 | -4,969774193 | 1,869583 | -2,65823 | 0,007855 | 1 | 0 | Promoter (<=1kb) | 18 | 3267572 | 3281769 | 14198 | 2 | 12916 | ENSMUST00000147138.8 | -730 | ENSMUSG00000063889 | Crem | cAMP responsive element modulator |
| chr14 | 1,04E+08 | 1,04E+08 | 860 | * | 5,903132 | -4,969608075 | 1,819965 | -2,73061 | 0,006322 | 1 | 0 | Distal Intergenic | 14 | 1,04E+08 | 1,04E+08 | 6460 | 2 | 70898 | ENSMUST00000160117.2 | -48266 | ENSMUSG00000089707 | Slain1os | SLAIN motif family, member 1, opposite strand |
| chr2 | 1,15E+08 | 1,15E+08 | 832 | * | 6,229574 | -4,963760521 | 1,871468 | -2,65233 | 0,007994 | 1 | 0 | Intron (ENSMUST0000( | 2 | 1,15E+08 | 1,15E+08 | 15017 | 2 | 78193 | ENSMUST00000152815.2 | 232889 | ENSMUSG00000087309 | 4930528P14Rik | RIKEN cDNA 4930528P14 gene |
| chr3 | 1,47E+08 | 1,47E+08 | 863 | * | 8,847834 | -4,962374586 | 1,687672 | -2,94037 | 0,003278 | 0,686838043 | 1,449706 | Distal Intergenic | 3 | 1,47E+08 | 1,47E+08 | 4401 | 1 | 1,05E+08 | ENSMUST00000195873.2 | 283003 | | Gm37041 | predicted gene, 37041 |
| chr1 | 1,49E+08 | 1,49E+08 | 807 | * | 7,489211 | -4,962358133 | 1,829263 | -2,71276 | 0,006672 | 1 | 0 | Distal Intergenic | 1 | 1,49E+08 | 1,49E+08 | 36562 | 2 | 1,05E+08 | ENSMUST00000245751.1 | 380756 | | Gm29398 | predicted gene 29398 |
| chr7 | 91068353 | 91069558 | 1206 | * | 9,188384 | -4,961928877 | 1,725129 | -2,87627 | 0,004024 | 0,75508254 | 1,211315 | Intron (ENSMUST0000( | 7 | 91192018 | 91521636 | 329619 | 1 | 23859 | ENSMUST00000239136.2 | -122460 | ENSMUSG00000052572 | Dlg2 | discs large MAGUK scaffold protein 2 |
| chr9 | 67608455 | 67609738 | 1284 | * | 8,55515 | -4,961875716 | 1,695199 | -2,92702 | 0,003422 | 0,627785088 | 2,237403 | Distal Intergenic | 9 | 67634999 | 67635062 | 64 | 2 | 1,02E+08 | ENSMUST00000184532.3 | 25324 | ENSMUSG00000098937 | Mir7241 | microRNA 7241 |
| chrX | 1,29E+08 | 1,29E+08 | 927 | * | 6,656673 | -4,960600728 | 1,899645 | -2,61133 | 0,009019 | 1 | 0 | Distal Intergenic | 20 | 1,29E+08 | 1,29E+08 | 430730 | 1 | 54004 | ENSMUST00000127577.8 | -77899 | ENSMUSG00000034480 | Diaph2 | diaphanous related formin 2 |
| chr13 | 32144589 | 32145268 | 680 | * | 7,286383 | -4,958528281 | 1,862863 | -2,66178 | 0,007773 | 1 | 0 | Intron (ENSMUST0000( | 13 | 32155069 | 32156312 | 1244 | 1 | 1,03E+08 | ENSMUST00000221737.2 | -9801 | | Gm35732 | predicted gene, 35732 |
| chr11 | 1,06E+08 | 1,06E+08 | 1281 | * | 7,377737 | -4,956081536 | 1,799551 | -2,75407 | 0,005886 | 1 | 0 | Intron (ENSMUST0000( | 11 | 1,06E+08 | 1,06E+08 | 1290 | 1 | 77097 | ENSMUST00000167695.2 | 10735 | ENSMUSG00000053580 | Tanc2 | tetratricopeptide repeat, ankyrin repeat and coiled-coil containing 2 |
| chr5 | 8341377 | 8342072 | 696 | * | 6,569828 | -4,955701719 | 1,853091 | -2,67429 | 0,007489 | 1 | 0 | Intron (ENSMUST0000( | 5 | 8292723 | 8292866 | 144 | 1 | 1,15E+08 | ENSMUST00000157478.3 | 48654 | ENSMUSG00000088103 | Gm23993 | predicted gene, 23993 |
| chr14 | 1,07E+08 | 1,07E+08 | 1029 | * | 6,233677 | -4,953084406 | 1,913563 | -2,58841 | 0,009642 | 1 | 0 | Distal Intergenic | 14 | 1,07E+08 | 1,07E+08 | 127 | 1 | 1,15E+08 | ENSMUST00000104036.3 | 108395 | ENSMUSG00000077224 | Gm23437 | predicted gene, 23437 |
| chr5 | 1,29E+08 | 1,29E+08 | 928 | * | 6,658684 | -4,950727468 | 1,844442 | -2,68413 | 0,007272 | 1 | 0 | Distal Intergenic | 5 | 1,29E+08 | 1,29E+08 | 106924 | 1 | 243277 | ENSMUST00000156437.3 | 108999 | ENSMUSG00000044017 | Adgrd1 | adhesion G protein-coupled receptor D1 |
| chr7 | 46020554 | 46021176 | 623 | * | 6,357927 | -4,947357004 | 1,90967 | -2,59069 | 0,009578 | 1 | 0 | Distal Intergenic | 7 | 46025898 | 46028523 | 2626 | 1 | 17927 | ENSMUST00000072514.3 | -4722 | ENSMUSG00000009471 | Myod1 | myogenic differentiation 1 |
| chr15 | 1,02E+08 | 1,02E+08 | 1218 | * | 14,73064 | -4,946783857 | 1,60073 | -3,09033 | 0,001999 | 0,641187423 | 1,214904 | Distal Intergenic | 15 | 1,02E+08 | 1,02E+08 | 8570 | 2 | 77055 | ENSMUST00000100179.2 | -9085 | ENSMUSG00000075402 | Krt76 | keratin 76 |
| chr16 | 28709414 | 28710194 | 781 | * | 7,903132 | -4,945685055 | 1,746818 | -2,83125 | 0,004637 | 1 | 0 | Intron (ENSMUST0000( | 16 | 28644928 | 28748425 | 103498 | 2 | 239796 | ENSMUST00000100023.3 | 38231 | ENSMUSG00000051065 | Mb21d2 | Mab-21 domain containing 2 |
| chr11 | 19786732 | 19787940 | 1209 | * | 7,742358 | -4,945233848 | 1,895245 | -2,60928 | 0,009073 | 0,881405079 | 1,360622 | Distal Intergenic | 11 | 19874375 | 19974026 | 99652 | 1 | 114716 | ENSMUST00000093299.13 | -86435 | ENSMUSG00000045671 | Spred2 | sprouty-related EVH1 domain containing 2 |
| chr12 | 66679569 | 66680310 | 742 | * | 6,933624 | -4,939317749 | 1,84071 | -2,68338 | 0,007288 | 0,841453364 | 1,360622 | Promoter (2-3kb) | 12 | 66666376 | 66676887 | 10512 | 2 | 320772 | ENSMUST00000179729.2 | -2682 | ENSMUSG00000034912 | Mdga2 | MAM domain containing glycosylphosphatidylinositol anchor 2 |
| chr8 | 1,26E+08 | 1,26E+08 | 938 | * | 11,19616 | -4,938463925 | 1,730751 | -2,85337 | 0,004326 | 0,819720472 | 0,918255 | Distal Intergenic | 8 | 1,26E+08 | 1,26E+08 | 2543 | 1 | 1,03E+08 | ENSMUST00000245925.1 | -91373 | | Gm30713 | predicted gene, 30713 |
| chr11 | 1,22E+08 | 1,22E+08 | 1164 | * | 7,251523 | -4,936627393 | 1,750568 | -2,82002 | 0,004802 | 1 | 0 | Intron (ENSMUST0000( | 11 | 1,22E+08 | 1,22E+08 | 56957 | 2 | 210004 | ENSMUST00000067399.14 | 3575 | ENSMUSG00000046605 | B3gntl1 | UDP-GlcNAc:betaGal beta-1,3-N-acetylglucosaminyltransferase-like 1 |
| chr14 | 14154638 | 14155367 | 730 | * | 7,941665 | -4,936159875 | 1,818821 | -2,71393 | 0,006649 | 1 | 0 | Promoter (2-3kb) | 14 | 14151750 | 14172217 | 20468 | 1 | 268709 | ENSMUST00000120411.9 | 2888 | ENSMUSG00000021750 | Fam107a | family with sequence similarity 107, member A |
| chr11 | 58286746 | 58287493 | 748 | * | 6,886065 | -4,936136551 | 1,824623 | -2,70529 | 0,006824 | 1 | 0 | Exon (ENSMUST00000: | 11 | 58269869 | 58281554 | 11686 | 1 | 70163 | ENSMUST00000108826.3 | 16877 | ENSMUSG00000013643 | Lypd8 | LY6/PLAUR domain containing 8 |
| chr18 | 63530099 | 63530982 | 884 | * | 6,45843 | -4,936012477 | 1,858196 | -2,65635 | 0,007899 | 1 | 0 | Promoter (1-2kb) | 18 | 63532285 | 63554268 | 21984 | 1 | 1,05E+08 | ENSMUST00000183218.3 | -1303 | | Gm26972 | predicted gene, 26972 |
| chr3 | 1,41E+08 | 1,41E+08 | 795 | * | 7,306206 | -4,936010921 | 1,858562 | -2,65582 | 0,007912 | 1 | 0 | Distal Intergenic | 3 | 1,41E+08 | 1,41E+08 | 1372 | 1 | 75852 | ENSMUST00000196030.2 | 47026 | | 4930590L14Rik | RIKEN cDNA 4930590L14 gene |
| chr2 | 1,54E+08 | 1,54E+08 | 961 | * | 6,261095 | -4,935997347 | 1,862479 | -2,65023 | 0,008044 | 1 | 0 | Distal Intergenic | 2 | 1,54E+08 | 1,54E+08 | 13420 | 1 | 433492 | ENSMUST00000088921.6 | 18382 | ENSMUSG00000067996 | Bpifb9b | BPI fold containing family B, member 9B |
| chrX | 40901672 | 40902827 | 1156 | * | 5,472802 | -4,935958151 | 1,850206 | -2,66779 | 0,007635 | 1 | 0 | Promoter (<=1kb) | 20 | 40892393 | 40902592 | 10200 | 2 | 331401 | ENSMUST00000134084.2 | 0 | ENSMUSG00000037475 | Thoc2 | THO complex 2 |
| chr11 | 57877636 | 57878385 | 750 | * | 6,143243 | -4,935941618 | 1,878764 | -2,62723 | 0,008608 | 1 | 0 | Distal Intergenic | 11 | 57899890 | 57952857 | 52968 | 1 | 73158 | ENSMUST00000071487.13 | -21505 | ENSMUSG00000037331 | Larp1 | La ribonucleoprotein domain family, member 1 |
| chr12 | 75872430 | 75873526 | 1097 | * | 5,775818 | -4,935912496 | 1,889162 | -2,61275 | 0,008982 | 1 | 0 | Intron (ENSMUST0000( | 12 | 75865124 | 75952677 | 87554 | 1 | 319565 | ENSMUST00000157033.2 | 7306 | ENSMUSG00000063450 | Syne2 | spectrin repeat containing, nuclear envelope 2 |
| chr2 | 74466185 | 74467220 | 1036 | * | 11,54884 | -4,93560443 | 1,758433 | -2,80682 | 0,005003 | 0,882702232 | 0,744375 | Distal Intergenic | 2 | 74454986 | 74455092 | 107 | 1 | 1,15E+08 | ENSMUST00000230528.3 | 11199 | ENSMUSG00000119655 | Gm22975 | predicted gene, 22975 |
| chr14 | 1,14E+08 | 1,14E+08 | 1170 | * | 6,913489 | -4,93506323 | 1,781598 | -2,77002 | 0,005605 | 1 | 0 | Distal Intergenic | 14 | 1,15E+08 | 1,15E+08 | 29363 | 1 | 75822 | ENSMUST00000226121.2 | -1230499 | ENSMUSG00000115320 | 4930524C18Rik | RIKEN cDNA 4930524C18 gene |
| chr13 | 1,04E+08 | 1,04E+08 | 972 | * | 6,103598 | -4,934036765 | 1,909659 | -2,58373 | 0,009774 | 1 | 0 | Intron (ENSMUST0000( | 13 | 1,04E+08 | 1,04E+08 | 78335 | 2 | 59079 | ENSMUST00000189323.2 | -44571 | ENSMUSG00000021709 | Erbin | Erbb2 interacting protein |
| chr19 | 55356754 | 55357587 | 834 | * | 6,625113 | -4,933262457 | 1,878613 | -2,62601 | 0,008639 | 1 | 0 | Intron (ENSMUST0000( | 19 | 55369446 | 55435411 | 65966 | 1 | 53611 | ENSMUST00000224396.2 | -11859 | ENSMUSG00000024983 | Vti1a | vesicle transport through interaction with t-SNAREs 1A |
| chr5 | 39414398 | 39415198 | 801 | * | 7,430139 | -4,933260651 | 1,766189 | -2,79317 | 0,005219 | 0,783821651 | 1,360622 | Intron (ENSMUST0000( | 5 | 39411293 | 39464427 | 53135 | 2 | 1,05E+08 | ENSMUST00000202596.2 | 49229 | | Gm40293 | predicted gene, 40293 |
| chr7 | 10026547 | 10027671 | 1125 | * | 5,429088 | -4,930312285 | 1,853129 | -2,66053 | 0,007802 | 1 | 0 | Distal Intergenic | 7 | 10007755 | 10009278 | 1524 | 2 | 171264 | ENSMUST00000060374.5 | -17269 | ENSMUSG00000043066 | Vmn1r66 | vomeronasal 1 receptor 66 |
| chr3 | 85925700 | 85926285 | 586 | * | 5,829866 | -4,924161659 | 1,886113 | -2,61075 | 0,009035 | 1 | 0 | Intron (ENSMUST0000( | 3 | 85901117 | 85909798 | 8682 | 2 | 368202 | ENSMUST00000061343.4 | -15902 | ENSMUSG00000049013 | Prss48 | protease, serine 48 |
| chr10 | 39377985 | 39378899 | 915 | * | 7,70497 | -4,924161503 | 1,725676 | -2,85347 | 0,004324 | 1 | 0 | Intron (ENSMUST0000( | 10 | 39387735 | 39440418 | 52684 | 1 | 14360 | ENSMUST00000136659.2 | -8836 | ENSMUSG00000019843 | Fyn | Fyn proto-oncogene |
| chrX | 79268788 | 79269555 | 768 | * | 5,396946 | -4,923032444 | 1,893035 | -2,6006 | 0,009306 | 1 | 0 | Distal Intergenic | 20 | 78666837 | 78715041 | 48205 | 2 | 71967 | ENSMUST00000052283.7 | -553747 | ENSMUSG00000046942 | Mageb16 | MAGE family member B16 |
| chr3 | 26754549 | 26755385 | 837 | * | 7,342633 | -4,922344893 | 1,768766 | -2,78293 | 0,005387 | 1 | 0 | Intron (ENSMUST0000( | 3 | 26691799 | 26981628 | 289830 | 1 | 70862 | ENSMUST00000108305.2 | 62750 | ENSMUSG00000039335 | Spata16 | spermatogenesis associated 16 |
| chr3 | 63805838 | 63806956 | 1119 | * | 7,558041 | -4,920721027 | 1,837829 | -2,67746 | 0,007418 | 1 | 0 | Promoter (<=1kb) | 3 | 63604152 | 63806893 | 202742 | 2 | 269437 | ENSMUST00000177143.8 | 0 | ENSMUSG00000036834 | Plch1 | phospholipase C, eta 1 |
| chr4 | 1,2E+08 | 1,2E+08 | 1159 | * | 7,765235 | -4,917940418 | 1,732266 | -2,83902 | 0,004525 | 1 | 0 | Intron (ENSMUST0000( | 4 | 1,2E+08 | 1,2E+08 | 323551 | 1 | 16656 | ENSMUST00000106307.9 | 67992 | ENSMUSG00000028634 | Hivep3 | human immunodeficiency virus type I enhancer binding protein 3 |
| chr19 | 36803141 | 36803916 | 776 | * | 7,65965 | -4,914113574 | 1,805205 | -2,72219 | 0,006485 | 1 | 0 | Distal Intergenic | 19 | 36811632 | 36870877 | 59246 | 1 | 74493 | ENSMUST00000025729.12 | -7716 | ENSMUSG00000024811 | Tnks2 | tankyrase, TRF1-interacting ankyrin-related ADP-ribose polymerase 2 |
| chr3 | 96855662 | 96856535 | 874 | * | 8,125282 | -4,913600415 | 1,836584 | -2,6754 | 0,007464 | 0,755390167 | 2,237403 | Intron (ENSMUST0000( | 3 | 96820882 | 96833336 | 12455 | 2 | 14616 | ENSMUST00000062944.7 | -22326 | ENSMUSG00000049908 | Gja8 | gap junction protein, alpha 8 |
| chr18 | 77151918 | 77152745 | 828 | * | 6,636319 | -4,913076917 | 1,874131 | -2,62152 | 0,008754 | 1 | 0 | Promoter (<=1kb) | 18 | 77152904 | 77241496 | 88593 | 1 | 17344 | ENSMUST00000114777.10 | -159 | ENSMUSG00000025423 | Pias2 | protein inhibitor of activated STAT 2 |
| chr8 | 50199615 | 50200545 | 931 | * | 5,724636 | -4,912691619 | 1,905977 | -2,57752 | 0,009951 | 1 | 0 | Distal Intergenic | 8 | 49938516 | 49974130 | 35615 | 1 | 75359 | ENSMUST00000126285.2 | 261099 | ENSMUSG00000031559 | 4930555F03Rik | RIKEN cDNA 4930555F03 gene |
| chr18 | 50130000 | 50131116 | 1117 | * | 7,8432 | -4,910100455 | 1,753645 | -2,79994 | 0,005111 | 1 | 0 | Intron (ENSMUST0000( | 18 | 50112602 | 50240240 | 127639 | 1 | 106869 | ENSMUST00000153545.2 | 17398 | ENSMUSG00000062210 | Tnfaip8 | tumor necrosis factor, alpha-induced protein 8 |
| chr6 | 7755131 | 7756401 | 1271 | * | 13,19136 | -4,909175091 | 1,642605 | -2,98865 | 0,002802 | 0,78061212 | 0,738441 | Distal Intergenic | 6 | 7766394 | 7777681 | 11288 | 1 | 75487 | ENSMUST00000203458.2 | -9993 | ENSMUSG00000107394 | 1700012J15Rik | RIKEN cDNA 1700012J15 gene |
| chrX | 1,35E+08 | 1,35E+08 | 1600 | * | 9,266869 | -4,906573732 | 1,696613 | -2,89198 | 0,003828 | 0,748971518 | 1,211315 | Distal Intergenic | 20 | 1,35E+08 | 1,35E+08 | 1146 | 1 | 12070 | ENSMUST00000113113.2 | 14780 | ENSMUSG00000046432 | Bex3 | brain expressed X-linked 3 |
| chrX | 61406697 | 61407528 | 832 | * | 5,755356 | -4,904375796 | 1,903076 | -2,57708 | 0,009964 | 1 | 0 | Distal Intergenic | 20 | 61330748 | 61335684 | 4937 | 1 | 74437 | ENSMUST00000051414.2 | 75949 | ENSMUSG00000045330 | Magec2 | MAGE family member C2 |
| chr3 | 93151019 | 93152136 | 1118 | * | 8,167548 | -4,903669821 | 1,734544 | -2,82707 | 0,004698 | 0,753191087 | 1,425929 | Distal Intergenic | 3 | 93180848 | 93200993 | 20146 | 1 | 14246 | ENSMUST00000214837.2 | -28712 | | Flg | filaggrin |
| chr6 | 33320584 | 33321796 | 1213 | * | 9,773746 | -4,903146874 | 1,754747 | -2,79422 | 0,005203 | 0,770673161 | 1,449706 | Intron (ENSMUST0000( | 6 | 33310352 | 33325069 | 14718 | 1 | 20336 | ENSMUST00000156207.2 | 10232 | ENSMUSG00000029763 | Exoc4 | exocyst complex component 4 |
| chr12 | 15190494 | 15191638 | 1145 | * | 8,201011 | -4,902294798 | 1,83535 | -2,67104 | 0,007562 | 0,757706088 | 2,237403 | Distal Intergenic | 12 | 15118118 | 15187519 | 69402 | 1 | 1,03E+08 | ENSMUST00000222816.2 | 72376 | | Gm36128 | predicted gene, 36128 |

| | | | | | | | | | | | | | | | | | | | | | | | |
|---|---|---|---|---|---|---|---|---|---|---|---|---|---|---|---|---|---|---|---|---|---|---|---|
| chr3 | 82358913 | 82359718 | 806 | * | 5,732194 | -4,902080911 | 1,8355 | -2,67071 | 0,007569 | 1 | 0 | Distal Intergenic | 3 | 82287380 | 82290617 | 3238 | 1 | 213582 | ENSMUST00000195793.2 | 71533 | ENSMUSG00000033900 | Map9 | microtubule-associated protein 9 |
| chr4 | 65040557 | 65041892 | 1336 | * | 8,855257 | -4,902078562 | 1,813441 | -2,70319 | 0,006868 | 0,831521609 | 1,360622 | Promoter (<=1kb) | 4 | 65042411 | 65275746 | 233336 | 1 | 18491 | ENSMUST00000084501.4 | -519 | ENSMUSG00000028370 | Pappa | pregnancy-associated plasma protein A |
| chr6 | 62026142 | 62026868 | 727 | * | 8,623026 | -4,900428365 | 1,738454 | -2,81884 | 0,00482 | 0,767135014 | 1,360622 | Intron (ENSMUST0000C | 6 | 61933673 | 62359849 | 426177 | 1 | 232035 | ENSMUST00000232648.2 | 92469 | ENSMUSG00000039578 | Ccser1 | coiled-coil serine rich 1 |
| chr16 | 27698896 | 27699713 | 818 | * | 8,668361 | -4,899089956 | 1,745656 | -2,80645 | 0,005009 | 0,68526372 | 2,237403 | Intron (ENSMUST0000C | 16 | 27668682 | 27744880 | 76199 | 1 | 1E+08 | ENSMUST00000231532.2 | 30214 | ENSMUSG00000116654 | Gm10823 | predicted gene 10823 |
| chr13 | 1,1E+08 | 1,1E+08 | 882 | * | 8,489051 | -4,898570845 | 1,734457 | -2,82427 | 0,004739 | 0,682206193 | 2,237403 | Intron (ENSMUST0000C | 13 | 1,1E+08 | 1,1E+08 | 225964 | 2 | 67295 | ENSMUST00000167824.3 | 53553 | ENSMUSG00000021700 | Rab3c | RAB3C, member RAS oncogene family |
| chr2 | 78763153 | 78763864 | 712 | * | 7,599288 | -4,897566342 | 1,773984 | -2,76077 | 0,005766 | 1 | 0 | Distal Intergenic | 2 | 78748737 | 78750396 | 1660 | 1 | 22193 | ENSMUST00000153266.2 | 14416 | ENSMUSG00000027011 | Ube2e3 | ubiquitin-conjugating enzyme E2E 3 |
| chr14 | 22413558 | 22414656 | 1099 | * | 7,614243 | -4,897489923 | 1,816885 | -2,69554 | 0,007027 | 1 | 0 | Intron (ENSMUST0000C | 14 | 22360233 | 22634507 | 274275 | 1 | 76633 | ENSMUST00000159107.2 | 53325 | ENSMUSG00000063458 | Lrmda | leucine rich melanocyte differentiation associated |
| chr6 | 60849801 | 60850339 | 539 | * | 5,533467 | -4,896579123 | 1,882902 | -2,60055 | 0,009307 | 1 | 0 | Distal Intergenic | 6 | 60708559 | 60806839 | 98281 | 2 | 20617 | ENSMUST00000163779.8 | -42962 | ENSMUSG00000025889 | Snca | synuclein, alpha |
| chr5 | 62898224 | 62898622 | 399 | * | 5,664325 | -4,895190997 | 1,826549 | -2,68002 | 0,007362 | 1 | 0 | Intron (ENSMUST0000C | 5 | 62863960 | 62888164 | 24205 | 2 | 212285 | ENSMUST00000161911.2 | -10060 | ENSMUSG00000037999 | Arap2 | ArfGAP with RhoGAP domain, ankyrin repeat and PH domain 2 |
| chr14 | 33559543 | 33560279 | 737 | * | 6,104914 | -4,889427997 | 1,897665 | -2,57655 | 0,009979 | 1 | 0 | Distal Intergenic | 14 | 33531716 | 33533240 | 1525 | 2 | 218902 | ENSMUST00000226156.2 | -26303 | ENSMUSG00000115751 | 4930503F20Rik | RIKEN cDNA 4930503F20 gene |
| chr14 | 1,16E+08 | 1,16E+08 | 778 | * | 6,047603 | -4,884733147 | 1,800029 | -2,7137 | 0,006654 | 1 | 0 | Intron (ENSMUST0000C | 14 | 1,16E+08 | 1,16E+08 | 13002 | 2 | 75088 | ENSMUST00000176582.2 | -204673 | ENSMUSG00000093551 | 4930505G20Rik | RIKEN cDNA 4930505G20 gene |
| chrX | 52418311 | 52419667 | 1357 | * | 8,875428 | -4,883082545 | 1,764079 | -2,76806 | 0,005639 | 0,817544002 | 1,211315 | Intron (ENSMUST0000C | 20 | 52433475 | 52459379 | 25905 | 2 | 70380 | ENSMUST00000152743.8 | 39712 | ENSMUSG00000023074 | Mospd1 | motile sperm domain containing 1 |
| chrX | 49195470 | 49197107 | 1638 | * | 8,243786 | -4,882898307 | 1,751717 | -2,78749 | 0,005312 | 0,775744126 | 1,449706 | Distal Intergenic | 20 | 49098183 | 49099112 | 930 | 2 | 258385 | ENSMUST00000057071.2 | -96358 | ENSMUSG00000050504 | Olfr1323 | olfactory receptor 1323 |
| chr4 | 44456557 | 44457611 | 1055 | * | 9,516976 | -4,881692661 | 1,747692 | -2,79322 | 0,005219 | 0,783815908 | 1,360622 | Distal Intergenic | 4 | 44412469 | 44412595 | 127 | 2 | 1,15E+08 | ENSMUST00000104399.2 | -43962 | ENSMUSG00000077587 | Gm24376 | predicted gene, 24376 |
| chr6 | 1,16E+08 | 1,16E+08 | 815 | * | 9,380624 | -4,880644723 | 1,673767 | -2,91596 | 0,003546 | 0,704364304 | 1,449706 | Promoter (1-2kb) | 6 | 1,16E+08 | 1,16E+08 | 2517 | 2 | 11689 | ENSMUST00000169625.2 | 1026 | ENSMUSG00000025701 | Alox5 | arachidonate 5-lipoxygenase |
| chr18 | 82276176 | 82276765 | 590 | * | 7,040921 | -4,880250054 | 1,750013 | -2,78869 | 0,005292 | 0,786955198 | 1,360622 | Exon (ENSMUST000002 | 18 | 82410505 | 82424902 | 14398 | 2 | 14427 | ENSMUST00000065224.8 | 148137 | ENSMUSG00000024553 | Galr1 | galanin receptor 1 |
| chr10 | 82396036 | 82397356 | 1321 | * | 8,74693 | -4,877881412 | 1,833847 | -2,65992 | 0,007816 | 0,842444373 | 1,449706 | Distal Intergenic | 10 | 82455685 | 82458760 | 3076 | 2 | 544717 | ENSMUST00000176200.8 | 61404 | ENSMUSG00000063320 | 1190007I07Rik | RIKEN cDNA 1190007I07 gene |
| chr14 | 30122371 | 30123817 | 1447 | * | 10,81556 | -4,876654485 | 1,657128 | -2,94283 | 0,003252 | 0,767135014 | 0,918255 | Intron (ENSMUST0000C | 14 | 29763465 | 30075484 | 312020 | 2 | 12289 | ENSMUST00000224785.3 | -46887 | ENSMUSG00000015968 | Cacna1d | calcium channel, voltage-dependent, L type, alpha 1D subunit |
| chr14 | 53351856 | 53353325 | 1470 | * | 9,555265 | -4,876047439 | 1,689897 | -2,88541 | 0,003909 | 0,721416593 | 1,449706 | Distal Intergenic | 14 | 53269874 | 53276923 | 7050 | 2 | 1,03E+08 | ENSMUST00000199271.2 | -74933 | ENSMUSG00000104598 | Gm30214 | predicted gene, 30214 |
| chr11 | 50981519 | 50982141 | 623 | * | 7,468788 | -4,874788436 | 1,851689 | -2,63262 | 0,008473 | 1 | 0 | Exon (ENSMUST00000 | 11 | 50975914 | 50982660 | 6747 | 1 | 619293 | ENSMUST00000101255.2 | 5605 | ENSMUSG00000072982 | 9230009I02Rik | RIKEN cDNA 9230009I02 gene |
| chrX | 1,58E+08 | 1,58E+08 | 1426 | * | 7,575755 | -4,874562439 | 1,804899 | -2,70074 | 0,006919 | 1 | 0 | Intron (ENSMUST0000C | 20 | 1,58E+08 | 1,58E+08 | 55744 | 1 | 110651 | ENSMUST00000126686.8 | -12910 | ENSMUSG00000031309 | Rps6ka3 | ribosomal protein S6 kinase polypeptide 3 |
| chr10 | 77746637 | 77748114 | 1478 | * | 11,17334 | -4,871103603 | 1,663515 | -2,9282 | 0,003409 | 0,814558 | 0,744375 | Promoter (<=1kb) | 10 | 77748470 | 77748545 | 76 | 2 | 1,15E+08 | ENSMUST00000177209.3 | 431 | ENSMUSG00000093784 | Gm22394 | predicted gene, 22394 |
| chr2 | 1,75E+08 | 1,75E+08 | 603 | * | 7,67099 | -4,870774919 | 1,734218 | -2,80863 | 0,004975 | 1 | 0 | Distal Intergenic | 2 | 1,75E+08 | 1,75E+08 | 10243 | 1 | 1,03E+08 | ENSMUST00000131690.2 | 61007 | | Gm14616 | predicted gene 14616 |
| chr16 | 91182474 | 91184037 | 1564 | * | 11,82856 | -4,869297422 | 1,639335 | -2,97029 | 0,002975 | 0,711373629 | 1,173325 | Promoter (<=1kb) | 16 | 91184677 | 91188664 | 3988 | 1 | 15976 | ENSMUST00000139503.2 | -640 | ENSMUSG00000022971 | Ifnar2 | interferon (alpha and beta) receptor 2 |
| chr5 | 46883854 | 46884733 | 880 | * | 6,669115 | -4,869017133 | 1,777568 | -2,73915 | 0,00616 | 1 | 0 | Distal Intergenic | 5 | 46084160 | 46089751 | 5592 | 2 | 78106 | ENSMUST00000198088.2 | -794103 | ENSMUSG00000106602 | 4930405L22Rik | RIKEN cDNA 4930405L22 gene |
| chrX | 60122433 | 60123340 | 908 | * | 6,115769 | -4,86771933 | 1,866847 | -2,60745 | 0,009122 | 1 | 0 | Distal Intergenic | 20 | 60159984 | 60336090 | 176107 | 1 | 245440 | ENSMUST00000232361.2 | -36644 | ENSMUSG00000118608 | Gm4988 | predicted gene 4988 |
| chr3 | 40182570 | 40183748 | 1179 | * | 8,233876 | -4,866412249 | 1,773361 | -2,74417 | 0,006066 | 0,722439117 | 2,237403 | Distal Intergenic | 3 | 40562579 | 40563923 | 1345 | 2 | 1E+08 | ENSMUST00000202778.2 | 380175 | ENSMUSG00000037884 | 1700017G19Rik | RIKEN cDNA 1700017G19 gene |
| chr12 | 1,02E+08 | 1,02E+08 | 988 | * | 7,714544 | -4,863835556 | 1,716148 | -2,83416 | 0,004595 | 1 | 0 | Distal Intergenic | 12 | 1,02E+08 | 1,02E+08 | 8963 | 2 | 1,03E+08 | ENSMUST00000221227.2 | -57199 | ENSMUSG00000113902 | Ndufb1 | NADH:ubiquinone oxidoreductase subunit B1 |
| chr7 | 1,21E+08 | 1,21E+08 | 839 | * | 8,472003 | -4,863119821 | 1,810076 | -2,68669 | 0,007216 | 0,749931768 | 2,237403 | Intron (ENSMUST0000C | 7 | 1,21E+08 | 1,21E+08 | 44702 | 1 | 246190 | ENSMUST00000163275.2 | 30207 | ENSMUSG00000034990 | Otoa | otoancorin |
| chrX | 98996911 | 98997626 | 716 | * | 6,455186 | -4,863041193 | 1,864722 | -2,60792 | 0,00911 | 1 | 0 | Distal Intergenic | 20 | 99019212 | 99354905 | 335694 | 1 | 13607 | ENSMUST00000134582.2 | -21586 | ENSMUSG00000059327 | Eda | ectodysplasin-A |
| chr12 | 29633330 | 29634392 | 1063 | * | 8,992865 | -4,86264895 | 1,706421 | -2,84962 | 0,004377 | 0,739716578 | 1,449706 | Intron (ENSMUST0000C | 12 | 29616699 | 29835277 | 218579 | 1 | 17933 | ENSMUST00000217961.2 | 16631 | ENSMUSG00000061911 | Myt1l | myelin transcription factor 1-like |
| chr17 | 47407527 | 47409202 | 1676 | * | 11,80338 | -4,862485317 | 1,662622 | -2,92459 | 0,003449 | 0,818130284 | 0,738441 | Distal Intergenic | 17 | 47451801 | 47640462 | 188662 | 1 | 224829 | ENSMUST00000188947.7 | -42599 | ENSMUSG00000064043 | Trerf1 | transcriptional regulating factor 1 |
| chr2 | 43848206 | 43848727 | 522 | * | 7,100105 | -4,861073939 | 1,866042 | -2,60502 | 0,009187 | 1 | 0 | Intron (ENSMUST0000C | 2 | 43955053 | 44032468 | 77416 | 1 | 76117 | ENSMUST00000128630.2 | -106326 | ENSMUSG00000049744 | Arhgap15 | Rho GTPase activating protein 15 |
| chr16 | 79232286 | 79233140 | 855 | * | 7,3871 | -4,857983711 | 1,881634 | -2,58179 | 0,009829 | 1 | 0 | Distal Intergenic | 16 | 78750352 | 78887985 | 137634 | 2 | 19146 | ENSMUST00000060402.6 | -344301 | ENSMUSG00000022857 | Tmprss15 | transmembrane protease, serine 15 |
| chr17 | 15046262 | 15047385 | 1124 | * | 10,84111 | -4,85610313 | 1,650396 | -2,94239 | 0,003257 | 0,767439474 | 0,918255 | Intron (ENSMUST0000C | 17 | 15012203 | 15012266 | 64 | 2 | 1,15E+08 | ENSMUST00000158428.3 | -33996 | ENSMUSG00000089053 | Gm23352 | predicted gene, 23352 |
| chr3 | 1,04E+08 | 1,04E+08 | 838 | * | 6,696573 | -4,851327364 | 1,843308 | -2,63186 | 0,008492 | 1 | 0 | Promoter (1-2kb) | 3 | 1,04E+08 | 1,04E+08 | 47514 | 2 | 15257 | ENSMUST00000106845.9 | 1800 | ENSMUSG00000008730 | Hipk1 | homeodomain interacting protein kinase 1 |
| chr7 | 43830292 | 43830857 | 566 | * | 6,73084 | -4,85107727 | 1,880749 | -2,57933 | 0,009899 | 1 | 0 | Distal Intergenic | 7 | 43837660 | 43841879 | 4220 | 1 | 16617 | ENSMUST00000073713.8 | -6803 | ENSMUSG00000063713 | Klk1b24 | kallikrein 1-related peptidase b24 |
| chrX | 1,62E+08 | 1,62E+08 | 1517 | * | 12,84814 | -4,848784665 | 1,746128 | -2,77688 | 0,005488 | 0,858623476 | 0,918255 | Distal Intergenic | 20 | 1,62E+08 | 1,62E+08 | 35774 | 2 | 14829 | ENSMUST00000033730.3 | 77169 | ENSMUSG00000031364 | Grpr | gastrin releasing peptide receptor |
| chr16 | 46709730 | 46711149 | 1420 | * | 8,327581 | -4,848207201 | 1,736978 | -2,79117 | 0,005252 | 0,785681303 | 1,360622 | Distal Intergenic | 16 | 46279215 | 46318888 | 39674 | 2 | 58998 | ENSMUST00000133935.2 | -390842 | ENSMUSG00000022656 | Nectin3 | nectin cell adhesion molecule 3 |
| chr18 | 21646207 | 21647178 | 972 | * | 12,22607 | -4,846650562 | 1,635712 | -2,96302 | 0,003046 | 0,755058193 | 0,918255 | Distal Intergenic | 18 | 21513882 | 21513988 | 107 | 2 | 1,15E+08 | ENSMUST00000240423.1 | -132219 | ENSMUSG00000119907 | Gm22886 | predicted gene, 22886 |
| chr16 | 29592401 | 29593535 | 1135 | * | 9,257215 | -4,846076652 | 1,76994 | -2,73799 | 0,006182 | 0,833333099 | 1,211315 | Distal Intergenic | 16 | 29629876 | 29656632 | 26757 | 1 | 1,03E+08 | ENSMUST00000181815.2 | -36341 | | Gm32740 | predicted gene, 32740 |
| chr11 | 95496145 | 95497195 | 1051 | * | 6,938164 | -4,845470292 | 1,83433 | -2,64155 | 0,008253 | 0,860883941 | 1,360622 | Distal Intergenic | 11 | 95459644 | 95478561 | 18918 | 2 | 18053 | ENSMUST00000000122.7 | -17584 | ENSMUSG00000000120 | Ngfr | nerve growth factor receptor (TNFR superfamily, member 16) |
| chr10 | 49997368 | 49998542 | 1175 | * | 8,863406 | -4,843049524 | 1,689721 | -2,86618 | 0,004155 | 0,730469267 | 1,449706 | Distal Intergenic | 10 | 48970929 | 49664862 | 693934 | 2 | 14806 | ENSMUST00000218823.2 | -332506 | ENSMUSG00000056073 | Grik2 | glutamate receptor, ionotropic, kainate 2 (beta 2) |
| chr18 | 35744884 | 35745808 | 925 | * | 8,88268 | -4,842631673 | 1,77396 | -2,72984 | 0,006336 | 0,808893772 | 1,425929 | Promoter (<=1kb) | 18 | 35744038 | 35746927 | 2890 | 1 | 67869 | ENSMUST00000237730.2 | 846 | ENSMUSG00000037058 | Paip2 | polyadenylate-binding protein-interacting protein 2 |
| chr8 | 25112822 | 25114184 | 1363 | * | 14,12611 | -4,837070027 | 1,639989 | -2,94945 | 0,003183 | 0,802323085 | 0,738441 | Intron (ENSMUST0000C | 8 | 25092266 | 25118270 | 26005 | 2 | 13524 | ENSMUST00000138232.2 | 4086 | ENSMUSG00000031552 | Adam18 | a disintegrin and metallopeptidase domain 18 |
| chr2 | 91259846 | 91260623 | 778 | * | 6,113026 | -4,835560753 | 1,800004 | -2,68642 | 0,007222 | 1 | 0 | Intron (ENSMUST0000C | 2 | 91105413 | 91274967 | 169555 | 2 | 228356 | ENSMUST00000064652.14 | 14344 | ENSMUSG00000040591 | Cstpp1 | centriolar satellite-associated tubulin polyglutamylase complex regulator 1 |
| chr2 | 1,22E+08 | 1,22E+08 | 1354 | * | 11,48225 | -4,832233841 | 1,642673 | -2,94169 | 0,003264 | 0,713633527 | 1,2763 | Distal Intergenic | 2 | 1,21E+08 | 1,22E+08 | 261559 | 2 | 228564 | ENSMUST00000138157.8 | -10100 | ENSMUSG00000027238 | Frmd5 | FERM domain containing 5 |
| chr6 | 97979884 | 97980629 | 746 | * | 7,649441 | -4,83183988 | 1,789447 | -2,70019 | 0,00693 | 0,832960417 | 1,360622 | Intron (ENSMUST0000C | 6 | 97968882 | 97995136 | 26255 | 1 | 17342 | ENSMUST00000203938.2 | 11002 | ENSMUSG00000035158 | Mitf | melanogenesis associated transcription factor |
| chr9 | 1,15E+08 | 1,15E+08 | 763 | * | 7,561289 | -4,831773915 | 1,748511 | -2,76337 | 0,005721 | 1 | 0 | Distal Intergenic | 9 | 1,15E+08 | 1,15E+08 | 54808 | 2 | 70031 | ENSMUST00000047013.4 | -7392 | ENSMUSG00000041012 | Cmtm8 | CKLF-like MARVEL transmembrane domain containing 8 |
| chr13 | 41446152 | 41446887 | 736 | * | 7,337871 | -4,827049035 | 1,805946 | -2,67286 | 0,007521 | 1 | 0 | Distal Intergenic | 13 | 41403320 | 41430053 | 26734 | 1 | 108934 | ENSMUST00000165561.4 | 42832 | ENSMUSG00000091264 | Smim13 | small integral membrane protein 13 |
| chr1 | 73634064 | 73634718 | 655 | * | 6,849338 | -4,824356385 | 1,779584 | -2,71095 | 0,006709 | 1 | 0 | Distal Intergenic | 1 | 73562360 | 73568949 | 6590 | 2 | 1,03E+08 | ENSMUST00000186453.2 | -65115 | | C530043A13Rik | RIKEN cDNA C530043A13 gene |
| chr18 | 64573644 | 64574561 | 918 | * | 6,883044 | -4,823819026 | 1,829063 | -2,63732 | 0,008356 | 1 | 0 | Distal Intergenic | 18 | 64590965 | 64597397 | 6433 | 2 | 14151 | ENSMUST00000238082.2 | 22836 | ENSMUSG00000024588 | Fech | ferrochelatase |
| chr4 | 13540798 | 13541596 | 799 | * | 8,226558 | -4,823195681 | 1,751049 | -2,75446 | 0,005879 | 0,802490409 | 1,360622 | Distal Intergenic | 4 | 13743436 | 13893649 | 150214 | 1 | 12395 | ENSMUST00000006761.10 | -201840 | ENSMUSG00000006586 | Runx1t1 | RUNX1 translocation partner 1 |
| chr3 | 1,31E+08 | 1,31E+08 | 792 | * | 8,278667 | -4,820122566 | 1,772298 | -2,7197 | 0,006534 | 0,732489153 | 2,237403 | Intron (ENSMUST0000C | 3 | 1,31E+08 | 1,31E+08 | 18266 | 1 | 23971 | ENSMUST00000196638.2 | 58917 | ENSMUSG00000028032 | Papss1 | 3'-phosphoadenosine 5'-phosphosulfate synthase 1 |
| chr13 | 64774820 | 64775772 | 953 | * | 8,991332 | -4,818184888 | 1,705128 | -2,8257 | 0,004718 | 0,681459438 | 2,237403 | Distal Intergenic | 13 | 64629060 | 64645606 | 16547 | 2 | 68880 | ENSMUST00000180282.2 | -129214 | ENSMUSG00000096537 | Fam240b | family with sequence similarity 240 member B |
| chr1 | 1,41E+08 | 1,41E+08 | 828 | * | 6,801126 | -4,815181102 | 1,78654 | -2,69526 | 0,007033 | 1 | 0 | Distal Intergenic | 1 | 1,4E+08 | 1,41E+08 | 86229 | 2 | 75910 | ENSMUST00000193820.2 | -244582 | ENSMUSG00000102368 | 4930590L20Rik | RIKEN cDNA 4930590L20 gene |
| chr1 | 1,67E+08 | 1,67E+08 | 1244 | * | 7,511899 | -4,812265069 | 1,807572 | -2,66228 | 0,007761 | 1 | 0 | Distal Intergenic | 1 | 1,67E+08 | 1,67E+08 | 21112 | 2 | 240899 | ENSMUST00000036643.6 | -63306 | ENSMUSG00000040485 | Lrrc52 | leucine rich repeat containing 52 |
| chr13 | 63703425 | 63704170 | 746 | * | 6,210463 | -4,811969443 | 1,832838 | -2,62542 | 0,008654 | 1 | 0 | Intron (ENSMUST0000C | 13 | 63656144 | 63711629 | 55486 | 2 | 19206 | ENSMUST00000195258.7 | 7459 | ENSMUSG00000021466 | Ptch1 | patched 1 |
| chr8 | 64520714 | 64521434 | 721 | * | 7,312825 | -4,810256645 | 1,746385 | -2,75441 | 0,00588 | 1 | 0 | Intron (ENSMUST0000C | 8 | 64377728 | 64405204 | 27477 | 2 | 244495 | ENSMUST00000179944.2 | -115510 | ENSMUSG00000094443 | Sgo2b | shugoshin 2B |
| chr5 | 52917988 | 52918757 | 770 | * | 7,143872 | -4,808257264 | 1,797199 | -2,67542 | 0,007464 | 0,846832214 | 1,360622 | Exon (ENSMUST00000 | 5 | 52911944 | 52914825 | 2882 | 1 | 67073 | ENSMUST00000145825.2 | 6044 | ENSMUSG00000029186 | Pi4k2b | phosphatidylinositol 4-kinase type 2 beta |
| chr3 | 96973569 | 96974833 | 1265 | * | 10,47379 | -4,806117307 | 1,685239 | -2,85189 | 0,004346 | 0,820868587 | 0,918255 | Intron (ENSMUST0000C | 3 | 96953972 | 97000101 | 46130 | 2 | 75888 | ENSMUST00000197696.2 | 25268 | ENSMUSG00000104766 | 4930573H18Rik | RIKEN cDNA 4930573H18 gene |
| chr2 | 1,41E+08 | 1,41E+08 | 1036 | * | 5,674833 | -4,805816869 | 1,829906 | -2,62626 | 0,008633 | 1 | 0 | Intron (ENSMUST0000C | 2 | 1,41E+08 | 1,41E+08 | 56561 | 2 | 1,03E+08 | ENSMUST00000151538.2 | 45663 | | Macrod2os1 | mono-ADP ribosylhydrolase 2, opposite strand 1 |
| chr5 | 68859364 | 68860259 | 896 | * | 7,410287 | -4,802705414 | 1,803074 | -2,66362 | 0,00773 | 1 | 0 | Distal Intergenic | 5 | 68723513 | 68723639 | 127 | 2 | 1,15E+08 | ENSMUST00000157255.3 | -135725 | ENSMUSG00000087880 | Gm23605 | predicted gene, 23605 |
| chr4 | 70495742 | 70496829 | 1088 | * | 10,85287 | -4,801302 | 1,752191 | -2,74017 | 0,006141 | 0,913059589 | 0,744375 | Distal Intergenic | 4 | 70380063 | 70453232 | 73170 | 2 | 230316 | ENSMUST00000037881.3 | -42510 | ENSMUSG00000039270 | Megf9 | multiple EGF-like-domains 9 |
| chr2 | 1,2E+08 | 1,2E+08 | 1337 | * | 8,037632 | -4,800361611 | 1,733834 | -2,76864 | 0,005629 | 0,793995086 | 1,360622 | Distal Intergenic | 2 | 1,2E+08 | 1,2E+08 | 70522 | 1 | 29808 | ENSMUST00000046717.13 | -42417 | ENSMUSG00000033943 | Mga | MAX gene associated |
| chr7 | 1,42E+08 | 1,42E+08 | 1013 | * | 9,445989 | -4,798775843 | 1,768653 | -2,71324 | 0,006663 | 0,847020973 | 1,211315 | Intron (ENSMUST0000C | 7 | 1,42E+08 | 1,42E+08 | 7263 | 2 | 21823 | ENSMUST00000000219.10 | -3734 | ENSMUSG00000000214 | Th | tyrosine hydroxylase |
| chr7 | 65232271 | 65233301 | 1031 | * | 7,150008 | -4,798228314 | 1,835304 | -2,61441 | 0,008938 | 1 | 0 | Promoter (<=1kb) | 7 | 65231818 | 65268632 | 36815 | 1 | 407810 | ENSMUST00000154632.2 | 453 | ENSMUSG00000086662 | BC046251 | cDNA sequence BC046251 |
| chr9 | 14734679 | 14735469 | 791 | * | 7,081575 | -4,797948968 | 1,779427 | -2,69635 | 0,00701 | 1 | 0 | Intron (ENSMUST0000C | 9 | 14708462 | 14715149 | 6688 | 1 | 17535 | ENSMUST00000136568.2 | 26217 | ENSMUSG00000031928 | Mre11a | MRE11A homolog A, double strand break repair nuclease |
| chr10 | 46656654 | 46657741 | 1088 | * | 5,959062 | -4,796646787 | 1,818913 | -2,6371 | 0,008362 | 1 | 0 | Distal Intergenic | 10 | 46714442 | 46714603 | 162 | 2 | 1,15E+08 | ENSMUST00000083784.3 | 56862 | ENSMUSG00000065718 | Gm25650 | predicted gene, 25650 |
| chr11 | 36163734 | 36164863 | 1130 | * | 12,38231 | -4,795727301 | 1,308914 | -3,6639 | 0,000248 | 0,282235674 | 1,2763 | Exon (ENSMUST00000 | 11 | 36275950 | 36323103 | 47154 | 2 | 23964 | ENSMUST00000128501.2 | 158240 | ENSMUSG00000049336 | Tenm2 | teneurin transmembrane protein 2 |
| chr16 | 19767408 | 19768381 | 974 | * | 8,746055 | -4,795502359 | 1,733606 | -2,7662 | 0,005671 | 0,787852124 | 1,449706 | Exon (ENSMUST00000 | 16 | 19775700 | 19785069 | 9370 | 2 | 239743 | ENSMUST00000171910.2 | 16688 | ENSMUSG00000043008 | Klhl6 | kelch-like 6 |
| chr11 | 1,2E+08 | 1,2E+08 | 1325 | * | 10,48937 | -4,79259131 | 1,686945 | -2,84099 | 0,004497 | 0,860573043 | 0,744375 | Exon (ENSMUST00000 | 11 | 1,2E+08 | 1,2E+08 | 209867 | 1 | 74370 | ENSMUST00000127899.8 | 69115 | ENSMUSG00000025583 | Rptor | regulatory associated protein of MTOR, complex 1 |
| chr9 | 1,04E+08 | 1,04E+08 | 754 | * | 7,77798 | -4,791622492 | 1,734553 | -2,76245 | 0,005737 | 1 | 0 | Intron (ENSMUST0000C | 9 | 1,04E+08 | 1,05E+08 | 407846 | 1 | 74020 | ENSMUST00000213120.2 | 29275 | ENSMUSG00000032564 | Cpne4 | copine IV |
| chr4 | 66734453 | 66735482 | 1030 | * | 9,280205 | -4,785809085 | 1,682886 | -2,84381 | 0,004458 | 0,671251031 | 2,237403 | Distal Intergenic | 4 | 66745821 | 66761037 | 15217 | 1 | 21898 | ENSMUST00000048096.12 | -10339 | ENSMUSG00000039005 | Tlr4 | toll-like receptor 4 |
| chr16 | 23708302 | 23709140 | 839 | * | 6,15575 | -4,781113274 | 1,845761 | -2,59032 | 0,009589 | 1 | 0 | Promoter (<=1kb) | 16 | 23708323 | 23709708 | 1386 | 2 | 20604 | ENSMUST00000004480.5 | 568 | ENSMUSG00000004366 | Sst | somatostatin |
| chr13 | 33437169 | 33438256 | 1088 | * | 7,879878 | -4,779036703 | 1,749993 | -2,73089 | 0,006316 | 1 | 0 | Exon (ENSMUST00000 | 13 | 33433595 | 33444833 | 11239 | 1 | 20710 | ENSMUST00000071873.4 | 3574 | ENSMUSG00000062342 | Serpinb9e | serine (or cysteine) peptidase inhibitor, clade B, member 9e |
| chr10 | 74233843 | 74234822 | 980 | * | 9,910577 | -4,778512768 | 1,669216 | -2,86273 | 0,0042 | 0,770104803 | 1,173325 | Intron (ENSMUST0000C | 10 | 74221705 | 74232556 | 10852 | 1 | 11994 | ENSMUST00000139106.3 | 12138 | ENSMUSG00000052613 | Pcdh15 | protocadherin 15 |
| chr4 | 1,17E+08 | 1,17E+08 | 736 | * | 7,079016 | -4,77673315 | 1,790362 | -2,66803 | 0,00763 | 1 | 0 | Intron (ENSMUST0000C | 4 | 1,17E+08 | 1,17E+08 | 14140 | 2 | 66743 | ENSMUST00000150148.2 | -37210 | ENSMUSG00000028677 | Rnf220 | ring finger protein 220 |
| chr9 | 55425933 | 55427153 | 1221 | * | 9,769149 | -4,774451542 | 1,744511 | -2,73684 | 0,006203 | 0,802925002 | 1,425929 | Exon (ENSMUST000002 | 9 | 55361792 | 55419527 | 57736 | 2 | 110842 | ENSMUST00000034866.9 | -6406 | ENSMUSG00000032314 | Etfa | electron transferring flavoprotein, alpha polypeptide |
| chr13 | 78347589 | 78348884 | 1296 | * | 7,900631 | -4,773985437 | 1,810914 | -2,63623 | 0,008383 | 1 | 0 | Promoter (<=1kb) | 13 | 78337092 | 78347876 | 10785 | 2 | 13865 | ENSMUST00000091458.13 | 0 | ENSMUSG00000069171 | Nr2f1 | nuclear receptor subfamily 2, group F, member 1 |
| chrX | 1,56E+08 | 1,56E+08 | 1182 | * | 8,000012 | -4,770759171 | 1,747573 | -2,72993 | 0,006335 | 1 | 0 | Intron (ENSMUST0000C | 20 | 1,56E+08 | 1,56E+08 | 4868 | 2 | 20603 | ENSMUST00000146026.2 | -13931 | ENSMUSG00000071708 | Sms | spermine synthase |
| chr7 | 1,36E+08 | 1,36E+08 | 800 | * | 11,06558 | -4,76963154 | 1,726438 | -2,7627 | 0,005733 | 0,810897439 | 1,2763 | Distal Intergenic | 7 | 1,36E+08 | 1,36E+08 | 42650 | 2 | 621677 | ENSMUST00000211142.2 | -28989 | ENSMUSG00000110101 | Gm6249 | predicted gene 6249 |
| chr1 | 44373215 | 44373873 | 659 | * | 7,420107 | -4,766484315 | 1,753577 | -2,71815 | 0,006565 | 1 | 0 | Distal Intergenic | 1 | 44431969 | 44433607 | 1639 | 1 | 1,05E+08 | ENSMUST00000190951.2 | -58096 | ENSMUSG00000100769 | Gm28893 | predicted gene 28893 |
| chr7 | 1,19E+08 | 1,19E+08 | 1658 | * | 11,49116 | -4,766165428 | 1,63916 | -2,90769 | 0,003641 | 0,826676469 | 0,738441 | Promoter (1-2kb) | 7 | 1,19E+08 | 1,19E+08 | 18526 | 1 | 20216 | ENSMUST00000106526.2 | -1205 | ENSMUSG00000030935 | Acsm3 | acyl-CoA synthetase medium-chain family member 3 |
| chr7 | 17682474 | 17683276 | 803 | * | 10,42771 | -4,764852145 | 1,711578 | -2,78389 | 0,005371 | 0,85617852 | 0,918255 | Distal Intergenic | 7 | 17706049 | 17712481 | 6433 | 1 | 66996 | ENSMUST00000094799.3 | -22773 | ENSMUSG00000030368 | Ceacam11 | carcinoembryonic antigen-related cell adhesion molecule 11 |
| chr5 | 1,5E+08 | 1,5E+08 | 1312 | * | 11,295 | -4,762486109 | 1,675785 | -2,84194 | 0,004484 | 0,782451902 | 1,173325 | Distal Intergenic | 5 | 1,5E+08 | 1,5E+08 | 6666 | 1 | 381694 | ENSMUST00000201088.2 | 25197 | ENSMUSG00000051950 | B3glct | beta-3-glucosyltransferase |
| chrX | 11296659 | 11298442 | 1784 | * | 9,382165 | -4,762135588 | 1,717222 | -2,77316 | 0,005551 | 0,786955198 | 1,425929 | Distal Intergenic | 20 | 11194061 | 11194390 | 330 | 1 | 236663 | ENSMUST00000091180.3 | 102598 | ENSMUSG00000069038 | H2al1j | H2A histone family member L1J |
| chr13 | 51987546 | 51989103 | 1558 | * | 10,01129 | -4,761632265 | 1,710768 | -2,78333 | 0,00538 | 0,777172375 | 1,449706 | Distal Intergenic | 13 | 52000714 | 52002504 | 1791 | 1 | 23882 | ENSMUST00000021903.3 | -11611 | ENSMUSG00000021453 | Gadd45g | growth arrest and DNA-damage-inducible 45 gamma |
| chr11 | 69860629 | 69861747 | 1119 | * | 6,993122 | -4,761438538 | 1,846225 | -2,57901 | 0,009908 | 1 | 0 | Promoter (<=1kb) | 11 | 69859058 | 69860229 | 1172 | 2 | 54351 | ENSMUST00000151766.2 | -400 | ENSMUSG00000018565 | Elp5 | elongator acetyltransferase complex subunit 5 |
| chr11 | 1,09E+08 | 1,09E+08 | 1014 | * | 11,54588 | -4,760430855 | 1,6495 | -2,88598 | 0,003902 | 0,800381675 | 0,918255 | Intron (ENSMUST0000C | 11 | 1,09E+08 | 1,09E+08 | 6610 | 1 | 629945 | ENSMUST00000143781.2 | -7521 | ENSMUSG00000085811 | Cep112it | centrosomal protein 112, intronic transcript |
| chr2 | 99080534 | 99081726 | 1193 | * | 10,0684 | -4,758083936 | 1,674882 | -2,84085 | 0,004499 | 0,793995086 | 1,086118 | Distal Intergenic | 2 | 97298434 | 97459408 | 160975 | 1 | 241568 | ENSMUST00000170144.2 | 1782100 | ENSMUSG00000050587 | Lrrc4c | leucine rich repeat containing 4C |
| chr9 | 32915841 | 32916668 | 828 | * | 8,098215 | -4,755969059 | 1,708468 | -2,78376 | 0,005373 | 1 | 0 | Distal Intergenic | 9 | 32943087 | 33002095 | 59009 | 1 | 1,03E+08 | ENSMUST00000184476.2 | -26419 | ENSMUSG00000098519 | Gm27166 | predicted gene 27166 |
| chr10 | 30773108 | 30774256 | 1149 | * | 10,39615 | -4,755740043 | 1,647055 | -2,88742 | 0,003884 | 0,79897927 | 0,918255 | Distal Intergenic | 10 | 30818375 | 30818481 | 107 | 2 | 1,15E+08 | ENSMUST00000240080.1 | 44225 | ENSMUSG00000119396 | Gm24168 | predicted gene, 24168 |
| chr5 | 67423825 | 67425000 | 1176 | * | 11,85662 | -4,754227807 | 1,663152 | -2,85856 | 0,004256 | 0,816911501 | 0,918255 | 3' UTR | 5 | 67418239 | 67426349 | 8111 | 1 | 67878 | ENSMUST00000163028.3 | 5586 | ENSMUSG00000037720 | Tmem33 | transmembrane protein 33 |
| chr11 | 71745268 | 71746028 | 761 | * | 7,621588 | -4,75339027 | 1,743034 | -2,72708 | 0,00639 | 1 | 0 | Distal Intergenic | 11 | 71738902 | 71739032 | 131 | 2 | 1,15E+08 | ENSMUST00000104268.3 | -6236 | ENSMUSG00000077456 | Gm22927 | predicted gene, 22927 |
| chr19 | 20166310 | 20167432 | 1123 | * | 9,631891 | -4,753120997 | 1,687928 | -2,81595 | 0,004863 | 0,810132289 | 1,086118 | Distal Intergenic | 19 | 19983353 | 20038122 | 54770 | 2 | 1E+08 | ENSMUST00000237242.2 | -128188 | | Gm3728 | predicted gene 3728 |
| chr3 | 1,04E+08 | 1,04E+08 | 971 | * | 9,046978 | -4,750596611 | 1,765671 | -2,69053 | 0,007134 | 0,826357927 | 1,449706 | Intron (ENSMUST0000C | 3 | 1,04E+08 | 1,04E+08 | 1146 | 1 | 18685 | ENSMUST00000130385.2 | 47346 | ENSMUSG00000058388 | Phtf1 | putative homeodomain transcription factor 1 |
| chr14 | 4970911 | 4971779 | 869 | * | 6,857218 | -4,747617672 | 1,770612 | -2,68134 | 0,007333 | 1 | 0 | Distal Intergenic | 14 | 4506571 | 4809435 | 302865 | 2 | 21834 | ENSMUST00000224470.2 | -161476 | ENSMUSG00000021779 | Thrb | thyroid hormone receptor beta |
| chr1 | 52122042 | 52122927 | 886 | * | 6,765859 | -4,743653092 | 1,773584 | -2,67461 | 0,007482 | 1 | 0 | Exon (ENSMUST00000 | 1 | 52116420 | 52195246 | 78827 | 1 | 20846 | ENSMUST00000185516.2 | 5622 | ENSMUSG00000026104 | Stat1 | signal transducer and activator of transcription 1 |
| chr16 | 43997228 | 43998133 | 906 | * | 6,401512 | -4,740847103 | 1,794829 | -2,64139 | 0,008257 | 1 | 0 | Intron (ENSMUST0000C | 16 | 43993760 | 44047826 | 54067 | 1 | 207806 | ENSMUST00000169582.3 | 3468 | ENSMUSG00000068284 | Usf3 | upstream transcription factor family member 3 |
| chr6 | 88771283 | 88772417 | 1135 | * | 13,51529 | -4,740120851 | 1,603553 | -2,95601 | 0,003116 | 0,758890329 | 0,918255 | Intron (ENSMUST0000C | 6 | 88800112 | 88802815 | 2704 | 1 | 23945 | ENSMUST00000205045.2 | -27695 | ENSMUSG00000033174 | Mgll | monoglyceride lipase |

| | | | | | | | | | | | | | | | | | | | | | | | |
|---|---|---|---|---|---|---|---|---|---|---|---|---|---|---|---|---|---|---|---|---|---|---|---|
| chr19 | 7731391 | 7732026 | 636 | * | 7,488956 | -4,735338665 | 1,805779 | -2,62233 | 0,008733 | 1 | 0 | Distal Intergenic | 19 | 7684047 | 7688675 | 4629 | 2 | 207151 | ENSMUST00000141771.2 | -42716 | ENSMUSG00000024757 | Slc22a19 | solute carrier family 22 (organic anion transporter), member 19 |
| chr5 | 12875689 | 12876711 | 1023 | * | 8,529745 | -4,732590359 | 1,786442 | -2,64917 | 0,008069 | 0,771248365 | 2,237403 | Distal Intergenic | 5 | 12625761 | 12638719 | 12959 | 1 | 108151 | ENSMUST00000195923.2 | 249928 | ENSMUSG00000040254 | Sema3d | sema domain, immunoglobulin domain (Ig), short basic domain, secreted, (semaphorin) 3D |
| chr14 | 68344535 | 68345697 | 1163 | * | 7,227708 | -4,728745433 | 1,82217 | -2,59512 | 0,009456 | 1 | 0 | Intron (ENSMUST0000( | 14 | 68320039 | 68362295 | 42257 | 2 | 18040 | ENSMUST00000111089.8 | 16598 | ENSMUSG00000022054 | Nefm | neurofilament, medium polypeptide |
| chrX | 1,36E+08 | 1,36E+08 | 1125 | * | 9,274899 | -4,72226611 | 1,690146 | -2,794 | 0,005206 | 0,693003605 | 2,237403 | Distal Intergenic | 20 | 1,36E+08 | 1,38E+08 | 1276339 | 1 | 60367 | ENSMUST00000113063.8 | -133397 | ENSMUSG00000059203 | Il1rapl2 | interleukin 1 receptor accessory protein-like 2 |
| chr18 | 8298951 | 8299666 | 716 | * | 6,131707 | -4,72134429 | 1,827025 | -2,58417 | 0,009761 | 1 | 0 | Distal Intergenic | 18 | 7910619 | 7918503 | 7885 | 1 | 225131 | ENSMUST00000167542.2 | 388332 | ENSMUSG00000024283 | Wac | WW domain containing adaptor with coiled-coil |
| chr14 | 1,21E+08 | 1,21E+08 | 776 | * | 6,495844 | -4,718827998 | 1,784346 | -2,64457 | 0,008179 | 1 | 0 | Intron (ENSMUST0000( | 14 | 1,21E+08 | 1,22E+08 | 13014 | 1 | 223254 | ENSMUST00000137971.2 | -43482 | ENSMUSG00000025555 | Farp1 | FERM, RhoGEF (Arhgef) and pleckstrin domain protein 1 (chondrocyte-derived) |
| chr11 | 20909295 | 20910107 | 813 | * | 8,521285 | -4,717173877 | 1,697258 | -2,77929 | 0,005448 | 0,791068542 | 1,360622 | Distal Intergenic | 11 | 20797753 | 20797881 | 129 | 2 | 1,15E+08 | ENSMUST00000082737.3 | -111414 | ENSMUSG00000064671 | Gm22807 | predicted gene, 22807 |
| chr3 | 1,45E+08 | 1,45E+08 | 956 | * | 7,339165 | -4,717110386 | 1,75014 | -2,69528 | 0,007033 | 1 | 0 | Intron (ENSMUST0000( | 3 | 1,45E+08 | 1,45E+08 | 21521 | 1 | 71355 | ENSMUST00000139001.2 | 109685 | ENSMUSG00000028197 | Col24a1 | collagen, type XXIV, alpha 1 |
| chr2 | 1,32E+08 | 1,32E+08 | 1077 | * | 8,211963 | -4,711472822 | 1,714897 | -2,74738 | 0,006007 | 0,793995086 | 1,449706 | Distal Intergenic | 2 | 1,32E+08 | 1,32E+08 | 5094 | 1 | 26434 | ENSMUST00000110169.2 | 17348 | ENSMUSG00000027338 | Prnd | prion like protein doppel |
| chr11 | 82678028 | 82679211 | 1184 | * | 8,608482 | -4,707379364 | 1,699686 | -2,76956 | 0,005613 | 0,709779237 | 2,237403 | Exon (ENSMUST00000( | 11 | 82672029 | 82695024 | 22996 | 1 | 16882 | ENSMUST00000173722.8 | 5999 | ENSMUSG00000020697 | Lig3 | ligase III, DNA, ATP-dependent |
| chr3 | 40552658 | 40553554 | 897 | * | 7,599502 | -4,705896006 | 1,80423 | -2,60826 | 0,0091 | 1 | 0 | Distal Intergenic | 3 | 40562579 | 40563923 | 1345 | 2 | 1E+08 | ENSMUST00000202778.2 | 10369 | ENSMUSG00000037884 | 1700017G19Rik | RIKEN cDNA 1700017G19 gene |
| chr6 | 12010332 | 12011124 | 793 | * | 8,355513 | -4,701448223 | 1,784219 | -2,63502 | 0,008413 | 1 | 0 | Intron (ENSMUST0000( | 6 | 12006819 | 12081099 | 74281 | 1 | 75725 | ENSMUST00000203045.2 | 3513 | ENSMUSG00000029629 | Phf14 | PHD finger protein 14 |
| chr19 | 13569978 | 13570759 | 782 | * | 7,653025 | -4,701025603 | 1,790257 | -2,62589 | 0,008642 | 0,872043312 | 1,360622 | Distal Intergenic | 19 | 13562670 | 13563617 | 948 | 1 | 258288 | ENSMUST00000074180.2 | 7308 | ENSMUSG00000096289 | Olfr1484 | olfactory receptor 1484 |
| chr12 | 76863275 | 76864089 | 815 | * | 6,93119 | -4,70057826 | 1,77718 | -2,64496 | 0,00817 | 1 | 0 | Promoter (1-2kb) | 12 | 76846896 | 76865148 | 18253 | 2 | 104886 | ENSMUST00000118604.8 | 1059 | ENSMUSG00000021062 | Rab15 | RAB15, member RAS oncogene family |
| chr1 | 1,69E+08 | 1,69E+08 | 1900 | * | 17,23568 | -4,696522013 | 1,593313 | -2,94765 | 0,003202 | 0,594430593 | 2,507013 | Distal Intergenic | 1 | 1,69E+08 | 1,7E+08 | 40313 | 1 | 19737 | ENSMUST00000027997.9 | -18494 | ENSMUSG00000026678 | Rgs5 | regulator of G-protein signaling 5 |
| chr9 | 58997669 | 58999142 | 1474 | * | 9,360852 | -4,69599683 | 1,782286 | -2,63482 | 0,008418 | 0,855226996 | 1,449706 | Distal Intergenic | 9 | 58781970 | 58943724 | 161755 | 2 | 18007 | ENSMUST00000068664.7 | -53945 | ENSMUSG00000032340 | Neo1 | neogenin |
| chr11 | 56242378 | 56243170 | 793 | * | 7,44551 | -4,692662762 | 1,79232 | -2,61821 | 0,008839 | 1 | 0 | Distal Intergenic | 11 | 56153991 | 56154066 | 76 | 2 | 1,15E+08 | ENSMUST00000083701.3 | -88312 | ENSMUSG00000065635 | Gm24253 | predicted gene, 24253 |
| chr9 | 22682205 | 22683158 | 954 | * | 8,627704 | -4,691327506 | 1,732892 | -2,70723 | 0,006785 | 0,818408612 | 1,449706 | Intron (ENSMUST0000( | 9 | 22590046 | 22799570 | 209525 | 1 | 319845 | ENSMUST00000124076.2 | 92159 | ENSMUSG00000035919 | Bbs9 | Bardet-Biedl syndrome 9 (human) |
| chr9 | 36500501 | 36501182 | 682 | * | 8,956367 | -4,690767959 | 1,714566 | -2,73583 | 0,006222 | 0,834909202 | 1,211315 | Distal Intergenic | 9 | 36492574 | 36493931 | 1358 | 2 | 1E+08 | ENSMUST00000170030.2 | -6570 | ENSMUSG00000091248 | Pate8 | prostate and testis expressed 8 |
| chr8 | 54120887 | 54121737 | 851 | * | 7,512054 | -4,690024225 | 1,819148 | -2,57814 | 0,009933 | 1 | 0 | Distal Intergenic | 8 | 54039902 | 54092100 | 52199 | 2 | 234258 | ENSMUST00000047768.11 | -28787 | ENSMUSG00000039396 | Neil3 | nei like 3 (E. coli) |
| chr1 | 47831450 | 47832303 | 854 | * | 7,450873 | -4,688116091 | 1,767226 | -2,65281 | 0,007982 | 1 | 0 | Distal Intergenic | 1 | 47506180 | 47506284 | 105 | 2 | 1,02E+08 | ENSMUST00000183639.3 | -325166 | ENSMUSG00000098876 | Mir6350 | microRNA 6350 |
| chrX | 81499902 | 81501144 | 1243 | * | 7,603322 | -4,681964105 | 1,811897 | -2,58401 | 0,009766 | 1 | 0 | Distal Intergenic | 20 | 81444697 | 81444833 | 137 | 2 | 1,15E+08 | ENSMUST00000083047.3 | -55069 | ENSMUSG00000119611 | Gm24706 | predicted gene, 24706 |
| chr4 | 31145369 | 31146018 | 650 | * | 6,553297 | -4,681339356 | 1,813046 | -2,58203 | 0,009822 | 1 | 0 | Distal Intergenic | 4 | 31174909 | 31175012 | 104 | 2 | 1,15E+08 | ENSMUST00000083119.2 | 28994 | ENSMUSG00000065053 | Gm25705 | predicted gene, 25705 |
| chr7 | 65078817 | 65080665 | 1849 | * | 11,43379 | -4,681179827 | 1,636005 | -2,86135 | 0,004218 | 0,753976089 | 1,2763 | Intron (ENSMUST0000( | 7 | 64945913 | 65020987 | 75075 | 2 | 21872 | ENSMUST00000102592.10 | -57830 | ENSMUSG00000030516 | Tjp1 | tight junction protein 1 |
| chrX | 77624148 | 77625498 | 1351 | * | 6,302035 | -4,681126806 | 1,792291 | -2,61181 | 0,009006 | 1 | 0 | Promoter (1-2kb) | 20 | 77493219 | 77627388 | 134170 | 2 | 546336 | ENSMUST00000177904.8 | 1890 | ENSMUSG00000047996 | Prrg1 | proline rich Gla (G-carboxyglutamic acid) 1 |
| chr16 | 34016457 | 34017318 | 862 | * | 8,346681 | -4,679386882 | 1,793897 | -2,6085 | 0,009094 | 0,867904199 | 1,449706 | Intron (ENSMUST0000( | 16 | 33833502 | 33983649 | 150148 | 2 | 545156 | ENSMUST00000114963.8 | -32808 | ENSMUSG00000061751 | Kalrn | kalirin, RhoGEF kinase |
| chr5 | 22340215 | 22341172 | 958 | * | 10,29691 | -4,67339464 | 1,66753 | -2,80259 | 0,005069 | 0,768389035 | 1,425929 | Promoter (<=1kb) | 5 | 22265311 | 22339667 | 74357 | 2 | 19699 | ENSMUST00000162876.2 | -548 | ENSMUSG00000042453 | Reln | reelin |
| chr9 | 27224648 | 27225730 | 1083 | * | 9,548298 | -4,673219124 | 1,676646 | -2,78724 | 0,005316 | 0,812305378 | 1,173325 | Intron (ENSMUST0000( | 9 | 27238236 | 27246059 | 7824 | 1 | 235086 | ENSMUST00000214187.2 | -12506 | ENSMUSG00000034275 | Igsf9b | immunoglobulin superfamily, member 9B |
| chr12 | 1E+08 | 1E+08 | 1007 | * | 7,15911 | -4,672498941 | 1,762242 | -2,65145 | 0,008015 | 1 | 0 | Promoter (<=1kb) | 12 | 1E+08 | 1E+08 | 3036 | 2 | 1,03E+08 | ENSMUST00000245285.1 | 745 | | Gm36263 | predicted gene, 36263 |
| chr18 | 48494174 | 48494953 | 780 | * | 9,38093 | -4,671569027 | 1,775574 | -2,63102 | 0,008513 | 0,889260882 | 1,211315 | Distal Intergenic | 18 | 48178402 | 48181479 | 3078 | 1 | 433182 | ENSMUST00000076155.6 | 315772 | ENSMUSG00000059040 | Eno1b | enolase 1B, retrotransposed |
| chr13 | 4723970 | 4724961 | 992 | * | 8,578913 | -4,671557326 | 1,709737 | -2,73233 | 0,006289 | 0,80696549 | 1,425929 | Intron (ENSMUST0000( | 13 | 4693906 | 4799063 | 105158 | 1 | 1,05E+08 | ENSMUST00000221900.2 | 30064 | | Gm40658 | predicted gene, 40658 |
| chr4 | 26376197 | 26376907 | 711 | * | 6,976596 | -4,666497982 | 1,76578 | -2,64274 | 0,008224 | 1 | 0 | Distal Intergenic | 4 | 26340751 | 26346891 | 6141 | 2 | 242362 | ENSMUST00000153813.2 | -29306 | ENSMUSG00000040520 | Manea | mannosidase, endo-alpha |
| chr1 | 48243845 | 48244910 | 1066 | * | 7,509717 | -4,662609793 | 1,768251 | -2,63685 | 0,008368 | 1 | 0 | Distal Intergenic | 1 | 47506180 | 47506284 | 105 | 2 | 1,02E+08 | ENSMUST00000183639.3 | -737561 | ENSMUSG00000098876 | Mir6350 | microRNA 6350 |
| chr14 | 1,17E+08 | 1,17E+08 | 1713 | * | 9,69503 | -4,661726514 | 1,499717 | -3,1084 | 0,001881 | 0,597812173 | 1,425929 | Intron (ENSMUST0000( | 14 | 1,17E+08 | 1,18E+08 | 701884 | 1 | 23888 | ENSMUST00000123239.2 | -466354 | ENSMUSG00000058571 | Gpc6 | glypican 6 |
| chr15 | 27382571 | 27383439 | 869 | * | 7,355465 | -4,660918383 | 1,758161 | -2,65102 | 0,008025 | 0,857326444 | 1,360622 | Distal Intergenic | 15 | 27466763 | 27594995 | 128233 | 1 | 11732 | ENSMUST00000022875.7 | -83324 | ENSMUSG00000022265 | Ank | progressive ankylosis |
| chr7 | 1,34E+08 | 1,34E+08 | 772 | * | 7,863604 | -4,657907822 | 1,757587 | -2,65017 | 0,008045 | 1 | 0 | Intron (ENSMUST0000( | 7 | 1,34E+08 | 1,34E+08 | 5729 | 1 | 330662 | ENSMUST00000210121.2 | -24346 | ENSMUSG00000058325 | Dock1 | dedicator of cytokinesis 1 |
| chr2 | 1,03E+08 | 1,03E+08 | 1070 | * | 8,873399 | -4,656665845 | 1,799852 | -2,58725 | 0,009675 | 0,908065779 | 1,211315 | Distal Intergenic | 2 | 1,03E+08 | 1,03E+08 | 39846 | 2 | 13661 | ENSMUST00000090475.10 | -6984 | ENSMUSG00000012350 | Ehf | ets homologous factor |
| chr1 | 1,92E+08 | 1,92E+08 | 842 | * | 8,579281 | -4,656046199 | 1,703325 | -2,73351 | 0,006266 | 1 | 0 | Intron (ENSMUST0000( | 1 | 1,92E+08 | 1,92E+08 | 319386 | 1 | 16510 | ENSMUST00000110844.3 | 49739 | ENSMUSG00000058248 | Kcnh1 | potassium voltage-gated channel, subfamily H (eag-related), member 1 |
| chr17 | 7745218 | 7746828 | 1611 | * | 15,37887 | -4,650791971 | 1,601091 | -2,90476 | 0,003675 | 0,730689299 | 1,2763 | Distal Intergenic | 17 | 7795022 | 7797221 | 2200 | 1 | 1,03E+08 | ENSMUST00000232610.2 | -48194 | ENSMUSG00000116585 | Gm33086 | predicted gene, 33086 |
| chr1 | 1,29E+08 | 1,29E+08 | 807 | * | 6,978562 | -4,650769853 | 1,765382 | -2,63443 | 0,008428 | 0,864595549 | 1,360622 | Distal Intergenic | 1 | 1,29E+08 | 1,29E+08 | 111 | 2 | 1,15E+08 | ENSMUST00000082803.3 | 119321 | ENSMUSG00000064737 | Gm23056 | predicted gene, 23056 |
| chr8 | 1,1E+08 | 1,1E+08 | 555 | * | 8,478351 | -4,64618563 | 1,741654 | -2,66769 | 0,007638 | 0,851512567 | 1,360622 | Distal Intergenic | 8 | 1,1E+08 | 1,1E+08 | 178418 | 1 | 628686 | ENSMUST00000180524.8 | -41779 | ENSMUSG00000097393 | D030068K23Rik | RIKEN cDNA D030068K23 gene |
| chr4 | 63804587 | 63805193 | 607 | * | 6,55347 | -4,645182356 | 1,785881 | -2,60106 | 0,009294 | 1 | 0 | Distal Intergenic | 4 | 63749545 | 63779584 | 30040 | 2 | 21949 | ENSMUST00000030047.3 | -25003 | ENSMUSG00000028362 | Tnfsf8 | tumor necrosis factor (ligand) superfamily, member 8 |
| chr5 | 45165014 | 45165765 | 752 | * | 7,916961 | -4,63985762 | 1,752288 | -2,64789 | 0,0081 | 1 | 0 | Distal Intergenic | 5 | 45123925 | 45125209 | 1285 | 1 | 75783 | ENSMUST00000079389.6 | 41089 | ENSMUSG00000062496 | 4930431F12Rik | RIKEN cDNA 4930431F12 gene |
| chr11 | 21131947 | 21133077 | 1131 | * | 7,667313 | -4,639482787 | 1,756559 | -2,64123 | 0,00826 | 1 | 0 | Distal Intergenic | 11 | 21085469 | 21100323 | 14855 | 1 | 67245 | ENSMUST00000101477.2 | 46478 | ENSMUSG00000020134 | Peli1 | pellino 1 |
| chr7 | 45815116 | 45816957 | 1842 | * | 18,13001 | -4,638897598 | 1,616541 | -2,86964 | 0,004109 | 0,644704154 | 2,423594 | Exon (ENSMUST00000( | 7 | 45824835 | 45829410 | 4576 | 2 | 20927 | ENSMUST00000210511.2 | 12453 | ENSMUSG00000040136 | Abcc8 | ATP-binding cassette, sub-family C (CFTR/MRP), member 8 |
| chr1 | 11136632 | 11137738 | 1107 | * | 8,062448 | -4,638496703 | 1,715202 | -2,70434 | 0,006844 | 0,830950621 | 1,360622 | Intron (ENSMUST0000( | 1 | 11168754 | 11220295 | 51542 | 1 | 109294 | ENSMUST00000189822.2 | -31016 | ENSMUSG00000048960 | Prex2 | phosphatidylinositol-3,4,5-trisphosphate-dependent Rac exchange factor 2 |
| chr7 | 78753617 | 78754831 | 1215 | * | 7,984041 | -4,63776836 | 1,735392 | -2,67246 | 0,00753 | 0,848821558 | 1,360622 | Intron (ENSMUST0000( | 7 | 78749499 | 78764375 | 14877 | 1 | 11595 | ENSMUST00000206779.2 | 4118 | ENSMUSG00000030607 | Acan | aggrecan |
| chrX | 1,37E+08 | 1,37E+08 | 1193 | * | 8,609163 | -4,637522693 | 1,727157 | -2,68506 | 0,007252 | 0,829907213 | 1,449706 | Intron (ENSMUST0000( | 20 | 1,37E+08 | 1,37E+08 | 1416 | 2 | 67944 | ENSMUST00000096314.3 | 129932 | ENSMUSG00000071686 | Tex13a | testis expressed 13A |
| chr13 | 36181414 | 36182385 | 972 | * | 7,445257 | -4,637345199 | 1,781414 | -2,60318 | 0,009236 | 1 | 0 | Intron (ENSMUST0000( | 13 | 36174844 | 36175503 | 660 | 2 | 1E+08 | ENSMUST00000224867.2 | -5911 | | Tmed10-ps | transmembrane p24 trafficking protein 10, pseudogene |
| chr2 | 1,39E+08 | 1,39E+08 | 727 | * | 7,980549 | -4,633985605 | 1,723096 | -2,68934 | 0,007159 | 0,838625535 | 1,360622 | Intron (ENSMUST0000( | 2 | 1,39E+08 | 1,39E+08 | 143144 | 1 | 228677 | ENSMUST00000110083.8 | -97503 | ENSMUSG00000039092 | Sptlc3 | serine palmitoyltransferase, long chain base subunit 3 |
| chr1 | 14730908 | 14731796 | 889 | * | 7,989447 | -4,630820581 | 1,75656 | -2,6363 | 0,008382 | 1 | 0 | Distal Intergenic | 1 | 14823161 | 14847155 | 23995 | 1 | 1,05E+08 | ENSMUST00000187401.2 | -91365 | | Gm9947 | predicted gene 9947 |
| chr4 | 21380469 | 21381128 | 660 | * | 11,74925 | -4,628715085 | 1,703495 | -2,71719 | 0,006584 | 0,891097601 | 0,918255 | Distal Intergenic | 4 | 21679747 | 21685781 | 6035 | 2 | 230025 | ENSMUST00000149516.8 | 304653 | ENSMUSG00000040478 | Prdm13 | PR domain containing 13 |
| chr7 | 3123043 | 3123993 | 951 | * | 9,059733 | -4,624581617 | 1,699612 | -2,72096 | 0,006509 | 0,813839948 | 1,425929 | Distal Intergenic | 7 | 3178146 | 3194832 | 16687 | 1 | 73949 | ENSMUST00000203447.2 | -54153 | ENSMUSG00000107956 | Speer9-ps1 | spermatogenesis associated glutamate (E)-rich protein 9, pseudogene 1 |
| chr12 | 67666496 | 67667559 | 1064 | * | 8,486495 | -4,623588986 | 1,715319 | -2,69547 | 0,007029 | 0,748382289 | 2,237403 | Distal Intergenic | 12 | 67659070 | 67659255 | 186 | 1 | 1,15E+08 | ENSMUST00000122499.3 | 7426 | ENSMUSG00000084448 | Gm26256 | predicted gene, 26256 |
| chr14 | 56276625 | 56277556 | 932 | * | 7,71546 | -4,622061242 | 1,737064 | -2,66085 | 0,007794 | 1 | 0 | Promoter (1-2kb) | 14 | 56279498 | 56282099 | 2602 | 1 | 17225 | ENSMUST00000015576.6 | -1942 | ENSMUSG00000022226 | Mcpt2 | mast cell protease 2 |
| chr6 | 1,45E+08 | 1,45E+08 | 1062 | * | 14,07614 | -4,620154598 | 1,599088 | -2,88924 | 0,003862 | 0,740176831 | 1,2763 | Intron (ENSMUST0000( | 6 | 1,45E+08 | 1,45E+08 | 741 | 1 | 381827 | ENSMUST00000148424.2 | -5674 | | 1700073E17Rik | ribosomal protein L7 pseudogene |
| chr10 | 1,1E+08 | 1,1E+08 | 1038 | * | 7,578531 | -4,616244226 | 1,743998 | -2,64693 | 0,008123 | 1 | 0 | Intron (ENSMUST0000( | 10 | 1,1E+08 | 1,1E+08 | 23391 | 2 | 260315 | ENSMUST00000161798.2 | -7475 | ENSMUSG00000020181 | Nav3 | neuron navigator 3 |
| chr4 | 59410805 | 59411528 | 724 | * | 6,910532 | -4,615901749 | 1,790876 | -2,57745 | 0,009953 | 1 | 0 | Exon (ENSMUST00000( | 4 | 59314683 | 59438633 | 123951 | 2 | 634731 | ENSMUST00000040166.14 | 27105 | ENSMUSG00000038578 | Susd1 | sushi domain containing 1 |
| chr11 | 85640194 | 85641471 | 1278 | * | 9,44585 | -4,612022778 | 1,724762 | -2,67401 | 0,007495 | 0,836351745 | 1,449706 | Intron (ENSMUST0000( | 11 | 85651442 | 85651528 | 87 | 2 | 1,01E+08 | ENSMUST00000175537.3 | 10057 | ENSMUSG00000093278 | Mir5110 | microRNA 5110 |
| chr10 | 22879911 | 22880685 | 775 | * | 8,699856 | -4,611722437 | 1,711664 | -2,69429 | 0,007054 | 0,826716028 | 1,425929 | Distal Intergenic | 10 | 23042763 | 23042869 | 107 | 2 | 1,15E+08 | ENSMUST00000239688.1 | 162184 | ENSMUSG00000119293 | Gm24655 | predicted gene, 24655 |
| chr11 | 60672415 | 60673477 | 1063 | * | 7,90598 | -4,607020391 | 1,717408 | -2,68254 | 0,007306 | 1 | 0 | Intron (ENSMUST0000( | 11 | 60668351 | 60679113 | 10763 | 1 | 237782 | ENSMUST00000056907.7 | 4064 | ENSMUSG00000049323 | Smcr8 | Smith-Magenis syndrome chromosome region, candidate 8 homolog (human) |
| chr4 | 92946865 | 92949216 | 2352 | * | 16,74185 | -4,604484414 | 1,318063 | -3,49337 | 0,000477 | 0,272208984 | 2,663619 | Distal Intergenic | 4 | 92754712 | 92754819 | 108 | 2 | 1,15E+08 | ENSMUST00000158289.3 | -192046 | ENSMUSG00000088914 | Gm22605 | predicted gene, 22605 |
| chr14 | 1,15E+08 | 1,15E+08 | 1640 | * | 7,403433 | -4,599865549 | 1,782865 | -2,58004 | 0,009879 | 1 | 0 | Distal Intergenic | 14 | 1,15E+08 | 1,15E+08 | 84 | 1 | 723905 | ENSMUST00000083574.3 | -7027 | ENSMUSG00000065508 | Mir17 | microRNA 17 |
| chr2 | 1,14E+08 | 1,14E+08 | 667 | * | 8,010138 | -4,596363902 | 1,712096 | -2,68464 | 0,007261 | 1 | 0 | Exon (ENSMUST00001 | 2 | 1,14E+08 | 1,14E+08 | 17498 | 2 | 75166 | ENSMUST00000123377.2 | 9103 | ENSMUSG00000084798 | 4930533B01Rik | RIKEN cDNA 4930533B01 gene |
| chr3 | 76486340 | 76487214 | 875 | * | 6,893357 | -4,590570983 | 1,775043 | -2,58618 | 0,009705 | 1 | 0 | Intron (ENSMUST0000( | 3 | 76500857 | 76614856 | 114000 | 1 | 213262 | ENSMUST00000162471.2 | -13643 | ENSMUSG00000034098 | Fstl5 | follistatin-like 5 |
| chr4 | 48111677 | 48113199 | 1523 | * | 9,959385 | -4,588356902 | 1,726022 | -2,65834 | 0,007853 | 0,881672947 | 1,173325 | Distal Intergenic | 4 | 48124915 | 48185911 | 60997 | 1 | 67727 | ENSMUST00000107721.8 | -11716 | ENSMUSG00000061455 | Stx17 | syntaxin 17 |
| chr2 | 91398001 | 91398979 | 979 | * | 6,930194 | -4,583785232 | 1,758851 | -2,60613 | 0,009157 | 1 | 0 | Exon (ENSMUST00001 | 2 | 91385585 | 91395516 | 9932 | 1 | 75786 | ENSMUST00000133633.2 | 12416 | ENSMUSG00000040549 | Ckap5 | cytoskeleton associated protein 5 |
| chr11 | 1,08E+08 | 1,08E+08 | 920 | * | 11,74677 | -4,58269068 | 1,6744 | -2,73692 | 0,006202 | 0,824134241 | 1,2763 | Intron (ENSMUST0000( | 11 | 1,08E+08 | 1,08E+08 | 6883 | 2 | 54377 | ENSMUST00000134076.2 | -10830 | ENSMUSG00000020723 | Cacng4 | calcium channel, voltage-dependent, gamma subunit 4 |
| chr8 | 1,13E+08 | 1,13E+08 | 497 | * | 6,658169 | -4,581661523 | 1,775919 | -2,57988 | 0,009883 | 1 | 0 | 3' UTR | 8 | 1,13E+08 | 1,13E+08 | 1056 | 2 | 627828 | ENSMUST00000071732.7 | -3554 | ENSMUSG00000069014 | Gm6793 | heterogeneous nuclear ribonucleoprotein A3 pseudogene |
| chr16 | 34234718 | 34235722 | 1005 | * | 8,100271 | -4,579335438 | 1,738379 | -2,63426 | 0,008432 | 1 | 0 | Intron (ENSMUST0000( | 16 | 33833507 | 34212479 | 378973 | 2 | 545156 | ENSMUST00000142817.8 | -22239 | ENSMUSG00000061751 | Kalrn | kalirin, RhoGEF kinase |
| chr1 | 87999903 | 88000713 | 811 | * | 8,221894 | -4,574754815 | 1,704268 | -2,68429 | 0,007268 | 0,832984033 | 1,425929 | Promoter (1-2kb) | 1 | 87998522 | 88146719 | 148198 | 1 | 394434 | ENSMUST00000073772.5 | 1381 | ENSMUSG00000090175 | Ugt1a9 | UDP glucuronosyltransferase 1 family, polypeptide A9 |
| chr18 | 71430230 | 71431565 | 1336 | * | 7,170077 | -4,571969476 | 1,750895 | -2,61122 | 0,009022 | 1 | 0 | Intron (ENSMUST0000( | 18 | 70750844 | 70758848 | 8005 | 1 | 17191 | ENSMUST00000128555.2 | 679386 | ENSMUSG00000024513 | Mbd2 | methyl-CpG binding domain protein 2 |
| chr11 | 43001277 | 43002038 | 762 | * | 7,543328 | -4,571645013 | 1,742993 | -2,62287 | 0,008719 | 1 | 0 | Intron (ENSMUST0000( | 11 | 43040704 | 43050302 | 9599 | 1 | 319767 | ENSMUST00000148911.2 | -38666 | ENSMUSG00000055415 | Atp10b | ATPase, class V, type 10B |
| chr2 | 1,74E+08 | 1,74E+08 | 911 | * | 11,75176 | -4,56062765 | 1,671113 | -2,7291 | 0,006351 | 0,887238309 | 0,918255 | Distal Intergenic | 2 | 1,74E+08 | 1,74E+08 | 2767 | 1 | 1,03E+08 | ENSMUST00000241747.1 | -5917 | | Gm30143 | predicted gene, 30143 |
| chr11 | 41492573 | 41493859 | 1287 | * | 9,798939 | -4,555333334 | 1,671449 | -2,72538 | 0,006423 | 0,839085757 | 1,211315 | Distal Intergenic | 11 | 41601169 | 41859514 | 258346 | 1 | 1,03E+08 | ENSMUST00000155032.8 | -107310 | ENSMUSG00000087306 | A230004M16Rik | RIKEN cDNA A230004M16 gene |
| chr2 | 1,05E+08 | 1,05E+08 | 1449 | * | 14,04786 | -4,548243325 | 1,328967 | -3,42239 | 0,000621 | 0,44905823 | 1,075735 | Exon (ENSMUST00001 | 2 | 1,05E+08 | 1,05E+08 | 13280 | 2 | 329480 | ENSMUST00000135153.8 | 12751 | ENSMUSG00000074987 | Wt1os | WT1 transcription factor, opposite strand |
| chr1 | 70387936 | 70389202 | 1267 | * | 14,68923 | -4,547957242 | 1,263125 | -3,60056 | 0,000318 | 0,300106667 | 1,41696 | Intron (ENSMUST0000( | 1 | 70635974 | 70636080 | 107 | 1 | 1,15E+08 | ENSMUST00000102000.3 | -246772 | ENSMUSG00000075952 | Gm23422 | predicted gene, 23422 |
| chr14 | 27617641 | 27618989 | 1349 | * | 9,868604 | -4,540951447 | 1,756979 | -2,58452 | 0,009751 | 0,924754734 | 1,086118 | Intron (ENSMUST0000( | 14 | 27598371 | 27601820 | 3450 | 1 | 1,03E+08 | ENSMUST00000210524.2 | 19270 | | Gm34996 | predicted gene, 34996 |
| chr9 | 58198715 | 58199679 | 965 | * | 8,558737 | -4,537460346 | 1,752592 | -2,589 | 0,009626 | 0,890233764 | 1,360622 | Intron (ENSMUST0000( | 9 | 58195021 | 58220469 | 25449 | 2 | 16949 | ENSMUST00000061799.10 | 20790 | ENSMUSG00000032334 | Loxl1 | lysyl oxidase-like 1 |
| chr3 | 41001997 | 41003047 | 1051 | * | 9,968433 | -4,535504483 | 1,710131 | -2,65214 | 0,007998 | 0,879010025 | 1,211315 | Distal Intergenic | 3 | 40991168 | 40993048 | 1881 | 1 | 214048 | ENSMUST00000199797.2 | 10829 | ENSMUSG00000025762 | Larp1b | La ribonucleoprotein domain family, member 1B |
| chr5 | 77979338 | 77980467 | 1130 | * | 10,67713 | -4,534762158 | 1,639125 | -2,76657 | 0,005665 | 0,823736512 | 1,173325 | Distal Intergenic | 5 | 77613335 | 77616698 | 3364 | 1 | 69370 | ENSMUST00000201654.2 | 366003 | ENSMUSG00000107232 | 1700017L05Rik | RIKEN cDNA 1700017L05 gene |
| chr1 | 1,31E+08 | 1,31E+08 | 1282 | * | 10,43957 | -4,534327901 | 1,645327 | -2,75588 | 0,005853 | 0,905781613 | 0,744375 | Promoter (1-2kb) | 1 | 1,31E+08 | 1,31E+08 | 67692 | 2 | 14270 | ENSMUST00000187273.3 | -1135 | ENSMUSG00000026425 | Srgap2 | SLIT-ROBO Rho GTPase activating protein 2 |
| chr1 | 34437555 | 34438378 | 824 | * | 9,197172 | -4,526856383 | 1,740229 | -2,6013 | 0,009287 | 0,902187657 | 1,211315 | Distal Intergenic | 1 | 34454844 | 34457067 | 2224 | 2 | 1E+08 | ENSMUST00000188254.3 | 18689 | | Gm3052 | predicted gene 3052 |
| chr8 | 53539858 | 53540451 | 594 | * | 7,414891 | -4,525895204 | 1,737618 | -2,60465 | 0,009197 | 1 | 0 | Distal Intergenic | 8 | 53889135 | 53933533 | 44399 | 1 | 1,05E+08 | ENSMUST00000210964.2 | -348684 | | Gm39180 | predicted gene, 39180 |
| chr13 | 46514593 | 46515626 | 1034 | * | 8,214885 | -4,524885911 | 1,753151 | -2,581 | 0,009851 | 0,884485418 | 1,449706 | Distal Intergenic | 13 | 46571910 | 46584571 | 12662 | 1 | 666794 | ENSMUST00000037923.5 | -56284 | ENSMUSG00000038132 | Rbm24 | RNA binding motif protein 24 |
| chr10 | 16283697 | 16284379 | 683 | * | 7,961614 | -4,524200736 | 1,475024 | -3,06721 | 0,002161 | 1 | 0 | Distal Intergenic | 10 | 16827043 | 16838928 | 11886 | 2 | 1,01E+08 | ENSMUST00000208191.2 | 554549 | ENSMUSG00000101969 | Gm20125 | predicted gene, 20125 |
| chr18 | 14742711 | 14744117 | 1407 | * | 11,37257 | -4,52090974 | 1,676632 | -2,69642 | 0,007009 | 0,934396727 | 0,744375 | Distal Intergenic | 18 | 14766599 | 14770463 | 3865 | 2 | 268996 | ENSMUST00000238844.2 | 26346 | ENSMUSG00000037013 | Ss18 | SS18, nBAF chromatin remodeling complex subunit |
| chr8 | 11373158 | 11374989 | 1832 | * | 17,39212 | -4,51649514 | 1,584244 | -2,85088 | 0,00436 | 0,658336602 | 2,386008 | Intron (ENSMUST0000( | 8 | 11248423 | 11362826 | 114404 | 2 | 12826 | ENSMUST00000033898.10 | -10332 | ENSMUSG00000031502 | Col4a1 | collagen, type IV, alpha 1 |
| chr2 | 1,49E+08 | 1,49E+08 | 1114 | * | 9,991102 | -4,509635741 | 1,391979 | -3,23973 | 0,001196 | 0,563032042 | 1,086118 | Distal Intergenic | 2 | 1,49E+08 | 1,49E+08 | 12909 | 2 | 1,15E+08 | ENSMUST00000109942.3 | -61261 | | Gm1330 | predicted gene 1330 |
| chr1 | 1,69E+08 | 1,69E+08 | 1318 | * | 9,830094 | -4,50215929 | 1,693413 | -2,65863 | 0,007846 | 0,875818484 | 1,211315 | Distal Intergenic | 1 | 1,69E+08 | 1,69E+08 | 116 | 2 | 1,02E+08 | ENSMUST00000184979.3 | 104298 | ENSMUSG00000098829 | Mir6354 | microRNA 6354 |
| chr16 | 9941649 | 9942394 | 746 | * | 7,623592 | -4,493160754 | 1,730571 | -2,59635 | 0,009422 | 1 | 0 | Distal Intergenic | 16 | 9988090 | 9992494 | 4405 | 1 | 68172 | ENSMUST00000121292.8 | -45696 | ENSMUSG00000039209 | Rpl39l | ribosomal protein L39-like |
| chr16 | 56883943 | 56884734 | 792 | * | 9,495302 | -4,483337277 | 1,364686 | -3,28525 | 0,001019 | 0,410415129 | 2,237403 | Intron (ENSMUST0000( | 16 | 56857330 | 56891715 | 34386 | 2 | 69457 | ENSMUST00000067173.8 | 6981 | ENSMUSG00000046748 | Tmem45a2 | transmembrane protein 45A2 |
| chr2 | 42445021 | 42446066 | 1046 | * | 8,415271 | -4,4823486 | 1,723659 | -2,60049 | 0,009309 | 0,794811783 | 2,237403 | Intron (ENSMUST0000( | 2 | 41944131 | 42542938 | 598808 | 2 | 94217 | ENSMUST00000204204.2 | 96872 | ENSMUSG00000049252 | Lrp1b | low density lipoprotein-related protein 1B |
| chrX | 1,06E+08 | 1,06E+08 | 1168 | * | 9,161952 | -4,479069124 | 1,688687 | -2,6524 | 0,007992 | 0,769692849 | 2,237403 | Distal Intergenic | 20 | 1,06E+08 | 1,06E+08 | 6278 | 2 | 16431 | ENSMUST00000033591.6 | 55575 | ENSMUSG00000031239 | Itm2a | integral membrane protein 2A |
| chr15 | 4998012 | 4998867 | 856 | * | 8,016784 | -4,470792206 | 1,724147 | -2,59305 | 0,009513 | 1 | 0 | Distal Intergenic | 15 | 4977964 | 4991554 | 13591 | 1 | 223825 | ENSMUST00000226849.2 | 20048 | ENSMUSG00000022155 | Mroh2b | maestro heat-like repeat family member 2B |
| chr15 | 55746779 | 55747352 | 574 | * | 10,25085 | -4,447117083 | 1,71209 | -2,59748 | 0,009391 | 0,875818484 | 1,449706 | Intron (ENSMUST0000( | 15 | 55655334 | 55769706 | 114373 | 2 | 20649 | ENSMUST00000140574.2 | 22354 | ENSMUSG00000060429 | Sntb1 | syntrophin, basic 1 |
| chr11 | 21734475 | 21735637 | 1163 | * | 8,906487 | -4,435318406 | 1,710566 | -2,5929 | 0,009517 | 0,88870234 | 1,360622 | Intron (ENSMUST0000( | 11 | 21763399 | 21767341 | 3943 | 1 | 216560 | ENSMUST00000128794.2 | -27762 | ENSMUSG00000020319 | Wdpcp | WD repeat containing planar cell polarity effector |
| chr14 | 29186290 | 29187714 | 1425 | * | 11,6551 | -4,433377 | 1,334669 | -3,32171 | 0,000895 | 0,582111135 | 0,738441 | Intron (ENSMUST0000( | 14 | 28786153 | 29082791 | 296639 | 2 | 12294 | ENSMUST00000225863.2 | -103499 | ENSMUSG00000021991 | Cacna2d3 | calcium channel, voltage-dependent, alpha2/delta subunit 3 |
| chr5 | 99860544 | 99861828 | 1285 | * | 8,9454 | -4,429416485 | 1,709586 | -2,59093 | 0,009572 | 0,879010025 | 1,449706 | Intron (ENSMUST0000( | 5 | 99365285 | 99876671 | 511387 | 2 | 320292 | ENSMUST00000209346.2 | 14843 | ENSMUSG00000089809 | Rasgef1b | RasGEF domain family, member 1B |
| chr5 | 1,1E+08 | 1,1E+08 | 1602 | * | 11,30018 | -4,401722334 | 1,655745 | -2,65845 | 0,00785 | 0,918200407 | 0,918255 | Distal Intergenic | 5 | 1,1E+08 | 1,1E+08 | 16497 | 2 | 77200 | ENSMUST00000092720.4 | 30871 | ENSMUSG00000072763 | 5430403G16Rik | RIKEN cDNA 5430403G16 gene |
| chr16 | 6889015 | 6889984 | 970 | * | 8,444254 | -4,398718371 | 1,707306 | -2,57641 | 0,009983 | 0,88871498 | 1,425929 | Promoter (1-2kb) | 16 | 6887710 | 7227677 | 339968 | 1 | 268859 | ENSMUST00000229741.2 | 1305 | ENSMUSG00000008658 | Rbfox1 | RNA binding protein, fox-1 homolog (C. elegans) 1 |

| | | | | | | | | | | | | | | | | | | | | | | | |
|---|---|---|---|---|---|---|---|---|---|---|---|---|---|---|---|---|---|---|---|---|---|---|---|
| chr1 | 1,95E+08 | 1,95E+08 | 995 | * | 9,726117 | -4,396267509 | 1,661309 | -2,64627 | 0,008139 | 0,851512567 | 1,449706 | Distal Intergenic | 1 | 1,95E+08 | 1,95E+08 | 39906 | 2 | 12902 | ENSMUST00000195120.6 | -188316 | ENSMUSG00000026616 | Cr2 | complement receptor 2 |
| chr5 | 26778367 | 26779301 | 935 | * | 8,682409 | -4,388893937 | 1,699993 | -2,58171 | 0,009831 | 0,805916009 | 2,237403 | Distal Intergenic | 5 | 26665070 | 26698312 | 33243 | 2 | 75942 | ENSMUST00000197091.5 | -80055 | ENSMUSG00000029044 | 4930584F24Rik | RIKEN cDNA 4930584F24 gene |
| chr16 | 52233595 | 52234493 | 899 | * | 8,259755 | -4,383820052 | 1,468604 | -2,98503 | 0,002836 | 0,680476059 | 1,360622 | Intron (ENSMUST00000 | 16 | 52071830 | 52272828 | 200999 | 2 | 11658 | ENSMUST00000170035.8 | 38335 | ENSMUSG00000022636 | Alcam | activated leukocyte cell adhesion molecule |
| chr10 | 74120259 | 74121545 | 1287 | * | 10,33276 | -4,383636603 | 1,66559 | -2,63188 | 0,008491 | 0,856202617 | 1,449706 | Intron (ENSMUST00000 | 10 | 74138983 | 74166882 | 27900 | 1 | 11994 | ENSMUST00000147455.2 | -17438 | ENSMUSG00000052613 | Pcdh15 | protocadherin 15 |
| chr13 | 1,01E+08 | 1,01E+08 | 1339 | * | 10,65049 | -4,377266626 | 1,654761 | -2,64526 | 0,008163 | 0,888042352 | 1,173325 | Intron (ENSMUST00000 | 13 | 1,01E+08 | 1,01E+08 | 39116 | 1 | 71174 | ENSMUST00000209771.2 | 102364 | ENSMUSG00000100733 | 4933416M06Rik | RIKEN cDNA 4933416M06 gene |
| chr3 | 1,08E+08 | 1,08E+08 | 1034 | * | 8,0123 | -4,371256428 | 1,491389 | -2,931 | 0,003379 | 0,707651812 | 1,360622 | Distal Intergenic | 3 | 1,08E+08 | 1,08E+08 | 2200 | 2 | 12977 | ENSMUST00000156820.2 | 22423 | ENSMUSG00000014599 | Csf1 | colony stimulating factor 1 (macrophage) |
| chr6 | 84270426 | 84271328 | 903 | * | 14,72719 | -4,364386505 | 1,587473 | -2,74927 | 0,005973 | 0,818408612 | 1,2763 | Distal Intergenic | 6 | 84187466 | 84188023 | 558 | 1 | 26903 | ENSMUST00000128965.2 | 82960 | ENSMUSG00000033788 | Dysf | dysferlin |
| chr13 | 45044237 | 45045452 | 1216 | * | 10,15948 | -4,361135373 | 1,346883 | -3,23795 | 0,001204 | 0,546791007 | 1,173325 | Intron (ENSMUST00000 | 13 | 44994259 | 45049957 | 55699 | 1 | 16468 | ENSMUST00000173003.2 | 49978 | ENSMUSG00000038518 | Jarid2 | jumonji, AT rich interactive domain 2 |
| chr11 | 85251881 | 85253476 | 1596 | * | 12,46814 | -4,355889406 | 1,659173 | -2,62534 | 0,008656 | 0,884058067 | 1,2763 | Exon (ENSMUST000001 | 11 | 85244038 | 85716176 | 472139 | 1 | 192197 | ENSMUST00000092821.10 | 7843 | ENSMUSG00000059439 | Bcas3 | BCAS3 microtubule associated cell migration factor |
| chr16 | 54506657 | 54508211 | 1555 | * | 14,75005 | -4,352333364 | 1,336899 | -3,25554 | 0,001132 | 0,504335604 | 1,41696 | Distal Intergenic | 16 | 54792794 | 54792900 | 107 | 1 | 1,15E+08 | ENSMUST00000082615.3 | -284583 | ENSMUSG00000064549 | Gm22977 | predicted gene, 22977 |
| chr18 | 57249683 | 57251657 | 1975 | * | 13,50976 | -4,347125355 | 1,323354 | -3,28493 | 0,00102 | 0,534793902 | 1,075735 | Exon (ENSMUST000002 | 18 | 57266162 | 57430539 | 164378 | 1 | 70417 | ENSMUST00000075770.13 | -14505 | ENSMUSG00000024593 | Megf10 | multiple EGF-like-domains 10 |
| chr7 | 3518461 | 3519746 | 1286 | * | 11,24368 | -4,345302157 | 1,214528 | -3,57777 | 0,000347 | 0,322750682 | 1,2763 | Intron (ENSMUST00000 | 7 | 3499873 | 3523536 | 23664 | 1 | 1,03E+08 | ENSMUST00000203144.2 | 18588 | | Gm35037 | predicted gene, 35037 |
| chr12 | 31292295 | 31293346 | 1052 | * | 10,14476 | -4,34402658 | 1,667098 | -2,60574 | 0,009168 | 0,900637308 | 1,211315 | Promoter (1-2kb) | 12 | 31290769 | 31290978 | 210 | 1 | 1,03E+08 | ENSMUST00000219110.2 | 1526 | ENSMUSG00000112187 | Gm6993 | predicted gene 6993 |
| chr12 | 62315691 | 62316910 | 1220 | * | 8,677876 | -4,334802811 | 1,425935 | -3,03997 | 0,002366 | 0,640427178 | 1,449706 | Distal Intergenic | 12 | 62734608 | 62735031 | 424 | 1 | 73309 | ENSMUST00000172433.3 | -417698 | | Spanxn4 | SPANX family, member N4 |
| chr2 | 1,65E+08 | 1,65E+08 | 1221 | * | 12,2004 | -4,333933442 | 1,312519 | -3,302 | 0,00096 | 0,507398906 | 1,173325 | Intron (ENSMUST00000 | 2 | 1,65E+08 | 1,65E+08 | 12058 | 2 | 72211 | ENSMUST00000109304.2 | -34207 | ENSMUSG00000078935 | 1700025C18Rik | RIKEN cDNA 1700025C18 gene |
| chr3 | 56567501 | 56568434 | 934 | * | 8,130399 | -4,332812495 | 1,546772 | -2,8012 | 0,005091 | 0,769692849 | 1,425929 | Distal Intergenic | 3 | 56352703 | 56352833 | 131 | 2 | 1,15E+08 | ENSMUST00000104522.3 | -214668 | ENSMUSG00000119391 | Gm25727 | predicted gene, 25727 |
| chr17 | 55778299 | 55779278 | 980 | * | 9,037115 | -4,331693727 | 1,454162 | -2,97883 | 0,002894 | 0,671793638 | 1,449706 | Intron (ENSMUST00000 | 17 | 55753245 | 55789369 | 36125 | 1 | 240119 | ENSMUST00000153220.2 | 25054 | ENSMUSG00000024172 | St6gal2 | beta galactoside alpha 2,6 sialyltransferase 2 |
| chr16 | 35423686 | 35424221 | 536 | * | 8,012139 | -4,329503123 | 1,523663 | -2,84151 | 0,00449 | 1 | 0 | Intron (ENSMUST00000 | 16 | 35478465 | 35478857 | 393 | 1 | 20357 | ENSMUST00000139805.2 | -54244 | ENSMUSG00000052133 | Sema5b | sema domain, seven thrombospondin repeats (type 1 and type 1-like), transmembrane domain (TM) and short cytoplasmic domain, (semaphorin) 5B |
| chr2 | 88048252 | 88048992 | 741 | * | 6,867031 | -4,324560516 | 1,610359 | -2,68546 | 0,007243 | 1 | 0 | Distal Intergenic | 2 | 88054424 | 88055374 | 951 | 2 | 258525 | ENSMUST00000099831.2 | 6382 | ENSMUSG00000075133 | Olfr1170 | olfactory receptor 1170 |
| chr11 | 99404908 | 99406195 | 1288 | * | 11,84506 | -4,323570471 | 1,319444 | -3,27681 | 0,00105 | 0,613809578 | 0,738441 | 3' UTR | 11 | 99405450 | 99412084 | 6635 | 2 | 237934 | ENSMUST00000076948.2 | 5889 | ENSMUSG00000064165 | Krt39 | keratin 39 |
| chr3 | 74724581 | 74725276 | 696 | * | 10,5151 | -4,316650812 | 1,380297 | -3,12733 | 0,001764 | 0,697824004 | 0,738441 | Distal Intergenic | 3 | 74968668 | 75051010 | 82343 | 2 | 213234 | ENSMUST00000107776.8 | 325734 | ENSMUSG00000034151 | Zbbx | zinc finger, B-box domain containing |
| chr10 | 1,13E+08 | 1,13E+08 | 721 | * | 9,923138 | -4,315629029 | 1,501155 | -2,87487 | 0,004042 | 0,7299084 | 1,425929 | Distal Intergenic | 10 | 1,13E+08 | 1,13E+08 | 11133 | 2 | 1,05E+08 | ENSMUST00000218612.2 | 77703 | | Gm40765 | predicted gene, 40765 |
| chr11 | 80198601 | 80199203 | 603 | * | 6,400519 | -4,313570404 | 1,599306 | -2,69715 | 0,006994 | 1 | 0 | Intron (ENSMUST00000 | 11 | 80191738 | 80246781 | 55044 | 1 | 246104 | ENSMUST00000017836.8 | 6863 | ENSMUSG00000017692 | Rhbdl3 | rhomboid like 3 |
| chr4 | 73397295 | 73398667 | 1373 | * | 13,23733 | -4,297803152 | 1,188825 | -3,61517 | 0,0003 | 0,350503854 | 0,918255 | Distal Intergenic | 4 | 73319269 | 73323309 | 4041 | 2 | 433719 | ENSMUST00000084480.6 | -73986 | ENSMUSG00000066137 | Gm11487 | predicted gene 11487 |
| chr5 | 83620100 | 83621729 | 1630 | * | 13,17384 | -4,292370666 | 1,333521 | -3,21882 | 0,001287 | 0,649854434 | 0,738441 | Distal Intergenic | 5 | 83425992 | 83503042 | 77051 | 2 | 243078 | ENSMUST00000053543.11 | -117058 | ENSMUSG00000049537 | Tecrl | trans-2,3-enoyl-CoA reductase-like |
| chr8 | 44881619 | 44882661 | 1043 | * | 9,614515 | -4,288035569 | 1,411903 | -3,03706 | 0,002389 | 0,643106401 | 1,425929 | Distal Intergenic | 8 | 45388484 | 45505294 | 116811 | 1 | 14107 | ENSMUST00000191428.7 | -505823 | ENSMUSG00000070047 | Fat1 | FAT atypical cadherin 1 |
| chr7 | 9835753 | 9837250 | 1498 | * | 10,70873 | -4,285036963 | 1,342865 | -3,19097 | 0,001418 | 0,581323192 | 1,173325 | Promoter (2-3kb) | 7 | 9821125 | 9839668 | 18544 | 2 | 1E+08 | ENSMUST00000094863.6 | 2418 | ENSMUSG00000058685 | Vmn2r51 | vomeronasal 2, receptor 51 |
| chr1 | 15293211 | 15294666 | 1456 | * | 11,49025 | -4,277538527 | 1,36123 | -3,14241 | 0,001676 | 0,686589525 | 0,744375 | Distal Intergenic | 1 | 15357478 | 15782772 | 425295 | 1 | 98741 | ENSMUST00000175681.3 | -62812 | ENSMUSG00000092083 | Kcnb2 | potassium voltage gated channel, Shab-related subfamily, member 2 |
| chr18 | 46194237 | 46195160 | 924 | * | 9,747864 | -4,272421903 | 1,391917 | -3,06945 | 0,002145 | 0,624579927 | 1,425929 | Distal Intergenic | 18 | 45818260 | 46178327 | 360068 | 2 | 107003 | ENSMUST00000237850.2 | -15910 | | A330093E20Rik | RIKEN cDNA A330093E20 gene |
| chr13 | 97680962 | 97682127 | 1166 | * | 7,685786 | -4,27039829 | 1,540133 | -2,77275 | 0,005559 | 1 | 0 | Distal Intergenic | 13 | 97696405 | 97713926 | 17522 | 1 | 328314 | ENSMUST00000210897.3 | -14278 | ENSMUSG00000110235 | Gm5086 | predicted gene 5086 |
| chr2 | 89788846 | 89789894 | 1049 | * | 9,399001 | -4,262916455 | 1,386118 | -3,07544 | 0,002102 | 0,617500523 | 1,449706 | Distal Intergenic | 2 | 89773528 | 89774457 | 930 | 2 | 258338 | ENSMUST00000090695.3 | -14389 | ENSMUSG00000068806 | Olfr1259 | olfactory receptor 1259 |
| chr4 | 1,15E+08 | 1,15E+08 | 1072 | * | 9,161512 | -4,239783196 | 1,400017 | -3,02838 | 0,002459 | 0,644704154 | 1,449706 | Intron (ENSMUST00000 | 4 | 1,15E+08 | 1,15E+08 | 17796 | 2 | 66588 | ENSMUST00000129317.2 | 18055 | ENSMUSG00000028719 | Cmpk1 | cytidine monophosphate (UMP-CMP) kinase 1 |
| chr5 | 1,44E+08 | 1,44E+08 | 953 | * | 9,847532 | -4,235049117 | 1,395354 | -3,03511 | 0,002404 | 0,671156916 | 1,211315 | Distal Intergenic | 5 | 1,44E+08 | 1,44E+08 | 11577 | 1 | 53324 | ENSMUST00000071782.8 | -85415 | ENSMUSG00000059991 | Nptx2 | neuronal pentraxin 2 |
| chr3 | 1,32E+08 | 1,32E+08 | 856 | * | 10,45176 | -4,233386439 | 1,400215 | -3,02338 | 0,0025 | 0,72253719 | 0,918255 | Distal Intergenic | 3 | 1,32E+08 | 1,32E+08 | 15529 | 1 | 433653 | ENSMUST00000212804.2 | -111804 | ENSMUSG00000091721 | Gimd1 | GIMAP family P-loop NTPase domain containing 1 |
| chr18 | 58284833 | 58286022 | 1190 | * | 8,522441 | -4,225535447 | 1,313172 | -3,21781 | 0,001292 | 0,526973991 | 1,425929 | Intron (ENSMUST00000 | 18 | 58318633 | 58333777 | 15145 | 2 | 14119 | ENSMUST00000235779.2 | 47755 | ENSMUSG00000024598 | Fbn2 | fibrillin 2 |
| chrX | 92284474 | 92285903 | 1430 | * | 8,132849 | -4,223516844 | 1,48888 | -2,83671 | 0,004558 | 0,756018836 | 1,360622 | Distal Intergenic | 20 | 92295080 | 92295212 | 133 | 1 | 1,15E+08 | ENSMUST00000083332.3 | -9177 | ENSMUSG00000065266 | Gm23454 | predicted gene, 23454 |
| chr4 | 48809735 | 48810612 | 878 | * | 11,56032 | -4,218793203 | 1,356565 | -3,10991 | 0,001871 | 0,710166078 | 0,744375 | Distal Intergenic | 4 | 48898593 | 48963237 | 64645 | 1 | 1,03E+08 | ENSMUST00000131099.2 | -87981 | ENSMUSG00000084787 | Gm12436 | predicted gene 12436 |
| chr9 | 39194477 | 39195246 | 770 | * | 8,027698 | -4,212909879 | 1,596498 | -2,63884 | 0,008319 | 1 | 0 | Distal Intergenic | 9 | 39199220 | 39210777 | 11558 | 2 | 257924 | ENSMUST00000215471.2 | 15531 | ENSMUSG00000111190 | Olfr947-ps1 | olfactory receptor 947, pseudogene 1 |
| chr6 | 43522459 | 43523276 | 818 | * | 8,61765 | -4,206883032 | 1,523186 | -2,7619 | 0,005747 | 1 | 0 | Intron (ENSMUST00000 | 6 | 43323776 | 43453454 | 129679 | 2 | 29807 | ENSMUST00000149424.2 | -69005 | ENSMUSG00000029735 | Tpk1 | thiamine pyrophosphokinase |
| chr7 | 83149281 | 83150952 | 1672 | * | 13,19402 | -4,205513394 | 1,309877 | -3,21062 | 0,001325 | 0,566892297 | 1,173325 | Distal Intergenic | 7 | 83192078 | 83199464 | 7387 | 2 | 1E+08 | ENSMUST00000208877.2 | 48512 | | Gm10610 | predicted gene 10610 |
| chr5 | 1,51E+08 | 1,51E+08 | 1180 | * | 9,121861 | -4,202505104 | 1,528785 | -2,74892 | 0,005979 | 0,793703679 | 1,449706 | Intron (ENSMUST00000 | 5 | 1,51E+08 | 1,51E+08 | 43743 | 1 | 1,03E+08 | ENSMUST00000201101.2 | 9612 | | Gm36447 | predicted gene, 36447 |
| chr14 | 1,2E+08 | 1,2E+08 | 1409 | * | 9,349726 | -4,194392316 | 1,419621 | -2,95459 | 0,003131 | 0,71419937 | 1,211315 | Intron (ENSMUST00000 | 14 | 1,2E+08 | 1,2E+08 | 18597 | 1 | 74845 | ENSMUST00000228885.2 | 15948 | | 4930404K13Rik | RIKEN cDNA 4930404K13 gene |
| chr18 | 43769898 | 43770707 | 810 | * | 7,009111 | -4,193561664 | 1,619533 | -2,58936 | 0,009615 | 1 | 0 | Intron (ENSMUST00000 | 18 | 43723582 | 43820690 | 97109 | 2 | 76217 | ENSMUST00000237069.2 | 49983 | ENSMUSG00000024502 | Jakmip2 | janus kinase and microtubule interacting protein 2 |
| chr1 | 81580054 | 81580997 | 944 | * | 10,59693 | -4,18760583 | 1,378442 | -3,03793 | 0,002382 | 0,714956493 | 0,918255 | Distal Intergenic | 1 | 81055300 | 81319479 | 264180 | 1 | 241134 | ENSMUST00000123285.2 | 524754 | ENSMUSG00000054976 | Nyap2 | neuronal tyrosine-phophorylated phosphoinositide 3-kinase adaptor 2 |
| chr18 | 77419711 | 77420894 | 1184 | * | 14,5278 | -4,184506505 | 1,312788 | -3,1875 | 0,001435 | 0,47475665 | 2,141612 | Exon (ENSMUST00000 | 18 | 77395446 | 77470024 | 74579 | 1 | 240411 | ENSMUST00000035501.11 | 24265 | ENSMUSG00000032818 | Loxhd1 | lipoxygenase homology domains 1 |
| chr16 | 47135841 | 47137945 | 2105 | * | 16,05951 | -4,177122795 | 1,057382 | -3,95044 | 7,8E-05 | 0,107571107 | 2,141612 | Distal Intergenic | 16 | 47326682 | 47326788 | 107 | 2 | 1,15E+08 | ENSMUST00000157107.4 | 188843 | ENSMUSG00000119721 | Gm24651 | predicted gene, 24651 |
| chr11 | 71703818 | 71704801 | 984 | * | 8,131248 | -4,174318572 | 1,536782 | -2,71627 | 0,006602 | 0,816751204 | 1,425929 | Distal Intergenic | 11 | 71738902 | 71739032 | 131 | 2 | 1,15E+08 | ENSMUST00000104268.3 | 34231 | ENSMUSG00000077456 | Gm22927 | predicted gene, 22927 |
| chr17 | 27578863 | 27579979 | 1117 | * | 14,36233 | -4,17328013 | 1,299003 | -3,21268 | 0,001315 | 0,547497234 | 1,2763 | Promoter (1-2kb) | 17 | 27577071 | 27577129 | 59 | 1 | 1,02E+08 | ENSMUST00000200050.3 | 1792 | ENSMUSG00000106215 | Mir7218 | microRNA 7218 |
| chr12 | 52510763 | 52512596 | 1834 | * | 14,96348 | -4,173047679 | 1,59882 | -2,61008 | 0,009052 | 0,877892722 | 1,37683 | Distal Intergenic | 12 | 52522522 | 52527392 | 4871 | 2 | 1,03E+08 | ENSMUST00000219885.2 | 14796 | ENSMUSG00000111982 | Gm35135 | predicted gene, 35135 |
| chr12 | 82934687 | 82936749 | 2063 | * | 15,33251 | -4,168979177 | 1,242524 | -3,35525 | 0,000793 | 0,445547689 | 1,41696 | Intron (ENSMUST00000 | 12 | 82979294 | 82985929 | 6636 | 2 | 74283 | ENSMUST00000187123.2 | 49180 | ENSMUSG00000100890 | 1700085C21Rik | RIKEN cDNA 1700085C21 gene |
| chr13 | 83629842 | 83630819 | 978 | * | 9,240636 | -4,16817767 | 1,387955 | -3,00311 | 0,002672 | 0,684274806 | 1,211315 | Distal Intergenic | 13 | 83652153 | 83811462 | 159310 | 1 | 17260 | ENSMUST00000198199.5 | -21334 | ENSMUSG00000005583 | Mef2c | myocyte enhancer factor 2C |
| chr8 | 45767737 | 45768387 | 651 | * | 8,175587 | -4,168172931 | 1,506743 | -2,76635 | 0,005669 | 0,787701468 | 1,449706 | Intron (ENSMUST00000 | 8 | 45722708 | 45761503 | 38796 | 2 | 16621 | ENSMUST00000116473.5 | -6234 | ENSMUSG00000109764 | Klkb1 | kallikrein B, plasma 1 |
| chr16 | 18753598 | 18754855 | 1258 | * | 10,8081 | -4,1633817 | 1,360246 | -3,06076 | 0,002208 | 0,699276165 | 0,918255 | Promoter (1-2kb) | 16 | 18751763 | 18767914 | 16152 | 1 | 15260 | ENSMUST00000150799.3 | 1835 | ENSMUSG00000022702 | Hira | histone cell cycle regulator |
| chr7 | 1,42E+08 | 1,42E+08 | 1740 | * | 14,69362 | -4,154245818 | 1,319638 | -3,14802 | 0,001644 | 0,57879681 | 1,37683 | Distal Intergenic | 7 | 1,42E+08 | 1,42E+08 | 12186 | 2 | 320802 | ENSMUST00000131791.2 | 13729 | ENSMUSG00000045777 | Ifitm10 | interferon induced transmembrane protein 10 |
| chr7 | 67258059 | 67259331 | 1273 | * | 10,80008 | -4,151394212 | 1,392496 | -2,98126 | 0,002871 | 0,747644399 | 0,918255 | Promoter (1-2kb) | 7 | 67181494 | 67257010 | 75517 | 2 | 67867 | ENSMUST00000191062.7 | -1049 | ENSMUSG00000030556 | Lrrc28 | leucine rich repeat containing 28 |
| chr3 | 52977492 | 52978734 | 1243 | * | 9,370207 | -4,151004758 | 1,367565 | -3,03533 | 0,002403 | 0,671156916 | 1,211315 | Intron (ENSMUST00000 | 3 | 52949004 | 53169100 | 220097 | 1 | 108927 | ENSMUST00000147139.2 | 28488 | ENSMUSG00000048332 | Lhfp | lipoma HMGIC fusion partner |
| chr15 | 11085746 | 11087498 | 1753 | * | 12,14657 | -4,150374617 | 1,382425 | -3,00224 | 0,00268 | 0,732373293 | 0,918255 | Intron (ENSMUST00000 | 15 | 11064904 | 11217207 | 152304 | 1 | 239337 | ENSMUST00000228940.2 | 20842 | ENSMUSG00000047497 | Adamts12 | a disintegrin-like and metallopeptidase (reprolysin type) with thrombospondin type 1 motif, 12 |
| chrX | 89332985 | 89333940 | 956 | * | 6,41791 | -4,149736229 | 1,609153 | -2,57883 | 0,009913 | 1 | 0 | Distal Intergenic | 20 | 88973704 | 88976458 | 2755 | 2 | 245511 | ENSMUST00000088146.3 | -356527 | ENSMUSG00000035387 | 4930415L06Rik | RIKEN cDNA 4930415L06 gene |
| chr3 | 1,01E+08 | 1,01E+08 | 1358 | * | 10,53144 | -4,149710209 | 1,347218 | -3,08021 | 0,002069 | 0,686762073 | 0,918255 | Distal Intergenic | 3 | 1,01E+08 | 1,01E+08 | 71464 | 1 | 242122 | ENSMUST00000054791.9 | -33212 | ENSMUSG00000051076 | Vtcn1 | V-set domain containing T cell activation inhibitor 1 |
| chr17 | 18004568 | 18005699 | 1132 | * | 7,76683 | -4,148778927 | 1,517569 | -2,73383 | 0,00626 | 1 | 0 | Distal Intergenic | 17 | 18047420 | 18059333 | 11914 | 1 | 75202 | ENSMUST00000226899.2 | -41721 | ENSMUSG00000080316 | Spaca6 | sperm acrosome associated 6 |
| chr10 | 1,28E+08 | 1,28E+08 | 916 | * | 6,539518 | -4,147221879 | 1,596422 | -2,59782 | 0,009382 | 1 | 0 | Promoter (<=1kb) | 10 | 1,28E+08 | 1,28E+08 | 828 | 1 | 56530 | ENSMUST00000219199.2 | 385 | ENSMUSG00000025381 | Cnpy2 | canopy FGF signaling regulator 2 |
| chr5 | 42290655 | 42291336 | 682 | * | 8,075308 | -4,140846056 | 1,512592 | -2,73758 | 0,006189 | 1 | 0 | Intron (ENSMUST00000 | 5 | 42225303 | 42374141 | 148839 | 1 | 433882 | ENSMUST00000087332.5 | 65352 | ENSMUSG00000067285 | Gm16223 | predicted gene 16223 |
| chr16 | 40788750 | 40790433 | 1684 | * | 13,79966 | -4,140173372 | 1,2813 | -3,23123 | 0,001233 | 0,596684816 | 0,938682 | Intron (ENSMUST00000 | 16 | 41353220 | 41966576 | 613357 | 1 | 268890 | ENSMUST00000078873.11 | -562787 | ENSMUSG00000061080 | Lsamp | limbic system-associated membrane protein |
| chr18 | 18555103 | 18556344 | 1242 | * | 8,958317 | -4,138806747 | 1,459218 | -2,83632 | 0,004564 | 0,749550311 | 1,425929 | Distal Intergenic | 18 | 19266274 | 19334773 | 68500 | 1 | 1,03E+08 | ENSMUST00000234128.2 | -709930 | | Gm31086 | predicted gene, 31086 |
| chr11 | 1,11E+08 | 1,11E+08 | 789 | * | 9,507075 | -4,130822615 | 1,26767 | -3,2586 | 0,00112 | 0,501362189 | 1,425929 | Distal Intergenic | 11 | 1,11E+08 | 1,11E+08 | 2269 | 2 | 1,03E+08 | ENSMUST00000247083.1 | 209981 | | Gm30158 | predicted gene, 30158 |
| chr16 | 49497112 | 49498091 | 980 | * | 10,57377 | -4,126618849 | 1,360111 | -3,03403 | 0,002413 | 0,715249603 | 0,918255 | Distal Intergenic | 16 | 49519596 | 49585489 | 65894 | 1 | 73916 | ENSMUST00000046777.11 | -21505 | ENSMUSG00000032965 | Ift57 | intraflagellar transport 57 |
| chr5 | 1,34E+08 | 1,34E+08 | 737 | * | 8,050716 | -4,125738154 | 1,433463 | -2,87816 | 0,004 | 0,734161328 | 1,360622 | Distal Intergenic | 5 | 1,34E+08 | 1,34E+08 | 63817 | 2 | 1,03E+08 | ENSMUST00000196125.2 | -47556 | | Gm36667 | predicted gene, 36667 |
| chr3 | 1,34E+08 | 1,34E+08 | 662 | * | 8,917852 | -4,125375888 | 1,278443 | -3,22688 | 0,001251 | 0,518149475 | 1,449706 | Distal Intergenic | 3 | 1,35E+08 | 1,35E+08 | 105573 | 1 | 21338 | ENSMUST00000029822.6 | -35057 | ENSMUSG00000028172 | Tacr3 | tachykinin receptor 3 |
| chr8 | 1,15E+08 | 1,15E+08 | 697 | * | 7,216563 | -4,125183436 | 1,530343 | -2,69559 | 0,007026 | 1 | 0 | Intron (ENSMUST00000 | 8 | 1,15E+08 | 1,15E+08 | 5955 | 1 | 270097 | ENSMUST00000124143.2 | -65511 | ENSMUSG00000046844 | Vat1l | vesicle amine transport protein 1 like |
| chr1 | 1,91E+08 | 1,91E+08 | 1115 | * | 8,437137 | -4,122212595 | 1,460278 | -2,8229 | 0,004759 | 0,683067851 | 2,237403 | Distal Intergenic | 1 | 1,91E+08 | 1,91E+08 | 1565 | 1 | 226856 | ENSMUST00000133076.7 | -53007 | ENSMUSG00000026623 | Lpgat1 | lysophosphatidylglycerol acyltransferase 1 |
| chr2 | 1,7E+08 | 1,7E+08 | 1531 | * | 13,88139 | -4,120489615 | 1,341322 | -3,07196 | 0,002127 | 0,686950995 | 0,938682 | Intron (ENSMUST00000 | 2 | 1,7E+08 | 1,7E+08 | 27259 | 1 | 228911 | ENSMUST00000140699.2 | 15545 | ENSMUSG00000047907 | Tshz2 | teashirt zinc finger family member 2 |
| chr19 | 43324106 | 43325215 | 1110 | * | 11,59657 | -4,11705875 | 1,495135 | -2,75364 | 0,005894 | 0,906138253 | 0,744375 | Intron (ENSMUST00000 | 19 | 43282640 | 43376693 | 94054 | 2 | 545291 | ENSMUST00000236823.2 | 51478 | ENSMUSG00000074852 | Hpse2 | heparanase 2 |
| chrX | 1,57E+08 | 1,57E+08 | 878 | * | 8,388813 | -4,110778078 | 1,495242 | -2,74924 | 0,005973 | 0,718208008 | 2,237403 | Intron (ENSMUST00000 | 20 | 1,57E+08 | 1,57E+08 | 221859 | 2 | 245684 | ENSMUST00000026750.15 | 26714 | ENSMUSG00000025658 | Cnksr2 | connector enhancer of kinase suppressor of Ras 2 |
| chr1 | 95544981 | 95545759 | 779 | * | 8,75384 | -4,105998371 | 1,410746 | -2,91052 | 0,003608 | 1 | 0 | Intron (ENSMUST00000 | 1 | 95546276 | 95595260 | 48985 | 2 | 20452 | ENSMUST00000189556.2 | 49501 | ENSMUSG00000040710 | St8sia4 | ST8 alpha-N-acetyl-neuraminide alpha-2,8-sialyltransferase 4 |
| chr17 | 80221527 | 80222501 | 975 | * | 8,730875 | -4,101781559 | 1,410359 | -2,90833 | 0,003634 | 0,639383182 | 2,237403 | Distal Intergenic | 17 | 80155821 | 80203552 | 47732 | 2 | 56298 | ENSMUST00000112437.8 | -17975 | ENSMUSG00000059811 | Atl2 | atlastin GTPase 2 |
| chr7 | 1,2E+08 | 1,2E+08 | 1255 | * | 13,78341 | -4,10112398 | 1,303035 | -3,14736 | 0,001648 | 0,650529916 | 0,938682 | Exon (ENSMUST00000 | 7 | 1,2E+08 | 1,2E+08 | 15732 | 2 | 12971 | ENSMUST00000033198.6 | 3413 | ENSMUSG00000030905 | Crym | crystallin, mu |
| chr1 | 57901851 | 57903415 | 1565 | * | 13,06727 | -4,099840579 | 1,157529 | -3,54189 | 0,000397 | 0,339611164 | 1,2763 | Intron (ENSMUST00000 | 1 | 57884730 | 57986124 | 101395 | 1 | 67198 | ENSMUST00000169772.3 | 17121 | ENSMUSG00000038305 | Spats2l | spermatogenesis associated, serine-rich 2-like |
| chr7 | 1,23E+08 | 1,23E+08 | 1589 | * | 16,72824 | -4,098797439 | 1,064828 | -3,84926 | 0,000118 | 0,13101642 | 2,386008 | Distal Intergenic | 7 | 1,23E+08 | 1,23E+08 | 6236 | 2 | 210162 | ENSMUST00000128217.2 | -101106 | ENSMUSG00000030757 | Zkscan2 | zinc finger with KRAB and SCAN domains 2 |
| chr8 | 51855770 | 51856524 | 755 | * | 11,34074 | -4,097785157 | 1,235753 | -3,31602 | 0,000913 | 0,484531896 | 1,2763 | Distal Intergenic | 8 | 51251821 | 51370032 | 118212 | 2 | 23965 | ENSMUST00000211313.2 | -485738 | ENSMUSG00000031561 | Tenm3 | teneurin transmembrane protein 3 |
| chrX | 1,1E+08 | 1,1E+08 | 1392 | * | 10,9887 | -4,096915363 | 1,371425 | -2,98734 | 0,002814 | 0,742868694 | 0,918255 | Distal Intergenic | 20 | 1,1E+08 | 1,1E+08 | 2928 | 1 | 18994 | ENSMUST00000078229.5 | 58741 | ENSMUSG00000056854 | Pou3f4 | POU domain, class 3, transcription factor 4 |
| chrX | 1,26E+08 | 1,26E+08 | 888 | * | 7,942723 | -4,090925277 | 1,579683 | -2,58971 | 0,009606 | 1 | 0 | Distal Intergenic | 20 | 1,26E+08 | 1,26E+08 | 2037 | 2 | 1,19E+08 | ENSMUST00000117443.2 | 35424 | ENSMUSG00000082772 | LOC118568449 | vigilin-like |
| chr8 | 39484302 | 39484952 | 651 | * | 7,567578 | -4,090039374 | 1,51551 | -2,69879 | 0,006959 | 1 | 0 | Intron (ENSMUST00000 | 8 | 39473064 | 39540476 | 67413 | 1 | 80286 | ENSMUST00000239511.1 | 11238 | ENSMUSG00000118664 | Tusc3 | tumor suppressor candidate 3 |
| chr8 | 19431052 | 19431812 | 761 | * | 7,674699 | -4,08454951 | 1,475072 | -2,76905 | 0,005622 | 1 | 0 | Distal Intergenic | 8 | 19495785 | 19497622 | 1838 | 2 | 244334 | ENSMUST00000033854.4 | 65810 | ENSMUSG00000031471 | Defb8 | defensin beta 8 |
| chr13 | 14563639 | 14564654 | 1016 | * | 11,71364 | -4,084304092 | 1,301605 | -3,1379 | 0,001702 | 0,688853937 | 0,744375 | Intron (ENSMUST00000 | 13 | 14420046 | 14493542 | 73497 | 2 | 94253 | ENSMUST00000222241.2 | -70097 | ENSMUSG00000021301 | Hecw1 | HECT, C2 and WW domain containing E3 ubiquitin protein ligase 1 |
| chr18 | 40579760 | 40580608 | 849 | * | 7,859651 | -4,083059254 | 1,508106 | -2,70741 | 0,006781 | 1 | 0 | Intron (ENSMUST00000 | 18 | 40541138 | 40664683 | 123546 | 1 | 383348 | ENSMUST00000236301.2 | 38622 | ENSMUSG00000051401 | Kctd16 | potassium channel tetramerisation domain containing 16 |
| chr2 | 1,79E+08 | 1,79E+08 | 691 | * | 9,355773 | -4,081079793 | 1,328687 | -3,07151 | 0,00213 | 0,650999838 | 1,211315 | Distal Intergenic | 2 | 1,79E+08 | 1,79E+08 | 1953 | 1 | 68175 | ENSMUST00000041998.3 | -397890 | ENSMUSG00000039138 | 4930591A17Rik | RIKEN cDNA 4930591A17 gene |
| chr9 | 69688896 | 69689805 | 910 | * | 7,595934 | -4,077122545 | 1,45509 | -2,80197 | 0,005079 | 1 | 0 | Intron (ENSMUST00000 | 9 | 69674895 | 69676274 | 1380 | 1 | 1,03E+08 | ENSMUST00000243201.1 | 14001 | | Gm31530 | predicted gene, 31530 |
| chr19 | 54179976 | 54181487 | 1512 | * | 13,38059 | -4,072395327 | 1,330312 | -3,06123 | 0,002204 | 0,676726988 | 1,075735 | Distal Intergenic | 19 | 54032582 | 54037413 | 4832 | 1 | 11551 | ENSMUST00000036700.7 | 147394 | ENSMUSG00000033717 | Adra2a | adrenergic receptor, alpha 2a |
| chr7 | 90163032 | 90164211 | 1180 | * | 9,742747 | -4,064126629 | 1,412081 | -2,87811 | 0,004001 | 0,728037066 | 1,425929 | Intron (ENSMUST00000 | 7 | 90126439 | 92098455 | 1972017 | 1 | 23859 | ENSMUST00000231777.3 | 36593 | ENSMUSG00000052572 | Dlg2 | discs large MAGUK scaffold protein 2 |
| chr7 | 1,09E+08 | 1,09E+08 | 955 | * | 11,24598 | -4,062575502 | 1,189805 | -3,41449 | 0,000639 | 0,426043749 | 1,2763 | Distal Intergenic | 7 | 1,08E+08 | 1,08E+08 | 960 | 2 | 277935 | ENSMUST00000084752.2 | -9809 | ENSMUSG00000066239 | Olfr519 | olfactory receptor 519 |
| chr13 | 44261506 | 44262885 | 1380 | * | 11,43849 | -4,062211923 | 1,317073 | -3,08427 | 0,002041 | 0,649854434 | 1,173325 | Distal Intergenic | 13 | 44245504 | 44253436 | 7933 | 2 | 1,03E+08 | ENSMUST00000223038.2 | -8070 | ENSMUSG00000113276 | Gm33630 | predicted gene, 33630 |
| chr7 | 1,19E+08 | 1,19E+08 | 798 | * | 9,504923 | -4,061276153 | 1,462715 | -2,77653 | 0,005494 | 0,792225394 | 1,360622 | Distal Intergenic | 7 | 1,19E+08 | 1,19E+08 | 23165 | 2 | 64297 | ENSMUST00000008878.10 | -88243 | ENSMUSG00000008734 | Gprc5b | G protein-coupled receptor, family C, group 5, member B |
| chr19 | 54816078 | 54817115 | 1038 | * | 8,686965 | -4,060521743 | 1,440394 | -2,81904 | 0,004817 | 0,683210259 | 2,237403 | Distal Intergenic | 19 | 55058166 | 55087883 | 29718 | 2 | 14732 | ENSMUST00000061856.6 | 270768 | ENSMUSG00000024978 | Gpam | glycerol-3-phosphate acyltransferase, mitochondrial |
| chr5 | 1,41E+08 | 1,41E+08 | 1249 | * | 11,4635 | -4,052583267 | 1,367274 | -2,96399 | 0,003037 | 0,793220117 | 0,738441 | Intron (ENSMUST00000 | 5 | 1,41E+08 | 1,41E+08 | 7374 | 2 | 1,03E+08 | ENSMUST00000244363.1 | -42241 | | Gm16035 | predicted gene 16035 |
| chr7 | 1,13E+08 | 1,13E+08 | 1056 | * | 10,49965 | -4,052252641 | 1,45095 | -2,79283 | 0,005225 | 0,888802661 | 0,744375 | Distal Intergenic | 7 | 1,13E+08 | 1,13E+08 | 26990 | 1 | 1,03E+08 | ENSMUST00000210768.2 | -12007 | | Gm33586 | predicted gene, 33586 |
| chr1 | 1,7E+08 | 1,7E+08 | 2032 | * | 18,62759 | -4,051194017 | 1,090822 | -3,71389 | 0,000204 | 0,170809454 | 2,663619 | Exon (ENSMUST000001 | 1 | 1,7E+08 | 1,7E+08 | 24987 | 2 | 70729 | ENSMUST00000161221.8 | -62133 | ENSMUSG00000038473 | Nos1ap | nitric oxide synthase 1 (neuronal) adaptor protein |
| chr12 | 41348132 | 41349122 | 991 | * | 9,738593 | -4,049604796 | 1,364234 | -2,96841 | 0,002993 | 0,724018326 | 1,086118 | Intron (ENSMUST00000 | 12 | 41398440 | 41398571 | 132 | 2 | 1,15E+08 | ENSMUST00000104425.3 | 49449 | ENSMUSG00000119635 | Gm25497 | predicted gene, 25497 |
| chrX | 75744146 | 75745082 | 937 | * | 6,99439 | -4,044982024 | 1,489208 | -2,7162 | 0,006604 | 1 | 0 | Distal Intergenic | 20 | 75626214 | 75646530 | 20317 | 2 | 74987 | ENSMUST00000114054.3 | -97616 | ENSMUSG00000079525 | Cldn34d | claudin 34D |
| chr3 | 1,24E+08 | 1,24E+08 | 906 | * | 8,713354 | -4,04323178 | 1,465947 | -2,7581 | 0,005814 | 0,791147121 | 1,449706 | Distal Intergenic | 3 | 1,24E+08 | 1,24E+08 | 7885 | 2 | 384775 | ENSMUST00000180033.8 | -27336 | ENSMUSG00000039174 | 1700003H04Rik | RIKEN cDNA 1700003H04 gene |
| chrX | 80648820 | 80649860 | 1041 | * | 6,537732 | -4,042976695 | 1,531406 | -2,64004 | 0,00829 | 1 | 0 | Distal Intergenic | 20 | 80463181 | 80501727 | 38547 | 2 | 245492 | ENSMUST00000080083.3 | -147093 | ENSMUSG00000060673 | 4930595M18Rik | RIKEN cDNA 4930595M18 gene |
| chr7 | 1,34E+08 | 1,34E+08 | 941 | * | 7,273296 | -4,04126681 | 1,51998 | -2,65876 | 0,007843 | 1 | 0 | Promoter (<=1kb) | 7 | 1,33E+08 | 1,34E+08 | 75070 | 2 | 11489 | ENSMUST00000138363.2 | 0 | ENSMUSG00000054555 | Adam12 | a disintegrin and metallopeptidase domain 12 (meltrin alpha) |

| | | | | | | | | | | | | | | | | | | | | | | | |
|---|---|---|---|---|---|---|---|---|---|---|---|---|---|---|---|---|---|---|---|---|---|---|---|
| chr1 | 29021634 | 29023598 | 1965 | * | 10,46138 | -4,040233846 | 1,37227 | -2,9442 | 0,003238 | 0,711373629 | 1,2763 | Distal Intergenic | 1 | 28977117 | 28977420 | 304 | 1 | 623924 | ENSMUST00000081488.3 | 44517 | ENSMUSG00000058590 | Gm6462 | predicted gene 6462 |
| chr4 | 1,5E+08 | 1,5E+08 | 1059 | * | 9,553522 | -4,039661433 | 1,437139 | -2,81091 | 0,00494 | 0,813191811 | 1,086118 | Distal Intergenic | 4 | 1,5E+08 | 1,5E+08 | 691 | 2 | 1,05E+08 | ENSMUST00000134059.2 | -5793 | | Gm13073 | predicted gene 13073 |
| chr1 | 1,46E+08 | 1,46E+08 | 1171 | * | 8,030092 | -4,036142062 | 1,469825 | -2,746 | 0,006033 | 1 | 0 | Intron (ENSMUST0000 | 1 | 1,46E+08 | 1,46E+08 | 19817 | 1 | 215378 | ENSMUST00000125331.2 | 67590 | ENSMUSG00000035131 | Brinp3 | bone morphogenetic protein/retinoic acid inducible neural specific 3 |
| chr3 | 65816201 | 65817875 | 1675 | * | 15,56734 | -4,03396058 | 1,325562 | -3,04321 | 0,002341 | 0,641470306 | 1,41696 | Distal Intergenic | 3 | 65854203 | 65855419 | 1217 | 2 | 56706 | ENSMUST00000149729.2 | 37544 | ENSMUSG00000027829 | Ccnl1 | cyclin L1 |
| chr19 | 37492456 | 37493608 | 1153 | * | 10,31096 | -4,03278583 | 1,423132 | -2,83374 | 0,004601 | 0,749931768 | 1,425929 | Distal Intergenic | 19 | 37509327 | 37672507 | 163181 | 1 | 107371 | ENSMUST00000238954.2 | -15719 | ENSMUSG00000053799 | Exoc6 | exocyst complex component 6 |
| chr11 | 28325267 | 28326471 | 1205 | * | 9,02018 | -4,031011329 | 1,403804 | -2,87149 | 0,004085 | 0,728748169 | 1,449706 | Distal Intergenic | 11 | 28338817 | 28384110 | 45294 | 2 | 216613 | ENSMUST00000160016.8 | 57639 | ENSMUSG00000032878 | Ccdc85a | coiled-coil domain containing 85A |
| chr7 | 1,17E+08 | 1,17E+08 | 935 | * | 9,658643 | -4,030690098 | 1,372811 | -2,93609 | 0,003324 | 0,723239852 | 1,211315 | Intron (ENSMUST0000 | 7 | 1,17E+08 | 1,17E+08 | 73868 | 1 | 233781 | ENSMUST00000161889.2 | 152138 | ENSMUSG00000030657 | Xylt1 | xylosyltransferase 1 |
| chr14 | 3307281 | 3308728 | 1448 | * | 9,743223 | -4,030485533 | 1,40197 | -2,87487 | 0,004042 | 0,72659841 | 1,449706 | Exon (ENSMUST00000 | 14 | 3575429 | 3839361 | 263933 | 1 | 218793 | ENSMUST00000124353.2 | -266701 | ENSMUSG00000058317 | Ube2e2 | ubiquitin-conjugating enzyme E2E 2 |
| chr11 | 90932109 | 90933035 | 927 | * | 6,707234 | -4,030418488 | 1,539258 | -2,61842 | 0,008834 | 1 | 0 | Distal Intergenic | 11 | 90774171 | 90785825 | 11655 | 1 | 73922 | ENSMUST00000132218.2 | 157938 | ENSMUSG00000085940 | 4930405D11Rik | RIKEN cDNA 4930405D11 gene |
| chr10 | 1,1E+08 | 1,1E+08 | 560 | * | 7,694846 | -4,023503723 | 1,488839 | -2,70244 | 0,006883 | 1 | 0 | Distal Intergenic | 10 | 1,1E+08 | 1,1E+08 | 557102 | 2 | 260315 | ENSMUST00000160159.3 | -55567 | ENSMUSG00000020181 | Nav3 | neuron navigator 3 |
| chr3 | 14746026 | 14747105 | 1080 | * | 11,83031 | -4,02300185 | 1,214893 | -3,31141 | 0,000928 | 0,488376383 | 1,2763 | Distal Intergenic | 3 | 14706787 | 14728062 | 21276 | 1 | 71934 | ENSMUST00000029071.9 | 39239 | ENSMUSG00000027555 | Car13 | carbonic anhydrase 13 |
| chr1 | 63502457 | 63503325 | 869 | * | 9,77168 | -4,018529183 | 1,415524 | -2,8389 | 0,004527 | 0,74907225 | 1,425929 | Intron (ENSMUST0000 | 1 | 63509778 | 63584717 | 74940 | 1 | 23792 | ENSMUST00000190658.7 | -6453 | ENSMUSG00000025964 | Adam23 | a disintegrin and metallopeptidase domain 23 |
| chr4 | 8157286 | 8158086 | 801 | * | 9,936743 | -4,014371922 | 1,42837 | -2,81046 | 0,004947 | 0,759970924 | 1,449706 | Intron (ENSMUST0000 | 4 | 8143362 | 8239041 | 95680 | 2 | 12319 | ENSMUST00000066674.8 | 80955 | ENSMUSG00000041261 | Car8 | carbonic anhydrase 8 |
| chr2 | 1,41E+08 | 1,41E+08 | 1423 | * | 8,07696 | -4,010509435 | 1,5461 | -2,59395 | 0,009488 | 1 | 0 | Intron (ENSMUST0000 | 2 | 1,41E+08 | 1,42E+08 | 837291 | 1 | 72899 | ENSMUST00000138786.2 | -11971 | ENSMUSG00000068205 | Macrod2 | mono-ADP ribosylhydrolase 2 |
| chr14 | 68239473 | 68240574 | 1102 | * | 9,224926 | -4,010477144 | 1,404194 | -2,85607 | 0,004289 | 0,767135014 | 1,211315 | Distal Intergenic | 14 | 68281877 | 68288779 | 6903 | 2 | 1,03E+08 | ENSMUST00000223853.2 | 48205 | | Gm31107 | predicted gene, 31107 |
| chrX | 19273439 | 19274518 | 1080 | * | 6,439206 | -4,006830797 | 1,515832 | -2,64332 | 0,00821 | 1 | 0 | Distal Intergenic | 20 | 19221240 | 19221346 | 107 | 1 | 1,15E+08 | ENSMUST00000104089.3 | 52199 | ENSMUSG00000077277 | Gm22783 | predicted gene, 22783 |
| chr5 | 78354826 | 78355798 | 973 | * | 7,302639 | -4,005777784 | 1,503535 | -2,66424 | 0,007716 | 0,852340527 | 1,360622 | Distal Intergenic | 5 | 78927706 | 78927841 | 136 | 1 | 1,15E+08 | ENSMUST00000103990.3 | -571908 | ENSMUSG00000077178 | Gm25756 | predicted gene, 25756 |
| chr13 | 47031111 | 47032580 | 1470 | * | 12,79554 | -4,005568783 | 1,323259 | -3,02705 | 0,00247 | 0,755814512 | 0,738441 | Intron (ENSMUST0000 | 13 | 46932380 | 47049181 | 116802 | 2 | 16553 | ENSMUST00000225591.2 | 16601 | ENSMUSG00000021375 | Kif13a | kinesin family member 13A |
| chr12 | 41393918 | 41394664 | 747 | * | 6,923972 | -4,003343766 | 1,550141 | -2,58257 | 0,009807 | 1 | 0 | Intron (ENSMUST0000 | 12 | 41398440 | 41398571 | 132 | 2 | 1,15E+08 | ENSMUST00000104425.3 | 3907 | ENSMUSG00000119635 | Gm25497 | predicted gene, 25497 |
| chrX | 1,09E+08 | 1,09E+08 | 1404 | * | 8,544678 | -4,003028145 | 1,450765 | -2,75925 | 0,005793 | 0,790570581 | 1,449706 | Distal Intergenic | 20 | 1,09E+08 | 1,09E+08 | 2406 | 1 | 634340 | ENSMUST00000122296.2 | -66750 | ENSMUSG00000082853 | Gm7134 | kinesin family member 18B pseudogene |
| chr9 | 84087800 | 84088516 | 717 | * | 6,392815 | -4,00160051 | 1,544252 | -2,59129 | 0,009562 | 1 | 0 | Distal Intergenic | 9 | 84070408 | 84070592 | 185 | 1 | 1,15E+08 | ENSMUST00000104478.4 | 17392 | ENSMUSG00000077666 | Gm26146 | predicted gene, 26146 |
| chr5 | 51131488 | 51132389 | 902 | * | 8,771585 | -3,999950286 | 1,549687 | -2,58113 | 0,009848 | 0,884485418 | 1,449706 | Distal Intergenic | 5 | 50606004 | 50701833 | 95830 | 2 | 1,03E+08 | ENSMUST00000197457.5 | -429655 | | Gm36814 | predicted gene, 36814 |
| chr8 | 1,14E+08 | 1,14E+08 | 925 | * | 8,555023 | -3,993753319 | 1,438367 | -2,77659 | 0,005493 | 0,780196848 | 1,449706 | Intron (ENSMUST0000 | 8 | 1,14E+08 | 1,14E+08 | 124538 | 2 | 1,15E+08 | ENSMUST00000213082.2 | 11132 | | Gm45904 | predicted gene 45904 |
| chr4 | 37544853 | 37545933 | 1081 | * | 8,011995 | -3,991143001 | 1,510114 | -2,64294 | 0,008219 | 0,8605359 | 1,360622 | Distal Intergenic | 4 | 37249403 | 37332515 | 83113 | 1 | 1,03E+08 | ENSMUST00000147108.2 | 295450 | ENSMUSG00000086026 | Gm12374 | predicted gene 12374 |
| chr11 | 67966063 | 67967710 | 1648 | * | 11,92083 | -3,988820215 | 1,216909 | -3,27783 | 0,001046 | 0,611385361 | 0,744375 | Intron (ENSMUST0000 | 11 | 67857332 | 67864467 | 7136 | 1 | 55943 | ENSMUST00000142649.2 | 108731 | ENSMUSG00000020903 | Stx8 | syntaxin 8 |
| chr7 | 58600940 | 58601964 | 1025 | * | 9,695212 | -3,98430821 | 1,385237 | -2,87627 | 0,004024 | 0,777172375 | 1,086118 | Exon (ENSMUST00000 | 7 | 58878498 | 58939567 | 61070 | 1 | 22215 | ENSMUST00000202945.4 | -276534 | ENSMUSG00000025326 | Ube3a | ubiquitin protein ligase E3A |
| chr6 | 48244303 | 48245356 | 1054 | * | 14,48104 | -3,984149839 | 1,122999 | -3,54778 | 0,000388 | 0,327375387 | 1,37683 | Distal Intergenic | 6 | 48107840 | 48136214 | 28375 | 2 | 1,01E+08 | ENSMUST00000180741.3 | -108089 | ENSMUSG00000107875 | Gm16630 | predicted gene, 16630 |
| chr5 | 51194563 | 51196162 | 1600 | * | 12,42253 | -3,98070778 | 1,305572 | -3,04901 | 0,002296 | 0,745788938 | 0,738441 | Distal Intergenic | 5 | 50606004 | 50701833 | 95830 | 2 | 1,03E+08 | ENSMUST00000197457.5 | -492730 | | Gm36814 | predicted gene, 36814 |
| chr1 | 1,21E+08 | 1,21E+08 | 1148 | * | 10,93404 | -3,978723084 | 1,391277 | -2,85976 | 0,00424 | 0,75508254 | 1,2763 | Promoter (2-3kb) | 1 | 1,21E+08 | 1,21E+08 | 68664 | 1 | 69904 | ENSMUST00000186982.3 | 2042 | | 2610027F03Rik | RIKEN cDNA 2610027F03 gene |
| chr11 | 76296690 | 76298367 | 1678 | * | 13,7858 | -3,978031653 | 1,275317 | -3,11925 | 0,001813 | 0,644368101 | 1,075735 | Promoter (<=1kb) | 11 | 76297778 | 76305155 | 7378 | 1 | 56322 | ENSMUST00000120699.8 | 0 | ENSMUSG00000020843 | Timm22 | translocase of inner mitochondrial membrane 22 |
| chr14 | 36859577 | 36860794 | 1218 | * | 9,295627 | -3,97754708 | 1,364763 | -2,91446 | 0,003563 | 0,70513707 | 1,449706 | Promoter (2-3kb) | 14 | 36853355 | 36857229 | 3875 | 2 | 66092 | ENSMUST00000223921.2 | -2348 | ENSMUSG00000041028 | Ghitm | growth hormone inducible transmembrane protein |
| chr8 | 17724118 | 17725207 | 1090 | * | 9,10765 | -3,971802987 | 1,524089 | -2,60602 | 0,00916 | 0,792545355 | 2,237403 | Distal Intergenic | 8 | 16394877 | 17585602 | 1190726 | 2 | 94109 | ENSMUST00000137947.2 | -138516 | ENSMUSG00000060924 | Csmd1 | CUB and Sushi multiple domains 1 |
| chr1 | 13236474 | 13237424 | 951 | * | 8,959597 | -3,971640872 | 1,397099 | -2,84278 | 0,004472 | 0,752939853 | 1,360622 | Exon (ENSMUST00000 | 1 | 13218653 | 13221242 | 2590 | 2 | 17978 | ENSMUST00000133232.2 | -15232 | ENSMUSG00000005886 | Ncoa2 | nuclear receptor coactivator 2 |
| chr12 | 1,04E+08 | 1,04E+08 | 2048 | * | 20,88335 | -3,969905746 | 1,057529 | -3,75394 | 0,000174 | 0,147378366 | 2,951933 | Promoter (<=1kb) | 12 | 1,04E+08 | 1,04E+08 | 2122 | 2 | 14836 | ENSMUST00000021513.6 | -687 | ENSMUSG00000021095 | Gsc | goosecoid homeobox |
| chr11 | 1,01E+08 | 1,01E+08 | 1536 | * | 12,29627 | -3,968979246 | 1,302694 | -3,04675 | 0,002313 | 0,670234202 | 1,173325 | Promoter (<=1kb) | 11 | 1,01E+08 | 1,01E+08 | 1860 | 2 | 15171 | ENSMUST00000107360.3 | 0 | ENSMUSG00000045471 | Hcrt | hypocretin |
| chr7 | 60160448 | 60162397 | 1950 | * | 11,23726 | -3,966624137 | 1,488276 | -2,66525 | 0,007693 | 0,86031879 | 1,2763 | Distal Intergenic | 7 | 59805357 | 59805848 | 492 | 1 | 664849 | ENSMUST00000057611.6 | 355091 | | Gm7367 | 1110014K08Rik pseudogene |
| chr14 | 1E+08 | 1E+08 | 1772 | * | 11,11644 | -3,965573442 | 1,235036 | -3,2109 | 0,001323 | 0,566578624 | 1,173325 | Distal Intergenic | 14 | 1E+08 | 1E+08 | 297 | 2 | 546623 | ENSMUST00000147698.2 | -25008 | | Gm5958 | predicted pseudogene 5958 |
| chrX | 1,62E+08 | 1,62E+08 | 1287 | * | 10,48932 | -3,963723501 | 1,360383 | -2,91368 | 0,003572 | 0,8237677 | 0,738441 | Distal Intergenic | 20 | 1,62E+08 | 1,62E+08 | 107 | 2 | 1,15E+08 | ENSMUST00000082494.3 | 12824 | ENSMUSG00000118912 | Gm22746 | predicted gene, 22746 |
| chr4 | 35831515 | 35832652 | 1138 | * | 6,978903 | -3,962135395 | 1,466217 | -2,70229 | 0,006886 | 0,83238039 | 1,360622 | Intron (ENSMUST0000 | 4 | 35706647 | 35845204 | 138558 | 2 | 242384 | ENSMUST00000164772.8 | 12552 | ENSMUSG00000045083 | Lingo2 | leucine rich repeat and Ig domain containing 2 |
| chr2 | 65438336 | 65439123 | 788 | * | 6,581495 | -3,95925665 | 1,530508 | -2,58689 | 0,009685 | 1 | 0 | Distal Intergenic | 2 | 65451115 | 65520519 | 69405 | 1 | 110876 | ENSMUST00000144254.6 | -11992 | ENSMUSG00000075318 | Scn2a | sodium channel, voltage-gated, type II, alpha |
| chr15 | 47395837 | 47396431 | 595 | * | 7,295627 | -3,957409648 | 1,477328 | -2,67876 | 0,007389 | 0,844723341 | 1,360622 | Distal Intergenic | 15 | 47445122 | 47478597 | 33476 | 2 | 239420 | ENSMUST00000162006.2 | 82166 | ENSMUSG00000022311 | Csmd3 | CUB and Sushi multiple domains 3 |
| chr13 | 14205942 | 14207136 | 1195 | * | 8,177198 | -3,955970424 | 1,443163 | -2,74118 | 0,006122 | 1 | 0 | Intron (ENSMUST0000 | 13 | 14172534 | 14214164 | 41631 | 2 | 70430 | ENSMUST00000159893.8 | 7028 | ENSMUSG00000039233 | Tbce | tubulin-specific chaperone E |
| chr19 | 37730955 | 37732726 | 1772 | * | 12,95899 | -3,953934749 | 1,293622 | -3,05648 | 0,002239 | 0,650529916 | 1,2763 | Distal Intergenic | 19 | 37686533 | 37687456 | 924 | 1 | 13082 | ENSMUST00000236500.2 | 44422 | ENSMUSG00000024987 | Cyp26a1 | cytochrome P450, family 26, subfamily a, polypeptide 1 |
| chr7 | 85378282 | 85379179 | 898 | * | 8,057508 | -3,953488977 | 1,451821 | -2,72312 | 0,006467 | 1 | 0 | Distal Intergenic | 7 | 85385148 | 85404305 | 19158 | 2 | 244114 | ENSMUST00000063425.9 | 25126 | ENSMUSG00000051877 | Vmn2r72 | vomeronasal 2, receptor 72 |
| chr12 | 1,03E+08 | 1,03E+08 | 1177 | * | 10,41625 | -3,945954641 | 1,339302 | -2,94628 | 0,003216 | 0,710505631 | 1,2763 | Intron (ENSMUST0000 | 12 | 1,03E+08 | 1,03E+08 | 16180 | 1 | 217843 | ENSMUST00000177984.2 | 6328 | ENSMUSG00000021198 | Unc79 | unc-79 homolog |
| chrX | 83537276 | 83538219 | 944 | * | 7,361862 | -3,937053277 | 1,473724 | -2,6715 | 0,007551 | 1 | 0 | Intron (ENSMUST0000 | 20 | 83642807 | 84188019 | 545213 | 1 | 13405 | ENSMUST00000146331.8 | -104588 | ENSMUSG00000045103 | Dmd | dystrophin, muscular dystrophy |
| chr17 | 71355553 | 71356728 | 1176 | * | 12,00397 | -3,936893193 | 1,196179 | -3,29122 | 0,000998 | 0,500381023 | 1,2763 | Intron (ENSMUST0000 | 17 | 71326586 | 71390722 | 64137 | 1 | 17929 | ENSMUST00000232851.2 | 28967 | ENSMUSG00000024049 | Myom1 | myomesin 1 |
| chr4 | 1,54E+08 | 1,54E+08 | 1111 | * | 13,73947 | -3,935444674 | 1,182091 | -3,32922 | 0,000871 | 0,534793902 | 0,918255 | Exon (ENSMUST00000 | 4 | 1,54E+08 | 1,54E+08 | 34215 | 1 | 230971 | ENSMUST00000152159.8 | 4539 | ENSMUSG00000057751 | Megf6 | multiple EGF-like-domains 6 |
| chr9 | 42054619 | 42055734 | 1116 | * | 8,053744 | -3,934688801 | 1,428039 | -2,75531 | 0,005864 | 1 | 0 | Promoter (<=1kb) | 9 | 42053657 | 42055560 | 1904 | 1 | 1,05E+08 | ENSMUST00000217090.2 | 962 | | Gm39321 | predicted gene, 39321 |
| chr17 | 4518275 | 4519248 | 974 | * | 9,815172 | -3,932496639 | 1,433816 | -2,74268 | 0,006094 | 0,837036698 | 1,173325 | Distal Intergenic | 17 | 4685253 | 4687746 | 2494 | 1 | 75117 | ENSMUST00000232624.2 | -166005 | ENSMUSG00000116767 | 4930517M08Rik | RIKEN cDNA 4930517M08 gene |
| chr8 | 4740383 | 4741595 | 1213 | * | 11,4801 | -3,929192947 | 1,24052 | -3,16738 | 0,001538 | 0,678079528 | 0,744375 | Distal Intergenic | 8 | 4728431 | 4738497 | 10067 | 1 | 623223 | ENSMUST00000181337.3 | 11952 | | Gm6410 | predicted gene 6410 |
| chr3 | 31507877 | 31509170 | 1294 | * | 13,20036 | -3,928754256 | 1,186949 | -3,30996 | 0,000933 | 0,490086417 | 1,2763 | Distal Intergenic | 3 | 31365072 | 31369157 | 4086 | 1 | 1,03E+08 | ENSMUST00000195091.2 | 142805 | | Gm15496 | predicted gene 15496 |
| chr8 | 10041727 | 10042845 | 1119 | * | 8,121704 | -3,925670366 | 1,461698 | -2,68569 | 0,007238 | 0,749931768 | 2,237403 | 3' UTR | 8 | 10056467 | 10085996 | 29530 | 1 | 24099 | ENSMUST00000208683.2 | -13622 | ENSMUSG00000031497 | Tnfsf13b | tumor necrosis factor (ligand) superfamily, member 13b |
| chr2 | 1,46E+08 | 1,46E+08 | 1608 | * | 11,38112 | -3,923105309 | 1,32541 | -2,95992 | 0,003077 | 0,795341563 | 0,738441 | Promoter (2-3kb) | 2 | 1,46E+08 | 1,46E+08 | 3178 | 1 | 74065 | ENSMUST00000147212.2 | 2927 | ENSMUSG00000085796 | 4933406D12Rik | RIKEN cDNA 4933406D12 gene |
| chr7 | 61552935 | 61554623 | 1689 | * | 12,66044 | -3,92225113 | 1,332715 | -2,94305 | 0,00325 | 0,767135014 | 0,918255 | Intron (ENSMUST0000 | 7 | 61357487 | 61561150 | 203664 | 2 | 319893 | ENSMUST00000208961.3 | 6527 | ENSMUSG00000109394 | A230057D06Rik | RIKEN cDNA A230057D06 gene |
| chr5 | 1,05E+08 | 1,05E+08 | 1609 | * | 13,64175 | -3,920891766 | 1,130784 | -3,46741 | 0,000526 | 0,483145429 | 0,738441 | 3' UTR | 5 | 1,05E+08 | 1,05E+08 | 2628 | 2 | 236573 | ENSMUST00000199453.2 | 4110 | ENSMUSG00000029298 | Gbp9 | guanylate-binding protein 9 |
| chr10 | 30818401 | 30819903 | 1503 | * | 10,44075 | -3,919184496 | 1,397305 | -2,80482 | 0,005035 | 0,801688106 | 1,173325 | Promoter (<=1kb) | 10 | 30818375 | 30818481 | 107 | 2 | 1,15E+08 | ENSMUST00000240080.1 | 0 | ENSMUSG00000119396 | Gm24168 | predicted gene, 24168 |
| chr4 | 95760321 | 95761866 | 1546 | * | 11,51403 | -3,918948118 | 1,346811 | -2,9098 | 0,003617 | 0,825384969 | 0,738441 | Intron (ENSMUST0000 | 4 | 95737862 | 95815171 | 77310 | 1 | 75578 | ENSMUST00000142384.2 | 22459 | ENSMUSG00000028573 | Fggy | FGGY carbohydrate kinase domain containing |
| chr4 | 91725009 | 91726176 | 1168 | * | 7,909156 | -3,917199633 | 1,468986 | -2,6666 | 0,007662 | 1 | 0 | Distal Intergenic | 4 | 91693784 | 91694792 | 1009 | 2 | 620016 | ENSMUST00000122162.2 | -30217 | ENSMUSG00000100863 | Gm12669 | glutaredoxin 3 pseudogene |
| chr13 | 46504242 | 46505265 | 1024 | * | 10,83269 | -3,906979315 | 1,403745 | -2,78325 | 0,005382 | 0,892445563 | 0,744375 | Distal Intergenic | 13 | 46571910 | 46584571 | 12662 | 1 | 666794 | ENSMUST00000037923.5 | -66645 | ENSMUSG00000038132 | Rbm24 | RNA binding motif protein 24 |
| chr8 | 1,26E+08 | 1,26E+08 | 724 | * | 10,87782 | -3,903366875 | 1,420898 | -2,74711 | 0,006012 | 0,819720472 | 1,2763 | Distal Intergenic | 8 | 1,26E+08 | 1,26E+08 | 3542 | 1 | 74393 | ENSMUST00000053078.5 | -28104 | ENSMUSG00000050930 | Map10 | microtubule-associated protein 10 |
| chr12 | 99885266 | 99886308 | 1043 | * | 10,78992 | -3,902442694 | 1,411623 | -2,76451 | 0,005701 | 0,903101391 | 0,738441 | Exon (ENSMUST00000 | 12 | 99881725 | 99885563 | 3839 | 1 | 104884 | ENSMUST00000220685.2 | 3541 | ENSMUSG00000021177 | Tdp1 | tyrosyl-DNA phosphodiesterase 1 |
| chr3 | 18844733 | 18845610 | 878 | * | 8,319788 | -3,902405764 | 1,397936 | -2,79155 | 0,005246 | 1 | 0 | Distal Intergenic | 3 | 18522451 | 18534641 | 12191 | 2 | 73959 | ENSMUST00000192010.2 | -310092 | ENSMUSG00000102699 | 4930433B08Rik | RIKEN cDNA 4930433B08 gene |
| chr4 | 95103002 | 95104457 | 1456 | * | 13,24955 | -3,898457213 | 1,160199 | -3,36016 | 0,000779 | 0,514210241 | 0,918255 | Distal Intergenic | 4 | 95081599 | 95094485 | 12887 | 1 | 1,05E+08 | ENSMUST00000126143.2 | 21403 | | Gm12708 | predicted gene 12708 |
| chr16 | 89799260 | 89801132 | 1873 | * | 17,75719 | -3,895615829 | 1,081125 | -3,6033 | 0,000314 | 0,245270361 | 2,154504 | Intron (ENSMUST0000 | 16 | 89752853 | 89776968 | 24116 | 2 | 21844 | ENSMUST00000136760.2 | -22292 | ENSMUSG00000002489 | Tiam1 | T cell lymphoma invasion and metastasis 1 |
| chr5 | 46861920 | 46862815 | 896 | * | 8,110372 | -3,894539682 | 1,476914 | -2,63694 | 0,008366 | 0,856353593 | 1,425929 | Distal Intergenic | 5 | 46084160 | 46089751 | 5592 | 2 | 78106 | ENSMUST00000198088.2 | -772169 | ENSMUSG00000106602 | 4930405L22Rik | RIKEN cDNA 4930405L22 gene |
| chr14 | 89257138 | 89258048 | 911 | * | 7,57228 | -3,89254093 | 1,464329 | -2,65824 | 0,007855 | 0,853945114 | 1,360622 | Distal Intergenic | 14 | 88873469 | 89001197 | 127729 | 1 | 1,05E+08 | ENSMUST00000227074.2 | 383669 | | Gm41225 | predicted gene, 41225 |
| chrX | 1,27E+08 | 1,27E+08 | 2198 | * | 11,62771 | -3,892332693 | 1,25504 | -3,10136 | 0,001926 | 0,715249603 | 0,738441 | Distal Intergenic | 20 | 1,27E+08 | 1,27E+08 | 107 | 2 | 1,15E+08 | ENSMUST00000240076.1 | -106318 | ENSMUSG00000119321 | Gm25061 | predicted gene, 25061 |
| chr12 | 75054764 | 75055721 | 958 | * | 10,7325 | -3,890830475 | 1,414085 | -2,75148 | 0,005933 | 0,908873708 | 0,738441 | Intron (ENSMUST0000 | 12 | 74943994 | 75224106 | 280113 | 2 | 238271 | ENSMUST00000042299.4 | 168385 | ENSMUSG00000034402 | Kcnh5 | potassium voltage-gated channel, subfamily H (eag-related), member 5 |
| chr19 | 41763098 | 41764093 | 996 | * | 7,990009 | -3,887489765 | 1,48948 | -2,60996 | 0,009055 | 0,88052247 | 1,360622 | Intron (ENSMUST0000 | 19 | 41755027 | 41790486 | 35460 | 2 | 71085 | ENSMUST00000026150.15 | 26393 | ENSMUSG00000025154 | Arhgap19 | Rho GTPase activating protein 19 |
| chr19 | 30974973 | 30977484 | 2512 | * | 18,80561 | -3,887164622 | 1,251176 | -3,10681 | 0,001891 | 0,479521813 | 2,713306 | Intron (ENSMUST0000 | 19 | 31060237 | 31064469 | 4233 | 1 | 83410 | ENSMUST00000066039.8 | -82753 | ENSMUSG00000053536 | Cstf2t | cleavage stimulation factor, 3' pre-RNA subunit 2, tau |
| chrX | 1,37E+08 | 1,37E+08 | 871 | * | 7,753841 | -3,883945433 | 1,502094 | -2,58569 | 0,009719 | 1 | 0 | Intron (ENSMUST0000 | 20 | 1,37E+08 | 1,38E+08 | 1194905 | 1 | 60367 | ENSMUST00000075471.4 | -36224 | ENSMUSG00000059203 | Il1rapl2 | interleukin 1 receptor accessory protein-like 2 |
| chr3 | 1,07E+08 | 1,07E+08 | 1184 | * | 7,971833 | -3,878186359 | 1,424782 | -2,72195 | 0,00649 | 1 | 0 | Distal Intergenic | 3 | 1,07E+08 | 1,07E+08 | 52055 | 2 | 1,03E+08 | ENSMUST00000197932.2 | -21324 | ENSMUSG00000105102 | Gm35507 | predicted gene, 35507 |
| chr17 | 23687092 | 23687790 | 699 | * | 8,537059 | -3,877987508 | 1,468884 | -2,64009 | 0,008288 | 0,777172375 | 2,237403 | Intron (ENSMUST0000 | 17 | 23674147 | 23698673 | 24527 | 2 | 619788 | ENSMUST00000171996.3 | 10883 | ENSMUSG00000091407 | Vmn2r117 | vomeronasal 2, receptor 117 |
| chr14 | 85444539 | 85446300 | 1762 | * | 12,09295 | -3,876461447 | 1,262195 | -3,07121 | 0,002132 | 0,657635782 | 1,173325 | Distal Intergenic | 14 | 84696164 | 84713864 | 17701 | 2 | 319903 | ENSMUST00000228125.2 | -730675 | ENSMUSG00000115529 | 9630013A20Rik | RIKEN cDNA 9630013A20 gene |
| chr16 | 59239757 | 59241283 | 1527 | * | 13,365 | -3,875442268 | 1,277407 | -3,03384 | 0,002415 | 0,715387412 | 0,918255 | Intron (ENSMUST0000 | 16 | 59227695 | 59284867 | 57173 | 1 | 328699 | ENSMUST00000114341.4 | 12062 | ENSMUSG00000074991 | Gabrr3 | gamma-aminobutyric acid (GABA) receptor, rho 3 |
| chr16 | 18330699 | 18331447 | 749 | * | 7,311779 | -3,872878393 | 1,462811 | -2,64756 | 0,008108 | 1 | 0 | Intron (ENSMUST0000 | 16 | 18317690 | 18383132 | 65443 | 1 | 13972 | ENSMUST00000232235.2 | 13009 | ENSMUSG00000000884 | Gnb1l | guanine nucleotide binding protein (G protein), beta polypeptide 1-like |
| chr15 | 98478977 | 98480240 | 1264 | * | 11,29874 | -3,869339521 | 1,409672 | -2,74485 | 0,006054 | 0,910279554 | 0,744375 | Intron (ENSMUST0000 | 15 | 98468885 | 98487461 | 18577 | 1 | 73863 | ENSMUST00000023728.8 | 10092 | ENSMUSG00000022993 | Tex49 | testis expressed 49 |
| chr5 | 7977103 | 7978150 | 1048 | * | 7,902446 | -3,868969447 | 1,458112 | -2,65341 | 0,007968 | 0,856202617 | 1,360622 | Intron (ENSMUST0000 | 5 | 7730108 | 8010346 | 280239 | 2 | 1,19E+08 | ENSMUST00000198825.2 | 32196 | | LOC118568749 | uncharacterized LOC118568749 |
| chr5 | 1,04E+08 | 1,04E+08 | 748 | * | 8,407059 | -3,864406748 | 1,418766 | -2,72378 | 0,006454 | 0,809369408 | 1,449706 | Exon (ENSMUST00000 | 5 | 1,04E+08 | 1,04E+08 | 61 | 1 | 1,01E+08 | ENSMUST00000176065.3 | 11009 | ENSMUSG00000093690 | Mir5619 | microRNA 5619 |
| chr16 | 30110771 | 30111682 | 912 | * | 8,177001 | -3,861530896 | 1,459773 | -2,6453 | 0,008162 | 0,852537034 | 1,425929 | Distal Intergenic | 16 | 30088120 | 30102074 | 13955 | 2 | 74488 | ENSMUST00000064606.8 | -8697 | ENSMUSG00000052316 | Lrrc15 | leucine rich repeat containing 15 |
| chr5 | 39989666 | 39991042 | 1377 | * | 10,58605 | -3,859577081 | 1,435993 | -2,68774 | 0,007194 | 0,85188813 | 1,2763 | Distal Intergenic | 5 | 39771278 | 39912818 | 141541 | 2 | 15476 | ENSMUST00000117944.2 | -76848 | ENSMUSG00000051022 | Hs3st1 | heparan sulfate (glucosamine) 3-O-sulfotransferase 1 |
| chrX | 52696992 | 52698735 | 1744 | * | 14,51579 | -3,859494251 | 1,186134 | -3,25384 | 0,001139 | 0,53261455 | 1,214904 | Exon (ENSMUST00000 | 20 | 52698835 | 52712448 | 13614 | 2 | 331578 | ENSMUST00000187290.7 | 13713 | ENSMUSG00000085998 | AW822252 | expressed sequence AW822252 |
| chr5 | 8002169 | 8003216 | 1048 | * | 8,926978 | -3,858811588 | 1,399396 | -2,75748 | 0,005825 | 0,715249603 | 2,237403 | Intron (ENSMUST0000 | 5 | 7730108 | 8010346 | 280239 | 2 | 1,19E+08 | ENSMUST00000198825.2 | 7130 | | LOC118568749 | uncharacterized LOC118568749 |
| chr18 | 24958056 | 24959005 | 950 | * | 8,704512 | -3,858642686 | 1,430533 | -2,69735 | 0,006989 | 1 | 0 | Intron (ENSMUST0000 | 18 | 24842502 | 25266558 | 424057 | 1 | 225288 | ENSMUST00000234834.2 | 115554 | ENSMUSG00000034295 | Fhod3 | formin homology 2 domain containing 3 |
| chr14 | 9991682 | 9993845 | 2164 | * | 13,34183 | -3,857947422 | 1,299082 | -2,96975 | 0,00298 | 0,792225394 | 0,738441 | Intron (ENSMUST0000 | 14 | 9980246 | 10093085 | 112840 | 1 | 27062 | ENSMUST00000224581.2 | 11436 | ENSMUSG00000054423 | Cadps | Ca2+-dependent secretion activator |
| chr3 | 28201136 | 28202838 | 1703 | * | 11,33796 | -3,856354171 | 1,338358 | -2,88141 | 0,003959 | 0,841622493 | 0,738441 | Distal Intergenic | 3 | 28230702 | 28266358 | 35657 | 1 | 1,05E+08 | ENSMUST00000209618.2 | -27864 | | Gm42196 | predicted gene, 42196 |
| chr6 | 73541604 | 73542237 | 634 | * | 7,829466 | -3,855377637 | 1,470357 | -2,62207 | 0,00874 | 1 | 0 | Distal Intergenic | 6 | 73563092 | 73585656 | 22565 | 2 | 1,03E+08 | ENSMUST00000204638.2 | 43419 | ENSMUSG00000107441 | Gm31747 | predicted gene, 31747 |
| chr13 | 1,09E+08 | 1,09E+08 | 1063 | * | 10,1062 | -3,855112887 | 1,421583 | -2,71185 | 0,006691 | 0,852340527 | 1,173325 | Intron (ENSMUST0000 | 13 | 1,09E+08 | 1,09E+08 | 81 | 1 | 1E+08 | ENSMUST00000103758.4 | -11608 | ENSMUSG00000076946 | Mir582 | microRNA 582 |
| chr12 | 3741631 | 3742757 | 1127 | * | 12,66711 | -3,853390422 | 1,362151 | -2,8289 | 0,004671 | 0,832960417 | 0,918255 | Intron (ENSMUST0000 | 12 | 3736756 | 3768968 | 32213 | 1 | 13528 | ENSMUST00000173074.2 | 4875 | ENSMUSG00000071454 | Dtnb | dystrobrevin, beta |
| chr11 | 36295977 | 36296886 | 910 | * | 8,228461 | -3,852815389 | 1,451745 | -2,65392 | 0,007956 | 1 | 0 | Intron (ENSMUST0000 | 11 | 36275950 | 36323103 | 47154 | 2 | 23964 | ENSMUST00000128501.2 | 26217 | ENSMUSG00000049336 | Tenm2 | teneurin transmembrane protein 2 |
| chr14 | 1,14E+08 | 1,14E+08 | 1882 | * | 9,766931 | -3,849016612 | 1,214246 | -3,16988 | 0,001525 | 0,595550945 | 1,173325 | Distal Intergenic | 14 | 1,15E+08 | 1,15E+08 | 29363 | 1 | 75822 | ENSMUST00000226121.2 | -1491035 | ENSMUSG00000115320 | 4930524C18Rik | RIKEN cDNA 4930524C18 gene |
| chr11 | 18584502 | 18585222 | 721 | * | 8,175827 | -3,846653065 | 1,484844 | -2,59061 | 0,009581 | 0,889425844 | 1,360622 | Intron (ENSMUST0000 | 11 | 18688695 | 18688796 | 102 | 1 | 1,15E+08 | ENSMUST00000083450.4 | -103473 | ENSMUSG00000065384 | Gm23093 | predicted gene, 23093 |
| chrX | 1,68E+08 | 1,68E+08 | 1118 | * | 8,102201 | -3,841620048 | 1,430208 | -2,68606 | 0,00723 | 1 | 0 | Intron (ENSMUST0000 | 20 | 1,68E+08 | 1,68E+08 | 509337 | 1 | 11856 | ENSMUST00000033721.13 | 93278 | ENSMUSG00000031355 | Arhgap6 | Rho GTPase activating protein 6 |
| chr12 | 54036083 | 54037363 | 1281 | * | 9,958751 | -3,841085526 | 1,384937 | -2,77347 | 0,005546 | 0,782813788 | 1,449706 | Intron (ENSMUST0000 | 12 | 54138213 | 54153668 | 15456 | 2 | 1,03E+08 | ENSMUST00000219480.2 | 116305 | | Gm35818 | predicted gene, 35818 |
| chr2 | 73269418 | 73270593 | 1176 | * | 9,827624 | -3,838642569 | 1,396377 | -2,749 | 0,005978 | 0,832960417 | 1,173325 | Exon (ENSMUST00000 | 2 | 73274796 | 73283054 | 8259 | 2 | 215280 | ENSMUST00000151939.2 | 12461 | ENSMUSG00000075284 | Wipf1 | WAS/WASL interacting protein family, member 1 |
| chr11 | 79556194 | 79557477 | 1284 | * | 12,35054 | -3,831707381 | 1,442213 | -2,65682 | 0,007888 | 0,91880156 | 0,918255 | Intron (ENSMUST0000 | 11 | 79551388 | 79574324 | 22937 | 1 | 268451 | ENSMUST00000155381.2 | 4806 | ENSMUSG00000017639 | Rab11fip4 | RAB11 family interacting protein 4 (class II) |
| chr11 | 55605441 | 55606601 | 1161 | * | 10,93601 | -3,831157313 | 1,37968 | -2,77685 | 0,005489 | 0,818408612 | 1,173325 | Distal Intergenic | 11 | 55662914 | 55961674 | 298761 | 1 | 1,15E+08 | ENSMUST00000134285.2 | -56313 | | Gm12239 | predicted gene 12239 |
| chrX | 1,46E+08 | 1,46E+08 | 1548 | * | 7,315135 | -3,831143041 | 1,447338 | -2,64703 | 0,00812 | 0,858508307 | 1,360622 | Distal Intergenic | 20 | 1,46E+08 | 1,46E+08 | 26816 | 1 | 15560 | ENSMUST00000149260.8 | -3130 | ENSMUSG00000041380 | Htr2c | 5-hydroxytryptamine (serotonin) receptor 2C |
| chr6 | 1,05E+08 | 1,05E+08 | 819 | * | 9,855193 | -3,828635382 | 1,407138 | -2,72087 | 0,006511 | 0,810897439 | 1,449706 | Distal Intergenic | 6 | 1,05E+08 | 1,05E+08 | 11403 | 2 | 1,01E+08 | ENSMUST00000205113.2 | 38257 | ENSMUSG00000108237 | Gm19757 | predicted gene, 19757 |
| chr12 | 92988382 | 92989187 | 806 | * | 9,189684 | -3,826872424 | 1,339984 | -2,85591 | 0,004291 | 0,667187627 | 2,237403 | Distal Intergenic | 12 | 93383232 | 93411597 | 28366 | 2 | 75295 | ENSMUST00000223202.2 | 422410 | ENSMUSG00000113164 | 4930559C10Rik | RIKEN cDNA 4930559C10 gene |

| | | | | | | | | | | | | | | | | | | | | | | | |
|---|---|---|---|---|---|---|---|---|---|---|---|---|---|---|---|---|---|---|---|---|---|---|---|
| chr16 | 52309776 | 52311018 | 1243 | * | 11,30135 | -3,826679946 | 1,468389 | -2,60604 | 0,00916 | 0,974006322 | 0,738441 | Distal Intergenic | 16 | 52069359 | 52274437 | 205079 | 2 | 11658 | ENSMUST00000023312.14 | -35339 | ENSMUSG00000022636 | Alcam | activated leukocyte cell adhesion molecule |
| chr15 | 1,03E+08 | 1,03E+08 | 1107 | * | 9,521876 | -3,825148668 | 1,388419 | -2,75504 | 0,005869 | 0,793243838 | 1,425929 | Promoter (1-2kb) | 15 | 1,03E+08 | 1,03E+08 | 19662 | 1 | 105855 | ENSMUST00000230276.2 | 1384 | ENSMUSG00000022488 | Nckap1l | NCK associated protein 1 like |
| chr15 | 84342514 | 84343448 | 935 | * | 12,0663 | -3,824931571 | 1,337511 | -2,85974 | 0,00424 | 0,853363379 | 0,738441 | Distal Intergenic | 15 | 84263406 | 84331298 | 67893 | 2 | 72301 | ENSMUST00000239449.2 | -11216 | ENSMUSG00000062760 | Shisal1 | shisa like 1 |
| chr11 | 19569455 | 19570721 | 1267 | * | 13,2259 | -3,824344164 | 1,166679 | -3,27797 | 0,001046 | 0,568412386 | 0,918255 | Distal Intergenic | 11 | 19643155 | 19643298 | 144 | 2 | 1,15E+08 | ENSMUST00000158585.3 | 72577 | ENSMUSG00000089210 | Gm23386 | predicted gene, 23386 |
| chr9 | 76872161 | 76873533 | 1373 | * | 9,059583 | -3,822004408 | 1,247669 | -3,06332 | 0,002189 | 0,625646014 | 1,449706 | Intron (ENSMUST0000 | 9 | 76858975 | 76953031 | 94057 | 2 | 26944 | ENSMUST00000184897.8 | 79498 | ENSMUSG00000032357 | Tinag | tubulointerstitial nephritis antigen |
| chr15 | 29071024 | 29072095 | 1072 | * | 11,25037 | -3,821408434 | 1,228436 | -3,11079 | 0,001866 | 0,675576354 | 0,918255 | Distal Intergenic | 15 | 29713477 | 29713550 | 74 | 2 | 1,01E+08 | ENSMUST00000174898.3 | 641455 | ENSMUSG00000092639 | Mir3964 | microRNA 3964 |
| chrX | 85103826 | 85105377 | 1552 | * | 10,37193 | -3,821179234 | 1,214674 | -3,14585 | 0,001656 | 0,611772917 | 1,173325 | Distal Intergenic | 20 | 85087805 | 85090919 | 3115 | 1 | 73495 | ENSMUST00000047945.2 | 16021 | ENSMUSG00000035454 | Samt3 | spermatogenesis associated multipass transmembrane protein 3 |
| chr11 | 4169972 | 4171920 | 1949 | * | 16,83262 | -3,820421365 | 1,282984 | -2,97776 | 0,002904 | 0,603645287 | 2,141612 | Promoter (1-2kb) | 11 | 4168225 | 4172409 | 4185 | 1 | 71962 | ENSMUST00000020699.4 | 1747 | ENSMUSG00000020424 | Castor1 | cytosolic arginine sensor for mTORC1 subunit 1 |
| chr16 | 45583615 | 45584905 | 1291 | * | 10,99392 | -3,819948935 | 1,248363 | -3,05997 | 0,002214 | 0,736639439 | 0,744375 | Intron (ENSMUST0000 | 16 | 45583063 | 45592646 | 9584 | 2 | 208177 | ENSMUST00000151300.2 | 7741 | ENSMUSG00000033149 | Phldb2 | pleckstrin homology like domain, family B, member 2 |
| chr11 | 91209443 | 91210706 | 1264 | * | 10,36469 | -3,81847365 | 1,271742 | -3,00255 | 0,002677 | 0,732200941 | 0,918255 | Distal Intergenic | 11 | 91466141 | 91468384 | 2244 | 2 | 73470 | ENSMUST00000061019.6 | 257678 | ENSMUSG00000046755 | Kif2b | kinesin family member 2B |
| chr12 | 90216703 | 90217716 | 1014 | * | 8,510355 | -3,816493496 | 1,478588 | -2,58117 | 0,009846 | 0,892743851 | 1,360622 | Intron (ENSMUST0000 | 12 | 90366441 | 90375598 | 9158 | 2 | 1,05E+08 | ENSMUST00000220610.2 | 157882 | ENSMUSG00000113001 | Gm40538 | predicted gene, 40538 |
| chr14 | 78282601 | 78284592 | 1992 | * | 11,51071 | -3,816088778 | 1,320668 | -2,88951 | 0,003858 | 0,836571183 | 0,744375 | Intron (ENSMUST0000 | 14 | 78320183 | 78326421 | 6239 | 2 | 219170 | ENSMUST00000173826.2 | 41829 | ENSMUSG00000045655 | Fam216b | family with sequence similarity 216, member B |
| chr8 | 1,09E+08 | 1,09E+08 | 1018 | * | 13,91516 | -3,816051919 | 1,140621 | -3,34559 | 0,000821 | 0,496560937 | 1,075735 | Intron (ENSMUST0000 | 8 | 1,09E+08 | 1,09E+08 | 9318 | 2 | 1,05E+08 | ENSMUST00000210837.2 | 61435 | | Gm39244 | predicted gene, 39244 |
| chr8 | 95419585 | 95420556 | 972 | * | 8,486029 | -3,813007015 | 1,442221 | -2,64384 | 0,008197 | 1 | 0 | Promoter (2-3kb) | 8 | 95401522 | 95422906 | 21385 | 2 | 67801 | ENSMUST00000034227.6 | 2350 | ENSMUSG00000031775 | Pllp | plasma membrane proteolipid |
| chr10 | 29357766 | 29358908 | 1143 | * | 9,125211 | -3,812552995 | 1,275639 | -2,98874 | 0,002801 | 0,669639313 | 1,425929 | Intron (ENSMUST0000 | 10 | 29328412 | 29380726 | 52315 | 2 | 72780 | ENSMUST00000215256.2 | 21818 | ENSMUSG00000019880 | Rspo3 | R-spondin 3 |
| chr10 | 99249379 | 99250788 | 1410 | * | 11,82775 | -3,81084214 | 1,45389 | -2,62114 | 0,008764 | 0,899471409 | 1,173325 | Intron (ENSMUST0000 | 10 | 99238652 | 99249134 | 10483 | 1 | 1,19E+08 | ENSMUST00000218725.2 | 10727 | | LOC118567817 | uncharacterized LOC118567817 |
| chr10 | 44302068 | 44303383 | 1316 | * | 8,955365 | -3,807336872 | 1,461321 | -2,60541 | 0,009177 | 0,900637308 | 1,211315 | Promoter (1-2kb) | 10 | 44294396 | 44300320 | 5925 | 2 | 1,03E+08 | ENSMUST00000215400.2 | -1748 | | Gm35028 | predicted gene, 35028 |
| chr5 | 66969592 | 66970404 | 813 | * | 11,23504 | -3,806914781 | 1,447106 | -2,63071 | 0,008521 | 0,931010269 | 0,918255 | Intron (ENSMUST0000 | 5 | 66903232 | 67111021 | 207790 | 1 | 77569 | ENSMUST00000130228.2 | 66360 | ENSMUSG00000037736 | Limch1 | LIM and calponin homology domains 1 |
| chr7 | 41319606 | 41320613 | 1008 | * | 11,30907 | -3,805165424 | 1,347933 | -2,82296 | 0,004758 | 0,792545355 | 1,173325 | Promoter (2-3kb) | 7 | 41317438 | 41376617 | 59180 | 1 | 75835 | ENSMUST00000150147.8 | 2168 | ENSMUSG00000074164 | Vmn2r-ps54 | vomeronasal 2, receptor, pseudogene 54 |
| chr11 | 73154106 | 73155074 | 969 | * | 9,481214 | -3,804511905 | 1,439446 | -2,64304 | 0,008217 | 0,852340527 | 1,449706 | Distal Intergenic | 11 | 73158214 | 73191189 | 32976 | 1 | 246788 | ENSMUST00000049676.3 | -3140 | ENSMUSG00000043029 | Trpv3 | transient receptor potential cation channel, subfamily V, member 3 |
| chr11 | 1,1E+08 | 1,1E+08 | 923 | * | 12,08382 | -3,803964457 | 1,173899 | -3,24045 | 0,001193 | 0,595550945 | 0,918255 | Intron (ENSMUST0000 | 11 | 1,1E+08 | 1,1E+08 | 67448 | 2 | 217262 | ENSMUST00000044850.4 | -17767 | ENSMUSG00000041797 | Abca9 | ATP-binding cassette, sub-family A (ABC1), member 9 |
| chr13 | 71745980 | 71747083 | 1104 | * | 9,418306 | -3,803509969 | 1,407067 | -2,70315 | 0,006869 | 0,8315492 | 1,360622 | Intron (ENSMUST0000 | 13 | 71635726 | 71760828 | 125103 | 1 | 1,03E+08 | ENSMUST00000221719.2 | 110254 | | 1700112M02Rik | RIKEN cDNA 1700112M02 gene |
| chr18 | 58193003 | 58194087 | 1085 | * | 10,21934 | -3,802857012 | 1,360524 | -2,79514 | 0,005188 | 0,801562884 | 1,211315 | Intron (ENSMUST0000 | 18 | 58160598 | 58169050 | 8453 | 2 | 14119 | ENSMUST00000235281.2 | -23953 | ENSMUSG00000024598 | Fbn2 | fibrillin 2 |
| chr2 | 87113232 | 87114289 | 1058 | * | 7,026898 | -3,800612585 | 1,466147 | -2,59225 | 0,009535 | 0,88871498 | 1,360622 | Exon (ENSMUST00000 | 2 | 87099190 | 87100113 | 924 | 1 | 257875 | ENSMUST00000090717.5 | 14042 | ENSMUSG00000070855 | Olfr1116 | olfactory receptor 1116 |
| chr18 | 65009157 | 65010831 | 1675 | * | 12,42242 | -3,797128532 | 1,33539 | -2,84346 | 0,004463 | 0,781219703 | 1,173325 | Distal Intergenic | 18 | 65020776 | 65346475 | 325700 | 1 | 83814 | ENSMUST00000235343.2 | -9945 | ENSMUSG00000024589 | Nedd4l | neural precursor cell expressed, developmentally down-regulated gene 4-like |
| chr2 | 97640415 | 97641535 | 1121 | * | 11,16527 | -3,795799581 | 1,209633 | -3,13798 | 0,001701 | 0,660698266 | 0,918255 | Distal Intergenic | 2 | 97298434 | 97459408 | 160975 | 1 | 241568 | ENSMUST00000170144.2 | 341981 | ENSMUSG00000050587 | Lrrc4c | leucine rich repeat containing 4C |
| chr7 | 16405165 | 16406568 | 1404 | * | 14,7866 | -3,793045678 | 1,276812 | -2,97071 | 0,002971 | 0,704364304 | 1,214904 | 3' UTR | 7 | 16405256 | 16409630 | 4375 | 2 | 101434 | ENSMUST00000108498.3 | 3062 | ENSMUSG00000078795 | Ceacam15 | carcinoembryonic antigen-related cell adhesion molecule 15 |
| chr9 | 82517437 | 82518091 | 655 | * | 7,747476 | -3,792169612 | 1,432877 | -2,64654 | 0,008132 | 0,858623476 | 1,360622 | Distal Intergenic | 9 | 82711593 | 82729740 | 18148 | 1 | 65099 | ENSMUST00000113245.9 | -193502 | ENSMUSG00000032251 | Irak1bp1 | interleukin-1 receptor-associated kinase 1 binding protein 1 |
| chr1 | 1,57E+08 | 1,57E+08 | 1371 | * | 10,90535 | -3,78766644 | 1,440488 | -2,62943 | 0,008553 | 0,962965811 | 0,744375 | Distal Intergenic | 1 | 1,57E+08 | 1,57E+08 | 30052 | 1 | 226527 | ENSMUST00000193791.6 | -13061 | ENSMUSG00000033488 | Cryzl2 | crystallin zeta like 2 |
| chr5 | 41052400 | 41053632 | 1233 | * | 9,268522 | -3,785693342 | 1,4298 | -2,64771 | 0,008104 | 0,881662676 | 1,211315 | Distal Intergenic | 5 | 41783076 | 41784845 | 1770 | 2 | 100972 | ENSMUST00000200702.2 | 731213 | ENSMUSG00000029128 | Rab28 | RAB28, member RAS oncogene family |
| chr3 | 57614399 | 57615605 | 1207 | * | 10,21682 | -3,784914654 | 1,395442 | -2,71234 | 0,006681 | 0,852340527 | 1,173325 | Distal Intergenic | 3 | 57564814 | 57600220 | 35407 | 2 | 242037 | ENSMUST00000200665.2 | -14179 | ENSMUSG00000074591 | Ankub1 | ankyrin repeat and ubiquitin domain containing 1 |
| chr8 | 1,09E+08 | 1,09E+08 | 1055 | * | 11,29907 | -3,783220661 | 1,463143 | -2,58568 | 0,009719 | 0,913502848 | 1,173325 | Intron (ENSMUST0000 | 8 | 1,09E+08 | 1,09E+08 | 396257 | 1 | 11906 | ENSMUST00000221632.2 | 223565 | ENSMUSG00000038872 | Zfhx3 | zinc finger homeobox 3 |
| chr14 | 48378262 | 48379368 | 1107 | * | 9,054333 | -3,777625384 | 1,374175 | -2,74901 | 0,005977 | 0,805916009 | 1,360622 | Intron (ENSMUST0000 | 14 | 48359486 | 48477769 | 118284 | 1 | 93834 | ENSMUST00000226260.2 | 18776 | ENSMUSG00000021846 | Peli2 | pellino 2 |
| chr11 | 44773728 | 44774991 | 1264 | * | 10,76528 | -3,776254592 | 1,193886 | -3,16299 | 0,001562 | 0,681307794 | 0,744375 | Exon (ENSMUST00000 | 11 | 44802965 | 44863741 | 60777 | 1 | 13591 | ENSMUST00000140822.2 | -27974 | ENSMUSG00000057098 | Ebf1 | early B cell factor 1 |
| chr16 | 7185328 | 7186738 | 1411 | * | 11,07576 | -3,774564946 | 1,366527 | -2,76216 | 0,005742 | 0,903101391 | 0,744375 | Intron (ENSMUST0000 | 16 | 7170834 | 7227833 | 57000 | 1 | 268859 | ENSMUST00000231031.2 | 14494 | ENSMUSG00000008658 | Rbfox1 | RNA binding protein, fox-1 homolog (C. elegans) 1 |
| chr13 | 27213930 | 27215274 | 1345 | * | 11,22214 | -3,773124084 | 1,243628 | -3,03397 | 0,002414 | 0,750475959 | 0,744375 | Intron (ENSMUST0000 | 13 | 27241553 | 27249187 | 7635 | 1 | 19109 | ENSMUST00000110369.10 | -26279 | ENSMUSG00000021342 | Prl | prolactin |
| chr13 | 26143399 | 26144461 | 1063 | * | 8,993784 | -3,772078308 | 1,403017 | -2,68855 | 0,007176 | 0,838625535 | 1,360622 | Intron (ENSMUST0000 | 13 | 26105016 | 26150299 | 45284 | 2 | 1,01E+08 | ENSMUST00000139335.3 | 5838 | ENSMUSG00000086531 | Gm11351 | predicted gene 11351 |
| chr2 | 82247738 | 82249084 | 1347 | * | 9,241637 | -3,771824334 | 1,39176 | -2,71011 | 0,006726 | 0,849859156 | 1,211315 | Distal Intergenic | 2 | 82205716 | 82222546 | 16831 | 2 | 228116 | ENSMUST00000128608.2 | -25192 | ENSMUSG00000087061 | A130030D18Rik | RIKEN cDNA A130030D18 gene |
| chr9 | 97132759 | 97133725 | 967 | * | 10,22614 | -3,771518895 | 1,383427 | -2,72621 | 0,006407 | 0,845747925 | 1,173325 | Distal Intergenic | 9 | 97157765 | 97163946 | 6182 | 1 | 1,03E+08 | ENSMUST00000245996.1 | -24040 | | Gm33302 | predicted gene, 33302 |
| chr7 | 1,41E+08 | 1,41E+08 | 1733 | * | 14,99648 | -3,770497553 | 1,241042 | -3,03817 | 0,00238 | 0,564733506 | 2,141612 | Promoter (<=1kb) | 7 | 1,41E+08 | 1,41E+08 | 1090 | 2 | 71345 | ENSMUST00000210927.2 | 0 | ENSMUSG00000054662 | Ano9 | anoctamin 9 |
| chr12 | 42181052 | 42181802 | 751 | * | 8,653357 | -3,769564846 | 1,416826 | -2,66057 | 0,007801 | 0,852732156 | 1,360622 | Intron (ENSMUST0000 | 12 | 41965491 | 42063539 | 98049 | 2 | 1,08E+08 | ENSMUST00000222917.2 | -117513 | | Gm46348 | predicted gene, 46348 |
| chrX | 75438887 | 75440434 | 1548 | * | 9,470036 | -3,767538581 | 1,396414 | -2,69801 | 0,006976 | 0,853738252 | 1,211315 | Promoter (1-2kb) | 20 | 75436956 | 75441585 | 4630 | 1 | 619294 | ENSMUST00000101419.2 | 1931 | ENSMUSG00000073085 | Cldn34b4 | claudin 34B4 |
| chr1 | 98290998 | 98291547 | 550 | * | 8,542131 | -3,764309652 | 1,451013 | -2,59426 | 0,009479 | 1 | 0 | Distal Intergenic | 1 | 98348849 | 98437105 | 88257 | 1 | 70866 | ENSMUST00000027575.13 | -57302 | ENSMUSG00000026336 | Slco6d1 | solute carrier organic anion transporter family, member 6d1 |
| chr4 | 46107066 | 46108064 | 999 | * | 8,596024 | -3,76274277 | 1,398497 | -2,69056 | 0,007133 | 0,749550311 | 2,237403 | Intron (ENSMUST0000 | 4 | 46100243 | 46114439 | 14197 | 1 | 21916 | ENSMUST00000155730.2 | 6823 | ENSMUSG00000028328 | Tmod1 | tropomodulin 1 |
| chr10 | 20306485 | 20307883 | 1399 | * | 15,53092 | -3,760335675 | 1,11735 | -3,36541 | 0,000764 | 0,365634218 | 2,141612 | Intron (ENSMUST0000 | 10 | 20223753 | 20236641 | 12889 | 1 | 71804 | ENSMUST00000213374.2 | 82732 | ENSMUSG00000019992 | Mtfr2 | mitochondrial fission regulator 2 |
| chr13 | 39442051 | 39443403 | 1353 | * | 11,37252 | -3,759805008 | 1,259435 | -2,98531 | 0,002833 | 0,700182401 | 1,173325 | Distal Intergenic | 13 | 39116112 | 39144851 | 28740 | 2 | 108652 | ENSMUST00000021870.12 | -297200 | ENSMUSG00000021432 | Slc35b3 | solute carrier family 35, member B3 |
| chr12 | 53820490 | 53821829 | 1340 | * | 11,50906 | -3,75705236 | 1,275703 | -2,94508 | 0,003229 | 0,72352563 | 1,173325 | Intron (ENSMUST0000 | 12 | 54138213 | 54153668 | 15456 | 2 | 1,03E+08 | ENSMUST00000219480.2 | 331839 | | Gm35818 | predicted gene, 35818 |
| chr7 | 68022512 | 68023710 | 1199 | * | 10,21448 | -3,756555852 | 1,356054 | -2,77021 | 0,005602 | 0,788774532 | 1,425929 | Distal Intergenic | 7 | 67988416 | 68002949 | 14534 | 2 | 1,05E+08 | ENSMUST00000129965.2 | -19563 | | Gm16158 | predicted gene 16158 |
| chrX | 1,54E+08 | 1,54E+08 | 1930 | * | 12,37212 | -3,752719351 | 1,322289 | -2,83805 | 0,004539 | 0,798887389 | 1,075735 | Intron (ENSMUST0000 | 20 | 1,54E+08 | 1,55E+08 | 721113 | 1 | 1,03E+08 | ENSMUST00000112542.2 | 36953 | | Gm15155 | predicted gene 15155 |
| chr19 | 42615364 | 42616255 | 892 | * | 9,372983 | -3,75200106 | 1,451356 | -2,58517 | 0,009733 | 0,908873708 | 1,211315 | Distal Intergenic | 19 | 42583545 | 42601252 | 17708 | 2 | 67573 | ENSMUST00000164786.8 | -14112 | ENSMUSG00000025185 | Loxl4 | lysyl oxidase-like 4 |
| chr5 | 1,28E+08 | 1,28E+08 | 827 | * | 7,067973 | -3,750492042 | 1,395402 | -2,68775 | 0,007194 | 1 | 0 | Intron (ENSMUST0000 | 5 | 1,28E+08 | 1,28E+08 | 32839 | 1 | 319804 | ENSMUST00000134529.2 | 281965 | ENSMUSG00000049971 | Glt1d1 | glycosyltransferase 1 domain containing 1 |
| chr1 | 1,91E+08 | 1,91E+08 | 1339 | * | 10,55992 | -3,749390411 | 1,341214 | -2,79552 | 0,005182 | 0,851959528 | 0,918255 | Promoter (1-2kb) | 1 | 1,91E+08 | 1,91E+08 | 41664 | 1 | 226856 | ENSMUST00000130876.2 | 1709 | ENSMUSG00000026623 | Lpgat1 | lysophosphatidylglycerol acyltransferase 1 |
| chr4 | 27361324 | 27362119 | 796 | * | 7,383626 | -3,749126414 | 1,401596 | -2,6749 | 0,007475 | 1 | 0 | Distal Intergenic | 4 | 27700671 | 27700777 | 107 | 1 | 1,15E+08 | ENSMUST00000179794.2 | -338552 | ENSMUSG00000095003 | Gm25173 | predicted gene, 25173 |
| chr3 | 31018828 | 31020258 | 1431 | * | 12,57374 | -3,747533589 | 1,173386 | -3,19378 | 0,001404 | 0,563032042 | 1,2763 | Promoter (<=1kb) | 3 | 31012536 | 31020084 | 7549 | 2 | 241915 | ENSMUST00000150939.3 | 0 | ENSMUSG00000037652 | Phc3 | polyhomeotic 3 |
| chr1 | 1,23E+08 | 1,23E+08 | 907 | * | 8,9467 | -3,743851382 | 1,327947 | -2,81928 | 0,004813 | 0,79031971 | 1,211315 | Distal Intergenic | 1 | 1,22E+08 | 1,22E+08 | 144 | 1 | 1,15E+08 | ENSMUST00000158565.3 | 81450 | ENSMUSG00000089190 | Gm25771 | predicted gene, 25771 |
| chr4 | 1,56E+08 | 1,56E+08 | 1335 | * | 9,39893 | -3,743596016 | 1,412985 | -2,64942 | 0,008063 | 0,770994815 | 2,237403 | 3' UTR | 4 | 1,56E+08 | 1,56E+08 | 4430 | 1 | 14688 | ENSMUST00000176308.2 | 3208 | ENSMUSG00000029064 | Gnb1 | guanine nucleotide binding protein (G protein), beta 1 |
| chr5 | 1,44E+08 | 1,44E+08 | 1395 | * | 10,0114 | -3,742437448 | 1,449349 | -2,58215 | 0,009819 | 0,910106437 | 1,211315 | Intron (ENSMUST0000 | 5 | 1,44E+08 | 1,44E+08 | 3126 | 1 | 1,01E+08 | ENSMUST00000197311.2 | -9300 | ENSMUSG00000104600 | Dmrt1i | Dmrt1 interacting ncRNA |
| chr1 | 80222915 | 80224210 | 1296 | * | 9,615339 | -3,741597413 | 1,393663 | -2,68472 | 0,007259 | 0,832960417 | 1,425929 | Distal Intergenic | 1 | 80202761 | 80207332 | 4572 | 1 | 69565 | ENSMUST00000191452.3 | 20154 | | 2310015K22Rik | RIKEN cDNA 2310015K22 gene |
| chr5 | 1,01E+08 | 1,01E+08 | 1615 | * | 10,22196 | -3,739167348 | 1,377132 | -2,71519 | 0,006624 | 0,861872447 | 1,086118 | Distal Intergenic | 5 | 1,01E+08 | 1,01E+08 | 3707 | 2 | 1,03E+08 | ENSMUST00000199211.2 | -334851 | ENSMUSG00000106255 | Gm36793 | predicted gene, 36793 |
| chr14 | 6813474 | 6815170 | 1697 | * | 8,189403 | -3,739153211 | 1,430835 | -2,61327 | 0,008968 | 0,878632224 | 1,360622 | Distal Intergenic | 14 | 6685105 | 6699664 | 14560 | 2 | 1,03E+08 | ENSMUST00000224585.2 | -113810 | | Gm31804 | predicted gene, 31804 |
| chr5 | 1,19E+08 | 1,19E+08 | 1346 | * | 12,09483 | -3,737972042 | 1,274035 | -2,93396 | 0,003347 | 0,715249603 | 1,2763 | Distal Intergenic | 5 | 1,19E+08 | 1,19E+08 | 5535 | 2 | 665194 | ENSMUST00000200918.2 | -149533 | ENSMUSG00000107334 | Gm7538 | predicted gene 7538 |
| chr7 | 53116446 | 53117645 | 1200 | * | 10,59026 | -3,737818593 | 1,11649 | -3,34783 | 0,000814 | 0,479393375 | 1,173325 | Distal Intergenic | 7 | 53541509 | 53541637 | 129 | 2 | 1,02E+08 | ENSMUST00000184140.3 | 423992 | ENSMUSG00000099176 | Mir6238 | microRNA 6238 |
| chr7 | 81260327 | 81261767 | 1441 | * | 13,38347 | -3,734596964 | 1,213854 | -3,07664 | 0,002093 | 0,685026746 | 0,938682 | 3' UTR | 7 | 81264100 | 81268653 | 4554 | 2 | 26557 | ENSMUST00000207991.2 | 6886 | ENSMUSG00000025813 | Homer2 | homer scaffolding protein 2 |
| chr12 | 43060443 | 43062045 | 1603 | * | 15,50912 | -3,731523224 | 1,100769 | -3,38992 | 0,000699 | 0,428016132 | 1,37683 | Promoter (<=1kb) | 12 | 43013667 | 43062242 | 48576 | 2 | 1,03E+08 | ENSMUST00000220044.2 | 197 | ENSMUSG00000112407 | Gm33340 | predicted gene, 33340 |
| chr4 | 82413361 | 82415605 | 2245 | * | 16,51062 | -3,730293676 | 1,251928 | -2,97964 | 0,002886 | 0,581453237 | 2,386008 | Promoter (1-2kb) | 4 | 82416781 | 82417553 | 773 | 2 | 18028 | ENSMUST00000152588.2 | 1948 | ENSMUSG00000008575 | Nfib | nuclear factor I/B |
| chr11 | 83962677 | 83964448 | 1772 | * | 11,8183 | -3,728887362 | 1,335329 | -2,79249 | 0,00523 | 0,890233764 | 0,738441 | Promoter (2-3kb) | 11 | 83939806 | 83960087 | 20282 | 2 | 56405 | ENSMUST00000138208.2 | -2590 | ENSMUSG00000018648 | Dusp14 | dual specificity phosphatase 14 |
| chr14 | 54159706 | 54160877 | 1172 | * | 14,35036 | -3,727388396 | 1,0341 | -3,60448 | 0,000313 | 0,332694435 | 1,075735 | Intron (ENSMUST0000 | 14 | 54155784 | 54198972 | 43189 | 2 | 1,03E+08 | ENSMUST00000198142.2 | 38095 | | Gm30392 | predicted gene, 30392 |
| chr4 | 99373892 | 99375239 | 1348 | * | 12,45707 | -3,723672329 | 1,170169 | -3,18217 | 0,001462 | 0,569556737 | 1,2763 | Distal Intergenic | 4 | 99544536 | 99546859 | 2324 | 1 | 15221 | ENSMUST00000087285.5 | -169297 | ENSMUSG00000067261 | Foxd3 | forkhead box D3 |
| chr4 | 22557976 | 22558775 | 800 | * | 8,70538 | -3,720259161 | 1,431482 | -2,59889 | 0,009353 | 0,886332554 | 1,360622 | Distal Intergenic | 4 | 22490548 | 22493126 | 2579 | 1 | 1,06E+08 | ENSMUST00000203909.2 | 67428 | ENSMUSG00000107859 | Gm30731 | predicted gene, 30731 |
| chr18 | 79892061 | 79893547 | 1487 | * | 11,45481 | -3,718326843 | 1,26246 | -2,9453 | 0,003226 | 0,767061106 | 0,918255 | Distal Intergenic | 18 | 80059500 | 80059745 | 246 | 1 | 1,03E+08 | ENSMUST00000237241.2 | -165953 | ENSMUSG00000118357 | Gm36627 | predicted gene, 36627 |
| chr2 | 1,57E+08 | 1,57E+08 | 2142 | * | 19,2132 | -3,716317708 | 1,009626 | -3,68088 | 0,000232 | 0,182130938 | 2,713306 | 3' UTR | 2 | 1,57E+08 | 1,57E+08 | 8919 | 2 | 56045 | ENSMUST00000160213.2 | -4240 | ENSMUSG00000027639 | Samhd1 | SAM domain and HD domain, 1 |
| chr17 | 83414308 | 83415103 | 796 | * | 10,15503 | -3,715125208 | 1,373588 | -2,70469 | 0,006837 | 0,868766096 | 1,086118 | Intron (ENSMUST0000 | 17 | 83405695 | 83423064 | 17370 | 2 | 1,08E+08 | ENSMUST00000234289.2 | 7961 | | Gm46606 | predicted gene, 46606 |
| chr19 | 15650699 | 15652268 | 1570 | * | 13,71032 | -3,712573573 | 1,214678 | -3,05643 | 0,00224 | 0,650529916 | 1,2763 | Distal Intergenic | 19 | 15657368 | 15657474 | 107 | 1 | 1,15E+08 | ENSMUST00000083389.3 | -5100 | ENSMUSG00000118757 | Gm24319 | predicted gene, 24319 |
| chr9 | 1,06E+08 | 1,06E+08 | 836 | * | 9,491111 | -3,71177023 | 1,421776 | -2,61066 | 0,009037 | 0,865559856 | 1,449706 | Promoter (<=1kb) | 9 | 1,06E+08 | 1,06E+08 | 2193 | 1 | 75669 | ENSMUST00000188385.2 | -695 | ENSMUSG00000032571 | Pik3r4 | phosphoinositide-3-kinase regulatory subunit 4 |
| chr14 | 13087365 | 13088821 | 1457 | * | 13,86985 | -3,710643102 | 1,172265 | -3,16536 | 0,001549 | 0,616178941 | 1,075735 | Distal Intergenic | 14 | 12583005 | 12585578 | 2574 | 2 | 1E+08 | ENSMUST00000224435.2 | -501787 | ENSMUSG00000089686 | Mnd1-ps | Mnd1 retrotransposed pseudogene |
| chr2 | 1,16E+08 | 1,16E+08 | 1073 | * | 10,52272 | -3,709747125 | 1,376204 | -2,69564 | 0,007025 | 0,93486226 | 0,738441 | Distal Intergenic | 2 | 1,16E+08 | 1,16E+08 | 932 | 2 | 69974 | ENSMUST00000195368.2 | -349071 | | 2810405F15Rik | RIKEN cDNA 2810405F15 gene |
| chr16 | 18555961 | 18557264 | 1304 | * | 8,278526 | -3,706707413 | 1,428245 | -2,59529 | 0,009451 | 0,877294452 | 1,449706 | Distal Intergenic | 16 | 18595597 | 18597012 | 1416 | 1 | 12741 | ENSMUST00000043577.3 | -38333 | ENSMUSG00000041378 | Cldn5 | claudin 5 |
| chr17 | 43146535 | 43148288 | 1754 | * | 15,16534 | -3,702244073 | 1,049629 | -3,52719 | 0,00042 | 0,282588188 | 2,141612 | Intron (ENSMUST0000 | 17 | 43103843 | 43110779 | 6937 | 2 | 12488 | ENSMUST00000233350.2 | -35756 | ENSMUSG00000061665 | Cd2ap | CD2-associated protein |
| chr11 | 81541268 | 81542591 | 1324 | * | 13,14458 | -3,701228982 | 1,15657 | -3,20018 | 0,001373 | 0,594430593 | 1,075735 | Intron (ENSMUST0000 | 11 | 81478187 | 81484304 | 6118 | 1 | 1,03E+08 | ENSMUST00000156193.2 | 63081 | | Gm11418 | predicted gene 11418 |
| chr12 | 81443593 | 81444984 | 1392 | * | 8,708841 | -3,699624097 | 1,408592 | -2,62647 | 0,008628 | 0,78560377 | 2,237403 | Intron (ENSMUST0000 | 12 | 81423765 | 81426238 | 2474 | 2 | 214321 | ENSMUST00000062182.8 | -17355 | ENSMUSG00000072974 | Gm4787 | predicted gene 4787 |
| chr14 | 59777737 | 59778974 | 1238 | * | 11,95293 | -3,697561399 | 1,393819 | -2,65283 | 0,007982 | 0,954588896 | 0,744375 | Intron (ENSMUST0000 | 14 | 59798345 | 59835285 | 36941 | 2 | 71891 | ENSMUST00000022555.11 | 56311 | ENSMUSG00000021982 | Cdadc1 | cytidine and dCMP deaminase domain containing 1 |
| chr7 | 1,07E+08 | 1,07E+08 | 843 | * | 7,554379 | -3,697379897 | 1,393173 | -2,65393 | 0,007956 | 0,85617852 | 1,360622 | Promoter (2-3kb) | 7 | 1,07E+08 | 1,07E+08 | 22777 | 1 | 244198 | ENSMUST00000239087.2 | 2214 | ENSMUSG00000051041 | Olfml1 | olfactomedin-like 1 |
| chr8 | 4524417 | 4525061 | 645 | * | 8,084022 | -3,691631722 | 1,379309 | -2,67644 | 0,007441 | 1 | 0 | Distal Intergenic | 8 | 4543026 | 4568641 | 25616 | 1 | 67260 | ENSMUST00000177010.10 | -17965 | ENSMUSG00000008206 | Cers4 | ceramide synthase 4 |
| chr19 | 26891535 | 26893283 | 1749 | * | 13,04173 | -3,690774027 | 1,203662 | -3,06629 | 0,002167 | 0,695892636 | 0,918255 | Distal Intergenic | 19 | 26863325 | 26866047 | 2723 | 1 | 329047 | ENSMUST00000099536.4 | 28210 | ENSMUSG00000074913 | Gm815 | predicted gene 815 |
| chr14 | 1,01E+08 | 1,01E+08 | 1889 | * | 14,2864 | -3,682912604 | 1,055492 | -3,48929 | 0,000484 | 0,430943288 | 0,938682 | Distal Intergenic | 14 | 1,01E+08 | 1,01E+08 | 2380 | 2 | 76627 | ENSMUST00000057718.11 | 136663 | ENSMUSG00000042888 | Prr30 | proline rich 30 |
| chr13 | 53823714 | 53825160 | 1447 | * | 11,20747 | -3,682592809 | 1,330297 | -2,76825 | 0,005636 | 0,900637308 | 0,744375 | Distal Intergenic | 13 | 53802621 | 53804116 | 1496 | 1 | 75252 | ENSMUST00000221718.2 | 21093 | ENSMUSG00000113329 | 4930555G21Rik | RIKEN cDNA 4930555G21 gene |
| chr9 | 1,22E+08 | 1,22E+08 | 1207 | * | 9,815865 | -3,681935246 | 1,401236 | -2,62763 | 0,008598 | 0,860455002 | 1,425929 | Promoter (<=1kb) | 9 | 1,22E+08 | 1,22E+08 | 12894 | 2 | 1,08E+08 | ENSMUST00000216452.2 | 41 | ENSMUSG00000097572 | Gm26797 | predicted gene, 26797 |
| chr4 | 1,06E+08 | 1,06E+08 | 1756 | * | 12,67698 | -3,679414028 | 1,349359 | -2,72679 | 0,006395 | 0,921073486 | 0,738441 | Distal Intergenic | 4 | 1,06E+08 | 1,06E+08 | 113 | 2 | 1,15E+08 | ENSMUST00000157308.3 | 225046 | ENSMUSG00000087933 | Gm26047 | predicted gene, 26047 |
| chr9 | 51690124 | 51691909 | 1786 | * | 14,83165 | -3,677546656 | 1,326728 | -2,77189 | 0,005573 | 0,814686254 | 1,214904 | Intron (ENSMUST0000 | 9 | 51676960 | 51761951 | 84992 | 1 | 244867 | ENSMUST00000130405.2 | 13164 | ENSMUSG00000053199 | Arhgap20 | Rho GTPase activating protein 20 |
| chr7 | 1,39E+08 | 1,39E+08 | 796 | * | 10,41948 | -3,677094764 | 1,422984 | -2,58407 | 0,009764 | 0,984317753 | 0,738441 | Exon (ENSMUST00000 | 7 | 1,39E+08 | 1,39E+08 | 4339 | 1 | 74004 | ENSMUST00000238324.2 | -14721 | ENSMUSG00000056856 | Jakmip3 | janus kinase and microtubule interacting protein 3 |
| chr14 | 60022447 | 60023398 | 952 | * | 8,940106 | -3,676527501 | 1,40334 | -2,61984 | 0,008797 | 0,860889191 | 1,449706 | Intron (ENSMUST0000 | 14 | 59864638 | 59867126 | 2489 | 1 | 219134 | ENSMUST00000224155.2 | 157809 | ENSMUSG00000044461 | Shisa2 | shisa family member 2 |
| chr4 | 54643774 | 54645067 | 1294 | * | 13,88845 | -3,67596603 | 1,378504 | -2,66663 | 0,007662 | 0,859692746 | 1,2763 | Distal Intergenic | 4 | 54945048 | 55008314 | 63267 | 1 | 242466 | ENSMUST00000133895.8 | -299981 | ENSMUSG00000060206 | Zfp462 | zinc finger protein 462 |
| chr14 | 70118146 | 70119590 | 1445 | * | 7,914141 | -3,675782282 | 1,420291 | -2,58805 | 0,009652 | 1 | 0 | Intron (ENSMUST0000 | 14 | 70077869 | 70297367 | 219499 | 1 | 73523 | ENSMUST00000022678.5 | 40277 | ENSMUSG00000022085 | Pebp4 | phosphatidylethanolamine binding protein 4 |
| chr14 | 28703616 | 28705422 | 1807 | * | 15,2693 | -3,674791293 | 1,2684 | -2,89719 | 0,003765 | 0,744477896 | 1,214904 | Exon (ENSMUST00000 | 14 | 28627204 | 28691428 | 64225 | 2 | 12294 | ENSMUST00000225985.2 | -12188 | ENSMUSG00000021991 | Cacna2d3 | calcium channel, voltage-dependent, alpha2/delta subunit 3 |
| chr3 | 1,47E+08 | 1,47E+08 | 982 | * | 8,101249 | -3,674520887 | 1,380973 | -2,66082 | 0,007795 | 0,852732156 | 1,360622 | Exon (ENSMUST00000 | 3 | 1,47E+08 | 1,47E+08 | 4401 | 1 | 1,05E+08 | ENSMUST00000195873.2 | 190950 | | Gm37041 | predicted gene, 37041 |
| chr4 | 93117505 | 93119089 | 1585 | * | 11,50265 | -3,673538969 | 1,21445 | -3,02486 | 0,002487 | 0,756849896 | 0,738441 | Distal Intergenic | 4 | 93222375 | 93223748 | 1374 | 2 | 69136 | ENSMUST00000066774.6 | 104659 | ENSMUSG00000054000 | Tusc1 | tumor suppressor candidate 1 |
| chr19 | 51882615 | 51883945 | 1331 | * | 9,665956 | -3,673248314 | 1,397318 | -2,62879 | 0,008569 | 0,857624805 | 1,449706 | Intron (ENSMUST0000 | 19 | 52252735 | 52253914 | 1180 | 1 | 16333 | ENSMUST00000039652.6 | -368790 | ENSMUSG00000035804 | Ins1 | insulin I |
| chr3 | 1,02E+08 | 1,02E+08 | 1262 | * | 11,62154 | -3,670871407 | 1,204827 | -3,0468 | 0,002313 | 0,747833008 | 0,738441 | Distal Intergenic | 3 | 1,01E+08 | 1,02E+08 | 28466 | 2 | 11928 | ENSMUST00000036493.8 | -58673 | ENSMUSG00000033161 | Atp1a1 | ATPase, Na+/K+ transporting, alpha 1 polypeptide |
| chr7 | 79506295 | 79507814 | 1520 | * | 12,52526 | -3,670433138 | 1,236989 | -2,96723 | 0,003005 | 0,792545355 | 0,738441 | Promoter (2-3kb) | 7 | 79491804 | 79510807 | 19004 | 2 | 16790 | ENSMUST00000206235.2 | 2993 | ENSMUSG00000039062 | Anpep | alanyl (membrane) aminopeptidase |
| chrX | 1,27E+08 | 1,27E+08 | 1495 | * | 10,21053 | -3,66960669 | 1,255654 | -2,92247 | 0,003473 | 0,702848616 | 1,425929 | Distal Intergenic | 20 | 1,28E+08 | 1,28E+08 | 108 | 1 | 1,15E+08 | ENSMUST00000083964.3 | -117177 | ENSMUSG00000065898 | Gm24522 | predicted gene, 24522 |
| chr16 | 95091741 | 95093289 | 1549 | * | 14,21252 | -3,668446811 | 1,203813 | -3,04736 | 0,002309 | 0,705187732 | 0,938682 | 5' UTR | 16 | 95080052 | 95092756 | 12705 | 1 | 16516 | ENSMUST00000138329.2 | 11689 | ENSMUSG00000062609 | Kcnj15 | potassium inwardly-rectifying channel, subfamily J, member 15 |

| | | | | | | | | | | | | | | | | | | | | | | | |
|---|---|---|---|---|---|---|---|---|---|---|---|---|---|---|---|---|---|---|---|---|---|---|---|
| chr16 | 82225156 | 82226004 | 849 | * | 8,853916 | -3,668169181 | 1,258509 | -2,91469 | 0,00356 | 0,705098751 | 1,449706 | Distal Intergenic | 16 | 82340117 | 82340223 | 107 | 1 | 1,15E+08 | ENSMUST00000240414.1 | -114113 | ENSMUSG00000119122 | Gm23355 | predicted gene, 23355 |
| chr9 | 1,1E+08 | 1,1E+08 | 1287 | * | 9,808261 | -3,662512879 | 1,399061 | -2,61784 | 0,008849 | 0,90060438 | 1,173325 | Distal Intergenic | 9 | 1,1E+08 | 1,1E+08 | 8321 | 1 | 78709 | ENSMUST00000200555.5 | -5065 | ENSMUSG00000050074 | Spink8 | serine peptidase inhibitor, Kazal type 8 |
| chr6 | 76734719 | 76735592 | 874 | * | 10,65091 | -3,662312982 | 1,363818 | -2,68534 | 0,007246 | 0,852340527 | 1,2763 | Distal Intergenic | 6 | 76859739 | 76879871 | 20133 | 2 | 12386 | ENSMUST00000162991.2 | 144279 | ENSMUSG00000063063 | Ctnna2 | catenin (cadherin associated protein), alpha 2 |
| chr6 | 47207879 | 47208478 | 600 | * | 9,394496 | -3,657823233 | 1,415142 | -2,58477 | 0,009744 | 0,882702232 | 1,449706 | Intron (ENSMUST0000 | 6 | 47221321 | 47281143 | 59823 | 1 | 66797 | ENSMUST00000199100.5 | -12843 | ENSMUSG00000039419 | Cntnap2 | contactin associated protein-like 2 |
| chr3 | 69186281 | 69187615 | 1335 | * | 9,553828 | -3,654829031 | 1,406294 | -2,59891 | 0,009352 | 0,877892722 | 1,425929 | Distal Intergenic | 3 | 69224194 | 69468135 | 243942 | 1 | 242083 | ENSMUST00000029355.9 | -36579 | ENSMUSG00000027784 | Ppm1l | protein phosphatase 1 (formerly 2C)-like |
| chr10 | 96569666 | 96571192 | 1527 | * | 16,79064 | -3,654603856 | 1,154519 | -3,16548 | 0,001548 | 0,474532331 | 2,318434 | Distal Intergenic | 10 | 96636901 | 96720570 | 83670 | 1 | 1,08E+08 | ENSMUST00000218446.2 | -65709 | | Gm46240 | predicted gene, 46240 |
| chr5 | 6413309 | 6414296 | 988 | * | 8,832827 | -3,652719129 | 1,318113 | -2,77117 | 0,005585 | 0,708539289 | 2,237403 | Distal Intergenic | 5 | 6064863 | 6065024 | 162 | 2 | 1,15E+08 | ENSMUST00000158067.3 | -348285 | ENSMUSG00000088692 | Gm23838 | predicted gene, 23838 |
| chr5 | 1,48E+08 | 1,48E+08 | 911 | * | 8,446398 | -3,650798588 | 1,403201 | -2,60176 | 0,009275 | 0,885263498 | 1,360622 | Intron (ENSMUST0000 | 5 | 1,48E+08 | 1,48E+08 | 487 | 2 | 14254 | ENSMUST00000142477.2 | 12919 | ENSMUSG00000029648 | Flt1 | FMS-like tyrosine kinase 1 |
| chr13 | 14969903 | 14971319 | 1417 | * | 10,75194 | -3,649692508 | 1,345696 | -2,71212 | 0,006685 | 0,852340527 | 1,173325 | Distal Intergenic | 13 | 14804862 | 14811499 | 6638 | 1 | 105351 | ENSMUST00000222052.2 | 165041 | ENSMUSG00000039182 | AW209491 | expressed sequence AW209491 |
| chr7 | 83045419 | 83046612 | 1194 | * | 9,076392 | -3,649268884 | 1,379812 | -2,64476 | 0,008175 | 0,775111864 | 2,237403 | Distal Intergenic | 7 | 82990141 | 83018637 | 28497 | 2 | 1,03E+08 | ENSMUST00000208430.2 | -26782 | | Gm29976 | predicted gene, 29976 |
| chr19 | 56059033 | 56060183 | 1151 | * | 9,354203 | -3,648959979 | 1,410609 | -2,5868 | 0,009687 | 0,881662676 | 1,449706 | Distal Intergenic | 19 | 56094680 | 56129039 | 34360 | 1 | 1,03E+08 | ENSMUST00000235362.2 | -34497 | | Gm31912 | predicted gene, 31912 |
| chr2 | 1,41E+08 | 1,41E+08 | 2163 | * | 12,25388 | -3,648473185 | 1,21665 | -2,99879 | 0,002711 | 0,682691026 | 1,2763 | Intron (ENSMUST0000 | 2 | 1,41E+08 | 1,42E+08 | 837291 | 1 | 72899 | ENSMUST00000138786.2 | -19801 | ENSMUSG00000068205 | Macrod2 | mono-ADP ribosylhydrolase 2 |
| chr12 | 79392167 | 79393167 | 1001 | * | 8,397925 | -3,647402342 | 1,330525 | -2,74133 | 0,006119 | 0,798887389 | 1,449706 | Intron (ENSMUST0000 | 12 | 79374127 | 79748630 | 374504 | 1 | 19363 | ENSMUST00000221257.2 | 18040 | ENSMUSG00000059060 | Rad51b | RAD51 paralog B |
| chr3 | 1,42E+08 | 1,42E+08 | 1032 | * | 9,501593 | -3,646242124 | 1,399071 | -2,60619 | 0,009156 | 0,873955012 | 1,425929 | Intron (ENSMUST0000 | 3 | 1,42E+08 | 1,42E+08 | 145129 | 2 | 12167 | ENSMUST00000029948.15 | 14245 | ENSMUSG00000052430 | Bmpr1b | bone morphogenetic protein receptor, type 1B |
| chr7 | 1,2E+08 | 1,2E+08 | 880 | * | 10,59579 | -3,644626134 | 1,226681 | -2,97113 | 0,002967 | 0,790460916 | 0,744375 | Intron (ENSMUST0000 | 7 | 1,2E+08 | 1,2E+08 | 11729 | 1 | 72074 | ENSMUST00000033201.7 | 3321 | ENSMUSG00000030909 | Anks4b | ankyrin repeat and sterile alpha motif domain containing 4B |
| chr13 | 98242908 | 98244334 | 1427 | * | 11,31771 | -3,642505859 | 1,253062 | -2,90688 | 0,00365 | 0,826716028 | 0,738441 | Intron (ENSMUST0000 | 13 | 98282060 | 98312495 | 30436 | 2 | 110596 | ENSMUST00000224866.2 | 68161 | ENSMUSG00000021662 | Arhgef28 | Rho guanine nucleotide exchange factor (GEF) 28 |
| chr4 | 1,19E+08 | 1,19E+08 | 911 | * | 9,779476 | -3,639613763 | 1,382599 | -2,63244 | 0,008477 | 0,904389377 | 1,086118 | Promoter (2-3kb) | 4 | 1,19E+08 | 1,19E+08 | 4442 | 1 | 242653 | ENSMUST00000084309.12 | -2731 | ENSMUSG00000066058 | Cldn19 | claudin 19 |
| chr11 | 16890491 | 16892026 | 1536 | * | 12,97884 | -3,638193171 | 1,326356 | -2,743 | 0,006088 | 0,836575905 | 1,173325 | Intron (ENSMUST0000 | 11 | 16885154 | 16901072 | 15919 | 2 | 72735 | ENSMUST00000245314.1 | 9046 | | Eldr | Egfr long non-coding downstream RNA |
| chr3 | 1,2E+08 | 1,2E+08 | 717 | * | 6,453636 | -3,636745519 | 1,409907 | -2,57942 | 0,009897 | 1 | 0 | Distal Intergenic | 3 | 1,2E+08 | 1,2E+08 | 128 | 2 | 1,15E+08 | ENSMUST00000158402.3 | 54321 | ENSMUSG00000089027 | Gm25013 | predicted gene, 25013 |
| chr2 | 45101755 | 45102667 | 913 | * | 9,947002 | -3,635444368 | 1,40486 | -2,58776 | 0,00966 | 0,912474253 | 1,173325 | Distal Intergenic | 2 | 45129338 | 45169052 | 39715 | 1 | 110153 | ENSMUST00000244809.1 | -26671 | | Vis1 | viral integration site 1 |
| chr12 | 33640631 | 33641751 | 1121 | * | 11,61947 | -3,634190608 | 1,202533 | -3,02211 | 0,00251 | 0,722855505 | 0,918255 | Distal Intergenic | 12 | 33502711 | 33502842 | 132 | 2 | 1,15E+08 | ENSMUST00000239981.1 | -137789 | ENSMUSG00000119573 | Gm23529 | predicted gene, 23529 |
| chr15 | 80486148 | 80486658 | 511 | * | 6,848617 | -3,633510887 | 1,401163 | -2,59321 | 0,009508 | 1 | 0 | Intron (ENSMUST0000 | 15 | 80507684 | 80537055 | 29372 | 1 | 17444 | ENSMUST00000043149.9 | -21026 | ENSMUSG00000042351 | Grap2 | GRB2-related adaptor protein 2 |
| chr5 | 58116044 | 58117424 | 1381 | * | 9,823438 | -3,631956238 | 1,237236 | -2,93554 | 0,00333 | 0,742884189 | 1,086118 | Intron (ENSMUST0000 | 5 | 57879551 | 58286796 | 407246 | 1 | 54216 | ENSMUST00000192048.2 | 236493 | ENSMUSG00000029108 | Pcdh7 | protocadherin 7 |
| chr17 | 52700622 | 52701692 | 1071 | * | 10,70316 | -3,630147063 | 1,37001 | -2,64972 | 0,008056 | 0,955124605 | 0,744375 | Distal Intergenic | 17 | 52611595 | 52633610 | 22016 | 1 | 1,05E+08 | ENSMUST00000232754.2 | 89027 | ENSMUSG00000117076 | Gm41600 | predicted gene, 41600 |
| chr2 | 22239239 | 22240204 | 966 | * | 12,07632 | -3,627885941 | 1,205066 | -3,01053 | 0,002608 | 0,685026746 | 1,173325 | Intron (ENSMUST0000 | 2 | 22245868 | 22412265 | 166398 | 1 | 667663 | ENSMUST00000153002.3 | -5664 | ENSMUSG00000025716 | Myo3a | myosin IIIA |
| chr14 | 79978172 | 79981071 | 2900 | * | 18,3011 | -3,627349123 | 1,007597 | -3,6 | 0,000318 | 0,229205408 | 2,423594 | Promoter (2-3kb) | 14 | 79975871 | 79975968 | 98 | 1 | 791080 | ENSMUST00000103254.3 | 2301 | ENSMUSG00000076453 | Mir759 | microRNA 759 |
| chr10 | 1E+08 | 1E+08 | 1111 | * | 8,761223 | -3,624597951 | 1,28761 | -2,81498 | 0,004878 | 0,760613407 | 1,425929 | Intron (ENSMUST0000 | 10 | 1E+08 | 1E+08 | 3620 | 1 | 216274 | ENSMUST00000220331.2 | 7808 | ENSMUSG00000019971 | Cep290 | centrosomal protein 290 |
| chr5 | 1,48E+08 | 1,48E+08 | 918 | * | 7,934616 | -3,623704772 | 1,351787 | -2,68068 | 0,007347 | 1 | 0 | Promoter (2-3kb) | 5 | 1,48E+08 | 1,48E+08 | 15996 | 1 | 66537 | ENSMUST00000031654.10 | -2691 | ENSMUSG00000029649 | Pomp | proteasome maturation protein |
| chr5 | 42145720 | 42146788 | 1069 | * | 8,898148 | -3,622455643 | 1,281022 | -2,82779 | 0,004687 | 0,681119785 | 2,237403 | Distal Intergenic | 5 | 42225303 | 42374141 | 148839 | 1 | 433882 | ENSMUST00000087332.5 | -78515 | ENSMUSG00000067285 | Gm16223 | predicted gene 16223 |
| chr3 | 1,2E+08 | 1,2E+08 | 1054 | * | 8,945062 | -3,620957503 | 1,396863 | -2,59221 | 0,009536 | 0,88871498 | 1,360622 | Exon (ENSMUST00000 | 3 | 1,2E+08 | 1,2E+08 | 128 | 2 | 1,15E+08 | ENSMUST00000158402.3 | -4160 | ENSMUSG00000089027 | Gm25013 | predicted gene, 25013 |
| chr11 | 42892195 | 42894500 | 2306 | * | 16,424 | -3,619563541 | 1,005808 | -3,59866 | 0,00032 | 0,236730442 | 2,318434 | Distal Intergenic | 11 | 43040704 | 43050302 | 9599 | 1 | 319767 | ENSMUST00000148911.2 | -146204 | ENSMUSG00000055415 | Atp10b | ATPase, class V, type 10B |
| chr10 | 20339920 | 20341701 | 1782 | * | 16,76271 | -3,618731518 | 1,057836 | -3,42088 | 0,000624 | 0,332694435 | 2,141612 | Intron (ENSMUST0000 | 10 | 20274674 | 20424074 | 149401 | 2 | 29863 | ENSMUST00000169016.8 | 82373 | ENSMUSG00000019990 | Pde7b | phosphodiesterase 7B |
| chrX | 76809215 | 76810031 | 817 | * | 9,02166 | -3,618145206 | 1,286643 | -2,81208 | 0,004922 | 0,759000138 | 1,449706 | Intron (ENSMUST0000 | 20 | 76815061 | 76839578 | 24518 | 2 | 19108 | ENSMUST00000114044.2 | 29547 | ENSMUSG00000035725 | Prkx | protein kinase, X-linked |
| chr11 | 66331667 | 66332917 | 1251 | * | 10,58282 | -3,618087059 | 1,192229 | -3,03472 | 0,002408 | 0,663408464 | 1,2763 | Intron (ENSMUST0000 | 11 | 66265959 | 66393640 | 127682 | 2 | 380702 | ENSMUST00000134562.2 | 60723 | ENSMUSG00000053930 | Shisa6 | shisa family member 6 |
| chr16 | 95450912 | 95452418 | 1507 | * | 13,41372 | -3,615285918 | 1,166195 | -3,10007 | 0,001935 | 0,681459438 | 0,918255 | Distal Intergenic | 16 | 95502942 | 95522095 | 19154 | 1 | 23872 | ENSMUST00000023612.17 | -50524 | ENSMUSG00000022895 | Ets2 | E26 avian leukemia oncogene 2, 3' domain |
| chr11 | 46450654 | 46451395 | 742 | * | 9,010962 | -3,614786058 | 1,348011 | -2,68157 | 0,007328 | 0,834936846 | 1,425929 | Distal Intergenic | 11 | 46462363 | 46480059 | 17697 | 1 | 407814 | ENSMUST00000050937.7 | -10968 | ENSMUSG00000046974 | BC053393 | cDNA sequence BC053393 |
| chr15 | 22411172 | 22411968 | 797 | * | 9,771162 | -3,61466074 | 1,371266 | -2,636 | 0,008389 | 0,854157826 | 1,449706 | Distal Intergenic | 15 | 22549108 | 23400653 | 851546 | 1 | 320865 | ENSMUST00000163361.8 | -137140 | ENSMUSG00000040420 | Cdh18 | cadherin 18 |
| chr11 | 1,17E+08 | 1,17E+08 | 811 | * | 11,05802 | -3,611669498 | 1,283351 | -2,81425 | 0,004889 | 0,839831097 | 0,918255 | Promoter (<=1kb) | 11 | 1,17E+08 | 1,17E+08 | 13868 | 1 | 268510 | ENSMUST00000126757.2 | -941 | ENSMUSG00000043857 | Mgat5b | mannoside acetylglucosaminyltransferase 5, isoenzyme B |
| chr17 | 69548758 | 69550486 | 1729 | * | 15,84603 | -3,609744567 | 1,014671 | -3,55755 | 0,000374 | 0,333467627 | 1,2763 | Promoter (<=1kb) | 17 | 69550631 | 69596980 | 46350 | 1 | 13823 | ENSMUST00000223703.2 | -145 | ENSMUSG00000024044 | Epb41l3 | erythrocyte membrane protein band 4.1 like 3 |
| chrX | 86442302 | 86443640 | 1339 | * | 8,072223 | -3,608283426 | 1,39548 | -2,58569 | 0,009718 | 1 | 0 | Intron (ENSMUST0000 | 20 | 85790848 | 86933841 | 1142994 | 2 | 331461 | ENSMUST00000078875.8 | 490201 | ENSMUSG00000052372 | Il1rapl1 | interleukin 1 receptor accessory protein-like 1 |
| chr5 | 1,42E+08 | 1,42E+08 | 1327 | * | 11,75156 | -3,607174484 | 1,265595 | -2,85018 | 0,004369 | 0,777604125 | 1,173325 | Intron (ENSMUST0000 | 5 | 1,42E+08 | 1,42E+08 | 107 | 2 | 1,15E+08 | ENSMUST00000158539.3 | -36255 | ENSMUSG00000089164 | Gm24131 | predicted gene, 24131 |
| chr10 | 74035874 | 74036949 | 1076 | * | 8,981843 | -3,60684335 | 1,38566 | -2,60298 | 0,009242 | 0,872427545 | 1,449706 | Intron (ENSMUST0000 | 10 | 74138983 | 74166882 | 27900 | 1 | 11994 | ENSMUST00000147455.2 | -102034 | ENSMUSG00000052613 | Pcdh15 | protocadherin 15 |
| chr1 | 1,8E+08 | 1,8E+08 | 1223 | * | 7,829442 | -3,603875412 | 1,337423 | -2,69464 | 0,007046 | 1 | 0 | Intron (ENSMUST0000 | 1 | 1,8E+08 | 1,8E+08 | 47647 | 1 | 226751 | ENSMUST00000132894.3 | 24434 | ENSMUSG00000026490 | Cdc42bpa | CDC42 binding protein kinase alpha |
| chr8 | 1,3E+08 | 1,3E+08 | 1358 | * | 16,77201 | -3,602905532 | 0,970311 | -3,71315 | 0,000205 | 0,185760459 | 2,318434 | Exon (ENSMUST00000 | 8 | 1,3E+08 | 1,3E+08 | 132 | 1 | 1,15E+08 | ENSMUST00000104350.3 | -99446 | ENSMUSG00000077538 | Gm22545 | predicted gene, 22545 |
| chr14 | 84940233 | 84941700 | 1468 | * | 7,038704 | -3,601672224 | 1,383404 | -2,60348 | 0,009228 | 1 | 0 | Distal Intergenic | 14 | 84696164 | 84713864 | 17701 | 2 | 319903 | ENSMUST00000228125.2 | -226369 | ENSMUSG00000115529 | 9630013A20Rik | RIKEN cDNA 9630013A20 gene |
| chr8 | 1,08E+08 | 1,08E+08 | 996 | * | 10,95207 | -3,601667439 | 1,343431 | -2,68095 | 0,007341 | 0,853738252 | 1,2763 | Distal Intergenic | 8 | 1,08E+08 | 1,08E+08 | 8084 | 2 | 17463 | ENSMUST00000044106.6 | 11222 | ENSMUSG00000039067 | Psmd7 | proteasome (prosome, macropain) 26S subunit, non-ATPase, 7 |
| chr10 | 10186202 | 10187125 | 924 | * | 8,737911 | -3,597777923 | 1,343142 | -2,67863 | 0,007392 | 0,836571183 | 1,425929 | Distal Intergenic | 10 | 10111205 | 10131220 | 20016 | 1 | 1,03E+08 | ENSMUST00000208935.2 | 74997 | ENSMUSG00000109160 | 4930598N05Rik | RIKEN cDNA 4930598N05 gene |
| chr12 | 70346271 | 70347806 | 1536 | * | 11,16012 | -3,596570057 | 1,336334 | -2,69137 | 0,007116 | 0,903294867 | 0,918255 | Intron (ENSMUST0000 | 12 | 70339107 | 70363029 | 23923 | 2 | 94090 | ENSMUST00000220557.2 | 15223 | ENSMUSG00000021071 | Trim9 | tripartite motif-containing 9 |
| chr5 | 68152782 | 68153827 | 1046 | * | 8,355426 | -3,593739816 | 1,29827 | -2,7681 | 0,005638 | 0,710505631 | 2,237403 | Distal Intergenic | 5 | 68189178 | 68323741 | 134564 | 1 | 433899 | ENSMUST00000094715.9 | -35351 | ENSMUSG00000068082 | Grxcr1 | glutaredoxin, cysteine rich 1 |
| chr9 | 29903181 | 29904832 | 1652 | * | 12,10916 | -3,592572491 | 1,316837 | -2,72818 | 0,006368 | 0,919921445 | 0,738441 | Distal Intergenic | 9 | 29852844 | 29874437 | 21594 | 2 | 235106 | ENSMUST00000151977.2 | -28744 | ENSMUSG00000059974 | Ntm | neurotrimin |
| chr7 | 87626845 | 87627976 | 1132 | * | 9,420089 | -3,589808572 | 1,250966 | -2,86963 | 0,00411 | 0,7299084 | 1,449706 | Intron (ENSMUST0000 | 7 | 87736396 | 87736459 | 64 | 2 | 1,15E+08 | ENSMUST00000157322.3 | 108483 | ENSMUSG00000087947 | Gm24249 | predicted gene, 24249 |
| chrX | 1,58E+08 | 1,58E+08 | 1546 | * | 12,09497 | -3,589300754 | 1,27072 | -2,82462 | 0,004734 | 0,872184583 | 0,744375 | Distal Intergenic | 20 | 1,58E+08 | 1,58E+08 | 154196 | 1 | 110651 | ENSMUST00000112493.8 | -401351 | ENSMUSG00000031309 | Rps6ka3 | ribosomal protein S6 kinase polypeptide 3 |
| chrX | 65336489 | 65338758 | 2270 | * | 13,20439 | -3,587236142 | 1,064997 | -3,36831 | 0,000756 | 0,480916943 | 1,075735 | Distal Intergenic | 20 | 65692924 | 65704999 | 12076 | 1 | 245450 | ENSMUST00000166241.2 | -354166 | ENSMUSG00000036790 | Slitrk2 | SLIT and NTRK-like family, member 2 |
| chr18 | 41703323 | 41704856 | 1534 | * | 10,83493 | -3,583596545 | 1,316009 | -2,72308 | 0,006468 | 0,83238039 | 1,2763 | Distal Intergenic | 18 | 41880299 | 42007899 | 127601 | 2 | 1,15E+08 | ENSMUST00000235677.2 | 303043 | | Gm50410 | predicted gene, 50410 |
| chr7 | 60376054 | 60378357 | 2304 | * | 18,14702 | -3,581771294 | 0,99839 | -3,58755 | 0,000334 | 0,231796186 | 2,507013 | Distal Intergenic | 7 | 60565298 | 60572651 | 7354 | 2 | 1,01E+08 | ENSMUST00000208825.2 | 194294 | | Gm29729 | predicted gene, 29729 |
| chr2 | 43734635 | 43735260 | 626 | * | 9,186158 | -3,580924382 | 1,293601 | -2,76818 | 0,005637 | 0,789027175 | 1,425929 | Exon (ENSMUST00000 | 2 | 43638882 | 44277635 | 638754 | 1 | 76117 | ENSMUST00000055776.8 | 95753 | ENSMUSG00000049744 | Arhgap15 | Rho GTPase activating protein 15 |
| chr2 | 76180064 | 76181266 | 1203 | * | 12,26703 | -3,580533153 | 1,256506 | -2,84959 | 0,004378 | 0,857955756 | 0,738441 | Distal Intergenic | 2 | 76184286 | 76190794 | 6509 | 2 | 13067 | ENSMUST00000028430.6 | 9528 | ENSMUSG00000056436 | Cyct | cytochrome c, testis |
| chr17 | 11021044 | 11022516 | 1473 | * | 12,97265 | -3,580374695 | 1,160438 | -3,08537 | 0,002033 | 0,663569499 | 1,075735 | Intron (ENSMUST0000 | 17 | 11026721 | 11045512 | 18792 | 1 | 1,03E+08 | ENSMUST00000162432.2 | -4205 | | Gm16169 | predicted gene 16169 |
| chr9 | 1,08E+08 | 1,08E+08 | 1227 | * | 12,79098 | -3,577597531 | 1,303708 | -2,74417 | 0,006066 | 0,879131976 | 0,918255 | Intron (ENSMUST0000 | 9 | 1,08E+08 | 1,08E+08 | 3019 | 1 | 22258 | ENSMUST00000194034.2 | 3765 | ENSMUSG00000032612 | Usp4 | ubiquitin specific peptidase 4 (proto-oncogene) |
| chr4 | 70179401 | 70180849 | 1449 | * | 16,13576 | -3,577406087 | 1,099293 | -3,25428 | 0,001137 | 0,400771934 | 2,663619 | Promoter (2-3kb) | 4 | 70156587 | 70182917 | 26331 | 2 | 214444 | ENSMUST00000138561.8 | 2068 | ENSMUSG00000039298 | Cdk5rap2 | CDK5 regulatory subunit associated protein 2 |
| chr1 | 1,3E+08 | 1,3E+08 | 1148 | * | 10,3662 | -3,574457318 | 1,225788 | -2,91605 | 0,003545 | 0,725842761 | 1,2763 | Intron (ENSMUST0000 | 1 | 1,3E+08 | 1,3E+08 | 267643 | 1 | 210417 | ENSMUST00000151700.8 | -41440 | ENSMUSG00000042581 | Thsd7b | thrombospondin, type I, domain containing 7B |
| chr13 | 56531037 | 56531949 | 913 | * | 10,9714 | -3,573787048 | 1,362298 | -2,62335 | 0,008707 | 0,966603024 | 0,738441 | Distal Intergenic | 13 | 56510714 | 56513582 | 2869 | 1 | 1E+08 | ENSMUST00000099476.4 | 20323 | ENSMUSG00000074885 | Gm10782 | predicted gene 10782 |
| chr9 | 70021850 | 70023123 | 1274 | * | 11,74556 | -3,572740556 | 1,182608 | -3,02107 | 0,002519 | 0,759970924 | 0,738441 | Intron (ENSMUST0000 | 9 | 70009832 | 70010358 | 527 | 2 | 76886 | ENSMUST00000145427.2 | -11492 | ENSMUSG00000032224 | Fam81a | family with sequence similarity 81, member A |
| chr18 | 3152449 | 3153489 | 1041 | * | 11,65506 | -3,571333314 | 1,212109 | -2,94638 | 0,003215 | 0,723183077 | 1,173325 | Distal Intergenic | 18 | 3122492 | 3123412 | 921 | 2 | 1E+08 | ENSMUST00000165255.3 | -29037 | ENSMUSG00000091539 | Vmn1r238 | vomeronasal 1 receptor, 238 |
| chr8 | 99007349 | 99008660 | 1312 | * | 14,38246 | -3,569563923 | 1,027042 | -3,47558 | 0,00051 | 0,3894426 | 1,2763 | Distal Intergenic | 8 | 99401709 | 99401834 | 126 | 2 | 1,15E+08 | ENSMUST00000116830.3 | 393174 | ENSMUSG00000080480 | Gm23494 | predicted gene, 23494 |
| chr14 | 64389548 | 64390975 | 1428 | * | 9,143834 | -3,568137173 | 1,383594 | -2,57889 | 0,009912 | 0,88797938 | 1,425929 | Intron (ENSMUST0000 | 14 | 64353763 | 64365054 | 11292 | 1 | 78569 | ENSMUST00000225122.2 | 35785 | ENSMUSG00000114284 | 9630015K15Rik | RIKEN cDNA 9630015K15 gene |
| chr17 | 45722164 | 45723295 | 1132 | * | 10,05182 | -3,568123182 | 1,372552 | -2,59963 | 0,009333 | 0,902886974 | 1,211315 | Exon (ENSMUST00000 | 17 | 45719298 | 45726610 | 7313 | 2 | 71702 | ENSMUST00000232900.2 | 3315 | ENSMUSG00000023932 | Cdc5l | cell division cycle 5-like (S. pombe) |
| chr1 | 1,05E+08 | 1,05E+08 | 1572 | * | 10,90786 | -3,561922675 | 1,199697 | -2,96902 | 0,002988 | 0,792545355 | 0,738441 | Intron (ENSMUST0000 | 1 | 1,05E+08 | 1,05E+08 | 226953 | 1 | 23836 | ENSMUST00000062528.9 | 131759 | ENSMUSG00000050840 | Cdh20 | cadherin 20 |
| chr16 | 38550360 | 38551278 | 919 | * | 8,888743 | -3,561097755 | 1,315005 | -2,70805 | 0,006768 | 0,740176831 | 2,237403 | Distal Intergenic | 16 | 38562626 | 38589411 | 26786 | 1 | 56375 | ENSMUST00000023482.13 | -11348 | ENSMUSG00000022793 | B4galt4 | UDP-Gal:betaGlcNAc beta 1,4-galactosyltransferase, polypeptide 4 |
| chr13 | 26612433 | 26613935 | 1503 | * | 10,50563 | -3,560522113 | 1,256024 | -2,83476 | 0,004586 | 0,86525631 | 0,738441 | Distal Intergenic | 13 | 26467133 | 26496389 | 29257 | 2 | 73576 | ENSMUST00000154703.3 | -116044 | ENSMUSG00000084811 | 1700092E19Rik | RIKEN cDNA 1700092E19 gene |
| chr7 | 1,16E+08 | 1,16E+08 | 1156 | * | 11,08736 | -3,558585135 | 1,19606 | -2,97526 | 0,002927 | 0,689688276 | 1,2763 | Intron (ENSMUST0000 | 7 | 1,15E+08 | 1,16E+08 | 523743 | 2 | 20679 | ENSMUST00000072804.11 | 29501 | ENSMUSG00000051910 | Sox6 | SRY (sex determining region Y)-box 6 |
| chr1 | 1,73E+08 | 1,73E+08 | 1395 | * | 10,79013 | -3,55693055 | 1,282196 | -2,77409 | 0,005536 | 0,860322035 | 0,918255 | Intron (ENSMUST0000 | 1 | 1,73E+08 | 1,73E+08 | 105 | 2 | 1,15E+08 | ENSMUST00000093672.3 | 31222 | ENSMUSG00000070182 | Gm24817 | predicted gene, 24817 |
| chr5 | 59382623 | 59384422 | 1800 | * | 7,722332 | -3,55639198 | 1,308577 | -2,71776 | 0,006573 | 1 | 0 | Distal Intergenic | 5 | 59372289 | 59372395 | 107 | 2 | 1,15E+08 | ENSMUST00000239835.1 | -10228 | ENSMUSG00000119859 | Gm22308 | predicted gene, 22308 |
| chr11 | 73050450 | 73052467 | 2018 | * | 21,20343 | -3,555598489 | 1,050441 | -3,38486 | 0,000712 | 0,302204121 | 3,104559 | Promoter (<=1kb) | 11 | 73051247 | 73063511 | 12265 | 1 | 94045 | ENSMUST00000006104.10 | 0 | ENSMUSG00000005950 | P2rx5 | purinergic receptor P2X, ligand-gated ion channel, 5 |
| chr4 | 44667445 | 44668693 | 1249 | * | 15,36801 | -3,553412532 | 1,12333 | -3,16328 | 0,00156 | 0,563032042 | 1,41696 | Intron (ENSMUST0000 | 4 | 44664415 | 44685347 | 20933 | 1 | 1,03E+08 | ENSMUST00000140826.2 | 3030 | | Gm12462 | predicted gene 12462 |
| chr14 | 32958436 | 32960300 | 1865 | * | 18,39105 | -3,552451981 | 0,936723 | -3,79243 | 0,000149 | 0,148120959 | 2,507013 | Intron (ENSMUST0000 | 14 | 32972347 | 33065260 | 92914 | 1 | 239027 | ENSMUST00000131086.3 | -12047 | ENSMUSG00000063506 | Arhgap22 | Rho GTPase activating protein 22 |
| chr2 | 1,01E+08 | 1,01E+08 | 1008 | * | 12,13859 | -3,552109751 | 1,100609 | -3,2274 | 0,001249 | 0,644704154 | 0,738441 | Distal Intergenic | 2 | 1,01E+08 | 1,01E+08 | 14001 | 2 | 68170 | ENSMUST00000177152.2 | 277425 | ENSMUSG00000027165 | Iftap | intraflagellar transport associated protein |
| chr6 | 23455116 | 23455792 | 677 | * | 11,32627 | -3,551846822 | 1,090589 | -3,25682 | 0,001127 | 0,627785088 | 0,738441 | Intron (ENSMUST0000 | 6 | 23262842 | 23355918 | 93077 | 2 | 320405 | ENSMUST00000156986.8 | -99198 | ENSMUSG00000017978 | Cadps2 | Ca2+-dependent activator protein for secretion 2 |
| chr7 | 1,42E+08 | 1,42E+08 | 1404 | * | 12,65167 | -3,550615097 | 1,077286 | -3,29589 | 0,000981 | 0,497688602 | 1,2763 | Distal Intergenic | 7 | 1,42E+08 | 1,42E+08 | 12128 | 2 | 13033 | ENSMUST00000151120.9 | -16737 | ENSMUSG00000007891 | Ctsd | cathepsin D |
| chr17 | 36609699 | 36611263 | 1565 | * | 11,71139 | -3,550475853 | 1,263307 | -2,81046 | 0,004947 | 0,842187815 | 0,918255 | Distal Intergenic | 17 | 36595119 | 36597382 | 2264 | 2 | 333715 | ENSMUST00000023845.7 | -12317 | ENSMUSG00000023083 | H2-M10.2 | histocompatibility 2, M region locus 10.2 |
| chr11 | 32281162 | 32282547 | 1386 | * | 10,74419 | -3,54930195 | 1,344375 | -2,64011 | 0,008288 | 0,876148358 | 1,2763 | Distal Intergenic | 11 | 32297820 | 32378173 | 80354 | 1 | 268396 | ENSMUST00000038753.6 | -15273 | ENSMUSG00000040711 | Sh3pxd2b | SH3 and PX domains 2B |
| chrX | 1,51E+08 | 1,51E+08 | 1882 | * | 14,94111 | -3,548955679 | 1,013495 | -3,5017 | 0,000462 | 0,377927525 | 1,214904 | Intron (ENSMUST0000 | 20 | 1,51E+08 | 1,51E+08 | 636 | 1 | 635396 | ENSMUST00000164764.2 | -3106 | ENSMUSG00000091863 | Cldn34a | claudin 34A |
| chr16 | 76853503 | 76854612 | 1110 | * | 9,421249 | -3,548539447 | 1,320012 | -2,68826 | 0,007182 | 0,858079007 | 1,211315 | Intron (ENSMUST0000 | 16 | 76878390 | 76890816 | 12427 | 1 | 30940 | ENSMUST00000232538.2 | -23778 | ENSMUSG00000022867 | Usp25 | ubiquitin specific peptidase 25 |
| chr6 | 16357491 | 16358143 | 653 | * | 8,85747 | -3,547596341 | 1,271873 | -2,78927 | 0,005283 | 0,786955198 | 1,360622 | Distal Intergenic | 6 | 16468235 | 16469903 | 1669 | 2 | 1,03E+08 | ENSMUST00000203052.2 | 111760 | | Gm36406 | predicted gene, 36406 |
| chr10 | 36358911 | 36361743 | 2833 | * | 27,911721 | -3,547011384 | 0,8007477 | -4,4296244 | 9,44E-06 | 0,020676456 | 2,9510621 | Distal Intergenic | 10 | 36382810 | 36709769 | 326960 | 1 | 319415 | ENSMUST00000167191.8 | -21067 | ENSMUSG00000044499 | Hs3st5 | heparan sulfate (glucosamine) 3-O-sulfotransferase 5 |
| chr19 | 42520150 | 42521516 | 1367 | * | 11,61733 | -3,546393646 | 1,373929 | -2,58121 | 0,009846 | 0,915421656 | 1,173325 | Intron (ENSMUST0000 | 19 | 42507261 | 42564539 | 57279 | 1 | 52013 | ENSMUST00000160893.2 | 12889 | ENSMUSG00000025184 | R3hcc1l | R3H domain and coiled-coil containing 1 like |
| chr16 | 82769487 | 82770572 | 1086 | * | 9,940539 | -3,54634566 | 1,244262 | -2,85016 | 0,00437 | 0,771248365 | 1,211315 | Distal Intergenic | 16 | 82340117 | 82340223 | 107 | 1 | 1,15E+08 | ENSMUST00000240414.1 | 429370 | ENSMUSG00000119122 | Gm23355 | predicted gene, 23355 |
| chr17 | 64628066 | 64629734 | 1669 | * | 14,03665 | -3,54584996 | 1,123343 | -3,15652 | 0,001597 | 0,645537369 | 0,938682 | Exon (ENSMUST00020 | 17 | 64588001 | 64620026 | 32026 | 2 | 224938 | ENSMUST00000024889.14 | -8040 | ENSMUSG00000024083 | Pja2 | praja ring finger ubiquitin ligase 2 |
| chr4 | 27184726 | 27185826 | 1101 | * | 8,173236 | -3,545600578 | 1,315719 | -2,6948 | 0,007043 | 0,748382289 | 2,237403 | Distal Intergenic | 4 | 26635820 | 26705449 | 69630 | 2 | 75360 | ENSMUST00000154459.2 | -479277 | ENSMUSG00000086257 | 4930548K13Rik | RIKEN cDNA 4930548K13 gene |
| chr15 | 1,03E+08 | 1,03E+08 | 1125 | * | 14,70451 | -3,53861357 | 1,156415 | -3,05999 | 0,002213 | 0,649217039 | 1,2763 | Distal Intergenic | 15 | 1,03E+08 | 1,03E+08 | 6987 | 2 | 1,03E+08 | ENSMUST00000188137.3 | -10558 | | Gm28876 | predicted gene 28876 |
| chr19 | 28264191 | 28265497 | 1307 | * | 10,7687 | -3,537039836 | 1,352261 | -2,61565 | 0,008906 | 0,888080529 | 1,2763 | Promoter (2-3kb) | 19 | 28241351 | 28261306 | 19956 | 2 | 226075 | ENSMUST00000159639.2 | -2885 | ENSMUSG00000052942 | Glis3 | GLIS family zinc finger 3 |
| chr8 | 69741816 | 69743071 | 1256 | * | 11,84047 | -3,535855129 | 1,36016 | -2,59959 | 0,009334 | 0,974266082 | 0,744375 | Intron (ENSMUST0000 | 8 | 69753428 | 69755814 | 2387 | 1 | 211135 | ENSMUST00000078257.7 | -10357 | ENSMUSG00000079038 | D130040H23Rik | RIKEN cDNA D130040H23 gene |
| chr13 | 46326627 | 46328003 | 1377 | * | 10,41191 | -3,534197817 | 1,368164 | -2,58317 | 0,00979 | 0,914590236 | 1,173325 | Distal Intergenic | 13 | 46427197 | 46453591 | 26395 | 1 | 380842 | ENSMUST00000076622.4 | -99194 | ENSMUSG00000063529 | Stmnd1 | stathmin domain containing 1 |
| chr7 | 52656190 | 52657409 | 1220 | * | 12,76696 | -3,533945429 | 1,178237 | -2,99935 | 0,002706 | 0,775744126 | 0,738441 | Distal Intergenic | 7 | 52576818 | 52576925 | 108 | 1 | 1,15E+08 | ENSMUST00000093693.3 | 79372 | ENSMUSG00000070203 | Gm22444 | predicted gene, 22444 |
| chr5 | 1,03E+08 | 1,03E+08 | 1138 | * | 9,7489 | -3,533317024 | 1,244197 | -2,83984 | 0,004514 | 0,74907225 | 1,425929 | Intron (ENSMUST0000 | 5 | 1,03E+08 | 1,03E+08 | 39122 | 2 | 26414 | ENSMUST00000128869.8 | 14547 | ENSMUSG00000046709 | Mapk10 | mitogen-activated protein kinase 10 |
| chr2 | 85717000 | 85718158 | 1159 | * | 10,47542 | -3,532033372 | 1,302892 | -2,71092 | 0,00671 | 0,838625535 | 1,2763 | Promoter (<=1kb) | 2 | 85717146 | 85718081 | 936 | 1 | 258580 | ENSMUST00000056408.2 | 0 | ENSMUSG00000050128 | Olfr1023 | olfactory receptor 1023 |
| chr3 | 1,44E+08 | 1,44E+08 | 1904 | * | 13,83194 | -3,531239033 | 1,004281 | -3,51619 | 0,000438 | 0,453798789 | 0,738441 | Intron (ENSMUST0000 | 3 | 1,44E+08 | 1,44E+08 | 33851 | 2 | 1,03E+08 | ENSMUST00000198202.2 | 6716 | | Gm34078 | predicted gene, 34078 |
| chr11 | 51929109 | 51930692 | 1584 | * | 8,465265 | -3,529652339 | 1,319495 | -2,675 | 0,007473 | 0,836008094 | 1,449706 | Exon (ENSMUST00000 | 11 | 51924357 | 51980605 | 56249 | 1 | 213084 | ENSMUST00000207272.2 | 4752 | ENSMUSG00000020389 | Cdkl3 | cyclin-dependent kinase-like 3 |
| chr2 | 1,64E+08 | 1,64E+08 | 1523 | * | 11,38654 | -3,527356184 | 1,231268 | -2,86481 | 0,004173 | 0,813191811 | 0,918255 | Intron (ENSMUST0000 | 2 | 1,64E+08 | 1,64E+08 | 33501 | 2 | 20568 | ENSMUST00000167427.2 | 8713 | ENSMUSG00000017002 | Slpi | secretory leukocyte peptidase inhibitor |
| chr10 | 1,24E+08 | 1,24E+08 | 1188 | * | 8,110805 | -3,526503715 | 1,353215 | -2,60602 | 0,00916 | 0,883012141 | 1,360622 | Distal Intergenic | 10 | 1,24E+08 | 1,24E+08 | 55415 | 1 | 75030 | ENSMUST00000219167.2 | -20545 | ENSMUSG00000112364 | 4930503E24Rik | eukaryotic translation initiation factor 3, subunit I pseudogene |
| chr3 | 77089841 | 77091093 | 1253 | * | 11,15596 | -3,526348662 | 1,178941 | -2,99111 | 0,00278 | 0,777919825 | 0,744375 | Distal Intergenic | 3 | 77012516 | 77012592 | 77 | 2 | 1,15E+08 | ENSMUST00000157856.4 | -77249 | | Gm23644 | predicted gene, 23644 |

| | | | | | | | | | | | | | | | | | | | | | | | |
|---|---|---|---|---|---|---|---|---|---|---|---|---|---|---|---|---|---|---|---|---|---|---|---|
| chr14 | 12944697 | 12945509 | 813 | * | 11,18201 | -3,52623851 | 1,168109 | -3,01876 | 0,002538 | 0,761677776 | 0,738441 | Distal Intergenic | 14 | 12583005 | 12585578 | 2574 | 2 | 1E+08 | ENSMUST00000224435.2 | -359119 | ENSMUSG00000089686 | Mnd1-ps | Mnd1 retrotransposed pseudogene |
| chr11 | 43763062 | 43764312 | 1251 | * | 12,78721 | -3,525480864 | 1,125419 | -3,1326 | 0,001733 | 0,663408464 | 0,918255 | Intron (ENSMUST0000( | 11 | 43665433 | 43792037 | 126605 | 2 | 11548 | ENSMUST00000067258.9 | 27725 | ENSMUSG00000050541 | Adra1b | adrenergic receptor, alpha 1b |
| chr7 | 69076686 | 69078401 | 1716 | * | 14,15455 | -3,523071647 | 1,087757 | -3,23884 | 0,0012 | 0,596107452 | 0,918255 | Distal Intergenic | 7 | 69004120 | 69046680 | 42561 | 2 | 1,03E+08 | ENSMUST00000209231.2 | -30006 | | Gm34838 | predicted gene, 34838 |
| chr17 | 63108317 | 63109992 | 1676 | * | 18,14902 | -3,518861615 | 1,024686 | -3,43409 | 0,000595 | 0,306095127 | 2,507013 | Intron (ENSMUST0000( | 17 | 62914314 | 63188139 | 273826 | 2 | 13640 | ENSMUST00000078839.5 | 78147 | ENSMUSG00000048915 | Efna5 | ephrin A5 |
| chr16 | 45002226 | 45003304 | 1079 | * | 9,910851 | -3,518341276 | 1,248997 | -2,81693 | 0,004848 | 0,755814512 | 1,449706 | Exon (ENSMUST00000( | 16 | 44986246 | 45008257 | 22012 | 1 | 67841 | ENSMUST00000150390.2 | 15980 | ENSMUSG00000022663 | Atg3 | autophagy related 3 |
| chr11 | 88764349 | 88766197 | 1849 | * | 13,06986 | -3,513105049 | 1,264913 | -2,77735 | 0,00548 | 0,898584511 | 0,738441 | Intron (ENSMUST0000( | 11 | 88721618 | 88755412 | 33795 | 2 | 11640 | ENSMUST00000018572.11 | -8937 | ENSMUSG00000018428 | Akap1 | A kinase (PRKA) anchor protein 1 |
| chr15 | 60718750 | 60719911 | 1162 | * | 16,65671 | -3,510899788 | 1,104268 | -3,17939 | 0,001476 | 0,443457296 | 2,663619 | Intron (ENSMUST0000( | 15 | 60690843 | 60696929 | 6087 | 2 | 399603 | ENSMUST00000100635.5 | -21821 | ENSMUSG00000072568 | Lratd2 | LRAT domain containing 1 |
| chr11 | 44781173 | 44782555 | 1383 | * | 14,06576 | -3,51078965 | 1,119636 | -3,13565 | 0,001715 | 0,661775851 | 0,918255 | Intron (ENSMUST0000( | 11 | 44802965 | 44863741 | 60777 | 1 | 13591 | ENSMUST00000140822.2 | -20410 | ENSMUSG00000057098 | Ebf1 | early B cell factor 1 |
| chr10 | 1,09E+08 | 1,09E+08 | 1012 | * | 10,10932 | -3,510561453 | 1,218401 | -2,88128 | 0,003961 | 0,751314174 | 1,211315 | Distal Intergenic | 10 | 1,09E+08 | 1,09E+08 | 106 | 2 | 1,15E+08 | ENSMUST00000083868.3 | -119801 | ENSMUSG00000065802 | Gm23041 | predicted gene, 23041 |
| chr3 | 1,42E+08 | 1,42E+08 | 1163 | * | 9,256015 | -3,510195017 | 1,329057 | -2,64112 | 0,008263 | 0,852340527 | 1,449706 | Exon (ENSMUST00000( | 3 | 1,42E+08 | 1,42E+08 | 54612 | 2 | 56376 | ENSMUST00000200650.5 | 55644 | ENSMUSG00000028273 | Pdlim5 | PDZ and LIM domain 5 |
| chr9 | 1,07E+08 | 1,07E+08 | 2057 | * | 15,06149 | -3,507208891 | 1,082637 | -3,2395 | 0,001197 | 0,532631848 | 1,2763 | Distal Intergenic | 9 | 1,07E+08 | 1,07E+08 | 58570 | 1 | 321006 | ENSMUST00000159645.8 | -11651 | ENSMUSG00000040325 | Dcaf1 | DDB1 and CUL4 associated factor 1 |
| chr4 | 3478126 | 3479273 | 1148 | * | 10,97571 | -3,506752457 | 1,270561 | -2,76 | 0,00578 | 0,905015433 | 0,738441 | Distal Intergenic | 4 | 3496978 | 3516515 | 19538 | 2 | 666513 | ENSMUST00000121996.2 | 37242 | | Gm11787 | predicted gene 11787 |
| chr1 | 43003248 | 43004287 | 1040 | * | 8,249844 | -3,50542573 | 1,327224 | -2,64117 | 0,008262 | 0,861009039 | 1,360622 | Intron (ENSMUST0000( | 1 | 42992299 | 43069941 | 77643 | 1 | 93690 | ENSMUST00000179655.2 | 10949 | ENSMUSG00000041907 | Gpr45 | G protein-coupled receptor 45 |
| chr14 | 28280876 | 28282248 | 1373 | * | 13,80153 | -3,504120779 | 1,010586 | -3,46741 | 0,000525 | 0,483145429 | 0,738441 | Distal Intergenic | 14 | 28233408 | 28233864 | 457 | 1 | 22418 | ENSMUST00000146192.2 | 47468 | ENSMUSG00000021994 | Wnt5a | wingless-type MMTV integration site family, member 5A |
| chrX | 85671195 | 85672407 | 1213 | * | 9,03068 | -3,5033714 | 1,270884 | -2,75664 | 0,00584 | 0,801688106 | 1,360622 | Distal Intergenic | 20 | 85293862 | 85348699 | 54838 | 2 | 434903 | ENSMUST00000096378.3 | -322496 | ENSMUSG00000035427 | Mageb4 | MAGE family member B4 |
| chr5 | 83204429 | 83205306 | 878 | * | 7,524956 | -3,5022823 | 1,349957 | -2,59437 | 0,009477 | 1 | 0 | Distal Intergenic | 5 | 82946791 | 82946912 | 122 | 2 | 1E+08 | ENSMUST00000116936.3 | -257517 | ENSMUSG00000080586 | Mir1187 | microRNA 1187 |
| chr1 | 1,87E+08 | 1,87E+08 | 1288 | * | 13,68545 | -3,497192662 | 1,263199 | -2,76852 | 0,005631 | 0,901463351 | 0,738441 | Exon (ENSMUST00000: | 1 | 1,87E+08 | 1,88E+08 | 184047 | 1 | 26381 | ENSMUST00000152927.2 | 16830 | ENSMUSG00000026610 | Esrrg | estrogen-related receptor gamma |
| chr2 | 98051814 | 98052750 | 937 | * | 7,848857 | -3,496979056 | 1,249986 | -2,79762 | 0,005148 | 1 | 0 | Distal Intergenic | 2 | 97298434 | 97459408 | 160975 | 1 | 241568 | ENSMUST00000170144.2 | 753380 | ENSMUSG00000050587 | Lrrc4c | leucine rich repeat containing 4C |
| chr1 | 17414951 | 17416326 | 1376 | * | 12,04283 | -3,496648783 | 0,991999 | -3,52485 | 0,000424 | 0,446770792 | 0,744375 | Distal Intergenic | 1 | 17238627 | 17243327 | 4701 | 1 | 1,03E+08 | ENSMUST00000187817.2 | 176324 | | Gm28784 | predicted gene 28784 |
| chr2 | 57117804 | 57118622 | 819 | * | 10,00583 | -3,496417982 | 1,265714 | -2,76241 | 0,005738 | 0,838878373 | 1,086118 | Distal Intergenic | 2 | 57127647 | 57257865 | 130219 | 1 | 14571 | ENSMUST00000141536.8 | -9025 | ENSMUSG00000026827 | Gpd2 | glycerol phosphate dehydrogenase 2, mitochondrial |
| chr2 | 1,23E+08 | 1,23E+08 | 1000 | * | 8,475154 | -3,496054613 | 1,340541 | -2,60794 | 0,009109 | 0,792545355 | 2,237403 | Distal Intergenic | 2 | 1,23E+08 | 1,23E+08 | 9798 | 1 | 59010 | ENSMUST00000176343.2 | 557140 | ENSMUSG00000005803 | Sqor | sulfide quinone oxidoreductase |
| chr1 | 29866979 | 29867770 | 792 | * | 7,921718 | -3,495741615 | 1,326422 | -2,63547 | 0,008402 | 1 | 0 | Distal Intergenic | 1 | 29460569 | 29460693 | 125 | 1 | 1,02E+08 | ENSMUST00000183870.3 | 406410 | ENSMUSG00000099257 | Mir6342 | microRNA 6342 |
| chrX | 55058069 | 55059948 | 1880 | * | 11,90649 | -3,493823845 | 1,090587 | -3,20362 | 0,001357 | 0,659240897 | 0,738441 | Distal Intergenic | 20 | 55181932 | 55198858 | 16927 | 1 | 70691 | ENSMUST00000033458.3 | -121984 | ENSMUSG00000031125 | 3830403N18Rik | RIKEN cDNA 3830403N18 gene |
| chr13 | 1,1E+08 | 1,1E+08 | 1094 | * | 9,959498 | -3,488431016 | 1,336864 | -2,60941 | 0,00907 | 0,899620932 | 1,211315 | Distal Intergenic | 13 | 1,1E+08 | 1,1E+08 | 4572 | 2 | 238875 | ENSMUST00000058806.7 | 6343 | ENSMUSG00000046006 | Gapt | Grb2-binding adaptor, transmembrane |
| chr2 | 1,04E+08 | 1,04E+08 | 864 | * | 8,366314 | -3,48610348 | 1,326539 | -2,62797 | 0,00859 | 0,86031879 | 1,425929 | Promoter (<=1kb) | 2 | 1,04E+08 | 1,04E+08 | 139 | 1 | 1,15E+08 | ENSMUST00000157114.3 | -385 | ENSMUSG00000087739 | Gm24644 | predicted gene, 24644 |
| chr10 | 46769879 | 46771198 | 1320 | * | 9,80738 | -3,484529588 | 1,262135 | -2,76082 | 0,005766 | 0,792545355 | 1,425929 | Distal Intergenic | 10 | 46714442 | 46714603 | 162 | 2 | 1,15E+08 | ENSMUST00000083784.3 | -55276 | ENSMUSG00000065718 | Gm25650 | predicted gene, 25650 |
| chr3 | 1,4E+08 | 1,4E+08 | 841 | * | 10,14055 | -3,481615851 | 1,201518 | -2,89768 | 0,003759 | 0,749550311 | 1,173325 | Distal Intergenic | 3 | 1,4E+08 | 1,4E+08 | 122 | 1 | 1,02E+08 | ENSMUST00000184427.3 | 171922 | ENSMUSG00000099169 | Mir7j | microRNA 7j |
| chrX | 44945086 | 44946606 | 1521 | * | 8,914751 | -3,480270531 | 1,276062 | -2,72735 | 0,006384 | 0,730469267 | 2,237403 | Distal Intergenic | 20 | 44884238 | 44884344 | 107 | 2 | 1,15E+08 | ENSMUST00000082530.3 | -60742 | ENSMUSG00000119236 | Gm25078 | predicted gene, 25078 |
| chrX | 68651509 | 68652930 | 1422 | * | 8,190174 | -3,477653983 | 1,320085 | -2,63442 | 0,008428 | 1 | 0 | Intron (ENSMUST0000( | 20 | 68404306 | 68828782 | 424477 | 1 | 14266 | ENSMUST00000143097.2 | 247203 | ENSMUSG00000031189 | Aff2 | AF4/FMR2 family, member 2 |
| chr9 | 58682836 | 58684110 | 1275 | * | 10,80567 | -3,470021363 | 1,228114 | -2,82549 | 0,004721 | 0,776899207 | 1,2763 | Distal Intergenic | 9 | 58573116 | 58651247 | 78132 | 2 | 73673 | ENSMUST00000186960.2 | -31589 | ENSMUSG00000074269 | Rec114 | REC114 meiotic recombination protein |
| chr1 | 79532943 | 79533984 | 1042 | * | 11,03303 | -3,464510243 | 1,177514 | -2,94222 | 0,003259 | 0,713406565 | 1,2763 | Distal Intergenic | 1 | 79491456 | 79491581 | 126 | 2 | 1,15E+08 | ENSMUST00000104594.3 | -41362 | ENSMUSG00000077782 | Gm24160 | predicted gene, 24160 |
| chrX | 20228025 | 20229776 | 1752 | * | 8,045537 | -3,46007851 | 1,31363 | -2,63398 | 0,008439 | 1 | 0 | Promoter (<=1kb) | 20 | 20230720 | 20267092 | 36373 | 1 | 19889 | ENSMUST00000033372.13 | -944 | ENSMUSG00000060090 | Rp2 | retinitis pigmentosa 2 homolog |
| chr1 | 41998395 | 41999592 | 1198 | * | 11,51361 | -3,457245592 | 1,168536 | -2,95861 | 0,00309 | 0,756849896 | 0,918255 | Distal Intergenic | 1 | 41957512 | 41959245 | 1734 | 1 | 1,03E+08 | ENSMUST00000187227.2 | 40883 | | Gm29260 | predicted gene 29260 |
| chr13 | 14683183 | 14683973 | 791 | * | 9,951602 | -3,452989823 | 1,260994 | -2,73831 | 0,006176 | 0,832984033 | 1,211315 | Intron (ENSMUST0000( | 13 | 14552309 | 14696160 | 143852 | 2 | 94253 | ENSMUST00000221274.2 | 12187 | ENSMUSG00000021301 | Hecw1 | HECT, C2 and WW domain containing E3 ubiquitin protein ligase 1 |
| chr1 | 93666633 | 93668147 | 1515 | * | 10,98465 | -3,451083628 | 1,202881 | -2,86902 | 0,004118 | 0,810451808 | 0,918255 | Intron (ENSMUST0000( | 1 | 93633109 | 93659648 | 26540 | 2 | 67026 | ENSMUST00000189728.7 | -6985 | ENSMUSG00000026279 | Thap4 | THAP domain containing 4 |
| chr15 | 15407877 | 15409206 | 1330 | * | 8,171513 | -3,449109016 | 1,32485 | -2,6034 | 0,009231 | 0,793703679 | 2,237403 | Distal Intergenic | 15 | 15780080 | 15823510 | 43431 | 1 | 1,03E+08 | ENSMUST00000226912.2 | -370874 | | 4930557F08Rik | RIKEN cDNA 4930557F08 gene |
| chr8 | 45670881 | 45672042 | 1162 | * | 14,03369 | -3,448741451 | 1,13609 | -3,03562 | 0,0024 | 0,685078899 | 1,075735 | Distal Intergenic | 8 | 45639778 | 45649578 | 9801 | 1 | 1,03E+08 | ENSMUST00000209762.2 | 31103 | | Gm30504 | predicted gene, 30504 |
| chr12 | 71454427 | 71455854 | 1428 | * | 12,38681 | -3,446138691 | 1,179292 | -2,92221 | 0,003476 | 0,81949316 | 0,738441 | Intron (ENSMUST0000( | 12 | 71429928 | 71430982 | 1055 | 2 | 67348 | ENSMUST00000190526.2 | -23445 | ENSMUSG00000100668 | 1700083H02Rik | RIKEN cDNA 1700083H02 gene |
| chr1 | 1,64E+08 | 1,64E+08 | 1983 | * | 14,99384 | -3,445801286 | 1,174607 | -2,93358 | 0,003351 | 0,702508842 | 1,37683 | Distal Intergenic | 1 | 1,64E+08 | 1,64E+08 | 22234 | 2 | 240879 | ENSMUST00000160495.2 | -17082 | ENSMUSG00000040113 | Mettl11b | methyltransferase like 11B |
| chr11 | 1,14E+08 | 1,14E+08 | 909 | * | 10,63272 | -3,444656518 | 1,213464 | -2,8387 | 0,00453 | 0,861411338 | 0,744375 | Distal Intergenic | 11 | 1,14E+08 | 1,14E+08 | 942 | 1 | 73536 | ENSMUST00000106604.2 | 11009 | ENSMUSG00000078600 | 1700092K14Rik | RIKEN cDNA 1700092K14 gene |
| chrX | 1,6E+08 | 1,6E+08 | 1111 | * | 9,450533 | -3,444057804 | 1,238948 | -2,77982 | 0,005439 | 0,778555725 | 1,449706 | Distal Intergenic | 20 | 1,6E+08 | 1,6E+08 | 67874 | 1 | 1,08E+08 | ENSMUST00000207320.2 | 84373 | | Gm15262 | predicted gene 15262 |
| chrX | 58321593 | 58323244 | 1652 | * | 11,254 | -3,443317522 | 1,087777 | -3,16546 | 0,001548 | 0,644495993 | 0,918255 | Intron (ENSMUST0000( | 20 | 58107505 | 58211524 | 104020 | 2 | 14168 | ENSMUST00000119306.2 | -110069 | ENSMUSG00000031137 | Fgf13 | fibroblast growth factor 13 |
| chr16 | 36217001 | 36217752 | 752 | * | 8,368133 | -3,441868375 | 1,309173 | -2,62904 | 0,008563 | 0,859725538 | 1,425929 | Distal Intergenic | 16 | 36208340 | 36211484 | 3145 | 2 | 639105 | ENSMUST00000114851.3 | -5517 | | Gm7260 | predicted gene 7260 |
| chr9 | 89819974 | 89821249 | 1276 | * | 9,099751 | -3,441446465 | 1,312158 | -2,62274 | 0,008723 | 0,891607598 | 1,211315 | Intron (ENSMUST0000( | 9 | 89791965 | 89808793 | 16829 | 1 | 19417 | ENSMUST00000190073.2 | 28009 | ENSMUSG00000032356 | Rasgrf1 | RAS protein-specific guanine nucleotide-releasing factor 1 |
| chr9 | 84008617 | 84009998 | 1382 | * | 11,20879 | -3,441024282 | 1,221016 | -2,81817 | 0,00483 | 0,838625535 | 0,918255 | Distal Intergenic | 9 | 83989560 | 84006293 | 16734 | 1 | 12040 | ENSMUST00000186136.2 | 19057 | ENSMUSG00000032263 | Bckdhb | branched chain ketoacid dehydrogenase E1, beta polypeptide |
| chr5 | 57831485 | 57832912 | 1428 | * | 8,959087 | -3,439387938 | 1,290217 | -2,66574 | 0,007682 | 0,838878373 | 1,449706 | Intron (ENSMUST0000( | 5 | 57727424 | 57875262 | 147839 | 2 | 319216 | ENSMUST00000180708.3 | 42350 | ENSMUSG00000097216 | 4932441J04Rik | RIKEN cDNA 4932441J04 gene |
| chrX | 8592928 | 8594607 | 1680 | * | 10,0996 | -3,435924616 | 1,305016 | -2,63286 | 0,008467 | 0,89184433 | 1,173325 | Distal Intergenic | 20 | 8614671 | 8620826 | 6156 | 1 | 382206 | ENSMUST00000103001.4 | -20064 | ENSMUSG00000035371 | Ssx9 | synovial sarcoma, X 9 |
| chr5 | 1,01E+08 | 1,01E+08 | 1373 | * | 9,897175 | -3,43406593 | 1,326321 | -2,58917 | 0,009621 | 0,923399738 | 1,086118 | Distal Intergenic | 5 | 1,01E+08 | 1,01E+08 | 18521 | 2 | 231507 | ENSMUST00000031264.12 | -33220 | ENSMUSG00000029322 | Plac8 | placenta-specific 8 |
| chr14 | 35354220 | 35355455 | 1236 | * | 13,16527 | -3,43363057 | 1,129188 | -3,0408 | 0,00236 | 0,713663698 | 0,918255 | Distal Intergenic | 14 | 35291951 | 35303071 | 11121 | 1 | 14803 | ENSMUST00000173307.2 | 62269 | ENSMUSG00000041078 | Grid1 | glutamate receptor, ionotropic, delta 1 |
| chr10 | 1,02E+08 | 1,02E+08 | 943 | * | 9,218485 | -3,433145172 | 1,249021 | -2,74867 | 0,005984 | 0,806376005 | 1,360622 | Intron (ENSMUST0000( | 10 | 1,02E+08 | 1,02E+08 | 81 | 1 | 1,01E+08 | ENSMUST00000175211.3 | -16145 | ENSMUSG00000092952 | Mir3059 | microRNA 3059 |
| chr8 | 52263117 | 52264713 | 1597 | * | 12,86685 | -3,432263048 | 1,191887 | -2,87969 | 0,003981 | 0,777021969 | 1,075735 | Distal Intergenic | 8 | 52789950 | 52790140 | 191 | 2 | 1,15E+08 | ENSMUST00000082748.3 | 525427 | ENSMUSG00000064682 | Gm25813 | predicted gene, 25813 |
| chr2 | 5781047 | 5782562 | 1516 | * | 14,64532 | -3,431222067 | 1,214806 | -2,8245 | 0,004735 | 0,777172375 | 1,2763 | Distal Intergenic | 2 | 5799476 | 5803409 | 3934 | 2 | 98828 | ENSMUST00000130142.2 | 20847 | ENSMUSG00000039128 | Cdc123 | cell division cycle 123 |
| chr2 | 1,61E+08 | 1,61E+08 | 1603 | * | 14,25165 | -3,430881282 | 1,029139 | -3,33374 | 0,000857 | 0,57149888 | 0,738441 | Intron (ENSMUST0000( | 2 | 1,62E+08 | 1,62E+08 | 102147 | 2 | 19281 | ENSMUST00000153770.2 | 155385 | ENSMUSG00000053141 | Ptprt | protein tyrosine phosphatase, receptor type, T |
| chr13 | 89544073 | 89545525 | 1453 | * | 9,625464 | -3,428543879 | 1,276814 | -2,68523 | 0,007248 | 0,864313266 | 1,173325 | Distal Intergenic | 13 | 89687915 | 89759771 | 71857 | 1 | 12950 | ENSMUST00000022108.9 | -142390 | ENSMUSG00000021613 | Hapln1 | hyaluronan and proteoglycan link protein 1 |
| chr2 | 18238358 | 18239388 | 1031 | * | 10,60068 | -3,424677437 | 1,318888 | -2,59664 | 0,009414 | 0,976107896 | 0,744375 | Intron (ENSMUST0000( | 2 | 18200465 | 18224368 | 23904 | 2 | 13418 | ENSMUST00000148401.8 | -13990 | ENSMUSG00000026740 | Dnajc1 | DnaJ heat shock protein family (Hsp40) member C1 |
| chr13 | 10353129 | 10354203 | 1075 | * | 9,604549 | -3,422234053 | 1,223934 | -2,79609 | 0,005172 | 0,801134483 | 1,211315 | Intron (ENSMUST0000( | 13 | 10407631 | 10708929 | 301299 | 1 | 1,08E+08 | ENSMUST00000180807.3 | -53428 | | Gm26861 | predicted gene, 26861 |
| chr8 | 51808599 | 51809629 | 1031 | * | 9,729797 | -3,422231942 | 1,248646 | -2,74075 | 0,00613 | 0,79897927 | 1,449706 | Distal Intergenic | 8 | 51251821 | 51370032 | 118212 | 2 | 23965 | ENSMUST00000211313.2 | -438567 | ENSMUSG00000031561 | Tenm3 | teneurin transmembrane protein 3 |
| chr4 | 1,17E+08 | 1,17E+08 | 1278 | * | 9,65301 | -3,422196802 | 1,231696 | -2,77844 | 0,005462 | 0,81148874 | 1,211315 | Intron (ENSMUST0000( | 4 | 1,17E+08 | 1,17E+08 | 449 | 2 | 381544 | ENSMUST00000141554.2 | -6530 | ENSMUSG00000060268 | Armh1 | armadillo-like helical domain containing 1 |
| chr10 | 17831699 | 17832978 | 1280 | * | 10,69246 | -3,422099352 | 1,311845 | -2,60862 | 0,009091 | 0,972507971 | 0,744375 | Distal Intergenic | 10 | 17774783 | 17823815 | 49033 | 2 | 380629 | ENSMUST00000037879.8 | -7884 | ENSMUSG00000039879 | Heca | hdc homolog, cell cycle regulator |
| chr15 | 34103757 | 34105705 | 1949 | * | 11,85333 | -3,419747306 | 1,195553 | -2,86039 | 0,004231 | 0,81623383 | 0,918255 | Intron (ENSMUST0000( | 15 | 34114233 | 34115348 | 1116 | 1 | 67154 | ENSMUST00000226331.2 | -8528 | ENSMUSG00000022255 | Mtdh | metadherin |
| chr3 | 56746933 | 56748603 | 1671 | * | 15,4367 | -3,415353968 | 1,171197 | -2,91612 | 0,003544 | 0,713797407 | 1,37683 | Distal Intergenic | 3 | 56961571 | 56961700 | 130 | 1 | 1,15E+08 | ENSMUST00000239689.1 | -212968 | ENSMUSG00000118960 | Gm22269 | predicted gene, 22269 |
| chr9 | 70970482 | 70972416 | 1935 | * | 13,93196 | -3,413430038 | 1,284373 | -2,65766 | 0,007868 | 0,915421656 | 0,938682 | Distal Intergenic | 9 | 70976432 | 70983809 | 7378 | 1 | 1,08E+08 | ENSMUST00000240854.1 | -4016 | | Gm46120 | predicted gene, 46120 |
| chr5 | 1,39E+08 | 1,39E+08 | 1107 | * | 9,033886 | -3,411605264 | 1,246802 | -2,73629 | 0,006214 | 0,725842761 | 2,237403 | Distal Intergenic | 5 | 1,39E+08 | 1,39E+08 | 55564 | 1 | 80752 | ENSMUST00000026972.8 | -34723 | ENSMUSG00000025854 | Fam20c | FAM20C, golgi associated secretory pathway kinase |
| chr14 | 28043281 | 28044917 | 1637 | * | 14,67747 | -3,405701644 | 1,169611 | -2,91182 | 0,003593 | 0,735602255 | 1,214904 | Intron (ENSMUST0000( | 14 | 28177478 | 28197960 | 20483 | 1 | 238988 | ENSMUST00000224045.2 | -132561 | ENSMUSG00000040640 | Erc2 | ELKS/RAB6-interacting/CAST family member 2 |
| chr1 | 72957832 | 72959900 | 2069 | * | 13,89423 | -3,404562513 | 1,283155 | -2,65327 | 0,007972 | 0,897709339 | 1,075735 | Distal Intergenic | 1 | 72897091 | 72914043 | 16953 | 2 | 16011 | ENSMUST00000027377.9 | -43789 | ENSMUSG00000026185 | Igfbp5 | insulin-like growth factor binding protein 5 |
| chr3 | 85144414 | 85145255 | 842 | * | 10,88275 | -3,403881312 | 1,316999 | -2,58457 | 0,00975 | 0,982231644 | 0,744375 | Intron (ENSMUST0000( | 3 | 85224826 | 85239337 | 14512 | 1 | 73283 | ENSMUST00000189046.2 | -79571 | ENSMUSG00000099995 | 1700036G14Rik | RIKEN cDNA 1700036G14 gene |
| chr16 | 26745471 | 26746959 | 1489 | * | 13,49556 | -3,40199909 | 1,165919 | -2,91787 | 0,003524 | 0,778555725 | 0,938682 | Intron (ENSMUST0000( | 16 | 26732303 | 26784356 | 52054 | 2 | 239789 | ENSMUST00000231417.2 | 37397 | ENSMUSG00000068428 | Gmnc | geminin coiled-coil domain containing |
| chr7 | 17350761 | 17352814 | 2054 | * | 16,78045 | -3,400985763 | 0,989615 | -3,43668 | 0,000589 | 0,309888367 | 2,386008 | Exon (ENSMUST00020: | 7 | 17371404 | 17384270 | 12867 | 2 | 671132 | ENSMUST00000206108.2 | 31456 | ENSMUSG00000108654 | Gm9519 | predicted gene 9519 |
| chr16 | 16933699 | 16935050 | 1352 | * | 10,25535 | -3,400074409 | 1,277311 | -2,6619 | 0,00777 | 0,845344544 | 1,425929 | Promoter (<=1kb) | 16 | 16931399 | 16935640 | 4242 | 2 | 70458 | ENSMUST00000231630.2 | 590 | ENSMUSG00000049916 | 2610318N02Rik | RIKEN cDNA 2610318N02 gene |
| chr10 | 75276147 | 75277218 | 1072 | * | 9,943514 | -3,396391704 | 1,25073 | -2,71553 | 0,006617 | 0,85188813 | 1,173325 | 3' UTR | 10 | 75264015 | 75266153 | 2139 | 1 | 103149 | ENSMUST00000129946.2 | 12132 | ENSMUSG00000033427 | Upb1 | ureidopropionase, beta |
| chr11 | 1,09E+08 | 1,09E+08 | 1140 | * | 12,30571 | -3,39589251 | 1,177559 | -2,88384 | 0,003929 | 0,839768694 | 0,738441 | Distal Intergenic | 11 | 1,09E+08 | 1,09E+08 | 7163 | 1 | 12006 | ENSMUST00000139682.2 | 59865 | ENSMUSG00000000142 | Axin2 | axin 2 |
| chr3 | 19397909 | 19399114 | 1206 | * | 9,363376 | -3,394610037 | 1,239388 | -2,73894 | 0,006164 | 0,802490409 | 1,425929 | Distal Intergenic | 3 | 19277272 | 19365486 | 88215 | 2 | 18583 | ENSMUST00000099195.10 | -32423 | ENSMUSG00000069094 | Pde7a | phosphodiesterase 7A |
| chr14 | 1,23E+08 | 1,23E+08 | 1405 | * | 9,516811 | -3,394405821 | 1,284893 | -2,64178 | 0,008247 | 0,884930392 | 1,211315 | Intron (ENSMUST0000( | 14 | 1,23E+08 | 1,23E+08 | 84606 | 1 | 380934 | ENSMUST00000226524.2 | 13978 | ENSMUSG00000115016 | AA536875 | expressed sequence AA536875 |
| chr6 | 65839755 | 65840669 | 915 | * | 10,74391 | -3,392534561 | 1,22704 | -2,76481 | 0,005696 | 0,824134241 | 1,173325 | Exon (ENSMUST00000( | 6 | 65756085 | 65913118 | 157034 | 1 | 70779 | ENSMUST00000172638.2 | 83670 | ENSMUSG00000029913 | Prdm5 | PR domain containing 5 |
| chr10 | 21439280 | 21440404 | 1125 | * | 11,96391 | -3,390882515 | 1,237995 | -2,73901 | 0,006162 | 0,823736512 | 1,2763 | Distal Intergenic | 10 | 21469044 | 21498275 | 29232 | 1 | 67692 | ENSMUST00000057341.6 | -28640 | ENSMUSG00000050844 | 1700020N01Rik | RIKEN cDNA 1700020N01 gene |
| chr9 | 1E+08 | 1E+08 | 1251 | * | 15,7891 | -3,388980071 | 1,052553 | -3,21977 | 0,001283 | 0,439076133 | 2,386008 | Distal Intergenic | 9 | 1E+08 | 1E+08 | 7470 | 2 | 1,03E+08 | ENSMUST00000214991.2 | 30697 | | Gm34397 | predicted gene, 34397 |
| chr2 | 83141400 | 83142186 | 787 | * | 11,53549 | -3,388194327 | 1,09577 | -3,09207 | 0,001988 | 0,644704154 | 1,173325 | Distal Intergenic | 2 | 82811140 | 82839281 | 28142 | 1 | 241516 | ENSMUST00000132967.2 | 330260 | ENSMUSG00000075249 | Fsip2 | fibrous sheath-interacting protein 2 |
| chrX | 1,16E+08 | 1,16E+08 | 1809 | * | 7,953262 | -3,385891813 | 1,296265 | -2,61204 | 0,009 | 1 | 0 | Distal Intergenic | 20 | 1,16E+08 | 1,16E+08 | 348 | 1 | 1E+08 | ENSMUST00000122446.2 | 206788 | ENSMUSG00000067441 | H2ab1 | H2A.B variant histone 1 |
| chr9 | 81062099 | 81063118 | 1020 | * | 12,96813 | -3,384154368 | 1,086229 | -3,11551 | 0,001836 | 0,671937722 | 0,918255 | Intron (ENSMUST0000( | 9 | 81513604 | 81528698 | 15095 | 1 | 319405 | ENSMUST00000181893.9 | -450486 | ENSMUSG00000097466 | D430036J16Rik | RIKEN cDNA D430036J16 gene |
| chr1 | 1E+08 | 1E+08 | 863 | * | 8,795051 | -3,383684053 | 1,279263 | -2,64503 | 0,008168 | 1 | 0 | Intron (ENSMUST0000( | 1 | 1E+08 | 1E+08 | 490 | 1 | 1,15E+08 | ENSMUST00000190817.2 | 104228 | | Gm29667 | predicted gene 29667 |
| chr6 | 1,06E+08 | 1,06E+08 | 729 | * | 11,82041 | -3,381712429 | 1,189169 | -2,84376 | 0,004458 | 0,860530858 | 0,738441 | Intron (ENSMUST0000( | 6 | 1,06E+08 | 1,06E+08 | 226685 | 1 | 269784 | ENSMUST00000130040.2 | 24500 | ENSMUSG00000064293 | Cntn4 | contactin 4 |
| chr4 | 58147779 | 58148966 | 1188 | * | 11,25014 | -3,376664014 | 1,226656 | -2,75274 | 0,00591 | 0,874247416 | 0,918255 | Intron (ENSMUST0000( | 4 | 58042442 | 58206596 | 164155 | 2 | 64817 | ENSMUST00000042850.9 | 57630 | ENSMUSG00000028369 | Svep1 | sushi, von Willebrand factor type A, EGF and pentraxin domain containing 1 |
| chr2 | 1,58E+08 | 1,58E+08 | 1151 | * | 10,01002 | -3,374659835 | 1,256349 | -2,68609 | 0,007229 | 0,829772916 | 1,449706 | Exon (ENSMUST00000( | 2 | 1,58E+08 | 1,58E+08 | 46712 | 1 | 66642 | ENSMUST00000156300.2 | 15942 | ENSMUSG00000027649 | Ctnnbl1 | catenin, beta like 1 |
| chr7 | 15888276 | 15889437 | 1162 | * | 9,749709 | -3,372839605 | 1,275194 | -2,64496 | 0,00817 | 0,88319889 | 1,211315 | Intron (ENSMUST0000( | 7 | 15909015 | 15912718 | 3704 | 1 | 17537 | ENSMUST00000176342.8 | -19578 | ENSMUSG00000041420 | Meis3 | Meis homeobox 3 |
| chr12 | 68642607 | 68643683 | 1077 | * | 9,792165 | -3,369391244 | 1,257147 | -2,68019 | 0,007358 | 0,83238039 | 1,449706 | Distal Intergenic | 12 | 69176610 | 69191850 | 15241 | 1 | 1,03E+08 | ENSMUST00000222345.2 | -532927 | ENSMUSG00000113158 | Gm31447 | predicted gene, 31447 |
| chr2 | 1,4E+08 | 1,4E+08 | 1236 | * | 12,01656 | -3,363728413 | 1,136685 | -2,95925 | 0,003084 | 0,700182401 | 1,2763 | Intron (ENSMUST0000( | 2 | 1,4E+08 | 1,4E+08 | 65683 | 2 | 75812 | ENSMUST00000124949.2 | -7144 | ENSMUSG00000039033 | Tasp1 | taspase, threonine aspartase 1 |
| chr6 | 20124239 | 20125092 | 854 | * | 10,9652 | -3,36296891 | 1,200155 | -2,80211 | 0,005077 | 0,886332554 | 0,738441 | Distal Intergenic | 6 | 19624608 | 19624714 | 107 | 1 | 1,15E+08 | ENSMUST00000083768.3 | 499631 | ENSMUSG00000065702 | Gm22828 | predicted gene, 22828 |
| chr3 | 14257191 | 14258892 | 1702 | * | 16,48972 | -3,36203625 | 0,987808 | -3,40353 | 0,000665 | 0,339611164 | 2,141612 | Distal Intergenic | 3 | 14428177 | 14442647 | 14471 | 1 | 622229 | ENSMUST00000192393.2 | -169285 | | Gm6300 | solute carrier family 7 (cationic amino acid transporter, y+ system), member pseudogene |
| chrX | 1,08E+08 | 1,08E+08 | 1336 | * | 9,270744 | -3,36196118 | 1,237868 | -2,71593 | 0,006609 | 0,73516474 | 2,237403 | Distal Intergenic | 20 | 1,08E+08 | 1,08E+08 | 3682 | 1 | 622976 | ENSMUST00000060013.4 | 18411 | ENSMUSG00000048621 | Gm6377 | predicted gene 6377 |
| chr11 | 66228548 | 66229855 | 1308 | * | 10,50067 | -3,360331813 | 1,219094 | -2,75642 | 0,005844 | 0,906138253 | 0,738441 | Intron (ENSMUST0000( | 11 | 66265959 | 66393640 | 127682 | 2 | 380702 | ENSMUST00000134562.2 | 163785 | ENSMUSG00000053930 | Shisa6 | shisa family member 6 |
| chr1 | 96625268 | 96626143 | 876 | * | 9,545217 | -3,360054039 | 1,147967 | -2,92696 | 0,003423 | 0,748641341 | 1,086118 | Distal Intergenic | 1 | 96539782 | 96590298 | 50517 | 2 | 75159 | ENSMUST00000189056.2 | -34970 | ENSMUSG00000100589 | 4930533P14Rik | RIKEN cDNA 4930533P14 gene |
| chr4 | 68112962 | 68113838 | 877 | * | 11,87247 | -3,360014967 | 1,137482 | -2,95391 | 0,003138 | 0,705187732 | 1,2763 | Distal Intergenic | 4 | 68100769 | 68100939 | 171 | 1 | 1,15E+08 | ENSMUST00000082828.3 | 12193 | ENSMUSG00000064762 | Gm23738 | predicted gene, 23738 |
| chr11 | 24904405 | 24905617 | 1213 | * | 10,72689 | -3,359803866 | 1,152956 | -2,91408 | 0,003567 | 0,740669062 | 1,173325 | Distal Intergenic | 11 | 24853983 | 24876058 | 22076 | 2 | 1,03E+08 | ENSMUST00000126063.8 | -28347 | ENSMUSG00000087145 | Gm12665 | predicted gene 12665 |
| chr9 | 16453106 | 16453988 | 883 | * | 9,531131 | -3,35933868 | 1,173087 | -2,86367 | 0,004188 | 0,661775851 | 2,237403 | Distal Intergenic | 9 | 15821485 | 16412581 | 591097 | 2 | 270120 | ENSMUST00000217308.3 | -40525 | ENSMUSG00000074505 | Fat3 | FAT atypical cadherin 3 |
| chr5 | 1,29E+08 | 1,29E+08 | 1360 | * | 14,43399 | -3,357855893 | 1,138869 | -2,94841 | 0,003194 | 0,70944879 | 1,2763 | Distal Intergenic | 5 | 1,29E+08 | 1,29E+08 | 3250 | 1 | 93897 | ENSMUST00000117102.4 | 38245 | ENSMUSG00000081683 | Fzd10 | frizzled class receptor 10 |
| chr10 | 23580711 | 23581583 | 873 | * | 10,92878 | -3,357268647 | 1,115066 | -3,01082 | 0,002605 | 0,767135014 | 0,738441 | Distal Intergenic | 10 | 23661245 | 23661373 | 129 | 2 | 1,01E+08 | ENSMUST00000093556.3 | 79790 | ENSMUSG00000070063 | Snora33 | small nucleolar RNA, H/ACA box 33 |
| chr10 | 10267216 | 10268651 | 1436 | * | 9,938101 | -3,354705097 | 1,248986 | -2,68594 | 0,007233 | 0,83238039 | 1,425929 | 3' UTR | 10 | 10211453 | 10287488 | 76036 | 2 | 215772 | ENSMUST00000148816.8 | 18837 | ENSMUSG00000050994 | Adgb | androglobin |
| chr3 | 33016317 | 33017349 | 1033 | * | 10,05412 | -3,353700647 | 1,299045 | -2,58167 | 0,009832 | 0,884058067 | 1,449706 | Intron (ENSMUST0000( | 3 | 33004779 | 33039228 | 34450 | 2 | 58869 | ENSMUST00000108221.2 | 21879 | ENSMUSG00000027674 | Pex5l | peroxisomal biogenesis factor 5-like |
| chr1 | 41009144 | 41009859 | 716 | * | 9,713253 | -3,350169793 | 1,298323 | -2,58038 | 0,009869 | 0,887238309 | 1,425929 | Distal Intergenic | 1 | 40844761 | 40896047 | 51287 | 1 | 381339 | ENSMUST00000114765.4 | 164383 | ENSMUSG00000079588 | Tmem182 | transmembrane protein 182 |
| chr14 | 95336408 | 95338342 | 1935 | * | 8,915839 | -3,349337733 | 1,258344 | -2,6617 | 0,007775 | 0,852340527 | 1,360622 | Distal Intergenic | 14 | 95167137 | 95167262 | 126 | 1 | 1,15E+08 | ENSMUST00000158759.3 | 169271 | ENSMUSG00000089384 | Gm24680 | predicted gene, 24680 |
| chr15 | 64389347 | 64390895 | 1549 | * | 10,74307 | -3,348201 | 1,219548 | -2,74544 | 0,006043 | 0,878272222 | 0,918255 | Distal Intergenic | 15 | 64499258 | 64505357 | 6100 | 2 | 1,03E+08 | ENSMUST00000227641.2 | 114462 | | Gm30691 | predicted gene, 30691 |

| | | | | | | | | | | | | | | | | | | | | | | | |
|---|---|---|---|---|---|---|---|---|---|---|---|---|---|---|---|---|---|---|---|---|---|---|---|
| chr2 | 1,61E+08 | 1,61E+08 | 1473 | * | 12,88826 | -3,343029791 | 1,104252 | -3,02742 | 0,002467 | 0,681331037 | 1,173325 | Promoter (1-2kb) | 2 | 1,61E+08 | 1,61E+08 | 22740 | 2 | 71389 | ENSMUST00000130265.2 | 1764 | ENSMUSG00000057133 | Chd6 | chromodomain helicase DNA binding protein 6 |
| chr7 | 72026488 | 72027383 | 896 | * | 9,339605 | -3,342718956 | 1,277158 | -2,61731 | 0,008863 | 0,862146257 | 1,449706 | Distal Intergenic | 7 | 71956657 | 71958449 | 1793 | 1 | 1,05E+08 | ENSMUST00000244156.1 | 69831 | | Gm39035 | predicted gene, 39035 |
| chr14 | 23224110 | 23226188 | 2079 | * | 15,186 | -3,341868971 | 1,076487 | -3,10442 | 0,001907 | 0,602555268 | 1,41696 | Distal Intergenic | 14 | 23088262 | 23144639 | 56378 | 2 | 791307 | ENSMUST00000090409.2 | -79471 | ENSMUSG00000068680 | Gm10248 | predicted gene 10248 |
| chr1 | 88326207 | 88327604 | 1398 | * | 11,59703 | -3,340298587 | 1,215685 | -2,74767 | 0,006002 | 0,909192827 | 0,744375 | Distal Intergenic | 1 | 88334683 | 88354160 | 19478 | 1 | 75396 | ENSMUST00000027518.12 | -7079 | ENSMUSG00000026295 | Spp2 | secreted phosphoprotein 2 |
| chr9 | 1,12E+08 | 1,12E+08 | 1055 | * | 9,459979 | -3,339915384 | 1,29612 | -2,57686 | 0,00997 | 0,886332554 | 1,449706 | Distal Intergenic | 9 | 1,12E+08 | 1,12E+08 | 132 | 1 | 1,15E+08 | ENSMUST00000239605.1 | -19068 | ENSMUSG00000119291 | Gm24957 | predicted gene, 24957 |
| chr10 | 25982821 | 25984283 | 1463 | * | 12,29523 | -3,339744418 | 1,223022 | -2,73073 | 0,006319 | 0,886332554 | 0,918255 | Exon (ENSMUST000002 | 10 | 25867084 | 25954950 | 87867 | 2 | 77220 | ENSMUST00000066049.7 | -27871 | ENSMUSG00000049420 | Tmem200a | transmembrane protein 200A |
| chr3 | 22235899 | 22238095 | 2197 | * | 17,87578 | -3,338689423 | 1,049813 | -3,18027 | 0,001471 | 0,459933416 | 2,386008 | Intron (ENSMUST00000 | 3 | 22257356 | 22264743 | 7388 | 1 | 81004 | ENSMUST00000201467.2 | -19261 | ENSMUSG00000027630 | Tbl1xr1 | transducin (beta)-like 1X-linked receptor 1 |
| chr19 | 59607045 | 59608088 | 1044 | * | 11,08703 | -3,334350829 | 1,246056 | -2,67592 | 0,007452 | 0,85617852 | 1,2763 | Distal Intergenic | 19 | 59649755 | 59669638 | 19884 | 1 | 1,01E+08 | ENSMUST00000236333.2 | -41667 | | Gm19956 | predicted gene, 19956 |
| chr17 | 43940810 | 43942113 | 1304 | * | 11,90277 | -3,331557331 | 1,234296 | -2,69916 | 0,006952 | 0,900637308 | 0,918255 | Promoter (<=1kb) | 17 | 43926226 | 43941190 | 14965 | 2 | 210510 | ENSMUST00000168073.9 | 0 | ENSMUSG00000040140 | Tdrd6 | tudor domain containing 6 |
| chr16 | 97796208 | 97797266 | 1059 | * | 9,940229 | -3,331552658 | 1,28694 | -2,58874 | 0,009633 | 0,912096919 | 1,173325 | Promoter (1-2kb) | 16 | 97795799 | 97798403 | 2605 | 2 | 1,08E+08 | ENSMUST00000244430.1 | 1137 | | C030010L15Rik | RIKEN cDNA C030010L15 gene |
| chr13 | 1,16E+08 | 1,16E+08 | 1012 | * | 11,87532 | -3,33061365 | 1,133815 | -2,93753 | 0,003308 | 0,728793055 | 1,173325 | Distal Intergenic | 13 | 1,16E+08 | 1,16E+08 | 3957 | 2 | 16392 | ENSMUST00000176444.2 | 73308 | ENSMUSG00000042258 | Isl1 | ISL1 transcription factor, LIM/homeodomain |
| chrX | 1,6E+08 | 1,6E+08 | 1518 | * | 10,04367 | -3,330057485 | 1,266008 | -2,63036 | 0,008529 | 0,858706485 | 1,425929 | Distal Intergenic | 20 | 1,61E+08 | 1,61E+08 | 62426 | 1 | 24004 | ENSMUST00000112338.2 | -34030 | ENSMUSG00000043518 | Rai2 | retinoic acid induced 2 |
| chr2 | 28908657 | 28909722 | 1066 | * | 12,28638 | -3,329721589 | 1,186814 | -2,8056 | 0,005022 | 0,789027175 | 1,2763 | Intron (ENSMUST00000 | 2 | 28817846 | 28929372 | 111527 | 2 | 329375 | ENSMUST00000190049.2 | 19650 | ENSMUSG00000079502 | Cfap77 | cilia and flagella associated protein 77 |
| chr6 | 22916290 | 22917568 | 1279 | * | 10,69989 | -3,329711137 | 1,228124 | -2,71122 | 0,006704 | 0,926527938 | 0,738441 | Intron (ENSMUST00000 | 6 | 22875776 | 23052616 | 176841 | 1 | 19283 | ENSMUST00000202102.2 | 40514 | ENSMUSG00000068748 | Ptprz1 | protein tyrosine phosphatase, receptor type Z, polypeptide 1 |
| chr2 | 4297684 | 4298645 | 962 | * | 9,378591 | -3,32748848 | 1,263975 | -2,63256 | 0,008474 | 0,85617852 | 1,449706 | Intron (ENSMUST00000 | 2 | 4305329 | 4577250 | 271922 | 1 | 209630 | ENSMUST00000175669.8 | -6684 | ENSMUSG00000026657 | Frmd4a | FERM domain containing 4A |
| chr1 | 13330873 | 13332124 | 1252 | * | 10,51106 | -3,326469865 | 1,159374 | -2,86919 | 0,004115 | 0,749931768 | 1,2763 | Intron (ENSMUST00000 | 1 | 13260330 | 13369946 | 109617 | 2 | 17978 | ENSMUST00000145280.2 | 37822 | ENSMUSG00000005886 | Ncoa2 | nuclear receptor coactivator 2 |
| chr9 | 1,11E+08 | 1,11E+08 | 2663 | * | 21,30062 | -3,326203513 | 0,90212 | -3,6871 | 0,000227 | 0,173497184 | 2,880017 | Promoter (2-3kb) | 9 | 1,11E+08 | 1,11E+08 | 2020 | 1 | 791282 | ENSMUST00000080872.5 | 2380 | ENSMUSG00000057802 | Gm10030 | predicted gene 10030 |
| chr6 | 74979549 | 74980395 | 847 | * | 9,591327 | -3,324963125 | 1,249031 | -2,66203 | 0,007767 | 0,84525842 | 1,425929 | Distal Intergenic | 6 | 74428780 | 74428887 | 108 | 2 | 1,15E+08 | ENSMUST00000082474.3 | -550662 | ENSMUSG00000064408 | Gm23924 | predicted gene, 23924 |
| chr1 | 40556031 | 40558067 | 2037 | * | 13,32497 | -3,32424454 | 1,047173 | -3,17449 | 0,001501 | 0,61075929 | 1,075735 | Promoter (1-2kb) | 1 | 40554608 | 40566077 | 11470 | 1 | 16174 | ENSMUST00000159724.2 | 1423 | ENSMUSG00000026068 | Il18rap | interleukin 18 receptor accessory protein |
| chr1 | 1,94E+08 | 1,94E+08 | 1398 | * | 10,02592 | -3,323858944 | 1,085872 | -3,06101 | 0,002206 | 0,65808656 | 1,211315 | Distal Intergenic | 1 | 1,94E+08 | 1,94E+08 | 10573 | 2 | 74942 | ENSMUST00000193395.2 | -69692 | ENSMUSG00000103692 | 4930503O07Rik | RIKEN cDNA 4930503O07 gene |
| chrX | 1,21E+08 | 1,21E+08 | 1540 | * | 9,311207 | -3,316588717 | 1,17608 | -2,82004 | 0,004802 | 0,754906705 | 1,449706 | Distal Intergenic | 20 | 1,21E+08 | 1,21E+08 | 2837 | 2 | 54561 | ENSMUST00000079490.6 | -189330 | ENSMUSG00000055733 | Nap1l3 | nucleosome assembly protein 1-like 3 |
| chrX | 39899665 | 39901191 | 1527 | * | 9,337986 | -3,314552231 | 1,205723 | -2,74902 | 0,005977 | 0,718390388 | 2,237403 | Distal Intergenic | 20 | 40461136 | 40461391 | 256 | 1 | 1,08E+08 | ENSMUST00000121134.2 | -559945 | ENSMUSG00000083515 | Gm14577 | predicted gene 14577 |
| chr6 | 16127265 | 16128145 | 881 | * | 9,70977 | -3,313885713 | 1,273731 | -2,60172 | 0,009276 | 0,906138253 | 1,173325 | Distal Intergenic | 6 | 16468235 | 16469903 | 1669 | 2 | 1,03E+08 | ENSMUST00000203052.2 | 341758 | | Gm36406 | predicted gene, 36406 |
| chr1 | 1,19E+08 | 1,19E+08 | 1677 | * | 15,78787 | -3,312005151 | 0,995888 | -3,32568 | 0,000882 | 0,352029189 | 2,663619 | Distal Intergenic | 1 | 1,19E+08 | 1,19E+08 | 219488 | 2 | 14633 | ENSMUST00000062483.15 | -14743 | ENSMUSG00000048402 | Gli2 | GLI-Kruppel family member GLI2 |
| chr18 | 58147102 | 58148493 | 1392 | * | 11,1013 | -3,311799858 | 1,155155 | -2,86697 | 0,004144 | 0,81148874 | 0,918255 | Intron (ENSMUST00000 | 18 | 58158173 | 58159952 | 1780 | 2 | 14119 | ENSMUST00000236932.2 | 11459 | ENSMUSG00000024598 | Fbn2 | fibrillin 2 |
| chr7 | 1,29E+08 | 1,29E+08 | 949 | * | 9,738818 | -3,308570366 | 1,279876 | -2,58507 | 0,009736 | 0,885395021 | 1,425929 | Intron (ENSMUST00000 | 7 | 1,29E+08 | 1,29E+08 | 25976 | 1 | 1,03E+08 | ENSMUST00000205836.2 | 18030 | | Gm33248 | predicted gene, 33248 |
| chrX | 1,33E+08 | 1,33E+08 | 1449 | * | 10,67278 | -3,304665971 | 1,253972 | -2,63536 | 0,008405 | 0,962000766 | 0,738441 | Exon (ENSMUST000000 | 20 | 1,33E+08 | 1,33E+08 | 20528 | 1 | 74448 | ENSMUST00000064476.5 | 15270 | ENSMUSG00000052549 | Arl13a | ADP-ribosylation factor-like 13A |
| chr1 | 84370155 | 84371568 | 1414 | * | 11,17544 | -3,303309673 | 1,157341 | -2,85422 | 0,004314 | 0,85617852 | 0,738441 | Exon (ENSMUST000001 | 1 | 84085515 | 84341901 | 256387 | 2 | 98496 | ENSMUST00000175996.8 | -28254 | ENSMUSG00000045658 | Pid1 | phosphotyrosine interaction domain containing 1 |
| chr2 | 77033760 | 77035657 | 1898 | * | 11,18464 | -3,29970074 | 1,20601 | -2,73605 | 0,006218 | 0,824134241 | 1,2763 | Intron (ENSMUST00000 | 2 | 77027229 | 77029771 | 2543 | 2 | 228071 | ENSMUST00000145366.2 | -3989 | ENSMUSG00000042272 | Sestd1 | SEC14 and spectrin domains 1 |
| chr8 | 3860965 | 3862903 | 1939 | * | 14,44534 | -3,297742859 | 1,16652 | -2,82699 | 0,004699 | 0,775744126 | 1,2763 | Distal Intergenic | 8 | 3853069 | 3853358 | 290 | 2 | 19785 | ENSMUST00000157400.3 | -7607 | ENSMUSG00000088025 | Rprl3 | ribonuclease P RNA-like 3 |
| chr15 | 53858281 | 53859241 | 961 | * | 13,71688 | -3,292921232 | 1,17351 | -2,80604 | 0,005015 | 0,845747925 | 0,918255 | Distal Intergenic | 15 | 53682190 | 53765933 | 83744 | 2 | 320679 | ENSMUST00000132362.2 | -92348 | ENSMUSG00000058656 | Samd12 | sterile alpha motif domain containing 12 |
| chr10 | 87462503 | 87464107 | 1605 | * | 13,32809 | -3,292563549 | 1,11631 | -2,94951 | 0,003183 | 0,758498689 | 0,938682 | Distal Intergenic | 10 | 87382535 | 87406194 | 23660 | 1 | 18478 | ENSMUST00000217864.2 | 79968 | ENSMUSG00000020051 | Pah | phenylalanine hydroxylase |
| chr1 | 55402766 | 55404283 | 1518 | * | 13,80226 | -3,292002145 | 1,138869 | -2,89059 | 0,003845 | 0,792545355 | 0,938682 | Promoter (<=1kb) | 1 | 55385416 | 55402628 | 17213 | 2 | 75388 | ENSMUST00000159398.2 | -138 | ENSMUSG00000025977 | Boll | boule homolog, RNA binding protein |
| chr7 | 1,37E+08 | 1,37E+08 | 907 | * | 9,628683 | -3,289811817 | 1,261984 | -2,60686 | 0,009138 | 0,90060438 | 1,211315 | Intron (ENSMUST00000 | 7 | 1,36E+08 | 1,37E+08 | 235653 | 1 | 17314 | ENSMUST00000081510.4 | 159239 | ENSMUSG00000054612 | Mgmt | O-6-methylguanine-DNA methyltransferase |
| chr11 | 59364550 | 59366388 | 1839 | * | 11,6891 | -3,284798353 | 1,150482 | -2,85515 | 0,004302 | 0,775744126 | 1,173325 | Promoter (1-2kb) | 11 | 59352023 | 59363300 | 11278 | 2 | 237775 | ENSMUST00000057799.8 | -1250 | ENSMUSG00000054519 | Zfp867 | zinc finger protein 867 |
| chr5 | 4470818 | 4472330 | 1513 | * | 9,76866 | -3,284346416 | 1,226436 | -2,67796 | 0,007407 | 0,837092547 | 1,425929 | Distal Intergenic | 5 | 4242367 | 4247651 | 5285 | 1 | 208595 | ENSMUST00000177258.2 | 228451 | ENSMUSG00000053178 | Mterf1b | mitochondrial transcription termination factor 1b |
| chr5 | 49943486 | 49944822 | 1337 | * | 13,38064 | -3,283526274 | 1,107794 | -2,96402 | 0,003036 | 0,793220117 | 0,738441 | Distal Intergenic | 5 | 50123452 | 50129417 | 5966 | 2 | 70693 | ENSMUST00000196177.2 | 184595 | ENSMUSG00000029090 | Adgra3 | adhesion G protein-coupled receptor A3 |
| chr11 | 52664895 | 52666305 | 1411 | * | 13,03389 | -3,281673031 | 1,106795 | -2,96502 | 0,003027 | 0,71419937 | 1,173325 | Intron (ENSMUST00000 | 11 | 52655461 | 53079365 | 423905 | 1 | 320027 | ENSMUST00000036796.8 | 9434 | ENSMUSG00000036264 | Fstl4 | follistatin-like 4 |
| chr11 | 72304295 | 72305112 | 818 | * | 9,363648 | -3,28007197 | 1,269823 | -2,58309 | 0,009792 | 0,883696596 | 1,449706 | Promoter (1-2kb) | 11 | 72280940 | 72302539 | 21600 | 2 | 276829 | ENSMUST00000050226.7 | -1756 | ENSMUSG00000045667 | Smtnl2 | smoothelin-like 2 |
| chr1 | 1,54E+08 | 1,54E+08 | 914 | * | 10,28928 | -3,279070846 | 1,220908 | -2,68576 | 0,007236 | 0,829907213 | 1,449706 | Exon (ENSMUST000001 | 1 | 1,54E+08 | 1,54E+08 | 7180 | 2 | 12290 | ENSMUST00000226546.2 | 4453 | ENSMUSG00000004110 | Cacna1e | calcium channel, voltage-dependent, R type, alpha 1E subunit |
| chr3 | 1,19E+08 | 1,19E+08 | 1766 | * | 15,47967 | -3,278553974 | 0,98544 | -3,32699 | 0,000878 | 0,477622452 | 1,2763 | Distal Intergenic | 3 | 1,19E+08 | 1,19E+08 | 131 | 2 | 1,15E+08 | ENSMUST00000157181.3 | -33316 | ENSMUSG00000087806 | Gm23432 | predicted gene, 23432 |
| chr4 | 20700606 | 20701981 | 1376 | * | 9,984179 | -3,274090219 | 1,210933 | -2,70377 | 0,006856 | 0,820725534 | 1,449706 | Intron (ENSMUST00000 | 4 | 20118874 | 20778567 | 659694 | 2 | 269513 | ENSMUST00000119374.8 | 76586 | ENSMUSG00000055761 | Nkain3 | Na+/K+ transporting ATPase interacting 3 |
| chr16 | 28458989 | 28460009 | 1021 | * | 12,73177 | -3,271226705 | 1,0371 | -3,15421 | 0,001609 | 0,650529916 | 0,918255 | Intron (ENSMUST00000 | 16 | 28418687 | 28418795 | 109 | 2 | 751543 | ENSMUST00000102110.3 | -40194 | ENSMUSG00000076064 | Mir690 | microRNA 690 |
| chrX | 1,58E+08 | 1,58E+08 | 2070 | * | 10,64004 | -3,270962485 | 1,230838 | -2,65751 | 0,007872 | 0,953262301 | 0,738441 | Exon (ENSMUST000001 | 20 | 1,58E+08 | 1,58E+08 | 10135 | 1 | 382252 | ENSMUST00000131623.2 | 10737 | ENSMUSG00000044150 | Bclaf3 | Bclaf1 and Thrap3 family member 3 |
| chr14 | 30143831 | 30144440 | 610 | * | 9,913513 | -3,268477216 | 1,26418 | -2,58545 | 0,009725 | 0,908787948 | 1,211315 | Intron (ENSMUST00000 | 14 | 30201492 | 30232451 | 30960 | 1 | 75901 | ENSMUST00000224453.2 | -57052 | ENSMUSG00000021962 | Dcp1a | decapping mRNA 1A |
| chr7 | 1,19E+08 | 1,19E+08 | 1267 | * | 10,3835 | -3,268271405 | 1,188526 | -2,74985 | 0,005962 | 0,832746263 | 1,173325 | Distal Intergenic | 7 | 1,19E+08 | 1,19E+08 | 43857 | 2 | 209776 | ENSMUST00000084650.6 | 86287 | ENSMUSG00000066197 | Gpr139 | G protein-coupled receptor 139 |
| chr14 | 60601756 | 60603192 | 1437 | * | 13,43966 | -3,267360804 | 1,190177 | -2,74527 | 0,006046 | 0,911967721 | 0,738441 | Distal Intergenic | 14 | 60610708 | 60618104 | 7397 | 1 | 72125 | ENSMUST00000225247.3 | -7516 | ENSMUSG00000021986 | Amer2 | APC membrane recruitment 2 |
| chr9 | 99594190 | 99595367 | 1178 | * | 9,981278 | -3,264485489 | 1,213475 | -2,6902 | 0,007141 | 0,826626424 | 1,449706 | Promoter (2-3kb) | 9 | 99571514 | 99592116 | 20603 | 2 | 56492 | ENSMUST00000035048.12 | -2074 | ENSMUSG00000032473 | Cldn18 | claudin 18 |
| chr1 | 1,77E+08 | 1,77E+08 | 1564 | * | 13,19744 | -3,263257821 | 1,257561 | -2,59491 | 0,009462 | 0,977736121 | 0,738441 | Promoter (2-3kb) | 1 | 1,77E+08 | 1,77E+08 | 108 | 1 | 1,15E+08 | ENSMUST00000157250.4 | -2931 | ENSMUSG00000087875 | Gm26104 | predicted gene, 26104 |
| chrX | 99118476 | 99120370 | 1895 | * | 10,42552 | -3,261228511 | 1,204298 | -2,70799 | 0,006769 | 0,928382841 | 0,738441 | Intron (ENSMUST00000 | 20 | 99019387 | 99440762 | 421376 | 1 | 13607 | ENSMUST00000071453.3 | 99089 | ENSMUSG00000059327 | Eda | ectodysplasin-A |
| chr9 | 38206420 | 38207847 | 1428 | * | 9,19018 | -3,259468706 | 1,151139 | -2,83152 | 0,004633 | 0,75938015 | 1,360622 | Distal Intergenic | 9 | 38179811 | 38180761 | 951 | 1 | 258875 | ENSMUST00000093864.2 | 26609 | ENSMUSG00000096427 | Olfr895 | olfactory receptor 895 |
| chr2 | 1,46E+08 | 1,46E+08 | 875 | * | 9,321325 | -3,259252207 | 1,24136 | -2,62555 | 0,008651 | 0,890735468 | 1,211315 | Intron (ENSMUST00000 | 2 | 1,46E+08 | 1,46E+08 | 16592 | 2 | 241694 | ENSMUST00000124583.2 | -37555 | ENSMUSG00000037110 | Ralgapa2 | Ral GTPase activating protein, alpha subunit 2 (catalytic) |
| chr17 | 11193956 | 11196623 | 2668 | * | 25,24295 | -3,258099973 | 1,248148 | -2,61035 | 0,009045 | 0,748382289 | 2,880017 | Intron (ENSMUST00000 | 17 | 11286012 | 11712060 | 426049 | 1 | 50873 | ENSMUST00000168593.9 | -89389 | ENSMUSG00000023826 | Prkn | parkin RBR E3 ubiquitin protein ligase |
| chr2 | 1,3E+08 | 1,3E+08 | 1209 | * | 10,44749 | -3,254040749 | 1,174384 | -2,77085 | 0,005591 | 0,900637308 | 0,738441 | Distal Intergenic | 2 | 1,3E+08 | 1,3E+08 | 743 | 2 | 77006 | ENSMUST00000153271.2 | 3295 | ENSMUSG00000068290 | Ddrgk1 | DDRGK domain containing 1 |
| chrX | 50355025 | 50356419 | 1395 | * | 8,984537 | -3,253574606 | 1,18052 | -2,75605 | 0,00585 | 0,802195585 | 1,360622 | Distal Intergenic | 20 | 50253294 | 50295409 | 42116 | 2 | 171170 | ENSMUST00000156028.2 | -59616 | ENSMUSG00000036109 | Mbnl3 | muscleblind like splicing factor 3 |
| chr9 | 1,13E+08 | 1,13E+08 | 1472 | * | 10,35159 | -3,252957549 | 1,220249 | -2,66582 | 0,00768 | 0,842187815 | 1,425929 | Distal Intergenic | 9 | 1,13E+08 | 1,13E+08 | 31649 | 1 | 1,08E+08 | ENSMUST00000242854.1 | -274469 | | Gm46133 | predicted gene, 46133 |
| chr8 | 38847162 | 38848177 | 1016 | * | 10,11997 | -3,250072046 | 1,140103 | -2,85068 | 0,004363 | 0,770994815 | 1,211315 | Intron (ENSMUST00000 | 8 | 38719287 | 38719356 | 70 | 2 | 723860 | ENSMUST00000083523.5 | -127806 | ENSMUSG00000065457 | Mir383 | microRNA 383 |
| chr6 | 43813539 | 43814263 | 725 | * | 9,743095 | -3,249670891 | 1,256152 | -2,587 | 0,009681 | 0,923399738 | 1,086118 | Distal Intergenic | 6 | 43864128 | 43864258 | 131 | 1 | 1,15E+08 | ENSMUST00000103958.3 | -49865 | ENSMUSG00000077146 | Gm22939 | predicted gene, 22939 |
| chr12 | 28349603 | 28351397 | 1795 | * | 14,98363 | -3,246820942 | 1,15543 | -2,81005 | 0,004953 | 0,770673161 | 1,37683 | Exon (ENSMUST000002 | 12 | 28400262 | 28400393 | 132 | 1 | 1,15E+08 | ENSMUST00000240083.1 | -48865 | ENSMUSG00000118901 | Gm25564 | predicted gene, 25564 |
| chr9 | 69854557 | 69856092 | 1536 | * | 15,14898 | -3,244744981 | 1,005064 | -3,2284 | 0,001245 | 0,526851572 | 1,37683 | Distal Intergenic | 9 | 69896748 | 69915599 | 18852 | 1 | 12175 | ENSMUST00000034754.12 | -40656 | ENSMUSG00000011958 | Bnip2 | BCL2/adenovirus E1B interacting protein 2 |
| chr11 | 96316796 | 96318215 | 1420 | * | 11,8885 | -3,244621068 | 1,225552 | -2,64748 | 0,008109 | 0,955649401 | 0,744375 | Distal Intergenic | 11 | 96339149 | 96355373 | 16225 | 2 | 1E+08 | ENSMUST00000153012.2 | 37158 | | Gm11529 | predicted gene 11529 |
| chr11 | 31988069 | 31989643 | 1575 | * | 18,88037 | -3,24437471 | 0,933009 | -3,47733 | 0,000506 | 0,302171532 | 2,221611 | Intron (ENSMUST00000 | 11 | 31950502 | 31983414 | 32913 | 1 | 18197 | ENSMUST00000093219.4 | 37567 | ENSMUSG00000020297 | Nsg2 | neuron specific gene family member 2 |
| chr9 | 9080839 | 9082177 | 1339 | * | 11,51208 | -3,244132542 | 1,142344 | -2,83989 | 0,004513 | 0,862146257 | 0,738441 | Intron (ENSMUST00000 | 9 | 9236293 | 9260888 | 24596 | 1 | 1,01E+08 | ENSMUST00000189204.2 | -154116 | ENSMUSG00000085253 | Gm16833 | predicted gene, 16833 |
| chr10 | 17981576 | 17983282 | 1707 | * | 11,23099 | -3,241586449 | 1,212478 | -2,67352 | 0,007506 | 0,911482541 | 0,918255 | Promoter (1-2kb) | 10 | 17979960 | 17983217 | 3258 | 1 | 19707 | ENSMUST00000216576.2 | 1616 | ENSMUSG00000019854 | Reps1 | RalBP1 associated Eps domain containing protein |
| chr11 | 63058360 | 63059326 | 967 | * | 10,5428 | -3,239951945 | 1,253086 | -2,58558 | 0,009722 | 0,913502848 | 1,173325 | Distal Intergenic | 11 | 63023894 | 63049392 | 25499 | 1 | 18858 | ENSMUST00000108700.2 | 34466 | ENSMUSG00000018217 | Pmp22 | peripheral myelin protein 22 |
| chr16 | 32563793 | 32565961 | 2169 | * | 13,71864 | -3,239535575 | 1,213113 | -2,67043 | 0,007575 | 0,909280719 | 0,938682 | Intron (ENSMUST00000 | 16 | 32555015 | 32602596 | 47582 | 1 | 140474 | ENSMUST00000239554.1 | 8778 | ENSMUSG00000118672 | Muc4 | mucin 4 |
| chr19 | 17803450 | 17804681 | 1232 | * | 10,74895 | -3,238689781 | 1,224314 | -2,64531 | 0,008162 | 0,956533297 | 0,738441 | Intron (ENSMUST00000 | 19 | 17650816 | 17814981 | 164166 | 2 | 18552 | ENSMUST00000235505.2 | 10300 | ENSMUSG00000024713 | Pcsk5 | proprotein convertase subtilisin/kexin type 5 |
| chr12 | 99038767 | 99040507 | 1741 | * | 19,08171 | -3,238324319 | 0,907819 | -3,56715 | 0,000361 | 0,243374613 | 2,507013 | Distal Intergenic | 12 | 99127536 | 99129067 | 1532 | 2 | 67639 | ENSMUST00000223511.2 | 88560 | ENSMUSG00000113210 | 4930474N09Rik | RIKEN cDNA 4930474N09 gene |
| chr16 | 49125881 | 49127028 | 1148 | * | 10,79138 | -3,236465322 | 1,21606 | -2,66144 | 0,007781 | 0,880128915 | 1,173325 | Distal Intergenic | 16 | 49031031 | 49076135 | 45105 | 2 | 73255 | ENSMUST00000098226.5 | -49746 | ENSMUSG00000073976 | 1700026J12Rik | RIKEN cDNA 1700026J12 gene |
| chr3 | 29588765 | 29589692 | 928 | * | 8,567812 | -3,236015483 | 1,202436 | -2,69122 | 0,007119 | 0,749550311 | 2,237403 | Intron (ENSMUST00000 | 3 | 29568055 | 29724883 | 156829 | 1 | 75740 | ENSMUST00000140288.2 | 20710 | ENSMUSG00000063600 | Egfem1 | EGF-like and EMI domain containing 1 |
| chr2 | 1,12E+08 | 1,12E+08 | 986 | * | 10,51259 | -3,235230983 | 1,209889 | -2,67399 | 0,007495 | 0,872628491 | 1,173325 | Intron (ENSMUST00000 | 2 | 1,11E+08 | 1,12E+08 | 25883 | 1 | 258199 | ENSMUST00000208175.2 | 16781 | ENSMUSG00000109389 | Olfr1300-ps1 | olfactory receptor 1300, pseudogene 1 |
| chr3 | 17357601 | 17358895 | 1295 | * | 12,36263 | -3,235210447 | 1,077919 | -3,00135 | 0,002688 | 0,681421811 | 1,2763 | Distal Intergenic | 3 | 17382858 | 17388589 | 5732 | 2 | 1,03E+08 | ENSMUST00000191898.2 | 29694 | | Gm30340 | predicted gene, 30340 |
| chr3 | 82573042 | 82573901 | 860 | * | 10,35156 | -3,233150446 | 1,233382 | -2,62137 | 0,008758 | 0,892445563 | 1,211315 | Promoter (<=1kb) | 3 | 82567828 | 82572207 | 4380 | 2 | 1,03E+08 | ENSMUST00000193554.2 | -835 | ENSMUSG00000102629 | Gm29999 | predicted gene, 29999 |
| chr9 | 49023354 | 49024827 | 1474 | * | 13,09637 | -3,232722256 | 1,215228 | -2,66018 | 0,00781 | 0,951821 | 0,738441 | 3' UTR | 9 | 49020302 | 49028885 | 8584 | 1 | 80893 | ENSMUST00000170426.2 | 3052 | ENSMUSG00000032268 | Tmprss5 | transmembrane protease, serine 5 (spinesin) |
| chr6 | 65072060 | 65072806 | 747 | * | 11,5136 | -3,231674021 | 1,201604 | -2,68947 | 0,007157 | 0,937460407 | 0,738441 | Exon (ENSMUST000002 | 6 | 65052007 | 65071174 | 19168 | 1 | 13990 | ENSMUST00000203411.2 | 20053 | ENSMUSG00000029920 | Smarcad1 | SWI/SNF-related, matrix-associated actin-dependent regulator of chromatin, subfamily a, containing DEAD/H box 1 |
| chr5 | 1,06E+08 | 1,06E+08 | 1056 | * | 10,83022 | -3,231079484 | 1,194105 | -2,70586 | 0,006813 | 0,929295662 | 0,738441 | Distal Intergenic | 5 | 1,06E+08 | 1,06E+08 | 1878 | 1 | 1,03E+08 | ENSMUST00000200592.2 | -52153 | ENSMUSG00000105508 | Gm32921 | predicted gene, 32921 |
| chr8 | 97471667 | 97472920 | 1254 | * | 11,95257 | -3,23003154 | 1,215035 | -2,65839 | 0,007852 | 0,863495249 | 1,2763 | Distal Intergenic | 8 | 97098355 | 97130490 | 32136 | 2 | 1,03E+08 | ENSMUST00000212119.2 | -341177 | ENSMUSG00000110633 | Gm32122 | predicted gene, 32122 |
| chr1 | 1,5E+08 | 1,5E+08 | 1220 | * | 10,38186 | -3,226177147 | 1,133703 | -2,8457 | 0,004431 | 0,764063654 | 1,2763 | Distal Intergenic | 1 | 1,5E+08 | 1,5E+08 | 14489 | 1 | 20028 | ENSMUST00000165062.8 | -10165 | ENSMUSG00000006007 | Pdc | phosducin |
| chr1 | 85645358 | 85647085 | 1728 | * | 20,4286 | -3,222002738 | 0,942634 | -3,41808 | 0,000631 | 0,303890196 | 2,713306 | 3' UTR | 1 | 85631163 | 85635716 | 4554 | 1 | 20684 | ENSMUST00000141709.2 | 14195 | ENSMUSG00000026222 | Sp100 | nuclear antigen Sp100 |
| chr16 | 71810746 | 71811951 | 1206 | * | 9,923165 | -3,221426221 | 1,248912 | -2,57939 | 0,009898 | 0,915629184 | 1,173325 | Distal Intergenic | 16 | 72023271 | 72031190 | 7920 | 2 | 319533 | ENSMUST00000231223.2 | 219239 | ENSMUSG00000116769 | 8030451O07Rik | RIKEN cDNA 8030451O07 gene |
| chr19 | 58236792 | 58238409 | 1618 | * | 16,25228 | -3,218542594 | 0,923476 | -3,48525 | 0,000492 | 0,275349208 | 2,663619 | Intron (ENSMUST00000 | 19 | 58080246 | 58084463 | 4218 | 2 | 1,15E+08 | ENSMUST00000235250.2 | -152329 | | Gm50287 | predicted gene, 50287 |
| chr1 | 1,18E+08 | 1,18E+08 | 1383 | * | 10,69116 | -3,218287154 | 1,235125 | -2,60564 | 0,00917 | 0,973894282 | 0,744375 | Promoter (<=1kb) | 1 | 1,18E+08 | 1,18E+08 | 6138 | 1 | 1,03E+08 | ENSMUST00000190194.2 | -110 | | Gm28867 | predicted gene 28867 |
| chr14 | 29643707 | 29645484 | 1778 | * | 13,70294 | -3,217565936 | 1,063425 | -3,02566 | 0,002481 | 0,669377201 | 1,2763 | Distal Intergenic | 14 | 29690265 | 29697619 | 7355 | 1 | 80795 | ENSMUST00000112268.4 | -44781 | ENSMUSG00000042682 | Selenok | selenoprotein K |
| chr19 | 36276021 | 36277237 | 1217 | * | 11,35978 | -3,21628611 | 1,170527 | -2,74772 | 0,006001 | 0,833926809 | 1,173325 | Distal Intergenic | 19 | 36216285 | 36241837 | 25553 | 2 | 1,15E+08 | ENSMUST00000236548.3 | -34184 | | LOC115489009 | uncharacterized LOC115489009 |
| chr1 | 90789317 | 90791065 | 1749 | * | 13,58087 | -3,215975051 | 1,170105 | -2,74845 | 0,005988 | 0,819338417 | 1,2763 | Distal Intergenic | 1 | 90749993 | 90771693 | 21701 | 2 | 12835 | ENSMUST00000187753.2 | -17624 | ENSMUSG00000048126 | Col6a3 | collagen, type VI, alpha 3 |
| chr13 | 44442479 | 44443656 | 1178 | * | 10,44138 | -3,215486597 | 1,20813 | -2,66154 | 0,007778 | 0,916531797 | 0,918255 | Distal Intergenic | 13 | 44404438 | 44433925 | 29488 | 2 | 432754 | ENSMUST00000222409.2 | -8554 | | Gm29676 | predicted gene, 29676 |
| chr15 | 19199815 | 19200468 | 654 | * | 8,38651 | -3,213875276 | 1,149479 | -2,79594 | 0,005175 | 0,769692849 | 1,449706 | Intron (ENSMUST00000 | 15 | 19185459 | 19203415 | 17957 | 1 | 1,03E+08 | ENSMUST00000226343.2 | 14356 | ENSMUSG00000115803 | Gm35496 | predicted gene, 35496 |
| chr12 | 35614466 | 35615973 | 1508 | * | 11,36152 | -3,212950312 | 1,097628 | -2,92718 | 0,003421 | 0,718585444 | 1,2763 | Exon (ENSMUST000002 | 12 | 35547973 | 35585037 | 37065 | 2 | 11622 | ENSMUST00000116436.9 | -29429 | ENSMUSG00000019256 | Ahr | aryl-hydrocarbon receptor |
| chr3 | 62474747 | 62475587 | 841 | * | 9,441924 | -3,209368359 | 1,236884 | -2,59472 | 0,009467 | 0,87765772 | 1,449706 | Intron (ENSMUST00000 | 3 | 62502229 | 62510195 | 7967 | 2 | 229357 | ENSMUST00000149007.2 | 34608 | ENSMUSG00000043441 | Gpr149 | G protein-coupled receptor 149 |
| chr12 | 97192439 | 97193216 | 778 | * | 10,5042 | -3,209107615 | 1,221153 | -2,62793 | 0,008591 | 0,883012141 | 1,2763 | Distal Intergenic | 12 | 97279623 | 97282627 | 3005 | 1 | 1,05E+08 | ENSMUST00000218233.2 | -86407 | | Gm40552 | predicted gene, 40552 |
| chr7 | 1,2E+08 | 1,2E+08 | 1105 | * | 10,75513 | -3,208695261 | 1,197816 | -2,67879 | 0,007389 | 0,941748492 | 0,744375 | Intron (ENSMUST00000 | 7 | 1,2E+08 | 1,2E+08 | 6912 | 2 | 381917 | ENSMUST00000208910.2 | -49483 | ENSMUSG00000052273 | Dnah3 | dynein, axonemal, heavy chain 3 |
| chr2 | 25815837 | 25817332 | 1496 | * | 16,04242 | -3,208471898 | 1,040709 | -3,08297 | 0,002049 | 0,524368688 | 2,318434 | 3' UTR | 2 | 25819955 | 25824626 | 4672 | 2 | 227634 | ENSMUST00000148146.2 | 7294 | ENSMUSG00000026933 | Camsap1 | calmodulin regulated spectrin-associated protein 1 |
| chr16 | 57037128 | 57038398 | 1271 | * | 10,44789 | -3,20713038 | 1,200267 | -2,67201 | 0,00754 | 0,946556802 | 0,744375 | Intron (ENSMUST00000 | 16 | 56989710 | 57051729 | 62020 | 2 | 67581 | ENSMUST00000226586.2 | 13331 | ENSMUSG00000022749 | Tbc1d23 | TBC1 domain family, member 23 |
| chr10 | 74152709 | 74153585 | 877 | * | 9,667121 | -3,205555812 | 1,179628 | -2,71743 | 0,006579 | 0,81655291 | 1,425929 | 5' UTR | 10 | 74138983 | 74166882 | 27900 | 1 | 11994 | ENSMUST00000147455.2 | 13726 | ENSMUSG00000052613 | Pcdh15 | protocadherin 15 |
| chr3 | 24791506 | 24792595 | 1090 | * | 12,81875 | -3,204696034 | 1,001747 | -3,19911 | 0,001379 | 0,574952042 | 1,173325 | Intron (ENSMUST00000 | 3 | 23852266 | 24837786 | 985521 | 2 | 635702 | ENSMUST00000203414.2 | 45191 | ENSMUSG00000102758 | Naaladl2 | N-acetylated alpha-linked acidic dipeptidase-like 2 |
| chr10 | 20157384 | 20159159 | 1776 | * | 14,34188 | -3,202917778 | 1,142841 | -2,80259 | 0,005069 | 0,842722325 | 0,938682 | Intron (ENSMUST00000 | 10 | 20151640 | 20162372 | 10733 | 2 | 1,15E+08 | ENSMUST00000215889.2 | 3213 | | Gm48249 | predicted gene, 48249 |
| chr1 | 14251425 | 14252966 | 1542 | * | 12,96197 | -3,201525589 | 1,14944 | -2,78529 | 0,005348 | 0,797327098 | 1,2763 | Exon (ENSMUST000201 | 1 | 14278150 | 14301716 | 23567 | 2 | 14048 | ENSMUST00000189526.2 | 48750 | ENSMUSG00000025932 | Eya1 | EYA transcriptional coactivator and phosphatase 1 |
| chr4 | 92341496 | 92343206 | 1711 | * | 11,74694 | -3,199545059 | 1,156806 | -2,76585 | 0,005678 | 0,901463351 | 0,744375 | Distal Intergenic | 4 | 92416788 | 92436541 | 19754 | 1 | 1,08E+08 | ENSMUST00000130501.2 | -73582 | | Gm12637 | predicted gene 12637 |
| chr9 | 1,21E+08 | 1,21E+08 | 1768 | * | 12,40901 | -3,198837669 | 1,241528 | -2,57653 | 0,00998 | 0,955649401 | 0,918255 | Promoter (<=1kb) | 9 | 1,21E+08 | 1,21E+08 | 148258 | 2 | 209012 | ENSMUST00000170406.8 | 0 | ENSMUSG00000040936 | Ulk4 | unc-51-like kinase 4 |
| chr14 | 1,03E+08 | 1,03E+08 | 2050 | * | 12,7299 | -3,197448501 | 1,070716 | -2,98627 | 0,002824 | 0,684274806 | 1,2763 | Distal Intergenic | 14 | 1,03E+08 | 1,03E+08 | 132 | 2 | 1,15E+08 | ENSMUST00000240023.1 | -214630 | ENSMUSG00000119010 | Gm22347 | predicted gene, 22347 |
| chr15 | 89394366 | 89397301 | 2936 | * | 30,208201 | -3,196316893 | 0,7592008 | -4,2101072 | 2,552E-05 | 0,04457732 | 2,8800171 | Intron (ENSMUST000001 | 15 | 89385486 | 89444462 | 58977 | 1 | 58234 | ENSMUST00000066545.14 | 8880 | ENSMUSG00000022623 | Shank3 | SH3 and multiple ankyrin repeat domains 3 |

| | | | | | | | | | | | | | | | | | | | | | | | |
|---|---|---|---|---|---|---|---|---|---|---|---|---|---|---|---|---|---|---|---|---|---|---|---|
| chr18 | 28919660 | 28920735 | 1076 | * | 10,53059 | -3,19566601 | 1,139477 | -2,8045 | 0,005039 | 0,789027175 | 1,2763 | Distal Intergenic | 18 | 28844914 | 28859900 | 14987 | 1 | 408061 | ENSMUST00000234187.2 | 74746 | ENSMUSG00000117478 | BC051408 | cDNA sequence BC051408 |
| chr14 | 1,05E+08 | 1,05E+08 | 2083 | * | 11,99637 | -3,19236459 | 1,031865 | -3,09378 | 0,001976 | 0,644222245 | 1,173325 | Distal Intergenic | 14 | 1,05E+08 | 1,05E+08 | 134 | 1 | 1,15E+08 | ENSMUST00000104237.3 | 23460 | ENSMUSG00000077425 | Gm24280 | predicted gene, 24280 |
| chr17 | 70142396 | 70144192 | 1797 | * | 18,28043 | -3,188283166 | 0,940718 | -3,3892 | 0,000701 | 0,326832856 | 2,507013 | Distal Intergenic | 17 | 70200755 | 70230990 | 30236 | 2 | 1,03E+08 | ENSMUST00000232745.2 | 86798 | | Gm36594 | predicted gene, 36594 |
| chr10 | 44322152 | 44323275 | 1124 | * | 11,22044 | -3,185055373 | 1,197391 | -2,66 | 0,007814 | 0,951979183 | 0,738441 | Exon (ENSMUST000000 | 10 | 44315663 | 44333206 | 17544 | 2 | 12142 | ENSMUST00000218369.2 | 9931 | ENSMUSG00000038151 | Prdm1 | PR domain containing 1, with ZNF domain |
| chrX | 79189883 | 79191709 | 1827 | * | 10,77063 | -3,184411432 | 1,230642 | -2,5876 | 0,009665 | 0,95067847 | 0,918255 | Distal Intergenic | 20 | 78666837 | 78715041 | 48205 | 2 | 71967 | ENSMUST00000052283.7 | -474842 | ENSMUSG00000046942 | Mageb16 | MAGE family member B16 |
| chr14 | 8296411 | 8298117 | 1707 | * | 10,12296 | -3,183677052 | 1,128308 | -2,82164 | 0,004778 | 0,806442491 | 1,086118 | Promoter (2-3kb) | 14 | 8293646 | 8294668 | 1023 | 1 | 258387 | ENSMUST00000035250.6 | 2765 | ENSMUSG00000052417 | Olfr720 | olfactory receptor 720 |
| chr5 | 1,49E+08 | 1,49E+08 | 1821 | * | 15,71566 | -3,183339605 | 1,043117 | -3,05176 | 0,002275 | 0,541111532 | 2,318434 | Distal Intergenic | 5 | 1,49E+08 | 1,49E+08 | 3397 | 2 | 231912 | ENSMUST00000139269.2 | 12634 | ENSMUSG00000041298 | Katnal1 | katanin p60 subunit A-like 1 |
| chr6 | 1,12E+08 | 1,12E+08 | 1006 | * | 12,91262 | -3,183296662 | 1,000049 | -3,18314 | 0,001457 | 0,603552508 | 1,075735 | Exon (ENSMUST000000 | 6 | 1,12E+08 | 1,12E+08 | 2981 | 2 | 1,01E+08 | ENSMUST00000053559.5 | -40055 | | 5031434C07Rik | RIKEN cDNA 5031434C07 gene |
| chr13 | 6585881 | 6587106 | 1226 | * | 11,00754 | -3,181309566 | 1,229035 | -2,58846 | 0,009641 | 0,912096919 | 1,173325 | Distal Intergenic | 13 | 6598185 | 6630078 | 31894 | 1 | 69617 | ENSMUST00000021611.10 | -11079 | ENSMUSG00000021193 | Pitrm1 | pitrilysin metallepetidase 1 |
| chr5 | 58591459 | 58593367 | 1909 | * | 8,571879 | -3,180874492 | 1,218916 | -2,60959 | 0,009065 | 0,792545355 | 2,237403 | Distal Intergenic | 5 | 58600582 | 58610342 | 9761 | 2 | 78163 | ENSMUST00000199751.2 | 16975 | ENSMUSG00000105234 | 4930459L07Rik | RIKEN cDNA 4930459L07 gene |
| chr12 | 95179065 | 95180662 | 1598 | * | 13,64479 | -3,178742164 | 1,085805 | -2,92754 | 0,003417 | 0,77749164 | 0,918255 | Distal Intergenic | 12 | 95345619 | 95345725 | 107 | 2 | 1,15E+08 | ENSMUST00000240345.1 | 165063 | ENSMUSG00001118669 | Gm26055 | predicted gene, 26055 |
| chr18 | 48705068 | 48705857 | 790 | * | 10,13897 | -3,177659865 | 1,148555 | -2,76666 | 0,005663 | 0,823736512 | 1,173325 | Distal Intergenic | 18 | 48178402 | 48181479 | 3078 | 1 | 433182 | ENSMUST00000076155.6 | 526666 | ENSMUSG00000059040 | Eno1b | enolase 1B, retrotransposed |
| chr14 | 56289549 | 56290378 | 830 | * | 10,46999 | -3,177333533 | 1,069303 | -2,97141 | 0,002964 | 0,692794047 | 1,2763 | Distal Intergenic | 14 | 56297086 | 56299770 | 2685 | 2 | 17227 | ENSMUST00000043249.10 | 9392 | ENSMUSG00000061068 | Mcpt4 | mast cell protease 4 |
| chr11 | 65730136 | 65730947 | 812 | * | 9,557622 | -3,177261133 | 1,221602 | -2,6009 | 0,009298 | 0,918554156 | 1,086118 | Intron (ENSMUST000000 | 11 | 65742011 | 65746533 | 4523 | 2 | 237806 | ENSMUST00000136124.2 | 15586 | ENSMUSG00000056752 | Dnah9 | dynein, axonemal, heavy chain 9 |
| chr17 | 36845709 | 36846693 | 985 | * | 11,30063 | -3,17693675 | 1,174335 | -2,70531 | 0,006824 | 0,898643449 | 0,918255 | Distal Intergenic | 17 | 36857967 | 36860142 | 2176 | 1 | 224754 | ENSMUST00000041964.7 | -11274 | ENSMUSG00000037537 | H2-M11 | histocompatibility 2, M region locus 11 |
| chr10 | 1,16E+08 | 1,16E+08 | 1128 | * | 11,39526 | -3,175249268 | 1,096849 | -2,89488 | 0,003793 | 0,832960417 | 0,744375 | Distal Intergenic | 10 | 1,16E+08 | 1,16E+08 | 16481 | 1 | 216350 | ENSMUST00000217900.2 | -75321 | ENSMUSG00000034127 | Tspan8 | tetraspanin 8 |
| chr14 | 38197511 | 38198882 | 1372 | * | 11,8235 | -3,174933469 | 1,200121 | -2,64551 | 0,008157 | 0,956450448 | 0,744375 | Intron (ENSMUST000000 | 14 | 38039861 | 38059083 | 19223 | 1 | 1,03E+08 | ENSMUST00000225614.2 | 157650 | | Gm31456 | predicted gene, 31456 |
| chr2 | 1,64E+08 | 1,64E+08 | 3073 | * | 25,0414 | -3,173066918 | 0,798426 | -3,97415 | 7,06E-05 | 0,08392487 | 2,951062 | Promoter (1-2kb) | 2 | 1,64E+08 | 1,64E+08 | 2438 | 2 | 20568 | ENSMUST00000109367.10 | -1485 | ENSMUSG00000017002 | Slpi | secretory leukocyte peptidase inhibitor |
| chr1 | 25819644 | 25820707 | 1064 | * | 11,13807 | -3,17200596 | 1,188424 | -2,66909 | 0,007606 | 0,858257847 | 1,2763 | Intron (ENSMUST000000 | 1 | 25106631 | 25865220 | 758590 | 2 | 210933 | ENSMUST00000146592.8 | 44513 | ENSMUSG00000033569 | Adgrb3 | adhesion G protein-coupled receptor B3 |
| chr3 | 38148575 | 38150085 | 1511 | * | 13,56202 | -3,170780248 | 1,089998 | -2,90898 | 0,003626 | 0,789027175 | 0,918255 | Exon (ENSMUST000001 | 3 | 38049254 | 38049384 | 131 | 2 | 1,15E+08 | ENSMUST00000104540.3 | -99191 | ENSMUSG00000118777 | Gm22899 | predicted gene, 22899 |
| chr2 | 16718237 | 16720005 | 1769 | * | 12,06127 | -3,170137663 | 1,175445 | -2,69697 | 0,006997 | 0,934391503 | 0,744375 | Exon (ENSMUST000000 | 2 | 16643507 | 16756501 | 112995 | 1 | 67448 | ENSMUST00000126173.2 | 74730 | ENSMUSG00000026748 | Plxdc2 | plexin domain containing 2 |
| chr6 | 55384423 | 55385218 | 796 | * | 10,01883 | -3,169990254 | 1,140716 | -2,77895 | 0,005454 | 0,817544002 | 1,173325 | Promoter (2-3kb) | 6 | 55382297 | 55382585 | 289 | 1 | 330324 | ENSMUST00000209841.2 | 2126 | ENSMUSG00000108228 | 6430584L05Rik | RIKEN cDNA 6430584L05 gene |
| chr13 | 1,11E+08 | 1,11E+08 | 1121 | * | 12,09 | -3,167983605 | 1,142571 | -2,77268 | 0,00556 | 0,860883941 | 0,918255 | Distal Intergenic | 13 | 1,11E+08 | 1,11E+08 | 1239 | 1 | 20620 | ENSMUST00000225156.2 | 41407 | ENSMUSG00000021701 | Plk2 | polo like kinase 2 |
| chr7 | 1,2E+08 | 1,2E+08 | 1870 | * | 14,21279 | -3,167030623 | 1,045236 | -3,02997 | 0,002446 | 0,717609615 | 0,918255 | Promoter (<=1kb) | 7 | 1,2E+08 | 1,2E+08 | 9886 | 1 | 75073 | ENSMUST00000208641.2 | -624 | ENSMUSG00000109414 | 4930505K13Rik | RIKEN cDNA 4930505K13 gene |
| chr8 | 1,09E+08 | 1,09E+08 | 1271 | * | 14,10707 | -3,163167888 | 1,081578 | -2,92459 | 0,003449 | 0,818130284 | 0,738441 | Intron (ENSMUST000000 | 8 | 1,09E+08 | 1,09E+08 | 396257 | 1 | 11906 | ENSMUST00000221632.2 | 147976 | ENSMUSG00000038872 | Zfhx3 | zinc finger homeobox 3 |
| chr17 | 43889205 | 43889676 | 472 | * | 12,83421 | -3,161091437 | 1,100287 | -2,87297 | 0,004066 | 0,749550311 | 1,2763 | Intron (ENSMUST000000 | 17 | 43879606 | 43918527 | 38922 | 1 | 27226 | ENSMUST00000163489.2 | 9599 | ENSMUSG00000023913 | Pla2g7 | phospholipase A2, group VII (platelet-activating factor acetylhydrolase, plasma) |
| chr1 | 1,66E+08 | 1,66E+08 | 1423 | * | 12,20177 | -3,160829298 | 1,136953 | -2,78009 | 0,005434 | 0,894522042 | 0,744375 | Distal Intergenic | 1 | 1,66E+08 | 1,66E+08 | 29744 | 2 | 240892 | ENSMUST00000085992.4 | 50583 | ENSMUSG00000026564 | Dusp27 | dual specificity phosphatase 27 (putative) |
| chr13 | 80998035 | 80999253 | 1219 | * | 10,48647 | -3,160413012 | 1,173776 | -2,69252 | 0,007091 | 0,849977165 | 1,2763 | Distal Intergenic | 13 | 81031503 | 81037762 | 6260 | 1 | 105171 | ENSMUST00000159090.2 | -32250 | ENSMUSG00000074794 | Arrdc3 | arrestin domain containing 3 |
| chr12 | 1,03E+08 | 1,03E+08 | 2099 | * | 16,27873 | -3,159452408 | 1,208825 | -2,61366 | 0,008958 | 0,785796708 | 2,318434 | Intron (ENSMUST000000 | 12 | 1,03E+08 | 1,03E+08 | 37560 | 2 | 238386 | ENSMUST00000220596.2 | 15947 | ENSMUSG00000041702 | Btbd7 | BTB (POZ) domain containing 7 |
| chr9 | 50025807 | 50027823 | 2017 | * | 18,74235 | -3,158742205 | 0,935242 | -3,37746 | 0,000732 | 0,355694809 | 2,154504 | Distal Intergenic | 9 | 50143823 | 50144482 | 660 | 2 | 71873 | ENSMUST00000215655.2 | 116659 | ENSMUSG00000110916 | 2310003N18Rik | RIKEN cDNA 2310003N18 gene |
| chr13 | 59828132 | 59829590 | 1459 | * | 13,56969 | -3,158371282 | 1,081109 | -2,92142 | 0,003484 | 0,722798324 | 1,2763 | Distal Intergenic | 13 | 59782440 | 59823625 | 41186 | 2 | 105348 | ENSMUST00000022039.7 | -4507 | ENSMUSG00000021556 | Golm1 | golgi membrane protein 1 |
| chrX | 1,66E+08 | 1,66E+08 | 2230 | * | 16,75197 | -3,158104521 | 1,0106 | -3,12498 | 0,001778 | 0,61075929 | 1,2763 | Distal Intergenic | 20 | 1,66E+08 | 1,66E+08 | 36428 | 2 | 110639 | ENSMUST00000026839.5 | -24825 | ENSMUSG00000025742 | Prps2 | phosphoribosyl pyrophosphate synthetase 2 |
| chr11 | 92390889 | 92392128 | 1240 | * | 8,780171 | -3,156681168 | 1,154826 | -2,73347 | 0,006267 | 0,802490409 | 1,449706 | Distal Intergenic | 11 | 92801123 | 92801265 | 143 | 1 | 1,15E+08 | ENSMUST00000157185.3 | -408995 | ENSMUSG00000087810 | Gm25113 | predicted gene, 25113 |
| chr2 | 1,13E+08 | 1,13E+08 | 1241 | * | 11,26979 | -3,155827193 | 1,147789 | -2,74948 | 0,005969 | 0,832960417 | 1,173325 | Promoter (<=1kb) | 2 | 1,13E+08 | 1,13E+08 | 166 | 1 | 1,15E+08 | ENSMUST00000176208.3 | 0 | ENSMUSG00000093660 | Gm26505 | predicted gene, 26505 |
| chr10 | 44912029 | 44913809 | 1781 | * | 14,65825 | -3,15475178 | 1,060855 | -2,97378 | 0,002942 | 0,683140223 | 1,37683 | Distal Intergenic | 10 | 44900398 | 44904248 | 3851 | 1 | 1,05E+08 | ENSMUST00000214642.2 | 11631 | | Gm40639 | predicted gene, 40639 |
| chr7 | 39176547 | 39177905 | 1359 | * | 11,90854 | -3,154405607 | 1,180991 | -2,67098 | 0,007563 | 0,947196831 | 0,744375 | Intron (ENSMUST000000 | 7 | 39167190 | 39189844 | 22655 | 1 | 70227 | ENSMUST00000108015.4 | 9357 | ENSMUSG00000068959 | Zfp619 | zinc finger protein 619 |
| chr1 | 55627084 | 55628993 | 1910 | * | 15,12627 | -3,154235651 | 0,922957 | -3,41753 | 0,000632 | 0,334207335 | 2,141612 | Intron (ENSMUST000000 | 1 | 55741184 | 55774386 | 33203 | 1 | 227120 | ENSMUST00000187059.2 | -112191 | ENSMUSG00000038349 | Plcl1 | phospholipase C-like 1 |
| chr9 | 1,07E+08 | 1,07E+08 | 1214 | * | 13,57089 | -3,153558899 | 1,067832 | -2,95324 | 0,003145 | 0,732824791 | 1,075735 | Promoter (1-2kb) | 9 | 1,07E+08 | 1,07E+08 | 966 | 2 | 81000 | ENSMUST00000190363.2 | 1036 | ENSMUSG00000040661 | Rad54l2 | RAD54 like 2 (S. cerevisiae) |
| chr3 | 66151640 | 66153167 | 1528 | * | 10,97104 | -3,152767566 | 1,21959 | -2,58511 | 0,009735 | 0,952402626 | 0,918255 | Exon (ENSMUST000000 | 3 | 66127331 | 66133226 | 5896 | 1 | 19288 | ENSMUST00000029421.6 | 24309 | ENSMUSG00000027832 | Ptx3 | pentraxin related gene |
| chr1 | 1,77E+08 | 1,77E+08 | 1949 | * | 13,61239 | -3,152638842 | 1,157331 | -2,72406 | 0,006449 | 0,831753076 | 1,2763 | Intron (ENSMUST000000 | 1 | 1,77E+08 | 1,77E+08 | 226653 | 2 | 23797 | ENSMUST00000111160.9 | 54186 | ENSMUSG00000019699 | Akt3 | thymoma viral proto-oncogene 3 |
| chr10 | 67338241 | 67341149 | 2909 | * | 26,18729 | -3,151167505 | 0,754331 | -4,17743 | 2,95E-05 | 0,05090669 | 2,725491 | Distal Intergenic | 10 | 67371305 | 67376076 | 4772 | 1 | 13654 | ENSMUST00000145754.8 | -30156 | ENSMUSG00000037868 | Egr2 | early growth response 2 |
| chr2 | 1,04E+08 | 1,04E+08 | 1182 | * | 12,74805 | -3,150808402 | 1,021101 | -3,0857 | 0,002031 | 0,663408464 | 1,075735 | Exon (ENSMUST000000 | 2 | 1,04E+08 | 1,04E+08 | 7899 | 1 | 228410 | ENSMUST00000124944.2 | -7158 | ENSMUSG00000027176 | Cstf3 | cleavage stimulation factor, 3' pre-RNA, subunit 3 |
| chr7 | 1,29E+08 | 1,29E+08 | 1659 | * | 13,73053 | -3,15071586 | 1,085766 | -2,90184 | 0,00371 | 0,731734453 | 1,2763 | Distal Intergenic | 7 | 1,29E+08 | 1,29E+08 | 72720 | 1 | 1,05E+08 | ENSMUST00000206679.2 | -12929 | | Gm39094 | predicted gene, 39094 |
| chr2 | 69358209 | 69360118 | 1910 | * | 16,25428 | -3,15055296 | 0,962285 | -3,27403 | 0,00106 | 0,410415129 | 2,318434 | Intron (ENSMUST000000 | 2 | 69337967 | 69339489 | 1523 | 2 | 14725 | ENSMUST00000128742.2 | -18720 | ENSMUSG00000027070 | Lrp2 | low density lipoprotein receptor-related protein 2 |
| chr12 | 98015597 | 98016243 | 647 | * | 9,733686 | -3,147619093 | 1,22012 | -2,57976 | 0,009887 | 0,887433944 | 1,425929 | Distal Intergenic | 12 | 98042614 | 98074820 | 32207 | 2 | 1,03E+08 | ENSMUST00000219545.2 | 58577 | ENSMUSG00000112812 | Gm47415 | predicted gene, 47415 |
| chr10 | 14926317 | 14927462 | 1146 | * | 12,44028 | -3,147115044 | 1,050015 | -2,99721 | 0,002725 | 0,73648081 | 0,918255 | Distal Intergenic | 10 | 14860414 | 14860543 | 130 | 1 | 1,15E+08 | ENSMUST00000158105.3 | 65903 | ENSMUSG00000088730 | Gm23892 | predicted gene, 23892 |
| chr7 | 41399958 | 41400843 | 886 | * | 10,42645 | -3,146083954 | 1,195642 | -2,63129 | 0,008506 | 0,930105186 | 0,918255 | Intron (ENSMUST000000 | 7 | 41336885 | 41351731 | 14847 | 1 | 75835 | ENSMUST00000146087.2 | 63073 | ENSMUSG00000074164 | Vmn2r-ps54 | vomeronasal 2, receptor, pseudogene 54 |
| chr12 | 97173382 | 97175037 | 1656 | * | 11,70176 | -3,142441491 | 1,099842 | -2,85718 | 0,004274 | 0,817544002 | 0,918255 | Distal Intergenic | 12 | 97279623 | 97282627 | 3005 | 1 | 1,05E+08 | ENSMUST00000218233.2 | -104586 | | Gm40552 | predicted gene, 40552 |
| chr15 | 67819170 | 67820196 | 1027 | * | 10,97616 | -3,141896374 | 1,136394 | -2,7648 | 0,005696 | 0,809505441 | 1,2763 | Distal Intergenic | 15 | 67956170 | 68130636 | 174467 | 2 | 380993 | ENSMUST00000162173.8 | 310440 | ENSMUSG00000022335 | Zfat | zinc finger and AT hook domain containing |
| chr15 | 70491163 | 70493264 | 2102 | * | 17,73437 | -3,141645278 | 1,009902 | -3,11084 | 0,001866 | 0,497895643 | 2,423594 | Distal Intergenic | 15 | 69883472 | 69949690 | 66219 | 1 | 1,01E+08 | ENSMUST00000230802.2 | 607691 | ENSMUSG00000116483 | Gm19782 | predicted gene, 19782 |
| chr3 | 85406609 | 85408374 | 1766 | * | 11,88783 | -3,139607102 | 1,212672 | -2,589 | 0,009626 | 0,90060438 | 1,2763 | Intron (ENSMUST000000 | 3 | 85374684 | 85374805 | 122 | 1 | 1E+08 | ENSMUST00000116774.3 | 31925 | ENSMUSG00000080424 | Mir466k | microRNA 466k |
| chr1 | 1,31E+08 | 1,31E+08 | 1126 | * | 11,20978 | -3,138358229 | 1,19779 | -2,62012 | 0,00879 | 0,966603024 | 0,744375 | 3' UTR | 1 | 1,31E+08 | 1,31E+08 | 5132 | 1 | 16865 | ENSMUST00000153651.2 | 5914 | ENSMUSG00000026427 | Eif2d | eukaryotic translation initiation factor 2D |
| chr19 | 19982104 | 19984900 | 2797 | * | 19,34511 | -3,138136158 | 0,937667 | -3,34675 | 0,000818 | 0,337054778 | 2,713306 | Promoter (2-3kb) | 19 | 19983304 | 19987508 | 4205 | 2 | 1E+08 | ENSMUST00000243433.1 | 2608 | | Gm3728 | predicted gene 3728 |
| chr4 | 73681637 | 73683046 | 1410 | * | 13,11496 | -3,137507108 | 0,993337 | -3,15855 | 0,001586 | 0,649217039 | 0,918255 | Intron (ENSMUST000000 | 4 | 73632816 | 73708839 | 76024 | 2 | 242505 | ENSMUST00000058292.13 | 25793 | ENSMUSG00000043003 | Rasef | RAS and EF hand domain containing |
| chr14 | 14045675 | 14047051 | 1377 | * | 13,83555 | -3,137378568 | 1,004255 | -3,12409 | 0,001784 | 0,61075929 | 1,2763 | Distal Intergenic | 14 | 14091019 | 14120836 | 29818 | 1 | 18300 | ENSMUST00000022269.7 | -43968 | ENSMUSG00000021749 | Fam3d | FAM3 metabolism regulating signaling molecule D |
| chr1 | 1,58E+08 | 1,58E+08 | 890 | * | 13,12962 | -3,137206549 | 1,105044 | -2,83899 | 0,004526 | 0,798502641 | 1,075735 | Intron (ENSMUST000000 | 1 | 1,58E+08 | 1,58E+08 | 69268 | 2 | 1,15E+08 | ENSMUST00000192868.2 | -19295 | | Gm37679 | predicted gene, 37679 |
| chr2 | 1,09E+08 | 1,09E+08 | 650 | * | 10,46903 | -3,13635725 | 1,139912 | -2,7514 | 0,005934 | 0,87537569 | 0,918255 | Distal Intergenic | 2 | 1,1E+08 | 1,1E+08 | 52308 | 1 | 12064 | ENSMUST00000053317.12 | -96411 | ENSMUSG00000048482 | Bdnf | brain derived neurotrophic factor |
| chr14 | 98913630 | 98914987 | 1358 | * | 12,19565 | -3,135794525 | 1,160876 | -2,70123 | 0,006908 | 0,931030181 | 0,744375 | Distal Intergenic | 14 | 98861626 | 98861732 | 107 | 1 | 1,15E+08 | ENSMUST00000180098.3 | 52004 | ENSMUSG00000119462 | Gm26139 | predicted gene, 26139 |
| chr2 | 1,25E+08 | 1,25E+08 | 1249 | * | 10,90999 | -3,135566947 | 1,126282 | -2,784 | 0,005369 | 0,891774732 | 0,744375 | Exon (ENSMUST000000 | 2 | 1,25E+08 | 1,25E+08 | 62026 | 2 | 99100 | ENSMUST00000089776.3 | 3930 | ENSMUSG00000068394 | Cep152 | centrosomal protein 152 |
| chr3 | 1,18E+08 | 1,18E+08 | 998 | * | 9,763202 | -3,135137118 | 1,202034 | -2,60819 | 0,009102 | 0,903444531 | 1,173325 | Distal Intergenic | 3 | 1,18E+08 | 1,18E+08 | 64536 | 2 | 76561 | ENSMUST00000198499.5 | 98338 | ENSMUSG00000028007 | Snx7 | sorting nexin 7 |
| chr11 | 65913936 | 65915174 | 1239 | * | 14,98154 | -3,13326978 | 1,013993 | -3,09003 | 0,002001 | 0,641208222 | 1,214904 | Intron (ENSMUST000000 | 11 | 65722229 | 65896021 | 173793 | 2 | 237806 | ENSMUST00000152386.8 | -17915 | ENSMUSG00000056752 | Dnah9 | dynein, axonemal, heavy chain 9 |
| chr12 | 73631956 | 73633419 | 1464 | * | 12,06597 | -3,132054368 | 1,162121 | -2,69512 | 0,007036 | 0,902063278 | 0,918255 | Promoter (<=1kb) | 12 | 73631617 | 73808316 | 176700 | 1 | 18755 | ENSMUST00000119092.2 | 339 | ENSMUSG00000021108 | Prkch | protein kinase C, eta |
| chr5 | 50765996 | 50768069 | 2074 | * | 15,42258 | -3,131636219 | 1,031502 | -3,036 | 0,002397 | 0,670234202 | 1,214904 | Intron (ENSMUST000000 | 5 | 50606004 | 50701833 | 95830 | 2 | 1,03E+08 | ENSMUST00000197457.5 | -64163 | | Gm36814 | predicted gene, 36814 |
| chr7 | 18278276 | 18279388 | 1113 | * | 12,41671 | -3,131266 | 1,175548 | -2,66366 | 0,007729 | 0,878632224 | 1,173325 | Distal Intergenic | 7 | 18283010 | 18283116 | 107 | 1 | 1,15E+08 | ENSMUST00000240048.1 | -3622 | ENSMUSG00000119780 | Gm26062 | predicted gene, 26062 |
| chr9 | 46575444 | 46577006 | 1563 | * | 12,90934 | -3,129292326 | 1,079733 | -2,89821 | 0,003753 | 0,749550311 | 1,173325 | Promoter (<=1kb) | 9 | 46577571 | 46586066 | 8496 | 1 | 1,03E+08 | ENSMUST00000213995.2 | -565 | ENSMUSG00000111800 | Gm31557 | predicted gene, 31557 |
| chr19 | 58774089 | 58775505 | 1417 | * | 14,12891 | -3,126145987 | 1,021677 | -3,05982 | 0,002215 | 0,649217039 | 1,2763 | 3' UTR | 19 | 58774428 | 58781715 | 7288 | 2 | 67507 | ENSMUST00000166692.2 | 6210 | ENSMUSG00000026931 | 1700019N19Rik | RIKEN cDNA 1700019N19 gene |
| chr17 | 20251428 | 20253205 | 1778 | * | 10,26761 | -3,125549544 | 1,176015 | -2,65775 | 0,007867 | 0,876149777 | 1,211315 | Exon (ENSMUST000002 | 17 | 20248680 | 20262880 | 14201 | 2 | 22313 | ENSMUST00000233647.2 | 9675 | ENSMUSG00000090315 | Vmn2r104 | vomeronasal 2, receptor 104 |
| chr1 | 1,55E+08 | 1,55E+08 | 950 | * | 10,50224 | -3,123905392 | 1,158626 | -2,69621 | 0,007013 | 0,847020973 | 1,2763 | Distal Intergenic | 1 | 1,55E+08 | 1,55E+08 | 13232 | 2 | 1,05E+08 | ENSMUST00000243429.1 | 61297 | | Gm41979 | predicted gene, 41979 |
| chr9 | 51738046 | 51739585 | 1540 | * | 15,45324 | -3,122986672 | 0,984596 | -3,17185 | 0,001515 | 0,588308481 | 1,214904 | Exon (ENSMUST000000 | 9 | 51743072 | 51759949 | 16878 | 1 | 244867 | ENSMUST00000149754.2 | -3487 | ENSMUSG00000053199 | Arhgap20 | Rho GTPase activating protein 20 |
| chr3 | 1,41E+08 | 1,41E+08 | 1476 | * | 14,40497 | -3,122300142 | 1,122862 | -2,78066 | 0,005425 | 0,854052567 | 0,938682 | Distal Intergenic | 3 | 1,41E+08 | 1,41E+08 | 3470 | 2 | 1,03E+08 | ENSMUST00000199504.2 | 31215 | | Gm32754 | predicted gene, 32754 |
| chr16 | 65458013 | 65459892 | 1880 | * | 12,94801 | -3,121509692 | 1,112554 | -2,80572 | 0,005021 | 0,801266925 | 1,173325 | Promoter (2-3kb) | 16 | 65461988 | 65551514 | 89527 | 1 | 1,05E+08 | ENSMUST00000241896.1 | -2096 | | Gm41469 | predicted gene, 41469 |
| chr9 | 52416877 | 52417925 | 1049 | * | 9,613464 | -3,120596592 | 1,199194 | -2,60225 | 0,009262 | 0,906138253 | 1,173325 | Distal Intergenic | 9 | 52313396 | 52381646 | 68251 | 1 | 1,03E+08 | ENSMUST00000215193.2 | 103481 | | Gm33125 | predicted gene, 33125 |
| chr18 | 55497297 | 55498724 | 1428 | * | 10,06008 | -3,11954685 | 1,167898 | -2,67108 | 0,007561 | 0,874247416 | 1,173325 | Distal Intergenic | 18 | 55190657 | 55212486 | 21830 | 1 | 1E+08 | ENSMUST00000181765.2 | 306640 | | Gm4221 | predicted gene 4221 |
| chr4 | 1,11E+08 | 1,11E+08 | 1339 | * | 14,70098 | -3,118824924 | 1,038059 | -3,00448 | 0,00266 | 0,668684091 | 1,37683 | Intron (ENSMUST000000 | 4 | 1,11E+08 | 1,11E+08 | 140199 | 1 | 78933 | ENSMUST00000136433.2 | -350119 | ENSMUSG00000061298 | Agbl4 | ATP/GTP binding protein-like 4 |
| chr9 | 46452752 | 46455764 | 3013 | * | 24,03753 | -3,116539841 | 0,771609 | -4,03901 | 5,37E-05 | 0,070521466 | 2,951933 | Exon (ENSMUST000002 | 9 | 46444055 | 46454674 | 10620 | 1 | 1,03E+08 | ENSMUST00000214727.2 | 8697 | | Gm31432 | predicted gene, 31432 |
| chr12 | 31791338 | 31793019 | 1682 | * | 13,1801 | -3,113630529 | 1,159327 | -2,68572 | 0,007237 | 0,906013031 | 0,918255 | Intron (ENSMUST000000 | 12 | 31758711 | 31763946 | 5236 | 2 | 73010 | ENSMUST00000176710.2 | -27392 | ENSMUSG00000044067 | Gpr22 | G protein-coupled receptor 22 |
| chrX | 1,58E+08 | 1,58E+08 | 1427 | * | 11,18753 | -3,112965564 | 1,114456 | -2,79326 | 0,005218 | 0,792545355 | 1,2763 | Intron (ENSMUST000000 | 20 | 1,58E+08 | 1,59E+08 | 168085 | 1 | 58194 | ENSMUST00000123433.8 | -9752 | ENSMUSG00000040990 | Sh3kbp1 | SH3-domain kinase binding protein 1 |
| chr13 | 15468751 | 15470191 | 1441 | * | 15,97949 | -3,112397574 | 1,05466 | -2,95109 | 0,003167 | 0,605351622 | 2,318434 | Distal Intergenic | 13 | 15550209 | 15587685 | 37477 | 1 | 1,01E+08 | ENSMUST00000222267.2 | -80018 | ENSMUSG00000113410 | Gm20043 | predicted gene, 20043 |
| chr8 | 89418613 | 89419626 | 1014 | * | 13,71997 | -3,112092527 | 1,050687 | -2,96196 | 0,003057 | 0,75508254 | 0,918255 | Distal Intergenic | 8 | 89423656 | 89478573 | 54918 | 1 | 74256 | ENSMUST00000043526.15 | -4030 | ENSMUSG00000036712 | Cyld | CYLD lysine 63 deubiquitinase |
| chr1 | 68736323 | 68737699 | 1377 | * | 10,96543 | -3,11142308 | 1,139696 | -2,73005 | 0,006333 | 0,918664394 | 0,738441 | Intron (ENSMUST000000 | 1 | 68337309 | 68369375 | 32067 | 2 | 13869 | ENSMUST00000126682.2 | -366948 | ENSMUSG00000062209 | Erbb4 | erb-b2 receptor tyrosine kinase 4 |
| chr17 | 15381787 | 15383767 | 1981 | * | 23,41525 | -3,107437427 | 1,004266 | -3,09424 | 0,001973 | 0,474715691 | 2,951062 | Intron (ENSMUST000000 | 17 | 15372421 | 15393320 | 20900 | 1 | 1,03E+08 | ENSMUST00000231330.2 | 9366 | | Gm35455 | predicted gene, 35455 |
| chrX | 1,39E+08 | 1,39E+08 | 703 | * | 9,106062 | -3,106282754 | 1,183547 | -2,62455 | 0,008676 | 0,858755542 | 1,449706 | 3' UTR | 20 | 1,39E+08 | 1,39E+08 | 26502 | 2 | 237073 | ENSMUST00000138635.8 | 49940 | ENSMUSG00000031433 | Rbm41 | RNA binding motif protein 41 |
| chr10 | 45204506 | 45205605 | 1100 | * | 10,28125 | -3,10610937 | 1,202633 | -2,58276 | 0,009801 | 0,886332554 | 1,425929 | Distal Intergenic | 10 | 45211868 | 45248575 | 36708 | 1 | 23828 | ENSMUST00000095715.5 | -6263 | ENSMUSG00000071317 | Bves | blood vessel epicardial substance |
| chr11 | 1,13E+08 | 1,13E+08 | 1785 | * | 18,65805 | -3,105727083 | 0,946932 | -3,27978 | 0,001039 | 0,401822115 | 2,423594 | Intron (ENSMUST000000 | 11 | 1,13E+08 | 1,13E+08 | 320904 | 2 | 69806 | ENSMUST00000071539.10 | 33036 | ENSMUSG00000041654 | Slc39a11 | solute carrier family 39 (metal ion transporter), member 11 |
| chr12 | 1,19E+08 | 1,19E+08 | 1160 | * | 10,8609 | -3,105593692 | 1,186988 | -2,61637 | 0,008887 | 0,966989334 | 0,744375 | Distal Intergenic | 12 | 1,19E+08 | 1,19E+08 | 37308 | 2 | 320910 | ENSMUST00000137804.2 | 122652 | ENSMUSG00000025321 | Itgb8 | integrin beta 8 |
| chr16 | 36967009 | 36968671 | 1663 | * | 11,32521 | -3,105255062 | 1,134544 | -2,73701 | 0,0062 | 0,915421656 | 0,738441 | Intron (ENSMUST000000 | 16 | 36977114 | 37024938 | 47825 | 2 | 207227 | ENSMUST00000149984.2 | 56267 | ENSMUSG00000022829 | Stxbp5l | syntaxin binding protein 5-like |
| chr6 | 1,46E+08 | 1,46E+08 | 663 | * | 13,03075 | -3,105242652 | 1,086134 | -2,85899 | 0,00425 | 0,853738252 | 0,738441 | Distal Intergenic | 6 | 1,46E+08 | 1,46E+08 | 5723 | 1 | 22147 | ENSMUST00000087445.7 | 45316 | ENSMUSG00000067338 | Tuba3b | tubulin, alpha 3B |
| chr7 | 61908831 | 61910332 | 1502 | * | 11,72477 | -3,103523501 | 1,107652 | -2,80189 | 0,00508 | 0,802490409 | 1,173325 | Distal Intergenic | 7 | 61996317 | 62000010 | 3694 | 1 | 17984 | ENSMUST00000038775.6 | -85985 | ENSMUSG00000033585 | Ndn | necdin, MAGE family member |
| chr10 | 77504171 | 77506114 | 1944 | * | 18,38796 | -3,103121517 | 0,961178 | -3,22846 | 0,001245 | 0,430115816 | 2,423594 | Distal Intergenic | 10 | 77522403 | 77722603 | 200201 | 1 | 252974 | ENSMUST00000092366.4 | -16289 | ENSMUSG00000069581 | Tspear | thrombospondin type laminin G domain and EAR repeats |
| chr10 | 26961653 | 26963163 | 1511 | * | 11,43542 | -3,100745443 | 1,194913 | -2,59496 | 0,00946 | 0,976466474 | 0,744375 | Intron (ENSMUST000000 | 10 | 26857491 | 26932641 | 75151 | 2 | 16773 | ENSMUST00000219763.2 | -29012 | ENSMUSG00000019899 | Lama2 | laminin, alpha 2 |
| chr5 | 84630036 | 84631095 | 1060 | * | 12,90536 | -3,100330298 | 1,194094 | -2,59639 | 0,009421 | 0,908806502 | 1,173325 | Distal Intergenic | 5 | 84478634 | 84565241 | 86608 | 2 | 13839 | ENSMUST00000136775.2 | -64795 | ENSMUSG00000029245 | Epha5 | Eph receptor A5 |
| chr1 | 55552414 | 55553492 | 1079 | * | 9,900069 | -3,099632553 | 1,137534 | -2,72487 | 0,006433 | 0,83918951 | 1,211315 | Intron (ENSMUST000000 | 1 | 55483233 | 55484183 | 951 | 2 | 626009 | ENSMUST00000186507.2 | -68231 | | Gm6644 | Akr1b3 pseudogene |
| chr18 | 73935482 | 73936907 | 1426 | * | 11,59935 | -3,098530966 | 1,195276 | -2,59232 | 0,009533 | 0,899471409 | 1,2763 | Intron (ENSMUST000000 | 18 | 73902974 | 73948509 | 45536 | 2 | 107029 | ENSMUST00000025439.5 | 11602 | ENSMUSG00000024556 | Me2 | malic enzyme 2, NAD(+)-dependent, mitochondrial |
| chr5 | 20405958 | 20407028 | 1071 | * | 10,48377 | -3,097103089 | 1,174557 | -2,63683 | 0,008369 | 0,877892722 | 1,2763 | Promoter (<=1kb) | 5 | 20407567 | 20670611 | 263045 | 1 | 50791 | ENSMUST00000197822.5 | -539 | ENSMUSG00000040003 | Magi2 | membrane associated guanylate kinase, WW and PDZ domain containing 2 |
| chrX | 1,23E+08 | 1,23E+08 | 1431 | * | 8,289803 | -3,09709021 | 1,194234 | -2,59337 | 0,009504 | 0,877892722 | 1,449706 | Distal Intergenic | 20 | 1,23E+08 | 1,23E+08 | 10992 | 2 | 1E+08 | ENSMUST00000094491.5 | 30341 | ENSMUSG00000072049 | Vmn2r121 | vomeronasal 2, receptor 121 |
| chr12 | 78701110 | 78703356 | 2247 | * | 19,62821 | -3,093770146 | 0,964021 | -3,20923 | 0,001331 | 0,425952262 | 2,663619 | Exon (ENSMUST000000 | 12 | 78729421 | 78730867 | 1447 | 1 | 268566 | ENSMUST00000219629.2 | -26065 | ENSMUSG00000047454 | Gphn | gephyrin |
| chr3 | 16140789 | 16141908 | 1120 | * | 11,69521 | -3,093068556 | 1,144837 | -2,70175 | 0,006897 | 0,856964735 | 1,173325 | Distal Intergenic | 3 | 16001034 | 16060601 | 59568 | 2 | 381484 | ENSMUST00000194367.6 | -80188 | ENSMUSG00000078780 | Gm5150 | predicted gene 5150 |
| chr12 | 52110160 | 52111839 | 1680 | * | 11,62502 | -3,092840403 | 1,198991 | -2,57954 | 0,009893 | 0,903294867 | 1,2763 | Distal Intergenic | 12 | 52144520 | 52359527 | 215008 | 1 | 76826 | ENSMUST00000040090.16 | -32681 | ENSMUSG00000035142 | Nubpl | nucleotide binding protein-like |
| chr18 | 3230417 | 3231484 | 1068 | * | 11,42342 | -3,092313397 | 1,107213 | -2,79288 | 0,005224 | 0,888740228 | 0,744375 | Distal Intergenic | 18 | 3267546 | 3281078 | 13533 | 2 | 12916 | ENSMUST00000049942.13 | 49594 | ENSMUSG00000063889 | Crem | cAMP responsive element modulator |
| chr19 | 26181709 | 26182984 | 1276 | * | 10,66497 | -3,090529981 | 1,161704 | -2,66034 | 0,007806 | 0,917631345 | 0,918255 | Distal Intergenic | 19 | 26015816 | 26023831 | 8016 | 2 | 73433 | ENSMUST00000237184.2 | -157878 | ENSMUSG00000118016 | 1700048O14Rik | RIKEN cDNA 1700048O14 gene |
| chr7 | 1,13E+08 | 1,13E+08 | 1212 | * | 11,01538 | -3,090416033 | 1,132939 | -2,72779 | 0,006376 | 0,918554156 | 0,744375 | Distal Intergenic | 7 | 1,13E+08 | 1,13E+08 | 7553 | 2 | 1,08E+08 | ENSMUST00000211589.2 | 28012 | | Gm5600 | predicted gene 5600 |

| | | | | | | | | | | | | | | | | | | | | | | | |
|---|---|---|---|---|---|---|---|---|---|---|---|---|---|---|---|---|---|---|---|---|---|---|---|
| chr14 | 28083012 | 28084558 | 1547 | * | 13,00127 | -3,089006142 | 1,19238 | -2,59062 | 0,00958 | 0,900603852 | 1,2763 | Intron (ENSMUST0000( | 14 | 28177478 | 28197960 | 20483 | 1 | 238988 | ENSMUST00000224045.2 | -92920 | ENSMUSG00000040640 | Erc2 | ELKS/RAB6-interacting/CAST family member 2 |
| chr2 | 50924584 | 50925453 | 870 | * | 12,03396 | -3,088266945 | 1,110599 | -2,78072 | 0,005424 | 0,895765172 | 0,738441 | Distal Intergenic | 2 | 51027081 | 51039106 | 12026 | 2 | 74194 | ENSMUST00000140864.2 | 113653 | ENSMUSG00000017144 | Rnd3 | Rho family GTPase 3 |
| chr4 | 49452328 | 49453495 | 1168 | * | 13,97687 | -3,088055211 | 1,144817 | -2,69742 | 0,006988 | 0,846674939 | 1,2763 | Promoter (1-2kb) | 4 | 49447109 | 49451137 | 4029 | 2 | 230161 | ENSMUST00000095086.3 | -1191 | ENSMUSG00000070985 | Acnat1 | acyl-coenzyme A amino acid N-acyltransferase 1 |
| chr11 | 85216825 | 85218301 | 1477 | * | 12,26266 | -3,0874396 | 1,133952 | -2,72273 | 0,006475 | 0,860218139 | 1,075735 | Exon (ENSMUST00000( | 11 | 85202990 | 85228029 | 25040 | 1 | 53892 | ENSMUST00000127717.2 | 13835 | ENSMUSG00000020525 | Ppm1d | protein phosphatase 1D magnesium-dependent, delta isoform |
| chr18 | 18557000 | 18559096 | 2097 | * | 17,50236 | -3,087018951 | 1,03323 | -2,98774 | 0,002811 | 0,551447786 | 2,663619 | Distal Intergenic | 18 | 19266274 | 19334773 | 68500 | 1 | 1,03E+08 | ENSMUST00000234128.2 | -707178 | | Gm31086 | predicted gene, 31086 |
| chr1 | 1,31E+08 | 1,31E+08 | 808 | * | 10,70983 | -3,087001673 | 1,158359 | -2,66498 | 0,007699 | 0,915421656 | 0,918255 | Distal Intergenic | 1 | 1,32E+08 | 1,32E+08 | 12762 | 1 | 26361 | ENSMUST00000027690.7 | -29449 | ENSMUSG00000026432 | Avpr1b | arginine vasopressin receptor 1B |
| chr7 | 33631976 | 33633255 | 1280 | * | 10,87894 | -3,083972899 | 1,182358 | -2,60832 | 0,009099 | 0,941444936 | 0,918255 | Distal Intergenic | 7 | 33642427 | 33644465 | 2039 | 2 | 110187 | ENSMUST00000085623.4 | 11210 | ENSMUSG00000066586 | Scgb2b26 | secretoglobin, family 2B, member 26 |
| chr8 | 1,17E+08 | 1,17E+08 | 1267 | * | 13,95469 | -3,08375732 | 0,914793 | -3,37099 | 0,000749 | 0,507398906 | 0,918255 | Distal Intergenic | 8 | 1,17E+08 | 1,17E+08 | 11426 | 1 | 1,03E+08 | ENSMUST00000241934.1 | 160758 | | Gm30925 | predicted gene, 30925 |
| chr10 | 61254761 | 61256158 | 1398 | * | 13,62495 | -3,083752574 | 1,164197 | -2,64882 | 0,008077 | 0,923078468 | 0,918255 | Promoter (1-2kb) | 10 | 61253751 | 61261117 | 7367 | 1 | 18119 | ENSMUST00000049339.7 | 1010 | ENSMUSG00000037171 | Nodal | nodal |
| chr3 | 13873612 | 13875009 | 1398 | * | 14,00973 | -3,082914436 | 1,044315 | -2,95209 | 0,003156 | 0,801134483 | 0,738441 | Intron (ENSMUST0000( | 3 | 13841908 | 13842367 | 460 | 1 | 76897 | ENSMUST00000108373.4 | 31704 | ENSMUSG00000039717 | Ralyl | RALY RNA binding protein-like |
| chr14 | 89844815 | 89846321 | 1507 | * | 9,343779 | -3,082038919 | 1,143637 | -2,69494 | 0,00704 | 0,836508023 | 1,360622 | Distal Intergenic | 14 | 89944241 | 89944372 | 132 | 1 | 1,15E+08 | ENSMUST00000104220.3 | -97920 | ENSMUSG00000077408 | Gm25415 | predicted gene, 25415 |
| chr8 | 1,21E+08 | 1,21E+08 | 1255 | * | 15,3828 | -3,079917667 | 0,976495 | -3,15406 | 0,00161 | 0,591609342 | 1,2763 | Promoter (<=1kb) | 8 | 1,21E+08 | 1,21E+08 | 4665 | 2 | 272551 | ENSMUST00000155609.2 | 837 | ENSMUSG00000031821 | Gins2 | GINS complex subunit 2 (Psf2 homolog) |
| chr15 | 4134964 | 4136325 | 1362 | * | 11,02813 | -3,078227479 | 1,124386 | -2,7377 | 0,006187 | 0,915421656 | 0,738441 | Intron (ENSMUST0000( | 15 | 4172005 | 4183462 | 11458 | 1 | 67041 | ENSMUST00000124591.8 | -35680 | ENSMUSG00000022186 | Oxct1 | 3-oxoacid CoA transferase 1 |
| chr10 | 1,06E+08 | 1,06E+08 | 1327 | * | 11,35162 | -3,07803124 | 1,154897 | -2,6652 | 0,007694 | 0,915421656 | 0,918255 | Distal Intergenic | 10 | 1,05E+08 | 1,05E+08 | 9283 | 1 | 327812 | ENSMUST00000243515.1 | 91633 | ENSMUSG00000085282 | Gm15663 | predicted gene 15663 |
| chr4 | 71600195 | 71601327 | 1133 | * | 13,27869 | -3,074574286 | 1,061306 | -2,89697 | 0,003768 | 0,791242424 | 0,938682 | Distal Intergenic | 4 | 71673880 | 71676202 | 2323 | 2 | 1,03E+08 | ENSMUST00000084489.3 | 74875 | ENSMUSG00000066141 | Gm11232 | predicted gene 11232 |
| chr11 | 65105665 | 65106852 | 1188 | * | 11,78925 | -3,072977972 | 1,137881 | -2,70061 | 0,006921 | 0,844192936 | 1,2763 | Promoter (2-3kb) | 11 | 65069254 | 65109385 | 40132 | 2 | 214384 | ENSMUST00000101042.9 | 2533 | ENSMUSG00000020542 | Myocd | myocardin |
| chr9 | 1,06E+08 | 1,06E+08 | 1312 | * | 13,13705 | -3,072579088 | 1,105803 | -2,7786 | 0,005459 | 0,817544002 | 1,173325 | Distal Intergenic | 9 | 1,06E+08 | 1,06E+08 | 11019 | 1 | 654798 | ENSMUST00000143956.2 | -15101 | | D030055H07Rik | RIKEN cDNA D030055H07 gene |
| chr2 | 1,59E+08 | 1,59E+08 | 839 | * | 9,862478 | -3,072225874 | 1,181699 | -2,59984 | 0,009327 | 0,918758168 | 1,086118 | Intron (ENSMUST0000( | 2 | 1,59E+08 | 1,59E+08 | 7531 | 1 | 71715 | ENSMUST00000157070.2 | 10903 | ENSMUSG00000027655 | Dhx35 | DEAH (Asp-Glu-Ala-His) box polypeptide 35 |
| chr2 | 3903585 | 3904221 | 637 | * | 9,939994 | -3,071841628 | 1,127629 | -2,72416 | 0,006447 | 0,858257847 | 1,086118 | Distal Intergenic | 2 | 3856067 | 3878612 | 22546 | 1 | 1,03E+08 | ENSMUST00000141222.2 | 47518 | | Gm36031 | predicted gene, 36031 |
| chr1 | 1,77E+08 | 1,77E+08 | 1281 | * | 15,02937 | -3,070591898 | 1,042193 | -2,94628 | 0,003216 | 0,693610666 | 1,37683 | Distal Intergenic | 1 | 1,77E+08 | 1,77E+08 | 108 | 1 | 1,15E+08 | ENSMUST00000157250.4 | -5881 | ENSMUSG00000087875 | Gm26104 | predicted gene, 26104 |
| chr7 | 84359609 | 84360768 | 1160 | * | 14,09059 | -3,070435317 | 1,062583 | -2,8896 | 0,003857 | 0,793028263 | 0,938682 | Intron (ENSMUST0000( | 7 | 84283190 | 84339167 | 55978 | 2 | 65098 | ENSMUST00000207975.2 | -20442 | ENSMUSG00000030629 | Zfand6 | zinc finger, AN1-type domain 6 |
| chr7 | 1,3E+08 | 1,3E+08 | 2372 | * | 15,44929 | -3,069911692 | 1,132776 | -2,71008 | 0,006727 | 0,838625535 | 1,2763 | Promoter (<=1kb) | 7 | 1,3E+08 | 1,3E+08 | 39349 | 2 | 67872 | ENSMUST00000159694.3 | 0 | ENSMUSG00000040331 | Nsmce4a | NSE4 homolog A, SMC5-SMC6 complex component |
| chr13 | 20405724 | 20406776 | 1053 | * | 10,88932 | -3,068964552 | 1,08837 | -2,81978 | 0,004806 | 0,792545355 | 1,173325 | Intron (ENSMUST0000( | 13 | 20435240 | 20500623 | 65384 | 1 | 140580 | ENSMUST00000220751.2 | -28464 | ENSMUSG00000041112 | Elmo1 | engulfment and cell motility 1 |
| chr7 | 71712260 | 71713407 | 1148 | * | 11,81606 | -3,068462643 | 1,085513 | -2,82674 | 0,004702 | 0,870290757 | 0,744375 | Distal Intergenic | 7 | 71730161 | 71760331 | 30171 | 2 | 244049 | ENSMUST00000206466.2 | 46924 | ENSMUSG00000032776 | Mctp2 | multiple C2 domains, transmembrane 2 |
| chr13 | 41776854 | 41779136 | 2283 | * | 17,06499 | -3,068124394 | 0,952283 | -3,22186 | 0,001274 | 0,437520575 | 2,386008 | Intron (ENSMUST0000( | 13 | 41759692 | 41794831 | 35140 | 1 | 621976 | ENSMUST00000129449.3 | 17162 | ENSMUSG00000087370 | Tmem170b | transmembrane protein 170B |
| chr19 | 54735158 | 54736455 | 1298 | * | 11,74292 | -3,067936884 | 1,102328 | -2,78314 | 0,005383 | 0,798887389 | 1,2763 | Distal Intergenic | 19 | 55058166 | 55087883 | 29718 | 2 | 14732 | ENSMUST00000061856.6 | 351428 | ENSMUSG00000024978 | Gpam | glycerol-3-phosphate acyltransferase, mitochondrial |
| chr14 | 85735214 | 85736360 | 1147 | * | 10,43452 | -3,066145578 | 1,078003 | -2,84428 | 0,004451 | 0,86031879 | 0,738441 | Distal Intergenic | 14 | 86410363 | 86417021 | 6659 | 1 | 1,03E+08 | ENSMUST00000227116.2 | -674003 | | Gm32729 | predicted gene, 32729 |
| chr13 | 33777367 | 33778767 | 1401 | * | 14,22449 | -3,065529987 | 1,093039 | -2,80459 | 0,005038 | 0,841702416 | 0,938682 | Distal Intergenic | 13 | 33746469 | 33753300 | 6832 | 2 | 619715 | ENSMUST00000222963.2 | -24067 | ENSMUSG00000112963 | Gm6093 | predicted gene 6093 |
| chr9 | 61207472 | 61209645 | 2174 | * | 20,83946 | -3,065421074 | 1,015522 | -3,01857 | 0,00254 | 0,524368688 | 2,880017 | Distal Intergenic | 9 | 61166509 | 61201091 | 34583 | 2 | 622037 | ENSMUST00000213377.2 | -6381 | ENSMUSG00000111658 | B930092H01Rik | RIKEN cDNA B930092H01 gene |
| chr17 | 11260599 | 11262283 | 1685 | * | 13,8661 | -3,063493116 | 1,166614 | -2,62597 | 0,00864 | 0,964229994 | 0,738441 | Intron (ENSMUST0000( | 17 | 11286012 | 11712060 | 426049 | 1 | 50873 | ENSMUST00000168593.9 | -23729 | ENSMUSG00000023826 | Prkn | parkin RBR E3 ubiquitin protein ligase |
| chr11 | 40254415 | 40255790 | 1376 | * | 11,63509 | -3,06179913 | 1,026062 | -2,98403 | 0,002845 | 0,744477896 | 0,918255 | Distal Intergenic | 11 | 40365545 | 40366504 | 960 | 2 | 78343 | ENSMUST00000123904.2 | 110714 | ENSMUSG00000085588 | 3110004A20Rik | RIKEN cDNA 3110004A20 gene |
| chr11 | 36305181 | 36307585 | 2405 | * | 17,0333 | -3,061570864 | 0,978463 | -3,12896 | 0,001754 | 0,507398906 | 2,141612 | Intron (ENSMUST0000( | 11 | 36275950 | 36323103 | 47154 | 2 | 23964 | ENSMUST00000128501.2 | 15518 | ENSMUSG00000049336 | Tenm2 | teneurin transmembrane protein 2 |
| chr1 | 1,67E+08 | 1,67E+08 | 1643 | * | 12,52856 | -3,057967024 | 1,160947 | -2,63403 | 0,008438 | 0,905322447 | 1,075735 | Distal Intergenic | 1 | 1,67E+08 | 1,67E+08 | 41240 | 1 | 20183 | ENSMUST00000015987.10 | -35540 | ENSMUSG00000015843 | Rxrg | retinoid X receptor gamma |
| chr18 | 82632280 | 82633261 | 982 | * | 10,83324 | -3,057798705 | 1,110535 | -2,75345 | 0,005897 | 0,908065779 | 0,738441 | Promoter (2-3kb) | 18 | 82613601 | 82635875 | 22275 | 2 | 329002 | ENSMUST00000183324.8 | 2614 | ENSMUSG00000041258 | Zfp236 | zinc finger protein 236 |
| chr9 | 85733263 | 85734345 | 1083 | * | 11,47687 | -3,056711769 | 1,054889 | -2,89766 | 0,00376 | 0,792545355 | 0,918255 | Distal Intergenic | 9 | 85702355 | 85726376 | 24022 | 2 | 442799 | ENSMUST00000185960.2 | -6887 | | 9330154J02Rik | RIKEN cDNA 9330154J02 gene |
| chr9 | 55797466 | 55799357 | 1892 | * | 14,31864 | -3,056555694 | 0,99594 | -3,06901 | 0,002148 | 0,671821937 | 1,075735 | Intron (ENSMUST0000( | 9 | 55457167 | 55826889 | 369723 | 2 | 244891 | ENSMUST00000037408.9 | 27532 | ENSMUSG00000034007 | Scaper | S phase cyclin A-associated protein in the ER |
| chr5 | 30124817 | 30126903 | 2087 | * | 13,07256 | -3,056039299 | 1,142237 | -2,67549 | 0,007462 | 0,85617852 | 1,2763 | Distal Intergenic | 5 | 30209471 | 30212759 | 3289 | 2 | 97209 | ENSMUST00000199600.2 | 85856 | | A230098N10Rik | RIKEN cDNA A230098N10 gene |
| chr9 | 25517930 | 25520139 | 2210 | * | 22,17283 | -3,054428233 | 0,82178 | -3,71685 | 0,000202 | 0,155627448 | 3,104559 | Distal Intergenic | 9 | 25480974 | 25481101 | 128 | 2 | 1,15E+08 | ENSMUST00000104509.3 | -36829 | ENSMUSG00000077697 | Gm25346 | predicted gene, 25346 |
| chr4 | 94343179 | 94344392 | 1214 | * | 13,35322 | -3,053700767 | 1,079567 | -2,82863 | 0,004675 | 0,832960417 | 0,918255 | Distal Intergenic | 4 | 94349166 | 94365417 | 16252 | 2 | 1,15E+08 | ENSMUST00000243092.1 | 21025 | | Gm12655 | predicted gene 12655 |
| chr5 | 1,42E+08 | 1,42E+08 | 1445 | * | 16,00764 | -3,052627524 | 0,924145 | -3,30319 | 0,000956 | 0,408648517 | 2,141612 | Intron (ENSMUST0000( | 5 | 1,42E+08 | 1,42E+08 | 18124 | 1 | 17425 | ENSMUST00000198422.2 | -245179 | ENSMUSG00000056493 | Foxk1 | forkhead box K1 |
| chr13 | 40183089 | 40183988 | 900 | * | 12,13863 | -3,052139829 | 1,099734 | -2,77534 | 0,005514 | 0,897633667 | 0,744375 | Intron (ENSMUST0000( | 13 | 40012339 | 40012503 | 165 | 2 | 1,15E+08 | ENSMUST00000083322.3 | -170586 | ENSMUSG00000065256 | Gm23972 | predicted gene, 23972 |
| chr8 | 1,03E+08 | 1,03E+08 | 1465 | * | 12,49629 | -3,052068402 | 1,070455 | -2,85119 | 0,004356 | 0,857326444 | 0,738441 | Distal Intergenic | 8 | 1,03E+08 | 1,03E+08 | 2716 | 2 | 12552 | ENSMUST00000210578.2 | 482299 | ENSMUSG00000031673 | Cdh11 | cadherin 11 |
| chr12 | 59913032 | 59914121 | 1090 | * | 9,344608 | -3,051744491 | 1,133556 | -2,69219 | 0,007099 | 0,828987261 | 1,425929 | Distal Intergenic | 12 | 59719776 | 59719906 | 131 | 1 | 1,15E+08 | ENSMUST00000104415.3 | 193256 | ENSMUSG00000077603 | Gm22673 | predicted gene, 22673 |
| chr17 | 90999359 | 91000294 | 936 | * | 12,09358 | -3,051581988 | 1,155768 | -2,64031 | 0,008283 | 0,959532911 | 0,738441 | Intron (ENSMUST0000( | 17 | 90897622 | 91008202 | 110581 | 2 | 18189 | ENSMUST00000197224.2 | 7908 | ENSMUSG00000024109 | Nrxn1 | neurexin I |
| chr18 | 20325648 | 20326742 | 1095 | * | 12,98583 | -3,051276043 | 1,079516 | -2,82652 | 0,004706 | 0,791147121 | 1,173325 | Distal Intergenic | 18 | 20380397 | 20418088 | 37692 | 1 | 211924 | ENSMUST00000054128.7 | -53655 | ENSMUSG00000034774 | Dsg1c | desmoglein 1 gamma |
| chr9 | 3240266 | 3241244 | 979 | * | 10,11413 | -3,051194031 | 1,076962 | -2,83315 | 0,004609 | 0,788774532 | 1,173325 | Distal Intergenic | 9 | 3199894 | 3215230 | 15337 | 1 | 1,03E+08 | ENSMUST00000212296.2 | 40372 | | Gm31138 | predicted gene, 31138 |
| chr16 | 84645714 | 84646684 | 971 | * | 10,10336 | -3,050425128 | 1,154045 | -2,64325 | 0,008212 | 0,884058067 | 1,211315 | Promoter (2-3kb) | 16 | 84649471 | 84657317 | 7847 | 1 | 14390 | ENSMUST00000126031.8 | -2787 | ENSMUSG00000008976 | Gabpa | GA repeat binding protein, alpha |
| chr13 | 75781932 | 75783278 | 1347 | * | 13,6925 | -3,049812255 | 1,064402 | -2,86528 | 0,004166 | 0,751231966 | 1,2763 | Distal Intergenic | 13 | 75793015 | 75793290 | 276 | 1 | 751556 | ENSMUST00000099371.5 | -9737 | ENSMUSG00000076236 | Mir682 | microRNA 682 |
| chr1 | 1,08E+08 | 1,08E+08 | 1442 | * | 14,86657 | -3,048693629 | 1,007363 | -3,02641 | 0,002475 | 0,669377201 | 1,2763 | Promoter (2-3kb) | 1 | 1,08E+08 | 1,08E+08 | 18904 | 1 | 20725 | ENSMUST00000000514.11 | -2066 | ENSMUSG00000026315 | Serpinb8 | serine (or cysteine) peptidase inhibitor, clade B, member 8 |
| chr3 | 1,09E+08 | 1,09E+08 | 1611 | * | 14,56941 | -3,047991225 | 1,061924 | -2,87025 | 0,004101 | 0,731244872 | 1,41696 | Distal Intergenic | 3 | 1,09E+08 | 1,1E+08 | 345046 | 1 | 57257 | ENSMUST00000046864.14 | -38548 | ENSMUSG00000033721 | Vav3 | vav 3 oncogene |
| chrX | 1,38E+08 | 1,38E+08 | 1510 | * | 10,30363 | -3,047968249 | 1,082978 | -2,81443 | 0,004886 | 0,760759863 | 1,425929 | Distal Intergenic | 20 | 1,38E+08 | 1,38E+08 | 1782 | 1 | 71248 | ENSMUST00000113046.2 | -38915 | ENSMUSG00000042525 | 4933428M09Rik | RIKEN cDNA 4933428M09 gene |
| chr5 | 81189724 | 81191243 | 1520 | * | 11,01896 | -3,047432495 | 1,164802 | -2,61627 | 0,00889 | 0,888080529 | 1,2763 | Intron (ENSMUST0000( | 5 | 81169657 | 81708898 | 539242 | 1 | 319387 | ENSMUST00000127046.2 | 20067 | ENSMUSG00000037605 | Adgrl3 | adhesion G protein-coupled receptor L3 |
| chr6 | 86743595 | 86745028 | 1434 | * | 16,11016 | -3,047376868 | 1,101 | -2,76783 | 0,005643 | 0,695892636 | 2,386008 | Promoter (<=1kb) | 6 | 86724453 | 86742892 | 18440 | 2 | 11746 | ENSMUST00000129772.2 | -703 | ENSMUSG00000029994 | Anxa4 | annexin A4 |
| chr4 | 48698115 | 48699914 | 1800 | * | 14,14244 | -3,045432013 | 1,090037 | -2,79388 | 0,005208 | 0,792545355 | 1,2763 | Distal Intergenic | 4 | 48663514 | 48673502 | 9989 | 1 | 68016 | ENSMUST00000030033.5 | 34601 | ENSMUSG00000028348 | Cavin4 | caveolae associated 4 |
| chr1 | 1,39E+08 | 1,39E+08 | 1685 | * | 12,29643 | -3,044040166 | 1,155583 | -2,6342 | 0,008434 | 0,879169754 | 1,2763 | Intron (ENSMUST0000( | 1 | 1,39E+08 | 1,39E+08 | 1591 | 1 | 78229 | ENSMUST00000200494.2 | 42827 | ENSMUSG00000104900 | 4930596I21Rik | RIKEN cDNA 4930596I21 gene |
| chr4 | 29085087 | 29086168 | 1082 | * | 12,77451 | -3,043771099 | 1,150356 | -2,64594 | 0,008146 | 0,923399738 | 0,918255 | Intron (ENSMUST0000( | 4 | 29068079 | 29088811 | 20733 | 1 | 1,03E+08 | ENSMUST00000154813.2 | 17008 | | Gm11917 | predicted gene 11917 |
| chr9 | 76319967 | 76321846 | 1880 | * | 12,60911 | -3,04365918 | 1,015919 | -2,99597 | 0,002736 | 0,777172375 | 0,738441 | Distal Intergenic | 9 | 76137236 | 76231138 | 93903 | 2 | 387285 | ENSMUST00000184757.2 | -88829 | ENSMUSG00000032360 | Hcrtr2 | hypocretin (orexin) receptor 2 |
| chr6 | 1,21E+08 | 1,21E+08 | 953 | * | 12,93186 | -3,042961022 | 1,078926 | -2,82036 | 0,004797 | 0,778555725 | 1,2763 | Distal Intergenic | 6 | 1,21E+08 | 1,21E+08 | 8084 | 1 | 73505 | ENSMUST00000204672.2 | 53314 | | 1700072O05Rik | RIKEN cDNA 1700072O05 gene |
| chr17 | 21295201 | 21296676 | 1476 | * | 10,40433 | -3,042156971 | 1,13389 | -2,68294 | 0,007298 | 0,939041504 | 0,744375 | Distal Intergenic | 17 | 21311470 | 21323609 | 12140 | 1 | 328766 | ENSMUST00000188185.2 | -14794 | ENSMUSG00000102080 | Gm5092 | predicted gene 5092 |
| chr2 | 64701427 | 64702225 | 799 | * | 11,28251 | -3,041970809 | 1,146865 | -2,65242 | 0,007992 | 0,955124605 | 0,738441 | Distal Intergenic | 2 | 64742820 | 64745363 | 2544 | 2 | 50915 | ENSMUST00000134020.2 | 43138 | ENSMUSG00000026888 | Grb14 | growth factor receptor bound protein 14 |
| chr8 | 47497869 | 47499704 | 1836 | * | 21,63131 | -3,038569272 | 0,823677 | -3,68903 | 0,000225 | 0,173046115 | 2,880017 | Intron (ENSMUST0000( | 8 | 47521966 | 47536511 | 14546 | 1 | 320981 | ENSMUST00000149593.2 | -22262 | ENSMUSG00000038173 | Enpp6 | ectonucleotide pyrophosphatase/phosphodiesterase 6 |
| chr16 | 62356418 | 62357630 | 1213 | * | 11,25245 | -3,038015446 | 1,161705 | -2,61514 | 0,008919 | 0,937836704 | 0,918255 | Distal Intergenic | 16 | 62391399 | 62391502 | 104 | 1 | 1,15E+08 | ENSMUST00000093663.3 | -33769 | ENSMUSG00000070173 | Gm24164 | predicted gene, 24164 |
| chr12 | 21956877 | 21958879 | 2003 | * | 11,06419 | -3,03370978 | 1,097365 | -2,76454 | 0,0057 | 0,903101391 | 0,738441 | Distal Intergenic | 12 | 21826160 | 21826708 | 549 | 2 | 434693 | ENSMUST00000164169.2 | -130169 | | Mrto4-ps2 | mRNA turnover 4, pseudogene 2 |
| chr14 | 61543028 | 61544432 | 1405 | * | 14,03493 | -3,032814943 | 1,082977 | -2,80044 | 0,005103 | 0,887107752 | 0,738441 | Promoter (<=1kb) | 14 | 61530429 | 61542783 | 12355 | 2 | 67355 | ENSMUST00000190737.2 | -245 | ENSMUSG00000101505 | 1700109G14Rik | RIKEN cDNA 1700109G14 gene |
| chr1 | 49641765 | 49642507 | 743 | * | 9,279303 | -3,03187049 | 1,140963 | -2,65729 | 0,007877 | 0,844744375 | 1,449706 | Distal Intergenic | 1 | 49283775 | 49379602 | 95828 | 1 | 442837 | ENSMUST00000188673.7 | 357990 | ENSMUSG00000051616 | C230029F24Rik | RIKEN cDNA C230029F24 gene |
| chrX | 1,44E+08 | 1,44E+08 | 1596 | * | 12,47466 | -3,031400131 | 1,134278 | -2,67254 | 0,007528 | 0,947196831 | 0,738441 | Distal Intergenic | 20 | 1,44E+08 | 1,44E+08 | 145 | 1 | 1,15E+08 | ENSMUST00000104042.3 | -26397 | ENSMUSG00000077230 | Gm25097 | predicted gene, 25097 |
| chr7 | 64950987 | 64952727 | 1741 | * | 15,1061 | -3,027051394 | 1,000247 | -3,0263 | 0,002476 | 0,654833721 | 1,37683 | Promoter (<=1kb) | 7 | 64947252 | 64952053 | 4802 | 2 | 21872 | ENSMUST00000132036.2 | 0 | ENSMUSG00000030516 | Tjp1 | tight junction protein 1 |
| chr5 | 27879796 | 27881108 | 1313 | * | 13,96645 | -3,02681201 | 1,031378 | -2,93473 | 0,003338 | 0,745178802 | 1,075735 | Intron (ENSMUST0000( | 5 | 27949083 | 27956293 | 7211 | 2 | 55982 | ENSMUST00000197625.2 | 75185 | ENSMUSG00000002221 | Paxip1 | PAX interacting (with transcription-activation domain) protein 1 |
| chr17 | 44587152 | 44588501 | 1350 | * | 13,64909 | -3,025547062 | 1,021079 | -2,96309 | 0,003046 | 0,697700764 | 1,2763 | 3' UTR | 17 | 44547993 | 44562606 | 14614 | 1 | 224796 | ENSMUST00000233193.2 | 39159 | ENSMUSG00000023959 | Clic5 | chloride intracellular channel 5 |
| chr5 | 1,4E+08 | 1,4E+08 | 1628 | * | 14,03104 | -3,024449603 | 1,107361 | -2,73122 | 0,00631 | 0,856353593 | 1,075735 | Promoter (2-3kb) | 5 | 1,4E+08 | 1,4E+08 | 95583 | 2 | 17120 | ENSMUST00000132832.2 | 2691 | ENSMUSG00000029554 | Mad1l1 | MAD1 mitotic arrest deficient 1-like 1 |
| chr10 | 37405483 | 37406931 | 1449 | * | 14,91512 | -3,024031697 | 0,92939 | -3,25378 | 0,001139 | 0,50985457 | 1,37683 | Distal Intergenic | 10 | 37213482 | 37255648 | 42167 | 1 | 1,03E+08 | ENSMUST00000181602.2 | 192001 | | Gm26535 | predicted gene, 26535 |
| chr2 | 1,59E+08 | 1,59E+08 | 1674 | * | 14,10916 | -3,023243048 | 1,030022 | -2,93512 | 0,003334 | 0,744477896 | 1,075735 | Distal Intergenic | 2 | 1,6E+08 | 1,6E+08 | 33948 | 1 | 71081 | ENSMUST00000133091.2 | -367749 | ENSMUSG00000085198 | 4933416E03Rik | RIKEN cDNA 4933416E03 gene |
| chr18 | 55931618 | 55933122 | 1505 | * | 10,69853 | -3,021011954 | 1,148556 | -2,63027 | 0,008532 | 0,931442063 | 0,918255 | Distal Intergenic | 18 | 56326071 | 56326205 | 135 | 1 | 1,15E+08 | ENSMUST00000157815.3 | -392949 | ENSMUSG00000088440 | Gm23555 | predicted gene, 23555 |
| chr13 | 34587857 | 34590613 | 2757 | * | 22,37893 | -3,01759375 | 0,869311 | -3,47125 | 0,000518 | 0,281843592 | 2,67243 | Distal Intergenic | 13 | 34363141 | 34529165 | 166025 | 2 | 73102 | ENSMUST00000040336.12 | -58692 | ENSMUSG00000038267 | Slc22a23 | solute carrier family 22, member 23 |
| chr15 | 67687077 | 67688124 | 1048 | * | 11,75249 | -3,016908549 | 1,103393 | -2,73421 | 0,006253 | 0,915421656 | 0,744375 | Distal Intergenic | 15 | 67956170 | 68130636 | 174467 | 2 | 380993 | ENSMUST00000162173.8 | 442512 | ENSMUSG00000022335 | Zfat | zinc finger and AT hook domain containing |
| chr15 | 61463266 | 61464165 | 900 | * | 12,97401 | -3,015251903 | 1,049022 | -2,87435 | 0,004049 | 0,762980415 | 1,173325 | Intron (ENSMUST0000( | 15 | 61406525 | 61617033 | 210509 | 2 | 432956 | ENSMUST00000229538.2 | 152868 | ENSMUSG00000116031 | D030024E09Rik | RIKEN cDNA D030024E09 gene |
| chr7 | 10381299 | 10382435 | 1137 | * | 9,35864 | -3,013902906 | 1,139328 | -2,64533 | 0,008161 | 0,88303073 | 1,211315 | Distal Intergenic | 7 | 10367514 | 10368410 | 897 | 1 | 171262 | ENSMUST00000055847.4 | 13785 | ENSMUSG00000045340 | Vmn1r70 | vomeronasal 1 receptor 70 |
| chr1 | 1,88E+08 | 1,88E+08 | 1215 | * | 13,06898 | -3,013694839 | 1,085317 | -2,77679 | 0,00549 | 0,89873691 | 0,738441 | Promoter (2-3kb) | 1 | 1,88E+08 | 1,88E+08 | 217059 | 1 | 26381 | ENSMUST00000027906.13 | -2049 | ENSMUSG00000026610 | Esrrg | estrogen-related receptor gamma |
| chr12 | 40002936 | 40004769 | 1834 | * | 18,38429 | -3,011823012 | 0,91905 | -3,2771 | 0,001049 | 0,421090004 | 2,154504 | Promoter (2-3kb) | 12 | 39998628 | 40000631 | 2004 | 2 | 1,05E+08 | ENSMUST00000222820.2 | -2305 | | Gm40399 | predicted gene, 40399 |
| chr14 | 69009448 | 69010990 | 1543 | * | 10,10609 | -3,011165769 | 1,138767 | -2,64423 | 0,008188 | 0,888080529 | 1,173325 | Distal Intergenic | 14 | 68845183 | 68893291 | 48109 | 2 | 13522 | ENSMUST00000224039.2 | -116157 | ENSMUSG00000014725 | Adam28 | a disintegrin and metallopeptidase domain 28 |
| chr18 | 57789359 | 57790829 | 1471 | * | 14,90165 | -3,010199653 | 0,928275 | -3,24279 | 0,001184 | 0,531203841 | 1,2763 | Intron (ENSMUST0000( | 18 | 57802525 | 57802630 | 106 | 2 | 1,15E+08 | ENSMUST00000158151.3 | 11801 | ENSMUSG00000088776 | Gm26038 | predicted gene, 26038 |
| chr11 | 88993693 | 88995560 | 1868 | * | 17,23718 | -3,00971259 | 0,873444 | -3,4458 | 0,000569 | 0,304413931 | 2,423594 | Distal Intergenic | 11 | 88965835 | 88983791 | 17957 | 1 | 217071 | ENSMUST00000127186.2 | 27858 | ENSMUSG00000072553 | Gm525 | predicted gene 525 |
| chrX | 1,02E+08 | 1,02E+08 | 2109 | * | 11,95884 | -3,009499274 | 0,992982 | -3,03077 | 0,002439 | 0,717133806 | 0,918255 | Intron (ENSMUST0000( | 20 | 1,02E+08 | 1,02E+08 | 26587 | 2 | 75106 | ENSMUST00000183292.8 | -9382 | ENSMUSG00000031325 | 4930519F16Rik | RIKEN cDNA 4930519F16 gene |
| chr4 | 35516423 | 35517880 | 1458 | * | 13,68386 | -3,008708476 | 1,104853 | -2,72317 | 0,006466 | 0,922186553 | 0,738441 | Distal Intergenic | 4 | 35212423 | 35226175 | 13753 | 2 | 73205 | ENSMUST00000156472.2 | -290248 | ENSMUSG00000028300 | C9orf72 | C9orf72, member of C9orf72-SMCR8 complex |
| chr7 | 1,29E+08 | 1,29E+08 | 1823 | * | 15,80799 | -3,008012191 | 1,072611 | -2,80438 | 0,005041 | 0,681633938 | 2,386008 | Distal Intergenic | 7 | 1,28E+08 | 1,29E+08 | 132820 | 1 | 1,03E+08 | ENSMUST00000205528.2 | 210618 | | Gm32884 | predicted gene, 32884 |
| chr1 | 98160833 | 98161979 | 1147 | * | 10,62028 | -3,007431899 | 1,089611 | -2,7601 | 0,005778 | 0,812199284 | 1,2763 | Distal Intergenic | 1 | 98101373 | 98118423 | 17051 | 1 | 1,03E+08 | ENSMUST00000189232.2 | 59460 | ENSMUSG00000100711 | Gm29461 | predicted gene 29461 |
| chr13 | 19317257 | 19318539 | 1283 | * | 10,80664 | -3,005951255 | 1,083992 | -2,77304 | 0,005554 | 0,899471409 | 0,744375 | Intron (ENSMUST0000( | 13 | 19303959 | 19326182 | 22224 | 1 | 218038 | ENSMUST00000197545.2 | 13298 | ENSMUSG00000021314 | Amph | amphiphysin |
| chr7 | 1,35E+08 | 1,35E+08 | 1802 | * | 18,80882 | -3,005908468 | 1,055466 | -2,84794 | 0,0044 | 0,641470306 | 2,663619 | Distal Intergenic | 7 | 1,35E+08 | 1,35E+08 | 3315 | 1 | 270004 | ENSMUST00000060356.8 | -46965 | ENSMUSG00000048377 | Foxi2 | forkhead box I2 |
| chr10 | 64050557 | 64051597 | 1041 | * | 9,446094 | -3,005277594 | 1,12591 | -2,6692 | 0,007603 | 0,868022856 | 1,211315 | Intron (ENSMUST0000( | 10 | 63765669 | 63926056 | 160388 | 2 | 216028 | ENSMUST00000131945.2 | -124501 | ENSMUSG00000042846 | Lrrtm3 | leucine rich repeat transmembrane neuronal 3 |
| chr3 | 41930662 | 41931669 | 1008 | * | 12,21865 | -3,005140677 | 1,064706 | -2,82251 | 0,004765 | 0,836575905 | 0,918255 | Exon (ENSMUST00001 | 3 | 41705631 | 41710927 | 5297 | 1 | 73852 | ENSMUST00000195030.2 | 225031 | ENSMUSG00000025766 | D3Ertd751e | DNA segment, Chr 3, ERATO Doi 751, expressed |
| chr16 | 11964364 | 11966152 | 1789 | * | 15,73718 | -2,999772135 | 1,136835 | -2,63871 | 0,008322 | 0,877294452 | 1,2763 | Intron (ENSMUST0000( | 16 | 11802584 | 11813260 | 10677 | 1 | 72555 | ENSMUST00000229804.2 | 161780 | ENSMUSG00000022494 | Shisa9 | shisa family member 9 |
| chr1 | 61857624 | 61859253 | 1630 | * | 13,09851 | -2,999504764 | 1,007539 | -2,97706 | 0,00291 | 0,789027175 | 0,738441 | Intron (ENSMUST0000( | 1 | 61806574 | 61890621 | 84048 | 2 | 74661 | ENSMUST00000138535.8 | 31368 | | Pard3bos1 | par-3 family cell polarity regulator beta, opposite strand 1 |
| chr11 | 39001964 | 39002910 | 947 | * | 10,00014 | -2,998275244 | 1,153492 | -2,5993 | 0,009341 | 0,874247416 | 1,449706 | Distal Intergenic | 11 | 38688998 | 38689104 | 107 | 2 | 1,15E+08 | ENSMUST00000240209.1 | -312860 | ENSMUSG00000119681 | Gm23520 | predicted gene, 23520 |
| chr8 | 82859777 | 82860914 | 1138 | * | 12,76931 | -2,997838987 | 1,101475 | -2,72166 | 0,006495 | 0,860530858 | 1,075735 | Distal Intergenic | 8 | 82879349 | 82879411 | 63 | 2 | 1,15E+08 | ENSMUST00000158522.3 | 18497 | ENSMUSG00000089147 | Gm25267 | predicted gene, 25267 |
| chr8 | 8894435 | 8895485 | 1051 | * | 13,71726 | -2,996596908 | 1,12816 | -2,65618 | 0,007903 | 0,864595549 | 1,2763 | Intron (ENSMUST0000( | 8 | 8785970 | 8805337 | 19368 | 1 | 1,05E+08 | ENSMUST00000209128.2 | 108465 | | Gm39127 | predicted gene, 39127 |
| chr4 | 15653829 | 15654893 | 1065 | * | 10,99384 | -2,995405656 | 1,137622 | -2,63304 | 0,008462 | 0,880128915 | 1,2763 | Distal Intergenic | 4 | 15881264 | 15908064 | 26801 | 1 | 12307 | ENSMUST00000029876.2 | -226371 | ENSMUSG00000028222 | Calb1 | calbindin 1 |
| chr13 | 71265026 | 71266380 | 1355 | * | 15,68977 | -2,993971936 | 1,059954 | -2,82462 | 0,004734 | 0,684274806 | 2,141612 | Distal Intergenic | 13 | 71255208 | 71255328 | 121 | 1 | 1E+08 | ENSMUST00000116930.3 | 9818 | ENSMUSG00000080580 | Mir466f-4 | microRNA 466f-4 |
| chr13 | 79988021 | 79989603 | 1583 | * | 10,3437 | -2,993614926 | 1,148027 | -2,60762 | 0,009118 | 0,915421656 | 1,086118 | Distal Intergenic | 13 | 80306782 | 80306909 | 128 | 1 | 1,15E+08 | ENSMUST00000122712.4 | -317179 | ENSMUSG00000084661 | Gm24635 | predicted gene, 24635 |
| chr9 | 55106411 | 55107770 | 1360 | * | 10,45399 | -2,991147713 | 1,135785 | -2,63355 | 0,00845 | 0,96198404 | 0,744375 | Intron (ENSMUST0000( | 9 | 55102129 | 55113494 | 11366 | 1 | 109161 | ENSMUST00000148468.2 | 4282 | ENSMUSG00000032307 | Ube2q2 | ubiquitin-conjugating enzyme E2Q family member 2 |
| chr9 | 69781697 | 69784006 | 2310 | * | 20,05312 | -2,990956321 | 0,902015 | -3,31586 | 0,000914 | 0,369592988 | 2,507013 | Distal Intergenic | 9 | 69674895 | 69676274 | 1380 | 1 | 1,03E+08 | ENSMUST00000243201.1 | 106802 | | Gm31530 | predicted gene, 31530 |
| chr5 | 1,3E+08 | 1,3E+08 | 1631 | * | 14,60321 | -2,988932646 | 1,011652 | -2,95451 | 0,003132 | 0,705098751 | 1,2763 | Distal Intergenic | 5 | 1,3E+08 | 1,3E+08 | 70164 | 1 | 231769 | ENSMUST00000053737.9 | -42260 | ENSMUSG00000029439 | Sfswap | splicing factor SWAP |

| | | | | | | | | | | | | | | | | | | | | | | | |
|---|---|---|---|---|---|---|---|---|---|---|---|---|---|---|---|---|---|---|---|---|---|---|---|
| chr13 | 54819918 | 54821245 | 1328 | * | 12,6964 | -2,988365042 | 1,036504 | -2,88312 | 0,003938 | 0,801688106 | 0,918255 | Distal Intergenic | 13 | 54830460 | 54831885 | 1426 | 2 | 105239 | ENSMUST00000145215.2 | 10640 | ENSMUSG00000034928 | Rnf44 | ring finger protein 44 |
| chr1 | 70795148 | 70796268 | 1121 | * | 10,26306 | -2,988168836 | 1,121444 | -2,66457 | 0,007709 | 0,877892722 | 1,173325 | Intron (ENSMUST0000( | 1 | 70765082 | 70921648 | 156567 | 1 | 320460 | ENSMUST00000162182.2 | 30066 | ENSMUSG00000045648 | Vwc2l | von Willebrand factor C domain-containing protein 2-like |
| chr2 | 78966423 | 78967504 | 1082 | * | 12,08923 | -2,986296996 | 1,096602 | -2,72323 | 0,006465 | 0,83238039 | 1,2763 | Distal Intergenic | 2 | 78953071 | 78953706 | 636 | 1 | 1,03E+08 | ENSMUST00000246949.1 | 13352 | | Gm35748 | predicted gene, 35748 |
| chrX | 1,28E+08 | 1,28E+08 | 1432 | * | 10,55681 | -2,985662457 | 1,05728 | -2,82391 | 0,004744 | 0,872427545 | 0,744375 | Distal Intergenic | 20 | 1,28E+08 | 1,28E+08 | 132 | 2 | 1,15E+08 | ENSMUST00000158659.3 | -119087 | ENSMUSG00000089284 | Gm26029 | predicted gene, 26029 |
| chr11 | 22023436 | 22025259 | 1824 | * | 14,94921 | -2,985401865 | 1,052589 | -2,83625 | 0,004565 | 0,769939394 | 1,2763 | Intron (ENSMUST0000( | 11 | 21946858 | 21952897 | 6040 | 2 | 18423 | ENSMUST00000147486.3 | -70539 | ENSMUSG00000005917 | Otx1 | orthodenticle homeobox 1 |
| chr2 | 1,65E+08 | 1,65E+08 | 1615 | * | 14,96973 | -2,98456108 | 1,030923 | -2,89504 | 0,003791 | 0,73648081 | 1,2763 | Intron (ENSMUST0000( | 2 | 1,65E+08 | 1,65E+08 | 17033 | 1 | 67538 | ENSMUST00000052107.5 | 7452 | ENSMUSG00000045822 | Zswim3 | zinc finger SWIM-type containing 3 |
| chr2 | 80186064 | 80187220 | 1157 | * | 11,83442 | -2,983300949 | 1,11967 | -2,66445 | 0,007712 | 0,948006995 | 0,744375 | Distal Intergenic | 2 | 80177998 | 80181286 | 3289 | 1 | 66861 | ENSMUST00000154912.2 | 8066 | ENSMUSG00000027006 | Dnajc10 | DnaJ heat shock protein family (Hsp40) member C10 |
| chr3 | 99132506 | 99133755 | 1250 | * | 15,61207 | -2,980111249 | 1,055326 | -2,82388 | 0,004745 | 0,77746544 | 1,2763 | Intron (ENSMUST0000( | 3 | 99147697 | 99220381 | 72685 | 1 | 21384 | ENSMUST00000151606.8 | -13942 | ENSMUSG00000027868 | Tbx15 | T-box 15 |
| chr14 | 8392151 | 8393928 | 1778 | * | 11,61623 | -2,978817938 | 1,124302 | -2,64948 | 0,008062 | 0,870290757 | 1,2763 | Intron (ENSMUST0000( | 14 | 8379295 | 8382294 | 3000 | 2 | 246103 | ENSMUST00000224616.2 | -9857 | ENSMUSG00000021738 | Atxn7 | ataxin 7 |
| chr7 | 1,15E+08 | 1,15E+08 | 1189 | * | 13,3922 | -2,978688751 | 1,106053 | -2,69308 | 0,00708 | 0,90060438 | 0,938682 | Intron (ENSMUST0000( | 7 | 1,15E+08 | 1,15E+08 | 81768 | 2 | 20679 | ENSMUST00000205479.2 | 16395 | ENSMUSG00000051910 | Sox6 | SRY (sex determining region Y)-box 6 |
| chr15 | 69030183 | 69031086 | 904 | * | 10,68252 | -2,977564694 | 1,142202 | -2,60686 | 0,009138 | 0,974006322 | 0,738441 | Distal Intergenic | 15 | 69105780 | 69106848 | 1069 | 2 | 1,01E+08 | ENSMUST00000229754.2 | 75762 | ENSMUSG00000115935 | 4930573C08Rik | RIKEN cDNA 4930573C08 gene |
| chr1 | 61863927 | 61865062 | 1136 | * | 11,36136 | -2,976891519 | 1,154826 | -2,57778 | 0,009944 | 0,986468657 | 0,738441 | Intron (ENSMUST0000( | 1 | 61806574 | 61890621 | 84048 | 2 | 74661 | ENSMUST00000138535.8 | 25559 | | Pard3bos1 | par-3 family cell polarity regulator beta, opposite strand 1 |
| chr9 | 91236342 | 91237541 | 1200 | * | 12,09809 | -2,975287073 | 1,088811 | -2,7326 | 0,006284 | 0,82646024 | 1,2763 | Distal Intergenic | 9 | 91244466 | 91260968 | 16503 | 1 | 22774 | ENSMUST00000173342.8 | -6925 | ENSMUSG00000036972 | Zic4 | zinc finger protein of the cerebellum 4 |
| chr10 | 95270016 | 95271335 | 1320 | * | 14,52291 | -2,970581053 | 1,02142 | -2,90829 | 0,003634 | 0,713633527 | 1,41696 | Intron (ENSMUST0000( | 10 | 95253242 | 95279547 | 26306 | 1 | 70523 | ENSMUST00000244592.1 | 16774 | ENSMUSG00000097766 | 5730420D15Rik | RIKEN cDNA 5730420D15 gene |
| chr18 | 44043049 | 44044360 | 1312 | * | 11,11584 | -2,964777495 | 1,066839 | -2,77903 | 0,005452 | 0,801134483 | 1,2763 | Intron (ENSMUST0000( | 18 | 44096302 | 44155568 | 59267 | 1 | 72432 | ENSMUST00000069245.8 | -51942 | ENSMUSG00000055561 | Spink5 | serine peptidase inhibitor, Kazal type 5 |
| chr12 | 18093540 | 18094411 | 872 | * | 10,29453 | -2,964663019 | 1,150198 | -2,57752 | 0,009951 | 0,886332554 | 1,449706 | Intron (ENSMUST0000( | 12 | 18111968 | 18114868 | 2901 | 1 | 245297 | ENSMUST00000223186.2 | -17557 | | Gm4983 | predicted gene 4983 |
| chr12 | 1,07E+08 | 1,07E+08 | 1642 | * | 15,43771 | -2,962843608 | 1,148639 | -2,57944 | 0,009896 | 0,888080529 | 1,41696 | Distal Intergenic | 12 | 1,07E+08 | 1,07E+08 | 13044 | 2 | 1,19E+08 | ENSMUST00000148122.2 | -256612 | | Gm16084 | predicted gene 16084 |
| chr2 | 95311457 | 95312406 | 950 | * | 9,725698 | -2,962442676 | 1,11283 | -2,66208 | 0,007766 | 0,841453364 | 1,449706 | Distal Intergenic | 2 | 95224587 | 95227996 | 3410 | 1 | 1,03E+08 | ENSMUST00000148637.2 | 86870 | | Gm13794 | predicted gene 13794 |
| chr9 | 8676384 | 8677586 | 1203 | * | 12,10866 | -2,962392027 | 1,070551 | -2,76717 | 0,005655 | 0,808330792 | 1,2763 | Intron (ENSMUST0000( | 9 | 8715323 | 8740943 | 25621 | 2 | 1,03E+08 | ENSMUST00000214779.2 | 63357 | ENSMUSG00000111602 | Gm32014 | predicted gene, 32014 |
| chr13 | 1,05E+08 | 1,05E+08 | 1894 | * | 12,10996 | -2,96186368 | 1,102577 | -2,68631 | 0,007225 | 0,863571204 | 1,173325 | 3' UTR | 13 | 1,04E+08 | 1,05E+08 | 68399 | 1 | 108154 | ENSMUST00000224504.2 | 67703 | ENSMUSG00000046169 | Adamts6 | a disintegrin-like and metallopeptidase (reprolysin type) with thrombospondin type 1 motif, 6 |
| chr1 | 8550761 | 8552044 | 1284 | * | 11,29889 | -2,961345981 | 1,140723 | -2,59602 | 0,009431 | 0,947773602 | 0,918255 | Intron (ENSMUST0000( | 1 | 8433407 | 8540744 | 107338 | 2 | 71096 | ENSMUST00000115484.9 | -10017 | ENSMUSG00000025909 | Sntg1 | syntrophin, gamma 1 |
| chr9 | 25697400 | 25698869 | 1470 | * | 12,82828 | -2,958772895 | 1,0507 | -2,816 | 0,004863 | 0,81200135 | 1,075735 | Distal Intergenic | 9 | 25480974 | 25481101 | 128 | 2 | 1,15E+08 | ENSMUST00000104509.3 | -216299 | ENSMUSG00000077697 | Gm25346 | predicted gene, 25346 |
| chr14 | 14007310 | 14009376 | 2067 | * | 24,16179 | -2,955628205 | 0,814537 | -3,6286 | 0,000285 | 0,199811501 | 2,858511 | Intron (ENSMUST0000( | 14 | 14091019 | 14120836 | 29818 | 1 | 18300 | ENSMUST00000022269.7 | -81643 | ENSMUSG00000021749 | Fam3d | FAM3 metabolism regulating signaling molecule D |
| chr7 | 1,08E+08 | 1,08E+08 | 971 | * | 12,74269 | -2,952917514 | 1,133455 | -2,60524 | 0,009181 | 0,974006322 | 0,738441 | Distal Intergenic | 7 | 1,08E+08 | 1,08E+08 | 1440 | 1 | 258361 | ENSMUST00000210990.2 | -5617 | ENSMUSG00000110253 | Olfr495 | olfactory receptor 495 |
| chr13 | 19723364 | 19725119 | 1756 | * | 14,79788 | -2,950196174 | 1,042271 | -2,83055 | 0,004647 | 0,683966322 | 2,141612 | Distal Intergenic | 13 | 19647017 | 19688537 | 41521 | 2 | 319466 | ENSMUST00000221607.2 | -34827 | ENSMUSG00000114189 | 9330199G10Rik | RIKEN cDNA 9330199G10 gene |
| chrX | 1,36E+08 | 1,36E+08 | 1591 | * | 14,25105 | -2,950138039 | 0,991473 | -2,97551 | 0,002925 | 0,722439117 | 1,075735 | Distal Intergenic | 20 | 1,36E+08 | 1,38E+08 | 1276339 | 1 | 60367 | ENSMUST00000113063.8 | -152108 | ENSMUSG00000059203 | Il1rapl2 | interleukin 1 receptor accessory protein-like 2 |
| chr19 | 36243456 | 36245300 | 1845 | * | 20,6142 | -2,948842077 | 0,798236 | -3,6942 | 0,000221 | 0,177369126 | 2,67243 | Promoter (1-2kb) | 19 | 36216285 | 36241837 | 25553 | 2 | 1,15E+08 | ENSMUST00000236548.3 | -1619 | | LOC115489009 | uncharacterized LOC115489009 |
| chr14 | 21404071 | 21405904 | 1834 | * | 13,04735 | -2,947786089 | 1,065946 | -2,76542 | 0,005685 | 0,809369408 | 1,2763 | Intron (ENSMUST0000( | 14 | 21419138 | 21420402 | 1265 | 1 | 11534 | ENSMUST00000224974.2 | -13234 | ENSMUSG00000039197 | Adk | adenosine kinase |
| chr7 | 87328324 | 87330240 | 1917 | * | 15,89324 | -2,947586946 | 0,990911 | -2,97462 | 0,002933 | 0,563032042 | 2,663619 | Intron (ENSMUST0000( | 7 | 87252297 | 87458828 | 206532 | 1 | 108071 | ENSMUST00000138732.3 | 76027 | ENSMUSG00000049583 | Grm5 | glutamate receptor, metabotropic 5 |
| chr11 | 66737022 | 66739067 | 2046 | * | 15,22099 | -2,946628051 | 0,971206 | -3,03399 | 0,002413 | 0,64586006 | 1,41696 | Distal Intergenic | 11 | 66802807 | 66820702 | 17896 | 1 | 193003 | ENSMUST00000123434.3 | -63740 | ENSMUSG00000048070 | Pirt | phosphoinositide-interacting regulator of transient receptor potential channels |
| chr6 | 85711679 | 85713480 | 1802 | * | 14,11546 | -2,942892113 | 1,055894 | -2,78711 | 0,005318 | 0,891636336 | 0,738441 | Intron (ENSMUST0000( | 6 | 85683709 | 85690187 | 6479 | 2 | 1E+08 | ENSMUST00000160534.2 | -21492 | ENSMUSG00000089694 | Nat8f7 | N-acetyltransferase 8 (GCN5-related) family member 7 |
| chr3 | 1,29E+08 | 1,29E+08 | 1989 | * | 22,55952 | -2,941894702 | 0,811452 | -3,62547 | 0,000288 | 0,190625259 | 3,104559 | Distal Intergenic | 3 | 1,29E+08 | 1,29E+08 | 1043 | 1 | 435755 | ENSMUST00000199748.2 | 10412 | ENSMUSG00000105712 | Pou5f1-rs4 | POU domain, class 5, transcription factor 1, related sequence 4 |
| chr12 | 1,1E+08 | 1,1E+08 | 991 | * | 11,87451 | -2,941578208 | 1,055192 | -2,78772 | 0,005308 | 0,854052567 | 0,918255 | Distal Intergenic | 12 | 1,1E+08 | 1,1E+08 | 1495 | 1 | 1E+08 | ENSMUST00000182300.2 | 22682 | ENSMUSG00000097391 | Mirg | miRNA containing gene |
| chr16 | 35023771 | 35025094 | 1324 | * | 12,21093 | -2,940893793 | 1,137999 | -2,58427 | 0,009759 | 0,984310536 | 0,738441 | Intron (ENSMUST0000( | 16 | 34975247 | 35126108 | 150862 | 1 | 224129 | ENSMUST00000114913.3 | 48524 | ENSMUSG00000022840 | Adcy5 | adenylate cyclase 5 |
| chr18 | 37334364 | 37335991 | 1628 | * | 12,85827 | -2,933863558 | 1,04705 | -2,80203 | 0,005078 | 0,820023113 | 1,075735 | Intron (ENSMUST0000( | 18 | 37276556 | 37320710 | 44155 | 1 | 353237 | ENSMUST00000047479.3 | 57808 | ENSMUSG00000102697 | Pcdhac2 | protocadherin alpha subfamily C, 2 |
| chr13 | 95565729 | 95568267 | 2539 | * | 18,07553 | -2,93276113 | 0,935868 | -3,13373 | 0,001726 | 0,477622452 | 2,507013 | 3' UTR | 13 | 95567885 | 95581339 | 13455 | 2 | 12919 | ENSMUST00000045583.9 | 13072 | ENSMUSG00000021680 | Crhbp | corticotropin releasing hormone binding protein |
| chr1 | 1,45E+08 | 1,45E+08 | 2230 | * | 13,58627 | -2,93192995 | 0,960798 | -3,05156 | 0,002277 | 0,744054256 | 0,738441 | Distal Intergenic | 1 | 1,45E+08 | 1,45E+08 | 127 | 2 | 1,15E+08 | ENSMUST00000157954.3 | -132222 | ENSMUSG00000088579 | Gm25460 | predicted gene, 25460 |
| chr1 | 42313422 | 42315201 | 1780 | * | 14,7887 | -2,931074453 | 1,017962 | -2,87936 | 0,003985 | 0,747027154 | 1,2763 | Intron (ENSMUST0000( | 1 | 42301088 | 42343568 | 42481 | 1 | 1,03E+08 | ENSMUST00000244990.1 | 12334 | | Gm28175 | predicted gene 28175 |
| chr4 | 8782823 | 8784192 | 1370 | * | 14,68886 | -2,930301426 | 1,073442 | -2,72982 | 0,006337 | 0,83712454 | 1,214904 | Intron (ENSMUST0000( | 4 | 8751438 | 8867224 | 115787 | 1 | 320790 | ENSMUST00000170391.2 | 31385 | ENSMUSG00000041235 | Chd7 | chromodomain helicase DNA binding protein 7 |
| chr2 | 1,48E+08 | 1,48E+08 | 1370 | * | 12,00517 | -2,930234955 | 1,103662 | -2,65501 | 0,007931 | 0,953233027 | 0,744375 | Distal Intergenic | 2 | 1,48E+08 | 1,48E+08 | 313 | 1 | 1,15E+08 | ENSMUST00000158833.4 | -14714 | ENSMUSG00000089458 | Gm24221 | predicted gene, 24221 |
| chr12 | 95755962 | 95757498 | 1537 | * | 12,95674 | -2,928975951 | 1,034107 | -2,83237 | 0,00462 | 0,801688106 | 1,075735 | Distal Intergenic | 12 | 95662131 | 95747881 | 85751 | 1 | 399558 | ENSMUST00000110117.2 | 93831 | ENSMUSG00000047414 | Flrt2 | fibronectin leucine rich transmembrane protein 2 |
| chr3 | 52214786 | 52216101 | 1316 | * | 15,00286 | -2,928005997 | 1,118836 | -2,61701 | 0,00887 | 0,874247416 | 1,37683 | Intron (ENSMUST0000( | 3 | 52175757 | 52260642 | 84886 | 1 | 56458 | ENSMUST00000053764.7 | 39029 | ENSMUSG00000044167 | Foxo1 | forkhead box O1 |
| chr5 | 1,24E+08 | 1,24E+08 | 1536 | * | 12,44282 | -2,925700893 | 1,074408 | -2,72308 | 0,006468 | 0,889472689 | 0,918255 | Promoter (<=1kb) | 5 | 1,24E+08 | 1,24E+08 | 8139 | 1 | 208908 | ENSMUST00000169898.3 | 377 | ENSMUSG00000061882 | Ccdc62 | coiled-coil domain containing 62 |
| chr13 | 91631504 | 91632961 | 1458 | * | 12,42481 | -2,925478532 | 1,062306 | -2,75389 | 0,005889 | 0,907729723 | 0,738441 | Intron (ENSMUST0000( | 13 | 91609300 | 91672985 | 63686 | 1 | 66970 | ENSMUST00000141381.2 | 22204 | ENSMUSG00000003992 | Ssbp2 | single-stranded DNA binding protein 2 |
| chrX | 1,26E+08 | 1,26E+08 | 2598 | * | 12,71704 | -2,92486292 | 1,069173 | -2,73563 | 0,006226 | 0,854157826 | 1,075735 | Distal Intergenic | 20 | 1,26E+08 | 1,26E+08 | 105 | 1 | 1,15E+08 | ENSMUST00000158747.3 | 4347 | ENSMUSG00000089372 | Gm22139 | predicted gene, 22139 |
| chr14 | 1,23E+08 | 1,23E+08 | 2037 | * | 11,91946 | -2,924276259 | 1,083971 | -2,69774 | 0,006981 | 0,901314414 | 0,918255 | Intron (ENSMUST0000( | 14 | 1,23E+08 | 1,23E+08 | 6306 | 1 | 380934 | ENSMUST00000226639.2 | -20770 | ENSMUSG00000115016 | AA536875 | expressed sequence AA536875 |
| chr12 | 77638272 | 77640111 | 1840 | * | 16,20642 | -2,923934608 | 0,886764 | -3,29731 | 0,000976 | 0,370253435 | 2,663619 | Distal Intergenic | 12 | 77600121 | 77616483 | 16363 | 1 | 1,03E+08 | ENSMUST00000218335.2 | 38151 | | Gm35240 | predicted gene, 35240 |
| chr4 | 56885012 | 56886594 | 1583 | * | 13,30738 | -2,922236636 | 1,123234 | -2,60163 | 0,009278 | 0,89325945 | 1,2763 | Exon (ENSMUST00000( | 4 | 56895229 | 56895298 | 70 | 2 | 723837 | ENSMUST00000083610.2 | 8704 | ENSMUSG00000065544 | Mir32 | microRNA 32 |
| chr4 | 1,4E+08 | 1,4E+08 | 1357 | * | 13,15205 | -2,922090656 | 1,062629 | -2,74987 | 0,005962 | 0,849352323 | 1,075735 | Promoter (2-3kb) | 4 | 1,4E+08 | 1,4E+08 | 9822 | 1 | 1,03E+08 | ENSMUST00000128563.2 | -2862 | | Gm1667 | predicted gene 1667 |
| chr1 | 39492101 | 39493573 | 1473 | * | 14,04193 | -2,919304084 | 1,087647 | -2,68405 | 0,007274 | 0,906138253 | 0,918255 | Intron (ENSMUST0000( | 1 | 39411377 | 39517742 | 106366 | 2 | 54610 | ENSMUST00000192531.6 | 24169 | ENSMUSG00000003134 | Tbc1d8 | TBC1 domain family, member 8 |
| chr4 | 1,25E+08 | 1,25E+08 | 2275 | * | 22,62489 | -2,917300822 | 0,907964 | -3,21301 | 0,001314 | 0,400053238 | 3,104559 | Intron (ENSMUST0000( | 4 | 1,25E+08 | 1,25E+08 | 22670 | 1 | 192199 | ENSMUST00000030687.8 | -13922 | ENSMUSG00000028871 | Rspo1 | R-spondin 1 |
| chr19 | 23639430 | 23640740 | 1311 | * | 13,21784 | -2,917194103 | 1,082185 | -2,69565 | 0,007025 | 0,87765772 | 1,075735 | Distal Intergenic | 19 | 23536117 | 23630176 | 94060 | 2 | 67483 | ENSMUST00000035849.5 | -9254 | ENSMUSG00000033053 | Cfap95 | cilia and flagella associated protein 95 |
| chr7 | 1,17E+08 | 1,17E+08 | 2266 | * | 16,14188 | -2,916981998 | 0,97589 | -2,98905 | 0,002798 | 0,684274806 | 1,2763 | Distal Intergenic | 7 | 1,17E+08 | 1,17E+08 | 73868 | 1 | 233781 | ENSMUST00000161889.2 | 308394 | ENSMUSG00000030657 | Xylt1 | xylosyltransferase 1 |
| chr6 | 54494365 | 54495385 | 1021 | * | 11,61857 | -2,916090512 | 1,10765 | -2,63268 | 0,008471 | 0,929295662 | 0,918255 | Promoter (2-3kb) | 6 | 54483250 | 54497838 | 14589 | 2 | 69938 | ENSMUST00000203800.2 | 2453 | ENSMUSG00000019124 | Scrn1 | secernin 1 |
| chr12 | 1,12E+08 | 1,12E+08 | 1970 | * | 18,4171 | -2,915366658 | 1,017264 | -2,86589 | 0,004158 | 0,64236196 | 2,507013 | Intron (ENSMUST0000( | 12 | 1,12E+08 | 1,12E+08 | 35540 | 1 | 668303 | ENSMUST00000128402.3 | 6985 | ENSMUSG00000021294 | Kif26a | kinesin family member 26A |
| chr13 | 19214309 | 19215959 | 1651 | * | 13,86413 | -2,910036702 | 1,093939 | -2,66014 | 0,007811 | 0,951821 | 0,738441 | Exon (ENSMUST000002 | 13 | 19248987 | 19335082 | 86096 | 1 | 218038 | ENSMUST00000222698.2 | -33028 | ENSMUSG00000021314 | Amph | amphiphysin |
| chr17 | 3851621 | 3853474 | 1854 | * | 15,80139 | -2,90962958 | 0,925378 | -3,14426 | 0,001665 | 0,479521813 | 2,386008 | Distal Intergenic | 17 | 3685515 | 3746536 | 61022 | 2 | 224480 | ENSMUST00000115800.2 | -105085 | ENSMUSG00000023802 | Nox3 | NADPH oxidase 3 |
| chr1 | 1,01E+08 | 1,01E+08 | 1391 | * | 12,86716 | -2,907610689 | 1,079531 | -2,6934 | 0,007073 | 0,877892722 | 1,075735 | Distal Intergenic | 1 | 1E+08 | 1E+08 | 490 | 1 | 1,15E+08 | ENSMUST00000190817.2 | 389199 | | Gm29667 | predicted gene 29667 |
| chr19 | 54597596 | 54598791 | 1196 | * | 12,66033 | -2,9049643 | 1,121402 | -2,59048 | 0,009584 | 0,980429836 | 0,738441 | Distal Intergenic | 19 | 55058166 | 55087883 | 29718 | 2 | 14732 | ENSMUST00000061856.6 | 489092 | ENSMUSG00000024978 | Gpam | glycerol-3-phosphate acyltransferase, mitochondrial |
| chr13 | 85126811 | 85128242 | 1432 | * | 11,08299 | -2,904665337 | 1,085982 | -2,67469 | 0,00748 | 0,872427545 | 1,173325 | Distal Intergenic | 13 | 85337527 | 85361842 | 24316 | 1 | 66671 | ENSMUST00000022030.11 | -209285 | ENSMUSG00000021548 | Ccnh | cyclin H |
| chr1 | 1,23E+08 | 1,23E+08 | 1771 | * | 15,71503 | -2,903011371 | 0,958097 | -3,02998 | 0,002446 | 0,530563721 | 2,663619 | Distal Intergenic | 1 | 1,22E+08 | 1,22E+08 | 144 | 1 | 1,15E+08 | ENSMUST00000158565.3 | 194712 | ENSMUSG00000089190 | Gm25771 | predicted gene, 25771 |
| chr10 | 96584317 | 96586494 | 2178 | * | 19,00592 | -2,902630174 | 0,952947 | -3,04595 | 0,002319 | 0,549506271 | 2,221611 | Distal Intergenic | 10 | 96636901 | 96720570 | 83670 | 1 | 1,08E+08 | ENSMUST00000218446.2 | -50407 | | Gm46240 | predicted gene, 46240 |
| chr11 | 49686389 | 49687283 | 895 | * | 10,45013 | -2,902464559 | 1,088425 | -2,66666 | 0,007661 | 0,947971763 | 0,744375 | Promoter (1-2kb) | 11 | 49685014 | 49729440 | 44427 | 1 | 14584 | ENSMUST00000020629.5 | 1375 | ENSMUSG00000020363 | Gfpt2 | glutamine fructose-6-phosphate transaminase 2 |
| chr4 | 12127385 | 12129173 | 1789 | * | 16,00122 | -2,902289877 | 1,011826 | -2,86837 | 0,004126 | 0,627995376 | 2,663619 | Distal Intergenic | 4 | 12140264 | 12144688 | 4425 | 1 | 72397 | ENSMUST00000146416.2 | -11091 | ENSMUSG00000046667 | Rbm12b1 | RNA binding motif protein 12 B1 |
| chr6 | 36162450 | 36163294 | 845 | * | 12,93909 | -2,90227121 | 0,99863 | -2,90625 | 0,003658 | 0,827303855 | 0,738441 | Distal Intergenic | 6 | 36186555 | 36364786 | 178232 | 2 | 319472 | ENSMUST00000101535.9 | 201492 | ENSMUSG00000073154 | 9330158H04Rik | RIKEN cDNA 9330158H04 gene |
| chr17 | 25563795 | 25565540 | 1746 | * | 14,58971 | -2,901778333 | 1,103384 | -2,62989 | 0,008541 | 0,865002963 | 1,37683 | Promoter (<=1kb) | 17 | 25562219 | 25564501 | 2283 | 2 | 1,01E+08 | ENSMUST00000153118.2 | 0 | ENSMUSG00000024173 | Tpsab1 | tryptase alpha/beta 1 |
| chr9 | 30919826 | 30922231 | 2406 | * | 19,65939 | -2,901228114 | 0,88668 | -3,27201 | 0,001068 | 0,390164541 | 2,663619 | Promoter (<=1kb) | 9 | 30897263 | 30921359 | 24097 | 2 | 1,01E+08 | ENSMUST00000216803.2 | 0 | | Gm29724 | predicted gene, 29724 |
| chr10 | 1,14E+08 | 1,14E+08 | 1440 | * | 12,98081 | -2,901173109 | 1,097614 | -2,64316 | 0,008214 | 0,901529883 | 1,075735 | Distal Intergenic | 10 | 1,13E+08 | 1,13E+08 | 11133 | 2 | 1,05E+08 | ENSMUST00000218612.2 | -340524 | | Gm40765 | predicted gene, 40765 |
| chr9 | 28613916 | 28615710 | 1795 | * | 16,21105 | -2,900095991 | 0,991126 | -2,92606 | 0,003433 | 0,616178941 | 2,386008 | Intron (ENSMUST0000( | 9 | 28572925 | 28586536 | 13612 | 1 | 330908 | ENSMUST00000126845.2 | 40991 | ENSMUSG00000062257 | Opcml | opioid binding protein/cell adhesion molecule-like |
| chr16 | 84826060 | 84827070 | 1011 | * | 12,76788 | -2,897133262 | 1,046363 | -2,76877 | 0,005627 | 0,901463351 | 0,738441 | Promoter (1-2kb) | 16 | 84811193 | 84824513 | 13321 | 2 | 11820 | ENSMUST00000228509.2 | -1547 | ENSMUSG00000022892 | App | amyloid beta (A4) precursor protein |
| chr7 | 36897877 | 36899227 | 1351 | * | 14,66559 | -2,895854932 | 0,995654 | -2,9085 | 0,003632 | 0,644704154 | 2,141612 | Distal Intergenic | 7 | 36839966 | 36841136 | 1171 | 1 | 1,03E+08 | ENSMUST00000189179.2 | 57911 | | Gm28075 | predicted gene 28075 |
| chr10 | 74033992 | 74035384 | 1393 | * | 12,76483 | -2,894291834 | 1,029569 | -2,81117 | 0,004936 | 0,814686254 | 1,075735 | Intron (ENSMUST0000( | 10 | 74138983 | 74166882 | 27900 | 1 | 11994 | ENSMUST00000147455.2 | -103599 | ENSMUSG00000052613 | Pcdh15 | protocadherin 15 |
| chr13 | 40374218 | 40375820 | 1603 | * | 16,48647 | -2,893592057 | 0,925752 | -3,12567 | 0,001774 | 0,610248187 | 1,2763 | Intron (ENSMUST0000( | 13 | 40155358 | 40441487 | 286130 | 2 | 218165 | ENSMUST00000054635.8 | 65667 | ENSMUSG00000047094 | Ofcc1 | orofacial cleft 1 candidate 1 |
| chr10 | 65355343 | 65356996 | 1654 | * | 13,49224 | -2,892805005 | 1,016118 | -2,84692 | 0,004414 | 0,858625916 | 0,738441 | Distal Intergenic | 10 | 65370433 | 65371056 | 624 | 1 | 216036 | ENSMUST00000219802.2 | -13437 | ENSMUSG00000112341 | Gm4796 | predicted gene 4796 |
| chr13 | 63508100 | 63509570 | 1471 | * | 13,13147 | -2,892726148 | 0,998654 | -2,89663 | 0,003772 | 0,793028263 | 0,918255 | Exon (ENSMUST000002 | 13 | 63478353 | 63499986 | 21634 | 2 | 14088 | ENSMUST00000161656.2 | -8114 | ENSMUSG00000021461 | Fancc | Fanconi anemia, complementation group C |
| chr2 | 80074461 | 80075779 | 1319 | * | 16,31264 | -2,892238901 | 0,966142 | -2,9936 | 0,002757 | 0,577565003 | 2,318434 | Distal Intergenic | 2 | 80122816 | 80125700 | 2885 | 1 | 320769 | ENSMUST00000057072.6 | -47037 | ENSMUSG00000050114 | Prdx6b | peroxiredoxin 6B |
| chr13 | 78602406 | 78606050 | 3645 | * | 26,18384 | -2,888530098 | 0,74841 | -3,85956 | 0,000114 | 0,117479709 | 2,725491 | Distal Intergenic | 13 | 78631473 | 78710689 | 79217 | 1 | 1,03E+08 | ENSMUST00000224561.2 | -25423 | | Gm32067 | predicted gene, 32067 |
| chr2 | 1,04E+08 | 1,04E+08 | 1435 | * | 13,42821 | -2,886453957 | 0,989863 | -2,91601 | 0,003545 | 0,785709102 | 0,918255 | Intron (ENSMUST0000( | 2 | 1,04E+08 | 1,04E+08 | 63910 | 2 | 15259 | ENSMUST00000111124.8 | 20039 | ENSMUSG00000027177 | Hipk3 | homeodomain interacting protein kinase 3 |
| chr3 | 36248689 | 36250666 | 1978 | * | 14,27178 | -2,885810819 | 1,045406 | -2,76047 | 0,005772 | 0,81200135 | 1,2763 | Intron (ENSMUST0000( | 3 | 36241954 | 36276329 | 34376 | 2 | 229214 | ENSMUST00000197275.2 | 25663 | ENSMUSG00000058400 | Qrfpr | pyroglutamylated RFamide peptide receptor |
| chr3 | 1,21E+08 | 1,21E+08 | 1394 | * | 14,56862 | -2,884953716 | 1,050717 | -2,7457 | 0,006038 | 0,728793055 | 2,141612 | Intron (ENSMUST0000( | 3 | 1,2E+08 | 1,2E+08 | 128 | 2 | 1,15E+08 | ENSMUST00000158402.3 | -135879 | ENSMUSG00000089027 | Gm25013 | predicted gene, 25013 |
| chr11 | 28061323 | 28062481 | 1159 | * | 12,64181 | -2,884822633 | 1,08869 | -2,64981 | 0,008054 | 0,899620932 | 1,075735 | Distal Intergenic | 11 | 28338817 | 28384110 | 45294 | 2 | 216613 | ENSMUST00000160016.8 | 321629 | ENSMUSG00000032878 | Ccdc85a | coiled-coil domain containing 85A |
| chr2 | 81227358 | 81228359 | 1002 | * | 10,4187 | -2,884264404 | 1,069948 | -2,69571 | 0,007024 | 0,847111497 | 1,2763 | Distal Intergenic | 2 | 80814427 | 80814533 | 107 | 2 | 1,15E+08 | ENSMUST00000240007.1 | -412825 | ENSMUSG00000119241 | Gm23452 | predicted gene, 23452 |
| chr17 | 11548961 | 11550212 | 1252 | * | 11,58722 | -2,883678491 | 1,071425 | -2,69144 | 0,007114 | 0,935585339 | 0,744375 | Intron (ENSMUST0000( | 17 | 11522534 | 11829832 | 307299 | 1 | 50873 | ENSMUST00000186406.2 | 26427 | ENSMUSG00000023826 | Prkn | parkin RBR E3 ubiquitin protein ligase |
| chr1 | 1,64E+08 | 1,64E+08 | 2377 | * | 14,39967 | -2,882963373 | 1,045119 | -2,7585 | 0,005807 | 0,905781613 | 0,738441 | Distal Intergenic | 1 | 1,64E+08 | 1,64E+08 | 11574 | 1 | 1,03E+08 | ENSMUST00000181831.3 | -10763 | ENSMUSG00000097281 | Gm26685 | predicted gene, 26685 |
| chr19 | 57574815 | 57576330 | 1516 | * | 13,62093 | -2,882298884 | 0,991855 | -2,90597 | 0,003661 | 0,827707075 | 0,738441 | Distal Intergenic | 19 | 57599466 | 58121775 | 522310 | 1 | 226255 | ENSMUST00000077282.7 | -23136 | ENSMUSG00000054843 | Atrnl1 | attractin like 1 |
| chr5 | 1,13E+08 | 1,13E+08 | 1334 | * | 13,69734 | -2,881993598 | 1,038427 | -2,77534 | 0,005514 | 0,83481911 | 1,075735 | Intron (ENSMUST0000( | 5 | 1,13E+08 | 1,13E+08 | 98701 | 2 | 74376 | ENSMUST00000183029.8 | 9908 | ENSMUSG00000072720 | Myo18b | myosin XVIIIb |
| chr9 | 71054883 | 71056137 | 1255 | * | 11,96188 | -2,878702913 | 1,114084 | -2,58392 | 0,009768 | 0,982552203 | 0,744375 | Intron (ENSMUST0000( | 9 | 71017944 | 71069915 | 51972 | 2 | 64008 | ENSMUST00000060917.12 | 13778 | ENSMUSG00000032204 | Aqp9 | aquaporin 9 |
| chr8 | 1,11E+08 | 1,11E+08 | 1181 | * | 11,56854 | -2,878669896 | 1,117503 | -2,57598 | 0,009996 | 0,905015433 | 1,2763 | Distal Intergenic | 8 | 1,11E+08 | 1,11E+08 | 5550 | 1 | 234728 | ENSMUST00000189685.2 | -20693 | ENSMUSG00000046441 | Cmtr2 | cap methyltransferase 2 |
| chr19 | 39652576 | 39653666 | 1091 | * | 10,85622 | -2,874444829 | 1,103919 | -2,60385 | 0,009218 | 0,905781613 | 1,173325 | Distal Intergenic | 19 | 39597286 | 39637495 | 40210 | 2 | 545288 | ENSMUST00000067328.7 | -15081 | ENSMUSG00000062624 | Cyp2c67 | cytochrome P450, family 2, subfamily c, polypeptide 67 |
| chr11 | 38554368 | 38555678 | 1311 | * | 10,77295 | -2,874433924 | 0,988949 | -2,90655 | 0,003654 | 0,730469267 | 1,2763 | Distal Intergenic | 11 | 38688998 | 38689104 | 107 | 2 | 1,15E+08 | ENSMUST00000240209.1 | 133426 | ENSMUSG00000119681 | Gm23520 | predicted gene, 23520 |
| chr7 | 1,24E+08 | 1,24E+08 | 1679 | * | 16,2897 | -2,873523528 | 0,915543 | -3,1386 | 0,001698 | 0,59912059 | 1,2763 | Intron (ENSMUST0000( | 7 | 1,24E+08 | 1,24E+08 | 83266 | 2 | 73076 | ENSMUST00000130240.2 | 15594 | ENSMUSG00000086254 | 3100003L05Rik | RIKEN cDNA 3100003L05 gene |
| chr13 | 1,11E+08 | 1,11E+08 | 1802 | * | 14,13414 | -2,868784435 | 1,015417 | -2,82523 | 0,004725 | 0,805916009 | 1,075735 | Distal Intergenic | 13 | 1,11E+08 | 1,11E+08 | 17965 | 2 | 73129 | ENSMUST00000225326.2 | 148582 | ENSMUSG00000114824 | 3110015C05Rik | RIKEN cDNA 3110015C05 gene |
| chr1 | 1,21E+08 | 1,21E+08 | 1720 | * | 17,62171 | -2,868464016 | 0,98711 | -2,90592 | 0,003662 | 0,645382905 | 2,154504 | Distal Intergenic | 1 | 1,21E+08 | 1,21E+08 | 33046 | 1 | 98452 | ENSMUST00000245225.1 | -60211 | ENSMUSG00000097881 | Celrr | cerebellum expressed regulatory RNA |
| chr4 | 27929526 | 27931620 | 2095 | * | 15,12965 | -2,86778122 | 1,02569 | -2,79595 | 0,005175 | 0,778555725 | 1,37683 | Distal Intergenic | 4 | 27700671 | 27700777 | 107 | 1 | 1,15E+08 | ENSMUST00000179794.2 | 228855 | ENSMUSG00000095003 | Gm25173 | predicted gene, 25173 |
| chr7 | 27051342 | 27052478 | 1137 | * | 13,47969 | -2,867432524 | 0,946074 | -3,03088 | 0,002438 | 0,717133806 | 0,918255 | Promoter (1-2kb) | 7 | 27047525 | 27050034 | 2510 | 2 | 192192 | ENSMUST00000152053.2 | -1308 | ENSMUSG00000089832 | Shkbp1 | Sh3kbp1 binding protein 1 |
| chr2 | 27753911 | 27755364 | 1454 | * | 16,26179 | -2,867291684 | 0,915764 | -3,13104 | 0,001742 | 0,605351622 | 1,2763 | Distal Intergenic | 2 | 27776437 | 27929526 | 153090 | 1 | 12831 | ENSMUST00000028280.14 | -21073 | ENSMUSG00000026837 | Col5a1 | collagen, type V, alpha 1 |
| chr11 | 71507968 | 71509277 | 1310 | * | 11,14257 | -2,865106873 | 1,07604 | -2,66264 | 0,007753 | 0,879169754 | 1,173325 | Distal Intergenic | 11 | 71444234 | 71465421 | 21188 | 2 | 1,03E+08 | ENSMUST00000213352.2 | -42547 | | Gm33351 | predicted gene, 33351 |
| chr10 | 14031090 | 14032782 | 1693 | * | 18,00614 | -2,864904582 | 0,91043 | -3,14676 | 0,001651 | 0,478779213 | 2,386008 | Distal Intergenic | 10 | 14172665 | 14200194 | 27530 | 1 | 1,05E+08 | ENSMUST00000218016.2 | -139883 | | Gm40603 | predicted gene, 40603 |
| chr5 | 25418591 | 25420449 | 1859 | * | 14,66932 | -2,863332311 | 0,956584 | -2,99329 | 0,00276 | 0,594676756 | 2,141612 | Intron (ENSMUST0000( | 5 | 25427880 | 25470916 | 43037 | 1 | 231050 | ENSMUST00000045737.14 | -7431 | ENSMUSG00000038072 | Galnt11 | polypeptide N-acetylgalactosaminyltransferase 11 |
| chr12 | 1,2E+08 | 1,2E+08 | 1172 | * | 13,39527 | -2,863259039 | 0,988494 | -2,89659 | 0,003772 | 0,766579572 | 1,075735 | Exon (ENSMUST00020: | 12 | 1,2E+08 | 1,2E+08 | 73255 | 1 | 217951 | ENSMUST00000058644.15 | -25961 | ENSMUSG00000048004 | Tmem196 | transmembrane protein 196 |
| chr18 | 68672912 | 68673848 | 937 | * | 9,460849 | -2,859802196 | 1,105415 | -2,58708 | 0,009679 | 0,802490409 | 2,237403 | Intron (ENSMUST0000( | 18 | 68692493 | 68699212 | 6720 | 1 | 433197 | ENSMUST00000238121.2 | -18645 | | Gm29677 | predicted gene, 29677 |

| | | | | | | | | | | | | | | | | | | | | | | | |
|---|---|---|---|---|---|---|---|---|---|---|---|---|---|---|---|---|---|---|---|---|---|---|---|
| chr5 | 1,01E+08 | 1,01E+08 | 1288 | * | 14,94703 | -2,857192084 | 1,002166 | -2,85102 | 0,004358 | 0,675594321 | 2,141612 | Distal Intergenic | 5 | 1,01E+08 | 1,01E+08 | 3288 | 2 | 191578 | ENSMUST00000139203.2 | 11094 | ENSMUSG00000035266 | Helq | helicase, POLQ-like |
| chrX | 1,56E+08 | 1,56E+08 | 2004 | * | 15,2411 | -2,856001786 | 0,974272 | -2,93142 | 0,003374 | 0,725842761 | 1,214904 | Distal Intergenic | 20 | 1,56E+08 | 1,56E+08 | 132 | 2 | 1,15E+08 | ENSMUST00000104421.3 | 332071 | ENSMUSG00000077609 | Gm25795 | predicted gene, 25795 |
| chr4 | 1,53E+08 | 1,53E+08 | 1631 | * | 19,55255 | -2,853343315 | 0,829874 | -3,43828 | 0,000585 | 0,304322242 | 2,507013 | Distal Intergenic | 4 | 1,53E+08 | 1,54E+08 | 107949 | 2 | 230959 | ENSMUST00000149177.2 | 313674 | ENSMUSG00000039546 | Ajap1 | adherens junction associated protein 1 |
| chr7 | 71045856 | 71047855 | 2000 | * | 19,16979 | -2,853284196 | 0,91692 | -3,11181 | 0,001859 | 0,477594358 | 2,713306 | Distal Intergenic | 7 | 71020081 | 71029561 | 9481 | 1 | 1,08E+08 | ENSMUST00000189063.2 | 25775 | | Gm29328 | predicted gene 29328 |
| chr18 | 12519813 | 12521313 | 1501 | * | 14,37845 | -2,853090604 | 1,080576 | -2,64034 | 0,008282 | 0,876007548 | 1,2763 | Intron (ENSMUST00000 | 18 | 12466876 | 12716070 | 249195 | 1 | 16774 | ENSMUST00000092070.13 | 52937 | ENSMUSG00000024421 | Lama3 | laminin, alpha 3 |
| chr5 | 27221789 | 27222820 | 1032 | * | 16,16982 | -2,851945834 | 0,928678 | -3,07097 | 0,002134 | 0,541358161 | 2,141612 | Intron (ENSMUST00000 | 5 | 27254197 | 27932499 | 678303 | 1 | 13483 | ENSMUST00000122171.8 | -31377 | ENSMUSG00000061576 | Dpp6 | dipeptidylpeptidase 6 |
| chr10 | 1,21E+08 | 1,21E+08 | 1565 | * | 14,67016 | -2,851105451 | 1,084496 | -2,62897 | 0,008564 | 0,86525631 | 1,37683 | Distal Intergenic | 10 | 1,2E+08 | 1,21E+08 | 66532 | 2 | 74912 | ENSMUST00000219150.2 | -23117 | ENSMUSG00000112716 | 4930471E19Rik | RIKEN cDNA 4930471E19 gene |
| chrX | 78147662 | 78149477 | 1816 | * | 12,33293 | -2,851022473 | 0,992322 | -2,87308 | 0,004065 | 0,779936001 | 1,075735 | Distal Intergenic | 20 | 78374119 | 78393916 | 19798 | 2 | 636104 | ENSMUST00000199433.2 | 244439 | ENSMUSG00000073077 | Cfap47 | cilia and flagella associated protein 47 |
| chr7 | 1,29E+08 | 1,29E+08 | 2710 | * | 23,28153 | -2,850275963 | 0,861428 | -3,30878 | 0,000937 | 0,349574405 | 2,880017 | Promoter (<=1kb) | 7 | 1,29E+08 | 1,29E+08 | 10679 | 1 | 207425 | ENSMUST00000140877.8 | 0 | ENSMUSG00000042055 | Wdr11 | WD repeat domain 11 |
| chr12 | 24098261 | 24099748 | 1488 | * | 19,04571 | -2,847216437 | 0,948794 | -3,00088 | 0,002692 | 0,542967057 | 2,663619 | Intron (ENSMUST00000 | 12 | 24063355 | 24066998 | 3644 | 2 | 1,05E+08 | ENSMUST00000222563.2 | -31263 | | Gm5954 | predicted gene 5954 |
| chr16 | 6610162 | 6612342 | 2181 | * | 17,53948 | -2,846422195 | 0,994258 | -2,86286 | 0,004198 | 0,631820259 | 2,663619 | Intron (ENSMUST00000 | 16 | 6627086 | 7227677 | 600592 | 1 | 268859 | ENSMUST00000115841.10 | -14744 | ENSMUSG00000008658 | Rbfox1 | RNA binding protein, fox-1 homolog (C. elegans) 1 |
| chr16 | 67280168 | 67281329 | 1162 | * | 11,58493 | -2,843467234 | 1,028235 | -2,76539 | 0,005686 | 0,864595549 | 0,918255 | Intron (ENSMUST00000 | 16 | 66581676 | 67417381 | 835706 | 2 | 239857 | ENSMUST00000123266.2 | 136052 | ENSMUSG00000064115 | Cadm2 | cell adhesion molecule 2 |
| chr11 | 51340392 | 51342341 | 1950 | * | 17,86139 | -2,843464455 | 0,898564 | -3,16445 | 0,001554 | 0,470097239 | 2,386008 | Intron (ENSMUST00000 | 11 | 51465259 | 51471126 | 5868 | 1 | 237759 | ENSMUST00000151098.2 | -122918 | ENSMUSG00000063564 | Col23a1 | collagen, type XXIII, alpha 1 |
| chr7 | 23538020 | 23539254 | 1235 | * | 12,6668 | -2,842808585 | 1,02641 | -2,76966 | 0,005611 | 0,901178841 | 0,738441 | Promoter (2-3kb) | 7 | 23534237 | 23535151 | 915 | 2 | 546943 | ENSMUST00000171073.2 | -2869 | ENSMUSG00000096859 | Vmn1r176 | vomeronasal 1 receptor 176 |
| chr8 | 4556035 | 4557845 | 1811 | * | 19,64502 | -2,842449506 | 0,960073 | -2,96066 | 0,00307 | 0,570876293 | 2,663619 | Intron (ENSMUST00000 | 8 | 4543576 | 4573692 | 30117 | 1 | 67260 | ENSMUST00000176130.10 | 12459 | ENSMUSG00000008206 | Cers4 | ceramide synthase 4 |
| chr8 | 1,22E+08 | 1,22E+08 | 1507 | * | 17,76453 | -2,840663704 | 1,040914 | -2,72901 | 0,006353 | 0,715249603 | 2,423594 | Distal Intergenic | 8 | 1,22E+08 | 1,22E+08 | 6806 | 1 | 1,03E+08 | ENSMUST00000180597.3 | -5607 | | Gm26747 | predicted gene, 26747 |
| chrX | 21242288 | 21243811 | 1524 | * | 12,24533 | -2,840058429 | 1,095028 | -2,5936 | 0,009498 | 0,948006995 | 0,918255 | Distal Intergenic | 20 | 21227104 | 21230861 | 3758 | 1 | 331392 | ENSMUST00000127554.2 | 15184 | ENSMUSG00000051537 | Gm5124 | nucleolar and coiled-body phosphoprotein 1 pseudogene |
| chrX | 1,42E+08 | 1,42E+08 | 1559 | * | 14,33121 | -2,839978702 | 1,065825 | -2,66458 | 0,007708 | 0,891097601 | 1,075735 | Distal Intergenic | 20 | 1,42E+08 | 1,42E+08 | 4704 | 1 | 209540 | ENSMUST00000165829.3 | 90661 | ENSMUSG00000085584 | Rtl9 | retrotransposon Gag like 9 |
| chrX | 95575673 | 95577798 | 2126 | * | 12,69433 | -2,837175084 | 1,058383 | -2,68067 | 0,007348 | 0,885416414 | 1,075735 | Intron (ENSMUST00000 | 20 | 95528465 | 95529085 | 621 | 1 | 1,03E+08 | ENSMUST00000118924.2 | 47208 | | Rpl17-ps8 | ribosomal protein L17, pseudogene 8 |
| chr14 | 1,15E+08 | 1,15E+08 | 1722 | * | 11,34465 | -2,830906664 | 1,033913 | -2,73805 | 0,00618 | 0,915235483 | 0,738441 | Distal Intergenic | 14 | 1,15E+08 | 1,15E+08 | 84 | 1 | 723905 | ENSMUST00000083574.3 | -35558 | ENSMUSG00000065508 | Mir17 | microRNA 17 |
| chr1 | 37631186 | 37632822 | 1637 | * | 15,14457 | -2,830626975 | 0,97916 | -2,89087 | 0,003842 | 0,722439117 | 1,41696 | Distal Intergenic | 1 | 37653340 | 37654453 | 1114 | 1 | 1,03E+08 | ENSMUST00000160659.2 | -20518 | | 4930556I23Rik | RIKEN cDNA 4930556I23 gene |
| chr1 | 56569857 | 56572049 | 2193 | * | 17,64662 | -2,82730501 | 0,864276 | -3,2713 | 0,001071 | 0,425046984 | 2,154504 | Exon (ENSMUST000001 | 1 | 56675203 | 56676594 | 1392 | 2 | 71066 | ENSMUST00000062085.6 | 104545 | ENSMUSG00000045336 | Hsfy2 | heat shock transcription factor, Y-linked 2 |
| chr10 | 1,03E+08 | 1,03E+08 | 1070 | * | 12,66485 | -2,826981422 | 0,969413 | -2,91618 | 0,003543 | 0,78560377 | 0,918255 | Distal Intergenic | 10 | 1,03E+08 | 1,03E+08 | 3709 | 1 | 103098 | ENSMUST00000219936.2 | 92385 | ENSMUSG00000019894 | Slc6a15 | solute carrier family 6 (neurotransmitter transporter), member 15 |
| chr3 | 1,14E+08 | 1,14E+08 | 2190 | * | 16,97738 | -2,826594633 | 0,949962 | -2,97548 | 0,002925 | 0,605351622 | 2,141612 | Distal Intergenic | 3 | 1,15E+08 | 1,15E+08 | 132 | 2 | 1,15E+08 | ENSMUST00000104222.3 | 241954 | ENSMUSG00000077410 | Gm23750 | predicted gene, 23750 |
| chr2 | 70184637 | 70186328 | 1692 | * | 13,0867 | -2,825101773 | 0,972829 | -2,90401 | 0,003684 | 0,748382289 | 1,173325 | Intron (ENSMUST00000 | 2 | 70305267 | 70308073 | 2807 | 1 | 64406 | ENSMUST00000100043.3 | -118939 | ENSMUSG00000075304 | Sp5 | trans-acting transcription factor 5 |
| chr14 | 99350154 | 99352329 | 2176 | * | 11,63315 | -2,824703843 | 1,046013 | -2,70045 | 0,006925 | 0,90060438 | 0,918255 | Exon (ENSMUST00000 | 14 | 99337394 | 99351653 | 14260 | 1 | 52023 | ENSMUST00000228279.2 | 12760 | ENSMUSG00000022064 | Pibf1 | progesterone immunomodulatory binding factor 1 |
| chr5 | 55588115 | 55590316 | 2202 | * | 17,76484 | -2,824379661 | 0,87453 | -3,2296 | 0,00124 | 0,448852574 | 2,154504 | Distal Intergenic | 5 | 55460762 | 55460923 | 162 | 2 | 1,15E+08 | ENSMUST00000083861.3 | -127192 | ENSMUSG00000065795 | Gm25513 | predicted gene, 25513 |
| chr1 | 65298115 | 65299657 | 1543 | * | 11,97941 | -2,823350542 | 1,01394 | -2,78453 | 0,00536 | 0,797915302 | 1,2763 | Exon (ENSMUST00000 | 1 | 65321215 | 65384500 | 63286 | 1 | 213527 | ENSMUST00000140190.8 | -21558 | ENSMUSG00000025946 | Pth2r | parathyroid hormone 2 receptor |
| chr10 | 93668537 | 93670094 | 1558 | * | 13,70798 | -2,820801188 | 1,000866 | -2,81836 | 0,004827 | 0,810618681 | 1,075735 | Promoter (<=1kb) | 10 | 93669680 | 93693950 | 24271 | 1 | 327799 | ENSMUST00000216224.2 | 0 | ENSMUSG00000020020 | Usp44 | ubiquitin specific peptidase 44 |
| chr18 | 11205676 | 11206941 | 1266 | * | 13,55333 | -2,820597591 | 1,053966 | -2,67617 | 0,007447 | 0,910222792 | 0,918255 | Distal Intergenic | 18 | 11059047 | 11063461 | 4415 | 1 | 14465 | ENSMUST00000234971.2 | 146629 | ENSMUSG00000005836 | Gata6 | GATA binding protein 6 |
| chrX | 1,61E+08 | 1,61E+08 | 1584 | * | 13,5405 | -2,820113825 | 1,040649 | -2,70996 | 0,006729 | 0,8666955 | 1,075735 | Intron (ENSMUST00000 | 20 | 1,61E+08 | 1,61E+08 | 6360 | 2 | 194590 | ENSMUST00000155043.2 | 84736 | ENSMUSG00000040855 | Reps2 | RALBP1 associated Eps domain containing protein 2 |
| chr15 | 57533282 | 57534200 | 919 | * | 11,38362 | -2,820005401 | 1,065232 | -2,64732 | 0,008113 | 0,872043312 | 1,2763 | Distal Intergenic | 15 | 57558061 | 57703228 | 145168 | 1 | 387609 | ENSMUST00000096430.11 | -23861 | ENSMUSG00000071757 | Zhx2 | zinc fingers and homeoboxes 2 |
| chr18 | 43551927 | 43552916 | 990 | * | 10,65177 | -2,819461541 | 1,093756 | -2,57778 | 0,009944 | 0,985737745 | 0,744375 | Intron (ENSMUST00000 | 18 | 43454049 | 43571351 | 117303 | 2 | 22240 | ENSMUST00000121805.9 | 18435 | ENSMUSG00000024501 | Dpysl3 | dihydropyrimidinase-like 3 |
| chr8 | 1,03E+08 | 1,03E+08 | 1504 | * | 14,00235 | -2,815097397 | 0,973996 | -2,89026 | 0,003849 | 0,797327226 | 0,918255 | Distal Intergenic | 8 | 1,03E+08 | 1,03E+08 | 2716 | 2 | 12552 | ENSMUST00000210578.2 | 773445 | ENSMUSG00000031673 | Cdh11 | cadherin 11 |
| chr1 | 76153964 | 76155114 | 1151 | * | 11,39621 | -2,814810381 | 1,071732 | -2,62641 | 0,008629 | 0,963466134 | 0,744375 | Distal Intergenic | 1 | 75993295 | 75995393 | 2099 | 2 | 1,03E+08 | ENSMUST00000188786.2 | -158571 | | Gm29065 | predicted gene 29065 |
| chr10 | 14143827 | 14145828 | 2002 | * | 18,28344 | -2,814035191 | 0,837188 | -3,3613 | 0,000776 | 0,332694435 | 2,663619 | Distal Intergenic | 10 | 14172665 | 14200194 | 27530 | 1 | 1,05E+08 | ENSMUST00000218016.2 | -26837 | | Gm40603 | predicted gene, 40603 |
| chr1 | 27357755 | 27359303 | 1549 | * | 15,90553 | -2,81083593 | 0,996917 | -2,81953 | 0,004809 | 0,673641551 | 2,386008 | Distal Intergenic | 1 | 27923059 | 27941903 | 18845 | 1 | 433281 | ENSMUST00000185409.3 | -563756 | ENSMUSG00000101514 | Gm5524 | predicted gene 5524 |
| chr17 | 45968343 | 45970541 | 2199 | * | 19,34884 | -2,808171127 | 1,026842 | -2,73476 | 0,006243 | 0,694289896 | 2,663619 | Promoter (<=1kb) | 17 | 45941130 | 45970251 | 29122 | 2 | 268958 | ENSMUST00000120717.8 | 0 | ENSMUSG00000058626 | Capn11 | calpain 11 |
| chr16 | 36270938 | 36272403 | 1466 | * | 10,64047 | -2,807469702 | 1,033954 | -2,71527 | 0,006622 | 0,836575905 | 1,2763 | 3' UTR | 16 | 36270899 | 36275754 | 4856 | 2 | 20863 | ENSMUST00000068182.3 | 3351 | ENSMUSG00000054905 | Stfa3 | stefin A3 |
| chr12 | 1,13E+08 | 1,13E+08 | 1780 | * | 13,16774 | -2,807201735 | 1,083762 | -2,59024 | 0,009591 | 0,980607198 | 0,738441 | Promoter (<=1kb) | 12 | 1,13E+08 | 1,13E+08 | 5416 | 2 | 72308 | ENSMUST00000221409.2 | 0 | ENSMUSG00000011158 | Brf1 | BRF1, RNA polymerase III transcription initiation factor 90 kDa subunit |
| chrX | 76802624 | 76804617 | 1994 | * | 15,02471 | -2,804613175 | 0,971544 | -2,88676 | 0,003892 | 0,749550311 | 1,214904 | Distal Intergenic | 20 | 76815061 | 76839578 | 24518 | 2 | 19108 | ENSMUST00000114044.2 | 34961 | ENSMUSG00000035725 | Prkx | protein kinase, X-linked |
| chr17 | 23453006 | 23454315 | 1310 | * | 12,55716 | -2,802712562 | 1,079772 | -2,59565 | 0,009441 | 0,977574845 | 0,738441 | Distal Intergenic | 17 | 23410320 | 23412226 | 1907 | 2 | 22700 | ENSMUST00000151447.2 | -40780 | ENSMUSG00000002617 | Zfp40 | zinc finger protein 40 |
| chr6 | 1,37E+08 | 1,37E+08 | 1875 | * | 21,30404 | -2,80221478 | 0,839556 | -3,33774 | 0,000845 | 0,333467627 | 2,880017 | Distal Intergenic | 6 | 1,37E+08 | 1,37E+08 | 126102 | 1 | 19277 | ENSMUST00000203010.2 | -29009 | ENSMUSG00000030223 | Ptpro | protein tyrosine phosphatase, receptor type, O |
| chr17 | 92106221 | 92107543 | 1323 | * | 15,3149 | -2,801013826 | 0,857132 | -3,26789 | 0,001084 | 0,496713645 | 1,41696 | Distal Intergenic | 17 | 92119919 | 92120048 | 130 | 2 | 1,15E+08 | ENSMUST00000157317.3 | 12505 | ENSMUSG00000087942 | Gm24244 | predicted gene, 24244 |
| chr18 | 19885226 | 19886898 | 1673 | * | 16,78613 | -2,798496393 | 0,81626 | -3,42844 | 0,000607 | 0,313797801 | 2,386008 | Distal Intergenic | 18 | 20096027 | 20135012 | 38986 | 2 | 13507 | ENSMUST00000225110.2 | 248114 | ENSMUSG00000059898 | Dsc3 | desmocollin 3 |
| chr6 | 36665640 | 36666866 | 1227 | * | 14,63104 | -2,797734164 | 1,035203 | -2,70259 | 0,00688 | 0,749550311 | 2,141612 | Distal Intergenic | 6 | 36716394 | 36787114 | 70721 | 2 | 19242 | ENSMUST00000201321.2 | 120248 | ENSMUSG00000029838 | Ptn | pleiotrophin |
| chr16 | 15857727 | 15859124 | 1398 | * | 14,09061 | -2,795444083 | 0,967166 | -2,89034 | 0,003848 | 0,837112103 | 0,738441 | Intron (ENSMUST00000 | 16 | 15711249 | 15964668 | 253420 | 2 | 224008 | ENSMUST00000229846.2 | 105544 | ENSMUSG00000041974 | Spidr | scaffolding protein involved in DNA repair |
| chr1 | 1,8E+08 | 1,8E+08 | 1863 | * | 16,7577 | -2,795351078 | 0,968046 | -2,88762 | 0,003882 | 0,642983105 | 2,318434 | Intron (ENSMUST00000 | 1 | 1,8E+08 | 1,8E+08 | 10034 | 2 | 1,08E+08 | ENSMUST00000192794.2 | 4066 | | LOC108167750 | uncharacterized LOC108167750 |
| chr14 | 31016550 | 31017976 | 1427 | * | 15,7547 | -2,794942505 | 1,077685 | -2,59347 | 0,009501 | 0,792604439 | 2,318434 | Exon (ENSMUST00000 | 14 | 30983949 | 30991468 | 7520 | 2 | 110084 | ENSMUST00000228429.2 | -25082 | ENSMUSG00000019027 | Dnah1 | dynein, axonemal, heavy chain 1 |
| chr10 | 1,1E+08 | 1,1E+08 | 967 | * | 11,65461 | -2,794552031 | 1,075589 | -2,59816 | 0,009372 | 0,895765172 | 1,2763 | Intron (ENSMUST00000 | 10 | 1,1E+08 | 1,1E+08 | 318961 | 2 | 260315 | ENSMUST00000032719.15 | -95505 | ENSMUSG00000020181 | Nav3 | neuron navigator 3 |
| chr7 | 74476389 | 74478045 | 1657 | * | 14,12482 | -2,793906173 | 1,01137 | -2,7625 | 0,005736 | 0,903470246 | 0,738441 | Intron (ENSMUST00000 | 7 | 74473363 | 74503558 | 30196 | 2 | 76253 | ENSMUST00000205326.2 | 25513 | | 0610006L08Rik | RIKEN cDNA 0610006L08 gene |
| chr7 | 53848955 | 53850694 | 1740 | * | 13,07827 | -2,793555977 | 0,996344 | -2,80381 | 0,00505 | 0,885719815 | 0,738441 | Distal Intergenic | 7 | 53541509 | 53541637 | 129 | 2 | 1,02E+08 | ENSMUST00000184140.3 | -307318 | ENSMUSG00000099176 | Mir6238 | microRNA 6238 |
| chr7 | 1,24E+08 | 1,24E+08 | 1277 | * | 15,19998 | -2,790039935 | 0,913038 | -3,05578 | 0,002245 | 0,632712345 | 1,41696 | Intron (ENSMUST00000 | 7 | 1,24E+08 | 1,24E+08 | 83266 | 2 | 73076 | ENSMUST00000130240.2 | 67014 | ENSMUSG00000086254 | 3100003L05Rik | RIKEN cDNA 3100003L05 gene |
| chr5 | 1,27E+08 | 1,27E+08 | 1916 | * | 19,38066 | -2,789394821 | 0,911949 | -3,05872 | 0,002223 | 0,524368688 | 2,507013 | Distal Intergenic | 5 | 1,27E+08 | 1,27E+08 | 32560 | 1 | 1,05E+08 | ENSMUST00000198430.2 | 38725 | | Gm40331 | predicted gene, 40331 |
| chr3 | 1,58E+08 | 1,58E+08 | 770 | * | 11,64987 | -2,788896593 | 1,000176 | -2,78841 | 0,005297 | 0,853837252 | 0,918255 | Distal Intergenic | 3 | 1,58E+08 | 1,58E+08 | 477783 | 2 | 242274 | ENSMUST00000106044.6 | -171292 | ENSMUSG00000028176 | Lrrc7 | leucine rich repeat containing 7 |
| chr3 | 63352895 | 63354186 | 1292 | * | 14,16104 | -2,787740842 | 0,98108 | -2,8415 | 0,00449 | 0,767061106 | 1,2763 | Exon (ENSMUST000001 | 3 | 63346672 | 63365318 | 18647 | 1 | 1,03E+08 | ENSMUST00000192500.2 | 6223 | ENSMUSG00000104336 | Gm34240 | predicted gene, 34240 |
| chrX | 58156965 | 58158402 | 1438 | * | 10,63669 | -2,787598404 | 1,076813 | -2,58875 | 0,009633 | 0,912096919 | 1,173325 | Intron (ENSMUST00000 | 20 | 58143018 | 58143096 | 79 | 2 | 1E+08 | ENSMUST00000093602.4 | -13869 | ENSMUSG00000070110 | Mir504 | microRNA 504 |
| chr3 | 55652027 | 55653154 | 1128 | * | 10,23256 | -2,78757334 | 1,032234 | -2,70052 | 0,006923 | 0,857326444 | 1,173325 | Intron (ENSMUST00000 | 3 | 55603781 | 55631368 | 27588 | 2 | 26422 | ENSMUST00000198259.2 | -20659 | ENSMUSG00000027799 | Nbea | neurobeachin |
| chr12 | 1,08E+08 | 1,08E+08 | 1142 | * | 13,67231 | -2,786838493 | 1,025317 | -2,71803 | 0,006567 | 0,88871498 | 0,938682 | Distal Intergenic | 12 | 1,08E+08 | 1,08E+08 | 11749 | 2 | 74968 | ENSMUST00000130564.2 | 116389 | ENSMUSG00000085641 | 4930465M20Rik | RIKEN cDNA 4930465M20 gene |
| chr1 | 1,27E+08 | 1,27E+08 | 1627 | * | 15,43849 | -2,786424401 | 1,073776 | -2,59498 | 0,00946 | 0,807119269 | 2,141612 | Distal Intergenic | 1 | 1,27E+08 | 1,27E+08 | 183198 | 1 | 107895 | ENSMUST00000189427.7 | -68037 | ENSMUSG00000036155 | Mgat5 | mannoside acetylglucosaminyltransferase 5 |
| chr5 | 22176463 | 22178145 | 1683 | * | 14,67927 | -2,785463181 | 0,920552 | -3,02586 | 0,002479 | 0,571379912 | 2,141612 | Exon (ENSMUST000001 | 5 | 22181036 | 22215262 | 34227 | 2 | 19699 | ENSMUST00000161353.2 | 37117 | ENSMUSG00000042453 | Reln | reelin |
| chr3 | 1,58E+08 | 1,58E+08 | 1275 | * | 18,11445 | -2,783383142 | 0,871388 | -3,19419 | 0,001402 | 0,470097239 | 2,154504 | Distal Intergenic | 3 | 1,58E+08 | 1,58E+08 | 477783 | 2 | 242274 | ENSMUST00000106044.6 | -55947 | ENSMUSG00000028176 | Lrrc7 | leucine rich repeat containing 7 |
| chr13 | 1,06E+08 | 1,06E+08 | 1588 | * | 12,45465 | -2,780419009 | 0,946109 | -2,93879 | 0,003295 | 0,742868694 | 1,075735 | Distal Intergenic | 13 | 1,07E+08 | 1,07E+08 | 14510 | 2 | 1,03E+08 | ENSMUST00000225713.2 | 559247 | | Gm30411 | predicted gene, 30411 |
| chr14 | 1,05E+08 | 1,05E+08 | 2651 | * | 16,40457 | -2,779218388 | 0,812636 | -3,42 | 0,000626 | 0,304413931 | 2,663619 | Distal Intergenic | 14 | 1,05E+08 | 1,05E+08 | 134 | 1 | 1,15E+08 | ENSMUST00000104237.3 | 38252 | ENSMUSG00000077425 | Gm24280 | predicted gene, 24280 |
| chr11 | 35925875 | 35927957 | 2083 | * | 14,63845 | -2,778364163 | 0,935967 | -2,96844 | 0,002993 | 0,61075929 | 2,141612 | Intron (ENSMUST00000 | 11 | 35729227 | 35871354 | 142128 | 2 | 211652 | ENSMUST00000018993.7 | -54521 | ENSMUSG00000018849 | Wwc1 | WW, C2 and coiled-coil domain containing 1 |
| chr18 | 69325364 | 69326983 | 1620 | * | 15,73813 | -2,77722716 | 0,98071 | -2,83185 | 0,004628 | 0,64945399 | 2,663619 | Promoter (1-2kb) | 18 | 69326407 | 69328815 | 2409 | 2 | 1,03E+08 | ENSMUST00000211216.2 | 1832 | | Gm31294 | predicted gene, 31294 |
| chr10 | 6805191 | 6806350 | 1160 | * | 12,78058 | -2,777099823 | 1,028445 | -2,70029 | 0,006928 | 0,933631108 | 0,738441 | Intron (ENSMUST00000 | 10 | 6739099 | 6799987 | 60889 | 1 | 18390 | ENSMUST00000148625.2 | 66092 | ENSMUSG00000000766 | Oprm1 | opioid receptor, mu 1 |
| chr3 | 1,28E+08 | 1,28E+08 | 1479 | * | 11,93453 | -2,777025354 | 1,000781 | -2,77486 | 0,005523 | 0,860218139 | 0,918255 | Distal Intergenic | 3 | 1,28E+08 | 1,28E+08 | 2173 | 1 | 1,03E+08 | ENSMUST00000199122.2 | 24562 | | Gm35667 | predicted gene, 35667 |
| chr9 | 36966150 | 36967968 | 1819 | * | 18,26819 | -2,775686479 | 0,859439 | -3,22965 | 0,001239 | 0,414202366 | 2,663619 | Intron (ENSMUST00000 | 9 | 36803938 | 36994575 | 190638 | 2 | 208076 | ENSMUST00000177080.8 | 26607 | ENSMUSG00000035934 | Pknox2 | Pbx/knotted 1 homeobox 2 |
| chr1 | 1,76E+08 | 1,76E+08 | 1879 | * | 22,37214 | -2,775638066 | 0,730931 | -3,7974 | 0,000146 | 0,125710024 | 3,104559 | Distal Intergenic | 1 | 1,76E+08 | 1,76E+08 | 97163 | 2 | 1,03E+08 | ENSMUST00000194241.2 | -29751 | ENSMUSG00000104021 | Gm36536 | predicted gene, 36536 |
| chr14 | 23342561 | 23345173 | 2613 | * | 22,43801 | -2,773569382 | 0,907135 | -3,05751 | 0,002232 | 0,467440127 | 3,471678 | 3' UTR | 14 | 23088262 | 23144639 | 56378 | 2 | 791307 | ENSMUST00000090409.2 | -197922 | ENSMUSG00000068680 | Gm10248 | predicted gene 10248 |
| chr3 | 32099884 | 32101384 | 1501 | * | 17,17692 | -2,773005713 | 0,94826 | -2,92431 | 0,003452 | 0,614021623 | 2,423594 | Intron (ENSMUST00000 | 3 | 32210263 | 32252505 | 42243 | 1 | 72413 | ENSMUST00000194796.2 | -108879 | ENSMUSG00000037610 | Kcnmb2 | potassium large conductance calcium-activated channel, subfamily M, beta member 2 |
| chr16 | 9926720 | 9928718 | 1999 | * | 13,39037 | -2,772893069 | 0,961455 | -2,88406 | 0,003926 | 0,839562284 | 0,738441 | Distal Intergenic | 16 | 9988090 | 9992494 | 4405 | 1 | 68172 | ENSMUST00000121292.8 | -59372 | ENSMUSG00000039209 | Rpl39l | ribosomal protein L39-like |
| chr9 | 47136628 | 47138109 | 1482 | * | 13,5458 | -2,770317951 | 1,028547 | -2,69343 | 0,007072 | 0,90060438 | 0,938682 | Promoter (<=1kb) | 9 | 47093162 | 47138393 | 45232 | 2 | 1,03E+08 | ENSMUST00000217115.2 | 284 | | Gm31698 | predicted gene, 31698 |
| chr15 | 28485384 | 28486542 | 1159 | * | 13,84377 | -2,77030275 | 1,034117 | -2,67891 | 0,007386 | 0,886332554 | 1,075735 | Distal Intergenic | 15 | 28203898 | 28472198 | 268301 | 1 | 110082 | ENSMUST00000067048.8 | 281486 | ENSMUSG00000022262 | Dnah5 | dynein, axonemal, heavy chain 5 |
| chr1 | 65984225 | 65985634 | 1410 | * | 11,01117 | -2,769664718 | 1,029575 | -2,6901 | 0,007143 | 0,9371986 | 0,738441 | Distal Intergenic | 1 | 65965697 | 65967465 | 1769 | 1 | 12969 | ENSMUST00000114027.3 | 18528 | ENSMUSG00000025945 | Crygf | crystallin, gamma F |
| chr13 | 26695262 | 26697030 | 1769 | * | 15,4517 | -2,769652317 | 0,90246 | -3,069 | 0,002148 | 0,643955341 | 1,2763 | Distal Intergenic | 13 | 26467133 | 26496389 | 29257 | 2 | 73576 | ENSMUST00000154703.3 | -198873 | ENSMUSG00000084811 | 1700092E19Rik | RIKEN cDNA 1700092E19 gene |
| chr17 | 15038566 | 15040438 | 1873 | * | 14,98809 | -2,76945156 | 0,919011 | -3,01351 | 0,002582 | 0,662610953 | 1,37683 | 3' UTR | 17 | 15012203 | 15012266 | 64 | 2 | 1,15E+08 | ENSMUST00000158428.3 | -26300 | ENSMUSG00000089053 | Gm23352 | predicted gene, 23352 |
| chr5 | 5028774 | 5030848 | 2075 | * | 17,12278 | -2,769233053 | 1,024394 | -2,70329 | 0,006866 | 0,730689299 | 2,386008 | Intron (ENSMUST00000 | 5 | 5035111 | 5035237 | 127 | 2 | 1,15E+08 | ENSMUST00020182726.1 | 4389 | ENSMUSG00000105400 | Gm22897 | predicted gene, 22897 |
| chr7 | 1,44E+08 | 1,44E+08 | 2387 | * | 20,53322 | -2,768233877 | 0,933981 | -2,96391 | 0,003038 | 0,565876168 | 2,713306 | Intron (ENSMUST00000 | 7 | 1,44E+08 | 1,44E+08 | 248966 | 1 | 210274 | ENSMUST00000105900.9 | 17884 | ENSMUSG00000037541 | Shank2 | SH3 and multiple ankyrin repeat domains 2 |
| chr1 | 13163326 | 13165155 | 1830 | * | 13,55463 | -2,767986781 | 0,987501 | -2,80302 | 0,005063 | 0,81949316 | 1,075735 | Distal Intergenic | 1 | 13183681 | 13197387 | 13707 | 2 | 383491 | ENSMUST00000047577.7 | 32232 | ENSMUSG00000042414 | Prdm14 | PR domain containing 14 |
| chr10 | 27448619 | 27450080 | 1462 | * | 14,15855 | -2,766264992 | 1,007191 | -2,74652 | 0,006023 | 0,877892722 | 0,918255 | Intron (ENSMUST00000 | 10 | 27063544 | 27492792 | 429249 | 2 | 16773 | ENSMUST00000189575.2 | 42712 | ENSMUSG00000019899 | Lama2 | laminin, alpha 2 |
| chr1 | 1,1E+08 | 1,1E+08 | 1587 | * | 10,08019 | -2,765621312 | 1,064151 | -2,5989 | 0,009352 | 0,919447249 | 1,086118 | Distal Intergenic | 1 | 1,1E+08 | 1,1E+08 | 130 | 2 | 1,15E+08 | ENSMUST00000158412.3 | -152476 | ENSMUSG00000089037 | Gm22159 | predicted gene, 22159 |
| chr10 | 94941101 | 94943298 | 2198 | * | 16,29294 | -2,761087038 | 0,884669 | -3,12104 | 0,001802 | 0,498636665 | 2,318434 | Distal Intergenic | 10 | 94956444 | 94981687 | 25244 | 2 | 628062 | ENSMUST00000220248.2 | 38389 | ENSMUSG00000112481 | Gm29684 | predicted gene, 29684 |
| chr11 | 90177983 | 90178978 | 996 | * | 13,37975 | -2,759027678 | 0,977061 | -2,8238 | 0,004746 | 0,836351745 | 0,918255 | Distal Intergenic | 11 | 90140302 | 90169415 | 29114 | 1 | 67468 | ENSMUST00000004050.7 | 37681 | ENSMUSG00000003948 | Mmd | monocyte to macrophage differentiation-associated |
| chr10 | 79132701 | 79134724 | 2024 | * | 18,59142 | -2,758685501 | 0,831548 | -3,31753 | 0,000908 | 0,377927525 | 2,386008 | Distal Intergenic | 10 | 79083526 | 79130369 | 46844 | 1 | 216144 | ENSMUST00000020547.10 | 49175 | ENSMUSG00000055515 | Vmn2r81 | vomeronasal 2, receptor 81 |
| chr12 | 1,1E+08 | 1,1E+08 | 1901 | * | 23,39903 | -2,758186136 | 0,855451 | -3,22425 | 0,001263 | 0,401315146 | 2,951933 | Distal Intergenic | 12 | 1,1E+08 | 1,1E+08 | 10848 | 2 | 1,05E+08 | ENSMUST00000221773.2 | -18419 | | Gm40576 | predicted gene, 40576 |
| chr16 | 15165985 | 15167646 | 1662 | * | 15,32897 | -2,757943365 | 0,944409 | -2,92029 | 0,003497 | 0,730689299 | 1,214904 | Distal Intergenic | 16 | 15373745 | 15412336 | 38592 | 2 | 70620 | ENSMUST00000118236.8 | 244690 | ENSMUSG00000022674 | Ube2v2 | ubiquitin-conjugating enzyme E2 variant 2 |
| chr3 | 1,24E+08 | 1,24E+08 | 1762 | * | 14,55828 | -2,757110335 | 0,912974 | -3,01992 | 0,002528 | 0,670853401 | 1,2763 | Distal Intergenic | 3 | 1,24E+08 | 1,24E+08 | 430 | 1 | 668809 | ENSMUST00000168454.4 | -251456 | ENSMUSG00000090474 | Gm9372 | predicted gene 9372 |
| chr5 | 23406789 | 23408265 | 1477 | * | 15,64482 | -2,756904561 | 0,977607 | -2,82006 | 0,004802 | 0,789027175 | 1,214904 | Intron (ENSMUST00000 | 5 | 23305550 | 23480586 | 175037 | 1 | 269629 | ENSMUST00000196406.2 | 101239 | ENSMUSG00000106379 | Lhfpl3 | lipoma HMGIC fusion partner-like 3 |
| chr13 | 1,09E+08 | 1,09E+08 | 1500 | * | 15,96338 | -2,7565698 | 1,063193 | -2,59273 | 0,009522 | 0,793028263 | 2,318434 | Intron (ENSMUST00000 | 13 | 1,09E+08 | 1,1E+08 | 1091179 | 1 | 238871 | ENSMUST00000177907.8 | 96891 | ENSMUSG00000021699 | Pde4d | phosphodiesterase 4D, cAMP specific |
| chr16 | 46184649 | 46186126 | 1478 | * | 12,16352 | -2,755884304 | 0,981478 | -2,80789 | 0,004987 | 0,881706354 | 0,744375 | Distal Intergenic | 16 | 46215221 | 46231532 | 16312 | 2 | 58998 | ENSMUST00000121803.8 | 45406 | ENSMUSG00000022656 | Nectin3 | nectin cell adhesion molecule 3 |
| chr3 | 1,54E+08 | 1,54E+08 | 817 | * | 15,4604 | -2,754750123 | 0,978119 | -2,81637 | 0,004857 | 0,790570581 | 1,214904 | Promoter (2-3kb) | 3 | 1,54E+08 | 1,54E+08 | 103932 | 1 | 209601 | ENSMUST00000098496.9 | 2965 | ENSMUSG00000078161 | Erich3 | glutamate rich 3 |
| chr8 | 63327966 | 63329074 | 1109 | * | 10,83192 | -2,751993443 | 1,020571 | -2,69652 | 0,007007 | 0,934846269 | 0,738441 | Distal Intergenic | 8 | 63404043 | 63809886 | 405844 | 1 | 72902 | ENSMUST00000117377.8 | -74969 | ENSMUSG00000054162 | Spock3 | sparc/osteonectin, cwcv and kazal-like domains proteoglycan 3 |
| chr1 | 89077548 | 89079201 | 1654 | * | 17,07202 | -2,751222497 | 1,019206 | -2,69938 | 0,006947 | 0,845014042 | 1,2763 | Intron (ENSMUST00000 | 1 | 89029828 | 89030640 | 813 | 2 | 1,03E+08 | ENSMUST00000244652.1 | -46908 | | Gm34331 | predicted gene, 34331 |
| chr4 | 57341021 | 57342155 | 1135 | * | 10,32572 | -2,749841615 | 1,006117 | -2,73312 | 0,006274 | 0,840920027 | 1,173325 | Distal Intergenic | 4 | 57361294 | 57364252 | 2959 | 2 | 73349 | ENSMUST00000143574.2 | 22097 | | 1700042G15Rik | RIKEN cDNA 1700042G15 gene |
| chr13 | 1,14E+08 | 1,14E+08 | 1808 | * | 15,04605 | -2,749498092 | 0,957816 | -2,87059 | 0,004097 | 0,749550311 | 1,2763 | Exon (ENSMUST000002 | 13 | 1,14E+08 | 1,14E+08 | 56557 | 2 | 70917 | ENSMUST00000223835.2 | 36764 | ENSMUSG00000114790 | 4921509O07Rik | RIKEN cDNA 4921509O07 gene |
| chr18 | 3073687 | 3075057 | 1371 | * | 14,54005 | -2,74753449 | 0,927872 | -2,96111 | 0,003065 | 0,685026746 | 1,37683 | Distal Intergenic | 18 | 3122492 | 3123412 | 921 | 2 | 1E+08 | ENSMUST00000165255.3 | 48355 | ENSMUSG00000091539 | Vmn1r238 | vomeronasal 1 receptor, 238 |
| chr12 | 43201507 | 43203195 | 1689 | * | 16,96779 | -2,746325853 | 0,969415 | -2,83297 | 0,004612 | 0,671156916 | 2,318434 | Distal Intergenic | 12 | 43013861 | 43081000 | 67140 | 2 | 1,03E+08 | ENSMUST00000219475.2 | -120507 | ENSMUSG00000112407 | Gm33340 | predicted gene, 33340 |
| chr3 | 3638144 | 3639376 | 1233 | * | 14,24703 | -2,746264405 | 0,975475 | -2,81531 | 0,004873 | 0,839117124 | 0,918255 | Intron (ENSMUST00000 | 3 | 3699205 | 3724860 | 25656 | 1 | 30942 | ENSMUST00000108394.4 | -59829 | ENSMUSG00000017688 | Hnf4g | hepatocyte nuclear factor 4, gamma |
| chr7 | 1,38E+08 | 1,38E+08 | 840 | * | 13,07584 | -2,746220683 | 0,968372 | -2,83591 | 0,004569 | 0,770104803 | 1,2763 | Distal Intergenic | 7 | 1,38E+08 | 1,38E+08 | 13480 | 1 | 75187 | ENSMUST00000209357.2 | -56395 | ENSMUSG00000110037 | 4930543N07Rik | RIKEN cDNA 4930543N07 gene |

| | | | | | | | | | | | | | | | | | | | | | | | |
|---|---|---|---|---|---|---|---|---|---|---|---|---|---|---|---|---|---|---|---|---|---|---|---|
| chr8 | 26700752 | 26702094 | 1343 | * | 17,01211 | -2,745673691 | 1,037805 | -2,64566 | 0,008153 | 0,767061106 | 2,318434 | Distal Intergenic | 8 | 26648821 | 26650840 | 2020 | 1 | 73754 | ENSMUST00000209926.2 | 51931 | ENSMUSG00000037214 | Thap1 | THAP domain containing, apoptosis associated protein 1 |
| chr13 | 97597097 | 97598601 | 1505 | * | 14,84075 | -2,744552938 | 1,059157 | -2,59126 | 0,009562 | 0,883511731 | 1,41696 | Intron (ENSMUST0000 | 13 | 97619209 | 97633515 | 14307 | 1 | 1E+08 | ENSMUST00000179324.2 | -20608 | ENSMUSG00000078952 | Lncenc1 | long non-coding RNA, embryonic stem cells expressed 1 |
| chr7 | 1,4E+08 | 1,4E+08 | 1453 | * | 14,02828 | -2,743336551 | 0,984205 | -2,78736 | 0,005314 | 0,854157826 | 0,918255 | Promoter (1-2kb) | 7 | 1,4E+08 | 1,4E+08 | 10624 | 2 | 74075 | ENSMUST00000026553.6 | -1079 | ENSMUSG00000025480 | Syce1 | synaptonemal complex central element protein 1 |
| chr14 | 1,13E+08 | 1,13E+08 | 3063 | * | 13,34795 | -2,742332008 | 1,013579 | -2,70559 | 0,006818 | 0,893189326 | 0,938682 | Distal Intergenic | 14 | 1,12E+08 | 1,12E+08 | 3494 | 1 | 75409 | ENSMUST00000227891.2 | 696370 | ENSMUSG00000033214 | Slitrk5 | SLIT and NTRK-like family, member 5 |
| chr14 | 15996676 | 15998104 | 1429 | * | 17,23347 | -2,738880955 | 0,826982 | -3,3119 | 0,000927 | 0,402652506 | 2,154504 | Distal Intergenic | 14 | 16025162 | 16033976 | 8815 | 1 | 1E+08 | ENSMUST00000169152.8 | -27058 | ENSMUSG00000095360 | Gm3594 | predicted gene 3594 |
| chr2 | 1,34E+08 | 1,34E+08 | 1203 | * | 17,27217 | -2,738629925 | 0,904814 | -3,02673 | 0,002472 | 0,545426979 | 2,423594 | Distal Intergenic | 2 | 1,34E+08 | 1,34E+08 | 23548 | 2 | 15112 | ENSMUST00000140794.2 | 105115 | ENSMUSG00000027261 | Hao1 | hydroxyacid oxidase 1, liver |
| chr1 | 9648422 | 9649795 | 1374 | * | 12,98712 | -2,737742428 | 0,998238 | -2,74258 | 0,006096 | 0,880247401 | 0,918255 | Intron (ENSMUST0000 | 1 | 9671424 | 9697368 | 25945 | 1 | 76982 | ENSMUST00000088666.4 | -21629 | ENSMUSG00000067879 | Vxn | vexin |
| chr9 | 86690849 | 86692768 | 1920 | * | 19,31886 | -2,727867733 | 0,918919 | -2,96856 | 0,002992 | 0,578742906 | 2,507013 | Promoter (<=1kb) | 9 | 86647986 | 86690794 | 42809 | 2 | 20616 | ENSMUST00000190388.7 | -55 | ENSMUSG00000033419 | Snap91 | synaptosomal-associated protein 91 |
| chr8 | 55112161 | 55113025 | 865 | * | 10,45453 | -2,727088491 | 1,037127 | -2,62946 | 0,008552 | 0,963466134 | 0,738441 | Exon (ENSMUST00000 | 8 | 55102075 | 55107033 | 4959 | 2 | 244484 | ENSMUST00000153074.2 | -5128 | ENSMUSG00000039375 | Wdr17 | WD repeat domain 17 |
| chr14 | 61526048 | 61527742 | 1695 | * | 19,77812 | -2,726485482 | 0,82083 | -3,32162 | 0,000895 | 0,375922902 | 2,386008 | Distal Intergenic | 14 | 61530429 | 61542783 | 12355 | 2 | 67355 | ENSMUST00000190737.2 | 15041 | ENSMUSG00000101505 | 1700109G14Rik | RIKEN cDNA 1700109G14 gene |
| chr12 | 1,17E+08 | 1,17E+08 | 1791 | * | 19,99556 | -2,725631184 | 0,897744 | -3,03609 | 0,002397 | 0,536870492 | 2,507013 | Intron (ENSMUST0000 | 12 | 1,16E+08 | 1,16E+08 | 268 | 2 | 627375 | ENSMUST00000189273.2 | -318340 | | Gm6750 | predicted gene 6750 |
| chr13 | 31260117 | 31261554 | 1438 | * | 13,66211 | -2,724665962 | 1,038842 | -2,62279 | 0,008721 | 0,931442063 | 0,938682 | Exon (ENSMUST00000 | 13 | 31107639 | 31158076 | 50438 | 2 | 66482 | ENSMUST00000222133.2 | -102041 | ENSMUSG00000021357 | Exoc2 | exocyst complex component 2 |
| chr6 | 1,43E+08 | 1,43E+08 | 1028 | * | 11,52756 | -2,721421087 | 1,020552 | -2,66662 | 0,007662 | 0,87765772 | 1,173325 | Intron (ENSMUST0000 | 6 | 1,43E+08 | 1,43E+08 | 21777 | 2 | 74741 | ENSMUST00000204140.2 | -17284 | ENSMUSG00000030279 | C2cd5 | C2 calcium-dependent domain containing 5 |
| chr1 | 1,52E+08 | 1,52E+08 | 1717 | * | 14,55473 | -2,717854401 | 0,978811 | -2,77669 | 0,005492 | 0,802323085 | 1,2763 | Intron (ENSMUST0000 | 1 | 1,52E+08 | 1,52E+08 | 110866 | 1 | 269132 | ENSMUST00000044311.9 | 42574 | ENSMUSG00000032649 | Colgalt2 | collagen beta(1-O)galactosyltransferase 2 |
| chr8 | 57361445 | 57362891 | 1447 | * | 14,42204 | -2,714179946 | 1,034584 | -2,62345 | 0,008704 | 0,870290757 | 1,37683 | Distal Intergenic | 8 | 57313603 | 57341928 | 28326 | 2 | 1,03E+08 | ENSMUST00000209257.2 | -19517 | ENSMUSG00000110000 | Gm33501 | predicted gene, 33501 |
| chr8 | 3529322 | 3530585 | 1264 | * | 13,90857 | -2,714135719 | 0,942768 | -2,8789 | 0,003991 | 0,777172375 | 1,075735 | Intron (ENSMUST0000 | 8 | 3525849 | 3537181 | 11333 | 1 | 76406 | ENSMUST00000161680.2 | 3473 | ENSMUSG00000040340 | Tex45 | testis expressed 45 |
| chr8 | 25885726 | 25888233 | 2508 | * | 31,43505 | -2,711682676 | 0,767956 | -3,53104 | 0,000414 | 0,250313148 | 2,725491 | Promoter (<=1kb) | 8 | 25889036 | 25917452 | 28417 | 1 | 1,05E+08 | ENSMUST00000211730.2 | -803 | ENSMUSG00000110172 | Gm39147 | predicted gene, 39147 |
| chr2 | 1,24E+08 | 1,24E+08 | 1184 | * | 11,63878 | -2,710888604 | 1,019669 | -2,6586 | 0,007847 | 0,918200407 | 0,918255 | Distal Intergenic | 2 | 1,24E+08 | 1,24E+08 | 53385 | 1 | 214968 | ENSMUST00000137172.2 | -414339 | ENSMUSG00000027200 | Sema6d | sema domain, transmembrane domain (TM), and cytoplasmic domain, (semaphorin) 6D |
| chr18 | 14591931 | 14593407 | 1477 | * | 13,76505 | -2,71046783 | 1,023519 | -2,64819 | 0,008092 | 0,90060438 | 1,075735 | Distal Intergenic | 18 | 14446723 | 14464733 | 18011 | 2 | 1,03E+08 | ENSMUST00000234914.2 | -127198 | | Gm29992 | predicted gene, 29992 |
| chr3 | 1,12E+08 | 1,12E+08 | 1000 | * | 10,75618 | -2,708386829 | 1,021086 | -2,65246 | 0,007991 | 0,955124605 | 0,738441 | Distal Intergenic | 3 | 1,12E+08 | 1,12E+08 | 200099 | 1 | 625585 | ENSMUST00000198910.2 | -50792 | ENSMUSG00000104543 | Gm6602 | predicted gene 6602 |
| chr13 | 3415040 | 3416850 | 1811 | * | 17,99765 | -2,707416133 | 0,894593 | -3,02642 | 0,002475 | 0,531203841 | 2,663619 | Distal Intergenic | 13 | 3323289 | 3332998 | 9710 | 1 | 1,05E+08 | ENSMUST00000222272.2 | 91751 | | Gm40653 | predicted gene, 40653 |
| chr3 | 1,44E+08 | 1,44E+08 | 1349 | * | 14,73852 | -2,704749762 | 1,043439 | -2,59215 | 0,009538 | 0,809369408 | 2,141612 | Distal Intergenic | 3 | 1,44E+08 | 1,44E+08 | 76382 | 2 | 1,19E+08 | ENSMUST00000196668.2 | -86883 | | Gm42707 | predicted gene 42707 |
| chr1 | 93553427 | 93554274 | 848 | * | 9,818654 | -2,704734067 | 1,033806 | -2,61629 | 0,008889 | 0,911821655 | 1,086118 | Promoter (<=1kb) | 1 | 93551118 | 93553510 | 2393 | 2 | 59041 | ENSMUST00000125778.2 | 0 | ENSMUSG00000026277 | Stk25 | serine/threonine kinase 25 (yeast) |
| chr9 | 1,04E+08 | 1,04E+08 | 2299 | * | 17,97435 | -2,704546188 | 0,984768 | -2,74638 | 0,006026 | 0,686838043 | 2,663619 | Distal Intergenic | 9 | 1,04E+08 | 1,04E+08 | 23233 | 1 | 1,03E+08 | ENSMUST00000191599.3 | -10454 | ENSMUSG00000099599 | Gm28548 | predicted gene 28548 |
| chr1 | 16045089 | 16046700 | 1612 | * | 14,85992 | -2,703609673 | 0,981004 | -2,75596 | 0,005852 | 0,814686254 | 1,2763 | Distal Intergenic | 1 | 15924086 | 15962946 | 38861 | 2 | 226866 | ENSMUST00000040695.5 | -82143 | ENSMUSG00000032719 | Sbspon | somatomedin B and thrombospondin, type 1 domain containing |
| chr14 | 67531101 | 67533186 | 2086 | * | 13,47302 | -2,703546865 | 0,948096 | -2,85155 | 0,004351 | 0,857011814 | 0,738441 | Intron (ENSMUST0000 | 14 | 67542337 | 67552160 | 9824 | 2 | 628416 | ENSMUST00000100448.3 | 18974 | ENSMUSG00000075549 | Gm6878 | predicted gene 6878 |
| chr5 | 27102879 | 27104653 | 1775 | * | 16,54779 | -2,703511112 | 1,042785 | -2,59259 | 0,009526 | 0,769939394 | 2,663619 | Intron (ENSMUST0000 | 5 | 27109693 | 27932503 | 822811 | 1 | 13483 | ENSMUST00000120555.8 | -5040 | ENSMUSG00000061576 | Dpp6 | dipeptidylpeptidase 6 |
| chr14 | 12932329 | 12934207 | 1879 | * | 18,1767 | -2,702835817 | 0,840692 | -3,21501 | 0,001304 | 0,457049514 | 2,154504 | Distal Intergenic | 14 | 12583005 | 12585578 | 2574 | 2 | 1E+08 | ENSMUST00000224435.2 | -346751 | ENSMUSG00000089686 | Mnd1-ps | Mnd1 retrotransposed pseudogene |
| chr4 | 55545946 | 55548722 | 2777 | * | 17,82075 | -2,701570451 | 0,963692 | -2,80336 | 0,005057 | 0,664099998 | 2,663619 | Distal Intergenic | 4 | 55532736 | 55541745 | 9010 | 1 | 1,03E+08 | ENSMUST00000244104.1 | 13210 | | Gm34343 | predicted gene, 34343 |
| chr7 | 1,31E+08 | 1,31E+08 | 1741 | * | 18,26596 | -2,701421582 | 0,893368 | -3,02386 | 0,002496 | 0,549632826 | 2,386008 | Exon (ENSMUST00000 | 7 | 1,31E+08 | 1,31E+08 | 132031 | 1 | 71395 | ENSMUST00000208921.4 | 12903 | ENSMUSG00000006204 | Cdcp3 | CUB domain containing protein 3 |
| chr4 | 63708338 | 63710039 | 1702 | * | 12,89534 | -2,700065472 | 1,012833 | -2,66585 | 0,007679 | 0,914972668 | 0,918255 | Distal Intergenic | 4 | 63645321 | 63663350 | 18030 | 2 | 326623 | ENSMUST00000062246.8 | -44988 | ENSMUSG00000050395 | Tnfsf15 | tumor necrosis factor (ligand) superfamily, member 15 |
| chr7 | 92336057 | 92338043 | 1987 | * | 18,14221 | -2,699400107 | 0,965526 | -2,79578 | 0,005177 | 0,699034389 | 2,154504 | Distal Intergenic | 7 | 92292751 | 92319142 | 26392 | 2 | 74737 | ENSMUST00000119954.9 | -16915 | ENSMUSG00000041328 | Pcf11 | PCF11 cleavage and polyadenylation factor subunit |
| chr16 | 40645675 | 40647257 | 1583 | * | 11,32418 | -2,699044353 | 1,032541 | -2,61398 | 0,008949 | 0,901463351 | 1,173325 | Intron (ENSMUST0000 | 16 | 41353220 | 41966576 | 613357 | 1 | 268890 | ENSMUST00000078873.11 | -705963 | ENSMUSG00000061080 | Lsamp | limbic system-associated membrane protein |
| chr9 | 26437887 | 26439227 | 1341 | * | 14,20247 | -2,698435251 | 1,009864 | -2,67208 | 0,007538 | 0,88871498 | 1,075735 | Intron (ENSMUST0000 | 9 | 26430781 | 26461249 | 30469 | 2 | 1,03E+08 | ENSMUST00000216806.2 | 22022 | | Gm30313 | predicted gene, 30313 |
| chr16 | 54707585 | 54709436 | 1852 | * | 12,13898 | -2,698357978 | 0,975949 | -2,76486 | 0,005695 | 0,901762077 | 0,744375 | Distal Intergenic | 16 | 54792794 | 54792900 | 107 | 1 | 1,15E+08 | ENSMUST00000082615.3 | -83358 | ENSMUSG00000064549 | Gm22977 | predicted gene, 22977 |
| chr11 | 17346186 | 17347770 | 1585 | * | 14,7069 | -2,698205594 | 0,982968 | -2,74496 | 0,006052 | 0,806043807 | 1,37683 | Distal Intergenic | 11 | 17213621 | 17216560 | 2940 | 1 | 57316 | ENSMUST00000130330.2 | 132565 | ENSMUSG00000000581 | C1d | C1D nuclear receptor co-repressor |
| chr11 | 18146213 | 18148609 | 2397 | * | 17,38324 | -2,697342563 | 0,960851 | -2,80724 | 0,004997 | 0,692527675 | 2,154504 | Distal Intergenic | 11 | 18045986 | 18053384 | 7399 | 1 | 327872 | ENSMUST00000150694.2 | 100227 | ENSMUSG00000085453 | Gm12018 | predicted gene 12018 |
| chr10 | 86815932 | 86817772 | 1841 | * | 17,90679 | -2,697035499 | 0,827851 | -3,25788 | 0,001122 | 0,416497853 | 2,386008 | Exon (ENSMUST00000 | 10 | 86808080 | 86863879 | 55800 | 1 | 1,15E+08 | ENSMUST00000162837.2 | 7852 | | Gm16271 | predicted gene 16271 |
| chr10 | 23228727 | 23230639 | 1913 | * | 16,18946 | -2,694154952 | 0,96116 | -2,80302 | 0,005063 | 0,68259716 | 2,386008 | Promoter (2-3kb) | 10 | 22980662 | 23226684 | 246023 | 2 | 14051 | ENSMUST00000220299.2 | -2043 | ENSMUSG00000010461 | Eya4 | EYA transcriptional coactivator and phosphatase 4 |
| chr15 | 94117716 | 94118955 | 1240 | * | 19,65177 | -2,693015691 | 0,899259 | -2,99471 | 0,002747 | 0,570876293 | 2,386008 | Distal Intergenic | 15 | 94168044 | 94302139 | 134096 | 2 | 223838 | ENSMUST00000035342.11 | 183184 | ENSMUSG00000022449 | Adamts20 | a disintegrin-like and metallopeptidase (reprolysin type) with thrombospondin type 1 motif, 20 |
| chr10 | 48338900 | 48341674 | 2775 | * | 17,50363 | -2,691221735 | 0,845391 | -3,18341 | 0,001456 | 0,455547114 | 2,423594 | Distal Intergenic | 10 | 48495408 | 48495491 | 84 | 2 | 1,15E+08 | ENSMUST00000083696.4 | 153817 | | Gm24256 | predicted gene, 24256 |
| chr3 | 32184247 | 32185889 | 1643 | * | 16,38361 | -2,690884343 | 0,956181 | -2,8142 | 0,00489 | 0,659240897 | 2,663619 | Intron (ENSMUST0000 | 3 | 32210263 | 32252505 | 42243 | 1 | 72413 | ENSMUST00000194796.2 | -24374 | ENSMUSG00000037610 | Kcnmb2 | potassium large conductance calcium-activated channel, subfamily M, beta member 2 |
| chr7 | 18324097 | 18325274 | 1178 | * | 11,29502 | -2,690784619 | 1,012147 | -2,65849 | 0,007849 | 0,918200407 | 0,918255 | Distal Intergenic | 7 | 18290439 | 18301230 | 10792 | 2 | 545925 | ENSMUST00000094794.5 | -22867 | ENSMUSG00000070797 | Psg27 | pregnancy-specific glycoprotein 27 |
| chr12 | 74478990 | 74480180 | 1191 | * | 12,82557 | -2,689355316 | 1,006021 | -2,67326 | 0,007512 | 0,911626845 | 0,918255 | Distal Intergenic | 12 | 74360975 | 74363936 | 2962 | 2 | 1,03E+08 | ENSMUST00000223381.2 | -115054 | | Gm34552 | predicted gene, 34552 |
| chr1 | 1,57E+08 | 1,57E+08 | 1687 | * | 15,00457 | -2,688095049 | 1,03923 | -2,58662 | 0,009692 | 0,900910297 | 1,2763 | Intron (ENSMUST0000 | 1 | 1,57E+08 | 1,57E+08 | 80511 | 2 | 226525 | ENSMUST00000134543.8 | -24821 | ENSMUSG00000070565 | Rasal2 | RAS protein activator like 2 |
| chr16 | 75681781 | 75683032 | 1252 | * | 13,00612 | -2,687790507 | 0,977998 | -2,74826 | 0,005991 | 0,819350294 | 1,2763 | Intron (ENSMUST0000 | 16 | 75685140 | 75706167 | 21028 | 2 | 67742 | ENSMUST00000150248.2 | 23135 | ENSMUSG00000022876 | Samsn1 | SAM domain, SH3 domain and nuclear localization signals, 1 |
| chr9 | 1,17E+08 | 1,17E+08 | 858 | * | 10,35182 | -2,687781828 | 1,014591 | -2,64913 | 0,00807 | 0,886332554 | 1,173325 | Intron (ENSMUST0000 | 9 | 1,17E+08 | 1,17E+08 | 308475 | 2 | 207181 | ENSMUST00000084824.7 | 38436 | ENSMUSG00000039607 | Rbms3 | RNA binding motif, single stranded interacting protein |
| chr13 | 45206535 | 45208803 | 2269 | * | 18,51152 | -2,684130647 | 0,866502 | -3,09766 | 0,001951 | 0,490274805 | 2,663619 | Distal Intergenic | 13 | 45179728 | 45179831 | 104 | 2 | 1,15E+08 | ENSMUST00000083722.4 | -26704 | ENSMUSG00000065656 | Gm23104 | predicted gene, 23104 |
| chr1 | 1,72E+08 | 1,72E+08 | 1590 | * | 18,02374 | -2,681657488 | 0,968652 | -2,76844 | 0,005633 | 0,695892636 | 2,386008 | Exon (ENSMUST00000 | 1 | 1,72E+08 | 1,72E+08 | 6276 | 1 | 71870 | ENSMUST00000159395.2 | 3448 | ENSMUSG00000026546 | Cfap45 | cilia and flagella associated protein 45 |
| chr7 | 33688811 | 33691388 | 2578 | * | 23,38361 | -2,679779226 | 0,72321 | -3,70539 | 0,000211 | 0,167380886 | 2,858511 | Distal Intergenic | 7 | 33711395 | 33713292 | 1898 | 2 | 233099 | ENSMUST00000185399.2 | 21904 | ENSMUSG00000066584 | Scgb2b27 | secretoglobin, family 2B, member 27 |
| chr17 | 78263016 | 78263906 | 891 | * | 12,57908 | -2,67962473 | 1,028876 | -2,60442 | 0,009203 | 0,905322447 | 1,173325 | Distal Intergenic | 17 | 78506685 | 78507454 | 770 | 2 | 1,03E+08 | ENSMUST00000234450.2 | 243548 | ENSMUSG00000117648 | Gm32282 | predicted gene, 32282 |
| chr17 | 60083202 | 60084687 | 1486 | * | 14,20488 | -2,67763873 | 0,952849 | -2,81014 | 0,004952 | 0,815986445 | 1,075735 | Distal Intergenic | 17 | 59876554 | 59876660 | 107 | 2 | 1,15E+08 | ENSMUST00000104374.3 | -206542 | ENSMUSG00000077562 | Gm23203 | predicted gene, 23203 |
| chr1 | 1,06E+08 | 1,06E+08 | 1490 | * | 13,85004 | -2,676927586 | 0,974667 | -2,7465 | 0,006023 | 0,877892722 | 0,918255 | Intron (ENSMUST0000 | 1 | 1,06E+08 | 1,06E+08 | 5998 | 1 | 98432 | ENSMUST00000135810.3 | -21250 | ENSMUSG00000044340 | Phlpp1 | PH domain and leucine rich repeat protein phosphatase 1 |
| chr3 | 93250135 | 93251966 | 1832 | * | 11,53048 | -2,676109308 | 0,985541 | -2,71537 | 0,00662 | 0,923399738 | 0,744375 | Intron (ENSMUST0000 | 3 | 93227056 | 93240877 | 13822 | 1 | 68723 | ENSMUST00000093774.4 | 23079 | ENSMUSG00000041991 | Hrnr | hornerin |
| chr9 | 1,17E+08 | 1,17E+08 | 1899 | * | 19,22811 | -2,675523964 | 0,818839 | -3,26746 | 0,001085 | 0,393356861 | 2,663619 | Intron (ENSMUST0000 | 9 | 1,16E+08 | 1,17E+08 | 245225 | 2 | 207181 | ENSMUST00000173429.8 | 67038 | ENSMUSG00000039607 | Rbms3 | RNA binding motif, single stranded interacting protein |
| chr16 | 71690554 | 71692548 | 1995 | * | 21,86442 | -2,67541799 | 0,750456 | -3,56506 | 0,000364 | 0,225888909 | 2,880017 | Distal Intergenic | 16 | 71460676 | 71460806 | 131 | 1 | 1,15E+08 | ENSMUST00000083990.3 | 229878 | ENSMUSG00000065924 | Gm22797 | predicted gene, 22797 |
| chr9 | 1,22E+08 | 1,22E+08 | 2121 | * | 16,55799 | -2,671876632 | 1,011161 | -2,64239 | 0,008232 | 0,7843572 | 2,141612 | Intron (ENSMUST0000 | 9 | 1,22E+08 | 1,22E+08 | 11727 | 2 | 102566 | ENSMUST00000216738.2 | -31211 | ENSMUSG00000037949 | Ano10 | anoctamin 10 |
| chr13 | 44766050 | 44768069 | 2020 | * | 18,57854 | -2,671580663 | 0,990157 | -2,69814 | 0,006973 | 0,749931768 | 2,154504 | Distal Intergenic | 13 | 44690785 | 44722144 | 31360 | 2 | 1,03E+08 | ENSMUST00000222558.2 | -43906 | | Gm34276 | predicted gene, 34276 |
| chr13 | 1,1E+08 | 1,1E+08 | 1546 | * | 13,84191 | -2,669501314 | 0,93478 | -2,85575 | 0,004293 | 0,818408612 | 0,918255 | Intron (ENSMUST0000 | 13 | 1,1E+08 | 1,1E+08 | 369940 | 1 | 238871 | ENSMUST00000079975.11 | -6940 | ENSMUSG00000021699 | Pde4d | phosphodiesterase 4D, cAMP specific |
| chr3 | 1,44E+08 | 1,44E+08 | 2048 | * | 24,45083 | -2,669356665 | 0,733279 | -3,6403 | 0,000272 | 0,190625259 | 2,951062 | Promoter (1-2kb) | 3 | 1,44E+08 | 1,44E+08 | 76382 | 2 | 1,19E+08 | ENSMUST00000196668.2 | 1737 | | Gm42707 | predicted gene 42707 |
| chr7 | 54081376 | 54083312 | 1937 | * | 17,36612 | -2,668653622 | 0,930746 | -2,86722 | 0,004141 | 0,667992748 | 2,154504 | Distal Intergenic | 7 | 54485246 | 54899837 | 414592 | 1 | 233271 | ENSMUST00000206923.2 | -401934 | ENSMUSG00000063297 | Luzp2 | leucine zipper protein 2 |
| chr16 | 90674925 | 90676439 | 1515 | * | 18,15478 | -2,668506546 | 0,841998 | -3,16925 | 0,001528 | 0,457782113 | 2,507013 | Intron (ENSMUST0000 | 16 | 90633484 | 90647851 | 14368 | 2 | 1,03E+08 | ENSMUST00000232254.2 | -27074 | | Gm36363 | predicted gene, 36363 |
| chr19 | 40587888 | 40589461 | 1574 | * | 14,71991 | -2,662483384 | 0,994064 | -2,67838 | 0,007398 | 0,763224986 | 2,141612 | Intron (ENSMUST0000 | 19 | 40577879 | 40583306 | 5428 | 1 | 1,03E+08 | ENSMUST00000181109.3 | 10009 | | Gm27042 | predicted gene, 27042 |
| chr9 | 1,01E+08 | 1,01E+08 | 1807 | * | 15,47722 | -2,662364965 | 0,904005 | -2,94508 | 0,003229 | 0,711199341 | 1,2763 | Distal Intergenic | 9 | 1,01E+08 | 1,01E+08 | 853 | 2 | 382099 | ENSMUST00000075243.8 | -34658 | ENSMUSG00000061486 | Gm5161 | predicted pseudogene 5161 |
| chr13 | 38516342 | 38517640 | 1299 | * | 14,64357 | -2,660719679 | 0,909481 | -2,92554 | 0,003439 | 0,729588924 | 1,214904 | Exon (ENSMUST00000 | 13 | 38529083 | 38684278 | 155196 | 1 | 12161 | ENSMUST00000171970.3 | -11443 | ENSMUSG00000039004 | Bmp6 | bone morphogenetic protein 6 |
| chr17 | 55749174 | 55751300 | 2127 | * | 16,34415 | -2,660715781 | 0,896951 | -2,9664 | 0,003013 | 0,611385361 | 2,141612 | Promoter (1-2kb) | 17 | 55752383 | 55790033 | 37651 | 1 | 240119 | ENSMUST00000133899.2 | -1083 | ENSMUSG00000024172 | St6gal2 | beta galactoside alpha 2,6 sialyltransferase 2 |
| chr6 | 1,05E+08 | 1,05E+08 | 1451 | * | 18,49934 | -2,654688373 | 0,953885 | -2,78303 | 0,005385 | 0,683210259 | 2,507013 | Intron (ENSMUST0000 | 6 | 1,04E+08 | 1,05E+08 | 370154 | 1 | 53870 | ENSMUST00000162872.2 | 33274 | ENSMUSG00000030092 | Cntn6 | contactin 6 |
| chr5 | 26761157 | 26762530 | 1374 | * | 13,55242 | -2,653692331 | 1,02791 | -2,58164 | 0,009833 | 0,954456537 | 0,918255 | Distal Intergenic | 5 | 26665070 | 26698312 | 33243 | 2 | 75942 | ENSMUST00000197091.5 | -62845 | ENSMUSG00000029044 | 4930584F24Rik | RIKEN cDNA 4930584F24 gene |
| chr3 | 98988301 | 98990142 | 1842 | * | 19,41718 | -2,652265994 | 0,844805 | -3,1395 | 0,001692 | 0,476080922 | 2,507013 | Distal Intergenic | 3 | 99020471 | 99020599 | 129 | 2 | 1,15E+08 | ENSMUST00000083775.3 | 30457 | ENSMUSG00000065709 | Gm22341 | predicted gene, 22341 |
| chr1 | 1,39E+08 | 1,39E+08 | 1767 | * | 16,10145 | -2,651161415 | 0,969469 | -2,73465 | 0,006245 | 0,715249603 | 2,386008 | Intron (ENSMUST0000 | 1 | 1,38E+08 | 1,39E+08 | 136305 | 2 | 59125 | ENSMUST00000187407.7 | -21895 | ENSMUSG00000026393 | Nek7 | NIMA (never in mitosis gene a)-related expressed kinase 7 |
| chr19 | 43416392 | 43418425 | 2034 | * | 12,95133 | -2,65109864 | 1,006854 | -2,63305 | 0,008462 | 0,891774732 | 1,173325 | Distal Intergenic | 19 | 43428875 | 43483480 | 54606 | 1 | 83674 | ENSMUST00000223787.2 | -10450 | ENSMUSG00000025189 | Cnnm1 | cyclin M1 |
| chr19 | 51776518 | 51777858 | 1341 | * | 12,58595 | -2,650939503 | 1,015801 | -2,6097 | 0,009062 | 0,972947889 | 0,738441 | Distal Intergenic | 19 | 52252735 | 52253914 | 1180 | 1 | 16333 | ENSMUST00000039652.6 | -474877 | ENSMUSG00000035804 | Ins1 | insulin I |
| chr1 | 17190743 | 17192687 | 1945 | * | 18,83825 | -2,647972921 | 0,808076 | -3,27688 | 0,00105 | 0,406160411 | 2,386008 | Promoter (2-3kb) | 1 | 17194971 | 17199180 | 4210 | 1 | 1,08E+08 | ENSMUST00000188872.2 | -2284 | | Gm46085 | predicted gene, 46085 |
| chr8 | 77550892 | 77552232 | 1341 | * | 15,63863 | -2,6474586 | 1,025899 | -2,58062 | 0,009862 | 0,910268063 | 1,214904 | Intron (ENSMUST0000 | 8 | 77626070 | 77635042 | 8973 | 1 | 110784 | ENSMUST00000143284.2 | -73838 | ENSMUSG00000031618 | Nr3c2 | nuclear receptor subfamily 3, group C, member 2 |
| chr1 | 1,78E+08 | 1,78E+08 | 1514 | * | 13,15771 | -2,647013429 | 0,962407 | -2,75041 | 0,005952 | 0,875820727 | 0,918255 | Intron (ENSMUST0000 | 1 | 1,78E+08 | 1,78E+08 | 61492 | 1 | 78825 | ENSMUST00000159284.2 | 17207 | ENSMUSG00000026502 | Desi2 | desumoylating isopeptidase 2 |
| chrX | 1,55E+08 | 1,55E+08 | 1440 | * | 15,67929 | -2,644569896 | 0,91143 | -2,90156 | 0,003713 | 0,649217039 | 2,141612 | Intron (ENSMUST0000 | 20 | 1,54E+08 | 1,55E+08 | 721113 | 1 | 1,03E+08 | ENSMUST00000112542.2 | 104845 | | Gm15155 | predicted gene 15155 |
| chr1 | 1,92E+08 | 1,92E+08 | 1688 | * | 13,62611 | -2,64394457 | 0,967145 | -2,73376 | 0,006262 | 0,916652644 | 0,738441 | Intron (ENSMUST0000 | 1 | 1,92E+08 | 1,92E+08 | 319386 | 1 | 16510 | ENSMUST00000110844.3 | 87161 | ENSMUSG00000058248 | Kcnh1 | potassium voltage-gated channel, subfamily H (eag-related), member 1 |
| chr15 | 54581998 | 54583731 | 1734 | * | 19,97722 | -2,643664651 | 0,835604 | -3,16378 | 0,001557 | 0,451585113 | 2,663619 | Distal Intergenic | 15 | 54609098 | 54617435 | 8338 | 1 | 18133 | ENSMUST00000050027.9 | -25367 | ENSMUSG00000037362 | Ccn3 | cellular communication network factor 3 |
| chr7 | 47458070 | 47460797 | 2728 | * | 26,73485 | -2,641906683 | 0,686168 | -3,85023 | 0,000118 | 0,1203567 | 2,725491 | Exon (ENSMUST00000 | 7 | 47630585 | 47631843 | 1259 | 2 | 235854 | ENSMUST00000087092.4 | 171046 | ENSMUSG00000067173 | Mrgpra4 | MAS-related GPR, member A4 |
| chr6 | 30760058 | 30761317 | 1260 | * | 20,28936 | -2,641589076 | 0,917299 | -2,87975 | 0,00398 | 0,654833721 | 2,221611 | Intron (ENSMUST0000 | 6 | 30742088 | 30743149 | 1062 | 1 | 17294 | ENSMUST00000149496.2 | 17970 | ENSMUSG00000051855 | Mest | mesoderm specific transcript |
| chr13 | 88173035 | 88175890 | 2856 | * | 19,37645 | -2,640633153 | 0,890958 | -2,96381 | 0,003039 | 0,568612328 | 2,663619 | Distal Intergenic | 13 | 88669309 | 88726287 | 56979 | 1 | 1,03E+08 | ENSMUST00000224910.2 | -493419 | ENSMUSG00000114273 | Gm34585 | predicted gene, 34585 |
| chr7 | 27813615 | 27815580 | 1966 | * | 18,45255 | -2,640218193 | 0,873086 | -3,02401 | 0,002495 | 0,549632826 | 2,386008 | Promoter (<=1kb) | 7 | 27815039 | 27820286 | 5248 | 1 | 215384 | ENSMUST00000131090.2 | 0 | ENSMUSG00000047730 | Fcgbp | Fc fragment of IgG binding protein |
| chr12 | 1,15E+08 | 1,15E+08 | 1276 | * | 21,49387 | -2,639286431 | 0,80245 | -3,28903 | 0,001005 | 0,355694809 | 2,951933 | Distal Intergenic | 12 | 1,15E+08 | 1,15E+08 | 25225 | 1 | 1,03E+08 | ENSMUST00000195683.2 | 187033 | ENSMUSG00000103168 | Gm30948 | predicted gene, 30948 |
| chr9 | 23071300 | 23073225 | 1926 | * | 18,6475 | -2,63829122 | 0,914966 | -2,88348 | 0,003933 | 0,639383182 | 2,423594 | Distal Intergenic | 9 | 23134372 | 23396496 | 262125 | 1 | 73230 | ENSMUST00000071982.7 | -61147 | ENSMUSG00000031963 | Bmper | BMP-binding endothelial regulator |
| chr11 | 65030729 | 65031936 | 1208 | * | 11,91 | -2,637682981 | 1,015665 | -2,597 | 0,009404 | 0,976995956 | 0,738441 | Intron (ENSMUST0000 | 11 | 65042736 | 65050717 | 7982 | 2 | 1,15E+08 | ENSMUST00000108696.7 | 18781 | | 1700086D15Rik | RIKEN cDNA 1700086D15 gene |
| chr13 | 44412437 | 44413713 | 1277 | * | 14,08294 | -2,635866081 | 0,922149 | -2,85839 | 0,004258 | 0,789037101 | 1,075735 | Intron (ENSMUST0000 | 13 | 44402936 | 44406146 | 3211 | 2 | 432754 | ENSMUST00000222396.2 | -6291 | | Gm29676 | predicted gene, 29676 |
| chr7 | 13083497 | 13085399 | 1903 | * | 15,363 | -2,634053084 | 0,859982 | -3,06292 | 0,002192 | 0,655645354 | 1,214904 | Intron (ENSMUST0000 | 7 | 13063243 | 13094597 | 31355 | 1 | 232889 | ENSMUST00000043612.10 | 20254 | ENSMUSG00000033847 | Pla2g4c | phospholipase A2, group IVC (cytosolic, calcium-independent) |
| chr5 | 29410037 | 29412329 | 2293 | * | 17,67145 | -2,631203091 | 0,899777 | -2,92428 | 0,003453 | 0,607561104 | 2,507013 | 3' UTR | 5 | 29403669 | 29430299 | 26631 | 1 | 56874 | ENSMUST00000162975.2 | 6368 | ENSMUSG00000029130 | Rnf32 | ring finger protein 32 |
| chr1 | 1,54E+08 | 1,54E+08 | 2050 | * | 17,30137 | -2,631086475 | 0,929899 | -2,82943 | 0,004663 | 0,683761897 | 2,154504 | Exon (ENSMUST00000 | 1 | 1,54E+08 | 1,54E+08 | 16590 | 2 | 329271 | ENSMUST00000190663.2 | 14340 | ENSMUSG00000100334 | C230024C17Rik | RIKEN cDNA C230024C17 gene |
| chr1 | 84148313 | 84150088 | 1776 | * | 16,68941 | -2,630583342 | 1,001072 | -2,62777 | 0,008595 | 0,791242424 | 2,141612 | Intron (ENSMUST0000 | 1 | 84196570 | 84196682 | 113 | 2 | 1,02E+08 | ENSMUST00000183596.3 | 46594 | ENSMUSG00000098851 | Mir6353 | microRNA 6353 |
| chr10 | 4027246 | 4029246 | 2001 | * | 21,53569 | -2,625934043 | 0,781396 | -3,36057 | 0,000778 | 0,318448436 | 2,951933 | Intron (ENSMUST0000 | 10 | 4054313 | 4054469 | 157 | 1 | 1,15E+08 | ENSMUST00000083858.3 | -25067 | ENSMUSG00000065792 | Gm25515 | predicted gene, 25515 |
| chr10 | 1,09E+08 | 1,09E+08 | 1953 | * | 15,92828 | -2,625524773 | 0,795905 | -3,29879 | 0,000971 | 0,496560937 | 1,2763 | Distal Intergenic | 10 | 1,1E+08 | 1,1E+08 | 9500 | 2 | 260315 | ENSMUST00000161223.2 | 60658 | ENSMUSG00000020181 | Nav3 | neuron navigator 3 |
| chr15 | 7293361 | 7295256 | 1896 | * | 14,49552 | -2,625366384 | 0,999375 | -2,62701 | 0,008614 | 0,890237282 | 1,214904 | Intron (ENSMUST0000 | 15 | 7282992 | 7284249 | 1258 | 2 | 268780 | ENSMUST00000162105.2 | -9112 | ENSMUSG00000042961 | Egflam | EGF-like, fibronectin type III and laminin G domains |
| chr5 | 1,01E+08 | 1,01E+08 | 1933 | * | 16,15662 | -2,621746033 | 0,841252 | -3,11648 | 0,00183 | 0,61616728 | 1,2763 | Distal Intergenic | 5 | 1,01E+08 | 1,01E+08 | 18521 | 2 | 231507 | ENSMUST00000031264.12 | -22422 | ENSMUSG00000029322 | Plac8 | placenta-specific 8 |
| chr11 | 1,09E+08 | 1,09E+08 | 1220 | * | 16,16241 | -2,619308374 | 0,880824 | -2,9737 | 0,002942 | 0,691182187 | 1,2763 | Intron (ENSMUST0000 | 11 | 1,09E+08 | 1,09E+08 | 29577 | 1 | 74008 | ENSMUST00000106696.2 | 7647 | ENSMUSG00000020604 | Arsg | arylsulfatase G |
| chr10 | 31670028 | 31672119 | 2092 | * | 22,33568 | -2,618965324 | 0,794056 | -3,29821 | 0,000973 | 0,327363151 | 3,471678 | Intron (ENSMUST0000 | 10 | 31369189 | 31485180 | 115992 | 2 | 268291 | ENSMUST00000081989.8 | -184848 | ENSMUSG00000063760 | Rnf217 | ring finger protein 217 |
| chr7 | 25324906 | 25326891 | 1986 | * | 18,17644 | -2,61829768 | 0,978183 | -2,6767 | 0,007435 | 0,736412045 | 2,507013 | Promoter (<=1kb) | 7 | 25326079 | 25327850 | 1772 | 1 | 232984 | ENSMUST00000205281.2 | 0 | ENSMUSG00000059479 | B3gnt8 | UDP-GlcNAc:betaGal beta-1,3-N-acetylglucosaminyltransferase 8 |
| chr7 | 96688752 | 96689759 | 1008 | * | 14,11971 | -2,61792261 | 0,989822 | -2,64484 | 0,008173 | 0,956661559 | 0,738441 | Promoter (<=1kb) | 7 | 96689211 | 96712179 | 22969 | 1 | 244141 | ENSMUST00000150681.3 | 0 | ENSMUSG00000018995 | Nars2 | asparaginyl-tRNA synthetase 2 (mitochondrial)(putative) |
| chr4 | 31840384 | 31841455 | 1072 | * | 13,22602 | -2,61759876 | 1,005825 | -2,60244 | 0,009256 | 0,892968148 | 1,2763 | Intron (ENSMUST0000 | 4 | 31825469 | 31877549 | 52081 | 2 | 1,03E+08 | ENSMUST00000243650.1 | 36094 | | Gm32412 | predicted gene, 32412 |
| chr15 | 33981812 | 33983644 | 1833 | * | 17,43429 | -2,617246913 | 0,90269 | -2,89939 | 0,003739 | 0,627785088 | 2,423594 | Distal Intergenic | 15 | 34022141 | 34037926 | 15786 | 2 | 1,05E+08 | ENSMUST00000228182.2 | 54282 | ENSMUSG00000115428 | Gm41290 | predicted gene, 41290 |
| chr17 | 72408439 | 72409463 | 1025 | * | 14,00664 | -2,616609717 | 0,961671 | -2,7209 | 0,00651 | 0,923078468 | 0,738441 | Intron (ENSMUST0000 | 17 | 72163281 | 72171268 | 7988 | 1 | 78785 | ENSMUST00000230290.2 | 245158 | ENSMUSG00000024059 | Clip4 | CAP-GLY domain containing linker protein family, member 4 |

| | | | | | | | | | | | | | | | | | | | | | | | |
|---|---|---|---|---|---|---|---|---|---|---|---|---|---|---|---|---|---|---|---|---|---|---|---|
| chr12 | 68179067 | 68179993 | 927 | * | 13,2351 | -2,616095098 | 0,956056 | -2,73634 | 0,006213 | 0,854050055 | 1,075735 | Distal Intergenic | 12 | 67659070 | 67659255 | 186 | 1 | 1,15E+08 | ENSMUST00000122499.3 | 519997 | ENSMUSG00000084448 | Gm26256 | predicted gene, 26256 |
| chr1 | 33212110 | 33213391 | 1282 | * | 13,57343 | -2,614508961 | 0,994733 | -2,62835 | 0,00858 | 0,928298336 | 0,938682 | Distal Intergenic | 1 | 33231074 | 33231183 | 110 | 1 | 1,15E+08 | ENSMUST00000158125.3 | -17683 | ENSMUSG00000088750 | Gm24901 | predicted gene, 24901 |
| chr10 | 1,2E+08 | 1,2E+08 | 1641 | * | 14,0633 | -2,614003151 | 0,971061 | -2,6919 | 0,007105 | 0,850457798 | 1,2763 | Distal Intergenic | 10 | 1,2E+08 | 1,2E+08 | 115195 | 2 | 15364 | ENSMUST00000159699.2 | -63753 | ENSMUSG00000056758 | Hmga2 | high mobility group AT-hook 2 |
| chr7 | 1,02E+08 | 1,02E+08 | 3263 | * | 33,07294 | -2,613403438 | 0,639054 | -4,08949 | 4,32E-05 | 0,096492906 | 1,405709 | Intron (ENSMUST00000 | 7 | 1,02E+08 | 1,02E+08 | 2276 | 1 | 20866 | ENSMUST00000210834.2 | -59356 | ENSMUSG00000030987 | Stim1 | stromal interaction molecule 1 |
| chr4 | 51034097 | 51036087 | 1991 | * | 12,0277 | -2,613298163 | 1,004102 | -2,60262 | 0,009251 | 0,974006322 | 0,744375 | Distal Intergenic | 4 | 51216678 | 51229928 | 13251 | 1 | 74914 | ENSMUST00000166749.2 | -180591 | ENSMUSG00000039555 | Cylc2 | cylicin, basic protein of sperm head cytoskeleton 2 |
| chr18 | 5790167 | 5792434 | 2268 | * | 15,70449 | -2,611306367 | 0,904196 | -2,88799 | 0,003877 | 0,618136403 | 2,663619 | Distal Intergenic | 18 | 5731081 | 5772804 | 41724 | 1 | 21417 | ENSMUST00000177030.2 | 59086 | ENSMUSG00000024238 | Zeb1 | zinc finger E-box binding homeobox 1 |
| chr13 | 39379769 | 39381163 | 1395 | * | 13,17298 | -2,610597007 | 1,006649 | -2,59335 | 0,009504 | 0,910106437 | 1,173325 | Distal Intergenic | 13 | 39116112 | 39144851 | 28740 | 2 | 108652 | ENSMUST00000021870.12 | -234918 | ENSMUSG00000021432 | Slc35b3 | solute carrier family 35, member B3 |
| chr12 | 55694624 | 55696040 | 1417 | * | 14,37432 | -2,610119644 | 0,979891 | -2,66368 | 0,007729 | 0,915421656 | 0,918255 | Intron (ENSMUST00000 | 12 | 55650645 | 55689233 | 38589 | 2 | 56784 | ENSMUST00000219451.2 | -5391 | ENSMUSG00000021027 | Ralgapa1 | Ral GTPase activating protein, alpha subunit 1 |
| chr7 | 1,33E+08 | 1,33E+08 | 1660 | * | 18,03519 | -2,610005084 | 0,844402 | -3,09095 | 0,001995 | 0,531203841 | 2,154504 | Distal Intergenic | 7 | 1,33E+08 | 1,33E+08 | 14207 | 1 | 74970 | ENSMUST00000209850.2 | 224932 | ENSMUSG00000109816 | 4930483O08Rik | RIKEN cDNA 4930483O08 gene |
| chr2 | 75018612 | 75020547 | 1936 | * | 13,66319 | -2,60939005 | 0,987762 | -2,64172 | 0,008249 | 0,922828293 | 0,938682 | Promoter (1-2kb) | 2 | 75021417 | 75022166 | 750 | 2 | 1,05E+08 | ENSMUST00000118499.2 | 1619 | ENSMUSG00000081406 | Rps6-ps4 | ribosomal protein S6, pseudogene 4 |
| chr15 | 65007265 | 65008337 | 1073 | * | 14,01901 | -2,608670922 | 0,956844 | -2,72633 | 0,006404 | 0,885309127 | 0,938682 | Distal Intergenic | 15 | 64568933 | 64794145 | 225213 | 2 | 11514 | ENSMUST00000228014.2 | -213120 | ENSMUSG00000022376 | Adcy8 | adenylate cyclase 8 |
| chr16 | 45469611 | 45470968 | 1358 | * | 11,78274 | -2,608204192 | 0,985594 | -2,64633 | 0,008137 | 0,872427545 | 1,2763 | Intron (ENSMUST00000 | 16 | 45447211 | 45474431 | 27221 | 2 | 212998 | ENSMUST00000036732.9 | 3463 | ENSMUSG00000033187 | BC016579 | cDNA sequence, BC016579 |
| chr15 | 65767380 | 65768923 | 1544 | * | 22,23456 | -2,606456113 | 0,812652 | -3,20735 | 0,00134 | 0,402858015 | 3,104559 | Intron (ENSMUST00000 | 15 | 65748241 | 65764287 | 16047 | 2 | 18256 | ENSMUST00000135442.8 | -3093 | ENSMUSG00000015001 | Oc90 | otoconin 90 |
| chr1 | 25652963 | 25654612 | 1650 | * | 11,10692 | -2,605517845 | 0,980262 | -2,65798 | 0,007861 | 0,918554156 | 0,918255 | Exon (ENSMUST000001 | 1 | 25490152 | 25490255 | 104 | 1 | 1,15E+08 | ENSMUST00000122791.3 | 162811 | ENSMUSG00000084740 | Gm25294 | predicted gene, 25294 |
| chr18 | 70495153 | 70496602 | 1450 | * | 14,22524 | -2,605173659 | 0,949203 | -2,74459 | 0,006059 | 0,874820482 | 0,938682 | Distal Intergenic | 18 | 70585283 | 70597221 | 11939 | 2 | 269033 | ENSMUST00000213066.2 | 100619 | ENSMUSG00000044906 | 4930503L19Rik | RIKEN cDNA 4930503L19 gene |
| chr11 | 31958970 | 31961541 | 2572 | * | 20,21699 | -2,604386515 | 0,832444 | -3,1286 | 0,001756 | 0,48037412 | 2,507013 | Intron (ENSMUST00000 | 11 | 31950502 | 31983414 | 32913 | 1 | 18197 | ENSMUST00000093219.4 | 8468 | ENSMUSG00000020297 | Nsg2 | neuron specific gene family member 2 |
| chr1 | 93299625 | 93302242 | 2618 | * | 27,38581 | -2,60344354 | 0,747045 | -3,48499 | 0,000492 | 0,264977345 | 2,880017 | Promoter (<=1kb) | 1 | 93301652 | 93331124 | 29473 | 1 | 404545 | ENSMUST00000058682.11 | 0 | ENSMUSG00000034107 | Ano7 | anoctamin 7 |
| chr13 | 57123422 | 57124864 | 1443 | * | 17,55883 | -2,595201655 | 0,816357 | -3,179 | 0,001478 | 0,477594358 | 2,154504 | Distal Intergenic | 13 | 57078633 | 57078761 | 129 | 2 | 1,15E+08 | ENSMUST00000104091.3 | -44661 | ENSMUSG00000077279 | Gm22777 | predicted gene, 22777 |
| chr10 | 39893731 | 39895264 | 1534 | * | 11,9133 | -2,594691925 | 0,992813 | -2,61347 | 0,008963 | 0,938147218 | 0,918255 | Intron (ENSMUST00000 | 10 | 39883042 | 39901155 | 18114 | 2 | 215929 | ENSMUST00000176975.2 | 5891 | ENSMUSG00000038522 | Mfsd4b1 | major facilitator superfamily domain containing 4B1 |
| chr13 | 63671536 | 63673611 | 2076 | * | 20,08789 | -2,594655772 | 0,886995 | -2,92522 | 0,003442 | 0,593612587 | 2,713306 | 3' UTR | 13 | 63692673 | 63693130 | 458 | 2 | 19206 | ENSMUST00000220588.2 | 19519 | ENSMUSG00000021466 | Ptch1 | patched 1 |
| chr1 | 1,19E+08 | 1,19E+08 | 1824 | * | 20,65415 | -2,594421422 | 0,846184 | -3,06603 | 0,002169 | 0,463877619 | 3,471678 | Distal Intergenic | 1 | 1,19E+08 | 1,19E+08 | 219488 | 2 | 14633 | ENSMUST00000062483.15 | -9718 | ENSMUSG00000048402 | Gli2 | GLI-Kruppel family member GLI2 |
| chr1 | 96331576 | 96332920 | 1345 | * | 14,22659 | -2,591329231 | 0,982578 | -2,63728 | 0,008357 | 0,877892722 | 1,2763 | Distal Intergenic | 1 | 96544824 | 96544930 | 107 | 1 | 1,15E+08 | ENSMUST00001239562.1 | -211904 | ENSMUSG00000119190 | Gm26245 | predicted gene, 26245 |
| chr7 | 42055793 | 42057052 | 1260 | * | 10,57655 | -2,590236423 | 1,004977 | -2,57741 | 0,009954 | 0,904389377 | 1,2763 | Distal Intergenic | 7 | 42079252 | 42080148 | 897 | 2 | 628490 | ENSMUST00000173296.2 | 23096 | | Vmn2r-ps60 | vomeronasal 2, receptor, pseudogene 60 |
| chr1 | 16346407 | 16348017 | 1611 | * | 18,86748 | -2,589770156 | 0,788533 | -3,28429 | 0,001022 | 0,411735506 | 2,221611 | Intron (ENSMUST00000 | 1 | 16323853 | 16329081 | 5229 | 1 | 1,03E+08 | ENSMUST00000159190.2 | 22554 | ENSMUSG00000089982 | Gm7568 | predicted gene 7568 |
| chr18 | 66008681 | 66010324 | 1644 | * | 13,59709 | -2,589704987 | 0,98032 | -2,64169 | 0,008249 | 0,902200314 | 1,075735 | Promoter (2-3kb) | 18 | 66006575 | 66013329 | 6755 | 1 | 225642 | ENSMUST00000173530.4 | 2106 | ENSMUSG00000024517 | Grp | gastrin releasing peptide |
| chr6 | 1,21E+08 | 1,21E+08 | 1110 | * | 14,61725 | -2,588600593 | 0,958537 | -2,70057 | 0,006922 | 0,830950621 | 1,37683 | Distal Intergenic | 6 | 1,21E+08 | 1,21E+08 | 9282 | 1 | 1,03E+08 | ENSMUST00000244553.1 | 19736 | | Gm30842 | predicted gene, 30842 |
| chr7 | 11756340 | 11757213 | 874 | * | 17,66088 | -2,587155559 | 0,910583 | -2,84121 | 0,004494 | 0,654833721 | 2,507013 | Distal Intergenic | 7 | 11775226 | 11776146 | 921 | 1 | 628946 | ENSMUST00000164446.2 | -18013 | ENSMUSG00000095864 | Vmn1r77 | vomeronasal 1 receptor 77 |
| chr17 | 60192166 | 60193707 | 1542 | * | 15,50983 | -2,585018966 | 0,902886 | -2,86306 | 0,004196 | 0,669586355 | 2,141612 | Intron (ENSMUST00000 | 17 | 59876554 | 59876660 | 107 | 2 | 1,15E+08 | ENSMUST00000104374.3 | -315506 | ENSMUSG00000077562 | Gm23203 | predicted gene, 23203 |
| chr17 | 60711149 | 60712741 | 1593 | * | 13,96309 | -2,583988029 | 0,923346 | -2,79851 | 0,005134 | 0,791416391 | 1,2763 | Distal Intergenic | 17 | 60793002 | 60793164 | 163 | 1 | 1,15E+08 | ENSMUST00000082798.3 | -80261 | ENSMUSG00000064732 | Gm23447 | predicted gene, 23447 |
| chr6 | 1,04E+08 | 1,04E+08 | 1954 | * | 24,23538 | -2,58316768 | 0,688003 | -3,75459 | 0,000174 | 0,149049497 | 2,880017 | Intron (ENSMUST00000 | 6 | 1,04E+08 | 1,04E+08 | 62 | 1 | 1,15E+08 | ENSMUST00000157640.3 | 8248 | ENSMUSG00000088265 | Gm24784 | predicted gene, 24784 |
| chr18 | 85614863 | 85616499 | 1637 | * | 15,61301 | -2,583028955 | 0,914357 | -2,82497 | 0,004729 | 0,777172375 | 1,2763 | Distal Intergenic | 18 | 84983635 | 84999597 | 15963 | 1 | 50764 | ENSMUST00000225015.2 | 631228 | ENSMUSG00000034391 | Fbxo15 | F-box protein 15 |
| chr10 | 72466001 | 72468098 | 2098 | * | 14,9024 | -2,581042639 | 0,95551 | -2,70122 | 0,006909 | 0,852340527 | 1,214904 | Distal Intergenic | 10 | 72490677 | 72510796 | 20120 | 1 | 52696 | ENSMUST00000020081.11 | -22579 | ENSMUSG00000019923 | Zwint | ZW10 interactor |
| chr19 | 28832734 | 28834338 | 1605 | * | 13,30328 | -2,580890893 | 0,919082 | -2,80812 | 0,004983 | 0,816751204 | 1,075735 | Intron (ENSMUST00000 | 19 | 28812535 | 28883392 | 70858 | 1 | 20510 | ENSMUST00000161119.8 | 20199 | ENSMUSG00000024935 | Slc1a1 | solute carrier family 1 (neuronal/epithelial high affinity glutamate transporter, system Xag), member 1 |
| chr1 | 1,2E+08 | 1,2E+08 | 1012 | * | 14,98098 | -2,580802365 | 0,946801 | -2,72581 | 0,006414 | 0,812199284 | 1,41696 | Distal Intergenic | 1 | 1,2E+08 | 1,2E+08 | 13844 | 2 | 17167 | ENSMUST00000186432.3 | 75430 | ENSMUSG00000026390 | Marco | macrophage receptor with collagenous structure |
| chr2 | 1,6E+08 | 1,6E+08 | 1740 | * | 22,03975 | -2,576998979 | 0,771599 | -3,33982 | 0,000838 | 0,307730759 | 3,471678 | Distal Intergenic | 2 | 1,6E+08 | 1,6E+08 | 23967 | 1 | 71613 | ENSMUST00000140319.2 | -4462 | ENSMUSG00000086653 | 9130015L21Rik | RIKEN cDNA 9130015L21 gene |
| chr12 | 37813527 | 37816297 | 2771 | * | 23,4273 | -2,576778142 | 0,728701 | -3,53613 | 0,000406 | 0,236345598 | 2,951062 | Distal Intergenic | 12 | 37867725 | 38032009 | 164285 | 1 | 217480 | ENSMUST00000222337.2 | -51428 | ENSMUSG00000036095 | Dgkb | diacylglycerol kinase, beta |
| chr13 | 34942607 | 34945014 | 2408 | * | 19,20711 | -2,57655046 | 0,834809 | -3,0864 | 0,002026 | 0,505909454 | 2,507013 | Distal Intergenic | 13 | 34923595 | 34931605 | 8011 | 1 | 108161 | ENSMUST00000221037.2 | 19012 | ENSMUSG00000038246 | Fam50b | family with sequence similarity 50, member B |
| chr5 | 26431725 | 26433918 | 2194 | * | 19,41251 | -2,575983603 | 0,85572 | -3,01031 | 0,00261 | 0,537550416 | 2,713306 | Distal Intergenic | 5 | 26462689 | 26469306 | 6618 | 1 | 664837 | ENSMUST00000074148.7 | -28771 | ENSMUSG00000059645 | Gm7361 | predicted gene 7361 |
| chr6 | 59284772 | 59285930 | 1159 | * | 16,74096 | -2,575321406 | 0,953579 | -2,70069 | 0,00692 | 0,731734453 | 2,386008 | Distal Intergenic | 6 | 59185855 | 59189018 | 3164 | 1 | 68140 | ENSMUST00000062626.4 | 98917 | ENSMUSG00000049232 | Tigd2 | tigger transposable element derived 2 |
| chr9 | 86754344 | 86755894 | 1551 | * | 15,28888 | -2,575089307 | 0,973138 | -2,64617 | 0,008141 | 0,872533632 | 1,2763 | Intron (ENSMUST00000 | 9 | 86717162 | 86762450 | 45289 | 2 | 20616 | ENSMUST00000191290.3 | 6556 | ENSMUSG00000033419 | Snap91 | synaptosomal-associated protein 91 |
| chr10 | 33824750 | 33827597 | 2848 | * | 19,16949 | -2,574967477 | 0,913837 | -2,81776 | 0,004836 | 0,668446397 | 2,507013 | Promoter (<=1kb) | 10 | 33795138 | 33827255 | 32118 | 2 | 72580 | ENSMUST00000218055.2 | 0 | ENSMUSG00000039531 | Zup1 | zinc finger containing ubiquitin peptidase 1 |
| chr19 | 11377375 | 11379751 | 2377 | * | 17,69513 | -2,574185726 | 0,866305 | -2,97145 | 0,002964 | 0,576448911 | 2,507013 | Promoter (2-3kb) | 19 | 11382134 | 11393783 | 11650 | 1 | 64380 | ENSMUST00000153546.8 | -2383 | ENSMUSG00000024675 | Ms4a4c | membrane-spanning 4-domains, subfamily A, member 4C |
| chr18 | 30317045 | 30318884 | 1840 | * | 16,96513 | -2,573038751 | 0,871277 | -2,95318 | 0,003145 | 0,576083201 | 2,663619 | Distal Intergenic | 18 | 30400302 | 30405186 | 4885 | 1 | 225326 | ENSMUST00000234467.2 | -81418 | ENSMUSG00000033628 | Pik3c3 | phosphatidylinositol 3-kinase catalytic subunit type 3 |
| chr16 | 95073200 | 95075294 | 2095 | * | 26,18951 | -2,57174582 | 0,758386 | -3,39108 | 0,000696 | 0,3078074 | 2,880017 | Intron (ENSMUST00000 | 16 | 95080052 | 95092756 | 12705 | 1 | 16516 | ENSMUST00000138329.2 | -4758 | ENSMUSG00000062609 | Kcnj15 | potassium inwardly-rectifying channel, subfamily J, member 15 |
| chr8 | 18920759 | 18922544 | 1786 | * | 17,81842 | -2,571266546 | 0,958259 | -2,68327 | 0,007291 | 0,731734453 | 2,507013 | Exon (ENSMUST000001 | 8 | 18896295 | 18933437 | 37143 | 1 | 52123 | ENSMUST00000033847.5 | 24464 | ENSMUSG00000031467 | Agpat5 | 1-acylglycerol-3-phosphate O-acyltransferase 5 (lysophosphatidic acid acyltransferase, epsilon) |
| chr12 | 32937794 | 32940019 | 2226 | * | 17,98389 | -2,570982019 | 0,860848 | -2,98657 | 0,002821 | 0,566484927 | 2,507013 | Distal Intergenic | 12 | 32974954 | 33003719 | 28766 | 2 | 74076 | ENSMUST00000245098.1 | 63700 | ENSMUSG00000097494 | 4933406C10Rik | RIKEN cDNA 4933406C10 gene |
| chr2 | 52829681 | 52831567 | 1887 | * | 20,07227 | -2,570116469 | 0,929586 | -2,7648 | 0,005696 | 0,683140223 | 2,663619 | Intron (ENSMUST00000 | 2 | 52747880 | 53021890 | 274011 | 1 | 71409 | ENSMUST00000050719.13 | 81801 | ENSMUSG00000036053 | Fmnl2 | formin-like 2 |
| chr11 | 1,06E+08 | 1,06E+08 | 2321 | * | 20,29449 | -2,569537377 | 0,893902 | -2,87452 | 0,004046 | 0,637823949 | 2,507013 | Promoter (<=1kb) | 11 | 1,06E+08 | 1,06E+08 | 8509 | 1 | 11421 | ENSMUST00000151657.2 | 0 | ENSMUSG00000020681 | Ace | angiotensin I converting enzyme (peptidyl-dipeptidase A) 1 |
| chr1 | 1,31E+08 | 1,31E+08 | 2189 | * | 23,06927 | -2,569478068 | 0,806948 | -3,18419 | 0,001452 | 0,440188707 | 2,67243 | Promoter (<=1kb) | 1 | 1,31E+08 | 1,31E+08 | 3923 | 2 | 93672 | ENSMUST00000187650.7 | 0 | ENSMUSG00000026420 | Il24 | interleukin 24 |
| chr12 | 97327337 | 97329868 | 2532 | * | 18,8883 | -2,567969172 | 0,925057 | -2,77601 | 0,005503 | 0,675437713 | 2,713306 | Distal Intergenic | 12 | 97279623 | 97282627 | 3005 | 1 | 1,05E+08 | ENSMUST00000218233.2 | 47714 | | Gm40552 | predicted gene, 40552 |
| chr4 | 21277287 | 21279345 | 2059 | * | 21,77351 | -2,567241835 | 0,723865 | -3,54657 | 0,00039 | 0,225397758 | 3,104559 | Distal Intergenic | 4 | 21679747 | 21685781 | 6035 | 2 | 230025 | ENSMUST00000149516.8 | 406436 | ENSMUSG00000040478 | Prdm13 | PR domain containing 13 |
| chr11 | 63703181 | 63705013 | 1833 | * | 17,38554 | -2,563333386 | 0,844323 | -3,03596 | 0,002398 | 0,564864958 | 2,154504 | Distal Intergenic | 11 | 63776618 | 63813116 | 36499 | 2 | 54710 | ENSMUST00000094103.4 | 108103 | ENSMUSG00000070407 | Hs3st3b1 | heparan sulfate (glucosamine) 3-O-sulfotransferase 3B1 |
| chr10 | 96335761 | 96337175 | 1415 | * | 12,27588 | -2,562309732 | 0,982022 | -2,60922 | 0,009075 | 0,940922799 | 0,918255 | Intron (ENSMUST00000 | 10 | 96359946 | 96431468 | 71523 | 2 | 1,03E+08 | ENSMUST00000244443.1 | 94293 | | Gm33843 | predicted gene, 33843 |
| chr14 | 16612832 | 16614475 | 1644 | * | 14,08977 | -2,560710128 | 0,911927 | -2,80802 | 0,004985 | 0,83918951 | 0,938682 | Distal Intergenic | 14 | 16589391 | 16611855 | 22465 | 1 | 1E+08 | ENSMUST00000164484.8 | 23441 | | Gm3373 | predicted gene 3373 |
| chr10 | 9841401 | 9842745 | 1345 | * | 13,10459 | -2,559625345 | 0,956499 | -2,67603 | 0,00745 | 0,945372047 | 0,738441 | Distal Intergenic | 10 | 9852434 | 10083725 | 231292 | 1 | 1,03E+08 | ENSMUST00000219328.2 | -9689 | | Gm28905 | predicted gene 28905 |
| chr1 | 1,02E+08 | 1,02E+08 | 2058 | * | 18,64375 | -2,557793211 | 0,759028 | -3,36983 | 0,000752 | 0,339611164 | 2,423594 | Distal Intergenic | 1 | 1,01E+08 | 1,01E+08 | 19394 | 1 | 1,01E+08 | ENSMUST00000187932.2 | 451839 | ENSMUSG00000100684 | Gm20268 | predicted gene, 20268 |
| chr6 | 1,12E+08 | 1,12E+08 | 772 | * | 14,6234 | -2,55770872 | 0,922413 | -2,77285 | 0,005557 | 0,803086765 | 1,2763 | Distal Intergenic | 6 | 1,12E+08 | 1,12E+08 | 1812 | 1 | 67336 | ENSMUST00000186102.2 | -158535 | ENSMUSG00000100219 | 1700054K19Rik | RIKEN cDNA 1700054K19 gene |
| chr12 | 1,07E+08 | 1,07E+08 | 1925 | * | 17,2035 | -2,557191702 | 0,922919 | -2,77077 | 0,005592 | 0,691182187 | 2,423594 | Distal Intergenic | 12 | 1,07E+08 | 1,07E+08 | 10314 | 2 | 71033 | ENSMUST00000149382.2 | -49306 | ENSMUSG00000085820 | 4933406K04Rik | RIKEN cDNA 4933406K04 gene |
| chr7 | 1,24E+08 | 1,24E+08 | 1316 | * | 16,06503 | -2,554942823 | 0,899966 | -2,83893 | 0,004526 | 0,669586355 | 2,318434 | Distal Intergenic | 7 | 1,24E+08 | 1,24E+08 | 83266 | 2 | 73076 | ENSMUST00000130240.2 | -22450 | ENSMUSG00000086254 | 3100003L05Rik | RIKEN cDNA 3100003L05 gene |
| chr7 | 1,19E+08 | 1,19E+08 | 3200 | * | 37,28074 | -2,554901039 | 0,620139 | -4,11988 | 3,79E-05 | 0,065127074 | 2,377674 | Distal Intergenic | 7 | 1,19E+08 | 1,19E+08 | 126 | 2 | 1,15E+08 | ENSMUST00000104465.3 | 46460 | ENSMUSG00000077653 | Gm23321 | predicted gene, 23321 |
| chr17 | 13604718 | 13605839 | 1122 | * | 13,26698 | -2,551784945 | 0,979459 | -2,6053 | 0,009179 | 0,91804783 | 1,075735 | Exon (ENSMUST000002 | 17 | 13619304 | 13619468 | 165 | 2 | 1,15E+08 | ENSMUST00020183776.1 | 13629 | ENSMUSG00002075210 | Gm25730 | predicted gene, 25730 |
| chr5 | 1,3E+08 | 1,3E+08 | 1865 | * | 16,64775 | -2,549228597 | 0,869665 | -2,93128 | 0,003376 | 0,612704358 | 2,386008 | Promoter (<=1kb) | 5 | 1,3E+08 | 1,3E+08 | 1756 | 1 | 22021 | ENSMUST00000128335.2 | 0 | ENSMUSG00000034118 | Tpst1 | protein-tyrosine sulfotransferase 1 |
| chr2 | 1,22E+08 | 1,22E+08 | 1861 | * | 19,47854 | -2,549163751 | 0,850505 | -2,99724 | 0,002724 | 0,542857998 | 2,713306 | Distal Intergenic | 2 | 1,22E+08 | 1,22E+08 | 69236 | 1 | 319996 | ENSMUST00000110586.10 | -16142 | ENSMUSG00000060227 | Golm2 | golgi membrane protein 2 |
| chr12 | 1,02E+08 | 1,02E+08 | 1199 | * | 13,80512 | -2,548607123 | 0,980058 | -2,60047 | 0,00931 | 0,941631683 | 0,938682 | Intron (ENSMUST00000 | 12 | 1,02E+08 | 1,02E+08 | 60504 | 2 | 74413 | ENSMUST00000160715.8 | 87942 | ENSMUSG00000021187 | Tc2n | tandem C2 domains, nuclear |
| chr6 | 24408616 | 24409564 | 949 | * | 13,85786 | -2,548443553 | 0,971934 | -2,62203 | 0,008741 | 0,910268063 | 1,075735 | Distal Intergenic | 6 | 24444864 | 24515066 | 70203 | 2 | 214704 | ENSMUST00000052277.5 | 105502 | ENSMUSG00000046192 | Iqub | IQ motif and ubiquitin domain containing |
| chr9 | 49202877 | 49204355 | 1479 | * | 17,07917 | -2,547955754 | 0,788833 | -3,23003 | 0,001238 | 0,437520575 | 2,318434 | Distal Intergenic | 9 | 49161831 | 49191894 | 30064 | 1 | 235327 | ENSMUST00000216428.2 | 41046 | ENSMUSG00000054156 | Gm4894 | predicted gene 4894 |
| chr12 | 1,15E+08 | 1,15E+08 | 556 | * | 15,64176 | -2,54006798 | 0,949826 | -2,67425 | 0,00749 | 0,864017288 | 1,214904 | Distal Intergenic | 12 | 1,15E+08 | 1,15E+08 | 25225 | 1 | 1,03E+08 | ENSMUST00000195683.2 | 117305 | ENSMUSG00000103168 | Gm30948 | predicted gene, 30948 |
| chr4 | 1,02E+08 | 1,02E+08 | 1306 | * | 17,58822 | -2,539118145 | 0,873378 | -2,90724 | 0,003646 | 0,618136403 | 2,507013 | Intron (ENSMUST00000 | 4 | 1,02E+08 | 1,02E+08 | 301183 | 1 | 18578 | ENSMUST00000106907.9 | -31360 | ENSMUSG00000028525 | Pde4b | phosphodiesterase 4B, cAMP specific |
| chr5 | 76918223 | 76919820 | 1598 | * | 15,15552 | -2,537776052 | 0,936613 | -2,70952 | 0,006738 | 0,849437958 | 1,214904 | Intron (ENSMUST00000 | 5 | 76900215 | 77021177 | 120963 | 1 | 320827 | ENSMUST00000121160.8 | 18008 | ENSMUSG00000036377 | Cracd | capping protein inhibiting regulator of actin |
| chr13 | 77316801 | 77319129 | 2329 | * | 16,19842 | -2,537068134 | 0,897586 | -2,82655 | 0,004705 | 0,65166214 | 2,663619 | Exon (ENSMUST000002 | 13 | 77341153 | 77410088 | 68936 | 1 | 72371 | ENSMUST00000225760.2 | -22024 | ENSMUSG00000071252 | 2210408I21Rik | RIKEN cDNA 2210408I21 gene |
| chr5 | 1,04E+08 | 1,04E+08 | 2492 | * | 15,8981 | -2,53316572 | 0,946163 | -2,6773 | 0,007422 | 0,764024179 | 2,141612 | Intron (ENSMUST00000 | 5 | 1,04E+08 | 1,04E+08 | 27237 | 1 | 17355 | ENSMUST00000152145.2 | 4290 | ENSMUSG00000029313 | Aff1 | AF4/FMR2 family, member 1 |
| chr2 | 1,48E+08 | 1,48E+08 | 2138 | * | 17,43898 | -2,531394253 | 0,860164 | -2,94292 | 0,003251 | 0,595365039 | 2,507013 | Intron (ENSMUST00000 | 2 | 1,48E+08 | 1,48E+08 | 6924 | 2 | 17064 | ENSMUST00000099269.4 | -28349 | ENSMUSG00000027435 | Cd93 | CD93 antigen |
| chr3 | 74892892 | 74894652 | 1761 | * | 16,91813 | -2,530873674 | 0,866166 | -2,92193 | 0,003479 | 0,639383182 | 2,141612 | Distal Intergenic | 3 | 74968668 | 75051010 | 82343 | 2 | 213234 | ENSMUST00000107776.8 | 156358 | ENSMUSG00000034151 | Zbbx | zinc finger, B-box domain containing |
| chr7 | 1,39E+08 | 1,39E+08 | 1961 | * | 19,41788 | -2,52836567 | 0,861593 | -2,93453 | 0,003341 | 0,590570804 | 2,663619 | Intron (ENSMUST00000 | 7 | 1,39E+08 | 1,39E+08 | 19614 | 1 | 76484 | ENSMUST00000121839.2 | 7286 | ENSMUSG00000066129 | Kndc1 | kinase non-catalytic C-lobe domain (KIND) containing 1 |
| chr7 | 138027152 | 138032124 | 4973 | * | 58,824701 | -2,527213503 | 0,5685975 | -4,444644 | 8,804E-06 | 0,017605583 | 3,439501 | Distal Intergenic | 7 | 137860316 | 137999459 | 139144 | 2 | 70571 | ENSMUST00000162222.2 | -27693 | ENSMUSG00000091002 | Tcerg1l | transcription elongation regulator 1-like |
| chr15 | 82420947 | 82422576 | 1630 | * | 18,42099 | -2,526307818 | 0,8517 | -2,9662 | 0,003015 | 0,61075929 | 2,154504 | Exon (ENSMUST000002 | 15 | 82439273 | 82442035 | 2763 | 1 | 380997 | ENSMUST00000229103.2 | -16697 | ENSMUSG00000096852 | Cyp2d12 | cytochrome P450, family 2, subfamily d, polypeptide 12 |
| chr15 | 1,04E+08 | 1,04E+08 | 1790 | * | 21,60484 | -2,526122013 | 0,775356 | -3,25802 | 0,001122 | 0,348387064 | 3,471678 | Distal Intergenic | 15 | 1,04E+08 | 1,04E+08 | 3377 | 2 | 20771 | ENSMUST00000078842.11 | 37074 | ENSMUSG00000060311 | Mucl1 | mucin-like 1 |
| chr9 | 9799176 | 9800528 | 1353 | * | 16,31831 | -2,526036013 | 0,9354 | -2,70049 | 0,006924 | 0,715171447 | 2,663619 | Intron (ENSMUST00000 | 9 | 9894436 | 9894526 | 91 | 1 | 1,02E+08 | ENSMUST00000183808.3 | -93908 | ENSMUSG00000098408 | Mir6237 | microRNA 6237 |
| chr16 | 16299567 | 16301147 | 1581 | * | 13,5465 | -2,522137931 | 0,887328 | -2,8424 | 0,004478 | 0,860883941 | 0,738441 | Exon (ENSMUST000001 | 16 | 16253570 | 16322086 | 68517 | 2 | 224014 | ENSMUST00000161188.9 | 20939 | ENSMUSG00000022788 | Fgd4 | FYVE, RhoGEF and PH domain containing 4 |
| chr6 | 17409481 | 17411376 | 1896 | * | 21,32946 | -2,519595168 | 0,824657 | -3,05532 | 0,002248 | 0,487260898 | 3,104559 | Distal Intergenic | 6 | 17436305 | 17436411 | 107 | 1 | 1,15E+08 | ENSMUST00000240512.1 | -24929 | ENSMUSG00000119734 | Gm25021 | predicted gene, 25021 |
| chr13 | 53801216 | 53803320 | 2105 | * | 19,92073 | -2,518941637 | 0,848391 | -2,96908 | 0,002987 | 0,563032042 | 2,713306 | Promoter (<=1kb) | 13 | 53802621 | 53804116 | 1496 | 1 | 75252 | ENSMUST00000221718.2 | 0 | ENSMUSG00000113329 | 4930555G21Rik | RIKEN cDNA 4930555G21 gene |
| chr7 | 97671755 | 97674642 | 2888 | * | 27,02264 | -2,518165922 | 0,675988 | -3,72516 | 0,000195 | 0,156945882 | 2,951062 | Intron (ENSMUST00000 | 7 | 97711850 | 97714409 | 2560 | 2 | 17921 | ENSMUST00000124787.2 | 39767 | ENSMUSG00000030761 | Myo7a | myosin VIIA |
| chr7 | 1,17E+08 | 1,17E+08 | 2341 | * | 17,54175 | -2,51699423 | 0,868963 | -2,89655 | 0,003773 | 0,650529916 | 2,154504 | Distal Intergenic | 7 | 1,17E+08 | 1,17E+08 | 107 | 2 | 1,15E+08 | ENSMUST00000240319.1 | -33311 | ENSMUSG00000119801 | Gm26054 | predicted gene, 26054 |
| chr8 | 1,19E+08 | 1,19E+08 | 1567 | * | 24,44788 | -2,514758754 | 0,73408 | -3,42573 | 0,000613 | 0,302204121 | 2,67243 | Distal Intergenic | 8 | 1,19E+08 | 1,19E+08 | 10299 | 2 | 68533 | ENSMUST00000034303.3 | -148589 | ENSMUSG00000031843 | Mphosph6 | M phase phosphoprotein 6 |
| chr17 | 52868893 | 52870606 | 1714 | * | 15,31241 | -2,51426147 | 0,873116 | -2,87964 | 0,003981 | 0,752089169 | 1,214904 | Distal Intergenic | 17 | 52900009 | 52909688 | 9680 | 2 | 1,08E+08 | ENSMUST00000183438.2 | 39082 | ENSMUSG00000098552 | Gm27217 | predicted gene 27217 |
| chr16 | 35752685 | 35754500 | 1816 | * | 16,26236 | -2,513508805 | 0,850217 | -2,95631 | 0,003113 | 0,601894237 | 2,318434 | Exon (ENSMUST000000 | 16 | 35758840 | 35792975 | 34136 | 1 | 80285 | ENSMUST00000114878.8 | -4340 | ENSMUSG00000022906 | Parp9 | poly (ADP-ribose) polymerase family, member 9 |
| chr18 | 51038312 | 51040808 | 2497 | * | 21,56416 | -2,511182198 | 0,729007 | -3,44466 | 0,000572 | 0,281843592 | 2,951933 | Distal Intergenic | 18 | 51250805 | 51438262 | 187458 | 1 | 71373 | ENSMUST00000116639.4 | -209997 | ENSMUSG00000073565 | Prr16 | proline rich 16 |
| chr14 | 45529320 | 45530891 | 1572 | * | 13,08538 | -2,510497996 | 0,961183 | -2,61188 | 0,009004 | 0,902325717 | 1,173325 | Exon (ENSMUST000000 | 14 | 45525194 | 45536522 | 11329 | 2 | 50527 | ENSMUST00000227147.2 | 5631 | ENSMUSG00000021831 | Ero1a | endoplasmic reticulum oxidoreductase 1 alpha |
| chr16 | 7822888 | 7824212 | 1325 | * | 15,64044 | -2,510454224 | 0,89845 | -2,7942 | 0,005203 | 0,775744126 | 1,41696 | Distal Intergenic | 16 | 8154857 | 8154960 | 104 | 1 | 1,15E+08 | ENSMUST00000158968.3 | -330645 | ENSMUSG00000089593 | Gm26159 | predicted gene, 26159 |
| chr17 | 60239974 | 60241675 | 1702 | * | 14,40903 | -2,510252397 | 0,948301 | -2,6471 | 0,008118 | 0,872185848 | 1,2763 | Intron (ENSMUST00000 | 17 | 59876554 | 59876660 | 107 | 2 | 1,15E+08 | ENSMUST00000104374.3 | -363314 | ENSMUSG00000077562 | Gm23203 | predicted gene, 23203 |
| chr13 | 43018877 | 43021493 | 2617 | * | 19,75242 | -2,507907304 | 0,901574 | -2,7817 | 0,005408 | 0,68353261 | 2,507013 | Promoter (<=1kb) | 13 | 43019723 | 43288780 | 269058 | 1 | 218194 | ENSMUST00000131942.2 | 0 | ENSMUSG00000054728 | Phactr1 | phosphatase and actin regulator 1 |
| chr17 | 44569680 | 44571387 | 1708 | * | 18,67272 | -2,50685703 | 0,844307 | -2,96913 | 0,002986 | 0,565876168 | 2,663619 | Intron (ENSMUST00000 | 17 | 44547993 | 44562606 | 14614 | 1 | 224796 | ENSMUST00000233193.2 | 21687 | ENSMUSG00000023959 | Clic5 | chloride intracellular channel 5 |
| chr5 | 27836223 | 27839304 | 3082 | * | 32,52027 | -2,506469828 | 0,787365 | -3,18336 | 0,001456 | 0,547949061 | 1,405709 | Exon (ENSMUST000000 | 5 | 27757549 | 27758873 | 1325 | 2 | 1,05E+08 | ENSMUST00000124392.2 | -77350 | | Gm16058 | predicted gene 16058 |
| chr14 | 36201031 | 36202325 | 1295 | * | 12,19834 | -2,505526149 | 0,965435 | -2,59523 | 0,009453 | 0,977736121 | 0,738441 | Distal Intergenic | 14 | 36489382 | 36489487 | 106 | 1 | 1,15E+08 | ENSMUST00000082843.3 | -287057 | ENSMUSG00000064777 | Gm25204 | predicted gene, 25204 |
| chr1 | 1,5E+08 | 1,5E+08 | 1687 | * | 18,08624 | -2,504280213 | 0,857783 | -2,91948 | 0,003506 | 0,617289329 | 2,423594 | Distal Intergenic | 1 | 1,5E+08 | 1,5E+08 | 465 | 2 | 1E+08 | ENSMUST00000121422.2 | 14847 | ENSMUSG00000031167 | Rbm3-ps | RNA binding motif (RNP1, RRM) protein 3 |
| chr4 | 66296029 | 66297667 | 1639 | * | 16,33635 | -2,50383142 | 0,858327 | -2,91711 | 0,003533 | 0,641208222 | 2,141612 | Intron (ENSMUST00000 | 4 | 65299040 | 66322720 | 1023681 | 2 | 56079 | ENSMUST00000084496.3 | 25053 | ENSMUSG00000028373 | Astn2 | astrotactin 2 |
| chr1 | 8348446 | 8349680 | 1235 | * | 12,94926 | -2,503461081 | 0,935984 | -2,67468 | 0,00748 | 0,872427545 | 1,173325 | Distal Intergenic | 1 | 8433649 | 8540703 | 107055 | 2 | 71096 | ENSMUST00000135046.4 | 191023 | ENSMUSG00000025909 | Sntg1 | syntrophin, gamma 1 |
| chr1 | 78040844 | 78042494 | 1651 | * | 13,04002 | -2,503400838 | 0,968619 | -2,5845 | 0,009752 | 0,914358307 | 1,173325 | Distal Intergenic | 1 | 78077904 | 78173475 | 95572 | 2 | 18505 | ENSMUST00000004994.16 | 130981 | ENSMUSG00000004872 | Pax3 | paired box 3 |
| chr6 | 1,16E+08 | 1,16E+08 | 1457 | * | 16,10344 | -2,503209161 | 0,933219 | -2,68234 | 0,007311 | 0,742868694 | 2,386008 | Promoter (2-3kb) | 6 | 1,16E+08 | 1,16E+08 | 40033 | 2 | 330401 | ENSMUST00000173031.2 | -2832 | ENSMUSG00000030126 | Tmcc1 | transmembrane and coiled coil domains 1 |
| chr18 | 20340386 | 20341718 | 1333 | * | 12,52638 | -2,502778143 | 0,937402 | -2,66991 | 0,007587 | 0,947773602 | 0,738441 | Distal Intergenic | 18 | 20380397 | 20418088 | 37692 | 1 | 211924 | ENSMUST00000054128.7 | -38679 | ENSMUSG00000034774 | Dsg1c | desmoglein 1 gamma |
| chr16 | 65657193 | 65658869 | 1677 | * | 14,37209 | -2,502140675 | 0,883331 | -2,83262 | 0,004617 | 0,801688106 | 1,075735 | 3' UTR | 16 | 65625499 | 65658781 | 33283 | 1 | 73569 | ENSMUST00000228708.2 | 31694 | ENSMUSG00000091243 | Vgll3 | vestigial like family member 3 |

| | | | | | | | | | | | | | | | | | | | | | | | |
|---|---|---|---|---|---|---|---|---|---|---|---|---|---|---|---|---|---|---|---|---|---|---|---|
| chr1 | 1,05E+08 | 1,05E+08 | 3152 | * | 33,9743 | -2,498944244 | 0,606083 | -4,1231 | 3,74E-05 | 0,065047955 | 2,377674 | Distal Intergenic | 1 | 1,05E+08 | 1,05E+08 | 10913 | 1 | 1,03E+08 | ENSMUST00000187882.2 | 81630 | | Gm29012 | predicted gene 29012 |
| chr7 | 93311462 | 93312830 | 1369 | * | 14,8892 | -2,498761106 | 0,935463 | -2,67115 | 0,007559 | 0,86525631 | 1,214904 | Distal Intergenic | 7 | 93691898 | 93998607 | 306710 | 1 | 1,03E+08 | ENSMUST00000206116.2 | -379068 | | Gm32647 | predicted gene, 32647 |
| chr3 | 79297561 | 79299201 | 1641 | * | 19,26978 | -2,498172324 | 0,792142 | -3,15369 | 0,001612 | 0,467440127 | 2,507013 | Intron (ENSMUST0000( | 3 | 79252686 | 79338593 | 85908 | 1 | 1E+08 | ENSMUST00000199658.2 | 44875 | ENSMUSG00000091685 | Gm17359 | predicted gene, 17359 |
| chr6 | 89191282 | 89193144 | 1863 | * | 18,44773 | -2,49767033 | 0,81323 | -3,0713 | 0,002131 | 0,514210241 | 2,507013 | Promoter (<=1kb) | 6 | 89188147 | 89193219 | 5073 | 2 | 330379 | ENSMUST00000079186.3 | 75 | | Gm839 | predicted gene 839 |
| chr1 | 40672094 | 40673328 | 1235 | * | 13,65064 | -2,495782173 | 0,940387 | -2,654 | 0,007954 | 0,954881543 | 0,738441 | Distal Intergenic | 1 | 40639762 | 40642063 | 2302 | 1 | 110895 | ENSMUST00000193371.2 | 32332 | ENSMUSG00000026065 | Slc9a4 | solute carrier family 9 (sodium/hydrogen exchanger), member 4 |
| chr2 | 1,27E+08 | 1,27E+08 | 1945 | * | 17,26925 | -2,495420385 | 0,917942 | -2,71849 | 0,006558 | 0,722249331 | 2,423594 | Distal Intergenic | 2 | 1,27E+08 | 1,27E+08 | 28628 | 1 | 108011 | ENSMUST00000149254.9 | -25204 | ENSMUSG00000001998 | Ap4e1 | adaptor-related protein complex AP-4, epsilon 1 |
| chr1 | 1,06E+08 | 1,06E+08 | 2288 | * | 22,29288 | -2,494290618 | 0,796633 | -3,13104 | 0,001742 | 0,456947326 | 2,880017 | Distal Intergenic | 1 | 1,06E+08 | 1,06E+08 | 136 | 2 | 1,02E+08 | ENSMUST00000183743.3 | 98728 | ENSMUSG00000098344 | Mir3473f | microRNA 3473f |
| chr4 | 55858812 | 55860321 | 1510 | * | 18,13176 | -2,493131694 | 0,960266 | -2,59629 | 0,009424 | 0,786955198 | 2,423594 | Distal Intergenic | 4 | 55969023 | 55993739 | 24717 | 2 | 1,03E+08 | ENSMUST00000134099.2 | 133418 | | Gm12519 | predicted gene 12519 |
| chr11 | 46970539 | 46972648 | 2110 | * | 19,51974 | -2,492744854 | 0,923861 | -2,69818 | 0,006972 | 0,726036755 | 2,507013 | Intron (ENSMUST0000( | 11 | 46701627 | 46735159 | 33533 | 1 | 276891 | ENSMUST00000068877.7 | 268912 | ENSMUSG00000055546 | Timd4 | T cell immunoglobulin and mucin domain containing 4 |
| chr17 | 65216296 | 65217851 | 1556 | * | 17,59367 | -2,490832262 | 0,95203 | -2,61634 | 0,008888 | 0,767253638 | 2,507013 | Distal Intergenic | 17 | 65251716 | 65251847 | 132 | 2 | 1,15E+08 | ENSMUST00000104503.4 | 33996 | ENSMUSG00000077691 | Gm25344 | predicted gene, 25344 |
| chr5 | 1,26E+08 | 1,26E+08 | 1558 | * | 20,92113 | -2,490511837 | 0,874894 | -2,84665 | 0,004418 | 0,61384098 | 3,104559 | Distal Intergenic | 5 | 1,26E+08 | 1,26E+08 | 107 | 2 | 1,15E+08 | ENSMUST00000239978.1 | 171691 | ENSMUSG00000119346 | Gm23151 | predicted gene, 23151 |
| chr7 | 5378774 | 5380182 | 1409 | * | 13,78706 | -2,490478159 | 0,963241 | -2,58552 | 0,009723 | 0,901463351 | 1,2763 | Distal Intergenic | 7 | 5354207 | 5362964 | 8758 | 1 | 272350 | ENSMUST00000207531.2 | 24567 | ENSMUSG00000048398 | Gm5065 | predicted gene 5065 |
| chr11 | 14971783 | 14974488 | 2706 | * | 16,64115 | -2,486703157 | 0,909478 | -2,73421 | 0,006253 | 0,825384969 | 1,2763 | Distal Intergenic | 11 | 15362300 | 15362406 | 107 | 1 | 1,15E+08 | ENSMUST00000240290.1 | -387812 | ENSMUSG00000119754 | Gm24707 | predicted gene, 24707 |
| chr5 | 99845732 | 99848027 | 2296 | * | 22,15603 | -2,484890126 | 0,871712 | -2,85059 | 0,004364 | 0,621406209 | 2,951933 | Intron (ENSMUST0000( | 5 | 99830298 | 99863629 | 33332 | 1 | 1,03E+08 | ENSMUST00000161197.2 | 15434 | ENSMUSG00000089743 | Gm16226 | predicted gene 16226 |
| chr1 | 1,03E+08 | 1,03E+08 | 1786 | * | 16,97962 | -2,483466138 | 0,852443 | -2,91335 | 0,003576 | 0,627785088 | 2,318434 | Intron (ENSMUST0000( | 1 | 1,03E+08 | 1,03E+08 | 134 | 2 | 1,15E+08 | ENSMUST00000157283.3 | -265174 | ENSMUSG00000087908 | Gm22034 | predicted gene, 22034 |
| chr9 | 89677199 | 89679347 | 2149 | * | 19,81347 | -2,482969312 | 0,842635 | -2,94667 | 0,003212 | 0,576696438 | 2,713306 | Distal Intergenic | 9 | 89684313 | 89684381 | 69 | 2 | 387179 | ENSMUST00000083662.3 | 5034 | ENSMUSG00000065596 | Mir184 | microRNA 184 |
| chr11 | 54234879 | 54236211 | 1333 | * | 13,65358 | -2,48242033 | 0,943647 | -2,63067 | 0,008522 | 0,881662676 | 1,2763 | Promoter (<=1kb) | 11 | 54236714 | 54252354 | 15641 | 1 | 216739 | ENSMUST00000139484.2 | -503 | ENSMUSG00000020333 | Acsl6 | acyl-CoA synthetase long-chain family member 6 |
| chr9 | 1,21E+08 | 1,21E+08 | 1919 | * | 17,23037 | -2,481755837 | 0,837268 | -2,96411 | 0,003036 | 0,61140987 | 2,154504 | 3' UTR | 9 | 1,21E+08 | 1,21E+08 | 32682 | 2 | 209012 | ENSMUST00000164969.2 | -14809 | ENSMUSG00000040936 | Ulk4 | unc-51-like kinase 4 |
| chr3 | 99699168 | 99701371 | 2204 | * | 20,70098 | -2,481511741 | 0,878463 | -2,82483 | 0,00473 | 0,641187423 | 2,880017 | Distal Intergenic | 3 | 99615056 | 99615163 | 108 | 2 | 1,15E+08 | ENSMUST00000082956.3 | -84005 | ENSMUSG00000064890 | Gm22505 | predicted gene, 22505 |
| chr10 | 35031703 | 35032742 | 1040 | * | 13,60102 | -2,479576017 | 0,910522 | -2,72325 | 0,006464 | 0,889425844 | 0,918255 | Distal Intergenic | 10 | 34988194 | 34988327 | 134 | 1 | 1,15E+08 | ENSMUST00000104561.3 | 43509 | ENSMUSG00000077749 | Gm26341 | predicted gene, 26341 |
| chr14 | 28646306 | 28648811 | 2506 | * | 19,42485 | -2,478475733 | 0,892134 | -2,77814 | 0,005467 | 0,683988405 | 2,507013 | Intron (ENSMUST0000( | 14 | 28630009 | 28691370 | 61362 | 2 | 12294 | ENSMUST00000225668.2 | 42559 | ENSMUSG00000021991 | Cacna2d3 | calcium channel, voltage-dependent, alpha2/delta subunit 3 |
| chr11 | 91607085 | 91608512 | 1428 | * | 14,9296 | -2,477203433 | 0,863536 | -2,86867 | 0,004122 | 0,749931768 | 1,2763 | Distal Intergenic | 11 | 91466141 | 91468384 | 2244 | 2 | 73470 | ENSMUST00000061019.6 | -138701 | ENSMUSG00000046755 | Kif2b | kinesin family member 2B |
| chr17 | 42671758 | 42673922 | 2165 | * | 18,28467 | -2,477133727 | 0,818314 | -3,02712 | 0,002469 | 0,531203841 | 2,663619 | Intron (ENSMUST0000( | 17 | 42627491 | 42815968 | 188478 | 1 | 627626 | ENSMUST00000232968.2 | 44267 | ENSMUSG00000042256 | Ptchd4 | patched domain containing 4 |
| chr2 | 1,73E+08 | 1,73E+08 | 1205 | * | 13,59824 | -2,476396195 | 0,930721 | -2,66073 | 0,007797 | 0,892811906 | 1,075735 | Distal Intergenic | 2 | 1,73E+08 | 1,73E+08 | 1155 | 1 | 1,03E+08 | ENSMUST00000147357.2 | 14927 | ENSMUSG00000086114 | Gm14642 | predicted gene 14642 |
| chr5 | 1,06E+08 | 1,06E+08 | 2758 | * | 22,34185 | -2,476259918 | 0,898803 | -2,75506 | 0,005868 | 0,671156916 | 2,951933 | Distal Intergenic | 5 | 1,06E+08 | 1,06E+08 | 8457 | 1 | 1,03E+08 | ENSMUST00000200427.2 | 89048 | | Gm32736 | predicted gene, 32736 |
| chr2 | 1,51E+08 | 1,51E+08 | 2053 | * | 17,01386 | -2,476251456 | 0,960677 | -2,57761 | 0,009949 | 0,777919825 | 2,663619 | Exon (ENSMUST00000( | 2 | 1,51E+08 | 1,52E+08 | 23130 | 2 | 668929 | ENSMUST00000096439.4 | 13667 | ENSMUSG00000074704 | Rad21l | RAD21-like (S. pombe) |
| chr4 | 9560198 | 9562737 | 2540 | * | 21,51826 | -2,476088677 | 0,900556 | -2,74951 | 0,005968 | 0,647880877 | 3,471678 | Intron (ENSMUST0000( | 4 | 9574872 | 9595689 | 20818 | 2 | 65973 | ENSMUST00000124408.2 | 32952 | ENSMUSG00000028207 | Asph | aspartate-beta-hydroxylase |
| chr2 | 1,18E+08 | 1,18E+08 | 1557 | * | 14,51371 | -2,475388351 | 0,927252 | -2,6696 | 0,007594 | 0,841453364 | 1,41696 | Intron (ENSMUST0000( | 2 | 1,18E+08 | 1,18E+08 | 3253 | 1 | 73358 | ENSMUST00000125654.2 | 7651 | ENSMUSG00000085889 | 1700054M17Rik | RIKEN cDNA 1700054M17 gene |
| chr4 | 81484501 | 81487296 | 2796 | * | 23,36291 | -2,474034806 | 0,83499 | -2,96295 | 0,003047 | 0,55292773 | 2,858511 | Distal Intergenic | 4 | 81334777 | 81361052 | 26276 | 2 | 17475 | ENSMUST00000142273.2 | -123449 | ENSMUSG00000028402 | Mpdz | multiple PDZ domain crumbs cell polarity complex component |
| chr15 | 27694777 | 27696364 | 1588 | * | 16,0253 | -2,47344922 | 0,91694 | -2,6975 | 0,006986 | 0,715249603 | 2,663619 | Distal Intergenic | 15 | 27655157 | 27681665 | 26509 | 2 | 223433 | ENSMUST00000100739.5 | -13112 | ENSMUSG00000056069 | Otulinl | OTU deubiquitinase with linear linkage specificity like |
| chr10 | 54343733 | 54345583 | 1851 | * | 13,36136 | -2,47277668 | 0,923855 | -2,67659 | 0,007438 | 0,910106437 | 0,918255 | Distal Intergenic | 10 | 54187321 | 54187427 | 107 | 2 | 1,15E+08 | ENSMUST00000239711.1 | -156306 | ENSMUSG00000119760 | Gm26177 | predicted gene, 26177 |
| chr1 | 1,19E+08 | 1,19E+08 | 1784 | * | 17,38817 | -2,472332347 | 0,907086 | -2,72558 | 0,006419 | 0,713633527 | 2,507013 | Distal Intergenic | 1 | 1,19E+08 | 1,19E+08 | 9822 | 1 | 81879 | ENSMUST00000190652.2 | -12523 | ENSMUSG00000026380 | Tfcp2l1 | transcription factor CP2-like 1 |
| chr6 | 87418432 | 87419887 | 1456 | * | 16,66635 | -2,471765938 | 0,94042 | -2,62836 | 0,00858 | 0,791068542 | 2,141612 | Distal Intergenic | 6 | 87405976 | 87414659 | 8684 | 1 | 12154 | ENSMUST00000032125.7 | 12456 | ENSMUSG00000030046 | Bmp10 | bone morphogenetic protein 10 |
| chr12 | 43950531 | 43952384 | 1854 | * | 25,45296 | -2,471704005 | 0,760223 | -3,25129 | 0,001149 | 0,383459188 | 2,951062 | Distal Intergenic | 12 | 44002462 | 44007880 | 5419 | 2 | 1,08E+08 | ENSMUST00000219678.2 | 55496 | | Gm46349 | predicted gene, 46349 |
| chr12 | 77677763 | 77680108 | 2346 | * | 19,05249 | -2,470549931 | 0,909886 | -2,71523 | 0,006623 | 0,703687259 | 2,713306 | Distal Intergenic | 12 | 77600121 | 77616483 | 16363 | 1 | 1,03E+08 | ENSMUST00000218335.2 | 77642 | | Gm35240 | predicted gene, 35240 |
| chr6 | 61421113 | 61422147 | 1035 | * | 14,34907 | -2,467367 | 0,904835 | -2,72687 | 0,006394 | 0,830133334 | 1,2763 | Intron (ENSMUST0000( | 6 | 61351024 | 61447249 | 96226 | 1 | 232035 | ENSMUST00000203112.2 | 70089 | ENSMUSG00000039578 | Ccser1 | coiled-coil serine rich 1 |
| chr12 | 70179235 | 70181060 | 1826 | * | 15,40776 | -2,466458374 | 0,936377 | -2,63404 | 0,008437 | 0,879289589 | 1,2763 | Distal Intergenic | 12 | 70058209 | 70160491 | 102283 | 2 | 18080 | ENSMUST00000222237.2 | -18744 | ENSMUSG00000021068 | Nin | ninein |
| chr10 | 15599065 | 15600956 | 1892 | * | 16,03873 | -2,466012348 | 0,853374 | -2,88972 | 0,003856 | 0,739716578 | 1,2763 | Intron (ENSMUST0000( | 10 | 15610660 | 15614570 | 3911 | 2 | 1,03E+08 | ENSMUST00000218861.2 | 13614 | | Gm32283 | predicted gene, 32283 |
| chr4 | 17761456 | 17762805 | 1350 | * | 11,96614 | -2,463985974 | 0,949714 | -2,59445 | 0,009474 | 0,948006995 | 0,918255 | Distal Intergenic | 4 | 17852893 | 18117457 | 264565 | 1 | 17389 | ENSMUST00000183662.8 | -90088 | ENSMUSG00000028226 | Mmp16 | matrix metallopeptidase 16 |
| chr14 | 1,05E+08 | 1,05E+08 | 2814 | * | 17,22781 | -2,462864188 | 0,934487 | -2,63553 | 0,008401 | 0,756025543 | 2,507013 | Distal Intergenic | 14 | 1,05E+08 | 1,05E+08 | 134 | 1 | 1,15E+08 | ENSMUST00000104237.3 | 58898 | ENSMUSG00000077425 | Gm24280 | predicted gene, 24280 |
| chr16 | 92874479 | 92876169 | 1691 | * | 21,28987 | -2,462773913 | 0,793572 | -3,1034 | 0,001913 | 0,46944092 | 2,951933 | Distal Intergenic | 16 | 92398354 | 92623037 | 224684 | 2 | 12394 | ENSMUST00000023673.14 | -251442 | ENSMUSG00000022952 | Runx1 | runt related transcription factor 1 |
| chr12 | 35880622 | 35882734 | 2113 | * | 20,0519 | -2,461521788 | 0,859914 | -2,86252 | 0,004203 | 0,663630601 | 2,221611 | Distal Intergenic | 12 | 35975619 | 35999735 | 24117 | 1 | 403205 | ENSMUST00000042101.5 | -92885 | ENSMUSG00000036231 | Agr3 | anterior gradient 3 |
| chr11 | 82471914 | 82474253 | 2340 | * | 22,26645 | -2,46016426 | 0,876351 | -2,80728 | 0,004996 | 0,637143933 | 3,104559 | Distal Intergenic | 11 | 82486899 | 82487005 | 107 | 2 | 1,15E+08 | ENSMUST00000083077.3 | 12752 | ENSMUSG00000119381 | Gm24612 | predicted gene, 24612 |
| chr17 | 73283309 | 73284868 | 1560 | * | 13,33518 | -2,459716671 | 0,932146 | -2,63877 | 0,008321 | 0,923399738 | 0,938682 | Distal Intergenic | 17 | 73294793 | 73294899 | 107 | 1 | 1,15E+08 | ENSMUST00000093636.4 | -9925 | ENSMUSG00000118930 | Gm23649 | predicted gene, 23649 |
| chr17 | 37544063 | 37545542 | 1480 | * | 17,24959 | -2,459143752 | 0,948348 | -2,59308 | 0,009512 | 0,807119269 | 2,154504 | Promoter (<=1kb) | 17 | 37542327 | 37543259 | 933 | 2 | 258505 | ENSMUST00000207414.3 | -804 | ENSMUSG00000058802 | Olfr97 | olfactory receptor 97 |
| chr1 | 22018978 | 22020152 | 1175 | * | 12,59554 | -2,457945617 | 0,928815 | -2,64633 | 0,008137 | 0,887433944 | 1,173325 | Intron (ENSMUST0000( | 1 | 21468627 | 22031805 | 563179 | 2 | 226922 | ENSMUST00000029667.13 | 11653 | ENSMUSG00000028033 | Kcnq5 | potassium voltage-gated channel, subfamily Q, member 5 |
| chr9 | 1,15E+08 | 1,15E+08 | 1454 | * | 16,00687 | -2,455758156 | 0,838253 | -2,92962 | 0,003394 | 0,717133806 | 1,2763 | Distal Intergenic | 9 | 1,15E+08 | 1,15E+08 | 253655 | 1 | 74486 | ENSMUST00000183104.8 | -18071 | ENSMUSG00000040875 | Osbpl10 | oxysterol binding protein-like 10 |
| chr10 | 91373467 | 91374806 | 1340 | * | 16,49654 | -2,455537865 | 0,916162 | -2,68024 | 0,007357 | 0,853738252 | 1,2763 | Distal Intergenic | 10 | 91044120 | 91056504 | 12385 | 2 | 1,03E+08 | ENSMUST00000216311.2 | -316963 | ENSMUSG00000111544 | 4930534H03Rik | RIKEN cDNA 4930534H03 gene |
| chr13 | 69052994 | 69054409 | 1416 | * | 19,29002 | -2,453459636 | 0,924094 | -2,65499 | 0,007931 | 0,769692849 | 2,221611 | Intron (ENSMUST0000( | 13 | 69146912 | 69179593 | 32682 | 1 | 1,03E+08 | ENSMUST00000221765.2 | -92503 | | Gm35161 | predicted gene, 35161 |
| chr14 | 40719959 | 40721218 | 1260 | * | 15,66825 | -2,451972113 | 0,841517 | -2,91375 | 0,003571 | 0,642870233 | 2,141612 | Exon (ENSMUST00000( | 14 | 40720830 | 40725061 | 4232 | 2 | 70564 | ENSMUST00000135797.2 | 3843 | ENSMUSG00000021792 | Prxl2a | peroxiredoxin like 2A |
| chr5 | 61286284 | 61288913 | 2630 | * | 16,29997 | -2,450023775 | 0,925544 | -2,64712 | 0,008118 | 0,766298356 | 2,318434 | Distal Intergenic | 5 | 60658176 | 60658283 | 108 | 2 | 1,15E+08 | ENSMUST00000157423.3 | -628001 | ENSMUSG00000088048 | Gm23331 | predicted gene, 23331 |
| chr11 | 28819941 | 28821829 | 1889 | * | 16,5788 | -2,449842336 | 0,873599 | -2,80431 | 0,005042 | 0,789027175 | 1,2763 | Intron (ENSMUST0000( | 11 | 28803244 | 28871526 | 68283 | 1 | 216616 | ENSMUST00000124103.2 | 16697 | ENSMUSG00000020467 | Efemp1 | epidermal growth factor-containing fibulin-like extracellular matrix protein 1 |
| chr9 | 1,23E+08 | 1,23E+08 | 1772 | * | 15,44595 | -2,449697386 | 0,919254 | -2,66487 | 0,007702 | 0,770673161 | 2,141612 | Intron (ENSMUST0000( | 9 | 1,23E+08 | 1,23E+08 | 4899 | 1 | 83493 | ENSMUST00000217089.2 | -10468 | ENSMUSG00000025240 | Sacm1l | SAC1 suppressor of actin mutations 1-like (yeast) |
| chr9 | 22745764 | 22748381 | 2618 | * | 19,99472 | -2,449377517 | 0,750623 | -3,26313 | 0,001102 | 0,406160411 | 2,507013 | Intron (ENSMUST0000( | 9 | 22590046 | 22799570 | 209525 | 1 | 319845 | ENSMUST00000124076.2 | 155718 | ENSMUSG00000035919 | Bbs9 | Bardet-Biedl syndrome 9 (human) |
| chr12 | 1,05E+08 | 1,05E+08 | 1263 | * | 13,65445 | -2,448599033 | 0,937158 | -2,61279 | 0,008981 | 0,935723116 | 0,938682 | Distal Intergenic | 12 | 1,04E+08 | 1,04E+08 | 6010 | 1 | 791311 | ENSMUST00000222329.2 | 58108 | | Gm10000 | predicted gene 10000 |
| chr14 | 80017908 | 80020138 | 2231 | * | 19,82766 | -2,443424146 | 0,875929 | -2,78952 | 0,005279 | 0,669491826 | 2,713306 | Distal Intergenic | 14 | 80004220 | 80008752 | 4533 | 2 | 18530 | ENSMUST00000195355.2 | -9156 | ENSMUSG00000036422 | Pcdh8 | protocadherin 8 |
| chr9 | 18272894 | 18274425 | 1532 | * | 15,49411 | -2,441503276 | 0,846573 | -2,88399 | 0,003927 | 0,725737145 | 1,41696 | Intron (ENSMUST0000( | 9 | 18292065 | 18297218 | 5154 | 2 | 72560 | ENSMUST00000168934.2 | 22793 | ENSMUSG00000043943 | Naalad2 | N-acetylated alpha-linked acidic dipeptidase 2 |
| chr16 | 12913548 | 12915485 | 1938 | * | 16,3294 | -2,441092527 | 0,922576 | -2,64595 | 0,008146 | 0,767061106 | 2,318434 | Distal Intergenic | 16 | 12927548 | 12966598 | 39051 | 1 | 50505 | ENSMUST00000023206.14 | -12063 | ENSMUSG00000022545 | Ercc4 | excision repair cross-complementing rodent repair deficiency, complementation group 4 |
| chr4 | 38718936 | 38720867 | 1932 | * | 20,271 | -2,44089585 | 0,843854 | -2,89256 | 0,003821 | 0,61616728 | 2,663619 | Intron (ENSMUST0000( | 4 | 38645427 | 38855463 | 210037 | 2 | 1,03E+08 | ENSMUST00000244574.1 | 134596 | ENSMUSG00000085632 | Gm12381 | predicted gene 12381 |
| chr17 | 41658133 | 41660080 | 1948 | * | 13,60792 | -2,439149668 | 0,898912 | -2,71345 | 0,006659 | 0,890237282 | 0,938682 | Distal Intergenic | 17 | 41266079 | 41269872 | 3794 | 1 | 17850 | ENSMUST00000233222.2 | 392054 | ENSMUSG00000023921 | Mmut | methylmalonyl-Coenzyme A mutase |
| chr7 | 1,18E+08 | 1,18E+08 | 2293 | * | 19,1843 | -2,437177051 | 0,791257 | -3,08013 | 0,002069 | 0,498636665 | 2,663619 | Distal Intergenic | 7 | 1,18E+08 | 1,18E+08 | 636 | 1 | 1,03E+08 | ENSMUST00000208337.2 | 41420 | | Gm35309 | predicted gene, 35309 |
| chr17 | 37451580 | 37453837 | 2258 | * | 21,49627 | -2,437068035 | 0,718639 | -3,39123 | 0,000696 | 0,300106667 | 3,104559 | Intron (ENSMUST0000( | 17 | 37461898 | 37462942 | 1045 | 2 | 258051 | ENSMUST00000171679.2 | 9105 | ENSMUSG00000091601 | Olfr93 | olfactory receptor 93 |
| chr17 | 37312490 | 37314362 | 1873 | * | 17,70462 | -2,43664763 | 0,922446 | -2,64151 | 0,008254 | 0,744477896 | 2,663619 | Promoter (<=1kb) | 17 | 37313374 | 37320334 | 6961 | 1 | 22715 | ENSMUST00000172527.8 | 0 | ENSMUSG00000036036 | Zfp57 | zinc finger protein 57 |
| chr1 | 1,76E+08 | 1,76E+08 | 1565 | * | 17,09998 | -2,436335729 | 0,943942 | -2,58102 | 0,009851 | 0,903101391 | 1,2763 | Intron (ENSMUST0000( | 1 | 1,76E+08 | 1,76E+08 | 14535 | 2 | 319455 | ENSMUST00000111166.2 | 27761 | ENSMUSG00000055214 | Pld5 | phospholipase D family, member 5 |
| chr15 | 63546083 | 63547190 | 1108 | * | 15,59104 | -2,436334181 | 0,925277 | -2,63309 | 0,008461 | 0,863495249 | 1,37683 | Intron (ENSMUST0000( | 15 | 63589667 | 63609529 | 19863 | 1 | 73994 | ENSMUST00000226508.2 | -42477 | | 4930449C09Rik | RIKEN cDNA 4930449C09 gene |
| chr13 | 53316015 | 53317759 | 1745 | * | 14,95537 | -2,434107032 | 0,935434 | -2,60212 | 0,009265 | 0,802490409 | 2,141612 | Intron (ENSMUST0000( | 13 | 53285996 | 53429694 | 143699 | 2 | 26564 | ENSMUST00000149299.2 | 111935 | ENSMUSG00000021464 | Ror2 | receptor tyrosine kinase-like orphan receptor 2 |
| chr5 | 1,51E+08 | 1,51E+08 | 1133 | * | 13,58557 | -2,432866449 | 0,921626 | -2,63976 | 0,008297 | 0,923203349 | 0,938682 | Intron (ENSMUST0000( | 5 | 1,51E+08 | 1,51E+08 | 6137 | 1 | 100710 | ENSMUST00000201920.2 | -5910 | ENSMUSG00000034021 | Pds5b | PDS5 cohesin associated factor B |
| chr10 | 86010574 | 86013513 | 2940 | * | 27,08241 | -2,432311162 | 0,770348 | -3,15742 | 0,001592 | 0,454149353 | 2,67243 | Intron (ENSMUST0000( | 10 | 85931723 | 85949450 | 17728 | 1 | 78170 | ENSMUST00000218077.2 | 78851 | ENSMUSG00000112493 | 4930486F22Rik | RIKEN cDNA 4930486F22 gene |
| chr8 | 4035408 | 4037350 | 1943 | * | 16,98806 | -2,431925063 | 0,810067 | -3,00213 | 0,002681 | 0,589834336 | 2,141612 | Distal Intergenic | 8 | 3990294 | 4004746 | 14453 | 2 | 170776 | ENSMUST00000127592.2 | -30662 | ENSMUSG00000040165 | Cd209c | CD209c antigen |
| chr2 | 54233191 | 54234102 | 912 | * | 13,83151 | -2,431848826 | 0,908582 | -2,67653 | 0,007439 | 0,906348091 | 0,938682 | Distal Intergenic | 2 | 54326329 | 55008321 | 681993 | 1 | 271786 | ENSMUST00000112636.8 | -92227 | ENSMUSG00000060988 | Galnt13 | polypeptide N-acetylgalactosaminyltransferase 13 |
| chr2 | 1,33E+08 | 1,33E+08 | 2040 | * | 16,27181 | -2,430577108 | 0,941678 | -2,58111 | 0,009848 | 0,903101391 | 1,2763 | Distal Intergenic | 2 | 1,33E+08 | 1,33E+08 | 21268 | 1 | 381426 | ENSMUST00000149965.2 | -35874 | | A430048G15Rik | RIKEN cDNA A430048G15 gene |
| chr12 | 37824323 | 37825839 | 1517 | * | 14,03783 | -2,429559178 | 0,890428 | -2,72853 | 0,006362 | 0,919610766 | 0,738441 | Distal Intergenic | 12 | 37867725 | 38032009 | 164285 | 1 | 217480 | ENSMUST00000222337.2 | -41886 | ENSMUSG00000036095 | Dgkb | diacylglycerol kinase, beta |
| chr8 | 9984157 | 9986105 | 1949 | * | 16,88234 | -2,42752859 | 0,877119 | -2,76762 | 0,005647 | 0,695892636 | 2,386008 | Distal Intergenic | 8 | 10019049 | 10026299 | 7251 | 2 | 319583 | ENSMUST00000095476.6 | 40194 | ENSMUSG00000049717 | Lig4 | ligase IV, DNA, ATP-dependent |
| chr15 | 27078299 | 27079773 | 1475 | * | 15,90599 | -2,427495695 | 0,829062 | -2,928 | 0,003411 | 0,594430593 | 2,663619 | Distal Intergenic | 15 | 27061822 | 27061952 | 131 | 2 | 1,15E+08 | ENSMUST00000157566.3 | -16347 | ENSMUSG00000088191 | Gm23903 | predicted gene, 23903 |
| chr15 | 30761952 | 30763590 | 1639 | * | 19,21361 | -2,426687659 | 0,835188 | -2,90556 | 0,003666 | 0,607561104 | 2,663619 | Intron (ENSMUST0000( | 15 | 30683305 | 30807046 | 123742 | 1 | 18163 | ENSMUST00000227065.2 | 78647 | ENSMUSG00000022240 | Ctnnd2 | catenin (cadherin associated protein), delta 2 |
| chr1 | 41528936 | 41530612 | 1677 | * | 16,32186 | -2,426515528 | 0,82973 | -2,92447 | 0,00345 | 0,637442941 | 2,141612 | Distal Intergenic | 1 | 41782848 | 41978749 | 195902 | 1 | 1,03E+08 | ENSMUST00000247631.1 | -252236 | | Gm29260 | predicted gene 29260 |
| chr3 | 1,47E+08 | 1,47E+08 | 1844 | * | 18,30183 | -2,425962121 | 0,893201 | -2,71603 | 0,006607 | 0,722798324 | 2,423594 | Distal Intergenic | 3 | 1,47E+08 | 1,47E+08 | 11284 | 1 | 1,03E+08 | ENSMUST00000196485.2 | -118220 | | Gm36831 | predicted gene, 36831 |
| chr12 | 64128023 | 64130301 | 2279 | * | 18,07047 | -2,425064871 | 0,797765 | -3,03982 | 0,002367 | 0,533987851 | 2,507013 | Distal Intergenic | 12 | 64518104 | 64521464 | 3361 | 2 | 623046 | ENSMUST00000059833.8 | 391163 | ENSMUSG00000043060 | Fscb | fibrous sheath CABYR binding protein |
| chr2 | 81529302 | 81531592 | 2291 | * | 15,19708 | -2,422890735 | 0,892135 | -2,71584 | 0,006611 | 0,836571183 | 1,2763 | Distal Intergenic | 2 | 81883566 | 82090223 | 206658 | 1 | 241514 | ENSMUST00000047527.8 | -351974 | ENSMUSG00000070866 | Zfp804a | zinc finger protein 804A |
| chr18 | 71443058 | 71445124 | 2067 | * | 16,43082 | -2,421046973 | 0,902835 | -2,68161 | 0,007327 | 0,74323496 | 2,386008 | Intron (ENSMUST0000( | 18 | 70750844 | 70758848 | 8005 | 1 | 17191 | ENSMUST00000128555.2 | 692214 | ENSMUSG00000024513 | Mbd2 | methyl-CpG binding domain protein 2 |
| chr6 | 36390538 | 36391859 | 1322 | * | 21,34449 | -2,420283863 | 0,833353 | -2,90427 | 0,003681 | 0,607561104 | 2,67243 | Intron (ENSMUST0000( | 6 | 36398677 | 36398760 | 84 | 1 | 735279 | ENSMUST00000093567.3 | -6818 | ENSMUSG00000070075 | Mir490 | microRNA 490 |
| chr1 | 1,74E+08 | 1,74E+08 | 2186 | * | 16,93099 | -2,419540099 | 0,867253 | -2,78989 | 0,005273 | 0,705458567 | 2,141612 | Promoter (<=1kb) | 1 | 1,74E+08 | 1,74E+08 | 19758 | 2 | 226695 | ENSMUST00000059226.7 | 0 | ENSMUSG00000054203 | Ifi205 | interferon activated gene 205 |
| chr2 | 1,52E+08 | 1,52E+08 | 1782 | * | 17,59384 | -2,419203423 | 0,847701 | -2,85384 | 0,004319 | 0,656950206 | 2,386008 | Promoter (2-3kb) | 2 | 1,52E+08 | 1,52E+08 | 5303 | 1 | 108832 | ENSMUST00000060196.3 | -2750 | ENSMUSG00000044364 | Tmem74b | transmembrane protein 74B |
| chr6 | 1,37E+08 | 1,37E+08 | 1572 | * | 20,20366 | -2,419004697 | 0,791061 | -3,05792 | 0,002229 | 0,511485635 | 2,663619 | Exon (ENSMUST000002 | 6 | 1,37E+08 | 1,37E+08 | 52513 | 1 | 19277 | ENSMUST00000167002.4 | -32497 | ENSMUSG00000030223 | Ptpro | protein tyrosine phosphatase, receptor type, O |
| chr1 | 1,73E+08 | 1,73E+08 | 2341 | * | 21,1774 | -2,41818319 | 0,847413 | -2,85361 | 0,004323 | 0,61941203 | 2,951933 | Promoter (1-2kb) | 1 | 1,73E+08 | 1,73E+08 | 21110 | 1 | 1,03E+08 | ENSMUST00000245598.1 | 1295 | | Gm35686 | predicted gene, 35686 |
| chr14 | 37629534 | 37631136 | 1603 | * | 15,72937 | -2,417958094 | 0,917795 | -2,63453 | 0,008425 | 0,748971518 | 2,663619 | Distal Intergenic | 14 | 37323064 | 37328745 | 5682 | 2 | 1,03E+08 | ENSMUST00000224767.2 | -300789 | ENSMUSG00000114334 | Gm31393 | predicted gene, 31393 |
| chr18 | 70243198 | 70245878 | 2681 | * | 24,48519 | -2,417456121 | 0,717512 | -3,36922 | 0,000754 | 0,318448436 | 2,858511 | Intron (ENSMUST0000( | 18 | 70271235 | 70274645 | 3411 | 2 | 80718 | ENSMUST00000127217.2 | 28767 | ENSMUSG00000024511 | Rab27b | RAB27B, member RAS oncogene family |
| chr1 | 45783618 | 45785579 | 1962 | * | 18,52344 | -2,417398813 | 0,817632 | -2,95658 | 0,003111 | 0,586565597 | 2,507013 | Promoter (<=1kb) | 1 | 45786055 | 45786185 | 131 | 1 | 1,15E+08 | ENSMUST00000083321.3 | -476 | ENSMUSG00000065255 | Gm23974 | predicted gene, 23974 |
| chr2 | 1,05E+08 | 1,05E+08 | 2041 | * | 18,34487 | -2,416790007 | 0,881679 | -2,74112 | 0,006123 | 0,703968308 | 2,507013 | Distal Intergenic | 2 | 1,05E+08 | 1,05E+08 | 857 | 1 | 1,05E+08 | ENSMUST00000130677.2 | -26408 | | 4930527A07Rik | RIKEN cDNA 4930527A07 gene |
| chr14 | 38323002 | 38324470 | 1469 | * | 15,11593 | -2,415810418 | 0,905337 | -2,66841 | 0,007621 | 0,858623476 | 1,2763 | Intron (ENSMUST0000( | 14 | 38039861 | 38059083 | 19223 | 1 | 1,03E+08 | ENSMUST00000225614.2 | 283141 | | Gm31456 | predicted gene, 31456 |
| chr12 | 1,04E+08 | 1,04E+08 | 1656 | * | 16,64688 | -2,414957715 | 0,830794 | -2,90681 | 0,003651 | 0,60631654 | 2,663619 | Exon (ENSMUST000001 | 12 | 1,04E+08 | 1,04E+08 | 5480 | 2 | 71907 | ENSMUST00000164023.2 | -10550 | ENSMUSG00000058260 | Serpina9 | serine (or cysteine) peptidase inhibitor, clade A (alpha-1 antiproteinase, antitrypsin), member 9 |
| chr14 | 1,02E+08 | 1,02E+08 | 1763 | * | 14,5073 | -2,413949148 | 0,90889 | -2,65593 | 0,007909 | 0,851014755 | 1,41696 | Distal Intergenic | 14 | 1,02E+08 | 1,02E+08 | 20856 | 2 | 210789 | ENSMUST00000161304.2 | 90949 | ENSMUSG00000033083 | Tbc1d4 | TBC1 domain family, member 4 |
| chr17 | 50625606 | 50627069 | 1464 | * | 18,52326 | -2,413185448 | 0,814673 | -2,96215 | 0,003055 | 0,613331296 | 2,154504 | Distal Intergenic | 17 | 50646280 | 50705503 | 59224 | 1 | 1,03E+08 | ENSMUST00000233175.2 | -19211 | | Gm31015 | predicted gene, 31015 |
| chr18 | 30261123 | 30262752 | 1630 | * | 19,34524 | -2,412657782 | 0,879826 | -2,7422 | 0,006103 | 0,686297586 | 2,713306 | Distal Intergenic | 18 | 30400302 | 30405186 | 4885 | 1 | 225326 | ENSMUST00000234467.2 | -137550 | ENSMUSG00000033628 | Pik3c3 | phosphatidylinositol 3-kinase catalytic subunit type 3 |
| chr4 | 62961204 | 62963566 | 2363 | * | 20,86483 | -2,410304085 | 0,785293 | -3,0693 | 0,002146 | 0,492834344 | 2,880017 | Intron (ENSMUST0000( | 4 | 63038637 | 63039665 | 1029 | 1 | 72701 | ENSMUST00000151324.2 | -75071 | ENSMUSG00000028358 | Zfp618 | zinc finger protein 618 |
| chr19 | 55381653 | 55383961 | 2309 | * | 18,77026 | -2,40942578 | 0,86058 | -2,79977 | 0,005114 | 0,690933518 | 2,221611 | Intron (ENSMUST0000( | 19 | 55369446 | 55435411 | 65966 | 1 | 53611 | ENSMUST00000224396.2 | 12207 | ENSMUSG00000024983 | Vti1a | vesicle transport through interaction with t-SNAREs 1A |
| chr13 | 8755822 | 8757143 | 1322 | * | 15,40767 | -2,409350107 | 0,88142 | -2,73349 | 0,006267 | 0,836074214 | 1,214904 | Intron (ENSMUST0000( | 13 | 8885425 | 8887270 | 1846 | 2 | 207615 | ENSMUST00000177409.2 | 130127 | ENSMUSG00000021147 | Wdr37 | WD repeat domain 37 |
| chr5 | 1,33E+08 | 1,33E+08 | 1713 | * | 15,82711 | -2,409242402 | 0,90069 | -2,67488 | 0,007476 | 0,856202617 | 1,2763 | Distal Intergenic | 5 | 1,34E+08 | 1,34E+08 | 63817 | 2 | 1,03E+08 | ENSMUST00000196125.2 | 138576 | | Gm36667 | predicted gene, 36667 |
| chr9 | 54071026 | 54073144 | 2119 | * | 18,01008 | -2,408454767 | 0,933402 | -2,5803 | 0,009872 | 0,777172375 | 2,663619 | Distal Intergenic | 9 | 54073221 | 54100726 | 27506 | 2 | 13075 | ENSMUST00000034811.8 | 27582 | ENSMUSG00000032274 | Cyp19a1 | cytochrome P450, family 19, subfamily a, polypeptide 1 |
| chrX | 1,33E+08 | 1,33E+08 | 1574 | * | 20,28555 | -2,408397976 | 0,889886 | -2,70641 | 0,006801 | 0,713406565 | 2,663619 | Promoter (1-2kb) | 20 | 1,33E+08 | 1,33E+08 | 14598 | 1 | 64103 | ENSMUST00000033602.9 | -1514 | ENSMUSG00000031250 | Tnmd | tenomodulin |
| chr12 | 59685097 | 59686527 | 1431 | * | 15,35477 | -2,407144886 | 0,813998 | -2,95719 | 0,003105 | 0,702508842 | 1,2763 | Distal Intergenic | 12 | 59712440 | 59712570 | 131 | 2 | 1,15E+08 | ENSMUST00000104398.3 | 26043 | ENSMUSG00000077586 | Gm24377 | predicted gene, 24377 |

| | | | | | | | | | | | | | | | | | | | | | | |
|---|---|---|---|---|---|---|---|---|---|---|---|---|---|---|---|---|---|---|---|---|---|---|
| chr6 | 93213622 | 93216108 | 2487 | * | 19,86034 | -2,406336081 | 0,820249 | -2,93366 | 0,00335 | 0,625491631 | 2,221611 | Exon (ENSMUST00000 | 6 | 93312439 | 93312569 | 131 | 2 | 1,15E+08 | ENSMUST00000103992.3 | 96461 | ENSMUSG00000077180 | Gm25094 | predicted gene, 25094 |
| chr8 | 12160389 | 12162486 | 2098 | * | 19,04077 | -2,403711844 | 0,851688 | -2,82229 | 0,004768 | 0,683210259 | 2,221611 | Distal Intergenic | 8 | 12328819 | 12330657 | 1839 | 1 | 619290 | ENSMUST00000098935.2 | -166333 | ENSMUSG00000074473 | A230072I06Rik | RIKEN cDNA A230072I06 gene |
| chr7 | 69482785 | 69484324 | 1540 | * | 13,82858 | -2,402810902 | 0,883129 | -2,72079 | 0,006513 | 0,923078468 | 0,738441 | Distal Intergenic | 7 | 69607524 | 69607652 | 129 | 2 | 1,15E+08 | ENSMUST00000103978.3 | 123328 | ENSMUSG00000077166 | Gm24120 | predicted gene, 24120 |
| chr7 | 90315817 | 90317311 | 1495 | * | 14,25433 | -2,40064509 | 0,848708 | -2,82859 | 0,004675 | 0,775744126 | 1,2763 | Intron (ENSMUST00000 | 7 | 90126439 | 92098455 | 1972017 | 1 | 23859 | ENSMUST00000231777.3 | 189378 | ENSMUSG00000052572 | Dlg2 | discs large MAGUK scaffold protein 2 |
| chr11 | 1,15E+08 | 1,15E+08 | 1565 | * | 18,66457 | -2,399651669 | 0,869205 | -2,76074 | 0,005767 | 0,700182401 | 2,386008 | Downstream (<=300bp | 11 | 1,15E+08 | 1,15E+08 | 4487 | 1 | 432611 | ENSMUST00000130692.2 | 5299 | ENSMUSG00000034706 | Dnai2 | dynein axonemal intermediate chain 2 |
| chr8 | 69868732 | 69870190 | 1459 | * | 16,23826 | -2,398556379 | 0,868939 | -2,76033 | 0,005774 | 0,700691674 | 2,386008 | Distal Intergenic | 8 | 69819162 | 69848191 | 29030 | 2 | 378466 | ENSMUST00000239456.2 | -20541 | ENSMUSG00000110444 | Gm10033 | predicted gene 10033 |
| chr16 | 17125606 | 17126598 | 993 | * | 13,38853 | -2,398198914 | 0,914972 | -2,62106 | 0,008766 | 0,886332554 | 1,2763 | Promoter (<=1kb) | 16 | 17115448 | 17125594 | 10147 | 2 | 224020 | ENSMUST00000231683.2 | -12 | ENSMUSG00000041720 | Pi4ka | phosphatidylinositol 4-kinase alpha |
| chr10 | 27162328 | 27164219 | 1892 | * | 16,99119 | -2,398190902 | 0,832857 | -2,87947 | 0,003983 | 0,74685607 | 1,2763 | Intron (ENSMUST00000 | 10 | 27080797 | 27088250 | 7454 | 2 | 16773 | ENSMUST00000239098.2 | -74078 | ENSMUSG00000019899 | Lama2 | laminin, alpha 2 |
| chr1 | 98146166 | 98148145 | 1980 | * | 21,7045 | -2,395236537 | 0,780973 | -3,06699 | 0,002162 | 0,494205337 | 2,880017 | Distal Intergenic | 1 | 98101373 | 98118423 | 17051 | 1 | 1,03E+08 | ENSMUST00000189232.2 | 44793 | ENSMUSG00000100711 | Gm29461 | predicted gene 29461 |
| chr6 | 13133523 | 13134842 | 1320 | * | 20,73278 | -2,390096636 | 0,831994 | -2,87273 | 0,004069 | 0,574310558 | 3,471678 | Distal Intergenic | 6 | 13111298 | 13111426 | 129 | 1 | 1,15E+08 | ENSMUST00000104455.3 | 22225 | ENSMUSG00000077643 | Gm24985 | predicted gene, 24985 |
| chr12 | 1,02E+08 | 1,02E+08 | 1706 | * | 19,89061 | -2,38979139 | 0,750888 | -3,18262 | 0,001459 | 0,471617287 | 2,221611 | Distal Intergenic | 12 | 1,02E+08 | 1,02E+08 | 60504 | 2 | 74413 | ENSMUST00000160715.8 | 80563 | ENSMUSG00000021187 | Tc2n | tandem C2 domains, nuclear |
| chr1 | 95209942 | 95211873 | 1932 | * | 15,54698 | -2,389594285 | 0,831619 | -2,87342 | 0,00406 | 0,663630601 | 2,141612 | Distal Intergenic | 1 | 95241348 | 95263009 | 21662 | 1 | 67698 | ENSMUST00000059975.8 | -29475 | ENSMUSG00000051185 | Fam174a | family with sequence similarity 174, member A |
| chr10 | 15982556 | 15985132 | 2577 | * | 20,10328 | -2,38915416 | 0,788743 | -3,02907 | 0,002453 | 0,531203841 | 2,663619 | Distal Intergenic | 10 | 15667525 | 15682813 | 15289 | 1 | 77538 | ENSMUST00000218833.2 | 315031 | ENSMUSG00000112227 | C330004P14Rik | RIKEN cDNA C330004P14 gene |
| chr10 | 37130098 | 37131535 | 1438 | * | 13,63175 | -2,388370599 | 0,883667 | -2,7028 | 0,006876 | 0,932174684 | 0,738441 | Distal Intergenic | 10 | 37213482 | 37255648 | 42167 | 1 | 1,03E+08 | ENSMUST00000181602.2 | -81947 | | Gm26535 | predicted gene, 26535 |
| chr10 | 78519525 | 78521399 | 1875 | * | 19,75697 | -2,387356709 | 0,831753 | -2,87027 | 0,004101 | 0,627785088 | 2,663619 | Distal Intergenic | 10 | 78547831 | 78553240 | 5410 | 2 | 12365 | ENSMUST00000219237.2 | 31841 | ENSMUSG00000005355 | Casp14 | caspase 14 |
| chr7 | 1,13E+08 | 1,13E+08 | 2044 | * | 19,24971 | -2,387013748 | 0,788822 | -3,02605 | 0,002478 | 0,53146022 | 2,663619 | Distal Intergenic | 7 | 1,13E+08 | 1,13E+08 | 26990 | 1 | 1,03E+08 | ENSMUST00000210768.2 | -13176 | | Gm33586 | predicted gene, 33586 |
| chr11 | 81878199 | 81880446 | 2248 | * | 17,13716 | -2,38645244 | 0,899675 | -2,65257 | 0,007988 | 0,749550311 | 2,507013 | Promoter (1-2kb) | 11 | 81876232 | 81879595 | 3364 | 1 | 1,03E+08 | ENSMUST00000150772.2 | 1967 | | Gm11419 | predicted gene 11419 |
| chr18 | 60320454 | 60322585 | 2132 | * | 17,02317 | -2,383685339 | 0,924635 | -2,57797 | 0,009938 | 0,795181342 | 2,386008 | Distal Intergenic | 18 | 60345152 | 60380892 | 35741 | 1 | 240327 | ENSMUST00000031549.6 | -22567 | ENSMUSG00000073555 | Gm4951 | predicted gene 4951 |
| chr17 | 49525330 | 49526883 | 1554 | * | 15,69057 | -2,382631802 | 0,908769 | -2,62182 | 0,008746 | 0,891431005 | 1,214904 | Distal Intergenic | 17 | 49599770 | 49599929 | 160 | 1 | 1,15E+08 | ENSMUST00000082607.3 | -72887 | ENSMUSG00000064541 | Gm22978 | predicted gene, 22978 |
| chr9 | 1,17E+08 | 1,17E+08 | 1539 | * | 16,62835 | -2,380793656 | 0,906088 | -2,62755 | 0,0086 | 0,791416391 | 2,141612 | Intron (ENSMUST00000 | 9 | 1,17E+08 | 1,17E+08 | 2182 | 1 | 1,03E+08 | ENSMUST00000172075.3 | 186615 | ENSMUSG00000079669 | Gm17396 | predicted gene, 17396 |
| chr8 | 1,27E+08 | 1,27E+08 | 2201 | * | 22,13364 | -2,374547876 | 0,904985 | -2,62385 | 0,008694 | 0,709502177 | 3,471678 | Promoter (<=1kb) | 8 | 1,27E+08 | 1,27E+08 | 49737 | 2 | 212728 | ENSMUST00000170518.3 | 510 | ENSMUSG00000090290 | Tarbp1 | TAR RNA binding protein 1 |
| chr4 | 1,03E+08 | 1,03E+08 | 2021 | * | 16,84468 | -2,373879348 | 0,897837 | -2,644 | 0,008193 | 0,782813788 | 2,141612 | Distal Intergenic | 4 | 1,03E+08 | 1,03E+08 | 128 | 1 | 1,15E+08 | ENSMUST00000104072.3 | -43830 | ENSMUSG00000077260 | Gm24869 | predicted gene, 24869 |
| chr9 | 99911770 | 99913053 | 1284 | * | 14,82366 | -2,371765839 | 0,89024 | -2,66419 | 0,007717 | 0,770994815 | 2,141612 | Intron (ENSMUST00000 | 9 | 99905701 | 99916703 | 11003 | 1 | 108970 | ENSMUST00000215896.2 | 6069 | ENSMUSG00000101584 | 4930519F24Rik | RIKEN cDNA 4930519F24 gene |
| chr9 | 94373459 | 94374636 | 1178 | * | 13,23977 | -2,370371525 | 0,919375 | -2,57824 | 0,00993 | 0,916457354 | 1,173325 | Distal Intergenic | 9 | 94399917 | 94420134 | 20218 | 2 | 68861 | ENSMUST00000113028.2 | 45498 | ENSMUSG00000045414 | Dipk2a | divergent protein kinase domain 2A |
| chr10 | 11978745 | 11983394 | 4650 | * | 39,12053 | -2,369876108 | 0,617876 | -3,83552 | 0,000125 | 0,13754731 | 2,338387 | Distal Intergenic | 10 | 12239154 | 12239317 | 164 | 1 | 1,15E+08 | ENSMUST00000083162.3 | -255760 | ENSMUSG00000065096 | Gm23680 | predicted gene, 23680 |
| chr11 | 23594934 | 23597232 | 2299 | * | 16,58908 | -2,369483729 | 0,877098 | -2,7015 | 0,006903 | 0,749931768 | 2,141612 | 3' UTR | 11 | 23523788 | 23583639 | 59852 | 2 | 71675 | ENSMUST00000109532.9 | -11295 | ENSMUSG00000042208 | Sanbr | SANT and BTB domain regulator of CSR |
| chr8 | 94289315 | 94290280 | 966 | * | 13,64984 | -2,368146657 | 0,916404 | -2,58417 | 0,009761 | 0,948538535 | 0,938682 | Distal Intergenic | 8 | 94292749 | 94302071 | 9323 | 2 | 1,03E+08 | ENSMUST00000212213.2 | 11791 | | Gm26843 | predicted gene, 26843 |
| chr5 | 90345937 | 90347258 | 1322 | * | 13,74389 | -2,365853756 | 0,905245 | -2,6135 | 0,008962 | 0,888934304 | 1,2763 | Distal Intergenic | 5 | 90362586 | 90363440 | 855 | 2 | 231430 | ENSMUST00000133905.6 | 16182 | ENSMUSG00000035505 | Cox18 | cytochrome c oxidase assembly protein 18 |
| chr3 | 23017617 | 23019140 | 1524 | * | 13,78833 | -2,365738489 | 0,87712 | -2,69717 | 0,006993 | 0,899299554 | 0,938682 | Distal Intergenic | 3 | 23515214 | 23575562 | 60349 | 1 | 1E+08 | ENSMUST00000194267.2 | -496074 | ENSMUSG00000102203 | 4930412E21Rik | RIKEN cDNA 4930412E21 gene |
| chr8 | 46748847 | 46750472 | 1626 | * | 17,29757 | -2,365638192 | 0,86481 | -2,73544 | 0,00623 | 0,715249603 | 2,386008 | Promoter (1-2kb) | 8 | 46745076 | 46747708 | 2633 | 2 | 234219 | ENSMUST00000058636.9 | -1139 | ENSMUSG00000047171 | Helt | helt bHLH transcription factor |
| chr8 | 52405663 | 52407305 | 1643 | * | 14,68164 | -2,365377645 | 0,911008 | -2,59644 | 0,00942 | 0,903294867 | 1,214904 | Distal Intergenic | 8 | 52789950 | 52790140 | 191 | 2 | 1,15E+08 | ENSMUST00000082748.3 | 382835 | ENSMUSG00000064682 | Gm25813 | predicted gene, 25813 |
| chr3 | 1,12E+08 | 1,12E+08 | 2064 | * | 21,29149 | -2,365018623 | 0,744724 | -3,1757 | 0,001495 | 0,43148287 | 2,880017 | Distal Intergenic | 3 | 1,12E+08 | 1,12E+08 | 127 | 2 | 1,15E+08 | ENSMUST00000158392.3 | 53513 | ENSMUSG00000089017 | Gm25519 | predicted gene, 25519 |
| chr5 | 7036722 | 7038761 | 2040 | * | 18,03961 | -2,364618707 | 0,834561 | -2,83337 | 0,004606 | 0,683140223 | 2,154504 | Intron (ENSMUST00000 | 5 | 6819030 | 6926523 | 107494 | 2 | 207618 | ENSMUST00000164784.2 | -110199 | ENSMUSG00000092094 | Zfp804b | zinc finger protein 804B |
| chr12 | 43544548 | 43545733 | 1186 | * | 14,51805 | -2,362826469 | 0,886996 | -2,66385 | 0,007725 | 0,851512567 | 1,37683 | Distal Intergenic | 12 | 43789820 | 43789948 | 129 | 1 | 1,15E+08 | ENSMUST00000083070.3 | -244087 | ENSMUSG00000065004 | Gm26326 | predicted gene, 26326 |
| chr7 | 1,19E+08 | 1,19E+08 | 1865 | * | 15,68518 | -2,362332077 | 0,895117 | -2,63913 | 0,008312 | 0,876996413 | 1,2763 | Promoter (<=1kb) | 7 | 1,19E+08 | 1,19E+08 | 25068 | 1 | 434234 | ENSMUST00000207042.2 | -804 | ENSMUSG00000030924 | Rexo5 | RNA exonuclease 5 |
| chr4 | 1,35E+08 | 1,35E+08 | 2725 | * | 19,72053 | -2,362235804 | 0,843495 | -2,80053 | 0,005102 | 0,662924926 | 2,713306 | Distal Intergenic | 4 | 1,35E+08 | 1,35E+08 | 3544 | 1 | 73937 | ENSMUST00000153347.2 | -18988 | ENSMUSG00000086788 | 1700029M20Rik | RIKEN cDNA 1700029M20 gene |
| chr15 | 61226207 | 61227723 | 1517 | * | 18,41664 | -2,361257034 | 0,831474 | -2,83985 | 0,004514 | 0,663630601 | 2,386008 | Promoter (2-3kb) | 15 | 61229720 | 61229853 | 134 | 2 | 1,15E+08 | ENSMUST00000083878.3 | 2130 | ENSMUSG00000065812 | Gm24696 | predicted gene, 24696 |
| chr14 | 28274710 | 28276452 | 1743 | * | 16,66325 | -2,360546026 | 0,89485 | -2,63792 | 0,008342 | 0,770673161 | 2,318434 | Distal Intergenic | 14 | 28233408 | 28233864 | 457 | 1 | 22418 | ENSMUST00000146192.2 | 41302 | ENSMUSG00000021994 | Wnt5a | wingless-type MMTV integration site family, member 5A |
| chrX | 1,63E+08 | 1,63E+08 | 2252 | * | 19,90768 | -2,358688317 | 0,733143 | -3,21723 | 0,001294 | 0,421090004 | 2,663619 | Intron (ENSMUST00000 | 20 | 1,63E+08 | 1,63E+08 | 16097 | 2 | 56078 | ENSMUST00000126650.2 | 30742 | ENSMUSG00000031373 | Car5b | carbonic anhydrase 5b, mitochondrial |
| chr17 | 37574873 | 37576925 | 2053 | * | 17,81523 | -2,356739023 | 0,842098 | -2,79865 | 0,005132 | 0,668427355 | 2,663619 | Promoter (<=1kb) | 17 | 37573602 | 37574640 | 1039 | 2 | 258503 | ENSMUST00000080759.5 | -233 | ENSMUSG00000063660 | Olfr98 | olfactory receptor 98 |
| chr5 | 88639824 | 88642029 | 2206 | * | 19,01795 | -2,356619677 | 0,822232 | -2,86613 | 0,004155 | 0,642322948 | 2,507013 | Exon (ENSMUST00000 | 5 | 88635841 | 88653908 | 18068 | 1 | 13801 | ENSMUST00000031222.9 | 3983 | ENSMUSG00000029286 | Enam | enamelin |
| chr4 | 93266761 | 93268848 | 2088 | * | 16,8625 | -2,355699766 | 0,909305 | -2,59066 | 0,009579 | 0,793995086 | 2,318434 | Distal Intergenic | 4 | 93222375 | 93223748 | 1374 | 2 | 69136 | ENSMUST00000066774.6 | -43013 | ENSMUSG00000054000 | Tusc1 | tumor suppressor candidate 1 |
| chr14 | 29584604 | 29586230 | 1627 | * | 17,35981 | -2,35387156 | 0,845957 | -2,7825 | 0,005394 | 0,685026746 | 2,423594 | Distal Intergenic | 14 | 29508785 | 29574301 | 65517 | 2 | 1,03E+08 | ENSMUST00000223916.2 | -10303 | | Gm35281 | predicted gene, 35281 |
| chr1 | 1,31E+08 | 1,31E+08 | 1645 | * | 15,03046 | -2,351984667 | 0,814879 | -2,8863 | 0,003898 | 0,742014238 | 1,2763 | Distal Intergenic | 1 | 1,31E+08 | 1,31E+08 | 13807 | 1 | 64435 | ENSMUST00000112477.9 | -4952 | ENSMUSG00000026415 | Fcamr | Fc receptor, IgA, IgM, high affinity |
| chr1 | 1,11E+08 | 1,11E+08 | 1381 | * | 13,66638 | -2,350126595 | 0,910985 | -2,57976 | 0,009887 | 0,986185663 | 0,738441 | Intron (ENSMUST00000 | 1 | 1,11E+08 | 1,11E+08 | 89038 | 2 | 227485 | ENSMUST00000094626.5 | 31598 | ENSMUSG00000047216 | Cdh19 | cadherin 19, type 2 |
| chr4 | 1,1E+08 | 1,1E+08 | 2247 | * | 20,52814 | -2,348775055 | 0,730245 | -3,21642 | 0,001298 | 0,440188707 | 2,386008 | Exon (ENSMUST00000 | 4 | 1,1E+08 | 1,1E+08 | 16790 | 2 | 15572 | ENSMUST00000153200.2 | -3485 | ENSMUSG00000028546 | Elavl4 | ELAV like RNA binding protein 4 |
| chr1 | 15733868 | 15735788 | 1921 | * | 18,11585 | -2,347899266 | 0,82888 | -2,83262 | 0,004617 | 0,649217039 | 2,663619 | Intron (ENSMUST00000 | 1 | 15828056 | 15828187 | 132 | 1 | 1,15E+08 | ENSMUST00000104189.3 | -92268 | ENSMUSG00000077377 | Gm25168 | predicted gene, 25168 |
| chr14 | 59885489 | 59888251 | 2763 | * | 32,16706 | -2,347234477 | 0,692885 | -3,38763 | 0,000705 | 0,33490115 | 2,377674 | Intron (ENSMUST00000 | 14 | 59864638 | 59867126 | 2489 | 1 | 219134 | ENSMUST00000224155.2 | 20851 | ENSMUSG00000044461 | Shisa2 | shisa family member 2 |
| chr8 | 29469738 | 29471698 | 1961 | * | 27,87775 | -2,347010513 | 0,64936 | -3,61435 | 0,000301 | 0,2135993 | 2,67243 | Intron (ENSMUST00000 | 8 | 29142453 | 29709455 | 567003 | 2 | 210801 | ENSMUST00000209401.2 | 237757 | ENSMUSG00000063626 | Unc5d | unc-5 netrin receptor D |
| chr9 | 37950402 | 37951762 | 1361 | * | 13,43698 | -2,344969388 | 0,885266 | -2,64889 | 0,008076 | 0,923078468 | 0,918255 | Distal Intergenic | 9 | 37958520 | 37959449 | 930 | 1 | 257996 | ENSMUST00000077732.2 | -6758 | ENSMUSG00000095893 | Olfr884 | olfactory receptor 884 |
| chr14 | 33019278 | 33021425 | 2148 | * | 22,58626 | -2,343411105 | 0,863612 | -2,7135 | 0,006658 | 0,683210259 | 3,104559 | Exon (ENSMUST00000 | 14 | 33028075 | 33091889 | 63815 | 1 | 239027 | ENSMUST00000137590.8 | -6650 | ENSMUSG00000063506 | Arhgap22 | Rho GTPase activating protein 22 |
| chr3 | 45200112 | 45201961 | 1850 | * | 18,10783 | -2,342927984 | 0,823111 | -2,84643 | 0,004421 | 0,660698266 | 2,386008 | Distal Intergenic | 3 | 45281480 | 45332640 | 51161 | 2 | 72511 | ENSMUST00000191843.2 | 130679 | ENSMUSG00000097040 | 2610316D01Rik | RIKEN cDNA 2610316D01 gene |
| chr3 | 1,18E+08 | 1,18E+08 | 1748 | * | 20,10519 | -2,341766599 | 0,86662 | -2,70218 | 0,006889 | 0,71419937 | 2,663619 | Distal Intergenic | 3 | 1,18E+08 | 1,18E+08 | 87290 | 2 | 76561 | ENSMUST00000029639.12 | -71098 | ENSMUSG00000028007 | Snx7 | sorting nexin 7 |
| chr12 | 1,06E+08 | 1,06E+08 | 1953 | * | 14,63912 | -2,341651838 | 0,893424 | -2,62098 | 0,008768 | 0,89184433 | 1,214904 | Distal Intergenic | 12 | 1,06E+08 | 1,06E+08 | 5014 | 2 | 76639 | ENSMUST00000167922.2 | -82556 | ENSMUSG00000092066 | 1700121N20Rik | RIKEN cDNA 1700121N20 gene |
| chr19 | 43085048 | 43088260 | 3213 | * | 28,62892 | -2,34090663 | 0,720652 | -3,24832 | 0,001161 | 0,404612944 | 2,67243 | Intron (ENSMUST00000 | 19 | 42897023 | 43212128 | 315106 | 2 | 545291 | ENSMUST00000238153.2 | 123868 | ENSMUSG00000074852 | Hpse2 | heparanase 2 |
| chr11 | 66158998 | 66160322 | 1325 | * | 18,75798 | -2,338329068 | 0,78076 | -2,99494 | 0,002745 | 0,570876293 | 2,386008 | Intron (ENSMUST00000 | 11 | 66049578 | 66059377 | 9800 | 2 | 237806 | ENSMUST00000142836.2 | -99621 | ENSMUSG00000056752 | Dnah9 | dynein, axonemal, heavy chain 9 |
| chr6 | 3774306 | 3776089 | 1784 | * | 22,96379 | -2,338197114 | 0,770122 | -3,03614 | 0,002396 | 0,524856133 | 2,67243 | Intron (ENSMUST00000 | 6 | 3685720 | 3764714 | 78995 | 2 | 12311 | ENSMUST00000115622.8 | -9592 | ENSMUSG00000023964 | Calcr | calcitonin receptor |
| chr9 | 18609961 | 18612108 | 2148 | * | 18,19449 | -2,33793025 | 0,771539 | -3,03022 | 0,002444 | 0,54658246 | 2,386008 | Distal Intergenic | 9 | 18406814 | 18585826 | 179013 | 2 | 73732 | ENSMUST00000208663.2 | -24135 | ENSMUSG00000109564 | Muc16 | mucin 16 |
| chr10 | 97645095 | 97646526 | 1432 | * | 14,98665 | -2,337607707 | 0,84765 | -2,75775 | 0,00582 | 0,813839948 | 1,2763 | Intron (ENSMUST00000 | 10 | 97580619 | 97580746 | 128 | 2 | 1,15E+08 | ENSMUST00000158472.3 | -64349 | ENSMUSG00000089097 | Gm25658 | predicted gene, 25658 |
| chr2 | 94400139 | 94401536 | 1398 | * | 20,52096 | -2,337041258 | 0,774144 | -3,01887 | 0,002537 | 0,542967057 | 2,507013 | Distal Intergenic | 2 | 94452303 | 94452433 | 131 | 1 | 1,15E+08 | ENSMUST00000104490.3 | -50767 | ENSMUSG00000077678 | Gm24318 | predicted gene, 24318 |
| chr11 | 48470391 | 48472514 | 2124 | * | 22,41157 | -2,336953025 | 0,876606 | -2,66591 | 0,007678 | 0,715249603 | 2,951933 | Distal Intergenic | 11 | 48691159 | 48695801 | 4643 | 1 | 14694 | ENSMUST00000142269.2 | -218645 | ENSMUSG00000020372 | Rack1 | receptor for activated C kinase 1 |
| chrX | 1,68E+08 | 1,68E+08 | 1701 | * | 16,0813 | -2,336063687 | 0,89044 | -2,62349 | 0,008703 | 0,778555725 | 2,318434 | Distal Intergenic | 20 | 1,68E+08 | 1,68E+08 | 509337 | 1 | 11856 | ENSMUST00000033721.13 | -40214 | ENSMUSG00000031355 | Arhgap6 | Rho GTPase activating protein 6 |
| chr18 | 66742636 | 66743892 | 1257 | * | 15,15784 | -2,335874379 | 0,846291 | -2,76013 | 0,005778 | 0,792545355 | 1,41696 | Distal Intergenic | 18 | 66769899 | 66807573 | 37675 | 2 | 1,03E+08 | ENSMUST00000237763.2 | 63681 | | Gm30190 | predicted gene, 30190 |
| chr13 | 1,02E+08 | 1,02E+08 | 2439 | * | 19,9212 | -2,331601502 | 0,846665 | -2,75387 | 0,00589 | 0,683966322 | 2,713306 | Distal Intergenic | 13 | 1,02E+08 | 1,02E+08 | 8582 | 1 | 1,03E+08 | ENSMUST00000243228.1 | -258162 | | Gm36703 | predicted gene, 36703 |
| chr17 | 42828220 | 42829951 | 1732 | * | 15,53112 | -2,330023051 | 0,894183 | -2,60576 | 0,009167 | 0,90060438 | 1,214904 | Distal Intergenic | 17 | 42867674 | 42922286 | 54613 | 2 | 353344 | ENSMUST00000068355.8 | 92335 | ENSMUSG00000043972 | Opn5 | opsin 5 |
| chr17 | 78761950 | 78763501 | 1552 | * | 19,69682 | -2,328194112 | 0,852288 | -2,7317 | 0,006301 | 0,730024514 | 2,221611 | Distal Intergenic | 17 | 78685314 | 78725581 | 40268 | 2 | 225020 | ENSMUST00000112487.3 | -36369 | ENSMUSG00000056121 | Fez2 | fasciculation and elongation protein zeta 2 (zygin II) |
| chr10 | 69567534 | 69569589 | 2056 | * | 20,30392 | -2,327019985 | 0,814083 | -2,85846 | 0,004257 | 0,653987087 | 2,386008 | Intron (ENSMUST00000 | 10 | 69542308 | 69658465 | 116158 | 1 | 11735 | ENSMUST00000182162.8 | 25226 | ENSMUSG00000069601 | Ank3 | ankyrin 3, epithelial |
| chr14 | 50981741 | 50984176 | 2436 | * | 28,83382 | -2,325882334 | 0,629258 | -3,69623 | 0,000219 | 0,170809454 | 2,858511 | Promoter (<=1kb) | 14 | 50973628 | 50982551 | 8924 | 2 | 56858 | ENSMUST00000214290.2 | 0 | ENSMUSG00000059069 | Olfr749 | olfactory receptor 749 |
| chr2 | 1,25E+08 | 1,25E+08 | 2365 | * | 18,31051 | -2,324698183 | 0,802182 | -2,89797 | 0,003756 | 0,627995982 | 2,423594 | Distal Intergenic | 2 | 1,25E+08 | 1,25E+08 | 11127 | 1 | 110074 | ENSMUST00000051605.9 | 17723 | ENSMUSG00000027203 | Dut | deoxyuridine triphosphatase |
| chr19 | 40715763 | 40717514 | 1752 | * | 17,77895 | -2,324446744 | 0,86097 | -2,6998 | 0,006938 | 0,72508691 | 2,507013 | Promoter (1-2kb) | 19 | 40717881 | 40718748 | 868 | 2 | 1,05E+08 | ENSMUST00000245142.1 | 1234 | | Gm41846 | predicted gene, 41846 |
| chr9 | 1,19E+08 | 1,19E+08 | 1310 | * | 15,61904 | -2,323830877 | 0,853104 | -2,72397 | 0,00645 | 0,83918951 | 1,214904 | Exon (ENSMUST00000 | 9 | 1,19E+08 | 1,19E+08 | 79271 | 2 | 20271 | ENSMUST00000065196.13 | 18755 | ENSMUSG00000032511 | Scn5a | sodium channel, voltage-gated, type V, alpha |
| chr6 | 57697954 | 57699344 | 1391 | * | 19,02886 | -2,32101412 | 0,785081 | -2,9564 | 0,003113 | 0,595849045 | 2,386008 | Intron (ENSMUST00000 | 6 | 57680072 | 57716423 | 36352 | 1 | 71835 | ENSMUST00000153777.2 | 17882 | ENSMUSG00000062190 | Lancl2 | LanC (bacterial lantibiotic synthetase component C)-like 2 |
| chrX | 1,68E+08 | 1,68E+08 | 1975 | * | 16,07641 | -2,320780841 | 0,875937 | -2,64948 | 0,008061 | 0,757694328 | 2,386008 | Intron (ENSMUST00000 | 20 | 1,68E+08 | 1,68E+08 | 267825 | 1 | 11856 | ENSMUST00000139146.8 | 30873 | ENSMUSG00000031355 | Arhgap6 | Rho GTPase activating protein 6 |
| chr19 | 17457676 | 17459854 | 2179 | * | 19,60118 | -2,320432861 | 0,895391 | -2,59153 | 0,009555 | 0,802320835 | 2,221611 | Intron (ENSMUST00000 | 19 | 17374801 | 17377325 | 2525 | 1 | 54391 | ENSMUST00000146171.2 | 82875 | ENSMUSG00000024712 | Rfk | riboflavin kinase |
| chr13 | 77371654 | 77373228 | 1575 | * | 19,68277 | -2,320396454 | 0,837487 | -2,77066 | 0,005594 | 0,685078899 | 2,507013 | Intron (ENSMUST00000 | 13 | 77341153 | 77410088 | 68936 | 1 | 72371 | ENSMUST00000225760.2 | 30501 | ENSMUSG00000071252 | 2210408I21Rik | RIKEN cDNA 2210408I21 gene |
| chr14 | 29873432 | 29875199 | 1768 | * | 18,29273 | -2,319709942 | 0,877836 | -2,64253 | 0,008229 | 0,744224125 | 2,663619 | Intron (ENSMUST00000 | 14 | 29851796 | 29864538 | 12743 | 2 | 12289 | ENSMUST00000238336.2 | -8894 | ENSMUSG00000015968 | Cacna1d | calcium channel, voltage-dependent, L type, alpha 1D subunit |
| chr1 | 1,62E+08 | 1,62E+08 | 1792 | * | 18,18944 | -2,318920366 | 0,868947 | -2,66865 | 0,007616 | 0,748382289 | 2,423594 | Intron (ENSMUST00000 | 1 | 1,62E+08 | 1,62E+08 | 38567 | 2 | 103967 | ENSMUST00000161894.2 | 11256 | ENSMUSG00000040265 | Dnm3 | dynamin 3 |
| chr1 | 61629408 | 61631431 | 2024 | * | 16,92025 | -2,315867958 | 0,865942 | -2,67439 | 0,007487 | 0,728215953 | 2,663619 | Distal Intergenic | 1 | 61677983 | 62117418 | 439436 | 1 | 72823 | ENSMUST00000138768.8 | -46552 | ENSMUSG00000052062 | Pard3b | par-3 family cell polarity regulator beta |
| chr17 | 16337227 | 16339032 | 1806 | * | 17,9344 | -2,315171496 | 0,843215 | -2,74565 | 0,006039 | 0,699278451 | 2,507013 | Distal Intergenic | 17 | 16057811 | 16057917 | 107 | 2 | 1,15E+08 | ENSMUST00000239713.1 | -279310 | ENSMUSG00000119921 | Gm25598 | predicted gene, 25598 |
| chr8 | 1,09E+08 | 1,09E+08 | 1868 | * | 24,81586 | -2,312969052 | 0,764706 | -3,02465 | 0,002489 | 0,518149475 | 2,880017 | Intron (ENSMUST00000 | 8 | 1,09E+08 | 1,09E+08 | 9318 | 2 | 1,05E+08 | ENSMUST00000210837.2 | 201350 | | Gm39244 | predicted gene, 39244 |
| chr11 | 65150798 | 65152876 | 2079 | * | 17,75043 | -2,312054582 | 0,795532 | -2,9063 | 0,003657 | 0,625646014 | 2,423594 | Intron (ENSMUST00000 | 11 | 65114560 | 65160680 | 46121 | 2 | 214384 | ENSMUST00000150823.2 | 7804 | ENSMUSG00000020542 | Myocd | myocardin |
| chr2 | 1,48E+08 | 1,48E+08 | 1320 | * | 16,52874 | -2,311211705 | 0,865079 | -2,67168 | 0,007547 | 0,751210692 | 2,318434 | Distal Intergenic | 2 | 1,48E+08 | 1,48E+08 | 107 | 2 | 1,15E+08 | ENSMUST00000240448.1 | 62716 | ENSMUSG00000119413 | Gm25516 | predicted gene, 25516 |
| chr1 | 1,58E+08 | 1,58E+08 | 2384 | * | 21,55691 | -2,306655482 | 0,772038 | -2,98775 | 0,00281 | 0,526946456 | 3,104559 | Distal Intergenic | 1 | 1,58E+08 | 1,58E+08 | 761 | 2 | 240843 | ENSMUST00000194578.2 | 51802 | ENSMUSG00000004031 | Brinp2 | bone morphogenic protein/retinoic acid inducible neural-specific 2 |
| chr12 | 97155689 | 97158157 | 2469 | * | 24,35365 | -2,306507286 | 0,686641 | -3,35912 | 0,000782 | 0,322750682 | 2,880017 | Distal Intergenic | 12 | 97279623 | 97282627 | 3005 | 1 | 1,05E+08 | ENSMUST00000218233.2 | -121466 | | Gm40552 | predicted gene, 40552 |
| chr4 | 29876455 | 29878323 | 1869 | * | 14,73723 | -2,304667168 | 0,867658 | -2,65619 | 0,007903 | 0,853297048 | 1,37683 | Distal Intergenic | 4 | 29068079 | 29088811 | 20733 | 1 | 1,03E+08 | ENSMUST00000154813.2 | 808376 | | Gm11917 | predicted gene 11917 |
| chr16 | 26231939 | 26233751 | 1813 | * | 18,0604 | -2,301886804 | 0,83308 | -2,76311 | 0,005725 | 0,688853937 | 2,507013 | Distal Intergenic | 16 | 26175392 | 26190591 | 15200 | 2 | 12737 | ENSMUST00000023154.3 | -41348 | ENSMUSG00000022512 | Cldn1 | claudin 1 |
| chr15 | 86545925 | 86547528 | 1604 | * | 20,16747 | -2,299456267 | 0,760176 | -3,0249 | 0,002487 | 0,532146359 | 2,663619 | Distal Intergenic | 15 | 86588492 | 86595746 | 7255 | 1 | 75160 | ENSMUST00000229409.2 | -40964 | | 4930543I03Rik | RIKEN cDNA 4930543I03 gene |
| chr3 | 27576870 | 27579318 | 2449 | * | 25,7812 | -2,299160924 | 0,757942 | -3,03342 | 0,002418 | 0,524368688 | 2,725491 | Intron (ENSMUST00000 | 3 | 27639049 | 27639144 | 96 | 2 | 1,01E+08 | ENSMUST00000174954.3 | 59826 | ENSMUSG00000092695 | Mir3092 | microRNA 3092 |
| chr15 | 70699715 | 70700994 | 1280 | * | 13,75709 | -2,298800925 | 0,873352 | -2,63216 | 0,008484 | 0,880247401 | 1,2763 | Distal Intergenic | 15 | 71317525 | 71334249 | 16725 | 2 | 70363 | ENSMUST00000229634.2 | 633255 | ENSMUSG00000036800 | Fam135b | family with sequence similarity 135, member B |
| chr12 | 27330822 | 27333313 | 2492 | * | 23,83937 | -2,298656477 | 0,709527 | -3,2397 | 0,001197 | 0,392614537 | 2,951933 | Distal Intergenic | 12 | 27384263 | 27392573 | 8311 | 2 | 20666 | ENSMUST00000079063.7 | 59260 | ENSMUSG00000063632 | Sox11 | SRY (sex determining region Y)-box 11 |
| chr1 | 1,71E+08 | 1,71E+08 | 1948 | * | 15,11905 | -2,298097481 | 0,886285 | -2,59296 | 0,009515 | 0,808876909 | 2,141612 | Intron (ENSMUST00000 | 1 | 1,71E+08 | 1,71E+08 | 14182 | 2 | 1,03E+08 | ENSMUST00000244344.1 | 4127 | | Gm26641 | predicted gene, 26641 |
| chr1 | 83073828 | 83075208 | 1381 | * | 24,20211 | -2,295825485 | 0,690454 | -3,32509 | 0,000884 | 0,352029189 | 2,67243 | Distal Intergenic | 1 | 83094487 | 83096887 | 2401 | 1 | 20297 | ENSMUST00000027351.13 | -19279 | ENSMUSG00000026166 | Ccl20 | chemokine (C-C motif) ligand 20 |
| chr7 | 1,42E+08 | 1,42E+08 | 2360 | * | 22,33101 | -2,295769081 | 0,789284 | -2,90867 | 0,00363 | 0,591387753 | 2,880017 | Promoter (<=1kb) | 7 | 1,42E+08 | 1,42E+08 | 15698 | 2 | 677289 | ENSMUST00000054910.9 | 0 | ENSMUSG00000043795 | Prr33 | proline rich 33 |
| chr2 | 85238895 | 85240547 | 1653 | * | 18,62681 | -2,295000589 | 0,837301 | -2,74095 | 0,006126 | 0,704211591 | 2,507013 | 5' UTR | 2 | 85244277 | 85245221 | 945 | 2 | 258427 | ENSMUST00000099926.2 | 4674 | ENSMUSG00000075220 | Olfr993 | olfactory receptor 993 |
| chrX | 1,04E+08 | 1,04E+08 | 2181 | * | 15,54427 | -2,293048896 | 0,859632 | -2,66748 | 0,007642 | 0,769692849 | 2,141612 | Intron (ENSMUST00000 | 20 | 1,04E+08 | 1,04E+08 | 25680 | 1 | 76643 | ENSMUST00000189166.2 | 11680 | ENSMUSG00000100364 | 1700121L16Rik | RIKEN cDNA 1700121L16 gene |
| chr2 | 1,5E+08 | 1,5E+08 | 1662 | * | 17,92918 | -2,292981615 | 0,86386 | -2,65434 | 0,007946 | 0,755177881 | 2,386008 | Distal Intergenic | 2 | 1,5E+08 | 1,5E+08 | 19487 | 2 | 104348 | ENSMUST00000094538.6 | 78529 | ENSMUSG00000068134 | Zfp120 | zinc finger protein 120 |
| chr17 | 43887327 | 43888943 | 1617 | * | 19,38905 | -2,292844604 | 0,74618 | -3,07278 | 0,002121 | 0,536282961 | 2,221611 | Intron (ENSMUST00000 | 17 | 43879606 | 43918527 | 38922 | 1 | 27226 | ENSMUST00000163489.2 | 7721 | ENSMUSG00000023913 | Pla2g7 | phospholipase A2, group VII (platelet-activating factor acetylhydrolase, plasma) |
| chr4 | 69946518 | 69949295 | 2778 | * | 34,24662 | -2,290332324 | 0,585076 | -3,91459 | 9,06E-05 | 0,155022149 | 1,405709 | Distal Intergenic | 4 | 70160708 | 70162089 | 1382 | 2 | 214444 | ENSMUST00000124251.2 | 212794 | ENSMUSG00000039298 | Cdk5rap2 | CDK5 regulatory subunit associated protein 2 |
| chr9 | 1,13E+08 | 1,13E+08 | 1967 | * | 17,72405 | -2,289601093 | 0,797135 | -2,87229 | 0,004075 | 0,646274978 | 2,386008 | Distal Intergenic | 9 | 1,13E+08 | 1,13E+08 | 13892 | 2 | 1,03E+08 | ENSMUST00000214286.2 | -9403 | | Gm36485 | predicted gene, 36485 |
| chr9 | 47547688 | 47549759 | 2072 | * | 17,41 | -2,28836143 | 0,84676 | -2,70249 | 0,006882 | 0,749550311 | 2,154504 | Intron (ENSMUST00000 | 9 | 47441671 | 47761688 | 320018 | 1 | 54725 | ENSMUST00000034581.4 | 106017 | ENSMUSG00000032076 | Cadm1 | cell adhesion molecule 1 |
| chr14 | 19611466 | 19613119 | 1654 | * | 17,97458 | -2,28817923 | 0,867448 | -2,63783 | 0,008344 | 0,747833008 | 2,663619 | Distal Intergenic | 14 | 19614974 | 19619621 | 4648 | 2 | 1E+08 | ENSMUST00000112595.3 | 6502 | ENSMUSG00000093979 | Gm2237 | predicted gene 2237 |

| | | | | | | | | | | | | | | | | | | | | | | | |
|---|---|---|---|---|---|---|---|---|---|---|---|---|---|---|---|---|---|---|---|---|---|---|---|
| chr17 | 3910113 | 3912410 | 2298 | * | 21,15595 | -2,287824722 | 0,759801 | -3,01108 | 0,002603 | 0,494045114 | 3,471678 | Distal Intergenic | 17 | 3685515 | 3746536 | 61022 | 2 | 224480 | ENSMUST00000115800.2 | -163577 | ENSMUSG00000023802 | Nox3 | NADPH oxidase 3 |
| chr2 | 1,04E+08 | 1,04E+08 | 1880 | * | 15,71882 | -2,286071007 | 0,886439 | -2,57894 | 0,00991 | 0,77749164 | 2,663619 | Intron (ENSMUST00000 | 2 | 1,04E+08 | 1,04E+08 | 50319 | 2 | 241589 | ENSMUST00000149165.2 | -53974 | ENSMUSG00000068373 | D430041D05Rik | RIKEN cDNA D430041D05 gene |
| chr18 | 85183255 | 85185109 | 1855 | * | 16,58674 | -2,284476682 | 0,869165 | -2,62836 | 0,00858 | 0,791068542 | 2,141612 | Distal Intergenic | 18 | 84983635 | 84999597 | 15963 | 1 | 50764 | ENSMUST00000225015.2 | 199620 | ENSMUSG00000034391 | Fbxo15 | F-box protein 15 |
| chr19 | 21768053 | 21769776 | 1724 | * | 15,27354 | -2,28417122 | 0,880714 | -2,59354 | 0,009499 | 0,808557537 | 2,141612 | Intron (ENSMUST00000 | 19 | 21755752 | 21785266 | 29515 | 1 | 83921 | ENSMUST00000238079.2 | 12301 | ENSMUSG00000024754 | Cemip2 | cell migration inducing hyaluronidase 2 |
| chr7 | 1,34E+08 | 1,34E+08 | 2625 | * | 18,09148 | -2,28303066 | 0,820986 | -2,78084 | 0,005422 | 0,685188007 | 2,423594 | Exon (ENSMUST000002 | 7 | 1,34E+08 | 1,34E+08 | 206290 | 1 | 330662 | ENSMUST00000211570.2 | -8264 | ENSMUSG00000058325 | Dock1 | dedicator of cytokinesis 1 |
| chr11 | 72996805 | 72998553 | 1749 | * | 18,16208 | -2,282938761 | 0,855366 | -2,66896 | 0,007609 | 0,767135014 | 2,154504 | Intron (ENSMUST00000 | 11 | 72981409 | 73038272 | 56864 | 1 | 16407 | ENSMUST00000006101.4 | 15396 | ENSMUSG00000005947 | Itgae | integrin alpha E, epithelial-associated |
| chr9 | 85864162 | 85865691 | 1530 | * | 16,48373 | -2,282134113 | 0,883701 | -2,58247 | 0,00981 | 0,775744126 | 2,663619 | Distal Intergenic | 9 | 85702355 | 85726376 | 24022 | 2 | 442799 | ENSMUST00000185960.2 | -137786 | | 9330154J02Rik | RIKEN cDNA 9330154J02 gene |
| chr5 | 1,47E+08 | 1,47E+08 | 2113 | * | 22,99788 | -2,281168508 | 0,800087 | -2,85115 | 0,004356 | 0,621084104 | 2,951062 | Distal Intergenic | 5 | 1,47E+08 | 1,47E+08 | 875 | 1 | 75949 | ENSMUST00000197707.2 | -45806 | | 4930573C15Rik | RIKEN cDNA 4930573C15 gene |
| chr9 | 67798084 | 67800365 | 2282 | * | 16,88428 | -2,280813398 | 0,881024 | -2,58882 | 0,009631 | 0,90060438 | 1,2763 | Exon (ENSMUST000000 | 9 | 67747694 | 67813092 | 65399 | 1 | 320528 | ENSMUST00000217386.2 | 50390 | ENSMUSG00000035284 | Vps13c | vacuolar protein sorting 13C |
| chr7 | 1,28E+08 | 1,28E+08 | 1485 | * | 18,02029 | -2,278029309 | 0,868972 | -2,62152 | 0,008754 | 0,792545355 | 2,154504 | Intron (ENSMUST00000 | 7 | 1,28E+08 | 1,29E+08 | 132820 | 1 | 1,03E+08 | ENSMUST00000205528.2 | 13684 | | Gm32884 | predicted gene, 32884 |
| chr11 | 73852056 | 73853751 | 1696 | * | 17,73284 | -2,277729997 | 0,774581 | -2,9406 | 0,003276 | 0,585155013 | 2,663619 | Promoter (1-2kb) | 11 | 73855356 | 73856446 | 1091 | 1 | 258343 | ENSMUST00000121280.3 | -1605 | ENSMUSG00000057050 | Olfr397 | olfactory receptor 397 |
| chr5 | 4329766 | 4331190 | 1425 | * | 15,86245 | -2,277462306 | 0,868891 | -2,62111 | 0,008764 | 0,752939853 | 2,663619 | Distal Intergenic | 5 | 4242367 | 4247651 | 5285 | 1 | 208595 | ENSMUST00000177258.2 | 87399 | ENSMUSG00000053178 | Mterf1b | mitochondrial transcription termination factor 1b |
| chr10 | 1,16E+08 | 1,16E+08 | 2028 | * | 22,30358 | -2,274467187 | 0,723344 | -3,14438 | 0,001664 | 0,44621096 | 2,951933 | Distal Intergenic | 10 | 1,16E+08 | 1,16E+08 | 16481 | 1 | 216350 | ENSMUST00000217900.2 | -41883 | ENSMUSG00000034127 | Tspan8 | tetraspanin 8 |
| chr4 | 34168960 | 34170554 | 1595 | * | 18,18952 | -2,274065705 | 0,804522 | -2,82661 | 0,004704 | 0,669377201 | 2,423594 | Distal Intergenic | 4 | 34086126 | 34086233 | 108 | 1 | 1,15E+08 | ENSMUST00000157216.2 | 82834 | ENSMUSG00000087841 | Gm25623 | predicted gene, 25623 |
| chr1 | 83340997 | 83342485 | 1489 | * | 16,01024 | -2,273484747 | 0,858371 | -2,6486 | 0,008083 | 0,740451144 | 2,663619 | Intron (ENSMUST00000 | 1 | 83316867 | 83385860 | 68994 | 2 | 77629 | ENSMUST00000160237.2 | 43375 | ENSMUSG00000026163 | Sphkap | SPHK1 interactor, AKAP domain containing |
| chr16 | 56427582 | 56429748 | 2167 | * | 18,17224 | -2,273262668 | 0,745454 | -3,0495 | 0,002292 | 0,537550416 | 2,386008 | Intron (ENSMUST00000 | 16 | 56511240 | 56526057 | 14818 | 2 | 21787 | ENSMUST00000127994.8 | 96309 | ENSMUSG00000022757 | Tfg | Trk-fused gene |
| chr4 | 63717923 | 63719406 | 1484 | * | 20,08528 | -2,272908934 | 0,771434 | -2,94634 | 0,003216 | 0,617289329 | 2,221611 | Distal Intergenic | 4 | 63645321 | 63663350 | 18030 | 2 | 326623 | ENSMUST00000062246.8 | -54573 | ENSMUSG00000050395 | Tnfsf15 | tumor necrosis factor (ligand) superfamily, member 15 |
| chr17 | 90093094 | 90094885 | 1792 | * | 26,5216 | -2,271056908 | 0,739249 | -3,07211 | 0,002125 | 0,486196597 | 2,951062 | Distal Intergenic | 17 | 90263016 | 90284213 | 21198 | 2 | 1,05E+08 | ENSMUST00000234603.2 | 189328 | | Gm41653 | predicted gene, 41653 |
| chr2 | 1,41E+08 | 1,41E+08 | 1798 | * | 15,11574 | -2,269965335 | 0,869324 | -2,61118 | 0,009023 | 0,890147477 | 1,2763 | Intron (ENSMUST00000 | 2 | 1,41E+08 | 1,41E+08 | 13272 | 2 | 71436 | ENSMUST00000110057.3 | -97723 | ENSMUSG00000051379 | Flrt3 | fibronectin leucine rich transmembrane protein 3 |
| chr5 | 62739983 | 62742061 | 2079 | * | 17,30809 | -2,269698063 | 0,784043 | -2,89487 | 0,003793 | 0,630714459 | 2,423594 | Distal Intergenic | 5 | 62759788 | 62769717 | 9930 | 2 | 212285 | ENSMUST00000160620.3 | 27656 | ENSMUSG00000037999 | Arap2 | ArfGAP with RhoGAP domain, ankyrin repeat and PH domain 2 |
| chr7 | 31147674 | 31149741 | 2068 | * | 17,36282 | -2,268633886 | 0,867257 | -2,61587 | 0,0089 | 0,775744126 | 2,423594 | Distal Intergenic | 7 | 31100459 | 31102601 | 2143 | 2 | 1,03E+08 | ENSMUST00000186705.2 | -45073 | ENSMUSG00000101325 | Scgb2b4-ps | secretoglobin, family 2B, member 4, pseudogene |
| chr13 | 1,11E+08 | 1,11E+08 | 2370 | * | 21,73437 | -2,268479483 | 0,718893 | -3,15552 | 0,001602 | 0,440073451 | 2,951933 | Distal Intergenic | 13 | 1,11E+08 | 1,11E+08 | 3014 | 1 | 1,03E+08 | ENSMUST00000225669.2 | 39950 | | Gm33172 | predicted gene, 33172 |
| chr8 | 98407031 | 98408069 | 1039 | * | 17,32743 | -2,268255921 | 0,79941 | -2,83741 | 0,004548 | 0,662625192 | 2,423594 | Distal Intergenic | 8 | 98566680 | 98573899 | 7220 | 1 | 1,05E+08 | ENSMUST00000209813.2 | -158611 | | Gm39231 | predicted gene, 39231 |
| chr12 | 15188349 | 15190424 | 2076 | * | 18,07947 | -2,267960634 | 0,799365 | -2,8372 | 0,004551 | 0,681608406 | 2,154504 | Distal Intergenic | 12 | 15118118 | 15187519 | 69402 | 1 | 1,03E+08 | ENSMUST00000222816.2 | 70231 | | Gm36128 | predicted gene, 36128 |
| chr9 | 97594343 | 97597362 | 3020 | * | 26,55486 | -2,26758951 | 0,643495 | -3,52386 | 0,000425 | 0,247743734 | 2,858511 | Intron (ENSMUST00000 | 9 | 97321554 | 97321680 | 127 | 2 | 1,15E+08 | ENSMUST00000104020.3 | -272663 | ENSMUSG00000077208 | Gm25607 | predicted gene, 25607 |
| chr2 | 1,48E+08 | 1,48E+08 | 2158 | * | 18,32223 | -2,267088301 | 0,834164 | -2,7178 | 0,006572 | 0,722439117 | 2,423594 | Distal Intergenic | 2 | 1,49E+08 | 1,49E+08 | 3427 | 1 | 56488 | ENSMUST00000109961.8 | -36559 | ENSMUSG00000036992 | Nxt1 | NTF2-related export protein 1 |
| chr15 | 97183294 | 97184311 | 1018 | * | 19,83586 | -2,266887671 | 0,807265 | -2,80811 | 0,004983 | 0,671156916 | 2,507013 | Intron (ENSMUST00000 | 15 | 97146954 | 97175219 | 28266 | 1 | 239647 | ENSMUST00000226615.2 | 36340 | ENSMUSG00000044250 | Pced1b | PC-esterase domain containing 1B |
| chr9 | 68576070 | 68578397 | 2328 | * | 22,67909 | -2,26606711 | 0,783853 | -2,89093 | 0,003841 | 0,595637142 | 2,951933 | Intron (ENSMUST00000 | 9 | 68562585 | 68571462 | 8878 | 1 | 19883 | ENSMUST00000143507.2 | 13485 | ENSMUSG00000032238 | Rora | RAR-related orphan receptor alpha |
| chr11 | 56783072 | 56784775 | 1704 | * | 14,91835 | -2,266010989 | 0,846792 | -2,67599 | 0,007451 | 0,863136415 | 1,214904 | Distal Intergenic | 11 | 56902213 | 56906117 | 3905 | 1 | 14799 | ENSMUST00000125292.2 | -117438 | ENSMUSG00000020524 | Gria1 | glutamate receptor, ionotropic, AMPA1 (alpha 1) |
| chr12 | 60636454 | 60638336 | 1883 | * | 20,69088 | -2,265428274 | 0,745499 | -3,03881 | 0,002375 | 0,509154993 | 2,880017 | Distal Intergenic | 12 | 60782284 | 60782390 | 107 | 1 | 1,15E+08 | ENSMUST00000158978.3 | -143948 | ENSMUSG00000119332 | Gm23059 | predicted gene, 23059 |
| chr2 | 1,44E+08 | 1,44E+08 | 2187 | * | 22,10844 | -2,264843571 | 0,829728 | -2,72962 | 0,006341 | 0,675966474 | 3,104559 | Exon (ENSMUST000000 | 2 | 1,44E+08 | 1,44E+08 | 41604 | 2 | 378775 | ENSMUST00000153878.8 | 13426 | ENSMUSG00000087115 | Pcsk2os2 | proprotein convertase subtilisin/kexin type 2, opposite strand 2 |
| chr16 | 88175046 | 88177434 | 2389 | * | 25,37602 | -2,263190677 | 0,674896 | -3,35339 | 0,000798 | 0,336438515 | 2,67243 | Promoter (1-2kb) | 16 | 88164790 | 88178584 | 13795 | 2 | 1,03E+08 | ENSMUST00000231384.2 | 1150 | ENSMUSG00000116754 | Gm33255 | predicted gene, 33255 |
| chr10 | 1,11E+08 | 1,11E+08 | 2019 | * | 21,64859 | -2,263019304 | 0,708818 | -3,19267 | 0,00141 | 0,410415129 | 3,104559 | Exon (ENSMUST000001 | 10 | 1,11E+08 | 1,11E+08 | 128 | 1 | 1,15E+08 | ENSMUST00000158836.3 | 3421 | ENSMUSG00000089461 | Gm23698 | predicted gene, 23698 |
| chr13 | 1,03E+08 | 1,03E+08 | 1657 | * | 16,39772 | -2,263000425 | 0,86102 | -2,62828 | 0,008582 | 0,749931768 | 2,663619 | Intron (ENSMUST00000 | 13 | 1,03E+08 | 1,03E+08 | 302701 | 2 | 328329 | ENSMUST00000166336.9 | 13194 | ENSMUSG00000034751 | Mast4 | microtubule associated serine/threonine kinase family member 4 |
| chr7 | 17465051 | 17467097 | 2047 | * | 16,58413 | -2,262761579 | 0,80666 | -2,8051 | 0,00503 | 0,663496633 | 2,663619 | Intron (ENSMUST00000 | 7 | 17447163 | 17495057 | 47895 | 1 | 73250 | ENSMUST00000081907.8 | 17888 | ENSMUSG00000008789 | Ceacam5 | carcinoembryonic antigen-related cell adhesion molecule 5 |
| chr1 | 1,44E+08 | 1,44E+08 | 2023 | * | 18,25451 | -2,262366033 | 0,778247 | -2,907 | 0,003649 | 0,624905941 | 2,423594 | Distal Intergenic | 1 | 1,44E+08 | 1,44E+08 | 2150 | 2 | 19735 | ENSMUST00000153527.3 | 44332 | ENSMUSG00000026360 | Rgs2 | regulator of G-protein signaling 2 |
| chr13 | 56411612 | 56414358 | 2747 | * | 33,3467 | -2,260070104 | 0,699147 | -3,23261 | 0,001227 | 0,399560242 | 2,925497 | Distal Intergenic | 13 | 56398291 | 56399976 | 1686 | 2 | 18014 | ENSMUST00000058475.6 | -11636 | ENSMUSG00000048904 | Neurog1 | neurogenin 1 |
| chr1 | 1,58E+08 | 1,58E+08 | 1899 | * | 16,37262 | -2,258991282 | 0,870253 | -2,59579 | 0,009437 | 0,789027175 | 2,386008 | Distal Intergenic | 1 | 1,57E+08 | 1,57E+08 | 5913 | 1 | 89867 | ENSMUST00000155665.2 | 247662 | ENSMUSG00000026589 | Sec16b | SEC16 homolog B (S. cerevisiae) |
| chr10 | 40831316 | 40834304 | 2989 | * | 24,13425 | -2,258229259 | 0,833993 | -2,70773 | 0,006775 | 0,696194408 | 2,858511 | Distal Intergenic | 10 | 40759823 | 40814566 | 54744 | 1 | 83767 | ENSMUST00000105509.2 | 71493 | ENSMUSG00000019831 | Wasf1 | WASP family, member 1 |
| chr7 | 1,32E+08 | 1,32E+08 | 2383 | * | 22,88744 | -2,256992764 | 0,749415 | -3,01167 | 0,002598 | 0,539604518 | 2,67243 | Promoter (<=1kb) | 7 | 1,32E+08 | 1,32E+08 | 97558 | 2 | 55987 | ENSMUST00000127039.8 | 0 | ENSMUSG00000030862 | Cpxm2 | carboxypeptidase X 2 (M14 family) |
| chr14 | 28366703 | 28368746 | 2044 | * | 20,23354 | -2,256555984 | 0,79423 | -2,84119 | 0,004495 | 0,644557958 | 2,663619 | Promoter (2-3kb) | 14 | 28371608 | 28426438 | 54831 | 1 | 1,03E+08 | ENSMUST00000245373.1 | -2862 | | Gm35164 | predicted gene, 35164 |
| chr13 | 21044084 | 21045853 | 1770 | * | 16,57168 | -2,256534293 | 0,812358 | -2,77776 | 0,005474 | 0,689688276 | 2,386008 | Intron (ENSMUST00000 | 13 | 20978305 | 21036082 | 57778 | 1 | 27052 | ENSMUST00000221982.2 | 65779 | ENSMUSG00000021322 | Aoah | acyloxyacyl hydrolase |
| chr7 | 1,31E+08 | 1,31E+08 | 1738 | * | 19,24612 | -2,254784986 | 0,741827 | -3,0395 | 0,00237 | 0,54122637 | 2,386008 | Intron (ENSMUST00000 | 7 | 1,31E+08 | 1,31E+08 | 10455 | 1 | 12237 | ENSMUST00000207442.2 | 68808 | ENSMUSG00000066979 | Bub3 | BUB3 mitotic checkpoint protein |
| chr10 | 51385375 | 51386775 | 1401 | * | 18,11843 | -2,253638036 | 0,845734 | -2,66471 | 0,007705 | 0,743536224 | 2,507013 | Intron (ENSMUST00000 | 10 | 51365626 | 51398543 | 32918 | 2 | 1,05E+08 | ENSMUST00000218413.2 | 11768 | | Gm40645 | predicted gene, 40645 |
| chr3 | 4769434 | 4771855 | 2422 | * | 26,7968 | -2,252696407 | 0,714366 | -3,15342 | 0,001614 | 0,440933973 | 2,951062 | Distal Intergenic | 3 | 4862388 | 4879967 | 17580 | 1 | 1,01E+08 | ENSMUST00000245577.1 | -90533 | ENSMUSG00000098659 | 1110015O18Rik | RIKEN cDNA 1110015O18 gene |
| chr7 | 63387665 | 63389440 | 1776 | * | 19,89001 | -2,251566538 | 0,804585 | -2,79842 | 0,005135 | 0,691532203 | 2,221611 | Intron (ENSMUST00000 | 7 | 63404460 | 63407673 | 3214 | 1 | 170711 | ENSMUST00000177511.2 | -15020 | ENSMUSG00000033510 | Otud7a | OTU domain containing 7A |
| chr12 | 37403569 | 37406324 | 2756 | * | 24,27283 | -2,250824547 | 0,681827 | -3,30117 | 0,000963 | 0,349985669 | 2,951933 | Intron (ENSMUST00000 | 12 | 37495184 | 37632199 | 137016 | 1 | 319660 | ENSMUST00000221949.2 | -88860 | ENSMUSG00000050103 | Agmo | alkylglycerol monooxygenase |
| chr6 | 1,08E+08 | 1,08E+08 | 1016 | * | 17,86152 | -2,250397727 | 0,807907 | -2,78546 | 0,005345 | 0,68526372 | 2,386008 | Distal Intergenic | 6 | 1,08E+08 | 1,08E+08 | 12078 | 1 | 74729 | ENSMUST00000049246.7 | -219192 | ENSMUSG00000034639 | Setmar | SET domain without mariner transposase fusion |
| chr5 | 1,12E+08 | 1,12E+08 | 1475 | * | 17,17955 | -2,250205412 | 0,868388 | -2,59125 | 0,009563 | 0,770673161 | 2,663619 | Distal Intergenic | 5 | 1,12E+08 | 1,12E+08 | 4432 | 2 | 1,03E+08 | ENSMUST00000197648.2 | 74649 | | Gm36794 | predicted gene, 36794 |
| chr17 | 67120356 | 67122647 | 2292 | * | 27,79814 | -2,250028388 | 0,672737 | -3,34459 | 0,000824 | 0,328946709 | 2,880017 | Promoter (2-3kb) | 17 | 67069174 | 67124870 | 55697 | 2 | 19274 | ENSMUST00000225554.2 | 2223 | ENSMUSG00000033278 | Ptprm | protein tyrosine phosphatase, receptor type, M |
| chr2 | 13014735 | 13018876 | 4142 | * | 35,8505 | -2,249313659 | 0,571354 | -3,93682 | 8,26E-05 | 0,147030992 | 1,405709 | Promoter (<=1kb) | 2 | 13008268 | 13016617 | 8350 | 2 | 227580 | ENSMUST00000061545.7 | 0 | ENSMUSG00000049630 | C1ql3 | C1q-like 3 |
| chr10 | 31002357 | 31003979 | 1623 | * | 19,85659 | -2,247866066 | 0,801912 | -2,80313 | 0,005061 | 0,68259716 | 2,386008 | Distal Intergenic | 10 | 31039522 | 31039628 | 107 | 2 | 1,15E+08 | ENSMUST00000240069.1 | 35649 | ENSMUSG00000118948 | Gm23790 | predicted gene, 23790 |
| chr11 | 1,09E+08 | 1,09E+08 | 2083 | * | 14,88349 | -2,246971882 | 0,861408 | -2,60849 | 0,009094 | 0,873937167 | 1,41696 | Distal Intergenic | 11 | 1,09E+08 | 1,09E+08 | 107 | 2 | 1,15E+08 | ENSMUST00000158564.3 | 11746 | ENSMUSG00000119031 | Gm24149 | predicted gene, 24149 |
| chr14 | 1,17E+08 | 1,17E+08 | 2331 | * | 16,5817 | -2,2461682 | 0,841417 | -2,66951 | 0,007596 | 0,730469267 | 2,663619 | Distal Intergenic | 14 | 1,17E+08 | 1,18E+08 | 701884 | 1 | 23888 | ENSMUST00000123239.2 | -145631 | ENSMUSG00000058571 | Gpc6 | glypican 6 |
| chr9 | 30357141 | 30359045 | 1905 | * | 18,42597 | -2,245747839 | 0,798966 | -2,81082 | 0,004942 | 0,678869598 | 2,386008 | Intron (ENSMUST00000 | 9 | 30347153 | 30375969 | 28817 | 1 | 102607 | ENSMUST00000216545.2 | 9988 | ENSMUSG00000031993 | Snx19 | sorting nexin 19 |
| chr14 | 1,02E+08 | 1,02E+08 | 2729 | * | 13,88958 | -2,245255147 | 0,855233 | -2,62531 | 0,008657 | 0,934396727 | 0,918255 | Distal Intergenic | 14 | 1,02E+08 | 1,02E+08 | 20856 | 2 | 210789 | ENSMUST00000161304.2 | 120725 | ENSMUSG00000033083 | Tbc1d4 | TBC1 domain family, member 4 |
| chr4 | 57011445 | 57013518 | 2074 | * | 17,71494 | -2,244335778 | 0,797002 | -2,81597 | 0,004863 | 0,68680243 | 2,154504 | Intron (ENSMUST00000 | 4 | 56995678 | 57027676 | 31999 | 2 | 54357 | ENSMUST00000044022.5 | 14158 | ENSMUSG00000028434 | Epb41l4b | erythrocyte membrane protein band 4.1 like 4b |
| chr14 | 34745407 | 34747097 | 1691 | * | 18,19343 | -2,244076891 | 0,781282 | -2,8723 | 0,004075 | 0,663630601 | 2,154504 | Intron (ENSMUST00000 | 14 | 34616566 | 34616663 | 98 | 1 | 723847 | ENSMUST00000083547.3 | 128841 | ENSMUSG00000065481 | Mir346 | microRNA 346 |
| chr2 | 5333677 | 5335773 | 2097 | * | 19,84716 | -2,243352267 | 0,798758 | -2,80855 | 0,004977 | 0,680780911 | 2,386008 | Intron (ENSMUST00000 | 2 | 5264740 | 5264926 | 187 | 2 | 1,15E+08 | ENSMUST00000122657.3 | -68751 | ENSMUSG00000084606 | Gm23118 | predicted gene, 23118 |
| chr10 | 61646506 | 61648431 | 1926 | * | 22,6649 | -2,243172367 | 0,832808 | -2,69351 | 0,00707 | 0,688021707 | 3,104559 | Distal Intergenic | 10 | 61574444 | 61619926 | 45483 | 2 | 404634 | ENSMUST00000020283.5 | -26580 | ENSMUSG00000020086 | Macroh2a2 | macroH2A.2 histone |
| chr9 | 1,16E+08 | 1,16E+08 | 2095 | * | 20,60202 | -2,243165961 | 0,779575 | -2,87742 | 0,004009 | 0,594676756 | 3,104559 | Distal Intergenic | 9 | 1,16E+08 | 1,16E+08 | 8341 | 1 | 78580 | ENSMUST00000217456.2 | 10100 | | D730003K21Rik | RIKEN cDNA D730003K21 gene |
| chr14 | 21659992 | 21662260 | 2269 | * | 18,31247 | -2,242715804 | 0,861466 | -2,60337 | 0,009231 | 0,775744126 | 2,507013 | Intron (ENSMUST00000 | 14 | 21672610 | 21684657 | 12048 | 1 | 54169 | ENSMUST00000182732.2 | -10350 | ENSMUSG00000021767 | Kat6b | K(lysine) acetyltransferase 6B |
| chr13 | 69828019 | 69829443 | 1425 | * | 26,58858 | -2,242456098 | 0,699136 | -3,20747 | 0,001339 | 0,410415129 | 2,951062 | Exon (ENSMUST000002 | 13 | 69828708 | 69845795 | 17088 | 2 | 1,03E+08 | ENSMUST00000223164.2 | 16352 | | Gm35618 | predicted gene, 35618 |
| chr9 | 24335964 | 24338994 | 3031 | * | 29,10951 | -2,24204092 | 0,666243 | -3,3652 | 0,000765 | 0,321880003 | 2,858511 | Promoter (<=1kb) | 9 | 24333585 | 24337714 | 4130 | 2 | 244745 | ENSMUST00000149001.2 | 0 | ENSMUSG00000043067 | Dpy19l1 | dpy-19-like 1 (C. elegans) |
| chr3 | 62365878 | 62367885 | 2008 | * | 17,63247 | -2,234857758 | 0,793106 | -2,81786 | 0,004835 | 0,675197144 | 2,386008 | 3' UTR | 3 | 62378066 | 62380008 | 1943 | 2 | 72162 | ENSMUST00000160858.2 | 12123 | ENSMUSG00000027770 | Dhx36 | DEAH (Asp-Glu-Ala-His) box polypeptide 36 |
| chrX | 1,39E+08 | 1,39E+08 | 2143 | * | 18,93938 | -2,234053058 | 0,815462 | -2,73961 | 0,006151 | 0,705098751 | 2,507013 | Intron (ENSMUST00000 | 20 | 1,39E+08 | 1,39E+08 | 65 | 1 | 1E+08 | ENSMUST00000175361.3 | -114700 | ENSMUSG00000093102 | Mir3475 | microRNA 3475 |
| chr2 | 17079069 | 17080695 | 1627 | * | 14,52024 | -2,233978801 | 0,861206 | -2,59401 | 0,009486 | 0,881675712 | 1,41696 | Distal Intergenic | 2 | 17353505 | 17365897 | 12393 | 2 | 74103 | ENSMUST00000138241.2 | 285202 | ENSMUSG00000053702 | Nebl | nebulette |
| chr3 | 1,36E+08 | 1,36E+08 | 1730 | * | 16,06528 | -2,232827848 | 0,846765 | -2,63689 | 0,008367 | 0,766579572 | 2,386008 | Distal Intergenic | 3 | 1,36E+08 | 1,36E+08 | 14080 | 2 | 70055 | ENSMUST00000189233.2 | 16965 | ENSMUSG00000099508 | 1700030L20Rik | RIKEN cDNA 1700030L20 gene |
| chr3 | 1,19E+08 | 1,19E+08 | 1578 | * | 17,94506 | -2,232118287 | 0,803314 | -2,77864 | 0,005459 | 0,689042734 | 2,386008 | Distal Intergenic | 3 | 1,19E+08 | 1,19E+08 | 131 | 2 | 1,15E+08 | ENSMUST00000157181.3 | 23975 | ENSMUSG00000087806 | Gm23432 | predicted gene, 23432 |
| chr12 | 82847518 | 82849128 | 1611 | * | 17,44122 | -2,231853379 | 0,832233 | -2,68177 | 0,007323 | 0,7591629 | 2,154504 | Intron (ENSMUST00000 | 12 | 82819095 | 82819230 | 136 | 2 | 1,15E+08 | ENSMUST00000104442.3 | -28288 | ENSMUSG00000077630 | Gm22149 | predicted gene, 22149 |
| chrX | 1,31E+08 | 1,31E+08 | 2378 | * | 18,30124 | -2,231834413 | 0,813965 | -2,74193 | 0,006108 | 0,710505631 | 2,423594 | Distal Intergenic | 20 | 1,31E+08 | 1,31E+08 | 131 | 2 | 1,15E+08 | ENSMUST00000104440.3 | 29223 | ENSMUSG00000077628 | Gm23768 | predicted gene, 23768 |
| chr19 | 17902858 | 17905570 | 2713 | * | 21,19925 | -2,231784941 | 0,802089 | -2,78246 | 0,005395 | 0,658336602 | 2,951933 | Exon (ENSMUST000002 | 19 | 17842681 | 17842783 | 103 | 2 | 1,15E+08 | ENSMUST00000157745.3 | -60075 | ENSMUSG00000088370 | Gm22880 | predicted gene, 22880 |
| chr5 | 1,51E+08 | 1,51E+08 | 2168 | * | 19,29644 | -2,231433626 | 0,761622 | -2,92985 | 0,003391 | 0,594430593 | 2,663619 | Distal Intergenic | 5 | 1,51E+08 | 1,51E+08 | 25720 | 2 | 243362 | ENSMUST00000202385.4 | 52250 | ENSMUSG00000016128 | Stard13 | StAR-related lipid transfer (START) domain containing 13 |
| chr16 | 44217650 | 44220404 | 2755 | * | 24,88474 | -2,23081037 | 0,732761 | -3,04439 | 0,002332 | 0,506156746 | 2,880017 | Promoter (1-2kb) | 16 | 44221784 | 44302457 | 80674 | 1 | 212517 | ENSMUST00000120049.8 | -1380 | ENSMUSG00000071550 | Cfap44 | cilia and flagella associated protein 44 |
| chr6 | 1,15E+08 | 1,15E+08 | 1266 | * | 18,31741 | -2,229440408 | 0,821277 | -2,7146 | 0,006636 | 0,716908183 | 2,507013 | Intron (ENSMUST00000 | 6 | 1,15E+08 | 1,15E+08 | 3072 | 1 | 74244 | ENSMUST00000182934.2 | -58133 | ENSMUSG00000030314 | Atg7 | autophagy related 7 |
| chr10 | 8410073 | 8412409 | 2337 | * | 22,40171 | -2,229175588 | 0,768697 | -2,89994 | 0,003732 | 0,580030873 | 3,104559 | Distal Intergenic | 10 | 8080573 | 8394589 | 314017 | 2 | 338362 | ENSMUST00000061601.9 | -15484 | ENSMUSG00000047712 | Ust | uronyl-2-sulfotransferase |
| chr17 | 39076105 | 39078341 | 2237 | * | 22,0403 | -2,22852138 | 0,735758 | -3,02888 | 0,002455 | 0,502272318 | 3,104559 | Distal Intergenic | 17 | 38950338 | 38956544 | 6207 | 1 | 1E+08 | ENSMUST00000174742.3 | 125767 | ENSMUSG00000092342 | Esp31 | exocrine gland secreted peptide 31 |
| chr5 | 96524904 | 96527718 | 2815 | * | 23,84521 | -2,227517453 | 0,725223 | -3,07149 | 0,00213 | 0,492273754 | 2,880017 | Promoter (2-3kb) | 5 | 96522294 | 96535493 | 13200 | 1 | 231470 | ENSMUST00000196751.2 | 2610 | ENSMUSG00000034687 | Fras1 | Fraser extracellular matrix complex subunit 1 |
| chr1 | 1,67E+08 | 1,67E+08 | 2512 | * | 20,73507 | -2,227015315 | 0,790756 | -2,81631 | 0,004858 | 0,61075929 | 3,471678 | Distal Intergenic | 1 | 1,67E+08 | 1,67E+08 | 25706 | 2 | 1,01E+08 | ENSMUST00000180995.3 | 50536 | ENSMUSG00000102548 | Gm16701 | predicted gene, 16701 |
| chr10 | 1,03E+08 | 1,03E+08 | 2293 | * | 18,47237 | -2,225569844 | 0,769975 | -2,89045 | 0,003847 | 0,627785088 | 2,507013 | Distal Intergenic | 10 | 1,03E+08 | 1,03E+08 | 14424 | 2 | 216285 | ENSMUST00000164226.3 | 33098 | ENSMUSG00000036602 | Alx1 | ALX homeobox 1 |
| chr15 | 84591487 | 84592675 | 1189 | * | 15,83241 | -2,225565137 | 0,827437 | -2,68971 | 0,007151 | 0,755814512 | 2,141612 | Distal Intergenic | 15 | 84604253 | 84656408 | 52156 | 1 | 73167 | ENSMUST00000006029.11 | -11578 | ENSMUSG00000078954 | Arhgap8 | Rho GTPase activating protein 8 |
| chr11 | 1,05E+08 | 1,05E+08 | 2155 | * | 18,33344 | -2,225206281 | 0,807357 | -2,75616 | 0,005848 | 0,684274806 | 2,663619 | Promoter (1-2kb) | 11 | 1,05E+08 | 1,05E+08 | 105 | 1 | 1,15E+08 | ENSMUST00000158854.3 | -1308 | ENSMUSG00000089479 | Gm25362 | predicted gene, 25362 |
| chr6 | 43315874 | 43317144 | 1271 | * | 16,40456 | -2,221966115 | 0,795225 | -2,79414 | 0,005204 | 0,792545355 | 1,2763 | Distal Intergenic | 6 | 43280608 | 43286488 | 5881 | 2 | 18291 | ENSMUST00000031749.16 | -29386 | ENSMUSG00000029736 | Nobox | NOBOX oogenesis homeobox |
| chr13 | 90488137 | 90490088 | 1952 | * | 15,26155 | -2,220173961 | 0,822424 | -2,69955 | 0,006943 | 0,825384969 | 1,41696 | Distal Intergenic | 13 | 90247100 | 90247233 | 134 | 1 | 1,15E+08 | ENSMUST00000157683.3 | 241037 | ENSMUSG00000088308 | Gm24507 | predicted gene, 24507 |
| chr16 | 81166301 | 81168354 | 2054 | * | 17,97494 | -2,219380083 | 0,715148 | -3,10338 | 0,001913 | 0,496560937 | 2,507013 | Intron (ENSMUST00000 | 16 | 81262511 | 81314918 | 52408 | 1 | 17968 | ENSMUST00000232550.2 | -94157 | ENSMUSG00000022762 | Ncam2 | neural cell adhesion molecule 2 |
| chr15 | 44555250 | 44556997 | 1748 | * | 26,1259 | -2,218097072 | 0,76291 | -2,90741 | 0,003644 | 0,600307614 | 2,725491 | Intron (ENSMUST00000 | 15 | 44535531 | 44611184 | 75654 | 2 | 319613 | ENSMUST00000226825.2 | 54187 | ENSMUSG00000022340 | Sybu | syntabulin (syntaxin-interacting) |
| chr18 | 48759732 | 48761216 | 1485 | * | 17,44027 | -2,217146708 | 0,799857 | -2,77193 | 0,005573 | 0,693610666 | 2,386008 | Exon (ENSMUST000001 | 18 | 48178402 | 48181479 | 3078 | 1 | 433182 | ENSMUST00000076155.6 | 581330 | ENSMUSG00000059040 | Eno1b | enolase 1B, retrotransposed |
| chr2 | 1,42E+08 | 1,42E+08 | 1969 | * | 16,78167 | -2,216434202 | 0,806486 | -2,74826 | 0,005991 | 0,727652793 | 2,141612 | Intron (ENSMUST00000 | 2 | 1,42E+08 | 1,42E+08 | 57769 | 1 | 72899 | ENSMUST00000156619.2 | -243022 | ENSMUSG00000068205 | Macrod2 | mono-ADP ribosylhydrolase 2 |
| chr17 | 44137994 | 44140644 | 2651 | * | 22,25737 | -2,216357318 | 0,707657 | -3,13196 | 0,001736 | 0,426043749 | 3,471678 | Intron (ENSMUST00000 | 17 | 44114894 | 44350403 | 235510 | 1 | 53901 | ENSMUST00000044895.13 | 23100 | ENSMUSG00000039601 | Rcan2 | regulator of calcineurin 2 |
| chr13 | 1,06E+08 | 1,06E+08 | 1926 | * | 18,45547 | -2,214859507 | 0,834151 | -2,65523 | 0,007925 | 0,75508254 | 2,386008 | Distal Intergenic | 13 | 1,06E+08 | 1,06E+08 | 4484 | 1 | 15550 | ENSMUST00000022235.6 | 74916 | ENSMUSG00000021721 | Htr1a | 5-hydroxytryptamine (serotonin) receptor 1A |
| chr17 | 55743368 | 55744921 | 1554 | * | 14,68908 | -2,214618299 | 0,834897 | -2,65257 | 0,007988 | 0,877892722 | 1,214904 | Distal Intergenic | 17 | 55752383 | 55790033 | 37651 | 1 | 240119 | ENSMUST00000133899.2 | -7462 | ENSMUSG00000024172 | St6gal2 | beta galactoside alpha 2,6 sialyltransferase 2 |
| chr3 | 1,26E+08 | 1,26E+08 | 1781 | * | 17,68078 | -2,213365246 | 0,833109 | -2,65675 | 0,00789 | 0,736169201 | 2,663619 | Distal Intergenic | 3 | 1,26E+08 | 1,26E+08 | 50380 | 2 | 22239 | ENSMUST00000198610.2 | 61269 | ENSMUSG00000032854 | Ugt8a | UDP galactosyltransferase 8A |
| chr16 | 96021427 | 96024373 | 2947 | * | 31,86818 | -2,213024993 | 0,730131 | -3,031 | 0,002437 | 0,509737802 | 2,951062 | Promoter (<=1kb) | 16 | 96021699 | 96030125 | 8427 | 1 | 50795 | ENSMUST00000145069.3 | 0 | ENSMUSG00000040666 | Sh3bgr | SH3-binding domain glutamic acid-rich protein |
| chr15 | 25253919 | 25255618 | 1700 | * | 21,60717 | -2,211696013 | 0,719713 | -3,07303 | 0,002119 | 0,490966492 | 2,880017 | Distal Intergenic | 15 | 25137617 | 25238916 | 101300 | 2 | 1,03E+08 | ENSMUST00000228644.2 | -15003 | | Gm35996 | predicted gene, 35996 |
| chr3 | 1,15E+08 | 1,15E+08 | 2886 | * | 31,33236 | -2,211625327 | 0,621051 | -3,5611 | 0,000369 | 0,22800456 | 2,858511 | Distal Intergenic | 3 | 1,16E+08 | 1,16E+08 | 4640 | 2 | 13609 | ENSMUST00000055676.4 | 11188 | ENSMUSG00000045092 | S1pr1 | sphingosine-1-phosphate receptor 1 |
| chr2 | 17186274 | 17188849 | 2576 | * | 28,00106 | -2,209530399 | 0,642126 | -3,44096 | 0,00058 | 0,296099576 | 2,67243 | Exon (ENSMUST000001 | 2 | 17353505 | 17365897 | 12393 | 2 | 74103 | ENSMUST00000138241.2 | 177048 | ENSMUSG00000053702 | Nebl | nebulette |
| chr18 | 24483621 | 24485594 | 1974 | * | 17,67249 | -2,209196765 | 0,803998 | -2,74776 | 0,006 | 0,706976265 | 2,423594 | Distal Intergenic | 18 | 24472423 | 24477687 | 5265 | 1 | 328919 | ENSMUST00000234938.2 | 11198 | ENSMUSG00000117570 | Gm38410 | predicted gene, 38410 |
| chr15 | 85057282 | 85059275 | 1994 | * | 29,47467 | -2,207259201 | 0,74654 | -2,95665 | 0,00311 | 0,551069862 | 2,951062 | Distal Intergenic | 15 | 85033237 | 85040345 | 7109 | 2 | 1,01E+08 | ENSMUST00000229478.3 | -16937 | ENSMUSG00000115924 | Gm19277 | predicted gene, 19277 |
| chr16 | 84320985 | 84323342 | 2358 | * | 24,54393 | -2,206492076 | 0,800884 | -2,75507 | 0,005868 | 0,684274806 | 2,67243 | Downstream (<=300bp | 16 | 84312257 | 84319200 | 6944 | 1 | 402727 | ENSMUST00000210306.2 | 8728 | | A730009L09Rik | RIKEN cDNA A730009L09 gene |
| chr11 | 81422877 | 81425649 | 2773 | * | 22,83047 | -2,20598305 | 0,844518 | -2,61212 | 0,008998 | 0,756849896 | 2,67243 | Intron (ENSMUST00000 | 11 | 81463481 | 81464299 | 819 | 2 | 237880 | ENSMUST00000117522.2 | 38650 | ENSMUSG00000080980 | 1700071K01Rik | RIKEN cDNA 1700071K01 gene |
| chr9 | 10027552 | 10029588 | 2037 | * | 17,61175 | -2,205332459 | 0,843503 | -2,61449 | 0,008936 | 0,77580395 | 2,423594 | Intron (ENSMUST00000 | 9 | 9894436 | 9894526 | 91 | 1 | 1,02E+08 | ENSMUST00000183808.3 | 133116 | ENSMUSG00000098408 | Mir6237 | microRNA 6237 |

| | | | | | | | | | | | | | | | | | | | | | | | |
|---|---|---|---|---|---|---|---|---|---|---|---|---|---|---|---|---|---|---|---|---|---|---|---|
| chr12 | 63078780 | 63080711 | 1932 | * | 16,14936 | -2,205266057 | 0,815155 | -2,70533 | 0,006824 | 0,713633527 | 2,663619 | Distal Intergenic | 12 | 62734608 | 62735031 | 424 | 1 | 73309 | ENSMUST00000172433.3 | 344172 | | Spanxn4 | SPANX family, member N4 |
| chr2 | 84297469 | 84300315 | 2847 | * | 25,45448 | -2,205198303 | 0,663154 | -3,32532 | 0,000883 | 0,348632365 | 2,725491 | Intron (ENSMUST0000( | 2 | 84263199 | 84304128 | 40930 | 2 | 21788 | ENSMUST00000111722.8 | 3813 | ENSMUSG00000027082 | Tfpi | tissue factor pathway inhibitor |
| chr16 | 94612714 | 94614360 | 1647 | * | 21,47242 | -2,204933494 | 0,825245 | -2,67185 | 0,007543 | 0,715249603 | 2,880017 | Intron (ENSMUST0000( | 16 | 94562512 | 94657531 | 95020 | 2 | 16522 | ENSMUST00000232562.2 | 43171 | ENSMUSG00000043301 | Kcnj6 | potassium inwardly-rectifying channel, subfamily J, member 6 |
| chr2 | 1,47E+08 | 1,47E+08 | 2333 | * | 22,58873 | -2,204850176 | 0,827347 | -2,66497 | 0,0077 | 0,718390388 | 2,880017 | Distal Intergenic | 2 | 1,48E+08 | 1,48E+08 | 6940 | 1 | 381384 | ENSMUST00000152542.2 | -86571 | ENSMUSG00000051986 | A530006G24Rik | RIKEN cDNA A530006G24 gene |
| chr7 | 64807589 | 64809829 | 2241 | * | 22,63555 | -2,204748107 | 0,831415 | -2,6518 | 0,008006 | 0,71419937 | 3,104559 | Promoter (1-2kb) | 7 | 64405839 | 64806318 | 400480 | 2 | 70638 | ENSMUST00000119118.8 | -1271 | ENSMUSG00000030518 | Fam189a1 | family with sequence similarity 189, member A1 |
| chr10 | 91457580 | 91459154 | 1575 | * | 19,90999 | -2,202997611 | 0,795172 | -2,77047 | 0,005598 | 0,677933849 | 2,713306 | Distal Intergenic | 10 | 91724691 | 91789442 | 64752 | 1 | 1,03E+08 | ENSMUST00000217529.2 | -265537 | ENSMUSG00000111657 | Gm31592 | predicted gene, 31592 |
| chr12 | 42134675 | 42137197 | 2523 | * | 20,88586 | -2,202067995 | 0,712898 | -3,08889 | 0,002009 | 0,494205337 | 2,67243 | Exon (ENSMUST00000: | 12 | 41965491 | 42063539 | 98049 | 2 | 1,08E+08 | ENSMUST00000222917.2 | -71136 | | Gm46348 | predicted gene, 46348 |
| chr2 | 1,35E+08 | 1,35E+08 | 1632 | * | 18,49644 | -2,201376287 | 0,742871 | -2,96334 | 0,003043 | 0,590939713 | 2,423594 | Intron (ENSMUST0000( | 2 | 1,35E+08 | 1,35E+08 | 6341 | 1 | 18795 | ENSMUST00000201524.2 | 31163 | ENSMUSG00000051177 | Plcb1 | phospholipase C, beta 1 |
| chr17 | 17091583 | 17093956 | 2374 | * | 22,26421 | -2,198291486 | 0,810555 | -2,71208 | 0,006686 | 0,686950995 | 2,951933 | Distal Intergenic | 17 | 17171801 | 17194425 | 22625 | 1 | 213811 | ENSMUST00000190415.3 | -77845 | ENSMUSG00000060149 | BC002059 | cDNA sequence BC002059 |
| chr14 | 24447030 | 24449354 | 2325 | * | 21,43282 | -2,198090664 | 0,826063 | -2,66092 | 0,007793 | 0,716908183 | 2,951933 | Distal Intergenic | 14 | 24489055 | 24496471 | 7417 | 2 | 1,15E+08 | ENSMUST00000224174.2 | 47117 | | Gm47814 | predicted gene, 47814 |
| chr16 | 51927108 | 51929567 | 2460 | * | 19,52158 | -2,196678553 | 0,725792 | -3,02659 | 0,002473 | 0,531203841 | 2,663619 | Promoter (1-2kb) | 16 | 51925378 | 51925510 | 133 | 2 | 1,15E+08 | ENSMUST00000158181.4 | -1598 | ENSMUSG00000119052 | Gm23513 | predicted gene, 23513 |
| chr2 | 12507009 | 12509274 | 2266 | * | 19,80309 | -2,194112297 | 0,834786 | -2,62835 | 0,00858 | 0,770673161 | 2,386008 | Distal Intergenic | 2 | 12352074 | 12424281 | 72208 | 2 | 66960 | ENSMUST00000028105.13 | -82728 | ENSMUSG00000026767 | Mindy3 | MINDY lysine 48 deubiquitinase 3 |
| chr7 | 58815450 | 58817520 | 2071 | * | 19,65608 | -2,193796366 | 0,785411 | -2,79318 | 0,005219 | 0,684274806 | 2,386008 | Distal Intergenic | 7 | 58878498 | 58939567 | 61070 | 1 | 22215 | ENSMUST00000202945.4 | -60978 | ENSMUSG00000025326 | Ube3a | ubiquitin protein ligase E3A |
| chr6 | 8665430 | 8667356 | 1927 | * | 27,47884 | -2,191340574 | 0,708764 | -3,09178 | 0,00199 | 0,492834344 | 2,67243 | 3' UTR | 6 | 8658991 | 8672317 | 13327 | 2 | 15893 | ENSMUST00000127398.8 | 4961 | ENSMUSG00000062995 | Ica1 | islet cell autoantigen 1 |
| chr6 | 32218798 | 32219817 | 1020 | * | 16,70589 | -2,191318073 | 0,828828 | -2,64387 | 0,008196 | 0,767135014 | 2,318434 | Intron (ENSMUST0000( | 6 | 32089078 | 32090161 | 1084 | 1 | 1,05E+08 | ENSMUST00000140465.2 | 129720 | | Gm13850 | predicted gene 13850 |
| chr10 | 52420511 | 52422646 | 2136 | * | 22,62613 | -2,190917431 | 0,765101 | -2,86357 | 0,004189 | 0,618136403 | 2,880017 | Promoter (<=1kb) | 10 | 52418416 | 52421835 | 3420 | 2 | 22639 | ENSMUST00000180673.2 | 0 | ENSMUSG00000049576 | Zfa-ps | zinc finger protein, autosomal, pseudogene |
| chr16 | 6753834 | 6756583 | 2750 | * | 22,57964 | -2,190017474 | 0,834163 | -2,62541 | 0,008655 | 0,708539289 | 3,471678 | Intron (ENSMUST0000( | 16 | 6627086 | 7227677 | 600592 | 1 | 268859 | ENSMUST00000115841.10 | 126748 | ENSMUSG00000008658 | Rbfox1 | RNA binding protein, fox-1 homolog (C. elegans) 1 |
| chr11 | 64375786 | 64377884 | 2099 | * | 20,54714 | -2,189250705 | 0,787741 | -2,77915 | 0,00545 | 0,676476319 | 2,663619 | Intron (ENSMUST0000( | 11 | 64326158 | 64413667 | 87510 | 1 | 15478 | ENSMUST00000058652.6 | 49628 | ENSMUSG00000047759 | Hs3st3a1 | heparan sulfate (glucosamine) 3-O-sulfotransferase 3A1 |
| chr11 | 1,09E+08 | 1,09E+08 | 2181 | * | 21,20691 | -2,188580017 | 0,733731 | -2,98281 | 0,002856 | 0,508113534 | 3,471678 | Intron (ENSMUST0000( | 11 | 1,09E+08 | 1,09E+08 | 98635 | 1 | 76380 | ENSMUST00000151755.2 | -11289 | ENSMUSG00000020728 | Cep112 | centrosomal protein 112 |
| chr18 | 9626249 | 9627990 | 1742 | * | 17,18403 | -2,188174076 | 0,786476 | -2,78225 | 0,005398 | 0,685026746 | 2,423594 | Distal Intergenic | 18 | 9618419 | 9619469 | 1051 | 2 | 26369 | ENSMUST00000234003.2 | -6780 | ENSMUSG00000050996 | Cetn1 | centrin 1 |
| chr10 | 96177659 | 96179398 | 1740 | * | 17,62371 | -2,186929518 | 0,782383 | -2,79522 | 0,005187 | 0,699278451 | 2,154504 | Intron (ENSMUST0000( | 10 | 96145993 | 96198229 | 52237 | 1 | 1,03E+08 | ENSMUST00000218518.2 | 31666 | | Gm33782 | predicted gene, 33782 |
| chr1 | 20523073 | 20524444 | 1372 | * | 15,39566 | -2,18687189 | 0,815738 | -2,68085 | 0,007344 | 0,760759863 | 2,141612 | Intron (ENSMUST0000( | 1 | 20515035 | 20515136 | 102 | 2 | 1,15E+08 | ENSMUST00000093665.3 | -7937 | ENSMUSG00000070175 | Gm24162 | predicted gene, 24162 |
| chr2 | 58717698 | 58719610 | 1913 | * | 24,1631 | -2,186682091 | 0,793945 | -2,7542 | 0,005884 | 0,671421781 | 2,951062 | Exon (ENSMUST00000: | 2 | 58655337 | 58680314 | 24978 | 1 | 76654 | ENSMUST00000128864.2 | 62361 | ENSMUSG00000026839 | Upp2 | uridine phosphorylase 2 |
| chr14 | 42037426 | 42039066 | 1641 | * | 20,90538 | -2,186598392 | 0,810429 | -2,69807 | 0,006974 | 0,700999262 | 2,880017 | Distal Intergenic | 14 | 42076616 | 42082005 | 5390 | 2 | 1,01E+08 | ENSMUST00000166674.2 | 42939 | ENSMUSG00000092152 | Gm17026 | predicted gene 17026 |
| chr9 | 1,11E+08 | 1,11E+08 | 1952 | * | 18,31058 | -2,186386718 | 0,838666 | -2,60698 | 0,009134 | 0,779303858 | 2,423594 | Distal Intergenic | 9 | 1,11E+08 | 1,11E+08 | 50038 | 1 | 245038 | ENSMUST00000111879.5 | 56509 | ENSMUSG00000032500 | Dclk3 | doublecortin-like kinase 3 |
| chr10 | 23481357 | 23484903 | 3547 | * | 29,90466 | -2,184968529 | 0,653735 | -3,34228 | 0,000831 | 0,337811311 | 2,725491 | Promoter (<=1kb) | 10 | 23448918 | 23485329 | 36412 | 2 | 1,15E+08 | ENSMUST00000219823.2 | 426 | | Gm47843 | predicted gene, 47843 |
| chr5 | 77718985 | 77720616 | 1632 | * | 18,34138 | -2,18427552 | 0,781567 | -2,79474 | 0,005194 | 0,683988405 | 2,386008 | Distal Intergenic | 5 | 77613335 | 77616698 | 3364 | 1 | 69370 | ENSMUST00000201654.2 | 105650 | ENSMUSG00000107232 | 1700017L05Rik | RIKEN cDNA 1700017L05 gene |
| chr7 | 76293425 | 76294886 | 1462 | * | 13,9111 | -2,184013443 | 0,828131 | -2,63728 | 0,008357 | 0,903470246 | 1,075735 | Intron (ENSMUST0000( | 7 | 76060810 | 76416356 | 355547 | 1 | 244071 | ENSMUST00000107442.3 | 232615 | ENSMUSG00000025754 | Agbl1 | ATP/GTP binding protein-like 1 |
| chr10 | 86923435 | 86926595 | 3161 | * | 25,44061 | -2,183885949 | 0,680343 | -3,20998 | 0,001327 | 0,409763611 | 2,951062 | Intron (ENSMUST0000( | 10 | 86893910 | 87066450 | 172541 | 1 | 76640 | ENSMUST00000169849.3 | 29525 | ENSMUSG00000047129 | 1700113H08Rik | RIKEN cDNA 1700113H08 gene |
| chr6 | 1,48E+08 | 1,48E+08 | 1409 | * | 19,85966 | -2,183756917 | 0,791236 | -2,75993 | 0,005781 | 0,683067851 | 2,713306 | Distal Intergenic | 6 | 1,48E+08 | 1,48E+08 | 129591 | 1 | 330450 | ENSMUST00000032443.14 | -111298 | ENSMUSG00000030303 | Far2 | fatty acyl CoA reductase 2 |
| chr17 | 6965244 | 6966601 | 1358 | * | 20,0589 | -2,18363957 | 0,773359 | -2,82358 | 0,004749 | 0,663630601 | 2,507013 | Intron (ENSMUST0000( | 17 | 6973229 | 7005235 | 32007 | 1 | 83672 | ENSMUST00000161118.8 | -6628 | ENSMUSG00000041831 | Sytl3 | synaptotagmin-like 3 |
| chr16 | 14539471 | 14541971 | 2501 | * | 23,88863 | -2,183025917 | 0,662868 | -3,2933 | 0,00099 | 0,359463243 | 2,858511 | Distal Intergenic | 16 | 14523716 | 14527249 | 3534 | 1 | 20583 | ENSMUST00000023356.8 | 15755 | ENSMUSG00000022676 | Snai2 | snail family zinc finger 2 |
| chr7 | 1,42E+08 | 1,42E+08 | 2420 | * | 23,42987 | -2,182851422 | 0,777325 | -2,80816 | 0,004983 | 0,649854434 | 2,858511 | Promoter (1-2kb) | 7 | 1,42E+08 | 1,42E+08 | 20855 | 2 | 16334 | ENSMUST00000105934.8 | 1960 | ENSMUSG00000000215 | Ins2 | insulin II |
| chr15 | 44350774 | 44352801 | 2028 | * | 20,50907 | -2,180843765 | 0,759466 | -2,87155 | 0,004085 | 0,639997962 | 2,507013 | Intron (ENSMUST0000( | 15 | 44320949 | 44464765 | 143817 | 1 | 192190 | ENSMUST00000209244.2 | 29825 | ENSMUSG00000038725 | Pkhd1l1 | polycystic kidney and hepatic disease 1-like 1 |
| chr4 | 77129342 | 77131846 | 2505 | * | 22,38792 | -2,179912668 | 0,716463 | -3,0426 | 0,002345 | 0,476080922 | 3,471678 | Intron (ENSMUST0000( | 4 | 75863043 | 76512536 | 649494 | 2 | 19266 | ENSMUST00000173376.8 | -616806 | ENSMUSG00000028399 | Ptprd | protein tyrosine phosphatase, receptor type, D |
| chr3 | 31298437 | 31301928 | 3492 | * | 32,01075 | -2,179781634 | 0,635196 | -3,43167 | 0,0006 | 0,297590871 | 2,725491 | Intron (ENSMUST0000( | 3 | 31276869 | 31312543 | 35675 | 2 | 241919 | ENSMUST00000108245.2 | 10615 | ENSMUSG00000069072 | Slc7a14 | solute carrier family 7 (cationic amino acid transporter, y+ system), member 14 |
| chr19 | 48424494 | 48426621 | 2128 | * | 18,47459 | -2,179716473 | 0,792629 | -2,74998 | 0,00596 | 0,705187732 | 2,423594 | Intron (ENSMUST0000( | 19 | 48418399 | 48643749 | 225351 | 1 | 66673 | ENSMUST00000236094.2 | 6095 | ENSMUSG00000063434 | Sorcs3 | sortilin-related VPS10 domain containing receptor 3 |
| chr12 | 24517101 | 24520981 | 3881 | * | 34,08061 | -2,178113162 | 0,707375 | -3,07915 | 0,002076 | 0,492834344 | 2,787255 | Distal Intergenic | 12 | 24548358 | 24608538 | 60181 | 1 | 21340 | ENSMUST00000075954.9 | -27377 | ENSMUSG00000059669 | Taf1b | TATA-box binding protein associated factor, RNA polymerase I, B |
| chr11 | 41382100 | 41384183 | 2084 | * | 20,47715 | -2,177846397 | 0,775684 | -2,80765 | 0,004991 | 0,68605621 | 2,221611 | Distal Intergenic | 11 | 41601169 | 41859514 | 258346 | 1 | 1,03E+08 | ENSMUST00000155032.8 | -216986 | ENSMUSG00000087306 | A230004M16Rik | RIKEN cDNA A230004M16 gene |
| chr2 | 1,78E+08 | 1,78E+08 | 1478 | * | 20,13724 | -2,176964371 | 0,836095 | -2,60373 | 0,009222 | 0,7591629 | 2,713306 | Exon (ENSMUST00000: | 2 | 1,78E+08 | 1,78E+08 | 8700 | 1 | 74189 | ENSMUST00000133267.3 | 4878 | ENSMUSG00000027525 | Phactr3 | phosphatase and actin regulator 3 |
| chrX | 1,31E+08 | 1,31E+08 | 2098 | * | 16,28385 | -2,175783195 | 0,826132 | -2,6337 | 0,008446 | 0,879615137 | 1,2763 | Distal Intergenic | 20 | 1,31E+08 | 1,31E+08 | 131 | 2 | 1,15E+08 | ENSMUST00000104440.3 | 350378 | ENSMUSG00000077628 | Gm23768 | predicted gene, 23768 |
| chr10 | 55200391 | 55203030 | 2640 | * | 23,05924 | -2,174576466 | 0,721377 | -3,01448 | 0,002574 | 0,521453391 | 2,951933 | Distal Intergenic | 10 | 54589072 | 54589133 | 62 | 2 | 1,15E+08 | ENSMUST00000158362.3 | -611258 | ENSMUSG00000088987 | Gm25602 | predicted gene, 25602 |
| chr7 | 38014182 | 38015719 | 1538 | * | 22,06272 | -2,172572317 | 0,6903 | -3,14729 | 0,001648 | 0,44771988 | 2,880017 | Exon (ENSMUST00000: | 7 | 37961420 | 37970847 | 9428 | 2 | 66161 | ENSMUST00000032585.8 | -43335 | ENSMUSG00000030423 | Pop4 | processing of precursor 4, ribonuclease P/MRP family, (S. cerevisiae) |
| chr3 | 46794129 | 46795692 | 1564 | * | 15,69178 | -2,172498391 | 0,805968 | -2,69552 | 0,007028 | 0,854157826 | 1,214904 | Distal Intergenic | 3 | 46855103 | 46855217 | 115 | 2 | 1,02E+08 | ENSMUST00000183392.3 | 59525 | ENSMUSG00000098417 | Mir6379 | microRNA 6379 |
| chr10 | 91413303 | 91414730 | 1428 | * | 16,66495 | -2,172246686 | 0,763441 | -2,84534 | 0,004436 | 0,661358703 | 2,386008 | Distal Intergenic | 10 | 91724691 | 91789442 | 64752 | 1 | 1,03E+08 | ENSMUST00000217529.2 | -309961 | ENSMUSG00000111657 | Gm31592 | predicted gene, 31592 |
| chr11 | 52494211 | 52497790 | 3580 | * | 30,63493 | -2,171277271 | 0,66723 | -3,25417 | 0,001137 | 0,388793511 | 2,858511 | Distal Intergenic | 11 | 52655461 | 53079365 | 423905 | 1 | 320027 | ENSMUST00000036796.8 | -157671 | ENSMUSG00000036264 | Fstl4 | follistatin-like 4 |
| chr3 | 13204524 | 13206418 | 1895 | * | 16,63291 | -2,171076104 | 0,808909 | -2,68396 | 0,007276 | 0,748382289 | 2,318434 | Distal Intergenic | 3 | 13536715 | 13842163 | 305449 | 1 | 76897 | ENSMUST00000191806.3 | -330297 | ENSMUSG00000039717 | Ralyl | RALY RNA binding protein-like |
| chr5 | 45139380 | 45141811 | 2432 | * | 23,00742 | -2,170873115 | 0,792913 | -2,73784 | 0,006184 | 0,681119785 | 2,951933 | Exon (ENSMUST00000( | 5 | 45123925 | 45125209 | 1285 | 1 | 75783 | ENSMUST00000079389.6 | 15455 | ENSMUSG00000062496 | 4930431F12Rik | RIKEN cDNA 4930431F12 gene |
| chr1 | 1,67E+08 | 1,67E+08 | 2511 | * | 21,44438 | -2,170278456 | 0,767935 | -2,82612 | 0,004712 | 0,626725358 | 3,104559 | Promoter (<=1kb) | 1 | 1,67E+08 | 1,67E+08 | 21313 | 1 | 20183 | ENSMUST00000111380.2 | -902 | ENSMUSG00000015843 | Rxrg | retinoid X receptor gamma |
| chr17 | 44015564 | 44017808 | 2245 | * | 19,74985 | -2,166960037 | 0,749474 | -2,89131 | 0,003836 | 0,627785088 | 2,507013 | Intron (ENSMUST0000( | 17 | 44009254 | 44009443 | 190 | 1 | 1,15E+08 | ENSMUST00000122525.4 | 6310 | ENSMUSG00000084474 | Gm25783 | predicted gene, 25783 |
| chr13 | 44288895 | 44290909 | 2015 | * | 16,59508 | -2,166842807 | 0,804035 | -2,69496 | 0,00704 | 0,740451144 | 2,318434 | Distal Intergenic | 13 | 44304071 | 44305429 | 1359 | 1 | 1,03E+08 | ENSMUST00000221119.2 | -13162 | | Gm33684 | predicted gene, 33684 |
| chr1 | 1,88E+08 | 1,88E+08 | 2240 | * | 17,20954 | -2,166185736 | 0,79871 | -2,71211 | 0,006686 | 0,717868719 | 2,507013 | Intron (ENSMUST0000( | 1 | 1,88E+08 | 1,88E+08 | 95498 | 1 | 22283 | ENSMUST00000142189.8 | -52210 | ENSMUSG00000026609 | Ush2a | usherin |
| chr17 | 95037606 | 95038674 | 1069 | * | 23,24586 | -2,166008941 | 0,706923 | -3,06399 | 0,002184 | 0,495652322 | 2,880017 | Intron (ENSMUST0000( | 17 | 95030018 | 95057320 | 27303 | 2 | 76781 | ENSMUST00000234990.2 | 18646 | ENSMUSG00000055660 | Mettl4 | methyltransferase like 4 |
| chr5 | 1,48E+08 | 1,48E+08 | 1849 | * | 19,29198 | -2,165145851 | 0,769495 | -2,81372 | 0,004897 | 0,669586355 | 2,507013 | Intron (ENSMUST0000( | 5 | 1,48E+08 | 1,48E+08 | 48694 | 1 | 77521 | ENSMUST00000110514.8 | 7010 | ENSMUSG00000029651 | Mtus2 | microtubule associated tumor suppressor candidate 2 |
| chr2 | 68313607 | 68316315 | 2709 | * | 27,42433 | -2,164586799 | 0,682263 | -3,17266 | 0,001511 | 0,44621096 | 2,67243 | Exon (ENSMUST00020: | 2 | 68040789 | 68302612 | 261824 | 2 | 53416 | ENSMUST00000102715.4 | -10995 | ENSMUSG00000027030 | Stk39 | serine/threonine kinase 39 |
| chr14 | 36284417 | 36286631 | 2215 | * | 22,80826 | -2,163868718 | 0,687557 | -3,14718 | 0,001649 | 0,444967102 | 2,951933 | Distal Intergenic | 14 | 36489382 | 36489487 | 106 | 1 | 1,15E+08 | ENSMUST00000082843.3 | -202751 | ENSMUSG00000064777 | Gm25204 | predicted gene, 25204 |
| chr10 | 1,02E+08 | 1,02E+08 | 1600 | * | 19,94871 | -2,163706715 | 0,78207 | -2,76664 | 0,005664 | 0,686838043 | 2,507013 | Intron (ENSMUST0000( | 10 | 1,02E+08 | 1,02E+08 | 81 | 1 | 1,01E+08 | ENSMUST00000175211.3 | -39388 | ENSMUSG00000092952 | Mir3059 | microRNA 3059 |
| chr8 | 37788238 | 37790627 | 2390 | * | 24,72966 | -2,163665706 | 0,727836 | -2,97274 | 0,002952 | 0,54190677 | 2,951933 | Exon (ENSMUST00000: | 8 | 37784631 | 37784732 | 102 | 2 | 1,15E+08 | ENSMUST00000083343.3 | -3506 | ENSMUSG00000065277 | Gm25126 | predicted gene, 25126 |
| chr6 | 52543679 | 52545303 | 1625 | * | 20,57657 | -2,163012156 | 0,83432 | -2,59255 | 0,009527 | 0,801688106 | 2,221611 | Intron (ENSMUST0000( | 6 | 52525837 | 52533900 | 8064 | 2 | 58875 | ENSMUST00000149471.2 | -9779 | ENSMUSG00000029776 | Hibadh | 3-hydroxyisobutyrate dehydrogenase |
| chr6 | 1,24E+08 | 1,24E+08 | 1379 | * | 22,24283 | -2,161302401 | 0,766604 | -2,81932 | 0,004813 | 0,639997962 | 2,951933 | Intron (ENSMUST0000( | 6 | 1,24E+08 | 1,24E+08 | 46034 | 1 | 243628 | ENSMUST00000233759.2 | 8830 | ENSMUSG00000072780 | Vmn2r24 | vomeronasal 2, receptor 24 |
| chr10 | 1,08E+08 | 1,08E+08 | 1628 | * | 16,14735 | -2,160874217 | 0,830032 | -2,60336 | 0,009231 | 0,892478752 | 1,2763 | Exon (ENSMUST00000: | 10 | 1,08E+08 | 1,08E+08 | 151604 | 2 | 628870 | ENSMUST00000165341.5 | 57440 | ENSMUSG00000091455 | Otogl | otogelin-like |
| chr10 | 95961621 | 95964232 | 2612 | * | 25,82515 | -2,160785988 | 0,632984 | -3,41365 | 0,000641 | 0,307219482 | 2,67243 | Exon (ENSMUST00000: | 10 | 95883490 | 95931867 | 48378 | 2 | 1,19E+08 | ENSMUST00000233655.2 | -29754 | | Gm49817 | predicted gene, 49817 |
| chr13 | 57018405 | 57020008 | 1604 | * | 19,19807 | -2,160166278 | 0,819051 | -2,6374 | 0,008354 | 0,778618001 | 2,221611 | Intron (ENSMUST0000( | 13 | 56921577 | 57035929 | 114353 | 2 | 26946 | ENSMUST00000173067.9 | 15921 | ENSMUSG00000021541 | Trpc7 | transient receptor potential cation channel, subfamily C, member 7 |
| chr8 | 92899668 | 92901468 | 1801 | * | 26,90675 | -2,157959341 | 0,686084 | -3,14533 | 0,001659 | 0,450394359 | 2,858511 | Intron (ENSMUST0000( | 8 | 92766545 | 92766648 | 104 | 1 | 1,15E+08 | ENSMUST00000083702.3 | 133123 | ENSMUSG00000065636 | Gm24254 | predicted gene, 24254 |
| chr13 | 16390021 | 16392121 | 2101 | * | 17,60621 | -2,156928901 | 0,836132 | -2,57965 | 0,00989 | 0,777172375 | 2,663619 | Distal Intergenic | 13 | 16215631 | 16215737 | 107 | 2 | 1,15E+08 | ENSMUST00000239694.1 | -174284 | ENSMUSG00000119058 | Gm22628 | predicted gene, 22628 |
| chr3 | 1,36E+08 | 1,36E+08 | 1882 | * | 22,86195 | -2,15511549 | 0,784232 | -2,74806 | 0,005995 | 0,680148888 | 2,880017 | Intron (ENSMUST0000( | 3 | 1,36E+08 | 1,36E+08 | 795 | 2 | 242248 | ENSMUST00000196131.2 | 6603 | ENSMUSG00000037922 | Bank1 | B cell scaffold protein with ankyrin repeats 1 |
| chr9 | 1,04E+08 | 1,04E+08 | 2821 | * | 23,24878 | -2,154551732 | 0,825114 | -2,61122 | 0,009022 | 0,747833008 | 2,880017 | Promoter (2-3kb) | 9 | 1,04E+08 | 1,05E+08 | 306146 | 1 | 74020 | ENSMUST00000157006.8 | -2536 | ENSMUSG00000032564 | Cpne4 | copine IV |
| chr11 | 26782208 | 26784331 | 2124 | * | 18,06179 | -2,153196323 | 0,756749 | -2,84532 | 0,004437 | 0,658408085 | 2,423594 | Exon (ENSMUST00000: | 11 | 26497848 | 26543999 | 46152 | 2 | 69922 | ENSMUST00000156264.2 | -238209 | ENSMUSG00000064090 | Vrk2 | vaccinia related kinase 2 |
| chr16 | 12745219 | 12748312 | 3094 | * | 26,68345 | -2,152740477 | 0,707164 | -3,04419 | 0,002333 | 0,50615802 | 2,880017 | Intron (ENSMUST0000( | 16 | 12446948 | 12763799 | 316852 | 2 | 1,05E+08 | ENSMUST00000230696.2 | 15487 | | Gm45919 | predicted gene, 45919 |
| chr6 | 81885296 | 81886351 | 1056 | * | 17,46181 | -2,150845258 | 0,804709 | -2,67282 | 0,007522 | 0,765231686 | 2,154504 | Promoter (1-2kb) | 6 | 81877445 | 81887781 | 10337 | 2 | 74220 | ENSMUST00000191101.2 | 1430 | ENSMUSG00000101404 | 1700009C05Rik | RIKEN cDNA 1700009C05 gene |
| chr13 | 50316921 | 50320393 | 3473 | * | 26,83548 | -2,150195743 | 0,684221 | -3,14255 | 0,001675 | 0,459179243 | 2,725491 | Exon (ENSMUST00000: | 13 | 50291105 | 50300706 | 9602 | 1 | 71252 | ENSMUST00000222134.2 | 25816 | | 4933433N18Rik | RIKEN cDNA 4933433N18 gene |
| chr16 | 12225539 | 12227930 | 2392 | * | 25,02511 | -2,149252516 | 0,768496 | -2,7967 | 0,005163 | 0,654833721 | 2,880017 | Exon (ENSMUST00000: | 16 | 12522232 | 12524604 | 2373 | 1 | 1,03E+08 | ENSMUST00000230690.2 | -294302 | | Gm38619 | predicted gene, 38619 |
| chr2 | 83168073 | 83170028 | 1956 | * | 19,99995 | -2,147410191 | 0,784263 | -2,73812 | 0,006179 | 0,714462637 | 2,386008 | Distal Intergenic | 2 | 83474779 | 83494966 | 20188 | 1 | 69082 | ENSMUST00000081591.7 | -304751 | ENSMUSG00000027091 | Zc3h15 | zinc finger CCCH-type containing 15 |
| chr9 | 92311352 | 92313053 | 1702 | * | 15,9565 | -2,14603136 | 0,819176 | -2,61974 | 0,0088 | 0,77580395 | 2,386008 | Distal Intergenic | 9 | 92339426 | 92374509 | 35084 | 1 | 235527 | ENSMUST00000034941.9 | -26373 | ENSMUSG00000032377 | Plscr4 | phospholipid scramblase 4 |
| chr18 | 39225438 | 39228523 | 3086 | * | 31,63001 | -2,143163199 | 0,647166 | -3,31161 | 0,000928 | 0,355694809 | 2,725491 | Intron (ENSMUST0000( | 18 | 39232364 | 39259986 | 27623 | 1 | 71302 | ENSMUST00000123820.2 | -3841 | ENSMUSG00000036452 | Arhgap26 | Rho GTPase activating protein 26 |
| chr11 | 81273914 | 81276274 | 2361 | * | 23,95972 | -2,141476725 | 0,738187 | -2,90099 | 0,00372 | 0,591387753 | 2,951933 | Exon (ENSMUST00000: | 11 | 81463481 | 81464299 | 819 | 2 | 237880 | ENSMUST00000117522.2 | 188025 | ENSMUSG00000080980 | 1700071K01Rik | RIKEN cDNA 1700071K01 gene |
| chr5 | 32394416 | 32396885 | 2470 | * | 25,14054 | -2,141474905 | 0,820488 | -2,61 | 0,009054 | 0,758346111 | 2,67243 | Promoter (<=1kb) | 5 | 32396428 | 32479588 | 83161 | 1 | 665270 | ENSMUST00000202201.4 | 0 | ENSMUSG00000029134 | Plb1 | phospholipase B1 |
| chr2 | 1,6E+08 | 1,6E+08 | 2010 | * | 23,63877 | -2,140347293 | 0,774708 | -2,76278 | 0,005731 | 0,672298194 | 2,858511 | Distal Intergenic | 2 | 1,6E+08 | 1,6E+08 | 23967 | 1 | 71613 | ENSMUST00000140319.2 | 146694 | ENSMUSG00000086653 | 9130015L21Rik | RIKEN cDNA 9130015L21 gene |
| chr10 | 4543008 | 4544791 | 1784 | * | 22,21953 | -2,138878948 | 0,759739 | -2,81528 | 0,004873 | 0,658200216 | 2,67243 | Distal Intergenic | 10 | 4561593 | 4843083 | 281491 | 1 | 13982 | ENSMUST00000149195.8 | -16802 | ENSMUSG00000019768 | Esr1 | estrogen receptor 1 (alpha) |
| chr18 | 26603180 | 26605095 | 1916 | * | 15,94957 | -2,13867317 | 0,817614 | -2,61575 | 0,008903 | 0,888080529 | 1,2763 | Distal Intergenic | 18 | 26628488 | 26628618 | 131 | 1 | 1,15E+08 | ENSMUST00000157473.3 | -23393 | ENSMUSG00000088098 | Gm23674 | predicted gene, 23674 |
| chr7 | 76922394 | 76924166 | 1773 | * | 16,91478 | -2,13778915 | 0,796949 | -2,68247 | 0,007308 | 0,723239852 | 2,663619 | Distal Intergenic | 7 | 76060810 | 76416356 | 355547 | 1 | 244071 | ENSMUST00000107442.3 | 861584 | ENSMUSG00000025754 | Agbl1 | ATP/GTP binding protein-like 1 |
| chr10 | 1,05E+08 | 1,05E+08 | 2284 | * | 17,81873 | -2,136575069 | 0,787247 | -2,71398 | 0,006648 | 0,708539289 | 2,663619 | Distal Intergenic | 10 | 1,05E+08 | 1,05E+08 | 105 | 2 | 1,15E+08 | ENSMUST00000158394.3 | 167058 | ENSMUSG00000089019 | Gm25522 | predicted gene, 25522 |
| chr8 | 1,21E+08 | 1,21E+08 | 2165 | * | 27,18344 | -2,13631678 | 0,673585 | -3,17156 | 0,001516 | 0,430115816 | 2,951062 | Distal Intergenic | 8 | 1,21E+08 | 1,21E+08 | 1503 | 1 | 15900 | ENSMUST00000160594.2 | 9650 | ENSMUSG00000041515 | Irf8 | interferon regulatory factor 8 |
| chr2 | 6928015 | 6929791 | 1777 | * | 17,51777 | -2,135860578 | 0,81418 | -2,62333 | 0,008708 | 0,751314174 | 2,663619 | Intron (ENSMUST0000( | 2 | 6546680 | 6889807 | 343128 | 2 | 14007 | ENSMUST00000114934.11 | -38208 | ENSMUSG00000002107 | Celf2 | CUGBP, Elav-like family member 2 |
| chr5 | 127738217 | 127742249 | 4033 | * | 47,501616 | -2,133155011 | 0,4886587 | -4,3653266 | 1,269E-05 | 0,030043467 | 2,3776742 | Exon (ENSMUST0000013 | 5 | 127734097 | 127766935 | 32839 | 1 | 319804 | ENSMUST00000134529.2 | 4120 | ENSMUSG00000049971 | Glt1d1 | glycosyltransferase 1 domain containing 1 |
| chr11 | 94355526 | 94357528 | 2003 | * | 20,20479 | -2,132098738 | 0,7536 | -2,82922 | 0,004666 | 0,669586355 | 2,386008 | Exon (ENSMUST00000: | 11 | 94317778 | 94364467 | 46690 | 2 | 12291 | ENSMUST00000152811.2 | 6939 | ENSMUSG00000020866 | Cacna1g | calcium channel, voltage-dependent, T type, alpha 1G subunit |
| chr3 | 49900697 | 49902780 | 2084 | * | 16,26118 | -2,131209481 | 0,811544 | -2,62612 | 0,008637 | 0,79206913 | 2,141612 | Intron (ENSMUST0000( | 3 | 49699720 | 49711774 | 12055 | 2 | 73173 | ENSMUST00000193600.2 | -188923 | ENSMUSG00000037892 | Pcdh18 | protocadherin 18 |
| chr9 | 76490896 | 76494507 | 3612 | * | 29,57325 | -2,127088822 | 0,63071 | -3,37253 | 0,000745 | 0,315367183 | 2,880017 | Distal Intergenic | 9 | 76397336 | 76474398 | 77063 | 2 | 208994 | ENSMUST00000183437.8 | -16498 | ENSMUSG00000032358 | Fam83b | family with sequence similarity 83, member B |
| chrX | 57196069 | 57198674 | 2606 | * | 16,44777 | -2,125307192 | 0,794332 | -2,67559 | 0,00746 | 0,764849314 | 2,141612 | Distal Intergenic | 20 | 57076370 | 57087096 | 10727 | 1 | 22773 | ENSMUST00000088629.4 | 119699 | ENSMUSG00000067860 | Zic3 | zinc finger protein of the cerebellum 3 |
| chr11 | 39393422 | 39395607 | 2186 | * | 18,66493 | -2,123711618 | 0,801419 | -2,64994 | 0,008051 | 0,75508254 | 2,423594 | Distal Intergenic | 11 | 39721483 | 39721627 | 145 | 1 | 1,15E+08 | ENSMUST00000157978.4 | -325876 | ENSMUSG00000088603 | Gm24192 | predicted gene, 24192 |
| chr14 | 46418719 | 46420909 | 2191 | * | 21,72134 | -2,123708523 | 0,793094 | -2,67775 | 0,007412 | 0,725842761 | 2,67243 | Distal Intergenic | 14 | 46458549 | 46466805 | 8257 | 1 | 1,03E+08 | ENSMUST00000242583.1 | -37640 | | Gm34499 | predicted gene, 34499 |
| chr5 | 22246692 | 22248985 | 2294 | * | 20,43112 | -2,123514554 | 0,795503 | -2,6694 | 0,007599 | 0,740451144 | 2,507013 | Intron (ENSMUST0000( | 5 | 22252522 | 22253751 | 1230 | 2 | 19699 | ENSMUST00000159741.2 | 4766 | ENSMUSG00000042453 | Reln | reelin |
| chr18 | 61463163 | 61465522 | 2360 | * | 20,40381 | -2,123342742 | 0,797555 | -2,66231 | 0,007761 | 0,744477896 | 2,507013 | Intron (ENSMUST0000( | 18 | 61408094 | 61422995 | 14902 | 1 | 225600 | ENSMUST00000135688.2 | 55069 | ENSMUSG00000024575 | Pde6a | phosphodiesterase 6A, cGMP-specific, rod, alpha |
| chr2 | 1,43E+08 | 1,43E+08 | 2414 | * | 27,10531 | -2,12110097 | 0,632404 | -3,35403 | 0,000796 | 0,335900794 | 2,67243 | Intron (ENSMUST0000( | 2 | 1,43E+08 | 1,43E+08 | 202516 | 2 | 16558 | ENSMUST00000211861.2 | 85177 | ENSMUSG00000038844 | Kif16b | kinesin family member 16B |
| chr16 | 3153797 | 3154120 | 324 | * | 18,46175 | -2,120706873 | 0,775248 | -2,73552 | 0,006228 | 0,713663698 | 2,423594 | Distal Intergenic | 16 | 3286324 | 3298915 | 12592 | 1 | 1,03E+08 | ENSMUST00000229453.2 | -132204 | ENSMUSG00000116455 | Gm35974 | predicted gene, 35974 |
| chr17 | 65485021 | 65487364 | 2344 | * | 26,869 | -2,119869104 | 0,68126 | -3,11169 | 0,00186 | 0,464338132 | 2,951062 | Distal Intergenic | 17 | 65251716 | 65251847 | 132 | 2 | 1,15E+08 | ENSMUST00000104503.4 | -233174 | ENSMUSG00000077691 | Gm25344 | predicted gene, 25344 |
| chr10 | 95284100 | 95286448 | 2349 | * | 19,51291 | -2,119387056 | 0,81573 | -2,59815 | 0,009373 | 0,777938745 | 2,507013 | Intron (ENSMUST0000( | 10 | 95253242 | 95279547 | 26306 | 1 | 70523 | ENSMUST00000244592.1 | 30858 | ENSMUSG00000097766 | 5730420D15Rik | RIKEN cDNA 5730420D15 gene |
| chr16 | 69860278 | 69862587 | 2310 | * | 20,51455 | -2,118827011 | 0,740848 | -2,86 | 0,004236 | 0,644704154 | 2,507013 | Exon (ENSMUST00000: | 16 | 69653762 | 69660632 | 6871 | 2 | 224318 | ENSMUST00000076500.14 | -199646 | ENSMUSG00000063163 | Speer2 | spermatogenesis associated glutamate (E)-rich protein 2 |
| chr4 | 1,49E+08 | 1,49E+08 | 1840 | * | 18,08864 | -2,118186519 | 0,779449 | -2,71754 | 0,006577 | 0,705941691 | 2,663619 | Promoter (<=1kb) | 4 | 1,49E+08 | 1,49E+08 | 28256 | 1 | 230909 | ENSMUST00000105698.3 | 0 | ENSMUSG00000070577 | Gm572 | predicted gene 572 |
| chr9 | 49565530 | 49568448 | 2919 | * | 27,17123 | -2,116053304 | 0,751451 | -2,81596 | 0,004863 | 0,641208222 | 2,951062 | Intron (ENSMUST0000( | 9 | 49413448 | 49479849 | 66402 | 2 | 17967 | ENSMUST00000194252.6 | -85681 | ENSMUSG00000039542 | Ncam1 | neural cell adhesion molecule 1 |
| chr17 | 72728853 | 72731113 | 2261 | * | 24,1729 | -2,115769586 | 0,742365 | -2,85004 | 0,004371 | 0,626725358 | 2,880017 | Intron (ENSMUST0000( | 17 | 72175967 | 72910822 | 734856 | 2 | 11682 | ENSMUST00000086639.6 | 179709 | ENSMUSG00000055471 | Alk | anaplastic lymphoma kinase |

| | | | | | | | | | | | | | | | | | | | | | | | |
|---|---|---|---|---|---|---|---|---|---|---|---|---|---|---|---|---|---|---|---|---|---|---|---|
| chr13 | 1,08E+08 | 1,08E+08 | 2446 | * | 26,4874 | -2,114929804 | 0,659885 | -3,205 | 0,001351 | 0,424708674 | 2,725491 | Exon (ENSMUST00002 | 13 | 1,08E+08 | 1,08E+08 | 12897 | 1 | 105120 | ENSMUST00000223617.2 | 10333 | ENSMUSG00000114780 | AI197445 | expressed sequence AI197445 |
| chr1 | 96760350 | 96762109 | 1760 | * | 15,8025 | -2,111681284 | 0,784708 | -2,69104 | 0,007123 | 0,75508254 | 2,141612 | Intron (ENSMUST00000 | 1 | 96798851 | 96835521 | 36671 | 1 | 329217 | ENSMUST00000188631.5 | -36742 | ENSMUSG00000097814 | Panct2 | pluripotency-associated noncoding transcript 2 |
| chr11 | 74437590 | 74440484 | 2895 | * | 29,24225 | -2,110691206 | 0,670666 | -3,14716 | 0,001649 | 0,44905823 | 2,858511 | Intron (ENSMUST00000 | 11 | 74327750 | 74413577 | 85828 | 2 | 380711 | ENSMUST00000208896.2 | -24013 | ENSMUSG00000038807 | Rap1gap2 | RAP1 GTPase activating protein 2 |
| chr9 | 34721238 | 34723682 | 2445 | * | 26,50357 | -2,1102613 | 0,731896 | -2,88328 | 0,003936 | 0,607561104 | 2,858511 | Intron (ENSMUST00000 | 9 | 34829529 | 34840805 | 11277 | 2 | 442804 | ENSMUST00000066604.4 | 117123 | | Kirrel3os | kirre like nephrin family adhesion molecule 3, opposite strand |
| chr11 | 17026556 | 17029153 | 2598 | * | 22,62797 | -2,108133542 | 0,788573 | -2,67335 | 0,00751 | 0,715249603 | 2,880017 | 3' UTR | 11 | 16921464 | 17002381 | 80918 | 2 | 56193 | ENSMUST00000020321.13 | -24175 | ENSMUSG00000020120 | Plek | pleckstrin |
| chr1 | 1,87E+08 | 1,87E+08 | 1915 | * | 20,0039 | -2,107856414 | 0,784344 | -2,68741 | 0,007201 | 0,722180633 | 2,663619 | Intron (ENSMUST00000 | 1 | 1,87E+08 | 1,88E+08 | 184047 | 1 | 26381 | ENSMUST00000152927.2 | 122718 | ENSMUSG00000026610 | Esrrg | estrogen-related receptor gamma |
| chr1 | 1,3E+08 | 1,3E+08 | 2855 | * | 25,83337 | -2,107794449 | 0,651673 | -3,23444 | 0,001219 | 0,396928896 | 2,951062 | Intron (ENSMUST00000 | 1 | 1,3E+08 | 1,3E+08 | 267643 | 1 | 210417 | ENSMUST00000151700.8 | 68402 | ENSMUSG00000042581 | Thsd7b | thrombospondin, type I, domain containing 7B |
| chr18 | 70045210 | 70046869 | 1660 | * | 16,45623 | -2,107446282 | 0,807725 | -2,60911 | 0,009078 | 0,799514271 | 2,141612 | Distal Intergenic | 18 | 70058497 | 70077115 | 18619 | 1 | 381175 | ENSMUST00000151773.2 | -11628 | ENSMUSG00000038903 | Ccdc68 | coiled-coil domain containing 68 |
| chr7 | 1,21E+08 | 1,21E+08 | 2554 | * | 26,62267 | -2,10666033 | 0,64388 | -3,27182 | 0,001069 | 0,37346973 | 2,880017 | 3' UTR | 7 | 1,21E+08 | 1,21E+08 | 44702 | 1 | 246190 | ENSMUST00000163275.2 | 23943 | ENSMUSG00000034990 | Otoa | otoancorin |
| chr10 | 98601004 | 98603186 | 2183 | * | 16,40209 | -2,105954485 | 0,786684 | -2,677 | 0,007428 | 0,72659841 | 2,663619 | Distal Intergenic | 10 | 98750268 | 98822984 | 72717 | 1 | 67972 | ENSMUST00000219557.2 | -147082 | ENSMUSG00000019943 | Atp2b1 | ATPase, Ca++ transporting, plasma membrane 1 |
| chr17 | 40637863 | 40640284 | 2422 | * | 19,25737 | -2,105505949 | 0,799015 | -2,63513 | 0,008411 | 0,779936001 | 2,221611 | Distal Intergenic | 17 | 40604649 | 40630098 | 25450 | 2 | 11571 | ENSMUST00000026498.5 | -7765 | ENSMUSG00000025431 | Crisp1 | cysteine-rich secretory protein 1 |
| chr16 | 26212863 | 26214224 | 1362 | * | 17,73813 | -2,103983345 | 0,758755 | -2,77294 | 0,005555 | 0,689688276 | 2,423594 | Distal Intergenic | 16 | 26175392 | 26190591 | 15200 | 2 | 12737 | ENSMUST00000023154.3 | -22272 | ENSMUSG00000022512 | Cldn1 | claudin 1 |
| chr7 | 27274625 | 27276944 | 2320 | * | 21,52864 | -2,103801809 | 0,78383 | -2,684 | 0,007275 | 0,72253719 | 2,67243 | Promoter (<=1kb) | 7 | 27273892 | 27279845 | 5954 | 1 | 66367 | ENSMUST00000186501.2 | 733 | ENSMUSG00000049643 | 2310022A10Rik | RIKEN cDNA 2310022A10 gene |
| chr1 | 1,06E+08 | 1,06E+08 | 2415 | * | 23,9115 | -2,103092305 | 0,736361 | -2,85606 | 0,004289 | 0,618136403 | 2,951062 | Intron (ENSMUST00000 | 1 | 1,06E+08 | 1,06E+08 | 222499 | 1 | 98432 | ENSMUST00000061047.7 | 38856 | ENSMUSG00000044340 | Phlpp1 | PH domain and leucine rich repeat protein phosphatase 1 |
| chr15 | 20672694 | 20673939 | 1246 | * | 16,69783 | -2,09804839 | 0,796271 | -2,63484 | 0,008418 | 0,748971518 | 2,663619 | Intron (ENSMUST00000 | 15 | 20665300 | 20666836 | 1537 | 2 | 64833 | ENSMUST00000052910.6 | -5858 | ENSMUSG00000047565 | Acot10 | acyl-CoA thioesterase 10 |
| chr16 | 20041219 | 20043629 | 2411 | * | 23,70727 | -2,097688728 | 0,683194 | -3,07042 | 0,002138 | 0,504427013 | 2,67243 | Promoter (<=1kb) | 16 | 20041506 | 20047747 | 6242 | 1 | 208146 | ENSMUST00000232671.2 | 0 | ENSMUSG00000041215 | Yeats2 | YEATS domain containing 2 |
| chr12 | 74321320 | 74325042 | 3723 | * | 31,68533 | -2,096750628 | 0,609148 | -3,4421 | 0,000577 | 0,282588188 | 2,951062 | Distal Intergenic | 12 | 74331016 | 74342701 | 11686 | 1 | 74284 | ENSMUST00000095617.3 | -5974 | ENSMUSG00000071265 | 1700086L19Rik | RIKEN cDNA 1700086L19 gene |
| chr3 | 39857297 | 39859496 | 2200 | * | 15,6543 | -2,09623168 | 0,810531 | -2,58624 | 0,009703 | 0,889546768 | 1,37683 | Distal Intergenic | 3 | 39720437 | 39720596 | 160 | 1 | 1,15E+08 | ENSMUST00000084011.4 | 136860 | ENSMUSG00000065945 | Gm23796 | predicted gene, 23796 |
| chr10 | 97926589 | 97931196 | 4608 | * | 38,01789 | -2,09578963 | 0,597735 | -3,50622 | 0,000455 | 0,261608871 | 2,725491 | Distal Intergenic | 10 | 97517276 | 97802952 | 285677 | 2 | 1E+08 | ENSMUST00000220233.2 | -123637 | ENSMUSG00000074776 | Gm10754 | predicted gene 10754 |
| chr17 | 7059772 | 7061753 | 1982 | * | 16,81353 | -2,0954219 | 0,805773 | -2,60051 | 0,009308 | 0,787452947 | 2,386008 | Distal Intergenic | 17 | 7005440 | 7050183 | 44744 | 2 | 22350 | ENSMUST00000064234.7 | -9589 | ENSMUSG00000052397 | Ezr | ezrin |
| chr1 | 1,63E+08 | 1,63E+08 | 2283 | * | 21,16616 | -2,095292379 | 0,773894 | -2,70747 | 0,00678 | 0,695401133 | 2,880017 | Exon (ENSMUST00001 | 1 | 1,63E+08 | 1,63E+08 | 4452 | 2 | 226562 | ENSMUST00000182535.2 | 10667 | ENSMUSG00000040225 | Prrc2c | proline-rich coiled-coil 2C |
| chr4 | 87043797 | 87045784 | 1988 | * | 23,32576 | -2,094158538 | 0,670912 | -3,12136 | 0,0018 | 0,47561414 | 2,67243 | Intron (ENSMUST00000 | 4 | 86946520 | 87094522 | 148003 | 2 | 76376 | ENSMUST00000146815.2 | 48738 | ENSMUSG00000037996 | Slc24a2 | solute carrier family 24 (sodium/potassium/calcium exchanger), member 2 |
| chr11 | 43971348 | 43973775 | 2428 | * | 25,24614 | -2,091634726 | 0,736175 | -2,84122 | 0,004494 | 0,629778374 | 2,880017 | Intron (ENSMUST00000 | 11 | 43849916 | 44025814 | 175899 | 1 | 1,03E+08 | ENSMUST00000148504.2 | 121432 | ENSMUSG00000087172 | Gm12153 | predicted gene 12153 |
| chr10 | 1,07E+08 | 1,07E+08 | 2507 | * | 23,97387 | -2,091587363 | 0,699408 | -2,99051 | 0,002785 | 0,548877304 | 2,67243 | Intron (ENSMUST00000 | 10 | 1,07E+08 | 1,07E+08 | 1885 | 2 | 17878 | ENSMUST00000044210.5 | -53906 | ENSMUSG00000035923 | Myf6 | myogenic factor 6 |
| chr8 | 60554172 | 60555865 | 1694 | * | 17,85475 | -2,091120046 | 0,80903 | -2,58472 | 0,009746 | 0,792545355 | 2,386008 | Distal Intergenic | 8 | 60532624 | 60532727 | 104 | 1 | 1,15E+08 | ENSMUST00000083793.3 | 21548 | ENSMUSG00000065727 | Gm26164 | predicted gene, 26164 |
| chr2 | 1,6E+08 | 1,6E+08 | 1446 | * | 18,62993 | -2,091042166 | 0,766188 | -2,72915 | 0,00635 | 0,734572036 | 2,154504 | Distal Intergenic | 2 | 1,6E+08 | 1,6E+08 | 33948 | 1 | 71081 | ENSMUST00000133091.2 | -89988 | ENSMUSG00000085198 | 4933416E03Rik | RIKEN cDNA 4933416E03 gene |
| chr12 | 25670342 | 25673437 | 3096 | * | 24,72134 | -2,091038114 | 0,670428 | -3,11896 | 0,001815 | 0,459192069 | 2,951933 | Distal Intergenic | 12 | 25444754 | 25521978 | 77225 | 1 | 328066 | ENSMUST00000221270.2 | 225588 | | Gm36723 | predicted gene, 36723 |
| chr3 | 58866796 | 58869089 | 2294 | * | 24,68949 | -2,089062432 | 0,742261 | -2,81446 | 0,004886 | 0,646275731 | 2,858511 | Distal Intergenic | 3 | 58883051 | 58883152 | 102 | 1 | 1,15E+08 | ENSMUST00000157190.3 | -13962 | ENSMUSG00000087815 | Gm22491 | predicted gene, 22491 |
| chr13 | 88855469 | 88857749 | 2281 | * | 21,35923 | -2,088722001 | 0,78162 | -2,6723 | 0,007533 | 0,701807025 | 3,104559 | Distal Intergenic | 13 | 88969591 | 89348657 | 379067 | 1 | 13612 | ENSMUST00000118731.8 | -111842 | ENSMUSG00000034488 | Edil3 | EGF-like repeats and discoidin I-like domains 3 |
| chr11 | 45073102 | 45075031 | 1930 | * | 25,94943 | -2,087477854 | 0,735286 | -2,839 | 0,004526 | 0,631320778 | 2,880017 | Intron (ENSMUST00000 | 11 | 44993693 | 45007755 | 14063 | 2 | 1,01E+08 | ENSMUST00000143869.2 | -65347 | ENSMUSG00000084773 | Gm12159 | predicted gene 12159 |
| chr10 | 85352821 | 85354809 | 1989 | * | 17,96644 | -2,086807018 | 0,806555 | -2,58731 | 0,009673 | 0,773870972 | 2,663619 | Intron (ENSMUST00000 | 10 | 85434265 | 85496156 | 61892 | 1 | 74007 | ENSMUST00000105306.3 | -79456 | ENSMUSG00000020042 | Btbd11 | BTB (POZ) domain containing 11 |
| chr4 | 1,31E+08 | 1,31E+08 | 2953 | * | 30,71283 | -2,0843634 | 0,652178 | -3,196 | 0,001393 | 0,421372944 | 2,858511 | Distal Intergenic | 4 | 1,31E+08 | 1,31E+08 | 9643 | 2 | 77835 | ENSMUST00000153810.2 | -77951 | ENSMUSG00000086262 | A930031H19Rik | RIKEN cDNA A930031H19 gene |
| chr10 | 7024021 | 7026017 | 1997 | * | 19,95899 | -2,082838199 | 0,761003 | -2,73697 | 0,006201 | 0,726927364 | 2,221611 | Distal Intergenic | 10 | 6861543 | 7002478 | 140936 | 2 | 320495 | ENSMUST00000086896.10 | -21543 | ENSMUSG00000064065 | Ipcef1 | interaction protein for cytohesin exchange factors 1 |
| chr17 | 89134902 | 89136404 | 1503 | * | 22,71268 | -2,082531821 | 0,693219 | -3,00415 | 0,002663 | 0,531203841 | 2,880017 | Distal Intergenic | 17 | 89047134 | 89099418 | 52285 | 2 | 16867 | ENSMUST00000234368.2 | -35484 | ENSMUSG00000024107 | Lhcgr | luteinizing hormone/choriogonadotropin receptor |
| chr14 | 19735896 | 19737472 | 1577 | * | 21,59856 | -2,082194589 | 0,750736 | -2,77354 | 0,005545 | 0,668290775 | 2,880017 | Distal Intergenic | 14 | 19758074 | 19775083 | 17010 | 1 | 1,03E+08 | ENSMUST00000241873.1 | -20602 | | Gm29899 | predicted gene, 29899 |
| chr6 | 16419928 | 16421591 | 1664 | * | 17,18864 | -2,08193714 | 0,785592 | -2,65015 | 0,008046 | 0,757118792 | 2,386008 | Distal Intergenic | 6 | 16468235 | 16469903 | 1669 | 2 | 1,03E+08 | ENSMUST00000203052.2 | 48312 | | Gm36406 | predicted gene, 36406 |
| chr2 | 1,73E+08 | 1,73E+08 | 3580 | * | 36,55807 | -2,081115712 | 0,570829 | -3,64578 | 0,000267 | 0,189492453 | 2,925497 | Intron (ENSMUST00000 | 2 | 1,73E+08 | 1,73E+08 | 23975 | 2 | 12162 | ENSMUST00000137247.2 | 53857 | ENSMUSG00000008999 | Bmp7 | bone morphogenetic protein 7 |
| chr7 | 5198175 | 5200530 | 2356 | * | 28,03544 | -2,081079785 | 0,66277 | -3,13997 | 0,00169 | 0,453464543 | 2,858511 | Promoter (<=1kb) | 7 | 5197915 | 5199746 | 1832 | 2 | 81015 | ENSMUST00000169556.3 | 0 | ENSMUSG00000091874 | Vmn1r56 | vomeronasal 1 receptor 56 |
| chr6 | 23272438 | 23274273 | 1836 | * | 23,31086 | -2,079617488 | 0,674769 | -3,08197 | 0,002056 | 0,484153011 | 2,880017 | Intron (ENSMUST00000 | 6 | 23245043 | 23248361 | 3319 | 2 | 73191 | ENSMUST00000031709.7 | -24077 | ENSMUSG00000029697 | Fezf1 | Fez family zinc finger 1 |
| chr10 | 1,23E+08 | 1,23E+08 | 2618 | * | 26,2869 | -2,07924526 | 0,65043 | -3,19672 | 0,00139 | 0,420798623 | 2,880017 | Intron (ENSMUST00000 | 10 | 1,23E+08 | 1,24E+08 | 475963 | 1 | 268354 | ENSMUST00000219723.2 | 278165 | ENSMUSG00000044071 | Tafa2 | TAFA chemokine like family member 2 |
| chr11 | 21092317 | 21095098 | 2782 | * | 23,0681 | -2,079047866 | 0,789108 | -2,63468 | 0,008422 | 0,730469267 | 2,951062 | Exon (ENSMUST00001 | 11 | 21085469 | 21100323 | 14855 | 1 | 67245 | ENSMUST00000101477.2 | 6848 | ENSMUSG00000020134 | Peli1 | pellino 1 |
| chr10 | 1,16E+08 | 1,16E+08 | 1698 | * | 18,4338 | -2,075855609 | 0,773658 | -2,68317 | 0,007293 | 0,731734453 | 2,507013 | Distal Intergenic | 10 | 1,16E+08 | 1,16E+08 | 256720 | 1 | 19279 | ENSMUST00000063470.11 | -26339 | ENSMUSG00000020151 | Ptprr | protein tyrosine phosphatase, receptor type, R |
| chr19 | 55680851 | 55683600 | 2750 | * | 23,724 | -2,075680329 | 0,696598 | -2,97974 | 0,002885 | 0,542967057 | 2,858511 | Distal Intergenic | 19 | 55730252 | 55921661 | 191410 | 1 | 21416 | ENSMUST00000111662.11 | -46652 | ENSMUSG00000024985 | Tcf7l2 | transcription factor 7 like 2, T cell specific, HMG box |
| chr1 | 62152690 | 62156082 | 3393 | * | 27,64408 | -2,074637865 | 0,635851 | -3,26277 | 0,001103 | 0,392614537 | 2,725491 | Intron (ENSMUST00000 | 1 | 62170354 | 62175268 | 4915 | 2 | 1,05E+08 | ENSMUST00000127498.3 | 19186 | ENSMUSG00000085019 | Pard3bos2 | par-3 family cell polarity regulator beta, opposite strand 2 |
| chr3 | 37326744 | 37329046 | 2303 | * | 20,97463 | -2,073942236 | 0,752652 | -2,75551 | 0,00586 | 0,644557958 | 3,471678 | Distal Intergenic | 3 | 37362909 | 37366569 | 3661 | 2 | 207175 | ENSMUST00000075537.7 | 37523 | ENSMUSG00000045031 | Cetn4 | centrin 4 |
| chr10 | 50583807 | 50586250 | 2444 | * | 20,00952 | -2,072948977 | 0,802733 | -2,58236 | 0,009813 | 0,807119269 | 2,221611 | Exon (ENSMUST00000 | 10 | 50629937 | 50643603 | 13667 | 1 | 77987 | ENSMUST00000219172.2 | -43687 | ENSMUSG00000038774 | Ascc3 | activating signal cointegrator 1 complex subunit 3 |
| chr19 | 56626968 | 56629672 | 2705 | * | 25,96598 | -2,072579871 | 0,683304 | -3,03318 | 0,00242 | 0,524368688 | 2,725491 | Distal Intergenic | 19 | 56644943 | 56653043 | 8101 | 2 | 1,03E+08 | ENSMUST00000237615.2 | 23371 | | Gm32441 | predicted gene, 32441 |
| chr10 | 68405265 | 68407545 | 2281 | * | 21,4338 | -2,072399868 | 0,798206 | -2,59632 | 0,009423 | 0,740451144 | 3,104559 | Distal Intergenic | 10 | 68266783 | 68377726 | 110944 | 2 | 73287 | ENSMUST00000020103.9 | -27539 | ENSMUSG00000019945 | Cabcoco1 | ciliary associated calcium binding coiled-coil 1 |
| chr6 | 67791772 | 67793583 | 1812 | * | 25,47425 | -2,072315334 | 0,688936 | -3,00799 | 0,00263 | 0,524368688 | 2,951062 | Distal Intergenic | 6 | 67781359 | 67781494 | 136 | 2 | 1,15E+08 | ENSMUST00000157292.3 | -10278 | ENSMUSG00000087917 | Gm24911 | predicted gene, 24911 |
| chr1 | 51423540 | 51426566 | 3027 | * | 32,9159 | -2,071054749 | 0,59966 | -3,45371 | 0,000553 | 0,385677228 | 1,405709 | Distal Intergenic | 1 | 51257545 | 51372938 | 115394 | 2 | 442821 | ENSMUST00000189867.2 | -50602 | ENSMUSG00000101799 | 9330175M20Rik | RIKEN cDNA 9330175M20 gene |
| chr15 | 41925724 | 41928529 | 2806 | * | 32,46598 | -2,070803886 | 0,619587 | -3,34223 | 0,000831 | 0,361786359 | 2,377674 | Intron (ENSMUST00000 | 15 | 41727472 | 41733116 | 5645 | 2 | 223513 | ENSMUST00000054742.7 | -192608 | ENSMUSG00000042895 | Abra | actin-binding Rho activating protein |
| chr1 | 78345292 | 78347514 | 2223 | * | 22,70795 | -2,070540525 | 0,693085 | -2,98743 | 0,002813 | 0,535609098 | 2,951933 | Intron (ENSMUST00000 | 1 | 78286982 | 78396926 | 109945 | 1 | 433323 | ENSMUST00000036172.10 | 58310 | ENSMUSG00000032908 | Sgpp2 | sphingosine-1-phosphate phosphatase 2 |
| chr7 | 72843824 | 72845747 | 1924 | * | 24,87156 | -2,068358827 | 0,645018 | -3,20667 | 0,001343 | 0,416350185 | 2,858511 | Distal Intergenic | 7 | 72835551 | 72839194 | 3644 | 1 | 1,01E+08 | ENSMUST00000206663.2 | 8273 | | Gm20083 | predicted gene, 20083 |
| chr6 | 95651869 | 95654626 | 2758 | * | 29,53398 | -2,068232895 | 0,717335 | -2,88322 | 0,003936 | 0,605351622 | 2,880017 | Intron (ENSMUST00000 | 6 | 95451123 | 95669863 | 218741 | 2 | 20917 | ENSMUST00000079847.8 | 15237 | ENSMUSG00000061838 | Suclg2 | succinate-Coenzyme A ligase, GDP-forming, beta subunit |
| chr5 | 54346235 | 54349114 | 2880 | * | 33,66301 | -2,067554058 | 0,610486 | -3,38673 | 0,000707 | 0,426043749 | 1,405709 | Distal Intergenic | 5 | 54273199 | 54275929 | 2731 | 1 | 116873 | ENSMUST00000202342.2 | 73036 | ENSMUSG00000039156 | Stim2 | stromal interaction molecule 2 |
| chr11 | 24536961 | 24538893 | 1933 | * | 21,35893 | -2,066713639 | 0,732941 | -2,81975 | 0,004806 | 0,642870233 | 2,880017 | Intron (ENSMUST00000 | 11 | 24431909 | 24519952 | 88044 | 1 | 71316 | ENSMUST00000152726.2 | 105052 | ENSMUSG00000086892 | 4933430M04Rik | RIKEN cDNA 4933430M04 gene |
| chr12 | 5955534 | 5958107 | 2574 | * | 25,72116 | -2,066490728 | 0,652824 | -3,16546 | 0,001548 | 0,439829468 | 2,858511 | Intron (ENSMUST00000 | 12 | 5858234 | 5906142 | 47909 | 1 | 1,03E+08 | ENSMUST00000219472.2 | 97300 | | Gm32442 | predicted gene, 32442 |
| chr11 | 19969776 | 19972679 | 2904 | * | 23,33952 | -2,065337911 | 0,766799 | -2,69346 | 0,007072 | 0,698318058 | 2,951933 | 3' UTR | 11 | 19904570 | 19968133 | 63564 | 1 | 114716 | ENSMUST00000123036.2 | 65206 | ENSMUSG00000045671 | Spred2 | sprouty-related EVH1 domain containing 2 |
| chr2 | 1,01E+08 | 1,01E+08 | 2604 | * | 22,21573 | -2,064634605 | 0,671797 | -3,0733 | 0,002117 | 0,457782113 | 3,471678 | Distal Intergenic | 2 | 1,01E+08 | 1,01E+08 | 14001 | 2 | 68170 | ENSMUST00000177152.2 | 606397 | ENSMUSG00000027165 | Iftap | intraflagellar transport associated protein |
| chr3 | 33113910 | 33115990 | 2081 | * | 20,27968 | -2,06400771 | 0,765862 | -2,69501 | 0,007039 | 0,728037066 | 2,507013 | Intron (ENSMUST00000 | 3 | 33003557 | 33136217 | 132661 | 2 | 58869 | ENSMUST00000108225.10 | 20227 | ENSMUSG00000027674 | Pex5l | peroxisomal biogenesis factor 5-like |
| chr5 | 29341928 | 29344721 | 2794 | * | 24,6397 | -2,061407547 | 0,739379 | -2,78803 | 0,005303 | 0,671156916 | 2,67243 | Distal Intergenic | 5 | 29400990 | 29408133 | 7144 | 1 | 56874 | ENSMUST00000160888.8 | -56269 | ENSMUSG00000029130 | Rnf32 | ring finger protein 32 |
| chr9 | 31795211 | 31797264 | 2054 | * | 18,98796 | -2,057098863 | 0,756319 | -2,71988 | 0,006531 | 0,734161328 | 2,221611 | Intron (ENSMUST00000 | 9 | 31817289 | 31822494 | 5206 | 1 | 1,03E+08 | ENSMUST00000216981.2 | -20025 | ENSMUSG00000110813 | Gm31497 | predicted gene, 31497 |
| chr1 | 79606905 | 79610095 | 3191 | * | 33,2969 | -2,056708452 | 0,613075 | -3,35474 | 0,000794 | 0,44621096 | 1,405709 | Intron (ENSMUST00000 | 1 | 79586562 | 79649682 | 63121 | 2 | 252903 | ENSMUST00000159312.2 | 39587 | ENSMUSG00000054702 | Ap1s3 | adaptor-related protein complex AP-1, sigma 3 |
| chr18 | 7079491 | 7081695 | 2205 | * | 18,42451 | -2,05568902 | 0,795428 | -2,58438 | 0,009755 | 0,786955198 | 2,507013 | Distal Intergenic | 18 | 6934518 | 7004780 | 70263 | 2 | 210719 | ENSMUST00000079788.7 | -74711 | ENSMUSG00000061013 | Mkx | mohawk homeobox |
| chr14 | 33884295 | 33887338 | 3044 | * | 26,45004 | -2,055096063 | 0,788643 | -2,60586 | 0,009164 | 0,749550311 | 2,858511 | Distal Intergenic | 14 | 33865949 | 33874446 | 8498 | 2 | 19065 | ENSMUST00000052164.4 | -9849 | ENSMUSG00000048337 | Npy4r | neuropeptide Y receptor Y4 |
| chr18 | 81048795 | 81050751 | 1957 | * | 21,13408 | -2,055062933 | 0,778497 | -2,63978 | 0,008296 | 0,730689299 | 2,880017 | Distal Intergenic | 18 | 81030671 | 81031790 | 1120 | 1 | 1,03E+08 | ENSMUST00000184638.2 | 18124 | ENSMUSG00000098720 | Gm27239 | predicted gene 27239 |
| chr16 | 67489471 | 67492338 | 2868 | * | 31,81849 | -2,053340306 | 0,633492 | -3,24131 | 0,00119 | 0,398211274 | 2,858511 | Distal Intergenic | 16 | 66459076 | 67417796 | 958721 | 2 | 239857 | ENSMUST00000120898.8 | -71675 | ENSMUSG00000064115 | Cadm2 | cell adhesion molecule 2 |
| chr5 | 22998998 | 23001004 | 2007 | * | 21,46384 | -2,051715852 | 0,750494 | -2,73382 | 0,00626 | 0,683988405 | 2,880017 | Intron (ENSMUST00000 | 5 | 22943908 | 22951885 | 7978 | 2 | 330052 | ENSMUST00000199056.2 | -47113 | ENSMUSG00000105076 | A930003O13Rik | RIKEN cDNA A930003O13 gene |
| chr12 | 7863229 | 7865375 | 2147 | * | 20,54486 | -2,051671645 | 0,756732 | -2,71122 | 0,006704 | 0,705941691 | 2,713306 | Intron (ENSMUST00000 | 12 | 7574002 | 7909691 | 335690 | 2 | 1,03E+08 | ENSMUST00000219198.2 | 44316 | ENSMUSG00000112323 | Gm32828 | predicted gene, 32828 |
| chr2 | 14300240 | 14302010 | 1771 | * | 20,80345 | -2,051188383 | 0,775562 | -2,64478 | 0,008174 | 0,715706132 | 3,104559 | Intron (ENSMUST00000 | 2 | 14333372 | 14336738 | 3367 | 1 | 17533 | ENSMUST00000146168.3 | -31362 | ENSMUSG00000026712 | Mrc1 | mannose receptor, C type 1 |
| chr3 | 1,42E+08 | 1,42E+08 | 3191 | * | 33,02392 | -2,051135278 | 0,573389 | -3,57721 | 0,000347 | 0,219863416 | 2,925497 | Exon (ENSMUST00001 | 3 | 1,41E+08 | 1,41E+08 | 2135 | 1 | 22253 | ENSMUST00000123631.2 | 13474 | ENSMUSG00000059921 | Unc5c | unc-5 netrin receptor C |
| chr18 | 29939720 | 29941691 | 1972 | * | 17,44452 | -2,049769329 | 0,759998 | -2,69707 | 0,006995 | 0,730832709 | 2,423594 | Distal Intergenic | 18 | 29580877 | 29629622 | 48746 | 2 | 78058 | ENSMUST00000234290.2 | -310098 | ENSMUSG00000117618 | 4930578E11Rik | RIKEN cDNA 4930578E11 gene |
| chr6 | 1,17E+08 | 1,17E+08 | 2046 | * | 21,28322 | -2,049355523 | 0,750134 | -2,73199 | 0,006295 | 0,695401133 | 2,67243 | Distal Intergenic | 6 | 1,17E+08 | 1,17E+08 | 37851 | 2 | 319776 | ENSMUST00000056623.13 | -196849 | ENSMUSG00000048108 | Tmem72 | transmembrane protein 72 |
| chr16 | 80387110 | 80389069 | 1960 | * | 21,74627 | -2,048651231 | 0,71608 | -2,86092 | 0,004224 | 0,620083984 | 2,880017 | Distal Intergenic | 16 | 79796264 | 79797434 | 1171 | 1 | 78654 | ENSMUST00000191029.2 | 590846 | ENSMUSG00000101810 | 1700066C05Rik | RIKEN cDNA 1700066C05 gene |
| chr17 | 85772757 | 85774418 | 1662 | * | 26,00298 | -2,047353554 | 0,700644 | -2,9221 | 0,003477 | 0,582425528 | 2,858511 | Distal Intergenic | 17 | 85921036 | 85932650 | 11615 | 1 | 20473 | ENSMUST00000175898.4 | -146618 | ENSMUSG00000038805 | Six3 | sine oculis-related homeobox 3 |
| chr2 | 1,07E+08 | 1,07E+08 | 1657 | * | 19,28176 | -2,046765879 | 0,724253 | -2,82604 | 0,004713 | 0,651804778 | 2,663619 | Distal Intergenic | 2 | 1,07E+08 | 1,07E+08 | 7864 | 1 | 16492 | ENSMUST00000037012.3 | -14857 | ENSMUSG00000042604 | Kcna4 | potassium voltage-gated channel, shaker-related subfamily, member 4 |
| chr6 | 34952778 | 34955676 | 2899 | * | 34,74528 | -2,046732559 | 0,620795 | -3,29695 | 0,000977 | 0,479521813 | 1,405709 | Promoter (<=1kb) | 6 | 34922213 | 34954934 | 32722 | 2 | 243753 | ENSMUST00000044387.14 | 0 | ENSMUSG00000029847 | Slc23a4 | solute carrier family 23 member 4 |
| chr2 | 1,2E+08 | 1,2E+08 | 3035 | * | 26,02668 | -2,04609457 | 0,643278 | -3,18073 | 0,001469 | 0,425046984 | 2,951062 | Intron (ENSMUST00000 | 2 | 1,2E+08 | 1,2E+08 | 1408 | 1 | 26390 | ENSMUST00000137760.2 | -4229 | ENSMUSG00000033902 | Mapkbp1 | mitogen-activated protein kinase binding protein 1 |
| chr3 | 84363098 | 84365111 | 2014 | * | 21,09379 | -2,044582261 | 0,750406 | -2,72463 | 0,006437 | 0,683988405 | 2,951933 | Promoter (2-3kb) | 3 | 84356116 | 84361070 | 4955 | 2 | 229474 | ENSMUST00000125643.2 | -2028 | ENSMUSG00000041842 | Fhdc1 | FH2 domain containing 1 |
| chr9 | 1,01E+08 | 1,01E+08 | 2251 | * | 20,58634 | -2,043654222 | 0,725285 | -2,81773 | 0,004837 | 0,64068479 | 2,951933 | Distal Intergenic | 9 | 1,01E+08 | 1,01E+08 | 853 | 2 | 382099 | ENSMUST00000075243.8 | -165306 | ENSMUSG00000061486 | Gm5161 | predicted pseudogene 5161 |
| chr17 | 91223741 | 91224632 | 892 | * | 18,02203 | -2,039954058 | 0,786522 | -2,59364 | 0,009497 | 0,769360374 | 2,663619 | Intron (ENSMUST00000 | 17 | 91168597 | 91174530 | 5934 | 1 | 78800 | ENSMUST00000162121.2 | 55144 | ENSMUSG00000090160 | 4930480K15Rik | RIKEN cDNA 4930480K15 gene |
| chr10 | 52093486 | 52095562 | 2077 | * | 16,81599 | -2,039937154 | 0,779483 | -2,61704 | 0,00887 | 0,755177881 | 2,663619 | Distal Intergenic | 10 | 52109715 | 52197472 | 87758 | 1 | 66686 | ENSMUST00000069004.14 | -14153 | ENSMUSG00000019891 | Dcbld1 | discoidin, CUB and LCCL domain containing 1 |
| chr6 | 1,26E+08 | 1,26E+08 | 2975 | * | 39,65668 | -2,039850044 | 0,604713 | -3,37325 | 0,000743 | 0,313797801 | 2,925497 | Distal Intergenic | 6 | 1,26E+08 | 1,26E+08 | 65499 | 2 | 18205 | ENSMUST00000050484.9 | -78939 | ENSMUSG00000049107 | Ntf3 | neurotrophin 3 |
| chr3 | 54774878 | 54777473 | 2596 | * | 23,98658 | -2,039619649 | 0,690773 | -2,95266 | 0,00315 | 0,56314244 | 2,858511 | Distal Intergenic | 3 | 54804489 | 54822452 | 17964 | 2 | 329641 | ENSMUST00000162201.3 | 44979 | ENSMUSG00000056306 | Sertm1 | serine rich and transmembrane domain containing 1 |
| chr1 | 76308262 | 76310230 | 1969 | * | 21,06257 | -2,038333557 | 0,782095 | -2,60625 | 0,009154 | 0,749236309 | 2,880017 | Distal Intergenic | 1 | 76397329 | 76505059 | 107731 | 1 | 1,19E+08 | ENSMUST00000185857.2 | -87099 | | Gm29069 | predicted gene 29069 |
| chr1 | 1,31E+08 | 1,31E+08 | 2258 | * | 24,61881 | -2,034900908 | 0,677085 | -3,00538 | 0,002652 | 0,54122637 | 2,67243 | Distal Intergenic | 1 | 1,31E+08 | 1,31E+08 | 7300 | 2 | 329244 | ENSMUST00000187916.7 | -16201 | ENSMUSG00000016524 | Il19 | interleukin 19 |
| chr2 | 4184488 | 4186753 | 2266 | * | 20,6952 | -2,033207542 | 0,726121 | -2,8001 | 0,005109 | 0,641187423 | 3,104559 | Exon (ENSMUST00001 | 2 | 4157683 | 4618753 | 461071 | 1 | 209630 | ENSMUST00000091497.11 | 26805 | ENSMUSG00000026657 | Frmd4a | FERM domain containing 4A |
| chrX | 1,57E+08 | 1,57E+08 | 2785 | * | 23,17713 | -2,03313258 | 0,771749 | -2,63445 | 0,008427 | 0,732824791 | 2,880017 | Intron (ENSMUST00000 | 20 | 1,57E+08 | 1,57E+08 | 221859 | 2 | 245684 | ENSMUST00000026750.15 | 62405 | ENSMUSG00000025658 | Cnksr2 | connector enhancer of kinase suppressor of Ras 2 |
| chr15 | 57371606 | 57374282 | 2677 | * | 38,79872 | -2,033131891 | 0,637488 | -3,18929 | 0,001426 | 0,429625689 | 2,787255 | Distal Intergenic | 15 | 57107167 | 57341021 | 233855 | 2 | 210463 | ENSMUST00000110196.8 | -30585 | ENSMUSG00000022366 | Slc22a22 | solute carrier family 22 (organic cation transporter), member 22 |
| chr12 | 1,02E+08 | 1,02E+08 | 2138 | * | 22,2088 | -2,032678613 | 0,754821 | -2,69293 | 0,007083 | 0,716908183 | 2,67243 | Intron (ENSMUST00000 | 12 | 1,02E+08 | 1,02E+08 | 137331 | 1 | 238384 | ENSMUST00000235018.2 | 49749 | ENSMUSG00000041771 | Slc24a4 | solute carrier family 24 (sodium/potassium/calcium exchanger), member 4 |
| chr3 | 26455437 | 26457822 | 2386 | * | 21,63694 | -2,031162303 | 0,733302 | -2,76989 | 0,005608 | 0,681307794 | 2,67243 | Distal Intergenic | 3 | 25480379 | 26386609 | 906231 | 2 | 192167 | ENSMUST00000193603.6 | -68828 | ENSMUSG00000063887 | Nlgn1 | neuroligin 1 |
| chr13 | 75042068 | 75044002 | 1935 | * | 19,63903 | -2,030928038 | 0,748671 | -2,71271 | 0,006674 | 0,738600147 | 2,221611 | Distal Intergenic | 13 | 74840487 | 74956929 | 116443 | 2 | 12380 | ENSMUST00000223206.2 | -85139 | ENSMUSG00000021585 | Cast | calpastatin |
| chr18 | 8567076 | 8569586 | 2511 | * | 22,68422 | -2,030780391 | 0,749452 | -2,70969 | 0,006735 | 0,683966322 | 3,104559 | Distal Intergenic | 18 | 9212163 | 9218136 | 5974 | 1 | 14370 | ENSMUST00000041080.7 | -642577 | ENSMUSG00000036904 | Fzd8 | frizzled class receptor 8 |
| chr8 | 1,17E+08 | 1,17E+08 | 2486 | * | 27,47653 | -2,030363281 | 0,665374 | -3,05146 | 0,002277 | 0,511342349 | 2,725491 | Distal Intergenic | 8 | 1,17E+08 | 1,17E+08 | 11426 | 1 | 1,03E+08 | ENSMUST00000241934.1 | 36310 | | Gm30925 | predicted gene, 30925 |
| chr7 | 29067698 | 29070397 | 2700 | * | 25,32213 | -2,029324242 | 0,71197 | -2,85029 | 0,004368 | 0,635792383 | 2,725491 | Promoter (<=1kb) | 7 | 29066129 | 29067251 | 1123 | 2 | 74206 | ENSMUST00000183275.2 | -447 | ENSMUSG00000030583 | Sipa1l3 | signal-induced proliferation-associated 1 like 3 |
| chr3 | 53171306 | 53175575 | 4270 | * | 40,4028 | -2,028554992 | 0,574059 | -3,5337 | 0,00041 | 0,269133964 | 2,338387 | Distal Intergenic | 3 | 53360040 | 53365218 | 5179 | 1 | 1,03E+08 | ENSMUST00000132485.2 | -184465 | | Gm16206 | predicted gene 16206 |
| chr13 | 1,18E+08 | 1,18E+08 | 2639 | * | 22,20251 | -2,027966448 | 0,768776 | -2,63791 | 0,008342 | 0,729319353 | 2,951933 | Distal Intergenic | 13 | 1,18E+08 | 1,18E+08 | 130 | 1 | 1,15E+08 | ENSMUST00000157530.3 | -29189 | ENSMUSG00000088155 | Gm26349 | predicted gene, 26349 |
| chr6 | 1,35E+08 | 1,35E+08 | 1888 | * | 27,51636 | -2,027950724 | 0,736692 | -2,75278 | 0,005909 | 0,676922067 | 2,880017 | Intron (ENSMUST00000 | 6 | 1,35E+08 | 1,35E+08 | 6481 | 1 | 1,01E+08 | ENSMUST00000204330.2 | -8400 | ENSMUSG00000108239 | Gm19434 | predicted gene, 19434 |
| chr17 | 18895116 | 18896709 | 1594 | * | 17,27805 | -2,027687746 | 0,776922 | -2,6099 | 0,009057 | 0,7591629 | 2,663619 | Distal Intergenic | 17 | 18801989 | 18818419 | 16431 | 1 | 433070 | ENSMUST00000165692.8 | 93127 | ENSMUSG00000091679 | Vmn2r96 | vomeronasal 2, receptor 96 |
| chr1 | 38416506 | 38418511 | 2006 | * | 29,75737 | -2,026704278 | 0,62809 | -3,22677 | 0,001252 | 0,406085428 | 2,858511 | Intron (ENSMUST00000 | 1 | 38424030 | 38433055 | 9026 | 1 | 1,03E+08 | ENSMUST00000143261.8 | -5519 | | Gm16152 | predicted gene 16152 |
| chr8 | 88194080 | 88196160 | 2081 | * | 22,64964 | -2,026330632 | 0,711098 | -2,84958 | 0,004378 | 0,622205297 | 2,951933 | Promoter (1-2kb) | 8 | 88195495 | 88197696 | 2202 | 2 | 12404 | ENSMUST00000211279.2 | 1536 | ENSMUSG00000031654 | Cbln1 | cerebellin 1 precursor protein |

| | | | | | | | | | | | | | | | | | | | | | | |
|---|---|---|---|---|---|---|---|---|---|---|---|---|---|---|---|---|---|---|---|---|---|---|
| chr5 | 40176840 | 40178885 | 2046 | * | 20,98635 | -2,025965856 | 0,695102 | -2,91463 | 0,003561 | 0,546791007 | 3,471678 Distal Intergenic | 5 | 40133014 | 40142971 | 9958 | 2 | 1,03E+08 | ENSMUST00000200925.2 | -33869 | ENSMUSG00000107070 | Gm35191 | predicted gene, 35191 |
| chr2 | 1,6E+08 | 1,6E+08 | 1794 | * | 22,85727 | -2,024486542 | 0,743072 | -2,72448 | 0,00644 | 0,686762073 | 2,858511 Distal Intergenic | 2 | 1,6E+08 | 1,6E+08 | 51593 | 2 | 1,05E+08 | ENSMUST00000109473.2 | 115891 | | Gm14221 | predicted gene 14221 |
| chr2 | 1,74E+08 | 1,74E+08 | 2426 | * | 19,43194 | -2,023821377 | 0,784737 | -2,57898 | 0,009909 | 0,809505441 | 2,221611 Intron (ENSMUST0000( | 2 | 1,74E+08 | 1,74E+08 | 48886 | 1 | 14683 | ENSMUST00000087876.11 | 13311 | ENSMUSG00000027523 | Gnas | GNAS (guanine nucleotide binding protein, alpha stimulating) complex locus |
| chr10 | 87451829 | 87454563 | 2735 | * | 25,38537 | -2,020397393 | 0,722807 | -2,79521 | 0,005187 | 0,656950206 | 2,858511 Distal Intergenic | 10 | 87382535 | 87406194 | 23660 | 1 | 18478 | ENSMUST00000217864.2 | 69294 | ENSMUSG00000020051 | Pah | phenylalanine hydroxylase |
| chr9 | 32235644 | 32238606 | 2963 | * | 31,71593 | -2,018707406 | 0,640701 | -3,15078 | 0,001628 | 0,457782113 | 2,67243 Intron (ENSMUST0000( | 9 | 32227196 | 32255533 | 28338 | 2 | 16521 | ENSMUST00000216033.2 | 16927 | ENSMUSG00000032034 | Kcnj5 | potassium inwardly-rectifying channel, subfamily J, member 5 |
| chr12 | 80715955 | 80718530 | 2576 | * | 21,72087 | -2,016961527 | 0,752127 | -2,68168 | 0,007325 | 0,707552275 | 2,951933 Promoter (2-3kb) | 12 | 80713365 | 80726649 | 13285 | 1 | 328133 | ENSMUST00000219689.2 | 2590 | ENSMUSG00000048833 | Slc39a9 | solute carrier family 39 (zinc transporter), member 9 |
| chr5 | 1,01E+08 | 1,01E+08 | 1555 | * | 17,22419 | -2,016270575 | 0,780593 | -2,583 | 0,009795 | 0,792465459 | 2,423594 Distal Intergenic | 5 | 1,01E+08 | 1,01E+08 | 3707 | 2 | 1,03E+08 | ENSMUST00000199211.2 | -123621 | ENSMUSG00000106255 | Gm36793 | predicted gene, 36793 |
| chr8 | 25398436 | 25400780 | 2345 | * | 25,21629 | -2,015671698 | 0,723771 | -2,78496 | 0,005353 | 0,672273099 | 2,67243 Intron (ENSMUST0000( | 8 | 25326162 | 25374611 | 48450 | 2 | 353188 | ENSMUST00000140573.8 | -23825 | ENSMUSG00000037437 | Adam32 | a disintegrin and metallopeptidase domain 32 |
| chr11 | 43839747 | 43841859 | 2113 | * | 18,51064 | -2,011958382 | 0,759013 | -2,65075 | 0,008031 | 0,739435642 | 2,663619 Distal Intergenic | 11 | 43849916 | 44025814 | 175899 | 1 | 1,03E+08 | ENSMUST00000148504.2 | -8057 | ENSMUSG00000087172 | Gm12153 | predicted gene 12153 |
| chrX | 1,32E+08 | 1,32E+08 | 3093 | * | 22,87819 | -2,0102147 | 0,714851 | -2,81207 | 0,004922 | 0,642870233 | 2,951062 Exon (ENSMUST000001 | 20 | 1,31E+08 | 1,31E+08 | 131 | 2 | 1,15E+08 | ENSMUST00000104440.3 | -539818 | ENSMUSG00000077628 | Gm23768 | predicted gene, 23768 |
| chr16 | 47102592 | 47104329 | 1738 | * | 16,11289 | -2,009585767 | 0,772228 | -2,60232 | 0,00926 | 0,790570581 | 2,318434 Distal Intergenic | 16 | 47326682 | 47326788 | 107 | 2 | 1,15E+08 | ENSMUST00000157107.4 | 222459 | ENSMUSG00000119721 | Gm24651 | predicted gene, 24651 |
| chr1 | 1,24E+08 | 1,24E+08 | 2628 | * | 27,56888 | -2,009365972 | 0,630662 | -3,18612 | 0,001442 | 0,439573569 | 2,67243 Intron (ENSMUST0000( | 1 | 1,24E+08 | 1,24E+08 | 104 | 1 | 1,15E+08 | ENSMUST00000083406.3 | -122932 | ENSMUSG00000065340 | Gm25578 | predicted gene, 25578 |
| chr13 | 65145649 | 65147218 | 1570 | * | 20,02416 | -2,009149775 | 0,754678 | -2,66226 | 0,007762 | 0,730469267 | 2,713306 Exon (ENSMUST000002 | 13 | 65180872 | 65185818 | 4947 | 1 | 238683 | ENSMUST00000099427.3 | -33654 | ENSMUSG00000074849 | Spata31d1c | spermatogenesis associated 31 subfamily D, member 1C |
| chr8 | 69814376 | 69816202 | 1827 | * | 20,63329 | -2,008905158 | 0,705869 | -2,846 | 0,004427 | 0,594430593 | 3,471678 Distal Intergenic | 8 | 69819162 | 69848174 | 29013 | 2 | 378466 | ENSMUST00000072427.7 | 31972 | ENSMUSG00000110444 | Gm10033 | predicted gene 10033 |
| chr3 | 76328583 | 76330586 | 2004 | * | 20,21712 | -2,008530979 | 0,752657 | -2,66859 | 0,007617 | 0,730469267 | 2,663619 Intron (ENSMUST0000( | 3 | 76500857 | 76614856 | 114000 | 1 | 213262 | ENSMUST00000162471.2 | -170271 | ENSMUSG00000034098 | Fstl5 | follistatin-like 5 |
| chr6 | 35571125 | 35573470 | 2346 | * | 27,97459 | -2,007823796 | 0,712293 | -2,81882 | 0,00482 | 0,640437129 | 2,951062 Distal Intergenic | 6 | 35485841 | 35516823 | 30983 | 2 | 14489 | ENSMUST00000031866.9 | -54302 | ENSMUSG00000029840 | Mtpn | myotrophin |
| chr8 | 1,1E+08 | 1,1E+08 | 2064 | * | 23,82145 | -2,00533249 | 0,702102 | -2,85619 | 0,004288 | 0,623120685 | 2,880017 Promoter (2-3kb) | 8 | 1,1E+08 | 1,1E+08 | 47597 | 1 | 56523 | ENSMUST00000212003.2 | 2027 | ENSMUSG00000031727 | Pmfbp1 | polyamine modulated factor 1 binding protein 1 |
| chr16 | 19675388 | 19679929 | 4542 | * | 43,45729 | -2,004659322 | 0,58864 | -3,40558 | 0,00066 | 0,327375387 | 2,338387 Distal Intergenic | 16 | 19695317 | 19703014 | 7698 | 2 | 68162 | ENSMUST00000100083.5 | 23085 | ENSMUSG00000075330 | A930003A15Rik | RIKEN cDNA A930003A15 gene |
| chr5 | 63868249 | 63870372 | 2124 | * | 17,50505 | -2,003569352 | 0,774757 | -2,58606 | 0,009708 | 0,792545355 | 2,386008 Intron (ENSMUST0000( | 5 | 63806676 | 63957990 | 151315 | 1 | 319807 | ENSMUST00000162166.3 | 61573 | ENSMUSG00000090061 | Nwd2 | NACHT and WD repeat domain containing 2 |
| chr2 | 1,73E+08 | 1,73E+08 | 2162 | * | 27,91378 | -2,003494319 | 0,741244 | -2,70288 | 0,006874 | 0,693003605 | 2,951062 Distal Intergenic | 2 | 1,72E+08 | 1,72E+08 | 1403 | 1 | 21420 | ENSMUST00000142633.2 | 179268 | ENSMUSG00000028640 | Tfap2c | transcription factor AP-2, gamma |
| chr18 | 25577268 | 25581259 | 3992 | * | 34,40235 | -2,00314926 | 0,712994 | -2,80949 | 0,004962 | 0,653585323 | 2,787255 Intron (ENSMUST0000( | 18 | 25612545 | 25619954 | 7410 | 2 | 108013 | ENSMUST00000225528.2 | 38695 | ENSMUSG00000024268 | Celf4 | CUGBP, Elav-like family member 4 |
| chr13 | 1,14E+08 | 1,14E+08 | 1442 | * | 17,02997 | -2,00001681 | 0,760411 | -2,63018 | 0,008534 | 0,775744126 | 2,318434 Distal Intergenic | 13 | 1,14E+08 | 1,14E+08 | 27245 | 2 | 319803 | ENSMUST00000225044.2 | -51800 | ENSMUSG00000114213 | A430090L17Rik | RIKEN cDNA A430090L17 gene |
| chr15 | 88618536 | 88621261 | 2726 | * | 25,1875 | 2,000087912 | 0,700833 | 2,853871 | 0,004319 | 0,625704414 | 2,858511 Promoter (<=1kb) | 15 | 88571237 | 88618436 | 47200 | 2 | 223770 | ENSMUST00000109381.9 | -100 | ENSMUSG00000022387 | Brd1 | bromodomain containing 1 |
| chr14 | 21736379 | 21740607 | 4229 | * | 37,02004 | 2,000227209 | 0,542439 | 3,68747 | 0,000226 | 0,17530799 | 2,787255 Exon (ENSMUST00000( | 14 | 21735014 | 21753170 | 18157 | 2 | 435391 | ENSMUST00000224735.2 | 12563 | ENSMUSG00000063821 | Dusp29 | dual specificity phosphatase 29 |
| chr2 | 29972447 | 29974284 | 1838 | * | 24,77731 | 2,000365732 | 0,714133 | 2,80111 | 0,005093 | 0,665162479 | 2,67243 Promoter (<=1kb) | 2 | 29974665 | 29977048 | 2384 | 1 | 263803 | ENSMUST00000156197.2 | -381 | ENSMUSG00000026785 | Pkn3 | protein kinase N3 |
| chr5 | 1,36E+08 | 1,36E+08 | 2764 | * | 37,43914 | 2,001021344 | 0,619831 | 3,228334 | 0,001245 | 0,410415129 | 2,725491 Promoter (<=1kb) | 5 | 1,36E+08 | 1,36E+08 | 2104 | 2 | 109267 | ENSMUST00000054895.4 | 0 | ENSMUSG00000029699 | Ssc4d | scavenger receptor cysteine rich family, 4 domains |
| chr11 | 92144725 | 92146857 | 2133 | * | 19,20926 | 2,001053993 | 0,701864 | 2,851056 | 0,004357 | 0,636647377 | 2,713306 Distal Intergenic | 11 | 92801123 | 92801265 | 143 | 1 | 1,15E+08 | ENSMUST00000157185.3 | -654266 | ENSMUSG00000087810 | Gm25113 | predicted gene, 25113 |
| chr2 | 1,67E+08 | 1,67E+08 | 1389 | * | 31,36543 | 2,001646118 | 0,588307 | 3,402384 | 0,000668 | 0,301441602 | 2,951062 Distal Intergenic | 2 | 1,67E+08 | 1,67E+08 | 50740 | 2 | 56336 | ENSMUST00000109221.9 | -34350 | ENSMUSG00000017929 | B4galt5 | UDP-Gal:betaGlcNAc beta 1,4-galactosyltransferase, polypeptide 5 |
| chr2 | 31544594 | 31547590 | 2997 | * | 37,057 | 2,001736702 | 0,573097 | 3,492842 | 0,000478 | 0,259309257 | 2,925497 3' UTR | 2 | 31560727 | 31571361 | 10635 | 1 | 227715 | ENSMUST00000038474.14 | -13137 | ENSMUSG00000039356 | Exosc2 | exosome component 2 |
| chr8 | 46688391 | 46690209 | 1819 | * | 17,7074 | 2,001794594 | 0,767737 | 2,607395 | 0,009123 | 0,773936378 | 2,507013 Distal Intergenic | 8 | 46659834 | 46664321 | 4488 | 2 | 11739 | ENSMUST00000034049.5 | -24070 | ENSMUSG00000031633 | Slc25a4 | solute carrier family 25 (mitochondrial carrier, adenine nucleotide translocator), member 4 |
| chr19 | 6107074 | 6109036 | 1963 | * | 22,96586 | 2,002802734 | 0,707658 | 2,830184 | 0,004652 | 0,639997962 | 2,858511 Promoter (<=1kb) | 19 | 6107874 | 6109549 | 1676 | 1 | 14109 | ENSMUST00000178310.9 | 0 | ENSMUSG00000038274 | Fau | Finkel-Biskis-Reilly murine sarcoma virus (FBR-MuSV) ubiquitously expressed (fox derived) |
| chr12 | 40639824 | 40641821 | 1998 | * | 21,6375 | 2,00302927 | 0,728544 | 2,749359 | 0,005971 | 0,674343184 | 2,951933 Intron (ENSMUST0000( | 12 | 40496335 | 40896179 | 399845 | 1 | 238130 | ENSMUST00000220912.2 | 143489 | ENSMUSG00000035954 | Dock4 | dedicator of cytokinesis 4 |
| chr11 | 83190580 | 83194319 | 3740 | * | 28,769 | 2,003163023 | 0,706722 | 2,834442 | 0,004591 | 0,647632102 | 2,67243 Promoter (<=1kb) | 11 | 83188856 | 83193412 | 4557 | 2 | 103737 | ENSMUST00000176374.2 | 0 | ENSMUSG00000018733 | Pex12 | peroxisomal biogenesis factor 12 |
| chr7 | 1,35E+08 | 1,35E+08 | 3225 | * | 30,1089 | 2,003393143 | 0,608808 | 3,290684 | 0,000999 | 0,370253435 | 2,725491 Promoter (2-3kb) | 7 | 1,35E+08 | 1,35E+08 | 89968 | 1 | 19267 | ENSMUST00000211788.2 | 2218 | ENSMUSG00000041836 | Ptpre | protein tyrosine phosphatase, receptor type, E |
| chr6 | 1,27E+08 | 1,27E+08 | 2279 | * | 25,42659 | 2,00371533 | 0,672863 | 2,977894 | 0,002902 | 0,541180509 | 2,951062 Distal Intergenic | 6 | 1,27E+08 | 1,27E+08 | 6301 | 2 | 1,03E+08 | ENSMUST00000210335.2 | -12536 | | Gm33320 | predicted gene, 33320 |
| chr7 | 16584267 | 16586729 | 2463 | * | 22,92365 | 2,004537629 | 0,749462 | 2,674636 | 0,007481 | 0,715156227 | 2,880017 Promoter (1-2kb) | 7 | 16582634 | 16587907 | 5274 | 1 | 101540 | ENSMUST00000205456.2 | 1633 | ENSMUSG00000041187 | Prkd2 | protein kinase D2 |
| chr2 | 1,72E+08 | 1,72E+08 | 1903 | * | 28,27937 | 2,004931213 | 0,683962 | 2,931347 | 0,003375 | 0,569348679 | 2,951062 Promoter (2-3kb) | 2 | 1,72E+08 | 1,72E+08 | 9030 | 1 | 21420 | ENSMUST00000030391.9 | -2110 | ENSMUSG00000028640 | Tfap2c | transcription factor AP-2, gamma |
| chr11 | 88385219 | 88388619 | 3401 | * | 35,57725 | 2,00515411 | 0,576065 | 3,480778 | 0,0005 | 0,275159027 | 2,725491 Intron (ENSMUST0000( | 11 | 88256341 | 88480973 | 224633 | 2 | 76626 | ENSMUST00000148013.8 | 92354 | ENSMUSG00000069769 | Msi2 | musashi RNA-binding protein 2 |
| chr9 | 85056595 | 85058482 | 1888 | * | 20,89912 | 2,005318905 | 0,737602 | 2,718701 | 0,006554 | 0,681459438 | 3,104559 Distal Intergenic | 9 | 85202492 | 85209177 | 6686 | 2 | 212943 | ENSMUST00000034802.15 | 150695 | ENSMUSG00000032265 | Tent5a | terminal nucleotidyltransferase 5A |
| chr8 | 70996063 | 71000042 | 3980 | * | 57,48413 | 2,00550905 | 0,52011 | 3,855934 | 0,000115 | 0,091686978 | 4,055758 Intron (ENSMUST0000( | 8 | 70992338 | 71045508 | 53171 | 1 | 13716 | ENSMUST00000093454.8 | 3725 | ENSMUSG00000070002 | Ell | elongation factor RNA polymerase II |
| chr2 | 71794186 | 71795530 | 1345 | * | 19,14368 | 2,005885927 | 0,737043 | 2,721532 | 0,006498 | 0,714462637 | 2,507013 Distal Intergenic | 2 | 71811584 | 72087818 | 276235 | 1 | 56508 | ENSMUST00000090826.12 | -16054 | ENSMUSG00000049044 | Rapgef4 | Rap guanine nucleotide exchange factor (GEF) 4 |
| chr1 | 4938469 | 4940812 | 2344 | * | 19,14015 | 2,006000085 | 0,719457 | 2,788213 | 0,0053 | 0,669586355 | 2,713306 Intron (ENSMUST0000( | 1 | 4928261 | 4968132 | 39872 | 1 | 21399 | ENSMUST00000165720.3 | 10208 | ENSMUSG00000033813 | Tcea1 | transcription elongation factor A (SII) 1 |
| chr4 | 1,08E+08 | 1,08E+08 | 2504 | * | 23,96428 | 2,006376366 | 0,665577 | 3,014492 | 0,002574 | 0,521453391 | 2,951933 Distal Intergenic | 4 | 1,08E+08 | 1,08E+08 | 6572 | 2 | 67305 | ENSMUST00000030332.7 | -9454 | ENSMUSG00000028597 | Gpx7 | glutathione peroxidase 7 |
| chr7 | 74442926 | 74445316 | 2391 | * | 19,75916 | 2,006671419 | 0,679666 | 2,952437 | 0,003153 | 0,591141332 | 2,507013 Distal Intergenic | 7 | 74415735 | 74420991 | 5257 | 1 | 1,03E+08 | ENSMUST00000206203.2 | 27191 | ENSMUSG00000108468 | Gm30459 | predicted gene, 30459 |
| chr2 | 29441573 | 29443486 | 1914 | * | 22,26006 | 2,006889707 | 0,763762 | 2,627638 | 0,008598 | 0,737081993 | 2,880017 Distal Intergenic | 2 | 29394883 | 29414693 | 19811 | 1 | 68975 | ENSMUST00000147547.2 | 46690 | ENSMUSG00000026799 | Med27 | mediator complex subunit 27 |
| chr4 | 81966621 | 81968548 | 1928 | * | 16,90861 | 2,00739756 | 0,732437 | 2,74071 | 0,006131 | 0,716908183 | 2,318434 Exon (ENSMUST000001 | 4 | 81983668 | 82021046 | 37379 | 1 | 545641 | ENSMUST00000137810.4 | -15120 | ENSMUSG00000085022 | Gm5860 | predicted gene 5860 |
| chr6 | 86528510 | 86530814 | 2305 | * | 26,68247 | 2,007402801 | 0,770001 | 2,607013 | 0,009134 | 0,756025543 | 2,725491 Intron (ENSMUST0000( | 6 | 86524942 | 86541432 | 16491 | 1 | 72012 | ENSMUST00000240938.1 | 3568 | ENSMUSG00000097048 | 1600020E01Rik | RIKEN cDNA 1600020E01 gene |
| chr8 | 24928236 | 24931108 | 2873 | * | 25,68185 | 2,0075238 | 0,760162 | 2,640915 | 0,008268 | 0,740451144 | 2,725491 Promoter (<=1kb) | 8 | 24927196 | 24929000 | 1805 | 2 | 69068 | ENSMUST00000052622.6 | 0 | ENSMUSG00000056313 | Tcim | transcriptional and immune response regulator |
| chr2 | 1,53E+08 | 1,53E+08 | 4469 | * | 38,92729 | 2,007866696 | 0,644666 | 3,114582 | 0,001842 | 0,498011371 | 2,377674 Promoter (<=1kb) | 2 | 1,53E+08 | 1,53E+08 | 39210 | 1 | 14950 | ENSMUST00000125366.8 | 0 | ENSMUSG00000019188 | H13 | histocompatibility 13 |
| chr8 | 1,05E+08 | 1,05E+08 | 2217 | * | 22,98932 | 2,007900809 | 0,704435 | 2,85037 | 0,004367 | 0,627785088 | 2,858511 Promoter (1-2kb) | 8 | 1,05E+08 | 1,05E+08 | 47617 | 2 | 97487 | ENSMUST00000179802.2 | 1474 | ENSMUSG00000096188 | Cmtm4 | CKLF-like MARVEL transmembrane domain containing 4 |
| chr17 | 88270474 | 88274307 | 3834 | * | 37,12712 | 2,008133508 | 0,548521 | 3,660994 | 0,000251 | 0,190625259 | 2,725491 Distal Intergenic | 17 | 88282490 | 88298320 | 15831 | 1 | 17688 | ENSMUST00000005503.5 | -8183 | ENSMUSG00000005370 | Msh6 | mutS homolog 6 |
| chr5 | 21128979 | 21131034 | 2056 | * | 19,41115 | 2,009240258 | 0,767837 | 2,616755 | 0,008877 | 0,755289138 | 2,663619 Intron (ENSMUST0000( | 5 | 21125140 | 21150502 | 25363 | 2 | 242860 | ENSMUST00000197089.2 | 19468 | ENSMUSG00000039968 | Rsbn1l | round spermatid basic protein 1-like |
| chr14 | 31669084 | 31670837 | 1754 | * | 20,32804 | 2,00978469 | 0,772632 | 2,601219 | 0,009289 | 0,760759863 | 2,713306 Distal Intergenic | 14 | 31651545 | 31653675 | 2131 | 1 | 1,03E+08 | ENSMUST00000242479.1 | 17539 | | Gm53172 | predicted gene, 53172 |
| chr16 | 90531863 | 90536269 | 4407 | * | 79,42139 | 2,009970335 | 0,720297 | 2,790474 | 0,005263 | 0,595550945 | 4,055758 Promoter (<=1kb) | 16 | 90535095 | 90546673 | 11579 | 1 | 77037 | ENSMUST00000125429.3 | 0 | ENSMUSG00000039956 | Mrap | melanocortin 2 receptor accessory protein |
| chr8 | 50751343 | 50753803 | 2461 | * | 21,67381 | 2,010313707 | 0,671721 | 2,992783 | 0,002764 | 0,532496361 | 2,951933 Intron (ENSMUST0000( | 8 | 50821147 | 50869123 | 47977 | 1 | 1E+08 | ENSMUST00000210011.2 | -67344 | ENSMUSG00000110281 | Gm2516 | predicted gene 2516 |
| chr10 | 21664934 | 21667535 | 2602 | * | 29,86981 | 2,01080963 | 0,657505 | 3,058242 | 0,002226 | 0,499240204 | 2,858511 Distal Intergenic | 10 | 21706398 | 21720750 | 14353 | 2 | 628133 | ENSMUST00000180731.2 | 53215 | | Platr6 | pluripotency associated transcript 6 |
| chr13 | 108827992 | 108832739 | 4748 | * | 53,774099 | 2,011208706 | 0,4612175 | 4,360651 | 1,297E-05 | 0,030165351 | 2,3776742 Intron (ENSMUST000001 | 13 | 108790711 | 110088008 | 1297298 | 1 | 238871 | ENSMUST00000122041.8 | 37281 | ENSMUSG00000021699 | Pde4d | phosphodiesterase 4D, cAMP specific |
| chr9 | 40832129 | 40834400 | 2272 | * | 22,00362 | 2,011662446 | 0,736607 | 2,730983 | 0,006315 | 0,658015022 | 3,471678 Exon (ENSMUST00000( | 9 | 40806147 | 40808914 | 2768 | 2 | 70989 | ENSMUST00000162122.2 | -23215 | ENSMUSG00000032023 | Jhy | junctional cadherin complex regulator |
| chr7 | 29137535 | 29141080 | 3546 | * | 35,18038 | 2,011900864 | 0,566269 | 3,552909 | 0,000381 | 0,322750682 | 1,405709 Intron (ENSMUST0000( | 7 | 29070213 | 29077170 | 6958 | 2 | 74206 | ENSMUST00000182529.2 | -60365 | ENSMUSG00000030583 | Sipa1l3 | signal-induced proliferation-associated 1 like 3 |
| chr4 | 46038073 | 46043294 | 5222 | * | 49,8099 | 2,012030554 | 0,540502 | 3,722523 | 0,000197 | 0,165924532 | 2,725491 Promoter (<=1kb) | 4 | 46038935 | 46116032 | 77098 | 1 | 21916 | ENSMUST00000107773.3 | 0 | ENSMUSG00000028328 | Tmod1 | tropomodulin 1 |
| chr12 | 1,08E+08 | 1,08E+08 | 2225 | * | 20,33433 | 2,012684145 | 0,687011 | 2,929626 | 0,003394 | 0,627785088 | 2,221611 Distal Intergenic | 12 | 1,08E+08 | 1,08E+08 | 11749 | 2 | 74968 | ENSMUST00000130564.2 | 28271 | ENSMUSG00000085641 | 4930465M20Rik | RIKEN cDNA 4930465M20 gene |
| chr4 | 1,33E+08 | 1,33E+08 | 2206 | * | 21,82515 | 2,012749101 | 0,719644 | 2,796866 | 0,00516 | 0,650270742 | 2,951933 Promoter (1-2kb) | 4 | 1,33E+08 | 1,33E+08 | 10732 | 2 | 52174 | ENSMUST00000134781.8 | 1663 | ENSMUSG00000028857 | Tmem222 | transmembrane protein 222 |
| chr7 | 78979620 | 78981657 | 2038 | * | 20,55266 | 2,013195739 | 0,771586 | 2,609167 | 0,009076 | 0,755814512 | 2,713306 Intron (ENSMUST0000( | 7 | 79025276 | 79027451 | 2176 | 2 | 19771 | ENSMUST00000206624.2 | 45794 | ENSMUSG00000039194 | Rlbp1 | retinaldehyde binding protein 1 |
| chr15 | 83555101 | 83556930 | 1830 | * | 21,16032 | 2,013351707 | 0,738575 | 2,725997 | 0,006411 | 0,685026746 | 2,880017 Intron (ENSMUST0000( | 15 | 83491524 | 83609127 | 117604 | 2 | 64706 | ENSMUST00000171496.9 | 52197 | ENSMUSG00000016763 | Scube1 | signal peptide, CUB domain, EGF-like 1 |
| chr1 | 59378543 | 59380770 | 2228 | * | 19,21159 | 2,014801452 | 0,724969 | 2,779155 | 0,00545 | 0,67266775 | 2,713306 Intron (ENSMUST0000( | 1 | 59338095 | 59391607 | 53513 | 1 | 271697 | ENSMUST00000162906.2 | 40448 | ENSMUSG00000026023 | Cdk15 | cyclin-dependent kinase 15 |
| chr8 | 85176883 | 85179316 | 2434 | * | 29,45558 | 2,015036107 | 0,633131 | 3,182654 | 0,001459 | 0,428853558 | 2,858511 Intron (ENSMUST0000( | 8 | 85142413 | 85366875 | 224463 | 1 | 12286 | ENSMUST00000215756.2 | 34470 | ENSMUSG00000034656 | Cacna1a | calcium channel, voltage-dependent, P/Q type, alpha 1A subunit |
| chr18 | 75092861 | 75097030 | 4170 | * | 33,60581 | 2,015087876 | 0,669882 | 3,008124 | 0,002629 | 0,661775851 | 1,405709 Promoter (<=1kb) | 18 | 75072393 | 75094334 | 21942 | 2 | 16891 | ENSMUST00000066532.5 | 0 | ENSMUSG00000053846 | Lipg | lipase, endothelial |
| chr7 | 68526354 | 68528530 | 2177 | * | 22,84878 | 2,015128589 | 0,695691 | 2,896584 | 0,003772 | 0,594430593 | 2,951933 Intron (ENSMUST0000( | 7 | 68559158 | 68564346 | 5189 | 2 | 1,03E+08 | ENSMUST00000208180.2 | 35816 | ENSMUSG00000109407 | Gm34664 | predicted gene, 34664 |
| chr5 | 65592263 | 65595864 | 3602 | * | 41,97065 | 2,015591141 | 0,523943 | 3,84697 | 0,00012 | 0,133396034 | 2,338387 Promoter (<=1kb) | 5 | 65570564 | 65593292 | 22729 | 2 | 22235 | ENSMUST00000031103.14 | 0 | ENSMUSG00000029201 | Ugdh | UDP-glucose dehydrogenase |
| chr1 | 1,21E+08 | 1,21E+08 | 2057 | * | 20,54431 | 2,015960501 | 0,761129 | 2,648645 | 0,008082 | 0,740451144 | 2,663619 Exon (ENSMUST00000( | 1 | 1,21E+08 | 1,21E+08 | 70479 | 1 | 70829 | ENSMUST00000112621.2 | 5816 | ENSMUSG00000026339 | Ccdc93 | coiled-coil domain containing 93 |
| chr2 | 93054589 | 93056530 | 1942 | * | 20,1051 | 2,016124402 | 0,744183 | 2,709179 | 0,006745 | 0,740451144 | 2,221611 Promoter (1-2kb) | 2 | 93034592 | 93058131 | 23540 | 2 | 241556 | ENSMUST00000028646.4 | 1601 | ENSMUSG00000027217 | Tspan18 | tetraspanin 18 |
| chr6 | 1,13E+08 | 1,13E+08 | 2288 | * | 26,30887 | 2,017476434 | 0,721622 | 2,795753 | 0,005178 | 0,650529916 | 2,951062 Exon (ENSMUST00000( | 6 | 1,13E+08 | 1,13E+08 | 9831 | 1 | 72895 | ENSMUST00000156048.2 | 13489 | ENSMUSG00000034269 | Setd5 | SET domain containing 5 |
| chr15 | 77150498 | 77153222 | 2725 | * | 22,4522 | 2,01753585 | 0,729506 | 2,765621 | 0,005681 | 0,64068479 | 3,471678 Intron (ENSMUST0000( | 15 | 76982266 | 77129786 | 147521 | 2 | 93686 | ENSMUST00000228558.2 | -20712 | ENSMUSG00000033565 | Rbfox2 | RNA binding protein, fox-1 homolog (C. elegans) 2 |
| chr4 | 66853482 | 66855509 | 2028 | * | 18,013 | 2,017578995 | 0,741048 | 2,722604 | 0,006477 | 0,702396314 | 2,663619 Distal Intergenic | 4 | 66746439 | 66757855 | 11417 | 1 | 21898 | ENSMUST00000147008.2 | 107043 | ENSMUSG00000039005 | Tlr4 | toll-like receptor 4 |
| chr13 | 1,03E+08 | 1,03E+08 | 3425 | * | 39,81365 | 2,018089959 | 0,594272 | 3,395902 | 0,000684 | 0,311687208 | 2,725491 Intron (ENSMUST0000( | 13 | 1,03E+08 | 1,03E+08 | 163818 | 2 | 328329 | ENSMUST00000167462.8 | 4272 | ENSMUSG00000034751 | Mast4 | microtubule associated serine/threonine kinase family member 4 |
| chr8 | 72049779 | 72052414 | 2636 | * | 29,41652 | 2,018805634 | 0,603042 | 3,347703 | 0,000815 | 0,327375387 | 2,880017 Promoter (<=1kb) | 8 | 72050292 | 72060580 | 10289 | 1 | 1E+08 | ENSMUST00000143662.8 | 0 | ENSMUSG00000043243 | Niban3 | niban apoptosis regulator 3 |
| chr12 | 80183605 | 80185953 | 2349 | * | 36,19734 | 2,019630624 | 0,66996 | 3,014554 | 0,002574 | 0,531203841 | 2,787255 Distal Intergenic | 12 | 80170745 | 80179640 | 8896 | 2 | 69548 | ENSMUST00000180643.4 | -3965 | | 2310015A10Rik | RIKEN cDNA 2310015A10 gene |
| chr17 | 36087144 | 36089150 | 2007 | * | 20,20804 | 2,01967941 | 0,767775 | 2,630561 | 0,008524 | 0,7591629 | 2,507013 Exon (ENSMUST000001 | 17 | 36083342 | 36091579 | 8238 | 1 | 74601 | ENSMUST00000172874.2 | 3802 | ENSMUSG00000092171 | 4833427F10Rik | RIKEN cDNA 4833427F10 gene |
| chr1 | 7156854 | 7159217 | 2364 | * | 22,43595 | 2,021140201 | 0,688062 | 2,93744 | 0,003309 | 0,565876168 | 2,951933 Promoter (<=1kb) | 1 | 7159144 | 7243852 | 84709 | 1 | 319263 | ENSMUST00000061280.17 | 0 | ENSMUSG00000051285 | Pcmtd1 | protein-L-isoaspartate (D-aspartate) O-methyltransferase domain containing 1 |
| chr15 | 81809977 | 81813036 | 3060 | * | 35,47052 | 2,021656582 | 0,699802 | 2,888896 | 0,003866 | 0,610143359 | 2,787255 Promoter (<=1kb) | 15 | 81772429 | 81810365 | 37937 | 2 | 78929 | ENSMUST00000230669.2 | 0 | ENSMUSG00000022476 | Polr3h | polymerase (RNA) III (DNA directed) polypeptide H |
| chr10 | 39902434 | 39904874 | 2441 | * | 19,3451 | 2,021674899 | 0,739362 | 2,734352 | 0,00625 | 0,691026033 | 2,713306 Promoter (1-2kb) | 10 | 39877571 | 39901264 | 23694 | 2 | 215929 | ENSMUST00000163705.3 | -1170 | ENSMUSG00000038522 | Mfsd4b1 | major facilitator superfamily domain containing 4B1 |
| chr18 | 36459876 | 36462423 | 2548 | * | 44,46172 | 2,02195277 | 0,534553 | 3,782509 | 0,000155 | 0,156161007 | 2,377674 Distal Intergenic | 18 | 36475573 | 36526430 | 50858 | 1 | 66060 | ENSMUST00000139727.3 | -13150 | ENSMUSG00000046727 | Cystm1 | cysteine-rich transmembrane module containing 1 |
| chr9 | 13008094 | 13009983 | 1890 | * | 19,45283 | 2,022567972 | 0,778966 | 2,596478 | 0,009419 | 0,767135014 | 2,663619 Distal Intergenic | 9 | 13091272 | 13105860 | 14589 | 2 | 78219 | ENSMUST00000216694.2 | 95877 | ENSMUSG00000111830 | 4930568E12Rik | RIKEN cDNA 4930568E12 gene |
| chr11 | 48741237 | 48744309 | 3073 | * | 28,67372 | 2,022700491 | 0,570668 | 3,544443 | 0,000393 | 0,243803137 | 2,725491 Promoter (2-3kb) | 11 | 48738729 | 48743036 | 4308 | 1 | 94089 | ENSMUST00000131270.2 | 2508 | ENSMUSG00000040350 | Trim7 | tripartite motif-containing 7 |
| chr4 | 45486704 | 45490130 | 3427 | * | 43,82362 | 2,022793411 | 0,570668 | 3,544608 | 0,000393 | 0,262754801 | 2,338387 Promoter (<=1kb) | 4 | 45447506 | 45489403 | 41898 | 2 | 230126 | ENSMUST00000125599.2 | 0 | ENSMUSG00000044813 | Shb | src homology 2 domain-containing transforming protein B |
| chr9 | 9110696 | 9113612 | 2917 | * | 20,91597 | 2,022806865 | 0,712249 | 2,840026 | 0,004511 | 0,644557958 | 2,67243 Intron (ENSMUST0000( | 9 | 9236293 | 9260888 | 24596 | 1 | 1,01E+08 | ENSMUST00000189204.2 | -122681 | ENSMUSG00000085253 | Gm16833 | predicted gene, 16833 |
| chr10 | 31814630 | 31816680 | 2051 | * | 22,35313 | 2,022857024 | 0,772957 | 2,617039 | 0,00887 | 0,75508254 | 2,67243 Intron (ENSMUST0000( | 10 | 31369189 | 31485180 | 115992 | 2 | 268291 | ENSMUST00000081989.8 | -329450 | ENSMUSG00000063760 | Rnf217 | ring finger protein 217 |
| chr11 | 69226584 | 69228974 | 2391 | * | 20,62106 | 2,023877694 | 0,769498 | 2,630127 | 0,008535 | 0,723275538 | 3,104559 Promoter (2-3kb) | 11 | 69231287 | 69233467 | 2181 | 1 | 80515 | ENSMUST00000129321.2 | -2313 | ENSMUSG00000043419 | Rnf227 | ring finger protein 227 |
| chr6 | 50542051 | 50545273 | 3223 | * | 40,09177 | 2,02423926 | 0,506302 | 3,998083 | 6,39E-05 | 0,082076621 | 2,787255 Promoter (<=1kb) | 6 | 50539543 | 50543518 | 3976 | 2 | 13063 | ENSMUST00000161401.2 | 0 | ENSMUSG00000063694 | Cycs | cytochrome c, somatic |
| chr6 | 51843540 | 51846328 | 2789 | * | 29,56683 | 2,025059468 | 0,624345 | 3,243496 | 0,001181 | 0,40679506 | 2,67243 Promoter (2-3kb) | 6 | 51836084 | 51848909 | 12826 | 2 | 54353 | ENSMUST00000205174.2 | 2581 | ENSMUSG00000059182 | Skap2 | src family associated phosphoprotein 2 |
| chr4 | 1,26E+08 | 1,26E+08 | 2408 | * | 23,06566 | 2,025631676 | 0,731816 | 2,767952 | 0,005641 | 0,669586355 | 2,880017 Promoter (<=1kb) | 4 | 1,26E+08 | 1,26E+08 | 43932 | 2 | 76850 | ENSMUST00000084289.5 | -224 | ENSMUSG00000042500 | Ago4 | argonaute RISC catalytic subunit 4 |
| chr3 | 94343855 | 94345343 | 1489 | * | 18,08394 | 2,025897502 | 0,783966 | 2,584167 | 0,009761 | 0,775744126 | 2,663619 Promoter (<=1kb) | 3 | 94340473 | 94343923 | 3451 | 2 | 53814 | ENSMUST00000191506.8 | 0 | ENSMUSG00000028141 | Oaz3 | ornithine decarboxylase antizyme 3 |
| chr10 | 13708101 | 13709898 | 1798 | * | 18,73728 | 2,0259407 | 0,782665 | 2,588517 | 0,009639 | 0,79031971 | 2,423594 Intron (ENSMUST0000( | 10 | 13529332 | 13720030 | 190699 | 2 | 66253 | ENSMUST00000162174.8 | 10132 | ENSMUSG00000019806 | Aig1 | androgen-induced 1 |
| chr7 | 1,32E+08 | 1,32E+08 | 2735 | * | 26,66256 | 2,026724062 | 0,702044 | 2,88689 | 0,003891 | 0,618136403 | 2,67243 3' UTR | 7 | 1,32E+08 | 1,32E+08 | 63552 | 2 | 77938 | ENSMUST00000106168.2 | 15057 | ENSMUSG00000030956 | Fam53b | family with sequence similarity 53, member B |
| chr15 | 3572020 | 3574170 | 2151 | * | 21,73986 | 2,027817735 | 0,705279 | 2,875201 | 0,004038 | 0,571756101 | 3,471678 Exon (ENSMUST000001 | 15 | 3348491 | 3611324 | 262834 | 2 | 14600 | ENSMUST00000161561.8 | 37154 | ENSMUSG00000055737 | Ghr | growth hormone receptor |
| chr17 | 57106365 | 57108297 | 1933 | * | 17,63683 | 2,027988729 | 0,753768 | 2,690468 | 0,007135 | 0,734194343 | 2,423594 Exon (ENSMUST000002 | 17 | 57100882 | 57102378 | 1497 | 1 | 1,05E+08 | ENSMUST00000247457.1 | 5483 | | Gm41604 | predicted gene, 41604 |
| chr8 | 81300000 | 81301769 | 1770 | * | 18,45174 | 2,028006471 | 0,786725 | 2,577784 | 0,009944 | 0,792873671 | 2,423594 Distal Intergenic | 8 | 81337709 | 81421985 | 84277 | 1 | 333315 | ENSMUST00000039695.7 | -35940 | ENSMUSG00000042353 | Frem3 | Fras1 related extracellular matrix protein 3 |
| chr6 | 5370734 | 5372776 | 2043 | * | 23,28364 | 2,028350793 | 0,727567 | 2,787853 | 0,005306 | 0,659642963 | 2,880017 Distal Intergenic | 6 | 5383386 | 5433022 | 49637 | 1 | 65255 | ENSMUST00000043294.12 | -10610 | ENSMUSG00000042607 | Asb4 | ankyrin repeat and SOCS box-containing 4 |
| chr5 | 1,39E+08 | 1,39E+08 | 2827 | * | 24,73149 | 2,029033381 | 0,687248 | 2,952403 | 0,003153 | 0,553866303 | 2,951933 Promoter (2-3kb) | 5 | 1,39E+08 | 1,39E+08 | 25920 | 1 | 77053 | ENSMUST00000146715.8 | 2502 | ENSMUSG00000036817 | Sun1 | Sad1 and UNC84 domain containing 1 |
| chr12 | 59068090 | 59069989 | 1900 | * | 17,42068 | 2,031684342 | 0,737792 | 2,753737 | 0,005892 | 0,705941691 | 2,386008 Promoter (1-2kb) | 12 | 59071452 | 59074673 | 3222 | 1 | 66603 | ENSMUST00000182710.2 | -1463 | ENSMUSG00000060121 | Gemin2 | gem nuclear organelle associated protein 2 |
| chr5 | 31204953 | 31206895 | 1943 | * | 26,58366 | 2,032371096 | 0,726138 | 2,798876 | 0,005128 | 0,654833721 | 2,858511 Promoter (<=1kb) | 5 | 31193380 | 31205346 | 11967 | 2 | 330064 | ENSMUST00000202520.4 | 0 | ENSMUSG00000006641 | Slc5a6 | solute carrier family 5 (sodium-dependent vitamin transporter), member 6 |

| | | | | | | | | | | | | | | | | | | | | | | | |
|---|---|---|---|---|---|---|---|---|---|---|---|---|---|---|---|---|---|---|---|---|---|---|---|
| chr6 | 1,25E+08 | 1,25E+08 | 1861 | * | 21,38675 | 2,032416782 | 0,731954 | 2,776699 | 0,005491 | 0,6318891 | 3,471678 | Promoter (1-2kb) | 6 | 1,25E+08 | 1,25E+08 | 1278 | 2 | 14791 | ENSMUST00000125795.2 | -1862 | ENSMUSG00000004268 | Emg1 | EMG1 N1-specific pseudouridine methyltransferase |
| chr8 | 96225929 | 96228684 | 2756 | * | 30,29619 | 2,032591737 | 0,662299 | 3,068995 | 0,002148 | 0,492839239 | 2,880017 | Promoter (2-3kb) | 8 | 96223075 | 96227576 | 4502 | 1 | 71040 | ENSMUST00000212927.2 | 2854 | ENSMUSG00000110427 | 4933406B17Rik | RIKEN cDNA 4933406B17 gene |
| chr18 | 42397688 | 42400509 | 2822 | * | 24,05774 | 2,032810525 | 0,668714 | 3,039882 | 0,002367 | 0,504427013 | 2,951062 | Promoter (2-3kb) | 18 | 42335363 | 42395259 | 59897 | 2 | 107045 | ENSMUST00000097590.5 | -2429 | ENSMUSG00000024493 | Lars | leucyl-tRNA synthetase |
| chr8 | 71727533 | 71730093 | 2561 | * | 25,81583 | 2,033151989 | 0,668728 | 3,040325 | 0,002363 | 0,524368688 | 2,67243 | Promoter (1-2kb) | 8 | 71703769 | 71725578 | 21810 | 2 | 76478 | ENSMUST00000035960.13 | -1955 | ENSMUSG00000035439 | Haus8 | 4HAUS augmin-like complex, subunit 8 |
| chr1 | 1,23E+08 | 1,23E+08 | 1632 | * | 19,80163 | 2,033371146 | 0,760003 | 2,675479 | 0,007462 | 0,727652793 | 2,663619 | Distal Intergenic | 1 | 1,22E+08 | 1,22E+08 | 144 | 1 | 1,15E+08 | ENSMUST00000158565.3 | 40027 | ENSMUSG00000089190 | Gm25771 | predicted gene, 25771 |
| chr4 | 46418737 | 46421615 | 2879 | * | 26,34264 | 2,034572874 | 0,683491 | 2,976737 | 0,002913 | 0,542967057 | 2,880017 | Exon (ENSMUST00000 | 4 | 46393989 | 46413506 | 19518 | 2 | 93966 | ENSMUST00000071096.3 | -5231 | ENSMUSG00000028332 | Hemgn | hemogen |
| chr19 | 5460226 | 5462965 | 2740 | * | 24,43482 | 2,034630454 | 0,723234 | 2,813238 | 0,004905 | 0,659240897 | 2,67243 | Promoter (1-2kb) | 19 | 5464905 | 5467645 | 2741 | 1 | 74931 | ENSMUST00000241938.1 | -1940 | ENSMUSG00000086938 | 4930481A15Rik | RIKEN cDNA 4930481A15 gene |
| chr12 | 77813249 | 77815451 | 2203 | * | 24,22122 | 2,035256964 | 0,733047 | 2,776434 | 0,005496 | 0,676922067 | 2,67243 | Distal Intergenic | 12 | 77861454 | 77861583 | 130 | 2 | 1,15E+08 | ENSMUST00000104519.3 | 46132 | ENSMUSG00000077707 | Gm24070 | predicted gene, 24070 |
| chr19 | 41902786 | 41905071 | 2286 | * | 22,37982 | 2,035286651 | 0,747747 | 2,721893 | 0,006491 | 0,686762073 | 2,880017 | Promoter (2-3kb) | 19 | 41900515 | 41904525 | 4011 | 1 | 18648 | ENSMUST00000235468.2 | 2271 | ENSMUSG00000011752 | Pgam1 | phosphoglycerate mutase 1 |
| chr1 | 59190568 | 59193596 | 3029 | * | 25,83656 | 2,035638339 | 0,652949 | 3,117607 | 0,001823 | 0,466212802 | 2,858511 | Promoter (<=1kb) | 1 | 59178317 | 59191506 | 13190 | 2 | 227157 | ENSMUST00000189884.7 | 0 | ENSMUSG00000079550 | Mpp4 | membrane protein, palmitoylated 4 (MAGUK p55 subfamily member 4) |
| chr4 | 41418363 | 41420348 | 1986 | * | 18,29068 | 2,03567967 | 0,786889 | 2,586998 | 0,009682 | 0,811102052 | 2,154504 | Intron (ENSMUST0000 | 4 | 41395185 | 41409778 | 14594 | 2 | 109242 | ENSMUST00000141951.2 | -8585 | ENSMUSG00000028438 | Kif24 | kinesin family member 24 |
| chr5 | 1,42E+08 | 1,42E+08 | 2013 | * | 21,26145 | 2,03582544 | 0,716172 | 2,842649 | 0,004474 | 0,616346987 | 3,104559 | 3' UTR | 5 | 1,42E+08 | 1,42E+08 | 2611 | 1 | 231855 | ENSMUST00000198967.2 | -5910 | ENSMUSG00000039623 | Ap5z1 | adaptor-related protein complex 5, zeta 1 subunit |
| chr4 | 40720650 | 40722591 | 1942 | * | 20,01323 | 2,035864397 | 0,746195 | 2,728329 | 0,006366 | 0,69472367 | 2,713306 | Promoter (<=1kb) | 4 | 40722150 | 40723795 | 1646 | 1 | 15502 | ENSMUST00000139406.2 | 0 | ENSMUSG00000028410 | Dnaja1 | DnaJ heat shock protein family (Hsp40) member A1 |
| chr2 | 1,02E+08 | 1,02E+08 | 3958 | * | 32,63352 | 2,035996126 | 0,64863 | 3,138915 | 0,001696 | 0,450044524 | 2,925497 | Intron (ENSMUST0000 | 2 | 1,02E+08 | 1,02E+08 | 27097 | 1 | 1,03E+08 | ENSMUST00000140238.2 | -56188 | | Gm13919 | predicted gene 13919 |
| chr19 | 40383216 | 40385485 | 2270 | * | 26,80948 | 2,036430254 | 0,706286 | 2,883294 | 0,003935 | 0,621084104 | 2,67243 | Exon (ENSMUST00000 | 19 | 40283493 | 40390711 | 107219 | 2 | 20411 | ENSMUST00000099466.10 | 5226 | ENSMUSG00000025006 | Sorbs1 | sorbin and SH3 domain containing 1 |
| chr5 | 91027178 | 91030503 | 3326 | * | 28,92277 | 2,036734423 | 0,746288 | 2,729154 | 0,00635 | 0,683210259 | 2,951062 | Distal Intergenic | 5 | 91039100 | 91040974 | 1875 | 1 | 14825 | ENSMUST00000031327.9 | -8597 | ENSMUSG00000029380 | Cxcl1 | chemokine (C-X-C motif) ligand 1 |
| chr1 | 37924486 | 37927118 | 2633 | * | 25,65 | 2,036747174 | 0,729542 | 2,791815 | 0,005241 | 0,669637528 | 2,67243 | Promoter (2-3kb) | 1 | 37921010 | 37929357 | 8348 | 2 | 69028 | ENSMUST00000138806.2 | 2239 | ENSMUSG00000026088 | Mitd1 | MIT, microtubule interacting and transport, domain containing 1 |
| chr13 | 21992864 | 21995891 | 3028 | * | 26,54111 | 2,037075271 | 0,674004 | 3,022351 | 0,002508 | 0,514210241 | 2,951062 | Promoter (<=1kb) | 13 | 21994635 | 21995114 | 480 | 1 | 665433 | ENSMUST00000091745.6 | 0 | ENSMUSG00000094248 | H2ac23 | H2A clustered histone 23 |
| chr2 | 1,57E+08 | 1,57E+08 | 2912 | * | 26,3991 | 2,03720976 | 0,619243 | 3,289838 | 0,001002 | 0,355694809 | 2,951062 | Promoter (<=1kb) | 2 | 1,57E+08 | 1,57E+08 | 10436 | 1 | 67388 | ENSMUST00000029165.4 | 0 | ENSMUSG00000027637 | Rab5if | RAB5 interacting factor |
| chr9 | 1,21E+08 | 1,21E+08 | 1668 | * | 17,00265 | 2,038392662 | 0,781994 | 2,60666 | 0,009143 | 0,891097601 | 1,2763 | Promoter (<=1kb) | 9 | 1,21E+08 | 1,21E+08 | 43320 | 1 | 67095 | ENSMUST00000210351.2 | 0 | ENSMUSG00000032536 | Trak1 | trafficking protein, kinesin binding 1 |
| chr14 | 45731008 | 45734344 | 3337 | * | 31,02809 | 2,038809499 | 0,642379 | 3,173844 | 0,001504 | 0,434822663 | 2,858511 | Intron (ENSMUST0000 | 14 | 45706771 | 45715318 | 8548 | 2 | 218952 | ENSMUST00000141424.2 | -15690 | ENSMUSG00000037712 | Fermt2 | fermitin family member 2 |
| chr5 | 33986894 | 33989313 | 2420 | * | 16,36006 | 2,039144126 | 0,765128 | 2,665101 | 0,007697 | 0,730832709 | 2,663619 | Intron (ENSMUST0000 | 5 | 33978173 | 34003526 | 25354 | 1 | 107823 | ENSMUST00000155880.8 | 8721 | ENSMUSG00000057406 | Nsd2 | nuclear receptor binding SET domain protein 2 |
| chr5 | 38719156 | 38721615 | 2460 | * | 18,76391 | 2,039217081 | 0,772953 | 2,638217 | 0,008334 | 0,75508254 | 2,507013 | Promoter (<=1kb) | 5 | 38703018 | 38720564 | 17547 | 2 | 22388 | ENSMUST00000202766.4 | 0 | ENSMUSG00000005103 | Wdr1 | WD repeat domain 1 |
| chr13 | 55697915 | 55699705 | 1791 | * | 19,13731 | 2,039399535 | 0,770031 | 2,648465 | 0,008086 | 0,749931768 | 2,507013 | Promoter (<=1kb) | 13 | 55687132 | 55700142 | 13011 | 2 | 212483 | ENSMUST00000224486.2 | 437 | ENSMUSG00000021495 | Fam193b | family with sequence similarity 193, member B |
| chr3 | 51608247 | 51610954 | 2708 | * | 23,65634 | 2,040135698 | 0,696361 | 2,929709 | 0,003393 | 0,575442234 | 2,880017 | Intron (ENSMUST0000 | 3 | 51589043 | 51590098 | 1056 | 1 | 211666 | ENSMUST00000195423.2 | 19204 | ENSMUSG00000074604 | Mgst2 | microsomal glutathione S-transferase 2 |
| chr9 | 1,2E+08 | 1,2E+08 | 2926 | * | 26,34996 | 2,040724127 | 0,69533 | 2,934899 | 0,003337 | 0,573118536 | 2,858511 | Promoter (2-3kb) | 9 | 1,2E+08 | 1,2E+08 | 9844 | 2 | 22437 | ENSMUST00000111635.4 | -2371 | ENSMUSG00000079243 | Xirp1 | xin actin-binding repeat containing 1 |
| chr5 | 34347467 | 34350447 | 2981 | * | 28,96713 | 2,041505479 | 0,651236 | 3,134815 | 0,00172 | 0,464338132 | 2,725491 | Promoter (2-3kb) | 5 | 34333923 | 34345064 | 11142 | 2 | 17122 | ENSMUST00000119171.2 | -2403 | ENSMUSG00000037235 | Mxd4 | Max dimerization protein 4 |
| chr11 | 84708975 | 84711129 | 2155 | * | 26,32107 | 2,041702108 | 0,687875 | 2,968129 | 0,002996 | 0,549912148 | 2,858511 | Promoter (<=1kb) | 11 | 84703887 | 84710341 | 6455 | 2 | 217038 | ENSMUST00000018549.8 | 0 | ENSMUSG00000018405 | Mrm1 | mitochondrial rRNA methyltransferase 1 |
| chr10 | 37252031 | 37253505 | 1475 | * | 15,62697 | 2,041835144 | 0,78912 | 2,587483 | 0,009668 | 0,888934304 | 1,37683 | Exon (ENSMUST00000 | 10 | 37213482 | 37255648 | 42167 | 1 | 1,03E+08 | ENSMUST00000181602.2 | 38549 | | Gm26535 | predicted gene, 26535 |
| chr10 | 1,02E+08 | 1,02E+08 | 1877 | * | 20,91464 | 2,041990947 | 0,667956 | 3,057075 | 0,002235 | 0,486196597 | 3,104559 | Intron (ENSMUST0000 | 10 | 1,02E+08 | 1,02E+08 | 232391 | 1 | 67569 | ENSMUST00000020039.13 | 42568 | ENSMUSG00000019888 | Mgat4c | MGAT4 family, member C |
| chr4 | 1,27E+08 | 1,27E+08 | 2101 | * | 18,58335 | 2,043247146 | 0,758716 | 2,693033 | 0,007081 | 0,73648081 | 2,386008 | Promoter (<=1kb) | 4 | 1,27E+08 | 1,27E+08 | 15836 | 1 | 100317 | ENSMUST00000141918.2 | -388 | ENSMUSG00000028830 | AU040320 | expressed sequence AU040320 |
| chr5 | 64423399 | 64425714 | 2316 | * | 19,79779 | 2,043507238 | 0,753993 | 2,710245 | 0,006723 | 0,728748169 | 2,386008 | Intron (ENSMUST0000 | 5 | 64432763 | 64442082 | 9320 | 1 | 57915 | ENSMUST00000147348.6 | -7049 | ENSMUSG00000029174 | Tbc1d1 | TBC1 domain family, member 1 |
| chr3 | 90589198 | 90591357 | 2160 | * | 21,10249 | 2,04359675 | 0,714001 | 2,862175 | 0,004207 | 0,614207576 | 2,951933 | Distal Intergenic | 3 | 90599939 | 90603018 | 3080 | 2 | 20202 | ENSMUST00000069960.12 | 11661 | ENSMUSG00000056071 | S100a9 | S100 calcium binding protein A9 (calgranulin B) |
| chr10 | 67020287 | 67022189 | 1903 | * | 15,41451 | 2,043711015 | 0,78417 | 2,60621 | 0,009155 | 0,891097601 | 1,2763 | Promoter (<=1kb) | 10 | 67021528 | 67051950 | 30423 | 1 | 108829 | ENSMUST00000173661.8 | 0 | ENSMUSG00000037876 | Jmjd1c | jumonji domain containing 1C |
| chr6 | 146747538 | 146749036 | 1499 | * | 1011,4745 | 2,043820497 | 0,3312032 | 6,1708966 | 6,79E-10 | 8,16586E-06 | 4,0557576 | Promoter (2-3kb) | 6 | 146751602 | 146772904 | 21303 | 1 | 69371 | ENSMUST00000032433.9 | -2566 | ENSMUSG00000030292 | Smco2 | single-pass membrane protein with coiled-coil domains 2 |
| chr4 | 45011532 | 45014448 | 2917 | * | 31,2101 | 2,044233357 | 0,604518 | 3,381592 | 0,000721 | 0,322703004 | 2,67243 | Promoter (<=1kb) | 4 | 44991242 | 45012394 | 21153 | 2 | 230119 | ENSMUST00000055028.9 | 0 | ENSMUSG00000049657 | Zbtb5 | zinc finger and BTB domain containing 5 |
| chr3 | 28412592 | 28415122 | 2531 | * | 17,59067 | 2,04468001 | 0,757134 | 2,700551 | 0,006922 | 0,715156227 | 2,663619 | Exon (ENSMUST00000 | 3 | 28463580 | 28650232 | 186653 | 1 | 665113 | ENSMUST00000161423.2 | -48458 | ENSMUSG00000027692 | Tnik | TRAF2 and NCK interacting kinase |
| chr7 | 1,45E+08 | 1,45E+08 | 1665 | * | 15,71807 | 2,045795722 | 0,789007 | 2,592873 | 0,009518 | 0,769699963 | 2,663619 | Distal Intergenic | 7 | 1,45E+08 | 1,45E+08 | 9252 | 1 | 211578 | ENSMUST00000062163.8 | 19335 | ENSMUSG00000051207 | Mrgprd | MAS-related GPR, member D |
| chr7 | 1,01E+08 | 1,01E+08 | 3264 | * | 27,76376 | 2,046073395 | 0,62155 | 3,291889 | 0,000995 | 0,369614065 | 2,725491 | 3' UTR | 7 | 1,01E+08 | 1,01E+08 | 2891 | 2 | 207212 | ENSMUST00000208482.2 | 5544 | ENSMUSG00000032875 | Arhgef17 | Rho guanine nucleotide exchange factor (GEF) 17 |
| chr1 | 1,77E+08 | 1,77E+08 | 1565 | * | 31,6473 | 2,04745641 | 0,663104 | 3,087685 | 0,002017 | 0,494741046 | 2,67243 | Promoter (<=1kb) | 1 | 1,77E+08 | 1,77E+08 | 80406 | 2 | 545389 | ENSMUST00000195717.6 | 0 | ENSMUSG00000057335 | Cep170 | centrosomal protein 170 |
| chr13 | 24749198 | 24751416 | 2219 | * | 23,29456 | 2,047700073 | 0,712908 | 2,872322 | 0,004075 | 0,627785088 | 2,67243 | Intron (ENSMUST0000 | 13 | 24766172 | 24917799 | 151628 | 1 | 193385 | ENSMUST00000110383.8 | -14756 | ENSMUSG00000036006 | Ripor2 | RHO family interacting cell polarization regulator 2 |
| chr5 | 8228754 | 8230382 | 1629 | * | 17,41883 | 2,047894973 | 0,706127 | 2,900178 | 0,00373 | 0,649217039 | 2,154504 | Exon (ENSMUST00000 | 5 | 8127839 | 8171095 | 43257 | 2 | 11496 | ENSMUST00000124121.8 | -57659 | ENSMUSG00000040537 | Adam22 | a disintegrin and metallopeptidase domain 22 |
| chr4 | 1,5E+08 | 1,5E+08 | 2933 | * | 21,24972 | 2,048128496 | 0,737018 | 2,77894 | 0,005454 | 0,631320778 | 3,471678 | Promoter (<=1kb) | 4 | 1,5E+08 | 1,5E+08 | 1161 | 1 | 13806 | ENSMUST00000133789.2 | 0 | ENSMUSG00000063524 | Eno1 | enolase 1, alpha non-neuron |
| chr8 | 1,07E+08 | 1,07E+08 | 3873 | * | 26,78998 | 2,048429076 | 0,757596 | 2,703853 | 0,006854 | 0,697016988 | 2,880017 | Exon (ENSMUST00000 | 8 | 1,07E+08 | 1,07E+08 | 348 | 2 | 77411 | ENSMUST00000142050.2 | 28261 | ENSMUSG00000084128 | Esrp2 | epithelial splicing regulatory protein 2 |
| chr2 | 1,3E+08 | 1,3E+08 | 2375 | * | 21,10247 | 2,049007172 | 0,725518 | 2,824198 | 0,00474 | 0,605351622 | 3,471678 | Promoter (1-2kb) | 2 | 1,3E+08 | 1,3E+08 | 1425 | 1 | 66766 | ENSMUST00000055421.6 | 1239 | ENSMUSG00000049692 | Tmem239 | transmembrane 239 |
| chr7 | 1,45E+08 | 1,45E+08 | 5142 | * | 52,99645 | 2,049672702 | 0,538079 | 3,809239 | 0,000139 | 0,133396034 | 2,725491 | Distal Intergenic | 7 | 1,45E+08 | 1,45E+08 | 794 | 2 | 1E+08 | ENSMUST00000207703.2 | 93465 | ENSMUSG00000109305 | Smim38 | small integral membrane protein 38 |
| chr6 | 72338083 | 72340789 | 2707 | * | 41,70546 | 2,04984505 | 0,597997 | 3,427853 | 0,000608 | 0,29714458 | 2,787255 | Promoter (<=1kb) | 6 | 72336697 | 72339376 | 2680 | 2 | 66510 | ENSMUST00000069595.13 | 0 | ENSMUSG00000055850 | Rnf181 | ring finger protein 181 |
| chr6 | 1,2E+08 | 1,2E+08 | 2216 | * | 26,21477 | 2,049893351 | 0,692846 | 2,958657 | 0,00309 | 0,54964001 | 2,951062 | Promoter (<=1kb) | 6 | 1,2E+08 | 1,2E+08 | 6512 | 1 | 29862 | ENSMUST00000035244.6 | 698 | ENSMUSG00000041377 | Ninj2 | ninjurin 2 |
| chr1 | 1,8E+08 | 1,8E+08 | 3268 | * | 27,10077 | 2,049927306 | 0,639928 | 3,203372 | 0,001358 | 0,418252864 | 2,858511 | Intron (ENSMUST0000 | 1 | 1,8E+08 | 1,8E+08 | 47647 | 1 | 226751 | ENSMUST00000132894.3 | 37279 | ENSMUSG00000026490 | Cdc42bpa | CDC42 binding protein kinase alpha |
| chr9 | 60157557 | 60159969 | 2413 | * | 19,22365 | 2,049991557 | 0,779728 | 2,629111 | 0,008561 | 0,770673161 | 2,386008 | Exon (ENSMUST00000 | 9 | 60241425 | 60242251 | 827 | 1 | 1,15E+08 | ENSMUST00000214235.2 | -81456 | ENSMUSG00000111296 | 2010001M07Rik | RIKEN cDNA 2010001M07 gene |
| chr4 | 1,56E+08 | 1,56E+08 | 1609 | * | 14,57006 | 2,050158357 | 0,791001 | 2,591852 | 0,009546 | 0,809505441 | 2,141612 | Distal Intergenic | 4 | 1,56E+08 | 1,56E+08 | 5881 | 2 | 231004 | ENSMUST00000219393.2 | -71277 | ENSMUSG00000096351 | Samd11 | sterile alpha motif domain containing 11 |
| chr11 | 8570042 | 8574145 | 4104 | * | 34,65756 | 2,050330047 | 0,660372 | 3,104813 | 0,001904 | 0,603552508 | 1,405709 | Promoter (<=1kb) | 11 | 8381652 | 8574887 | 193236 | 2 | 319939 | ENSMUST00000239091.2 | 742 | ENSMUSG00000020422 | Tns3 | tensin 3 |
| chr12 | 21432219 | 21434574 | 2356 | * | 22,60786 | 2,050381873 | 0,758324 | 2,703833 | 0,006854 | 0,697016988 | 2,880017 | Distal Intergenic | 12 | 21373518 | 21423633 | 50116 | 2 | 11491 | ENSMUST00000064536.13 | -8586 | ENSMUSG00000052593 | Adam17 | a disintegrin and metallopeptidase domain 17 |
| chr7 | 73267022 | 73269294 | 2273 | * | 32,32792 | 2,051248089 | 0,653246 | 3,140083 | 0,001689 | 0,579872171 | 1,405709 | Distal Intergenic | 7 | 73295733 | 73295837 | 105 | 2 | 1,15E+08 | ENSMUST00000083795.3 | 26543 | ENSMUSG00000065729 | Gm26176 | predicted gene, 26176 |
| chr7 | 1,3E+08 | 1,3E+08 | 3719 | * | 32,83517 | 2,051272828 | 0,655166 | 3,130923 | 0,001743 | 0,454149353 | 2,925497 | Intron (ENSMUST0000 | 7 | 1,3E+08 | 1,3E+08 | 120522 | 1 | 57752 | ENSMUST00000207999.2 | 5745 | ENSMUSG00000030852 | Tacc2 | transforming, acidic coiled-coil containing protein 2 |
| chr5 | 1,36E+08 | 1,36E+08 | 3546 | * | 27,82378 | 2,051642383 | 0,631524 | 3,248714 | 0,001159 | 0,391231126 | 2,880017 | Promoter (<=1kb) | 5 | 1,36E+08 | 1,36E+08 | 6093 | 1 | 20022 | ENSMUST00000111127.8 | 0 | ENSMUSG00000039771 | Polr2j | polymerase (RNA) II (DNA directed) polypeptide J |
| chr12 | 74010378 | 74012696 | 2319 | * | 23,15275 | 2,051771667 | 0,708709 | 2,895083 | 0,003791 | 0,59835171 | 2,858511 | Promoter (<=1kb) | 12 | 74011255 | 74014773 | 3519 | 1 | 75627 | ENSMUST00000222025.2 | 0 | ENSMUSG00000021113 | Snapc1 | small nuclear RNA activating complex, polypeptide 1 |
| chr2 | 1,31E+08 | 1,31E+08 | 4113 | * | 26,49379 | 2,051799952 | 0,690432 | 2,971762 | 0,002961 | 0,542827503 | 2,951062 | Promoter (<=1kb) | 2 | 1,31E+08 | 1,31E+08 | 20943 | 1 | 74450 | ENSMUST00000131779.8 | 0 | ENSMUSG00000037514 | Pank2 | pantothenate kinase 2 |
| chr2 | 30390500 | 30392204 | 1705 | * | 25,09316 | 2,051887461 | 0,717877 | 2,85827 | 0,00426 | 0,616699946 | 2,951933 | Distal Intergenic | 2 | 30361549 | 30364231 | 2683 | 2 | 72500 | ENSMUST00000065134.4 | -26269 | ENSMUSG00000089762 | Ier5l | immediate early response 5-like |
| chr18 | 29412320 | 29414535 | 2216 | * | 24,6232 | 2,051890931 | 0,764694 | 2,683283 | 0,00729 | 0,705941691 | 2,951933 | Distal Intergenic | 18 | 29575253 | 29599160 | 23908 | 1 | 78199 | ENSMUST00000234621.2 | -160718 | ENSMUSG00000117554 | 4930527G23Rik | RIKEN cDNA 4930527G23 gene |
| chr14 | 14052564 | 14053856 | 1293 | * | 18,22891 | 2,051909539 | 0,752513 | 2,726742 | 0,006396 | 0,713406565 | 2,507013 | Distal Intergenic | 14 | 14091019 | 14120836 | 29818 | 1 | 18300 | ENSMUST00000022269.7 | -37163 | ENSMUSG00000021749 | Fam3d | FAM3 metabolism regulating signaling molecule D |
| chr16 | 25906510 | 25908558 | 2049 | * | 20,38594 | 2,051929113 | 0,79617 | 2,577251 | 0,009959 | 0,775744126 | 2,713306 | Intron (ENSMUST0000 | 16 | 25778038 | 25924534 | 146497 | 2 | 210530 | ENSMUST00000039990.6 | 15976 | ENSMUSG00000038168 | P3h2 | prolyl 3-hydroxylase 2 |
| chr15 | 82249804 | 82251277 | 1474 | * | 18,56022 | 2,052377721 | 0,753914 | 2,722296 | 0,006483 | 0,722439117 | 2,386008 | Exon (ENSMUST00000 | 15 | 82256023 | 82257647 | 1625 | 2 | 56448 | ENSMUST00000229599.2 | 6370 | ENSMUSG00000061740 | Cyp2d22 | cytochrome P450, family 2, subfamily d, polypeptide 22 |
| chr7 | 1,01E+08 | 1,01E+08 | 2418 | * | 18,71829 | 2,053145373 | 0,740817 | 2,771461 | 0,005581 | 0,694030854 | 2,386008 | Promoter (<=1kb) | 7 | 1,01E+08 | 1,01E+08 | 8413 | 2 | 11871 | ENSMUST00000173420.2 | -267 | ENSMUSG00000092517 | Art2a | ADP-ribosyltransferase 2a |
| chr2 | 1,29E+08 | 1,29E+08 | 2669 | * | 20,31095 | 2,053360951 | 0,780839 | 2,629684 | 0,008546 | 0,749550311 | 2,663619 | Exon (ENSMUST00000 | 2 | 1,29E+08 | 1,29E+08 | 23100 | 2 | 668894 | ENSMUST00000145798.4 | 5698 | ENSMUSG00000079051 | Vinac1 | vinculin/alpha-catenin family member 1 |
| chr9 | 44007574 | 44010062 | 2489 | * | 22,42517 | 2,053805114 | 0,727631 | 2,822593 | 0,004764 | 0,627785088 | 3,104559 | Promoter (2-3kb) | 9 | 44013026 | 44020484 | 7459 | 1 | 235312 | ENSMUST00000034654.9 | -2964 | ENSMUSG00000079592 | C1qtnf5 | C1q and tumor necrosis factor related protein 5 |
| chr2 | 1,54E+08 | 1,54E+08 | 1729 | * | 19,73131 | 2,05413405 | 0,786872 | 2,610507 | 0,009041 | 0,792545355 | 2,221611 | Exon (ENSMUST00000 | 2 | 1,54E+08 | 1,54E+08 | 1311 | 1 | 66557 | ENSMUST00000129776.2 | 4427 | ENSMUSG00000027481 | Bpifb2 | BPI fold containing family B, member 2 |
| chr5 | 1,39E+08 | 1,39E+08 | 2155 | * | 19,21784 | 2,054304579 | 0,731481 | 2,808419 | 0,004979 | 0,659240897 | 2,713306 | Promoter (1-2kb) | 5 | 1,39E+08 | 1,39E+08 | 35800 | 1 | 433956 | ENSMUST00000196441.2 | 1720 | ENSMUSG00000025857 | Dnaaf5 | dynein, axonemal assembly factor 5 |
| chr9 | 1,09E+08 | 1,09E+08 | 2757 | * | 26,49523 | 2,055433994 | 0,672667 | 3,055648 | 0,002246 | 0,496560937 | 2,951062 | Intron (ENSMUST0000 | 9 | 1,09E+08 | 1,09E+08 | 16820 | 2 | 320083 | ENSMUST00000196410.2 | -7358 | ENSMUSG00000074062 | Fbxw16 | F-box and WD-40 domain protein 16 |
| chr11 | 95789725 | 95792457 | 2733 | * | 20,93761 | 2,055507851 | 0,783842 | 2,622351 | 0,008733 | 0,751314174 | 2,67243 | Intron (ENSMUST0000 | 11 | 95756769 | 95805717 | 48949 | 2 | 14422 | ENSMUST00000038343.7 | 13260 | ENSMUSG00000013418 | B4galnt2 | beta-1,4-N-acetyl-galactosaminyl transferase 2 |
| chr10 | 1,27E+08 | 1,27E+08 | 2835 | * | 28,09592 | 2,055990789 | 0,65573 | 3,135421 | 0,001716 | 0,451060501 | 2,951062 | 3' UTR | 10 | 1,27E+08 | 1,27E+08 | 34031 | 2 | 16572 | ENSMUST00000217895.2 | 23675 | ENSMUSG00000074657 | Kif5a | kinesin family member 5A |
| chr7 | 109886126 | 109889465 | 3340 | * | 53,918717 | 2,056154954 | 0,4460686 | 4,6095042 | 4,036E-06 | 0,012861975 | 2,3383871 | Distal Intergenic | 7 | 109919948 | 109939492 | 19545 | 2 | 319934 | ENSMUST00000165992.2 | 50027 | ENSMUSG00000038371 | Sbf2 | SET binding factor 2 |
| chr1 | 86286666 | 86289074 | 2409 | * | 25,46859 | 2,05654808 | 0,648552 | 3,170986 | 0,001519 | 0,444359691 | 2,725491 | Promoter (<=1kb) | 1 | 86272441 | 86287122 | 14682 | 2 | 17975 | ENSMUST00000027438.8 | 0 | ENSMUSG00000026234 | Ncl | nucleolin |
| chr9 | 1,22E+08 | 1,22E+08 | 1815 | * | 24,51741 | 2,056766944 | 0,775756 | 2,651308 | 0,008018 | 0,738379885 | 2,67243 | Promoter (<=1kb) | 9 | 1,22E+08 | 1,22E+08 | 2593 | 1 | 26901 | ENSMUST00000035113.11 | 0 | ENSMUSG00000032526 | Ss18l2 | SS18, nBAF chromatin remodeling complex subunit like 2 |
| chr7 | 1,3E+08 | 1,3E+08 | 3081 | * | 23,96916 | 2,056796801 | 0,71717 | 2,867933 | 0,004132 | 0,61616728 | 2,880017 | Distal Intergenic | 7 | 1,3E+08 | 1,3E+08 | 110 | 1 | 1,15E+08 | ENSMUST00000083829.3 | 41218 | ENSMUSG00000065763 | Gm23847 | predicted gene, 23847 |
| chr11 | 69678723 | 69681307 | 2585 | * | 25,00763 | 2,05750253 | 0,792143 | 2,597386 | 0,009394 | 0,749550311 | 2,951933 | Intron (ENSMUST0000 | 11 | 69683605 | 69684646 | 1042 | 2 | 11443 | ENSMUST00000129381.2 | 3339 | ENSMUSG00000041189 | Chrnb1 | cholinergic receptor, nicotinic, beta polypeptide 1 (muscle) |
| chr9 | 62339826 | 62341605 | 1780 | * | 19,70365 | 2,057659192 | 0,752909 | 2,732946 | 0,006277 | 0,695401133 | 2,663619 | Intron (ENSMUST0000 | 9 | 62335545 | 62336532 | 988 | 2 | 235431 | ENSMUST00000151604.2 | -3294 | ENSMUSG00000041729 | Coro2b | coronin, actin binding protein, 2B |
| chr10 | 67700724 | 67703454 | 2731 | * | 21,20842 | 2,058027112 | 0,776446 | 2,650575 | 0,008035 | 0,72253719 | 2,951933 | Distal Intergenic | 10 | 67725783 | 67734837 | 9055 | 2 | 216049 | ENSMUST00000127869.2 | 31383 | ENSMUSG00000037855 | Zfp365 | zinc finger protein 365 |
| chr19 | 47293535 | 47296518 | 2984 | * | 25,60849 | 2,058926893 | 0,757703 | 2,717327 | 0,006581 | 0,690933518 | 2,858511 | Intron (ENSMUST0000 | 19 | 47255352 | 47303190 | 47839 | 2 | 14218 | ENSMUST00000237796.2 | 6672 | ENSMUSG00000053617 | Sh3pxd2a | SH3 and PX domains 2A |
| chr19 | 4337452 | 4339137 | 1686 | * | 17,51512 | 2,06030673 | 0,799341 | 2,577507 | 0,009952 | 0,777919825 | 2,663619 | Promoter (<=1kb) | 19 | 4336647 | 4337877 | 1231 | 2 | 110355 | ENSMUST00000235681.2 | 0 | ENSMUSG00000024858 | Grk2 | G protein-coupled receptor kinase 2 |
| chr8 | 1,25E+08 | 1,25E+08 | 2799 | * | 27,85939 | 2,061079773 | 0,713308 | 2,889467 | 0,003859 | 0,596145184 | 2,951062 | 3' UTR | 8 | 1,25E+08 | 1,25E+08 | 25112 | 1 | 382038 | ENSMUST00000034457.9 | 14780 | ENSMUSG00000031976 | Urb2 | URB2 ribosome biogenesis 2 homolog (S. cerevisiae) |
| chr8 | 85596839 | 85598780 | 1942 | * | 26,62498 | 2,061218237 | 0,70114 | 2,93981 | 0,003284 | 0,568574173 | 2,880017 | Promoter (<=1kb) | 8 | 85598740 | 85614075 | 15336 | 1 | 71846 | ENSMUST00000170296.2 | 0 | ENSMUSG00000003824 | Syce2 | synaptonemal complex central element protein 2 |
| chr4 | 156114078 | 156118076 | 3999 | * | 72,622911 | 2,061613998 | 0,4144048 | 4,9748802 | 6,529E-07 | 0,00252152 | 3,177995 | Promoter (<=1kb) | 4 | 156106465 | 156116104 | 9640 | 2 | 100038526 | ENSMUST00000181389.3 | 0 | | Gm16008 | predicted gene 16008 |
| chr11 | 57886806 | 57889899 | 3094 | * | 28,58393 | 2,062893646 | 0,672862 | 3,065851 | 0,002171 | 0,490966492 | 2,951062 | Distal Intergenic | 11 | 57899890 | 57952857 | 52968 | 1 | 73158 | ENSMUST00000071487.13 | -9991 | ENSMUSG00000037331 | Larp1 | La ribonucleoprotein domain family, member 1 |
| chr7 | 1E+08 | 1E+08 | 4045 | * | 37,27135 | 2,06314894 | 0,619703 | 3,329252 | 0,000871 | 0,370253435 | 2,377674 | Exon (ENSMUST00000 | 7 | 1E+08 | 1E+08 | 49823 | 1 | 319604 | ENSMUST00000216021.2 | 22412 | ENSMUSG00000029461 | Fam168a | family with sequence similarity 168, member A |
| chr7 | 1,27E+08 | 1,27E+08 | 1929 | * | 18,40162 | 2,063353295 | 0,748772 | 2,755648 | 0,005858 | 0,700182401 | 2,423594 | Promoter (<=1kb) | 7 | 1,27E+08 | 1,27E+08 | 27762 | 2 | 78388 | ENSMUST00000165096.9 | 113 | ENSMUSG00000030681 | Mvp | major vault protein |
| chr6 | 86016511 | 86017918 | 1408 | * | 19,78966 | 2,063460633 | 0,766068 | 2,693572 | 0,007069 | 0,717133806 | 2,663619 | Exon (ENSMUST00000 | 6 | 86005729 | 86086449 | 80721 | 1 | 11519 | ENSMUST00000203366.3 | 10782 | ENSMUSG00000030000 | Add2 | adducin 2 (beta) |
| chr1 | 37285789 | 37288139 | 2351 | * | 22,33562 | 2,06441951 | 0,697184 | 2,961084 | 0,003066 | 0,55292773 | 2,880017 | Intron (ENSMUST0000 | 1 | 37258286 | 37302465 | 44180 | 1 | 12790 | ENSMUST00000027288.10 | 27503 | ENSMUSG00000026114 | Cnga3 | cyclic nucleotide gated channel alpha 3 |
| chr12 | 69620408 | 69624900 | 4493 | * | 30,19474 | 2,064458619 | 0,694021 | 2,974635 | 0,002933 | 0,54122637 | 2,951062 | 3' UTR | 12 | 69629050 | 69629709 | 660 | 2 | 207965 | ENSMUST00000238563.2 | 4809 | ENSMUSG00000049882 | Vcpkmt | valosin containing protein lysine (K) methyltransferase |
| chr14 | 31336479 | 31339244 | 2766 | * | 23,18609 | 2,064630908 | 0,769755 | 2,682193 | 0,007314 | 0,723027638 | 2,67243 | Promoter (<=1kb) | 14 | 31329622 | 31336268 | 6647 | 2 | 56794 | ENSMUST00000140078.9 | -211 | ENSMUSG00000021884 | Hacl1 | 2-hydroxyacyl-CoA lyase 1 |
| chr9 | 70039833 | 70042176 | 2344 | * | 18,48095 | 2,065022193 | 0,778619 | 2,652161 | 0,007998 | 0,756030966 | 2,386008 | Intron (ENSMUST0000 | 9 | 69995793 | 70048860 | 53068 | 2 | 76886 | ENSMUST00000034749.16 | 6684 | ENSMUSG00000032224 | Fam81a | family with sequence similarity 81, member A |
| chr19 | 5550277 | 5552447 | 2171 | * | 17,67228 | 2,065202908 | 0,727836 | 2,837455 | 0,004547 | 0,681459438 | 2,154504 | Promoter (1-2kb) | 19 | 5552795 | 5553815 | 1021 | 2 | 75555 | ENSMUST00000189704.2 | 1368 | ENSMUSG00000100937 | Nscme3l | NSE3 homolog, SMC5-SMC6 complex component like |
| chr2 | 31559393 | 31561569 | 2177 | * | 32,74717 | 2,065249612 | 0,581658 | 3,550623 | 0,000384 | 0,323558126 | 1,405709 | Promoter (<=1kb) | 2 | 31560727 | 31571361 | 10635 | 1 | 227715 | ENSMUST00000038474.14 | 0 | ENSMUSG00000039356 | Exosc2 | exosome component 2 |
| chr3 | 83747837 | 83751437 | 3601 | * | 27,62046 | 2,065488876 | 0,742439 | 2,782032 | 0,005402 | 0,663569499 | 2,858511 | Promoter (<=1kb) | 3 | 83743579 | 83749074 | 5496 | 2 | 24088 | ENSMUST00000029623.11 | 0 | ENSMUSG00000027995 | Tlr2 | toll-like receptor 2 |
| chr1 | 75143371 | 75146120 | 2750 | * | 27,98712 | 2,065603925 | 0,599005 | 3,44839 | 0,000564 | 0,282588188 | 2,880017 | Promoter (<=1kb) | 1 | 75145290 | 75147396 | 2107 | 1 | 68818 | ENSMUST00000162768.8 | 0 | ENSMUSG00000026197 | Zfand2b | zinc finger, AN1 type domain 2B |
| chr17 | 32595440 | 32597872 | 2433 | * | 20,72034 | 2,06603635 | 0,709788 | 2,91078 | 0,003605 | 0,589834336 | 2,880017 | 3' UTR | 17 | 32580788 | 32588426 | 7639 | 2 | 22404 | ENSMUST00000169741.2 | -7014 | ENSMUSG00000024050 | Wiz | widely-interspaced zinc finger motifs |
| chr6 | 86503155 | 86505121 | 1967 | * | 26,00294 | 2,066095955 | 0,68217 | 3,028711 | 0,002456 | 0,514468591 | 2,880017 | Promoter (<=1kb) | 6 | 86501474 | 86503303 | 1830 | 2 | 23983 | ENSMUST00000053015.7 | 0 | ENSMUSG00000051695 | Pcbp1 | poly(rC) binding protein 1 |
| chr5 | 98573343 | 98575010 | 1668 | * | 17,09268 | 2,066216495 | 0,799309 | 2,585004 | 0,009738 | 0,90148771 | 1,2763 | Intron (ENSMUST0000 | 5 | 98623415 | 98623523 | 109 | 2 | 735265 | ENSMUST00000197105.3 | 48513 | ENSMUSG00000105814 | Mir703 | microRNA 703 |
| chr10 | 1,27E+08 | 1,27E+08 | 2790 | * | 42,49232 | 2,066438025 | 0,493429 | 4,187913 | 2,82E-05 | 0,053529752 | 2,338387 | Intron (ENSMUST0000 | 10 | 1,27E+08 | 1,27E+08 | 23107 | 2 | 216440 | ENSMUST00000218798.2 | 11411 | ENSMUSG00000040462 | Os9 | amplified in osteosarcoma |
| chr2 | 61626061 | 61628254 | 2194 | * | 19,76787 | 2,068096128 | 0,714479 | 2,894553 | 0,003797 | 0,626412475 | 2,507013 | Exon (ENSMUST00000 | 2 | 61633274 | 61634902 | 1629 | 1 | 21375 | ENSMUST00000136867.2 | -5020 | ENSMUSG00000035033 | Tbr1 | T-box brain transcription factor 1 |
| chr11 | 1,17E+08 | 1,17E+08 | 3180 | * | 41,13499 | 2,068871643 | 0,510377 | 4,053617 | 5,04E-05 | 0,078945816 | 2,377674 | Promoter (<=1kb) | 11 | 1,17E+08 | 1,17E+08 | 5850 | 2 | 107817 | ENSMUST00000141735.8 | 0 | ENSMUSG00000056962 | Jmjd6 | jumonji domain containing 6 |

| | | | | | | | | | | | | | | | | | | | | | | | |
|---|---|---|---|---|---|---|---|---|---|---|---|---|---|---|---|---|---|---|---|---|---|---|---|
| chr15 | 78727248 | 78730684 | 3437 | * | 47,1129 | 2,06890735 | 0,614524 | 3,366685 | 0,000761 | 0,346327963 | 2,377674 | Promoter (<=1kb) | 15 | 78726824 | 78735097 | 8274 | 1 | 104445 | ENSMUST00000059619.3 | 424 | ENSMUSG00000049521 | Cdc42ep1 | CDC42 effector protein (Rho GTPase binding) 1 |
| chr7 | 28792739 | 28795902 | 3164 | * | 29,1051 | 2,068964647 | 0,605246 | 3,418387 | 0,00063 | 0,303366229 | 2,725491 | Exon (ENSMUST00000 | 7 | 28783061 | 28783119 | 59 | 2 | 1E+08 | ENSMUST00000158594.3 | -9620 | ENSMUSG00000089219 | Mir1963 | microRNA 1963 |
| chr3 | 81731069 | 81733824 | 2756 | * | 17,14529 | 2,069272858 | 0,776651 | 2,664353 | 0,007714 | 0,770104803 | 2,154504 | Distal Intergenic | 3 | 81709332 | 81712468 | 3137 | 1 | 320566 | ENSMUST00000193672.2 | 21737 | ENSMUSG00000103695 | A830029E22Rik | RIKEN cDNA A830029E22 gene |
| chr7 | 80860423 | 80862920 | 2498 | * | 29,95847 | 2,070612462 | 0,600874 | 3,446001 | 0,000569 | 0,284503918 | 2,858511 | Promoter (<=1kb) | 7 | 80863344 | 80984281 | 120938 | 1 | 18584 | ENSMUST00000026672.8 | -424 | ENSMUSG00000025584 | Pde8a | phosphodiesterase 8A |
| chr8 | 85811782 | 85813872 | 2091 | * | 22,92966 | 2,070955843 | 0,758273 | 2,731147 | 0,006311 | 0,684274806 | 2,880017 | Promoter (<=1kb) | 8 | 85813087 | 85815663 | 2577 | 1 | 17159 | ENSMUST00000211223.2 | 0 | ENSMUSG00000005142 | Man2b1 | mannosidase 2, alpha B1 |
| chr15 | 79142093 | 79145166 | 3074 | * | 29,11547 | 2,071230354 | 0,654811 | 3,163096 | 0,001561 | 0,451585113 | 2,67243 | Promoter (<=1kb) | 15 | 79142531 | 79143551 | 1021 | 2 | 207495 | ENSMUST00000170254.2 | 0 | ENSMUSG00000018126 | Baiap2l2 | BAI1-associated protein 2-like 2 |
| chr18 | 33313327 | 33316021 | 2695 | * | 25,71925 | 2,071650962 | 0,614353 | 3,372085 | 0,000746 | 0,326832856 | 2,67243 | Exon (ENSMUST00000 | 18 | 33336458 | 33346819 | 10362 | 2 | 170459 | ENSMUST00000119991.8 | 30798 | ENSMUSG00000024378 | Stard4 | StAR-related lipid transfer (START) domain containing 4 |
| chr3 | 95166339 | 95169177 | 2839 | * | 25,46102 | 2,071900762 | 0,684453 | 3,027088 | 0,002469 | 0,518149475 | 2,858511 | Exon (ENSMUST00000 | 3 | 95172469 | 95176255 | 3787 | 2 | 229589 | ENSMUST00000148959.2 | 7078 | ENSMUSG00000015711 | Prune1 | prune exopolyphosphatase |
| chr13 | 73084639 | 73086111 | 1473 | * | 18,68758 | 2,072376582 | 0,754797 | 2,745609 | 0,00604 | 0,711199341 | 2,386008 | Distal Intergenic | 13 | 72957720 | 72964882 | 7163 | 2 | 78446 | ENSMUST00000223149.2 | -119757 | ENSMUSG00000113082 | D730050B12Rik | RIKEN cDNA D730050B12 gene |
| chr17 | 25488863 | 25490830 | 1968 | * | 15,82128 | 2,072521205 | 0,802609 | 2,582229 | 0,009816 | 0,793220117 | 2,386008 | Promoter (1-2kb) | 17 | 25488376 | 25492129 | 3754 | 2 | 22196 | ENSMUST00000173231.2 | 1299 | ENSMUSG00000015120 | Ube2i | ubiquitin-conjugating enzyme E2I |
| chr8 | 1,2E+08 | 1,2E+08 | 1594 | * | 33,91269 | 2,073126836 | 0,641422 | 3,232079 | 0,001229 | 0,431115599 | 2,377674 | Promoter (<=1kb) | 8 | 1,2E+08 | 1,2E+08 | 11413 | 2 | 1,03E+08 | ENSMUST00000243033.1 | 0 | | Gm32006 | predicted gene, 32006 |
| chr11 | 69424961 | 69429074 | 4114 | * | 25,27431 | 2,073322629 | 0,77108 | 2,688854 | 0,00717 | 0,718881903 | 2,67243 | Intron (ENSMUST0000 | 11 | 69311635 | 69435382 | 123748 | 2 | 327954 | ENSMUST00000035539.12 | 6308 | ENSMUSG00000005237 | Dnah2 | dynein, axonemal, heavy chain 2 |
| chr9 | 53447571 | 53449518 | 1948 | * | 19,72543 | 2,073542436 | 0,803686 | 2,58004 | 0,009879 | 0,789027175 | 2,507013 | Promoter (<=1kb) | 9 | 53350668 | 53448040 | 97373 | 2 | 11920 | ENSMUST00000232179.2 | 0 | ENSMUSG00000034218 | Atm | ataxia telangiectasia mutated |
| chr7 | 99158321 | 99160837 | 2517 | * | 20,48143 | 2,073628845 | 0,718228 | 2,887146 | 0,003888 | 0,615933766 | 2,713306 | Distal Intergenic | 7 | 99184673 | 99255978 | 71306 | 1 | 109689 | ENSMUST00000098266.9 | -23836 | ENSMUSG00000018909 | Arrb1 | arrestin, beta 1 |
| chr16 | 21928461 | 21931899 | 3439 | * | 28,03409 | 2,074065937 | 0,657042 | 3,156673 | 0,001596 | 0,454419039 | 2,67243 | Intron (ENSMUST0000 | 16 | 21906278 | 21939986 | 33709 | 2 | 319765 | ENSMUST00000232504.2 | 8087 | ENSMUSG00000033581 | Igf2bp2 | insulin-like growth factor 2 mRNA binding protein 2 |
| chr12 | 65217401 | 65221264 | 3864 | * | 30,32085 | 2,074341639 | 0,600334 | 3,455314 | 0,00055 | 0,284660899 | 2,725491 | Promoter (<=1kb) | 12 | 65179508 | 65219354 | 39847 | 2 | 217653 | ENSMUST00000052201.9 | 0 | ENSMUSG00000047534 | Mis18bp1 | MIS18 binding protein 1 |
| chr8 | 1,21E+08 | 1,21E+08 | 2790 | * | 36,52724 | 2,075057015 | 0,683562 | 3,035652 | 0,0024 | 0,517350387 | 2,787255 | Intron (ENSMUST0000 | 8 | 1,21E+08 | 1,21E+08 | 325179 | 1 | 382034 | ENSMUST00000180448.3 | 94668 | ENSMUSG00000031822 | Gse1 | genetic suppressor element 1, coiled-coil protein |
| chr13 | 21342081 | 21345005 | 2925 | * | 22,61675 | 2,075064342 | 0,687109 | 3,019992 | 0,002528 | 0,534363468 | 2,67243 | Distal Intergenic | 13 | 21326188 | 21327251 | 1064 | 2 | 258527 | ENSMUST00000055298.6 | -14830 | ENSMUSG00000045474 | Olfr1368 | olfactory receptor 1368 |
| chr12 | 1,1E+08 | 1,1E+08 | 2795 | * | 23,32984 | 2,07541194 | 0,74083 | 2,801468 | 0,005087 | 0,65166214 | 2,880017 | Distal Intergenic | 12 | 1,1E+08 | 1,1E+08 | 3880 | 1 | 26931 | ENSMUST00000222276.2 | -46621 | ENSMUSG00000017843 | Ppp2r5c | protein phosphatase 2, regulatory subunit B', gamma |
| chr10 | 81440805 | 81443515 | 2711 | * | 34,98156 | 2,075829748 | 0,574136 | 3,615573 | 0,0003 | 0,206624632 | 2,787255 | Promoter (<=1kb) | 10 | 81442104 | 81456537 | 14434 | 1 | 268319 | ENSMUST00000126940.2 | 0 | ENSMUSG00000074862 | BC025920 | zinc finger protein pseudogene |
| chr6 | 91178153 | 91180820 | 2668 | * | 38,66201 | 2,07783088 | 0,553177 | 3,756177 | 0,000173 | 0,16596695 | 2,377674 | Distal Intergenic | 6 | 91189437 | 91249521 | 60085 | 1 | 14115 | ENSMUST00000113498.9 | -8617 | ENSMUSG00000064080 | Fbln2 | fibulin 2 |
| chr8 | 86159397 | 86163008 | 3612 | * | 33,21731 | 2,078115445 | 0,63924 | 3,250917 | 0,00115 | 0,507398906 | 1,405709 | Promoter (<=1kb) | 8 | 86135388 | 86159462 | 24075 | 2 | 66714 | ENSMUST00000047749.7 | 0 | ENSMUSG00000036934 | 4921524J17Rik | RIKEN cDNA 4921524J17 gene |
| chr17 | 84505485 | 84506568 | 1084 | * | 19,73269 | 2,078448005 | 0,719698 | 2,887944 | 0,003878 | 0,639496036 | 2,386008 | Intron (ENSMUST0000 | 17 | 84491359 | 84495375 | 4017 | 2 | 12193 | ENSMUST00000060366.7 | -10110 | ENSMUSG00000045817 | Zfp36l2 | zinc finger protein 36, C3H type-like 2 |
| chr12 | 79950627 | 79953215 | 2589 | * | 21,44355 | 2,078994499 | 0,756892 | 2,746753 | 0,006019 | 0,686762073 | 2,67243 | Distal Intergenic | 12 | 79938062 | 79938677 | 616 | 1 | 1,08E+08 | ENSMUST00000243367.1 | 12565 | | Gm46362 | predicted gene, 46362 |
| chr19 | 45269629 | 45272397 | 2769 | * | 24,47162 | 2,079528088 | 0,779297 | 2,668467 | 0,00762 | 0,716908183 | 2,880017 | Distal Intergenic | 19 | 45288909 | 45291776 | 2868 | 2 | 1,03E+08 | ENSMUST00000187954.3 | 19379 | ENSMUSG00000100417 | Gm29595 | predicted gene 29595 |
| chr11 | 78970538 | 78975029 | 4492 | * | 40,14412 | 2,079528152 | 0,639891 | 3,249815 | 0,001155 | 0,396931054 | 2,787255 | Intron (ENSMUST0000 | 11 | 78921451 | 78965312 | 43862 | 2 | 16706 | ENSMUST00000208969.2 | -5226 | ENSMUSG00000018334 | Ksr1 | kinase suppressor of ras 1 |
| chr11 | 1,07E+08 | 1,07E+08 | 3033 | * | 23,45586 | 2,079565987 | 0,723018 | 2,876231 | 0,004025 | 0,61075929 | 2,880017 | Promoter (1-2kb) | 11 | 1,07E+08 | 1,07E+08 | 8711 | 2 | 66313 | ENSMUST00000144027.2 | 1549 | ENSMUSG00000018363 | Smurf2 | SMAD specific E3 ubiquitin protein ligase 2 |
| chr3 | 1,54E+08 | 1,54E+08 | 1622 | * | 17,45513 | 2,079676034 | 0,753643 | 2,759498 | 0,005789 | 0,701807025 | 2,386008 | Intron (ENSMUST0000 | 3 | 1,54E+08 | 1,54E+08 | 2219 | 1 | 209601 | ENSMUST00000192912.2 | 38529 | ENSMUSG00000078161 | Erich3 | glutamate rich 3 |
| chr11 | 19960694 | 19962882 | 2189 | * | 25,79814 | 2,07979488 | 0,650922 | 3,195153 | 0,001398 | 0,422040194 | 2,858511 | Intron (ENSMUST0000 | 11 | 19904570 | 19968133 | 63564 | 1 | 114716 | ENSMUST00000123036.2 | 56124 | ENSMUSG00000045671 | Spred2 | sprouty-related EVH1 domain containing 2 |
| chr13 | 18330526 | 18331915 | 1390 | * | 15,85163 | 2,08019753 | 0,783747 | 2,654168 | 0,00795 | 0,777172375 | 2,141612 | Intron (ENSMUST0000 | 13 | 18299709 | 18347120 | 47412 | 2 | 218030 | ENSMUST00000176795.8 | 15205 | ENSMUSG00000009734 | Pou6f2 | POU domain, class 6, transcription factor 2 |
| chr12 | 1,09E+08 | 1,09E+08 | 2326 | * | 24,04877 | 2,080239569 | 0,781364 | 2,662317 | 0,00776 | 0,731734453 | 2,67243 | Intron (ENSMUST0000 | 12 | 1,09E+08 | 1,09E+08 | 9411 | 2 | 22375 | ENSMUST00000162952.2 | 3994 | ENSMUSG00000021266 | Wars | tryptophanyl-tRNA synthetase |
| chr15 | 27782654 | 27789342 | 6689 | * | 80,078316 | 2,082241923 | 0,4566233 | 4,5600867 | 5,113E-06 | 0,010095388 | 4,0557576 | Promoter (<=1kb) | 15 | 27773974 | 27788724 | 14751 | 2 | 223435 | ENSMUST00000226287.2 | 0 | ENSMUSG00000022263 | Trio | triple functional domain (PTPRF interacting) |
| chr5 | 23764502 | 23767701 | 3200 | * | 36,61624 | 2,083817387 | 0,542146 | 3,843645 | 0,000121 | 0,133396034 | 2,377674 | Intron (ENSMUST0000 | 5 | 23730729 | 23747116 | 16388 | 2 | 20817 | ENSMUST00000200173.2 | -17386 | ENSMUSG00000062604 | Srpk2 | serine/arginine-rich protein specific kinase 2 |
| chr5 | 28945662 | 28947752 | 2091 | * | 18,11637 | 2,084269766 | 0,776302 | 2,684868 | 0,007256 | 0,756849896 | 2,154504 | Exon (ENSMUST00000 | 5 | 28661813 | 28672254 | 10442 | 2 | 20423 | ENSMUST00000002708.5 | -273408 | ENSMUSG00000002633 | Shh | sonic hedgehog |
| chr10 | 82501568 | 82503770 | 2203 | * | 19,59498 | 2,084522131 | 0,794237 | 2,62456 | 0,008676 | 0,787687627 | 2,221611 | Promoter (2-3kb) | 10 | 82500471 | 82506302 | 5832 | 2 | 74782 | ENSMUST00000155529.2 | 2532 | ENSMUSG00000020251 | Glt8d2 | glycosyltransferase 8 domain containing 2 |
| chr3 | 95139602 | 95141796 | 2195 | * | 21,93055 | 2,084523336 | 0,795016 | 2,62199 | 0,008742 | 0,751314174 | 2,67243 | Promoter (<=1kb) | 3 | 95139604 | 95141973 | 2370 | 1 | 57912 | ENSMUST00000107199.2 | 0 | ENSMUSG00000046722 | Cdc42se1 | CDC42 small effector 1 |
| chr2 | 1,68E+08 | 1,68E+08 | 3022 | * | 25,25477 | 2,084553194 | 0,672637 | 3,099074 | 0,001941 | 0,488164756 | 2,67243 | Intron (ENSMUST0000 | 2 | 1,68E+08 | 1,68E+08 | 1504 | 1 | 12608 | ENSMUST00000070642.4 | -5815 | ENSMUSG00000056501 | Cebpb | CCAAT/enhancer binding protein (C/EBP), beta |
| chr2 | 1,22E+08 | 1,22E+08 | 1974 | * | 18,61742 | 2,084596976 | 0,69052 | 3,01888 | 0,002537 | 0,55292773 | 2,386008 | Promoter (<=1kb) | 2 | 1,22E+08 | 1,22E+08 | 18328 | 1 | 70928 | ENSMUST00000036089.8 | 0 | ENSMUSG00000033368 | Trim69 | tripartite motif-containing 69 |
| chrX | 57679901 | 57682560 | 2660 | * | 17,93668 | 2,0848299 | 0,764151 | 2,728297 | 0,006366 | 0,697903123 | 2,663619 | Distal Intergenic | 20 | 57954720 | 57965664 | 10945 | 1 | 75352 | ENSMUST00000118305.8 | -272160 | ENSMUSG00000046180 | Magea13 | MAGE family member A13 |
| chr7 | 1,39E+08 | 1,39E+08 | 1790 | * | 18,15593 | 2,085883227 | 0,746746 | 2,793298 | 0,005217 | 0,683210259 | 2,423594 | Promoter (<=1kb) | 7 | 1,39E+08 | 1,39E+08 | 121434 | 1 | 74004 | ENSMUST00000106111.9 | -79 | ENSMUSG00000056856 | Jakmip3 | janus kinase and microtubule interacting protein 3 |
| chr4 | 1,34E+08 | 1,34E+08 | 3182 | * | 25,65637 | 2,086084856 | 0,722109 | 2,888877 | 0,003866 | 0,613419803 | 2,725491 | Promoter (<=1kb) | 4 | 1,34E+08 | 1,34E+08 | 22596 | 1 | 67586 | ENSMUST00000131329.8 | -326 | ENSMUSG00000012126 | Ubxn11 | UBX domain protein 11 |
| chr17 | 47221874 | 47224438 | 2565 | * | 34,15658 | 2,08615221 | 0,59255 | 3,520636 | 0,000431 | 0,273126507 | 2,377674 | Promoter (<=1kb) | 17 | 47221404 | 47231273 | 9870 | 1 | 19133 | ENSMUST00000162469.2 | 470 | ENSMUSG00000023978 | Prph2 | peripherin 2 |
| chr15 | 68102447 | 68103988 | 1542 | * | 19,3802 | 2,08648127 | 0,786829 | 2,651761 | 0,008007 | 0,75632407 | 2,386008 | Intron (ENSMUST0000 | 15 | 67956170 | 68130636 | 174467 | 2 | 380993 | ENSMUST00000162173.8 | 26648 | ENSMUSG00000022335 | Zfat | zinc finger and AT hook domain containing |
| chr7 | 1,05E+08 | 1,05E+08 | 2187 | * | 22,35664 | 2,086493681 | 0,672863 | 3,100918 | 0,001929 | 0,461973573 | 3,104559 | Distal Intergenic | 7 | 1,05E+08 | 1,05E+08 | 2218 | 2 | 109042 | ENSMUST00000047040.4 | 5777 | ENSMUSG00000037060 | Cavin3 | caveolae associated 3 |
| chr11 | 59302785 | 59306479 | 3695 | * | 34,01539 | 2,086506857 | 0,562683 | 3,708141 | 0,000209 | 0,184990586 | 2,377674 | Intron (ENSMUST0000 | 11 | 59329082 | 59338317 | 9236 | 2 | 67826 | ENSMUST00000145550.2 | 31838 | ENSMUSG00000009894 | Snap47 | synaptosomal-associated protein, 47 |
| chr3 | 1,29E+08 | 1,29E+08 | 2009 | * | 21,10687 | 2,087306513 | 0,737585 | 2,829922 | 0,004656 | 0,649854434 | 2,67243 | Distal Intergenic | 3 | 1,29E+08 | 1,29E+08 | 19713 | 1 | 18741 | ENSMUST00000174661.9 | -264459 | ENSMUSG00000028023 | Pitx2 | paired-like homeodomain transcription factor 2 |
| chr18 | 53399742 | 53401848 | 2107 | * | 23,91482 | 2,088239264 | 0,663843 | 3,145684 | 0,001657 | 0,445842166 | 2,951062 | Intron (ENSMUST0000 | 18 | 53378841 | 53523895 | 145055 | 1 | 69226 | ENSMUST00000025417.9 | 20901 | ENSMUSG00000024535 | Snx24 | sorting nexing 24 |
| chr9 | 21222962 | 21226321 | 3360 | * | 33,44524 | 2,088741602 | 0,672853 | 3,104307 | 0,001907 | 0,504427013 | 2,377674 | Promoter (<=1kb) | 9 | 21205571 | 21223612 | 18042 | 2 | 11768 | ENSMUST00000213762.2 | 0 | ENSMUSG00000003309 | Ap1m2 | adaptor protein complex AP-1, mu 2 subunit |
| chr17 | 5708396 | 5709866 | 1471 | * | 17,06201 | 2,088956475 | 0,767184 | 2,72289 | 0,006471 | 0,702222274 | 2,663619 | Intron (ENSMUST0000 | 17 | 5848932 | 5853168 | 4237 | 1 | 78512 | ENSMUST00000180924.8 | -139066 | ENSMUSG00000096965 | 3300005D01Rik | RIKEN cDNA 3300005D01 gene |
| chr5 | 1,29E+08 | 1,29E+08 | 1749 | * | 20,8659 | 2,089232534 | 0,768282 | 2,719356 | 0,006541 | 0,684391766 | 2,951933 | Distal Intergenic | 5 | 1,29E+08 | 1,29E+08 | 3004 | 2 | 13852 | ENSMUST00000128448.2 | 4726 | ENSMUSG00000029428 | Stx2 | syntaxin 2 |
| chr12 | 1,11E+08 | 1,11E+08 | 2307 | * | 23,97967 | 2,089473844 | 0,765777 | 2,728567 | 0,006361 | 0,683337777 | 2,951062 | 3' UTR | 12 | 1,11E+08 | 1,11E+08 | 9926 | 1 | 69641 | ENSMUST00000194118.2 | 13652 | ENSMUSG00000037957 | Wdr20 | WD repeat domain 20 |
| chr13 | 55611867 | 55614659 | 2793 | * | 22,85571 | 2,08961349 | 0,707274 | 2,954461 | 0,003132 | 0,56254855 | 2,858511 | Promoter (<=1kb) | 13 | 55612080 | 55620967 | 8888 | 1 | 432763 | ENSMUST00000046533.9 | 0 | ENSMUSG00000034686 | Prr7 | proline rich 7 (synaptic) |
| chr17 | 57338420 | 57341861 | 3442 | * | 64,76454 | 2,089729685 | 0,517845 | 4,035433 | 5,45E-05 | 0,068345002 | 3,177995 | Promoter (<=1kb) | 17 | 57328051 | 57338488 | 10438 | 2 | 16549 | ENSMUST00000007814.10 | 0 | ENSMUSG00000007670 | Khsrp | KH-type splicing regulatory protein |
| chr2 | 59959430 | 59960425 | 996 | * | 16,67879 | 2,089949934 | 0,809308 | 2,58239 | 0,009812 | 0,775744126 | 2,663619 | Intron (ENSMUST0000 | 2 | 59809019 | 59956084 | 147066 | 2 | 407823 | ENSMUST00000137480.8 | -3346 | ENSMUSG00000026987 | Baz2b | bromodomain adjacent to zinc finger domain, 2B |
| chr11 | 1,21E+08 | 1,21E+08 | 3901 | * | 35,56226 | 2,090043618 | 0,678694 | 3,079507 | 0,002073 | 0,483145429 | 2,925497 | Promoter (<=1kb) | 11 | 1,21E+08 | 1,21E+08 | 2558 | 2 | 209011 | ENSMUST00000146809.2 | 0 | ENSMUSG00000025138 | Sirt7 | sirtuin 7 |
| chr8 | 1,12E+08 | 1,12E+08 | 1428 | * | 30,35993 | 2,090162063 | 0,58947 | 3,545831 | 0,000391 | 0,243032283 | 2,725491 | Promoter (1-2kb) | 8 | 1,12E+08 | 1,12E+08 | 12714 | 2 | 12927 | ENSMUST00000212147.2 | 1315 | ENSMUSG00000031955 | Bcar1 | breast cancer anti-estrogen resistance 1 |
| chr8 | 95716210 | 95719240 | 3031 | * | 37,39954 | 2,090545773 | 0,622195 | 3,359952 | 0,00078 | 0,328946709 | 2,725491 | Promoter (1-2kb) | 8 | 95720902 | 95732528 | 11627 | 1 | 14766 | ENSMUST00000212141.2 | -1662 | ENSMUSG00000031785 | Adgrg1 | adhesion G protein-coupled receptor G1 |
| chr2 | 92580560 | 92582976 | 2417 | * | 31,4485 | 2,091928279 | 0,633697 | 3,301149 | 0,000963 | 0,367735927 | 2,67243 | Intron (ENSMUST0000 | 2 | 92572575 | 92589553 | 16979 | 1 | 1,05E+08 | ENSMUST00000152230.2 | 7985 | | Gm13816 | predicted gene 13816 |
| chr14 | 37530650 | 37531922 | 1273 | * | 16,27881 | 2,092335878 | 0,737524 | 2,836973 | 0,004554 | 0,665572106 | 2,386008 | Distal Intergenic | 14 | 37323064 | 37328745 | 5682 | 2 | 1,03E+08 | ENSMUST00000224767.2 | -201905 | ENSMUSG00000114334 | Gm31393 | predicted gene, 31393 |
| chr5 | 24589498 | 24591342 | 1845 | * | 17,72334 | 2,093003949 | 0,789704 | 2,650364 | 0,008041 | 0,749550311 | 2,507013 | Promoter (1-2kb) | 5 | 24592103 | 24592964 | 862 | 2 | 213948 | ENSMUST00000128831.2 | 1622 | ENSMUSG00000038295 | Atg9b | autophagy related 9B |
| chr5 | 1,37E+08 | 1,37E+08 | 3726 | * | 32,75857 | 2,093348416 | 0,561941 | 3,725207 | 0,000195 | 0,157661852 | 2,925497 | Distal Intergenic | 5 | 1,37E+08 | 1,37E+08 | 10564 | 2 | 384309 | ENSMUST00000054384.6 | 12025 | ENSMUSG00000043279 | Trim56 | tripartite motif-containing 56 |
| chr17 | 74847326 | 74849434 | 2109 | * | 20,73918 | 2,093364588 | 0,685119 | 3,055474 | 0,002247 | 0,496560937 | 2,951933 | Intron (ENSMUST0000 | 17 | 74853193 | 74856103 | 2911 | 1 | 12211 | ENSMUST00000182631.2 | -3759 | ENSMUSG00000024073 | Birc6 | baculoviral IAP repeat-containing 6 |
| chr11 | 1,15E+08 | 1,15E+08 | 2956 | * | 26,94716 | 2,093487913 | 0,756476 | 2,767422 | 0,00565 | 0,678869598 | 2,725491 | Promoter (1-2kb) | 11 | 1,15E+08 | 1,15E+08 | 749 | 1 | 68572 | ENSMUST00000134386.2 | 1116 | ENSMUSG00000018858 | Mrpl58 | mitochondrial ribosomal protein L58 |
| chr6 | 64850469 | 64851607 | 1139 | * | 16,92506 | 2,093620165 | 0,803146 | 2,606774 | 0,00914 | 0,761148741 | 2,663619 | Distal Intergenic | 6 | 64706109 | 64708229 | 2121 | 1 | 11921 | ENSMUST00000101351.6 | 144360 | ENSMUSG00000073043 | Atoh1 | atonal bHLH transcription factor 1 |
| chr19 | 6101525 | 6104342 | 2818 | * | 26,72595 | 2,093670498 | 0,795179 | 2,632955 | 0,008465 | 0,745401733 | 2,725491 | Promoter (<=1kb) | 19 | 6101364 | 6101422 | 59 | 1 | 1,02E+08 | ENSMUST00000184338.3 | 161 | ENSMUSG00000098928 | Mir6988 | microRNA 6988 |
| chr4 | 1,39E+08 | 1,39E+08 | 3536 | * | 26,60872 | 2,093842958 | 0,684106 | 3,0607 | 0,002208 | 0,497406263 | 2,880017 | Exon (ENSMUST00000 | 4 | 1,39E+08 | 1,39E+08 | 18567 | 2 | 73162 | ENSMUST00000097830.4 | 12467 | ENSMUSG00000041161 | Otud3 | OTU domain containing 3 |
| chr4 | 1,41E+08 | 1,41E+08 | 3142 | * | 24,46068 | 2,093880863 | 0,676272 | 3,09621 | 0,00196 | 0,474289497 | 2,951062 | Promoter (<=1kb) | 4 | 1,41E+08 | 1,41E+08 | 8629 | 1 | 18600 | ENSMUST00000148160.2 | 0 | ENSMUSG00000028927 | Padi2 | peptidyl arginine deiminase, type II |
| chr18 | 33924276 | 33926522 | 2247 | * | 14,67032 | 2,094281644 | 0,801055 | 2,614404 | 0,008938 | 0,86955927 | 1,41696 | Promoter (1-2kb) | 18 | 33927889 | 33929050 | 1162 | 1 | 69749 | ENSMUST00000146010.4 | -1367 | ENSMUSG00000087590 | Epb41l4aos | erythrocyte membrane protein band 4.1 like 4a, opposite strand |
| chr4 | 1,18E+08 | 1,18E+08 | 2284 | * | 22,80931 | 2,09462634 | 0,786447 | 2,663405 | 0,007735 | 0,715249603 | 2,951933 | Promoter (<=1kb) | 4 | 1,18E+08 | 1,18E+08 | 20514 | 2 | 230673 | ENSMUST00000036156.6 | 0 | ENSMUSG00000033365 | Ipo13 | importin 13 |
| chr8 | 41591378 | 41594021 | 2644 | * | 27,38099 | 2,0948371 | 0,682965 | 3,067269 | 0,00216 | 0,502454322 | 2,725491 | Distal Intergenic | 8 | 41503137 | 41586763 | 83627 | 2 | 102103 | ENSMUST00000145860.2 | -4615 | ENSMUSG00000045636 | Mtus1 | mitochondrial tumor suppressor 1 |
| chr5 | 1,41E+08 | 1,41E+08 | 2140 | * | 25,24774 | 2,094908646 | 0,770925 | 2,717395 | 0,00658 | 0,685026746 | 2,951062 | 3' UTR | 5 | 1,41E+08 | 1,41E+08 | 12157 | 1 | 231842 | ENSMUST00000176035.2 | 9689 | ENSMUSG00000050022 | Amz1 | archaelysin family metallopeptidase 1 |
| chr16 | 94028254 | 94030574 | 2321 | * | 28,99014 | 2,094921456 | 0,61133 | 3,426825 | 0,000611 | 0,293348204 | 2,880017 | Exon (ENSMUST00000 | 16 | 93935526 | 94023976 | 88451 | 2 | 110948 | ENSMUST00000227698.2 | -4278 | ENSMUSG00000040820 | Hlcs | holocarboxylase synthetase (biotin- [propriony-Coenzyme A-carboxylase (ATP-hydrolysing)] ligase) |
| chr15 | 15935823 | 15937246 | 1424 | * | 16,87883 | 2,095173747 | 0,747472 | 2,803012 | 0,005063 | 0,664469307 | 2,663619 | Distal Intergenic | 15 | 15819313 | 15819440 | 128 | 1 | 1,15E+08 | ENSMUST00000158079.4 | 116510 | ENSMUSG00000088704 | Gm25550 | predicted gene, 25550 |
| chr2 | 29502066 | 29504269 | 2204 | * | 18,927 | 2,095628834 | 0,770195 | 2,720907 | 0,00651 | 0,698702076 | 2,713306 | Distal Intergenic | 2 | 29509732 | 29630365 | 120634 | 1 | 107746 | ENSMUST00000095087.11 | -5463 | ENSMUSG00000039844 | Rapgef1 | Rap guanine nucleotide exchange factor (GEF) 1 |
| chr7 | 1,14E+08 | 1,14E+08 | 3100 | * | 42,29428 | 2,095714595 | 0,513521 | 4,08107 | 4,48E-05 | 0,063587778 | 2,925497 | Intron (ENSMUST0000 | 7 | 1,14E+08 | 1,14E+08 | 19871 | 1 | 233752 | ENSMUST00000136347.2 | 6270 | ENSMUSG00000048782 | Insc | INSC spindle orientation adaptor protein |
| chr19 | 59321631 | 59324801 | 3171 | * | 19,35215 | 2,09641974 | 0,805249 | 2,603441 | 0,009229 | 0,785866008 | 2,386008 | Intron (ENSMUST0000 | 19 | 59285610 | 59334212 | 48603 | 2 | 107368 | ENSMUST00000099274.4 | 9411 | ENSMUSG00000074746 | Pdzd8 | PDZ domain containing 8 |
| chr16 | 92348261 | 92350507 | 2247 | * | 24,8287 | 2,096488445 | 0,637204 | 3,290139 | 0,001001 | 0,36185416 | 2,858511 | Exon (ENSMUST00000 | 16 | 92295022 | 92338131 | 43110 | 1 | 209195 | ENSMUST00000023670.4 | 53239 | ENSMUSG00000022949 | Clic6 | chloride intracellular channel 6 |
| chr19 | 36158806 | 36161650 | 2845 | * | 26,24516 | 2,09655094 | 0,607284 | 3,452337 | 0,000556 | 0,278016366 | 2,951062 | Promoter (<=1kb) | 19 | 36159980 | 36199018 | 39039 | 1 | 1,03E+08 | ENSMUST00000236660.2 | 0 | ENSMUSG00000117998 | Gm32081 | predicted gene, 32081 |
| chr2 | 57005109 | 57007974 | 2866 | * | 33,93474 | 2,096986395 | 0,621194 | 3,375734 | 0,000736 | 0,312110535 | 2,925497 | Promoter (<=1kb) | 2 | 57002015 | 57005050 | 3036 | 2 | 18227 | ENSMUST00000140165.2 | -59 | ENSMUSG00000026826 | Nr4a2 | nuclear receptor subfamily 4, group A, member 2 |
| chr18 | 65329364 | 65330600 | 1237 | * | 17,74876 | 2,097054592 | 0,776893 | 2,699284 | 0,006949 | 0,730469267 | 2,423594 | Intron (ENSMUST0000 | 18 | 65336936 | 65345920 | 8985 | 1 | 83814 | ENSMUST00000237818.2 | -6336 | ENSMUSG00000024589 | Nedd4l | neural precursor cell expressed, developmentally down-regulated gene 4-like |
| chr18 | 32277954 | 32281064 | 3111 | * | 28,32147 | 2,097075411 | 0,628837 | 3,334845 | 0,000853 | 0,331767406 | 2,951062 | Distal Intergenic | 18 | 32266200 | 32272623 | 6424 | 2 | 19123 | ENSMUST00000234375.2 | -5331 | ENSMUSG00000024386 | Proc | protein C |
| chr17 | 36129567 | 36131638 | 2072 | * | 22,87102 | 2,097591861 | 0,652074 | 3,216803 | 0,001296 | 0,421090004 | 2,67243 | Promoter (<=1kb) | 17 | 36132576 | 36133815 | 1240 | 1 | 15937 | ENSMUST00000003635.7 | -938 | ENSMUSG00000003541 | Ier3 | immediate early response 3 |
| chr18 | 8422743 | 8425199 | 2457 | * | 18,45121 | 2,098647678 | 0,734751 | 2,85627 | 0,004287 | 0,651710641 | 2,423594 | Distal Intergenic | 18 | 7910619 | 7918503 | 7885 | 1 | 225131 | ENSMUST00000167542.2 | 512124 | ENSMUSG00000024283 | Wac | WW domain containing adaptor with coiled-coil |
| chr19 | 26822229 | 26824168 | 1940 | * | 22,43313 | 2,099124237 | 0,65196 | 3,219715 | 0,001283 | 0,372510154 | 3,471678 | Distal Intergenic | 19 | 26746201 | 26801307 | 55107 | 2 | 70942 | ENSMUST00000177048.2 | -20922 | ENSMUSG00000093772 | 4931403E22Rik | RIKEN cDNA 4931403E22 gene |
| chr19 | 4559627 | 4560792 | 1166 | * | 30,70128 | 2,100131038 | 0,669731 | 3,135784 | 0,001714 | 0,454149353 | 2,880017 | Promoter (<=1kb) | 19 | 4560500 | 4671780 | 111281 | 1 | 18563 | ENSMUST00000068004.13 | 0 | ENSMUSG00000024892 | Pcx | pyruvate carboxylase |
| chr18 | 75489777 | 75491538 | 1762 | * | 17,27501 | 2,100156227 | 0,794077 | 2,644775 | 0,008175 | 0,757118792 | 2,423594 | Distal Intergenic | 18 | 75491711 | 75500160 | 8450 | 2 | 1,03E+08 | ENSMUST00000174338.3 | 8622 | | Gm20544 | predicted gene 20544 |
| chr7 | 29664574 | 29667121 | 2548 | * | 20,10628 | 2,100613399 | 0,808045 | 2,599623 | 0,009333 | 0,766329882 | 2,663619 | Exon (ENSMUST00000 | 7 | 29656427 | 29669046 | 12620 | 1 | 625421 | ENSMUST00000238888.2 | 8147 | | C230062I16Rik | RIKEN cDNA C230062I16 gene |
| chr7 | 1,32E+08 | 1,32E+08 | 2064 | * | 20,16554 | 2,100734482 | 0,761185 | 2,75982 | 0,005783 | 0,683117834 | 2,713306 | Promoter (<=1kb) | 7 | 1,32E+08 | 1,32E+08 | 23445 | 2 | 77938 | ENSMUST00000148303.2 | 540 | ENSMUSG00000030956 | Fam53b | family with sequence similarity 53, member B |
| chr7 | 1,26E+08 | 1,26E+08 | 2485 | * | 30,56881 | 2,100989352 | 0,698575 | 3,007535 | 0,002634 | 0,54122637 | 2,67243 | Promoter (<=1kb) | 7 | 1,26E+08 | 1,26E+08 | 2428 | 2 | 56312 | ENSMUST00000032961.4 | 0 | ENSMUSG00000030717 | Nupr1 | nuclear protein transcription regulator 1 |
| chr1 | 33095484 | 33097308 | 1825 | * | 14,56564 | 2,101247569 | 0,804867 | 2,610678 | 0,009036 | 0,798887389 | 2,141612 | Distal Intergenic | 1 | 33231074 | 33231183 | 110 | 1 | 1,15E+08 | ENSMUST00000158125.3 | -133766 | ENSMUSG00000088750 | Gm24901 | predicted gene, 24901 |
| chr3 | 84554505 | 84556049 | 1545 | * | 16,42968 | 2,101282747 | 0,801251 | 2,622504 | 0,008729 | 0,885719815 | 1,2763 | Intron (ENSMUST0000 | 3 | 84573499 | 84611882 | 38384 | 1 | 320782 | ENSMUST00000107682.2 | -17450 | ENSMUSG00000056498 | Tmem154 | transmembrane protein 154 |
| chr15 | 7810659 | 7812061 | 1403 | * | 16,04044 | 2,102423079 | 0,796967 | 2,638029 | 0,008339 | 0,770673161 | 2,318434 | Distal Intergenic | 15 | 7840327 | 7867056 | 26730 | 1 | 14573 | ENSMUST00000022744.5 | -28266 | ENSMUSG00000022144 | Gdnf | glial cell line derived neurotrophic factor |
| chr2 | 59811714 | 59813589 | 1876 | * | 21,51672 | 2,10280655 | 0,772407 | 2,722409 | 0,006481 | 0,661775851 | 3,471678 | Intron (ENSMUST0000 | 2 | 59783806 | 59790387 | 6582 | 2 | 407823 | ENSMUST00000135722.2 | -21327 | ENSMUSG00000026987 | Baz2b | bromodomain adjacent to zinc finger domain, 2B |
| chr8 | 1,23E+08 | 1,23E+08 | 3352 | * | 47,39442 | 2,102836079 | 0,526027 | 3,997578 | 6,4E-05 | 0,082076621 | 2,787255 | Distal Intergenic | 8 | 1,23E+08 | 1,23E+08 | 7803 | 1 | 195209 | ENSMUST00000247413.1 | 45061 | ENSMUSG00000043903 | Zfp469 | zinc finger protein 469 |
| chr15 | 1,02E+08 | 1,02E+08 | 1960 | * | 16,63909 | 2,103161029 | 0,784064 | 2,682386 | 0,00731 | 0,759970924 | 2,141612 | 3' UTR | 15 | 1,02E+08 | 1,02E+08 | 4313 | 1 | 1,03E+08 | ENSMUST00000229977.2 | 4049 | ENSMUSG00000116402 | Gm36246 | predicted gene, 36246 |
| chr12 | 1,11E+08 | 1,11E+08 | 2230 | * | 20,2662 | 2,103440929 | 0,762191 | 2,759728 | 0,005785 | 0,691182187 | 2,507013 | Promoter (2-3kb) | 12 | 1,11E+08 | 1,11E+08 | 13762 | 2 | 66323 | ENSMUST00000222915.2 | 2398 | ENSMUSG00000056508 | 1700001K19Rik | RIKEN cDNA 1700001K19 gene |
| chr10 | 41028125 | 41030487 | 2363 | * | 21,73122 | 2,103652711 | 0,773219 | 2,720643 | 0,006516 | 0,684274806 | 2,951933 | Distal Intergenic | 10 | 40979000 | 41014192 | 35193 | 2 | 1,03E+08 | ENSMUST00000214352.2 | -13933 | | Gm32338 | predicted gene, 32338 |
| chr4 | 1,25E+08 | 1,25E+08 | 2954 | * | 23,61819 | 2,103681332 | 0,7926 | 2,654153 | 0,007951 | 0,736703271 | 2,67243 | Promoter (2-3kb) | 4 | 1,25E+08 | 1,25E+08 | 2230 | 1 | 230737 | ENSMUST00000147550.2 | -2999 | ENSMUSG00000028869 | Gnl2 | guanine nucleotide binding protein-like 2 (nucleolar) |
| chr2 | 1,55E+08 | 1,55E+08 | 2337 | * | 32,69383 | 2,103971007 | 0,673923 | 3,121975 | 0,001796 | 0,594430593 | 1,405709 | Promoter (<=1kb) | 2 | 1,55E+08 | 1,55E+08 | 13465 | 2 | 207182 | ENSMUST00000147601.2 | 0 | ENSMUSG00000027603 | Ggt7 | gamma-glutamyltransferase 7 |
| chr2 | 27874542 | 27875911 | 1370 | * | 25,77492 | 2,104116172 | 0,737056 | 2,854759 | 0,004307 | 0,625178494 | 2,858511 | Exon (ENSMUST00000 | 2 | 27907453 | 27927027 | 19575 | 1 | 12831 | ENSMUST00000145423.2 | -31542 | ENSMUSG00000026837 | Col5a1 | collagen, type V, alpha 1 |
| chr2 | 25429113 | 25433112 | 4000 | * | 76,355228 | 2,104569737 | 0,4224202 | 4,9821712 | 6,287E-07 | 0,002468468 | 3,177995 | Promoter (<=1kb) | 2 | 25408878 | 25429152 | 20275 | 2 | 22030 | ENSMUST00000151742.2 | 0 | ENSMUSG00000026942 | Traf2 | TNF receptor-associated factor 2 |

| | | | | | | | | | | | | | | | | | | | | | | | |
|---|---|---|---|---|---|---|---|---|---|---|---|---|---|---|---|---|---|---|---|---|---|---|---|
| chr5 | 21349467 | 21351354 | 1888 | * | 19,20268 | 2,104632409 | 0,794191 | 2,650033 | 0,008048 | 0,749550311 | 2,507013 | Distal Intergenic | 5 | 21322081 | 21322199 | 119 | 2 | 1,15E+08 | ENSMUST00000157188.3 | -27268 | ENSMUSG00000087813 | Gm22490 | predicted gene, 22490 |
| chr2 | 25423825 | 25425830 | 2006 | * | 25,51746 | 2,105094317 | 0,622085 | 3,383935 | 0,000715 | 0,30807497 | 2,951062 | Intron (ENSMUST0000 | 2 | 25408878 | 25429152 | 20275 | 2 | 22030 | ENSMUST00000151742.2 | 3322 | ENSMUSG00000026942 | Traf2 | TNF receptor-associated factor 2 |
| chr5 | 91081876 | 91083792 | 1917 | * | 15,68127 | 2,105679982 | 0,756515 | 2,783395 | 0,005379 | 0,798887389 | 1,2763 | Promoter (2-3kb) | 5 | 91079139 | 91169227 | 90089 | 1 | 665563 | ENSMUST00000071652.6 | 2737 | ENSMUSG00000029376 | Mthfd2l | methylenetetrahydrofolate dehydrogenase (NADP+ dependent) 2-like |
| chr1 | 20918037 | 20919679 | 1643 | * | 17,58961 | 2,105738218 | 0,798484 | 2,637169 | 0,00836 | 0,766329882 | 2,386008 | Distal Intergenic | 1 | 20912759 | 20912890 | 132 | 2 | 1,15E+08 | ENSMUST00000158233.3 | -5147 | ENSMUSG00000088858 | Gm24723 | predicted gene, 24723 |
| chr6 | 90724183 | 90726322 | 2140 | * | 22,16281 | 2,105758829 | 0,767108 | 2,745061 | 0,00605 | 0,686839406 | 2,67243 | Intron (ENSMUST0000 | 6 | 90671581 | 90734532 | 62952 | 2 | 232227 | ENSMUST00000146977.2 | 8210 | ENSMUSG00000034312 | Iqsec1 | IQ motif and Sec7 domain 1 |
| chr12 | 76883770 | 76885279 | 1510 | * | 20,50133 | 2,106023272 | 0,779342 | 2,702309 | 0,006886 | 0,744477896 | 2,221611 | Promoter (<=1kb) | 12 | 76884188 | 76968186 | 83999 | 1 | 110606 | ENSMUST00000041008.10 | 0 | ENSMUSG00000033373 | Fntb | farnesyltransferase, CAAX box, beta |
| chr7 | 1,17E+08 | 1,17E+08 | 3309 | * | 26,18999 | 2,106052424 | 0,712238 | 2,95695 | 0,003107 | 0,571756101 | 2,67243 | Distal Intergenic | 7 | 1,18E+08 | 1,18E+08 | 5566 | 2 | 267019 | ENSMUST00000134270.2 | 251922 | ENSMUSG00000008683 | Rps15a | ribosomal protein S15A |
| chr5 | 57431484 | 57433618 | 2135 | * | 17,59997 | 2,106221481 | 0,758171 | 2,77803 | 0,005469 | 0,68680243 | 2,423594 | Distal Intergenic | 5 | 57285987 | 57286114 | 128 | 2 | 1,15E+08 | ENSMUST00000104145.3 | -145370 | ENSMUSG00000077333 | Gm23015 | predicted gene, 23015 |
| chr16 | 32465714 | 32468511 | 2798 | * | 22,97645 | 2,106382905 | 0,774703 | 2,718957 | 0,006549 | 0,6845989 | 2,951933 | Promoter (1-2kb) | 16 | 32463777 | 32487112 | 23336 | 1 | 51789 | ENSMUST00000164358.3 | 1937 | ENSMUSG00000022791 | Tnk2 | tyrosine kinase, non-receptor, 2 |
| chr9 | 1,08E+08 | 1,08E+08 | 1828 | * | 23,23918 | 2,106403953 | 0,799175 | 2,635723 | 0,008396 | 0,732200941 | 2,880017 | Intron (ENSMUST0000 | 9 | 1,08E+08 | 1,08E+08 | 388 | 1 | 76873 | ENSMUST00000195536.2 | 6379 | ENSMUSG00000102224 | 4930447F24Rik | RIKEN cDNA 4930447F24 gene |
| chr16 | 10102440 | 10105266 | 2827 | * | 35,39609 | 2,106494261 | 0,560622 | 3,757424 | 0,000172 | 0,222881372 | 1,405709 | Exon (ENSMUST00000 | 16 | 10099613 | 10131832 | 32220 | 2 | 13731 | ENSMUST00000078357.5 | 26566 | ENSMUSG00000022505 | Emp2 | epithelial membrane protein 2 |
| chr12 | 79646916 | 79649069 | 2154 | * | 22,62444 | 2,106517869 | 0,697142 | 3,021649 | 0,002514 | 0,506156746 | 3,104559 | Intron (ENSMUST0000 | 12 | 79374127 | 79748630 | 374504 | 1 | 19363 | ENSMUST00000221257.2 | 272789 | ENSMUSG00000059060 | Rad51b | RAD51 paralog B |
| chr8 | 1,26E+08 | 1,26E+08 | 2935 | * | 42,99944 | 2,106877188 | 0,587885 | 3,583825 | 0,000339 | 0,216236188 | 2,925497 | Intron (ENSMUST0000 | 8 | 1,26E+08 | 1,26E+08 | 40738 | 2 | 112405 | ENSMUST00000034469.7 | 20610 | ENSMUSG00000031987 | Egln1 | egl-9 family hypoxia-inducible factor 1 |
| chr6 | 81250981 | 81252418 | 1438 | * | 19,6986 | 2,107263075 | 0,745273 | 2,827505 | 0,004691 | 0,649217039 | 2,713306 | Distal Intergenic | 6 | 81615292 | 81615450 | 159 | 1 | 1,15E+08 | ENSMUST00000083278.3 | -362874 | ENSMUSG00000065212 | Gm26264 | predicted gene, 26264 |
| chr7 | 1,11E+08 | 1,11E+08 | 3021 | * | 22,39937 | 2,10731537 | 0,708079 | 2,976103 | 0,002919 | 0,533665288 | 3,104559 | Intron (ENSMUST0000 | 7 | 1,11E+08 | 1,11E+08 | 5049 | 2 | 75479 | ENSMUST00000210788.2 | -245772 | ENSMUSG00000110424 | 1700012D14Rik | RIKEN cDNA 1700012D14 gene |
| chr12 | 85911079 | 85913521 | 2443 | * | 19,56783 | 2,108401314 | 0,708271 | 2,976828 | 0,002912 | 0,583673368 | 2,386008 | Exon (ENSMUST00000 | 12 | 85925760 | 85938973 | 13214 | 1 | 320244 | ENSMUST00000175899.2 | -12239 | ENSMUSG00000012609 | Ttll5 | tubulin tyrosine ligase-like family, member 5 |
| chr18 | 34222379 | 34224772 | 2394 | * | 20,31701 | 2,108763758 | 0,755948 | 2,789562 | 0,005278 | 0,670853401 | 2,663619 | Distal Intergenic | 18 | 34178641 | 34186087 | 7447 | 2 | 1,05E+08 | ENSMUST00000235466.2 | -36292 | ENSMUSG00000117961 | Gm50265 | predicted gene, 50265 |
| chr6 | 1,37E+08 | 1,37E+08 | 2593 | * | 40,18841 | 2,108904319 | 0,598033 | 3,526403 | 0,000421 | 0,252389304 | 2,725491 | Intron (ENSMUST0000 | 6 | 1,37E+08 | 1,37E+08 | 8595 | 1 | 19277 | ENSMUST00000203127.2 | 3777 | ENSMUSG00000030223 | Ptpro | protein tyrosine phosphatase, receptor type, O |
| chr8 | 71206887 | 71208078 | 1192 | * | 25,41671 | 2,109209423 | 0,811858 | 2,598002 | 0,009377 | 0,751210692 | 2,880017 | Promoter (<=1kb) | 8 | 71207328 | 71211323 | 3996 | 1 | 19339 | ENSMUST00000110093.9 | 0 | ENSMUSG00000031840 | Rab3a | RAB3A, member RAS oncogene family |
| chr6 | 71419708 | 71420896 | 1189 | * | 14,35182 | 2,109620307 | 0,817723 | 2,579872 | 0,009884 | 0,954881543 | 0,918255 | Promoter (2-3kb) | 6 | 71365618 | 71417621 | 52004 | 2 | 68477 | ENSMUST00000002292.15 | -2087 | ENSMUSG00000002222 | Rmnd5a | required for meiotic nuclear division 5 homolog A |
| chr8 | 127575688 | 127577957 | 2270 | * | 56,966693 | 2,109628143 | 0,4395625 | 4,7993812 | 1,592E-06 | 0,00399751 | 4,0557576 | Exon (ENSMUST0000021 | 8 | 127565983 | 127578223 | 12241 | 1 | 414091 | ENSMUST00000212914.2 | 9705 | | A630001O12Rik | RIKEN cDNA A630001O12 gene |
| chr5 | 1,25E+08 | 1,25E+08 | 1972 | * | 20,98692 | 2,109820849 | 0,775691 | 2,719925 | 0,00653 | 0,684274806 | 2,951933 | Promoter (<=1kb) | 5 | 1,25E+08 | 1,25E+08 | 162067 | 2 | 20602 | ENSMUST00000111398.8 | -22 | ENSMUSG00000029478 | Ncor2 | nuclear receptor co-repressor 2 |
| chr8 | 41554178 | 41555400 | 1223 | * | 16,79579 | 2,11036111 | 0,815808 | 2,586837 | 0,009686 | 0,773936378 | 2,663619 | Intron (ENSMUST0000 | 8 | 41443955 | 41567704 | 123750 | 2 | 102103 | ENSMUST00000118835.8 | 12304 | ENSMUSG00000045636 | Mtus1 | mitochondrial tumor suppressor 1 |
| chr7 | 34311286 | 34314370 | 3085 | * | 27,52379 | 2,110457683 | 0,803059 | 2,628023 | 0,008588 | 0,749550311 | 2,67243 | Distal Intergenic | 7 | 34339607 | 34348169 | 8563 | 2 | 233107 | ENSMUST00000205419.2 | 33799 | ENSMUSG00000030499 | Kctd15 | potassium channel tetramerisation domain containing 15 |
| chr17 | 81084043 | 81085767 | 1725 | * | 19,14967 | 2,111450452 | 0,730936 | 2,888695 | 0,003868 | 0,627785088 | 2,507013 | Distal Intergenic | 17 | 80961290 | 81035914 | 74625 | 2 | 225028 | ENSMUST00000234249.2 | -48129 | ENSMUSG00000024242 | Map4k3 | mitogen-activated protein kinase kinase kinase kinase 3 |
| chr1 | 88121236 | 88123358 | 2123 | * | 21,24189 | 2,11179255 | 0,711317 | 2,96885 | 0,002989 | 0,537712701 | 3,104559 | Intron (ENSMUST0000 | 1 | 88128323 | 88146719 | 18397 | 1 | 22236 | ENSMUST00000049289.9 | -4965 | ENSMUSG00000090171 | Ugt1a2 | UDP glucuronosyltransferase 1 family, polypeptide A2 |
| chr2 | 30126186 | 30127880 | 1695 | * | 29,68325 | 2,111900713 | 0,625673 | 3,375405 | 0,000737 | 0,322750682 | 2,725491 | Promoter (<=1kb) | 2 | 30127727 | 30153797 | 26071 | 1 | 241296 | ENSMUST00000113654.8 | 0 | ENSMUSG00000007476 | Lrrc8a | leucine rich repeat containing 8A VRAC subunit A |
| chr8 | 1,24E+08 | 1,24E+08 | 2906 | * | 28,03997 | 2,112078646 | 0,723392 | 2,919689 | 0,003504 | 0,594676756 | 2,725491 | Distal Intergenic | 8 | 1,24E+08 | 1,24E+08 | 2057 | 1 | 69581 | ENSMUST00000136615.2 | -11057 | ENSMUSG00000039960 | Rhou | ras homolog family member U |
| chr6 | 108634740 | 108640023 | 5284 | * | 84,577525 | 2,112305251 | 0,3794113 | 5,5673227 | 2,587E-08 | 0,000164635 | 4,2003037 | Promoter (<=1kb) | 6 | 108637590 | 108643886 | 6297 | 1 | 20893 | ENSMUST00000032194.11 | 0 | ENSMUSG00000030103 | Bhlhe40 | basic helix-loop-helix family, member e40 |
| chr1 | 53732903 | 53734566 | 1664 | * | 17,33344 | 2,112416463 | 0,799381 | 2,642564 | 0,008228 | 0,744224125 | 2,663619 | Intron (ENSMUST0000 | 1 | 53436165 | 53745920 | 309756 | 2 | 627872 | ENSMUST00000094964.7 | 11354 | ENSMUSG00000096141 | Dnah7a | dynein, axonemal, heavy chain 7A |
| chr19 | 44941309 | 44943197 | 1889 | * | 17,12875 | 2,112607464 | 0,75228 | 2,808271 | 0,004981 | 0,693003605 | 2,141612 | Intron (ENSMUST0000 | 19 | 44933461 | 44933567 | 107 | 1 | 1,15E+08 | ENSMUST00000083225.3 | 7848 | ENSMUSG00000065159 | Gm25482 | predicted gene, 25482 |
| chr19 | 43654462 | 43657118 | 2657 | * | 20,51413 | 2,11282425 | 0,72736 | 2,904785 | 0,003675 | 0,627785088 | 2,386008 | Promoter (<=1kb) | 19 | 43652240 | 43655582 | 3343 | 2 | 246696 | ENSMUST00000237364.2 | 0 | ENSMUSG00000040414 | Slc25a28 | solute carrier family 25, member 28 |
| chr4 | 1,41E+08 | 1,41E+08 | 1595 | * | 21,00723 | 2,112880774 | 0,773527 | 2,731488 | 0,006305 | 0,675437713 | 3,104559 | Promoter (2-3kb) | 4 | 1,41E+08 | 1,41E+08 | 1430 | 1 | 1,01E+08 | ENSMUST00000142429.3 | -2596 | ENSMUSG00000085395 | Gm13056 | predicted gene 13056 |
| chr8 | 1,12E+08 | 1,12E+08 | 3260 | * | 32,48242 | 2,112922799 | 0,5902 | 3,580011 | 0,000344 | 0,308866996 | 1,405709 | Distal Intergenic | 8 | 1,12E+08 | 1,12E+08 | 101319 | 2 | 20340 | ENSMUST00000003404.9 | -5302 | ENSMUSG00000003316 | Glg1 | golgi apparatus protein 1 |
| chr6 | 83089862 | 83092021 | 2160 | * | 21,29891 | 2,113058567 | 0,707923 | 2,98487 | 0,002837 | 0,507398906 | 3,471678 | Promoter (<=1kb) | 6 | 83092477 | 83095879 | 3403 | 1 | 57377 | ENSMUST00000032114.8 | -456 | ENSMUSG00000030036 | Mogs | mannosyl-oligosaccharide glucosidase |
| chr5 | 35950842 | 35954619 | 3778 | * | 40,59429 | 2,11340293 | 0,647111 | 3,265905 | 0,001091 | 0,414938321 | 2,338387 | Intron (ENSMUST0000 | 5 | 35971703 | 36000548 | 28846 | 1 | 231148 | ENSMUST00000130233.8 | -17084 | ENSMUSG00000029095 | Ablim2 | actin-binding LIM protein 2 |
| chr6 | 1,29E+08 | 1,29E+08 | 2522 | * | 34,87113 | 2,113443058 | 0,628361 | 3,363423 | 0,00077 | 0,327375387 | 2,725491 | Promoter (2-3kb) | 6 | 1,29E+08 | 1,29E+08 | 614 | 2 | 1,15E+08 | ENSMUST00000203263.2 | 2430 | | Gm44066 | predicted gene, 44066 |
| chr9 | 22385384 | 22387873 | 2490 | * | 26,10506 | 2,113570665 | 0,718567 | 2,94137 | 0,003268 | 0,563858922 | 2,951062 | Promoter (<=1kb) | 9 | 22387011 | 22799576 | 412566 | 1 | 319845 | ENSMUST00000039798.16 | 0 | ENSMUSG00000035919 | Bbs9 | Bardet-Biedl syndrome 9 (human) |
| chr4 | 1,39E+08 | 1,39E+08 | 2402 | * | 24,92171 | 2,113889956 | 0,73891 | 2,860821 | 0,004225 | 0,615591422 | 2,951933 | Promoter (<=1kb) | 4 | 1,39E+08 | 1,39E+08 | 17833 | 1 | 77056 | ENSMUST00000123636.8 | 0 | ENSMUSG00000041143 | Tmco4 | transmembrane and coiled-coil domains 4 |
| chr15 | 75480141 | 75483363 | 3223 | * | 38,10505 | 2,114727901 | 0,569508 | 3,713251 | 0,000205 | 0,166885401 | 2,787255 | Distal Intergenic | 15 | 75488528 | 75497149 | 8622 | 1 | 22701 | ENSMUST00000161785.8 | -5165 | ENSMUSG00000047003 | Zfp41 | zinc finger protein 41 |
| chrY | 90736063 | 90736866 | 804 | * | 21,21566 | 2,115489203 | 0,747903 | 2,828561 | 0,004676 | 0,65050992 | 2,67243 | Distal Intergenic | 21 | 90763696 | 90766736 | 3041 | 2 | 654820 | ENSMUST00000238676.2 | 29870 | ENSMUSG00000072844 | G530011O06Rik | RIKEN cDNA G530011O06 gene |
| chr1 | 4854230 | 4856281 | 2052 | * | 18,02209 | 2,11593405 | 0,749366 | 2,823634 | 0,004748 | 0,663630601 | 2,507013 | Promoter (<=1kb) | 1 | 4843429 | 4855933 | 12505 | 2 | 27395 | ENSMUST00000130201.8 | 0 | ENSMUSG00000033845 | Mrpl15 | mitochondrial ribosomal protein L15 |
| chr13 | 99576298 | 99579923 | 3626 | * | 29,29893 | 2,116136303 | 0,712139 | 2,971521 | 0,002963 | 0,563858922 | 2,67243 | Promoter (<=1kb) | 13 | 99570809 | 99580452 | 9644 | 2 | 17755 | ENSMUST00000223693.2 | 529 | ENSMUSG00000052727 | Map1b | microtubule-associated protein 1B |
| chr8 | 1,06E+08 | 1,06E+08 | 1665 | * | 18,32338 | 2,116707483 | 0,775457 | 2,729626 | 0,006341 | 0,715249603 | 2,423594 | Distal Intergenic | 8 | 1,06E+08 | 1,06E+08 | 4505 | 1 | 13909 | ENSMUST00000173088.2 | 17710 | ENSMUSG00000062181 | Ces3b | carboxylesterase 3B |
| chr10 | 1,28E+08 | 1,28E+08 | 2421 | * | 23,4611 | 2,11741257 | 0,782474 | 2,70605 | 0,006809 | 0,691182187 | 2,951062 | Promoter (1-2kb) | 10 | 1,28E+08 | 1,28E+08 | 4457 | 2 | 11536 | ENSMUST00000079692.6 | -1138 | ENSMUSG00000058396 | Gpr182 | G protein-coupled receptor 182 |
| chr5 | 39671585 | 39674728 | 3144 | * | 25,28305 | 2,11782918 | 0,730348 | 2,899755 | 0,003735 | 0,60631654 | 2,725491 | Intron (ENSMUST0000 | 5 | 39678319 | 39678447 | 129 | 2 | 1,15E+08 | ENSMUST00000104334.3 | 3719 | ENSMUSG00000077522 | Gm25377 | predicted gene, 25377 |
| chr18 | 5190904 | 5193712 | 2809 | * | 25,26073 | 2,117978073 | 0,793201 | 2,670165 | 0,007581 | 0,73009546 | 2,67243 | Distal Intergenic | 18 | 5162839 | 5165731 | 2893 | 2 | 1,03E+08 | ENSMUST00000180850.3 | -25173 | ENSMUSG00000097888 | Gm26682 | predicted gene, 26682 |
| chr13 | 32111796 | 32114140 | 2345 | * | 20,86734 | 2,118855023 | 0,731408 | 2,896953 | 0,003768 | 0,559013874 | 3,471678 | Intron (ENSMUST0000 | 13 | 32155069 | 32156312 | 1244 | 1 | 1,03E+08 | ENSMUST00000221737.2 | -40929 | | Gm35732 | predicted gene, 35732 |
| chr5 | 45881647 | 45883757 | 2111 | * | 18,51242 | 2,118906096 | 0,74313 | 2,851326 | 0,004354 | 0,658200216 | 2,386008 | Promoter (1-2kb) | 5 | 45877009 | 45884920 | 7912 | 2 | 1E+08 | ENSMUST00000200142.2 | 1163 | ENSMUSG00000015882 | Gm3414 | predicted gene 3414 |
| chr15 | 81962823 | 81965934 | 3112 | * | 31,20309 | 2,118956628 | 0,735115 | 2,882482 | 0,003946 | 0,621084104 | 2,67243 | Promoter (<=1kb) | 15 | 81966665 | 82011010 | 44346 | 1 | 74369 | ENSMUST00000189540.7 | -731 | ENSMUSG00000068117 | Mei1 | meiotic double-stranded break formation protein 1 |
| chr16 | 93218212 | 93219717 | 1506 | * | 16,50278 | 2,119459682 | 0,775018 | 2,734725 | 0,006243 | 0,825197108 | 1,2763 | Intron (ENSMUST0000 | 16 | 93188565 | 93222798 | 34234 | 2 | 72264 | ENSMUST00000189879.7 | 3081 | ENSMUSG00000099968 | 1700029J03Rik | RIKEN cDNA 1700029J03 gene |
| chr4 | 1,38E+08 | 1,38E+08 | 1778 | * | 14,06497 | 2,120376212 | 0,812966 | 2,608198 | 0,009102 | 0,890735468 | 1,2763 | Promoter (2-3kb) | 4 | 1,38E+08 | 1,38E+08 | 29465 | 2 | 72269 | ENSMUST00000030535.4 | 2305 | ENSMUSG00000028755 | Cda | cytidine deaminase |
| chr10 | 80456369 | 80458536 | 2168 | * | 40,420176 | 2,120532031 | 0,4857419 | 4,3655533 | 1,268E-05 | 0,030043467 | 2,3776742 | Promoter (<=1kb) | 10 | 80458672 | 80476575 | 17904 | 1 | 103236 | ENSMUST00000079773.14 | -136 | ENSMUSG00000003345 | Csnk1g2 | casein kinase 1, gamma 2 |
| chr4 | 1,5E+08 | 1,5E+08 | 2702 | * | 31,49418 | 2,120915058 | 0,577304 | 3,673829 | 0,000239 | 0,178303794 | 2,858511 | Promoter (<=1kb) | 4 | 1,5E+08 | 1,5E+08 | 691 | 2 | 1,05E+08 | ENSMUST00000134059.2 | 0 | | Gm13073 | predicted gene 13073 |
| chr11 | 97551256 | 97553852 | 2597 | * | 32,51661 | 2,12091959 | 0,621266 | 3,413867 | 0,00064 | 0,304413931 | 2,725491 | Promoter (<=1kb) | 11 | 97535010 | 97551193 | 16184 | 2 | 1,08E+08 | ENSMUST00000134427.2 | -63 | | Gm11613 | predicted gene 11613 |
| chr7 | 1,26E+08 | 1,26E+08 | 1897 | * | 18,40024 | 2,121268355 | 0,721186 | 2,941363 | 0,003268 | 0,60312831 | 2,423594 | Promoter (<=1kb) | 7 | 1,26E+08 | 1,26E+08 | 27764 | 2 | 74204 | ENSMUST00000166538.2 | 5 | ENSMUSG00000000131 | Xpo6 | exportin 6 |
| chr1 | 1,28E+08 | 1,28E+08 | 2994 | * | 31,43538 | 2,122225526 | 0,677218 | 3,133739 | 0,001726 | 0,451585113 | 2,951062 | Promoter (<=1kb) | 1 | 1,28E+08 | 1,28E+08 | 53653 | 2 | 226414 | ENSMUST00000027602.15 | 0 | ENSMUSG00000026356 | Dars | aspartyl-tRNA synthetase |
| chr10 | 24009755 | 24011797 | 2043 | * | 19,21264 | 2,122346091 | 0,810073 | 2,619943 | 0,008794 | 0,766607481 | 2,507013 | Promoter (1-2kb) | 10 | 24001721 | 24008385 | 6665 | 2 | 1,05E+08 | ENSMUST00000243923.1 | -1370 | | Gm40614 | predicted gene, 40614 |
| chr3 | 63872961 | 63875606 | 2646 | * | 17,15024 | 2,122401121 | 0,7668 | 2,767867 | 0,005642 | 0,682206193 | 2,663619 | Promoter (<=1kb) | 3 | 63849751 | 63872189 | 22439 | 2 | 11416 | ENSMUST00000029402.15 | -772 | ENSMUSG00000027822 | Slc33a1 | solute carrier family 33 (acetyl-CoA transporter), member 1 |
| chr8 | 41691197 | 41694218 | 3022 | * | 34,72353 | 2,122522749 | 0,636816 | 3,333024 | 0,000859 | 0,457838671 | 1,405709 | Promoter (<=1kb) | 8 | 41692789 | 41728131 | 35343 | 1 | 18536 | ENSMUST00000211090.2 | 0 | ENSMUSG00000031592 | Pcm1 | pericentriolar material 1 |
| chr6 | 19250202 | 19251149 | 948 | * | 17,05304 | 2,12266459 | 0,766592 | 2,768962 | 0,005624 | 0,695401133 | 2,386008 | Intron (ENSMUST0000 | 6 | 19247898 | 19278842 | 30945 | 2 | 74725 | ENSMUST00000200755.2 | 27693 | ENSMUSG00000107217 | 4930524B17Rik | RIKEN cDNA 4930524B17 gene |
| chr5 | 1,18E+08 | 1,18E+08 | 2266 | * | 24,46992 | 2,123396763 | 0,721598 | 2,942631 | 0,003254 | 0,568412386 | 2,858511 | Promoter (<=1kb) | 5 | 1,18E+08 | 1,18E+08 | 54126 | 2 | 269695 | ENSMUST00000117177.8 | 0 | ENSMUSG00000032850 | Rnft2 | ring finger protein, transmembrane 2 |
| chr5 | 74648853 | 74650830 | 1978 | * | 20,28504 | 2,124558734 | 0,727951 | 2,918544 | 0,003517 | 0,595178563 | 2,713306 | Intron (ENSMUST0000 | 5 | 74619034 | 74639787 | 20754 | 2 | 212986 | ENSMUST00000142216.2 | -9066 | ENSMUSG00000062110 | Scfd2 | Sec1 family domain containing 2 |
| chr6 | 75265655 | 75266596 | 942 | * | 16,20791 | 2,124690192 | 0,801934 | 2,649457 | 0,008062 | 0,77899407 | 2,141612 | Distal Intergenic | 6 | 74428780 | 74428887 | 108 | 2 | 1,15E+08 | ENSMUST00000082474.3 | -836768 | ENSMUSG00000064408 | Gm23924 | predicted gene, 23924 |
| chr10 | 80408450 | 80410812 | 2363 | * | 22,50822 | 2,125389171 | 0,761804 | 2,789943 | 0,005272 | 0,670234202 | 2,67243 | Promoter (2-3kb) | 10 | 80402958 | 80413155 | 10198 | 2 | 118445 | ENSMUST00000038558.9 | 2343 | ENSMUSG00000035397 | Klf16 | Kruppel-like factor 16 |
| chr4 | 72749697 | 72750927 | 1231 | * | 16,41887 | 2,125742899 | 0,75401 | 2,819249 | 0,004814 | 0,679450705 | 2,318434 | Distal Intergenic | 4 | 72768820 | 72770869 | 2050 | 2 | 353204 | ENSMUST00000179234.2 | 19942 | ENSMUSG00000059343 | Aldoart1 | aldolase 1 A, retrogene 1 |
| chr8 | 25954768 | 25956745 | 1978 | * | 32,91838 | 2,12641066 | 0,66539 | 3,195737 | 0,001395 | 0,451739959 | 2,377674 | Exon (ENSMUST000002 | 8 | 25933684 | 25944123 | 10440 | 2 | 1,03E+08 | ENSMUST00000210204.2 | -10645 | | Gm30978 | predicted gene, 30978 |
| chr7 | 26859878 | 26862785 | 2908 | * | 29,09879 | 2,126517872 | 0,659156 | 3,22612 | 0,001255 | 0,406085428 | 2,858511 | Promoter (2-3kb) | 7 | 26864958 | 26865707 | 750 | 2 | 112406 | ENSMUST00000134906.2 | 2922 | ENSMUSG00000058709 | Egln2 | egl-9 family hypoxia-inducible factor 2 |
| chr17 | 6242528 | 6244563 | 2036 | * | 20,98274 | 2,126535488 | 0,6895 | 3,084172 | 0,002041 | 0,483145429 | 2,880017 | Intron (ENSMUST0000 | 17 | 6282927 | 6301403 | 18477 | 1 | 68842 | ENSMUST00000137063.2 | -38364 | ENSMUSG00000034377 | Tulp4 | tubby like protein 4 |
| chr5 | 66699783 | 66702127 | 2345 | * | 22,00007 | 2,126537973 | 0,693361 | 3,067001 | 0,002162 | 0,494205337 | 2,880017 | Promoter (2-3kb) | 5 | 66609610 | 66704300 | 94691 | 2 | 11787 | ENSMUST00000161716.8 | 2173 | ENSMUSG00000029207 | Apbb2 | amyloid beta (A4) precursor protein-binding, family B, member 2 |
| chr13 | 14128616 | 14130435 | 1820 | * | 23,11426 | 2,126594232 | 0,727441 | 2,923389 | 0,003462 | 0,582111135 | 2,858511 | Promoter (<=1kb) | 13 | 14129054 | 14173659 | 44606 | 1 | 97884 | ENSMUST00000099747.5 | 0 | ENSMUSG00000039242 | B3galnt2 | UDP-GalNAc:betaGlcNAc beta 1,3-galactosaminyltransferase, polypeptide 2 |
| chr14 | 1,22E+08 | 1,22E+08 | 2769 | * | 27,50276 | 2,126713362 | 0,58996 | 3,604841 | 0,000312 | 0,205744841 | 2,951062 | Intron (ENSMUST0000 | 14 | 1,22E+08 | 1,22E+08 | 195666 | 2 | 105445 | ENSMUST00000239077.2 | -13020 | ENSMUSG00000025558 | Dock9 | dedicator of cytokinesis 9 |
| chr15 | 98411789 | 98414326 | 2538 | * | 34,67022 | 2,126742299 | 0,553935 | 3,839337 | 0,000123 | 0,184990586 | 1,405709 | Distal Intergenic | 15 | 98417597 | 98417734 | 138 | 2 | 1E+08 | ENSMUST00020182350.1 | 3408 | ENSMUSG00002076288 | Snora34 | small nucleolar RNA, H/ACA box 34 |
| chr8 | 105034235 | 105037940 | 3706 | * | 69,351601 | 2,12688909 | 0,4573953 | 4,650002 | 3,319E-06 | 0,00813231 | 3,439501 | Promoter (<=1kb) | 8 | 105007979 | 105036713 | 28735 | 2 | 100504164 | ENSMUST00000162616.8 | 0 | ENSMUSG00000110430 | Cmtm1 | CKLF-like MARVEL transmembrane domain containing 1 |
| chr2 | 1,29E+08 | 1,29E+08 | 2508 | * | 21,79642 | 2,126903275 | 0,674372 | 3,153903 | 0,001611 | 0,440518878 | 2,951933 | 3' UTR | 2 | 1,29E+08 | 1,29E+08 | 36144 | 1 | 72477 | ENSMUST00000110325.8 | -14851 | ENSMUSG00000014353 | Tmem87b | transmembrane protein 87B |
| chr1 | 24255251 | 24256750 | 1500 | * | 14,30538 | 2,127980459 | 0,792161 | 2,686298 | 0,007225 | 0,905781613 | 0,918255 | Exon (ENSMUST00000 | 1 | 24268598 | 24270036 | 1439 | 1 | 12839 | ENSMUST00000147902.2 | -11848 | ENSMUSG00000026147 | Col9a1 | collagen, type IX, alpha 1 |
| chr16 | 87292254 | 87294489 | 2236 | * | 18,09818 | 2,12866838 | 0,825681 | 2,578075 | 0,009935 | 0,792742583 | 2,423594 | Promoter (<=1kb) | 16 | 87280214 | 87292761 | 12548 | 2 | 12469 | ENSMUST00000026704.14 | 0 | ENSMUSG00000025613 | Cct8 | chaperonin containing Tcp1, subunit 8 (theta) |
| chr14 | 22976732 | 22978447 | 1716 | * | 14,5192 | 2,128926363 | 0,813142 | 2,618147 | 0,008841 | 0,793708774 | 2,141612 | Exon (ENSMUST000002 | 14 | 23088262 | 23144639 | 56378 | 2 | 791307 | ENSMUST00000090409.2 | 166192 | ENSMUSG00000068680 | Gm10248 | predicted gene 10248 |
| chr4 | 12087129 | 12090531 | 3403 | * | 24,46456 | 2,129246383 | 0,740876 | 2,873958 | 0,004054 | 0,611385361 | 2,880017 | Promoter (<=1kb) | 4 | 12039355 | 12087957 | 48603 | 2 | 329795 | ENSMUST00000050686.10 | 0 | ENSMUSG00000049488 | Tmem67 | transmembrane protein 67 |
| chr2 | 11320019 | 11321327 | 1309 | * | 18,33113 | 2,12958317 | 0,799746 | 2,662826 | 0,007749 | 0,770673161 | 2,154504 | Exon (ENSMUST000001 | 2 | 11330251 | 11337393 | 7143 | 2 | 1,19E+08 | ENSMUST00000247076.1 | 16066 | | Gm38171 | predicted gene, 38171 |
| chr1 | 64268434 | 64269959 | 1526 | * | 16,34893 | 2,130261553 | 0,75505 | 2,821353 | 0,004782 | 0,685078899 | 2,141612 | Intron (ENSMUST0000 | 1 | 64118262 | 64161441 | 43180 | 2 | 93691 | ENSMUST00000135075.2 | -106993 | ENSMUSG00000025959 | Klf7 | Kruppel-like factor 7 (ubiquitous) |
| chr4 | 1,4E+08 | 1,4E+08 | 2151 | * | 22,41708 | 2,130446908 | 0,764424 | 2,786995 | 0,00532 | 0,646818417 | 3,104559 | Distal Intergenic | 4 | 1,4E+08 | 1,4E+08 | 9408 | 1 | 1,03E+08 | ENSMUST00000135716.3 | 51687 | ENSMUSG00000086437 | Gm13021 | predicted gene 13021 |
| chr8 | 1,28E+08 | 1,28E+08 | 3631 | * | 31,32672 | 2,130997468 | 0,62182 | 3,427031 | 0,00061 | 0,293348204 | 2,880017 | Distal Intergenic | 8 | 1,27E+08 | 1,27E+08 | 2764 | 2 | 1,15E+08 | ENSMUST00000212695.2 | -27334 | | Gm45805 | predicted gene 45805 |
| chr8 | 13331720 | 13333949 | 2230 | * | 21,82846 | 2,131106136 | 0,71757 | 2,969891 | 0,002979 | 0,547830407 | 2,880017 | Promoter (<=1kb) | 8 | 13328529 | 13331592 | 3064 | 2 | 102323 | ENSMUST00000204840.2 | -128 | ENSMUSG00000038506 | Dcun1d2 | DCN1, defective in cullin neddylation 1, domain containing 2 (S. cerevisiae) |
| chr10 | 80381400 | 80383861 | 2462 | * | 23,13535 | 2,131538965 | 0,731347 | 2,91454 | 0,003562 | 0,580633053 | 2,951933 | Promoter (<=1kb) | 10 | 80376756 | 80381670 | 4915 | 2 | 66932 | ENSMUST00000182682.8 | 0 | ENSMUSG00000047417 | Rexo1 | REX1, RNA exonuclease 1 |
| chr2 | 75594680 | 75597205 | 2526 | * | 20,23622 | 2,132609461 | 0,816079 | 2,613238 | 0,008969 | 0,792225394 | 2,221611 | Distal Intergenic | 2 | 75607495 | 75609446 | 1952 | 1 | 1,03E+08 | ENSMUST00000242477.1 | -10290 | | Gm13657 | predicted gene 13657 |
| chr6 | 87436777 | 87437716 | 940 | * | 17,62869 | 2,132746681 | 0,794416 | 2,684673 | 0,00726 | 0,737937087 | 2,423594 | Promoter (2-3kb) | 6 | 87437097 | 87440629 | 3533 | 2 | 232201 | ENSMUST00000203559.2 | 2913 | ENSMUSG00000030047 | Arhgap25 | Rho GTPase activating protein 25 |
| chr10 | 77034578 | 77035950 | 1373 | * | 16,72358 | 2,133068947 | 0,826302 | 2,581464 | 0,009838 | 0,902917968 | 1,2763 | Distal Intergenic | 10 | 76888012 | 77002382 | 114371 | 2 | 12822 | ENSMUST00000081654.13 | -32196 | ENSMUSG00000001435 | Col18a1 | collagen, type XVIII, alpha 1 |
| chr18 | 60781075 | 60782663 | 1589 | * | 16,71155 | 2,133579768 | 0,752373 | 2,835802 | 0,004571 | 0,666986161 | 2,386008 | Promoter (<=1kb) | 18 | 60733281 | 60781391 | 48111 | 2 | 104027 | ENSMUST00000143275.3 | 0 | ENSMUSG00000043079 | Synpo | synaptopodin |
| chr2 | 30407931 | 30409916 | 1986 | * | 26,64304 | 2,134046385 | 0,802842 | 2,658116 | 0,007858 | 0,723275538 | 2,858511 | Distal Intergenic | 2 | 30424741 | 30429034 | 4294 | 1 | 1,03E+08 | ENSMUST00000145413.2 | -14825 | ENSMUSG00000085275 | Gm14487 | predicted gene 14487 |
| chr17 | 46314892 | 46316938 | 2047 | * | 34,6986 | 2,134644992 | 0,56011 | 3,811117 | 0,000138 | 0,13101642 | 2,787255 | Distal Intergenic | 17 | 46329638 | 46332298 | 2661 | 2 | 22339 | ENSMUST00000143985.2 | 15360 | ENSMUSG00000023951 | Vegfa | vascular endothelial growth factor A |
| chr10 | 80592764 | 80595245 | 2482 | * | 30,03667 | 2,135169867 | 0,672069 | 3,177009 | 0,001488 | 0,444201987 | 2,67243 | Promoter (1-2kb) | 10 | 80591297 | 80602421 | 11125 | 1 | 208266 | ENSMUST00000147579.2 | 1467 | ENSMUSG00000061589 | Dot1l | DOT1-like, histone H3 methyltransferase (S. cerevisiae) |
| chr11 | 62492283 | 62495593 | 3311 | * | 41,67249 | 2,135275416 | 0,550099 | 3,881617 | 0,000104 | 0,120486853 | 2,377674 | Promoter (<=1kb) | 11 | 62493702 | 62495635 | 1934 | 1 | 1E+08 | ENSMUST00000245938.1 | 0 | ENSMUSG00000064437 | Snord49b | small nucleolar RNA, C/D box 49B |
| chr10 | 83478747 | 83482598 | 3852 | * | 62,486102 | 2,135505483 | 0,4744251 | 4,5012485 | 6,756E-06 | 0,015138847 | 3,177995 | Promoter (1-2kb) | 10 | 83456577 | 83484467 | 27891 | 2 | 216190 | ENSMUST00000150685.8 | 1869 | ENSMUSG00000020263 | Appl2 | adaptor protein, phosphotyrosine interaction, PH domain and leucine zipper containing 2 |
| chr5 | 66636574 | 66640086 | 3513 | * | 28,57391 | 2,135752675 | 0,645471 | 3,308828 | 0,000937 | 0,36185416 | 2,67243 | Intron (ENSMUST0000 | 5 | 66459548 | 66649514 | 189967 | 2 | 11787 | ENSMUST00000160063.5 | 9428 | ENSMUSG00000029207 | Apbb2 | amyloid beta (A4) precursor protein-binding, family B, member 2 |
| chr18 | 75617040 | 75620407 | 3368 | * | 31,22772 | 2,136850285 | 0,586693 | 3,642196 | 0,00027 | 0,201373005 | 2,67243 | Intron (ENSMUST0000 | 18 | 75647716 | 75655877 | 8162 | 1 | 1E+08 | ENSMUST00000181913.2 | -27309 | ENSMUSG00000097712 | Gm10532 | predicted gene 10532 |
| chr15 | 80101939 | 80104950 | 3012 | * | 64,93057 | 2,137506387 | 0,4466043 | 4,7861303 | 1,7E-06 | 0,004911375 | 3,439501 | Distal Intergenic | 15 | 80118219 | 80137572 | 19354 | 1 | 239555 | ENSMUST00000023048.12 | -13269 | ENSMUSG00000022412 | Mief1 | mitochondrial elongation factor 1 |
| chr7 | 44170139 | 44174323 | 4185 | * | 32,73439 | 2,137624372 | 0,62425 | 3,424309 | 0,000616 | 0,301441602 | 2,725491 | Promoter (<=1kb) | 7 | 44151123 | 44174080 | 22958 | 2 | 233199 | ENSMUST00000165208.4 | 0 | ENSMUSG00000038670 | Mybpc2 | myosin binding protein C, fast-type |
| chr6 | 53743172 | 53744912 | 1741 | * | 19,48251 | 2,137790327 | 0,807072 | 2,648821 | 0,008077 | 0,758230902 | 2,386008 | Distal Intergenic | 6 | 53795728 | 53845168 | 49441 | 1 | 1E+08 | ENSMUST00000204648.2 | -50816 | | Gm16499 | predicted gene 16499 |
| chr12 | 70585760 | 70589345 | 3586 | * | 41,32439 | 2,137961682 | 0,5005277 | 4,2714157 | 1,942E-05 | 0,041477448 | 2,3776742 | Exon (ENSMUST0000022 | 12 | 70569372 | 70591361 | 21990 | 1 | 102634692 | ENSMUST00000221234.2 | 16388 | | Gm32219 | predicted gene, 32219 |
| chr12 | 8406245 | 8410129 | 3885 | * | 36,05684 | 2,138040017 | 0,548561 | 3,897544 | 9,72E-05 | 0,16124834 | 1,405709 | Distal Intergenic | 12 | 8385460 | 8386717 | 1258 | 2 | 1,19E+08 | ENSMUST00000217669.2 | -19528 | | Gm48071 | predicted gene, 48071 |

| | | | | | | | | | | | | | | | | | | | | | | | |
|---|---|---|---|---|---|---|---|---|---|---|---|---|---|---|---|---|---|---|---|---|---|---|---|
| chr2 | 70963512 | 70966482 | 2971 | * | 24,81953 | 2,139132867 | 0,769694 | 2,7792 | 0,005449 | 0,675709769 | 2,67243 | Intron (ENSMUST0000( | 2 | 70948267 | 70973270 | 25004 | 1 | 73649 | ENSMUST00000028403.3 | 15245 | ENSMUSG00000027015 | Cybrd1 | cytochrome b reductase 1 |
| chr16 | 93358222 | 93360361 | 2140 | * | 16,13282 | 2,139228346 | 0,80355 | 2,66222 | 0,007763 | 0,773432032 | 2,141612 | Distal Intergenic | 16 | 93380450 | 93380896 | 447 | 2 | 224440 | ENSMUST00000233912.2 | 20535 | ENSMUSG00000022948 | Setd4 | SET domain containing 4 |
| chr7 | 80436376 | 80438590 | 2215 | * | 25,15008 | 2,139724984 | 0,70862 | 3,019565 | 0,002531 | 0,516158525 | 2,951933 | Intron (ENSMUST0000( | 7 | 80399125 | 80453011 | 53887 | 2 | 29875 | ENSMUST00000206149.2 | 14421 | ENSMUSG00000030536 | Iqgap1 | IQ motif containing GTPase activating protein 1 |
| chr6 | 47857035 | 47858660 | 1626 | * | 20,02959 | 2,139765982 | 0,780617 | 2,74112 | 0,006123 | 0,68680243 | 2,713306 | Promoter (2-3kb) | 6 | 47854590 | 47869747 | 15158 | 1 | 101095 | ENSMUST00000125957.2 | 2445 | ENSMUSG00000025821 | Zfp282 | zinc finger protein 282 |
| chr7 | 76742852 | 76744658 | 1807 | * | 21,28016 | 2,139953816 | 0,711758 | 3,006576 | 0,002642 | 0,530557026 | 2,880017 | Intron (ENSMUST0000( | 7 | 76060810 | 76416356 | 355547 | 1 | 244071 | ENSMUST00000107442.3 | 682042 | ENSMUSG00000025754 | Agbl1 | ATP/GTP binding protein-like 1 |
| chr8 | 1,26E+08 | 1,26E+08 | 2519 | * | 25,00878 | 2,14012739 | 0,686289 | 3,118407 | 0,001818 | 0,45991801 | 2,951062 | Exon (ENSMUST00000: | 8 | 1,26E+08 | 1,26E+08 | 2543 | 1 | 1,03E+08 | ENSMUST00000245925.1 | 34854 | | Gm30713 | predicted gene, 30713 |
| chr15 | 40796762 | 40798638 | 1877 | * | 21,124 | 2,140268661 | 0,709927 | 3,014773 | 0,002572 | 0,524454869 | 2,880017 | Intron (ENSMUST0000( | 15 | 41036883 | 41037264 | 382 | 1 | 1E+08 | ENSMUST00000060066.6 | -238245 | | 4930555K19Rik | leucine rich repeat and sterile alpha motif containing 1 pseudogene |
| chr2 | 167489961 | 167493834 | 3874 | * | 70,025178 | 2,140896461 | 0,4334837 | 4,9388165 | 7,86E-07 | 0,002823289 | 3,2706213 | Promoter (2-3kb) | 2 | 167485656 | 167487097 | 1442 | 2 | 407243 | ENSMUST00000142148.2 | -2864 | ENSMUSG00000090213 | Peds1 | plasmanylethanolamine desaturase 1 |
| chr5 | 31732673 | 31734613 | 1941 | * | 26,7223 | 2,141113159 | 0,701151 | 3,05371 | 0,00226 | 0,50985457 | 2,725491 | Distal Intergenic | 5 | 31726932 | 31728817 | 1886 | 2 | 100740 | ENSMUST00000200738.3 | -3856 | | AI839979 | expressed sequence AI839979 |
| chr15 | 77687030 | 77689763 | 2734 | * | 33,39465 | 2,141223659 | 0,629202 | 3,40308 | 0,000666 | 0,418252864 | 1,405709 | Exon (ENSMUST00000: | 15 | 77678852 | 77697431 | 18580 | 2 | 17886 | ENSMUST00000124844.8 | 7668 | ENSMUSG00000022443 | Myh9 | myosin, heavy polypeptide 9, non-muscle |
| chr1 | 1,78E+08 | 1,78E+08 | 2558 | * | 17,28982 | 2,141376433 | 0,751126 | 2,85089 | 0,00436 | 0,64068479 | 2,663619 | Intron (ENSMUST0000( | 1 | 1,78E+08 | 1,78E+08 | 78411 | 1 | 68226 | ENSMUST00000027775.9 | 12970 | ENSMUSG00000026495 | Efcab2 | EF-hand calcium binding domain 2 |
| chr5 | 78157407 | 78159233 | 1827 | * | 17,37433 | 2,141786115 | 0,774516 | 2,765322 | 0,005687 | 0,69472367 | 2,423594 | Distal Intergenic | 5 | 77613335 | 77616698 | 3364 | 1 | 69370 | ENSMUST00000201654.2 | 544072 | ENSMUSG00000107232 | 1700017L05Rik | RIKEN cDNA 1700017L05 gene |
| chr19 | 57050620 | 57053474 | 2855 | * | 24,43076 | 2,141982084 | 0,697628 | 3,070377 | 0,002138 | 0,493950739 | 2,858511 | Exon (ENSMUST00000( | 19 | 57027879 | 57035670 | 7792 | 2 | 226251 | ENSMUST00000137389.8 | -14950 | ENSMUSG00000025085 | Ablim1 | actin-binding LIM protein 1 |
| chr7 | 27184929 | 27187047 | 2119 | * | 38,70297 | 2,142014089 | 0,640881 | 3,342294 | 0,000831 | 0,337811311 | 2,725491 | Promoter (<=1kb) | 7 | 27186335 | 27189741 | 3407 | 1 | 55942 | ENSMUST00000008528.8 | 0 | ENSMUSG00000008384 | Sertad1 | SERTA domain containing 1 |
| chr10 | 60652517 | 60653964 | 1448 | * | 23,58251 | 2,142294288 | 0,782758 | 2,736854 | 0,006203 | 0,681331037 | 2,951933 | Promoter (2-3kb) | 10 | 60598389 | 60656420 | 58032 | 2 | 107449 | ENSMUST00000218316.2 | 2456 | ENSMUSG00000020099 | Unc5b | unc-5 netrin receptor B |
| chr6 | 5443066 | 5445313 | 2248 | * | 26,89002 | 2,14230106 | 0,75701 | 2,829952 | 0,004655 | 0,646274978 | 2,725491 | Distal Intergenic | 6 | 5491591 | 5496261 | 4671 | 2 | 27273 | ENSMUST00000203347.3 | 50948 | ENSMUSG00000019577 | Pdk4 | pyruvate dehydrogenase kinase, isoenzyme 4 |
| chr11 | 61156412 | 61158585 | 2174 | * | 19,20765 | 2,142659555 | 0,740357 | 2,894089 | 0,003803 | 0,626725358 | 2,507013 | Promoter (<=1kb) | 11 | 61114243 | 61158011 | 43769 | 2 | 11671 | ENSMUST00000066277.10 | 0 | ENSMUSG00000010025 | Aldh3a2 | aldehyde dehydrogenase family 3, subfamily A2 |
| chr17 | 35285943 | 35287891 | 1949 | * | 21,99754 | 2,142933699 | 0,745693 | 2,873747 | 0,004056 | 0,611385361 | 2,880017 | Promoter (<=1kb) | 17 | 35286301 | 35289026 | 2726 | 1 | 68468 | ENSMUST00000173478.2 | 0 | ENSMUSG00000092586 | Ly6g6c | lymphocyte antigen 6 complex, locus G6C |
| chr19 | 56867930 | 56869650 | 1721 | * | 20,08361 | 2,14333685 | 0,774862 | 2,766087 | 0,005673 | 0,686838043 | 2,507013 | 5' UTR | 19 | 56862681 | 56900510 | 37830 | 1 | 240675 | ENSMUST00000026068.8 | 5249 | ENSMUSG00000025082 | Vwa2 | von Willebrand factor A domain containing 2 |
| chr5 | 76465321 | 76467043 | 1723 | * | 17,61958 | 2,143352267 | 0,830181 | 2,581789 | 0,009829 | 0,793567638 | 2,386008 | Exon (ENSMUST00000: | 5 | 76459962 | 76478980 | 19019 | 2 | 79455 | ENSMUST00000122213.8 | 11937 | ENSMUSG00000029235 | Pdcl2 | phosducin-like 2 |
| chr13 | 38718387 | 38720711 | 2325 | * | 18,62015 | 2,143951827 | 0,801386 | 2,675304 | 0,007466 | 0,727657783 | 2,663619 | Exon (ENSMUST00000: | 13 | 38684249 | 38712800 | 28552 | 2 | 105245 | ENSMUST00000162075.8 | -5587 | ENSMUSG00000038991 | Txndc5 | thioredoxin domain containing 5 |
| chr3 | 56459537 | 56463257 | 3721 | * | 32,19373 | 2,144614194 | 0,558122 | 3,842553 | 0,000122 | 0,120486853 | 2,787255 | Distal Intergenic | 3 | 56352703 | 56352833 | 131 | 2 | 1,15E+08 | ENSMUST00000104522.3 | -106704 | ENSMUSG00000119391 | Gm25727 | predicted gene, 25727 |
| chr2 | 38954827 | 38956621 | 1795 | * | 22,38018 | 2,145349867 | 0,707144 | 3,033821 | 0,002415 | 0,507398906 | 2,951933 | Promoter (<=1kb) | 2 | 38906167 | 38955553 | 49387 | 2 | 76899 | ENSMUST00000039165.15 | 0 | ENSMUSG00000026754 | Golga1 | golgi autoantigen, golgin subfamily a, 1 |
| chr9 | 71632166 | 71635754 | 3589 | * | 29,67056 | 2,14576864 | 0,678421 | 3,162886 | 0,001562 | 0,44002295 | 2,880017 | 5' UTR | 9 | 71536493 | 71678842 | 142350 | 2 | 68178 | ENSMUST00000122065.2 | 43088 | ENSMUSG00000032232 | Cgnl1 | cingulin-like 1 |
| chr15 | 96606779 | 96609474 | 2696 | * | 42,06738 | 2,147090504 | 0,538995 | 3,983508 | 6,79E-05 | 0,092920819 | 2,338387 | Exon (ENSMUST00000: | 15 | 96585273 | 96597611 | 12339 | 2 | 67760 | ENSMUST00000023099.8 | -9168 | ENSMUSG00000022462 | Slc38a2 | solute carrier family 38, member 2 |
| chr6 | 30890074 | 30891827 | 1754 | * | 19,61513 | 2,147128739 | 0,69947 | 3,06965 | 0,002143 | 0,502272318 | 2,713306 | Promoter (<=1kb) | 6 | 30873916 | 30892508 | 18593 | 2 | 116732 | ENSMUST00000048580.2 | 681 | ENSMUSG00000039032 | Tsga13 | testis specific gene A13 |
| chr17 | 77497500 | 77499081 | 1582 | * | 17,3162 | 2,147902889 | 0,809639 | 2,652913 | 0,00798 | 0,749550311 | 2,507013 | Distal Intergenic | 17 | 78506685 | 78507454 | 770 | 2 | 1,03E+08 | ENSMUST00000234450.2 | 1008373 | ENSMUSG00000117648 | Gm32282 | predicted gene, 32282 |
| chr1 | 1,32E+08 | 1,32E+08 | 3041 | * | 24,16262 | 2,148984097 | 0,779714 | 2,756118 | 0,005849 | 0,670853401 | 2,951933 | Promoter (<=1kb) | 1 | 1,32E+08 | 1,32E+08 | 27448 | 2 | 18557 | ENSMUST00000027697.12 | 0 | ENSMUSG00000026437 | Cdk18 | cyclin-dependent kinase 18 |
| chr5 | 1,41E+08 | 1,41E+08 | 2015 | * | 22,40538 | 2,149150967 | 0,736975 | 2,916179 | 0,003543 | 0,545597917 | 3,471678 | Promoter (1-2kb) | 5 | 1,41E+08 | 1,41E+08 | 12157 | 1 | 231842 | ENSMUST00000176035.2 | -1166 | ENSMUSG00000050022 | Amz1 | archaelysin family metallopeptidase 1 |
| chr12 | 55509595 | 55511616 | 2022 | * | 19,73399 | 2,149732422 | 0,736676 | 2,918151 | 0,003521 | 0,597812173 | 2,663619 | Distal Intergenic | 12 | 55537192 | 55538595 | 1404 | 2 | 18035 | ENSMUST00000219869.2 | 26979 | ENSMUSG00000021025 | Nfkbia | nuclear factor of kappa light polypeptide gene enhancer in B cells inhibitor, alpha |
| chr4 | 97644834 | 97647360 | 2527 | * | 24,83079 | 2,15045037 | 0,69802 | 3,080787 | 0,002065 | 0,497895643 | 2,67243 | Promoter (<=1kb) | 4 | 97642405 | 97645626 | 3222 | 2 | 1,05E+08 | ENSMUST00000149078.2 | 0 | | Gm38709 | predicted gene, 38709 |
| chr18 | 61807129 | 61810865 | 3737 | * | 34,68039 | 2,150468236 | 0,630108 | 3,412859 | 0,000643 | 0,410415129 | 1,405709 | Distal Intergenic | 18 | 61817958 | 61825609 | 7652 | 1 | 56069 | ENSMUST00000237575.2 | -7093 | ENSMUSG00000024578 | Il17b | interleukin 17B |
| chr11 | 1,15E+08 | 1,15E+08 | 2182 | * | 24,04427 | 2,150746266 | 0,75172 | 2,861101 | 0,004222 | 0,621135422 | 2,858511 | Distal Intergenic | 11 | 1,15E+08 | 1,15E+08 | 3808 | 1 | 67671 | ENSMUST00000106602.10 | -30585 | ENSMUSG00000057322 | Rpl38 | ribosomal protein L38 |
| chr11 | 1,15E+08 | 1,15E+08 | 1638 | * | 16,54584 | 2,150977654 | 0,834165 | 2,578598 | 0,00992 | 0,903476157 | 1,2763 | Distal Intergenic | 11 | 1,15E+08 | 1,15E+08 | 17751 | 1 | 432611 | ENSMUST00000151053.8 | -3370 | ENSMUSG00000034706 | Dnai2 | dynein axonemal intermediate chain 2 |
| chr7 | 1,01E+08 | 1,01E+08 | 2941 | * | 33,99004 | 2,15158974 | 0,680536 | 3,16161 | 0,001569 | 0,44621096 | 2,787255 | Intron (ENSMUST0000( | 7 | 1,01E+08 | 1,01E+08 | 22030 | 1 | 20480 | ENSMUST00000150832.2 | 18411 | ENSMUSG00000001829 | Clpb | ClpB caseinolytic peptidase B |
| chr16 | 17632852 | 17635078 | 2227 | * | 27,56914 | 2,15169306 | 0,780394 | 2,75719 | 0,00583 | 0,67040149 | 2,951062 | Exon (ENSMUST00000: | 16 | 17622673 | 17624825 | 2153 | 1 | 224024 | ENSMUST00000231301.2 | 10179 | ENSMUSG00000012017 | Scarf2 | scavenger receptor class F, member 2 |
| chr4 | 1,51E+08 | 1,51E+08 | 2070 | * | 18,03345 | 2,151729648 | 0,798477 | 2,694791 | 0,007043 | 0,734804995 | 2,386008 | Intron (ENSMUST0000( | 4 | 1,51E+08 | 1,51E+08 | 42413 | 1 | 68703 | ENSMUST00000137112.2 | 11182 | ENSMUSG00000039852 | Rere | arginine glutamic acid dipeptide (RE) repeats |
| chr15 | 79230896 | 79233571 | 2676 | * | 40,19921 | 2,152480956 | 0,602829 | 3,570631 | 0,000356 | 0,222720036 | 2,925497 | Promoter (<=1kb) | 15 | 79231720 | 79243276 | 11557 | 1 | 17133 | ENSMUST00000096350.11 | 0 | ENSMUSG00000042622 | Maff | v-maf musculoaponeurotic fibrosarcoma oncogene family, protein F (avian) |
| chr11 | 1,13E+08 | 1,13E+08 | 1270 | * | 21,14899 | 2,152681015 | 0,790723 | 2,722422 | 0,006481 | 0,684274806 | 2,951933 | Intron (ENSMUST0000( | 11 | 1,13E+08 | 1,13E+08 | 129057 | 2 | 72386 | ENSMUST00000244333.1 | 111488 | ENSMUSG00000087259 | 2610035D17Rik | RIKEN cDNA 2610035D17 gene |
| chr13 | 36693077 | 36695070 | 1994 | * | 17,51449 | 2,152711358 | 0,766992 | 2,806694 | 0,005005 | 0,681307794 | 2,386008 | Intron (ENSMUST0000( | 13 | 36796790 | 36804413 | 7624 | 1 | 1,08E+08 | ENSMUST00000225239.2 | -101720 | | Gm46410 | predicted gene, 46410 |
| chr2 | 1,65E+08 | 1,65E+08 | 2095 | * | 30,42761 | 2,152884989 | 0,642029 | 3,353255 | 0,000799 | 0,322750682 | 2,951062 | Promoter (<=1kb) | 2 | 1,65E+08 | 1,65E+08 | 7147 | 1 | 19025 | ENSMUST00000103093.10 | 0 | ENSMUSG00000017760 | Ctsa | cathepsin A |
| chr17 | 6188742 | 6191347 | 2606 | * | 23,61959 | 2,153585114 | 0,812909 | 2,649232 | 0,008068 | 0,722855505 | 2,951062 | Promoter (<=1kb) | 17 | 6188423 | 6290903 | 102481 | 1 | 68842 | ENSMUST00000039655.3 | 319 | ENSMUSG00000034377 | Tulp4 | tubby like protein 4 |
| chr7 | 37702111 | 37704510 | 2400 | * | 22,31527 | 2,153804207 | 0,760448 | 2,832283 | 0,004622 | 0,599349201 | 3,471678 | Intron (ENSMUST0000( | 7 | 37669030 | 37718947 | 49918 | 2 | 19777 | ENSMUST00000206169.2 | 14437 | ENSMUSG00000030421 | Uri1 | URI1, prefoldin-like chaperone |
| chr11 | 76455728 | 76457387 | 1660 | * | 16,88792 | 2,1538043 | 0,835615 | 2,577509 | 0,009952 | 0,777919825 | 2,663619 | Intron (ENSMUST0000( | 11 | 76307560 | 76462361 | 154802 | 2 | 109934 | ENSMUST00000072740.13 | 4974 | ENSMUSG00000017631 | Abr | active BCR-related gene |
| chr11 | 1,19E+08 | 1,19E+08 | 1585 | * | 16,44938 | 2,154945548 | 0,807424 | 2,668915 | 0,00761 | 0,858273023 | 1,2763 | 3' UTR | 11 | 1,19E+08 | 1,19E+08 | 62 | 2 | 1,02E+08 | ENSMUST00000183965.3 | 6923 | ENSMUSG00000098704 | Mir6934 | microRNA 6934 |
| chr1 | 1,82E+08 | 1,82E+08 | 1843 | * | 20,39355 | 2,155485165 | 0,782244 | 2,755513 | 0,00586 | 0,683710007 | 2,713306 | Distal Intergenic | 1 | 1,82E+08 | 1,82E+08 | 33704 | 2 | 12334 | ENSMUST00000192483.2 | -9933 | ENSMUSG00000026509 | Capn2 | calpain 2 |
| chr4 | 43441325 | 43443075 | 1751 | * | 22,01659 | 2,155595696 | 0,748563 | 2,879647 | 0,003981 | 0,623120685 | 2,67243 | Promoter (<=1kb) | 4 | 43441939 | 43448064 | 6126 | 1 | 21754 | ENSMUST00000060864.13 | 0 | ENSMUSG00000028458 | Tesk1 | testis specific protein kinase 1 |
| chr15 | 25983173 | 25984501 | 1329 | * | 14,39073 | 2,156205437 | 0,828867 | 2,601389 | 0,009285 | 0,940994296 | 0,938682 | Promoter (<=1kb) | 15 | 25984452 | 25990550 | 6099 | 1 | 52521 | ENSMUST00000227912.2 | 0 | ENSMUSG00000052253 | Zfp622 | zinc finger protein 622 |
| chr10 | 81099606 | 81100883 | 1278 | * | 16,65917 | 2,156557994 | 0,806703 | 2,6733 | 0,007511 | 0,728793055 | 2,663619 | Promoter (1-2kb) | 10 | 81101264 | 81102758 | 1495 | 2 | 66047 | ENSMUST00000145452.2 | 1875 | ENSMUSG00000034932 | Mrpl54 | mitochondrial ribosomal protein L54 |
| chr17 | 23884271 | 23886020 | 1750 | * | 17,70599 | 2,156558555 | 0,822846 | 2,620854 | 0,008771 | 0,792545355 | 2,154504 | Promoter (<=1kb) | 17 | 23885100 | 23886066 | 967 | 1 | 212733 | ENSMUST00000233930.2 | 0 | ENSMUSG00000043782 | Bicdl2 | BICD family like cargo adaptor 2 |
| chr1 | 75411477 | 75414653 | 3177 | * | 35,44351 | 2,156734943 | 0,6732 | 3,203708 | 0,001357 | 0,538066226 | 1,405709 | Promoter (<=1kb) | 1 | 75412574 | 75419818 | 7245 | 1 | 69080 | ENSMUST00000037796.14 | 0 | ENSMUSG00000033021 | Gmppa | GDP-mannose pyrophosphorylase A |
| chr8 | 37125577 | 37127713 | 2137 | * | 18,73478 | 2,157786077 | 0,772211 | 2,794295 | 0,005201 | 0,683988405 | 2,386008 | Intron (ENSMUST0000( | 8 | 37034905 | 37081097 | 46193 | 2 | 50768 | ENSMUST00000033923.14 | -44480 | ENSMUSG00000031523 | Dlc1 | deleted in liver cancer 1 |
| chr15 | 83357997 | 83359763 | 1767 | * | 21,57233 | 2,158021894 | 0,804746 | 2,68162 | 0,007327 | 0,723215019 | 2,67243 | Distal Intergenic | 15 | 83259808 | 83348807 | 89000 | 2 | 23970 | ENSMUST00000171436.8 | -9190 | ENSMUSG00000016664 | Pacsin2 | protein kinase C and casein kinase substrate in neurons 2 |
| chr15 | 1,02E+08 | 1,02E+08 | 1603 | * | 16,8117 | 2,158231253 | 0,767897 | 2,810574 | 0,004945 | 0,786235538 | 1,2763 | Distal Intergenic | 15 | 1,02E+08 | 1,02E+08 | 1753 | 1 | 432986 | ENSMUST00000231005.2 | -8781 | | Gm5477 | type II keratin Kb16P |
| chr1 | 1,65E+08 | 1,65E+08 | 2106 | * | 16,94258 | 2,158342232 | 0,811218 | 2,660619 | 0,0078 | 0,756849896 | 2,318434 | Promoter (<=1kb) | 1 | 1,65E+08 | 1,65E+08 | 95497 | 2 | 1,03E+08 | ENSMUST00000193955.2 | 0 | | Gm32569 | predicted gene, 32569 |
| chr8 | 26208985 | 26211619 | 2635 | * | 27,58698 | 2,15846287 | 0,778735 | 2,771756 | 0,005575 | 0,676476319 | 2,725491 | Promoter (<=1kb) | 8 | 26210064 | 26214914 | 4851 | 1 | 71910 | ENSMUST00000068916.16 | 0 | ENSMUSG00000031570 | Plpp5 | phospholipid phosphatase 5 |
| chr5 | 34705707 | 34707852 | 2146 | * | 19,13169 | 2,158689155 | 0,806144 | 2,677795 | 0,007411 | 0,735590291 | 2,507013 | Promoter (<=1kb) | 5 | 34706920 | 34720969 | 14050 | 1 | 24055 | ENSMUST00000118545.8 | 0 | ENSMUSG00000054520 | Sh3bp2 | SH3-domain binding protein 2 |
| chr2 | 13414277 | 13416276 | 2000 | * | 20,35421 | 2,15880236 | 0,764295 | 2,824567 | 0,004734 | 0,653331264 | 2,663619 | Intron (ENSMUST0000( | 2 | 13426840 | 13460438 | 33599 | 2 | 65969 | ENSMUST00000195447.2 | 44162 | ENSMUSG00000026726 | Cubn | cubilin (intrinsic factor-cobalamin receptor) |
| chr19 | 37322740 | 37324848 | 2109 | * | 17,70194 | 2,159001419 | 0,794436 | 2,717653 | 0,006575 | 0,705941691 | 2,663619 | Intron (ENSMUST0000( | 19 | 37281049 | 37315251 | 34203 | 2 | 15925 | ENSMUST00000154308.2 | -7489 | ENSMUSG00000056999 | Ide | insulin degrading enzyme |
| chr2 | 1,69E+08 | 1,69E+08 | 1804 | * | 16,36948 | 2,159047523 | 0,793216 | 2,72189 | 0,006491 | 0,727652793 | 2,318434 | Exon (ENSMUST00000: | 2 | 1,69E+08 | 1,69E+08 | 4692 | 2 | 22722 | ENSMUST00000136242.2 | 18207 | ENSMUSG00000027551 | Zfp64 | zinc finger protein 64 |
| chr4 | 57789690 | 57791616 | 1927 | * | 19,26344 | 2,159282435 | 0,728038 | 2,965893 | 0,003018 | 0,580348965 | 2,507013 | Intron (ENSMUST0000( | 4 | 57782180 | 57855202 | 73023 | 1 | 677884 | ENSMUST00000124581.2 | 7510 | ENSMUSG00000038729 | Pakap | paralemmin A kinase anchor protein |
| chr16 | 5284413 | 5287575 | 3163 | * | 27,74422 | 2,159791132 | 0,733901 | 2,94289 | 0,003252 | 0,578728206 | 2,725491 | Distal Intergenic | 16 | 5249345 | 5268689 | 19345 | 1 | 1,05E+08 | ENSMUST00000242230.1 | 35068 | | Gm41414 | predicted gene, 41414 |
| chr4 | 1,05E+08 | 1,05E+08 | 4005 | * | 32,81216 | 2,15981638 | 0,605193 | 3,568803 | 0,000359 | 0,227550076 | 2,787255 | Intron (ENSMUST0000( | 4 | 1,05E+08 | 1,05E+08 | 75418 | 1 | 67916 | ENSMUST00000064139.8 | 57929 | ENSMUSG00000028517 | Plpp3 | phospholipid phosphatase 3 |
| chr2 | 1,56E+08 | 1,56E+08 | 1860 | * | 19,59245 | 2,16016181 | 0,792093 | 2,727157 | 0,006388 | 0,695401133 | 2,713306 | Intron (ENSMUST0000( | 2 | 1,56E+08 | 1,57E+08 | 150659 | 1 | 228836 | ENSMUST00000169464.9 | 7813 | ENSMUSG00000061689 | Dlgap4 | DLG associated protein 4 |
| chr19 | 5065624 | 5069236 | 3613 | * | 31,48723 | 2,160606546 | 0,636838 | 3,392712 | 0,000692 | 0,315367183 | 2,67243 | Distal Intergenic | 19 | 5073888 | 5082000 | 8113 | 1 | 13340 | ENSMUST00000236152.2 | -4652 | ENSMUSG00000024891 | Slc29a2 | solute carrier family 29 (nucleoside transporters), member 2 |
| chr2 | 5981850 | 5983788 | 1939 | * | 19,62096 | 2,161403957 | 0,825281 | 2,618993 | 0,008819 | 0,776632464 | 2,386008 | Intron (ENSMUST0000( | 2 | 5956280 | 6061514 | 105235 | 1 | 326622 | ENSMUST00000060092.13 | 25570 | ENSMUSG00000043241 | Upf2 | UPF2 regulator of nonsense transcripts homolog (yeast) |
| chr10 | 80701430 | 80703236 | 1807 | * | 20,6511 | 2,162105907 | 0,811978 | 2,662763 | 0,00775 | 0,685078899 | 3,471678 | Promoter (2-3kb) | 10 | 80699315 | 80701580 | 2266 | 1 | 73218 | ENSMUST00000219136.2 | 2115 | ENSMUSG00000035206 | Sppl2b | signal peptide peptidase like 2B |
| chr9 | 96746485 | 96749074 | 2590 | * | 21,06224 | 2,162777845 | 0,724331 | 2,985899 | 0,002827 | 0,541111532 | 2,880017 | Promoter (1-2kb) | 9 | 96738523 | 96745035 | 6513 | 2 | 235534 | ENSMUST00000141127.2 | -1450 | ENSMUSG00000043587 | Pxylp1 | 2-phosphoxylose phosphatase 1 |
| chr18 | 4452772 | 4454801 | 2030 | * | 17,63534 | 2,163608945 | 0,815786 | 2,652177 | 0,007997 | 0,738600147 | 2,663619 | Distal Intergenic | 18 | 4383246 | 4397330 | 14085 | 1 | 67440 | ENSMUST00000234924.2 | 69526 | ENSMUSG00000024234 | Mtpap | mitochondrial poly(A) polymerase |
| chr2 | 90507546 | 90510851 | 3306 | * | 25,1615 | 2,1641424 | 0,69092 | 3,132263 | 0,001735 | 0,456474583 | 2,880017 | Promoter (<=1kb) | 2 | 90507559 | 90566672 | 59114 | 1 | 59015 | ENSMUST00000057481.7 | 0 | ENSMUSG00000051329 | Nup160 | nucleoporin 160 |
| chr5 | 24112582 | 24114898 | 2317 | * | 27,66239 | 2,164193128 | 0,648718 | 3,33611 | 0,00085 | 0,334207335 | 2,880017 | Distal Intergenic | 5 | 24120274 | 24127902 | 7629 | 2 | 84652 | ENSMUST00000126828.2 | 13004 | ENSMUSG00000028995 | Fam126a | family with sequence similarity 126, member A |
| chr15 | 37782597 | 37784841 | 2245 | * | 27,38496 | 2,164386021 | 0,67621 | 3,200758 | 0,001371 | 0,426043749 | 2,725491 | Intron (ENSMUST0000( | 15 | 37366419 | 37792244 | 425826 | 2 | 52589 | ENSMUST00000168992.8 | 7403 | ENSMUSG00000051359 | Ncald | neurocalcin delta |
| chr12 | 6726604 | 6728331 | 1728 | * | 13,95167 | 2,164582937 | 0,835041 | 2,592189 | 0,009537 | 0,979538317 | 0,738441 | Distal Intergenic | 12 | 6548124 | 6548615 | 492 | 2 | 1E+08 | ENSMUST00000218888.2 | -177989 | ENSMUSG00000111967 | Gm3678 | predicted gene 3678 |
| chr13 | 94590587 | 94592318 | 1732 | * | 18,03741 | 2,165008625 | 0,759977 | 2,848783 | 0,004389 | 0,675594321 | 2,154504 | Exon (ENSMUST00000( | 13 | 94609371 | 94702821 | 93451 | 1 | 11774 | ENSMUST00000231916.2 | -17053 | ENSMUSG00000021686 | Ap3b1 | adaptor-related protein complex 3, beta 1 subunit |
| chr16 | 7987098 | 7989418 | 2321 | * | 20,09762 | 2,165223857 | 0,761403 | 2,84373 | 0,004459 | 0,653416641 | 2,507013 | Distal Intergenic | 16 | 8154857 | 8154960 | 104 | 1 | 1,15E+08 | ENSMUST00000158968.3 | -165439 | ENSMUSG00000089593 | Gm26159 | predicted gene, 26159 |
| chr17 | 24692819 | 24695233 | 2415 | * | 29,37833 | 2,165354784 | 0,656143 | 3,300127 | 0,000966 | 0,364434126 | 2,725491 | Promoter (<=1kb) | 17 | 24692858 | 24694443 | 1586 | 1 | 319259 | ENSMUST00000054946.10 | 0 | ENSMUSG00000045744 | Bricd5 | BRICHOS domain containing 5 |
| chr7 | 76000979 | 76002742 | 1764 | * | 17,2251 | 2,166206044 | 0,823233 | 2,631341 | 0,008505 | 0,758671033 | 2,507013 | Intron (ENSMUST0000( | 7 | 76060810 | 76349871 | 289062 | 1 | 244071 | ENSMUST00000026854.10 | -58068 | ENSMUSG00000025754 | Agbl1 | ATP/GTP binding protein-like 1 |
| chr4 | 1,07E+08 | 1,07E+08 | 2313 | * | 20,19442 | 2,166566558 | 0,719743 | 3,010193 | 0,002611 | 0,562310527 | 2,386008 | Promoter (<=1kb) | 4 | 1,07E+08 | 1,07E+08 | 107837 | 1 | 72475 | ENSMUST00000148842.8 | 0 | ENSMUSG00000061887 | Ssbp3 | single-stranded DNA binding protein 3 |
| chr4 | 46154678 | 46157920 | 3243 | * | 26,20137 | 2,167167075 | 0,753243 | 2,877117 | 0,004013 | 0,621084104 | 2,725491 | Promoter (<=1kb) | 4 | 46157901 | 46160099 | 2199 | 1 | 433702 | ENSMUST00000131187.2 | 0 | ENSMUSG00000028330 | Ncbp1 | nuclear cap binding protein subunit 1 |
| chr1 | 33803136 | 33805619 | 2484 | * | 19,32841 | 2,167555475 | 0,827058 | 2,620803 | 0,008772 | 0,753158126 | 2,663619 | 3' UTR | 1 | 33784565 | 33796876 | 12312 | 2 | 213539 | ENSMUST00000044691.9 | -6260 | ENSMUSG00000042215 | Bag2 | BCL2-associated athanogene 2 |
| chr5 | 1,43E+08 | 1,43E+08 | 2073 | * | 25,41211 | 2,168401841 | 0,764565 | 2,836125 | 0,004566 | 0,63293289 | 2,880017 | Distal Intergenic | 5 | 1,43E+08 | 1,43E+08 | 1905 | 1 | 14086 | ENSMUST00000135293.2 | 18431 | ENSMUSG00000029581 | Fscn1 | fascin actin-bundling protein 1 |
| chr1 | 1,43E+08 | 1,43E+08 | 3373 | * | 27,0024 | 2,170157566 | 0,753988 | 2,878239 | 0,003999 | 0,624283749 | 2,67243 | Distal Intergenic | 1 | 1,43E+08 | 1,44E+08 | 34421 | 2 | 214498 | ENSMUST00000162638.2 | 695652 | ENSMUSG00000026361 | Cdc73 | cell division cycle 73, Paf1/RNA polymerase II complex component |
| chr12 | 1,1E+08 | 1,1E+08 | 3003 | * | 27,91852 | 2,170515959 | 0,764344 | 2,83971 | 0,004515 | 0,631028885 | 2,880017 | Distal Intergenic | 12 | 1,1E+08 | 1,1E+08 | 3880 | 1 | 26931 | ENSMUST00000222276.2 | -43324 | ENSMUSG00000017843 | Ppp2r5c | protein phosphatase 2, regulatory subunit B', gamma |
| chr8 | 127277437 | 127280485 | 3049 | * | 47,966356 | 2,171519337 | 0,4636542 | 4,6834888 | 2,82E-06 | 0,008394893 | 2,7872553 | Distal Intergenic | 8 | 127315035 | 127320725 | 5691 | 2 | 270110 | ENSMUST00000054960.9 | 40240 | ENSMUSG00000051495 | Irf2bp2 | interferon regulatory factor 2 binding protein 2 |
| chr10 | 43413949 | 43416090 | 2142 | * | 21,56897 | 2,172111433 | 0,708346 | 3,066457 | 0,002166 | 0,494205337 | 2,880017 | Promoter (<=1kb) | 10 | 43401129 | 43416964 | 15836 | 2 | 67851 | ENSMUST00000147196.3 | 874 | ENSMUSG00000019797 | Mtres1 | mitochondrial transcription rescue factor 1 |
| chr16 | 38318886 | 38320410 | 1525 | * | 15,59185 | 2,172432215 | 0,775397 | 2,801705 | 0,005083 | 0,770673161 | 1,41696 | 3' UTR | 16 | 38337022 | 38343003 | 5982 | 2 | 76916 | ENSMUST00000147543.2 | 22593 | ENSMUSG00000002846 | Timmdc1 | translocase of inner mitochondrial membrane domain containing 1 |
| chr3 | 26599897 | 26601731 | 1835 | * | 17,60885 | 2,172597173 | 0,781804 | 2,778955 | 0,005453 | 0,688853937 | 2,386008 | Distal Intergenic | 3 | 26691769 | 27037361 | 345593 | 1 | 70862 | ENSMUST00000047005.11 | -90038 | ENSMUSG00000039335 | Spata16 | spermatogenesis associated 16 |
| chr12 | 16898068 | 16899836 | 1769 | * | 16,66684 | 2,173141911 | 0,800483 | 2,714787 | 0,006632 | 0,730469267 | 2,318434 | Promoter (<=1kb) | 12 | 16894917 | 16900782 | 5866 | 2 | 1,03E+08 | ENSMUST00000245045.1 | 946 | | Gm36495 | predicted gene, 36495 |
| chr10 | 80458883 | 80460434 | 1552 | * | 20,28543 | 2,17348756 | 0,794414 | 2,735962 | 0,00622 | 0,715249603 | 2,386008 | Promoter (<=1kb) | 10 | 80458672 | 80476575 | 17904 | 1 | 103236 | ENSMUST00000079773.14 | 211 | ENSMUSG00000003345 | Csnk1g2 | casein kinase 1, gamma 2 |
| chr2 | 1,54E+08 | 1,54E+08 | 2506 | * | 21,40731 | 2,173644877 | 0,747173 | 2,909158 | 0,003624 | 0,549506271 | 3,471678 | 3' UTR | 2 | 1,54E+08 | 1,54E+08 | 37631 | 2 | 66971 | ENSMUST00000109731.8 | -3156 | ENSMUSG00000027487 | Cdk5rap1 | CDK5 regulatory subunit associated protein 1 |
| chr5 | 1,09E+08 | 1,09E+08 | 1285 | * | 21,24949 | 2,17368459 | 0,813881 | 2,670763 | 0,007568 | 0,715249603 | 2,880017 | Promoter (2-3kb) | 5 | 1,09E+08 | 1,09E+08 | 799 | 1 | 116701 | ENSMUST00000199802.2 | -2115 | ENSMUSG00000008090 | Fgfrl1 | fibroblast growth factor receptor-like 1 |
| chr10 | 99688265 | 99690221 | 1957 | * | 20,60194 | 2,173818534 | 0,826853 | 2,629028 | 0,008563 | 0,724313953 | 3,104559 | Intron (ENSMUST0000( | 10 | 99593568 | 99595520 | 1953 | 2 | 71832 | ENSMUST00000056085.6 | -92745 | ENSMUSG00000046934 | Csl | citrate synthase like |
| chr11 | 36195647 | 36197237 | 1591 | * | 18,55731 | 2,174094695 | 0,771277 | 2,818824 | 0,00482 | 0,671156916 | 2,423594 | Intron (ENSMUST0000( | 11 | 36275950 | 36323103 | 47154 | 2 | 23964 | ENSMUST00000128501.2 | 125866 | ENSMUSG00000049336 | Tenm2 | teneurin transmembrane protein 2 |
| chr13 | 55467340 | 55469891 | 2552 | * | 26,29731 | 2,174499263 | 0,773228 | 2,812236 | 0,00492 | 0,642870233 | 2,951062 | Promoter (<=1kb) | 13 | 55467556 | 55469759 | 2204 | 2 | 19336 | ENSMUST00000225289.2 | 0 | ENSMUSG00000034789 | Rab24 | RAB24, member RAS oncogene family |
| chr5 | 23977481 | 23980750 | 3270 | * | 28,57048 | 2,174991798 | 0,634207 | 3,429467 | 0,000605 | 0,298861419 | 2,725491 | Promoter (2-3kb) | 5 | 23965264 | 23983376 | 18113 | 2 | 78697 | ENSMUST00000151449.2 | 2626 | ENSMUSG00000057541 | Pus7 | pseudouridylate synthase 7 |
| chr5 | 1,45E+08 | 1,45E+08 | 1905 | * | 21,44875 | 2,175779712 | 0,761129 | 2,85862 | 0,004255 | 0,616346987 | 2,951933 | Distal Intergenic | 5 | 1,45E+08 | 1,45E+08 | 171 | 2 | 1,08E+08 | ENSMUST00000121256.2 | 33225 | | Gm6272 | predicted pseudogene 6272 |
| chr12 | 52950875 | 52952612 | 1738 | * | 15,57818 | 2,175809834 | 0,838209 | 2,595784 | 0,009438 | 0,886275091 | 1,37683 | Intron (ENSMUST0000( | 12 | 52842619 | 53195546 | 352928 | 1 | 238161 | ENSMUST00000219786.2 | 108256 | ENSMUSG00000061603 | Akap6 | A kinase (PRKA) anchor protein 6 |
| chr4 | 1,33E+08 | 1,33E+08 | 1420 | * | 14,7898 | 2,176003925 | 0,826043 | 2,63425 | 0,008432 | 0,858257847 | 1,41696 | Promoter (<=1kb) | 4 | 1,33E+08 | 1,33E+08 | 69252 | 1 | 242687 | ENSMUST00000084241.12 | 0 | ENSMUSG00000028868 | Wasf2 | WASP family, member 2 |
| chr1 | 93721778 | 93725432 | 3655 | * | 34,34123 | 2,176263259 | 0,663642 | 3,279275 | 0,001041 | 0,406085428 | 2,377674 | Promoter (1-2kb) | 1 | 93720446 | 93726723 | 6278 | 2 | 21915 | ENSMUST00000112893.2 | 1291 | ENSMUSG00000026281 | Dtymk | deoxythymidylate kinase |
| chr19 | 57192392 | 57193808 | 1417 | * | 20,60052 | 2,176416867 | 0,762307 | 2,855039 | 0,004303 | 0,586565597 | 3,471678 | Intron (ENSMUST0000( | 19 | 57021706 | 57186063 | 164358 | 2 | 226251 | ENSMUST00000099294.9 | -6329 | ENSMUSG00000025085 | Ablim1 | actin-binding LIM protein 1 |
| chr3 | 1,02E+08 | 1,02E+08 | 1734 | * | 19,84083 | 2,176431813 | 0,831422 | 2,617723 | 0,008852 | 0,767135014 | 2,507013 | Intron (ENSMUST0000( | 3 | 1,02E+08 | 1,02E+08 | 23148 | 2 | 242125 | ENSMUST00000061831.11 | 3485 | ENSMUSG00000044313 | Mab21l3 | mab-21-like 3 |

| | | | | | | | | | | | | | | | | | | | | | | | |
|---|---|---|---|---|---|---|---|---|---|---|---|---|---|---|---|---|---|---|---|---|---|---|---|
| chr1 | 1,66E+08 | 1,66E+08 | 2086 | * | 23,06522 | 2,17651678 | 0,7259 | 2,998369 | 0,002714 | 0,543993049 | 2,67243 | Distal Intergenic | 1 | 1,66E+08 | 1,66E+08 | 14525 | 1 | 27878 | ENSMUST00000027846.8 | -41104 | ENSMUSG00000026563 | Tada1 | transcriptional adaptor 1 |
| chr9 | 78353297 | 78355309 | 2013 | * | 14,31556 | 2,176663 | 0,819431 | 2,65631 | 0,0079 | 0,895765172 | 1,075735 | Promoter (<=1kb) | 9 | 78355490 | 78381434 | 25945 | 1 | 68291 | ENSMUST00000034896.13 | -181 | ENSMUSG00000032342 | Mto1 | mitochondrial tRNA translation optimization 1 |
| chr1 | 1,03E+08 | 1,03E+08 | 1733 | * | 17,49594 | 2,177254851 | 0,770116 | 2,827176 | 0,004696 | 0,651384755 | 2,663619 | Distal Intergenic | 1 | 1,03E+08 | 1,03E+08 | 134 | 2 | 1,15E+08 | ENSMUST00000157283.3 | 66548 | ENSMUSG00000087908 | Gm22034 | predicted gene, 22034 |
| chr7 | 64095599 | 64097195 | 1597 | * | 18,19616 | 2,177942586 | 0,752158 | 2,895591 | 0,003784 | 0,63298934 | 2,386008 | Distal Intergenic | 7 | 64042523 | 64061869 | 19347 | 1 | 73724 | ENSMUST00000206194.2 | 53076 | ENSMUSG00000033429 | Mcee | methylmalonyl CoA epimerase |
| chr10 | 48104095 | 48106092 | 1998 | * | 19,48444 | 2,177978711 | 0,775073 | 2,810031 | 0,004954 | 0,685026746 | 2,221611 | Distal Intergenic | 10 | 48495408 | 48495491 | 84 | 2 | 1,15E+08 | ENSMUST00000083696.4 | 389399 | | Gm24256 | predicted gene, 24256 |
| chr7 | 83140390 | 83142444 | 2055 | * | 18,05637 | 2,17800728 | 0,800517 | 2,720752 | 0,006513 | 0,715156227 | 2,507013 | Distal Intergenic | 7 | 83192078 | 83199464 | 7387 | 2 | 1E+08 | ENSMUST00000208877.2 | 57020 | | Gm10610 | predicted gene 10610 |
| chr10 | 31809450 | 31811295 | 1846 | * | 20,51016 | 2,178966406 | 0,844553 | 2,580022 | 0,009879 | 0,793995086 | 2,386008 | Intron (ENSMUST0000 | 10 | 31369189 | 31485180 | 115992 | 2 | 268291 | ENSMUST00000081989.8 | -324270 | ENSMUSG00000063760 | Rnf217 | ring finger protein 217 |
| chr3 | 85132868 | 85135335 | 2468 | * | 19,13089 | 2,179019924 | 0,827761 | 2,632426 | 0,008478 | 0,757937851 | 2,507013 | Intron (ENSMUST0000 | 3 | 85224826 | 85239337 | 14512 | 1 | 73283 | ENSMUST00000189046.2 | -89491 | ENSMUSG00000099995 | 1700036G14Rik | RIKEN cDNA 1700036G14 gene |
| chr8 | 1,06E+08 | 1,06E+08 | 3100 | * | 27,1303 | 2,179785424 | 0,682372 | 3,194422 | 0,001401 | 0,418252864 | 2,951062 | Intron (ENSMUST0000 | 8 | 1,06E+08 | 1,06E+08 | 50452 | 1 | 270091 | ENSMUST00000216765.2 | 5250 | ENSMUSG00000054320 | Lrrc36 | leucine rich repeat containing 36 |
| chr12 | 1,12E+08 | 1,12E+08 | 2603 | * | 21,36113 | 2,180512382 | 0,811435 | 2,687229 | 0,007205 | 0,692527675 | 3,104559 | Promoter (<=1kb) | 12 | 1,12E+08 | 1,12E+08 | 2130 | 2 | 1,01E+08 | ENSMUST00000221597.3 | 0 | ENSMUSG00000096995 | 2810029C07Rik | RIKEN cDNA 2810029C07 gene |
| chr17 | 26800521 | 26802495 | 1975 | * | 20,07649 | 2,180711423 | 0,736148 | 2,962327 | 0,003053 | 0,569763832 | 2,663619 | Intron (ENSMUST0000 | 17 | 26813850 | 26874326 | 60477 | 1 | 67458 | ENSMUST00000235915.2 | -11355 | ENSMUSG00000001576 | Ergic1 | endoplasmic reticulum-golgi intermediate compartment (ERGIC) 1 |
| chr1 | 16182941 | 16185220 | 2280 | * | 19,1999 | 2,180961879 | 0,804573 | 2,710708 | 0,006714 | 0,710505631 | 2,663619 | Intron (ENSMUST0000 | 1 | 16175998 | 16203958 | 27961 | 1 | 98711 | ENSMUST00000027053.8 | 6943 | ENSMUSG00000025921 | Rdh10 | retinol dehydrogenase 10 (all-trans) |
| chr10 | 62376056 | 62378865 | 2810 | * | 25,05244 | 2,181023939 | 0,704911 | 3,094044 | 0,001974 | 0,492203416 | 2,67243 | Intron (ENSMUST0000 | 10 | 62329612 | 62363230 | 33619 | 2 | 19073 | ENSMUST00000160987.8 | -12826 | ENSMUSG00000020077 | Srgn | serglycin |
| chr11 | 1,18E+08 | 1,18E+08 | 2190 | * | 29,44755 | 2,18138481 | 0,650795 | 3,351878 | 0,000803 | 0,327375387 | 2,858511 | Promoter (<=1kb) | 11 | 1,18E+08 | 1,18E+08 | 30581 | 2 | 72344 | ENSMUST00000106296.9 | 0 | ENSMUSG00000033909 | Usp36 | ubiquitin specific peptidase 36 |
| chr4 | 43382228 | 43385187 | 2960 | * | 27,88142 | 2,181432601 | 0,759524 | 2,872105 | 0,004077 | 0,624283749 | 2,725491 | Promoter (<=1kb) | 4 | 43383449 | 43414840 | 31392 | 1 | 100213 | ENSMUST00000135216.2 | 0 | ENSMUSG00000035969 | Rusc2 | RUN and SH3 domain containing 2 |
| chr8 | 1,24E+08 | 1,24E+08 | 2912 | * | 31,90447 | 2,181939346 | 0,638765 | 3,41587 | 0,000636 | 0,295868206 | 2,951062 | Distal Intergenic | 8 | 1,24E+08 | 1,24E+08 | 2057 | 1 | 69581 | ENSMUST00000136615.2 | -3012 | ENSMUSG00000039960 | Rhou | ras homolog family member U |
| chr14 | 52423535 | 52425664 | 2130 | * | 16,82288 | 2,182240352 | 0,802513 | 2,719258 | 0,006543 | 0,723215019 | 2,386008 | Intron (ENSMUST0000 | 14 | 52397871 | 52434873 | 37003 | 2 | 114741 | ENSMUST00000046709.9 | 9209 | ENSMUSG00000035726 | Supt16 | SPT16, facilitates chromatin remodeling subunit |
| chr1 | 1,03E+08 | 1,03E+08 | 1551 | * | 16,47857 | 2,18235577 | 0,838793 | 2,601782 | 0,009274 | 0,89325945 | 1,2763 | Distal Intergenic | 1 | 1,03E+08 | 1,03E+08 | 134 | 2 | 1,15E+08 | ENSMUST00000157283.3 | 33469 | ENSMUSG00000087908 | Gm22034 | predicted gene, 22034 |
| chr5 | 53020440 | 53024501 | 4062 | * | 42,11531 | 2,182525311 | 0,536793 | 4,06586 | 4,79E-05 | 0,068345002 | 2,787255 | Promoter (<=1kb) | 5 | 53022955 | 53024076 | 1122 | 1 | 52206 | ENSMUST00000138629.2 | 0 | ENSMUSG00000029176 | Anapc4 | anaphase promoting complex subunit 4 |
| chr15 | 53555566 | 53557160 | 1595 | * | 20,11646 | 2,182533517 | 0,80476 | 2,71203 | 0,006687 | 0,739106282 | 2,221611 | Intron (ENSMUST0000 | 15 | 53477601 | 53477675 | 75 | 2 | 1,01E+08 | ENSMUST00000200096.3 | -77891 | ENSMUSG00000104794 | Mir28c | microRNA 28c |
| chr1 | 86397988 | 86399278 | 1291 | * | 21,66987 | 2,182631089 | 0,784312 | 2,78286 | 0,005388 | 0,658200216 | 2,951933 | Distal Intergenic | 1 | 86354051 | 86355774 | 1724 | 1 | 71863 | ENSMUST00000046004.6 | 43937 | ENSMUSG00000036574 | Tex44 | testis expressed 44 |
| chr19 | 47905999 | 47909227 | 3229 | * | 29,36983 | 2,183524537 | 0,741226 | 2,94583 | 0,003221 | 0,565507616 | 2,880017 | Promoter (<=1kb) | 19 | 47775019 | 47907726 | 132708 | 2 | 1E+08 | ENSMUST00000026048.10 | 0 | ENSMUSG00000044948 | Cfap43 | cilia and flagella associated protein 43 |
| chr4 | 1,06E+08 | 1,06E+08 | 2198 | * | 23,12235 | 2,183962046 | 0,758768 | 2,8783 | 0,003998 | 0,61075929 | 2,858511 | Exon (ENSMUST00000 | 4 | 1,06E+08 | 1,06E+08 | 2360 | 1 | 329908 | ENSMUST00000150521.2 | -39722 | ENSMUSG00000028514 | Usp24 | ubiquitin specific peptidase 24 |
| chr10 | 89278598 | 89280520 | 1923 | * | 23,91022 | 2,184430169 | 0,708598 | 3,082752 | 0,002051 | 0,479521813 | 2,951933 | Promoter (<=1kb) | 10 | 89244685 | 89279829 | 35145 | 2 | 237436 | ENSMUST00000099374.9 | 0 | ENSMUSG00000074802 | Gas2l3 | growth arrest-specific 2 like 3 |
| chr4 | 1,26E+08 | 1,26E+08 | 2915 | * | 18,44102 | 2,184886141 | 0,798359 | 2,736723 | 0,006205 | 0,713633527 | 2,423594 | Exon (ENSMUST00000 | 4 | 1,26E+08 | 1,26E+08 | 5326 | 2 | 214150 | ENSMUST00000127831.2 | -20337 | ENSMUSG00000028842 | Ago3 | argonaute RISC catalytic subunit 3 |
| chr7 | 4139471 | 4140963 | 1493 | * | 22,50005 | 2,185051855 | 0,784391 | 2,785666 | 0,005342 | 0,671923639 | 2,67243 | Promoter (<=1kb) | 7 | 4129339 | 4139925 | 10587 | 2 | 1E+08 | ENSMUST00000148318.2 | 0 | ENSMUSG00000085886 | D030047H15Rik | RIKEN cDNA D030047H15 gene |
| chr7 | 1,39E+08 | 1,39E+08 | 1808 | * | 17,33921 | 2,185411175 | 0,783151 | 2,790538 | 0,005262 | 0,683966322 | 2,423594 | Promoter (<=1kb) | 7 | 1,39E+08 | 1,39E+08 | 5307 | 1 | 76612 | ENSMUST00000130248.2 | 328 | ENSMUSG00000015980 | Lrrc27 | leucine rich repeat containing 27 |
| chr3 | 45477403 | 45479414 | 2012 | * | 16,34797 | 2,185953694 | 0,79747 | 2,741111 | 0,006123 | 0,730469267 | 2,141612 | Distal Intergenic | 3 | 45371019 | 45393745 | 22727 | 2 | 72274 | ENSMUST00000186818.3 | -83658 | ENSMUSG00000100799 | 1700027H10Rik | RIKEN cDNA 1700027H10 gene |
| chr19 | 24569659 | 24571839 | 2181 | * | 22,85696 | 2,187236708 | 0,697959 | 3,133762 | 0,001726 | 0,468195777 | 2,67243 | Distal Intergenic | 19 | 24533668 | 24534594 | 927 | 1 | 1,15E+08 | ENSMUST00000235982.3 | 35991 | | Gm50341 | predicted gene, 50341 |
| chr15 | 83365520 | 83367860 | 2341 | * | 24,84505 | 2,188169182 | 0,786033 | 2,783812 | 0,005372 | 0,67266775 | 2,67243 | Distal Intergenic | 15 | 83259808 | 83348807 | 89000 | 2 | 23970 | ENSMUST00000171436.8 | -16713 | ENSMUSG00000016664 | Pacsin2 | protein kinase C and casein kinase substrate in neurons 2 |
| chr2 | 1,66E+08 | 1,66E+08 | 2305 | * | 19,8257 | 2,188237309 | 0,804574 | 2,719746 | 0,006533 | 0,723027638 | 2,386008 | Promoter (<=1kb) | 2 | 1,66E+08 | 1,66E+08 | 24823 | 2 | 228880 | ENSMUST00000149369.8 | -465 | ENSMUSG00000039671 | Zmynd8 | zinc finger, MYND-type containing 8 |
| chr19 | 5859729 | 5861945 | 2217 | * | 61,28677 | 2,188333754 | 0,768002 | 2,849384 | 0,00438 | 0,55735526 | 4,055758 | Distal Intergenic | 19 | 5845717 | 5852704 | 6988 | 2 | 72289 | ENSMUST00000172812.4 | -7025 | ENSMUSG00000092341 | Malat1 | metastasis associated lung adenocarcinoma transcript 1 (non-coding RNA) |
| chr5 | 4840359 | 4842509 | 2151 | * | 21,18984 | 2,188696834 | 0,796021 | 2,749546 | 0,005968 | 0,678869598 | 2,880017 | Intron (ENSMUST0000 | 5 | 4803839 | 4808035 | 4197 | 2 | 14362 | ENSMUST00000054294.7 | -32324 | ENSMUSG00000044674 | Fzd1 | frizzled class receptor 1 |
| chr6 | 1,35E+08 | 1,35E+08 | 2246 | * | 26,51748 | 2,18876748 | 0,624814 | 3,503071 | 0,00046 | 0,263182818 | 2,725491 | Promoter (<=1kb) | 6 | 1,35E+08 | 1,35E+08 | 6481 | 1 | 1,01E+08 | ENSMUST00000204330.2 | -646 | ENSMUSG00000108239 | Gm19434 | predicted gene, 19434 |
| chr7 | 1,01E+08 | 1,01E+08 | 2206 | * | 19,50507 | 2,189373418 | 0,801313 | 2,732232 | 0,006291 | 0,709779237 | 2,507013 | Intron (ENSMUST0000 | 7 | 1,01E+08 | 1,01E+08 | 68126 | 1 | 20480 | ENSMUST00000153473.2 | 33049 | ENSMUSG00000001829 | Clpb | ClpB caseinolytic peptidase B |
| chr18 | 23571205 | 23573110 | 1906 | * | 23,25836 | 2,189750795 | 0,6697 | 3,269751 | 0,001076 | 0,369614065 | 2,951062 | Intron (ENSMUST0000 | 18 | 23548567 | 23737288 | 188722 | 1 | 13527 | ENSMUST00000047954.15 | 22638 | ENSMUSG00000024302 | Dtna | dystrobrevin alpha |
| chr15 | 37128504 | 37132147 | 3644 | * | 36,52821 | 2,190122033 | 0,525086 | 4,17098 | 3,03E-05 | 0,051012164 | 2,787255 | Distal Intergenic | 15 | 37233280 | 37333891 | 100612 | 1 | 252973 | ENSMUST00000161405.8 | -101133 | ENSMUSG00000022286 | Grhl2 | grainyhead like transcription factor 2 |
| chr15 | 83188468 | 83191432 | 2965 | * | 44,51194 | 2,190511582 | 0,605998 | 3,614718 | 0,000301 | 0,206624632 | 2,787255 | Promoter (<=1kb) | 15 | 83183941 | 83189542 | 5602 | 2 | 66251 | ENSMUST00000226816.2 | 0 | ENSMUSG00000054277 | Arfgap3 | ADP-ribosylation factor GTPase activating protein 3 |
| chr4 | 64254641 | 64256892 | 2252 | * | 15,84715 | 2,191047631 | 0,83163 | 2,634642 | 0,008423 | 0,767135014 | 2,386008 | Distal Intergenic | 4 | 64172290 | 64194840 | 22551 | 2 | 1,03E+08 | ENSMUST00000141162.2 | -59801 | ENSMUSG00000085032 | Gm11217 | predicted gene 11217 |
| chr16 | 22551935 | 22556568 | 4634 | * | 51,536397 | 2,191620801 | 0,5038089 | 4,350103 | 1,361E-05 | 0,02869889 | 2,7254908 | Distal Intergenic | 16 | 22623953 | 22630327 | 6375 | 2 | 12970 | ENSMUST00000040592.6 | 73759 | ENSMUSG00000033501 | Crygs | crystallin, gamma S |
| chr11 | 1,1E+08 | 1,1E+08 | 1430 | * | 15,38078 | 2,192032362 | 0,803742 | 2,727283 | 0,006386 | 0,829907213 | 1,2763 | Promoter (<=1kb) | 11 | 1,1E+08 | 1,1E+08 | 52531 | 2 | 208659 | ENSMUST00000155559.8 | 0 | ENSMUSG00000020614 | Fam20a | FAM20A, golgi associated secretory pathway pseudokinase |
| chr6 | 47925101 | 47927202 | 2102 | * | 28,29559 | 2,192108093 | 0,608967 | 3,599718 | 0,000319 | 0,206624632 | 2,951062 | Promoter (1-2kb) | 6 | 47923605 | 47930508 | 6904 | 1 | 232785 | ENSMUST00000132381.2 | 1496 | ENSMUSG00000072653 | Zfp783 | zinc finger protein 783 |
| chr14 | 31064189 | 31066193 | 2005 | * | 16,75357 | 2,192399835 | 0,831881 | 2,635472 | 0,008402 | 0,767061106 | 2,386008 | Intron (ENSMUST0000 | 14 | 31058664 | 31093941 | 35278 | 1 | 12339 | ENSMUST00000152182.2 | 5525 | ENSMUSG00000021893 | Capn7 | calpain 7 |
| chr9 | 89935676 | 89937217 | 1542 | * | 15,21897 | 2,193055873 | 0,811911 | 2,701104 | 0,006911 | 0,843602846 | 1,2763 | Promoter (<=1kb) | 9 | 89936205 | 89947475 | 11271 | 1 | 13036 | ENSMUST00000132718.8 | 0 | ENSMUSG00000032359 | Ctsh | cathepsin H |
| chr10 | 81364885 | 81367642 | 2758 | * | 30,01655 | 2,193073088 | 0,638526 | 3,434588 | 0,000593 | 0,284660899 | 2,951062 | Promoter (<=1kb) | 10 | 81366121 | 81367849 | 1729 | 2 | 14672 | ENSMUST00000146984.2 | 207 | ENSMUSG00000034781 | Gna11 | guanine nucleotide binding protein, alpha 11 |
| chr15 | 81123403 | 81125395 | 1993 | * | 23,39776 | 2,193166284 | 0,832117 | 2,635645 | 0,008398 | 0,730469267 | 2,951062 | Downstream (<=300bp | 15 | 81119700 | 81123165 | 3466 | 1 | 207911 | ENSMUST00000166855.3 | 3703 | ENSMUSG00000050164 | Mchr1 | melanin-concentrating hormone receptor 1 |
| chr15 | 98490065 | 98493854 | 3790 | * | 76,556859 | 2,194092247 | 0,4399817 | 4,9867811 | 6,139E-07 | 0,002378936 | 3,2706213 | Promoter (<=1kb) | 15 | 98490588 | 98492115 | 1528 | 2 | 11512 | ENSMUST00000228596.2 | 0 | ENSMUSG00000022994 | Adcy6 | adenylate cyclase 6 |
| chr11 | 72430410 | 72434140 | 3731 | * | 35,54154 | 2,194228569 | 0,53404 | 4,108732 | 3,98E-05 | 0,061989207 | 2,725491 | Exon (ENSMUST00000 | 11 | 72388986 | 72441127 | 52142 | 2 | 77577 | ENSMUST00000021154.7 | 6987 | ENSMUSG00000020798 | Spns3 | spinster homolog 3 |
| chr7 | 97891327 | 97893292 | 1966 | * | 21,02977 | 2,194282461 | 0,720839 | 3,044066 | 0,002334 | 0,475225141 | 3,471678 | Exon (ENSMUST00000 | 7 | 97841624 | 97848688 | 7065 | 2 | 272411 | ENSMUST00000098278.4 | -42639 | ENSMUSG00000074004 | B3gnt6 | UDP-GlcNAc:betaGal beta-1,3-N-acetylglucosaminyltransferase 6 (core 3 synthase) |
| chr4 | 1,36E+08 | 1,36E+08 | 2471 | * | 34,28407 | 2,194315532 | 0,673472 | 3,258215 | 0,001121 | 0,417228227 | 2,377674 | Promoter (<=1kb) | 4 | 1,36E+08 | 1,36E+08 | 5645 | 1 | 22704 | ENSMUST00000144217.8 | 0 | ENSMUSG00000051351 | Zfp46 | zinc finger protein 46 |
| chr4 | 46030762 | 46032822 | 2061 | * | 18,45934 | 2,194580158 | 0,811408 | 2,704658 | 0,006837 | 0,730469267 | 2,386008 | Intron (ENSMUST0000 | 4 | 46026101 | 46034761 | 8661 | 1 | 100121 | ENSMUST00000155936.2 | 4661 | ENSMUSG00000035517 | Tdrd7 | tudor domain containing 7 |
| chr18 | 12773651 | 12776687 | 3037 | * | 25,54649 | 2,195725625 | 0,729436 | 3,010168 | 0,002611 | 0,528836551 | 2,858511 | Promoter (<=1kb) | 18 | 12776401 | 12822502 | 46102 | 1 | 72747 | ENSMUST00000141060.2 | 0 | ENSMUSG00000024424 | Ttc39c | tetratricopeptide repeat domain 39C |
| chr7 | 1,27E+08 | 1,27E+08 | 2427 | * | 16,70512 | 2,196006646 | 0,817646 | 2,685768 | 0,007236 | 0,722439117 | 2,663619 | Promoter (<=1kb) | 7 | 1,27E+08 | 1,27E+08 | 15058 | 1 | 1E+08 | ENSMUST00000185401.7 | -675 | ENSMUSG00000053877 | Srcap | Snf2-related CREBBP activator protein |
| chr10 | 96294344 | 96297456 | 3113 | * | 36,03663 | 2,196838684 | 0,536491 | 4,094826 | 4,22E-05 | 0,061899193 | 2,925497 | Distal Intergenic | 10 | 96359946 | 96431468 | 71523 | 2 | 1,03E+08 | ENSMUST00000244443.1 | 134012 | | Gm33843 | predicted gene, 33843 |
| chr3 | 1,23E+08 | 1,23E+08 | 2729 | * | 26,30381 | 2,196870264 | 0,679056 | 3,235184 | 0,001216 | 0,399951667 | 2,880017 | Promoter (<=1kb) | 3 | 1,23E+08 | 1,23E+08 | 53018 | 2 | 99526 | ENSMUST00000090379.7 | 0 | ENSMUSG00000039701 | Usp53 | ubiquitin specific peptidase 53 |
| chr7 | 24412791 | 24414980 | 2190 | * | 22,67394 | 2,197176023 | 0,723331 | 3,037581 | 0,002385 | 0,505909454 | 2,951933 | 3' UTR | 7 | 24408684 | 24413960 | 5277 | 1 | 232972 | ENSMUST00000051714.9 | 4107 | ENSMUSG00000045587 | Lypd10 | Ly6/PLAUR domain containing 10 |
| chr4 | 47310102 | 47311729 | 1628 | * | 17,87223 | 2,197506173 | 0,736327 | 2,984417 | 0,002841 | 0,568205102 | 2,507013 | Exon (ENSMUST00000 | 4 | 47303722 | 47313162 | 9441 | 1 | 12819 | ENSMUST00000107730.3 | 6380 | ENSMUSG00000028339 | Col15a1 | collagen, type XV, alpha 1 |
| chr5 | 1,25E+08 | 1,25E+08 | 2749 | * | 24,33803 | 2,197703169 | 0,802569 | 2,738335 | 0,006175 | 0,681119785 | 2,951062 | Promoter (<=1kb) | 5 | 1,25E+08 | 1,25E+08 | 34502 | 2 | 1,03E+08 | ENSMUST00000198586.2 | 0 | | Gm32585 | predicted gene, 32585 |
| chr10 | 1,19E+08 | 1,19E+08 | 1655 | * | 23,55771 | 2,198768469 | 0,675428 | 3,25537 | 0,001132 | 0,400053238 | 2,67243 | Distal Intergenic | 10 | 1,19E+08 | 1,19E+08 | 31222 | 2 | 1,03E+08 | ENSMUST00000217700.2 | -9026 | | Gm34045 | predicted gene, 34045 |
| chr13 | 37721218 | 37724111 | 2894 | * | 22,84494 | 2,198832724 | 0,728991 | 3,016269 | 0,002559 | 0,524368688 | 2,880017 | Promoter (1-2kb) | 13 | 37725333 | 37732419 | 7087 | 1 | 1,03E+08 | ENSMUST00000224508.3 | -1222 | | Lncbate6 | brown adipose tissue enriched long noncoding RNA 6 |
| chr5 | 1,16E+08 | 1,16E+08 | 1796 | * | 14,03716 | 2,199528095 | 0,845424 | 2,601686 | 0,009277 | 0,89325945 | 1,2763 | Intron (ENSMUST0000 | 5 | 1,16E+08 | 1,16E+08 | 26100 | 1 | 17690 | ENSMUST00000150779.8 | -16201 | ENSMUSG00000054256 | Msi1 | musashi RNA-binding protein 1 |
| chr4 | 1,01E+08 | 1,01E+08 | 2332 | * | 21,6469 | 2,199534997 | 0,749803 | 2,933483 | 0,003352 | 0,571963524 | 2,880017 | Promoter (<=1kb) | 4 | 1,01E+08 | 1,01E+08 | 9725 | 2 | 77866 | ENSMUST00000144604.3 | -607 | ENSMUSG00000086782 | E130102H24Rik | RIKEN cDNA E130102H24 gene |
| chr4 | 46507355 | 46508823 | 1469 | * | 21,62197 | 2,199627693 | 0,681302 | 3,228566 | 0,001244 | 0,414202366 | 2,67243 | Exon (ENSMUST00000 | 4 | 46495155 | 46495285 | 131 | 1 | 1,15E+08 | ENSMUST00000082476.2 | 12200 | ENSMUSG00000064410 | Gm22247 | predicted gene, 22247 |
| chr8 | 71248726 | 71251630 | 2905 | * | 86,09975 | 2,199829279 | 0,67793 | 3,244923 | 0,001175 | 0,367076136 | 3,270621 | Promoter (<=1kb) | 8 | 71250779 | 71250836 | 58 | 1 | 1,02E+08 | ENSMUST00000184074.3 | 0 | ENSMUSG00000099031 | Mir7240 | microRNA 7240 |
| chr12 | 55492616 | 55494470 | 1855 | * | 23,06735 | 2,200158581 | 0,719465 | 3,058048 | 0,002228 | 0,498011371 | 2,880017 | Distal Intergenic | 12 | 55454168 | 55459072 | 4905 | 1 | 26443 | ENSMUST00000162972.2 | 38448 | ENSMUSG00000021024 | Psma6 | proteasome subunit alpha 6 |
| chr7 | 46261087 | 46264202 | 3116 | * | 25,46662 | 2,200452523 | 0,699542 | 3,145564 | 0,001658 | 0,448852574 | 2,880017 | Exon (ENSMUST00000 | 7 | 46092645 | 46252666 | 160022 | 2 | 27414 | ENSMUST00000156147.8 | -8421 | ENSMUSG00000030839 | Sergef | secretion regulating guanine nucleotide exchange factor |
| chr4 | 1,36E+08 | 1,36E+08 | 2539 | * | 21,05418 | 2,201280831 | 0,803195 | 2,740654 | 0,006132 | 0,689688276 | 2,67243 | Intron (ENSMUST0000 | 4 | 1,36E+08 | 1,36E+08 | 1966 | 2 | 13844 | ENSMUST00000151502.2 | -4334 | ENSMUSG00000028664 | Ephb2 | Eph receptor B2 |
| chr3 | 87007846 | 87009786 | 1941 | * | 19,07449 | 2,201448372 | 0,756075 | 2,911681 | 0,003595 | 0,59811801 | 2,713306 | Intron (ENSMUST0000 | 3 | 86974200 | 86974331 | 132 | 1 | 1,15E+08 | ENSMUST00000104484.3 | 33646 | ENSMUSG00000077672 | Gm24472 | predicted gene, 24472 |
| chr8 | 1,28E+08 | 1,28E+08 | 1804 | * | 20,189 | 2,201532442 | 0,830407 | 2,651148 | 0,008022 | 0,739258701 | 2,663619 | Distal Intergenic | 8 | 1,28E+08 | 1,28E+08 | 31125 | 1 | 93742 | ENSMUST00000159940.2 | 184292 | ENSMUSG00000025812 | Pard3 | par-3 family cell polarity regulator |
| chr3 | 62678840 | 62680612 | 1773 | * | 16,29362 | 2,201710098 | 0,777039 | 2,833463 | 0,004605 | 0,649217039 | 2,663619 | Distal Intergenic | 3 | 62436498 | 62512561 | 76064 | 2 | 229357 | ENSMUST00000058535.6 | -166279 | ENSMUSG00000043441 | Gpr149 | G protein-coupled receptor 149 |
| chr7 | 98132821 | 98134765 | 1945 | * | 20,16768 | 2,201734999 | 0,787053 | 2,797442 | 0,005151 | 0,668772562 | 2,663619 | Distal Intergenic | 7 | 98138490 | 98147461 | 8972 | 1 | 434215 | ENSMUST00000205937.2 | -3725 | ENSMUSG00000090958 | Lrrc32 | leucine rich repeat containing 32 |
| chr3 | 1,22E+08 | 1,22E+08 | 2819 | * | 29,03894 | 2,203837684 | 0,691176 | 3,188535 | 0,00143 | 0,421090004 | 2,951062 | Intron (ENSMUST0000 | 3 | 1,22E+08 | 1,22E+08 | 133013 | 1 | 29815 | ENSMUST00000196134.2 | 22516 | ENSMUSG00000028121 | Bcar3 | breast cancer anti-estrogen resistance 3 |
| chr11 | 78302377 | 78305057 | 2681 | * | 33,5164 | 2,203939446 | 0,61509 | 3,583115 | 0,00034 | 0,224776719 | 2,725491 | Intron (ENSMUST0000 | 11 | 78287913 | 78313043 | 25131 | 2 | 20500 | ENSMUST00000001122.6 | 7986 | ENSMUSG00000001095 | Slc13a2 | solute carrier family 13 (sodium-dependent dicarboxylate transporter), member 2 |
| chr1 | 56826573 | 56828128 | 1556 | * | 15,24985 | 2,204593968 | 0,851697 | 2,588471 | 0,00964 | 0,906489483 | 1,214904 | Distal Intergenic | 1 | 56675203 | 56676594 | 1392 | 2 | 71066 | ENSMUST00000062085.6 | -149979 | ENSMUSG00000045336 | Hsfy2 | heat shock transcription factor, Y-linked 2 |
| chr15 | 6070024 | 6072556 | 2533 | * | 18,59326 | 2,204769522 | 0,836201 | 2,636651 | 0,008373 | 0,748382289 | 2,663619 | Distal Intergenic | 15 | 6180698 | 6180829 | 132 | 2 | 1,15E+08 | ENSMUST00000240204.1 | 108273 | ENSMUSG00000118903 | Gm23139 | predicted gene, 23139 |
| chr12 | 10377469 | 10379881 | 2413 | * | 17,52713 | 2,204946925 | 0,805753 | 2,736504 | 0,00621 | 0,713633527 | 2,423594 | Distal Intergenic | 12 | 10419973 | 10425935 | 5963 | 1 | 70881 | ENSMUST00000143739.9 | -40092 | ENSMUSG00000020622 | Nt5c1b | 5'-nucleotidase, cytosolic IB |
| chr9 | 1,21E+08 | 1,21E+08 | 1389 | * | 18,95503 | 2,205826675 | 0,806314 | 2,735692 | 0,006225 | 0,693610666 | 2,663619 | Exon (ENSMUST00000 | 9 | 1,21E+08 | 1,21E+08 | 5142 | 2 | 1,05E+08 | ENSMUST00000214714.2 | 12926 | | Gm19385 | predicted gene, 19385 |
| chr7 | 78934977 | 78936302 | 1326 | * | 20,37108 | 2,205859496 | 0,734702 | 3,002385 | 0,002679 | 0,580348965 | 2,221611 | Intron (ENSMUST0000 | 7 | 78923002 | 78995119 | 72118 | 1 | 54608 | ENSMUST00000135333.3 | 11975 | ENSMUSG00000039202 | Abhd2 | abhydrolase domain containing 2 |
| chr1 | 53523678 | 53525092 | 1415 | * | 17,50568 | 2,205987091 | 0,763638 | 2,888788 | 0,003867 | 0,634611828 | 2,423594 | Intron (ENSMUST0000 | 1 | 53386990 | 53391911 | 4922 | 2 | 70396 | ENSMUST00000147021.9 | -131767 | ENSMUSG00000026095 | Asnsd1 | asparagine synthetase domain containing 1 |
| chr13 | 1,15E+08 | 1,15E+08 | 2318 | * | 21,69896 | 2,206000038 | 0,741196 | 2,976271 | 0,002918 | 0,512326648 | 3,471678 | Intron (ENSMUST0000 | 13 | 1,15E+08 | 1,15E+08 | 90586 | 1 | 75161 | ENSMUST00000181553.2 | 5889 | ENSMUSG00000051758 | 4930544M13Rik | RIKEN cDNA 4930544M13 gene |
| chr13 | 23070110 | 23072098 | 1989 | * | 17,45958 | 2,206057593 | 0,836333 | 2,637775 | 0,008345 | 0,75508254 | 2,507013 | Promoter (1-2kb) | 13 | 23067212 | 23068375 | 1164 | 2 | 171275 | ENSMUST00000095960.5 | -1735 | ENSMUSG00000071490 | Vmn1r212 | vomeronasal 1 receptor 212 |
| chr4 | 56836305 | 56839086 | 2782 | * | 31,73437 | 2,206812059 | 0,59839 | 3,687913 | 0,000226 | 0,173329714 | 2,880017 | Promoter (<=1kb) | 4 | 56829525 | 56838584 | 9060 | 2 | 54366 | ENSMUST00000154511.2 | 0 | ENSMUSG00000038816 | Ctnnal1 | catenin (cadherin associated protein), alpha-like 1 |
| chr17 | 45063592 | 45065951 | 2360 | * | 17,9568 | 2,206994894 | 0,718125 | 3,073274 | 0,002117 | 0,524368688 | 2,386008 | Intron (ENSMUST0000 | 17 | 44914888 | 45047535 | 132648 | 2 | 12393 | ENSMUST00000113572.9 | -16057 | ENSMUSG00000039153 | Runx2 | runt related transcription factor 2 |
| chr11 | 1,16E+08 | 1,16E+08 | 4381 | * | 43,86146 | 2,207028774 | 0,560203 | 3,939697 | 8,16E-05 | 0,090744111 | 2,925497 | Promoter (<=1kb) | 11 | 1,16E+08 | 1,16E+08 | 18734 | 2 | 53413 | ENSMUST00000021147.14 | 0 | ENSMUSG00000020792 | Exoc7 | exocyst complex component 7 |
| chr5 | 65513393 | 65515137 | 1745 | * | 18,87219 | 2,207570086 | 0,803779 | 2,746488 | 0,006024 | 0,722439117 | 2,221611 | Intron (ENSMUST0000 | 5 | 65505751 | 65536871 | 31121 | 1 | 83379 | ENSMUST00000205084.2 | 7642 | ENSMUSG00000029195 | Klb | klotho beta |
| chr8 | 15044324 | 15046845 | 2522 | * | 22,4714 | 2,208006716 | 0,717689 | 3,07655 | 0,002094 | 0,488376383 | 2,880017 | Promoter (<=1kb) | 8 | 15044121 | 15046042 | 1922 | 2 | 1,03E+08 | ENSMUST00000246690.1 | 0 | | Gm35149 | predicted gene, 35149 |
| chr3 | 1,03E+08 | 1,03E+08 | 1489 | * | 23,34888 | 2,20840223 | 0,791901 | 2,788734 | 0,005291 | 0,659240897 | 2,880017 | Promoter (<=1kb) | 3 | 1,03E+08 | 1,03E+08 | 37761 | 1 | 229663 | ENSMUST00000029446.13 | 0 | ENSMUSG00000068823 | Csde1 | cold shock domain containing E1, RNA binding |
| chr15 | 73057153 | 73059891 | 2739 | * | 31,94479 | 2,208710855 | 0,642922 | 3,435424 | 0,000592 | 0,288385947 | 2,880017 | Promoter (<=1kb) | 15 | 72967693 | 73056784 | 89092 | 2 | 239528 | ENSMUST00000044113.12 | -369 | ENSMUSG00000036698 | Ago2 | argonaute RISC catalytic subunit 2 |
| chr6 | 1,16E+08 | 1,16E+08 | 1636 | * | 21,43422 | 2,209128946 | 0,694823 | 3,179414 | 0,001476 | 0,429988769 | 2,880017 | Promoter (2-3kb) | 6 | 1,16E+08 | 1,16E+08 | 63 | 1 | 1,02E+08 | ENSMUST00000183825.3 | 2378 | ENSMUSG00000099163 | Mir7660 | microRNA 7660 |
| chr9 | 21801787 | 21803426 | 1640 | * | 16,05411 | 2,209613482 | 0,8491 | 2,602302 | 0,00926 | 0,764402529 | 2,663619 | Exon (ENSMUST00000 | 9 | 21826188 | 21829385 | 3198 | 2 | 19340 | ENSMUST00000128442.2 | 25959 | ENSMUSG00000019066 | Rab3d | RAB3D, member RAS oncogene family |
| chr7 | 67410495 | 67415558 | 5064 | * | 50,684366 | 2,210026161 | 0,5143059 | 4,2971042 | 1,73E-05 | 0,033161239 | 2,7872553 | Promoter (1-2kb) | 7 | 67379916 | 67409490 | 29575 | 2 | 233335 | ENSMUST00000051389.10 | -1005 | ENSMUSG00000030554 | Synm | synemin, intermediate filament protein |
| chr5 | 36942621 | 36944861 | 2241 | * | 25,27662 | 2,210031053 | 0,710252 | 3,111616 | 0,001861 | 0,479521813 | 2,67243 | Distal Intergenic | 5 | 36952211 | 36954116 | 1906 | 2 | 67568 | ENSMUST00000068795.4 | 9255 | ENSMUSG00000055302 | Mrfap1 | Morf4 family associated protein 1 |
| chr16 | 24313917 | 24316710 | 2794 | * | 26,37015 | 2,210844549 | 0,710966 | 3,109633 | 0,001873 | 0,470097239 | 2,880017 | Intron (ENSMUST0000 | 16 | 24348144 | 24349112 | 969 | 1 | 624582 | ENSMUST00000231737.2 | -31434 | | Morf4l1-ps1 | mortality factor 4 like 1, pseudogene 1 |
| chr6 | 1,22E+08 | 1,22E+08 | 822 | * | 14,94337 | 2,211281016 | 0,804143 | 2,749861 | 0,005962 | 0,825778563 | 1,214904 | Distal Intergenic | 6 | 1,22E+08 | 1,22E+08 | 3013 | 1 | 232345 | ENSMUST00000204850.2 | -32375 | ENSMUSG00000030111 | A2m | alpha-2-macroglobulin |
| chr4 | 1,29E+08 | 1,29E+08 | 2335 | * | 18,16954 | 2,211629931 | 0,762234 | 2,901509 | 0,003714 | 0,648791129 | 2,154504 | 3' UTR | 4 | 1,29E+08 | 1,29E+08 | 19733 | 1 | 68828 | ENSMUST00000146448.2 | 19672 | ENSMUSG00000001333 | Sync | syncoilin |
| chr4 | 59328663 | 59330818 | 2156 | * | 19,44128 | 2,211703505 | 0,73243 | 3,01968 | 0,00253 | 0,542967057 | 2,507013 | Promoter (2-3kb) | 4 | 59329210 | 59333022 | 3813 | 2 | 634731 | ENSMUST00000136562.2 | 2204 | ENSMUSG00000038578 | Susd1 | sushi domain containing 1 |
| chr12 | 1,05E+08 | 1,05E+08 | 1718 | * | 21,28785 | 2,211912049 | 0,762981 | 2,89904 | 0,003743 | 0,580348965 | 3,104559 | Promoter (<=1kb) | 12 | 1,05E+08 | 1,05E+08 | 1680 | 2 | 1E+08 | ENSMUST00000245499.1 | 0 | ENSMUSG00002076601 | Scarna13 | small Cajal body-specific RNA 1 |
| chr13 | 23867739 | 23870688 | 2950 | * | 31,29747 | 2,212486435 | 0,650647 | 3,400439 | 0,000673 | 0,309888367 | 2,725491 | Promoter (<=1kb) | 13 | 23865450 | 23867931 | 2482 | 2 | 319164 | ENSMUST00000171127.4 | 0 | ENSMUSG00000069270 | H2ac6 | H2A clustered histone 6 |
| chr8 | 41451443 | 41453291 | 1849 | * | 18,63039 | 2,212487198 | 0,803098 | 2,754941 | 0,00587 | 0,684274806 | 2,663619 | Promoter (<=1kb) | 8 | 41451807 | 41453242 | 1436 | 2 | 102103 | ENSMUST00000135194.2 | 0 | ENSMUSG00000045636 | Mtus1 | mitochondrial tumor suppressor 1 |
| chr11 | 61459199 | 61461139 | 1941 | * | 17,6487 | 2,212513457 | 0,79882 | 2,769728 | 0,00561 | 0,681421811 | 2,663619 | Intron (ENSMUST0000 | 11 | 61437332 | 61470484 | 33153 | 2 | 13855 | ENSMUST00000147501.8 | 9345 | ENSMUSG00000001036 | Epn2 | epsin 2 |
| chr19 | 22688802 | 22691322 | 2521 | * | 22,80677 | 2,212986888 | 0,704486 | 3,141278 | 0,001682 | 0,451585113 | 2,880017 | Exon (ENSMUST00000 | 19 | 22670186 | 22746744 | 76559 | 1 | 226025 | ENSMUST00000236009.2 | 18616 | ENSMUSG00000052387 | Trpm3 | transient receptor potential cation channel, subfamily M, member 3 |

| | | | | | | | | | | | | | | | | | | | | | | | |
|---|---|---|---|---|---|---|---|---|---|---|---|---|---|---|---|---|---|---|---|---|---|---|---|
| chr4 | 1,29E+08 | 1,29E+08 | 2751 | * | 26,60315 | 2,213321792 | 0,642226 | 3,44633 | 0,000568 | 0,280986007 | 2,951062 | Intron (ENSMUST0000 | 4 | 1,29E+08 | 1,29E+08 | 19733 | 1 | 68828 | ENSMUST00000146448.2 | 14413 | ENSMUSG00000001333 | Sync | syncoilin |
| chr6 | 1,27E+08 | 1,27E+08 | 2032 | * | 20,24851 | 2,213457078 | 0,794259 | 2,786821 | 0,005323 | 0,682691026 | 2,507013 | Distal Intergenic | 6 | 1,27E+08 | 1,27E+08 | 11782 | 2 | 232386 | ENSMUST00000202005.2 | -20612 | ENSMUSG00000107153 | Gm38404 | predicted gene, 38404 |
| chr19 | 6886431 | 6889385 | 2955 | * | 48,383058 | 2,213867659 | 0,5168989 | 4,28298 | 1,844E-05 | 0,04029432 | 2,3383871 | Promoter (<=1kb) | 19 | 6884065 | 6887346 | 3282 | 2 | 54683 | ENSMUST00000145300.2 | 0 | ENSMUSG00000024953 | Prdx5 | peroxiredoxin 5 |
| chr16 | 32044894 | 32047593 | 2700 | * | 24,09946 | 2,21404038 | 0,719684 | 3,076408 | 0,002095 | 0,483145429 | 2,951062 | Distal Intergenic | 16 | 32062530 | 32062673 | 144 | 2 | 1,15E+08 | ENSMUST00000116790.3 | 15080 | ENSMUSG00000080440 | Gm25848 | predicted gene, 25848 |
| chr15 | 23056508 | 23057794 | 1287 | * | 13,5279 | 2,214275476 | 0,845978 | 2,617414 | 0,00886 | 0,934396727 | 0,938682 | Intron (ENSMUST0000 | 15 | 23036549 | 23474504 | 437956 | 1 | 320865 | ENSMUST00000164787.8 | 19959 | ENSMUSG00000040420 | Cdh18 | cadherin 18 |
| chr13 | 98422033 | 98424561 | 2529 | * | 16,8838 | 2,214540567 | 0,842653 | 2,628059 | 0,008587 | 0,791147121 | 2,141612 | Exon (ENSMUST00000 | 13 | 98416400 | 98430278 | 13879 | 1 | 1,03E+08 | ENSMUST00000178658.8 | 5633 | ENSMUSG00000096330 | Gm21976 | predicted gene 21976 |
| chr11 | 1,03E+08 | 1,03E+08 | 2271 | * | 22,18695 | 2,215290396 | 0,856408 | 2,586723 | 0,009689 | 0,756849896 | 2,880017 | Distal Intergenic | 11 | 1,03E+08 | 1,03E+08 | 167 | 2 | 544817 | ENSMUST00000163250.2 | 16288 | ENSMUSG00000034255 | Arhgap27 | Rho GTPase activating protein 27 |
| chr2 | 78405511 | 78408106 | 2596 | * | 24,10826 | 2,215338848 | 0,73441 | 3,016488 | 0,002557 | 0,524482861 | 2,858511 | Distal Intergenic | 2 | 78123516 | 78132734 | 9219 | 1 | 329436 | ENSMUST00000240588.1 | 281995 | ENSMUSG00000054510 | Gm14461 | predicted gene 14461 |
| chr7 | 76692890 | 76694484 | 1595 | * | 17,07313 | 2,215769417 | 0,786723 | 2,816455 | 0,004856 | 0,687090886 | 2,141612 | Intron (ENSMUST0000 | 7 | 76060810 | 76416356 | 355547 | 1 | 244071 | ENSMUST00000107442.3 | 632080 | ENSMUSG00000025754 | Agbl1 | ATP/GTP binding protein-like 1 |
| chr7 | 80602581 | 80604840 | 2260 | * | 21,67532 | 2,21638568 | 0,717355 | 3,089662 | 0,002004 | 0,494045114 | 2,67243 | Distal Intergenic | 7 | 80565837 | 80597528 | 31692 | 2 | 56529 | ENSMUST00000119980.2 | -5053 | ENSMUSG00000025724 | Sec11a | SEC11 homolog A, signal peptidase complex subunit |
| chr11 | 1,06E+08 | 1,06E+08 | 2643 | * | 31,71447 | 2,217022146 | 0,661175 | 3,353155 | 0,000799 | 0,322750682 | 2,951062 | Distal Intergenic | 11 | 1,06E+08 | 1,06E+08 | 17538 | 2 | 13056 | ENSMUST00000143251.8 | -5228 | ENSMUSG00000019590 | Cyb561 | cytochrome b-561 |
| chr5 | 1,37E+08 | 1,37E+08 | 3477 | * | 38,18152 | 2,217205617 | 0,584187 | 3,795369 | 0,000147 | 0,138409996 | 2,725491 | Promoter (<=1kb) | 5 | 1,37E+08 | 1,37E+08 | 9630 | 1 | 26433 | ENSMUST00000004968.11 | 0 | ENSMUSG00000004846 | Plod3 | procollagen-lysine, 2-oxoglutarate 5-dioxygenase 3 |
| chr8 | 57963195 | 57965266 | 2072 | * | 30,86129 | 2,217579796 | 0,566335 | 3,915665 | 9,02E-05 | 0,102151284 | 2,67243 | Promoter (<=1kb) | 8 | 57964941 | 57969033 | 4093 | 1 | 97165 | ENSMUST00000067925.8 | 0 | ENSMUSG00000054717 | Hmgb2 | high mobility group box 2 |
| chr16 | 14133475 | 14137059 | 3585 | * | 29,83055 | 2,217654139 | 0,601141 | 3,689073 | 0,000225 | 0,179303674 | 2,67243 | Promoter (<=1kb) | 16 | 14117108 | 14135211 | 18104 | 2 | 66086 | ENSMUST00000120707.2 | 0 | ENSMUSG00000022677 | Cep20 | centrosomal protein 20 |
| chrX | 1,34E+08 | 1,34E+08 | 1790 | * | 17,71583 | 2,217754675 | 0,860802 | 2,576382 | 0,009984 | 0,790460916 | 2,507013 | Promoter (<=1kb) | 20 | 1,34E+08 | 1,34E+08 | 2955 | 2 | 278097 | ENSMUST00000052431.12 | -734 | ENSMUSG00000050394 | Armcx6 | armadillo repeat containing, X-linked 6 |
| chr12 | 1,02E+08 | 1,02E+08 | 3938 | * | 41,83931 | 2,218211162 | 0,548935 | 4,040934 | 5,32E-05 | 0,080985792 | 2,377674 | Promoter (<=1kb) | 12 | 1,02E+08 | 1,02E+08 | 45716 | 2 | 19141 | ENSMUST00000021607.9 | 0 | ENSMUSG00000021190 | Lgmn | legumain |
| chr10 | 59662245 | 59665760 | 3516 | * | 28,73658 | 2,218353816 | 0,663176 | 3,345048 | 0,000823 | 0,328946709 | 2,880017 | Promoter (<=1kb) | 10 | 59663043 | 59676189 | 13147 | 1 | 216001 | ENSMUST00000169842.2 | 0 | ENSMUSG00000020111 | Micu1 | mitochondrial calcium uptake 1 |
| chr5 | 1,48E+08 | 1,48E+08 | 1501 | * | 14,04127 | 2,218369167 | 0,828761 | 2,676729 | 0,007434 | 0,855969265 | 1,2763 | Promoter (<=1kb) | 5 | 1,48E+08 | 1,48E+08 | 14894 | 1 | 66537 | ENSMUST00000201376.4 | 0 | ENSMUSG00000029649 | Pomp | proteasome maturation protein |
| chr4 | 6125660 | 6127336 | 1677 | * | 18,07105 | 2,218574488 | 0,860934 | 2,57694 | 0,009968 | 0,816590739 | 2,154504 | Distal Intergenic | 4 | 6191098 | 6221688 | 30591 | 1 | 68053 | ENSMUST00000029907.6 | -63762 | ENSMUSG00000028243 | Ubxn2b | UBX domain protein 2B |
| chr18 | 73948562 | 73950582 | 2021 | * | 20,86438 | 2,218764861 | 0,741293 | 2,993101 | 0,002762 | 0,547647187 | 2,67243 | Promoter (<=1kb) | 18 | 73930733 | 73948520 | 17788 | 2 | 107029 | ENSMUST00000237619.2 | -42 | ENSMUSG00000024556 | Me2 | malic enzyme 2, NAD(+)-dependent, mitochondrial |
| chr12 | 69404729 | 69407156 | 2428 | * | 14,81991 | 2,218915866 | 0,830418 | 2,672047 | 0,007539 | 0,857338919 | 1,2763 | Promoter (<=1kb) | 12 | 69357296 | 69403939 | 46644 | 2 | 66244 | ENSMUST00000021368.10 | -790 | ENSMUSG00000020982 | Nemf | nuclear export mediator factor |
| chr17 | 13082067 | 13083515 | 1449 | * | 16,19032 | 2,218937129 | 0,848404 | 2,615425 | 0,008912 | 0,888080529 | 1,2763 | Intron (ENSMUST0000 | 17 | 13059966 | 13087030 | 27065 | 2 | 17171 | ENSMUST00000089015.10 | 3515 | ENSMUSG00000068037 | Mas1 | MAS1 oncogene |
| chr16 | 21241524 | 21243562 | 2039 | * | 28,79732 | 2,218971683 | 0,644784 | 3,441418 | 0,000579 | 0,293348204 | 2,725491 | Promoter (<=1kb) | 16 | 21241868 | 21397997 | 156130 | 1 | 209018 | ENSMUST00000096191.11 | 0 | ENSMUSG00000033653 | Vps8 | VPS8 CORVET complex subunit |
| chr17 | 32705887 | 32708263 | 2377 | * | 23,83458 | 2,219094553 | 0,756729 | 2,932483 | 0,003363 | 0,568412386 | 2,951933 | Exon (ENSMUST00000 | 17 | 32697225 | 32702392 | 5168 | 1 | 320997 | ENSMUST00000236005.2 | 8662 | ENSMUSG00000061126 | Cyp4f39 | cytochrome P450, family 4, subfamily f, polypeptide 39 |
| chr10 | 75035712 | 75037697 | 1986 | * | 16,99441 | 2,219800045 | 0,823035 | 2,69709 | 0,006995 | 0,715249603 | 2,663619 | Distal Intergenic | 10 | 75047905 | 75146897 | 98993 | 1 | 74392 | ENSMUST00000218766.2 | -10208 | ENSMUSG00000033444 | Specc1l | sperm antigen with calponin homology and coiled-coil domains 1-like |
| chr11 | 41103830 | 41105754 | 1925 | * | 15,48341 | 2,220347957 | 0,778567 | 2,851838 | 0,004347 | 0,743137506 | 1,41696 | Distal Intergenic | 11 | 40639379 | 40646138 | 6760 | 2 | 12450 | ENSMUST00000020576.8 | -457692 | ENSMUSG00000020326 | Ccng1 | cyclin G1 |
| chr16 | 52406031 | 52407722 | 1692 | * | 15,65207 | 2,220547868 | 0,816782 | 2,718653 | 0,006555 | 0,834909202 | 1,2763 | Distal Intergenic | 16 | 52069359 | 52274437 | 205079 | 2 | 11658 | ENSMUST00000023312.14 | -131594 | ENSMUSG00000022636 | Alcam | activated leukocyte cell adhesion molecule |
| chr8 | 1,18E+08 | 1,18E+08 | 2944 | * | 40,34468 | 2,220753553 | 0,547808 | 4,053892 | 5,04E-05 | 0,070291925 | 2,787255 | Intron (ENSMUST0000 | 8 | 1,18E+08 | 1,18E+08 | 189072 | 1 | 74440 | ENSMUST00000211913.2 | 29823 | ENSMUSG00000034390 | Cmip | c-Maf inducing protein |
| chr10 | 83017392 | 83019779 | 2388 | * | 22,66423 | 2,221659469 | 0,685952 | 3,238797 | 0,0012 | 0,393315816 | 2,951933 | Intron (ENSMUST0000 | 10 | 82974984 | 82977872 | 2889 | 2 | 1,15E+08 | ENSMUST00000160997.2 | -39520 | | 1700025N21Rik | RIKEN cDNA 1700025N21 gene |
| chr9 | 66896883 | 66899014 | 2132 | * | 16,72692 | 2,221790492 | 0,836594 | 2,655758 | 0,007913 | 0,755058193 | 2,386008 | Distal Intergenic | 9 | 66862675 | 66882766 | 20092 | 2 | 80907 | ENSMUST00000034929.7 | -14117 | ENSMUSG00000032370 | Lactb | lactamase, beta |
| chr9 | 59566468 | 59568091 | 1624 | * | 31,15795 | 2,222003705 | 0,69132 | 3,214145 | 0,001308 | 0,407158022 | 2,951062 | Promoter (<=1kb) | 9 | 59565526 | 59577881 | 12356 | 1 | 18746 | ENSMUST00000213930.2 | 942 | ENSMUSG00000032294 | Pkm | pyruvate kinase, muscle |
| chr4 | 45217014 | 45218725 | 1712 | * | 15,69522 | 2,222025622 | 0,849393 | 2,616016 | 0,008896 | 0,868136045 | 1,41696 | Intron (ENSMUST0000 | 4 | 45203925 | 45285936 | 82012 | 1 | 666060 | ENSMUST00000107804.2 | 13089 | ENSMUSG00000035615 | Frmpd1 | FERM and PDZ domain containing 1 |
| chr7 | 28648061 | 28650299 | 2239 | * | 20,25659 | 2,222418287 | 0,802752 | 2,768499 | 0,005632 | 0,695892636 | 2,386008 | Promoter (<=1kb) | 7 | 28612803 | 28649184 | 36382 | 2 | 60595 | ENSMUST00000148196.3 | 0 | ENSMUSG00000054808 | Actn4 | actinin alpha 4 |
| chr1 | 1,56E+08 | 1,56E+08 | 2247 | * | 21,88702 | 2,223307636 | 0,778392 | 2,856282 | 0,004286 | 0,623120685 | 2,880017 | Promoter (<=1kb) | 1 | 1,56E+08 | 1,56E+08 | 128292 | 2 | 74081 | ENSMUST00000138762.8 | -52 | ENSMUSG00000033671 | Cep350 | centrosomal protein 350 |
| chr9 | 65719858 | 65722066 | 2209 | * | 19,16421 | 2,223996977 | 0,834575 | 2,664827 | 0,007703 | 0,749931768 | 2,386008 | Intron (ENSMUST0000 | 9 | 65599673 | 65734846 | 135174 | 2 | 214812 | ENSMUST00000159109.2 | 12780 | ENSMUSG00000040524 | Zfp609 | zinc finger protein 609 |
| chr1 | 75221103 | 75222862 | 1760 | * | 21,08475 | 2,224591964 | 0,835449 | 2,66275 | 0,00775 | 0,708539289 | 3,104559 | 3' UTR | 1 | 75218053 | 75221571 | 3519 | 1 | 56812 | ENSMUST00000190445.2 | 3050 | ENSMUSG00000026203 | Dnajb2 | DnaJ heat shock protein family (Hsp40) member B2 |
| chr8 | 86362909 | 86365643 | 2735 | * | 19,28285 | 2,225450239 | 0,852032 | 2,611934 | 0,009003 | 0,757706088 | 2,663619 | 3' UTR | 8 | 86363217 | 86417602 | 54386 | 2 | 74513 | ENSMUST00000109686.4 | 51959 | ENSMUSG00000036902 | Neto2 | neuropilin (NRP) and tolloid (TLL)-like 2 |
| chr11 | 1,09E+08 | 1,09E+08 | 1667 | * | 22,71773 | 2,226100121 | 0,706371 | 3,151461 | 0,001625 | 0,457337755 | 2,67243 | Exon (ENSMUST00000 | 11 | 1,09E+08 | 1,09E+08 | 29577 | 1 | 74008 | ENSMUST00000106696.2 | -7608 | ENSMUSG00000020604 | Arsg | arylsulfatase G |
| chr7 | 73181902 | 73184528 | 2627 | * | 21,59017 | 2,226105107 | 0,680613 | 3,270737 | 0,001073 | 0,339457681 | 3,471678 | Promoter (2-3kb) | 7 | 73157608 | 73187369 | 29762 | 2 | 244059 | ENSMUST00000172704.6 | 2841 | ENSMUSG00000078671 | Chd2 | chromodomain helicase DNA binding protein 2 |
| chr3 | 42470134 | 42471964 | 1831 | * | 15,93552 | 2,226135023 | 0,826657 | 2,692935 | 0,007083 | 0,73648081 | 2,386008 | Distal Intergenic | 3 | 41705631 | 41710927 | 5297 | 1 | 73852 | ENSMUST00000195030.2 | 764503 | ENSMUSG00000025766 | D3Ertd751e | DNA segment, Chr 3, ERATO Doi 751, expressed |
| chr8 | 1,27E+08 | 1,27E+08 | 1838 | * | 22,60834 | 2,226456501 | 0,714639 | 3,115497 | 0,001836 | 0,466212802 | 2,880017 | Distal Intergenic | 8 | 1,27E+08 | 1,27E+08 | 2764 | 2 | 1,15E+08 | ENSMUST00000212695.2 | -7986 | | Gm45805 | predicted gene 45805 |
| chr19 | 44994278 | 44996673 | 2396 | * | 30,45301 | 2,22660706 | 0,675507 | 3,296201 | 0,00098 | 0,352527836 | 2,951062 | Promoter (<=1kb) | 19 | 44994997 | 45001201 | 6205 | 1 | 226153 | ENSMUST00000026227.3 | 0 | ENSMUSG00000025209 | Twnk | twinkle mtDNA helicase |
| chr12 | 1,11E+08 | 1,11E+08 | 2247 | * | 20,56025 | 2,226749042 | 0,77292 | 2,880955 | 0,003965 | 0,6318891 | 2,507013 | Distal Intergenic | 12 | 1,11E+08 | 1,11E+08 | 100784 | 1 | 22031 | ENSMUST00000117269.8 | -7389 | ENSMUSG00000021277 | Traf3 | TNF receptor-associated factor 3 |
| chr17 | 7093574 | 7096067 | 2494 | * | 20,77947 | 2,226785875 | 0,836336 | 2,662549 | 0,007755 | 0,731734453 | 2,67243 | Promoter (<=1kb) | 17 | 7065782 | 7095310 | 29529 | 2 | 1,03E+08 | ENSMUST00000246841.1 | 0 | | Gm2885 | predicted gene 2885 |
| chr12 | 1,11E+08 | 1,11E+08 | 1959 | * | 16,88465 | 2,226818383 | 0,81687 | 2,726036 | 0,00641 | 0,72508691 | 2,318434 | Promoter (<=1kb) | 12 | 1,11E+08 | 1,11E+08 | 24642 | 2 | 78062 | ENSMUST00000121608.2 | 0 | ENSMUSG00000083193 | 4930595D18Rik | RIKEN cDNA 4930595D18 gene |
| chr11 | 5801114 | 5803182 | 2069 | * | 21,46918 | 2,228371796 | 0,793287 | 2,809037 | 0,004969 | 0,661358703 | 2,67243 | Distal Intergenic | 11 | 5811947 | 5822088 | 10142 | 1 | 11568 | ENSMUST00000102923.10 | -8765 | ENSMUSG00000020473 | Aebp1 | AE binding protein 1 |
| chr19 | 36768968 | 36770694 | 1727 | * | 22,29461 | 2,228526672 | 0,726157 | 3,068933 | 0,002148 | 0,492839239 | 2,880017 | Intron (ENSMUST0000 | 19 | 36811632 | 36870877 | 59246 | 1 | 74493 | ENSMUST00000025729.12 | -40938 | ENSMUSG00000024811 | Tnks2 | tankyrase, TRF1-interacting ankyrin-related ADP-ribose polymerase 2 |
| chr6 | 86134129 | 86135672 | 1544 | * | 24,23638 | 2,229004139 | 0,753087 | 2,959822 | 0,003078 | 0,570876293 | 2,67243 | Distal Intergenic | 6 | 86172205 | 86252701 | 80497 | 1 | 21802 | ENSMUST00000032066.13 | -36533 | ENSMUSG00000029999 | Tgfa | transforming growth factor alpha |
| chr18 | 34793008 | 34795622 | 2615 | * | 18,05768 | 2,229157975 | 0,848137 | 2,6283 | 0,008581 | 0,770673161 | 2,386008 | Distal Intergenic | 18 | 34764736 | 34784788 | 20053 | 2 | 52563 | ENSMUST00000133181.2 | -8220 | ENSMUSG00000024370 | Cdc23 | CDC23 cell division cycle 23 |
| chr14 | 60494102 | 60496014 | 1913 | * | 16,90629 | 2,229360243 | 0,807734 | 2,760018 | 0,00578 | 0,683966322 | 2,663619 | Distal Intergenic | 14 | 60455527 | 60488956 | 33430 | 2 | 71844 | ENSMUST00000041905.8 | -5146 | ENSMUSG00000114797 | Nupl1 | nucleoporin like 1 |
| chr5 | 12382316 | 12383688 | 1373 | * | 16,95868 | 2,229616161 | 0,847947 | 2,629428 | 0,008553 | 0,790570581 | 2,141612 | Distal Intergenic | 5 | 12433352 | 12638915 | 205564 | 1 | 108151 | ENSMUST00000030868.11 | -49664 | ENSMUSG00000040254 | Sema3d | sema domain, immunoglobulin domain (Ig), short basic domain, secreted, (semaphorin) 3D |
| chr5 | 31787283 | 31788672 | 1390 | * | 27,27697 | 2,22998899 | 0,616278 | 3,618479 | 0,000296 | 0,203673596 | 2,880017 | Intron (ENSMUST0000 | 5 | 31781783 | 31797882 | 16100 | 2 | 71336 | ENSMUST00000201744.2 | 9210 | ENSMUSG00000029136 | Rbks | ribokinase |
| chr11 | 11758682 | 11760451 | 1770 | * | 21,50697 | 2,230389399 | 0,732534 | 3,044757 | 0,002329 | 0,505985803 | 2,880017 | Promoter (<=1kb) | 11 | 11737431 | 11758925 | 21495 | 2 | 60530 | ENSMUST00000152345.8 | 0 | ENSMUSG00000035455 | Fignl1 | fidgetin-like 1 |
| chr12 | 85232350 | 85236293 | 3944 | * | 38,99239 | 2,230642626 | 0,560812 | 3,977522 | 6,96E-05 | 0,083814446 | 2,925497 | Exon (ENSMUST00000 | 12 | 85213409 | 85224564 | 11156 | 2 | 18654 | ENSMUST00000223220.2 | -7786 | ENSMUSG00000004791 | Pgf | placental growth factor |
| chr19 | 52744291 | 52745758 | 1468 | * | 15,68554 | 2,230866043 | 0,82275 | 2,711474 | 0,006698 | 0,747833008 | 2,141612 | Distal Intergenic | 19 | 52911612 | 52915362 | 3751 | 1 | 1E+08 | ENSMUST00000236393.2 | -165854 | ENSMUSG00000117896 | AA387883 | expressed sequence AA387883 |
| chr7 | 65451258 | 65453237 | 1980 | * | 17,03274 | 2,231056604 | 0,815186 | 2,736868 | 0,006203 | 0,731263337 | 2,141612 | Promoter (<=1kb) | 7 | 65452758 | 65455998 | 3241 | 1 | 1,01E+08 | ENSMUST00000243649.1 | 0 | ENSMUSG00000100094 | 1810008I18Rik | RIKEN cDNA 1810008I18 gene |
| chr17 | 46041969 | 46044769 | 2801 | * | 36,11314 | 2,231081149 | 0,613492 | 3,636689 | 0,000276 | 0,192569014 | 2,925497 | Promoter (<=1kb) | 17 | 46043790 | 46044770 | 981 | 2 | 72240 | ENSMUST00000178179.2 | 1 | ENSMUSG00000094690 | 1600014C23Rik | RIKEN cDNA 1600014C23 gene |
| chr19 | 42111392 | 42113466 | 2075 | * | 16,00556 | 2,231195849 | 0,84137 | 2,651861 | 0,008005 | 0,75628971 | 2,386008 | 5' UTR | 19 | 42113218 | 42117428 | 4211 | 2 | 69534 | ENSMUST00000161873.2 | 3962 | ENSMUSG00000018821 | Avpi1 | arginine vasopressin-induced 1 |
| chr8 | 1,21E+08 | 1,21E+08 | 2079 | * | 27,0796 | 2,231939554 | 0,770661 | 2,896136 | 0,003778 | 0,597812173 | 2,858511 | Exon (ENSMUST00000 | 8 | 1,21E+08 | 1,21E+08 | 23797 | 2 | 434353 | ENSMUST00000183235.2 | 23669 | ENSMUSG00000097960 | A330074K22Rik | RIKEN cDNA A330074K22 gene |
| chr11 | 79086311 | 79088703 | 2393 | * | 18,92616 | 2,232071794 | 0,799476 | 2,79192 | 0,00524 | 0,684274806 | 2,386008 | Distal Intergenic | 11 | 79131133 | 79133775 | 2643 | 2 | 78889 | ENSMUST00000153585.2 | 45072 | ENSMUSG00000017677 | Wsb1 | WD repeat and SOCS box-containing 1 |
| chr3 | 1,46E+08 | 1,46E+08 | 1588 | * | 17,0923 | 2,232359635 | 0,808993 | 2,759429 | 0,00579 | 0,683988405 | 2,663619 | Distal Intergenic | 3 | 1,46E+08 | 1,46E+08 | 10095 | 1 | 12042 | ENSMUST00000029842.9 | -16185 | ENSMUSG00000028191 | Bcl10 | B cell leukemia/lymphoma 10 |
| chr8 | 72989747 | 72991446 | 1700 | * | 28,11538 | 2,232693628 | 0,596244 | 3,744595 | 0,000181 | 0,152532781 | 2,880017 | Promoter (2-3kb) | 8 | 72993862 | 73011229 | 17368 | 1 | 11767 | ENSMUST00000003117.15 | -2416 | ENSMUSG00000003033 | Ap1m1 | adaptor-related protein complex AP-1, mu subunit 1 |
| chr19 | 46580475 | 46583142 | 2668 | * | 23,83929 | 2,233156877 | 0,864804 | 2,582269 | 0,009815 | 0,775744126 | 2,67243 | Promoter (<=1kb) | 19 | 46579560 | 46585337 | 5778 | 1 | 94279 | ENSMUST00000237001.2 | 915 | ENSMUSG00000025036 | Sfxn2 | sideroflexin 2 |
| chr6 | 13676826 | 13677722 | 897 | * | 12,2818 | 2,233224144 | 0,863942 | 2,584922 | 0,00974 | 0,952447946 | 0,918255 | Promoter (<=1kb) | 6 | 13626814 | 13677937 | 51124 | 2 | 101148 | ENSMUST00000203078.2 | 215 | ENSMUSG00000042742 | Bmt2 | base methyltransferase of 25S rRNA 2 |
| chr1 | 89407104 | 89408382 | 1279 | * | 14,33986 | 2,233245615 | 0,862968 | 2,587867 | 0,009657 | 0,95061795 | 0,918255 | Intron (ENSMUST0000 | 1 | 89382596 | 89817723 | 435128 | 1 | 347722 | ENSMUST00000190096.7 | 24508 | ENSMUSG00000055013 | Agap1 | ArfGAP with GTPase domain, ankyrin repeat and PH domain 1 |
| chr13 | 23804200 | 23806641 | 2442 | * | 23,10472 | 2,233522992 | 0,723121 | 3,088728 | 0,00201 | 0,479521813 | 2,880017 | Promoter (<=1kb) | 13 | 23767653 | 23805077 | 37425 | 2 | 319179 | ENSMUST00000091704.7 | 0 | ENSMUSG00000047246 | H2bc6 | H2B clustered histone 6 |
| chr11 | 97250988 | 97253096 | 2109 | * | 14,66243 | 2,233854207 | 0,836035 | 2,671963 | 0,007541 | 0,865252447 | 1,214904 | Promoter (<=1kb) | 11 | 97253261 | 97289368 | 36108 | 1 | 192157 | ENSMUST00000238729.2 | -165 | ENSMUSG00000038485 | Socs7 | suppressor of cytokine signaling 7 |
| chr5 | 35792708 | 35795072 | 2365 | * | 28,77562 | 2,23388342 | 0,657535 | 3,39736 | 0,00068 | 0,305414759 | 2,880017 | Distal Intergenic | 5 | 35764303 | 35767263 | 2961 | 1 | 80911 | ENSMUST00000154796.2 | 28405 | ENSMUSG00000029098 | Acox3 | acyl-Coenzyme A oxidase 3, pristanoyl |
| chrX | 1,3E+08 | 1,3E+08 | 1617 | * | 12,95211 | 2,23460591 | 0,85203 | 2,622685 | 0,008724 | 0,966603024 | 0,738441 | Distal Intergenic | 20 | 1,3E+08 | 1,3E+08 | 129 | 2 | 1,15E+08 | ENSMUST00000104191.3 | 257845 | ENSMUSG00000077379 | Gm25170 | predicted gene, 25170 |
| chrX | 73436404 | 73438027 | 1624 | * | 13,85821 | 2,234638787 | 0,812839 | 2,749179 | 0,005974 | 0,818408612 | 1,2763 | Promoter (<=1kb) | 20 | 73436896 | 73477599 | 40704 | 1 | 16151 | ENSMUST00000144947.8 | 0 | ENSMUSG00000004221 | Ikbkg | inhibitor of kappaB kinase gamma |
| chr15 | 36792346 | 36793091 | 746 | * | 15,15755 | 2,234816357 | 0,859682 | 2,599585 | 0,009334 | 0,878632224 | 1,41696 | Promoter (<=1kb) | 15 | 36790956 | 36792658 | 1703 | 2 | 22631 | ENSMUST00000126184.2 | 0 | ENSMUSG00000022285 | Ywhaz | tyrosine 3-monooxygenase/tryptophan 5-monooxygenase activation protein, zeta polypeptide |
| chr2 | 31089116 | 31092538 | 3423 | * | 40,09769 | 2,23559863 | 0,603431 | 3,704812 | 0,000212 | 0,188932071 | 2,338387 | Promoter (2-3kb) | 2 | 31086908 | 31097719 | 10812 | 1 | 277463 | ENSMUST00000149622.2 | 2208 | ENSMUSG00000000194 | Gpr107 | G protein-coupled receptor 107 |
| chr5 | 82729937 | 82732894 | 2958 | * | 21,0731 | 2,235876733 | 0,756047 | 2,957327 | 0,003103 | 0,524454869 | 3,471678 | Distal Intergenic | 5 | 82946791 | 82946912 | 122 | 2 | 1E+08 | ENSMUST00000116936.3 | 214018 | ENSMUSG00000080586 | Mir1187 | microRNA 1187 |
| chr18 | 30527324 | 30529919 | 2596 | * | 25,59443 | 2,23592956 | 0,78086 | 2,86342 | 0,004191 | 0,627785088 | 2,725491 | Distal Intergenic | 18 | 30405980 | 30481173 | 75194 | 1 | 225326 | ENSMUST00000131405.2 | 121344 | ENSMUSG00000033628 | Pik3c3 | phosphatidylinositol 3-kinase catalytic subunit type 3 |
| chr7 | 1,28E+08 | 1,28E+08 | 2004 | * | 15,58144 | 2,236037943 | 0,8426 | 2,653737 | 0,007961 | 0,8656457 | 1,2763 | Intron (ENSMUST0000 | 7 | 1,28E+08 | 1,28E+08 | 10781 | 1 | 16409 | ENSMUST00000134694.2 | -10774 | ENSMUSG00000030786 | Itgam | integrin alpha M |
| chr4 | 1,14E+08 | 1,14E+08 | 1672 | * | 18,13726 | 2,236042975 | 0,717976 | 3,114369 | 0,001843 | 0,497895643 | 2,386008 | Intron (ENSMUST0000 | 4 | 1,14E+08 | 1,14E+08 | 208375 | 1 | 666048 | ENSMUST00000094894.4 | 21688 | ENSMUSG00000070867 | Trabd2b | TraB domain containing 2B |
| chr18 | 5607346 | 5610618 | 3273 | * | 28,69913 | 2,23643055 | 0,650795 | 3,436461 | 0,000589 | 0,297781562 | 2,67243 | Intron (ENSMUST0000 | 18 | 5603744 | 5642873 | 39130 | 1 | 21417 | ENSMUST00000159477.2 | 3602 | ENSMUSG00000024238 | Zeb1 | zinc finger E-box binding homeobox 1 |
| chr7 | 1,26E+08 | 1,26E+08 | 1767 | * | 18,95827 | 2,236736409 | 0,868351 | 2,575842 | 0,01 | 0,790460916 | 2,507013 | Promoter (<=1kb) | 7 | 1,26E+08 | 1,26E+08 | 10700 | 2 | 233871 | ENSMUST00000167759.8 | 0 | ENSMUSG00000032637 | Atxn2l | ataxin 2-like |
| chr9 | 63215863 | 63218467 | 2605 | * | 25,0305 | 2,237184036 | 0,688171 | 3,250912 | 0,00115 | 0,402652506 | 2,67243 | Intron (ENSMUST0000 | 9 | 63123553 | 63210425 | 86873 | 2 | 23938 | ENSMUST00000213604.2 | -5438 | ENSMUSG00000058444 | Map2k5 | mitogen-activated protein kinase kinase 5 |
| chr2 | 1,32E+08 | 1,32E+08 | 2670 | * | 25,11437 | 2,237212813 | 0,640118 | 3,494999 | 0,000474 | 0,256539258 | 2,951933 | Promoter (<=1kb) | 2 | 1,31E+08 | 1,32E+08 | 68942 | 2 | 1,19E+08 | ENSMUST00000138080.2 | 0 | | Gm14280 | predicted gene 14280 |
| chr3 | 89858455 | 89860073 | 1619 | * | 16,2737 | 2,237242592 | 0,848034 | 2,638153 | 0,008336 | 0,747833008 | 2,663619 | Promoter (<=1kb) | 3 | 89846796 | 89860672 | 13877 | 2 | 54667 | ENSMUST00000163354.8 | 599 | ENSMUSG00000060671 | Atp8b2 | ATPase, class I, type 8B, member 2 |
| chr7 | 72668356 | 72670597 | 2242 | * | 24,79824 | 2,237525673 | 0,695543 | 3,216946 | 0,001296 | 0,410415129 | 2,858511 | Distal Intergenic | 7 | 72747497 | 72751816 | 4320 | 2 | 625243 | ENSMUST00000205529.2 | 81219 | ENSMUSG00000108383 | Gm6567 | predicted gene 6567 |
| chr19 | 13814735 | 13816822 | 2088 | * | 19,1353 | 2,237837904 | 0,746032 | 2,999653 | 0,002703 | 0,543902653 | 2,663619 | Promoter (<=1kb) | 19 | 13815588 | 13816535 | 948 | 2 | 258626 | ENSMUST00000073966.2 | 0 | ENSMUSG00000057270 | Olfr1501 | olfactory receptor 1501 |
| chr15 | 43663774 | 43665447 | 1674 | * | 24,1929 | 2,238138183 | 0,754276 | 2,967267 | 0,003005 | 0,566484927 | 2,67243 | Distal Intergenic | 15 | 43728042 | 43733432 | 5391 | 2 | 239408 | ENSMUST00000067469.6 | 67985 | ENSMUSG00000054409 | Tmem74 | transmembrane protein 74 |
| chr5 | 1,47E+08 | 1,47E+08 | 2929 | * | 27,15496 | 2,238270275 | 0,631419 | 3,544824 | 0,000393 | 0,24621716 | 2,67243 | Distal Intergenic | 5 | 1,47E+08 | 1,47E+08 | 13458 | 2 | 1,03E+08 | ENSMUST00000200363.2 | -40047 | ENSMUSG00000105907 | Gm34333 | predicted gene, 34333 |
| chr19 | 44271005 | 44273500 | 2496 | * | 56,926447 | 2,238656524 | 0,4456024 | 5,0238874 | 5,064E-07 | 0,002004214 | 3,2706213 | Distal Intergenic | 19 | 44282113 | 44295303 | 13191 | 1 | 20250 | ENSMUST00000026221.7 | -8613 | ENSMUSG00000025203 | Scd2 | stearoyl-Coenzyme A desaturase 2 |
| chr1 | 25386795 | 25388023 | 1229 | * | 13,00972 | 2,238957953 | 0,865032 | 2,588295 | 0,009645 | 0,981411547 | 0,738441 | Exon (ENSMUST00000 | 1 | 25318526 | 25318632 | 107 | 2 | 1,15E+08 | ENSMUST00000240417.1 | -68163 | ENSMUSG00000119153 | Gm22260 | predicted gene, 22260 |
| chr10 | 82986600 | 82989571 | 2972 | * | 29,3825 | 2,239428789 | 0,70603 | 3,171859 | 0,001515 | 0,443906448 | 2,725491 | Intron (ENSMUST0000 | 10 | 82974984 | 82977872 | 2889 | 2 | 1,15E+08 | ENSMUST00000160997.2 | -8728 | | 1700025N21Rik | RIKEN cDNA 1700025N21 gene |
| chr13 | 17979809 | 17981566 | 1758 | * | 21,82883 | 2,241132288 | 0,85606 | 2,617963 | 0,008846 | 0,754485841 | 2,67243 | Promoter (<=1kb) | 13 | 17884900 | 17979682 | 94783 | 2 | 69562 | ENSMUST00000223490.2 | -127 | ENSMUSG00000041297 | Cdk13 | cyclin-dependent kinase 13 |
| chr8 | 1,25E+08 | 1,25E+08 | 2633 | * | 26,11186 | 2,241365499 | 0,77048 | 2,909051 | 0,003625 | 0,591141332 | 2,880017 | Intron (ENSMUST0000 | 8 | 1,25E+08 | 1,25E+08 | 102803 | 1 | 108148 | ENSMUST00000142578.8 | 20322 | ENSMUSG00000089704 | Galnt2 | polypeptide N-acetylgalactosaminyltransferase 2 |
| chr9 | 23595265 | 23596788 | 1524 | * | 14,58834 | 2,241524763 | 0,856402 | 2,617374 | 0,008861 | 0,793995086 | 2,141612 | Distal Intergenic | 9 | 23285228 | 23318127 | 32900 | 1 | 73230 | ENSMUST00000214050.2 | 310037 | ENSMUSG00000031963 | Bmper | BMP-binding endothelial regulator |
| chr4 | 1,42E+08 | 1,42E+08 | 1982 | * | 18,56431 | 2,241716983 | 0,782472 | 2,864915 | 0,004171 | 0,630824695 | 2,663619 | Intron (ENSMUST0000 | 4 | 1,42E+08 | 1,42E+08 | 40822 | 2 | 71529 | ENSMUST00000173068.8 | -13471 | ENSMUSG00000040606 | Kazn | kazrin, periplakin interacting protein |
| chr8 | 1,26E+08 | 1,26E+08 | 1838 | * | 20,56337 | 2,241993728 | 0,784871 | 2,856512 | 0,004283 | 0,654833721 | 2,386008 | Intron (ENSMUST0000 | 8 | 1,26E+08 | 1,26E+08 | 206919 | 1 | 244667 | ENSMUST00000121953.2 | 13389 | ENSMUSG00000043051 | Disc1 | disrupted in schizophrenia 1 |
| chr3 | 88074975 | 88077457 | 2483 | * | 23,18912 | 2,242130078 | 0,84053 | 2,667519 | 0,007641 | 0,714462637 | 2,951933 | Promoter (<=1kb) | 3 | 88075559 | 88076213 | 655 | 1 | 17261 | ENSMUST00000152861.2 | 0 | ENSMUSG00000001419 | Mef2d | myocyte enhancer factor 2D |
| chr2 | 27334317 | 27335572 | 1256 | * | 20,87894 | 2,24231636 | 0,722926 | 3,101725 | 0,001924 | 0,486196597 | 2,67243 | Promoter (1-2kb) | 2 | 27332515 | 27333116 | 602 | 1 | 1,01E+08 | ENSMUST00000209262.2 | 1802 | ENSMUSG00000109946 | Brd3os | bromodomain containing 3, opposite strand |
| chr12 | 76457485 | 76459459 | 1975 | * | 16,09238 | 2,242534286 | 0,86459 | 2,593754 | 0,009493 | 0,792545355 | 2,318434 | Distal Intergenic | 12 | 76464312 | 76486266 | 21955 | 1 | 210762 | ENSMUST00000063977.9 | -4853 | ENSMUSG00000052221 | Ppp1r36 | protein phosphatase 1, regulatory subunit 36 |
| chr17 | 26998892 | 27002682 | 3791 | * | 28,28366 | 2,242943925 | 0,73728 | 3,042186 | 0,002349 | 0,507398906 | 2,880017 | Promoter (<=1kb) | 17 | 27000040 | 27011495 | 11456 | 1 | 224630 | ENSMUST00000015725.16 | 0 | ENSMUSG00000024191 | Bnip1 | BCL2/adenovirus E1B interacting protein 1 |
| chr17 | 47304449 | 47306639 | 2191 | * | 19,17974 | 2,243383455 | 0,801157 | 2,800179 | 0,005107 | 0,675594321 | 2,507013 | Intron (ENSMUST0000 | 17 | 47241174 | 47321405 | 80232 | 2 | 224826 | ENSMUST00000224759.2 | 14766 | ENSMUSG00000023977 | Ubr2 | ubiquitin protein ligase E3 component n-recognin 2 |
| chr5 | 1,36E+08 | 1,36E+08 | 1089 | * | 13,88651 | 2,243462882 | 0,852528 | 2,631541 | 0,0085 | 0,926147584 | 0,938682 | Intron (ENSMUST0000 | 5 | 1,36E+08 | 1,36E+08 | 28029 | 2 | 13047 | ENSMUST00000163906.2 | -27246 | ENSMUSG00000029705 | Cux1 | cut-like homeobox 1 |
| chr16 | 24996647 | 24998696 | 2050 | * | 19,79931 | 2,243850941 | 0,797121 | 2,814944 | 0,004879 | 0,654833721 | 2,713306 | Promoter (<=1kb) | 16 | 24963558 | 24996835 | 33278 | 2 | 1,03E+08 | ENSMUST00000241640.1 | 0 | | Gm32196 | predicted gene, 32196 |
| chr4 | 1,3E+08 | 1,3E+08 | 4636 | * | 59,42867 | 2,244509571 | 0,783424 | 2,864999 | 0,00417 | 0,54122637 | 4,200304 | Promoter (<=1kb) | 4 | 1,3E+08 | 1,3E+08 | 10349 | 2 | 94242 | ENSMUST00000030560.9 | 0 | ENSMUSG00000028776 | Tinagl1 | tubulointerstitial nephritis antigen-like 1 |

| | | | | | | | | | | | | | | | | | | | | | | |
|---|---|---|---|---|---|---|---|---|---|---|---|---|---|---|---|---|---|---|---|---|---|---|
| chr18 | 66430606 | 66433416 | 2811 | * | 24,13469 | 2,244811271 | 0,666688 | 3,367109 | 0,00076 | 0,315049828 | 2,951062 | Promoter (2-3kb) | 18 | 66402092 | 66435812 | 33721 | 2 | 320924 | ENSMUST00000151343.3 | 2396 | ENSMUSG00000046318 | Ccbe1 | collagen and calcium binding EGF domains 1 |
| chr5 | 1,4E+08 | 1,4E+08 | 1945 | * | 22,78337 | 2,244845775 | 0,803473 | 2,793929 | 0,005207 | 0,656750434 | 2,880017 | Promoter (<=1kb) | 5 | 1,4E+08 | 1,4E+08 | 11141 | 1 | 17135 | ENSMUST00000018287.10 | 0 | ENSMUSG00000018143 | Mafk | v-maf musculoaponeurotic fibrosarcoma oncogene family, protein K (avian) |
| chr18 | 9211142 | 9213412 | 2271 | * | 22,63045 | 2,245322937 | 0,824839 | 2,722134 | 0,006486 | 0,661775851 | 3,471678 | Promoter (<=1kb) | 18 | 9212163 | 9218136 | 5974 | 1 | 14370 | ENSMUST00000041080.7 | 0 | ENSMUSG00000036904 | Fzd8 | frizzled class receptor 8 |
| chr9 | 88599527 | 88602825 | 3299 | * | 32,3558 | 2,24597869 | 0,676079 | 3,322065 | 0,000894 | 0,375922902 | 2,377674 | Promoter (<=1kb) | 9 | 88601682 | 88688317 | 86636 | 1 | 1E+08 | ENSMUST00000188891.7 | 0 | ENSMUSG00000074149 | Gm10634 | predicted gene 10634 |
| chr13 | 56654319 | 56656461 | 2143 | * | 16,34465 | 2,245998709 | 0,792762 | 2,833131 | 0,004609 | 0,649217039 | 2,663619 | Distal Intergenic | 13 | 56670285 | 56675280 | 4996 | 1 | 213311 | ENSMUST00000121095.8 | -13824 | ENSMUSG00000035509 | Fbxl21 | F-box and leucine-rich repeat protein 21 |
| chr17 | 88160940 | 88162043 | 1104 | * | 16,01307 | 2,246024387 | 0,800583 | 2,805485 | 0,005024 | 0,663408464 | 2,663619 | Distal Intergenic | 17 | 88053229 | 88105422 | 52194 | 2 | 210741 | ENSMUST00000055221.9 | -55518 | ENSMUSG00000050138 | Kcnk12 | potassium channel, subfamily K, member 12 |
| chr4 | 46049128 | 46051746 | 2619 | * | 25,04412 | 2,246060479 | 0,663292 | 3,386234 | 0,000709 | 0,307219482 | 2,951062 | Intron (ENSMUST00000 | 4 | 46039209 | 46090897 | 51689 | 1 | 21916 | ENSMUST00000156200.8 | 9919 | ENSMUSG00000028328 | Tmod1 | tropomodulin 1 |
| chr19 | 4512844 | 4514191 | 1348 | * | 13,05921 | 2,247037012 | 0,865215 | 2,597086 | 0,009402 | 0,908738779 | 1,173325 | Intron (ENSMUST00000 | 19 | 4497745 | 4502179 | 4435 | 2 | 171180 | ENSMUST00000128114.2 | -10665 | ENSMUSG00000049303 | Syt12 | synaptotagmin XII |
| chr8 | 1,12E+08 | 1,12E+08 | 1118 | * | 16,56885 | 2,247175383 | 0,808026 | 2,781069 | 0,005418 | 0,675594321 | 2,663619 | Intron (ENSMUST00000 | 8 | 1,12E+08 | 1,12E+08 | 12714 | 2 | 12927 | ENSMUST00000212147.2 | 6547 | ENSMUSG00000031955 | Bcar1 | breast cancer anti-estrogen resistance 1 |
| chr1 | 1,32E+08 | 1,32E+08 | 1971 | * | 23,64723 | 2,24729919 | 0,67517 | 3,328492 | 0,000873 | 0,334753716 | 2,951062 | Distal Intergenic | 1 | 1,32E+08 | 1,32E+08 | 2784 | 1 | 98396 | ENSMUST00000132585.2 | -3266 | ENSMUSG00000013275 | Slc41a1 | solute carrier family 41, member 1 |
| chr7 | 80454464 | 80457216 | 2753 | * | 19,68926 | 2,247460844 | 0,740725 | 3,034135 | 0,002412 | 0,537550416 | 2,507013 | Promoter (1-2kb) | 7 | 80361331 | 80453164 | 91834 | 2 | 29875 | ENSMUST00000167377.3 | -1300 | ENSMUSG00000030536 | Iqgap1 | IQ motif containing GTPase activating protein 1 |
| chr11 | 76091892 | 76093213 | 1322 | * | 16,80115 | 2,248782185 | 0,734397 | 3,062081 | 0,002198 | 0,531203841 | 2,386008 | Promoter (<=1kb) | 11 | 76092841 | 76100242 | 7402 | 1 | 116972 | ENSMUST00000094014.10 | 0 | ENSMUSG00000069808 | Tlcd3a | TLC domain containing 3A |
| chr13 | 65640884 | 65641780 | 897 | * | 24,88058 | 2,249350535 | 0,747461 | 3,009321 | 0,002618 | 0,524368688 | 2,951933 | Exon (ENSMUST000002 | 13 | 65660492 | 65660662 | 171 | 1 | 1,03E+08 | ENSMUST00000081471.3 | -18712 | | Gm10139 | predicted gene 10139 |
| chr19 | 42744609 | 42746836 | 2228 | * | 38,58339 | 2,24992755 | 0,559006 | 4,024875 | 5,7E-05 | 0,084745052 | 2,338387 | Promoter (<=1kb) | 19 | 42744022 | 42746268 | 2247 | 2 | 192236 | ENSMUST00000161252.2 | 0 | ENSMUSG00000025188 | Hps1 | HPS1, biogenesis of lysosomal organelles complex 3 subunit 1 |
| chr15 | 25985090 | 25986890 | 1801 | * | 20,06011 | 2,250603561 | 0,675709 | 3,330731 | 0,000866 | 0,369395048 | 2,386008 | Promoter (<=1kb) | 15 | 25986881 | 25997372 | 10492 | 1 | 52521 | ENSMUST00000226884.2 | 0 | ENSMUSG00000052253 | Zfp622 | zinc finger protein 622 |
| chr10 | 60870258 | 60871889 | 1632 | * | 15,88761 | 2,25069457 | 0,868988 | 2,590018 | 0,009597 | 0,770994815 | 2,663619 | Intron (ENSMUST00000 | 10 | 60867082 | 60873859 | 6778 | 1 | 327760 | ENSMUST00000177492.2 | 3176 | ENSMUSG00000093568 | Gm20611 | predicted gene 20611 |
| chr13 | 55753556 | 55756092 | 2537 | * | 21,80195 | 2,251304712 | 0,793003 | 2,83896 | 0,004526 | 0,627785088 | 2,951933 | Exon (ENSMUST000001 | 13 | 55747878 | 55758255 | 10378 | 1 | 218271 | ENSMUST00000064701.8 | 5678 | ENSMUSG00000021504 | B4galt7 | xylosylprotein beta1,4-galactosyltransferase, polypeptide 7 (galactosyltransferase I) |
| chr7 | 45479665 | 45482304 | 2640 | * | 24,77461 | 2,251306885 | 0,721399 | 3,120753 | 0,001804 | 0,457838671 | 2,951062 | Promoter (<=1kb) | 7 | 45474647 | 45480368 | 5722 | 2 | 101612 | ENSMUST00000131384.3 | 0 | ENSMUSG00000053801 | Grwd1 | glutamate-rich WD repeat containing 1 |
| chr8 | 1,2E+08 | 1,2E+08 | 2172 | * | 23,13961 | 2,25136984 | 0,768688 | 2,928846 | 0,003402 | 0,570876293 | 2,951933 | Promoter (<=1kb) | 8 | 1,2E+08 | 1,2E+08 | 23220 | 1 | 68270 | ENSMUST00000093100.3 | 14 | ENSMUSG00000031831 | Dnaaf1 | dynein, axonemal assembly factor 1 |
| chr10 | 1,14E+08 | 1,14E+08 | 1541 | * | 14,39824 | 2,251857183 | 0,872848 | 2,579896 | 0,009883 | 0,903294867 | 1,2763 | Distal Intergenic | 10 | 1,13E+08 | 1,13E+08 | 11133 | 2 | 1,05E+08 | ENSMUST00000218612.2 | -241213 | | Gm40765 | predicted gene, 40765 |
| chr11 | 86746463 | 86748295 | 1833 | * | 19,99917 | 2,252313105 | 0,851523 | 2,64504 | 0,008168 | 0,751231966 | 2,507013 | Distal Intergenic | 11 | 86659672 | 86698572 | 38901 | 2 | 67487 | ENSMUST00000018569.14 | -47891 | ENSMUSG00000018425 | Dhx40 | DEAH (Asp-Glu-Ala-His) box polypeptide 40 |
| chr15 | 71761003 | 71763538 | 2536 | * | 25,62915 | 2,252579032 | 0,788729 | 2,85596 | 0,004291 | 0,623120685 | 2,880017 | Exon (ENSMUST000002 | 15 | 71740730 | 71769465 | 28736 | 2 | 69700 | ENSMUST00000160513.2 | 5927 | ENSMUSG00000079022 | Col22a1 | collagen, type XXII, alpha 1 |
| chr2 | 1,69E+08 | 1,69E+08 | 1443 | * | 15,51364 | 2,253069416 | 0,767127 | 2,937021 | 0,003314 | 0,699278451 | 1,37683 | Distal Intergenic | 2 | 1,69E+08 | 1,69E+08 | 6444 | 1 | 1,05E+08 | ENSMUST00000148458.2 | -192164 | | Gm14250 | predicted gene 14250 |
| chr19 | 57690262 | 57692908 | 2647 | * | 23,59902 | 2,253402382 | 0,757548 | 2,974599 | 0,002934 | 0,546791007 | 2,858511 | Exon (ENSMUST000000 | 19 | 57599466 | 58121775 | 522310 | 1 | 226255 | ENSMUST00000077282.7 | 90796 | ENSMUSG00000054843 | Atrnl1 | attractin like 1 |
| chr4 | 1,49E+08 | 1,49E+08 | 3025 | * | 22,59767 | 2,255095862 | 0,853548 | 2,642027 | 0,008241 | 0,717133806 | 3,104559 | Promoter (<=1kb) | 4 | 1,49E+08 | 1,49E+08 | 3803 | 2 | 69928 | ENSMUST00000150150.3 | 0 | ENSMUSG00000073705 | Cenps | centromere protein S |
| chr13 | 21898751 | 21900666 | 1916 | * | 20,18434 | 2,255570492 | 0,727122 | 3,10205 | 0,001922 | 0,505078837 | 2,386008 | Promoter (<=1kb) | 13 | 21899933 | 21900313 | 381 | 2 | 319185 | ENSMUST00000091756.2 | 0 | ENSMUSG00000094338 | H2bc13 | H2B clustered histone 13 |
| chr10 | 1,25E+08 | 1,25E+08 | 1336 | * | 13,78649 | 2,255897635 | 0,84927 | 2,656279 | 0,007901 | 0,895765172 | 1,075735 | Distal Intergenic | 10 | 1,25E+08 | 1,25E+08 | 23819 | 1 | 1,03E+08 | ENSMUST00000219518.2 | -97944 | | Gm36719 | predicted gene, 36719 |
| chr15 | 52770792 | 52771902 | 1111 | * | 14,62279 | 2,256416383 | 0,85354 | 2,643596 | 0,008203 | 0,783315811 | 2,141612 | Distal Intergenic | 15 | 52646777 | 52649663 | 2887 | 2 | 1,03E+08 | ENSMUST00000228167.2 | -121129 | | Gm34794 | predicted gene, 34794 |
| chr2 | 69959787 | 69961991 | 2205 | * | 16,84962 | 2,256427732 | 0,801216 | 2,816256 | 0,004859 | 0,687150104 | 2,141612 | Intron (ENSMUST00000 | 2 | 69926642 | 70013611 | 86970 | 1 | 329421 | ENSMUST00000112241.2 | 33145 | ENSMUSG00000042064 | Myo3b | myosin IIIB |
| chr8 | 1E+08 | 1E+08 | 1571 | * | 17,27022 | 2,25696251 | 0,799943 | 2,821404 | 0,004781 | 0,670234202 | 2,423594 | Intron (ENSMUST00000 | 8 | 1E+08 | 1E+08 | 316765 | 1 | 234624 | ENSMUST00000153366.2 | 287431 | ENSMUSG00000052479 | A330008L17Rik | RIKEN cDNA A330008L17 gene |
| chr2 | 1,32E+08 | 1,32E+08 | 2279 | * | 15,61077 | 2,257094284 | 0,851872 | 2,64957 | 0,008059 | 0,856202617 | 1,37683 | Distal Intergenic | 2 | 1,31E+08 | 1,32E+08 | 68942 | 2 | 1,19E+08 | ENSMUST00000138080.2 | -26456 | | Gm14280 | predicted gene 14280 |
| chr4 | 65452921 | 65454846 | 1926 | * | 16,84428 | 2,257295563 | 0,781003 | 2,890254 | 0,003849 | 0,64236196 | 2,318434 | Intron (ENSMUST00000 | 4 | 65523223 | 65534475 | 11253 | 1 | 69807 | ENSMUST00000050850.14 | -68377 | ENSMUSG00000051675 | Trim32 | tripartite motif-containing 32 |
| chr12 | 1,02E+08 | 1,02E+08 | 1875 | * | 15,27135 | 2,257509292 | 0,875031 | 2,579921 | 0,009882 | 0,903294867 | 1,2763 | Intron (ENSMUST00000 | 12 | 1,02E+08 | 1,02E+08 | 45716 | 2 | 19141 | ENSMUST00000021607.9 | 6372 | ENSMUSG00000021190 | Lgmn | legumain |
| chr17 | 49113087 | 49114880 | 1794 | * | 20,18735 | 2,257775337 | 0,67611 | 3,339361 | 0,00084 | 0,377305803 | 2,221611 | Distal Intergenic | 17 | 49239407 | 49404616 | 165210 | 1 | 70530 | ENSMUST00000046254.3 | -124527 | ENSMUSG00000040490 | Lrfn2 | leucine rich repeat and fibronectin type III domain containing 2 |
| chr17 | 83048311 | 83050464 | 2154 | * | 23,4209 | 2,25829837 | 0,760861 | 2,968083 | 0,002997 | 0,543902653 | 2,951933 | Distal Intergenic | 17 | 83341023 | 83385654 | 44632 | 2 | 1,01E+08 | ENSMUST00000234063.2 | 335190 | ENSMUSG00000117399 | Gm19689 | predicted gene, 19689 |
| chr3 | 1,08E+08 | 1,08E+08 | 1570 | * | 19,28637 | 2,258397302 | 0,86987 | 2,596248 | 0,009425 | 0,764280266 | 2,713306 | Promoter (<=1kb) | 3 | 1,08E+08 | 1,08E+08 | 1056 | 2 | 14862 | ENSMUST00000198532.2 | 0 | ENSMUSG00000058135 | Gstm1 | glutathione S-transferase, mu 1 |
| chr5 | 1,38E+08 | 1,38E+08 | 2401 | * | 26,70571 | 2,258481929 | 0,681052 | 3,316166 | 0,000913 | 0,339523086 | 2,951062 | Promoter (<=1kb) | 5 | 1,38E+08 | 1,38E+08 | 17640 | 1 | 50765 | ENSMUST00000198866.5 | 0 | ENSMUSG00000029716 | Tfr2 | transferrin receptor 2 |
| chr4 | 1,35E+08 | 1,35E+08 | 3035 | * | 26,76868 | 2,259092628 | 0,645776 | 3,498259 | 0,000468 | 0,254650393 | 2,951062 | 3' UTR | 4 | 1,35E+08 | 1,35E+08 | 8455 | 2 | 66146 | ENSMUST00000150031.2 | 4017 | ENSMUSG00000028826 | Maco1 | macoilin 1 |
| chr5 | 74976983 | 74979314 | 2332 | * | 21,20607 | 2,260237145 | 0,76395 | 2,95862 | 0,00309 | 0,54122637 | 3,104559 | Distal Intergenic | 5 | 74892177 | 74895112 | 2936 | 2 | 1,19E+08 | ENSMUST00000137731.3 | -81871 | | Gm15985 | predicted gene 15985 |
| chr4 | 1,25E+08 | 1,25E+08 | 2222 | * | 29,04571 | 2,260307426 | 0,709307 | 3,186642 | 0,001439 | 0,439336412 | 2,67243 | Promoter (<=1kb) | 4 | 1,25E+08 | 1,25E+08 | 2427 | 2 | 230738 | ENSMUST00000131685.2 | 0 | ENSMUSG00000042677 | Zc3h12a | zinc finger CCCH type containing 12A |
| chr16 | 8648581 | 8651841 | 3261 | * | 22,41172 | 2,26038238 | 0,772774 | 2,925025 | 0,003444 | 0,578908869 | 2,880017 | Promoter (<=1kb) | 16 | 8649185 | 8661995 | 12811 | 1 | 69053 | ENSMUST00000229318.2 | 0 | ENSMUSG00000022507 | 1810013L24Rik | RIKEN cDNA 1810013L24 gene |
| chr17 | 56583435 | 56585183 | 1749 | * | 22,75825 | 2,26055198 | 0,748589 | 3,019749 | 0,00253 | 0,516158525 | 2,951062 | Promoter (<=1kb) | 17 | 56576319 | 56583786 | 7468 | 2 | 106759 | ENSMUST00000058136.9 | 0 | ENSMUSG00000047123 | Ticam1 | toll-like receptor adaptor molecule 1 |
| chr16 | 5902949 | 5906888 | 3940 | * | 32,5778 | 2,260745246 | 0,596546 | 3,789727 | 0,000151 | 0,133396034 | 2,925497 | Intron (ENSMUST00000 | 16 | 5793449 | 5816360 | 22912 | 1 | 68238 | ENSMUST00000190515.2 | 109500 | ENSMUSG00000101481 | 1700123O21Rik | RIKEN cDNA 1700123O21 gene |
| chr16 | 92375569 | 92377078 | 1510 | * | 14,84462 | 2,26085259 | 0,850294 | 2,658908 | 0,007839 | 0,848799361 | 1,41696 | Distal Intergenic | 16 | 92295022 | 92338131 | 43110 | 1 | 209195 | ENSMUST00000023670.4 | 80547 | ENSMUSG00000022949 | Clic6 | chloride intracellular channel 6 |
| chr3 | 1,08E+08 | 1,08E+08 | 2778 | * | 25,03427 | 2,261078724 | 0,721215 | 3,135099 | 0,001718 | 0,467440127 | 2,67243 | Promoter (<=1kb) | 3 | 1,08E+08 | 1,08E+08 | 38867 | 2 | 14679 | ENSMUST00000000001.5 | 0 | ENSMUSG00000000001 | Gnai3 | guanine nucleotide binding protein (G protein), alpha inhibiting 3 |
| chr5 | 1,16E+08 | 1,16E+08 | 1412 | * | 15,07598 | 2,261361408 | 0,82547 | 2,739482 | 0,006154 | 0,83238039 | 1,214904 | 3' UTR | 5 | 1,16E+08 | 1,16E+08 | 3617 | 1 | 12704 | ENSMUST00000134609.2 | -7750 | ENSMUSG00000029516 | Cit | citron |
| chr15 | 52791408 | 52794017 | 2610 | * | 31,74576 | 2,261578621 | 0,605771 | 3,733388 | 0,000189 | 0,154711074 | 2,951062 | Distal Intergenic | 15 | 52646777 | 52649663 | 2887 | 2 | 1,03E+08 | ENSMUST00000228167.2 | -141745 | | Gm34794 | predicted gene, 34794 |
| chr2 | 1,67E+08 | 1,67E+08 | 3439 | * | 32,86487 | 2,261611529 | 0,607005 | 3,725853 | 0,000195 | 0,23710373 | 1,405709 | Promoter (1-2kb) | 2 | 1,67E+08 | 1,67E+08 | 14115 | 2 | 1,03E+08 | ENSMUST00000245024.1 | 1816 | | Gm31619 | predicted gene, 31619 |
| chr4 | 1,5E+08 | 1,5E+08 | 2795 | * | 25,07296 | 2,261877947 | 0,772483 | 2,928061 | 0,003411 | 0,594430593 | 2,67243 | Promoter (<=1kb) | 4 | 1,5E+08 | 1,5E+08 | 5388 | 1 | 13806 | ENSMUST00000080149.6 | -170 | ENSMUSG00000063524 | Eno1 | enolase 1, alpha non-neuron |
| chr2 | 1,66E+08 | 1,66E+08 | 1962 | * | 18,35674 | 2,262132568 | 0,788867 | 2,86757 | 0,004136 | 0,646266933 | 2,423594 | Distal Intergenic | 2 | 1,66E+08 | 1,66E+08 | 20924 | 1 | 670775 | ENSMUST00000152987.2 | 49898 | ENSMUSG00000084840 | Gm11468 | predicted gene 11468 |
| chr4 | 43458099 | 43459823 | 1725 | * | 18,48353 | 2,262326238 | 0,816871 | 2,769504 | 0,005614 | 0,695401133 | 2,386008 | Distal Intergenic | 4 | 43447724 | 43454628 | 6905 | 2 | 12517 | ENSMUST00000030179.11 | -3471 | ENSMUSG00000028459 | Cd72 | CD72 antigen |
| chr2 | 1,63E+08 | 1,63E+08 | 1923 | * | 14,74771 | 2,262345463 | 0,844067 | 2,680291 | 0,007356 | 0,853738252 | 1,2763 | Exon (ENSMUST000000 | 2 | 1,63E+08 | 1,63E+08 | 3907 | 1 | 76080 | ENSMUST00000132904.2 | 14255 | ENSMUSG00000017679 | Ttpal | tocopherol (alpha) transfer protein-like |
| chr11 | 76975809 | 76979186 | 3378 | * | 26,11765 | 2,262673906 | 0,715462 | 3,162537 | 0,001564 | 0,436322355 | 2,951062 | Distal Intergenic | 11 | 76945320 | 76969261 | 23942 | 2 | 237859 | ENSMUST00000134685.2 | -6548 | ENSMUSG00000037958 | Nsrp1 | nuclear speckle regulatory protein 1 |
| chr15 | 71745834 | 71747245 | 1412 | * | 16,83723 | 2,26299327 | 0,75636 | 2,991951 | 0,002772 | 0,595051702 | 2,141612 | Intron (ENSMUST00000 | 15 | 71740730 | 71769465 | 28736 | 2 | 69700 | ENSMUST00000160513.2 | 22220 | ENSMUSG00000079022 | Col22a1 | collagen, type XXII, alpha 1 |
| chr8 | 65095956 | 65097750 | 1795 | * | 23,30268 | 2,263264099 | 0,696641 | 3,248824 | 0,001159 | 0,404488619 | 2,67243 | Intron (ENSMUST00000 | 8 | 65070635 | 65075031 | 4397 | 2 | 12876 | ENSMUST00000211312.2 | -20925 | ENSMUSG00000037852 | Cpe | carboxypeptidase E |
| chr12 | 91540874 | 91542587 | 1714 | * | 15,97854 | 2,263284471 | 0,821533 | 2,754953 | 0,00587 | 0,684274806 | 2,663619 | Promoter (<=1kb) | 12 | 91540204 | 91540346 | 143 | 2 | 1,15E+08 | ENSMUST00000116715.3 | -528 | ENSMUSG00000080365 | Gm25776 | predicted gene, 25776 |
| chr1 | 1,64E+08 | 1,64E+08 | 2958 | * | 22,69477 | 2,263369301 | 0,745005 | 3,03806 | 0,002381 | 0,477594358 | 3,471678 | Promoter (1-2kb) | 1 | 1,64E+08 | 1,64E+08 | 1099 | 1 | 116914 | ENSMUST00000160773.2 | -1473 | ENSMUSG00000040918 | Slc19a2 | solute carrier family 19 (thiamine transporter), member 2 |
| chr15 | 62306157 | 62308311 | 2155 | * | 28,30697 | 2,263395564 | 0,658908 | 3,435069 | 0,000592 | 0,288385947 | 2,880017 | Distal Intergenic | 15 | 62439824 | 62654636 | 214813 | 1 | 1,05E+08 | ENSMUST00000230029.2 | -131513 | | Gm41333 | predicted gene, 41333 |
| chr12 | 13303250 | 13305655 | 2406 | * | 17,56321 | 2,263471845 | 0,742645 | 3,047852 | 0,002305 | 0,518149475 | 2,663619 | Distal Intergenic | 12 | 13266974 | 13299214 | 32241 | 2 | 104721 | ENSMUST00000071103.10 | -4036 | ENSMUSG00000037149 | Ddx1 | DEAD box helicase 1 |
| chr13 | 23738132 | 23740890 | 2759 | * | 23,90332 | 2,263473937 | 0,819135 | 2,763248 | 0,005723 | 0,669377201 | 2,951933 | Promoter (<=1kb) | 13 | 23737436 | 23744860 | 7425 | 1 | 14957 | ENSMUST00000045301.9 | 696 | ENSMUSG00000052565 | H1f3 | H1.3 linker histone, cluster member |
| chr11 | 1,17E+08 | 1,17E+08 | 2118 | * | 24,52725 | 2,263716967 | 0,690771 | 3,277089 | 0,001049 | 0,370253435 | 2,858511 | Intron (ENSMUST00000 | 11 | 1,16E+08 | 1,17E+08 | 26036 | 2 | 217344 | ENSMUST00000103029.10 | 4322 | ENSMUSG00000020806 | Rhbdf2 | rhomboid 5 homolog 2 |
| chr5 | 70574262 | 70575663 | 1402 | * | 14,46661 | 2,264612149 | 0,826613 | 2,739629 | 0,006151 | 0,802671652 | 1,41696 | Distal Intergenic | 5 | 70708184 | 70708290 | 107 | 2 | 1,15E+08 | ENSMUST00000240250.2 | 132627 | ENSMUSG00000118789 | Gm26072 | predicted gene, 26072 |
| chr3 | 95595187 | 95597138 | 1952 | * | 18,91592 | 2,2650658 | 0,723635 | 3,130123 | 0,001747 | 0,471171835 | 2,663619 | Promoter (<=1kb) | 3 | 95583511 | 95595228 | 11718 | 2 | 229595 | ENSMUST00000117782.8 | 0 | ENSMUSG00000015850 | Adamtsl4 | ADAMTS-like 4 |
| chr11 | 87527771 | 87530430 | 2660 | * | 16,09891 | 2,265305197 | 0,860174 | 2,633544 | 0,00845 | 0,74907225 | 2,663619 | Exon (ENSMUST000000 | 11 | 87507990 | 87550368 | 42379 | 1 | 327992 | ENSMUST00000093956.4 | 19781 | ENSMUSG00000070345 | Hsf5 | heat shock transcription factor family member 5 |
| chr12 | 70041145 | 70043896 | 2752 | * | 19,83862 | 2,265563899 | 0,783914 | 2,890068 | 0,003852 | 0,617289329 | 2,663619 | Promoter (<=1kb) | 12 | 70041564 | 70041666 | 103 | 2 | 1,15E+08 | ENSMUST00000158528.4 | 0 | ENSMUSG00002075853 | Gm23579 | predicted gene, 23579 |
| chr6 | 50236940 | 50239957 | 3018 | * | 43,87895 | 2,265735272 | 0,574318 | 3,945087 | 7,98E-05 | 0,090117734 | 2,925497 | Promoter (<=1kb) | 6 | 50166349 | 50238735 | 72387 | 2 | 54722 | ENSMUST00000165099.8 | 0 | ENSMUSG00000029821 | Gsdme | gasdermin E |
| chr11 | 1,16E+08 | 1,16E+08 | 2301 | * | 17,70244 | 2,266662876 | 0,732941 | 3,09256 | 0,001984 | 0,531203841 | 2,154504 | Exon (ENSMUST000001 | 11 | 1,16E+08 | 1,16E+08 | 31862 | 1 | 57230 | ENSMUST00000021101.4 | 3204 | ENSMUSG00000020755 | Sap30bp | SAP30 binding protein |
| chr5 | 28401629 | 28403161 | 1533 | * | 14,06266 | 2,266894263 | 0,867183 | 2,61409 | 0,008947 | 0,970928491 | 0,738441 | Distal Intergenic | 5 | 28407447 | 28415020 | 7574 | 2 | 269637 | ENSMUST00000118882.2 | 11859 | ENSMUSG00000044681 | Cnpy1 | canopy FGF signaling regulator 1 |
| chr7 | 44428358 | 44429840 | 1483 | * | 14,1557 | 2,266935356 | 0,843359 | 2,687984 | 0,007188 | 0,937845932 | 0,738441 | Distal Intergenic | 7 | 44439631 | 44440072 | 442 | 2 | 1,01E+08 | ENSMUST00000207888.2 | 10232 | | Gm29791 | predicted gene, 29791 |
| chr7 | 1,32E+08 | 1,32E+08 | 2765 | * | 26,33073 | 2,267173039 | 0,734013 | 3,088736 | 0,00201 | 0,477253929 | 2,951062 | Promoter (1-2kb) | 7 | 1,32E+08 | 1,32E+08 | 53828 | 2 | 77938 | ENSMUST00000106165.8 | 1454 | ENSMUSG00000030956 | Fam53b | family with sequence similarity 53, member B |
| chr15 | 81482632 | 81484837 | 2206 | * | 15,09959 | 2,267403286 | 0,80821 | 2,805464 | 0,005024 | 0,774782871 | 1,37683 | Intron (ENSMUST00000 | 15 | 81476294 | 81476400 | 107 | 1 | 1,15E+08 | ENSMUST00000122790.4 | 6338 | ENSMUSG00000084739 | Gm22067 | predicted gene, 22067 |
| chr15 | 36878953 | 36881762 | 2810 | * | 31,71932 | 2,267530355 | 0,595539 | 3,807528 | 0,00014 | 0,129229882 | 2,880017 | Promoter (<=1kb) | 15 | 36870744 | 36880121 | 9378 | 2 | 1E+08 | ENSMUST00000100713.4 | 0 | ENSMUSG00000072618 | Gm10384 | predicted gene 10384 |
| chr7 | 1,22E+08 | 1,22E+08 | 1877 | * | 17,95166 | 2,267797966 | 0,803331 | 2,822994 | 0,004758 | 0,671421781 | 2,386008 | Distal Intergenic | 7 | 1,22E+08 | 1,22E+08 | 611 | 1 | 70261 | ENSMUST00000160828.2 | 18685 | ENSMUSG00000030865 | Chp2 | calcineurin-like EF hand protein 2 |
| chr5 | 34670390 | 34672935 | 2546 | * | 21,29694 | 2,268887602 | 0,725757 | 3,126234 | 0,001771 | 0,455547114 | 2,951933 | Promoter (<=1kb) | 5 | 34653431 | 34671307 | 17877 | 2 | 231130 | ENSMUST00000030991.14 | 0 | ENSMUSG00000059866 | Tnip2 | TNFAIP3 interacting protein 2 |
| chr17 | 24910328 | 24912326 | 1999 | * | 17,986 | 2,269261051 | 0,796787 | 2,848015 | 0,004399 | 0,650529916 | 2,507013 | Promoter (1-2kb) | 17 | 24904066 | 24908929 | 4864 | 2 | 20974 | ENSMUST00000007236.5 | -1399 | ENSMUSG00000007021 | Syngr3 | synaptogyrin 3 |
| chr15 | 66854995 | 66857888 | 2894 | * | 58,143219 | 2,269284721 | 0,4582438 | 4,9521337 | 7,34E-07 | 0,002656105 | 3,2706213 | Intron (ENSMUST000001 | 15 | 66818386 | 66841489 | 23104 | 2 | 17988 | ENSMUST00000164070.2 | -13506 | ENSMUSG00000005125 | Ndrg1 | N-myc downstream regulated gene 1 |
| chr19 | 37935373 | 37937381 | 2009 | * | 20,65197 | 2,269716878 | 0,745385 | 3,045025 | 0,002327 | 0,494045114 | 3,104559 | Exon (ENSMUST000002 | 19 | 37907791 | 37928429 | 20639 | 2 | 226101 | ENSMUST00000224518.2 | -6944 | ENSMUSG00000048612 | Myof | myoferlin |
| chr7 | 1,1E+08 | 1,1E+08 | 2825 | * | 24,67577 | 2,270345845 | 0,682393 | 3,327035 | 0,000878 | 0,335478401 | 2,951062 | Exon (ENSMUST000000 | 7 | 1,1E+08 | 1,1E+08 | 184 | 1 | 1E+08 | ENSMUST00000082517.3 | -8293 | ENSMUSG00000064451 | Snora23 | small nucleolar RNA, H/ACA box 23 |
| chr1 | 1,71E+08 | 1,71E+08 | 2965 | * | 24,4255 | 2,27118651 | 0,864207 | 2,62806 | 0,008587 | 0,738600147 | 2,858511 | Distal Intergenic | 1 | 1,71E+08 | 1,71E+08 | 165 | 1 | 1,15E+08 | ENSMUST00000083971.3 | 6157 | ENSMUSG00000065905 | Gm26110 | predicted gene, 26110 |
| chr12 | 84234713 | 84238401 | 3689 | * | 28,02538 | 2,271721446 | 0,632269 | 3,592964 | 0,000327 | 0,214556109 | 2,858511 | Promoter (2-3kb) | 12 | 84196052 | 84240789 | 44738 | 2 | 238317 | ENSMUST00000110294.2 | 2388 | ENSMUSG00000042507 | Mideas | mitotic deacetylase associated SANT domain protein |
| chr8 | 71557704 | 71559028 | 1325 | * | 19,74563 | 2,273812891 | 0,855202 | 2,658802 | 0,007842 | 0,730832709 | 2,713306 | Intron (ENSMUST00000 | 8 | 71459372 | 71538402 | 79031 | 2 | 69419 | ENSMUST00000212266.2 | -19302 | ENSMUSG00000101854 | 1700026F02Rik | RIKEN cDNA 1700026F02 gene |
| chr11 | 1,06E+08 | 1,06E+08 | 1904 | * | 16,5003 | 2,273839495 | 0,843345 | 2,696214 | 0,007013 | 0,847020973 | 1,2763 | Exon (ENSMUST000001 | 11 | 1,06E+08 | 1,06E+08 | 2210 | 2 | 67803 | ENSMUST00000106875.2 | -6916 | ENSMUSG00000040699 | Limd2 | LIM domain containing 2 |
| chr2 | 1,53E+08 | 1,53E+08 | 2150 | * | 22,62854 | 2,274960442 | 0,769097 | 2,957965 | 0,003097 | 0,57149888 | 2,67243 | Promoter (<=1kb) | 2 | 1,53E+08 | 1,53E+08 | 1052 | 1 | 78151 | ENSMUST00000109783.2 | 400 | | 4930404H24Rik | RIKEN cDNA 4930404H24 gene |
| chr14 | 55098996 | 55104067 | 5072 | * | 39,83112 | 2,275229244 | 0,566153 | 4,018756 | 5,85E-05 | 0,08586327 | 2,338387 | Promoter (<=1kb) | 14 | 55091244 | 55101615 | 10372 | 2 | 239099 | ENSMUST00000142283.4 | 0 | ENSMUSG00000057156 | Homez | homeodomain leucine zipper-encoding gene |
| chr13 | 6633554 | 6635504 | 1951 | * | 18,14165 | 2,275288202 | 0,770226 | 2,954054 | 0,003136 | 0,595051702 | 2,423594 | Promoter (1-2kb) | 13 | 6631504 | 6637131 | 5628 | 2 | 56421 | ENSMUST00000142972.2 | 1627 | ENSMUSG00000021196 | Pfkp | phosphofructokinase, platelet |
| chr17 | 44999527 | 45000796 | 1270 | * | 12,00347 | 2,27552285 | 0,882564 | 2,578309 | 0,009929 | 0,985419174 | 0,744375 | Intron (ENSMUST00000 | 17 | 44950568 | 45045580 | 95013 | 2 | 12393 | ENSMUST00000160199.2 | 44784 | ENSMUSG00000039153 | Runx2 | runt related transcription factor 2 |
| chr9 | 62188297 | 62193341 | 5045 | * | 57,138013 | 2,275940004 | 0,5362158 | 4,2444475 | 2,191E-05 | 0,030165351 | 4,0557576 | Promoter (<=1kb) | 9 | 62178011 | 62189490 | 11480 | 2 | 66712 | ENSMUST00000056949.5 | 0 | ENSMUSG00000046846 | Spesp1 | sperm equatorial segment protein 1 |
| chr14 | 77023154 | 77026513 | 3360 | * | 30,83957 | 2,2761303 | 0,579103 | 3,930442 | 8,48E-05 | 0,096870739 | 2,725491 | Distal Intergenic | 14 | 77018267 | 77021652 | 3386 | 1 | 1,03E+08 | ENSMUST00000226490.2 | 4887 | | Gm30246 | predicted gene, 30246 |
| chr11 | 69753420 | 69755660 | 2241 | * | 25,16157 | 2,276571235 | 0,660438 | 3,447064 | 0,000567 | 0,280905021 | 2,951933 | Distal Intergenic | 11 | 69745306 | 69749556 | 4251 | 2 | 83813 | ENSMUST00000108626.8 | -3864 | ENSMUSG00000001583 | Tnk1 | tyrosine kinase, non-receptor, 1 |
| chr6 | 54078904 | 54079966 | 1063 | * | 16,15068 | 2,276664908 | 0,843271 | 2,699801 | 0,006938 | 0,73764377 | 2,318434 | Intron (ENSMUST00000 | 6 | 54074705 | 54197522 | 122818 | 1 | 69993 | ENSMUST00000204410.2 | 4199 | ENSMUSG00000004633 | Chn2 | chimerin 2 |
| chr12 | 85531272 | 85536089 | 4818 | * | 43,5983 | 2,276825548 | 0,566439 | 4,019539 | 5,83E-05 | 0,075415099 | 2,925497 | Exon (ENSMUST000002 | 12 | 85521685 | 85522837 | 1153 | 1 | 14281 | ENSMUST00000134311.2 | 9587 | ENSMUSG00000021250 | Fos | FBJ osteosarcoma oncogene |
| chr9 | 41287065 | 41290378 | 3314 | * | 28,95042 | 2,277181407 | 0,665217 | 3,423217 | 0,000619 | 0,296046677 | 2,858511 | Promoter (<=1kb) | 9 | 41287605 | 41440221 | 152617 | 1 | 73144 | ENSMUST00000233800.2 | 0 | ENSMUSG00000074415 | Mir100hg | Mir100 Mirlet7a-2 Mir125b-1 cluster host gene |
| chr13 | 64395815 | 64397641 | 1827 | * | 16,30631 | 2,277261201 | 0,86883 | 2,621065 | 0,008766 | 0,775744126 | 2,386008 | Promoter (<=1kb) | 13 | 64343172 | 64396456 | 53285 | 2 | 218294 | ENSMUST00000109769.10 | 0 | ENSMUSG00000033102 | Cdc14b | CDC14 cell division cycle 14B |
| chr8 | 70220330 | 70221679 | 1350 | * | 13,52862 | 2,277621596 | 0,869505 | 2,619448 | 0,008807 | 0,936214558 | 0,918255 | Promoter (1-2kb) | 8 | 70195834 | 70219329 | 23496 | 2 | 620419 | ENSMUST00000126915.2 | -1001 | ENSMUSG00000092260 | Zfp963 | zinc finger protein 963 |
| chr5 | 1,02E+08 | 1,02E+08 | 1303 | * | 16,11658 | 2,27770679 | 0,872213 | 2,61141 | 0,009017 | 0,779936001 | 2,386008 | Distal Intergenic | 5 | 1,02E+08 | 1,02E+08 | 6858 | 2 | 18096 | ENSMUST00000044125.11 | 104486 | ENSMUSG00000035187 | Nkx6-1 | NK6 homeobox 1 |
| chr9 | 88226251 | 88228631 | 2381 | * | 21,88194 | 2,277878197 | 0,682727 | 3,336442 | 0,000849 | 0,344155199 | 2,67243 | Intron (ENSMUST00000 | 9 | 88237740 | 88240056 | 2317 | 1 | 23959 | ENSMUST00000217134.2 | -9109 | ENSMUSG00000032420 | Nt5e | 5' nucleotidase, ecto |
| chr11 | 76365365 | 76368046 | 2682 | * | 19,02795 | 2,278244998 | 0,827481 | 2,753229 | 0,005901 | 0,684391766 | 2,663619 | Intron (ENSMUST00000 | 11 | 76307577 | 76359291 | 51715 | 2 | 109934 | ENSMUST00000065028.14 | -6074 | ENSMUSG00000017631 | Abr | active BCR-related gene |
| chr8 | 1,06E+08 | 1,06E+08 | 1246 | * | 15,36366 | 2,278315471 | 0,867587 | 2,626036 | 0,008639 | 0,883970984 | 1,2763 | Promoter (1-2kb) | 8 | 1,06E+08 | 1,06E+08 | 3080 | 1 | 234695 | ENSMUST00000212643.2 | -1283 | ENSMUSG00000050357 | Carmil2 | capping protein regulator and myosin 1 linker 2 |
| chr11 | 68743729 | 68745637 | 1909 | * | 21,42719 | 2,278449228 | 0,735627 | 3,097287 | 0,001953 | 0,47348041 | 2,951933 | Promoter (<=1kb) | 11 | 68712260 | 68743961 | 31702 | 2 | 83431 | ENSMUST00000018880.14 | 0 | ENSMUSG00000018736 | Ndel1 | nudE neurodevelopment protein 1 like 1 |
| chr12 | 10219312 | 10222079 | 2768 | * | 17,86293 | 2,278455052 | 0,728834 | 3,126163 | 0,001771 | 0,48280639 | 2,507013 | Distal Intergenic | 12 | 10419973 | 10425935 | 5963 | 1 | 70881 | ENSMUST00000143739.9 | -197894 | ENSMUSG00000020622 | Nt5c1b | 5'-nucleotidase, cytosolic IB |
| chr5 | 76690362 | 76693584 | 3223 | * | 29,06166 | 2,278579242 | 0,617913 | 3,687542 | 0,000226 | 0,170809454 | 2,951062 | Promoter (<=1kb) | 5 | 76689791 | 76697362 | 7572 | 1 | 69940 | ENSMUST00000141884.2 | 571 | ENSMUSG00000036435 | Exoc1 | exocyst complex component 1 |

| | | | | | | | | | | | | | | | | | | | | | | | |
|---|---|---|---|---|---|---|---|---|---|---|---|---|---|---|---|---|---|---|---|---|---|---|---|
| chr8 | 54982752 | 54985521 | 2770 | * | 26,56996 | 2,279084029 | 0,713971 | 3,192125 | 0,001412 | 0,419066905 | 2,951062 | Promoter (<=1kb) | 8 | 54973468 | 54983033 | 9566 | 2 | 76687 | ENSMUST00000067476.9 | 0 | ENSMUSG00000054408 | Spcs3 | signal peptidase complex subunit 3 homolog (S. cerevisiae) |
| chr11 | 94215310 | 94218923 | 3614 | * | 26,18855 | 2,280202768 | 0,728396 | 3,130444 | 0,001745 | 0,457244879 | 2,880017 | Promoter (<=1kb) | 11 | 94218827 | 94232667 | 13841 | 1 | 71452 | ENSMUST00000107818.9 | 0 | ENSMUSG00000020864 | Ankrd40 | ankyrin repeat domain 40 |
| chr6 | 68687013 | 68688117 | 1105 | * | 15,29693 | 2,280482186 | 0,832325 | 2,739895 | 0,006146 | 0,802671652 | 1,41696 | Distal Intergenic | 6 | 68857162 | 68857658 | 497 | 2 | 1,08E+08 | ENSMUST00000198756.2 | 169541 | | Gm53190 | predicted gene, 53190 |
| chr6 | 1,23E+08 | 1,23E+08 | 1760 | * | 23,43419 | 2,28064648 | 0,691479 | 3,298215 | 0,000973 | 0,355694809 | 2,858511 | Intron (ENSMUST0000( | 6 | 1,23E+08 | 1,23E+08 | 3029 | 2 | 20527 | ENSMUST00000168801.2 | -17336 | ENSMUSG00000003153 | Slc2a3 | solute carrier family 2 (facilitated glucose transporter), member 3 |
| chr10 | 71181860 | 71183643 | 1784 | * | 22,92676 | 2,2807537 | 0,664285 | 3,433397 | 0,000596 | 0,284660899 | 2,951933 | Promoter (<=1kb) | 10 | 71183593 | 71221715 | 38123 | 1 | 69718 | ENSMUST00000079252.13 | 0 | ENSMUSG00000060733 | Ipmk | inositol polyphosphate multikinase |
| chr9 | 21834225 | 21837280 | 3056 | * | 30,59461 | 2,281549993 | 0,588099 | 3,879536 | 0,000105 | 0,113364373 | 2,67243 | Promoter (<=1kb) | 9 | 21832304 | 21837615 | 5312 | 2 | 235043 | ENSMUST00000217263.2 | 335 | ENSMUSG00000040883 | Tmem205 | transmembrane protein 205 |
| chr19 | 4622112 | 4624326 | 2215 | * | 17,29725 | 2,281555724 | 0,845767 | 2,697617 | 0,006984 | 0,750475959 | 2,154504 | Intron (ENSMUST0000( | 19 | 4608838 | 4652049 | 43212 | 1 | 18563 | ENSMUST00000236457.2 | 13274 | ENSMUSG00000024892 | Pcx | pyruvate carboxylase |
| chr11 | 20195454 | 20197079 | 1626 | * | 18,06462 | 2,281579577 | 0,838769 | 2,720154 | 0,006525 | 0,722999566 | 2,386008 | Promoter (2-3kb) | 11 | 20191523 | 20199383 | 7861 | 2 | 216543 | ENSMUST00000161218.2 | 2304 | ENSMUSG00000044066 | Cep68 | centrosomal protein 68 |
| chr2 | 1,59E+08 | 1,59E+08 | 2011 | * | 16,77942 | 2,282198336 | 0,873361 | 2,613122 | 0,008972 | 0,756849896 | 2,663619 | Intron (ENSMUST0000( | 2 | 1,59E+08 | 1,59E+08 | 70200 | 1 | 228852 | ENSMUST00000145073.2 | -12609 | ENSMUSG00000037754 | Ppp1r16b | protein phosphatase 1, regulatory subunit 16B |
| chr15 | 89238793 | 89242192 | 3400 | * | 49,434998 | 2,282271033 | 0,4926925 | 4,6322419 | 3,617E-06 | 0,009977325 | 2,9254973 | Promoter (<=1kb) | 15 | 89235207 | 89239862 | 4656 | 2 | 105847 | ENSMUST00000023283.6 | 0 | ENSMUSG00000022614 | Lmf2 | lipase maturation factor 2 |
| chr4 | 1,51E+08 | 1,51E+08 | 2202 | * | 15,46626 | 2,282914386 | 0,844671 | 2,702727 | 0,006877 | 0,749550311 | 2,141612 | Promoter (<=1kb) | 4 | 1,51E+08 | 1,51E+08 | 41014 | 2 | 18628 | ENSMUST00000103204.11 | 0 | ENSMUSG00000028957 | Per3 | period circadian clock 3 |
| chr18 | 74644028 | 74646353 | 2326 | * | 18,93157 | 2,282991935 | 0,743422 | 3,070924 | 0,002134 | 0,504427013 | 2,663619 | Exon (ENSMUST00000 | 18 | 74658324 | 74664177 | 5854 | 2 | 73606 | ENSMUST00000137034.2 | 17824 | | 1700120E14Rik | RIKEN cDNA 1700120E14 gene |
| chr14 | 47336375 | 47338840 | 2466 | * | 27,07016 | 2,283207386 | 0,646829 | 3,529847 | 0,000416 | 0,244895089 | 2,858511 | Intron (ENSMUST0000( | 14 | 47324592 | 47326558 | 1967 | 1 | 74480 | ENSMUST00000130046.2 | 11783 | ENSMUSG00000021838 | Samd4 | sterile alpha motif domain containing 4 |
| chr8 | 4040421 | 4042412 | 1992 | * | 23,20823 | 2,283505752 | 0,709553 | 3,218229 | 0,00129 | 0,408869321 | 2,880017 | Distal Intergenic | 8 | 3990294 | 4004746 | 14453 | 2 | 170776 | ENSMUST00000127592.2 | -35675 | ENSMUSG00000040165 | Cd209c | CD209c antigen |
| chr10 | 78105509 | 78107189 | 1681 | * | 16,8836 | 2,283773909 | 0,870403 | 2,623813 | 0,008695 | 0,751231966 | 2,663619 | 3' UTR | 10 | 78084714 | 78099973 | 15260 | 2 | 1,03E+08 | ENSMUST00000218662.2 | -5536 | | Gm30122 | predicted gene, 30122 |
| chr15 | 84136821 | 84138907 | 2087 | * | 24,8554 | 2,283775909 | 0,741439 | 3,080193 | 0,002069 | 0,48037412 | 2,951933 | Promoter (2-3kb) | 15 | 84141341 | 84174635 | 33295 | 1 | 170736 | ENSMUST00000146020.2 | -2434 | ENSMUSG00000022438 | Parvb | parvin, beta |
| chr7 | 76606065 | 76607424 | 1360 | * | 15,5551 | 2,284755308 | 0,828797 | 2,756713 | 0,005839 | 0,823446343 | 1,214904 | Intron (ENSMUST0000( | 7 | 76060810 | 76416356 | 355547 | 1 | 244071 | ENSMUST00000107442.3 | 545255 | ENSMUSG00000025754 | Agbl1 | ATP/GTP binding protein-like 1 |
| chr17 | 44954657 | 44957190 | 2534 | * | 19,6359 | 2,285907405 | 0,760043 | 3,007604 | 0,002633 | 0,54122637 | 2,663619 | Intron (ENSMUST0000( | 17 | 44872877 | 44874059 | 1183 | 1 | 1,05E+08 | ENSMUST00000160203.2 | 81780 | | Runx2os3 | runt related transcription factor 2, opposite strand 3 |
| chr14 | 21895948 | 21898531 | 2584 | * | 22,70087 | 2,285962475 | 0,758729 | 3,012882 | 0,002588 | 0,538977324 | 2,67243 | Promoter (<=1kb) | 14 | 21897646 | 21901108 | 3463 | 1 | 319486 | ENSMUST00000170150.9 | 0 | ENSMUSG00000090564 | A430057M04Rik | RIKEN cDNA A430057M04 gene |
| chr16 | 31656573 | 31658058 | 1486 | * | 13,75856 | 2,286127666 | 0,859885 | 2,658643 | 0,007846 | 0,914972668 | 0,938682 | Promoter (<=1kb) | 16 | 31656977 | 31670851 | 13875 | 1 | 13383 | ENSMUST00000147382.2 | 0 | ENSMUSG00000022770 | Dlg1 | discs large MAGUK scaffold protein 1 |
| chr1 | 1,62E+08 | 1,62E+08 | 1228 | * | 18,32337 | 2,286284208 | 0,840697 | 2,719512 | 0,006538 | 0,715249603 | 2,507013 | Intron (ENSMUST0000( | 1 | 1,62E+08 | 1,62E+08 | 31156 | 2 | 103967 | ENSMUST00000160173.2 | -50981 | ENSMUSG00000040265 | Dnm3 | dynamin 3 |
| chr15 | 73024726 | 73031211 | 6486 | * | 131,56198 | 2,286418145 | 0,3630552 | 6,2977141 | 3,021E-10 | 5,12589E-06 | 3,2706213 | Intron (ENSMUST000000 | 15 | 73000271 | 73049696 | 49426 | 2 | 239528 | ENSMUST00000230464.2 | 18485 | ENSMUSG00000036698 | Ago2 | argonaute RISC catalytic subunit 2 |
| chr6 | 86506380 | 86508406 | 2027 | * | 20,65452 | 2,286544229 | 0,783359 | 2,918896 | 0,003513 | 0,542967057 | 3,471678 | Promoter (1-2kb) | 6 | 86504456 | 86540487 | 36032 | 1 | 72012 | ENSMUST00000205028.2 | 1924 | ENSMUSG00000097048 | 1600020E01Rik | RIKEN cDNA 1600020E01 gene |
| chr9 | 1,04E+08 | 1,04E+08 | 2982 | * | 38,02642 | 2,287335275 | 0,613448 | 3,728657 | 0,000193 | 0,17530799 | 2,377674 | Promoter (<=1kb) | 9 | 1,04E+08 | 1,04E+08 | 624 | 2 | 1,03E+08 | ENSMUST00000216588.3 | 0 | ENSMUSG00000111035 | Gm33054 | predicted gene, 33054 |
| chr2 | 6736520 | 6739069 | 2550 | * | 20,17183 | 2,28734207 | 0,716256 | 3,19347 | 0,001406 | 0,431374403 | 2,713306 | Intron (ENSMUST0000( | 2 | 6628339 | 6727027 | 98689 | 2 | 14007 | ENSMUST00000126742.2 | -9493 | ENSMUSG00000002107 | Celf2 | CUGBP, Elav-like family member 2 |
| chr4 | 1,37E+08 | 1,37E+08 | 1451 | * | 19,60947 | 2,287514842 | 0,760964 | 3,006074 | 0,002646 | 0,539997763 | 2,713306 | Promoter (<=1kb) | 4 | 1,37E+08 | 1,37E+08 | 101862 | 1 | 15530 | ENSMUST00000030547.15 | -94 | ENSMUSG00000028763 | Hspg2 | perlecan (heparan sulfate proteoglycan 2) |
| chr15 | 36977896 | 36979756 | 1861 | * | 21,20339 | 2,287701111 | 0,761809 | 3,002985 | 0,002673 | 0,531203841 | 2,880017 | Distal Intergenic | 15 | 37000872 | 37002279 | 1408 | 2 | 68036 | ENSMUST00000226257.2 | 22523 | ENSMUSG00000062397 | Zfp706 | zinc finger protein 706 |
| chr9 | 1,03E+08 | 1,03E+08 | 2231 | * | 13,73466 | 2,288040403 | 0,825699 | 2,771033 | 0,005588 | 0,858257847 | 0,938682 | Distal Intergenic | 9 | 1,03E+08 | 1,03E+08 | 7564 | 1 | 69010 | ENSMUST00000186693.2 | 9193 | ENSMUSG00000035048 | Anapc13 | anaphase promoting complex subunit 13 |
| chr2 | 24796555 | 24800357 | 3803 | * | 41,01862 | 2,288127102 | 0,620777 | 3,68591 | 0,000228 | 0,175928278 | 2,787255 | Intron (ENSMUST0000( | 2 | 24681155 | 24809586 | 128432 | 2 | 77683 | ENSMUST00000046227.12 | 9229 | ENSMUSG00000036893 | Ehmt1 | euchromatic histone methyltransferase 1 |
| chr12 | 1,11E+08 | 1,11E+08 | 2611 | * | 24,25473 | 2,288333239 | 0,793471 | 2,883952 | 0,003927 | 0,59912059 | 2,951933 | Intron (ENSMUST0000( | 12 | 1,11E+08 | 1,11E+08 | 26086 | 2 | 26448 | ENSMUST00000177224.2 | 16934 | ENSMUSG00000056458 | Mok | MOK protein kinase |
| chr4 | 43834322 | 43836086 | 1765 | * | 14,35027 | 2,288432291 | 0,831185 | 2,753216 | 0,005901 | 0,81655291 | 1,2763 | Promoter (<=1kb) | 4 | 43834752 | 43836517 | 1766 | 2 | 1E+08 | ENSMUST00000215442.2 | 431 | ENSMUSG00000111611 | Olfr157 | olfactory receptor 157 |
| chr12 | 50122001 | 50123634 | 1634 | * | 15,30903 | 2,288504081 | 0,814498 | 2,809711 | 0,004959 | 0,766212249 | 1,41696 | Distal Intergenic | 12 | 50167092 | 50192019 | 24928 | 1 | 1,05E+08 | ENSMUST00000218806.2 | -43458 | ENSMUSG00000112353 | Gm40418 | predicted gene, 40418 |
| chr10 | 77136304 | 77137842 | 1539 | * | 17,25588 | 2,288930904 | 0,865207 | 2,645531 | 0,008156 | 0,751210692 | 2,507013 | Intron (ENSMUST0000( | 10 | 77102751 | 77103398 | 648 | 1 | 80294 | ENSMUST00000218117.2 | 33553 | ENSMUSG00000020260 | Pofut2 | protein O-fucosyltransferase 2 |
| chr2 | 74587181 | 74588790 | 1610 | * | 16,05224 | 2,289393131 | 0,850505 | 2,691805 | 0,007107 | 0,850457798 | 1,2763 | Promoter (2-3kb) | 2 | 74580206 | 74584386 | 4181 | 2 | 1,03E+08 | ENSMUST00000230704.2 | -2795 | | Haglr | Hoxd antisense growth associated long non-coding RNA |
| chr11 | 73594667 | 73596835 | 2169 | * | 16,53571 | 2,290195751 | 0,815137 | 2,809583 | 0,004961 | 0,692179368 | 2,141612 | Intron (ENSMUST0000( | 11 | 73584749 | 73601589 | 16841 | 1 | 76390 | ENSMUST00000145996.2 | 9918 | ENSMUSG00000060630 | Zfp735 | zinc finger protein 735 |
| chr16 | 95501018 | 95503352 | 2335 | * | 24,85731 | 2,290718206 | 0,710774 | 3,222851 | 0,001269 | 0,407436312 | 2,858511 | Promoter (<=1kb) | 16 | 95502942 | 95522095 | 19154 | 1 | 23872 | ENSMUST00000023612.17 | 0 | ENSMUSG00000022895 | Ets2 | E26 avian leukemia oncogene 2, 3' domain |
| chr17 | 26795227 | 26796964 | 1738 | * | 16,25839 | 2,291186262 | 0,831242 | 2,756341 | 0,005845 | 0,722798324 | 2,141612 | Intron (ENSMUST0000( | 17 | 26724565 | 26781102 | 56538 | 2 | 19252 | ENSMUST00000235532.2 | -14125 | ENSMUSG00000024190 | Dusp1 | dual specificity phosphatase 1 |
| chr5 | 70387739 | 70389505 | 1767 | * | 16,14992 | 2,291290129 | 0,834755 | 2,744867 | 0,006054 | 0,729319353 | 2,141612 | Distal Intergenic | 5 | 70344128 | 70344189 | 62 | 2 | 1,15E+08 | ENSMUST00000158238.3 | -43550 | ENSMUSG00000088863 | Gm23067 | predicted gene, 23067 |
| chr11 | 69531562 | 69534482 | 2921 | * | 21,45459 | 2,291393509 | 0,687912 | 3,330939 | 0,000866 | 0,348632365 | 2,67243 | Promoter (<=1kb) | 11 | 69534374 | 69540627 | 6254 | 1 | 23879 | ENSMUST00000139605.2 | 0 | ENSMUSG00000018765 | Fxr2 | fragile X mental retardation, autosomal homolog 2 |
| chr13 | 64234662 | 64236100 | 1439 | * | 15,3803 | 2,291816762 | 0,799371 | 2,867025 | 0,004144 | 0,732824791 | 1,41696 | Promoter (<=1kb) | 13 | 64206088 | 64237000 | 30913 | 2 | 15487 | ENSMUST00000039832.7 | 900 | ENSMUSG00000033122 | Hsd17b3 | hydroxysteroid (17-beta) dehydrogenase 3 |
| chr7 | 1,27E+08 | 1,27E+08 | 1713 | * | 13,40122 | 2,292302858 | 0,884849 | 2,590614 | 0,00958 | 0,947196831 | 0,938682 | Promoter (<=1kb) | 7 | 1,27E+08 | 1,27E+08 | 8296 | 1 | 233905 | ENSMUST00000050383.8 | 599 | ENSMUSG00000049739 | Zfp646 | zinc finger protein 646 |
| chr12 | 32671514 | 32673298 | 1785 | * | 20,53468 | 2,292641881 | 0,748109 | 3,064581 | 0,00218 | 0,507398906 | 2,663619 | Distal Intergenic | 12 | 32440767 | 32481922 | 41156 | 2 | 1,05E+08 | ENSMUST00000218751.2 | -189592 | | Gm40375 | predicted gene, 40375 |
| chr5 | 32112819 | 32114696 | 1878 | * | 17,18624 | 2,2926576 | 0,858963 | 2,6691 | 0,007605 | 0,749550311 | 2,386008 | Intron (ENSMUST0000( | 5 | 32203322 | 32242097 | 38776 | 1 | 107976 | ENSMUST00000202080.2 | -88626 | ENSMUSG00000052139 | Babam2 | BRISC and BRCA1 A complex member 2 |
| chr9 | 65367586 | 65369904 | 2319 | * | 20,36149 | 2,29327541 | 0,798537 | 2,871846 | 0,004081 | 0,659240897 | 2,221611 | Promoter (<=1kb) | 9 | 65368229 | 65395752 | 27524 | 1 | 27965 | ENSMUST00000034955.8 | 0 | ENSMUSG00000032388 | Spg21 | SPG21, maspardin |
| chr11 | 45908890 | 45911587 | 2698 | * | 19,78643 | 2,293325732 | 0,722628 | 3,17359 | 0,001506 | 0,44621096 | 2,663619 | Downstream (<=300bp | 11 | 45946819 | 46038170 | 91352 | 1 | 11492 | ENSMUST00000011400.8 | -35232 | ENSMUSG00000011256 | Adam19 | a disintegrin and metallopeptidase domain 19 (meltrin beta) |
| chr10 | 80790542 | 80792203 | 1662 | * | 21,01804 | 2,294001058 | 0,711411 | 3,224579 | 0,001262 | 0,406085428 | 2,880017 | Exon (ENSMUST00000 | 10 | 80766646 | 80767593 | 948 | 1 | 17873 | ENSMUST00000219449.2 | 23896 | ENSMUSG00000015312 | Gadd45b | growth arrest and DNA-damage-inducible 45 beta |
| chr16 | 49715488 | 49717573 | 2086 | * | 21,19646 | 2,294119935 | 0,725317 | 3,16292 | 0,001562 | 0,451585113 | 2,67243 | Exon (ENSMUST00000 | 16 | 49687196 | 49731454 | 44259 | 1 | 16423 | ENSMUST00000229104.2 | 28292 | ENSMUSG00000055447 | Cd47 | CD47 antigen (Rh-related antigen, integrin-associated signal transducer) |
| chr4 | 1E+08 | 1E+08 | 3010 | * | 25,98891 | 2,29449315 | 0,689648 | 3,327051 | 0,000878 | 0,338260507 | 2,858511 | Intron (ENSMUST0000( | 4 | 99976818 | 99978038 | 1221 | 1 | 1,05E+08 | ENSMUST00000134206.2 | 101933 | | Gm12701 | predicted gene 12701 |
| chr16 | 93089330 | 93090676 | 1347 | * | 15,0276 | 2,294753567 | 0,861291 | 2,664319 | 0,007714 | 0,871920697 | 1,214904 | Distal Intergenic | 16 | 93140612 | 93150077 | 9466 | 2 | 69857 | ENSMUST00000185971.7 | 59401 | ENSMUSG00000100277 | 1810053B23Rik | RIKEN cDNA 1810053B23 gene |
| chr1 | 33522724 | 33525472 | 2749 | * | 19,86032 | 2,29525183 | 0,859757 | 2,669651 | 0,007593 | 0,727652793 | 2,713306 | Exon (ENSMUST00000( | 1 | 33461814 | 33461919 | 106 | 1 | 1,15E+08 | ENSMUST00000083326.3 | 60910 | ENSMUSG00000065260 | Gm23453 | predicted gene, 23453 |
| chr8 | 27351266 | 27352705 | 1440 | * | 16,02936 | 2,295285447 | 0,793821 | 2,891439 | 0,003835 | 0,654756507 | 2,141612 | Intron (ENSMUST0000( | 8 | 27304621 | 27370610 | 65990 | 2 | 72316 | ENSMUST00000210732.2 | 17905 | ENSMUSG00000100335 | 2310008N11Rik | RIKEN cDNA 2310008N11 gene |
| chr17 | 46828947 | 46830984 | 2038 | * | 16,25352 | 2,295417978 | 0,854036 | 2,687729 | 0,007194 | 0,756538269 | 2,141612 | Promoter (<=1kb) | 17 | 46830386 | 46831497 | 1112 | 2 | 78309 | ENSMUST00000183016.8 | 513 | ENSMUSG00000040327 | Cul9 | cullin 9 |
| chr18 | 16786585 | 16788666 | 2082 | * | 17,2098 | 2,295835154 | 0,851194 | 2,697193 | 0,006993 | 0,733134639 | 2,386008 | Intron (ENSMUST0000( | 18 | 16722080 | 16803126 | 81047 | 2 | 12558 | ENSMUST00000115850.2 | 14460 | ENSMUSG00000024304 | Cdh2 | cadherin 2 |
| chr11 | 120346076 | 120349945 | 3870 | * | 47,583362 | 2,296908685 | 0,5452243 | 4,2127778 | 2,522E-05 | 0,045976292 | 2,7254908 | Promoter (<=1kb) | 11 | 120347432 | 120348894 | 1463 | 2 | 66431 | ENSMUST00000044007.3 | 0 | ENSMUSG00000039670 | Oxld1 | oxidoreductase like domain containing 1 |
| chr19 | 57348206 | 57349787 | 1582 | * | 14,50105 | 2,297466843 | 0,873695 | 2,629598 | 0,008549 | 0,860530858 | 1,41696 | Promoter (<=1kb) | 19 | 57349112 | 57378018 | 28907 | 1 | 226252 | ENSMUST00000036407.6 | 0 | ENSMUSG00000033478 | Fhip2a | FHF complex subunit HOOK interacting protein 2A |
| chr17 | 31904101 | 31908446 | 4346 | * | 54,303948 | 2,297481315 | 0,4683343 | 4,9056437 | 9,312E-07 | 0,003690733 | 2,7254908 | Exon (ENSMUST0000023 | 17 | 31898454 | 31900479 | 2026 | 1 | 12954 | ENSMUST00000228305.2 | 5647 | ENSMUSG00000024041 | Cryaa | crystallin, alpha A |
| chr8 | 1,07E+08 | 1,07E+08 | 2606 | * | 26,31617 | 2,297620057 | 0,656442 | 3,500112 | 0,000465 | 0,268627242 | 2,67243 | Intron (ENSMUST0000( | 8 | 1,07E+08 | 1,07E+08 | 23117 | 1 | 272538 | ENSMUST00000211979.2 | 26254 | ENSMUSG00000041949 | Tango6 | transport and golgi organization 6 |
| chr2 | 31385940 | 31387700 | 1761 | * | 21,337 | 2,297854389 | 0,839608 | 2,736817 | 0,006204 | 0,671902369 | 3,104559 | Promoter (<=1kb) | 2 | 31387784 | 31400167 | 12384 | 1 | 11898 | ENSMUST00000126474.7 | -84 | ENSMUSG00000076441 | Ass1 | argininosuccinate synthetase 1 |
| chr12 | 55126481 | 55127559 | 1079 | * | 13,81876 | 2,298147314 | 0,856875 | 2,682011 | 0,007318 | 0,884651431 | 1,075735 | Promoter (<=1kb) | 12 | 55127047 | 55162152 | 35106 | 1 | 24067 | ENSMUST00000220578.2 | 0 | ENSMUSG00000073079 | Srp54a | signal recognition particle 54A |
| chr10 | 79716055 | 79718067 | 2013 | * | 21,46595 | 2,298573823 | 0,756234 | 3,039501 | 0,00237 | 0,524368688 | 2,67243 | Promoter (<=1kb) | 10 | 79716814 | 79719008 | 2195 | 1 | 19152 | ENSMUST00000165982.2 | 0 | ENSMUSG00000057729 | Prtn3 | proteinase 3 |
| chr17 | 14711970 | 14713694 | 1725 | * | 17,04171 | 2,298856578 | 0,842095 | 2,729926 | 0,006335 | 0,717133806 | 2,386008 | Exon (ENSMUST00000 | 17 | 14669833 | 14716053 | 46221 | 1 | 1,03E+08 | ENSMUST00000232609.2 | 42137 | | Gm38551 | predicted gene, 38551 |
| chr17 | 46397855 | 46400213 | 2359 | * | 39,59971 | 2,299040858 | 0,553816 | 4,151274 | 3,31E-05 | 0,053034304 | 2,787255 | Distal Intergenic | 17 | 46421912 | 46439836 | 17925 | 1 | 68565 | ENSMUST00000024763.10 | -21699 | ENSMUSG00000023967 | Mrps18a | mitochondrial ribosomal protein S18A |
| chr19 | 46950815 | 46952935 | 2121 | * | 18,89568 | 2,299056698 | 0,81808 | 2,810306 | 0,004949 | 0,670293874 | 2,507013 | Promoter (<=1kb) | 19 | 46892317 | 46950981 | 58665 | 2 | 76952 | ENSMUST00000237639.2 | 0 | ENSMUSG00000025041 | Nt5c2 | 5'-nucleotidase, cytosolic II |
| chr5 | 1,08E+08 | 1,08E+08 | 1697 | * | 13,44982 | 2,299676776 | 0,856747 | 2,684197 | 0,00727 | 0,852340527 | 1,2763 | Promoter (<=1kb) | 5 | 1,08E+08 | 1,08E+08 | 79033 | 2 | 67266 | ENSMUST00000031198.11 | -90 | ENSMUSG00000029270 | Dipk1a | divergent protein kinase domain 1A |
| chr9 | 21510595 | 21513591 | 2997 | * | 25,31459 | 2,299762984 | 0,720262 | 3,192951 | 0,001408 | 0,422040194 | 2,880017 | Distal Intergenic | 9 | 21519159 | 21519623 | 465 | 2 | 666043 | ENSMUST00000217049.2 | 6032 | | Gm7904 | predicted gene 7904 |
| chr16 | 37700042 | 37701698 | 1657 | * | 17,10127 | 2,300048961 | 0,767657 | 2,996194 | 0,002734 | 0,594430593 | 2,141612 | Exon (ENSMUST00000( | 16 | 37688751 | 37709220 | 20470 | 1 | 320184 | ENSMUST00000078717.7 | 11291 | ENSMUSG00000034158 | Lrrc58 | leucine rich repeat containing 58 |
| chr18 | 66437291 | 66439528 | 2238 | * | 21,22779 | 2,301103394 | 0,794547 | 2,896118 | 0,003778 | 0,582425528 | 3,104559 | Promoter (1-2kb) | 18 | 66402092 | 66435812 | 33721 | 2 | 320924 | ENSMUST00000151343.3 | -1479 | ENSMUSG00000046318 | Ccbe1 | collagen and calcium binding EGF domains 1 |
| chr12 | 1E+08 | 1E+08 | 2244 | * | 21,66428 | 2,301353547 | 0,766708 | 3,001604 | 0,002686 | 0,527676616 | 2,951933 | 3' UTR | 12 | 1E+08 | 1E+08 | 12031 | 1 | 1E+08 | ENSMUST00000220721.2 | 10890 | ENSMUSG00000113679 | Gm10432 | predicted gene 10432 |
| chr5 | 7282745 | 7284352 | 1608 | * | 16,71322 | 2,301570818 | 0,777731 | 2,959342 | 0,003083 | 0,594696308 | 2,386008 | Exon (ENSMUST00000 | 5 | 7227706 | 7227856 | 151 | 2 | 1,15E+08 | ENSMUST00000082917.3 | -54889 | ENSMUSG00000064851 | Gm24525 | predicted gene, 24525 |
| chr4 | 1,16E+08 | 1,16E+08 | 2713 | * | 24,81401 | 2,302317897 | 0,758366 | 3,035894 | 0,002398 | 0,512077677 | 2,858511 | Promoter (<=1kb) | 4 | 1,16E+08 | 1,16E+08 | 30687 | 1 | 16351 | ENSMUST00000106479.8 | 0 | ENSMUSG00000028696 | Ipp | IAP promoted placental gene |
| chr12 | 83008678 | 83011597 | 2920 | * | 27,59383 | 2,302769056 | 0,675289 | 3,41005 | 0,00065 | 0,308275655 | 2,67243 | Intron (ENSMUST0000( | 12 | 82979294 | 82985929 | 6636 | 2 | 74283 | ENSMUST00000187123.2 | -22749 | ENSMUSG00000100890 | 1700085C21Rik | RIKEN cDNA 1700085C21 gene |
| chr14 | 25859719 | 25863391 | 3673 | * | 39,440292 | 2,302787128 | 0,5442621 | 4,2310258 | 2,326E-05 | 0,048228233 | 2,3383871 | Intron (ENSMUST000000 | 14 | 25842607 | 25887228 | 44622 | 1 | 11744 | ENSMUST00000133547.2 | 17112 | ENSMUSG00000021866 | Anxa11 | annexin A11 |
| chr4 | 1,56E+08 | 1,56E+08 | 2603 | * | 24,43943 | 2,302979831 | 0,817616 | 2,816701 | 0,004852 | 0,641187423 | 2,951933 | Promoter (<=1kb) | 4 | 1,56E+08 | 1,56E+08 | 2730 | 2 | 26561 | ENSMUST00000030937.2 | 0 | ENSMUSG00000029061 | Mmp23 | matrix metallopeptidase 23 |
| chr2 | 1,19E+08 | 1,19E+08 | 2750 | * | 23,9967 | 2,3032227 | 0,664653 | 3,4653 | 0,00053 | 0,275349208 | 2,880017 | Promoter (<=1kb) | 2 | 1,19E+08 | 1,19E+08 | 3153 | 1 | 69065 | ENSMUST00000028780.4 | 0 | ENSMUSG00000027313 | Chac1 | ChaC, cation transport regulator 1 |
| chr7 | 73261025 | 73262287 | 1263 | * | 13,1925 | 2,303606555 | 0,873637 | 2,6368 | 0,008369 | 0,877892722 | 1,2763 | Intron (ENSMUST0000( | 7 | 73295733 | 73295837 | 105 | 2 | 1,15E+08 | ENSMUST00000083795.3 | 33550 | ENSMUSG00000065729 | Gm26176 | predicted gene, 26176 |
| chr15 | 73609719 | 73612322 | 2604 | * | 61,700851 | 2,303659556 | 0,4743526 | 4,8564287 | 1,195E-06 | 0,003692163 | 3,439501 | Intron (ENSMUST0000023 | 15 | 73619483 | 73623876 | 4394 | 1 | 19245 | ENSMUST00000230307.2 | -7161 | ENSMUSG00000059895 | Ptp4a3 | protein tyrosine phosphatase 4a3 |
| chr8 | 25174032 | 25176108 | 2077 | * | 16,32303 | 2,304439596 | 0,802741 | 2,870715 | 0,004095 | 0,66649935 | 2,141612 | Promoter (1-2kb) | 8 | 25167241 | 25177979 | 10739 | 2 | 11497 | ENSMUST00000167431.2 | 1871 | ENSMUSG00000031553 | Adam3 | a disintegrin and metallopeptidase domain 3 (cyritestin) |
| chr18 | 47493396 | 47495424 | 2029 | * | 23,01555 | 2,304922199 | 0,713735 | 3,229382 | 0,001241 | 0,402652506 | 2,880017 | Intron (ENSMUST0000( | 18 | 47436741 | 47501335 | 64595 | 2 | 20358 | ENSMUST00000144223.2 | 5911 | ENSMUSG00000019647 | Sema6a | sema domain, transmembrane domain (TM), and cytoplasmic domain, (semaphorin) 6A |
| chr12 | 45099946 | 45101509 | 1564 | * | 15,24367 | 2,305028106 | 0,871351 | 2,64535 | 0,008161 | 0,87277522 | 1,2763 | Intron (ENSMUST0000( | 12 | 44899271 | 45120895 | 221625 | 2 | 217517 | ENSMUST00000120531.8 | 19386 | ENSMUSG00000046314 | Stxbp6 | syntaxin binding protein 6 (amisyn) |
| chr5 | 1,37E+08 | 1,37E+08 | 1719 | * | 13,56955 | 2,305387132 | 0,884734 | 2,60574 | 0,009168 | 0,942900113 | 0,918255 | Distal Intergenic | 5 | 1,37E+08 | 1,37E+08 | 2186 | 2 | 384309 | ENSMUST00000152207.2 | -4502 | ENSMUSG00000043279 | Trim56 | tripartite motif-containing 56 |
| chr8 | 1,1E+08 | 1,1E+08 | 2122 | * | 24,03935 | 2,305519037 | 0,625397 | 3,686488 | 0,000227 | 0,173939409 | 2,858511 | Intron (ENSMUST0000( | 8 | 1,09E+08 | 1,1E+08 | 246987 | 1 | 11906 | ENSMUST00000043896.10 | 80600 | ENSMUSG00000038872 | Zfhx3 | zinc finger homeobox 3 |
| chr8 | 70774088 | 70775664 | 1577 | * | 19,09263 | 2,306232902 | 0,812986 | 2,836745 | 0,004558 | 0,676922067 | 2,221611 | 5' UTR | 8 | 70771000 | 70775339 | 4340 | 1 | 93898 | ENSMUST00000136257.3 | 3088 | ENSMUSG00000087408 | Cers1 | ceramide synthase 1 |
| chr3 | 145408008 | 145411839 | 3832 | * | 39,927124 | 2,306310474 | 0,5135711 | 4,4907323 | 7,098E-06 | 0,017590954 | 2,7872553 | Distal Intergenic | 3 | 145357824 | 145383764 | 25941 | 1 | 100216343 | ENSMUST00000197148.3 | 50184 | | Gm17501 | predicted gene, 17501 |
| chr10 | 77843501 | 77846048 | 2548 | * | 41,68754 | 2,306968192 | 0,585794 | 3,938188 | 8,21E-05 | 0,103768001 | 2,377674 | Promoter (<=1kb) | 10 | 77822781 | 77845641 | 22861 | 2 | 18641 | ENSMUST00000020522.9 | 0 | ENSMUSG00000020277 | Pfkl | phosphofructokinase, liver, B-type |
| chr16 | 24485432 | 24487528 | 2097 | * | 16,51496 | 2,30742972 | 0,845161 | 2,730165 | 0,00633 | 0,722798324 | 2,318434 | Intron (ENSMUST0000( | 16 | 24540605 | 24798280 | 257676 | 1 | 210126 | ENSMUST00000115314.4 | -53077 | ENSMUSG00000033306 | Lpp | LIM domain containing preferred translocation partner in lipoma |
| chr18 | 9136036 | 9137539 | 1504 | * | 13,74759 | 2,307460456 | 0,872677 | 2,644118 | 0,00819 | 0,923399738 | 0,918255 | Distal Intergenic | 18 | 9212163 | 9218136 | 5974 | 1 | 14370 | ENSMUST00000041080.7 | -74624 | ENSMUSG00000036904 | Fzd8 | frizzled class receptor 8 |
| chr5 | 39150938 | 39152518 | 1581 | * | 13,16258 | 2,307940502 | 0,864906 | 2,668428 | 0,007621 | 0,948006995 | 0,738441 | Distal Intergenic | 5 | 38863805 | 39034155 | 170351 | 2 | 27278 | ENSMUST00000171633.5 | -116783 | ENSMUSG00000039315 | Clnk | cytokine-dependent hematopoietic cell linker |
| chr4 | 11704925 | 11706801 | 1877 | * | 16,74847 | 2,308008547 | 0,843839 | 2,735129 | 0,006236 | 0,731734453 | 2,141612 | Promoter (<=1kb) | 4 | 11705807 | 11714752 | 8946 | 1 | 14579 | ENSMUST00000029868.7 | 0 | ENSMUSG00000028214 | Gem | GTP binding protein (gene overexpressed in skeletal muscle) |
| chr2 | 1,65E+08 | 1,65E+08 | 1524 | * | 13,19016 | 2,308198999 | 0,893162 | 2,5843 | 0,009758 | 0,984310536 | 0,738441 | Promoter (1-2kb) | 2 | 1,65E+08 | 1,65E+08 | 16131 | 1 | 170441 | ENSMUST00000029196.5 | 1261 | ENSMUSG00000027661 | Slc2a10 | solute carrier family 2 (facilitated glucose transporter), member 10 |
| chr3 | 1,23E+08 | 1,23E+08 | 1366 | * | 18,48417 | 2,308398517 | 0,859719 | 2,685062 | 0,007252 | 0,731089925 | 2,507013 | Distal Intergenic | 3 | 1,23E+08 | 1,23E+08 | 28810 | 2 | 59006 | ENSMUST00000029761.14 | -27544 | ENSMUSG00000028116 | Myoz2 | myozenin 2 |
| chr2 | 35158571 | 35160869 | 2299 | * | 41,407 | 2,308809668 | 0,634871 | 3,636662 | 0,000276 | 0,201373005 | 2,725491 | Intron (ENSMUST0000( | 2 | 35172392 | 35197904 | 25513 | 1 | 227753 | ENSMUST00000028239.8 | -11523 | ENSMUSG00000026879 | Gsn | gelsolin |
| chr10 | 1,21E+08 | 1,21E+08 | 3674 | * | 34,27678 | 2,308899666 | 0,655025 | 3,524903 | 0,000424 | 0,244286346 | 2,925497 | Intron (ENSMUST0000( | 10 | 1,21E+08 | 1,21E+08 | 47827 | 2 | 192678 | ENSMUST00000219500.2 | 28755 | ENSMUSG00000025795 | Rassf3 | Ras association (RalGDS/AF-6) domain family member 3 |
| chr7 | 45822813 | 45824873 | 2061 | * | 22,57021 | 2,308927732 | 0,723756 | 3,1902 | 0,001422 | 0,437316709 | 2,67243 | 3' UTR | 7 | 45824835 | 45829410 | 4576 | 2 | 20927 | ENSMUST00000210511.2 | 4537 | ENSMUSG00000040136 | Abcc8 | ATP-binding cassette, sub-family C (CFTR/MRP), member 8 |
| chr5 | 34742015 | 34745376 | 3362 | * | 25,15125 | 2,309038623 | 0,69565 | 3,319256 | 0,000903 | 0,340936969 | 2,880017 | Intron (ENSMUST0000( | 5 | 34731495 | 34731557 | 63 | 1 | 1,02E+08 | ENSMUST00000184791.3 | 10520 | ENSMUSG00000099082 | Mir7036b | microRNA 7036b |
| chr3 | 95774252 | 95777222 | 2971 | * | 20,83424 | 2,309111995 | 0,755733 | 3,055462 | 0,002247 | 0,469069098 | 3,471678 | Promoter (<=1kb) | 3 | 95769966 | 95777931 | 7966 | 2 | 66292 | ENSMUST00000072587.5 | 709 | ENSMUSG00000054312 | Mrps21 | mitochondrial ribosomal protein S21 |
| chr6 | 1,28E+08 | 1,28E+08 | 2708 | * | 26,79599 | 2,309660945 | 0,772025 | 2,991692 | 0,002774 | 0,537794802 | 2,880017 | Promoter (1-2kb) | 6 | 1,28E+08 | 1,28E+08 | 68846 | 2 | 109246 | ENSMUST00000123786.9 | 1717 | ENSMUSG00000030352 | Tspan9 | tetraspanin 9 |
| chr11 | 94163833 | 94165725 | 1893 | * | 15,20786 | 2,310401016 | 0,865729 | 2,668735 | 0,007614 | 0,841775205 | 1,41696 | Distal Intergenic | 11 | 94172699 | 94177079 | 4381 | 1 | 1,03E+08 | ENSMUST00000208510.3 | -6974 | ENSMUSG00000094091 | Ankrd40cl | ANKRD40 C-terminal like |
| chr11 | 60146244 | 60147658 | 1415 | * | 18,42857 | 2,310898085 | 0,891063 | 2,593417 | 0,009503 | 0,790460916 | 2,386008 | Exon (ENSMUST00000 | 11 | 60120728 | 60136495 | 15768 | 2 | 216810 | ENSMUST00000143124.8 | -9749 | ENSMUSG00000000538 | Tom1l2 | target of myb1-like 2 (chicken) |
| chr1 | 87478039 | 87480651 | 2613 | * | 28,36345 | 2,311406712 | 0,643564 | 3,591574 | 0,000329 | 0,221323359 | 2,725491 | Intron (ENSMUST0000( | 1 | 87404556 | 87501592 | 97037 | 2 | 53972 | ENSMUST00000068681.12 | 20941 | ENSMUSG00000026259 | Ngef | neuronal guanine nucleotide exchange factor |
| chr11 | 69863842 | 69866078 | 2237 | * | 16,13324 | 2,312174733 | 0,834769 | 2,769837 | 0,005608 | 0,681421811 | 2,663619 | Exon (ENSMUST00000 | 11 | 69859058 | 69860229 | 1172 | 2 | 54351 | ENSMUST00000151766.2 | -3613 | ENSMUSG00000018565 | Elp5 | elongator acetyltransferase complex subunit 5 |
| chr2 | 1,56E+08 | 1,56E+08 | 2806 | * | 24,11978 | 2,3124553 | 0,671079 | 3,445875 | 0,000569 | 0,283414546 | 2,880017 | Promoter (2-3kb) | 2 | 1,56E+08 | 1,56E+08 | 3356 | 1 | 17391 | ENSMUST00000135945.2 | 2187 | ENSMUSG00000027612 | Mmp24 | matrix metallopeptidase 24 |

| | | | | | | | | | | | | | | | | | | | | | | | |
|---|---|---|---|---|---|---|---|---|---|---|---|---|---|---|---|---|---|---|---|---|---|---|---|
| chr12 | 75992681 | 75993854 | 1174 | * | 15,14444 | 2,312624917 | 0,795531 | 2,90702 | 0,003649 | 0,730469267 | 1,2763 | Exon (ENSMUST00000( | 12 | 75955992 | 75960661 | 4670 | 1 | 319565 | ENSMUST00000124685.2 | 36689 | ENSMUSG00000063450 | Syne2 | spectrin repeat containing, nuclear envelope 2 |
| chr10 | 60025720 | 60028770 | 3051 | * | 27,36613 | 2,312700647 | 0,781114 | 2,960772 | 0,003069 | 0,566484927 | 2,725491 | Promoter (1-2kb) | 10 | 60017356 | 60024529 | 7174 | 2 | 53374 | ENSMUST00000068690.7 | -1191 | ENSMUSG00000057337 | Chst3 | carbohydrate sulfotransferase 3 |
| chr8 | 36631312 | 36634248 | 2937 | * | 45,636181 | 2,312765281 | 0,4957251 | 4,665419 | 3,08E-06 | 0,010321083 | 2,3776742 | Distal Intergenic | 8 | 36626633 | 36628742 | 2110 | 1 | 330738 | ENSMUST00000211253.2 | 4679 | ENSMUSG00000109704 | Gm38414 | predicted gene, 38414 |
| chr4 | 91583133 | 91584779 | 1647 | * | 14,51755 | 2,312801664 | 0,856452 | 2,700444 | 0,006925 | 0,830950621 | 1,37683 | Distal Intergenic | 4 | 91693784 | 91694792 | 1009 | 2 | 620016 | ENSMUST00000122162.2 | 110013 | ENSMUSG00000100863 | Gm12669 | glutaredoxin 3 pseudogene |
| chr1 | 70626120 | 70627796 | 1677 | * | 11,8207 | 2,312816374 | 0,875577 | 2,641476 | 0,008255 | 0,924754734 | 0,918255 | Intron (ENSMUST00000( | 1 | 70635974 | 70636080 | 107 | 1 | 1,15E+08 | ENSMUST00000102000.3 | -8178 | ENSMUSG00000075952 | Gm23422 | predicted gene, 23422 |
| chr5 | 1,35E+08 | 1,35E+08 | 2046 | * | 18,83933 | 2,313064609 | 0,880904 | 2,625784 | 0,008645 | 0,749550311 | 2,713306 | Promoter (<=1kb) | 5 | 1,35E+08 | 1,35E+08 | 28614 | 2 | 215114 | ENSMUST00000202643.2 | 621 | ENSMUSG00000039959 | Hip1 | huntingtin interacting protein 1 |
| chr2 | 26452999 | 26454949 | 1951 | * | 25,67261 | 2,313435271 | 0,702455 | 3,293359 | 0,00099 | 0,353824371 | 2,951062 | Distal Intergenic | 2 | 26470026 | 26479396 | 9371 | 1 | 353156 | ENSMUST00000152713.9 | -15077 | ENSMUSG00000026921 | Egfl7 | EGF-like domain 7 |
| chr8 | 1,16E+08 | 1,16E+08 | 2517 | * | 27,39051 | 2,313469541 | 0,711961 | 3,249433 | 0,001156 | 0,400277096 | 2,725491 | Intron (ENSMUST00000( | 8 | 1,15E+08 | 1,16E+08 | 639976 | 1 | 80707 | ENSMUST00000162052.8 | 392152 | ENSMUSG00000004637 | Wwox | WW domain-containing oxidoreductase |
| chr4 | 1,36E+08 | 1,36E+08 | 1651 | * | 13,03774 | 2,31354766 | 0,89002 | 2,599433 | 0,009338 | 0,947196831 | 0,918255 | Distal Intergenic | 4 | 1,36E+08 | 1,36E+08 | 853 | 1 | 15903 | ENSMUST00000151001.2 | 9842 | ENSMUSG00000007872 | Id3 | inhibitor of DNA binding 3 |
| chr18 | 90418506 | 90419785 | 1280 | * | 16,53505 | 2,313981664 | 0,876884 | 2,638869 | 0,008318 | 0,770673161 | 2,318434 | Distal Intergenic | 18 | 90478118 | 90501584 | 23467 | 2 | 1,15E+08 | ENSMUST00000237124.2 | 81799 | | Gm18280 | predicted gene, 18280 |
| chr7 | 80497496 | 80499339 | 1844 | * | 17,51226 | 2,314026494 | 0,834294 | 2,773635 | 0,005543 | 0,713797407 | 2,154504 | Distal Intergenic | 7 | 80510668 | 80524717 | 14050 | 1 | 22691 | ENSMUST00000132163.8 | -11329 | ENSMUSG00000038797 | Zscan2 | zinc finger and SCAN domain containing 2 |
| chr18 | 73708188 | 73709924 | 1737 | * | 16,22668 | 2,314334153 | 0,85727 | 2,699656 | 0,006941 | 0,750475959 | 2,141612 | Promoter (2-3kb) | 18 | 73706115 | 73725646 | 19532 | 1 | 240396 | ENSMUST00000091852.5 | 2073 | ENSMUSG00000037253 | Mex3c | mex3 RNA binding family member C |
| chr4 | 1,5E+08 | 1,5E+08 | 1376 | * | 14,45178 | 2,315394994 | 0,839752 | 2,757237 | 0,005829 | 0,813839948 | 1,2763 | Distal Intergenic | 4 | 1,5E+08 | 1,5E+08 | 14318 | 2 | 12353 | ENSMUST00000030817.5 | -15267 | ENSMUSG00000028972 | Car6 | carbonic anhydrase 6 |
| chr2 | 1,54E+08 | 1,54E+08 | 2117 | * | 20,23045 | 2,316016968 | 0,772535 | 2,997946 | 0,002718 | 0,568412386 | 2,386008 | Promoter (1-2kb) | 2 | 1,54E+08 | 1,54E+08 | 2612 | 2 | 56846 | ENSMUST00000149725.2 | 1056 | ENSMUSG00000027489 | Necab3 | N-terminal EF-hand calcium binding protein 3 |
| chr18 | 60748612 | 60751819 | 3208 | * | 40,03147 | 2,31629941 | 0,595994 | 3,886446 | 0,000102 | 0,109316219 | 2,725491 | Intron (ENSMUST00000( | 18 | 60737181 | 60756397 | 19217 | 2 | 104027 | ENSMUST00000123288.2 | 4578 | ENSMUSG00000043079 | Synpo | synaptopodin |
| chr1 | 8410397 | 8412220 | 1824 | * | 20,26299 | 2,317687649 | 0,772444 | 3,000461 | 0,002696 | 0,542967057 | 2,663619 | Distal Intergenic | 1 | 8433649 | 8540703 | 107055 | 2 | 71096 | ENSMUST00000135046.4 | 128483 | ENSMUSG00000025909 | Sntg1 | syntrophin, gamma 1 |
| chr1 | 1,7E+08 | 1,7E+08 | 1734 | * | 14,26385 | 2,318403957 | 0,87892 | 2,637787 | 0,008345 | 0,961259828 | 0,738441 | Promoter (2-3kb) | 1 | 1,7E+08 | 1,7E+08 | 74989 | 2 | 18214 | ENSMUST00000195867.6 | -2460 | ENSMUSG00000026674 | Ddr2 | discoidin domain receptor family, member 2 |
| chr1 | 11014859 | 11016475 | 1617 | * | 17,98111 | 2,318493352 | 0,835605 | 2,774628 | 0,005526 | 0,713633527 | 2,154504 | Distal Intergenic | 1 | 11063689 | 11373905 | 310217 | 1 | 109294 | ENSMUST00000027056.12 | -47214 | ENSMUSG00000048960 | Prex2 | phosphatidylinositol-3,4,5-trisphosphate-dependent Rac exchange factor 2 |
| chr11 | 10907241 | 10908687 | 1447 | * | 15,94174 | 2,319049428 | 0,767081 | 3,023211 | 0,002501 | 0,549912148 | 2,386008 | Distal Intergenic | 11 | 11064223 | 11213526 | 149304 | 1 | 319922 | ENSMUST00000109681.8 | -155536 | ENSMUSG00000050830 | Vwc2 | von Willebrand factor C domain containing 2 |
| chr9 | 1,06E+08 | 1,06E+08 | 2076 | * | 16,9352 | 2,319205513 | 0,821892 | 2,821789 | 0,004776 | 0,654833721 | 2,663619 | Exon (ENSMUST000001 | 9 | 1,06E+08 | 1,06E+08 | 1267 | 2 | 68553 | ENSMUST00000137847.2 | 14731 | ENSMUSG00000032572 | Col6a4 | collagen, type VI, alpha 4 |
| chr9 | 96834666 | 96836192 | 1527 | * | 23,32087 | 2,319532253 | 0,770592 | 3,010066 | 0,002612 | 0,527189687 | 2,880017 | Intron (ENSMUST00000( | 9 | 96826025 | 96854028 | 28004 | 2 | 211949 | ENSMUST00000159298.2 | 17836 | ENSMUSG00000046997 | Spsb4 | splA/ryanodine receptor domain and SOCS box containing 4 |
| chr6 | 29427190 | 29428535 | 1346 | * | 22,0412 | 2,320103142 | 0,670399 | 3,460781 | 0,000539 | 0,284660899 | 2,67243 | Downstream (<=300bp | 6 | 29433255 | 29439055 | 5801 | 1 | 68794 | ENSMUST00000148404.2 | -4720 | ENSMUSG00000068699 | Flnc | filamin C, gamma |
| chr14 | 16307308 | 16308858 | 1551 | * | 17,5771 | 2,320111734 | 0,881183 | 2,632951 | 0,008465 | 0,765231686 | 2,423594 | Intron (ENSMUST00000( | 14 | 16297772 | 16315771 | 18000 | 1 | 1,05E+08 | ENSMUST00000180950.2 | 9536 | | Gm26991 | predicted gene, 26991 |
| chr15 | 78819494 | 78821054 | 1561 | * | 14,40564 | 2,320306939 | 0,875918 | 2,649001 | 0,008073 | 0,870441665 | 1,2763 | Promoter (<=1kb) | 15 | 78819408 | 78820781 | 1374 | 1 | 97961 | ENSMUST00000123013.2 | 86 | ENSMUSG00000033099 | Nol12 | nucleolar protein 12 |
| chr4 | 57174198 | 57176445 | 2248 | * | 17,47225 | 2,320345696 | 0,894692 | 2,593457 | 0,009502 | 0,788675375 | 2,423594 | Promoter (2-3kb) | 4 | 57172071 | 57176554 | 4484 | 1 | 329851 | ENSMUST00000152074.2 | 2127 | ENSMUSG00000085967 | Gm12530 | predicted gene 12530 |
| chr8 | 35219701 | 35221967 | 2267 | * | 33,59237 | 2,320612109 | 0,561345 | 4,134019 | 3,56E-05 | 0,054027489 | 2,925497 | Distal Intergenic | 8 | 35274451 | 35287048 | 12598 | 1 | 319520 | ENSMUST00000033930.5 | -52484 | ENSMUSG00000031530 | Dusp4 | dual specificity phosphatase 4 |
| chr8 | 70226790 | 70228093 | 1304 | * | 16,14487 | 2,321107364 | 0,875572 | 2,65096 | 0,008026 | 0,778321623 | 2,141612 | Promoter (<=1kb) | 8 | 70216493 | 70227561 | 11069 | 2 | 330788 | ENSMUST00000137573.2 | 0 | ENSMUSG00000043090 | Zfp866 | zinc finger protein 866 |
| chr2 | 86879643 | 86881544 | 1902 | * | 16,98582 | 2,321548627 | 0,805333 | 2,88272 | 0,003943 | 0,744224125 | 1,2763 | Promoter (<=1kb) | 2 | 86878640 | 86879578 | 939 | 2 | 258747 | ENSMUST00000099867.2 | -65 | ENSMUSG00000075164 | Olfr1106 | olfactory receptor 1106 |
| chr4 | 1,4E+08 | 1,4E+08 | 2179 | * | 22,46753 | 2,321650021 | 0,724968 | 3,202417 | 0,001363 | 0,406085428 | 3,104559 | Intron (ENSMUST00000( | 4 | 1,4E+08 | 1,4E+08 | 57429 | 2 | 72754 | ENSMUST00000143614.8 | -4377 | ENSMUSG00000040964 | Arhgef10l | Rho guanine nucleotide exchange factor (GEF) 10-like |
| chr4 | 1,39E+08 | 1,39E+08 | 1317 | * | 25,29947 | 2,321690669 | 0,815478 | 2,84703 | 0,004413 | 0,627785088 | 2,858511 | Promoter (<=1kb) | 4 | 1,39E+08 | 1,39E+08 | 45663 | 1 | 212632 | ENSMUST00000174078.2 | -10 | ENSMUSG00000041025 | Iffo2 | intermediate filament family orphan 2 |
| chr2 | 26284300 | 26286206 | 1907 | * | 19,7107 | 2,322208137 | 0,790715 | 2,936845 | 0,003316 | 0,623611487 | 2,221611 | 3' UTR | 2 | 26286528 | 26289483 | 2956 | 2 | 64436 | ENSMUST00000131109.8 | 3277 | ENSMUSG00000026925 | Inpp5e | inositol polyphosphate-5-phosphatase E |
| chr8 | 72182830 | 72186007 | 3178 | * | 60,703633 | 2,322225752 | 0,4203762 | 5,5241614 | 3,311E-08 | 0,000243164 | 3,2706213 | Distal Intergenic | 8 | 72161038 | 72178360 | 17323 | 2 | 74015 | ENSMUST00000146100.8 | -4470 | ENSMUSG00000070000 | Fcho1 | FCH domain only 1 |
| chr16 | 9369434 | 9370809 | 1376 | * | 14,85294 | 2,322937859 | 0,880026 | 2,639623 | 0,0083 | 0,876338958 | 1,2763 | Distal Intergenic | 16 | 9579363 | 9615197 | 35835 | 2 | 14811 | ENSMUST00000199267.2 | 244388 | ENSMUSG00000059003 | Grin2a | glutamate receptor, ionotropic, NMDA2A (epsilon 1) |
| chr5 | 1,07E+08 | 1,07E+08 | 2711 | * | 22,26074 | 2,3229885 | 0,743697 | 3,123569 | 0,001787 | 0,46163452 | 2,880017 | Exon (ENSMUST00000( | 5 | 1,07E+08 | 1,07E+08 | 14727 | 2 | 21814 | ENSMUST00000136882.2 | -7414 | ENSMUSG00000029287 | Tgfbr3 | transforming growth factor, beta receptor III |
| chr2 | 1,66E+08 | 1,66E+08 | 2567 | * | 24,01975 | 2,323277146 | 0,843191 | 2,755339 | 0,005863 | 0,675594321 | 2,880017 | Distal Intergenic | 2 | 1,66E+08 | 1,66E+08 | 11188 | 1 | 319684 | ENSMUST00000245642.1 | 18193 | | 5031425F14Rik | RIKEN cDNA 5031425F14 gene |
| chr11 | 79144070 | 79146249 | 2180 | * | 28,34249 | 2,323295378 | 0,671848 | 3,458065 | 0,000544 | 0,279693823 | 2,858511 | Promoter (<=1kb) | 11 | 79130198 | 79145497 | 15300 | 2 | 78889 | ENSMUST00000017821.12 | 0 | ENSMUSG00000017677 | Wsb1 | WD repeat and SOCS box-containing 1 |
| chr5 | 29627340 | 29629775 | 2436 | * | 32,9453 | 2,323311121 | 0,567033 | 4,097311 | 4,18E-05 | 0,069348015 | 2,377674 | Distal Intergenic | 5 | 29639662 | 29662841 | 23180 | 1 | 433864 | ENSMUST00000001611.11 | -9887 | ENSMUSG00000001569 | Nom1 | nucleolar protein with MIF4G domain 1 |
| chr16 | 10853400 | 10855756 | 2357 | * | 20,31221 | 2,323923414 | 0,824335 | 2,819149 | 0,004815 | 0,673983361 | 2,386008 | Exon (ENSMUST000002 | 16 | 10849820 | 10854284 | 4465 | 1 | 1E+08 | ENSMUST00000229764.2 | 3580 | | Gm10832 | predicted gene 10832 |
| chr10 | 6954848 | 6956825 | 1978 | * | 19,56617 | 2,3239954 | 0,727817 | 3,193102 | 0,001408 | 0,46544192 | 2,221611 | Intron (ENSMUST00000( | 10 | 6906062 | 6950444 | 44383 | 2 | 320495 | ENSMUST00000151960.2 | -4404 | ENSMUSG00000064065 | Ipcef1 | interaction protein for cytohesin exchange factors 1 |
| chr2 | 92261168 | 92263499 | 2332 | * | 20,65942 | 2,324318075 | 0,887328 | 2,619456 | 0,008807 | 0,737047928 | 2,951933 | Promoter (<=1kb) | 2 | 92257249 | 92262737 | 5489 | 2 | 12953 | ENSMUST00000144842.2 | 0 | ENSMUSG00000068742 | Cry2 | cryptochrome 2 (photolyase-like) |
| chr7 | 1,05E+08 | 1,05E+08 | 1392 | * | 16,05231 | 2,324863146 | 0,761772 | 3,051914 | 0,002274 | 0,65364112 | 1,2763 | 3' UTR | 7 | 1,05E+08 | 1,05E+08 | 4225 | 1 | 77505 | ENSMUST00000128388.2 | -4472 | ENSMUSG00000030882 | Dnhd1 | dynein heavy chain domain 1 |
| chr15 | 4054410 | 4056495 | 2086 | * | 17,74505 | 2,325021174 | 0,849581 | 2,736669 | 0,006206 | 0,706976265 | 2,507013 | Promoter (<=1kb) | 15 | 4055865 | 4184826 | 128962 | 1 | 67041 | ENSMUST00000110690.9 | 0 | ENSMUSG00000022186 | Oxct1 | 3-oxoacid CoA transferase 1 |
| chr3 | 1,09E+08 | 1,09E+08 | 2993 | * | 43,57834 | 2,325149605 | 0,582018 | 3,994981 | 6,47E-05 | 0,082292767 | 2,787255 | Promoter (<=1kb) | 3 | 1,09E+08 | 1,09E+08 | 12460 | 2 | 320181 | ENSMUST00000139137.3 | 0 | ENSMUSG00000045326 | Fndc7 | fibronectin type III domain containing 7 |
| chr2 | 1,03E+08 | 1,03E+08 | 1401 | * | 13,37077 | 2,32673473 | 0,885092 | 2,628805 | 0,008569 | 0,927963368 | 0,938682 | Distal Intergenic | 2 | 1,03E+08 | 1,03E+08 | 39297 | 1 | 13711 | ENSMUST00000239273.2 | -3309 | ENSMUSG00000027186 | Elf5 | E74-like factor 5 |
| chr19 | 43662929 | 43664497 | 1569 | * | 22,3243 | 2,326866698 | 0,706914 | 3,291584 | 0,000996 | 0,354474598 | 2,951933 | Promoter (<=1kb) | 19 | 43663572 | 43665615 | 2044 | 1 | 1,01E+08 | ENSMUST00000238165.3 | 0 | ENSMUSG00000117698 | BC037704 | cDNA sequence BC037704 |
| chr14 | 1,06E+08 | 1,06E+08 | 3112 | * | 16,67698 | 2,327456921 | 0,853994 | 2,725378 | 0,006423 | 0,738247573 | 2,141612 | Distal Intergenic | 14 | 1,06E+08 | 1,06E+08 | 18317 | 2 | 620376 | ENSMUST00000211689.2 | -30949 | ENSMUSG00000110080 | Gm6145 | predicted gene 6145 |
| chr13 | 51615952 | 51617230 | 1279 | * | 12,3394 | 2,328612696 | 0,883787 | 2,634811 | 0,008418 | 0,928382841 | 0,918255 | Intron (ENSMUST00000( | 13 | 51532172 | 51572722 | 40551 | 2 | 1,03E+08 | ENSMUST00000222417.2 | -43230 | ENSMUSG00000112959 | Gm32834 | predicted gene, 32834 |
| chr18 | 64178307 | 64180393 | 2087 | * | 18,28848 | 2,328678484 | 0,8093 | 2,877397 | 0,00401 | 0,643386705 | 2,386008 | Distal Intergenic | 18 | 64300797 | 64301845 | 1049 | 2 | 73588 | ENSMUST00000236604.2 | 121452 | ENSMUSG00000086128 | St8sia3os | ST8 alpha-N-acetyl-neuraminide alpha-2,8-sialyltransferase 3, opposite strand |
| chr15 | 81371966 | 81374820 | 2855 | * | 24,15446 | 2,328716947 | 0,762126 | 3,055555 | 0,002246 | 0,50153942 | 2,858511 | Distal Intergenic | 15 | 81357823 | 81359773 | 1951 | 1 | 56438 | ENSMUST00000229991.2 | 14143 | ENSMUSG00000022400 | Rbx1 | ring-box 1 |
| chr3 | 1,35E+08 | 1,35E+08 | 3067 | * | 26,26619 | 2,329668817 | 0,709747 | 3,282395 | 0,001029 | 0,381011816 | 2,67243 | Intron (ENSMUST00000( | 3 | 1,35E+08 | 1,35E+08 | 27205 | 1 | 110173 | ENSMUST00000144654.2 | 12577 | ENSMUSG00000028164 | Manba | mannosidase, beta A, lysosomal |
| chr6 | 64946238 | 64947758 | 1521 | * | 19,21133 | 2,329953781 | 0,77628 | 3,001434 | 0,002687 | 0,542967057 | 2,663619 | Distal Intergenic | 6 | 65019567 | 65033422 | 13856 | 1 | 13990 | ENSMUST00000204801.3 | -71809 | ENSMUSG00000029920 | Smarcad1 | SWI/SNF-related, matrix-associated actin-dependent regulator of chromatin, subfamily a, containing DEAD/H box 1 |
| chr10 | 6806815 | 6808711 | 1897 | * | 13,17385 | 2,329991297 | 0,888384 | 2,62273 | 0,008723 | 0,93486226 | 0,918255 | Intron (ENSMUST00000( | 10 | 6739099 | 6799987 | 60889 | 1 | 18390 | ENSMUST00000148625.2 | 67716 | ENSMUSG00000000766 | Oprm1 | opioid receptor, mu 1 |
| chr11 | 35734478 | 35737085 | 2608 | * | 31,336665 | 2,329998504 | 0,5414522 | 4,3032396 | 1,683E-05 | 0,031671937 | 2,8800171 | Exon (ENSMUST0000001 | 11 | 35743022 | 35744381 | 1360 | 2 | 211652 | ENSMUST00000131001.2 | 7296 | ENSMUSG00000018849 | Wwc1 | WW, C2 and coiled-coil domain containing 1 |
| chr9 | 36681594 | 36683490 | 1897 | * | 14,25392 | 2,330442746 | 0,839646 | 2,775507 | 0,005512 | 0,859692746 | 0,918255 | Promoter (2-3kb) | 9 | 36679157 | 36693973 | 14817 | 1 | 1,03E+08 | ENSMUST00000180474.5 | 2437 | ENSMUSG00000096993 | Gm26787 | predicted gene, 26787 |
| chr6 | 1,21E+08 | 1,21E+08 | 1611 | * | 19,37941 | 2,33087386 | 0,75528 | 3,086105 | 0,002028 | 0,514210241 | 2,386008 | Promoter (1-2kb) | 6 | 1,21E+08 | 1,21E+08 | 56605 | 1 | 94044 | ENSMUST00000009256.4 | 1547 | ENSMUSG00000009112 | Bcl2l13 | BCL2-like 13 (apoptosis facilitator) |
| chr15 | 84827349 | 84828804 | 1456 | * | 18,93092 | 2,331328835 | 0,797753 | 2,92237 | 0,003474 | 0,618136403 | 2,386008 | 3' UTR | 15 | 84835727 | 84835824 | 98 | 2 | 1,01E+08 | ENSMUST00000116791.3 | 7020 | ENSMUSG00000080441 | Mir1249 | microRNA 1249 |
| chr18 | 7799220 | 7801292 | 2073 | * | 18,39177 | 2,331503557 | 0,901746 | 2,585545 | 0,009723 | 0,81148874 | 2,154504 | Distal Intergenic | 18 | 7835177 | 7835283 | 107 | 1 | 1,15E+08 | ENSMUST00000082888.2 | -33885 | ENSMUSG00000064822 | Gm24292 | predicted gene, 24292 |
| chr17 | 15600552 | 15602837 | 2286 | * | 19,39361 | 2,331719908 | 0,754186 | 3,091706 | 0,00199 | 0,490652834 | 2,713306 | Distal Intergenic | 17 | 15595056 | 15597134 | 2079 | 2 | 13388 | ENSMUST00000143460.2 | -3418 | ENSMUSG00000014773 | Dll1 | delta like canonical Notch ligand 1 |
| chr10 | 19329134 | 19331058 | 1925 | * | 19,38357 | 2,331745215 | 0,743828 | 3,134789 | 0,00172 | 0,46544192 | 2,713306 | Distal Intergenic | 10 | 19362463 | 19383059 | 20597 | 1 | 1,03E+08 | ENSMUST00000214178.2 | -31405 | ENSMUSG00000110902 | Gm33104 | predicted gene, 33104 |
| chr6 | 91266694 | 91268341 | 1648 | * | 22,60272 | 2,331773253 | 0,863095 | 2,701643 | 0,0069 | 0,698318058 | 2,880017 | Distal Intergenic | 6 | 91242861 | 91249413 | 6553 | 1 | 14115 | ENSMUST00000137029.2 | 23833 | ENSMUSG00000064080 | Fbln2 | fibulin 2 |
| chr12 | 1,12E+08 | 1,12E+08 | 2158 | * | 23,26313 | 2,333049924 | 0,763461 | 3,055885 | 0,002244 | 0,501362189 | 2,858511 | Promoter (<=1kb) | 12 | 1,12E+08 | 1,12E+08 | 629 | 1 | 68520 | ENSMUST00000221313.2 | 0 | ENSMUSG00000021286 | Zfyve21 | zinc finger, FYVE domain containing 21 |
| chr12 | 87033785 | 87036003 | 2219 | * | 18,37176 | 2,333114518 | 0,815608 | 2,860583 | 0,004229 | 0,649854434 | 2,423594 | Promoter (<=1kb) | 12 | 87030155 | 87035450 | 5296 | 2 | 238331 | ENSMUST00000095521.3 | 0 | ENSMUSG00000048483 | Zdhhc22 | zinc finger, DHHC-type containing 22 |
| chr2 | 41373577 | 41374833 | 1257 | * | 13,50393 | 2,333424147 | 0,853122 | 2,735158 | 0,006235 | 0,91615004 | 0,738441 | Intron (ENSMUST00000( | 2 | 40691405 | 41679090 | 987686 | 2 | 94217 | ENSMUST00000167270.5 | 304257 | ENSMUSG00000049252 | Lrp1b | low density lipoprotein-related protein 1B |
| chr19 | 28966647 | 28969452 | 2806 | * | 24,10794 | 2,333459254 | 0,68517 | 3,405665 | 0,00066 | 0,300106667 | 2,951933 | Promoter (<=1kb) | 19 | 28967752 | 28993171 | 25420 | 1 | 67072 | ENSMUST00000225310.2 | 0 | ENSMUSG00000024780 | Cdc37l1 | cell division cycle 37-like 1 |
| chr3 | 65836630 | 65838289 | 1660 | * | 22,68118 | 2,333765513 | 0,771259 | 3,025917 | 0,002479 | 0,504335604 | 3,104559 | Distal Intergenic | 3 | 65854203 | 65855419 | 1217 | 2 | 56706 | ENSMUST00000149729.2 | 17130 | ENSMUSG00000027829 | Ccnl1 | cyclin L1 |
| chr17 | 74534510 | 74535808 | 1299 | * | 12,68096 | 2,33399407 | 0,887126 | 2,63096 | 0,008514 | 0,906138253 | 1,075735 | Promoter (1-2kb) | 17 | 74508111 | 74537496 | 29386 | 2 | 76890 | ENSMUST00000233495.2 | 1688 | ENSMUSG00000058704 | Memo1 | mediator of cell motility 1 |
| chr8 | 72180455 | 72182598 | 2144 | * | 29,98147 | 2,334233724 | 0,707765 | 3,298035 | 0,000974 | 0,352029189 | 2,951062 | Promoter (2-3kb) | 8 | 72161038 | 72178360 | 17323 | 2 | 74015 | ENSMUST00000146100.8 | -2095 | ENSMUSG00000070000 | Fcho1 | FCH domain only 1 |
| chr15 | 36860664 | 36862971 | 2308 | * | 22,22155 | 2,335267139 | 0,69387 | 3,365567 | 0,000764 | 0,309527539 | 3,104559 | Distal Intergenic | 15 | 36870744 | 36880121 | 9378 | 2 | 1E+08 | ENSMUST00000100713.4 | 17150 | ENSMUSG00000072618 | Gm10384 | predicted gene 10384 |
| chr5 | 65649956 | 65652279 | 2324 | * | 19,78134 | 2,335453685 | 0,730289 | 3,197984 | 0,001384 | 0,428853558 | 2,713306 | Promoter (<=1kb) | 5 | 65584770 | 65650176 | 65407 | 2 | 68552 | ENSMUST00000196121.2 | 0 | ENSMUSG00000037822 | Smim14 | small integral membrane protein 14 |
| chr4 | 1,39E+08 | 1,39E+08 | 2147 | * | 25,26417 | 2,335504125 | 0,65615 | 3,559406 | 0,000372 | 0,225497398 | 2,951062 | Exon (ENSMUST00000( | 4 | 1,39E+08 | 1,39E+08 | 16307 | 1 | 83770 | ENSMUST00000166773.2 | 12468 | ENSMUSG00000028738 | Tas1r2 | taste receptor, type 1, member 2 |
| chr9 | 69337161 | 69339036 | 1876 | * | 19,33533 | 2,33619162 | 0,838111 | 2,787448 | 0,005312 | 0,669586355 | 2,713306 | Intron (ENSMUST00000( | 9 | 69324155 | 69334306 | 10152 | 1 | 93697 | ENSMUST00000147558.2 | 13006 | ENSMUSG00000032235 | Ice2 | interactor of little elongation complex ELL subunit 2 |
| chr5 | 56989518 | 56991273 | 1756 | * | 14,49427 | 2,336577598 | 0,869011 | 2,688777 | 0,007171 | 0,83238039 | 1,41696 | Distal Intergenic | 5 | 56893432 | 56893561 | 130 | 1 | 1,15E+08 | ENSMUST00000158109.3 | 96086 | ENSMUSG00000088734 | Gm23896 | predicted gene, 23896 |
| chr3 | 56868670 | 56870453 | 1784 | * | 15,60108 | 2,336771698 | 0,888312 | 2,630574 | 0,008524 | 0,888740228 | 1,214904 | Distal Intergenic | 3 | 56961571 | 56961700 | 130 | 1 | 1,15E+08 | ENSMUST00000239689.1 | -91118 | ENSMUSG00000118960 | Gm22269 | predicted gene, 22269 |
| chr7 | 84851879 | 84854158 | 2280 | * | 21,38092 | 2,33695658 | 0,803011 | 2,910241 | 0,003611 | 0,548877304 | 3,471678 | Distal Intergenic | 7 | 84870346 | 84886991 | 16646 | 2 | 620697 | ENSMUST00000061074.10 | 32833 | ENSMUSG00000096861 | Vmn2r68 | vomeronasal 2, receptor 68 |
| chr5 | 34046663 | 34048730 | 2068 | * | 17,36292 | 2,337042744 | 0,886988 | 2,634807 | 0,008419 | 0,787687627 | 2,154504 | Promoter (<=1kb) | 5 | 34049249 | 34052439 | 3191 | 1 | 107823 | ENSMUST00000201572.2 | -519 | ENSMUSG00000057406 | Nsd2 | nuclear receptor binding SET domain protein 2 |
| chr8 | 72125972 | 72127352 | 1381 | * | 16,68075 | 2,337211648 | 0,778513 | 3,00215 | 0,002681 | 0,681316955 | 1,2763 | Promoter (1-2kb) | 8 | 72077061 | 72124401 | 47341 | 2 | 382018 | ENSMUST00000030170.15 | -1571 | ENSMUSG00000034799 | Unc13a | unc-13 homolog A |
| chr7 | 33932261 | 33934732 | 2472 | * | 65,24386 | 2,337412993 | 0,771356 | 3,030264 | 0,002443 | 0,456474583 | 4,055758 | Promoter (<=1kb) | 7 | 33933802 | 33939632 | 5831 | 1 | 1E+08 | ENSMUST00000127010.2 | 0 | ENSMUSG00000085105 | Gm12758 | predicted gene 12758 |
| chr2 | 25473824 | 25474962 | 1139 | * | 13,41129 | 2,337457009 | 0,9061 | 2,579691 | 0,009889 | 0,954881543 | 0,918255 | Promoter (<=1kb) | 2 | 25474147 | 25475072 | 926 | 2 | 227624 | ENSMUST00000128942.2 | 110 | ENSMUSG00000015087 | Rabl6 | RAB, member RAS oncogene family-like 6 |
| chr7 | 25013254 | 25014298 | 1045 | * | 14,52231 | 2,338217663 | 0,901118 | 2,594795 | 0,009465 | 0,886332554 | 1,37683 | Promoter (2-3kb) | 7 | 25016589 | 25065342 | 48754 | 1 | 269878 | ENSMUST00000128119.2 | -2291 | ENSMUSG00000045039 | Megf8 | multiple EGF-like-domains 8 |
| chr13 | 1,03E+08 | 1,03E+08 | 3093 | * | 22,04727 | 2,338650098 | 0,726667 | 3,218324 | 0,001289 | 0,420798623 | 2,67243 | Intron (ENSMUST00000( | 13 | 1,03E+08 | 1,03E+08 | 45894 | 1 | 17079 | ENSMUST00000172138.2 | 31550 | ENSMUSG00000021624 | Cd180 | CD180 antigen |
| chr10 | 42152938 | 42154237 | 1300 | * | 18,61781 | 2,339113436 | 0,788736 | 2,965649 | 0,00302 | 0,568221164 | 2,663619 | Promoter (<=1kb) | 10 | 42123427 | 42152751 | 29325 | 2 | 56484 | ENSMUST00000105501.2 | -187 | ENSMUSG00000048756 | Foxo3 | forkhead box O3 |
| chr9 | 94869050 | 94870381 | 1332 | * | 11,85147 | 2,339149614 | 0,907659 | 2,577124 | 0,009963 | 0,904389377 | 1,2763 | Intron (ENSMUST00000( | 9 | 95003739 | 95006604 | 2866 | 1 | 331004 | ENSMUST00000162329.2 | -133358 | ENSMUSG00000031129 | Slc9a9 | solute carrier family 9 (sodium/hydrogen exchanger), member 9 |
| chr6 | 94616894 | 94620114 | 3221 | * | 40,57418 | 2,339504741 | 0,5313354 | 4,4030657 | 1,067E-05 | 0,026684056 | 2,3776742 | Intron (ENSMUST0000002 | 6 | 94603431 | 94632506 | 29076 | 2 | 16206 | ENSMUST00000150811.8 | 12392 | ENSMUSG00000030029 | Lrig1 | leucine-rich repeats and immunoglobulin-like domains 1 |
| chr5 | 1,49E+08 | 1,49E+08 | 1536 | * | 17,13624 | 2,339589114 | 0,801496 | 2,919027 | 0,003511 | 0,610904497 | 2,507013 | Exon (ENSMUST000001 | 5 | 1,5E+08 | 1,5E+08 | 6439 | 1 | 381693 | ENSMUST00000202635.2 | -7320 | ENSMUSG00000029658 | Wdr95 | WD40 repeat domain 95 |
| chr2 | 1,11E+08 | 1,11E+08 | 1019 | * | 12,25941 | 2,341125154 | 0,905422 | 2,585672 | 0,009719 | 0,913502848 | 1,173325 | Distal Intergenic | 2 | 1,11E+08 | 1,11E+08 | 1064 | 1 | 258379 | ENSMUST00000209096.2 | 9985 | ENSMUSG00000108931 | Olfr1284 | olfactory receptor 1284 |
| chr17 | 74390058 | 74393350 | 3293 | * | 20,52321 | 2,342286108 | 0,903284 | 2,593077 | 0,009512 | 0,78061212 | 2,507013 | Promoter (2-3kb) | 17 | 74385744 | 74387288 | 1545 | 2 | 1,03E+08 | ENSMUST00000233356.2 | -2770 | ENSMUSG00000117226 | Gm31235 | predicted gene, 31235 |
| chr7 | 95871092 | 95873156 | 2065 | * | 23,42985 | 2,3424771 | 0,635774 | 3,684447 | 0,000229 | 0,173939409 | 2,880017 | Intron (ENSMUST00000( | 7 | 95860878 | 96557761 | 696884 | 1 | 23966 | ENSMUST00000107165.8 | 10214 | ENSMUSG00000048078 | Tenm4 | teneurin transmembrane protein 4 |
| chr11 | 72012682 | 72014881 | 2200 | * | 20,91981 | 2,342533594 | 0,86446 | 2,709822 | 0,006732 | 0,683966322 | 3,104559 | Intron (ENSMUST00000( | 11 | 71938354 | 72026547 | 88194 | 2 | 327958 | ENSMUST00000108508.3 | 11666 | ENSMUSG00000040543 | Pitpnm3 | PITPNM family member 3 |
| chr15 | 62110621 | 62112354 | 1734 | * | 25,42079 | 2,342943784 | 0,685714 | 3,416793 | 0,000634 | 0,297590871 | 2,880017 | Intron (ENSMUST00000( | 15 | 62094452 | 62114354 | 19903 | 1 | 19296 | ENSMUST00000182141.2 | 16169 | ENSMUSG00000097039 | Pvt1 | Pvt1 oncogene |
| chr7 | 84339046 | 84340859 | 1814 | * | 19,00801 | 2,343112846 | 0,727852 | 3,219214 | 0,001285 | 0,418252864 | 2,713306 | Promoter (<=1kb) | 7 | 84262974 | 84339165 | 76192 | 2 | 65098 | ENSMUST00000209117.2 | 0 | ENSMUSG00000030629 | Zfand6 | zinc finger, AN1-type domain 6 |
| chr5 | 37838510 | 37840764 | 2255 | * | 21,26542 | 2,343505106 | 0,716439 | 3,271045 | 0,001072 | 0,390164541 | 2,67243 | Intron (ENSMUST00000( | 5 | 37604169 | 37874515 | 270347 | 2 | 64293 | ENSMUST00000094836.6 | 33751 | ENSMUSG00000029123 | Stk32b | serine/threonine kinase 32B |
| chr10 | 80936564 | 80939335 | 2772 | * | 24,57929 | 2,343816554 | 0,629227 | 3,724915 | 0,000195 | 0,159633916 | 2,880017 | Promoter (1-2kb) | 10 | 80940450 | 80940509 | 60 | 1 | 1,01E+08 | ENSMUST00000176537.3 | -1115 | ENSMUSG00000093498 | Mir5615-2 | microRNA 5615-2 |
| chr10 | 49179762 | 49182417 | 2656 | * | 20,95961 | 2,344479539 | 0,822053 | 2,851981 | 0,004345 | 0,61075929 | 3,104559 | Intron (ENSMUST00000( | 10 | 48976933 | 49148901 | 171969 | 2 | 14806 | ENSMUST00000219509.2 | -30861 | ENSMUSG00000056073 | Grik2 | glutamate receptor, ionotropic, kainate 2 (beta 2) |
| chr3 | 1,52E+08 | 1,52E+08 | 1448 | * | 16,26381 | 2,344810234 | 0,872117 | 2,688642 | 0,007174 | 0,738747729 | 2,386008 | Promoter (<=1kb) | 3 | 1,52E+08 | 1,52E+08 | 9891 | 2 | 67035 | ENSMUST00000050073.13 | 0 | ENSMUSG00000028035 | Dnajb4 | DnaJ heat shock protein family (Hsp40) member B4 |
| chr1 | 64653336 | 64656979 | 3644 | * | 43,18316 | 2,344825464 | 0,560773 | 4,181413 | 2,9E-05 | 0,050236155 | 2,725491 | Promoter (<=1kb) | 1 | 64645632 | 64656401 | 10770 | 2 | 67099 | ENSMUST00000114079.9 | 0 | ENSMUSG00000025956 | Mettl21a | methyltransferase like 21A |
| chr14 | 52263557 | 52265796 | 2240 | * | 21,4678 | 2,344841816 | 0,806205 | 2,908495 | 0,003632 | 0,585155013 | 2,951933 | Distal Intergenic | 14 | 52243539 | 52258190 | 14652 | 2 | 69890 | ENSMUST00000067549.15 | -5367 | ENSMUSG00000049295 | Zfp219 | zinc finger protein 219 |
| chr10 | 84573429 | 84575047 | 1619 | * | 21,67278 | 2,345062986 | 0,7139 | 3,28486 | 0,00102 | 0,35147835 | 3,104559 | Distal Intergenic | 10 | 84591926 | 84742008 | 150083 | 1 | 71137 | ENSMUST00000060397.13 | -16879 | ENSMUSG00000020037 | Rfx4 | regulatory factor X, 4 (influences HLA class II expression) |
| chr4 | 1,25E+08 | 1,25E+08 | 2139 | * | 20,62342 | 2,345143699 | 0,769843 | 3,046262 | 0,002317 | 0,474532331 | 3,471678 | Intron (ENSMUST00000( | 4 | 1,24E+08 | 1,24E+08 | 5456 | 1 | 108809 | ENSMUST00000133499.2 | 39788 | | 4933407E24Rik | RIKEN cDNA 4933407E24 gene |
| chr2 | 41387684 | 41389116 | 1433 | * | 14,98523 | 2,345306813 | 0,806797 | 2,906936 | 0,00365 | 0,738600147 | 1,214904 | Exon (ENSMUST00000( | 2 | 40691405 | 41679090 | 987686 | 2 | 94217 | ENSMUST00000167270.5 | 289974 | ENSMUSG00000049252 | Lrp1b | low density lipoprotein-related protein 1B |
| chr4 | 1,27E+08 | 1,27E+08 | 1929 | * | 14,56644 | 2,345525042 | 0,867221 | 2,704645 | 0,006838 | 0,823736512 | 1,41696 | Intron (ENSMUST00000( | 4 | 1,27E+08 | 1,27E+08 | 1811 | 2 | 67785 | ENSMUST00000150467.2 | -4796 | ENSMUSG00000042446 | Zmym4 | zinc finger, MYM-type 4 |
| chr9 | 1,19E+08 | 1,19E+08 | 1702 | * | 13,98537 | 2,345739409 | 0,875094 | 2,680556 | 0,00735 | 0,885572768 | 1,075735 | 3' UTR | 9 | 1,19E+08 | 1,19E+08 | 2218 | 2 | 108737 | ENSMUST00000136902.2 | -7272 | ENSMUSG00000036737 | Oxsr1 | oxidative-stress responsive 1 |

| | | | | | | | | | | | | | | | | | | | | | | | |
|---|---|---|---|---|---|---|---|---|---|---|---|---|---|---|---|---|---|---|---|---|---|---|---|
| chr11 | 57732913 | 57735091 | 2179 | * | 14,66236 | 2,34637725 | 0,862271 | 2,721159 | 0,006505 | 0,740451144 | 2,141612 | Distal Intergenic | 11 | 57719531 | 57723644 | 4114 | 2 | 15110 | ENSMUST00000160392.9 | -9269 | ENSMUSG00000037335 | Hand1 | heart and neural crest derivatives expressed 1 |
| chr6 | 12126182 | 12127476 | 1295 | * | 14,06739 | 2,347083098 | 0,89028 | 2,636342 | 0,008381 | 0,96198404 | 0,738441 | Distal Intergenic | 6 | 12099517 | 12109582 | 10066 | 2 | 625347 | ENSMUST00000080891.6 | -16600 | ENSMUSG00000057223 | Gm6578 | mitochondrial ribosomal protein L32 pseudogene |
| chr9 | 1,1E+08 | 1,1E+08 | 1865 | * | 24,37522 | 2,347249067 | 0,765314 | 3,067039 | 0,002162 | 0,506040436 | 2,67243 | Promoter (1-2kb) | 9 | 1,1E+08 | 1,1E+08 | 484 | 1 | 235623 | ENSMUST00000196424.2 | 1741 | ENSMUSG00000032485 | Scap | SREBF chaperone |
| chr5 | 1,24E+08 | 1,24E+08 | 1538 | * | 19,39564 | 2,347645409 | 0,713865 | 3,288642 | 0,001007 | 0,37346973 | 2,713306 | Exon (ENSMUST00000 | 5 | 1,24E+08 | 1,24E+08 | 5578 | 2 | 330192 | ENSMUST00000199306.2 | 3881 | ENSMUSG00000066278 | Vps37b | vacuolar protein sorting 37B |
| chr16 | 3895269 | 3897259 | 1991 | * | 18,20205 | 2,347727909 | 0,791092 | 2,967704 | 0,003 | 0,591141332 | 2,386008 | Promoter (<=1kb) | 16 | 3857835 | 3895691 | 37857 | 2 | 68015 | ENSMUST00000006137.9 | 0 | ENSMUSG00000005981 | Trap1 | TNF receptor-associated protein 1 |
| chr14 | 26795257 | 26797443 | 2187 | * | 20,06611 | 2,347756048 | 0,90899 | 2,582818 | 0,0098 | 0,80696549 | 2,221611 | Intron (ENSMUST00000 | 14 | 26789354 | 26822933 | 33580 | 1 | 171463 | ENSMUST00000226105.2 | 5903 | ENSMUSG00000040717 | Il17rd | interleukin 17 receptor D |
| chr8 | 1,07E+08 | 1,07E+08 | 1352 | * | 16,03893 | 2,348350629 | 0,891241 | 2,634923 | 0,008416 | 0,773819636 | 2,318434 | Intron (ENSMUST00000 | 8 | 1,07E+08 | 1,07E+08 | 26089 | 2 | 58994 | ENSMUST00000212896.2 | -5726 | ENSMUSG00000031906 | Smpd3 | sphingomyelin phosphodiesterase 3, neutral |
| chr9 | 89599480 | 89600692 | 1213 | * | 13,73546 | 2,348386835 | 0,884329 | 2,655557 | 0,007918 | 0,954588896 | 0,738441 | Distal Intergenic | 9 | 89581259 | 89587121 | 5863 | 2 | 66111 | ENSMUST00000058488.9 | -12359 | ENSMUSG00000032353 | Tmed3 | transmembrane p24 trafficking protein 3 |
| chr13 | 6226142 | 6228522 | 2381 | * | 17,83906 | 2,348670702 | 0,875279 | 2,683339 | 0,007289 | 0,742014238 | 2,386008 | Distal Intergenic | 13 | 6330414 | 6344750 | 14337 | 2 | 1,03E+08 | ENSMUST00000221255.2 | 116228 | ENSMUSG00000114097 | Gm35615 | predicted gene, 35615 |
| chr18 | 36803121 | 36805203 | 2083 | * | 19,0834 | 2,34885419 | 0,824904 | 2,847428 | 0,004407 | 0,639383182 | 2,713306 | Promoter (<=1kb) | 18 | 36799734 | 36803868 | 4135 | 2 | 24068 | ENSMUST00000173875.9 | 0 | ENSMUSG00000006050 | Sra1 | steroid receptor RNA activator 1 |
| chr18 | 32518380 | 32520220 | 1841 | * | 17,63355 | 2,349877147 | 0,83351 | 2,819256 | 0,004814 | 0,685078899 | 2,154504 | Intron (ENSMUST00000 | 18 | 32504580 | 32511331 | 6752 | 2 | 402767 | ENSMUST00000234978.2 | -7049 | | A830052D11Rik | RIKEN cDNA A830052D11 gene |
| chr6 | 87866444 | 87867834 | 1391 | * | 18,67839 | 2,350211364 | 0,843258 | 2,78706 | 0,005319 | 0,68259716 | 2,507013 | Promoter (1-2kb) | 6 | 87864939 | 87871944 | 7006 | 1 | 54161 | ENSMUST00000137717.2 | 1505 | ENSMUSG00000030058 | Copg1 | coatomer protein complex, subunit gamma 1 |
| chr4 | 1,39E+08 | 1,39E+08 | 1834 | * | 21,05481 | 2,350292509 | 0,755391 | 3,111357 | 0,001862 | 0,455547114 | 3,104559 | Exon (ENSMUST00000 | 4 | 1,39E+08 | 1,39E+08 | 3133 | 1 | 69116 | ENSMUST00000129779.2 | 6756 | ENSMUSG00000066036 | Ubr4 | ubiquitin protein ligase E3 component n-recognin 4 |
| chr11 | 54165496 | 54167727 | 2232 | * | 20,44827 | 2,350764057 | 0,718472 | 3,271895 | 0,001068 | 0,40037769 | 2,507013 | Intron (ENSMUST00000 | 11 | 54156129 | 54158103 | 1975 | 2 | 16187 | ENSMUST00000019058.6 | -7393 | ENSMUSG00000018914 | Il3 | interleukin 3 |
| chr16 | 20542718 | 20546468 | 3751 | * | 25,8192 | 2,351547239 | 0,65128 | 3,610653 | 0,000305 | 0,206624632 | 2,858511 | Promoter (2-3kb) | 16 | 20544337 | 20549301 | 4965 | 2 | 21832 | ENSMUST00000231826.2 | 2833 | ENSMUSG00000022847 | Thpo | thrombopoietin |
| chr11 | 75404461 | 75406805 | 2345 | * | 18,78533 | 2,351767761 | 0,889064 | 2,645219 | 0,008164 | 0,751210692 | 2,507013 | Promoter (<=1kb) | 11 | 75404401 | 75417404 | 13004 | 1 | 380713 | ENSMUST00000123819.2 | 60 | ENSMUSG00000038188 | Scarf1 | scavenger receptor class F, member 1 |
| chr13 | 52608157 | 52610328 | 2172 | * | 20,87446 | 2,352051608 | 0,747568 | 3,146272 | 0,001654 | 0,445695953 | 2,951933 | Distal Intergenic | 13 | 52658416 | 52685315 | 26900 | 2 | 68203 | ENSMUST00000057442.8 | 74987 | ENSMUSG00000047842 | Diras2 | DIRAS family, GTP-binding RAS-like 2 |
| chr8 | 1,11E+08 | 1,11E+08 | 1755 | * | 19,70667 | 2,352098364 | 0,760909 | 3,091167 | 0,001994 | 0,524856133 | 2,221611 | Intron (ENSMUST00000 | 8 | 1,11E+08 | 1,11E+08 | 19925 | 1 | 244654 | ENSMUST00000052457.15 | -19313 | ENSMUSG00000033763 | Mtss2 | MTSS I-BAR domain containing 2 |
| chr10 | 1,21E+08 | 1,21E+08 | 1533 | * | 12,06845 | 2,352220018 | 0,899808 | 2,614134 | 0,008945 | 0,901463351 | 1,173325 | Promoter (1-2kb) | 10 | 1,21E+08 | 1,21E+08 | 1927 | 2 | 56480 | ENSMUST00000218460.2 | 1248 | ENSMUSG00000020115 | Tbk1 | TANK-binding kinase 1 |
| chr11 | 36941674 | 36943035 | 1362 | * | 12,80626 | 2,352674085 | 0,898487 | 2,618483 | 0,008832 | 0,90060438 | 1,173325 | Intron (ENSMUST00000 | 11 | 36030307 | 36834993 | 804687 | 2 | 23964 | ENSMUST00000146113.8 | -106681 | ENSMUSG00000049336 | Tenm2 | teneurin transmembrane protein 2 |
| chr16 | 94291250 | 94292807 | 1558 | * | 15,25183 | 2,352919504 | 0,858185 | 2,741738 | 0,006112 | 0,822789084 | 1,2763 | Promoter (1-2kb) | 16 | 94289636 | 94322976 | 33341 | 1 | 1,03E+08 | ENSMUST00000210205.2 | 1614 | | Gm31323 | predicted gene, 31323 |
| chr10 | 59536933 | 59539705 | 2773 | * | 20,57064 | 2,352954854 | 0,735287 | 3,200048 | 0,001374 | 0,448852574 | 2,386008 | Promoter (<=1kb) | 10 | 59538299 | 59699954 | 161656 | 1 | 216001 | ENSMUST00000165563.8 | 0 | ENSMUSG00000020111 | Micu1 | mitochondrial calcium uptake 1 |
| chr4 | 71155822 | 71157420 | 1599 | * | 17,11565 | 2,353137121 | 0,727206 | 3,235861 | 0,001213 | 0,410415129 | 2,663619 | Distal Intergenic | 4 | 70993346 | 71049370 | 56025 | 2 | 1,03E+08 | ENSMUST00000143796.2 | -106452 | ENSMUSG00000084937 | Gm11228 | predicted gene 11228 |
| chr7 | 99207013 | 99208565 | 1553 | * | 17,08539 | 2,353276609 | 0,83746 | 2,810017 | 0,004954 | 0,786955198 | 1,2763 | Intron (ENSMUST00000 | 7 | 99201476 | 99201570 | 95 | 1 | 723840 | ENSMUST00000083637.3 | 5537 | ENSMUSG00000065571 | Mir326 | microRNA 326 |
| chr3 | 36548923 | 36550575 | 1653 | * | 17,96482 | 2,353313306 | 0,832633 | 2,826351 | 0,004708 | 0,651710641 | 2,663619 | Distal Intergenic | 3 | 36553953 | 36560958 | 7006 | 2 | 329623 | ENSMUST00000140203.2 | 10383 | ENSMUSG00000085320 | Gm11548 | predicted gene 11548 |
| chr2 | 1,58E+08 | 1,58E+08 | 1424 | * | 15,25773 | 2,353563142 | 0,893535 | 2,633992 | 0,008439 | 0,887885738 | 1,214904 | Distal Intergenic | 2 | 1,58E+08 | 1,58E+08 | 134 | 2 | 1,15E+08 | ENSMUST00000104102.3 | 23052 | ENSMUSG00000077290 | Gm25407 | predicted gene, 25407 |
| chr2 | 14060566 | 14062093 | 1528 | * | 17,93474 | 2,353918554 | 0,77115 | 3,05248 | 0,00227 | 0,535120883 | 2,386008 | Promoter (<=1kb) | 2 | 13855093 | 14060847 | 205755 | 2 | 30963 | ENSMUST00000131730.7 | 0 | ENSMUSG00000063275 | Hacd1 | 3-hydroxyacyl-CoA dehydratase 1 |
| chr17 | 47449711 | 47451373 | 1663 | * | 15,30098 | 2,35405588 | 0,858941 | 2,740649 | 0,006132 | 0,831521609 | 1,214904 | Promoter (<=1kb) | 17 | 47451801 | 47640462 | 188662 | 1 | 224829 | ENSMUST00000188947.7 | -428 | ENSMUSG00000064043 | Trerf1 | transcriptional regulating factor 1 |
| chr1 | 57752989 | 57754063 | 1075 | * | 14,26625 | 2,354549585 | 0,826935 | 2,84732 | 0,004409 | 0,819350294 | 0,938682 | Distal Intergenic | 1 | 57813321 | 57924927 | 111607 | 1 | 67198 | ENSMUST00000172287.8 | -59258 | ENSMUSG00000038305 | Spats2l | spermatogenesis associated, serine-rich 2-like |
| chr13 | 62905158 | 62907091 | 1934 | * | 20,0965 | 2,355199634 | 0,747171 | 3,152155 | 0,001621 | 0,477253929 | 2,386008 | Promoter (1-2kb) | 13 | 62886357 | 62908506 | 22150 | 2 | 435366 | ENSMUST00000220864.2 | 1415 | ENSMUSG00000074863 | Platr25 | pluripotency associated transcript 25 |
| chr19 | 53261085 | 53262915 | 1831 | * | 26,34988 | 2,355664918 | 0,61786 | 3,81262 | 0,000138 | 0,133396034 | 2,67243 | Distal Intergenic | 19 | 53237316 | 53253836 | 16521 | 2 | 69319 | ENSMUST00000235503.2 | -7249 | ENSMUSG00000118314 | 1700001K23Rik | RIKEN cDNA 1700001K23 gene |
| chr14 | 58407042 | 58408310 | 1269 | * | 16,78331 | 2,355693308 | 0,89152 | 2,642333 | 0,008234 | 0,874458545 | 1,2763 | Distal Intergenic | 14 | 58313964 | 58346849 | 32886 | 1 | 14180 | ENSMUST00000166770.2 | 93078 | ENSMUSG00000021974 | Fgf9 | fibroblast growth factor 9 |
| chr4 | 46388712 | 46391010 | 2299 | * | 17,91851 | 2,355851052 | 0,875752 | 2,690089 | 0,007143 | 0,734572036 | 2,423594 | Promoter (<=1kb) | 4 | 46376505 | 46389418 | 12914 | 2 | 74753 | ENSMUST00000151903.8 | 0 | ENSMUSG00000028331 | Trmo | tRNA methyltransferase O |
| chr10 | 81121418 | 81122765 | 1348 | * | 12,69658 | 2,356438223 | 0,877938 | 2,68406 | 0,007273 | 0,906138253 | 0,918255 | Promoter (<=1kb) | 10 | 81117021 | 81121603 | 4583 | 2 | 27375 | ENSMUST00000218484.2 | 0 | ENSMUSG00000034917 | Tjp3 | tight junction protein 3 |
| chrX | 1,63E+08 | 1,63E+08 | 2336 | * | 23,01909 | 2,35678839 | 0,650079 | 3,625386 | 0,000289 | 0,206624632 | 2,67243 | Promoter (<=1kb) | 20 | 1,63E+08 | 1,63E+08 | 12050 | 1 | 18700 | ENSMUST00000208741.2 | 0 | ENSMUSG00000031381 | Piga | phosphatidylinositol glycan anchor biosynthesis, class A |
| chr7 | 1,26E+08 | 1,26E+08 | 2212 | * | 17,59688 | 2,357206432 | 0,783809 | 3,007375 | 0,002635 | 0,563591278 | 2,386008 | Promoter (2-3kb) | 7 | 1,26E+08 | 1,26E+08 | 983 | 2 | 16797 | ENSMUST00000205426.2 | 2458 | ENSMUSG00000030742 | Lat | linker for activation of T cells |
| chr9 | 40872673 | 40874325 | 1653 | * | 17,51857 | 2,357365044 | 0,838004 | 2,813072 | 0,004907 | 0,669586355 | 2,507013 | Promoter (<=1kb) | 9 | 40872182 | 40873629 | 1448 | 2 | 70989 | ENSMUST00000160120.2 | 0 | ENSMUSG00000032023 | Jhy | junctional cadherin complex regulator |
| chr13 | 31357801 | 31360547 | 2747 | * | 20,31756 | 2,358354572 | 0,840355 | 2,806379 | 0,00501 | 0,660150434 | 2,713306 | Distal Intergenic | 13 | 31397393 | 31425069 | 27677 | 2 | 73924 | ENSMUST00000153175.3 | 64522 | ENSMUSG00000087460 | 4930401O12Rik | RIKEN cDNA 4930401O12 gene |
| chr16 | 91310429 | 91312661 | 2233 | * | 17,14293 | 2,358791125 | 0,876925 | 2,689845 | 0,007149 | 0,718881903 | 2,663619 | Exon (ENSMUST00000 | 16 | 91302192 | 91303554 | 1363 | 1 | 15975 | ENSMUST00000232453.2 | 8237 | ENSMUSG00000022967 | Ifnar1 | interferon (alpha and beta) receptor 1 |
| chr13 | 1,05E+08 | 1,05E+08 | 1533 | * | 15,82013 | 2,358884364 | 0,816905 | 2,887588 | 0,003882 | 0,656950206 | 2,141612 | Distal Intergenic | 13 | 1,05E+08 | 1,05E+08 | 18611 | 2 | 77803 | ENSMUST00000043061.5 | -80292 | ENSMUSG00000042655 | Shisal2b | shisa like 2B |
| chr2 | 127597129 | 127604379 | 7251 | * | 58,917413 | 2,359073387 | 0,515767 | 4,5739126 | 4,787E-06 | 0,01138298 | 3,2706213 | Promoter (1-2kb) | 2 | 127587542 | 127595990 | 8449 | 2 | 53885 | ENSMUST00000148033.2 | -1139 | ENSMUSG00000027378 | Nphp1 | nephronophthisis 1 (juvenile) homolog (human) |
| chr6 | 1,25E+08 | 1,25E+08 | 4163 | * | 169,2427 | 2,360555259 | 0,611869 | 3,857941 | 0,000114 | 0,101149739 | 3,439501 | Promoter (<=1kb) | 6 | 1,25E+08 | 1,25E+08 | 4615 | 2 | 14433 | ENSMUST00000117757.9 | 0 | ENSMUSG00000057666 | Gapdh | glyceraldehyde-3-phosphate dehydrogenase |
| chr16 | 15765162 | 15767585 | 2424 | * | 34,634278 | 2,360864765 | 0,5185632 | 4,5527039 | 5,296E-06 | 0,013744849 | 2,7872553 | Intron (ENSMUST000000 | 16 | 15705150 | 15708895 | 3746 | 1 | 12609 | ENSMUST00000096232.6 | 60012 | ENSMUSG00000071637 | Cebpd | CCAAT/enhancer binding protein (C/EBP), delta |
| chr9 | 44319494 | 44321386 | 1893 | * | 13,0283 | 2,361316082 | 0,911229 | 2,591352 | 0,00956 | 0,899620932 | 1,2763 | Promoter (<=1kb) | 9 | 44319308 | 44320652 | 1345 | 1 | 75617 | ENSMUST00000217620.2 | 186 | ENSMUSG00000009927 | Rps25 | ribosomal protein S25 |
| chr2 | 1,32E+08 | 1,32E+08 | 2334 | * | 32,07044 | 2,361420882 | 0,620699 | 3,804454 | 0,000142 | 0,201373005 | 1,405709 | Intron (ENSMUST00000 | 2 | 1,32E+08 | 1,32E+08 | 70275 | 2 | 54338 | ENSMUST00000127724.8 | -7160 | ENSMUSG00000027340 | Slc23a2 | solute carrier family 23 (nucleobase transporters), member 2 |
| chr19 | 11985837 | 11988634 | 2798 | * | 21,00222 | 2,36145298 | 0,852472 | 2,770123 | 0,005604 | 0,637823949 | 3,471678 | Distal Intergenic | 19 | 12013172 | 12014104 | 933 | 2 | 258675 | ENSMUST00000087831.2 | 25470 | ENSMUSG00000067529 | Olfr1423 | olfactory receptor 1423 |
| chr14 | 30183432 | 30184940 | 1509 | * | 17,95472 | 2,361544803 | 0,826412 | 2,857587 | 0,004269 | 0,670853401 | 2,154504 | Intron (ENSMUST00000 | 14 | 30201492 | 30232451 | 30960 | 1 | 75901 | ENSMUST00000224453.2 | -16552 | ENSMUSG00000021962 | Dcp1a | decapping mRNA 1A |
| chr12 | 34243626 | 34245916 | 2291 | * | 16,88258 | 2,361853819 | 0,814595 | 2,899421 | 0,003739 | 0,630824695 | 2,386008 | Intron (ENSMUST00000 | 12 | 34007670 | 34009828 | 2159 | 1 | 22160 | ENSMUST00000049089.7 | 235956 | ENSMUSG00000035799 | Twist1 | twist basic helix-loop-helix transcription factor 1 |
| chr17 | 68024025 | 68025675 | 1651 | * | 16,76581 | 2,361972946 | 0,853322 | 2,767974 | 0,005641 | 0,695892636 | 2,386008 | Exon (ENSMUST00000 | 17 | 68004260 | 68063907 | 59648 | 1 | 16772 | ENSMUST00000233821.2 | 19765 | ENSMUSG00000032796 | Lama1 | laminin, alpha 1 |
| chr8 | 1,18E+08 | 1,18E+08 | 1539 | * | 18,47319 | 2,362768015 | 0,814382 | 2,901303 | 0,003716 | 0,61075929 | 2,663619 | Promoter (1-2kb) | 8 | 1,18E+08 | 1,18E+08 | 202367 | 1 | 74440 | ENSMUST00000166750.9 | -1598 | ENSMUSG00000034390 | Cmip | c-Maf inducing protein |
| chr8 | 1,23E+08 | 1,23E+08 | 1687 | * | 16,37881 | 2,363037257 | 0,829575 | 2,848492 | 0,004393 | 0,676922067 | 2,141612 | Promoter (<=1kb) | 8 | 1,23E+08 | 1,23E+08 | 2141 | 2 | 50917 | ENSMUST00000211906.2 | 333 | ENSMUSG00000015027 | Galns | galactosamine (N-acetyl)-6-sulfate sulfatase |
| chr6 | 34660237 | 34661918 | 1682 | * | 18,94597 | 2,363280435 | 0,774837 | 3,050034 | 0,002288 | 0,537244646 | 2,386008 | Intron (ENSMUST00000 | 6 | 34686377 | 34752408 | 66032 | 1 | 109624 | ENSMUST00000115021.8 | -24459 | ENSMUSG00000029761 | Cald1 | caldesmon 1 |
| chr16 | 22272823 | 22274483 | 1661 | * | 12,7074 | 2,363418595 | 0,87973 | 2,686529 | 0,00722 | 0,905781613 | 0,918255 | Distal Intergenic | 16 | 22200059 | 22258469 | 58411 | 2 | 104156 | ENSMUST00000079601.13 | -14354 | ENSMUSG00000013089 | Etv5 | ets variant 5 |
| chr4 | 1,51E+08 | 1,51E+08 | 2758 | * | 28,86353 | 2,363640731 | 0,771545 | 3,063518 | 0,002188 | 0,504427013 | 2,725491 | Promoter (<=1kb) | 4 | 1,51E+08 | 1,51E+08 | 19850 | 1 | 21942 | ENSMUST00000135169.8 | 0 | ENSMUSG00000028965 | Tnfrsf9 | tumor necrosis factor receptor superfamily, member 9 |
| chr9 | 1,06E+08 | 1,06E+08 | 1610 | * | 15,91842 | 2,364412244 | 0,86697 | 2,727214 | 0,006387 | 0,718444962 | 2,386008 | Intron (ENSMUST00000 | 9 | 1,06E+08 | 1,06E+08 | 5857 | 1 | 75669 | ENSMUST00000214254.2 | 21217 | ENSMUSG00000032571 | Pik3r4 | phosphoinositide-3-kinase regulatory subunit 4 |
| chr12 | 33364786 | 33366722 | 1937 | * | 18,61534 | 2,364664631 | 0,849173 | 2,784666 | 0,005358 | 0,707972935 | 2,154504 | Promoter (<=1kb) | 12 | 33365396 | 33422356 | 56961 | 1 | 380753 | ENSMUST00000154742.2 | 0 | ENSMUSG00000020564 | Atxn7l1 | ataxin 7-like 1 |
| chr4 | 57605807 | 57607578 | 1772 | * | 23,39476 | 2,36515711 | 0,658626 | 3,591047 | 0,000329 | 0,212327236 | 2,951062 | Intron (ENSMUST00000 | 4 | 57637823 | 57894718 | 256896 | 1 | 677884 | ENSMUST00000150412.4 | -30245 | ENSMUSG00000038729 | Pakap | paralemmin A kinase anchor protein |
| chr8 | 1,15E+08 | 1,15E+08 | 1284 | * | 15,47121 | 2,365386857 | 0,908438 | 2,603796 | 0,00922 | 0,900890887 | 1,214904 | Distal Intergenic | 8 | 1,14E+08 | 1,15E+08 | 84214 | 2 | 208936 | ENSMUST00000213061.2 | -136194 | ENSMUSG00000053399 | Adamts18 | a disintegrin-like and metallopeptidase (reprolysin type) with thrombospondin type 1 motif, 18 |
| chr15 | 98681868 | 98682946 | 1079 | * | 11,48114 | 2,365500236 | 0,906973 | 2,608127 | 0,009104 | 0,903444531 | 1,173325 | Distal Intergenic | 15 | 98687738 | 98691718 | 3981 | 1 | 22408 | ENSMUST00000023734.8 | -4792 | ENSMUSG00000022997 | Wnt1 | wingless-type MMTV integration site family, member 1 |
| chr9 | 41162542 | 41167187 | 4646 | * | 38,16204 | 2,365828082 | 0,596313 | 3,967427 | 7,27E-05 | 0,089013319 | 2,725491 | Distal Intergenic | 9 | 41128462 | 41129167 | 706 | 1 | 78500 | ENSMUST00000245921.1 | 34080 | ENSMUSG00000100837 | 1700063D05Rik | RIKEN cDNA 1700063D05 gene |
| chr7 | 1,28E+08 | 1,28E+08 | 2347 | * | 18,67733 | 2,365941002 | 0,861234 | 2,747152 | 0,006012 | 0,726797252 | 2,154504 | Distal Intergenic | 7 | 1,28E+08 | 1,28E+08 | 2697 | 2 | 21843 | ENSMUST00000133444.8 | 11283 | ENSMUSG00000030846 | Tial1 | Tia1 cytotoxic granule-associated RNA binding protein-like 1 |
| chr7 | 29606936 | 29610020 | 3085 | * | 25,92752 | 2,366081011 | 0,706013 | 3,351326 | 0,000804 | 0,327375387 | 2,858511 | Promoter (<=1kb) | 7 | 29607648 | 29616238 | 8591 | 1 | 73729 | ENSMUST00000187028.7 | 0 | ENSMUSG00000099689 | Zfp383 | zinc finger protein 383 |
| chr10 | 41910519 | 41912438 | 1920 | * | 15,61014 | 2,366329307 | 0,871535 | 2,715129 | 0,006625 | 0,823446343 | 1,37683 | Intron (ENSMUST00000 | 10 | 41792803 | 41894438 | 101636 | 2 | 213402 | ENSMUST00000161927.8 | -16081 | ENSMUSG00000071324 | Armc2 | armadillo repeat containing 2 |
| chr8 | 56007703 | 56009847 | 2145 | * | 24,72322 | 2,367426319 | 0,76029 | 3,113845 | 0,001847 | 0,478766182 | 2,67243 | Distal Intergenic | 8 | 56323947 | 56359983 | 36037 | 2 | 244486 | ENSMUST00000053441.5 | 350136 | ENSMUSG00000046258 | Adam29 | a disintegrin and metallopeptidase domain 29 |
| chr18 | 67906867 | 67909691 | 2825 | * | 30,58192 | 2,367467904 | 0,569362 | 4,15811 | 3,21E-05 | 0,05191976 | 2,858511 | Promoter (<=1kb) | 18 | 67907946 | 67928557 | 20612 | 1 | 72124 | ENSMUST00000025421.9 | 0 | ENSMUSG00000079614 | Seh1l | SEH1-like (S. cerevisiae |
| chr3 | 86250419 | 86251784 | 1366 | * | 13,17101 | 2,368099537 | 0,887004 | 2,669774 | 0,00759 | 0,913241432 | 0,918255 | Intron (ENSMUST00000 | 3 | 86271764 | 86271932 | 169 | 1 | 1,15E+08 | ENSMUST00000083830.3 | -19980 | ENSMUSG00000065764 | Gm23850 | predicted gene, 23850 |
| chr1 | 22418572 | 22420411 | 1840 | * | 17,84493 | 2,368261505 | 0,781958 | 3,028629 | 0,002457 | 0,547497234 | 2,386008 | Intron (ENSMUST00000 | 1 | 22357139 | 22386029 | 28891 | 2 | 116837 | ENSMUST00000164877.8 | -32543 | ENSMUSG00000041670 | Rims1 | regulating synaptic membrane exocytosis 1 |
| chr15 | 81717677 | 81720605 | 2929 | * | 26,50725 | 2,369797303 | 0,654918 | 3,618466 | 0,000296 | 0,204368914 | 2,858511 | Exon (ENSMUST00000 | 15 | 81705144 | 81707980 | 2837 | 1 | 21685 | ENSMUST00000168632.2 | 12533 | ENSMUSG00000022389 | Tef | thyrotroph embryonic factor |
| chr15 | 25649237 | 25651914 | 2678 | * | 23,4661 | 2,370112831 | 0,781672 | 3,032108 | 0,002429 | 0,527638486 | 2,67243 | Intron (ENSMUST00000 | 15 | 25674596 | 25724036 | 49441 | 1 | 17909 | ENSMUST00000130517.8 | -22682 | ENSMUSG00000022272 | Myo10 | myosin X |
| chr17 | 35227383 | 35229161 | 1779 | * | 16,97625 | 2,370927974 | 0,818782 | 2,895677 | 0,003783 | 0,613712777 | 2,663619 | Promoter (1-2kb) | 17 | 35230191 | 35231375 | 1185 | 1 | 22321 | ENSMUST00000173911.2 | -1030 | ENSMUSG00000007029 | Vars | valyl-tRNA synthetase |
| chr11 | 1,21E+08 | 1,21E+08 | 2120 | * | 24,4966 | 2,371096642 | 0,746595 | 3,175881 | 0,001494 | 0,433562155 | 2,858511 | Promoter (<=1kb) | 11 | 1,21E+08 | 1,21E+08 | 1059 | 2 | 14104 | ENSMUST00000155276.3 | 0 | ENSMUSG00000025153 | Fasn | fatty acid synthase |
| chr16 | 8173864 | 8176322 | 2459 | * | 23,02073 | 2,371718016 | 0,735457 | 3,224823 | 0,001261 | 0,400863018 | 2,951062 | Distal Intergenic | 16 | 8154857 | 8154960 | 104 | 1 | 1,15E+08 | ENSMUST00000158968.3 | 19007 | ENSMUSG00000089593 | Gm26159 | predicted gene, 26159 |
| chr17 | 25652828 | 25654366 | 1539 | * | 39,00277 | 2,371736453 | 0,571124 | 4,152752 | 3,29E-05 | 0,059457361 | 2,377674 | Promoter (<=1kb) | 17 | 25650813 | 25652757 | 1945 | 2 | 58226 | ENSMUST00000159052.2 | -71 | ENSMUSG00000024112 | Cacna1h | calcium channel, voltage-dependent, T type, alpha 1H subunit |
| chr5 | 34920828 | 34923046 | 2219 | * | 17,88339 | 2,371761751 | 0,84129 | 2,819196 | 0,004814 | 0,685078899 | 2,154504 | Promoter (1-2kb) | 5 | 34919105 | 34972053 | 52949 | 1 | 15194 | ENSMUST00000123755.8 | 1723 | ENSMUSG00000029104 | Htt | huntingtin |
| chr2 | 1,23E+08 | 1,23E+08 | 2034 | * | 19,19271 | 2,37236073 | 0,811277 | 2,924231 | 0,003453 | 0,62816697 | 2,221611 | Distal Intergenic | 2 | 1,22E+08 | 1,22E+08 | 79 | 1 | 387165 | ENSMUST00000102354.4 | 17960 | ENSMUSG00000076309 | Mir147 | microRNA 147 |
| chr9 | 22352091 | 22354442 | 2352 | * | 27,7575 | 2,372487229 | 0,63399 | 3,742152 | 0,000182 | 0,153389938 | 2,880017 | Intron (ENSMUST00000 | 9 | 22341561 | 22355977 | 14417 | 1 | 68283 | ENSMUST00000213133.2 | 10530 | ENSMUSG00000036411 | 9530077C05Rik | RIKEN cDNA 9530077C05 gene |
| chr8 | 23019844 | 23022059 | 2216 | * | 19,94011 | 2,372520906 | 0,731496 | 3,243382 | 0,001181 | 0,407158022 | 2,663619 | Intron (ENSMUST00000 | 8 | 22996242 | 23025329 | 29088 | 1 | 20516 | ENSMUST00000210854.2 | 23602 | ENSMUSG00000037656 | Slc20a2 | solute carrier family 20, member 2 |
| chr11 | 79406065 | 79408042 | 1978 | * | 16,18662 | 2,373260861 | 0,912444 | 2,600992 | 0,009295 | 0,787155588 | 2,386008 | 5' UTR | 11 | 79405882 | 79414588 | 8707 | 2 | 216984 | ENSMUST00000179322.2 | 6546 | ENSMUSG00000093938 | Evi2b | ecotropic viral integration site 2b |
| chr4 | 1,36E+08 | 1,36E+08 | 3947 | * | 31,80107 | 2,373757776 | 0,701318 | 3,384708 | 0,000713 | 0,30988873 | 2,880017 | Promoter (1-2kb) | 4 | 1,36E+08 | 1,36E+08 | 10220 | 1 | 15356 | ENSMUST00000151879.2 | 1157 | ENSMUSG00000028672 | Hmgcl | 3-hydroxy-3-methylglutaryl-Coenzyme A lyase |
| chr13 | 82026938 | 82029212 | 2275 | * | 20,78533 | 2,373851799 | 0,801974 | 2,960012 | 0,003076 | 0,54122637 | 3,104559 | Distal Intergenic | 13 | 81940096 | 81941787 | 1692 | 1 | 12626 | ENSMUST00000224168.2 | 86842 | ENSMUSG00000021537 | Cetn3 | centrin 3 |
| chr13 | 1,04E+08 | 1,04E+08 | 3319 | * | 26,06574 | 2,374472217 | 0,706773 | 3,359595 | 0,000781 | 0,323558126 | 2,858511 | Promoter (<=1kb) | 13 | 1,04E+08 | 1,04E+08 | 38633 | 2 | 75805 | ENSMUST00000225324.2 | 0 | ENSMUSG00000021710 | Nln | neurolysin (metallopeptidase M3 family) |
| chr11 | 115181885 | 115185635 | 3751 | * | 58,683049 | 2,37488343 | 0,5374311 | 4,4189541 | 9,918E-06 | 0,02002598 | 3,2706213 | Promoter (2-3kb) | 11 | 115170448 | 115188340 | 17893 | 2 | 328035 | ENSMUST00000056153.8 | 2705 | ENSMUSG00000044788 | Fads6 | fatty acid desaturase domain family, member 6 |
| chr3 | 1,05E+08 | 1,05E+08 | 2703 | * | 25,13145 | 2,375948022 | 0,647753 | 3,667983 | 0,000244 | 0,177779799 | 2,951062 | Distal Intergenic | 3 | 1,05E+08 | 1,05E+08 | 16650 | 2 | 22414 | ENSMUST00000029429.6 | 26800 | ENSMUSG00000027840 | Wnt2b | wingless-type MMTV integration site family, member 2B |
| chr13 | 5610161 | 5612482 | 2322 | * | 16,89436 | 2,375960651 | 0,898426 | 2,644581 | 0,008179 | 0,782362632 | 2,141612 | Exon (ENSMUST00000 | 13 | 5504908 | 5610328 | 105421 | 1 | 1,05E+08 | ENSMUST00000223343.2 | 105253 | | Gm40661 | predicted gene, 40661 |
| chr9 | 1,2E+08 | 1,2E+08 | 2525 | * | 29,75351 | 2,376265204 | 0,707719 | 3,357641 | 0,000786 | 0,333881695 | 2,67243 | Distal Intergenic | 9 | 1,2E+08 | 1,2E+08 | 2409 | 1 | 68969 | ENSMUST00000217553.2 | -3825 | ENSMUSG00000006941 | Eif1b | eukaryotic translation initiation factor 1B |
| chr15 | 99691320 | 99693538 | 2219 | * | 24,09135 | 2,376599486 | 0,72406 | 3,282322 | 0,00103 | 0,361567622 | 2,951062 | Exon (ENSMUST00000 | 15 | 99694299 | 99700014 | 5716 | 2 | 65970 | ENSMUST00000230033.2 | 6476 | ENSMUSG00000023022 | Lima1 | LIM domain and actin binding 1 |
| chr5 | 75746918 | 75749007 | 2090 | * | 14,73415 | 2,376740171 | 0,900252 | 2,640085 | 0,008289 | 0,85617852 | 1,41696 | Intron (ENSMUST00000 | 5 | 75735615 | 75817382 | 81768 | 1 | 16590 | ENSMUST00000005815.7 | 11303 | ENSMUSG00000005672 | Kit | KIT proto-oncogene receptor tyrosine kinase |
| chr8 | 44789132 | 44790913 | 1782 | * | 25,33853 | 2,377226759 | 0,816984 | 2,909758 | 0,003617 | 0,59839358 | 2,725491 | Distal Intergenic | 8 | 45388484 | 45505294 | 116811 | 1 | 14107 | ENSMUST00000191428.7 | -597571 | ENSMUSG00000070047 | Fat1 | FAT atypical cadherin 1 |
| chr2 | 13519237 | 13521053 | 1817 | * | 14,42167 | 2,377648652 | 0,873064 | 2,723338 | 0,006463 | 0,819350294 | 1,37683 | Exon (ENSMUST00000 | 2 | 13281149 | 13496624 | 215476 | 2 | 65969 | ENSMUST00000091436.7 | -22613 | ENSMUSG00000026726 | Cubn | cubilin (intrinsic factor-cobalamin receptor) |
| chr6 | 1,25E+08 | 1,25E+08 | 1617 | * | 22,63316 | 2,378103744 | 0,796439 | 2,985922 | 0,002827 | 0,528005996 | 3,104559 | Distal Intergenic | 6 | 1,25E+08 | 1,25E+08 | 26176 | 1 | 1,03E+08 | ENSMUST00000243273.1 | -4763 | | Gm32673 | predicted gene, 32673 |
| chr9 | 95902987 | 95904607 | 1621 | * | 20,51226 | 2,378579538 | 0,756417 | 3,144534 | 0,001664 | 0,474289497 | 2,507013 | Exon (ENSMUST00000 | 9 | 95855570 | 95860386 | 4817 | 1 | 24127 | ENSMUST00000189806.2 | 47417 | ENSMUSG00000032410 | Xrn1 | 5'-3' exoribonuclease 1 |
| chr16 | 3067687 | 3068101 | 415 | * | 12,07158 | 2,378767542 | 0,884832 | 2,688383 | 0,00718 | 0,936915369 | 0,744375 | Exon (ENSMUST00020 | 16 | 3286324 | 3298915 | 12592 | 1 | 1,03E+08 | ENSMUST00000229453.2 | -218223 | ENSMUSG00000116455 | Gm35974 | predicted gene, 35974 |
| chr9 | 64158283 | 64161398 | 3116 | * | 30,94827 | 2,379237134 | 0,638703 | 3,725109 | 0,000195 | 0,156945882 | 2,951062 | Promoter (<=1kb) | 9 | 64093052 | 64160913 | 67862 | 2 | 26395 | ENSMUST00000005066.9 | 0 | ENSMUSG00000004936 | Map2k1 | mitogen-activated protein kinase kinase 1 |
| chr1 | 1,41E+08 | 1,41E+08 | 1926 | * | 17,16804 | 2,379242441 | 0,823141 | 2,890444 | 0,003847 | 0,627785088 | 2,507013 | Intron (ENSMUST00000 | 1 | 1,4E+08 | 1,41E+08 | 86229 | 2 | 75910 | ENSMUST00000193820.2 | 6085 | ENSMUSG00000102368 | 4930590L20Rik | RIKEN cDNA 4930590L20 gene |
| chr19 | 41268180 | 41270024 | 1845 | * | 16,21873 | 2,379796026 | 0,833305 | 2,855851 | 0,004292 | 0,660398635 | 2,318434 | Intron (ENSMUST00000 | 19 | 41199283 | 41252436 | 53154 | 2 | 107358 | ENSMUST00000025989.10 | -15744 | ENSMUSG00000025016 | Tm9sf3 | transmembrane 9 superfamily member 3 |
| chr18 | 86604962 | 86606081 | 1120 | * | 15,35149 | 2,379987937 | 0,922966 | 2,578629 | 0,009919 | 0,910915024 | 1,214904 | Distal Intergenic | 18 | 86517081 | 86523561 | 6481 | 1 | 246317 | ENSMUST00000235167.2 | 87881 | ENSMUSG00000050321 | Neto1 | neuropilin (NRP) and tolloid (TLL)-like 1 |
| chr13 | 81029856 | 81031196 | 1341 | * | 14,98772 | 2,380087947 | 0,844075 | 2,819757 | 0,004806 | 0,789027175 | 1,214904 | Promoter (<=1kb) | 13 | 81031503 | 81037762 | 6260 | 1 | 105171 | ENSMUST00000159090.2 | -307 | ENSMUSG00000074794 | Arrdc3 | arrestin domain containing 3 |
| chr14 | 52339722 | 52342187 | 2466 | * | 21,40526 | 2,380408073 | 0,893666 | 2,663643 | 0,00773 | 0,685026746 | 3,471678 | Promoter (<=1kb) | 14 | 52310834 | 52341485 | 30652 | 2 | 15381 | ENSMUST00000111610.12 | 0 | ENSMUSG00000060373 | Hnrnpc | heterogeneous nuclear ribonucleoprotein C |
| chr1 | 71328878 | 71330853 | 1976 | * | 16,71911 | 2,380478284 | 0,853214 | 2,790013 | 0,005271 | 0,670853401 | 2,663619 | Exon (ENSMUST00000 | 1 | 71329172 | 71346246 | 17075 | 2 | 74591 | ENSMUST00000188752.2 | 15393 | ENSMUSG00000050296 | Abca12 | ATP-binding cassette, sub-family A (ABC1), member 12 |
| chr10 | 61265373 | 61267663 | 2291 | * | 20,84211 | 2,380940372 | 0,848122 | 2,807309 | 0,004996 | 0,649217039 | 2,880017 | 3' UTR | 10 | 61253751 | 61261117 | 7367 | 1 | 18119 | ENSMUST00000049339.7 | 11622 | ENSMUSG00000037171 | Nodal | nodal |
| chr19 | 47201987 | 47203302 | 1316 | * | 11,75459 | 2,381807369 | 0,879395 | 2,708463 | 0,00676 | 0,838878373 | 1,2763 | Intron (ENSMUST00000 | 19 | 47186680 | 47190471 | 3792 | 2 | 1,03E+08 | ENSMUST00000236668.2 | -11516 | | Gm36693 | predicted gene, 36693 |

| | | | | | | | | | | | | | | | | | | | | | | |
|---|---|---|---|---|---|---|---|---|---|---|---|---|---|---|---|---|---|---|---|---|---|---|
| chr3 | 1,31E+08 | 1,31E+08 | 1408 | * | 13,15832 | 2,382069457 | 0,905685 | 2,63013 | 0,008535 | 0,881662676 | 1,2763 | Distal Intergenic | 3 | 1,31E+08 | 1,32E+08 | 270939 | 1 | 1,03E+08 | ENSMUST00000197726.2 | -90181 | | Gm29865 | predicted gene, 29865 |
| chr6 | 3499061 | 3501582 | 2522 | * | 24,78829 | 2,382644035 | 0,815902 | 2,920257 | 0,003497 | 0,582232389 | 2,880017 | Promoter (<=1kb) | 6 | 3498426 | 3521345 | 22920 | 1 | 73288 | ENSMUST00000184780.8 | 635 | ENSMUSG00000001376 | Vps50 | VPS50 EARP/GARPII complex subunit |
| chr7 | 41055161 | 41056469 | 1309 | * | 13,95534 | 2,38287191 | 0,837572 | 2,844977 | 0,004441 | 0,820796659 | 0,938682 | Intron (ENSMUST0000( | 7 | 41043122 | 41046871 | 3750 | 1 | 1,03E+08 | ENSMUST00000170929.2 | 12039 | ENSMUSG00000091864 | Gm17102 | predicted gene 17102 |
| chr11 | 68968231 | 68970147 | 1917 | * | 18,79021 | 2,383026746 | 0,783409 | 3,041869 | 0,002351 | 0,551991925 | 2,221611 | Distal Intergenic | 11 | 68964253 | 68964387 | 135 | 1 | 1E+08 | ENSMUST00000082965.3 | 3978 | ENSMUSG00000064899 | Snord118 | small nucleolar RNA, C/D box 118 |
| chr13 | 1,14E+08 | 1,14E+08 | 1917 | * | 16,9961 | 2,383218291 | 0,917842 | 2,596545 | 0,009417 | 0,896435577 | 1,2763 | Exon (ENSMUST00000( | 13 | 1,13E+08 | 1,14E+08 | 28109 | 1 | 408066 | ENSMUST00000135096.2 | 15469 | ENSMUSG00000021763 | Cspg4b | chondroitin sulfate proteoglycan 4B |
| chr15 | 9970186 | 9971510 | 1325 | * | 11,87701 | 2,383474449 | 0,908103 | 2,624674 | 0,008673 | 0,884485418 | 1,2763 | Distal Intergenic | 15 | 10177324 | 10329474 | 152151 | 1 | 19116 | ENSMUST00000005400.13 | -205814 | ENSMUSG00000005268 | Prlr | prolactin receptor |
| chr11 | 62766974 | 62768945 | 1972 | * | 16,40239 | 2,383651029 | 0,853871 | 2,791583 | 0,005245 | 0,684274806 | 2,386008 | Promoter (1-2kb) | 11 | 62765685 | 62767579 | 1895 | 1 | 213980 | ENSMUST00000127646.2 | 1289 | ENSMUSG00000090173 | Fbxw10 | F-box and WD-40 domain protein 10 |
| chr4 | 1,06E+08 | 1,07E+08 | 2646 | * | 22,34863 | 2,384382269 | 0,789691 | 3,019386 | 0,002533 | 0,516270433 | 2,951933 | Distal Intergenic | 4 | 1,06E+08 | 1,06E+08 | 1652 | 1 | 230576 | ENSMUST00000136505.2 | 4862 | ENSMUSG00000034919 | Ttc22 | tetratricopeptide repeat domain 22 |
| chr11 | 17895610 | 17897375 | 1766 | * | 13,49946 | 2,384831141 | 0,867776 | 2,748211 | 0,005992 | 0,819350294 | 1,2763 | Exon (ENSMUST00000( | 11 | 17888756 | 17903875 | 15120 | 2 | 68145 | ENSMUST00000076661.7 | 6500 | ENSMUSG00000016984 | Etaa1 | Ewing tumor-associated antigen 1 |
| chr11 | 1,15E+08 | 1,15E+08 | 2645 | * | 27,40347 | 2,385375113 | 0,764414 | 3,120529 | 0,001805 | 0,463890057 | 2,880017 | Promoter (<=1kb) | 11 | 1,15E+08 | 1,15E+08 | 17893 | 2 | 328035 | ENSMUST00000056153.8 | 0 | ENSMUSG00000044788 | Fads6 | fatty acid desaturase domain family, member 6 |
| chr1 | 32935979 | 32937297 | 1319 | * | 18,56378 | 2,386432589 | 0,813573 | 2,933276 | 0,003354 | 0,611385361 | 2,386008 | Distal Intergenic | 1 | 33231074 | 33231183 | 110 | 1 | 1,15E+08 | ENSMUST00000158125.3 | -293777 | ENSMUSG00000088750 | Gm24901 | predicted gene, 24901 |
| chr5 | 1,2E+08 | 1,2E+08 | 3281 | * | 26,5117 | 2,386875306 | 0,73278 | 3,25729 | 0,001125 | 0,399754181 | 2,67243 | Distal Intergenic | 5 | 1,2E+08 | 1,2E+08 | 6144 | 2 | 433944 | ENSMUST00000135385.3 | -19981 | ENSMUSG00000086847 | Tbx3os2 | T-box 3, opposite strand 2 |
| chr3 | 1,32E+08 | 1,32E+08 | 1304 | * | 14,84089 | 2,387187341 | 0,900392 | 2,651277 | 0,008019 | 0,778321623 | 2,141612 | Distal Intergenic | 3 | 1,32E+08 | 1,32E+08 | 95013 | 1 | 56811 | ENSMUST00000029665.7 | 213972 | ENSMUSG00000028031 | Dkk2 | dickkopf WNT signaling pathway inhibitor 2 |
| chr7 | 6364851 | 6366883 | 2033 | * | 55,29647 | 2,387722269 | 0,922829 | 2,587395 | 0,00967 | 0,764024179 | 2,787255 | Promoter (<=1kb) | 7 | 6366279 | 6384708 | 18430 | 1 | 330463 | ENSMUST00000207347.2 | 0 | ENSMUSG00000055150 | Zfp78 | zinc finger protein 78 |
| chr1 | 84858285 | 84860964 | 2680 | * | 31,96941 | 2,387867858 | 0,63535 | 3,75835 | 0,000171 | 0,145302235 | 2,951062 | Exon (ENSMUST00000( | 1 | 84846872 | 84878208 | 31337 | 1 | 66153 | ENSMUST00000151737.2 | 11413 | ENSMUSG00000073633 | Fbxo36 | F-box protein 36 |
| chr14 | 25841628 | 25843765 | 2138 | * | 19,97991 | 2,388230669 | 0,756709 | 3,156076 | 0,001599 | 0,452555282 | 2,713306 | Promoter (<=1kb) | 14 | 25842580 | 25887228 | 44649 | 1 | 11744 | ENSMUST00000022416.15 | 0 | ENSMUSG00000021866 | Anxa11 | annexin A11 |
| chr10 | 80524723 | 80526976 | 2254 | * | 24,34496 | 2,3882631 | 0,750152 | 3,183707 | 0,001454 | 0,428190429 | 2,858511 | Promoter (2-3kb) | 10 | 80527052 | 80529570 | 2519 | 2 | 208228 | ENSMUST00000220138.2 | 2594 | ENSMUSG00000003348 | Mob3a | MOB kinase activator 3A |
| chr11 | 95960877 | 95962455 | 1579 | * | 13,27918 | 2,388299511 | 0,860462 | 2,775601 | 0,00551 | 0,834498178 | 1,075735 | 3' UTR | 11 | 95959693 | 95965863 | 6171 | 2 | 11951 | ENSMUST00000178611.8 | 3408 | ENSMUSG00000006057 | Atp5g1 | ATP synthase, H+ transporting, mitochondrial F0 complex, subunit C1 (subunit 9) |
| chr3 | 88412720 | 88414130 | 1411 | * | 16,27052 | 2,388327327 | 0,794698 | 3,005329 | 0,002653 | 0,679990945 | 1,2763 | Promoter (2-3kb) | 3 | 88389153 | 88410644 | 21492 | 2 | 16905 | ENSMUST00000135494.4 | -2076 | ENSMUSG00000028063 | Lmna | lamin A |
| chr6 | 30283836 | 30284971 | 1136 | * | 16,50112 | 2,388610607 | 0,852488 | 2,801928 | 0,00508 | 0,696406045 | 2,141612 | Intron (ENSMUST0000( | 6 | 30214860 | 30304521 | 89662 | 2 | 22214 | ENSMUST00000094543.3 | 19550 | ENSMUSG00000039159 | Ube2h | ubiquitin-conjugating enzyme E2H |
| chr11 | 5736405 | 5737976 | 1572 | * | 15,24724 | 2,388768116 | 0,849121 | 2,813225 | 0,004905 | 0,792225394 | 1,214904 | Promoter (<=1kb) | 11 | 5738488 | 5750629 | 12142 | 1 | 13169 | ENSMUST00000109845.8 | -512 | ENSMUSG00000020476 | Dbnl | drebrin-like |
| chr11 | 97362859 | 97364179 | 1321 | * | 11,84695 | 2,39036302 | 0,914615 | 2,61352 | 0,008961 | 0,971104542 | 0,738441 | Intron (ENSMUST0000( | 11 | 97381970 | 97393224 | 11255 | 1 | 58996 | ENSMUST00000093940.4 | -17791 | ENSMUSG00000049807 | Arhgap23 | Rho GTPase activating protein 23 |
| chr1 | 1,14E+08 | 1,14E+08 | 2555 | * | 21,80624 | 2,391139015 | 0,862098 | 2,773629 | 0,005543 | 0,668267959 | 2,880017 | Distal Intergenic | 1 | 1,16E+08 | 1,16E+08 | 15632 | 2 | 1,08E+08 | ENSMUST00000186137.7 | 1590163 | | Gm50605 | predicted gene, 50605 |
| chr6 | 86769748 | 86771846 | 2099 | * | 24,70903 | 2,391529084 | 0,771996 | 3,097851 | 0,001949 | 0,477253929 | 2,858511 | Promoter (<=1kb) | 6 | 86713823 | 86770566 | 56744 | 2 | 11746 | ENSMUST00000001187.15 | 0 | ENSMUSG00000029994 | Anxa4 | annexin A4 |
| chr6 | 29280180 | 29282246 | 2067 | * | 22,8736 | 2,391684789 | 0,71633 | 3,338803 | 0,000841 | 0,328946709 | 2,951062 | Promoter (<=1kb) | 6 | 29281140 | 29290679 | 9540 | 1 | 245884 | ENSMUST00000115286.4 | 0 | ENSMUSG00000079652 | Garin1a | golgi associated RAB2 interactor 1A |
| chr16 | 20219951 | 20221524 | 1574 | * | 15,17426 | 2,39213522 | 0,829317 | 2,884466 | 0,003921 | 0,743408116 | 1,2763 | Promoter (<=1kb) | 16 | 20217567 | 20219036 | 1470 | 2 | 27416 | ENSMUST00000128978.2 | -915 | ENSMUSG00000022822 | Abcc5 | ATP-binding cassette, sub-family C (CFTR/MRP), member 5 |
| chr12 | 25147591 | 25150584 | 2994 | * | 38,260489 | 2,392269541 | 0,5661718 | 4,2253422 | 2,386E-05 | 0,048727933 | 2,3383871 | Promoter (<=1kb) | 12 | 25143798 | 25147139 | 3342 | 2 | 15902 | ENSMUST00000221761.2 | -452 | ENSMUSG00000020644 | Id2 | inhibitor of DNA binding 2 |
| chr5 | 1,23E+08 | 1,23E+08 | 3241 | * | 31,05049 | 2,393465638 | 0,600118 | 3,988325 | 6,65E-05 | 0,082272052 | 2,880017 | Distal Intergenic | 5 | 1,23E+08 | 1,23E+08 | 45886 | 2 | 11938 | ENSMUST00000031423.10 | -21119 | ENSMUSG00000029467 | Atp2a2 | ATPase, Ca++ transporting, cardiac muscle, slow twitch 2 |
| chr13 | 47433376 | 47435212 | 1837 | * | 18,80952 | 2,394263023 | 0,847593 | 2,824779 | 0,004731 | 0,663408464 | 2,507013 | Distal Intergenic | 13 | 47514862 | 47593030 | 78169 | 1 | 1,03E+08 | ENSMUST00000224000.2 | -79650 | | Gm35733 | predicted gene, 35733 |
| chr1 | 59551085 | 59553072 | 1988 | * | 19,67861 | 2,394501412 | 0,757865 | 3,159536 | 0,00158 | 0,453798789 | 2,663619 | Promoter (2-3kb) | 1 | 59555423 | 59673668 | 118246 | 1 | 381260 | ENSMUST00000114243.8 | -2351 | ENSMUSG00000047361 | Gm973 | predicted gene 973 |
| chr5 | 1,48E+08 | 1,48E+08 | 2431 | * | 22,44975 | 2,394732239 | 0,700329 | 3,419438 | 0,000628 | 0,273126507 | 3,471678 | Promoter (<=1kb) | 5 | 1,48E+08 | 1,48E+08 | 72495 | 2 | 11987 | ENSMUST00000048116.15 | 0 | ENSMUSG00000041313 | Slc7a1 | solute carrier family 7 (cationic amino acid transporter, y+ system), member 1 |
| chr3 | 57408274 | 57410353 | 2080 | * | 21,03079 | 2,394988545 | 0,893382 | 2,680811 | 0,007344 | 0,723275538 | 2,67243 | Exon (ENSMUST00000( | 3 | 57435653 | 57436202 | 550 | 1 | 1,15E+08 | ENSMUST00000120831.2 | -25300 | | Gm16016 | NEDD8-conjugating enzyme UBE2F pseudogene |
| chr9 | 1,17E+08 | 1,17E+08 | 1266 | * | 15,18216 | 2,395520585 | 0,926863 | 2,584547 | 0,009751 | 0,908787948 | 1,214904 | Intron (ENSMUST0000( | 9 | 1,16E+08 | 1,17E+08 | 239998 | 2 | 207181 | ENSMUST00000172469.3 | -37983 | ENSMUSG00000039607 | Rbms3 | RNA binding motif, single stranded interacting protein |
| chr18 | 62148334 | 62149979 | 1646 | * | 15,00503 | 2,396009657 | 0,878962 | 2,725953 | 0,006412 | 0,81218711 | 1,41696 | 3' UTR | 18 | 62111030 | 62148790 | 37761 | 1 | 225608 | ENSMUST00000236759.2 | 37304 | ENSMUSG00000045629 | Sh3tc2 | SH3 domain and tetratricopeptide repeats 2 |
| chr17 | 46062529 | 46066328 | 3800 | * | 65,2245 | 2,396166527 | 0,4936758 | 4,8537251 | 1,212E-06 | 0,00336247 | 4,0557576 | Promoter (1-2kb) | 17 | 46067636 | 46099520 | 31885 | 1 | 433111 | ENSMUST00000180519.2 | -1308 | ENSMUSG00000097727 | F630040K05Rik | RIKEN cDNA F630040K05 gene |
| chr7 | 19241839 | 19243513 | 1675 | * | 19,73968 | 2,397385003 | 0,788272 | 3,041315 | 0,002355 | 0,524368688 | 2,663619 | Promoter (<=1kb) | 7 | 19238411 | 19242292 | 3882 | 2 | 232946 | ENSMUST00000077408.8 | 0 | ENSMUSG00000057667 | Bloc1s3 | biogenesis of lysosomal organelles complex-1, subunit 3 |
| chr17 | 25722280 | 25725875 | 3596 | * | 30,78498 | 2,397539881 | 0,650475 | 3,685829 | 0,000228 | 0,178303794 | 2,725491 | Distal Intergenic | 17 | 25708849 | 25716262 | 7414 | 2 | 20609 | ENSMUST00000051864.5 | -6018 | ENSMUSG00000050824 | Sstr5 | somatostatin receptor 5 |
| chr10 | 61120908 | 61123060 | 2153 | * | 27,56587 | 2,397610549 | 0,775098 | 3,0933 | 0,001979 | 0,474900193 | 2,951062 | Distal Intergenic | 10 | 61133612 | 61140459 | 6848 | 1 | 18646 | ENSMUST00000219375.2 | -10552 | ENSMUSG00000037202 | Prf1 | perforin 1 (pore forming protein) |
| chr16 | 73817928 | 73819873 | 1946 | * | 16,9815 | 2,397866633 | 0,836963 | 2,864962 | 0,004171 | 0,669586355 | 2,141612 | Intron (ENSMUST0000( | 16 | 73778989 | 73824777 | 45789 | 2 | 268902 | ENSMUST00000147408.3 | 4904 | ENSMUSG00000052516 | Robo2 | roundabout guidance receptor 2 |
| chr3 | 1,57E+08 | 1,57E+08 | 865 | * | 15,14464 | 2,398175578 | 0,89101 | 2,691524 | 0,007113 | 0,850518759 | 1,2763 | Distal Intergenic | 3 | 1,57E+08 | 1,57E+08 | 107 | 2 | 1,15E+08 | ENSMUST00000240136.1 | -40840 | ENSMUSG00000118829 | Gm23379 | predicted gene, 23379 |
| chr8 | 15042256 | 15044220 | 1965 | * | 31,21821 | 2,398233154 | 0,628233 | 3,817427 | 0,000135 | 0,125710024 | 2,858511 | Promoter (1-2kb) | 8 | 15044121 | 15046042 | 1922 | 2 | 1,03E+08 | ENSMUST00000246690.1 | 1822 | | Gm35149 | predicted gene, 35149 |
| chr18 | 31999095 | 32001450 | 2356 | * | 21,81706 | 2,398349999 | 0,739171 | 3,244647 | 0,001176 | 0,377581283 | 3,104559 | Intron (ENSMUST0000( | 18 | 32015195 | 32019423 | 4229 | 1 | 74320 | ENSMUST00000234137.2 | -13745 | ENSMUSG00000024400 | Wdr33 | WD repeat domain 33 |
| chr7 | 80390647 | 80392478 | 1832 | * | 17,46752 | 2,398723275 | 0,868294 | 2,76257 | 0,005735 | 0,683210259 | 2,663619 | Exon (ENSMUST00000( | 7 | 80376527 | 80379996 | 3470 | 2 | 29875 | ENSMUST00000205606.2 | -10651 | ENSMUSG00000030536 | Iqgap1 | IQ motif containing GTPase activating protein 1 |
| chr6 | 84213163 | 84214312 | 1150 | * | 14,84256 | 2,39886567 | 0,844705 | 2,839887 | 0,004513 | 0,777172375 | 1,214904 | Intron (ENSMUST0000( | 6 | 84187466 | 84188023 | 558 | 1 | 26903 | ENSMUST00000128965.2 | 25697 | ENSMUSG00000033788 | Dysf | dysferlin |
| chr3 | 1,08E+08 | 1,08E+08 | 1339 | * | 16,20754 | 2,399740459 | 0,82815 | 2,897711 | 0,003759 | 0,734252598 | 1,2763 | Distal Intergenic | 3 | 1,08E+08 | 1,08E+08 | 33441 | 2 | 229709 | ENSMUST00000029490.15 | -6621 | ENSMUSG00000027893 | Ahcyl1 | S-adenosylhomocysteine hydrolase-like 1 |
| chr6 | 40166230 | 40167914 | 1685 | * | 24,77479 | 2,399761202 | 0,702394 | 3,416546 | 0,000634 | 0,297590871 | 2,880017 | Intron (ENSMUST0000( | 6 | 40087185 | 40225371 | 138187 | 1 | 434008 | ENSMUST00000061740.8 | 79045 | ENSMUSG00000057716 | Tmem178b | transmembrane protein 178B |
| chr7 | 42930680 | 42933246 | 2567 | * | 28,75081 | 2,400241163 | 0,726491 | 3,303883 | 0,000954 | 0,352029189 | 2,880017 | Distal Intergenic | 7 | 42903163 | 42915072 | 11910 | 1 | 1,03E+08 | ENSMUST00000243584.1 | 27517 | | Gm35588 | predicted gene, 35588 |
| chr6 | 54796135 | 54800274 | 4140 | * | 49,732533 | 2,400380469 | 0,4855776 | 4,9433507 | 7,679E-07 | 0,003573187 | 2,3383871 | Promoter (1-2kb) | 6 | 54794433 | 54862096 | 67664 | 1 | 387524 | ENSMUST00000127331.2 | 1702 | ENSMUSG00000058446 | Znrf2 | zinc and ring finger 2 |
| chr10 | 75669716 | 75671297 | 1582 | * | 16,67988 | 2,400662202 | 0,783185 | 3,065256 | 0,002175 | 0,532631848 | 2,318434 | Promoter (<=1kb) | 10 | 75667679 | 75670715 | 3037 | 2 | 14872 | ENSMUST00000038257.7 | 0 | ENSMUSG00000033318 | Gstt2 | glutathione S-transferase, theta 2 |
| chr12 | 83609697 | 83611731 | 2035 | * | 19,54274 | 2,4014164 | 0,925075 | 2,595914 | 0,009434 | 0,764402529 | 2,713306 | Promoter (<=1kb) | 12 | 83599186 | 83609234 | 10049 | 2 | 217695 | ENSMUST00000222448.2 | -463 | ENSMUSG00000042628 | Zfyve1 | zinc finger, FYVE domain containing 1 |
| chr4 | 1,5E+08 | 1,5E+08 | 1944 | * | 18,05022 | 2,401595047 | 0,895537 | 2,681737 | 0,007324 | 0,7591629 | 2,154504 | Distal Intergenic | 4 | 1,5E+08 | 1,5E+08 | 102 | 1 | 723848 | ENSMUST00000083559.3 | -6668 | ENSMUSG00000065493 | Mir34a | microRNA 34a |
| chr15 | 27877430 | 27878492 | 1063 | * | 15,49634 | 2,402365578 | 0,889754 | 2,700033 | 0,006933 | 0,844744375 | 1,2763 | Intron (ENSMUST0000( | 15 | 27746986 | 27841827 | 94842 | 2 | 223435 | ENSMUST00000226644.2 | -35603 | ENSMUSG00000022263 | Trio | triple functional domain (PTPRF interacting) |
| chr4 | 1,43E+08 | 1,43E+08 | 1516 | * | 15,32841 | 2,402894735 | 0,864621 | 2,77913 | 0,00545 | 0,801134483 | 1,2763 | Intron (ENSMUST0000( | 4 | 1,43E+08 | 1,43E+08 | 63 | 2 | 1,02E+08 | ENSMUST00000197026.3 | -28369 | ENSMUSG00000106597 | Mir7021 | microRNA 7021 |
| chr5 | 1,07E+08 | 1,07E+08 | 2508 | * | 21,45278 | 2,402909547 | 0,840035 | 2,860487 | 0,00423 | 0,605265376 | 3,104559 | Distal Intergenic | 5 | 1,07E+08 | 1,07E+08 | 4086 | 1 | 12545 | ENSMUST00000123546.2 | 14746 | ENSMUSG00000029283 | Cdc7 | cell division cycle 7 (S. cerevisiae) |
| chr10 | 1,17E+08 | 1,17E+08 | 1402 | * | 13,24355 | 2,40311708 | 0,909759 | 2,641487 | 0,008254 | 0,924754734 | 0,918255 | Intron (ENSMUST0000( | 10 | 1,17E+08 | 1,17E+08 | 12814 | 2 | 12461 | ENSMUST00000047672.9 | -28023 | ENSMUSG00000034024 | Cct2 | chaperonin containing Tcp1, subunit 2 (beta) |
| chr6 | 38408194 | 38411030 | 2837 | * | 39,528897 | 2,403356623 | 0,5088238 | 4,7233571 | 2,32E-06 | 0,007486076 | 2,7254908 | Promoter (<=1kb) | 6 | 38410885 | 38489698 | 78814 | 1 | 320538 | ENSMUST00000160583.8 | 0 | ENSMUSG00000038538 | Ubn2 | ubinuclein 2 |
| chr8 | 1,08E+08 | 1,08E+08 | 1984 | * | 16,36082 | 2,403867596 | 0,839055 | 2,864971 | 0,00417 | 0,650270742 | 2,386008 | Promoter (<=1kb) | 8 | 1,08E+08 | 1,08E+08 | 82 | 2 | 1,02E+08 | ENSMUST00000184187.3 | 354 | ENSMUSG00000098671 | Mir7075 | microRNA 7075 |
| chr15 | 78483788 | 78486158 | 2371 | * | 22,64844 | 2,403967899 | 0,797764 | 3,013383 | 0,002584 | 0,524856133 | 2,880017 | Promoter (2-3kb) | 15 | 78481349 | 78503318 | 21970 | 1 | 72318 | ENSMUST00000229248.2 | 2439 | ENSMUSG00000018008 | Cyth4 | cytohesin 4 |
| chr3 | 83976308 | 83977848 | 1541 | * | 15,74597 | 2,406004943 | 0,932567 | 2,579981 | 0,009881 | 0,777172375 | 2,663619 | Distal Intergenic | 3 | 83805516 | 83947482 | 141967 | 2 | 229473 | ENSMUST00000052342.9 | -28826 | ENSMUSG00000033767 | Tmem131l | transmembrane 131 like |
| chr15 | 72961636 | 72963571 | 1936 | * | 25,60057 | 2,406039431 | 0,720486 | 3,339469 | 0,000839 | 0,333539548 | 2,858511 | Promoter (<=1kb) | 15 | 72962241 | 72965924 | 3684 | 1 | 93696 | ENSMUST00000089765.9 | 0 | ENSMUSG00000068391 | Chrac1 | chromatin accessibility complex 1 |
| chr13 | 49366707 | 49368039 | 1333 | * | 25,21168 | 2,406286585 | 0,744509 | 3,232046 | 0,001229 | 0,402652506 | 2,858511 | Promoter (1-2kb) | 13 | 49356426 | 49369336 | 12911 | 2 | 68480 | ENSMUST00000221443.2 | 1297 | ENSMUSG00000037960 | Card19 | caspase recruitment domain family, member 19 |
| chr2 | 31843402 | 31846869 | 3468 | * | 60,863879 | 2,406402439 | 0,5013255 | 4,8000799 | 1,586E-06 | 0,00399751 | 4,0557576 | Promoter (2-3kb) | 2 | 31840529 | 31861350 | 20822 | 1 | 108897 | ENSMUST00000138833.8 | 2873 | ENSMUSG00000001864 | Aif1l | allograft inflammatory factor 1-like |
| chr6 | 86169253 | 86170885 | 1633 | * | 18,29794 | 2,40702791 | 0,833565 | 2,887629 | 0,003882 | 0,618282231 | 2,663619 | Promoter (1-2kb) | 6 | 86172205 | 86252701 | 80497 | 1 | 21802 | ENSMUST00000032066.13 | -1320 | ENSMUSG00000029999 | Tgfa | transforming growth factor alpha |
| chr16 | 20592607 | 20594572 | 1966 | * | 15,75349 | 2,407043886 | 0,867347 | 2,775179 | 0,005517 | 0,697016988 | 2,318434 | Distal Intergenic | 16 | 20564058 | 20564222 | 165 | 1 | 1,15E+08 | ENSMUST00000083438.3 | 28549 | ENSMUSG00000065372 | Gm22741 | predicted gene, 22741 |
| chr4 | 1,24E+08 | 1,24E+08 | 1373 | * | 16,82174 | 2,407240711 | 0,906339 | 2,656006 | 0,007907 | 0,755058193 | 2,386008 | Distal Intergenic | 4 | 1,24E+08 | 1,24E+08 | 14265 | 1 | 54170 | ENSMUST00000155454.2 | 100563 | ENSMUSG00000028646 | Rragc | Ras-related GTP binding C |
| chr19 | 46956513 | 46958348 | 1836 | * | 17,94346 | 2,407392043 | 0,874288 | 2,753545 | 0,005895 | 0,684274806 | 2,663619 | Promoter (<=1kb) | 19 | 46871756 | 46958007 | 86252 | 2 | 76952 | ENSMUST00000235234.2 | 0 | ENSMUSG00000025041 | Nt5c2 | 5'-nucleotidase, cytosolic II |
| chr11 | 4737624 | 4739473 | 1850 | * | 18,33144 | 2,407516095 | 0,835468 | 2,881636 | 0,003956 | 0,659240897 | 2,154504 | Promoter (2-3kb) | 11 | 4717953 | 4735107 | 17155 | 2 | 18016 | ENSMUST00000141342.3 | -2517 | ENSMUSG00000009073 | Nf2 | neurofibromin 2 |
| chr2 | 1,21E+08 | 1,21E+08 | 1804 | * | 16,7533 | 2,407611427 | 0,902651 | 2,667267 | 0,007647 | 0,859319705 | 1,2763 | Intron (ENSMUST0000( | 2 | 1,21E+08 | 1,21E+08 | 14997 | 2 | 68968 | ENSMUST00000110700.2 | -29812 | ENSMUSG00000027284 | Cdan1 | congenital dyserythropoietic anemia, type I (human) |
| chr9 | 65193422 | 65196285 | 2864 | * | 21,48794 | 2,408260236 | 0,897657 | 2,682828 | 0,0073 | 0,722832903 | 2,67243 | Distal Intergenic | 9 | 65201542 | 65237939 | 36398 | 1 | 270166 | ENSMUST00000015501.11 | -5257 | ENSMUSG00000015357 | Clpx | caseinolytic mitochondrial matrix peptidase chaperone subunit |
| chr12 | 56464634 | 56465821 | 1188 | * | 13,07198 | 2,408394703 | 0,884873 | 2,72174 | 0,006494 | 0,890232174 | 0,918255 | Distal Intergenic | 12 | 56375088 | 56392679 | 17592 | 2 | 217588 | ENSMUST00000021416.9 | -71955 | ENSMUSG00000021028 | Mbip | MAP3K12 binding inhibitory protein 1 |
| chr6 | 29851791 | 29854548 | 2758 | * | 33,26703 | 2,409014745 | 0,637059 | 3,781464 | 0,000156 | 0,143892051 | 2,725491 | Promoter (<=1kb) | 6 | 29853759 | 29909214 | 55456 | 1 | 74340 | ENSMUST00000064872.13 | 0 | ENSMUSG00000029772 | Ahcyl2 | S-adenosylhomocysteine hydrolase-like 2 |
| chr1 | 1,63E+08 | 1,63E+08 | 1598 | * | 14,78715 | 2,409183769 | 0,876333 | 2,749165 | 0,005975 | 0,798887389 | 1,41696 | Distal Intergenic | 1 | 1,63E+08 | 1,63E+08 | 18762 | 2 | 98376 | ENSMUST00000045138.6 | -121360 | ENSMUSG00000040124 | Gorab | golgin, RAB6-interacting |
| chr13 | 111933698 | 111936118 | 2421 | * | 37,839535 | 2,409470761 | 0,5545774 | 4,3446974 | 1,395E-05 | 0,029206134 | 2,7254908 | Intron (ENSMUST000001( | 13 | 111904518 | 111944849 | 40332 | 2 | 26401 | ENSMUST00000145055.2 | 8731 | ENSMUSG00000021754 | Map3k1 | mitogen-activated protein kinase kinase kinase 1 |
| chr19 | 25485504 | 25487044 | 1541 | * | 14,65157 | 2,409537538 | 0,926072 | 2,601889 | 0,009271 | 0,87775273 | 1,41696 | Promoter (2-3kb) | 19 | 25483071 | 25537629 | 54559 | 1 | 50796 | ENSMUST00000160814.2 | 2433 | ENSMUSG00000024837 | Dmrt1 | doublesex and mab-3 related transcription factor 1 |
| chr5 | 66793123 | 66795292 | 2170 | * | 19,24355 | 2,409739098 | 0,706554 | 3,410553 | 0,000648 | 0,322750682 | 2,386008 | Intron (ENSMUST0000( | 5 | 66456071 | 66776127 | 320057 | 2 | 11787 | ENSMUST00000159512.8 | -16996 | ENSMUSG00000029207 | Apbb2 | amyloid beta (A4) precursor protein-binding, family B, member 2 |
| chr14 | 47770154 | 47771788 | 1635 | * | 14,80107 | 2,409870617 | 0,837604 | 2,877101 | 0,004013 | 0,7299084 | 1,41696 | Distal Intergenic | 14 | 47763536 | 47766733 | 3198 | 1 | 78938 | ENSMUST00000228634.2 | 6618 | ENSMUSG00000037536 | Fbxo34 | F-box protein 34 |
| chr5 | 1,05E+08 | 1,05E+08 | 2240 | * | 19,25035 | 2,410041793 | 0,866231 | 2,782217 | 0,005399 | 0,686838043 | 2,386008 | Promoter (2-3kb) | 5 | 1,05E+08 | 1,05E+08 | 24074 | 2 | 626391 | ENSMUST00000191135.2 | 2126 | ENSMUSG00000072774 | Zfp951 | zinc finger protein 951 |
| chr10 | 82949500 | 82951954 | 2455 | * | 18,00384 | 2,410618175 | 0,791021 | 3,047478 | 0,002308 | 0,531203841 | 2,507013 | Intron (ENSMUST0000( | 10 | 82974984 | 82977872 | 2889 | 2 | 1,15E+08 | ENSMUST00000160997.2 | 25918 | | 1700025N21Rik | RIKEN cDNA 1700025N21 gene |
| chr19 | 5836065 | 5838815 | 2751 | * | 39,235837 | 2,41068203 | 0,5203386 | 4,6329099 | 3,606E-06 | 0,010488962 | 2,7254908 | Distal Intergenic | 19 | 5849028 | 5849113 | 86 | 2 | 72289 | ENSMUST00020182077.1 | 10298 | ENSMUSG00000092341 | Malat1 | metastasis associated lung adenocarcinoma transcript 1 (non-coding RNA) |
| chr3 | 5337806 | 5340050 | 2245 | * | 13,83194 | 2,410740799 | 0,898637 | 2,682665 | 0,007304 | 0,852537034 | 1,2763 | Intron (ENSMUST0000( | 3 | 5306732 | 5480917 | 174186 | 1 | 80892 | ENSMUST00000176383.3 | 31074 | ENSMUSG00000025255 | Zfhx4 | zinc finger homeodomain 4 |
| chr19 | 5735602 | 5738066 | 2465 | * | 26,76786 | 2,412061465 | 0,79357 | 3,039509 | 0,00237 | 0,50985457 | 2,858511 | Promoter (<=1kb) | 19 | 5714668 | 5738177 | 23510 | 2 | 104401 | ENSMUST00000068169.12 | 111 | ENSMUSG00000054874 | Pcnx3 | pecanex homolog 3 |
| chr2 | 1,55E+08 | 1,55E+08 | 1898 | * | 17,29879 | 2,412199244 | 0,793337 | 3,040572 | 0,002361 | 0,539620856 | 2,423594 | Distal Intergenic | 2 | 1,55E+08 | 1,55E+08 | 3212 | 1 | 75608 | ENSMUST00000136788.2 | 10354 | ENSMUSG00000038467 | Chmp4b | charged multivesicular body protein 4B |
| chr19 | 43711095 | 43713185 | 2091 | * | 18,43716 | 2,412697818 | 0,857375 | 2,814052 | 0,004892 | 0,687319907 | 2,154504 | Promoter (1-2kb) | 19 | 43714806 | 43716575 | 1770 | 1 | 93685 | ENSMUST00000131293.2 | -1621 | ENSMUSG00000025192 | Entpd7 | ectonucleoside triphosphate diphosphohydrolase 7 |
| chr11 | 76822911 | 76824840 | 1930 | * | 15,88567 | 2,413277465 | 0,927692 | 2,601378 | 0,009285 | 0,802671652 | 2,141612 | Intron (ENSMUST0000( | 11 | 76815635 | 76841962 | 26328 | 1 | 104184 | ENSMUST00000140781.9 | 7276 | ENSMUSG00000020840 | Blmh | bleomycin hydrolase |
| chr1 | 1,53E+08 | 1,53E+08 | 1727 | * | 18,27593 | 2,414220377 | 0,864621 | 2,79223 | 0,005235 | 0,702222274 | 2,154504 | Intron (ENSMUST0000( | 1 | 1,53E+08 | 1,53E+08 | 40874 | 1 | 226518 | ENSMUST00000187569.2 | -19508 | ENSMUSG00000042751 | Nmnat2 | nicotinamide nucleotide adenylyltransferase 2 |
| chr14 | 60892939 | 60894967 | 2029 | * | 22,92015 | 2,415837212 | 0,774654 | 3,118601 | 0,001817 | 0,477253929 | 2,67243 | Intron (ENSMUST0000( | 14 | 60872204 | 60928342 | 56139 | 1 | 219140 | ENSMUST00000159135.2 | 20735 | ENSMUSG00000021990 | Spata13 | spermatogenesis associated 13 |
| chr12 | 46211526 | 46212934 | 1409 | * | 13,46638 | 2,416765066 | 0,911337 | 2,65189 | 0,008004 | 0,92207189 | 0,918255 | Distal Intergenic | 12 | 46031126 | 46031332 | 207 | 1 | 1,15E+08 | ENSMUST00000082866.3 | 180400 | ENSMUSG00000064800 | Gm24948 | predicted gene, 24948 |
| chr18 | 68670405 | 68672402 | 1998 | * | 16,15318 | 2,416949232 | 0,865817 | 2,791524 | 0,005246 | 0,705025276 | 2,141612 | Intron (ENSMUST0000( | 18 | 68692493 | 68699212 | 6720 | 1 | 433197 | ENSMUST00000238121.2 | -20091 | | Gm29677 | predicted gene, 29677 |
| chr12 | 40848714 | 40850352 | 1639 | * | 14,39879 | 2,417523875 | 0,830353 | 2,911442 | 0,003598 | 0,824134241 | 0,738441 | Exon (ENSMUST00000( | 12 | 40844233 | 40848081 | 3849 | 1 | 238130 | ENSMUST00000220850.2 | 4481 | ENSMUSG00000035954 | Dock4 | dedicator of cytokinesis 4 |
| chr8 | 1,07E+08 | 1,07E+08 | 2703 | * | 31,70729 | 2,417806508 | 0,675635 | 3,578571 | 0,000345 | 0,226269722 | 2,725491 | Distal Intergenic | 8 | 1,07E+08 | 1,07E+08 | 11272 | 1 | 22751 | ENSMUST00000213045.2 | -26592 | ENSMUSG00000031907 | Zfp90 | zinc finger protein 90 |
| chr18 | 42632920 | 42634741 | 1822 | * | 13,68415 | 2,41793266 | 0,905127 | 2,671375 | 0,007554 | 0,909022873 | 0,938682 | Distal Intergenic | 18 | 42644552 | 42708858 | 64307 | 1 | 56070 | ENSMUST00000237602.2 | -9811 | ENSMUSG00000024498 | Tcerg1 | transcription elongation regulator 1 (CA150) |
| chr8 | 84310220 | 84311480 | 1261 | * | 12,0817 | 2,418035683 | 0,904144 | 2,674393 | 0,007486 | 0,946556802 | 0,738441 | Exon (ENSMUST00000( | 8 | 84302673 | 84312548 | 9876 | 1 | 1,05E+08 | ENSMUST00000212132.2 | 7547 | | D830024N08Rik | RIKEN cDNA D830024N08 gene |
| chr3 | 27748598 | 27750357 | 1760 | * | 15,57094 | 2,418402597 | 0,86715 | 2,788911 | 0,005289 | 0,793995086 | 1,2763 | Intron (ENSMUST0000( | 3 | 27470311 | 27764547 | 294237 | 2 | 72007 | ENSMUST00000195008.6 | 14190 | ENSMUSG00000039286 | Fndc3b | fibronectin type III domain containing 3B |
| chr12 | 102671582 | 102675129 | 3548 | * | 61,775623 | 2,418996549 | 0,4960325 | 4,8766895 | 1,079E-06 | 0,003457727 | 3,439501 | Promoter (<=1kb) | 12 | 102534841 | 102671189 | 136349 | 2 | 217837 | ENSMUST00000046518.12 | -393 | ENSMUSG00000057963 | Itpk1 | inositol 1,3,4-triphosphate 5/6 kinase |
| chr4 | 1,34E+08 | 1,34E+08 | 1699 | * | 15,97294 | 2,419021222 | 0,810188 | 2,985754 | 0,002829 | 0,55292773 | 2,663619 | Promoter (2-3kb) | 4 | 1,34E+08 | 1,34E+08 | 12117 | 1 | 71904 | ENSMUST00000081525.4 | -2048 | ENSMUSG00000037348 | Paqr7 | progestin and adipoQ receptor family member VII |
| chr3 | 96168756 | 96172414 | 3659 | * | 41,52894 | 2,419221309 | 0,594158 | 4,071678 | 4,67E-05 | 0,075415099 | 2,338387 | Promoter (<=1kb) | 3 | 96168998 | 96170627 | 1630 | 2 | 97122 | ENSMUST00000171473.3 | 0 | ENSMUSG00000091405 | H4c14 | H4 clustered histone 14 |
| chr10 | 73923923 | 73925051 | 1129 | * | 15,43921 | 2,419964783 | 0,903789 | 2,677577 | 0,007416 | 0,837650061 | 1,41696 | Intron (ENSMUST0000( | 10 | 73782888 | 74482000 | 699113 | 1 | 11994 | ENSMUST00000195531.6 | 141035 | ENSMUSG00000052613 | Pcdh15 | protocadherin 15 |
| chr15 | 1,03E+08 | 1,03E+08 | 1560 | * | 22,23133 | 2,420045954 | 0,900599 | 2,687152 | 0,007206 | 0,721416593 | 2,67243 | Intron (ENSMUST0000( | 15 | 1,03E+08 | 1,03E+08 | 2309 | 1 | 15425 | ENSMUST00000001711.6 | -3264 | ENSMUSG00000001661 | Hoxc6 | homeobox C6 |
| chr5 | 1,45E+08 | 1,45E+08 | 1208 | * | 14,93455 | 2,420162105 | 0,85193 | 2,840799 | 0,0045 | 0,767135014 | 1,2763 | Promoter (<=1kb) | 5 | 1,45E+08 | 1,45E+08 | 13403 | 2 | 75788 | ENSMUST00000198621.2 | -181 | ENSMUSG00000038780 | Smurf1 | SMAD specific E3 ubiquitin protein ligase 1 |
| chr15 | 16239661 | 16241578 | 1918 | * | 20,50414 | 2,420256519 | 0,846163 | 2,860272 | 0,004233 | 0,665162479 | 2,221611 | Distal Intergenic | 15 | 16489101 | 16496898 | 7798 | 2 | 1,05E+08 | ENSMUST00000227529.2 | 255320 | ENSMUSG00000115614 | Gm41276 | predicted gene, 41276 |
| chr7 | 33928565 | 33930355 | 1791 | * | 20,87363 | 2,420339551 | 0,755059 | 3,205495 | 0,001348 | 0,426343337 | 2,67243 | Promoter (<=1kb) | 7 | 33900755 | 33929761 | 29007 | 2 | 14751 | ENSMUST00000038027.6 | 0 | ENSMUSG00000036427 | Gpi1 | glucose-6-phosphate isomerase 1 |

| | | | | | | | | | | | | | | | | | | | | | | | |
|---|---|---|---|---|---|---|---|---|---|---|---|---|---|---|---|---|---|---|---|---|---|---|---|
| chr19 | 3372458 | 3375076 | 2619 | * | 23,77285 | 2,42066592 | 0,736653 | 3,286033 | 0,001016 | 0,377927525 | 2,67243 | Promoter (<=1kb) | 19 | 3373285 | 3435733 | 62449 | 1 | 12894 | ENSMUST00000025835.6 | 0 | ENSMUSG00000024900 | Cpt1a | carnitine palmitoyltransferase 1a, liver |
| chr3 | 83352861 | 83355706 | 2846 | * | 26,86923 | 2,422298498 | 0,6154 | 3,936134 | 8,28E-05 | 0,092441431 | 2,880017 | Distal Intergenic | 3 | 83462670 | 83481429 | 18760 | 2 | 69460 | ENSMUST00000185938.2 | 125723 | ENSMUSG00000101334 | 1700028M03Rik | RIKEN cDNA 1700028M03 gene |
| chr5 | 92232310 | 92234387 | 2078 | * | 17,16325 | 2,422407205 | 0,898511 | 2,696024 | 0,007017 | 0,727652793 | 2,507013 | Promoter (<=1kb) | 5 | 92200023 | 92231578 | 31556 | 2 | 23881 | ENSMUST00000113127.7 | -732 | ENSMUSG00000029405 | G3bp2 | GTPase activating protein (SH3 domain) binding protein 2 |
| chr2 | 71702517 | 71705912 | 3396 | * | 62,70053 | 2,422437523 | 0,4861225 | 4,9831835 | 6,255E-07 | 0,002468468 | 3,177995 | Promoter (<=1kb) | 2 | 71703568 | 71734202 | 30635 | 1 | 228026 | ENSMUST00000006669.6 | 0 | ENSMUSG00000006494 | Pdk1 | pyruvate dehydrogenase kinase, isoenzyme 1 |
| chr13 | 98163367 | 98165787 | 2421 | * | 28,08281 | 2,42265639 | 0,615431 | 3,936519 | 8,27E-05 | 0,096868676 | 2,67243 | Intron (ENSMUST0000 | 13 | 98065138 | 98169187 | 104050 | 2 | 110596 | ENSMUST00000225663.2 | 3400 | ENSMUSG00000021662 | Arhgef28 | Rho guanine nucleotide exchange factor (GEF) 28 |
| chr16 | 55851104 | 55852235 | 1132 | * | 12,37964 | 2,423470568 | 0,932942 | 2,597665 | 0,009386 | 0,89593761 | 1,2763 | Promoter (1-2kb) | 16 | 55835869 | 55850102 | 14234 | 2 | 76302 | ENSMUST00000114444.9 | -1002 | ENSMUSG00000071533 | Pcnp | PEST proteolytic signal containing nuclear protein |
| chr11 | 54670877 | 54672280 | 1404 | * | 14,01505 | 2,423516696 | 0,877385 | 2,762205 | 0,005741 | 0,862146257 | 0,938682 | Intron (ENSMUST0000 | 11 | 54631136 | 54678451 | 47316 | 2 | 72729 | ENSMUST00000135903.8 | 6171 | ENSMUSG00000052298 | Cdc42se2 | CDC42 small effector 2 |
| chr8 | 34106818 | 34109431 | 2614 | * | 25,78931 | 2,423545641 | 0,652445 | 3,714558 | 0,000204 | 0,163971232 | 2,858511 | 3' UTR | 8 | 34116352 | 34126282 | 9931 | 2 | 108159 | ENSMUST00000210807.2 | 16851 | ENSMUSG00000052906 | Ubxn8 | UBX domain protein 8 |
| chr4 | 34979840 | 34982403 | 2564 | * | 21,46302 | 2,424960194 | 0,773358 | 3,135626 | 0,001715 | 0,424243345 | 3,471678 | Intron (ENSMUST0000 | 4 | 34909789 | 34911481 | 1693 | 2 | 73696 | ENSMUST00000125820.2 | -68359 | ENSMUSG00000085824 | Platr9 | pluripotency associated transcript 9 |
| chr8 | 65087918 | 65089803 | 1886 | * | 17,0573 | 2,42511969 | 0,803051 | 3,019883 | 0,002529 | 0,670853401 | 1,2763 | Intron (ENSMUST0000 | 8 | 65070635 | 65075031 | 4397 | 2 | 12876 | ENSMUST00000211312.2 | -12887 | ENSMUSG00000037852 | Cpe | carboxypeptidase E |
| chr17 | 71035879 | 71037242 | 1364 | * | 13,49071 | 2,425228609 | 0,934509 | 2,59519 | 0,009454 | 0,897633667 | 1,2763 | Intron (ENSMUST0000 | 17 | 71072834 | 71112992 | 40159 | 2 | 1,05E+08 | ENSMUST00000233559.2 | 75750 | ENSMUSG00000117231 | Gm41609 | predicted gene, 41609 |
| chr16 | 33031182 | 33033052 | 1871 | * | 14,9436 | 2,425758741 | 0,93026 | 2,607613 | 0,009118 | 0,80080207 | 2,141612 | Intron (ENSMUST0000 | 16 | 33044968 | 33057348 | 12381 | 1 | 106326 | ENSMUST00000231617.2 | -11916 | ENSMUSG00000022807 | Osbpl11 | oxysterol binding protein-like 11 |
| chr8 | 27748441 | 27751700 | 3260 | * | 47,360916 | 2,425768577 | 0,4923468 | 4,9269513 | 8,352E-07 | 0,003425236 | 2,7254908 | Promoter (<=1kb) | 8 | 27750357 | 27766702 | 16346 | 1 | 13685 | ENSMUST00000033880.7 | 0 | ENSMUSG00000031490 | Eif4ebp1 | eukaryotic translation initiation factor 4E binding protein 1 |
| chr4 | 82326374 | 82328167 | 1794 | * | 15,54716 | 2,425899125 | 0,894633 | 2,711613 | 0,006696 | 0,847307764 | 1,214904 | Intron (ENSMUST0000 | 4 | 82416781 | 82417553 | 773 | 2 | 18028 | ENSMUST00000152588.2 | 89386 | ENSMUSG00000008575 | Nfib | nuclear factor I/B |
| chr12 | 4891667 | 4894848 | 3182 | * | 37,122995 | 2,426082874 | 0,5650871 | 4,2932904 | 1,76E-05 | 0,038073228 | 2,3776742 | Promoter (<=1kb) | 12 | 4893303 | 4906967 | 13665 | 1 | 238037 | ENSMUST00000220215.2 | 0 | ENSMUSG00000051721 | Wdcp | WD repeat and coiled coil containing |
| chr8 | 127461227 | 127464450 | 3224 | * | 53,927022 | 2,426162529 | 0,4675152 | 5,1894838 | 2,109E-07 | 0,001297716 | 2,3776742 | Distal Intergenic | 8 | 127482321 | 127485084 | 2764 | 2 | 115486981 | ENSMUST00000212695.2 | 20634 | | Gm45805 | predicted gene 45805 |
| chr6 | 38319837 | 38321202 | 1366 | * | 16,62827 | 2,426633781 | 0,920332 | 2,636694 | 0,008372 | 0,787687702 | 2,141612 | Intron (ENSMUST0000 | 6 | 38282221 | 38331208 | 48988 | 2 | 78781 | ENSMUST00000114900.8 | 10006 | ENSMUSG00000029826 | Zc3hav1 | zinc finger CCCH type, antiviral 1 |
| chr1 | 75168416 | 75170398 | 1983 | * | 23,97335 | 2,427556629 | 0,659876 | 3,678807 | 0,000234 | 0,17530799 | 2,880017 | Promoter (<=1kb) | 1 | 75157504 | 75168570 | 11067 | 2 | 245860 | ENSMUST00000189702.7 | 0 | ENSMUSG00000033124 | Atg9a | autophagy related 9A |
| chr11 | 50341899 | 50343755 | 1857 | * | 15,38571 | 2,427664513 | 0,884054 | 2,746059 | 0,006032 | 0,728748169 | 2,141612 | Distal Intergenic | 11 | 50280113 | 50321952 | 41840 | 2 | 216724 | ENSMUST00000020643.4 | -19947 | ENSMUSG00000020375 | Rufy1 | RUN and FYVE domain containing 1 |
| chr8 | 89084883 | 89086651 | 1769 | * | 13,58141 | 2,427751009 | 0,895784 | 2,710197 | 0,006724 | 0,891097601 | 0,938682 | Promoter (1-2kb) | 8 | 89084160 | 89088642 | 4483 | 2 | 26992 | ENSMUST00000135471.2 | 1991 | ENSMUSG00000031660 | Brd7 | bromodomain containing 7 |
| chr4 | 54006956 | 54008453 | 1498 | * | 14,3089 | 2,427785492 | 0,914175 | 2,655712 | 0,007914 | 0,915629184 | 0,938682 | Promoter (2-3kb) | 4 | 54004057 | 54004187 | 131 | 1 | 1,15E+08 | ENSMUST00000158927.2 | 2899 | ENSMUSG00000089552 | Gm24177 | predicted gene, 24177 |
| chr17 | 15946997 | 15948655 | 1659 | * | 15,98346 | 2,428139441 | 0,810269 | 2,996707 | 0,002729 | 0,683140223 | 1,2763 | Exon (ENSMUST00000 | 17 | 15958797 | 15975610 | 16814 | 1 | 12648 | ENSMUST00000174461.2 | -10142 | ENSMUSG00000023852 | Chd1 | chromodomain helicase DNA binding protein 1 |
| chr3 | 5254378 | 5256299 | 1922 | * | 17,04065 | 2,428563358 | 0,925957 | 2,622759 | 0,008722 | 0,792545355 | 2,141612 | Intron (ENSMUST0000 | 3 | 5283586 | 5478225 | 194640 | 1 | 80892 | ENSMUST00000175866.8 | -27287 | ENSMUSG00000025255 | Zfhx4 | zinc finger homeodomain 4 |
| chr10 | 1,08E+08 | 1,08E+08 | 1561 | * | 16,12991 | 2,428637768 | 0,807762 | 3,006624 | 0,002642 | 0,568412386 | 2,318434 | Intron (ENSMUST0000 | 10 | 1,08E+08 | 1,08E+08 | 134 | 1 | 1,15E+08 | ENSMUST00000083719.3 | 13667 | ENSMUSG00000065653 | Gm23105 | predicted gene, 23105 |
| chr14 | 50986935 | 50988377 | 1443 | * | 15,34079 | 2,429014846 | 0,886626 | 2,739617 | 0,006151 | 0,730469267 | 2,141612 | Distal Intergenic | 14 | 50973628 | 50982551 | 8924 | 2 | 56858 | ENSMUST00000214290.2 | -4384 | ENSMUSG00000059069 | Olfr749 | olfactory receptor 749 |
| chr1 | 1,57E+08 | 1,57E+08 | 1514 | * | 13,20507 | 2,429405009 | 0,883093 | 2,751017 | 0,005941 | 0,909022873 | 0,738441 | Intron (ENSMUST0000 | 1 | 1,57E+08 | 1,57E+08 | 80511 | 2 | 226525 | ENSMUST00000134543.8 | -66218 | ENSMUSG00000070565 | Rasal2 | RAS protein activator like 2 |
| chr10 | 32068967 | 32070340 | 1374 | * | 15,47041 | 2,430157229 | 0,79009 | 3,075799 | 0,002099 | 0,649217039 | 1,214904 | Intron (ENSMUST0000 | 10 | 32205761 | 32286331 | 80571 | 2 | 432450 | ENSMUST00000092603.5 | 215991 | ENSMUSG00000069670 | Nkain2 | Na+/K+ transporting ATPase interacting 2 |
| chr7 | 1,1E+08 | 1,1E+08 | 2042 | * | 14,80672 | 2,430864614 | 0,869081 | 2,797052 | 0,005157 | 0,773936378 | 1,41696 | Intron (ENSMUST0000 | 7 | 1,1E+08 | 1,1E+08 | 21241 | 1 | 22390 | ENSMUST00000033326.10 | 9445 | ENSMUSG00000031016 | Wee1 | WEE 1 homolog 1 (S. pombe) |
| chr5 | 129266397 | 129269050 | 2654 | * | 45,900856 | 2,431640692 | 0,4674361 | 5,2020813 | 1,971E-07 | 0,001237282 | 2,3776742 | Exon (ENSMUST0000005 | 5 | 129173821 | 129280744 | 106924 | 1 | 243277 | ENSMUST00000156437.3 | 92576 | ENSMUSG00000044017 | Adgrd1 | adhesion G protein-coupled receptor D1 |
| chr7 | 34020180 | 34021930 | 1751 | * | 14,85108 | 2,431674472 | 0,841531 | 2,889584 | 0,003858 | 0,72253719 | 1,41696 | Distal Intergenic | 7 | 33936132 | 34012976 | 76845 | 2 | 233103 | ENSMUST00000108074.8 | -7204 | ENSMUSG00000066571 | Garre1 | granule associated Rac and RHOG effector 1 |
| chr8 | 1,26E+08 | 1,26E+08 | 2923 | * | 28,37836 | 2,431817305 | 0,745777 | 3,260782 | 0,001111 | 0,383091533 | 2,858511 | Promoter (<=1kb) | 8 | 1,26E+08 | 1,26E+08 | 21196 | 1 | 53424 | ENSMUST00000075896.7 | 0 | ENSMUSG00000056820 | Tsnax | translin-associated factor X |
| chr2 | 1,67E+08 | 1,67E+08 | 2494 | * | 17,37044 | 2,433009124 | 0,929687 | 2,617018 | 0,00887 | 0,775744126 | 2,423594 | Distal Intergenic | 2 | 1,67E+08 | 1,67E+08 | 1858 | 1 | 76367 | ENSMUST00000065753.2 | -3912 | ENSMUSG00000042854 | Trp53rkb | transformation related protein 53 regulating kinase B |
| chr8 | 1,3E+08 | 1,3E+08 | 985 | * | 16,97499 | 2,433168393 | 0,756948 | 3,214445 | 0,001307 | 0,457838671 | 2,141612 | Intron (ENSMUST0000 | 8 | 1,3E+08 | 1,3E+08 | 132 | 1 | 1,15E+08 | ENSMUST00000104350.3 | 7552 | ENSMUSG00000077538 | Gm22545 | predicted gene, 22545 |
| chr8 | 84938945 | 84940434 | 1490 | * | 18,47356 | 2,433215971 | 0,89186 | 2,728248 | 0,006367 | 0,712027511 | 2,507013 | Exon (ENSMUST00000 | 8 | 84935549 | 84936238 | 690 | 1 | 723960 | ENSMUST00000180630.2 | 3396 | ENSMUSG00000065541 | Mir24-2 | microRNA 24-2 |
| chr11 | 64388559 | 64389592 | 1034 | * | 13,08979 | 2,433742495 | 0,911561 | 2,669861 | 0,007588 | 0,913191717 | 0,918255 | Intron (ENSMUST0000 | 11 | 64326158 | 64413667 | 87510 | 1 | 15478 | ENSMUST00000058652.6 | 62401 | ENSMUSG00000047759 | Hs3st3a1 | heparan sulfate (glucosamine) 3-O-sulfotransferase 3A1 |
| chr15 | 83445434 | 83447650 | 2217 | * | 26,42142 | 2,434816655 | 0,709478 | 3,431841 | 0,000599 | 0,290624195 | 2,880017 | Promoter (<=1kb) | 15 | 83447793 | 83458404 | 10612 | 1 | 12257 | ENSMUST00000047419.8 | -143 | ENSMUSG00000041736 | Tspo | translocator protein |
| chr5 | 33394443 | 33396985 | 2543 | * | 26,24624 | 2,434856757 | 0,688758 | 3,535143 | 0,000408 | 0,247743734 | 2,725491 | Distal Intergenic | 5 | 33405512 | 33406594 | 1083 | 2 | 13016 | ENSMUST00000202135.2 | 9609 | ENSMUSG00000037373 | Ctbp1 | C-terminal binding protein 1 |
| chr8 | 1,28E+08 | 1,28E+08 | 1567 | * | 16,22308 | 2,43501293 | 0,793062 | 3,070393 | 0,002138 | 0,504427013 | 2,663619 | Distal Intergenic | 8 | 1,28E+08 | 1,28E+08 | 31125 | 1 | 93742 | ENSMUST00000159940.2 | 145938 | ENSMUSG00000025812 | Pard3 | par-3 family cell polarity regulator |
| chr2 | 93424594 | 93426929 | 2336 | * | 22,35162 | 2,435098914 | 0,736273 | 3,307333 | 0,000942 | 0,337696055 | 3,104559 | Distal Intergenic | 2 | 93472729 | 93511684 | 38956 | 1 | 11695 | ENSMUST00000042078.10 | -45800 | ENSMUSG00000040310 | Alx4 | aristaless-like homeobox 4 |
| chr13 | 31165903 | 31167879 | 1977 | * | 16,10732 | 2,435909505 | 0,800176 | 3,044217 | 0,002333 | 0,540220995 | 2,386008 | Distal Intergenic | 13 | 31107639 | 31158076 | 50438 | 2 | 66482 | ENSMUST00000222133.2 | -7827 | ENSMUSG00000021357 | Exoc2 | exocyst complex component 2 |
| chr11 | 61810125 | 61812139 | 2015 | * | 15,88721 | 2,437493147 | 0,911177 | 2,675104 | 0,007471 | 0,747833008 | 2,386008 | Intron (ENSMUST0000 | 11 | 61780616 | 61807044 | 26429 | 2 | 56697 | ENSMUST00000058173.12 | -3081 | ENSMUSG00000047804 | Akap10 | A kinase (PRKA) anchor protein 10 |
| chr15 | 7891143 | 7893385 | 2243 | * | 16,12891 | 2,438288617 | 0,908037 | 2,685232 | 0,007248 | 0,740451144 | 2,386008 | Distal Intergenic | 15 | 7902536 | 7915858 | 13323 | 2 | 545085 | ENSMUST00000226928.2 | 22473 | ENSMUSG00000039828 | Wdr70 | WD repeat domain 70 |
| chr3 | 1,23E+08 | 1,23E+08 | 1951 | * | 16,41842 | 2,438407469 | 0,801558 | 3,042086 | 0,002349 | 0,562910619 | 2,141612 | Distal Intergenic | 3 | 1,23E+08 | 1,23E+08 | 130 | 2 | 1,15E+08 | ENSMUST00000116895.3 | -24906 | ENSMUSG00000080545 | Gm22713 | predicted gene, 22713 |
| chr8 | 1,16E+08 | 1,16E+08 | 2083 | * | 21,34902 | 2,439394419 | 0,841961 | 2,897278 | 0,003764 | 0,594430593 | 2,951933 | Intron (ENSMUST0000 | 8 | 1,16E+08 | 1,16E+08 | 104 | 2 | 1,15E+08 | ENSMUST00000083156.3 | 257583 | ENSMUSG00000065090 | Gm23677 | predicted gene, 23677 |
| chr16 | 8417147 | 8418512 | 1366 | * | 13,22597 | 2,439473017 | 0,876577 | 2,782953 | 0,005387 | 0,798887389 | 1,2763 | Intron (ENSMUST0000 | 16 | 8430742 | 8436441 | 5700 | 1 | 268860 | ENSMUST00000128459.2 | -12230 | ENSMUSG00000057880 | Abat | 4-aminobutyrate aminotransferase |
| chr10 | 62173790 | 62174785 | 996 | * | 13,8924 | 2,439902211 | 0,862633 | 2,828437 | 0,004678 | 0,832960417 | 0,918255 | Promoter (1-2kb) | 10 | 62175980 | 62185650 | 9671 | 1 | 70162 | ENSMUST00000123457.3 | -1195 | | Hk1os | hexokinase 1, opposite strand |
| chr1 | 28276707 | 28278079 | 1373 | * | 13,66128 | 2,440786381 | 0,942337 | 2,590143 | 0,009594 | 0,923399738 | 1,075735 | Distal Intergenic | 1 | 27923059 | 27941903 | 18845 | 1 | 433281 | ENSMUST00000185409.3 | 353648 | ENSMUSG00000101514 | Gm5524 | predicted gene 5524 |
| chr17 | 26916695 | 26920200 | 3506 | * | 26,3771 | 2,441182604 | 0,716892 | 3,405229 | 0,000661 | 0,303366229 | 2,858511 | 3' UTR | 17 | 26934624 | 26995609 | 60986 | 1 | 77128 | ENSMUST00000062519.14 | -14424 | ENSMUSG00000048249 | Crebrf | CREB3 regulatory factor |
| chr12 | 65119930 | 65121655 | 1726 | * | 18,0745 | 2,441194252 | 0,924843 | 2,639578 | 0,008301 | 0,760759863 | 2,423594 | Promoter (<=1kb) | 12 | 65109198 | 65120736 | 11539 | 2 | 30795 | ENSMUST00000221166.2 | 0 | ENSMUSG00000020949 | Fkbp3 | FK506 binding protein 3 |
| chr5 | 53229083 | 53230147 | 1065 | * | 14,52647 | 2,441368138 | 0,843597 | 2,893998 | 0,003804 | 0,653106633 | 2,141612 | Downstream (<=300bp | 5 | 53206695 | 53223642 | 16948 | 1 | 20531 | ENSMUST00000147243.4 | 22388 | ENSMUSG00000029188 | Slc34a2 | solute carrier family 34 (sodium phosphate), member 2 |
| chr9 | 65070918 | 65074107 | 3190 | * | 25,491 | 2,441583498 | 0,693564 | 3,520342 | 0,000431 | 0,254009046 | 2,725491 | Intron (ENSMUST0000 | 9 | 65088020 | 65089004 | 985 | 1 | 19289 | ENSMUST00000217135.2 | -13913 | ENSMUSG00000032394 | Igdcc3 | immunoglobulin superfamily, DCC subclass, member 3 |
| chr3 | 88941418 | 88944102 | 2685 | * | 19,24416 | 2,442017866 | 0,87285 | 2,79775 | 0,005146 | 0,663630601 | 2,713306 | Exon (ENSMUST00001 | 3 | 88930490 | 88935575 | 5086 | 1 | 192195 | ENSMUST00000198846.2 | 10928 | ENSMUSG00000028053 | Ash1l | ASH1 like histone lysine methyltransferase |
| chr4 | 1,48E+08 | 1,48E+08 | 692 | * | 21,12715 | 2,442101625 | 0,71414 | 3,419642 | 0,000627 | 0,293348204 | 2,951933 | Distal Intergenic | 4 | 1,48E+08 | 1,48E+08 | 21070 | 1 | 195531 | ENSMUST00000105721.9 | -18722 | ENSMUSG00000078496 | Zfp982 | zinc finger protein 982 |
| chr11 | 70910081 | 70912871 | 2791 | * | 20,50712 | 2,442563216 | 0,788729 | 3,096833 | 0,001956 | 0,490833296 | 2,663619 | Promoter (<=1kb) | 11 | 70898268 | 70910129 | 11862 | 2 | 116891 | ENSMUST00000143850.8 | 0 | ENSMUSG00000018442 | Derl2 | Der1-like domain family, member 2 |
| chr6 | 61346360 | 61347560 | 1201 | * | 12,13845 | 2,442689427 | 0,882932 | 2,766566 | 0,005665 | 0,823736512 | 1,173325 | Intron (ENSMUST0000 | 6 | 61350981 | 61788702 | 437722 | 1 | 232035 | ENSMUST00000067307.11 | -3421 | ENSMUSG00000039578 | Ccser1 | coiled-coil serine rich 1 |
| chr10 | 77404262 | 77407360 | 3099 | * | 32,30543 | 2,442695406 | 0,667154 | 3,661366 | 0,000251 | 0,188876298 | 2,787255 | Distal Intergenic | 10 | 77417554 | 77434566 | 17013 | 1 | 108705 | ENSMUST00000009435.12 | -10194 | ENSMUSG00000009291 | Pttg1ip | pituitary tumor-transforming 1 interacting protein |
| chr15 | 95267689 | 95269890 | 2202 | * | 40,607452 | 2,44345678 | 0,5127411 | 4,7654786 | 1,884E-06 | 0,00609733 | 2,9254973 | Intron (ENSMUST000002 | 15 | 95084587 | 95426059 | 341473 | 2 | 54003 | ENSMUST00000229981.2 | 156169 | ENSMUSG00000022454 | Nell2 | NEL-like 2 |
| chr15 | 8183505 | 8185448 | 1944 | * | 17,46136 | 2,443688986 | 0,858006 | 2,848104 | 0,004398 | 0,656950206 | 2,423594 | Promoter (2-3kb) | 15 | 8180567 | 8188308 | 7742 | 1 | 170762 | ENSMUST00000230925.2 | 2938 | ENSMUSG00000022142 | Nup155 | nucleoporin 155 |
| chr12 | 27615794 | 27618051 | 2258 | * | 18,42506 | 2,44425674 | 0,884218 | 2,764315 | 0,005704 | 0,697963122 | 2,386008 | Distal Intergenic | 12 | 27648200 | 27665361 | 17162 | 1 | 328069 | ENSMUST00000220893.2 | -30149 | | Gm4166 | predicted gene 4166 |
| chr1 | 74065940 | 74069251 | 3312 | * | 57,55431 | 2,444991395 | 0,4568309 | 5,3520709 | 8,695E-08 | 0,000528281 | 3,2706213 | Intron (ENSMUST000001 | 1 | 74032322 | 74076292 | 43971 | 2 | 21961 | ENSMUST00000187281.7 | 7041 | ENSMUSG00000055322 | Tns1 | tensin 1 |
| chr15 | 84609344 | 84610482 | 1139 | * | 13,23053 | 2,445363194 | 0,889157 | 2,750205 | 0,005956 | 0,909245362 | 0,738441 | Intron (ENSMUST0000 | 15 | 84598830 | 84604467 | 5638 | 2 | 328576 | ENSMUST00000229841.2 | -4877 | ENSMUSG00000116396 | Gm20556 | predicted gene, 20556 |
| chr2 | 1,04E+08 | 1,04E+08 | 1841 | * | 19,19709 | 2,446671716 | 0,743285 | 3,2917 | 0,000996 | 0,388415573 | 2,507013 | Exon (ENSMUST00000 | 2 | 1,04E+08 | 1,04E+08 | 2183 | 1 | 99382 | ENSMUST00000138580.2 | -31018 | ENSMUSG00000032724 | Abtb2 | ankyrin repeat and BTB (POZ) domain containing 2 |
| chr15 | 95747160 | 95748261 | 1102 | * | 15,26408 | 2,446983185 | 0,935087 | 2,616852 | 0,008874 | 0,793995086 | 2,141612 | Intron (ENSMUST0000 | 15 | 95722339 | 95727953 | 5615 | 2 | 1,03E+08 | ENSMUST00000168960.3 | -19207 | ENSMUSG00000078648 | Gm17546 | predicted gene, 17546 |
| chr9 | 40452558 | 40454256 | 1699 | * | 13,43556 | 2,447309679 | 0,936082 | 2,614418 | 0,008938 | 0,970928491 | 0,738441 | Distal Intergenic | 9 | 40458656 | 40472689 | 14034 | 1 | 1,03E+08 | ENSMUST00000216525.2 | -4400 | ENSMUSG00000110922 | Gm35177 | predicted gene, 35177 |
| chr2 | 1,32E+08 | 1,32E+08 | 1896 | * | 16,25288 | 2,447663216 | 0,911029 | 2,686701 | 0,007216 | 0,740176831 | 2,386008 | Distal Intergenic | 2 | 1,32E+08 | 1,32E+08 | 3772 | 2 | 414081 | ENSMUST00000140443.2 | 47309 | ENSMUSG00000086822 | 5330413P13Rik | RIKEN cDNA 5330413P13 gene |
| chr7 | 1,44E+08 | 1,44E+08 | 1289 | * | 15,26614 | 2,448404782 | 0,84679 | 2,891397 | 0,003835 | 0,72659841 | 1,37683 | Promoter (<=1kb) | 7 | 1,44E+08 | 1,44E+08 | 5146 | 2 | 14082 | ENSMUST00000033394.8 | 400 | ENSMUSG00000031077 | Fadd | Fas (TNFRSF6)-associated via death domain |
| chr10 | 1,3E+08 | 1,3E+08 | 1110 | * | 12,00327 | 2,44843073 | 0,911918 | 2,684925 | 0,007255 | 0,93974765 | 0,738441 | Distal Intergenic | 10 | 1,3E+08 | 1,3E+08 | 1058 | 2 | 258297 | ENSMUST00000058123.6 | -18174 | ENSMUSG00000045559 | Olfr827 | olfactory receptor 827 |
| chr3 | 1,13E+08 | 1,13E+08 | 2696 | * | 24,28589 | 2,448904727 | 0,864064 | 2,83417 | 0,004594 | 0,647880877 | 2,67243 | Intron (ENSMUST0000 | 3 | 1,13E+08 | 1,13E+08 | 41713 | 2 | 11722 | ENSMUST00000174147.2 | 5797 | ENSMUSG00000074264 | Amy1 | amylase 1, salivary |
| chr2 | 1,64E+08 | 1,64E+08 | 2143 | * | 18,43981 | 2,448944511 | 0,844553 | 2,899693 | 0,003735 | 0,611385361 | 2,663619 | Promoter (<=1kb) | 2 | 1,64E+08 | 1,64E+08 | 6443 | 1 | 52840 | ENSMUST00000109349.9 | 0 | ENSMUSG00000017734 | Dbndd2 | dysbindin (dystrobrevin binding protein 1) domain containing 2 |
| chr19 | 56800328 | 56802267 | 1940 | * | 14,62728 | 2,449208971 | 0,900921 | 2,71856 | 0,006557 | 0,834909202 | 1,2763 | Promoter (<=1kb) | 19 | 56779395 | 56802115 | 22721 | 2 | 213993 | ENSMUST00000076085.4 | 0 | ENSMUSG00000035173 | Ccdc186 | coiled-coil domain containing 186 |
| chr1 | 1,31E+08 | 1,31E+08 | 2067 | * | 16,21013 | 2,449879085 | 0,844758 | 2,900094 | 0,003731 | 0,630453518 | 2,386008 | Distal Intergenic | 1 | 1,31E+08 | 1,31E+08 | 11970 | 1 | 108900 | ENSMUST00000068613.5 | 27495 | ENSMUSG00000055184 | Fam72a | family with sequence similarity 72, member A |
| chr2 | 89006466 | 89007723 | 1258 | * | 11,07753 | 2,450022628 | 0,950558 | 2,577457 | 0,009953 | 0,985829131 | 0,744375 | Distal Intergenic | 2 | 89000563 | 89001589 | 1027 | 2 | 258893 | ENSMUST00000188861.3 | -4877 | ENSMUSG00000101918 | Olfr1225 | olfactory receptor 1225 |
| chr11 | 1,03E+08 | 1,03E+08 | 2125 | * | 25,54151 | 2,450174637 | 0,723846 | 3,38494 | 0,000712 | 0,310869445 | 2,858511 | Exon (ENSMUST00001 | 11 | 1,03E+08 | 1,03E+08 | 18824 | 1 | 11488 | ENSMUST00000068150.7 | -10583 | ENSMUSG00000020926 | Adam11 | a disintegrin and metallopeptidase domain 11 |
| chr3 | 58167285 | 58168942 | 1658 | * | 12,84659 | 2,450833254 | 0,932995 | 2,626845 | 0,008618 | 0,908689178 | 1,075735 | Distal Intergenic | 3 | 58197672 | 58197778 | 107 | 2 | 1,15E+08 | ENSMUST00000083788.3 | 28836 | ENSMUSG00000065722 | Gm26166 | predicted gene, 26166 |
| chr8 | 1,15E+08 | 1,15E+08 | 1918 | * | 22,01816 | 2,451097536 | 0,817377 | 2,998737 | 0,002711 | 0,530563721 | 2,951933 | Promoter (<=1kb) | 8 | 1,15E+08 | 1,16E+08 | 913053 | 1 | 80707 | ENSMUST00000004756.14 | 0 | ENSMUSG00000004637 | Wwox | WW domain-containing oxidoreductase |
| chr12 | 93646591 | 93647643 | 1053 | * | 12,84523 | 2,451737041 | 0,913413 | 2,68415 | 0,007271 | 0,906138253 | 0,918255 | Distal Intergenic | 12 | 93713276 | 93713436 | 161 | 1 | 1,15E+08 | ENSMUST00000083369.4 | -65633 | ENSMUSG00000065303 | Gm23249 | predicted gene, 23249 |
| chr14 | 39089627 | 39091237 | 1611 | * | 16,32444 | 2,45280929 | 0,867571 | 2,827215 | 0,004695 | 0,670064583 | 2,386008 | Intron (ENSMUST0000 | 14 | 38731144 | 39194623 | 463480 | 2 | 18183 | ENSMUST00000176122.2 | 103386 | ENSMUSG00000041014 | Nrg3 | neuregulin 3 |
| chr10 | 1,18E+08 | 1,18E+08 | 3110 | * | 30,62111 | 2,453647794 | 0,629272 | 3,899186 | 9,65E-05 | 0,102151284 | 2,858511 | Distal Intergenic | 10 | 1,18E+08 | 1,18E+08 | 21884 | 2 | 17246 | ENSMUST00000105263.9 | -9930 | ENSMUSG00000020184 | Mdm2 | transformed mouse 3T3 cell double minute 2 |
| chr17 | 21693702 | 21698456 | 4755 | * | 58,980809 | 2,45448784 | 0,4636587 | 5,2937387 | 1,198E-07 | 0,00056002 | 4,2003037 | Exon (ENSMUST000002 | 17 | 21709260 | 21730735 | 21476 | 1 | 24132 | ENSMUST00000233158.2 | -10804 | ENSMUSG00000057409 | Zfp53 | zinc finger protein 53 |
| chr6 | 1,18E+08 | 1,18E+08 | 1929 | * | 19,6893 | 2,454870335 | 0,775212 | 3,166706 | 0,001542 | 0,446344042 | 2,713306 | Promoter (2-3kb) | 6 | 1,18E+08 | 1,18E+08 | 7402 | 1 | 98758 | ENSMUST00000179478.8 | -2664 | ENSMUSG00000042079 | Hnrnpf | heterogeneous nuclear ribonucleoprotein F |
| chr8 | 27465480 | 27466995 | 1516 | * | 24,2845 | 2,454974716 | 0,728898 | 3,368064 | 0,000757 | 0,314438111 | 2,951062 | Promoter (<=1kb) | 8 | 27458699 | 27466955 | 8257 | 2 | 1,05E+08 | ENSMUST00000241140.1 | 0 | | Gm39150 | predicted gene, 39150 |
| chr11 | 61273724 | 61276175 | 2452 | * | 25,326 | 2,455292108 | 0,685719 | 3,580612 | 0,000343 | 0,216386913 | 2,951062 | Distal Intergenic | 11 | 61253712 | 61269171 | 15460 | 2 | 67473 | ENSMUST00000148671.2 | -4553 | ENSMUSG00000010122 | Slc47a1 | solute carrier family 47, member 1 |
| chr15 | 81438371 | 81440420 | 2050 | * | 31,39037 | 2,456768346 | 0,708422 | 3,467944 | 0,000524 | 0,274176375 | 2,880017 | Distal Intergenic | 15 | 81468375 | 81469229 | 855 | 2 | 68635 | ENSMUST00000192082.2 | 28809 | | 1110025M09Rik | RIKEN cDNA 1110025M09 gene |
| chr15 | 98462287 | 98464473 | 2187 | * | 21,99176 | 2,456769888 | 0,696923 | 3,525169 | 0,000423 | 0,242896956 | 2,951933 | Promoter (<=1kb) | 15 | 98442448 | 98463020 | 20573 | 2 | 12455 | ENSMUST00000164294.8 | 0 | ENSMUSG00000011960 | Ccnt1 | cyclin T1 |
| chr4 | 1,55E+08 | 1,55E+08 | 1823 | * | 20,80062 | 2,457343399 | 0,809749 | 3,034697 | 0,002408 | 0,511439669 | 2,880017 | Intron (ENSMUST0000 | 4 | 1,55E+08 | 1,55E+08 | 68461 | 2 | 20481 | ENSMUST00000084103.10 | 12472 | ENSMUSG00000029050 | Ski | ski sarcoma viral oncogene homolog (avian) |
| chr3 | 1,17E+08 | 1,17E+08 | 2336 | * | 18,51518 | 2,457472127 | 0,796834 | 3,084044 | 0,002042 | 0,497406263 | 2,663619 | Promoter (<=1kb) | 3 | 1,17E+08 | 1,17E+08 | 68168 | 2 | 77559 | ENSMUST00000040603.14 | 0 | ENSMUSG00000033400 | Agl | amylo-1,6-glucosidase, 4-alpha-glucanotransferase |
| chr17 | 28039621 | 28042749 | 3129 | * | 31,78897 | 2,458638807 | 0,634837 | 3,872866 | 0,000108 | 0,113460495 | 2,725491 | Promoter (<=1kb) | 17 | 27972181 | 28039622 | 67442 | 2 | 224647 | ENSMUST00000233131.2 | 0 | ENSMUSG00000056692 | Ilrun | inflammation and lipid regulator with UBA-like and NBR1-like domains |
| chr13 | 1,11E+08 | 1,11E+08 | 2168 | * | 20,17561 | 2,459555796 | 0,768233 | 3,201575 | 0,001367 | 0,440320279 | 2,507013 | Distal Intergenic | 13 | 1,11E+08 | 1,11E+08 | 1239 | 1 | 20620 | ENSMUST00000225156.2 | 28186 | ENSMUSG00000021701 | Plk2 | polo like kinase 2 |
| chr16 | 18035632 | 18037265 | 1634 | * | 17,68375 | 2,460841061 | 0,892999 | 2,755703 | 0,005857 | 0,722439117 | 2,154504 | Promoter (<=1kb) | 16 | 18035743 | 18038205 | 2463 | 1 | 1,03E+08 | ENSMUST00000231860.2 | 0 | ENSMUSG00000090777 | Ccdc188 | coiled-coil domain containing 188 |
| chr7 | 1,1E+08 | 1,1E+08 | 1898 | * | 14,54142 | 2,461840355 | 0,853839 | 2,883261 | 0,003936 | 0,659240897 | 2,141612 | Intron (ENSMUST0000 | 7 | 1,1E+08 | 1,1E+08 | 19545 | 2 | 319934 | ENSMUST00000165992.2 | -4299 | ENSMUSG00000038371 | Sbf2 | SET binding factor 2 |
| chr12 | 75292429 | 75293559 | 1131 | * | 13,39597 | 2,462574914 | 0,860315 | 2,86241 | 0,004204 | 0,753191087 | 1,2763 | Distal Intergenic | 12 | 75355096 | 75442591 | 87496 | 1 | 80837 | ENSMUST00000118602.8 | -61537 | ENSMUSG00000046768 | Rhoj | ras homolog family member J |
| chr16 | 7563380 | 7565539 | 2160 | * | 18,1639 | 2,462817355 | 0,899866 | 2,736872 | 0,006203 | 0,693003605 | 2,663619 | Distal Intergenic | 16 | 7170834 | 7227833 | 57000 | 1 | 268859 | ENSMUST00000231031.2 | 392546 | ENSMUSG00000008658 | Rbfox1 | RNA binding protein, fox-1 homolog (C. elegans) 1 |
| chrX | 7706969 | 7708153 | 1185 | * | 14,38384 | 2,463105555 | 0,909822 | 2,707239 | 0,006785 | 0,839557728 | 1,2763 | Promoter (<=1kb) | 20 | 7707603 | 7708247 | 645 | 1 | 54644 | ENSMUST00000152899.2 | 0 | ENSMUSG00000031154 | Otud5 | OTU domain containing 5 |
| chr10 | 26326439 | 26328454 | 2016 | * | 17,93952 | 2,463585444 | 0,72806 | 3,383765 | 0,000715 | 0,334207335 | 2,423594 | Distal Intergenic | 10 | 26369094 | 26374743 | 5650 | 1 | 1,03E+08 | ENSMUST00000219589.2 | -40640 | | Gm30034 | predicted gene, 30034 |
| chr8 | 1,07E+08 | 1,07E+08 | 1108 | * | 13,71625 | 2,463756118 | 0,895801 | 2,750339 | 0,005953 | 0,875821625 | 0,918255 | Exon (ENSMUST00001 | 8 | 1,07E+08 | 1,07E+08 | 10538 | 2 | 20498 | ENSMUST00000132231.2 | 3669 | ENSMUSG00000017765 | Slc12a4 | solute carrier family 12, member 4 |
| chr19 | 42490656 | 42493631 | 2976 | * | 26,5913 | 2,463843319 | 0,762333 | 3,231979 | 0,001229 | 0,411735506 | 2,67243 | Distal Intergenic | 19 | 42507198 | 42580782 | 73585 | 1 | 52013 | ENSMUST00000026188.10 | -13567 | ENSMUSG00000025184 | R3hcc1l | R3H domain and coiled-coil containing 1 like |
| chr3 | 86049872 | 86051221 | 1350 | * | 19,80321 | 2,463872351 | 0,778732 | 3,163955 | 0,001556 | 0,47038366 | 2,386008 | Promoter (<=1kb) | 3 | 86045247 | 86050009 | 4763 | 2 | 20091 | ENSMUST00000029722.7 | 0 | ENSMUSG00000028081 | Rps3a1 | ribosomal protein S3A1 |
| chr5 | 51895188 | 51896650 | 1463 | * | 14,02216 | 2,46406893 | 0,851009 | 2,895467 | 0,003786 | 0,834870998 | 0,738441 | Distal Intergenic | 5 | 51653122 | 51725068 | 71947 | 2 | 19017 | ENSMUST00000127135.3 | -170120 | ENSMUSG00000029167 | Ppargc1a | peroxisome proliferative activated receptor, gamma, coactivator 1 alpha |
| chr7 | 98560046 | 98561898 | 1853 | * | 13,58837 | 2,465271051 | 0,939176 | 2,62493 | 0,008667 | 0,965053147 | 0,738441 | Intron (ENSMUST0000 | 7 | 98554294 | 98557262 | 2969 | 1 | 666529 | ENSMUST00000207697.2 | 5752 | | Gm8149 | predicted gene 8149 |
| chr12 | 34586947 | 34588047 | 1101 | * | 12,01488 | 2,46534409 | 0,951831 | 2,590108 | 0,009595 | 0,978800182 | 0,744375 | Intron (ENSMUST0000 | 12 | 34421496 | 34578873 | 157378 | 2 | 79221 | ENSMUST00000110819.4 | -8074 | ENSMUSG00000004698 | Hdac9 | histone deacetylase 9 |
| chrX | 92874432 | 92877387 | 2956 | * | 14,92393 | 2,466410282 | 0,945286 | 2,609169 | 0,009076 | 0,890608507 | 1,2763 | Promoter (<=1kb) | 20 | 92808213 | 92875807 | 67595 | 2 | 236900 | ENSMUST00000045748.7 | 0 | ENSMUSG00000035232 | Pdk3 | pyruvate dehydrogenase kinase, isoenzyme 3 |

| | | | | | | | | | | | | | | | | | | | | | | | |
|---|---|---|---|---|---|---|---|---|---|---|---|---|---|---|---|---|---|---|---|---|---|---|---|
| chr14 | 65580133 | 65582869 | 2737 | * | 39,17328 | 2,466451208 | 0,644534 | 3,826718 | 0,00013 | 0,126759383 | 2,725491 | Distal Intergenic | 14 | 65595838 | 65623837 | 28000 | 1 | 380912 | ENSMUST00000224623.2 | -12969 | ENSMUSG00000034522 | Zfp395 | zinc finger protein 395 |
| chr18 | 64557575 | 64559078 | 1504 | * | 12,20461 | 2,466756293 | 0,883973 | 2,790533 | 0,005262 | 0,852732156 | 0,918255 | Distal Intergenic | 18 | 64590965 | 64597397 | 6433 | 2 | 14151 | ENSMUST00000238082.2 | 38319 | ENSMUSG00000024588 | Fech | ferrochelatase |
| chr18 | 38056092 | 38057717 | 1626 | * | 12,92644 | 2,466813091 | 0,907372 | 2,718636 | 0,006555 | 0,890735468 | 0,918255 | Intron (ENSMUST00000 | 18 | 37976656 | 38068434 | 91779 | 2 | 13367 | ENSMUST00000025337.14 | 10717 | ENSMUSG00000024456 | Diaph1 | diaphanous related formin 1 |
| chr7 | 34041192 | 34042572 | 1381 | * | 20,1841 | 2,46684924 | 0,793909 | 3,10722 | 0,001889 | 0,514210241 | 2,221611 | Distal Intergenic | 7 | 34044088 | 34052920 | 8833 | 2 | 67070 | ENSMUST00000206830.2 | 10348 | ENSMUSG00000066568 | Lsm14a | LSM14A mRNA processing body assembly factor |
| chr4 | 52909338 | 52911196 | 1859 | * | 13,95682 | 2,467538519 | 0,904663 | 2,727577 | 0,00638 | 0,920164459 | 0,738441 | Promoter (<=1kb) | 4 | 52910833 | 52911794 | 962 | 2 | 258836 | ENSMUST00000051600.3 | 598 | ENSMUSG00000051593 | Olfr272 | olfactory receptor 272 |
| chr17 | 34157760 | 34160066 | 2307 | * | 21,75321 | 2,467786672 | 0,886357 | 2,784191 | 0,005366 | 0,661775851 | 2,880017 | Promoter (<=1kb) | 17 | 34157795 | 34159317 | 1523 | 2 | 14976 | ENSMUST00000025163.14 | 0 | ENSMUSG00000024309 | Pfdn6 | prefoldin subunit 6 |
| chr6 | 66441626 | 66443626 | 2001 | * | 19,45087 | 2,467946198 | 0,770147 | 3,204512 | 0,001353 | 0,428016132 | 2,663619 | Promoter (<=1kb) | 6 | 66428933 | 66442683 | 13751 | 2 | 72484 | ENSMUST00000203716.2 | 0 | ENSMUSG00000107479 | 2610300M13Rik | RIKEN cDNA 2610300M13 gene |
| chr7 | 44935557 | 44939043 | 3487 | * | 32,71002 | 2,467968476 | 0,734212 | 3,361382 | 0,000776 | 0,442614296 | 1,405709 | Promoter (<=1kb) | 7 | 44936560 | 44938422 | 1863 | 1 | 76713 | ENSMUST00000210733.2 | 0 | ENSMUSG00000070568 | Slc6a21 | solute carrier family 6 member 21 |
| chr2 | 1,56E+08 | 1,56E+08 | 2122 | * | 13,86073 | 2,467970843 | 0,945495 | 2,610242 | 0,009048 | 0,915421656 | 1,075735 | Intron (ENSMUST00000 | 2 | 1,56E+08 | 1,57E+08 | 150659 | 1 | 228836 | ENSMUST00000169464.9 | 20171 | ENSMUSG00000061689 | Dlgap4 | DLG associated protein 4 |
| chr6 | 83252245 | 83253756 | 1512 | * | 16,50724 | 2,468644857 | 0,787491 | 3,134824 | 0,00172 | 0,486196597 | 2,386008 | Promoter (<=1kb) | 6 | 83254375 | 83265433 | 11059 | 1 | 232156 | ENSMUST00000141863.8 | -619 | ENSMUSG00000068323 | Slc4a5 | solute carrier family 4, sodium bicarbonate cotransporter, member 5 |
| chr11 | 32387462 | 32389264 | 1803 | * | 15,11885 | 2,469016508 | 0,813815 | 3,03388 | 0,002414 | 0,671156916 | 1,214904 | Distal Intergenic | 11 | 32405370 | 32466687 | 61318 | 1 | 327900 | ENSMUST00000051053.5 | -16106 | ENSMUSG00000044949 | Ubtd2 | ubiquitin domain containing 2 |
| chr10 | 79936956 | 79937805 | 850 | * | 11,34199 | 2,470668859 | 0,949282 | 2,602671 | 0,00925 | 0,974266082 | 0,738441 | Promoter (<=1kb) | 10 | 79893605 | 79937021 | 43417 | 2 | 216161 | ENSMUST00000218630.2 | 0 | ENSMUSG00000035673 | Sbno2 | strawberry notch 2 |
| chr18 | 76121691 | 76123090 | 1400 | * | 17,52417 | 2,470853231 | 0,909276 | 2,717387 | 0,00658 | 0,715249603 | 2,507013 | Intron (ENSMUST00000 | 18 | 76192529 | 76279514 | 86986 | 1 | 207259 | ENSMUST00000167921.2 | -69439 | ENSMUSG00000044646 | Zbtb7c | zinc finger and BTB domain containing 7C |
| chr10 | 42009048 | 42010922 | 1875 | * | 16,8022 | 2,470915968 | 0,814172 | 3,034883 | 0,002406 | 0,542967057 | 2,386008 | Distal Intergenic | 10 | 41917587 | 41944651 | 27065 | 2 | 1,05E+08 | ENSMUST00000241848.1 | -64397 | | Gm40630 | predicted gene, 40630 |
| chr10 | 26860284 | 26861785 | 1502 | * | 11,57714 | 2,47158194 | 0,956668 | 2,583532 | 0,009779 | 0,914539344 | 1,173325 | Promoter (2-3kb) | 10 | 26856032 | 26864510 | 8479 | 2 | 16773 | ENSMUST00000186279.2 | 2725 | ENSMUSG00000019899 | Lama2 | laminin, alpha 2 |
| chr12 | 86836025 | 86838499 | 2475 | * | 34,7552 | 2,471623077 | 0,601149 | 4,111499 | 3,93E-05 | 0,090744111 | 1,405709 | Distal Intergenic | 12 | 86787781 | 86801168 | 13388 | 1 | 627607 | ENSMUST00000223197.2 | 48244 | ENSMUSG00000059114 | Lrrc74a | leucine rich repeat containing 74A |
| chr17 | 56813418 | 56816057 | 2640 | * | 34,20513 | 2,472197605 | 0,608947 | 4,059794 | 4,91E-05 | 0,067555009 | 2,925497 | Distal Intergenic | 17 | 56818248 | 56819431 | 1184 | 2 | 20834 | ENSMUST00000052211.4 | 3374 | ENSMUSG00000044526 | Znrf4 | zinc and ring finger 4 |
| chr16 | 20440359 | 20442170 | 1812 | * | 14,829 | 2,472816647 | 0,844862 | 2,926887 | 0,003424 | 0,707972935 | 1,37683 | Promoter (<=1kb) | 16 | 20437965 | 20440037 | 2073 | 2 | 73047 | ENSMUST00000052939.4 | -322 | ENSMUSG00000051146 | Camk2n2 | calcium/calmodulin-dependent protein kinase II inhibitor 2 |
| chr1 | 27972835 | 27974926 | 2092 | * | 18,67998 | 2,472956984 | 0,803895 | 3,076218 | 0,002096 | 0,518149475 | 2,423594 | Distal Intergenic | 1 | 27923059 | 27941903 | 18845 | 1 | 433281 | ENSMUST00000185409.3 | 49776 | ENSMUSG00000101514 | Gm5524 | predicted gene 5524 |
| chr2 | 59712513 | 59713965 | 1453 | * | 17,49176 | 2,473692082 | 0,876761 | 2,821398 | 0,004781 | 0,670234202 | 2,423594 | Promoter (<=1kb) | 2 | 59682708 | 59712928 | 30221 | 2 | 72137 | ENSMUST00000028368.14 | 0 | ENSMUSG00000026988 | Wdsub1 | WD repeat, SAM and U-box domain containing 1 |
| chr18 | 75429842 | 75432596 | 2755 | * | 36,11815 | 2,473846287 | 0,611967 | 4,04245 | 5,29E-05 | 0,080985792 | 2,377674 | Promoter (<=1kb) | 18 | 75421726 | 75430388 | 8663 | 2 | 1,01E+08 | ENSMUST00000190465.2 | 0 | ENSMUSG00000102047 | 2010010A06Rik | RIKEN cDNA 2010010A06 gene |
| chr19 | 6349733 | 6352365 | 2633 | * | 25,13373 | 2,47391485 | 0,694799 | 3,56062 | 0,00037 | 0,225394068 | 2,951933 | Promoter (1-2kb) | 19 | 6347944 | 6348390 | 447 | 1 | 13660 | ENSMUST00000148547.2 | 1789 | ENSMUSG00000024772 | Ehd1 | EH-domain containing 1 |
| chrX | 1,67E+08 | 1,67E+08 | 2302 | * | 16,57047 | 2,473925887 | 0,858729 | 2,880915 | 0,003965 | 0,64236196 | 2,386008 | Intron (ENSMUST00000 | 20 | 1,66E+08 | 1,67E+08 | 648854 | 2 | 333605 | ENSMUST00000112146.9 | 102882 | ENSMUSG00000049176 | Frmpd4 | FERM and PDZ domain containing 4 |
| chr15 | 83050452 | 83052307 | 1856 | * | 31,80608 | 2,474084171 | 0,675252 | 3,66394 | 0,000248 | 0,190625259 | 2,67243 | Promoter (2-3kb) | 15 | 83037695 | 83054378 | 16684 | 2 | 109754 | ENSMUST00000162834.3 | 2071 | ENSMUSG00000018042 | Cyb5r3 | cytochrome b5 reductase 3 |
| chr9 | 1,1E+08 | 1,1E+08 | 1923 | * | 22,56637 | 2,474127659 | 0,854676 | 2,894814 | 0,003794 | 0,613712777 | 2,67243 | Promoter (<=1kb) | 9 | 1,1E+08 | 1,1E+08 | 16151 | 1 | 72341 | ENSMUST00000199592.5 | 0 | ENSMUSG00000054836 | Elp6 | elongator acetyltransferase complex subunit 6 |
| chr7 | 1,31E+08 | 1,31E+08 | 1335 | * | 14,3358 | 2,475303729 | 0,90016 | 2,74985 | 0,005962 | 0,81815673 | 1,2763 | Intron (ENSMUST00000 | 7 | 1,31E+08 | 1,31E+08 | 774 | 1 | 56213 | ENSMUST00000140741.2 | -12753 | ENSMUSG00000006205 | Htra1 | HtrA serine peptidase 1 |
| chr5 | 1,25E+08 | 1,25E+08 | 2045 | * | 15,45072 | 2,475351995 | 0,861544 | 2,873159 | 0,004064 | 0,755814512 | 1,214904 | Distal Intergenic | 5 | 1,25E+08 | 1,25E+08 | 1528 | 2 | 22190 | ENSMUST00000108707.3 | 9552 | ENSMUSG00000008348 | Ubc | ubiquitin C |
| chr7 | 15768243 | 15771172 | 2930 | * | 25,5136 | 2,475627609 | 0,844975 | 2,929825 | 0,003392 | 0,570555504 | 2,951062 | Intron (ENSMUST00000 | 7 | 15704597 | 15781846 | 77250 | 2 | 243842 | ENSMUST00000210781.2 | 10674 | ENSMUSG00000070808 | Bicra | BRD4 interacting chromatin remodeling complex associated protein |
| chr9 | 61950129 | 61952204 | 2076 | * | 16,49534 | 2,475653123 | 0,788025 | 3,141592 | 0,00168 | 0,464338132 | 2,663619 | Distal Intergenic | 9 | 61861020 | 61934138 | 73119 | 2 | 74090 | ENSMUST00000034817.11 | -15991 | ENSMUSG00000032278 | Paqr5 | progestin and adipoQ receptor family member V |
| chr7 | 45139087 | 45142518 | 3432 | * | 30,24888 | 2,475871352 | 0,615253 | 4,024148 | 5,72E-05 | 0,079351667 | 2,67243 | Promoter (<=1kb) | 7 | 45120411 | 45138229 | 17819 | 2 | 71755 | ENSMUST00000011526.7 | -858 | ENSMUSG00000011382 | Dhdh | dihydrodiol dehydrogenase (dimeric) |
| chr1 | 73187389 | 73188354 | 966 | * | 11,50574 | 2,476594243 | 0,957246 | 2,587207 | 0,009676 | 0,900637308 | 1,2763 | Distal Intergenic | 1 | 73308223 | 73345519 | 37297 | 1 | 1,05E+08 | ENSMUST00000190578.2 | -119869 | | Gm29185 | predicted gene 29185 |
| chr2 | 75891829 | 75893496 | 1668 | * | 12,07593 | 2,476991534 | 0,949646 | 2,608331 | 0,009098 | 0,973898515 | 0,738441 | Intron (ENSMUST00000 | 2 | 75819485 | 75836387 | 16903 | 2 | 241489 | ENSMUST00000124825.2 | -55442 | ENSMUSG00000075270 | Pde11a | phosphodiesterase 11A |
| chrX | 1,54E+08 | 1,54E+08 | 1838 | * | 17,0971 | 2,477207887 | 0,758291 | 3,266831 | 0,001088 | 0,393994242 | 2,663619 | Distal Intergenic | 20 | 1,53E+08 | 1,53E+08 | 108 | 1 | 1,15E+08 | ENSMUST00000082884.3 | 54590 | ENSMUSG00000064818 | Gm22359 | predicted gene, 22359 |
| chr1 | 1,91E+08 | 1,91E+08 | 1699 | * | 12,78366 | 2,477329164 | 0,954958 | 2,594176 | 0,009482 | 0,978028525 | 0,738441 | Distal Intergenic | 1 | 1,91E+08 | 1,91E+08 | 28709 | 2 | 1,03E+08 | ENSMUST00000192778.2 | -17274 | ENSMUSG00000103114 | Gm32200 | predicted gene, 32200 |
| chr2 | 25114402 | 25116613 | 2212 | * | 61,129619 | 2,477385192 | 0,5338277 | 4,6407957 | 3,471E-06 | 0,008394893 | 3,439501 | Promoter (<=1kb) | 2 | 25112172 | 25114714 | 2543 | 2 | 227613 | ENSMUST00000043584.5 | 0 | ENSMUSG00000036752 | Tubb4b | tubulin, beta 4B class IVB |
| chr5 | 39078508 | 39079804 | 1297 | * | 12,77147 | 2,477423129 | 0,944613 | 2,622685 | 0,008724 | 0,93486226 | 0,918255 | Distal Intergenic | 5 | 38863805 | 39034155 | 170351 | 2 | 27278 | ENSMUST00000171633.5 | -44353 | ENSMUSG00000039315 | Clnk | cytokine-dependent hematopoietic cell linker |
| chr9 | 16980569 | 16981862 | 1294 | * | 13,31311 | 2,478028675 | 0,928209 | 2,669689 | 0,007592 | 0,858257847 | 1,2763 | Distal Intergenic | 9 | 15821485 | 16412581 | 591097 | 2 | 270120 | ENSMUST00000217308.3 | -567988 | ENSMUSG00000074505 | Fat3 | FAT atypical cadherin 3 |
| chr5 | 34738326 | 34740313 | 1988 | * | 17,99754 | 2,478633113 | 0,890916 | 2,782117 | 0,005401 | 0,683210259 | 2,507013 | Intron (ENSMUST00000 | 5 | 34731495 | 34731557 | 63 | 1 | 1,02E+08 | ENSMUST00000184791.3 | 6831 | ENSMUSG00000099082 | Mir7036b | microRNA 7036b |
| chr3 | 89887967 | 89890185 | 2219 | * | 18,74357 | 2,478852743 | 0,940886 | 2,634594 | 0,008424 | 0,767135014 | 2,386008 | Distal Intergenic | 3 | 89875306 | 89875470 | 165 | 2 | 1,15E+08 | ENSMUST00000083111.3 | -12497 | ENSMUSG00000065045 | Gm24046 | predicted gene, 24046 |
| chr19 | 6177494 | 6178951 | 1458 | * | 38,661476 | 2,479134059 | 0,5733419 | 4,3240067 | 1,532E-05 | 0,0307765 | 2,7254908 | Promoter (<=1kb) | 19 | 6169429 | 6178334 | 8906 | 2 | 69024 | ENSMUST00000025702.14 | 0 | ENSMUSG00000024787 | Snx15 | sorting nexin 15 |
| chr14 | 25638093 | 25639669 | 1577 | * | 16,00692 | 2,479169311 | 0,913483 | 2,713974 | 0,006648 | 0,72659841 | 2,386008 | Intron (ENSMUST00000 | 14 | 25657254 | 25658829 | 1576 | 1 | 328365 | ENSMUST00000162546.2 | -17585 | ENSMUSG00000007817 | Zmiz1 | zinc finger, MIZ-type containing 1 |
| chr8 | 1,05E+08 | 1,05E+08 | 1121 | * | 12,78049 | 2,479413361 | 0,874201 | 2,836207 | 0,004565 | 0,864414194 | 0,738441 | Promoter (2-3kb) | 8 | 1,05E+08 | 1,05E+08 | 1859 | 2 | 68523 | ENSMUST00000163140.2 | 2237 | ENSMUSG00000031879 | Ciao2b | cytosolic iron-sulfur assembly component 2B |
| chr6 | 72199542 | 72200451 | 910 | * | 16,69189 | 2,479659523 | 0,888471 | 2,79093 | 0,005256 | 0,684274806 | 2,386008 | Intron (ENSMUST00000 | 6 | 72183161 | 72211521 | 28361 | 2 | 71093 | ENSMUST00000206425.2 | 11070 | ENSMUSG00000037621 | Atoh8 | atonal bHLH transcription factor 8 |
| chr14 | 30332719 | 30333763 | 1045 | * | 12,58695 | 2,480584549 | 0,958494 | 2,588001 | 0,009653 | 0,912372166 | 1,173325 | Promoter (<=1kb) | 14 | 30317315 | 30332259 | 14945 | 2 | 18753 | ENSMUST00000022521.13 | -460 | ENSMUSG00000021948 | Prkcd | protein kinase C, delta |
| chr9 | 62719019 | 62722593 | 3575 | * | 35,206609 | 2,480707615 | 0,5504911 | 4,5063534 | 6,595E-06 | 0,016860099 | 2,7254908 | Promoter (<=1kb) | 9 | 62699103 | 62719216 | 20114 | 2 | 14155 | ENSMUST00000034775.10 | 0 | ENSMUSG00000032244 | Fem1b | fem 1 homolog b |
| chr4 | 91276057 | 91277041 | 985 | * | 10,61597 | 2,480720891 | 0,940219 | 2,63845 | 0,008329 | 0,890237282 | 1,173325 | Intron (ENSMUST00000 | 4 | 91268669 | 91288213 | 19545 | 2 | 15569 | ENSMUST00000144769.2 | 11172 | ENSMUSG00000008489 | Elavl2 | ELAV like RNA binding protein 1 |
| chr12 | 84582171 | 84583899 | 1729 | * | 18,79111 | 2,481433798 | 0,849791 | 2,92005 | 0,0035 | 0,61075929 | 2,507013 | Intron (ENSMUST00000 | 12 | 84616536 | 84642231 | 25696 | 1 | 12677 | ENSMUST00000021665.12 | -32637 | ENSMUSG00000021239 | Vsx2 | visual system homeobox 2 |
| chr6 | 13677960 | 13680138 | 2179 | * | 23,95663 | 2,481444905 | 0,772253 | 3,213252 | 0,001312 | 0,407736912 | 2,951933 | Promoter (<=1kb) | 6 | 13625672 | 13677965 | 52294 | 2 | 101148 | ENSMUST00000045235.8 | 0 | ENSMUSG00000042742 | Bmt2 | base methyltransferase of 25S rRNA 2 |
| chr6 | 86877185 | 86879119 | 1935 | * | 20,34706 | 2,481758176 | 0,822788 | 3,01628 | 0,002559 | 0,554723541 | 2,386008 | Intron (ENSMUST00000 | 6 | 86828167 | 86912489 | 84323 | 1 | 269774 | ENSMUST00000156885.2 | 49018 | ENSMUSG00000057230 | Aak1 | AP2 associated kinase 1 |
| chr4 | 24094032 | 24096438 | 2407 | * | 17,30871 | 2,481762322 | 0,893862 | 2,776449 | 0,005496 | 0,677933849 | 2,663619 | Distal Intergenic | 4 | 24496451 | 24581338 | 84888 | 1 | 212377 | ENSMUST00000173079.8 | -400013 | ENSMUSG00000045751 | Mms22l | MMS22-like, DNA repair protein |
| chr14 | 24904800 | 24906495 | 1696 | * | 18,61687 | 2,482352862 | 0,864833 | 2,870327 | 0,0041 | 0,665053167 | 2,154504 | Distal Intergenic | 14 | 25109808 | 25113350 | 3543 | 1 | 1,03E+08 | ENSMUST00000225462.2 | -203313 | ENSMUSG00000114288 | Gm32123 | predicted gene, 32123 |
| chr12 | 37834387 | 37836598 | 2212 | * | 15,39189 | 2,482516168 | 0,946773 | 2,622081 | 0,008739 | 0,864017288 | 1,41696 | Distal Intergenic | 12 | 37867725 | 38032009 | 164285 | 1 | 217480 | ENSMUST00000222337.2 | -31127 | ENSMUSG00000036095 | Dgkb | diacylglycerol kinase, beta |
| chr11 | 13404374 | 13405873 | 1500 | * | 12,20653 | 2,482709727 | 0,907589 | 2,7355 | 0,006229 | 0,824314641 | 1,2763 | Distal Intergenic | 11 | 14022103 | 14129041 | 106939 | 2 | 75280 | ENSMUST00000126260.2 | 723168 | ENSMUSG00000085452 | 4930554G24Rik | RIKEN cDNA 4930554G24 gene |
| chr18 | 67872645 | 67874560 | 1916 | * | 18,21132 | 2,482781286 | 0,864897 | 2,87061 | 0,004097 | 0,640566274 | 2,507013 | Distal Intergenic | 18 | 67804282 | 67857665 | 53384 | 2 | 19255 | ENSMUST00000122412.2 | -14980 | ENSMUSG00000024539 | Ptpn2 | protein tyrosine phosphatase, non-receptor type 2 |
| chr9 | 23375658 | 23377762 | 2105 | * | 24,24516 | 2,48348907 | 0,666696 | 3,72507 | 0,000195 | 0,156945882 | 2,951062 | Intron (ENSMUST00000 | 9 | 23285228 | 23318127 | 32900 | 1 | 73230 | ENSMUST00000214050.2 | 90430 | ENSMUSG00000031963 | Bmper | BMP-binding endothelial regulator |
| chr19 | 47259376 | 47260607 | 1232 | * | 13,09775 | 2,484164125 | 0,933871 | 2,660072 | 0,007812 | 0,893166641 | 1,075735 | Promoter (1-2kb) | 19 | 47261958 | 47262028 | 71 | 2 | 1,02E+08 | ENSMUST00000183658.3 | 1421 | ENSMUSG00000098786 | Mir6995 | microRNA 6995 |
| chr11 | 1,15E+08 | 1,15E+08 | 1879 | * | 17,97778 | 2,484309202 | 0,89949 | 2,761909 | 0,005746 | 0,717868719 | 2,154504 | Promoter (<=1kb) | 11 | 1,15E+08 | 1,15E+08 | 2837 | 1 | 286942 | ENSMUST00000138340.2 | 0 | ENSMUSG00000010021 | Kif19a | kinesin family member 19A |
| chr14 | 65643733 | 65645487 | 1755 | * | 10,75747 | 2,484480212 | 0,942188 | 2,636927 | 0,008366 | 0,96198404 | 0,738441 | Intron (ENSMUST00000 | 14 | 65633129 | 65634199 | 1071 | 1 | 380912 | ENSMUST00000225512.2 | 10604 | ENSMUSG00000034522 | Zfp395 | zinc finger protein 395 |
| chr10 | 84244932 | 84248309 | 3378 | * | 29,84056 | 2,484829644 | 0,721421 | 3,444355 | 0,000572 | 0,284660899 | 2,858511 | Exon (ENSMUST00000 | 10 | 84211160 | 84228252 | 17093 | 2 | 77976 | ENSMUST00000129355.2 | -16680 | ENSMUSG00000020032 | Nuak1 | NUAK family, SNF1-like kinase, 1 |
| chr11 | 1,11E+08 | 1,11E+08 | 1248 | * | 17,94792 | 2,484992493 | 0,819412 | 3,032651 | 0,002424 | 0,543993049 | 2,386008 | Distal Intergenic | 11 | 1,1E+08 | 1,1E+08 | 1893 | 1 | 1,03E+08 | ENSMUST00000144689.2 | 168136 | | Gm11682 | predicted gene 11682 |
| chr5 | 1,35E+08 | 1,35E+08 | 1158 | * | 11,46164 | 2,486534582 | 0,930414 | 2,672505 | 0,007529 | 0,857326444 | 1,2763 | Promoter (1-2kb) | 5 | 1,35E+08 | 1,35E+08 | 771 | 1 | 79565 | ENSMUST00000148286.2 | 1710 | ENSMUSG00000040557 | Mettl27 | methyltransferase like 27 |
| chr15 | 73213421 | 73214795 | 1375 | * | 16,60209 | 2,486790465 | 0,802551 | 3,098608 | 0,001944 | 0,489733767 | 2,663619 | Promoter (<=1kb) | 15 | 73087406 | 73215176 | 127771 | 2 | 14083 | ENSMUST00000228628.2 | 381 | ENSMUSG00000022607 | Ptk2 | PTK2 protein tyrosine kinase 2 |
| chr15 | 55009821 | 55011593 | 1773 | * | 17,79289 | 2,48698814 | 0,805705 | 3,086723 | 0,002024 | 0,505746397 | 2,507013 | Intron (ENSMUST00000 | 15 | 54996849 | 55072281 | 75433 | 1 | 97998 | ENSMUST00000226687.2 | 12972 | ENSMUSG00000022419 | Deptor | DEP domain containing MTOR-interacting protein |
| chr8 | 1,18E+08 | 1,18E+08 | 2442 | * | 32,31393 | 2,487133384 | 0,711202 | 3,497085 | 0,00047 | 0,283414546 | 2,377674 | Distal Intergenic | 8 | 1,18E+08 | 1,18E+08 | 136852 | 1 | 234779 | ENSMUST00000081232.9 | -11200 | ENSMUSG00000034330 | Plcg2 | phospholipase C, gamma 2 |
| chr11 | 22679589 | 22680713 | 1125 | * | 13,8764 | 2,48779267 | 0,881574 | 2,82199 | 0,004773 | 0,837097674 | 0,918255 | Exon (ENSMUST00000 | 11 | 22567408 | 22568487 | 1080 | 1 | 1E+08 | ENSMUST00000142151.2 | 112181 | | Gm12052 | predicted gene 12052 |
| chr8 | 95977193 | 95979603 | 2411 | * | 26,4837 | 2,488106565 | 0,733959 | 3,389981 | 0,000699 | 0,314438111 | 2,725491 | Exon (ENSMUST00000 | 8 | 95967609 | 96008218 | 40610 | 2 | 333329 | ENSMUST00000120044.9 | 28615 | ENSMUSG00000031789 | Cngb1 | cyclic nucleotide gated channel beta 1 |
| chr17 | 86452841 | 86454096 | 1256 | * | 13,30585 | 2,48866991 | 0,958831 | 2,595525 | 0,009445 | 0,944637293 | 0,938682 | Promoter (<=1kb) | 17 | 86292093 | 86452603 | 160511 | 2 | 78586 | ENSMUST00000095187.4 | -238 | ENSMUSG00000024135 | Srbd1 | S1 RNA binding domain 1 |
| chr5 | 1,19E+08 | 1,19E+08 | 2275 | * | 18,4801 | 2,489107537 | 0,884987 | 2,812593 | 0,004914 | 0,675437713 | 2,423594 | Distal Intergenic | 5 | 1,19E+08 | 1,19E+08 | 13530 | 1 | 76199 | ENSMUST00000202787.2 | 91748 | ENSMUSG00000018076 | Med13l | mediator complex subunit 13-like |
| chr14 | 1,22E+08 | 1,22E+08 | 2688 | * | 31,46629 | 2,491363926 | 0,662245 | 3,761998 | 0,000169 | 0,15128023 | 2,725491 | Distal Intergenic | 14 | 1,22E+08 | 1,22E+08 | 4033 | 2 | 1,03E+08 | ENSMUST00000227451.2 | 18961 | | Gm33299 | predicted gene, 33299 |
| chr15 | 77597810 | 77599086 | 1277 | * | 13,00363 | 2,491380159 | 0,936628 | 2,659947 | 0,007815 | 0,862488992 | 1,2763 | Exon (ENSMUST00000 | 15 | 77601836 | 77607049 | 5214 | 2 | 1,03E+08 | ENSMUST00000230236.2 | 7963 | | Gm36329 | predicted gene, 36329 |
| chr2 | 31830677 | 31832331 | 1655 | * | 17,72835 | 2,491476518 | 0,823839 | 3,024228 | 0,002493 | 0,547497234 | 2,423594 | Exon (ENSMUST00000 | 2 | 31840151 | 31863454 | 23304 | 1 | 108897 | ENSMUST00000001920.13 | -7820 | ENSMUSG00000001864 | Aif1l | allograft inflammatory factor 1-like |
| chr4 | 1,38E+08 | 1,38E+08 | 1397 | * | 12,37198 | 2,491666398 | 0,940478 | 2,649361 | 0,008064 | 0,886332554 | 1,173325 | Distal Intergenic | 4 | 1,38E+08 | 1,38E+08 | 7094 | 2 | 26971 | ENSMUST00000030526.7 | -6875 | ENSMUSG00000028749 | Pla2g2f | phospholipase A2, group IIF |
| chr19 | 60094966 | 60097270 | 2305 | * | 15,21667 | 2,492686857 | 0,87353 | 2,853578 | 0,004323 | 0,758328549 | 1,2763 | Distal Intergenic | 19 | 60132997 | 60160005 | 27009 | 1 | 107376 | ENSMUST00000244672.1 | -35727 | ENSMUSG00000053117 | E330013P04Rik | RIKEN cDNA E330013P04 gene |
| chr8 | 1,12E+08 | 1,12E+08 | 1642 | * | 19,71477 | 2,493170457 | 0,725342 | 3,437233 | 0,000588 | 0,304413931 | 2,507013 | Intron (ENSMUST00000 | 8 | 1,12E+08 | 1,12E+08 | 12714 | 2 | 12927 | ENSMUST00000212147.2 | 7838 | ENSMUSG00000031955 | Bcar1 | breast cancer anti-estrogen resistance 1 |
| chr5 | 1,2E+08 | 1,2E+08 | 2801 | * | 21,43506 | 2,493214747 | 0,840604 | 2,965978 | 0,003017 | 0,539604518 | 3,104559 | Distal Intergenic | 5 | 1,2E+08 | 1,2E+08 | 9160 | 2 | 1,03E+08 | ENSMUST00000201435.2 | 44376 | ENSMUSG00000106938 | Gm31314 | predicted gene, 31314 |
| chr5 | 96980117 | 96982245 | 2129 | * | 19,04448 | 2,493338308 | 0,899026 | 2,773377 | 0,005548 | 0,684274806 | 2,507013 | Promoter (2-3kb) | 5 | 96977942 | 96993541 | 15600 | 1 | 11745 | ENSMUST00000199390.3 | 2175 | ENSMUSG00000029484 | Anxa3 | annexin A3 |
| chr10 | 66700295 | 66701493 | 1199 | * | 13,03712 | 2,493358274 | 0,941925 | 2,647089 | 0,008119 | 0,887238309 | 1,173325 | Intron (ENSMUST00000 | 10 | 66725450 | 66727391 | 1942 | 2 | 1,19E+08 | ENSMUST00000217953.2 | 25898 | | Gm47903 | predicted gene, 47903 |
| chr17 | 24634922 | 24636751 | 1830 | * | 14,01582 | 2,494253605 | 0,84414 | 2,954786 | 0,003129 | 0,759484484 | 0,918255 | Promoter (<=1kb) | 17 | 24637144 | 24639305 | 2162 | 1 | 19826 | ENSMUST00000234089.2 | -393 | ENSMUSG00000034681 | Rnps1 | RNA binding protein with serine rich domain 1 |
| chr4 | 1,31E+08 | 1,31E+08 | 1218 | * | 14,4473 | 2,494486416 | 0,879827 | 2,835202 | 0,00458 | 0,683140223 | 2,141612 | Distal Intergenic | 4 | 1,31E+08 | 1,31E+08 | 134 | 1 | 1,15E+08 | ENSMUST00000157416.3 | 104142 | ENSMUSG00000088041 | Gm25261 | predicted gene, 25261 |
| chr18 | 12757983 | 12759950 | 1968 | * | 17,62388 | 2,495305661 | 0,832774 | 2,996378 | 0,002732 | 0,560563432 | 2,507013 | Intron (ENSMUST00000 | 18 | 12776401 | 12822502 | 46102 | 1 | 72747 | ENSMUST00000141060.2 | -16451 | ENSMUSG00000024424 | Ttc39c | tetratricopeptide repeat domain 39C |
| chr1 | 86465551 | 86467266 | 1716 | * | 13,80709 | 2,496193452 | 0,911216 | 2,739408 | 0,006155 | 0,877892722 | 0,938682 | Distal Intergenic | 1 | 86454529 | 86457931 | 3403 | 1 | 19231 | ENSMUST00000188533.2 | 11022 | ENSMUSG00000026238 | Ptma | prothymosin alpha |
| chr18 | 76575438 | 76576606 | 1169 | * | 14,27033 | 2,496259634 | 0,965586 | 2,585229 | 0,009731 | 0,948006995 | 0,938682 | Distal Intergenic | 18 | 76704983 | 76720565 | 15583 | 2 | 1,03E+08 | ENSMUST00000235949.2 | 143959 | ENSMUSG00000117960 | Gm31933 | predicted gene, 31933 |
| chr15 | 99645709 | 99647405 | 1697 | * | 21,46935 | 2,498284558 | 0,837846 | 2,981796 | 0,002866 | 0,539604518 | 2,951933 | Promoter (<=1kb) | 15 | 99635216 | 99644951 | 9736 | 2 | 71949 | ENSMUST00000176970.8 | -758 | ENSMUSG00000023021 | Cers5 | ceramide synthase 5 |
| chr9 | 1,01E+08 | 1,01E+08 | 2874 | * | 20,59435 | 2,498467789 | 0,725915 | 3,441819 | 0,000578 | 0,284660899 | 2,880017 | Promoter (<=1kb) | 9 | 1,01E+08 | 1,01E+08 | 146712 | 2 | 235542 | ENSMUST00000075941.12 | 441 | ENSMUSG00000043154 | Ppp2r3a | protein phosphatase 2, regulatory subunit B'', alpha |
| chr11 | 1,01E+08 | 1,01E+08 | 2745 | * | 21,60406 | 2,499177642 | 0,830897 | 3,007807 | 0,002631 | 0,54122637 | 2,67243 | Promoter (2-3kb) | 11 | 1,01E+08 | 1,01E+08 | 3614 | 2 | 19183 | ENSMUST00000019447.15 | -2304 | ENSMUSG00000019303 | Psmc3ip | proteasome (prosome, macropain) 26S subunit, ATPase 3, interacting protein |
| chr13 | 18939112 | 18941642 | 2531 | * | 19,37599 | 2,499209618 | 0,884982 | 2,824024 | 0,004742 | 0,683140223 | 2,221611 | Intron (ENSMUST00000 | 13 | 18964013 | 19011825 | 47813 | 1 | 218035 | ENSMUST00000222813.2 | -22371 | ENSMUSG00000041236 | Vps41 | VPS41 HOPS complex subunit |
| chr11 | 83156830 | 83159232 | 2403 | * | 17,83046 | 2,499228903 | 0,78746 | 3,173785 | 0,001505 | 0,455547114 | 2,507013 | Distal Intergenic | 11 | 83162324 | 83162514 | 191 | 1 | 1,15E+08 | ENSMUST00000084010.3 | -3092 | ENSMUSG00000119838 | Gm23444 | predicted gene, 23444 |
| chr1 | 3269124 | 3270570 | 1447 | * | 11,50371 | 2,499779664 | 0,957353 | 2,611137 | 0,009024 | 0,939804939 | 0,918255 | Distal Intergenic | 1 | 3276746 | 3285855 | 9110 | 2 | 497097 | ENSMUST00000159265.2 | 15285 | ENSMUSG00000051951 | Xkr4 | X-linked Kx blood group related 4 |
| chr5 | 1,15E+08 | 1,15E+08 | 1552 | * | 17,67434 | 2,49999347 | 0,828491 | 3,017527 | 0,002548 | 0,576475101 | 2,154504 | 3' UTR | 5 | 1,15E+08 | 1,15E+08 | 72 | 1 | 1,02E+08 | ENSMUST00000183408.3 | 15056 | ENSMUSG00000098731 | Mir7027 | microRNA 7027 |
| chr7 | 12121307 | 12122854 | 1548 | * | 21,99902 | 2,500556107 | 0,653696 | 3,825257 | 0,000131 | 0,123146349 | 2,880017 | Distal Intergenic | 7 | 12094240 | 12096691 | 2452 | 2 | 171267 | ENSMUST00000086228.3 | -24616 | ENSMUSG00000066803 | Vmn1r84 | vomeronasal 1 receptor 84 |
| chr9 | 69322152 | 69323583 | 1432 | * | 12,66408 | 2,500577123 | 0,938632 | 2,664066 | 0,00772 | 0,891097601 | 1,075735 | Promoter (<=1kb) | 9 | 69323997 | 69332987 | 8991 | 1 | 93697 | ENSMUST00000146368.8 | -414 | ENSMUSG00000032235 | Ice2 | interactor of little elongation complex ELL subunit 2 |
| chr7 | 1,37E+08 | 1,37E+08 | 1114 | * | 14,9382 | 2,500809074 | 0,944286 | 2,648361 | 0,008088 | 0,852340527 | 1,41696 | Distal Intergenic | 7 | 1,37E+08 | 1,37E+08 | 4481 | 1 | 30926 | ENSMUST00000211598.2 | 281929 | ENSMUSG00000031068 | Glrx3 | glutaredoxin 3 |
| chr19 | 42314372 | 42317362 | 2991 | * | 22,76171 | 2,50086388 | 0,716188 | 3,49191 | 0,00048 | 0,262421328 | 2,858511 | Intron (ENSMUST00000 | 19 | 42271971 | 42276393 | 4423 | 2 | 72832 | ENSMUST00000238182.2 | -37979 | ENSMUSG00000042401 | Crtac1 | cartilage acidic protein 1 |
| chr14 | 31870569 | 31873140 | 2572 | * | 20,52239 | 2,500931966 | 0,790282 | 3,164607 | 0,001553 | 0,470097239 | 2,386008 | Promoter (1-2kb) | 14 | 31869544 | 31880327 | 10784 | 1 | 17695 | ENSMUST00000130397.2 | 1025 | ENSMUSG00000021907 | Msmb | beta-microseminoprotein |
| chr5 | 13707218 | 13709181 | 1964 | * | 20,71206 | 2,501189945 | 0,894936 | 2,794824 | 0,005193 | 0,669377201 | 2,67243 | Distal Intergenic | 5 | 13818676 | 13821901 | 3226 | 1 | 626053 | ENSMUST00000170514.2 | -109495 | | Gm6647 | predicted gene 6647 |
| chr7 | 75046265 | 75048479 | 2215 | * | 30,70731 | 2,501358888 | 0,652673 | 3,832486 | 0,000127 | 0,121267917 | 2,858511 | Distal Intergenic | 7 | 75105282 | 75404357 | 299076 | 1 | 75547 | ENSMUST00000166315.7 | -56803 | ENSMUSG00000066406 | Akap13 | A kinase (PRKA) anchor protein 13 |
| chr10 | 59705206 | 59706774 | 1569 | * | 16,58693 | 2,501733069 | 0,894412 | 2,79707 | 0,005157 | 0,685026746 | 2,318434 | Distal Intergenic | 10 | 59697340 | 59699225 | 1886 | 1 | 216001 | ENSMUST00000166565.2 | 7866 | ENSMUSG00000020111 | Micu1 | mitochondrial calcium uptake 1 |
| chr4 | 1,39E+08 | 1,39E+08 | 3984 | * | 37,10813 | 2,502279946 | 0,629777 | 3,97328 | 7,09E-05 | 0,095764633 | 2,338387 | Promoter (<=1kb) | 4 | 1,39E+08 | 1,39E+08 | 58401 | 1 | 12345 | ENSMUST00000042675.9 | 0 | ENSMUSG00000028745 | Capzb | capping protein (actin filament) muscle Z-line, beta |
| chr4 | 43645910 | 43648376 | 2467 | * | 23,71584 | 2,502338268 | 0,668571 | 3,742816 | 0,000182 | 0,159757724 | 2,67243 | Promoter (<=1kb) | 4 | 43646560 | 43650309 | 3750 | 1 | 230103 | ENSMUST00000130093.8 | 0 | ENSMUSG00000028469 | Npr2 | natriuretic peptide receptor 2 |
| chr5 | 1,24E+08 | 1,24E+08 | 2139 | * | 22,26006 | 2,502607576 | 0,725247 | 3,450696 | 0,000559 | 0,256659223 | 3,471678 | Promoter (2-3kb) | 5 | 1,24E+08 | 1,24E+08 | 4496 | 2 | 330192 | ENSMUST00000199741.2 | 2909 | ENSMUSG00000066278 | Vps37b | vacuolar protein sorting 37B |
| chr1 | 46892670 | 46894235 | 1566 | * | 15,02695 | 2,503831622 | 0,81104 | 3,087188 | 0,002021 | 0,620292045 | 1,37683 | Promoter (<=1kb) | 1 | 46846704 | 46893206 | 46503 | 2 | 227059 | ENSMUST00000027131.6 | 0 | ENSMUSG00000025986 | Slc39a10 | solute carrier family 39 (zinc transporter), member 10 |

| | | | | | | | | | | | | | | | | | | | | | | | |
|---|---|---|---|---|---|---|---|---|---|---|---|---|---|---|---|---|---|---|---|---|---|---|---|
| chr10 | 83516374 | 83518146 | 1773 | * | 18,08214 | 2,503874125 | 0,952791 | 2,627937 | 0,00859 | 0,760759863 | 2,507013 | Distal Intergenic | 10 | 83435897 | 83484602 | 48706 | 2 | 216190 | ENSMUST00000020500.14 | -31772 | ENSMUSG00000020263 | Appl2 | adaptor protein, phosphotyrosine interaction, PH domain and leucine zipper containing 2 |
| chr18 | 75319952 | 75322056 | 2105 | * | 21,48101 | 2,504052858 | 0,820984 | 3,050062 | 0,002288 | 0,503036965 | 2,880017 | Intron (ENSMUST0000( | 18 | 75258496 | 75284099 | 25604 | 1 | 69190 | ENSMUST00000236840.2 | 61456 | ENSMUSG00000035765 | Dym | dymeclin |
| chr3 | 1,24E+08 | 1,24E+08 | 2119 | * | 16,00659 | 2,50416103 | 0,955047 | 2,62203 | 0,008741 | 0,886041774 | 1,2763 | Distal Intergenic | 3 | 1,24E+08 | 1,24E+08 | 7885 | 2 | 384775 | ENSMUST00000180033.8 | -3554 | ENSMUSG00000039174 | 1700003H04Rik | RIKEN cDNA 1700003H04 gene |
| chr19 | 42106462 | 42108001 | 1540 | * | 13,56826 | 2,504232725 | 0,867555 | 2,886542 | 0,003895 | 0,838625535 | 0,738441 | Exon (ENSMUST00000 | 19 | 42103117 | 42106985 | 3869 | 1 | 84095 | ENSMUST00000238085.2 | 3345 | ENSMUSG00000025178 | Pi4k2a | phosphatidylinositol 4-kinase type 2 alpha |
| chr15 | 64176921 | 64178092 | 1172 | * | 12,35482 | 2,504441035 | 0,938805 | 2,66769 | 0,007637 | 0,858755542 | 1,2763 | Intron (ENSMUST0000( | 15 | 64063386 | 64183538 | 120153 | 2 | 13196 | ENSMUST00000176516.2 | 5446 | ENSMUSG00000022377 | Asap1 | ArfGAP with SH3 domain, ankyrin repeat and PH domain1 |
| chr17 | 56931849 | 56933599 | 1751 | * | 21,06275 | 2,504897239 | 0,829682 | 3,019104 | 0,002535 | 0,524362003 | 2,880017 | Promoter (<=1kb) | 17 | 56921297 | 56933875 | 12579 | 2 | 74142 | ENSMUST00000233732.2 | 276 | ENSMUSG00000041168 | Lonp1 | lon peptidase 1, mitochondrial |
| chr5 | 1,36E+08 | 1,36E+08 | 1948 | * | 20,57257 | 2,505504735 | 0,782215 | 3,203089 | 0,00136 | 0,428853558 | 2,663619 | Promoter (1-2kb) | 5 | 1,36E+08 | 1,36E+08 | 9833 | 1 | 69665 | ENSMUST00000139999.8 | -1349 | ENSMUSG00000006143 | Upk3bl | uroplakin 3B-like |
| chr15 | 78917613 | 78919171 | 1559 | * | 13,71407 | 2,505764125 | 0,891686 | 2,810143 | 0,004952 | 0,838625535 | 0,938682 | Promoter (<=1kb) | 15 | 78917698 | 78920513 | 2816 | 1 | 26912 | ENSMUST00000229276.2 | 0 | ENSMUSG00000006378 | Gcat | glycine C-acetyltransferase (2-amino-3-ketobutyrate-coenzyme A ligase) |
| chr11 | 94212895 | 94214835 | 1941 | * | 16,47832 | 2,505900554 | 0,878539 | 2,852349 | 0,00434 | 0,639997962 | 2,663619 | Promoter (<=1kb) | 11 | 94181965 | 94212814 | 30850 | 2 | 67684 | ENSMUST00000107820.2 | -81 | ENSMUSG00000020863 | Luc7l3 | LUC7-like 3 (S. cerevisiae) |
| chr12 | 3627468 | 3629318 | 1851 | * | 14,95184 | 2,50651013 | 0,86919 | 2,88373 | 0,00393 | 0,730469267 | 1,37683 | Intron (ENSMUST0000( | 12 | 3622984 | 3831391 | 208408 | 1 | 13528 | ENSMUST00000077930.13 | 4484 | ENSMUSG00000071454 | Dtnb | dystrobrevin, beta |
| chr9 | 62256066 | 62257822 | 1757 | * | 26,77375 | 2,506936309 | 0,703949 | 3,561249 | 0,000369 | 0,236733567 | 2,67243 | Intron (ENSMUST0000( | 9 | 62252081 | 62281997 | 29917 | 1 | 11737 | ENSMUST00000126451.8 | 3985 | ENSMUSG00000032249 | Anp32a | acidic (leucine-rich) nuclear phosphoprotein 32 family, member A |
| chr8 | 25505307 | 25507793 | 2487 | * | 20,39156 | 2,507063817 | 0,739546 | 3,390003 | 0,000699 | 0,326832856 | 2,507013 | Promoter (<=1kb) | 8 | 25439627 | 25506938 | 67312 | 2 | 11502 | ENSMUST00000084035.12 | 0 | ENSMUSG00000031555 | Adam9 | a disintegrin and metallopeptidase domain 9 (meltrin gamma) |
| chr19 | 44202915 | 44204379 | 1465 | * | 11,72867 | 2,50726508 | 0,966871 | 2,593174 | 0,009509 | 0,948006995 | 0,918255 | Exon (ENSMUST00000( | 19 | 44191708 | 44232455 | 40748 | 1 | 30049 | ENSMUST00000026220.7 | 11207 | ENSMUSG00000025202 | Scd3 | stearoyl-coenzyme A desaturase 3 |
| chr3 | 1,39E+08 | 1,39E+08 | 2074 | * | 16,91425 | 2,50782095 | 0,877657 | 2,857405 | 0,004271 | 0,654756507 | 2,386008 | Intron (ENSMUST0000( | 3 | 1,39E+08 | 1,39E+08 | 42165 | 1 | 56224 | ENSMUST00000121826.3 | -4991 | ENSMUSG00000028152 | Tspan5 | tetraspanin 5 |
| chr1 | 1,56E+08 | 1,56E+08 | 1699 | * | 16,93849 | 2,508045528 | 0,806364 | 3,110314 | 0,001869 | 0,480547165 | 2,663619 | Downstream (<=300bp | 1 | 1,56E+08 | 1,56E+08 | 1630 | 1 | 240832 | ENSMUST00000125371.2 | 5814 | ENSMUSG00000050565 | Tor1aip2 | torsin A interacting protein 2 |
| chr6 | 1,2E+08 | 1,2E+08 | 1358 | * | 12,70318 | 2,50808432 | 0,965618 | 2,597388 | 0,009394 | 0,92207189 | 1,075735 | Intron (ENSMUST0000( | 6 | 1,2E+08 | 1,2E+08 | 22976 | 2 | 232341 | ENSMUST00000161392.8 | -16545 | ENSMUSG00000045962 | Wnk1 | WNK lysine deficient protein kinase 1 |
| chr5 | 1,25E+08 | 1,25E+08 | 1439 | * | 14,36516 | 2,508350584 | 0,871576 | 2,877947 | 0,004003 | 0,801054956 | 0,938682 | Distal Intergenic | 5 | 1,25E+08 | 1,25E+08 | 34502 | 2 | 1,03E+08 | ENSMUST00000198586.2 | 41915 | | Gm32585 | predicted gene, 32585 |
| chr19 | 14643067 | 14644518 | 1452 | * | 11,88187 | 2,50883917 | 0,87226 | 2,87625 | 0,004024 | 0,843085444 | 0,744375 | Distal Intergenic | 19 | 14425514 | 14575415 | 149902 | 2 | 21888 | ENSMUST00000167776.3 | -67652 | ENSMUSG00000024642 | Tle4 | transducin-like enhancer of split 4 |
| chr9 | 42929427 | 42931724 | 2298 | * | 19,13709 | 2,510325112 | 0,805076 | 3,118123 | 0,00182 | 0,507398906 | 2,221611 | 3' UTR | 9 | 42937300 | 42939114 | 1815 | 2 | 69632 | ENSMUST00000214175.2 | 7390 | ENSMUSG00000059495 | Arhgef12 | Rho guanine nucleotide exchange factor (GEF) 12 |
| chr6 | 52881932 | 52883246 | 1315 | * | 16,67025 | 2,510372141 | 0,960741 | 2,612955 | 0,008976 | 0,889364623 | 1,2763 | Intron (ENSMUST0000( | 6 | 52745782 | 52886605 | 140824 | 2 | 231986 | ENSMUST00000136250.6 | 3359 | ENSMUSG00000063568 | Jazf1 | JAZF zinc finger 1 |
| chr4 | 1,29E+08 | 1,29E+08 | 1616 | * | 15,13417 | 2,511043992 | 0,843209 | 2,977961 | 0,002902 | 0,681459438 | 1,37683 | Promoter (2-3kb) | 4 | 1,29E+08 | 1,29E+08 | 10300 | 1 | 11637 | ENSMUST00000141982.2 | -2417 | ENSMUSG00000028792 | Ak2 | adenylate kinase 2 |
| chr14 | 57984531 | 57985793 | 1263 | * | 11,89409 | 2,511564047 | 0,891342 | 2,817734 | 0,004836 | 0,876195292 | 0,744375 | Promoter (<=1kb) | 14 | 57971710 | 57984035 | 12326 | 2 | 50523 | ENSMUST00000173077.2 | -496 | ENSMUSG00000021959 | Lats2 | large tumor suppressor 2 |
| chr7 | 44773537 | 44775381 | 1845 | * | 22,39873 | 2,511740226 | 0,646807 | 3,883293 | 0,000103 | 0,104080853 | 2,951933 | Promoter (<=1kb) | 7 | 44771803 | 44773863 | 2061 | 2 | 27207 | ENSMUST00000003521.10 | 0 | ENSMUSG00000003429 | Rps11 | ribosomal protein S11 |
| chr16 | 11729006 | 11730743 | 1738 | * | 15,51685 | 2,512076082 | 0,94177 | 2,6674 | 0,007644 | 0,849859156 | 1,37683 | Promoter (1-2kb) | 16 | 11702087 | 11727309 | 25223 | 2 | 223978 | ENSMUST00000073371.7 | -1697 | ENSMUSG00000065979 | Cpped1 | calcineurin-like phosphoesterase domain containing 1 |
| chr2 | 16811479 | 16813177 | 1699 | * | 16,91948 | 2,512356619 | 0,842863 | 2,980743 | 0,002876 | 0,55735526 | 2,663619 | Distal Intergenic | 2 | 16643507 | 16756501 | 112995 | 1 | 67448 | ENSMUST00000126173.2 | 167972 | ENSMUSG00000026748 | Plxdc2 | plexin domain containing 2 |
| chr14 | 76070962 | 76072479 | 1518 | * | 13,76764 | 2,512568565 | 0,935221 | 2,686603 | 0,007218 | 0,902600349 | 0,938682 | Distal Intergenic | 14 | 76080819 | 76082404 | 1586 | 2 | 1E+08 | ENSMUST00000131222.2 | 9925 | | Gm4285 | predicted gene 4285 |
| chr1 | 93394702 | 93395940 | 1239 | * | 12,57672 | 2,512751131 | 0,946298 | 2,655348 | 0,007923 | 0,883601315 | 1,173325 | Intron (ENSMUST0000( | 1 | 93368410 | 93406099 | 37690 | 2 | 110611 | ENSMUST00000188165.2 | 10159 | ENSMUSG00000034088 | Hdlbp | high density lipoprotein (HDL) binding protein |
| chr19 | 38272669 | 38274357 | 1689 | * | 14,20106 | 2,513655596 | 0,919619 | 2,733367 | 0,006269 | 0,855969265 | 1,075735 | Intron (ENSMUST0000( | 19 | 38253858 | 38289717 | 35860 | 1 | 56839 | ENSMUST00000130039.3 | 18811 | ENSMUSG00000067242 | Lgi1 | leucine-rich repeat LGI family, member 1 |
| chr9 | 69985271 | 69988654 | 3384 | * | 30,14747 | 2,513967488 | 0,696444 | 3,609721 | 0,000307 | 0,214706231 | 2,67243 | Distal Intergenic | 9 | 70009832 | 70010358 | 527 | 2 | 76886 | ENSMUST00000145427.2 | 21704 | ENSMUSG00000032224 | Fam81a | family with sequence similarity 81, member A |
| chr10 | 62322385 | 62325564 | 3180 | * | 23,54284 | 2,515239384 | 0,70601 | 3,56261 | 0,000367 | 0,224762615 | 2,951062 | Promoter (<=1kb) | 10 | 62291051 | 62322511 | 31461 | 2 | 30930 | ENSMUST00000219574.2 | 0 | ENSMUSG00000020078 | Vps26a | VPS26 retromer complex component A |
| chr19 | 50828171 | 50829922 | 1752 | * | 15,37813 | 2,51564311 | 0,86878 | 2,895604 | 0,003784 | 0,651709303 | 2,141612 | Distal Intergenic | 19 | 50136098 | 50667084 | 530987 | 2 | 58178 | ENSMUST00000164039.9 | -161087 | ENSMUSG00000043531 | Sorcs1 | sortilin-related VPS10 domain containing receptor 1 |
| chr13 | 63233336 | 63234971 | 1636 | * | 12,74738 | 2,515936597 | 0,903977 | 2,783187 | 0,005383 | 0,829907213 | 1,075735 | Intron (ENSMUST0000( | 13 | 63215929 | 63388125 | 172197 | 1 | 72061 | ENSMUST00000221676.2 | 17407 | ENSMUSG00000021458 | Aopep | aminopeptidase O |
| chr17 | 34100813 | 34102782 | 1970 | * | 16,07561 | 2,516560256 | 0,83367 | 3,018655 | 0,002539 | 0,577007376 | 2,141612 | Promoter (<=1kb) | 17 | 34094642 | 34102200 | 7559 | 2 | 1,01E+08 | ENSMUST00000173982.2 | 0 | ENSMUSG00000079553 | Kifc1 | kinesin family member C1 |
| chr10 | 80240359 | 80241652 | 1294 | * | 15,36399 | 2,516935856 | 0,965753 | 2,606191 | 0,009156 | 0,801134483 | 2,141612 | Promoter (<=1kb) | 10 | 80238831 | 80242600 | 3770 | 2 | 66594 | ENSMUST00000138428.8 | 948 | ENSMUSG00000020163 | Uqcr11 | ubiquinol-cytochrome c reductase, complex III subunit XI |
| chr12 | 69434242 | 69435951 | 1710 | * | 20,68033 | 2,5169939 | 0,75652 | 3,327069 | 0,000878 | 0,337811311 | 2,880017 | Distal Intergenic | 12 | 69418924 | 69420557 | 1634 | 1 | 11845 | ENSMUST00000050063.9 | 15318 | ENSMUSG00000044147 | Arf6 | ADP-ribosylation factor 6 |
| chr9 | 21293917 | 21295990 | 2074 | * | 21,08349 | 2,517063246 | 0,778415 | 3,233574 | 0,001223 | 0,385754389 | 3,104559 | Exon (ENSMUST00000 | 9 | 21299827 | 21306696 | 6870 | 1 | 16201 | ENSMUST00000214821.2 | -3837 | ENSMUSG00000032178 | Ilf3 | interleukin enhancer binding factor 3 |
| chr1 | 78296417 | 78298075 | 1659 | * | 14,86483 | 2,517503786 | 0,945239 | 2,663352 | 0,007737 | 0,845747925 | 1,41696 | Intron (ENSMUST0000( | 1 | 78286982 | 78396926 | 109945 | 1 | 433323 | ENSMUST00000036172.10 | 9435 | ENSMUSG00000032908 | Sgpp2 | sphingosine-1-phosphate phosphatase 2 |
| chr5 | 88911661 | 88914638 | 2978 | * | 40,850416 | 2,517636007 | 0,4963071 | 5,0727385 | 3,921E-07 | 0,001777866 | 2,9254973 | Promoter (<=1kb) | 5 | 88912855 | 88931140 | 18286 | 1 | 13178 | ENSMUST00000031311.10 | 0 | ENSMUSG00000029366 | Dck | deoxycytidine kinase |
| chr15 | 93940081 | 93941119 | 1039 | * | 14,91166 | 2,517705757 | 0,94816 | 2,655359 | 0,007922 | 0,853738252 | 1,37683 | Distal Intergenic | 15 | 93802737 | 93824992 | 22256 | 2 | 77248 | ENSMUST00000230109.2 | -115089 | ENSMUSG00000116360 | 9430014N10Rik | RIKEN cDNA 9430014N10 gene |
| chr6 | 125137010 | 125139957 | 2948 | * | 48,947738 | 2,518271624 | 0,5462392 | 4,6101993 | 4,023E-06 | 0,011411459 | 2,7254908 | Promoter (<=1kb) | 6 | 125138816 | 125139490 | 675 | 2 | 14433 | ENSMUST00000147954.2 | 0 | ENSMUSG00000057666 | Gapdh | glyceraldehyde-3-phosphate dehydrogenase |
| chr9 | 22113235 | 22114733 | 1499 | * | 13,45044 | 2,5188225 | 0,865464 | 2,910373 | 0,00361 | 0,824944942 | 0,738441 | 3' UTR | 9 | 22108363 | 22111920 | 3558 | 1 | 619310 | ENSMUST00000091508.5 | 4872 | ENSMUSG00000074472 | Zfp872 | zinc finger protein 872 |
| chr13 | 1,02E+08 | 1,02E+08 | 1807 | * | 16,76036 | 2,519004022 | 0,875485 | 2,877268 | 0,004011 | 0,748382289 | 1,2763 | Distal Intergenic | 13 | 1,02E+08 | 1,02E+08 | 8582 | 1 | 1,03E+08 | ENSMUST00000243228.1 | -147334 | | Gm36703 | predicted gene, 36703 |
| chr4 | 1,29E+08 | 1,29E+08 | 1602 | * | 14,98192 | 2,51924086 | 0,891773 | 2,824979 | 0,004728 | 0,786955198 | 1,214904 | Promoter (<=1kb) | 4 | 1,29E+08 | 1,29E+08 | 10300 | 1 | 11637 | ENSMUST00000141982.2 | -32 | ENSMUSG00000028792 | Ak2 | adenylate kinase 2 |
| chr9 | 35048838 | 35050779 | 1942 | * | 18,6808 | 2,519368682 | 0,872396 | 2,887872 | 0,003879 | 0,62786102 | 2,507013 | Intron (ENSMUST0000( | 9 | 35055932 | 35056075 | 144 | 1 | 1,15E+08 | ENSMUST00000240399.1 | -5153 | ENSMUSG00000119783 | Gm24262 | predicted gene, 24262 |
| chr7 | 67462055 | 67463280 | 1226 | * | 14,79255 | 2,519551245 | 0,826845 | 3,047189 | 0,00231 | 0,55735526 | 2,141612 | Distal Intergenic | 7 | 67468998 | 67479366 | 10369 | 1 | 1,03E+08 | ENSMUST00000208898.2 | -5718 | | Gm34350 | predicted gene, 34350 |
| chr10 | 1,27E+08 | 1,27E+08 | 1546 | * | 13,84253 | 2,519814862 | 0,969768 | 2,598368 | 0,009367 | 0,942793663 | 0,938682 | Promoter (<=1kb) | 10 | 1,27E+08 | 1,27E+08 | 15565 | 2 | 216443 | ENSMUST00000171564.8 | -966 | ENSMUSG00000040354 | Mars1 | methionine-tRNA synthetase 1 |
| chr7 | 1,11E+08 | 1,11E+08 | 1079 | * | 13,59978 | 2,520070715 | 0,885061 | 2,847341 | 0,004409 | 0,823446343 | 0,918255 | Intron (ENSMUST0000( | 7 | 1,11E+08 | 1,11E+08 | 308317 | 2 | 233733 | ENSMUST00000049430.15 | 202700 | ENSMUSG00000038296 | Galnt18 | polypeptide N-acetylgalactosaminyltransferase 18 |
| chr1 | 1,36E+08 | 1,36E+08 | 905 | * | 19,54701 | 2,521644066 | 0,798626 | 3,15748 | 0,001591 | 0,45444759 | 2,663619 | Intron (ENSMUST0000( | 1 | 1,36E+08 | 1,36E+08 | 18540 | 1 | 16565 | ENSMUST00000122892.2 | 8383 | ENSMUSG00000041642 | Kif21b | kinesin family member 21B |
| chr7 | 66002209 | 66004409 | 2201 | * | 18,74087 | 2,521994173 | 0,856566 | 2,944306 | 0,003237 | 0,624579927 | 2,154504 | Intron (ENSMUST0000( | 7 | 65990428 | 65993013 | 2586 | 2 | 233328 | ENSMUST00000130438.3 | -9196 | ENSMUSG00000015133 | Lrrk1 | leucine-rich repeat kinase 1 |
| chr12 | 97767426 | 97768909 | 1484 | * | 13,61191 | 2,522076381 | 0,824008 | 3,060743 | 0,002208 | 0,649217039 | 1,2763 | Distal Intergenic | 12 | 97701700 | 97706503 | 4804 | 2 | 1,03E+08 | ENSMUST00000217709.2 | -60923 | ENSMUSG00000112524 | Gm35274 | predicted gene, 35274 |
| chr10 | 39490292 | 39491466 | 1175 | * | 12,15151 | 2,522155836 | 0,945914 | 2,666368 | 0,007668 | 0,87775273 | 1,173325 | Promoter (<=1kb) | 10 | 39471845 | 39489678 | 17834 | 2 | 1,01E+08 | ENSMUST00000143877.3 | -614 | ENSMUSG00000086915 | Gm16364 | predicted gene 16364 |
| chr11 | 3913640 | 3915179 | 1540 | * | 13,05413 | 2,522746633 | 0,938914 | 2,686878 | 0,007212 | 0,863495249 | 1,173325 | Promoter (<=1kb) | 11 | 3913975 | 3930004 | 16030 | 1 | 64934 | ENSMUST00000020705.5 | 0 | ENSMUSG00000020430 | Pes1 | pescadillo ribosomal biogenesis factor 1 |
| chr11 | 87356398 | 87357870 | 1473 | * | 13,75903 | 2,522858378 | 0,854605 | 2,952076 | 0,003156 | 0,756849896 | 0,938682 | Intron (ENSMUST0000( | 11 | 87353112 | 87353327 | 216 | 1 | 19861 | ENSMUST00020183227.1 | 3286 | ENSMUSG00002074955 | Snord3b4 | small nucleolar RNA, C/D box 3B4 |
| chr8 | 25763824 | 25765808 | 1985 | * | 48,257651 | 2,522887895 | 0,5385897 | 4,6842487 | 2,81E-06 | 0,008394893 | 2,7872553 | Distal Intergenic | 8 | 25672775 | 25746604 | 73830 | 2 | 320165 | ENSMUST00000209827.2 | -17220 | ENSMUSG00000065954 | Tacc1 | transforming, acidic coiled-coil containing protein 1 |
| chr7 | 35968958 | 35969960 | 1003 | * | 12,34203 | 2,523750104 | 0,975765 | 2,586432 | 0,009698 | 0,901178841 | 1,2763 | Distal Intergenic | 7 | 35796803 | 35799943 | 3141 | 1 | 1,05E+08 | ENSMUST00000205504.2 | 172155 | | Gm38991 | predicted gene, 38991 |
| chr5 | 1,21E+08 | 1,21E+08 | 1036 | * | 12,78205 | 2,52418492 | 0,976454 | 2,585051 | 0,009736 | 0,983764849 | 0,738441 | Exon (ENSMUST00000 | 5 | 1,21E+08 | 1,21E+08 | 9796 | 2 | 19247 | ENSMUST00000148871.2 | -20770 | ENSMUSG00000043733 | Ptpn11 | protein tyrosine phosphatase, non-receptor type 11 |
| chr1 | 1,83E+08 | 1,83E+08 | 1956 | * | 22,51329 | 2,524512309 | 0,696762 | 3,623204 | 0,000291 | 0,198315008 | 2,951933 | Intron (ENSMUST0000( | 1 | 1,83E+08 | 1,83E+08 | 131697 | 1 | 96935 | ENSMUST00000085724.5 | 38238 | ENSMUSG00000038576 | Susd4 | sushi domain containing 4 |
| chr12 | 24863146 | 24864188 | 1043 | * | 12,60329 | 2,526492722 | 0,945137 | 2,673149 | 0,007514 | 0,857011814 | 1,2763 | Intron (ENSMUST0000( | 12 | 24880878 | 24989301 | 108424 | 1 | 67216 | ENSMUST00000222198.2 | -16690 | ENSMUSG00000020646 | Mboat2 | membrane bound O-acyltransferase domain containing 2 |
| chr2 | 11781862 | 11784004 | 2143 | * | 18,48028 | 2,527002689 | 0,911123 | 2,773504 | 0,005546 | 0,684274806 | 2,507013 | Promoter (<=1kb) | 2 | 11783302 | 11795136 | 11835 | 1 | 320816 | ENSMUST00000056108.12 | 0 | ENSMUSG00000047909 | Ankrd16 | ankyrin repeat domain 16 |
| chr10 | 24121765 | 24123202 | 1438 | * | 11,89146 | 2,527313131 | 0,966295 | 2,615466 | 0,008911 | 0,888080529 | 1,2763 | Intron (ENSMUST0000( | 10 | 24099443 | 24107097 | 7655 | 1 | 59012 | ENSMUST00000154916.2 | 22322 | ENSMUSG00000020000 | Moxd1 | monooxygenase, DBH-like 1 |
| chr4 | 46852478 | 46854794 | 2317 | * | 26,62131 | 2,527672546 | 0,654639 | 3,861173 | 0,000113 | 0,113460495 | 2,858511 | Exon (ENSMUST00000 | 4 | 46798561 | 46859902 | 61342 | 2 | 242425 | ENSMUST00000206773.2 | 5108 | ENSMUSG00000039809 | Gabbr2 | gamma-aminobutyric acid (GABA) B receptor, 2 |
| chr15 | 62152472 | 62153783 | 1312 | * | 22,43434 | 2,527966841 | 0,845922 | 2,988418 | 0,002804 | 0,534851486 | 2,951933 | Intron (ENSMUST0000( | 15 | 62094452 | 62114354 | 19903 | 1 | 19296 | ENSMUST00000182141.2 | 58020 | ENSMUSG00000097039 | Pvt1 | Pvt1 oncogene |
| chr9 | 81929717 | 81930780 | 1064 | * | 12,43628 | 2,528616659 | 0,950974 | 2,658975 | 0,007838 | 0,863457117 | 1,2763 | Intron (ENSMUST0000( | 9 | 81745762 | 81908511 | 162750 | 1 | 75033 | ENSMUST00000189391.2 | 183955 | ENSMUSG00000043289 | Mei4 | meiotic double-stranded break formation protein 4 |
| chr15 | 5144203 | 5147788 | 3586 | * | 24,18805 | 2,529184885 | 0,852477 | 2,966865 | 0,003009 | 0,545154698 | 2,951933 | Promoter (<=1kb) | 15 | 5146127 | 5148622 | 2496 | 1 | 67281 | ENSMUST00000045356.9 | 0 | ENSMUSG00000041841 | Rpl37 | ribosomal protein L37 |
| chr9 | 30224853 | 30226775 | 1923 | * | 18,81751 | 2,529824716 | 0,797728 | 3,171288 | 0,001518 | 0,465974207 | 2,386008 | Distal Intergenic | 9 | 30203164 | 30203270 | 107 | 2 | 1,15E+08 | ENSMUST00000239738.1 | -21583 | ENSMUSG00000119318 | Gm26435 | predicted gene, 26435 |
| chr8 | 127635445 | 127637984 | 2540 | * | 41,035807 | 2,530003982 | 0,520316 | 4,8624374 | 1,159E-06 | 0,004200523 | 2,7872553 | Promoter (<=1kb) | 8 | 127541240 | 127634965 | 93726 | 2 | 102634034 | ENSMUST00000212639.2 | -480 | | Gm31718 | predicted gene, 31718 |
| chr5 | 53745762 | 53747807 | 2046 | * | 22,1564 | 2,530893666 | 0,720148 | 3,514407 | 0,000441 | 0,247743734 | 2,951933 | Promoter (<=1kb) | 5 | 53747556 | 53814704 | 67149 | 1 | 19664 | ENSMUST00000037618.13 | 0 | ENSMUSG00000039191 | Rbpj | recombination signal binding protein for immunoglobulin kappa J region |
| chr5 | 1,15E+08 | 1,15E+08 | 2147 | * | 18,80722 | 2,531581031 | 0,7899 | 3,20494 | 0,001351 | 0,457337755 | 2,221611 | Intron (ENSMUST0000( | 5 | 1,15E+08 | 1,15E+08 | 530 | 1 | 1,03E+08 | ENSMUST00000148916.2 | 11373 | | Gm13830 | predicted gene 13830 |
| chr11 | 64867715 | 64869930 | 2216 | * | 14,48716 | 2,531860308 | 0,92816 | 2,727828 | 0,006375 | 0,838625535 | 1,214904 | Promoter (<=1kb) | 11 | 64869864 | 64892895 | 23032 | 1 | 68626 | ENSMUST00000071891.12 | 0 | ENSMUSG00000020549 | Elac2 | elaC ribonuclease Z 2 |
| chr10 | 80770380 | 80771333 | 954 | * | 15,85088 | 2,532065263 | 0,96404 | 2,626514 | 0,008626 | 0,883917268 | 1,2763 | Distal Intergenic | 10 | 80766646 | 80767593 | 948 | 1 | 17873 | ENSMUST00000219449.2 | 3734 | ENSMUSG00000015312 | Gadd45b | growth arrest and DNA-damage-inducible 45 beta |
| chr8 | 38833259 | 38835008 | 1750 | * | 14,58085 | 2,532396106 | 0,937085 | 2,702419 | 0,006884 | 0,829907213 | 1,37683 | Intron (ENSMUST0000( | 8 | 38719287 | 38719356 | 70 | 2 | 723860 | ENSMUST00000083523.5 | -113903 | ENSMUSG00000065457 | Mir383 | microRNA 383 |
| chr6 | 1,47E+08 | 1,47E+08 | 1360 | * | 14,92013 | 2,532580139 | 0,826981 | 3,062439 | 0,002195 | 0,54670137 | 2,141612 | Promoter (<=1kb) | 6 | 1,47E+08 | 1,47E+08 | 5808 | 1 | 67533 | ENSMUST00000126774.2 | 124 | ENSMUSG00000016487 | Ppfibp1 | PTPRF interacting protein, binding protein 1 (liprin beta 1) |
| chr4 | 71842712 | 71844147 | 1436 | * | 13,9341 | 2,533042918 | 0,958379 | 2,643049 | 0,008216 | 0,957878022 | 0,738441 | Distal Intergenic | 4 | 71673880 | 71676202 | 2323 | 2 | 1,03E+08 | ENSMUST00000084489.3 | -166510 | ENSMUSG00000066141 | Gm11232 | predicted gene 11232 |
| chr6 | 94677018 | 94678177 | 1160 | * | 17,40274 | 2,533213834 | 0,913504 | 2,773075 | 0,005553 | 0,693003605 | 2,386008 | Promoter (<=1kb) | 6 | 94581510 | 94677139 | 95630 | 2 | 16206 | ENSMUST00000204645.3 | 0 | ENSMUSG00000030029 | Lrig1 | leucine-rich repeats and immunoglobulin-like domains 1 |
| chr14 | 36911995 | 36913513 | 1519 | * | 14,09429 | 2,533294184 | 0,935634 | 2,70757 | 0,006778 | 0,83918951 | 1,2763 | Distal Intergenic | 14 | 36946826 | 36946958 | 133 | 1 | 1,15E+08 | ENSMUST00000104118.3 | -33313 | ENSMUSG00000077306 | Gm22469 | predicted gene, 22469 |
| chr12 | 74000095 | 74001896 | 1802 | * | 19,64187 | 2,533597343 | 0,791705 | 3,200177 | 0,001373 | 0,448852574 | 2,386008 | Distal Intergenic | 12 | 73889813 | 73996588 | 106776 | 2 | 1,03E+08 | ENSMUST00000221427.2 | -3507 | | Gm15283 | predicted gene 15283 |
| chr11 | 83188448 | 83190417 | 1970 | * | 15,51408 | 2,533633505 | 0,966187 | 2,622302 | 0,008734 | 0,891097601 | 1,214904 | Promoter (<=1kb) | 11 | 83185468 | 83189803 | 4336 | 2 | 103737 | ENSMUST00000018877.9 | 0 | ENSMUSG00000018733 | Pex12 | peroxisomal biogenesis factor 12 |
| chr18 | 81448997 | 81450215 | 1219 | * | 12,12437 | 2,53391854 | 0,929581 | 2,725873 | 0,006413 | 0,830950621 | 1,2763 | Distal Intergenic | 18 | 81477398 | 81497187 | 19790 | 2 | 1,03E+08 | ENSMUST00000235899.2 | 46972 | | Gm30288 | predicted gene, 30288 |
| chr7 | 65728516 | 65729990 | 1475 | * | 16,63566 | 2,534755026 | 0,819208 | 3,094155 | 0,001974 | 0,62786102 | 1,2763 | Promoter (<=1kb) | 7 | 65729397 | 65739153 | 9757 | 1 | 109815 | ENSMUST00000101801.8 | 0 | ENSMUSG00000075701 | Selenos | selenoprotein S |
| chr7 | 30878274 | 30879721 | 1448 | * | 13,60999 | 2,534902933 | 0,972814 | 2,605741 | 0,009168 | 0,891097601 | 1,2763 | Distal Intergenic | 7 | 30861783 | 30870066 | 8284 | 1 | 1E+08 | ENSMUST00000190947.2 | 16491 | ENSMUSG00000100985 | Gm10640 | predicted gene 10640 |
| chr3 | 1,03E+08 | 1,03E+08 | 1859 | * | 18,79636 | 2,534958325 | 0,8545 | 2,966598 | 0,003011 | 0,579872171 | 2,507013 | Distal Intergenic | 3 | 1,03E+08 | 1,03E+08 | 79483 | 1 | 94093 | ENSMUST00000029444.13 | -47872 | ENSMUSG00000033014 | Trim33 | tripartite motif-containing 33 |
| chr4 | 50362940 | 50364178 | 1239 | * | 14,46845 | 2,535300353 | 0,935537 | 2,709995 | 0,006728 | 0,838625535 | 1,2763 | Distal Intergenic | 4 | 50286746 | 50286877 | 132 | 1 | 1,15E+08 | ENSMUST00000104120.2 | 76194 | ENSMUSG00000095394 | Gm25485 | predicted gene, 25485 |
| chr17 | 13134399 | 13136117 | 1719 | * | 23,39305 | 2,536226577 | 0,72281 | 3,508845 | 0,00045 | 0,253278065 | 2,858511 | Promoter (<=1kb) | 17 | 13134588 | 13137001 | 2414 | 1 | 21454 | ENSMUST00000147724.8 | 0 | ENSMUSG00000068039 | Tcp1 | t-complex protein 1 |
| chr11 | 70805655 | 70807663 | 2009 | * | 15,79997 | 2,536331593 | 0,861363 | 2,944555 | 0,003234 | 0,625646014 | 2,141612 | Intron (ENSMUST0000( | 11 | 70834764 | 70835314 | 551 | 2 | 19069 | ENSMUST00000128193.2 | 27651 | ENSMUSG00000040667 | Nup88 | nucleoporin 88 |
| chr4 | 1,34E+08 | 1,34E+08 | 1459 | * | 12,68825 | 2,536507291 | 0,916935 | 2,76629 | 0,00567 | 0,823736512 | 1,173325 | 3' UTR | 4 | 1,34E+08 | 1,34E+08 | 5796 | 2 | 76824 | ENSMUST00000129836.2 | 8843 | ENSMUSG00000046671 | Mtfr1l | mitochondrial fission regulator 1-like |
| chr19 | 45990116 | 45993618 | 3503 | * | 28,93754 | 2,537332812 | 0,634976 | 3,995951 | 6,44E-05 | 0,084344578 | 2,67243 | Promoter (<=1kb) | 19 | 45991947 | 45994612 | 2666 | 1 | 20170 | ENSMUST00000099393.4 | 0 | ENSMUSG00000074811 | Hps6 | HPS6, biogenesis of lysosomal organelles complex 2 subunit 3 |
| chr7 | 15990035 | 15991612 | 1578 | * | 11,59406 | 2,537383215 | 0,919005 | 2,761011 | 0,005762 | 0,868022856 | 0,918255 | Promoter (1-2kb) | 7 | 15980670 | 15993263 | 12594 | 2 | 12273 | ENSMUST00000168818.2 | 1651 | ENSMUSG00000049130 | C5ar1 | complement component 5a receptor 1 |
| chr5 | 1,37E+08 | 1,37E+08 | 2194 | * | 27,19133 | 2,537766413 | 0,655144 | 3,873603 | 0,000107 | 0,114310976 | 2,67243 | Distal Intergenic | 5 | 1,37E+08 | 1,37E+08 | 2186 | 2 | 384309 | ENSMUST00000152207.2 | -6367 | ENSMUSG00000043279 | Trim56 | tripartite motif-containing 56 |
| chr12 | 76909756 | 76911100 | 1345 | * | 14,63526 | 2,53812504 | 0,894291 | 2,838141 | 0,004538 | 0,768389035 | 1,2763 | Intron (ENSMUST0000( | 12 | 76887069 | 76934579 | 47511 | 1 | 110606 | ENSMUST00000123887.2 | 22687 | ENSMUSG00000033373 | Fntb | farnesyltransferase, CAAX box, beta |
| chr17 | 73973076 | 73974698 | 1623 | * | 20,65248 | 2,538423755 | 0,811588 | 3,127725 | 0,001762 | 0,454361911 | 2,951933 | Intron (ENSMUST0000( | 17 | 73826620 | 74017061 | 190442 | 2 | 71685 | ENSMUST00000233326.2 | 42363 | ENSMUSG00000024064 | Galnt14 | polypeptide N-acetylgalactosaminyltransferase 14 |
| chr2 | 1,26E+08 | 1,26E+08 | 1327 | * | 17,2628 | 2,538806098 | 0,963329 | 2,63545 | 0,008403 | 0,764024179 | 2,423594 | 3' UTR | 2 | 1,26E+08 | 1,27E+08 | 9599 | 2 | 14391 | ENSMUST00000137335.2 | 13331 | ENSMUSG00000027361 | Gabpb1 | GA repeat binding protein, beta 1 |
| chr2 | 72151107 | 72152564 | 1458 | * | 14,87749 | 2,539689188 | 0,852336 | 2,97968 | 0,002885 | 0,675594321 | 1,41696 | Intron (ENSMUST0000( | 2 | 72128245 | 72185966 | 57722 | 1 | 65964 | ENSMUST00000144111.2 | 22862 | ENSMUSG00000004085 | Map3k20 | mitogen-activated protein kinase kinase kinase 20 |
| chr4 | 19842365 | 19843710 | 1346 | * | 13,93842 | 2,540407572 | 0,899195 | 2,825201 | 0,004725 | 0,777172375 | 1,2763 | Downstream (<=300bp | 4 | 19818725 | 19842218 | 23494 | 1 | 74087 | ENSMUST00000035890.8 | 23640 | ENSMUSG00000041052 | Slc7a13 | solute carrier family 7, (cationic amino acid transporter, y+ system) member 13 |
| chr14 | 31127601 | 31129015 | 1415 | * | 12,04363 | 2,540780913 | 0,953506 | 2,664671 | 0,007706 | 0,915421656 | 0,918255 | Promoter (<=1kb) | 14 | 31099552 | 31129342 | 29791 | 2 | 24056 | ENSMUST00000147586.2 | 327 | ENSMUSG00000021892 | Sh3bp5 | SH3-domain binding protein 5 (BTK-associated) |
| chr13 | 1,12E+08 | 1,12E+08 | 1273 | * | 16,77827 | 2,540859078 | 0,890546 | 2,853148 | 0,004329 | 0,661775851 | 2,318434 | Promoter (<=1kb) | 13 | 1,12E+08 | 1,12E+08 | 1783 | 1 | 1,01E+08 | ENSMUST00000149252.2 | 256 | | Gm15327 | predicted gene 15327 |
| chr6 | 125167261 | 125170376 | 3116 | * | 42,558018 | 2,541071769 | 0,5548818 | 4,5794826 | 4,661E-06 | 0,012861975 | 2,7254908 | Promoter (<=1kb) | 6 | 125168764 | 125170521 | 1758 | 1 | 66493 | ENSMUST00000136804.2 | 0 | ENSMUSG00000030335 | Mrpl51 | mitochondrial ribosomal protein L51 |
| chr19 | 46262415 | 46263886 | 1472 | * | 12,63437 | 2,541499005 | 0,871876 | 2,914976 | 0,003557 | 0,786234479 | 0,918255 | Intron (ENSMUST0000( | 19 | 46258157 | 46261833 | 3677 | 1 | 107338 | ENSMUST00000176422.2 | 4258 | ENSMUSG00000025224 | Gbf1 | golgi-specific brefeldin A-resistance factor 1 |
| chr7 | 45673936 | 45675182 | 1247 | * | 13,36556 | 2,542169566 | 0,965774 | 2,632262 | 0,008482 | 0,880247401 | 1,2763 | Intron (ENSMUST0000( | 7 | 45625804 | 45679726 | 53923 | 2 | 27421 | ENSMUST00000002850.8 | 4544 | ENSMUSG00000030834 | Abcc6 | ATP-binding cassette, sub-family C (CFTR/MRP), member 6 |
| chrX | 20146295 | 20149494 | 3200 | * | 18,03884 | 2,542514726 | 0,879336 | 2,891402 | 0,003835 | 0,616178941 | 2,663619 | Intron (ENSMUST0000( | 20 | 20018691 | 20158009 | 139319 | 2 | 236727 | ENSMUST00000115393.3 | 8515 | ENSMUSG00000037341 | Slc9a7 | solute carrier family 9 (sodium/hydrogen exchanger), member 7 |
| chr1 | 51811784 | 51812766 | 983 | * | 12,78723 | 2,542555006 | 0,924553 | 2,750038 | 0,005959 | 0,832641105 | 1,173325 | Promoter (1-2kb) | 1 | 51801681 | 51810038 | 8358 | 2 | 17912 | ENSMUST00000140731.8 | -1746 | ENSMUSG00000018417 | Myo1b | myosin IB |
| chr5 | 1,23E+08 | 1,23E+08 | 2767 | * | 26,70241 | 2,542657362 | 0,736999 | 3,450016 | 0,000561 | 0,288385947 | 2,725491 | Promoter (<=1kb) | 5 | 1,23E+08 | 1,23E+08 | 487 | 2 | 1,03E+08 | ENSMUST00000246777.1 | 0 | | Gm33044 | predicted gene, 33044 |

| | | | | | | | | | | | | | | | | | | | | | | | |
|---|---|---|---|---|---|---|---|---|---|---|---|---|---|---|---|---|---|---|---|---|---|---|---|
| chr4 | 1,56E+08 | 1,56E+08 | 2089 | * | 18,54902 | 2,542683667 | 0,779606 | 3,2615 | 0,001108 | 0,430115816 | 2,154504 | Promoter (<=1kb) | 4 | 1,56E+08 | 1,56E+08 | 19140 | 1 | 71957 | ENSMUST00000120794.8 | 0 | ENSMUSG00000029034 | Ints11 | integrator complex subunit 11 |
| chr3 | 93444335 | 93445958 | 1624 | * | 15,14515 | 2,542705649 | 0,83799 | 3,034293 | 0,002411 | 0,663565385 | 1,2763 | Distal Intergenic | 3 | 93462387 | 93471950 | 9564 | 1 | 20194 | ENSMUST00000045756.14 | -16429 | ENSMUSG00000041959 | S100a10 | S100 calcium binding protein A10 (calpactin) |
| chr12 | 84033124 | 84034933 | 1810 | * | 12,1148 | 2,542977441 | 0,940019 | 2,705241 | 0,006825 | 0,855636327 | 1,173325 | Promoter (<=1kb) | 12 | 84034635 | 84040647 | 6013 | 1 | 171210 | ENSMUST00000021649.8 | 0 | ENSMUSG00000021226 | Acot2 | acyl-CoA thioesterase 2 |
| chr13 | 1,15E+08 | 1,15E+08 | 1884 | * | 16,0333 | 2,543080212 | 0,855859 | 2,971376 | 0,002965 | 0,594430593 | 2,318434 | Distal Intergenic | 13 | 1,15E+08 | 1,15E+08 | 6662 | 2 | 14313 | ENSMUST00000223640.3 | -36196 | ENSMUSG00000021765 | Fst | follistatin |
| chr5 | 32239194 | 32241652 | 2459 | * | 21,26418 | 2,543915513 | 0,731303 | 3,478608 | 0,000504 | 0,264977345 | 2,951933 | Promoter (2-3kb) | 5 | 32237041 | 32242090 | 5050 | 1 | 107976 | ENSMUST00000202087.2 | 2153 | ENSMUSG00000052139 | Babam2 | BRISC and BRCA1 A complex member 2 |
| chr4 | 1,39E+08 | 1,39E+08 | 2055 | * | 22,09083 | 2,544010324 | 0,74055 | 3,435301 | 0,000592 | 0,264977345 | 3,471678 | Distal Intergenic | 4 | 1,39E+08 | 1,39E+08 | 16307 | 1 | 83770 | ENSMUST00000166773.2 | 33535 | ENSMUSG00000028738 | Tas1r2 | taste receptor, type 1, member 2 |
| chr7 | 40326374 | 40327629 | 1256 | * | 13,49688 | 2,545109494 | 0,961934 | 2,645825 | 0,008149 | 0,923399738 | 0,918255 | Promoter (2-3kb) | 7 | 40324161 | 40340697 | 16537 | 1 | 71029 | ENSMUST00000207559.2 | 2213 | ENSMUSG00000102004 | 4933402C06Rik | RIKEN cDNA 4933402C06 gene |
| chr18 | 78172735 | 78175160 | 2426 | * | 30,209955 | 2,54538917 | 0,5762949 | 4,4168168 | 1,002E-05 | 0,021507905 | 2,9510621 | Intron (ENSMUST000001 | 18 | 78143306 | 78166641 | 23336 | 2 | 108052 | ENSMUST00000091813.12 | -6094 | ENSMUSG00000059336 | Slc14a1 | solute carrier family 14 (urea transporter), member 1 |
| chr10 | 79534601 | 79541440 | 6840 | * | 89,47774 | 2,545567432 | 0,670246 | 3,797961 | 0,000146 | 0,123779239 | 3,177995 | Promoter (<=1kb) | 10 | 79539318 | 79539424 | 107 | 2 | 1,15E+08 | ENSMUST00000083286.3 | 0 | ENSMUSG00000065220 | Gm25794 | predicted gene, 25794 |
| chr3 | 51543985 | 51546066 | 2082 | * | 18,18009 | 2,545635448 | 0,930474 | 2,735847 | 0,006222 | 0,715249603 | 2,386008 | Exon (ENSMUST00000 | 3 | 51567781 | 51589228 | 21448 | 1 | 211666 | ENSMUST00000160753.8 | -21715 | ENSMUSG00000074604 | Mgst2 | microsomal glutathione S-transferase 2 |
| chr13 | 67901833 | 67903474 | 1642 | * | 15,45826 | 2,545709749 | 0,971461 | 2,620495 | 0,00878 | 0,886332554 | 1,2763 | Promoter (<=1kb) | 13 | 67895919 | 67903347 | 7429 | 2 | 22746 | ENSMUST00000144183.3 | 0 | ENSMUSG00000058331 | Zfp85 | zinc finger protein 85 |
| chr19 | 10098938 | 10100422 | 1485 | * | 14,19029 | 2,545849211 | 0,847499 | 3,003957 | 0,002665 | 0,680780911 | 1,2763 | Distal Intergenic | 19 | 10040129 | 10079110 | 38982 | 2 | 56473 | ENSMUST00000025567.9 | -19828 | ENSMUSG00000024665 | Fads2 | fatty acid desaturase 2 |
| chr9 | 121685512 | 121689542 | 4031 | * | 45,573181 | 2,546064863 | 0,541161 | 4,7048196 | 2,541E-06 | 0,00896874 | 2,3383871 | Promoter (<=1kb) | 9 | 121686746 | 121688275 | 1530 | 1 | 115487203 | ENSMUST00000217491.2 | 0 | ENSMUSG00000111079 | Gm47112 | predicted gene, 47112 |
| chr16 | 30004885 | 30008513 | 3629 | * | 42,969851 | 2,546184999 | 0,5382609 | 4,730392 | 2,241E-06 | 0,007032136 | 2,9254973 | Distal Intergenic | 16 | 30013889 | 30019117 | 5229 | 1 | 75531 | ENSMUST00000187479.2 | -5376 | | 1700025H01Rik | RIKEN cDNA 1700025H01 gene |
| chr12 | 79522094 | 79524324 | 2231 | * | 16,9437 | 2,546495402 | 0,951097 | 2,677428 | 0,007419 | 0,72659841 | 2,663619 | Intron (ENSMUST0000 | 12 | 79374127 | 79748630 | 374504 | 1 | 19363 | ENSMUST00000221257.2 | 147967 | ENSMUSG00000059060 | Rad51b | RAD51 paralog B |
| chr10 | 1,18E+08 | 1,18E+08 | 1604 | * | 12,29092 | 2,547435888 | 0,839496 | 3,034484 | 0,002409 | 0,751231966 | 0,738441 | Distal Intergenic | 10 | 1,18E+08 | 1,18E+08 | 22834 | 1 | 71067 | ENSMUST00000217950.2 | 54729 | ENSMUSG00000112500 | 4933411E08Rik | RIKEN cDNA 4933411E08 gene |
| chr17 | 35073665 | 35077608 | 3944 | * | 33,47002 | 2,547576572 | 0,738114 | 3,451466 | 0,000558 | 0,388075979 | 1,405709 | Promoter (<=1kb) | 17 | 35076545 | 35076608 | 64 | 2 | 1,02E+08 | ENSMUST00000184774.3 | 0 | ENSMUSG00000098445 | Mir6972 | microRNA 6972 |
| chr9 | 43861279 | 43864489 | 3211 | * | 23,45912 | 2,547801985 | 0,73424 | 3,469987 | 0,00052 | 0,270514576 | 2,951062 | Intron (ENSMUST0000 | 9 | 43873156 | 43881468 | 8313 | 2 | 1,03E+08 | ENSMUST00000217507.2 | 16979 | | Gm30373 | predicted gene, 30373 |
| chr5 | 1,43E+08 | 1,43E+08 | 1787 | * | 13,34153 | 2,548133313 | 0,911271 | 2,796241 | 0,00517 | 0,823446343 | 1,075735 | Promoter (<=1kb) | 5 | 1,43E+08 | 1,43E+08 | 18064 | 1 | 66913 | ENSMUST00000110731.4 | 0 | ENSMUSG00000079111 | Kdelr2 | KDEL (Lys-Asp-Glu-Leu) endoplasmic reticulum protein retention receptor 2 |
| chr13 | 60379842 | 60380834 | 993 | * | 11,98398 | 2,549077099 | 0,945337 | 2,696474 | 0,007008 | 0,934846269 | 0,738441 | Distal Intergenic | 13 | 60353285 | 60363837 | 10553 | 1 | 1,01E+08 | ENSMUST00000224064.2 | 26557 | ENSMUSG00000050765 | Gm5084 | predicted gene 5084 |
| chr3 | 97673877 | 97676454 | 2578 | * | 49,5197 | 2,549456511 | 0,57022 | 4,4710048 | 7,785E-06 | 0,018663151 | 2,7872553 | Promoter (<=1kb) | 3 | 97646170 | 97675325 | 29156 | 2 | 83679 | ENSMUST00000045243.15 | 0 | ENSMUSG00000038170 | Pde4dip | phosphodiesterase 4D interacting protein (myomegalin) |
| chr17 | 67931899 | 67933407 | 1509 | * | 13,78829 | 2,549891767 | 0,89536 | 2,847895 | 0,004401 | 0,762875156 | 1,2763 | Promoter (2-3kb) | 17 | 67866743 | 67936327 | 69585 | 2 | 1,03E+08 | ENSMUST00000233130.2 | 2920 | ENSMUSG00000117029 | Gm36201 | predicted gene, 36201 |
| chr15 | 85304790 | 85306208 | 1419 | * | 11,59617 | 2,549965346 | 0,947499 | 2,69126 | 0,007118 | 0,860883941 | 1,173325 | Intron (ENSMUST0000 | 15 | 85235264 | 85235455 | 192 | 2 | 1,15E+08 | ENSMUST00000082830.3 | -69335 | ENSMUSG00000064764 | Gm23517 | predicted gene, 23517 |
| chr2 | 1,02E+08 | 1,02E+08 | 1318 | * | 12,32154 | 2,550093709 | 0,951812 | 2,679199 | 0,00738 | 0,886332554 | 1,075735 | Distal Intergenic | 2 | 1,02E+08 | 1,02E+08 | 3134 | 2 | 14221 | ENSMUST00000099678.5 | -14499 | ENSMUSG00000075012 | Fjx1 | four jointed box 1 |
| chr18 | 56904541 | 56905868 | 1328 | * | 15,1329 | 2,550507458 | 0,928497 | 2,74692 | 0,006016 | 0,728037066 | 2,141612 | Promoter (1-2kb) | 18 | 56902789 | 56934760 | 31972 | 1 | 1,05E+08 | ENSMUST00000128240.2 | 1752 | | Gm15345 | predicted gene 15345 |
| chr17 | 35357351 | 35359362 | 2012 | * | 16,51669 | 2,550720806 | 0,835065 | 3,054518 | 0,002254 | 0,534110533 | 2,386008 | Promoter (<=1kb) | 17 | 35357549 | 35358826 | 1278 | 1 | 224727 | ENSMUST00000172925.2 | 0 | ENSMUSG00000024392 | Bag6 | BCL2-associated athanogene 6 |
| chr11 | 5106005 | 5107366 | 1362 | * | 15,16062 | 2,551027252 | 0,937775 | 2,720298 | 0,006522 | 0,841717562 | 1,214904 | Distal Intergenic | 11 | 5056265 | 5102257 | 45993 | 2 | 140703 | ENSMUST00000062821.13 | -3748 | ENSMUSG00000034164 | Emid1 | EMI domain containing 1 |
| chr8 | 1,16E+08 | 1,16E+08 | 1103 | * | 15,84046 | 2,551705105 | 0,792147 | 3,221251 | 0,001276 | 0,54190677 | 1,2763 | Intron (ENSMUST0000 | 8 | 1,16E+08 | 1,16E+08 | 104 | 2 | 1,15E+08 | ENSMUST00000083156.3 | 395419 | ENSMUSG00000065090 | Gm23677 | predicted gene, 23677 |
| chr7 | 27877647 | 27879452 | 1806 | * | 14,8894 | 2,551725254 | 0,91733 | 2,781688 | 0,005408 | 0,788774532 | 1,37683 | Promoter (<=1kb) | 7 | 27878894 | 27886719 | 7826 | 1 | 13549 | ENSMUST00000085901.13 | 0 | ENSMUSG00000002409 | Dyrk1b | dual-specificity tyrosine-(Y)-phosphorylation regulated kinase 1b |
| chr17 | 25074009 | 25075271 | 1263 | * | 13,39769 | 2,551855945 | 0,970349 | 2,629834 | 0,008543 | 0,881672947 | 1,2763 | Promoter (1-2kb) | 17 | 25077175 | 25083420 | 6246 | 1 | 14651 | ENSMUST00000163132.2 | -1904 | ENSMUSG00000024158 | Hagh | hydroxyacyl glutathione hydrolase |
| chr5 | 137479190 | 137481055 | 1866 | * | 31,931259 | 2,552406931 | 0,5601138 | 4,5569435 | 5,19E-06 | 0,013744849 | 2,7254908 | Distal Intergenic | 5 | 137481282 | 137484078 | 2797 | 2 | 13856 | ENSMUST00000031723.14 | 3023 | ENSMUSG00000029711 | Epo | erythropoietin |
| chr15 | 82914211 | 82915290 | 1080 | * | 13,04635 | 2,552598774 | 0,913068 | 2,795628 | 0,00518 | 0,823446343 | 1,075735 | Promoter (<=1kb) | 15 | 82881927 | 82914151 | 32225 | 2 | 74039 | ENSMUST00000023076.6 | -60 | ENSMUSG00000058099 | Nfam1 | Nfat activating molecule with ITAM motif 1 |
| chr4 | 1,35E+08 | 1,35E+08 | 1666 | * | 13,33057 | 2,553089727 | 0,93936 | 2,717905 | 0,00657 | 0,86208821 | 1,075735 | Distal Intergenic | 4 | 1,35E+08 | 1,35E+08 | 2171 | 1 | 1,03E+08 | ENSMUST00000145711.2 | -4406 | | Gm12984 | predicted gene 12984 |
| chr14 | 27543959 | 27546722 | 2764 | * | 24,71287 | 2,553476715 | 0,817834 | 3,122245 | 0,001795 | 0,475134502 | 2,67243 | Intron (ENSMUST0000 | 14 | 27520515 | 27634859 | 114345 | 1 | 238988 | ENSMUST00000211636.2 | 23444 | ENSMUSG00000040640 | Erc2 | ELKS/RAB6-interacting/CAST family member 2 |
| chr9 | 51222722 | 51224021 | 1300 | * | 14,39329 | 2,553876772 | 0,943185 | 2,707715 | 0,006775 | 0,896435577 | 0,918255 | Promoter (<=1kb) | 9 | 51200986 | 51224015 | 23030 | 2 | 69809 | ENSMUST00000181366.3 | 0 | ENSMUSG00000036027 | 1810046K07Rik | RIKEN cDNA 1810046K07 gene |
| chr8 | 56985435 | 56986341 | 907 | * | 12,43402 | 2,55404593 | 0,962574 | 2,65335 | 0,00797 | 0,865873364 | 1,2763 | Exon (ENSMUST00000 | 8 | 56974407 | 56974538 | 132 | 2 | 1,15E+08 | ENSMUST00000104397.3 | -10897 | ENSMUSG00000077585 | Gm24375 | predicted gene, 24375 |
| chr18 | 61476322 | 61478432 | 2111 | * | 20,50445 | 2,554476097 | 0,761256 | 3,355609 | 0,000792 | 0,332921339 | 2,713306 | Intron (ENSMUST0000 | 18 | 61530906 | 61530971 | 66 | 2 | 723889 | ENSMUST00000198300.3 | 52539 | ENSMUSG00000105200 | Mir378a | microRNA 378a |
| chr18 | 48927307 | 48929456 | 2150 | * | 17,40783 | 2,554485264 | 0,844197 | 3,025935 | 0,002479 | 0,546144416 | 2,423594 | Distal Intergenic | 18 | 49656356 | 49657507 | 1152 | 1 | 67321 | ENSMUST00000190201.2 | -726900 | ENSMUSG00000100297 | 1700044K03Rik | RIKEN cDNA 1700044K03 gene |
| chr16 | 4982124 | 4984977 | 2854 | * | 18,99068 | 2,55461214 | 0,940622 | 2,715876 | 0,00661 | 0,736703271 | 2,221611 | Exon (ENSMUST00000 | 16 | 4964973 | 5005135 | 40163 | 1 | 665119 | ENSMUST00000230616.2 | 17151 | ENSMUSG00000091712 | Sec14l5 | SEC14-like lipid binding 5 |
| chr10 | 48529827 | 48531777 | 1951 | * | 14,70827 | 2,554701358 | 0,882441 | 2,895039 | 0,003791 | 0,723677315 | 1,37683 | Distal Intergenic | 10 | 48495408 | 48495491 | 84 | 2 | 1,15E+08 | ENSMUST00000083696.4 | -34336 | | Gm24256 | predicted gene, 24256 |
| chr8 | 41577440 | 41580526 | 3087 | * | 28,54985 | 2,555694037 | 0,747197 | 3,420375 | 0,000625 | 0,302204121 | 2,725491 | Intron (ENSMUST0000 | 8 | 41537600 | 41586755 | 49156 | 2 | 102103 | ENSMUST00000155055.2 | 6229 | ENSMUSG00000045636 | Mtus1 | mitochondrial tumor suppressor 1 |
| chr14 | 31312706 | 31314216 | 1511 | * | 15,67837 | 2,556183297 | 0,971897 | 2,630097 | 0,008536 | 0,881662676 | 1,2763 | Promoter (<=1kb) | 14 | 31274866 | 31313300 | 38435 | 2 | 1,03E+08 | ENSMUST00000124014.8 | 0 | ENSMUSG00000057606 | LOC102639719 | uncharacterized LOC102639719 |
| chr4 | 1,49E+08 | 1,49E+08 | 2973 | * | 25,26984 | 2,556587202 | 0,693186 | 3,688169 | 0,000226 | 0,170809454 | 2,951062 | Distal Intergenic | 4 | 1,49E+08 | 1,49E+08 | 2334 | 2 | 63958 | ENSMUST00000155968.8 | 5236 | ENSMUSG00000028960 | Ube4b | ubiquitination factor E4B |
| chr9 | 83102888 | 83104794 | 1907 | * | 17,37515 | 2,55666341 | 0,924531 | 2,765361 | 0,005686 | 0,716907932 | 2,154504 | Distal Intergenic | 9 | 83122534 | 83136075 | 13542 | 2 | 1,05E+08 | ENSMUST00000184605.8 | 31281 | ENSMUSG00000098747 | Gm27216 | predicted gene 27216 |
| chr5 | 1,43E+08 | 1,43E+08 | 1580 | * | 21,83827 | 2,556992862 | 0,854875 | 2,991071 | 0,00278 | 0,538066226 | 2,880017 | Distal Intergenic | 5 | 1,43E+08 | 1,43E+08 | 13003 | 2 | 231863 | ENSMUST00000156725.2 | -3315 | ENSMUSG00000066640 | Fbxl18 | F-box and leucine-rich repeat protein 18 |
| chr16 | 56323798 | 56324487 | 690 | * | 10,37318 | 2,55871347 | 0,975776 | 2,622233 | 0,008736 | 0,93486226 | 0,918255 | Intron (ENSMUST0000 | 16 | 56298263 | 56510498 | 212236 | 1 | 320712 | ENSMUST00000096012.11 | 25535 | ENSMUSG00000035258 | Abi3bp | ABI family member 3 binding protein |
| chr6 | 31056601 | 31060507 | 3907 | * | 89,413475 | 2,55911083 | 0,4361017 | 5,8681509 | 4,407E-09 | 4,6497E-05 | 3,177995 | Exon (ENSMUST0000023 | 6 | 31040027 | 31040430 | 404 | 2 | 387223 | ENSMUST00000183056.2 | -16171 | ENSMUSG00000065604 | Mir29b-1 | microRNA 29b-1 |
| chr5 | 3461914 | 3463399 | 1486 | * | 13,2326 | 2,559242943 | 0,977891 | 2,617105 | 0,008868 | 0,968558269 | 0,738441 | Intron (ENSMUST0000 | 5 | 3394312 | 3572225 | 177914 | 1 | 12571 | ENSMUST00000042410.5 | 67602 | ENSMUSG00000040274 | Cdk6 | cyclin-dependent kinase 6 |
| chr11 | 1,01E+08 | 1,01E+08 | 2055 | * | 20,00706 | 2,559510787 | 0,850566 | 3,009184 | 0,00262 | 0,549632826 | 2,507013 | Exon (ENSMUST00000 | 11 | 1,01E+08 | 1,01E+08 | 23205 | 2 | 19345 | ENSMUST00000019317.12 | -4288 | ENSMUSG00000019173 | Rab5c | RAB5C, member RAS oncogene family |
| chr4 | 1,29E+08 | 1,29E+08 | 1316 | * | 12,52725 | 2,559715865 | 0,930713 | 2,750275 | 0,005955 | 0,83238039 | 1,173325 | 3' UTR | 4 | 1,29E+08 | 1,29E+08 | 2991 | 1 | 67106 | ENSMUST00000140272.2 | 5820 | ENSMUSG00000057572 | Zbtb8os | zinc finger and BTB domain containing 8 opposite strand |
| chr17 | 34917572 | 34919274 | 1703 | * | 14,66526 | 2,560027279 | 0,890369 | 2,875245 | 0,004037 | 0,75508254 | 1,214904 | Exon (ENSMUST000002 | 17 | 34906670 | 34906776 | 107 | 2 | 1,15E+08 | ENSMUST00000239676.1 | -10796 | ENSMUSG00000118746 | Gm24389 | predicted gene, 24389 |
| chr18 | 47901232 | 47903244 | 2013 | * | 10,61683 | 2,560137121 | 0,965211 | 2,652413 | 0,007992 | 0,954706987 | 0,744375 | Intron (ENSMUST0000 | 18 | 47894156 | 47922600 | 28445 | 2 | 1E+08 | ENSMUST00000236862.2 | 19356 | ENSMUSG00000118113 | Gm4146 | predicted gene 4146 |
| chr7 | 73255493 | 73257051 | 1559 | * | 12,54931 | 2,561444547 | 0,984105 | 2,602815 | 0,009246 | 0,892811906 | 1,2763 | Distal Intergenic | 7 | 73295733 | 73295837 | 105 | 2 | 1,15E+08 | ENSMUST00000083795.3 | 38786 | ENSMUSG00000065729 | Gm26176 | predicted gene, 26176 |
| chr15 | 71581041 | 71582538 | 1498 | * | 13,90043 | 2,561847395 | 0,853821 | 3,000452 | 0,002696 | 0,709612585 | 1,075735 | Intron (ENSMUST0000 | 15 | 71397582 | 71598582 | 201001 | 2 | 70363 | ENSMUST00000229189.2 | 16044 | ENSMUSG00000036800 | Fam135b | family with sequence similarity 135, member B |
| chr11 | 62302050 | 62304152 | 2103 | * | 17,74609 | 2,562407581 | 0,838438 | 3,056167 | 0,002242 | 0,548877304 | 2,154504 | Intron (ENSMUST0000 | 11 | 62286012 | 62292376 | 6365 | 2 | 20185 | ENSMUST00000124027.3 | -9674 | ENSMUSG00000018501 | Ncor1 | nuclear receptor co-repressor 1 |
| chr9 | 55015958 | 55018540 | 2583 | * | 17,02831 | 2,562538849 | 0,839979 | 3,050717 | 0,002283 | 0,55292773 | 2,141612 | Distal Intergenic | 9 | 55056138 | 55114813 | 58676 | 1 | 109161 | ENSMUST00000059555.15 | -37598 | ENSMUSG00000032307 | Ube2q2 | ubiquitin-conjugating enzyme E2Q family member 2 |
| chr1 | 1,19E+08 | 1,19E+08 | 1446 | * | 13,3734 | 2,562871995 | 0,983635 | 2,605512 | 0,009174 | 0,91804783 | 1,075735 | Promoter (1-2kb) | 1 | 1,19E+08 | 1,19E+08 | 2824 | 1 | 226351 | ENSMUST00000183952.2 | 1384 | ENSMUSG00000098923 | Tmem185b | transmembrane protein 185B |
| chr6 | 86250179 | 86252944 | 2766 | * | 40,036668 | 2,563101011 | 0,5927482 | 4,3240973 | 1,532E-05 | 0,030265668 | 2,7872553 | 3' UTR | 6 | 86188576 | 86249201 | 60626 | 1 | 21802 | ENSMUST00000194449.3 | 61603 | ENSMUSG00000029999 | Tgfa | transforming growth factor alpha |
| chr6 | 1,04E+08 | 1,04E+08 | 1081 | * | 13,29185 | 2,563190641 | 0,947058 | 2,706478 | 0,0068 | 0,892811906 | 0,938682 | Intron (ENSMUST0000 | 6 | 1,04E+08 | 1,04E+08 | 62 | 1 | 1,15E+08 | ENSMUST00000157640.3 | 3318 | ENSMUSG00000088265 | Gm24784 | predicted gene, 24784 |
| chr10 | 3351697 | 3353145 | 1449 | * | 12,2108 | 2,56331654 | 0,975103 | 2,628764 | 0,00857 | 0,963466134 | 0,738441 | Intron (ENSMUST0000 | 10 | 3316544 | 3414975 | 98432 | 1 | 76142 | ENSMUST00000217573.2 | 35153 | ENSMUSG00000040653 | Ppp1r14c | protein phosphatase 1, regulatory inhibitor subunit 14C |
| chr9 | 59712605 | 59714592 | 1988 | * | 17,18742 | 2,563603656 | 0,880364 | 2,911981 | 0,003591 | 0,62192524 | 2,423594 | Intron (ENSMUST0000 | 9 | 59658179 | 59836149 | 177971 | 1 | 270163 | ENSMUST00000136740.8 | 54426 | ENSMUSG00000039585 | Myo9a | myosin IXa |
| chr2 | 32251474 | 32253782 | 2309 | * | 26,60152 | 2,56371581 | 0,751673 | 3,410681 | 0,000648 | 0,300782365 | 2,880017 | Promoter (<=1kb) | 2 | 32253022 | 32257419 | 4398 | 1 | 68379 | ENSMUST00000125769.8 | 0 | ENSMUSG00000039205 | Ciz1 | CDKN1A interacting zinc finger protein 1 |
| chr10 | 62134624 | 62137946 | 3323 | * | 37,79457 | 2,56395774 | 0,633552 | 4,046956 | 5,19E-05 | 0,080010047 | 2,377674 | Exon (ENSMUST00000 | 10 | 62120388 | 62126108 | 5721 | 2 | 15275 | ENSMUST00000161160.2 | -8516 | ENSMUSG00000037012 | Hk1 | hexokinase 1 |
| chr17 | 32074977 | 32077026 | 2050 | * | 22,58357 | 2,564133146 | 0,751561 | 3,411743 | 0,000645 | 0,290624195 | 3,104559 | Promoter (<=1kb) | 17 | 32063224 | 32074778 | 11555 | 2 | 17691 | ENSMUST00000024839.6 | -199 | ENSMUSG00000024042 | Sik1 | salt inducible kinase 1 |
| chr12 | 35971198 | 35972642 | 1445 | * | 14,88281 | 2,56435396 | 0,94306 | 2,719185 | 0,006544 | 0,842187815 | 1,214904 | Promoter (2-3kb) | 12 | 35975619 | 35999735 | 24117 | 1 | 403205 | ENSMUST00000042101.5 | -2977 | ENSMUSG00000036231 | Agr3 | anterior gradient 3 |
| chr18 | 16352478 | 16353522 | 1045 | * | 14,42031 | 2,564609115 | 0,825565 | 3,10649 | 0,001893 | 0,622205297 | 1,2763 | Distal Intergenic | 18 | 16170186 | 16248454 | 78269 | 2 | 1,05E+08 | ENSMUST00000242720.1 | -104024 | | Gm41673 | predicted gene, 41673 |
| chr1 | 84389357 | 84391317 | 1961 | * | 16,54908 | 2,564923078 | 0,86523 | 2,964442 | 0,003032 | 0,568412386 | 2,663619 | Intron (ENSMUST0000 | 1 | 84085515 | 84341901 | 256387 | 2 | 98496 | ENSMUST00000175996.8 | -47456 | ENSMUSG00000045658 | Pid1 | phosphotyrosine interaction domain containing 1 |
| chr7 | 79909710 | 79912386 | 2677 | * | 19,13295 | 2,565060407 | 0,816156 | 3,142857 | 0,001673 | 0,474532331 | 2,507013 | Promoter (<=1kb) | 7 | 79910963 | 79915122 | 4160 | 1 | 83485 | ENSMUST00000117989.2 | 0 | ENSMUSG00000047084 | Ngrn | neugrin, neurite outgrowth associated |
| chr2 | 1,56E+08 | 1,56E+08 | 2956 | * | 25,47172 | 2,565257194 | 0,695063 | 3,690684 | 0,000224 | 0,172398224 | 2,880017 | Intron (ENSMUST0000 | 2 | 1,56E+08 | 1,56E+08 | 43025 | 1 | 17391 | ENSMUST00000029141.6 | 10997 | ENSMUSG00000027612 | Mmp24 | matrix metallopeptidase 24 |
| chr5 | 1,45E+08 | 1,45E+08 | 1862 | * | 15,81632 | 2,565547552 | 0,850976 | 3,014831 | 0,002571 | 0,538066226 | 2,663619 | 3' UTR | 5 | 1,45E+08 | 1,45E+08 | 5526 | 1 | 72611 | ENSMUST00000085661.9 | 9775 | ENSMUSG00000007812 | Zfp655 | zinc finger protein 655 |
| chr16 | 95880435 | 95882698 | 2264 | * | 20,09308 | 2,565925949 | 0,797548 | 3,217269 | 0,001294 | 0,421090004 | 2,663619 | Promoter (<=1kb) | 16 | 95793292 | 95883628 | 90337 | 2 | 93871 | ENSMUST00000099502.9 | 930 | ENSMUSG00000022914 | Brwd1 | bromodomain and WD repeat domain containing 1 |
| chr12 | 1,05E+08 | 1,05E+08 | 1955 | * | 17,02787 | 2,565939148 | 0,81018 | 3,167123 | 0,00154 | 0,474289497 | 2,318434 | Distal Intergenic | 12 | 1,05E+08 | 1,05E+08 | 79877 | 2 | 212073 | ENSMUST00000067005.11 | -6168 | ENSMUSG00000054150 | Syne3 | spectrin repeat containing, nuclear envelope family member 3 |
| chr9 | 1,21E+08 | 1,21E+08 | 2310 | * | 14,09664 | 2,566030005 | 0,957176 | 2,680834 | 0,007344 | 0,908237331 | 0,918255 | Intron (ENSMUST0000 | 9 | 1,21E+08 | 1,21E+08 | 100558 | 2 | 209012 | ENSMUST00000170237.8 | 16722 | ENSMUSG00000040936 | Ulk4 | unc-51-like kinase 4 |
| chr11 | 96536474 | 96538625 | 2152 | * | 12,80584 | 2,566845385 | 0,990424 | 2,591663 | 0,009551 | 0,910299316 | 1,173325 | Intron (ENSMUST0000 | 11 | 96599061 | 96604028 | 4968 | 1 | 78473 | ENSMUST00000123814.2 | -60436 | ENSMUSG00000057058 | Skap1 | src family associated phosphoprotein 1 |
| chr18 | 36537132 | 36539255 | 2124 | * | 38,36226 | 2,567057834 | 0,5102203 | 5,0312734 | 4,872E-07 | 0,00252152 | 2,3383871 | 3' UTR | 18 | 36498834 | 36535593 | 36760 | 1 | 66060 | ENSMUST00000144158.2 | 38298 | ENSMUSG00000046727 | Cystm1 | cysteine-rich transmembrane module containing 1 |
| chr9 | 77939507 | 77940687 | 1181 | * | 13,10808 | 2,568064403 | 0,913491 | 2,811265 | 0,004935 | 0,881662676 | 0,738441 | Distal Intergenic | 9 | 77922300 | 77927985 | 5686 | 1 | 1,05E+08 | ENSMUST00000217068.2 | 17207 | | Gm39375 | predicted gene, 39375 |
| chr6 | 84294087 | 84295235 | 1149 | * | 14,15429 | 2,568285896 | 0,923589 | 2,780767 | 0,005423 | 0,800381675 | 1,2763 | Distal Intergenic | 6 | 84389924 | 84400820 | 10897 | 2 | 1E+08 | ENSMUST00000203521.2 | 105585 | ENSMUSG00000107797 | Gm10445 | predicted gene 10445 |
| chr3 | 1,01E+08 | 1,01E+08 | 1554 | * | 15,95746 | 2,568301468 | 0,915826 | 2,804356 | 0,005042 | 0,663630601 | 2,663619 | Distal Intergenic | 3 | 1,01E+08 | 1,01E+08 | 9501 | 1 | 229644 | ENSMUST00000094048.10 | -3354 | ENSMUSG00000033233 | Trim45 | tripartite motif-containing 45 |
| chr5 | 1,15E+08 | 1,15E+08 | 1657 | * | 15,89995 | 2,568492739 | 0,991532 | 2,590428 | 0,009586 | 0,793995086 | 2,318434 | Promoter (<=1kb) | 5 | 1,15E+08 | 1,15E+08 | 6437 | 2 | 330171 | ENSMUST00000134173.8 | 0 | ENSMUSG00000001098 | Kctd10 | potassium channel tetramerisation domain containing 10 |
| chr10 | 80099195 | 80100773 | 1579 | * | 14,82819 | 2,568719642 | 0,771995 | 3,327378 | 0,000877 | 0,486479832 | 1,214904 | Promoter (<=1kb) | 10 | 80100818 | 80124242 | 23425 | 1 | 70248 | ENSMUST00000105362.8 | -45 | ENSMUSG00000069565 | Dazap1 | DAZ associated protein 1 |
| chr14 | 51556573 | 51557337 | 765 | * | 11,59656 | 2,568730775 | 0,990875 | 2,592386 | 0,009531 | 0,948199575 | 0,918255 | Intron (ENSMUST0000 | 14 | 51554792 | 51582946 | 28155 | 2 | 75377 | ENSMUST00000228344.2 | 25609 | ENSMUSG00000115540 | 4930597G03Rik | RIKEN cDNA 4930597G03 gene |
| chr8 | 85710107 | 85713733 | 3627 | * | 85,925117 | 2,569145013 | 0,4335386 | 5,9259891 | 3,104E-09 | 3,72592E-05 | 3,2706213 | Exon (ENSMUST000002 | 8 | 85717232 | 85729972 | 12741 | 1 | 170833 | ENSMUST00000209764.2 | -3499 | ENSMUSG00000052566 | Hook2 | hook microtubule tethering protein 2 |
| chr9 | 59444581 | 59446957 | 2377 | * | 29,79527 | 2,569221714 | 0,747698 | 3,436173 | 0,00059 | 0,296046677 | 2,725491 | Promoter (<=1kb) | 9 | 59446823 | 59472392 | 25570 | 1 | 15211 | ENSMUST00000026262.8 | 0 | ENSMUSG00000025232 | Hexa | hexosaminidase A |
| chr14 | 24037017 | 24039037 | 2021 | * | 17,18464 | 2,569710478 | 0,984921 | 2,609052 | 0,009079 | 0,759660077 | 2,663619 | Intron (ENSMUST0000 | 14 | 24053912 | 24058403 | 4492 | 1 | 78019 | ENSMUST00000223609.2 | -14875 | ENSMUSG00000114953 | 4930519K11Rik | RIKEN cDNA 4930519K11 gene |
| chr15 | 85559344 | 85560484 | 1141 | * | 14,02179 | 2,569967139 | 0,982927 | 2,614606 | 0,008933 | 0,937845932 | 0,918255 | Intron (ENSMUST0000 | 15 | 85563644 | 85591725 | 28082 | 1 | 1,03E+08 | ENSMUST00000245865.1 | -3160 | | Lncppara | long noncoding RNA near Ppara |
| chr3 | 94349815 | 94351820 | 2006 | * | 16,8279 | 2,570852136 | 0,832612 | 3,087696 | 0,002017 | 0,631836405 | 1,2763 | Promoter (<=1kb) | 3 | 94340694 | 94351140 | 10447 | 2 | 53814 | ENSMUST00000203883.3 | 0 | ENSMUSG00000028141 | Oaz3 | ornithine decarboxylase antizyme 3 |
| chr11 | 59052811 | 59055957 | 3147 | * | 47,175443 | 2,571379499 | 0,5890167 | 4,3655461 | 1,268E-05 | 0,030043467 | 2,3776742 | Promoter (<=1kb) | 11 | 59046195 | 59054565 | 8371 | 2 | 216792 | ENSMUST00000054523.6 | 0 | ENSMUSG00000049287 | Iba57 | IBA57 homolog, iron-sulfur cluster assembly |
| chr13 | 56520103 | 56521379 | 1277 | * | 17,22425 | 2,57140347 | 0,838406 | 3,067015 | 0,002162 | 0,506770495 | 2,663619 | Distal Intergenic | 13 | 56510714 | 56513582 | 2869 | 1 | 1E+08 | ENSMUST00000099476.4 | 9389 | ENSMUSG00000074885 | Gm10782 | predicted gene 10782 |
| chr2 | 31382532 | 31385474 | 2943 | * | 42,87894 | 2,571666984 | 0,5474893 | 4,6972 | 2,638E-06 | 0,008107936 | 2,7872553 | Promoter (2-3kb) | 2 | 31387784 | 31400167 | 12384 | 1 | 11898 | ENSMUST00000126474.7 | -2310 | ENSMUSG00000076441 | Ass1 | argininosuccinate synthetase 1 |
| chr12 | 58621287 | 58622439 | 1153 | * | 11,47324 | 2,571940327 | 0,986491 | 2,60716 | 0,00913 | 0,941899824 | 0,918255 | Distal Intergenic | 12 | 58311506 | 58316076 | 4571 | 2 | 66864 | ENSMUST00000062254.4 | -305211 | ENSMUSG00000045930 | Clec14a | C-type lectin domain family 14, member a |
| chr12 | 99598004 | 99600402 | 2399 | * | 22,93708 | 2,572069101 | 0,688102 | 3,737921 | 0,000186 | 0,155627448 | 2,858511 | Distal Intergenic | 12 | 99592312 | 99594233 | 1922 | 2 | 73457 | ENSMUST00000185726.2 | -3771 | ENSMUSG00000100211 | 1700064M15Rik | RIKEN cDNA 1700064M15 gene |
| chr19 | 29618725 | 29620405 | 1681 | * | 12,189 | 2,572399839 | 0,96714 | 2,6598 | 0,007819 | 0,950586732 | 0,744375 | Promoter (1-2kb) | 19 | 29604448 | 29616963 | 12516 | 2 | 226090 | ENSMUST00000161124.8 | -1762 | ENSMUSG00000046324 | Ermp1 | endoplasmic reticulum metallopeptidase 1 |
| chr6 | 33776537 | 33778229 | 1693 | * | 17,89369 | 2,572479973 | 0,971726 | 2,647331 | 0,008113 | 0,749931768 | 2,507013 | Intron (ENSMUST0000 | 6 | 33839035 | 33851116 | 12082 | 1 | 20336 | ENSMUST00000134653.2 | -60806 | ENSMUSG00000029763 | Exoc4 | exocyst complex component 4 |
| chr5 | 1,21E+08 | 1,21E+08 | 1483 | * | 16,11094 | 2,572841074 | 0,846121 | 3,04075 | 0,00236 | 0,660341484 | 1,2763 | Promoter (<=1kb) | 5 | 1,21E+08 | 1,21E+08 | 18355 | 2 | 71772 | ENSMUST00000131831.8 | 459 | ENSMUSG00000029598 | Plbd2 | phospholipase B domain containing 2 |
| chr11 | 61906549 | 61908509 | 1961 | * | 16,06753 | 2,572858981 | 0,824079 | 3,122101 | 0,001796 | 0,498011371 | 2,318434 | Promoter (1-2kb) | 11 | 61909674 | 62032552 | 122879 | 1 | 432572 | ENSMUST00000049836.14 | -1165 | ENSMUSG00000042331 | Specc1 | sperm antigen with calponin homology and coiled-coil domains 1 |
| chr2 | 32523598 | 32525124 | 1527 | * | 20,70511 | 2,573079444 | 0,865453 | 2,9731 | 0,002948 | 0,534851486 | 3,104559 | 3' UTR | 2 | 32519533 | 32523412 | 3880 | 1 | 11636 | ENSMUST00000135392.2 | 4065 | ENSMUSG00000026817 | Ak1 | adenylate kinase 1 |
| chr15 | 88958600 | 88960167 | 1568 | * | 30,39855 | 2,573197682 | 0,661624 | 3,889214 | 0,000101 | 0,104080853 | 2,880017 | Promoter (<=1kb) | 15 | 88959327 | 88969512 | 10186 | 1 | 67976 | ENSMUST00000168863.8 | 0 | ENSMUSG00000015363 | Trabd | TraB domain containing |
| chr10 | 1,09E+08 | 1,09E+08 | 1902 | * | 15,38138 | 2,573711632 | 0,842546 | 3,054682 | 0,002253 | 0,660932959 | 1,214904 | Distal Intergenic | 10 | 1,08E+08 | 1,09E+08 | 513331 | 2 | 20979 | ENSMUST00000064054.14 | -156810 | ENSMUSG00000035864 | Syt1 | synaptotagmin I |
| chr4 | 21975989 | 21977227 | 1239 | * | 14,73905 | 2,574930187 | 0,937148 | 2,747625 | 0,006003 | 0,826676469 | 1,214904 | Intron (ENSMUST0000 | 4 | 21958456 | 21960807 | 2352 | 1 | 76132 | ENSMUST00000155220.2 | 17533 | ENSMUSG00000028246 | Faxc | failed axon connections homolog |
| chr8 | 1,27E+08 | 1,27E+08 | 2843 | * | 31,51072 | 2,575166893 | 0,67176 | 3,833464 | 0,000126 | 0,119411163 | 2,951062 | Distal Intergenic | 8 | 1,27E+08 | 1,27E+08 | 49737 | 2 | 212728 | ENSMUST00000170518.3 | -49196 | ENSMUSG00000090290 | Tarbp1 | TAR RNA binding protein 1 |
| chr17 | 32254653 | 32256471 | 1819 | * | 15,6924 | 2,575582522 | 0,939749 | 2,740714 | 0,006131 | 0,809369408 | 1,37683 | Promoter (<=1kb) | 17 | 32255074 | 32281839 | 26766 | 1 | 72462 | ENSMUST00000081339.13 | 0 | ENSMUSG00000058392 | Rrp1b | ribosomal RNA processing 1B |

| | | | | | | | | | | | | | | | | | | | | | | | |
|---|---|---|---|---|---|---|---|---|---|---|---|---|---|---|---|---|---|---|---|---|---|---|---|
| chr6 | 59172664 | 59174180 | 1517 | * | 22,43347 | 2,576078032 | 0,691115 | 3,727422 | 0,000193 | 0,142131669 | 3,471678 | Distal Intergenic | 6 | 59185855 | 59189018 | 3164 | 1 | 68140 | ENSMUST00000062626.4 | -11675 | ENSMUSG00000049232 | Tigd2 | tigger transposable element derived 2 |
| chr5 | 1,29E+08 | 1,29E+08 | 1666 | * | 15,70681 | 2,576243964 | 0,9164 | 2,811265 | 0,004935 | 0,683067851 | 2,318434 | Distal Intergenic | 5 | 1,29E+08 | 1,29E+08 | 107850 | 1 | 243277 | ENSMUST00000056617.14 | -29270 | ENSMUSG00000044017 | Adgrd1 | adhesion G protein-coupled receptor D1 |
| chr2 | 1,03E+08 | 1,03E+08 | 1916 | * | 17,11206 | 2,576249287 | 0,818052 | 3,149248 | 0,001637 | 0,477594358 | 2,386008 | Exon (ENSMUST00000 | 2 | 1,03E+08 | 1,03E+08 | 107 | 1 | 1,15E+08 | ENSMUST00000239821.1 | -44217 | ENSMUSG00000119260 | Gm23731 | predicted gene, 23731 |
| chr18 | 62108779 | 62110834 | 2056 | * | 12,17299 | 2,576890926 | 0,917023 | 2,810062 | 0,004953 | 0,842458878 | 0,918255 | Promoter (<=1kb) | 18 | 62111030 | 62148790 | 37761 | 1 | 225608 | ENSMUST00000236759.2 | -196 | ENSMUSG00000045629 | Sh3tc2 | SH3 domain and tetratricopeptide repeats 2 |
| chr2 | 1,18E+08 | 1,18E+08 | 1751 | * | 16,268 | 2,577892533 | 0,842561 | 3,059592 | 0,002216 | 0,531203841 | 2,386008 | Promoter (1-2kb) | 2 | 1,18E+08 | 1,18E+08 | 126533 | 2 | 71313 | ENSMUST00000231133.2 | 1892 | ENSMUSG00000027344 | Fsip1 | fibrous sheath-interacting protein 1 |
| chr1 | 1,69E+08 | 1,69E+08 | 1611 | * | 17,44157 | 2,578571271 | 0,968241 | 2,663151 | 0,007741 | 0,770673161 | 2,154504 | Distal Intergenic | 1 | 1,69E+08 | 1,69E+08 | 116 | 2 | 1,02E+08 | ENSMUST00000184979.3 | 152267 | ENSMUSG00000098829 | Mir6354 | microRNA 6354 |
| chr15 | 5468218 | 5469424 | 1207 | * | 12,97542 | 2,57886353 | 0,973926 | 2,647905 | 0,008099 | 0,923078468 | 0,918255 | Distal Intergenic | 15 | 5525747 | 5624465 | 98719 | 1 | 71432 | ENSMUST00000211750.2 | -56323 | ENSMUSG00000110018 | 5430437J10Rik | RIKEN cDNA 5430437J10 gene |
| chr11 | 20822813 | 20823997 | 1185 | * | 11,8251 | 2,579162812 | 0,985299 | 2,617644 | 0,008854 | 0,90060438 | 1,173325 | Distal Intergenic | 11 | 20797753 | 20797881 | 129 | 2 | 1,15E+08 | ENSMUST00000082737.3 | -24932 | ENSMUSG00000064671 | Gm22807 | predicted gene, 22807 |
| chr10 | 19062549 | 19065656 | 3108 | * | 20,58558 | 2,580507612 | 0,889207 | 2,902032 | 0,003708 | 0,605351622 | 2,713306 | Intron (ENSMUST00000 | 10 | 19042986 | 19082450 | 39465 | 1 | 1,01E+08 | ENSMUST00000216683.2 | 19563 | | Gm20139 | predicted gene, 20139 |
| chr15 | 1,02E+08 | 1,02E+08 | 1435 | * | 14,08749 | 2,580652981 | 0,928938 | 2,778068 | 0,005468 | 0,858257847 | 0,918255 | Distal Intergenic | 15 | 1,02E+08 | 1,02E+08 | 5231 | 1 | 1,03E+08 | ENSMUST00000230247.2 | -8505 | | Gm36026 | predicted gene, 36026 |
| chr19 | 12289814 | 12291560 | 1747 | * | 14,20631 | 2,581178183 | 0,914388 | 2,822846 | 0,00476 | 0,83238039 | 0,938682 | Distal Intergenic | 19 | 12296571 | 12302465 | 5895 | 2 | 258117 | ENSMUST00000207241.3 | 10905 | ENSMUSG00000096436 | Olfr1437 | olfactory receptor 1437 |
| chr5 | 59328975 | 59331011 | 2037 | * | 10,3225 | 2,581202384 | 0,965289 | 2,674019 | 0,007495 | 0,864414194 | 1,211315 | Distal Intergenic | 5 | 59372289 | 59372395 | 107 | 2 | 1,15E+08 | ENSMUST00000239835.1 | 41384 | ENSMUSG00000119859 | Gm22308 | predicted gene, 22308 |
| chr1 | 1,81E+08 | 1,81E+08 | 1711 | * | 17,83848 | 2,58141951 | 0,837094 | 3,083787 | 0,002044 | 0,534793902 | 2,154504 | Promoter (<=1kb) | 1 | 1,81E+08 | 1,81E+08 | 9829 | 1 | 208795 | ENSMUST00000161847.8 | 0 | ENSMUSG00000026519 | Tmem63a | transmembrane protein 63a |
| chr4 | 43924729 | 43928183 | 3455 | * | 31,80887 | 2,581460495 | 0,699879 | 3,688437 | 0,000226 | 0,173211012 | 2,880017 | Exon (ENSMUST00000 | 4 | 43940398 | 43942382 | 1985 | 1 | 53614 | ENSMUST00000130415.2 | -12215 | ENSMUSG00000028476 | Reck | reversion-inducing-cysteine-rich protein with kazal motifs |
| chr4 | 1,5E+08 | 1,5E+08 | 1181 | * | 12,46052 | 2,581491744 | 0,911678 | 2,831583 | 0,004632 | 0,831521609 | 0,918255 | Distal Intergenic | 4 | 1,5E+08 | 1,5E+08 | 7569 | 1 | 13806 | ENSMUST00000150175.8 | -5096 | ENSMUSG00000063524 | Eno1 | enolase 1, alpha non-neuron |
| chr19 | 13337807 | 13339497 | 1691 | * | 12,28452 | 2,581799885 | 0,958214 | 2,694387 | 0,007052 | 0,848477082 | 1,2763 | Promoter (2-3kb) | 19 | 13341994 | 13342920 | 927 | 1 | 258686 | ENSMUST00000054687.3 | -2497 | ENSMUSG00000049015 | Olfr1467 | olfactory receptor 1467 |
| chr8 | 70974886 | 70976678 | 1793 | * | 21,38387 | 2,582018567 | 0,718917 | 3,591539 | 0,000329 | 0,223919137 | 2,67243 | Promoter (<=1kb) | 8 | 70960922 | 70975813 | 14892 | 2 | 22186 | ENSMUST00000121623.8 | 0 | ENSMUSG00000090137 | Uba52 | ubiquitin A-52 residue ribosomal protein fusion product 1 |
| chr15 | 77755028 | 77756571 | 1544 | * | 16,00098 | 2,58223827 | 0,833095 | 3,099574 | 0,001938 | 0,526946456 | 2,141612 | Distal Intergenic | 15 | 77644787 | 77726375 | 81589 | 2 | 17886 | ENSMUST00000016771.13 | -28653 | ENSMUSG00000022443 | Myh9 | myosin, heavy polypeptide 9, non-muscle |
| chr19 | 12829321 | 12830594 | 1274 | * | 18,25079 | 2,58234313 | 0,893007 | 2,89174 | 0,003831 | 0,616178941 | 2,663619 | Distal Intergenic | 19 | 12839106 | 12840234 | 1129 | 1 | 258697 | ENSMUST00000059675.4 | -8512 | ENSMUSG00000046272 | Olfr1444 | olfactory receptor 1444 |
| chrX | 1,5E+08 | 1,5E+08 | 1883 | * | 21,08002 | 2,583230979 | 0,823381 | 3,137346 | 0,001705 | 0,44905823 | 2,951933 | Exon (ENSMUST00000 | 20 | 1,5E+08 | 1,5E+08 | 6101 | 2 | 237107 | ENSMUST00000146742.2 | 67663 | ENSMUSG00000025266 | Gnl3l | guanine nucleotide binding protein-like 3 (nucleolar)-like |
| chr3 | 51230391 | 51231955 | 1565 | * | 14,68076 | 2,58414734 | 0,917801 | 2,815585 | 0,004869 | 0,761670222 | 1,41696 | Promoter (1-2kb) | 3 | 51163106 | 51233408 | 70303 | 2 | 69257 | ENSMUST00000163748.8 | 1453 | ENSMUSG00000037174 | Elf2 | E74-like factor 2 |
| chr9 | 1,07E+08 | 1,07E+08 | 1467 | * | 12,47217 | 2,584281048 | 0,880217 | 2,93596 | 0,003325 | 0,773092266 | 0,918255 | Promoter (1-2kb) | 9 | 1,07E+08 | 1,07E+08 | 23270 | 2 | 208869 | ENSMUST00000165075.2 | -1557 | ENSMUSG00000039716 | Dock3 | dedicator of cyto-kinesis 3 |
| chr15 | 100673000 | 100675737 | 2738 | * | 79,268161 | 2,584483479 | 0,4683483 | 5,5182936 | 3,423E-08 | 0,000244135 | 3,2706213 | Intron (ENSMUST000000 | 15 | 100679135 | 100713816 | 34682 | 1 | 59033 | ENSMUST00000162049.2 | -3398 | ENSMUSG00000023032 | Slc4a8 | solute carrier family 4 (anion exchanger), member 8 |
| chr8 | 1,08E+08 | 1,08E+08 | 1108 | * | 14,60715 | 2,58453192 | 0,874916 | 2,954036 | 0,003136 | 0,688021707 | 1,37683 | Distal Intergenic | 8 | 1,08E+08 | 1,08E+08 | 12347 | 1 | 67644 | ENSMUST00000241277.1 | 38566 | | 4930517G24Rik | RIKEN cDNA 4930517G24 gene |
| chr10 | 40283071 | 40284809 | 1739 | * | 15,81376 | 2,584798179 | 0,882566 | 2,92873 | 0,003403 | 0,717868719 | 1,2763 | Intron (ENSMUST00000 | 10 | 40225522 | 40358097 | 132576 | 1 | 78334 | ENSMUST00000214659.2 | 57549 | ENSMUSG00000038481 | Cdk19 | cyclin-dependent kinase 19 |
| chr6 | 1,28E+08 | 1,28E+08 | 1363 | * | 20,38063 | 2,585012683 | 0,810679 | 3,188701 | 0,001429 | 0,467440127 | 2,221611 | Promoter (<=1kb) | 6 | 1,28E+08 | 1,28E+08 | 15488 | 2 | 101142 | ENSMUST00000001559.11 | 0 | ENSMUSG00000001518 | Itfg2 | integrin alpha FG-GAP repeat containing 2 |
| chr19 | 46656710 | 46657879 | 1170 | * | 14,16528 | 2,585245671 | 0,957993 | 2,698606 | 0,006963 | 0,875818484 | 1,075735 | Promoter (<=1kb) | 19 | 46656526 | 46657952 | 1427 | 2 | 13074 | ENSMUST00000131142.2 | 73 | ENSMUSG00000003555 | Cyp17a1 | cytochrome P450, family 17, subfamily a, polypeptide 1 |
| chr18 | 15024806 | 15026540 | 1735 | * | 11,96946 | 2,585742596 | 0,975713 | 2,650107 | 0,008047 | 0,955649401 | 0,738441 | Intron (ENSMUST00000 | 18 | 15122052 | 15181044 | 58993 | 1 | 74456 | ENSMUST00000234445.2 | -95512 | ENSMUSG00000117450 | 4933424G05Rik | RIKEN cDNA 4933424G05 gene |
| chr11 | 54005276 | 54006837 | 1562 | * | 15,51321 | 2,586368999 | 0,973624 | 2,656435 | 0,007897 | 0,864414194 | 1,2763 | Promoter (1-2kb) | 11 | 54004029 | 54005100 | 1072 | 1 | 18452 | ENSMUST00000155299.2 | 1247 | ENSMUSG00000018906 | P4ha2 | procollagen-proline, 2-oxoglutarate 4-dioxygenase (proline 4-hydroxylase), alpha II polypeptide |
| chr2 | 1,56E+08 | 1,56E+08 | 1390 | * | 15,49704 | 2,586808268 | 0,869399 | 2,975397 | 0,002926 | 0,682889642 | 1,37683 | Promoter (<=1kb) | 2 | 1,56E+08 | 1,56E+08 | 1172 | 2 | 266692 | ENSMUST00000183353.2 | -253 | ENSMUSG00000074643 | Cpne1 | copine I |
| chr2 | 132052473 | 132054885 | 2413 | * | 40,341032 | 2,586936829 | 0,4961076 | 5,2144669 | 1,843E-07 | 0,001087015 | 2,7872553 | Intron (ENSMUST000001 | 2 | 132081412 | 132089574 | 8163 | 2 | 70612 | ENSMUST00000110164.8 | 34689 | ENSMUSG00000027341 | Tmem230 | transmembrane protein 230 |
| chr1 | 61516117 | 61517231 | 1115 | * | 13,64714 | 2,587773058 | 0,889848 | 2,908108 | 0,003636 | 0,730469267 | 1,2763 | Distal Intergenic | 1 | 61417591 | 61435063 | 17473 | 1 | 77341 | ENSMUST00000141000.2 | 98526 | ENSMUSG00000085447 | 9530026F06Rik | RIKEN cDNA 9530026F06 gene |
| chr16 | 32051369 | 32052585 | 1217 | * | 10,59876 | 2,587812527 | 0,975617 | 2,652488 | 0,00799 | 0,866942328 | 1,2763 | 3' UTR | 16 | 32062530 | 32062673 | 144 | 2 | 1,15E+08 | ENSMUST00000116790.3 | 10088 | ENSMUSG00000080440 | Gm25848 | predicted gene, 25848 |
| chr13 | 53742638 | 53744371 | 1734 | * | 14,35592 | 2,587829032 | 0,94645 | 2,734249 | 0,006252 | 0,880965195 | 0,938682 | Distal Intergenic | 13 | 53789930 | 53793229 | 3300 | 2 | 77398 | ENSMUST00000221882.2 | 48858 | ENSMUSG00000113599 | 9530014B07Rik | RIKEN cDNA 9530014B07 gene |
| chr5 | 74277994 | 74279067 | 1074 | * | 10,27635 | 2,588508017 | 0,998033 | 2,593609 | 0,009497 | 0,880965195 | 1,425929 | Distal Intergenic | 5 | 74254191 | 74254311 | 121 | 1 | 1E+08 | ENSMUST00000175614.3 | 23803 | ENSMUSG00000093355 | Snora26 | small nucleolar RNA, H/ACA box 26 |
| chr3 | 95522741 | 95524853 | 2113 | * | 17,49923 | 2,588740943 | 0,901838 | 2,870516 | 0,004098 | 0,664801704 | 2,154504 | 5' UTR | 3 | 95532322 | 95539413 | 7092 | 1 | 56205 | ENSMUST00000058230.13 | -7469 | ENSMUSG00000038619 | Ensa | endosulfine alpha |
| chr7 | 66103879 | 66106442 | 2564 | * | 16,70919 | 2,589228781 | 0,831331 | 3,114559 | 0,001842 | 0,497895643 | 2,386008 | Distal Intergenic | 7 | 66040638 | 66077265 | 36628 | 2 | 56847 | ENSMUST00000015278.15 | -26614 | ENSMUSG00000015134 | Aldh1a3 | aldehyde dehydrogenase family 1, subfamily A3 |
| chr13 | 6081761 | 6084553 | 2793 | * | 20,39639 | 2,589730793 | 0,752267 | 3,442567 | 0,000576 | 0,315383621 | 2,221611 | Distal Intergenic | 13 | 6027784 | 6064284 | 36501 | 1 | 1,08E+08 | ENSMUST00000221291.2 | 53977 | | Gm46401 | predicted gene, 46401 |
| chr3 | 99047970 | 99049427 | 1458 | * | 14,89468 | 2,589959869 | 0,956347 | 2,708181 | 0,006765 | 0,838976992 | 1,2763 | Promoter (<=1kb) | 3 | 99048384 | 99127519 | 79136 | 1 | 70560 | ENSMUST00000004343.7 | 0 | ENSMUSG00000004233 | Wars2 | tryptophanyl tRNA synthetase 2 (mitochondrial) |
| chr15 | 82548896 | 82551116 | 2221 | * | 18,08539 | 2,590267121 | 0,862451 | 3,003379 | 0,00267 | 0,55292773 | 2,507013 | Distal Intergenic | 15 | 82520388 | 82526247 | 5860 | 2 | 68444 | ENSMUST00000238885.2 | -22649 | ENSMUSG00000118516 | Cyp2d13 | cytochrome P450, family 2, subfamily d, polypeptide 13 |
| chr15 | 38075522 | 38076790 | 1269 | * | 12,48534 | 2,590300668 | 0,992581 | 2,609663 | 0,009063 | 0,903101391 | 1,173325 | Promoter (1-2kb) | 15 | 37968537 | 38078669 | 110133 | 2 | 70790 | ENSMUST00000226414.2 | 1879 | ENSMUSG00000037487 | Ubr5 | ubiquitin protein ligase E3 component n-recognin 5 |
| chr18 | 59959413 | 59960967 | 1555 | * | 14,78654 | 2,590501987 | 0,943128 | 2,746714 | 0,00602 | 0,827392612 | 1,214904 | Intron (ENSMUST00000 | 18 | 59958503 | 59988534 | 30032 | 2 | 1,03E+08 | ENSMUST00000237574.2 | 27567 | ENSMUSG00000117701 | Gm35234 | predicted gene, 35234 |
| chr3 | 1,17E+08 | 1,17E+08 | 1036 | * | 11,3287 | 2,590959939 | 0,968337 | 2,67568 | 0,007458 | 0,85617852 | 1,2763 | Distal Intergenic | 3 | 1,16E+08 | 1,17E+08 | 18402 | 2 | 229782 | ENSMUST00000169530.2 | -23260 | ENSMUSG00000027957 | Slc35a3 | solute carrier family 35 (UDP-N-acetylglucosamine (UDP-GlcNAc) transporter), member 3 |
| chr8 | 72688244 | 72689697 | 1454 | * | 12,53972 | 2,591276455 | 0,930112 | 2,785984 | 0,005337 | 0,855495103 | 0,918255 | 3' UTR | 8 | 72676696 | 72688473 | 11778 | 1 | 170938 | ENSMUST00000119003.2 | 11548 | ENSMUSG00000066880 | Zfp617 | zinc finger protein 617 |
| chr15 | 99936939 | 99938151 | 1213 | * | 13,11941 | 2,592060687 | 0,915479 | 2,831372 | 0,004635 | 0,802323085 | 1,075735 | Promoter (<=1kb) | 15 | 99936545 | 1E+08 | 180810 | 1 | 239667 | ENSMUST00000100203.10 | 394 | ENSMUSG00000023026 | Dip2b | disco interacting protein 2 homolog B |
| chr15 | 37982693 | 37984556 | 1864 | * | 15,22932 | 2,592704094 | 0,986735 | 2,627559 | 0,0086 | 0,890233764 | 1,214904 | Promoter (<=1kb) | 15 | 37968103 | 37984611 | 16509 | 2 | 70790 | ENSMUST00000228029.2 | 55 | ENSMUSG00000037487 | Ubr5 | ubiquitin protein ligase E3 component n-recognin 5 |
| chr2 | 34553608 | 34554598 | 991 | * | 11,08733 | 2,593064344 | 1,001602 | 2,588916 | 0,009628 | 0,912096919 | 1,173325 | Distal Intergenic | 2 | 34567906 | 34574305 | 6400 | 2 | 66691 | ENSMUST00000138203.2 | 19707 | ENSMUSG00000026867 | Gapvd1 | GTPase activating protein and VPS9 domains 1 |
| chr14 | 25832302 | 25833696 | 1395 | * | 14,74058 | 2,593130207 | 1,00195 | 2,588083 | 0,009651 | 0,900637308 | 1,2763 | Distal Intergenic | 14 | 25842580 | 25887228 | 44649 | 1 | 11744 | ENSMUST00000022416.15 | -8884 | ENSMUSG00000021866 | Anxa11 | annexin A11 |
| chr12 | 55489340 | 55491570 | 2231 | * | 45,098057 | 2,593242029 | 0,5153074 | 5,0324171 | 4,843E-07 | 0,00252152 | 2,3383871 | Distal Intergenic | 12 | 55454168 | 55459072 | 4905 | 1 | 26443 | ENSMUST00000162972.2 | 35172 | ENSMUSG00000021024 | Psma6 | proteasome subunit alpha 6 |
| chr17 | 80989445 | 80991202 | 1758 | * | 22,02418 | 2,593553926 | 0,878745 | 2,95143 | 0,003163 | 0,562958709 | 2,880017 | Intron (ENSMUST00000 | 17 | 80887946 | 81035453 | 147508 | 2 | 225028 | ENSMUST00000234133.2 | 44251 | ENSMUSG00000024242 | Map4k3 | mitogen-activated protein kinase kinase kinase kinase 3 |
| chr12 | 1,12E+08 | 1,12E+08 | 1282 | * | 14,85668 | 2,593842876 | 0,985184 | 2,632852 | 0,008467 | 0,863495249 | 1,37683 | Promoter (<=1kb) | 12 | 1,12E+08 | 1,12E+08 | 1310 | 1 | 104816 | ENSMUST00000222645.2 | 0 | ENSMUSG00000037686 | Aspg | asparaginase |
| chr1 | 5761013 | 5762405 | 1393 | * | 13,18959 | 2,593904248 | 0,890609 | 2,912505 | 0,003585 | 0,742014238 | 1,173325 | Distal Intergenic | 1 | 5666741 | 5673376 | 6636 | 1 | 18387 | ENSMUST00000159083.2 | 94272 | ENSMUSG00000025905 | Oprk1 | opioid receptor, kappa 1 |
| chr5 | 27630608 | 27631871 | 1264 | * | 12,85071 | 2,593923225 | 0,978964 | 2,64966 | 0,008057 | 0,922788862 | 0,918255 | Intron (ENSMUST00000 | 5 | 27702535 | 27706360 | 3826 | 2 | 73526 | ENSMUST00000155721.2 | 74489 | ENSMUSG00000048703 | Speer4b | spermatogenesis associated glutamate (E)-rich protein 4B |
| chr12 | 94335916 | 94336775 | 860 | * | 10,83633 | 2,59494236 | 1,000092 | 2,594702 | 0,009467 | 0,898273833 | 1,2763 | Distal Intergenic | 12 | 93894112 | 93894238 | 127 | 1 | 1,02E+08 | ENSMUST00000183568.3 | 441804 | ENSMUSG00000098931 | Mir8099-2 | microRNA 8099-2 |
| chr3 | 1,35E+08 | 1,35E+08 | 1409 | * | 14,49116 | 2,595109951 | 0,929939 | 2,790625 | 0,005261 | 0,782403314 | 1,37683 | Intron (ENSMUST00000 | 3 | 1,35E+08 | 1,35E+08 | 105573 | 1 | 21338 | ENSMUST00000029822.6 | 14200 | ENSMUSG00000028172 | Tacr3 | tachykinin receptor 3 |
| chr2 | 44857902 | 44860188 | 2287 | * | 25,07953 | 2,595568457 | 0,680326 | 3,815183 | 0,000136 | 0,124112216 | 2,951062 | Distal Intergenic | 2 | 44525276 | 44817669 | 292394 | 2 | 227835 | ENSMUST00000148279.8 | -40233 | ENSMUSG00000036890 | Gtdc1 | glycosyltransferase-like domain containing 1 |
| chr11 | 52188123 | 52190217 | 2095 | * | 12,01705 | 2,595854365 | 0,979788 | 2,649404 | 0,008063 | 0,922828293 | 0,918255 | Distal Intergenic | 11 | 52144063 | 52174158 | 30096 | 2 | 21414 | ENSMUST00000109071.3 | -13965 | ENSMUSG00000000782 | Tcf7 | transcription factor 7, T cell specific |
| chr6 | 54791967 | 54793794 | 1828 | * | 22,50769 | 2,595952591 | 0,828559 | 3,133095 | 0,00173 | 0,455547114 | 2,880017 | Promoter (<=1kb) | 6 | 54793901 | 54870485 | 76585 | 1 | 387524 | ENSMUST00000079869.13 | -107 | ENSMUSG00000058446 | Znrf2 | zinc and ring finger 2 |
| chr2 | 96968559 | 96970424 | 1866 | * | 13,71123 | 2,596508001 | 0,908449 | 2,858178 | 0,004261 | 0,817520905 | 0,918255 | Intron (ENSMUST00000 | 2 | 97068163 | 97068291 | 129 | 1 | 1,15E+08 | ENSMUST00000158008.3 | -97739 | ENSMUSG00000088633 | Gm22313 | predicted gene, 22313 |
| chr11 | 92921676 | 92923132 | 1457 | * | 13,69831 | 2,597136847 | 0,922521 | 2,815259 | 0,004874 | 0,782360218 | 1,2763 | Distal Intergenic | 11 | 92988854 | 92990692 | 1839 | 1 | 72605 | ENSMUST00000149611.2 | -65722 | ENSMUSG00000056158 | Car10 | carbonic anhydrase 10 |
| chr4 | 93154560 | 93155652 | 1093 | * | 13,93925 | 2,597526704 | 0,911825 | 2,848713 | 0,00439 | 0,858257847 | 0,738441 | Distal Intergenic | 4 | 93222375 | 93223748 | 1374 | 2 | 69136 | ENSMUST00000066774.6 | 68096 | ENSMUSG00000054000 | Tusc1 | tumor suppressor candidate 1 |
| chr10 | 79948896 | 79951950 | 3055 | * | 42,305259 | 2,59766053 | 0,6026084 | 4,3106944 | 1,627E-05 | 0,03581197 | 2,3776742 | Promoter (<=1kb) | 10 | 79951637 | 79966303 | 14667 | 1 | 20869 | ENSMUST00000144883.8 | 0 | ENSMUSG00000003068 | Stk11 | serine/threonine kinase 11 |
| chr6 | 78707928 | 78708997 | 1070 | * | 14,72473 | 2,597730572 | 0,880855 | 2,949102 | 0,003187 | 0,715249603 | 1,214904 | Distal Intergenic | 6 | 78888150 | 78888313 | 164 | 1 | 1,15E+08 | ENSMUST00000240322.1 | -179153 | ENSMUSG00000118890 | Gm23727 | predicted gene, 23727 |
| chr18 | 73884774 | 73885805 | 1032 | * | 10,70542 | 2,598217929 | 0,999066 | 2,600648 | 0,009305 | 0,974266082 | 0,744375 | Promoter (1-2kb) | 18 | 73868109 | 73887550 | 19442 | 2 | 114615 | ENSMUST00000041138.3 | 1745 | ENSMUSG00000036941 | Elac1 | elaC ribonuclease Z 1 |
| chr8 | 79156948 | 79158879 | 1932 | * | 15,31115 | 2,599222643 | 0,939607 | 2,766286 | 0,00567 | 0,791068542 | 1,41696 | Distal Intergenic | 8 | 79159639 | 79163274 | 3636 | 2 | 18997 | ENSMUST00000034115.5 | 4395 | ENSMUSG00000031688 | Pou4f2 | POU domain, class 4, transcription factor 2 |
| chr7 | 1,21E+08 | 1,21E+08 | 1871 | * | 14,62326 | 2,599728372 | 0,955075 | 2,722014 | 0,006489 | 0,819720472 | 1,37683 | Promoter (<=1kb) | 7 | 1,21E+08 | 1,21E+08 | 65233 | 2 | 76179 | ENSMUST00000046929.7 | -338 | ENSMUSG00000063317 | Usp31 | ubiquitin specific peptidase 31 |
| chr11 | 6643572 | 6645916 | 2345 | * | 28,70821 | 2,599844787 | 0,637047 | 4,081085 | 4,48E-05 | 0,063501557 | 2,951062 | Distal Intergenic | 11 | 6666001 | 6670037 | 4037 | 2 | 626419 | ENSMUST00000139769.2 | 24121 | ENSMUSG00000086808 | Gm11981 | predicted gene 11981 |
| chr11 | 76462571 | 76464858 | 2288 | * | 19,72546 | 2,600024135 | 0,854284 | 3,043514 | 0,002338 | 0,523637575 | 2,663619 | Promoter (<=1kb) | 11 | 76307560 | 76462361 | 154802 | 2 | 109934 | ENSMUST00000072740.13 | -210 | ENSMUSG00000017631 | Abr | active BCR-related gene |
| chr15 | 66910462 | 66913224 | 2763 | * | 61,86348 | 2,600027952 | 0,4404439 | 5,9031991 | 3,565E-09 | 3,98744E-05 | 3,2706213 | Promoter (<=1kb) | 15 | 66911907 | 66912673 | 767 | 1 | 102633543 | ENSMUST00000230911.2 | 0 | | Gm31342 | predicted gene, 31342 |
| chrX | 1,1E+08 | 1,1E+08 | 2246 | * | 15,42407 | 2,600231257 | 0,868548 | 2,99377 | 0,002756 | 0,594676756 | 2,141612 | Intron (ENSMUST00000 | 20 | 1,1E+08 | 1,1E+08 | 29097 | 2 | 67071 | ENSMUST00000123213.8 | -4707 | ENSMUSG00000025665 | Rps6ka6 | ribosomal protein S6 kinase polypeptide 6 |
| chr16 | 31797584 | 31799703 | 2120 | * | 18,50435 | 2,600417584 | 0,724501 | 3,589251 | 0,000332 | 0,236050623 | 2,423594 | Intron (ENSMUST00000 | 16 | 31805649 | 31805941 | 293 | 2 | 1,02E+08 | ENSMUST00000232628.2 | 6238 | | Gm15694 | NADH dehydrogenase (ubiquinone) 1 alpha subcomplex, 7 (B14.5a), pseudogene |
| chr1 | 1,71E+08 | 1,71E+08 | 3111 | * | 25,15116 | 2,601278241 | 0,79888 | 3,256155 | 0,001129 | 0,379156186 | 2,951933 | Promoter (<=1kb) | 1 | 1,71E+08 | 1,71E+08 | 107 | 2 | 1,15E+08 | ENSMUST00000179933.3 | 0 | ENSMUSG00000119049 | Gm25235 | predicted gene, 25235 |
| chr10 | 92512867 | 92514500 | 1634 | * | 16,07502 | 2,601321456 | 0,863352 | 3,013049 | 0,002586 | 0,565091829 | 2,318434 | Distal Intergenic | 10 | 92524726 | 92524870 | 145 | 2 | 1,15E+08 | ENSMUST00000157787.3 | 10370 | ENSMUSG00000088412 | Gm24241 | predicted gene, 24241 |
| chr17 | 26822830 | 26825458 | 2629 | * | 25,38341 | 2,601583806 | 0,684433 | 3,80108 | 0,000144 | 0,138163056 | 2,67243 | Promoter (1-2kb) | 17 | 26826805 | 26874322 | 47518 | 1 | 67458 | ENSMUST00000235197.2 | -1347 | ENSMUSG00000001576 | Ergic1 | endoplasmic reticulum-golgi intermediate compartment (ERGIC) 1 |
| chr11 | 52546254 | 52547705 | 1452 | * | 15,55808 | 2,602092363 | 0,847515 | 3,07026 | 0,002139 | 0,54190677 | 2,141612 | Distal Intergenic | 11 | 52655461 | 53079365 | 423905 | 1 | 320027 | ENSMUST00000036796.8 | -107756 | ENSMUSG00000036264 | Fstl4 | follistatin-like 4 |
| chr14 | 21870990 | 21872171 | 1182 | * | 14,79679 | 2,602205766 | 0,885836 | 2,93757 | 0,003308 | 0,714462637 | 1,2763 | Distal Intergenic | 14 | 21875306 | 21890637 | 15332 | 1 | 22334 | ENSMUST00000224373.2 | -3135 | ENSMUSG00000021771 | Vdac2 | voltage-dependent anion channel 2 |
| chr14 | 25722681 | 25724909 | 2229 | * | 16,26578 | 2,602418467 | 0,965194 | 2,696264 | 0,007012 | 0,847020973 | 1,2763 | Promoter (2-3kb) | 14 | 25712068 | 25720216 | 8149 | 2 | 71918 | ENSMUST00000184642.2 | -2465 | ENSMUSG00000055538 | Zcchc24 | zinc finger, CCHC domain containing 24 |
| chr8 | 79818538 | 79820356 | 1819 | * | 17,95368 | 2,603351358 | 0,723416 | 3,59869 | 0,00032 | 0,221323359 | 2,663619 | Exon (ENSMUST00000 | 8 | 79844956 | 79920395 | 75440 | 1 | 622675 | ENSMUST00000148713.8 | -24600 | ENSMUSG00000071064 | Zfp827 | zinc finger protein 827 |
| chr10 | 81031184 | 81032734 | 1551 | * | 14,98197 | 2,603462898 | 1,009267 | 2,579558 | 0,009893 | 0,910299316 | 1,214904 | Promoter (<=1kb) | 10 | 81030517 | 81045257 | 14741 | 1 | 69678 | ENSMUST00000129950.3 | 667 | | Atcayos | ataxia, cerebellar, Cayman type, opposite strand |
| chr5 | 1,02E+08 | 1,02E+08 | 934 | * | 10,57071 | 2,60364104 | 0,959695 | 2,712989 | 0,006668 | 0,893166641 | 0,918255 | Promoter (<=1kb) | 5 | 1,02E+08 | 1,02E+08 | 14900 | 1 | 74596 | ENSMUST00000132213.2 | -213 | ENSMUSG00000029330 | Cds1 | CDP-diacylglycerol synthase 1 |
| chr15 | 8507279 | 8509581 | 2303 | * | 19,77408 | 2,603858716 | 0,765805 | 3,400158 | 0,000673 | 0,322225015 | 2,507013 | Promoter (2-3kb) | 15 | 8511603 | 8529827 | 18225 | 1 | 1,03E+08 | ENSMUST00000227472.2 | -2022 | | Gm31282 | predicted gene, 31282 |
| chr15 | 84419515 | 84421396 | 1882 | * | 23,02053 | 2,604334577 | 0,737947 | 3,529164 | 0,000417 | 0,252389304 | 2,67243 | Distal Intergenic | 15 | 84437599 | 84442024 | 4426 | 2 | 223732 | ENSMUST00000069476.5 | 20628 | ENSMUSG00000055745 | Rtl6 | retrotransposon Gag like 6 |
| chr11 | 5804066 | 5806945 | 2880 | * | 24,20153 | 2,60458719 | 0,7641 | 3,408698 | 0,000653 | 0,301441602 | 2,880017 | Distal Intergenic | 11 | 5811947 | 5822088 | 10142 | 1 | 11568 | ENSMUST00000102923.10 | -5002 | ENSMUSG00000020473 | Aebp1 | AE binding protein 1 |
| chr3 | 21574052 | 21575677 | 1626 | * | 14,93857 | 2,604962241 | 0,860749 | 3,02639 | 0,002475 | 0,675594321 | 1,214904 | Distal Intergenic | 3 | 21756979 | 21766624 | 9646 | 1 | 1,03E+08 | ENSMUST00000200150.2 | -181302 | ENSMUSG00000105440 | Gm31693 | predicted gene, 31693 |
| chr14 | 22671966 | 22673589 | 1624 | * | 13,07792 | 2,605004921 | 0,950447 | 2,740821 | 0,006129 | 0,852340527 | 1,075735 | Intron (ENSMUST00000 | 14 | 22646581 | 22649336 | 2756 | 1 | 76633 | ENSMUST00000226003.2 | 25385 | ENSMUSG00000063458 | Lrmda | leucine rich melanocyte differentiation associated |
| chr6 | 83142021 | 83142911 | 891 | * | 15,03569 | 2,605032647 | 0,829593 | 3,140133 | 0,001689 | 0,584572307 | 1,37683 | Promoter (<=1kb) | 6 | 83142902 | 83177098 | 34197 | 1 | 13191 | ENSMUST00000077407.12 | 0 | ENSMUSG00000031865 | Dctn1 | dynactin 1 |
| chr7 | 139549922 | 139553564 | 3643 | * | 42,003068 | 2,605042363 | 0,5891491 | 4,4217034 | 9,793E-06 | 0,022589498 | 2,7254908 | 3' UTR | 7 | 139552355 | 139558632 | 6278 | 2 | 381933 | ENSMUST00000209335.2 | 5068 | ENSMUSG00000073795 | Spef1l | sperm flagellar 1 like |
| chr7 | 12513174 | 12514540 | 1367 | * | 17,54867 | 2,605436055 | 0,863464 | 3,017424 | 0,002549 | 0,544107745 | 2,507013 | Promoter (<=1kb) | 7 | 12502019 | 12514581 | 12563 | 2 | 232875 | ENSMUST00000211392.2 | 41 | ENSMUSG00000070822 | Zscan18 | zinc finger and SCAN domain containing 18 |
| chr4 | 1,42E+08 | 1,42E+08 | 896 | * | 12,70298 | 2,605555329 | 0,877753 | 2,968439 | 0,002993 | 0,751231966 | 0,918255 | Intron (ENSMUST00000 | 4 | 1,42E+08 | 1,42E+08 | 3806 | 1 | 545691 | ENSMUST00000146094.2 | 8479 | ENSMUSG00000085424 | Fhad1os1 | forkhead-associated (FHA) phosphopeptide binding domain 1, opposite strand 1 |
| chr15 | 81544310 | 81548104 | 3795 | * | 30,19825 | 2,605585536 | 0,788599 | 3,304068 | 0,000953 | 0,352774915 | 2,858511 | Promoter (<=1kb) | 15 | 81548090 | 81572516 | 24427 | 1 | 214669 | ENSMUST00000023029.15 | 0 | ENSMUSG00000022394 | L3mbtl2 | L3MBTL2 polycomb repressive complex 1 subunit |
| chr7 | 98657987 | 98659681 | 1695 | * | 10,97441 | 2,60654411 | 0,941515 | 2,768457 | 0,005632 | 0,807032197 | 1,2763 | Intron (ENSMUST00000 | 7 | 98535957 | 98665851 | 129895 | 2 | 78610 | ENSMUST00000209123.2 | 6170 | ENSMUSG00000035354 | Uvrag | UV radiation resistance associated gene |
| chr2 | 72313464 | 72315291 | 1828 | * | 14,96529 | 2,606762202 | 0,88918 | 2,931646 | 0,003372 | 0,631820259 | 2,141612 | Promoter (<=1kb) | 2 | 72313908 | 72315087 | 1180 | 1 | 66953 | ENSMUST00000130730.2 | 0 | ENSMUSG00000055612 | Cdca7 | cell division cycle associated 7 |
| chr15 | 39888690 | 39889263 | 574 | * | 11,33626 | 2,607665927 | 0,957924 | 2,722206 | 0,006485 | 0,922828293 | 0,738441 | Distal Intergenic | 15 | 39908587 | 39992378 | 83792 | 1 | 1,03E+08 | ENSMUST00000227360.2 | -19324 | ENSMUSG00000115142 | Gm33251 | predicted gene, 33251 |
| chr7 | 1,26E+08 | 1,26E+08 | 3155 | * | 20,53044 | 2,60770823 | 0,868738 | 3,001721 | 0,002685 | 0,54122637 | 2,713306 | Exon (ENSMUST00000 | 7 | 1,26E+08 | 1,26E+08 | 107 | 2 | 1,15E+08 | ENSMUST00000122504.3 | 9975 | ENSMUSG00000084453 | Gm24596 | predicted gene, 24596 |
| chr8 | 1,24E+08 | 1,24E+08 | 1264 | * | 13,26838 | 2,607914634 | 0,909809 | 2,866443 | 0,004151 | 0,78560377 | 1,075735 | 3' UTR | 8 | 1,24E+08 | 1,24E+08 | 10109 | 2 | 57429 | ENSMUST00000146007.2 | 8315 | ENSMUSG00000000739 | Sult5a1 | sulfotransferase family 5A, member 1 |
| chr7 | 80477819 | 80479353 | 1535 | * | 22,45709 | 2,608769955 | 0,677403 | 3,851135 | 0,000118 | 0,120888649 | 2,67243 | Promoter (2-3kb) | 7 | 80418107 | 80475722 | 57616 | 2 | 29875 | ENSMUST00000205304.2 | -2097 | ENSMUSG00000030536 | Iqgap1 | IQ motif containing GTPase activating protein 1 |
| chr14 | 58134589 | 58136069 | 1481 | * | 16,09617 | 2,609148499 | 0,827909 | 3,151493 | 0,001624 | 0,495652322 | 2,141612 | Distal Intergenic | 14 | 58070160 | 58127733 | 57574 | 2 | 75965 | ENSMUST00000089473.5 | -6856 | ENSMUSG00000021969 | Zdhhc20 | zinc finger, DHHC domain containing 20 |
| chr13 | 1,16E+08 | 1,16E+08 | 1254 | * | 13,66031 | 2,609168053 | 0,93223 | 2,798845 | 0,005129 | 0,821456993 | 1,075735 | Distal Intergenic | 13 | 1,16E+08 | 1,16E+08 | 3957 | 2 | 16392 | ENSMUST00000176444.2 | 121248 | ENSMUSG00000042258 | Isl1 | ISL1 transcription factor, LIM/homeodomain |
| chr8 | 1,28E+08 | 1,28E+08 | 1114 | * | 20,09735 | 2,609744466 | 0,705413 | 3,699597 | 0,000216 | 0,188932071 | 2,386008 | Intron (ENSMUST00000 | 8 | 1,28E+08 | 1,28E+08 | 12241 | 1 | 414091 | ENSMUST00000212914.2 | 12816 | | A630001O12Rik | RIKEN cDNA A630001O12 gene |
| chr16 | 13714303 | 13716788 | 2486 | * | 16,53747 | 2,610343373 | 0,844614 | 3,090575 | 0,001998 | 0,517479776 | 2,318434 | Intron (ENSMUST00000 | 16 | 13699473 | 13720839 | 21367 | 2 | 94184 | ENSMUST00000148551.2 | 4051 | ENSMUSG00000022680 | Pdxdc1 | pyridoxal-dependent decarboxylase domain containing 1 |
| chr5 | 1,37E+08 | 1,37E+08 | 1678 | * | 13,65816 | 2,610664284 | 0,937664 | 2,784222 | 0,005366 | 0,85617852 | 0,918255 | Promoter (<=1kb) | 5 | 1,37E+08 | 1,37E+08 | 6960 | 1 | 381677 | ENSMUST00000190827.7 | -437 | ENSMUSG00000037428 | Vgf | VGF nerve growth factor inducible |

| | | | | | | | | | | | | | | | | | | | | | | | |
|---|---|---|---|---|---|---|---|---|---|---|---|---|---|---|---|---|---|---|---|---|---|---|---|
| chr15 | 99191635 | 99192855 | 1221 | * | 16,03265 | 2,610866533 | 0,897649 | 2,90856 | 0,003631 | 0,644704154 | 2,141612 | Promoter (<=1kb) | 15 | 99192968 | 99203846 | 10879 | 1 | 54614 | ENSMUST00000128352.8 | -113 | ENSMUSG00000023007 | Prpf40b | pre-mRNA processing factor 40B |
| chr12 | 46115796 | 46117988 | 2193 | * | 17,35478 | 2,610869741 | 0,791016 | 3,300652 | 0,000965 | 0,408869321 | 2,154504 | Distal Intergenic | 12 | 46031126 | 46031332 | 207 | 1 | 1,15E+08 | ENSMUST00000082866.3 | 84670 | ENSMUSG00000064800 | Gm24948 | predicted gene, 24948 |
| chr11 | 69587303 | 69588664 | 1362 | * | 11,93675 | 2,610957125 | 0,94424 | 2,76514 | 0,00569 | 0,824134241 | 1,173325 | Promoter (<=1kb) | 11 | 69573403 | 69586924 | 13522 | 2 | 619441 | ENSMUST00000180587.8 | -379 | ENSMUSG00000018752 | Tnfsfm13 | tumor necrosis factor (ligand) superfamily, membrane-bound member 13 |
| chrX | 1,51E+08 | 1,51E+08 | 2750 | * | 15,11061 | 2,611039011 | 0,868674 | 3,005776 | 0,002649 | 0,661775851 | 1,41696 | 3' UTR | 20 | 1,51E+08 | 1,51E+08 | 3919 | 1 | 59026 | ENSMUST00000239245.2 | -5110 | ENSMUSG00000025261 | Huwe1 | HECT, UBA and WWE domain containing 1 |
| chr3 | 63080780 | 63082204 | 1425 | * | 14,6265 | 2,611562557 | 0,82052 | 3,182812 | 0,001459 | 0,477253929 | 2,141612 | Exon (ENSMUST00000 | 3 | 63059638 | 63059779 | 142 | 1 | 1,15E+08 | ENSMUST00000158550.3 | 21142 | ENSMUSG00000089175 | Gm22433 | predicted gene, 22433 |
| chr1 | 64834411 | 64835381 | 971 | * | 11,91406 | 2,61186614 | 1,005777 | 2,596863 | 0,009408 | 0,976076191 | 0,744375 | Intron (ENSMUST00000 | 1 | 64832968 | 64875923 | 42956 | 2 | 241075 | ENSMUST00000187188.2 | 40542 | ENSMUSG00000051344 | Plekhm3 | pleckstrin homology domain containing, family M, member 3 |
| chr11 | 15535523 | 15536787 | 1265 | * | 12,51034 | 2,612112965 | 0,860029 | 3,037238 | 0,002388 | 0,661775851 | 1,2763 | Distal Intergenic | 11 | 15402422 | 15402551 | 130 | 1 | 1,15E+08 | ENSMUST00000158734.3 | 133101 | ENSMUSG00000089359 | Gm25490 | predicted gene, 25490 |
| chr10 | 94647234 | 94648891 | 1658 | * | 16,21879 | 2,612605251 | 0,868386 | 3,008576 | 0,002625 | 0,584572307 | 2,141612 | Intron (ENSMUST00000 | 10 | 94663084 | 94665408 | 2325 | 2 | 54712 | ENSMUST00000218839.2 | 16517 | ENSMUSG00000074785 | Plxnc1 | plexin C1 |
| chr12 | 63654148 | 63655381 | 1234 | * | 10,94139 | 2,612646234 | 0,999419 | 2,614164 | 0,008945 | 0,970928491 | 0,738441 | Promoter (1-2kb) | 12 | 63656642 | 63656748 | 107 | 1 | 1,15E+08 | ENSMUST00000083024.4 | -1261 | ENSMUSG00000118736 | Gm24921 | predicted gene, 24921 |
| chr3 | 72494674 | 72495907 | 1234 | * | 12,37686 | 2,612651224 | 0,947106 | 2,758562 | 0,005806 | 0,842680506 | 1,075735 | Distal Intergenic | 3 | 72795893 | 72874199 | 78307 | 2 | 69983 | ENSMUST00000094190.5 | 378292 | ENSMUSG00000027790 | Sis | sucrase isomaltase (alpha-glucosidase) |
| chr3 | 88957662 | 88959852 | 2191 | * | 16,7511 | 2,613038756 | 0,994756 | 2,626815 | 0,008619 | 0,883601315 | 1,2763 | Promoter (<=1kb) | 3 | 88960109 | 88963087 | 2979 | 1 | 192195 | ENSMUST00000189824.2 | -257 | ENSMUSG00000028053 | Ash1l | ASH1 like histone lysine methyltransferase |
| chr9 | 92957636 | 92959636 | 2001 | * | 15,95259 | 2,613189689 | 0,998206 | 2,617885 | 0,008848 | 0,777172375 | 2,386008 | Distal Intergenic | 9 | 92862048 | 92887792 | 25745 | 2 | 78349 | ENSMUST00000190562.7 | -69844 | ENSMUSG00000100294 | 1700034K08Rik | RIKEN cDNA 1700034K08 gene |
| chr5 | 81616558 | 81617961 | 1404 | * | 10,73894 | 2,614466979 | 0,957404 | 2,730788 | 0,006318 | 0,916926803 | 0,744375 | Intron (ENSMUST00000 | 5 | 81457872 | 81944481 | 486610 | 1 | 319387 | ENSMUST00000072521.11 | 158686 | ENSMUSG00000037605 | Adgrl3 | adhesion G protein-coupled receptor L3 |
| chr18 | 61777054 | 61779668 | 2615 | * | 22,44983 | 2,614519494 | 0,821019 | 3,184481 | 0,00145 | 0,426043749 | 2,880017 | Promoter (<=1kb) | 18 | 61777201 | 61778564 | 1364 | 1 | 545261 | ENSMUST00000240555.1 | 0 | ENSMUSG00000098098 | Bvht | braveheart long non-coding RNA |
| chr2 | 1,67E+08 | 1,67E+08 | 2507 | * | 30,26445 | 2,6148054 | 0,659803 | 3,963007 | 7,4E-05 | 0,089931799 | 2,725491 | Distal Intergenic | 2 | 1,67E+08 | 1,67E+08 | 55289 | 1 | 77031 | ENSMUST00000047815.13 | -33438 | ENSMUSG00000039463 | Slc9a8 | solute carrier family 9 (sodium/hydrogen exchanger), member 8 |
| chr2 | 25081009 | 25083027 | 2019 | * | 24,57733 | 2,615327868 | 0,675455 | 3,871952 | 0,000108 | 0,107816957 | 2,951933 | 3' UTR | 2 | 25085069 | 25086771 | 1703 | 2 | 227612 | ENSMUST00000142857.2 | 3744 | ENSMUSG00000059555 | Tor4a | torsin family 4, member A |
| chr2 | 97222411 | 97223936 | 1526 | * | 13,50604 | 2,615573758 | 0,904026 | 2,893252 | 0,003813 | 0,737185115 | 1,2763 | Intron (ENSMUST00000 | 2 | 97298002 | 97462009 | 164008 | 1 | 241568 | ENSMUST00000059049.8 | -74066 | ENSMUSG00000050587 | Lrrc4c | leucine rich repeat containing 4C |
| chr6 | 1,24E+08 | 1,24E+08 | 1932 | * | 21,97135 | 2,61561013 | 0,833342 | 3,138699 | 0,001697 | 0,448852574 | 2,951933 | Distal Intergenic | 6 | 1,24E+08 | 1,24E+08 | 17531 | 1 | 232371 | ENSMUST00000049124.10 | -6813 | ENSMUSG00000038527 | C1rl | complement component 1, r subcomponent-like |
| chr11 | 94814133 | 94816188 | 2056 | * | 13,68659 | 2,615666 | 0,974016 | 2,685445 | 0,007243 | 0,883012141 | 1,075735 | Distal Intergenic | 11 | 94827050 | 94843868 | 16819 | 1 | 12842 | ENSMUST00000001547.8 | -10862 | ENSMUSG00000001506 | Col1a1 | collagen, type I, alpha 1 |
| chr12 | 1,09E+08 | 1,09E+08 | 1638 | * | 13,99534 | 2,616174113 | 0,872047 | 3,000037 | 0,002699 | 0,734161328 | 0,918255 | Promoter (2-3kb) | 12 | 1,09E+08 | 1,09E+08 | 5687 | 1 | 104910 | ENSMUST00000057026.10 | -2838 | ENSMUSG00000048856 | Slc25a47 | solute carrier family 25, member 47 |
| chr6 | 7569856 | 7570473 | 618 | * | 12,35025 | 2,616399305 | 0,96231 | 2,718873 | 0,00655 | 0,851014755 | 1,173325 | Distal Intergenic | 6 | 7556091 | 7559114 | 3024 | 1 | 21333 | ENSMUST00000185130.2 | 13765 | ENSMUSG00000061762 | Tac1 | tachykinin 1 |
| chr1 | 1,1E+08 | 1,1E+08 | 901 | * | 11,02771 | 2,61675784 | 0,962694 | 2,718161 | 0,006565 | 0,891057096 | 0,918255 | Intron (ENSMUST00000 | 1 | 1,1E+08 | 1,1E+08 | 145208 | 1 | 241201 | ENSMUST00000027542.13 | 40323 | ENSMUSG00000026312 | Cdh7 | cadherin 7, type 2 |
| chr17 | 46109146 | 46110962 | 1817 | * | 39,354923 | 2,616764404 | 0,5769265 | 4,5356984 | 5,741E-06 | 0,014734867 | 2,7872553 | Distal Intergenic | 17 | 46117130 | 46133249 | 16120 | 2 | 102639361 | ENSMUST00000233237.2 | 22287 | | Gm35692 | predicted gene, 35692 |
| chr7 | 45545681 | 45548053 | 2373 | * | 21,60285 | 2,616897347 | 0,786087 | 3,329017 | 0,000872 | 0,349985669 | 2,67243 | Promoter (<=1kb) | 7 | 45534646 | 45546123 | 11478 | 2 | 58867 | ENSMUST00000211254.2 | 0 | ENSMUSG00000040231 | Syngr4 | synaptogyrin 4 |
| chr11 | 21812768 | 21814080 | 1313 | * | 12,85124 | 2,617091057 | 0,886575 | 2,951913 | 0,003158 | 0,720584502 | 1,173325 | Intron (ENSMUST00000 | 11 | 21763399 | 21767341 | 3943 | 1 | 216560 | ENSMUST00000128794.2 | 49369 | ENSMUSG00000020319 | Wdpcp | WD repeat containing planar cell polarity effector |
| chr18 | 75548498 | 75550741 | 2244 | * | 20,61666 | 2,617592252 | 0,776899 | 3,369285 | 0,000754 | 0,308275655 | 3,104559 | Distal Intergenic | 18 | 75507985 | 75527072 | 19088 | 1 | 17131 | ENSMUST00000172718.2 | 40513 | ENSMUSG00000025880 | Smad7 | SMAD family member 7 |
| chr2 | 38266676 | 38267576 | 901 | * | 14,04642 | 2,617594989 | 0,929313 | 2,816698 | 0,004852 | 0,877892722 | 0,738441 | Distal Intergenic | 2 | 38249592 | 38253992 | 4401 | 2 | 1,03E+08 | ENSMUST00000137134.2 | -12684 | | Gm13584 | predicted gene 13584 |
| chr10 | 13718610 | 13720369 | 1760 | * | 12,88382 | 2,617776147 | 0,924893 | 2,830357 | 0,00465 | 0,802490409 | 1,075735 | Promoter (<=1kb) | 10 | 13529332 | 13720030 | 190699 | 2 | 66253 | ENSMUST00000162174.8 | 0 | ENSMUSG00000019806 | Aig1 | androgen-induced 1 |
| chr2 | 83636763 | 83638619 | 1857 | * | 12,13318 | 2,61833324 | 0,959781 | 2,728052 | 0,006371 | 0,887593795 | 0,918255 | 3' UTR | 2 | 83633265 | 83637260 | 3996 | 1 | 16410 | ENSMUST00000131192.2 | 3498 | ENSMUSG00000027087 | Itgav | integrin alpha V |
| chr8 | 88720687 | 88722234 | 1548 | * | 18,49596 | 2,618572804 | 0,822192 | 3,18487 | 0,001448 | 0,4572736 | 2,386008 | Distal Intergenic | 8 | 88388438 | 88686223 | 297786 | 2 | 94187 | ENSMUST00000109655.9 | -34464 | ENSMUSG00000045333 | Zfp423 | zinc finger protein 423 |
| chr7 | 25032717 | 25034621 | 1905 | * | 20,16144 | 2,619071394 | 0,793263 | 3,301645 | 0,000961 | 0,364328215 | 2,713306 | Exon (ENSMUST00000 | 7 | 25045406 | 25065339 | 19934 | 1 | 269878 | ENSMUST00000153077.2 | -10785 | ENSMUSG00000045039 | Megf8 | multiple EGF-like-domains 8 |
| chr8 | 1,23E+08 | 1,23E+08 | 1683 | * | 21,20132 | 2,619410785 | 0,737955 | 3,549553 | 0,000386 | 0,228944295 | 2,951933 | Promoter (1-2kb) | 8 | 1,23E+08 | 1,23E+08 | 3780 | 2 | 170732 | ENSMUST00000044123.2 | 1243 | ENSMUSG00000039079 | Trhr2 | thyrotropin releasing hormone receptor 2 |
| chr15 | 77448743 | 77450708 | 1966 | * | 21,38826 | 2,619515701 | 0,835494 | 3,13529 | 0,001717 | 0,442939362 | 3,104559 | Promoter (2-3kb) | 15 | 77442439 | 77446092 | 3654 | 2 | 76409 | ENSMUST00000190233.2 | -2651 | ENSMUSG00000101113 | 1700025B11Rik | RIKEN cDNA 1700025B11 gene |
| chr11 | 75652141 | 75654896 | 2756 | * | 17,98434 | 2,619720916 | 0,922032 | 2,841248 | 0,004494 | 0,644557958 | 2,663619 | Intron (ENSMUST00000 | 11 | 75642430 | 75650141 | 7712 | 1 | 22627 | ENSMUST00000155057.2 | 9711 | ENSMUSG00000020849 | Ywhae | tyrosine 3-monooxygenase/tryptophan 5-monooxygenase activation protein, epsilon polypeptide |
| chr5 | 1,22E+08 | 1,22E+08 | 2311 | * | 23,54142 | 2,619968697 | 0,703198 | 3,725792 | 0,000195 | 0,15931259 | 2,880017 | Promoter (<=1kb) | 5 | 1,22E+08 | 1,22E+08 | 26569 | 1 | 320717 | ENSMUST00000119015.2 | 0 | ENSMUSG00000038582 | Pptc7 | PTC7 protein phosphatase homolog |
| chr13 | 50164315 | 50166038 | 1724 | * | 16,76465 | 2,620236962 | 0,912498 | 2,8715 | 0,004085 | 0,646594296 | 2,386008 | Distal Intergenic | 13 | 50291105 | 50300706 | 9602 | 1 | 71252 | ENSMUST00000222134.2 | -125067 | | 4933433N18Rik | RIKEN cDNA 4933433N18 gene |
| chr15 | 98605830 | 98608885 | 3056 | * | 31,922344 | 2,622451717 | 0,6199026 | 4,2304253 | 2,332E-05 | 0,043315528 | 2,7254908 | Promoter (<=1kb) | 15 | 98606088 | 98621217 | 15130 | 1 | 105833 | ENSMUST00000003444.6 | 0 | ENSMUSG00000003354 | Ccdc65 | coiled-coil domain containing 65 |
| chr10 | 98330133 | 98332624 | 2492 | * | 16,93997 | 2,622531777 | 0,959679 | 2,732718 | 0,006281 | 0,722439117 | 2,318434 | Distal Intergenic | 10 | 98373340 | 98404898 | 31559 | 1 | 1,03E+08 | ENSMUST00000219170.2 | -40716 | | Gm34297 | predicted gene, 34297 |
| chr8 | 11302211 | 11303808 | 1598 | * | 13,01613 | 2,62549524 | 0,959253 | 2,737022 | 0,0062 | 0,838878373 | 1,173325 | Intron (ENSMUST00000 | 8 | 11268035 | 11291148 | 23114 | 2 | 12826 | ENSMUST00000209661.2 | -11063 | ENSMUSG00000031502 | Col4a1 | collagen, type IV, alpha 1 |
| chr2 | 90793653 | 90794666 | 1014 | * | 9,808612 | 2,626262724 | 0,987822 | 2,65864 | 0,007846 | 0,881662676 | 1,173325 | Promoter (<=1kb) | 2 | 90795437 | 90818161 | 22725 | 1 | 13046 | ENSMUST00000133657.2 | -771 | ENSMUSG00000005506 | Celf1 | CUGBP, Elav-like family member 1 |
| chr10 | 79547479 | 79550359 | 2881 | * | 31,94905 | 2,62635529 | 0,5865587 | 4,4775662 | 7,55E-06 | 0,018654836 | 2,7254908 | Promoter (1-2kb) | 10 | 79545939 | 79547544 | 1606 | 1 | 12215 | ENSMUST00000180235.2 | 1540 | ENSMUSG00000023175 | Bsg | basigin |
| chr15 | 40079347 | 40080740 | 1394 | * | 12,50967 | 2,62646163 | 0,956075 | 2,747129 | 0,006012 | 0,851183656 | 1,075735 | Distal Intergenic | 15 | 40026003 | 40031472 | 5470 | 1 | 1,03E+08 | ENSMUST00000228503.2 | 53344 | ENSMUSG00000115367 | Gm33301 | predicted gene, 33301 |
| chr4 | 1,49E+08 | 1,49E+08 | 1283 | * | 10,67801 | 2,626692426 | 1,00102 | 2,624017 | 0,00869 | 0,934396727 | 0,918255 | Promoter (<=1kb) | 4 | 1,49E+08 | 1,49E+08 | 43224 | 2 | 16561 | ENSMUST00000150230.2 | -254 | ENSMUSG00000063077 | Kif1b | kinesin family member 1B |
| chr19 | 29555576 | 29556805 | 1230 | * | 12,32593 | 2,627780244 | 0,986019 | 2,66504 | 0,007698 | 0,891057096 | 1,075735 | Intron (ENSMUST00000 | 19 | 29572928 | 29582585 | 9658 | 1 | 226089 | ENSMUST00000160452.2 | -16123 | ENSMUSG00000038658 | Ric1 | RAB6A GEF complex partner 1 |
| chr1 | 1,32E+08 | 1,32E+08 | 2980 | * | 22,20041 | 2,628747578 | 0,719292 | 3,654633 | 0,000258 | 0,184528934 | 2,951933 | Promoter (2-3kb) | 1 | 1,32E+08 | 1,32E+08 | 17168 | 1 | 74137 | ENSMUST00000072177.14 | -2594 | ENSMUSG00000009772 | Nuak2 | NUAK family, SNF1-like kinase, 2 |
| chr11 | 63961236 | 63962373 | 1138 | * | 14,33892 | 2,628968496 | 0,977056 | 2,690704 | 0,00713 | 0,880128915 | 1,075735 | Exon (ENSMUST00000 | 11 | 63970270 | 63974091 | 3822 | 1 | 66456 | ENSMUST00000135037.9 | -7897 | ENSMUSG00000087497 | 2810001G20Rik | RIKEN cDNA 2810001G20 gene |
| chr12 | 9161879 | 9165030 | 3152 | * | 44,816686 | 2,629257871 | 0,5080816 | 5,1748737 | 2,281E-07 | 0,001389179 | 2,3383871 | Distal Intergenic | 12 | 9080028 | 9086397 | 6370 | 1 | 75516 | ENSMUST00000219470.2 | 81851 | ENSMUSG00000066637 | Ttc32 | tetratricopeptide repeat domain 32 |
| chr8 | 23027278 | 23028774 | 1497 | * | 14,2252 | 2,629629846 | 0,949912 | 2,768288 | 0,005635 | 0,863172166 | 0,918255 | Intron (ENSMUST00000 | 8 | 22996242 | 23025329 | 29088 | 1 | 20516 | ENSMUST00000210854.2 | 31036 | ENSMUSG00000037656 | Slc20a2 | solute carrier family 20, member 2 |
| chr10 | 82566022 | 82567505 | 1484 | * | 17,65483 | 2,630008919 | 0,791862 | 3,321298 | 0,000896 | 0,365691948 | 2,507013 | Promoter (1-2kb) | 10 | 82564183 | 82577229 | 13047 | 1 | 67933 | ENSMUST00000160681.2 | 1839 | ENSMUSG00000020246 | Hcfc2 | host cell factor C2 |
| chr5 | 75205931 | 75208122 | 2192 | * | 20,01882 | 2,630298867 | 0,890636 | 2,95328 | 0,003144 | 0,612704358 | 2,221611 | Promoter (<=1kb) | 5 | 75158649 | 75205435 | 46787 | 2 | 74277 | ENSMUST00000075452.7 | -496 | ENSMUSG00000029229 | Chic2 | cysteine-rich hydrophobic domain 2 |
| chr4 | 1,39E+08 | 1,39E+08 | 1953 | * | 16,72205 | 2,630492775 | 0,897756 | 2,930076 | 0,003389 | 0,619070779 | 2,318434 | Distal Intergenic | 4 | 1,39E+08 | 1,39E+08 | 2303 | 1 | 69116 | ENSMUST00000135534.2 | 14114 | ENSMUSG00000066036 | Ubr4 | ubiquitin protein ligase E3 component n-recognin 4 |
| chr8 | 122870281 | 122872889 | 2609 | * | 93,281707 | 2,63095413 | 0,4033715 | 6,5224088 | 6,919E-11 | 1,54955E-06 | 3,2706213 | Exon (ENSMUST0000024 | 8 | 122866751 | 122874553 | 7803 | 1 | 195209 | ENSMUST00000247413.1 | 3530 | ENSMUSG00000043903 | Zfp469 | zinc finger protein 469 |
| chr17 | 79253717 | 79254695 | 979 | * | 14,42381 | 2,631446285 | 0,99936 | 2,633131 | 0,00846 | 0,888080529 | 1,214904 | Promoter (<=1kb) | 17 | 79253457 | 79254602 | 1146 | 1 | 73694 | ENSMUST00000232821.2 | 260 | ENSMUSG00000024082 | Ndufaf7 | NADH:ubiquinone oxidoreductase complex assembly factor 7 |
| chr6 | 88020511 | 88021657 | 1147 | * | 15,32484 | 2,631787657 | 0,987599 | 2,664834 | 0,007703 | 0,844777257 | 1,41696 | Promoter (<=1kb) | 6 | 87976099 | 88021999 | 45901 | 2 | 19349 | ENSMUST00000113598.8 | 342 | ENSMUSG00000079477 | Rab7 | RAB7, member RAS oncogene family |
| chr18 | 35503715 | 35505894 | 2180 | * | 19,08708 | 2,631792588 | 0,737112 | 3,570412 | 0,000356 | 0,247743734 | 2,386008 | Intron (ENSMUST00000 | 18 | 35399449 | 35476120 | 76672 | 2 | 81500 | ENSMUST00000236387.2 | -27595 | ENSMUSG00000024357 | Sil1 | endoplasmic reticulum chaperone SIL1 homolog (S. cerevisiae) |
| chr15 | 77207430 | 77210051 | 2622 | * | 22,28913 | 2,632854598 | 0,738729 | 3,564031 | 0,000365 | 0,226269722 | 2,880017 | Distal Intergenic | 15 | 76997806 | 77191204 | 193399 | 2 | 93686 | ENSMUST00000228582.2 | -16226 | ENSMUSG00000033565 | Rbfox2 | RNA binding protein, fox-1 homolog (C. elegans) 2 |
| chr5 | 35739713 | 35741516 | 1804 | * | 20,94932 | 2,633259217 | 0,835963 | 3,14997 | 0,001633 | 0,442939362 | 2,951933 | Promoter (<=1kb) | 5 | 35740384 | 35771145 | 30762 | 1 | 80911 | ENSMUST00000068947.14 | 0 | ENSMUSG00000029098 | Acox3 | acyl-Coenzyme A oxidase 3, pristanoyl |
| chr9 | 63039212 | 63042465 | 3254 | * | 28,02988 | 2,633563484 | 0,77887 | 3,381263 | 0,000722 | 0,312753404 | 2,858511 | Promoter (<=1kb) | 9 | 63040565 | 63040671 | 107 | 1 | 1,15E+08 | ENSMUST00000240511.1 | 0 | ENSMUSG00000118764 | Gm25064 | predicted gene, 25064 |
| chr7 | 41201178 | 41202678 | 1501 | * | 11,27103 | 2,633671009 | 1,005713 | 2,61871 | 0,008826 | 0,90060438 | 1,173325 | Intron (ENSMUST00000 | 7 | 41195836 | 41210219 | 14384 | 2 | 1,03E+08 | ENSMUST00000164677.2 | 7541 | ENSMUSG00000090744 | Gm6871 | predicted gene 6871 |
| chr1 | 1,56E+08 | 1,56E+08 | 2790 | * | 24,45083 | 2,633817259 | 0,741556 | 3,551745 | 0,000383 | 0,242864164 | 2,67243 | Exon (ENSMUST00000 | 1 | 1,56E+08 | 1,56E+08 | 37012 | 2 | 20652 | ENSMUST00000051396.13 | 10216 | ENSMUSG00000026600 | Soat1 | sterol O-acyltransferase 1 |
| chr6 | 81627819 | 81628350 | 532 | * | 10,19817 | 2,633962225 | 0,998265 | 2,638539 | 0,008326 | 0,886332554 | 1,211315 | Distal Intergenic | 6 | 81615292 | 81615450 | 159 | 1 | 1,15E+08 | ENSMUST00000083278.3 | 12527 | ENSMUSG00000065212 | Gm26264 | predicted gene, 26264 |
| chr13 | 52734547 | 52736320 | 1774 | * | 17,01493 | 2,635692937 | 0,86822 | 3,035742 | 0,002399 | 0,525510069 | 2,663619 | Promoter (<=1kb) | 13 | 52737209 | 52802828 | 65620 | 1 | 20963 | ENSMUST00000120135.8 | -889 | ENSMUSG00000021457 | Syk | spleen tyrosine kinase |
| chr14 | 20473214 | 20474465 | 1252 | * | 12,6058 | 2,636192688 | 0,95487 | 2,760787 | 0,005766 | 0,826310388 | 1,173325 | Exon (ENSMUST00000 | 14 | 20475477 | 20480056 | 4580 | 2 | 76670 | ENSMUST00000148692.2 | 5591 | ENSMUSG00000039543 | Cfap70 | cilia and flagella associated protein 70 |
| chr19 | 53300772 | 53303479 | 2708 | * | 27,66441 | 2,636265881 | 0,689951 | 3,820947 | 0,000133 | 0,124780314 | 2,858511 | Promoter (<=1kb) | 19 | 53301497 | 53360891 | 59395 | 1 | 17859 | ENSMUST00000237480.2 | 0 | ENSMUSG00000025025 | Mxi1 | MAX interactor 1, dimerization protein |
| chr12 | 84909482 | 84911319 | 1838 | * | 18,15277 | 2,636813162 | 0,824487 | 3,198127 | 0,001383 | 0,467362358 | 2,154504 | Intron (ENSMUST00000 | 12 | 84922543 | 84924906 | 2364 | 1 | 1E+08 | ENSMUST00000171853.2 | -11224 | ENSMUSG00000092130 | D030025P21Rik | RIKEN cDNA D030025P21 gene |
| chr3 | 50208770 | 50210439 | 1670 | * | 19,68357 | 2,637645213 | 0,850416 | 3,101596 | 0,001925 | 0,486847323 | 2,663619 | Exon (ENSMUST00000 | 3 | 50319387 | 50398060 | 78674 | 2 | 26570 | ENSMUST00000029297.6 | 187621 | ENSMUSG00000027737 | Slc7a11 | solute carrier family 7 (cationic amino acid transporter, y+ system), member 11 |
| chr14 | 14342858 | 14347363 | 4506 | * | 40,412884 | 2,639115395 | 0,5765132 | 4,5777189 | 4,701E-06 | 0,012199007 | 2,9254973 | Exon (ENSMUST0000022 | 14 | 14329077 | 14334419 | 5343 | 2 | 218699 | ENSMUST00000224462.2 | -8439 | ENSMUSG00000033885 | Pxk | PX domain containing serine/threonine kinase |
| chr6 | 1,28E+08 | 1,28E+08 | 1715 | * | 23,63576 | 2,639455303 | 0,809492 | 3,260632 | 0,001112 | 0,38136622 | 2,880017 | Intron (ENSMUST00000 | 6 | 1,28E+08 | 1,28E+08 | 36110 | 1 | 68498 | ENSMUST00000149801.2 | 27482 | ENSMUSG00000030351 | Tspan11 | tetraspanin 11 |
| chr13 | 54257410 | 54258563 | 1154 | * | 10,87119 | 2,639539072 | 1,016554 | 2,596555 | 0,009416 | 0,976107896 | 0,744375 | Intron (ENSMUST00000 | 13 | 54225888 | 54262361 | 36474 | 1 | 14057 | ENSMUST00000021930.10 | 31522 | ENSMUSG00000021474 | Sfxn1 | sideroflexin 1 |
| chr13 | 95904239 | 95905967 | 1729 | * | 15,08942 | 2,639913521 | 0,942732 | 2,800281 | 0,005106 | 0,770994815 | 1,41696 | Promoter (2-3kb) | 13 | 95901635 | 95912143 | 10509 | 1 | 1,03E+08 | ENSMUST00000247030.1 | 2604 | | Gm32702 | predicted gene, 32702 |
| chr15 | 71754734 | 71755803 | 1070 | * | 11,68445 | 2,640708494 | 1,023723 | 2,579514 | 0,009894 | 0,984933022 | 0,744375 | Intron (ENSMUST00000 | 15 | 71740730 | 71769465 | 28736 | 2 | 69700 | ENSMUST00000160513.2 | 13662 | ENSMUSG00000079022 | Col22a1 | collagen, type XXII, alpha 1 |
| chr6 | 1,25E+08 | 1,25E+08 | 1166 | * | 22,11276 | 2,641377686 | 0,762655 | 3,463397 | 0,000533 | 0,265946 | 3,104559 | Promoter (<=1kb) | 6 | 1,25E+08 | 1,25E+08 | 1278 | 2 | 68298 | ENSMUST00000188119.2 | -120 | ENSMUSG00000038252 | Ncapd2 | non-SMC condensin I complex, subunit D2 |
| chr5 | 44551736 | 44552894 | 1159 | * | 11,97439 | 2,641419096 | 1,009242 | 2,61723 | 0,008865 | 0,900637308 | 1,173325 | Distal Intergenic | 5 | 44570135 | 44584934 | 14800 | 2 | 1,03E+08 | ENSMUST00000199005.2 | 32040 | ENSMUSG00000104800 | Gm36017 | predicted gene, 36017 |
| chr1 | 17572424 | 17573910 | 1487 | * | 12,49571 | 2,642025144 | 1,002384 | 2,635741 | 0,008395 | 0,891057096 | 1,173325 | Distal Intergenic | 1 | 17672125 | 17701163 | 29039 | 1 | 94227 | ENSMUST00000088476.4 | -98215 | ENSMUSG00000067780 | Pi15 | peptidase inhibitor 15 |
| chr5 | 1,4E+08 | 1,4E+08 | 1658 | * | 13,02749 | 2,642359378 | 0,947702 | 2,788175 | 0,005301 | 0,826676469 | 1,075735 | Distal Intergenic | 5 | 1,4E+08 | 1,4E+08 | 3294 | 2 | 66506 | ENSMUST00000031531.14 | -3543 | ENSMUSG00000029551 | Psmg3 | proteasome (prosome, macropain) assembly chaperone 3 |
| chr12 | 86833344 | 86835116 | 1773 | * | 28,891551 | 2,64258494 | 0,6107654 | 4,3266778 | 1,514E-05 | 0,030904397 | 2,6724296 | Distal Intergenic | 12 | 86787781 | 86801168 | 13388 | 1 | 627607 | ENSMUST00000223197.2 | 45563 | ENSMUSG00000059114 | Lrrc74a | leucine rich repeat containing 74A |
| chr19 | 53128976 | 53131458 | 2483 | * | 45,143175 | 2,643041015 | 0,5969891 | 4,4272855 | 9,543E-06 | 0,024692459 | 2,3383871 | Promoter (<=1kb) | 19 | 53130005 | 53205366 | 75362 | 1 | 27360 | ENSMUST00000235133.2 | 0 | ENSMUSG00000025026 | Add3 | adducin 3 (gamma) |
| chr11 | 32406637 | 32408415 | 1779 | * | 11,52775 | 2,643953054 | 1,026034 | 2,576866 | 0,00997 | 0,955649401 | 0,918255 | Promoter (1-2kb) | 11 | 32405370 | 32466687 | 61318 | 1 | 327900 | ENSMUST00000051053.5 | 1267 | ENSMUSG00000044949 | Ubtd2 | ubiquitin domain containing 2 |
| chr14 | 52153391 | 52154303 | 913 | * | 9,971085 | 2,644004476 | 0,9638 | 2,743311 | 0,006082 | 0,801054956 | 1,425929 | Promoter (1-2kb) | 14 | 52155882 | 52158157 | 2276 | 1 | 219038 | ENSMUST00000008957.13 | -1579 | ENSMUSG00000008813 | Tppp2 | tubulin polymerization-promoting protein family member 2 |
| chr5 | 1,24E+08 | 1,24E+08 | 2046 | * | 19,55286 | 2,644325505 | 0,769813 | 3,435024 | 0,000593 | 0,321298166 | 2,221611 | Intron (ENSMUST00000 | 5 | 1,24E+08 | 1,24E+08 | 41513 | 2 | 19679 | ENSMUST00000161644.3 | 10295 | ENSMUSG00000029406 | Pitpnm2 | phosphatidylinositol transfer protein, membrane-associated 2 |
| chr17 | 78703969 | 78705451 | 1483 | * | 18,09561 | 2,644711711 | 0,838422 | 3,154393 | 0,001608 | 0,456907656 | 2,663619 | Intron (ENSMUST00000 | 17 | 78685315 | 78708859 | 23545 | 2 | 225020 | ENSMUST00000234601.2 | 3408 | ENSMUSG00000056121 | Fez2 | fasciculation and elongation protein zeta 2 (zygin II) |
| chr9 | 61864461 | 61867573 | 3113 | * | 23,24504 | 2,644753983 | 0,758554 | 3,486575 | 0,000489 | 0,26105284 | 2,951933 | Intron (ENSMUST00000 | 9 | 61824562 | 61854056 | 29495 | 2 | 71819 | ENSMUST00000034815.9 | -10405 | ENSMUSG00000032254 | Kif23 | kinesin family member 23 |
| chr5 | 18287244 | 18288608 | 1365 | * | 18,0588 | 2,644972374 | 0,846342 | 3,125183 | 0,001777 | 0,492834344 | 2,386008 | Distal Intergenic | 5 | 18167547 | 18224832 | 57286 | 1 | 242851 | ENSMUST00000030561.9 | 119697 | ENSMUSG00000028777 | Gnat3 | guanine nucleotide binding protein, alpha transducing 3 |
| chr12 | 1,12E+08 | 1,12E+08 | 1435 | * | 12,53479 | 2,645316991 | 0,963322 | 2,746035 | 0,006032 | 0,835026365 | 1,173325 | Intron (ENSMUST00000 | 12 | 1,12E+08 | 1,12E+08 | 41122 | 2 | 21981 | ENSMUST00000220927.2 | 30318 | ENSMUSG00000021285 | Ppp1r13b | protein phosphatase 1, regulatory subunit 13B |
| chr13 | 60431135 | 60432620 | 1486 | * | 14,14465 | 2,645427153 | 0,979232 | 2,701533 | 0,006902 | 0,843167315 | 1,2763 | Distal Intergenic | 13 | 60353285 | 60363837 | 10553 | 1 | 1,01E+08 | ENSMUST00000224064.2 | 77850 | ENSMUSG00000050765 | Gm5084 | predicted gene 5084 |
| chr7 | 98698284 | 98699391 | 1108 | * | 11,34262 | 2,646407152 | 1,008007 | 2,625385 | 0,008655 | 0,934396727 | 0,918255 | Intron (ENSMUST00000 | 7 | 98535957 | 98665851 | 129895 | 2 | 78610 | ENSMUST00000209123.2 | -32433 | ENSMUSG00000035354 | Uvrag | UV radiation resistance associated gene |
| chr10 | 1,18E+08 | 1,18E+08 | 1039 | * | 12,30925 | 2,646667681 | 0,9775 | 2,707588 | 0,006777 | 0,869016875 | 1,075735 | Intron (ENSMUST00000 | 10 | 1,18E+08 | 1,18E+08 | 53318 | 2 | 103214 | ENSMUST00000239416.2 | 47016 | ENSMUSG00000112230 | Ifngas1 | Ifng antisense RNA 1 |
| chr4 | 41347610 | 41350024 | 2415 | * | 22,39153 | 2,64674879 | 0,807441 | 3,277945 | 0,001046 | 0,337054778 | 3,471678 | Promoter (<=1kb) | 4 | 41348996 | 41389766 | 40771 | 1 | 67123 | ENSMUST00000108060.10 | 0 | ENSMUSG00000028437 | Ubap1 | ubiquitin-associated protein 1 |
| chr4 | 58715116 | 58717115 | 2000 | * | 15,83925 | 2,646822694 | 0,988049 | 2,678838 | 0,007388 | 0,749550311 | 2,318434 | Distal Intergenic | 4 | 58784779 | 58785722 | 944 | 2 | 258922 | ENSMUST00000059608.5 | 68607 | ENSMUSG00000043385 | Olfr267 | olfactory receptor 267 |
| chr19 | 14556381 | 14557821 | 1441 | * | 15,86179 | 2,6475007 | 1,009958 | 2,621396 | 0,008757 | 0,775744126 | 2,386008 | Intron (ENSMUST00000 | 19 | 14521090 | 14572903 | 51814 | 2 | 21888 | ENSMUST00000235975.2 | 15082 | ENSMUSG00000024642 | Tle4 | transducin-like enhancer of split 4 |
| chr18 | 35086757 | 35089363 | 2607 | * | 29,15038 | 2,648263333 | 0,664295 | 3,986578 | 6,7E-05 | 0,084947995 | 2,725491 | Promoter (<=1kb) | 18 | 35070467 | 35087410 | 16944 | 2 | 15526 | ENSMUST00000025217.11 | 0 | ENSMUSG00000024359 | Hspa9 | heat shock protein 9 |
| chr13 | 45838124 | 45840820 | 2697 | * | 23,69312 | 2,648435496 | 0,780433 | 3,393546 | 0,00069 | 0,304355254 | 2,951933 | Intron (ENSMUST00000 | 13 | 45692285 | 45696026 | 3742 | 1 | 66355 | ENSMUST00000129176.2 | 145839 | ENSMUSG00000000253 | Gmpr | guanosine monophosphate reductase |
| chr3 | 1,09E+08 | 1,09E+08 | 2023 | * | 21,66638 | 2,648466202 | 0,730518 | 3,625461 | 0,000288 | 0,197827068 | 2,951933 | 3' UTR | 3 | 1,09E+08 | 1,09E+08 | 14789 | 1 | 329738 | ENSMUST00000147916.2 | 11032 | ENSMUSG00000049565 | Aknad1 | AKNA domain containing 1 |
| chr12 | 45118628 | 45120830 | 2203 | * | 17,38193 | 2,648599394 | 0,838893 | 3,157255 | 0,001593 | 0,465630924 | 2,507013 | Promoter (<=1kb) | 12 | 44899271 | 45120895 | 221625 | 2 | 217517 | ENSMUST00000120531.8 | 65 | ENSMUSG00000046314 | Stxbp6 | syntaxin binding protein 6 (amisyn) |
| chr15 | 31223668 | 31224918 | 1251 | * | 17,66425 | 2,649082884 | 0,763791 | 3,468335 | 0,000524 | 0,297583251 | 2,386008 | Promoter (<=1kb) | 15 | 31224460 | 31274487 | 50028 | 1 | 223453 | ENSMUST00000044524.16 | 0 | ENSMUSG00000039168 | Dap | death-associated protein |
| chr8 | 1,06E+08 | 1,06E+08 | 1778 | * | 16,2239 | 2,649649357 | 0,941676 | 2,81376 | 0,004897 | 0,783821651 | 1,2763 | Promoter (<=1kb) | 8 | 1,06E+08 | 1,06E+08 | 1601 | 2 | 11604 | ENSMUST00000005849.11 | -416 | ENSMUSG00000005705 | Agrp | agouti related neuropeptide |
| chr15 | 75584865 | 75586410 | 1546 | * | 18,59028 | 2,649998429 | 0,97413 | 2,720375 | 0,006521 | 0,715249603 | 2,507013 | Promoter (<=1kb) | 15 | 75584437 | 75585323 | 887 | 1 | 14787 | ENSMUST00000229670.2 | 428 | ENSMUSG00000022580 | Rhpn1 | rhophilin, Rho GTPase binding protein 1 |
| chr7 | 63588407 | 63590843 | 2437 | * | 52,30779 | 2,651026928 | 0,991144 | 2,674714 | 0,007479 | 0,723239852 | 2,725491 | Promoter (<=1kb) | 7 | 63536099 | 63588663 | 52565 | 2 | 50794 | ENSMUST00000063694.10 | 0 | ENSMUSG00000052040 | Klf13 | Kruppel-like factor 13 |
| chr4 | 62398547 | 62400294 | 1748 | * | 17,21243 | 2,651251445 | 0,832529 | 3,184574 | 0,00145 | 0,440188707 | 2,663619 | Promoter (<=1kb) | 4 | 62398326 | 62404656 | 6331 | 1 | 192120 | ENSMUST00000132237.2 | 221 | ENSMUSG00000028392 | Bspry | B-box and SPRY domain containing |
| chr15 | 42374367 | 42375488 | 1122 | * | 13,72216 | 2,651663476 | 0,891177 | 2,975462 | 0,002925 | 0,749550311 | 0,918255 | Intron (ENSMUST00000 | 15 | 42496702 | 42540145 | 43444 | 2 | 1,05E+08 | ENSMUST00000227738.2 | 164657 | | Gm41313 | predicted gene, 41313 |
| chr13 | 97062190 | 97063714 | 1525 | * | 15,47917 | 2,652309636 | 0,831924 | 3,188163 | 0,001432 | 0,542967057 | 1,41696 | Promoter (<=1kb) | 13 | 97061197 | 97087414 | 26218 | 1 | 218476 | ENSMUST00000171324.3 | 993 | ENSMUSG00000091387 | Gcnt4 | glucosaminyl (N-acetyl) transferase 4, core 2 (beta-1,6-N-acetylglucosaminyltransferase) |

| | | | | | | | | | | | | | | | | | | | | | | | |
|---|---|---|---|---|---|---|---|---|---|---|---|---|---|---|---|---|---|---|---|---|---|---|---|
| chr11 | 77332044 | 77333895 | 1852 | * | 11,73926 | 2,652329095 | 0,985417 | 2,69158 | 0,007111 | 0,903294867 | 0,918255 | Exon (ENSMUST00000 | 11 | 77353237 | 77361310 | 8074 | 1 | 216961 | ENSMUST00000102493.8 | -19342 | ENSMUSG00000020836 | Coro6 | coronin 6 |
| chr4 | 1,34E+08 | 1,34E+08 | 1316 | * | 11,85828 | 2,65261695 | 1,015112 | 2,613128 | 0,008972 | 0,901762077 | 1,173325 | Promoter (2-3kb) | 4 | 1,34E+08 | 1,34E+08 | 6211 | 2 | 76824 | ENSMUST00000146808.2 | -2027 | ENSMUSG00000046671 | Mtfr1l | mitochondrial fission regulator 1-like |
| chr11 | 56504360 | 56506412 | 2053 | * | 15,85598 | 2,653445647 | 0,80519 | 3,29543 | 0,000983 | 0,497895643 | 1,2763 | Distal Intergenic | 11 | 56153991 | 56154066 | 76 | 2 | 1,15E+08 | ENSMUST00000083701.3 | -350294 | ENSMUSG00000065635 | Gm24253 | predicted gene, 24253 |
| chr13 | 9238500 | 9239840 | 1341 | * | 10,93235 | 2,654209136 | 1,024676 | 2,59029 | 0,00959 | 0,911412811 | 1,173325 | Distal Intergenic | 13 | 9218397 | 9223157 | 4761 | 1 | 217980 | ENSMUST00000189330.2 | 20103 | ENSMUSG00000033499 | Larp4b | La ribonucleoprotein domain family, member 4B |
| chr17 | 67935835 | 67937458 | 1624 | * | 13,31565 | 2,655072262 | 0,977155 | 2,717146 | 0,006585 | 0,891097601 | 0,918255 | Promoter (<=1kb) | 17 | 67866743 | 67936327 | 69585 | 2 | 1,03E+08 | ENSMUST00000233130.2 | 0 | ENSMUSG00000117029 | Gm36201 | predicted gene, 36201 |
| chr17 | 48716562 | 48717370 | 809 | * | 13,94047 | 2,655620343 | 0,878207 | 3,02391 | 0,002495 | 0,722439117 | 0,918255 | Promoter (<=1kb) | 17 | 48717042 | 48724282 | 7241 | 1 | 106821 | ENSMUST00000167180.8 | 0 | ENSMUSG00000040771 | Oard1 | O-acyl-ADP-ribose deacylase 1 |
| chr2 | 44817430 | 44818776 | 1347 | * | 10,88961 | 2,655707597 | 1,005843 | 2,640282 | 0,008284 | 0,924988593 | 0,918255 | Promoter (<=1kb) | 2 | 44525276 | 44817669 | 292394 | 2 | 227835 | ENSMUST00000148279.8 | 0 | ENSMUSG00000036890 | Gtdc1 | glycosyltransferase-like domain containing 1 |
| chr8 | 27701466 | 27703516 | 2051 | * | 20,01428 | 2,65589086 | 0,964777 | 2,752855 | 0,005908 | 0,706976265 | 2,386008 | Intron (ENSMUST0000 | 8 | 27687487 | 27692575 | 5089 | 2 | 76615 | ENSMUST00000038174.8 | -8891 | ENSMUSG00000039720 | Got1l1 | glutamic-oxaloacetic transaminase 1-like 1 |
| chr1 | 41336356 | 41337515 | 1160 | * | 10,53081 | 2,65590351 | 0,983565 | 2,700283 | 0,006928 | 0,933631108 | 0,738441 | Distal Intergenic | 1 | 41206747 | 41220412 | 13666 | 2 | 74897 | ENSMUST00000194002.2 | -115944 | ENSMUSG00000104072 | 4930448I06Rik | RIKEN cDNA 4930448I06 gene |
| chr15 | 96183103 | 96184758 | 1656 | * | 17,54985 | 2,656394871 | 0,851064 | 3,121263 | 0,001801 | 0,476080922 | 2,663619 | Promoter (<=1kb) | 15 | 96185399 | 96249919 | 64521 | 1 | 77044 | ENSMUST00000134985.9 | -641 | ENSMUSG00000033237 | Arid2 | AT rich interactive domain 2 (ARID, RFX-like) |
| chr1 | 1,82E+08 | 1,82E+08 | 1199 | * | 8,942458 | 2,656768036 | 1,025351 | 2,591081 | 0,009567 | 0,878992922 | 1,449706 | Intron (ENSMUST0000 | 1 | 1,82E+08 | 1,82E+08 | 3744 | 1 | 209456 | ENSMUST00000191626.2 | 11678 | ENSMUSG00000026510 | Trp53bp2 | transformation related protein 53 binding protein 2 |
| chr17 | 25147026 | 25148846 | 1821 | * | 15,01902 | 2,656935503 | 0,93739 | 2,834398 | 0,004591 | 0,770673161 | 1,2763 | Intron (ENSMUST0000 | 17 | 25134328 | 25143185 | 8858 | 2 | 30957 | ENSMUST00000126280.2 | -3841 | ENSMUSG00000024163 | Mapk8ip3 | mitogen-activated protein kinase 8 interacting protein 3 |
| chr9 | 20984562 | 20986531 | 1970 | * | 15,94294 | 2,657180063 | 0,94762 | 2,804056 | 0,005046 | 0,683988405 | 2,318434 | Promoter (<=1kb) | 9 | 20978816 | 20984840 | 6025 | 2 | 68165 | ENSMUST00000010348.7 | 0 | ENSMUSG00000079677 | Fdx2 | ferredoxin 2 |
| chr5 | 75227741 | 75229760 | 2020 | * | 19,68378 | 2,658094994 | 0,795304 | 3,342236 | 0,000831 | 0,338031787 | 2,713306 | Distal Intergenic | 5 | 75236262 | 75238554 | 2293 | 1 | 14843 | ENSMUST00000040477.4 | -6502 | ENSMUSG00000035946 | Gsx2 | GS homeobox 2 |
| chr9 | 1,2E+08 | 1,2E+08 | 2047 | * | 17,61356 | 2,658158428 | 0,899947 | 2,953684 | 0,00314 | 0,618282231 | 2,154504 | Distal Intergenic | 9 | 1,2E+08 | 1,2E+08 | 727 | 1 | 76100 | ENSMUST00000064646.5 | 10745 | ENSMUSG00000052658 | 5830454E08Rik | RIKEN cDNA 5830454E08 gene |
| chr5 | 1,41E+08 | 1,41E+08 | 2080 | * | 19,26923 | 2,658404229 | 0,855118 | 3,108814 | 0,001878 | 0,51411469 | 2,221611 | Promoter (<=1kb) | 5 | 1,41E+08 | 1,41E+08 | 12157 | 1 | 231842 | ENSMUST00000176035.2 | 0 | ENSMUSG00000050022 | Amz1 | archaelysin family metallopeptidase 1 |
| chr5 | 1,21E+08 | 1,21E+08 | 1777 | * | 14,73099 | 2,658578874 | 0,938367 | 2,833197 | 0,004608 | 0,683210259 | 2,141612 | Distal Intergenic | 5 | 1,21E+08 | 1,21E+08 | 12114 | 2 | 1,03E+08 | ENSMUST00000185026.4 | -15104 | ENSMUSG00000098649 | Gm27199 | predicted gene 27199 |
| chr5 | 1,15E+08 | 1,15E+08 | 2732 | * | 28,03101 | 2,65868676 | 0,688645 | 3,860753 | 0,000113 | 0,117111673 | 2,725491 | Promoter (<=1kb) | 5 | 1,15E+08 | 1,15E+08 | 18110 | 1 | 1,03E+08 | ENSMUST00000149609.3 | 0 | | Gm13830 | predicted gene 13830 |
| chr5 | 1,21E+08 | 1,21E+08 | 2224 | * | 18,12006 | 2,659043247 | 0,870171 | 3,05577 | 0,002245 | 0,524482861 | 2,507013 | Exon (ENSMUST000001 | 5 | 1,21E+08 | 1,21E+08 | 9796 | 2 | 19247 | ENSMUST00000148871.2 | -22308 | ENSMUSG00000043733 | Ptpn11 | protein tyrosine phosphatase, non-receptor type 11 |
| chr1 | 1,52E+08 | 1,52E+08 | 1647 | * | 15,49905 | 2,65985001 | 0,876243 | 3,035518 | 0,002401 | 0,644704154 | 1,41696 | Promoter (<=1kb) | 1 | 1,52E+08 | 1,52E+08 | 42176 | 1 | 63913 | ENSMUST00000086267.6 | -807 | ENSMUSG00000026483 | Niban1 | niban apoptosis regulator 1 |
| chr11 | 51141483 | 51142658 | 1176 | * | 12,22164 | 2,659884439 | 1,003727 | 2,650008 | 0,008049 | 0,886332554 | 1,173325 | Distal Intergenic | 11 | 51144620 | 51147091 | 2472 | 2 | 193286 | ENSMUST00000231433.2 | 4433 | ENSMUSG00000045942 | BC049762 | cDNA sequence BC049762 |
| chr7 | 45237999 | 45239404 | 1406 | * | 17,55035 | 2,65988516 | 0,815626 | 3,261156 | 0,00111 | 0,414202366 | 2,386008 | 3' UTR | 7 | 45234845 | 45239130 | 4286 | 1 | 12036 | ENSMUST00000209410.2 | 3154 | ENSMUSG00000030826 | Bcat2 | branched chain aminotransferase 2, mitochondrial |
| chr5 | 1,49E+08 | 1,49E+08 | 2928 | * | 23,38507 | 2,66029755 | 0,698456 | 3,808824 | 0,00014 | 0,134807641 | 2,67243 | Intron (ENSMUST0000 | 5 | 1,49E+08 | 1,49E+08 | 822 | 1 | 1,05E+08 | ENSMUST00000202340.2 | 10532 | | BC028471 | cDNA sequence BC028471 |
| chr12 | 63786390 | 63787826 | 1437 | * | 14,04557 | 2,660674724 | 0,911718 | 2,918308 | 0,003519 | 0,724246685 | 1,2763 | Distal Intergenic | 12 | 63656642 | 63656748 | 107 | 1 | 1,15E+08 | ENSMUST00000083024.4 | 129748 | ENSMUSG00000118736 | Gm24921 | predicted gene, 24921 |
| chr11 | 51526316 | 51528983 | 2668 | * | 22,10234 | 2,661742153 | 0,7009 | 3,797606 | 0,000146 | 0,139668493 | 2,67243 | Promoter (<=1kb) | 11 | 51514498 | 51526474 | 11977 | 2 | 66089 | ENSMUST00000136020.8 | 0 | ENSMUSG00000001054 | Rmnd5b | required for meiotic nuclear division 5 homolog B |
| chr6 | 1,19E+08 | 1,19E+08 | 1016 | * | 11,1834 | 2,662382225 | 1,026263 | 2,59425 | 0,00948 | 0,909806909 | 1,173325 | Promoter (<=1kb) | 6 | 1,19E+08 | 1,19E+08 | 34922 | 2 | 68465 | ENSMUST00000169744.8 | -105 | ENSMUSG00000030168 | Adipor2 | adiponectin receptor 2 |
| chr2 | 1,73E+08 | 1,73E+08 | 1296 | * | 15,37583 | 2,662447453 | 0,837633 | 3,178538 | 0,00148 | 0,57149888 | 1,2763 | Distal Intergenic | 2 | 1,73E+08 | 1,73E+08 | 2102 | 2 | 1,03E+08 | ENSMUST00000241106.1 | 16659 | | Gm36462 | predicted gene, 36462 |
| chr10 | 90958470 | 90961051 | 2582 | * | 22,75492 | 2,662573172 | 0,851535 | 3,126791 | 0,001767 | 0,455296541 | 2,951062 | Promoter (<=1kb) | 10 | 90952436 | 90959921 | 7486 | 2 | 18674 | ENSMUST00000076694.13 | 0 | ENSMUSG00000061904 | Slc25a3 | solute carrier family 25 (mitochondrial carrier, phosphate carrier), member 3 |
| chrX | 1,03E+08 | 1,03E+08 | 1889 | * | 13,83501 | 2,662896911 | 1,024036 | 2,600395 | 0,009312 | 0,975728078 | 0,738441 | Promoter (<=1kb) | 20 | 1,03E+08 | 1,03E+08 | 54243 | 2 | 78878 | ENSMUST00000236107.2 | 0 | ENSMUSG00000086370 | Ftx | Ftx transcript, Xist regulator (non-protein coding) |
| chr15 | 1,01E+08 | 1,01E+08 | 1442 | * | 23,47524 | 2,662933924 | 0,720291 | 3,697024 | 0,000218 | 0,170809454 | 2,858511 | Promoter (<=1kb) | 15 | 1,01E+08 | 1,01E+08 | 62222 | 1 | 59033 | ENSMUST00000023776.13 | 0 | ENSMUSG00000023032 | Slc4a8 | solute carrier family 4 (anion exchanger), member 8 |
| chr9 | 1,11E+08 | 1,11E+08 | 1441 | * | 13,85049 | 2,662983401 | 0,919545 | 2,895979 | 0,00378 | 0,793243838 | 0,918255 | Distal Intergenic | 9 | 1,11E+08 | 1,11E+08 | 7149 | 1 | 77781 | ENSMUST00000060711.8 | 11389 | ENSMUSG00000046785 | Epm2aip1 | EPM2A (laforin) interacting protein 1 |
| chr1 | 1,46E+08 | 1,46E+08 | 1916 | * | 15,79956 | 2,66354136 | 0,96503 | 2,760062 | 0,005779 | 0,721416593 | 2,141612 | Intron (ENSMUST0000 | 1 | 1,46E+08 | 1,46E+08 | 19817 | 1 | 215378 | ENSMUST00000125331.2 | 8712 | ENSMUSG00000035131 | Brinp3 | bone morphogenetic protein/retinoic acid inducible neural specific 3 |
| chr11 | 75837477 | 75838729 | 1253 | * | 14,22352 | 2,663618179 | 1,010165 | 2,636815 | 0,008369 | 0,927820586 | 0,918255 | Distal Intergenic | 11 | 75797398 | 75829255 | 31858 | 2 | 380714 | ENSMUST00000152035.2 | -8222 | ENSMUSG00000020847 | Rph3al | rabphilin 3A-like (without C2 domains) |
| chr15 | 36774817 | 36776482 | 1666 | * | 16,19497 | 2,663718188 | 0,919578 | 2,896674 | 0,003771 | 0,651089638 | 2,141612 | Exon (ENSMUST00000 | 15 | 36790956 | 36792658 | 1703 | 2 | 22631 | ENSMUST00000126184.2 | 16176 | ENSMUSG00000022285 | Ywhaz | tyrosine 3-monooxygenase/tryptophan 5-monooxygenase activation protein, zeta polypeptide |
| chr7 | 1,26E+08 | 1,26E+08 | 2219 | * | 19,15142 | 2,663759611 | 0,866529 | 3,074059 | 0,002112 | 0,534918345 | 2,221611 | Promoter (<=1kb) | 7 | 1,26E+08 | 1,26E+08 | 621 | 2 | 12752 | ENSMUST00000139766.2 | 0 | ENSMUSG00000030720 | Cln3 | ceroid lipofuscinosis, neuronal 3, juvenile (Batten, Spielmeyer-Vogt disease) |
| chr7 | 99616301 | 99618114 | 1814 | * | 17,37671 | 2,663911907 | 0,87811 | 3,033688 | 0,002416 | 0,527189687 | 2,663619 | Promoter (1-2kb) | 7 | 99584485 | 99614941 | 30457 | 2 | 108937 | ENSMUST00000174264.2 | -1360 | ENSMUSG00000058761 | Rnf169 | ring finger protein 169 |
| chr18 | 36651245 | 36653234 | 1990 | * | 17,46507 | 2,66395999 | 0,782484 | 3,404491 | 0,000663 | 0,319016519 | 2,507013 | Promoter (2-3kb) | 18 | 36637980 | 36648858 | 10879 | 2 | 15200 | ENSMUST00000025363.7 | -2387 | ENSMUSG00000024486 | Hbegf | heparin-binding EGF-like growth factor |
| chr12 | 1,05E+08 | 1,05E+08 | 2234 | * | 20,56851 | 2,664349035 | 0,864914 | 3,080478 | 0,002067 | 0,498526197 | 2,663619 | Exon (ENSMUST000002 | 12 | 1,05E+08 | 1,05E+08 | 65 | 2 | 1,01E+08 | ENSMUST00000198802.3 | 9304 | ENSMUSG00000106366 | Mir3069 | microRNA 3069 |
| chr6 | 85779259 | 85780938 | 1680 | * | 12,81482 | 2,664492346 | 0,977904 | 2,724697 | 0,006436 | 0,831521609 | 1,2763 | Exon (ENSMUST000001 | 6 | 85785467 | 85786845 | 1379 | 2 | 1,01E+08 | ENSMUST00000087656.5 | 5907 | ENSMUSG00000079495 | Nat8f6 | N-acetyltransferase 8 (GCN5-related) family member 6 |
| chr14 | 26356470 | 26358441 | 1972 | * | 12,59093 | 2,664892277 | 0,964221 | 2,763778 | 0,005714 | 0,810132289 | 1,2763 | Promoter (<=1kb) | 14 | 26359229 | 26378413 | 19185 | 1 | 11843 | ENSMUST00000022429.9 | -788 | ENSMUSG00000021877 | Arf4 | ADP-ribosylation factor 4 |
| chr6 | 86621723 | 86623275 | 1553 | * | 22,59336 | 2,666096816 | 0,84022 | 3,173095 | 0,001508 | 0,421090004 | 3,104559 | Distal Intergenic | 6 | 86605146 | 86606692 | 1547 | 1 | 67855 | ENSMUST00000043400.9 | 16577 | ENSMUSG00000033508 | Asprv1 | aspartic peptidase, retroviral-like 1 |
| chr4 | 15244816 | 15246185 | 1370 | * | 16,59809 | 2,666551854 | 0,983163 | 2,712218 | 0,006683 | 0,74685607 | 2,141612 | Distal Intergenic | 4 | 15265831 | 15286753 | 20923 | 1 | 100201 | ENSMUST00000062684.9 | -19646 | ENSMUSG00000043252 | Tmem64 | transmembrane protein 64 |
| chr3 | 69582101 | 69584034 | 1934 | * | 19,63755 | 2,666664459 | 0,775006 | 3,440829 | 0,00058 | 0,308275655 | 2,386008 | Distal Intergenic | 3 | 69629318 | 69631794 | 2477 | 1 | 97112 | ENSMUST00000149680.8 | -45284 | ENSMUSG00000027787 | Nmd3 | NMD3 ribosome export adaptor |
| chr11 | 1,21E+08 | 1,21E+08 | 1913 | * | 19,01877 | 2,667506857 | 0,852362 | 3,129546 | 0,001751 | 0,479521813 | 2,507013 | Promoter (<=1kb) | 11 | 1,21E+08 | 1,21E+08 | 27222 | 2 | 66840 | ENSMUST00000026173.13 | 0 | ENSMUSG00000025173 | Wdr45b | WD repeat domain 45B |
| chr7 | 27889645 | 27891613 | 1969 | * | 14,71445 | 2,668284475 | 0,973901 | 2,739791 | 0,006148 | 0,809505441 | 1,37683 | Distal Intergenic | 7 | 27880934 | 27882192 | 1259 | 1 | 13549 | ENSMUST00000249248.1 | 8711 | ENSMUSG00000002409 | Dyrk1b | dual-specificity tyrosine-(Y)-phosphorylation regulated kinase 1b |
| chr13 | 54430801 | 54432674 | 1874 | * | 16,86285 | 2,668340746 | 0,893376 | 2,986807 | 0,002819 | 0,551991925 | 2,663619 | Distal Intergenic | 13 | 54455978 | 54466816 | 10839 | 1 | 12890 | ENSMUST00000221369.2 | -23304 | ENSMUSG00000025867 | Cplx2 | complexin 2 |
| chr4 | 1,39E+08 | 1,39E+08 | 1568 | * | 13,92841 | 2,668544429 | 0,874582 | 3,051222 | 0,002279 | 0,654404145 | 1,2763 | Exon (ENSMUST000001 | 4 | 1,39E+08 | 1,39E+08 | 37504 | 1 | 69116 | ENSMUST00000129949.8 | -10603 | ENSMUSG00000066036 | Ubr4 | ubiquitin protein ligase E3 component n-recognin 4 |
| chr13 | 18642078 | 18643741 | 1664 | * | 14,30192 | 2,668552018 | 0,923054 | 2,891002 | 0,00384 | 0,739058507 | 1,2763 | Distal Intergenic | 13 | 18520810 | 18572271 | 51462 | 2 | 218030 | ENSMUST00000223058.2 | -69807 | ENSMUSG00000009734 | Pou6f2 | POU domain, class 6, transcription factor 2 |
| chr1 | 1,82E+08 | 1,82E+08 | 1819 | * | 14,52007 | 2,668662369 | 0,922232 | 2,893699 | 0,003807 | 0,653331264 | 2,141612 | Distal Intergenic | 1 | 1,82E+08 | 1,82E+08 | 24386 | 2 | 67948 | ENSMUST00000194213.2 | -29379 | ENSMUSG00000047539 | Fbxo28 | F-box protein 28 |
| chr2 | 1,2E+08 | 1,2E+08 | 2386 | * | 19,55173 | 2,670025255 | 0,834169 | 3,200821 | 0,00137 | 0,426489623 | 2,713306 | Exon (ENSMUST00000 | 2 | 1,2E+08 | 1,2E+08 | 969 | 1 | 76246 | ENSMUST00000126984.2 | -5745 | ENSMUSG00000027304 | Rtf1 | RTF1, Paf1/RNA polymerase II complex component |
| chr10 | 1,28E+08 | 1,28E+08 | 2469 | * | 20,56627 | 2,670289428 | 0,882672 | 3,025235 | 0,002484 | 0,54122637 | 2,507013 | Promoter (<=1kb) | 10 | 1,28E+08 | 1,28E+08 | 18319 | 1 | 56351 | ENSMUST00000052798.14 | 0 | ENSMUSG00000071072 | Ptges3 | prostaglandin E synthase 3 |
| chr1 | 92754441 | 92755913 | 1473 | * | 13,65319 | 2,670597663 | 0,949149 | 2,813675 | 0,004898 | 0,813409596 | 1,075735 | Intron (ENSMUST0000 | 1 | 92759367 | 92788501 | 29135 | 1 | 14733 | ENSMUST00000045970.8 | -3454 | ENSMUSG00000034220 | Gpc1 | glypican 1 |
| chr10 | 3921195 | 3924828 | 3634 | * | 41,061648 | 2,671124446 | 0,6025617 | 4,4329478 | 9,295E-06 | 0,024016264 | 2,3776742 | Promoter (<=1kb) | 10 | 3923118 | 4106111 | 182994 | 1 | 270685 | ENSMUST00000117291.8 | 0 | ENSMUSG00000040675 | Mthfd1l | methylenetetrahydrofolate dehydrogenase (NADP+ dependent) 1-like |
| chr5 | 1,39E+08 | 1,39E+08 | 2072 | * | 20,50046 | 2,67148339 | 0,92906 | 2,87547 | 0,004034 | 0,622809134 | 2,713306 | Intron (ENSMUST0000 | 5 | 1,39E+08 | 1,39E+08 | 10342 | 1 | 319772 | ENSMUST00000163940.2 | -4193 | ENSMUSG00000044092 | C130050O18Rik | RIKEN cDNA C130050O18 gene |
| chr9 | 56348748 | 56351169 | 2422 | * | 18,07097 | 2,671951869 | 0,883558 | 3,024082 | 0,002494 | 0,54122637 | 2,507013 | Intron (ENSMUST0000 | 9 | 56326114 | 56383401 | 57288 | 1 | 66867 | ENSMUST00000214869.2 | 22634 | ENSMUSG00000032329 | Hmg20a | high mobility group 20A |
| chr3 | 90172936 | 90173734 | 799 | * | 11,56825 | 2,67200882 | 0,937038 | 2,851547 | 0,004351 | 0,857011814 | 0,738441 | Promoter (<=1kb) | 3 | 90173821 | 90187976 | 14156 | 1 | 229541 | ENSMUST00000098914.10 | -87 | ENSMUSG00000042404 | Dennd4b | DENN/MADD domain containing 4B |
| chr11 | 1,19E+08 | 1,19E+08 | 1557 | * | 15,22201 | 2,67223443 | 0,893259 | 2,991555 | 0,002776 | 0,688853937 | 1,214904 | Promoter (<=1kb) | 11 | 1,19E+08 | 1,19E+08 | 5098 | 2 | 1,05E+08 | ENSMUST00000156576.2 | 0 | | Gm11754 | predicted gene 11754 |
| chr13 | 37366109 | 37367868 | 1760 | * | 11,57463 | 2,672967588 | 1,037538 | 2,576259 | 0,009988 | 0,955649401 | 0,918255 | Distal Intergenic | 13 | 37421625 | 37428883 | 7259 | 2 | 1,03E+08 | ENSMUST00000225067.2 | 61015 | | Gm30600 | predicted gene, 30600 |
| chr1 | 1,48E+08 | 1,48E+08 | 1180 | * | 11,76364 | 2,673662206 | 0,93521 | 2,858891 | 0,004251 | 0,853738252 | 0,738441 | Distal Intergenic | 1 | 1,48E+08 | 1,48E+08 | 129 | 2 | 1,15E+08 | ENSMUST00000157957.3 | 509924 | ENSMUSG00000088582 | Gm22966 | predicted gene, 22966 |
| chr6 | 29733433 | 29735228 | 1796 | * | 17,42058 | 2,673769839 | 1,011576 | 2,643173 | 0,008213 | 0,744054256 | 2,663619 | Promoter (<=1kb) | 6 | 29735693 | 29761364 | 25672 | 1 | 319757 | ENSMUST00000001812.5 | -465 | ENSMUSG00000001761 | Smo | smoothened, frizzled class receptor |
| chr7 | 75076153 | 75078148 | 1996 | * | 22,29162 | 2,673910786 | 0,682019 | 3,92058 | 8,83E-05 | 0,096470298 | 2,880017 | Distal Intergenic | 7 | 75105282 | 75404357 | 299076 | 1 | 75547 | ENSMUST00000166315.7 | -27134 | ENSMUSG00000066406 | Akap13 | A kinase (PRKA) anchor protein 13 |
| chr8 | 95741013 | 95742472 | 1460 | * | 17,69719 | 2,674305782 | 0,960379 | 2,784637 | 0,005359 | 0,684274806 | 2,423594 | Promoter (1-2kb) | 8 | 95744320 | 95771878 | 27559 | 1 | 54672 | ENSMUST00000051259.10 | -1848 | ENSMUSG00000060470 | Adgrg3 | adhesion G protein-coupled receptor G3 |
| chr11 | 60426888 | 60428807 | 1920 | * | 27,41302 | 2,674826236 | 0,647006 | 4,134159 | 3,56E-05 | 0,057325307 | 2,725491 | Promoter (<=1kb) | 11 | 60427207 | 60445193 | 17987 | 1 | 268420 | ENSMUST00000134770.2 | 0 | ENSMUSG00000042650 | Alkbh5 | alkB homolog 5, RNA demethylase |
| chr2 | 1,58E+08 | 1,58E+08 | 2353 | * | 18,81954 | 2,675069314 | 0,799231 | 3,347055 | 0,000817 | 0,338031787 | 2,663619 | 3' UTR | 2 | 1,58E+08 | 1,58E+08 | 3742 | 1 | 73399 | ENSMUST00000109506.3 | -4423 | ENSMUSG00000078957 | 1700060C20Rik | RIKEN cDNA 1700060C20 gene |
| chr19 | 47658296 | 47659283 | 988 | * | 16,58879 | 2,675686019 | 0,87377 | 3,062233 | 0,002197 | 0,648067121 | 1,2763 | Promoter (<=1kb) | 19 | 47657722 | 47659487 | 1766 | 2 | 12821 | ENSMUST00000145254.2 | 204 | ENSMUSG00000025064 | Col17a1 | collagen, type XVII, alpha 1 |
| chr5 | 1,35E+08 | 1,35E+08 | 1418 | * | 12,41801 | 2,675994391 | 1,011792 | 2,644808 | 0,008174 | 0,923399738 | 0,918255 | Distal Intergenic | 5 | 1,35E+08 | 1,35E+08 | 2293 | 2 | 14371 | ENSMUST00000062572.3 | -8388 | ENSMUSG00000049551 | Fzd9 | frizzled class receptor 9 |
| chr10 | 59782936 | 59783951 | 1016 | * | 17,31854 | 2,676024948 | 0,74409 | 3,596371 | 0,000323 | 0,234308661 | 2,386008 | Distal Intergenic | 10 | 59785491 | 59787656 | 2166 | 2 | 74747 | ENSMUST00000020308.5 | 3705 | ENSMUSG00000020108 | Ddit4 | DNA-damage-inducible transcript 4 |
| chr13 | 55853716 | 55855710 | 1995 | * | 14,53983 | 2,676277927 | 0,921365 | 2,904688 | 0,003676 | 0,647512418 | 2,141612 | Distal Intergenic | 13 | 55862463 | 55874040 | 11578 | 1 | 69672 | ENSMUST00000021959.11 | -6753 | ENSMUSG00000021497 | Txndc15 | thioredoxin domain containing 15 |
| chr12 | 84941008 | 84942506 | 1499 | * | 14,0769 | 2,676503854 | 0,978036 | 2,736609 | 0,006208 | 0,879511593 | 0,938682 | Distal Intergenic | 12 | 84829986 | 84923306 | 93321 | 2 | 16997 | ENSMUST00000002073.13 | -17702 | ENSMUSG00000002020 | Ltbp2 | latent transforming growth factor beta binding protein 2 |
| chr18 | 87757532 | 87759199 | 1668 | * | 13,79893 | 2,677186219 | 0,894356 | 2,993423 | 0,002759 | 0,738600147 | 0,918255 | Distal Intergenic | 18 | 87943555 | 87943688 | 134 | 2 | 1,15E+08 | ENSMUST00000104453.3 | 184489 | ENSMUSG00000077641 | Gm24987 | predicted gene, 24987 |
| chr1 | 173739070 | 173742829 | 3760 | * | 50,976311 | 2,677435503 | 0,5748078 | 4,6579664 | 3,193E-06 | 0,010596832 | 2,3776742 | Promoter (<=1kb) | 1 | 173723911 | 173740612 | 16702 | 2 | 381308 | ENSMUST00000009340.10 | 0 | ENSMUSG00000026536 | Ifi211 | interferon activated gene 211 |
| chr9 | 1,09E+08 | 1,09E+08 | 1742 | * | 16,21011 | 2,678228409 | 1,010571 | 2,650212 | 0,008044 | 0,739716578 | 2,663619 | Promoter (<=1kb) | 9 | 1,09E+08 | 1,09E+08 | 1155 | 1 | 171530 | ENSMUST00000052724.5 | -674 | ENSMUSG00000049699 | Ucn2 | urocortin 2 |
| chr15 | 12367168 | 12368930 | 1763 | * | 12,30053 | 2,678261298 | 1,002637 | 2,671219 | 0,007558 | 0,912452206 | 0,918255 | Exon (ENSMUST00000 | 15 | 12405018 | 12412466 | 7449 | 2 | 68070 | ENSMUST00000191339.2 | 43536 | ENSMUSG00000022197 | Pdzd2 | PDZ domain containing 2 |
| chr8 | 34205681 | 34207377 | 1697 | * | 17,95947 | 2,678735615 | 0,795498 | 3,367371 | 0,000759 | 0,344492479 | 2,386008 | Distal Intergenic | 8 | 34220562 | 34221842 | 1281 | 2 | 1,15E+08 | ENSMUST00000079376.12 | 14465 | | 1700104B16Rik | RIKEN cDNA 1700104B16 gene |
| chr1 | 1,34E+08 | 1,34E+08 | 2357 | * | 23,5314 | 2,680136519 | 0,834942 | 3,209966 | 0,001328 | 0,414107862 | 2,858511 | Promoter (<=1kb) | 1 | 1,34E+08 | 1,34E+08 | 36207 | 2 | 11539 | ENSMUST00000086465.6 | -94 | ENSMUSG00000042429 | Adora1 | adenosine A1 receptor |
| chr2 | 1,3E+08 | 1,3E+08 | 1161 | * | 11,44448 | 2,680474022 | 1,033906 | 2,592571 | 0,009526 | 0,899471409 | 1,2763 | Exon (ENSMUST000001 | 2 | 1,3E+08 | 1,3E+08 | 74543 | 1 | 228598 | ENSMUST00000140169.8 | 10633 | ENSMUSG00000053552 | Ebf4 | early B cell factor 4 |
| chr1 | 6654142 | 6655412 | 1271 | * | 14,88759 | 2,680552173 | 0,915937 | 2,926569 | 0,003427 | 0,635193806 | 2,141612 | Intron (ENSMUST0000 | 1 | 6557555 | 6931164 | 373610 | 1 | 240690 | ENSMUST00000131494.8 | 96587 | ENSMUSG00000033740 | St18 | suppression of tumorigenicity 18 |
| chr8 | 69329224 | 69329971 | 748 | * | 9,504462 | 2,6807131 | 1,008046 | 2,659317 | 0,00783 | 0,853738252 | 1,360622 | Distal Intergenic | 8 | 69333143 | 69359586 | 26444 | 1 | 16956 | ENSMUST00000015712.15 | -3172 | ENSMUSG00000015568 | Lpl | lipoprotein lipase |
| chr12 | 1,19E+08 | 1,19E+08 | 1456 | * | 18,05716 | 2,681000806 | 0,898634 | 2,983416 | 0,00285 | 0,578908869 | 2,386008 | Distal Intergenic | 12 | 1,19E+08 | 1,19E+08 | 37308 | 2 | 320910 | ENSMUST00000137804.2 | 102171 | ENSMUSG00000025321 | Itgb8 | integrin beta 8 |
| chr18 | 36745831 | 36746935 | 1105 | * | 11,39527 | 2,6811254 | 0,925229 | 2,897797 | 0,003758 | 0,749550311 | 1,173325 | Promoter (<=1kb) | 18 | 36746962 | 36767410 | 20449 | 1 | 108857 | ENSMUST00000136030.2 | -27 | ENSMUSG00000024483 | Ankhd1 | ankyrin repeat and KH domain containing 1 |
| chr15 | 31685030 | 31685855 | 826 | * | 10,69294 | 2,681781615 | 1,017631 | 2,635318 | 0,008406 | 0,961052412 | 0,744375 | Distal Intergenic | 15 | 31602264 | 31618755 | 16492 | 1 | 68073 | ENSMUST00000161061.3 | 82766 | ENSMUSG00000039065 | Atpsckmt | ATP synthase C subunit lysine N-methyltransferase |
| chr14 | 55130291 | 55131656 | 1366 | * | 14,0649 | 2,682448052 | 0,940653 | 2,851687 | 0,004349 | 0,857011814 | 0,738441 | Promoter (<=1kb) | 14 | 55131605 | 55135613 | 4009 | 1 | 54196 | ENSMUST00000116476.9 | 0 | ENSMUSG00000022194 | Pabpn1 | poly(A) binding protein, nuclear 1 |
| chr7 | 67675109 | 67677483 | 2375 | * | 14,42638 | 2,682600881 | 1,013251 | 2,647519 | 0,008108 | 0,852537034 | 1,41696 | Intron (ENSMUST0000 | 7 | 67602575 | 67883416 | 280842 | 1 | 16001 | ENSMUST00000005671.10 | 72534 | ENSMUSG00000005533 | Igf1r | insulin-like growth factor I receptor |
| chr9 | 41555756 | 41557757 | 2002 | * | 18,97556 | 2,683047505 | 0,903978 | 2,968046 | 0,002997 | 0,578908869 | 2,507013 | Intron (ENSMUST0000 | 9 | 41570460 | 41586350 | 15891 | 2 | 1,03E+08 | ENSMUST00000214294.2 | 28593 | | Gm35835 | predicted gene, 35835 |
| chr5 | 1,49E+08 | 1,49E+08 | 1687 | * | 14,13732 | 2,68311144 | 0,862523 | 3,110772 | 0,001866 | 0,649217039 | 1,075735 | Intron (ENSMUST0000 | 5 | 1,49E+08 | 1,49E+08 | 3199 | 2 | 15289 | ENSMUST00000125605.2 | -10869 | ENSMUSG00000066551 | Hmgb1 | high mobility group box 1 |
| chr11 | 5304368 | 5306418 | 2051 | * | 16,52614 | 2,683216654 | 0,916227 | 2,928551 | 0,003405 | 0,614881386 | 2,386008 | Intron (ENSMUST0000 | 11 | 5228602 | 5331740 | 103139 | 2 | 407821 | ENSMUST00000172492.8 | 25322 | ENSMUSG00000041961 | Znrf3 | zinc and ring finger 3 |
| chr5 | 1,19E+08 | 1,19E+08 | 1792 | * | 12,95999 | 2,68324645 | 1,003067 | 2,675042 | 0,007472 | 0,946228625 | 0,738441 | Distal Intergenic | 5 | 1,19E+08 | 1,19E+08 | 6310 | 1 | 73477 | ENSMUST00000190550.2 | 16789 | ENSMUSG00000100573 | 1700081H04Rik | RIKEN cDNA 1700081H04 gene |
| chr7 | 1E+08 | 1E+08 | 2079 | * | 16,83401 | 2,683695132 | 0,943956 | 2,843029 | 0,004469 | 0,766298356 | 1,2763 | Intron (ENSMUST0000 | 7 | 1E+08 | 1E+08 | 8988 | 2 | 27276 | ENSMUST00000135255.2 | 4300 | ENSMUSG00000030701 | Plekhb1 | pleckstrin homology domain containing, family B (evectins) member 1 |
| chr11 | 1,18E+08 | 1,18E+08 | 1073 | * | 11,86475 | 2,683965447 | 1,00575 | 2,66862 | 0,007616 | 0,948006995 | 0,738441 | Exon (ENSMUST00000 | 11 | 1,18E+08 | 1,18E+08 | 11669 | 1 | 74451 | ENSMUST00000155507.2 | 20234 | ENSMUSG00000017715 | Pgs1 | phosphatidylglycerophosphate synthase 1 |
| chr5 | 66874495 | 66875591 | 1097 | * | 12,59303 | 2,684628723 | 0,940843 | 2,853429 | 0,004325 | 0,856202617 | 0,738441 | Distal Intergenic | 5 | 66903170 | 67132005 | 228836 | 1 | 77569 | ENSMUST00000201852.4 | -27579 | ENSMUSG00000037736 | Limch1 | LIM and calponin homology domains 1 |
| chr19 | 8746516 | 8748341 | 1826 | * | 14,0519 | 2,685619898 | 1,026272 | 2,616868 | 0,008874 | 0,888019398 | 1,2763 | Promoter (<=1kb) | 19 | 8747456 | 8748285 | 830 | 1 | 53319 | ENSMUST00000183898.2 | 0 | ENSMUSG00000010097 | Nxf1 | nuclear RNA export factor 1 |
| chr12 | 79817092 | 79818832 | 1741 | * | 14,0863 | 2,686970472 | 0,963659 | 2,788299 | 0,005299 | 0,853837252 | 0,918255 | Intron (ENSMUST0000 | 12 | 79938062 | 79938677 | 616 | 1 | 1,08E+08 | ENSMUST00000243367.1 | -119230 | | Gm46362 | predicted gene, 46362 |
| chr15 | 54599446 | 54601043 | 1598 | * | 16,60547 | 2,687250903 | 0,91397 | 2,940195 | 0,00328 | 0,713663698 | 1,2763 | Distal Intergenic | 15 | 54609098 | 54617435 | 8338 | 1 | 18133 | ENSMUST00000050027.9 | -8055 | ENSMUSG00000037362 | Ccn3 | cellular communication network factor 3 |
| chr11 | 1,15E+08 | 1,15E+08 | 1626 | * | 14,24455 | 2,687402409 | 0,991279 | 2,711044 | 0,006707 | 0,894566572 | 0,918255 | Intron (ENSMUST0000 | 11 | 1,15E+08 | 1,15E+08 | 17751 | 1 | 432611 | ENSMUST00000151053.8 | -12604 | ENSMUSG00000034706 | Dnai2 | dynein axonemal intermediate chain 2 |
| chr17 | 47358897 | 47360548 | 1652 | * | 15,10423 | 2,687631229 | 0,914352 | 2,939384 | 0,003289 | 0,714056355 | 1,2763 | Distal Intergenic | 17 | 47267551 | 47321482 | 53932 | 2 | 224826 | ENSMUST00000225599.2 | -37415 | ENSMUSG00000023977 | Ubr2 | ubiquitin protein ligase E3 component n-recognin 2 |
| chr12 | 6300665 | 6301498 | 834 | * | 12,28081 | 2,688108599 | 0,964098 | 2,788212 | 0,0053 | 0,853870296 | 0,918255 | Distal Intergenic | 12 | 6548124 | 6548615 | 492 | 2 | 1E+08 | ENSMUST00000218888.2 | 247117 | ENSMUSG00000111967 | Gm3678 | predicted gene 3678 |
| chr18 | 47935642 | 47938029 | 2388 | * | 20,39712 | 2,688971274 | 0,807001 | 3,332053 | 0,000862 | 0,348632365 | 2,663619 | Distal Intergenic | 18 | 47894156 | 47922600 | 28445 | 2 | 1E+08 | ENSMUST00000236862.2 | -13042 | ENSMUSG00000118113 | Gm4146 | predicted gene 4146 |
| chr9 | 36879514 | 36880844 | 1331 | * | 11,49806 | 2,689015935 | 0,989974 | 2,716249 | 0,006603 | 0,836487307 | 1,2763 | 5' UTR | 9 | 36803774 | 36898568 | 94795 | 2 | 208076 | ENSMUST00000175938.9 | 17724 | ENSMUSG00000035934 | Pknox2 | Pbx/knotted 1 homeobox 2 |
| chr1 | 65231302 | 65232751 | 1450 | * | 10,79168 | 2,689050527 | 0,992377 | 2,709707 | 0,006734 | 0,927945139 | 0,738441 | 3' UTR | 1 | 65235919 | 65255421 | 19503 | 1 | 18711 | ENSMUST00000190847.2 | -3168 | ENSMUSG00000025949 | Pikfyve | phosphoinositide kinase, FYVE type zinc finger containing |
| chr4 | 1,17E+08 | 1,17E+08 | 1789 | * | 15,40888 | 2,689498219 | 0,937786 | 2,867923 | 0,004132 | 0,75003142 | 1,2763 | Distal Intergenic | 4 | 1,17E+08 | 1,17E+08 | 5309 | 2 | 12795 | ENSMUST00000076859.12 | -8248 | ENSMUSG00000028680 | Plk3 | polo like kinase 3 |
| chr1 | 1,31E+08 | 1,31E+08 | 2346 | * | 15,28812 | 2,689895257 | 0,910588 | 2,95402 | 0,003137 | 0,71419937 | 1,214904 | 3' UTR | 1 | 1,31E+08 | 1,31E+08 | 38976 | 2 | 18640 | ENSMUST00000189534.7 | 4062 | ENSMUSG00000026409 | Pfkfb2 | 6-phosphofructo-2-kinase/fructose-2,6-biphosphatase 2 |

| | | | | | | | | | | | | | | | | | | | | | | | |
|---|---|---|---|---|---|---|---|---|---|---|---|---|---|---|---|---|---|---|---|---|---|---|---|
| chr18 | 12776954 | 12780056 | 3103 | * | 25,42601 | 2,690168766 | 0,781435 | 3,442603 | 0,000576 | 0,293307161 | 2,725491 | Promoter (<=1kb) | 18 | 12776419 | 12870113 | 93695 | 1 | 72747 | ENSMUST00000025294.9 | 535 | ENSMUSG00000024424 | Ttc39c | tetratricopeptide repeat domain 39C |
| chr9 | 88365821 | 88367157 | 1337 | * | 12,75125 | 2,690352819 | 0,895592 | 3,003995 | 0,002665 | 0,687319907 | 1,173325 | Promoter (1-2kb) | 9 | 88361715 | 88364627 | 2913 | 2 | 56403 | ENSMUST00000172508.2 | -1194 | ENSMUSG00000032423 | Syncrip | synaptotagmin binding, cytoplasmic RNA interacting protein |
| chr19 | 37996552 | 37998171 | 1620 | * | 16,59682 | 2,690390616 | 0,998207 | 2,695224 | 0,007034 | 0,752939853 | 2,141612 | Intron (ENSMUST0000( | 19 | 37887758 | 38031799 | 144042 | 2 | 226101 | ENSMUST00000226068.2 | 33628 | ENSMUSG00000048612 | Myof | myoferlin |
| chr5 | 31854850 | 31856435 | 1586 | * | 10,11318 | 2,690601113 | 1,008174 | 2,668787 | 0,007613 | 0,88871498 | 1,086118 | Promoter (<=1kb) | 5 | 31855028 | 32242306 | 387279 | 1 | 107976 | ENSMUST00000201352.4 | 0 | ENSMUSG00000052139 | Babam2 | BRISC and BRCA1 A complex member 2 |
| chr16 | 32839154 | 32840477 | 1324 | * | 11,66846 | 2,691239413 | 1,007298 | 2,671742 | 0,007546 | 0,947196831 | 0,738441 | Exon (ENSMUST00000: | 16 | 32837320 | 32844307 | 6988 | 2 | 69707 | ENSMUST00000133219.2 | 3830 | ENSMUSG00000035578 | Iqcg | IQ motif containing G |
| chr18 | 67744047 | 67745949 | 1903 | * | 16,96356 | 2,691299625 | 0,966175 | 2,785521 | 0,005344 | 0,797091634 | 1,2763 | Promoter (<=1kb) | 18 | 67621279 | 67743860 | 122582 | 2 | 68166 | ENSMUST00000115050.10 | -187 | ENSMUSG00000024533 | Spire1 | spire type actin nucleation factor 1 |
| chr7 | 66461456 | 66463266 | 1811 | * | 21,18575 | 2,692213473 | 0,831742 | 3,236836 | 0,001209 | 0,394841966 | 2,951933 | Intron (ENSMUST0000( | 7 | 66489483 | 66619303 | 129821 | 1 | 233332 | ENSMUST00000156396.8 | -26217 | ENSMUSG00000058145 | Adamts17 | a disintegrin-like and metallopeptidase (reprolysin type) with thrombospondin type 1 motif, 17 |
| chr4 | 68613037 | 68614369 | 1333 | * | 10,40926 | 2,693081036 | 1,039649 | 2,590374 | 0,009587 | 0,949251015 | 0,918255 | Distal Intergenic | 4 | 68744818 | 68747595 | 2778 | 2 | 56710 | ENSMUST00000140572.2 | 133226 | ENSMUSG00000028351 | Brinp1 | bone morphogenic protein/retinoic acid inducible neural specific 1 |
| chr19 | 44302598 | 44304535 | 1938 | * | 23,57191 | 2,693176785 | 0,850864 | 3,165225 | 0,00155 | 0,439157569 | 2,880017 | Exon (ENSMUST000002 | 19 | 44291610 | 44291670 | 61 | 1 | 1,01E+08 | ENSMUST00000175574.3 | 10988 | ENSMUSG00000093315 | Mir5114 | microRNA 5114 |
| chr11 | 1,01E+08 | 1,01E+08 | 1686 | * | 18,64928 | 2,693792739 | 0,867732 | 3,104408 | 0,001907 | 0,484531896 | 2,663619 | Exon (ENSMUST00000: | 11 | 1,01E+08 | 1,01E+08 | 10343 | 1 | 14377 | ENSMUST00000019469.3 | -14814 | ENSMUSG00000078650 | G6pc | glucose-6-phosphatase, catalytic |
| chr9 | 1,03E+08 | 1,03E+08 | 1888 | * | 16,13947 | 2,69403154 | 0,858089 | 3,139571 | 0,001692 | 0,465630924 | 2,663619 | Distal Intergenic | 9 | 1,03E+08 | 1,03E+08 | 32064 | 1 | 24059 | ENSMUST00000185215.2 | -16767 | ENSMUSG00000032548 | Slco2a1 | solute carrier organic anion transporter family, member 2a1 |
| chr4 | 82155019 | 82156338 | 1320 | * | 14,34845 | 2,694763325 | 0,888882 | 3,031631 | 0,002432 | 0,665162479 | 1,2763 | Distal Intergenic | 4 | 81952907 | 81984130 | 31224 | 2 | 1,03E+08 | ENSMUST00000243987.1 | -170889 | ENSMUSG00000087110 | Gm11264 | predicted gene 11264 |
| chr16 | 39951979 | 39953188 | 1210 | * | 12,26614 | 2,695376645 | 1,022109 | 2,637074 | 0,008362 | 0,877892722 | 1,2763 | Intron (ENSMUST0000( | 16 | 39804957 | 40478248 | 673292 | 1 | 268890 | ENSMUST00000189829.2 | 147022 | ENSMUSG00000061080 | Lsamp | limbic system-associated membrane protein |
| chr7 | 26930838 | 26932971 | 2134 | * | 31,74543 | 2,695441701 | 0,728177 | 3,701628 | 0,000214 | 0,174628563 | 2,67243 | Promoter (<=1kb) | 7 | 26932448 | 26957375 | 24928 | 1 | 76889 | ENSMUST00000003860.13 | 0 | ENSMUSG00000003762 | Coq8b | coenzyme Q8B |
| chr7 | 79930642 | 79932293 | 1652 | * | 15,6549 | 2,696000114 | 0,830886 | 3,24473 | 0,001176 | 0,509880389 | 1,41696 | 3' UTR | 7 | 79924042 | 79933823 | 9782 | 1 | 233405 | ENSMUST00000145594.3 | 6600 | ENSMUSG00000030534 | Vps33b | vacuolar protein sorting 33B |
| chr10 | 79938246 | 79941434 | 3189 | * | 43,92791 | 2,696366091 | 0,934907 | 2,884101 | 0,003925 | 0,616178941 | 2,725491 | Promoter (<=1kb) | 10 | 79892826 | 79940204 | 47379 | 2 | 216161 | ENSMUST00000219260.2 | 0 | ENSMUSG00000035673 | Sbno2 | strawberry notch 2 |
| chr11 | 1,03E+08 | 1,03E+08 | 2095 | * | 19,98465 | 2,696415882 | 0,861482 | 3,129973 | 0,001748 | 0,501659463 | 2,221611 | Distal Intergenic | 11 | 1,03E+08 | 1,03E+08 | 167 | 2 | 544817 | ENSMUST00000163250.2 | 20621 | ENSMUSG00000034255 | Arhgap27 | Rho GTPase activating protein 27 |
| chr12 | 32045272 | 32047466 | 2195 | * | 14,9996 | 2,696491877 | 0,818038 | 3,296293 | 0,00098 | 0,410415129 | 2,141612 | Intron (ENSMUST0000( | 12 | 31976253 | 32000534 | 24282 | 2 | 73389 | ENSMUST00000172314.9 | -44738 | ENSMUSG00000002996 | Hbp1 | high mobility group box transcription factor 1 |
| chr15 | 29778353 | 29779476 | 1124 | * | 12,19918 | 2,697337993 | 1,010645 | 2,668927 | 0,007609 | 0,947971763 | 0,738441 | Distal Intergenic | 15 | 29713477 | 29713550 | 74 | 2 | 1,01E+08 | ENSMUST00000174898.3 | -64803 | ENSMUSG00000092639 | Mir3964 | microRNA 3964 |
| chr5 | 32290146 | 32293915 | 3770 | * | 50,602029 | 2,6990564 | 0,5156404 | 5,2343778 | 1,655E-07 | 0,001120595 | 2,3776742 | Promoter (<=1kb) | 5 | 32293145 | 32315186 | 22042 | 1 | 14284 | ENSMUST00000031017.11 | 0 | ENSMUSG00000029135 | Fosl2 | fos-like antigen 2 |
| chr17 | 10654170 | 10655609 | 1440 | * | 10,71006 | 2,699555972 | 0,997474 | 2,706393 | 0,006802 | 0,928277543 | 0,744375 | Intron (ENSMUST0000( | 17 | 10675145 | 10693052 | 17908 | 1 | 319750 | ENSMUST00000160763.2 | -19536 | ENSMUSG00000089633 | A230009B12Rik | RIKEN cDNA A230009B12 gene |
| chr7 | 1,01E+08 | 1,01E+08 | 1894 | * | 16,45379 | 2,69978465 | 0,908269 | 2,97245 | 0,002954 | 0,563858922 | 2,663619 | Promoter (2-3kb) | 7 | 1,01E+08 | 1,01E+08 | 2891 | 2 | 207212 | ENSMUST00000208482.2 | 2302 | ENSMUSG00000032875 | Arhgef17 | Rho guanine nucleotide exchange factor (GEF) 17 |
| chr4 | 93982550 | 93984120 | 1571 | * | 13,59368 | 2,700318214 | 0,902021 | 2,993633 | 0,002757 | 0,683710007 | 1,2763 | Intron (ENSMUST0000( | 4 | 93977813 | 94313825 | 336013 | 2 | 1,08E+08 | ENSMUST00000126270.2 | 329705 | | Gm12648 | predicted gene 12648 |
| chr8 | 87004519 | 87005674 | 1156 | * | 11,30772 | 2,700587646 | 1,046103 | 2,581568 | 0,009835 | 0,954496109 | 0,918255 | Distal Intergenic | 8 | 86762275 | 86787270 | 24996 | 1 | 102093 | ENSMUST00000162305.2 | 242244 | ENSMUSG00000036879 | Phkb | phosphorylase kinase beta |
| chr1 | 1,82E+08 | 1,82E+08 | 1486 | * | 13,14076 | 2,700953116 | 0,890615 | 3,032682 | 0,002424 | 0,678018213 | 1,173325 | Distal Intergenic | 1 | 1,82E+08 | 1,82E+08 | 115591 | 2 | 13800 | ENSMUST00000111024.10 | -41594 | ENSMUSG00000022995 | Enah | ENAH actin regulator |
| chr12 | 1,12E+08 | 1,12E+08 | 1422 | * | 14,04225 | 2,701361555 | 1,036125 | 2,607178 | 0,009129 | 0,974006322 | 0,738441 | Promoter (1-2kb) | 12 | 1,12E+08 | 1,12E+08 | 2494 | 2 | 77675 | ENSMUST00000221314.2 | -1579 | ENSMUSG00000113769 | 5033406O09Rik | RIKEN cDNA 5033406O09 gene |
| chr19 | 13248765 | 13249814 | 1050 | * | 11,59665 | 2,701669189 | 1,033472 | 2,614169 | 0,008944 | 0,970928491 | 0,738441 | Distal Intergenic | 19 | 13278365 | 13278471 | 107 | 2 | 1,15E+08 | ENSMUST00000082890.3 | 28657 | ENSMUSG00000119251 | Gm22443 | predicted gene, 22443 |
| chr2 | 18578669 | 18579529 | 861 | * | 10,19285 | 2,701673641 | 1,041202 | 2,594764 | 0,009466 | 0,909280719 | 1,173325 | Distal Intergenic | 2 | 18677195 | 18681041 | 3847 | 1 | 12238 | ENSMUST00000171845.8 | -97666 | ENSMUSG00000051154 | Commd3 | COMM domain containing 3 |
| chr13 | 36807144 | 36808678 | 1535 | * | 14,5156 | 2,702051293 | 1,041426 | 2,594569 | 0,009471 | 0,904215312 | 1,214904 | Intron (ENSMUST0000( | 13 | 36796946 | 36837047 | 40102 | 1 | 1,08E+08 | ENSMUST00000226032.2 | 10198 | | Gm46410 | predicted gene, 46410 |
| chr8 | 89639555 | 89640163 | 609 | * | 11,19397 | 2,70219369 | 0,986092 | 2,740305 | 0,006138 | 0,914358307 | 0,738441 | Distal Intergenic | 8 | 89751363 | 89751459 | 97 | 1 | 1,02E+08 | ENSMUST00000183771.3 | -111200 | ENSMUSG00000099078 | Mir8110 | microRNA 8110 |
| chr17 | 12943122 | 12944445 | 1324 | * | 14,21978 | 2,702484362 | 1,019106 | 2,651819 | 0,008006 | 0,91804783 | 0,938682 | Exon (ENSMUST000000 | 17 | 12960198 | 13079023 | 118826 | 1 | 104103 | ENSMUST00000160932.3 | -15753 | ENSMUSG00000078247 | Airn | antisense Igf2r RNA |
| chr12 | 63374643 | 63376462 | 1820 | * | 13,95152 | 2,70313935 | 0,987713 | 2,736765 | 0,006205 | 0,915421656 | 0,738441 | Distal Intergenic | 12 | 63656642 | 63656748 | 107 | 1 | 1,15E+08 | ENSMUST00000083024.4 | -280180 | ENSMUSG00000118736 | Gm24921 | predicted gene, 24921 |
| chr18 | 46730576 | 46732035 | 1460 | * | 15,26884 | 2,703427197 | 0,936917 | 2,885449 | 0,003909 | 0,730215586 | 1,37683 | Promoter (<=1kb) | 18 | 46730768 | 46742316 | 11549 | 1 | 13664 | ENSMUST00000238168.2 | 0 | ENSMUSG00000057561 | Eif1a | eukaryotic translation initiation factor 1A |
| chr5 | 56101637 | 56103086 | 1450 | * | 13,9975 | 2,703441923 | 0,998273 | 2,708119 | 0,006767 | 0,896435577 | 0,918255 | Distal Intergenic | 5 | 55460762 | 55460923 | 162 | 2 | 1,15E+08 | ENSMUST00000083861.3 | -640714 | ENSMUSG00000065795 | Gm25513 | predicted gene, 25513 |
| chr7 | 80992637 | 80994688 | 2052 | * | 14,46878 | 2,70366331 | 0,938666 | 2,880325 | 0,003973 | 0,751314174 | 1,214904 | Promoter (<=1kb) | 7 | 80992480 | 80994279 | 1800 | 2 | 20068 | ENSMUST00000130487.8 | 0 | ENSMUSG00000061787 | Rps17 | ribosomal protein S17 |
| chrX | 1,12E+08 | 1,12E+08 | 2382 | * | 16,62394 | 2,704074126 | 0,856233 | 3,158106 | 0,001588 | 0,477594358 | 2,318434 | Promoter (<=1kb) | 20 | 1,12E+08 | 1,13E+08 | 538877 | 1 | 93837 | ENSMUST00000113380.8 | 0 | ENSMUSG00000025592 | Dach2 | dachshund family transcription factor 2 |
| chr16 | 44060738 | 44062662 | 1925 | * | 17,75623 | 2,704094764 | 0,874898 | 3,090754 | 0,001996 | 0,531203841 | 2,154504 | 3' UTR | 16 | 44060543 | 44089895 | 29353 | 2 | 320007 | ENSMUST00000127567.8 | 27233 | ENSMUSG00000022696 | Sidt1 | SID1 transmembrane family, member 1 |
| chr11 | 4258123 | 4260246 | 2124 | * | 29,73424 | 2,704536688 | 0,671466 | 4,02781 | 5,63E-05 | 0,074939782 | 2,858511 | Distal Intergenic | 11 | 4274149 | 4274255 | 107 | 2 | 1,15E+08 | ENSMUST00000240522.1 | 14009 | ENSMUSG00000119807 | Gm24803 | predicted gene, 24803 |
| chr14 | 25138645 | 25140250 | 1606 | * | 19,9651 | 2,705148796 | 0,815254 | 3,318168 | 0,000906 | 0,355694809 | 2,663619 | Exon (ENSMUST000002 | 14 | 25140219 | 25143665 | 3447 | 2 | 75862 | ENSMUST00000225429.2 | 3415 | ENSMUSG00000114916 | 4930572O13Rik | RIKEN cDNA 4930572O13 gene |
| chr13 | 99104060 | 99105639 | 1580 | * | 11,60854 | 2,705188968 | 0,983578 | 2,750356 | 0,005953 | 0,83238039 | 1,173325 | Distal Intergenic | 13 | 99120593 | 99121073 | 481 | 2 | 69382 | ENSMUST00000054425.7 | 15434 | ENSMUSG00000045022 | H2bl1 | H2B.L histone variant 1 |
| chr11 | 88609312 | 88610600 | 1289 | * | 18,55904 | 2,705258434 | 0,78046 | 3,466234 | 0,000528 | 0,296794689 | 2,423594 | Promoter (<=1kb) | 11 | 88609354 | 88619986 | 10633 | 1 | 193280 | ENSMUST00000136201.3 | 0 | | C030037D09Rik | RIKEN cDNA C030037D09 gene |
| chr19 | 43512339 | 43514183 | 1845 | * | 21,95477 | 2,705412953 | 0,688031 | 3,932109 | 8,42E-05 | 0,089931799 | 3,104559 | Promoter (<=1kb) | 19 | 43488191 | 43513044 | 24854 | 2 | 14718 | ENSMUST00000026196.14 | 0 | ENSMUSG00000025190 | Got1 | glutamic-oxaloacetic transaminase 1, soluble |
| chr9 | 20788651 | 20790396 | 1746 | * | 29,32782 | 2,705799428 | 0,66172 | 4,089042 | 4,33E-05 | 0,065127074 | 2,725491 | Promoter (<=1kb) | 9 | 20785104 | 20791023 | 5920 | 2 | 70726 | ENSMUST00000043726.8 | 627 | ENSMUSG00000038742 | Angptl6 | angiopoietin-like 6 |
| chr9 | 34893409 | 34894862 | 1454 | * | 14,88427 | 2,706656245 | 0,929063 | 2,913317 | 0,003576 | 0,642870233 | 2,141612 | Intron (ENSMUST0000( | 9 | 34927748 | 34942172 | 14425 | 1 | 67703 | ENSMUST00000188658.2 | -32886 | ENSMUSG00000032036 | Kirrel3 | kirre like nephrin family adhesion molecule 3 |
| chr19 | 47210236 | 47212035 | 1800 | * | 17,17578 | 2,706779405 | 0,847504 | 3,193824 | 0,001404 | 0,434848027 | 2,663619 | Exon (ENSMUST000002 | 19 | 47217350 | 47247874 | 30525 | 1 | 18011 | ENSMUST00000111807.5 | -5315 | ENSMUSG00000006435 | Neurl1a | neuralized E3 ubiquitin protein ligase 1A |
| chr8 | 1,07E+08 | 1,07E+08 | 3400 | * | 38,40251 | 2,707552356 | 0,661818 | 4,091083 | 4,29E-05 | 0,062163559 | 2,925497 | Distal Intergenic | 8 | 1,07E+08 | 1,07E+08 | 11272 | 1 | 22751 | ENSMUST00000213045.2 | -33804 | ENSMUSG00000031907 | Zfp90 | zinc finger protein 90 |
| chr3 | 5518317 | 5519696 | 1380 | * | 12,01062 | 2,708713944 | 1,040381 | 2,60358 | 0,009226 | 0,974006322 | 0,744375 | Distal Intergenic | 3 | 5578055 | 5580393 | 2339 | 2 | 78039 | ENSMUST00000176666.2 | 60697 | ENSMUSG00000093557 | 4930555M17Rik | RIKEN cDNA 4930555M17 gene |
| chr6 | 49071672 | 49072555 | 884 | * | 13,82544 | 2,708861976 | 0,945794 | 2,864114 | 0,004182 | 0,852340527 | 0,738441 | Promoter (<=1kb) | 6 | 49065062 | 49071145 | 6084 | 2 | 140488 | ENSMUST00000205057.2 | -527 | ENSMUSG00000029814 | Igf2bp3 | insulin-like growth factor 2 mRNA binding protein 3 |
| chr8 | 94822694 | 94824081 | 1388 | * | 16,02466 | 2,709665348 | 0,968433 | 2,797988 | 0,005142 | 0,698838646 | 2,141612 | Promoter (1-2kb) | 8 | 94794582 | 94825556 | 30975 | 2 | 67378 | ENSMUST00000034206.6 | 1475 | ENSMUSG00000031755 | Bbs2 | Bardet-Biedl syndrome 2 (human) |
| chr7 | 43226254 | 43228008 | 1755 | * | 12,72265 | 2,709717261 | 0,979877 | 2,765364 | 0,005686 | 0,809369408 | 1,2763 | Promoter (1-2kb) | 7 | 43229034 | 43242659 | 13626 | 1 | 210105 | ENSMUST00000058104.8 | -1026 | ENSMUSG00000030469 | Zfp719 | zinc finger protein 719 |
| chr10 | 79602595 | 79604365 | 1771 | * | 45,02008 | 2,710124549 | 0,958174 | 2,828426 | 0,004678 | 0,643386705 | 2,787255 | Promoter (<=1kb) | 10 | 79594349 | 79602786 | 8438 | 2 | 70294 | ENSMUST00000047203.9 | 0 | ENSMUSG00000035890 | Rnf126 | ring finger protein 126 |
| chr10 | 67065969 | 67067347 | 1379 | * | 11,91437 | 2,710628038 | 1,006914 | 2,692014 | 0,007102 | 0,850402396 | 1,2763 | Intron (ENSMUST0000( | 10 | 67073927 | 67088451 | 14525 | 1 | 108829 | ENSMUST00000173762.2 | -6580 | ENSMUSG00000037876 | Jmjd1c | jumonji domain containing 1C |
| chr2 | 148520809 | 148524158 | 3350 | * | 44,198258 | 2,710666888 | 0,5455934 | 4,9682916 | 6,755E-07 | 0,003219946 | 2,3776742 | Promoter (<=1kb) | 2 | 148522943 | 148534869 | 11927 | 1 | 74533 | ENSMUST00000028928.8 | 0 | ENSMUSG00000027439 | Gzf1 | GDNF-inducible zinc finger protein 1 |
| chr11 | 79713245 | 79715149 | 1905 | * | 17,06394 | 2,710700617 | 0,876162 | 3,093834 | 0,001976 | 0,514468591 | 2,318434 | Intron (ENSMUST0000( | 11 | 79687019 | 79695998 | 8980 | 2 | 1,05E+08 | ENSMUST00000162268.2 | -17247 | | 9130204K15Rik | RIKEN cDNA 9130204K15 gene |
| chr4 | 62139014 | 62139710 | 697 | * | 10,44159 | 2,710887959 | 1,022572 | 2,65105 | 0,008024 | 0,885719815 | 1,173325 | Intron (ENSMUST0000( | 4 | 62126557 | 62149284 | 22728 | 2 | 1,08E+08 | ENSMUST00000212325.2 | 9574 | | Gm30511 | predicted gene, 30511 |
| chr12 | 35630715 | 35632557 | 1843 | * | 14,77887 | 2,710926229 | 0,919423 | 2,948507 | 0,003193 | 0,709438124 | 1,2763 | Exon (ENSMUST000002 | 12 | 35547973 | 35585037 | 37065 | 2 | 11622 | ENSMUST00000116436.9 | -45678 | ENSMUSG00000019256 | Ahr | aryl-hydrocarbon receptor |
| chr8 | 1,03E+08 | 1,03E+08 | 1013 | * | 11,48565 | 2,711116209 | 1,040608 | 2,605318 | 0,009179 | 0,891097601 | 1,2763 | Distal Intergenic | 8 | 1,03E+08 | 1,03E+08 | 2716 | 2 | 12552 | ENSMUST00000210578.2 | 548266 | ENSMUSG00000031673 | Cdh11 | cadherin 11 |
| chr17 | 77774996 | 77776240 | 1245 | * | 22,11166 | 2,712055735 | 0,773281 | 3,507207 | 0,000453 | 0,253278065 | 2,880017 | Distal Intergenic | 17 | 78506685 | 78507454 | 770 | 2 | 1,03E+08 | ENSMUST00000234450.2 | 731214 | ENSMUSG00000117648 | Gm32282 | predicted gene, 32282 |
| chr15 | 16057051 | 16057973 | 923 | * | 10,25935 | 2,712609607 | 0,971204 | 2,793039 | 0,005222 | 0,802323085 | 1,211315 | Distal Intergenic | 15 | 15819313 | 15819440 | 128 | 1 | 1,15E+08 | ENSMUST00000158079.4 | 237738 | ENSMUSG00000088704 | Gm25550 | predicted gene, 25550 |
| chr18 | 34445455 | 34446453 | 999 | * | 11,86035 | 2,712856131 | 1,023787 | 2,649825 | 0,008053 | 0,86971946 | 1,2763 | 3' UTR | 18 | 34434314 | 34443790 | 9477 | 1 | 11789 | ENSMUST00000167136.2 | 11141 | ENSMUSG00000005871 | Apc | APC, WNT signaling pathway regulator |
| chr7 | 44197882 | 44199637 | 1756 | * | 16,42398 | 2,712885641 | 0,976637 | 2,777782 | 0,005473 | 0,689688276 | 2,386008 | Promoter (<=1kb) | 7 | 44182170 | 44198271 | 16102 | 2 | 18971 | ENSMUST00000049343.15 | 0 | ENSMUSG00000038644 | Pold1 | polymerase (DNA directed), delta 1, catalytic subunit |
| chr5 | 1,24E+08 | 1,24E+08 | 1253 | * | 11,04207 | 2,713600256 | 1,017276 | 2,667517 | 0,007641 | 0,947773602 | 0,744375 | Distal Intergenic | 5 | 1,24E+08 | 1,24E+08 | 52099 | 1 | 208104 | ENSMUST00000111596.8 | -5252 | ENSMUSG00000038342 | Mlxip | MLX interacting protein |
| chr10 | 63093526 | 63095782 | 2257 | * | 18,01973 | 2,714158914 | 0,99883 | 2,717339 | 0,006581 | 0,705941691 | 2,663619 | Intron (ENSMUST0000( | 10 | 63081601 | 63109346 | 27746 | 1 | 67345 | ENSMUST00000218533.2 | 11925 | ENSMUSG00000020064 | Herc4 | hect domain and RLD 4 |
| chr9 | 22067009 | 22069800 | 2792 | * | 26,51118 | 2,714652017 | 0,80369 | 3,377734 | 0,000731 | 0,313895232 | 2,858511 | Promoter (<=1kb) | 9 | 22068142 | 22069654 | 1513 | 1 | 66268 | ENSMUST00000123680.2 | 0 | ENSMUSG00000010607 | Pigyl | phosphatidylinositol glycan anchor biosynthesis, class Y-like |
| chr1 | 1,57E+08 | 1,57E+08 | 1669 | * | 18,63762 | 2,714676608 | 0,795551 | 3,412324 | 0,000644 | 0,320417818 | 2,423594 | Intron (ENSMUST0000( | 1 | 1,57E+08 | 1,57E+08 | 38184 | 2 | 78255 | ENSMUST00000189648.2 | 39825 | ENSMUSG00000026594 | Ralgps2 | Ral GEF with PH domain and SH3 binding motif 2 |
| chr15 | 66378871 | 66380492 | 1622 | * | 19,68823 | 2,714970225 | 0,722905 | 3,755639 | 0,000173 | 0,161106071 | 2,507013 | Distal Intergenic | 15 | 66251707 | 66372759 | 121053 | 2 | 54562 | ENSMUST00000023006.7 | -6112 | ENSMUSG00000022375 | Lrrc6 | leucine rich repeat containing 6 (testis) |
| chr8 | 1,06E+08 | 1,06E+08 | 883 | * | 11,24011 | 2,715033174 | 1,038779 | 2,613678 | 0,008957 | 0,888740228 | 1,2763 | Intron (ENSMUST0000( | 8 | 1,06E+08 | 1,06E+08 | 2476 | 2 | 279499 | ENSMUST00000168196.2 | -11456 | ENSMUSG00000051648 | Kctd19 | potassium channel tetramerisation domain containing 19 |
| chr11 | 33476007 | 33477134 | 1128 | * | 12,41808 | 2,715716344 | 1,008381 | 2,693144 | 0,007078 | 0,849437958 | 1,2763 | Distal Intergenic | 11 | 33161795 | 33463746 | 301952 | 2 | 66011 | ENSMUST00000102815.10 | -12261 | ENSMUSG00000040594 | Ranbp17 | RAN binding protein 17 |
| chr7 | 1,39E+08 | 1,39E+08 | 1667 | * | 13,6613 | 2,716496052 | 0,962173 | 2,823291 | 0,004753 | 0,874247416 | 0,738441 | Intron (ENSMUST0000( | 7 | 1,39E+08 | 1,39E+08 | 57423 | 1 | 212111 | ENSMUST00000138183.2 | 10360 | ENSMUSG00000025477 | Inpp5a | inositol polyphosphate-5-phosphatase A |
| chr2 | 93588168 | 93589111 | 944 | * | 11,79654 | 2,717063787 | 1,008893 | 2,693113 | 0,007079 | 0,93486226 | 0,744375 | Intron (ENSMUST0000( | 2 | 93592906 | 93626334 | 33429 | 2 | 14043 | ENSMUST00000152344.2 | 37223 | ENSMUSG00000027198 | Ext2 | exostosin glycosyltransferase 2 |
| chr16 | 18551776 | 18552876 | 1101 | * | 14,29873 | 2,717998026 | 0,873126 | 3,11295 | 0,001852 | 0,670234202 | 0,938682 | Distal Intergenic | 16 | 18595597 | 18597012 | 1416 | 1 | 12741 | ENSMUST00000043577.3 | -42721 | ENSMUSG00000041378 | Cldn5 | claudin 5 |
| chr7 | 1,18E+08 | 1,18E+08 | 1605 | * | 14,98774 | 2,718088322 | 0,884751 | 3,07215 | 0,002125 | 0,65050992 | 1,214904 | 3' UTR | 7 | 1,18E+08 | 1,18E+08 | 5566 | 2 | 267019 | ENSMUST00000134270.2 | 7745 | ENSMUSG00000008683 | Rps15a | ribosomal protein S15A |
| chr19 | 4458463 | 4460513 | 2051 | * | 16,0953 | 2,718210679 | 0,881455 | 3,083775 | 0,002044 | 0,515999383 | 2,386008 | Distal Intergenic | 19 | 4364447 | 4448313 | 83867 | 2 | 225876 | ENSMUST00000047898.14 | -10150 | ENSMUSG00000054611 | Kdm2a | lysine (K)-specific demethylase 2A |
| chr3 | 1,4E+08 | 1,4E+08 | 1615 | * | 10,77242 | 2,71825404 | 1,044233 | 2,603111 | 0,009238 | 0,892743851 | 1,2763 | Distal Intergenic | 3 | 1,4E+08 | 1,4E+08 | 122 | 1 | 1,02E+08 | ENSMUST00000184427.3 | 366582 | ENSMUSG00000099169 | Mir7j | microRNA 7j |
| chr6 | 1,46E+08 | 1,46E+08 | 755 | * | 11,42254 | 2,718838104 | 0,987719 | 2,752642 | 0,005912 | 0,90673102 | 0,744375 | Distal Intergenic | 6 | 1,46E+08 | 1,46E+08 | 7581 | 2 | 1,03E+08 | ENSMUST00000144757.2 | 63642 | | Gm15704 | predicted gene 15704 |
| chr18 | 36894893 | 36896398 | 1506 | * | 13,93411 | 2,720007002 | 0,884042 | 3,076785 | 0,002092 | 0,640889834 | 1,2763 | Promoter (<=1kb) | 18 | 36895027 | 36895992 | 966 | 1 | 67936 | ENSMUST00000236324.2 | 0 | ENSMUSG00000042660 | Wdr55 | WD repeat domain 55 |
| chr1 | 1,35E+08 | 1,35E+08 | 1653 | * | 15,09636 | 2,72013303 | 1,039499 | 2,616774 | 0,008877 | 0,874247416 | 1,37683 | Exon (ENSMUST000000 | 1 | 1,35E+08 | 1,35E+08 | 81 | 2 | 1,01E+08 | ENSMUST00000175639.3 | -5432 | ENSMUSG00000093515 | Mir1231 | microRNA 1231 |
| chr7 | 1,3E+08 | 1,3E+08 | 2302 | * | 18,46215 | 2,720321784 | 0,949942 | 2,863672 | 0,004188 | 0,642870233 | 2,507013 | Promoter (1-2kb) | 7 | 1,3E+08 | 1,3E+08 | 115207 | 1 | 57752 | ENSMUST00000209108.2 | 1745 | ENSMUSG00000030852 | Tacc2 | transforming, acidic coiled-coil containing protein 2 |
| chr6 | 51358508 | 51361591 | 3084 | * | 30,87146 | 2,720752498 | 0,746625 | 3,644066 | 0,000268 | 0,198315008 | 2,725491 | Promoter (1-2kb) | 6 | 51356690 | 51369771 | 13082 | 1 | 625167 | ENSMUST00000204678.2 | 1818 | ENSMUSG00000107660 | Gm6559 | predicted gene 6559 |
| chr4 | 69923731 | 69924383 | 653 | * | 9,338491 | 2,721353513 | 1,021431 | 2,664256 | 0,007716 | 0,852340527 | 1,360622 | Distal Intergenic | 4 | 70160708 | 70162089 | 1382 | 2 | 214444 | ENSMUST00000124251.2 | 237706 | ENSMUSG00000039298 | Cdk5rap2 | CDK5 regulatory subunit associated protein 2 |
| chr5 | 35100858 | 35102930 | 2073 | * | 15,62857 | 2,721357509 | 0,911724 | 2,984847 | 0,002837 | 0,59811801 | 2,141612 | Distal Intergenic | 5 | 35106789 | 35190939 | 84151 | 1 | 71729 | ENSMUST00000030984.14 | -3859 | ENSMUSG00000029101 | Rgs12 | regulator of G-protein signaling 12 |
| chr15 | 63515028 | 63516384 | 1357 | * | 11,87631 | 2,721436791 | 0,975639 | 2,789389 | 0,005281 | 0,811413964 | 1,173325 | Intron (ENSMUST0000( | 15 | 63589667 | 63609529 | 19863 | 1 | 73994 | ENSMUST00000226508.2 | -73283 | | 4930449C09Rik | RIKEN cDNA 4930449C09 gene |
| chr7 | 1,31E+08 | 1,31E+08 | 997 | * | 9,914268 | 2,721992729 | 1,052811 | 2,585453 | 0,009725 | 0,885309127 | 1,425929 | Exon (ENSMUST000000 | 7 | 1,31E+08 | 1,31E+08 | 13739 | 2 | 16433 | ENSMUST00000046611.9 | 10454 | ENSMUSG00000040205 | Cuzd1 | CUB and zona pellucida-like domains 1 |
| chr16 | 17538326 | 17541483 | 3158 | * | 21,06516 | 2,722305466 | 0,784181 | 3,471528 | 0,000518 | 0,247743734 | 3,471678 | Promoter (<=1kb) | 16 | 17469072 | 17540764 | 71693 | 2 | 94112 | ENSMUST00000232645.2 | 0 | ENSMUSG00000012114 | Med15 | mediator complex subunit 15 |
| chr11 | 29881664 | 29882662 | 999 | * | 13,42164 | 2,722532883 | 1,03046 | 2,642056 | 0,00824 | 0,922828293 | 0,938682 | Intron (ENSMUST0000( | 11 | 29799461 | 29799600 | 140 | 1 | 1,15E+08 | ENSMUST00000157510.3 | 82203 | ENSMUSG00000088135 | Gm22588 | predicted gene, 22588 |
| chr5 | 97537041 | 97538830 | 1790 | * | 13,31932 | 2,722654622 | 1,006867 | 2,704086 | 0,006849 | 0,894365639 | 0,938682 | Promoter (1-2kb) | 5 | 97530057 | 97540238 | 10182 | 2 | 97243 | ENSMUST00000060265.6 | 1408 | ENSMUSG00000046000 | Naa11 | N(alpha)-acetyltransferase 11, NatA catalytic subunit |
| chr4 | 1,2E+08 | 1,2E+08 | 2373 | * | 20,0727 | 2,723618396 | 0,926093 | 2,940976 | 0,003272 | 0,621084104 | 2,221611 | Intron (ENSMUST0000( | 4 | 1,21E+08 | 1,21E+08 | 1249 | 1 | 56222 | ENSMUST00000094814.6 | -37173 | ENSMUSG00000070803 | Cited4 | Cbp/p300-interacting transactivator, with Glu/Asp-rich carboxy-terminal domain, 4 |
| chr13 | 53691270 | 53692719 | 1450 | * | 16,22717 | 2,724896549 | 0,933861 | 2,917883 | 0,003524 | 0,724997822 | 1,2763 | Distal Intergenic | 13 | 53657642 | 53687293 | 29652 | 2 | 1,03E+08 | ENSMUST00000223156.2 | -3977 | | Gm34022 | predicted gene, 34022 |
| chr4 | 1,41E+08 | 1,41E+08 | 1929 | * | 17,42479 | 2,724992587 | 1,015928 | 2,682271 | 0,007312 | 0,723275538 | 2,663619 | Promoter (<=1kb) | 4 | 1,41E+08 | 1,41E+08 | 5258 | 2 | 213989 | ENSMUST00000143154.8 | -250 | ENSMUSG00000043085 | Tmem82 | transmembrane protein 82 |
| chr2 | 94129601 | 94131103 | 1503 | * | 17,42523 | 2,725453126 | 0,804405 | 3,388159 | 0,000704 | 0,327375387 | 2,507013 | Distal Intergenic | 2 | 94082220 | 94095374 | 13155 | 2 | 414123 | ENSMUST00000246180.1 | -34227 | | Mir670hg | MIR670 host gene (non-protein coding) |
| chr3 | 1,56E+08 | 1,56E+08 | 774 | * | 9,500258 | 2,725577037 | 1,048419 | 2,599703 | 0,00933 | 0,902886632 | 1,211315 | Intron (ENSMUST0000( | 3 | 1,56E+08 | 1,56E+08 | 14822 | 2 | 329782 | ENSMUST00000148224.2 | 41013 | ENSMUSG00000085084 | 4930570G19Rik | RIKEN cDNA 4930570G19 gene |
| chr10 | 75609061 | 75610215 | 1155 | * | 13,35146 | 2,725630122 | 1,046412 | 2,604739 | 0,009194 | 0,891636336 | 1,2763 | Promoter (<=1kb) | 10 | 75607067 | 75609248 | 2182 | 2 | 13202 | ENSMUST00000001716.8 | 0 | ENSMUSG00000001666 | Ddt | D-dopachrome tautomerase |
| chr11 | 26390504 | 26391998 | 1495 | * | 10,72702 | 2,725884081 | 1,058197 | 2,575971 | 0,009996 | 0,955649401 | 0,918255 | Intron (ENSMUST0000( | 11 | 26418335 | 26421430 | 3096 | 1 | 67030 | ENSMUST00000134445.2 | -26337 | ENSMUSG00000004018 | Fancl | Fanconi anemia, complementation group L |
| chr4 | 1,3E+08 | 1,3E+08 | 1972 | * | 14,65269 | 2,726011339 | 1,016311 | 2,682262 | 0,007313 | 0,836236014 | 1,41696 | Exon (ENSMUST000000 | 4 | 1,3E+08 | 1,3E+08 | 4674 | 2 | 66264 | ENSMUST00000030586.15 | -6481 | ENSMUSG00000028795 | Ccdc28b | coiled coil domain containing 28B |
| chr11 | 38108196 | 38109535 | 1340 | * | 9,821822 | 2,726149634 | 1,039895 | 2,621562 | 0,008753 | 0,892357983 | 1,211315 | Distal Intergenic | 11 | 38384157 | 38410840 | 26684 | 2 | 1,01E+08 | ENSMUST00000133599.2 | 301305 | | Gm12130 | predicted gene 12130 |
| chr12 | 75831501 | 75832976 | 1476 | * | 13,62848 | 2,726433845 | 1,038187 | 2,626149 | 0,008636 | 0,964229994 | 0,738441 | Distal Intergenic | 12 | 75864908 | 75927280 | 62373 | 1 | 319565 | ENSMUST00000142864.8 | -31932 | ENSMUSG00000063450 | Syne2 | spectrin repeat containing, nuclear envelope 2 |
| chr11 | 51803398 | 51805128 | 1731 | * | 14,93084 | 2,727169095 | 0,955465 | 2,854284 | 0,004313 | 0,767860254 | 1,214904 | Distal Intergenic | 11 | 51789793 | 51800451 | 10659 | 1 | 1,05E+08 | ENSMUST00000212253.2 | 13605 | ENSMUSG00000110618 | Gm39822 | predicted gene, 39822 |
| chr9 | 30117674 | 30118448 | 775 | * | 12,77944 | 2,727445197 | 0,909482 | 2,9989 | 0,00271 | 0,691249627 | 1,173325 | Distal Intergenic | 9 | 30203164 | 30203270 | 107 | 2 | 1,15E+08 | ENSMUST00000239738.1 | 84822 | ENSMUSG00000119318 | Gm26435 | predicted gene, 26435 |
| chr2 | 1,31E+08 | 1,31E+08 | 2143 | * | 15,10697 | 2,727468339 | 0,878864 | 3,103403 | 0,001913 | 0,61020361 | 1,37683 | Intron (ENSMUST0000( | 2 | 1,31E+08 | 1,31E+08 | 26658 | 2 | 613258 | ENSMUST00000128176.2 | 12514 | | A730017L22Rik | RIKEN cDNA A730017L22 gene |
| chr3 | 16276312 | 16278263 | 1952 | * | 14,37752 | 2,727583317 | 0,910057 | 2,997158 | 0,002725 | 0,71122523 | 1,075735 | Distal Intergenic | 3 | 16237400 | 16271019 | 33620 | 1 | 229096 | ENSMUST00000108346.5 | 38912 | ENSMUSG00000047213 | Ythdf3 | YTH N6-methyladenosine RNA binding protein 3 |
| chr12 | 15867595 | 15869288 | 1694 | * | 13,20651 | 2,727641613 | 0,996925 | 2,736055 | 0,006218 | 0,915421656 | 0,738441 | Promoter (<=1kb) | 12 | 15867004 | 15938156 | 71153 | 1 | 432636 | ENSMUST00000180759.2 | 591 | | Gm5432 | predicted gene 5432 |

| | | | | | | | | | | | | | | | | | | | | | | | |
|---|---|---|---|---|---|---|---|---|---|---|---|---|---|---|---|---|---|---|---|---|---|---|---|
| chr3 | 101017717 | 101020402 | 2686 | * | 32,832019 | 2,728191825 | 0,5684094 | 4,7996952 | 1,589E-06 | 0,005619226 | 2,7254908 | Promoter (<=1kb) | 3 | 100947548 | 101017594 | 70047 | 2 | 19221 | ENSMUST00000102694.4 | -123 | ENSMUSG00000027864 | Ptgfrn | prostaglandin F2 receptor negative regulator |
| chr8 | 39610273 | 39611838 | 1566 | * | 19,76625 | 2,728202273 | 0,825482 | 3,304981 | 0,00095 | 0,377581283 | 2,507013 | 3' UTR | 8 | 39473064 | 39540476 | 67413 | 1 | 80286 | ENSMUST00000239511.1 | 137209 | ENSMUSG00000118664 | Tusc3 | tumor suppressor candidate 3 |
| chr16 | 31325176 | 31327026 | 1851 | * | 25,46305 | 2,728563728 | 0,709111 | 3,847863 | 0,000119 | 0,120486853 | 2,725491 | Promoter (<=1kb) | 16 | 31327205 | 31330595 | 3391 | 1 | 1,08E+08 | ENSMUST00000231805.2 | -179 | | Gm46560 | predicted gene, 46560 |
| chr6 | 54460995 | 54462599 | 1605 | * | 15,94396 | 2,730015037 | 0,966577 | 2,824415 | 0,004737 | 0,676053959 | 2,318434 | Promoter (2-3kb) | 6 | 54464891 | 54468527 | 3637 | 1 | 330319 | ENSMUST00000131043.2 | -2292 | ENSMUSG00000086040 | Wipf3 | WAS/WASL interacting protein family, member 3 |
| chr11 | 43478018 | 43479430 | 1413 | * | 11,63681 | 2,730466549 | 1,044961 | 2,612984 | 0,008976 | 0,889364623 | 1,2763 | Exon (ENSMUST00000 | 11 | 43486876 | 43492367 | 5492 | 2 | 16204 | ENSMUST00000020672.5 | 12937 | ENSMUSG00000020405 | Fabp6 | fatty acid binding protein 6 |
| chr11 | 1,01E+08 | 1,01E+08 | 1384 | * | 17,33475 | 2,730557213 | 0,891335 | 3,063446 | 0,002188 | 0,530541227 | 2,386008 | Distal Intergenic | 11 | 1,01E+08 | 1,01E+08 | 10870 | 1 | 11754 | ENSMUST00000017316.7 | 16978 | ENSMUSG00000019326 | Aoc3 | amine oxidase, copper containing 3 |
| chr3 | 95557259 | 95560024 | 2766 | * | 59,031812 | 2,73112487 | 0,4675201 | 5,8417267 | 5,166E-09 | 4,45697E-05 | 4,0557576 | Distal Intergenic | 3 | 95566099 | 95570487 | 4389 | 1 | 17210 | ENSMUST00000037947.15 | -6075 | ENSMUSG00000038612 | Mcl1 | myeloid cell leukemia sequence 1 |
| chr18 | 52513459 | 52514666 | 1208 | * | 10,92883 | 2,731616171 | 1,038617 | 2,63005 | 0,008537 | 0,881662676 | 1,2763 | Distal Intergenic | 18 | 52464621 | 52466068 | 1448 | 1 | 67634 | ENSMUST00000025388.7 | 48838 | ENSMUSG00000024510 | Ftmt | ferritin mitochondrial |
| chr18 | 65879139 | 65881057 | 1919 | * | 18,11329 | 2,732237714 | 0,903182 | 3,025125 | 0,002485 | 0,548877304 | 2,386008 | Promoter (1-2kb) | 18 | 65882085 | 65883461 | 1377 | 1 | 319888 | ENSMUST00000134343.2 | -1028 | ENSMUSG00000046610 | Oacyl | O-acyltransferase like |
| chr11 | 97073205 | 97074618 | 1414 | * | 10,80788 | 2,732399239 | 1,015567 | 2,690517 | 0,007134 | 0,936950943 | 0,738441 | Intron (ENSMUST0000 | 11 | 97050540 | 97078707 | 28168 | 2 | 16211 | ENSMUST00000001479.5 | 4089 | ENSMUSG00000001440 | Kpnb1 | karyopherin (importin) beta 1 |
| chr17 | 81078811 | 81081230 | 2420 | * | 27,00538 | 2,732529264 | 0,736794 | 3,708677 | 0,000208 | 0,165924532 | 2,880017 | Distal Intergenic | 17 | 80961290 | 81035914 | 74625 | 2 | 225028 | ENSMUST00000234249.2 | -42897 | ENSMUSG00000024242 | Map4k3 | mitogen-activated protein kinase kinase kinase kinase 3 |
| chr8 | 127320846 | 127323811 | 2966 | * | 83,192765 | 2,732530315 | 0,3823428 | 7,146807 | 8,882E-13 | 3,09407E-08 | 4,2003037 | Promoter (<=1kb) | 8 | 127315035 | 127320725 | 5691 | 2 | 270110 | ENSMUST00000054960.9 | -121 | ENSMUSG00000051495 | Irf2bp2 | interferon regulatory factor 2 binding protein 2 |
| chr15 | 11368793 | 11371157 | 2365 | * | 15,56422 | 2,732623706 | 0,828723 | 3,297391 | 0,000976 | 0,504427013 | 1,214904 | Distal Intergenic | 15 | 11361697 | 11361823 | 127 | 2 | 1,15E+08 | ENSMUST00000158679.3 | -6970 | ENSMUSG00000089304 | Gm24844 | predicted gene, 24844 |
| chr13 | 94508371 | 94509965 | 1595 | * | 12,32354 | 2,732645127 | 1,01438 | 2,693908 | 0,007062 | 0,902600349 | 0,918255 | Intron (ENSMUST0000 | 13 | 94495468 | 94702825 | 207358 | 1 | 11774 | ENSMUST00000022196.5 | 12903 | ENSMUSG00000021686 | Ap3b1 | adaptor-related protein complex 3, beta 1 subunit |
| chr4 | 1,26E+08 | 1,26E+08 | 2502 | * | 16,22803 | 2,7329825 | 0,952131 | 2,870384 | 0,0041 | 0,651710641 | 2,318434 | Promoter (<=1kb) | 4 | 1,26E+08 | 1,26E+08 | 1360 | 2 | 230753 | ENSMUST00000169855.2 | -271 | ENSMUSG00000043962 | Thrap3 | thyroid hormone receptor associated protein 3 |
| chr5 | 66704031 | 66705166 | 1136 | * | 18,31917 | 2,733611114 | 0,770794 | 3,546487 | 0,00039 | 0,245757594 | 2,663619 | Promoter (<=1kb) | 5 | 66609610 | 66704300 | 94691 | 2 | 11787 | ENSMUST00000161716.8 | 0 | ENSMUSG00000029207 | Apbb2 | amyloid beta (A4) precursor protein-binding, family B, member 2 |
| chr3 | 51900869 | 51902062 | 1194 | * | 15,32612 | 2,733778095 | 0,874381 | 3,126529 | 0,001769 | 0,618136403 | 1,214904 | Intron (ENSMUST0000 | 3 | 51593328 | 52012497 | 419170 | 2 | 433586 | ENSMUST00000121440.4 | 110435 | ENSMUSG00000061143 | Maml3 | mastermind like transcriptional coactivator 3 |
| chr8 | 1,12E+08 | 1,12E+08 | 1384 | * | 13,01888 | 2,734071512 | 1,054324 | 2,5932 | 0,009509 | 0,899299554 | 1,2763 | Intron (ENSMUST0000 | 8 | 1,12E+08 | 1,12E+08 | 20252 | 2 | 20340 | ENSMUST00000168741.8 | -7687 | ENSMUSG00000003316 | Glg1 | golgi apparatus protein 1 |
| chr17 | 56631585 | 56633105 | 1521 | * | 42,91118 | 2,734313473 | 0,890009 | 3,07223 | 0,002125 | 0,496560937 | 2,787255 | Promoter (<=1kb) | 17 | 56633062 | 56663056 | 29995 | 1 | 193796 | ENSMUST00000139679.8 | 0 | ENSMUSG00000024201 | Kdm4b | lysine (K)-specific demethylase 4B |
| chr9 | 72625511 | 72627489 | 1979 | * | 14,01568 | 2,734565821 | 0,88598 | 3,086487 | 0,002025 | 0,723215019 | 0,738441 | 3' UTR | 9 | 72646754 | 72655440 | 8687 | 1 | 17999 | ENSMUST00000184020.8 | -19265 | ENSMUSG00000032216 | Nedd4 | neural precursor cell expressed, developmentally down-regulated 4 |
| chr5 | 1,14E+08 | 1,14E+08 | 1406 | * | 12,44184 | 2,734856733 | 1,049821 | 2,605069 | 0,009186 | 0,905015433 | 1,173325 | Exon (ENSMUST00000 | 5 | 1,14E+08 | 1,15E+08 | 49636 | 1 | 231646 | ENSMUST00000202006.4 | 7955 | ENSMUSG00000066952 | Myo1h | myosin 1H |
| chr7 | 1,26E+08 | 1,26E+08 | 1682 | * | 12,33696 | 2,735275853 | 0,970736 | 2,817734 | 0,004836 | 0,838625535 | 0,918255 | Promoter (1-2kb) | 7 | 1,26E+08 | 1,26E+08 | 5563 | 1 | 68952 | ENSMUST00000207020.2 | 1673 | ENSMUSG00000058966 | Tlcd3b | TLC domain containing 3B |
| chr19 | 6380818 | 6382652 | 1835 | * | 16,66282 | 2,735429881 | 0,792135 | 3,453238 | 0,000554 | 0,404488619 | 1,2763 | Promoter (2-3kb) | 19 | 6385009 | 6390910 | 5902 | 1 | 17283 | ENSMUST00000113502.10 | -2357 | ENSMUSG00000024947 | Men1 | multiple endocrine neoplasia 1 |
| chr19 | 43120818 | 43122236 | 1419 | * | 13,79364 | 2,735931002 | 0,860514 | 3,179415 | 0,001476 | 0,631326702 | 0,938682 | Intron (ENSMUST0000 | 19 | 42897023 | 43212128 | 315106 | 2 | 545291 | ENSMUST00000238153.2 | 89892 | ENSMUSG00000074852 | Hpse2 | heparanase 2 |
| chr5 | 1,44E+08 | 1,44E+08 | 1508 | * | 13,36206 | 2,737226928 | 0,983557 | 2,782987 | 0,005386 | 0,798887389 | 1,2763 | Intron (ENSMUST0000 | 5 | 1,44E+08 | 1,44E+08 | 87769 | 1 | 231876 | ENSMUST00000041804.8 | 7717 | ENSMUSG00000038970 | Lmtk2 | lemur tyrosine kinase 2 |
| chr9 | 62254331 | 62254927 | 597 | * | 10,17908 | 2,737931511 | 0,956943 | 2,861124 | 0,004221 | 0,770673161 | 1,173325 | Promoter (2-3kb) | 9 | 62252081 | 62281997 | 29917 | 1 | 11737 | ENSMUST00000126451.8 | 2250 | ENSMUSG00000032249 | Anp32a | acidic (leucine-rich) nuclear phosphoprotein 32 family, member A |
| chr14 | 25882078 | 25883878 | 1801 | * | 17,87283 | 2,738039945 | 0,773332 | 3,540577 | 0,000399 | 0,260200653 | 2,423594 | Exon (ENSMUST00000 | 14 | 25888828 | 25902898 | 14071 | 2 | 211623 | ENSMUST00000183431.8 | 19020 | ENSMUSG00000095304 | Plac9 | placenta specific 9 |
| chr6 | 21795417 | 21796941 | 1525 | * | 17,94591 | 2,738285659 | 0,92762 | 2,951948 | 0,003158 | 0,597840795 | 2,386008 | Exon (ENSMUST00000 | 6 | 21799949 | 21851913 | 51965 | 2 | 269831 | ENSMUST00000134635.2 | 54972 | ENSMUSG00000029669 | Tspan12 | tetraspanin 12 |
| chr3 | 94319600 | 94321142 | 1543 | * | 14,87861 | 2,738905529 | 0,960983 | 2,850109 | 0,00437 | 0,744054256 | 1,41696 | Promoter (<=1kb) | 3 | 94320580 | 94341975 | 21396 | 1 | 72634 | ENSMUST00000166032.8 | 0 | ENSMUSG00000041912 | Tdrkh | tudor and KH domain containing protein |
| chr7 | 81570725 | 81572377 | 1653 | * | 13,13501 | 2,73930033 | 0,95304 | 2,874275 | 0,00405 | 0,80696549 | 0,918255 | Intron (ENSMUST0000 | 7 | 81547559 | 81583771 | 36213 | 2 | 29877 | ENSMUST00000149288.2 | 11394 | ENSMUSG00000025104 | Hdgfl3 | HDGF like 3 |
| chr5 | 1,2E+08 | 1,2E+08 | 1391 | * | 14,02502 | 2,739532469 | 0,972839 | 2,816018 | 0,004862 | 0,838878373 | 0,918255 | Exon (ENSMUST00000 | 5 | 1,2E+08 | 1,2E+08 | 8956 | 1 | 21388 | ENSMUST00000202504.3 | 5065 | ENSMUSG00000018263 | Tbx5 | T-box 5 |
| chr14 | 70440441 | 70442549 | 2109 | * | 20,80435 | 2,739560739 | 0,892354 | 3,070039 | 0,00214 | 0,492834344 | 2,880017 | Promoter (<=1kb) | 14 | 70418381 | 70442072 | 23692 | 2 | 20410 | ENSMUST00000227259.2 | 0 | ENSMUSG00000022091 | Sorbs3 | sorbin and SH3 domain containing 3 |
| chr9 | 1,08E+08 | 1,08E+08 | 1712 | * | 15,12589 | 2,740821815 | 1,047367 | 2,616868 | 0,008874 | 0,894566572 | 1,214904 | Intron (ENSMUST0000 | 9 | 1,08E+08 | 1,08E+08 | 3351 | 1 | 69232 | ENSMUST00000194139.2 | -7832 | ENSMUSG00000006673 | Qrich1 | glutamine-rich 1 |
| chr17 | 13226342 | 13228442 | 2101 | * | 16,42756 | 2,741417075 | 0,851829 | 3,218272 | 0,00129 | 0,439591476 | 2,386008 | Promoter (<=1kb) | 17 | 13226883 | 13258950 | 32068 | 1 | 1,03E+08 | ENSMUST00000232726.2 | 0 | | Gm36117 | predicted gene, 36117 |
| chr4 | 1,01E+08 | 1,01E+08 | 2430 | * | 22,32587 | 2,741661167 | 0,703138 | 3,899179 | 9,65E-05 | 0,096870739 | 3,104559 | Intron (ENSMUST0000 | 4 | 1,01E+08 | 1,01E+08 | 104 | 1 | 1,15E+08 | ENSMUST00000104489.3 | 15327 | ENSMUSG00000077677 | Gm24468 | predicted gene, 24468 |
| chr4 | 57797981 | 57799573 | 1593 | * | 27,07196 | 2,742226055 | 0,674825 | 4,063612 | 4,83E-05 | 0,070291925 | 2,67243 | Intron (ENSMUST0000 | 4 | 57782180 | 57855202 | 73023 | 1 | 677884 | ENSMUST00000124581.2 | 15801 | ENSMUSG00000038729 | Pakap | paralemmin A kinase anchor protein |
| chr2 | 1,28E+08 | 1,28E+08 | 2149 | * | 12,93222 | 2,742321385 | 0,985707 | 2,782087 | 0,005401 | 0,815986445 | 1,173325 | 3' UTR | 2 | 1,28E+08 | 1,28E+08 | 35232 | 1 | 1,03E+08 | ENSMUST00000243378.1 | -6219 | | Gm14009 | predicted gene 14009 |
| chr5 | 1,4E+08 | 1,4E+08 | 1392 | * | 16,10984 | 2,742546205 | 0,886128 | 3,094978 | 0,001968 | 0,627785088 | 1,2763 | Intron (ENSMUST0000 | 5 | 1,4E+08 | 1,4E+08 | 166225 | 2 | 17120 | ENSMUST00000199237.5 | -23351 | ENSMUSG00000029554 | Mad1l1 | MAD1 mitotic arrest deficient 1-like 1 |
| chr1 | 80614454 | 80615703 | 1250 | * | 11,84264 | 2,742768856 | 1,015565 | 2,700731 | 0,006919 | 0,933541856 | 0,738441 | Intron (ENSMUST0000 | 1 | 80479430 | 80642992 | 163563 | 2 | 210293 | ENSMUST00000190983.7 | 27289 | ENSMUSG00000038608 | Dock10 | dedicator of cytokinesis 10 |
| chr2 | 1,1E+08 | 1,1E+08 | 2008 | * | 14,99653 | 2,742773343 | 0,863168 | 3,177566 | 0,001485 | 0,584366651 | 1,214904 | Promoter (1-2kb) | 2 | 1,1E+08 | 1,1E+08 | 933 | 1 | 107515 | ENSMUST00000142869.2 | -1416 | ENSMUSG00000050199 | Lgr4 | leucine-rich repeat-containing G protein-coupled receptor 4 |
| chr7 | 46672014 | 46672892 | 879 | * | 13,32493 | 2,74431686 | 0,95915 | 2,861196 | 0,00422 | 0,852340527 | 0,738441 | Distal Intergenic | 7 | 46658272 | 46659323 | 1052 | 1 | 1,03E+08 | ENSMUST00000241262.1 | 13742 | ENSMUSG00000109118 | Gm32031 | predicted gene, 32031 |
| chr7 | 44195854 | 44197873 | 2020 | * | 17,6847 | 2,744410257 | 0,812363 | 3,378306 | 0,000729 | 0,337811311 | 2,386008 | Promoter (<=1kb) | 7 | 44191526 | 44198157 | 6632 | 2 | 18971 | ENSMUST00000145956.2 | 284 | ENSMUSG00000038644 | Pold1 | polymerase (DNA directed), delta 1, catalytic subunit |
| chr11 | 87583976 | 87585782 | 1807 | * | 11,92703 | 2,744556565 | 0,998941 | 2,747466 | 0,006006 | 0,834186259 | 1,173325 | Exon (ENSMUST00000 | 11 | 87590723 | 87626365 | 35643 | 1 | 207742 | ENSMUST00000040089.5 | -4941 | ENSMUSG00000034177 | Rnf43 | ring finger protein 43 |
| chr12 | 3576044 | 3577039 | 996 | * | 11,05471 | 2,744662474 | 0,960177 | 2,858497 | 0,004257 | 0,853738252 | 0,738441 | Distal Intergenic | 12 | 3543761 | 3554748 | 10988 | 1 | 75302 | ENSMUST00000152196.2 | 32283 | ENSMUSG00000037486 | Asxl2 | ASXL transcriptional regulator 2 |
| chr8 | 13804223 | 13805922 | 1700 | * | 13,04119 | 2,746393181 | 0,999065 | 2,748965 | 0,005978 | 0,818587238 | 1,2763 | Promoter (1-2kb) | 8 | 13807676 | 13816497 | 8822 | 1 | 69957 | ENSMUST00000130173.9 | -1754 | ENSMUSG00000038416 | Cdc16 | CDC16 cell division cycle 16 |
| chr11 | 85740679 | 85743005 | 2327 | * | 23,37951 | 2,74736064 | 0,806765 | 3,405402 | 0,000661 | 0,309888367 | 2,67243 | Exon (ENSMUST00000 | 11 | 85723377 | 85732774 | 9398 | 1 | 21385 | ENSMUST00000000095.7 | 17302 | ENSMUSG00000000093 | Tbx2 | T-box 2 |
| chr7 | 39914712 | 39916698 | 1987 | * | 21,24678 | 2,747638891 | 0,6992 | 3,92969 | 8,51E-05 | 0,097946479 | 2,67243 | Intron (ENSMUST0000 | 7 | 40324161 | 40332685 | 8525 | 1 | 71029 | ENSMUST00000206289.2 | -407463 | ENSMUSG00000102004 | 4933402C06Rik | RIKEN cDNA 4933402C06 gene |
| chr15 | 76653657 | 76655937 | 2281 | * | 32,49079 | 2,748166442 | 0,709809 | 3,8717 | 0,000108 | 0,108693506 | 2,925497 | Intron (ENSMUST0000 | 15 | 76635877 | 76679025 | 43149 | 2 | 223666 | ENSMUST00000176855.9 | 23088 | ENSMUSG00000033697 | Arhgap39 | Rho GTPase activating protein 39 |
| chr14 | 22777319 | 22779148 | 1830 | * | 15,896 | 2,749123526 | 0,837529 | 3,28242 | 0,001029 | 0,505412507 | 1,2763 | Intron (ENSMUST0000 | 14 | 22646581 | 22649336 | 2756 | 1 | 76633 | ENSMUST00000226003.2 | 130738 | ENSMUSG00000063458 | Lrmda | leucine rich melanocyte differentiation associated |
| chr4 | 1,37E+08 | 1,37E+08 | 1895 | * | 20,10406 | 2,749187326 | 0,960633 | 2,86185 | 0,004212 | 0,6516284 | 2,386008 | Exon (ENSMUST00000 | 4 | 1,37E+08 | 1,37E+08 | 6798 | 2 | 230848 | ENSMUST00000218160.2 | 6166 | ENSMUSG00000060862 | Zbtb40 | zinc finger and BTB domain containing 40 |
| chr13 | 60926202 | 60927395 | 1194 | * | 11,30865 | 2,749542868 | 1,066307 | 2,578565 | 0,009921 | 0,954902749 | 0,918255 | Distal Intergenic | 13 | 60929701 | 60934178 | 4478 | 2 | 218275 | ENSMUST00000026078.10 | 6783 | ENSMUSG00000042243 | BC051665 | cDNA sequence BC051665 |
| chr16 | 75061124 | 75062325 | 1202 | * | 16,47121 | 2,749847756 | 0,818221 | 3,360765 | 0,000777 | 0,369395048 | 2,141612 | Distal Intergenic | 16 | 75179597 | 75244082 | 64486 | 2 | 268902 | ENSMUST00000243278.1 | 181757 | ENSMUSG00000052516 | Robo2 | roundabout guidance receptor 2 |
| chr6 | 1,16E+08 | 1,16E+08 | 1577 | * | 18,78614 | 2,750346035 | 0,955243 | 2,879212 | 0,003987 | 0,634319754 | 2,507013 | Promoter (<=1kb) | 6 | 1,16E+08 | 1,16E+08 | 174876 | 2 | 330401 | ENSMUST00000088896.10 | -118 | ENSMUSG00000030126 | Tmcc1 | transmembrane and coiled coil domains 1 |
| chr6 | 1,16E+08 | 1,16E+08 | 1522 | * | 14,17446 | 2,750860767 | 0,887138 | 3,100825 | 0,00193 | 0,681331037 | 0,918255 | Intron (ENSMUST0000 | 6 | 1,16E+08 | 1,16E+08 | 35244 | 2 | 330401 | ENSMUST00000173140.4 | 9320 | ENSMUSG00000030126 | Tmcc1 | transmembrane and coiled coil domains 1 |
| chr8 | 70212493 | 70213602 | 1110 | * | 13,37012 | 2,751021848 | 0,927667 | 2,965527 | 0,003022 | 0,749781925 | 0,938682 | Intron (ENSMUST0000 | 8 | 70195834 | 70219329 | 23496 | 2 | 620419 | ENSMUST00000126915.2 | 5727 | ENSMUSG00000092260 | Zfp963 | zinc finger protein 963 |
| chr5 | 1E+08 | 1E+08 | 1632 | * | 15,68849 | 2,751064424 | 0,91269 | 3,014237 | 0,002576 | 0,661775851 | 1,37683 | Intron (ENSMUST0000 | 5 | 1,01E+08 | 1,01E+08 | 2125 | 2 | 69162 | ENSMUST00000182664.2 | 37113 | ENSMUSG00000035325 | Sec31a | Sec31 homolog A (S. cerevisiae) |
| chr7 | 79579432 | 79580805 | 1374 | * | 10,18723 | 2,751376989 | 1,053608 | 2,611385 | 0,009018 | 0,902481521 | 1,173325 | Exon (ENSMUST00000 | 7 | 79575109 | 79585107 | 9999 | 2 | 70420 | ENSMUST00000048731.6 | 4302 | ENSMUSG00000039043 | Arpin | actin-related protein 2/3 complex inhibitor |
| chr5 | 1,49E+08 | 1,49E+08 | 1585 | * | 13,1659 | 2,75174076 | 1,060788 | 2,594053 | 0,009485 | 0,923078468 | 1,075735 | Promoter (2-3kb) | 5 | 1,49E+08 | 1,49E+08 | 30814 | 1 | 75860 | ENSMUST00000201610.2 | 2122 | ENSMUSG00000029660 | Tex26 | testis expressed 26 |
| chr8 | 37198212 | 37200178 | 1967 | * | 15,90174 | 2,752016127 | 1,042555 | 2,639686 | 0,008298 | 0,764402529 | 2,386008 | Promoter (<=1kb) | 8 | 37034905 | 37200208 | 165304 | 2 | 50768 | ENSMUST00000098826.10 | 30 | ENSMUSG00000031523 | Dlc1 | deleted in liver cancer 1 |
| chr1 | 1,65E+08 | 1,65E+08 | 1580 | * | 15,6646 | 2,752292448 | 0,973765 | 2,826443 | 0,004707 | 0,684274806 | 2,141612 | Distal Intergenic | 1 | 1,65E+08 | 1,65E+08 | 1524 | 2 | 68481 | ENSMUST00000192716.2 | 17038 | ENSMUSG00000026566 | Mpzl1 | myelin protein zero-like 1 |
| chr13 | 1,03E+08 | 1,03E+08 | 1150 | * | 10,3812 | 2,752335443 | 1,063608 | 2,587734 | 0,009661 | 0,900637308 | 1,2763 | Promoter (1-2kb) | 13 | 1,03E+08 | 1,03E+08 | 163818 | 2 | 328329 | ENSMUST00000167462.8 | 1155 | ENSMUSG00000034751 | Mast4 | microtubule associated serine/threonine kinase family member 4 |
| chr15 | 93449180 | 93450766 | 1587 | * | 24,75501 | 2,752582726 | 0,757432 | 3,634097 | 0,000279 | 0,19659001 | 2,880017 | Intron (ENSMUST0000 | 15 | 93396995 | 93417404 | 20410 | 2 | 106042 | ENSMUST00000109255.3 | -31776 | ENSMUSG00000036158 | Prickle1 | prickle planar cell polarity protein 1 |
| chr18 | 34937820 | 34939643 | 1824 | * | 14,21255 | 2,752861405 | 0,904146 | 3,044708 | 0,002329 | 0,658200216 | 1,2763 | Intron (ENSMUST0000 | 18 | 34946610 | 34952837 | 6228 | 1 | 277250 | ENSMUST00000224065.2 | -6967 | ENSMUSG00000038773 | Kdm3b | KDM3B lysine (K)-specific demethylase 3B |
| chr11 | 1,02E+08 | 1,02E+08 | 2192 | * | 15,06939 | 2,753455312 | 1,019843 | 2,699882 | 0,006936 | 0,844744745 | 1,2763 | Distal Intergenic | 11 | 1,02E+08 | 1,02E+08 | 3965 | 1 | 217214 | ENSMUST00000129799.8 | -7480 | ENSMUSG00000048217 | Nags | N-acetylglutamate synthase |
| chr12 | 44574759 | 44576665 | 1907 | * | 19,17029 | 2,753662593 | 0,88962 | 3,095324 | 0,001966 | 0,524368688 | 2,221611 | Promoter (2-3kb) | 12 | 44579211 | 44591654 | 12444 | 1 | 319504 | ENSMUST00000218431.2 | -2546 | ENSMUSG00000020598 | Nrcam | neuronal cell adhesion molecule |
| chr4 | 1,39E+08 | 1,39E+08 | 1286 | * | 12,57905 | 2,754067572 | 0,943626 | 2,918602 | 0,003516 | 0,821417341 | 0,738441 | Promoter (<=1kb) | 4 | 1,39E+08 | 1,39E+08 | 4876 | 1 | 26970 | ENSMUST00000030531.14 | 0 | ENSMUSG00000028751 | Pla2g2e | phospholipase A2, group IIE |
| chr8 | 1,26E+08 | 1,26E+08 | 1156 | * | 12,82509 | 2,7544626 | 0,98579 | 2,794168 | 0,005203 | 0,889546768 | 0,738441 | Promoter (<=1kb) | 8 | 1,26E+08 | 1,26E+08 | 151733 | 2 | 244668 | ENSMUST00000212987.2 | -210 | ENSMUSG00000001995 | Sipa1l2 | signal-induced proliferation-associated 1 like 2 |
| chr1 | 78673679 | 78675311 | 1633 | * | 12,8559 | 2,754571562 | 0,88398 | 3,1161 | 0,001833 | 0,63007937 | 1,173325 | Promoter (<=1kb) | 1 | 78674373 | 78679006 | 4634 | 1 | 74205 | ENSMUST00000148608.2 | 0 | ENSMUSG00000032883 | Acsl3 | acyl-CoA synthetase long-chain family member 3 |
| chr4 | 1,4E+08 | 1,4E+08 | 1150 | * | 10,79167 | 2,755109489 | 1,032868 | 2,667437 | 0,007643 | 0,948006995 | 0,738441 | Intron (ENSMUST0000 | 4 | 1,4E+08 | 1,4E+08 | 9408 | 1 | 1,03E+08 | ENSMUST00000135716.3 | 7733 | ENSMUSG00000086437 | Gm13021 | predicted gene 13021 |
| chr5 | 1,15E+08 | 1,15E+08 | 812 | * | 11,16105 | 2,755334605 | 1,033428 | 2,666209 | 0,007671 | 0,859983798 | 1,2763 | Promoter (2-3kb) | 5 | 1,15E+08 | 1,15E+08 | 4857 | 2 | 1,05E+08 | ENSMUST00000245927.1 | 2365 | | Gm13822 | predicted gene 13822 |
| chr4 | 1,37E+08 | 1,37E+08 | 1911 | * | 23,12662 | 2,755353094 | 0,788982 | 3,492288 | 0,000479 | 0,272208984 | 2,67243 | Exon (ENSMUST00000 | 4 | 1,37E+08 | 1,37E+08 | 6439 | 1 | 15530 | ENSMUST00000238544.2 | -20415 | ENSMUSG00000028763 | Hspg2 | perlecan (heparan sulfate proteoglycan 2) |
| chr5 | 1,4E+08 | 1,4E+08 | 1413 | * | 15,47377 | 2,755871717 | 0,918789 | 2,999461 | 0,002705 | 0,685026746 | 1,214904 | Intron (ENSMUST0000 | 5 | 1,4E+08 | 1,4E+08 | 26139 | 2 | 231830 | ENSMUST00000170773.6 | 5304 | ENSMUSG00000036718 | Micall2 | MICAL-like 2 |
| chr8 | 71725701 | 71727254 | 1554 | * | 17,03895 | 2,756545748 | 0,955803 | 2,884012 | 0,003926 | 0,644704154 | 2,318434 | Promoter (<=1kb) | 8 | 71703769 | 71725578 | 21810 | 2 | 76478 | ENSMUST00000035960.13 | -123 | ENSMUSG00000035439 | Haus8 | 4HAUS augmin-like complex, subunit 8 |
| chr19 | 3573818 | 3576401 | 2584 | * | 20,54913 | 2,757182236 | 0,808785 | 3,409044 | 0,000652 | 0,323558126 | 2,386008 | Promoter (2-3kb) | 19 | 3561495 | 3571747 | 10253 | 2 | 52036 | ENSMUST00000225339.2 | -2071 | ENSMUSG00000024908 | Ppp6r3 | protein phosphatase 6, regulatory subunit 3 |
| chr4 | 1,41E+08 | 1,41E+08 | 2186 | * | 17,69816 | 2,757815588 | 0,912097 | 3,0236 | 0,002498 | 0,54964001 | 2,386008 | Promoter (2-3kb) | 4 | 1,41E+08 | 1,41E+08 | 1444 | 2 | 66147 | ENSMUST00000132704.2 | 2473 | ENSMUSG00000028923 | Necap2 | NECAP endocytosis associated 2 |
| chr12 | 82347342 | 82349031 | 1690 | * | 12,63803 | 2,758125615 | 1,029557 | 2,678943 | 0,007385 | 0,886332554 | 1,075735 | Intron (ENSMUST0000 | 12 | 82358123 | 82498556 | 140434 | 1 | 217692 | ENSMUST00000053969.7 | -9092 | ENSMUSG00000042700 | Sipa1l1 | signal-induced proliferation-associated 1 like 1 |
| chr13 | 55626923 | 55629091 | 2169 | * | 16,89282 | 2,758143878 | 1,006768 | 2,739601 | 0,006151 | 0,730469267 | 2,141612 | Promoter (<=1kb) | 13 | 55629191 | 55629957 | 767 | 2 | 56320 | ENSMUST00000139516.2 | 866 | ENSMUSG00000034675 | Dbn1 | drebrin 1 |
| chr9 | 1,08E+08 | 1,08E+08 | 1763 | * | 16,92242 | 2,758286749 | 0,857029 | 3,218429 | 0,001289 | 0,439573569 | 2,386008 | Promoter (<=1kb) | 9 | 1,08E+08 | 1,08E+08 | 902 | 1 | 23918 | ENSMUST00000195456.2 | 21 | ENSMUSG00000062867 | Impdh2 | inosine monophosphate dehydrogenase 2 |
| chr13 | 21960681 | 21961751 | 1071 | * | 13,66574 | 2,759303004 | 0,930687 | 2,964803 | 0,003029 | 0,749931768 | 0,938682 | Distal Intergenic | 13 | 21964053 | 21964795 | 743 | 2 | 56702 | ENSMUST00000080511.3 | 3044 | ENSMUSG00000058773 | H1f5 | H1.5 linker histone, cluster member |
| chr7 | 7123199 | 7125087 | 1889 | * | 19,27965 | 2,759628933 | 0,763163 | 3,61604 | 0,000299 | 0,210093832 | 2,713306 | Promoter (<=1kb) | 7 | 7117689 | 7124487 | 6799 | 2 | 232853 | ENSMUST00000056246.8 | 0 | ENSMUSG00000062116 | Zfp954 | zinc finger protein 954 |
| chr16 | 32147434 | 32149733 | 2300 | * | 17,12755 | 2,760100353 | 0,878824 | 3,140676 | 0,001686 | 0,597812173 | 1,2763 | Promoter (1-2kb) | 16 | 32151075 | 32171881 | 20807 | 1 | 224111 | ENSMUST00000232016.2 | -1342 | ENSMUSG00000053774 | Ubxn7 | UBX domain protein 7 |
| chr4 | 1,3E+08 | 1,3E+08 | 1583 | * | 11,86469 | 2,760502377 | 1,041602 | 2,650248 | 0,008043 | 0,955649401 | 0,738441 | Promoter (1-2kb) | 4 | 1,3E+08 | 1,31E+08 | 116849 | 1 | 80912 | ENSMUST00000097862.3 | 1576 | ENSMUSG00000028580 | Pum1 | pumilio RNA-binding family member 1 |
| chr2 | 1,54E+08 | 1,54E+08 | 1371 | * | 20,76426 | 2,76125771 | 0,933036 | 2,959432 | 0,003082 | 0,554161661 | 2,880017 | Promoter (2-3kb) | 2 | 1,54E+08 | 1,54E+08 | 14600 | 1 | 378700 | ENSMUST00000109760.2 | 2812 | ENSMUSG00000068008 | Bpifb3 | BPI fold containing family B, member 3 |
| chr8 | 53159693 | 53160519 | 827 | * | 13,32668 | 2,761659375 | 0,856937 | 3,222708 | 0,00127 | 0,603552508 | 0,938682 | Distal Intergenic | 8 | 52809646 | 52856374 | 46729 | 1 | 330767 | ENSMUST00000210977.2 | 350047 | ENSMUSG00000110013 | 1190028D05Rik | RIKEN cDNA 1190028D05 gene |
| chr19 | 38075223 | 38076991 | 1769 | * | 24,59212 | 2,761919069 | 0,828958 | 3,331796 | 0,000863 | 0,337396172 | 2,858511 | Distal Intergenic | 19 | 38085519 | 38102711 | 17193 | 1 | 107221 | ENSMUST00000067098.8 | -8528 | ENSMUSG00000054200 | Ffar4 | free fatty acid receptor 4 |
| chr15 | 77064718 | 77066621 | 1904 | * | 13,1295 | 2,762564326 | 0,980611 | 2,817187 | 0,004845 | 0,838625535 | 0,918255 | Intron (ENSMUST0000 | 15 | 77001151 | 77087636 | 86486 | 2 | 93686 | ENSMUST00000226877.2 | 21015 | ENSMUSG00000033565 | Rbfox2 | RNA binding protein, fox-1 homolog (C. elegans) 2 |
| chr2 | 1,18E+08 | 1,18E+08 | 2346 | * | 16,34168 | 2,762680667 | 0,862351 | 3,203661 | 0,001357 | 0,451585113 | 2,318434 | Intron (ENSMUST0000 | 2 | 1,18E+08 | 1,18E+08 | 96310 | 2 | 381413 | ENSMUST00000039160.3 | 30264 | ENSMUSG00000040133 | Gpr176 | G protein-coupled receptor 176 |
| chr2 | 1,32E+08 | 1,32E+08 | 2015 | * | 17,88732 | 2,763033197 | 0,788836 | 3,502673 | 0,000461 | 0,294606173 | 2,154504 | Intron (ENSMUST0000 | 2 | 1,31E+08 | 1,32E+08 | 68942 | 2 | 1,19E+08 | ENSMUST00000138080.2 | 20359 | | Gm14280 | predicted gene 14280 |
| chr2 | 1,7E+08 | 1,7E+08 | 1216 | * | 10,6894 | 2,7637172 | 1,062157 | 2,601986 | 0,009269 | 0,893166641 | 1,2763 | Promoter (<=1kb) | 2 | 1,7E+08 | 1,7E+08 | 24075 | 1 | 73303 | ENSMUST00000131931.2 | -323 | ENSMUSG00000086999 | Bcas1os2 | brain enriched myelin associated protein 1, opposite strand 2 |
| chr18 | 75939290 | 75941145 | 1856 | * | 20,28419 | 2,764277656 | 0,956722 | 2,889322 | 0,003861 | 0,617489886 | 2,663619 | Distal Intergenic | 18 | 75953249 | 76281635 | 328387 | 1 | 207259 | ENSMUST00000058997.15 | -12104 | ENSMUSG00000044646 | Zbtb7c | zinc finger and BTB domain containing 7C |
| chr3 | 34431301 | 34432618 | 1318 | * | 10,69163 | 2,76438264 | 0,999803 | 2,764927 | 0,005694 | 0,902917968 | 0,738441 | Intron (ENSMUST0000 | 3 | 34352321 | 34405861 | 53541 | 2 | 1,03E+08 | ENSMUST00000174005.3 | -25440 | ENSMUSG00000092196 | Gm38505 | predicted gene, 38505 |
| chr13 | 97718089 | 97720376 | 2288 | * | 14,865 | 2,76458377 | 0,973896 | 2,838684 | 0,00453 | 0,752939853 | 1,37683 | Exon (ENSMUST00000 | 13 | 97696479 | 97720504 | 24026 | 1 | 328314 | ENSMUST00000210827.2 | 21610 | ENSMUSG00000110235 | Gm5086 | predicted gene 5086 |
| chr15 | 55702319 | 55703838 | 1520 | * | 14,84236 | 2,764692694 | 0,98714 | 2,80071 | 0,005099 | 0,797736349 | 1,214904 | Intron (ENSMUST0000 | 15 | 55655334 | 55769706 | 114373 | 2 | 20649 | ENSMUST00000140574.2 | 65868 | ENSMUSG00000060429 | Sntb1 | syntrophin, basic 1 |
| chr5 | 53666207 | 53667582 | 1376 | * | 22,28939 | 2,764706478 | 0,749701 | 3,687746 | 0,000226 | 0,166376752 | 3,104559 | Intron (ENSMUST0000 | 5 | 53703693 | 53704664 | 972 | 2 | 1,05E+08 | ENSMUST00000202154.2 | 37082 | | Gm40304 | predicted gene, 40304 |
| chr11 | 96284978 | 96286499 | 1522 | * | 12,08112 | 2,764722102 | 1,02044 | 2,709344 | 0,006742 | 0,927963368 | 0,738441 | Distal Intergenic | 11 | 96257084 | 96258047 | 964 | 1 | 15407 | ENSMUST00000123805.2 | 27894 | ENSMUSG00000018973 | Hoxb1 | homeobox B1 |
| chr6 | 30869238 | 30870588 | 1351 | * | 12,03164 | 2,764811848 | 0,968248 | 2,85548 | 0,004297 | 0,756849896 | 1,2763 | Promoter (2-3kb) | 6 | 30835738 | 30873361 | 37624 | 2 | 54160 | ENSMUST00000122833.8 | 2773 | ENSMUSG00000025607 | Copg2 | coatomer protein complex, subunit gamma 2 |
| chr4 | 1,35E+08 | 1,35E+08 | 1522 | * | 22,64568 | 2,764990844 | 0,754985 | 3,662312 | 0,00025 | 0,191521485 | 2,67243 | Promoter (<=1kb) | 4 | 1,35E+08 | 1,35E+08 | 50587 | 2 | 66146 | ENSMUST00000030628.15 | -17 | ENSMUSG00000028826 | Maco1 | macoilin 1 |
| chr15 | 5920544 | 5922526 | 1983 | * | 17,20078 | 2,765480628 | 0,88991 | 3,107597 | 0,001886 | 0,502272318 | 2,386008 | Distal Intergenic | 15 | 6180698 | 6180829 | 132 | 2 | 1,15E+08 | ENSMUST00000240204.1 | 258303 | ENSMUSG00000118903 | Gm23139 | predicted gene, 23139 |
| chr4 | 9668532 | 9671182 | 2651 | * | 46,729379 | 2,765716755 | 0,5110486 | 5,4118465 | 6,238E-08 | 0,000527818 | 2,3776742 | Promoter (<=1kb) | 4 | 9449079 | 9669162 | 220084 | 2 | 65973 | ENSMUST00000078139.13 | 0 | ENSMUSG00000028207 | Asph | aspartate-beta-hydroxylase |
| chr2 | 167615206 | 167617552 | 2347 | * | 40,123286 | 2,766002416 | 0,5595327 | 4,9434149 | 7,677E-07 | 0,003016716 | 2,9254973 | Promoter (1-2kb) | 2 | 167619387 | 167624489 | 5103 | 1 | 100048985 | ENSMUST00000139927.2 | -1835 | | Gm14321 | predicted gene 14321 |
| chr13 | 38267709 | 38269123 | 1415 | * | 10,03995 | 2,766422195 | 1,015525 | 2,724131 | 0,006447 | 0,847020973 | 1,173325 | Distal Intergenic | 13 | 38241191 | 38245289 | 4099 | 1 | 71340 | ENSMUST00000225174.2 | 26518 | ENSMUSG00000021428 | Riok1 | RIO kinase 1 |

| | | | | | | | | | | | | | | | | | | | | | | | |
|---|---|---|---|---|---|---|---|---|---|---|---|---|---|---|---|---|---|---|---|---|---|---|---|
| chr7 | 1,27E+08 | 1,27E+08 | 1867 | * | 15,60227 | 2,766508597 | 0,834908 | 3,313547 | 0,000921 | 0,495652322 | 1,214904 | Distal Intergenic | 7 | 1,27E+08 | 1,27E+08 | 983 | 2 | 70233 | ENSMUST00000205355.2 | 24142 | ENSMUSG00000042502 | Cd2bp2 | CD2 cytoplasmic tail binding protein 2 |
| chr11 | 94568233 | 94570352 | 2120 | * | 16,50646 | 2,766602488 | 0,91829 | 3,012776 | 0,002589 | 0,675437713 | 1,2763 | Promoter (<=1kb) | 11 | 94554677 | 94568341 | 13665 | 2 | 217119 | ENSMUST00000116349.9 | 0 | ENSMUSG00000020868 | Xylt2 | xylosyltransferase II |
| chr16 | 68483841 | 68484683 | 843 | * | 11,08586 | 2,766755974 | 1,013606 | 2,729616 | 0,006341 | 0,917631345 | 0,744375 | Distal Intergenic | 16 | 66459076 | 67417796 | 958721 | 2 | 239857 | ENSMUST00000120898.8 | -1066045 | ENSMUSG00000064115 | Cadm2 | cell adhesion molecule 2 |
| chr5 | 49462197 | 49463647 | 1451 | * | 12,36865 | 2,767571567 | 0,945157 | 2,928162 | 0,00341 | 0,74907225 | 1,075735 | Intron (ENSMUST0000( | 5 | 48546854 | 49443001 | 896148 | 2 | 80334 | ENSMUST00000175660.5 | -19196 | ENSMUSG00000029088 | Kcnip4 | Kv channel interacting protein 4 |
| chr4 | 1,19E+08 | 1,19E+08 | 1942 | * | 17,20966 | 2,76797566 | 0,877125 | 3,155738 | 0,001601 | 0,474900193 | 2,386008 | Intron (ENSMUST0000( | 4 | 1,19E+08 | 1,19E+08 | 33224 | 1 | 230700 | ENSMUST00000160219.2 | 5648 | ENSMUSG00000032998 | Foxj3 | forkhead box J3 |
| chr19 | 35591932 | 35593445 | 1514 | * | 15,40428 | 2,768147538 | 0,846167 | 3,271398 | 0,00107 | 0,498636665 | 1,37683 | Distal Intergenic | 19 | 35737333 | 35737465 | 133 | 2 | 1,15E+08 | ENSMUST00000104017.3 | 144020 | ENSMUSG00000077205 | Gm25611 | predicted gene, 25611 |
| chr7 | 1,12E+08 | 1,12E+08 | 1035 | * | 9,284761 | 2,769194142 | 1,066624 | 2,596222 | 0,009426 | 0,903608597 | 1,211315 | Distal Intergenic | 7 | 1,12E+08 | 1,12E+08 | 87881 | 1 | 74996 | ENSMUST00000106653.4 | -3353 | ENSMUSG00000059263 | Usp47 | ubiquitin specific peptidase 47 |
| chr3 | 1,23E+08 | 1,23E+08 | 1290 | * | 10,7053 | 2,769476331 | 0,979533 | 2,827344 | 0,004694 | 0,872043312 | 0,738441 | Intron (ENSMUST0000( | 3 | 1,23E+08 | 1,23E+08 | 32523 | 2 | 71647 | ENSMUST00000200530.2 | -39057 | ENSMUSG00000106583 | 4930447N08Rik | RIKEN cDNA 4930447N08 gene |
| chr4 | 56393740 | 56395040 | 1301 | * | 17,26231 | 2,769890677 | 0,797112 | 3,474909 | 0,000511 | 0,294632081 | 2,386008 | Distal Intergenic | 4 | 56678904 | 56679035 | 132 | 2 | 1,15E+08 | ENSMUST00000104475.2 | 283995 | ENSMUSG00000077663 | Gm26144 | predicted gene, 26144 |
| chr17 | 47596779 | 47598606 | 1828 | * | 15,4926 | 2,769913075 | 0,959586 | 2,88657 | 0,003895 | 0,658015022 | 2,141612 | 5' UTR | 17 | 47604995 | 47670388 | 65394 | 1 | 224829 | ENSMUST00000190020.4 | -6389 | ENSMUSG00000064043 | Trerf1 | transcriptional regulating factor 1 |
| chr13 | 35056971 | 35059304 | 2334 | * | 21,05751 | 2,770020237 | 0,932739 | 2,969771 | 0,00298 | 0,565091829 | 2,67243 | Promoter (<=1kb) | 13 | 35059285 | 35090047 | 30763 | 1 | 19134 | ENSMUST00000077853.5 | 0 | ENSMUSG00000021413 | Prpf4b | pre-mRNA processing factor 4B |
| chr5 | 45056908 | 45059056 | 2149 | * | 21,43103 | 2,770565036 | 0,782199 | 3,54202 | 0,000397 | 0,214556109 | 3,471678 | Distal Intergenic | 5 | 45123542 | 45136147 | 12606 | 1 | 75783 | ENSMUST00000199309.5 | -64486 | ENSMUSG00000062496 | 4930431F12Rik | RIKEN cDNA 4930431F12 gene |
| chr19 | 8949603 | 8951174 | 1572 | * | 10,85646 | 2,770883896 | 1,075683 | 2,575929 | 0,009997 | 0,986692175 | 0,738441 | Promoter (<=1kb) | 19 | 8949768 | 8955843 | 6076 | 1 | 67160 | ENSMUST00000135025.2 | 0 | ENSMUSG00000071644 | Eef1g | eukaryotic translation elongation factor 1 gamma |
| chr9 | 61843749 | 61845774 | 2026 | * | 20,48861 | 2,771109321 | 0,81439 | 3,40268 | 0,000667 | 0,311505128 | 2,663619 | Promoter (<=1kb) | 9 | 61830133 | 61844058 | 13926 | 2 | 71819 | ENSMUST00000216717.2 | 0 | ENSMUSG00000032254 | Kif23 | kinesin family member 23 |
| chr1 | 1,53E+08 | 1,53E+08 | 1006 | * | 11,04624 | 2,772743073 | 1,064573 | 2,604558 | 0,009199 | 0,974006322 | 0,744375 | Promoter (<=1kb) | 1 | 1,53E+08 | 1,53E+08 | 63777 | 2 | 226517 | ENSMUST00000111836.4 | -417 | ENSMUSG00000042772 | Smg7 | SMG7 nonsense mediated mRNA decay factor |
| chr1 | 56693636 | 56694781 | 1146 | * | 10,97159 | 2,77328014 | 1,004499 | 2,760858 | 0,005765 | 0,903294867 | 0,744375 | Distal Intergenic | 1 | 56675203 | 56676594 | 1392 | 2 | 71066 | ENSMUST00000062085.6 | -17042 | ENSMUSG00000045336 | Hsfy2 | heat shock transcription factor, Y-linked 2 |
| chr5 | 1,45E+08 | 1,45E+08 | 1447 | * | 17,12927 | 2,7737711 | 0,842378 | 3,292786 | 0,000992 | 0,413326369 | 2,141612 | Promoter (<=1kb) | 5 | 1,45E+08 | 1,45E+08 | 3313 | 1 | 100683 | ENSMUST00000143357.2 | -834 | ENSMUSG00000045482 | Trrap | transformation/transcription domain-associated protein |
| chr11 | 16457626 | 16459266 | 1641 | * | 15,93026 | 2,774750703 | 0,971213 | 2,856996 | 0,004277 | 0,671850201 | 2,141612 | Promoter (<=1kb) | 11 | 16450530 | 16458484 | 7955 | 2 | 20335 | ENSMUST00000109643.8 | 0 | ENSMUSG00000078974 | Sec61g | SEC61, gamma subunit |
| chr1 | 74125112 | 74126884 | 1773 | * | 23,41038 | 2,774780068 | 0,755548 | 3,672538 | 0,00024 | 0,17823463 | 2,880017 | Intron (ENSMUST0000( | 1 | 73949391 | 74130510 | 181120 | 2 | 21961 | ENSMUST00000187584.7 | 3626 | ENSMUSG00000055322 | Tns1 | tensin 1 |
| chr11 | 1,02E+08 | 1,02E+08 | 1443 | * | 13,806 | 2,774858833 | 0,981511 | 2,827131 | 0,004697 | 0,833926809 | 0,918255 | Promoter (1-2kb) | 11 | 1,02E+08 | 1,02E+08 | 8352 | 1 | 51799 | ENSMUST00000107102.8 | -1230 | ENSMUSG00000006575 | Rundc3a | RUN domain containing 3A |
| chr11 | 86374644 | 86376180 | 1537 | * | 23,18324 | 2,775462866 | 0,742385 | 3,738574 | 0,000185 | 0,152532781 | 2,951062 | Promoter (<=1kb) | 11 | 86375483 | 86389851 | 14369 | 1 | 76892 | ENSMUST00000020827.7 | 0 | ENSMUSG00000020521 | Rnft1 | ring finger protein, transmembrane 1 |
| chr7 | 44759101 | 44760637 | 1537 | * | 11,7696 | 2,775483662 | 1,03378 | 2,684791 | 0,007258 | 0,937850254 | 0,744375 | Distal Intergenic | 7 | 44742414 | 44753275 | 10862 | 2 | 14132 | ENSMUST00000003512.9 | -5826 | ENSMUSG00000003420 | Fcgrt | Fc fragment of IgG receptor and transporter |
| chr4 | 1,3E+08 | 1,3E+08 | 1447 | * | 12,22588 | 2,775651 | 1,068842 | 2,596876 | 0,009408 | 0,92207189 | 1,075735 | Distal Intergenic | 4 | 1,3E+08 | 1,3E+08 | 9404 | 2 | 94242 | ENSMUST00000105998.8 | -4274 | ENSMUSG00000028776 | Tinagl1 | tubulointerstitial nephritis antigen-like 1 |
| chr1 | 1,36E+08 | 1,36E+08 | 1056 | * | 9,658214 | 2,775815005 | 1,075003 | 2,582147 | 0,009819 | 0,883970984 | 1,449706 | Intron (ENSMUST0000( | 1 | 1,36E+08 | 1,36E+08 | 5036 | 1 | 13007 | ENSMUST00000246868.1 | -12023 | ENSMUSG00000026421 | Csrp1 | cysteine and glycine-rich protein 1 |
| chr19 | 49779103 | 49780812 | 1710 | * | 10,78569 | 2,776693616 | 0,955544 | 2,905878 | 0,003662 | 0,790460916 | 0,918255 | Distal Intergenic | 19 | 50139405 | 50241116 | 101712 | 2 | 58178 | ENSMUST00000168357.9 | 460304 | ENSMUSG00000043531 | Sorcs1 | sortilin-related VPS10 domain containing receptor 1 |
| chr1 | 38582921 | 38584287 | 1367 | * | 13,55694 | 2,778279236 | 1,021795 | 2,719019 | 0,006548 | 0,888080529 | 0,938682 | Intron (ENSMUST0000( | 1 | 38216407 | 38666266 | 449860 | 2 | 16764 | ENSMUST00000095027.9 | 81979 | ENSMUSG00000037138 | Aff3 | AF4/FMR2 family, member 3 |
| chr14 | 58131888 | 58134310 | 2423 | * | 18,32109 | 2,77841804 | 0,821245 | 3,383178 | 0,000717 | 0,328946709 | 2,507013 | Distal Intergenic | 14 | 58070160 | 58127733 | 57574 | 2 | 75965 | ENSMUST00000089473.5 | -4155 | ENSMUSG00000021969 | Zdhhc20 | zinc finger, DHHC domain containing 20 |
| chr13 | 54718837 | 54720969 | 2133 | * | 17,82924 | 2,778870961 | 0,810197 | 3,429871 | 0,000604 | 0,307682443 | 2,507013 | Promoter (1-2kb) | 13 | 54722828 | 54728941 | 6114 | 1 | 56795 | ENSMUST00000026988.11 | -1859 | ENSMUSG00000025870 | Arl10 | ADP-ribosylation factor-like 10 |
| chr14 | 26692523 | 26694450 | 1928 | * | 20,07837 | 2,779117449 | 0,771451 | 3,602454 | 0,000315 | 0,230379597 | 2,386008 | Promoter (<=1kb) | 14 | 26640945 | 26693189 | 52245 | 2 | 72993 | ENSMUST00000036570.5 | 0 | ENSMUSG00000040760 | Appl1 | adaptor protein, phosphotyrosine interaction, PH domain and leucine zipper containing 1 |
| chr4 | 1,35E+08 | 1,35E+08 | 1680 | * | 20,24277 | 2,779347714 | 0,79408 | 3,500087 | 0,000465 | 0,268684353 | 2,663619 | Distal Intergenic | 4 | 1,35E+08 | 1,35E+08 | 5069 | 1 | 68592 | ENSMUST00000144037.2 | 54822 | ENSMUSG00000028821 | Syf2 | SYF2 homolog, RNA splicing factor (S. cerevisiae) |
| chr2 | 71474170 | 71475725 | 1556 | * | 22,9382 | 2,779757191 | 0,761682 | 3,649497 | 0,000263 | 0,190625259 | 2,858511 | Distal Intergenic | 2 | 71549682 | 71561320 | 11639 | 1 | 381371 | ENSMUST00000245331.1 | -73957 | ENSMUSG00000064263 | Platr26 | pluripotency associated transcript 26 |
| chr5 | 56142996 | 56144774 | 1779 | * | 14,3912 | 2,780426094 | 0,896267 | 3,102228 | 0,001921 | 0,681119785 | 0,918255 | Distal Intergenic | 5 | 55460762 | 55460923 | 162 | 2 | 1,15E+08 | ENSMUST00000083861.3 | -682073 | ENSMUSG00000065795 | Gm25513 | predicted gene, 25513 |
| chr13 | 96806373 | 96808065 | 1693 | * | 12,23582 | 2,780497996 | 1,004086 | 2,769184 | 0,00562 | 0,806557336 | 1,2763 | Promoter (<=1kb) | 13 | 96785475 | 96807444 | 21970 | 2 | 15357 | ENSMUST00000022176.15 | 0 | ENSMUSG00000021670 | Hmgcr | 3-hydroxy-3-methylglutaryl-Coenzyme A reductase |
| chr10 | 80765193 | 80767644 | 2452 | * | 25,06345 | 2,781225917 | 0,809949 | 3,433827 | 0,000595 | 0,290624195 | 2,858511 | Promoter (<=1kb) | 10 | 80765907 | 80768038 | 2132 | 1 | 17873 | ENSMUST00000015456.10 | 0 | ENSMUSG00000015312 | Gadd45b | growth arrest and DNA-damage-inducible 45 beta |
| chr10 | 30602398 | 30604111 | 1714 | * | 12,40446 | 2,78123883 | 1,02074 | 2,724729 | 0,006435 | 0,846832214 | 1,173325 | Intron (ENSMUST0000( | 10 | 30570161 | 30598701 | 28541 | 2 | 211329 | ENSMUST00000217398.2 | -3697 | ENSMUSG00000039697 | Ncoa7 | nuclear receptor coactivator 7 |
| chr11 | 1,01E+08 | 1,01E+08 | 2693 | * | 21,71719 | 2,781504654 | 0,766516 | 3,628762 | 0,000285 | 0,198692711 | 2,880017 | 3' UTR | 11 | 1,01E+08 | 1,01E+08 | 4926 | 2 | 80860 | ENSMUST00000017891.14 | -9378 | ENSMUSG00000017747 | Ghdc | GH3 domain containing |
| chrX | 35818101 | 35819192 | 1092 | * | 12,1641 | 2,781514615 | 0,959438 | 2,899109 | 0,003742 | 0,830950621 | 0,744375 | Distal Intergenic | 20 | 35861859 | 35869732 | 7874 | 1 | 53328 | ENSMUST00000073339.7 | -42667 | ENSMUSG00000006373 | Pgrmc1 | progesterone receptor membrane component 1 |
| chr9 | 61927069 | 61929220 | 2152 | * | 22,79283 | 2,781822431 | 0,814179 | 3,41672 | 0,000634 | 0,306095127 | 2,67243 | Exon (ENSMUST00020: | 9 | 61860763 | 61934083 | 73321 | 2 | 74090 | ENSMUST00000135050.2 | 4863 | ENSMUSG00000032278 | Paqr5 | progestin and adipoQ receptor family member V |
| chr10 | 13713110 | 13714373 | 1264 | * | 15,99101 | 2,782112861 | 0,84069 | 3,309321 | 0,000935 | 0,406085428 | 2,141612 | Intron (ENSMUST0000( | 10 | 13529332 | 13720030 | 190699 | 2 | 66253 | ENSMUST00000162174.8 | 5657 | ENSMUSG00000019806 | Aig1 | androgen-induced 1 |
| chr2 | 5746814 | 5747668 | 855 | * | 10,61563 | 2,783113937 | 1,070533 | 2,599746 | 0,009329 | 0,974266082 | 0,744375 | Distal Intergenic | 2 | 5298268 | 5719326 | 421059 | 2 | 227541 | ENSMUST00000044009.14 | -27488 | ENSMUSG00000039145 | Camk1d | calcium/calmodulin-dependent protein kinase ID |
| chr4 | 1,31E+08 | 1,31E+08 | 1888 | * | 15,8322 | 2,783330177 | 0,863839 | 3,222048 | 0,001273 | 0,454149353 | 2,141612 | Distal Intergenic | 4 | 1,31E+08 | 1,31E+08 | 5406 | 1 | 20970 | ENSMUST00000140623.2 | 16256 | ENSMUSG00000025743 | Sdc3 | syndecan 3 |
| chr9 | 58765455 | 58766553 | 1099 | * | 15,71824 | 2,783371384 | 0,974664 | 2,855724 | 0,004294 | 0,672298194 | 2,141612 | Exon (ENSMUST00000( | 9 | 58795769 | 58801217 | 5449 | 2 | 18007 | ENSMUST00000215165.2 | 34664 | ENSMUSG00000032340 | Neo1 | neogenin |
| chr13 | 63713804 | 63718472 | 4669 | * | 42,903468 | 2,784379374 | 0,5509833 | 5,0534732 | 4,338E-07 | 0,002004214 | 2,7872553 | Promoter (<=1kb) | 13 | 63656142 | 63713549 | 57408 | 2 | 19206 | ENSMUST00000021921.11 | -255 | ENSMUSG00000021466 | Ptch1 | patched 1 |
| chr4 | 1,36E+08 | 1,36E+08 | 1035 | * | 10,22677 | 2,784689651 | 1,039459 | 2,67898 | 0,007385 | 0,868766096 | 1,173325 | Promoter (<=1kb) | 4 | 1,36E+08 | 1,36E+08 | 3663 | 2 | 67025 | ENSMUST00000102536.11 | 0 | ENSMUSG00000059291 | Rpl11 | ribosomal protein L11 |
| chr11 | 30161807 | 30163456 | 1650 | * | 12,43069 | 2,784862541 | 1,072805 | 2,595871 | 0,009435 | 0,947773602 | 0,918255 | Intron (ENSMUST0000( | 11 | 30049396 | 30169772 | 120377 | 2 | 20742 | ENSMUST00000011877.13 | 6316 | ENSMUSG00000020315 | Sptbn1 | spectrin beta, non-erythrocytic 1 |
| chr7 | 100019547 | 100022665 | 3119 | * | 42,588017 | 2,785313896 | 0,5390754 | 5,1668351 | 2,381E-07 | 0,001425027 | 2,3383871 | Promoter (<=1kb) | 7 | 100021440 | 100118668 | 97229 | 1 | 277939 | ENSMUST00000098259.11 | 0 | ENSMUSG00000047248 | C2cd3 | C2 calcium-dependent domain containing 3 |
| chr16 | 23338323 | 23339862 | 1540 | * | 13,67364 | 2,785669088 | 0,938062 | 2,969601 | 0,002982 | 0,693610666 | 1,2763 | Promoter (<=1kb) | 16 | 23268167 | 23339565 | 71399 | 2 | 17174 | ENSMUST00000089883.7 | 0 | ENSMUSG00000022887 | Masp1 | mannan-binding lectin serine peptidase 1 |
| chr16 | 4866853 | 4868312 | 1460 | * | 12,09101 | 2,785677103 | 1,023178 | 2,722574 | 0,006478 | 0,889546768 | 0,918255 | Promoter (<=1kb) | 16 | 4867921 | 4904153 | 36233 | 1 | 170644 | ENSMUST00000230703.2 | 0 | ENSMUSG00000039473 | Ubn1 | ubinuclein 1 |
| chr2 | 13067289 | 13068755 | 1467 | * | 9,984329 | 2,785949904 | 1,023624 | 2,721652 | 0,006496 | 0,813409596 | 1,425929 | Distal Intergenic | 2 | 13008268 | 13016617 | 8350 | 2 | 227580 | ENSMUST00000061545.7 | -50672 | ENSMUSG00000049630 | C1ql3 | C1q-like 3 |
| chr8 | 1,23E+08 | 1,23E+08 | 1010 | * | 13,12847 | 2,786240164 | 1,00744 | 2,765664 | 0,005681 | 0,902613747 | 0,738441 | Promoter (<=1kb) | 8 | 1,23E+08 | 1,23E+08 | 1366 | 2 | 68718 | ENSMUST00000136356.2 | 0 | ENSMUSG00000014470 | Rnf166 | ring finger protein 166 |
| chr9 | 54322524 | 54324128 | 1605 | * | 11,88579 | 2,78639182 | 1,048453 | 2,657622 | 0,007869 | 0,953262301 | 0,738441 | Exon (ENSMUST00000: | 9 | 54272544 | 54352704 | 80161 | 2 | 235380 | ENSMUST00000123709.8 | 28576 | ENSMUSG00000041268 | Dmxl2 | Dmx-like 2 |
| chr11 | 6198452 | 6200281 | 1830 | * | 15,20274 | 2,786431936 | 1,031309 | 2,701841 | 0,006896 | 0,852340527 | 1,214904 | Distal Intergenic | 11 | 6215307 | 6216392 | 1086 | 2 | 52513 | ENSMUST00000139073.2 | 16111 | ENSMUSG00000004393 | Ddx56 | DEAD box helicase 56 |
| chr11 | 30899503 | 30901452 | 1950 | * | 11,42394 | 2,786516772 | 0,933734 | 2,984272 | 0,002843 | 0,685026746 | 1,2763 | Promoter (1-2kb) | 11 | 30881751 | 30898469 | 16719 | 2 | 66753 | ENSMUST00000129593.9 | -1034 | ENSMUSG00000020311 | Erlec1 | endoplasmic reticulum lectin 1 |
| chr6 | 50216938 | 50217759 | 822 | * | 12,62175 | 2,786699628 | 1,069927 | 2,60457 | 0,009199 | 0,905214371 | 1,173325 | Intron (ENSMUST0000( | 6 | 50177204 | 50238727 | 61524 | 2 | 54722 | ENSMUST00000101405.10 | 20968 | ENSMUSG00000029821 | Gsdme | gasdermin E |
| chr12 | 10286831 | 10288494 | 1664 | * | 13,46868 | 2,787290287 | 1,015404 | 2,745005 | 0,006051 | 0,878632224 | 0,918255 | Distal Intergenic | 12 | 10419973 | 10425935 | 5963 | 1 | 70881 | ENSMUST00000143739.9 | -131479 | ENSMUSG00000020622 | Nt5c1b | 5'-nucleotidase, cytosolic IB |
| chr1 | 1,9E+08 | 1,9E+08 | 2029 | * | 11,82839 | 2,787558901 | 0,977046 | 2,853047 | 0,00433 | 0,855636327 | 0,744375 | Exon (ENSMUST00000( | 1 | 1,9E+08 | 1,9E+08 | 775 | 1 | 19250 | ENSMUST00000148129.2 | -4761 | ENSMUSG00000026604 | Ptpn14 | protein tyrosine phosphatase, non-receptor type 14 |
| chr15 | 1,03E+08 | 1,03E+08 | 1455 | * | 15,94532 | 2,787562836 | 0,896979 | 3,107723 | 0,001885 | 0,502272318 | 2,386008 | Distal Intergenic | 15 | 1,03E+08 | 1,03E+08 | 2931 | 2 | 67488 | ENSMUST00000168112.8 | 10323 | ENSMUSG00000023055 | Calcoco1 | calcium binding and coiled coil domain 1 |
| chr19 | 57038286 | 57039599 | 1314 | * | 12,63603 | 2,788585485 | 1,023242 | 2,725246 | 0,006425 | 0,846686317 | 1,173325 | Promoter (2-3kb) | 19 | 57027879 | 57035670 | 7792 | 2 | 226251 | ENSMUST00000137389.8 | -2616 | ENSMUSG00000025085 | Ablim1 | actin-binding LIM protein 1 |
| chr18 | 26299078 | 26300101 | 1024 | * | 11,15097 | 2,789383624 | 0,974874 | 2,861275 | 0,004219 | 0,770673161 | 1,173325 | Intron (ENSMUST0000( | 18 | 26145360 | 26348706 | 203347 | 1 | 1,03E+08 | ENSMUST00000234272.2 | 153718 | | Gm33228 | predicted gene, 33228 |
| chr12 | 88652019 | 88652843 | 825 | * | 10,06285 | 2,789565071 | 1,04401 | 2,671972 | 0,007541 | 0,865306051 | 1,211315 | Intron (ENSMUST0000( | 12 | 88689646 | 89228341 | 538696 | 1 | 18191 | ENSMUST00000167734.8 | -36803 | ENSMUSG00000066392 | Nrxn3 | neurexin III |
| chr5 | 23307943 | 23308982 | 1040 | * | 10,10779 | 2,789964 | 1,063378 | 2,623681 | 0,008699 | 0,908469938 | 1,086118 | Promoter (2-3kb) | 5 | 23305550 | 23480586 | 175037 | 1 | 269629 | ENSMUST00000196406.2 | 2393 | ENSMUSG00000106379 | Lhfpl3 | lipoma HMGIC fusion partner-like 3 |
| chr11 | 75385651 | 75387622 | 1972 | * | 24,01089 | 2,790094786 | 0,704303 | 3,961496 | 7,45E-05 | 0,086053637 | 2,951062 | Promoter (2-3kb) | 11 | 75383072 | 75384761 | 1690 | 1 | 192159 | ENSMUST00000133995.2 | 2579 | ENSMUSG00000020850 | Prpf8 | pre-mRNA processing factor 8 |
| chr17 | 46346641 | 46348862 | 2222 | * | 39,809033 | 2,790238494 | 0,6118966 | 4,5599839 | 5,116E-06 | 0,012970123 | 2,9254973 | Distal Intergenic | 17 | 46329469 | 46343295 | 13827 | 2 | 22339 | ENSMUST00000142351.9 | -3346 | ENSMUSG00000023951 | Vegfa | vascular endothelial growth factor A |
| chr5 | 89063867 | 89064819 | 953 | * | 9,256924 | 2,790684139 | 1,077331 | 2,590369 | 0,009587 | 0,801671497 | 2,237403 | Intron (ENSMUST0000( | 5 | 89082391 | 89102806 | 20416 | 1 | 54403 | ENSMUST00000144713.2 | -17572 | ENSMUSG00000060961 | Slc4a4 | solute carrier family 4 (anion exchanger), member 4 |
| chr5 | 8672660 | 8673780 | 1121 | * | 11,06059 | 2,790722884 | 1,06064 | 2,63117 | 0,008509 | 0,881370225 | 1,2763 | Promoter (<=1kb) | 5 | 8540334 | 8672952 | 132619 | 2 | 242819 | ENSMUST00000047485.15 | 0 | ENSMUSG00000040570 | Rundc3b | RUN domain containing 3B |
| chr17 | 8820417 | 8821455 | 1039 | * | 11,80309 | 2,791175059 | 0,972605 | 2,869793 | 0,004107 | 0,848080889 | 0,744375 | Intron (ENSMUST0000( | 17 | 8745858 | 9017563 | 271706 | 1 | 23984 | ENSMUST00000233016.2 | 74559 | ENSMUSG00000023868 | Pde10a | phosphodiesterase 10A |
| chr12 | 59266283 | 59267503 | 1221 | * | 16,40106 | 2,792113613 | 0,903761 | 3,089437 | 0,002005 | 0,532496361 | 2,141612 | Promoter (<=1kb) | 12 | 59247441 | 59266511 | 19071 | 2 | 70611 | ENSMUST00000043204.8 | 0 | ENSMUSG00000035329 | Fbxo33 | F-box protein 33 |
| chr13 | 23611179 | 23613184 | 2006 | * | 14,56362 | 2,792179709 | 0,894125 | 3,122808 | 0,001791 | 0,621084104 | 1,214904 | Promoter (1-2kb) | 13 | 23612145 | 23614898 | 2754 | 2 | 320842 | ENSMUST00000241463.1 | 1714 | ENSMUSG00000085024 | C230035I16Rik | RIKEN cDNA C230035I16 gene |
| chr15 | 99889941 | 99891493 | 1553 | * | 16,54398 | 2,792186624 | 0,893872 | 3,123699 | 0,001786 | 0,61075929 | 1,2763 | 3' UTR | 15 | 99898435 | 99903183 | 4749 | 1 | 207214 | ENSMUST00000229426.2 | -6942 | ENSMUSG00000023025 | Larp4 | La ribonucleoprotein domain family, member 4 |
| chr15 | 88748100 | 88750031 | 1932 | * | 21,26277 | 2,792609113 | 0,779617 | 3,582026 | 0,000341 | 0,226640038 | 2,67243 | Promoter (1-2kb) | 15 | 88746389 | 88749929 | 3541 | 1 | 223775 | ENSMUST00000042818.11 | 1711 | ENSMUSG00000035828 | Pim3 | proviral integration site 3 |
| chr8 | 31861771 | 31866311 | 4541 | * | 62,099212 | 2,793580989 | 0,4609606 | 6,0603471 | 1,358E-09 | 1,76557E-05 | 3,439501 | Promoter (2-3kb) | 8 | 31868841 | 31872467 | 3627 | 1 | 74517 | ENSMUST00000209524.2 | -2530 | ENSMUSG00000109915 | 7420700N18Rik | RIKEN cDNA 7420700N18 gene |
| chr10 | 72777826 | 72779237 | 1412 | * | 10,18839 | 2,794884586 | 0,972153 | 2,874943 | 0,004041 | 0,7299084 | 1,425929 | Intron (ENSMUST0000( | 10 | 72776070 | 72785624 | 9555 | 2 | 1,03E+08 | ENSMUST00000219415.2 | 6387 | | Gm34776 | predicted gene, 34776 |
| chr5 | 1,25E+08 | 1,25E+08 | 1232 | * | 14,02924 | 2,7952835 | 0,987247 | 2,831392 | 0,004635 | 0,831521609 | 0,918255 | Exon (ENSMUST00000: | 5 | 1,25E+08 | 1,25E+08 | 28788 | 1 | 56087 | ENSMUST00000196708.2 | 18456 | ENSMUSG00000038011 | Dnah10 | dynein, axonemal, heavy chain 10 |
| chr10 | 77359356 | 77361499 | 2144 | * | 15,63655 | 2,795546008 | 1,043583 | 2,678796 | 0,007389 | 0,841689315 | 1,37683 | Distal Intergenic | 10 | 77366086 | 77401520 | 35435 | 1 | 16414 | ENSMUST00000000299.14 | -4587 | ENSMUSG00000000290 | Itgb2 | integrin beta 2 |
| chr1 | 1,57E+08 | 1,57E+08 | 880 | * | 10,11699 | 2,795584307 | 1,036933 | 2,696012 | 0,007018 | 0,858706485 | 1,173325 | Intron (ENSMUST0000( | 1 | 1,57E+08 | 1,57E+08 | 126546 | 2 | 78255 | ENSMUST00000185198.7 | -57836 | ENSMUSG00000026594 | Ralgps2 | Ral GEF with PH domain and SH3 binding motif 2 |
| chr10 | 79611991 | 79612991 | 1001 | * | 9,676179 | 2,79609238 | 1,054796 | 2,650837 | 0,008029 | 0,849121913 | 1,449706 | Promoter (<=1kb) | 10 | 79613106 | 79618464 | 5359 | 1 | 83554 | ENSMUST00000020575.5 | -115 | ENSMUSG00000020325 | Fstl3 | follistatin-like 3 |
| chr5 | 35854613 | 35856396 | 1784 | * | 15,43944 | 2,796132306 | 0,89473 | 3,125114 | 0,001777 | 0,619158066 | 1,214904 | 3' UTR | 5 | 35854527 | 35873595 | 19069 | 2 | 231147 | ENSMUST00000129664.8 | 17199 | ENSMUSG00000036553 | Sh3tc1 | SH3 domain and tetratricopeptide repeats 1 |
| chr5 | 1,4E+08 | 1,4E+08 | 1413 | * | 14,34181 | 2,79660389 | 1,007533 | 2,775695 | 0,005508 | 0,899471409 | 0,738441 | Promoter (<=1kb) | 5 | 1,4E+08 | 1,4E+08 | 24033 | 1 | 27979 | ENSMUST00000100507.8 | 0 | ENSMUSG00000056076 | Eif3b | eukaryotic translation initiation factor 3, subunit B |
| chr13 | 1,09E+08 | 1,09E+08 | 1632 | * | 11,6351 | 2,796906436 | 1,070763 | 2,612068 | 0,009 | 0,939041504 | 0,918255 | Intron (ENSMUST0000( | 13 | 1,08E+08 | 1,09E+08 | 16531 | 1 | 218581 | ENSMUST00000172402.2 | 32324 | ENSMUSG00000021697 | Depdc1b | DEP domain containing 1B |
| chrX | 1,38E+08 | 1,38E+08 | 950 | * | 9,661292 | 2,797367981 | 1,079873 | 2,590459 | 0,009585 | 0,923078468 | 1,086118 | Distal Intergenic | 20 | 1,38E+08 | 1,38E+08 | 7163 | 1 | 75125 | ENSMUST00000033625.2 | 13388 | ENSMUSG00000031270 | 4930513O06Rik | RIKEN cDNA 4930513O06 gene |
| chr4 | 1,17E+08 | 1,17E+08 | 1938 | * | 17,35225 | 2,797601825 | 1,019502 | 2,744087 | 0,006068 | 0,688021707 | 2,663619 | Promoter (<=1kb) | 4 | 1,17E+08 | 1,17E+08 | 1743 | 1 | 70603 | ENSMUST00000135449.2 | 0 | ENSMUSG00000028687 | Mutyh | mutY DNA glycosylase |
| chr16 | 46755139 | 46756549 | 1411 | * | 10,28927 | 2,798150008 | 1,082812 | 2,584152 | 0,009762 | 0,909192827 | 1,211315 | Distal Intergenic | 16 | 46279215 | 46318888 | 39674 | 2 | 58998 | ENSMUST00000133935.2 | -436251 | ENSMUSG00000022656 | Nectin3 | nectin cell adhesion molecule 3 |
| chr5 | 1,43E+08 | 1,43E+08 | 1277 | * | 10,9574 | 2,79887993 | 1,046419 | 2,674722 | 0,007479 | 0,94432823 | 0,744375 | Exon (ENSMUST00000: | 5 | 1,43E+08 | 1,43E+08 | 18064 | 1 | 66913 | ENSMUST00000110731.4 | 13329 | ENSMUSG00000079111 | Kdelr2 | KDEL (Lys-Asp-Glu-Leu) endoplasmic reticulum protein retention receptor 2 |
| chr7 | 16520528 | 16521731 | 1204 | * | 10,83064 | 2,799142276 | 1,065007 | 2,628286 | 0,008582 | 0,932975916 | 0,918255 | Promoter (<=1kb) | 7 | 16520296 | 16527721 | 7426 | 1 | 20514 | ENSMUST00000147814.2 | 232 | ENSMUSG00000001918 | Slc1a5 | solute carrier family 1 (neutral amino acid transporter), member 5 |
| chr18 | 23936211 | 23938370 | 2160 | * | 20,20335 | 2,800352733 | 0,98681 | 2,837784 | 0,004543 | 0,656950206 | 2,507013 | Promoter (<=1kb) | 18 | 23937041 | 24026914 | 89874 | 1 | 212307 | ENSMUST00000025127.5 | 0 | ENSMUSG00000024277 | Mapre2 | microtubule-associated protein, RP/EB family, member 2 |
| chrX | 35815192 | 35817645 | 2454 | * | 13,31337 | 2,80043113 | 0,865787 | 3,234551 | 0,001218 | 0,641208222 | 0,738441 | Distal Intergenic | 20 | 35861859 | 35869732 | 7874 | 1 | 53328 | ENSMUST00000073339.7 | -44214 | ENSMUSG00000006373 | Pgrmc1 | progesterone receptor membrane component 1 |
| chr9 | 65172752 | 65174500 | 1749 | * | 15,98196 | 2,801384973 | 0,824766 | 3,396581 | 0,000682 | 0,328946709 | 2,386008 | Promoter (<=1kb) | 9 | 65173105 | 65187817 | 14713 | 1 | 214425 | ENSMUST00000141382.2 | 0 | ENSMUSG00000042254 | Cilp | cartilage intermediate layer protein, nucleotide pyrophosphohydrolase |
| chr7 | 91507168 | 91508070 | 903 | * | 9,845628 | 2,801395839 | 1,025278 | 2,732328 | 0,006289 | 0,80696549 | 1,425929 | Intron (ENSMUST0000( | 7 | 91360628 | 92096211 | 735584 | 1 | 23859 | ENSMUST00000107193.8 | 146540 | ENSMUSG00000052572 | Dlg2 | discs large MAGUK scaffold protein 2 |
| chr6 | 1,47E+08 | 1,47E+08 | 1189 | * | 14,17291 | 2,803276039 | 0,95635 | 2,931224 | 0,003376 | 0,775744126 | 0,918255 | Exon (ENSMUST00000: | 6 | 1,47E+08 | 1,47E+08 | 11273 | 1 | 232536 | ENSMUST00000137556.3 | 4749 | ENSMUSG00000040112 | Mrps35 | mitochondrial ribosomal protein S35 |
| chr8 | 1,28E+08 | 1,28E+08 | 1626 | * | 32,75616 | 2,803288433 | 0,682342 | 4,108335 | 3,99E-05 | 0,09162632 | 1,405709 | Promoter (<=1kb) | 8 | 1,28E+08 | 1,28E+08 | 12241 | 1 | 414091 | ENSMUST00000212914.2 | 0 | | A630001O12Rik | RIKEN cDNA A630001O12 gene |
| chr15 | 79022128 | 79023424 | 1297 | * | 12,64838 | 2,803903058 | 1,032164 | 2,716529 | 0,006597 | 0,863136415 | 1,075735 | Promoter (<=1kb) | 15 | 79021620 | 79023475 | 1856 | 2 | 27660 | ENSMUST00000191006.2 | 51 | ENSMUSG00000033029 | 1700088E04Rik | RIKEN cDNA 1700088E04 gene |
| chr16 | 38163114 | 38164844 | 1731 | * | 12,90613 | 2,804316482 | 0,94957 | 2,953248 | 0,003144 | 0,705594569 | 1,2763 | Promoter (<=1kb) | 16 | 38118116 | 38162505 | 44390 | 2 | 320214 | ENSMUST00000023501.16 | -609 | ENSMUSG00000022805 | Cfap91 | cilia and flagella associated protein 91 |
| chr17 | 32538784 | 32541155 | 2372 | * | 21,69158 | 2,804699019 | 0,772502 | 3,630667 | 0,000283 | 0,189492453 | 3,104559 | Promoter (<=1kb) | 17 | 32522646 | 32540159 | 17514 | 2 | 56399 | ENSMUST00000002699.7 | 0 | ENSMUSG00000024045 | Akap8 | A kinase (PRKA) anchor protein 8 |
| chr3 | 1,46E+08 | 1,46E+08 | 1230 | * | 12,83856 | 2,804812886 | 1,053863 | 2,661458 | 0,00778 | 0,916531797 | 0,918255 | Promoter (2-3kb) | 3 | 1,46E+08 | 1,46E+08 | 12879 | 2 | 70285 | ENSMUST00000196909.2 | 2292 | ENSMUSG00000028187 | Rpf1 | ribosome production factor 1 homolog |
| chr1 | 1,35E+08 | 1,35E+08 | 2260 | * | 18,40192 | 2,805944709 | 0,836129 | 3,355874 | 0,000791 | 0,335241819 | 2,663619 | Exon (ENSMUST00000( | 1 | 1,35E+08 | 1,35E+08 | 87354 | 2 | 329251 | ENSMUST00000156348.8 | -31323 | ENSMUSG00000073557 | Ppp1r12b | protein phosphatase 1, regulatory subunit 12B |
| chr13 | 52032361 | 52034031 | 1671 | * | 14,86663 | 2,805963559 | 1,057766 | 2,652727 | 0,007984 | 0,854774997 | 1,37683 | Distal Intergenic | 13 | 52000780 | 52002260 | 1481 | 1 | 23882 | ENSMUST00000132480.2 | 31581 | ENSMUSG00000021453 | Gadd45g | growth arrest and DNA-damage-inducible 45 gamma |
| chr4 | 1,31E+08 | 1,31E+08 | 2638 | * | 21,37263 | 2,806119141 | 0,781706 | 3,589739 | 0,000331 | 0,213080468 | 2,951933 | Intron (ENSMUST0000( | 4 | 1,31E+08 | 1,31E+08 | 7122 | 1 | 20970 | ENSMUST00000152591.8 | -4891 | ENSMUSG00000025743 | Sdc3 | syndecan 3 |
| chr18 | 76372970 | 76374451 | 1482 | * | 12,66672 | 2,806579983 | 0,952907 | 2,945283 | 0,003227 | 0,711174281 | 1,2763 | Promoter (<=1kb) | 18 | 76374651 | 76444034 | 69384 | 1 | 17126 | ENSMUST00000168423.9 | -200 | ENSMUSG00000024563 | Smad2 | SMAD family member 2 |
| chr5 | 98301072 | 98302080 | 1009 | * | 10,07118 | 2,80738016 | 1,081203 | 2,596535 | 0,009417 | 0,908787948 | 1,173325 | Exon (ENSMUST00000: | 5 | 98315057 | 98335313 | 20257 | 1 | 77630 | ENSMUST00000210477.2 | -12977 | ENSMUSG00000035456 | Prdm8 | PR domain containing 8 |
| chr12 | 85293735 | 85295398 | 1664 | * | 16,60822 | 2,807693095 | 0,913986 | 3,071922 | 0,002127 | 0,524368688 | 2,386008 | Promoter (<=1kb) | 12 | 85292697 | 85292759 | 63 | 2 | 1,02E+08 | ENSMUST00000184173.3 | -976 | ENSMUSG00000098263 | Mir6938 | microRNA 6938 |
| chr8 | 1,27E+08 | 1,27E+08 | 1010 | * | 16,53013 | 2,807952583 | 0,809644 | 3,468134 | 0,000524 | 0,297583251 | 2,386008 | Intron (ENSMUST0000( | 8 | 1,27E+08 | 1,27E+08 | 5691 | 2 | 270110 | ENSMUST00000054960.9 | 8921 | ENSMUSG00000051495 | Irf2bp2 | interferon regulatory factor 2 binding protein 2 |

| | | | | | | | | | | | | | | | | | | | | | | | |
|---|---|---|---|---|---|---|---|---|---|---|---|---|---|---|---|---|---|---|---|---|---|---|---|
| chr5 | 34345042 | 34347286 | 2245 | * | 30,88237 | 2,808073141 | 0,675358 | 4,157901 | 3,21E-05 | 0,05285139 | 2,725491 | Promoter (<=1kb) | 5 | 34331231 | 34345054 | 13824 | 2 | 17122 | ENSMUST00000042701.13 | 0 | ENSMUSG00000037235 | Mxd4 | Max dimerization protein 4 |
| chr10 | 83173480 | 83174790 | 1311 | * | 14,34903 | 2,808221943 | 0,83664 | 3,356549 | 0,000789 | 0,512326648 | 0,938682 | Promoter (<=1kb) | 10 | 83066712 | 83173746 | 107035 | 2 | 338365 | ENSMUST00000039956.6 | 0 | ENSMUSG00000034591 | Slc41a2 | solute carrier family 41, member 2 |
| chr17 | 45367375 | 45368957 | 1583 | * | 15,20678 | 2,808426975 | 0,913565 | 3,07414 | 0,002111 | 0,54122637 | 2,141612 | Intron (ENSMUST00000 | 17 | 45190657 | 45191182 | 526 | 2 | 75341 | ENSMUST00000233907.2 | -176193 | | 4930564C03Rik | RIKEN cDNA 4930564C03 gene |
| chr15 | 76167053 | 76169175 | 2123 | * | 21,6975 | 2,810790129 | 0,839941 | 3,346415 | 0,000819 | 0,326089819 | 2,951933 | 3' UTR | 15 | 76175079 | 76175168 | 90 | 2 | 1,01E+08 | ENSMUST00000175428.3 | 5993 | ENSMUSG00000093169 | Mir3079 | microRNA 3079 |
| chr4 | 1,09E+08 | 1,09E+08 | 1612 | * | 10,69014 | 2,811032829 | 0,978788 | 2,871953 | 0,004079 | 0,749550311 | 1,2763 | Intron (ENSMUST00000 | 4 | 1,09E+08 | 1,09E+08 | 69679 | 2 | 100273 | ENSMUST00000139248.8 | -10237 | ENSMUSG00000028559 | Osbpl9 | oxysterol binding protein-like 9 |
| chr5 | 22542152 | 22543482 | 1331 | * | 15,27789 | 2,812175717 | 1,003524 | 2,8023 | 0,005074 | 0,79031971 | 1,2763 | Intron (ENSMUST00000 | 5 | 22090248 | 22549688 | 459441 | 2 | 19699 | ENSMUST00000062372.14 | 6206 | ENSMUSG00000042453 | Reln | reelin |
| chr7 | 97480859 | 97482022 | 1164 | * | 11,20493 | 2,812606166 | 1,052171 | 2,673145 | 0,007514 | 0,873025333 | 1,173325 | Promoter (<=1kb) | 7 | 97480135 | 97515333 | 35199 | 1 | 18479 | ENSMUST00000206351.2 | 724 | ENSMUSG00000030774 | Pak1 | p21 (RAC1) activated kinase 1 |
| chr3 | 1,04E+08 | 1,04E+08 | 2500 | * | 21,09595 | 2,814735836 | 0,734497 | 3,832196 | 0,000127 | 0,120915146 | 2,880017 | Distal Intergenic | 3 | 1,04E+08 | 1,04E+08 | 207116 | 2 | 99470 | ENSMUST00000064371.14 | -18237 | ENSMUSG00000052539 | Magi3 | membrane associated guanylate kinase, WW and PDZ domain containing 3 |
| chr10 | 81028788 | 81029981 | 1194 | * | 15,02843 | 2,815008151 | 0,871839 | 3,228816 | 0,001243 | 0,523637575 | 1,41696 | Promoter (<=1kb) | 10 | 81030443 | 81034275 | 3833 | 1 | 69678 | ENSMUST00000129834.3 | -462 | | Atcayos | ataxia, cerebellar, Cayman type, opposite strand |
| chr11 | 16783136 | 16784179 | 1044 | * | 12,1216 | 2,815434587 | 1,008255 | 2,792384 | 0,005232 | 0,852340527 | 0,918255 | Promoter (2-3kb) | 11 | 16765431 | 16780702 | 15272 | 2 | 319954 | ENSMUST00000150329.2 | -2434 | | Egfros | epidermal growth factor receptor, opposite strand |
| chr19 | 27125790 | 27127586 | 1797 | * | 14,11788 | 2,815503223 | 1,002292 | 2,809065 | 0,004969 | 0,81655291 | 1,075735 | Intron (ENSMUST00000 | 19 | 27079351 | 27092885 | 13535 | 2 | 1,03E+08 | ENSMUST00000235490.2 | -32905 | | Gm35438 | predicted gene, 35438 |
| chr4 | 1,36E+08 | 1,36E+08 | 2588 | * | 20,23569 | 2,81552605 | 0,726529 | 3,875309 | 0,000106 | 0,119105693 | 2,507013 | Distal Intergenic | 4 | 1,36E+08 | 1,36E+08 | 1896 | 1 | 15903 | ENSMUST00000008016.3 | -10457 | ENSMUSG00000007872 | Id3 | inhibitor of DNA binding 3 |
| chr11 | 49082762 | 49084937 | 2176 | * | 13,7632 | 2,815604095 | 0,944363 | 2,981484 | 0,002869 | 0,742014238 | 0,938682 | Distal Intergenic | 11 | 49056412 | 49077986 | 21575 | 2 | 237754 | ENSMUST00000046522.13 | -4776 | ENSMUSG00000040283 | Btnl9 | butyrophilin-like 9 |
| chr3 | 1,09E+08 | 1,09E+08 | 1559 | * | 13,63889 | 2,816346982 | 0,980308 | 2,87292 | 0,004067 | 0,802490409 | 0,938682 | Promoter (<=1kb) | 3 | 1,09E+08 | 1,09E+08 | 2675 | 1 | 66715 | ENSMUST00000197427.5 | 0 | ENSMUSG00000045662 | Henmt1 | HEN1 methyltransferase homolog 1 (Arabidopsis) |
| chr4 | 1,5E+08 | 1,5E+08 | 1452 | * | 23,31769 | 2,816517476 | 0,770486 | 3,655509 | 0,000257 | 0,184384686 | 2,951933 | 3' UTR | 4 | 1,5E+08 | 1,5E+08 | 5418 | 2 | 100198 | ENSMUST00000153394.2 | 20622 | ENSMUSG00000028980 | H6pd | hexose-6-phosphate dehydrogenase (glucose 1-dehydrogenase) |
| chr5 | 34029233 | 34030460 | 1228 | * | 11,26716 | 2,816637504 | 0,987061 | 2,853561 | 0,004323 | 0,775744126 | 1,173325 | 3' UTR | 5 | 34035559 | 34037424 | 1866 | 1 | 107823 | ENSMUST00000202912.2 | -5099 | ENSMUSG00000057406 | Nsd2 | nuclear receptor binding SET domain protein 2 |
| chr13 | 52498461 | 52500430 | 1970 | * | 14,62456 | 2,816844016 | 0,933314 | 3,018109 | 0,002544 | 0,681119785 | 1,214904 | Distal Intergenic | 13 | 52334447 | 52335673 | 1227 | 1 | 1,19E+08 | ENSMUST00000222876.2 | 164014 | | Gm48199 | predicted gene, 48199 |
| chr1 | 1,66E+08 | 1,66E+08 | 1487 | * | 11,88249 | 2,817150428 | 1,080224 | 2,607932 | 0,009109 | 0,974006322 | 0,738441 | Distal Intergenic | 1 | 1,66E+08 | 1,66E+08 | 11563 | 1 | 433375 | ENSMUST00000111432.10 | -18338 | ENSMUSG00000040713 | Creg1 | cellular repressor of E1A-stimulated genes 1 |
| chr15 | 36848393 | 36851441 | 3049 | * | 27,00206 | 2,817488927 | 0,6822 | 4,130004 | 3,63E-05 | 0,058184583 | 2,725491 | Distal Intergenic | 15 | 36870744 | 36880121 | 9378 | 2 | 1E+08 | ENSMUST00000100713.4 | 28680 | ENSMUSG00000072618 | Gm10384 | predicted gene 10384 |
| chr14 | 31889146 | 31891099 | 1954 | * | 16,88763 | 2,817975652 | 0,941619 | 2,992693 | 0,002765 | 0,594676756 | 2,141612 | Promoter (1-2kb) | 14 | 31888078 | 31894579 | 6502 | 1 | 27057 | ENSMUST00000164341.2 | 1068 | ENSMUSG00000056234 | Ncoa4 | nuclear receptor coactivator 4 |
| chr2 | 31575305 | 31576681 | 1377 | * | 13,30921 | 2,818023451 | 0,974746 | 2,891034 | 0,00384 | 0,792545355 | 0,938682 | Promoter (1-2kb) | 2 | 31578388 | 31588348 | 9961 | 1 | 11350 | ENSMUST00000146537.2 | -1707 | ENSMUSG00000026842 | Abl1 | c-abl oncogene 1, non-receptor tyrosine kinase |
| chr11 | 54104222 | 54107441 | 3220 | * | 26,54833 | 2,818378728 | 0,702708 | 4,010742 | 6,05E-05 | 0,080985792 | 2,725491 | Promoter (<=1kb) | 11 | 54068598 | 54104111 | 35514 | 2 | 71056 | ENSMUST00000131019.2 | -111 | ENSMUSG00000086209 | 4933405E24Rik | RIKEN cDNA 4933405E24 gene |
| chr5 | 91036877 | 91037968 | 1092 | * | 11,15869 | 2,818413151 | 0,959164 | 2,938406 | 0,003299 | 0,71419937 | 1,2763 | Promoter (1-2kb) | 5 | 91039100 | 91040974 | 1875 | 1 | 14825 | ENSMUST00000031327.9 | -1132 | ENSMUSG00000029380 | Cxcl1 | chemokine (C-X-C motif) ligand 1 |
| chr5 | 1,22E+08 | 1,22E+08 | 2647 | * | 22,32914 | 2,818982159 | 0,777897 | 3,62385 | 0,00029 | 0,198315008 | 2,951933 | Promoter (<=1kb) | 5 | 1,22E+08 | 1,22E+08 | 9156 | 1 | 20239 | ENSMUST00000159928.2 | 0 | ENSMUSG00000042605 | Atxn2 | ataxin 2 |
| chr14 | 45562975 | 45564157 | 1183 | * | 15,0022 | 2,819387017 | 0,842656 | 3,345835 | 0,00082 | 0,449809164 | 1,41696 | Distal Intergenic | 14 | 45567245 | 45587162 | 19918 | 1 | 67089 | ENSMUST00000022380.9 | -3088 | ENSMUSG00000021832 | Psmc6 | proteasome (prosome, macropain) 26S subunit, ATPase, 6 |
| chr12 | 85155931 | 85157803 | 1873 | * | 15,47127 | 2,819700038 | 0,991997 | 2,842448 | 0,004477 | 0,748971518 | 1,41696 | Promoter (<=1kb) | 12 | 85153089 | 85157533 | 4445 | 2 | 73422 | ENSMUST00000221003.2 | 0 | ENSMUSG00000042320 | Prox2 | prospero homeobox 2 |
| chr5 | 1,43E+08 | 1,43E+08 | 756 | * | 13,3037 | 2,820275515 | 1,003028 | 2,811761 | 0,004927 | 0,838625535 | 0,938682 | Intron (ENSMUST00000 | 5 | 1,43E+08 | 1,43E+08 | 12129 | 2 | 231861 | ENSMUST00000155461.2 | 3521 | ENSMUSG00000039477 | Tnrc18 | trinucleotide repeat containing 18 |
| chr10 | 89329834 | 89331183 | 1350 | * | 15,47192 | 2,821250356 | 0,881392 | 3,200902 | 0,00137 | 0,554978012 | 1,2763 | Intron (ENSMUST00000 | 10 | 89290096 | 89342511 | 52416 | 2 | 20186 | ENSMUST00000058126.15 | 11328 | ENSMUSG00000047638 | Nr1h4 | nuclear receptor subfamily 1, group H, member 4 |
| chr6 | 29979795 | 29981586 | 1792 | * | 24,02928 | 2,821930551 | 0,689377 | 4,093448 | 4,25E-05 | 0,065127074 | 2,67243 | Distal Intergenic | 6 | 29985234 | 29993777 | 8544 | 1 | 76419 | ENSMUST00000141679.3 | -3648 | | Smkr-ps | smal lysine rich protein 1, pseudogene |
| chr11 | 1,01E+08 | 1,01E+08 | 1864 | * | 13,37045 | 2,823175887 | 1,056115 | 2,673171 | 0,007514 | 0,908469938 | 0,938682 | 3' UTR | 11 | 1,01E+08 | 1,01E+08 | 6145 | 2 | 19285 | ENSMUST00000132934.2 | 10442 | ENSMUSG00000004044 | Cavin1 | caveolae associated 1 |
| chr1 | 1,33E+08 | 1,33E+08 | 2526 | * | 18,41695 | 2,823424215 | 0,847485 | 3,331534 | 0,000864 | 0,365335255 | 2,423594 | Downstream (<=300bp | 1 | 1,33E+08 | 1,33E+08 | 13227 | 1 | 68338 | ENSMUST00000094557.7 | -3679 | ENSMUSG00000103421 | Golt1a | golgi transport 1A |
| chr17 | 24686915 | 24689825 | 2911 | * | 49,773102 | 2,823487908 | 0,5291172 | 5,3362236 | 9,49E-08 | 0,000759549 | 2,3383871 | Promoter (<=1kb) | 17 | 24662753 | 24689287 | 26535 | 2 | 13560 | ENSMUST00000226654.2 | 0 | ENSMUSG00000024137 | E4f1 | E4F transcription factor 1 |
| chr3 | 36552385 | 36556090 | 3706 | * | 39,241819 | 2,823616721 | 0,5489983 | 5,1432161 | 2,701E-07 | 0,001523506 | 2,3776742 | Exon (ENSMUST0000014 | 3 | 36553953 | 36560958 | 7006 | 2 | 329623 | ENSMUST00000140203.2 | 4868 | ENSMUSG00000085320 | Gm11548 | predicted gene 11548 |
| chr4 | 1,3E+08 | 1,3E+08 | 1659 | * | 17,52511 | 2,824054195 | 0,825674 | 3,4203 | 0,000626 | 0,317496979 | 2,386008 | Distal Intergenic | 4 | 1,3E+08 | 1,3E+08 | 27867 | 1 | 622480 | ENSMUST00000084263.6 | -4145 | ENSMUSG00000028784 | Spocd1 | SPOC domain containing 1 |
| chr16 | 55651839 | 55653492 | 1654 | * | 12,60061 | 2,824277658 | 0,914346 | 3,08885 | 0,002009 | 0,72253719 | 0,738441 | Intron (ENSMUST00000 | 16 | 55631741 | 55659217 | 27477 | 2 | 80859 | ENSMUST00000096026.9 | 5725 | ENSMUSG00000035356 | Nfkbiz | nuclear factor of kappa light polypeptide gene enhancer in B cells inhibitor, zeta |
| chr2 | 34414592 | 34416124 | 1533 | * | 13,91752 | 2,824695535 | 1,093701 | 2,582693 | 0,009803 | 0,902481521 | 1,2763 | Intron (ENSMUST00000 | 2 | 34334359 | 34387885 | 53527 | 1 | 227743 | ENSMUST00000137021.2 | 80233 | ENSMUSG00000038696 | Mapkap1 | mitogen-activated protein kinase associated protein 1 |
| chr12 | 70730609 | 70732410 | 1802 | * | 17,94155 | 2,825389253 | 0,876368 | 3,223977 | 0,001264 | 0,418252864 | 2,663619 | Promoter (1-2kb) | 12 | 70722621 | 70728624 | 6004 | 2 | 1,05E+08 | ENSMUST00000222630.2 | -1985 | | Gm32369 | predicted gene, 32369 |
| chr9 | 78262411 | 78264840 | 2430 | * | 19,61338 | 2,825749469 | 0,924297 | 3,057186 | 0,002234 | 0,542967057 | 2,221611 | Promoter (<=1kb) | 9 | 78239369 | 78263070 | 23702 | 2 | 14858 | ENSMUST00000125479.8 | 0 | ENSMUSG00000057933 | Gsta2 | glutathione S-transferase, alpha 2 (Yc2) |
| chr19 | 16793718 | 16795659 | 1942 | * | 12,77583 | 2,825837582 | 1,06547 | 2,652199 | 0,007997 | 0,955202728 | 0,738441 | Distal Intergenic | 19 | 16729684 | 16758297 | 28614 | 2 | 271564 | ENSMUST00000225764.2 | -35421 | ENSMUSG00000046230 | Vps13a | vacuolar protein sorting 13A |
| chr13 | 49733947 | 49735874 | 1928 | * | 12,86722 | 2,826395242 | 1,062848 | 2,659266 | 0,007831 | 0,863172166 | 1,2763 | Promoter (<=1kb) | 13 | 49735938 | 49746298 | 10361 | 1 | 27047 | ENSMUST00000221170.2 | -64 | ENSMUSG00000048368 | Omd | osteomodulin |
| chr19 | 48950629 | 48952282 | 1654 | * | 16,04305 | 2,826758101 | 0,930364 | 3,038334 | 0,002379 | 0,541401776 | 2,386008 | Distal Intergenic | 19 | 48692655 | 48792976 | 100322 | 1 | 66673 | ENSMUST00000235967.2 | 257974 | ENSMUSG00000063434 | Sorcs3 | sortilin-related VPS10 domain containing receptor 3 |
| chr3 | 37554081 | 37555749 | 1669 | * | 11,4881 | 2,827091858 | 1,090118 | 2,593383 | 0,009504 | 0,977532477 | 0,744375 | Intron (ENSMUST00000 | 3 | 37547337 | 37633158 | 85822 | 1 | 57815 | ENSMUST00000200093.2 | 6744 | ENSMUSG00000027722 | Spata5 | spermatogenesis associated 5 |
| chr7 | 1,16E+08 | 1,16E+08 | 1258 | * | 12,09983 | 2,827193971 | 1,095442 | 2,580871 | 0,009855 | 0,954706987 | 0,918255 | Intron (ENSMUST00000 | 7 | 1,15E+08 | 1,16E+08 | 523743 | 2 | 20679 | ENSMUST00000072804.11 | 8190 | ENSMUSG00000051910 | Sox6 | SRY (sex determining region Y)-box 6 |
| chr7 | 16655549 | 16657754 | 2206 | * | 21,86195 | 2,827786802 | 0,811522 | 3,484548 | 0,000493 | 0,26172012 | 2,951933 | Promoter (<=1kb) | 7 | 16651069 | 16657754 | 6686 | 2 | 12315 | ENSMUST00000173557.2 | 0 | ENSMUSG00000019370 | Calm3 | calmodulin 3 |
| chr5 | 19980242 | 19981881 | 1640 | * | 14,52997 | 2,827815691 | 1,059246 | 2,669649 | 0,007593 | 0,858257847 | 1,2763 | Intron (ENSMUST00000 | 5 | 20112516 | 20909790 | 797275 | 1 | 50791 | ENSMUST00000101558.10 | -130635 | ENSMUSG00000040003 | Magi2 | membrane associated guanylate kinase, WW and PDZ domain containing 2 |
| chr9 | 57026117 | 57027523 | 1407 | * | 12,91585 | 2,827833346 | 0,968172 | 2,920796 | 0,003491 | 0,736703271 | 1,173325 | Exon (ENSMUST000001 | 9 | 57020745 | 57032475 | 11731 | 1 | 20466 | ENSMUST00000167963.2 | 5372 | ENSMUSG00000042557 | Sin3a | transcriptional regulator, SIN3A (yeast) |
| chr15 | 54039204 | 54040068 | 865 | * | 11,60132 | 2,82833586 | 1,096077 | 2,580417 | 0,009868 | 0,984504123 | 0,744375 | Distal Intergenic | 15 | 54114015 | 54141880 | 27866 | 2 | 18383 | ENSMUST00000079772.4 | 101812 | ENSMUSG00000063727 | Tnfrsf11b | tumor necrosis factor receptor superfamily, member 11b (osteoprotegerin) |
| chr3 | 30647159 | 30648633 | 1475 | * | 13,98088 | 2,828631208 | 1,003159 | 2,819724 | 0,004806 | 0,834323723 | 0,938682 | Distal Intergenic | 3 | 30651222 | 30654088 | 2867 | 2 | 76652 | ENSMUST00000047630.7 | 5455 | ENSMUSG00000037737 | Actrt3 | actin related protein T3 |
| chr2 | 33918903 | 33923245 | 4343 | * | 74,166763 | 2,828657026 | 0,4409888 | 6,4143506 | 1,414E-10 | 2,45309E-06 | 3,439501 | Exon (ENSMUST0000013 | 2 | 33868258 | 33870336 | 2079 | 1 | 105244124 | ENSMUST00000156633.2 | 50645 | | Gm13404 | predicted gene 13404 |
| chr7 | 79541482 | 79542916 | 1435 | * | 12,98512 | 2,828822093 | 0,997789 | 2,835089 | 0,004581 | 0,80080207 | 1,075735 | Intron (ENSMUST00000 | 7 | 79532259 | 79570344 | 38086 | 2 | 11778 | ENSMUST00000206725.2 | 27428 | ENSMUSG00000063801 | Ap3s2 | adaptor-related protein complex 3, sigma 2 subunit |
| chrX | 1,52E+08 | 1,52E+08 | 1268 | * | 11,05064 | 2,828825252 | 1,016688 | 2,782392 | 0,005396 | 0,894789713 | 0,738441 | Distal Intergenic | 20 | 1,52E+08 | 1,52E+08 | 3344 | 1 | 54609 | ENSMUST00000060714.10 | 6869 | ENSMUSG00000050148 | Ubqln2 | ubiquilin 2 |
| chr6 | 84992019 | 84993885 | 1867 | * | 17,43919 | 2,829166158 | 0,941261 | 3,00572 | 0,00265 | 0,585155013 | 2,154504 | Intron (ENSMUST00000 | 6 | 84811188 | 85046439 | 235252 | 2 | 75914 | ENSMUST00000046334.15 | 52554 | ENSMUSG00000033769 | Exoc6b | exocyst complex component 6B |
| chr13 | 23726328 | 23729351 | 3024 | * | 32,941312 | 2,829425096 | 0,6609477 | 4,280861 | 1,862E-05 | 0,03581197 | 2,7254908 | Promoter (<=1kb) | 13 | 23728222 | 23729482 | 1261 | 1 | 260423 | ENSMUST00000075558.5 | 0 | ENSMUSG00000100210 | H3c7 | H3 clustered histone 7 |
| chr8 | 70340009 | 70341993 | 1985 | * | 27,61077 | 2,82951361 | 0,815081 | 3,471452 | 0,000518 | 0,279427465 | 2,725491 | Promoter (<=1kb) | 8 | 70340440 | 70340835 | 396 | 2 | 234365 | ENSMUST00000152456.2 | 0 | ENSMUSG00000048967 | Yjefn3 | YjeF N-terminal domain containing 3 |
| chr11 | 8627230 | 8630047 | 2818 | * | 36,414682 | 2,829696953 | 0,5386322 | 5,2534867 | 1,492E-07 | 0,001081418 | 2,3776742 | Distal Intergenic | 11 | 8381652 | 8614681 | 233030 | 2 | 319939 | ENSMUST00000239111.2 | -12549 | ENSMUSG00000020422 | Tns3 | tensin 3 |
| chr11 | 68548748 | 68549563 | 816 | * | 9,907671 | 2,829810267 | 1,095363 | 2,583445 | 0,009782 | 0,924781491 | 1,086118 | Distal Intergenic | 11 | 68582385 | 68590085 | 7701 | 1 | 77579 | ENSMUST00000141558.2 | -32822 | ENSMUSG00000020900 | Myh10 | myosin, heavy polypeptide 10, non-muscle |
| chr18 | 44726851 | 44727849 | 999 | * | 9,836267 | 2,831005722 | 1,059192 | 2,672798 | 0,007522 | 0,86525631 | 1,211315 | Intron (ENSMUST00000 | 18 | 44794732 | 44809338 | 14607 | 1 | 626275 | ENSMUST00000181094.2 | -66883 | ENSMUSG00000097610 | A930012L18Rik | RIKEN cDNA A930012L18 gene |
| chr7 | 79960037 | 79961902 | 1866 | * | 13,8569 | 2,8324599 | 0,941836 | 3,007381 | 0,002635 | 0,728037066 | 0,938682 | Promoter (<=1kb) | 7 | 79960597 | 79963242 | 2646 | 1 | 233406 | ENSMUST00000172781.8 | 0 | ENSMUSG00000038943 | Prc1 | protein regulator of cytokinesis 1 |
| chr17 | 3321301 | 3324132 | 2832 | * | 30,96288 | 2,833345071 | 0,702409 | 4,033755 | 5,49E-05 | 0,077423487 | 2,67243 | Exon (ENSMUST000002 | 17 | 3320956 | 3333482 | 12527 | 2 | 1,03E+08 | ENSMUST00000232068.2 | 9350 | ENSMUSG00000116874 | Gm34035 | predicted gene, 34035 |
| chr3 | 84800644 | 84801943 | 1300 | * | 11,7547 | 2,833371432 | 1,023185 | 2,769167 | 0,00562 | 0,822931039 | 1,173325 | Intron (ENSMUST00000 | 3 | 84771452 | 84886505 | 115054 | 1 | 50754 | ENSMUST00000107678.8 | 29192 | ENSMUSG00000028086 | Fbxw7 | F-box and WD-40 domain protein 7 |
| chr8 | 61444821 | 61447939 | 3119 | * | 68,022632 | 2,834122859 | 0,4294907 | 6,5987995 | 4,145E-11 | 1,02087E-06 | 3,439501 | Promoter (<=1kb) | 8 | 61446229 | 61584373 | 138145 | 1 | 18004 | ENSMUST00000120689.8 | 0 | ENSMUSG00000031644 | Nek1 | NIMA (never in mitosis gene a)-related expressed kinase 1 |
| chr9 | 21131012 | 21132655 | 1644 | * | 17,3054 | 2,834336076 | 0,962232 | 2,945586 | 0,003223 | 0,59912059 | 2,423594 | Distal Intergenic | 9 | 21147419 | 21148489 | 1071 | 2 | 50868 | ENSMUST00000193247.2 | 15834 | ENSMUSG00000003308 | Keap1 | kelch-like ECH-associated protein 1 |
| chr10 | 79516487 | 79517812 | 1326 | * | 15,81961 | 2,834668676 | 1,0024 | 2,827882 | 0,004686 | 0,684274806 | 2,141612 | Promoter (<=1kb) | 10 | 79518029 | 79524232 | 6204 | 1 | 216150 | ENSMUST00000020550.13 | -217 | ENSMUSG00000020307 | Cdc34 | cell division cycle 34 |
| chr11 | 1,17E+08 | 1,17E+08 | 1349 | * | 12,97756 | 2,83506168 | 0,947328 | 2,992692 | 0,002765 | 0,778321623 | 0,738441 | Intron (ENSMUST00000 | 11 | 1,17E+08 | 1,17E+08 | 96068 | 1 | 53860 | ENSMUST00000019038.15 | -13168 | ENSMUSG00000059248 | Septin9 | septin 9 |
| chr8 | 22964855 | 22966489 | 1635 | * | 12,74982 | 2,835322837 | 1,086426 | 2,609771 | 0,00906 | 0,890237282 | 1,2763 | Promoter (<=1kb) | 8 | 22966804 | 23059628 | 92825 | 1 | 20516 | ENSMUST00000067786.9 | -315 | ENSMUSG00000037656 | Slc20a2 | solute carrier family 20, member 2 |
| chr1 | 1,71E+08 | 1,71E+08 | 3325 | * | 21,7752 | 2,836086671 | 0,759818 | 3,732589 | 0,00019 | 0,154924569 | 2,951933 | Promoter (<=1kb) | 1 | 1,71E+08 | 1,71E+08 | 1326 | 1 | 11807 | ENSMUST00000111321.8 | 0 | ENSMUSG00000005681 | Apoa2 | apolipoprotein A-II |
| chr1 | 1,15E+08 | 1,15E+08 | 958 | * | 9,190402 | 2,836805064 | 1,064602 | 2,664663 | 0,007707 | 0,852340527 | 1,360622 | Distal Intergenic | 1 | 1,16E+08 | 1,16E+08 | 15632 | 2 | 1,08E+08 | ENSMUST00000186137.7 | 936337 | | Gm50605 | predicted gene, 50605 |
| chr3 | 79770266 | 79771024 | 759 | * | 8,066806 | 2,837073199 | 1,101118 | 2,57654 | 0,009979 | 1 | 0 | Intron (ENSMUST00000 | 3 | 79739369 | 79760080 | 20712 | 2 | 70652 | ENSMUST00000193410.3 | -10186 | ENSMUSG00000027956 | Tmem144 | transmembrane protein 144 |
| chr4 | 1,49E+08 | 1,49E+08 | 1910 | * | 16,70776 | 2,837158905 | 0,810414 | 3,500875 | 0,000464 | 0,268627242 | 2,663619 | Promoter (1-2kb) | 4 | 1,49E+08 | 1,49E+08 | 8088 | 1 | 50912 | ENSMUST00000126897.2 | 1589 | ENSMUSG00000017264 | Exosc10 | exosome component 10 |
| chr9 | 123683363 | 123686326 | 2964 | * | 26,747338 | 2,837563439 | 0,6344788 | 4,4722748 | 7,739E-06 | 0,018326614 | 2,8800171 | Promoter (2-3kb) | 9 | 123618575 | 123680964 | 62390 | 2 | 17281 | ENSMUST00000167595.9 | -2399 | ENSMUSG00000025241 | Fyco1 | FYVE and coiled-coil domain containing 1 |
| chr13 | 55860684 | 55862397 | 1714 | * | 15,06273 | 2,837855259 | 1,003432 | 2,82815 | 0,004682 | 0,684274806 | 2,141612 | Promoter (<=1kb) | 13 | 55862463 | 55874040 | 11578 | 1 | 69672 | ENSMUST00000021959.11 | -66 | ENSMUSG00000021497 | Txndc15 | thioredoxin domain containing 15 |
| chr16 | 32427156 | 32428205 | 1050 | * | 15,53514 | 2,837935199 | 1,046292 | 2,712373 | 0,00668 | 0,847020973 | 1,214904 | Promoter (<=1kb) | 16 | 32427738 | 32451612 | 23875 | 1 | 22042 | ENSMUST00000023486.15 | 0 | ENSMUSG00000022797 | Tfrc | transferrin receptor |
| chr19 | 38586327 | 38589003 | 2677 | * | 20,83766 | 2,83833957 | 0,936097 | 3,032099 | 0,002429 | 0,500623832 | 3,104559 | Intron (ENSMUST00000 | 19 | 38571812 | 38640451 | 68640 | 1 | 74055 | ENSMUST00000182999.2 | 14515 | ENSMUSG00000024998 | Plce1 | phospholipase C, epsilon 1 |
| chr1 | 1,68E+08 | 1,68E+08 | 3459 | * | 28,6091 | 2,838404828 | 0,778869 | 3,644266 | 0,000268 | 0,200622128 | 2,67243 | Promoter (2-3kb) | 1 | 1,68E+08 | 1,68E+08 | 272773 | 2 | 18514 | ENSMUST00000064438.11 | 2837 | ENSMUSG00000052534 | Pbx1 | pre B cell leukemia homeobox 1 |
| chr13 | 44563896 | 44565129 | 1234 | * | 13,41143 | 2,838976102 | 0,964943 | 2,942117 | 0,00326 | 0,740451144 | 1,075735 | Intron (ENSMUST00000 | 13 | 44517312 | 44544593 | 27282 | 2 | 1,03E+08 | ENSMUST00000222879.2 | -19303 | | Gm33958 | predicted gene, 33958 |
| chr18 | 34639964 | 34643460 | 3497 | * | 46,469143 | 2,839269932 | 0,5965225 | 4,7597027 | 1,939E-06 | 0,007372728 | 2,3383871 | Promoter (<=1kb) | 18 | 34627926 | 34639884 | 11959 | 2 | 225358 | ENSMUST00000236598.2 | -80 | ENSMUSG00000036501 | Fam13b | family with sequence similarity 13, member B |
| chr19 | 41899594 | 41902661 | 3068 | * | 53,87945 | 2,840878169 | 1,009518 | 2,814095 | 0,004891 | 0,654833721 | 2,725491 | Promoter (<=1kb) | 19 | 41900362 | 41907099 | 6738 | 1 | 18648 | ENSMUST00000011896.8 | 0 | ENSMUSG00000011752 | Pgam1 | phosphoglycerate mutase 1 |
| chr4 | 1,07E+08 | 1,07E+08 | 1874 | * | 18,30165 | 2,840988468 | 0,914453 | 3,106763 | 0,001891 | 0,483145429 | 2,663619 | Intron (ENSMUST00000 | 4 | 1,07E+08 | 1,07E+08 | 196064 | 1 | 230587 | ENSMUST00000106738.2 | 23512 | ENSMUSG00000034762 | Glis1 | GLIS family zinc finger 1 |
| chr1 | 1,11E+08 | 1,11E+08 | 1220 | * | 8,948148 | 2,841933305 | 1,092598 | 2,601079 | 0,009293 | 0,885416414 | 1,360622 | Distal Intergenic | 1 | 1,12E+08 | 1,12E+08 | 6067 | 2 | 319901 | ENSMUST00000186365.2 | 376647 | ENSMUSG00000038702 | Dsel | dermatan sulfate epimerase-like |
| chr6 | 20491786 | 20492638 | 853 | * | 12,93849 | 2,841981931 | 0,93527 | 3,038674 | 0,002376 | 0,749781925 | 0,738441 | Distal Intergenic | 6 | 21013111 | 21013241 | 131 | 2 | 1,15E+08 | ENSMUST00000157206.3 | 520603 | ENSMUSG00000087831 | Gm23960 | predicted gene, 23960 |
| chr1 | 75279761 | 75280903 | 1143 | * | 12,59893 | 2,842192378 | 0,982726 | 2,892152 | 0,003826 | 0,738140609 | 1,2763 | Distal Intergenic | 1 | 75285827 | 75290403 | 4577 | 2 | 13437 | ENSMUST00000186278.2 | 9500 | ENSMUSG00000026209 | Dnpep | aspartyl aminopeptidase |
| chr10 | 28017264 | 28018613 | 1350 | * | 13,42928 | 2,842313992 | 0,952798 | 2,983124 | 0,002853 | 0,716994918 | 1,075735 | Intron (ENSMUST00000 | 10 | 27951006 | 28140669 | 189664 | 1 | 19272 | ENSMUST00000219761.2 | 66258 | ENSMUSG00000019889 | Ptprk | protein tyrosine phosphatase, receptor type, K |
| chr4 | 1,09E+08 | 1,09E+08 | 1149 | * | 11,13868 | 2,842504769 | 1,092339 | 2,602218 | 0,009262 | 0,945372047 | 0,918255 | Intron (ENSMUST00000 | 4 | 1,08E+08 | 1,09E+08 | 84618 | 2 | 230597 | ENSMUST00000042185.8 | 9703 | ENSMUSG00000034557 | Zfyve9 | zinc finger, FYVE domain containing 9 |
| chr11 | 9705688 | 9706941 | 1254 | * | 11,1818 | 2,84269701 | 1,025793 | 2,771219 | 0,005585 | 0,804972409 | 1,2763 | Distal Intergenic | 11 | 9143451 | 9206178 | 62728 | 1 | 268379 | ENSMUST00000151522.2 | 562237 | ENSMUSG00000004668 | Abca13 | ATP-binding cassette, sub-family A (ABC1), member 13 |
| chr3 | 87981128 | 87983249 | 2122 | * | 15,51456 | 2,843192208 | 0,975852 | 2,913547 | 0,003573 | 0,642870233 | 2,141612 | Promoter (<=1kb) | 3 | 87977334 | 87982824 | 5491 | 2 | 214191 | ENSMUST00000238508.2 | 0 | ENSMUSG00000051036 | Ttc24 | tetratricopeptide repeat domain 24 |
| chr17 | 73089537 | 73090892 | 1356 | * | 13,62448 | 2,843577272 | 0,985919 | 2,88419 | 0,003924 | 0,801134483 | 0,918255 | Distal Intergenic | 17 | 73143448 | 73158190 | 14743 | 1 | 383295 | ENSMUST00000045174.7 | -52556 | ENSMUSG00000039770 | Ypel5 | yippee like 5 |
| chr10 | 20792616 | 20794420 | 1805 | * | 11,00455 | 2,843800084 | 1,0395 | 2,73574 | 0,006224 | 0,914972668 | 0,744375 | Distal Intergenic | 10 | 20742000 | 20767448 | 25449 | 2 | 1,19E+08 | ENSMUST00000241754.1 | -25168 | | Gm53617 | predicted gene, 53617 |
| chr1 | 1,51E+08 | 1,51E+08 | 870 | * | 9,651235 | 2,844177286 | 1,050224 | 2,708162 | 0,006766 | 0,853827002 | 1,173325 | Exon (ENSMUST00000 | 1 | 1,5E+08 | 1,5E+08 | 19529 | 2 | 545370 | ENSMUST00000177036.2 | -160464 | ENSMUSG00000066842 | Hmcn1 | hemicentin 1 |
| chr13 | 41554077 | 41556413 | 2337 | * | 24,26148 | 2,84461577 | 0,721103 | 3,944811 | 7,99E-05 | 0,090530822 | 2,880017 | Exon (ENSMUST000002 | 13 | 41471396 | 41513215 | 41820 | 2 | 18003 | ENSMUST00000224803.2 | -40862 | ENSMUSG00000021365 | Nedd9 | neural precursor cell expressed, developmentally down-regulated gene 9 |
| chr8 | 16203147 | 16203794 | 648 | * | 8,361794 | 2,844628363 | 1,101123 | 2,583388 | 0,009784 | 1 | 0 | Intron (ENSMUST00000 | 8 | 16218771 | 16218852 | 82 | 2 | 1,01E+08 | ENSMUST00000175401.3 | 15058 | ENSMUSG00000093142 | Mir3106 | microRNA 3106 |
| chr10 | 59540544 | 59542282 | 1739 | * | 12,88844 | 2,844716484 | 1,096632 | 2,594049 | 0,009485 | 0,948006995 | 0,918255 | Promoter (2-3kb) | 10 | 59538480 | 59587527 | 49048 | 1 | 216001 | ENSMUST00000165405.2 | 2064 | ENSMUSG00000020111 | Micu1 | mitochondrial calcium uptake 1 |
| chr5 | 1,01E+08 | 1,01E+08 | 2039 | * | 16,36276 | 2,844785109 | 1,020269 | 2,78827 | 0,005299 | 0,671156916 | 2,663619 | Promoter (<=1kb) | 5 | 1,01E+08 | 1,01E+08 | 9071 | 2 | 1,01E+08 | ENSMUST00000181873.9 | 0 | ENSMUSG00000097772 | 5430416N02Rik | RIKEN cDNA 5430416N02 gene |
| chr13 | 8551395 | 8553323 | 1929 | * | 17,75989 | 2,844805568 | 0,774919 | 3,671099 | 0,000242 | 0,189033898 | 2,663619 | Intron (ENSMUST00000 | 13 | 8252958 | 8808908 | 555951 | 1 | 94191 | ENSMUST00000123187.2 | 298437 | ENSMUSG00000052551 | Adarb2 | adenosine deaminase, RNA-specific, B2 |
| chr1 | 1,52E+08 | 1,52E+08 | 1275 | * | 13,62361 | 2,844815533 | 1,022458 | 2,782329 | 0,005397 | 0,79897927 | 1,2763 | Intron (ENSMUST00000 | 1 | 1,52E+08 | 1,52E+08 | 42176 | 1 | 63913 | ENSMUST00000086267.6 | -18214 | ENSMUSG00000026483 | Niban1 | niban apoptosis regulator 1 |
| chr4 | 59331174 | 59332455 | 1282 | * | 14,05193 | 2,845639001 | 0,873988 | 3,255925 | 0,00113 | 0,580348965 | 0,938682 | Promoter (<=1kb) | 4 | 59329210 | 59333022 | 3813 | 2 | 634731 | ENSMUST00000136562.2 | 567 | ENSMUSG00000038578 | Susd1 | sushi domain containing 1 |
| chr18 | 42723808 | 42725617 | 1810 | * | 13,14822 | 2,846261236 | 1,017516 | 2,797265 | 0,005154 | 0,888080529 | 0,738441 | Intron (ENSMUST00000 | 18 | 42710946 | 42712717 | 1772 | 2 | 240239 | ENSMUST00000054738.5 | -11091 | ENSMUSG00000042816 | Gpr151 | G protein-coupled receptor 151 |
| chr15 | 78393861 | 78395369 | 1509 | * | 17,36096 | 2,846380404 | 0,893114 | 3,187028 | 0,001437 | 0,453798789 | 2,423594 | Distal Intergenic | 15 | 78363456 | 78379471 | 16016 | 2 | 16185 | ENSMUST00000089398.9 | -14390 | ENSMUSG00000068227 | Il2rb | interleukin 2 receptor, beta chain |
| chr15 | 62166546 | 62168042 | 1497 | * | 20,44982 | 2,847844817 | 0,894382 | 3,18415 | 0,001452 | 0,440320279 | 2,663619 | Intron (ENSMUST00000 | 15 | 62094452 | 62114354 | 19903 | 1 | 19296 | ENSMUST00000182141.2 | 72094 | ENSMUSG00000097039 | Pvt1 | Pvt1 oncogene |
| chr1 | 74406520 | 74409107 | 2588 | * | 19,37683 | 2,847852501 | 0,857778 | 3,320035 | 0,0009 | 0,367076136 | 2,507013 | Promoter (2-3kb) | 1 | 74403690 | 74405901 | 2212 | 1 | 241112 | ENSMUST00000186688.2 | 2830 | ENSMUSG00000073650 | Catip | ciliogenesis associated TTC17 interacting protein |
| chr4 | 1,32E+08 | 1,32E+08 | 2296 | * | 16,85659 | 2,84790928 | 0,975883 | 2,918288 | 0,00352 | 0,641187423 | 2,141612 | Exon (ENSMUST000001 | 4 | 1,32E+08 | 1,32E+08 | 44433 | 2 | 269587 | ENSMUST00000146443.8 | 3749 | ENSMUSG00000028906 | Epb41 | erythrocyte membrane protein band 4.1 |
| chr18 | 39672532 | 39674803 | 2272 | * | 16,09168 | 2,84851143 | 0,946731 | 3,008788 | 0,002623 | 0,584572307 | 2,141612 | Promoter (<=1kb) | 18 | 39672413 | 39781584 | 109172 | 1 | 1,03E+08 | ENSMUST00000241486.1 | 119 | | Gm30961 | predicted gene, 30961 |

| | | | | | | | | | | | | | | | | | | | | | | |
|---|---|---|---|---|---|---|---|---|---|---|---|---|---|---|---|---|---|---|---|---|---|---|
| chr5 | 36881331 | 36884333 | 3003 | * | 40,687793 | 2,84859816 | 0,5120045 | 5,5636197 | 2,642E-08 | 0,000240064 | 2,7254908 Distal Intergenic | 5 | 36904722 | 36905994 | 1273 | 2 | 117197 | ENSMUST00000071949.5 | 21661 | ENSMUSG00000060708 | Bloc1s4 | biogenesis of lysosomal organelles complex-1, subunit 4, cappuccino |
| chr12 | 1,11E+08 | 1,11E+08 | 1592 | * | 13,10762 | 2,849471793 | 0,997431 | 2,856812 | 0,004279 | 0,756113531 | 1,2763 Promoter (<=1kb) | 12 | 1,11E+08 | 1,11E+08 | 84743 | 2 | 217866 | ENSMUST00000041965.5 | -127 | ENSMUSG00000021279 | Cdc42bpb | CDC42 binding protein kinase beta |
| chr6 | 43558426 | 43559256 | 831 | * | 11,66822 | 2,849915537 | 1,023259 | 2,785137 | 0,005351 | 0,855969265 | 0,918255 Intron (ENSMUST0000 | 6 | 43322007 | 43602900 | 280894 | 2 | 29807 | ENSMUST00000203649.3 | 43644 | ENSMUSG00000029735 | Tpk1 | thiamine pyrophosphokinase |
| chr5 | 31997348 | 31999229 | 1882 | * | 13,34297 | 2,849946011 | 1,099053 | 2,593093 | 0,009512 | 0,899308755 | 1,2763 Intron (ENSMUST0000 | 5 | 31942727 | 31977987 | 35261 | 1 | 107976 | ENSMUST00000202270.2 | 54621 | ENSMUSG00000052139 | Babam2 | BRISC and BRCA1 A complex member 2 |
| chr9 | 64266343 | 64267738 | 1396 | * | 14,67586 | 2,850255768 | 0,971112 | 2,935043 | 0,003335 | 0,695401133 | 1,41696 Distal Intergenic | 9 | 64214039 | 64248570 | 34532 | 2 | 213550 | ENSMUST00000068367.14 | -17773 | ENSMUSG00000032396 | Dis3l | DIS3 like exosome 3'-5' exoribonuclease |
| chr14 | 31894399 | 31896807 | 2409 | * | 16,66132 | 2,850418573 | 1,009638 | 2,82321 | 0,004755 | 0,671251031 | 2,386008 Promoter (1-2kb) | 14 | 31892815 | 31899885 | 7071 | 1 | 27057 | ENSMUST00000227112.2 | 1584 | ENSMUSG00000056234 | Ncoa4 | nuclear receptor coactivator 4 |
| chr19 | 37098183 | 37099862 | 1680 | * | 10,99583 | 2,851129208 | 1,048461 | 2,719348 | 0,006541 | 0,922828293 | 0,744375 Intron (ENSMUST0000 | 19 | 37151243 | 37173940 | 22698 | 1 | 319931 | ENSMUST00000180609.2 | -51381 | ENSMUSG00000097591 | A330032B11Rik | RIKEN cDNA A330032B11 gene |
| chr1 | 62772885 | 62774552 | 1668 | * | 10,17802 | 2,851309329 | 1,045278 | 2,727801 | 0,006376 | 0,838625535 | 1,211315 Intron (ENSMUST0000 | 1 | 62753882 | 62757232 | 3351 | 2 | 1,05E+08 | ENSMUST00000189205.2 | -15653 | | Gm29083 | predicted gene 29083 |
| chr2 | 1,21E+08 | 1,21E+08 | 1361 | * | 9,848667 | 2,851590705 | 1,026106 | 2,779042 | 0,005452 | 0,778618001 | 1,449706 Distal Intergenic | 2 | 1,21E+08 | 1,21E+08 | 13833 | 2 | 1E+08 | ENSMUST00000056732.4 | -4037 | ENSMUSG00000048222 | Mfap1b | microfibrillar-associated protein 1B |
| chr17 | 41051032 | 41052198 | 1167 | * | 13,44004 | 2,852966281 | 0,893181 | 3,194165 | 0,001402 | 0,595849045 | 1,075735 Distal Intergenic | 17 | 41038011 | 41042673 | 4663 | 1 | 619517 | ENSMUST00000233513.2 | 13021 | ENSMUSG00000073396 | Esp1 | exocrine gland secreted peptide 1 |
| chr2 | 41247446 | 41248705 | 1260 | * | 14,69795 | 2,853769215 | 0,918459 | 3,107129 | 0,001889 | 0,60653653 | 1,37683 Intron (ENSMUST0000 | 2 | 40691405 | 41679090 | 987686 | 2 | 94217 | ENSMUST00000167270.5 | 430385 | ENSMUSG00000049252 | Lrp1b | low density lipoprotein-related protein 1B |
| chr4 | 9050419 | 9052616 | 2198 | * | 17,0384 | 2,853779866 | 0,976798 | 2,921567 | 0,003483 | 0,624579927 | 2,318434 Distal Intergenic | 4 | 9210003 | 9210108 | 106 | 1 | 1,15E+08 | ENSMUST00000082782.2 | -157387 | ENSMUSG00000075958 | Gm23423 | predicted gene, 23423 |
| chr7 | 95925562 | 95926530 | 969 | * | 11,92276 | 2,853873516 | 1,050132 | 2,717634 | 0,006575 | 0,923078468 | 0,744375 Intron (ENSMUST0000 | 7 | 95898631 | 95898739 | 109 | 1 | 735284 | ENSMUST00000102189.3 | 26931 | ENSMUSG00000076143 | Mir708 | microRNA 708 |
| chr12 | 81504275 | 81505230 | 956 | * | 11,95066 | 2,85441808 | 0,999852 | 2,854841 | 0,004306 | 0,854157826 | 0,744375 Intron (ENSMUST0000 | 12 | 81518867 | 81531603 | 12737 | 2 | 66272 | ENSMUST00000171237.8 | 26373 | ENSMUSG00000091803 | Cox16 | cytochrome c oxidase assembly protein 16 |
| chr9 | 1,19E+08 | 1,19E+08 | 1566 | * | 20,098 | 2,854449428 | 0,804753 | 3,546989 | 0,00039 | 0,252389304 | 2,507013 Promoter (<=1kb) | 9 | 1,19E+08 | 1,19E+08 | 83996 | 2 | 108737 | ENSMUST00000040853.11 | 0 | ENSMUSG00000036737 | Oxsr1 | oxidative-stress responsive 1 |
| chr13 | 54614189 | 54616268 | 2080 | * | 14,0636 | 2,85516945 | 0,960591 | 2,972305 | 0,002956 | 0,723275538 | 1,075735 Promoter (<=1kb) | 13 | 54611495 | 54616601 | 5107 | 2 | 73666 | ENSMUST00000151158.2 | 333 | ENSMUSG00000025872 | Thoc3 | THO complex 3 |
| chr10 | 74940729 | 74941848 | 1120 | * | 9,853182 | 2,856322367 | 1,079994 | 2,644756 | 0,008175 | 0,90060438 | 1,086118 Intron (ENSMUST0000 | 10 | 74896424 | 75020753 | 124330 | 1 | 110279 | ENSMUST00000164107.3 | 44305 | ENSMUSG00000009681 | Bcr | BCR activator of RhoGEF and GTPase |
| chrX | 84385584 | 84387199 | 1616 | * | 10,7664 | 2,856660027 | 1,058388 | 2,699068 | 0,006953 | 0,900637308 | 0,918255 Distal Intergenic | 20 | 84293285 | 84313894 | 20610 | 1 | 71996 | ENSMUST00000048382.5 | 92299 | ENSMUSG00000035491 | Fthl17a | ferritin, heavy polypeptide-like 17, member A |
| chr8 | 73369324 | 73370176 | 853 | * | 11,65936 | 2,856985874 | 1,081293 | 2,642195 | 0,008237 | 0,958290471 | 0,738441 Distal Intergenic | 8 | 73373339 | 73420341 | 47003 | 1 | 319555 | ENSMUST00000161386.8 | -3163 | ENSMUSG00000048148 | Nwd1 | NACHT and WD repeat domain containing 1 |
| chr16 | 33470633 | 33472494 | 1862 | * | 14,17058 | 2,857154187 | 0,941083 | 3,036027 | 0,002397 | 0,713406565 | 0,938682 Intron (ENSMUST0000 | 16 | 33504740 | 33587183 | 82444 | 1 | 77446 | ENSMUST00000232568.2 | -32246 | ENSMUSG00000075254 | Heg1 | heart development protein with EGF-like domains 1 |
| chr1 | 1,43E+08 | 1,43E+08 | 830 | * | 9,994516 | 2,857155408 | 1,029351 | 2,775687 | 0,005509 | 0,781165481 | 1,449706 Distal Intergenic | 1 | 1,43E+08 | 1,44E+08 | 34421 | 2 | 214498 | ENSMUST00000162638.2 | 989122 | ENSMUSG00000026361 | Cdc73 | cell division cycle 73, Paf1/RNA polymerase II complex component |
| chr12 | 1,12E+08 | 1,12E+08 | 1600 | * | 12,11888 | 2,857388436 | 1,098243 | 2,601781 | 0,009274 | 0,974801712 | 0,738441 Promoter (<=1kb) | 12 | 1,12E+08 | 1,12E+08 | 12176 | 2 | 1,01E+08 | ENSMUST00000221848.2 | 298 | ENSMUSG00000047828 | A730018C14Rik | RIKEN cDNA A730018C14 gene |
| chr8 | 84902066 | 84903291 | 1226 | * | 15,6416 | 2,85763946 | 0,903014 | 3,164556 | 0,001553 | 0,487591669 | 2,141612 Promoter (<=1kb) | 8 | 84900362 | 84903181 | 2820 | 2 | 244551 | ENSMUST00000070102.6 | 0 | ENSMUSG00000056155 | Nanos3 | nanos C2HC-type zinc finger 3 |
| chr2 | 65429543 | 65430708 | 1166 | * | 13,08148 | 2,85816047 | 1,002649 | 2,850608 | 0,004364 | 0,777468971 | 1,173325 Distal Intergenic | 2 | 65451115 | 65520519 | 69405 | 1 | 110876 | ENSMUST00000144254.6 | -20407 | ENSMUSG00000075318 | Scn2a | sodium channel, voltage-gated, type II, alpha |
| chr18 | 17086291 | 17087104 | 814 | * | 10,06967 | 2,858173762 | 1,090453 | 2,621089 | 0,008765 | 0,860855381 | 1,449706 Distal Intergenic | 18 | 17152108 | 17270616 | 118509 | 1 | 75433 | ENSMUST00000147119.8 | -65004 | ENSMUSG00000087170 | 1700001G01Rik | RIKEN cDNA 1700001G01 gene |
| chr12 | 80457112 | 80459783 | 2672 | * | 20,15112 | 2,858256891 | 0,787877 | 3,627794 | 0,000286 | 0,227550076 | 2,221611 Intron (ENSMUST0000 | 12 | 80438387 | 80438525 | 139 | 2 | 1E+08 | ENSMUST00020182423.1 | -18587 | ENSMUSG00000112285 | Scarna3b | small Cajal body-specific RNA 3B |
| chr2 | 38823470 | 38824518 | 1049 | * | 12,63653 | 2,858980258 | 1,095674 | 2,609335 | 0,009072 | 0,903294867 | 1,173325 Promoter (1-2kb) | 2 | 38821990 | 38853765 | 31776 | 1 | 241327 | ENSMUST00000057279.6 | 1480 | ENSMUSG00000046618 | Olfml2a | olfactomedin-like 2A |
| chr15 | 99337857 | 99338537 | 681 | * | 9,886292 | 2,859278482 | 1,062532 | 2,691005 | 0,007124 | 0,860965195 | 1,173325 Intron (ENSMUST0000 | 15 | 99325582 | 99333194 | 7613 | 1 | 1,03E+08 | ENSMUST00000159738.2 | 12275 | | Nckap5los | NCK-associated protein 5-like, opposite strand |
| chr15 | 36285542 | 36287168 | 1627 | * | 12,66095 | 2,859867299 | 1,06447 | 2,686658 | 0,007217 | 0,863495249 | 1,173325 Promoter (2-3kb) | 15 | 36240079 | 36283293 | 43215 | 2 | 30945 | ENSMUST00000222890.10 | -2249 | ENSMUSG00000022280 | Rnf19a | ring finger protein 19A |
| chr9 | 59516070 | 59518471 | 2402 | * | 23,09894 | 2,86034101 | 0,802212 | 3,565568 | 0,000363 | 0,223919137 | 2,951062 Distal Intergenic | 9 | 59511316 | 59513384 | 2069 | 1 | 76183 | ENSMUST00000129728.2 | 4754 | ENSMUSG00000032297 | Celf6 | CUGBP, Elav-like family member 6 |
| chrX | 1,62E+08 | 1,62E+08 | 1194 | * | 9,37042 | 2,860891659 | 1,005209 | 2,846067 | 0,004426 | 0,670853401 | 2,237403 Distal Intergenic | 20 | 1,62E+08 | 1,62E+08 | 120 | 2 | 1,15E+08 | ENSMUST00000158798.3 | 79687 | ENSMUSG00000089423 | Gm26007 | predicted gene, 26007 |
| chr2 | 156294994 | 156299631 | 4638 | * | 49,430733 | 2,86223113 | 0,5074887 | 5,6399903 | 1,701E-08 | 0,000164635 | 2,7254908 Exon (ENSMUST000001 | 2 | 156317454 | 156385132 | 67679 | 1 | 13821 | ENSMUST00000029155.16 | -17823 | ENSMUSG00000027624 | Epb41l1 | erythrocyte membrane protein band 4.1 like 1 |
| chr13 | 34226009 | 34227179 | 1171 | * | 11,17974 | 2,863343059 | 1,054957 | 2,71418 | 0,006644 | 0,892655056 | 0,918255 Intron (ENSMUST0000 | 13 | 34221640 | 34234306 | 12667 | 1 | 68021 | ENSMUST00000222691.2 | 4369 | ENSMUSG00000038286 | Bphl | biphenyl hydrolase-like (serine hydrolase, breast epithelial mucin-associated antigen) |
| chr9 | 41726831 | 41728194 | 1364 | * | 17,39336 | 2,86412893 | 0,917733 | 3,120872 | 0,001803 | 0,494741046 | 2,386008 Promoter (<=1kb) | 9 | 41726900 | 41749714 | 22815 | 1 | 1,03E+08 | ENSMUST00000213355.2 | 0 | | Gm35940 | predicted gene, 35940 |
| chr16 | 7464286 | 7465243 | 958 | * | 9,03934 | 2,864366482 | 1,109551 | 2,581554 | 0,009836 | 0,892445563 | 1,360622 Distal Intergenic | 16 | 7170834 | 7227833 | 57000 | 1 | 268859 | ENSMUST00000231031.2 | 293452 | ENSMUSG00000008658 | Rbfox1 | RNA binding protein, fox-1 homolog (C. elegans) 1 |
| chr6 | 1,23E+08 | 1,23E+08 | 1215 | * | 16,63815 | 2,864510773 | 0,8736 | 3,278972 | 0,001042 | 0,421090004 | 2,141612 Promoter (<=1kb) | 6 | 1,23E+08 | 1,23E+08 | 13149 | 1 | 17474 | ENSMUST00000032240.4 | 17 | ENSMUSG00000030144 | Clec4d | C-type lectin domain family 4, member d |
| chrX | 56755427 | 56757107 | 1681 | * | 9,936584 | 2,864861692 | 1,102088 | 2,599484 | 0,009336 | 0,874247416 | 1,449706 Distal Intergenic | 20 | 56542028 | 56549117 | 7090 | 2 | 245424 | ENSMUST00000057645.6 | -206310 | ENSMUSG00000036357 | Gpr101 | G protein-coupled receptor 101 |
| chr1 | 55066435 | 55067636 | 1202 | * | 13,98842 | 2,866974877 | 1,093532 | 2,621757 | 0,008748 | 0,935170595 | 0,918255 Promoter (<=1kb) | 1 | 55024328 | 55066640 | 42313 | 2 | 81898 | ENSMUST00000027127.14 | 0 | ENSMUSG00000025982 | Sf3b1 | splicing factor 3b, subunit 1 |
| chr2 | 1,31E+08 | 1,31E+08 | 1102 | * | 11,49177 | 2,867096347 | 1,069937 | 2,679687 | 0,007369 | 0,942793663 | 0,738441 Promoter (<=1kb) | 2 | 1,31E+08 | 1,31E+08 | 29002 | 1 | 11990 | ENSMUST00000156982.2 | 30 | ENSMUSG00000027312 | Atrn | attractin |
| chr1 | 62789147 | 62792302 | 3156 | * | 21,55296 | 2,868039635 | 0,829006 | 3,459614 | 0,000541 | 0,268627242 | 3,104559 Intron (ENSMUST0000 | 1 | 62753882 | 62757232 | 3351 | 2 | 1,05E+08 | ENSMUST00000189205.2 | -31915 | | Gm29083 | predicted gene 29083 |
| chr3 | 94446418 | 94448477 | 2060 | * | 16,4249 | 2,868534757 | 0,818762 | 3,503503 | 0,000459 | 0,280986007 | 2,386008 Intron (ENSMUST0000 | 3 | 94408348 | 94431598 | 23251 | 2 | 76742 | ENSMUST00000196675.5 | -14820 | ENSMUSG00000028136 | Snx27 | sorting nexin family member 27 |
| chr2 | 1,48E+08 | 1,48E+08 | 1047 | * | 12,64809 | 2,868766201 | 1,007365 | 2,847792 | 0,004402 | 0,792545355 | 1,075735 Distal Intergenic | 2 | 1,48E+08 | 1,48E+08 | 313 | 1 | 1,15E+08 | ENSMUST00000158833.4 | 78699 | ENSMUSG00000089458 | Gm24221 | predicted gene, 24221 |
| chr1 | 62461929 | 62463278 | 1350 | * | 11,40003 | 2,869354943 | 0,991861 | 2,892902 | 0,003817 | 0,794811783 | 0,918255 Intron (ENSMUST0000 | 1 | 62742444 | 62779091 | 36648 | 1 | 18187 | ENSMUST00000126344.2 | -279166 | ENSMUSG00000025969 | Nrp2 | neuropilin 2 |
| chr6 | 9307319 | 9308143 | 825 | * | 12,71828 | 2,869604239 | 1,102202 | 2,603519 | 0,009227 | 0,892445563 | 1,2763 Distal Intergenic | 6 | 8949676 | 9249030 | 299355 | 1 | 18231 | ENSMUST00000060369.4 | 357643 | ENSMUSG00000046178 | Nxph1 | neurexophilin 1 |
| chr7 | 1,01E+08 | 1,01E+08 | 1963 | * | 17,8989 | 2,869639701 | 0,892842 | 3,214051 | 0,001309 | 0,457298447 | 2,154504 Promoter (1-2kb) | 7 | 1,01E+08 | 1,01E+08 | 26762 | 2 | 233571 | ENSMUST00000060174.6 | 1606 | ENSMUSG00000048779 | P2ry6 | pyrimidinergic receptor P2Y, G-protein coupled, 6 |
| chr14 | 31376412 | 31377971 | 1560 | * | 14,80012 | 2,869716287 | 0,970605 | 2,956625 | 0,00311 | 0,618056482 | 2,141612 Intron (ENSMUST0000 | 14 | 31357605 | 31363243 | 5639 | 2 | 56794 | ENSMUST00000135751.2 | -13169 | ENSMUSG00000021884 | Hacl1 | 2-hydroxyacyl-CoA lyase 1 |
| chr7 | 1,26E+08 | 1,26E+08 | 1277 | * | 13,29921 | 2,870125684 | 0,990214 | 2,898491 | 0,00375 | 0,765163361 | 1,075735 Promoter (<=1kb) | 7 | 1,26E+08 | 1,26E+08 | 4392 | 1 | 70314 | ENSMUST00000148536.2 | 0 | ENSMUSG00000030727 | Rabep2 | rabaptin, RAB GTPase binding effector protein 2 |
| chr11 | 17164073 | 17165361 | 1289 | * | 11,38394 | 2,870430221 | 1,02283 | 2,806361 | 0,00501 | 0,883012141 | 0,744375 Promoter (2-3kb) | 11 | 17161918 | 17183796 | 21879 | 1 | 103784 | ENSMUST00000046955.7 | 2155 | ENSMUSG00000078970 | Dnaaf10 | dynein axonemal assembly factor 10 |
| chr3 | 48898332 | 48899604 | 1273 | * | 9,511023 | 2,870444414 | 1,082627 | 2,65137 | 0,008017 | 0,879615137 | 1,211315 Distal Intergenic | 3 | 48909171 | 48909334 | 164 | 1 | 1,15E+08 | ENSMUST00000239931.1 | -9567 | ENSMUSG00000118871 | Gm23511 | predicted gene, 23511 |
| chr3 | 65865525 | 65868256 | 2732 | * | 27,97886 | 2,870747227 | 0,711172 | 4,036642 | 5,42E-05 | 0,072320047 | 2,880017 Promoter (<=1kb) | 3 | 65853572 | 65865583 | 12012 | 2 | 56706 | ENSMUST00000129002.8 | 0 | ENSMUSG00000027829 | Ccnl1 | cyclin L1 |
| chr13 | 30201360 | 30202642 | 1283 | * | 10,10549 | 2,87078781 | 1,051807 | 2,729387 | 0,006345 | 0,809369408 | 1,425929 Distal Intergenic | 13 | 30090558 | 30170046 | 79489 | 2 | 13557 | ENSMUST00000102948.11 | -31314 | ENSMUSG00000016477 | E2f3 | E2F transcription factor 3 |
| chr15 | 91631392 | 91632812 | 1421 | * | 18,03598 | 2,871064519 | 0,878692 | 3,267429 | 0,001085 | 0,426043749 | 2,154504 Exon (ENSMUST000000 | 15 | 91623151 | 91673313 | 50163 | 1 | 66725 | ENSMUST00000140734.8 | 8241 | ENSMUSG00000036273 | Lrrk2 | leucine-rich repeat kinase 2 |
| chr3 | 95124745 | 95127067 | 2323 | * | 29,342114 | 2,871594715 | 0,5979807 | 4,802153 | 1,57E-06 | 0,005348945 | 2,8800171 Promoter (<=1kb) | 3 | 95124768 | 95133969 | 9202 | 1 | 100504029 | ENSMUST00000181819.2 | 0 | ENSMUSG00000097167 | Gm16740 | predicted gene, 16740 |
| chr4 | 34333750 | 34334882 | 1133 | * | 12,31371 | 2,873692006 | 1,086949 | 2,643815 | 0,008198 | 0,874025796 | 1,2763 Distal Intergenic | 4 | 34550937 | 34566908 | 15972 | 1 | 433693 | ENSMUST00000084299.6 | -216055 | ENSMUSG00000028291 | Akirin2 | akirin 2 |
| chr4 | 150320172 | 150322864 | 2693 | * | 85,768248 | 2,87371679 | 0,429886 | 6,6848352 | 2,312E-11 | 6,36368E-07 | 3,439501 Promoter (<=1kb) | 4 | 150321178 | 150328746 | 7569 | 1 | 13806 | ENSMUST00000150175.8 | 0 | ENSMUSG00000063524 | Eno1 | enolase 1, alpha non-neuron |
| chr14 | 31471097 | 31472885 | 1789 | * | 11,73621 | 2,875129677 | 1,104601 | 2,602867 | 0,009245 | 0,974266082 | 0,738441 Promoter (1-2kb) | 14 | 31467262 | 31469407 | 2146 | 2 | 105522 | ENSMUST00000226963.2 | -1690 | ENSMUSG00000014496 | Ankrd28 | ankyrin repeat domain 28 |
| chr11 | 1,16E+08 | 1,16E+08 | 1823 | * | 15,31275 | 2,875400468 | 0,98788 | 2,910679 | 0,003606 | 0,729319353 | 1,2763 Promoter (2-3kb) | 11 | 1,16E+08 | 1,16E+08 | 10647 | 1 | 71947 | ENSMUST00000134948.2 | 2162 | ENSMUSG00000020747 | Tmem94 | transmembrane protein 94 |
| chr16 | 33334889 | 33336604 | 1716 | * | 12,53671 | 2,875635219 | 0,955442 | 3,009744 | 0,002615 | 0,702222274 | 1,075735 Promoter (1-2kb) | 16 | 33337698 | 33483778 | 146081 | 1 | 171286 | ENSMUST00000121925.8 | -1094 | ENSMUSG00000035506 | Slc12a8 | solute carrier family 12 (potassium/chloride transporters), member 8 |
| chr7 | 51968729 | 51969896 | 1168 | * | 9,37348 | 2,876368976 | 1,102614 | 2,608683 | 0,009089 | 0,872427545 | 1,425929 Distal Intergenic | 7 | 51661431 | 51665476 | 4046 | 2 | 1,01E+08 | ENSMUST00000180038.8 | -303253 | ENSMUSG00000094445 | 1700015G11Rik | RIKEN cDNA 1700015G11 gene |
| chr1 | 1,5E+08 | 1,5E+08 | 2215 | * | 24,96191 | 2,876482757 | 0,704681 | 4,081963 | 4,47E-05 | 0,063407977 | 2,951933 Promoter (<=1kb) | 1 | 1,5E+08 | 1,5E+08 | 56978 | 1 | 108989 | ENSMUST00000124973.9 | 0 | ENSMUSG00000006005 | Tpr | translocated promoter region, nuclear basket protein |
| chr9 | 1,21E+08 | 1,21E+08 | 1262 | * | 18,70216 | 2,876618004 | 0,886453 | 3,245089 | 0,001174 | 0,423546149 | 2,386008 3' UTR | 9 | 1,21E+08 | 1,21E+08 | 18020 | 1 | 67095 | ENSMUST00000209446.2 | 14532 | ENSMUSG00000032536 | Trak1 | trafficking protein, kinesin binding 1 |
| chr18 | 25996260 | 25997555 | 1296 | * | 14,02318 | 2,876991674 | 0,97844 | 2,940388 | 0,003278 | 0,765231686 | 0,938682 Distal Intergenic | 18 | 25972134 | 25992477 | 20344 | 1 | 1,03E+08 | ENSMUST00000234817.2 | 24126 | | Gm33228 | predicted gene, 33228 |
| chr4 | 1,55E+08 | 1,55E+08 | 1393 | * | 13,73924 | 2,877050098 | 0,921576 | 3,121882 | 0,001797 | 0,642870233 | 1,075735 Downstream (<=300bp | 4 | 1,55E+08 | 1,55E+08 | 18089 | 1 | 76866 | ENSMUST00000129415.2 | 34867 | ENSMUSG00000029049 | Morn1 | MORN repeat containing 1 |
| chr17 | 36009988 | 36011315 | 1328 | * | 18,70999 | 2,877931867 | 0,765629 | 3,758912 | 0,000171 | 0,162641545 | 2,423594 Promoter (<=1kb) | 17 | 36010095 | 36010201 | 107 | 2 | 1,15E+08 | ENSMUST00001239549.1 | 0 | ENSMUSG00001118668 | Gm23864 | predicted gene, 23864 |
| chr2 | 5952050 | 5953002 | 953 | * | 8,208273 | 2,878157667 | 1,110437 | 2,591914 | 0,009544 | 0,881672947 | 1,425929 Distal Intergenic | 2 | 5956280 | 6061514 | 105235 | 1 | 326622 | ENSMUST00000060092.13 | -3278 | ENSMUSG00000043241 | Upf2 | UPF2 regulator of nonsense transcripts homolog (yeast) |
| chr4 | 1,34E+08 | 1,34E+08 | 1779 | * | 12,68148 | 2,878440819 | 1,04764 | 2,747548 | 0,006004 | 0,851014755 | 1,075735 Intron (ENSMUST0000 | 4 | 1,34E+08 | 1,34E+08 | 135879 | 2 | 230815 | ENSMUST00000176606.2 | 44888 | ENSMUSG00000037306 | Man1c1 | mannosidase, alpha, class 1C, member 1 |
| chr17 | 35900668 | 35902447 | 1780 | * | 15,48776 | 2,878531595 | 0,786964 | 3,657769 | 0,000254 | 0,275385512 | 1,37683 Distal Intergenic | 17 | 35912441 | 35961459 | 49019 | 1 | 433102 | ENSMUST00000174521.2 | -9994 | ENSMUSG00000090509 | Sfta2 | surfactant associated 2 |
| chr10 | 67897679 | 67899157 | 1479 | * | 12,66105 | 2,879726528 | 1,043413 | 2,75991 | 0,005782 | 0,905015433 | 0,738441 Distal Intergenic | 10 | 67879798 | 67886876 | 7079 | 1 | 170799 | ENSMUST00000123147.2 | 17881 | ENSMUSG00000037846 | Rtkn2 | rhotekin 2 |
| chr8 | 71938753 | 71940049 | 1297 | * | 10,85124 | 2,879824924 | 1,00927 | 2,853375 | 0,004326 | 0,758346111 | 1,2763 Promoter (<=1kb) | 8 | 71928663 | 71938607 | 9945 | 2 | 382014 | ENSMUST00000213382.2 | -146 | ENSMUSG00000034863 | Ano8 | anoctamin 8 |
| chr11 | 22280528 | 22281905 | 1378 | * | 12,92872 | 2,880254355 | 0,945366 | 3,04671 | 0,002314 | 0,710505631 | 0,918255 Intron (ENSMUST0000 | 11 | 22182002 | 22292292 | 110291 | 2 | 216565 | ENSMUST00000137594.2 | 10387 | ENSMUSG00000042302 | Ehbp1 | EH domain binding protein 1 |
| chr11 | 1,03E+08 | 1,03E+08 | 2110 | * | 19,92502 | 2,880292944 | 0,840733 | 3,425931 | 0,000613 | 0,302471858 | 2,663619 Intron (ENSMUST0000 | 11 | 1,03E+08 | 1,03E+08 | 9273 | 1 | 1,05E+08 | ENSMUST00000212331.3 | 3761 | | Gm39397 | predicted gene, 39397 |
| chr6 | 1,42E+08 | 1,42E+08 | 1575 | * | 11,49631 | 2,88066083 | 0,953405 | 3,021444 | 0,002516 | 0,683210259 | 1,173325 Distal Intergenic | 6 | 1,42E+08 | 1,42E+08 | 47474 | 2 | 28254 | ENSMUST00000174455.2 | -15946 | ENSMUSG00000079262 | Slco1a6 | solute carrier organic anion transporter family, member 1a6 |
| chr9 | 84160905 | 84162582 | 1678 | * | 12,55437 | 2,880913848 | 1,0869 | 2,650579 | 0,008035 | 0,868766096 | 1,2763 Distal Intergenic | 9 | 84070408 | 84070592 | 185 | 1 | 1,15E+08 | ENSMUST00000104478.4 | 90497 | ENSMUSG00000077666 | Gm26146 | predicted gene, 26146 |
| chr2 | 30660403 | 30661424 | 1022 | * | 11,72487 | 2,880984903 | 0,922734 | 3,122226 | 0,001795 | 0,611385361 | 1,2763 Distal Intergenic | 2 | 30685480 | 30686676 | 1197 | 2 | 73598 | ENSMUST00000130396.8 | 25252 | ENSMUSG00000044320 | 1700001O22Rik | RIKEN cDNA 1700001O22 gene |
| chr9 | 53433788 | 53435256 | 1469 | * | 13,61412 | 2,881080202 | 1,034175 | 2,785873 | 0,005338 | 0,827954389 | 1,075735 Intron (ENSMUST0000 | 9 | 53350449 | 53445266 | 94818 | 2 | 11920 | ENSMUST00000118282.9 | 10010 | ENSMUSG00000034218 | Atm | ataxia telangiectasia mutated |
| chr4 | 1,36E+08 | 1,36E+08 | 1643 | * | 20,92687 | 2,88108348 | 0,730407 | 3,944493 | 8E-05 | 0,086915529 | 3,104559 Exon (ENSMUST000000 | 4 | 1,36E+08 | 1,36E+08 | 23576 | 1 | 21401 | ENSMUST00000136812.2 | -17746 | ENSMUSG00000001604 | Tcea3 | transcription elongation factor A (SII), 3 |
| chr15 | 79055492 | 79056381 | 890 | * | 9,400751 | 2,881175722 | 1,114381 | 2,58545 | 0,009725 | 0,802671652 | 2,237403 Promoter (<=1kb) | 15 | 79056920 | 79100697 | 43778 | 1 | 1E+08 | ENSMUST00000229207.2 | -539 | ENSMUSG00000075555 | Gm10863 | predicted gene 10863 |
| chr14 | 30928841 | 30930064 | 1224 | * | 17,4296 | 2,881207875 | 0,959436 | 3,003021 | 0,002673 | 0,586565597 | 2,154504 Promoter (<=1kb) | 14 | 30905270 | 30928783 | 23514 | 2 | 64652 | ENSMUST00000172142.8 | -58 | ENSMUSG00000021910 | Nisch | nischarin |
| chr3 | 1,22E+08 | 1,22E+08 | 2481 | * | 19,08941 | 2,881417755 | 0,936925 | 3,075398 | 0,002102 | 0,498011371 | 2,713306 Promoter (1-2kb) | 3 | 1,22E+08 | 1,22E+08 | 11357 | 1 | 14630 | ENSMUST00000149983.4 | -1128 | ENSMUSG00000028124 | Gclm | glutamate-cysteine ligase, modifier subunit |
| chr9 | 62061146 | 62062683 | 1538 | * | 13,60382 | 2,88173136 | 1,06429 | 2,707655 | 0,006776 | 0,869013335 | 1,075735 Distal Intergenic | 9 | 61977750 | 62029937 | 52188 | 2 | 93683 | ENSMUST00000185873.2 | -31209 | ENSMUSG00000032252 | Glce | glucuronyl C5-epimerase |
| chr11 | 8506791 | 8508720 | 1930 | * | 18,07832 | 2,88177288 | 0,80659 | 3,572786 | 0,000353 | 0,259958999 | 2,154504 Intron (ENSMUST0000 | 11 | 8487539 | 8495351 | 7813 | 2 | 319939 | ENSMUST00000131941.2 | -11440 | ENSMUSG00000020422 | Tns3 | tensin 3 |
| chr19 | 56583098 | 56584449 | 1352 | * | 13,69025 | 2,881927502 | 0,977744 | 2,947527 | 0,003203 | 0,802671652 | 0,738441 Exon (ENSMUST000000 | 19 | 56559229 | 56580076 | 20848 | 1 | 66866 | ENSMUST00000236979.2 | 23869 | ENSMUSG00000025078 | Nhlrc2 | NHL repeat containing 2 |
| chr14 | 23636842 | 23638547 | 1706 | * | 15,5632 | 2,882461233 | 0,91013 | 3,167086 | 0,00154 | 0,559413722 | 1,41696 Intron (ENSMUST0000 | 14 | 23551193 | 23604866 | 53674 | 2 | 16531 | ENSMUST00000212576.2 | -31976 | ENSMUSG00000063142 | Kcnma1 | potassium large conductance calcium-activated channel, subfamily M, alpha member 1 |
| chr16 | 4695317 | 4699926 | 4610 | * | 64,190616 | 2,882705474 | 0,4867736 | 5,9220657 | 3,179E-09 | 3,04561E-05 | 4,2003037 Promoter (<=1kb) | 16 | 4692642 | 4698179 | 5538 | 2 | 207740 | ENSMUST00000037843.7 | 0 | ENSMUSG00000039568 | Ubald1 | UBA-like domain containing 1 |
| chrX | 98271823 | 98274009 | 2187 | * | 13,85828 | 2,883232602 | 0,936509 | 3,078703 | 0,002079 | 0,728651153 | 0,738441 Distal Intergenic | 20 | 98181737 | 98190836 | 9100 | 1 | 13641 | ENSMUST00000151750.2 | 90086 | ENSMUSG00000031217 | Efnb1 | ephrin B1 |
| chr5 | 14035643 | 14036783 | 1141 | * | 10,64097 | 2,883777324 | 0,99837 | 2,888484 | 0,003871 | 0,753418617 | 1,173325 Distal Intergenic | 5 | 14075290 | 14304328 | 229039 | 1 | 20349 | ENSMUST00000130116.2 | -38507 | ENSMUSG00000063531 | Sema3e | sema domain, immunoglobulin domain (Ig), short basic domain, secreted, (semaphorin) 3E |
| chr13 | 52155661 | 52156878 | 1218 | * | 11,07923 | 2,883944069 | 1,029572 | 2,80111 | 0,005093 | 0,849437958 | 0,918255 Distal Intergenic | 13 | 52250777 | 52278197 | 27421 | 1 | 74050 | ENSMUST00000220691.3 | -93899 | ENSMUSG00000113211 | 4921525O09Rik | RIKEN cDNA 4921525O09 gene |
| chr5 | 77514854 | 77515704 | 851 | * | 9,175298 | 2,884599102 | 1,090995 | 2,644008 | 0,008193 | 0,852104582 | 1,449706 Intron (ENSMUST0000 | 5 | 77534446 | 77541530 | 7085 | 1 | 1,03E+08 | ENSMUST00000202460.2 | -18742 | | Gm34648 | predicted gene, 34648 |
| chr4 | 1,36E+08 | 1,36E+08 | 1194 | * | 15,35848 | 2,884963859 | 0,86535 | 3,333869 | 0,000856 | 0,462239985 | 1,37683 Promoter (2-3kb) | 4 | 1,36E+08 | 1,36E+08 | 16490 | 1 | 242707 | ENSMUST00000178843.3 | -2887 | ENSMUSG00000070683 | Lactbl1 | lactamase, beta-like 1 |
| chr19 | 36817791 | 36820770 | 2980 | * | 20,33091 | 2,88516396 | 0,890277 | 3,240747 | 0,001192 | 0,408869321 | 2,663619 Intron (ENSMUST0000 | 19 | 36811804 | 36833337 | 21534 | 1 | 74493 | ENSMUST00000168656.8 | 5987 | ENSMUSG00000024811 | Tnks2 | tankyrase, TRF1-interacting ankyrin-related ADP-ribose polymerase 2 |
| chr14 | 20794952 | 20796649 | 1698 | * | 13,12636 | 2,885815048 | 1,108548 | 2,603238 | 0,009235 | 0,944611132 | 0,918255 Exon (ENSMUST000000 | 14 | 20786677 | 20789465 | 2789 | 2 | 12325 | ENSMUST00000223863.2 | -5487 | ENSMUSG00000021820 | Camk2g | calcium/calmodulin-dependent protein kinase II gamma |
| chr4 | 1,36E+08 | 1,36E+08 | 1216 | * | 11,00149 | 2,88586148 | 0,987681 | 2,921854 | 0,00348 | 0,736412045 | 1,173325 Promoter (<=1kb) | 4 | 1,36E+08 | 1,36E+08 | 580 | 2 | 67025 | ENSMUST00000133613.2 | 0 | ENSMUSG00000059291 | Rpl11 | ribosomal protein L11 |
| chr19 | 23982045 | 23983863 | 1819 | * | 14,61394 | 2,886487197 | 1,029042 | 2,805023 | 0,005031 | 0,793995086 | 1,214904 Exon (ENSMUST00000 | 19 | 23950114 | 24008383 | 58270 | 2 | 381217 | ENSMUST00000096164.6 | 24520 | ENSMUSG00000071604 | Fam189a2 | family with sequence similarity 189, member A2 |
| chr5 | 139184652 | 139187679 | 3028 | * | 31,454189 | 2,886868956 | 0,6104251 | 4,7292766 | 2,253E-06 | 0,007087666 | 2,8585108 Promoter (<=1kb) | 5 | 139186392 | 139235595 | 49204 | 1 | 77053 | ENSMUST00000058716.14 | 0 | ENSMUSG00000036817 | Sun1 | Sad1 and UNC84 domain containing 1 |
| chr12 | 84226384 | 84228464 | 2081 | * | 19,7577 | 2,887777353 | 0,799512 | 3,611925 | 0,000304 | 0,214556109 | 2,663619 Intron (ENSMUST0000 | 12 | 84196052 | 84240789 | 44738 | 2 | 238317 | ENSMUST00000110294.2 | 12325 | ENSMUSG00000042507 | Mideas | mitotic deacetylase associated SANT domain protein |
| chr4 | 53409640 | 53411162 | 1523 | * | 12,73262 | 2,887869061 | 1,068643 | 2,702371 | 0,006885 | 0,932399283 | 0,738441 Distal Intergenic | 4 | 53401948 | 53404030 | 2083 | 2 | 1,03E+08 | ENSMUST00000154377.2 | -5610 | | Gm12496 | predicted gene 12496 |
| chr12 | 19783854 | 19785446 | 1593 | * | 9,766051 | 2,88795338 | 1,105796 | 2,611651 | 0,009011 | 0,898985075 | 1,211315 Distal Intergenic | 12 | 19874892 | 19880373 | 5482 | 2 | 1,05E+08 | ENSMUST00000223533.2 | 94927 | | Gm38702 | predicted gene, 38702 |
| chr8 | 39472505 | 39474261 | 1757 | * | 18,63936 | 2,888201765 | 0,776259 | 3,720669 | 0,000199 | 0,190625259 | 2,154504 Promoter (<=1kb) | 8 | 39472999 | 39619367 | 146369 | 1 | 80286 | ENSMUST00000239508.1 | 0 | ENSMUSG00000118664 | Tusc3 | tumor suppressor candidate 3 |
| chr1 | 1,81E+08 | 1,81E+08 | 1924 | * | 12,79491 | 2,889505176 | 0,939381 | 3,075966 | 0,002098 | 0,641187423 | 1,2763 3' UTR | 1 | 1,81E+08 | 1,81E+08 | 4154 | 1 | 208768 | ENSMUST00000194948.2 | 5420 | ENSMUSG00000038806 | Sde2 | SDE2 telomere maintenance homolog (S. pombe) |
| chr1 | 1,38E+08 | 1,38E+08 | 1670 | * | 15,65314 | 2,889638933 | 0,817104 | 3,536438 | 0,000406 | 0,352867589 | 1,214904 Distal Intergenic | 1 | 1,38E+08 | 1,38E+08 | 18362 | 1 | 1E+08 | ENSMUST00000245018.1 | -115146 | | Gm4258 | predicted gene 4258 |
| chr11 | 1,17E+08 | 1,17E+08 | 1697 | * | 15,31332 | 2,889967655 | 0,936377 | 3,086329 | 0,002026 | 0,615459792 | 1,41696 Promoter (<=1kb) | 11 | 1,17E+08 | 1,17E+08 | 4130 | 1 | 1E+08 | ENSMUST00000247579.1 | 0 | ENSMUSG00000065640 | Snord1c | small nucleolar RNA, C/D box 1C |

| | | | | | | | | | | | | | | | | | | | | | | | |
|---|---|---|---|---|---|---|---|---|---|---|---|---|---|---|---|---|---|---|---|---|---|---|---|
| chr18 | 80229320 | 80230849 | 1530 | * | 16,05962 | 2,890192362 | 0,851344 | 3,394859 | 0,000687 | 0,315279198 | 2,663619 | Promoter (<=1kb) | 18 | 80229454 | 80236782 | 7329 | 1 | 1,05E+08 | ENSMUST00000243491.1 | 0 | ENSMUSG00000118061 | Rbfaos | ribosome binding factor A, opposite strand |
| chr5 | 1,04E+08 | 1,04E+08 | 1497 | * | 12,50873 | 2,890795225 | 0,97649 | 2,960393 | 0,003072 | 0,794811783 | 0,738441 | Promoter (<=1kb) | 5 | 1,04E+08 | 1,04E+08 | 13866 | 2 | 243168 | ENSMUST00000120320.8 | 495 | ENSMUSG00000034528 | Hsd17b13 | hydroxysteroid (17-beta) dehydrogenase 13 |
| chr11 | 77703590 | 77705083 | 1494 | * | 12,29957 | 2,891422349 | 1,082857 | 2,67018 | 0,007581 | 0,947773602 | 0,738441 | Promoter (1-2kb) | 11 | 77702341 | 77756806 | 54466 | 1 | 360013 | ENSMUST00000167856.8 | 1249 | ENSMUSG00000000631 | Myo18a | myosin XVIIIA |
| chr18 | 61817403 | 61819173 | 1771 | * | 14,83758 | 2,89158352 | 1,07981 | 2,677864 | 0,007409 | 0,85518169 | 1,2763 | Promoter (<=1kb) | 18 | 61817958 | 61825609 | 7652 | 1 | 56069 | ENSMUST00000237575.2 | 0 | ENSMUSG00000024578 | Il17b | interleukin 17B |
| chr4 | 28045787 | 28046953 | 1167 | * | 15,03216 | 2,891603749 | 0,898947 | 3,216656 | 0,001297 | 0,533665288 | 1,37683 | Distal Intergenic | 4 | 28365316 | 28365444 | 129 | 1 | 1,15E+08 | ENSMUST00000104333.2 | -318363 | ENSMUSG00000077521 | Gm25378 | predicted gene, 25378 |
| chr7 | 1,25E+08 | 1,25E+08 | 1373 | * | 12,38094 | 2,893135744 | 0,974509 | 2,968813 | 0,00299 | 0,725884027 | 1,075735 | Distal Intergenic | 7 | 1,25E+08 | 1,25E+08 | 27355 | 1 | 16190 | ENSMUST00000033004.8 | -24532 | ENSMUSG00000030748 | Il4ra | interleukin 4 receptor, alpha |
| chr11 | 20124935 | 20126123 | 1189 | * | 9,566579 | 2,894111982 | 1,116881 | 2,591245 | 0,009563 | 0,906138253 | 1,211315 | Distal Intergenic | 11 | 20151432 | 20176856 | 25425 | 1 | 19324 | ENSMUST00000020358.12 | -25309 | ENSMUSG00000020149 | Rab1a | RAB1A, member RAS oncogene family |
| chr17 | 11539043 | 11540018 | 976 | * | 8,912408 | 2,894140371 | 1,108358 | 2,611197 | 0,009023 | 0,865306051 | 1,449706 | Intron (ENSMUST0000 | 17 | 11522534 | 11829832 | 307299 | 1 | 50873 | ENSMUST00000186406.2 | 16509 | ENSMUSG00000023826 | Prkn | parkin RBR E3 ubiquitin protein ligase |
| chr15 | 38382190 | 38385631 | 3442 | * | 57,064698 | 2,89451162 | 0,4634872 | 6,2450734 | 4,236E-10 | 5,60348E-06 | 4,0557576 | Distal Intergenic | 15 | 38385878 | 38404398 | 18521 | 2 | 105245929 | ENSMUST00000227422.2 | 18767 | | Gm41307 | predicted gene, 41307 |
| chr4 | 1,29E+08 | 1,29E+08 | 1703 | * | 16,59857 | 2,895136835 | 1,014736 | 2,853094 | 0,00433 | 0,639496036 | 2,663619 | Promoter (2-3kb) | 4 | 1,29E+08 | 1,29E+08 | 28866 | 2 | 97130 | ENSMUST00000052602.6 | -2271 | ENSMUSG00000050390 | C77080 | expressed sequence C77080 |
| chr5 | 1,49E+08 | 1,49E+08 | 925 | * | 10,0317 | 2,895178378 | 0,98856 | 2,928681 | 0,003404 | 0,697709534 | 1,425929 | Promoter (<=1kb) | 5 | 1,49E+08 | 1,49E+08 | 1409 | 1 | 71473 | ENSMUST00000110507.2 | 0 | ENSMUSG00000079069 | 8430423G03Rik | RIKEN cDNA 8430423G03 gene |
| chr13 | 21934045 | 21935654 | 1610 | * | 15,70307 | 2,895831992 | 0,964899 | 3,001177 | 0,002689 | 0,684274806 | 1,214904 | Promoter (<=1kb) | 13 | 21934364 | 21934675 | 312 | 2 | 319160 | ENSMUST00000102983.2 | 0 | ENSMUSG00000064288 | H4c12 | H4 clustered histone 12 |
| chr15 | 20138859 | 20140248 | 1390 | * | 10,12962 | 2,896481854 | 1,105188 | 2,620806 | 0,008772 | 0,892763628 | 1,211315 | Distal Intergenic | 15 | 20184918 | 20185047 | 130 | 1 | 1,15E+08 | ENSMUST00000158695.3 | -44670 | ENSMUSG00000089320 | Gm25976 | predicted gene, 25976 |
| chr19 | 42108447 | 42109899 | 1453 | * | 12,23731 | 2,896632652 | 1,0846 | 2,670692 | 0,007569 | 0,947639325 | 0,738441 | 3' UTR | 19 | 42103117 | 42106985 | 3869 | 1 | 84095 | ENSMUST00000238085.2 | 5330 | ENSMUSG00000025178 | Pi4k2a | phosphatidylinositol 4-kinase type 2 alpha |
| chr2 | 1,53E+08 | 1,53E+08 | 1642 | * | 16,47622 | 2,896898687 | 1,077896 | 2,687549 | 0,007198 | 0,756677862 | 2,141612 | Promoter (1-2kb) | 2 | 1,53E+08 | 1,53E+08 | 1311 | 2 | 14239 | ENSMUST00000099200.3 | 1427 | ENSMUSG00000074676 | Foxs1 | forkhead box S1 |
| chr16 | 24269237 | 24270051 | 815 | * | 8,681871 | 2,897800915 | 1,098571 | 2,63779 | 0,008345 | 0,777919825 | 2,237403 | Promoter (2-3kb) | 16 | 24266841 | 24799780 | 532940 | 1 | 210126 | ENSMUST00000078988.10 | 2396 | ENSMUSG00000033306 | Lpp | LIM domain containing preferred translocation partner in lipoma |
| chr4 | 1,42E+08 | 1,42E+08 | 1410 | * | 13,57907 | 2,898195113 | 0,902539 | 3,211158 | 0,001322 | 0,611385361 | 0,938682 | Intron (ENSMUST0000 | 4 | 1,42E+08 | 1,42E+08 | 100162 | 2 | 71529 | ENSMUST00000036476.10 | -20107 | ENSMUSG00000040606 | Kazn | kazrin, periplakin interacting protein |
| chr16 | 50148633 | 50149556 | 924 | * | 8,096549 | 2,899441611 | 1,121473 | 2,585388 | 0,009727 | 1 | 0 | Promoter (1-2kb) | 16 | 50017522 | 50151356 | 133835 | 2 | 70508 | ENSMUST00000114488.8 | 1800 | ENSMUSG00000022641 | Bbx | bobby sox HMG box containing |
| chr15 | 86410232 | 86411446 | 1215 | * | 12,91555 | 2,899587065 | 0,952492 | 3,044212 | 0,002333 | 0,671156916 | 1,173325 | Distal Intergenic | 15 | 86588492 | 86595746 | 7255 | 1 | 75160 | ENSMUST00000229409.2 | -177046 | | 4930543I03Rik | RIKEN cDNA 4930543I03 gene |
| chr1 | 92672099 | 92673774 | 1676 | * | 15,14118 | 2,899642063 | 1,114047 | 2,602802 | 0,009247 | 0,892811906 | 1,2763 | Distal Intergenic | 1 | 92653363 | 92658482 | 5120 | 2 | 1,03E+08 | ENSMUST00000243412.1 | -13617 | | Gm29483 | predicted gene 29483 |
| chr16 | 32793533 | 32795124 | 1592 | * | 13,47944 | 2,899833928 | 1,042203 | 2,782408 | 0,005396 | 0,830474929 | 1,075735 | 3' UTR | 16 | 32799767 | 32818948 | 19182 | 1 | 70144 | ENSMUST00000163428.8 | -4643 | ENSMUSG00000022801 | Lrch3 | leucine-rich repeats and calponin homology (CH) domain containing 3 |
| chr7 | 46505958 | 46507398 | 1441 | * | 14,09323 | 2,900463458 | 0,883312 | 3,283621 | 0,001025 | 0,565091829 | 0,918255 | Distal Intergenic | 7 | 46510627 | 46527566 | 16940 | 1 | 16833 | ENSMUST00000014545.11 | -3229 | ENSMUSG00000030851 | Ldhc | lactate dehydrogenase C |
| chr7 | 1,26E+08 | 1,26E+08 | 1940 | * | 22,02525 | 2,901630538 | 0,733751 | 3,954516 | 7,67E-05 | 0,085136329 | 3,104559 | Promoter (<=1kb) | 7 | 1,25E+08 | 1,26E+08 | 203992 | 2 | 269994 | ENSMUST00000073935.7 | -153 | ENSMUSG00000046182 | Gsg1l | GSG1-like |
| chr5 | 34707961 | 34709782 | 1822 | * | 17,29139 | 2,901743496 | 1,045693 | 2,774948 | 0,005521 | 0,678869598 | 2,663619 | Promoter (<=1kb) | 5 | 34708889 | 34710136 | 1248 | 1 | 24055 | ENSMUST00000202745.2 | 0 | ENSMUSG00000054520 | Sh3bp2 | SH3-domain binding protein 2 |
| chr11 | 40617736 | 40618794 | 1059 | * | 10,05125 | 2,901897902 | 1,078747 | 2,690064 | 0,007144 | 0,857326444 | 1,211315 | Intron (ENSMUST0000 | 11 | 40615458 | 40624218 | 8761 | 2 | 15366 | ENSMUST00000152749.2 | 5424 | ENSMUSG00000020330 | Hmmr | hyaluronan mediated motility receptor (RHAMM) |
| chr18 | 39850511 | 39852116 | 1606 | * | 12,78959 | 2,90197705 | 0,928343 | 3,125976 | 0,001772 | 0,610143359 | 1,2763 | Intron (ENSMUST0000 | 18 | 39906550 | 39909135 | 2586 | 1 | 18459 | ENSMUST00000063219.3 | -54434 | ENSMUSG00000051732 | Pabpc2 | poly(A) binding protein, cytoplasmic 2 |
| chr7 | 15940789 | 15943893 | 3105 | * | 38,875827 | 2,902084948 | 0,5944755 | 4,8817572 | 1,051E-06 | 0,00373207 | 2,9254973 | Promoter (2-3kb) | 7 | 15931072 | 15938126 | 7055 | 2 | 71723 | ENSMUST00000128452.2 | -2663 | ENSMUSG00000006019 | Dhx34 | DEAH (Asp-Glu-Ala-His) box polypeptide 34 |
| chr7 | 1,12E+08 | 1,12E+08 | 1394 | * | 11,29747 | 2,903308951 | 1,026885 | 2,827297 | 0,004694 | 0,872043312 | 0,738441 | Intron (ENSMUST0000 | 7 | 1,12E+08 | 1,12E+08 | 69270 | 1 | 57342 | ENSMUST00000106640.2 | -28743 | ENSMUSG00000030770 | Parva | parvin, alpha |
| chr17 | 47812606 | 47814230 | 1625 | * | 17,5838 | 2,903919253 | 0,822788 | 3,529366 | 0,000417 | 0,261079505 | 2,507013 | Promoter (<=1kb) | 17 | 47794289 | 47813212 | 18924 | 2 | 63856 | ENSMUST00000233174.2 | 0 | ENSMUSG00000023980 | Taf8 | TATA-box binding protein associated factor 8 |
| chr5 | 1,44E+08 | 1,44E+08 | 2563 | * | 20,4132 | 2,904951303 | 0,767338 | 3,785752 | 0,000153 | 0,144460057 | 2,663619 | Promoter (1-2kb) | 5 | 1,43E+08 | 1,44E+08 | 24403 | 2 | 19353 | ENSMUST00000080537.14 | -1726 | ENSMUSG00000001847 | Rac1 | Rac family small GTPase 1 |
| chr11 | 92880609 | 92882187 | 1579 | * | 11,67182 | 2,905157311 | 1,089126 | 2,667421 | 0,007644 | 0,947773602 | 0,744375 | Distal Intergenic | 11 | 92801123 | 92801265 | 143 | 1 | 1,15E+08 | ENSMUST00000157185.3 | 79486 | ENSMUSG00000087810 | Gm25113 | predicted gene, 25113 |
| chr2 | 1,58E+08 | 1,58E+08 | 1622 | * | 11,51726 | 2,905910605 | 1,096817 | 2,649404 | 0,008063 | 0,955787258 | 0,738441 | Promoter (2-3kb) | 2 | 1,58E+08 | 1,58E+08 | 30035 | 2 | 21817 | ENSMUST00000103122.10 | -2699 | ENSMUSG00000037820 | Tgm2 | transglutaminase 2, C polypeptide |
| chr11 | 64177118 | 64178633 | 1516 | * | 11,69465 | 2,90610977 | 1,11916 | 2,596689 | 0,009413 | 0,97723982 | 0,738441 | Distal Intergenic | 11 | 64055399 | 64072997 | 17599 | 1 | 1,03E+08 | ENSMUST00000145095.2 | 121719 | ENSMUSG00000087622 | Gm12290 | predicted gene 12290 |
| chr3 | 28526549 | 28527820 | 1272 | * | 11,84461 | 2,907410656 | 1,114014 | 2,609852 | 0,009058 | 0,971329855 | 0,744375 | Intron (ENSMUST0000 | 3 | 28474092 | 28474167 | 76 | 1 | 1,01E+08 | ENSMUST00000175249.3 | 52457 | ENSMUSG00000092990 | Mir466q | microRNA 466q |
| chr1 | 1,64E+08 | 1,64E+08 | 1299 | * | 9,703781 | 2,908283655 | 1,049596 | 2,770859 | 0,005591 | 0,81655291 | 1,211315 | Intron (ENSMUST0000 | 1 | 1,64E+08 | 1,64E+08 | 12263 | 1 | 74895 | ENSMUST00000027867.7 | 8859 | ENSMUSG00000026578 | Ccdc181 | coiled-coil domain containing 181 |
| chr10 | 81093731 | 81095559 | 1829 | * | 16,07887 | 2,908294725 | 1,033859 | 2,813048 | 0,004907 | 0,676922067 | 2,386008 | Promoter (<=1kb) | 10 | 81093752 | 81098809 | 5058 | 1 | 17179 | ENSMUST00000120265.2 | 0 | ENSMUSG00000004933 | Matk | megakaryocyte-associated tyrosine kinase |
| chr4 | 1,34E+08 | 1,34E+08 | 1640 | * | 18,58952 | 2,909154462 | 0,811261 | 3,585965 | 0,000336 | 0,239591648 | 2,386008 | Intron (ENSMUST0000 | 4 | 1,34E+08 | 1,34E+08 | 129 | 2 | 1,02E+08 | ENSMUST00000185007.3 | -10887 | ENSMUSG00000099289 | Mir6403 | microRNA 6403 |
| chr18 | 69769794 | 69771581 | 1788 | * | 17,14477 | 2,909180112 | 0,809694 | 3,592937 | 0,000327 | 0,228721288 | 2,507013 | Exon (ENSMUST00000 | 18 | 69745515 | 69816269 | 70755 | 1 | 21413 | ENSMUST00000202474.4 | 24279 | ENSMUSG00000053477 | Tcf4 | transcription factor 4 |
| chr17 | 24384916 | 24387579 | 2664 | * | 17,61884 | 2,9099458 | 0,806508 | 3,608081 | 0,000308 | 0,216236188 | 2,663619 | Promoter (<=1kb) | 17 | 24382840 | 24386003 | 3164 | 2 | 11984 | ENSMUST00000151061.2 | 0 | ENSMUSG00000024121 | Atp6v0c | ATPase, H+ transporting, lysosomal V0 subunit C |
| chr2 | 1,03E+08 | 1,03E+08 | 1023 | * | 10,19554 | 2,909999513 | 1,097886 | 2,650547 | 0,008036 | 0,85188813 | 1,425929 | Intron (ENSMUST0000 | 2 | 1,03E+08 | 1,03E+08 | 40059 | 2 | 27402 | ENSMUST00000132449.8 | 23544 | ENSMUSG00000010914 | Pdhx | pyruvate dehydrogenase complex, component X |
| chr3 | 1,05E+08 | 1,05E+08 | 1721 | * | 33,68478 | 2,910735798 | 0,700697 | 4,154059 | 3,27E-05 | 0,05285139 | 2,787255 | Promoter (1-2kb) | 3 | 1,05E+08 | 1,05E+08 | 51232 | 2 | 80281 | ENSMUST00000077548.12 | -1413 | ENSMUSG00000062127 | Cttnbp2nl | CTTNBP2 N-terminal like |
| chr14 | 23238103 | 23238991 | 889 | * | 8,238303 | 2,911096392 | 1,101741 | 2,64227 | 0,008235 | 0,854050055 | 1,425929 | Distal Intergenic | 14 | 23088262 | 23144639 | 56378 | 2 | 791307 | ENSMUST00000090409.2 | -93464 | ENSMUSG00000068680 | Gm10248 | predicted gene 10248 |
| chr15 | 36283069 | 36285203 | 2135 | * | 37,346377 | 2,91128791 | 0,5373311 | 5,4180519 | 6,025E-08 | 0,000451853 | 2,7872553 | Promoter (<=1kb) | 15 | 36240079 | 36283293 | 43215 | 2 | 30945 | ENSMUST00000022890.10 | 0 | ENSMUSG00000022280 | Rnf19a | ring finger protein 19A |
| chr13 | 69758417 | 69760138 | 1722 | * | 34,045922 | 2,912478529 | 0,675683 | 4,3104216 | 1,629E-05 | 0,03581197 | 2,3776742 | Promoter (<=1kb) | 13 | 69760135 | 69782453 | 22319 | 1 | 28114 | ENSMUST00000109699.11 | 0 | ENSMUSG00000021595 | Nsun2 | NOL1/NOP2/Sun domain family member 2 |
| chr9 | 53761555 | 53763222 | 1668 | * | 11,59244 | 2,913850916 | 1,059609 | 2,74993 | 0,005961 | 0,832746263 | 1,173325 | 3' UTR | 9 | 53757448 | 53761844 | 4397 | 1 | 66402 | ENSMUST00000048485.7 | 4107 | ENSMUSG00000042045 | Sln | sarcolipin |
| chr11 | 79904766 | 79906537 | 1772 | * | 12,22826 | 2,91479678 | 1,003454 | 2,904763 | 0,003675 | 0,730689299 | 1,2763 | Exon (ENSMUST00000 | 11 | 79916021 | 79922827 | 6807 | 1 | 52615 | ENSMUST00000144188.2 | -9484 | ENSMUSG00000017548 | Suz12 | SUZ12 polycomb repressive complex 2 subunit |
| chr1 | 161704262 | 161706010 | 1749 | * | 45,217664 | 2,914816915 | 0,5736791 | 5,0809186 | 3,756E-07 | 0,001777866 | 2,7872553 | Promoter (<=1kb) | 1 | 161643683 | 161704251 | 60569 | 2 | 226551 | ENSMUST00000048377.11 | -11 | ENSMUSG00000040297 | Suco | SUN domain containing ossification factor |
| chr6 | 1,42E+08 | 1,42E+08 | 1978 | * | 19,01292 | 2,915075998 | 0,927386 | 3,143324 | 0,00167 | 0,463890057 | 2,663619 | Intron (ENSMUST0000 | 6 | 1,42E+08 | 1,42E+08 | 31254 | 2 | 108096 | ENSMUST00000111822.8 | -6058 | ENSMUSG00000063975 | Slco1a5 | solute carrier organic anion transporter family, member 1a5 |
| chr11 | 1,04E+08 | 1,04E+08 | 1366 | * | 12,56866 | 2,915141081 | 1,041455 | 2,799103 | 0,005124 | 0,851014755 | 0,918255 | Intron (ENSMUST0000 | 11 | 1,04E+08 | 1,04E+08 | 4930 | 2 | 217232 | ENSMUST00000127506.2 | -10938 | ENSMUSG00000020687 | Cdc27 | cell division cycle 27 |
| chr11 | 54930750 | 54933336 | 2587 | * | 21,10689 | 2,91545088 | 0,874334 | 3,334482 | 0,000855 | 0,309527539 | 3,471678 | Distal Intergenic | 11 | 54869934 | 54924271 | 54338 | 2 | 11749 | ENSMUST00000108883.10 | -6479 | ENSMUSG00000018340 | Anxa6 | annexin A6 |
| chr2 | 1,56E+08 | 1,56E+08 | 1649 | * | 16,28483 | 2,915547895 | 0,890253 | 3,274964 | 0,001057 | 0,424243345 | 2,141612 | Intron (ENSMUST0000 | 2 | 1,56E+08 | 1,57E+08 | 150659 | 1 | 228836 | ENSMUST00000169464.9 | 4174 | ENSMUSG00000061689 | Dlgap4 | DLG associated protein 4 |
| chr7 | 110228549 | 110232431 | 3883 | * | 62,538436 | 2,91563421 | 0,5322871 | 5,477559 | 4,312E-08 | 0,000298894 | 3,2706213 | Promoter (1-2kb) | 7 | 110227235 | 110227758 | 524 | 1 | 11535 | ENSMUST00000185766.2 | 1314 | ENSMUSG00000030790 | Adm | adrenomedullin |
| chr2 | 82424599 | 82425521 | 923 | * | 12,22859 | 2,916173687 | 0,899345 | 3,242554 | 0,001185 | 0,563032042 | 1,075735 | Distal Intergenic | 2 | 82205716 | 82222546 | 16831 | 2 | 228116 | ENSMUST00000128608.2 | -202053 | ENSMUSG00000087061 | A130030D18Rik | RIKEN cDNA A130030D18 gene |
| chr1 | 53656560 | 53657773 | 1214 | * | 10,51163 | 2,916495361 | 1,061072 | 2,74863 | 0,005984 | 0,908970978 | 0,744375 | Exon (ENSMUST00000 | 1 | 53436165 | 53745920 | 309756 | 2 | 627872 | ENSMUST00000094964.7 | 88147 | ENSMUSG00000096141 | Dnah7a | dynein, axonemal, heavy chain 7A |
| chr9 | 44091820 | 44092643 | 824 | * | 9,867031 | 2,916859446 | 1,014195 | 2,876035 | 0,004027 | 0,777172375 | 1,086118 | Exon (ENSMUST00000 | 9 | 44080317 | 44097262 | 16946 | 2 | 12402 | ENSMUST00000206629.2 | 4619 | ENSMUSG00000034342 | Cbl | Casitas B-lineage lymphoma |
| chr4 | 1,08E+08 | 1,08E+08 | 1052 | * | 10,08566 | 2,916960618 | 1,088004 | 2,68102 | 0,00734 | 0,8666955 | 1,173325 | Promoter (<=1kb) | 4 | 1,08E+08 | 1,08E+08 | 69285 | 1 | 1,03E+08 | ENSMUST00000143064.8 | 0 | | 1700047F07Rik | RIKEN cDNA 1700047F07 gene |
| chr19 | 30715997 | 30717335 | 1339 | * | 11,35718 | 2,918323922 | 1,132657 | 2,576529 | 0,00998 | 0,986301782 | 0,744375 | Intron (ENSMUST0000 | 19 | 30545427 | 30707982 | 162556 | 2 | 19091 | ENSMUST00000239061.2 | -8015 | ENSMUSG00000052920 | Prkg1 | protein kinase, cGMP-dependent, type I |
| chr1 | 75445354 | 75446908 | 1555 | * | 16,13123 | 2,918330572 | 0,924913 | 3,155249 | 0,001604 | 0,591135276 | 1,2763 | Exon (ENSMUST00000 | 1 | 75451220 | 75455390 | 4171 | 2 | 74241 | ENSMUST00000143496.8 | 8482 | ENSMUSG00000032997 | Chpf | chondroitin polymerizing factor |
| chr12 | 32869589 | 32870394 | 806 | * | 14,1289 | 2,918585858 | 0,824407 | 3,540223 | 0,0004 | 0,40037769 | 0,938682 | Promoter (<=1kb) | 12 | 32870334 | 32903348 | 33015 | 1 | 59027 | ENSMUST00000020886.9 | 0 | ENSMUSG00000020572 | Nampt | nicotinamide phosphoribosyltransferase |
| chr14 | 31844580 | 31846097 | 1518 | * | 12,82317 | 2,91864991 | 0,98202 | 2,972088 | 0,002958 | 0,791068542 | 0,738441 | Distal Intergenic | 14 | 31863980 | 31880324 | 16345 | 1 | 17695 | ENSMUST00000022464.14 | -17883 | ENSMUSG00000021907 | Msmb | beta-microseminoprotein |
| chr4 | 1,34E+08 | 1,34E+08 | 2155 | * | 20,05382 | 2,919020675 | 0,973709 | 2,997838 | 0,002719 | 0,584533232 | 2,221611 | Promoter (<=1kb) | 4 | 1,34E+08 | 1,34E+08 | 10359 | 2 | 194231 | ENSMUST00000030645.9 | 0 | ENSMUSG00000028841 | Cnksr1 | connector enhancer of kinase suppressor of Ras 1 |
| chr11 | 1,12E+08 | 1,12E+08 | 1383 | * | 9,18443 | 2,919679656 | 1,092421 | 2,67267 | 0,007525 | 0,86525631 | 1,211315 | Distal Intergenic | 11 | 1,11E+08 | 1,11E+08 | 2269 | 2 | 1,03E+08 | ENSMUST00000247083.1 | -483596 | | Gm30158 | predicted gene, 30158 |
| chr11 | 75238188 | 75239872 | 1685 | * | 13,7123 | 2,920447863 | 1,098014 | 2,659755 | 0,00782 | 0,89325945 | 1,075735 | Promoter (<=1kb) | 11 | 75239259 | 75296531 | 57273 | 1 | 319822 | ENSMUST00000044530.3 | 0 | ENSMUSG00000018809 | Smyd4 | SET and MYND domain containing 4 |
| chr17 | 26206924 | 26208206 | 1283 | * | 13,22217 | 2,922213232 | 1,095521 | 2,667418 | 0,007644 | 0,890436263 | 1,075735 | Promoter (2-3kb) | 17 | 26204856 | 26210004 | 5149 | 1 | 69431 | ENSMUST00000235322.2 | 2068 | ENSMUSG00000097523 | 1700022N22Rik | RIKEN cDNA 1700022N22 gene |
| chr4 | 107914119 | 107917697 | 3579 | * | 38,188941 | 2,922282383 | 0,6266328 | 4,6634683 | 3,109E-06 | 0,00932046 | 2,7254908 | Intron (ENSMUST000000 | 4 | 107901465 | 107905076 | 3612 | 2 | 20280 | ENSMUST00000145721.2 | -9043 | ENSMUSG00000028603 | Scp2 | sterol carrier protein 2, liver |
| chr3 | 1,28E+08 | 1,28E+08 | 591 | * | 10,72943 | 2,922837728 | 1,067908 | 2,736975 | 0,006201 | 0,838878373 | 1,173325 | Promoter (1-2kb) | 3 | 1,28E+08 | 1,28E+08 | 635 | 2 | 211556 | ENSMUST00000197450.2 | 1649 | ENSMUSG00000074238 | Ap1ar | adaptor-related protein complex 1 associated regulatory protein |
| chr1 | 1,06E+08 | 1,06E+08 | 1609 | * | 13,63846 | 2,923030042 | 1,069553 | 2,732944 | 0,006277 | 0,917631345 | 0,738441 | 3' UTR | 1 | 1,06E+08 | 1,06E+08 | 10010 | 1 | 227446 | ENSMUST00000188588.2 | 3581 | ENSMUSG00000026319 | Relch | RAB11 binding and LisH domain, coiled-coil and HEAT repeat containing |
| chr6 | 13829695 | 13830157 | 463 | * | 7,918396 | 2,923095044 | 1,127448 | 2,592666 | 0,009524 | 0,88871498 | 1,360622 | Distal Intergenic | 6 | 13835067 | 13838453 | 3387 | 2 | 64450 | ENSMUST00000115492.2 | 8296 | ENSMUSG00000048216 | Gpr85 | G protein-coupled receptor 85 |
| chr1 | 181251574 | 181253797 | 2224 | * | 43,389053 | 2,924157417 | 0,5171471 | 5,6544018 | 1,564E-08 | 0,000159232 | 2,7254908 | Intron (ENSMUST000001 | 1 | 181180991 | 181286817 | 105827 | 1 | 72978 | ENSMUST00000209607.2 | 70583 | ENSMUSG00000026514 | Cnih3 | cornichon family AMPA receptor auxiliary protein 3 |
| chr2 | 1,74E+08 | 1,74E+08 | 634 | * | 11,28234 | 2,924323797 | 1,12053 | 2,609769 | 0,00906 | 0,972947889 | 0,738441 | Distal Intergenic | 2 | 1,74E+08 | 1,74E+08 | 58881 | 2 | 1E+08 | ENSMUST00000131346.2 | -20225 | ENSMUSG00000074555 | Gm10714 | predicted gene 10714 |
| chr13 | 67977120 | 67978332 | 1213 | * | 11,55905 | 2,924955787 | 1,033606 | 2,829856 | 0,004657 | 0,869491026 | 0,738441 | Promoter (<=1kb) | 13 | 67944713 | 67979104 | 34392 | 2 | 320163 | ENSMUST00000181912.2 | 772 | ENSMUSG00000097276 | 4930525G20Rik | RIKEN cDNA 4930525G20 gene |
| chr11 | 38932034 | 38933659 | 1626 | * | 12,58353 | 2,925256122 | 0,94116 | 3,108138 | 0,001883 | 0,621084104 | 1,2763 | Distal Intergenic | 11 | 38688998 | 38689104 | 107 | 2 | 1,15E+08 | ENSMUST00000240209.1 | -242930 | ENSMUSG00000119681 | Gm23520 | predicted gene, 23520 |
| chr16 | 74297740 | 74298633 | 894 | * | 9,721276 | 2,926545129 | 1,087765 | 2,690419 | 0,007136 | 0,861411338 | 1,173325 | Distal Intergenic | 16 | 74146683 | 74208713 | 62031 | 2 | 268902 | ENSMUST00000156010.2 | -89027 | ENSMUSG00000052516 | Robo2 | roundabout guidance receptor 2 |
| chr9 | 69356404 | 69357702 | 1299 | * | 15,37021 | 2,926824257 | 0,873153 | 3,352018 | 0,000802 | 0,375922902 | 2,141612 | Exon (ENSMUST00000 | 9 | 69360902 | 69399077 | 38176 | 1 | 12306 | ENSMUST00000034756.15 | -3200 | ENSMUSG00000032231 | Anxa2 | annexin A2 |
| chr7 | 65743835 | 65745253 | 1419 | * | 14,11778 | 2,927228643 | 1,037306 | 2,821953 | 0,004773 | 0,808608011 | 1,075735 | Distal Intergenic | 7 | 65759263 | 65823546 | 64284 | 1 | 269941 | ENSMUST00000036372.8 | -14010 | ENSMUSG00000032640 | Chsy1 | chondroitin sulfate synthase 1 |
| chr2 | 31587283 | 31588993 | 1711 | * | 18,90898 | 2,927642824 | 0,941641 | 3,109084 | 0,001877 | 0,501659463 | 2,386008 | Exon (ENSMUST00000 | 2 | 31579623 | 31669036 | 89414 | 1 | 11350 | ENSMUST00000135233.3 | 7660 | ENSMUSG00000026842 | Abl1 | c-abl oncogene 1, non-receptor tyrosine kinase |
| chr9 | 61837865 | 61839846 | 1982 | * | 16,37858 | 2,92807176 | 0,968241 | 3,024114 | 0,002494 | 0,532496361 | 2,663619 | Promoter (<=1kb) | 9 | 61834834 | 61839820 | 4987 | 2 | 71819 | ENSMUST00000215965.2 | 0 | ENSMUSG00000032254 | Kif23 | kinesin family member 23 |
| chr12 | 46108341 | 46110394 | 2054 | * | 16,55002 | 2,928859245 | 1,005645 | 2,912419 | 0,003586 | 0,728181287 | 1,2763 | Distal Intergenic | 12 | 46031126 | 46031332 | 207 | 1 | 1,15E+08 | ENSMUST00000082866.3 | 77215 | ENSMUSG00000064800 | Gm24948 | predicted gene, 24948 |
| chr2 | 70959109 | 70960753 | 1645 | * | 11,87138 | 2,929565765 | 1,020022 | 2,872061 | 0,004078 | 0,846796411 | 0,744375 | Exon (ENSMUST00000 | 2 | 70948267 | 70973270 | 25004 | 1 | 73649 | ENSMUST00000028403.3 | 10842 | ENSMUSG00000027015 | Cybrd1 | cytochrome b reductase 1 |
| chr7 | 97598204 | 97599697 | 1494 | * | 12,63479 | 2,930185458 | 1,087068 | 2,695493 | 0,007028 | 0,87775273 | 1,075735 | Intron (ENSMUST0000 | 7 | 97607028 | 97697818 | 90791 | 1 | 233537 | ENSMUST00000170049.2 | -7331 | ENSMUSG00000035582 | Gdpd4 | glycerophosphodiester phosphodiesterase domain containing 4 |
| chr13 | 59916573 | 59918627 | 2055 | * | 28,405959 | 2,93026255 | 0,6143019 | 4,7700695 | 1,842E-06 | 0,006356998 | 2,7254908 | Promoter (<=1kb) | 13 | 59917311 | 59919222 | 1912 | 1 | 13996 | ENSMUST00000225619.2 | 0 | | Etohd2 | ethanol decreased 2 |
| chr8 | 82905474 | 82906949 | 1476 | * | 14,49749 | 2,93087524 | 1,012727 | 2,894043 | 0,003803 | 0,747833008 | 1,214904 | Distal Intergenic | 8 | 82879349 | 82879411 | 63 | 2 | 1,15E+08 | ENSMUST00000158522.3 | -26063 | ENSMUSG00000089147 | Gm25267 | predicted gene, 25267 |
| chr1 | 37288390 | 37290124 | 1735 | * | 16,62572 | 2,930994973 | 0,860086 | 3,407792 | 0,000655 | 0,337811311 | 2,141612 | Intron (ENSMUST0000 | 1 | 37258286 | 37302465 | 44180 | 1 | 12790 | ENSMUST00000027288.10 | 30104 | ENSMUSG00000026114 | Cnga3 | cyclic nucleotide gated channel alpha 3 |
| chr4 | 1,41E+08 | 1,41E+08 | 1677 | * | 17,69142 | 2,931673423 | 0,866652 | 3,382758 | 0,000718 | 0,337054778 | 2,386008 | Intron (ENSMUST0000 | 4 | 1,41E+08 | 1,41E+08 | 12539 | 2 | 230872 | ENSMUST00000151455.8 | -9541 | ENSMUSG00000040860 | Crocc | ciliary rootlet coiled-coil, rootletin |
| chr4 | 1,25E+08 | 1,25E+08 | 1906 | * | 17,23263 | 2,932892922 | 0,932575 | 3,144941 | 0,001661 | 0,497895643 | 2,154504 | Promoter (<=1kb) | 4 | 1,25E+08 | 1,25E+08 | 10366 | 2 | 75563 | ENSMUST00000036383.4 | 0 | ENSMUSG00000042707 | Dnali1 | dynein, axonemal, light intermediate polypeptide 1 |
| chr17 | 67313819 | 67314969 | 1151 | * | 14,66544 | 2,933925441 | 0,992404 | 2,956382 | 0,003113 | 0,703368984 | 1,2763 | Intron (ENSMUST0000 | 17 | 67333142 | 67333230 | 89 | 2 | 1,01E+08 | ENSMUST00000176783.3 | 18261 | ENSMUSG00000093508 | Mir5709 | microRNA 5709 |
| chr12 | 1,13E+08 | 1,13E+08 | 1644 | * | 15,85588 | 2,934153043 | 0,964898 | 3,040895 | 0,002359 | 0,563032042 | 2,141612 | Intron (ENSMUST0000 | 12 | 1,13E+08 | 1,13E+08 | 38269 | 1 | 116870 | ENSMUST00000130926.8 | 10035 | ENSMUSG00000021144 | Mta1 | metastasis associated 1 |
| chr9 | 21974821 | 21976439 | 1619 | * | 12,26784 | 2,934291908 | 1,074493 | 2,730861 | 0,006317 | 0,842187815 | 1,173325 | Exon (ENSMUST00000 | 9 | 21982298 | 21996476 | 14179 | 1 | 1,01E+08 | ENSMUST00000216270.2 | -5859 | ENSMUSG00000096981 | Gm16845 | predicted gene, 16845 |
| chr7 | 97233326 | 97235853 | 2528 | * | 16,97575 | 2,934468427 | 1,003251 | 2,92496 | 0,003445 | 0,595046521 | 2,663619 | Intron (ENSMUST0000 | 7 | 97229114 | 97288670 | 59557 | 1 | 233532 | ENSMUST00000127891.3 | 4212 | ENSMUSG00000035623 | Rsf1 | remodeling and spacing factor 1 |
| chr1 | 1,65E+08 | 1,65E+08 | 2476 | * | 15,63163 | 2,934986285 | 1,007907 | 2,911962 | 0,003592 | 0,728544502 | 1,2763 | 3' UTR | 1 | 1,65E+08 | 1,65E+08 | 1524 | 2 | 68481 | ENSMUST00000192716.2 | 26623 | ENSMUSG00000026566 | Mpzl1 | myelin protein zero-like 1 |
| chr8 | 61656904 | 61658285 | 1382 | * | 10,9458 | 2,935740289 | 1,091033 | 2,69079 | 0,007128 | 0,903470246 | 0,918255 | Distal Intergenic | 8 | 61676906 | 61849105 | 172200 | 1 | 59009 | ENSMUST00000034060.7 | -18621 | ENSMUSG00000031642 | Sh3rf1 | SH3 domain containing ring finger 1 |
| chr7 | 1,43E+08 | 1,43E+08 | 1353 | * | 14,15461 | 2,936538225 | 0,968022 | 3,033543 | 0,002417 | 0,713797407 | 0,938682 | Intron (ENSMUST0000 | 7 | 1,43E+08 | 1,43E+08 | 2066 | 2 | 79196 | ENSMUST00000149736.2 | -15272 | ENSMUSG00000037606 | Osbpl5 | oxysterol binding protein-like 5 |
| chr2 | 41998804 | 42000236 | 1433 | * | 12,44862 | 2,937387923 | 0,987503 | 2,974561 | 0,002934 | 0,79031971 | 0,738441 | Intron (ENSMUST0000 | 2 | 40691405 | 41679090 | 987686 | 2 | 94217 | ENSMUST00000167270.5 | -319714 | ENSMUSG00000049252 | Lrp1b | low density lipoprotein-related protein 1B |
| chr13 | 24803477 | 24806018 | 2542 | * | 23,50272 | 2,938305494 | 0,872646 | 3,367121 | 0,00076 | 0,318448436 | 2,880017 | Intron (ENSMUST0000 | 13 | 24798610 | 24889880 | 91271 | 1 | 193385 | ENSMUST00000132689.8 | 4867 | ENSMUSG00000036006 | Ripor2 | RHO family interacting cell polarization regulator 2 |
| chr13 | 26884185 | 26885386 | 1202 | * | 11,22538 | 2,938311949 | 1,014149 | 2,897317 | 0,003764 | 0,734721579 | 1,2763 | Distal Intergenic | 13 | 26952156 | 26954148 | 1993 | 2 | 15192 | ENSMUST00000055915.6 | 68762 | ENSMUSG00000045835 | Hdgfl1 | HDGF like 1 |
| chr6 | 43985858 | 43986513 | 656 | * | 10,19747 | 2,938693968 | 1,140787 | 2,576023 | 0,009994 | 0,917631345 | 1,173325 | Distal Intergenic | 6 | 44057422 | 44060289 | 2868 | 2 | 1,03E+08 | ENSMUST00000203777.2 | 73776 | ENSMUSG00000107598 | Gm35216 | predicted gene, 35216 |
| chr14 | 65821129 | 65822860 | 1732 | * | 10,20547 | 2,938873229 | 0,98919 | 2,970989 | 0,002968 | 0,705083711 | 1,211315 | Intron (ENSMUST0000 | 14 | 65767904 | 65830499 | 62596 | 2 | 74195 | ENSMUST00000224743.2 | 7639 | ENSMUSG00000022031 | Elp3 | elongator acetyltransferase complex subunit 3 |
| chr4 | 1,5E+08 | 1,5E+08 | 2198 | * | 18,42695 | 2,939067952 | 0,817263 | 3,596232 | 0,000323 | 0,231796186 | 2,423594 | Exon (ENSMUST00000 | 4 | 1,5E+08 | 1,5E+08 | 24887 | 1 | 56485 | ENSMUST00000030826.4 | -5766 | ENSMUSG00000028976 | Slc2a5 | solute carrier family 2 (facilitated glucose transporter), member 5 |
| chr4 | 1,23E+08 | 1,23E+08 | 1153 | * | 10,64125 | 2,939294458 | 1,004363 | 2,926527 | 0,003428 | 0,817520905 | 0,738441 | Distal Intergenic | 4 | 1,23E+08 | 1,23E+08 | 11475 | 1 | 170638 | ENSMUST00000059667.9 | -7649 | ENSMUSG00000046093 | Hpcal4 | hippocalcin-like 4 |
| chr14 | 31489240 | 31491109 | 1870 | * | 13,32847 | 2,939308058 | 0,924161 | 3,180515 | 0,00147 | 0,605351622 | 1,075735 | Intron (ENSMUST0000 | 14 | 31420725 | 31502350 | 81626 | 2 | 105522 | ENSMUST00000227863.2 | 11241 | ENSMUSG00000014496 | Ankrd28 | ankyrin repeat domain 28 |
| chr7 | 4077640 | 4078454 | 815 | * | 8,774875 | 2,939708525 | 1,081237 | 2,718837 | 0,006551 | 0,823446343 | 1,360622 | Exon (ENSMUST00000 | 7 | 4010072 | 4066203 | 56132 | 2 | 52855 | ENSMUST00000086400.13 | -11437 | ENSMUSG00000055541 | Lair1 | leukocyte-associated Ig-like receptor 1 |

| | | | | | | | | | | | | | | | | | | | | | | | |
|---|---|---|---|---|---|---|---|---|---|---|---|---|---|---|---|---|---|---|---|---|---|---|---|
| chr4 | 45823091 | 45824477 | 1387 | * | 13,60763 | 2,939766955 | 0,882417 | 3,331495 | 0,000864 | 0,504427013 | 1,075735 | Promoter (2-3kb) | 4 | 45809468 | 45826923 | 17456 | 2 | 75426 | ENSMUST00000044297.7 | 2446 | ENSMUSG00000035551 | Igfbpl1 | insulin-like growth factor binding protein-like 1 |
| chr11 | 96997958 | 96999911 | 1954 | * | 13,26997 | 2,94028656 | 0,907137 | 3,241282 | 0,00119 | 0,563858922 | 1,075735 | Intron (ENSMUST0000 | 11 | 96988897 | 97006157 | 17261 | 2 | 57765 | ENSMUST00000001484.3 | 6246 | ENSMUSG00000001444 | Tbx21 | T-box 21 |
| chr11 | 8551330 | 8552916 | 1587 | * | 12,94881 | 2,940465376 | 1,112074 | 2,644126 | 0,00819 | 0,923399738 | 0,918255 | Intron (ENSMUST0000 | 11 | 8381652 | 8574887 | 193236 | 2 | 319939 | ENSMUST00000239091.2 | 21971 | ENSMUSG00000020422 | Tns3 | tensin 3 |
| chr2 | 1,29E+08 | 1,29E+08 | 1715 | * | 12,35163 | 2,940799117 | 1,108593 | 2,652731 | 0,007984 | 0,954902749 | 0,738441 | Distal Intergenic | 2 | 1,29E+08 | 1,29E+08 | 36144 | 1 | 72477 | ENSMUST00000110325.8 | -5270 | ENSMUSG00000014353 | Tmem87b | transmembrane protein 87B |
| chr1 | 6334695 | 6335867 | 1173 | * | 10,47656 | 2,940873523 | 1,103988 | 2,663863 | 0,007725 | 0,948066884 | 0,744375 | Promoter (<=1kb) | 1 | 6335830 | 6342989 | 7160 | 1 | 12421 | ENSMUST00000162418.2 | 0 | ENSMUSG00000025907 | Rb1cc1 | RB1-inducible coiled-coil 1 |
| chr12 | 85500072 | 85502759 | 2688 | * | 37,325509 | 2,942043402 | 0,6253742 | 4,704453 | 2,545E-06 | 0,007611064 | 2,9254973 | Distal Intergenic | 12 | 85475884 | 85483023 | 7140 | 2 | 105244959 | ENSMUST00000221505.2 | -17049 | | Gm40477 | predicted gene, 40477 |
| chr1 | 34147293 | 34149306 | 2014 | * | 19,72866 | 2,942241304 | 0,938699 | 3,134384 | 0,001722 | 0,477622452 | 2,507013 | Intron (ENSMUST0000 | 1 | 34156440 | 34156512 | 73 | 1 | 1,02E+08 | ENSMUST00000198033.3 | -7134 | ENSMUSG00000105982 | Mir6896 | microRNA 6896 |
| chr16 | 32599002 | 32600424 | 1423 | * | 11,8561 | 2,942662818 | 1,081195 | 2,721677 | 0,006495 | 0,92207189 | 0,744375 | Promoter (2-3kb) | 16 | 32596657 | 32602754 | 6098 | 1 | 140474 | ENSMUST00000239560.1 | 2345 | ENSMUSG00000118672 | Muc4 | mucin 4 |
| chr17 | 85398693 | 85400284 | 1592 | * | 16,45557 | 2,94267173 | 0,955173 | 3,080772 | 0,002065 | 0,524368688 | 2,318434 | Promoter (<=1kb) | 17 | 85399511 | 85473936 | 74426 | 1 | 1,03E+08 | ENSMUST00000234501.2 | 0 | | Gm36826 | predicted gene, 36826 |
| chr15 | 79295019 | 79297135 | 2117 | * | 17,85097 | 2,942832414 | 0,993171 | 2,963066 | 0,003046 | 0,612498827 | 2,154504 | Distal Intergenic | 15 | 79244884 | 79287769 | 42886 | 2 | 223693 | ENSMUST00000074991.10 | -7250 | ENSMUSG00000009035 | Tmem184b | transmembrane protein 184b |
| chr3 | 1,44E+08 | 1,44E+08 | 1236 | * | 9,48124 | 2,943347593 | 1,090606 | 2,698818 | 0,006959 | 0,823446343 | 1,449706 | Promoter (1-2kb) | 3 | 1,44E+08 | 1,44E+08 | 26969 | 1 | 93684 | ENSMUST00000106211.3 | 1533 | ENSMUSG00000037072 | Selenof | selenoprotein F |
| chr4 | 1,29E+08 | 1,29E+08 | 1700 | * | 11,98921 | 2,943954317 | 1,092858 | 2,693812 | 0,007064 | 0,902613747 | 0,918255 | Promoter (<=1kb) | 4 | 1,29E+08 | 1,3E+08 | 5321 | 1 | 76799 | ENSMUST00000172635.2 | 712 | ENSMUSG00000028797 | Tmem234 | transmembrane protein 234 |
| chr15 | 35370803 | 35372990 | 2188 | * | 15,42267 | 2,944687186 | 0,886476 | 3,321791 | 0,000894 | 0,490999135 | 1,214904 | Promoter (<=1kb) | 15 | 35371306 | 35931375 | 560070 | 1 | 666173 | ENSMUST00000048646.9 | 0 | ENSMUSG00000037646 | Vps13b | vacuolar protein sorting 13B |
| chr14 | 71140683 | 71143490 | 2808 | * | 15,27993 | 2,945773011 | 1,012136 | 2,910451 | 0,003609 | 0,73648081 | 1,214904 | Intron (ENSMUST0000 | 14 | 71127968 | 71215936 | 87969 | 1 | 14586 | ENSMUST00000227697.2 | 12715 | ENSMUSG00000022103 | Gfra2 | glial cell line derived neurotrophic factor family receptor alpha 2 |
| chr1 | 1,24E+08 | 1,24E+08 | 811 | * | 8,55387 | 2,946516509 | 1,129126 | 2,609555 | 0,009066 | 0,792545355 | 2,237403 | Intron (ENSMUST0000 | 1 | 1,24E+08 | 1,24E+08 | 104 | 1 | 1,15E+08 | ENSMUST00000083406.3 | 59684 | ENSMUSG00000065340 | Gm25578 | predicted gene, 25578 |
| chr16 | 6941265 | 6942167 | 903 | * | 9,799201 | 2,94751805 | 1,099081 | 2,681802 | 0,007323 | 0,865825879 | 1,173325 | Intron (ENSMUST0000 | 16 | 6887710 | 7227677 | 339968 | 1 | 268859 | ENSMUST00000229741.2 | 53555 | ENSMUSG00000008658 | Rbfox1 | RNA binding protein, fox-1 homolog (C. elegans) 1 |
| chr13 | 30583412 | 30585009 | 1598 | * | 14,06818 | 2,947587365 | 0,943712 | 3,123396 | 0,001788 | 0,669586355 | 0,918255 | Exon (ENSMUST00000 | 13 | 30529524 | 30565089 | 35566 | 1 | 11607 | ENSMUST00000221743.2 | 53888 | ENSMUSG00000049115 | Agtr1a | angiotensin II receptor, type 1a |
| chr8 | 1,24E+08 | 1,24E+08 | 1006 | * | 10,34209 | 2,948059369 | 0,990136 | 2,977429 | 0,002907 | 0,7183981 | 1,086118 | Promoter (<=1kb) | 8 | 1,24E+08 | 1,24E+08 | 1996 | 2 | 14087 | ENSMUST00000213090.2 | 0 | ENSMUSG00000032815 | Fanca | Fanconi anemia, complementation group A |
| chr4 | 1,29E+08 | 1,29E+08 | 1817 | * | 13,97454 | 2,948245369 | 0,948798 | 3,107348 | 0,001888 | 0,650529916 | 1,075735 | Promoter (1-2kb) | 4 | 1,29E+08 | 1,29E+08 | 14177 | 2 | 433759 | ENSMUST00000139305.2 | 1806 | ENSMUSG00000028800 | Hdac1 | histone deacetylase 1 |
| chr2 | 1,46E+08 | 1,46E+08 | 941 | * | 9,571427 | 2,948333574 | 1,138858 | 2,588852 | 0,00963 | 0,923399738 | 1,086118 | Intron (ENSMUST0000 | 2 | 1,46E+08 | 1,46E+08 | 16592 | 2 | 241694 | ENSMUST00000124583.2 | -62209 | ENSMUSG00000037110 | Ralgapa2 | Ral GTPase activating protein, alpha subunit 2 (catalytic) |
| chr8 | 4660221 | 4661520 | 1300 | * | 11,07869 | 2,94848895 | 1,125331 | 2,620108 | 0,00879 | 0,899673069 | 1,173325 | Promoter (1-2kb) | 8 | 4663167 | 4680231 | 17065 | 1 | 233987 | ENSMUST00000202692.5 | -1647 | ENSMUSG00000058748 | Zfp958 | zinc finger protein 958 |
| chr11 | 1,01E+08 | 1,01E+08 | 1777 | * | 16,42913 | 2,948903684 | 0,989125 | 2,981326 | 0,00287 | 0,686261859 | 1,2763 | Exon (ENSMUST00000 | 11 | 1,01E+08 | 1,01E+08 | 1012 | 1 | 11975 | ENSMUST00000127810.2 | 9987 | ENSMUSG00000019302 | Atp6v0a1 | ATPase, H+ transporting, lysosomal V0 subunit A1 |
| chr9 | 66210046 | 66211495 | 1450 | * | 12,95297 | 2,949199314 | 1,002607 | 2,941532 | 0,003266 | 0,768389035 | 0,918255 | Distal Intergenic | 9 | 66175656 | 66178997 | 3342 | 1 | 13143 | ENSMUST00000129778.2 | 34390 | ENSMUSG00000032380 | Dapk2 | death-associated protein kinase 2 |
| chr19 | 36713547 | 36716977 | 3431 | * | 38,862025 | 2,949765423 | 0,5318173 | 5,5465761 | 2,913E-08 | 0,000244135 | 2,7872553 | Promoter (<=1kb) | 19 | 36709131 | 36714053 | 4923 | 2 | 53412 | ENSMUST00000087321.4 | 0 | ENSMUSG00000067279 | Ppp1r3c | protein phosphatase 1, regulatory subunit 3C |
| chr16 | 85909440 | 85910176 | 737 | * | 9,79463 | 2,949974355 | 1,143902 | 2,578871 | 0,009912 | 0,927963368 | 1,086118 | Distal Intergenic | 16 | 86033059 | 86066038 | 32980 | 1 | 1,03E+08 | ENSMUST00000231827.2 | -122883 | | Gm32357 | predicted gene, 32357 |
| chr11 | 13153618 | 13154803 | 1186 | * | 10,38353 | 2,950017955 | 1,124688 | 2,622967 | 0,008717 | 0,964323611 | 0,744375 | Distal Intergenic | 11 | 12642813 | 12642942 | 130 | 1 | 1,15E+08 | ENSMUST00000157884.3 | 510805 | ENSMUSG00000088509 | Gm22789 | predicted gene, 22789 |
| chr12 | 35195505 | 35196964 | 1460 | * | 14,09066 | 2,950083297 | 1,095442 | 2,693053 | 0,00708 | 0,878304325 | 1,075735 | 3' UTR | 12 | 35244810 | 35244896 | 87 | 2 | 751520 | ENSMUST00000102298.3 | 47932 | ENSMUSG00000076253 | Mir680-3 | microRNA 680-3 |
| chr12 | 69216067 | 69217445 | 1379 | * | 10,49092 | 2,950462681 | 1,102278 | 2,676696 | 0,007435 | 0,871154332 | 1,173325 | Promoter (<=1kb) | 12 | 69215616 | 69221778 | 6163 | 1 | 69706 | ENSMUST00000222382.2 | 451 | ENSMUSG00000034883 | Lrr1 | leucine rich repeat protein 1 |
| chr13 | 23713915 | 23716150 | 2236 | * | 27,99837 | 2,951067245 | 0,735921 | 4,010033 | 6,07E-05 | 0,078945816 | 2,858511 | Promoter (<=1kb) | 13 | 23711167 | 23715104 | 3938 | 2 | 1E+08 | ENSMUST00000244140.1 | 0 | | Gm11335 | predicted gene 11335 |
| chr4 | 1,39E+08 | 1,39E+08 | 2011 | * | 14,4765 | 2,951073297 | 1,01933 | 2,89511 | 0,00379 | 0,718390388 | 1,41696 | Intron (ENSMUST0000 | 4 | 1,39E+08 | 1,39E+08 | 9328 | 2 | 212555 | ENSMUST00000141007.2 | 4652 | ENSMUSG00000028744 | Slc66a1 | solute carrier family 66 member 1 |
| chr10 | 59156920 | 59160624 | 3705 | * | 44,275751 | 2,951374298 | 0,5766897 | 5,117786 | 3,091E-07 | 0,001466209 | 2,9254973 | Promoter (<=1kb) | 10 | 59159118 | 59209126 | 50009 | 1 | 18451 | ENSMUST00000009789.15 | 0 | ENSMUSG00000019916 | P4ha1 | procollagen-proline, 2-oxoglutarate 4-dioxygenase (proline 4-hydroxylase), alpha 1 polypeptide |
| chr17 | 45617972 | 45618976 | 1005 | * | 11,56745 | 2,952304118 | 1,02333 | 2,884997 | 0,003914 | 0,838976992 | 0,738441 | Distal Intergenic | 17 | 45609026 | 45610144 | 1119 | 1 | 1,03E+08 | ENSMUST00000247429.1 | 8946 | | Gm35204 | predicted gene, 35204 |
| chr11 | 1,03E+08 | 1,03E+08 | 2087 | * | 21,07253 | 2,952466453 | 0,929715 | 3,17567 | 0,001495 | 0,420289001 | 3,104559 | Promoter (<=1kb) | 11 | 1,03E+08 | 1,03E+08 | 3495 | 1 | 192231 | ENSMUST00000053063.7 | 0 | ENSMUSG00000048878 | Hexim1 | hexamethylene bis-acetamide inducible 1 |
| chr5 | 75625505 | 75628019 | 2515 | * | 18,36111 | 2,952621044 | 0,901449 | 3,275418 | 0,001055 | 0,40679506 | 2,386008 | Distal Intergenic | 5 | 75721706 | 75721810 | 105 | 2 | 1,15E+08 | ENSMUST00000158453.3 | 93791 | ENSMUSG00000089078 | Gm24502 | predicted gene, 24502 |
| chr4 | 59067807 | 59069156 | 1350 | * | 13,9493 | 2,952630844 | 0,972909 | 3,034847 | 0,002407 | 0,751231966 | 0,738441 | Intron (ENSMUST0000 | 4 | 59035772 | 59041903 | 6132 | 1 | 14700 | ENSMUST00000126507.2 | 32035 | ENSMUSG00000038607 | Gng10 | guanine nucleotide binding protein (G protein), gamma 10 |
| chr2 | 73731724 | 73733212 | 1489 | * | 9,257135 | 2,952658901 | 1,087879 | 2,714143 | 0,006645 | 0,846832214 | 1,211315 | Distal Intergenic | 2 | 73739527 | 73740803 | 1277 | 2 | 228033 | ENSMUST00000131045.2 | 7591 | ENSMUSG00000018770 | Atp5g3 | ATP synthase, H+ transporting, mitochondrial F0 complex, subunit C3 (subunit 9) |
| chr6 | 91639058 | 91641295 | 2238 | * | 37,951007 | 2,952703922 | 0,5167645 | 5,7138287 | 1,105E-08 | 0,000118124 | 2,7872553 | Distal Intergenic | 6 | 91661034 | 91736047 | 75014 | 1 | 21366 | ENSMUST00000032185.9 | -19739 | ENSMUSG00000030096 | Slc6a6 | solute carrier family 6 (neurotransmitter transporter, taurine), member 6 |
| chr7 | 34924273 | 34925965 | 1693 | * | 17,85279 | 2,95354074 | 0,83316 | 3,544984 | 0,000393 | 0,260174751 | 2,386008 | Distal Intergenic | 7 | 34916660 | 34916785 | 126 | 1 | 1,15E+08 | ENSMUST00000157682.3 | 7613 | ENSMUSG00000088307 | Gm24505 | predicted gene, 24505 |
| chr19 | 5090393 | 5091973 | 1581 | * | 14,74366 | 2,95449415 | 1,009542 | 2,926569 | 0,003427 | 0,708350412 | 1,37683 | Promoter (<=1kb) | 19 | 5091382 | 5099945 | 8564 | 1 | 107392 | ENSMUST00000116567.4 | 0 | ENSMUSG00000080268 | Brms1 | breast cancer metastasis-suppressor 1 |
| chr4 | 1,56E+08 | 1,56E+08 | 1316 | * | 11,5301 | 2,95502372 | 1,109897 | 2,662432 | 0,007758 | 0,916242407 | 0,918255 | Promoter (<=1kb) | 4 | 1,56E+08 | 1,56E+08 | 31711 | 2 | 11603 | ENSMUST00000180572.2 | -452 | ENSMUSG00000041936 | Agrn | agrin |
| chr7 | 80396944 | 80398667 | 1724 | * | 13,00299 | 2,955392963 | 0,952792 | 3,101824 | 0,001923 | 0,654756507 | 1,075735 | Exon (ENSMUST00000 | 7 | 80376527 | 80379996 | 3470 | 2 | 29875 | ENSMUST00000205606.2 | -16948 | ENSMUSG00000030536 | Iqgap1 | IQ motif containing GTPase activating protein 1 |
| chr11 | 16315306 | 16316329 | 1024 | * | 9,500161 | 2,955445245 | 1,010027 | 2,926105 | 0,003432 | 0,695892636 | 1,449706 | Intron (ENSMUST0000 | 11 | 16213218 | 16234551 | 21334 | 1 | 211739 | ENSMUST00000207256.2 | 102088 | ENSMUSG00000048834 | Vstm2a | V-set and transmembrane domain containing 2A |
| chr14 | 81292324 | 81293899 | 1576 | * | 14,18236 | 2,956256844 | 0,994409 | 2,972879 | 0,00295 | 0,723215019 | 1,075735 | Distal Intergenic | 14 | 81923250 | 81958568 | 35319 | 1 | 1,05E+08 | ENSMUST00000227374.2 | -629351 | | Gm41219 | predicted gene, 41219 |
| chr2 | 14624864 | 14625993 | 1130 | * | 11,88516 | 2,956903721 | 1,011188 | 2,924189 | 0,003454 | 0,818408612 | 0,738441 | Intron (ENSMUST0000 | 2 | 14609933 | 14965668 | 355736 | 1 | 12296 | ENSMUST00000156027.8 | 14931 | ENSMUSG00000057914 | Cacnb2 | calcium channel, voltage-dependent, beta 2 subunit |
| chr6 | 1,01E+08 | 1,01E+08 | 667 | * | 9,962064 | 2,957347659 | 1,109012 | 2,66665 | 0,007661 | 0,841767996 | 1,425929 | Distal Intergenic | 6 | 1,01E+08 | 1,02E+08 | 17638 | 1 | 1,03E+08 | ENSMUST00000205133.2 | -19891 | | Gm34335 | predicted gene, 34335 |
| chr16 | 15800604 | 15801938 | 1335 | * | 12,50828 | 2,958200625 | 1,124115 | 2,631582 | 0,008499 | 0,906138253 | 1,075735 | Intron (ENSMUST0000 | 16 | 15705150 | 15708895 | 3746 | 1 | 12609 | ENSMUST00000096232.6 | 95454 | ENSMUSG00000071637 | Cebpd | CCAAT/enhancer binding protein (C/EBP), delta |
| chr15 | 54973981 | 54975041 | 1061 | * | 10,87113 | 2,958247316 | 1,110732 | 2,66333 | 0,007737 | 0,949733311 | 0,738441 | Promoter (<=1kb) | 15 | 54975713 | 55122667 | 146955 | 1 | 97998 | ENSMUST00000096433.10 | -672 | ENSMUSG00000022419 | Deptor | DEP domain containing MTOR-interacting protein |
| chr13 | 90944678 | 90946144 | 1467 | * | 11,95961 | 2,959726079 | 1,08953 | 2,716516 | 0,006597 | 0,851704515 | 1,173325 | Distal Intergenic | 13 | 91031572 | 91046943 | 15372 | 2 | 435376 | ENSMUST00000182104.2 | 100799 | ENSMUSG00000078958 | Atp6ap1l | ATPase, H+ transporting, lysosomal accessory protein 1-like |
| chr19 | 30263378 | 30264915 | 1538 | * | 13,52446 | 2,960459863 | 0,934624 | 3,167541 | 0,001537 | 0,580348965 | 1,2763 | Distal Intergenic | 19 | 30210342 | 30217087 | 6746 | 1 | 17195 | ENSMUST00000025797.7 | 53036 | ENSMUSG00000024863 | Mbl2 | mannose-binding lectin (protein C) 2 |
| chr10 | 62442918 | 62444202 | 1285 | * | 11,13377 | 2,960905202 | 1,109529 | 2,668614 | 0,007616 | 0,948006995 | 0,738441 | Distal Intergenic | 10 | 62416030 | 62438060 | 22031 | 2 | 56200 | ENSMUST00000045866.9 | -4858 | ENSMUSG00000020075 | Ddx21 | DExD box helicase 21 |
| chr7 | 98305003 | 98306830 | 1828 | * | 23,62001 | 2,961154314 | 0,75768 | 3,908184 | 9,3E-05 | 0,099733893 | 2,858511 | Promoter (<=1kb) | 7 | 98236344 | 98305986 | 69643 | 2 | 233545 | ENSMUST00000205276.2 | 0 | ENSMUSG00000035401 | Emsy | EMSY, BRCA2-interacting transcriptional repressor |
| chr3 | 55728297 | 55729585 | 1289 | * | 10,12395 | 2,961288016 | 1,111388 | 2,664494 | 0,00771 | 0,890237282 | 1,086118 | Intron (ENSMUST0000 | 3 | 55754526 | 55755489 | 964 | 1 | 71182 | ENSMUST00000199732.2 | -24941 | ENSMUSG00000106572 | 4933417G07Rik | SEC13 homolog pseudogene |
| chr19 | 28866269 | 28867510 | 1242 | * | 14,09142 | 2,961548854 | 0,956209 | 3,097176 | 0,001954 | 0,680148888 | 0,938682 | Promoter (2-3kb) | 19 | 28870413 | 28889277 | 18865 | 1 | 20510 | ENSMUST00000160702.8 | -2903 | ENSMUSG00000024935 | Slc1a1 | solute carrier family 1 (neuronal/epithelial high affinity glutamate transporter, system Xag), member 1 |
| chr9 | 63648938 | 63651581 | 2644 | * | 28,645159 | 2,962473896 | 0,6496511 | 4,5601002 | 5,113E-06 | 0,013096578 | 2,8800171 | Promoter (<=1kb) | 9 | 63575095 | 63650547 | 75453 | 2 | 17127 | ENSMUST00000137065.2 | 0 | ENSMUSG00000032402 | Smad3 | SMAD family member 3 |
| chr9 | 31703533 | 31705011 | 1479 | * | 14,84757 | 2,962754076 | 0,908867 | 3,259831 | 0,001115 | 0,432339608 | 2,141612 | Distal Intergenic | 9 | 31817289 | 31822494 | 5206 | 1 | 1,03E+08 | ENSMUST00000216981.2 | -112278 | ENSMUSG00000110813 | Gm31497 | predicted gene, 31497 |
| chr16 | 29029383 | 29030643 | 1261 | * | 11,34093 | 2,964209559 | 1,08782 | 2,724908 | 0,006432 | 0,920164459 | 0,744375 | Promoter (<=1kb) | 16 | 29028922 | 29039091 | 10170 | 1 | 27281 | ENSMUST00000161294.2 | 461 | ENSMUSG00000022525 | Plaat1 | phospholipase A and acyltransferase 1 |
| chr8 | 122951091 | 122955437 | 4347 | * | 86,004906 | 2,964242264 | 0,4523309 | 6,5532602 | 5,629E-11 | 1,07857E-06 | 4,2003037 | Distal Intergenic | 8 | 122985359 | 122999389 | 14031 | 1 | 195209 | ENSMUST00000187142.3 | -29922 | ENSMUSG00000043903 | Zfp469 | zinc finger protein 469 |
| chr11 | 1,19E+08 | 1,19E+08 | 1984 | * | 17,28181 | 2,964878667 | 1,05399 | 2,813004 | 0,004908 | 0,676922067 | 2,386008 | Exon (ENSMUST00000 | 11 | 1,19E+08 | 1,19E+08 | 94319 | 1 | 672511 | ENSMUST00000131035.10 | 32647 | ENSMUSG00000070327 | Rnf213 | ring finger protein 213 |
| chr11 | 82515888 | 82517627 | 1740 | * | 15,78035 | 2,96545378 | 0,998489 | 2,969941 | 0,002979 | 0,565507616 | 2,663619 | Distal Intergenic | 11 | 82524179 | 82527135 | 2957 | 1 | 791422 | ENSMUST00000125600.2 | -6552 | ENSMUSG00000069796 | Gm11426 | predicted gene 11426 |
| chr19 | 47864224 | 47865721 | 1498 | * | 15,50866 | 2,965787276 | 1,121899 | 2,643543 | 0,008204 | 0,858257847 | 1,37683 | Exon (ENSMUST00000 | 19 | 47858701 | 47874737 | 16037 | 1 | 68214 | ENSMUST00000135016.3 | 5523 | ENSMUSG00000025069 | Gsto2 | glutathione S-transferase omega 2 |
| chr10 | 81214105 | 81215624 | 1520 | * | 47,58735 | 2,966896257 | 1,080561 | 2,7457 | 0,006038 | 0,71419937 | 2,338387 | Promoter (<=1kb) | 10 | 81202056 | 81214350 | 12295 | 2 | 56371 | ENSMUST00000140901.8 | 0 | ENSMUSG00000020235 | Fzr1 | fizzy and cell division cycle 20 related 1 |
| chr13 | 99654191 | 99655726 | 1536 | * | 11,4242 | 2,968098784 | 0,968455 | 3,064778 | 0,002178 | 0,734804995 | 0,738441 | Promoter (1-2kb) | 13 | 99653137 | 99658268 | 5132 | 1 | 1,03E+08 | ENSMUST00000181939.3 | 1054 | | Gm26559 | predicted gene, 26559 |
| chr10 | 75401756 | 75402994 | 1239 | * | 15,68652 | 2,968128731 | 0,878525 | 3,378536 | 0,000729 | 0,430115816 | 1,41696 | Promoter (<=1kb) | 10 | 75402127 | 75414751 | 12625 | 1 | 14598 | ENSMUST00000129232.8 | 0 | ENSMUSG00000006345 | Ggt1 | gamma-glutamyltransferase 1 |
| chr19 | 5488598 | 5490526 | 1929 | * | 11,5754 | 2,968283651 | 1,106849 | 2,681743 | 0,007324 | 0,865825879 | 1,173325 | Exon (ENSMUST00000 | 19 | 5497575 | 5505974 | 8400 | 1 | 14283 | ENSMUST00000025850.7 | -7049 | ENSMUSG00000024912 | Fosl1 | fos-like antigen 1 |
| chr7 | 79415749 | 79417514 | 1766 | * | 15,48089 | 2,968491471 | 1,017482 | 2,917487 | 0,003529 | 0,708529912 | 1,41696 | Exon (ENSMUST00000 | 7 | 79392911 | 79435698 | 42788 | 1 | 626359 | ENSMUST00000035622.8 | 22838 | ENSMUSG00000039099 | Wdr93 | WD repeat domain 93 |
| chr5 | 1,25E+08 | 1,25E+08 | 2041 | * | 21,13767 | 2,968543858 | 0,942416 | 3,149928 | 0,001633 | 0,446292698 | 2,880017 | Distal Intergenic | 5 | 1,25E+08 | 1,25E+08 | 4062 | 1 | 269704 | ENSMUST00000134473.2 | 55774 | ENSMUSG00000079215 | Zfp664 | zinc finger protein 664 |
| chr11 | 3709118 | 3710765 | 1648 | * | 18,92838 | 2,96855229 | 0,831379 | 3,570635 | 0,000356 | 0,229205408 | 2,713306 | Intron (ENSMUST0000 | 11 | 3667852 | 3723789 | 55938 | 2 | 74309 | ENSMUST00000127371.2 | 13024 | ENSMUSG00000020435 | Osbp2 | oxysterol binding protein 2 |
| chr13 | 23944091 | 23947345 | 3255 | * | 37,476822 | 2,968785646 | 0,6071253 | 4,8899057 | 1,009E-06 | 0,003692163 | 2,9254973 | Promoter (<=1kb) | 13 | 23944675 | 23945213 | 539 | 2 | 326619 | ENSMUST00000102964.4 | 0 | ENSMUSG00000060093 | H4c1 | H4 clustered histone 1 |
| chr12 | 32515123 | 32516309 | 1187 | * | 11,06504 | 2,96961893 | 1,113471 | 2,666993 | 0,007653 | 0,948006995 | 0,738441 | Distal Intergenic | 12 | 32440767 | 32481922 | 41156 | 2 | 1,05E+08 | ENSMUST00000218751.2 | -33201 | | Gm40375 | predicted gene, 40375 |
| chr5 | 1,21E+08 | 1,21E+08 | 2286 | * | 16,39142 | 2,970262501 | 0,903132 | 3,288846 | 0,001006 | 0,501831163 | 1,2763 | Intron (ENSMUST0000 | 5 | 1,21E+08 | 1,22E+08 | 148359 | 1 | 269700 | ENSMUST00000042614.13 | 28039 | ENSMUSG00000042744 | Hectd4 | HECT domain E3 ubiquitin protein ligase 4 |
| chr2 | 1,19E+08 | 1,19E+08 | 2918 | * | 22,50569 | 2,970543012 | 0,773915 | 3,838333 | 0,000124 | 0,105761771 | 3,471678 | Promoter (1-2kb) | 2 | 1,19E+08 | 1,19E+08 | 20922 | 2 | 18796 | ENSMUST00000102524.8 | -1440 | ENSMUSG00000040061 | Plcb2 | phospholipase C, beta 2 |
| chr2 | 13561251 | 13563116 | 1866 | * | 30,7927 | 2,970616314 | 0,74644 | 3,979713 | 6,9E-05 | 0,086053637 | 2,725491 | Distal Intergenic | 2 | 13513825 | 13549479 | 35655 | 2 | 13434 | ENSMUST00000124488.7 | -11772 | ENSMUSG00000026723 | Trdmt1 | tRNA aspartic acid methyltransferase 1 |
| chr4 | 99648244 | 99650002 | 1759 | * | 15,25583 | 2,970779694 | 0,921182 | 3,224964 | 0,00126 | 0,524368688 | 1,41696 | Intron (ENSMUST0000 | 4 | 99607389 | 99627710 | 20322 | 1 | 320438 | ENSMUST00000124147.2 | 40855 | ENSMUSG00000073792 | Alg6 | asparagine-linked glycosylation 6 (alpha-1,3,-glucosyltransferase) |
| chr9 | 57876389 | 57877570 | 1182 | * | 10,55812 | 2,971026374 | 1,1062 | 2,685795 | 0,007236 | 0,939041504 | 0,738441 | Distal Intergenic | 9 | 57905324 | 57914734 | 9411 | 1 | 13070 | ENSMUST00000181289.2 | -27754 | ENSMUSG00000032323 | Cyp11a1 | cytochrome P450, family 11, subfamily a, polypeptide 1 |
| chr8 | 13810670 | 13812011 | 1342 | * | 11,65032 | 2,971239107 | 1,06442 | 2,791415 | 0,005248 | 0,890475095 | 0,738441 | Promoter (1-2kb) | 8 | 13808767 | 13813010 | 4244 | 1 | 69957 | ENSMUST00000129872.2 | 1903 | ENSMUSG00000038416 | Cdc16 | CDC16 cell division cycle 16 |
| chr6 | 93547568 | 93548187 | 620 | * | 8,974143 | 2,971386521 | 1,052535 | 2,823076 | 0,004757 | 0,788774532 | 1,211315 | Distal Intergenic | 6 | 93486554 | 93486683 | 130 | 1 | 1,15E+08 | ENSMUST00000158005.4 | 61014 | ENSMUSG00000088630 | Gm22312 | predicted gene, 22312 |
| chr7 | 24179958 | 24181478 | 1521 | * | 13,83932 | 2,971627469 | 1,101293 | 2,698308 | 0,006969 | 0,934396727 | 0,738441 | Promoter (<=1kb) | 7 | 24181448 | 24203964 | 22517 | 1 | 260299 | ENSMUST00000068023.8 | 0 | ENSMUSG00000054793 | Cadm4 | cell adhesion molecule 4 |
| chr5 | 28222174 | 28223239 | 1066 | * | 9,975218 | 2,971756937 | 1,13175 | 2,625806 | 0,008644 | 0,890735468 | 1,211315 | Distal Intergenic | 5 | 28258922 | 28260456 | 1535 | 2 | 1,03E+08 | ENSMUST00000246890.1 | 37217 | | Gm26608 | predicted gene, 26608 |
| chr11 | 1,21E+08 | 1,21E+08 | 1429 | * | 21,07118 | 2,972582418 | 0,889886 | 3,34041 | 0,000837 | 0,322690573 | 3,104559 | Promoter (<=1kb) | 11 | 1,21E+08 | 1,21E+08 | 2791 | 2 | 20892 | ENSMUST00000106154.8 | 0 | ENSMUSG00000025144 | Cenpx | centromere protein X |
| chr13 | 53067745 | 53068778 | 1034 | * | 8,155209 | 2,972745439 | 1,125379 | 2,64155 | 0,008253 | 0,854157826 | 1,425929 | Promoter (<=1kb) | 13 | 53064359 | 53066982 | 2624 | 2 | 11992 | ENSMUST00000230563.2 | -763 | ENSMUSG00000021460 | Auh | AU RNA binding protein/enoyl-coenzyme A hydratase |
| chr17 | 27333519 | 27334642 | 1124 | * | 11,63907 | 2,973048427 | 1,111799 | 2,674089 | 0,007493 | 0,872576097 | 1,173325 | Exon (ENSMUST00000 | 17 | 27325151 | 27326323 | 1173 | 1 | 16440 | ENSMUST00000235521.2 | 8368 | ENSMUSG00000042644 | Itpr3 | inositol 1,4,5-triphosphate receptor 3 |
| chrX | 1,32E+08 | 1,32E+08 | 1269 | * | 11,71134 | 2,973508242 | 0,938707 | 3,167664 | 0,001537 | 0,677933849 | 0,744375 | Distal Intergenic | 20 | 1,33E+08 | 1,33E+08 | 3672 | 2 | 279653 | ENSMUST00000193485.2 | 295888 | ENSMUSG00000051323 | Pcdh19 | protocadherin 19 |
| chr13 | 22219154 | 22222240 | 3087 | * | 38,992028 | 2,973788805 | 0,6090788 | 4,8824371 | 1,048E-06 | 0,00373207 | 2,9254973 | Promoter (<=1kb) | 13 | 22219334 | 22219738 | 405 | 2 | 319168 | ENSMUST00000091742.6 | 0 | ENSMUSG00000069302 | H2ac12 | H2A clustered histone 12 |
| chr17 | 28098819 | 28101497 | 2679 | * | 21,81814 | 2,973937221 | 0,88242 | 3,370205 | 0,000751 | 0,316567501 | 2,880017 | Exon (ENSMUST00000 | 17 | 28109079 | 28114340 | 5262 | 1 | 224648 | ENSMUST00000137825.2 | -7582 | ENSMUSG00000039512 | Uhrf1bp1 | UHRF1 (ICBP90) binding protein 1 |
| chr1 | 1,77E+08 | 1,77E+08 | 997 | * | 13,17847 | 2,974301225 | 1,123063 | 2,648383 | 0,008088 | 0,956450448 | 0,738441 | Promoter (<=1kb) | 1 | 1,77E+08 | 1,77E+08 | 80406 | 2 | 545389 | ENSMUST00000195717.6 | 368 | ENSMUSG00000057335 | Cep170 | centrosomal protein 170 |
| chr7 | 18943260 | 18944876 | 1617 | * | 16,19854 | 2,974483458 | 0,922458 | 3,224518 | 0,001262 | 0,418252864 | 2,663619 | Promoter (1-2kb) | 7 | 18946463 | 18958099 | 11637 | 1 | 319197 | ENSMUST00000060225.6 | -1587 | ENSMUSG00000044317 | Gpr4 | G protein-coupled receptor 4 |
| chr13 | 5321894 | 5323100 | 1207 | * | 9,682745 | 2,975489368 | 0,996773 | 2,985121 | 0,002835 | 0,694157242 | 1,211315 | Distal Intergenic | 13 | 5239802 | 5239933 | 132 | 2 | 1,15E+08 | ENSMUST00000104285.3 | -81961 | ENSMUSG00000077473 | Gm24790 | predicted gene, 24790 |
| chr19 | 23796236 | 23797711 | 1476 | * | 11,74949 | 2,975977959 | 1,115496 | 2,667853 | 0,007634 | 0,876507886 | 1,173325 | Intron (ENSMUST0000 | 19 | 23736251 | 23926960 | 190710 | 1 | 319924 | ENSMUST00000025830.9 | 59985 | ENSMUSG00000024897 | Apba1 | amyloid beta (A4) precursor protein binding, family A, member 1 |
| chr8 | 31323453 | 31324181 | 729 | * | 10,1757 | 2,976104967 | 1,129465 | 2,634969 | 0,008414 | 0,85518169 | 1,449706 | Distal Intergenic | 8 | 31579499 | 31587075 | 7577 | 1 | 66959 | ENSMUST00000036631.14 | -255318 | ENSMUSG00000039661 | Dusp26 | dual specificity phosphatase 26 (putative) |
| chr12 | 42582122 | 42583105 | 984 | * | 10,3678 | 2,977187919 | 1,140265 | 2,610961 | 0,009029 | 0,971049273 | 0,744375 | Distal Intergenic | 12 | 43013667 | 43062242 | 48576 | 2 | 1,03E+08 | ENSMUST00000220044.2 | 479137 | ENSMUSG00000112407 | Gm33340 | predicted gene, 33340 |
| chr15 | 38421628 | 38422229 | 602 | * | 9,422924 | 2,977411547 | 1,033393 | 2,881199 | 0,003962 | 0,652050642 | 2,237403 | Distal Intergenic | 15 | 38385878 | 38404398 | 18521 | 2 | 1,05E+08 | ENSMUST00000227422.2 | -17230 | | Gm41307 | predicted gene, 41307 |
| chr8 | 1,23E+08 | 1,23E+08 | 1292 | * | 18,89485 | 2,977524364 | 0,923805 | 3,22311 | 0,001268 | 0,418252864 | 2,663619 | Distal Intergenic | 8 | 1,23E+08 | 1,23E+08 | 14031 | 1 | 195209 | ENSMUST00000187142.3 | -21845 | ENSMUSG00000043903 | Zfp469 | zinc finger protein 469 |
| chr18 | 61711488 | 61712911 | 1424 | * | 11,85191 | 2,977885756 | 1,135666 | 2,622149 | 0,008738 | 0,885980681 | 1,2763 | Promoter (2-3kb) | 18 | 61708588 | 61715189 | 6602 | 1 | 93687 | ENSMUST00000164253.2 | 2900 | ENSMUSG00000024576 | Csnk1a1 | casein kinase 1, alpha 1 |
| chr3 | 1,08E+08 | 1,08E+08 | 1326 | * | 11,39167 | 2,978396135 | 1,151163 | 2,587293 | 0,009673 | 0,912942638 | 1,173325 | Promoter (<=1kb) | 3 | 1,08E+08 | 1,08E+08 | 24816 | 2 | 229709 | ENSMUST00000153623.2 | 0 | ENSMUSG00000027893 | Ahcyl1 | S-adenosylhomocysteine hydrolase-like 1 |
| chr4 | 1,09E+08 | 1,09E+08 | 1559 | * | 13,91151 | 2,978548102 | 1,040994 | 2,861255 | 0,00422 | 0,852340527 | 0,738441 | Promoter (<=1kb) | 4 | 1,09E+08 | 1,09E+08 | 140620 | 2 | 100273 | ENSMUST00000194478.6 | 0 | ENSMUSG00000028559 | Osbpl9 | oxysterol binding protein-like 9 |
| chr12 | 1,13E+08 | 1,13E+08 | 2058 | * | 21,2051 | 2,978732627 | 0,829109 | 3,59269 | 0,000327 | 0,214423576 | 2,880017 | Exon (ENSMUST00000 | 12 | 1,13E+08 | 1,13E+08 | 2049 | 1 | 116870 | ENSMUST00000134488.2 | -3682 | ENSMUSG00000021144 | Mta1 | metastasis associated 1 |
| chr19 | 15943298 | 15944495 | 1198 | * | 12,47871 | 2,978791203 | 0,99167 | 3,003812 | 0,002666 | 0,731734453 | 0,918255 | Intron (ENSMUST0000 | 19 | 15933345 | 15962057 | 28713 | 2 | 208518 | ENSMUST00000236749.2 | 17562 | ENSMUSG00000041491 | Cep78 | centrosomal protein 78 |
| chr3 | 95132877 | 95134217 | 1341 | * | 9,699256 | 2,979426116 | 1,076242 | 2,768361 | 0,005634 | 0,789027175 | 1,425929 | Promoter (1-2kb) | 3 | 95126446 | 95135988 | 9543 | 2 | 56772 | ENSMUST00000065482.6 | 1771 | ENSMUSG00000053192 | Mllt11 | myeloid/lymphoid or mixed-lineage leukemia; translocated to, 11 |
| chr16 | 30469414 | 30470669 | 1256 | * | 13,07545 | 2,979717447 | 1,063319 | 2,80228 | 0,005074 | 0,79031971 | 1,2763 | Distal Intergenic | 16 | 30418541 | 30421615 | 3075 | 1 | 224093 | ENSMUST00000059078.4 | 50873 | ENSMUSG00000046546 | Fam43a | family with sequence similarity 43, member A |
| chr11 | 1,2E+08 | 1,2E+08 | 1412 | * | 13,60131 | 2,979884044 | 1,038397 | 2,869697 | 0,004109 | 0,805038542 | 0,938682 | Promoter (<=1kb) | 11 | 1,2E+08 | 1,2E+08 | 1898 | 1 | 208092 | ENSMUST00000137055.2 | 0 | ENSMUSG00000025371 | Chmp6 | charged multivesicular body protein 6 |
| chr17 | 46592942 | 46594273 | 1332 | * | 11,55423 | 2,980291887 | 1,15429 | 2,581927 | 0,009825 | 0,95413859 | 0,918255 | Promoter (<=1kb) | 17 | 46568777 | 46593952 | 25176 | 2 | 74094 | ENSMUST00000180283.2 | 0 | ENSMUSG00000012296 | Tjap1 | tight junction associated protein 1 |

| | | | | | | | | | | | | | | | | | | | | | | | |
|---|---|---|---|---|---|---|---|---|---|---|---|---|---|---|---|---|---|---|---|---|---|---|---|
| chr8 | 35283961 | 35285320 | 1360 | * | 10,30251 | 2,980398824 | 1,080297 | 2,75887 | 0,0058 | 0,792545355 | 1,425929 | Exon (ENSMUST00000 | 8 | 35274451 | 35287048 | 12598 | 1 | 319520 | ENSMUST00000033930.5 | 9510 | ENSMUSG00000031530 | Dusp4 | dual specificity phosphatase 4 |
| chr8 | 45253259 | 45254253 | 995 | * | 9,858358 | 2,98042734 | 1,122624 | 2,654876 | 0,007934 | 0,883918379 | 1,173325 | Distal Intergenic | 8 | 45388484 | 45505294 | 116811 | 1 | 14107 | ENSMUST00000191428.7 | -134231 | ENSMUSG00000070047 | Fat1 | FAT atypical cadherin 1 |
| chr10 | 86619635 | 86621146 | 1512 | * | 12,92093 | 2,980722983 | 0,912468 | 3,266661 | 0,001088 | 0,577007376 | 0,918255 | Intron (ENSMUST00000 | 10 | 86614869 | 86674253 | 59385 | 1 | 103466 | ENSMUST00000099396.3 | 4766 | ENSMUSG00000054027 | Nt5dc3 | 5'-nucleotidase domain containing 3 |
| chr10 | 13923029 | 13925702 | 2674 | * | 21,07009 | 2,981014213 | 0,901803 | 3,305615 | 0,000948 | 0,351510656 | 2,880017 | Intron (ENSMUST00000 | 10 | 13842768 | 13955419 | 112652 | 1 | 15273 | ENSMUST00000191464.2 | 80261 | ENSMUSG00000015501 | Hivep2 | human immunodeficiency virus type I enhancer binding protein 2 |
| chr3 | 145627784 | 145630391 | 2608 | * | 32,186216 | 2,981153706 | 0,6771689 | 4,4023783 | 1,071E-05 | 0,024041634 | 2,7254908 | Promoter (<=1kb) | 3 | 145630017 | 145640111 | 10095 | 1 | 12042 | ENSMUST00000029842.9 | 0 | ENSMUSG00000028191 | Bcl10 | B cell leukemia/lymphoma 10 |
| chr11 | 5319295 | 5320609 | 1315 | * | 10,42068 | 2,981326363 | 1,153562 | 2,584453 | 0,009753 | 0,98239328 | 0,744375 | 5' UTR | 11 | 5228602 | 5331740 | 103139 | 2 | 407821 | ENSMUST00000172492.8 | 11131 | ENSMUSG00000041961 | Znrf3 | zinc and ring finger 3 |
| chr10 | 8116692 | 8117670 | 979 | * | 10,12415 | 2,982092679 | 1,154065 | 2,58399 | 0,009766 | 0,914482087 | 1,173325 | Intron (ENSMUST00000 | 10 | 8080520 | 8178006 | 97487 | 2 | 338362 | ENSMUST00000219116.2 | 60336 | ENSMUSG00000047712 | Ust | uronyl-2-sulfotransferase |
| chr4 | 1,36E+08 | 1,36E+08 | 1372 | * | 12,90025 | 2,982423497 | 0,872694 | 3,417489 | 0,000632 | 0,477622452 | 0,918255 | Distal Intergenic | 4 | 1,36E+08 | 1,36E+08 | 1896 | 1 | 15903 | ENSMUST00000008016.3 | -24245 | ENSMUSG00000007872 | Id3 | inhibitor of DNA binding 3 |
| chr15 | 9070204 | 9071517 | 1314 | * | 14,83713 | 2,982735585 | 0,921773 | 3,235869 | 0,001213 | 0,44621096 | 2,141612 | Promoter (<=1kb) | 15 | 9071340 | 9110979 | 39640 | 1 | 68646 | ENSMUST00000190591.10 | 0 | ENSMUSG00000022253 | Nadk2 | NAD kinase 2, mitochondrial |
| chr13 | 21546794 | 21548390 | 1597 | * | 13,99075 | 2,982960313 | 1,052026 | 2,835445 | 0,004576 | 0,824314641 | 0,938682 | Promoter (<=1kb) | 13 | 21546990 | 21556459 | 9470 | 1 | 22758 | ENSMUST00000225545.2 | 0 | ENSMUSG00000036721 | Zscan12 | zinc finger and SCAN domain containing 12 |
| chr18 | 21906390 | 21907432 | 1043 | * | 10,6735 | 2,983220598 | 1,046078 | 2,851814 | 0,004347 | 0,75938015 | 1,2763 | Distal Intergenic | 18 | 21683436 | 21787775 | 104340 | 2 | 225266 | ENSMUST00000122333.2 | -118615 | ENSMUSG00000042514 | Klhl14 | kelch-like 14 |
| chr13 | 99548910 | 99550463 | 1554 | * | 13,1658 | 2,983361042 | 1,027408 | 2,903773 | 0,003687 | 0,730832709 | 1,2763 | Promoter (<=1kb) | 13 | 99533432 | 99549129 | 15698 | 2 | 320893 | ENSMUST00000180491.2 | 0 | | 6430562O15Rik | RIKEN cDNA 6430562O15 gene |
| chr18 | 71802098 | 71803104 | 1007 | * | 10,63886 | 2,983552158 | 1,041676 | 2,864184 | 0,004181 | 0,851512567 | 0,744375 | Intron (ENSMUST00000 | 18 | 71392061 | 72484088 | 1092028 | 2 | 13176 | ENSMUST00000073379.6 | 680984 | ENSMUSG00000060534 | Dcc | deleted in colorectal carcinoma |
| chr3 | 65778053 | 65779234 | 1182 | * | 16,51646 | 2,984953986 | 0,929182 | 3,212455 | 0,001316 | 0,459179243 | 2,141612 | Intron (ENSMUST00000 | 3 | 65854203 | 65855419 | 1217 | 2 | 56706 | ENSMUST00000149729.2 | 76185 | ENSMUSG00000027829 | Ccnl1 | cyclin L1 |
| chr8 | 25005958 | 25006932 | 975 | * | 10,40517 | 2,985040435 | 1,149236 | 2,597412 | 0,009393 | 0,908469938 | 1,173325 | Distal Intergenic | 8 | 25023472 | 25030812 | 7341 | 2 | 209176 | ENSMUST00000138335.2 | 23880 | ENSMUSG00000031549 | Ido2 | indoleamine 2,3-dioxygenase 2 |
| chr4 | 1,36E+08 | 1,36E+08 | 1542 | * | 14,61274 | 2,985374059 | 1,12098 | 2,663183 | 0,007741 | 0,851704515 | 1,37683 | Exon (ENSMUST00000 | 4 | 1,36E+08 | 1,36E+08 | 17375 | 1 | 242702 | ENSMUST00000145304.8 | -14762 | ENSMUSG00000037139 | Myom3 | myomesin family, member 3 |
| chr4 | 1,05E+08 | 1,05E+08 | 1393 | * | 11,09694 | 2,985380211 | 1,139437 | 2,62005 | 0,008792 | 0,966989334 | 0,738441 | Promoter (1-2kb) | 4 | 1,05E+08 | 1,05E+08 | 75418 | 1 | 67916 | ENSMUST00000064139.8 | -1921 | ENSMUSG00000028517 | Plpp3 | phospholipid phosphatase 3 |
| chr7 | 71248808 | 71250610 | 1803 | * | 12,22954 | 2,986124241 | 1,071598 | 2,786609 | 0,005326 | 0,813025434 | 1,173325 | Distal Intergenic | 7 | 71331523 | 71338195 | 6673 | 2 | 1,15E+08 | ENSMUST00000208786.2 | 87585 | | Gm44689 | predicted gene 44689 |
| chr19 | 57615456 | 57616419 | 964 | * | 10,41518 | 2,986755033 | 1,118009 | 2,671495 | 0,007551 | 0,857842921 | 1,2763 | Intron (ENSMUST00000 | 19 | 57599466 | 58121775 | 522310 | 1 | 226255 | ENSMUST00000077282.7 | 15990 | ENSMUSG00000054843 | Atrnl1 | attractin like 1 |
| chr2 | 30603793 | 30604786 | 994 | * | 13,23445 | 2,986878383 | 1,15027 | 2,596675 | 0,009413 | 0,908787948 | 1,173325 | Promoter (2-3kb) | 2 | 30603162 | 30607055 | 3894 | 2 | 1E+08 | ENSMUST00000131207.2 | 2269 | | Gm14488 | predicted gene 14488 |
| chr17 | 18000427 | 18002433 | 2007 | * | 14,90333 | 2,987680966 | 0,966089 | 3,092551 | 0,001984 | 0,640437129 | 1,214904 | Distal Intergenic | 17 | 18047420 | 18059333 | 11914 | 1 | 75202 | ENSMUST00000226899.2 | -44987 | ENSMUSG00000080316 | Spaca6 | sperm acrosome associated 6 |
| chr9 | 14170553 | 14172211 | 1659 | * | 18,94173 | 2,987854475 | 0,76564 | 3,902426 | 9,52E-05 | 0,119411163 | 2,221611 | Promoter (2-3kb) | 9 | 14170337 | 14175099 | 4763 | 2 | 1,03E+08 | ENSMUST00000214060.2 | 2888 | | Gm34069 | predicted gene, 34069 |
| chr5 | 24954153 | 24955135 | 983 | * | 10,14639 | 2,988014408 | 1,124435 | 2,657347 | 0,007876 | 0,882294105 | 1,173325 | Promoter (1-2kb) | 5 | 24956001 | 24956322 | 322 | 2 | 214301 | ENSMUST00000123386.2 | 1187 | ENSMUSG00000038135 | Crygn | crystallin, gamma N |
| chr2 | 1,65E+08 | 1,65E+08 | 1560 | * | 14,95092 | 2,988157227 | 1,054622 | 2,833392 | 0,004606 | 0,755814512 | 1,37683 | Distal Intergenic | 2 | 1,65E+08 | 1,65E+08 | 1288 | 2 | 228875 | ENSMUST00000125550.2 | 8130 | ENSMUSG00000017664 | Slc35c2 | solute carrier family 35, member C2 |
| chr11 | 68989171 | 68990274 | 1104 | * | 17,34285 | 2,988518442 | 0,864839 | 3,455577 | 0,000549 | 0,301441602 | 2,423594 | Promoter (<=1kb) | 11 | 68989774 | 69000786 | 11013 | 1 | 18626 | ENSMUST00000021271.14 | 0 | ENSMUSG00000020893 | Per1 | period circadian clock 1 |
| chr1 | 53226200 | 53227624 | 1425 | * | 16,44551 | 2,988572345 | 0,911694 | 3,278044 | 0,001045 | 0,409763611 | 2,318434 | Promoter (<=1kb) | 1 | 53197736 | 53226795 | 29060 | 2 | 69397 | ENSMUST00000072235.10 | 0 | ENSMUSG00000060715 | 1700019A02Rik | RIKEN cDNA 1700019A02 gene |
| chr13 | 22212988 | 22214607 | 1620 | * | 11,17303 | 2,988794941 | 1,03104 | 2,898817 | 0,003746 | 0,749550311 | 1,173325 | Intron (ENSMUST00000 | 13 | 22219334 | 22219738 | 405 | 2 | 319168 | ENSMUST00000091742.6 | 5131 | ENSMUSG00000069302 | H2ac12 | H2A clustered histone 12 |
| chr19 | 46963444 | 46964736 | 1293 | * | 12,90976 | 2,988945146 | 0,937435 | 3,188428 | 0,00143 | 0,565876168 | 1,2763 | Intron (ENSMUST00000 | 19 | 46871756 | 46958007 | 86252 | 2 | 76952 | ENSMUST00000235234.2 | -5437 | ENSMUSG00000025041 | Nt5c2 | 5'-nucleotidase, cytosolic II |
| chr3 | 1,55E+08 | 1,55E+08 | 744 | * | 13,0163 | 2,989160674 | 0,94902 | 3,149734 | 0,001634 | 0,61075929 | 1,173325 | Intron (ENSMUST00000 | 3 | 1,55E+08 | 1,55E+08 | 44161 | 1 | 74435 | ENSMUST00000194376.2 | 67049 | ENSMUSG00000028182 | Lrriq3 | leucine-rich repeats and IQ motif containing 3 |
| chr1 | 89539403 | 89540930 | 1528 | * | 13,52132 | 2,989625629 | 1,157389 | 2,583077 | 0,009792 | 0,984611743 | 0,738441 | Intron (ENSMUST00000 | 1 | 89507958 | 89822999 | 315042 | 1 | 347722 | ENSMUST00000074945.8 | 31445 | ENSMUSG00000055013 | Agap1 | ArfGAP with GTPase domain, ankyrin repeat and PH domain 1 |
| chr3 | 57761797 | 57762848 | 1052 | * | 10,03134 | 2,989659229 | 1,080314 | 2,767399 | 0,005651 | 0,817940051 | 1,211315 | Distal Intergenic | 3 | 57749316 | 57755500 | 6185 | 2 | 18645 | ENSMUST00000066882.10 | -6297 | ENSMUSG00000027805 | Pfn2 | profilin 2 |
| chr9 | 58084694 | 58087502 | 2809 | * | 21,68077 | 2,989976072 | 0,800072 | 3,737133 | 0,000186 | 0,155253659 | 2,880017 | Intron (ENSMUST00000 | 9 | 58063551 | 58066504 | 2954 | 2 | 26968 | ENSMUST00000041477.15 | -18190 | ENSMUSG00000037206 | Islr | immunoglobulin superfamily containing leucine-rich repeat |
| chr9 | 1,14E+08 | 1,14E+08 | 1780 | * | 10,59613 | 2,990180829 | 1,147355 | 2,606152 | 0,009157 | 0,891097601 | 1,2763 | Promoter (1-2kb) | 9 | 1,14E+08 | 1,14E+08 | 10065 | 2 | 18571 | ENSMUST00000156425.2 | 1597 | ENSMUSG00000032504 | Pdcd6ip | programmed cell death 6 interacting protein |
| chr13 | 52151758 | 52155085 | 3328 | * | 28,36832 | 2,990261844 | 0,746997 | 4,003046 | 6,25E-05 | 0,080010047 | 2,880017 | Distal Intergenic | 13 | 52250777 | 52278197 | 27421 | 1 | 74050 | ENSMUST00000220691.3 | -95692 | ENSMUSG00000113211 | 4921525O09Rik | RIKEN cDNA 4921525O09 gene |
| chr3 | 1,09E+08 | 1,09E+08 | 1316 | * | 14,85295 | 2,991063989 | 1,005449 | 2,974854 | 0,002931 | 0,689914111 | 1,2763 | Distal Intergenic | 3 | 1,09E+08 | 1,1E+08 | 345046 | 1 | 57257 | ENSMUST00000046864.14 | -87800 | ENSMUSG00000033721 | Vav3 | vav 3 oncogene |
| chr11 | 1,2E+08 | 1,2E+08 | 2302 | * | 49,97641 | 2,991658066 | 1,033168 | 2,895617 | 0,003784 | 0,639171992 | 2,338387 | Promoter (<=1kb) | 11 | 1,2E+08 | 1,2E+08 | 12956 | 2 | 18453 | ENSMUST00000026122.11 | 0 | ENSMUSG00000025130 | P4hb | prolyl 4-hydroxylase, beta polypeptide |
| chr12 | 13264893 | 13266216 | 1324 | * | 11,10799 | 2,991719219 | 1,004905 | 2,977117 | 0,00291 | 0,688182394 | 1,2763 | Distal Intergenic | 12 | 13275162 | 13279380 | 4219 | 2 | 104721 | ENSMUST00000221028.2 | 13164 | ENSMUSG00000037149 | Ddx1 | DEAD box helicase 1 |
| chr9 | 61892108 | 61894235 | 2128 | * | 17,66862 | 2,991990865 | 0,825665 | 3,623737 | 0,00029 | 0,233755249 | 2,154504 | Intron (ENSMUST00000 | 9 | 61863328 | 61883855 | 20528 | 2 | 74090 | ENSMUST00000113990.2 | -8253 | ENSMUSG00000032278 | Paqr5 | progestin and adipoQ receptor family member V |
| chr5 | 1,35E+08 | 1,35E+08 | 2231 | * | 22,42115 | 2,99205273 | 0,762285 | 3,925111 | 8,67E-05 | 0,090744111 | 3,104559 | Promoter (<=1kb) | 5 | 1,35E+08 | 1,35E+08 | 13573 | 1 | 12054 | ENSMUST00000031692.12 | 0 | ENSMUSG00000029681 | Bcl7b | B cell CLL/lymphoma 7B |
| chr19 | 9013441 | 9014914 | 1474 | * | 13,21116 | 2,992118625 | 1,100885 | 2,717921 | 0,006569 | 0,891057096 | 0,918255 | Exon (ENSMUST00000 | 19 | 8977024 | 9054278 | 77255 | 1 | 66395 | ENSMUST00000237033.2 | 36417 | ENSMUSG00000069833 | Ahnak | AHNAK nucleoprotein (desmoyokin) |
| chr4 | 1,39E+08 | 1,39E+08 | 1160 | * | 11,86042 | 2,992395626 | 1,08246 | 2,76444 | 0,005702 | 0,86525631 | 0,918255 | Intron (ENSMUST00000 | 4 | 1,39E+08 | 1,39E+08 | 63726 | 1 | 77056 | ENSMUST00000124835.3 | 26115 | ENSMUSG00000041143 | Tmco4 | transmembrane and coiled-coil domains 4 |
| chr1 | 1,36E+08 | 1,36E+08 | 1503 | * | 15,20419 | 2,992433983 | 1,054008 | 2,839101 | 0,004524 | 0,749550311 | 1,41696 | Distal Intergenic | 1 | 1,36E+08 | 1,36E+08 | 14745 | 1 | 16763 | ENSMUST00000038760.10 | -3483 | ENSMUSG00000041782 | Lad1 | ladinin |
| chr8 | 1,23E+08 | 1,23E+08 | 1675 | * | 17,34855 | 2,992821244 | 1,038375 | 2,882217 | 0,003949 | 0,642024518 | 2,386008 | Exon (ENSMUST00000 | 8 | 1,23E+08 | 1,23E+08 | 13526 | 2 | 319463 | ENSMUST00000212334.2 | 12228 | | C230057M02Rik | RIKEN cDNA C230057M02 gene |
| chr1 | 1,71E+08 | 1,71E+08 | 2263 | * | 25,79206 | 2,993245289 | 0,742186 | 4,03301 | 5,51E-05 | 0,07748799 | 2,67243 | Promoter (<=1kb) | 1 | 1,71E+08 | 1,71E+08 | 12912 | 1 | 75472 | ENSMUST00000129651.8 | 0 | ENSMUSG00000026649 | Cfap126 | cilia and flagella associated protein 126 |
| chr8 | 1,23E+08 | 1,23E+08 | 1204 | * | 12,89767 | 2,99415557 | 1,133124 | 2,64239 | 0,008232 | 0,874399847 | 1,2763 | Intron (ENSMUST00000 | 8 | 1,23E+08 | 1,23E+08 | 1085 | 2 | 234825 | ENSMUST00000174206.2 | 15511 | ENSMUSG00000040263 | Klhdc4 | kelch domain containing 4 |
| chr3 | 76299361 | 76300622 | 1262 | * | 8,547158 | 2,99434612 | 1,142737 | 2,620328 | 0,008785 | 0,863794395 | 1,425929 | Intron (ENSMUST00000 | 3 | 76500857 | 76614856 | 114000 | 1 | 213262 | ENSMUST00000162471.2 | -200235 | ENSMUSG00000034098 | Fstl5 | follistatin-like 5 |
| chr19 | 42125478 | 42126989 | 1512 | * | 11,94955 | 2,994643914 | 1,093031 | 2,739762 | 0,006148 | 0,881683465 | 0,918255 | Distal Intergenic | 19 | 42111712 | 42117498 | 5787 | 2 | 69534 | ENSMUST00000118965.4 | -7980 | ENSMUSG00000018821 | Avpi1 | arginine vasopressin-induced 1 |
| chr18 | 82971800 | 82973074 | 1275 | * | 18,09317 | 2,995158515 | 0,871987 | 3,434865 | 0,000593 | 0,308866996 | 2,423594 | Intron (ENSMUST00000 | 18 | 83005933 | 83019825 | 13893 | 1 | 329003 | ENSMUST00000236468.2 | -32859 | ENSMUSG00000058881 | Zfp516 | zinc finger protein 516 |
| chr19 | 46887846 | 46889973 | 2128 | * | 15,48053 | 2,995496228 | 0,962306 | 3,11283 | 0,001853 | 0,602555268 | 1,37683 | Intron (ENSMUST00000 | 19 | 46877447 | 46898014 | 20568 | 2 | 76952 | ENSMUST00000174731.3 | 8041 | ENSMUSG00000025041 | Nt5c2 | 5'-nucleotidase, cytosolic II |
| chr11 | 78149991 | 78150985 | 995 | * | 8,501951 | 2,995516648 | 1,145938 | 2,61403 | 0,008948 | 1 | 0 | Promoter (1-2kb) | 11 | 78152578 | 78154858 | 2281 | 1 | 72503 | ENSMUST00000125459.2 | -1593 | ENSMUSG00000010277 | 2610507B11Rik | RIKEN cDNA 2610507B11 gene |
| chr3 | 1,46E+08 | 1,46E+08 | 1017 | * | 12,37387 | 2,995555181 | 1,054311 | 2,841244 | 0,004494 | 0,767061106 | 1,2763 | Promoter (1-2kb) | 3 | 1,46E+08 | 1,46E+08 | 55456 | 1 | 22262 | ENSMUST00000199489.5 | -1973 | ENSMUSG00000028186 | Uox | urate oxidase |
| chr19 | 3757643 | 3760505 | 2863 | * | 24,977245 | 2,995653502 | 0,7052962 | 4,2473692 | 2,163E-05 | 0,041223772 | 2,6724296 | Promoter (<=1kb) | 19 | 3739686 | 3758176 | 18491 | 2 | 102640578 | ENSMUST00000236650.2 | 0 | ENSMUSG00000118249 | Gm36608 | predicted gene, 36608 |
| chr11 | 1,04E+08 | 1,04E+08 | 1171 | * | 10,97863 | 2,995815205 | 1,046169 | 2,863605 | 0,004188 | 0,752744767 | 1,2763 | Promoter (<=1kb) | 11 | 1,04E+08 | 1,04E+08 | 18467 | 1 | 11841 | ENSMUST00000063347.12 | 865 | ENSMUSG00000062421 | Arf2 | ADP-ribosylation factor 2 |
| chr11 | 76967994 | 76970584 | 2591 | * | 18,74539 | 2,99589543 | 0,990361 | 3,025055 | 0,002486 | 0,570876293 | 2,154504 | Promoter (<=1kb) | 11 | 76935118 | 76969261 | 34144 | 2 | 237859 | ENSMUST00000102494.8 | 0 | ENSMUSG00000037958 | Nsrp1 | nuclear speckle regulatory protein 1 |
| chr4 | 28291463 | 28292560 | 1098 | * | 10,93649 | 2,995974909 | 1,107821 | 2,704386 | 0,006843 | 0,929109173 | 0,744375 | Distal Intergenic | 4 | 28365316 | 28365444 | 129 | 1 | 1,15E+08 | ENSMUST00000104333.2 | -72756 | ENSMUSG00000077521 | Gm25378 | predicted gene, 25378 |
| chr4 | 88154057 | 88155326 | 1270 | * | 10,27217 | 2,996879676 | 1,067616 | 2,807077 | 0,004999 | 0,815986445 | 1,086118 | Intron (ENSMUST00000 | 4 | 88205872 | 88221867 | 15996 | 1 | 230393 | ENSMUST00000144388.2 | -50546 | ENSMUSG00000038368 | Focad | focadhesin |
| chr5 | 1,08E+08 | 1,08E+08 | 2040 | * | 10,91441 | 2,997007471 | 1,078639 | 2,778509 | 0,005461 | 0,817544002 | 1,173325 | Intron (ENSMUST00000 | 5 | 1,08E+08 | 1,08E+08 | 48729 | 2 | 14020 | ENSMUST00000155955.2 | 11043 | ENSMUSG00000011831 | Evi5 | ecotropic viral integration site 5 |
| chr10 | 1,01E+08 | 1,01E+08 | 1612 | * | 15,19071 | 2,997473734 | 0,959084 | 3,12535 | 0,001776 | 0,50985457 | 2,141612 | Distal Intergenic | 10 | 1,01E+08 | 1,01E+08 | 34101 | 2 | 1,03E+08 | ENSMUST00000218270.2 | -202035 | ENSMUSG00000112104 | Gm35722 | predicted gene, 35722 |
| chr12 | 43793589 | 43794831 | 1243 | * | 11,9973 | 2,998172559 | 1,018228 | 2,9445 | 0,003235 | 0,802671652 | 0,744375 | Distal Intergenic | 12 | 43789820 | 43789948 | 129 | 1 | 1,15E+08 | ENSMUST00000083070.3 | 3769 | ENSMUSG00000065004 | Gm26326 | predicted gene, 26326 |
| chr13 | 34330751 | 34332464 | 1714 | * | 12,8548 | 2,998369532 | 1,08921 | 2,752792 | 0,005909 | 0,908374536 | 0,738441 | Distal Intergenic | 13 | 34317843 | 34321637 | 3795 | 1 | 1,03E+08 | ENSMUST00000221821.2 | 12908 | | Gm36500 | predicted gene, 36500 |
| chr7 | 1E+08 | 1E+08 | 1316 | * | 12,08789 | 2,998564826 | 1,138225 | 2,634421 | 0,008428 | 0,96198404 | 0,744375 | Intron (ENSMUST00000 | 7 | 1E+08 | 1E+08 | 49823 | 1 | 319604 | ENSMUST00000216021.2 | -3010 | ENSMUSG00000029461 | Fam168a | family with sequence similarity 168, member A |
| chr10 | 1,11E+08 | 1,11E+08 | 1748 | * | 19,66884 | 3,000309121 | 0,852251 | 3,520451 | 0,000431 | 0,265946 | 2,507013 | Promoter (<=1kb) | 10 | 1,11E+08 | 1,11E+08 | 132498 | 1 | 237542 | ENSMUST00000105275.9 | 0 | ENSMUSG00000020189 | Osbpl8 | oxysterol binding protein-like 8 |
| chr19 | 16138517 | 16140344 | 1828 | * | 17,72838 | 3,001203033 | 0,880009 | 3,410421 | 0,000649 | 0,315367183 | 2,507013 | Promoter (1-2kb) | 19 | 16141676 | 16142169 | 494 | 2 | 1,18E+08 | ENSMUST00000075293.5 | 1825 | | Gm53055 | peptidylprolyl isomerase A, pseudogene 8 |
| chr2 | 1,03E+08 | 1,03E+08 | 1133 | * | 10,82791 | 3,002089042 | 1,138089 | 2,637834 | 0,008344 | 0,87775273 | 1,2763 | Promoter (<=1kb) | 2 | 1,03E+08 | 1,03E+08 | 26928 | 1 | 13711 | ENSMUST00000028609.14 | 0 | ENSMUSG00000027186 | Elf5 | E74-like factor 5 |
| chr9 | 108013411 | 108016419 | 3009 | * | 65,542312 | 3,003179621 | 0,4420171 | 6,7942614 | 1,089E-11 | 2,60733E-07 | 4,2003037 | Promoter (<=1kb) | 9 | 108014054 | 108015766 | 1713 | 1 | 76873 | ENSMUST00000194140.2 | 0 | ENSMUSG00000102224 | 4930447F24Rik | RIKEN cDNA 4930447F24 gene |
| chr1 | 1,82E+08 | 1,82E+08 | 1518 | * | 14,78815 | 3,003255411 | 0,894986 | 3,355644 | 0,000792 | 0,470187393 | 1,214904 | Intron (ENSMUST00000 | 1 | 1,82E+08 | 1,82E+08 | 4475 | 1 | 240960 | ENSMUST00000160793.2 | 5223 | ENSMUSG00000047369 | Dnah14 | dynein, axonemal, heavy chain 14 |
| chr7 | 99998198 | 99999685 | 1488 | * | 12,92777 | 3,003696776 | 1,063549 | 2,824221 | 0,00474 | 0,777172375 | 1,2763 | Intron (ENSMUST00000 | 7 | 99975944 | 99994257 | 18314 | 2 | 72590 | ENSMUST00000207634.2 | -3941 | ENSMUSG00000030718 | Ppme1 | protein phosphatase methylesterase 1 |
| chr13 | 81110371 | 81112559 | 2189 | * | 26,974601 | 3,004097392 | 0,6643169 | 4,5220849 | 6,123E-06 | 0,016104274 | 2,6724296 | Promoter (<=1kb) | 13 | 81105784 | 81111849 | 6066 | 2 | 100043074 | ENSMUST00000181103.2 | 0 | | Gm4211 | predicted gene 4211 |
| chr15 | 25664232 | 25665441 | 1210 | * | 13,44309 | 3,005165885 | 0,979414 | 3,068332 | 0,002153 | 0,689688276 | 0,938682 | Intron (ENSMUST00000 | 15 | 25674596 | 25724036 | 49441 | 1 | 17909 | ENSMUST00000130517.8 | -9155 | ENSMUSG00000022272 | Myo10 | myosin X |
| chr1 | 75480005 | 75482535 | 2531 | * | 34,353057 | 3,005678333 | 0,5867965 | 5,1221821 | 3,02E-07 | 0,00149878 | 2,7254908 | Promoter (<=1kb) | 1 | 75467178 | 75482285 | 15108 | 2 | 98733 | ENSMUST00000155084.8 | 0 | ENSMUSG00000026211 | Obsl1 | obscurin-like 1 |
| chr7 | 29682136 | 29683835 | 1700 | * | 18,7555 | 3,006140965 | 0,770008 | 3,90404 | 9,46E-05 | 0,119192643 | 2,221611 | Promoter (<=1kb) | 7 | 29683380 | 29724536 | 41157 | 1 | 243905 | ENSMUST00000148442.8 | 0 | ENSMUSG00000074221 | Zfp568 | zinc finger protein 568 |
| chr12 | 23193446 | 23193904 | 459 | * | 8,292974 | 3,007249164 | 1,16259 | 2,586681 | 0,009691 | 1 | 0 | Distal Intergenic | 12 | 23162536 | 23165585 | 3050 | 2 | 668662 | ENSMUST00000222008.2 | -27861 | | Gm9292 | predicted gene 9292 |
| chr12 | 1,03E+08 | 1,03E+08 | 1360 | * | 10,78613 | 3,007432192 | 1,050219 | 2,863624 | 0,004188 | 0,752744767 | 1,2763 | Distal Intergenic | 12 | 1,03E+08 | 1,03E+08 | 3408 | 1 | 12652 | ENSMUST00000223100.2 | 7634 | ENSMUSG00000021194 | Chga | chromogranin A |
| chr5 | 71932118 | 71933701 | 1584 | * | 16,22335 | 3,007451033 | 0,839398 | 3,582867 | 0,00034 | 0,245757594 | 2,318434 | Intron (ENSMUST00000 | 5 | 71857275 | 72306380 | 449106 | 1 | 14400 | ENSMUST00000031122.9 | 74843 | ENSMUSG00000029212 | Gabrb1 | gamma-aminobutyric acid (GABA) A receptor, subunit beta 1 |
| chr13 | 62038623 | 62039791 | 1169 | * | 8,581293 | 3,007558635 | 1,116091 | 2,694726 | 0,007045 | 1 | 0 | Distal Intergenic | 13 | 61952112 | 61974330 | 22219 | 1 | 638502 | ENSMUST00000244075.1 | 86511 | | Gm7240 | predicted gene 7240 |
| chr2 | 98606953 | 98607878 | 926 | * | 10,18995 | 3,007747633 | 1,023867 | 2,937635 | 0,003307 | 0,741363157 | 1,086118 | Distal Intergenic | 2 | 97298434 | 97459408 | 160975 | 1 | 241568 | ENSMUST00000170144.2 | 1308519 | ENSMUSG00000050587 | Lrrc4c | leucine rich repeat containing 4C |
| chr3 | 94986356 | 94988178 | 1823 | * | 10,59975 | 3,00818885 | 1,07897 | 2,788018 | 0,005303 | 0,890608507 | 0,744375 | Promoter (<=1kb) | 3 | 94978137 | 94985827 | 7691 | 2 | 18720 | ENSMUST00000107231.2 | -529 | ENSMUSG00000028126 | Pip5k1a | phosphatidylinositol-4-phosphate 5-kinase, type 1 alpha |
| chr8 | 95913357 | 95914305 | 949 | * | 10,686 | 3,008266379 | 1,099295 | 2,736543 | 0,006209 | 0,824134241 | 1,2763 | Intron (ENSMUST00000 | 8 | 95836877 | 95909419 | 72543 | 2 | 16582 | ENSMUST00000212424.2 | -3938 | ENSMUSG00000031788 | Kifc3 | kinesin family member C3 |
| chr6 | 1,01E+08 | 1,01E+08 | 1121 | * | 17,46952 | 3,008679139 | 0,780045 | 3,857059 | 0,000115 | 0,137154233 | 2,154504 | Intron (ENSMUST00000 | 6 | 1,01E+08 | 1,01E+08 | 13503 | 1 | 232314 | ENSMUST00000204001.2 | -8247 | ENSMUSG00000052144 | Ppp4r2 | protein phosphatase 4, regulatory subunit 2 |
| chr5 | 1,25E+08 | 1,25E+08 | 2281 | * | 15,75605 | 3,008911814 | 0,84245 | 3,57162 | 0,000355 | 0,247743734 | 2,386008 | Promoter (2-3kb) | 5 | 1,25E+08 | 1,25E+08 | 16797 | 1 | 67848 | ENSMUST00000071057.14 | -2453 | ENSMUSG00000029389 | Ddx55 | DEAD box helicase 55 |
| chr6 | 71683758 | 71684656 | 899 | * | 13,90663 | 3,009028819 | 1,036081 | 2,904241 | 0,003681 | 0,730765405 | 1,2763 | Promoter (<=1kb) | 6 | 71684545 | 71787694 | 103150 | 1 | 52250 | ENSMUST00000121469.2 | 0 | ENSMUSG00000052852 | Reep1 | receptor accessory protein 1 |
| chr15 | 83255860 | 83257647 | 1788 | * | 19,67007 | 3,009979823 | 0,939159 | 3,204975 | 0,001351 | 0,44621096 | 2,386008 | Distal Intergenic | 15 | 83238048 | 83251483 | 13436 | 2 | 69291 | ENSMUST00000178628.9 | -4377 | | 1700001L05Rik | RIKEN cDNA 1700001L05 gene |
| chr1 | 86078181 | 86079496 | 1316 | * | 11,20235 | 3,01015356 | 0,987335 | 3,048767 | 0,002298 | 0,669586355 | 1,173325 | Intron (ENSMUST00000 | 1 | 86082502 | 86174048 | 91547 | 1 | 78795 | ENSMUST00000123618.8 | -3006 | ENSMUSG00000062590 | Armc9 | armadillo repeat containing 9 |
| chr6 | 1,34E+08 | 1,34E+08 | 711 | * | 12,2826 | 3,010703922 | 1,151604 | 2,614357 | 0,00894 | 0,914358307 | 1,075735 | Intron (ENSMUST00000 | 6 | 1,34E+08 | 1,34E+08 | 3129 | 2 | 1,15E+08 | ENSMUST00000164616.2 | -13340 | | Gm52920 | predicted gene, 52920 |
| chr4 | 68073631 | 68074605 | 975 | * | 8,648504 | 3,010971775 | 1,164729 | 2,585128 | 0,009734 | 0,891057096 | 1,360622 | Distal Intergenic | 4 | 68100769 | 68100939 | 171 | 1 | 1,15E+08 | ENSMUST00000082828.3 | -26164 | ENSMUSG00000064762 | Gm23738 | predicted gene, 23738 |
| chr16 | 22763828 | 22765170 | 1343 | * | 14,17548 | 3,011514562 | 0,961512 | 3,13206 | 0,001736 | 0,695401133 | 0,738441 | Distal Intergenic | 16 | 22769822 | 22780406 | 10585 | 1 | 94175 | ENSMUST00000023590.9 | -4652 | ENSMUSG00000022877 | Hrg | histidine-rich glycoprotein |
| chr13 | 42745915 | 42748022 | 2108 | * | 14,54411 | 3,011587025 | 1,000531 | 3,00999 | 0,002613 | 0,683210259 | 1,214904 | Distal Intergenic | 13 | 42834099 | 43292000 | 457902 | 1 | 218194 | ENSMUST00000128646.8 | -86077 | ENSMUSG00000054728 | Phactr1 | phosphatase and actin regulator 1 |
| chr15 | 76107209 | 76108255 | 1047 | * | 10,84908 | 3,011598133 | 1,160776 | 2,594469 | 0,009474 | 0,977962962 | 0,738441 | Intron (ENSMUST00000 | 15 | 76055175 | 76113697 | 58523 | 2 | 18810 | ENSMUST00000074834.12 | 5442 | ENSMUSG00000022565 | Plec | plectin |
| chr18 | 24414669 | 24416324 | 1656 | * | 11,94394 | 3,011651918 | 1,030828 | 2,921585 | 0,003483 | 0,722798324 | 1,2763 | Promoter (1-2kb) | 18 | 24413090 | 24416248 | 3159 | 1 | 14423 | ENSMUST00000170191.2 | 1579 | ENSMUSG00000000420 | Galnt1 | polypeptide N-acetylgalactosaminyltransferase 1 |
| chr11 | 3529706 | 3531566 | 1861 | * | 26,09649 | 3,011698537 | 0,6846101 | 4,3991444 | 1,087E-05 | 0,023535556 | 2,8585108 | Distal Intergenic | 11 | 3482460 | 3490612 | 8153 | 2 | 29856 | ENSMUST00000156201.8 | -39094 | ENSMUSG00000020439 | Smtn | smoothelin |
| chr9 | 1,09E+08 | 1,09E+08 | 1406 | * | 10,56826 | 3,012130589 | 1,148539 | 2,622576 | 0,008727 | 0,964625216 | 0,744375 | Distal Intergenic | 9 | 1,09E+08 | 1,09E+08 | 22595 | 2 | 320082 | ENSMUST00000054925.13 | -15190 | ENSMUSG00000047237 | Fbxw21 | F-box and WD-40 domain protein 21 |
| chr9 | 16578724 | 16579382 | 659 | * | 9,912265 | 3,01373369 | 1,109791 | 2,715588 | 0,006616 | 0,861581225 | 1,086118 | Distal Intergenic | 9 | 15821485 | 16412581 | 591097 | 2 | 270120 | ENSMUST00000217308.3 | -166143 | ENSMUSG00000074505 | Fat3 | FAT atypical cadherin 3 |
| chr4 | 1,33E+08 | 1,33E+08 | 2651 | * | 19,07144 | 3,013770412 | 0,835825 | 3,605742 | 0,000311 | 0,228721288 | 2,386008 | Promoter (<=1kb) | 4 | 1,33E+08 | 1,33E+08 | 11748 | 2 | 52174 | ENSMUST00000105907.9 | 0 | ENSMUSG00000028857 | Tmem222 | transmembrane protein 222 |
| chr4 | 1,08E+08 | 1,08E+08 | 1234 | * | 10,28346 | 3,014040375 | 1,145506 | 2,631187 | 0,008509 | 0,905015433 | 1,086118 | Promoter (2-3kb) | 4 | 1,08E+08 | 1,08E+08 | 69949 | 1 | 16975 | ENSMUST00000146552.9 | 2559 | ENSMUSG00000028613 | Lrp8 | low density lipoprotein receptor-related protein 8, apolipoprotein e receptor |
| chr7 | 14362633 | 14363944 | 1312 | * | 9,30265 | 3,014914866 | 1,160374 | 2,598226 | 0,009371 | 0,877892722 | 1,425929 | Distal Intergenic | 7 | 14264577 | 14356404 | 91828 | 2 | 1E+08 | ENSMUST00000044683.12 | -6229 | ENSMUSG00000041596 | Nlrp5-ps | NLR family, pyrin domain containing 5, pseudogene |
| chr11 | 80150508 | 80151747 | 1240 | * | 10,44465 | 3,014980065 | 1,11285 | 2,709243 | 0,006744 | 0,926256234 | 0,744375 | Intron (ENSMUST00000 | 11 | 80154877 | 80157577 | 2701 | 1 | 59040 | ENSMUST00000236882.2 | -3130 | ENSMUSG00000017686 | Rhot1 | ras homolog family member T1 |
| chr18 | 60638608 | 60639639 | 1032 | * | 9,747409 | 3,015118857 | 1,100459 | 2,739874 | 0,006146 | 0,802323085 | 1,425929 | Distal Intergenic | 18 | 60607263 | 60635059 | 27797 | 2 | 106878 | ENSMUST00000042710.8 | -3549 | ENSMUSG00000038059 | Smim3 | small integral membrane protein 3 |
| chr1 | 1,94E+08 | 1,94E+08 | 1457 | * | 11,59378 | 3,015513226 | 1,116189 | 2,701614 | 0,0069 | 0,932975916 | 0,738441 | Intron (ENSMUST00000 | 1 | 1,94E+08 | 1,94E+08 | 10573 | 2 | 74942 | ENSMUST00000193395.2 | 5181 | ENSMUSG00000103692 | 4930503O07Rik | RIKEN cDNA 4930503O07 gene |
| chr5 | 1,1E+08 | 1,1E+08 | 1138 | * | 10,4426 | 3,016117365 | 1,151572 | 2,61913 | 0,008815 | 0,966901185 | 0,744375 | Distal Intergenic | 5 | 1,1E+08 | 1,1E+08 | 38772 | 1 | 384221 | ENSMUST00000232722.2 | -29968 | ENSMUSG00000091879 | Vmn2r17 | vomeronasal 2, receptor 17 |
| chr10 | 30438954 | 30440331 | 1378 | * | 13,95328 | 3,016613801 | 0,887734 | 3,398104 | 0,000679 | 0,530664257 | 0,738441 | Intron (ENSMUST00000 | 10 | 30435117 | 30466105 | 30989 | 2 | 73681 | ENSMUST00000215041.2 | 25774 | ENSMUSG00000019792 | Trmt11 | tRNA methyltransferase 11 |
| chr6 | 1,28E+08 | 1,28E+08 | 950 | * | 13,00305 | 3,01662867 | 0,979472 | 3,079853 | 0,002071 | 0,68680243 | 0,918255 | Promoter (<=1kb) | 6 | 1,28E+08 | 1,28E+08 | 428 | 2 | 101142 | ENSMUST00000204362.2 | -162 | ENSMUSG00000001518 | Itfg2 | integrin alpha FG-GAP repeat containing 2 |

| | | | | | | | | | | | | | | | | | | | | | | | |
|---|---|---|---|---|---|---|---|---|---|---|---|---|---|---|---|---|---|---|---|---|---|---|---|
| chr9 | 15129528 | 15130458 | 931 | * | 10,34576 | 3,01747024 | 0,990728 | 3,04571 | 0,002321 | 0,667043202 | 1,211315 | Distal Intergenic | 9 | 15150341 | 15170712 | 20372 | 1 | 69137 | ENSMUST00000034413.8 | -19883 | ENSMUSG00000031937 | Vstm5 | V-set and transmembrane domain containing 5 |
| chr15 | 97726199 | 97726700 | 502 | * | 10,03503 | 3,018196242 | 1,109519 | 2,720275 | 0,006523 | 0,842187815 | 1,211315 | Promoter (2-3kb) | 15 | 97695680 | 97729350 | 33671 | 2 | 56233 | ENSMUST00000156045.8 | 2650 | ENSMUSG00000022475 | Hdac7 | histone deacetylase 7 |
| chr19 | 6294049 | 6295543 | 1495 | * | 13,38019 | 3,018399092 | 0,991415 | 3,044537 | 0,00233 | 0,683938614 | 1,075735 | Promoter (<=1kb) | 19 | 6295495 | 6312325 | 16831 | 1 | 329015 | ENSMUST00000145600.2 | 0 | ENSMUSG00000024773 | Atg2a | autophagy related 2A |
| chr16 | 20223858 | 20226061 | 2204 | * | 20,45924 | 3,018511298 | 0,75562 | 3,994748 | 6,48E-05 | 0,084704777 | 2,663619 | Exon (ENSMUST00000 | 16 | 20217567 | 20219036 | 1470 | 2 | 27416 | ENSMUST00000128978.2 | -4822 | ENSMUSG00000022822 | Abcc5 | ATP-binding cassette, sub-family C (CFTR/MRP), member 5 |
| chr13 | 21564731 | 21566106 | 1376 | * | 12,78144 | 3,018536419 | 0,993768 | 3,037465 | 0,002386 | 0,661775851 | 1,2763 | Exon (ENSMUST00000 | 13 | 21572775 | 21578933 | 6159 | 2 | 72739 | ENSMUST00000145631.9 | 12827 | ENSMUSG00000021327 | Zkscan3 | zinc finger with KRAB and SCAN domains 3 |
| chr4 | 48099486 | 48100921 | 1436 | * | 12,94816 | 3,019131132 | 0,98241 | 3,073189 | 0,002118 | 0,670185906 | 1,075735 | Distal Intergenic | 4 | 48124915 | 48185911 | 60997 | 1 | 67727 | ENSMUST00000107721.8 | -23994 | ENSMUSG00000061455 | Stx17 | syntaxin 17 |
| chr13 | 99569083 | 99570613 | 1531 | * | 9,286702 | 3,019240599 | 1,139394 | 2,649865 | 0,008052 | 0,85203509 | 1,425929 | Exon (ENSMUST00000 | 13 | 99570809 | 99580452 | 9644 | 2 | 17755 | ENSMUST00000223693.2 | 9839 | ENSMUSG00000052727 | Map1b | microtubule-associated protein 1B |
| chr5 | 1,24E+08 | 1,24E+08 | 1610 | * | 12,33196 | 3,019344596 | 1,107651 | 2,725898 | 0,006413 | 0,888080529 | 0,918255 | Promoter (2-3kb) | 5 | 1,24E+08 | 1,24E+08 | 106824 | 2 | 56430 | ENSMUST00000111566.9 | -2466 | ENSMUSG00000049550 | Clip1 | CAP-GLY domain containing linker protein 1 |
| chr19 | 60786085 | 60787989 | 1905 | * | 11,31591 | 3,019864032 | 1,057662 | 2,855225 | 0,004301 | 0,81856079 | 0,918255 | Distal Intergenic | 19 | 60761433 | 60779096 | 17664 | 2 | 13669 | ENSMUST00000235706.2 | -6989 | ENSMUSG00000024991 | Eif3a | eukaryotic translation initiation factor 3, subunit A |
| chr9 | 30596912 | 30597905 | 994 | * | 8,746335 | 3,020371973 | 1,167877 | 2,586207 | 0,009704 | 1 | 0 | Distal Intergenic | 9 | 30633331 | 30692664 | 59334 | 2 | 1,03E+08 | ENSMUST00000241822.1 | 94759 | ENSMUSG00000111867 | Gm31013 | predicted gene, 31013 |
| chr1 | 1,55E+08 | 1,55E+08 | 1760 | * | 12,52876 | 3,020528726 | 1,098 | 2,750937 | 0,005942 | 0,909022873 | 0,738441 | Promoter (1-2kb) | 1 | 1,55E+08 | 1,55E+08 | 19161 | 1 | 58244 | ENSMUST00000191769.2 | 1568 | ENSMUSG00000026470 | Stx6 | syntaxin 6 |
| chr13 | 81805151 | 81806620 | 1470 | * | 11,68118 | 3,020704459 | 1,120625 | 2,695554 | 0,007027 | 0,847125405 | 1,2763 | Promoter (<=1kb) | 13 | 81805782 | 81820990 | 15209 | 1 | 80289 | ENSMUST00000224300.2 | 0 | ENSMUSG00000035840 | Lysmd3 | LysM, putative peptidoglycan-binding, domain containing 3 |
| chr14 | 55327982 | 55329224 | 1243 | * | 10,73215 | 3,020822904 | 1,099573 | 2,747269 | 0,006009 | 0,910299316 | 0,738441 | Promoter (<=1kb) | 14 | 55297719 | 55329781 | 32063 | 2 | 239102 | ENSMUST00000036328.9 | 557 | ENSMUSG00000040721 | Zfhx2 | zinc finger homeobox 2 |
| chr12 | 55756932 | 55758754 | 1823 | * | 16,78375 | 3,021894441 | 0,840386 | 3,59584 | 0,000323 | 0,249239 | 2,141612 | Intron (ENSMUST0000 | 12 | 55723875 | 55744684 | 20810 | 2 | 56784 | ENSMUST00000219566.2 | -12248 | ENSMUSG00000021027 | Ralgapa1 | Ral GTPase activating protein, alpha subunit 1 |
| chr6 | 86781741 | 86782576 | 836 | * | 10,78817 | 3,022793413 | 1,127062 | 2,682011 | 0,007318 | 0,8656457 | 1,173325 | Distal Intergenic | 6 | 86713823 | 86770566 | 56744 | 2 | 11746 | ENSMUST00000001187.15 | -11175 | ENSMUSG00000029994 | Anxa4 | annexin A4 |
| chr17 | 25632683 | 25634468 | 1786 | * | 16,47557 | 3,022929903 | 0,819058 | 3,690739 | 0,000224 | 0,204595855 | 2,141612 | Intron (ENSMUST0000 | 17 | 25616599 | 25652252 | 35654 | 2 | 58226 | ENSMUST00000159623.2 | 17784 | ENSMUSG00000024112 | Cacna1h | calcium channel, voltage-dependent, T type, alpha 1H subunit |
| chr7 | 1,09E+08 | 1,09E+08 | 1562 | * | 11,65828 | 3,024090076 | 1,121074 | 2,697494 | 0,006986 | 0,858257847 | 1,173325 | Intron (ENSMUST0000 | 7 | 1,09E+08 | 1,09E+08 | 6893 | 2 | 56786 | ENSMUST00000128043.2 | 4821 | ENSMUSG00000031021 | Tmem9b | TMEM9 domain family, member B |
| chr7 | 1,12E+08 | 1,12E+08 | 1941 | * | 15,12492 | 3,025613043 | 1,07658 | 2,810393 | 0,004948 | 0,765231686 | 1,41696 | Intron (ENSMUST0000 | 7 | 1,12E+08 | 1,12E+08 | 220142 | 1 | 21676 | ENSMUST00000084705.13 | -18975 | ENSMUSG00000055320 | Tead1 | TEA domain family member 1 |
| chr13 | 91668217 | 91670929 | 2713 | * | 24,53789 | 3,026310544 | 0,875104 | 3,458228 | 0,000544 | 0,278016366 | 2,880017 | Intron (ENSMUST0000 | 13 | 91687462 | 91805895 | 118434 | 1 | 66970 | ENSMUST00000133984.2 | -16533 | ENSMUSG00000003992 | Ssbp2 | single-stranded DNA binding protein 2 |
| chr16 | 21916117 | 21917359 | 1243 | * | 8,824383 | 3,026629623 | 1,156377 | 2,617337 | 0,008862 | 0,7893429 | 2,237403 | Intron (ENSMUST0000 | 16 | 21877824 | 21907868 | 30045 | 2 | 319765 | ENSMUST00000134188.8 | -8249 | ENSMUSG00000033581 | Igf2bp2 | insulin-like growth factor 2 mRNA binding protein 2 |
| chr5 | 93413047 | 93415550 | 2504 | * | 39,129055 | 3,026672681 | 0,6176422 | 4,9003658 | 9,566E-07 | 0,003692163 | 2,7254908 | Promoter (<=1kb) | 5 | 93415116 | 93424090 | 8975 | 1 | 12452 | ENSMUST00000031331.13 | 0 | ENSMUSG00000029385 | Ccng2 | cyclin G2 |
| chr11 | 56253162 | 56254252 | 1091 | * | 8,693103 | 3,026761856 | 1,137658 | 2,66052 | 0,007802 | 1 | 0 | Distal Intergenic | 11 | 56153991 | 56154066 | 76 | 2 | 1,15E+08 | ENSMUST00000083701.3 | -99096 | ENSMUSG00000065635 | Gm24253 | predicted gene, 24253 |
| chr13 | 38831442 | 38832295 | 854 | * | 7,761768 | 3,026857066 | 1,169599 | 2,587944 | 0,009655 | 1 | 0 | Intron (ENSMUST0000 | 13 | 38838496 | 38842931 | 4436 | 2 | 66143 | ENSMUST00000223823.2 | 10636 | ENSMUSG00000001707 | Eef1e1 | eukaryotic translation elongation factor 1 epsilon 1 |
| chr9 | 1,15E+08 | 1,15E+08 | 1110 | * | 10,07348 | 3,026935291 | 1,1228 | 2,695882 | 0,00702 | 0,875473843 | 1,086118 | Intron (ENSMUST0000 | 9 | 1,15E+08 | 1,15E+08 | 96274 | 1 | 74486 | ENSMUST00000182413.2 | -11326 | ENSMUSG00000040875 | Osbpl10 | oxysterol binding protein-like 10 |
| chr15 | 63860830 | 63862546 | 1717 | * | 13,93412 | 3,027545085 | 1,020401 | 2,967014 | 0,003007 | 0,695401133 | 1,2763 | Intron (ENSMUST0000 | 15 | 63802791 | 63869818 | 67028 | 2 | 223601 | ENSMUST00000164532.3 | 7272 | ENSMUSG00000022378 | Cyrib | CYFIP related Rac1 interactor B |
| chr13 | 51671135 | 51673571 | 2437 | * | 19,26488 | 3,027552207 | 0,929563 | 3,256964 | 0,001126 | 0,39738041 | 2,713306 | Intron (ENSMUST0000 | 13 | 51585077 | 51721120 | 136044 | 2 | 20418 | ENSMUST00000021898.6 | 47549 | ENSMUSG00000021448 | Shc3 | src homology 2 domain-containing transforming protein C3 |
| chr17 | 21450581 | 21451350 | 770 | * | 9,469148 | 3,028149149 | 1,081932 | 2,798836 | 0,005129 | 0,779303858 | 1,360622 | Promoter (1-2kb) | 17 | 21449088 | 21450077 | 990 | 1 | 171232 | ENSMUST00000079633.3 | 1493 | ENSMUSG00000057203 | Vmn1r234 | vomeronasal 1 receptor 234 |
| chr9 | 72615651 | 72617027 | 1377 | * | 11,80002 | 3,028150411 | 1,101459 | 2,749218 | 0,005974 | 0,910076379 | 0,738441 | Exon (ENSMUST00000 | 9 | 72600721 | 72657134 | 56414 | 1 | 17999 | ENSMUST00000238315.2 | 14930 | ENSMUSG00000032216 | Nedd4 | neural precursor cell expressed, developmentally down-regulated 4 |
| chr13 | 44382235 | 44383674 | 1440 | * | 11,15568 | 3,02872398 | 1,147646 | 2,639074 | 0,008313 | 0,960268559 | 0,738441 | Distal Intergenic | 13 | 44347065 | 44369963 | 22899 | 2 | 1,01E+08 | ENSMUST00000223513.2 | -12272 | ENSMUSG00000113186 | A330076C08Rik | RIKEN cDNA A330076C08 gene |
| chr4 | 80934840 | 80935945 | 1106 | * | 10,24854 | 3,029066892 | 1,023677 | 2,959006 | 0,003086 | 0,730215586 | 1,086118 | Distal Intergenic | 4 | 80828883 | 80873865 | 44983 | 1 | 52829 | ENSMUST00000055922.4 | 105957 | ENSMUSG00000048706 | Lurap1l | leucine rich adaptor protein 1-like |
| chr2 | 1,19E+08 | 1,19E+08 | 980 | * | 11,17238 | 3,029618279 | 1,044333 | 2,901009 | 0,00372 | 0,792545355 | 0,918255 | Exon (ENSMUST00000 | 2 | 1,19E+08 | 1,19E+08 | 5128 | 2 | 214239 | ENSMUST00000151270.2 | 3440 | ENSMUSG00000045838 | Ccdc9b | coiled-coil domain containing 9B |
| chr8 | 25496262 | 25497941 | 1680 | * | 15,57655 | 3,03038859 | 1,013559 | 2,98985 | 0,002791 | 0,670185906 | 1,41696 | Exon (ENSMUST00000 | 8 | 25480288 | 25487806 | 7519 | 2 | 11502 | ENSMUST00000211319.2 | -8456 | ENSMUSG00000031555 | Adam9 | a disintegrin and metallopeptidase domain 9 (meltrin gamma) |
| chr15 | 8158181 | 8159902 | 1722 | * | 16,77977 | 3,03038994 | 0,839078 | 3,611573 | 0,000304 | 0,241529234 | 2,141612 | Promoter (<=1kb) | 15 | 8159582 | 8164418 | 4837 | 1 | 170762 | ENSMUST00000229466.2 | 0 | ENSMUSG00000022142 | Nup155 | nucleoporin 155 |
| chr4 | 1,39E+08 | 1,39E+08 | 1710 | * | 13,56277 | 3,031005559 | 1,083545 | 2,797304 | 0,005153 | 0,846985985 | 0,938682 | Promoter (<=1kb) | 4 | 1,39E+08 | 1,39E+08 | 3657 | 1 | 69116 | ENSMUST00000145659.2 | 0 | ENSMUSG00000066036 | Ubr4 | ubiquitin protein ligase E3 component n-recognin 4 |
| chr14 | 65568282 | 65571255 | 2974 | * | 25,767695 | 3,031276426 | 0,6757243 | 4,4859666 | 7,258E-06 | 0,017063542 | 2,9510621 | Intron (ENSMUST000002 | 14 | 65595838 | 65623837 | 28000 | 1 | 380912 | ENSMUST00000224623.2 | -24583 | ENSMUSG00000034522 | Zfp395 | zinc finger protein 395 |
| chr10 | 76351995 | 76353161 | 1167 | * | 10,78582 | 3,032039557 | 1,012947 | 2,993285 | 0,00276 | 0,777938745 | 0,738441 | Distal Intergenic | 10 | 76346728 | 76348299 | 1572 | 1 | 54387 | ENSMUST00000218904.2 | 5267 | ENSMUSG00000001150 | Mcm3ap | minichromosome maintenance complex component 3 associated protein |
| chr7 | 1,27E+08 | 1,27E+08 | 3050 | * | 25,26117 | 3,032834043 | 0,768187 | 3,948041 | 7,88E-05 | 0,092690398 | 2,725491 | Promoter (<=1kb) | 7 | 1,27E+08 | 1,27E+08 | 2528 | 1 | 233878 | ENSMUST00000134471.2 | 0 | ENSMUSG00000030683 | Sez6l2 | seizure related 6 homolog like 2 |
| chrX | 13396914 | 13398538 | 1625 | * | 13,94243 | 3,033038545 | 0,999544 | 3,034422 | 0,00241 | 0,751231966 | 0,738441 | Intron (ENSMUST0000 | 20 | 13388021 | 13393007 | 4987 | 2 | 12361 | ENSMUST00000115435.9 | -3907 | ENSMUSG00000031012 | Cask | calcium/calmodulin-dependent serine protein kinase (MAGUK family) |
| chr16 | 24829501 | 24831375 | 1875 | * | 14,67495 | 3,034324064 | 1,02042 | 2,973603 | 0,002943 | 0,691182187 | 1,2763 | Distal Intergenic | 16 | 24878389 | 24887808 | 9420 | 1 | 319487 | ENSMUST00000181655.2 | -47014 | ENSMUSG00000096960 | A230028O05Rik | RIKEN cDNA A230028O05 gene |
| chr2 | 1,67E+08 | 1,67E+08 | 1392 | * | 13,01031 | 3,034671737 | 0,998161 | 3,040263 | 0,002364 | 0,749550311 | 0,738441 | Promoter (<=1kb) | 2 | 1,67E+08 | 1,67E+08 | 1662 | 1 | 228889 | ENSMUST00000126409.2 | 0 | ENSMUSG00000017999 | Ddx27 | DEAD box helicase 27 |
| chr5 | 1,49E+08 | 1,49E+08 | 1400 | * | 11,58277 | 3,034938016 | 1,009091 | 3,007596 | 0,002633 | 0,769360374 | 0,738441 | Promoter (<=1kb) | 5 | 1,49E+08 | 1,49E+08 | 5856 | 1 | 1,01E+08 | ENSMUST00000244958.1 | 0 | ENSMUSG00000086610 | Gm15408 | predicted gene 15408 |
| chr12 | 62640017 | 62641281 | 1265 | * | 8,74129 | 3,035148553 | 1,146731 | 2,646782 | 0,008126 | 0,852340527 | 1,425929 | Distal Intergenic | 12 | 62734608 | 62735031 | 424 | 1 | 73309 | ENSMUST00000172433.3 | -93327 | | Spanxn4 | SPANX family, member N4 |
| chr3 | 1,23E+08 | 1,23E+08 | 1398 | * | 11,82533 | 3,035598476 | 1,106298 | 2,743924 | 0,006071 | 0,910447564 | 0,744375 | Intron (ENSMUST0000 | 3 | 1,23E+08 | 1,23E+08 | 11893 | 1 | 242202 | ENSMUST00000199253.2 | -8035 | ENSMUSG00000053965 | Pde5a | phosphodiesterase 5A, cGMP-specific |
| chr9 | 89855694 | 89857143 | 1450 | * | 15,23892 | 3,035607682 | 0,944664 | 3,213425 | 0,001312 | 0,531203841 | 1,41696 | Exon (ENSMUST00000 | 9 | 89791965 | 89808793 | 16829 | 1 | 19417 | ENSMUST00000190073.2 | 63729 | ENSMUSG00000032356 | Rasgrf1 | RAS protein-specific guanine nucleotide-releasing factor 1 |
| chr4 | 30031337 | 30032649 | 1313 | * | 10,30701 | 3,03581761 | 1,041952 | 2,913586 | 0,003573 | 0,752939853 | 1,086118 | Distal Intergenic | 4 | 29068079 | 29088811 | 20733 | 1 | 1,03E+08 | ENSMUST00000154813.2 | 963258 | | Gm11917 | predicted gene 11917 |
| chr13 | 1,15E+08 | 1,15E+08 | 1548 | * | 11,30806 | 3,037804563 | 1,140507 | 2,663556 | 0,007732 | 0,878632224 | 1,173325 | Distal Intergenic | 13 | 1,15E+08 | 1,15E+08 | 10488 | 1 | 17434 | ENSMUST00000166104.9 | -54801 | ENSMUSG00000015536 | Mocs2 | molybdenum cofactor synthesis 2 |
| chr16 | 15646842 | 15648282 | 1441 | * | 12,48593 | 3,03829724 | 0,88574 | 3,430236 | 0,000603 | 0,445796299 | 1,075735 | Exon (ENSMUST00000 | 16 | 15666307 | 15680586 | 14280 | 2 | 72083 | ENSMUST00000137864.2 | 32304 | ENSMUSG00000022671 | Mzt2 | mitotic spindle organizing protein 2 |
| chr11 | 84230666 | 84232176 | 1511 | * | 10,60719 | 3,03841116 | 1,145013 | 2,653605 | 0,007964 | 0,884323351 | 1,173325 | Intron (ENSMUST0000 | 11 | 84171281 | 84202227 | 30947 | 1 | 107476 | ENSMUST00000183887.2 | 59385 | ENSMUSG00000020532 | Acaca | acetyl-Coenzyme A carboxylase alpha |
| chr7 | 1,13E+08 | 1,13E+08 | 1573 | * | 17,63847 | 3,039683674 | 0,982618 | 3,093455 | 0,001978 | 0,530563721 | 2,154504 | Distal Intergenic | 7 | 1,13E+08 | 1,13E+08 | 41856 | 2 | 75239 | ENSMUST00000209589.2 | -9286 | ENSMUSG00000110009 | 4930543E12Rik | RIKEN cDNA 4930543E12 gene |
| chr4 | 74254438 | 74256061 | 1624 | * | 11,60247 | 3,039816035 | 0,974172 | 3,120411 | 0,001806 | 0,701807025 | 0,744375 | Exon (ENSMUST00000 | 4 | 74181047 | 74191896 | 10850 | 1 | 76804 | ENSMUST00000150439.2 | 73391 | ENSMUSG00000028397 | Kdm4c | lysine (K)-specific demethylase 4C |
| chr18 | 61681207 | 61682402 | 1196 | * | 15,59717 | 3,040115886 | 1,001004 | 3,037066 | 0,002389 | 0,565091829 | 2,141612 | Distal Intergenic | 18 | 61688345 | 61723132 | 34788 | 1 | 93687 | ENSMUST00000165123.8 | -5943 | ENSMUSG00000024576 | Csnk1a1 | casein kinase 1, alpha 1 |
| chr9 | 1,03E+08 | 1,03E+08 | 1667 | * | 14,68708 | 3,040660194 | 0,985228 | 3,086251 | 0,002027 | 0,642870233 | 1,214904 | Distal Intergenic | 9 | 1,03E+08 | 1,03E+08 | 50871 | 1 | 270192 | ENSMUST00000189134.2 | -13051 | ENSMUSG00000032549 | Rab6b | RAB6B, member RAS oncogene family |
| chr9 | 59305987 | 59307008 | 1022 | * | 8,920297 | 3,040844234 | 1,119647 | 2,715895 | 0,00661 | 0,845724506 | 1,211315 | Intron (ENSMUST0000 | 9 | 59302168 | 59313077 | 10910 | 2 | 23806 | ENSMUST00000171856.2 | 6069 | ENSMUSG00000025234 | Arih1 | ariadne RBR E3 ubiquitin protein ligase 1 |
| chr7 | 60690979 | 60692074 | 1096 | * | 7,849988 | 3,041313934 | 1,148747 | 2,647506 | 0,008109 | 0,858257847 | 1,360622 | Intron (ENSMUST0000 | 7 | 60662432 | 60705129 | 42698 | 2 | 27493 | ENSMUST00000245926.1 | 13055 | | A230006K03Rik | RIKEN cDNA A230006K03 gene |
| chr6 | 59185695 | 59187374 | 1680 | * | 14,85302 | 3,041375865 | 1,026157 | 2,963851 | 0,003038 | 0,709360512 | 1,214904 | Promoter (<=1kb) | 6 | 59185855 | 59189018 | 3164 | 1 | 68140 | ENSMUST00000062626.4 | 0 | ENSMUSG00000049232 | Tigd2 | tigger transposable element derived 2 |
| chr9 | 82701222 | 82702206 | 985 | * | 11,81192 | 3,041807715 | 1,016193 | 2,993337 | 0,002759 | 0,738600147 | 0,918255 | Distal Intergenic | 9 | 82711593 | 82729740 | 18148 | 1 | 65099 | ENSMUST00000113245.9 | -9387 | ENSMUSG00000032251 | Irak1bp1 | interleukin-1 receptor-associated kinase 1 binding protein 1 |
| chr15 | 22479958 | 22480805 | 848 | * | 10,76773 | 3,042552655 | 1,165446 | 2,610634 | 0,009037 | 0,890233764 | 1,2763 | Distal Intergenic | 15 | 22549108 | 23400653 | 851546 | 1 | 320865 | ENSMUST00000163361.8 | -68303 | ENSMUSG00000040420 | Cdh18 | cadherin 18 |
| chr11 | 35658945 | 35660299 | 1355 | * | 13,15849 | 3,042625325 | 1,03013 | 2,953633 | 0,003141 | 0,732373293 | 1,075735 | Promoter (<=1kb) | 11 | 35660311 | 35682112 | 21802 | 1 | 211347 | ENSMUST00000018990.8 | -12 | ENSMUSG00000018846 | Pank3 | pantothenate kinase 3 |
| chr13 | 23600334 | 23602530 | 2197 | * | 21,7385 | 3,043176272 | 0,872152 | 3,489273 | 0,000484 | 0,273126507 | 2,67243 | Distal Intergenic | 13 | 23602531 | 23608036 | 5506 | 2 | 30946 | ENSMUST00000041782.4 | 5506 | ENSMUSG00000036376 | Abt1 | activator of basal transcription 1 |
| chr17 | 6220628 | 6224498 | 3871 | * | 25,6608 | 3,043282813 | 0,811197 | 3,751594 | 0,000176 | 0,148120959 | 2,951062 | Intron (ENSMUST0000 | 17 | 6188423 | 6290903 | 102481 | 1 | 68842 | ENSMUST00000039655.3 | 32205 | ENSMUSG00000034377 | Tulp4 | tubby like protein 4 |
| chr15 | 75786127 | 75789195 | 3069 | * | 27,9067 | 3,043307548 | 0,819194 | 3,715003 | 0,000203 | 0,168300715 | 2,725491 | 3' UTR | 15 | 75788326 | 75793409 | 5084 | 2 | 66194 | ENSMUST00000053918.9 | 4214 | ENSMUSG00000022571 | Pycrl | pyrroline-5-carboxylate reductase-like |
| chr4 | 1,5E+08 | 1,5E+08 | 1659 | * | 13,94241 | 3,043528344 | 0,909886 | 3,344955 | 0,000823 | 0,523637575 | 0,938682 | Distal Intergenic | 4 | 1,5E+08 | 1,5E+08 | 3738 | 2 | 63954 | ENSMUST00000148596.8 | 7210 | ENSMUSG00000028996 | Rbp7 | retinol binding protein 7, cellular |
| chr10 | 1,28E+08 | 1,28E+08 | 888 | * | 8,755745 | 3,043802908 | 1,144829 | 2,658741 | 0,007843 | 0,847020973 | 1,425929 | Exon (ENSMUST000002 | 10 | 1,28E+08 | 1,28E+08 | 131 | 2 | 1,15E+08 | ENSMUST00000158062.3 | -8239 | ENSMUSG00000088687 | Gm23996 | predicted gene, 23996 |
| chr15 | 82030517 | 82031670 | 1154 | * | 14,35941 | 3,043930496 | 0,973039 | 3,128272 | 0,001758 | 0,666864737 | 0,918255 | Promoter (<=1kb) | 15 | 82031382 | 82089580 | 58199 | 1 | 20788 | ENSMUST00000023100.8 | 0 | ENSMUSG00000022463 | Srebf2 | sterol regulatory element binding factor 2 |
| chr11 | 84069137 | 84070344 | 1208 | * | 11,21791 | 3,044309567 | 1,075131 | 2,83157 | 0,004632 | 0,773819636 | 1,2763 | Promoter (<=1kb) | 11 | 84070657 | 84105021 | 34365 | 1 | 107476 | ENSMUST00000137500.9 | -313 | ENSMUSG00000020532 | Acaca | acetyl-Coenzyme A carboxylase alpha |
| chr2 | 1,32E+08 | 1,32E+08 | 1927 | * | 13,81197 | 3,044493858 | 0,99954 | 3,045896 | 0,00232 | 0,748382289 | 0,738441 | 3' UTR | 2 | 1,32E+08 | 1,32E+08 | 20252 | 2 | 215653 | ENSMUST00000155829.2 | 31583 | ENSMUSG00000027339 | Rassf2 | Ras association (RalGDS/AF-6) domain family member 2 |
| chr3 | 57210318 | 57212490 | 2173 | * | 18,11731 | 3,04450077 | 0,960629 | 3,169277 | 0,001528 | 0,483145429 | 2,154504 | Promoter (<=1kb) | 3 | 57194905 | 57209409 | 14505 | 2 | 17112 | ENSMUST00000029376.13 | -909 | ENSMUSG00000027800 | Tm4sf1 | transmembrane 4 superfamily member 1 |
| chr2 | 60263402 | 60264022 | 621 | * | 11,95558 | 3,04515647 | 0,982366 | 3,09982 | 0,001936 | 0,626725358 | 1,2763 | Exon (ENSMUST000001 | 2 | 60250369 | 60254651 | 4283 | 2 | 18779 | ENSMUST00000126327.2 | -8751 | ENSMUSG00000054580 | Pla2r1 | phospholipase A2 receptor 1 |
| chr2 | 44497058 | 44497970 | 913 | * | 11,50922 | 3,045432385 | 1,098663 | 2,771945 | 0,005572 | 0,821417341 | 1,173325 | Intron (ENSMUST0000 | 2 | 44460382 | 44461726 | 1345 | 2 | 227835 | ENSMUST00000143333.2 | -35332 | ENSMUSG00000036890 | Gtdc1 | glycosyltransferase-like domain containing 1 |
| chr4 | 1,56E+08 | 1,56E+08 | 968 | * | 9,529253 | 3,045537589 | 1,055723 | 2,884789 | 0,003917 | 0,723275538 | 1,425929 | Promoter (1-2kb) | 4 | 1,56E+08 | 1,56E+08 | 1848 | 1 | 100382 | ENSMUST00000105140.3 | 1192 | ENSMUSG00000078349 | AW011738 | expressed sequence AW011738 |
| chr1 | 36266161 | 36268077 | 1917 | * | 13,17737 | 3,045632564 | 1,12102 | 2,716841 | 0,006591 | 0,851512567 | 1,173325 | Exon (ENSMUST00000 | 1 | 36217686 | 36283188 | 65503 | 2 | 320011 | ENSMUST00000174716.8 | 15111 | ENSMUSG00000037470 | Uggt1 | UDP-glucose glycoprotein glucosyltransferase 1 |
| chr5 | 1,36E+08 | 1,36E+08 | 962 | * | 9,479998 | 3,046272189 | 1,125908 | 2,705615 | 0,006818 | 0,741363157 | 2,237403 | Distal Intergenic | 5 | 1,36E+08 | 1,36E+08 | 13759 | 1 | 215160 | ENSMUST00000043707.7 | -12012 | ENSMUSG00000039917 | Rhbdd2 | rhomboid domain containing 2 |
| chr2 | 1,21E+08 | 1,21E+08 | 792 | * | 8,162122 | 3,046797774 | 1,150505 | 2,648227 | 0,008092 | 0,858257847 | 1,360622 | Promoter (1-2kb) | 2 | 1,21E+08 | 1,21E+08 | 16204 | 2 | 640543 | ENSMUST00000238428.2 | 1100 | ENSMUSG00000079103 | Tgm7 | transglutaminase 7 |
| chr5 | 1,35E+08 | 1,35E+08 | 2559 | * | 19,56504 | 3,046880394 | 1,123727 | 2,711406 | 0,0067 | 0,705941691 | 2,713306 | Promoter (<=1kb) | 5 | 1,35E+08 | 1,35E+08 | 27619 | 1 | 20907 | ENSMUST00000005509.11 | 0 | ENSMUSG00000007207 | Stx1a | syntaxin 1A (brain) |
| chr4 | 86490280 | 86491911 | 1632 | * | 14,63494 | 3,047009397 | 0,967826 | 3,148304 | 0,001642 | 0,578728206 | 1,37683 | Promoter (1-2kb) | 4 | 86493905 | 86495522 | 1618 | 1 | 68441 | ENSMUST00000091064.8 | -1994 | ENSMUSG00000070934 | Rraga | Ras-related GTP binding A |
| chr5 | 1,21E+08 | 1,21E+08 | 2162 | * | 18,34074 | 3,048129452 | 0,984708 | 3,095466 | 0,001965 | 0,50121588 | 2,507013 | Exon (ENSMUST00000 | 5 | 1,21E+08 | 1,21E+08 | 13724 | 2 | 246727 | ENSMUST00000166703.2 | 5850 | ENSMUSG00000032661 | Oas3 | 2'-5' oligoadenylate synthetase 3 |
| chr2 | 26556010 | 26557197 | 1188 | * | 16,91369 | 3,049720793 | 0,976723 | 3,122402 | 0,001794 | 0,494205337 | 2,386008 | Distal Intergenic | 2 | 26557686 | 26561294 | 3609 | 2 | 16821 | ENSMUST00000028283.2 | 4097 | ENSMUSG00000026919 | Lcn4 | lipocalin 4 |
| chr2 | 34487814 | 34488836 | 1023 | * | 10,41234 | 3,04992766 | 1,131422 | 2,69566 | 0,007025 | 0,93486226 | 0,738441 | Intron (ENSMUST0000 | 2 | 34509594 | 34513931 | 4338 | 1 | 227743 | ENSMUST00000129633.2 | -20758 | ENSMUSG00000038696 | Mapkap1 | mitogen-activated protein kinase associated protein 1 |
| chr17 | 24253530 | 24255217 | 1688 | * | 25,56518 | 3,049955666 | 0,6678508 | 4,5668219 | 4,952E-06 | 0,012861975 | 2,8800171 | Promoter (1-2kb) | 17 | 24257118 | 24264928 | 7811 | 1 | 213171 | ENSMUST00000059482.6 | -1901 | ENSMUSG00000050762 | Prss27 | protease, serine 27 |
| chr18 | 13569103 | 13570402 | 1300 | * | 11,41063 | 3,050500072 | 1,059781 | 2,878426 | 0,003997 | 0,843548266 | 0,738441 | Distal Intergenic | 18 | 13595473 | 13734457 | 138985 | 2 | 1,05E+08 | ENSMUST00000234018.2 | 164055 | | Gm41670 | predicted gene, 41670 |
| chr10 | 83578981 | 83580839 | 1859 | * | 16,06479 | 3,050854117 | 1,019119 | 2,993618 | 0,002757 | 0,683710007 | 1,2763 | Intron (ENSMUST0000 | 10 | 83573773 | 83598027 | 24255 | 1 | 69784 | ENSMUST00000218623.2 | 5208 | ENSMUSG00000087651 | 1500009L16Rik | RIKEN cDNA 1500009L16 gene |
| chr12 | 59677124 | 59678158 | 1035 | * | 10,29285 | 3,050927473 | 1,151957 | 2,648473 | 0,008086 | 0,886373014 | 1,173325 | Distal Intergenic | 12 | 59712440 | 59712570 | 131 | 2 | 1,15E+08 | ENSMUST00000104398.3 | 34412 | ENSMUSG00000077586 | Gm24377 | predicted gene, 24377 |
| chr8 | 127249391 | 127250620 | 1230 | * | 28,823924 | 3,051049503 | 0,6759924 | 4,5134373 | 6,379E-06 | 0,015325349 | 2,9510621 | Distal Intergenic | 8 | 127152068 | 127201804 | 49737 | 2 | 212728 | ENSMUST00000170518.3 | -47587 | ENSMUSG00000090290 | Tarbp1 | TAR RNA binding protein 1 |
| chr18 | 23919301 | 23921263 | 1963 | * | 16,01435 | 3,051175157 | 0,967362 | 3,154121 | 0,00161 | 0,494205337 | 2,141612 | Intron (ENSMUST0000 | 18 | 23907921 | 23908046 | 126 | 2 | 1,02E+08 | ENSMUST00000184278.3 | -11255 | ENSMUSG00000098607 | Mir6360 | microRNA 6360 |
| chr9 | 62724859 | 62726441 | 1583 | * | 10,55104 | 3,052152169 | 1,152319 | 2,648704 | 0,00808 | 0,886332554 | 1,173325 | Distal Intergenic | 9 | 62699103 | 62719216 | 20114 | 2 | 14155 | ENSMUST00000034775.10 | -5643 | ENSMUSG00000032244 | Fem1b | fem 1 homolog b |
| chr11 | 44588000 | 44589293 | 1294 | * | 15,11991 | 3,052272888 | 1,014888 | 3,007496 | 0,002634 | 0,585155013 | 2,141612 | Intron (ENSMUST0000 | 11 | 44512588 | 44534286 | 21699 | 1 | 13591 | ENSMUST00000138452.2 | 75412 | ENSMUSG00000057098 | Ebf1 | early B cell factor 1 |
| chr5 | 22753994 | 22755846 | 1853 | * | 16,473 | 3,053270636 | 0,903445 | 3,379586 | 0,000726 | 0,355428041 | 2,141612 | Promoter (<=1kb) | 5 | 22691483 | 22755302 | 63820 | 2 | 26429 | ENSMUST00000135906.8 | 0 | ENSMUSG00000029012 | Orc5 | origin recognition complex, subunit 5 |
| chr16 | 6909058 | 6910401 | 1344 | * | 10,7614 | 3,053603188 | 1,114904 | 2,738893 | 0,006165 | 0,882355787 | 0,918255 | Intron (ENSMUST0000 | 16 | 6887710 | 7227677 | 339968 | 1 | 268859 | ENSMUST00000229741.2 | 21348 | ENSMUSG00000008658 | Rbfox1 | RNA binding protein, fox-1 homolog (C. elegans) 1 |
| chr11 | 97199634 | 97204117 | 4484 | * | 54,039169 | 3,053641089 | 0,5438361 | 5,615003 | 1,966E-08 | 0,000171651 | 2,9254973 | Promoter (<=1kb) | 11 | 97192965 | 97201105 | 8141 | 2 | 102633095 | ENSMUST00000240797.1 | 0 | | Gm31005 | predicted gene, 31005 |
| chr13 | 74354570 | 74356354 | 1785 | * | 12,1102 | 3,053759612 | 0,949834 | 3,215046 | 0,001304 | 0,546062568 | 1,2763 | Promoter (<=1kb) | 13 | 74341389 | 74355212 | 13824 | 2 | 211446 | ENSMUST00000222213.2 | 0 | ENSMUSG00000034152 | Exoc3 | exocyst complex component 3 |
| chr3 | 51302330 | 51306167 | 3838 | * | 42,766374 | 3,054069501 | 0,5688204 | 5,3691277 | 7,912E-08 | 0,000645269 | 2,3776742 | Promoter (<=1kb) | 3 | 51295833 | 51303947 | 8115 | 2 | 67749 | ENSMUST00000108046.8 | 0 | ENSMUSG00000037161 | Mgarp | mitochondria localized glutamic acid rich protein |
| chr19 | 15850831 | 15851701 | 871 | * | 10,71859 | 3,054481314 | 1,058213 | 2,886453 | 0,003896 | 0,799782439 | 0,918255 | Distal Intergenic | 19 | 15882494 | 15887076 | 4583 | 2 | 107272 | ENSMUST00000161707.2 | 35375 | ENSMUSG00000024640 | Psat1 | phosphoserine aminotransferase 1 |
| chr15 | 66867914 | 66868851 | 938 | * | 11,3545 | 3,055426096 | 1,003845 | 3,043723 | 0,002337 | 0,71242838 | 0,918255 | Intron (ENSMUST0000 | 15 | 66818386 | 66841489 | 23104 | 2 | 17988 | ENSMUST00000164070.2 | -26425 | ENSMUSG00000005125 | Ndrg1 | N-myc downstream regulated gene 1 |
| chr8 | 1,26E+08 | 1,26E+08 | 2314 | * | 21,47881 | 3,056861525 | 0,771946 | 3,959942 | 7,5E-05 | 0,086053637 | 2,951933 | Promoter (<=1kb) | 8 | 1,26E+08 | 1,26E+08 | 52772 | 2 | 244668 | ENSMUST00000212604.2 | 403 | ENSMUSG00000001995 | Sipa1l2 | signal-induced proliferation-associated 1 like 2 |
| chr5 | 1E+08 | 1E+08 | 786 | * | 8,709635 | 3,058394989 | 1,173883 | 2,605367 | 0,009178 | 1 | 0 | Intron (ENSMUST0000 | 5 | 1E+08 | 1E+08 | 38850 | 2 | 231503 | ENSMUST00000124145.8 | 5375 | ENSMUSG00000050640 | Tmem150c | transmembrane protein 150C |
| chr18 | 34697222 | 34700001 | 2780 | * | 21,25362 | 3,058504453 | 0,744424 | 4,108552 | 3,98E-05 | 0,0522907 | 3,471678 | Intron (ENSMUST0000 | 18 | 34700091 | 34712136 | 12046 | 2 | 75533 | ENSMUST00000154342.2 | 12135 | ENSMUSG00000035984 | Nme5 | NME/NM23 family member 5 |
| chr19 | 57612715 | 57613621 | 907 | * | 10,11253 | 3,058525436 | 1,144926 | 2,671374 | 0,007554 | 0,888042352 | 1,086118 | Intron (ENSMUST0000 | 19 | 57599466 | 58121775 | 522310 | 1 | 226255 | ENSMUST00000077282.7 | 13249 | ENSMUSG00000054843 | Atrnl1 | attractin like 1 |
| chr5 | 1,23E+08 | 1,23E+08 | 1589 | * | 11,23893 | 3,05861146 | 1,102304 | 2,774744 | 0,005525 | 0,899471409 | 0,738441 | Promoter (1-2kb) | 5 | 1,23E+08 | 1,23E+08 | 727 | 1 | 208043 | ENSMUST00000162839.3 | 1814 | ENSMUSG00000038384 | Setd1b | SET domain containing 1B |
| chr14 | 61409717 | 61410832 | 1116 | * | 24,99504 | 3,058645327 | 0,781347 | 3,914578 | 9,06E-05 | 0,096470298 | 2,951062 | Promoter (<=1kb) | 14 | 61410439 | 61433112 | 22674 | 1 | 50720 | ENSMUST00000121091.2 | 0 | ENSMUSG00000048279 | Sacs | sacsin |
| chr10 | 61456145 | 61457647 | 1503 | * | 15,42932 | 3,058838043 | 0,965234 | 3,169011 | 0,00153 | 0,591141332 | 1,214904 | Intron (ENSMUST0000 | 10 | 61431271 | 61464344 | 33074 | 1 | 237362 | ENSMUST00000020287.8 | 24874 | ENSMUSG00000020090 | Npffr1 | neuropeptide FF receptor 1 |
| chr17 | 33999759 | 34001705 | 1947 | * | 14,29631 | 3,059329704 | 1,144002 | 2,674234 | 0,00749 | 0,856353593 | 1,2763 | Promoter (<=1kb) | 17 | 33992724 | 34000804 | 8081 | 2 | 57875 | ENSMUST00000002360.17 | 0 | ENSMUSG00000002289 | Angptl4 | angiopoietin-like 4 |

| | | | | | | | | | | | | | | | | | | | | | | | |
|---|---|---|---|---|---|---|---|---|---|---|---|---|---|---|---|---|---|---|---|---|---|---|---|
| chr11 | 1,15E+08 | 1,15E+08 | 1728 | * | 14,07496 | 3,060137638 | 0,983637 | 3,111042 | 0,001864 | 0,710786992 | 0,738441 | Distal Intergenic | 11 | 1,15E+08 | 1,15E+08 | 8923 | 1 | 68572 | ENSMUST00000153983.8 | -5109 | ENSMUSG00000018858 | Mrpl58 | mitochondrial ribosomal protein L58 |
| chr4 | 1,39E+08 | 1,39E+08 | 1436 | * | 12,10043 | 3,060416738 | 1,092485 | 2,801335 | 0,005089 | 0,886373014 | 0,738441 | Promoter (<=1kb) | 4 | 1,39E+08 | 1,39E+08 | 7683 | 1 | 110198 | ENSMUST00000073787.7 | 0 | ENSMUSG00000028743 | Akr7a5 | aldo-keto reductase family 7, member A5 (aflatoxin aldehyde reductase) |
| chr2 | 34258030 | 34259289 | 1260 | * | 11,66169 | 3,060452915 | 0,998453 | 3,065196 | 0,002175 | 0,646033227 | 1,2763 | Promoter (1-2kb) | 2 | 34066871 | 34260858 | 193988 | 2 | 18516 | ENSMUST00000175855.3 | 1569 | ENSMUSG00000038718 | Pbx3 | pre B cell leukemia homeobox 3 |
| chr11 | 97301913 | 97303069 | 1157 | * | 10,80848 | 3,062637915 | 1,168034 | 2,622045 | 0,00874 | 0,965053522 | 0,744375 | Distal Intergenic | 11 | 97306359 | 97393228 | 86870 | 1 | 58996 | ENSMUST00000121799.8 | -3290 | ENSMUSG00000049807 | Arhgap23 | Rho GTPase activating protein 23 |
| chr8 | 63315262 | 63316140 | 879 | * | 12,04598 | 3,063321133 | 1,100114 | 2,78455 | 0,00536 | 0,893069436 | 0,738441 | Distal Intergenic | 8 | 63404043 | 63809886 | 405844 | 1 | 72902 | ENSMUST00000117377.8 | -87903 | ENSMUSG00000054162 | Spock3 | sparc/osteonectin, cwcv and kazal-like domains proteoglycan 3 |
| chr8 | 1,25E+08 | 1,25E+08 | 977 | * | 10,01587 | 3,063562064 | 1,080442 | 2,835471 | 0,004576 | 0,787366762 | 1,173325 | Distal Intergenic | 8 | 1,25E+08 | 1,25E+08 | 31242 | 1 | 76332 | ENSMUST00000034460.11 | -5529 | ENSMUSG00000031979 | Cog2 | component of oligomeric golgi complex 2 |
| chr10 | 80611554 | 80613494 | 1941 | * | 17,15059 | 3,063659789 | 0,881855 | 3,474108 | 0,000513 | 0,280986007 | 2,663619 | Promoter (<=1kb) | 10 | 80613390 | 80630225 | 16836 | 1 | 208266 | ENSMUST00000150338.8 | 0 | ENSMUSG00000061589 | Dot1l | DOT1-like, histone H3 methyltransferase (S. cerevisiae) |
| chr15 | 57846822 | 57847926 | 1105 | * | 15,25968 | 3,064182487 | 1,022181 | 2,997691 | 0,00272 | 0,68526372 | 1,214904 | Promoter (<=1kb) | 15 | 57848815 | 57874405 | 25591 | 1 | 239463 | ENSMUST00000160942.8 | -889 | ENSMUSG00000051225 | Fam83a | family with sequence similarity 83, member A |
| chr18 | 36536117 | 36536631 | 515 | * | 7,534973 | 3,064525026 | 1,162113 | 2,637029 | 0,008364 | 1 | 0 | 3' UTR | 18 | 36498834 | 36535593 | 36760 | 1 | 66060 | ENSMUST00000144158.2 | 37283 | ENSMUSG00000046727 | Cystm1 | cysteine-rich transmembrane module containing 1 |
| chr7 | 75095297 | 75096389 | 1093 | * | 8,764324 | 3,066858209 | 1,146389 | 2,675233 | 0,007468 | 0,837814951 | 1,425929 | Distal Intergenic | 7 | 75105282 | 75404357 | 299076 | 1 | 75547 | ENSMUST00000166315.7 | -8893 | ENSMUSG00000066406 | Akap13 | A kinase (PRKA) anchor protein 13 |
| chr15 | 99048227 | 99048996 | 770 | * | 8,001044 | 3,067039301 | 1,165798 | 2,630849 | 0,008517 | 1 | 0 | Intron (ENSMUST0000 | 15 | 99030948 | 99031053 | 106 | 2 | 1,19E+08 | ENSMUST00000157752.3 | -17174 | | LOC118568219 | 60S ribosomal protein L35a-like |
| chr6 | 1,34E+08 | 1,34E+08 | 1434 | * | 17,32115 | 3,067095863 | 0,989674 | 3,099098 | 0,001941 | 0,498011371 | 2,507013 | Intron (ENSMUST0000 | 6 | 1,34E+08 | 1,34E+08 | 3129 | 2 | 1,15E+08 | ENSMUST00000164616.2 | 47783 | | Gm52920 | predicted gene, 52920 |
| chr9 | 59205472 | 59206616 | 1145 | * | 11,1275 | 3,06722876 | 1,01579 | 3,01955 | 0,002532 | 0,760759863 | 0,738441 | Exon (ENSMUST00000 | 9 | 59198866 | 59223483 | 24618 | 1 | 72141 | ENSMUST00000214361.2 | 6606 | ENSMUSG00000025236 | Adpgk | ADP-dependent glucokinase |
| chrX | 90970975 | 90972091 | 1117 | * | 10,53515 | 3,06751176 | 1,107031 | 2,770937 | 0,00559 | 0,805225994 | 1,2763 | Distal Intergenic | 20 | 90975552 | 90976586 | 1035 | 2 | 278087 | ENSMUST00000104922.2 | 4495 | ENSMUSG00000078123 | Gm5071 | predicted pseudogene 5071 |
| chr1 | 1,52E+08 | 1,52E+08 | 1297 | * | 11,06543 | 3,068332172 | 1,131072 | 2,712765 | 0,006672 | 0,89325945 | 0,918255 | Exon (ENSMUST00000 | 1 | 1,52E+08 | 1,52E+08 | 35217 | 2 | 19731 | ENSMUST00000128251.8 | 67750 | ENSMUSG00000026482 | Rgl1 | ral guanine nucleotide dissociation stimulator,-like 1 |
| chr19 | 21066175 | 21067088 | 914 | * | 8,354435 | 3,069304402 | 1,169966 | 2,623413 | 0,008705 | 1 | 0 | Distal Intergenic | 19 | 21081866 | 21084521 | 2656 | 1 | 75354 | ENSMUST00000236278.2 | -14778 | ENSMUSG00000118047 | 4930554I06Rik | RIKEN cDNA 4930554I06 gene |
| chr4 | 1,36E+08 | 1,36E+08 | 1474 | * | 12,06898 | 3,06934118 | 1,139357 | 2,693924 | 0,007062 | 0,860218139 | 1,173325 | Distal Intergenic | 4 | 1,36E+08 | 1,36E+08 | 1896 | 1 | 15903 | ENSMUST00000008016.3 | -16804 | ENSMUSG00000007872 | Id3 | inhibitor of DNA binding 3 |
| chr17 | 55062874 | 55063942 | 1069 | * | 10,32988 | 3,069594075 | 1,101602 | 2,786483 | 0,005328 | 0,813025434 | 1,173325 | Distal Intergenic | 17 | 55159563 | 55167473 | 7911 | 1 | 1,03E+08 | ENSMUST00000233856.2 | -95621 | | Gm32055 | predicted gene, 32055 |
| chr1 | 33194370 | 33195735 | 1366 | * | 12,31592 | 3,069777128 | 1,009007 | 3,042375 | 0,002347 | 0,659533991 | 1,2763 | Distal Intergenic | 1 | 33231074 | 33231183 | 110 | 1 | 1,15E+08 | ENSMUST00000158125.3 | -35339 | ENSMUSG00000088750 | Gm24901 | predicted gene, 24901 |
| chr14 | 23632443 | 23633821 | 1379 | * | 10,97901 | 3,070070299 | 1,148617 | 2,672842 | 0,007521 | 0,947196831 | 0,738441 | Intron (ENSMUST0000 | 14 | 23551193 | 23604866 | 53674 | 2 | 16531 | ENSMUST00000212576.2 | -27577 | ENSMUSG00000063142 | Kcnma1 | potassium large conductance calcium-activated channel, subfamily M, alpha member 1 |
| chr1 | 31266417 | 31267272 | 856 | * | 9,123177 | 3,0703274 | 1,108288 | 2,770334 | 0,0056 | 0,788774532 | 1,425929 | Distal Intergenic | 1 | 31261919 | 31263369 | 1451 | 1 | 74708 | ENSMUST00000027230.3 | 4498 | ENSMUSG00000026063 | Dnaaf6 | dynein axonemal assembly factor 6 |
| chr5 | 1,4E+08 | 1,4E+08 | 1492 | * | 16,66544 | 3,070896472 | 0,922389 | 3,329286 | 0,000871 | 0,477253929 | 1,2763 | Intron (ENSMUST0000 | 5 | 1,4E+08 | 1,4E+08 | 26796 | 2 | 17120 | ENSMUST00000128579.2 | 37122 | ENSMUSG00000029554 | Mad1l1 | MAD1 mitotic arrest deficient 1-like 1 |
| chr10 | 80091032 | 80092233 | 1202 | * | 13,97319 | 3,071028878 | 0,991246 | 3,09815 | 0,001947 | 0,679069545 | 0,938682 | Promoter (1-2kb) | 10 | 80089149 | 80090974 | 1826 | 1 | 75406 | ENSMUST00000155336.2 | 1883 | ENSMUSG00000020153 | Ndufs7 | NADH:ubiquinone oxidoreductase core subunit S7 |
| chr6 | 34616514 | 34617744 | 1231 | * | 12,5244 | 3,071297454 | 1,180678 | 2,6013 | 0,009287 | 0,975155433 | 0,738441 | Intron (ENSMUST0000 | 6 | 34628289 | 34630829 | 2541 | 2 | 1E+08 | ENSMUST00000137136.2 | 13085 | ENSMUSG00000085382 | Gm13861 | predicted gene 13861 |
| chr1 | 90979646 | 90981014 | 1369 | * | 11,96173 | 3,071402434 | 1,169137 | 2,627069 | 0,008612 | 0,964160506 | 0,738441 | Promoter (<=1kb) | 1 | 90981166 | 91056666 | 75501 | 1 | 16978 | ENSMUST00000068167.13 | -152 | ENSMUSG00000026305 | Lrrfip1 | leucine rich repeat (in FLII) interacting protein 1 |
| chr11 | 50816266 | 50817839 | 1574 | * | 13,87632 | 3,071661578 | 0,975371 | 3,149223 | 0,001637 | 0,594430593 | 1,2763 | Intron (ENSMUST0000 | 11 | 50819391 | 50821476 | 2086 | 2 | 27274 | ENSMUST00000152296.2 | 3637 | ENSMUSG00000020335 | Zfp354b | zinc finger protein 354B |
| chr12 | 72739364 | 72741515 | 2152 | * | 14,48564 | 3,07189228 | 1,166736 | 2,632894 | 0,008466 | 0,880145259 | 1,2763 | Distal Intergenic | 12 | 72697127 | 72711683 | 14557 | 2 | 66375 | ENSMUST00000021512.11 | -27681 | ENSMUSG00000021094 | Dhrs7 | dehydrogenase/reductase (SDR family) member 7 |
| chr5 | 34010772 | 34011981 | 1210 | * | 8,855436 | 3,072672895 | 1,117939 | 2,748515 | 0,005987 | 0,793708774 | 1,449706 | Exon (ENSMUST00000 | 5 | 34018413 | 34034634 | 16222 | 1 | 107823 | ENSMUST00000142080.2 | -6432 | ENSMUSG00000057406 | Nsd2 | nuclear receptor binding SET domain protein 2 |
| chr12 | 79498467 | 79500063 | 1597 | * | 12,16259 | 3,073025633 | 1,043833 | 2,943981 | 0,00324 | 0,805745288 | 0,738441 | Intron (ENSMUST0000 | 12 | 79374127 | 79748630 | 374504 | 1 | 19363 | ENSMUST00000221257.2 | 124340 | ENSMUSG00000059060 | Rad51b | RAD51 paralog B |
| chr15 | 68701184 | 68701960 | 777 | * | 10,53442 | 3,073426204 | 1,107316 | 2,775564 | 0,005511 | 0,897433455 | 0,744375 | Intron (ENSMUST0000 | 15 | 68750164 | 68755425 | 5262 | 1 | 1,05E+08 | ENSMUST00000230367.2 | -48204 | ENSMUSG00000069996 | Gm41341 | predicted gene, 41341 |
| chr4 | 11696720 | 11698580 | 1861 | * | 16,56972 | 3,073449815 | 0,901904 | 3,407735 | 0,000655 | 0,323988916 | 2,386008 | Distal Intergenic | 4 | 11704457 | 11714752 | 10296 | 1 | 14579 | ENSMUST00000108304.9 | -5877 | ENSMUSG00000028214 | Gem | GTP binding protein (gene overexpressed in skeletal muscle) |
| chr5 | 44959652 | 44961637 | 1986 | * | 12,87632 | 3,073497832 | 1,055691 | 2,911362 | 0,003599 | 0,755814512 | 1,075735 | Promoter (2-3kb) | 5 | 44629474 | 44957022 | 327549 | 2 | 16826 | ENSMUST00000199256.5 | -2630 | ENSMUSG00000039706 | Ldb2 | LIM domain binding 2 |
| chr1 | 1,66E+08 | 1,66E+08 | 1569 | * | 10,15006 | 3,073715238 | 1,139961 | 2,696334 | 0,007011 | 0,854157826 | 1,211315 | Distal Intergenic | 1 | 1,66E+08 | 1,66E+08 | 13976 | 2 | 226601 | ENSMUST00000143922.2 | -73723 | ENSMUSG00000086056 | Gm4846 | predicted gene 4846 |
| chr1 | 173594268 | 173595979 | 1712 | * | 29,587862 | 3,073848742 | 0,5932685 | 5,1812104 | 2,205E-07 | 0,00116151 | 2,9510621 | Promoter (<=1kb) | 1 | 173574859 | 173594485 | 19627 | 2 | 15951 | ENSMUST00000111214.4 | 0 | ENSMUSG00000073489 | Ifi204 | interferon activated gene 204 |
| chr10 | 80771750 | 80773829 | 2080 | * | 35,526315 | 3,074684739 | 0,6479568 | 4,7452005 | 2,083E-06 | 0,011411459 | 1,4057095 | Distal Intergenic | 10 | 80766646 | 80767593 | 948 | 1 | 17873 | ENSMUST00000219449.2 | 5104 | ENSMUSG00000015312 | Gadd45b | growth arrest and DNA-damage-inducible 45 beta |
| chr7 | 29149267 | 29150234 | 968 | * | 12,43707 | 3,074825486 | 1,134565 | 2,710135 | 0,006726 | 0,838625535 | 1,2763 | Intron (ENSMUST0000 | 7 | 29083681 | 29204877 | 121197 | 2 | 74206 | ENSMUST00000182011.8 | 54643 | ENSMUSG00000030583 | Sipa1l3 | signal-induced proliferation-associated 1 like 3 |
| chr11 | 69327457 | 69328755 | 1299 | * | 10,58694 | 3,075910047 | 1,025478 | 2,99949 | 0,002704 | 0,682206193 | 1,2763 | Exon (ENSMUST00000 | 11 | 69289334 | 69304501 | 15168 | 2 | 216850 | ENSMUST00000094077.5 | -22956 | ENSMUSG00000018476 | Kdm6b | KDM1 lysine (K)-specific demethylase 6B |
| chr3 | 1,03E+08 | 1,03E+08 | 1567 | * | 10,20468 | 3,077013477 | 1,146448 | 2,683954 | 0,007276 | 0,830691256 | 1,449706 | Distal Intergenic | 3 | 1,03E+08 | 1,03E+08 | 37761 | 1 | 229663 | ENSMUST00000029446.13 | -4375 | ENSMUSG00000068823 | Csde1 | cold shock domain containing E1, RNA binding |
| chr7 | 1,01E+08 | 1,01E+08 | 1030 | * | 9,052999 | 3,078530618 | 1,165873 | 2,640538 | 0,008277 | 0,852340527 | 1,449706 | Promoter (2-3kb) | 7 | 1,01E+08 | 1,01E+08 | 68126 | 1 | 20480 | ENSMUST00000153473.2 | 2742 | ENSMUSG00000001829 | Clpb | ClpB caseinolytic peptidase B |
| chr17 | 32820149 | 32821763 | 1615 | * | 20,01084 | 3,078593634 | 0,77202 | 3,987714 | 6,67E-05 | 0,084947995 | 2,713306 | Exon (ENSMUST00000 | 17 | 32840283 | 32855158 | 14876 | 1 | 677156 | ENSMUST00000077639.7 | -18520 | ENSMUSG00000062464 | Cyp4f37 | cytochrome P450, family 4, subfamily f, polypeptide 37 |
| chr5 | 1,07E+08 | 1,07E+08 | 1897 | * | 13,5624 | 3,079203809 | 0,925487 | 3,327116 | 0,000877 | 0,477622452 | 1,2763 | Promoter (<=1kb) | 5 | 1,07E+08 | 1,07E+08 | 20110 | 1 | 12545 | ENSMUST00000117196.9 | 0 | ENSMUSG00000029283 | Cdc7 | cell division cycle 7 (S. cerevisiae) |
| chr17 | 88410256 | 88411418 | 1163 | * | 14,38622 | 3,0794405 | 0,968966 | 3,178067 | 0,001483 | 0,607577254 | 1,075735 | Distal Intergenic | 17 | 88432940 | 88438434 | 5495 | 1 | 225058 | ENSMUST00000183295.2 | -21522 | ENSMUSG00000058869 | Gm4832 | predicted gene 4832 |
| chr11 | 42921238 | 42922485 | 1248 | * | 11,55479 | 3,079921164 | 1,135516 | 2,712354 | 0,006681 | 0,852340527 | 1,173325 | Distal Intergenic | 11 | 43040704 | 43050302 | 9599 | 1 | 319767 | ENSMUST00000148911.2 | -118219 | ENSMUSG00000055415 | Atp10b | ATPase, class V, type 10B |
| chr11 | 60363821 | 60365203 | 1383 | * | 10,71234 | 3,080055708 | 1,092881 | 2,81829 | 0,004828 | 0,793243838 | 1,173325 | Intron (ENSMUST0000 | 11 | 60360165 | 60419195 | 59031 | 1 | 17910 | ENSMUST00000094135.9 | 3656 | ENSMUSG00000042678 | Myo15 | myosin XV |
| chr5 | 124153757 | 124156619 | 2863 | * | 38,164305 | 3,080545208 | 0,5956016 | 5,1721575 | 2,314E-07 | 0,001237282 | 2,7872553 | Intron (ENSMUST000000 | 5 | 124144199 | 124149776 | 5578 | 2 | 330192 | ENSMUST00000199306.2 | -3981 | ENSMUSG00000066278 | Vps37b | vacuolar protein sorting 37B |
| chr5 | 1,37E+08 | 1,37E+08 | 1083 | * | 9,081777 | 3,081792689 | 1,161128 | 2,654136 | 0,007951 | 0,85617852 | 1,360622 | Distal Intergenic | 5 | 1,37E+08 | 1,37E+08 | 2092 | 2 | 18787 | ENSMUST00000199832.2 | -21028 | ENSMUSG00000037411 | Serpine1 | serine (or cysteine) peptidase inhibitor, clade E, member 1 |
| chr15 | 1,03E+08 | 1,03E+08 | 1106 | * | 10,1791 | 3,081893049 | 1,09329 | 2,818917 | 0,004819 | 0,79031971 | 1,211315 | Promoter (1-2kb) | 15 | 1,03E+08 | 1,03E+08 | 26954 | 1 | 56447 | ENSMUST00000230893.2 | 1477 | ENSMUSG00000060992 | Copz1 | coatomer protein complex, subunit zeta 1 |
| chr6 | 30616912 | 30618065 | 1154 | * | 16,24114 | 3,082524112 | 0,94207 | 3,272077 | 0,001068 | 0,425424592 | 2,141612 | Intron (ENSMUST0000 | 6 | 30612554 | 30631435 | 18882 | 1 | 74649 | ENSMUST00000165949.2 | 4358 | ENSMUSG00000029788 | Cpa5 | carboxypeptidase A5 |
| chr8 | 1,07E+08 | 1,07E+08 | 1127 | * | 10,21028 | 3,083423359 | 1,065221 | 2,894633 | 0,003796 | 0,765659517 | 1,086118 | Promoter (2-3kb) | 8 | 1,07E+08 | 1,07E+08 | 63 | 2 | 1,02E+08 | ENSMUST00000183884.3 | -2119 | ENSMUSG00000099131 | Mir7074 | microRNA 7074 |
| chr6 | 79411240 | 79411980 | 741 | * | 7,858293 | 3,083669397 | 1,101064 | 2,800626 | 0,0051 | 1 | 0 | Distal Intergenic | 6 | 79794899 | 79795347 | 449 | 1 | 1E+08 | ENSMUST00000179797.3 | -382919 | ENSMUSG00000096887 | Gm20594 | predicted gene, 20594 |
| chr2 | 56212471 | 56213286 | 816 | * | 8,652383 | 3,083823142 | 1,186148 | 2,599863 | 0,009326 | 0,874247416 | 1,449706 | Distal Intergenic | 2 | 56675823 | 56675919 | 97 | 1 | 1,02E+08 | ENSMUST00000184814.3 | -462537 | ENSMUSG00000098437 | Mir195b | microRNA 195b |
| chr10 | 1,28E+08 | 1,28E+08 | 2046 | * | 16,92656 | 3,083826914 | 1,033785 | 2,983045 | 0,002854 | 0,685148714 | 1,2763 | Promoter (<=1kb) | 10 | 1,28E+08 | 1,28E+08 | 43373 | 2 | 56516 | ENSMUST00000092033.4 | 0 | ENSMUSG00000040043 | Rbms2 | RNA binding motif, single stranded interacting protein 2 |
| chr3 | 87067356 | 87068597 | 1242 | * | 8,792571 | 3,084529913 | 1,138684 | 2,708855 | 0,006752 | 0,820889278 | 1,425929 | Exon (ENSMUST00000 | 3 | 86991935 | 87081857 | 89923 | 2 | 170643 | ENSMUST00000041732.9 | 13260 | ENSMUSG00000041734 | Kirrel | kirre like nephrin family adhesion molecule 1 |
| chr2 | 13409478 | 13410887 | 1410 | * | 15,43472 | 3,084582122 | 0,930652 | 3,314432 | 0,000918 | 0,474532331 | 1,37683 | Intron (ENSMUST0000 | 2 | 13426840 | 13460438 | 33599 | 2 | 65969 | ENSMUST00000195447.2 | 49551 | ENSMUSG00000026726 | Cubn | cubilin (intrinsic factor-cobalamin receptor) |
| chr17 | 46045139 | 46047575 | 2437 | * | 68,232562 | 3,085867804 | 0,4106532 | 7,5145349 | 5,711E-14 | 4,16979E-09 | 3,177995 | Promoter (<=1kb) | 17 | 46045432 | 46092178 | 46747 | 1 | 433111 | ENSMUST00000180558.9 | 0 | ENSMUSG00000097727 | F630040K05Rik | RIKEN cDNA F630040K05 gene |
| chr6 | 51568945 | 51570973 | 2029 | * | 16,53947 | 3,086927525 | 0,875444 | 3,526127 | 0,000422 | 0,352029189 | 1,2763 | Distal Intergenic | 6 | 51521546 | 51566023 | 44478 | 1 | 71982 | ENSMUST00000149024.2 | 47399 | ENSMUSG00000038301 | Snx10 | sorting nexin 10 |
| chr18 | 19284314 | 19285895 | 1582 | * | 13,19762 | 3,087182039 | 0,908233 | 3,399107 | 0,000676 | 0,464166555 | 1,075735 | Intron (ENSMUST0000 | 18 | 19266274 | 19334773 | 68500 | 1 | 1,03E+08 | ENSMUST00000234128.2 | 18040 | | Gm31086 | predicted gene, 31086 |
| chr5 | 1,25E+08 | 1,25E+08 | 1689 | * | 14,89567 | 3,088041446 | 1,050298 | 2,940159 | 0,00328 | 0,697241323 | 1,37683 | Distal Intergenic | 5 | 1,25E+08 | 1,25E+08 | 9586 | 1 | 1,05E+08 | ENSMUST00000200649.2 | -7173 | | Gm40323 | predicted gene, 40323 |
| chr3 | 27673898 | 27674786 | 889 | * | 9,994464 | 3,089062412 | 1,171273 | 2,637354 | 0,008356 | 0,902613747 | 1,086118 | Exon (ENSMUST00000 | 3 | 27639049 | 27639144 | 96 | 2 | 1,01E+08 | ENSMUST00000174954.3 | -34754 | ENSMUSG00000092695 | Mir3092 | microRNA 3092 |
| chr7 | 87449248 | 87450785 | 1538 | * | 11,93141 | 3,090441994 | 1,141791 | 2,706663 | 0,006796 | 0,929178509 | 0,738441 | Intron (ENSMUST0000 | 7 | 87252297 | 87458828 | 206532 | 1 | 108071 | ENSMUST00000138732.3 | 196951 | ENSMUSG00000049583 | Grm5 | glutamate receptor, metabotropic 5 |
| chr17 | 33748256 | 33749490 | 1235 | * | 11,15411 | 3,090703439 | 1,041691 | 2,967006 | 0,003007 | 0,792545355 | 0,738441 | Promoter (<=1kb) | 17 | 33747288 | 33772422 | 25135 | 1 | 224697 | ENSMUST00000173013.8 | 968 | ENSMUSG00000024299 | Adamts10 | a disintegrin-like and metallopeptidase (reprolysin type) with thrombospondin type 1 motif, 10 |
| chr17 | 43733196 | 43734127 | 932 | * | 12,35568 | 3,091168348 | 1,063017 | 2,90792 | 0,003638 | 0,757975613 | 1,075735 | Exon (ENSMUST00000 | 17 | 43737903 | 43767662 | 29760 | 1 | 224792 | ENSMUST00000225962.2 | -3776 | ENSMUSG00000056492 | Adgrf5 | adhesion G protein-coupled receptor F5 |
| chr10 | 62174998 | 62179835 | 4838 | * | 87,716451 | 3,091367682 | 0,7138113 | 4,330791 | 1,486E-05 | 0,022826798 | 4,0557576 | Promoter (<=1kb) | 10 | 62104634 | 62176321 | 71688 | 2 | 15275 | ENSMUST00000099691.11 | 0 | ENSMUSG00000037012 | Hk1 | hexokinase 1 |
| chr7 | 46496316 | 46497358 | 1043 | * | 11,07469 | 3,091501578 | 1,023644 | 3,020094 | 0,002527 | 0,670853401 | 1,2763 | Promoter (<=1kb) | 7 | 46496506 | 46505050 | 8545 | 1 | 16828 | ENSMUST00000209984.2 | 0 | ENSMUSG00000063229 | Ldha | lactate dehydrogenase A |
| chr13 | 36298975 | 36300243 | 1269 | * | 9,782091 | 3,091904525 | 1,084212 | 2,851753 | 0,004348 | 0,770673161 | 1,211315 | Promoter (1-2kb) | 13 | 36301395 | 36721567 | 420173 | 1 | 69955 | ENSMUST00000021857.13 | -1152 | ENSMUSG00000021420 | Fars2 | phenylalanine-tRNA synthetase 2 (mitochondrial) |
| chr19 | 13284411 | 13286064 | 1654 | * | 11,56855 | 3,092012543 | 1,160882 | 2,663503 | 0,007733 | 0,915421656 | 0,918255 | Distal Intergenic | 19 | 13290724 | 13291647 | 924 | 2 | 258121 | ENSMUST00000080142.2 | 5583 | ENSMUSG00000062199 | Olfr1465 | olfactory receptor 1465 |
| chr1 | 90949659 | 90952882 | 3224 | * | 53,405721 | 3,092723803 | 0,4802975 | 6,439184 | 1,201E-10 | 2,45309E-06 | 2,9254973 | Intron (ENSMUST000001 | 1 | 90926501 | 91056662 | 130162 | 1 | 16978 | ENSMUST00000068116.13 | 23158 | ENSMUSG00000026305 | Lrrfip1 | leucine rich repeat (in FLII) interacting protein 1 |
| chr9 | 62107480 | 62109014 | 1535 | * | 10,85813 | 3,095018534 | 1,164187 | 2,658523 | 0,007848 | 0,918200407 | 0,918255 | Distal Intergenic | 9 | 62149878 | 62151382 | 1505 | 2 | 330958 | ENSMUST00000215490.2 | 42368 | ENSMUSG00000111232 | A730043L09Rik | RIKEN cDNA A730043L09 gene |
| chr4 | 1,33E+08 | 1,33E+08 | 1840 | * | 20,15913 | 3,095152916 | 0,772496 | 4,006692 | 6,16E-05 | 0,090530822 | 2,221611 | Distal Intergenic | 4 | 1,33E+08 | 1,33E+08 | 2349 | 1 | 20544 | ENSMUST00000141658.2 | 28563 | ENSMUSG00000028854 | Slc9a1 | solute carrier family 9 (sodium/hydrogen exchanger), member 1 |
| chr5 | 28379163 | 28380905 | 1743 | * | 15,76201 | 3,096289483 | 0,963715 | 3,212869 | 0,001314 | 0,446292698 | 2,318434 | Distal Intergenic | 5 | 28370692 | 28377164 | 6473 | 1 | 13799 | ENSMUST00000036177.9 | 8471 | ENSMUSG00000039095 | En2 | engrailed 2 |
| chr12 | 91163602 | 91164848 | 1247 | * | 13,4041 | 3,097104791 | 0,903164 | 3,429171 | 0,000605 | 0,468032694 | 0,938682 | Intron (ENSMUST0000 | 12 | 90989174 | 91150546 | 161373 | 2 | 75216 | ENSMUST00000133329.2 | -13056 | ENSMUSG00000061533 | Cep128 | centrosomal protein 128 |
| chr4 | 1,5E+08 | 1,5E+08 | 1675 | * | 15,42794 | 3,09750852 | 0,899244 | 3,444569 | 0,000572 | 0,321510378 | 2,141612 | Exon (ENSMUST00000 | 4 | 1,5E+08 | 1,5E+08 | 8912 | 2 | 12353 | ENSMUST00000134648.8 | 5974 | ENSMUSG00000028972 | Car6 | carbonic anhydrase 6 |
| chr11 | 74700848 | 74702673 | 1826 | * | 24,001279 | 3,098166077 | 0,7028089 | 4,4082627 | 1,042E-05 | 0,022659302 | 2,8800171 | Promoter (1-2kb) | 11 | 74699355 | 74708337 | 8983 | 1 | 67493 | ENSMUST00000134571.2 | 1493 | ENSMUSG00000010554 | Mettl16 | methyltransferase like 16 |
| chr9 | 36677703 | 36679852 | 2150 | * | 24,172229 | 3,098551261 | 0,7381133 | 4,1979343 | 2,694E-05 | 0,045482826 | 2,9510621 | Promoter (<=1kb) | 9 | 36679157 | 36693973 | 14817 | 1 | 102637672 | ENSMUST00000180474.5 | 0 | ENSMUSG00000096993 | Gm26787 | predicted gene, 26787 |
| chr9 | 1,09E+08 | 1,09E+08 | 1912 | * | 17,73465 | 3,098723239 | 0,908737 | 3,409922 | 0,00065 | 0,308866996 | 2,663619 | Promoter (<=1kb) | 9 | 1,09E+08 | 1,09E+08 | 4701 | 2 | 66167 | ENSMUST00000167504.3 | 0 | ENSMUSG00000091537 | Tma7 | translational machinery associated 7 |
| chr7 | 27498962 | 27500793 | 1832 | * | 12,44383 | 3,100090781 | 1,085831 | 2,855041 | 0,004303 | 0,855969265 | 0,738441 | Distal Intergenic | 7 | 27506621 | 27522341 | 15721 | 1 | 71163 | ENSMUST00000205671.2 | -5828 | ENSMUSG00000030604 | Zfp626 | zinc finger protein 626 |
| chr9 | 89713150 | 89714479 | 1330 | * | 14,06677 | 3,100250172 | 1,071823 | 2,892502 | 0,003822 | 0,737853731 | 1,2763 | Intron (ENSMUST0000 | 9 | 89709035 | 89720373 | 11339 | 1 | 74700 | ENSMUST00000184759.2 | 4115 | ENSMUSG00000098627 | 4930524O08Rik | RIKEN cDNA 4930524O08 gene |
| chr16 | 39581281 | 39582346 | 1066 | * | 10,50681 | 3,100314258 | 1,012074 | 3,063329 | 0,002189 | 0,734161328 | 0,744375 | Distal Intergenic | 16 | 39804723 | 40800071 | 995349 | 1 | 268890 | ENSMUST00000189229.7 | -222377 | ENSMUSG00000061080 | Lsamp | limbic system-associated membrane protein |
| chr9 | 62261745 | 62263228 | 1484 | * | 11,8509 | 3,10068861 | 1,067522 | 2,904568 | 0,003678 | 0,826676469 | 0,744375 | Intron (ENSMUST0000 | 9 | 62252081 | 62281997 | 29917 | 1 | 11737 | ENSMUST00000126451.8 | 9664 | ENSMUSG00000032249 | Anp32a | acidic (leucine-rich) nuclear phosphoprotein 32 family, member A |
| chr11 | 78056063 | 78057256 | 1194 | * | 11,22194 | 3,101953082 | 1,149806 | 2,697805 | 0,00698 | 0,858257847 | 1,173325 | Promoter (<=1kb) | 11 | 78049325 | 78056415 | 7091 | 2 | 22032 | ENSMUST00000149437.8 | 0 | ENSMUSG00000017386 | Traf4 | TNF receptor associated factor 4 |
| chr3 | 1,02E+08 | 1,02E+08 | 1419 | * | 10,67379 | 3,102251793 | 1,147826 | 2,702719 | 0,006877 | 0,856317799 | 1,173325 | 3' UTR | 3 | 1,02E+08 | 1,02E+08 | 35277 | 2 | 242125 | ENSMUST00000118411.2 | -7565 | ENSMUSG00000044313 | Mab21l3 | mab-21-like 3 |
| chr5 | 27996467 | 27997629 | 1163 | * | 11,98244 | 3,102889438 | 1,091703 | 2,842246 | 0,00448 | 0,860883941 | 0,738441 | Promoter (<=1kb) | 5 | 27945078 | 27996548 | 51471 | 2 | 55982 | ENSMUST00000002291.12 | 0 | ENSMUSG00000002221 | Paxip1 | PAX interacting (with transcription-activation domain) protein 1 |
| chr1 | 88388122 | 88389469 | 1348 | * | 13,89212 | 3,103274849 | 1,037414 | 2,991356 | 0,002777 | 0,735602255 | 0,938682 | Promoter (<=1kb) | 1 | 88371691 | 88390330 | 18640 | 2 | 1,03E+08 | ENSMUST00000245480.1 | 861 | ENSMUSG00000099553 | Gm29538 | predicted gene 29538 |
| chr5 | 1,06E+08 | 1,06E+08 | 1387 | * | 12,11233 | 3,103312087 | 1,104964 | 2,808518 | 0,004977 | 0,881662676 | 0,744375 | Promoter (<=1kb) | 5 | 1,06E+08 | 1,06E+08 | 5562 | 1 | 54367 | ENSMUST00000135374.2 | 298 | ENSMUSG00000029290 | Zfp326 | zinc finger protein 326 |
| chr19 | 60297916 | 60298852 | 937 | * | 8,156471 | 3,103390902 | 1,142077 | 2,717321 | 0,006581 | 1 | 0 | Distal Intergenic | 19 | 60187867 | 60215133 | 27267 | 2 | 76539 | ENSMUST00000237339.2 | -82783 | ENSMUSG00000057858 | Fam204a | family with sequence similarity 204, member A |
| chr10 | 74155668 | 74156862 | 1195 | * | 11,96258 | 3,10423265 | 1,053263 | 2,947254 | 0,003206 | 0,710067561 | 1,2763 | Intron (ENSMUST0000 | 10 | 74138983 | 74166882 | 27900 | 1 | 11994 | ENSMUST00000147455.2 | 16685 | ENSMUSG00000052613 | Pcdh15 | protocadherin 15 |
| chr8 | 38175213 | 38176001 | 789 | * | 12,9175 | 3,104722149 | 0,970382 | 3,199483 | 0,001377 | 0,594430593 | 1,075735 | Intron (ENSMUST0000 | 8 | 37784631 | 37784732 | 102 | 2 | 1,15E+08 | ENSMUST00000083343.3 | -390481 | ENSMUSG00000065277 | Gm25126 | predicted gene, 25126 |
| chr13 | 90036963 | 90038202 | 1240 | * | 10,75 | 3,105492877 | 0,999973 | 3,105575 | 0,001899 | 0,678498769 | 0,918255 | Intron (ENSMUST0000 | 13 | 89803431 | 89890628 | 87198 | 2 | 13003 | ENSMUST00000109546.9 | -146335 | ENSMUSG00000021614 | Vcan | versican |
| chr14 | 58032093 | 58035743 | 3651 | * | 42,424276 | 3,1055456 | 0,5708681 | 5,4400403 | 5,327E-08 | 0,00047006 | 2,3383871 | Promoter (<=1kb) | 14 | 58035637 | 58044095 | 8459 | 1 | 20220 | ENSMUST00000128764.9 | 0 | ENSMUSG00000021963 | Sap18 | Sin3-associated polypeptide 18 |
| chr16 | 55641501 | 55644218 | 2718 | * | 22,711115 | 3,105762546 | 0,6781203 | 4,5799584 | 4,651E-06 | 0,012021261 | 2,9510621 | Promoter (<=1kb) | 16 | 55631738 | 55642501 | 10764 | 2 | 80859 | ENSMUST00000114458.8 | 0 | ENSMUSG00000035356 | Nfkbiz | nuclear factor of kappa light polypeptide gene enhancer in B cells inhibitor, zeta |
| chr4 | 24173499 | 24174704 | 1206 | * | 12,26078 | 3,106367673 | 1,01453 | 3,061878 | 0,0022 | 0,676450861 | 1,075735 | Distal Intergenic | 4 | 24496451 | 24581338 | 84888 | 1 | 212377 | ENSMUST00000173079.8 | -321747 | ENSMUSG00000045751 | Mms22l | MMS22-like, DNA repair protein |
| chr6 | 31066731 | 31068667 | 1937 | * | 44,442323 | 3,107303872 | 0,5199384 | 5,9762929 | 2,283E-09 | 3,74095E-05 | 2,3383871 | Intron (ENSMUST000002 | 6 | 31040027 | 31040430 | 404 | 2 | 387223 | ENSMUST00000183056.2 | -26301 | ENSMUSG00000065604 | Mir29b-1 | microRNA 29b-1 |
| chr17 | 65283278 | 65284101 | 824 | * | 9,122754 | 3,107998223 | 1,164771 | 2,668335 | 0,007623 | 0,851504891 | 1,360622 | Distal Intergenic | 17 | 65251716 | 65251847 | 132 | 2 | 1,15E+08 | ENSMUST00000104503.4 | -31431 | ENSMUSG00000077691 | Gm25344 | predicted gene, 25344 |
| chr10 | 1,28E+08 | 1,28E+08 | 1961 | * | 17,37137 | 3,108177659 | 0,847987 | 3,665361 | 0,000247 | 0,190625259 | 2,663619 | Promoter (1-2kb) | 10 | 1,28E+08 | 1,28E+08 | 7504 | 1 | 68094 | ENSMUST00000217751.2 | 1157 | ENSMUSG00000025369 | Smarcc2 | SWI/SNF related, matrix associated, actin dependent regulator of chromatin, subfamily c, member 2 |
| chr19 | 41257705 | 41259248 | 1544 | * | 13,86832 | 3,108473434 | 1,033443 | 3,007881 | 0,002631 | 0,704364304 | 1,075735 | Intron (ENSMUST0000 | 19 | 41199283 | 41252436 | 53154 | 2 | 107358 | ENSMUST00000025989.10 | -5269 | ENSMUSG00000025016 | Tm9sf3 | transmembrane 9 superfamily member 3 |
| chr18 | 29521270 | 29522547 | 1278 | * | 10,03879 | 3,10894539 | 1,058782 | 2,936342 | 0,003321 | 0,742014238 | 1,086118 | Distal Intergenic | 18 | 29575253 | 29599160 | 23908 | 1 | 78199 | ENSMUST00000234621.2 | -52706 | ENSMUSG00000117554 | 4930527G23Rik | RIKEN cDNA 4930527G23 gene |
| chr4 | 1,11E+08 | 1,11E+08 | 789 | * | 10,85964 | 3,109867671 | 1,074147 | 2,895198 | 0,003789 | 0,832960417 | 0,744375 | Intron (ENSMUST0000 | 4 | 1,11E+08 | 1,11E+08 | 501580 | 1 | 78933 | ENSMUST00000148038.2 | 85329 | ENSMUSG00000061298 | Agbl4 | ATP/GTP binding protein-like 4 |
| chr7 | 80183427 | 80186041 | 2615 | * | 28,02113 | 3,110100124 | 0,752552 | 4,132738 | 3,58E-05 | 0,058289444 | 2,67243 | Promoter (<=1kb) | 7 | 80104741 | 80184867 | 80127 | 2 | 12144 | ENSMUST00000081314.11 | 0 | ENSMUSG00000030528 | Blm | Bloom syndrome, RecQ like helicase |
| chr12 | 79561443 | 79562352 | 910 | * | 9,555181 | 3,111073245 | 1,113072 | 2,795034 | 0,005189 | 0,773936378 | 1,425929 | Intron (ENSMUST0000 | 12 | 79374127 | 79748630 | 374504 | 1 | 19363 | ENSMUST00000221257.2 | 187316 | ENSMUSG00000059060 | Rad51b | RAD51 paralog B |
| chr7 | 110295849 | 110298692 | 2844 | * | 23,833754 | 3,111086031 | 0,7202287 | 4,3195805 | 1,563E-05 | 0,030043467 | 2,9519335 | Promoter (<=1kb) | 7 | 110289370 | 110298941 | 9572 | 2 | 100861673 | ENSMUST00000210822.2 | 249 | ENSMUSG00000109700 | Gm21123 | predicted gene, 21123 |
| chr4 | 33294423 | 33295900 | 1478 | * | 11,43483 | 3,111579644 | 1,18214 | 2,632158 | 0,008484 | 0,962965811 | 0,738441 | Distal Intergenic | 4 | 33246012 | 33290163 | 44152 | 2 | 108767 | ENSMUST00000137589.2 | -4260 | ENSMUSG00000040128 | Pnrc1 | proline-rich nuclear receptor coactivator 1 |

| | | | | | | | | | | | | | | | | | | | | | | | |
|---|---|---|---|---|---|---|---|---|---|---|---|---|---|---|---|---|---|---|---|---|---|---|---|
| chr1 | 98410713 | 98412194 | 1482 | * | 12,21945 | 3,11194615 | 0,971545 | 3,203091 | 0,00136 | 0,659642963 | 0,738441 | Intron (ENSMUST0000( | 1 | 98427614 | 98437108 | 9495 | 1 | 70866 | ENSMUST00000162782.2 | -15420 | ENSMUSG00000026336 | Slco6d1 | solute carrier organic anion transporter family, member 6d1 |
| chr5 | 14561472 | 14562978 | 1507 | * | 15,79253 | 3,112241599 | 0,891169 | 3,492314 | 0,000479 | 0,272716235 | 2,663619 | Promoter (1-2kb) | 5 | 14564932 | 14846007 | 281076 | 1 | 26875 | ENSMUST00000182407.8 | -1954 | ENSMUSG00000061601 | Pclo | piccolo (presynaptic cytomatrix protein) |
| chr6 | 72514773 | 72516364 | 1592 | * | 39,793713 | 3,112444118 | 0,6631374 | 4,6935132 | 2,686E-06 | 0,009275024 | 2,3776742 | Distal Intergenic | 6 | 72521374 | 72539966 | 18593 | 1 | 12332 | ENSMUST00000071044.13 | -5010 | ENSMUSG00000056737 | Capg | capping protein (actin filament), gelsolin-like |
| chr17 | 79399758 | 79401307 | 1550 | * | 15,74206 | 3,112582389 | 0,981029 | 3,172774 | 0,00151 | 0,576475101 | 1,2763 | Distal Intergenic | 17 | 79384431 | 79389282 | 4852 | 1 | 70536 | ENSMUST00000233899.2 | 15327 | ENSMUSG00000024084 | Qpct | glutaminyl-peptide cyclotransferase (glutaminyl cyclase) |
| chr9 | 98959033 | 98959555 | 523 | * | 8,487263 | 3,112643312 | 1,193422 | 2,608166 | 0,009103 | 0,792545355 | 2,237403 | Intron (ENSMUST0000( | 9 | 98888843 | 98917768 | 28926 | 2 | 382097 | ENSMUST00000076730.8 | -41265 | ENSMUSG00000044860 | Gm1123 | predicted gene 1123 |
| chr4 | 1,3E+08 | 1,3E+08 | 1116 | * | 10,17327 | 3,112976013 | 1,026028 | 3,034006 | 0,002413 | 0,644704154 | 1,425929 | Intron (ENSMUST0000( | 4 | 1,3E+08 | 1,3E+08 | 73 | 1 | 1E+08 | ENSMUST00000083262.3 | -36411 | ENSMUSG00000065196 | Snord85 | small nucleolar RNA, C/D box 85 |
| chr5 | 30140650 | 30141835 | 1186 | * | 8,717938 | 3,113160067 | 1,202602 | 2,588687 | 0,009634 | 1 | 0 | Distal Intergenic | 5 | 30209471 | 30212759 | 3289 | 2 | 97209 | ENSMUST00000199600.2 | 70924 | | A230098N10Rik | RIKEN cDNA A230098N10 gene |
| chr11 | 23024414 | 23025988 | 1575 | * | 15,8901 | 3,11338071 | 0,898122 | 3,466547 | 0,000527 | 0,283414546 | 2,663619 | Exon (ENSMUST00000: | 11 | 23017998 | 23025574 | 7577 | 1 | 1E+08 | ENSMUST00000128996.2 | 6416 | | Gm12059 | predicted gene 12059 |
| chr7 | 31359869 | 31361274 | 1406 | * | 11,82958 | 3,113468645 | 1,032291 | 3,016076 | 0,002561 | 0,683988405 | 1,173325 | Distal Intergenic | 7 | 31317168 | 31318951 | 1784 | 2 | 1,01E+08 | ENSMUST00000188132.2 | -40918 | ENSMUSG00000102002 | Scgb2b6 | secretoglobin, family 2B, member 6 |
| chr1 | 1,38E+08 | 1,38E+08 | 1336 | * | 11,83019 | 3,114129871 | 0,94341 | 3,300929 | 0,000964 | 0,494741046 | 1,2763 | Distal Intergenic | 1 | 1,38E+08 | 1,38E+08 | 15725 | 1 | 338375 | ENSMUST00000027643.6 | 27242 | ENSMUSG00000026394 | Atp6v1g3 | ATPase, H+ transporting, lysosomal V1 subunit G3 |
| chr16 | 43180625 | 43181676 | 1052 | * | 9,473712 | 3,114671001 | 1,139819 | 2,732601 | 0,006284 | 0,728037066 | 2,237403 | Promoter (1-2kb) | 16 | 43183411 | 43390124 | 206714 | 1 | 56490 | ENSMUST00000141941.8 | -1735 | ENSMUSG00000022708 | Zbtb20 | zinc finger and BTB domain containing 20 |
| chr5 | 1,36E+08 | 1,36E+08 | 1025 | * | 10,06355 | 3,114763709 | 1,168398 | 2,66584 | 0,00768 | 0,838878373 | 1,449706 | Exon (ENSMUST00000( | 5 | 1,36E+08 | 1,36E+08 | 5031 | 1 | 17448 | ENSMUST00000143747.2 | 4032 | ENSMUSG00000019179 | Mdh2 | malate dehydrogenase 2, NAD (mitochondrial) |
| chr5 | 1,26E+08 | 1,26E+08 | 2188 | * | 23,63025 | 3,114868376 | 0,772135 | 4,034097 | 5,48E-05 | 0,071862525 | 2,951062 | Promoter (<=1kb) | 5 | 1,25E+08 | 1,26E+08 | 20264 | 2 | 208144 | ENSMUST00000169485.6 | 0 | ENSMUSG00000029480 | Dhx37 | DEAH (Asp-Glu-Ala-His) box polypeptide 37 |
| chr19 | 36975431 | 36977141 | 1711 | * | 11,42121 | 3,115255493 | 1,07784 | 2,890275 | 0,003849 | 0,836351745 | 0,744375 | Intron (ENSMUST0000( | 19 | 36938715 | 36943232 | 4518 | 1 | 107182 | ENSMUST00000236343.2 | 36716 | ENSMUSG00000040565 | Btaf1 | B-TFIID TATA-box binding protein associated factor 1 |
| chr2 | 27357848 | 27359108 | 1261 | * | 14,58743 | 3,115510608 | 0,990714 | 3,144714 | 0,001662 | 0,498577247 | 2,141612 | Intron (ENSMUST0000( | 2 | 27353904 | 27354594 | 691 | 2 | 67382 | ENSMUST00000144758.2 | -3254 | ENSMUSG00000026918 | Brd3 | bromodomain containing 3 |
| chr6 | 47762694 | 47764621 | 1928 | * | 14,68617 | 3,116308782 | 1,042438 | 2,989442 | 0,002795 | 0,684274806 | 1,2763 | Promoter (<=1kb) | 6 | 47765003 | 47765114 | 112 | 2 | 19872 | ENSMUST00000083767.3 | 493 | ENSMUSG00000065701 | Rny1 | RNA, Y1 small cytoplasmic, Ro-associated |
| chr14 | 89166954 | 89167838 | 885 | * | 7,317443 | 3,11671832 | 1,196282 | 2,605337 | 0,009178 | 1 | 0 | Distal Intergenic | 14 | 88873469 | 89001197 | 127729 | 1 | 1,05E+08 | ENSMUST00000227074.2 | 293485 | | Gm41225 | predicted gene, 41225 |
| chr17 | 27406282 | 27407900 | 1619 | * | 16,0845 | 3,117090433 | 0,970432 | 3,212064 | 0,001318 | 0,424477229 | 2,663619 | Distal Intergenic | 17 | 27409086 | 27412454 | 3369 | 2 | 224640 | ENSMUST00000237625.2 | 4554 | ENSMUSG00000044857 | Lemd2 | LEM domain containing 2 |
| chr11 | 1,02E+08 | 1,02E+08 | 3183 | * | 59,31171 | 3,118306117 | 0,809202 | 3,853558 | 0,000116 | 0,090530822 | 4,200304 | Promoter (1-2kb) | 11 | 1,02E+08 | 1,02E+08 | 16306 | 1 | 217216 | ENSMUST00000100392.5 | -1789 | ENSMUSG00000034773 | Hrob | homologous recombination factor with OB-fold |
| chr10 | 58281540 | 58282591 | 1052 | * | 13,26439 | 3,118635041 | 1,017538 | 3,064884 | 0,002178 | 0,696366028 | 0,918255 | Promoter (<=1kb) | 10 | 58282742 | 58294821 | 12080 | 1 | 19386 | ENSMUST00000219332.2 | -151 | ENSMUSG00000003226 | Ranbp2 | RAN binding protein 2 |
| chr12 | 21590794 | 21591625 | 832 | * | 8,655543 | 3,118899302 | 1,13922 | 2,73775 | 0,006186 | 0,802671652 | 1,425929 | Promoter (1-2kb) | 12 | 21593258 | 21604692 | 11435 | 1 | 1,15E+08 | ENSMUST00000211164.3 | -1633 | | LOC115487949 | uncharacterized LOC115487949 |
| chr6 | 30453671 | 30454816 | 1146 | * | 13,58348 | 3,119105785 | 1,088374 | 2,86584 | 0,004159 | 0,851741608 | 0,738441 | 3' UTR | 6 | 30460127 | 30463260 | 3134 | 1 | 69337 | ENSMUST00000143687.2 | -5311 | ENSMUSG00000085470 | 1700012C14Rik | RIKEN cDNA 1700012C14 gene |
| chr13 | 42451920 | 42453196 | 1277 | * | 9,235223 | 3,120587634 | 1,209213 | 2,580677 | 0,009861 | 0,884588199 | 1,449706 | Promoter (1-2kb) | 13 | 42454952 | 42461466 | 6515 | 1 | 13614 | ENSMUST00000021796.9 | -1756 | ENSMUSG00000021367 | Edn1 | endothelin 1 |
| chr3 | 1,31E+08 | 1,31E+08 | 1903 | * | 20,63803 | 3,120879017 | 0,808953 | 3,857922 | 0,000114 | 0,11391768 | 2,880017 | Distal Intergenic | 3 | 1,31E+08 | 1,31E+08 | 2752 | 1 | 1,19E+08 | ENSMUST00000198737.2 | 75936 | | Gm43351 | predicted gene 43351 |
| chr19 | 42340345 | 42341585 | 1241 | * | 11,0592 | 3,121093955 | 1,198256 | 2,604696 | 0,009196 | 0,891636336 | 1,2763 | Intron (ENSMUST0000( | 19 | 42398833 | 42402685 | 3853 | 1 | 1,03E+08 | ENSMUST00000235253.2 | -57248 | | Gm34299 | predicted gene, 34299 |
| chr1 | 1,64E+08 | 1,64E+08 | 1669 | * | 13,1468 | 3,121416135 | 1,111952 | 2,80715 | 0,004998 | 0,80080207 | 1,173325 | Exon (ENSMUST00000( | 1 | 1,64E+08 | 1,64E+08 | 133605 | 1 | 171567 | ENSMUST00000193808.6 | -4941 | ENSMUSG00000026575 | Nme7 | NME/NM23 family member 7 |
| chrX | 90874480 | 90875531 | 1052 | * | 8,050434 | 3,124005887 | 1,15999 | 2,693133 | 0,007078 | 1 | 0 | Distal Intergenic | 20 | 90823150 | 90826624 | 3475 | 2 | 74271 | ENSMUST00000082048.3 | -47856 | ENSMUSG00000061392 | Mageb5 | MAGE family member B5 |
| chr1 | 85963135 | 85963992 | 858 | * | 7,408367 | 3,124178733 | 1,19088 | 2,623421 | 0,008705 | 1 | 0 | Distal Intergenic | 1 | 85973585 | 85981075 | 7491 | 1 | 72792 | ENSMUST00000027429.11 | -9593 | ENSMUSG00000026227 | 2810459M11Rik | RIKEN cDNA 2810459M11 gene |
| chr7 | 1,33E+08 | 1,33E+08 | 1884 | * | 15,65799 | 3,124232822 | 0,927395 | 3,368828 | 0,000755 | 0,441389356 | 1,37683 | Distal Intergenic | 7 | 1,33E+08 | 1,33E+08 | 14207 | 1 | 74970 | ENSMUST00000209850.2 | 182306 | ENSMUSG00000109816 | 4930483O08Rik | RIKEN cDNA 4930483O08 gene |
| chr19 | 15684076 | 15684984 | 909 | * | 9,567386 | 3,124468002 | 1,117089 | 2,796974 | 0,005158 | 0,80080207 | 1,211315 | Distal Intergenic | 19 | 15657368 | 15657474 | 107 | 1 | 1,15E+08 | ENSMUST00000083389.3 | 26708 | ENSMUSG00000118757 | Gm24319 | predicted gene, 24319 |
| chr7 | 96966499 | 96968596 | 2098 | * | 20,41563 | 3,125012072 | 0,775104 | 4,031735 | 5,54E-05 | 0,082292767 | 2,386008 | Promoter (1-2kb) | 7 | 96961795 | 96965202 | 3408 | 2 | 244144 | ENSMUST00000180476.3 | -1297 | ENSMUSG00000035713 | Usp35 | ubiquitin specific peptidase 35 |
| chr2 | 70011632 | 70013100 | 1469 | * | 12,37652 | 3,125086152 | 0,959994 | 3,255317 | 0,001133 | 0,524368688 | 1,2763 | 3' UTR | 2 | 69926642 | 70013611 | 86970 | 1 | 329421 | ENSMUST00000112241.2 | 84990 | ENSMUSG00000042064 | Myo3b | myosin IIIB |
| chr9 | 57852990 | 57854605 | 1616 | * | 15,93272 | 3,126221961 | 1,081128 | 2,891631 | 0,003832 | 0,636647377 | 2,386008 | Intron (ENSMUST0000( | 9 | 57847396 | 57870148 | 22753 | 1 | 20361 | ENSMUST00000043059.9 | 5594 | ENSMUSG00000038264 | Sema7a | sema domain, immunoglobulin domain (Ig), and GPI membrane anchor, (semaphorin) 7A |
| chr12 | 82865552 | 82868064 | 2513 | * | 14,90752 | 3,127104109 | 0,976931 | 3,200946 | 0,00137 | 0,54122637 | 1,37683 | Intron (ENSMUST0000( | 12 | 82819095 | 82819230 | 136 | 2 | 1,15E+08 | ENSMUST00000104442.3 | -46322 | ENSMUSG00000077630 | Gm22149 | predicted gene, 22149 |
| chr11 | 6306859 | 6308044 | 1186 | * | 11,63396 | 3,127242571 | 1,143855 | 2,733951 | 0,006258 | 0,825493135 | 1,2763 | Downstream (<=300bp | 11 | 6339074 | 6356158 | 17085 | 1 | 52915 | ENSMUST00000109787.8 | -31030 | ENSMUSG00000041164 | Zmiz2 | zinc finger, MIZ-type containing 2 |
| chr16 | 20457724 | 20459072 | 1349 | * | 11,38045 | 3,127650041 | 1,210379 | 2,584026 | 0,009765 | 0,982552203 | 0,744375 | Promoter (<=1kb) | 16 | 20457956 | 20462445 | 4490 | 1 | 107522 | ENSMUST00000231739.2 | 0 | ENSMUSG00000022842 | Ece2 | endothelin converting enzyme 2 |
| chr12 | 10306386 | 10307647 | 1262 | * | 10,49579 | 3,127761858 | 1,078011 | 2,901419 | 0,003715 | 0,829907213 | 0,744375 | Distal Intergenic | 12 | 10419973 | 10425935 | 5963 | 1 | 70881 | ENSMUST00000143739.9 | -112326 | ENSMUSG00000020622 | Nt5c1b | 5'-nucleotidase, cytosolic IB |
| chr3 | 1,05E+08 | 1,05E+08 | 871 | * | 12,41982 | 3,127788446 | 1,158091 | 2,700814 | 0,006917 | 0,90060438 | 0,918255 | Distal Intergenic | 3 | 1,05E+08 | 1,05E+08 | 10133 | 2 | 78635 | ENSMUST00000143893.2 | 106456 | ENSMUSG00000074346 | Kcnd3os | potassium voltage-gated channel, Shal-related family, member 3, opposite strand |
| chr4 | 33509751 | 33510645 | 895 | * | 10,32397 | 3,1280885 | 1,067088 | 2,931424 | 0,003374 | 0,745502963 | 1,086118 | Distal Intergenic | 4 | 33438081 | 33438209 | 129 | 1 | 1,02E+08 | ENSMUST00000183962.2 | 71670 | ENSMUSG00000098310 | Mir8118 | microRNA 8118 |
| chr15 | 98848843 | 98850229 | 1387 | * | 15,46716 | 3,128195106 | 1,00628 | 3,108672 | 0,001879 | 0,523637575 | 2,141612 | Promoter (1-2kb) | 15 | 98830018 | 98851432 | 21415 | 2 | 22143 | ENSMUST00000134214.3 | 1203 | ENSMUSG00000023004 | Tuba1b | tubulin, alpha 1B |
| chr16 | 10951085 | 10952188 | 1104 | * | 10,24897 | 3,128969534 | 1,142521 | 2,738653 | 0,006169 | 0,852340527 | 1,086118 | Promoter (<=1kb) | 16 | 10922475 | 10952417 | 29943 | 2 | 106200 | ENSMUST00000155221.8 | 229 | ENSMUSG00000022498 | Txndc11 | thioredoxin domain containing 11 |
| chr2 | 1,65E+08 | 1,65E+08 | 1119 | * | 10,94729 | 3,129175514 | 1,173081 | 2,667485 | 0,007642 | 0,914417016 | 0,918255 | 5' UTR | 2 | 1,65E+08 | 1,65E+08 | 1983 | 2 | 228869 | ENSMUST00000123347.2 | -4262 | ENSMUSG00000039804 | Ncoa5 | nuclear receptor coactivator 5 |
| chr2 | 1,36E+08 | 1,36E+08 | 854 | * | 12,20391 | 3,129978059 | 0,994763 | 3,146456 | 0,001653 | 0,59511204 | 1,2763 | Intron (ENSMUST0000( | 2 | 1,36E+08 | 1,36E+08 | 306835 | 2 | 241656 | ENSMUST00000035264.9 | 72471 | ENSMUSG00000039913 | Pak5 | p21 (RAC1) activated kinase 5 |
| chr4 | 1,24E+08 | 1,24E+08 | 1428 | * | 13,48259 | 3,130226567 | 0,995357 | 3,144828 | 0,001662 | 0,656950206 | 0,918255 | Promoter (<=1kb) | 4 | 1,24E+08 | 1,24E+08 | 5477 | 2 | 595136 | ENSMUST00000106206.8 | 0 | ENSMUSG00000028648 | Ndufs5 | NADH:ubiquinone oxidoreductase core subunit S5 |
| chr16 | 24182545 | 24183454 | 910 | * | 8,443846 | 3,131318961 | 1,178403 | 2,657257 | 0,007878 | 0,844744375 | 1,449706 | Intron (ENSMUST0000( | 16 | 24089588 | 24198874 | 109287 | 2 | 1,03E+08 | ENSMUST00000231672.2 | 15420 | | Gm38456 | predicted gene, 38456 |
| chr11 | 1,01E+08 | 1,01E+08 | 1527 | * | 22,8823 | 3,132348453 | 0,835773 | 3,747844 | 0,000178 | 0,152163457 | 2,858511 | Promoter (<=1kb) | 11 | 1,01E+08 | 1,01E+08 | 8673 | 2 | 217198 | ENSMUST00000043397.14 | 0 | ENSMUSG00000035172 | Plekhh3 | pleckstrin homology domain containing, family H (with MyTH4 domain) member 3 |
| chr5 | 1,37E+08 | 1,37E+08 | 907 | * | 8,614309 | 3,133832269 | 1,209526 | 2,590959 | 0,009571 | 0,889364747 | 1,360622 | Distal Intergenic | 5 | 1,37E+08 | 1,37E+08 | 7949 | 1 | 59310 | ENSMUST00000196068.5 | -48337 | ENSMUSG00000005474 | Myl10 | myosin, light chain 10, regulatory |
| chr17 | 66430460 | 66432868 | 2409 | * | 18,21309 | 3,134825005 | 0,916593 | 3,420083 | 0,000626 | 0,317511466 | 2,386008 | Promoter (<=1kb) | 17 | 66430515 | 66437779 | 7265 | 1 | 320209 | ENSMUST00000225956.2 | 0 | ENSMUSG00000035842 | Ddx11 | DEAD/H box helicase 11 |
| chr5 | 137284468 | 137286363 | 1896 | * | 21,203773 | 3,134996618 | 0,7543128 | 4,1560965 | 3,237E-05 | 0,04965487 | 3,104559 | Promoter (<=1kb) | 5 | 137285781 | 137288416 | 2636 | 1 | 11423 | ENSMUST00000132191.8 | 0 | ENSMUSG00000023328 | Ache | acetylcholinesterase |
| chr4 | 41314768 | 41316247 | 1480 | * | 12,33011 | 3,135744205 | 1,068227 | 2,935467 | 0,00333 | 0,81148874 | 0,738441 | Promoter (<=1kb) | 4 | 41314960 | 41320630 | 5671 | 1 | 1,08E+08 | ENSMUST00000245488.1 | 0 | | Gm46891 | predicted gene, 46891 |
| chr6 | 112671660 | 112675688 | 4029 | * | 60,768273 | 3,13597884 | 0,4889405 | 6,4138251 | 1,419E-10 | 2,40889E-06 | 4,0557576 | Promoter (<=1kb) | 6 | 112596811 | 112673647 | 76837 | 2 | 58186 | ENSMUST00000077088.11 | 0 | ENSMUSG00000030254 | Rad18 | RAD18 E3 ubiquitin protein ligase |
| chr5 | 1,5E+08 | 1,5E+08 | 1056 | * | 11,47704 | 3,136062767 | 1,031996 | 3,038832 | 0,002375 | 0,71419937 | 0,918255 | Exon (ENSMUST00000: | 5 | 1,5E+08 | 1,5E+08 | 3486 | 1 | 140498 | ENSMUST00000138901.2 | -15302 | ENSMUSG00000053368 | Rxfp2 | relaxin/insulin-like family peptide receptor 2 |
| chr7 | 19375264 | 19375940 | 677 | * | 8,866616 | 3,136792734 | 1,164339 | 2,694054 | 0,007059 | 0,824134241 | 1,449706 | Intron (ENSMUST0000( | 7 | 19368881 | 19371671 | 2791 | 2 | 56457 | ENSMUST00000208868.2 | -3593 | ENSMUSG00000002981 | Clptm1 | cleft lip and palate associated transmembrane protein 1 |
| chr14 | 59692199 | 59693242 | 1044 | * | 9,262291 | 3,137426696 | 1,202041 | 2,610083 | 0,009052 | 0,899471409 | 1,211315 | Intron (ENSMUST0000( | 14 | 59696587 | 59737118 | 40532 | 1 | 69008 | ENSMUST00000223678.2 | -3345 | ENSMUSG00000021981 | Cab39l | calcium binding protein 39-like |
| chr12 | 1,11E+08 | 1,11E+08 | 2167 | * | 12,85522 | 3,139257048 | 0,97022 | 3,235615 | 0,001214 | 0,597812173 | 0,918255 | Promoter (1-2kb) | 12 | 1,11E+08 | 1,11E+08 | 66295 | 1 | 69641 | ENSMUST00000193053.6 | -1913 | ENSMUSG00000037957 | Wdr20 | WD repeat domain 20 |
| chr8 | 1,06E+08 | 1,06E+08 | 2542 | * | 17,42315 | 3,139952535 | 0,862961 | 3,63858 | 0,000274 | 0,213937229 | 2,423594 | Promoter (1-2kb) | 8 | 1,06E+08 | 1,06E+08 | 107 | 1 | 1,15E+08 | ENSMUST00000240010.1 | 1005 | ENSMUSG00000119598 | Gm22063 | predicted gene, 22063 |
| chr11 | 48159943 | 48161110 | 1168 | * | 10,64189 | 3,140097162 | 1,109847 | 2,829308 | 0,004665 | 0,870290757 | 0,738441 | Distal Intergenic | 11 | 47469632 | 47879796 | 410165 | 2 | 24052 | ENSMUST00000128558.2 | -280147 | ENSMUSG00000020354 | Sgcd | sarcoglycan, delta (dystrophin-associated glycoprotein) |
| chr4 | 1,3E+08 | 1,3E+08 | 1358 | * | 8,731775 | 3,140444262 | 1,194983 | 2,628024 | 0,008588 | 0,869872986 | 1,360622 | Promoter (2-3kb) | 4 | 1,3E+08 | 1,3E+08 | 83 | 1 | 1,15E+08 | ENSMUST00000083856.3 | 2018 | ENSMUSG00000119649 | Gm24091 | predicted gene, 24091 |
| chr3 | 71861909 | 71863571 | 1663 | * | 11,29994 | 3,140730212 | 0,96701 | 3,247877 | 0,001163 | 0,63007937 | 0,744375 | Distal Intergenic | 3 | 72795893 | 72874199 | 78307 | 2 | 69983 | ENSMUST00000094190.5 | 1010628 | ENSMUSG00000027790 | Sis | sucrase isomaltase (alpha-glucosidase) |
| chr12 | 81508468 | 81509869 | 1402 | * | 13,44807 | 3,140784977 | 1,091678 | 2,877025 | 0,004014 | 0,801134483 | 0,938682 | Intron (ENSMUST0000( | 12 | 81518867 | 81531603 | 12737 | 2 | 66272 | ENSMUST00000171237.8 | 21734 | ENSMUSG00000091803 | Cox16 | cytochrome c oxidase assembly protein 16 |
| chr3 | 1,38E+08 | 1,38E+08 | 1033 | * | 9,634719 | 3,140936923 | 1,206918 | 2,602445 | 0,009256 | 0,872576097 | 1,449706 | Distal Intergenic | 3 | 1,38E+08 | 1,38E+08 | 15450 | 2 | 74635 | ENSMUST00000197761.2 | 83970 | ENSMUSG00000105838 | 4930425O10Rik | RIKEN cDNA 4930425O10 gene |
| chr11 | 1,15E+08 | 1,15E+08 | 2200 | * | 15,44135 | 3,141436513 | 0,994577 | 3,158567 | 0,001585 | 0,595849045 | 1,214904 | Intron (ENSMUST0000( | 11 | 1,15E+08 | 1,15E+08 | 12716 | 2 | 170930 | ENSMUST00000117589.8 | 7244 | ENSMUSG00000020738 | Sumo2 | small ubiquitin-like modifier 2 |
| chr16 | 73407067 | 73408578 | 1512 | * | 13,03086 | 3,142465146 | 0,929823 | 3,379638 | 0,000726 | 0,541133622 | 0,738441 | Distal Intergenic | 16 | 73219681 | 73219847 | 167 | 2 | 1,15E+08 | ENSMUST00000082896.3 | -187220 | ENSMUSG00000064830 | Gm25699 | predicted gene, 25699 |
| chr4 | 1,29E+08 | 1,29E+08 | 1300 | * | 10,05431 | 3,142557174 | 1,166984 | 2,692889 | 0,007084 | 0,824883149 | 1,449706 | Intron (ENSMUST0000( | 4 | 1,29E+08 | 1,29E+08 | 19986 | 1 | 67525 | ENSMUST00000147852.2 | -30272 | ENSMUSG00000041000 | Trim62 | tripartite motif-containing 62 |
| chr11 | 97863517 | 97864841 | 1325 | * | 10,86237 | 3,142747079 | 0,999593 | 3,144028 | 0,001666 | 0,68680243 | 0,738441 | Intron (ENSMUST0000( | 11 | 97859618 | 97860061 | 444 | 1 | 1,03E+08 | ENSMUST00000118176.2 | 3899 | | Gm11633 | predicted gene 11633 |
| chr2 | 31371887 | 31372991 | 1105 | * | 16,72987 | 3,143294347 | 0,85705 | 3,667573 | 0,000245 | 0,280905021 | 1,2763 | Intron (ENSMUST0000( | 2 | 31360219 | 31410684 | 50466 | 1 | 11898 | ENSMUST00000102840.5 | 11668 | ENSMUSG00000076441 | Ass1 | argininosuccinate synthetase 1 |
| chr15 | 99598246 | 99599375 | 1130 | * | 12,57533 | 3,143360328 | 1,031455 | 3,047501 | 0,002308 | 0,669774031 | 1,173325 | Promoter (<=1kb) | 15 | 99600010 | 99611872 | 11863 | 1 | 83797 | ENSMUST00000023759.6 | -635 | ENSMUSG00000023018 | Smarcd1 | SWI/SNF related, matrix associated, actin dependent regulator of chromatin, subfamily d, member 1 |
| chr17 | 71042589 | 71043825 | 1237 | * | 12,16777 | 3,144036031 | 1,004977 | 3,128466 | 0,001757 | 0,697016988 | 0,738441 | Intron (ENSMUST0000( | 17 | 71072834 | 71112992 | 40159 | 2 | 1,05E+08 | ENSMUST00000233559.2 | 69167 | ENSMUSG00000117231 | Gm41609 | predicted gene, 41609 |
| chr11 | 6058592 | 6060129 | 1538 | * | 14,02065 | 3,144090938 | 1,06139 | 2,962239 | 0,003054 | 0,730215586 | 1,075735 | Intron (ENSMUST0000( | 11 | 6055986 | 6069409 | 13424 | 2 | 209586 | ENSMUST00000109805.8 | 9280 | ENSMUSG00000053838 | Nudcd3 | NudC domain containing 3 |
| chr8 | 96623431 | 96624848 | 1418 | * | 18,68552 | 3,144112049 | 0,806513 | 3,898403 | 9,68E-05 | 0,114217481 | 2,423594 | Distal Intergenic | 8 | 96590762 | 96615175 | 24414 | 2 | 14719 | ENSMUST00000034097.8 | -8256 | ENSMUSG00000031672 | Got2 | glutamatic-oxaloacetic transaminase 2, mitochondrial |
| chr7 | 34614472 | 34615524 | 1053 | * | 10,71743 | 3,144125804 | 1,176037 | 2,673493 | 0,007507 | 0,872808323 | 1,173325 | Promoter (2-3kb) | 7 | 34611804 | 34744133 | 132330 | 1 | 18624 | ENSMUST00000075068.14 | 2668 | ENSMUSG00000063931 | Pepd | peptidase D |
| chr11 | 1,01E+08 | 1,01E+08 | 1126 | * | 9,929554 | 3,144430968 | 1,051899 | 2,98929 | 0,002796 | 0,668446397 | 1,449706 | Promoter (<=1kb) | 11 | 1,01E+08 | 1,01E+08 | 1200 | 2 | 80861 | ENSMUST00000150656.2 | 0 | ENSMUSG00000017830 | Dhx58 | DEXH (Asp-Glu-X-His) box polypeptide 58 |
| chr5 | 31211232 | 31213114 | 1883 | * | 20,72972 | 3,144586645 | 0,808347 | 3,890145 | 0,0001 | 0,10248494 | 2,951933 | Promoter (<=1kb) | 5 | 31212124 | 31235823 | 23700 | 1 | 69719 | ENSMUST00000013773.12 | 0 | ENSMUSG00000013629 | Cad | carbamoyl-phosphate synthetase 2, aspartate transcarbamylase, and dihydroorotase |
| chr4 | 1E+08 | 1E+08 | 1322 | * | 12,0988 | 3,144839558 | 1,169419 | 2,689233 | 0,007162 | 0,86195649 | 1,173325 | Intron (ENSMUST0000( | 4 | 99976818 | 99978038 | 1221 | 1 | 1,05E+08 | ENSMUST00000134206.2 | 61793 | | Gm12701 | predicted gene 12701 |
| chr5 | 1,37E+08 | 1,37E+08 | 1527 | * | 11,91343 | 3,145620251 | 1,138578 | 2,762762 | 0,005731 | 0,902886632 | 0,744375 | Intron (ENSMUST0000( | 5 | 1,36E+08 | 1,37E+08 | 290947 | 2 | 13047 | ENSMUST00000176172.8 | 75473 | ENSMUSG00000029705 | Cux1 | cut-like homeobox 1 |
| chr2 | 33124692 | 33125941 | 1250 | * | 13,94102 | 3,146310448 | 1,058387 | 2,972742 | 0,002952 | 0,749550311 | 0,918255 | Intron (ENSMUST0000( | 2 | 33106129 | 33132405 | 26277 | 1 | 26360 | ENSMUST00000193373.3 | 18563 | ENSMUSG00000004105 | Angptl2 | angiopoietin-like 2 |
| chr6 | 6792784 | 6793725 | 942 | * | 10,92699 | 3,146631096 | 1,169784 | 2,689924 | 0,007147 | 0,861581225 | 1,173325 | Intron (ENSMUST0000( | 6 | 6824188 | 6830791 | 6604 | 2 | 320038 | ENSMUST00000210702.2 | 37066 | ENSMUSG00000090063 | Dlx6os1 | distal-less homeobox 6, opposite strand 1 |
| chr11 | 1,02E+08 | 1,02E+08 | 813 | * | 8,269867 | 3,149026114 | 1,087161 | 2,89656 | 0,003773 | 0,715249603 | 1,449706 | Promoter (<=1kb) | 11 | 1,02E+08 | 1,02E+08 | 22733 | 2 | 268490 | ENSMUST00000107156.9 | 242 | ENSMUSG00000020922 | Lsm12 | LSM12 homolog |
| chr16 | 4523977 | 4524611 | 635 | * | 7,186767 | 3,149883364 | 1,176868 | 2,676498 | 0,00744 | 1 | 0 | 3' UTR | 16 | 4532373 | 4536952 | 4580 | 2 | 67824 | ENSMUST00000124977.2 | 12341 | ENSMUSG00000063445 | Nmral1 | NmrA-like family domain containing 1 |
| chr11 | 35872736 | 35873558 | 823 | * | 8,364773 | 3,150567659 | 1,204414 | 2,615851 | 0,008901 | 0,87775273 | 1,360622 | Promoter (1-2kb) | 11 | 35729227 | 35871354 | 142128 | 2 | 211652 | ENSMUST00000018993.7 | -1382 | ENSMUSG00000018849 | Wwc1 | WW, C2 and coiled-coil domain containing 1 |
| chr7 | 36515333 | 36516336 | 1004 | * | 8,392718 | 3,150572604 | 1,215918 | 2,591105 | 0,009567 | 1 | 0 | Distal Intergenic | 7 | 36397642 | 36401538 | 3897 | 1 | 243931 | ENSMUST00000176413.2 | 117691 | ENSMUSG00000021217 | Tshz3 | teashirt zinc finger family member 3 |
| chr1 | 74104338 | 74105655 | 1318 | * | 13,1358 | 3,151414487 | 1,051902 | 2,99592 | 0,002736 | 0,737047122 | 0,918255 | Intron (ENSMUST0000( | 1 | 73949391 | 74130510 | 181120 | 2 | 21961 | ENSMUST00000187584.7 | 24855 | ENSMUSG00000055322 | Tns1 | tensin 1 |
| chr4 | 1,16E+08 | 1,16E+08 | 1426 | * | 8,419545 | 3,153215291 | 1,165625 | 2,705172 | 0,006827 | 0,822789084 | 1,425929 | Intron (ENSMUST0000( | 4 | 1,16E+08 | 1,16E+08 | 6892 | 2 | 72181 | ENSMUST00000136655.3 | 7027 | ENSMUSG00000028706 | Nsun4 | NOL1/NOP2/Sun domain family, member 4 |
| chr11 | 96150379 | 96151788 | 1410 | * | 15,83711 | 3,153647694 | 1,128377 | 2,794853 | 0,005192 | 0,669586355 | 2,663619 | Promoter (<=1kb) | 11 | 96152436 | 96155694 | 3259 | 1 | 193022 | ENSMUST00000143887.2 | -648 | | Gm53 | predicted gene 53 |
| chr5 | 1,24E+08 | 1,24E+08 | 1295 | * | 11,04237 | 3,153956105 | 1,102815 | 2,859915 | 0,004238 | 0,81655291 | 0,918255 | Promoter (2-3kb) | 5 | 1,24E+08 | 1,24E+08 | 5957 | 1 | 208908 | ENSMUST00000166983.3 | 2967 | ENSMUSG00000061882 | Ccdc62 | coiled-coil domain containing 62 |
| chr17 | 51406777 | 51407803 | 1027 | * | 8,995151 | 3,154071217 | 1,108229 | 2,846046 | 0,004427 | 0,751210692 | 1,360622 | Intron (ENSMUST0000( | 17 | 51040160 | 51466406 | 426247 | 2 | 72238 | ENSMUST00000224977.2 | 58603 | ENSMUSG00000023923 | Tbc1d5 | TBC1 domain family, member 5 |
| chr6 | 50623175 | 50624105 | 931 | * | 12,37052 | 3,156951997 | 1,117215 | 2,825734 | 0,004717 | 0,791416391 | 1,173325 | Distal Intergenic | 6 | 50627652 | 50631419 | 3768 | 2 | 60531 | ENSMUST00000031853.8 | 7314 | ENSMUSG00000029831 | Npvf | neuropeptide VF precursor |
| chr12 | 86031802 | 86033507 | 1706 | * | 14,37755 | 3,157161738 | 0,896486 | 3,521708 | 0,000429 | 0,353999513 | 1,2763 | Intron (ENSMUST0000( | 12 | 86008775 | 86100296 | 91522 | 1 | 320244 | ENSMUST00000220906.2 | 23027 | ENSMUSG00000012609 | Ttll5 | tubulin tyrosine ligase-like family, member 5 |
| chr6 | 21984905 | 21986520 | 1616 | * | 20,12367 | 3,157760607 | 0,834527 | 3,783891 | 0,000154 | 0,161408969 | 2,221611 | Promoter (<=1kb) | 6 | 21985915 | 22256403 | 270489 | 1 | 214642 | ENSMUST00000115383.9 | 0 | ENSMUSG00000062980 | Cped1 | cadherin-like and PC-esterase domain containing 1 |
| chr5 | 31160245 | 31162120 | 1876 | * | 12,67968 | 3,157770159 | 1,000771 | 3,155338 | 0,001603 | 0,623120685 | 1,075735 | Distal Intergenic | 5 | 31194231 | 31195338 | 1108 | 2 | 330064 | ENSMUST00000201017.2 | 33218 | ENSMUSG00000006641 | Slc5a6 | solute carrier family 5 (sodium-dependent vitamin transporter), member 6 |
| chr12 | 84281568 | 84283695 | 2128 | * | 19,07311 | 3,157988839 | 0,838756 | 3,765088 | 0,000166 | 0,161606382 | 2,386008 | Distal Intergenic | 12 | 84199233 | 84265655 | 66423 | 2 | 238317 | ENSMUST00000220974.2 | -15913 | ENSMUSG00000042507 | Mideas | mitotic deacetylase associated SANT domain protein |
| chr11 | 50183030 | 50184614 | 1585 | * | 36,942721 | 3,158471631 | 0,5493616 | 5,7493488 | 8,959E-09 | 9,4785E-05 | 2,9254973 | Promoter (<=1kb) | 11 | 50146461 | 50183138 | 36678 | 2 | 103806 | ENSMUST00000059458.5 | 0 | ENSMUSG00000050567 | Maml1 | mastermind like transcriptional coactivator 1 |
| chr1 | 58931607 | 58932729 | 1123 | * | 10,72339 | 3,159055498 | 1,149367 | 2,748519 | 0,005987 | 0,833321855 | 1,173325 | Promoter (<=1kb) | 1 | 58889244 | 58931951 | 42708 | 2 | 1,01E+08 | ENSMUST00000188624.2 | 0 | | Gm28802 | predicted gene 28802 |
| chr14 | 84227612 | 84228974 | 1363 | * | 10,45743 | 3,159065229 | 1,043411 | 3,027632 | 0,002465 | 0,668684091 | 1,2763 | Distal Intergenic | 14 | 84283871 | 84312730 | 28860 | 2 | 1,03E+08 | ENSMUST00000228897.2 | 83756 | ENSMUSG00000115043 | Gm32455 | predicted gene, 32455 |
| chr17 | 6244898 | 6246457 | 1560 | * | 12,40988 | 3,1596388 | 1,144017 | 2,761882 | 0,005747 | 0,840850009 | 1,075735 | Intron (ENSMUST0000( | 17 | 6282927 | 6301403 | 18477 | 1 | 68842 | ENSMUST00000137063.2 | -36470 | ENSMUSG00000034377 | Tulp4 | tubby like protein 4 |
| chr6 | 65722488 | 65723362 | 875 | * | 10,48956 | 3,160282238 | 1,159785 | 2,724887 | 0,006432 | 0,922063891 | 0,738441 | Distal Intergenic | 6 | 65755972 | 65913360 | 157389 | 1 | 70779 | ENSMUST00000031976.14 | -32610 | ENSMUSG00000029913 | Prdm5 | PR domain containing 5 |
| chr6 | 39405691 | 39406777 | 1087 | * | 13,27424 | 3,160408798 | 1,125983 | 2,8068 | 0,005004 | 0,840073741 | 0,938682 | Distal Intergenic | 6 | 39399223 | 39399353 | 131 | 2 | 1,15E+08 | ENSMUST00000082966.3 | -6338 | ENSMUSG00000064900 | Gm25402 | predicted gene, 25402 |
| chr2 | 25436668 | 25437426 | 759 | * | 15,02927 | 3,160629724 | 1,041751 | 3,033958 | 0,002414 | 0,64586006 | 1,41696 | Promoter (<=1kb) | 2 | 25407994 | 25436952 | 28959 | 2 | 22030 | ENSMUST00000028311.13 | 0 | ENSMUSG00000026942 | Traf2 | TNF receptor-associated factor 2 |
| chr11 | 49729074 | 49731038 | 1965 | * | 15,33483 | 3,160949963 | 0,89148 | 3,545734 | 0,000392 | 0,347797317 | 1,214904 | 3' UTR | 11 | 49737578 | 49777248 | 39671 | 1 | 26420 | ENSMUST00000109179.9 | -6540 | ENSMUSG00000020366 | Mapk9 | mitogen-activated protein kinase 9 |
| chr12 | 36060121 | 36061294 | 1174 | * | 10,37764 | 3,162251011 | 1,106921 | 2,8568 | 0,004279 | 0,85473227 | 0,738441 | Distal Intergenic | 12 | 36042906 | 36054086 | 11181 | 1 | 23795 | ENSMUST00000020898.12 | 17215 | ENSMUSG00000020581 | Agr2 | anterior gradient 2 |
| chr3 | 99976745 | 99977899 | 1155 | * | 10,23373 | 3,162532933 | 1,133451 | 2,790179 | 0,005268 | 0,810751536 | 1,173325 | Intron (ENSMUST0000( | 3 | 99934924 | 99946267 | 11344 | 1 | 74362 | ENSMUST00000143050.2 | 41821 | ENSMUSG00000027867 | Spag17 | sperm associated antigen 17 |

| | | | | | | | | | | | | | | | | | | | | | | |
|---|---|---|---|---|---|---|---|---|---|---|---|---|---|---|---|---|---|---|---|---|---|---|
| chr11 | 31771327 | 31773058 | 1732 | * | 15,42236 | 3,162914273 | 1,100798 | 2,873291 | 0,004062 | 0,730469267 | 1,41696 | Promoter (1-2kb) | 11 | 31768110 | 31774473 | 6364 | 2 | 414116 | ENSMUST00000126265.2 | 1415 | ENSMUSG00000085772 | D630024D03Rik | RIKEN cDNA D630024D03 gene |
| chr10 | 79872610 | 79874177 | 1568 | * | 12,37911 | 3,163160549 | 1,1518 | 2,746275 | 0,006028 | 0,877892722 | 0,918255 | Promoter (<=1kb) | 10 | 79871783 | 79873019 | 1237 | 2 | 66420 | ENSMUST00000147507.8 | 0 | ENSMUSG00000004667 | Polr2e | polymerase (RNA) II (DNA directed) polypeptide E |
| chr5 | 1,15E+08 | 1,15E+08 | 1438 | * | 9,432188 | 3,163756555 | 1,13833 | 2,779296 | 0,005448 | 0,811407117 | 1,211315 | Promoter (1-2kb) | 5 | 1,15E+08 | 1,15E+08 | 1007 | 2 | 330173 | ENSMUST00000100850.6 | 1133 | ENSMUSG00000072694 | 2610524H06Rik | RIKEN cDNA 2610524H06 gene |
| chr19 | 24690895 | 24692789 | 1895 | * | 13,5309 | 3,164372877 | 0,97135 | 3,257705 | 0,001123 | 0,627067801 | 0,738441 | Intron (ENSMUST0000 | 19 | 24651372 | 24659597 | 8226 | 1 | 226040 | ENSMUST00000057243.6 | 39523 | ENSMUSG00000048572 | Tmem252 | transmembrane protein 252 |
| chr9 | 121613639 | 121615761 | 2123 | * | 31,205874 | 3,164666189 | 0,7221147 | 4,382498 | 1,173E-05 | 0,024692459 | 2,8800171 | Exon (ENSMUST000000 | 9 | 121618938 | 121620152 | 1215 | 2 | 74770 | ENSMUST00000215910.2 | 4391 | ENSMUSG00000032523 | Hhatl | hedgehog acyltransferase-like |
| chr13 | 45420237 | 45421148 | 912 | * | 9,040225 | 3,164699425 | 1,169258 | 2,706587 | 0,006798 | 0,821456993 | 1,425929 | Promoter (<=1kb) | 13 | 45400108 | 45421187 | 21080 | 2 | 1,03E+08 | ENSMUST00000222218.2 | 39 | ENSMUSG00000113919 | Gm34466 | predicted gene, 34466 |
| chr1 | 74044961 | 74046412 | 1452 | * | 11,37033 | 3,164727708 | 1,071144 | 2,95453 | 0,003131 | 0,718085364 | 1,173325 | Intron (ENSMUST0000 | 1 | 74031364 | 74040726 | 9363 | 2 | 21961 | ENSMUST00000190389.7 | -4235 | ENSMUSG00000055322 | Tns1 | tensin 1 |
| chrX | 37208600 | 37210521 | 1922 | * | 11,87171 | 3,164906571 | 1,07934 | 2,932261 | 0,003365 | 0,813584786 | 0,738441 | Promoter (<=1kb) | 20 | 37210096 | 37218844 | 8749 | 1 | 73614 | ENSMUST00000152730.4 | 0 | ENSMUSG00000050197 | Rhox13 | reproductive homeobox 13 |
| chr5 | 1,04E+08 | 1,04E+08 | 1794 | * | 13,69714 | 3,165123722 | 1,157847 | 2,733629 | 0,006264 | 0,885309127 | 0,918255 | Distal Intergenic | 5 | 1,04E+08 | 1,04E+08 | 34623 | 2 | 13602 | ENSMUST00000031249.8 | -18929 | ENSMUSG00000029309 | Sparcl1 | SPARC-like 1 |
| chr5 | 1,25E+08 | 1,25E+08 | 903 | * | 10,50849 | 3,165346796 | 1,169198 | 2,707279 | 0,006784 | 0,854052567 | 1,173325 | Distal Intergenic | 5 | 1,25E+08 | 1,25E+08 | 34502 | 2 | 1,03E+08 | ENSMUST00000198586.2 | 34724 | | Gm32585 | predicted gene, 32585 |
| chr2 | 26125783 | 26127128 | 1346 | * | 12,88396 | 3,16571368 | 1,038102 | 3,049521 | 0,002292 | 0,745196503 | 0,738441 | Promoter (<=1kb) | 2 | 26099137 | 26127537 | 28401 | 2 | 227638 | ENSMUST00000036187.9 | 409 | ENSMUSG00000036327 | Qsox2 | quiescin Q6 sulfhydryl oxidase 2 |
| chr11 | 76293380 | 76294376 | 997 | * | 9,093483 | 3,165861763 | 1,219397 | 2,596252 | 0,009425 | 0,876338958 | 1,449706 | Distal Intergenic | 11 | 76297778 | 76305155 | 7378 | 1 | 56322 | ENSMUST00000120699.8 | -3402 | ENSMUSG00000020843 | Timm22 | translocase of inner mitochondrial membrane 22 |
| chr12 | 24942201 | 24943345 | 1145 | * | 8,251445 | 3,166264542 | 1,225768 | 2,583086 | 0,009792 | 0,891333894 | 1,360622 | Intron (ENSMUST0000 | 12 | 24881630 | 25010285 | 128656 | 1 | 67216 | ENSMUST00000222994.2 | 60571 | ENSMUSG00000020646 | Mboat2 | membrane bound O-acyltransferase domain containing 2 |
| chr3 | 98981349 | 98982435 | 1087 | * | 11,64259 | 3,166384115 | 1,12884 | 2,804989 | 0,005032 | 0,846832214 | 0,918255 | Distal Intergenic | 3 | 98957138 | 98961060 | 3923 | 1 | 1,03E+08 | ENSMUST00000155224.3 | 24211 | ENSMUSG00000086833 | Gm12440 | predicted gene 12440 |
| chr18 | 35090356 | 35092600 | 2245 | * | 24,9626 | 3,166427768 | 0,7147822 | 4,4299196 | 9,427E-06 | 0,020676456 | 2,9519335 | Promoter (2-3kb) | 18 | 35070467 | 35087410 | 16944 | 2 | 15526 | ENSMUST00000025217.11 | -2946 | ENSMUSG00000024359 | Hspa9 | heat shock protein 9 |
| chr5 | 65909513 | 65910783 | 1271 | * | 10,20656 | 3,166494121 | 1,153665 | 2,744726 | 0,006056 | 0,850729797 | 1,086118 | Distal Intergenic | 5 | 65920864 | 65987451 | 66588 | 1 | 333789 | ENSMUST00000201489.4 | -10081 | ENSMUSG00000037795 | N4bp2 | NEDD4 binding protein 2 |
| chr12 | 77893850 | 77894654 | 805 | * | 8,804412 | 3,167930786 | 1,1694 | 2,709022 | 0,006748 | 0,817544002 | 1,449706 | Distal Intergenic | 12 | 77861454 | 77861583 | 130 | 2 | 1,15E+08 | ENSMUST00000104519.3 | -32267 | ENSMUSG00000077707 | Gm24070 | predicted gene, 24070 |
| chr3 | 38330808 | 38332344 | 1537 | * | 15,07005 | 3,16808658 | 1,038188 | 3,051553 | 0,002277 | 0,653981288 | 1,2763 | Distal Intergenic | 3 | 38257300 | 38262268 | 4969 | 2 | 71342 | ENSMUST00000197658.2 | -68540 | ENSMUSG00000106002 | 5430434I15Rik | RIKEN cDNA 5430434I15 gene |
| chr5 | 1,23E+08 | 1,23E+08 | 1572 | * | 10,57002 | 3,168225485 | 1,167474 | 2,713744 | 0,006653 | 0,852340527 | 1,173325 | Promoter (<=1kb) | 5 | 1,23E+08 | 1,23E+08 | 5228 | 1 | 56378 | ENSMUST00000141395.2 | 972 | ENSMUSG00000029465 | Arpc3 | actin related protein 2/3 complex, subunit 3 |
| chr5 | 99040103 | 99041223 | 1121 | * | 7,197985 | 3,168577244 | 1,217637 | 2,602236 | 0,009262 | 1 | 0 | Distal Intergenic | 5 | 99002310 | 99032255 | 29946 | 1 | 110075 | ENSMUST00000200388.2 | 37793 | ENSMUSG00000029335 | Bmp3 | bone morphogenetic protein 3 |
| chr14 | 67337806 | 67339378 | 1573 | * | 22,939914 | 3,169013851 | 0,6873131 | 4,6107281 | 4,013E-06 | 0,010675515 | 2,9510621 | Distal Intergenic | 14 | 67251505 | 67309893 | 58389 | 2 | 71978 | ENSMUST00000089230.7 | -27913 | ENSMUSG00000022052 | Ppp2r2a | protein phosphatase 2, regulatory subunit B, alpha |
| chr3 | 1,11E+08 | 1,11E+08 | 1081 | * | 9,563604 | 3,170396419 | 1,150621 | 2,755379 | 0,005862 | 0,829907213 | 1,173325 | Distal Intergenic | 3 | 1,1E+08 | 1,1E+08 | 105 | 2 | 1,15E+08 | ENSMUST00000104065.3 | -401211 | ENSMUSG00000119642 | Gm26076 | predicted gene, 26076 |
| chr14 | 56114326 | 56115481 | 1156 | * | 10,08349 | 3,170476419 | 1,17577 | 2,696511 | 0,007007 | 0,8237677 | 1,449706 | Distal Intergenic | 14 | 56119518 | 56121689 | 2172 | 2 | 56410 | ENSMUST00000172378.2 | 6208 | ENSMUSG00000040380 | Cbln3 | cerebellin 3 precursor protein |
| chr3 | 1,16E+08 | 1,16E+08 | 1018 | * | 11,95741 | 3,170581219 | 1,073855 | 2,952522 | 0,003152 | 0,705941691 | 1,2763 | Intron (ENSMUST0000 | 3 | 1,16E+08 | 1,16E+08 | 100440 | 2 | 229776 | ENSMUST00000197830.5 | 83363 | ENSMUSG00000033502 | Cdc14a | CDC14 cell division cycle 14A |
| chr15 | 12086845 | 12088219 | 1375 | * | 9,742801 | 3,171342746 | 1,043942 | 3,037854 | 0,002383 | 0,684274806 | 1,086118 | Promoter (1-2kb) | 15 | 12068334 | 12085283 | 16950 | 2 | 1,03E+08 | ENSMUST00000227908.2 | -1562 | | Gm34759 | predicted gene, 34759 |
| chr11 | 60425826 | 60426804 | 979 | * | 9,597065 | 3,171754862 | 1,218768 | 2,602427 | 0,009257 | 0,901729831 | 1,211315 | Promoter (<=1kb) | 11 | 60427207 | 60445193 | 17987 | 1 | 268420 | ENSMUST00000134770.2 | -403 | ENSMUSG00000042650 | Alkbh5 | alkB homolog 5, RNA demethylase |
| chr16 | 39990686 | 39991972 | 1287 | * | 13,15008 | 3,172263773 | 0,939547 | 3,376376 | 0,000734 | 0,463890057 | 1,173325 | Intron (ENSMUST0000 | 16 | 39804957 | 40478248 | 673292 | 1 | 268890 | ENSMUST00000189829.2 | 185729 | ENSMUSG00000061080 | Lsamp | limbic system-associated membrane protein |
| chr8 | 1,24E+08 | 1,24E+08 | 893 | * | 11,47122 | 3,172327491 | 1,075431 | 2,949818 | 0,00318 | 0,764024179 | 0,918255 | Promoter (2-3kb) | 8 | 1,24E+08 | 1,24E+08 | 42303 | 1 | 257633 | ENSMUST00000212790.2 | 2967 | ENSMUSG00000015016 | Acsf3 | acyl-CoA synthetase family member 3 |
| chr5 | 1,23E+08 | 1,23E+08 | 945 | * | 7,897579 | 3,173479728 | 1,210735 | 2,621118 | 0,008764 | 1 | 0 | Distal Intergenic | 5 | 1,23E+08 | 1,23E+08 | 48268 | 1 | 330189 | ENSMUST00000161660.8 | -4215 | ENSMUSG00000054434 | Tmem120b | transmembrane protein 120B |
| chr6 | 1,34E+08 | 1,34E+08 | 1410 | * | 12,90141 | 3,173863273 | 0,958906 | 3,309879 | 0,000933 | 0,544061719 | 0,918255 | 5' UTR | 6 | 1,34E+08 | 1,34E+08 | 28420 | 1 | 14011 | ENSMUST00000169529.2 | -10238 | ENSMUSG00000030199 | Etv6 | ets variant 6 |
| chr3 | 94854542 | 94857093 | 2552 | * | 18,36754 | 3,174055329 | 0,910117 | 3,487525 | 0,000488 | 0,284660899 | 2,423594 | Distal Intergenic | 3 | 94861386 | 94866807 | 5422 | 1 | 53970 | ENSMUST00000107260.9 | -4293 | ENSMUSG00000005774 | Rfx5 | regulatory factor X, 5 (influences HLA class II expression) |
| chr7 | 1,27E+08 | 1,27E+08 | 2399 | * | 22,29643 | 3,174381707 | 0,795405 | 3,990899 | 6,58E-05 | 0,080985792 | 2,951933 | Promoter (1-2kb) | 7 | 1,27E+08 | 1,27E+08 | 9654 | 2 | 244219 | ENSMUST00000106261.8 | 1847 | ENSMUSG00000049728 | Zfp668 | zinc finger protein 668 |
| chr6 | 38325749 | 38327158 | 1410 | * | 18,34155 | 3,175352769 | 0,882463 | 3,598283 | 0,00032 | 0,221323359 | 2,663619 | Intron (ENSMUST0000 | 6 | 38282221 | 38331208 | 48988 | 2 | 78781 | ENSMUST00000114900.8 | 4050 | ENSMUSG00000029826 | Zc3hav1 | zinc finger CCCH type, antiviral 1 |
| chr5 | 21937278 | 21939321 | 2044 | * | 16,31947 | 3,175639694 | 0,909154 | 3,49296 | 0,000478 | 0,284660899 | 2,386008 | Promoter (2-3kb) | 5 | 21942139 | 21962150 | 20012 | 1 | 73078 | ENSMUST00000030882.12 | -2818 | ENSMUSG00000029017 | Pmpcb | peptidase (mitochondrial processing) beta |
| chr1 | 71265281 | 71266682 | 1402 | * | 9,407515 | 3,175673823 | 1,126526 | 2,818997 | 0,004817 | 0,79031971 | 1,211315 | Distal Intergenic | 1 | 71329172 | 71346246 | 17075 | 2 | 74591 | ENSMUST00000188752.2 | 79564 | ENSMUSG00000050296 | Abca12 | ATP-binding cassette, sub-family A (ABC1), member 12 |
| chr15 | 99566540 | 99568526 | 1987 | * | 23,96222 | 3,175878856 | 0,888367 | 3,574963 | 0,00035 | 0,219863416 | 2,951933 | Promoter (<=1kb) | 15 | 99568249 | 99599009 | 30761 | 1 | 11419 | ENSMUST00000023758.9 | 0 | ENSMUSG00000023017 | Asic1 | acid-sensing (proton-gated) ion channel 1 |
| chr4 | 38702079 | 38703205 | 1127 | * | 11,26969 | 3,175885628 | 1,023957 | 3,10158 | 0,001925 | 0,71419937 | 0,744375 | Intron (ENSMUST0000 | 4 | 38645427 | 38855463 | 210037 | 2 | 1,03E+08 | ENSMUST00000244574.1 | 152258 | ENSMUSG00000085632 | Gm12381 | predicted gene 12381 |
| chr15 | 13461502 | 13462361 | 860 | * | 9,866706 | 3,176330422 | 1,080959 | 2,938437 | 0,003299 | 0,740451144 | 1,086118 | Distal Intergenic | 15 | 13059337 | 13173761 | 114425 | 2 | 12563 | ENSMUST00000226917.2 | -287741 | ENSMUSG00000039385 | Cdh6 | cadherin 6 |
| chr5 | 31447205 | 31449403 | 2199 | * | 23,587115 | 3,177506338 | 0,7365656 | 4,3139489 | 1,604E-05 | 0,030165351 | 2,9510621 | Promoter (<=1kb) | 5 | 31410621 | 31448451 | 37831 | 2 | 67661 | ENSMUST00000041565.11 | 0 | ENSMUSG00000038564 | Ift172 | intraflagellar transport 172 |
| chr9 | 1,18E+08 | 1,18E+08 | 1562 | * | 14,98155 | 3,178024575 | 0,887284 | 3,581745 | 0,000341 | 0,310253846 | 1,37683 | Intron (ENSMUST0000 | 9 | 1,16E+08 | 1,17E+08 | 1051889 | 2 | 207181 | ENSMUST00000164018.9 | -53126 | ENSMUSG00000039607 | Rbms3 | RNA binding motif, single stranded interacting protein |
| chr18 | 68099305 | 68100207 | 903 | * | 10,01364 | 3,178501815 | 1,16753 | 2,722415 | 0,006481 | 0,809718961 | 1,449706 | Intron (ENSMUST0000 | 18 | 68066328 | 68388622 | 322295 | 1 | 52662 | ENSMUST00000063775.5 | 32977 | ENSMUSG00000024544 | Ldlrad4 | low density lipoprotein receptor class A domain containing 4 |
| chr16 | 33096176 | 33097168 | 993 | * | 9,410468 | 3,178877203 | 1,227297 | 2,590146 | 0,009594 | 0,879615137 | 1,449706 | Intron (ENSMUST0000 | 16 | 33088128 | 33090215 | 2088 | 1 | 69150 | ENSMUST00000231242.2 | 8048 | ENSMUSG00000022808 | Snx4 | sorting nexin 4 |
| chr9 | 31946889 | 31948116 | 1228 | * | 11,33595 | 3,179584962 | 1,196227 | 2,658012 | 0,00786 | 0,881706354 | 1,173325 | Distal Intergenic | 9 | 31858314 | 31916756 | 58443 | 2 | 1,19E+08 | ENSMUST00000216368.2 | -30133 | | Gm47465 | predicted gene, 47465 |
| chr8 | 1,08E+08 | 1,08E+08 | 3405 | * | 22,16729 | 3,179714025 | 0,809565 | 3,927681 | 8,58E-05 | 0,094915397 | 2,880017 | Distal Intergenic | 8 | 1,08E+08 | 1,08E+08 | 83965 | 1 | 20650 | ENSMUST00000212524.2 | -3444 | ENSMUSG00000041308 | Sntb2 | syntrophin, basic 2 |
| chr7 | 1,39E+08 | 1,39E+08 | 977 | * | 10,11595 | 3,179926208 | 1,156584 | 2,749412 | 0,00597 | 0,832960417 | 1,173325 | Promoter (1-2kb) | 7 | 1,39E+08 | 1,39E+08 | 1571 | 2 | 14912 | ENSMUST00000106095.3 | 1683 | ENSMUSG00000041309 | Nkx6-2 | NK6 homeobox 2 |
| chr4 | 1,4E+08 | 1,4E+08 | 1369 | * | 11,11821 | 3,18027085 | 1,02804 | 3,093529 | 0,001978 | 0,718444962 | 0,738441 | Intron (ENSMUST0000 | 4 | 1,4E+08 | 1,4E+08 | 11176 | 2 | 72754 | ENSMUST00000125045.2 | -3518 | ENSMUSG00000040964 | Arhgef10l | Rho guanine nucleotide exchange factor (GEF) 10-like |
| chr5 | 77304633 | 77307283 | 2651 | * | 21,17978 | 3,180318665 | 0,7671 | 4,145898 | 3,38E-05 | 0,051012164 | 3,104559 | Distal Intergenic | 5 | 77264657 | 77270195 | 5539 | 2 | 1,05E+08 | ENSMUST00000156195.2 | -34438 | | Gm15831 | predicted gene 15831 |
| chr9 | 22262320 | 22263089 | 770 | * | 9,49348 | 3,180738752 | 1,097131 | 2,899141 | 0,003742 | 0,715249603 | 1,425929 | Promoter (<=1kb) | 9 | 22245449 | 22262552 | 17104 | 2 | 68743 | ENSMUST00000217010.2 | 0 | ENSMUSG00000036777 | Anln | anillin, actin binding protein |
| chr16 | 84968661 | 84970397 | 1737 | * | 12,6469 | 3,181437655 | 1,027407 | 3,096569 | 0,001958 | 0,716908183 | 0,738441 | Promoter (<=1kb) | 16 | 84791592 | 84970592 | 179001 | 2 | 11820 | ENSMUST00000227990.2 | 195 | ENSMUSG00000022892 | App | amyloid beta (A4) precursor protein |
| chr3 | 37367406 | 37368678 | 1273 | * | 13,17838 | 3,181620925 | 1,082706 | 2,938582 | 0,003297 | 0,770673161 | 0,918255 | Promoter (<=1kb) | 3 | 37363494 | 37366857 | 3364 | 2 | 207175 | ENSMUST00000125252.3 | -549 | ENSMUSG00000045031 | Cetn4 | centrin 4 |
| chr17 | 24388146 | 24390123 | 1978 | * | 20,14658 | 3,181654558 | 0,859984 | 3,699667 | 0,000216 | 0,188932071 | 2,386008 | Promoter (<=1kb) | 17 | 24382840 | 24388435 | 5596 | 2 | 11984 | ENSMUST00000024932.12 | 0 | ENSMUSG00000024121 | Atp6v0c | ATPase, H+ transporting, lysosomal V0 subunit C |
| chr19 | 38288891 | 38290307 | 1417 | * | 10,05554 | 3,181714943 | 1,161199 | 2,740026 | 0,006143 | 0,799514271 | 1,449706 | 3' UTR | 19 | 38288531 | 38303718 | 15188 | 2 | 1,05E+08 | ENSMUST00000200561.2 | 13411 | | Gm9886 | predicted gene 9886 |
| chr8 | 46763590 | 46764668 | 1079 | * | 9,859157 | 3,181878708 | 1,20583 | 2,638745 | 0,008321 | 0,853297048 | 1,449706 | Intron (ENSMUST0000 | 8 | 46752168 | 46778711 | 26544 | 1 | 1,05E+08 | ENSMUST00000210181.2 | 11422 | ENSMUSG00000110179 | Gm39168 | predicted gene, 39168 |
| chrX | 1,42E+08 | 1,42E+08 | 1806 | * | 13,37675 | 3,182242817 | 1,029585 | 3,090801 | 0,001996 | 0,661059054 | 1,075735 | Intron (ENSMUST0000 | 20 | 1,42E+08 | 1,42E+08 | 115583 | 2 | 56068 | ENSMUST00000041317.3 | 67793 | ENSMUSG00000042225 | Ammecr1 | Alport syndrome, mental retardation, midface hypoplasia and elliptocytosis chromosomal region gene 1 |
| chr5 | 22818865 | 22819988 | 1124 | * | 9,881717 | 3,182440033 | 1,086313 | 2,929579 | 0,003394 | 0,727652793 | 1,211315 | Intron (ENSMUST0000 | 5 | 22850117 | 22850181 | 65 | 2 | 1,15E+08 | ENSMUST00000174963.3 | 30193 | ENSMUSG00000092704 | Gm24512 | predicted gene, 24512 |
| chr8 | 94190874 | 94191876 | 1003 | * | 7,531024 | 3,182966849 | 1,21765 | 2,614025 | 0,008948 | 1 | 0 | Distal Intergenic | 8 | 94142165 | 94154256 | 12092 | 2 | 1,03E+08 | ENSMUST00000212816.2 | -36618 | | Gm30132 | predicted gene, 30132 |
| chr4 | 58393203 | 58394776 | 1574 | * | 16,9049 | 3,183327877 | 0,869841 | 3,659666 | 0,000253 | 0,205744841 | 2,386008 | Distal Intergenic | 4 | 58441012 | 58441098 | 87 | 2 | 1,01E+08 | ENSMUST00000175552.2 | 46322 | ENSMUSG00000093293 | Mir3095 | microRNA 3095 |
| chr2 | 1,26E+08 | 1,26E+08 | 849 | * | 9,762013 | 3,183480419 | 1,234907 | 2,577911 | 0,00994 | 0,886332554 | 1,449706 | Promoter (1-2kb) | 2 | 1,26E+08 | 1,26E+08 | 120015 | 2 | 241633 | ENSMUST00000134977.8 | 1457 | ENSMUSG00000060131 | Atp8b4 | ATPase, class I, type 8B, member 4 |
| chr2 | 75541495 | 75543237 | 1743 | * | 13,86292 | 3,184048297 | 1,052663 | 3,024757 | 0,002488 | 0,756849896 | 0,738441 | Intron (ENSMUST0000 | 2 | 75505857 | 75534985 | 29129 | 2 | 18024 | ENSMUST00000102672.5 | -6510 | ENSMUSG00000015839 | Nfe2l2 | nuclear factor, erythroid derived 2, like 2 |
| chr1 | 1,53E+08 | 1,53E+08 | 1646 | * | 13,1545 | 3,1842737 | 0,971993 | 3,276025 | 0,001053 | 0,540558173 | 1,075735 | Exon (ENSMUST00000 | 1 | 1,52E+08 | 1,53E+08 | 108267 | 2 | 19731 | ENSMUST00000027760.14 | -59832 | ENSMUSG00000026482 | Rgl1 | ral guanine nucleotide dissociation stimulator,-like 1 |
| chr19 | 56173747 | 56174839 | 1093 | * | 9,831409 | 3,184720424 | 1,124508 | 2,832101 | 0,004624 | 0,80080207 | 1,086118 | Distal Intergenic | 19 | 56094742 | 56099437 | 4696 | 1 | 1,03E+08 | ENSMUST00000238174.2 | 79005 | | Gm31912 | predicted gene, 31912 |
| chr5 | 1,15E+08 | 1,15E+08 | 1668 | * | 14,53616 | 3,184804318 | 0,893493 | 3,564441 | 0,000365 | 0,264977345 | 2,141612 | Intron (ENSMUST0000 | 5 | 1,15E+08 | 1,15E+08 | 22088 | 2 | 21405 | ENSMUST00000031535.12 | -11587 | ENSMUSG00000029556 | Hnf1a | HNF1 homeobox A |
| chr10 | 80870208 | 80871661 | 1454 | * | 15,58011 | 3,184937242 | 0,998748 | 3,188929 | 0,001428 | 0,542967057 | 1,41696 | Promoter (<=1kb) | 10 | 80864375 | 80869746 | 5372 | 2 | 106947 | ENSMUST00000168076.2 | -462 | ENSMUSG00000046822 | Slc39a3 | solute carrier family 39 (zinc transporter), member 3 |
| chr16 | 8956233 | 8957364 | 1132 | * | 11,58077 | 3,185348823 | 1,17277 | 2,71609 | 0,006606 | 0,851726441 | 1,173325 | Distal Intergenic | 16 | 8867836 | 8873262 | 5427 | 1 | 1,03E+08 | ENSMUST00000229201.2 | 88397 | | Gm30567 | predicted gene, 30567 |
| chr6 | 1,25E+08 | 1,25E+08 | 1653 | * | 16,83611 | 3,185799922 | 0,959792 | 3,319262 | 0,000903 | 0,483103348 | 1,2763 | Intron (ENSMUST0000 | 6 | 1,25E+08 | 1,25E+08 | 6340 | 2 | 69202 | ENSMUST00000204671.3 | -3259 | ENSMUSG00000030122 | Ptms | parathymosin |
| chr8 | 4725853 | 4729377 | 3525 | * | 50,417399 | 3,186416139 | 0,5834564 | 5,4612755 | 4,727E-08 | 0,000356271 | 2,9254973 | Promoter (<=1kb) | 8 | 4728431 | 4738497 | 10067 | 1 | 623223 | ENSMUST00000181337.3 | 0 | | Gm6410 | predicted gene 6410 |
| chr19 | 55048088 | 55050314 | 2227 | * | 20,82241 | 3,187691568 | 0,865347 | 3,683713 | 0,00023 | 0,156945882 | 3,471678 | Distal Intergenic | 19 | 55058166 | 55087883 | 29718 | 2 | 14732 | ENSMUST00000061856.6 | 37569 | ENSMUSG00000024978 | Gpam | glycerol-3-phosphate acyltransferase, mitochondrial |
| chr1 | 1,69E+08 | 1,69E+08 | 2498 | * | 15,21667 | 3,188549315 | 1,052498 | 3,029506 | 0,00245 | 0,673689499 | 1,214904 | Distal Intergenic | 1 | 1,69E+08 | 1,69E+08 | 16286 | 2 | 66977 | ENSMUST00000195342.6 | 141298 | ENSMUSG00000026683 | Nuf2 | NUF2, NDC80 kinetochore complex component |
| chr9 | 58548485 | 58550726 | 2242 | * | 15,60971 | 3,188909223 | 1,169273 | 2,727259 | 0,006386 | 0,838625535 | 1,214904 | Intron (ENSMUST0000 | 9 | 58557482 | 58559340 | 1859 | 1 | 20320 | ENSMUST00000177021.2 | -6756 | ENSMUSG00000032336 | Nptn | neuroplastin |
| chr18 | 77039111 | 77040421 | 1311 | * | 11,75216 | 3,189419054 | 1,176527 | 2,710875 | 0,006711 | 0,838625535 | 1,2763 | Intron (ENSMUST0000 | 18 | 77032133 | 77060598 | 28466 | 1 | 76987 | ENSMUST00000148955.3 | 6978 | ENSMUSG00000025421 | Hdhd2 | haloacid dehalogenase-like hydrolase domain containing 2 |
| chr1 | 1,89E+08 | 1,89E+08 | 1103 | * | 8,972447 | 3,190026583 | 1,211449 | 2,633233 | 0,008458 | 0,85617852 | 1,449706 | Promoter (<=1kb) | 1 | 1,89E+08 | 1,89E+08 | 5193 | 2 | 16526 | ENSMUST00000193686.2 | 0 | ENSMUSG00000037624 | Kcnk2 | potassium channel, subfamily K, member 2 |
| chr12 | 84624925 | 84626145 | 1221 | * | 12,64103 | 3,190483081 | 1,097465 | 2,907138 | 0,003648 | 0,826676469 | 0,738441 | Intron (ENSMUST0000 | 12 | 84616614 | 84642231 | 25618 | 1 | 12677 | ENSMUST00000169934.4 | 8311 | ENSMUSG00000021239 | Vsx2 | visual system homeobox 2 |
| chr15 | 10819790 | 10821000 | 1211 | * | 12,94952 | 3,190563769 | 1,13234 | 2,817673 | 0,004837 | 0,838625535 | 0,918255 | Intron (ENSMUST0000 | 15 | 10814913 | 10835300 | 20388 | 1 | 1,01E+08 | ENSMUST00000228418.2 | 4877 | | Gm19276 | predicted gene, 19276 |
| chr2 | 55378519 | 55379551 | 1033 | * | 10,04177 | 3,191011998 | 1,048358 | 3,04382 | 0,002336 | 0,64068479 | 1,425929 | Intron (ENSMUST0000 | 2 | 55327177 | 55488152 | 160976 | 1 | 16519 | ENSMUST00000112632.2 | 51342 | ENSMUSG00000026824 | Kcnj3 | potassium inwardly-rectifying channel, subfamily J, member 3 |
| chr4 | 86573647 | 86574642 | 996 | * | 12,60103 | 3,191337734 | 0,95608 | 3,337939 | 0,000844 | 0,486196597 | 1,173325 | Promoter (2-3kb) | 4 | 86575144 | 86576881 | 1738 | 2 | 11520 | ENSMUST00000138605.2 | 2239 | ENSMUSG00000028494 | Plin2 | perilipin 2 |
| chr19 | 3898352 | 3899726 | 1375 | * | 14,44446 | 3,191371485 | 0,952587 | 3,350214 | 0,000807 | 0,446770792 | 1,41696 | Promoter (2-3kb) | 19 | 3901773 | 3944365 | 42593 | 1 | 12660 | ENSMUST00000128694.8 | -2047 | ENSMUSG00000024843 | Chka | choline kinase alpha |
| chr6 | 72101378 | 72102420 | 1043 | * | 11,81468 | 3,192004024 | 1,056414 | 3,021547 | 0,002515 | 0,757694328 | 0,744375 | Promoter (1-2kb) | 6 | 72097197 | 72100181 | 2985 | 2 | 71265 | ENSMUST00000205571.2 | -1197 | ENSMUSG00000108709 | 4933431G14Rik | RIKEN cDNA 4933431G14 gene |
| chr6 | 15843360 | 15844197 | 838 | * | 10,39069 | 3,192499775 | 1,237935 | 2,578892 | 0,009912 | 0,903470246 | 1,2763 | Distal Intergenic | 6 | 15727798 | 15770348 | 42551 | 1 | 16543 | ENSMUST00000128849.3 | 115562 | ENSMUSG00000041390 | Mdfic | MyoD family inhibitor domain containing |
| chr2 | 33222522 | 33223427 | 906 | * | 11,64435 | 3,192732994 | 1,141974 | 2,795802 | 0,005177 | 0,888042352 | 0,744375 | Intron (ENSMUST0000 | 2 | 33023429 | 33261441 | 238013 | 2 | 241308 | ENSMUST00000131298.7 | 38014 | ENSMUSG00000038831 | Ralgps1 | Ral GEF with PH domain and SH3 binding motif 1 |
| chr2 | 15676707 | 15677875 | 1169 | * | 9,185427 | 3,192784004 | 1,140747 | 2,798854 | 0,005128 | 0,767135014 | 1,449706 | Intron (ENSMUST0000 | 2 | 15531290 | 16260366 | 729077 | 1 | 1,03E+08 | ENSMUST00000146205.3 | 145417 | ENSMUSG00000075520 | Malrd1 | MAM and LDL receptor class A domain containing 1 |
| chrX | 1,26E+08 | 1,26E+08 | 948 | * | 7,954799 | 3,19328084 | 1,150311 | 2,776015 | 0,005503 | 1 | 0 | Distal Intergenic | 20 | 1,26E+08 | 1,26E+08 | 2037 | 2 | 1,19E+08 | ENSMUST00000117443.2 | -10113 | ENSMUSG00000082772 | LOC118568449 | vigilin-like |
| chr8 | 1,22E+08 | 1,22E+08 | 1331 | * | 25,18508 | 3,19445648 | 1,230859 | 2,595306 | 0,009451 | 0,749550311 | 2,951062 | Distal Intergenic | 8 | 1,22E+08 | 1,22E+08 | 6806 | 1 | 1,03E+08 | ENSMUST00000180597.3 | 23613 | | Gm26747 | predicted gene, 26747 |
| chr15 | 67505516 | 67506532 | 1017 | * | 11,09148 | 3,194561952 | 1,048922 | 3,045567 | 0,002322 | 0,711174281 | 0,918255 | Distal Intergenic | 15 | 67098618 | 67248943 | 150326 | 1 | 69334 | ENSMUST00000187233.2 | 406898 | ENSMUSG00000102069 | 1700012I11Rik | RIKEN cDNA 1700012I11 gene |
| chr9 | 66458671 | 66460287 | 1617 | * | 11,63369 | 3,194807992 | 1,034075 | 3,089531 | 0,002005 | 0,646275731 | 1,173325 | Intron (ENSMUST0000 | 9 | 66434258 | 66447633 | 13376 | 2 | 235441 | ENSMUST00000124694.2 | -11038 | ENSMUSG00000032376 | Usp3 | ubiquitin specific peptidase 3 |
| chr9 | 1,11E+08 | 1,11E+08 | 1207 | * | 9,677472 | 3,195343321 | 1,165208 | 2,742295 | 0,006101 | 0,837097674 | 1,173325 | Intron (ENSMUST0000 | 9 | 1,11E+08 | 1,11E+08 | 18569 | 1 | 791282 | ENSMUST00000197321.6 | -11428 | ENSMUSG00000057802 | Gm10030 | predicted gene 10030 |
| chr15 | 77202837 | 77203974 | 1138 | * | 14,529 | 3,19703125 | 1,005902 | 3,178274 | 0,001482 | 0,549632826 | 1,41696 | Distal Intergenic | 15 | 76997806 | 77191204 | 193399 | 2 | 93686 | ENSMUST00000228582.2 | -11633 | ENSMUSG00000033565 | Rbfox2 | RNA binding protein, fox-1 homolog (C. elegans) 2 |
| chr12 | 70644949 | 70646577 | 1629 | * | 17,72247 | 3,197167139 | 1,003243 | 3,186833 | 0,001438 | 0,474532331 | 2,154504 | Distal Intergenic | 12 | 70696719 | 70697578 | 860 | 1 | 1,03E+08 | ENSMUST00000243885.1 | -50142 | | Gm24474 | predicted gene, 24474 |
| chr13 | 28335467 | 28336642 | 1176 | * | 8,181835 | 3,197867492 | 1,199895 | 2,665122 | 0,007696 | 0,83918951 | 1,449706 | 3' UTR | 13 | 28326467 | 28335594 | 9128 | 1 | 28078 | ENSMUST00000017208.5 | 9000 | ENSMUSG00000017064 | Prl5a1 | prolactin family 5, subfamily a, member 1 |
| chr9 | 1,22E+08 | 1,22E+08 | 1666 | * | 16,96025 | 3,198056372 | 0,831449 | 3,846364 | 0,00012 | 0,14233436 | 2,141612 | Promoter (1-2kb) | 9 | 1,22E+08 | 1,22E+08 | 2593 | 1 | 26901 | ENSMUST00000035113.11 | -1449 | ENSMUSG00000032526 | Ss18l2 | SS18, nBAF chromatin remodeling complex subunit like 2 |
| chr7 | 97141660 | 97142593 | 934 | * | 10,2413 | 3,198416623 | 1,200371 | 2,664524 | 0,00771 | 0,872204014 | 1,211315 | Intron (ENSMUST0000 | 7 | 97150986 | 97157076 | 6091 | 1 | 101861 | ENSMUST00000205876.2 | -8393 | ENSMUSG00000025133 | Ints4 | integrator complex subunit 4 |
| chr7 | 28641850 | 28643035 | 1186 | * | 9,536603 | 3,199306756 | 1,223491 | 2,614899 | 0,008925 | 0,896435577 | 1,211315 | Exon (ENSMUST00000 | 7 | 28612803 | 28649184 | 36382 | 2 | 60595 | ENSMUST00000148196.3 | 6149 | ENSMUSG00000054808 | Actn4 | actinin alpha 4 |
| chr9 | 56370141 | 56372210 | 2070 | * | 17,05398 | 3,199365089 | 0,900839 | 3,551538 | 0,000383 | 0,337054778 | 1,2763 | Intron (ENSMUST0000 | 9 | 56326114 | 56383401 | 57288 | 1 | 66867 | ENSMUST00000214869.2 | 44027 | ENSMUSG00000032329 | Hmg20a | high mobility group 20A |
| chr13 | 93434900 | 93436538 | 1639 | * | 11,23396 | 3,200015601 | 1,226219 | 2,609662 | 0,009063 | 0,972947889 | 0,738441 | Promoter (<=1kb) | 13 | 93436143 | 93478420 | 42278 | 1 | 26556 | ENSMUST00000220609.2 | 0 | ENSMUSG00000007617 | Homer1 | homer scaffolding protein 1 |
| chr3 | 1,33E+08 | 1,33E+08 | 1872 | * | 14,08987 | 3,200907433 | 1,072183 | 2,985412 | 0,002832 | 0,744054256 | 0,918255 | Intron (ENSMUST0000 | 3 | 1,33E+08 | 1,33E+08 | 36395 | 2 | 1,01E+08 | ENSMUST00000197386.2 | 6113 | ENSMUSG00000106230 | Gm29811 | predicted gene, 29811 |
| chrX | 1,68E+08 | 1,68E+08 | 1194 | * | 16,53865 | 3,201643168 | 0,862363 | 3,712641 | 0,000205 | 0,18252995 | 2,386008 | Promoter (<=1kb) | 20 | 1,68E+08 | 1,68E+08 | 40474 | 1 | 1,01E+08 | ENSMUST00000156210.8 | 0 | ENSMUSG00000087159 | Gm15246 | predicted gene 15246 |
| chr14 | 23341248 | 23342367 | 1120 | * | 11,43515 | 3,202136723 | 1,090909 | 2,935293 | 0,003332 | 0,81148874 | 0,738441 | 3' UTR | 14 | 23088262 | 23144639 | 56378 | 2 | 791307 | ENSMUST00000090409.2 | -196609 | ENSMUSG00000068680 | Gm10248 | predicted gene 10248 |
| chr9 | 95392530 | 95395483 | 2954 | * | 21,67839 | 3,202498761 | 0,784939 | 4,079935 | 4,5E-05 | 0,063587778 | 2,951933 | Promoter (<=1kb) | 9 | 95338951 | 95393947 | 54997 | 2 | 67958 | ENSMUST00000079659.12 | 0 | ENSMUSG00000032407 | U2surp | U2 snRNP-associated SURP domain containing |
| chr16 | 95911250 | 95912623 | 1374 | * | 13,27414 | 3,202578284 | 1,053349 | 3,040378 | 0,002363 | 0,710397763 | 0,938682 | Intron (ENSMUST0000 | 16 | 95923434 | 95926539 | 3106 | 2 | 15312 | ENSMUST00000233638.2 | 13916 | ENSMUSG00000040681 | Hmgn1 | high mobility group nucleosomal binding domain 1 |
| chr11 | 68945228 | 68946773 | 1546 | * | 12,09391 | 3,20307762 | 1,026161 | 3,121418 | 0,0018 | 0,669586355 | 0,918255 | Distal Intergenic | 11 | 68950543 | 68952404 | 1862 | 1 | 71923 | ENSMUST00000051888.4 | -3770 | ENSMUSG00000045176 | Borcs6 | BLOC-1 related complex subunit 6 |
| chr14 | 25606119 | 25607821 | 1703 | * | 15,64766 | 3,203084973 | 1,002366 | 3,195523 | 0,001396 | 0,54122637 | 1,41696 | Promoter (<=1kb) | 14 | 25607781 | 25644081 | 36301 | 1 | 328365 | ENSMUST00000160190.2 | 0 | ENSMUSG00000007817 | Zmiz1 | zinc finger, MIZ-type containing 1 |
| chr10 | 92846937 | 92847977 | 1041 | * | 9,586676 | 3,204300744 | 1,091368 | 2,936041 | 0,003324 | 0,689688276 | 1,449706 | 3' UTR | 10 | 92857573 | 92860979 | 3407 | 2 | 380654 | ENSMUST00000165090.2 | 13002 | ENSMUSG00000020014 | Cfap54 | cilia and flagella associated protein 54 |

| | | | | | | | | | | | | | | | | | | | | | | | |
|---|---|---|---|---|---|---|---|---|---|---|---|---|---|---|---|---|---|---|---|---|---|---|---|
| chr6 | 48367061 | 48368227 | 1167 | * | 10,48974 | 3,204671901 | 1,20488 | 2,659744 | 0,00782 | 0,950586732 | 0,744375 | Intron (ENSMUST0000( | 6 | 48351070 | 48372342 | 21273 | 2 | 1,01E+08 | ENSMUST00000181943.3 | 4115 | ENSMUSG00000107875 | Gm16630 | predicted gene, 16630 |
| chr18 | 67900109 | 67901427 | 1319 | * | 9,266229 | 3,205143644 | 1,150799 | 2,785147 | 0,00535 | 0,6980047 | 2,237403 | Exon (ENSMUST000002 | 18 | 67907946 | 67928557 | 20612 | 1 | 72124 | ENSMUST00000025421.9 | -6519 | ENSMUSG00000079614 | Seh1l | SEH1-like (S. cerevisiae |
| chr2 | 93121061 | 93122295 | 1235 | * | 11,89335 | 3,206066317 | 1,158007 | 2,768607 | 0,00563 | 0,80696549 | 1,2763 | Exon (ENSMUST000001 | 2 | 93141608 | 93142860 | 1253 | 2 | 241556 | ENSMUST00000181716.8 | 20565 | ENSMUSG00000027217 | Tspan18 | tetraspanin 18 |
| chr19 | 37379932 | 37381420 | 1489 | * | 9,959605 | 3,206717393 | 1,231463 | 2,603991 | 0,009215 | 0,872043312 | 1,449706 | Intron (ENSMUST0000( | 19 | 37390629 | 37394219 | 3591 | 1 | 16551 | ENSMUST00000236027.2 | -9209 | ENSMUSG00000012443 | Kif11 | kinesin family member 11 |
| chr17 | 31943199 | 31944317 | 1119 | * | 10,37103 | 3,206901109 | 1,240234 | 2,585723 | 0,009718 | 0,901463351 | 1,2763 | Distal Intergenic | 17 | 31898454 | 31900479 | 2026 | 1 | 12954 | ENSMUST00000228305.2 | 44745 | ENSMUSG00000024041 | Cryaa | crystallin, alpha A |
| chr15 | 55686308 | 55687015 | 708 | * | 9,630847 | 3,207601846 | 1,151775 | 2,78492 | 0,005354 | 0,826676469 | 1,086118 | Intron (ENSMUST0000( | 15 | 55655334 | 55769706 | 114373 | 2 | 20649 | ENSMUST00000140574.2 | 82691 | ENSMUSG00000060429 | Sntb1 | syntrophin, basic 1 |
| chr18 | 11478150 | 11479710 | 1561 | * | 12,86757 | 3,208418085 | 1,110995 | 2,887877 | 0,003879 | 0,838625535 | 0,738441 | Distal Intergenic | 18 | 11766333 | 11876264 | 109932 | 1 | 225182 | ENSMUST00000115861.9 | -286623 | ENSMUSG00000041238 | Rbbp8 | retinoblastoma binding protein 8, endonuclease |
| chr4 | 53831246 | 53832628 | 1383 | * | 11,28349 | 3,208615068 | 1,215684 | 2,639349 | 0,008307 | 0,960042286 | 0,738441 | Exon (ENSMUST000001 | 4 | 53826057 | 53847844 | 21788 | 1 | 52076 | ENSMUST00000151206.2 | 5189 | ENSMUSG00000028420 | Tmem38b | transmembrane protein 38B |
| chr8 | 111817209 | 111820259 | 3051 | * | 28,495515 | 3,209405932 | 0,7607391 | 4,2187998 | 2,456E-05 | 0,04517794 | 2,7254908 | Promoter (<=1kb) | 8 | 111796130 | 111819058 | 22929 | 2 | 353287 | ENSMUST00000190222.2 | 0 | ENSMUSG00000033633 | Clec18a | C-type lectin domain family 18, member A |
| chr14 | 55114609 | 55115451 | 843 | * | 10,92504 | 3,21010285 | 1,167524 | 2,749496 | 0,005969 | 0,832960417 | 1,173325 | Promoter (<=1kb) | 14 | 55110138 | 55115007 | 4870 | 2 | 105651 | ENSMUST00000177403.2 | 0 | ENSMUSG00000072494 | Ppp1r3e | protein phosphatase 1, regulatory subunit 3E |
| chr12 | 1,12E+08 | 1,12E+08 | 967 | * | 9,319802 | 3,210119944 | 1,159117 | 2,769453 | 0,005615 | 0,789027175 | 1,425929 | Distal Intergenic | 12 | 1,12E+08 | 1,12E+08 | 14269 | 2 | 320390 | ENSMUST00000223323.2 | 19583 | ENSMUSG00000113120 | E330035G20Rik | RIKEN cDNA E330035G20 gene |
| chr4 | 77795427 | 77796755 | 1329 | * | 9,74928 | 3,210129268 | 1,236056 | 2,597074 | 0,009402 | 0,908738779 | 1,173325 | Intron (ENSMUST0000( | 4 | 76511788 | 78129976 | 1618189 | 2 | 19266 | ENSMUST00000138955.8 | 333221 | ENSMUSG00000028399 | Ptprd | protein tyrosine phosphatase, receptor type, D |
| chr16 | 38255808 | 38256837 | 1030 | * | 10,00931 | 3,210604769 | 1,150976 | 2,789464 | 0,00528 | 0,824134241 | 1,086118 | Promoter (2-3kb) | 16 | 38216479 | 38253507 | 37029 | 2 | 85031 | ENSMUST00000002926.8 | -2301 | ENSMUSG00000002847 | Pla1a | phospholipase A1 member A |
| chr7 | 44137681 | 44138699 | 1019 | * | 12,49632 | 3,210956974 | 1,016268 | 3,159558 | 0,00158 | 0,586304784 | 1,2763 | Promoter (2-3kb) | 7 | 44140309 | 44141442 | 1134 | 2 | 69683 | ENSMUST00000147102.2 | 2743 | ENSMUSG00000008140 | Emc10 | ER membrane protein complex subunit 10 |
| chr12 | 84300568 | 84303151 | 2584 | * | 18,78114 | 3,211326902 | 1,041597 | 3,083081 | 0,002049 | 0,507398906 | 2,507013 | Distal Intergenic | 12 | 84332006 | 84362606 | 30601 | 1 | 77219 | ENSMUST00000123614.8 | -28855 | ENSMUSG00000072946 | Ptgr2 | prostaglandin reductase 2 |
| chr5 | 61013878 | 61015420 | 1543 | * | 11,05464 | 3,211382797 | 1,024994 | 3,133074 | 0,00173 | 0,694263044 | 0,738441 | Distal Intergenic | 5 | 60658176 | 60658283 | 108 | 2 | 1,15E+08 | ENSMUST00000157423.3 | -355595 | ENSMUSG00000088048 | Gm23331 | predicted gene, 23331 |
| chr5 | 37007930 | 37009029 | 1100 | * | 10,26174 | 3,211576231 | 1,196465 | 2,684221 | 0,00727 | 0,859692746 | 1,211315 | Distal Intergenic | 5 | 37025857 | 37112422 | 86566 | 1 | 269643 | ENSMUST00000031003.11 | -16828 | ENSMUSG00000029120 | Ppp2r2c | protein phosphatase 2, regulatory subunit B, gamma |
| chr19 | 18651486 | 18652935 | 1450 | * | 9,59159 | 3,212548749 | 1,151711 | 2,789371 | 0,005281 | 0,811413964 | 1,173325 | Promoter (2-3kb) | 19 | 18648693 | 18655159 | 6467 | 1 | 67383 | ENSMUST00000236401.2 | 2793 | ENSMUSG00000024726 | Carnmt1 | carnosine N-methyltransferase 1 |
| chr10 | 38458206 | 38459391 | 1186 | * | 9,565558 | 3,213274329 | 1,211691 | 2,651892 | 0,008004 | 0,896435577 | 1,086118 | Distal Intergenic | 10 | 38491371 | 38491477 | 107 | 2 | 1,15E+08 | ENSMUST00000240107.1 | 32086 | ENSMUSG00000119503 | Gm22911 | predicted gene, 22911 |
| chr1 | 75167010 | 75168299 | 1290 | * | 12,29301 | 3,213352628 | 1,063824 | 3,020567 | 0,002523 | 0,760397726 | 0,738441 | Promoter (<=1kb) | 1 | 75157509 | 75167026 | 9518 | 2 | 245860 | ENSMUST00000189820.7 | 0 | ENSMUSG00000033124 | Atg9a | autophagy related 9A |
| chr15 | 80406187 | 80407208 | 1022 | * | 9,535969 | 3,216003936 | 1,205182 | 2,66848 | 0,00762 | 0,75938015 | 2,237403 | Intron (ENSMUST0000( | 15 | 80336441 | 80444671 | 108231 | 2 | 383075 | ENSMUST00000096347.4 | 37463 | ENSMUSG00000050439 | Enthd1 | ENTH domain containing 1 |
| chr8 | 1,21E+08 | 1,21E+08 | 779 | * | 10,98105 | 3,216481334 | 1,1831 | 2,718688 | 0,006554 | 0,923078468 | 0,744375 | Distal Intergenic | 8 | 1,21E+08 | 1,21E+08 | 12535 | 2 | 72042 | ENSMUST00000212676.2 | 15315 | ENSMUSG00000031827 | Cotl1 | coactosin-like 1 (Dictyostelium) |
| chr2 | 72582616 | 72584829 | 2214 | * | 15,92072 | 3,216690856 | 0,998275 | 3,222251 | 0,001272 | 0,418823795 | 2,663619 | Promoter (1-2kb) | 2 | 72576849 | 72581197 | 4349 | 2 | 76218 | ENSMUST00000125382.2 | -1419 | ENSMUSG00000085427 | 6430710C18Rik | RIKEN cDNA 6430710C18 gene |
| chr3 | 1,34E+08 | 1,34E+08 | 1650 | * | 16,77238 | 3,217066957 | 0,880321 | 3,654424 | 0,000258 | 0,284660899 | 1,2763 | Intron (ENSMUST0000( | 3 | 1,34E+08 | 1,34E+08 | 2600 | 2 | 1,03E+08 | ENSMUST00000200327.2 | -117719 | | Gm30484 | predicted gene, 30484 |
| chr7 | 4149431 | 4150499 | 1069 | * | 12,4686 | 3,217245839 | 1,173054 | 2,742623 | 0,006095 | 0,91311537 | 0,738441 | Promoter (1-2kb) | 7 | 4147719 | 4148641 | 923 | 1 | 232798 | ENSMUST00000146434.2 | 1712 | ENSMUSG00000035545 | Leng8 | leukocyte receptor cluster (LRC) member 8 |
| chr17 | 89849556 | 89850697 | 1142 | * | 12,92565 | 3,217654815 | 1,020957 | 3,151605 | 0,001624 | 0,651710641 | 0,918255 | Distal Intergenic | 17 | 89686669 | 89692421 | 5753 | 1 | 1E+08 | ENSMUST00000234513.2 | 162887 | ENSMUSG00000117524 | Gm4719 | predicted gene 4719 |
| chr17 | 29313898 | 29314885 | 988 | * | 13,67112 | 3,218482101 | 1,004721 | 3,203359 | 0,001358 | 0,621585803 | 0,918255 | Promoter (1-2kb) | 17 | 29312755 | 29319701 | 6947 | 1 | 12575 | ENSMUST00000233296.2 | 1143 | ENSMUSG00000023067 | Cdkn1a | cyclin-dependent kinase inhibitor 1A (P21) |
| chr19 | 55366794 | 55367930 | 1137 | * | 9,451132 | 3,218871707 | 1,129213 | 2,850545 | 0,004364 | 0,749550311 | 1,360622 | Promoter (1-2kb) | 19 | 55369446 | 55435411 | 65966 | 1 | 53611 | ENSMUST00000224396.2 | -1516 | ENSMUSG00000024983 | Vti1a | vesicle transport through interaction with t-SNAREs 1A |
| chr11 | 84055480 | 84057178 | 1699 | * | 9,220783 | 3,219543816 | 1,244799 | 2,586397 | 0,009699 | 0,881662676 | 1,449706 | Promoter (1-2kb) | 11 | 84039177 | 84058302 | 19126 | 2 | 628813 | ENSMUST00000050771.8 | 1124 | ENSMUSG00000051452 | Gm11437 | predicted gene 11437 |
| chr13 | 45780471 | 45781709 | 1239 | * | 12,01188 | 3,219584068 | 1,173594 | 2,743355 | 0,006081 | 0,836571183 | 1,173325 | Intron (ENSMUST0000( | 13 | 45692285 | 45696026 | 3742 | 1 | 66355 | ENSMUST00000129176.2 | 88186 | ENSMUSG00000000253 | Gmpr | guanosine monophosphate reductase |
| chr3 | 58070808 | 58073016 | 2209 | * | 14,62431 | 3,220289017 | 1,17447 | 2,741909 | 0,006108 | 0,830950621 | 1,214904 | Promoter (<=1kb) | 3 | 58071353 | 58075037 | 3685 | 1 | 1,05E+08 | ENSMUST00000200072.2 | 0 | | Gm40056 | predicted gene, 40056 |
| chr4 | 8555450 | 8557144 | 1695 | * | 15,24292 | 3,222413726 | 1,056316 | 3,050616 | 0,002284 | 0,662903125 | 1,214904 | Intron (ENSMUST0000( | 4 | 8535644 | 8607778 | 72135 | 1 | 59021 | ENSMUST00000060232.8 | 19806 | ENSMUSG00000047187 | Rab2a | RAB2A, member RAS oncogene family |
| chr19 | 44233927 | 44237815 | 3889 | * | 99,968666 | 3,222756954 | 0,6392669 | 5,0413321 | 4,623E-07 | 0,00149878 | 4,2003037 | Promoter (<=1kb) | 19 | 44214235 | 44235009 | 20775 | 2 | 329064 | ENSMUST00000236516.2 | 0 | ENSMUSG00000037578 | Pkd2l1 | polycystic kidney disease 2-like 1 |
| chr14 | 20793236 | 20794679 | 1444 | * | 15,42612 | 3,222904128 | 0,910692 | 3,538963 | 0,000402 | 0,277776482 | 2,141612 | Exon (ENSMUST000001 | 14 | 20786677 | 20789465 | 2789 | 2 | 12325 | ENSMUST00000223863.2 | -3771 | ENSMUSG00000021820 | Camk2g | calcium/calmodulin-dependent protein kinase II gamma |
| chr9 | 58884961 | 58886507 | 1547 | * | 12,09692 | 3,224005308 | 1,069699 | 3,013937 | 0,002579 | 0,674235791 | 1,2763 | Promoter (2-3kb) | 9 | 58820119 | 58889391 | 69273 | 2 | 18007 | ENSMUST00000215026.2 | 2884 | ENSMUSG00000032340 | Neo1 | neogenin |
| chr9 | 61451391 | 61452485 | 1095 | * | 11,29552 | 3,224718442 | 1,126079 | 2,863669 | 0,004188 | 0,851704515 | 0,744375 | Distal Intergenic | 9 | 61417005 | 61441529 | 24525 | 1 | 1,03E+08 | ENSMUST00000214104.2 | 34386 | ENSMUSG00000111381 | Gm34424 | predicted gene, 34424 |
| chr17 | 68928371 | 68929742 | 1372 | * | 11,66311 | 3,22494436 | 1,126977 | 2,861587 | 0,004215 | 0,753831088 | 1,2763 | Intron (ENSMUST0000( | 17 | 69080815 | 69087081 | 6267 | 1 | 320858 | ENSMUST00000233696.2 | -151073 | ENSMUSG00000041565 | L3mbtl4 | L3MBTL4 histone methyl-lysine binding protein |
| chr1 | 1,34E+08 | 1,34E+08 | 1619 | * | 25,8923 | 3,224995246 | 0,776053 | 4,155636 | 3,24E-05 | 0,051943575 | 2,880017 | Exon (ENSMUST000002 | 1 | 1,34E+08 | 1,34E+08 | 3951 | 2 | 12227 | ENSMUST00000020692.7 | -3248 | ENSMUSG00000020423 | Btg2 | BTG anti-proliferation factor 2 |
| chr5 | 1,4E+08 | 1,4E+08 | 1451 | * | 11,62015 | 3,225158091 | 0,989446 | 3,25956 | 0,001116 | 0,625704414 | 0,738441 | Intron (ENSMUST0000( | 5 | 1,4E+08 | 1,4E+08 | 62577 | 1 | 243312 | ENSMUST00000198608.2 | 14734 | ENSMUSG00000048988 | Elfn1 | leucine rich repeat and fibronectin type III, extracellular 1 |
| chr10 | 82825108 | 82826280 | 1173 | * | 11,12412 | 3,225584577 | 1,19283 | 2,704144 | 0,006848 | 0,85617852 | 1,173325 | Intron (ENSMUST0000( | 10 | 82821332 | 83031734 | 210403 | 1 | 58250 | ENSMUST00000040110.8 | 3776 | ENSMUSG00000034612 | Chst11 | carbohydrate sulfotransferase 11 |
| chr2 | 1,8E+08 | 1,8E+08 | 1057 | * | 9,157784 | 3,226094144 | 1,120476 | 2,879216 | 0,003987 | 0,752975043 | 1,211315 | Promoter (2-3kb) | 2 | 1,8E+08 | 1,8E+08 | 6592 | 1 | 72075 | ENSMUST00000029087.4 | -2083 | ENSMUSG00000049401 | Ogfr | opioid growth factor receptor |
| chr5 | 35166070 | 35167032 | 963 | * | 13,05688 | 3,22656944 | 0,929275 | 3,472137 | 0,000516 | 0,406160411 | 1,173325 | Intron (ENSMUST0000( | 5 | 35171664 | 35176836 | 5173 | 1 | 71729 | ENSMUST00000225237.2 | -4632 | ENSMUSG00000029101 | Rgs12 | regulator of G-protein signaling 12 |
| chr3 | 1,16E+08 | 1,16E+08 | 1239 | * | 10,77979 | 3,226626702 | 1,184544 | 2,72394 | 0,006451 | 0,92121789 | 0,744375 | Distal Intergenic | 3 | 1,16E+08 | 1,16E+08 | 132 | 2 | 1,15E+08 | ENSMUST00000104553.3 | 17033 | ENSMUSG00000119256 | Gm26344 | predicted gene, 26344 |
| chr7 | 82143122 | 82144099 | 978 | * | 8,562947 | 3,227225155 | 1,127762 | 2,861619 | 0,004215 | 1 | 0 | Intron (ENSMUST0000( | 7 | 82057682 | 82115052 | 57371 | 1 | 269959 | ENSMUST00000174495.2 | 85440 | ENSMUSG00000070469 | Adamtsl3 | ADAMTS-like 3 |
| chr14 | 45587021 | 45588508 | 1488 | * | 10,09315 | 3,227387519 | 1,232955 | 2,617603 | 0,008855 | 0,910353918 | 1,086118 | Promoter (<=1kb) | 14 | 45588550 | 45622964 | 34415 | 1 | 56291 | ENSMUST00000226873.2 | -42 | ENSMUSG00000053205 | Styx | serine/threonine/tyrosine interaction protein |
| chr1 | 1,71E+08 | 1,71E+08 | 1929 | * | 13,50121 | 3,227456481 | 1,024665 | 3,149766 | 0,001634 | 0,626872733 | 1,075735 | Promoter (<=1kb) | 1 | 1,71E+08 | 1,71E+08 | 6462 | 2 | 66155 | ENSMUST00000080001.9 | -433 | ENSMUSG00000062963 | Ufc1 | ubiquitin-fold modifier conjugating enzyme 1 |
| chr14 | 45556118 | 45558365 | 2248 | * | 49,502428 | 3,227823556 | 0,4812687 | 6,7069048 | 1,988E-11 | 6,52205E-07 | 2,7254908 | Promoter (<=1kb) | 14 | 45539366 | 45556228 | 16863 | 2 | 50527 | ENSMUST00000227315.2 | 0 | ENSMUSG00000021831 | Ero1a | endoplasmic reticulum oxidoreductase 1 alpha |
| chr9 | 84095914 | 84096868 | 955 | * | 10,58919 | 3,229026727 | 1,156543 | 2,791963 | 0,005239 | 0,809505441 | 1,173325 | Distal Intergenic | 9 | 84070408 | 84070592 | 185 | 1 | 1,15E+08 | ENSMUST00000104478.4 | 25506 | ENSMUSG00000077666 | Gm26146 | predicted gene, 26146 |
| chr17 | 26361074 | 26362585 | 1512 | * | 12,2286 | 3,229163798 | 1,127076 | 2,865082 | 0,004169 | 0,852021046 | 0,738441 | Promoter (<=1kb) | 17 | 26361592 | 26409205 | 47614 | 1 | 12005 | ENSMUST00000163421.9 | 0 | ENSMUSG00000024182 | Axin1 | axin 1 |
| chr9 | 65125608 | 65129201 | 3594 | * | 27,676741 | 3,23047794 | 0,6754402 | 4,7827744 | 1,729E-06 | 0,006041404 | 2,7254908 | Intron (ENSMUST000002 | 9 | 65122427 | 65146506 | 24080 | 1 | 214424 | ENSMUST00000213396.2 | 3181 | ENSMUSG00000032392 | Parp16 | poly (ADP-ribose) polymerase family, member 16 |
| chr17 | 47174435 | 47175580 | 1146 | * | 12,07898 | 3,230558216 | 1,07634 | 3,00143 | 0,002687 | 0,773936378 | 0,738441 | Distal Intergenic | 17 | 47136721 | 47169408 | 32688 | 2 | 210982 | ENSMUST00000233455.2 | -5027 | ENSMUSG00000036568 | Bicral | BRD4 interacting chromatin remodeling complex associated protein like |
| chr7 | 29594587 | 29595969 | 1383 | * | 11,60685 | 3,230834529 | 1,239731 | 2,606077 | 0,009159 | 0,891097601 | 1,2763 | Promoter (2-3kb) | 7 | 29593504 | 29598288 | 4785 | 2 | 22689 | ENSMUST00000172448.2 | 2319 | ENSMUSG00000062040 | Zfp27 | zinc finger protein 27 |
| chr8 | 70662979 | 70664460 | 1482 | * | 18,19455 | 3,230973878 | 0,900478 | 3,588067 | 0,000333 | 0,252389304 | 2,154504 | Promoter (<=1kb) | 8 | 70636990 | 70664906 | 27917 | 2 | 73095 | ENSMUST00000063788.8 | 446 | ENSMUSG00000002346 | Slc25a42 | solute carrier family 25, member 42 |
| chr16 | 6986346 | 6987360 | 1015 | * | 9,808822 | 3,231683118 | 1,195668 | 2,702826 | 0,006875 | 0,821398149 | 1,449706 | Intron (ENSMUST0000( | 16 | 7011611 | 7227833 | 216223 | 1 | 268859 | ENSMUST00000231194.2 | -24251 | ENSMUSG00000008658 | Rbfox1 | RNA binding protein, fox-1 homolog (C. elegans) 1 |
| chr12 | 51857774 | 51859662 | 1889 | * | 13,7084 | 3,232300014 | 0,950349 | 3,401171 | 0,000671 | 0,434788971 | 1,2763 | Intron (ENSMUST0000( | 12 | 51872966 | 51876101 | 3136 | 2 | 207304 | ENSMUST00000217804.2 | 16439 | ENSMUSG00000035247 | Hectd1 | HECT domain E3 ubiquitin protein ligase 1 |
| chr10 | 61455012 | 61456041 | 1030 | * | 11,38704 | 3,232393373 | 1,113856 | 2,901985 | 0,003708 | 0,830858561 | 0,738441 | Intron (ENSMUST0000( | 10 | 61431271 | 61464344 | 33074 | 1 | 237362 | ENSMUST00000020287.8 | 23741 | ENSMUSG00000020090 | Npffr1 | neuropeptide FF receptor 1 |
| chr14 | 1,15E+08 | 1,15E+08 | 1169 | * | 6,64772 | 3,232834236 | 1,251083 | 2,584028 | 0,009765 | 1 | 0 | Intron (ENSMUST0000( | 14 | 1,15E+08 | 1,17E+08 | 1431792 | 1 | 103978 | ENSMUST00000022707.7 | 50707 | ENSMUSG00000022112 | Gpc5 | glypican 5 |
| chr1 | 1,8E+08 | 1,8E+08 | 1127 | * | 12,29719 | 3,232963569 | 1,136383 | 2,844959 | 0,004442 | 0,793708774 | 1,075735 | Intron (ENSMUST0000( | 1 | 1,8E+08 | 1,8E+08 | 47647 | 1 | 226751 | ENSMUST00000132894.3 | 28092 | ENSMUSG00000026490 | Cdc42bpa | CDC42 binding protein kinase alpha |
| chr19 | 58025980 | 58027621 | 1642 | * | 12,15914 | 3,233024995 | 1,155467 | 2,798024 | 0,005142 | 0,79185145 | 1,2763 | Intron (ENSMUST0000( | 19 | 58039599 | 58039668 | 70 | 1 | 1,01E+08 | ENSMUST00000176631.3 | -11978 | ENSMUSG00000093757 | Mir5623 | microRNA 5623 |
| chr18 | 31940405 | 31941167 | 763 | * | 8,704641 | 3,233573942 | 1,250722 | 2,585365 | 0,009728 | 0,802671652 | 2,237403 | Intron (ENSMUST0000( | 18 | 31937229 | 31968475 | 31247 | 1 | 74320 | ENSMUST00000234344.2 | 3176 | ENSMUSG00000024400 | Wdr33 | WD repeat domain 33 |
| chr1 | 1,46E+08 | 1,46E+08 | 1789 | * | 11,2761 | 3,233873299 | 1,034427 | 3,126247 | 0,001771 | 0,698472175 | 0,738441 | Distal Intergenic | 1 | 1,46E+08 | 1,47E+08 | 407712 | 1 | 215378 | ENSMUST00000074622.11 | -405289 | ENSMUSG00000035131 | Brinp3 | bone morphogenetic protein/retinoic acid inducible neural specific 3 |
| chr18 | 22493708 | 22494748 | 1041 | * | 11,33868 | 3,233904225 | 1,037934 | 3,115712 | 0,001835 | 0,671902369 | 0,918255 | Intron (ENSMUST0000( | 18 | 22478146 | 22663284 | 185139 | 1 | 211961 | ENSMUST00000097655.4 | 15562 | ENSMUSG00000045215 | Asxl3 | ASXL transcriptional regulator 3 |
| chr7 | 24439175 | 24440024 | 850 | * | 11,44964 | 3,233955251 | 1,114454 | 2,90183 | 0,00371 | 0,748971518 | 1,173325 | Distal Intergenic | 7 | 24444552 | 24450246 | 5695 | 2 | 68891 | ENSMUST00000206160.2 | 10222 | ENSMUSG00000052212 | Cd177 | CD177 antigen |
| chr2 | 1,8E+08 | 1,8E+08 | 1218 | * | 13,10895 | 3,234067286 | 0,989715 | 3,267674 | 0,001084 | 0,514210241 | 1,2763 | Distal Intergenic | 2 | 1,8E+08 | 1,8E+08 | 10475 | 1 | 241846 | ENSMUST00000055485.12 | -12172 | ENSMUSG00000039108 | Lsm14b | LSM family member 14B |
| chr11 | 1,14E+08 | 1,14E+08 | 1291 | * | 11,56221 | 3,234320872 | 1,154485 | 2,801528 | 0,005086 | 0,849121913 | 0,918255 | Intron (ENSMUST0000( | 11 | 1,14E+08 | 1,14E+08 | 1380 | 1 | 20606 | ENSMUST00000067591.3 | 3468 | ENSMUSG00000047904 | Sstr2 | somatostatin receptor 2 |
| chrX | 79991531 | 79992552 | 1022 | * | 7,275302 | 3,234674066 | 1,229522 | 2,630838 | 0,008517 | 1 | 0 | Distal Intergenic | 20 | 80114304 | 80138382 | 24079 | 1 | 192216 | ENSMUST00000150052.2 | -121752 | ENSMUSG00000025666 | Tmem47 | transmembrane protein 47 |
| chr17 | 18784577 | 18785622 | 1046 | * | 9,469584 | 3,23488085 | 1,061106 | 3,048594 | 0,002299 | 0,637630418 | 1,425929 | Distal Intergenic | 17 | 18793413 | 18821975 | 28563 | 1 | 433070 | ENSMUST00000177244.4 | -7791 | ENSMUSG00000091679 | Vmn2r96 | vomeronasal 2, receptor 96 |
| chr10 | 20776977 | 20778165 | 1189 | * | 10,29682 | 3,235100537 | 1,12792 | 2,8682 | 0,004128 | 0,781219703 | 1,086118 | Distal Intergenic | 10 | 20742000 | 20767448 | 25449 | 2 | 1,19E+08 | ENSMUST00000241754.1 | -9529 | | Gm53617 | predicted gene, 53617 |
| chr13 | 53529963 | 53531981 | 2019 | * | 13,25552 | 3,235424858 | 1,227625 | 2,635517 | 0,008401 | 0,962000766 | 0,738441 | Promoter (<=1kb) | 13 | 53486784 | 53531433 | 44650 | 2 | 268656 | ENSMUST00000021920.8 | 0 | ENSMUSG00000021468 | Sptlc1 | serine palmitoyltransferase, long chain base subunit 1 |
| chr2 | 1,57E+08 | 1,57E+08 | 1035 | * | 9,195616 | 3,235689622 | 1,25605 | 2,576083 | 0,009993 | 0,88871498 | 1,425929 | Promoter (1-2kb) | 2 | 1,57E+08 | 1,57E+08 | 2835 | 1 | 228836 | ENSMUST00000123730.2 | 1288 | ENSMUSG00000061689 | Dlgap4 | DLG associated protein 4 |
| chr19 | 48631651 | 48632824 | 1174 | * | 11,04019 | 3,235961413 | 1,150606 | 2,812399 | 0,004917 | 0,797327098 | 1,173325 | Intron (ENSMUST0000( | 19 | 48692655 | 48792976 | 100322 | 1 | 66673 | ENSMUST00000235967.2 | -59831 | ENSMUSG00000063434 | Sorcs3 | sortilin-related VPS10 domain containing receptor 3 |
| chr1 | 168150989 | 168152522 | 1534 | * | 18,808155 | 3,236304424 | 0,7350424 | 4,4028815 | 1,068E-05 | 0,027873477 | 2,2216115 | Intron (ENSMUST000001 | 1 | 167986301 | 168259073 | 272773 | 2 | 18514 | ENSMUST00000064438.11 | 106551 | ENSMUSG00000052534 | Pbx1 | pre B cell leukemia homeobox 1 |
| chr14 | 45360545 | 45361909 | 1365 | * | 14,29638 | 3,236678322 | 1,044534 | 3,098681 | 0,001944 | 0,678869598 | 0,938682 | Distal Intergenic | 14 | 45402278 | 45406782 | 4505 | 2 | 70561 | ENSMUST00000156600.2 | 44873 | ENSMUSG00000021830 | Txndc16 | thioredoxin domain containing 16 |
| chr11 | 1,21E+08 | 1,21E+08 | 2214 | * | 13,66272 | 3,236814042 | 1,02664 | 3,152823 | 0,001617 | 0,651089638 | 0,918255 | Promoter (2-3kb) | 11 | 1,21E+08 | 1,21E+08 | 4726 | 2 | 217370 | ENSMUST00000147490.6 | -2535 | ENSMUSG00000039294 | Cybc1 | cytochrome b 245 chaperone 1 |
| chr3 | 1,42E+08 | 1,42E+08 | 1021 | * | 9,369949 | 3,237273982 | 1,209131 | 2,677356 | 0,007421 | 0,837097674 | 1,425929 | Intron (ENSMUST0000( | 3 | 1,42E+08 | 1,42E+08 | 55016 | 2 | 56376 | ENSMUST00000197808.5 | -26525 | ENSMUSG00000028273 | Pdlim5 | PDZ and LIM domain 5 |
| chr17 | 28022435 | 28025827 | 3393 | * | 44,555207 | 3,237593446 | 0,6133423 | 5,2786078 | 1,302E-07 | 0,000844569 | 2,7872553 | Intron (ENSMUST000001 | 17 | 27972251 | 28039426 | 67176 | 2 | 224647 | ENSMUST00000233508.2 | 13599 | ENSMUSG00000056692 | Ilrun | inflammation and lipid regulator with UBA-like and NBR1-like domains |
| chr2 | 35328694 | 35329656 | 963 | * | 13,20131 | 3,237703835 | 1,066282 | 3,036442 | 0,002394 | 0,750475959 | 0,738441 | Promoter (2-3kb) | 2 | 35297957 | 35332113 | 34157 | 2 | 14594 | ENSMUST00000131745.8 | 2457 | ENSMUSG00000035778 | Ggta1 | glycoprotein galactosyltransferase alpha 1, 3 |
| chr10 | 1,23E+08 | 1,23E+08 | 1190 | * | 10,73016 | 3,238216025 | 1,252219 | 2,585983 | 0,00971 | 0,951821 | 0,918255 | Promoter (<=1kb) | 10 | 1,23E+08 | 1,23E+08 | 85 | 2 | 387251 | ENSMUST00000083472.5 | 621 | ENSMUSG00000065406 | Mirlet7i | microRNA let7i |
| chr15 | 57746049 | 57746620 | 572 | * | 7,127659 | 3,239105783 | 1,245413 | 2,600828 | 0,0093 | 0,885612383 | 1,360622 | Intron (ENSMUST0000( | 15 | 57735059 | 57753790 | 18732 | 2 | 67819 | ENSMUST00000226911.2 | 7170 | ENSMUSG00000022365 | Derl1 | Der1-like domain family, member 1 |
| chr8 | 13446311 | 13447442 | 1132 | * | 11,41645 | 3,239282311 | 1,190714 | 2,720453 | 0,006519 | 0,922205132 | 0,744375 | Promoter (<=1kb) | 8 | 13436205 | 13446825 | 10621 | 2 | 11945 | ENSMUST00000033826.4 | 0 | ENSMUSG00000031449 | Atp4b | ATPase, H+/K+ exchanging, beta polypeptide |
| chr2 | 1,2E+08 | 1,2E+08 | 1186 | * | 10,37833 | 3,240696594 | 1,106426 | 2,928976 | 0,003401 | 0,816165716 | 0,738441 | Exon (ENSMUST00000( | 2 | 1,2E+08 | 1,2E+08 | 2410 | 1 | 29808 | ENSMUST00000150742.2 | -3362 | ENSMUSG00000033943 | Mga | MAX gene associated |
| chr14 | 5272618 | 5273573 | 956 | * | 8,665358 | 3,240805136 | 1,207343 | 2,684246 | 0,007269 | 0,832984033 | 1,425929 | Distal Intergenic | 14 | 5363820 | 5364135 | 316 | 2 | 1E+08 | ENSMUST00000225472.2 | 90562 | | Pou5f1-rs6 | POU domain, class 5, transcription factor 1, related sequence 6 |
| chr19 | 9001155 | 9002916 | 1762 | * | 12,68579 | 3,241119908 | 1,080427 | 2,999851 | 0,002701 | 0,709889128 | 1,075735 | Intron (ENSMUST0000( | 19 | 8977024 | 9054278 | 77255 | 1 | 66395 | ENSMUST00000237033.2 | 24131 | ENSMUSG00000069833 | Ahnak | AHNAK nucleoprotein (desmoyokin) |
| chr17 | 27785893 | 27788147 | 2255 | * | 15,52746 | 3,24146461 | 1,027767 | 3,153891 | 0,001611 | 0,494205337 | 2,141612 | Promoter (1-2kb) | 17 | 27782547 | 27784730 | 2184 | 2 | 106672 | ENSMUST00000154473.9 | -1163 | ENSMUSG00000062753 | AI413582 | expressed sequence AI413582 |
| chr17 | 46197686 | 46198634 | 949 | * | 14,39582 | 3,241724588 | 0,969828 | 3,342575 | 0,00083 | 0,470097239 | 1,2763 | Promoter (1-2kb) | 17 | 46196272 | 46199556 | 3285 | 1 | 1,05E+08 | ENSMUST00000233085.2 | 1414 | ENSMUSG00000117048 | Gm41584 | predicted gene, 41584 |
| chr4 | 69300650 | 69301897 | 1248 | * | 11,59266 | 3,241776823 | 1,176219 | 2,756099 | 0,00585 | 0,829621864 | 1,173325 | Intron (ENSMUST0000( | 4 | 69028908 | 69029014 | 107 | 1 | 1,15E+08 | ENSMUST00000083881.3 | 271742 | ENSMUSG00000065815 | Gm24697 | predicted gene, 24697 |
| chr15 | 37377120 | 37377959 | 840 | * | 9,630732 | 3,241785379 | 1,205768 | 2,688564 | 0,007176 | 0,826891088 | 1,449706 | Intron (ENSMUST0000( | 15 | 37335981 | 37361184 | 25204 | 1 | 252973 | ENSMUST00000159221.2 | 41139 | ENSMUSG00000022286 | Grhl2 | grainyhead like transcription factor 2 |
| chr6 | 55022626 | 55023457 | 832 | * | 9,2044 | 3,243328888 | 1,209906 | 2,680644 | 0,007348 | 0,83238039 | 1,449706 | Promoter (1-2kb) | 6 | 55025048 | 55027951 | 2904 | 1 | 353172 | ENSMUST00000205258.2 | -1591 | ENSMUSG00000029777 | Gars | glycyl-tRNA synthetase |
| chr6 | 31076456 | 31079862 | 3407 | * | 64,86793 | 3,243999414 | 0,4583783 | 7,077123 | 1,472E-12 | 4,69975E-08 | 4,2003037 | Exon (ENSMUST000001 | 6 | 31040027 | 31040430 | 404 | 2 | 387223 | ENSMUST00000183056.2 | -36026 | ENSMUSG00000065604 | Mir29b-1 | microRNA 29b-1 |
| chr19 | 10035491 | 10036653 | 1163 | * | 11,40214 | 3,244763635 | 1,059257 | 3,063246 | 0,00219 | 0,736412045 | 0,738441 | Promoter (1-2kb) | 19 | 10033616 | 10037036 | 3421 | 1 | 60527 | ENSMUST00000237226.2 | 1875 | ENSMUSG00000024664 | Fads3 | fatty acid desaturase 3 |
| chr11 | 1,19E+08 | 1,19E+08 | 1113 | * | 15,08435 | 3,245217128 | 0,930785 | 3,486537 | 0,000489 | 0,302171532 | 2,141612 | 3' UTR | 11 | 1,19E+08 | 1,19E+08 | 15411 | 2 | 1,05E+08 | ENSMUST00000144096.2 | -5957 | | Gm11755 | predicted gene 11755 |
| chr9 | 92419506 | 92421508 | 2003 | * | 20,89539 | 3,245523572 | 0,861546 | 3,767091 | 0,000165 | 0,14233436 | 2,951933 | Promoter (<=1kb) | 9 | 92421989 | 92422136 | 148 | 2 | 78575 | ENSMUST00000186512.2 | 628 | ENSMUSG00000103622 | B430319G15Rik | RIKEN cDNA B430319G15 gene |
| chr4 | 1,16E+08 | 1,16E+08 | 2188 | * | 20,3057 | 3,245914438 | 0,916199 | 3,542804 | 0,000396 | 0,26105284 | 2,386008 | Promoter (<=1kb) | 4 | 1,16E+08 | 1,16E+08 | 36692 | 1 | 230648 | ENSMUST00000106525.9 | 0 | ENSMUSG00000034210 | Efcab14 | EF-hand calcium binding domain 14 |
| chr1 | 1,72E+08 | 1,72E+08 | 1331 | * | 13,26417 | 3,246156692 | 1,136134 | 2,857195 | 0,004274 | 0,756018836 | 1,2763 | Distal Intergenic | 1 | 1,72E+08 | 1,72E+08 | 5282 | 2 | 18071 | ENSMUST00000059794.4 | 10636 | ENSMUSG00000051251 | Nhlh1 | nescient helix loop helix 1 |
| chr5 | 1,19E+08 | 1,19E+08 | 2197 | * | 15,22187 | 3,246189648 | 0,99646 | 3,257721 | 0,001123 | 0,502454322 | 1,41696 | Distal Intergenic | 5 | 1,19E+08 | 1,19E+08 | 6310 | 1 | 73477 | ENSMUST00000190550.2 | -23328 | ENSMUSG00000100573 | 1700081H04Rik | RIKEN cDNA 1700081H04 gene |
| chr13 | 31438645 | 31440561 | 1917 | * | 17,64838 | 3,249508647 | 0,928908 | 3,498203 | 0,000468 | 0,282588188 | 2,386008 | Distal Intergenic | 13 | 31397393 | 31425069 | 27677 | 2 | 73924 | ENSMUST00000153175.3 | -13576 | ENSMUSG00000087460 | 4930401O12Rik | RIKEN cDNA 4930401O12 gene |
| chr14 | 70443028 | 70444235 | 1208 | * | 8,656743 | 3,249883514 | 1,23834 | 2,624388 | 0,00868 | 1 | 0 | Promoter (<=1kb) | 14 | 70417917 | 70443471 | 25555 | 2 | 20410 | ENSMUST00000227653.2 | 0 | ENSMUSG00000022091 | Sorbs3 | sorbin and SH3 domain containing 3 |
| chr18 | 31112916 | 31114050 | 1135 | * | 9,159502 | 3,250117416 | 1,17876 | 2,757234 | 0,005829 | 0,801625187 | 1,360622 | Intron (ENSMUST0000( | 18 | 31286324 | 31345785 | 59462 | 2 | 19762 | ENSMUST00000153196.2 | 231735 | ENSMUSG00000057455 | Rit2 | Ras-like without CAAX 2 |
| chr18 | 57359058 | 57360519 | 1462 | * | 12,41951 | 3,251021371 | 1,062679 | 3,05927 | 0,002219 | 0,700182401 | 0,918255 | Intron (ENSMUST0000( | 18 | 57275858 | 57425800 | 149943 | 1 | 70417 | ENSMUST00000139892.2 | 83200 | ENSMUSG00000024593 | Megf10 | multiple EGF-like-domains 10 |
| chr5 | 17643918 | 17644979 | 1062 | * | 7,699871 | 3,251119013 | 1,163306 | 2,794723 | 0,005194 | 1 | 0 | Distal Intergenic | 5 | 17681096 | 17685934 | 4839 | 1 | 70935 | ENSMUST00000076099.4 | -36117 | ENSMUSG00000058643 | Speer4f1 | spermatogenesis associated glutamate (E)-rich protein 4F1 |
| chr9 | 1,23E+08 | 1,23E+08 | 754 | * | 8,359139 | 3,251301358 | 1,222575 | 2,659387 | 0,007828 | 0,853738252 | 1,360622 | Distal Intergenic | 9 | 1,23E+08 | 1,23E+08 | 1712 | 2 | 72309 | ENSMUST00000068140.6 | -18231 | ENSMUSG00000054871 | Tmem158 | transmembrane protein 158 |

| | | | | | | | | | | | | | | | | | | | | | | | |
|---|---|---|---|---|---|---|---|---|---|---|---|---|---|---|---|---|---|---|---|---|---|---|---|
| chr4 | 39022923 | 39024118 | 1196 | * | 8,195282 | 3,251543847 | 1,160733 | 2,801286 | 0,00509 | 0,687937632 | 2,237403 | Distal Intergenic | 4 | 38571667 | 38855483 | 283817 | 2 | 1,03E+08 | ENSMUST00000126314.2 | -167440 | ENSMUSG00000085632 | Gm12381 | predicted gene 12381 |
| chr9 | 6667577 | 6669083 | 1507 | * | 15,70935 | 3,251736402 | 0,905148 | 3,592492 | 0,000328 | 0,236247134 | 2,386008 | Distal Intergenic | 9 | 6706342 | 6706446 | 105 | 2 | 1,15E+08 | ENSMUST00000083448.2 | 37363 | ENSMUSG00000065382 | Gm23097 | predicted gene, 23097 |
| chr3 | 1,28E+08 | 1,28E+08 | 1536 | * | 11,53049 | 3,251876811 | 1,221717 | 2,661727 | 0,007774 | 0,949251015 | 0,744375 | Distal Intergenic | 3 | 1,28E+08 | 1,28E+08 | 107025 | 2 | 402774 | ENSMUST00000196895.5 | -107032 | ENSMUSG00000105816 | D030025E07Rik | RIKEN cDNA D030025E07 gene |
| chr6 | 72069600 | 72070254 | 655 | * | 7,721797 | 3,252338458 | 1,17124 | 2,776834 | 0,005489 | 1 | 0 | Distal Intergenic | 6 | 72074576 | 72131555 | 56980 | 1 | 20454 | ENSMUST00000069994.11 | -4322 | ENSMUSG00000056091 | St3gal5 | ST3 beta-galactoside alpha-2,3-sialyltransferase 5 |
| chr2 | 1,64E+08 | 1,64E+08 | 1561 | * | 15,78149 | 3,252504191 | 0,915969 | 3,550889 | 0,000384 | 0,272208984 | 2,141612 | Distal Intergenic | 2 | 1,64E+08 | 1,64E+08 | 1724 | 1 | 78928 | ENSMUST00000131976.2 | 13130 | ENSMUSG00000017721 | Pigt | phosphatidylinositol glycan anchor biosynthesis, class T |
| chr5 | 87049995 | 87051046 | 1052 | * | 11,18574 | 3,252630351 | 1,235458 | 2,632731 | 0,00847 | 0,891860659 | 1,173325 | Intron (ENSMUST00000 | 5 | 87037626 | 87054796 | 17171 | 2 | 100727 | ENSMUST00000031181.16 | 3750 | ENSMUSG00000029260 | Ugt2b34 | UDP glucuronosyltransferase 2 family, polypeptide B34 |
| chr4 | 1,51E+08 | 1,51E+08 | 1813 | * | 16,70645 | 3,252634664 | 1,105902 | 2,941159 | 0,00327 | 0,584914035 | 2,663619 | Promoter (2-3kb) | 4 | 1,51E+08 | 1,51E+08 | 53546 | 2 | 100072 | ENSMUST00000143434.2 | -2726 | ENSMUSG00000014592 | Camta1 | calmodulin binding transcription activator 1 |
| chr9 | 1,19E+08 | 1,19E+08 | 1434 | * | 11,81774 | 3,252780499 | 1,120436 | 2,903137 | 0,003694 | 0,827954389 | 0,744375 | Distal Intergenic | 9 | 1,19E+08 | 1,19E+08 | 5278 | 1 | 22351 | ENSMUST00000135872.8 | -4399 | ENSMUSG00000038775 | Vill | villin-like |
| chr5 | 1,49E+08 | 1,49E+08 | 1269 | * | 13,21705 | 3,253233384 | 1,034072 | 3,146042 | 0,001655 | 0,627785088 | 1,075735 | Distal Intergenic | 5 | 1,49E+08 | 1,49E+08 | 1312 | 1 | 70138 | ENSMUST00000244021.1 | -36793 | ENSMUSG00000086494 | 2210417A02Rik | RIKEN cDNA 2210417A02 gene |
| chr13 | 51532966 | 51534159 | 1194 | * | 11,34729 | 3,253239314 | 1,14115 | 2,850843 | 0,00436 | 0,821417341 | 0,918255 | Intron (ENSMUST00000 | 13 | 51562675 | 51576833 | 14159 | 1 | 13610 | ENSMUST00000087978.5 | -28516 | ENSMUSG00000067586 | S1pr3 | sphingosine-1-phosphate receptor 3 |
| chr19 | 52388469 | 52389224 | 756 | * | 8,381098 | 3,2534074 | 1,225866 | 2,653967 | 0,007955 | 0,768389035 | 2,237403 | Distal Intergenic | 19 | 52252735 | 52253914 | 1180 | 1 | 16333 | ENSMUST00000039652.6 | 135734 | ENSMUSG00000035804 | Ins1 | insulin I |
| chr13 | 47294983 | 47296208 | 1226 | * | 9,191436 | 3,253436407 | 1,242178 | 2,619138 | 0,008815 | 0,861411338 | 1,449706 | Intron (ENSMUST00000 | 13 | 47297418 | 47302146 | 4729 | 2 | 319467 | ENSMUST00000224519.2 | 5938 | ENSMUSG00000114961 | A930002C04Rik | RIKEN cDNA A930002C04 gene |
| chr4 | 1,19E+08 | 1,19E+08 | 1179 | * | 13,9744 | 3,253681566 | 1,172653 | 2,774633 | 0,005526 | 0,860218139 | 0,918255 | Promoter (<=1kb) | 4 | 1,19E+08 | 1,19E+08 | 93056 | 2 | 73332 | ENSMUST00000238759.2 | 0 | ENSMUSG00000028637 | Ccdc30 | coiled-coil domain containing 30 |
| chr1 | 1,67E+08 | 1,67E+08 | 2136 | * | 15,60237 | 3,254572745 | 1,042161 | 3,122909 | 0,001791 | 0,595550945 | 1,37683 | Distal Intergenic | 1 | 1,67E+08 | 1,67E+08 | 25706 | 2 | 1,01E+08 | ENSMUST00000180995.3 | 93229 | ENSMUSG00000102548 | Gm16701 | predicted gene, 16701 |
| chr11 | 5575500 | 5577086 | 1587 | * | 10,73805 | 3,254930417 | 1,188892 | 2,737785 | 0,006185 | 0,915421656 | 0,738441 | Promoter (1-2kb) | 11 | 5578738 | 5595153 | 16416 | 1 | 76389 | ENSMUST00000137933.2 | -1652 | ENSMUSG00000020481 | Ankrd36 | ankyrin repeat domain 36 |
| chr4 | 1,52E+08 | 1,52E+08 | 1249 | * | 11,31608 | 3,255035613 | 1,175322 | 2,769483 | 0,005615 | 0,822557945 | 1,173325 | Intron (ENSMUST00000 | 4 | 1,52E+08 | 1,52E+08 | 391475 | 2 | 1,03E+08 | ENSMUST00000155685.8 | 166099 | | Gm38442 | predicted gene, 38442 |
| chr9 | 20863502 | 20865627 | 2126 | * | 23,44618 | 3,255040737 | 0,7302981 | 4,4571401 | 8,306E-06 | 0,020252769 | 2,6724296 | Promoter (<=1kb) | 9 | 20818505 | 20863919 | 45415 | 2 | 13433 | ENSMUST00000178110.3 | 0 | ENSMUSG00000004099 | Dnmt1 | DNA methyltransferase (cytosine-5) 1 |
| chr5 | 1,27E+08 | 1,27E+08 | 871 | * | 8,462489 | 3,255733104 | 1,242511 | 2,620286 | 0,008786 | 1 | 0 | Distal Intergenic | 5 | 1,27E+08 | 1,27E+08 | 13783 | 1 | 75916 | ENSMUST00000196995.5 | -49537 | ENSMUSG00000105308 | 4930572K03Rik | RIKEN cDNA 4930572K03 gene |
| chr15 | 26200967 | 26202063 | 1097 | * | 10,9395 | 3,256345637 | 1,009873 | 3,22451 | 0,001262 | 0,644704154 | 0,744375 | Distal Intergenic | 15 | 26309134 | 26312457 | 3324 | 1 | 211147 | ENSMUST00000126304.2 | -107071 | ENSMUSG00000022269 | Marchf11 | membrane associated ring-CH-type finger 11 |
| chr10 | 1,27E+08 | 1,27E+08 | 1299 | * | 8,380112 | 3,256403489 | 1,247918 | 2,609469 | 0,009068 | 0,8666955 | 1,449706 | Intron (ENSMUST00000 | 10 | 1,27E+08 | 1,27E+08 | 109 | 2 | 1,15E+08 | ENSMUST00000158744.3 | -6905 | ENSMUSG00000089369 | Gm23819 | predicted gene, 23819 |
| chr9 | 1,11E+08 | 1,11E+08 | 999 | * | 10,61155 | 3,256633232 | 1,15816 | 2,811902 | 0,004925 | 0,879615137 | 0,744375 | Exon (ENSMUST000000 | 9 | 1,11E+08 | 1,11E+08 | 31817 | 1 | 71268 | ENSMUST00000196703.6 | 5170 | ENSMUSG00000032497 | Lrrfip2 | leucine rich repeat (in FLII) interacting protein 2 |
| chr2 | 1,66E+08 | 1,66E+08 | 1482 | * | 13,51247 | 3,257760139 | 0,980675 | 3,321958 | 0,000894 | 0,48037412 | 1,2763 | Distal Intergenic | 2 | 1,66E+08 | 1,66E+08 | 81750 | 2 | 72043 | ENSMUST00000146497.9 | -15703 | ENSMUSG00000006800 | Sulf2 | sulfatase 2 |
| chr3 | 62932673 | 62933847 | 1175 | * | 11,62035 | 3,257873304 | 1,11422 | 2,923904 | 0,003457 | 0,818408612 | 0,738441 | Distal Intergenic | 3 | 63059638 | 63059779 | 142 | 1 | 1,15E+08 | ENSMUST00000158550.3 | -125791 | ENSMUSG00000089175 | Gm22433 | predicted gene, 22433 |
| chr11 | 84352597 | 84354020 | 1424 | * | 12,22161 | 3,259009595 | 1,000774 | 3,256489 | 0,001128 | 0,627785088 | 0,738441 | Intron (ENSMUST00000 | 11 | 84333335 | 84340359 | 7025 | 2 | 56321 | ENSMUST00000141093.2 | -12238 | ENSMUSG00000018697 | Aatf | apoptosis antagonizing transcription factor |
| chr4 | 43322228 | 43323412 | 1185 | * | 12,67824 | 3,259109049 | 1,117229 | 2,917135 | 0,003533 | 0,72508691 | 1,2763 | 3' UTR | 4 | 43336259 | 43340155 | 3897 | 2 | 1,08E+08 | ENSMUST00000245981.1 | 16743 | | Gm46851 | predicted gene, 46851 |
| chr1 | 39801172 | 39803740 | 2569 | * | 22,61645 | 3,26025366 | 0,950545 | 3,429879 | 0,000604 | 0,268627242 | 3,471678 | Distal Intergenic | 1 | 39843308 | 39844492 | 1185 | 2 | 1E+08 | ENSMUST00000171319.4 | 40752 | ENSMUSG00000091937 | Gm3646 | predicted gene 3646 |
| chr17 | 35245296 | 35246112 | 817 | * | 8,799657 | 3,260568647 | 1,124801 | 2,898797 | 0,003746 | 0,723677315 | 1,360622 | Promoter (<=1kb) | 17 | 35244935 | 35246635 | 1701 | 2 | 78376 | ENSMUST00000173211.8 | 523 | ENSMUSG00000036185 | Sapcd1 | suppressor APC domain containing 1 |
| chr3 | 90222552 | 90224375 | 1824 | * | 11,30778 | 3,260699166 | 1,115548 | 2,922958 | 0,003467 | 0,817552804 | 0,744375 | Exon (ENSMUST000002 | 3 | 90201390 | 90265680 | 64291 | 1 | 229542 | ENSMUST00000199607.5 | 21162 | ENSMUSG00000042390 | Gatad2b | GATA zinc finger domain containing 2B |
| chr4 | 1,3E+08 | 1,3E+08 | 1351 | * | 11,90679 | 3,261045553 | 1,123164 | 2,903445 | 0,003691 | 0,791416391 | 0,918255 | Exon (ENSMUST000001 | 4 | 1,3E+08 | 1,3E+08 | 18937 | 1 | 67149 | ENSMUST00000180498.2 | 17755 | ENSMUSG00000078532 | Nkain1 | Na+/K+ transporting ATPase interacting 1 |
| chr19 | 16733610 | 16736572 | 2963 | * | 18,68358 | 3,261052656 | 0,996293 | 3,273185 | 0,001063 | 0,406085428 | 2,423594 | Intron (ENSMUST00000 | 19 | 16699702 | 16728302 | 28601 | 2 | 271564 | ENSMUST00000223846.2 | -5308 | ENSMUSG00000046230 | Vps13a | vacuolar protein sorting 13A |
| chr12 | 82318475 | 82319511 | 1037 | * | 10,39115 | 3,261151036 | 1,231063 | 2,649053 | 0,008072 | 0,886332554 | 1,173325 | Intron (ENSMUST00000 | 12 | 82303359 | 82497045 | 193687 | 1 | 217692 | ENSMUST00000222714.2 | 15116 | ENSMUSG00000042700 | Sipa1l1 | signal-induced proliferation-associated 1 like 1 |
| chr12 | 84262322 | 84263571 | 1250 | * | 8,535724 | 3,261243855 | 1,210857 | 2,693334 | 0,007074 | 0,824314641 | 1,449706 | Promoter (2-3kb) | 12 | 84195950 | 84265655 | 69706 | 2 | 238317 | ENSMUST00000046266.13 | 2084 | ENSMUSG00000042507 | Mideas | mitotic deacetylase associated SANT domain protein |
| chr5 | 37131639 | 37132521 | 883 | * | 9,178116 | 3,261991294 | 1,20242 | 2,712855 | 0,006671 | 0,847197771 | 1,211315 | Promoter (1-2kb) | 5 | 37125845 | 37134300 | 8456 | 2 | 22393 | ENSMUST00000167937.2 | 1779 | ENSMUSG00000039474 | Wfs1 | wolframin ER transmembrane glycoprotein |
| chr14 | 24045765 | 24046776 | 1012 | * | 8,945991 | 3,262160672 | 1,236571 | 2,63807 | 0,008338 | 0,777852794 | 2,237403 | Intron (ENSMUST00000 | 14 | 24053912 | 24058403 | 4492 | 1 | 78019 | ENSMUST00000223609.2 | -7136 | ENSMUSG00000114953 | 4930519K11Rik | RIKEN cDNA 4930519K11 gene |
| chr8 | 1,11E+08 | 1,11E+08 | 1280 | * | 11,1712 | 3,262261096 | 1,094973 | 2,979307 | 0,002889 | 0,748382289 | 0,918255 | Promoter (2-3kb) | 8 | 1,11E+08 | 1,11E+08 | 9293 | 2 | 73608 | ENSMUST00000051430.7 | -2528 | ENSMUSG00000001672 | Marveld3 | MARVEL (membrane-associating) domain containing 3 |
| chr5 | 66298702 | 66299720 | 1019 | * | 8,724199 | 3,262430879 | 1,12111 | 2,910001 | 0,003614 | 0,717133806 | 1,360622 | Intron (ENSMUST00000 | 5 | 66182885 | 66308299 | 125415 | 2 | 245945 | ENSMUST00000202794.4 | 8579 | ENSMUSG00000070780 | Rbm47 | RNA binding motif protein 47 |
| chr10 | 1,15E+08 | 1,15E+08 | 1507 | * | 12,5704 | 3,262776572 | 0,990018 | 3,295676 | 0,000982 | 0,528623776 | 1,075735 | Distal Intergenic | 10 | 1,15E+08 | 1,15E+08 | 28508 | 2 | 216344 | ENSMUST00000020343.9 | -13766 | ENSMUSG00000020132 | Rab21 | RAB21, member RAS oncogene family |
| chr4 | 1,09E+08 | 1,09E+08 | 1841 | * | 11,35497 | 3,26296882 | 1,168802 | 2,79172 | 0,005243 | 0,852340527 | 0,918255 | Distal Intergenic | 4 | 1,09E+08 | 1,09E+08 | 419 | 2 | 1,03E+08 | ENSMUST00000154561.2 | 4408 | | Ttc39aos1 | Ttc39a opposite strand RNA 1 |
| chr13 | 29200844 | 29202584 | 1741 | * | 15,24015 | 3,263753533 | 0,920988 | 3,543752 | 0,000394 | 0,326827873 | 1,41696 | Promoter (<=1kb) | 13 | 29200691 | 29224319 | 23629 | 1 | 77920 | ENSMUST00000145801.2 | 153 | ENSMUSG00000086363 | A330102I10Rik | RIKEN cDNA A330102I10 gene |
| chr15 | 81158305 | 81160154 | 1850 | * | 15,90144 | 3,264782127 | 0,882776 | 3,698314 | 0,000217 | 0,201373005 | 2,141612 | Intron (ENSMUST00000 | 15 | 81126858 | 81136063 | 9206 | 2 | 78102 | ENSMUST00000230860.2 | -22242 | ENSMUSG00000115970 | 8430426J06Rik | RIKEN cDNA 8430426J06 gene |
| chr9 | 56127298 | 56128284 | 987 | * | 8,740113 | 3,266849989 | 1,203435 | 2,714604 | 0,006636 | 0,736412045 | 2,237403 | Intron (ENSMUST00000 | 9 | 56108410 | 56151349 | 42940 | 2 | 244895 | ENSMUST00000188142.7 | 23065 | ENSMUSG00000074305 | Peak1 | pseudopodium-enriched atypical kinase 1 |
| chr4 | 1,5E+08 | 1,5E+08 | 1049 | * | 10,22171 | 3,26707008 | 1,066741 | 3,062665 | 0,002194 | 0,627785088 | 1,425929 | Distal Intergenic | 4 | 1,5E+08 | 1,5E+08 | 104 | 2 | 1,15E+08 | ENSMUST00000122697.3 | 13193 | ENSMUSG00000084646 | Gm24381 | predicted gene, 24381 |
| chr12 | 99138827 | 99140494 | 1668 | * | 12,53549 | 3,267159413 | 1,028641 | 3,176191 | 0,001492 | 0,573678185 | 1,2763 | Distal Intergenic | 12 | 99127536 | 99129067 | 1532 | 2 | 67639 | ENSMUST00000223511.2 | -9760 | ENSMUSG00000113210 | 4930474N09Rik | RIKEN cDNA 4930474N09 gene |
| chr17 | 29733135 | 29735088 | 1954 | * | 14,7132 | 3,267167293 | 1,086811 | 3,006197 | 0,002645 | 0,586304784 | 2,141612 | Distal Intergenic | 17 | 29712381 | 29713938 | 1558 | 1 | 18712 | ENSMUST00000234270.2 | 20754 | ENSMUSG00000024014 | Pim1 | proviral integration site 1 |
| chr17 | 33191089 | 33192549 | 1461 | * | 9,747148 | 3,269204897 | 1,241808 | 2,632616 | 0,008473 | 0,85617852 | 1,449706 | Intron (ENSMUST00000 | 17 | 33184819 | 33195256 | 10438 | 1 | 224691 | ENSMUST00000234770.2 | 6270 | ENSMUSG00000053600 | Zfp472 | zinc finger protein 472 |
| chr3 | 1,08E+08 | 1,08E+08 | 1432 | * | 11,70104 | 3,270474628 | 1,165997 | 2,804873 | 0,005034 | 0,801688106 | 1,173325 | Promoter (<=1kb) | 3 | 1,08E+08 | 1,08E+08 | 5442 | 2 | 72023 | ENSMUST00000155099.2 | -864 | ENSMUSG00000048796 | Cyb561d1 | cytochrome b-561 domain containing 1 |
| chr18 | 50639940 | 50641074 | 1135 | * | 12,99241 | 3,270570544 | 1,156127 | 2,828903 | 0,004671 | 0,870441665 | 0,738441 | Distal Intergenic | 18 | 50700727 | 50701795 | 1069 | 2 | 67365 | ENSMUST00000056460.4 | 60721 | ENSMUSG00000048875 | Pudp | pseudouridine 5'-phosphatase |
| chr17 | 83167926 | 83168453 | 528 | * | 9,367313 | 3,271348937 | 1,18127 | 2,76935 | 0,005617 | 0,789027175 | 1,425929 | Distal Intergenic | 17 | 83341023 | 83385654 | 44632 | 2 | 1,01E+08 | ENSMUST00000234063.2 | 217201 | ENSMUSG00000117399 | Gm19689 | predicted gene, 19689 |
| chr2 | 1,2E+08 | 1,2E+08 | 1563 | * | 12,70025 | 3,271784065 | 1,042507 | 3,138381 | 0,001699 | 0,660341484 | 0,918255 | Distal Intergenic | 2 | 1,2E+08 | 1,2E+08 | 70522 | 1 | 29808 | ENSMUST00000046717.13 | -30101 | ENSMUSG00000033943 | Mga | MAX gene associated |
| chr5 | 1,48E+08 | 1,48E+08 | 1638 | * | 11,89749 | 3,27239998 | 1,13269 | 2,889052 | 0,003864 | 0,740193979 | 1,2763 | Intron (ENSMUST00000 | 5 | 1,48E+08 | 1,48E+08 | 19394 | 2 | 11987 | ENSMUST00000138596.2 | -5967 | ENSMUSG00000041313 | Slc7a1 | solute carrier family 7 (cationic amino acid transporter, y+ system), member 1 |
| chr11 | 1,04E+08 | 1,04E+08 | 1517 | * | 12,88052 | 3,272613686 | 1,095327 | 2,987796 | 0,00281 | 0,684274806 | 1,2763 | Intron (ENSMUST00000 | 11 | 1,04E+08 | 1,04E+08 | 4507 | 2 | 18195 | ENSMUST00000146036.2 | -9004 | ENSMUSG00000034187 | Nsf | N-ethylmaleimide sensitive fusion protein |
| chr6 | 1,3E+08 | 1,3E+08 | 729 | * | 10,80807 | 3,272956758 | 1,065224 | 3,072553 | 0,002122 | 0,642453589 | 1,2763 | Intron (ENSMUST00000 | 6 | 1,3E+08 | 1,3E+08 | 74 | 1 | 1,15E+08 | ENSMUST00000240055.2 | 3662 | | Gm24072 | predicted gene, 24072 |
| chr7 | 1,26E+08 | 1,26E+08 | 1399 | * | 12,62276 | 3,273334666 | 1,209192 | 2,707043 | 0,006789 | 0,929109173 | 0,738441 | Promoter (1-2kb) | 7 | 1,26E+08 | 1,26E+08 | 1111 | 2 | 56420 | ENSMUST00000206477.2 | 1564 | ENSMUSG00000030697 | Ppp4c | protein phosphatase 4, catalytic subunit |
| chr11 | 58829661 | 58830990 | 1330 | * | 11,21747 | 3,274191392 | 1,068116 | 3,065389 | 0,002174 | 0,734194343 | 0,738441 | Promoter (<=1kb) | 11 | 58823114 | 58829742 | 6629 | 2 | 108660 | ENSMUST00000094151.6 | 0 | ENSMUSG00000020496 | Rnf187 | ring finger protein 187 |
| chr2 | 1,53E+08 | 1,53E+08 | 1232 | * | 9,758948 | 3,274263262 | 1,265033 | 2,588284 | 0,009646 | 0,923399738 | 1,086118 | Distal Intergenic | 2 | 1,53E+08 | 1,53E+08 | 41978 | 1 | 16569 | ENSMUST00000028977.7 | -10829 | ENSMUSG00000027475 | Kif3b | kinesin family member 3B |
| chr16 | 24860585 | 24861303 | 719 | * | 8,902811 | 3,274455757 | 1,241416 | 2,637678 | 0,008348 | 0,886373014 | 1,211315 | Distal Intergenic | 16 | 24878389 | 24887808 | 9420 | 1 | 319487 | ENSMUST00000181655.2 | -17086 | ENSMUSG00000096960 | A230028O05Rik | RIKEN cDNA A230028O05 gene |
| chr6 | 8508348 | 8509519 | 1172 | * | 16,32432 | 3,275077929 | 0,871722 | 3,757023 | 0,000172 | 0,174628563 | 2,141612 | Promoter (<=1kb) | 6 | 8489941 | 8509172 | 19232 | 2 | 320312 | ENSMUST00000247340.1 | 0 | ENSMUSG00000087305 | A430035B10Rik | RIKEN cDNA A430035B10 gene |
| chr16 | 74593296 | 74594714 | 1419 | * | 9,102238 | 3,275867942 | 1,200686 | 2,728331 | 0,006366 | 0,730469267 | 2,237403 | Distal Intergenic | 16 | 74404645 | 74404770 | 126 | 1 | 1,15E+08 | ENSMUST00000157767.3 | 188651 | ENSMUSG00000088392 | Gm26307 | predicted gene, 26307 |
| chr19 | 59879633 | 59881754 | 2122 | * | 17,60128 | 3,276029851 | 1,026359 | 3,191894 | 0,001413 | 0,471171835 | 2,154504 | Distal Intergenic | 19 | 59895348 | 59931432 | 36085 | 2 | 74998 | ENSMUST00000170819.2 | 49678 | ENSMUSG00000040022 | Rab11fip2 | RAB11 family interacting protein 2 (class I) |
| chr5 | 1,22E+08 | 1,22E+08 | 2624 | * | 17,93889 | 3,276438262 | 0,936675 | 3,497946 | 0,000469 | 0,282588188 | 2,386008 | Promoter (2-3kb) | 5 | 1,22E+08 | 1,22E+08 | 26569 | 1 | 320717 | ENSMUST00000119015.2 | -2353 | ENSMUSG00000038582 | Pptc7 | PTC7 protein phosphatase homolog |
| chr11 | 79696043 | 79697063 | 1021 | * | 9,157908 | 3,277762575 | 1,235703 | 2,652548 | 0,007989 | 0,769692849 | 2,237403 | Promoter (<=1kb) | 11 | 79687019 | 79695998 | 8980 | 2 | 1,05E+08 | ENSMUST00000162268.2 | -45 | | 9130204K15Rik | RIKEN cDNA 9130204K15 gene |
| chr16 | 52278268 | 52279451 | 1184 | * | 10,03997 | 3,278064319 | 1,024171 | 3,2007 | 0,001371 | 0,591609342 | 1,086118 | Distal Intergenic | 16 | 52069359 | 52274437 | 205079 | 2 | 11658 | ENSMUST00000023312.14 | -3831 | ENSMUSG00000022636 | Alcam | activated leukocyte cell adhesion molecule |
| chr4 | 13279128 | 13280440 | 1313 | * | 12,32622 | 3,278077411 | 1,06767 | 3,070311 | 0,002138 | 0,693003605 | 0,918255 | Distal Intergenic | 4 | 13076140 | 13096806 | 20667 | 1 | 1,03E+08 | ENSMUST00000139614.2 | 202988 | | Gm11820 | predicted gene 11820 |
| chr9 | 22906473 | 22907970 | 1498 | * | 13,69071 | 3,279029615 | 1,041134 | 3,14948 | 0,001636 | 0,627067801 | 1,075735 | Distal Intergenic | 9 | 23134372 | 23396496 | 262125 | 1 | 73230 | ENSMUST00000071982.7 | -226402 | ENSMUSG00000031963 | Bmper | BMP-binding endothelial regulator |
| chr10 | 25258501 | 25259840 | 1340 | * | 11,8067 | 3,281189443 | 1,078577 | 3,042148 | 0,002349 | 0,713633527 | 0,918255 | Intron (ENSMUST00000 | 10 | 25274557 | 25360178 | 85622 | 1 | 13822 | ENSMUST00000219900.2 | -14717 | ENSMUSG00000019978 | Epb41l2 | erythrocyte membrane protein band 4.1 like 2 |
| chr10 | 1E+08 | 1E+08 | 1138 | * | 8,838673 | 3,281532659 | 1,228852 | 2,670405 | 0,007576 | 0,850457798 | 1,360622 | Distal Intergenic | 10 | 99965730 | 99979988 | 14259 | 2 | 1,03E+08 | ENSMUST00000218902.2 | -79582 | ENSMUSG00000112831 | Gm35533 | predicted gene, 35533 |
| chr4 | 74304293 | 74305375 | 1083 | * | 7,701796 | 3,283346218 | 1,2648 | 2,595941 | 0,009433 | 1 | 0 | Intron (ENSMUST00000 | 4 | 74181047 | 74191896 | 10850 | 1 | 76804 | ENSMUST00000150439.2 | 123246 | ENSMUSG00000028397 | Kdm4c | lysine (K)-specific demethylase 4C |
| chr9 | 83301099 | 83302670 | 1572 | * | 14,65617 | 3,283692208 | 0,904448 | 3,630602 | 0,000283 | 0,298645609 | 1,2763 | Intron (ENSMUST00000 | 9 | 83272346 | 83309014 | 36669 | 2 | 75782 | ENSMUST00000190514.7 | 6344 | ENSMUSG00000032258 | Lca5 | Leber congenital amaurosis 5 (human) |
| chr16 | 35589826 | 35591119 | 1294 | * | 9,026648 | 3,283877382 | 1,269834 | 2,586069 | 0,009708 | 0,908763428 | 1,211315 | Promoter (<=1kb) | 16 | 35590745 | 35645918 | 55174 | 1 | 66667 | ENSMUST00000231579.2 | 0 | ENSMUSG00000022849 | Hspbap1 | Hspb associated protein 1 |
| chr4 | 45006929 | 45008287 | 1359 | * | 12,6525 | 3,284725185 | 0,990027 | 3,317815 | 0,000907 | 0,54122637 | 0,918255 | Intron (ENSMUST00000 | 4 | 45011687 | 45015406 | 3720 | 1 | 73405 | ENSMUST00000186030.2 | -3400 | ENSMUSG00000070997 | 1700055D18Rik | RIKEN cDNA 1700055D18 gene |
| chr3 | 95777351 | 95779138 | 1788 | * | 13,65121 | 3,28538155 | 1,046996 | 3,137911 | 0,001702 | 0,691026033 | 0,738441 | Promoter (<=1kb) | 3 | 95778834 | 95795268 | 16435 | 1 | 1E+08 | ENSMUST00000142433.2 | 0 | ENSMUSG00000080727 | C920021L13Rik | RIKEN cDNA C920021L13 gene |
| chr8 | 1,08E+08 | 1,08E+08 | 1509 | * | 14,56588 | 3,286091944 | 1,135936 | 2,892849 | 0,003818 | 0,721176228 | 1,41696 | Promoter (1-2kb) | 8 | 1,08E+08 | 1,08E+08 | 219 | 2 | 1,15E+08 | ENSMUST00000212585.2 | 1460 | | Gm8940 | predicted gene 8940 |
| chr16 | 29907108 | 29910136 | 3029 | * | 41,607744 | 3,286410399 | 0,5801115 | 5,6651355 | 1,469E-08 | 0,000152202 | 2,7254908 | Distal Intergenic | 16 | 29885156 | 29886137 | 982 | 1 | 15205 | ENSMUST00000161839.2 | 21952 | ENSMUSG00000022528 | Hes1 | hes family bHLH transcription factor 1 |
| chr4 | 1,08E+08 | 1,08E+08 | 1701 | * | 24,51039 | 3,287233467 | 1,215157 | 2,705192 | 0,006826 | 0,713406565 | 2,67243 | Promoter (<=1kb) | 4 | 1,08E+08 | 1,08E+08 | 69949 | 1 | 16975 | ENSMUST00000146552.9 | 559 | ENSMUSG00000028613 | Lrp8 | low density lipoprotein receptor-related protein 8, apolipoprotein e receptor |
| chr18 | 12653762 | 12655019 | 1258 | * | 12,04953 | 3,287772945 | 1,084517 | 3,031554 | 0,002433 | 0,716908183 | 0,918255 | Intron (ENSMUST00000 | 18 | 12634247 | 12637483 | 3237 | 2 | 1,05E+08 | ENSMUST00000188118.2 | -16279 | | Gm29200 | predicted gene 29200 |
| chr14 | 58059174 | 58060728 | 1555 | * | 9,616537 | 3,287781399 | 1,187864 | 2,767809 | 0,005643 | 0,786955198 | 1,449706 | Promoter (2-3kb) | 14 | 58059344 | 58063565 | 4222 | 2 | 219114 | ENSMUST00000225753.2 | 2837 | ENSMUSG00000021965 | Ska3 | spindle and kinetochore associated complex subunit 3 |
| chr3 | 32450152 | 32452324 | 2173 | * | 12,42019 | 3,287860373 | 1,024231 | 3,210077 | 0,001327 | 0,56742763 | 1,173325 | Promoter (<=1kb) | 3 | 32451820 | 32520256 | 68437 | 1 | 18706 | ENSMUST00000108243.8 | 0 | ENSMUSG00000027665 | Pik3ca | phosphatidylinositol-4,5-bisphosphate 3-kinase catalytic subunit alpha |
| chr6 | 47294994 | 47297433 | 2440 | * | 37,303638 | 3,289781008 | 0,7007255 | 4,6948212 | 2,668E-06 | 0,008109885 | 2,7872553 | Distal Intergenic | 6 | 47221387 | 47276504 | 55118 | 1 | 66797 | ENSMUST00000060839.8 | 73607 | ENSMUSG00000039419 | Cntnap2 | contactin associated protein-like 2 |
| chr6 | 48554662 | 48557345 | 2684 | * | 33,851515 | 3,290469096 | 0,6299644 | 5,2232617 | 1,758E-07 | 0,001081418 | 2,7254908 | Distal Intergenic | 6 | 48546644 | 48549723 | 3080 | 2 | 71660 | ENSMUST00000204182.2 | -4939 | ENSMUSG00000009281 | Rarres2 | retinoic acid receptor responder (tazarotene induced) 2 |
| chr9 | 61963501 | 61964953 | 1453 | * | 11,65788 | 3,290653223 | 1,105402 | 2,976883 | 0,002912 | 0,68845579 | 1,2763 | 3' UTR | 9 | 61964530 | 61977888 | 13359 | 2 | 93683 | ENSMUST00000034785.8 | 12935 | ENSMUSG00000032252 | Glce | glucuronyl C5-epimerase |
| chr15 | 83280535 | 83281687 | 1153 | * | 9,252207 | 3,291637421 | 1,137546 | 2,89363 | 0,003808 | 0,727652793 | 1,360622 | Intron (ENSMUST00000 | 15 | 83261779 | 83272803 | 11025 | 2 | 23970 | ENSMUST00000230030.2 | -7732 | ENSMUSG00000016664 | Pacsin2 | protein kinase C and casein kinase substrate in neurons 2 |
| chr18 | 8552652 | 8554008 | 1357 | * | 9,998358 | 3,291972981 | 1,245404 | 2,643297 | 0,00821 | 0,852340527 | 1,449706 | Distal Intergenic | 18 | 7910619 | 7918503 | 7885 | 1 | 225131 | ENSMUST00000167542.2 | 642033 | ENSMUSG00000024283 | Wac | WW domain containing adaptor with coiled-coil |
| chr9 | 57302273 | 57304408 | 2136 | * | 20,27885 | 3,29225351 | 0,904747 | 3,638866 | 0,000274 | 0,223919137 | 2,221611 | Intron (ENSMUST00000 | 9 | 57322449 | 57328601 | 6153 | 2 | 66812 | ENSMUST00000217255.2 | 24193 | ENSMUSG00000063849 | Ppcdc | phosphopantothenoylcysteine decarboxylase |
| chr18 | 64616391 | 64618257 | 1867 | * | 15,00578 | 3,292837074 | 1,058973 | 3,109461 | 0,001874 | 0,62053486 | 1,2763 | 3' UTR | 18 | 64608218 | 64622012 | 13795 | 2 | 14151 | ENSMUST00000238149.2 | 3755 | ENSMUSG00000024588 | Fech | ferrochelatase |
| chr7 | 27094705 | 27095752 | 1048 | * | 16,30441 | 3,293073402 | 0,832698 | 3,954702 | 7,66E-05 | 0,148900155 | 1,2763 | Promoter (<=1kb) | 7 | 27055808 | 27095623 | 39816 | 2 | 80297 | ENSMUST00000108364.8 | 0 | ENSMUSG00000011751 | Sptbn4 | spectrin beta, non-erythrocytic 4 |
| chr5 | 59343715 | 59345083 | 1369 | * | 8,792763 | 3,293552208 | 1,133518 | 2,905601 | 0,003665 | 0,711174281 | 1,449706 | Distal Intergenic | 5 | 59372289 | 59372395 | 107 | 2 | 1,15E+08 | ENSMUST00000239835.1 | 27312 | ENSMUSG00000119859 | Gm22308 | predicted gene, 22308 |
| chr1 | 1,25E+08 | 1,25E+08 | 906 | * | 6,994302 | 3,293819697 | 1,273338 | 2,58676 | 0,009688 | 1 | 0 | Exon (ENSMUST000000 | 1 | 1,25E+08 | 1,26E+08 | 34805 | 1 | 74150 | ENSMUST00000027580.11 | 9031 | ENSMUSG00000026342 | Slc35f5 | solute carrier family 35, member F5 |
| chr18 | 75476892 | 75479067 | 2176 | * | 21,9813 | 3,294016811 | 0,811882 | 4,057261 | 4,97E-05 | 0,068345002 | 2,880017 | Distal Intergenic | 18 | 75491711 | 75500160 | 8450 | 2 | 1,03E+08 | ENSMUST00000174338.3 | 21093 | | Gm20544 | predicted gene 20544 |
| chr9 | 55339296 | 55340418 | 1123 | * | 15,11796 | 3,294122607 | 1,139908 | 2,889815 | 0,003855 | 0,739716578 | 1,2763 | Intron (ENSMUST00000 | 9 | 55381821 | 55394797 | 12977 | 2 | 110842 | ENSMUST00000139342.2 | 54379 | ENSMUSG00000032314 | Etfa | electron transferring flavoprotein, alpha polypeptide |
| chr18 | 41386450 | 41387028 | 579 | * | 9,807239 | 3,295070976 | 1,077754 | 3,057349 | 0,002233 | 0,627785088 | 1,449706 | Distal Intergenic | 18 | 40854557 | 40866926 | 12370 | 1 | 1,03E+08 | ENSMUST00000237626.2 | 531893 | ENSMUSG00000117847 | Gm31205 | predicted gene, 31205 |
| chr10 | 47673282 | 47674662 | 1381 | * | 9,273626 | 3,295404487 | 1,244924 | 2,647072 | 0,008119 | 0,773819636 | 2,237403 | Distal Intergenic | 10 | 46892675 | 46978928 | 86254 | 2 | 1,03E+08 | ENSMUST00000214355.3 | -694354 | | Gm35552 | predicted gene, 35552 |
| chr16 | 57519259 | 57520324 | 1066 | * | 9,814184 | 3,295498783 | 1,17354 | 2,808168 | 0,004982 | 0,793220117 | 1,211315 | Intron (ENSMUST00000 | 16 | 57444621 | 57575100 | 130480 | 2 | 12837 | ENSMUST00000089332.5 | 54776 | ENSMUSG00000068196 | Col8a1 | collagen, type VIII, alpha 1 |
| chr9 | 1,21E+08 | 1,21E+08 | 1551 | * | 11,07693 | 3,295674518 | 1,169973 | 2,816881 | 0,004849 | 0,838878373 | 0,918255 | Intron (ENSMUST00000 | 9 | 1,21E+08 | 1,21E+08 | 32682 | 2 | 209012 | ENSMUST00000164969.2 | -19823 | ENSMUSG00000040936 | Ulk4 | unc-51-like kinase 4 |
| chr11 | 50112626 | 50113943 | 1318 | * | 9,308908 | 3,296315013 | 1,234229 | 2,670749 | 0,007568 | 0,83726674 | 1,449706 | Promoter (2-3kb) | 11 | 50116162 | 50125930 | 9769 | 1 | 103534 | ENSMUST00000041725.14 | -2219 | ENSMUSG00000036620 | Mgat4b | mannoside acetylglucosaminyltransferase 4, isoenzyme B |
| chr7 | 28427047 | 28428684 | 1638 | * | 41,611849 | 3,296965538 | 0,5263413 | 6,2639308 | 3,754E-10 | 6,45401E-06 | 2,9254973 | Intron (ENSMUST000003 | 7 | 28431925 | 28436907 | 4983 | 1 | 50760 | ENSMUST00000167118.2 | -3241 | ENSMUSG00000030598 | Fbxo17 | F-box protein 17 |
| chr18 | 80495307 | 80496033 | 727 | * | 11,08658 | 3,297153994 | 0,977837 | 3,371886 | 0,000747 | 0,451585113 | 1,2763 | Exon (ENSMUST000000 | 18 | 80451180 | 80484074 | 32895 | 2 | 67655 | ENSMUST00000161003.2 | -11233 | ENSMUSG00000033323 | Ctdp1 | CTD (carboxy-terminal domain, RNA polymerase II, polypeptide A) phosphatase, subunit 1 |
| chr13 | 41803614 | 41805540 | 1927 | * | 15,92015 | 3,297805256 | 0,965254 | 3,416515 | 0,000634 | 0,334753716 | 2,141612 | Exon (ENSMUST000002 | 13 | 41759692 | 41794831 | 35140 | 1 | 621976 | ENSMUST00000129449.3 | 43922 | ENSMUSG00000087370 | Tmem170b | transmembrane protein 170B |
| chr5 | 1,3E+08 | 1,3E+08 | 1212 | * | 11,00623 | 3,298714722 | 1,222837 | 2,697593 | 0,006984 | 0,934396727 | 0,738441 | Promoter (<=1kb) | 5 | 1,3E+08 | 1,3E+08 | 21892 | 2 | 100678 | ENSMUST00000031399.13 | 0 | ENSMUSG00000029446 | Psph | phosphoserine phosphatase |
| chr5 | 1,19E+08 | 1,19E+08 | 2061 | * | 18,99313 | 3,300274225 | 1,017546 | 3,243365 | 0,001181 | 0,424527166 | 2,386008 | Distal Intergenic | 5 | 1,19E+08 | 1,19E+08 | 6310 | 1 | 73477 | ENSMUST00000190550.2 | -59798 | ENSMUSG00000100573 | 1700081H04Rik | RIKEN cDNA 1700081H04 gene |
| chr5 | 137879975 | 137882911 | 2937 | * | 37,993265 | 3,301178908 | 0,5960859 | 5,5380928 | 3,058E-08 | 0,000295704 | 2,3776742 | Exon (ENSMUST0000024 | 5 | 137869421 | 137870077 | 657 | 2 | 545812 | ENSMUST00000199773.2 | -9898 | ENSMUSG00000066682 | Pilrb2 | paired immunoglobin-like type 2 receptor beta 2 |
| chr3 | 31114533 | 31115780 | 1248 | * | 10,77374 | 3,301502619 | 1,041898 | 3,168739 | 0,001531 | 0,595849045 | 1,173325 | Exon (ENSMUST000002 | 3 | 31114951 | 31133794 | 18844 | 2 | 1,03E+08 | ENSMUST00000245305.1 | 18014 | | Gm33694 | predicted gene, 33694 |

| | | | | | | | | | | | | | | | | | | | | | | |
|---|---|---|---|---|---|---|---|---|---|---|---|---|---|---|---|---|---|---|---|---|---|---|
| chr7 | 1,21E+08 | 1,21E+08 | 1149 | * | 11,05797 | 3,301609728 | 1,083986 | 3,045806 | 0,002321 | 0,711174281 | 0,918255 | Promoter (<=1kb) | 7 | 1,21E+08 | 1,21E+08 | 492 | 2 | 668548 | ENSMUST00000077747.6 | -350 | | Gm9234 | predicted pseudogene 9234 |
| chr9 | 88318498 | 88320639 | 2142 | * | 13,29488 | 3,302138605 | 0,963731 | 3,426412 | 0,000612 | 0,419066905 | 1,2763 | Promoter (<=1kb) | 9 | 88258805 | 88320951 | 62147 | 2 | 244962 | ENSMUST00000173011.9 | 312 | ENSMUSG00000032422 | Snx14 | sorting nexin 14 |
| chr6 | 92750376 | 92751126 | 751 | * | 8,713706 | 3,303554615 | 1,240268 | 2,663581 | 0,007731 | 0,852340527 | 1,360622 | 3' UTR | 6 | 92793459 | 92820446 | 26988 | 1 | 74516 | ENSMUST00000057977.4 | -42333 | ENSMUSG00000048636 | A730049H05Rik | RIKEN cDNA A730049H05 gene |
| chr10 | 59515612 | 59516828 | 1217 | * | 10,89334 | 3,304167004 | 1,051741 | 3,141616 | 0,00168 | 0,68680243 | 0,744375 | Distal Intergenic | 10 | 59538299 | 59699954 | 161656 | 1 | 216001 | ENSMUST00000165563.8 | -21471 | ENSMUSG00000020111 | Micu1 | mitochondrial calcium uptake 1 |
| chr4 | 1,51E+08 | 1,51E+08 | 1679 | * | 14,34598 | 3,304787255 | 1,117541 | 2,957195 | 0,003105 | 0,75414571 | 0,938682 | Intron (ENSMUST00000 | 4 | 1,5E+08 | 1,51E+08 | 63988 | 1 | 68703 | ENSMUST00000131600.2 | 22437 | ENSMUSG00000039852 | Rere | arginine glutamic acid dipeptide (RE) repeats |
| chr8 | 30383517 | 30384444 | 928 | * | 10,28407 | 3,304987717 | 1,107226 | 2,984927 | 0,002836 | 0,671156916 | 1,425929 | Distal Intergenic | 8 | 30602338 | 30611242 | 8905 | 2 | 1,05E+08 | ENSMUST00000210257.2 | 226798 | | Gm39154 | predicted gene, 39154 |
| chr9 | 22974909 | 22976382 | 1474 | * | 17,18222 | 3,305301274 | 0,949257 | 3,48199 | 0,000498 | 0,27718041 | 2,663619 | Distal Intergenic | 9 | 23134372 | 23396496 | 262125 | 1 | 73230 | ENSMUST00000071982.7 | -157990 | ENSMUSG00000031963 | Bmper | BMP-binding endothelial regulator |
| chr7 | 66631297 | 66632464 | 1168 | * | 10,65357 | 3,305311981 | 1,272935 | 2,596606 | 0,009415 | 0,947711819 | 0,918255 | Intron (ENSMUST00000 | 7 | 66697219 | 66728052 | 30834 | 1 | 233332 | ENSMUST00000154103.2 | -64755 | ENSMUSG00000058145 | Adamts17 | a disintegrin-like and metallopeptidase (reprolysin type) with thrombospondin type 1 motif, 17 |
| chr10 | 59370946 | 59371915 | 970 | * | 9,043079 | 3,305321942 | 1,130707 | 2,923235 | 0,003464 | 0,627862701 | 2,237403 | Intron (ENSMUST00000 | 10 | 59284419 | 59293982 | 9564 | 2 | 215999 | ENSMUST00000009791.7 | -76964 | ENSMUSG00000009647 | Mcu | mitochondrial calcium uniporter |
| chr3 | 55101569 | 55103146 | 1578 | * | 11,73409 | 3,305650029 | 1,109167 | 2,980301 | 0,00288 | 0,748211534 | 0,918255 | Exon (ENSMUST000000 | 3 | 55089449 | 55117378 | 27930 | 1 | 74434 | ENSMUST00000029369.5 | 12120 | ENSMUSG00000027794 | Sohlh2 | spermatogenesis and oogenesis specific basic helix-loop-helix 2 |
| chr15 | 1E+08 | 1E+08 | 1046 | * | 14,53731 | 3,305805508 | 1,193743 | 2,769277 | 0,005618 | 0,806557336 | 1,2763 | Promoter (<=1kb) | 15 | 1E+08 | 1E+08 | 4911 | 2 | 19009 | ENSMUST00000177335.2 | -474 | ENSMUSG00000009739 | Pou6f1 | POU domain, class 6, transcription factor 1 |
| chr1 | 3969149 | 3970519 | 1371 | * | 10,6625 | 3,3058299 | 1,173027 | 2,818204 | 0,004829 | 0,876007548 | 0,744375 | Distal Intergenic | 1 | 3284705 | 3741721 | 457017 | 2 | 497097 | ENSMUST00000070533.5 | -227428 | ENSMUSG00000051951 | Xkr4 | X-linked Kx blood group related 4 |
| chrX | 1,53E+08 | 1,53E+08 | 1285 | * | 10,24272 | 3,305957784 | 1,136907 | 2,907851 | 0,003639 | 0,744866969 | 1,173325 | Distal Intergenic | 20 | 1,53E+08 | 1,53E+08 | 17675 | 1 | 70123 | ENSMUST00000140575.8 | -3519 | ENSMUSG00000086316 | Nbdy | negative regulator of P-body association |
| chr11 | 1,16E+08 | 1,16E+08 | 932 | * | 8,917251 | 3,307952814 | 1,199435 | 2,757926 | 0,005817 | 0,823446343 | 1,211315 | Promoter (<=1kb) | 11 | 1,16E+08 | 1,16E+08 | 3091 | 2 | 217342 | ENSMUST00000147851.2 | 0 | ENSMUSG00000020802 | Ube2o | ubiquitin-conjugating enzyme E2O |
| chr6 | 65822273 | 65823042 | 770 | * | 9,300701 | 3,30815255 | 1,232355 | 2,684415 | 0,007266 | 0,832960417 | 1,425929 | Intron (ENSMUST00000 | 6 | 65756085 | 65913118 | 157034 | 1 | 70779 | ENSMUST00000172638.2 | 66188 | ENSMUSG00000029913 | Prdm5 | PR domain containing 5 |
| chr17 | 36239474 | 36240552 | 1079 | * | 11,65276 | 3,308172469 | 1,17637 | 2,812186 | 0,004921 | 0,879289589 | 0,744375 | Promoter (2-3kb) | 17 | 36237386 | 36239221 | 1836 | 1 | 52040 | ENSMUST00000151375.2 | 2088 | ENSMUSG00000039220 | Ppp1r10 | protein phosphatase 1, regulatory subunit 10 |
| chr2 | 84379123 | 84381138 | 2016 | * | 19,43821 | 3,309226482 | 0,903352 | 3,663273 | 0,000249 | 0,204595855 | 2,386008 | Distal Intergenic | 2 | 84331089 | 84361718 | 30630 | 2 | 672763 | ENSMUST00000139492.3 | -17405 | ENSMUSG00000087362 | Gm13710 | predicted gene 13710 |
| chr5 | 97935920 | 97936890 | 971 | * | 9,016883 | 3,309450036 | 1,21151 | 2,731674 | 0,006301 | 0,802671652 | 1,449706 | Distal Intergenic | 5 | 97848713 | 97884671 | 35959 | 1 | 1,19E+08 | ENSMUST00000244134.1 | 87207 | | Gm54318 | predicted gene, 54318 |
| chr11 | 51967955 | 51968650 | 696 | * | 8,328247 | 3,311667312 | 1,161673 | 2,850775 | 0,004361 | 0,749550311 | 1,360622 | Intron (ENSMUST00000 | 11 | 51989508 | 52018605 | 29098 | 1 | 19052 | ENSMUST00000020608.3 | -20858 | ENSMUSG00000020349 | Ppp2ca | protein phosphatase 2 (formerly 2A), catalytic subunit, alpha isoform |
| chr6 | 87824821 | 87826125 | 1305 | * | 11,56965 | 3,312277442 | 1,283897 | 2,579862 | 0,009884 | 0,954881543 | 0,918255 | Promoter (1-2kb) | 6 | 87822512 | 87823299 | 788 | 2 | 12785 | ENSMUST00000204104.2 | -1522 | ENSMUSG00000030057 | Cnbp | cellular nucleic acid binding protein |
| chr18 | 50484590 | 50485450 | 861 | * | 7,842608 | 3,312362662 | 1,204733 | 2,749457 | 0,005969 | 1 | 0 | Distal Intergenic | 18 | 50411438 | 50416081 | 4644 | 1 | 225497 | ENSMUST00000238078.2 | 73152 | ENSMUSG00000035420 | Fam170a | family with sequence similarity 170, member A |
| chr16 | 82207392 | 82208141 | 750 | * | 9,350992 | 3,312517385 | 1,157337 | 2,862189 | 0,004207 | 0,734804995 | 1,425929 | Distal Intergenic | 16 | 82340117 | 82340223 | 107 | 1 | 1,15E+08 | ENSMUST00000240414.1 | -131976 | ENSMUSG00000119122 | Gm23355 | predicted gene, 23355 |
| chr4 | 135843083 | 135844927 | 1845 | * | 21,805139 | 3,31294573 | 0,7008984 | 4,7267131 | 2,282E-06 | 0,007032136 | 2,9519335 | Distal Intergenic | 4 | 135870808 | 135872703 | 1896 | 1 | 15903 | ENSMUST00000008016.3 | -25881 | ENSMUSG00000007872 | Id3 | inhibitor of DNA binding 3 |
| chr18 | 20745530 | 20746139 | 610 | * | 7,007375 | 3,313085608 | 1,224797 | 2,705007 | 0,00683 | 1 | 0 | Intron (ENSMUST00000 | 18 | 20731021 | 20798317 | 67297 | 2 | 1,03E+08 | ENSMUST00000130296.2 | 52178 | ENSMUSG00000085595 | Gm16090 | predicted gene 16090 |
| chr11 | 94091343 | 94093135 | 1793 | * | 14,91155 | 3,313196857 | 1,04947 | 3,157019 | 0,001594 | 0,492834344 | 2,141612 | Distal Intergenic | 11 | 94102280 | 94106321 | 4042 | 1 | 22057 | ENSMUST00000041589.6 | -9145 | ENSMUSG00000037573 | Tob1 | transducer of ErbB-2.1 |
| chr7 | 75430640 | 75433233 | 2594 | * | 31,630857 | 3,313533131 | 0,6570802 | 5,0428139 | 4,587E-07 | 0,002049962 | 2,8585108 | Promoter (<=1kb) | 7 | 75418786 | 75431869 | 13084 | 2 | 108167440 | ENSMUST00000181224.3 | 0 | | AU020206 | expressed sequence AU020206 |
| chr12 | 34090095 | 34091604 | 1510 | * | 15,19031 | 3,313544631 | 0,977761 | 3,388912 | 0,000702 | 0,351105383 | 2,141612 | Distal Intergenic | 12 | 34007670 | 34009828 | 2159 | 1 | 22160 | ENSMUST00000049089.7 | 82425 | ENSMUSG00000035799 | Twist1 | twist basic helix-loop-helix transcription factor 1 |
| chr8 | 74285031 | 74285759 | 729 | * | 8,214684 | 3,313720214 | 1,176347 | 2,816957 | 0,004848 | 1 | 0 | Distal Intergenic | 8 | 73541227 | 74080168 | 538942 | 2 | 16795 | ENSMUST00000212459.2 | -204863 | ENSMUSG00000004383 | Large1 | LARGE xylosyl- and glucuronyltransferase 1 |
| chr1 | 1,77E+08 | 1,77E+08 | 1205 | * | 9,765122 | 3,314447798 | 1,174253 | 2,8226 | 0,004764 | 0,788923377 | 1,211315 | Intron (ENSMUST00000 | 1 | 1,77E+08 | 1,77E+08 | 226653 | 2 | 23797 | ENSMUST00000111160.9 | 11070 | ENSMUSG00000019699 | Akt3 | thymoma viral proto-oncogene 3 |
| chr17 | 34122714 | 34123672 | 959 | * | 9,603125 | 3,314696896 | 1,159047 | 2,859847 | 0,004238 | 0,771766981 | 1,173325 | Promoter (<=1kb) | 17 | 34124109 | 34125649 | 1541 | 1 | 474160 | ENSMUST00000172817.2 | -437 | ENSMUSG00000092349 | Smim40 | small integral membrane protein 40 |
| chr15 | 101042841 | 101044527 | 1687 | * | 36,761223 | 3,315897201 | 0,5821151 | 5,6962913 | 1,224E-08 | 0,000126258 | 2,7872553 | 3' UTR | 15 | 101035481 | 101040505 | 5025 | 1 | 11482 | ENSMUST00000128525.2 | 7360 | ENSMUSG00000000530 | Acvrl1 | activin A receptor, type II-like 1 |
| chr17 | 25978646 | 25980315 | 1670 | * | 13,88909 | 3,31625622 | 0,954308 | 3,475038 | 0,000511 | 0,38991127 | 1,2763 | Exon (ENSMUST000002 | 17 | 25967589 | 25973352 | 5764 | 2 | 56047 | ENSMUST00000237359.2 | -5294 | ENSMUSG00000063011 | Msln | mesothelin |
| chr12 | 1,12E+08 | 1,12E+08 | 908 | * | 11,51084 | 3,317305618 | 1,227387 | 2,702738 | 0,006877 | 0,932174684 | 0,738441 | Promoter (2-3kb) | 12 | 1,12E+08 | 1,12E+08 | 14388 | 1 | 74691 | ENSMUST00000193586.2 | 2582 | ENSMUSG00000054003 | Tdrd9 | tudor domain containing 9 |
| chr10 | 74365111 | 74366004 | 894 | * | 8,953833 | 3,317371979 | 1,208573 | 2,744867 | 0,006054 | 0,829907213 | 1,211315 | Intron (ENSMUST00000 | 10 | 74420048 | 74460448 | 40401 | 1 | 11994 | ENSMUST00000128843.2 | -54044 | ENSMUSG00000052613 | Pcdh15 | protocadherin 15 |
| chr13 | 57211633 | 57212610 | 978 | * | 9,395851 | 3,31742733 | 1,184601 | 2,800461 | 0,005103 | 0,770104803 | 1,425929 | Distal Intergenic | 13 | 57078633 | 57078761 | 129 | 2 | 1,15E+08 | ENSMUST00000104091.3 | -132872 | ENSMUSG00000077279 | Gm22777 | predicted gene, 22777 |
| chr13 | 22017594 | 22019541 | 1948 | * | 19,026777 | 3,317878874 | 0,7735053 | 4,2894067 | 1,792E-05 | 0,035492893 | 2,6636191 | Promoter (<=1kb) | 13 | 22017196 | 22017677 | 482 | 2 | 319171 | ENSMUST00000081342.7 | 0 | ENSMUSG00000094777 | H2ac24 | H2A clustered histone 24 |
| chr11 | 77162548 | 77164241 | 1694 | * | 12,70252 | 3,318353601 | 1,05306 | 3,151153 | 0,001626 | 0,652050642 | 0,918255 | Intron (ENSMUST00000 | 11 | 77135754 | 77135841 | 88 | 2 | 1,15E+08 | ENSMUST00000104461.3 | -26707 | ENSMUSG00000077649 | Gm24989 | predicted gene, 24989 |
| chr10 | 4196091 | 4197240 | 1150 | * | 11,04727 | 3,318530266 | 1,237565 | 2,681501 | 0,007329 | 0,908065779 | 0,918255 | Distal Intergenic | 10 | 4216380 | 4309470 | 93091 | 1 | 83397 | ENSMUST00000045730.7 | -19140 | ENSMUSG00000038587 | Akap12 | A kinase (PRKA) anchor protein (gravin) 12 |
| chr13 | 54053729 | 54054944 | 1216 | * | 12,22398 | 3,31858949 | 1,103601 | 3,007056 | 0,002638 | 0,769692849 | 0,738441 | Distal Intergenic | 13 | 54064673 | 54069782 | 5110 | 2 | 78550 | ENSMUST00000243613.1 | 14838 | ENSMUSG00000114069 | E130119H09Rik | RIKEN cDNA E130119H09 gene |
| chr3 | 88202374 | 88203889 | 1516 | * | 13,36213 | 3,319332338 | 1,018368 | 3,259463 | 0,001116 | 0,582425528 | 0,918255 | Promoter (<=1kb) | 3 | 88190067 | 88204145 | 14079 | 2 | 76927 | ENSMUST00000010682.4 | 256 | ENSMUSG00000010538 | Tsacc | TSSK6 activating co-chaperone |
| chr1 | 27289643 | 27291300 | 1658 | * | 11,39131 | 3,319599306 | 1,064764 | 3,117684 | 0,001823 | 0,614953039 | 1,2763 | Distal Intergenic | 1 | 26720895 | 26726541 | 5647 | 2 | 210940 | ENSMUST00000097801.4 | -563102 | ENSMUSG00000073722 | 4931408C20Rik | RIKEN cDNA 4931408C20 gene |
| chr2 | 38533992 | 38535273 | 1282 | * | 16,65672 | 3,320887544 | 0,913928 | 3,633643 | 0,000279 | 0,297494857 | 1,2763 | Promoter (<=1kb) | 2 | 38478048 | 38534099 | 56052 | 2 | 19177 | ENSMUST00000028083.6 | 0 | ENSMUSG00000026750 | Psmb7 | proteasome (prosome, macropain) subunit, beta type 7 |
| chr19 | 16310836 | 16311909 | 1074 | * | 8,805724 | 3,322764274 | 1,212744 | 2,739872 | 0,006146 | 0,799539751 | 1,449706 | Intron (ENSMUST00000 | 19 | 16292256 | 16292368 | 113 | 2 | 1,02E+08 | ENSMUST00000184935.3 | -18468 | ENSMUSG00000098718 | Mir496b | microRNA 496b |
| chr2 | 1,18E+08 | 1,18E+08 | 909 | * | 8,588578 | 3,322772605 | 1,213039 | 2,739214 | 0,006159 | 0,812199284 | 1,360622 | Distal Intergenic | 2 | 1,18E+08 | 1,18E+08 | 6136 | 2 | 1,01E+08 | ENSMUST00000154409.2 | 80487 | | Gm13985 | predicted gene 13985 |
| chr17 | 34432408 | 34433341 | 934 | * | 9,451569 | 3,322924785 | 1,11912 | 2,96923 | 0,002985 | 0,681119785 | 1,425929 | Exon (ENSMUST000000 | 17 | 34426680 | 34428194 | 1515 | 1 | 21355 | ENSMUST00000145528.2 | 5728 | ENSMUSG00000024339 | Tap2 | transporter 2, ATP-binding cassette, sub-family B (MDR/TAP) |
| chr15 | 52936336 | 52937421 | 1086 | * | 15,37828 | 3,322950845 | 1,167307 | 2,84668 | 0,004418 | 0,773936378 | 1,214904 | 3' UTR | 15 | 52953327 | 52971264 | 17938 | 2 | 14042 | ENSMUST00000140798.2 | 33843 | ENSMUSG00000061731 | Ext1 | exostosin glycosyltransferase 1 |
| chr11 | 98819776 | 98825123 | 5348 | * | 110,6035 | 3,322951168 | 0,875602 | 3,795047 | 0,000148 | 0,123124216 | 3,270621 | Promoter (1-2kb) | 11 | 98818644 | 98864988 | 46345 | 1 | 19401 | ENSMUST00000107474.8 | 1132 | ENSMUSG00000037992 | Rara | retinoic acid receptor, alpha |
| chr1 | 11913485 | 11914739 | 1255 | * | 12,59076 | 3,324648229 | 1,000491 | 3,323016 | 0,00089 | 0,50985457 | 1,075735 | Intron (ENSMUST00000 | 1 | 11649530 | 11666855 | 17326 | 1 | 320492 | ENSMUST00000191437.2 | 263955 | ENSMUSG00000057715 | A830018L16Rik | RIKEN cDNA A830018L16 gene |
| chr1 | 36314753 | 36315569 | 817 | * | 8,544376 | 3,324692059 | 1,280529 | 2,596342 | 0,009422 | 0,876200449 | 1,449706 | Promoter (<=1kb) | 1 | 36307942 | 36313760 | 5819 | 2 | 214854 | ENSMUST00000188666.3 | -993 | ENSMUSG00000047180 | Neurl3 | neuralized E3 ubiquitin protein ligase 3 |
| chr3 | 36506994 | 36508638 | 1645 | * | 10,56599 | 3,325875004 | 1,039273 | 3,200195 | 0,001373 | 0,573832133 | 1,173325 | Promoter (<=1kb) | 3 | 36503313 | 36506477 | 3165 | 2 | 11747 | ENSMUST00000196097.2 | -517 | ENSMUSG00000027712 | Anxa5 | annexin A5 |
| chr10 | 78004892 | 78006278 | 1387 | * | 14,50378 | 3,326872029 | 0,950919 | 3,498585 | 0,000468 | 0,29714458 | 2,141612 | Promoter (<=1kb) | 10 | 77997900 | 78005616 | 7717 | 2 | 28295 | ENSMUST00000001242.9 | 0 | ENSMUSG00000053329 | Gatd3a | glutamine amidotransferase like class 1 domain containing 3A |
| chr7 | 24435330 | 24436262 | 933 | * | 11,09646 | 3,326994553 | 1,221352 | 2,724026 | 0,006449 | 0,92207189 | 0,738441 | Distal Intergenic | 7 | 24414448 | 24426333 | 11886 | 2 | 210155 | ENSMUST00000205376.2 | -8997 | ENSMUSG00000058717 | Lypd11 | Ly6/PLAUR domain containing 11 |
| chr4 | 60791963 | 60792809 | 847 | * | 11,33023 | 3,327270694 | 0,966632 | 3,442129 | 0,000577 | 0,464166555 | 0,918255 | Promoter (<=1kb) | 4 | 60792907 | 60795648 | 2742 | 1 | 1,03E+08 | ENSMUST00000098035.6 | -98 | | Gm21286 | predicted gene, 21286 |
| chr4 | 117871577 | 117874128 | 2552 | * | 37,561625 | 3,328149289 | 0,6415115 | 5,1879803 | 2,126E-07 | 0,00114665 | 2,9254973 | Exon (ENSMUST0000003 | 4 | 117797551 | 117889079 | 91529 | 2 | 20441 | ENSMUST00000138274.8 | 14951 | ENSMUSG00000028538 | St3gal3 | ST3 beta-galactoside alpha-2,3-sialyltransferase 3 |
| chr5 | 1,25E+08 | 1,25E+08 | 1630 | * | 9,680701 | 3,329533228 | 1,22585 | 2,716103 | 0,006606 | 0,816751204 | 1,425929 | Promoter (<=1kb) | 5 | 1,25E+08 | 1,25E+08 | 7417 | 1 | 56334 | ENSMUST00000135464.2 | 0 | ENSMUSG00000029390 | Tmed2 | transmembrane p24 trafficking protein 2 |
| chr12 | 46842608 | 46843709 | 1102 | * | 7,991197 | 3,329934164 | 1,232001 | 2,702866 | 0,006874 | 1 | 0 | Intron (ENSMUST00000 | 12 | 46758485 | 46863743 | 105259 | 2 | 664883 | ENSMUST00000219330.2 | 20034 | ENSMUSG00000021047 | Nova1 | NOVA alternative splicing regulator 1 |
| chr2 | 27909019 | 27909961 | 943 | * | 13,38033 | 3,330148543 | 1,094373 | 3,042974 | 0,002343 | 0,713406565 | 0,918255 | Promoter (1-2kb) | 2 | 27907453 | 27927027 | 19575 | 1 | 12831 | ENSMUST00000145423.2 | 1566 | ENSMUSG00000026837 | Col5a1 | collagen, type V, alpha 1 |
| chr9 | 71663436 | 71665141 | 1706 | * | 17,79085 | 3,330435458 | 0,838561 | 3,971606 | 7,14E-05 | 0,095171728 | 2,386008 | Intron (ENSMUST00000 | 9 | 71536493 | 71678842 | 142350 | 2 | 68178 | ENSMUST00000122065.2 | 13701 | ENSMUSG00000032232 | Cgnl1 | cingulin-like 1 |
| chrX | 1,58E+08 | 1,58E+08 | 826 | * | 9,885769 | 3,33065043 | 1,193476 | 2,790714 | 0,005259 | 0,824070388 | 1,086118 | Distal Intergenic | 20 | 1,58E+08 | 1,58E+08 | 154196 | 1 | 110651 | ENSMUST00000112493.8 | -83346 | ENSMUSG00000031309 | Rps6ka3 | ribosomal protein S6 kinase polypeptide 3 |
| chr4 | 47412947 | 47414348 | 1402 | * | 10,22093 | 3,331252565 | 1,168984 | 2,849698 | 0,004376 | 0,777919825 | 1,173325 | 3' UTR | 4 | 47386216 | 47411035 | 24820 | 1 | 21812 | ENSMUST00000107725.3 | 26731 | ENSMUSG00000007613 | Tgfbr1 | transforming growth factor, beta receptor I |
| chr2 | 1,16E+08 | 1,16E+08 | 1328 | * | 11,87144 | 3,331981736 | 1,008615 | 3,303523 | 0,000955 | 0,594430593 | 0,744375 | Exon (ENSMUST000001 | 2 | 1,16E+08 | 1,16E+08 | 932 | 2 | 69974 | ENSMUST00000195368.2 | -9161 | | 2810405F15Rik | RIKEN cDNA 2810405F15 gene |
| chr1 | 44185710 | 44187927 | 2218 | * | 17,71042 | 3,332256189 | 0,800793 | 4,161196 | 3,17E-05 | 0,055684611 | 2,507013 | Promoter (<=1kb) | 1 | 44187007 | 44220420 | 33414 | 1 | 22592 | ENSMUST00000027214.4 | 0 | ENSMUSG00000026048 | Ercc5 | excision repair cross-complementing rodent repair deficiency, complementation group 5 |
| chr8 | 48567337 | 48567965 | 629 | * | 9,356872 | 3,332478732 | 1,197974 | 2,781762 | 0,005406 | 0,700182401 | 2,237403 | Intron (ENSMUST00000 | 8 | 48563246 | 48606268 | 43023 | 1 | 320685 | ENSMUST00000174379.3 | 4091 | ENSMUSG00000031562 | Dctd | dCMP deaminase |
| chr15 | 78312163 | 78314182 | 2020 | * | 29,27194 | 3,332590863 | 1,135339 | 2,935327 | 0,003332 | 0,571379912 | 2,880017 | Promoter (<=1kb) | 15 | 78312764 | 78322697 | 9934 | 1 | 72844 | ENSMUST00000162517.9 | 0 | ENSMUSG00000033287 | Kctd17 | potassium channel tetramerisation domain containing 17 |
| chr12 | 75565908 | 75566731 | 824 | * | 11,02873 | 3,332651556 | 1,179777 | 2,824814 | 0,004731 | 0,835756183 | 0,918255 | Intron (ENSMUST00000 | 12 | 75540266 | 75596448 | 56183 | 2 | 26932 | ENSMUST00000218716.2 | 29717 | ENSMUSG00000021051 | Ppp2r5e | protein phosphatase 2, regulatory subunit B', epsilon |
| chr9 | 57525891 | 57526977 | 1087 | * | 15,2145 | 3,3336656 | 0,871503 | 3,82519 | 0,000131 | 0,206624632 | 1,214904 | Promoter (1-2kb) | 9 | 57514368 | 57528057 | 13690 | 2 | 235416 | ENSMUST00000044937.5 | 1080 | ENSMUSG00000056271 | Lman1l | lectin, mannose-binding 1 like |
| chr8 | 1,22E+08 | 1,22E+08 | 1248 | * | 14,9696 | 3,334659401 | 1,01436 | 3,287452 | 0,001011 | 0,510628937 | 1,214904 | Intron (ENSMUST00000 | 8 | 1,22E+08 | 1,22E+08 | 18737 | 2 | 76454 | ENSMUST00000212985.2 | 3052 | ENSMUSG00000052934 | Fbxo31 | F-box protein 31 |
| chr9 | 24926907 | 24928166 | 1260 | * | 10,12987 | 3,334713913 | 1,207275 | 2,762182 | 0,005742 | 0,7893429 | 1,449706 | Distal Intergenic | 9 | 24885504 | 24909716 | 24213 | 1 | 1,03E+08 | ENSMUST00000214469.2 | 41403 | ENSMUSG00000111866 | E130101E03Rik | RIKEN cDNA E130101E03 gene |
| chr10 | 80855059 | 80856533 | 1475 | * | 11,66287 | 3,334829285 | 1,177439 | 2,832274 | 0,004622 | 0,789027175 | 1,173325 | 3' UTR | 10 | 80860645 | 80862107 | 1463 | 1 | 1,19E+08 | ENSMUST00000151303.2 | -4112 | | Gm16099 | predicted gene 16099 |
| chr9 | 59383349 | 59385234 | 1886 | * | 11,58292 | 3,335561923 | 1,167012 | 2,858206 | 0,00426 | 0,853837252 | 0,738441 | Intron (ENSMUST00000 | 9 | 59333019 | 59393339 | 60321 | 2 | 23806 | ENSMUST00000168497.8 | 8105 | ENSMUSG00000025234 | Arih1 | ariadne RBR E3 ubiquitin protein ligase 1 |
| chr12 | 52386103 | 52387529 | 1427 | * | 10,60945 | 3,3356842 | 1,166567 | 2,859403 | 0,004244 | 0,772488766 | 1,173325 | Distal Intergenic | 12 | 52522522 | 52527392 | 4871 | 2 | 1,03E+08 | ENSMUST00000219885.2 | 139863 | ENSMUSG00000111982 | Gm35135 | predicted gene, 35135 |
| chrX | 1,64E+08 | 1,64E+08 | 1023 | * | 8,343134 | 3,335994759 | 1,235495 | 2,700129 | 0,006931 | 1 | 0 | Distal Intergenic | 20 | 1,64E+08 | 1,64E+08 | 198377 | 2 | 237213 | ENSMUST00000058787.9 | -17743 | ENSMUSG00000018589 | Glra2 | glycine receptor, alpha 2 subunit |
| chr5 | 73500018 | 73500832 | 815 | * | 12,90612 | 3,336439543 | 1,125201 | 2,965195 | 0,003025 | 0,728215953 | 1,075735 | Promoter (1-2kb) | 5 | 73485008 | 73498371 | 13364 | 2 | 433904 | ENSMUST00000200821.2 | -1647 | ENSMUSG00000029153 | Ociad2 | OCIA domain containing 2 |
| chr9 | 21655076 | 21655834 | 759 | * | 7,883192 | 3,336985901 | 1,254984 | 2,658986 | 0,007838 | 1 | 0 | Promoter (<=1kb) | 9 | 21656619 | 21661210 | 4592 | 1 | 16835 | ENSMUST00000214549.2 | -785 | ENSMUSG00000032193 | Ldlr | low density lipoprotein receptor |
| chr7 | 1,1E+08 | 1,1E+08 | 1212 | * | 10,11275 | 3,337047825 | 1,13869 | 2,930602 | 0,003383 | 0,746312988 | 1,086118 | Distal Intergenic | 7 | 1,1E+08 | 1,1E+08 | 9572 | 2 | 1,01E+08 | ENSMUST00000210822.2 | 12183 | ENSMUSG00000109700 | Gm21123 | predicted gene, 21123 |
| chr2 | 1,52E+08 | 1,52E+08 | 1652 | * | 15,03067 | 3,337115871 | 0,938628 | 3,555313 | 0,000378 | 0,323988916 | 1,37683 | Promoter (2-3kb) | 2 | 1,52E+08 | 1,52E+08 | 4453 | 2 | 20667 | ENSMUST00000182625.2 | 2872 | ENSMUSG00000051817 | Sox12 | SRY (sex determining region Y)-box 12 |
| chr11 | 77301944 | 77302685 | 742 | * | 7,53957 | 3,338692002 | 1,256334 | 2,657488 | 0,007873 | 1 | 0 | Intron (ENSMUST00000 | 11 | 77353237 | 77361310 | 8074 | 1 | 216961 | ENSMUST00000102493.8 | -50552 | ENSMUSG00000020836 | Coro6 | coronin 6 |
| chr5 | 143218822 | 143221911 | 3090 | * | 36,286501 | 3,338954862 | 0,6813098 | 4,9007882 | 9,545E-07 | 0,003692163 | 2,7254908 | Promoter (<=1kb) | 5 | 143220918 | 143233703 | 12786 | 1 | 231866 | ENSMUST00000077485.11 | 0 | ENSMUSG00000029587 | Zfp12 | zinc finger protein 12 |
| chr17 | 62147885 | 62148693 | 809 | * | 9,144187 | 3,339585816 | 1,29303 | 2,58276 | 0,009801 | 0,886332554 | 1,425929 | Distal Intergenic | 17 | 62763982 | 62764115 | 134 | 2 | 1,15E+08 | ENSMUST00000082636.3 | 615422 | ENSMUSG00000064570 | Gm25800 | predicted gene, 25800 |
| chr18 | 11827128 | 11828788 | 1661 | * | 12,17416 | 3,339710895 | 1,168106 | 2,859082 | 0,004249 | 0,75508254 | 1,2763 | Exon (ENSMUST000002 | 18 | 11855278 | 11856034 | 757 | 1 | 225182 | ENSMUST00000234161.2 | -26490 | ENSMUSG00000041238 | Rbbp8 | retinoblastoma binding protein 8, endonuclease |
| chr5 | 75288465 | 75289448 | 984 | * | 10,45429 | 3,339896136 | 1,082258 | 3,086043 | 0,002028 | 0,649217039 | 1,173325 | Exon (ENSMUST000002 | 5 | 75270982 | 75307768 | 36787 | 1 | 1,01E+08 | ENSMUST00000202993.2 | 17483 | ENSMUSG00000107192 | Gm19583 | predicted gene, 19583 |
| chr5 | 1,47E+08 | 1,47E+08 | 1476 | * | 16,23217 | 3,339937001 | 0,891983 | 3,744394 | 0,000181 | 0,172914801 | 2,318434 | Promoter (<=1kb) | 5 | 1,47E+08 | 1,47E+08 | 4143 | 1 | 19933 | ENSMUST00000035983.12 | 0 | ENSMUSG00000041453 | Rpl21 | ribosomal protein L21 |
| chr2 | 3237533 | 3238272 | 740 | * | 6,884372 | 3,340848562 | 1,245647 | 2,682018 | 0,007318 | 1 | 0 | Distal Intergenic | 2 | 3285249 | 3329914 | 44666 | 1 | 18108 | ENSMUST00000081932.13 | -46977 | ENSMUSG00000026643 | Nmt2 | N-myristoyltransferase 2 |
| chr2 | 154885233 | 154887396 | 2164 | * | 22,402203 | 3,340937951 | 0,7863897 | 4,2484507 | 2,153E-05 | 0,037616988 | 2,9519335 | Promoter (2-3kb) | 2 | 154889535 | 154892931 | 3397 | 1 | 50518 | ENSMUST00000148300.2 | -2139 | ENSMUSG00000027596 | a | nonagouti |
| chrX | 84010157 | 84011568 | 1412 | * | 12,83025 | 3,341213989 | 1,091895 | 3,060014 | 0,002213 | 0,649217039 | 1,2763 | Intron (ENSMUST00000 | 20 | 84091901 | 84099203 | 7303 | 1 | 13405 | ENSMUST00000128983.2 | -80333 | ENSMUSG00000045103 | Dmd | dystrophin, muscular dystrophy |
| chr5 | 44982017 | 44983661 | 1645 | * | 16,33085 | 3,34153245 | 0,867654 | 3,851227 | 0,000118 | 0,13288306 | 2,318434 | Distal Intergenic | 5 | 44629474 | 44957022 | 327549 | 2 | 16826 | ENSMUST00000199256.5 | -24995 | ENSMUSG00000039706 | Ldb2 | LIM domain binding 2 |
| chr13 | 61055324 | 61056151 | 828 | * | 9,950361 | 3,342026465 | 1,096295 | 3,048472 | 0,0023 | 0,664099998 | 1,211315 | Promoter (2-3kb) | 13 | 61049002 | 61052694 | 3693 | 2 | 116913 | ENSMUST00000080766.8 | -2630 | ENSMUSG00000062705 | Tpbpb | trophoblast specific protein beta |
| chr5 | 1,4E+08 | 1,4E+08 | 1789 | * | 13,21967 | 3,342151312 | 1,078397 | 3,099185 | 0,001941 | 0,715249603 | 0,738441 | Distal Intergenic | 5 | 1,4E+08 | 1,4E+08 | 1371 | 1 | 22255 | ENSMUST00000174792.2 | 24064 | ENSMUSG00000029546 | Uncx | UNC homeobox |
| chr11 | 87947654 | 87948623 | 970 | * | 7,150337 | 3,342575245 | 1,287136 | 2,596908 | 0,009407 | 0,887238309 | 1,360622 | Distal Intergenic | 11 | 87940238 | 87941877 | 1640 | 1 | 110809 | ENSMUST00000132983.3 | 7416 | ENSMUSG00000018379 | Srsf1 | serine and arginine-rich splicing factor 1 |
| chr18 | 40191225 | 40192241 | 1017 | * | 9,420332 | 3,342725198 | 1,222546 | 2,734233 | 0,006253 | 0,805745288 | 1,425929 | Distal Intergenic | 18 | 40039233 | 40062427 | 23195 | 2 | 1,05E+08 | ENSMUST00000237624.2 | -128798 | | Gm41708 | predicted gene, 41708 |
| chr3 | 34321666 | 34322989 | 1324 | * | 11,50679 | 3,343432035 | 1,241798 | 2,692413 | 0,007094 | 0,903101391 | 0,918255 | Intron (ENSMUST00000 | 3 | 34250900 | 34692632 | 441733 | 1 | 320478 | ENSMUST00000196058.5 | 70766 | ENSMUSG00000105265 | Sox2ot | SOX2 overlapping transcript (non-protein coding) |
| chr13 | 36327653 | 36328525 | 873 | * | 8,284576 | 3,343756113 | 1,289976 | 2,592107 | 0,009539 | 0,88871498 | 1,360622 | Intron (ENSMUST00000 | 13 | 36313883 | 36721567 | 407685 | 1 | 69955 | ENSMUST00000224241.2 | 13770 | ENSMUSG00000021420 | Fars2 | phenylalanine-tRNA synthetase 2 (mitochondrial) |
| chr19 | 8251108 | 8252304 | 1197 | * | 8,446318 | 3,343920911 | 1,294751 | 2,582675 | 0,009804 | 1 | 0 | Exon (ENSMUST000002 | 19 | 8229332 | 8294543 | 65212 | 1 | 1,19E+08 | ENSMUST00000235607.2 | 21776 | | LOC118568404 | uncharacterized LOC118568404 |
| chr9 | 1,12E+08 | 1,12E+08 | 729 | * | 8,783299 | 3,34481397 | 1,237258 | 2,703408 | 0,006863 | 0,831521609 | 1,360622 | Distal Intergenic | 9 | 1,12E+08 | 1,12E+08 | 6356 | 2 | 78028 | ENSMUST00000215688.2 | -45362 | ENSMUSG00000111054 | 4930545L08Rik | RIKEN cDNA 4930545L08 gene |
| chr4 | 27161468 | 27162738 | 1271 | * | 10,32387 | 3,347069558 | 1,147231 | 2,917521 | 0,003528 | 0,702508842 | 1,449706 | Distal Intergenic | 4 | 26635820 | 26705449 | 69630 | 2 | 75360 | ENSMUST00000154459.2 | -456019 | ENSMUSG00000086257 | 4930548K13Rik | RIKEN cDNA 4930548K13 gene |
| chr16 | 84868914 | 84870192 | 1279 | * | 11,26122 | 3,347161708 | 1,150928 | 2,908229 | 0,003635 | 0,744477896 | 1,173325 | Intron (ENSMUST00000 | 16 | 84876999 | 84880280 | 3282 | 1 | 1,19E+08 | ENSMUST00000227918.2 | -6807 | | Gm49227 | predicted gene, 49227 |
| chr1 | 1,35E+08 | 1,35E+08 | 1216 | * | 13,28286 | 3,347390791 | 1,08596 | 3,082426 | 0,002053 | 0,684274806 | 0,938682 | Intron (ENSMUST00000 | 1 | 1,35E+08 | 1,35E+08 | 902 | 1 | 67196 | ENSMUST00000139617.2 | 18839 | ENSMUSG00000026429 | Ube2t | ubiquitin-conjugating enzyme E2T |
| chr11 | 5785025 | 5786507 | 1483 | * | 11,95062 | 3,347688048 | 1,0547 | 3,174067 | 0,001503 | 0,575442234 | 1,2763 | Promoter (<=1kb) | 11 | 5784800 | 5786485 | 1686 | 2 | 54125 | ENSMUST00000156913.2 | 0 | ENSMUSG00000020474 | Polm | polymerase (DNA directed), mu |
| chr16 | 3253019 | 3254109 | 1091 | * | 8,439484 | 3,347973399 | 1,290297 | 2,594731 | 0,009467 | 0,880247401 | 1,425929 | Distal Intergenic | 16 | 3286324 | 3298915 | 12592 | 1 | 1,03E+08 | ENSMUST00000229453.2 | -32215 | ENSMUSG00000116455 | Gm35974 | predicted gene, 35974 |
| chr15 | 1,02E+08 | 1,02E+08 | 640 | * | 8,024936 | 3,34897784 | 1,210938 | 2,765607 | 0,005682 | 1 | 0 | Promoter (2-3kb) | 15 | 1,02E+08 | 1,02E+08 | 1079 | 2 | 16421 | ENSMUST00000229440.2 | -2376 | ENSMUSG00000001281 | Itgb7 | integrin beta 7 |
| chr4 | 1,1E+08 | 1,1E+08 | 1055 | * | 8,05029 | 3,350009346 | 1,270534 | 2,636695 | 0,008372 | 0,863495249 | 1,360622 | Intron (ENSMUST00000 | 4 | 1,1E+08 | 1,12E+08 | 1259720 | 1 | 78933 | ENSMUST00000097920.9 | 165552 | ENSMUSG00000061298 | Agbl4 | ATP/GTP binding protein-like 4 |

| | | | | | | | | | | | | | | | | | | | | | | | |
|---|---|---|---|---|---|---|---|---|---|---|---|---|---|---|---|---|---|---|---|---|---|---|---|
| chr15 | 71219152 | 71219749 | 598 | * | 7,060075 | 3,350654627 | 1,29446 | 2,588458 | 0,009641 | 1 | 0 | Distal Intergenic | 15 | 71317525 | 71334249 | 16725 | 2 | 70363 | ENSMUST00000229634.2 | 114500 | ENSMUSG00000036800 | Fam135b | family with sequence similarity 135, member B |
| chr11 | 90691720 | 90692903 | 1184 | * | 9,710355 | 3,350920549 | 1,213241 | 2,761957 | 0,005746 | 0,79185145 | 1,425929 | Promoter (2-3kb) | 11 | 90695518 | 90785975 | 90458 | 1 | 73922 | ENSMUST00000142521.2 | -2615 | ENSMUSG00000085940 | 4930405D11Rik | RIKEN cDNA 4930405D11 gene |
| chr8 | 24092490 | 24093533 | 1044 | * | 12,25695 | 3,351069045 | 0,982181 | 3,411866 | 0,000645 | 0,480916943 | 0,918255 | Distal Intergenic | 8 | 24126035 | 24392127 | 266093 | 1 | 320158 | ENSMUST00000131410.8 | -32502 | ENSMUSG00000037492 | Zmat4 | zinc finger, matrin type 4 |
| chr16 | 22468524 | 22469580 | 1057 | * | 9,849001 | 3,351661704 | 1,228662 | 2,727895 | 0,006374 | 0,838625535 | 1,211315 | Intron (ENSMUST00000 | 16 | 22414378 | 22475778 | 61401 | 2 | 110197 | ENSMUST00000150553.2 | 6198 | ENSMUSG00000022861 | Dgkg | diacylglycerol kinase, gamma |
| chr9 | 64072234 | 64073681 | 1448 | * | 10,50121 | 3,352031329 | 1,284105 | 2,610402 | 0,009044 | 0,939804939 | 0,918255 | 5' UTR | 9 | 64045275 | 64080161 | 34887 | 2 | 68014 | ENSMUST00000176299.8 | 6480 | ENSMUSG00000032400 | Zwilch | zwilch kinetochore protein |
| chr10 | 1,23E+08 | 1,23E+08 | 792 | * | 9,112955 | 3,352711596 | 1,280824 | 2,61762 | 0,008855 | 0,86525631 | 1,425929 | Promoter (<=1kb) | 10 | 1,23E+08 | 1,23E+08 | 91990 | 2 | 14479 | ENSMUST00000220377.2 | -867 | ENSMUSG00000020124 | Usp15 | ubiquitin specific peptidase 15 |
| chr2 | 59093201 | 59094082 | 882 | * | 10,81532 | 3,352882354 | 1,179518 | 2,842588 | 0,004475 | 0,766579572 | 1,2763 | Promoter (2-3kb) | 2 | 59096528 | 59138581 | 42054 | 1 | 227937 | ENSMUST00000124725.2 | -2446 | ENSMUSG00000026991 | Pkp4 | plakophilin 4 |
| chr3 | 96164786 | 96165707 | 922 | * | 8,00533 | 3,353188371 | 1,240752 | 2,702545 | 0,006881 | 1 | 0 | Promoter (1-2kb) | 3 | 96161935 | 96163593 | 1659 | 2 | 1,03E+08 | ENSMUST00000176275.2 | -1193 | | Gm20627 | predicted gene 20627 |
| chr2 | 51871430 | 51872426 | 997 | * | 9,044444 | 3,35325358 | 1,250595 | 2,681327 | 0,007333 | 0,752091016 | 2,237403 | Distal Intergenic | 2 | 51846056 | 51863506 | 17451 | 2 | 64685 | ENSMUST00000145656.2 | -7924 | ENSMUSG00000026946 | Nmi | N-myc (and STAT) interactor |
| chr10 | 1,2E+08 | 1,2E+08 | 1108 | * | 9,147933 | 3,353709407 | 1,226954 | 2,733363 | 0,006269 | 0,81623383 | 1,360622 | Promoter (1-2kb) | 10 | 1,2E+08 | 1,2E+08 | 16168 | 1 | 68212 | ENSMUST00000020446.11 | -1486 | ENSMUSG00000020225 | Tmbim4 | transmembrane BAX inhibitor motif containing 4 |
| chr13 | 41556421 | 41558995 | 2575 | * | 46,726368 | 3,354698949 | 0,6023403 | 5,5694411 | 2,556E-08 | 0,000230583 | 2,7872553 | Exon (ENSMUST0000022 | 13 | 41471396 | 41513215 | 41820 | 2 | 18003 | ENSMUST00000224803.2 | -43206 | ENSMUSG00000021365 | Nedd9 | neural precursor cell expressed, developmentally down-regulated gene 9 |
| chr12 | 77147289 | 77148205 | 917 | * | 9,879365 | 3,355816575 | 1,205518 | 2,783714 | 0,005374 | 0,826716028 | 1,086118 | Distal Intergenic | 12 | 77086371 | 77088975 | 2605 | 2 | 1,03E+08 | ENSMUST00000219319.2 | -58314 | | Gm35189 | predicted gene, 35189 |
| chr5 | 1,19E+08 | 1,19E+08 | 1210 | * | 9,426449 | 3,356986091 | 1,269645 | 2,644035 | 0,008192 | 0,883923857 | 1,211315 | Distal Intergenic | 5 | 1,19E+08 | 1,19E+08 | 159436 | 1 | 76199 | ENSMUST00000201614.2 | -50446 | ENSMUSG00000018076 | Med13l | mediator complex subunit 13-like |
| chr19 | 44777555 | 44779065 | 1511 | * | 14,27667 | 3,358580145 | 1,000691 | 3,35626 | 0,00079 | 0,554161661 | 0,738441 | Intron (ENSMUST00000 | 19 | 44796845 | 44796968 | 124 | 1 | 1,02E+08 | ENSMUST00000184584.3 | -17780 | ENSMUSG00000098952 | Mir6405 | microRNA 6405 |
| chr15 | 88960669 | 88961637 | 969 | * | 9,17116 | 3,35878008 | 1,279754 | 2,624551 | 0,008676 | 0,891097601 | 1,211315 | Promoter (<=1kb) | 15 | 88960327 | 88970336 | 10010 | 1 | 67976 | ENSMUST00000165690.2 | 342 | ENSMUSG00000015363 | Trabd | TraB domain containing |
| chr2 | 1,11E+08 | 1,11E+08 | 957 | * | 6,788192 | 3,359202287 | 1,274688 | 2,635314 | 0,008406 | 1 | 0 | Distal Intergenic | 2 | 1,11E+08 | 1,11E+08 | 939 | 2 | 258890 | ENSMUST00000099612.2 | 9099 | ENSMUSG00000094858 | Olfr1297 | olfactory receptor 1297 |
| chr10 | 3680604 | 3681764 | 1161 | * | 14,93309 | 3,360736192 | 0,943416 | 3,562305 | 0,000368 | 0,315383621 | 1,41696 | Exon (ENSMUST000002 | 10 | 3690364 | 3917303 | 226940 | 1 | 213783 | ENSMUST00000120274.8 | -8600 | ENSMUSG00000040624 | Plekhg1 | pleckstrin homology domain containing, family G (with RhoGef domain) member 1 |
| chr15 | 23079262 | 23080264 | 1003 | * | 8,175667 | 3,360790552 | 1,193854 | 2,815078 | 0,004877 | 0,756849896 | 1,449706 | Intron (ENSMUST00000 | 15 | 23036549 | 23474504 | 437956 | 1 | 320865 | ENSMUST00000164787.8 | 42713 | ENSMUSG00000040420 | Cdh18 | cadherin 18 |
| chr7 | 1,24E+08 | 1,24E+08 | 925 | * | 11,62576 | 3,362170642 | 1,105638 | 3,040932 | 0,002358 | 0,713663698 | 0,918255 | Intron (ENSMUST00000 | 7 | 1,24E+08 | 1,24E+08 | 103553 | 1 | 1,19E+08 | ENSMUST00000151884.2 | 62048 | | 4930551E15Rik | RIKEN cDNA 4930551E15 gene |
| chr11 | 60402886 | 60404361 | 1476 | * | 13,02869 | 3,36258053 | 1,074301 | 3,130017 | 0,001748 | 0,695892636 | 0,738441 | Promoter (2-3kb) | 11 | 60400360 | 60412812 | 12453 | 1 | 17910 | ENSMUST00000122825.2 | 2526 | ENSMUSG00000042678 | Myo15 | myosin XV |
| chr7 | 1,28E+08 | 1,28E+08 | 1457 | * | 13,65977 | 3,363584848 | 1,064421 | 3,160012 | 0,001578 | 0,586304784 | 1,2763 | Promoter (<=1kb) | 7 | 1,28E+08 | 1,28E+08 | 26792 | 2 | 233913 | ENSMUST00000033044.16 | 0 | ENSMUSG00000030780 | Rusf1 | RUS family member 1 |
| chr4 | 70771050 | 70772290 | 1241 | * | 10,95552 | 3,364139533 | 1,114865 | 3,017531 | 0,002548 | 0,725842761 | 0,918255 | Distal Intergenic | 4 | 70993346 | 71049370 | 56025 | 2 | 1,03E+08 | ENSMUST00000143796.2 | 277080 | ENSMUSG00000084937 | Gm11228 | predicted gene 11228 |
| chr1 | 84816755 | 84819071 | 2317 | * | 19,99582 | 3,364477175 | 0,82682 | 4,069178 | 4,72E-05 | 0,075059926 | 2,386008 | Promoter (<=1kb) | 1 | 84698910 | 84817025 | 118116 | 2 | 14897 | ENSMUST00000027421.13 | 0 | ENSMUSG00000026219 | Trip12 | thyroid hormone receptor interactor 12 |
| chr5 | 76408972 | 76410854 | 1883 | * | 11,63876 | 3,364703138 | 1,151705 | 2,921496 | 0,003484 | 0,78060962 | 0,918255 | Exon (ENSMUST000002 | 5 | 76361700 | 76421859 | 60160 | 2 | 12753 | ENSMUST00000202122.2 | 11005 | ENSMUSG00000029238 | Clock | circadian locomotor output cycles kaput |
| chr9 | 70316597 | 70317899 | 1303 | * | 12,14074 | 3,364801883 | 1,065277 | 3,158618 | 0,001585 | 0,683210259 | 0,738441 | Exon (ENSMUST00000 | 9 | 70312976 | 70313035 | 60 | 2 | 1,01E+08 | ENSMUST00000176002.3 | -3562 | ENSMUSG00000093763 | Mir5626 | microRNA 5626 |
| chr7 | 1,02E+08 | 1,02E+08 | 925 | * | 9,583567 | 3,365775273 | 1,23828 | 2,718104 | 0,006566 | 0,812305378 | 1,449706 | Promoter (1-2kb) | 7 | 1,02E+08 | 1,02E+08 | 1057 | 1 | 258358 | ENSMUST00000098218.3 | -1178 | ENSMUSG00000073967 | Olfr557 | olfactory receptor 557 |
| chr8 | 61435462 | 61438274 | 2813 | * | 69,732939 | 3,366221087 | 0,4482563 | 7,50959 | 5,931E-14 | 4,16979E-09 | 3,2706213 | Promoter (<=1kb) | 8 | 61363423 | 61436334 | 72912 | 2 | 12725 | ENSMUST00000093490.9 | 0 | ENSMUSG00000004319 | Clcn3 | chloride channel, voltage-sensitive 3 |
| chr15 | 28699523 | 28700384 | 862 | * | 11,24028 | 3,366393919 | 1,053018 | 3,196902 | 0,001389 | 0,661775851 | 0,744375 | Distal Intergenic | 15 | 28203898 | 28472198 | 268301 | 1 | 110082 | ENSMUST00000067048.8 | 495625 | ENSMUSG00000022262 | Dnah5 | dynein, axonemal, heavy chain 5 |
| chr7 | 47818172 | 47819070 | 899 | * | 8,323577 | 3,366636133 | 1,212189 | 2,777318 | 0,005481 | 1 | 0 | Promoter (<=1kb) | 7 | 47817731 | 47818767 | 1037 | 2 | 404239 | ENSMUST00000094389.4 | 0 | ENSMUSG00000070551 | Mrgprb5 | MAS-related GPR, member B5 |
| chr8 | 12670683 | 12671492 | 810 | * | 9,431968 | 3,366657827 | 1,216968 | 2,766431 | 0,005667 | 0,79031971 | 1,425929 | Intron (ENSMUST00000 | 8 | 12665167 | 12687955 | 22789 | 2 | 259279 | ENSMUST00000164611.8 | 16463 | ENSMUSG00000000759 | Tubgcp3 | tubulin, gamma complex associated protein 3 |
| chr9 | 56412683 | 56413935 | 1253 | * | 18,17744 | 3,367409497 | 0,839396 | 4,011706 | 6,03E-05 | 0,086053637 | 2,386008 | Exon (ENSMUST000002 | 9 | 56326114 | 56383401 | 57288 | 1 | 66867 | ENSMUST00000214869.2 | 86569 | ENSMUSG00000032329 | Hmg20a | high mobility group 20A |
| chr14 | 25332952 | 25333854 | 903 | * | 8,179985 | 3,36857899 | 1,19316 | 2,823241 | 0,004754 | 0,755289138 | 1,425929 | Intron (ENSMUST00000 | 14 | 25356225 | 25379059 | 22835 | 2 | 414118 | ENSMUST00000181582.2 | 45205 | | Zmiz1os1 | Zmiz1 opposite strand 1 |
| chr2 | 1,57E+08 | 1,57E+08 | 1341 | * | 11,49889 | 3,369343402 | 1,248586 | 2,698527 | 0,006965 | 0,845724506 | 1,2763 | Intron (ENSMUST00000 | 2 | 1,57E+08 | 1,57E+08 | 14818 | 2 | 29812 | ENSMUST00000140297.8 | -7711 | ENSMUSG00000027634 | Ndrg3 | N-myc downstream regulated gene 3 |
| chr10 | 22694975 | 22695795 | 821 | * | 9,357129 | 3,370661437 | 1,220795 | 2,761037 | 0,005762 | 0,799316832 | 1,360622 | Promoter (<=1kb) | 10 | 22693180 | 22696073 | 2894 | 2 | 21412 | ENSMUST00000049930.9 | 278 | ENSMUSG00000045680 | Tcf21 | transcription factor 21 |
| chr18 | 74981502 | 74982889 | 1388 | * | 29,14918 | 3,370893621 | 1,250275 | 2,696121 | 0,007015 | 0,702848616 | 2,880017 | Distal Intergenic | 18 | 74926454 | 74939279 | 12826 | 1 | 52538 | ENSMUST00000237306.2 | 55048 | ENSMUSG00000036880 | Acaa2 | acetyl-Coenzyme A acyltransferase 2 (mitochondrial 3-oxoacyl-Coenzyme A thiolase) |
| chr12 | 80728061 | 80729257 | 1197 | * | 10,69969 | 3,371296214 | 1,108771 | 3,040569 | 0,002361 | 0,660341484 | 1,2763 | 3' UTR | 12 | 80733144 | 80739240 | 6097 | 2 | 68354 | ENSMUST00000153297.2 | 9983 | ENSMUSG00000044062 | Plekhd1os | pleckstrin homology domain containing, family D (with coiled-coil domains) member 1, opposite strand |
| chr17 | 53995342 | 53997973 | 2632 | * | 20,71683 | 3,371769823 | 0,90385 | 3,730454 | 0,000191 | 0,151053691 | 3,104559 | Promoter (<=1kb) | 17 | 53981814 | 53996361 | 14548 | 2 | 72415 | ENSMUST00000024736.14 | 0 | ENSMUSG00000023940 | Sgo1 | shugoshin 1 |
| chr3 | 80529901 | 80531208 | 1308 | * | 11,45428 | 3,3717844 | 1,039942 | 3,242281 | 0,001186 | 0,594676756 | 0,918255 | Distal Intergenic | 3 | 80595347 | 80709886 | 114540 | 2 | 14800 | ENSMUST00000192463.2 | 178678 | ENSMUSG00000033981 | Gria2 | glutamate receptor, ionotropic, AMPA2 (alpha 2) |
| chr2 | 26802863 | 26804254 | 1392 | * | 10,78854 | 3,372637454 | 1,273097 | 2,649161 | 0,008069 | 0,955812308 | 0,738441 | Promoter (<=1kb) | 2 | 26802683 | 26803224 | 542 | 1 | 27176 | ENSMUST00000149339.2 | 180 | ENSMUSG00000062647 | Rpl7a | ribosomal protein L7A |
| chr16 | 23668102 | 23669525 | 1424 | * | 11,39507 | 3,372815156 | 1,085907 | 3,105988 | 0,001896 | 0,677966227 | 0,918255 | Distal Intergenic | 16 | 23708323 | 23709708 | 1386 | 2 | 20604 | ENSMUST00000004480.5 | 40183 | ENSMUSG00000004366 | Sst | somatostatin |
| chr16 | 7318179 | 7319150 | 972 | * | 8,803519 | 3,372901791 | 1,233919 | 2,733486 | 0,006267 | 0,727652793 | 2,237403 | Distal Intergenic | 16 | 7170834 | 7227833 | 57000 | 1 | 268859 | ENSMUST00000231031.2 | 147345 | ENSMUSG00000008658 | Rbfox1 | RNA binding protein, fox-1 homolog (C. elegans) 1 |
| chr18 | 48751570 | 48752322 | 753 | * | 8,37914 | 3,373007086 | 1,173014 | 2,875505 | 0,004034 | 0,729437292 | 1,425929 | Distal Intergenic | 18 | 48178402 | 48181479 | 3078 | 1 | 433182 | ENSMUST00000076155.6 | 573168 | ENSMUSG00000059040 | Eno1b | enolase 1B, retrotransposed |
| chr15 | 16730656 | 16731618 | 963 | * | 9,527634 | 3,37375784 | 1,154265 | 2,922861 | 0,003468 | 0,730469267 | 1,211315 | Promoter (1-2kb) | 15 | 16728842 | 16857180 | 128339 | 1 | 12565 | ENSMUST00000228307.2 | 1814 | ENSMUSG00000025370 | Cdh9 | cadherin 9 |
| chr8 | 22637808 | 22639290 | 1483 | * | 18,973605 | 3,374019062 | 0,7518123 | 4,4878476 | 7,195E-06 | 0,018343507 | 2,6636191 | Exon (ENSMUST0000003 | 8 | 22618302 | 22652510 | 34209 | 2 | 23954 | ENSMUST00000178324.2 | 13220 | ENSMUSG00000031478 | Nek3 | NIMA (never in mitosis gene a)-related expressed kinase 3 |
| chr5 | 137497776 | 137501696 | 3921 | * | 44,685936 | 3,374880195 | 0,6008801 | 5,616562 | 1,948E-08 | 0,000171651 | 2,9254973 | Promoter (<=1kb) | 5 | 137499701 | 137500780 | 1080 | 2 | 74097 | ENSMUST00000031728.5 | 0 | ENSMUSG00000029715 | Pop7 | processing of precursor 7, ribonuclease P family, (S. cerevisiae) |
| chr3 | 97832053 | 97833349 | 1297 | * | 8,381422 | 3,37552706 | 1,188933 | 2,839124 | 0,004524 | 0,74907225 | 1,425929 | Distal Intergenic | 3 | 97837489 | 97874334 | 36846 | 1 | 433632 | ENSMUST00000098828.2 | -4140 | ENSMUSG00000074388 | Gm5544 | predicted gene 5544 |
| chr9 | 22125322 | 22126664 | 1343 | * | 13,13498 | 3,377184596 | 1,012618 | 3,335103 | 0,000853 | 0,474532331 | 1,2763 | Promoter (<=1kb) | 9 | 22118077 | 22125156 | 7080 | 2 | 69862 | ENSMUST00000188595.2 | -166 | ENSMUSG00000090925 | 1810064F22Rik | RIKEN cDNA 1810064F22 gene |
| chr12 | 1,12E+08 | 1,12E+08 | 953 | * | 10,16465 | 3,377371659 | 1,255416 | 2,69024 | 0,00714 | 0,877892722 | 1,086118 | Promoter (<=1kb) | 12 | 1,12E+08 | 1,12E+08 | 35540 | 1 | 668303 | ENSMUST00000128402.3 | -928 | ENSMUSG00000021294 | Kif26a | kinesin family member 26A |
| chr4 | 1,51E+08 | 1,51E+08 | 1502 | * | 15,11932 | 3,377693864 | 0,861757 | 3,919543 | 8,87E-05 | 0,155022149 | 1,37683 | Promoter (<=1kb) | 4 | 1,51E+08 | 1,51E+08 | 160 | 1 | 1,15E+08 | ENSMUST00000082542.4 | 0 | ENSMUSG00000064476 | Gm23405 | predicted gene, 23405 |
| chr7 | 47366686 | 47368083 | 1398 | * | 14,75938 | 3,378216418 | 0,906702 | 3,725829 | 0,000195 | 0,251021298 | 1,2763 | Distal Intergenic | 7 | 47238698 | 47251120 | 12423 | 2 | 233222 | ENSMUST00000176369.2 | -115566 | ENSMUSG00000078698 | Mrgpra3 | MAS-related GPR, member A3 |
| chr11 | 57534892 | 57536199 | 1308 | * | 13,20056 | 3,378816309 | 1,026131 | 3,292771 | 0,000992 | 0,55735526 | 0,918255 | Promoter (<=1kb) | 11 | 57536268 | 57678340 | 142073 | 1 | 171212 | ENSMUST00000066987.14 | -69 | ENSMUSG00000020520 | Galnt10 | polypeptide N-acetylgalactosaminyltransferase 10 |
| chr8 | 1,24E+08 | 1,24E+08 | 1400 | * | 11,98234 | 3,378900696 | 1,062787 | 3,179284 | 0,001476 | 0,591141332 | 1,173325 | Promoter (<=1kb) | 8 | 1,24E+08 | 1,24E+08 | 2209 | 1 | 66855 | ENSMUST00000212741.2 | 212 | ENSMUSG00000001472 | Tcf25 | transcription factor 25 (basic helix-loop-helix) |
| chr5 | 1,36E+08 | 1,36E+08 | 970 | * | 9,732968 | 3,37907623 | 1,112453 | 3,0375 | 0,002385 | 0,684274806 | 1,086118 | Intron (ENSMUST00000 | 5 | 1,36E+08 | 1,36E+08 | 23255 | 2 | 269717 | ENSMUST00000041048.6 | -12607 | ENSMUSG00000039747 | Orai2 | ORAI calcium release-activated calcium modulator 2 |
| chr18 | 33345848 | 33347758 | 1911 | * | 15,6674 | 3,379506669 | 0,869266 | 3,88777 | 0,000101 | 0,173939409 | 1,2763 | Promoter (<=1kb) | 18 | 33332408 | 33346915 | 14508 | 2 | 170459 | ENSMUST00000025236.9 | 0 | ENSMUSG00000024378 | Stard4 | StAR-related lipid transfer (START) domain containing 4 |
| chr13 | 95213787 | 95214908 | 1122 | * | 9,427641 | 3,379645752 | 1,269043 | 2,663145 | 0,007741 | 0,872677147 | 1,211315 | Intron (ENSMUST00000 | 13 | 95179976 | 95189526 | 9551 | 2 | 218461 | ENSMUST00000160412.2 | -24261 | ENSMUSG00000021684 | Pde8b | phosphodiesterase 8B |
| chr3 | 1,43E+08 | 1,43E+08 | 1152 | * | 9,193976 | 3,381855685 | 1,264529 | 2,674399 | 0,007486 | 0,838625535 | 1,425929 | Intron (ENSMUST00000 | 3 | 1,43E+08 | 1,43E+08 | 17743 | 2 | 109333 | ENSMUST00000174680.2 | -18566 | ENSMUSG00000004591 | Pkn2 | protein kinase N2 |
| chrX | 1,5E+08 | 1,5E+08 | 967 | * | 7,710637 | 3,382273754 | 1,27648 | 2,649687 | 0,008057 | 1 | 0 | Distal Intergenic | 20 | 1,5E+08 | 1,5E+08 | 6101 | 2 | 237107 | ENSMUST00000146742.2 | 19228 | ENSMUSG00000025266 | Gnl3l | guanine nucleotide binding protein-like 3 (nucleolar)-like |
| chr6 | 1,43E+08 | 1,43E+08 | 1192 | * | 11,43834 | 3,382695133 | 1,026357 | 3,295827 | 0,000981 | 0,553963091 | 0,918255 | Distal Intergenic | 6 | 1,43E+08 | 1,43E+08 | 19029 | 1 | 12764 | ENSMUST00000032419.9 | -12615 | ENSMUSG00000030282 | Cmas | cytidine monophospho-N-acetylneuraminic acid synthetase |
| chr8 | 1,28E+08 | 1,28E+08 | 1136 | * | 12,84315 | 3,383378907 | 1,018152 | 3,323058 | 0,00089 | 0,50985457 | 1,075735 | Distal Intergenic | 8 | 1,27E+08 | 1,27E+08 | 2764 | 2 | 1,15E+08 | ENSMUST00000212695.2 | -19609 | | Gm45805 | predicted gene 45805 |
| chr11 | 55033590 | 55034588 | 999 | * | 7,860868 | 3,383422486 | 1,298062 | 2,606518 | 0,009147 | 1 | 0 | 3' UTR | 11 | 55027834 | 55031020 | 3187 | 1 | 1,03E+08 | ENSMUST00000155883.3 | 5756 | ENSMUSG00000085531 | Slc36a3os | solute carrier family 36 (proton/amino acid symporter), member 3, opposite strand |
| chr19 | 25959859 | 25960892 | 1034 | * | 8,493226 | 3,383483503 | 1,22608 | 2,759593 | 0,005787 | 0,792545355 | 1,425929 | Distal Intergenic | 19 | 26015816 | 26023831 | 8016 | 2 | 73433 | ENSMUST00000237184.2 | 62939 | ENSMUSG00000118016 | 1700048O14Rik | RIKEN cDNA 1700048O14 gene |
| chr5 | 24587821 | 24589442 | 1622 | * | 34,856337 | 3,383544006 | 0,6213927 | 5,4450977 | 5,178E-08 | 0,000407685 | 2,7254908 | 3' UTR | 5 | 24592103 | 24592964 | 862 | 2 | 213948 | ENSMUST00000128831.2 | 3522 | ENSMUSG00000038295 | Atg9b | autophagy related 9B |
| chr11 | 1,04E+08 | 1,04E+08 | 1484 | * | 12,67418 | 3,383727562 | 1,025173 | 3,300639 | 0,000965 | 0,494741046 | 1,2763 | Promoter (<=1kb) | 11 | 1,04E+08 | 1,04E+08 | 7952 | 1 | 399609 | ENSMUST00000246205.1 | 0 | | C130046K22Rik | RIKEN cDNA C130046K22 gene |
| chr7 | 30428166 | 30430417 | 2252 | * | 25,361594 | 3,384407487 | 0,7450662 | 4,542425 | 5,561E-06 | 0,01387426 | 2,8800171 | Promoter (<=1kb) | 7 | 30429004 | 30445354 | 16351 | 1 | 282619 | ENSMUST00000180820.8 | 0 | ENSMUSG00000046056 | Sbsn | suprabasin |
| chr16 | 10332376 | 10333569 | 1194 | * | 13,63997 | 3,384748307 | 1,081015 | 3,131084 | 0,001742 | 0,663630601 | 0,918255 | 3' UTR | 16 | 10348071 | 10349673 | 1603 | 2 | 58239 | ENSMUST00000183355.8 | 16104 | ENSMUSG00000038055 | Dexi | dexamethasone-induced transcript |
| chr8 | 78158982 | 78159808 | 827 | * | 9,645474 | 3,38533428 | 1,123364 | 3,013568 | 0,002582 | 0,684274806 | 1,173325 | Exon (ENSMUST00000 | 8 | 78243685 | 78245244 | 1560 | 1 | 70345 | ENSMUST00000184762.3 | -83877 | ENSMUSG00000097882 | 0610038B21Rik | RIKEN cDNA 0610038B21 gene |
| chr10 | 58183282 | 58184784 | 1503 | * | 10,84481 | 3,386001912 | 1,270188 | 2,665749 | 0,007682 | 0,860218139 | 1,2763 | Promoter (<=1kb) | 10 | 58185320 | 58185426 | 107 | 2 | 1,15E+08 | ENSMUST00000093703.3 | 642 | ENSMUSG00000070214 | Gm23058 | predicted gene, 23058 |
| chr7 | 1,06E+08 | 1,06E+08 | 978 | * | 9,874552 | 3,386622666 | 1,188181 | 2,850257 | 0,004368 | 0,742868694 | 1,425929 | Distal Intergenic | 7 | 1,06E+08 | 1,06E+08 | 136 | 1 | 1,15E+08 | ENSMUST00000104589.3 | 15610 | ENSMUSG00000077777 | Gm22372 | predicted gene, 22372 |
| chr11 | 53909758 | 53910847 | 1090 | * | 16,75651 | 3,38680524 | 0,860553 | 3,935617 | 8,3E-05 | 0,112690111 | 2,141612 | Intron (ENSMUST00000 | 11 | 53905086 | 53918778 | 13693 | 2 | 30805 | ENSMUST00000154369.2 | 7931 | ENSMUSG00000020334 | Slc22a4 | solute carrier family 22 (organic cation transporter), member 4 |
| chr3 | 79068533 | 79069743 | 1211 | * | 11,52502 | 3,387149746 | 1,2689 | 2,66936 | 0,0076 | 0,947196831 | 0,744375 | Intron (ENSMUST00000 | 3 | 79016373 | 79080505 | 64133 | 2 | 76089 | ENSMUST00000152275.8 | 10762 | ENSMUSG00000062232 | Rapgef2 | Rap guanine nucleotide exchange factor (GEF) 2 |
| chr7 | 34967100 | 34969985 | 2886 | * | 18,98478 | 3,387253021 | 0,887795 | 3,815356 | 0,000136 | 0,143779747 | 2,386008 | Intron (ENSMUST00000 | 7 | 34938216 | 34974958 | 36743 | 2 | 384605 | ENSMUST00000238407.2 | 4973 | ENSMUSG00000118454 | Wdr88 | WD repeat domain 88 |
| chr3 | 87601896 | 87603194 | 1299 | * | 11,44522 | 3,387443774 | 1,229781 | 2,754509 | 0,005878 | 0,906945156 | 0,738441 | Exon (ENSMUST000001 | 3 | 87646287 | 87648355 | 2069 | 2 | 74485 | ENSMUST00000174581.2 | 45161 | ENSMUSG00000023084 | Lrrc71 | leucine rich repeat containing 71 |
| chr2 | 1,56E+08 | 1,56E+08 | 2687 | * | 18,92088 | 3,387518965 | 0,933511 | 3,628793 | 0,000285 | 0,206624632 | 2,663619 | Promoter (<=1kb) | 2 | 1,56E+08 | 1,56E+08 | 17832 | 2 | 266692 | ENSMUST00000183972.2 | 0 | ENSMUSG00000074643 | Cpne1 | copine I |
| chr18 | 4866265 | 4867744 | 1480 | * | 14,1421 | 3,388076853 | 1,150726 | 2,944295 | 0,003237 | 0,739058507 | 1,075735 | Distal Intergenic | 18 | 4908916 | 4912273 | 3358 | 1 | 1,05E+08 | ENSMUST00000241253.1 | -41172 | | Gm41659 | predicted gene, 41659 |
| chr5 | 49615126 | 49616302 | 1177 | * | 10,42316 | 3,388823066 | 1,183528 | 2,863324 | 0,004192 | 0,851741608 | 0,744375 | Intron (ENSMUST00000 | 5 | 48546849 | 49682106 | 1135258 | 2 | 80334 | ENSMUST00000176191.8 | 65804 | ENSMUSG00000029088 | Kcnip4 | Kv channel interacting protein 4 |
| chr11 | 98527112 | 98527996 | 885 | * | 7,788394 | 3,388908792 | 1,306616 | 2,593654 | 0,009496 | 0,888080529 | 1,360622 | Intron (ENSMUST00000 | 11 | 98537585 | 98548790 | 11206 | 1 | 76758 | ENSMUST00000093938.10 | -9589 | ENSMUSG00000017211 | Gsdma2 | gasdermin A2 |
| chr8 | 94762714 | 94764861 | 2148 | * | 25,8613 | 3,389322251 | 0,7147948 | 4,7416716 | 2,12E-06 | 0,007032136 | 2,7254908 | Promoter (<=1kb) | 8 | 94763826 | 94794549 | 30724 | 1 | 270086 | ENSMUST00000109556.9 | 0 | ENSMUSG00000033009 | Ogfod1 | 2-oxoglutarate and iron-dependent oxygenase domain containing 1 |
| chr12 | 8886827 | 8888387 | 1561 | * | 12,61556 | 3,390190803 | 1,19886 | 2,827846 | 0,004686 | 0,775744126 | 1,2763 | Distal Intergenic | 12 | 8923856 | 8927382 | 3527 | 2 | 319503 | ENSMUST00000218928.2 | 38995 | | 9930038B18Rik | RIKEN cDNA 9930038B18 gene |
| chr14 | 20952779 | 20953777 | 999 | * | 9,156783 | 3,390476857 | 1,207033 | 2,808936 | 0,004971 | 0,792873671 | 1,211315 | Distal Intergenic | 14 | 20979466 | 21048849 | 69384 | 1 | 22330 | ENSMUST00000226113.2 | -25689 | ENSMUSG00000021823 | Vcl | vinculin |
| chr1 | 92812325 | 92813187 | 863 | * | 10,94447 | 3,390569471 | 1,232981 | 2,749896 | 0,005961 | 0,908469938 | 0,744375 | Exon (ENSMUST000001 | 1 | 92787525 | 92799442 | 11918 | 2 | 241158 | ENSMUST00000212504.2 | -12883 | ENSMUSG00000034212 | Ankmy1 | ankyrin repeat and MYND domain containing 1 |
| chr11 | 3712345 | 3713141 | 797 | * | 9,02316 | 3,392474097 | 1,278635 | 2,653201 | 0,007973 | 0,851183656 | 1,425929 | Intron (ENSMUST00000 | 11 | 3667852 | 3723789 | 55938 | 2 | 74309 | ENSMUST00000127371.2 | 10648 | ENSMUSG00000020435 | Osbp2 | oxysterol binding protein 2 |
| chr3 | 40847849 | 40849704 | 1856 | * | 30,11965 | 3,39292603 | 1,085222 | 3,126482 | 0,001769 | 0,459179243 | 2,880017 | Promoter (<=1kb) | 3 | 40848580 | 40892573 | 43994 | 1 | 269423 | ENSMUST00000159774.7 | 0 | ENSMUSG00000037818 | Abhd18 | abhydrolase domain containing 18 |
| chr5 | 22629591 | 22630468 | 878 | * | 9,522046 | 3,393123187 | 1,249637 | 2,715286 | 0,006622 | 0,817520905 | 1,425929 | Distal Intergenic | 5 | 22547069 | 22549700 | 2632 | 2 | 19699 | ENSMUST00000161782.2 | -79891 | ENSMUSG00000042453 | Reln | reelin |
| chr18 | 36933762 | 36936524 | 2763 | * | 19,98248 | 3,393849519 | 0,927657 | 3,658517 | 0,000254 | 0,214556109 | 2,221611 | Promoter (<=1kb) | 18 | 36934919 | 36935061 | 143 | 1 | 378472 | ENSMUST00020181660.1 | 0 | | Vaultrc5 | vault RNA component 5 |
| chr2 | 1,55E+08 | 1,55E+08 | 1294 | * | 12,65697 | 3,39473893 | 1,063872 | 3,190928 | 0,001418 | 0,627785088 | 0,918255 | Promoter (<=1kb) | 2 | 1,55E+08 | 1,55E+08 | 1180 | 1 | 66734 | ENSMUST00000154596.2 | 760 | ENSMUSG00000027602 | Map1lc3a | microtubule-associated protein 1 light chain 3 alpha |
| chr11 | 1,18E+08 | 1,18E+08 | 1508 | * | 11,39324 | 3,395239682 | 1,092711 | 3,107171 | 0,001889 | 0,676922067 | 0,918255 | 3' UTR | 11 | 1,18E+08 | 1,18E+08 | 3020 | 1 | 217364 | ENSMUST00000168689.2 | 5152 | ENSMUSG00000033857 | Engase | endo-beta-N-acetylglucosaminidase |
| chr15 | 77010736 | 77012245 | 1510 | * | 15,55379 | 3,395403493 | 1,086342 | 3,12554 | 0,001775 | 0,50985457 | 2,141612 | Intron (ENSMUST00000 | 15 | 77000570 | 77016980 | 16411 | 2 | 93686 | ENSMUST00000228190.2 | 4735 | ENSMUSG00000033565 | Rbfox2 | RNA binding protein, fox-1 homolog (C. elegans) 2 |
| chr1 | 74651356 | 74652182 | 827 | * | 8,175211 | 3,396066606 | 1,296586 | 2,619236 | 0,008813 | 1 | 0 | Exon (ENSMUST00000 | 1 | 74642369 | 74650415 | 8047 | 1 | 269209 | ENSMUST00000189830.2 | 8987 | ENSMUSG00000033276 | Stk36 | serine/threonine kinase 36 |
| chr11 | 28694056 | 28694795 | 740 | * | 7,566095 | 3,397290416 | 1,293798 | 2,625827 | 0,008644 | 1 | 0 | Promoter (1-2kb) | 11 | 28696191 | 28696276 | 86 | 1 | 735308 | ENSMUST00000102363.3 | -1396 | ENSMUSG00000076318 | Mir216b | microRNA 216b |
| chr17 | 74992845 | 74994071 | 1227 | * | 8,909034 | 3,397543051 | 1,177557 | 2,885246 | 0,003911 | 0,749931768 | 1,211315 | 3' UTR | 17 | 75004351 | 75010064 | 5714 | 1 | 12211 | ENSMUST00000182157.2 | -10280 | ENSMUSG00000024073 | Birc6 | baculoviral IAP repeat-containing 6 |
| chr2 | 71264199 | 71266291 | 2093 | * | 15,36765 | 3,397995323 | 0,883889 | 3,844367 | 0,000121 | 0,192725821 | 1,2763 | Exon (ENSMUST000001 | 2 | 71250436 | 71269156 | 18721 | 1 | 107435 | ENSMUST00000145147.2 | 13763 | ENSMUSG00000027018 | Hat1 | histone aminotransferase 1 |
| chr14 | 1,22E+08 | 1,22E+08 | 2132 | * | 14,51381 | 3,3982704 | 0,94977 | 3,577992 | 0,000346 | 0,308866996 | 1,41696 | Distal Intergenic | 14 | 1,22E+08 | 1,22E+08 | 4033 | 2 | 1,03E+08 | ENSMUST00000227451.2 | 11157 | | Gm33299 | predicted gene, 33299 |
| chr14 | 1,16E+08 | 1,16E+08 | 1466 | * | 8,595455 | 3,398961618 | 1,265348 | 2,686187 | 0,007227 | 0,749931768 | 2,237403 | Intron (ENSMUST00000 | 14 | 1,16E+08 | 1,16E+08 | 13002 | 2 | 75088 | ENSMUST00000176582.2 | -570124 | ENSMUSG00000093551 | 4930505G20Rik | RIKEN cDNA 4930505G20 gene |
| chr4 | 1,17E+08 | 1,17E+08 | 2254 | * | 14,19787 | 3,399125683 | 1,160355 | 2,929384 | 0,003396 | 0,815634049 | 0,738441 | Exon (ENSMUST000001 | 4 | 1,17E+08 | 1,17E+08 | 11875 | 1 | 18477 | ENSMUST00000030454.6 | 5410 | ENSMUSG00000028691 | Prdx1 | peroxiredoxin 1 |
| chr10 | 1,02E+08 | 1,02E+08 | 1817 | * | 12,9558 | 3,400108567 | 1,141304 | 2,979144 | 0,002891 | 0,748382289 | 0,918255 | Intron (ENSMUST00000 | 10 | 1,02E+08 | 1,02E+08 | 232391 | 1 | 67569 | ENSMUST00000020039.13 | 35857 | ENSMUSG00000019888 | Mgat4c | MGAT4 family, member C |
| chr10 | 86539563 | 86542444 | 2882 | * | 47,15557 | 3,400195907 | 0,930595 | 3,653787 | 0,000258 | 0,190625259 | 2,787255 | Promoter (<=1kb) | 10 | 86541675 | 86612708 | 71034 | 1 | 103220 | ENSMUST00000075632.14 | 0 | ENSMUSG00000044937 | Ttc41 | tetratricopeptide repeat domain 41 |
| chr3 | 56395468 | 56396493 | 1026 | * | 9,554335 | 3,401941102 | 1,242627 | 2,737701 | 0,006187 | 0,838625535 | 1,173325 | Distal Intergenic | 3 | 56352703 | 56352833 | 131 | 2 | 1,15E+08 | ENSMUST00000104522.3 | -42635 | ENSMUSG00000119391 | Gm25727 | predicted gene, 25727 |
| chr17 | 23173116 | 23173907 | 792 | * | 9,283993 | 3,402094307 | 1,272743 | 2,673041 | 0,007517 | 0,836571183 | 1,449706 | Exon (ENSMUST000002 | 17 | 23161958 | 23184632 | 22675 | 1 | 434701 | ENSMUST00000234875.2 | 11158 | ENSMUSG00000092111 | Vmn2r113 | vomeronasal 2, receptor 113 |
| chr15 | 78963733 | 78965418 | 1686 | * | 12,5403 | 3,403393067 | 1,09519 | 3,107582 | 0,001886 | 0,713406565 | 0,738441 | Promoter (1-2kb) | 15 | 78967249 | 78978605 | 11357 | 1 | 223691 | ENSMUST00000230032.2 | -1831 | ENSMUSG00000033047 | Eif3l | eukaryotic translation initiation factor 3, subunit L |
| chr8 | 88564223 | 88565542 | 1320 | * | 16,5175 | 3,403879073 | 0,93479 | 3,641331 | 0,000271 | 0,293348204 | 1,2763 | Intron (ENSMUST00000 | 8 | 88388438 | 88531067 | 142630 | 2 | 94187 | ENSMUST00000165770.9 | -33156 | ENSMUSG00000045333 | Zfp423 | zinc finger protein 423 |
| chr18 | 75498037 | 75500310 | 2274 | * | 20,819219 | 3,404734403 | 0,7848937 | 4,3378289 | 1,439E-05 | 0,024985923 | 3,4716776 | Promoter (<=1kb) | 18 | 75491711 | 75500160 | 8450 | 2 | 102636262 | ENSMUST00000174338.3 | 0 | | Gm20544 | predicted gene 20544 |

| | | | | | | | | | | | | | | | | | | | | | | | |
|---|---|---|---|---|---|---|---|---|---|---|---|---|---|---|---|---|---|---|---|---|---|---|---|
| chr1 | 53351582 | 53353760 | 2179 | * | 19,543631 | 3,405525067 | 0,7781157 | 4,3766305 | 1,205E-05 | 0,030165351 | 2,2216115 | Promoter (<=1kb) | 1 | 53352783 | 53365502 | 12720 | 1 | 72085 | ENSMUST00000027265.10 | 0 | ENSMUSG00000026096 | Osgepl1 | O-sialoglycoprotein endopeptidase-like 1 |
| chr17 | 3499201 | 3500844 | 1644 | * | 17,88314 | 3,405568902 | 0,925598 | 3,679316 | 0,000234 | 0,184990586 | 2,663619 | Exon (ENSMUST000001 | 17 | 3504521 | 3581619 | 77099 | 1 | 24001 | ENSMUST00000227405.2 | -3677 | ENSMUSG00000023800 | Tiam2 | T cell lymphoma invasion and metastasis 2 |
| chr3 | 1,43E+08 | 1,43E+08 | 1111 | * | 7,41126 | 3,40568963 | 1,304519 | 2,610686 | 0,009036 | 1 | 0 | Intron (ENSMUST00000 | 3 | 1,43E+08 | 1,43E+08 | 17743 | 2 | 109333 | ENSMUST00000174680.2 | -20463 | ENSMUSG00000004591 | Pkn2 | protein kinase N2 |
| chr18 | 63449464 | 63450493 | 1030 | * | 9,204621 | 3,406858866 | 1,118355 | 3,046313 | 0,002317 | 0,64586006 | 1,360622 | Intron (ENSMUST00000 | 18 | 63250737 | 63459720 | 208984 | 2 | 667742 | ENSMUST00000046860.7 | 9227 | ENSMUSG00000041482 | Piezo2 | piezo-type mechanosensitive ion channel component 2 |
| chr3 | 38806822 | 38808532 | 1711 | * | 10,56213 | 3,407634351 | 1,057549 | 3,2222 | 0,001272 | 0,60988361 | 0,918255 | Distal Intergenic | 3 | 38941089 | 39066134 | 125046 | 1 | 329628 | ENSMUST00000061260.8 | -132557 | ENSMUSG00000046743 | Fat4 | FAT atypical cadherin 4 |
| chr6 | 146859063 | 146862764 | 3702 | * | 40,45209 | 3,407756923 | 0,5777605 | 5,8982171 | 3,675E-09 | 4,51459E-05 | 2,7872553 | Intron (ENSMUST000000 | 6 | 146851827 | 146855920 | 4094 | 2 | 73344 | ENSMUST00000036592.15 | -3143 | ENSMUSG00000040163 | 1700034J05Rik | RIKEN cDNA 1700034J05 gene |
| chr4 | 4716762 | 4717911 | 1150 | * | 9,147047 | 3,407984372 | 1,215674 | 2,80337 | 0,005057 | 0,765231686 | 1,449706 | Distal Intergenic | 4 | 4729573 | 4729704 | 132 | 1 | 1,15E+08 | ENSMUST00000180325.2 | -11662 | ENSMUSG00000095971 | Gm26436 | predicted gene, 26436 |
| chr4 | 1,36E+08 | 1,36E+08 | 1164 | * | 14,15656 | 3,408139218 | 0,977449 | 3,486771 | 0,000489 | 0,473594154 | 0,738441 | Promoter (<=1kb) | 4 | 1,36E+08 | 1,36E+08 | 18396 | 2 | 27224 | ENSMUST00000030427.6 | 0 | ENSMUSG00000028668 | Eloa | elongin A |
| chr15 | 81111741 | 81113288 | 1548 | * | 13,20799 | 3,408294967 | 1,226862 | 2,778059 | 0,005468 | 0,858257847 | 0,918255 | Distal Intergenic | 15 | 81119700 | 81123165 | 3466 | 1 | 207911 | ENSMUST00000166855.3 | -6412 | ENSMUSG00000050164 | Mchr1 | melanin-concentrating hormone receptor 1 |
| chr5 | 77522978 | 77524512 | 1535 | * | 13,5085 | 3,409157717 | 1,068881 | 3,189464 | 0,001425 | 0,565251403 | 1,2763 | Intron (ENSMUST00000 | 5 | 77534446 | 77541530 | 7085 | 1 | 1,03E+08 | ENSMUST00000202460.2 | -9934 | | Gm34648 | predicted gene, 34648 |
| chr15 | 76503765 | 76504752 | 988 | * | 10,86876 | 3,40948717 | 1,108001 | 3,077153 | 0,00209 | 0,65364112 | 1,173325 | Promoter (2-3kb) | 15 | 76496591 | 76501584 | 4994 | 2 | 72027 | ENSMUST00000230977.2 | -2181 | ENSMUSG00000063354 | Slc39a4 | solute carrier family 39 (zinc transporter), member 4 |
| chr11 | 21518828 | 21520512 | 1685 | * | 14,33491 | 3,410227512 | 1,085636 | 3,141227 | 0,001682 | 0,688021707 | 0,738441 | Promoter (1-2kb) | 11 | 21506692 | 21521934 | 15243 | 2 | 17449 | ENSMUST00000239073.2 | 1422 | ENSMUSG00000020321 | Mdh1 | malate dehydrogenase 1, NAD (soluble) |
| chr6 | 52038856 | 52040619 | 1764 | * | 18,06296 | 3,410337137 | 1,009335 | 3,378796 | 0,000728 | 0,354548686 | 2,154504 | Distal Intergenic | 6 | 52079903 | 52091977 | 12075 | 1 | 1,03E+08 | ENSMUST00000189071.8 | -39284 | ENSMUSG00000085412 | Halr1 | Hoxa adjacent long noncoding RNA 1 |
| chr9 | 1,03E+08 | 1,03E+08 | 2269 | * | 17,1379 | 3,41051234 | 0,953311 | 3,577546 | 0,000347 | 0,242864164 | 2,423594 | Promoter (1-2kb) | 9 | 1,03E+08 | 1,03E+08 | 12747 | 2 | 321022 | ENSMUST00000035484.11 | -1478 | ENSMUSG00000032803 | Cdv3 | carnitine deficiency-associated gene expressed in ventricle 3 |
| chr2 | 98303105 | 98304069 | 965 | * | 9,47849 | 3,411322948 | 1,243767 | 2,742734 | 0,006093 | 0,798282642 | 1,449706 | Distal Intergenic | 2 | 97298434 | 97459408 | 160975 | 1 | 241568 | ENSMUST00000170144.2 | 1004671 | ENSMUSG00000050587 | Lrrc4c | leucine rich repeat containing 4C |
| chr16 | 32874201 | 32875838 | 1638 | * | 8,898692 | 3,411501884 | 1,275036 | 2,675613 | 0,007459 | 0,846832214 | 1,360622 | Promoter (<=1kb) | 16 | 32869877 | 32876555 | 6679 | 2 | 69707 | ENSMUST00000126793.2 | 717 | ENSMUSG00000035578 | Iqcg | IQ motif containing G |
| chr11 | 72687359 | 72688576 | 1218 | * | 11,16346 | 3,413371472 | 1,06355 | 3,209414 | 0,00133 | 0,568205102 | 1,173325 | Promoter (<=1kb) | 11 | 72687080 | 72817944 | 130865 | 1 | 195018 | ENSMUST00000207107.2 | 279 | ENSMUSG00000055670 | Zzef1 | zinc finger, ZZ-type with EF hand domain 1 |
| chr16 | 21171547 | 21173089 | 1543 | * | 13,39916 | 3,413377621 | 1,228908 | 2,777569 | 0,005477 | 0,85827362 | 0,918255 | Distal Intergenic | 16 | 21150643 | 21152139 | 1497 | 2 | 76222 | ENSMUST00000231504.3 | -19408 | ENSMUSG00000116632 | Magef1 | MAGE family member F1 |
| chr8 | 57940280 | 57943175 | 2896 | * | 47,533223 | 3,413432519 | 0,574724 | 5,9392554 | 2,863E-09 | 4,37941E-05 | 2,3383871 | Promoter (<=1kb) | 8 | 57935741 | 57940894 | 5154 | 2 | 60406 | ENSMUST00000034022.4 | 0 | ENSMUSG00000031609 | Sap30 | sin3 associated polypeptide |
| chr13 | 21621583 | 21622933 | 1351 | * | 8,197864 | 3,413986265 | 1,307801 | 2,610478 | 0,009042 | 0,880247401 | 1,360622 | Promoter (1-2kb) | 13 | 21624641 | 21630104 | 5464 | 1 | 1,05E+08 | ENSMUST00000122926.2 | -1708 | | 4930470G03Rik | RIKEN cDNA 4930470G03 gene |
| chr11 | 1,17E+08 | 1,17E+08 | 743 | * | 10,32223 | 3,414101866 | 1,164698 | 2,931319 | 0,003375 | 0,695892636 | 1,425929 | Promoter (<=1kb) | 11 | 1,17E+08 | 1,17E+08 | 13512 | 1 | 74136 | ENSMUST00000146585.2 | 942 | ENSMUSG00000020823 | Sec14l1 | SEC14-like lipid binding 1 |
| chrX | 1,18E+08 | 1,18E+08 | 1169 | * | 9,942495 | 3,415729293 | 1,177219 | 2,901524 | 0,003714 | 0,715249603 | 1,425929 | Distal Intergenic | 20 | 1,17E+08 | 1,17E+08 | 1030 | 2 | 245583 | ENSMUST00000073857.6 | -498036 | ENSMUSG00000100133 | Tgif2lx1 | TGFB-induced factor homeobox 2-like, X-linked 1 |
| chr16 | 85685600 | 85686673 | 1074 | * | 7,266072 | 3,416437967 | 1,288543 | 2,651397 | 0,008016 | 1 | 0 | Intron (ENSMUST00000 | 16 | 85653061 | 85698716 | 45656 | 2 | 23794 | ENSMUST00000023611.7 | 12043 | ENSMUSG00000022894 | Adamts5 | a disintegrin-like and metallopeptidase (reprolysin type) with thrombospondin type 1 motif, 5 (aggrecanase-2) |
| chr13 | 39001227 | 39002354 | 1128 | * | 13,36044 | 3,416854764 | 0,98656 | 3,463402 | 0,000533 | 0,39905285 | 1,2763 | Intron (ENSMUST00000 | 13 | 39045706 | 39092231 | 46526 | 1 | 1,08E+08 | ENSMUST00000225020.2 | -43352 | ENSMUSG00000114767 | Gm46392 | predicted gene, 46392 |
| chr10 | 31485544 | 31486348 | 805 | * | 21,661248 | 3,417099436 | 0,7262865 | 4,7048921 | 2,54E-06 | 0,007672933 | 2,8800171 | Promoter (<=1kb) | 10 | 31369189 | 31485180 | 115992 | 2 | 268291 | ENSMUST00000081989.8 | -364 | ENSMUSG00000063760 | Rnf217 | ring finger protein 217 |
| chr12 | 52681766 | 52683773 | 2008 | * | 17,95419 | 3,419610325 | 0,978206 | 3,495796 | 0,000473 | 0,278016366 | 2,507013 | Distal Intergenic | 12 | 52654453 | 52675595 | 21143 | 1 | 1,03E+08 | ENSMUST00000218346.2 | 27313 | | Gm35239 | predicted gene, 35239 |
| chr18 | 69683786 | 69685607 | 1822 | * | 15,36568 | 3,42050115 | 0,90296 | 3,7881 | 0,000152 | 0,206624632 | 1,41696 | Exon (ENSMUST00020 | 18 | 69696078 | 69776086 | 80009 | 1 | 21413 | ENSMUST00000201045.5 | -10471 | ENSMUSG00000053477 | Tcf4 | transcription factor 4 |
| chr3 | 1,38E+08 | 1,38E+08 | 1240 | * | 11,37277 | 3,421498104 | 1,138237 | 3,005962 | 0,002647 | 0,768389035 | 0,744375 | Promoter (<=1kb) | 3 | 1,38E+08 | 1,38E+08 | 12407 | 1 | 11532 | ENSMUST00000005964.7 | 0 | ENSMUSG00000028138 | Adh5 | alcohol dehydrogenase 5 (class III), chi polypeptide |
| chr14 | 70339191 | 70340720 | 1530 | * | 9,586972 | 3,421566302 | 1,305082 | 2,621726 | 0,008749 | 0,892066133 | 1,211315 | Promoter (1-2kb) | 14 | 70337614 | 70371975 | 34362 | 1 | 57784 | ENSMUST00000228049.2 | 1577 | ENSMUSG00000022089 | Bin3 | bridging integrator 3 |
| chr13 | 12041314 | 12043886 | 2573 | * | 18,23492 | 3,421755693 | 0,846352 | 4,042944 | 5,28E-05 | 0,080010047 | 2,423594 | Intron (ENSMUST00000 | 13 | 11932148 | 12121345 | 189198 | 2 | 20191 | ENSMUST00000221341.2 | 77459 | ENSMUSG00000021313 | Ryr2 | ryanodine receptor 2, cardiac |
| chr19 | 51877411 | 51878486 | 1076 | * | 9,899573 | 3,42184373 | 1,149313 | 2,977296 | 0,002908 | 0,672444708 | 1,449706 | Distal Intergenic | 19 | 52252735 | 52253914 | 1180 | 1 | 16333 | ENSMUST00000039652.6 | -374249 | ENSMUSG00000035804 | Ins1 | insulin I |
| chr13 | 7687164 | 7687941 | 778 | * | 8,340024 | 3,422328956 | 1,27811 | 2,677649 | 0,007414 | 1 | 0 | Distal Intergenic | 13 | 7496545 | 7515379 | 18835 | 1 | 1,03E+08 | ENSMUST00000221383.2 | 190619 | | Gm36074 | predicted gene, 36074 |
| chr17 | 65920527 | 65921800 | 1274 | * | 23,067039 | 3,422506207 | 0,8124305 | 4,2126757 | 2,524E-05 | 0,044544037 | 2,8585108 | Promoter (<=1kb) | 17 | 65885322 | 65920550 | 35229 | 2 | 30960 | ENSMUST00000024897.10 | 0 | ENSMUSG00000024091 | Vapa | vesicle-associated membrane protein, associated protein A |
| chr12 | 1,02E+08 | 1,02E+08 | 1194 | * | 7,796844 | 3,422537415 | 1,298131 | 2,636512 | 0,008376 | 1 | 0 | Promoter (1-2kb) | 12 | 1,02E+08 | 1,02E+08 | 25200 | 2 | 110616 | ENSMUST00000159883.2 | 1971 | ENSMUSG00000021189 | Atxn3 | ataxin 3 |
| chr5 | 51808644 | 51809238 | 595 | * | 7,61463 | 3,422909557 | 1,293364 | 2,646517 | 0,008133 | 1 | 0 | Distal Intergenic | 5 | 51653122 | 51725068 | 71947 | 2 | 19017 | ENSMUST00000127135.3 | -83576 | ENSMUSG00000029167 | Ppargc1a | peroxisome proliferative activated receptor, gamma, coactivator 1 alpha |
| chr2 | 1,32E+08 | 1,32E+08 | 1076 | * | 13,58565 | 3,422967382 | 0,923176 | 3,707815 | 0,000209 | 0,302119236 | 0,918255 | Intron (ENSMUST00000 | 2 | 1,32E+08 | 1,32E+08 | 8163 | 2 | 70612 | ENSMUST00000110164.8 | 40627 | ENSMUSG00000027341 | Tmem230 | transmembrane protein 230 |
| chr6 | 31061662 | 31062809 | 1148 | * | 22,0454 | 3,423009415 | 0,871034 | 3,929823 | 8,5E-05 | 0,09031013 | 3,104559 | Intron (ENSMUST00000 | 6 | 31040027 | 31040430 | 404 | 2 | 387223 | ENSMUST00000183056.2 | -21232 | ENSMUSG00000065604 | Mir29b-1 | microRNA 29b-1 |
| chr9 | 71265047 | 71266506 | 1460 | * | 12,74676 | 3,423264297 | 1,307867 | 2,617441 | 0,008859 | 0,912633142 | 1,075735 | Distal Intergenic | 9 | 71385753 | 71392240 | 6488 | 2 | 28015 | ENSMUST00000163972.2 | 125734 | ENSMUSG00000032199 | Polr2m | polymerase (RNA) II (DNA directed) polypeptide M |
| chr14 | 8601503 | 8603204 | 1702 | * | 8,781064 | 3,423543596 | 1,23214 | 2,778534 | 0,00546 | 0,791242424 | 1,360622 | Exon (ENSMUST000001 | 14 | 8555285 | 8655145 | 99861 | 1 | 238939 | ENSMUST00000144914.3 | 46218 | ENSMUSG00000084902 | Gm281 | predicted gene 281 |
| chr10 | 79821163 | 79822852 | 1690 | * | 19,9919 | 3,424402976 | 0,908878 | 3,767725 | 0,000165 | 0,156689151 | 2,507013 | Promoter (<=1kb) | 10 | 79821340 | 79821446 | 107 | 1 | 1,15E+08 | ENSMUST00000239704.1 | 0 | ENSMUSG00000118738 | Gm26243 | predicted gene, 26243 |
| chr15 | 79973241 | 79974562 | 1322 | * | 11,59287 | 3,424752165 | 1,097247 | 3,121221 | 0,001801 | 0,611772917 | 1,2763 | Promoter (<=1kb) | 15 | 79972648 | 79972754 | 107 | 2 | 1,15E+08 | ENSMUST00000082950.3 | -487 | ENSMUSG00000064884 | Gm24204 | predicted gene, 24204 |
| chr11 | 69597150 | 69599363 | 2214 | * | 15,75879 | 3,424830752 | 1,013547 | 3,379054 | 0,000727 | 0,337811311 | 2,386008 | Distal Intergenic | 11 | 69573403 | 69586924 | 13522 | 2 | 619441 | ENSMUST00000180587.8 | -10226 | ENSMUSG00000018752 | Tnfsfm13 | tumor necrosis factor (ligand) superfamily, membrane-bound member 13 |
| chr11 | 52171066 | 52172302 | 1237 | * | 11,92964 | 3,425069592 | 1,067202 | 3,209392 | 0,00133 | 0,654705303 | 0,744375 | Promoter (1-2kb) | 11 | 52143198 | 52173398 | 30201 | 2 | 21414 | ENSMUST00000086844.10 | 1096 | ENSMUSG00000000782 | Tcf7 | transcription factor 7, T cell specific |
| chr14 | 21037176 | 21038768 | 1593 | * | 9,220167 | 3,425146917 | 1,132272 | 3,02502 | 0,002486 | 0,649854434 | 1,425929 | Exon (ENSMUST00002 | 14 | 21072071 | 21077285 | 5215 | 1 | 22330 | ENSMUST00000224380.2 | -33303 | ENSMUSG00000021823 | Vcl | vinculin |
| chr16 | 33640009 | 33640885 | 877 | * | 7,581627 | 3,425300055 | 1,294211 | 2,646633 | 0,00813 | 1 | 0 | 3' UTR | 16 | 33650035 | 33769708 | 119674 | 1 | 16419 | ENSMUST00000115028.11 | -9150 | ENSMUSG00000022817 | Itgb5 | integrin beta 5 |
| chr11 | 69809024 | 69811001 | 1978 | * | 16,93157 | 3,426667752 | 1,010944 | 3,389571 | 0,0007 | 0,441389356 | 1,2763 | Promoter (<=1kb) | 11 | 69808008 | 69809130 | 1123 | 2 | 276770 | ENSMUST00000153652.8 | 0 | ENSMUSG00000078812 | Eif5a | eukaryotic translation initiation factor 5A |
| chr19 | 3668201 | 3670817 | 2617 | * | 20,84262 | 3,426683966 | 0,827791 | 4,13955 | 3,48E-05 | 0,05285139 | 2,951933 | Promoter (2-3kb) | 19 | 3634836 | 3665792 | 30957 | 2 | 16973 | ENSMUST00000177294.2 | -2409 | ENSMUSG00000024913 | Lrp5 | low density lipoprotein receptor-related protein 5 |
| chr4 | 14265911 | 14266844 | 934 | * | 11,63887 | 3,427206004 | 1,160137 | 2,954139 | 0,003135 | 0,718390388 | 1,173325 | Distal Intergenic | 4 | 14239618 | 14239744 | 127 | 1 | 1,15E+08 | ENSMUST00000157294.2 | 26293 | ENSMUSG00000087919 | Gm24908 | predicted gene, 24908 |
| chr16 | 50083659 | 50084305 | 647 | * | 7,109552 | 3,427325573 | 1,292363 | 2,651984 | 0,008002 | 1 | 0 | Intron (ENSMUST00000 | 16 | 50017522 | 50151356 | 133835 | 2 | 70508 | ENSMUST00000114488.8 | 67051 | ENSMUSG00000022641 | Bbx | bobby sox HMG box containing |
| chr3 | 95014978 | 95016584 | 1607 | * | 11,74925 | 3,427518957 | 1,066122 | 3,21494 | 0,001305 | 0,546082909 | 1,2763 | Promoter (<=1kb) | 3 | 94965854 | 95014241 | 48388 | 2 | 18720 | ENSMUST00000107233.9 | -737 | ENSMUSG00000028126 | Pip5k1a | phosphatidylinositol-4-phosphate 5-kinase, type 1 alpha |
| chr13 | 58865920 | 58866709 | 790 | * | 10,55069 | 3,42765227 | 1,076476 | 3,184142 | 0,001452 | 0,669774031 | 0,738441 | Distal Intergenic | 13 | 58888888 | 58910125 | 21238 | 2 | 67585 | ENSMUST00000225693.2 | 43416 | ENSMUSG00000114384 | 4930455J16Rik | RIKEN cDNA 4930455J16 gene |
| chr11 | 69773245 | 69775938 | 2694 | * | 55,160411 | 3,427780605 | 0,561549 | 6,1041524 | 1,033E-09 | 1,67231E-05 | 2,9254973 | Promoter (<=1kb) | 11 | 69772858 | 69772920 | 63 | 2 | 102466208 | ENSMUST00000198919.3 | -325 | ENSMUSG00000104996 | Mir6924 | microRNA 6924 |
| chr19 | 22495660 | 22496872 | 1213 | * | 8,778041 | 3,428143105 | 1,236368 | 2,772752 | 0,005558 | 0,787309858 | 1,425929 | Intron (ENSMUST00000 | 19 | 22415914 | 22426229 | 10316 | 2 | 1,03E+08 | ENSMUST00000185015.2 | -69431 | | Gm27151 | predicted gene 27151 |
| chr6 | 12105317 | 12105880 | 564 | * | 8,823871 | 3,428430938 | 1,309418 | 2,618286 | 0,008837 | 0,861872447 | 1,449706 | Intron (ENSMUST00000 | 6 | 12099517 | 12109582 | 10066 | 2 | 625347 | ENSMUST00000080891.6 | 3702 | ENSMUSG00000057223 | Gm6578 | mitochondrial ribosomal protein L32 pseudogene |
| chr6 | 37803223 | 37803881 | 659 | * | 7,341434 | 3,428745726 | 1,281445 | 2,675687 | 0,007458 | 0,846832214 | 1,360622 | Distal Intergenic | 6 | 37783021 | 37783485 | 465 | 1 | 1E+08 | ENSMUST00000122373.2 | 20202 | | Atp6v0c-ps2 | ATPase, H+ transporting, lysosomal V0 subunit C, pseudogene 2 |
| chr9 | 70061026 | 70062253 | 1228 | * | 9,735961 | 3,428976069 | 1,283809 | 2,67094 | 0,007564 | 0,839086232 | 1,425929 | Distal Intergenic | 9 | 70056088 | 70056218 | 131 | 2 | 1,15E+08 | ENSMUST00000104472.3 | -4808 | ENSMUSG00000077660 | Gm24257 | predicted gene, 24257 |
| chr10 | 79976649 | 79978320 | 1672 | * | 25,183092 | 3,42953782 | 0,7530615 | 4,5541273 | 5,26E-06 | 0,013096578 | 2,9519335 | Promoter (<=1kb) | 10 | 79973181 | 79976669 | 3489 | 2 | 100503659 | ENSMUST00000142853.3 | 0 | ENSMUSG00000035640 | Cbarp | calcium channel, voltage-dependent, beta subunit associated regulatory protein |
| chr15 | 9117568 | 9118544 | 977 | * | 9,101478 | 3,431052115 | 1,150741 | 2,981602 | 0,002867 | 0,670853401 | 1,449706 | Intron (ENSMUST00000 | 15 | 9103076 | 9106858 | 3783 | 1 | 68646 | ENSMUST00000186069.2 | 14492 | ENSMUSG00000022253 | Nadk2 | NAD kinase 2, mitochondrial |
| chr4 | 85575815 | 85577493 | 1679 | * | 15,22216 | 3,432659241 | 1,094025 | 3,137643 | 0,001703 | 0,586304784 | 1,37683 | Intron (ENSMUST00000 | 4 | 85432564 | 85701364 | 268801 | 1 | 77739 | ENSMUST00000147066.2 | 143251 | ENSMUSG00000066113 | Adamtsl1 | ADAMTS-like 1 |
| chr17 | 56343540 | 56345329 | 1790 | * | 24,008868 | 3,433032213 | 0,750027 | 4,5772117 | 4,712E-06 | 0,01245642 | 2,8585108 | Promoter (<=1kb) | 17 | 56323750 | 56343635 | 19886 | 2 | 20405 | ENSMUST00000003268.11 | 0 | ENSMUSG00000003200 | Sh3gl1 | SH3-domain GRB2-like 1 |
| chr19 | 42647730 | 42648896 | 1167 | * | 18,70702 | 3,433310715 | 0,958917 | 3,580404 | 0,000343 | 0,227550076 | 2,663619 | Distal Intergenic | 19 | 42688216 | 42688341 | 126 | 2 | 1,15E+08 | ENSMUST00000104580.3 | 39445 | ENSMUSG00000077768 | Gm25216 | predicted gene, 25216 |
| chr1 | 1,83E+08 | 1,83E+08 | 969 | * | 7,976352 | 3,433720425 | 1,321406 | 2,598535 | 0,009362 | 1 | 0 | Distal Intergenic | 1 | 1,83E+08 | 1,83E+08 | 17539 | 1 | 78772 | ENSMUST00000192527.2 | 25245 | ENSMUSG00000053461 | Hhipl2 | hedgehog interacting protein-like 2 |
| chr19 | 44709880 | 44711755 | 1876 | * | 18,39027 | 3,434466488 | 0,999237 | 3,437088 | 0,000588 | 0,323558126 | 2,154504 | Intron (ENSMUST00000 | 19 | 44728318 | 44731883 | 3566 | 1 | 1,03E+08 | ENSMUST00000236936.2 | -16563 | | Gm35610 | predicted gene, 35610 |
| chr11 | 97215755 | 97217075 | 1321 | * | 9,752165 | 3,43511006 | 1,255123 | 2,736872 | 0,006203 | 0,852340527 | 1,086118 | Exon (ENSMUST00000 | 11 | 97206734 | 97212551 | 5818 | 1 | 67036 | ENSMUST00000132168.2 | 9021 | ENSMUSG00000018882 | Mrpl45 | mitochondrial ribosomal protein L45 |
| chr9 | 31454217 | 31455862 | 1646 | * | 9,202488 | 3,435242795 | 1,227453 | 2,798676 | 0,005131 | 0,799397343 | 1,211315 | Distal Intergenic | 9 | 31470905 | 31490930 | 20026 | 1 | 1,03E+08 | ENSMUST00000216695.2 | -15043 | ENSMUSG00000111853 | Gm31408 | predicted gene, 31408 |
| chr15 | 94093775 | 94094437 | 663 | * | 7,197661 | 3,435538688 | 1,297845 | 2,647111 | 0,008118 | 1 | 0 | Distal Intergenic | 15 | 94168044 | 94302139 | 134096 | 2 | 223838 | ENSMUST00000035342.11 | 207702 | ENSMUSG00000022449 | Adamts20 | a disintegrin-like and metallopeptidase (reprolysin type) with thrombospondin type 1 motif, 20 |
| chr16 | 37692750 | 37694086 | 1337 | * | 10,5 | 3,435806289 | 1,15341 | 2,978826 | 0,002894 | 0,687090886 | 1,2763 | Intron (ENSMUST00000 | 16 | 37688751 | 37709220 | 20470 | 1 | 320184 | ENSMUST00000078717.7 | 3999 | ENSMUSG00000034158 | Lrrc58 | leucine rich repeat containing 58 |
| chr19 | 4874300 | 4876131 | 1832 | * | 17,12543 | 3,436977969 | 0,912491 | 3,766587 | 0,000165 | 0,151746892 | 2,663619 | Promoter (<=1kb) | 19 | 4875394 | 4876037 | 644 | 2 | 12460 | ENSMUST00000235216.2 | 0 | ENSMUSG00000034108 | Ccs | copper chaperone for superoxide dismutase |
| chr13 | 1,04E+08 | 1,04E+08 | 1603 | * | 14,15732 | 3,437291048 | 1,049059 | 3,276546 | 0,001051 | 0,61384098 | 0,738441 | Distal Intergenic | 13 | 1,03E+08 | 1,03E+08 | 1453 | 1 | 432800 | ENSMUST00000052030.6 | 132179 | | Gm5454 | predicted gene 5454 |
| chr16 | 61148730 | 61149498 | 769 | * | 10,68106 | 3,437850544 | 1,273687 | 2,699134 | 0,006952 | 0,84525842 | 1,2763 | Distal Intergenic | 16 | 60890381 | 60890457 | 77 | 1 | 1,15E+08 | ENSMUST00000158376.3 | 258349 | ENSMUSG00000089001 | Gm23852 | predicted gene, 23852 |
| chr8 | 46082632 | 46083724 | 1093 | * | 11,46233 | 3,438061548 | 1,123119 | 3,061173 | 0,002205 | 0,736169201 | 0,744375 | Promoter (1-2kb) | 8 | 46081238 | 46199669 | 118432 | 1 | 234214 | ENSMUST00000130850.8 | 1394 | ENSMUSG00000031626 | Sorbs2 | sorbin and SH3 domain containing 2 |
| chr12 | 69231265 | 69232582 | 1318 | * | 9,608455 | 3,438086141 | 1,330468 | 2,584118 | 0,009763 | 0,883012141 | 1,449706 | Promoter (<=1kb) | 12 | 69230931 | 69233544 | 2614 | 1 | 217664 | ENSMUST00000060579.10 | 334 | ENSMUSG00000043998 | Mgat2 | mannoside acetylglucosaminyltransferase 2 |
| chr10 | 1,18E+08 | 1,18E+08 | 1816 | * | 13,14024 | 3,43815864 | 1,125461 | 3,054889 | 0,002251 | 0,705098751 | 0,918255 | Promoter (<=1kb) | 10 | 1,18E+08 | 1,18E+08 | 32165 | 2 | 215449 | ENSMUST00000064667.9 | -60 | ENSMUSG00000052681 | Rap1b | RAS related protein 1b |
| chr12 | 83183036 | 83184459 | 1424 | * | 11,13358 | 3,43858161 | 1,116409 | 3,080038 | 0,00207 | 0,727657783 | 0,738441 | 3' UTR | 12 | 83162999 | 83184397 | 21399 | 1 | 50779 | ENSMUST00000201512.2 | 20037 | ENSMUSG00000021219 | Rgs6 | regulator of G-protein signaling 6 |
| chr19 | 46086205 | 46087664 | 1460 | * | 9,95605 | 3,439041438 | 1,158256 | 2,969154 | 0,002986 | 0,677439858 | 1,449706 | Distal Intergenic | 19 | 46064493 | 46068384 | 3892 | 1 | 70769 | ENSMUST00000224490.2 | 21712 | ENSMUSG00000015176 | Nolc1 | nucleolar and coiled-body phosphoprotein 1 |
| chr18 | 67722826 | 67724210 | 1385 | * | 12,78085 | 3,439251991 | 1,061198 | 3,240915 | 0,001191 | 0,563858922 | 1,075735 | Intron (ENSMUST00000 | 18 | 67621279 | 67743860 | 122582 | 2 | 68166 | ENSMUST00000115050.10 | 19650 | ENSMUSG00000024533 | Spire1 | spire type actin nucleation factor 1 |
| chr9 | 60523789 | 60524903 | 1115 | * | 17,06446 | 3,439379031 | 0,920685 | 3,735673 | 0,000187 | 0,17530799 | 2,318434 | Intron (ENSMUST00000 | 9 | 60495105 | 60505872 | 10768 | 2 | 102747 | ENSMUST00000136388.2 | -17917 | ENSMUSG00000047766 | Lrrc49 | leucine rich repeat containing 49 |
| chr7 | 12629554 | 12630908 | 1355 | * | 9,085575 | 3,439782772 | 1,241216 | 2,771302 | 0,005583 | 0,708539289 | 2,237403 | Promoter (<=1kb) | 7 | 12631742 | 12642166 | 10425 | 1 | 232878 | ENSMUST00000055528.11 | -834 | ENSMUSG00000054715 | Zscan22 | zinc finger and SCAN domain containing 22 |
| chr7 | 31335824 | 31336520 | 697 | * | 7,495875 | 3,439795192 | 1,332606 | 2,581255 | 0,009844 | 1 | 0 | Distal Intergenic | 7 | 31317168 | 31318951 | 1784 | 2 | 1,01E+08 | ENSMUST00000188132.2 | -16873 | ENSMUSG00000102002 | Scgb2b6 | secretoglobin, family 2B, member 6 |
| chr16 | 31436467 | 31437098 | 632 | * | 8,725056 | 3,439851774 | 1,245366 | 2,762121 | 0,005743 | 0,798887389 | 1,360622 | Promoter (<=1kb) | 16 | 31436553 | 31444110 | 7558 | 1 | 224098 | ENSMUST00000232340.2 | 0 | ENSMUSG00000116597 | Gm536 | predicted gene 536 |
| chr17 | 27851743 | 27852857 | 1115 | * | 9,501454 | 3,44116701 | 1,224323 | 2,810669 | 0,004944 | 0,764024179 | 1,425929 | Promoter (<=1kb) | 17 | 27850636 | 27853228 | 2593 | 2 | 67097 | ENSMUST00000150763.2 | 371 | ENSMUSG00000052146 | Rps10 | ribosomal protein S10 |
| chr15 | 53211555 | 53212711 | 1157 | * | 9,156758 | 3,441359216 | 1,259994 | 2,731251 | 0,006309 | 0,80763884 | 1,425929 | Promoter (1-2kb) | 15 | 52927434 | 53209555 | 282122 | 2 | 14042 | ENSMUST00000077273.9 | -2000 | ENSMUSG00000061731 | Ext1 | exostosin glycosyltransferase 1 |
| chr9 | 31297811 | 31299251 | 1441 | * | 11,57609 | 3,441465302 | 1,259818 | 2,731716 | 0,006301 | 0,826676469 | 1,2763 | Promoter (<=1kb) | 9 | 31297752 | 31310481 | 12730 | 1 | 235134 | ENSMUST00000215211.2 | 59 | ENSMUSG00000042185 | Nfrkb | nuclear factor related to kappa B binding protein |
| chr16 | 32944844 | 32945986 | 1143 | * | 10,3881 | 3,44198204 | 1,116088 | 3,08397 | 0,002043 | 0,72508691 | 0,738441 | Promoter (<=1kb) | 16 | 32944598 | 32945939 | 1342 | 1 | 239833 | ENSMUST00000155920.2 | 246 | ENSMUSG00000022802 | Lmln | leishmanolysin-like (metallopeptidase M8 family) |
| chr2 | 50823773 | 50824998 | 1226 | * | 14,5768 | 3,441985141 | 0,976042 | 3,526474 | 0,000421 | 0,282588188 | 2,141612 | Distal Intergenic | 2 | 50799696 | 50801861 | 2166 | 1 | 227885 | ENSMUST00000119159.2 | 24077 | ENSMUSG00000083270 | Gm13498 | heat shock protein 90 alpha (cytosolic), class B member 1 pseudogene |
| chr11 | 1,14E+08 | 1,14E+08 | 675 | * | 8,676068 | 3,442231615 | 1,244852 | 2,765173 | 0,005689 | 0,790561095 | 1,425929 | Intron (ENSMUST00000 | 11 | 1,14E+08 | 1,14E+08 | 19879 | 2 | 237979 | ENSMUST00000125879.2 | -56390 | ENSMUSG00000041592 | Sdk2 | sidekick cell adhesion molecule 2 |
| chr16 | 88837905 | 88838634 | 730 | * | 7,433671 | 3,44357568 | 1,26745 | 2,716932 | 0,006589 | 0,824070388 | 1,360622 | Promoter (<=1kb) | 16 | 88837811 | 88838360 | 550 | 2 | 1E+08 | ENSMUST00000089105.4 | 0 | ENSMUSG00000068074 | Gm10228 | predicted gene 10228 |
| chr19 | 3913953 | 3915020 | 1068 | * | 9,449754 | 3,443762726 | 1,217648 | 2,828208 | 0,004681 | 0,681119785 | 2,237403 | Exon (ENSMUST000001 | 19 | 3924664 | 3939934 | 15271 | 1 | 12660 | ENSMUST00000153343.2 | -9644 | ENSMUSG00000024843 | Chka | choline kinase alpha |
| chr2 | 38829260 | 38829900 | 641 | * | 8,27631 | 3,44384657 | 1,281096 | 2,688204 | 0,007184 | 1 | 0 | Exon (ENSMUST000001 | 2 | 38821990 | 38853765 | 31776 | 1 | 241327 | ENSMUST00000057279.6 | 7270 | ENSMUSG00000046618 | Olfml2a | olfactomedin-like 2A |
| chr8 | 1,07E+08 | 1,07E+08 | 1047 | * | 8,226106 | 3,444194707 | 1,304042 | 2,641168 | 0,008262 | 0,85473227 | 1,425929 | Intron (ENSMUST00000 | 8 | 1,07E+08 | 1,07E+08 | 17997 | 1 | 68051 | ENSMUST00000212042.2 | 3071 | ENSMUSG00000008450 | Nutf2 | nuclear transport factor 2 |
| chr14 | 3386184 | 3387113 | 930 | * | 7,241895 | 3,445529196 | 1,337339 | 2,576408 | 0,009983 | 0,895765172 | 1,360622 | Distal Intergenic | 14 | 3575429 | 3839361 | 263933 | 1 | 218793 | ENSMUST00000124353.2 | -188316 | ENSMUSG00000058317 | Ube2e2 | ubiquitin-conjugating enzyme E2E 2 |
| chr11 | 1,02E+08 | 1,02E+08 | 1086 | * | 11,36251 | 3,445589945 | 1,161524 | 2,96644 | 0,003013 | 0,792545355 | 0,744375 | Promoter (1-2kb) | 11 | 1,02E+08 | 1,02E+08 | 4042 | 1 | 72053 | ENSMUST00000156326.2 | 1858 | ENSMUSG00000034757 | Tmub2 | transmembrane and ubiquitin-like domain containing 2 |
| chr9 | 22065609 | 22066686 | 1078 | * | 9,280746 | 3,445918122 | 1,249689 | 2,757421 | 0,005826 | 0,715249603 | 2,237403 | Promoter (1-2kb) | 9 | 22068142 | 22069654 | 1513 | 1 | 66268 | ENSMUST00000123680.2 | -1456 | ENSMUSG00000010607 | Pigyl | phosphatidylinositol glycan anchor biosynthesis, class Y-like |
| chr2 | 1,31E+08 | 1,31E+08 | 1482 | * | 12,71965 | 3,446109374 | 1,152372 | 2,990448 | 0,002786 | 0,683988405 | 1,2763 | Intron (ENSMUST00000 | 2 | 1,31E+08 | 1,31E+08 | 34470 | 2 | 228602 | ENSMUST00000110243.8 | -24742 | ENSMUSG00000027309 | Dnaaf9 | dynein axonemal assembly factor 9 |
| chr13 | 21971261 | 21972344 | 1084 | * | 16,29366 | 3,446694544 | 0,989032 | 3,484919 | 0,000492 | 0,289014828 | 2,386008 | Promoter (<=1kb) | 13 | 21970996 | 21971388 | 393 | 2 | 319170 | ENSMUST00000091751.3 | 0 | ENSMUSG00000069309 | H2ac22 | H2A clustered histone 22 |
| chr10 | 76070796 | 76071912 | 1117 | * | 9,796013 | 3,447773611 | 1,25222 | 2,75333 | 0,005899 | 0,793995086 | 1,425929 | Promoter (1-2kb) | 10 | 76056911 | 76073646 | 16736 | 2 | 15468 | ENSMUST00000217726.2 | 1734 | ENSMUSG00000020230 | Prmt2 | protein arginine N-methyltransferase 2 |
| chr1 | 1,62E+08 | 1,62E+08 | 955 | * | 10,81652 | 3,448358509 | 1,198027 | 2,878364 | 0,003997 | 0,759484484 | 1,173325 | Intron (ENSMUST00000 | 1 | 1,62E+08 | 1,62E+08 | 27058 | 2 | 75368 | ENSMUST00000141397.2 | 22614 | ENSMUSG00000086277 | 4930558K02Rik | RIKEN cDNA 4930558K02 gene |
| chr10 | 80510696 | 80511664 | 969 | * | 12,06852 | 3,449228969 | 1,136222 | 3,035701 | 0,0024 | 0,676476319 | 1,173325 | Promoter (<=1kb) | 10 | 80501161 | 80512127 | 10967 | 2 | 17347 | ENSMUST00000200082.5 | 463 | ENSMUSG00000020190 | Mknk2 | MAP kinase-interacting serine/threonine kinase 2 |
| chr3 | 1,5E+08 | 1,5E+08 | 855 | * | 9,03139 | 3,449496865 | 1,225525 | 2,814709 | 0,004882 | 0,756849896 | 1,449706 | Distal Intergenic | 3 | 1,49E+08 | 1,49E+08 | 97515 | 2 | 1,03E+08 | ENSMUST00000197948.2 | -441491 | ENSMUSG00000104785 | Gm31121 | predicted gene, 31121 |
| chr14 | 31057438 | 31060087 | 2650 | * | 23,05244 | 3,449967075 | 0,830946 | 4,151854 | 3,3E-05 | 0,051835433 | 2,951062 | Promoter (<=1kb) | 14 | 31058595 | 31093941 | 35347 | 1 | 12339 | ENSMUST00000022451.14 | 0 | ENSMUSG00000021893 | Capn7 | calpain 7 |
| chr13 | 94099230 | 94100436 | 1207 | * | 10,91952 | 3,450178112 | 1,144817 | 3,013737 | 0,002581 | 0,674235791 | 1,2763 | Distal Intergenic | 13 | 94194304 | 94331917 | 137614 | 1 | 218454 | ENSMUST00000054274.8 | -93868 | ENSMUSG00000045312 | Lhfpl2 | lipoma HMGIC fusion partner-like 2 |

| | | | | | | | | | | | | | | | | | | | | | | | |
|---|---|---|---|---|---|---|---|---|---|---|---|---|---|---|---|---|---|---|---|---|---|---|---|
| chr4 | 1,55E+08 | 1,55E+08 | 1092 | * | 11,45912 | 3,450192174 | 1,18635 | 2,908242 | 0,003635 | 0,730469267 | 1,2763 | Promoter (<=1kb) | 4 | 1,55E+08 | 1,55E+08 | 4357 | 1 | 269614 | ENSMUST00000129386.2 | 96 | ENSMUSG00000029056 | Pank4 | pantothenate kinase 4 |
| chr1 | 36384513 | 36386234 | 1722 | * | 12,86118 | 3,450588803 | 1,121727 | 3,07614 | 0,002097 | 0,641187423 | 1,2763 | Promoter (1-2kb) | 1 | 36387763 | 36387839 | 77 | 2 | 1,02E+08 | ENSMUST00000184898.3 | 1605 | ENSMUSG00000098680 | Mir6898 | microRNA 6898 |
| chr6 | 91852723 | 91853544 | 822 | * | 10,2076 | 3,450996107 | 1,258876 | 2,741331 | 0,006119 | 0,831691385 | 1,211315 | Promoter (1-2kb) | 6 | 91855034 | 91861968 | 6935 | 1 | 232236 | ENSMUST00000138868.8 | -1490 | ENSMUSG00000034083 | Ccdc174 | coiled-coil domain containing 174 |
| chr13 | 53127503 | 53128854 | 1352 | * | 12,1061 | 3,452656992 | 1,031384 | 3,347596 | 0,000815 | 0,562508374 | 0,744375 | Intron (ENSMUST0000( | 13 | 53121245 | 53135109 | 13865 | 2 | 18030 | ENSMUST00000071065.8 | 6255 | ENSMUSG00000056749 | Nfil3 | nuclear factor, interleukin 3, regulated |
| chr13 | 52504323 | 52505573 | 1251 | * | 10,00632 | 3,452997443 | 1,113889 | 3,099948 | 0,001936 | 0,600125728 | 1,449706 | Distal Intergenic | 13 | 52334447 | 52335673 | 1227 | 1 | 1,19E+08 | ENSMUST00000222876.2 | 169876 | | Gm48199 | predicted gene, 48199 |
| chr5 | 27015474 | 27016824 | 1351 | * | 11,34431 | 3,45321122 | 1,298516 | 2,659352 | 0,007829 | 0,952402626 | 0,738441 | Distal Intergenic | 5 | 27022201 | 27055534 | 33334 | 1 | 13483 | ENSMUST00000149678.2 | -5377 | ENSMUSG00000061576 | Dpp6 | dipeptidylpeptidase 6 |
| chr16 | 88967421 | 88968354 | 934 | * | 8,638883 | 3,453434337 | 1,170641 | 2,950038 | 0,003177 | 0,684274806 | 1,449706 | Promoter (1-2kb) | 16 | 88966378 | 88966536 | 159 | 1 | 1,03E+08 | ENSMUST00000089098.2 | 1043 | ENSMUSG00000068068 | Gm7735 | predicted gene 7735 |
| chr3 | 74762741 | 74764388 | 1648 | * | 9,557412 | 3,453500051 | 1,211614 | 2,850331 | 0,004367 | 0,739258701 | 1,449706 | Distal Intergenic | 3 | 74968668 | 75051010 | 82343 | 2 | 213234 | ENSMUST00000107776.8 | 286622 | ENSMUSG00000034151 | Zbbx | zinc finger, B-box domain containing |
| chr4 | 1,17E+08 | 1,17E+08 | 1245 | * | 9,355308 | 3,455123707 | 1,307832 | 2,641871 | 0,008245 | 0,860883941 | 1,360622 | Intron (ENSMUST0000( | 4 | 1,17E+08 | 1,18E+08 | 109540 | 1 | 140546 | ENSMUST00000146384.8 | 5878 | ENSMUSG00000033423 | Eri3 | exoribonuclease 3 |
| chr17 | 3151299 | 3152155 | 857 | * | 7,749305 | 3,455759097 | 1,256371 | 2,750589 | 0,005949 | 1 | 0 | Exon (ENSMUST000002 | 17 | 3154825 | 3164752 | 9928 | 2 | 624086 | ENSMUST00000246188.1 | 12597 | | A230045G11Rik | RIKEN cDNA A230045G11 gene |
| chr7 | 46555539 | 46556989 | 1451 | * | 11,576 | 3,456121806 | 1,219805 | 2,83334 | 0,004606 | 0,864449641 | 0,744375 | Promoter (1-2kb) | 7 | 46539334 | 46558754 | 19421 | 2 | 22088 | ENSMUST00000209538.2 | 1765 | ENSMUSG00000014402 | Tsg101 | tumor susceptibility gene 101 |
| chr1 | 1,35E+08 | 1,35E+08 | 2134 | * | 17,19392 | 3,456711509 | 0,954712 | 3,620683 | 0,000294 | 0,223919137 | 2,386008 | Exon (ENSMUST00000( | 1 | 1,35E+08 | 1,35E+08 | 87354 | 2 | 329251 | ENSMUST00000156348.8 | -18479 | ENSMUSG00000073557 | Ppp1r12b | protein phosphatase 1, regulatory subunit 12B |
| chr3 | 36503074 | 36504635 | 1562 | * | 20,32928 | 3,457044059 | 0,843974 | 4,096152 | 4,2E-05 | 0,06508534 | 2,663619 | Promoter (1-2kb) | 3 | 36503313 | 36506477 | 3165 | 2 | 11747 | ENSMUST00000196097.2 | 1842 | ENSMUSG00000027712 | Anxa5 | annexin A5 |
| chr9 | 72300904 | 72302501 | 1598 | * | 12,56776 | 3,457329281 | 1,17884 | 2,932824 | 0,003359 | 0,813191811 | 0,738441 | Promoter (<=1kb) | 9 | 72303256 | 72311792 | 8537 | 1 | 1,03E+08 | ENSMUST00000184671.2 | -755 | | Gm27230 | predicted gene 27230 |
| chr9 | 1,18E+08 | 1,18E+08 | 952 | * | 9,332773 | 3,458099394 | 1,254693 | 2,756131 | 0,005849 | 0,823736512 | 1,211315 | Promoter (<=1kb) | 9 | 1,18E+08 | 1,18E+08 | 85069 | 2 | 67899 | ENSMUST00000215799.2 | 497 | ENSMUSG00000039163 | Cmc1 | COX assembly mitochondrial protein 1 |
| chr11 | 17290681 | 17291998 | 1318 | * | 12,28119 | 3,45898182 | 1,12176 | 3,083532 | 0,002046 | 0,649854434 | 1,173325 | Distal Intergenic | 11 | 17213621 | 17216560 | 2940 | 1 | 57316 | ENSMUST00000130330.2 | 77060 | ENSMUSG00000000581 | C1d | C1D nuclear receptor co-repressor |
| chr11 | 1,19E+08 | 1,19E+08 | 1260 | * | 10,15065 | 3,459861631 | 1,247023 | 2,774498 | 0,005529 | 0,782127811 | 1,449706 | Distal Intergenic | 11 | 1,19E+08 | 1,19E+08 | 4665 | 2 | 30951 | ENSMUST00000026663.8 | -3735 | ENSMUSG00000025578 | Cbx8 | chromobox 8 |
| chr7 | 88048114 | 88048946 | 833 | * | 8,246846 | 3,460248786 | 1,244759 | 2,779854 | 0,005438 | 1 | 0 | Distal Intergenic | 7 | 88079481 | 88140780 | 61300 | 1 | 72433 | ENSMUST00000107256.4 | -30535 | ENSMUSG00000030559 | Rab38 | RAB38, member RAS oncogene family |
| chr10 | 27253919 | 27255235 | 1317 | * | 12,2574 | 3,460623928 | 0,97332 | 3,555483 | 0,000377 | 0,36185416 | 1,075735 | Intron (ENSMUST0000( | 10 | 27265862 | 27273221 | 7360 | 1 | 75855 | ENSMUST00000187815.2 | -10627 | ENSMUSG00000101636 | 4930579H20Rik | RIKEN cDNA 4930579H20 gene |
| chr17 | 43823892 | 43824541 | 650 | * | 6,516715 | 3,461210352 | 1,297577 | 2,667442 | 0,007643 | 1 | 0 | Distal Intergenic | 17 | 43785215 | 43813703 | 28489 | 2 | 17287 | ENSMUST00000117137.8 | -10189 | ENSMUSG00000023914 | Mep1a | meprin 1 alpha |
| chr7 | 80981498 | 80982496 | 999 | * | 8,654133 | 3,4613173 | 1,245837 | 2,778306 | 0,005464 | 0,791416391 | 1,360622 | Intron (ENSMUST0000( | 7 | 80992480 | 80994279 | 1800 | 2 | 20068 | ENSMUST00000130487.8 | 11783 | ENSMUSG00000061787 | Rps17 | ribosomal protein S17 |
| chr2 | 32324093 | 32325035 | 943 | * | 7,451497 | 3,461705406 | 1,26145 | 2,744227 | 0,006065 | 1 | 0 | Promoter (2-3kb) | 2 | 32304499 | 32321131 | 16633 | 2 | 227731 | ENSMUST00000052119.14 | -2962 | ENSMUSG00000026819 | Slc25a25 | solute carrier family 25 (mitochondrial carrier, phosphate carrier), member 25 |
| chr5 | 97440783 | 97442304 | 1522 | * | 13,88482 | 3,461886038 | 1,031244 | 3,357 | 0,000788 | 0,512140351 | 0,938682 | Distal Intergenic | 5 | 97473763 | 97496298 | 22536 | 1 | 1,03E+08 | ENSMUST00000197615.2 | -31459 | ENSMUSG00000104717 | Gm33969 | predicted gene, 33969 |
| chr1 | 7755597 | 7756828 | 1232 | * | 8,406862 | 3,461906948 | 1,282591 | 2,699152 | 0,006952 | 0,824314641 | 1,425929 | Distal Intergenic | 1 | 7446112 | 7468093 | 21982 | 2 | 1,03E+08 | ENSMUST00000193732.2 | -287504 | | Gm26901 | predicted gene, 26901 |
| chr13 | 19647701 | 19648845 | 1145 | * | 10,6742 | 3,462103912 | 1,167401 | 2,96565 | 0,00302 | 0,695892636 | 1,2763 | Exon (ENSMUST000002 | 13 | 19649939 | 19688515 | 38577 | 2 | 319466 | ENSMUST00000221601.2 | 39670 | ENSMUSG00000114189 | 9330199G10Rik | RIKEN cDNA 9330199G10 gene |
| chr16 | 62465405 | 62466317 | 913 | * | 12,89847 | 3,462367596 | 1,131813 | 3,059134 | 0,00222 | 0,677933849 | 1,075735 | Distal Intergenic | 16 | 62391399 | 62391502 | 104 | 1 | 1,15E+08 | ENSMUST00000093663.3 | 74006 | ENSMUSG00000070173 | Gm24164 | predicted gene, 24164 |
| chr16 | 21936094 | 21938491 | 2398 | * | 40,058854 | 3,462814889 | 0,5349826 | 6,4727618 | 9,623E-11 | 2,45309E-06 | 2,3383871 | Promoter (1-2kb) | 16 | 21906278 | 21939986 | 33709 | 2 | 319765 | ENSMUST00000232504.2 | 1495 | ENSMUSG00000033581 | Igf2bp2 | insulin-like growth factor 2 mRNA binding protein 2 |
| chr14 | 1,25E+08 | 1,25E+08 | 906 | * | 8,343369 | 3,462817667 | 1,340141 | 2,58392 | 0,009768 | 0,891097601 | 1,360622 | Intron (ENSMUST0000( | 14 | 1,24E+08 | 1,25E+08 | 699221 | 2 | 14169 | ENSMUST00000095529.10 | 229903 | ENSMUSG00000025551 | Fgf14 | fibroblast growth factor 14 |
| chr5 | 8047455 | 8049142 | 1688 | * | 15,12468 | 3,463323078 | 1,115762 | 3,103999 | 0,001909 | 0,524368688 | 2,141612 | Distal Intergenic | 5 | 8010457 | 8032213 | 21757 | 1 | 117167 | ENSMUST00000115421.3 | 36998 | ENSMUSG00000012428 | Steap4 | STEAP family member 4 |
| chr2 | 154411779 | 154414473 | 2695 | * | 43,076538 | 3,463464996 | 0,5811842 | 5,9593243 | 2,533E-09 | 3,9873E-05 | 2,3776742 | Promoter (<=1kb) | 2 | 154401327 | 154411812 | 10486 | 2 | 13555 | ENSMUST00000103145.11 | 0 | ENSMUSG00000027490 | E2f1 | E2F transcription factor 1 |
| chr11 | 1,17E+08 | 1,17E+08 | 2071 | * | 11,99355 | 3,463542232 | 1,209068 | 2,864639 | 0,004175 | 0,851512567 | 0,744375 | Distal Intergenic | 11 | 1,17E+08 | 1,17E+08 | 980 | 1 | 1,06E+08 | ENSMUST00000124901.2 | -13097 | ENSMUSG00000086588 | Gm11729 | predicted gene 11729 |
| chr6 | 36928886 | 36929754 | 869 | * | 11,59029 | 3,463908817 | 1,084632 | 3,193625 | 0,001405 | 0,563032042 | 1,2763 | Intron (ENSMUST0000( | 6 | 36888402 | 37007963 | 119562 | 2 | 320127 | ENSMUST00000143046.2 | 78209 | ENSMUSG00000038665 | Dgki | diacylglycerol kinase, iota |
| chr5 | 31915618 | 31916365 | 748 | * | 7,896729 | 3,464198586 | 1,330824 | 2,603048 | 0,00924 | 1 | 0 | Intron (ENSMUST0000( | 5 | 31942727 | 31977987 | 35261 | 1 | 107976 | ENSMUST00000202270.2 | -26362 | ENSMUSG00000052139 | Babam2 | BRISC and BRCA1 A complex member 2 |
| chr14 | 54662790 | 54664671 | 1882 | * | 14,70165 | 3,464264235 | 1,003551 | 3,452008 | 0,000556 | 0,410544751 | 1,214904 | Promoter (<=1kb) | 14 | 54664366 | 54667213 | 2848 | 1 | 68836 | ENSMUST00000010550.12 | 0 | ENSMUSG00000010406 | Mrpl52 | mitochondrial ribosomal protein L52 |
| chr4 | 1,51E+08 | 1,51E+08 | 1397 | * | 9,013806 | 3,464362902 | 1,199987 | 2,887 | 0,003889 | 0,718881903 | 1,449706 | Promoter (<=1kb) | 4 | 1,51E+08 | 1,51E+08 | 19927 | 1 | 21942 | ENSMUST00000105671.8 | 284 | ENSMUSG00000028965 | Tnfrsf9 | tumor necrosis factor receptor superfamily, member 9 |
| chr5 | 61808151 | 61808969 | 819 | * | 6,770807 | 3,464453407 | 1,300059 | 2,664843 | 0,007702 | 1 | 0 | Distal Intergenic | 5 | 61966159 | 61968506 | 2348 | 1 | 14380 | ENSMUST00000053876.5 | -157190 | ENSMUSG00000089992 | G6pd2 | glucose-6-phosphate dehydrogenase 2 |
| chr7 | 99168152 | 99169038 | 887 | * | 9,07868 | 3,464607439 | 1,305656 | 2,653539 | 0,007965 | 0,856202617 | 1,360622 | Distal Intergenic | 7 | 99184673 | 99255978 | 71306 | 1 | 109689 | ENSMUST00000098266.9 | -15635 | ENSMUSG00000018909 | Arrb1 | arrestin, beta 1 |
| chr12 | 38168599 | 38169988 | 1390 | * | 11,16973 | 3,464930575 | 1,127242 | 3,073812 | 0,002113 | 0,730215586 | 0,744375 | Intron (ENSMUST0000( | 12 | 38069179 | 38224660 | 155482 | 1 | 217480 | ENSMUST00000220606.2 | 99420 | ENSMUSG00000036095 | Dgkb | diacylglycerol kinase, beta |
| chr5 | 1,11E+08 | 1,11E+08 | 1096 | * | 9,405702 | 3,465586271 | 1,230383 | 2,816672 | 0,004852 | 0,683966322 | 2,237403 | Promoter (<=1kb) | 5 | 1,11E+08 | 1,12E+08 | 57553 | 1 | 56305 | ENSMUST00000200298.2 | 735 | ENSMUSG00000050017 | Pitpnb | phosphatidylinositol transfer protein, beta |
| chr3 | 1,08E+08 | 1,08E+08 | 1341 | * | 10,22587 | 3,465893114 | 1,257956 | 2,755179 | 0,005866 | 0,793220117 | 1,425929 | Promoter (2-3kb) | 3 | 1,08E+08 | 1,08E+08 | 414 | 2 | 1E+08 | ENSMUST00000157560.3 | 2617 | | Scarna2 | small Cajal body-specific RNA 2 |
| chr18 | 62303523 | 62305501 | 1979 | * | 14,95209 | 3,466308799 | 1,08167 | 3,20459 | 0,001353 | 0,541111532 | 1,37683 | Distal Intergenic | 18 | 62310887 | 62313030 | 2144 | 2 | 11555 | ENSMUST00000053640.5 | 7529 | ENSMUSG00000045730 | Adrb2 | adrenergic receptor, beta 2 |
| chr10 | 13839348 | 13841965 | 2618 | * | 18,45064 | 3,46638362 | 0,858947 | 4,03562 | 5,45E-05 | 0,081998926 | 2,386008 | Promoter (<=1kb) | 10 | 13841819 | 14026333 | 184515 | 1 | 15273 | ENSMUST00000187083.7 | 0 | ENSMUSG00000015501 | Hivep2 | human immunodeficiency virus type I enhancer binding protein 2 |
| chr19 | 14605297 | 14606628 | 1332 | * | 10,62856 | 3,466550326 | 1,075667 | 3,222699 | 0,00127 | 0,609154465 | 0,918255 | Distal Intergenic | 19 | 14425514 | 14575415 | 149902 | 2 | 21888 | ENSMUST00000167776.3 | -29882 | ENSMUSG00000024642 | Tle4 | transducin-like enhancer of split 4 |
| chr9 | 1,08E+08 | 1,08E+08 | 820 | * | 8,32054 | 3,467035446 | 1,254237 | 2,764259 | 0,005705 | 0,791068542 | 1,425929 | Promoter (1-2kb) | 9 | 1,08E+08 | 1,08E+08 | 1487 | 1 | 22258 | ENSMUST00000192750.2 | -1488 | ENSMUSG00000032612 | Usp4 | ubiquitin specific peptidase 4 (proto-oncogene) |
| chrX | 85214836 | 85216109 | 1274 | * | 9,480237 | 3,467461523 | 1,338057 | 2,591416 | 0,009558 | 0,881927564 | 1,425929 | Distal Intergenic | 20 | 85235370 | 85239553 | 4184 | 1 | 11614 | ENSMUST00000026036.5 | -19261 | ENSMUSG00000025056 | Nr0b1 | nuclear receptor subfamily 0, group B, member 1 |
| chr1 | 10243056 | 10243946 | 891 | * | 7,450862 | 3,467831967 | 1,172637 | 2,957293 | 0,003104 | 1 | 0 | 3' UTR | 1 | 10230982 | 10231531 | 550 | 2 | 211673 | ENSMUST00000128834.2 | -11525 | ENSMUSG00000067851 | Arfgef1 | ADP-ribosylation factor guanine nucleotide-exchange factor 1(brefeldin A-inhibited) |
| chr11 | 49585683 | 49586561 | 879 | * | 7,337039 | 3,468438025 | 1,282542 | 2,704346 | 0,006844 | 1 | 0 | Intron (ENSMUST0000( | 11 | 49565525 | 49593459 | 27935 | 2 | 104625 | ENSMUST00000109183.3 | 6898 | ENSMUSG00000020362 | Cnot6 | CCR4-NOT transcription complex, subunit 6 |
| chr3 | 1,16E+08 | 1,16E+08 | 907 | * | 8,792336 | 3,468506879 | 1,342623 | 2,583382 | 0,009784 | 0,886332554 | 1,425929 | 3' UTR | 3 | 1,16E+08 | 1,16E+08 | 54 | 1 | 1E+08 | ENSMUST00000158636.3 | -32772 | ENSMUSG00000089261 | Mir669n | microRNA 669n |
| chr11 | 22667767 | 22670045 | 2279 | * | 29,364918 | 3,468633302 | 0,6546503 | 5,2984522 | 1,168E-07 | 0,00074937 | 2,9510621 | Distal Intergenic | 11 | 22567408 | 22568487 | 1080 | 1 | 100042827 | ENSMUST00000142151.2 | 100359 | | Gm12052 | predicted gene 12052 |
| chr3 | 1,38E+08 | 1,38E+08 | 842 | * | 7,677406 | 3,46982776 | 1,327936 | 2,612948 | 0,008976 | 1 | 0 | Distal Intergenic | 3 | 1,38E+08 | 1,38E+08 | 670 | 2 | 1,08E+08 | ENSMUST00000198830.3 | 8042 | | Gm43403 | predicted gene 43403 |
| chr13 | 55330980 | 55332241 | 1262 | * | 15,87786 | 3,470945666 | 0,952426 | 3,644321 | 0,000268 | 0,225376955 | 2,141612 | Distal Intergenic | 13 | 55357595 | 55463568 | 105974 | 1 | 18193 | ENSMUST00000224973.2 | -25354 | ENSMUSG00000021488 | Nsd1 | nuclear receptor-binding SET-domain protein 1 |
| chr10 | 63218136 | 63219296 | 1161 | * | 10,22813 | 3,471142425 | 1,130415 | 3,07068 | 0,002136 | 0,65808656 | 1,173325 | Promoter (<=1kb) | 10 | 63218222 | 63244059 | 25838 | 1 | 30045 | ENSMUST00000129625.8 | 0 | ENSMUSG00000036764 | Dnajc12 | DnaJ heat shock protein family (Hsp40) member C12 |
| chr6 | 1,24E+08 | 1,24E+08 | 1656 | * | 14,52525 | 3,472652704 | 1,033775 | 3,359196 | 0,000782 | 0,442614296 | 1,41696 | Distal Intergenic | 6 | 1,24E+08 | 1,24E+08 | 17531 | 1 | 232371 | ENSMUST00000049124.10 | -8971 | ENSMUSG00000038527 | C1rl | complement component 1, r subcomponent-like |
| chr13 | 89449715 | 89450859 | 1145 | * | 9,137342 | 3,473475853 | 1,202828 | 2,887758 | 0,00388 | 0,749550311 | 1,211315 | Intron (ENSMUST0000( | 13 | 89687915 | 89759771 | 71857 | 1 | 12950 | ENSMUST00000022108.9 | -237056 | ENSMUSG00000021613 | Hapln1 | hyaluronan and proteoglycan link protein 1 |
| chr9 | 49925708 | 49926559 | 852 | * | 8,05872 | 3,473543355 | 1,344613 | 2,583303 | 0,009786 | 1 | 0 | Distal Intergenic | 9 | 49737169 | 49737275 | 107 | 2 | 1,15E+08 | ENSMUST00000178565.3 | -188433 | ENSMUSG00000119677 | Gm25633 | predicted gene, 25633 |
| chr1 | 1,31E+08 | 1,31E+08 | 1126 | * | 8,894044 | 3,474287845 | 1,176363 | 2,953415 | 0,003143 | 0,611385361 | 2,237403 | Promoter (<=1kb) | 1 | 1,31E+08 | 1,31E+08 | 102433 | 2 | 14270 | ENSMUST00000185445.3 | -606 | ENSMUSG00000026425 | Srgap2 | SLIT-ROBO Rho GTPase activating protein 2 |
| chr11 | 96740432 | 96742206 | 1775 | * | 15,63688 | 3,475458145 | 1,010963 | 3,437769 | 0,000587 | 0,396931054 | 1,41696 | Promoter (<=1kb) | 11 | 96741219 | 96741291 | 73 | 1 | 387170 | ENSMUST00000083581.3 | 0 | ENSMUSG00000065515 | Mir152 | microRNA 152 |
| chr10 | 1,28E+08 | 1,28E+08 | 1226 | * | 11,6919 | 3,47622348 | 1,12038 | 3,102718 | 0,001918 | 0,680780911 | 0,918255 | 3' UTR | 10 | 1,28E+08 | 1,28E+08 | 912 | 2 | 56516 | ENSMUST00000217722.2 | 5846 | ENSMUSG00000040043 | Rbms2 | RNA binding motif, single stranded interacting protein 2 |
| chr5 | 31343261 | 31345082 | 1822 | * | 14,02461 | 3,477486926 | 1,082081 | 3,213704 | 0,00131 | 0,546805988 | 1,2763 | Promoter (2-3kb) | 5 | 31346913 | 31347260 | 348 | 2 | 13667 | ENSMUST00000200977.2 | 2178 | ENSMUSG00000029145 | Eif2b4 | eukaryotic translation initiation factor 2B, subunit 4 delta |
| chr12 | 67810181 | 67810792 | 612 | * | 6,661595 | 3,478022966 | 1,297429 | 2,680703 | 0,007347 | 1 | 0 | Distal Intergenic | 12 | 67659070 | 67659255 | 186 | 1 | 1,15E+08 | ENSMUST00000122499.3 | 151111 | ENSMUSG00000084448 | Gm26256 | predicted gene, 26256 |
| chr17 | 57335635 | 57336893 | 1259 | * | 12,98773 | 3,478735526 | 1,083852 | 3,209605 | 0,001329 | 0,617289329 | 0,918255 | Promoter (1-2kb) | 17 | 57328051 | 57338488 | 10438 | 2 | 16549 | ENSMUST00000007814.10 | 1595 | ENSMUSG00000007670 | Khsrp | KH-type splicing regulatory protein |
| chr11 | 68980003 | 68981209 | 1207 | * | 13,00385 | 3,478859552 | 1,044873 | 3,329456 | 0,00087 | 0,477253929 | 1,2763 | Promoter (<=1kb) | 11 | 68980432 | 68981002 | 571 | 1 | 22318 | ENSMUST00000146847.2 | 0 | ENSMUSG00000020894 | Vamp2 | vesicle-associated membrane protein 2 |
| chr18 | 5376356 | 5377280 | 925 | * | 10,02949 | 3,479592548 | 1,234218 | 2,819268 | 0,004813 | 0,808436696 | 1,086118 | Distal Intergenic | 18 | 5210124 | 5334807 | 124684 | 2 | 240186 | ENSMUST00000234241.2 | -41549 | ENSMUSG00000050945 | Zfp438 | zinc finger protein 438 |
| chr12 | 63412581 | 63413936 | 1356 | * | 9,468292 | 3,479859373 | 1,10776 | 3,141347 | 0,001682 | 0,610143359 | 1,211315 | Distal Intergenic | 12 | 63656642 | 63656748 | 107 | 1 | 1,15E+08 | ENSMUST00000083024.4 | -242706 | ENSMUSG00000118736 | Gm24921 | predicted gene, 24921 |
| chr10 | 66649669 | 66650894 | 1226 | * | 13,24368 | 3,480077279 | 1,204678 | 2,888803 | 0,003867 | 0,837814951 | 0,738441 | Intron (ENSMUST0000( | 10 | 66725450 | 66727391 | 1942 | 2 | 1,19E+08 | ENSMUST00000217953.2 | 76497 | | Gm47903 | predicted gene, 47903 |
| chr18 | 34762356 | 34763353 | 998 | * | 10,33475 | 3,480224964 | 1,197768 | 2,905593 | 0,003666 | 0,711174281 | 1,449706 | Promoter (<=1kb) | 18 | 34761976 | 34763682 | 1707 | 1 | 19348 | ENSMUST00000236114.2 | 380 | ENSMUSG00000003779 | Kif20a | kinesin family member 20A |
| chr4 | 1,54E+08 | 1,54E+08 | 520 | * | 7,5656 | 3,480469682 | 1,317154 | 2,642416 | 0,008232 | 0,860883941 | 1,360622 | Exon (ENSMUST000002 | 4 | 1,54E+08 | 1,54E+08 | 16039 | 2 | 381580 | ENSMUST00000047207.7 | -18791 | ENSMUSG00000039492 | Ccdc27 | coiled-coil domain containing 27 |
| chr1 | 76875808 | 76876960 | 1153 | * | 12,61773 | 3,48072174 | 1,051237 | 3,311073 | 0,000929 | 0,502454322 | 1,173325 | Intron (ENSMUST0000( | 1 | 76486197 | 76486280 | 84 | 1 | 1,02E+08 | ENSMUST00000183584.3 | 389611 | ENSMUSG00000098611 | Mir6343 | microRNA 6343 |
| chr8 | 1,21E+08 | 1,21E+08 | 1008 | * | 9,390978 | 3,480769341 | 1,27135 | 2,737853 | 0,006184 | 0,813191811 | 1,360622 | Intron (ENSMUST0000( | 8 | 1,21E+08 | 1,21E+08 | 325179 | 1 | 382034 | ENSMUST00000180448.3 | 39552 | ENSMUSG00000031822 | Gse1 | genetic suppressor element 1, coiled-coil protein |
| chr9 | 1,06E+08 | 1,06E+08 | 1306 | * | 11,85239 | 3,481072088 | 1,141595 | 3,049306 | 0,002294 | 0,669586355 | 1,173325 | Distal Intergenic | 9 | 1,06E+08 | 1,06E+08 | 5270 | 2 | 235582 | ENSMUST00000112543.9 | -4389 | ENSMUSG00000020258 | Glyctk | glycerate kinase |
| chr15 | 34853573 | 34854504 | 932 | * | 9,71831 | 3,481234365 | 1,141199 | 3,050505 | 0,002285 | 0,663408464 | 1,211315 | Distal Intergenic | 15 | 34838195 | 34843371 | 5177 | 1 | 16539 | ENSMUST00000228725.2 | 15378 | ENSMUSG00000050963 | Kcns2 | K+ voltage-gated channel, subfamily S, 2 |
| chr16 | 9542977 | 9543869 | 893 | * | 8,355505 | 3,482307785 | 1,332826 | 2,612726 | 0,008982 | 0,869013335 | 1,425929 | Intron (ENSMUST0000( | 16 | 9579363 | 9615197 | 35835 | 2 | 14811 | ENSMUST00000199267.2 | 71328 | ENSMUSG00000059003 | Grin2a | glutamate receptor, ionotropic, NMDA2A (epsilon 1) |
| chr3 | 49567890 | 49568629 | 740 | * | 6,645864 | 3,482788519 | 1,255898 | 2,773147 | 0,005552 | 1 | 0 | Distal Intergenic | 3 | 49613212 | 49613318 | 107 | 1 | 1,15E+08 | ENSMUST00000239579.1 | -44583 | ENSMUSG00000119731 | Gm23096 | predicted gene, 23096 |
| chr17 | 66263281 | 66264287 | 1007 | * | 8,267836 | 3,482946071 | 1,232978 | 2,824824 | 0,004731 | 0,755058193 | 1,425929 | Distal Intergenic | 17 | 66230060 | 66258221 | 28162 | 2 | 65960 | ENSMUST00000024906.6 | -5060 | ENSMUSG00000024098 | Twsg1 | twisted gastrulation BMP signaling modulator 1 |
| chr1 | 1,35E+08 | 1,35E+08 | 1432 | * | 9,493667 | 3,482959532 | 1,347352 | 2,585041 | 0,009737 | 0,891057096 | 1,360622 | Promoter (<=1kb) | 1 | 1,35E+08 | 1,35E+08 | 2881 | 1 | 67196 | ENSMUST00000188177.2 | -595 | ENSMUSG00000026429 | Ube2t | ubiquitin-conjugating enzyme E2T |
| chr2 | 1,3E+08 | 1,3E+08 | 1200 | * | 9,696298 | 3,482971704 | 1,32538 | 2,627904 | 0,008591 | 0,857955756 | 1,449706 | Intron (ENSMUST0000( | 2 | 1,3E+08 | 1,3E+08 | 1414 | 1 | 19262 | ENSMUST00000230516.2 | -32108 | ENSMUSG00000027303 | Ptpra | protein tyrosine phosphatase, receptor type, A |
| chr9 | 41519075 | 41520746 | 1672 | * | 11,94658 | 3,483577925 | 1,152144 | 3,023561 | 0,002498 | 0,75632407 | 0,744375 | Intron (ENSMUST0000( | 9 | 41511568 | 41524025 | 12458 | 1 | 73144 | ENSMUST00000188412.2 | 7507 | ENSMUSG00000074415 | Mir100hg | Mir100 Mirlet7a-2 Mir125b-1 cluster host gene |
| chr11 | 1,16E+08 | 1,16E+08 | 842 | * | 9,154882 | 3,484107347 | 1,343444 | 2,593415 | 0,009503 | 0,888080529 | 1,360622 | Promoter (2-3kb) | 11 | 1,16E+08 | 1,16E+08 | 4315 | 1 | 69535 | ENSMUST00000135460.2 | -2796 | ENSMUSG00000020778 | Ten1 | TEN1 telomerase capping complex subunit |
| chr7 | 97278442 | 97280467 | 2026 | * | 12,31123 | 3,484658277 | 1,104714 | 3,154353 | 0,001609 | 0,683988405 | 0,738441 | Intron (ENSMUST0000( | 7 | 97247556 | 97294319 | 46764 | 2 | 75448 | ENSMUST00000126902.2 | 13852 | ENSMUSG00000086433 | Rsf1os1 | remodeling and spacing factor 1, opposite strand 1 |
| chr19 | 24938093 | 24939407 | 1315 | * | 11,5061 | 3,484825995 | 1,093807 | 3,185961 | 0,001443 | 0,669586355 | 0,738441 | Promoter (<=1kb) | 19 | 24897280 | 24938965 | 41686 | 2 | 226043 | ENSMUST00000025815.10 | 0 | ENSMUSG00000024878 | Cbwd1 | COBW domain containing 1 |
| chr5 | 1,44E+08 | 1,44E+08 | 1124 | * | 8,26333 | 3,485381395 | 1,308635 | 2,663373 | 0,007736 | 0,84048153 | 1,449706 | Intron (ENSMUST0000( | 5 | 1,44E+08 | 1,44E+08 | 2481 | 1 | 1,01E+08 | ENSMUST00000196478.5 | 62393 | ENSMUSG00000104600 | Dmrt1i | Dmrt1 interacting ncRNA |
| chr9 | 66619554 | 66622057 | 2504 | * | 32,569453 | 3,486052602 | 0,6711099 | 5,1944586 | 2,053E-07 | 0,001275084 | 2,3776742 | Promoter (<=1kb) | 9 | 66620968 | 66674127 | 53160 | 1 | 76459 | ENSMUST00000071889.13 | 0 | ENSMUSG00000032373 | Car12 | carbonic anhydrase 12 |
| chr15 | 81804605 | 81806024 | 1420 | * | 14,11448 | 3,488651221 | 1,133881 | 3,076736 | 0,002093 | 0,640889834 | 1,2763 | Intron (ENSMUST0000( | 15 | 81801400 | 81809597 | 8198 | 2 | 78929 | ENSMUST00000230842.2 | 3573 | ENSMUSG00000022476 | Polr3h | polymerase (RNA) III (DNA directed) polypeptide H |
| chr9 | 1,08E+08 | 1,08E+08 | 1573 | * | 11,30201 | 3,489350278 | 1,19188 | 2,927602 | 0,003416 | 0,814686254 | 0,744375 | Promoter (1-2kb) | 9 | 1,08E+08 | 1,08E+08 | 15688 | 2 | 83486 | ENSMUST00000193342.6 | 1532 | ENSMUSG00000032580 | Rbm5 | RNA binding motif protein 5 |
| chr13 | 1,11E+08 | 1,11E+08 | 1121 | * | 8,292773 | 3,48940986 | 1,28177 | 2,722337 | 0,006482 | 0,809718961 | 1,449706 | Distal Intergenic | 13 | 1,11E+08 | 1,11E+08 | 4782 | 1 | 75139 | ENSMUST00000224654.2 | -43577 | ENSMUSG00000114472 | Gm38397 | predicted gene, 38397 |
| chr9 | 66851855 | 66853849 | 1995 | * | 15,4329 | 3,489540051 | 0,918909 | 3,797483 | 0,000146 | 0,221323359 | 1,214904 | Promoter (<=1kb) | 9 | 66853368 | 66856794 | 3427 | 1 | 67941 | ENSMUST00000040917.14 | 0 | ENSMUSG00000036781 | Rps27l | ribosomal protein S27-like |
| chr7 | 86526744 | 86528427 | 1684 | * | 21,78499 | 3,489837012 | 0,878115 | 3,974238 | 7,06E-05 | 0,088589416 | 2,67243 | Intron (ENSMUST0000( | 7 | 86564302 | 86576714 | 12413 | 1 | 637896 | ENSMUST00000233033.2 | -35875 | ENSMUSG00000091962 | Vmn2r78 | vomeronasal 2, receptor 78 |
| chr13 | 12396470 | 12398168 | 1699 | * | 10,20107 | 3,490828511 | 1,111128 | 3,141697 | 0,00168 | 0,615591422 | 1,173325 | Distal Intergenic | 13 | 12393039 | 12393518 | 480 | 1 | 432725 | ENSMUST00000180224.3 | 3431 | ENSMUSG00000094185 | Gm5445 | predicted gene 5445 |
| chr14 | 14375018 | 14376684 | 1667 | * | 13,48501 | 3,490877638 | 1,07984 | 3,232772 | 0,001226 | 0,568412386 | 1,075735 | Distal Intergenic | 14 | 14304656 | 14371562 | 66907 | 2 | 218699 | ENSMUST00000036682.9 | -3456 | ENSMUSG00000033885 | Pxk | PX domain containing serine/threonine kinase |
| chr17 | 84457719 | 84458433 | 715 | * | 9,385989 | 3,491253142 | 1,249905 | 2,793214 | 0,005219 | 0,783815908 | 1,360622 | Intron (ENSMUST0000( | 17 | 84441456 | 84463856 | 22401 | 2 | 1,01E+08 | ENSMUST00000234659.3 | 5423 | | Gm19696 | predicted gene, 19696 |
| chr14 | 25929201 | 25931163 | 1963 | * | 14,24475 | 3,491427336 | 1,134097 | 3,078596 | 0,00208 | 0,668772562 | 1,075735 | Promoter (1-2kb) | 14 | 25923724 | 25928096 | 4373 | 2 | 66039 | ENSMUST00000185006.9 | -1105 | ENSMUSG00000072676 | Tmem254 | transmembrane protein 254 |
| chr17 | 64201818 | 64202698 | 881 | * | 9,473718 | 3,492298419 | 1,274106 | 2,74098 | 0,006126 | 0,798887389 | 1,449706 | Promoter (<=1kb) | 17 | 64203017 | 64446458 | 243442 | 1 | 14158 | ENSMUST00000000129.14 | -319 | ENSMUSG00000000127 | Fer | fer (fms/fps related) protein kinase |
| chr9 | 88122210 | 88124449 | 2240 | * | 20,52691 | 3,493275503 | 0,882386 | 3,958898 | 7,53E-05 | 0,097525682 | 2,386008 | Distal Intergenic | 9 | 88141948 | 88145088 | 3141 | 1 | 1,03E+08 | ENSMUST00000190118.2 | -17499 | ENSMUSG00000100782 | Gm28231 | predicted gene 28231 |
| chr2 | 24855250 | 24856280 | 1031 | * | 12,17919 | 3,493293489 | 1,006618 | 3,470327 | 0,00052 | 0,407158022 | 1,173325 | Promoter (2-3kb) | 2 | 24852485 | 24859683 | 7199 | 1 | 67228 | ENSMUST00000124383.2 | 2765 | ENSMUSG00000026975 | Dph7 | diphthamine biosynethesis 7 |
| chr17 | 46826842 | 46828198 | 1357 | * | 11,05701 | 3,493510502 | 1,249725 | 2,795422 | 0,005183 | 0,85197474 | 0,918255 | Intron (ENSMUST0000( | 17 | 46830386 | 46831497 | 1112 | 2 | 78309 | ENSMUST00000183016.8 | 3299 | ENSMUSG00000040327 | Cul9 | cullin 9 |
| chr16 | 91914401 | 91915595 | 1195 | * | 9,700714 | 3,493688303 | 1,30298 | 2,681307 | 0,007334 | 0,83238039 | 1,449706 | Intron (ENSMUST0000( | 16 | 91923176 | 91983416 | 60241 | 1 | 1,05E+08 | ENSMUST00000231220.2 | -7581 | | Gm41496 | predicted gene, 41496 |
| chr19 | 3133826 | 3135636 | 1811 | * | 17,90693 | 3,493815196 | 0,868761 | 4,021605 | 5,78E-05 | 0,083814446 | 2,423594 | Intron (ENSMUST0000( | 19 | 3186984 | 3187090 | 107 | 2 | 1,15E+08 | ENSMUST00000101978.3 | 51454 | ENSMUSG00000118966 | Gm25431 | predicted gene, 25431 |
| chr4 | 1,32E+08 | 1,32E+08 | 2023 | * | 12,73937 | 3,495112517 | 1,168132 | 2,992054 | 0,002771 | 0,713978558 | 1,075735 | Promoter (2-3kb) | 4 | 1,32E+08 | 1,32E+08 | 66744 | 1 | 14050 | ENSMUST00000142301.8 | 2943 | ENSMUSG00000028886 | Eya3 | EYA transcriptional coactivator and phosphatase 3 |
| chr5 | 1,07E+08 | 1,07E+08 | 1447 | * | 19,74881 | 3,495116459 | 0,8431 | 4,145554 | 3,39E-05 | 0,058503909 | 2,507013 | Distal Intergenic | 5 | 1,07E+08 | 1,07E+08 | 20955 | 1 | 1,03E+08 | ENSMUST00000243509.1 | 31916 | | Gm33474 | predicted gene, 33474 |
| chr15 | 79360537 | 79361631 | 1095 | * | 12,93273 | 3,495253341 | 1,137878 | 3,071728 | 0,002128 | 0,642870233 | 1,2763 | Distal Intergenic | 15 | 79310459 | 79339767 | 29309 | 2 | 27373 | ENSMUST00000230599.2 | -20770 | ENSMUSG00000022433 | Csnk1e | casein kinase 1, epsilon |
| chr5 | 1,36E+08 | 1,36E+08 | 1142 | * | 10,31284 | 3,495600479 | 1,239117 | 2,821041 | 0,004787 | 0,789027175 | 1,211315 | Intron (ENSMUST0000( | 5 | 1,36E+08 | 1,36E+08 | 8964 | 2 | 215210 | ENSMUST00000043378.9 | -15654 | ENSMUSG00000039886 | Tmem120a | transmembrane protein 120A |

| | | | | | | | | | | | | | | | | | | | | | | |
|---|---|---|---|---|---|---|---|---|---|---|---|---|---|---|---|---|---|---|---|---|---|---|
| chr16 | 91156483 | 91157721 | 1239 | * | 13,88869 | 3,495697406 | 1,095819 | 3,190032 | 0,001423 | 0,597812173 | 1,075735 | Intron (ENSMUST0000 | 16 | 91162373 | 91164027 | 1655 | 2 | 1,03E+08 | ENSMUST00000244135.1 | 6306 | | Gm36587 | predicted gene, 36587 |
| chr14 | 25483359 | 25485031 | 1673 | * | 14,80872 | 3,496029491 | 1,045921 | 3,342537 | 0,00083 | 0,451739959 | 1,41696 | Intron (ENSMUST0000 | 14 | 25503079 | 25572454 | 69376 | 1 | 328365 | ENSMUST00000160680.3 | -18048 | ENSMUSG00000007817 | Zmiz1 | zinc finger, MIZ-type containing 1 |
| chr6 | 30003743 | 30004729 | 987 | * | 10,66592 | 3,496694587 | 1,210084 | 2,889629 | 0,003857 | 0,837306791 | 0,738441 | Distal Intergenic | 6 | 29986094 | 29993529 | 7436 | 1 | 76419 | ENSMUST00000238228.2 | 17649 | | Smkr-ps | smal lysine rich protein 1, pseudogene |
| chr6 | 59928984 | 59929956 | 973 | * | 10,22412 | 3,497263037 | 1,189178 | 2,940909 | 0,003273 | 0,68680243 | 1,449706 | Distal Intergenic | 6 | 60008952 | 60127030 | 118079 | 1 | 434019 | ENSMUST00000101355.9 | -78996 | ENSMUSG00000073045 | Gm5570 | predicted gene 5570 |
| chr9 | 22223255 | 22224273 | 1019 | * | 8,287419 | 3,498310372 | 1,329757 | 2,630789 | 0,008519 | 1 | 0 | Distal Intergenic | 9 | 22190363 | 22218944 | 28582 | 2 | 235050 | ENSMUST00000215202.2 | -4311 | ENSMUSG00000066829 | Zfp810 | zinc finger protein 810 |
| chr3 | 88571377 | 88573727 | 2351 | * | 19,14343 | 3,498991588 | 0,935351 | 3,740831 | 0,000183 | 0,15931259 | 2,713306 | Distal Intergenic | 3 | 88559205 | 88560449 | 1245 | 2 | 242093 | ENSMUST00000063119.5 | -10928 | ENSMUSG00000049741 | Rxfp4 | relaxin family peptide receptor 4 |
| chr10 | 97656755 | 97657723 | 969 | * | 9,940077 | 3,50057375 | 1,32143 | 2,649079 | 0,008071 | 0,886332554 | 1,173325 | Intron (ENSMUST0000 | 10 | 97580619 | 97580746 | 128 | 2 | 1,15E+08 | ENSMUST00000158472.3 | -76009 | ENSMUSG00000089097 | Gm25658 | predicted gene, 25658 |
| chr11 | 74788108 | 74788906 | 799 | * | 8,797824 | 3,501459543 | 1,323935 | 2,644737 | 0,008175 | 0,852732156 | 1,425929 | Promoter (<=1kb) | 11 | 74788897 | 74795766 | 6870 | 1 | 104662 | ENSMUST00000155702.8 | 0 | ENSMUSG00000038335 | Tsr1 | TSR1 20S rRNA accumulation |
| chr12 | 1,12E+08 | 1,12E+08 | 1239 | * | 12,25607 | 3,501476561 | 1,078382 | 3,246972 | 0,001166 | 0,593612587 | 0,918255 | Promoter (1-2kb) | 12 | 1,12E+08 | 1,12E+08 | 1334 | 2 | 70257 | ENSMUST00000223435.2 | 1902 | ENSMUSG00000021290 | Atp5mpl | ATP synthase membrane subunit 6.8PL |
| chrX | 38953124 | 38954535 | 1412 | * | 9,212575 | 3,502985228 | 1,27284 | 2,752101 | 0,005921 | 0,716908183 | 2,237403 | Promoter (<=1kb) | 20 | 38951791 | 38952481 | 691 | 2 | 1,01E+08 | ENSMUST00000249079.1 | -643 | ENSMUSG00000095240 | Cypt14 | cysteine-rich perinuclear theca 14 |
| chr2 | 1,65E+08 | 1,65E+08 | 958 | * | 8,942925 | 3,503024471 | 1,330181 | 2,633495 | 0,008451 | 0,888080529 | 1,211315 | Promoter (<=1kb) | 2 | 1,65E+08 | 1,65E+08 | 9984 | 2 | 18830 | ENSMUST00000142603.8 | 436 | ENSMUSG00000017754 | Pltp | phospholipid transfer protein |
| chr11 | 1,15E+08 | 1,15E+08 | 1728 | * | 14,21978 | 3,504147751 | 0,973941 | 3,597904 | 0,000321 | 0,356016677 | 0,938682 | Promoter (1-2kb) | 11 | 1,15E+08 | 1,15E+08 | 9379 | 1 | 237987 | ENSMUST00000106544.2 | 1029 | ENSMUSG00000050201 | Otop2 | otopetrin 2 |
| chr4 | 1,12E+08 | 1,12E+08 | 1121 | * | 9,188494 | 3,504176904 | 1,131438 | 3,097099 | 0,001954 | 0,60631654 | 1,425929 | Distal Intergenic | 4 | 1,12E+08 | 1,12E+08 | 68224 | 1 | 195564 | ENSMUST00000038455.12 | -6700 | ENSMUSG00000070868 | Skint3 | selection and upkeep of intraepithelial T cells 3 |
| chr7 | 28599039 | 28600496 | 1458 | * | 9,595978 | 3,504346889 | 1,330461 | 2,633935 | 0,00844 | 0,888080529 | 1,211315 | Promoter (1-2kb) | 7 | 28593136 | 28597618 | 4483 | 2 | 60595 | ENSMUST00000216863.2 | -1421 | ENSMUSG00000054808 | Actn4 | actinin alpha 4 |
| chr11 | 98859229 | 98860054 | 826 | * | 9,419605 | 3,504991242 | 1,317011 | 2,661322 | 0,007783 | 0,764492328 | 2,237403 | Intron (ENSMUST0000 | 11 | 98851238 | 98865768 | 14531 | 1 | 19401 | ENSMUST00000107473.3 | 7991 | ENSMUSG00000037992 | Rara | retinoic acid receptor, alpha |
| chr13 | 21905512 | 21906996 | 1485 | * | 12,18377 | 3,505126011 | 1,086976 | 3,224657 | 0,001261 | 0,644704154 | 0,744375 | Promoter (<=1kb) | 13 | 21906214 | 21906737 | 524 | 1 | 319186 | ENSMUST00000224651.2 | 0 | ENSMUSG00000114279 | H2bc14 | H2B clustered histone 14 |
| chr15 | 80667200 | 80668132 | 933 | * | 7,44978 | 3,505158592 | 1,314615 | 2,666301 | 0,007669 | 1 | 0 | Intron (ENSMUST0000 | 15 | 80682843 | 80765689 | 82847 | 1 | 213988 | ENSMUST00000226857.2 | -14711 | ENSMUSG00000047888 | Tnrc6b | trinucleotide repeat containing 6b |
| chr2 | 132004242 | 132006536 | 2295 | * | 32,24496 | 3,505885908 | 0,6636612 | 5,2826444 | 1,273E-07 | 0,000805219 | 2,9254973 | Intron (ENSMUST000001 | 2 | 131916754 | 131987028 | 70275 | 2 | 54338 | ENSMUST00000127724.8 | -17214 | ENSMUSG00000027340 | Slc23a2 | solute carrier family 23 (nucleobase transporters), member 2 |
| chr13 | 36766008 | 36768021 | 2014 | * | 14,33129 | 3,505963369 | 1,137005 | 3,083508 | 0,002046 | 0,725620806 | 0,738441 | Exon (ENSMUST000002 | 13 | 36796790 | 36804413 | 7624 | 1 | 1,08E+08 | ENSMUST00000225239.2 | -28769 | | Gm46410 | predicted gene, 46410 |
| chr1 | 1,53E+08 | 1,53E+08 | 977 | * | 8,992635 | 3,507993589 | 1,241536 | 2,825526 | 0,00472 | 0,764024179 | 1,360622 | Promoter (1-2kb) | 1 | 1,53E+08 | 1,53E+08 | 5897 | 1 | 67771 | ENSMUST00000143008.2 | 1963 | ENSMUSG00000008475 | Arpc5 | actin related protein 2/3 complex, subunit 5 |
| chr8 | 1,26E+08 | 1,26E+08 | 1500 | * | 14,73093 | 3,508618059 | 1,012768 | 3,464386 | 0,000531 | 0,398211274 | 1,2763 | Distal Intergenic | 8 | 1,26E+08 | 1,26E+08 | 40738 | 2 | 112405 | ENSMUST00000034469.7 | -8117 | ENSMUSG00000031987 | Egln1 | egl-9 family hypoxia-inducible factor 1 |
| chr8 | 1,23E+08 | 1,23E+08 | 1347 | * | 10,43565 | 3,508645268 | 1,198567 | 2,927368 | 0,003418 | 0,732009389 | 1,173325 | Intron (ENSMUST0000 | 8 | 1,23E+08 | 1,23E+08 | 2040 | 1 | 53325 | ENSMUST00000173753.2 | -9900 | ENSMUSG00000025316 | Banp | BTG3 associated nuclear protein |
| chr6 | 29909710 | 29910352 | 643 | * | 8,61035 | 3,510223176 | 1,178927 | 2,977473 | 0,002906 | 0,683140223 | 1,360622 | 3' UTR | 6 | 29903505 | 29909150 | 5646 | 1 | 74340 | ENSMUST00000128837.2 | 6205 | ENSMUSG00000029772 | Ahcyl2 | S-adenosylhomocysteine hydrolase-like 2 |
| chr7 | 1,13E+08 | 1,13E+08 | 1294 | * | 11,48338 | 3,510475259 | 1,092919 | 3,212018 | 0,001318 | 0,5663777 | 1,173325 | Distal Intergenic | 7 | 1,13E+08 | 1,13E+08 | 26990 | 1 | 1,03E+08 | ENSMUST00000210768.2 | 59562 | | Gm33586 | predicted gene, 33586 |
| chr3 | 1,45E+08 | 1,45E+08 | 2193 | * | 19,34514 | 3,510787515 | 0,849652 | 4,132031 | 3,6E-05 | 0,063329886 | 2,386008 | Promoter (<=1kb) | 3 | 1,45E+08 | 1,46E+08 | 132519 | 1 | 69219 | ENSMUST00000120310.2 | 0 | ENSMUSG00000028194 | Ddah1 | dimethylarginine dimethylaminohydrolase 1 |
| chr15 | 80005227 | 80006479 | 1253 | * | 10,57058 | 3,510829298 | 1,202589 | 2,919392 | 0,003507 | 0,737421635 | 1,173325 | Distal Intergenic | 15 | 79999674 | 80000466 | 793 | 1 | 20972 | ENSMUST00000135727.2 | 5553 | ENSMUSG00000022415 | Syngr1 | synaptogyrin 1 |
| chr3 | 49883112 | 49883788 | 677 | * | 7,262419 | 3,51149786 | 1,328088 | 2,644025 | 0,008193 | 1 | 0 | Intron (ENSMUST0000 | 3 | 49699720 | 49711774 | 12055 | 2 | 73173 | ENSMUST00000193600.2 | -171338 | ENSMUSG00000037892 | Pcdh18 | protocadherin 18 |
| chr2 | 1,43E+08 | 1,43E+08 | 1108 | * | 12,42389 | 3,511571841 | 1,044295 | 3,362625 | 0,000772 | 0,456947326 | 1,2763 | Distal Intergenic | 2 | 1,43E+08 | 1,43E+08 | 8008 | 2 | 378776 | ENSMUST00000156780.2 | 77323 | ENSMUSG00000086171 | Pcsk2os1 | proprotein convertase subtilisin/kexin type 2, opposite strand 1 |
| chr2 | 85945895 | 85947060 | 1166 | * | 10,8325 | 3,512358781 | 1,118383 | 3,140569 | 0,001686 | 0,659240897 | 0,918255 | Distal Intergenic | 2 | 85950567 | 85956700 | 6134 | 1 | 259015 | ENSMUST00000216665.3 | -3507 | | Olfr1038-ps | olfactory receptor 1038, pseudogene |
| chr11 | 51511243 | 51511799 | 557 | * | 6,160081 | 3,512521332 | 1,360416 | 2,581947 | 0,009824 | 1 | 0 | Promoter (<=1kb) | 11 | 51510598 | 51514256 | 3659 | 1 | 52530 | ENSMUST00000001083.4 | 645 | ENSMUSG00000001056 | Nhp2 | NHP2 ribonucleoprotein |
| chr12 | 31827830 | 31828941 | 1112 | * | 9,465719 | 3,513804117 | 1,28243 | 2,739959 | 0,006145 | 0,83238039 | 1,211315 | Intron (ENSMUST0000 | 12 | 31758711 | 31763946 | 5236 | 2 | 73010 | ENSMUST00000176710.2 | -63884 | ENSMUSG00000044067 | Gpr22 | G protein-coupled receptor 22 |
| chr15 | 35298457 | 35299629 | 1173 | * | 13,52303 | 3,51440546 | 1,028618 | 3,416627 | 0,000634 | 0,425046984 | 1,2763 | Promoter (<=1kb) | 15 | 35300446 | 35303450 | 3005 | 1 | 107587 | ENSMUST00000228152.2 | -817 | ENSMUSG00000022330 | Osr2 | odd-skipped related 2 |
| chrX | 98564149 | 98565882 | 1734 | * | 11,9502 | 3,514508232 | 1,076448 | 3,264912 | 0,001095 | 0,532073564 | 1,173325 | Distal Intergenic | 20 | 98509340 | 98514879 | 5540 | 2 | 18744 | ENSMUST00000113792.2 | -49270 | ENSMUSG00000034403 | Pja1 | praja ring finger ubiquitin ligase 1 |
| chr1 | 91618442 | 91619835 | 1394 | * | 10,41883 | 3,514683922 | 1,256249 | 2,797761 | 0,005146 | 0,887238309 | 0,744375 | Distal Intergenic | 1 | 91539128 | 91552222 | 13095 | 1 | 1,03E+08 | ENSMUST00000188800.2 | 79314 | | Gm28380 | predicted gene 28380 |
| chr6 | 59161638 | 59163079 | 1442 | * | 24,189529 | 3,514851957 | 0,73952 | 4,752883 | 2,005E-06 | 0,006356998 | 2,9519335 | Distal Intergenic | 6 | 59185855 | 59189018 | 3164 | 1 | 68140 | ENSMUST00000062626.4 | -22776 | ENSMUSG00000049232 | Tigd2 | tigger transposable element derived 2 |
| chr11 | 54100850 | 54101937 | 1088 | * | 9,039507 | 3,514893091 | 1,231136 | 2,854999 | 0,004304 | 0,748971518 | 1,360622 | Promoter (2-3kb) | 11 | 54068598 | 54104111 | 35514 | 2 | 71056 | ENSMUST00000131019.2 | 2174 | ENSMUSG00000086209 | 4933405E24Rik | RIKEN cDNA 4933405E24 gene |
| chr11 | 94114105 | 94115334 | 1230 | * | 12,78577 | 3,514901826 | 1,04501 | 3,36351 | 0,00077 | 0,456491176 | 1,2763 | Distal Intergenic | 11 | 94102280 | 94106321 | 4042 | 1 | 22057 | ENSMUST00000041589.6 | 11825 | ENSMUSG00000037573 | Tob1 | transducer of ErbB-2.1 |
| chr18 | 66978297 | 66979332 | 1036 | * | 8,322187 | 3,515061668 | 1,28599 | 2,733351 | 0,006269 | 1 | 0 | Distal Intergenic | 18 | 66990775 | 66993577 | 2803 | 2 | 17202 | ENSMUST00000057942.4 | 14245 | ENSMUSG00000047259 | Mc4r | melanocortin 4 receptor |
| chr11 | 55270380 | 55271690 | 1311 | * | 10,2164 | 3,515581425 | 1,287236 | 2,731108 | 0,006312 | 0,855495103 | 1,086118 | Distal Intergenic | 11 | 55286212 | 55290400 | 4189 | 2 | 20692 | ENSMUST00000123775.8 | 18710 | ENSMUSG00000018593 | Sparc | secreted acidic cysteine rich glycoprotein |
| chr7 | 1,26E+08 | 1,26E+08 | 839 | * | 12,73906 | 3,515853752 | 1,17457 | 2,993311 | 0,00276 | 0,683727424 | 1,2763 | Promoter (2-3kb) | 7 | 1,26E+08 | 1,26E+08 | 27764 | 2 | 74204 | ENSMUST00000166538.2 | 2236 | ENSMUSG00000000131 | Xpo6 | exportin 6 |
| chr17 | 67987111 | 67988252 | 1142 | * | 11,12504 | 3,516642708 | 1,131288 | 3,10853 | 0,00188 | 0,712714114 | 0,738441 | Distal Intergenic | 17 | 68004254 | 68129642 | 125389 | 1 | 16772 | ENSMUST00000035471.9 | -16002 | ENSMUSG00000032796 | Lama1 | laminin, alpha 1 |
| chr4 | 1,33E+08 | 1,33E+08 | 1314 | * | 10,15526 | 3,516757504 | 1,275118 | 2,757986 | 0,005816 | 0,841321009 | 1,086118 | Intron (ENSMUST0000 | 4 | 1,33E+08 | 1,33E+08 | 3101 | 2 | 93760 | ENSMUST00000139709.3 | -12141 | ENSMUSG00000007880 | Arid1a | AT rich interactive domain 1A (SWI-like) |
| chr3 | 86477527 | 86478632 | 1106 | * | 8,032501 | 3,516764655 | 1,361809 | 2,582421 | 0,009811 | 1 | 0 | Intron (ENSMUST0000 | 3 | 86453357 | 86455590 | 2234 | 2 | 23937 | ENSMUST00000191695.3 | -21937 | ENSMUSG00000057777 | Mab21l2 | mab-21-like 2 |
| chr4 | 1,05E+08 | 1,05E+08 | 1216 | * | 11,61499 | 3,518614261 | 1,269701 | 2,771214 | 0,005585 | 0,900637308 | 0,738441 | Intron (ENSMUST0000 | 4 | 1,05E+08 | 1,05E+08 | 75418 | 1 | 67916 | ENSMUST00000064139.8 | 69382 | ENSMUSG00000028517 | Plpp3 | phospholipid phosphatase 3 |
| chr7 | 1,26E+08 | 1,26E+08 | 2292 | * | 16,89586 | 3,520061082 | 1,091807 | 3,22407 | 0,001264 | 0,54122637 | 1,2763 | Promoter (<=1kb) | 7 | 1,26E+08 | 1,26E+08 | 6631 | 2 | 56420 | ENSMUST00000032936.8 | -423 | ENSMUSG00000030697 | Ppp4c | protein phosphatase 4, catalytic subunit |
| chr8 | 26259593 | 26260422 | 830 | * | 7,704757 | 3,52072976 | 1,346466 | 2,614793 | 0,008928 | 1 | 0 | Exon (ENSMUST000001 | 8 | 26259539 | 26267610 | 8072 | 2 | 67384 | ENSMUST00000210103.2 | 7188 | ENSMUSG00000037316 | Bag4 | BCL2-associated athanogene 4 |
| chr3 | 43100887 | 43101296 | 410 | * | 7,594875 | 3,521110495 | 1,337151 | 2,633293 | 0,008456 | 1 | 0 | Distal Intergenic | 3 | 41705631 | 41710927 | 5297 | 1 | 73852 | ENSMUST00000195030.2 | 1395256 | ENSMUSG00000025766 | D3Ertd751e | DNA segment, Chr 3, ERATO Doi 751, expressed |
| chr9 | 55441269 | 55441845 | 577 | * | 6,720203 | 3,521899795 | 1,313892 | 2,680509 | 0,007351 | 1 | 0 | Distal Intergenic | 9 | 55445956 | 55449854 | 3899 | 1 | 104360 | ENSMUST00000175950.8 | -4111 | ENSMUSG00000032318 | Isl2 | insulin related protein 2 (islet 2) |
| chr8 | 61680972 | 61682269 | 1298 | * | 12,7624 | 3,521929866 | 1,033726 | 3,407026 | 0,000657 | 0,445547689 | 1,173325 | Intron (ENSMUST0000 | 8 | 61677278 | 61839675 | 162398 | 1 | 59009 | ENSMUST00000209611.2 | 3694 | ENSMUSG00000031642 | Sh3rf1 | SH3 domain containing ring finger 1 |
| chr1 | 1,82E+08 | 1,82E+08 | 2111 | * | 13,68512 | 3,522398566 | 1,242639 | 2,834612 | 0,004588 | 0,824944942 | 0,938682 | Exon (ENSMUST000002 | 1 | 1,82E+08 | 1,82E+08 | 16457 | 1 | 240960 | ENSMUST00000097446.9 | -16828 | ENSMUSG00000047369 | Dnah14 | dynein, axonemal, heavy chain 14 |
| chr5 | 1,32E+08 | 1,32E+08 | 1550 | * | 10,02744 | 3,524374277 | 1,171697 | 3,007923 | 0,00263 | 0,659642963 | 1,425929 | Intron (ENSMUST0000 | 5 | 1,32E+08 | 1,32E+08 | 3500 | 1 | 75369 | ENSMUST00000198640.2 | -129049 | ENSMUSG00000105773 | 4930563F08Rik | RIKEN cDNA 4930563F08 gene |
| chr3 | 1,22E+08 | 1,22E+08 | 1555 | * | 12,9022 | 3,524862708 | 1,082801 | 3,255318 | 0,001133 | 0,550933624 | 1,075735 | Promoter (<=1kb) | 3 | 1,22E+08 | 1,22E+08 | 7110 | 1 | 29815 | ENSMUST00000200544.2 | 0 | ENSMUSG00000028121 | Bcar3 | breast cancer anti-estrogen resistance 3 |
| chr11 | 61537391 | 61538868 | 1478 | * | 16,48157 | 3,52544503 | 0,949275 | 3,713829 | 0,000204 | 0,193718476 | 2,141612 | Distal Intergenic | 11 | 61544091 | 61563610 | 19520 | 1 | 71520 | ENSMUST00000004959.3 | -5223 | ENSMUSG00000004837 | Grap | GRB2-related adaptor protein |
| chr6 | 87622562 | 87623678 | 1117 | * | 13,78266 | 3,526268648 | 1,147289 | 3,073567 | 0,002115 | 0,642322948 | 1,2763 | 3' UTR | 6 | 87634736 | 87640805 | 6070 | 1 | 1,15E+08 | ENSMUST00000204112.2 | -11058 | | Gm44097 | predicted gene, 44097 |
| chr12 | 21467293 | 21468785 | 1493 | * | 12,82125 | 3,527235717 | 1,007092 | 3,502398 | 0,000461 | 0,38567116 | 1,173325 | Promoter (<=1kb) | 12 | 21467324 | 21469390 | 2067 | 1 | 1E+08 | ENSMUST00000226239.2 | 0 | ENSMUSG00000097055 | Gm4419 | predicted gene 4419 |
| chr11 | 35728170 | 35730806 | 2637 | * | 35,991115 | 3,527550295 | 0,5528757 | 6,3803677 | 1,767E-10 | 6,45401E-06 | 1,4057095 | Promoter (2-3kb) | 11 | 35699208 | 35725333 | 26126 | 2 | 104458 | ENSMUST00000018992.4 | -2837 | ENSMUSG00000018848 | Rars | arginyl-tRNA synthetase |
| chr4 | 1,4E+08 | 1,4E+08 | 724 | * | 8,406714 | 3,528179011 | 1,356951 | 2,600079 | 0,00932 | 0,874247416 | 1,449706 | Intron (ENSMUST0000 | 4 | 1,4E+08 | 1,4E+08 | 219939 | 2 | 230868 | ENSMUST00000039331.9 | 13080 | ENSMUSG00000040972 | Igsf21 | immunoglobulin superfamily, member 21 |
| chr10 | 1,2E+08 | 1,2E+08 | 893 | * | 8,45401 | 3,530822791 | 1,360059 | 2,59608 | 0,009429 | 0,887433944 | 1,360622 | Distal Intergenic | 10 | 1,2E+08 | 1,2E+08 | 3271 | 1 | 320626 | ENSMUST00000161291.2 | -62919 | ENSMUSG00000090191 | 9230105E05Rik | RIKEN cDNA 9230105E05 gene |
| chr4 | 1,34E+08 | 1,34E+08 | 1403 | * | 11,04675 | 3,531133687 | 1,213111 | 2,910809 | 0,003605 | 0,824070388 | 0,744375 | Promoter (<=1kb) | 4 | 1,34E+08 | 1,34E+08 | 1412 | 1 | 77490 | ENSMUST00000135228.8 | -960 | ENSMUSG00000086322 | E130218I03Rik | RIKEN cDNA E130218I03 gene |
| chr10 | 26658972 | 26660191 | 1220 | * | 10,45026 | 3,531765821 | 1,149249 | 3,073108 | 0,002118 | 0,730469267 | 0,738441 | Intron (ENSMUST0000 | 10 | 26648473 | 26794644 | 146172 | 1 | 73910 | ENSMUST00000039557.9 | 10499 | ENSMUSG00000039031 | Arhgap18 | Rho GTPase activating protein 18 |
| chr10 | 91325616 | 91326354 | 739 | * | 8,349957 | 3,531811762 | 1,232062 | 2,866586 | 0,004149 | 0,741428016 | 1,360622 | Distal Intergenic | 10 | 91044120 | 91056504 | 12385 | 2 | 1,03E+08 | ENSMUST00000216311.2 | -269112 | ENSMUSG00000111544 | 4930534H03Rik | RIKEN cDNA 4930534H03 gene |
| chr4 | 68060181 | 68061348 | 1168 | * | 10,79807 | 3,532095831 | 1,2602 | 2,802806 | 0,005066 | 0,884930392 | 0,744375 | Distal Intergenic | 4 | 68100769 | 68100939 | 171 | 1 | 1,15E+08 | ENSMUST00000082828.3 | -39421 | ENSMUSG00000064762 | Gm23738 | predicted gene, 23738 |
| chr5 | 1,21E+08 | 1,21E+08 | 902 | * | 8,213458 | 3,532507668 | 1,291113 | 2,736018 | 0,006219 | 0,801688106 | 1,449706 | Intron (ENSMUST0000 | 5 | 1,21E+08 | 1,21E+08 | 5696 | 2 | 252972 | ENSMUST00000202072.2 | 5857 | ENSMUSG00000032741 | Tpcn1 | two pore channel 1 |
| chr2 | 31034628 | 31035830 | 1203 | * | 12,09547 | 3,532802059 | 1,286063 | 2,74699 | 0,006014 | 0,819720472 | 1,2763 | Promoter (2-3kb) | 2 | 30916219 | 31032020 | 115802 | 2 | 14269 | ENSMUST00000113560.8 | -2608 | ENSMUSG00000075415 | Fnbp1 | formin binding protein 1 |
| chr7 | 46024559 | 46025948 | 1390 | * | 13,31429 | 3,533627636 | 1,096131 | 3,223728 | 0,001265 | 0,607577254 | 0,918255 | Promoter (<=1kb) | 7 | 46025898 | 46028523 | 2626 | 1 | 17927 | ENSMUST00000072514.3 | 0 | ENSMUSG00000009471 | Myod1 | myogenic differentiation 1 |
| chrX | 1,02E+08 | 1,02E+08 | 1381 | * | 9,738137 | 3,534024354 | 1,248344 | 2,830969 | 0,004641 | 0,783315811 | 1,211315 | Distal Intergenic | 20 | 1,02E+08 | 1,02E+08 | 746 | 2 | 668339 | ENSMUST00000113610.3 | 3145 | ENSMUSG00000079479 | Gm9112 | predicted gene 9112 |
| chr1 | 92225975 | 92226627 | 653 | * | 8,514067 | 3,534068022 | 1,260634 | 2,803406 | 0,005057 | 0,777345889 | 1,360622 | Distal Intergenic | 1 | 92307317 | 92321355 | 14039 | 1 | 1,03E+08 | ENSMUST00000240981.1 | -80690 | | Gm35992 | predicted gene, 35992 |
| chr4 | 34275121 | 34276422 | 1302 | * | 10,67201 | 3,53408374 | 1,124437 | 3,142981 | 0,001672 | 0,596263346 | 1,2763 | Distal Intergenic | 4 | 34086126 | 34086233 | 108 | 1 | 1,15E+08 | ENSMUST00000157216.2 | 188995 | ENSMUSG00000087841 | Gm25623 | predicted gene, 25623 |
| chr10 | 37877870 | 37878721 | 852 | * | 8,172142 | 3,535014533 | 1,211894 | 2,916935 | 0,003535 | 0,631820259 | 2,237403 | Distal Intergenic | 10 | 37992241 | 37998057 | 5817 | 2 | 1,03E+08 | ENSMUST00000213538.2 | 119336 | | Gm31378 | predicted gene, 31378 |
| chr11 | 8591669 | 8592708 | 1040 | * | 9,275012 | 3,535078716 | 1,35333 | 2,612134 | 0,008998 | 0,898643449 | 1,211315 | Intron (ENSMUST0000 | 11 | 8381652 | 8574887 | 193236 | 2 | 319939 | ENSMUST00000239091.2 | -16782 | ENSMUSG00000020422 | Tns3 | tensin 3 |
| chr12 | 1,11E+08 | 1,11E+08 | 1359 | * | 13,02544 | 3,535144951 | 1,200041 | 2,945854 | 0,003221 | 0,803455139 | 0,738441 | Promoter (2-3kb) | 12 | 1,11E+08 | 1,11E+08 | 9926 | 1 | 69641 | ENSMUST00000194118.2 | 2914 | ENSMUSG00000037957 | Wdr20 | WD repeat domain 20 |
| chr12 | 33007115 | 33008116 | 1002 | * | 8,21697 | 3,535348384 | 1,291067 | 2,738316 | 0,006175 | 1 | 0 | Promoter (2-3kb) | 12 | 33004558 | 33017678 | 13121 | 1 | 19027 | ENSMUST00000129592.2 | 2557 | ENSMUSG00000020570 | Sypl | synaptophysin-like protein |
| chr10 | 45884090 | 45884884 | 795 | * | 8,078868 | 3,536436184 | 1,276907 | 2,769534 | 0,005614 | 1 | 0 | Distal Intergenic | 10 | 45713976 | 45728379 | 14404 | 1 | 320363 | ENSMUST00000215085.2 | 170114 | ENSMUSG00000110874 | D030045P18Rik | RIKEN cDNA D030045P18 gene |
| chr14 | 23689932 | 23691074 | 1143 | * | 12,52324 | 3,536441525 | 1,144711 | 3,089375 | 0,002006 | 0,646275731 | 1,173325 | Intron (ENSMUST0000 | 14 | 23349463 | 23700257 | 350795 | 2 | 16531 | ENSMUST00000112423.10 | 9183 | ENSMUSG00000063142 | Kcnma1 | potassium large conductance calcium-activated channel, subfamily M, alpha member 1 |
| chr13 | 52642459 | 52643383 | 925 | * | 8,332574 | 3,53708946 | 1,215654 | 2,909618 | 0,003619 | 0,717146731 | 1,360622 | Distal Intergenic | 13 | 52658416 | 52685315 | 26900 | 2 | 68203 | ENSMUST00000057442.8 | 41932 | ENSMUSG00000047842 | Diras2 | DIRAS family, GTP-binding RAS-like 2 |
| chr7 | 1,01E+08 | 1,01E+08 | 1556 | * | 15,80849 | 3,537298124 | 1,07138 | 3,301629 | 0,000961 | 0,391231126 | 2,386008 | Promoter (<=1kb) | 7 | 1,01E+08 | 1,01E+08 | 33555 | 1 | 69710 | ENSMUST00000084895.12 | -506 | ENSMUSG00000032812 | Arap1 | ArfGAP with RhoGAP domain, ankyrin repeat and PH domain 1 |
| chr17 | 31982427 | 31984110 | 1684 | * | 13,11302 | 3,53743988 | 1,029323 | 3,436666 | 0,000589 | 0,426043749 | 1,173325 | Distal Intergenic | 17 | 31898454 | 31900479 | 2026 | 1 | 12954 | ENSMUST00000228305.2 | 83973 | ENSMUSG00000024041 | Cryaa | crystallin, alpha A |
| chr9 | 33608653 | 33609462 | 810 | * | 8,305597 | 3,537450332 | 1,243435 | 2,844901 | 0,004443 | 0,746312988 | 1,425929 | Distal Intergenic | 9 | 33828788 | 33828918 | 131 | 1 | 1,15E+08 | ENSMUST00000158280.3 | -219326 | ENSMUSG00000088905 | Gm25439 | predicted gene, 25439 |
| chr9 | 1,1E+08 | 1,1E+08 | 1196 | * | 10,17479 | 3,538236359 | 1,331939 | 2,656455 | 0,007897 | 0,89325945 | 1,086118 | Distal Intergenic | 9 | 1,1E+08 | 1,1E+08 | 2239 | 2 | 12796 | ENSMUST00000112022.5 | -8148 | ENSMUSG00000038357 | Camp | cathelicidin antimicrobial peptide |
| chr19 | 41305704 | 41307175 | 1472 | * | 13,85826 | 3,538947497 | 1,150215 | 3,07677 | 0,002093 | 0,685026746 | 0,938682 | Intron (ENSMUST0000 | 19 | 41199283 | 41252436 | 53154 | 2 | 107358 | ENSMUST00000025989.10 | -53268 | ENSMUSG00000025016 | Tm9sf3 | transmembrane 9 superfamily member 3 |
| chr11 | 51838113 | 51839219 | 1107 | * | 9,31708 | 3,539267347 | 1,345969 | 2,629531 | 0,00855 | 0,857326444 | 1,449706 | Distal Intergenic | 11 | 51855209 | 51855316 | 108 | 1 | 1,15E+08 | ENSMUST00000175584.3 | -15990 | ENSMUSG00000093325 | Gm26061 | predicted gene, 26061 |
| chr4 | 59314658 | 59317144 | 2487 | * | 67,0299 | 3,539511855 | 0,4352529 | 8,1320803 | 4,22E-16 | 5,19159E-11 | 3,2706213 | Promoter (<=1kb) | 4 | 59314683 | 59315966 | 1284 | 2 | 634731 | ENSMUST00000136942.2 | 0 | ENSMUSG00000038578 | Susd1 | sushi domain containing 1 |
| chr4 | 62277551 | 62280098 | 2548 | * | 30,120183 | 3,539947498 | 0,7251619 | 4,881596 | 1,052E-06 | 0,003798148 | 2,8585108 | Promoter (<=1kb) | 4 | 62278964 | 62310006 | 31043 | 1 | 20529 | ENSMUST00000084526.12 | 0 | ENSMUSG00000066150 | Slc31a1 | solute carrier family 31, member 1 |
| chr17 | 25507422 | 25508640 | 1219 | * | 12,80182 | 3,540178495 | 1,091005 | 3,244879 | 0,001175 | 0,594430593 | 0,918255 | Distal Intergenic | 17 | 25517368 | 25519139 | 1772 | 1 | 328780 | ENSMUST00000037453.4 | -8728 | ENSMUSG00000056399 | Prss34 | protease, serine 34 |
| chr14 | 57150385 | 57151935 | 1551 | * | 9,278358 | 3,54037746 | 1,234908 | 2,866916 | 0,004145 | 0,731734453 | 1,425929 | Exon (ENSMUST00000 | 14 | 57158222 | 57197809 | 39588 | 1 | 76007 | ENSMUST00000224922.2 | -6287 | ENSMUSG00000021945 | Zmym2 | zinc finger, MYM-type 2 |
| chrX | 81231521 | 81233650 | 2130 | * | 12,85422 | 3,540668306 | 1,171306 | 3,022839 | 0,002504 | 0,722798324 | 0,918255 | Distal Intergenic | 20 | 81444697 | 81444833 | 137 | 2 | 1,15E+08 | ENSMUST00000083047.3 | 211183 | ENSMUSG00000119611 | Gm24706 | predicted gene, 24706 |
| chr4 | 1,16E+08 | 1,16E+08 | 984 | * | 8,243468 | 3,542641912 | 1,303066 | 2,718697 | 0,006554 | 0,815446267 | 1,425929 | Intron (ENSMUST0000 | 4 | 1,16E+08 | 1,16E+08 | 102354 | 2 | 17776 | ENSMUST00000151347.2 | -8161 | ENSMUSG00000003810 | Mast2 | microtubule associated serine/threonine kinase 2 |
| chr4 | 93876728 | 93877652 | 925 | * | 8,454614 | 3,543399657 | 1,29413 | 2,738056 | 0,00618 | 0,801054956 | 1,449706 | Distal Intergenic | 4 | 93341495 | 93559307 | 217813 | 2 | 1,03E+08 | ENSMUST00000137509.2 | -317421 | | Gm31828 | predicted gene, 31828 |
| chr11 | 27330738 | 27331959 | 1222 | * | 7,361707 | 3,543584762 | 1,32455 | 2,675313 | 0,007466 | 1 | 0 | Distal Intergenic | 11 | 27268847 | 27268978 | 132 | 2 | 1,15E+08 | ENSMUST00000157813.3 | -61760 | ENSMUSG00000119065 | Gm25387 | predicted gene, 25387 |
| chr4 | 41716221 | 41717133 | 913 | * | 8,80552 | 3,546655358 | 1,298751 | 2,73082 | 0,006318 | 0,817520905 | 1,360622 | Promoter (2-3kb) | 4 | 41712033 | 41713534 | 1502 | 2 | 69961 | ENSMUST00000038434.4 | -2687 | ENSMUSG00000036114 | Rpp25l | ribonuclease P/MRP 25 subunit-like |
| chr7 | 80505186 | 80506305 | 1120 | * | 9,917481 | 3,547601935 | 1,229636 | 2,885082 | 0,003913 | 0,755319554 | 1,173325 | Distal Intergenic | 7 | 80510668 | 80524717 | 14050 | 1 | 22691 | ENSMUST00000132163.8 | -4363 | ENSMUSG00000038797 | Zscan2 | zinc finger and SCAN domain containing 2 |
| chr13 | 1,03E+08 | 1,03E+08 | 1128 | * | 8,211674 | 3,54927094 | 1,268487 | 2,798035 | 0,005141 | 0,69016288 | 2,237403 | Intron (ENSMUST0000 | 13 | 1,03E+08 | 1,03E+08 | 3513 | 2 | 328329 | ENSMUST00000211607.2 | -66702 | ENSMUSG00000034751 | Mast4 | microtubule associated serine/threonine kinase family member 4 |
| chr8 | 1,05E+08 | 1,05E+08 | 562 | * | 8,773668 | 3,54955172 | 1,311851 | 2,705758 | 0,006815 | 0,83005113 | 1,360622 | Promoter (2-3kb) | 8 | 1,05E+08 | 1,05E+08 | 3063 | 2 | 234663 | ENSMUST00000212343.2 | 2987 | ENSMUSG00000035770 | Dync1li2 | dynein, cytoplasmic 1 light intermediate chain 2 |
| chr4 | 1,2E+08 | 1,2E+08 | 814 | * | 10,26439 | 3,550251073 | 1,16732 | 3,041369 | 0,002355 | 0,669377201 | 1,211315 | Distal Intergenic | 4 | 1,2E+08 | 1,2E+08 | 6155 | 1 | 13615 | ENSMUST00000030384.5 | -6422 | ENSMUSG00000028635 | Edn2 | endothelin 2 |
| chr15 | 7643629 | 7644465 | 837 | * | 8,405979 | 3,550827837 | 1,209123 | 2,936697 | 0,003317 | 0,703367735 | 1,360622 | Distal Intergenic | 15 | 7840327 | 7867056 | 26730 | 1 | 14573 | ENSMUST00000022744.5 | -195862 | ENSMUSG00000022144 | Gdnf | glial cell line derived neurotrophic factor |
| chr1 | 1,71E+08 | 1,71E+08 | 1318 | * | 11,54989 | 3,551049954 | 1,276467 | 2,781937 | 0,005404 | 0,894839376 | 0,738441 | Promoter (<=1kb) | 1 | 1,71E+08 | 1,71E+08 | 12270 | 2 | 226646 | ENSMUST00000013737.13 | 442 | ENSMUSG00000013593 | Ndufs2 | NADH:ubiquinone oxidoreductase core subunit S2 |
| chr8 | 75810068 | 75811683 | 1616 | * | 26,25962 | 3,551076331 | 0,6900965 | 5,1457681 | 2,664E-07 | 0,001329417 | 2,8800171 | Distal Intergenic | 8 | 75820249 | 75827217 | 6969 | 1 | 15368 | ENSMUST00000005548.8 | -8566 | ENSMUSG00000005413 | Hmox1 | heme oxygenase 1 |
| chr7 | 1,13E+08 | 1,13E+08 | 1141 | * | 8,882622 | 3,551193062 | 1,344416 | 2,641439 | 0,008255 | 0,860883941 | 1,360622 | Distal Intergenic | 7 | 1,13E+08 | 1,13E+08 | 53750 | 2 | 68815 | ENSMUST00000119278.8 | -9008 | ENSMUSG00000038187 | Btbd10 | BTB (POZ) domain containing 10 |
| chr4 | 1,41E+08 | 1,41E+08 | 1011 | * | 8,988001 | 3,55183546 | 1,271784 | 2,792798 | 0,005225 | 0,693610666 | 2,237403 | Intron (ENSMUST0000 | 4 | 1,41E+08 | 1,41E+08 | 6420 | 2 | 74202 | ENSMUST00000137491.2 | 9941 | ENSMUSG00000006219 | Fblim1 | filamin binding LIM protein 1 |
| chr4 | 37460979 | 37461858 | 880 | * | 7,765434 | 3,55231912 | 1,218784 | 2,914643 | 0,003561 | 1 | 0 | Distal Intergenic | 4 | 37249403 | 37332515 | 83113 | 1 | 1,03E+08 | ENSMUST00000147108.2 | 211576 | ENSMUSG00000086026 | Gm12374 | predicted gene 12374 |
| chr4 | 45433197 | 45433889 | 693 | * | 7,415242 | 3,553129771 | 1,370408 | 2,592754 | 0,009521 | 1 | 0 | Intron (ENSMUST0000 | 4 | 45395923 | 45408766 | 12844 | 2 | 230125 | ENSMUST00000116341.4 | -24431 | ENSMUSG00000045973 | Slc25a51 | solute carrier family 25, member 51 |

| | | | | | | | | | | | | | | | | | | | | | | | |
|---|---|---|---|---|---|---|---|---|---|---|---|---|---|---|---|---|---|---|---|---|---|---|---|
| chr1 | 45329172 | 45329921 | 750 | * | 7,205245 | 3,553349765 | 1,296109 | 2,741551 | 0,006115 | 1 | 0 | Distal Intergenic | 1 | 45350698 | 45364178 | 13481 | 1 | 12825 | ENSMUST00000129611.2 | -20777 | ENSMUSG00000026043 | Col3a1 | collagen, type III, alpha 1 |
| chr11 | 6218488 | 6220059 | 1572 | * | 11,01139 | 3,55382901 | 1,115974 | 3,184508 | 0,00145 | 0,669639313 | 0,738441 | Promoter (<=1kb) | 11 | 6215637 | 6217772 | 2136 | 2 | 52513 | ENSMUST00000151446.2 | -716 | ENSMUSG00000004393 | Ddx56 | DEAD box helicase 56 |
| chr10 | 79600850 | 79602253 | 1404 | * | 19,18472 | 3,553861062 | 0,882073 | 4,028988 | 5,6E-05 | 0,086053637 | 2,221611 | Promoter (<=1kb) | 10 | 79596874 | 79602744 | 5871 | 2 | 70294 | ENSMUST00000219189.2 | 491 | ENSMUSG00000035890 | Rnf126 | ring finger protein 126 |
| chr8 | 29312977 | 29313503 | 527 | * | 8,011123 | 3,553995385 | 1,348886 | 2,634762 | 0,00842 | 1 | 0 | Intron (ENSMUST0000( | 8 | 29083560 | 29084336 | 777 | 1 | 1,05E+08 | ENSMUST00000181267.2 | 229417 | | Gm26795 | predicted gene, 26795 |
| chr10 | 6457907 | 6458705 | 799 | * | 7,016997 | 3,554747889 | 1,348098 | 2,636862 | 0,008368 | 0,863495249 | 1,360622 | Distal Intergenic | 10 | 6708506 | 6929704 | 221199 | 1 | 18390 | ENSMUST00000147171.8 | -249801 | ENSMUSG00000000766 | Oprm1 | opioid receptor, mu 1 |
| chr6 | 29352396 | 29353428 | 1033 | * | 12,64213 | 3,554811152 | 1,267011 | 2,805667 | 0,005021 | 0,884588199 | 0,738441 | Intron (ENSMUST0000( | 6 | 29356470 | 29375024 | 18555 | 1 | 12321 | ENSMUST00000174096.7 | -3042 | ENSMUSG00000029767 | Calu | calumenin |
| chr11 | 98669871 | 98670587 | 717 | * | 8,531644 | 3,554811998 | 1,181532 | 3,008646 | 0,002624 | 0,574473041 | 2,237403 | Distal Intergenic | 11 | 98658758 | 98666159 | 7402 | 2 | 217166 | ENSMUST00000064941.7 | -3712 | ENSMUSG00000020889 | Nr1d1 | nuclear receptor subfamily 1, group D, member 1 |
| chr2 | 6688257 | 6689403 | 1147 | * | 10,72755 | 3,554835835 | 1,257104 | 2,827798 | 0,004687 | 0,871792855 | 0,738441 | Intron (ENSMUST0000( | 2 | 6551051 | 6726424 | 175374 | 2 | 14007 | ENSMUST00000142941.8 | 37021 | ENSMUSG00000002107 | Celf2 | CUGBP, Elav-like family member 2 |
| chr3 | 55802775 | 55804005 | 1231 | * | 9,93211 | 3,555658384 | 1,134524 | 3,134054 | 0,001724 | 0,61384098 | 1,211315 | Intron (ENSMUST0000( | 3 | 55797727 | 55844120 | 46394 | 1 | 1,03E+08 | ENSMUST00000199747.2 | 5048 | | Gm31914 | predicted gene, 31914 |
| chr5 | 66707442 | 66709264 | 1823 | * | 20,114272 | 3,555778249 | 0,790874 | 4,496011 | 6,924E-06 | 0,018654836 | 2,5070128 | Intron (ENSMUST000001( | 5 | 66609610 | 66704300 | 94691 | 2 | 11787 | ENSMUST00000161716.8 | -3142 | ENSMUSG00000029207 | Apbb2 | amyloid beta (A4) precursor protein-binding, family B, member 2 |
| chr5 | 81812685 | 81813496 | 812 | * | 7,094504 | 3,555963805 | 1,327348 | 2,678998 | 0,007384 | 1 | 0 | Intron (ENSMUST0000( | 5 | 81808150 | 81944481 | 136332 | 1 | 319387 | ENSMUST00000124117.2 | 4535 | ENSMUSG00000037605 | Adgrl3 | adhesion G protein-coupled receptor L3 |
| chr2 | 11690158 | 11690853 | 696 | * | 8,217004 | 3,556439345 | 1,288613 | 2,759898 | 0,005782 | 0,790460916 | 1,449706 | Intron (ENSMUST0000( | 2 | 11710101 | 11739128 | 29028 | 1 | 16169 | ENSMUST00000138349.8 | -19248 | ENSMUSG00000023206 | Il15ra | interleukin 15 receptor, alpha chain |
| chr2 | 1,58E+08 | 1,58E+08 | 1074 | * | 9,44116 | 3,556539366 | 1,226568 | 2,899586 | 0,003737 | 0,723275538 | 1,360622 | Exon (ENSMUST00000 | 2 | 1,58E+08 | 1,58E+08 | 26438 | 1 | 329547 | ENSMUST00000065039.3 | 23338 | ENSMUSG00000052922 | Bpi | bactericidal permeablility increasing protein |
| chr12 | 70498958 | 70500218 | 1261 | * | 12,23042 | 3,556914821 | 1,189261 | 2,990861 | 0,002782 | 0,71419937 | 1,075735 | Promoter (<=1kb) | 12 | 70499869 | 70514814 | 14946 | 1 | 72736 | ENSMUST00000021471.13 | 0 | ENSMUSG00000021072 | Tmx1 | thioredoxin-related transmembrane protein 1 |
| chr11 | 1,2E+08 | 1,2E+08 | 1247 | * | 10,46298 | 3,558237247 | 1,312813 | 2,710391 | 0,00672 | 0,838625535 | 1,2763 | Intron (ENSMUST0000( | 11 | 1,2E+08 | 1,2E+08 | 3676 | 2 | 72055 | ENSMUST00000152859.2 | 14595 | ENSMUSG00000061306 | Slc38a10 | solute carrier family 38, member 10 |
| chr2 | 1,81E+08 | 1,81E+08 | 1025 | * | 6,86722 | 3,559465968 | 1,377571 | 2,583872 | 0,00977 | 1 | 0 | Intron (ENSMUST0000( | 2 | 1,81E+08 | 1,81E+08 | 31157 | 2 | 229004 | ENSMUST00000130475.8 | 3308 | ENSMUSG00000038705 | Gmeb2 | glucocorticoid modulatory element binding protein 2 |
| chr17 | 25795176 | 25796210 | 1035 | * | 8,46776 | 3,560269437 | 1,296978 | 2,74505 | 0,00605 | 1 | 0 | Promoter (1-2kb) | 17 | 25798059 | 25875453 | 77395 | 1 | 76483 | ENSMUST00000141606.3 | -1849 | ENSMUSG00000002279 | Lmf1 | lipase maturation factor 1 |
| chr11 | 99465096 | 99466640 | 1545 | * | 12,62639 | 3,561980042 | 1,060747 | 3,357992 | 0,000785 | 0,55292773 | 0,738441 | Distal Intergenic | 11 | 99470802 | 99471842 | 1041 | 2 | 69664 | ENSMUST00000054532.5 | 5202 | ENSMUSG00000047253 | Krtap1-5 | keratin associated protein 1-5 |
| chr3 | 87565286 | 87566024 | 739 | * | 8,126455 | 3,562336926 | 1,230446 | 2,89516 | 0,00379 | 1 | 0 | Intron (ENSMUST0000( | 3 | 87526068 | 87596157 | 70090 | 1 | 213498 | ENSMUST00000212015.2 | 39218 | ENSMUSG00000041977 | Arhgef11 | Rho guanine nucleotide exchange factor (GEF) 11 |
| chr17 | 19277106 | 19277946 | 841 | * | 8,067245 | 3,563615929 | 1,284964 | 2,773319 | 0,005549 | 1 | 0 | Intron (ENSMUST0000( | 17 | 19273722 | 19302673 | 28952 | 1 | 224552 | ENSMUST00000170424.3 | 3384 | ENSMUSG00000096717 | Vmn2r98 | vomeronasal 2, receptor 98 |
| chr7 | 27153682 | 27155546 | 1865 | * | 14,49502 | 3,563801556 | 1,193963 | 2,984851 | 0,002837 | 0,693610666 | 1,214904 | Promoter (1-2kb) | 7 | 27151869 | 27162120 | 10252 | 1 | 233016 | ENSMUST00000153644.2 | 1813 | ENSMUSG00000040466 | Blvrb | biliverdin reductase B (flavin reductase (NADPH)) |
| chr1 | 54852848 | 54853984 | 1137 | * | 10,46881 | 3,563939992 | 1,146969 | 3,107267 | 0,001888 | 0,637143933 | 1,173325 | Promoter (<=1kb) | 1 | 54801527 | 54852602 | 51076 | 2 | 329154 | ENSMUST00000178905.2 | -246 | ENSMUSG00000052331 | Ankrd44 | ankyrin repeat domain 44 |
| chr2 | 1,68E+08 | 1,68E+08 | 1379 | * | 12,86569 | 3,565199475 | 1,239663 | 2,875944 | 0,004028 | 0,748971518 | 1,2763 | Distal Intergenic | 2 | 1,68E+08 | 1,68E+08 | 675 | 2 | 1,03E+08 | ENSMUST00000246946.1 | 32573 | | Gm32800 | predicted gene, 32800 |
| chr8 | 76536347 | 76536853 | 507 | * | 7,209284 | 3,565342411 | 1,325451 | 2,689908 | 0,007147 | 1 | 0 | Intron (ENSMUST0000( | 8 | 76748557 | 76748683 | 127 | 2 | 1,15E+08 | ENSMUST00000104349.3 | 211830 | ENSMUSG00000077537 | Gm22544 | predicted gene, 22544 |
| chr15 | 11475186 | 11476485 | 1300 | * | 8,069741 | 3,566292847 | 1,378427 | 2,587218 | 0,009675 | 1 | 0 | Distal Intergenic | 15 | 11382387 | 11399751 | 17365 | 2 | 110960 | ENSMUST00000022849.7 | -75435 | ENSMUSG00000022241 | Tars | threonyl-tRNA synthetase |
| chr11 | 1,17E+08 | 1,17E+08 | 1112 | * | 14,19392 | 3,567771595 | 1,017706 | 3,5057 | 0,000455 | 0,421090004 | 0,938682 | Promoter (<=1kb) | 11 | 1,17E+08 | 1,17E+08 | 3817 | 2 | 107817 | ENSMUST00000124831.2 | 0 | ENSMUSG00000056962 | Jmjd6 | jumonji domain containing 6 |
| chr2 | 1,41E+08 | 1,41E+08 | 1471 | * | 8,818827 | 3,568116337 | 1,359987 | 2,623639 | 0,0087 | 0,891097601 | 1,211315 | Intron (ENSMUST0000( | 2 | 1,41E+08 | 1,41E+08 | 56561 | 2 | 1,03E+08 | ENSMUST00000151538.2 | 33377 | | Macrod2os1 | mono-ADP ribosylhydrolase 2, opposite strand 1 |
| chr19 | 53739853 | 53741265 | 1413 | * | 12,8407 | 3,568397171 | 1,13282 | 3,150014 | 0,001633 | 0,684274806 | 0,738441 | Intron (ENSMUST0000( | 19 | 53781739 | 53802004 | 20266 | 1 | 73713 | ENSMUST00000162910.2 | -40474 | ENSMUSG00000043639 | Rbm20 | RNA binding motif protein 20 |
| chr9 | 55082552 | 55083738 | 1187 | * | 10,21052 | 3,56945226 | 1,31651 | 2,711298 | 0,006702 | 0,81655291 | 1,449706 | Exon (ENSMUST00000( | 9 | 55102129 | 55113494 | 11366 | 1 | 109161 | ENSMUST00000148468.2 | -18391 | ENSMUSG00000032307 | Ube2q2 | ubiquitin-conjugating enzyme E2Q family member 2 |
| chr4 | 59472686 | 59474195 | 1510 | * | 13,96756 | 3,570943204 | 1,041588 | 3,428365 | 0,000607 | 0,508068302 | 0,738441 | 3' UTR | 4 | 59476935 | 59485772 | 8838 | 2 | 230257 | ENSMUST00000173268.8 | 11577 | ENSMUSG00000028382 | Ptbp3 | polypyrimidine tract binding protein 3 |
| chr1 | 1,55E+08 | 1,55E+08 | 875 | * | 9,911271 | 3,571166078 | 1,374311 | 2,598515 | 0,009363 | 0,919753338 | 1,086118 | Promoter (2-3kb) | 1 | 1,55E+08 | 1,55E+08 | 7047 | 2 | 433367 | ENSMUST00000097531.4 | -2898 | ENSMUSG00000073535 | Gm5532 | predicted gene 5532 |
| chr1 | 90822910 | 90823708 | 799 | * | 8,955125 | 3,572497008 | 1,386747 | 2,57617 | 0,00999 | 0,809505441 | 2,237403 | Distal Intergenic | 1 | 90842807 | 90878864 | 36058 | 1 | 171531 | ENSMUST00000027528.7 | -19099 | ENSMUSG00000026303 | Mlph | melanophilin |
| chr18 | 67857447 | 67858318 | 872 | * | 8,244479 | 3,576116213 | 1,190708 | 3,003352 | 0,00267 | 0,659642963 | 1,449706 | Promoter (<=1kb) | 18 | 67798581 | 67857643 | 59063 | 2 | 19255 | ENSMUST00000120934.8 | 0 | ENSMUSG00000024539 | Ptpn2 | protein tyrosine phosphatase, non-receptor type 2 |
| chr3 | 74713752 | 74715094 | 1343 | * | 9,07917 | 3,57643833 | 1,317267 | 2,715045 | 0,006627 | 0,824134241 | 1,360622 | Distal Intergenic | 3 | 74968668 | 75051010 | 82343 | 2 | 213234 | ENSMUST00000107776.8 | 335916 | ENSMUSG00000034151 | Zbbx | zinc finger, B-box domain containing |
| chr19 | 16629588 | 16630574 | 987 | * | 8,462973 | 3,577299081 | 1,23179 | 2,904147 | 0,003683 | 0,71419937 | 1,425929 | 3' UTR | 19 | 16612571 | 16642012 | 29442 | 2 | 271564 | ENSMUST00000225233.2 | 11438 | ENSMUSG00000046230 | Vps13a | vacuolar protein sorting 13A |
| chr3 | 84219410 | 84221608 | 2199 | * | 12,44058 | 3,577595849 | 1,153151 | 3,102453 | 0,001919 | 0,624579927 | 1,2763 | Distal Intergenic | 3 | 84099518 | 84212753 | 113236 | 2 | 80890 | ENSMUST00000132283.8 | -6657 | ENSMUSG00000027993 | Trim2 | tripartite motif-containing 2 |
| chr8 | 54216831 | 54217806 | 976 | * | 8,095819 | 3,577763606 | 1,299961 | 2,752208 | 0,005919 | 0,802925002 | 1,360622 | Distal Intergenic | 8 | 54039902 | 54092100 | 52199 | 2 | 234258 | ENSMUST00000047768.11 | -124731 | ENSMUSG00000039396 | Neil3 | nei like 3 (E. coli) |
| chr9 | 38175592 | 38176923 | 1332 | * | 11,94395 | 3,578869551 | 1,115268 | 3,208977 | 0,001332 | 0,568412386 | 1,173325 | Promoter (2-3kb) | 9 | 38179147 | 38181031 | 1885 | 1 | 258875 | ENSMUST00000212156.3 | -2224 | ENSMUSG00000096427 | Olfr895 | olfactory receptor 895 |
| chr9 | 108151214 | 108153602 | 2389 | * | 20,047959 | 3,579171485 | 0,7946602 | 4,5040274 | 6,668E-06 | 0,018663151 | 2,3860078 | Distal Intergenic | 9 | 108081833 | 108141157 | 59325 | 2 | 13138 | ENSMUST00000191899.6 | -10057 | ENSMUSG00000039952 | Dag1 | dystroglycan 1 |
| chr3 | 1,05E+08 | 1,05E+08 | 2157 | * | 24,22537 | 3,579262385 | 0,865531 | 4,135339 | 3,54E-05 | 0,053529752 | 2,951933 | Distal Intergenic | 3 | 1,05E+08 | 1,05E+08 | 51232 | 2 | 80281 | ENSMUST00000077548.12 | -5465 | ENSMUSG00000062127 | Cttnbp2nl | CTTNBP2 N-terminal like |
| chr1 | 45978187 | 45979031 | 845 | * | 8,480075 | 3,580087436 | 1,168226 | 3,064551 | 0,00218 | 0,627785088 | 1,425929 | Distal Intergenic | 1 | 45960377 | 45965679 | 5303 | 2 | 53945 | ENSMUST00000187420.7 | -12508 | ENSMUSG00000025993 | Slc40a1 | solute carrier family 40 (iron-regulated transporter), member 1 |
| chr10 | 9825250 | 9827199 | 1950 | * | 9,787151 | 3,580163178 | 1,20404 | 2,973459 | 0,002945 | 0,722439117 | 1,086118 | Distal Intergenic | 10 | 9852434 | 10083725 | 231292 | 1 | 1,03E+08 | ENSMUST00000219328.2 | -25235 | | Gm28905 | predicted gene 28905 |
| chr2 | 29354636 | 29355513 | 878 | * | 11,29312 | 3,581774579 | 1,240215 | 2,888028 | 0,003877 | 0,838625535 | 0,738441 | Intron (ENSMUST0000( | 2 | 29365227 | 29365306 | 80 | 2 | 1,02E+08 | ENSMUST00000184068.3 | 9793 | ENSMUSG00000098539 | Mir133c | microRNA 133c |
| chr16 | 15214283 | 15215388 | 1106 | * | 11,62129 | 3,582335322 | 1,240127 | 2,888683 | 0,003869 | 0,753370437 | 1,173325 | Distal Intergenic | 16 | 15373745 | 15412336 | 38592 | 2 | 70620 | ENSMUST00000118236.8 | 196948 | ENSMUSG00000022674 | Ube2v2 | ubiquitin-conjugating enzyme E2 variant 2 |
| chr10 | 13845428 | 13846482 | 1055 | * | 8,477361 | 3,582851708 | 1,219289 | 2,938475 | 0,003298 | 0,687937632 | 1,449706 | Promoter (2-3kb) | 10 | 13842768 | 13955419 | 112652 | 1 | 15273 | ENSMUST00000191464.2 | 2660 | ENSMUSG00000015501 | Hivep2 | human immunodeficiency virus type I enhancer binding protein 2 |
| chr13 | 43920655 | 43921416 | 762 | * | 7,939163 | 3,583302327 | 1,358816 | 2,637076 | 0,008362 | 1 | 0 | Distal Intergenic | 13 | 43938251 | 43956608 | 18358 | 1 | 12522 | ENSMUST00000015540.4 | -16835 | ENSMUSG00000015396 | Cd83 | CD83 antigen |
| chr19 | 3844601 | 3845932 | 1332 | * | 11,33109 | 3,583924227 | 1,147255 | 3,123912 | 0,001785 | 0,669377201 | 0,918255 | Promoter (<=1kb) | 19 | 3846530 | 3855046 | 8517 | 1 | 225888 | ENSMUST00000236012.2 | -598 | ENSMUSG00000045098 | Kmt5b | lysine methyltransferase 5B |
| chr5 | 45818987 | 45820397 | 1411 | * | 12,66957 | 3,584445962 | 1,142798 | 3,136552 | 0,001709 | 0,634036279 | 1,075735 | Exon (ENSMUST00000 | 5 | 45826044 | 45827050 | 1007 | 2 | 69788 | ENSMUST00000196242.2 | 6653 | ENSMUSG00000104986 | 1600023N17Rik | RIKEN cDNA 1600023N17 gene |
| chr16 | 31327037 | 31328732 | 1696 | * | 13,52868 | 3,584817314 | 1,032149 | 3,473158 | 0,000514 | 0,391231126 | 1,2763 | Promoter (<=1kb) | 16 | 31327205 | 31330595 | 3391 | 1 | 1,08E+08 | ENSMUST00000231805.2 | 0 | | Gm46560 | predicted gene, 46560 |
| chr7 | 1,14E+08 | 1,14E+08 | 1296 | * | 10,87176 | 3,585389038 | 1,126166 | 3,183713 | 0,001454 | 0,586565597 | 1,173325 | Distal Intergenic | 7 | 1,14E+08 | 1,14E+08 | 106690 | 1 | 233752 | ENSMUST00000161800.3 | -5103 | ENSMUSG00000048782 | Insc | INSC spindle orientation adaptor protein |
| chr3 | 30810191 | 30811362 | 1172 | * | 9,883046 | 3,586467579 | 1,286627 | 2,787497 | 0,005312 | 0,805745288 | 1,211315 | 3' UTR | 3 | 30805194 | 30819644 | 14451 | 1 | 75953 | ENSMUST00000172593.2 | 4997 | ENSMUSG00000051860 | Samd7 | sterile alpha motif domain containing 7 |
| chr4 | 1,32E+08 | 1,32E+08 | 1195 | * | 8,202675 | 3,586478474 | 1,355015 | 2,646819 | 0,008125 | 0,773870972 | 2,237403 | Intron (ENSMUST0000( | 4 | 1,32E+08 | 1,32E+08 | 20739 | 2 | 100169 | ENSMUST00000136711.2 | 12446 | ENSMUSG00000066043 | Phactr4 | phosphatase and actin regulator 4 |
| chr5 | 1,2E+08 | 1,2E+08 | 1358 | * | 9,917366 | 3,586543361 | 1,358109 | 2,640836 | 0,00827 | 0,901463351 | 1,086118 | Distal Intergenic | 5 | 1,2E+08 | 1,2E+08 | 9160 | 2 | 1,03E+08 | ENSMUST00000201435.2 | 26859 | ENSMUSG00000106938 | Gm31314 | predicted gene, 31314 |
| chr12 | 79429636 | 79431141 | 1506 | * | 10,07429 | 3,588323738 | 1,339805 | 2,678244 | 0,007401 | 0,833355026 | 1,449706 | Intron (ENSMUST0000( | 12 | 79374127 | 79748630 | 374504 | 1 | 19363 | ENSMUST00000221257.2 | 55509 | ENSMUSG00000059060 | Rad51b | RAD51 paralog B |
| chr6 | 88750711 | 88751145 | 435 | * | 6,960884 | 3,589549132 | 1,369041 | 2,621945 | 0,008743 | 1 | 0 | Intron (ENSMUST0000( | 6 | 88693404 | 88702842 | 9439 | 2 | 1,03E+08 | ENSMUST00000181355.4 | -47869 | | Gm26588 | predicted gene, 26588 |
| chrX | 1,2E+08 | 1,2E+08 | 1363 | * | 8,879492 | 3,590456206 | 1,267387 | 2,83296 | 0,004612 | 0,758346111 | 1,360622 | Distal Intergenic | 20 | 1,2E+08 | 1,2E+08 | 128 | 1 | 1,15E+08 | ENSMUST00000104247.3 | 83635 | ENSMUSG00000077435 | Gm22590 | predicted gene, 22590 |
| chr10 | 42367153 | 42369151 | 1999 | * | 17,49991 | 3,591231148 | 0,928348 | 3,868411 | 0,00011 | 0,123779239 | 2,386008 | Distal Intergenic | 10 | 42378026 | 42410783 | 32758 | 1 | 54198 | ENSMUST00000105500.8 | -8875 | ENSMUSG00000019804 | Snx3 | sorting nexin 3 |
| chr5 | 1,21E+08 | 1,21E+08 | 839 | * | 9,170735 | 3,591798139 | 1,390815 | 2,582513 | 0,009808 | 0,883970984 | 1,449706 | Intron (ENSMUST0000( | 5 | 1,21E+08 | 1,21E+08 | 5696 | 2 | 252972 | ENSMUST00000202072.2 | 10048 | ENSMUSG00000032741 | Tpcn1 | two pore channel 1 |
| chr16 | 81556517 | 81557751 | 1235 | * | 8,883845 | 3,591937191 | 1,233777 | 2,911334 | 0,003599 | 0,73648081 | 1,211315 | Distal Intergenic | 16 | 81262511 | 81314918 | 52408 | 1 | 17968 | ENSMUST00000232550.2 | 294006 | ENSMUSG00000022762 | Ncam2 | neural cell adhesion molecule 2 |
| chr1 | 1,04E+08 | 1,04E+08 | 559 | * | 7,49641 | 3,592633552 | 1,360199 | 2,641256 | 0,00826 | 1 | 0 | Distal Intergenic | 1 | 1,03E+08 | 1,03E+08 | 134 | 2 | 1,15E+08 | ENSMUST00000157283.3 | -784475 | ENSMUSG00000087908 | Gm22034 | predicted gene, 22034 |
| chr3 | 95898873 | 95900230 | 1358 | * | 13,2894 | 3,592928336 | 1,119242 | 3,210145 | 0,001327 | 0,548877304 | 1,2763 | Promoter (2-3kb) | 3 | 95896131 | 95902980 | 6850 | 2 | 67220 | ENSMUST00000123006.8 | 2750 | ENSMUSG00000015745 | Plekho1 | pleckstrin homology domain containing, family O member 1 |
| chr2 | 1,27E+08 | 1,27E+08 | 663 | * | 6,576321 | 3,593634104 | 1,365702 | 2,631345 | 0,008505 | 1 | 0 | Distal Intergenic | 2 | 1,27E+08 | 1,27E+08 | 42845 | 2 | 66552 | ENSMUST00000028844.11 | -11917 | ENSMUSG00000027366 | Sppl2a | signal peptide peptidase like 2A |
| chr7 | 1,32E+08 | 1,32E+08 | 1463 | * | 10,98717 | 3,59364728 | 1,206305 | 2,979054 | 0,002891 | 0,786955198 | 0,744375 | Intron (ENSMUST0000( | 7 | 1,32E+08 | 1,32E+08 | 63552 | 2 | 77938 | ENSMUST00000106168.2 | 55492 | ENSMUSG00000030956 | Fam53b | family with sequence similarity 53, member B |
| chr13 | 48402474 | 48403800 | 1327 | * | 11,51126 | 3,593821276 | 1,166042 | 3,082068 | 0,002056 | 0,726422517 | 0,738441 | Intron (ENSMUST0000( | 13 | 48414704 | 48419502 | 4799 | 1 | 15904 | ENSMUST00000021810.3 | -10904 | ENSMUSG00000021379 | Id4 | inhibitor of DNA binding 4 |
| chr17 | 25233545 | 25234696 | 1152 | * | 10,94964 | 3,594103 | 1,040227 | 3,455114 | 0,00055 | 0,402652506 | 1,2763 | Promoter (<=1kb) | 17 | 25180200 | 25234204 | 54005 | 2 | 57354 | ENSMUST00000073337.8 | 0 | ENSMUSG00000038002 | Cramp1 | cramped chromatin regulator 1 |
| chr2 | 1,81E+08 | 1,81E+08 | 1381 | * | 17,17886 | 3,594175433 | 0,883321 | 4,068936 | 4,72E-05 | 0,069775067 | 2,663619 | Promoter (<=1kb) | 2 | 1,81E+08 | 1,81E+08 | 34649 | 1 | 13002 | ENSMUST00000072334.12 | 0 | ENSMUSG00000000826 | Dnajc5 | DnaJ heat shock protein family (Hsp40) member C5 |
| chr5 | 36986290 | 36988517 | 2228 | * | 19,70061 | 3,595161044 | 0,863509 | 4,163434 | 3,13E-05 | 0,05285139 | 2,663619 | Promoter (<=1kb) | 5 | 36964265 | 36987997 | 23733 | 2 | 17160 | ENSMUST00000031002.10 | 0 | ENSMUSG00000029119 | Man2b2 | mannosidase 2, alpha B2 |
| chr1 | 1,82E+08 | 1,82E+08 | 1166 | * | 8,290844 | 3,595398704 | 1,347512 | 2,668176 | 0,007626 | 0,840963129 | 1,425929 | Exon (ENSMUST00000 | 1 | 1,82E+08 | 1,82E+08 | 108899 | 2 | 13800 | ENSMUST00000193074.6 | 24803 | ENSMUSG00000022995 | Enah | ENAH actin regulator |
| chrX | 42496692 | 42497888 | 1197 | * | 7,457645 | 3,595501808 | 1,321697 | 2,720368 | 0,006521 | 1 | 0 | Intron (ENSMUST0000( | 20 | 41913235 | 42517880 | 604646 | 2 | 23963 | ENSMUST00000154433.8 | 19992 | ENSMUSG00000016150 | Tenm1 | teneurin transmembrane protein 1 |
| chr3 | 1,28E+08 | 1,28E+08 | 862 | * | 9,625914 | 3,596572927 | 1,202699 | 2,990417 | 0,002786 | 0,669586355 | 1,425929 | Distal Intergenic | 3 | 1,28E+08 | 1,28E+08 | 107025 | 2 | 402774 | ENSMUST00000196895.5 | -161018 | ENSMUSG00000105816 | D030025E07Rik | RIKEN cDNA D030025E07 gene |
| chr19 | 28602690 | 28603885 | 1196 | * | 8,246578 | 3,596763537 | 1,270811 | 2,830289 | 0,004651 | 0,749550311 | 1,449706 | Intron (ENSMUST0000( | 19 | 28509247 | 28644085 | 134839 | 2 | 226075 | ENSMUST00000159178.2 | 40200 | ENSMUSG00000052942 | Glis3 | GLIS family zinc finger 3 |
| chr14 | 1,17E+08 | 1,17E+08 | 688 | * | 6,938192 | 3,597010559 | 1,361955 | 2,641063 | 0,008265 | 1 | 0 | Exon (ENSMUST00000 | 14 | 1,17E+08 | 1,18E+08 | 701884 | 1 | 23888 | ENSMUST00000123239.2 | -174986 | ENSMUSG00000058571 | Gpc6 | glypican 6 |
| chr17 | 58730803 | 58731808 | 1006 | * | 7,291896 | 3,597541722 | 1,239526 | 2,902353 | 0,003704 | 1 | 0 | Distal Intergenic | 17 | 59185803 | 59191128 | 5326 | 2 | 69239 | ENSMUST00000177108.2 | 459320 | ENSMUSG00000024227 | Pdzph1 | PDZ and pleckstrin homology domains 1 |
| chr7 | 39167022 | 39168364 | 1343 | * | 8,009027 | 3,59756122 | 1,313055 | 2,739841 | 0,006147 | 1 | 0 | Promoter (<=1kb) | 7 | 39167190 | 39189844 | 22655 | 1 | 70227 | ENSMUST00000108015.4 | 0 | ENSMUSG00000068959 | Zfp619 | zinc finger protein 619 |
| chr9 | 46209105 | 46210551 | 1447 | * | 13,2063 | 3,598175497 | 1,156441 | 3,111423 | 0,001862 | 0,710505631 | 0,738441 | 3' UTR | 9 | 46202646 | 46210226 | 7581 | 1 | 215051 | ENSMUST00000213932.2 | 6459 | ENSMUSG00000032077 | Bud13 | BUD13 homolog |
| chr11 | 27067616 | 27069105 | 1490 | * | 12,2702 | 3,598783769 | 1,012377 | 3,554787 | 0,000378 | 0,347797317 | 1,173325 | Distal Intergenic | 11 | 27268847 | 27268978 | 132 | 2 | 1,15E+08 | ENSMUST00000157813.3 | 199873 | ENSMUSG00000119065 | Gm25387 | predicted gene, 25387 |
| chr5 | 1,23E+08 | 1,23E+08 | 2145 | * | 13,76197 | 3,600409217 | 0,998652 | 3,60527 | 0,000312 | 0,352527836 | 0,938682 | Intron (ENSMUST0000( | 5 | 1,23E+08 | 1,23E+08 | 14153 | 1 | 109305 | ENSMUST00000152213.3 | 4975 | ENSMUSG00000049686 | Orai1 | ORAI calcium release-activated calcium modulator 1 |
| chr1 | 34161921 | 34163957 | 2037 | * | 21,660774 | 3,600475969 | 0,7525051 | 4,7846534 | 1,713E-06 | 0,005619226 | 2,9519335 | Promoter (1-2kb) | 1 | 34160331 | 34193602 | 33272 | 1 | 13518 | ENSMUST00000183006.5 | 1590 | ENSMUSG00000026131 | Dst | dystonin |
| chr4 | 99785886 | 99788324 | 2439 | * | 57,335032 | 3,600520974 | 0,8677511 | 4,1492553 | 3,336E-05 | 0,049832569 | 3,177995 | Promoter (<=1kb) | 4 | 99654139 | 99787010 | 132872 | 2 | 67733 | ENSMUST00000123830.8 | 0 | ENSMUSG00000028549 | Itgb3bp | integrin beta 3 binding protein (beta3-endonexin) |
| chr5 | 1,16E+08 | 1,16E+08 | 819 | * | 8,13317 | 3,601105276 | 1,343927 | 2,67954 | 0,007372 | 0,836351745 | 1,425929 | Intron (ENSMUST0000( | 5 | 1,16E+08 | 1,16E+08 | 119739 | 2 | 269693 | ENSMUST00000086483.4 | 78351 | ENSMUSG00000043913 | Ccdc60 | coiled-coil domain containing 60 |
| chr5 | 32615121 | 32616568 | 1448 | * | 17,504362 | 3,602375878 | 0,8479117 | 4,248527 | 2,152E-05 | 0,04827315 | 2,1545044 | Promoter (<=1kb) | 5 | 32616187 | 32652867 | 36681 | 1 | 19046 | ENSMUST00000015100.15 | 0 | ENSMUSG00000014956 | Ppp1cb | protein phosphatase 1 catalytic subunit beta |
| chr5 | 1,49E+08 | 1,49E+08 | 1289 | * | 11,14042 | 3,602504993 | 1,165307 | 3,091466 | 0,001992 | 0,64497937 | 1,173325 | Intron (ENSMUST0000( | 5 | 1,49E+08 | 1,5E+08 | 83216 | 1 | 381693 | ENSMUST00000110502.7 | -22192 | ENSMUSG00000029658 | Wdr95 | WD40 repeat domain 95 |
| chr10 | 79672924 | 79674021 | 1098 | * | 9,806256 | 3,602626924 | 1,306899 | 2,756622 | 0,00584 | 0,792545355 | 1,425929 | Distal Intergenic | 10 | 79665120 | 79666114 | 995 | 1 | 78906 | ENSMUST00000218531.2 | 7804 | ENSMUSG00000035852 | Misp | mitotic spindle positioning |
| chr9 | 24333195 | 24334367 | 1173 | * | 8,119657 | 3,603449355 | 1,305254 | 2,760727 | 0,005767 | 0,714185185 | 2,237403 | Promoter (<=1kb) | 9 | 24323347 | 24335255 | 11909 | 2 | 244745 | ENSMUST00000152480.2 | 888 | ENSMUSG00000043067 | Dpy19l1 | dpy-19-like 1 (C. elegans) |
| chr17 | 46695157 | 46696468 | 1312 | * | 9,205708 | 3,60366538 | 1,302826 | 2,766036 | 0,005674 | 0,79031971 | 1,425929 | Promoter (<=1kb) | 17 | 46694677 | 46731846 | 37170 | 1 | 57908 | ENSMUST00000138127.8 | 480 | ENSMUSG00000015597 | Zfp318 | zinc finger protein 318 |
| chr4 | 1,37E+08 | 1,37E+08 | 1120 | * | 9,007348 | 3,60515489 | 1,198019 | 3,009263 | 0,002619 | 0,683727424 | 1,211315 | Distal Intergenic | 4 | 1,37E+08 | 1,37E+08 | 2178 | 1 | 69380 | ENSMUST00000048893.4 | -6609 | ENSMUSG00000041399 | 1700013G24Rik | RIKEN cDNA 1700013G24 gene |
| chr2 | 1,68E+08 | 1,68E+08 | 1206 | * | 9,579329 | 3,606516702 | 1,193172 | 3,022629 | 0,002506 | 0,683140223 | 1,173325 | Promoter (<=1kb) | 2 | 1,68E+08 | 1,68E+08 | 5103 | 1 | 1E+08 | ENSMUST00000139927.2 | -372 | | Gm14321 | predicted gene 14321 |
| chr2 | 1,66E+08 | 1,66E+08 | 1106 | * | 8,901497 | 3,607226815 | 1,307034 | 2,759856 | 0,005783 | 0,790460916 | 1,449706 | 3' UTR | 2 | 1,66E+08 | 1,66E+08 | 4661 | 2 | 277360 | ENSMUST00000136974.2 | 4563 | ENSMUSG00000039621 | Prex1 | phosphatidylinositol-3,4,5-trisphosphate-dependent Rac exchange factor 1 |
| chr6 | 134903576 | 134906054 | 2479 | * | 49,03799 | 3,607245604 | 0,6241989 | 5,7789999 | 7,515E-09 | 8,5085E-05 | 2,7872553 | Promoter (<=1kb) | 6 | 134906055 | 134928539 | 22485 | 1 | 381822 | ENSMUST00000247190.1 | -1 | ENSMUSG00000098318 | Lockd | lncRNA downstream of Cdkn1b |
| chr11 | 96135867 | 96137012 | 1146 | * | 9,408499 | 3,608326507 | 1,268353 | 2,844892 | 0,004443 | 0,751231966 | 1,360622 | Distal Intergenic | 11 | 96141924 | 96160073 | 18150 | 1 | 193022 | ENSMUST00000243907.1 | -4912 | | Gm53 | predicted gene 53 |
| chr8 | 1,21E+08 | 1,21E+08 | 1306 | * | 13,22687 | 3,608932009 | 1,060545 | 3,402902 | 0,000667 | 0,461626601 | 1,075735 | Intron (ENSMUST0000( | 8 | 1,21E+08 | 1,21E+08 | 4641 | 1 | 22224 | ENSMUST00000135210.2 | -8142 | ENSMUSG00000031826 | Usp10 | ubiquitin specific peptidase 10 |
| chrX | 1,66E+08 | 1,66E+08 | 965 | * | 9,616094 | 3,609366934 | 1,196909 | 3,015573 | 0,002565 | 0,682206193 | 1,211315 | Exon (ENSMUST00000 | 20 | 1,66E+08 | 1,66E+08 | 6781 | 2 | 1,03E+08 | ENSMUST00000153701.2 | 5806 | | Gm15230 | predicted gene 15230 |
| chr17 | 10271300 | 10272311 | 1012 | * | 11,61679 | 3,609630852 | 1,253708 | 2,879165 | 0,003987 | 0,802671652 | 0,918255 | Distal Intergenic | 17 | 10145522 | 10148696 | 3175 | 2 | 1,19E+08 | ENSMUST00000232911.2 | -122604 | | Gm34799 | predicted gene, 34799 |
| chr9 | 53141813 | 53143159 | 1347 | * | 11,86878 | 3,609836334 | 1,136993 | 3,174899 | 0,001499 | 0,574952042 | 1,2763 | Intron (ENSMUST0000( | 9 | 53009935 | 53159353 | 149419 | 2 | 77591 | ENSMUST00000065630.8 | 16194 | ENSMUSG00000053289 | Ddx10 | DEAD box helicase 10 |
| chr15 | 72915516 | 72916648 | 1133 | * | 9,64486 | 3,610872804 | 1,398966 | 2,581102 | 0,009849 | 0,910268063 | 1,211315 | Intron (ENSMUST0000( | 15 | 72461469 | 72927659 | 466191 | 2 | 76510 | ENSMUST00000023276.16 | 11011 | ENSMUSG00000047921 | Trappc9 | trafficking protein particle complex 9 |
| chr19 | 6673584 | 6674347 | 764 | * | 7,302286 | 3,610949973 | 1,381371 | 2,614034 | 0,008948 | 1 | 0 | Intron (ENSMUST0000( | 19 | 6585875 | 6593062 | 7188 | 2 | 20521 | ENSMUST00000113451.9 | -80522 | ENSMUSG00000061742 | Slc22a12 | solute carrier family 22 (organic anion/cation transporter), member 12 |
| chr14 | 1,09E+08 | 1,09E+08 | 1406 | * | 7,328428 | 3,611365937 | 1,287448 | 2,805058 | 0,005031 | 1 | 0 | Distal Intergenic | 14 | 1,09E+08 | 1,09E+08 | 5359 | 2 | 76965 | ENSMUST00000100322.4 | 104327 | ENSMUSG00000075478 | Slitrk1 | SLIT and NTRK-like family, member 1 |
| chr4 | 1,56E+08 | 1,56E+08 | 1053 | * | 8,269732 | 3,612812259 | 1,180572 | 3,060222 | 0,002212 | 0,627995982 | 1,425929 | Promoter (1-2kb) | 4 | 1,56E+08 | 1,56E+08 | 8460 | 2 | 1,01E+08 | ENSMUST00000154931.2 | 1681 | ENSMUSG00000086682 | Gm16023 | predicted gene 16023 |
| chr2 | 72891994 | 72892927 | 934 | * | 8,032319 | 3,613456717 | 1,354796 | 2,667159 | 0,00765 | 1 | 0 | Distal Intergenic | 2 | 72817585 | 72818951 | 1367 | 1 | 1,01E+08 | ENSMUST00000137941.2 | 74409 | | Sp3os | trans-acting transcription factor 3, opposite strand |
| chr11 | 79119509 | 79120655 | 1147 | * | 9,951314 | 3,613820968 | 1,350456 | 2,676 | 0,007451 | 0,863495249 | 1,211315 | Distal Intergenic | 11 | 79131133 | 79133775 | 2643 | 2 | 78889 | ENSMUST00000153585.2 | 13120 | ENSMUSG00000017677 | Wsb1 | WD repeat and SOCS box-containing 1 |
| chr18 | 12307546 | 12308789 | 1244 | * | 7,36953 | 3,61399031 | 1,396249 | 2,588356 | 0,009644 | 1 | 0 | Intron (ENSMUST0000( | 18 | 12312627 | 12317972 | 5346 | 1 | 76482 | ENSMUST00000134046.8 | -3838 | ENSMUSG00000024410 | Rmc1 | regulator of MON1-CCZ1 |

| | | | | | | | | | | | | | | | | | | | | | | | |
|---|---|---|---|---|---|---|---|---|---|---|---|---|---|---|---|---|---|---|---|---|---|---|---|
| chr1 | 1,48E+08 | 1,48E+08 | 1161 | * | 11,99312 | 3,614361821 | 1,066158 | 3,390081 | 0,000699 | 0,534363468 | 0,738441 | Distal Intergenic | 1 | 1,48E+08 | 1,48E+08 | 129 | 2 | 1,15E+08 | ENSMUST00000157957.3 | 399856 | ENSMUSG00000088582 | Gm22966 | predicted gene, 22966 |
| chr2 | 1,64E+08 | 1,64E+08 | 1613 | * | 18,11526 | 3,614978297 | 0,983554 | 3,675425 | 0,000237 | 0,192725821 | 2,507013 | Promoter (<=1kb) | 2 | 1,64E+08 | 1,64E+08 | 4221 | 1 | 66460 | ENSMUST00000155250.2 | 558 | ENSMUSG00000045503 | Sys1 | SYS1 Golgi-localized integral membrane protein homolog (S. cerevisiae) |
| chr12 | 1,05E+08 | 1,05E+08 | 949 | * | 8,556464 | 3,615059147 | 1,336034 | 2,705814 | 0,006814 | 1 | 0 | Intron (ENSMUST0000( | 12 | 1,05E+08 | 1,05E+08 | 6596 | 2 | 94040 | ENSMUST00000222323.2 | -21755 | ENSMUSG00000021097 | Clmn | calmin |
| chr18 | 75457553 | 75460580 | 3028 | * | 73,515081 | 3,615708005 | 0,4269254 | 8,46918 | 2,471E-17 | 4,90356E-12 | 4,0557576 | Distal Intergenic | 18 | 75428746 | 75430400 | 1655 | 2 | 100503117 | ENSMUST00000186356.2 | -27153 | ENSMUSG00000102047 | 2010010A06Rik | RIKEN cDNA 2010010A06 gene |
| chr16 | 49757761 | 49758771 | 1011 | * | 7,715092 | 3,616147573 | 1,380489 | 2,619468 | 0,008807 | 1 | 0 | Distal Intergenic | 16 | 49687196 | 49731454 | 44259 | 1 | 16423 | ENSMUST00000229104.2 | 70565 | ENSMUSG00000055447 | Cd47 | CD47 antigen (Rh-related antigen, integrin-associated signal transducer) |
| chr11 | 79037565 | 79039432 | 1868 | * | 17,65268 | 3,616360252 | 0,906109 | 3,991087 | 6,58E-05 | 0,089931799 | 2,423594 | Promoter (<=1kb) | 11 | 78917533 | 79037233 | 119701 | 2 | 16706 | ENSMUST00000108264.8 | -332 | ENSMUSG00000018334 | Ksr1 | kinase suppressor of ras 1 |
| chr11 | 42202821 | 42203713 | 893 | * | 6,906251 | 3,61649921 | 1,375397 | 2,629422 | 0,008553 | 1 | 0 | Intron (ENSMUST0000( | 11 | 42197264 | 42211876 | 14613 | 2 | 14399 | ENSMUST00000020703.13 | 8163 | ENSMUSG00000020428 | Gabra6 | gamma-aminobutyric acid (GABA) A receptor, subunit alpha 6 |
| chr4 | 46508909 | 46509814 | 906 | * | 6,668102 | 3,617235794 | 1,333714 | 2,712152 | 0,006685 | 1 | 0 | 3' UTR | 4 | 46495155 | 46495285 | 131 | 1 | 1,15E+08 | ENSMUST00000082476.2 | 13754 | ENSMUSG00000064410 | Gm22247 | predicted gene, 22247 |
| chr2 | 1,27E+08 | 1,27E+08 | 943 | * | 10,45226 | 3,617386292 | 1,246901 | 2,901101 | 0,003719 | 0,829907213 | 0,744375 | Distal Intergenic | 2 | 1,27E+08 | 1,27E+08 | 3936 | 1 | 11552 | ENSMUST00000071902.5 | 4668 | ENSMUSG00000058620 | Adra2b | adrenergic receptor, alpha 2b |
| chr15 | 77812471 | 77814865 | 2395 | * | 42,92214 | 3,61833827 | 1,154394 | 3,134405 | 0,001722 | 0,490999135 | 2,338387 | Promoter (<=1kb) | 15 | 77799247 | 77813206 | 13960 | 2 | 56551 | ENSMUST00000005487.12 | 0 | ENSMUSG00000005354 | Txn2 | thioredoxin 2 |
| chr4 | 124586408 | 124588732 | 2325 | * | 32,698562 | 3,618579692 | 0,6515516 | 5,5537886 | 2,795E-08 | 0,000243164 | 2,7872553 | Promoter (<=1kb) | 4 | 124571953 | 124587393 | 15441 | 2 | 67205 | ENSMUST00000030738.8 | 0 | ENSMUSG00000028907 | Utp11 | UTP11 small subunit processome component |
| chr3 | 65564706 | 65566842 | 2137 | * | 26,558135 | 3,618612265 | 0,7789392 | 4,6455646 | 3,391E-06 | 0,009572437 | 2,8800171 | Promoter (<=1kb) | 3 | 65566709 | 65566847 | 139 | 1 | 102465905 | ENSMUST00000185035.3 | 0 | ENSMUSG00000098944 | Mir8120 | microRNA 8120 |
| chr15 | 7648730 | 7649793 | 1064 | * | 7,799852 | 3,619637248 | 1,286095 | 2,81444 | 0,004886 | 1 | 0 | Distal Intergenic | 15 | 7840327 | 7867056 | 26730 | 1 | 14573 | ENSMUST00000022744.5 | -190534 | ENSMUSG00000022144 | Gdnf | glial cell line derived neurotrophic factor |
| chr4 | 1,2E+08 | 1,2E+08 | 1451 | * | 16,06809 | 3,619833862 | 0,930921 | 3,888443 | 0,000101 | 0,173810376 | 1,2763 | Intron (ENSMUST0000( | 4 | 1,2E+08 | 1,2E+08 | 83038 | 1 | 29871 | ENSMUST00000106301.8 | -20126 | ENSMUSG00000000085 | Scmh1 | sex comb on midleg homolog 1 |
| chr12 | 87757157 | 87758505 | 1349 | * | 8,97088 | 3,62033904 | 1,249553 | 2,897307 | 0,003764 | 0,72508691 | 1,360622 | Distal Intergenic | 12 | 87739195 | 87742525 | 3331 | 2 | 1,01E+08 | ENSMUST00000164517.3 | -14632 | ENSMUSG00000095724 | Eif1ad19 | eukaryotic translation initiation factor 1A domain containing 19 |
| chr10 | 80774440 | 80777185 | 2746 | * | 85,36086 | 3,62322225 | 0,943125 | 3,84172 | 0,000122 | 0,109316219 | 3,270621 | Distal Intergenic | 10 | 80766646 | 80767593 | 948 | 1 | 17873 | ENSMUST00000219449.2 | 7794 | ENSMUSG00000015312 | Gadd45b | growth arrest and DNA-damage-inducible 45 beta |
| chr3 | 1,18E+08 | 1,18E+08 | 932 | * | 10,81583 | 3,624076466 | 1,120815 | 3,23343 | 0,001223 | 0,641187423 | 0,744375 | Intron (ENSMUST0000( | 3 | 1,18E+08 | 1,18E+08 | 96057 | 1 | 1,03E+08 | ENSMUST00000181310.2 | 30889 | | Gm26871 | predicted gene, 26871 |
| chr9 | 56372235 | 56373443 | 1209 | * | 11,15608 | 3,624612124 | 1,217141 | 2,977972 | 0,002902 | 0,788774605 | 0,738441 | Intron (ENSMUST0000( | 9 | 56326114 | 56383401 | 57288 | 1 | 66867 | ENSMUST00000214869.2 | 46121 | ENSMUSG00000032329 | Hmg20a | high mobility group 20A |
| chr3 | 1,38E+08 | 1,38E+08 | 1272 | * | 10,55255 | 3,624863017 | 1,258739 | 2,879758 | 0,00398 | 0,802644208 | 0,918255 | Exon (ENSMUST00000( | 3 | 1,38E+08 | 1,38E+08 | 41524 | 2 | 17777 | ENSMUST00000029805.13 | 6704 | ENSMUSG00000028158 | Mttp | microsomal triglyceride transfer protein |
| chr1 | 180958517 | 180961595 | 3079 | * | 34,204833 | 3,624939461 | 0,7181523 | 5,0475917 | 4,474E-07 | 0,002001238 | 2,9254973 | Intron (ENSMUST000000: | 1 | 180914703 | 180971769 | 57067 | 2 | 67459 | ENSMUST00000027797.9 | 10174 | ENSMUSG00000026516 | Nvl | nuclear VCP-like |
| chr1 | 1,34E+08 | 1,34E+08 | 1550 | * | 13,46071 | 3,626366392 | 1,276185 | 2,841567 | 0,004489 | 0,822443391 | 0,938682 | Distal Intergenic | 1 | 1,34E+08 | 1,34E+08 | 3184 | 1 | 1,03E+08 | ENSMUST00000190927.2 | -11871 | ENSMUSG00000100830 | Gm28441 | predicted gene 28441 |
| chr15 | 81200140 | 81202112 | 1973 | * | 16,94578 | 3,627007399 | 0,97919 | 3,704091 | 0,000212 | 0,173939409 | 2,663619 | Distal Intergenic | 15 | 81134229 | 81197067 | 62839 | 2 | 78102 | ENSMUST00000230042.2 | -3073 | ENSMUSG00000115970 | 8430426J06Rik | RIKEN cDNA 8430426J06 gene |
| chr7 | 1,43E+08 | 1,43E+08 | 1185 | * | 9,776464 | 3,627374559 | 1,188513 | 3,052029 | 0,002273 | 0,669377201 | 1,173325 | Promoter (<=1kb) | 7 | 1,43E+08 | 1,43E+08 | 18259 | 2 | 79201 | ENSMUST00000152703.2 | -827 | ENSMUSG00000037613 | Tnfrsf23 | tumor necrosis factor receptor superfamily, member 23 |
| chr6 | 119369283 | 119371330 | 2048 | * | 32,290861 | 3,628293212 | 0,7391227 | 4,9089187 | 9,158E-07 | 0,003573187 | 2,7872553 | Promoter (2-3kb) | 6 | 119332427 | 119367209 | 34783 | 2 | 68465 | ENSMUST00000187699.7 | -2074 | ENSMUSG00000030168 | Adipor2 | adiponectin receptor 2 |
| chr11 | 96260553 | 96261792 | 1240 | * | 11,4329 | 3,629663708 | 1,298984 | 2,794233 | 0,005202 | 0,852340527 | 0,918255 | Distal Intergenic | 11 | 96257084 | 96258047 | 964 | 1 | 15407 | ENSMUST00000123805.2 | 3469 | ENSMUSG00000018973 | Hoxb1 | homeobox B1 |
| chr3 | 33533685 | 33534596 | 912 | * | 9,004085 | 3,62981828 | 1,31153 | 2,767621 | 0,005647 | 0,817671603 | 1,211315 | Distal Intergenic | 3 | 33447206 | 33447348 | 143 | 2 | 1,15E+08 | ENSMUST00000157635.3 | -86337 | ENSMUSG00000088260 | Gm24780 | predicted gene, 24780 |
| chr15 | 66868923 | 66871141 | 2219 | * | 54,068852 | 3,630415146 | 0,4696185 | 7,7305617 | 1,071E-14 | 1,26462E-09 | 2,7254908 | Intron (ENSMUST0000001 | 15 | 66818386 | 66841489 | 23104 | 2 | 17988 | ENSMUST00000164070.2 | -27434 | ENSMUSG00000005125 | Ndrg1 | N-myc downstream regulated gene 1 |
| chr15 | 76211278 | 76211962 | 685 | * | 8,643231 | 3,632411796 | 1,350792 | 2,689097 | 0,007165 | 0,749550311 | 2,237403 | Promoter (<=1kb) | 15 | 76211597 | 76214877 | 3281 | 1 | 109075 | ENSMUST00000059045.8 | 0 | ENSMUSG00000034259 | Exosc4 | exosome component 4 |
| chr2 | 1,58E+08 | 1,58E+08 | 659 | * | 6,513111 | 3,634470257 | 1,379853 | 2,633956 | 0,00844 | 1 | 0 | Promoter (2-3kb) | 2 | 1,58E+08 | 1,58E+08 | 15987 | 2 | 21817 | ENSMUST00000140923.8 | 2621 | ENSMUSG00000037820 | Tgm2 | transglutaminase 2, C polypeptide |
| chr13 | 69682952 | 69684464 | 1513 | * | 18,62457 | 3,634678099 | 0,97478 | 3,728716 | 0,000192 | 0,165924532 | 2,663619 | Promoter (<=1kb) | 13 | 69646078 | 69682736 | 36659 | 2 | 210106 | ENSMUST00000198607.5 | -216 | ENSMUSG00000034575 | Tent4a | terminal nucleotidyltransferase 4A |
| chr1 | 64254522 | 64255508 | 987 | * | 12,43078 | 3,634791284 | 1,147512 | 3,167542 | 0,001537 | 0,596263346 | 1,173325 | Intron (ENSMUST0000( | 1 | 64118262 | 64161441 | 43180 | 2 | 93691 | ENSMUST00000135075.2 | -93081 | ENSMUSG00000025959 | Klf7 | Kruppel-like factor 7 (ubiquitous) |
| chr7 | 1,45E+08 | 1,45E+08 | 952 | * | 7,236679 | 3,635955551 | 1,395212 | 2,606024 | 0,00916 | 1 | 0 | Distal Intergenic | 7 | 1,45E+08 | 1,45E+08 | 30089 | 2 | 233979 | ENSMUST00000058022.6 | -3534 | ENSMUSG00000048677 | Tpcn2 | two pore segment channel 2 |
| chr3 | 1,08E+08 | 1,08E+08 | 1722 | * | 13,62264 | 3,636179107 | 1,101959 | 3,29974 | 0,000968 | 0,547497234 | 0,938682 | Promoter (<=1kb) | 3 | 1,08E+08 | 1,08E+08 | 23049 | 1 | 26442 | ENSMUST00000090569.10 | 0 | ENSMUSG00000068749 | Psma5 | proteasome subunit alpha 5 |
| chr6 | 29765045 | 29766084 | 1040 | * | 8,027536 | 3,636678274 | 1,387283 | 2,62144 | 0,008756 | 1 | 0 | Promoter (2-3kb) | 6 | 29768442 | 29867418 | 98977 | 1 | 74340 | ENSMUST00000143707.2 | -2358 | ENSMUSG00000029772 | Ahcyl2 | S-adenosylhomocysteine hydrolase-like 2 |
| chr14 | 60926568 | 60927671 | 1104 | * | 8,795822 | 3,637326098 | 1,364786 | 2,665125 | 0,007696 | 0,83918951 | 1,449706 | Promoter (1-2kb) | 14 | 60929424 | 60947018 | 17595 | 1 | 219140 | ENSMUST00000160095.3 | -1753 | ENSMUSG00000021990 | Spata13 | spermatogenesis associated 13 |
| chr15 | 83033297 | 83034756 | 1460 | * | 24,642976 | 3,637346357 | 0,7000955 | 5,1955006 | 2,042E-07 | 0,001171089 | 2,6724296 | Promoter (<=1kb) | 15 | 83033845 | 83033995 | 151 | 1 | 104307 | ENSMUST00000083242.3 | 0 | ENSMUSG00000065176 | Rnu12 | RNA U12, small nuclear |
| chr4 | 1,48E+08 | 1,48E+08 | 622 | * | 6,308482 | 3,637769803 | 1,378666 | 2,638616 | 0,008325 | 1 | 0 | Distal Intergenic | 4 | 1,48E+08 | 1,48E+08 | 5804 | 1 | 230904 | ENSMUST00000047951.9 | 48712 | ENSMUSG00000041556 | Fbxo2 | F-box protein 2 |
| chr13 | 51553595 | 51555441 | 1847 | * | 15,28171 | 3,637940813 | 1,11855 | 3,252372 | 0,001144 | 0,533665288 | 1,214904 | Intron (ENSMUST0000( | 13 | 51562675 | 51576833 | 14159 | 1 | 13610 | ENSMUST00000087978.5 | -7234 | ENSMUSG00000067586 | S1pr3 | sphingosine-1-phosphate receptor 3 |
| chr2 | 1,55E+08 | 1,55E+08 | 1137 | * | 8,29454 | 3,638655109 | 1,346909 | 2,701485 | 0,006903 | 0,824070388 | 1,425929 | Promoter (2-3kb) | 2 | 1,55E+08 | 1,55E+08 | 3517 | 2 | 56406 | ENSMUST00000141928.2 | 2651 | ENSMUSG00000038369 | Ncoa6 | nuclear receptor coactivator 6 |
| chr12 | 52628684 | 52629673 | 990 | * | 8,86445 | 3,638779631 | 1,396491 | 2,605659 | 0,00917 | 0,88303073 | 1,360622 | Distal Intergenic | 12 | 52648378 | 52649101 | 724 | 2 | 73278 | ENSMUST00000191400.2 | 19428 | ENSMUSG00000100315 | 1700031P21Rik | RIKEN cDNA 1700031P21 gene |
| chr17 | 44995602 | 44996885 | 1284 | * | 9,425148 | 3,63919853 | 1,262148 | 2,883338 | 0,003935 | 0,731055602 | 1,360622 | Intron (ENSMUST0000( | 17 | 44950568 | 45045580 | 95013 | 2 | 12393 | ENSMUST00000160199.2 | 48695 | ENSMUSG00000039153 | Runx2 | runt related transcription factor 2 |
| chr19 | 27015197 | 27016462 | 1266 | * | 19,13684 | 3,639545855 | 0,879238 | 4,13943 | 3,48E-05 | 0,061989207 | 2,386008 | Distal Intergenic | 19 | 27079469 | 27080097 | 629 | 2 | 1,03E+08 | ENSMUST00000236708.2 | 63635 | | Gm35438 | predicted gene, 35438 |
| chr18 | 17636954 | 17637941 | 988 | * | 7,700911 | 3,639551432 | 1,30107 | 2,797353 | 0,005152 | 1 | 0 | Distal Intergenic | 18 | 17701803 | 17714047 | 12245 | 2 | 75162 | ENSMUST00000234987.2 | 76106 | ENSMUSG00000117615 | 4930545E07Rik | RIKEN cDNA 4930545E07 gene |
| chr13 | 23755122 | 23758629 | 3508 | * | 23,326899 | 3,639705494 | 0,8517974 | 4,2729711 | 1,929E-05 | 0,037473612 | 2,6724296 | Promoter (<=1kb) | 13 | 23758555 | 23759126 | 572 | 1 | 319165 | ENSMUST00000090776.7 | 0 | ENSMUSG00000071478 | H2ac7 | H2A clustered histone 7 |
| chr10 | 79991565 | 79992377 | 813 | * | 9,037697 | 3,639820854 | 1,21498 | 2,995787 | 0,002737 | 0,672782141 | 1,360622 | Exon (ENSMUST00000( | 10 | 79987072 | 79989530 | 2459 | 1 | 59090 | ENSMUST00000144526.2 | 4493 | ENSMUSG00000035621 | Midn | midnolin |
| chr10 | 55055996 | 55056814 | 819 | * | 9,144664 | 3,640158876 | 1,391865 | 2,615311 | 0,008915 | 0,863495249 | 1,449706 | Distal Intergenic | 10 | 54589072 | 54589133 | 62 | 2 | 1,15E+08 | ENSMUST00000158362.3 | -466863 | ENSMUSG00000088987 | Gm25602 | predicted gene, 25602 |
| chr13 | 95776528 | 95778714 | 2187 | * | 19,42521 | 3,641078343 | 0,921819 | 3,949885 | 7,82E-05 | 0,096906219 | 2,507013 | Intron (ENSMUST0000( | 13 | 95795083 | 95797606 | 2524 | 2 | 544963 | ENSMUST00000238404.2 | 18892 | ENSMUSG00000021676 | Iqgap2 | IQ motif containing GTPase activating protein 2 |
| chr4 | 1,23E+08 | 1,23E+08 | 1299 | * | 13,81462 | 3,641394386 | 1,101972 | 3,304435 | 0,000952 | 0,492839239 | 1,2763 | Exon (ENSMUST00000: | 4 | 1,23E+08 | 1,23E+08 | 8191 | 2 | 11426 | ENSMUST00000106213.9 | 25657 | ENSMUSG00000028649 | Macf1 | microtubule-actin crosslinking factor 1 |
| chr12 | 1,06E+08 | 1,06E+08 | 1212 | * | 8,850454 | 3,641494736 | 1,287045 | 2,829346 | 0,004664 | 0,760759863 | 1,360622 | Promoter (1-2kb) | 12 | 1,06E+08 | 1,06E+08 | 54114 | 1 | 18789 | ENSMUST00000109901.9 | -1488 | ENSMUSG00000021111 | Papola | poly (A) polymerase alpha |
| chr4 | 1,37E+08 | 1,37E+08 | 1422 | * | 11,89177 | 3,642058106 | 1,178168 | 3,091288 | 0,001993 | 0,645212515 | 1,173325 | Distal Intergenic | 4 | 1,37E+08 | 1,37E+08 | 1001 | 2 | 1,03E+08 | ENSMUST00000154285.2 | 20098 | | Gm13001 | predicted gene 13001 |
| chr18 | 89046806 | 89047862 | 1057 | * | 8,492926 | 3,643553916 | 1,282765 | 2,84039 | 0,004506 | 0,748971518 | 1,425929 | 3' UTR | 18 | 89053698 | 89093644 | 39947 | 1 | 246102 | ENSMUST00000235882.2 | -5836 | ENSMUSG00000023066 | Rttn | rotatin |
| chr7 | 91526697 | 91528161 | 1465 | * | 13,41097 | 3,643904251 | 1,005202 | 3,625045 | 0,000289 | 0,338590591 | 0,938682 | Intron (ENSMUST0000( | 7 | 91360628 | 92096211 | 735584 | 1 | 23859 | ENSMUST00000107193.8 | 166069 | ENSMUSG00000052572 | Dlg2 | discs large MAGUK scaffold protein 2 |
| chr18 | 63682849 | 63684488 | 1640 | * | 13,95338 | 3,64547052 | 1,095635 | 3,327267 | 0,000877 | 0,477622452 | 1,2763 | Distal Intergenic | 18 | 63666139 | 63666282 | 144 | 1 | 1,15E+08 | ENSMUST00000158101.3 | 16710 | ENSMUSG00000088726 | Gm22210 | predicted gene, 22210 |
| chr11 | 54421042 | 54421711 | 670 | * | 7,078839 | 3,645983604 | 1,381104 | 2,639905 | 0,008293 | 1 | 0 | Intron (ENSMUST0000( | 11 | 54413772 | 54587170 | 173399 | 1 | 192786 | ENSMUST00000207429.2 | 7270 | ENSMUSG00000037533 | Rapgef6 | Rap guanine nucleotide exchange factor (GEF) 6 |
| chr10 | 79878611 | 79879512 | 902 | * | 9,612063 | 3,646250776 | 1,235959 | 2,950139 | 0,003176 | 0,732824791 | 1,086118 | Promoter (2-3kb) | 10 | 79871783 | 79875629 | 3847 | 2 | 66420 | ENSMUST00000004786.10 | -2982 | ENSMUSG00000004667 | Polr2e | polymerase (RNA) II (DNA directed) polypeptide E |
| chr7 | 1,23E+08 | 1,23E+08 | 1525 | * | 11,78738 | 3,646379082 | 1,181286 | 3,086788 | 0,002023 | 0,722439117 | 0,744375 | Promoter (2-3kb) | 7 | 1,23E+08 | 1,23E+08 | 7699 | 1 | 19647 | ENSMUST00000206179.2 | 2542 | ENSMUSG00000030779 | Rbbp6 | retinoblastoma binding protein 6, ubiquitin ligase |
| chrX | 18577701 | 18578734 | 1034 | * | 6,290122 | 3,648666214 | 1,404329 | 2,598157 | 0,009373 | 1 | 0 | Distal Intergenic | 20 | 18281120 | 18327636 | 46517 | 2 | 75905 | ENSMUST00000147039.2 | -250065 | ENSMUSG00000037358 | Dipk2b | divergent protein kinase domain 2B |
| chr14 | 1,03E+08 | 1,03E+08 | 1347 | * | 8,688376 | 3,649965288 | 1,199638 | 3,042556 | 0,002346 | 0,648791129 | 1,360622 | Distal Intergenic | 14 | 1,03E+08 | 1,03E+08 | 16595 | 2 | 1,03E+08 | ENSMUST00000227348.2 | 223774 | ENSMUSG00000115816 | Gm34589 | predicted gene, 34589 |
| chr6 | 1,28E+08 | 1,28E+08 | 589 | * | 8,524262 | 3,650079415 | 1,363421 | 2,677147 | 0,007425 | 0,845724506 | 1,360622 | Promoter (2-3kb) | 6 | 1,28E+08 | 1,28E+08 | 10151 | 2 | 1,03E+08 | ENSMUST00000132657.2 | -2329 | | Gm15870 | predicted gene 15870 |
| chr13 | 34607403 | 34608518 | 1116 | * | 10,97841 | 3,650143 | 1,151317 | 3,170406 | 0,001522 | 0,642322948 | 0,918255 | Distal Intergenic | 13 | 34363141 | 34529165 | 166025 | 2 | 73102 | ENSMUST00000040336.12 | -78238 | ENSMUSG00000038267 | Slc22a23 | solute carrier family 22, member 23 |
| chr18 | 75625873 | 75629258 | 3386 | * | 88,948819 | 3,650371505 | 0,6996891 | 5,2171333 | 1,817E-07 | 0,000819457 | 4,0557576 | Intron (ENSMUST0000010 | 18 | 75647716 | 75655877 | 8162 | 1 | 100038353 | ENSMUST00000181913.2 | -18458 | ENSMUSG00000097712 | Gm10532 | predicted gene 10532 |
| chr17 | 48793148 | 48793906 | 759 | * | 8,845743 | 3,65090071 | 1,262955 | 2,89076 | 0,003843 | 0,716908183 | 1,449706 | Intron (ENSMUST0000( | 17 | 48769432 | 48846635 | 77204 | 1 | 76589 | ENSMUST00000162467.2 | 23716 | ENSMUSG00000043592 | Unc5cl | unc-5 family C-terminal like |
| chr11 | 84076004 | 84076953 | 950 | * | 7,141481 | 3,651219553 | 1,330667 | 2,743903 | 0,006071 | 0,809505441 | 1,360622 | Intron (ENSMUST0000( | 11 | 84070684 | 84105010 | 34327 | 1 | 107476 | ENSMUST00000136463.9 | 5320 | ENSMUSG00000020532 | Acaca | acetyl-Coenzyme A carboxylase alpha |
| chr4 | 55651697 | 55654427 | 2731 | * | 18,89087 | 3,651549121 | 1,074613 | 3,398015 | 0,000679 | 0,328694307 | 2,386008 | Distal Intergenic | 4 | 55605149 | 55636901 | 31753 | 1 | 1,03E+08 | ENSMUST00000123620.2 | 46548 | | Gm12506 | predicted gene 12506 |
| chr16 | 32886566 | 32887595 | 1030 | * | 8,655724 | 3,651787904 | 1,367509 | 2,670393 | 0,007576 | 0,758241749 | 2,237403 | Exon (ENSMUST00000( | 16 | 32882891 | 32948065 | 65175 | 1 | 239833 | ENSMUST00000023497.3 | 3675 | ENSMUSG00000022802 | Lmln | leishmanolysin-like (metallopeptidase M8 family) |
| chr17 | 26829505 | 26831125 | 1621 | * | 24,274233 | 3,652123834 | 0,7072322 | 5,1639669 | 2,418E-07 | 0,001311801 | 2,6724296 | Promoter (2-3kb) | 17 | 26826866 | 26868771 | 41906 | 1 | 67458 | ENSMUST00000170640.3 | 2639 | ENSMUSG00000001576 | Ergic1 | endoplasmic reticulum-golgi intermediate compartment (ERGIC) 1 |
| chr11 | 79989573 | 79990786 | 1214 | * | 8,666644 | 3,652348232 | 1,343666 | 2,718197 | 0,006564 | 0,823736512 | 1,360622 | Promoter (1-2kb) | 11 | 79992710 | 80026620 | 33911 | 1 | 237877 | ENSMUST00000151815.8 | -1924 | ENSMUSG00000017550 | Atad5 | ATPase family, AAA domain containing 5 |
| chr17 | 22801570 | 22802522 | 953 | * | 10,6383 | 3,652443039 | 1,122692 | 3,25329 | 0,001141 | 0,539604518 | 1,173325 | Distal Intergenic | 17 | 22763485 | 22792463 | 28979 | 2 | 210876 | ENSMUST00000234301.2 | -9107 | ENSMUSG00000095093 | Vmn2r111 | vomeronasal 2, receptor 111 |
| chr12 | 71260828 | 71261951 | 1124 | * | 9,562504 | 3,652527905 | 1,224284 | 2,9834 | 0,002851 | 0,669774031 | 1,449706 | Intron (ENSMUST0000( | 12 | 71197208 | 71201661 | 4454 | 1 | 76967 | ENSMUST00000134748.2 | 63620 | ENSMUSG00000034601 | 2700049A03Rik | RIKEN cDNA 2700049A03 gene |
| chr17 | 27425222 | 27426843 | 1622 | * | 11,9463 | 3,653058856 | 1,351485 | 2,702996 | 0,006872 | 0,929518563 | 0,744375 | Promoter (1-2kb) | 17 | 27408574 | 27423443 | 14870 | 2 | 224640 | ENSMUST00000055117.9 | -1779 | ENSMUSG00000044857 | Lemd2 | LEM domain containing 2 |
| chr5 | 32966849 | 32968454 | 1606 | * | 10,84442 | 3,653165636 | 1,144447 | 3,192079 | 0,001413 | 0,563858922 | 1,2763 | Intron (ENSMUST0000( | 5 | 32950983 | 32953448 | 2466 | 2 | 215476 | ENSMUST00000147752.2 | -13401 | ENSMUSG00000054280 | Prr14l | proline rich 14-like |
| chr19 | 45767849 | 45769668 | 1820 | * | 12,55428 | 3,653678078 | 1,355501 | 2,695445 | 0,007029 | 0,90199 | 0,918255 | Promoter (2-3kb) | 19 | 45738698 | 45771990 | 33293 | 2 | 76055 | ENSMUST00000026243.5 | 2322 | ENSMUSG00000025220 | Oga | O-GlcNAcase |
| chr13 | 50789681 | 50790450 | 770 | * | 8,518035 | 3,654044002 | 1,391339 | 2,626279 | 0,008632 | 0,78560377 | 2,237403 | Distal Intergenic | 13 | 50797268 | 50799710 | 2443 | 1 | 380845 | ENSMUST00000192039.6 | -6818 | ENSMUSG00000096641 | Gm904 | predicted gene 904 |
| chr14 | 30659888 | 30661038 | 1151 | * | 9,022744 | 3,654174909 | 1,238707 | 2,949991 | 0,003178 | 0,613712777 | 2,237403 | Exon (ENSMUST00000( | 14 | 30651137 | 30665234 | 14098 | 2 | 16424 | ENSMUST00000006704.17 | 4196 | ENSMUSG00000006529 | Itih1 | inter-alpha trypsin inhibitor, heavy chain 1 |
| chr10 | 67213349 | 67215466 | 2118 | * | 25,038759 | 3,654387101 | 0,7145849 | 5,1139997 | 3,154E-07 | 0,00149878 | 2,8585108 | Intron (ENSMUST000002: | 10 | 67259325 | 67260582 | 1258 | 1 | 102634442 | ENSMUST00000220168.2 | -43859 | | Gm32023 | predicted gene, 32023 |
| chr10 | 22546799 | 22547863 | 1065 | * | 11,47496 | 3,654522407 | 1,170102 | 3,123251 | 0,001789 | 0,699034389 | 0,744375 | Intron (ENSMUST0000( | 10 | 22524907 | 22540589 | 15683 | 1 | 353169 | ENSMUST00000159174.2 | 21892 | ENSMUSG00000037490 | Slc2a12 | solute carrier family 2 (facilitated glucose transporter), member 12 |
| chr1 | 26129253 | 26130560 | 1308 | * | 8,641494 | 3,654594931 | 1,254781 | 2,912536 | 0,003585 | 1 | 0 | Distal Intergenic | 1 | 25106631 | 25868788 | 762158 | 2 | 210933 | ENSMUST00000151309.8 | -260465 | ENSMUSG00000033569 | Adgrb3 | adhesion G protein-coupled receptor B3 |
| chr14 | 3280505 | 3281395 | 891 | * | 8,037181 | 3,654996044 | 1,389805 | 2,629862 | 0,008542 | 1 | 0 | Distal Intergenic | 14 | 3575429 | 3839361 | 263933 | 1 | 218793 | ENSMUST00000124353.2 | -294034 | ENSMUSG00000058317 | Ube2e2 | ubiquitin-conjugating enzyme E2E 2 |
| chr4 | 55532725 | 55534167 | 1443 | * | 13,04076 | 3,655846031 | 1,142801 | 3,199023 | 0,001379 | 0,55735526 | 1,2763 | Promoter (<=1kb) | 4 | 55532736 | 55541745 | 9010 | 1 | 1,03E+08 | ENSMUST00000244104.1 | 0 | | Gm34343 | predicted gene, 34343 |
| chr10 | 84227954 | 84228945 | 992 | * | 9,748288 | 3,655949683 | 1,27733 | 2,862181 | 0,004207 | 0,731734453 | 1,449706 | Promoter (<=1kb) | 10 | 84211160 | 84228252 | 17093 | 2 | 77976 | ENSMUST00000129355.2 | 0 | ENSMUSG00000020032 | Nuak1 | NUAK family, SNF1-like kinase, 1 |
| chr2 | 50931269 | 50931890 | 622 | * | 7,638245 | 3,656242254 | 1,37101 | 2,666823 | 0,007657 | 0,851704515 | 1,360622 | Distal Intergenic | 2 | 51027081 | 51039106 | 12026 | 2 | 74194 | ENSMUST00000140864.2 | 107216 | ENSMUSG00000017144 | Rnd3 | Rho family GTPase 3 |
| chr18 | 35113055 | 35114290 | 1236 | * | 10,03217 | 3,657649699 | 1,124914 | 3,251494 | 0,001148 | 0,505078837 | 1,425929 | Promoter (1-2kb) | 18 | 35115975 | 35242498 | 126524 | 1 | 1,03E+08 | ENSMUST00000237091.2 | -1685 | | Gm36037 | predicted gene, 36037 |
| chr9 | 86685762 | 86686728 | 967 | * | 7,773882 | 3,658630392 | 1,320108 | 2,771462 | 0,005581 | 1 | 0 | Exon (ENSMUST00000( | 9 | 86647986 | 86690794 | 42809 | 2 | 20616 | ENSMUST00000190388.7 | 4066 | ENSMUSG00000033419 | Snap91 | synaptosomal-associated protein 91 |
| chr11 | 21580666 | 21582008 | 1343 | * | 9,81006 | 3,658972315 | 1,291391 | 2,833357 | 0,004606 | 0,788774532 | 1,173325 | Intron (ENSMUST0000( | 11 | 21639583 | 21661910 | 22328 | 1 | 216560 | ENSMUST00000143608.2 | -57575 | ENSMUSG00000020319 | Wdpcp | WD repeat containing planar cell polarity effector |
| chr9 | 11146833 | 11147327 | 495 | * | 6,920095 | 3,659438495 | 1,37719 | 2,657178 | 0,00788 | 1 | 0 | Distal Intergenic | 9 | 9660974 | 10904780 | 1243807 | 2 | 244682 | ENSMUST00000160216.8 | -242053 | ENSMUSG00000039488 | Cntn5 | contactin 5 |
| chr10 | 34327115 | 34328755 | 1641 | * | 9,883331 | 3,660467435 | 1,329866 | 2,752508 | 0,005914 | 0,794354709 | 1,425929 | Distal Intergenic | 10 | 34359395 | 34487274 | 127880 | 1 | 14302 | ENSMUST00000019913.15 | -30640 | ENSMUSG00000019779 | Frk | fyn-related kinase |
| chr2 | 1,07E+08 | 1,07E+08 | 831 | * | 7,328517 | 3,660622657 | 1,355981 | 2,699612 | 0,006942 | 1 | 0 | Intron (ENSMUST0000( | 2 | 1,07E+08 | 1,07E+08 | 124 | 1 | 1,15E+08 | ENSMUST00000200346.3 | -21002 | ENSMUSG00000105886 | Gm22813 | predicted gene, 22813 |
| chr13 | 99896779 | 99897454 | 676 | * | 8,836866 | 3,661650208 | 1,247502 | 2,935186 | 0,003333 | 0,72352563 | 1,211315 | Distal Intergenic | 13 | 99828907 | 99829037 | 131 | 2 | 1,15E+08 | ENSMUST00000104485.3 | -67742 | ENSMUSG00000077673 | Gm24471 | predicted gene, 24471 |
| chr9 | 13351337 | 13353232 | 1896 | * | 13,59643 | 3,661866948 | 1,090522 | 3,357903 | 0,000785 | 0,511485635 | 0,938682 | Intron (ENSMUST0000( | 9 | 13342657 | 13344047 | 1391 | 1 | 18689 | ENSMUST00000216590.2 | 8680 | ENSMUSG00000031802 | Phxr4 | per-hexamer repeat gene 4 |
| chr19 | 44699912 | 44701018 | 1107 | * | 8,824377 | 3,664112122 | 1,366216 | 2,681942 | 0,00732 | 0,842187815 | 1,360622 | Intron (ENSMUST0000( | 19 | 44680796 | 44684888 | 4093 | 2 | 1,03E+08 | ENSMUST00000181365.4 | -15024 | ENSMUSG00000097683 | Gm26644 | predicted gene, 26644 |
| chr19 | 41997046 | 41997820 | 775 | * | 13,95141 | 3,664592814 | 1,087159 | 3,370798 | 0,00075 | 0,479521813 | 1,075735 | Intron (ENSMUST0000( | 19 | 42005588 | 42022134 | 16547 | 1 | 226122 | ENSMUST00000237817.2 | -7768 | ENSMUSG00000025171 | Ubtd1 | ubiquitin domain containing 1 |
| chr14 | 47225009 | 47226770 | 1762 | * | 12,00364 | 3,665768879 | 1,079684 | 3,395223 | 0,000686 | 0,451585113 | 1,173325 | Intron (ENSMUST0000( | 14 | 47238772 | 47339091 | 100320 | 1 | 74480 | ENSMUST00000125113.9 | -12002 | ENSMUSG00000021838 | Samd4 | sterile alpha motif domain containing 4 |
| chr10 | 77368754 | 77369789 | 1036 | * | 7,892613 | 3,665805515 | 1,323608 | 2,769556 | 0,005613 | 1 | 0 | Promoter (2-3kb) | 10 | 77366214 | 77401542 | 35329 | 1 | 16414 | ENSMUST00000153541.2 | 2540 | ENSMUSG00000000290 | Itgb2 | integrin beta 2 |
| chr5 | 1,46E+08 | 1,46E+08 | 1021 | * | 8,288829 | 3,66604297 | 1,384414 | 2,648082 | 0,008095 | 0,851014755 | 1,449706 | Intron (ENSMUST0000( | 5 | 1,46E+08 | 1,46E+08 | 31892 | 2 | 337924 | ENSMUST00000067479.6 | -22115 | ENSMUSG00000054417 | Cyp3a44 | cytochrome P450, family 3, subfamily a, polypeptide 44 |
| chr11 | 68960561 | 68962343 | 1783 | * | 19,75375 | 3,666106329 | 0,911454 | 4,022263 | 5,76E-05 | 0,084344578 | 2,386008 | Promoter (<=1kb) | 11 | 68961632 | 68963856 | 2225 | 1 | 66910 | ENSMUST00000075980.12 | 0 | ENSMUSG00000020895 | Tmem107 | transmembrane protein 107 |
| chr17 | 35941192 | 35943282 | 2091 | * | 14,94547 | 3,666239291 | 1,058666 | 3,463073 | 0,000534 | 0,376779196 | 1,41696 | Intron (ENSMUST0000( | 17 | 35928815 | 35937529 | 8715 | 2 | 672682 | ENSMUST00000164502.3 | -3663 | ENSMUSG00000090588 | Muc21 | mucin 21 |
| chr7 | 60396991 | 60397919 | 929 | * | 8,574497 | 3,666396682 | 1,362112 | 2,691699 | 0,007109 | 0,837306791 | 1,360622 | Distal Intergenic | 7 | 60565298 | 60572651 | 7354 | 2 | 1,01E+08 | ENSMUST00000208825.2 | 174732 | | Gm29729 | predicted gene, 29729 |
| chr15 | 8474338 | 8476555 | 2218 | * | 18,38055 | 3,666864194 | 1,089366 | 3,366053 | 0,000763 | 0,364217898 | 2,154504 | Promoter (<=1kb) | 15 | 8320101 | 8473947 | 153847 | 2 | 71175 | ENSMUST00000052965.8 | -391 | ENSMUSG00000022141 | Nipbl | NIPBL cohesin loading factor |
| chr4 | 1,09E+08 | 1,09E+08 | 2325 | * | 16,43976 | 3,667505094 | 1,029384 | 3,562816 | 0,000367 | 0,252371261 | 2,386008 | Promoter (<=1kb) | 4 | 1,09E+08 | 1,09E+08 | 1629 | 1 | 100087 | ENSMUST00000102738.4 | 0 | ENSMUSG00000073775 | Kti12 | KTI12 homolog, chromatin associated |
| chr1 | 1,51E+08 | 1,51E+08 | 1117 | * | 9,002081 | 3,669092168 | 1,225762 | 2,993315 | 0,00276 | 0,669159827 | 1,425929 | Distal Intergenic | 1 | 1,51E+08 | 1,52E+08 | 73713 | 1 | 63913 | ENSMUST00000134499.8 | -7861 | ENSMUSG00000026483 | Niban1 | niban apoptosis regulator 1 |
| chr13 | 39198924 | 39199980 | 1057 | * | 10,27422 | 3,669190919 | 1,245008 | 2,947121 | 0,003207 | 0,684274806 | 1,449706 | Distal Intergenic | 13 | 39116112 | 39144851 | 28740 | 2 | 108652 | ENSMUST00000021870.12 | -54073 | ENSMUSG00000021432 | Slc35b3 | solute carrier family 35, member B3 |

| | | | | | | | | | | | | | | | | | | | | | | | |
|---|---|---|---|---|---|---|---|---|---|---|---|---|---|---|---|---|---|---|---|---|---|---|---|
| chr4 | 25779871 | 25780992 | 1122 | * | 8,674459 | 3,669527288 | 1,308832 | 2,803665 | 0,005053 | 0,767208659 | 1,425929 | Intron (ENSMUST00000 | 4 | 25797703 | 25799855 | 2153 | 2 | 14348 | ENSMUST00000149875.2 | 18863 | ENSMUSG00000055373 | Fut9 | fucosyltransferase 9 |
| chr2 | 1,29E+08 | 1,29E+08 | 1032 | * | 7,869147 | 3,669799799 | 1,392995 | 2,634467 | 0,008427 | 1 | 0 | Promoter (1-2kb) | 2 | 1,29E+08 | 1,29E+08 | 2818 | 1 | 20017 | ENSMUST00000133345.2 | -1641 | ENSMUSG00000027395 | Polr1b | polymerase (RNA) I polypeptide B |
| chrX | 75054091 | 75055653 | 1563 | * | 8,033013 | 3,670082429 | 1,384617 | 2,650611 | 0,008035 | 1 | 0 | Distal Intergenic | 20 | 74929689 | 74929815 | 127 | 1 | 1,15E+08 | ENSMUST00000104462.3 | 124402 | ENSMUSG00000077650 | Gm23320 | predicted gene, 23320 |
| chr1 | 85663635 | 85665481 | 1847 | * | 17,51539 | 3,670112104 | 0,976833 | 3,757156 | 0,000172 | 0,163509282 | 2,423594 | Promoter (<=1kb) | 1 | 85644804 | 85664302 | 19499 | 2 | 319997 | ENSMUST00000159320.8 | 0 | ENSMUSG00000052760 | A630001G21Rik | RIKEN cDNA A630001G21 gene |
| chr6 | 1,25E+08 | 1,25E+08 | 700 | * | 12,23354 | 3,670435589 | 1,135045 | 3,233735 | 0,001222 | 0,548877304 | 1,173325 | Promoter (<=1kb) | 6 | 1,25E+08 | 1,25E+08 | 569 | 2 | 14433 | ENSMUST00000182464.2 | 0 | ENSMUSG00000057666 | Gapdh | glyceraldehyde-3-phosphate dehydrogenase |
| chrX | 1,58E+08 | 1,58E+08 | 1132 | * | 10,16991 | 3,67084141 | 1,221829 | 3,004381 | 0,002661 | 0,684274806 | 1,211315 | Distal Intergenic | 20 | 1,58E+08 | 1,59E+08 | 350798 | 1 | 58194 | ENSMUST00000112456.9 | -6436 | ENSMUSG00000040990 | Sh3kbp1 | SH3-domain kinase binding protein 1 |
| chr4 | 45589558 | 45590651 | 1094 | * | 8,319776 | 3,67087628 | 1,37162 | 2,676307 | 0,007444 | 1 | 0 | Distal Intergenic | 4 | 45548743 | 45548849 | 107 | 1 | 1,15E+08 | ENSMUST00000083388.2 | 40815 | ENSMUSG00000065322 | Gm22055 | predicted gene, 22055 |
| chr6 | 1,19E+08 | 1,19E+08 | 766 | * | 7,883207 | 3,671200111 | 1,353775 | 2,711825 | 0,006691 | 1 | 0 | Intron (ENSMUST00000 | 6 | 1,19E+08 | 1,19E+08 | 2963 | 1 | 101358 | ENSMUST00000032094.7 | 10281 | ENSMUSG00000030019 | Fbxl14 | F-box and leucine-rich repeat protein 14 |
| chr7 | 1,37E+08 | 1,37E+08 | 2676 | * | 20,10336 | 3,672485416 | 0,890322 | 4,124897 | 3,71E-05 | 0,067515008 | 2,221611 | Promoter (2-3kb) | 7 | 1,37E+08 | 1,37E+08 | 35189 | 2 | 77252 | ENSMUST00000068996.13 | -2351 | ENSMUSG00000040139 | 9430038I01Rik | RIKEN cDNA 9430038I01 gene |
| chr17 | 27112252 | 27114156 | 1905 | * | 16,20027 | 3,672913079 | 1,325132 | 2,771733 | 0,005576 | 0,699278451 | 2,318434 | Intron (ENSMUST00000 | 17 | 27079245 | 27105176 | 25932 | 2 | 1,03E+08 | ENSMUST00000236777.2 | -7076 | ENSMUSG00000078838 | Gm17382 | predicted gene, 17382 |
| chr7 | 87894646 | 87895608 | 963 | * | 7,438536 | 3,673676797 | 1,414765 | 2,596668 | 0,009413 | 1 | 0 | Distal Intergenic | 7 | 87927293 | 87960096 | 32804 | 1 | 13032 | ENSMUST00000032779.12 | -31685 | ENSMUSG00000030560 | Ctsc | cathepsin C |
| chr4 | 1,42E+08 | 1,42E+08 | 562 | * | 6,987937 | 3,673799342 | 1,420301 | 2,586635 | 0,009692 | 1 | 0 | Intron (ENSMUST00000 | 4 | 1,42E+08 | 1,42E+08 | 3806 | 1 | 545691 | ENSMUST00000146094.2 | -4613 | ENSMUSG00000085424 | Fhad1os1 | forkhead-associated (FHA) phosphopeptide binding domain 1, opposite strand 1 |
| chr4 | 82071809 | 82073263 | 1455 | * | 9,101076 | 3,674259496 | 1,226371 | 2,996043 | 0,002735 | 0,663408464 | 1,449706 | Distal Intergenic | 4 | 81952907 | 81984130 | 31224 | 2 | 1,03E+08 | ENSMUST00000243987.1 | -87679 | ENSMUSG00000087110 | Gm11264 | predicted gene 11264 |
| chr19 | 8955167 | 8956184 | 1018 | * | 8,847543 | 3,674289081 | 1,346609 | 2,72855 | 0,006361 | 0,838625535 | 1,211315 | 3' UTR | 19 | 8949768 | 8955843 | 6076 | 1 | 67160 | ENSMUST00000135025.2 | 5399 | ENSMUSG00000071644 | Eef1g | eukaryotic translation elongation factor 1 gamma |
| chr12 | 1,09E+08 | 1,09E+08 | 2179 | * | 20,18695 | 3,674785333 | 0,885776 | 4,148662 | 3,34E-05 | 0,058184583 | 2,507013 | Intron (ENSMUST00000 | 12 | 1,09E+08 | 1,09E+08 | 18282 | 1 | 104910 | ENSMUST00000221787.2 | 5233 | ENSMUSG00000048856 | Slc25a47 | solute carrier family 25, member 47 |
| chr7 | 29506434 | 29507177 | 744 | * | 7,353587 | 3,675252452 | 1,397967 | 2,628999 | 0,008564 | 1 | 0 | Distal Intergenic | 7 | 29515539 | 29530430 | 14892 | 1 | 233056 | ENSMUST00000178162.3 | -8362 | ENSMUSG00000011427 | Zfp790 | zinc finger protein 790 |
| chr11 | 4375377 | 4376738 | 1362 | * | 11,01678 | 3,675255826 | 1,220452 | 3,011388 | 0,002601 | 0,68443616 | 1,173325 | Promoter (<=1kb) | 11 | 4374068 | 4377262 | 3195 | 2 | 75828 | ENSMUST00000149443.2 | 524 | ENSMUSG00000020419 | Hormad2 | HORMA domain containing 2 |
| chr2 | 1,21E+08 | 1,21E+08 | 1940 | * | 14,39097 | 3,67527681 | 0,978728 | 3,755156 | 0,000173 | 0,278016366 | 0,918255 | Exon (ENSMUST00000 | 2 | 1,21E+08 | 1,21E+08 | 8908 | 2 | 22222 | ENSMUST00000154023.2 | -10763 | ENSMUSG00000027272 | Ubr1 | ubiquitin protein ligase E3 component n-recognin 1 |
| chr14 | 36706977 | 36708252 | 1276 | * | 10,99494 | 3,675342124 | 1,346943 | 2,728654 | 0,006359 | 0,887433944 | 0,918255 | Distal Intergenic | 14 | 36596893 | 36690734 | 93842 | 2 | 72972 | ENSMUST00000090024.11 | -16243 | ENSMUSG00000058690 | Ccser2 | coiled-coil serine rich 2 |
| chr11 | 32238219 | 32239251 | 1033 | * | 9,611589 | 3,675400334 | 1,279967 | 2,871481 | 0,004086 | 0,730469267 | 1,425929 | Promoter (1-2kb) | 11 | 32237001 | 32237784 | 784 | 1 | 544763 | ENSMUST00000151724.2 | 1218 | ENSMUSG00000073063 | Hbq1b | hemoglobin, theta 1B |
| chr10 | 1,01E+08 | 1,01E+08 | 518 | * | 9,139654 | 3,677268758 | 1,224846 | 3,002229 | 0,00268 | 0,684274806 | 1,211315 | Distal Intergenic | 10 | 1,01E+08 | 1,01E+08 | 34101 | 2 | 1,03E+08 | ENSMUST00000218270.2 | -385248 | ENSMUSG00000112104 | Gm35722 | predicted gene, 35722 |
| chr9 | 11791159 | 11791961 | 803 | * | 8,803074 | 3,67755102 | 1,249908 | 2,942258 | 0,003258 | 0,698318058 | 1,360622 | Distal Intergenic | 9 | 9660974 | 10904780 | 1243807 | 2 | 244682 | ENSMUST00000160216.8 | -886379 | ENSMUSG00000039488 | Cntn5 | contactin 5 |
| chr10 | 18849219 | 18850033 | 815 | * | 10,0513 | 3,678487232 | 1,302539 | 2,82409 | 0,004741 | 0,804027476 | 1,086118 | Distal Intergenic | 10 | 18883015 | 18884593 | 1579 | 2 | 21929 | ENSMUST00000154749.2 | 34560 | ENSMUSG00000019850 | Tnfaip3 | tumor necrosis factor, alpha-induced protein 3 |
| chr1 | 1,41E+08 | 1,41E+08 | 1017 | * | 7,658354 | 3,678832816 | 1,204717 | 3,05369 | 0,00226 | 0,64236196 | 1,360622 | Distal Intergenic | 1 | 1,4E+08 | 1,41E+08 | 86229 | 2 | 75910 | ENSMUST00000193820.2 | -274697 | ENSMUSG00000102368 | 4930590L20Rik | RIKEN cDNA 4930590L20 gene |
| chr17 | 27360800 | 27361803 | 1004 | * | 15,44785 | 3,678906839 | 0,95657 | 3,845937 | 0,00012 | 0,192431185 | 1,2763 | Distal Intergenic | 17 | 27341637 | 27352890 | 11254 | 2 | 67267 | ENSMUST00000025045.15 | -7910 | ENSMUSG00000024208 | Uqcc2 | ubiquinol-cytochrome c reductase complex assembly factor 2 |
| chr8 | 70987333 | 70988375 | 1043 | * | 12,29332 | 3,679045962 | 1,130415 | 3,254597 | 0,001136 | 0,538977324 | 1,173325 | Promoter (<=1kb) | 8 | 70986767 | 70987707 | 941 | 1 | 14232 | ENSMUST00000134893.2 | 566 | ENSMUSG00000019428 | Fkbp8 | FK506 binding protein 8 |
| chr8 | 88636444 | 88637036 | 593 | * | 7,869442 | 3,679834385 | 1,325368 | 2,776462 | 0,005495 | 0,792225394 | 1,360622 | Intron (ENSMUST00000 | 8 | 88388440 | 88622363 | 233924 | 2 | 94187 | ENSMUST00000052250.15 | -14081 | ENSMUSG00000045333 | Zfp423 | zinc finger protein 423 |
| chr7 | 1,08E+08 | 1,08E+08 | 975 | * | 8,433627 | 3,680208135 | 1,331692 | 2,763559 | 0,005717 | 1 | 0 | Promoter (<=1kb) | 7 | 1,08E+08 | 1,08E+08 | 945 | 2 | 258490 | ENSMUST00000079865.5 | -636 | ENSMUSG00000109497 | Olfr492 | olfactory receptor 492 |
| chr1 | 23489404 | 23490702 | 1299 | * | 11,75149 | 3,68095772 | 1,116035 | 3,298245 | 0,000973 | 0,596107452 | 0,744375 | Distal Intergenic | 1 | 23430830 | 23436853 | 6024 | 2 | 1,01E+08 | ENSMUST00000189537.2 | -52551 | | Gm19673 | predicted gene, 19673 |
| chr14 | 69675739 | 69677072 | 1334 | * | 9,920797 | 3,681085617 | 1,181389 | 3,115897 | 0,001834 | 0,62640387 | 1,211315 | Intron (ENSMUST00000 | 14 | 69656843 | 69663926 | 7084 | 2 | 1,01E+08 | ENSMUST00000184039.2 | -11813 | | Gm16867 | predicted gene, 16867 |
| chr1 | 89782733 | 89783468 | 736 | * | 8,94366 | 3,681136366 | 1,207968 | 3,047379 | 0,002308 | 0,634537746 | 1,449706 | Intron (ENSMUST00000 | 1 | 89716885 | 89762310 | 45426 | 1 | 347722 | ENSMUST00000212924.2 | 65848 | ENSMUSG00000055013 | Agap1 | ArfGAP with GTPase domain, ankyrin repeat and PH domain 1 |
| chr11 | 97305825 | 97306861 | 1037 | * | 10,3793 | 3,681637464 | 1,295898 | 2,840993 | 0,004497 | 0,861581225 | 0,738441 | Promoter (<=1kb) | 11 | 97306359 | 97393228 | 86870 | 1 | 58996 | ENSMUST00000121799.8 | 0 | ENSMUSG00000049807 | Arhgap23 | Rho GTPase activating protein 23 |
| chr10 | 42391189 | 42391938 | 750 | * | 8,165785 | 3,681670691 | 1,274731 | 2,888195 | 0,003875 | 0,649217039 | 2,237403 | Intron (ENSMUST00000 | 10 | 42378286 | 42411377 | 33092 | 1 | 54198 | ENSMUST00000105499.2 | 12903 | ENSMUSG00000019804 | Snx3 | sorting nexin 3 |
| chr3 | 1,09E+08 | 1,09E+08 | 743 | * | 6,948993 | 3,682312759 | 1,420898 | 2,591539 | 0,009555 | 1 | 0 | Intron (ENSMUST00000 | 3 | 1,09E+08 | 1,09E+08 | 14789 | 1 | 329738 | ENSMUST00000147916.2 | 10285 | ENSMUSG00000049565 | Aknad1 | AKNA domain containing 1 |
| chr19 | 50346574 | 50347838 | 1265 | * | 10,62981 | 3,682730771 | 1,185516 | 3,106437 | 0,001894 | 0,71238788 | 0,744375 | Intron (ENSMUST00000 | 19 | 50139405 | 50241116 | 101712 | 2 | 58178 | ENSMUST00000168357.9 | -105458 | ENSMUSG00000043531 | Sorcs1 | sortilin-related VPS10 domain containing receptor 1 |
| chr5 | 36617415 | 36618700 | 1286 | * | 11,6056 | 3,683954409 | 1,141183 | 3,22819 | 0,001246 | 0,642870233 | 0,744375 | Intron (ENSMUST00000 | 5 | 36613112 | 36622155 | 9044 | 2 | 76710 | ENSMUST00000044964.6 | 3455 | ENSMUSG00000039563 | 2210406O10Rik | RIKEN cDNA 2210406O10 gene |
| chr7 | 1,21E+08 | 1,21E+08 | 1399 | * | 9,372801 | 3,684251884 | 1,338318 | 2,752897 | 0,005907 | 0,802671652 | 1,360622 | Exon (ENSMUST00000 | 7 | 1,21E+08 | 1,21E+08 | 8034 | 2 | 76179 | ENSMUST00000205473.2 | -5856 | ENSMUSG00000063317 | Usp31 | ubiquitin specific peptidase 31 |
| chr2 | 83555434 | 83556036 | 603 | * | 6,770867 | 3,684586607 | 1,330098 | 2,770161 | 0,005603 | 1 | 0 | Promoter (<=1kb) | 2 | 83554899 | 83637260 | 82362 | 1 | 16410 | ENSMUST00000111740.9 | 535 | ENSMUSG00000027087 | Itgav | integrin alpha V |
| chr10 | 1,01E+08 | 1,01E+08 | 872 | * | 7,658511 | 3,685310013 | 1,272523 | 2,896066 | 0,003779 | 1 | 0 | Distal Intergenic | 10 | 1,02E+08 | 1,02E+08 | 331736 | 1 | 67569 | ENSMUST00000146230.8 | -20766 | ENSMUSG00000019888 | Mgat4c | MGAT4 family, member C |
| chr2 | 1,05E+08 | 1,05E+08 | 1054 | * | 8,632818 | 3,685562977 | 1,2596 | 2,925978 | 0,003434 | 1 | 0 | Promoter (1-2kb) | 2 | 1,05E+08 | 1,05E+08 | 17355 | 2 | 98221 | ENSMUST00000111110.3 | 1161 | ENSMUSG00000027170 | Eif3m | eukaryotic translation initiation factor 3, subunit M |
| chr11 | 1,04E+08 | 1,04E+08 | 1602 | * | 13,03921 | 3,685669814 | 1,335066 | 2,760666 | 0,005768 | 0,826313122 | 1,173325 | Distal Intergenic | 11 | 1,04E+08 | 1,04E+08 | 4039 | 1 | 11841 | ENSMUST00000154089.2 | 18068 | ENSMUSG00000062421 | Arf2 | ADP-ribosylation factor 2 |
| chr9 | 64920755 | 64921665 | 911 | * | 9,987254 | 3,685969387 | 1,259821 | 2,925788 | 0,003436 | 0,699278451 | 1,425929 | Intron (ENSMUST00000 | 9 | 64894265 | 64928975 | 34711 | 2 | 57874 | ENSMUST00000036615.7 | 7310 | ENSMUSG00000033629 | Hacd3 | 3-hydroxyacyl-CoA dehydratase 3 |
| chr17 | 27788154 | 27789201 | 1048 | * | 9,097884 | 3,685974008 | 1,296067 | 2,843969 | 0,004456 | 0,747386384 | 1,425929 | Distal Intergenic | 17 | 27782547 | 27784730 | 2184 | 2 | 106672 | ENSMUST00000154473.9 | -3424 | ENSMUSG00000062753 | AI413582 | expressed sequence AI413582 |
| chr17 | 34786216 | 34787757 | 1542 | * | 12,62262 | 3,686312499 | 1,257019 | 2,932583 | 0,003362 | 0,813409596 | 0,738441 | Promoter (2-3kb) | 17 | 34784110 | 34784551 | 442 | 1 | 18132 | ENSMUST00000152714.2 | 2106 | ENSMUSG00000015468 | Notch4 | notch 4 |
| chr16 | 71769718 | 71770593 | 876 | * | 6,808005 | 3,68717613 | 1,352961 | 2,725264 | 0,006425 | 1 | 0 | Distal Intergenic | 16 | 72023271 | 72031190 | 7920 | 2 | 319533 | ENSMUST00000231223.2 | 260597 | ENSMUSG00000116769 | 8030451O07Rik | RIKEN cDNA 8030451O07 gene |
| chr15 | 32610791 | 32611575 | 785 | * | 10,96308 | 3,687842091 | 1,225713 | 3,008732 | 0,002623 | 0,676922067 | 1,2763 | Intron (ENSMUST00000 | 15 | 32601821 | 32689782 | 87962 | 1 | 20356 | ENSMUST00000227429.2 | 8970 | ENSMUSG00000022231 | Sema5a | sema domain, seven thrombospondin repeats (type 1 and type 1-like), transmembrane domain (TM) and short cytoplasmic domain, (semaphorin) 5A |
| chr4 | 63713200 | 63714082 | 883 | * | 7,360132 | 3,68800216 | 1,404345 | 2,626136 | 0,008636 | 1 | 0 | Distal Intergenic | 4 | 63645321 | 63663350 | 18030 | 2 | 326623 | ENSMUST00000062246.8 | -49850 | ENSMUSG00000050395 | Tnfsf15 | tumor necrosis factor (ligand) superfamily, member 15 |
| chr11 | 83415807 | 83416915 | 1109 | * | 8,632882 | 3,688278949 | 1,280379 | 2,880614 | 0,003969 | 0,726422517 | 1,425929 | Promoter (2-3kb) | 11 | 83402562 | 83413001 | 10440 | 2 | 629303 | ENSMUST00000019069.4 | -2806 | ENSMUSG00000018925 | Heatr9 | HEAT repeat containing 9 |
| chr7 | 96797212 | 96799318 | 2107 | * | 17,32871 | 3,688641356 | 0,947018 | 3,895006 | 9,82E-05 | 0,123288945 | 2,154504 | Intron (ENSMUST00000 | 7 | 96730983 | 96954243 | 223261 | 1 | 14389 | ENSMUST00000206791.2 | 66229 | ENSMUSG00000004508 | Gab2 | growth factor receptor bound protein 2-associated protein 2 |
| chr15 | 42762050 | 42762863 | 814 | * | 10,61515 | 3,689197089 | 1,280103 | 2,881953 | 0,003952 | 0,744477896 | 1,2763 | Distal Intergenic | 15 | 42288119 | 42540373 | 252255 | 2 | 11600 | ENSMUST00000022921.7 | -221677 | ENSMUSG00000022309 | Angpt1 | angiopoietin 1 |
| chr13 | 94117744 | 94118354 | 611 | * | 6,483494 | 3,689601966 | 1,391964 | 2,650644 | 0,008034 | 1 | 0 | Distal Intergenic | 13 | 94194304 | 94331917 | 137614 | 1 | 218454 | ENSMUST00000054274.8 | -75950 | ENSMUSG00000045312 | Lhfpl2 | lipoma HMGIC fusion partner-like 2 |
| chr2 | 71940834 | 71941931 | 1098 | * | 8,727317 | 3,689894475 | 1,366258 | 2,70073 | 0,006919 | 0,82177581 | 1,449706 | Intron (ENSMUST00000 | 2 | 71971055 | 72026605 | 55551 | 1 | 56508 | ENSMUST00000122975.8 | -29124 | ENSMUSG00000049044 | Rapgef4 | Rap guanine nucleotide exchange factor (GEF) 4 |
| chr11 | 78422920 | 78423869 | 950 | * | 8,5826 | 3,69065171 | 1,385994 | 2,662819 | 0,007749 | 0,840920027 | 1,449706 | Intron (ENSMUST00000 | 11 | 78413676 | 78427097 | 13422 | 2 | 21927 | ENSMUST00000017759.9 | 3228 | ENSMUSG00000017615 | Tnfaip1 | tumor necrosis factor, alpha-induced protein 1 (endothelial) |
| chr6 | 29271995 | 29273729 | 1735 | * | 63,41019 | 3,691603197 | 1,114895 | 3,311168 | 0,000929 | 0,298416988 | 4,200304 | Promoter (<=1kb) | 6 | 29272487 | 29275185 | 2699 | 1 | 69573 | ENSMUST00000115289.2 | 0 | ENSMUSG00000043421 | Hilpda | hypoxia inducible lipid droplet associated |
| chr19 | 44298984 | 44300900 | 1917 | * | 27,48709 | 3,692130275 | 1,230603 | 3,000262 | 0,002697 | 0,532496361 | 2,880017 | Distal Intergenic | 19 | 44291610 | 44291670 | 61 | 1 | 1,01E+08 | ENSMUST00000175574.3 | 7374 | ENSMUSG00000093315 | Mir5114 | microRNA 5114 |
| chr9 | 59989252 | 59990767 | 1516 | * | 11,6114 | 3,693112947 | 1,152247 | 3,20514 | 0,00135 | 0,656950206 | 0,744375 | Intron (ENSMUST00000 | 9 | 59877002 | 60048503 | 171502 | 2 | 207596 | ENSMUST00000034829.6 | 57736 | ENSMUSG00000032289 | Thsd4 | thrombospondin, type I, domain containing 4 |
| chr4 | 1,07E+08 | 1,07E+08 | 1273 | * | 9,903642 | 3,693260806 | 1,35845 | 2,718731 | 0,006553 | 0,843429621 | 1,211315 | Promoter (2-3kb) | 4 | 1,07E+08 | 1,07E+08 | 15418 | 1 | 242603 | ENSMUST00000221740.2 | 2233 | ENSMUSG00000047636 | Cdcp2 | CUB domain containing protein 2 |
| chr5 | 1,06E+08 | 1,06E+08 | 1129 | * | 10,19321 | 3,694233611 | 1,338629 | 2,759714 | 0,005785 | 0,792545355 | 1,425929 | Distal Intergenic | 5 | 1,06E+08 | 1,06E+08 | 115247 | 1 | 231549 | ENSMUST00000060531.16 | -48928 | ENSMUSG00000046079 | Lrrc8d | leucine rich repeat containing 8D |
| chr5 | 45362087 | 45362808 | 722 | * | 8,009419 | 3,694371876 | 1,347194 | 2,742271 | 0,006102 | 1 | 0 | Intron (ENSMUST00000 | 5 | 45267971 | 45333035 | 65065 | 2 | 52401 | ENSMUST00000198493.2 | -29052 | ENSMUSG00000029092 | D5Ertd615e | DNA segment, Chr 5, ERATO Doi 615, expressed |
| chr4 | 87386081 | 87387229 | 1149 | * | 11,54576 | 3,694472002 | 1,142619 | 3,233337 | 0,001224 | 0,599552016 | 0,918255 | Distal Intergenic | 4 | 86901361 | 87148714 | 247354 | 2 | 76376 | ENSMUST00000107157.9 | -237367 | ENSMUSG00000037996 | Slc24a2 | solute carrier family 24 (sodium/potassium/calcium exchanger), member 2 |
| chr11 | 79981334 | 79983291 | 1958 | * | 11,0967 | 3,695105877 | 1,206955 | 3,061512 | 0,002202 | 0,735602255 | 0,744375 | Promoter (1-2kb) | 11 | 79980226 | 80026620 | 46395 | 1 | 237877 | ENSMUST00000108239.7 | 1108 | ENSMUSG00000017550 | Atad5 | ATPase family, AAA domain containing 5 |
| chr14 | 21150624 | 21152193 | 1570 | * | 12,54007 | 3,696039491 | 1,023367 | 3,611646 | 0,000304 | 0,327375387 | 1,075735 | Intron (ENSMUST00000 | 14 | 21126220 | 21498630 | 372411 | 1 | 11534 | ENSMUST00000224069.2 | 24404 | ENSMUSG00000039197 | Adk | adenosine kinase |
| chr3 | 83704385 | 83705613 | 1229 | * | 8,980848 | 3,696281628 | 1,328325 | 2,782664 | 0,005391 | 0,779936001 | 1,425929 | Distal Intergenic | 3 | 83674160 | 83679456 | 5297 | 1 | 20319 | ENSMUST00000154794.2 | 30225 | ENSMUSG00000027996 | Sfrp2 | secreted frizzled-related protein 2 |
| chr4 | 41267940 | 41269050 | 1111 | * | 9,006108 | 3,696672066 | 1,426159 | 2,592048 | 0,009541 | 0,88871498 | 1,360622 | Intron (ENSMUST00000 | 4 | 41221592 | 41275116 | 53525 | 2 | 68926 | ENSMUST00000132499.2 | 6066 | ENSMUSG00000028433 | Ubap2 | ubiquitin-associated protein 2 |
| chr5 | 1,48E+08 | 1,48E+08 | 929 | * | 9,078759 | 3,696893664 | 1,258881 | 2,936651 | 0,003318 | 0,693003605 | 1,425929 | Distal Intergenic | 5 | 1,48E+08 | 1,48E+08 | 72495 | 2 | 11987 | ENSMUST00000048116.15 | -19946 | ENSMUSG00000041313 | Slc7a1 | solute carrier family 7 (cationic amino acid transporter, y+ system), member 1 |
| chr4 | 96859811 | 96860892 | 1082 | * | 7,987036 | 3,696994713 | 1,379925 | 2,679127 | 0,007381 | 1 | 0 | Distal Intergenic | 4 | 96805257 | 96805320 | 64 | 1 | 1,15E+08 | ENSMUST00000184022.3 | 54554 | ENSMUSG00000098261 | Gm27521 | predicted gene, 27521 |
| chrX | 1,06E+08 | 1,06E+08 | 845 | * | 7,424506 | 3,697544913 | 1,374179 | 2,690729 | 0,00713 | 1 | 0 | Distal Intergenic | 20 | 1,06E+08 | 1,06E+08 | 6278 | 2 | 16431 | ENSMUST00000033591.6 | 33165 | ENSMUSG00000031239 | Itm2a | integral membrane protein 2A |
| chr6 | 50289305 | 50290263 | 959 | * | 9,281918 | 3,697854898 | 1,34291 | 2,753614 | 0,005894 | 0,824134241 | 1,211315 | Exon (ENSMUST00000 | 6 | 50272163 | 50277334 | 5172 | 2 | 71720 | ENSMUST00000141466.2 | -11971 | ENSMUSG00000029822 | Osbpl3 | oxysterol binding protein-like 3 |
| chr15 | 77336280 | 77338029 | 1750 | * | 14,88246 | 3,698459739 | 1,034712 | 3,574384 | 0,000351 | 0,324087962 | 1,2763 | Intron (ENSMUST00000 | 15 | 77309498 | 77331692 | 22195 | 2 | 278679 | ENSMUST00000149824.2 | -4588 | ENSMUSG00000068252 | Apol7b | apolipoprotein L 7b |
| chr12 | 28799144 | 28800328 | 1185 | * | 9,381993 | 3,698480817 | 1,307434 | 2,828809 | 0,004672 | 0,680780911 | 2,237403 | Promoter (<=1kb) | 12 | 28759506 | 28800376 | 40871 | 2 | 217449 | ENSMUST00000170994.3 | 48 | ENSMUSG00000020628 | Trappc12 | trafficking protein particle complex 12 |
| chr13 | 1,13E+08 | 1,13E+08 | 1927 | * | 12,9807 | 3,69880005 | 1,205227 | 3,068967 | 0,002148 | 0,731734453 | 0,738441 | Exon (ENSMUST00000 | 13 | 1,13E+08 | 1,13E+08 | 4862 | 2 | 72198 | ENSMUST00000225037.2 | -3763 | ENSMUSG00000016018 | Mtrex | Mtr4 exosome RNA helicase |
| chr10 | 28463451 | 28464264 | 814 | * | 11,02686 | 3,70130454 | 1,159534 | 3,192061 | 0,001413 | 0,627785088 | 0,918255 | Exon (ENSMUST00000 | 10 | 28456455 | 28467901 | 11447 | 1 | 19272 | ENSMUST00000220357.2 | 6996 | ENSMUSG00000019889 | Ptprk | protein tyrosine phosphatase, receptor type, K |
| chr14 | 30284985 | 30285793 | 809 | * | 6,997918 | 3,701341896 | 1,371252 | 2,699242 | 0,00695 | 1 | 0 | Intron (ENSMUST00000 | 14 | 30281647 | 30295581 | 13935 | 1 | 21881 | ENSMUST00000223633.2 | 3338 | ENSMUSG00000021957 | Tkt | transketolase |
| chr13 | 1,04E+08 | 1,04E+08 | 1281 | * | 9,346954 | 3,702204591 | 1,288754 | 2,8727 | 0,00407 | 0,73764377 | 1,360622 | Intron (ENSMUST00000 | 13 | 1,04E+08 | 1,04E+08 | 78335 | 2 | 59079 | ENSMUST00000189323.2 | -38576 | ENSMUSG00000021709 | Erbin | Erbb2 interacting protein |
| chr10 | 56257328 | 56258186 | 859 | * | 6,76402 | 3,70310933 | 1,373287 | 2,696529 | 0,007007 | 1 | 0 | Promoter (2-3kb) | 10 | 56255184 | 56266511 | 11328 | 1 | 14609 | ENSMUST00000220069.2 | 2144 | ENSMUSG00000050953 | Gja1 | gap junction protein, alpha 1 |
| chr4 | 19039056 | 19039909 | 854 | * | 8,643614 | 3,704234251 | 1,254218 | 2,953422 | 0,003143 | 0,611385361 | 2,237403 | Intron (ENSMUST00000 | 4 | 18767980 | 19122565 | 354586 | 2 | 435772 | ENSMUST00000137780.3 | 82656 | ENSMUSG00000073991 | Cnbd1 | cyclic nucleotide binding domain containing 1 |
| chr9 | 13355803 | 13356768 | 966 | * | 8,459847 | 3,705204835 | 1,19243 | 3,107274 | 0,001888 | 1 | 0 | Intron (ENSMUST00000 | 9 | 13342657 | 13344047 | 1391 | 1 | 18689 | ENSMUST00000216590.2 | 13146 | ENSMUSG00000031802 | Phxr4 | per-hexamer repeat gene 4 |
| chr13 | 68727088 | 68727640 | 553 | * | 5,880773 | 3,705422538 | 1,392697 | 2,66061 | 0,0078 | 1 | 0 | Promoter (2-3kb) | 13 | 68727591 | 68730013 | 2423 | 2 | 210009 | ENSMUST00000223055.2 | 2373 | ENSMUSG00000034617 | Mtrr | 5-methyltetrahydrofolate-homocysteine methyltransferase reductase |
| chr11 | 52250189 | 52251892 | 1704 | * | 56,84473 | 3,707743362 | 1,117706 | 3,31728 | 0,000909 | 0,326512715 | 3,270621 | Promoter (<=1kb) | 11 | 52251687 | 52280224 | 28538 | 1 | 22333 | ENSMUST00000102758.8 | 0 | ENSMUSG00000020402 | Vdac1 | voltage-dependent anion channel 1 |
| chr4 | 1,35E+08 | 1,35E+08 | 932 | * | 10,56782 | 3,708196286 | 1,109571 | 3,342008 | 0,000832 | 0,483145429 | 1,173325 | Distal Intergenic | 4 | 1,35E+08 | 1,35E+08 | 2227 | 2 | 1,08E+08 | ENSMUST00000146152.2 | 82960 | | Gm16225 | predicted gene 16225 |
| chr8 | 9397091 | 9397641 | 551 | * | 7,231132 | 3,70828279 | 1,379248 | 2,688626 | 0,007175 | 1 | 0 | Intron (ENSMUST00000 | 8 | 9257238 | 9373711 | 116474 | 2 | 270028 | ENSMUST00000207297.2 | -23380 | ENSMUSG00000079157 | Nalf1 | NALCN channel auxiliary factor 1 |
| chr10 | 1,15E+08 | 1,15E+08 | 647 | * | 7,492166 | 3,708353441 | 1,392924 | 2,66228 | 0,007761 | 1 | 0 | Intron (ENSMUST00000 | 10 | 1,15E+08 | 1,15E+08 | 12463 | 1 | 77798 | ENSMUST00000173620.2 | -13211 | ENSMUSG00000092210 | A930009A15Rik | RIKEN cDNA A930009A15 gene |
| chr1 | 1,21E+08 | 1,21E+08 | 969 | * | 9,821886 | 3,708813176 | 1,08248 | 3,426218 | 0,000612 | 0,399951667 | 1,449706 | Distal Intergenic | 1 | 1,21E+08 | 1,21E+08 | 4785 | 2 | 72365 | ENSMUST00000190690.2 | -24231 | | 2210011K15Rik | RIKEN cDNA 2210011K15 gene |
| chr2 | 1,03E+08 | 1,03E+08 | 823 | * | 6,98866 | 3,709277663 | 1,39987 | 2,64973 | 0,008056 | 1 | 0 | Exon (ENSMUST00000 | 2 | 1,03E+08 | 1,03E+08 | 30531 | 2 | 12359 | ENSMUST00000111168.4 | 17639 | ENSMUSG00000027187 | Cat | catalase |
| chr6 | 1,16E+08 | 1,16E+08 | 575 | * | 8,782099 | 3,711941064 | 1,323397 | 2,804859 | 0,005034 | 0,776966635 | 1,360622 | Promoter (<=1kb) | 6 | 1,16E+08 | 1,16E+08 | 9473 | 2 | 110157 | ENSMUST00000147979.3 | 0 | ENSMUSG00000000441 | Raf1 | v-raf-leukemia viral oncogene 1 |
| chr11 | 80307166 | 80308510 | 1345 | * | 12,40985 | 3,712133384 | 1,188489 | 3,123406 | 0,001788 | 0,64236196 | 1,075735 | Distal Intergenic | 11 | 80319441 | 80325364 | 5924 | 1 | 69077 | ENSMUST00000172641.2 | -10931 | ENSMUSG00000017428 | Psmd11 | proteasome (prosome, macropain) 26S subunit, non-ATPase, 11 |
| chr11 | 61543025 | 61543946 | 922 | * | 8,911237 | 3,713732409 | 1,200083 | 3,094564 | 0,001971 | 0,617307536 | 1,360622 | Promoter (<=1kb) | 11 | 61544091 | 61563610 | 19520 | 1 | 71520 | ENSMUST00000004959.3 | -145 | ENSMUSG00000004837 | Grap | GRB2-related adaptor protein |
| chr16 | 93044376 | 93045540 | 1165 | * | 7,869619 | 3,715517234 | 1,317275 | 2,820609 | 0,004793 | 1 | 0 | Distal Intergenic | 16 | 93140612 | 93150077 | 9466 | 2 | 69857 | ENSMUST00000185971.7 | 104537 | ENSMUSG00000100277 | 1810053B23Rik | RIKEN cDNA 1810053B23 gene |
| chr8 | 1,12E+08 | 1,12E+08 | 1291 | * | 11,18452 | 3,715609343 | 1,274878 | 2,914483 | 0,003563 | 0,823736512 | 0,738441 | Exon (ENSMUST00000 | 8 | 1,12E+08 | 1,12E+08 | 7115 | 2 | 101943 | ENSMUST00000212182.2 | 12179 | ENSMUSG00000033732 | Sf3b3 | splicing factor 3b, subunit 3 |
| chr5 | 1,16E+08 | 1,16E+08 | 1510 | * | 12,22299 | 3,715720623 | 1,069235 | 3,475122 | 0,000511 | 0,418252864 | 1,075735 | Promoter (2-3kb) | 5 | 1,16E+08 | 1,16E+08 | 1560 | 1 | 12704 | ENSMUST00000136780.2 | 2218 | ENSMUSG00000029516 | Cit | citron |
| chr19 | 23643909 | 23644995 | 1087 | * | 7,464774 | 3,717627529 | 1,400833 | 2,653868 | 0,007957 | 1 | 0 | Exon (ENSMUST00000 | 19 | 23536117 | 23630176 | 94060 | 2 | 67483 | ENSMUST00000035849.5 | -13733 | ENSMUSG00000033053 | Cfap95 | cilia and flagella associated protein 95 |
| chr9 | 96039756 | 96041110 | 1355 | * | 10,60715 | 3,718675055 | 1,253326 | 2,967044 | 0,003007 | 0,751712532 | 0,918255 | Intron (ENSMUST00000 | 9 | 96006273 | 96019873 | 13601 | 1 | 235533 | ENSMUST00000136496.2 | 33483 | ENSMUSG00000041440 | Gk5 | glycerol kinase 5 (putative) |
| chr16 | 80925305 | 80925987 | 683 | * | 8,233495 | 3,719491801 | 1,188567 | 3,129393 | 0,001752 | 1 | 0 | Distal Intergenic | 16 | 80997585 | 81394693 | 397109 | 1 | 17968 | ENSMUST00000037785.14 | -71598 | ENSMUSG00000022762 | Ncam2 | neural cell adhesion molecule 2 |
| chr14 | 54491198 | 54492285 | 1088 | * | 18,22727 | 3,719748552 | 1,029415 | 3,61346 | 0,000302 | 0,21428283 | 2,663619 | Promoter (<=1kb) | 14 | 54491645 | 54506532 | 14888 | 1 | 105501 | ENSMUST00000180359.8 | 0 | ENSMUSG00000040997 | Abhd4 | abhydrolase domain containing 4 |
| chr8 | 73436781 | 73437453 | 673 | * | 9,499041 | 3,721270113 | 1,357629 | 2,741007 | 0,006125 | 0,723027638 | 2,237403 | Intron (ENSMUST00000 | 8 | 73449913 | 73466534 | 16622 | 1 | 20467 | ENSMUST00000109950.5 | -12460 | ENSMUSG00000031622 | Sin3b | transcriptional regulator, SIN3B (yeast) |
| chr6 | 17583322 | 17584441 | 1120 | * | 14,9545 | 3,722432677 | 0,991221 | 3,755402 | 0,000173 | 0,222881372 | 1,41696 | Distal Intergenic | 6 | 17546972 | 17553429 | 6458 | 1 | 17295 | ENSMUST00000148903.2 | 36350 | ENSMUSG00000009376 | Met | met proto-oncogene |
| chr15 | 35800488 | 35801672 | 1185 | * | 10,82383 | 3,723277275 | 1,240559 | 3,001289 | 0,002688 | 0,773973551 | 0,738441 | Intron (ENSMUST00000 | 15 | 35787300 | 35790425 | 3126 | 1 | 74554 | ENSMUST00000228829.2 | 13188 | | 9130002K18Rik | RIKEN cDNA 9130002K18 gene |
| chr7 | 89278161 | 89279368 | 1208 | * | 8,603194 | 3,7233795 | 1,33658 | 2,785751 | 0,00534 | 0,697709534 | 2,237403 | Promoter (2-3kb) | 7 | 89281600 | 89459405 | 177806 | 1 | 109264 | ENSMUST00000159504.8 | -2232 | ENSMUSG00000030621 | Me3 | malic enzyme 3, NADP(+)-dependent, mitochondrial |
| chrX | 49465869 | 49466557 | 689 | * | 7,293242 | 3,723627081 | 1,43153 | 2,601152 | 0,009291 | 1 | 0 | Distal Intergenic | 20 | 49494690 | 49494796 | 107 | 1 | 1,15E+08 | ENSMUST00000240068.1 | -28133 | ENSMUSG00000119917 | Gm22288 | predicted gene, 22288 |
| chrX | 93736496 | 93737748 | 1253 | * | 12,26915 | 3,72389998 | 1,3283 | 2,803508 | 0,005055 | 0,819350294 | 1,075735 | Distal Intergenic | 20 | 93768175 | 93773793 | 5619 | 1 | 668166 | ENSMUST00000101388.4 | -30427 | ENSMUSG00000073062 | Zxdb | zinc finger, X-linked, duplicated B |
| chr5 | 14121610 | 14122967 | 1358 | * | 11,65864 | 3,724077881 | 1,166115 | 3,193577 | 0,001405 | 0,663408464 | 0,744375 | Intron (ENSMUST00000 | 5 | 14075290 | 14306703 | 231414 | 1 | 20349 | ENSMUST00000073957.8 | 46320 | ENSMUSG00000063531 | Sema3e | sema domain, immunoglobulin domain (Ig), short basic domain, secreted, (semaphorin) 3E |

| | | | | | | | | | | | | | | | | | | | | | | | |
|---|---|---|---|---|---|---|---|---|---|---|---|---|---|---|---|---|---|---|---|---|---|---|---|
| chr4 | 1,52E+08 | 1,52E+08 | 809 | * | 7,77975 | 3,724137049 | 1,371281 | 2,715809 | 0,006611 | 0,824134241 | 1,360622 | Promoter (<=1kb) | 4 | 1,52E+08 | 1,52E+08 | 3562 | 2 | 100090 | ENSMUST00000155441.2 | 96 | ENSMUSG00000028952 | Zbtb48 | zinc finger and BTB domain containing 48 |
| chr10 | 1,28E+08 | 1,28E+08 | 951 | * | 7,802012 | 3,724217277 | 1,366174 | 2,72602 | 0,00641 | 1 | 0 | Promoter (2-3kb) | 10 | 1,28E+08 | 1,28E+08 | 5898 | 2 | 210582 | ENSMUST00000220227.2 | -2118 | ENSMUSG00000039914 | Coq10a | coenzyme Q10A |
| chr3 | 63883124 | 63883982 | 859 | * | 7,299794 | 3,724631427 | 1,352552 | 2,75378 | 0,005891 | 1 | 0 | Promoter (<=1kb) | 3 | 63883527 | 63930000 | 46474 | 1 | 229363 | ENSMUST00000029405.8 | 0 | ENSMUSG00000027823 | Gmps | guanine monophosphate synthetase |
| chr12 | 16890052 | 16891451 | 1400 | * | 10,36463 | 3,725690041 | 1,232774 | 3,0222 | 0,002509 | 0,683140223 | 1,173325 | Distal Intergenic | 12 | 16894917 | 16900782 | 5866 | 2 | 1,03E+08 | ENSMUST00000245045.1 | 9331 | | Gm36495 | predicted gene, 36495 |
| chr1 | 75209513 | 75210923 | 1411 | * | 10,4406 | 3,725877937 | 1,155822 | 3,223573 | 0,001266 | 0,55735526 | 1,173325 | Promoter (<=1kb) | 1 | 75207507 | 75208746 | 1240 | 2 | 381272 | ENSMUST00000191271.2 | -767 | ENSMUSG00000096094 | A630095N17Rik | RIKEN cDNA A630095N17 gene |
| chr19 | 60523216 | 60524130 | 915 | * | 6,846265 | 3,726173485 | 1,35188 | 2,75629 | 0,005846 | 1 | 0 | Promoter (1-2kb) | 19 | 60522121 | 60525987 | 3867 | 2 | 78832 | ENSMUST00000238073.2 | 1857 | ENSMUSG00000033417 | Cacul1 | CDK2 associated, cullin domain 1 |
| chr3 | 41360431 | 41362055 | 1625 | * | 14,69738 | 3,726888486 | 1,132604 | 3,290548 | 0,001 | 0,482499612 | 1,41696 | Distal Intergenic | 3 | 41395778 | 41395902 | 125 | 2 | 1,15E+08 | ENSMUST00000158725.3 | 33847 | ENSMUSG00000089350 | Gm25487 | predicted gene, 25487 |
| chr11 | 1,01E+08 | 1,01E+08 | 1468 | * | 9,542503 | 3,729221999 | 1,256567 | 2,967787 | 0,003 | 0,681421811 | 1,425929 | Promoter (<=1kb) | 11 | 1,01E+08 | 1,01E+08 | 11864 | 2 | 12189 | ENSMUST00000156843.2 | 0 | ENSMUSG00000017146 | Brca1 | breast cancer 1, early onset |
| chr4 | 76363948 | 76364930 | 983 | * | 7,476859 | 3,731015797 | 1,317681 | 2,831501 | 0,004633 | 0,75938015 | 1,360622 | Intron (ENSMUST0000 | 4 | 75859475 | 76368217 | 508743 | 2 | 19266 | ENSMUST00000107289.9 | 3287 | ENSMUSG00000028399 | Ptprd | protein tyrosine phosphatase, receptor type, D |
| chr13 | 63378134 | 63380939 | 2806 | * | 28,248665 | 3,731723705 | 0,7197316 | 5,1848825 | 2,162E-07 | 0,00116151 | 2,8800171 | Intron (ENSMUST00000 | 13 | 63387843 | 63450096 | 62254 | 1 | 72061 | ENSMUST00000222929.2 | -6904 | ENSMUSG00000021458 | Aopep | aminopeptidase O |
| chr4 | 1,38E+08 | 1,38E+08 | 1096 | * | 8,918477 | 3,73193805 | 1,427845 | 2,613685 | 0,008957 | 0,791068542 | 2,237403 | Intron (ENSMUST0000 | 4 | 1,38E+08 | 1,38E+08 | 924 | 2 | 73563 | ENSMUST00000124620.2 | 19537 | ENSMUSG00000084804 | 1700095J12Rik | RIKEN cDNA 1700095J12 gene |
| chr16 | 90841399 | 90842195 | 797 | * | 9,123187 | 3,732401268 | 1,31755 | 2,832835 | 0,004614 | 0,749931768 | 1,425929 | Promoter (<=1kb) | 16 | 90810925 | 90841431 | 30507 | 2 | 67367 | ENSMUST00000118522.8 | 0 | ENSMUSG00000022974 | Paxbp1 | PAX3 and PAX7 binding protein 1 |
| chr3 | 129790814 | 129792911 | 2098 | * | 55,5319 | 3,73244974 | 0,5698825 | 6,5495077 | 5,773E-11 | 1,54955E-06 | 2,7254908 | Intron (ENSMUST00000 | 3 | 129782406 | 129784146 | 1741 | 2 | 99683 | ENSMUST00000172324.2 | -6668 | ENSMUSG00000001052 | Sec24b | Sec24 related gene family, member B (S. cerevisiae) |
| chr1 | 1,81E+08 | 1,81E+08 | 1217 | * | 11,33319 | 3,73251042 | 1,247107 | 2,992936 | 0,002763 | 0,683761897 | 1,2763 | Intron (ENSMUST0000 | 1 | 1,81E+08 | 1,81E+08 | 105827 | 1 | 72978 | ENSMUST00000209607.2 | 75712 | ENSMUSG00000026514 | Cnih3 | cornichon family AMPA receptor auxiliary protein 3 |
| chr9 | 96002126 | 96003390 | 1265 | * | 13,39241 | 3,732543985 | 0,994203 | 3,754308 | 0,000174 | 0,235194771 | 1,2763 | Promoter (<=1kb) | 9 | 96001485 | 96064863 | 63379 | 1 | 235533 | ENSMUST00000129774.2 | 641 | ENSMUSG00000041440 | Gk5 | glycerol kinase 5 (putative) |
| chr2 | 1,35E+08 | 1,35E+08 | 954 | * | 7,167279 | 3,736531429 | 1,392626 | 2,683082 | 0,007295 | 1 | 0 | Intron (ENSMUST0000 | 2 | 1,35E+08 | 1,35E+08 | 613146 | 1 | 18795 | ENSMUST00000070724.12 | 50162 | ENSMUSG00000051177 | Plcb1 | phospholipase C, beta 1 |
| chr1 | 63692114 | 63693497 | 1384 | * | 9,033803 | 3,737192476 | 1,168491 | 3,198307 | 0,001382 | 0,539352028 | 1,425929 | Intron (ENSMUST0000 | 1 | 63703379 | 63726069 | 22691 | 2 | 241073 | ENSMUST00000126954.2 | 32572 | ENSMUSG00000069085 | Dytn | dystrotelin |
| chr13 | 35588706 | 35589728 | 1023 | * | 6,995597 | 3,738183946 | 1,316967 | 2,838479 | 0,004533 | 1 | 0 | Intron (ENSMUST0000 | 13 | 35477130 | 35477260 | 131 | 2 | 1,15E+08 | ENSMUST00000104413.3 | -111446 | ENSMUSG00000077601 | Gm22674 | predicted gene, 22674 |
| chr7 | 1,39E+08 | 1,39E+08 | 1106 | * | 7,296809 | 3,738543592 | 1,35813 | 2,752715 | 0,00591 | 0,802671652 | 1,360622 | Distal Intergenic | 7 | 1,39E+08 | 1,4E+08 | 46839 | 1 | 76484 | ENSMUST00000053445.18 | -8316 | ENSMUSG00000066129 | Kndc1 | kinase non-catalytic C-lobe domain (KIND) containing 1 |
| chr19 | 47856274 | 47857278 | 1005 | * | 9,443646 | 3,739345388 | 1,203697 | 3,10655 | 0,001893 | 0,595849045 | 1,449706 | Promoter (<=1kb) | 19 | 47856080 | 47858749 | 2670 | 1 | 68214 | ENSMUST00000237607.2 | 194 | ENSMUSG00000025069 | Gsto2 | glutathione S-transferase omega 2 |
| chr10 | 86613517 | 86614916 | 1400 | * | 24,12345 | 3,740436802 | 1,317503 | 2,839035 | 0,004525 | 0,627785088 | 2,951062 | Promoter (<=1kb) | 10 | 86614869 | 86674253 | 59385 | 1 | 103466 | ENSMUST00000099396.3 | 0 | ENSMUSG00000054027 | Nt5dc3 | 5'-nucleotidase domain containing 3 |
| chr17 | 67492212 | 67493200 | 989 | * | 7,771042 | 3,740583644 | 1,25043 | 2,991439 | 0,002777 | 1 | 0 | Intron (ENSMUST0000 | 17 | 67263986 | 67354980 | 90995 | 2 | 19274 | ENSMUST00000224862.2 | -137232 | ENSMUSG00000033278 | Ptprm | protein tyrosine phosphatase, receptor type, M |
| chr12 | 76388441 | 76389975 | 1535 | * | 10,35213 | 3,741270907 | 1,253839 | 2,983853 | 0,002846 | 0,715249603 | 1,086118 | Distal Intergenic | 12 | 76394556 | 76403914 | 9359 | 2 | 109929 | ENSMUST00000176102.8 | 13939 | ENSMUSG00000056459 | Zbtb25 | zinc finger and BTB domain containing 25 |
| chr7 | 138511084 | 138512821 | 1738 | * | 34,068182 | 3,741962523 | 0,6443606 | 5,8072494 | 6,351E-09 | 8,5085E-05 | 2,3776742 | Promoter (<=1kb) | 7 | 138492565 | 138511235 | 18671 | 2 | 12176 | ENSMUST00000106112.2 | 0 | ENSMUSG00000078566 | Bnip3 | BCL2/adenovirus E1B interacting protein 3 |
| chr19 | 7501493 | 7502380 | 888 | * | 7,327636 | 3,742353767 | 1,416429 | 2,642104 | 0,008239 | 1 | 0 | Intron (ENSMUST0000 | 19 | 7471900 | 7511565 | 39666 | 1 | 109168 | ENSMUST00000236382.2 | 29593 | ENSMUSG00000024759 | Atl3 | atlastin GTPase 3 |
| chr19 | 45988204 | 45989905 | 1702 | * | 10,41354 | 3,743216213 | 1,296702 | 2,88672 | 0,003893 | 0,741447731 | 1,2763 | Promoter (1-2kb) | 19 | 45906883 | 45986927 | 80045 | 2 | 71617 | ENSMUST00000235131.2 | -1277 | ENSMUSG00000039901 | Armh3 | armadillo-like helical domain containing 3 |
| chr9 | 52725832 | 52727714 | 1883 | * | 9,867054 | 3,744522174 | 1,334354 | 2,806244 | 0,005012 | 0,81655291 | 1,086118 | Distal Intergenic | 9 | 52591163 | 52710074 | 118912 | 1 | 382139 | ENSMUST00000213681.2 | 134669 | ENSMUSG00000111020 | Gm1715 | predicted gene 1715 |
| chr10 | 1,22E+08 | 1,22E+08 | 837 | * | 6,830989 | 3,744835506 | 1,410909 | 2,6542 | 0,00795 | 1 | 0 | Intron (ENSMUST0000 | 10 | 1,22E+08 | 1,22E+08 | 16494 | 2 | 117600 | ENSMUST00000161156.2 | -95734 | ENSMUSG00000020121 | Srgap1 | SLIT-ROBO Rho GTPase activating protein 1 |
| chr13 | 24719582 | 24721800 | 2219 | * | 30,821614 | 3,745113683 | 0,7029101 | 5,3280121 | 9,929E-08 | 0,000698057 | 2,7254908 | Intron (ENSMUST000002 | 13 | 24685508 | 24889181 | 203674 | 1 | 193385 | ENSMUST00000238974.2 | 34074 | ENSMUSG00000036006 | Ripor2 | RHO family interacting cell polarization regulator 2 |
| chr10 | 80874629 | 80876558 | 1930 | * | 13,46543 | 3,745993848 | 1,136522 | 3,296015 | 0,000981 | 0,497663879 | 1,2763 | Promoter (1-2kb) | 10 | 80864372 | 80873260 | 8889 | 2 | 106947 | ENSMUST00000117276.11 | -1369 | ENSMUSG00000046822 | Slc39a3 | solute carrier family 39 (zinc transporter), member 3 |
| chr10 | 51635201 | 51636074 | 874 | * | 9,490196 | 3,746668236 | 1,118341 | 3,350201 | 0,000808 | 0,453798789 | 1,360622 | Promoter (1-2kb) | 10 | 51632270 | 51633290 | 1021 | 2 | 71090 | ENSMUST00000219766.2 | -1911 | ENSMUSG00000062224 | 4933411G06Rik | Sumo1 pseudogene |
| chr11 | 94394143 | 94395185 | 1043 | * | 9,25163 | 3,747005315 | 1,259341 | 2,97537 | 0,002926 | 0,676922067 | 1,425929 | Promoter (<=1kb) | 11 | 94392173 | 94394397 | 2225 | 2 | 104601 | ENSMUST00000139432.2 | 0 | ENSMUSG00000039110 | Mycbpap | MYCBP associated protein |
| chr4 | 1,36E+08 | 1,36E+08 | 1241 | * | 12,37882 | 3,747357903 | 1,189829 | 3,149494 | 0,001636 | 0,594430593 | 1,2763 | Promoter (<=1kb) | 4 | 1,36E+08 | 1,36E+08 | 23664 | 1 | 242705 | ENSMUST00000061721.6 | -661 | ENSMUSG00000018983 | E2f2 | E2F transcription factor 2 |
| chr14 | 67245712 | 67247985 | 2274 | * | 22,84488 | 3,747457778 | 1,280242 | 2,927147 | 0,003421 | 0,5770513 | 2,880017 | Promoter (<=1kb) | 14 | 67222688 | 67246305 | 23618 | 2 | 12177 | ENSMUST00000111115.8 | 0 | ENSMUSG00000022051 | Bnip3l | BCL2/adenovirus E1B interacting protein 3-like |
| chr4 | 1,35E+08 | 1,35E+08 | 1112 | * | 10,80854 | 3,747757151 | 1,208899 | 3,10014 | 0,001934 | 0,715156227 | 0,744375 | Distal Intergenic | 4 | 1,35E+08 | 1,35E+08 | 2171 | 1 | 1,03E+08 | ENSMUST00000145711.2 | -6613 | | Gm12984 | predicted gene 12984 |
| chr5 | 117500394 | 117503321 | 2928 | * | 28,88328 | 3,748960563 | 0,7962311 | 4,7083825 | 2,497E-06 | 0,007482484 | 2,9510621 | Promoter (<=1kb) | 5 | 117501587 | 117515872 | 14286 | 1 | 59043 | ENSMUST00000111959.2 | 0 | ENSMUSG00000029364 | Wsb2 | WD repeat and SOCS box-containing 2 |
| chr13 | 22198638 | 22200105 | 1468 | * | 9,148168 | 3,749548059 | 1,440674 | 2,602634 | 0,009251 | 0,872576097 | 1,449706 | Exon (ENSMUST00001 | 13 | 22191402 | 22193912 | 2511 | 2 | 54373 | ENSMUST00000130585.2 | -4726 | ENSMUSG00000006179 | Prss16 | protease, serine 16 (thymus) |
| chr5 | 35517253 | 35518576 | 1324 | * | 14,00458 | 3,75032792 | 1,0405 | 3,604352 | 0,000313 | 0,352774915 | 0,938682 | Promoter (<=1kb) | 5 | 35518885 | 35524785 | 5901 | 1 | 74429 | ENSMUST00000201379.4 | -309 | ENSMUSG00000106762 | 4930478P22Rik | RIKEN cDNA 4930478P22 gene |
| chr4 | 49441080 | 49441904 | 825 | * | 8,45754 | 3,750826038 | 1,30158 | 2,881748 | 0,003955 | 0,722798324 | 1,449706 | Distal Intergenic | 4 | 49447105 | 49451134 | 4030 | 2 | 230161 | ENSMUST00000107697.2 | 9230 | ENSMUSG00000070985 | Acnat1 | acyl-coenzyme A amino acid N-acyltransferase 1 |
| chr9 | 37269140 | 37270570 | 1431 | * | 13,25618 | 3,751767237 | 1,102516 | 3,402916 | 0,000667 | 0,446344042 | 1,173325 | Distal Intergenic | 9 | 37278647 | 37297871 | 19225 | 1 | 72927 | ENSMUST00000051839.9 | -8077 | ENSMUSG00000046240 | Hepacam | hepatocyte cell adhesion molecule |
| chr10 | 7841869 | 7842485 | 617 | * | 6,726052 | 3,752721407 | 1,383482 | 2,712519 | 0,006677 | 1 | 0 | Distal Intergenic | 10 | 7781417 | 7831994 | 50578 | 2 | 68652 | ENSMUST00000146444.8 | -9875 | ENSMUSG00000015755 | Tab2 | TGF-beta activated kinase 1/MAP3K7 binding protein 2 |
| chr3 | 95129857 | 95130814 | 958 | * | 9,34892 | 3,753026452 | 1,339486 | 2,80184 | 0,005081 | 0,797584062 | 1,211315 | Intron (ENSMUST0000 | 3 | 95106881 | 95125227 | 18347 | 2 | 213054 | ENSMUST00000138862.8 | -4630 | ENSMUSG00000038766 | Gabpb2 | GA repeat binding protein, beta 2 |
| chr1 | 64641626 | 64643008 | 1383 | * | 11,275 | 3,753646102 | 1,375751 | 2,728434 | 0,006364 | 0,887433944 | 0,918255 | 3' UTR | 1 | 64647115 | 64654790 | 7676 | 2 | 67099 | ENSMUST00000162806.8 | 11782 | ENSMUSG00000025956 | Mettl21a | methyltransferase like 21A |
| chr13 | 4290729 | 4291824 | 1096 | * | 9,902723 | 3,754501202 | 1,259756 | 2,98034 | 0,002879 | 0,705098751 | 1,173325 | Intron (ENSMUST0000 | 13 | 4295327 | 4298680 | 3354 | 1 | 432720 | ENSMUST00000129751.2 | -3503 | ENSMUSG00000071551 | Akr1c19 | aldo-keto reductase family 1, member C19 |
| chr11 | 50678261 | 50679262 | 1002 | * | 9,333758 | 3,755425262 | 1,277825 | 2,93892 | 0,003294 | 0,621084104 | 2,237403 | Intron (ENSMUST0000 | 11 | 50672597 | 50673625 | 1029 | 1 | 216725 | ENSMUST00000127534.2 | 5664 | ENSMUSG00000036545 | Adamts2 | a disintegrin-like and metallopeptidase (reprolysin type) with thrombospondin type 1 motif, 2 |
| chr10 | 1,04E+08 | 1,04E+08 | 1382 | * | 9,376812 | 3,755560889 | 1,304397 | 2,879155 | 0,003987 | 0,653696223 | 2,237403 | Distal Intergenic | 10 | 1,04E+08 | 1,04E+08 | 157 | 2 | 1,15E+08 | ENSMUST00000158557.3 | 68593 | ENSMUSG00000089182 | Gm22945 | predicted gene, 22945 |
| chr9 | 41431349 | 41432351 | 1003 | * | 9,195729 | 3,755997885 | 1,417728 | 2,649308 | 0,008066 | 0,771131827 | 2,237403 | Intron (ENSMUST0000 | 9 | 41442721 | 41442800 | 80 | 1 | 723892 | ENSMUST00000175270.3 | -10370 | ENSMUSG00000093011 | Mir100 | microRNA 100 |
| chr1 | 1,28E+08 | 1,28E+08 | 744 | * | 7,929269 | 3,756024005 | 1,388508 | 2,705078 | 0,006829 | 0,830691256 | 1,360622 | Promoter (<=1kb) | 1 | 1,28E+08 | 1,28E+08 | 75096 | 1 | 226407 | ENSMUST00000037649.6 | 0 | ENSMUSG00000036104 | Rab3gap1 | RAB3 GTPase activating protein subunit 1 |
| chr7 | 23782248 | 23783957 | 1710 | * | 17,67818 | 3,75603198 | 1,002479 | 3,746745 | 0,000179 | 0,178046761 | 2,154504 | Promoter (<=1kb) | 7 | 23781379 | 23807117 | 25739 | 1 | 210135 | ENSMUST00000203854.3 | 869 | ENSMUSG00000057101 | Zfp180 | zinc finger protein 180 |
| chr9 | 60408974 | 60410467 | 1494 | * | 15,59053 | 3,756940948 | 1,014991 | 3,701451 | 0,000214 | 0,252389304 | 1,37683 | Intron (ENSMUST0000 | 9 | 59874214 | 60404409 | 530196 | 2 | 207596 | ENSMUST00000171654.8 | -4565 | ENSMUSG00000032289 | Thsd4 | thrombospondin, type I, domain containing 4 |
| chr17 | 73596283 | 73597243 | 961 | * | 10,68167 | 3,757146985 | 1,249134 | 3,007801 | 0,002631 | 0,730469267 | 0,918255 | Distal Intergenic | 17 | 73559540 | 73567651 | 8112 | 1 | 328855 | ENSMUST00000233479.2 | 36743 | ENSMUSG00000117292 | E330032C10Rik | RIKEN cDNA E330032C10 gene |
| chr13 | 1,03E+08 | 1,03E+08 | 836 | * | 8,851297 | 3,758003937 | 1,157125 | 3,247707 | 0,001163 | 0,504427013 | 1,449706 | Distal Intergenic | 13 | 1,03E+08 | 1,03E+08 | 27099 | 2 | 1,03E+08 | ENSMUST00000225128.2 | -47394 | | Gm29927 | predicted gene, 29927 |
| chr11 | 21047929 | 21048810 | 882 | * | 8,091017 | 3,758193468 | 1,364415 | 2,754435 | 0,005879 | 1 | 0 | Promoter (<=1kb) | 11 | 21049534 | 21049827 | 294 | 1 | 1,08E+08 | ENSMUST00000121233.2 | -724 | | 2610024D14Rik | ATP synthase subunit g, mitochondrial pseudogene |
| chr13 | 76027650 | 76028487 | 838 | * | 8,609086 | 3,759352538 | 1,206403 | 3,116165 | 0,001832 | 0,591842599 | 1,449706 | Promoter (2-3kb) | 13 | 76020490 | 76025514 | 5025 | 2 | 73296 | ENSMUST00000222923.2 | -2136 | ENSMUSG00000021589 | Rhobtb3 | Rho-related BTB domain containing 3 |
| chr3 | 1,25E+08 | 1,25E+08 | 926 | * | 7,865655 | 3,760225715 | 1,38871 | 2,707712 | 0,006775 | 1 | 0 | Distal Intergenic | 3 | 1,25E+08 | 1,26E+08 | 324824 | 1 | 64580 | ENSMUST00000173932.8 | -48446 | ENSMUSG00000027971 | Ndst4 | N-deacetylase/N-sulfotransferase (heparin glucosaminyl) 4 |
| chr4 | 56430214 | 56431159 | 946 | * | 8,907492 | 3,760253015 | 1,346164 | 2,793309 | 0,005217 | 0,775744126 | 1,425929 | Distal Intergenic | 4 | 56678904 | 56679035 | 132 | 2 | 1,15E+08 | ENSMUST00000104475.2 | 247876 | ENSMUSG00000077663 | Gm26144 | predicted gene, 26144 |
| chr16 | 31820647 | 31823120 | 2474 | * | 18,28386 | 3,761118372 | 0,950444 | 3,957221 | 7,58E-05 | 0,097946479 | 2,386008 | Promoter (<=1kb) | 16 | 31778490 | 31822105 | 43616 | 2 | 320213 | ENSMUST00000231360.2 | 0 | ENSMUSG00000022772 | Senp5 | SUMO/sentrin specific peptidase 5 |
| chr1 | 92828421 | 92829336 | 916 | * | 7,786942 | 3,762078934 | 1,405056 | 2,677529 | 0,007417 | 1 | 0 | Promoter (1-2kb) | 1 | 92797851 | 92830628 | 32778 | 2 | 241158 | ENSMUST00000112998.2 | 1292 | ENSMUSG00000034212 | Ankmy1 | ankyrin repeat and MYND domain containing 1 |
| chrX | 6326027 | 6326919 | 893 | * | 6,318098 | 3,763106734 | 1,459904 | 2,57764 | 0,009948 | 1 | 0 | Promoter (<=1kb) | 20 | 6327796 | 6495594 | 167799 | 1 | 208431 | ENSMUST00000143641.4 | -877 | ENSMUSG00000068270 | Shroom4 | shroom family member 4 |
| chr9 | 43670579 | 43671857 | 1279 | * | 12,59153 | 3,763920335 | 1,196918 | 3,144676 | 0,001663 | 0,595849045 | 1,2763 | Intron (ENSMUST0000 | 9 | 43628138 | 43657397 | 29260 | 2 | 1,03E+08 | ENSMUST00000216604.2 | -13182 | | Gm30015 | predicted gene, 30015 |
| chr18 | 35845034 | 35845856 | 823 | * | 6,174733 | 3,764011559 | 1,459379 | 2,579187 | 0,009903 | 1 | 0 | Intron (ENSMUST0000 | 18 | 35816325 | 35841467 | 25143 | 2 | 76594 | ENSMUST00000237995.2 | -3567 | ENSMUSG00000024350 | Dnajc18 | DnaJ heat shock protein family (Hsp40) member C18 |
| chr6 | 43990733 | 43991339 | 607 | * | 6,365289 | 3,76403915 | 1,456733 | 2,58389 | 0,009769 | 1 | 0 | Distal Intergenic | 6 | 44057422 | 44060289 | 2868 | 2 | 1,03E+08 | ENSMUST00000203777.2 | 68950 | ENSMUSG00000107598 | Gm35216 | predicted gene, 35216 |
| chr11 | 1,04E+08 | 1,04E+08 | 1168 | * | 10,33452 | 3,764161632 | 1,140609 | 3,300134 | 0,000966 | 0,474532331 | 1,449706 | Intron (ENSMUST0000 | 11 | 1,04E+08 | 1,04E+08 | 6379 | 2 | 76719 | ENSMUST00000069188.7 | -11028 | ENSMUSG00000018412 | Kansl1 | KAT8 regulatory NSL complex subunit 1 |
| chr16 | 20472849 | 20474479 | 1631 | * | 9,21054 | 3,765156139 | 1,29387 | 2,909996 | 0,003614 | 0,737047928 | 1,211315 | Promoter (2-3kb) | 16 | 20470466 | 20471504 | 1039 | 1 | 21762 | ENSMUST00000169195.2 | 2383 | ENSMUSG00000006998 | Psmd2 | proteasome (prosome, macropain) 26S subunit, non-ATPase, 2 |
| chr1 | 1,85E+08 | 1,85E+08 | 1739 | * | 8,54868 | 3,765555517 | 1,318594 | 2,855736 | 0,004294 | 0,748971518 | 1,360622 | Promoter (<=1kb) | 1 | 1,85E+08 | 1,85E+08 | 47156 | 1 | 107508 | ENSMUST00000192284.6 | 0 | ENSMUSG00000026615 | Eprs | glutamyl-prolyl-tRNA synthetase |
| chr8 | 32708128 | 32710678 | 2551 | * | 35,462537 | 3,765982823 | 0,610795 | 6,1657071 | 7,017E-10 | 2,06005E-05 | 1,4057095 | Intron (ENSMUST000002 | 8 | 32676157 | 32676276 | 120 | 1 | 115487032 | ENSMUST00000158745.4 | 31971 | ENSMUSG00000089370 | Gm22140 | predicted gene, 22140 |
| chr11 | 1,19E+08 | 1,19E+08 | 1489 | * | 10,01478 | 3,766175683 | 1,169383 | 3,220653 | 0,001279 | 0,5515848 | 1,211315 | Intron (ENSMUST0000 | 11 | 1,19E+08 | 1,19E+08 | 2457 | 1 | 1,08E+08 | ENSMUST00000146016.2 | 14086 | | Gm11750 | predicted gene 11750 |
| chr16 | 32646388 | 32648665 | 2278 | * | 21,02372 | 3,766396223 | 0,976801 | 3,855849 | 0,000115 | 0,11427637 | 2,880017 | Promoter (1-2kb) | 16 | 32642077 | 32645290 | 3214 | 2 | 1,01E+08 | ENSMUST00000140229.2 | -1098 | ENSMUSG00000035629 | Rubcn | RUN domain and cysteine-rich domain containing, Beclin 1-interacting protein |
| chr2 | 1,27E+08 | 1,27E+08 | 1223 | * | 10,2903 | 3,766598914 | 1,319422 | 2,854735 | 0,004307 | 0,73648081 | 1,449706 | Promoter (<=1kb) | 2 | 1,27E+08 | 1,27E+08 | 14367 | 1 | 99138 | ENSMUST00000131196.2 | 0 | ENSMUSG00000027367 | Stard7 | START domain containing 7 |
| chr9 | 1,11E+08 | 1,11E+08 | 1063 | * | 11,77288 | 3,766946832 | 1,179465 | 3,193776 | 0,001404 | 0,663408464 | 0,744375 | Promoter (2-3kb) | 9 | 1,11E+08 | 1,11E+08 | 15564 | 2 | 19228 | ENSMUST00000199791.2 | -2055 | ENSMUSG00000032492 | Pth1r | parathyroid hormone 1 receptor |
| chr15 | 44481403 | 44482646 | 1244 | * | 16,60762 | 3,767040282 | 1,125527 | 3,346913 | 0,000817 | 0,466496079 | 1,2763 | Promoter (<=1kb) | 15 | 44482571 | 44504911 | 22341 | 1 | 55960 | ENSMUST00000227691.2 | 0 | ENSMUSG00000022339 | Ebag9 | estrogen receptor-binding fragment-associated gene 9 |
| chr12 | 1,12E+08 | 1,12E+08 | 1057 | * | 7,19143 | 3,767101056 | 1,420157 | 2,652595 | 0,007988 | 1 | 0 | Intron (ENSMUST0000 | 12 | 1,12E+08 | 1,12E+08 | 12594 | 1 | 74691 | ENSMUST00000194800.2 | 5235 | ENSMUSG00000054003 | Tdrd9 | tudor domain containing 9 |
| chr11 | 8576141 | 8577048 | 908 | * | 9,183948 | 3,76756155 | 1,382244 | 2,725685 | 0,006417 | 0,811413964 | 1,425929 | Promoter (1-2kb) | 11 | 8381652 | 8574887 | 193236 | 2 | 319939 | ENSMUST00000239091.2 | -1254 | ENSMUSG00000020422 | Tns3 | tensin 3 |
| chr4 | 1,24E+08 | 1,24E+08 | 551 | * | 8,801176 | 3,768933392 | 1,256171 | 3,000334 | 0,002697 | 0,580348965 | 2,237403 | Intron (ENSMUST0000 | 4 | 1,24E+08 | 1,24E+08 | 25892 | 2 | 1,05E+08 | ENSMUST00000137546.2 | 18222 | | 1700121C08Rik | RIKEN cDNA 1700121C08 gene |
| chr2 | 1,44E+08 | 1,44E+08 | 950 | * | 8,982738 | 3,769840783 | 1,21429 | 3,104564 | 0,001906 | 0,610904497 | 1,360622 | Promoter (<=1kb) | 2 | 1,44E+08 | 1,44E+08 | 6435 | 1 | 70408 | ENSMUST00000143292.8 | -54 | ENSMUSG00000027427 | Polr3f | polymerase (RNA) III (DNA directed) polypeptide F |
| chr5 | 24578764 | 24580066 | 1303 | * | 10,43691 | 3,770148754 | 1,248571 | 3,01957 | 0,002531 | 0,7591629 | 0,744375 | Intron (ENSMUST0000 | 5 | 24569808 | 24589472 | 19665 | 1 | 18127 | ENSMUST00000030834.7 | 8956 | ENSMUSG00000028978 | Nos3 | nitric oxide synthase 3, endothelial cell |
| chr2 | 29336363 | 29337288 | 926 | * | 10,48942 | 3,770343992 | 1,277604 | 2,951106 | 0,003166 | 0,80080207 | 0,744375 | Intron (ENSMUST0000 | 2 | 29365227 | 29365306 | 80 | 2 | 1,02E+08 | ENSMUST00000184068.3 | 28018 | ENSMUSG00000098539 | Mir133c | microRNA 133c |
| chr10 | 1,13E+08 | 1,13E+08 | 625 | * | 7,507525 | 3,771327153 | 1,361752 | 2,769467 | 0,005615 | 1 | 0 | Promoter (<=1kb) | 10 | 1,13E+08 | 1,13E+08 | 11133 | 2 | 1,05E+08 | ENSMUST00000218612.2 | -193 | | Gm40765 | predicted gene, 40765 |
| chr3 | 1,38E+08 | 1,38E+08 | 1974 | * | 17,52472 | 3,771862993 | 0,971191 | 3,883749 | 0,000103 | 0,126797776 | 2,154504 | Promoter (<=1kb) | 3 | 1,38E+08 | 1,39E+08 | 165711 | 1 | 56224 | ENSMUST00000029800.9 | -79 | ENSMUSG00000028152 | Tspan5 | tetraspanin 5 |
| chr18 | 32505581 | 32506470 | 890 | * | 8,432264 | 3,773282394 | 1,315874 | 2,86751 | 0,004137 | 0,731734453 | 1,425929 | Intron (ENSMUST0000 | 18 | 32492106 | 32510105 | 18000 | 2 | 402767 | ENSMUST00000234930.2 | 3635 | | A830052D11Rik | RIKEN cDNA A830052D11 gene |
| chr4 | 1,33E+08 | 1,33E+08 | 1106 | * | 8,885854 | 3,773679872 | 1,3919 | 2,711172 | 0,006705 | 0,819350294 | 1,425929 | Distal Intergenic | 4 | 1,33E+08 | 1,33E+08 | 3161 | 1 | 23957 | ENSMUST00000042706.3 | 3575 | ENSMUSG00000037583 | Nr0b2 | nuclear receptor subfamily 0, group B, member 2 |
| chr15 | 79340988 | 79342254 | 1267 | * | 12,26317 | 3,774334732 | 1,272605 | 2,965833 | 0,003019 | 0,728037066 | 1,075735 | Promoter (1-2kb) | 15 | 79310459 | 79339767 | 29309 | 2 | 27373 | ENSMUST00000230599.2 | -1221 | ENSMUSG00000022433 | Csnk1e | casein kinase 1, epsilon |
| chr12 | 1,08E+08 | 1,08E+08 | 1009 | * | 8,598617 | 3,774702497 | 1,388295 | 2,718948 | 0,006549 | 0,811757052 | 1,449706 | Intron (ENSMUST0000 | 12 | 1,08E+08 | 1,08E+08 | 6136 | 2 | 74968 | ENSMUST00000144556.2 | -40127 | ENSMUSG00000085641 | 4930465M20Rik | RIKEN cDNA 4930465M20 gene |
| chr7 | 18951926 | 18953035 | 1110 | * | 8,453841 | 3,774897747 | 1,380482 | 2,734478 | 0,006248 | 0,802323085 | 1,449706 | Intron (ENSMUST0000 | 7 | 18946463 | 18958099 | 11637 | 1 | 319197 | ENSMUST00000060225.6 | 5463 | ENSMUSG00000044317 | Gpr4 | G protein-coupled receptor 4 |
| chr2 | 1,03E+08 | 1,03E+08 | 1037 | * | 8,792961 | 3,774990263 | 1,33058 | 2,837101 | 0,004553 | 0,755814512 | 1,360622 | Exon (ENSMUST00000 | 2 | 1,03E+08 | 1,03E+08 | 107 | 1 | 1,15E+08 | ENSMUST00000239821.1 | -59025 | ENSMUSG00000119260 | Gm23731 | predicted gene, 23731 |
| chr7 | 1,02E+08 | 1,02E+08 | 1571 | * | 13,18558 | 3,775276361 | 1,205967 | 3,130498 | 0,001745 | 0,639383182 | 1,075735 | Intron (ENSMUST0000 | 7 | 1,02E+08 | 1,02E+08 | 45144 | 1 | 101706 | ENSMUST00000084852.13 | -4348 | ENSMUSG00000066306 | Numa1 | nuclear mitotic apparatus protein 1 |
| chr2 | 25426535 | 25427349 | 815 | * | 10,9708 | 3,775567772 | 1,092934 | 3,454525 | 0,000551 | 0,492478878 | 0,744375 | Promoter (1-2kb) | 2 | 25408878 | 25429152 | 20275 | 2 | 22030 | ENSMUST00000151742.2 | 1803 | ENSMUSG00000026942 | Traf2 | TNF receptor-associated factor 2 |
| chr8 | 4831065 | 4832464 | 1400 | * | 7,657433 | 3,775772731 | 1,339967 | 2,817811 | 0,004835 | 1 | 0 | Promoter (1-2kb) | 8 | 4813468 | 4829567 | 16100 | 2 | 20419 | ENSMUST00000208381.2 | -1498 | ENSMUSG00000022322 | Shcbp1 | Shc SH2-domain binding protein 1 |
| chr1 | 54768350 | 54768930 | 581 | * | 5,96945 | 3,776818826 | 1,453048 | 2,599238 | 0,009343 | 1 | 0 | Exon (ENSMUST00001 | 1 | 54768146 | 54774803 | 6658 | 2 | 329154 | ENSMUST00000178685.2 | 5873 | ENSMUSG00000052331 | Ankrd44 | ankyrin repeat domain 44 |
| chr5 | 12899797 | 12900523 | 727 | * | 8,494176 | 3,777808152 | 1,338544 | 2,822327 | 0,004768 | 0,683067851 | 2,237403 | Distal Intergenic | 5 | 12625761 | 12638719 | 12959 | 1 | 108151 | ENSMUST00000195923.2 | 274036 | ENSMUSG00000040254 | Sema3d | sema domain, immunoglobulin domain (Ig), short basic domain, secreted, (semaphorin) 3D |
| chr6 | 86379355 | 86380663 | 1309 | * | 10,94861 | 3,778252982 | 1,378946 | 2,739957 | 0,006145 | 0,91311537 | 0,744375 | Promoter (<=1kb) | 6 | 86381201 | 86408183 | 26983 | 1 | 21841 | ENSMUST00000095754.10 | -538 | ENSMUSG00000071337 | Tia1 | cytotoxic granule-associated RNA binding protein 1 |
| chr6 | 28589283 | 28589981 | 699 | * | 6,375203 | 3,778707238 | 1,424931 | 2,651853 | 0,008005 | 1 | 0 | Intron (ENSMUST0000 | 6 | 28561816 | 28724970 | 163155 | 1 | 56463 | ENSMUST00000171532.8 | 27467 | ENSMUSG00000001424 | Snd1 | staphylococcal nuclease and tudor domain containing 1 |
| chr12 | 96703059 | 96704061 | 1003 | * | 11,51774 | 3,780069837 | 1,120827 | 3,372572 | 0,000745 | 0,54190677 | 0,744375 | Distal Intergenic | 12 | 97279623 | 97282627 | 3005 | 1 | 1,05E+08 | ENSMUST00000218233.2 | -575562 | | Gm40552 | predicted gene, 40552 |
| chr5 | 1,22E+08 | 1,22E+08 | 1813 | * | 13,8554 | 3,780286686 | 1,187216 | 3,18416 | 0,001452 | 0,62786102 | 0,938682 | Promoter (<=1kb) | 5 | 1,22E+08 | 1,22E+08 | 3670 | 1 | 17906 | ENSMUST00000123913.8 | 0 | ENSMUSG00000013936 | Myl2 | myosin, light polypeptide 2, regulatory, cardiac, slow |
| chr19 | 43916435 | 43917343 | 909 | * | 7,22165 | 3,780303218 | 1,425447 | 2,652013 | 0,008001 | 1 | 0 | Intron (ENSMUST0000 | 19 | 43908912 | 43916367 | 7456 | 1 | 1,15E+08 | ENSMUST00000235838.2 | 7523 | | Gm50217 | predicted gene, 50217 |
| chr5 | 1,15E+08 | 1,15E+08 | 1724 | * | 9,373777 | 3,781047565 | 1,328581 | 2,845929 | 0,004428 | 0,751210692 | 1,360622 | Promoter (<=1kb) | 5 | 1,15E+08 | 1,15E+08 | 7940 | 2 | 384281 | ENSMUST00000139167.3 | 0 | ENSMUSG00000029536 | Gatc | glutamyl-tRNA(Gln) amidotransferase, subunit C |
| chr17 | 26772709 | 26773946 | 1238 | * | 13,147 | 3,782501952 | 1,171429 | 3,228965 | 0,001242 | 0,603552508 | 0,918255 | 5' UTR | 17 | 26780463 | 26875905 | 95443 | 1 | 67458 | ENSMUST00000167662.8 | -6517 | ENSMUSG00000001576 | Ergic1 | endoplasmic reticulum-golgi intermediate compartment (ERGIC) 1 |
| chrX | 81776315 | 81777228 | 914 | * | 6,601453 | 3,783009926 | 1,435856 | 2,634672 | 0,008422 | 1 | 0 | Distal Intergenic | 20 | 81735845 | 81735976 | 132 | 2 | 1,15E+08 | ENSMUST00000104486.3 | -40339 | ENSMUSG00000077674 | Gm24470 | predicted gene, 24470 |
| chr9 | 1,2E+08 | 1,2E+08 | 1344 | * | 9,97918 | 3,783156085 | 1,38884 | 2,723968 | 0,00645 | 0,809369408 | 1,449706 | Distal Intergenic | 9 | 1,2E+08 | 1,2E+08 | 120750 | 1 | 245049 | ENSMUST00000133173.3 | -8921 | ENSMUSG00000041794 | Myrip | myosin VIIA and Rab interacting protein |
| chr5 | 1,45E+08 | 1,45E+08 | 1080 | * | 7,484531 | 3,783163135 | 1,434845 | 2,636636 | 0,008373 | 1 | 0 | Intron (ENSMUST0000 | 5 | 1,45E+08 | 1,45E+08 | 13403 | 2 | 75788 | ENSMUST00000198621.2 | -16258 | ENSMUSG00000038780 | Smurf1 | SMAD specific E3 ubiquitin protein ligase 1 |
| chr1 | 33909621 | 33910383 | 763 | * | 7,783459 | 3,783507282 | 1,43148 | 2,643073 | 0,008216 | 1 | 0 | Intron (ENSMUST0000 | 1 | 33901776 | 33922561 | 20786 | 2 | 320705 | ENSMUST00000195350.6 | 12178 | ENSMUSG00000042182 | Bend6 | BEN domain containing 6 |

| | | | | | | | | | | | | | | | | | | | | | | | |
|---|---|---|---|---|---|---|---|---|---|---|---|---|---|---|---|---|---|---|---|---|---|---|---|
| chr11 | 55330187 | 55331592 | 1406 | * | 12,0772 | 3,783821779 | 1,118637 | 3,382529 | 0,000718 | 0,539618083 | 0,738441 | Distal Intergenic | 11 | 55286247 | 55314009 | 27763 | 2 | 20692 | ENSMUST00000214685.2 | -16178 | ENSMUSG00000018593 | Sparc | secreted acidic cysteine rich glycoprotein |
| chr11 | 54323008 | 54324016 | 1009 | * | 8,85738 | 3,784188936 | 1,365386 | 2,771515 | 0,00558 | 0,787934397 | 1,425929 | Distal Intergenic | 11 | 54329025 | 54409061 | 80037 | 1 | 216742 | ENSMUST00000046835.14 | -5009 | ENSMUSG00000035992 | Fnip1 | folliculin interacting protein 1 |
| chr9 | 66106487 | 66108384 | 1898 | * | 16,43841 | 3,784852785 | 1,074113 | 3,523701 | 0,000426 | 0,255794423 | 2,663619 | Intron (ENSMUST0000 | 9 | 66128029 | 66176232 | 48204 | 1 | 13143 | ENSMUST00000132987.8 | -19645 | ENSMUSG00000032380 | Dapk2 | death-associated protein kinase 2 |
| chr7 | 99164134 | 99165732 | 1599 | * | 11,51567 | 3,78574395 | 1,359273 | 2,785124 | 0,005351 | 0,891097601 | 0,744375 | Distal Intergenic | 7 | 99184673 | 99255978 | 71306 | 1 | 109689 | ENSMUST00000098266.9 | -18941 | ENSMUSG00000018909 | Arrb1 | arrestin, beta 1 |
| chr10 | 80400378 | 80401911 | 1534 | * | 10,45326 | 3,785972035 | 1,233476 | 3,069352 | 0,002145 | 0,730765405 | 0,744375 | Promoter (2-3kb) | 10 | 80376759 | 80397394 | 20636 | 2 | 66932 | ENSMUST00000057910.16 | -2984 | ENSMUSG00000047417 | Rexo1 | REX1, RNA exonuclease 1 |
| chr11 | 96825286 | 96826176 | 891 | * | 11,77273 | 3,78685267 | 1,20943 | 3,131106 | 0,001741 | 0,605351622 | 1,2763 | 5' UTR | 11 | 96822213 | 96826106 | 3894 | 1 | 217138 | ENSMUST00000107633.2 | 3073 | ENSMUSG00000047040 | Prr15l | proline rich 15-like |
| chr13 | 96053462 | 96054591 | 1130 | * | 9,446379 | 3,788008794 | 1,380175 | 2,744585 | 0,006059 | 0,722439117 | 2,237403 | Distal Intergenic | 13 | 96033470 | 96033628 | 159 | 2 | 1,15E+08 | ENSMUST00000083932.3 | -19834 | ENSMUSG00000065866 | Gm24032 | predicted gene, 24032 |
| chr4 | 1,34E+08 | 1,34E+08 | 1601 | * | 14,25267 | 3,788548528 | 1,104907 | 3,428839 | 0,000606 | 0,44621096 | 1,075735 | Promoter (<=1kb) | 4 | 1,34E+08 | 1,34E+08 | 12929 | 1 | 69885 | ENSMUST00000105866.3 | 0 | ENSMUSG00000078521 | Aunip | aurora kinase A and ninein interacting protein |
| chr10 | 25783969 | 25784790 | 822 | * | 8,825509 | 3,790027587 | 1,308679 | 2,896072 | 0,003779 | 0,716908183 | 1,425929 | Distal Intergenic | 10 | 25655212 | 25713335 | 58124 | 2 | 73803 | ENSMUST00000218300.2 | -70634 | ENSMUSG00000112444 | 4930401C15Rik | RIKEN cDNA 4930401C15 gene |
| chr1 | 72281500 | 72282106 | 607 | * | 6,838541 | 3,7916954 | 1,446416 | 2,621441 | 0,008756 | 1 | 0 | Promoter (1-2kb) | 1 | 72283372 | 72283562 | 191 | 1 | 1,15E+08 | ENSMUST00001239518.1 | -1266 | ENSMUSG00000119873 | Gm24497 | predicted gene, 24497 |
| chr18 | 67746324 | 67746896 | 573 | * | 6,521188 | 3,791995908 | 1,343274 | 2,822951 | 0,004758 | 1 | 0 | Promoter (2-3kb) | 18 | 67621279 | 67743860 | 122582 | 2 | 68166 | ENSMUST00000115050.10 | -2464 | ENSMUSG00000024533 | Spire1 | spire type actin nucleation factor 1 |
| chr17 | 45999334 | 46000614 | 1281 | * | 9,420425 | 3,793156735 | 1,378149 | 2,752356 | 0,005917 | 0,792545355 | 1,449706 | Promoter (1-2kb) | 17 | 45980940 | 45997831 | 16892 | 2 | 224807 | ENSMUST00000156254.8 | -1503 | ENSMUSG00000036026 | Tmem63b | transmembrane protein 63b |
| chr2 | 68757114 | 68757905 | 792 | * | 6,077437 | 3,793994599 | 1,469525 | 2,581783 | 0,009829 | 1 | 0 | Intron (ENSMUST0000 | 2 | 68777604 | 68881316 | 103713 | 1 | 241447 | ENSMUST00000155748.8 | -19699 | ENSMUSG00000027035 | Cers6 | ceramide synthase 6 |
| chr18 | 57007987 | 57008893 | 907 | * | 10,13996 | 3,794492116 | 1,178821 | 3,218887 | 0,001287 | 0,524368688 | 1,449706 | Intron (ENSMUST0000 | 18 | 56936741 | 57058587 | 121847 | 2 | 320253 | ENSMUST00000153044.2 | 49694 | ENSMUSG00000032656 | Marchf3 | membrane associated ring-CH-type finger 3 |
| chr17 | 26783814 | 26785026 | 1213 | * | 11,49398 | 3,794615026 | 1,327007 | 2,859529 | 0,004243 | 0,852340527 | 0,744375 | Promoter (2-3kb) | 17 | 26724565 | 26781102 | 56538 | 2 | 19252 | ENSMUST00000235532.2 | -2712 | ENSMUSG00000024190 | Dusp1 | dual specificity phosphatase 1 |
| chr17 | 27177145 | 27178082 | 938 | * | 8,124706 | 3,794753338 | 1,363084 | 2,783947 | 0,00537 | 0,698702076 | 2,237403 | Promoter (<=1kb) | 17 | 27176800 | 27179085 | 2286 | 1 | 240057 | ENSMUST00000202939.2 | 345 | ENSMUSG00000067629 | Syngap1 | synaptic Ras GTPase activating protein 1 homolog (rat) |
| chr7 | 55358277 | 55359308 | 1032 | * | 7,766027 | 3,795404446 | 1,371033 | 2,768281 | 0,005635 | 0,793995086 | 1,360622 | Distal Intergenic | 7 | 55351902 | 55351963 | 62 | 1 | 1,15E+08 | ENSMUST00000158015.3 | 6375 | ENSMUSG00000088640 | Gm26482 | predicted gene, 26482 |
| chr18 | 30413398 | 30415846 | 2449 | * | 22,012569 | 3,795532702 | 0,8621171 | 4,4025723 | 1,07E-05 | 0,02016051 | 3,4716776 | Exon (ENSMUST0000011 | 18 | 30405980 | 30481173 | 75194 | 1 | 225326 | ENSMUST00000131405.2 | 7418 | ENSMUSG00000033628 | Pik3c3 | phosphatidylinositol 3-kinase catalytic subunit type 3 |
| chr4 | 1,38E+08 | 1,38E+08 | 1569 | * | 10,98287 | 3,795597286 | 1,261918 | 3,007801 | 0,002631 | 0,685410602 | 1,173325 | Exon (ENSMUST00000 | 4 | 1,38E+08 | 1,38E+08 | 3018 | 2 | 68943 | ENSMUST00000105816.8 | -3855 | ENSMUSG00000028756 | Pink1 | PTEN induced putative kinase 1 |
| chr15 | 11738088 | 11738999 | 912 | * | 7,48595 | 3,797500697 | 1,293747 | 2,935273 | 0,003333 | 1 | 0 | Distal Intergenic | 15 | 11696896 | 11703789 | 6894 | 2 | 1,05E+08 | ENSMUST00000228108.2 | -34299 | ENSMUSG00000115585 | Gm41271 | predicted gene, 41271 |
| chr6 | 1,38E+08 | 1,38E+08 | 963 | * | 7,350407 | 3,797995048 | 1,250816 | 3,036415 | 0,002394 | 1 | 0 | Promoter (2-3kb) | 6 | 1,37E+08 | 1,38E+08 | 66165 | 2 | 13860 | ENSMUST00000146442.4 | 2770 | ENSMUSG00000015766 | Eps8 | epidermal growth factor receptor pathway substrate 8 |
| chr7 | 35436547 | 35437641 | 1095 | * | 13,82784 | 3,798547207 | 1,128561 | 3,365832 | 0,000763 | 0,507354269 | 0,938682 | Intron (ENSMUST0000 | 7 | 35424784 | 35429224 | 4441 | 2 | 233115 | ENSMUST00000155451.2 | -7323 | ENSMUSG00000043671 | Dpy19l3 | dpy-19-like 3 (C. elegans) |
| chr4 | 1,26E+08 | 1,26E+08 | 748 | * | 6,997929 | 3,798973927 | 1,468058 | 2,587755 | 0,00966 | 1 | 0 | 3' UTR | 4 | 1,26E+08 | 1,26E+08 | 762 | 2 | 236511 | ENSMUST00000156533.2 | 4456 | ENSMUSG00000041530 | Ago1 | argonaute RISC catalytic subunit 1 |
| chr4 | 76596977 | 76598060 | 1084 | * | 8,884428 | 3,799000419 | 1,271461 | 2,987901 | 0,002809 | 0,590939713 | 2,237403 | Intron (ENSMUST0000 | 4 | 75863043 | 76512536 | 649494 | 2 | 19266 | ENSMUST00000173376.8 | -84441 | ENSMUSG00000028399 | Ptprd | protein tyrosine phosphatase, receptor type, D |
| chr10 | 97736938 | 97737889 | 952 | * | 6,251892 | 3,799321437 | 1,463974 | 2,595212 | 0,009453 | 1 | 0 | Intron (ENSMUST0000 | 10 | 97517276 | 97802952 | 285677 | 2 | 1E+08 | ENSMUST00000220233.2 | 65063 | ENSMUSG00000074776 | Gm10754 | predicted gene 10754 |
| chr11 | 61481057 | 61482983 | 1927 | * | 12,32502 | 3,799969418 | 1,213512 | 3,131382 | 0,00174 | 0,605265376 | 1,2763 | Distal Intergenic | 11 | 61408076 | 61470513 | 62438 | 2 | 13855 | ENSMUST00000179936.8 | -10544 | ENSMUSG00000001036 | Epn2 | epsin 2 |
| chr2 | 1,56E+08 | 1,56E+08 | 1469 | * | 9,559105 | 3,801009886 | 1,402689 | 2,709803 | 0,006732 | 0,864595549 | 1,086118 | Promoter (1-2kb) | 2 | 1,56E+08 | 1,56E+08 | 27799 | 2 | 108687 | ENSMUST00000040833.5 | -1234 | ENSMUSG00000038312 | Edem2 | ER degradation enhancer, mannosidase alpha-like 2 |
| chr10 | 57045559 | 57046432 | 874 | * | 7,015792 | 3,801666299 | 1,475844 | 2,575926 | 0,009997 | 1 | 0 | Distal Intergenic | 10 | 57050126 | 57050225 | 100 | 1 | 1,15E+08 | ENSMUST00000157628.4 | -3694 | ENSMUSG00000088253 | Gm24285 | predicted gene, 24285 |
| chr18 | 7119308 | 7119940 | 633 | * | 7,349527 | 3,802484611 | 1,424915 | 2,668569 | 0,007618 | 0,851453105 | 1,360622 | Intron (ENSMUST0000 | 18 | 7129389 | 7184073 | 54685 | 2 | 74934 | ENSMUST00000235068.2 | 64133 | ENSMUSG00000061802 | Odad2 | outer dynein arm docking complex subunit 2 |
| chr11 | 16644988 | 16645669 | 682 | * | 7,766189 | 3,802773981 | 1,355484 | 2,805474 | 0,005024 | 0,776295651 | 1,360622 | Distal Intergenic | 11 | 16682605 | 16682711 | 107 | 2 | 1,15E+08 | ENSMUST00000240365.1 | 37042 | ENSMUSG00000119782 | Gm25698 | predicted gene, 25698 |
| chr15 | 64047586 | 64048754 | 1169 | * | 10,85381 | 3,80341834 | 1,369178 | 2,777885 | 0,005471 | 0,858257847 | 0,918255 | Intron (ENSMUST0000 | 15 | 64030788 | 64080340 | 49553 | 2 | 13196 | ENSMUST00000176358.2 | 31586 | ENSMUSG00000022377 | Asap1 | ArfGAP with SH3 domain, ankyrin repeat and PH domain1 |
| chr9 | 1,22E+08 | 1,22E+08 | 1058 | * | 10,60162 | 3,803874055 | 1,349359 | 2,819023 | 0,004817 | 0,779594379 | 1,2763 | Promoter (<=1kb) | 9 | 1,22E+08 | 1,22E+08 | 6128 | 1 | 72330 | ENSMUST00000098272.4 | 648 | ENSMUSG00000074001 | Klhl40 | kelch-like 40 |
| chr4 | 1,41E+08 | 1,41E+08 | 605 | * | 7,173221 | 3,804328008 | 1,318558 | 2,88522 | 0,003911 | 0,730607562 | 1,360622 | Distal Intergenic | 4 | 1,41E+08 | 1,41E+08 | 3434 | 2 | 277744 | ENSMUST00000094544.3 | 5289 | ENSMUSG00000070637 | Srarp | steroid receptor associated and regulated protein |
| chr19 | 41469552 | 41470734 | 1183 | * | 7,900896 | 3,804797446 | 1,400784 | 2,716191 | 0,006604 | 1 | 0 | Promoter (<=1kb) | 19 | 41471076 | 41550700 | 79625 | 1 | 212391 | ENSMUST00000067795.13 | -342 | ENSMUSG00000025019 | Lcor | ligand dependent nuclear receptor corepressor |
| chr18 | 74983164 | 74984237 | 1074 | * | 10,332 | 3,804812027 | 1,13835 | 3,342392 | 0,000831 | 0,496713645 | 1,086118 | Distal Intergenic | 18 | 74926454 | 74939279 | 12826 | 1 | 52538 | ENSMUST00000237306.2 | 56710 | ENSMUSG00000036880 | Acaa2 | acetyl-Coenzyme A acyltransferase 2 (mitochondrial 3-oxoacyl-Coenzyme A thiolase) |
| chrX | 1,53E+08 | 1,53E+08 | 815 | * | 7,024035 | 3,808453891 | 1,410559 | 2,69996 | 0,006935 | 1 | 0 | Distal Intergenic | 20 | 1,53E+08 | 1,53E+08 | 4168 | 2 | 75071 | ENSMUST00000082357.2 | 65455 | ENSMUSG00000059663 | Samt2b | spermatogenesis associated multipass transmembrane protein 2b |
| chr15 | 20331448 | 20332071 | 624 | * | 6,393819 | 3,80857523 | 1,469842 | 2,591146 | 0,009566 | 1 | 0 | Distal Intergenic | 15 | 20449205 | 20665946 | 216742 | 1 | 215654 | ENSMUST00000227439.2 | -117134 | ENSMUSG00000040452 | Cdh12 | cadherin 12 |
| chr9 | 69374248 | 69375570 | 1323 | * | 9,223451 | 3,80866885 | 1,355904 | 2,808952 | 0,00497 | 0,685026746 | 2,237403 | Exon (ENSMUST00000 | 9 | 69364226 | 69364313 | 88 | 1 | 1,01E+08 | ENSMUST00000175301.3 | 10022 | ENSMUSG00000093042 | Mir3109 | microRNA 3109 |
| chr12 | 27550176 | 27551222 | 1047 | * | 8,434237 | 3,808678859 | 1,457515 | 2,613132 | 0,008972 | 0,791147121 | 2,237403 | Distal Intergenic | 12 | 27648200 | 27665361 | 17162 | 1 | 328069 | ENSMUST00000220893.2 | -96978 | | Gm4166 | predicted gene 4166 |
| chr2 | 14955340 | 14956120 | 781 | * | 7,38468 | 3,808827148 | 1,34173 | 2,838742 | 0,004529 | 0,75508254 | 1,360622 | Intron (ENSMUST0000 | 2 | 14965587 | 14986183 | 20597 | 1 | 12296 | ENSMUST00000193522.2 | -9467 | ENSMUSG00000057914 | Cacnb2 | calcium channel, voltage-dependent, beta 2 subunit |
| chr12 | 81577852 | 81578829 | 978 | * | 8,66949 | 3,80914063 | 1,437375 | 2,650068 | 0,008048 | 0,770673161 | 2,237403 | Promoter (<=1kb) | 12 | 81521369 | 81579535 | 58167 | 2 | 1,06E+08 | ENSMUST00000170958.8 | 706 | ENSMUSG00000021139 | Gm20498 | predicted gene 20498 |
| chr19 | 43486942 | 43488378 | 1437 | * | 13,47358 | 3,809604265 | 1,025475 | 3,714967 | 0,000203 | 0,255268024 | 1,2763 | 3' UTR | 19 | 43488754 | 43493235 | 4482 | 2 | 14718 | ENSMUST00000132504.2 | 4857 | ENSMUSG00000025190 | Got1 | glutamic-oxaloacetic transaminase 1, soluble |
| chr9 | 19981212 | 19982129 | 918 | * | 11,68554 | 3,81247754 | 1,209061 | 3,153255 | 0,001615 | 0,607561104 | 1,173325 | Distal Intergenic | 9 | 19965762 | 19966757 | 996 | 2 | 257898 | ENSMUST00000060780.3 | -14455 | ENSMUSG00000044454 | Olfr867 | olfactory receptor 867 |
| chr12 | 81824344 | 81825625 | 1282 | * | 12,34043 | 3,812733032 | 1,330942 | 2,864687 | 0,004174 | 0,751314174 | 1,2763 | Promoter (2-3kb) | 12 | 81768813 | 81827648 | 58836 | 2 | 338372 | ENSMUST00000222322.2 | 2023 | ENSMUSG00000042724 | Map3k9 | mitogen-activated protein kinase kinase kinase 9 |
| chr17 | 28644567 | 28645895 | 1329 | * | 12,25576 | 3,813362397 | 1,202559 | 3,171039 | 0,001519 | 0,577829405 | 1,2763 | Promoter (1-2kb) | 17 | 28621670 | 28647603 | 25934 | 2 | 14229 | ENSMUST00000143685.3 | 1708 | ENSMUSG00000024222 | Fkbp5 | FK506 binding protein 5 |
| chr16 | 26923601 | 26924380 | 780 | * | 8,220578 | 3,813976161 | 1,374592 | 2,774623 | 0,005527 | 0,786347206 | 1,425929 | Distal Intergenic | 16 | 26777855 | 26810402 | 32548 | 2 | 239789 | ENSMUST00000231299.2 | -113199 | ENSMUSG00000068428 | Gmnc | geminin coiled-coil domain containing |
| chr8 | 1,08E+08 | 1,08E+08 | 1101 | * | 8,785227 | 3,814054525 | 1,325351 | 2,87777 | 0,004005 | 0,728037066 | 1,425929 | Intron (ENSMUST0000 | 8 | 1,08E+08 | 1,08E+08 | 110 | 2 | 1,15E+08 | ENSMUST00000158344.3 | 34328 | ENSMUSG00000088969 | Gm22085 | predicted gene, 22085 |
| chr17 | 29050314 | 29051803 | 1490 | * | 13,30987 | 3,815533349 | 1,34968 | 2,826991 | 0,004699 | 0,829907213 | 0,938682 | 3' UTR | 17 | 29047056 | 29055765 | 8710 | 1 | 268936 | ENSMUST00000127007.2 | 3258 | ENSMUSG00000063952 | Brpf3 | bromodomain and PHD finger containing, 3 |
| chr9 | 88132820 | 88134419 | 1600 | * | 8,910011 | 3,816014054 | 1,372538 | 2,780262 | 0,005432 | 0,778321623 | 1,449706 | Distal Intergenic | 9 | 88141948 | 88145088 | 3141 | 1 | 1,03E+08 | ENSMUST00000190118.2 | -7529 | ENSMUSG00000100782 | Gm28231 | predicted gene 28231 |
| chr9 | 38649983 | 38650878 | 896 | * | 8,113532 | 3,817416818 | 1,405642 | 2,715782 | 0,006612 | 0,817115778 | 1,425929 | Promoter (2-3kb) | 9 | 38647481 | 38653216 | 5736 | 1 | 67776 | ENSMUST00000137972.2 | 2502 | ENSMUSG00000023186 | Vwa5a | von Willebrand factor A domain containing 5A |
| chrX | 1,31E+08 | 1,31E+08 | 1037 | * | 6,815005 | 3,817683793 | 1,402481 | 2,722093 | 0,006487 | 1 | 0 | Distal Intergenic | 20 | 1,3E+08 | 1,3E+08 | 129 | 2 | 1,15E+08 | ENSMUST00000104191.3 | -718626 | ENSMUSG00000077379 | Gm25170 | predicted gene, 25170 |
| chr19 | 40724798 | 40725744 | 947 | * | 8,303592 | 3,819104897 | 1,314387 | 2,905617 | 0,003665 | 0,64068479 | 2,237403 | Promoter (<=1kb) | 19 | 40724628 | 40725309 | 682 | 1 | 12495 | ENSMUST00000151726.2 | 170 | ENSMUSG00000048120 | Entpd1 | ectonucleoside triphosphate diphosphohydrolase 1 |
| chrX | 1,61E+08 | 1,61E+08 | 972 | * | 7,163236 | 3,81935639 | 1,446981 | 2,639535 | 0,008302 | 1 | 0 | Distal Intergenic | 20 | 1,61E+08 | 1,61E+08 | 326435 | 2 | 195727 | ENSMUST00000087085.10 | -65065 | ENSMUSG00000059493 | Nhs | NHS actin remodeling regulator |
| chr8 | 44478733 | 44479486 | 754 | * | 7,359028 | 3,819371026 | 1,251589 | 3,051616 | 0,002276 | 1 | 0 | Distal Intergenic | 8 | 44103346 | 44163086 | 59741 | 2 | 252866 | ENSMUST00000110411.4 | -315647 | ENSMUSG00000079058 | Adam34 | a disintegrin and metallopeptidase domain 34 |
| chr13 | 41797144 | 41798427 | 1284 | * | 9,742642 | 3,819698513 | 1,223684 | 3,121475 | 0,001799 | 0,591609342 | 1,425929 | Distal Intergenic | 13 | 41759692 | 41794831 | 35140 | 1 | 621976 | ENSMUST00000129449.3 | 37452 | ENSMUSG00000087370 | Tmem170b | transmembrane protein 170B |
| chr18 | 5218221 | 5219028 | 808 | * | 6,880063 | 3,819747925 | 1,440928 | 2,650894 | 0,008028 | 1 | 0 | Intron (ENSMUST0000 | 18 | 5162839 | 5165731 | 2893 | 2 | 1,03E+08 | ENSMUST00000180850.3 | -52490 | ENSMUSG00000097888 | Gm26682 | predicted gene, 26682 |
| chrX | 1,4E+08 | 1,4E+08 | 853 | * | 6,449278 | 3,820620952 | 1,465362 | 2,607287 | 0,009126 | 1 | 0 | Intron (ENSMUST0000 | 20 | 1,4E+08 | 1,4E+08 | 21412 | 2 | 94216 | ENSMUST00000137051.8 | -37520 | ENSMUSG00000031273 | Col4a6 | collagen, type IV, alpha 6 |
| chr7 | 89837817 | 89838812 | 996 | * | 8,3603 | 3,820988514 | 1,461105 | 2,615136 | 0,008919 | 0,877892722 | 1,360622 | Promoter (<=1kb) | 7 | 89838011 | 89844922 | 6912 | 1 | 233489 | ENSMUST00000208891.2 | 0 | ENSMUSG00000039361 | Picalm | phosphatidylinositol binding clathrin assembly protein |
| chr12 | 80520459 | 80522028 | 1570 | * | 10,84982 | 3,823306052 | 1,186089 | 3,223456 | 0,001267 | 0,64558017 | 0,744375 | Promoter (<=1kb) | 12 | 80522854 | 80527943 | 5090 | 1 | 97827 | ENSMUST00000219477.2 | -826 | ENSMUSG00000032705 | Exd2 | exonuclease 3'-5' domain containing 2 |
| chr11 | 35724324 | 35725711 | 1388 | * | 14,05409 | 3,824396399 | 1,153205 | 3,316319 | 0,000912 | 0,514210241 | 1,075735 | Promoter (<=1kb) | 11 | 35699208 | 35725333 | 26126 | 2 | 104458 | ENSMUST00000018992.4 | 0 | ENSMUSG00000018848 | Rars | arginyl-tRNA synthetase |
| chr12 | 61287497 | 61288502 | 1006 | * | 7,984349 | 3,824453979 | 1,37107 | 2,789394 | 0,005281 | 1 | 0 | Distal Intergenic | 12 | 61535887 | 61568578 | 32692 | 2 | 1,01E+08 | ENSMUST00000244322.1 | 280076 | ENSMUSG00000113429 | Gm20063 | predicted gene, 20063 |
| chr3 | 36300065 | 36301474 | 1410 | * | 10,02868 | 3,824477613 | 1,299565 | 2,942892 | 0,003252 | 0,685410602 | 1,449706 | Distal Intergenic | 3 | 36233575 | 36276462 | 42888 | 2 | 229214 | ENSMUST00000091227.9 | -23603 | ENSMUSG00000058400 | Qrfpr | pyroglutamylated RFamide peptide receptor |
| chr4 | 119151606 | 119153154 | 1549 | * | 24,291496 | 3,825848184 | 0,7906406 | 4,8389218 | 1,305E-06 | 0,004421025 | 2,9519335 | Promoter (<=1kb) | 4 | 119135178 | 119151801 | 16624 | 2 | 22608 | ENSMUST00000079644.13 | 0 | ENSMUSG00000028639 | Ybx1 | Y box protein 1 |
| chr14 | 99512150 | 99512834 | 685 | * | 7,572792 | 3,826502655 | 1,377945 | 2,776964 | 0,005487 | 1 | 0 | Distal Intergenic | 14 | 99536127 | 99552472 | 16346 | 1 | 12224 | ENSMUST00000005279.8 | -23293 | ENSMUSG00000005148 | Klf5 | Kruppel-like factor 5 |
| chr11 | 3335151 | 3337683 | 2533 | * | 22,604347 | 3,826531433 | 0,818708 | 4,6738661 | 2,956E-06 | 0,008107936 | 3,104559 | Intron (ENSMUST000001 | 11 | 3294256 | 3321376 | 27121 | 2 | 16886 | ENSMUST00000134576.8 | -13775 | ENSMUSG00000020451 | Limk2 | LIM motif-containing protein kinase 2 |
| chr11 | 74676202 | 74677175 | 974 | * | 8,443846 | 3,826975838 | 1,348191 | 2,8386 | 0,004531 | 1 | 0 | Intron (ENSMUST0000 | 11 | 74661751 | 74708720 | 46970 | 1 | 67493 | ENSMUST00000092912.12 | 14451 | ENSMUSG00000010554 | Mettl16 | methyltransferase like 16 |
| chr10 | 13065849 | 13066970 | 1122 | * | 8,355477 | 3,827656102 | 1,388141 | 2,757397 | 0,005826 | 0,792545355 | 1,425929 | Promoter (1-2kb) | 10 | 13058156 | 13068172 | 10017 | 2 | 353258 | ENSMUST00000219824.2 | 1202 | ENSMUSG00000019814 | Ltv1 | LTV1 ribosome biogenesis factor |
| chr17 | 47818339 | 47819582 | 1244 | * | 10,75287 | 3,827966456 | 1,382304 | 2,769266 | 0,005618 | 0,901463351 | 0,738441 | Promoter (2-3kb) | 17 | 47816166 | 47909574 | 93409 | 1 | 12445 | ENSMUST00000182506.8 | 2173 | ENSMUSG00000034165 | Ccnd3 | cyclin D3 |
| chr1 | 1,81E+08 | 1,81E+08 | 982 | * | 6,814934 | 3,828200684 | 1,436959 | 2,664099 | 0,007719 | 1 | 0 | Distal Intergenic | 1 | 1,81E+08 | 1,81E+08 | 22705 | 1 | 1,03E+08 | ENSMUST00000240988.1 | 40119 | | Gm38664 | predicted gene, 38664 |
| chr2 | 84412295 | 84412959 | 665 | * | 7,464043 | 3,828586195 | 1,37877 | 2,776813 | 0,005489 | 1 | 0 | Intron (ENSMUST0000 | 2 | 84432877 | 84440021 | 7145 | 2 | 12388 | ENSMUST00000141151.8 | 27062 | ENSMUSG00000034101 | Ctnnd1 | catenin (cadherin associated protein), delta 1 |
| chr19 | 42652827 | 42654246 | 1420 | * | 9,70567 | 3,828934245 | 1,410791 | 2,714033 | 0,006647 | 0,846977404 | 1,211315 | Distal Intergenic | 19 | 42688216 | 42688341 | 126 | 2 | 1,15E+08 | ENSMUST00000104580.3 | 34095 | ENSMUSG00000077768 | Gm25216 | predicted gene, 25216 |
| chr11 | 1,16E+08 | 1,16E+08 | 870 | * | 8,723352 | 3,829645358 | 1,312868 | 2,917006 | 0,003534 | 0,715156227 | 1,360622 | Promoter (2-3kb) | 11 | 1,16E+08 | 1,16E+08 | 965 | 2 | 15081 | ENSMUST00000139378.2 | 2901 | ENSMUSG00000016559 | H3f3b | H3.3 histone B |
| chr8 | 80021637 | 80022434 | 798 | * | 11,24174 | 3,829868895 | 1,191146 | 3,215281 | 0,001303 | 0,650529916 | 0,744375 | Promoter (<=1kb) | 8 | 80021649 | 80035590 | 13942 | 1 | 234479 | ENSMUST00000181030.4 | 0 | | Gm4890 | predicted gene 4890 |
| chr17 | 74052606 | 74053428 | 823 | * | 8,058569 | 3,832195362 | 1,338943 | 2,862104 | 0,004208 | 1 | 0 | Distal Intergenic | 17 | 73879779 | 74017448 | 137670 | 2 | 71685 | ENSMUST00000146565.2 | -35158 | ENSMUSG00000024064 | Galnt14 | polypeptide N-acetylgalactosaminyltransferase 14 |
| chr2 | 1,09E+08 | 1,09E+08 | 844 | * | 7,588139 | 3,832765764 | 1,400762 | 2,7362 | 0,006215 | 1 | 0 | Intron (ENSMUST0000 | 2 | 1,09E+08 | 1,09E+08 | 185981 | 2 | 76894 | ENSMUST00000147376.8 | 5028 | ENSMUSG00000057234 | Mettl15 | methyltransferase like 15 |
| chr5 | 1,19E+08 | 1,19E+08 | 1181 | * | 10,02221 | 3,833185019 | 1,404687 | 2,728853 | 0,006356 | 0,843803477 | 1,173325 | Intron (ENSMUST0000 | 5 | 1,19E+08 | 1,19E+08 | 16598 | 1 | 76199 | ENSMUST00000202538.4 | -61230 | ENSMUSG00000018076 | Med13l | mediator complex subunit 13-like |
| chr11 | 1,07E+08 | 1,07E+08 | 769 | * | 10,35815 | 3,833804621 | 1,122943 | 3,414069 | 0,00064 | 0,40679506 | 1,449706 | Promoter (2-3kb) | 11 | 1,07E+08 | 1,07E+08 | 2052 | 1 | 320162 | ENSMUST00000124898.2 | 2984 | ENSMUSG00000018372 | Cep95 | centrosomal protein 95 |
| chr3 | 65831441 | 65832227 | 787 | * | 6,988108 | 3,834377944 | 1,420107 | 2,700063 | 0,006933 | 1 | 0 | Distal Intergenic | 3 | 65854203 | 65855419 | 1217 | 2 | 56706 | ENSMUST00000149729.2 | 23192 | ENSMUSG00000027829 | Ccnl1 | cyclin L1 |
| chr2 | 1,28E+08 | 1,28E+08 | 1307 | * | 12,23396 | 3,835340747 | 1,243944 | 3,083209 | 0,002048 | 0,649854434 | 1,173325 | Intron (ENSMUST0000 | 2 | 1,28E+08 | 1,28E+08 | 214451 | 1 | 74121 | ENSMUST00000110344.2 | 37576 | ENSMUSG00000027380 | Acoxl | acyl-Coenzyme A oxidase-like |
| chr1 | 1,64E+08 | 1,64E+08 | 755 | * | 7,209832 | 3,835722196 | 1,287859 | 2,97837 | 0,002898 | 0,683067851 | 1,360622 | Promoter (1-2kb) | 1 | 1,64E+08 | 1,64E+08 | 5208 | 2 | 381306 | ENSMUST00000161321.2 | -1298 | ENSMUSG00000041406 | BC055324 | cDNA sequence BC055324 |
| chr8 | 86366689 | 86367581 | 893 | * | 8,896134 | 3,835772886 | 1,231461 | 3,114815 | 0,001841 | 0,594676756 | 1,425929 | 3' UTR | 8 | 86363217 | 86417602 | 54386 | 2 | 74513 | ENSMUST00000109686.4 | 50021 | ENSMUSG00000036902 | Neto2 | neuropilin (NRP) and tolloid (TLL)-like 2 |
| chr5 | 35942773 | 35944508 | 1736 | * | 19,17389 | 3,836950571 | 1,05575 | 3,634336 | 0,000279 | 0,216838386 | 2,386008 | Intron (ENSMUST0000 | 5 | 35971703 | 36000548 | 28846 | 1 | 231148 | ENSMUST00000130233.8 | -27195 | ENSMUSG00000029095 | Ablim2 | actin-binding LIM protein 2 |
| chr7 | 66794612 | 66795256 | 645 | * | 6,928856 | 3,837237229 | 1,426019 | 2,690874 | 0,007127 | 1 | 0 | Intron (ENSMUST0000 | 7 | 66853830 | 66872206 | 18377 | 2 | 76628 | ENSMUST00000181631.2 | 76950 | ENSMUSG00000097527 | 1700112J16Rik | RIKEN cDNA 1700112J16 gene |
| chr5 | 16410467 | 16411943 | 1477 | * | 11,11221 | 3,837576841 | 1,370715 | 2,799691 | 0,005115 | 0,803455139 | 1,173325 | Intron (ENSMUST0000 | 5 | 16530983 | 16534881 | 3899 | 1 | 12293 | ENSMUST00000200158.2 | -119040 | ENSMUSG00000040118 | Cacna2d1 | calcium channel, voltage-dependent, alpha2/delta subunit 1 |
| chr2 | 155382179 | 155383786 | 1608 | * | 21,748519 | 3,839123144 | 0,8038737 | 4,775779 | 1,79E-06 | 0,005954891 | 2,8800171 | Intron (ENSMUST000000 | 2 | 155391141 | 155391778 | 638 | 1 | 60525 | ENSMUST00000143169.2 | -7355 | ENSMUSG00000027605 | Acss2 | acyl-CoA synthetase short-chain family member 2 |
| chr1 | 1,65E+08 | 1,65E+08 | 675 | * | 6,860492 | 3,839194931 | 1,404124 | 2,734228 | 0,006253 | 1 | 0 | Distal Intergenic | 1 | 1,65E+08 | 1,65E+08 | 20102 | 2 | 108735 | ENSMUST00000043338.10 | -13107 | ENSMUSG00000040848 | Sft2d2 | SFT2 domain containing 2 |
| chr9 | 44551463 | 44552717 | 1255 | * | 8,315317 | 3,840177393 | 1,402071 | 2,738933 | 0,006164 | 0,800767521 | 1,449706 | 3' UTR | 9 | 44581747 | 44583558 | 1812 | 2 | 1,15E+08 | ENSMUST00000244401.1 | 30841 | ENSMUSG00000110755 | BC049987 | cDNA sequence BC049987 |
| chr4 | 1,07E+08 | 1,07E+08 | 1214 | * | 12,33302 | 3,840477632 | 1,059856 | 3,623584 | 0,000291 | 0,385677228 | 0,738441 | Promoter (2-3kb) | 4 | 1,07E+08 | 1,07E+08 | 24166 | 2 | 329910 | ENSMUST00000140541.8 | -2906 | ENSMUSG00000034853 | Acot11 | acyl-CoA thioesterase 11 |
| chrX | 1,61E+08 | 1,61E+08 | 803 | * | 8,181559 | 3,841439043 | 1,436557 | 2,674059 | 0,007494 | 0,838625535 | 1,425929 | Intron (ENSMUST0000 | 20 | 1,61E+08 | 1,61E+08 | 17487 | 2 | 195726 | ENSMUST00000136860.3 | -36104 | ENSMUSG00000086540 | Scml1 | Scm polycomb group protein like 1 |
| chr2 | 1,53E+08 | 1,53E+08 | 1905 | * | 18,10535 | 3,842236017 | 0,95463 | 4,024843 | 5,7E-05 | 0,081885224 | 2,507013 | Distal Intergenic | 2 | 1,53E+08 | 1,53E+08 | 122510 | 2 | 329540 | ENSMUST00000035346.14 | -41110 | ENSMUSG00000061411 | Nol4l | nucleolar protein 4-like |
| chr12 | 1,09E+08 | 1,09E+08 | 930 | * | 8,113329 | 3,842825454 | 1,488054 | 2,582451 | 0,00981 | 1 | 0 | Promoter (<=1kb) | 12 | 1,09E+08 | 1,09E+08 | 33169 | 2 | 22375 | ENSMUST00000161410.8 | 0 | ENSMUSG00000021266 | Wars | tryptophanyl-tRNA synthetase |
| chr14 | 65351746 | 65352985 | 1240 | * | 7,047941 | 3,843914903 | 1,39945 | 2,746733 | 0,006019 | 1 | 0 | Intron (ENSMUST0000 | 14 | 65289509 | 65335555 | 46047 | 2 | 54616 | ENSMUST00000022550.8 | -16191 | ENSMUSG00000021978 | Extl3 | exostosin-like glycosyltransferase 3 |
| chr2 | 1,67E+08 | 1,67E+08 | 941 | * | 6,386616 | 3,846418639 | 1,401038 | 2,745407 | 0,006044 | 1 | 0 | Intron (ENSMUST0000 | 2 | 1,67E+08 | 1,67E+08 | 25845 | 1 | 77031 | ENSMUST00000148454.2 | 13204 | ENSMUSG00000039463 | Slc9a8 | solute carrier family 9 (sodium/hydrogen exchanger), member 8 |
| chr14 | 30094602 | 30095639 | 1038 | * | 9,776073 | 3,848038193 | 1,429308 | 2,692239 | 0,007097 | 0,856202617 | 1,211315 | Intron (ENSMUST0000 | 14 | 29763465 | 30075484 | 312020 | 2 | 12289 | ENSMUST00000224785.3 | -19118 | ENSMUSG00000015968 | Cacna1d | calcium channel, voltage-dependent, L type, alpha 1D subunit |
| chr15 | 74605878 | 74606803 | 926 | * | 10,59896 | 3,849844339 | 1,411783 | 2,726937 | 0,006393 | 0,9208979 | 0,738441 | Promoter (<=1kb) | 15 | 74604096 | 74606178 | 2083 | 2 | 68311 | ENSMUST00000023260.5 | 0 | ENSMUSG00000022595 | Lypd2 | Ly6/Plaur domain containing 2 |
| chr8 | 1,26E+08 | 1,26E+08 | 717 | * | 9,070623 | 3,850127566 | 1,220128 | 3,155511 | 0,001602 | 0,5663777 | 1,425929 | Promoter (<=1kb) | 8 | 1,26E+08 | 1,26E+08 | 40738 | 2 | 112405 | ENSMUST00000034469.7 | 769 | ENSMUSG00000031987 | Egln1 | egl-9 family hypoxia-inducible factor 1 |
| chr16 | 20168263 | 20169328 | 1066 | * | 7,782059 | 3,850775573 | 1,321224 | 2,914552 | 0,003562 | 1 | 0 | Intron (ENSMUST0000 | 16 | 20157679 | 20182756 | 25078 | 2 | 27416 | ENSMUST00000148003.8 | 13428 | ENSMUSG00000022822 | Abcc5 | ATP-binding cassette, sub-family C (CFTR/MRP), member 5 |
| chr9 | 1,1E+08 | 1,1E+08 | 949 | * | 6,771162 | 3,851977318 | 1,418789 | 2,714975 | 0,006628 | 1 | 0 | Promoter (<=1kb) | 9 | 1,1E+08 | 1,11E+08 | 44361 | 1 | 72654 | ENSMUST00000195979.2 | 0 | ENSMUSG00000019659 | Ccdc12 | coiled-coil domain containing 12 |
| chr4 | 1E+08 | 1E+08 | 1829 | * | 13,93772 | 3,852773301 | 1,045823 | 3,683963 | 0,00023 | 0,311294276 | 0,918255 | Intron (ENSMUST0000 | 4 | 99976818 | 99978038 | 1221 | 1 | 1,05E+08 | ENSMUST00000134206.2 | 108257 | | Gm12701 | predicted gene 12701 |
| chr18 | 49381360 | 49382616 | 1257 | * | 10,77076 | 3,853050462 | 1,271029 | 3,031441 | 0,002434 | 0,753219264 | 0,738441 | Distal Intergenic | 18 | 49656356 | 49657507 | 1152 | 1 | 67321 | ENSMUST00000190201.2 | -273740 | ENSMUSG00000100297 | 1700044K03Rik | RIKEN cDNA 1700044K03 gene |
| chr1 | 84481674 | 84482905 | 1232 | * | 9,206343 | 3,853132279 | 1,335405 | 2,885366 | 0,00391 | 0,721416593 | 1,449706 | Intron (ENSMUST0000 | 1 | 84560725 | 84601562 | 40838 | 2 | 227325 | ENSMUST00000191546.3 | 118657 | ENSMUSG00000036766 | Dner | delta/notch-like EGF repeat containing |

| | | | | | | | | | | | | | | | | | | | | | | | |
|---|---|---|---|---|---|---|---|---|---|---|---|---|---|---|---|---|---|---|---|---|---|---|---|
| chr11 | 6646073 | 6646984 | 912 | * | 7,991063 | 3,853895305 | 1,429893 | 2,695234 | 0,007034 | 1 | 0 | Distal Intergenic | 11 | 6666001 | 6670037 | 4037 | 2 | 626419 | ENSMUST00000139769.2 | 23053 | ENSMUSG00000086808 | Gm11981 | predicted gene 11981 |
| chr11 | 16900371 | 16901981 | 1611 | * | 15,31621 | 3,853931605 | 1,074215 | 3,587674 | 0,000334 | 0,252389304 | 2,141612 | Promoter (<=1kb) | 11 | 16901375 | 16904447 | 3073 | 1 | 319701 | ENSMUST00000109635.2 | 0 | ENSMUSG00000044966 | Fbxo48 | F-box protein 48 |
| chr16 | 31902122 | 31902789 | 668 | * | 5,870304 | 3,854155336 | 1,488137 | 2,58992 | 0,0096 | 1 | 0 | Distal Intergenic | 16 | 31835108 | 31898160 | 63053 | 2 | 224105 | ENSMUST00000023467.9 | -3962 | ENSMUSG00000022781 | Pak2 | p21 (RAC1) activated kinase 2 |
| chr9 | 32119185 | 32119955 | 771 | * | 7,043881 | 3,856071609 | 1,488558 | 2,590474 | 0,009584 | 1 | 0 | Promoter (<=1kb) | 9 | 32118744 | 32172514 | 53771 | 1 | 330914 | ENSMUST00000182802.8 | 441 | ENSMUSG00000041444 | Arhgap32 | Rho GTPase activating protein 32 |
| chrX | 1,11E+08 | 1,11E+08 | 1292 | * | 6,282495 | 3,858405169 | 1,363639 | 2,829492 | 0,004662 | 1 | 0 | Distal Intergenic | 20 | 1,1E+08 | 1,11E+08 | 127147 | 2 | 245596 | ENSMUST00000038472.7 | -73105 | ENSMUSG00000034551 | Hdx | highly divergent homeobox |
| chr16 | 48697175 | 48698140 | 966 | * | 11,75848 | 3,859145955 | 1,057359 | 3,649796 | 0,000262 | 0,328946709 | 0,918255 | Distal Intergenic | 16 | 48692984 | 48694859 | 1876 | 1 | 245195 | ENSMUST00000065666.6 | 4191 | ENSMUSG00000022651 | Retnlg | resistin like gamma |
| chr19 | 57303531 | 57305117 | 1587 | * | 11,64833 | 3,85915239 | 1,295573 | 2,978722 | 0,002895 | 0,786955198 | 0,744375 | Promoter (<=1kb) | 19 | 57025683 | 57303351 | 277669 | 2 | 226251 | ENSMUST00000156316.8 | -180 | ENSMUSG00000025085 | Ablim1 | actin-binding LIM protein 1 |
| chr18 | 67240987 | 67242218 | 1232 | * | 7,131386 | 3,859699693 | 1,384723 | 2,787345 | 0,005314 | 1 | 0 | Intron (ENSMUST0000( | 18 | 67221406 | 67325917 | 104512 | 1 | 14680 | ENSMUST00000236551.2 | 19581 | ENSMUSG00000024524 | Gnal | guanine nucleotide binding protein, alpha stimulating, olfactory type |
| chr10 | 11235234 | 11235858 | 625 | * | 7,627742 | 3,860211524 | 1,406148 | 2,745238 | 0,006047 | 1 | 0 | Intron (ENSMUST0000( | 10 | 11219197 | 11324442 | 105246 | 1 | 13853 | ENSMUST00000161438.2 | 16037 | ENSMUSG00000055493 | Epm2a | epilepsy, progressive myoclonic epilepsy, type 2 gene alpha |
| chr19 | 16718412 | 16719566 | 1155 | * | 7,631812 | 3,860760461 | 1,485663 | 2,598678 | 0,009358 | 1 | 0 | Promoter (<=1kb) | 19 | 16718344 | 16720402 | 2059 | 2 | 271564 | ENSMUST00000225079.2 | 836 | ENSMUSG00000046230 | Vps13a | vacuolar protein sorting 13A |
| chr6 | 1,44E+08 | 1,44E+08 | 877 | * | 7,366306 | 3,861177429 | 1,480757 | 2,60757 | 0,009119 | 0,881847489 | 1,360622 | Distal Intergenic | 6 | 1,44E+08 | 1,44E+08 | 57936 | 2 | 73425 | ENSMUST00000150559.2 | -97035 | ENSMUSG00000086887 | 1700060C16Rik | RIKEN cDNA 1700060C16 gene |
| chr2 | 97132294 | 97133093 | 800 | * | 7,970302 | 3,861369327 | 1,412008 | 2,734666 | 0,006244 | 1 | 0 | Intron (ENSMUST0000( | 2 | 97068163 | 97068291 | 129 | 1 | 1,15E+08 | ENSMUST00000158008.3 | 64131 | ENSMUSG00000088633 | Gm22313 | predicted gene, 22313 |
| chr7 | 76297599 | 76298776 | 1178 | * | 10,91505 | 3,861999079 | 1,158967 | 3,332276 | 0,000861 | 0,571379912 | 0,744375 | Intron (ENSMUST0000( | 7 | 76060810 | 76416356 | 355547 | 1 | 244071 | ENSMUST00000107442.3 | 236789 | ENSMUSG00000025754 | Agbl1 | ATP/GTP binding protein-like 1 |
| chr11 | 57350246 | 57351217 | 972 | * | 8,45966 | 3,862658567 | 1,460794 | 2,644219 | 0,008188 | 0,853363379 | 1,425929 | Distal Intergenic | 11 | 57396116 | 57398254 | 2139 | 2 | 67726 | ENSMUST00000129596.2 | 47037 | ENSMUSG00000020523 | Fam114a2 | family with sequence similarity 114, member A2 |
| chr3 | 65572456 | 65576154 | 3699 | * | 25,937984 | 3,863267323 | 0,8151859 | 4,7391243 | 2,146E-06 | 0,006985965 | 2,8585108 | Promoter (<=1kb) | 3 | 65573649 | 65593600 | 19952 | 1 | 624866 | ENSMUST00000099075.4 | 0 | ENSMUSG00000074579 | Lekr1 | leucine, glutamate and lysine rich 1 |
| chr13 | 55741581 | 55743937 | 2357 | * | 16,78517 | 3,86357197 | 1,095033 | 3,52827 | 0,000418 | 0,273126507 | 2,318434 | Promoter (<=1kb) | 13 | 55740989 | 55741924 | 936 | 1 | 67511 | ENSMUST00000225888.2 | 592 | ENSMUSG00000058569 | Tmed9 | transmembrane p24 trafficking protein 9 |
| chr10 | 94033160 | 94036431 | 3272 | * | 21,093255 | 3,864086632 | 0,8491743 | 4,5504047 | 5,354E-06 | 0,013269851 | 2,9519335 | Promoter (<=1kb) | 10 | 94034582 | 94057285 | 22704 | 1 | 66414 | ENSMUST00000135292.3 | 0 | ENSMUSG00000020022 | Ndufa12 | NADH:ubiquinone oxidoreductase subunit A12 |
| chr14 | 46389355 | 46390266 | 912 | * | 8,577427 | 3,864554216 | 1,198865 | 3,223509 | 0,001266 | 0,53146022 | 1,360622 | Distal Intergenic | 14 | 46458549 | 46466805 | 8257 | 1 | 1,03E+08 | ENSMUST00000242583.1 | -68283 | | Gm34499 | predicted gene, 34499 |
| chr13 | 96566120 | 96567783 | 1664 | * | 11,15123 | 3,86517151 | 1,187917 | 3,25374 | 0,001139 | 0,627785088 | 0,738441 | 3' UTR | 13 | 96553603 | 96572460 | 18858 | 2 | 271144 | ENSMUST00000181761.2 | 4677 | ENSMUSG00000047117 | Ankdd1b | ankyrin repeat and death domain containing 1B |
| chr10 | 27606981 | 27608170 | 1190 | * | 13,91201 | 3,866128402 | 1,085194 | 3,562614 | 0,000367 | 0,356016677 | 1,075735 | Distal Intergenic | 10 | 27522309 | 27574323 | 52015 | 2 | 1,03E+08 | ENSMUST00000218769.2 | -32658 | | Gm30165 | predicted gene, 30165 |
| chr17 | 24298926 | 24299945 | 1020 | * | 11,24518 | 3,866257806 | 1,285591 | 3,007379 | 0,002635 | 0,730469267 | 0,918255 | Exon (ENSMUST00000: | 17 | 24266708 | 24292459 | 25752 | 2 | 69259 | ENSMUST00000017090.6 | -6467 | ENSMUSG00000016946 | Kctd5 | potassium channel tetramerisation domain containing 5 |
| chr5 | 38407660 | 38409060 | 1401 | * | 8,166826 | 3,866372302 | 1,451033 | 2,664565 | 0,007709 | 1 | 0 | Distal Intergenic | 5 | 38417529 | 38426978 | 9450 | 1 | 66309 | ENSMUST00000087511.10 | -8469 | ENSMUSG00000067365 | Tmem128 | transmembrane protein 128 |
| chr18 | 60827230 | 60828011 | 782 | * | 7,940169 | 3,866703522 | 1,465995 | 2,637597 | 0,00835 | 1 | 0 | Promoter (2-3kb) | 18 | 60823129 | 60830144 | 7016 | 2 | 15531 | ENSMUST00000237572.2 | 2133 | ENSMUSG00000054008 | Ndst1 | N-deacetylase/N-sulfotransferase (heparan glucosaminyl) 1 |
| chr12 | 54896976 | 54897680 | 705 | * | 8,307029 | 3,867149279 | 1,36805 | 2,82676 | 0,004702 | 0,750518846 | 1,449706 | Distal Intergenic | 12 | 54907265 | 54908523 | 1259 | 2 | 12632 | ENSMUST00000223320.2 | 10843 | ENSMUSG00000062929 | Cfl2 | cofilin 2, muscle |
| chr12 | 73852701 | 73853992 | 1292 | * | 9,352172 | 3,867183688 | 1,46961 | 2,631436 | 0,008502 | 0,856353593 | 1,449706 | Intron (ENSMUST0000( | 12 | 73840262 | 73855185 | 14924 | 1 | 666385 | ENSMUST00000222997.2 | 12439 | | Gm8075 | predicted gene 8075 |
| chr6 | 83931173 | 83931838 | 666 | * | 10,37618 | 3,867431632 | 1,246395 | 3,102894 | 0,001916 | 0,680780911 | 0,918255 | Exon (ENSMUST00000: | 6 | 83924880 | 83946213 | 21334 | 1 | 18139 | ENSMUST00000205166.2 | 6293 | ENSMUSG00000030016 | Zfp638 | zinc finger protein 638 |
| chr11 | 79484288 | 79485193 | 906 | * | 9,397192 | 3,867440216 | 1,409675 | 2,743497 | 0,006079 | 0,722439117 | 2,237403 | Promoter (<=1kb) | 11 | 79485991 | 79486094 | 104 | 1 | 1,15E+08 | ENSMUST00000082552.4 | -798 | ENSMUSG00000064486 | Gm23293 | predicted gene, 23293 |
| chr14 | 45566292 | 45568160 | 1869 | * | 20,621197 | 3,867836623 | 0,8748434 | 4,4211762 | 9,817E-06 | 0,022826798 | 2,6724296 | Promoter (<=1kb) | 14 | 45567245 | 45587162 | 19918 | 1 | 67089 | ENSMUST00000022380.9 | 0 | ENSMUSG00000021832 | Psmc6 | proteasome (prosome, macropain) 26S subunit, ATPase, 6 |
| chr9 | 4687410 | 4688257 | 848 | * | 7,966135 | 3,869383748 | 1,28369 | 3,014266 | 0,002576 | 1 | 0 | Intron (ENSMUST0000( | 9 | 4417899 | 4795519 | 377621 | 2 | 14802 | ENSMUST00000212533.2 | 107262 | ENSMUSG00000025892 | Gria4 | glutamate receptor, ionotropic, AMPA4 (alpha 4) |
| chr2 | 1,29E+08 | 1,29E+08 | 2007 | * | 14,53532 | 3,869642715 | 1,297683 | 2,981962 | 0,002864 | 0,674235791 | 1,41696 | Distal Intergenic | 2 | 1,29E+08 | 1,29E+08 | 151 | 2 | 1,15E+08 | ENSMUST00000083123.3 | -14933 | ENSMUSG00000065057 | Gm25703 | predicted gene, 25703 |
| chr8 | 83816442 | 83817086 | 645 | * | 5,860044 | 3,872011808 | 1,48181 | 2,613029 | 0,008974 | 1 | 0 | 3' UTR | 8 | 83891981 | 83999051 | 107071 | 1 | 71310 | ENSMUST00000034145.11 | -74895 | ENSMUSG00000031709 | Tbc1d9 | TBC1 domain family, member 9 |
| chr8 | 95165110 | 95165767 | 658 | * | 5,921107 | 3,873461616 | 1,479544 | 2,61801 | 0,008844 | 1 | 0 | Intron (ENSMUST0000( | 8 | 95156441 | 95161731 | 5291 | 2 | 320377 | ENSMUST00000180445.3 | -3379 | | 9330175E14Rik | RIKEN cDNA 9330175E14 gene |
| chr1 | 91342450 | 91343806 | 1357 | * | 10,13092 | 3,874603184 | 1,335416 | 2,90142 | 0,003715 | 0,74907225 | 1,173325 | Promoter (<=1kb) | 1 | 91340070 | 91341760 | 1691 | 2 | 55927 | ENSMUST00000150520.2 | -690 | ENSMUSG00000067071 | Hes6 | hairy and enhancer of split 6 |
| chr12 | 72783482 | 72784345 | 864 | * | 7,596881 | 3,87624308 | 1,373122 | 2,822942 | 0,004759 | 1 | 0 | Distal Intergenic | 12 | 72804229 | 72841338 | 37110 | 1 | 19042 | ENSMUST00000221628.2 | -19884 | ENSMUSG00000021096 | Ppm1a | protein phosphatase 1A, magnesium dependent, alpha isoform |
| chr15 | 81924373 | 81925645 | 1273 | * | 10,29293 | 3,876963369 | 1,226909 | 3,159943 | 0,001578 | 0,618136403 | 1,086118 | 3' UTR | 15 | 81925935 | 81931720 | 5786 | 2 | 20826 | ENSMUST00000230953.2 | 6075 | ENSMUSG00000063480 | Snu13 | SNU13 homolog, small nuclear ribonucleoprotein (U4/U6.U5) |
| chr3 | 1,08E+08 | 1,08E+08 | 1239 | * | 8,801041 | 3,877103161 | 1,432949 | 2,705682 | 0,006816 | 0,851704515 | 1,211315 | Exon (ENSMUST00000( | 3 | 1,08E+08 | 1,08E+08 | 2829 | 2 | 329735 | ENSMUST00000146337.3 | 4505 | ENSMUSG00000086968 | 4933431E20Rik | RIKEN cDNA 4933431E20 gene |
| chr6 | 49130079 | 49130372 | 294 | * | 6,03994 | 3,877699487 | 1,474881 | 2,62916 | 0,00856 | 1 | 0 | Intron (ENSMUST0000( | 6 | 49084334 | 49088034 | 3701 | 2 | 140488 | ENSMUST00000203863.2 | -42045 | ENSMUSG00000029814 | Igf2bp3 | insulin-like growth factor 2 mRNA binding protein 3 |
| chrX | 70150475 | 70151684 | 1210 | * | 10,20693 | 3,877810358 | 1,247001 | 3,109708 | 0,001873 | 0,634611828 | 1,173325 | Intron (ENSMUST0000( | 20 | 70093862 | 70199662 | 105801 | 1 | 333639 | ENSMUST00000082088.10 | 56613 | ENSMUSG00000059401 | Mamld1 | mastermind-like domain containing 1 |
| chr2 | 1,07E+08 | 1,07E+08 | 912 | * | 6,919552 | 3,878037773 | 1,288987 | 3,008594 | 0,002625 | 1 | 0 | Distal Intergenic | 2 | 1,07E+08 | 1,07E+08 | 3078 | 1 | 16492 | ENSMUST00000173191.2 | 39376 | ENSMUSG00000042604 | Kcna4 | potassium voltage-gated channel, shaker-related subfamily, member 4 |
| chr7 | 18929527 | 18930633 | 1107 | * | 10,38515 | 3,878247969 | 1,391747 | 2,786603 | 0,005326 | 0,891774732 | 0,738441 | Exon (ENSMUST00000: | 7 | 18915531 | 18920296 | 4766 | 1 | 72205 | ENSMUST00000146095.2 | 13996 | ENSMUSG00000040811 | Eml2 | echinoderm microtubule associated protein like 2 |
| chr5 | 1,21E+08 | 1,21E+08 | 695 | * | 6,343395 | 3,878293412 | 1,494193 | 2,595577 | 0,009443 | 1 | 0 | Intron (ENSMUST0000( | 5 | 1,21E+08 | 1,21E+08 | 1442 | 1 | 170756 | ENSMUST00000123326.2 | 7133 | ENSMUSG00000032754 | Slc8b1 | solute carrier family 8 (sodium/lithium/calcium exchanger), member B1 |
| chr2 | 84284647 | 84285658 | 1012 | * | 7,470237 | 3,878658108 | 1,487892 | 2,606814 | 0,009139 | 0,882498405 | 1,360622 | Promoter (2-3kb) | 2 | 84270484 | 84288454 | 17971 | 2 | 21788 | ENSMUST00000150261.8 | 2796 | ENSMUSG00000027082 | Tfpi | tissue factor pathway inhibitor |
| chr19 | 35328837 | 35330309 | 1473 | * | 13,72146 | 3,879810736 | 1,043154 | 3,719307 | 0,0002 | 0,275988434 | 1,075735 | Distal Intergenic | 19 | 35180635 | 35180766 | 132 | 1 | 1,15E+08 | ENSMUST00000158462.3 | 148202 | ENSMUSG00000089087 | Gm24006 | predicted gene, 24006 |
| chr10 | 75376992 | 75378305 | 1314 | * | 10,17781 | 3,881086071 | 1,220374 | 3,180242 | 0,001472 | 0,603552508 | 1,086118 | Distal Intergenic | 10 | 75385959 | 75396164 | 10206 | 2 | 192734 | ENSMUST00000051129.10 | 17859 | ENSMUSG00000046807 | Lrrc75b | leucine rich repeat containing 75B |
| chr12 | 72196947 | 72198456 | 1510 | * | 11,53267 | 3,881096336 | 1,369042 | 2,8349 | 0,004584 | 0,787687627 | 1,173325 | Intron (ENSMUST0000( | 12 | 72148074 | 72231803 | 83730 | 2 | 73936 | ENSMUST00000021494.6 | 33347 | ENSMUSG00000021086 | Ccdc175 | coiled-coil domain containing 175 |
| chr3 | 51130771 | 51132293 | 1523 | * | 16,0603 | 3,881810219 | 0,975679 | 3,978572 | 6,93E-05 | 0,087643251 | 2,663619 | Promoter (<=1kb) | 3 | 51131868 | 51159065 | 27198 | 1 | 12457 | ENSMUST00000023849.15 | 0 | ENSMUSG00000023087 | Noct | nocturnin |
| chr6 | 1,15E+08 | 1,15E+08 | 1114 | * | 13,16429 | 3,882133616 | 1,218228 | 3,186704 | 0,001439 | 0,566578624 | 1,2763 | Distal Intergenic | 6 | 1,15E+08 | 1,15E+08 | 107231 | 2 | 232334 | ENSMUST00000139640.2 | -20044 | ENSMUSG00000030315 | Vgll4 | vestigial like family member 4 |
| chr4 | 1,25E+08 | 1,25E+08 | 1008 | * | 7,094049 | 3,882294544 | 1,507138 | 2,575939 | 0,009997 | 0,895870471 | 1,360622 | Intron (ENSMUST0000( | 4 | 1,25E+08 | 1,25E+08 | 2063 | 1 | 17764 | ENSMUST00000175875.2 | 19422 | ENSMUSG00000028890 | Mtf1 | metal response element binding transcription factor 1 |
| chr13 | 31315502 | 31316599 | 1098 | * | 9,533434 | 3,882884044 | 1,407305 | 2,759091 | 0,005796 | 0,714462637 | 2,237403 | Distal Intergenic | 13 | 31397393 | 31425069 | 27677 | 2 | 73924 | ENSMUST00000153175.3 | 108470 | ENSMUSG00000087460 | 4930401O12Rik | RIKEN cDNA 4930401O12 gene |
| chr19 | 43740862 | 43742140 | 1279 | * | 8,649066 | 3,886246878 | 1,333298 | 2,914761 | 0,00356 | 0,63445438 | 2,237403 | Promoter (<=1kb) | 19 | 43721693 | 43741439 | 19747 | 2 | 226139 | ENSMUST00000045562.6 | 0 | ENSMUSG00000040018 | Cox15 | cytochrome c oxidase assembly protein 15 |
| chr2 | 1,41E+08 | 1,41E+08 | 1246 | * | 7,344304 | 3,886700874 | 1,47892 | 2,628067 | 0,008587 | 1 | 0 | Intron (ENSMUST0000( | 2 | 1,41E+08 | 1,41E+08 | 56561 | 2 | 1,03E+08 | ENSMUST00000151538.2 | -77183 | | Macrod2os1 | mono-ADP ribosylhydrolase 2, opposite strand 1 |
| chr17 | 71038066 | 71038676 | 611 | * | 6,037846 | 3,886766012 | 1,482806 | 2,621224 | 0,008761 | 1 | 0 | Intron (ENSMUST0000( | 17 | 71072834 | 71112992 | 40159 | 2 | 1,05E+08 | ENSMUST00000233559.2 | 74316 | ENSMUSG00000117231 | Gm41609 | predicted gene, 41609 |
| chr5 | 1,09E+08 | 1,09E+08 | 1655 | * | 14,78484 | 3,886798143 | 1,191009 | 3,263451 | 0,001101 | 0,526973991 | 1,214904 | Intron (ENSMUST0000( | 5 | 1,09E+08 | 1,09E+08 | 39260 | 2 | 231589 | ENSMUST00000053253.10 | 31830 | ENSMUSG00000091635 | Vmn2r13 | vomeronasal 2, receptor 13 |
| chr9 | 12426890 | 12428149 | 1260 | * | 7,481978 | 3,888154482 | 1,436851 | 2,706025 | 0,006809 | 0,829907213 | 1,360622 | Distal Intergenic | 9 | 13091272 | 13105860 | 14589 | 2 | 78219 | ENSMUST00000216694.2 | 677711 | ENSMUSG00000111830 | 4930568E12Rik | RIKEN cDNA 4930568E12 gene |
| chr10 | 22459584 | 22460295 | 712 | * | 7,971192 | 3,888419976 | 1,339653 | 2,902558 | 0,003701 | 1 | 0 | Distal Intergenic | 10 | 22520910 | 22580184 | 59275 | 1 | 353169 | ENSMUST00000042261.5 | -60615 | ENSMUSG00000037490 | Slc2a12 | solute carrier family 2 (facilitated glucose transporter), member 12 |
| chr11 | 53358934 | 53361812 | 2879 | * | 20,9878 | 3,889908319 | 0,94527 | 4,115131 | 3,87E-05 | 0,058289444 | 2,880017 | Exon (ENSMUST00000( | 11 | 53367400 | 53371101 | 3702 | 2 | 237761 | ENSMUST00000104955.4 | 9289 | ENSMUSG00000044352 | Sowaha | sosondowah ankyrin repeat domain family member A |
| chr9 | 21704017 | 21705348 | 1332 | * | 12,87072 | 3,889938788 | 1,199315 | 3,243466 | 0,001181 | 0,594676756 | 0,918255 | Intron (ENSMUST0000( | 9 | 21679358 | 21709799 | 30442 | 2 | 235041 | ENSMUST00000216008.2 | 4451 | ENSMUSG00000032194 | Kank2 | KN motif and ankyrin repeat domains 2 |
| chr19 | 36678216 | 36680142 | 1927 | * | 14,60541 | 3,893602374 | 1,301062 | 2,992635 | 0,002766 | 0,594676756 | 2,141612 | Promoter (2-3kb) | 19 | 36591452 | 36683041 | 91590 | 2 | 668215 | ENSMUST00000149601.8 | 2899 | ENSMUSG00000087579 | Hectd2os | Hectd2, opposite strand |
| chr6 | 41669603 | 41670226 | 624 | * | 6,976772 | 3,893862147 | 1,338822 | 2,908424 | 0,003633 | 1 | 0 | Exon (ENSMUST00000: | 6 | 41665368 | 41665893 | 526 | 2 | 23925 | ENSMUST00000153760.2 | -3710 | ENSMUSG00000029866 | Kel | Kell blood group |
| chr9 | 24119337 | 24120056 | 720 | * | 6,565909 | 3,894832142 | 1,447688 | 2,690381 | 0,007137 | 1 | 0 | Intron (ENSMUST0000( | 9 | 24194729 | 24224639 | 29911 | 1 | 319239 | ENSMUST00000154644.2 | -74673 | ENSMUSG00000043659 | Npsr1 | neuropeptide S receptor 1 |
| chr7 | 19049102 | 19050025 | 924 | * | 7,891273 | 3,895353188 | 1,414175 | 2,754505 | 0,005878 | 0,802490409 | 1,360622 | Distal Intergenic | 7 | 19036621 | 19043976 | 7356 | 2 | 14282 | ENSMUST00000003640.4 | -5126 | ENSMUSG00000003545 | Fosb | FBJ osteosarcoma oncogene B |
| chr11 | 20495842 | 20497538 | 1697 | * | 13,58157 | 3,896759929 | 1,058526 | 3,681306 | 0,000232 | 0,295900293 | 1,075735 | Promoter (2-3kb) | 11 | 20493334 | 20603021 | 109688 | 1 | 58172 | ENSMUST00000093292.11 | 2508 | ENSMUSG00000049800 | Sertad2 | SERTA domain containing 2 |
| chr18 | 74315052 | 74316249 | 1198 | * | 12,141 | 3,897068581 | 1,152723 | 3,38075 | 0,000723 | 0,44621096 | 1,2763 | Distal Intergenic | 18 | 74329550 | 74340818 | 11269 | 2 | 66468 | ENSMUST00000237276.2 | 24569 | ENSMUSG00000036223 | Ska1 | spindle and kinetochore associated complex subunit 1 |
| chr6 | 5590958 | 5591785 | 828 | * | 10,64372 | 3,898981975 | 1,236055 | 3,154376 | 0,001608 | 0,683988405 | 0,738441 | Distal Intergenic | 6 | 5511171 | 5570319 | 59149 | 1 | 1,03E+08 | ENSMUST00000245058.1 | 79787 | | Gm34470 | predicted gene, 34470 |
| chrX | 1,38E+08 | 1,38E+08 | 750 | * | 8,525474 | 3,899084293 | 1,27434 | 3,05969 | 0,002216 | 0,627785088 | 1,449706 | Intron (ENSMUST0000( | 20 | 1,38E+08 | 1,38E+08 | 94663 | 1 | 27206 | ENSMUST00000064937.14 | -139371 | ENSMUSG00000052854 | Nrk | Nik related kinase |
| chr13 | 94892774 | 94894180 | 1407 | * | 11,72677 | 3,900676796 | 1,225012 | 3,184195 | 0,001452 | 0,568412386 | 1,2763 | Distal Intergenic | 13 | 94925418 | 94979430 | 54013 | 1 | 21371 | ENSMUST00000046644.7 | -31238 | ENSMUSG00000042043 | Tbca | tubulin cofactor A |
| chr13 | 22217827 | 22219031 | 1205 | * | 8,92618 | 3,900786929 | 1,284941 | 3,035771 | 0,002399 | 0,651384755 | 1,360622 | Promoter (<=1kb) | 13 | 22219334 | 22219738 | 405 | 2 | 319168 | ENSMUST00000091742.6 | 707 | ENSMUSG00000069302 | H2ac12 | H2A clustered histone 12 |
| chr15 | 80120103 | 80121587 | 1485 | * | 14,35748 | 3,900852546 | 1,106396 | 3,525728 | 0,000422 | 0,409763611 | 0,938682 | Promoter (1-2kb) | 15 | 80118326 | 80122510 | 4185 | 1 | 239555 | ENSMUST00000231110.2 | 1777 | ENSMUSG00000022412 | Mief1 | mitochondrial elongation factor 1 |
| chr1 | 59513798 | 59515165 | 1368 | * | 8,636823 | 3,900893146 | 1,439089 | 2,710668 | 0,006715 | 0,826676469 | 1,360622 | Distal Intergenic | 1 | 59521583 | 59526114 | 4532 | 1 | 14369 | ENSMUST00000114246.4 | -6418 | ENSMUSG00000041075 | Fzd7 | frizzled class receptor 7 |
| chr1 | 1,05E+08 | 1,05E+08 | 839 | * | 11,56654 | 3,901320025 | 1,150487 | 3,391017 | 0,000696 | 0,533665288 | 0,738441 | Intron (ENSMUST0000( | 1 | 1,05E+08 | 1,05E+08 | 226953 | 1 | 23836 | ENSMUST00000062528.9 | 133875 | ENSMUSG00000050840 | Cdh20 | cadherin 20 |
| chr10 | 77636362 | 77636894 | 533 | * | 5,47587 | 3,902647205 | 1,458581 | 2,675646 | 0,007459 | 1 | 0 | Promoter (1-2kb) | 10 | 77634221 | 77634967 | 747 | 2 | 1,01E+08 | ENSMUST00000178581.2 | -1395 | ENSMUSG00000096380 | Gm19668 | predicted gene, 19668 |
| chr4 | 79888410 | 79889160 | 751 | * | 6,858565 | 3,903086944 | 1,320435 | 2,95591 | 0,003117 | 1 | 0 | Distal Intergenic | 4 | 80752360 | 80755921 | 3562 | 1 | 22178 | ENSMUST00000133655.8 | -863200 | ENSMUSG00000005994 | Tyrp1 | tyrosinase-related protein 1 |
| chrX | 1,41E+08 | 1,41E+08 | 1003 | * | 7,383667 | 3,903148854 | 1,489168 | 2,621027 | 0,008767 | 1 | 0 | Distal Intergenic | 20 | 1,4E+08 | 1,41E+08 | 14266 | 2 | 16370 | ENSMUST00000067841.8 | -212852 | ENSMUSG00000054667 | Irs4 | insulin receptor substrate 4 |
| chr13 | 45818745 | 45819394 | 650 | * | 6,005956 | 3,90324937 | 1,441153 | 2,708421 | 0,00676 | 1 | 0 | Intron (ENSMUST0000( | 13 | 45692285 | 45696026 | 3742 | 1 | 66355 | ENSMUST00000129176.2 | 126460 | ENSMUSG00000000253 | Gmpr | guanosine monophosphate reductase |
| chrX | 7628096 | 7629255 | 1160 | * | 9,111333 | 3,903453494 | 1,406589 | 2,775119 | 0,005518 | 0,785866008 | 1,425929 | Promoter (<=1kb) | 20 | 7628799 | 7632310 | 3512 | 1 | 209446 | ENSMUST00000144715.2 | 0 | ENSMUSG00000000134 | Tfe3 | transcription factor E3 |
| chr2 | 70871376 | 70872940 | 1565 | * | 13,07733 | 3,90349414 | 1,155615 | 3,377851 | 0,000731 | 0,50314252 | 0,918255 | Intron (ENSMUST0000( | 2 | 70885672 | 70916962 | 31291 | 1 | 75763 | ENSMUST00000130292.8 | -12732 | ENSMUSG00000041966 | Dcaf17 | DDB1 and CUL4 associated factor 17 |
| chr10 | 88523756 | 88524780 | 1025 | * | 8,236955 | 3,905883714 | 1,326584 | 2,944316 | 0,003237 | 0,68731021 | 1,425929 | Promoter (2-3kb) | 10 | 88511752 | 88520877 | 9126 | 2 | 20728 | ENSMUST00000138734.2 | -2879 | ENSMUSG00000004359 | Spic | Spi-C transcription factor (Spi-1/PU.1 related) |
| chr5 | 1,1E+08 | 1,1E+08 | 1062 | * | 7,685295 | 3,905975375 | 1,275932 | 3,061273 | 0,002204 | 1 | 0 | Distal Intergenic | 5 | 1,1E+08 | 1,1E+08 | 16497 | 2 | 77200 | ENSMUST00000092720.4 | 29741 | ENSMUSG00000072763 | 5430403G16Rik | RIKEN cDNA 5430403G16 gene |
| chr11 | 97329999 | 97330978 | 980 | * | 8,392644 | 3,906852497 | 1,342326 | 2,91051 | 0,003608 | 0,711199341 | 1,425929 | Promoter (2-3kb) | 11 | 97327111 | 97335054 | 7944 | 1 | 58996 | ENSMUST00000152933.2 | 2888 | ENSMUSG00000049807 | Arhgap23 | Rho GTPase activating protein 23 |
| chr5 | 35848631 | 35849335 | 705 | * | 7,563661 | 3,907115558 | 1,515405 | 2,578266 | 0,00993 | 1 | 0 | Distal Intergenic | 5 | 35809367 | 35837126 | 27760 | 2 | 78558 | ENSMUST00000087629.10 | -11505 | ENSMUSG00000029096 | Htra3 | HtrA serine peptidase 3 |
| chr8 | 75814289 | 75814957 | 669 | * | 6,99842 | 3,907562077 | 1,307132 | 2,989415 | 0,002795 | 1 | 0 | Distal Intergenic | 8 | 75820249 | 75827217 | 6969 | 1 | 15368 | ENSMUST00000005548.8 | -5292 | ENSMUSG00000005413 | Hmox1 | heme oxygenase 1 |
| chr1 | 33780609 | 33781362 | 754 | * | 8,874216 | 3,908846522 | 1,379314 | 2,833907 | 0,004598 | 0,74907225 | 1,449706 | 3' UTR | 1 | 33777218 | 33779853 | 2636 | 1 | 19335 | ENSMUST00000132066.4 | 3391 | ENSMUSG00000004768 | Rab23 | RAB23, member RAS oncogene family |
| chrX | 47607330 | 47609861 | 2532 | * | 15,399894 | 3,90921514 | 0,8873606 | 4,4054416 | 1,056E-05 | 0,039302356 | 1,3768303 | Promoter (<=1kb) | 20 | 47608162 | 47619109 | 10948 | 1 | 19337 | ENSMUST00000033430.3 | 0 | ENSMUSG00000031104 | Rab33a | RAB33A, member RAS oncogene family |
| chr13 | 51057832 | 51059161 | 1330 | * | 8,327153 | 3,909251686 | 1,29265 | 3,024216 | 0,002493 | 0,650270742 | 1,425929 | Distal Intergenic | 13 | 50907427 | 50919284 | 11858 | 2 | 71654 | ENSMUST00000221281.2 | -138548 | ENSMUSG00000113652 | 4930518P08Rik | RIKEN cDNA 4930518P08 gene |
| chr13 | 50314692 | 50315797 | 1106 | * | 10,80429 | 3,909312649 | 1,133346 | 3,449354 | 0,000562 | 0,496560937 | 0,738441 | Distal Intergenic | 13 | 50291105 | 50300706 | 9602 | 1 | 71252 | ENSMUST00000222134.2 | 23587 | | 4933433N18Rik | RIKEN cDNA 4933433N18 gene |
| chr4 | 1,21E+08 | 1,21E+08 | 952 | * | 9,740302 | 3,910079242 | 1,271665 | 3,074771 | 0,002107 | 0,654833721 | 1,173325 | Intron (ENSMUST0000( | 4 | 1,21E+08 | 1,21E+08 | 58275 | 2 | 18046 | ENSMUST00000120779.8 | 6618 | ENSMUSG00000032897 | Nfyc | nuclear transcription factor-Y gamma |
| chr9 | 62296870 | 62298020 | 1151 | * | 8,981196 | 3,912594561 | 1,483522 | 2,637369 | 0,008355 | 0,777919825 | 2,237403 | Distal Intergenic | 9 | 62280721 | 62285063 | 4343 | 1 | 11737 | ENSMUST00000145679.2 | 16149 | ENSMUSG00000032249 | Anp32a | acidic (leucine-rich) nuclear phosphoprotein 32 family, member A |
| chr18 | 75342086 | 75342936 | 851 | * | 8,459425 | 3,913048046 | 1,347901 | 2,903068 | 0,003695 | 0,722439117 | 1,360622 | Intron (ENSMUST0000( | 18 | 75258496 | 75284099 | 25604 | 1 | 69190 | ENSMUST00000236840.2 | 83590 | ENSMUSG00000035765 | Dym | dymeclin |
| chr9 | 58499813 | 58501349 | 1537 | * | 10,93996 | 3,913628726 | 1,501444 | 2,606577 | 0,009145 | 0,974006322 | 0,738441 | Intron (ENSMUST0000( | 9 | 58489727 | 58550951 | 61225 | 1 | 20320 | ENSMUST00000176126.2 | 10086 | ENSMUSG00000032336 | Nptn | neuroplastin |
| chr17 | 11850875 | 11851649 | 775 | * | 7,709072 | 3,914151457 | 1,478027 | 2,648228 | 0,008091 | 1 | 0 | Intron (ENSMUST0000( | 17 | 12057489 | 12086448 | 28960 | 1 | 50873 | ENSMUST00000179624.8 | -205840 | ENSMUSG00000023826 | Prkn | parkin RBR E3 ubiquitin protein ligase |
| chr4 | 1,37E+08 | 1,37E+08 | 1072 | * | 10,11364 | 3,914539577 | 1,20398 | 3,251333 | 0,001149 | 0,505078837 | 1,425929 | Distal Intergenic | 4 | 1,36E+08 | 1,37E+08 | 188450 | 2 | 13844 | ENSMUST00000105845.9 | -12443 | ENSMUSG00000028664 | Ephb2 | Eph receptor B2 |
| chr12 | 1E+08 | 1E+08 | 759 | * | 7,008524 | 3,914719488 | 1,36617 | 2,86547 | 0,004164 | 1 | 0 | Distal Intergenic | 12 | 1E+08 | 1E+08 | 1093 | 1 | 12313 | ENSMUST00000127300.2 | 7825 | ENSMUSG00000001175 | Calm1 | calmodulin 1 |
| chr3 | 1,45E+08 | 1,45E+08 | 737 | * | 6,827672 | 3,915536452 | 1,408184 | 2,780557 | 0,005427 | 1 | 0 | Intron (ENSMUST0000( | 3 | 1,45E+08 | 1,45E+08 | 23256 | 1 | 71355 | ENSMUST00000127379.2 | -9832 | ENSMUSG00000028197 | Col24a1 | collagen, type XXIV, alpha 1 |
| chr16 | 4606633 | 4607404 | 772 | * | 9,482978 | 3,915559124 | 1,390265 | 2,816411 | 0,004856 | 0,756018836 | 1,449706 | Promoter (<=1kb) | 16 | 4585203 | 4607794 | 22592 | 2 | 66626 | ENSMUST00000117713.8 | 390 | ENSMUSG00000004071 | Cdip1 | cell death inducing Trp53 target 1 |
| chr9 | 14413625 | 14414772 | 1148 | * | 10,41527 | 3,915630888 | 1,255348 | 3,119159 | 0,001814 | 0,67039394 | 0,918255 | Promoter (1-2kb) | 9 | 14412182 | 14414216 | 2035 | 1 | 66070 | ENSMUST00000215143.2 | 1443 | ENSMUSG00000004096 | Cwc15 | CWC15 spliceosome-associated protein |
| chr9 | 88739411 | 88740132 | 722 | * | 6,770226 | 3,915736277 | 1,436979 | 2,724979 | 0,006431 | 1 | 0 | Promoter (<=1kb) | 9 | 88739746 | 88739817 | 72 | 2 | 1,15E+08 | ENSMUST00000240516.1 | 0 | ENSMUSG00000119491 | Gm26135 | predicted gene, 26135 |
| chr1 | 50442714 | 50443781 | 1068 | * | 6,875311 | 3,916673385 | 1,425547 | 2,747489 | 0,006005 | 1 | 0 | Distal Intergenic | 1 | 50939806 | 50954589 | 14784 | 1 | 56363 | ENSMUST00000139157.8 | -496025 | ENSMUSG00000026109 | Tmeff2 | transmembrane protein with EGF-like and two follistatin-like domains 2 |
| chr12 | 80308870 | 80309766 | 897 | * | 7,640714 | 3,917392158 | 1,332535 | 2,939805 | 0,003284 | 1 | 0 | Promoter (1-2kb) | 12 | 80214321 | 80307145 | 92825 | 2 | 109711 | ENSMUST00000021554.16 | -1725 | ENSMUSG00000015143 | Actn1 | actinin, alpha 1 |
| chr5 | 1,25E+08 | 1,25E+08 | 693 | * | 7,308574 | 3,917899218 | 1,29094 | 3,034919 | 0,002406 | 1 | 0 | Intron (ENSMUST0000( | 5 | 1,25E+08 | 1,25E+08 | 6373 | 2 | 20778 | ENSMUST00000124582.2 | 3633 | ENSMUSG00000037936 | Scarb1 | scavenger receptor class B, member 1 |
| chr4 | 45533918 | 45535559 | 1642 | * | 11,59351 | 3,91920448 | 1,231595 | 3,182218 | 0,001462 | 0,569556737 | 1,2763 | Promoter (1-2kb) | 4 | 45458320 | 45532470 | 74151 | 2 | 230126 | ENSMUST00000147448.2 | -1448 | ENSMUSG00000044813 | Shb | src homology 2 domain-containing transforming protein B |
| chr7 | 94329889 | 94331316 | 1428 | * | 13,14303 | 3,919796791 | 1,112613 | 3,523056 | 0,000427 | 0,388075979 | 1,075735 | Intron (ENSMUST0000( | 7 | 93846417 | 93846686 | 270 | 1 | 545982 | ENSMUST00000071770.4 | 483472 | | Gm5899 | predicted pseudogene 5899 |

| | | | | | | | | | | | | | | | | | | | | | | | |
|---|---|---|---|---|---|---|---|---|---|---|---|---|---|---|---|---|---|---|---|---|---|---|---|
| chr13 | 99502933 | 99505099 | 2167 | * | 19,14714 | 3,919967072 | 1,109735 | 3,532344 | 0,000412 | 0,275793374 | 2,221611 | Intron (ENSMUST00000 | 13 | 99482052 | 99540060 | 58009 | 1 | 218506 | ENSMUST00000225973.2 | 20881 | ENSMUSG00000041632 | Mrps27 | mitochondrial ribosomal protein S27 |
| chrX | 1,44E+08 | 1,44E+08 | 782 | * | 6,825721 | 3,920307108 | 1,518797 | 2,581192 | 0,009846 | 1 | 0 | Distal Intergenic | 20 | 1,44E+08 | 1,44E+08 | 29462 | 2 | 27494 | ENSMUST00000143610.3 | -49367 | ENSMUSG00000041688 | Amot | angiomotin |
| chr11 | 1,19E+08 | 1,19E+08 | 1447 | * | 11,56826 | 3,920425581 | 1,416512 | 2,767662 | 0,005646 | 0,900823955 | 0,744375 | Intron (ENSMUST00000 | 11 | 1,19E+08 | 1,19E+08 | 564 | 2 | 1,05E+08 | ENSMUST00000151091.2 | 9240 | | Gm11753 | predicted gene 11753 |
| chr4 | 1,35E+08 | 1,35E+08 | 682 | * | 6,803691 | 3,920569915 | 1,441472 | 2,719839 | 0,006531 | 1 | 0 | Promoter (<=1kb) | 4 | 1,35E+08 | 1,35E+08 | 1508 | 1 | 27981 | ENSMUST00000149003.2 | 114 | ENSMUSG00000037266 | Rsrp1 | arginine/serine rich protein 1 |
| chr18 | 4858913 | 4859607 | 695 | * | 7,819498 | 3,920645857 | 1,440385 | 2,721943 | 0,00649 | 0,821607359 | 1,360622 | Distal Intergenic | 18 | 4812486 | 4850959 | 38474 | 1 | 1E+08 | ENSMUST00000180904.2 | 46427 | ENSMUSG00000097043 | Gm10556 | predicted gene 10556 |
| chr11 | 1,13E+08 | 1,13E+08 | 1930 | * | 45,35771 | 3,920820118 | 1,508294 | 2,599506 | 0,009336 | 0,756018836 | 2,787255 | Intron (ENSMUST00000 | 11 | 1,13E+08 | 1,13E+08 | 129057 | 2 | 72386 | ENSMUST00000244333.1 | 108198 | ENSMUSG00000087259 | 2610035D17Rik | RIKEN cDNA 2610035D17 gene |
| chr12 | 67135548 | 67136177 | 630 | * | 6,039665 | 3,921035557 | 1,513648 | 2,590453 | 0,009585 | 1 | 0 | Intron (ENSMUST00000 | 12 | 66518289 | 67267816 | 749528 | 2 | 320772 | ENSMUST00000223141.2 | 131639 | ENSMUSG00000034912 | Mdga2 | MAM domain containing glycosylphosphatidylinositol anchor 2 |
| chr12 | 55038677 | 55039884 | 1208 | * | 8,247485 | 3,921226721 | 1,339772 | 2,926787 | 0,003425 | 0,695892636 | 1,449706 | Intron (ENSMUST00000 | 12 | 55043857 | 55043962 | 106 | 1 | 1,15E+08 | ENSMUST00000158297.3 | -3973 | ENSMUSG00000088922 | Gm24296 | predicted gene, 24296 |
| chr10 | 71180289 | 71181238 | 950 | * | 9,252798 | 3,921628223 | 1,328397 | 2,952152 | 0,003156 | 0,715249603 | 1,211315 | Promoter (<=1kb) | 10 | 71166310 | 71180784 | 14475 | 2 | 52637 | ENSMUST00000045887.9 | 0 | ENSMUSG00000037710 | Cisd1 | CDGSH iron sulfur domain 1 |
| chr17 | 32063184 | 32064338 | 1155 | * | 7,404775 | 3,921925497 | 1,518775 | 2,582296 | 0,009815 | 1 | 0 | 3' UTR | 17 | 32068728 | 32070549 | 1822 | 2 | 17691 | ENSMUST00000234867.2 | 6211 | ENSMUSG00000024042 | Sik1 | salt inducible kinase 1 |
| chr12 | 1,12E+08 | 1,12E+08 | 1337 | * | 11,40315 | 3,922269089 | 1,136531 | 3,451089 | 0,000558 | 0,418252864 | 1,173325 | Promoter (2-3kb) | 12 | 1,12E+08 | 1,12E+08 | 2047 | 2 | 1,01E+08 | ENSMUST00000245564.1 | 2363 | ENSMUSG00000096995 | 2810029C07Rik | RIKEN cDNA 2810029C07 gene |
| chr15 | 77924623 | 77925511 | 889 | * | 9,52115 | 3,922548131 | 1,456113 | 2,693848 | 0,007063 | 0,748971518 | 2,237403 | Intron (ENSMUST00000 | 15 | 77843198 | 77855013 | 11816 | 2 | 55944 | ENSMUST00000100484.6 | -69610 | ENSMUSG00000016554 | Eif3d | eukaryotic translation initiation factor 3, subunit D |
| chr9 | 21023577 | 21024476 | 900 | * | 8,383067 | 3,922744533 | 1,413692 | 2,774823 | 0,005523 | 0,786234479 | 1,425929 | Intron (ENSMUST00000 | 9 | 21015365 | 21038642 | 23278 | 2 | 54721 | ENSMUST00000001036.11 | 14166 | ENSMUSG00000032175 | Tyk2 | tyrosine kinase 2 |
| chr7 | 80866463 | 80867726 | 1264 | * | 11,38706 | 3,924434945 | 1,101586 | 3,562531 | 0,000367 | 0,425046984 | 0,738441 | Intron (ENSMUST00000 | 7 | 80863344 | 80984281 | 120938 | 1 | 18584 | ENSMUST00000026672.8 | 3119 | ENSMUSG00000025584 | Pde8a | phosphodiesterase 8A |
| chr14 | 89748319 | 89749304 | 986 | * | 5,737772 | 3,926098504 | 1,504438 | 2,609678 | 0,009063 | 1 | 0 | Distal Intergenic | 14 | 89944241 | 89944372 | 132 | 1 | 1,15E+08 | ENSMUST00000104220.3 | -194937 | ENSMUSG00000077408 | Gm25415 | predicted gene, 25415 |
| chr14 | 23055575 | 23057362 | 1788 | * | 13,00253 | 3,926950844 | 1,358921 | 2,889757 | 0,003855 | 0,770460067 | 1,075735 | Intron (ENSMUST00000 | 14 | 23088262 | 23144639 | 56378 | 2 | 791307 | ENSMUST00000090409.2 | 87277 | ENSMUSG00000068680 | Gm10248 | predicted gene 10248 |
| chr13 | 5174626 | 5175500 | 875 | * | 7,591653 | 3,927324623 | 1,355231 | 2,897901 | 0,003757 | 1 | 0 | Distal Intergenic | 13 | 5239802 | 5239933 | 132 | 2 | 1,15E+08 | ENSMUST00000104285.3 | 64433 | ENSMUSG00000077473 | Gm24790 | predicted gene, 24790 |
| chr11 | 88066097 | 88067375 | 1279 | * | 8,016893 | 3,927345722 | 1,5045 | 2,610399 | 0,009044 | 0,880247401 | 1,360622 | Intron (ENSMUST00000 | 11 | 88073129 | 88074106 | 978 | 1 | 103841 | ENSMUST00000132377.2 | -5754 | ENSMUSG00000018378 | Cuedc1 | CUE domain containing 1 |
| chr1 | 1,69E+08 | 1,69E+08 | 841 | * | 6,724748 | 3,927794114 | 1,441655 | 2,724504 | 0,00644 | 1 | 0 | Distal Intergenic | 1 | 1,68E+08 | 1,68E+08 | 120 | 1 | 1,02E+08 | ENSMUST00000183804.3 | 252529 | ENSMUSG00000098407 | Mir6348 | microRNA 6348 |
| chr5 | 19995255 | 19996286 | 1032 | * | 9,655716 | 3,927819477 | 1,223437 | 3,210478 | 0,001325 | 0,583571732 | 1,086118 | Intron (ENSMUST00000 | 5 | 20112516 | 20909790 | 797275 | 1 | 50791 | ENSMUST00000101558.10 | -116230 | ENSMUSG00000040003 | Magi2 | membrane associated guanylate kinase, WW and PDZ domain containing 2 |
| chr7 | 1,13E+08 | 1,13E+08 | 1662 | * | 9,420825 | 3,928221093 | 1,503125 | 2,613369 | 0,008965 | 0,868766096 | 1,425929 | Promoter (<=1kb) | 7 | 1,13E+08 | 1,13E+08 | 53714 | 2 | 68815 | ENSMUST00000047091.14 | 0 | ENSMUSG00000038187 | Btbd10 | BTB (POZ) domain containing 10 |
| chr7 | 80582605 | 80583938 | 1334 | * | 9,981232 | 3,928372577 | 1,515263 | 2,592535 | 0,009527 | 0,881662676 | 1,425929 | Intron (ENSMUST00000 | 7 | 80565125 | 80575398 | 10274 | 2 | 56529 | ENSMUST00000150049.8 | -7207 | ENSMUSG00000025724 | Sec11a | SEC11 homolog A, signal peptidase complex subunit |
| chr11 | 86514148 | 86515575 | 1428 | * | 12,92468 | 3,928480008 | 1,154526 | 3,402679 | 0,000667 | 0,526773513 | 0,738441 | Intron (ENSMUST00000 | 11 | 86521254 | 86521852 | 599 | 2 | 1,01E+08 | ENSMUST00000117851.2 | 6277 | ENSMUSG00000083992 | Gm11478 | predicted gene 11478 |
| chr18 | 66553911 | 66554781 | 871 | * | 7,461978 | 3,92982322 | 1,498125 | 2,623161 | 0,008712 | 1 | 0 | Distal Intergenic | 18 | 66591604 | 66598631 | 7028 | 1 | 58801 | ENSMUST00000025399.9 | -36823 | ENSMUSG00000024521 | Pmaip1 | phorbol-12-myristate-13-acetate-induced protein 1 |
| chr9 | 65005175 | 65006885 | 1711 | * | 13,04625 | 3,931040237 | 1,109747 | 3,542286 | 0,000397 | 0,438881512 | 0,738441 | Promoter (1-2kb) | 9 | 65008768 | 65045222 | 36455 | 1 | 56741 | ENSMUST00000035499.5 | -1883 | ENSMUSG00000032816 | Igdcc4 | immunoglobulin superfamily, DCC subclass, member 4 |
| chr10 | 59138823 | 59139631 | 809 | * | 7,720586 | 3,931169503 | 1,48273 | 2,651306 | 0,008018 | 0,857114361 | 1,360622 | Distal Intergenic | 10 | 59159118 | 59209126 | 50009 | 1 | 18451 | ENSMUST00000009789.15 | -19487 | ENSMUSG00000019916 | P4ha1 | procollagen-proline, 2-oxoglutarate 4-dioxygenase (proline 4-hydroxylase), alpha 1 polypeptide |
| chr13 | 99986120 | 99987146 | 1027 | * | 7,832816 | 3,931193724 | 1,482301 | 2,652088 | 0,008 | 1 | 0 | Distal Intergenic | 13 | 1E+08 | 1E+08 | 2153 | 2 | 27220 | ENSMUST00000022150.8 | 50003 | ENSMUSG00000021647 | Cartpt | CART prepropeptide |
| chr8 | 12163776 | 12164793 | 1018 | * | 11,44422 | 3,931201287 | 1,143909 | 3,436637 | 0,000589 | 0,426043749 | 1,173325 | Distal Intergenic | 8 | 12328819 | 12330657 | 1839 | 1 | 619290 | ENSMUST00000098935.2 | -164026 | ENSMUSG00000074473 | A230072I06Rik | RIKEN cDNA A230072I06 gene |
| chr1 | 89007926 | 89009736 | 1811 | * | 15,27436 | 3,93142912 | 1,093641 | 3,594807 | 0,000325 | 0,302204121 | 1,41696 | Intron (ENSMUST00000 | 1 | 88998175 | 89040582 | 42408 | 1 | 98402 | ENSMUST00000123745.2 | 9751 | ENSMUSG00000036206 | Sh3bp4 | SH3-domain binding protein 4 |
| chr14 | 31501597 | 31503932 | 2336 | * | 19,23958 | 3,931631035 | 1,123033 | 3,500904 | 0,000464 | 0,282235674 | 2,386008 | Promoter (<=1kb) | 14 | 31420725 | 31502350 | 81626 | 2 | 105522 | ENSMUST00000227863.2 | 0 | ENSMUSG00000014496 | Ankrd28 | ankyrin repeat domain 28 |
| chr1 | 93729186 | 93731710 | 2525 | * | 23,267036 | 3,931867988 | 0,8915206 | 4,4102941 | 1,032E-05 | 0,022589498 | 2,8800171 | Promoter (<=1kb) | 1 | 93720298 | 93729656 | 9359 | 2 | 21915 | ENSMUST00000027503.14 | 0 | ENSMUSG00000026281 | Dtymk | deoxythymidylate kinase |
| chr5 | 23743325 | 23744321 | 997 | * | 10,06382 | 3,933068366 | 1,383403 | 2,843039 | 0,004469 | 0,744054256 | 1,449706 | Promoter (2-3kb) | 5 | 23730729 | 23747116 | 16388 | 2 | 20817 | ENSMUST00000200173.2 | 2795 | ENSMUSG00000062604 | Srpk2 | serine/arginine-rich protein specific kinase 2 |
| chr12 | 69866495 | 69867364 | 870 | * | 7,231435 | 3,933311081 | 1,45143 | 2,709956 | 0,006729 | 1 | 0 | Promoter (<=1kb) | 12 | 69864556 | 69865631 | 1076 | 2 | 399510 | ENSMUST00000153550.2 | -864 | ENSMUSG00000034761 | Map4k5 | mitogen-activated protein kinase kinase kinase kinase 5 |
| chr6 | 27425175 | 27425728 | 554 | * | 7,146774 | 3,93479583 | 1,402535 | 2,805489 | 0,005024 | 1 | 0 | Intron (ENSMUST00000 | 6 | 27936654 | 27936749 | 96 | 2 | 735266 | ENSMUST00000102113.3 | 511021 | ENSMUSG00000076067 | Mir592 | microRNA 592 |
| chr1 | 1,36E+08 | 1,36E+08 | 635 | * | 6,520648 | 3,935261289 | 1,428052 | 2,755685 | 0,005857 | 1 | 0 | Intron (ENSMUST00000 | 1 | 1,36E+08 | 1,36E+08 | 81659 | 2 | 215690 | ENSMUST00000112103.2 | -6418 | ENSMUSG00000009418 | Nav1 | neuron navigator 1 |
| chr15 | 76044355 | 76046422 | 2068 | * | 14,82206 | 3,936982449 | 0,963966 | 4,08415 | 4,42E-05 | 0,10388 | 1,2763 | Distal Intergenic | 15 | 76059580 | 76072243 | 12664 | 2 | 18810 | ENSMUST00000165210.8 | 25821 | ENSMUSG00000022565 | Plec | plectin |
| chr15 | 6954463 | 6955593 | 1131 | * | 11,06898 | 3,937553606 | 1,285445 | 3,063183 | 0,00219 | 0,661775851 | 1,173325 | Distal Intergenic | 15 | 6844518 | 6904450 | 59933 | 2 | 18414 | ENSMUST00000176826.2 | -50013 | ENSMUSG00000022146 | Osmr | oncostatin M receptor |
| chr12 | 3475548 | 3477854 | 2307 | * | 25,849249 | 3,937612154 | 0,7221882 | 5,4523352 | 4,971E-08 | 0,000404597 | 2,6724296 | Promoter (<=1kb) | 12 | 3453878 | 3477011 | 23134 | 2 | 100043040 | ENSMUST00000168012.3 | 0 | ENSMUSG00000071456 | 1110002L01Rik | RIKEN cDNA 1110002L01 gene |
| chr4 | 1,33E+08 | 1,33E+08 | 1364 | * | 15,65349 | 3,937946276 | 1,143779 | 3,442927 | 0,000575 | 0,409763611 | 1,2763 | Distal Intergenic | 4 | 1,33E+08 | 1,33E+08 | 10266 | 1 | 14191 | ENSMUST00000124451.2 | -7658 | ENSMUSG00000028874 | Fgr | FGR proto-oncogene, Src family tyrosine kinase |
| chr18 | 64648665 | 64650018 | 1354 | * | 10,37461 | 3,938336992 | 1,210134 | 3,254464 | 0,001136 | 0,524368688 | 1,2763 | Promoter (<=1kb) | 18 | 64632718 | 64649723 | 17006 | 2 | 70223 | ENSMUST00000237400.2 | 0 | ENSMUSG00000024587 | Nars | asparaginyl-tRNA synthetase |
| chr2 | 96135764 | 96136679 | 916 | * | 7,299502 | 3,938947463 | 1,400085 | 2,813362 | 0,004903 | 1 | 0 | Distal Intergenic | 2 | 96148514 | 97462011 | 1313498 | 1 | 241568 | ENSMUST00000135431.8 | -11835 | ENSMUSG00000050587 | Lrrc4c | leucine rich repeat containing 4C |
| chr4 | 67456425 | 67457245 | 821 | * | 7,895028 | 3,939108776 | 1,36964 | 2,876017 | 0,004027 | 1 | 0 | Distal Intergenic | 4 | 68100769 | 68100939 | 171 | 1 | 1,15E+08 | ENSMUST00000082828.3 | -643524 | ENSMUSG00000064762 | Gm23738 | predicted gene, 23738 |
| chr15 | 97196840 | 97197373 | 534 | * | 7,941963 | 3,939496401 | 1,471784 | 2,676681 | 0,007436 | 1 | 0 | Intron (ENSMUST00000 | 15 | 97146954 | 97175219 | 28266 | 1 | 239647 | ENSMUST00000226615.2 | 49886 | ENSMUSG00000044250 | Pced1b | PC-esterase domain containing 1B |
| chr18 | 11771148 | 11772562 | 1415 | * | 9,321257 | 3,940221547 | 1,351363 | 2,915739 | 0,003548 | 0,715249603 | 1,360622 | Intron (ENSMUST00000 | 18 | 11766428 | 11876257 | 109830 | 1 | 225182 | ENSMUST00000235039.2 | 4720 | ENSMUSG00000041238 | Rbbp8 | retinoblastoma binding protein 8, endonuclease |
| chr6 | 85969959 | 85970316 | 358 | * | 8,488852 | 3,940726368 | 1,430436 | 2,754912 | 0,005871 | 1 | 0 | Intron (ENSMUST00000 | 6 | 85981362 | 85986337 | 4976 | 1 | 75355 | ENSMUST00000201164.2 | -11046 | ENSMUSG00000106791 | 4930553P18Rik | RIKEN cDNA 4930553P18 gene |
| chr5 | 1,4E+08 | 1,4E+08 | 1808 | * | 13,8862 | 3,942868049 | 1,021436 | 3,860122 | 0,000113 | 0,221323359 | 0,938682 | Intron (ENSMUST00000 | 5 | 1,4E+08 | 1,4E+08 | 62577 | 1 | 243312 | ENSMUST00000198608.2 | 4343 | ENSMUSG00000048988 | Elfn1 | leucine rich repeat and fibronectin type III, extracellular 1 |
| chr17 | 34695886 | 34696484 | 599 | * | 7,371659 | 3,943513851 | 1,443089 | 2,73269 | 0,006282 | 1 | 0 | Promoter (<=1kb) | 17 | 34685536 | 34696292 | 10757 | 2 | 632126 | ENSMUST00000236448.2 | 0 | ENSMUSG00000058435 | Btnl4 | butyrophilin-like 4 |
| chr14 | 14261982 | 14262908 | 927 | * | 10,2908 | 3,943960388 | 1,211693 | 3,254916 | 0,001134 | 0,50314252 | 1,425929 | Distal Intergenic | 14 | 14246212 | 14255878 | 9667 | 2 | 71393 | ENSMUST00000170111.3 | -6104 | ENSMUSG00000021752 | Kctd6 | potassium channel tetramerisation domain containing 6 |
| chr5 | 33602447 | 33603052 | 606 | * | 7,200486 | 3,944000436 | 1,359469 | 2,901134 | 0,003718 | 1 | 0 | Distal Intergenic | 5 | 33588078 | 33591320 | 3243 | 2 | 672284 | ENSMUST00000173348.2 | -11127 | ENSMUSG00000029112 | Nkx1-1 | NK1 homeobox 1 |
| chr8 | 8231919 | 8232525 | 607 | * | 5,902376 | 3,944558018 | 1,511203 | 2,610211 | 0,009049 | 1 | 0 | Intron (ENSMUST00000 | 8 | 8116666 | 8143355 | 26690 | 1 | 1,03E+08 | ENSMUST00000207352.2 | 115253 | | Gm31401 | predicted gene, 31401 |
| chr1 | 36141283 | 36142231 | 949 | * | 7,969806 | 3,944842195 | 1,486436 | 2,653894 | 0,007957 | 0,85617852 | 1,360622 | Exon (ENSMUST000001 | 1 | 36171099 | 36173635 | 2537 | 1 | 1,03E+08 | ENSMUST00000245804.1 | -28868 | | Gm33280 | predicted gene, 33280 |
| chr16 | 16804611 | 16806442 | 1832 | * | 16,73179 | 3,945229843 | 0,934889 | 4,219998 | 2,44E-05 | 0,052288507 | 2,141612 | Intron (ENSMUST00000 | 16 | 16801431 | 16856904 | 55474 | 1 | 26413 | ENSMUST00000115731.10 | 3180 | ENSMUSG00000063358 | Mapk1 | mitogen-activated protein kinase 1 |
| chr1 | 1,14E+08 | 1,14E+08 | 1075 | * | 9,293916 | 3,945395248 | 1,249517 | 3,157536 | 0,001591 | 0,573678185 | 1,360622 | Distal Intergenic | 1 | 1,16E+08 | 1,16E+08 | 15632 | 2 | 1,08E+08 | ENSMUST00000186137.7 | 1238382 | | Gm50605 | predicted gene, 50605 |
| chr6 | 54051877 | 54053783 | 1907 | * | 24,969496 | 3,946619851 | 0,8524231 | 4,6298835 | 3,659E-06 | 0,009977325 | 2,9510621 | Intron (ENSMUST000004 | 6 | 54074705 | 54197522 | 122818 | 1 | 69993 | ENSMUST00000204410.2 | -20922 | ENSMUSG00000004633 | Chn2 | chimerin 2 |
| chr9 | 1,23E+08 | 1,23E+08 | 889 | * | 7,75953 | 3,947046117 | 1,501188 | 2,629281 | 0,008557 | 0,868766096 | 1,360622 | Promoter (<=1kb) | 9 | 1,23E+08 | 1,23E+08 | 7957 | 1 | 22646 | ENSMUST00000051667.14 | -221 | ENSMUSG00000057895 | Zfp105 | zinc finger protein 105 |
| chr16 | 24289592 | 24290110 | 519 | * | 6,133815 | 3,950365451 | 1,438105 | 2,746925 | 0,006016 | 1 | 0 | Intron (ENSMUST00000 | 16 | 24266841 | 24799780 | 532940 | 1 | 210126 | ENSMUST00000078988.10 | 22751 | ENSMUSG00000033306 | Lpp | LIM domain containing preferred translocation partner in lipoma |
| chr11 | 1,16E+08 | 1,16E+08 | 1368 | * | 11,42104 | 3,950607197 | 1,263694 | 3,126236 | 0,001771 | 0,610143359 | 1,2763 | Intron (ENSMUST00000 | 11 | 1,16E+08 | 1,16E+08 | 1904 | 1 | 68528 | ENSMUST00000132961.2 | 6549 | ENSMUSG00000075420 | Smim6 | small integral membrane protein 6 |
| chr13 | 97159656 | 97160653 | 998 | * | 9,353259 | 3,950713515 | 1,250781 | 3,158597 | 0,001585 | 0,562139639 | 1,449706 | Intron (ENSMUST00000 | 13 | 97158643 | 97170841 | 12199 | 2 | 66479 | ENSMUST00000221277.2 | 10188 | ENSMUSG00000052075 | 1700029F12Rik | RIKEN cDNA 1700029F12 gene |
| chrX | 1,42E+08 | 1,42E+08 | 1315 | * | 8,260014 | 3,950951085 | 1,402828 | 2,816418 | 0,004856 | 0,75938015 | 1,425929 | Intron (ENSMUST00000 | 20 | 1,42E+08 | 1,42E+08 | 115583 | 2 | 56068 | ENSMUST00000041317.3 | 8524 | ENSMUSG00000042225 | Ammecr1 | Alport syndrome, mental retardation, midface hypoplasia and elliptocytosis chromosomal region gene 1 |
| chr5 | 1,4E+08 | 1,4E+08 | 727 | * | 6,740608 | 3,952993833 | 1,503632 | 2,628963 | 0,008565 | 1 | 0 | 3' UTR | 5 | 1,4E+08 | 1,4E+08 | 4818 | 1 | 17135 | ENSMUST00000110836.2 | 4378 | ENSMUSG00000018143 | Mafk | v-maf musculoaponeurotic fibrosarcoma oncogene family, protein K (avian) |
| chr19 | 11293824 | 11294932 | 1109 | * | 9,983848 | 3,95313201 | 1,433581 | 2,757522 | 0,005824 | 0,823446343 | 1,211315 | Promoter (2-3kb) | 19 | 11278613 | 11291818 | 13206 | 2 | 383435 | ENSMUST00000187467.2 | -2006 | ENSMUSG00000099398 | Ms4a14 | membrane-spanning 4-domains, subfamily A, member 14 |
| chr8 | 75760301 | 75761437 | 1137 | * | 29,193367 | 3,953174096 | 0,7490268 | 5,2777471 | 1,308E-07 | 0,00087517 | 2,6724296 | Promoter (<=1kb) | 8 | 75760333 | 75796749 | 36417 | 1 | 21968 | ENSMUST00000165630.3 | 0 | ENSMUSG00000042870 | Tom1 | target of myb1 trafficking protein |
| chr18 | 61500633 | 61501985 | 1353 | * | 10,78997 | 3,953795755 | 1,298228 | 3,045533 | 0,002323 | 0,670853401 | 1,173325 | Intron (ENSMUST00000 | 18 | 61530906 | 61530971 | 66 | 2 | 723889 | ENSMUST00000198300.3 | 28986 | ENSMUSG00000105200 | Mir378a | microRNA 378a |
| chr4 | 17671702 | 17672371 | 670 | * | 6,978585 | 3,955172556 | 1,50625 | 2,625841 | 0,008644 | 1 | 0 | Distal Intergenic | 4 | 17546447 | 17636191 | 89745 | 2 | 1,03E+08 | ENSMUST00000245354.1 | -35511 | | Gm30092 | predicted gene, 30092 |
| chr12 | 4779855 | 4780831 | 977 | * | 9,861681 | 3,955959817 | 1,387706 | 2,85072 | 0,004362 | 0,770994815 | 1,211315 | Exon (ENSMUST000002 | 12 | 4765549 | 4787771 | 22223 | 2 | 74855 | ENSMUST00000220978.2 | 6940 | ENSMUSG00000079177 | Fam228a | family with sequence similarity 228, member A |
| chr11 | 1,2E+08 | 1,2E+08 | 1043 | * | 8,448264 | 3,956281411 | 1,242275 | 3,184706 | 0,001449 | 0,55292773 | 1,360622 | Distal Intergenic | 11 | 1,2E+08 | 1,2E+08 | 8286 | 1 | 14527 | ENSMUST00000026119.8 | -14442 | ENSMUSG00000025127 | Gcgr | glucagon receptor |
| chr2 | 1,19E+08 | 1,19E+08 | 1389 | * | 10,36794 | 3,957366483 | 1,376727 | 2,874473 | 0,004047 | 0,762883845 | 1,173325 | Exon (ENSMUST00000 | 2 | 1,19E+08 | 1,19E+08 | 3153 | 1 | 69065 | ENSMUST00000028780.4 | 25605 | ENSMUSG00000027313 | Chac1 | ChaC, cation transport regulator 1 |
| chr7 | 1,26E+08 | 1,26E+08 | 848 | * | 7,849149 | 3,95822504 | 1,326226 | 2,984578 | 0,00284 | 1 | 0 | Intron (ENSMUST00000 | 7 | 1,25E+08 | 1,25E+08 | 2705 | 1 | 233865 | ENSMUST00000155059.3 | 103646 | ENSMUSG00000032743 | Katnip | katanin interacting protein |
| chr6 | 84144895 | 84146676 | 1782 | * | 18,26719 | 3,958713029 | 1,02992 | 3,843708 | 0,000121 | 0,123281796 | 2,663619 | Intron (ENSMUST00000 | 6 | 84123729 | 84129392 | 5664 | 1 | 26903 | ENSMUST00000142711.6 | 21166 | ENSMUSG00000033788 | Dysf | dysferlin |
| chr16 | 4108882 | 4110069 | 1188 | * | 9,719662 | 3,959062616 | 1,191389 | 3,323064 | 0,00089 | 0,507398906 | 1,086118 | Intron (ENSMUST00000 | 16 | 4048165 | 4048963 | 799 | 1 | 436332 | ENSMUST00000161633.2 | 60717 | ENSMUSG00000089645 | Gm5766 | ribosomal protein L7a pseudogene |
| chr12 | 3867067 | 3869391 | 2325 | * | 23,418335 | 3,960310432 | 0,9114781 | 4,3449319 | 1,393E-05 | 0,027906449 | 2,8800171 | Intron (ENSMUST000001 | 12 | 3857160 | 3963491 | 106332 | 1 | 13435 | ENSMUST00000174817.8 | 9907 | ENSMUSG00000020661 | Dnmt3a | DNA methyltransferase 3A |
| chr8 | 1,15E+08 | 1,15E+08 | 656 | * | 8,421808 | 3,960691508 | 1,461596 | 2,70984 | 0,006732 | 0,826716028 | 1,360622 | Intron (ENSMUST00000 | 8 | 1,15E+08 | 1,15E+08 | 9568 | 2 | 1,03E+08 | ENSMUST00000161914.2 | 16388 | ENSMUSG00000089797 | Gm16118 | predicted gene 16118 |
| chr6 | 1,01E+08 | 1,01E+08 | 1026 | * | 10,99283 | 3,960887247 | 1,236966 | 3,202098 | 0,001364 | 0,622584918 | 0,918255 | Promoter (1-2kb) | 6 | 1,01E+08 | 1,01E+08 | 36313 | 1 | 232314 | ENSMUST00000063854.7 | -1255 | ENSMUSG00000052144 | Ppp4r2 | protein phosphatase 4, regulatory subunit 2 |
| chr7 | 40577481 | 40578018 | 538 | * | 5,61414 | 3,96206642 | 1,451415 | 2,729796 | 0,006337 | 1 | 0 | Intron (ENSMUST00000 | 7 | 40467900 | 40550361 | 82462 | 2 | 320210 | ENSMUST00000189574.7 | -27120 | ENSMUSG00000100600 | A230077H06Rik | RIKEN cDNA A230077H06 gene |
| chr7 | 1,1E+08 | 1,1E+08 | 665 | * | 8,342202 | 3,962550656 | 1,328331 | 2,983104 | 0,002853 | 0,671937722 | 1,425929 | Intron (ENSMUST00000 | 7 | 1,1E+08 | 1,1E+08 | 133 | 1 | 1,15E+08 | ENSMUST00000082666.3 | 11080 | ENSMUSG00000064600 | Gm25636 | predicted gene, 25636 |
| chr11 | 82792038 | 82793346 | 1309 | * | 12,22368 | 3,963142077 | 1,252552 | 3,164055 | 0,001556 | 0,617289329 | 1,075735 | Promoter (<=1kb) | 11 | 82791598 | 82792642 | 1045 | 2 | 217011 | ENSMUST00000124109.2 | 0 | ENSMUSG00000020692 | Nle1 | notchless homolog 1 |
| chr11 | 101323471 | 101326279 | 2809 | * | 20,131492 | 3,963625751 | 0,833179 | 4,7572318 | 1,963E-06 | 0,007355195 | 2,3860078 | 3' UTR | 11 | 101333124 | 101336355 | 3232 | 1 | 19942 | ENSMUST00000077856.13 | -6845 | ENSMUSG00000063316 | Rpl27 | ribosomal protein L27 |
| chrX | 84300563 | 84301769 | 1207 | * | 7,784446 | 3,964664833 | 1,428179 | 2,776029 | 0,005503 | 1 | 0 | Intron (ENSMUST00000 | 20 | 84293285 | 84313894 | 20610 | 1 | 71996 | ENSMUST00000048382.5 | 7278 | ENSMUSG00000035491 | Fthl17a | ferritin, heavy polypeptide-like 17, member A |
| chr9 | 1,08E+08 | 1,08E+08 | 717 | * | 6,797005 | 3,965045932 | 1,442434 | 2,748858 | 0,00598 | 1 | 0 | Promoter (<=1kb) | 9 | 1,08E+08 | 1,08E+08 | 4187 | 1 | 76873 | ENSMUST00000194283.6 | 0 | ENSMUSG00000102224 | 4930447F24Rik | RIKEN cDNA 4930447F24 gene |
| chr14 | 59951253 | 59951766 | 514 | * | 7,032199 | 3,965339473 | 1,519693 | 2,609303 | 0,009073 | 1 | 0 | Intron (ENSMUST00000 | 14 | 59864638 | 59867126 | 2489 | 1 | 219134 | ENSMUST00000224155.2 | 86615 | ENSMUSG00000044461 | Shisa2 | shisa family member 2 |
| chr9 | 58816018 | 58817244 | 1227 | * | 9,107811 | 3,965839067 | 1,249506 | 3,173925 | 0,001504 | 0,475225141 | 2,237403 | Intron (ENSMUST00000 | 9 | 58795769 | 58801217 | 5449 | 2 | 18007 | ENSMUST00000215165.2 | -14801 | ENSMUSG00000032340 | Neo1 | neogenin |
| chr10 | 68097694 | 68098713 | 1020 | * | 8,433727 | 3,965984952 | 1,51265 | 2,621879 | 0,008745 | 0,787687702 | 2,237403 | Intron (ENSMUST00000 | 10 | 67931846 | 68114551 | 182706 | 2 | 71371 | ENSMUST00000020106.9 | 15838 | ENSMUSG00000019947 | Arid5b | AT rich interactive domain 5B (MRF1-like) |
| chr8 | 87045727 | 87047086 | 1360 | * | 12,64333 | 3,967435788 | 1,159906 | 3,42048 | 0,000625 | 0,437496229 | 1,173325 | Distal Intergenic | 8 | 87231212 | 87261376 | 30165 | 2 | 244562 | ENSMUST00000156066.8 | 214290 | ENSMUSG00000036872 | Abcc12 | ATP-binding cassette, sub-family C (CFTR/MRP), member 12 |
| chr9 | 76191026 | 76191950 | 925 | * | 6,905378 | 3,967781764 | 1,31161 | 3,025122 | 0,002485 | 1 | 0 | Intron (ENSMUST00000 | 9 | 76133162 | 76230893 | 97732 | 2 | 387285 | ENSMUST00000063140.15 | 38943 | ENSMUSG00000032360 | Hcrtr2 | hypocretin (orexin) receptor 2 |
| chr13 | 24833074 | 24833819 | 746 | * | 8,278754 | 3,968627144 | 1,441995 | 2,752178 | 0,00592 | 0,792545355 | 1,449706 | Intron (ENSMUST00000 | 13 | 24822696 | 24879708 | 57013 | 1 | 193385 | ENSMUST00000138547.8 | 10378 | ENSMUSG00000036006 | Ripor2 | RHO family interacting cell polarization regulator 2 |
| chr9 | 1,19E+08 | 1,19E+08 | 454 | * | 8,536737 | 3,968800992 | 1,225867 | 3,237546 | 0,001206 | 0,510628937 | 1,449706 | Promoter (1-2kb) | 9 | 1,19E+08 | 1,19E+08 | 31398 | 1 | 11481 | ENSMUST00000165044.3 | -1631 | ENSMUSG00000061393 | Acvr2b | activin receptor IIB |
| chr12 | 53019846 | 53021208 | 1363 | * | 10,4171 | 3,969037444 | 1,389975 | 2,855474 | 0,004297 | 0,855765563 | 0,738441 | Intron (ENSMUST00000 | 12 | 53093491 | 53093609 | 119 | 2 | 1,15E+08 | ENSMUST00000083299.3 | 72401 | ENSMUSG00000119139 | n-R5s58 | nuclear encoded rRNA 5S 58 |
| chr7 | 16664959 | 16666219 | 1261 | * | 10,17402 | 3,970082243 | 1,327298 | 2,991102 | 0,00278 | 0,689688276 | 1,211315 | Distal Intergenic | 7 | 16649304 | 16658039 | 8736 | 2 | 12315 | ENSMUST00000019514.10 | -6920 | ENSMUSG00000019370 | Calm3 | calmodulin 3 |
| chr13 | 15136708 | 15138057 | 1350 | * | 9,267561 | 3,970427476 | 1,265018 | 3,138634 | 0,001697 | 0,588341471 | 1,360622 | Distal Intergenic | 13 | 14804862 | 14811499 | 6638 | 1 | 105351 | ENSMUST00000222052.2 | 331846 | ENSMUSG00000039182 | AW209491 | expressed sequence AW209491 |
| chr1 | 86709183 | 86710686 | 1504 | * | 11,95868 | 3,970892181 | 1,181745 | 3,360193 | 0,000779 | 0,514210241 | 0,918255 | Intron (ENSMUST00000 | 1 | 86631565 | 86977814 | 346250 | 1 | 208718 | ENSMUST00000065694.8 | 77618 | ENSMUSG00000053333 | Dis3l2 | DIS3 like 3'-5' exoribonuclease 2 |
| chr15 | 79714471 | 79715011 | 541 | * | 6,18049 | 3,971221704 | 1,450578 | 2,737682 | 0,006187 | 1 | 0 | Promoter (2-3kb) | 15 | 79712837 | 79717999 | 5163 | 2 | 494448 | ENSMUST00000141345.2 | 2988 | ENSMUSG00000089715 | Cbx6 | chromobox 6 |
| chr7 | 1,42E+08 | 1,42E+08 | 1615 | * | 14,39026 | 3,97296277 | 1,030862 | 3,85402 | 0,000116 | 0,253278065 | 0,738441 | Promoter (2-3kb) | 7 | 1,42E+08 | 1,42E+08 | 1177 | 2 | 114666 | ENSMUST00000097942.3 | 2869 | ENSMUSG00000073785 | Krtap5-5 | keratin associated protein 5-5 |
| chr18 | 35924228 | 35925962 | 1735 | * | 12,54199 | 3,973049994 | 1,131881 | 3,51013 | 0,000448 | 0,457244879 | 0,738441 | Exon (ENSMUST000002 | 18 | 35920377 | 35940214 | 19838 | 1 | 56550 | ENSMUST00000236235.2 | 3851 | ENSMUSG00000091896 | Ube2d2a | ubiquitin-conjugating enzyme E2D 2A |
| chr14 | 26278084 | 26279045 | 962 | * | 9,279869 | 3,97361173 | 1,306499 | 3,041421 | 0,002355 | 0,639383182 | 1,449706 | Distal Intergenic | 14 | 26295011 | 26351738 | 56728 | 1 | 211922 | ENSMUST00000224248.2 | -15966 | ENSMUSG00000040818 | Dennd6a | DENN/MADD domain containing 6A |
| chr16 | 55449341 | 55450737 | 1397 | * | 8,722303 | 3,973951339 | 1,437205 | 2,765056 | 0,005691 | 0,796850747 | 1,360622 | Distal Intergenic | 16 | 55415145 | 55415251 | 107 | 2 | 1,15E+08 | ENSMUST00000240362.1 | -34090 | ENSMUSG00000119820 | Gm23003 | predicted gene, 23003 |
| chr8 | 1,25E+08 | 1,25E+08 | 735 | * | 8,747651 | 3,974747829 | 1,476713 | 2,691619 | 0,007111 | 0,825493135 | 1,449706 | Distal Intergenic | 8 | 1,24E+08 | 1,24E+08 | 10096 | 2 | 1E+08 | ENSMUST00000181805.3 | -21973 | ENSMUSG00000097203 | 4732419C18Rik | RIKEN cDNA 4732419C18 gene |
| chr10 | 1,27E+08 | 1,27E+08 | 1151 | * | 14,15136 | 3,975290459 | 1,168568 | 3,401848 | 0,000669 | 0,434635596 | 1,2763 | Promoter (2-3kb) | 10 | 1,27E+08 | 1,27E+08 | 17972 | 1 | 20852 | ENSMUST00000092074.12 | -2719 | ENSMUSG00000002147 | Stat6 | signal transducer and activator of transcription 6 |
| chrX | 1,69E+08 | 1,69E+08 | 699 | * | 5,898128 | 3,97531697 | 1,528699 | 2,600458 | 0,00931 | 1 | 0 | Intron (ENSMUST00000 | 20 | 1,69E+08 | 1,69E+08 | 11090 | 1 | 17318 | ENSMUST00000133857.8 | -11489 | ENSMUSG00000035299 | Mid1 | midline 1 |
| chr5 | 1,43E+08 | 1,43E+08 | 909 | * | 9,064085 | 3,975743199 | 1,306161 | 3,043838 | 0,002336 | 0,668446397 | 1,211315 | Exon (ENSMUST000001 | 5 | 1,43E+08 | 1,43E+08 | 23450 | 2 | 231863 | ENSMUST00000035985.8 | 16006 | ENSMUSG00000066640 | Fbxl18 | F-box and leucine-rich repeat protein 18 |
| chr10 | 36395890 | 36396447 | 558 | * | 6,62922 | 3,976111061 | 1,362791 | 2,917624 | 0,003527 | 1 | 0 | Intron (ENSMUST00000 | 10 | 36383403 | 36686034 | 302632 | 1 | 319415 | ENSMUST00000171521.2 | 12487 | ENSMUSG00000044499 | Hs3st5 | heparan sulfate (glucosamine) 3-O-sulfotransferase 5 |
| chr17 | 51034330 | 51034952 | 623 | * | 8,585139 | 3,977656625 | 1,497042 | 2,657011 | 0,007884 | 0,767061106 | 2,237403 | Distal Intergenic | 17 | 51005716 | 51006471 | 756 | 1 | 654432 | ENSMUST00000061681.9 | 28614 | | Gm7334 | B-cell translocation gene 3 pseudogene |

| | | | | | | | | | | | | | | | | | | | | | | | |
|---|---|---|---|---|---|---|---|---|---|---|---|---|---|---|---|---|---|---|---|---|---|---|---|
| chr1 | 1,68E+08 | 1,68E+08 | 993 | * | 7,87821 | 3,978464104 | 1,287409 | 3,090288 | 0,002 | 1 | 0 | Distal Intergenic | 1 | 1,68E+08 | 1,68E+08 | 6293 | 2 | 18514 | ENSMUST00000097471.4 | 216976 | ENSMUSG00000052534 | Pbx1 | pre B cell leukemia homeobox 1 |
| chr19 | 30924408 | 30926025 | 1618 | * | 13,30657 | 3,978671745 | 1,188941 | 3,3464 | 0,000819 | 0,521453391 | 0,938682 | Intron (ENSMUST0000( | 19 | 31060237 | 31064469 | 4233 | 1 | 83410 | ENSMUST00000066039.8 | -134212 | ENSMUSG00000053536 | Cstf2t | cleavage stimulation factor, 3' pre-RNA subunit 2, tau |
| chr8 | 1,29E+08 | 1,29E+08 | 766 | * | 6,967178 | 3,980039525 | 1,493905 | 2,664186 | 0,007717 | 1 | 0 | Intron (ENSMUST0000( | 8 | 1,29E+08 | 1,29E+08 | 102685 | 1 | 18186 | ENSMUST00000212933.2 | 8063 | ENSMUSG00000025810 | Nrp1 | neuropilin 1 |
| chr18 | 68604645 | 68606287 | 1643 | * | 17,01732 | 3,981771524 | 1,059714 | 3,757401 | 0,000172 | 0,23356078 | 1,2763 | Intron (ENSMUST0000( | 18 | 68539978 | 68562391 | 22414 | 2 | 17200 | ENSMUST00000052347.8 | -42254 | ENSMUSG00000045569 | Mc2r | melanocortin 2 receptor |
| chr16 | 64670399 | 64671684 | 1286 | * | 8,370307 | 3,982260087 | 1,339263 | 2,973471 | 0,002945 | 1 | 0 | Promoter (<=1kb) | 16 | 64600710 | 64672015 | 71306 | 2 | 72020 | ENSMUST00000207826.2 | 331 | ENSMUSG00000047141 | Zfp654 | zinc finger protein 654 |
| chr14 | 26251502 | 26253483 | 1982 | * | 13,87417 | 3,982407342 | 1,105507 | 3,602335 | 0,000315 | 0,309888367 | 1,2763 | Promoter (1-2kb) | 14 | 26134555 | 26254773 | 120219 | 2 | 83997 | ENSMUST00000090359.11 | 1290 | ENSMUSG00000021870 | Slmap | sarcolemma associated protein |
| chr11 | 94902362 | 94903513 | 1152 | * | 8,647643 | 3,982756344 | 1,530921 | 2,601543 | 0,009281 | 0,885309127 | 1,360622 | Promoter (1-2kb) | 11 | 94901107 | 94910942 | 9836 | 1 | 217125 | ENSMUST00000124735.2 | 1255 | ENSMUSG00000047181 | Samd14 | sterile alpha motif domain containing 14 |
| chr2 | 39194351 | 39194738 | 388 | * | 6,3616 | 3,982953237 | 1,450284 | 2,746327 | 0,006027 | 1 | 0 | Distal Intergenic | 2 | 39084366 | 39116463 | 32098 | 2 | 67857 | ENSMUST00000028087.6 | -77888 | ENSMUSG00000026753 | Ppp6c | protein phosphatase 6, catalytic subunit |
| chr17 | 45785114 | 45786338 | 1225 | * | 23,609707 | 3,983145246 | 0,694877 | 5,7321585 | 9,916E-09 | 0,000102012 | 2,9519335 | Promoter (<=1kb) | 17 | 45756803 | 45785790 | 28988 | 2 | 71020 | ENSMUST00000233769.2 | 0 | ENSMUSG00000023935 | Spats1 | spermatogenesis associated, serine-rich 1 |
| chr18 | 36458723 | 36459390 | 668 | * | 7,182704 | 3,983539724 | 1,518719 | 2,62296 | 0,008717 | 0,87311386 | 1,360622 | Distal Intergenic | 18 | 36475573 | 36526430 | 50858 | 1 | 66060 | ENSMUST00000139727.3 | -16183 | ENSMUSG00000046727 | Cystm1 | cysteine-rich transmembrane module containing 1 |
| chr5 | 125420655 | 125422258 | 1604 | * | 22,69909 | 3,983553565 | 0,8039415 | 4,9550293 | 7,232E-07 | 0,002928698 | 2,8800171 | Promoter (2-3kb) | 5 | 125354151 | 125418158 | 64008 | 2 | 20778 | ENSMUST00000086075.13 | -2497 | ENSMUSG00000037936 | Scarb1 | scavenger receptor class B, member 1 |
| chr11 | 95486016 | 95486966 | 951 | * | 8,770365 | 3,984102215 | 1,453242 | 2,741527 | 0,006115 | 0,798887389 | 1,449706 | Distal Intergenic | 11 | 95459644 | 95478561 | 18918 | 2 | 18053 | ENSMUST00000000122.7 | -7455 | ENSMUSG00000000120 | Ngfr | nerve growth factor receptor (TNFR superfamily, member 16) |
| chr10 | 94951139 | 94951875 | 737 | * | 8,475089 | 3,987274746 | 1,31287 | 3,037068 | 0,002389 | 0,650529916 | 1,360622 | Distal Intergenic | 10 | 94956444 | 94981687 | 25244 | 2 | 628062 | ENSMUST00000220248.2 | 29812 | ENSMUSG00000112481 | Gm29684 | predicted gene, 29684 |
| chr4 | 1,39E+08 | 1,39E+08 | 1455 | * | 7,658418 | 3,987640132 | 1,395273 | 2,857965 | 0,004264 | 1 | 0 | Distal Intergenic | 4 | 1,39E+08 | 1,39E+08 | 2169 | 2 | 69902 | ENSMUST00000131052.8 | 13760 | ENSMUSG00000028741 | Mrto4 | mRNA turnover 4, ribosome maturation factor |
| chr17 | 79407221 | 79408118 | 898 | * | 10,03986 | 3,98818571 | 1,237336 | 3,223203 | 0,001268 | 0,55735526 | 1,173325 | Distal Intergenic | 17 | 79384431 | 79389282 | 4852 | 1 | 70536 | ENSMUST00000233899.2 | 22790 | ENSMUSG00000024084 | Qpct | glutaminyl-peptide cyclotransferase (glutaminyl cyclase) |
| chr2 | 21342927 | 21343307 | 381 | * | 6,465395 | 3,988239101 | 1,545473 | 2,580595 | 0,009863 | 1 | 0 | Distal Intergenic | 2 | 21372353 | 21835358 | 463006 | 1 | 241263 | ENSMUST00000055946.8 | -29046 | ENSMUSG00000045967 | Gpr158 | G protein-coupled receptor 158 |
| chr1 | 59199905 | 59200853 | 949 | * | 6,774315 | 3,990150181 | 1,410863 | 2,828162 | 0,004682 | 1 | 0 | Promoter (<=1kb) | 1 | 59160094 | 59200795 | 40702 | 2 | 227157 | ENSMUST00000078874.14 | 0 | ENSMUSG00000079550 | Mpp4 | membrane protein, palmitoylated 4 (MAGUK p55 subfamily member 4) |
| chr3 | 95347135 | 95348540 | 1406 | * | 8,831035 | 3,990153943 | 1,368774 | 2,91513 | 0,003555 | 0,715249603 | 1,360622 | Intron (ENSMUST0000( | 3 | 95341830 | 95400001 | 58172 | 1 | 11863 | ENSMUST00000149051.8 | 5305 | ENSMUSG00000015522 | Arnt | aryl hydrocarbon receptor nuclear translocator |
| chr5 | 66130209 | 66131577 | 1369 | * | 12,12195 | 3,990266307 | 1,167249 | 3,418523 | 0,00063 | 0,438640586 | 1,173325 | Promoter (1-2kb) | 5 | 66133239 | 66134483 | 1245 | 1 | 231252 | ENSMUST00000201664.2 | -1662 | ENSMUSG00000029205 | Chrna9 | cholinergic receptor, nicotinic, alpha polypeptide 9 |
| chr13 | 53133797 | 53134757 | 961 | * | 6,613202 | 3,992942896 | 1,540238 | 2,592419 | 0,00953 | 1 | 0 | Promoter (<=1kb) | 13 | 53121245 | 53135109 | 13865 | 2 | 18030 | ENSMUST00000071065.8 | 352 | ENSMUSG00000056749 | Nfil3 | nuclear factor, interleukin 3, regulated |
| chr9 | 72492595 | 72493798 | 1204 | * | 9,662925 | 3,996129641 | 1,349104 | 2,962061 | 0,003056 | 0,683210259 | 1,425929 | Intron (ENSMUST0000( | 9 | 72540729 | 72540835 | 107 | 1 | 1,15E+08 | ENSMUST00000082801.3 | -46931 | ENSMUSG00000064735 | Gm23057 | predicted gene, 23057 |
| chr11 | 83603966 | 83605311 | 1346 | * | 10,06828 | 3,997022739 | 1,457115 | 2,743108 | 0,006086 | 0,836575905 | 1,173325 | Distal Intergenic | 11 | 83599951 | 83601524 | 1574 | 1 | 14038 | ENSMUST00000139762.2 | 4015 | ENSMUSG00000000983 | Wfdc18 | WAP four-disulfide core domain 18 |
| chr15 | 85791091 | 85792182 | 1092 | * | 10,78828 | 3,998755684 | 1,291773 | 3,095557 | 0,001964 | 0,627785088 | 1,2763 | Promoter (2-3kb) | 15 | 85785187 | 85788168 | 2982 | 2 | 12614 | ENSMUST00000226204.3 | -2923 | ENSMUSG00000016028 | Celsr1 | cadherin, EGF LAG seven-pass G-type receptor 1 |
| chr19 | 41332302 | 41333171 | 870 | * | 8,948954 | 3,999205184 | 1,39909 | 2,858433 | 0,004257 | 0,737081993 | 1,425929 | Intron (ENSMUST0000( | 19 | 41260816 | 41373535 | 112720 | 2 | 83490 | ENSMUST00000059672.9 | 40364 | ENSMUSG00000025017 | Pik3ap1 | phosphoinositide-3-kinase adaptor protein 1 |
| chrX | 46359687 | 46360988 | 1302 | * | 6,163902 | 3,999364773 | 1,550461 | 2,579468 | 0,009895 | 1 | 0 | Distal Intergenic | 20 | 46930807 | 46981273 | 50467 | 2 | 93761 | ENSMUST00000153548.9 | 620285 | ENSMUSG00000031099 | Smarca1 | SWI/SNF related, matrix associated, actin dependent regulator of chromatin, subfamily a, member 1 |
| chr13 | 52975306 | 52976131 | 826 | * | 8,22935 | 4,000959521 | 1,45002 | 2,759245 | 0,005794 | 0,790570581 | 1,449706 | Distal Intergenic | 13 | 52883958 | 52911241 | 27284 | 2 | 105404 | ENSMUST00000223209.2 | -64065 | | BB123696 | expressed sequence BB123696 |
| chr10 | 1,18E+08 | 1,18E+08 | 469 | * | 6,92868 | 4,001184513 | 1,368094 | 2,924642 | 0,003449 | 0,711307535 | 1,360622 | Distal Intergenic | 10 | 1,18E+08 | 1,18E+08 | 54539 | 2 | 103214 | ENSMUST00000241197.1 | -35236 | ENSMUSG00000112230 | Ifngas1 | Ifng antisense RNA 1 |
| chr4 | 91721495 | 91722485 | 991 | * | 6,560626 | 4,001379385 | 1,543344 | 2,592668 | 0,009523 | 1 | 0 | Distal Intergenic | 4 | 91693784 | 91694792 | 1009 | 2 | 620016 | ENSMUST00000122162.2 | -26703 | ENSMUSG00000100863 | Gm12669 | glutaredoxin 3 pseudogene |
| chr12 | 32074186 | 32074792 | 607 | * | 6,901523 | 4,00160455 | 1,547066 | 2,586577 | 0,009693 | 1 | 0 | Intron (ENSMUST0000( | 12 | 32008479 | 32110588 | 102110 | 2 | 19088 | ENSMUST00000146865.2 | 35796 | ENSMUSG00000002997 | Prkar2b | protein kinase, cAMP dependent regulatory, type II beta |
| chrX | 1,41E+08 | 1,41E+08 | 905 | * | 6,723104 | 4,003440601 | 1,500526 | 2,668026 | 0,00763 | 1 | 0 | Distal Intergenic | 20 | 1,41E+08 | 1,41E+08 | 432 | 2 | 1,05E+08 | ENSMUST00000117247.2 | -31389 | ENSMUSG00000081750 | Gm4995 | predicted pseudogene 4995 |
| chr2 | 63083778 | 63084677 | 900 | * | 9,060726 | 4,00347691 | 1,528147 | 2,619825 | 0,008797 | 0,788923377 | 2,237403 | Distal Intergenic | 2 | 62616625 | 63014631 | 398007 | 2 | 170738 | ENSMUST00000112454.8 | -69147 | ENSMUSG00000059742 | Kcnh7 | potassium voltage-gated channel, subfamily H (eag-related), member 7 |
| chrX | 1,28E+08 | 1,28E+08 | 819 | * | 5,616344 | 4,003541805 | 1,489688 | 2,687503 | 0,007199 | 1 | 0 | Distal Intergenic | 20 | 1,28E+08 | 1,28E+08 | 2850 | 1 | 75274 | ENSMUST00000122848.2 | -26810 | ENSMUSG00000086244 | 4930558G05Rik | RIKEN cDNA 4930558G05 gene |
| chr10 | 80995095 | 80996276 | 1182 | * | 12,25961 | 4,003783668 | 1,207221 | 3,31653 | 0,000911 | 0,484153011 | 1,2763 | Promoter (1-2kb) | 10 | 80991336 | 80993845 | 2510 | 2 | 59004 | ENSMUST00000147436.8 | -1250 | ENSMUSG00000004934 | Pias4 | protein inhibitor of activated STAT 4 |
| chr5 | 1,11E+08 | 1,11E+08 | 909 | * | 9,095866 | 4,003873412 | 1,41998 | 2,819668 | 0,004807 | 0,79015979 | 1,211315 | Intron (ENSMUST0000( | 5 | 1,11E+08 | 1,11E+08 | 17857 | 1 | 231605 | ENSMUST00000200404.5 | -28181 | ENSMUSG00000033316 | Galnt9 | polypeptide N-acetylgalactosaminyltransferase 9 |
| chr6 | 1,38E+08 | 1,38E+08 | 425 | * | 6,563136 | 4,004363814 | 1,473134 | 2,718262 | 0,006563 | 1 | 0 | Distal Intergenic | 6 | 1,38E+08 | 1,38E+08 | 16855 | 1 | 20901 | ENSMUST00000064910.7 | -11984 | ENSMUSG00000030224 | Strap | serine/threonine kinase receptor associated protein |
| chr19 | 28601210 | 28602672 | 1463 | * | 12,1675 | 4,005375185 | 1,276243 | 3,138411 | 0,001699 | 0,68845579 | 0,744375 | Intron (ENSMUST0000( | 19 | 28509247 | 28644085 | 134839 | 2 | 226075 | ENSMUST00000159178.2 | 41413 | ENSMUSG00000052942 | Glis3 | GLIS family zinc finger 3 |
| chr15 | 35539686 | 35540218 | 533 | * | 7,420019 | 4,005720466 | 1,546953 | 2,589426 | 0,009614 | 1 | 0 | Intron (ENSMUST0000( | 15 | 35487546 | 35487682 | 137 | 1 | 1,15E+08 | ENSMUST00000158508.4 | 52140 | ENSMUSG00000089133 | Gm24771 | predicted gene, 24771 |
| chr12 | 80748073 | 80749176 | 1104 | * | 11,31964 | 4,006908343 | 1,281903 | 3,125749 | 0,001774 | 0,625730806 | 1,173325 | Intron (ENSMUST0000( | 12 | 80753620 | 80767458 | 13839 | 1 | 217682 | ENSMUST00000152465.2 | -4444 | ENSMUSG00000066438 | Plekhd1 | pleckstrin homology domain containing, family D (with coiled-coil domains) member 1 |
| chr5 | 45470175 | 45471078 | 904 | * | 6,163651 | 4,007398268 | 1,473379 | 2,71987 | 0,006531 | 1 | 0 | Intron (ENSMUST0000( | 5 | 45166513 | 45581744 | 415232 | 2 | 52401 | ENSMUST00000197016.5 | 110666 | ENSMUSG00000029092 | D5Ertd615e | DNA segment, Chr 5, ERATO Doi 615, expressed |
| chr19 | 5578440 | 5580500 | 2061 | * | 30,447623 | 4,007766524 | 0,6580337 | 6,0905187 | 1,125E-09 | 1,76557E-05 | 2,8800171 | Intron (ENSMUST000002 | 19 | 5545302 | 5560566 | 15265 | 2 | 225861 | ENSMUST00000070172.6 | -17874 | ENSMUSG00000056185 | Snx32 | sorting nexin 32 |
| chr4 | 43093417 | 43094291 | 875 | * | 6,960187 | 4,008546552 | 1,520093 | 2,63704 | 0,008363 | 1 | 0 | Intron (ENSMUST0000( | 4 | 43115110 | 43237776 | 122667 | 1 | 22249 | ENSMUST00000132310.9 | -20819 | ENSMUSG00000028456 | Unc13b | unc-13 homolog B |
| chr9 | 21135935 | 21137844 | 1910 | * | 35,720557 | 4,008840028 | 0,6199135 | 6,4667735 | 1,001E-10 | 2,40889E-06 | 2,7872553 | Exon (ENSMUST000002 | 9 | 21147419 | 21148489 | 1071 | 2 | 50868 | ENSMUST00000193247.2 | 10645 | ENSMUSG00000003308 | Keap1 | kelch-like ECH-associated protein 1 |
| chr7 | 1,45E+08 | 1,45E+08 | 1093 | * | 9,493276 | 4,009592822 | 1,325273 | 3,025485 | 0,002482 | 0,676726988 | 1,211315 | Intron (ENSMUST0000( | 7 | 1,45E+08 | 1,45E+08 | 4998 | 2 | 233979 | ENSMUST00000208867.2 | 18599 | ENSMUSG00000048677 | Tpcn2 | two pore segment channel 2 |
| chr1 | 1,35E+08 | 1,35E+08 | 1182 | * | 12,35104 | 4,01031669 | 1,173133 | 3,418468 | 0,00063 | 0,514468591 | 0,738441 | Exon (ENSMUST00000( | 1 | 1,35E+08 | 1,35E+08 | 18842 | 2 | 329252 | ENSMUST00000139369.2 | -40476 | ENSMUSG00000042793 | Lgr6 | leucine-rich repeat-containing G protein-coupled receptor 6 |
| chr4 | 1,37E+08 | 1,37E+08 | 1350 | * | 11,44041 | 4,01131703 | 1,240459 | 3,233736 | 0,001222 | 0,548877304 | 1,173325 | Intron (ENSMUST0000( | 4 | 1,37E+08 | 1,37E+08 | 27398 | 2 | 13842 | ENSMUST00000030420.9 | 7503 | ENSMUSG00000028661 | Epha8 | Eph receptor A8 |
| chr16 | 92683291 | 92684231 | 941 | * | 8,56079 | 4,012821107 | 1,315032 | 3,0515 | 0,002277 | 1 | 0 | Distal Intergenic | 16 | 92398354 | 92623037 | 224684 | 2 | 12394 | ENSMUST00000023673.14 | -60254 | ENSMUSG00000022952 | Runx1 | runt related transcription factor 1 |
| chr1 | 1,31E+08 | 1,31E+08 | 914 | * | 6,746375 | 4,01431978 | 1,406851 | 2,853408 | 0,004325 | 1 | 0 | Distal Intergenic | 1 | 1,31E+08 | 1,31E+08 | 37605 | 2 | 54354 | ENSMUST00000124796.2 | -8136 | ENSMUSG00000026430 | Rassf5 | Ras association (RalGDS/AF-6) domain family member 5 |
| chr5 | 90375136 | 90376532 | 1397 | * | 10,20531 | 4,014414524 | 1,304262 | 3,077921 | 0,002085 | 0,648067121 | 1,211315 | 3' UTR | 5 | 90362583 | 90371860 | 9278 | 2 | 231430 | ENSMUST00000048363.10 | -3276 | ENSMUSG00000035505 | Cox18 | cytochrome c oxidase assembly protein 18 |
| chr19 | 11892453 | 11893542 | 1090 | * | 7,849048 | 4,01498058 | 1,353438 | 2,966506 | 0,003012 | 0,684274806 | 1,360622 | Promoter (2-3kb) | 19 | 11889807 | 11921367 | 31561 | 1 | 225929 | ENSMUST00000238190.2 | 2646 | ENSMUSG00000046139 | Patl1 | protein associated with topoisomerase II homolog 1 (yeast) |
| chr16 | 50153572 | 50154441 | 870 | * | 8,335456 | 4,016194149 | 1,340233 | 2,996638 | 0,00273 | 0,672444708 | 1,360622 | Promoter (2-3kb) | 16 | 50017522 | 50151356 | 133835 | 2 | 70508 | ENSMUST00000114488.8 | -2216 | ENSMUSG00000022641 | Bbx | bobby sox HMG box containing |
| chr19 | 23952444 | 23953326 | 883 | * | 8,059388 | 4,01772181 | 1,364033 | 2,945473 | 0,003225 | 1 | 0 | Promoter (<=1kb) | 19 | 23950576 | 23953799 | 3224 | 2 | 381217 | ENSMUST00000236123.2 | 473 | ENSMUSG00000071604 | Fam189a2 | family with sequence similarity 189, member A2 |
| chr5 | 43402053 | 43402988 | 936 | * | 7,008223 | 4,019503446 | 1,398848 | 2,873438 | 0,00406 | 1 | 0 | Intron (ENSMUST0000( | 5 | 43394658 | 43403802 | 9145 | 1 | 231207 | ENSMUST00000127599.3 | 7395 | ENSMUSG00000039782 | Cpeb2 | cytoplasmic polyadenylation element binding protein 2 |
| chr10 | 84549433 | 84550448 | 1016 | * | 9,241777 | 4,019774236 | 1,422447 | 2,825956 | 0,004714 | 0,764024179 | 1,360622 | Exon (ENSMUST00000( | 10 | 84591926 | 84742008 | 150083 | 1 | 71137 | ENSMUST00000060397.13 | -41478 | ENSMUSG00000020037 | Rfx4 | regulatory factor X, 4 (influences HLA class II expression) |
| chr19 | 5094562 | 5097069 | 2508 | * | 26,227737 | 4,021902948 | 0,8049697 | 4,9963409 | 5,843E-07 | 0,002511766 | 2,8800171 | Promoter (<=1kb) | 19 | 5096149 | 5099651 | 3503 | 1 | 107392 | ENSMUST00000237117.2 | 0 | ENSMUSG00000080268 | Brms1 | breast cancer metastasis-suppressor 1 |
| chr14 | 27317342 | 27319122 | 1781 | * | 11,12083 | 4,022177685 | 1,382668 | 2,908998 | 0,003626 | 0,826071267 | 0,738441 | Distal Intergenic | 14 | 27344385 | 28200492 | 856108 | 1 | 238988 | ENSMUST00000210135.2 | -25263 | ENSMUSG00000040640 | Erc2 | ELKS/RAB6-interacting/CAST family member 2 |
| chr3 | 95668706 | 95669713 | 1008 | * | 6,662454 | 4,022746086 | 1,48008 | 2,717925 | 0,006569 | 1 | 0 | Promoter (1-2kb) | 3 | 95662118 | 95667518 | 5401 | 2 | 71807 | ENSMUST00000197501.2 | -1188 | ENSMUSG00000028107 | Tars2 | threonyl-tRNA synthetase 2, mitochondrial (putative) |
| chr14 | 23182687 | 23183486 | 800 | * | 8,880447 | 4,022897169 | 1,485896 | 2,707388 | 0,006781 | 0,821417341 | 1,425929 | Distal Intergenic | 14 | 23088262 | 23144639 | 56378 | 2 | 791307 | ENSMUST00000090409.2 | -38048 | ENSMUSG00000068680 | Gm10248 | predicted gene 10248 |
| chrX | 1,2E+08 | 1,2E+08 | 687 | * | 5,986736 | 4,023035916 | 1,553212 | 2,59014 | 0,009594 | 1 | 0 | Distal Intergenic | 20 | 1,2E+08 | 1,2E+08 | 128 | 1 | 1,15E+08 | ENSMUST00000104247.3 | 345116 | ENSMUSG00000077435 | Gm22590 | predicted gene, 22590 |
| chr8 | 1,12E+08 | 1,12E+08 | 1534 | * | 19,20357 | 4,023458621 | 1,076517 | 3,737478 | 0,000186 | 0,161606382 | 2,663619 | Distal Intergenic | 8 | 1,12E+08 | 1,12E+08 | 74 | 2 | 1,02E+08 | ENSMUST00000183519.3 | 12559 | ENSMUSG00000099129 | Mir7076 | microRNA 7076 |
| chr4 | 83701818 | 83702939 | 1122 | * | 8,304594 | 4,024654261 | 1,4316 | 2,811298 | 0,004934 | 0,75938015 | 1,449706 | Intron (ENSMUST0000( | 4 | 83614192 | 83716913 | 102722 | 1 | 320226 | ENSMUST00000183729.2 | 87626 | ENSMUSG00000052407 | Ccdc171 | coiled-coil domain containing 171 |
| chr5 | 24904854 | 24905391 | 538 | * | 7,616967 | 4,024913439 | 1,329372 | 3,027681 | 0,002464 | 1 | 0 | Exon (ENSMUST000001 | 5 | 24901900 | 24909688 | 7789 | 2 | 269633 | ENSMUST00000181925.2 | 4297 | ENSMUSG00000055235 | Wdr86 | WD repeat domain 86 |
| chr13 | 3890923 | 3891384 | 462 | * | 5,992173 | 4,025169697 | 1,437314 | 2,80048 | 0,005103 | 1 | 0 | Intron (ENSMUST0000( | 13 | 3887757 | 3888673 | 917 | 1 | 80796 | ENSMUST00000042219.6 | 3166 | ENSMUSG00000033765 | Calm4 | calmodulin 4 |
| chr2 | 1,23E+08 | 1,23E+08 | 925 | * | 8,912905 | 4,02664671 | 1,256899 | 3,203637 | 0,001357 | 0,457244879 | 2,237403 | Distal Intergenic | 2 | 1,23E+08 | 1,23E+08 | 16072 | 1 | 414106 | ENSMUST00000142323.8 | -21726 | | 4930417H01Rik | RIKEN cDNA 4930417H01 gene |
| chr5 | 1,08E+08 | 1,08E+08 | 1607 | * | 13,80462 | 4,028775966 | 1,160932 | 3,470295 | 0,00052 | 0,393356861 | 1,2763 | Promoter (1-2kb) | 5 | 1,08E+08 | 1,08E+08 | 4431 | 2 | 14581 | ENSMUST00000159263.3 | -1264 | ENSMUSG00000029275 | Gfi1 | growth factor independent 1 transcription repressor |
| chr16 | 79938195 | 79938694 | 500 | * | 5,340224 | 4,028829518 | 1,532588 | 2,628775 | 0,008569 | 1 | 0 | Distal Intergenic | 16 | 79796264 | 79797434 | 1171 | 1 | 78654 | ENSMUST00000191029.2 | 141931 | ENSMUSG00000101810 | 1700066C05Rik | RIKEN cDNA 1700066C05 gene |
| chr10 | 62809543 | 62810556 | 1014 | * | 6,793562 | 4,030428191 | 1,546023 | 2,606966 | 0,009135 | 1 | 0 | 3' UTR | 10 | 62816002 | 62826072 | 10071 | 1 | 70432 | ENSMUST00000148259.2 | -5446 | ENSMUSG00000020070 | Rufy2 | RUN and FYVE domain-containing 2 |
| chr5 | 66357046 | 66358928 | 1883 | * | 13,07701 | 4,032768515 | 1,170464 | 3,445446 | 0,00057 | 0,436322355 | 1,075735 | Exon (ENSMUST000001 | 5 | 66323950 | 66348679 | 24730 | 2 | 73639 | ENSMUST00000201903.2 | -8367 | ENSMUSG00000107014 | 1700126H18Rik | RIKEN cDNA 1700126H18 gene |
| chr15 | 22302269 | 22302834 | 566 | * | 6,600625 | 4,034165976 | 1,477856 | 2,729743 | 0,006338 | 1 | 0 | Distal Intergenic | 15 | 22549108 | 23400653 | 851546 | 1 | 320865 | ENSMUST00000163361.8 | -246274 | ENSMUSG00000040420 | Cdh18 | cadherin 18 |
| chr7 | 1,15E+08 | 1,15E+08 | 632 | * | 6,696656 | 4,034466396 | 1,547991 | 2,606259 | 0,009154 | 1 | 0 | Intron (ENSMUST0000( | 7 | 1,15E+08 | 1,15E+08 | 85153 | 2 | 20679 | ENSMUST00000206775.2 | 100936 | ENSMUSG00000051910 | Sox6 | SRY (sex determining region Y)-box 6 |
| chr4 | 1,17E+08 | 1,17E+08 | 991 | * | 9,086592 | 4,03483636 | 1,295342 | 3,114881 | 0,00184 | 0,603552508 | 1,360622 | Intron (ENSMUST0000( | 4 | 1,17E+08 | 1,17E+08 | 8956 | 2 | 66743 | ENSMUST00000151829.2 | 3091 | ENSMUSG00000028677 | Rnf220 | ring finger protein 220 |
| chr14 | 8477347 | 8478068 | 722 | * | 6,18182 | 4,035131944 | 1,54992 | 2,603446 | 0,009229 | 1 | 0 | Promoter (2-3kb) | 14 | 8366256 | 8474756 | 108501 | 2 | 246103 | ENSMUST00000223714.2 | -2591 | ENSMUSG00000021738 | Atxn7 | ataxin 7 |
| chr4 | 31578839 | 31579725 | 887 | * | 10,52318 | 4,037327472 | 1,115968 | 3,617782 | 0,000297 | 0,304096805 | 1,2763 | Distal Intergenic | 4 | 31825469 | 31877549 | 52081 | 2 | 1,03E+08 | ENSMUST00000243650.1 | 297824 | | Gm32412 | predicted gene, 32412 |
| chr6 | 30206859 | 30207690 | 832 | * | 8,49917 | 4,037436829 | 1,388122 | 2,908561 | 0,003631 | 0,639383182 | 2,237403 | Distal Intergenic | 6 | 30214808 | 30243802 | 28995 | 2 | 22214 | ENSMUST00000128141.8 | 36112 | ENSMUSG00000039159 | Ube2h | ubiquitin-conjugating enzyme E2H |
| chr2 | 1,07E+08 | 1,07E+08 | 882 | * | 10,76329 | 4,038572669 | 1,127123 | 3,58308 | 0,00034 | 0,331767406 | 1,173325 | Intron (ENSMUST0000( | 2 | 1,07E+08 | 1,07E+08 | 53488 | 1 | 77015 | ENSMUST00000133401.2 | 37256 | ENSMUSG00000016386 | Mpped2 | metallophosphoesterase domain containing 2 |
| chr17 | 47413368 | 47414044 | 677 | * | 6,21534 | 4,039944001 | 1,487766 | 2,715444 | 0,006619 | 1 | 0 | Distal Intergenic | 17 | 47451801 | 47640462 | 188662 | 1 | 224829 | ENSMUST00000188947.7 | -37757 | ENSMUSG00000064043 | Trerf1 | transcriptional regulating factor 1 |
| chr12 | 72444304 | 72445159 | 856 | * | 6,555287 | 4,040474642 | 1,534558 | 2,632989 | 0,008464 | 1 | 0 | Intron (ENSMUST0000( | 12 | 72258526 | 72455828 | 197303 | 2 | 104001 | ENSMUST00000078505.14 | 10669 | ENSMUSG00000021087 | Rtn1 | reticulon 1 |
| chr7 | 30249786 | 30251142 | 1357 | * | 8,778866 | 4,04173234 | 1,34756 | 2,999296 | 0,002706 | 1 | 0 | Promoter (<=1kb) | 7 | 30251603 | 30254865 | 3263 | 1 | 243912 | ENSMUST00000126989.8 | -461 | ENSMUSG00000036854 | Hspb6 | heat shock protein, alpha-crystallin-related, B6 |
| chr3 | 43727175 | 43727790 | 616 | * | 5,327115 | 4,042303239 | 1,535107 | 2,633238 | 0,008457 | 1 | 0 | Distal Intergenic | 3 | 45281480 | 45332640 | 51161 | 2 | 72511 | ENSMUST00000191843.2 | 1604850 | ENSMUSG00000097040 | 2610316D01Rik | RIKEN cDNA 2610316D01 gene |
| chr9 | 61379495 | 61380340 | 846 | * | 6,919032 | 4,043206058 | 1,505517 | 2,685593 | 0,00724 | 1 | 0 | Exon (ENSMUST000002 | 9 | 61349873 | 61363229 | 13357 | 1 | 1,03E+08 | ENSMUST00000214224.2 | 29622 | ENSMUSG00000111033 | Gm34322 | predicted gene, 34322 |
| chr4 | 1,49E+08 | 1,49E+08 | 1252 | * | 8,700343 | 4,045101377 | 1,313579 | 3,079451 | 0,002074 | 0,531203841 | 2,237403 | Intron (ENSMUST0000( | 4 | 1,49E+08 | 1,49E+08 | 6639 | 1 | 1,03E+08 | ENSMUST00000153443.2 | 41333 | | Gm17029 | predicted gene 17029 |
| chr12 | 35361050 | 35361981 | 932 | * | 7,57115 | 4,045941327 | 1,477821 | 2,737774 | 0,006186 | 1 | 0 | Distal Intergenic | 12 | 35244810 | 35244896 | 87 | 2 | 751520 | ENSMUST00000102298.3 | -116154 | ENSMUSG00000076253 | Mir680-3 | microRNA 680-3 |
| chr7 | 94949138 | 94950092 | 955 | * | 6,604927 | 4,047772659 | 1,45892 | 2,774499 | 0,005529 | 1 | 0 | Distal Intergenic | 7 | 95091558 | 95091664 | 107 | 2 | 1,15E+08 | ENSMUST00000240171.1 | 141572 | ENSMUSG00000119638 | Gm25684 | predicted gene, 25684 |
| chr4 | 8547563 | 8548612 | 1050 | * | 7,129611 | 4,047858629 | 1,546657 | 2,617166 | 0,008866 | 1 | 0 | Intron (ENSMUST0000( | 4 | 8535644 | 8607778 | 72135 | 1 | 59021 | ENSMUST00000060232.8 | 11919 | ENSMUSG00000047187 | Rab2a | RAB2A, member RAS oncogene family |
| chr9 | 99924945 | 99925453 | 509 | * | 7,369503 | 4,047863165 | 1,532989 | 2,640504 | 0,008278 | 1 | 0 | Distal Intergenic | 9 | 99905701 | 99916703 | 11003 | 1 | 108970 | ENSMUST00000215896.2 | 19244 | ENSMUSG00000101584 | 4930519F24Rik | RIKEN cDNA 4930519F24 gene |
| chr2 | 1,04E+08 | 1,04E+08 | 1121 | * | 8,234659 | 4,048069918 | 1,457161 | 2,778053 | 0,005469 | 0,779366133 | 1,449706 | Promoter (<=1kb) | 2 | 1,04E+08 | 1,04E+08 | 1981 | 1 | 68243 | ENSMUST00000040374.6 | 0 | ENSMUSG00000032671 | A930018P22Rik | RIKEN cDNA A930018P22 gene |
| chr1 | 1,2E+08 | 1,2E+08 | 656 | * | 7,841221 | 4,048684934 | 1,343254 | 3,014088 | 0,002578 | 0,663630601 | 1,360622 | Distal Intergenic | 1 | 1,2E+08 | 1,2E+08 | 46053 | 2 | 68428 | ENSMUST00000112643.2 | -32280 | ENSMUSG00000026389 | Steap3 | STEAP family member 3 |
| chr13 | 19822839 | 19823829 | 991 | * | 7,589255 | 4,049667977 | 1,502436 | 2,695402 | 0,00703 | 1 | 0 | Distal Intergenic | 13 | 19808925 | 19816870 | 7946 | 1 | 20379 | ENSMUST00000222992.2 | 13914 | ENSMUSG00000021319 | Sfrp4 | secreted frizzled-related protein 4 |
| chr1 | 86265236 | 86265976 | 741 | * | 6,516058 | 4,050266877 | 1,562683 | 2,591867 | 0,009546 | 1 | 0 | Distal Intergenic | 1 | 86278218 | 86278933 | 716 | 2 | 17975 | ENSMUST00000188682.2 | 12957 | ENSMUSG00000026234 | Ncl | nucleolin |
| chr19 | 5652984 | 5654024 | 1041 | * | 11,76501 | 4,051287383 | 1,220732 | 3,318735 | 0,000904 | 0,483145429 | 1,2763 | Promoter (<=1kb) | 19 | 5653042 | 5653994 | 953 | 2 | 81601 | ENSMUST00000237072.2 | 0 | ENSMUSG00000024926 | Kat5 | K(lysine) acetyltransferase 5 |
| chr11 | 84775168 | 84776091 | 924 | * | 7,189862 | 4,051689817 | 1,549377 | 2,615044 | 0,008922 | 1 | 0 | Promoter (1-2kb) | 11 | 84773412 | 84801948 | 28537 | 1 | 66196 | ENSMUST00000141173.2 | 1756 | ENSMUSG00000020527 | Myo19 | myosin XIX |
| chr13 | 95744217 | 95746122 | 1906 | * | 15,95062 | 4,05203074 | 1,007672 | 4,021179 | 5,79E-05 | 0,123146349 | 1,2763 | Intron (ENSMUST0000( | 13 | 95738311 | 95754995 | 16685 | 2 | 14062 | ENSMUST00000059193.7 | 8873 | ENSMUSG00000048376 | F2r | coagulation factor II (thrombin) receptor |
| chr7 | 94914567 | 94916005 | 1439 | * | 7,154244 | 4,052080473 | 1,313382 | 3,085227 | 0,002034 | 1 | 0 | Distal Intergenic | 7 | 95091558 | 95091664 | 107 | 2 | 1,15E+08 | ENSMUST00000240171.1 | 175659 | ENSMUSG00000119638 | Gm25684 | predicted gene, 25684 |
| chr2 | 1,57E+08 | 1,57E+08 | 1056 | * | 8,822119 | 4,052388755 | 1,549943 | 2,614541 | 0,008935 | 0,896435577 | 1,211315 | Intron (ENSMUST0000( | 2 | 1,57E+08 | 1,57E+08 | 43667 | 2 | 29812 | ENSMUST00000156651.2 | 6457 | ENSMUSG00000027634 | Ndrg3 | N-myc downstream regulated gene 3 |
| chr17 | 16419386 | 16420724 | 1339 | * | 7,632087 | 4,05269595 | 1,250794 | 3,240099 | 0,001195 | 1 | 0 | Distal Intergenic | 17 | 16057811 | 16057917 | 107 | 2 | 1,15E+08 | ENSMUST00000239713.1 | -361469 | ENSMUSG00000119921 | Gm25598 | predicted gene, 25598 |
| chr17 | 27264234 | 27265443 | 1210 | * | 13,69708 | 4,053375901 | 1,098274 | 3,690678 | 0,000224 | 0,306095127 | 0,938682 | Distal Intergenic | 17 | 27276278 | 27341197 | 64920 | 1 | 16440 | ENSMUST00000049308.9 | -10835 | ENSMUSG00000042644 | Itpr3 | inositol 1,4,5-triphosphate receptor 3 |
| chr5 | 1,05E+08 | 1,05E+08 | 1430 | * | 8,357958 | 4,053584304 | 1,301463 | 3,114636 | 0,001842 | 0,593612587 | 1,449706 | Distal Intergenic | 5 | 1,05E+08 | 1,05E+08 | 14615 | 2 | 626391 | ENSMUST00000089906.7 | 113582 | ENSMUSG00000072774 | Zfp951 | zinc finger protein 951 |
| chr4 | 1,08E+08 | 1,08E+08 | 1276 | * | 9,482792 | 4,053995846 | 1,265092 | 3,204507 | 0,001353 | 0,54122637 | 1,360622 | Promoter (<=1kb) | 4 | 1,08E+08 | 1,08E+08 | 9572 | 1 | 74098 | ENSMUST00000135454.8 | -974 | ENSMUSG00000028608 | Czib | CXXC motif containing zinc binding protein |
| chr1 | 1,65E+08 | 1,65E+08 | 1482 | * | 9,485971 | 4,054211142 | 1,567948 | 2,58568 | 0,009719 | 0,885263498 | 1,425929 | Exon (ENSMUST000001 | 1 | 1,65E+08 | 1,65E+08 | 131768 | 2 | 74106 | ENSMUST00000027856.13 | 18447 | ENSMUSG00000026571 | Dcaf6 | DDB1 and CUL4 associated factor 6 |
| chr17 | 26828636 | 26829499 | 864 | * | 7,4554 | 4,055838812 | 1,433516 | 2,829293 | 0,004665 | 0,760759863 | 1,360622 | Promoter (1-2kb) | 17 | 26826866 | 26868771 | 41906 | 1 | 67458 | ENSMUST00000170640.3 | 1770 | ENSMUSG00000001576 | Ergic1 | endoplasmic reticulum-golgi intermediate compartment (ERGIC) 1 |
| chr3 | 61869979 | 61870522 | 544 | * | 6,143354 | 4,055941697 | 1,547089 | 2,62166 | 0,00875 | 1 | 0 | Distal Intergenic | 3 | 62245765 | 62369642 | 123878 | 1 | 622434 | ENSMUST00000079300.13 | -375243 | ENSMUSG00000036885 | Arhgef26 | Rho guanine nucleotide exchange factor (GEF) 26 |
| chr13 | 18149326 | 18151169 | 1844 | * | 14,83063 | 4,056236882 | 1,082025 | 3,748747 | 0,000178 | 0,245757594 | 1,214904 | Intron (ENSMUST0000( | 13 | 18135735 | 18167118 | 31384 | 2 | 67008 | ENSMUST00000223489.2 | 15949 | ENSMUSG00000075054 | Yae1d1 | Yae1 domain containing 1 |

| | | | | | | | | | | | | | | | | | | | | | | | |
|---|---|---|---|---|---|---|---|---|---|---|---|---|---|---|---|---|---|---|---|---|---|---|---|
| chr15 | 75606286 | 75606898 | 613 | * | 6,062533 | 4,057588684 | 1,563847 | 2,59462 | 0,00947 | 1 | 0 | Distal Intergenic | 15 | 75617339 | 75620077 | 2739 | 2 | 378435 | ENSMUST00000062002.6 | 13179 | ENSMUSG00000047591 | Mafa | v-maf musculoaponeurotic fibrosarcoma oncogene family, protein A (avian) |
| chr5 | 1,44E+08 | 1,44E+08 | 1049 | * | 7,78506 | 4,057668144 | 1,492337 | 2,719002 | 0,006548 | 0,823446343 | 1,360622 | Promoter (1-2kb) | 5 | 1,44E+08 | 1,44E+08 | 3093 | 2 | 70381 | ENSMUST00000147992.2 | 1278 | ENSMUSG00000066621 | Tecpr1 | tectonin beta-propeller repeat containing 1 |
| chr5 | 98669857 | 98670748 | 892 | * | 9,66836 | 4,058249277 | 1,286442 | 3,154631 | 0,001607 | 0,621585803 | 1,086118 | Intron (ENSMUST00000 | 5 | 98689343 | 98689473 | 131 | 1 | 1,15E+08 | ENSMUST00000104236.3 | -18595 | ENSMUSG00000077424 | Gm24279 | predicted gene, 24279 |
| chr9 | 60380179 | 60381257 | 1079 | * | 7,571938 | 4,058659589 | 1,51555 | 2,678011 | 0,007406 | 1 | 0 | Intron (ENSMUST00000 | 9 | 59874214 | 60404409 | 530196 | 2 | 207596 | ENSMUST00000171654.8 | 23152 | ENSMUSG00000032289 | Thsd4 | thrombospondin, type I, domain containing 4 |
| chrX | 1,5E+08 | 1,5E+08 | 1351 | * | 8,590294 | 4,059259501 | 1,330304 | 3,051377 | 0,002278 | 0,631320778 | 1,449706 | Distal Intergenic | 20 | 1,5E+08 | 1,5E+08 | 15070 | 2 | 237107 | ENSMUST00000156233.2 | -7260 | ENSMUSG00000025266 | Gnl3l | guanine nucleotide binding protein-like 3 (nucleolar)-like |
| chr8 | 1,05E+08 | 1,05E+08 | 563 | * | 5,632804 | 4,060106596 | 1,530281 | 2,653177 | 0,007974 | 1 | 0 | Distal Intergenic | 8 | 1,05E+08 | 1,05E+08 | 16604 | 2 | 1,01E+08 | ENSMUST00000159039.2 | -4880 | ENSMUSG00000110430 | Cmtm1 | CKLF-like MARVEL transmembrane domain containing 1 |
| chr10 | 83799277 | 83800298 | 1022 | * | 10,41519 | 4,06015917 | 1,347543 | 3,013009 | 0,002587 | 0,684274806 | 1,173325 | Distal Intergenic | 10 | 83928130 | 83932113 | 3984 | 1 | 1,03E+08 | ENSMUST00000213796.2 | -127832 | ENSMUSG00000111524 | Gm34113 | predicted gene, 34113 |
| chr17 | 29739331 | 29740255 | 925 | * | 9,475214 | 4,060293988 | 1,346419 | 3,015625 | 0,002565 | 0,682206193 | 1,211315 | Intron (ENSMUST00000 | 17 | 29712381 | 29713938 | 1558 | 1 | 18712 | ENSMUST00000234270.2 | 26950 | ENSMUSG00000024014 | Pim1 | proviral integration site 1 |
| chr1 | 1,78E+08 | 1,78E+08 | 1618 | * | 10,07843 | 4,061361994 | 1,279037 | 3,175328 | 0,001497 | 0,586304784 | 1,211315 | Distal Intergenic | 1 | 1,78E+08 | 1,78E+08 | 106 | 1 | 1,15E+08 | ENSMUST00000083020.3 | -18040 | ENSMUSG00000064954 | Gm24919 | predicted gene, 24919 |
| chr2 | 75469623 | 75471675 | 2053 | * | 21,306461 | 4,06469453 | 0,8232023 | 4,9376615 | 7,906E-07 | 0,002948921 | 3,104559 | Distal Intergenic | 2 | 75464379 | 75465121 | 743 | 1 | 105244209 | ENSMUST00000247464.1 | 5244 | | Gm39858 | predicted gene, 39858 |
| chr6 | 49231862 | 49233034 | 1173 | * | 8,603766 | 4,065207788 | 1,383508 | 2,938334 | 0,0033 | 0,691951235 | 1,425929 | Exon (ENSMUST000002 | 6 | 49220858 | 49240944 | 20087 | 2 | 101214 | ENSMUST00000204189.3 | 7910 | ENSMUSG00000029817 | Tra2a | transformer 2 alpha |
| chr10 | 16885287 | 16886668 | 1382 | * | 7,42989 | 4,066484017 | 1,361944 | 2,985793 | 0,002828 | 1 | 0 | Intron (ENSMUST00000 | 10 | 16851868 | 16872067 | 20200 | 2 | 1,01E+08 | ENSMUST00000208415.2 | -13220 | ENSMUSG00000101969 | Gm20125 | predicted gene, 20125 |
| chr2 | 71730700 | 71731946 | 1247 | * | 9,978993 | 4,067871474 | 1,251574 | 3,250204 | 0,001153 | 0,535402776 | 1,211315 | 3' UTR | 2 | 71717762 | 71726837 | 9076 | 1 | 228026 | ENSMUST00000129427.2 | 12938 | ENSMUSG00000006494 | Pdk1 | pyruvate dehydrogenase kinase, isoenzyme 1 |
| chr12 | 15302467 | 15303131 | 665 | * | 7,762981 | 4,068367143 | 1,311448 | 3,102195 | 0,001921 | 1 | 0 | Distal Intergenic | 12 | 15118118 | 15187519 | 69402 | 1 | 1,03E+08 | ENSMUST00000222816.2 | 184349 | | Gm36128 | predicted gene, 36128 |
| chr12 | 52838910 | 52840065 | 1156 | * | 7,635434 | 4,068924166 | 1,541343 | 2,639856 | 0,008294 | 1 | 0 | Promoter (2-3kb) | 12 | 52842619 | 53195546 | 352928 | 1 | 238161 | ENSMUST00000219786.2 | -2554 | ENSMUSG00000061603 | Akap6 | A kinase (PRKA) anchor protein 6 |
| chr13 | 63472527 | 63473705 | 1179 | * | 13,3678 | 4,068939387 | 1,108099 | 3,672 | 0,000241 | 0,317496979 | 0,918255 | Promoter (<=1kb) | 13 | 63460310 | 63472832 | 12523 | 2 | 14088 | ENSMUST00000160336.8 | 0 | ENSMUSG00000021461 | Fancc | Fanconi anemia, complementation group C |
| chr5 | 72527315 | 72528069 | 755 | * | 6,581202 | 4,069926945 | 1,579554 | 2,576631 | 0,009977 | 1 | 0 | Intron (ENSMUST00000 | 5 | 72510923 | 72518042 | 7120 | 2 | 53419 | ENSMUST00000176320.2 | -9273 | ENSMUSG00000005220 | Corin | corin, serine peptidase |
| chr10 | 81097465 | 81098341 | 877 | * | 9,336203 | 4,070696152 | 1,37249 | 2,965921 | 0,003018 | 0,682206193 | 1,425929 | Promoter (<=1kb) | 10 | 81097004 | 81098501 | 1498 | 1 | 17179 | ENSMUST00000148735.2 | 461 | ENSMUSG00000004933 | Matk | megakaryocyte-associated tyrosine kinase |
| chr5 | 1,29E+08 | 1,29E+08 | 787 | * | 6,885937 | 4,071993124 | 1,552617 | 2,622664 | 0,008725 | 1 | 0 | Distal Intergenic | 5 | 1,29E+08 | 1,29E+08 | 107850 | 1 | 243277 | ENSMUST00000056617.14 | -36248 | ENSMUSG00000044017 | Adgrd1 | adhesion G protein-coupled receptor D1 |
| chr11 | 85302901 | 85304107 | 1207 | * | 10,2428 | 4,073353102 | 1,367797 | 2,978038 | 0,002901 | 0,671937722 | 1,449706 | Intron (ENSMUST00000 | 11 | 85356985 | 85716884 | 359900 | 1 | 192197 | ENSMUST00000154396.8 | -52878 | ENSMUSG00000059439 | Bcas3 | BCAS3 microtubule associated cell migration factor |
| chr2 | 1,68E+08 | 1,68E+08 | 1501 | * | 12,59317 | 4,073379823 | 1,189299 | 3,425027 | 0,000615 | 0,447359401 | 1,075735 | Exon (ENSMUST000001 | 2 | 1,68E+08 | 1,68E+08 | 4591 | 1 | 1E+08 | ENSMUST00000127441.2 | 3052 | | Gm14321 | predicted gene 14321 |
| chr1 | 1,71E+08 | 1,71E+08 | 979 | * | 6,932518 | 4,074346384 | 1,487569 | 2,738929 | 0,006164 | 1 | 0 | Distal Intergenic | 1 | 1,71E+08 | 1,71E+08 | 897 | 2 | 14130 | ENSMUST00000161503.2 | 10239 | ENSMUSG00000026656 | Fcgr2b | Fc receptor, IgG, low affinity IIb |
| chr2 | 75490639 | 75491994 | 1356 | * | 8,828171 | 4,0762939 | 1,238773 | 3,290591 | 0,001 | 0,490931796 | 1,360622 | Promoter (<=1kb) | 2 | 75492833 | 75495799 | 2967 | 1 | 229279 | ENSMUST00000141974.2 | -839 | ENSMUSG00000059005 | Hnrnpa3 | heterogeneous nuclear ribonucleoprotein A3 |
| chr17 | 32323628 | 32325664 | 2037 | * | 19,43721 | 4,076909736 | 0,8581785 | 4,7506547 | 2,028E-06 | 0,007897566 | 2,2216115 | Promoter (<=1kb) | 17 | 32297771 | 32326324 | 28554 | 2 | 435518 | ENSMUST00000183934.2 | 660 | ENSMUSG00000002791 | Pdxk-ps | pyridoxal (pyridoxine, vitamin B6) kinase, pseudogene |
| chr4 | 1,16E+08 | 1,16E+08 | 1177 | * | 7,913181 | 4,077190132 | 1,535554 | 2,655192 | 0,007926 | 1 | 0 | Intron (ENSMUST00000 | 4 | 1,16E+08 | 1,16E+08 | 102354 | 2 | 17776 | ENSMUST00000151347.2 | -5777 | ENSMUSG00000003810 | Mast2 | microtubule associated serine/threonine kinase 2 |
| chr6 | 1,25E+08 | 1,25E+08 | 1080 | * | 8,762937 | 4,078240486 | 1,386864 | 2,94062 | 0,003276 | 0,699278451 | 1,360622 | Promoter (<=1kb) | 6 | 1,25E+08 | 1,25E+08 | 4186 | 2 | 50908 | ENSMUST00000159143.2 | 0 | ENSMUSG00000038521 | C1s1 | complement component 1, s subcomponent 1 |
| chr11 | 1,17E+08 | 1,17E+08 | 1312 | * | 12,94746 | 4,078449953 | 1,190733 | 3,425159 | 0,000614 | 0,47475665 | 0,918255 | Promoter (<=1kb) | 11 | 1,17E+08 | 1,17E+08 | 2723 | 2 | 1,03E+08 | ENSMUST00000212879.2 | 863 | | Gm34418 | predicted gene, 34418 |
| chr1 | 1,1E+08 | 1,1E+08 | 940 | * | 5,681512 | 4,078716369 | 1,445877 | 2,82093 | 0,004788 | 1 | 0 | Distal Intergenic | 1 | 1,1E+08 | 1,1E+08 | 130 | 2 | 1,15E+08 | ENSMUST00000158412.3 | -93590 | ENSMUSG00000089037 | Gm22159 | predicted gene, 22159 |
| chr7 | 34026162 | 34027740 | 1579 | * | 12,85791 | 4,079541923 | 1,045421 | 3,902296 | 9,53E-05 | 0,17530799 | 1,173325 | Distal Intergenic | 7 | 33936132 | 34012976 | 76845 | 2 | 233103 | ENSMUST00000108074.8 | -13186 | ENSMUSG00000066571 | Garre1 | granule associated Rac and RHOG effector 1 |
| chr11 | 81220826 | 81221391 | 566 | * | 6,441487 | 4,079687186 | 1,496151 | 2,726788 | 0,006395 | 1 | 0 | Intron (ENSMUST00000 | 11 | 80770995 | 81044147 | 273153 | 2 | 11418 | ENSMUST00000021045.13 | -176679 | ENSMUSG00000020704 | Asic2 | acid-sensing (proton-gated) ion channel 2 |
| chr19 | 6374284 | 6375399 | 1116 | * | 8,380124 | 4,08056934 | 1,552778 | 2,627916 | 0,008591 | 0,783872172 | 2,237403 | Promoter (2-3kb) | 19 | 6371838 | 6372262 | 425 | 1 | 240505 | ENSMUST00000141854.2 | 2446 | ENSMUSG00000024769 | Cdc42bpg | CDC42 binding protein kinase gamma (DMPK-like) |
| chr2 | 118165363 | 118167361 | 1999 | * | 19,267233 | 4,080864794 | 0,9375366 | 4,3527527 | 1,344E-05 | 0,028909459 | 2,6636191 | Intron (ENSMUST000000 | 2 | 118135395 | 118138647 | 3253 | 1 | 73358 | ENSMUST00000125654.2 | 29968 | ENSMUSG00000085889 | 1700054M17Rik | RIKEN cDNA 1700054M17 gene |
| chr14 | 99882550 | 99884023 | 1474 | * | 9,038721 | 4,081032187 | 1,542505 | 2,645718 | 0,008152 | 0,774037668 | 2,237403 | Distal Intergenic | 14 | 99759981 | 99814338 | 54358 | 1 | 1,05E+08 | ENSMUST00000228876.2 | 122569 | ENSMUSG00000115580 | Gm41230 | predicted gene, 41230 |
| chr3 | 87218030 | 87219632 | 1603 | * | 9,535889 | 4,082021411 | 1,433948 | 2,846701 | 0,004417 | 0,750831415 | 1,360622 | Distal Intergenic | 3 | 87265188 | 87278380 | 13193 | 1 | 11801 | ENSMUST00000015998.8 | -45556 | ENSMUSG00000015854 | Cd5l | CD5 antigen-like |
| chr3 | 1,25E+08 | 1,25E+08 | 1065 | * | 8,459701 | 4,082926222 | 1,42415 | 2,866922 | 0,004145 | 0,731734453 | 1,425929 | Distal Intergenic | 3 | 1,25E+08 | 1,26E+08 | 324824 | 1 | 64580 | ENSMUST00000173932.8 | -23944 | ENSMUSG00000027971 | Ndst4 | N-deacetylase/N-sulfotransferase (heparin glucosaminyl) 4 |
| chr1 | 65073054 | 65075056 | 2003 | * | 16,45993 | 4,083498684 | 1,284133 | 3,179967 | 0,001473 | 0,442939362 | 2,663619 | Promoter (1-2kb) | 1 | 65051848 | 65071918 | 20071 | 2 | 70861 | ENSMUST00000069142.12 | -1136 | ENSMUSG00000025955 | Akr1cl | aldo-keto reductase family 1, member C-like |
| chr7 | 77550609 | 77551394 | 786 | * | 8,021562 | 4,084307073 | 1,339981 | 3,048033 | 0,002303 | 1 | 0 | Distal Intergenic | 7 | 77830631 | 77842491 | 11861 | 2 | 18213 | ENSMUST00000205354.2 | 291097 | ENSMUSG00000059146 | Ntrk3 | neurotrophic tyrosine kinase, receptor, type 3 |
| chr7 | 1,16E+08 | 1,16E+08 | 609 | * | 6,191954 | 4,087521046 | 1,428318 | 2,861773 | 0,004213 | 1 | 0 | Distal Intergenic | 7 | 1,16E+08 | 1,16E+08 | 7222 | 1 | 53322 | ENSMUST00000183335.2 | 141614 | ENSMUSG00000030659 | Nucb2 | nucleobindin 2 |
| chr3 | 94779902 | 94780737 | 836 | * | 6,601297 | 4,088299422 | 1,505568 | 2,715453 | 0,006619 | 1 | 0 | Promoter (1-2kb) | 3 | 94778623 | 94781062 | 2440 | 1 | 229584 | ENSMUST00000125638.2 | 1279 | ENSMUSG00000038902 | Pogz | pogo transposable element with ZNF domain |
| chr17 | 35188635 | 35190790 | 2156 | * | 15,79978 | 4,088312206 | 1,17099 | 3,491329 | 0,000481 | 0,273126507 | 2,663619 | Promoter (<=1kb) | 17 | 35188166 | 35191132 | 2967 | 2 | 193740 | ENSMUST00000087328.4 | 342 | ENSMUSG00000091971 | Hspa1a | heat shock protein 1A |
| chr16 | 76349858 | 76351028 | 1171 | * | 9,592813 | 4,088983984 | 1,481411 | 2,760196 | 0,005777 | 0,790460916 | 1,449706 | Distal Intergenic | 16 | 76345683 | 76348112 | 2430 | 1 | 1,03E+08 | ENSMUST00000232268.2 | 4175 | ENSMUSG00000116701 | Gm30726 | predicted gene, 30726 |
| chr11 | 78656263 | 78657650 | 1388 | * | 15,13133 | 4,089965624 | 1,070694 | 3,81992 | 0,000133 | 0,192431185 | 1,41696 | Distal Intergenic | 11 | 78641332 | 78642555 | 1224 | 2 | 69109 | ENSMUST00000059468.6 | -13708 | ENSMUSG00000049489 | Ccnq | cyclin Q |
| chr10 | 48173955 | 48174693 | 739 | * | 8,187053 | 4,09027783 | 1,377927 | 2,968428 | 0,002993 | 0,677933849 | 1,449706 | Distal Intergenic | 10 | 48495408 | 48495491 | 84 | 2 | 1,15E+08 | ENSMUST00000083696.4 | 320798 | | Gm24256 | predicted gene, 24256 |
| chrX | 16484708 | 16486235 | 1528 | * | 10,40918 | 4,090400786 | 1,465957 | 2,790259 | 0,005267 | 0,890233764 | 0,744375 | Promoter (<=1kb) | 20 | 16485937 | 16554057 | 68121 | 1 | 17161 | ENSMUST00000026013.6 | 0 | ENSMUSG00000025037 | Maoa | monoamine oxidase A |
| chr11 | 1,18E+08 | 1,18E+08 | 1004 | * | 10,55082 | 4,090540443 | 1,181202 | 3,463031 | 0,000534 | 0,39905285 | 1,2763 | Intron (ENSMUST00000 | 11 | 1,18E+08 | 1,18E+08 | 67069 | 2 | 52897 | ENSMUST00000136551.3 | -36096 | ENSMUSG00000025576 | Rbfox3 | RNA binding protein, fox-1 homolog (C. elegans) 3 |
| chr19 | 58929462 | 58930808 | 1347 | * | 12,07214 | 4,090734817 | 1,230503 | 3,324442 | 0,000886 | 0,479393375 | 1,2763 | Promoter (1-2kb) | 19 | 58931857 | 58959851 | 27995 | 1 | 226265 | ENSMUST00000054280.10 | -1049 | ENSMUSG00000048029 | Eno4 | enolase 4 |
| chr19 | 15434318 | 15436582 | 2265 | * | 33,74368 | 4,090905054 | 1,354041 | 3,021257 | 0,002517 | 0,654756507 | 1,405709 | Distal Intergenic | 19 | 15657368 | 15657474 | 107 | 1 | 1,15E+08 | ENSMUST00000083389.3 | -220786 | ENSMUSG00000118757 | Gm24319 | predicted gene, 24319 |
| chr15 | 99786085 | 99786607 | 523 | * | 7,913671 | 4,09147037 | 1,510868 | 2,708027 | 0,006768 | 1 | 0 | Distal Intergenic | 15 | 99676351 | 99773337 | 96987 | 2 | 65970 | ENSMUST00000073691.5 | -12748 | ENSMUSG00000023022 | Lima1 | LIM domain and actin binding 1 |
| chr10 | 121461946 | 121463641 | 1696 | * | 20,936836 | 4,092234319 | 0,8402928 | 4,8700099 | 1,116E-06 | 0,004200523 | 2,6724296 | Promoter (<=1kb) | 10 | 121423285 | 121462221 | 38937 | 2 | 73192 | ENSMUST00000039810.8 | 0 | ENSMUSG00000034667 | Xpot | exportin, tRNA (nuclear export receptor for tRNAs) |
| chr1 | 1,42E+08 | 1,42E+08 | 1291 | * | 9,493042 | 4,092467569 | 1,326722 | 3,084647 | 0,002038 | 0,611385361 | 1,449706 | Distal Intergenic | 1 | 1,41E+08 | 1,41E+08 | 107 | 1 | 1,15E+08 | ENSMUST00000239951.1 | 294076 | ENSMUSG00000118759 | Gm25457 | predicted gene, 25457 |
| chr1 | 14886033 | 14887027 | 995 | * | 7,455852 | 4,092512771 | 1,497335 | 2,733198 | 0,006272 | 1 | 0 | Distal Intergenic | 1 | 14823570 | 14826214 | 2645 | 2 | 17681 | ENSMUST00000027062.7 | -59819 | ENSMUSG00000025930 | Msc | musculin |
| chr7 | 83626102 | 83628731 | 2630 | * | 22,276481 | 4,09290692 | 0,8673096 | 4,7190838 | 2,369E-06 | 0,007032136 | 3,104559 | Promoter (<=1kb) | 7 | 83599674 | 83625283 | 25610 | 2 | 80982 | ENSMUST00000142518.8 | -819 | ENSMUSG00000052353 | Cemip | cell migration inducing protein, hyaluronan binding |
| chr1 | 1,82E+08 | 1,82E+08 | 1481 | * | 11,54644 | 4,093279634 | 1,206798 | 3,391852 | 0,000694 | 0,453690896 | 1,173325 | Distal Intergenic | 1 | 1,82E+08 | 1,82E+08 | 6989 | 2 | 98386 | ENSMUST00000193030.6 | -18295 | ENSMUSG00000004880 | Lbr | lamin B receptor |
| chr8 | 1,3E+08 | 1,3E+08 | 975 | * | 10,0825 | 4,094709549 | 1,222698 | 3,348913 | 0,000811 | 0,478766182 | 1,173325 | Promoter (2-3kb) | 8 | 1,3E+08 | 1,3E+08 | 18833 | 1 | 1,03E+08 | ENSMUST00000245647.1 | 2155 | ENSMUSG00000110605 | Gm32856 | predicted gene, 32856 |
| chr2 | 1,08E+08 | 1,08E+08 | 770 | * | 5,741096 | 4,095003923 | 1,543483 | 2,653093 | 0,007976 | 1 | 0 | Intron (ENSMUST00000 | 2 | 1,08E+08 | 1,08E+08 | 107604 | 2 | 1,03E+08 | ENSMUST00000156541.2 | 74537 | ENSMUSG00000086758 | Gm13912 | predicted gene 13912 |
| chr4 | 1,24E+08 | 1,24E+08 | 1713 | * | 12,59669 | 4,096360223 | 1,042354 | 3,929914 | 8,5E-05 | 0,190625259 | 0,918255 | Promoter (<=1kb) | 4 | 1,24E+08 | 1,24E+08 | 2965 | 1 | 230726 | ENSMUST00000148797.2 | 0 | ENSMUSG00000043333 | Rhbdl2 | rhomboid like 2 |
| chr14 | 56591316 | 56592468 | 1153 | * | 10,96467 | 4,097336746 | 1,195887 | 3,426189 | 0,000612 | 0,419066905 | 1,2763 | Distal Intergenic | 14 | 56602525 | 56626007 | 23483 | 1 | 192113 | ENSMUST00000007340.4 | -10057 | ENSMUSG00000022229 | Atp12a | ATPase, H+/K+ transporting, nongastric, alpha polypeptide |
| chr7 | 19537043 | 19538388 | 1346 | * | 11,53734 | 4,098340678 | 1,360896 | 3,011502 | 0,0026 | 0,765231686 | 0,744375 | Distal Intergenic | 7 | 19542387 | 19543588 | 1202 | 2 | 12051 | ENSMUST00000135609.8 | 5200 | ENSMUSG00000053175 | Bcl3 | B cell leukemia/lymphoma 3 |
| chr1 | 166237416 | 166239316 | 1901 | * | 17,361655 | 4,099025919 | 0,976718 | 4,1967343 | 2,708E-05 | 0,048727933 | 2,6636191 | Promoter (<=1kb) | 1 | 166223738 | 166237432 | 13695 | 2 | 71592 | ENSMUST00000128861.3 | 0 | ENSMUSG00000040596 | Pogk | pogo transposable element with KRAB domain |
| chr9 | 42174515 | 42175205 | 691 | * | 7,246931 | 4,099285494 | 1,399444 | 2,929224 | 0,003398 | 1 | 0 | Promoter (<=1kb) | 9 | 42166029 | 42175496 | 9468 | 2 | 235293 | ENSMUST00000169609.2 | 291 | ENSMUSG00000032018 | Sc5d | sterol-C5-desaturase |
| chr14 | 57493417 | 57494980 | 1564 | * | 11,80332 | 4,101307757 | 1,363951 | 3,006931 | 0,002639 | 0,769699963 | 0,738441 | Distal Intergenic | 14 | 57570823 | 57570942 | 120 | 2 | 1,15E+08 | ENSMUST00000104309.3 | 75962 | ENSMUSG00000077497 | Gm23593 | predicted gene, 23593 |
| chrX | 1,39E+08 | 1,39E+08 | 860 | * | 6,893306 | 4,101362766 | 1,300548 | 3,153565 | 0,001613 | 1 | 0 | Intron (ENSMUST00000 | 20 | 1,39E+08 | 1,39E+08 | 18460 | 2 | 75746 | ENSMUST00000167664.2 | 7600 | ENSMUSG00000031434 | Morc4 | microrchidia 4 |
| chr1 | 88987800 | 88989417 | 1618 | * | 13,15177 | 4,101364452 | 1,372208 | 2,98888 | 0,0028 | 0,780381721 | 0,738441 | Distal Intergenic | 1 | 88998137 | 89082790 | 84654 | 1 | 98402 | ENSMUST00000066279.11 | -8720 | ENSMUSG00000036206 | Sh3bp4 | SH3-domain binding protein 4 |
| chr2 | 63090001 | 63090551 | 551 | * | 6,121312 | 4,102468111 | 1,573388 | 2,607411 | 0,009123 | 1 | 0 | Distal Intergenic | 2 | 62616625 | 63014631 | 398007 | 2 | 170738 | ENSMUST00000112454.8 | -75370 | ENSMUSG00000059742 | Kcnh7 | potassium voltage-gated channel, subfamily H (eag-related), member 7 |
| chr4 | 34877678 | 34878215 | 538 | * | 6,336638 | 4,104971248 | 1,541596 | 2,662806 | 0,007749 | 1 | 0 | Intron (ENSMUST00000 | 4 | 34803113 | 34882930 | 79818 | 2 | 30046 | ENSMUST00000098163.9 | 4715 | ENSMUSG00000039967 | Zfp292 | zinc finger protein 292 |
| chr16 | 33239111 | 33240157 | 1047 | * | 8,389151 | 4,106398996 | 1,458662 | 2,815183 | 0,004875 | 0,760549761 | 1,425929 | Intron (ENSMUST00000 | 16 | 33220815 | 33319013 | 98199 | 1 | 22661 | ENSMUST00000089677.7 | 18296 | ENSMUSG00000022811 | Zfp148 | zinc finger protein 148 |
| chr16 | 18100342 | 18102603 | 2262 | * | 18,1783 | 4,106867386 | 1,028464 | 3,993203 | 6,52E-05 | 0,095764633 | 2,154504 | Promoter (<=1kb) | 16 | 18102103 | 18102181 | 79 | 2 | 1E+08 | ENSMUST00000116802.5 | 0 | ENSMUSG00000080452 | Mir1306 | microRNA 1306 |
| chr2 | 1,09E+08 | 1,09E+08 | 668 | * | 6,100298 | 4,107476454 | 1,565818 | 2,623214 | 0,00871 | 1 | 0 | Intron (ENSMUST00000 | 2 | 1,09E+08 | 1,09E+08 | 185981 | 2 | 76894 | ENSMUST00000147376.8 | 80197 | ENSMUSG00000057234 | Mettl15 | methyltransferase like 15 |
| chr2 | 82347936 | 82348547 | 612 | * | 6,684807 | 4,107653902 | 1,371938 | 2,994052 | 0,002753 | 1 | 0 | Distal Intergenic | 2 | 82205716 | 82222546 | 16831 | 2 | 228116 | ENSMUST00000128608.2 | -125390 | ENSMUSG00000087061 | A130030D18Rik | RIKEN cDNA A130030D18 gene |
| chr19 | 6180532 | 6181000 | 469 | * | 6,41759 | 4,107675463 | 1,500616 | 2,737326 | 0,006194 | 1 | 0 | Promoter (2-3kb) | 19 | 6169429 | 6178334 | 8906 | 2 | 69024 | ENSMUST00000025702.14 | -2198 | ENSMUSG00000024787 | Snx15 | sorting nexin 15 |
| chr19 | 20326788 | 20328507 | 1720 | * | 12,50136 | 4,109294885 | 1,298734 | 3,164077 | 0,001556 | 0,617289329 | 1,075735 | Intron (ENSMUST00000 | 19 | 20350920 | 20352951 | 2032 | 2 | 16952 | ENSMUST00000163902.2 | 24444 | ENSMUSG00000024659 | Anxa1 | annexin A1 |
| chr19 | 37536200 | 37538552 | 2353 | * | 15,40629 | 4,111606859 | 1,276496 | 3,221011 | 0,001277 | 0,531203841 | 1,37683 | Promoter (<=1kb) | 19 | 37538866 | 37672507 | 133642 | 1 | 107371 | ENSMUST00000066439.8 | -314 | ENSMUSG00000053799 | Exoc6 | exocyst complex component 6 |
| chr7 | 1,12E+08 | 1,12E+08 | 966 | * | 8,484898 | 4,112172093 | 1,433573 | 2,868477 | 0,004125 | 0,659642963 | 2,237403 | Distal Intergenic | 7 | 1,12E+08 | 1,12E+08 | 130 | 2 | 1,15E+08 | ENSMUST00000175261.3 | 22648 | ENSMUSG00000093002 | Gm24154 | predicted gene, 24154 |
| chr18 | 55468067 | 55468665 | 599 | * | 5,582321 | 4,112818449 | 1,57795 | 2,606432 | 0,009149 | 1 | 0 | Distal Intergenic | 18 | 55190657 | 55212486 | 21830 | 1 | 1E+08 | ENSMUST00000181765.2 | 277410 | | Gm4221 | predicted gene 4221 |
| chr7 | 24829914 | 24831301 | 1388 | * | 10,69921 | 4,114378107 | 1,424782 | 2,887725 | 0,00388 | 0,740669062 | 1,2763 | Promoter (<=1kb) | 7 | 24791945 | 24831902 | 39958 | 2 | 18987 | ENSMUST00000108418.11 | 601 | ENSMUSG00000008496 | Pou2f2 | POU domain, class 2, transcription factor 2 |
| chr10 | 13843080 | 13844022 | 943 | * | 9,353701 | 4,114454869 | 1,462939 | 2,812458 | 0,004916 | 0,762875156 | 1,425929 | Promoter (<=1kb) | 10 | 13842768 | 13955419 | 112652 | 1 | 15273 | ENSMUST00000191464.2 | 312 | ENSMUSG00000015501 | Hivep2 | human immunodeficiency virus type I enhancer binding protein 2 |
| chrX | 26358391 | 26359546 | 1156 | * | 7,017292 | 4,114758857 | 1,478644 | 2,782791 | 0,005389 | 1 | 0 | Distal Intergenic | 20 | 26242277 | 26265344 | 23068 | 2 | 1E+08 | ENSMUST00000119590.2 | -93047 | ENSMUSG00000083628 | Gm2030 | predicted gene 2030 |
| chr11 | 69318930 | 69320747 | 1818 | * | 15,32418 | 4,116024464 | 1,084002 | 3,797063 | 0,000146 | 0,214706231 | 1,2763 | Exon (ENSMUST000000 | 11 | 69289334 | 69304501 | 15168 | 2 | 216850 | ENSMUST00000094077.5 | -14429 | ENSMUSG00000018476 | Kdm6b | KDM1 lysine (K)-specific demethylase 6B |
| chr17 | 63600547 | 63601267 | 721 | * | 7,15193 | 4,116567479 | 1,323086 | 3,111339 | 0,001862 | 1 | 0 | Intron (ENSMUST00000 | 17 | 63612441 | 63619966 | 7526 | 2 | 414079 | ENSMUST00000164605.8 | 18699 | ENSMUSG00000090565 | 4930405O22Rik | RIKEN cDNA 4930405O22 gene |
| chr4 | 1,11E+08 | 1,11E+08 | 850 | * | 6,200737 | 4,119126102 | 1,453781 | 2,833389 | 0,004606 | 1 | 0 | Intron (ENSMUST00000 | 4 | 1,1E+08 | 1,12E+08 | 1259720 | 1 | 78933 | ENSMUST00000097920.9 | 301392 | ENSMUSG00000061298 | Agbl4 | ATP/GTP binding protein-like 4 |
| chr7 | 98960660 | 98961526 | 867 | * | 8,432103 | 4,120074165 | 1,481657 | 2,780721 | 0,005424 | 0,778300587 | 1,449706 | Intron (ENSMUST00000 | 7 | 98970407 | 98971662 | 1256 | 1 | 17760 | ENSMUST00000129058.2 | -8881 | ENSMUSG00000055407 | Map6 | microtubule-associated protein 6 |
| chr11 | 97238429 | 97239445 | 1017 | * | 6,908018 | 4,121690087 | 1,537332 | 2,681067 | 0,007339 | 1 | 0 | Intron (ENSMUST00000 | 11 | 97222935 | 97242842 | 19908 | 2 | 217143 | ENSMUST00000093942.5 | 3397 | ENSMUSG00000070337 | Gpr179 | G protein-coupled receptor 179 |
| chr5 | 34682528 | 34684058 | 1531 | * | 14,57874 | 4,121881007 | 1,020121 | 4,040582 | 5,33E-05 | 0,117550795 | 1,2763 | Promoter (<=1kb) | 5 | 34683182 | 34709044 | 25863 | 1 | 24055 | ENSMUST00000125817.8 | 0 | ENSMUSG00000054520 | Sh3bp2 | SH3-domain binding protein 2 |
| chr18 | 7025529 | 7026445 | 917 | * | 9,190175 | 4,123056804 | 1,291723 | 3,191906 | 0,001413 | 0,548877304 | 1,360622 | Distal Intergenic | 18 | 6934518 | 7004780 | 70263 | 2 | 210719 | ENSMUST00000079788.7 | -20749 | ENSMUSG00000061013 | Mkx | mohawk homeobox |
| chr4 | 82317565 | 82319147 | 1583 | * | 14,07592 | 4,126957309 | 1,085502 | 3,801887 | 0,000144 | 0,213547685 | 1,2763 | Intron (ENSMUST00000 | 4 | 82416781 | 82417553 | 773 | 2 | 18028 | ENSMUST00000152588.2 | 98406 | ENSMUSG00000008575 | Nfib | nuclear factor I/B |
| chr12 | 73824868 | 73826725 | 1858 | * | 13,05244 | 4,128237689 | 1,111806 | 3,713093 | 0,000205 | 0,268684353 | 1,173325 | Promoter (1-2kb) | 12 | 73820368 | 73828279 | 7912 | 2 | 1,03E+08 | ENSMUST00000242536.1 | 1554 | | Gm34189 | predicted gene, 34189 |
| chr16 | 32084024 | 32084922 | 899 | * | 8,742501 | 4,128611244 | 1,422041 | 2,9033 | 0,003693 | 0,641208222 | 2,237403 | Promoter (1-2kb) | 16 | 32086268 | 32086401 | 134 | 2 | 1E+08 | ENSMUST00000158360.3 | 1479 | ENSMUSG00000088985 | Mir1946a | microRNA 1946a |
| chr13 | 55246263 | 55247781 | 1519 | * | 9,484552 | 4,128628418 | 1,421874 | 2,903652 | 0,003688 | 0,741363157 | 1,211315 | Promoter (<=1kb) | 13 | 55176008 | 55248067 | 72060 | 2 | 20184 | ENSMUST00000148702.8 | 286 | ENSMUSG00000025878 | Uimc1 | ubiquitin interaction motif containing 1 |
| chr13 | 83749948 | 83751374 | 1427 | * | 8,146177 | 4,129921149 | 1,533204 | 2,693654 | 0,007067 | 0,748971518 | 2,237403 | Intron (ENSMUST00000 | 13 | 83773722 | 83810855 | 37134 | 1 | 17260 | ENSMUST00000200138.5 | -22348 | ENSMUSG00000005583 | Mef2c | myocyte enhancer factor 2C |
| chr10 | 73830509 | 73831390 | 882 | * | 8,807729 | 4,130038196 | 1,190062 | 3,47044 | 0,00052 | 0,303890196 | 2,237403 | Intron (ENSMUST00000 | 10 | 73782888 | 74482000 | 699113 | 1 | 11994 | ENSMUST00000195531.6 | 47621 | ENSMUSG00000052613 | Pcdh15 | protocadherin 15 |
| chr15 | 1,02E+08 | 1,02E+08 | 1593 | * | 17,27934 | 4,130851154 | 1,136107 | 3,635969 | 0,000277 | 0,204574053 | 2,663619 | Promoter (1-2kb) | 15 | 1,02E+08 | 1,02E+08 | 517 | 2 | 16678 | ENSMUST00000230798.2 | 1815 | ENSMUSG00000046834 | Krt1 | keratin 1 |
| chr3 | 1,21E+08 | 1,21E+08 | 1136 | * | 6,812041 | 4,131609216 | 1,40685 | 2,93678 | 0,003316 | 1 | 0 | Distal Intergenic | 3 | 1,21E+08 | 1,21E+08 | 6360 | 2 | 1,01E+08 | ENSMUST00000050571.9 | -5155 | ENSMUSG00000044522 | A730020M07Rik | RIKEN cDNA A730020M07 gene |
| chrX | 58475246 | 58476557 | 1312 | * | 7,598194 | 4,132000727 | 1,459349 | 2,8314 | 0,004634 | 1 | 0 | Intron (ENSMUST00000 | 20 | 58177237 | 58612832 | 435596 | 2 | 14168 | ENSMUST00000124402.2 | 136275 | ENSMUSG00000031137 | Fgf13 | fibroblast growth factor 13 |
| chr10 | 1,27E+08 | 1,27E+08 | 879 | * | 7,612103 | 4,132236647 | 1,510163 | 2,736285 | 0,006214 | 1 | 0 | Intron (ENSMUST00000 | 10 | 1,27E+08 | 1,27E+08 | 1323 | 1 | 1,15E+08 | ENSMUST00000124182.2 | 6740 | | Gm16217 | predicted gene 16217 |
| chr19 | 41265032 | 41265831 | 800 | * | 7,630879 | 4,133736449 | 1,390023 | 2,973861 | 0,002941 | 1 | 0 | Intron (ENSMUST00000 | 19 | 41199283 | 41252436 | 53154 | 2 | 107358 | ENSMUST00000025989.10 | -12596 | ENSMUSG00000025016 | Tm9sf3 | transmembrane 9 superfamily member 3 |
| chr13 | 53135542 | 53136556 | 1015 | * | 7,545969 | 4,134064174 | 1,396547 | 2,960205 | 0,003074 | 0,686838043 | 1,360622 | Promoter (<=1kb) | 13 | 53121245 | 53135109 | 13865 | 2 | 18030 | ENSMUST00000071065.8 | -433 | ENSMUSG00000056749 | Nfil3 | nuclear factor, interleukin 3, regulated |
| chr2 | 15049119 | 15049872 | 754 | * | 10,85782 | 4,134531295 | 1,179062 | 3,506629 | 0,000454 | 0,424243345 | 0,918255 | Intron (ENSMUST00000 | 2 | 15052032 | 15054090 | 2059 | 2 | 74455 | ENSMUST00000192769.2 | 4218 | ENSMUSG00000026707 | Nsun6 | NOL1/NOP2/Sun domain family member 6 |
| chr14 | 65252963 | 65254341 | 1379 | * | 9,385759 | 4,134982648 | 1,518628 | 2,722841 | 0,006472 | 0,809505441 | 1,449706 | Exon (ENSMUST000000 | 14 | 65187762 | 65273009 | 85248 | 1 | 210925 | ENSMUST00000225790.2 | 65201 | ENSMUSG00000021975 | Ints9 | integrator complex subunit 9 |
| chr2 | 42998192 | 42998878 | 687 | * | 6,388095 | 4,135330355 | 1,422738 | 2,906601 | 0,003654 | 1 | 0 | Distal Intergenic | 2 | 43085690 | 43086577 | 888 | 1 | 1,19E+08 | ENSMUST00000247374.1 | -86812 | | Gm54112 | predicted gene, 54112 |
| chr5 | 74224508 | 74225995 | 1488 | * | 11,99285 | 4,136071531 | 1,218441 | 3,394561 | 0,000687 | 0,493992718 | 0,918255 | Promoter (<=1kb) | 5 | 74189977 | 74226409 | 36433 | 2 | 69727 | ENSMUST00000145016.3 | 414 | ENSMUSG00000054814 | Usp46 | ubiquitin specific peptidase 46 |
| chr7 | 77441208 | 77442409 | 1202 | * | 5,957467 | 4,136821931 | 1,544552 | 2,678331 | 0,007399 | 1 | 0 | Distal Intergenic | 7 | 77830631 | 77842491 | 11861 | 2 | 18213 | ENSMUST00000205354.2 | 400082 | ENSMUSG00000059146 | Ntrk3 | neurotrophic tyrosine kinase, receptor, type 3 |

| | | | | | | | | | | | | | | | | | | | | | | | |
|---|---|---|---|---|---|---|---|---|---|---|---|---|---|---|---|---|---|---|---|---|---|---|---|
| chr4 | 137520985 | 137522863 | 1879 | * | 19,197359 | 4,137070373 | 0,9243111 | 4,4758419 | 7,611E-06 | 0,02002598 | 2,5070128 | Promoter (<=1kb) | 4 | 137482941 | 137523541 | 40601 | 2 | 11647 | ENSMUST00000138463.2 | 678 | ENSMUSG00000028766 | Alpl | alkaline phosphatase, liver/bone/kidney |
| chr9 | 53436395 | 53437070 | 676 | * | 7,214954 | 4,137492769 | 1,53274 | 2,69941 | 0,006946 | 0,833355026 | 1,360622 | Intron (ENSMUST0000 | 9 | 53350449 | 53445266 | 94818 | 2 | 11920 | ENSMUST00000118282.9 | 8196 | ENSMUSG00000034218 | Atm | ataxia telangiectasia mutated |
| chr15 | 84170474 | 84172247 | 1774 | * | 18,282598 | 4,138184825 | 0,9601807 | 4,309798 | 1,634E-05 | 0,033022705 | 2,6636191 | Intron (ENSMUST000000 | 15 | 84155492 | 84158107 | 2616 | 1 | 170736 | ENSMUST00000131158.2 | 14982 | ENSMUSG00000022438 | Parvb | parvin, beta |
| chr5 | 32809916 | 32810960 | 1045 | * | 8,431876 | 4,139151739 | 1,470506 | 2,81478 | 0,004881 | 0,760759863 | 1,425929 | Promoter (1-2kb) | 5 | 32664528 | 32808409 | 143882 | 2 | 1,03E+08 | ENSMUST00000155976.2 | -1507 | ENSMUSG00000085234 | Gm15614 | predicted gene 15614 |
| chr5 | 31725053 | 31726195 | 1143 | * | 13,00263 | 4,139214003 | 1,132455 | 3,65508 | 0,000257 | 0,306649884 | 1,075735 | Promoter (2-3kb) | 5 | 31714397 | 31728698 | 14302 | 2 | 100740 | ENSMUST00000242635.1 | 2503 | | AI839979 | expressed sequence AI839979 |
| chr11 | 75633744 | 75634944 | 1201 | * | 7,826919 | 4,139428052 | 1,38743 | 2,983521 | 0,00285 | 1 | 0 | Intron (ENSMUST0000 | 11 | 75642430 | 75650141 | 7712 | 1 | 22627 | ENSMUST00000155057.2 | -7486 | ENSMUSG00000020849 | Ywhae | tyrosine 3-monooxygenase/tryptophan 5-monooxygenase activation protein, epsilon polypeptide |
| chr10 | 79604421 | 79605241 | 821 | * | 7,904929 | 4,139570787 | 1,503694 | 2,752935 | 0,005906 | 1 | 0 | Promoter (1-2kb) | 10 | 79594349 | 79602786 | 8438 | 2 | 70294 | ENSMUST00000047203.9 | -1635 | ENSMUSG00000035890 | Rnf126 | ring finger protein 126 |
| chr12 | 51393943 | 51395794 | 1852 | * | 33,265482 | 4,139709917 | 0,6825827 | 6,0647739 | 1,321E-09 | 1,92956E-05 | 2,9254973 | Promoter (<=1kb) | 12 | 51394844 | 51423769 | 28926 | 1 | 217558 | ENSMUST00000054308.13 | 0 | ENSMUSG00000035293 | G2e3 | G2/M-phase specific E3 ubiquitin ligase |
| chr4 | 94012405 | 94013391 | 987 | * | 6,716444 | 4,139758285 | 1,593073 | 2,598599 | 0,00936 | 1 | 0 | Intron (ENSMUST0000 | 4 | 93977813 | 94313825 | 336013 | 2 | 1,08E+08 | ENSMUST00000126270.2 | 300434 | | Gm12648 | predicted gene 12648 |
| chr3 | 51474329 | 51475388 | 1060 | * | 9,113427 | 4,144112581 | 1,427381 | 2,903299 | 0,003693 | 0,741428016 | 1,211315 | Exon (ENSMUST00000 | 3 | 51471062 | 51474532 | 3471 | 1 | 1E+08 | ENSMUST00000180616.4 | 3267 | | 5031434O11Rik | RIKEN cDNA 5031434O11 gene |
| chr3 | 86752915 | 86754960 | 2046 | * | 14,70617 | 4,144149149 | 1,053559 | 3,933475 | 8,37E-05 | 0,161106071 | 1,214904 | Intron (ENSMUST0000 | 3 | 86694656 | 86706467 | 11812 | 2 | 70762 | ENSMUST00000193400.6 | -46448 | ENSMUSG00000028078 | Dclk2 | doublecortin-like kinase 2 |
| chr11 | 1,17E+08 | 1,17E+08 | 950 | * | 7,00707 | 4,144552247 | 1,582345 | 2,619246 | 0,008812 | 1 | 0 | Exon (ENSMUST00020 | 11 | 1,17E+08 | 1,17E+08 | 13512 | 1 | 74136 | ENSMUST00000146585.2 | 16766 | ENSMUSG00000020823 | Sec14l1 | SEC14-like lipid binding 1 |
| chrX | 42635708 | 42636575 | 868 | * | 5,937099 | 4,144736551 | 1,538396 | 2,694193 | 0,007056 | 1 | 0 | Distal Intergenic | 20 | 42680037 | 42682409 | 2373 | 1 | 637093 | ENSMUST00000105113.4 | -43462 | ENSMUSG00000078320 | Tex13c1 | TEX13 family member C1 |
| chr13 | 55385602 | 55386562 | 961 | * | 9,177449 | 4,145521176 | 1,439918 | 2,878999 | 0,003989 | 0,733243707 | 1,360622 | Exon (ENSMUST00000 | 13 | 55411238 | 55425403 | 14166 | 1 | 18193 | ENSMUST00000223894.2 | -24676 | ENSMUSG00000021488 | Nsd1 | nuclear receptor-binding SET-domain protein 1 |
| chr1 | 56949229 | 56950017 | 789 | * | 6,941831 | 4,145749545 | 1,529405 | 2,710695 | 0,006714 | 1 | 0 | Intron (ENSMUST0000 | 1 | 56987293 | 57008999 | 21707 | 2 | 212712 | ENSMUST00000176759.2 | 58982 | ENSMUSG00000038331 | Satb2 | special AT-rich sequence binding protein 2 |
| chrX | 1,62E+08 | 1,62E+08 | 779 | * | 7,762324 | 4,147590357 | 1,42814 | 2,904191 | 0,003682 | 1 | 0 | Intron (ENSMUST0000 | 20 | 1,62E+08 | 1,62E+08 | 33563 | 2 | 353170 | ENSMUST00000131463.2 | 13679 | ENSMUSG00000038344 | Txlng | taxilin gamma |
| chr3 | 1,24E+08 | 1,24E+08 | 735 | * | 6,991473 | 4,148095443 | 1,596521 | 2,598208 | 0,009371 | 0,886332554 | 1,360622 | Distal Intergenic | 3 | 1,24E+08 | 1,24E+08 | 5284 | 2 | 384775 | ENSMUST00000177772.3 | 27823 | ENSMUSG00000039174 | 1700003H04Rik | RIKEN cDNA 1700003H04 gene |
| chr12 | 33881838 | 33882607 | 770 | * | 6,792812 | 4,148219923 | 1,5891 | 2,610421 | 0,009043 | 1 | 0 | Distal Intergenic | 12 | 33978424 | 33979309 | 886 | 1 | 114712 | ENSMUST00000061035.4 | -95817 | ENSMUSG00000046518 | Ferd3l | Fer3 like bHLH transcription factor |
| chr10 | 80613778 | 80614692 | 915 | * | 9,045001 | 4,149797241 | 1,27092 | 3,265193 | 0,001094 | 0,494741046 | 1,449706 | Promoter (<=1kb) | 10 | 80613390 | 80630225 | 16836 | 1 | 208266 | ENSMUST00000150338.8 | 388 | ENSMUSG00000061589 | Dot1l | DOT1-like, histone H3 methyltransferase (S. cerevisiae) |
| chr9 | 92422500 | 92423533 | 1034 | * | 7,940829 | 4,150822063 | 1,466356 | 2,830705 | 0,004645 | 1 | 0 | Promoter (<=1kb) | 9 | 92421989 | 92422136 | 148 | 2 | 78575 | ENSMUST00000186512.2 | -364 | ENSMUSG00000103622 | B430319G15Rik | RIKEN cDNA B430319G15 gene |
| chr15 | 38443096 | 38444169 | 1074 | * | 10,31572 | 4,152330117 | 1,171481 | 3,544515 | 0,000393 | 0,322750682 | 1,449706 | Intron (ENSMUST0000 | 15 | 38436469 | 38455378 | 18910 | 2 | 1E+08 | ENSMUST00000228650.2 | 11209 | | G930009F23Rik | RIKEN cDNA G930009F23 gene |
| chr5 | 82066517 | 82067177 | 661 | * | 6,758978 | 4,152942894 | 1,593287 | 2,606526 | 0,009147 | 1 | 0 | Distal Intergenic | 5 | 81919505 | 81972980 | 53476 | 1 | 319387 | ENSMUST00000148830.2 | 147012 | ENSMUSG00000037605 | Adgrl3 | adhesion G protein-coupled receptor L3 |
| chr15 | 78830529 | 78832633 | 2105 | * | 51,504781 | 4,152988572 | 0,5085841 | 8,1657859 | 3,193E-16 | 5,19159E-11 | 2,7872553 | Promoter (<=1kb) | 15 | 78831924 | 78853042 | 21119 | 1 | 110253 | ENSMUST00000140228.8 | 0 | ENSMUSG00000033088 | Triobp | TRIO and F-actin binding protein |
| chr2 | 25117012 | 25117424 | 413 | * | 8,768649 | 4,153338553 | 1,320594 | 3,145053 | 0,001661 | 1 | 0 | Promoter (2-3kb) | 2 | 25112172 | 25114714 | 2543 | 2 | 227613 | ENSMUST00000043584.5 | -2298 | ENSMUSG00000036752 | Tubb4b | tubulin, beta 4B class IVB |
| chr3 | 1,47E+08 | 1,47E+08 | 1166 | * | 8,959954 | 4,154503365 | 1,448878 | 2,867393 | 0,004139 | 0,730469267 | 1,449706 | Distal Intergenic | 3 | 1,47E+08 | 1,47E+08 | 4401 | 1 | 1,05E+08 | ENSMUST00000195873.2 | 320681 | | Gm37041 | predicted gene, 37041 |
| chr2 | 34107410 | 34108723 | 1314 | * | 13,39852 | 4,155186986 | 1,193882 | 3,480401 | 0,000501 | 0,440320279 | 0,918255 | Intron (ENSMUST0000 | 2 | 34062885 | 34178864 | 115980 | 2 | 18516 | ENSMUST00000176213.8 | 70141 | ENSMUSG00000038718 | Pbx3 | pre B cell leukemia homeobox 3 |
| chr12 | 85902218 | 85904184 | 1967 | * | 14,91428 | 4,155249211 | 1,144396 | 3,630955 | 0,000282 | 0,284602438 | 1,41696 | Exon (ENSMUST00000 | 12 | 85925760 | 85938973 | 13214 | 1 | 320244 | ENSMUST00000175899.2 | -21576 | ENSMUSG00000012609 | Ttll5 | tubulin tyrosine ligase-like family, member 5 |
| chr8 | 87550479 | 87551560 | 1082 | * | 8,12973 | 4,155482599 | 1,458441 | 2,849264 | 0,004382 | 1 | 0 | Intron (ENSMUST0000 | 8 | 87534788 | 87570673 | 35886 | 2 | 1,05E+08 | ENSMUST00000209389.2 | 19113 | | Gm39215 | predicted gene, 39215 |
| chr14 | 56047554 | 56048566 | 1013 | * | 9,567873 | 4,157873005 | 1,32844 | 3,129891 | 0,001749 | 0,584572307 | 1,425929 | Distal Intergenic | 14 | 56060601 | 56063689 | 3089 | 1 | 73181 | ENSMUST00000226979.2 | -12035 | ENSMUSG00000023411 | Nfatc4 | nuclear factor of activated T cells, cytoplasmic, calcineurin dependent 4 |
| chr2 | 1,67E+08 | 1,67E+08 | 612 | * | 9,096657 | 4,158111496 | 1,368939 | 3,03747 | 0,002386 | 0,642870233 | 1,425929 | Intron (ENSMUST0000 | 2 | 1,67E+08 | 1,67E+08 | 12719 | 1 | 81018 | ENSMUST00000127939.2 | 9398 | ENSMUSG00000006418 | Rnf114 | ring finger protein 114 |
| chr4 | 20998678 | 20999496 | 819 | * | 8,504124 | 4,158393193 | 1,484095 | 2,801973 | 0,005079 | 0,68731021 | 2,237403 | Distal Intergenic | 4 | 20243842 | 20778866 | 535025 | 2 | 269513 | ENSMUST00000102998.4 | -219812 | ENSMUSG00000055761 | Nkain3 | Na+/K+ transporting ATPase interacting 3 |
| chr4 | 1,07E+08 | 1,07E+08 | 808 | * | 10,08863 | 4,159426816 | 1,296612 | 3,207919 | 0,001337 | 0,568412386 | 1,173325 | Intron (ENSMUST0000 | 4 | 1,07E+08 | 1,07E+08 | 46269 | 1 | 230587 | ENSMUST00000138211.8 | -7727 | ENSMUSG00000034762 | Glis1 | GLIS family zinc finger 1 |
| chr1 | 1,5E+08 | 1,5E+08 | 950 | * | 6,883132 | 4,160105301 | 1,575185 | 2,641027 | 0,008266 | 0,861225402 | 1,360622 | Intron (ENSMUST0000 | 1 | 1,5E+08 | 1,5E+08 | 92804 | 2 | 18783 | ENSMUST00000111926.9 | -4805 | ENSMUSG00000056220 | Pla2g4a | phospholipase A2, group IVA (cytosolic, calcium-dependent) |
| chr14 | 121674229 | 121675657 | 1429 | * | 42,681261 | 4,161221729 | 0,5986582 | 6,9509143 | 3,629E-12 | 1,39715E-07 | 2,7872553 | Distal Intergenic | 14 | 121689142 | 121693174 | 4033 | 2 | 102636148 | ENSMUST00000227451.2 | 17517 | | Gm33299 | predicted gene, 33299 |
| chr9 | 41187767 | 41188583 | 817 | * | 7,522595 | 4,1616249 | 1,52528 | 2,728434 | 0,006364 | 1 | 0 | Distal Intergenic | 9 | 41238373 | 41440065 | 201693 | 1 | 73144 | ENSMUST00000233562.2 | -49790 | ENSMUSG00000074415 | Mir100hg | Mir100 Mirlet7a-2 Mir125b-1 cluster host gene |
| chr17 | 26390032 | 26390816 | 785 | * | 8,520626 | 4,161951572 | 1,464299 | 2,842283 | 0,004479 | 0,753013652 | 1,360622 | Intron (ENSMUST0000 | 17 | 26407127 | 26407769 | 643 | 1 | 12005 | ENSMUST00000169268.2 | -16311 | ENSMUSG00000024182 | Axin1 | axin 1 |
| chr15 | 36173362 | 36174617 | 1256 | * | 9,121602 | 4,161973785 | 1,26578 | 3,288069 | 0,001009 | 0,409763611 | 2,237403 | Promoter (<=1kb) | 15 | 36174156 | 36177153 | 2998 | 1 | 17749 | ENSMUST00000180159.8 | 0 | ENSMUSG00000045996 | Polr2k | polymerase (RNA) II (DNA directed) polypeptide K |
| chr17 | 53785669 | 53786297 | 629 | * | 10,3609 | 4,162042562 | 1,264642 | 3,291082 | 0,000998 | 0,477622452 | 1,449706 | Promoter (<=1kb) | 17 | 53786262 | 53814708 | 28447 | 1 | 271457 | ENSMUST00000017975.7 | 0 | ENSMUSG00000017831 | Rab5a | RAB5A, member RAS oncogene family |
| chr3 | 56545466 | 56546127 | 662 | * | 6,074864 | 4,163717913 | 1,614454 | 2,579026 | 0,009908 | 1 | 0 | Distal Intergenic | 3 | 56352703 | 56352833 | 131 | 2 | 1,15E+08 | ENSMUST00000104522.3 | -192633 | ENSMUSG00000119391 | Gm25727 | predicted gene, 25727 |
| chr7 | 1,3E+08 | 1,3E+08 | 1605 | * | 14,8852 | 4,167138618 | 1,17511 | 3,54617 | 0,000391 | 0,347771872 | 1,214904 | Intron (ENSMUST0000 | 7 | 1,3E+08 | 1,3E+08 | 103813 | 2 | 14183 | ENSMUST00000120187.9 | -11434 | ENSMUSG00000030849 | Fgfr2 | fibroblast growth factor receptor 2 |
| chr3 | 90087085 | 90087950 | 866 | * | 7,812046 | 4,168446396 | 1,490692 | 2,796317 | 0,005169 | 1 | 0 | Exon (ENSMUST00000 | 3 | 90105989 | 90106093 | 105 | 1 | 1,15E+08 | ENSMUST00000122578.3 | -18039 | ENSMUSG00000084527 | Gm24185 | predicted gene, 24185 |
| chr7 | 1,1E+08 | 1,1E+08 | 942 | * | 7,410227 | 4,169032092 | 1,479083 | 2,818661 | 0,004822 | 1 | 0 | Intron (ENSMUST0000 | 7 | 1,1E+08 | 1,1E+08 | 33673 | 1 | 20841 | ENSMUST00000169638.4 | 3329 | ENSMUSG00000061079 | Zfp143 | zinc finger protein 143 |
| chr3 | 37357664 | 37358372 | 709 | * | 6,323567 | 4,17212206 | 1,390124 | 3,00126 | 0,002689 | 1 | 0 | Distal Intergenic | 3 | 37362909 | 37366569 | 3661 | 2 | 207175 | ENSMUST00000075537.7 | 8197 | ENSMUSG00000045031 | Cetn4 | centrin 4 |
| chrX | 1,15E+08 | 1,15E+08 | 976 | * | 8,155142 | 4,172732936 | 1,348745 | 3,093789 | 0,001976 | 0,605265376 | 1,449706 | Distal Intergenic | 20 | 1,15E+08 | 1,15E+08 | 164 | 1 | 1,15E+08 | ENSMUST00000082617.3 | -33887 | ENSMUSG00000064551 | Gm24624 | predicted gene, 24624 |
| chr13 | 53260296 | 53261226 | 931 | * | 8,46898 | 4,173586776 | 1,364201 | 3,059364 | 0,002218 | 0,627785088 | 1,449706 | Distal Intergenic | 13 | 53155899 | 53172529 | 16631 | 1 | 1,03E+08 | ENSMUST00000222317.3 | 104397 | | Gm33424 | predicted gene, 33424 |
| chr3 | 38581820 | 38582690 | 871 | * | 7,707491 | 4,174950198 | 1,608424 | 2,595677 | 0,00944 | 0,887593795 | 1,360622 | Distal Intergenic | 3 | 38503408 | 38538993 | 35586 | 2 | 99696 | ENSMUST00000120875.8 | -42827 | ENSMUSG00000044864 | Ankrd50 | ankyrin repeat domain 50 |
| chr2 | 71557650 | 71558227 | 578 | * | 7,575113 | 4,177868418 | 1,348759 | 3,097565 | 0,001951 | 1 | 0 | Intron (ENSMUST0000 | 2 | 71549761 | 71561310 | 11550 | 1 | 381371 | ENSMUST00000079720.6 | 7889 | ENSMUSG00000064263 | Platr26 | pluripotency associated transcript 26 |
| chr2 | 1,29E+08 | 1,29E+08 | 771 | * | 7,506761 | 4,180581431 | 1,486449 | 2,812462 | 0,004916 | 1 | 0 | Promoter (<=1kb) | 2 | 1,29E+08 | 1,29E+08 | 4421 | 2 | 17222 | ENSMUST00000143007.2 | 0 | ENSMUSG00000014355 | Anapc1 | anaphase promoting complex subunit 1 |
| chr12 | 80030299 | 80031540 | 1242 | * | 11,09269 | 4,180940659 | 1,204429 | 3,471305 | 0,000518 | 0,44621096 | 0,918255 | Distal Intergenic | 12 | 79964619 | 79983038 | 18420 | 2 | 77290 | ENSMUST00000219857.2 | -47261 | ENSMUSG00000112765 | 9430078K24Rik | RIKEN cDNA 9430078K24 gene |
| chr7 | 1,35E+08 | 1,35E+08 | 868 | * | 7,576289 | 4,18186415 | 1,587046 | 2,634998 | 0,008414 | 1 | 0 | Intron (ENSMUST0000 | 7 | 1,34E+08 | 1,35E+08 | 54376 | 2 | 627214 | ENSMUST00000171394.3 | -71001 | ENSMUSG00000073805 | Insyn2a | inhibitory synaptic factor 2A |
| chr7 | 1,18E+08 | 1,18E+08 | 1301 | * | 14,86414 | 4,182864158 | 1,202187 | 3,479379 | 0,000503 | 0,369614065 | 1,37683 | Distal Intergenic | 7 | 1,18E+08 | 1,18E+08 | 1687 | 1 | 67745 | ENSMUST00000138180.2 | 8548 | ENSMUSG00000055159 | 4930583K01Rik | RIKEN cDNA 4930583K01 gene |
| chr5 | 21916876 | 21918333 | 1458 | * | 11,68064 | 4,182898804 | 1,399162 | 2,989574 | 0,002794 | 0,697016988 | 1,173325 | Distal Intergenic | 5 | 21867901 | 21906394 | 38494 | 2 | 242864 | ENSMUST00000060899.9 | -10482 | ENSMUSG00000044968 | Napepld | N-acyl phosphatidylethanolamine phospholipase D |
| chr13 | 1,04E+08 | 1,04E+08 | 755 | * | 6,414214 | 4,182950345 | 1,390054 | 3,0092 | 0,002619 | 1 | 0 | Exon (ENSMUST00000 | 13 | 1,04E+08 | 1,04E+08 | 21007 | 1 | 60411 | ENSMUST00000224500.2 | -3007 | ENSMUSG00000021714 | Cenpk | centromere protein K |
| chr2 | 1,3E+08 | 1,3E+08 | 1165 | * | 11,39731 | 4,183307352 | 1,331726 | 3,141267 | 0,001682 | 0,597812173 | 1,2763 | Distal Intergenic | 2 | 1,3E+08 | 1,3E+08 | 28458 | 1 | 67333 | ENSMUST00000165413.9 | -39184 | ENSMUSG00000037885 | Stk35 | serine/threonine kinase 35 |
| chr6 | 26132546 | 26133687 | 1142 | * | 15,47914 | 4,184776137 | 1,116437 | 3,748332 | 0,000178 | 0,225397758 | 1,41696 | Distal Intergenic | 6 | 25954484 | 25958158 | 3675 | 1 | 626904 | ENSMUST00000201080.2 | 178062 | ENSMUSG00000106799 | Gm20756 | predicted gene, 20756 |
| chr4 | 1,49E+08 | 1,49E+08 | 585 | * | 6,761304 | 4,185095889 | 1,602285 | 2,611955 | 0,009003 | 1 | 0 | Intron (ENSMUST0000 | 4 | 1,49E+08 | 1,49E+08 | 2094 | 2 | 230908 | ENSMUST00000186317.2 | 6528 | ENSMUSG00000041459 | Tardbp | TAR DNA binding protein |
| chr5 | 5587652 | 5588861 | 1210 | * | 9,405678 | 4,187407935 | 1,292527 | 3,239706 | 0,001197 | 0,54122637 | 1,211315 | 3' UTR | 5 | 5591922 | 5593446 | 1525 | 2 | 207704 | ENSMUST00000127670.2 | 4585 | ENSMUSG00000040464 | Gtpbp10 | GTP-binding protein 10 (putative) |
| chr13 | 95117782 | 95119229 | 1448 | * | 9,391167 | 4,188136689 | 1,350387 | 3,101434 | 0,001926 | 0,603552508 | 1,425929 | Intron (ENSMUST0000 | 13 | 95112946 | 95150985 | 38040 | 1 | 218460 | ENSMUST00000167155.2 | 4836 | ENSMUSG00000042015 | Wdr41 | WD repeat domain 41 |
| chr4 | 1,35E+08 | 1,35E+08 | 981 | * | 10,34994 | 4,188609964 | 1,247446 | 3,357748 | 0,000786 | 0,47475665 | 1,173325 | Distal Intergenic | 4 | 1,35E+08 | 1,35E+08 | 2171 | 1 | 1,03E+08 | ENSMUST00000145711.2 | -8476 | | Gm12984 | predicted gene 12984 |
| chr14 | 27092956 | 27093860 | 905 | * | 9,655759 | 4,18869324 | 1,229797 | 3,406003 | 0,000659 | 0,410415129 | 1,449706 | Intron (ENSMUST0000 | 14 | 27083346 | 27101604 | 18259 | 1 | 71704 | ENSMUST00000225494.2 | 9610 | ENSMUSG00000021895 | Arhgef3 | Rho guanine nucleotide exchange factor (GEF) 3 |
| chr5 | 1,08E+08 | 1,08E+08 | 790 | * | 8,326778 | 4,190773022 | 1,420571 | 2,950061 | 0,003177 | 0,685026746 | 1,425929 | Distal Intergenic | 5 | 1,08E+08 | 1,08E+08 | 79033 | 2 | 67266 | ENSMUST00000031198.11 | -23912 | ENSMUSG00000029270 | Dipk1a | divergent protein kinase domain 1A |
| chr10 | 53136265 | 53136986 | 722 | * | 6,663384 | 4,191121729 | 1,40522 | 2,982538 | 0,002859 | 1 | 0 | Exon (ENSMUST00002 | 10 | 53154141 | 53156244 | 2104 | 2 | 1E+08 | ENSMUST00000219673.2 | 19258 | ENSMUSG00000038594 | Cep85l | centrosomal protein 85-like |
| chr8 | 70768980 | 70769621 | 642 | * | 8,160898 | 4,191369598 | 1,351481 | 3,101316 | 0,001927 | 1 | 0 | Promoter (<=1kb) | 8 | 70768437 | 70784237 | 15801 | 1 | 93898 | ENSMUST00000140239.4 | 543 | ENSMUSG00000087408 | Cers1 | ceramide synthase 1 |
| chr15 | 97626048 | 97626427 | 380 | * | 5,881118 | 4,191509974 | 1,604443 | 2,612439 | 0,00899 | 1 | 0 | Promoter (2-3kb) | 15 | 97609772 | 97629220 | 19449 | 2 | 19011 | ENSMUST00000100249.10 | 2793 | ENSMUSG00000022468 | Endou | endonuclease, polyU-specific |
| chr1 | 1,35E+08 | 1,35E+08 | 787 | * | 7,025176 | 4,192552426 | 1,582691 | 2,649002 | 0,008073 | 1 | 0 | Exon (ENSMUST00020 | 1 | 1,35E+08 | 1,35E+08 | 2391 | 2 | 77552 | ENSMUST00000159918.2 | 19693 | ENSMUSG00000041889 | Shisa4 | shisa family member 4 |
| chr2 | 20143919 | 20144628 | 710 | * | 7,074295 | 4,193070419 | 1,578174 | 2,656913 | 0,007886 | 1 | 0 | Intron (ENSMUST0000 | 2 | 20285330 | 20285436 | 107 | 1 | 1,15E+08 | ENSMUST00000083319.3 | -140702 | ENSMUSG00000118880 | Gm23970 | predicted gene, 23970 |
| chrX | 44359814 | 44360951 | 1138 | * | 7,61162 | 4,194386203 | 1,342906 | 3,123366 | 0,001788 | 1 | 0 | Distal Intergenic | 20 | 44179781 | 44181668 | 1888 | 1 | 68800 | ENSMUST00000040002.4 | 180033 | ENSMUSG00000037086 | Prr32 | proline rich 32 |
| chr5 | 31768218 | 31769519 | 1302 | * | 12,61177 | 4,195305224 | 1,324297 | 3,16795 | 0,001535 | 0,596145184 | 1,173325 | Promoter (1-2kb) | 5 | 31771283 | 31779988 | 8706 | 1 | 66845 | ENSMUST00000031024.14 | -1764 | ENSMUSG00000106918 | Mrpl33 | mitochondrial ribosomal protein L33 |
| chr5 | 90502233 | 90503066 | 834 | * | 8,711813 | 4,195470031 | 1,334636 | 3,143531 | 0,001669 | 1 | 0 | Intron (ENSMUST0000 | 5 | 90401052 | 90514044 | 112993 | 2 | 81702 | ENSMUST00000218526.2 | 10978 | ENSMUSG00000055204 | Ankrd17 | ankyrin repeat domain 17 |
| chr11 | 32331606 | 32332420 | 815 | * | 8,996649 | 4,195865524 | 1,30838 | 3,206916 | 0,001342 | 0,563858922 | 1,211315 | Intron (ENSMUST0000 | 11 | 32297840 | 32340300 | 42461 | 1 | 268396 | ENSMUST00000138771.2 | 33766 | ENSMUSG00000040711 | Sh3pxd2b | SH3 and PX domains 2B |
| chr4 | 43923285 | 43924581 | 1297 | * | 10,08 | 4,195881205 | 1,281986 | 3,272955 | 0,001064 | 0,541053194 | 1,086118 | Intron (ENSMUST0000 | 4 | 43940398 | 43942382 | 1985 | 1 | 53614 | ENSMUST00000130415.2 | -15817 | ENSMUSG00000028476 | Reck | reversion-inducing-cysteine-rich protein with kazal motifs |
| chr11 | 63369691 | 63370531 | 841 | * | 7,086994 | 4,195911174 | 1,452596 | 2,888561 | 0,00387 | 1 | 0 | Distal Intergenic | 11 | 63023894 | 63049392 | 25499 | 1 | 18858 | ENSMUST00000108700.2 | 345797 | ENSMUSG00000018217 | Pmp22 | peripheral myelin protein 22 |
| chr18 | 88048830 | 88049501 | 672 | * | 6,138241 | 4,196328979 | 1,590317 | 2,638675 | 0,008323 | 1 | 0 | Distal Intergenic | 18 | 87943555 | 87943688 | 134 | 2 | 1,15E+08 | ENSMUST00000104453.3 | -105142 | ENSMUSG00000077641 | Gm24987 | predicted gene, 24987 |
| chr7 | 30072574 | 30073664 | 1091 | * | 10,42027 | 4,197257318 | 1,403702 | 2,990134 | 0,002789 | 0,684274806 | 1,2763 | Distal Intergenic | 7 | 30054914 | 30062197 | 7284 | 2 | 233067 | ENSMUST00000046351.7 | -10377 | ENSMUSG00000036957 | Lrfn3 | leucine rich repeat and fibronectin type III domain containing 3 |
| chr9 | 42914234 | 42915160 | 927 | * | 7,19315 | 4,198109492 | 1,496642 | 2,805019 | 0,005031 | 1 | 0 | Intron (ENSMUST0000 | 9 | 42937300 | 42939114 | 1815 | 2 | 69632 | ENSMUST00000214175.2 | 23954 | ENSMUSG00000059495 | Arhgef12 | Rho guanine nucleotide exchange factor (GEF) 12 |
| chr5 | 31889247 | 31890657 | 1411 | * | 9,022933 | 4,199126952 | 1,480753 | 2,835805 | 0,004571 | 0,756538269 | 1,360622 | Intron (ENSMUST0000 | 5 | 31859189 | 32215015 | 355827 | 1 | 107976 | ENSMUST00000200705.2 | 30058 | ENSMUSG00000052139 | Babam2 | BRISC and BRCA1 A complex member 2 |
| chr13 | 88434417 | 88435328 | 912 | * | 7,358457 | 4,201142299 | 1,402103 | 2,996315 | 0,002733 | 0,67266775 | 1,360622 | Distal Intergenic | 13 | 88669309 | 88726287 | 56979 | 1 | 1,03E+08 | ENSMUST00000224910.2 | -233981 | ENSMUSG00000114273 | Gm34585 | predicted gene, 34585 |
| chr11 | 58696719 | 58698039 | 1321 | * | 11,18617 | 4,201459626 | 1,315829 | 3,193015 | 0,001408 | 0,57981669 | 1,173325 | Distal Intergenic | 11 | 58683637 | 58692786 | 9150 | 2 | 75429 | ENSMUST00000060581.5 | -3933 | ENSMUSG00000049154 | Fam183b | family with sequence similarity 183, member B |
| chr12 | 80207604 | 80208851 | 1248 | * | 9,62059 | 4,20201693 | 1,418492 | 2,962312 | 0,003053 | 0,728215953 | 1,086118 | Distal Intergenic | 12 | 80215139 | 80216925 | 1787 | 2 | 109711 | ENSMUST00000219382.2 | 8074 | ENSMUSG00000015143 | Actn1 | actinin, alpha 1 |
| chr7 | 12687117 | 12688325 | 1209 | * | 8,60371 | 4,202239118 | 1,287191 | 3,264659 | 0,001096 | 0,505078837 | 1,360622 | Intron (ENSMUST0000 | 7 | 12682518 | 12695753 | 13236 | 2 | 69086 | ENSMUST00000181128.3 | 7428 | ENSMUSG00000097115 | 1810019N24Rik | RIKEN cDNA 1810019N24 gene |
| chr15 | 83368115 | 83368906 | 792 | * | 7,776884 | 4,204228446 | 1,389164 | 3,026446 | 0,002474 | 1 | 0 | 3' UTR | 15 | 83259808 | 83348807 | 89000 | 2 | 23970 | ENSMUST00000171436.8 | -19308 | ENSMUSG00000016664 | Pacsin2 | protein kinase C and casein kinase substrate in neurons 2 |
| chr5 | 80442618 | 80443615 | 998 | * | 8,264106 | 4,204646101 | 1,253079 | 3,355452 | 0,000792 | 0,440933973 | 1,449706 | Distal Intergenic | 5 | 80246755 | 80246861 | 107 | 1 | 1,15E+08 | ENSMUST00000239632.1 | 195863 | ENSMUSG00000119601 | Gm24304 | predicted gene, 24304 |
| chr2 | 1,3E+08 | 1,3E+08 | 2107 | * | 17,36563 | 4,204745365 | 1,096421 | 3,834973 | 0,000126 | 0,125710024 | 2,663619 | Intron (ENSMUST0000 | 2 | 1,3E+08 | 1,3E+08 | 103480 | 1 | 19262 | ENSMUST00000077303.4 | 24558 | ENSMUSG00000027303 | Ptpra | protein tyrosine phosphatase, receptor type, A |
| chr5 | 71640453 | 71641464 | 1012 | * | 9,60789 | 4,205354724 | 1,251031 | 3,361512 | 0,000775 | 0,473016631 | 1,173325 | Intron (ENSMUST0000 | 5 | 71600167 | 71705548 | 105382 | 2 | 78174 | ENSMUST00000050129.6 | 64084 | ENSMUSG00000049387 | Cox7b2 | cytochrome c oxidase subunit 7B2 |
| chr5 | 1,35E+08 | 1,35E+08 | 750 | * | 7,549465 | 4,205890847 | 1,416816 | 2,968552 | 0,002992 | 1 | 0 | Promoter (2-3kb) | 5 | 1,35E+08 | 1,35E+08 | 19770 | 2 | 22384 | ENSMUST00000202622.4 | -2896 | ENSMUSG00000040731 | Eif4h | eukaryotic translation initiation factor 4H |
| chr12 | 83712358 | 83713202 | 845 | * | 6,565261 | 4,206243217 | 1,538753 | 2,733541 | 0,006266 | 1 | 0 | Promoter (2-3kb) | 12 | 83709750 | 83710773 | 1024 | 1 | 67039 | ENSMUST00000183282.2 | 2608 | ENSMUSG00000010608 | Rbm25 | RNA binding motif protein 25 |
| chr3 | 88649796 | 88651346 | 1551 | * | 16,235687 | 4,207041152 | 0,9320499 | 4,5137512 | 6,369E-06 | 0,019779503 | 2,1416125 | Distal Intergenic | 3 | 88652007 | 88659358 | 7352 | 2 | 229521 | ENSMUST00000107503.8 | 8012 | ENSMUSG00000068923 | Syt11 | synaptotagmin XI |
| chr10 | 79550381 | 79551599 | 1219 | * | 10,08201 | 4,207053584 | 1,312705 | 3,204873 | 0,001351 | 0,565091829 | 1,211315 | Promoter (<=1kb) | 10 | 79552468 | 79571942 | 19475 | 1 | 15166 | ENSMUST00000020581.3 | -869 | ENSMUSG00000020331 | Hcn2 | hyperpolarization-activated, cyclic nucleotide-gated K+ 2 |
| chr2 | 30967335 | 30967843 | 509 | * | 5,564107 | 4,207070008 | 1,591909 | 2,642784 | 0,008223 | 1 | 0 | Intron (ENSMUST0000 | 2 | 30916219 | 30986512 | 70294 | 2 | 14269 | ENSMUST00000113559.10 | 18669 | ENSMUSG00000075415 | Fnbp1 | formin binding protein 1 |
| chr7 | 23515077 | 23515903 | 827 | * | 6,870124 | 4,207148073 | 1,594597 | 2,638376 | 0,00833 | 1 | 0 | Distal Intergenic | 7 | 23507711 | 23508625 | 915 | 2 | 622222 | ENSMUST00000166141.2 | -6452 | ENSMUSG00000095632 | Vmn1r175 | vomeronasal 1 receptor 175 |
| chr19 | 48057123 | 48057431 | 309 | * | 5,79824 | 4,207593914 | 1,564801 | 2,6889 | 0,007169 | 1 | 0 | Distal Intergenic | 19 | 47926151 | 48023826 | 97676 | 1 | 381229 | ENSMUST00000066308.9 | 130972 | ENSMUSG00000046585 | Cfap58 | cilia and flagella associated protein 58 |
| chr4 | 57717383 | 57718277 | 895 | * | 7,571202 | 4,207705433 | 1,520304 | 2,767674 | 0,005646 | 1 | 0 | Promoter (<=1kb) | 4 | 57717657 | 57856922 | 139266 | 1 | 677884 | ENSMUST00000098065.4 | 0 | ENSMUSG00000038729 | Pakap | paralemmin A kinase anchor protein |
| chr9 | 53331534 | 53333037 | 1504 | * | 16,72212 | 4,209031408 | 1,052497 | 3,999093 | 6,36E-05 | 0,131142082 | 1,2763 | 5' UTR | 9 | 53316598 | 53344020 | 27423 | 1 | 75311 | ENSMUST00000217318.2 | 14936 | ENSMUSG00000005131 | 4930550C14Rik | RIKEN cDNA 4930550C14 gene |
| chr11 | 3780445 | 3781336 | 892 | * | 8,179595 | 4,209228468 | 1,491194 | 2,822724 | 0,004762 | 0,752939853 | 1,449706 | Intron (ENSMUST0000 | 11 | 3653731 | 3813903 | 160173 | 2 | 74309 | ENSMUST00000070552.14 | 32567 | ENSMUSG00000020435 | Osbp2 | oxysterol binding protein 2 |
| chr8 | 28458318 | 28458838 | 521 | * | 6,608869 | 4,211186836 | 1,545866 | 2,72416 | 0,006447 | 1 | 0 | Distal Intergenic | 8 | 28572722 | 28574294 | 1573 | 2 | 666422 | ENSMUST00000210185.2 | 115456 | ENSMUSG00000110372 | Gm8096 | 3-phosphoglycerate dehydrogenase pseudogene |
| chr3 | 39695967 | 39696837 | 871 | * | 6,705087 | 4,211188423 | 1,530379 | 2,751729 | 0,005928 | 1 | 0 | Distal Intergenic | 3 | 39690113 | 39690231 | 119 | 2 | 1,15E+08 | ENSMUST00000083302.3 | -5736 | ENSMUSG00000118708 | n-R5s195 | nuclear encoded rRNA 5S 195 |
| chr7 | 1,28E+08 | 1,28E+08 | 929 | * | 5,636482 | 4,211436439 | 1,59906 | 2,633696 | 0,008446 | 1 | 0 | Promoter (<=1kb) | 7 | 1,28E+08 | 1,28E+08 | 1614 | 1 | 233908 | ENSMUST00000172755.2 | 100 | ENSMUSG00000030795 | Fus | fused in sarcoma |
| chrX | 24613680 | 24614695 | 1016 | * | 8,158461 | 4,211450169 | 1,547134 | 2,722097 | 0,006487 | 0,821456993 | 1,360622 | Distal Intergenic | 20 | 24619505 | 24639814 | 20310 | 2 | 546282 | ENSMUST00000178986.2 | 25119 | ENSMUSG00000093923 | Gm5935 | predicted gene 5935 |
| chr10 | 85018539 | 85020038 | 1500 | * | 8,864954 | 4,211533004 | 1,524471 | 2,762619 | 0,005734 | 0,791416391 | 1,425929 | Promoter (<=1kb) | 10 | 84967564 | 85020928 | 53365 | 2 | 12952 | ENSMUST00000020227.11 | 890 | ENSMUSG00000020038 | Cry1 | cryptochrome 1 (photolyase-like) |
| chr2 | 1,56E+08 | 1,56E+08 | 1520 | * | 12,995 | 4,211893055 | 1,234131 | 3,412841 | 0,000643 | 0,48037412 | 0,918255 | 3' UTR | 2 | 1,56E+08 | 1,56E+08 | 23486 | 2 | 56046 | ENSMUST00000162258.8 | 22718 | ENSMUSG00000005882 | Uqcc1 | ubiquinol-cytochrome c reductase complex assembly factor 1 |
| chr2 | 1,02E+08 | 1,02E+08 | 825 | * | 7,31419 | 4,212031145 | 1,47269 | 2,860093 | 0,004235 | 1 | 0 | Intron (ENSMUST0000 | 2 | 1,02E+08 | 1,02E+08 | 124696 | 2 | 72446 | ENSMUST00000163762.8 | 10942 | ENSMUSG00000032841 | Prr5l | proline rich 5 like |

| | | | | | | | | | | | | | | | | | | | | | | | |
|---|---|---|---|---|---|---|---|---|---|---|---|---|---|---|---|---|---|---|---|---|---|---|---|
| chrX | 47602362 | 47603530 | 1169 | * | 8,615878 | 4,212289132 | 1,293457 | 3,256613 | 0,001128 | 0,502272318 | 1,425929 | Promoter (<=1kb) | 20 | 47563821 | 47602436 | 38616 | 2 | 26926 | ENSMUST00000114945.9 | 0 | ENSMUSG00000036932 | Aifm1 | apoptosis-inducing factor, mitochondrion-associated 1 |
| chr4 | 80490254 | 80491416 | 1163 | * | 9,543529 | 4,212828787 | 1,268533 | 3,321024 | 0,000897 | 0,464166555 | 1,425929 | Distal Intergenic | 4 | 80752360 | 80755921 | 3562 | 1 | 22178 | ENSMUST00000133655.8 | -260944 | ENSMUSG00000005994 | Tyrp1 | tyrosinase-related protein 1 |
| chr18 | 39124654 | 39126558 | 1905 | * | 18,82583 | 4,214105711 | 1,081161 | 3,897761 | 9,71E-05 | 0,115300307 | 2,386008 | Promoter (<=1kb) | 18 | 39126198 | 39233202 | 107005 | 1 | 71302 | ENSMUST00000115574.2 | 0 | ENSMUSG00000036452 | Arhgap26 | Rho GTPase activating protein 26 |
| chr11 | 75613764 | 75614987 | 1224 | * | 8,812866 | 4,21665414 | 1,319817 | 3,194877 | 0,001399 | 0,571106708 | 1,211315 | Distal Intergenic | 11 | 75623695 | 75656671 | 32977 | 1 | 22627 | ENSMUST00000067664.10 | -8708 | ENSMUSG00000020849 | Ywhae | tyrosine 3-monooxygenase/tryptophan 5-monooxygenase activation protein, epsilon polypeptide |
| chr7 | 89911028 | 89912030 | 1003 | * | 8,525111 | 4,21767842 | 1,356841 | 3,108453 | 0,001881 | 0,512326648 | 2,237403 | Promoter (2-3kb) | 7 | 89873081 | 89914626 | 41546 | 2 | 75338 | ENSMUST00000107221.9 | 2596 | ENSMUSG00000030617 | Ccdc83 | coiled-coil domain containing 83 |
| chr15 | 99623158 | 99624461 | 1304 | * | 22,368944 | 4,217812588 | 0,9168307 | 4,6004269 | 4,216E-06 | 0,01138298 | 2,8800171 | Promoter (<=1kb) | 15 | 99623528 | 99627413 | 3886 | 1 | 66379 | ENSMUST00000023761.4 | 0 | ENSMUSG00000023020 | Cox14 | cytochrome c oxidase assembly protein 14 |
| chr13 | 45980517 | 45981974 | 1458 | * | 15,99374 | 4,218605219 | 1,02534 | 4,114349 | 3,88E-05 | 0,067555009 | 2,318434 | Intron (ENSMUST00000 | 13 | 45949019 | 46025764 | 76746 | 2 | 20238 | ENSMUST00000222610.2 | 43790 | ENSMUSG00000046876 | Atxn1 | ataxin 1 |
| chr11 | 1,13E+08 | 1,13E+08 | 897 | * | 9,987841 | 4,219292293 | 1,389587 | 3,036364 | 0,002394 | 0,670666828 | 1,211315 | Distal Intergenic | 11 | 1,13E+08 | 1,13E+08 | 129057 | 2 | 72386 | ENSMUST00000244333.1 | 182768 | ENSMUSG00000087259 | 2610035D17Rik | RIKEN cDNA 2610035D17 gene |
| chr5 | 70774263 | 70775474 | 1212 | * | 6,63073 | 4,219612127 | 1,516318 | 2,782802 | 0,005389 | 1 | 0 | Distal Intergenic | 5 | 70708184 | 70708290 | 107 | 2 | 1,15E+08 | ENSMUST00000240250.2 | -65973 | ENSMUSG00000118789 | Gm26072 | predicted gene, 26072 |
| chr5 | 68601013 | 68602106 | 1094 | * | 10,08168 | 4,219648278 | 1,437031 | 2,936366 | 0,003321 | 0,723215019 | 1,211315 | Distal Intergenic | 5 | 68723513 | 68723639 | 127 | 2 | 1,15E+08 | ENSMUST00000157255.3 | 121533 | ENSMUSG00000087880 | Gm23605 | predicted gene, 23605 |
| chr2 | 1,67E+08 | 1,67E+08 | 917 | * | 8,97558 | 4,220257638 | 1,445662 | 2,919255 | 0,003509 | 0,705327871 | 1,425929 | Intron (ENSMUST00000 | 2 | 1,67E+08 | 1,67E+08 | 3558 | 1 | 68941 | ENSMUST00000151217.8 | -27372 | | 1110018N20Rik | RIKEN cDNA 1110018N20 gene |
| chr14 | 88838303 | 88839275 | 973 | * | 7,064356 | 4,22037575 | 1,551516 | 2,720162 | 0,006525 | 1 | 0 | Distal Intergenic | 14 | 88873469 | 89001197 | 127729 | 1 | 1,05E+08 | ENSMUST00000227074.2 | -34194 | | Gm41225 | predicted gene, 41225 |
| chr12 | 85498688 | 85499370 | 683 | * | 10,38498 | 4,220395089 | 1,255468 | 3,361612 | 0,000775 | 0,4572736 | 1,2763 | Distal Intergenic | 12 | 85475884 | 85483023 | 7140 | 2 | 1,05E+08 | ENSMUST00000221505.2 | -15665 | | Gm40477 | predicted gene, 40477 |
| chr12 | 59107371 | 59108519 | 1149 | * | 8,698522 | 4,22151071 | 1,489363 | 2,83444 | 0,004591 | 1 | 0 | Promoter (<=1kb) | 12 | 59089878 | 59108247 | 18370 | 2 | 78232 | ENSMUST00000021380.10 | 0 | ENSMUSG00000020993 | Trappc6b | trafficking protein particle complex 6B |
| chr2 | 1,28E+08 | 1,28E+08 | 778 | * | 6,119889 | 4,22206803 | 1,548953 | 2,725757 | 0,006415 | 1 | 0 | Intron (ENSMUST00000 | 2 | 1,28E+08 | 1,28E+08 | 15009 | 2 | 1E+08 | ENSMUST00000239358.2 | 58440 | ENSMUSG00000074813 | Morrbid | myeloid RNA regulator of BCL2L11 induced cell death |
| chr11 | 43575159 | 43576243 | 1085 | * | 7,497111 | 4,222497487 | 1,47791 | 2,857073 | 0,004276 | 1 | 0 | Promoter (<=1kb) | 11 | 43574972 | 43598018 | 23047 | 1 | 70802 | ENSMUST00000109280.2 | 187 | ENSMUSG00000044950 | Pwwp2a | PWWP domain containing 2A |
| chr16 | 24561595 | 24562275 | 681 | * | 8,403773 | 4,223908718 | 1,345628 | 3,138988 | 0,001695 | 1 | 0 | Intron (ENSMUST00000 | 16 | 24540605 | 24798280 | 257676 | 1 | 210126 | ENSMUST00000115314.4 | 20990 | ENSMUSG00000033306 | Lpp | LIM domain containing preferred translocation partner in lipoma |
| chr7 | 79153776 | 79154509 | 734 | * | 9,991188 | 4,224284789 | 1,28315 | 3,29212 | 0,000994 | 0,479521813 | 1,425929 | Promoter (<=1kb) | 7 | 79155012 | 79155101 | 90 | 1 | 723968 | ENSMUST00000175497.3 | -503 | ENSMUSG00000093238 | Mir9-3 | microRNA 9-3 |
| chr19 | 15950557 | 15951652 | 1096 | * | 8,223301 | 4,22533596 | 1,486286 | 2,842881 | 0,004471 | 0,671902369 | 2,237403 | Intron (ENSMUST00000 | 19 | 15933345 | 15962057 | 28713 | 2 | 208518 | ENSMUST00000236749.2 | 10405 | ENSMUSG00000041491 | Cep78 | centrosomal protein 78 |
| chr12 | 84463645 | 84464802 | 1158 | * | 8,284429 | 4,225969658 | 1,614788 | 2,617043 | 0,00887 | 0,789538074 | 2,237403 | Promoter (<=1kb) | 12 | 84463970 | 84465352 | 1383 | 1 | 66381 | ENSMUST00000183146.2 | 0 | ENSMUSG00000098134 | Rnf113a2 | ring finger protein 113A2 |
| chr15 | 7270234 | 7271296 | 1063 | * | 9,239514 | 4,226872373 | 1,441181 | 2,932923 | 0,003358 | 0,692179368 | 1,449706 | Intron (ENSMUST00000 | 15 | 7282992 | 7284249 | 1258 | 2 | 268780 | ENSMUST00000162105.2 | 12953 | ENSMUSG00000042961 | Egflam | EGF-like, fibronectin type III and laminin G domains |
| chr11 | 1,2E+08 | 1,2E+08 | 1546 | * | 12,27201 | 4,228208807 | 1,286966 | 3,285408 | 0,001018 | 0,519493717 | 1,173325 | Distal Intergenic | 11 | 1,2E+08 | 1,2E+08 | 2034 | 2 | 1E+08 | ENSMUST00000243021.1 | -18954 | | 2810410L24Rik | RIKEN cDNA 2810410L24 gene |
| chr15 | 36166666 | 36168518 | 1853 | * | 13,41899 | 4,228531339 | 1,14892 | 3,680439 | 0,000233 | 0,342431125 | 0,738441 | Promoter (1-2kb) | 15 | 36169942 | 36170100 | 159 | 1 | 1,15E+08 | ENSMUST00000082609.3 | -1424 | ENSMUSG00000064543 | Gm22979 | predicted gene, 22979 |
| chr1 | 64448920 | 64449759 | 840 | * | 12,64839 | 4,231717462 | 1,270923 | 3,329641 | 0,00087 | 0,492478878 | 1,173325 | Distal Intergenic | 1 | 64571804 | 64636854 | 65051 | 1 | 12912 | ENSMUST00000185594.7 | -122045 | ENSMUSG00000025958 | Creb1 | cAMP responsive element binding protein 1 |
| chr7 | 1,22E+08 | 1,22E+08 | 1430 | * | 9,047015 | 4,231751102 | 1,538619 | 2,750357 | 0,005953 | 0,804972409 | 1,360622 | Exon (ENSMUST000002 | 7 | 1,22E+08 | 1,22E+08 | 41259 | 1 | 18751 | ENSMUST00000149583.2 | -10366 | ENSMUSG00000052889 | Prkcb | protein kinase C, beta |
| chr5 | 81522526 | 81523331 | 806 | * | 6,437483 | 4,232222635 | 1,401157 | 3,02052 | 0,002523 | 1 | 0 | Intron (ENSMUST00000 | 5 | 81457872 | 81944481 | 486610 | 1 | 319387 | ENSMUST00000072521.11 | 64654 | ENSMUSG00000037605 | Adgrl3 | adhesion G protein-coupled receptor L3 |
| chr10 | 128401385 | 128402819 | 1435 | * | 28,147801 | 4,233668395 | 0,7444795 | 5,6867493 | 1,295E-08 | 0,000126258 | 2,9510621 | Promoter (<=1kb) | 10 | 128393635 | 128401803 | 8169 | 2 | 18813 | ENSMUST00000026425.13 | 0 | ENSMUSG00000025364 | Pa2g4 | proliferation-associated 2G4 |
| chr6 | 11797920 | 11798655 | 736 | * | 6,168119 | 4,234573654 | 1,563591 | 2,708236 | 0,006764 | 1 | 0 | Distal Intergenic | 6 | 11900371 | 11907392 | 7022 | 2 | 17992 | ENSMUST00000204084.3 | 108737 | ENSMUSG00000029632 | Ndufa4 | Ndufa4, mitochondrial complex associated |
| chr11 | 1,2E+08 | 1,2E+08 | 1579 | * | 13,07776 | 4,234577383 | 1,075691 | 3,936613 | 8,26E-05 | 0,162641545 | 1,173325 | Exon (ENSMUST000002 | 11 | 1,2E+08 | 1,2E+08 | 5140 | 2 | 1,03E+08 | ENSMUST00000244839.1 | 3914 | | 2900052L18Rik | RIKEN cDNA 2900052L18 gene |
| chr2 | 1,2E+08 | 1,2E+08 | 1122 | * | 8,75016 | 4,235180079 | 1,50133 | 2,820953 | 0,004788 | 0,767061106 | 1,360622 | Intron (ENSMUST00000 | 2 | 1,2E+08 | 1,2E+08 | 35156 | 2 | 98878 | ENSMUST00000152366.8 | -6744 | ENSMUSG00000027293 | Ehd4 | EH-domain containing 4 |
| chr8 | 82188086 | 82188622 | 537 | * | 7,021547 | 4,235821342 | 1,494703 | 2,833888 | 0,004599 | 1 | 0 | Intron (ENSMUST00000 | 8 | 82069191 | 82851665 | 782475 | 1 | 234515 | ENSMUST00000215332.2 | 118895 | ENSMUSG00000037940 | Inpp4b | inositol polyphosphate-4-phosphatase, type II |
| chrX | 1,52E+08 | 1,52E+08 | 829 | * | 6,602073 | 4,23793875 | 1,579026 | 2,683894 | 0,007277 | 1 | 0 | Distal Intergenic | 20 | 1,52E+08 | 1,52E+08 | 3344 | 1 | 54609 | ENSMUST00000060714.10 | -20667 | ENSMUSG00000050148 | Ubqln2 | ubiquilin 2 |
| chr3 | 90151924 | 90153256 | 1333 | * | 8,939105 | 4,238985927 | 1,281849 | 3,306932 | 0,000943 | 0,39984309 | 2,237403 | Promoter (1-2kb) | 3 | 90144807 | 90150819 | 6013 | 2 | 78284 | ENSMUST00000029547.10 | -1105 | ENSMUSG00000027938 | Creb3l4 | cAMP responsive element binding protein 3-like 4 |
| chr1 | 1,35E+08 | 1,35E+08 | 923 | * | 8,324724 | 4,239467601 | 1,45785 | 2,908028 | 0,003637 | 0,709779237 | 1,449706 | Exon (ENSMUST000001 | 1 | 1,35E+08 | 1,35E+08 | 1758 | 2 | 329251 | ENSMUST00000146639.2 | 14231 | ENSMUSG00000073557 | Ppp1r12b | protein phosphatase 1, regulatory subunit 12B |
| chr8 | 47192789 | 47193659 | 871 | * | 7,780721 | 4,240716267 | 1,541009 | 2,751908 | 0,005925 | 1 | 0 | Promoter (<=1kb) | 8 | 47192796 | 47260410 | 67615 | 1 | 16363 | ENSMUST00000208507.2 | 0 | ENSMUSG00000031627 | Irf2 | interferon regulatory factor 2 |
| chr7 | 1,33E+08 | 1,33E+08 | 932 | * | 7,316322 | 4,241002897 | 1,641564 | 2,583513 | 0,00978 | 0,891097601 | 1,360622 | Distal Intergenic | 7 | 1,33E+08 | 1,33E+08 | 14207 | 1 | 74970 | ENSMUST00000209850.2 | 135814 | ENSMUSG00000109816 | 4930483O08Rik | RIKEN cDNA 4930483O08 gene |
| chr15 | 83230665 | 83231548 | 884 | * | 8,833642 | 4,242153899 | 1,355266 | 3,130125 | 0,001747 | 0,500183011 | 2,237403 | Promoter (2-3kb) | 15 | 83184022 | 83234375 | 50354 | 2 | 66251 | ENSMUST00000226124.2 | 2827 | ENSMUSG00000054277 | Arfgap3 | ADP-ribosylation factor GTPase activating protein 3 |
| chr2 | 1,48E+08 | 1,48E+08 | 908 | * | 8,19269 | 4,243211764 | 1,489655 | 2,848453 | 0,004393 | 0,669639313 | 2,237403 | Promoter (2-3kb) | 2 | 1,48E+08 | 1,48E+08 | 313 | 1 | 1,15E+08 | ENSMUST00000158833.4 | 2439 | ENSMUSG00000089458 | Gm24221 | predicted gene, 24221 |
| chr16 | 25235666 | 25238836 | 3171 | * | 22,66694 | 4,243260274 | 0,9573932 | 4,4320976 | 9,332E-06 | 0,020861092 | 2,8800171 | Intron (ENSMUST000000 | 16 | 25105571 | 25136985 | 31415 | 1 | 71338 | ENSMUST00000128595.2 | 130095 | ENSMUSG00000048399 | Tprg | transformation related protein 63 regulated |
| chr11 | 49993200 | 49993888 | 689 | * | 5,973147 | 4,244046508 | 1,568714 | 2,70543 | 0,006822 | 1 | 0 | Intron (ENSMUST00000 | 11 | 49984501 | 49995585 | 11085 | 1 | 59044 | ENSMUST00000143607.2 | 8699 | ENSMUSG00000020376 | Rnf130 | ring finger protein 130 |
| chr12 | 94902538 | 94903099 | 562 | * | 5,319574 | 4,244286798 | 1,627736 | 2,607479 | 0,009121 | 1 | 0 | Distal Intergenic | 12 | 95345619 | 95345725 | 107 | 2 | 1,15E+08 | ENSMUST00000240345.1 | 442626 | ENSMUSG00001118669 | Gm26055 | predicted gene, 26055 |
| chr5 | 74195402 | 74196096 | 695 | * | 8,272786 | 4,245922792 | 1,531881 | 2,771706 | 0,005576 | 0,708539289 | 2,237403 | Promoter (<=1kb) | 5 | 74162460 | 74196302 | 33843 | 2 | 69727 | ENSMUST00000128750.2 | 206 | ENSMUSG00000054814 | Usp46 | ubiquitin specific peptidase 46 |
| chr15 | 38054473 | 38055932 | 1460 | * | 11,12689 | 4,248355455 | 1,420979 | 2,989737 | 0,002792 | 0,697016988 | 1,173325 | Promoter (2-3kb) | 15 | 38058529 | 38058664 | 136 | 1 | 1,15E+08 | ENSMUST00000083918.3 | -2597 | ENSMUSG00000065852 | Gm22353 | predicted gene, 22353 |
| chr12 | 60289044 | 60290122 | 1079 | * | 7,573565 | 4,248358533 | 1,49168 | 2,848037 | 0,004399 | 1 | 0 | Distal Intergenic | 12 | 60364674 | 60364850 | 177 | 1 | 1,03E+08 | ENSMUST00000219590.2 | -74552 | ENSMUSG00000112092 | Gm32936 | predicted gene, 32936 |
| chr15 | 74844000 | 74844743 | 744 | * | 6,061742 | 4,248892027 | 1,622154 | 2,619291 | 0,008811 | 1 | 0 | Distal Intergenic | 15 | 74851383 | 74855279 | 3897 | 2 | 57248 | ENSMUST00000023250.11 | 10536 | ENSMUSG00000022586 | Ly6i | lymphocyte antigen 6 complex, locus I |
| chr3 | 21006582 | 21007203 | 622 | * | 5,493047 | 4,249564532 | 1,479973 | 2,87138 | 0,004087 | 1 | 0 | Distal Intergenic | 3 | 20557262 | 20560372 | 3111 | 2 | 1,08E+08 | ENSMUST00000195457.2 | -446210 | | Gm31466 | predicted gene, 31466 |
| chr7 | 90086328 | 90086950 | 623 | * | 5,986478 | 4,249867158 | 1,611094 | 2,637876 | 0,008343 | 1 | 0 | Distal Intergenic | 7 | 90091937 | 90095312 | 3376 | 1 | 233490 | ENSMUST00000061767.5 | -4987 | ENSMUSG00000051451 | Crebzf | CREB/ATF bZIP transcription factor |
| chr8 | 40954766 | 40955404 | 639 | * | 7,051628 | 4,251773493 | 1,516107 | 2,804402 | 0,005041 | 1 | 0 | Promoter (<=1kb) | 8 | 40947055 | 40954113 | 7059 | 2 | 18983 | ENSMUST00000142455.8 | -653 | ENSMUSG00000031601 | Cnot7 | CCR4-NOT transcription complex, subunit 7 |
| chr17 | 34919329 | 34920096 | 768 | * | 6,721776 | 4,252336941 | 1,416775 | 3,001421 | 0,002687 | 1 | 0 | Intron (ENSMUST00000 | 17 | 34906670 | 34906776 | 107 | 2 | 1,15E+08 | ENSMUST00000239676.1 | -12553 | ENSMUSG00000118746 | Gm24389 | predicted gene, 24389 |
| chr13 | 43501682 | 43503090 | 1409 | * | 8,779687 | 4,254145654 | 1,454427 | 2,924964 | 0,003445 | 0,627785088 | 2,237403 | Distal Intergenic | 13 | 43518972 | 43531780 | 12809 | 1 | 68346 | ENSMUST00000222489.2 | -15882 | ENSMUSG00000054021 | Sirt5 | sirtuin 5 |
| chr11 | 1,15E+08 | 1,15E+08 | 1503 | * | 15,36901 | 4,255269075 | 1,301492 | 3,269531 | 0,001077 | 0,426043749 | 2,141612 | Distal Intergenic | 11 | 1,15E+08 | 1,15E+08 | 12582 | 1 | 70355 | ENSMUST00000133245.2 | -9023 | ENSMUSG00000051043 | Gprc5c | G protein-coupled receptor, family C, group 5, member C |
| chr8 | 96160309 | 96161072 | 764 | * | 8,211553 | 4,255720934 | 1,438233 | 2,958992 | 0,003086 | 0,683761897 | 1,425929 | Promoter (<=1kb) | 8 | 96146877 | 96161385 | 14509 | 2 | 14894 | ENSMUST00000212731.2 | 313 | ENSMUSG00000031796 | Cfap20 | cilia and flagella associated protein 20 |
| chr10 | 27271705 | 27272279 | 575 | * | 8,100721 | 4,25684573 | 1,533566 | 2,775783 | 0,005507 | 1 | 0 | Exon (ENSMUST000001 | 10 | 27265862 | 27273221 | 7360 | 1 | 75855 | ENSMUST00000187815.2 | 5843 | ENSMUSG00000101636 | 4930579H20Rik | RIKEN cDNA 4930579H20 gene |
| chr7 | 4410360 | 4411111 | 752 | * | 6,149001 | 4,25883297 | 1,568991 | 2,714377 | 0,00664 | 1 | 0 | Distal Intergenic | 7 | 4366964 | 4400743 | 33780 | 2 | 243816 | ENSMUST00000206928.2 | -9617 | ENSMUSG00000078810 | Gp6 | glycoprotein 6 (platelet) |
| chr8 | 1,26E+08 | 1,26E+08 | 487 | * | 7,264019 | 4,258971703 | 1,515726 | 2,809855 | 0,004956 | 1 | 0 | Distal Intergenic | 8 | 1,26E+08 | 1,26E+08 | 2543 | 1 | 1,03E+08 | ENSMUST00000245925.1 | -31946 | | Gm30713 | predicted gene, 30713 |
| chr11 | 78699095 | 78700460 | 1366 | * | 9,228014 | 4,259106356 | 1,404892 | 3,031626 | 0,002432 | 0,643386705 | 1,449706 | Distal Intergenic | 11 | 78717410 | 78737696 | 20287 | 1 | 66274 | ENSMUST00000147875.9 | -16950 | ENSMUSG00000072640 | Lyrm9 | LYR motif containing 9 |
| chr13 | 1,12E+08 | 1,12E+08 | 1480 | * | 9,705068 | 4,260758062 | 1,389273 | 3,066899 | 0,002163 | 0,623564689 | 1,449706 | Distal Intergenic | 13 | 1,12E+08 | 1,12E+08 | 11831 | 1 | 1,03E+08 | ENSMUST00000156109.2 | 57742 | | Gm15322 | predicted gene 15322 |
| chr18 | 55059120 | 55060407 | 1288 | * | 7,448215 | 4,261767086 | 1,582591 | 2,692905 | 0,007083 | 1 | 0 | Intron (ENSMUST00000 | 18 | 55039980 | 55052486 | 12507 | 1 | 1,19E+08 | ENSMUST00000244785.1 | 19140 | | Gm54030 | predicted gene, 54030 |
| chr5 | 32110610 | 32111463 | 854 | * | 7,720477 | 4,261964856 | 1,48761 | 2,864974 | 0,00417 | 1 | 0 | Intron (ENSMUST00000 | 5 | 32203322 | 32242097 | 38776 | 1 | 107976 | ENSMUST00000202080.2 | -91859 | ENSMUSG00000052139 | Babam2 | BRISC and BRCA1 A complex member 2 |
| chr6 | 72718586 | 72719349 | 764 | * | 10,33211 | 4,262218654 | 1,460691 | 2,917947 | 0,003523 | 0,750518846 | 1,086118 | Promoter (1-2kb) | 6 | 72721003 | 72727005 | 6003 | 1 | 1,03E+08 | ENSMUST00000247353.1 | -1654 | | Gm31249 | predicted gene, 31249 |
| chr12 | 1,12E+08 | 1,12E+08 | 897 | * | 8,271116 | 4,262250813 | 1,601689 | 2,661098 | 0,007789 | 1 | 0 | Distal Intergenic | 12 | 1,12E+08 | 1,12E+08 | 10480 | 1 | 319942 | ENSMUST00000057465.7 | -9013 | ENSMUSG00000043122 | A530016L24Rik | RIKEN cDNA A530016L24 gene |
| chr9 | 7834454 | 7836086 | 1633 | * | 11,66414 | 4,263352572 | 1,395066 | 3,056023 | 0,002243 | 0,740451144 | 0,738441 | Promoter (<=1kb) | 9 | 7818237 | 7835256 | 17020 | 2 | 11797 | ENSMUST00000074246.7 | 0 | ENSMUSG00000057367 | Birc2 | baculoviral IAP repeat-containing 2 |
| chr11 | 68898554 | 68899930 | 1377 | * | 11,48041 | 4,263581298 | 1,429667 | 2,982219 | 0,002862 | 0,703368984 | 1,173325 | Promoter (<=1kb) | 11 | 68876523 | 68899286 | 22764 | 2 | 237823 | ENSMUST00000021282.12 | 0 | ENSMUSG00000020899 | Pfas | phosphoribosylformylglycinamidine synthase (FGAR amidotransferase) |
| chr1 | 1,09E+08 | 1,09E+08 | 762 | * | 7,923545 | 4,264544651 | 1,369569 | 3,113787 | 0,001847 | 0,604229357 | 1,360622 | Distal Intergenic | 1 | 1,1E+08 | 1,1E+08 | 130 | 2 | 1,15E+08 | ENSMUST00000158412.3 | 314032 | ENSMUSG00000089037 | Gm22159 | predicted gene, 22159 |
| chr2 | 1,32E+08 | 1,32E+08 | 1356 | * | 11,89451 | 4,26575898 | 1,286916 | 3,314713 | 0,000917 | 0,54190677 | 0,918255 | Exon (ENSMUST000001 | 2 | 1,32E+08 | 1,32E+08 | 8163 | 2 | 70612 | ENSMUST00000110164.8 | 26694 | ENSMUSG00000027341 | Tmem230 | transmembrane protein 230 |
| chr10 | 60773793 | 60774645 | 853 | * | 8,671073 | 4,266952158 | 1,349394 | 3,162124 | 0,001566 | 1 | 0 | Promoter (<=1kb) | 10 | 60764005 | 60773911 | 9907 | 2 | 73616 | ENSMUST00000036304.4 | 0 | ENSMUSG00000037247 | Pldi | polymorphic derived intron containing |
| chr12 | 34184377 | 34185413 | 1037 | * | 7,901948 | 4,26700177 | 1,469791 | 2,903134 | 0,003694 | 0,722439117 | 1,360622 | Intron (ENSMUST00000 | 12 | 34007670 | 34009828 | 2159 | 1 | 22160 | ENSMUST00000049089.7 | 176707 | ENSMUSG00000035799 | Twist1 | twist basic helix-loop-helix transcription factor 1 |
| chr12 | 31440622 | 31441485 | 864 | * | 7,768535 | 4,267442253 | 1,506955 | 2,831832 | 0,004628 | 1 | 0 | Promoter (<=1kb) | 12 | 31440870 | 31500971 | 60102 | 1 | 13487 | ENSMUST00000167432.8 | 0 | ENSMUSG00000001225 | Slc26a3 | solute carrier family 26, member 3 |
| chr1 | 97614762 | 97615556 | 795 | * | 8,159012 | 4,268001987 | 1,354153 | 3,151788 | 0,001623 | 0,578728206 | 1,360622 | Distal Intergenic | 1 | 97571628 | 97589799 | 18172 | 2 | 52392 | ENSMUST00000149927.2 | -24963 | ENSMUSG00000044768 | Macir | macrophage immunometabolism regulator |
| chr2 | 1,69E+08 | 1,69E+08 | 862 | * | 9,632506 | 4,268783222 | 1,449762 | 2,944471 | 0,003235 | 0,73688097 | 1,086118 | Distal Intergenic | 2 | 1,69E+08 | 1,69E+08 | 11330 | 2 | 99377 | ENSMUST00000125138.2 | -11366 | ENSMUSG00000027547 | Sall4 | spalt like transcription factor 4 |
| chrX | 58660292 | 58660793 | 502 | * | 4,913833 | 4,26919002 | 1,646154 | 2,593433 | 0,009502 | 1 | 0 | Distal Intergenic | 20 | 58296168 | 58613431 | 317264 | 2 | 14168 | ENSMUST00000131319.8 | -46861 | ENSMUSG00000031137 | Fgf13 | fibroblast growth factor 13 |
| chr5 | 75682730 | 75683738 | 1009 | * | 8,845453 | 4,269930547 | 1,437224 | 2,970957 | 0,002969 | 0,59839358 | 2,237403 | Distal Intergenic | 5 | 75721706 | 75721810 | 105 | 2 | 1,15E+08 | ENSMUST00000158453.3 | 38072 | ENSMUSG00000089078 | Gm24502 | predicted gene, 24502 |
| chr14 | 30908329 | 30909463 | 1135 | * | 10,48356 | 4,270566511 | 1,462005 | 2,921034 | 0,003489 | 0,736639439 | 1,173325 | Promoter (<=1kb) | 14 | 30893215 | 30909164 | 15950 | 2 | 64652 | ENSMUST00000170253.8 | 0 | ENSMUSG00000021910 | Nisch | nischarin |
| chr7 | 1,01E+08 | 1,01E+08 | 674 | * | 6,793208 | 4,270903972 | 1,50517 | 2,83749 | 0,004547 | 1 | 0 | Distal Intergenic | 7 | 1,01E+08 | 1,01E+08 | 4414 | 1 | 11859 | ENSMUST00000008090.11 | -3641 | ENSMUSG00000007946 | Phox2a | paired-like homeobox 2a |
| chr3 | 46602409 | 46603183 | 775 | * | 7,46847 | 4,271602129 | 1,533991 | 2,784634 | 0,005359 | 0,789027175 | 1,360622 | Distal Intergenic | 3 | 46396632 | 46402654 | 6023 | 2 | 241989 | ENSMUST00000192793.6 | -199755 | ENSMUSG00000090919 | Pabpc4l | poly(A) binding protein, cytoplasmic 4-like |
| chr7 | 63036570 | 63037270 | 701 | * | 8,091884 | 4,272525166 | 1,39822 | 3,055688 | 0,002245 | 1 | 0 | Intron (ENSMUST00000 | 7 | 62994438 | 63094277 | 99840 | 2 | 77631 | ENSMUST00000181327.2 | 57007 | ENSMUSG00000097682 | 4930554H23Rik | RIKEN cDNA 4930554H23 gene |
| chr4 | 21890277 | 21891138 | 862 | * | 6,182701 | 4,273602522 | 1,616797 | 2,643252 | 0,008211 | 1 | 0 | Intron (ENSMUST00000 | 4 | 21898712 | 21900445 | 1734 | 1 | 230027 | ENSMUST00000142244.2 | -7574 | ENSMUSG00000028247 | Coq3 | coenzyme Q3 methyltransferase |
| chr3 | 14374163 | 14375203 | 1041 | * | 9,57042 | 4,274612764 | 1,463869 | 2,920078 | 0,003499 | 0,705098751 | 1,425929 | Distal Intergenic | 3 | 14428177 | 14442647 | 14471 | 1 | 622229 | ENSMUST00000192393.2 | -52974 | | Gm6300 | solute carrier family 7 (cationic amino acid transporter, y+ system), member pseudogene |
| chr18 | 31028561 | 31029921 | 1361 | * | 15,3415 | 4,274690467 | 1,116031 | 3,83026 | 0,000128 | 0,190625259 | 1,37683 | Distal Intergenic | 18 | 31286324 | 31345785 | 59462 | 2 | 19762 | ENSMUST00000153196.2 | 315864 | ENSMUSG00000057455 | Rit2 | Ras-like without CAAX 2 |
| chr2 | 81423182 | 81424180 | 999 | * | 11,85193 | 4,274739303 | 1,097172 | 3,896142 | 9,77E-05 | 0,206624632 | 0,918255 | Distal Intergenic | 2 | 81883566 | 82090223 | 206658 | 1 | 241514 | ENSMUST00000047527.8 | -459386 | ENSMUSG00000070866 | Zfp804a | zinc finger protein 804A |
| chr18 | 16945128 | 16945605 | 478 | * | 6,521628 | 4,276649333 | 1,445435 | 2,958729 | 0,003089 | 1 | 0 | Promoter (2-3kb) | 18 | 16721934 | 16942303 | 220370 | 2 | 12558 | ENSMUST00000025166.14 | -2825 | ENSMUSG00000024304 | Cdh2 | cadherin 2 |
| chr14 | 58068600 | 58069531 | 932 | * | 5,457955 | 4,277342191 | 1,518483 | 2,816852 | 0,00485 | 1 | 0 | Exon (ENSMUST00000 | 14 | 58063691 | 58066753 | 3063 | 1 | 67840 | ENSMUST00000022538.5 | 4909 | ENSMUSG00000021967 | Mrpl57 | mitochondrial ribosomal protein L57 |
| chr11 | 1,03E+08 | 1,03E+08 | 543 | * | 5,210164 | 4,277804085 | 1,631161 | 2,622552 | 0,008727 | 1 | 0 | Exon (ENSMUST00000 | 11 | 1,03E+08 | 1,03E+08 | 11197 | 2 | 52715 | ENSMUST00000124869.8 | 7105 | ENSMUSG00000020925 | Ccdc43 | coiled-coil domain containing 43 |
| chr3 | 65370585 | 65371180 | 596 | * | 5,394934 | 4,278334926 | 1,618341 | 2,643654 | 0,008202 | 1 | 0 | Distal Intergenic | 3 | 65414678 | 65435605 | 20928 | 2 | 1,03E+08 | ENSMUST00000147424.8 | 64425 | | Gm38503 | predicted gene, 38503 |
| chr19 | 13557304 | 13557957 | 654 | * | 6,608725 | 4,278652849 | 1,46524 | 2,920103 | 0,003499 | 1 | 0 | Distal Intergenic | 19 | 13562670 | 13563617 | 948 | 1 | 258288 | ENSMUST00000074180.2 | -4713 | ENSMUSG00000096289 | Olfr1484 | olfactory receptor 1484 |
| chr2 | 1,46E+08 | 1,46E+08 | 975 | * | 7,147105 | 4,278842792 | 1,499313 | 2,853869 | 0,004319 | 0,74907225 | 1,360622 | Distal Intergenic | 2 | 1,46E+08 | 1,46E+08 | 269200 | 2 | 241694 | ENSMUST00000228797.2 | -11098 | ENSMUSG00000037110 | Ralgapa2 | Ral GTPase activating protein, alpha subunit 2 (catalytic) |
| chr7 | 1,43E+08 | 1,43E+08 | 1002 | * | 9,778263 | 4,279285675 | 1,449869 | 2,951499 | 0,003162 | 0,6845989 | 1,425929 | Promoter (2-3kb) | 7 | 1,43E+08 | 1,43E+08 | 18259 | 2 | 79201 | ENSMUST00000152703.2 | -2056 | ENSMUSG00000037613 | Tnfrsf23 | tumor necrosis factor receptor superfamily, member 23 |
| chr4 | 44183090 | 44183899 | 810 | * | 7,533594 | 4,280692308 | 1,416761 | 3,021464 | 0,002516 | 1 | 0 | Intron (ENSMUST00000 | 4 | 44126215 | 44168339 | 42125 | 2 | 73469 | ENSMUST00000045793.15 | -14751 | ENSMUSG00000035696 | Rnf38 | ring finger protein 38 |
| chr8 | 32173478 | 32174121 | 644 | * | 7,560376 | 4,282642649 | 1,589398 | 2,694506 | 0,007049 | 1 | 0 | Distal Intergenic | 8 | 32091163 | 32153856 | 62694 | 2 | 1,03E+08 | ENSMUST00000211064.2 | -19622 | ENSMUSG00000110329 | Gm38572 | predicted gene, 38572 |
| chr10 | 1,28E+08 | 1,28E+08 | 1082 | * | 8,368211 | 4,283446333 | 1,353782 | 3,164059 | 0,001556 | 0,479521813 | 2,237403 | Promoter (<=1kb) | 10 | 1,28E+08 | 1,28E+08 | 89 | 1 | 1,02E+08 | ENSMUST00000184718.3 | 0 | ENSMUSG00000098957 | Mir8105 | microRNA 8105 |
| chr19 | 36941211 | 36942155 | 945 | * | 10,40786 | 4,283671772 | 1,325806 | 3,230994 | 0,001234 | 0,642322948 | 0,744375 | Promoter (2-3kb) | 19 | 36938715 | 36943232 | 4518 | 1 | 107182 | ENSMUST00000236343.2 | 2496 | ENSMUSG00000040565 | Btaf1 | B-TFIID TATA-box binding protein associated factor 1 |
| chr3 | 1,03E+08 | 1,03E+08 | 949 | * | 9,078363 | 4,286244894 | 1,259506 | 3,403115 | 0,000666 | 0,411608797 | 1,449706 | Distal Intergenic | 3 | 1,03E+08 | 1,03E+08 | 13882 | 1 | 94093 | ENSMUST00000197365.2 | 26381 | ENSMUSG00000033014 | Trim33 | tripartite motif-containing 33 |
| chrX | 105228844 | 105231330 | 2487 | * | 44,532543 | 4,286953305 | 0,6125514 | 6,9985204 | 2,587E-12 | 1,27178E-07 | 2,3383871 | Promoter (<=1kb) | 20 | 105230706 | 105247305 | 16600 | 1 | 18655 | ENSMUST00000081593.13 | 0 | ENSMUSG00000062070 | Pgk1 | phosphoglycerate kinase 1 |
| chr15 | 1,01E+08 | 1,01E+08 | 871 | * | 9,327561 | 4,290894609 | 1,43007 | 3,000479 | 0,002696 | 0,580348965 | 2,237403 | Intron (ENSMUST00000 | 15 | 1,01E+08 | 1,01E+08 | 27395 | 1 | 1,03E+08 | ENSMUST00000229169.2 | 14227 | | Gm35853 | predicted gene, 35853 |
| chr4 | 1,26E+08 | 1,26E+08 | 1032 | * | 10,20646 | 4,291991581 | 1,303812 | 3,29188 | 0,000995 | 0,530305679 | 1,086118 | Intron (ENSMUST00000 | 4 | 1,26E+08 | 1,26E+08 | 9256 | 1 | 329941 | ENSMUST00000128435.2 | 5017 | ENSMUSG00000056174 | Col8a2 | collagen, type VIII, alpha 2 |
| chr11 | 62529193 | 62530218 | 1026 | * | 6,444868 | 4,292456811 | 1,586204 | 2,706119 | 0,006807 | 1 | 0 | Exon (ENSMUST000001 | 11 | 62525530 | 62529614 | 4085 | 1 | 78042 | ENSMUST00000142492.2 | 3663 | | Lrrc75aos1 | leucine rich repeat containing 75A, opposite strand 1 |
| chr1 | 60422301 | 60422818 | 518 | * | 5,942551 | 4,293014354 | 1,642326 | 2,613984 | 0,008949 | 1 | 0 | Intron (ENSMUST00000 | 1 | 60448778 | 60520317 | 71540 | 1 | 329165 | ENSMUST00000052332.15 | -25960 | ENSMUSG00000026782 | Abi2 | abl interactor 2 |
| chr6 | 1,19E+08 | 1,19E+08 | 1056 | * | 14,38429 | 4,294417921 | 1,345127 | 3,192575 | 0,00141 | 0,596263346 | 1,075735 | Promoter (2-3kb) | 6 | 1,19E+08 | 1,19E+08 | 14677 | 1 | 319734 | ENSMUST00000185965.2 | 2215 | ENSMUSG00000041460 | Cacna2d4 | calcium channel, voltage-dependent, alpha 2/delta subunit 4 |
| chr1 | 1,36E+08 | 1,36E+08 | 534 | * | 6,015733 | 4,294754403 | 1,601297 | 2,682047 | 0,007317 | 1 | 0 | Promoter (1-2kb) | 1 | 1,36E+08 | 1,36E+08 | 18540 | 1 | 16565 | ENSMUST00000122892.2 | 1769 | ENSMUSG00000041642 | Kif21b | kinesin family member 21B |
| chr8 | 76601975 | 76602910 | 936 | * | 12,5292 | 4,29492375 | 1,131687 | 3,795151 | 0,000148 | 0,225397758 | 1,173325 | Intron (ENSMUST00000 | 8 | 76748557 | 76748683 | 127 | 2 | 1,15E+08 | ENSMUST00000104349.3 | 145773 | ENSMUSG00000077537 | Gm22544 | predicted gene, 22544 |
| chr3 | 1,05E+08 | 1,05E+08 | 903 | * | 8,927212 | 4,296202754 | 1,41475 | 3,036723 | 0,002392 | 0,670293874 | 1,211315 | Distal Intergenic | 3 | 1,05E+08 | 1,05E+08 | 9974 | 1 | 1,19E+08 | ENSMUST00000198827.2 | -7621 | | LOC118568635 | uncharacterized LOC118568635 |

| | | | | | | | | | | | | | | | | | | | | | | | |
|---|---|---|---|---|---|---|---|---|---|---|---|---|---|---|---|---|---|---|---|---|---|---|---|
| chr19 | 60228203 | 60229418 | 1216 | * | 10,21269 | 4,296234771 | 1,361908 | 3,154571 | 0,001607 | 0,563858922 | 1,449706 | Distal Intergenic | 19 | 60187867 | 60215133 | 27267 | 2 | 76539 | ENSMUST00000237339.2 | -13070 | ENSMUSG00000057858 | Fam204a | family with sequence similarity 204, member A |
| chr1 | 1,7E+08 | 1,7E+08 | 671 | * | 6,229761 | 4,296893115 | 1,646836 | 2,609181 | 0,009076 | 1 | 0 | Promoter (<=1kb) | 1 | 1,7E+08 | 1,71E+08 | 38258 | 1 | 320078 | ENSMUST00000046792.9 | -737 | ENSMUSG00000038463 | Olfml2b | olfactomedin-like 2B |
| chr10 | 60446610 | 60448016 | 1407 | * | 10,89108 | 4,297127153 | 1,279506 | 3,358426 | 0,000784 | 0,459179243 | 1,2763 | Intron (ENSMUST0000( | 10 | 60464259 | 60488812 | 24554 | 2 | 22295 | ENSMUST00000153677.2 | 40796 | ENSMUSG00000012819 | Cdh23 | cadherin 23 (otocadherin) |
| chr3 | 95225833 | 95226731 | 899 | * | 8,040031 | 4,297570628 | 1,469262 | 2,924986 | 0,003445 | 1 | 0 | Promoter (<=1kb) | 3 | 95226103 | 95229266 | 3164 | 1 | 76893 | ENSMUST00000139866.2 | 0 | ENSMUSG00000015714 | Cers2 | ceramide synthase 2 |
| chr10 | 7730355 | 7730852 | 498 | * | 5,373579 | 4,299248901 | 1,626525 | 2,643211 | 0,008212 | 1 | 0 | Intron (ENSMUST0000( | 10 | 7724125 | 7724245 | 121 | 2 | 1,15E+08 | ENSMUST00000158232.3 | -6110 | ENSMUSG00000088857 | Gm24726 | predicted gene, 24726 |
| chr10 | 98747280 | 98748366 | 1087 | * | 5,587653 | 4,299330568 | 1,619952 | 2,653986 | 0,007955 | 1 | 0 | Promoter (1-2kb) | 10 | 98750268 | 98822984 | 72717 | 1 | 67972 | ENSMUST00000219557.2 | -1902 | ENSMUSG00000019943 | Atp2b1 | ATPase, Ca++ transporting, plasma membrane 1 |
| chr18 | 3399132 | 3399654 | 523 | * | 5,144063 | 4,300012418 | 1,51207 | 2,843792 | 0,004458 | 1 | 0 | Intron (ENSMUST0000( | 18 | 3383311 | 3417640 | 34330 | 1 | 71745 | ENSMUST00000234414.2 | 15821 | ENSMUSG00000024231 | Cul2 | cullin 2 |
| chrX | 35566472 | 35567856 | 1385 | * | 6,891632 | 4,300130623 | 1,639938 | 2,62213 | 0,008738 | 1 | 0 | Distal Intergenic | 20 | 35592006 | 35606264 | 14259 | 1 | 74365 | ENSMUST00000135758.2 | -24150 | ENSMUSG00000016239 | Lonrf3 | LON peptidase N-terminal domain and ring finger 3 |
| chr18 | 10561951 | 10563122 | 1172 | * | 9,662569 | 4,300609817 | 1,42976 | 3,007923 | 0,00263 | 0,659642963 | 1,425929 | 3' UTR | 18 | 10553786 | 10558837 | 5052 | 1 | 381157 | ENSMUST00000173261.2 | 8165 | ENSMUSG00000042942 | Greb1l | growth regulation by estrogen in breast cancer-like |
| chr2 | 1,31E+08 | 1,31E+08 | 1108 | * | 10,84709 | 4,300934833 | 1,331356 | 3,230491 | 0,001236 | 0,642870233 | 0,738441 | Promoter (<=1kb) | 2 | 1,31E+08 | 1,31E+08 | 17546 | 2 | 20612 | ENSMUST00000028794.10 | 405 | ENSMUSG00000027322 | Siglec1 | sialic acid binding Ig-like lectin 1, sialoadhesin |
| chr19 | 13193522 | 13194302 | 781 | * | 4,937668 | 4,302038464 | 1,645877 | 2,613827 | 0,008953 | 1 | 0 | Distal Intergenic | 19 | 13211616 | 13212548 | 933 | 1 | 258120 | ENSMUST00000064102.2 | -17314 | ENSMUSG00000096365 | Olfr1463 | olfactory receptor 1463 |
| chr13 | 23717767 | 23719645 | 1879 | * | 24,065944 | 4,302496843 | 0,9523054 | 4,5179803 | 6,243E-06 | 0,015418018 | 2,8585108 | Promoter (<=1kb) | 13 | 23718076 | 23718474 | 399 | 1 | 319173 | ENSMUST00000073261.3 | 0 | ENSMUSG00000061991 | H2ac10 | H2A clustered histone 10 |
| chr13 | 51750614 | 51751523 | 910 | * | 8,555352 | 4,303351033 | 1,40531 | 3,062207 | 0,002197 | 0,541111532 | 2,237403 | Distal Intergenic | 13 | 51585077 | 51723523 | 138447 | 2 | 20418 | ENSMUST00000239056.2 | -27091 | ENSMUSG00000021448 | Shc3 | src homology 2 domain-containing transforming protein C3 |
| chr2 | 1,2E+08 | 1,2E+08 | 747 | * | 10,26528 | 4,303743301 | 1,258455 | 3,419863 | 0,000627 | 0,431115599 | 1,211315 | 3' UTR | 2 | 1,2E+08 | 1,2E+08 | 2527 | 2 | 20402 | ENSMUST00000130303.2 | 3971 | ENSMUSG00000027288 | Zfp106 | zinc finger protein 106 |
| chr8 | 85367133 | 85368235 | 1103 | * | 11,03563 | 4,30420235 | 1,406821 | 3,059524 | 0,002217 | 0,64945399 | 1,2763 | Promoter (1-2kb) | 8 | 85365620 | 85366063 | 444 | 1 | 12286 | ENSMUST00000130507.2 | 1513 | ENSMUSG00000034656 | Cacna1a | calcium channel, voltage-dependent, P/Q type, alpha 1A subunit |
| chr4 | 1,32E+08 | 1,32E+08 | 1256 | * | 11,83867 | 4,30426885 | 1,44111 | 2,986773 | 0,002819 | 0,782360218 | 0,738441 | Promoter (1-2kb) | 4 | 1,32E+08 | 1,32E+08 | 24287 | 2 | 100340 | ENSMUST00000030709.9 | 1196 | ENSMUSG00000028885 | Smpdl3b | sphingomyelin phosphodiesterase, acid-like 3B |
| chr5 | 19175479 | 19176299 | 821 | * | 5,208752 | 4,304331832 | 1,669647 | 2,577989 | 0,009938 | 1 | 0 | Distal Intergenic | 5 | 19069575 | 19069702 | 128 | 1 | 1,15E+08 | ENSMUST00000158739.3 | 105904 | ENSMUSG00000089364 | Gm23822 | predicted gene, 23822 |
| chr19 | 57683444 | 57684918 | 1475 | * | 10,79232 | 4,304382617 | 1,207842 | 3,563697 | 0,000366 | 0,339611164 | 1,173325 | Exon (ENSMUST00000( | 19 | 57599466 | 58121775 | 522310 | 1 | 226255 | ENSMUST00000077282.7 | 83978 | ENSMUSG00000054843 | Atrnl1 | attractin like 1 |
| chr11 | 93853117 | 93854131 | 1015 | * | 8,177085 | 4,30673388 | 1,462182 | 2,945415 | 0,003225 | 0,686838043 | 1,425929 | Promoter (2-3kb) | 11 | 93850298 | 93856869 | 6572 | 2 | 18102 | ENSMUST00000021220.10 | 2738 | ENSMUSG00000037601 | Nme1 | NME/NM23 nucleoside diphosphate kinase 1 |
| chr10 | 75988348 | 75989242 | 895 | * | 6,746538 | 4,307475407 | 1,576851 | 2,731695 | 0,006301 | 1 | 0 | Intron (ENSMUST0000( | 10 | 75983285 | 76025099 | 41815 | 1 | 64452 | ENSMUST00000020450.4 | 5063 | ENSMUSG00000020229 | Slc5a4a | solute carrier family 5, member 4a |
| chr3 | 1,29E+08 | 1,29E+08 | 745 | * | 8,135682 | 4,308679579 | 1,443722 | 2,984425 | 0,002841 | 1 | 0 | Intron (ENSMUST0000( | 3 | 1,29E+08 | 1,29E+08 | 19713 | 1 | 18741 | ENSMUST00000174661.9 | -407341 | ENSMUSG00000028023 | Pitx2 | paired-like homeodomain transcription factor 2 |
| chr16 | 16922399 | 16923392 | 994 | * | 8,134048 | 4,311917645 | 1,444148 | 2,985786 | 0,002829 | 0,671156916 | 1,425929 | Promoter (<=1kb) | 16 | 16910711 | 16921491 | 10781 | 2 | 66053 | ENSMUST00000232169.2 | -908 | ENSMUSG00000022771 | Ppil2 | peptidylprolyl isomerase (cyclophilin)-like 2 |
| chr1 | 1,71E+08 | 1,71E+08 | 913 | * | 6,589154 | 4,312168146 | 1,648901 | 2,615177 | 0,008918 | 1 | 0 | Intron (ENSMUST0000( | 1 | 1,71E+08 | 1,71E+08 | 107 | 2 | 1,15E+08 | ENSMUST00000179933.3 | 32099 | ENSMUSG00000119049 | Gm25235 | predicted gene, 25235 |
| chr9 | 70076871 | 70077861 | 991 | * | 7,385416 | 4,313495465 | 1,568728 | 2,749678 | 0,005965 | 1 | 0 | Distal Intergenic | 9 | 70056088 | 70056218 | 131 | 2 | 1,15E+08 | ENSMUST00000104472.3 | -20653 | ENSMUSG00000077660 | Gm24257 | predicted gene, 24257 |
| chr10 | 58647988 | 58648940 | 953 | * | 9,636979 | 4,314520509 | 1,345489 | 3,206655 | 0,001343 | 0,563858922 | 1,211315 | Promoter (<=1kb) | 10 | 58649181 | 58885842 | 236662 | 1 | 237353 | ENSMUST00000135526.9 | -241 | ENSMUSG00000037990 | Sh3rf3 | SH3 domain containing ring finger 3 |
| chr4 | 62296691 | 62297468 | 778 | * | 6,957535 | 4,315120149 | 1,571725 | 2,745468 | 0,006042 | 1 | 0 | Intron (ENSMUST0000( | 4 | 62303738 | 62310006 | 6269 | 1 | 20529 | ENSMUST00000122092.2 | -6270 | ENSMUSG00000066150 | Slc31a1 | solute carrier family 31, member 1 |
| chr6 | 1,2E+08 | 1,2E+08 | 607 | * | 6,381018 | 4,315600293 | 1,639832 | 2,631734 | 0,008495 | 1 | 0 | Intron (ENSMUST0000( | 6 | 1,19E+08 | 1,2E+08 | 111817 | 2 | 22419 | ENSMUST00000178696.8 | -81203 | ENSMUSG00000030170 | Wnt5b | wingless-type MMTV integration site family, member 5B |
| chr13 | 58531745 | 58532726 | 982 | * | 8,353061 | 4,315818928 | 1,484916 | 2,906439 | 0,003656 | 0,710505631 | 1,449706 | Promoter (<=1kb) | 13 | 58527631 | 58533039 | 5409 | 2 | 70153 | ENSMUST00000022032.7 | 313 | ENSMUSG00000021550 | 2210016F16Rik | RIKEN cDNA 2210016F16 gene |
| chr13 | 46473998 | 46474723 | 726 | * | 7,545008 | 4,316407729 | 1,510899 | 2,856847 | 0,004279 | 1 | 0 | Distal Intergenic | 13 | 46427197 | 46453591 | 26395 | 1 | 380842 | ENSMUST00000076622.4 | 46801 | ENSMUSG00000063529 | Stmnd1 | stathmin domain containing 1 |
| chr12 | 1,01E+08 | 1,01E+08 | 1324 | * | 9,107914 | 4,320200638 | 1,440262 | 2,999594 | 0,002703 | 0,661775851 | 1,449706 | Intron (ENSMUST0000( | 12 | 1,01E+08 | 1,01E+08 | 107 | 2 | 1,15E+08 | ENSMUST00000240251.1 | -30863 | ENSMUSG00000119861 | Gm22749 | predicted gene, 22749 |
| chr4 | 51356692 | 51357649 | 958 | * | 7,988316 | 4,321989617 | 1,448098 | 2,984597 | 0,00284 | 0,680780911 | 1,360622 | Distal Intergenic | 4 | 51216678 | 51250622 | 33945 | 1 | 74914 | ENSMUST00000156384.4 | 140014 | ENSMUSG00000039555 | Cylc2 | cylicin, basic protein of sperm head cytoskeleton 2 |
| chr9 | 1,05E+08 | 1,05E+08 | 1034 | * | 8,455723 | 4,322115987 | 1,430887 | 3,020584 | 0,002523 | 0,566578624 | 2,237403 | Intron (ENSMUST0000( | 9 | 1,05E+08 | 1,05E+08 | 149743 | 1 | 74020 | ENSMUST00000213452.2 | 77529 | ENSMUSG00000032564 | Cpne4 | copine IV |
| chr1 | 1,33E+08 | 1,33E+08 | 1193 | * | 10,64587 | 4,322409609 | 1,422276 | 3,03908 | 0,002373 | 0,749550311 | 0,744375 | Distal Intergenic | 1 | 1,33E+08 | 1,33E+08 | 2511 | 1 | 280287 | ENSMUST00000178033.5 | 6640 | ENSMUSG00000116158 | Kiss1 | KiSS-1 metastasis-suppressor |
| chr14 | 34224908 | 34225634 | 727 | * | 7,045374 | 4,324235406 | 1,42249 | 3,039906 | 0,002367 | 1 | 0 | Promoter (<=1kb) | 14 | 34163063 | 34225298 | 62236 | 2 | 12166 | ENSMUST00000171343.9 | 0 | ENSMUSG00000021796 | Bmpr1a | bone morphogenetic protein receptor, type 1A |
| chr19 | 43664727 | 43666240 | 1514 | * | 11,32153 | 4,324498338 | 1,205994 | 3,585838 | 0,000336 | 0,409991363 | 0,744375 | Promoter (1-2kb) | 19 | 43663617 | 43665615 | 1999 | 1 | 1,01E+08 | ENSMUST00000236819.2 | 1110 | ENSMUSG00000117698 | BC037704 | cDNA sequence BC037704 |
| chr9 | 66941423 | 66943827 | 2405 | * | 15,36528 | 4,325637943 | 1,251341 | 3,456802 | 0,000547 | 0,313797801 | 2,141612 | Promoter (1-2kb) | 9 | 66937202 | 66940107 | 2906 | 2 | 22003 | ENSMUST00000156899.2 | -1316 | ENSMUSG00000032366 | Tpm1 | tropomyosin 1, alpha |
| chr13 | 53466046 | 53467225 | 1180 | * | 8,383258 | 4,326309875 | 1,509476 | 2,8661 | 0,004156 | 0,732113157 | 1,425929 | Distal Intergenic | 13 | 53263353 | 53440160 | 176808 | 2 | 26564 | ENSMUST00000021918.10 | -25886 | ENSMUSG00000021464 | Ror2 | receptor tyrosine kinase-like orphan receptor 2 |
| chrX | 70871838 | 70872978 | 1141 | * | 6,572712 | 4,327273161 | 1,590894 | 2,720027 | 0,006528 | 1 | 0 | Intron (ENSMUST0000( | 20 | 70860419 | 70864590 | 4172 | 1 | 67048 | ENSMUST00000138979.2 | 11419 | ENSMUSG00000073131 | Vma21 | VMA21 vacuolar H+-ATPase homolog (S. cerevisiae) |
| chr19 | 47204992 | 47205914 | 923 | * | 23,158344 | 4,327490271 | 0,8305098 | 5,2106434 | 1,882E-07 | 0,001081418 | 2,9519335 | Intron (ENSMUST000001 | 19 | 47217350 | 47247874 | 30525 | 1 | 18011 | ENSMUST00000111807.5 | -11436 | ENSMUSG00000006435 | Neurl1a | neuralized E3 ubiquitin protein ligase 1A |
| chr16 | 45710863 | 45712007 | 1145 | * | 9,443373 | 4,328088164 | 1,306823 | 3,311915 | 0,000927 | 0,396931054 | 2,237403 | Intron (ENSMUST0000( | 16 | 45566606 | 45664741 | 98136 | 2 | 208177 | ENSMUST00000036355.13 | -46122 | ENSMUSG00000033149 | Phldb2 | pleckstrin homology like domain, family B, member 2 |
| chr13 | 5626845 | 5628004 | 1160 | * | 7,765321 | 4,329573204 | 1,525971 | 2,837259 | 0,00455 | 1 | 0 | Intron (ENSMUST0000( | 13 | 5504908 | 5610328 | 105421 | 1 | 1,05E+08 | ENSMUST00000223343.2 | 121937 | | Gm40661 | predicted gene, 40661 |
| chrX | 92651958 | 92652730 | 773 | * | 6,066989 | 4,329821402 | 1,592681 | 2,718575 | 0,006556 | 1 | 0 | Intron (ENSMUST0000( | 20 | 92666904 | 92675721 | 8818 | 2 | 18968 | ENSMUST00000124236.2 | 22991 | ENSMUSG00000006678 | Pola1 | polymerase (DNA directed), alpha 1 |
| chrX | 13667860 | 13668802 | 943 | * | 7,528342 | 4,330466969 | 1,510042 | 2,86778 | 0,004134 | 1 | 0 | Intron (ENSMUST0000( | 20 | 13383319 | 13652269 | 268951 | 2 | 12361 | ENSMUST00000156096.8 | -15591 | ENSMUSG00000031012 | Cask | calcium/calmodulin-dependent serine protein kinase (MAGUK family) |
| chr8 | 16227578 | 16228386 | 809 | * | 7,404159 | 4,332112581 | 1,351267 | 3,205963 | 0,001346 | 1 | 0 | Intron (ENSMUST0000( | 8 | 16218771 | 16218852 | 82 | 2 | 1,01E+08 | ENSMUST00000175401.3 | -8726 | ENSMUSG00000093142 | Mir3106 | microRNA 3106 |
| chr9 | 87137522 | 87138877 | 1356 | * | 12,30067 | 4,332977813 | 1,424718 | 3,041289 | 0,002356 | 0,713633527 | 0,918255 | Promoter (<=1kb) | 9 | 87071630 | 87137589 | 65960 | 2 | 382090 | ENSMUST00000093802.6 | 0 | ENSMUSG00000056919 | Cep162 | centrosomal protein 162 |
| chr13 | 8152111 | 8153045 | 935 | * | 7,58397 | 4,334002551 | 1,526637 | 2,838921 | 0,004527 | 1 | 0 | Distal Intergenic | 13 | 8252902 | 8811566 | 558665 | 1 | 94191 | ENSMUST00000064473.13 | -99857 | ENSMUSG00000052551 | Adarb2 | adenosine deaminase, RNA-specific, B2 |
| chr5 | 90828418 | 90829236 | 819 | * | 7,921088 | 4,334484475 | 1,397219 | 3,102222 | 0,001921 | 1 | 0 | Distal Intergenic | 5 | 90751690 | 90788516 | 36827 | 2 | 73246 | ENSMUST00000202704.4 | -39902 | ENSMUSG00000029370 | Rassf6 | Ras association (RalGDS/AF-6) domain family member 6 |
| chr2 | 75426562 | 75427029 | 468 | * | 5,909508 | 4,334846578 | 1,567686 | 2,765124 | 0,00569 | 1 | 0 | Distal Intergenic | 2 | 75464373 | 75465119 | 747 | 1 | 1,05E+08 | ENSMUST00000247606.1 | -37344 | | Gm39858 | predicted gene, 39858 |
| chr11 | 60008438 | 60009005 | 568 | * | 6,022638 | 4,336731707 | 1,641464 | 2,64199 | 0,008242 | 1 | 0 | Promoter (2-3kb) | 11 | 60003043 | 60006083 | 3041 | 2 | 1,01E+08 | ENSMUST00000130746.3 | -2355 | ENSMUSG00000051008 | 4930412M03Rik | RIKEN cDNA 4930412M03 gene |
| chr1 | 95545886 | 95546719 | 834 | * | 6,790177 | 4,337171204 | 1,442837 | 3,006003 | 0,002647 | 1 | 0 | 3' UTR | 1 | 95546276 | 95595260 | 48985 | 2 | 20452 | ENSMUST00000189556.2 | 48541 | ENSMUSG00000040710 | St8sia4 | ST8 alpha-N-acetyl-neuraminide alpha-2,8-sialyltransferase 4 |
| chr11 | 60362723 | 60363816 | 1094 | * | 9,131801 | 4,33752905 | 1,389264 | 3,122178 | 0,001795 | 0,597812173 | 1,360622 | Promoter (2-3kb) | 11 | 60360165 | 60419195 | 59031 | 1 | 17910 | ENSMUST00000094135.9 | 2558 | ENSMUSG00000042678 | Myo15 | myosin XV |
| chr18 | 44605228 | 44606803 | 1576 | * | 25,31119 | 4,339814903 | 1,367362 | 3,17386 | 0,001504 | 0,442614296 | 2,725491 | Exon (ENSMUST00000( | 18 | 44513594 | 44558036 | 44443 | 1 | 70640 | ENSMUST00000025350.10 | 91634 | ENSMUSG00000024472 | Dcp2 | decapping mRNA 2 |
| chr19 | 37859371 | 37860027 | 657 | * | 6,334433 | 4,340059423 | 1,604422 | 2,705061 | 0,006829 | 1 | 0 | Exon (ENSMUST000002 | 19 | 37887916 | 37889951 | 2036 | 2 | 226101 | ENSMUST00000225435.2 | 29924 | ENSMUSG00000048612 | Myof | myoferlin |
| chr12 | 1,03E+08 | 1,03E+08 | 851 | * | 6,170985 | 4,340770299 | 1,655921 | 2,621364 | 0,008758 | 1 | 0 | Intron (ENSMUST0000( | 12 | 1,03E+08 | 1,03E+08 | 37560 | 2 | 238386 | ENSMUST00000220596.2 | 5971 | ENSMUSG00000041702 | Btbd7 | BTB (POZ) domain containing 7 |
| chr4 | 11007555 | 11009131 | 1577 | * | 12,2303 | 4,341975919 | 1,208698 | 3,592277 | 0,000328 | 0,326832856 | 1,173325 | Promoter (<=1kb) | 4 | 10988662 | 11007676 | 19015 | 2 | 71801 | ENSMUST00000054776.4 | 0 | ENSMUSG00000049969 | Plekhf2 | pleckstrin homology domain containing, family F (with FYVE domain) member 2 |
| chr10 | 80139570 | 80140504 | 935 | * | 7,167289 | 4,34200192 | 1,580787 | 2,746734 | 0,006019 | 1 | 0 | Promoter (1-2kb) | 10 | 80137654 | 80154090 | 16437 | 1 | 23805 | ENSMUST00000020349.7 | 1916 | ENSMUSG00000020135 | Apc2 | APC regulator of WNT signaling pathway 2 |
| chr6 | 1,29E+08 | 1,29E+08 | 639 | * | 5,401322 | 4,342224568 | 1,652227 | 2,628105 | 0,008586 | 1 | 0 | Promoter (1-2kb) | 6 | 1,29E+08 | 1,29E+08 | 614 | 2 | 1,15E+08 | ENSMUST00000203263.2 | 1719 | | Gm44066 | predicted gene, 44066 |
| chr19 | 43182937 | 43183538 | 602 | * | 6,346314 | 4,34286385 | 1,659673 | 2,616699 | 0,008878 | 1 | 0 | Intron (ENSMUST0000( | 19 | 42897023 | 43212128 | 315106 | 2 | 545291 | ENSMUST00000238153.2 | 28590 | ENSMUSG00000074852 | Hpse2 | heparanase 2 |
| chr2 | 1,03E+08 | 1,03E+08 | 937 | * | 7,842923 | 4,343031228 | 1,480095 | 2,934291 | 0,003343 | 1 | 0 | Promoter (<=1kb) | 2 | 1,03E+08 | 1,03E+08 | 3096 | 2 | 1,03E+08 | ENSMUST00000143995.2 | 701 | ENSMUSG00000087185 | Gm13872 | predicted gene 13872 |
| chr12 | 55053948 | 55054677 | 730 | * | 6,692563 | 4,343986667 | 1,593711 | 2,725705 | 0,006416 | 1 | 0 | Intron (ENSMUST0000( | 12 | 55001503 | 55061133 | 59631 | 2 | 217578 | ENSMUST00000172875.8 | 6456 | ENSMUSG00000035021 | Baz1a | bromodomain adjacent to zinc finger domain 1A |
| chr12 | 35868244 | 35869127 | 884 | * | 5,643017 | 4,345367894 | 1,627968 | 2,669198 | 0,007603 | 1 | 0 | Exon (ENSMUST000002 | 12 | 35975619 | 35999735 | 24117 | 1 | 403205 | ENSMUST00000042101.5 | -106492 | ENSMUSG00000036231 | Agr3 | anterior gradient 3 |
| chr7 | 29159803 | 29161352 | 1550 | * | 25,462415 | 4,346585716 | 0,7165587 | 6,0659171 | 1,312E-09 | 1,92956E-05 | 2,8800171 | Intron (ENSMUST000000 | 7 | 29083681 | 29204877 | 121197 | 2 | 74206 | ENSMUST00000182011.8 | 43525 | ENSMUSG00000030583 | Sipa1l3 | signal-induced proliferation-associated 1 like 3 |
| chr1 | 1,85E+08 | 1,85E+08 | 1395 | * | 7,37569 | 4,348250567 | 1,507052 | 2,88527 | 0,003911 | 1 | 0 | Distal Intergenic | 1 | 1,85E+08 | 1,85E+08 | 630 | 1 | 114668 | ENSMUST00000192616.2 | 181498 | ENSMUSG00000102185 | 5033404E19Rik | NSA2 ribosome biogenesis homolog pseudogene |
| chr8 | 67709557 | 67709988 | 432 | * | 5,508703 | 4,349497006 | 1,657884 | 2,623524 | 0,008703 | 1 | 0 | Distal Intergenic | 8 | 67731734 | 67731883 | 150 | 2 | 1,15E+08 | ENSMUST00000158755.4 | 21895 | ENSMUSG00000089380 | Gm24677 | predicted gene, 24677 |
| chr5 | 1,15E+08 | 1,15E+08 | 1024 | * | 6,482812 | 4,34971961 | 1,557145 | 2,793394 | 0,005216 | 1 | 0 | Promoter (2-3kb) | 5 | 1,15E+08 | 1,15E+08 | 5781 | 2 | 66236 | ENSMUST00000112160.4 | -2556 | ENSMUSG00000072694 | 1500011B03Rik | RIKEN cDNA 1500011B03 gene |
| chr2 | 72517275 | 72517888 | 614 | * | 7,124254 | 4,351654041 | 1,420631 | 3,063183 | 0,00219 | 1 | 0 | Distal Intergenic | 2 | 72533854 | 72559812 | 25959 | 1 | 74541 | ENSMUST00000124829.2 | -15966 | ENSMUSG00000085252 | 8430437L04Rik | RIKEN cDNA 8430437L04 gene |
| chr17 | 33790365 | 33791428 | 1064 | * | 9,235793 | 4,352095812 | 1,41582 | 3,073904 | 0,002113 | 0,627995982 | 1,360622 | Intron (ENSMUST0000( | 17 | 33801577 | 33826738 | 25162 | 1 | 17916 | ENSMUST00000173426.2 | -10149 | ENSMUSG00000024300 | Myo1f | myosin IF |
| chr6 | 59043444 | 59044566 | 1123 | * | 10,1213 | 4,352275011 | 1,395195 | 3,119474 | 0,001812 | 0,593612587 | 1,425929 | Distal Intergenic | 6 | 59076238 | 59076369 | 132 | 1 | 1,15E+08 | ENSMUST00000178930.3 | -31672 | ENSMUSG00000118703 | Gm22367 | predicted gene, 22367 |
| chr1 | 1,89E+08 | 1,89E+08 | 937 | * | 6,817717 | 4,355458241 | 1,472216 | 2,958437 | 0,003092 | 1 | 0 | Intron (ENSMUST0000( | 1 | 1,89E+08 | 1,89E+08 | 33373 | 2 | 16526 | ENSMUST00000192529.2 | -17684 | ENSMUSG00000037624 | Kcnk2 | potassium channel, subfamily K, member 2 |
| chr11 | 60681476 | 60682272 | 797 | * | 6,653434 | 4,359272729 | 1,548633 | 2,814917 | 0,004879 | 1 | 0 | Promoter (1-2kb) | 11 | 60680422 | 60683999 | 3578 | 2 | 20425 | ENSMUST00000135081.2 | 1727 | ENSMUSG00000020534 | Shmt1 | serine hydroxymethyltransferase 1 (soluble) |
| chr5 | 1,48E+08 | 1,48E+08 | 907 | * | 6,68696 | 4,359778523 | 1,561115 | 2,792733 | 0,005226 | 1 | 0 | Intron (ENSMUST0000( | 5 | 1,48E+08 | 1,48E+08 | 1311 | 2 | 71706 | ENSMUST00000152064.2 | -4488 | ENSMUSG00000029650 | Slc46a3 | solute carrier family 46, member 3 |
| chr13 | 84760389 | 84761296 | 908 | * | 9,104266 | 4,360865249 | 1,437945 | 3,032707 | 0,002424 | 0,65364112 | 1,360622 | Distal Intergenic | 13 | 84422963 | 84428936 | 5974 | 1 | 72745 | ENSMUST00000224744.2 | 337426 | ENSMUSG00000035762 | Tmem161b | transmembrane protein 161B |
| chr15 | 25646493 | 25647378 | 886 | * | 7,329136 | 4,360931287 | 1,471638 | 2,963317 | 0,003043 | 1 | 0 | Intron (ENSMUST0000( | 15 | 25622635 | 25702036 | 79402 | 1 | 17909 | ENSMUST00000127486.2 | 23858 | ENSMUSG00000022272 | Myo10 | myosin X |
| chr5 | 1,25E+08 | 1,25E+08 | 617 | * | 8,068916 | 4,361982871 | 1,578653 | 2,763104 | 0,005725 | 1 | 0 | Intron (ENSMUST0000( | 5 | 1,25E+08 | 1,25E+08 | 7472 | 2 | 20602 | ENSMUST00000199561.2 | -9615 | ENSMUSG00000029478 | Ncor2 | nuclear receptor co-repressor 2 |
| chr5 | 76288928 | 76289776 | 849 | * | 6,410468 | 4,362139631 | 1,691387 | 2,579032 | 0,009908 | 1 | 0 | Promoter (<=1kb) | 5 | 76288577 | 76297794 | 9218 | 1 | 57357 | ENSMUST00000127278.8 | 351 | ENSMUSG00000029233 | Srd5a3 | steroid 5 alpha-reductase 3 |
| chrX | 99702301 | 99703840 | 1540 | * | 7,41852 | 4,362470213 | 1,490072 | 2,927692 | 0,003415 | 1 | 0 | Intron (ENSMUST0000( | 20 | 99677566 | 99679913 | 2348 | 1 | 16571 | ENSMUST00000156386.2 | 24735 | ENSMUSG00000034311 | Kif4 | kinesin family member 4 |
| chr17 | 73231278 | 73232287 | 1010 | * | 7,380628 | 4,3627133 | 1,505106 | 2,898609 | 0,003748 | 1 | 0 | Intron (ENSMUST0000( | 17 | 73228183 | 73246340 | 18158 | 1 | 77889 | ENSMUST00000148556.2 | 3095 | ENSMUSG00000024063 | Lbh | limb-bud and heart |
| chr16 | 43254480 | 43255376 | 897 | * | 7,28717 | 4,364160377 | 1,549634 | 2,816252 | 0,004859 | 1 | 0 | Intron (ENSMUST0000( | 16 | 43230881 | 43240559 | 9679 | 2 | 626693 | ENSMUST00000155900.2 | -13921 | ENSMUSG00000085499 | Gm15713 | predicted gene 15713 |
| chrX | 60039422 | 60040146 | 725 | * | 6,1146 | 4,368528975 | 1,597625 | 2,73439 | 0,00625 | 1 | 0 | Distal Intergenic | 20 | 60085347 | 60119674 | 34328 | 1 | 1,03E+08 | ENSMUST00000142919.2 | -45201 | | Gm14664 | predicted gene 14664 |
| chr1 | 1,87E+08 | 1,87E+08 | 981 | * | 6,29555 | 4,369210951 | 1,623161 | 2,691791 | 0,007107 | 1 | 0 | Intron (ENSMUST0000( | 1 | 1,87E+08 | 1,87E+08 | 14788 | 1 | 329316 | ENSMUST00000195611.2 | -89791 | ENSMUSG00000103923 | Gm37896 | predicted gene, 37896 |
| chr18 | 56635782 | 56637212 | 1431 | * | 9,257408 | 4,369371222 | 1,316989 | 3,317698 | 0,000908 | 0,463890057 | 1,449706 | 3' UTR | 18 | 56602339 | 56625402 | 23064 | 1 | 107022 | ENSMUST00000237716.2 | 33443 | ENSMUSG00000001700 | Gramd3 | GRAM domain containing 3 |
| chr15 | 89004744 | 89005488 | 745 | * | 7,5405 | 4,371736359 | 1,562435 | 2,798027 | 0,005142 | 1 | 0 | Promoter (1-2kb) | 15 | 88984613 | 89007089 | 22477 | 2 | 328580 | ENSMUST00000041656.14 | 1601 | ENSMUSG00000051786 | Tubgcp6 | tubulin, gamma complex associated protein 6 |
| chr13 | 1,11E+08 | 1,11E+08 | 916 | * | 8,06985 | 4,37227078 | 1,333242 | 3,279428 | 0,00104 | 1 | 0 | Distal Intergenic | 13 | 1,11E+08 | 1,11E+08 | 3014 | 1 | 1,03E+08 | ENSMUST00000225669.2 | 11909 | | Gm33172 | predicted gene, 33172 |
| chr16 | 89553181 | 89554145 | 965 | * | 9,078413 | 4,373904668 | 1,417638 | 3,085347 | 0,002033 | 0,643386705 | 1,211315 | Distal Intergenic | 16 | 89585310 | 89592170 | 6861 | 2 | 21844 | ENSMUST00000178095.2 | 38025 | ENSMUSG00000002489 | Tiam1 | T cell lymphoma invasion and metastasis 1 |
| chr9 | 67570016 | 67571472 | 1457 | * | 9,418017 | 4,375089541 | 1,427374 | 3,065131 | 0,002176 | 0,654833721 | 1,211315 | Distal Intergenic | 9 | 67634999 | 67635062 | 64 | 2 | 1,02E+08 | ENSMUST00000184532.3 | 63590 | ENSMUSG00000098937 | Mir7241 | microRNA 7241 |
| chr13 | 9813020 | 9814137 | 1118 | * | 10,20507 | 4,375653025 | 1,217958 | 3,592615 | 0,000327 | 0,322750682 | 1,211315 | Promoter (<=1kb) | 13 | 9740409 | 9814386 | 73978 | 2 | 66505 | ENSMUST00000220996.2 | 249 | ENSMUSG00000021156 | Zmynd11 | zinc finger, MYND domain containing 11 |
| chr3 | 1,22E+08 | 1,22E+08 | 405 | * | 5,240512 | 4,375719162 | 1,601005 | 2,733108 | 0,006274 | 1 | 0 | Exon (ENSMUST000001 | 3 | 1,22E+08 | 1,22E+08 | 57 | 2 | 1,02E+08 | ENSMUST00000196051.3 | -8191 | ENSMUSG00000104773 | Mir7657 | microRNA 7657 |
| chr5 | 51668379 | 51669140 | 762 | * | 7,37551 | 4,375966778 | 1,517412 | 2,883836 | 0,003929 | 1 | 0 | Exon (ENSMUST000001 | 5 | 51614995 | 51711213 | 96219 | 2 | 19017 | ENSMUST00000031059.14 | 42073 | ENSMUSG00000029167 | Ppargc1a | peroxisome proliferative activated receptor, gamma, coactivator 1 alpha |
| chr6 | 31746576 | 31747394 | 819 | * | 8,171878 | 4,376939555 | 1,326834 | 3,298785 | 0,000971 | 1 | 0 | Intron (ENSMUST0000( | 6 | 31696316 | 31720029 | 23714 | 1 | 1,05E+08 | ENSMUST00000210985.2 | 50260 | | Gm13846 | predicted gene 13846 |
| chr2 | 1,72E+08 | 1,72E+08 | 1128 | * | 10,68343 | 4,377993342 | 1,398395 | 3,130728 | 0,001744 | 0,694263044 | 0,744375 | Promoter (<=1kb) | 2 | 1,72E+08 | 1,72E+08 | 2623 | 1 | 17201 | ENSMUST00000038532.2 | -258 | ENSMUSG00000038537 | Mc3r | melanocortin 3 receptor |
| chrX | 8052496 | 8053429 | 934 | * | 6,422086 | 4,378801474 | 1,640806 | 2,66869 | 0,007615 | 1 | 0 | Promoter (<=1kb) | 20 | 8051570 | 8053413 | 1844 | 2 | 13595 | ENSMUST00000140011.2 | 0 | ENSMUSG00000031168 | Ebp | phenylalkylamine Ca2+ antagonist (emopamil) binding protein |
| chr16 | 92086968 | 92087508 | 541 | * | 7,554329 | 4,383754643 | 1,536277 | 2,853492 | 0,004324 | 0,749236309 | 1,360622 | Promoter (1-2kb) | 16 | 92089277 | 92095017 | 5741 | 1 | 246133 | ENSMUST00000047383.10 | -1769 | ENSMUSG00000039672 | Kcne2 | potassium voltage-gated channel, Isk-related subfamily, gene 2 |
| chr4 | 3988766 | 3989425 | 660 | * | 6,123501 | 4,38493644 | 1,566618 | 2,798982 | 0,005126 | 1 | 0 | Distal Intergenic | 4 | 4012341 | 4016621 | 4281 | 2 | 242285 | ENSMUST00000126274.2 | 27196 | ENSMUSG00000028236 | Sdr16c5 | short chain dehydrogenase/reductase family 16C, member 5 |
| chr10 | 22602651 | 22603668 | 1018 | * | 7,571993 | 4,386211804 | 1,497379 | 2,92926 | 0,003398 | 1 | 0 | Promoter (1-2kb) | 10 | 22605271 | 22605331 | 61 | 2 | 1,02E+08 | ENSMUST00000183950.3 | 1663 | ENSMUSG00000098714 | Mir7663 | microRNA 7663 |
| chr10 | 1,22E+08 | 1,22E+08 | 1062 | * | 8,545592 | 4,386812747 | 1,428171 | 3,071629 | 0,002129 | 1 | 0 | Intron (ENSMUST0000( | 10 | 1,22E+08 | 1,22E+08 | 4869 | 2 | 319272 | ENSMUST00000219429.2 | 38817 | ENSMUSG00000112095 | Sros1 | non-coding RNA suppressor of Stat1 |
| chr17 | 32003831 | 32005590 | 1760 | * | 17,77695 | 4,386924388 | 0,9737397 | 4,5052334 | 6,63E-06 | 0,018112704 | 2,5070128 | Distal Intergenic | 17 | 32068728 | 32070549 | 1822 | 2 | 17691 | ENSMUST00000234867.2 | 64959 | ENSMUSG00000024042 | Sik1 | salt inducible kinase 1 |
| chr14 | 26368964 | 26370286 | 1323 | * | 8,748352 | 4,387935803 | 1,443634 | 3,039506 | 0,00237 | 0,640583529 | 1,449706 | Intron (ENSMUST0000( | 14 | 26374166 | 26375012 | 847 | 1 | 11843 | ENSMUST00000171207.2 | -3880 | ENSMUSG00000021877 | Arf4 | ADP-ribosylation factor 4 |
| chr2 | 30190857 | 30192024 | 1168 | * | 11,53285 | 4,388082963 | 1,284284 | 3,416753 | 0,000634 | 0,439573569 | 1,173325 | 3' UTR | 2 | 30176441 | 30216698 | 40258 | 1 | 227699 | ENSMUST00000133603.2 | 14416 | ENSMUSG00000052533 | Nup188 | nucleoporin 188 |
| chr5 | 29399256 | 29400927 | 1672 | * | 12,92701 | 4,38901747 | 1,05629 | 4,155124 | 3,25E-05 | 0,086132354 | 1,2763 | Promoter (<=1kb) | 5 | 29400990 | 29408133 | 7144 | 1 | 56874 | ENSMUST00000160888.8 | -63 | ENSMUSG00000029130 | Rnf32 | ring finger protein 32 |
| chr2 | 17846457 | 17847788 | 1332 | * | 12,61064 | 4,39170132 | 1,139853 | 3,852867 | 0,000117 | 0,226355781 | 0,918255 | Distal Intergenic | 2 | 17732197 | 17736275 | 4079 | 2 | 74103 | ENSMUST00000130725.2 | -110182 | ENSMUSG00000053702 | Nebl | nebulette |
| chr10 | 59974634 | 59976692 | 2059 | * | 20,011618 | 4,391724542 | 1,0434774 | 4,2087396 | 2,568E-05 | 0,046584446 | 2,713306 | Distal Intergenic | 10 | 59962598 | 59962655 | 58 | 1 | 102465546 | ENSMUST00000198998.3 | 12036 | ENSMUSG00000105468 | Mir6906 | microRNA 6906 |

| | | | | | | | | | | | | | | | | | | | | | | | |
|---|---|---|---|---|---|---|---|---|---|---|---|---|---|---|---|---|---|---|---|---|---|---|---|
| chr16 | 70110046 | 70111942 | 1897 | * | 33,50115 | 4,392053248 | 1,118052 | 3,928308 | 8,55E-05 | 0,093381457 | 2,925497 | Promoter (<=1kb) | 16 | 70110837 | 70366604 | 255768 | 1 | 74185 | ENSMUST00000163832.8 | 0 | ENSMUSG00000022707 | Gbe1 | glucan (1,4-alpha-), branching enzyme 1 |
| chr15 | 80258423 | 80259358 | 936 | * | 7,568325 | 4,392589441 | 1,53499 | 2,86164 | 0,004215 | 1 | 0 | Exon (ENSMUST00000: | 15 | 80273529 | 80275118 | 1590 | 1 | 239556 | ENSMUST00000161863.8 | -14171 | ENSMUSG00000022416 | Cacna1i | calcium channel, voltage-dependent, alpha 1I subunit |
| chr7 | 35029364 | 35030334 | 971 | * | 8,293806 | 4,392733472 | 1,273847 | 3,448401 | 0,000564 | 1 | 0 | Distal Intergenic | 7 | 35033595 | 35091714 | 58120 | 1 | 52428 | ENSMUST00000032705.13 | -3261 | ENSMUSG00000030494 | Rhpn2 | rhophilin, Rho GTPase binding protein 2 |
| chr14 | 76979713 | 76980641 | 929 | * | 6,133495 | 4,394039805 | 1,653947 | 2,656699 | 0,007891 | 1 | 0 | Distal Intergenic | 14 | 76997571 | 76999657 | 2087 | 1 | 73883 | ENSMUST00000228541.2 | -16930 | ENSMUSG00000115709 | 4930431P22Rik | RIKEN cDNA 4930431P22 gene |
| chr6 | 85098142 | 85098428 | 287 | * | 6,27188 | 4,394175452 | 1,620144 | 2,712213 | 0,006684 | 1 | 0 | Intron (ENSMUST0000( | 6 | 85088398 | 85103107 | 14710 | 2 | 545861 | ENSMUST00000203386.2 | 4679 | ENSMUSG00000072952 | Gm5878 | predicted gene 5878 |
| chr3 | 47219346 | 47220119 | 774 | * | 7,942109 | 4,394476699 | 1,44561 | 3,039878 | 0,002367 | 1 | 0 | Exon (ENSMUST00000: | 3 | 47155062 | 47155199 | 138 | 1 | 1,15E+08 | ENSMUST00000158114.3 | 64284 | ENSMUSG00000088739 | Gm23891 | predicted gene, 23891 |
| chr1 | 86746111 | 86747188 | 1078 | * | 8,403647 | 4,395343567 | 1,458262 | 3,014098 | 0,002577 | 1 | 0 | Intron (ENSMUST0000( | 1 | 86631565 | 86977814 | 346250 | 1 | 208718 | ENSMUST00000065694.8 | 114546 | ENSMUSG00000053333 | Dis3l2 | DIS3 like 3'-5' exoribonuclease 2 |
| chr3 | 21978317 | 21979041 | 725 | * | 7,107377 | 4,39535843 | 1,545108 | 2,844693 | 0,004445 | 1 | 0 | Distal Intergenic | 3 | 21807693 | 21854937 | 47245 | 2 | 319506 | ENSMUST00000191725.2 | -123380 | | 7530428D23Rik | RIKEN cDNA 7530428D23 gene |
| chr17 | 46355135 | 46355657 | 523 | * | 6,423739 | 4,396736674 | 1,681067 | 2,615443 | 0,008911 | 1 | 0 | Distal Intergenic | 17 | 46329469 | 46343295 | 13827 | 2 | 22339 | ENSMUST00000142351.9 | -11840 | ENSMUSG00000023951 | Vegfa | vascular endothelial growth factor A |
| chr7 | 94018524 | 94019307 | 784 | * | 7,27974 | 4,397951088 | 1,526102 | 2,88182 | 0,003954 | 0,731734453 | 1,360622 | Exon (ENSMUST00000: | 7 | 93846417 | 93846686 | 270 | 1 | 545982 | ENSMUST00000071770.4 | 172107 | | Gm5899 | predicted pseudogene 5899 |
| chr17 | 53795914 | 53796644 | 731 | * | 5,748011 | 4,398168302 | 1,658287 | 2,652236 | 0,007996 | 1 | 0 | Intron (ENSMUST0000( | 17 | 53786262 | 53814708 | 28447 | 1 | 271457 | ENSMUST00000017975.7 | 9652 | ENSMUSG00000017831 | Rab5a | RAB5A, member RAS oncogene family |
| chr12 | 70107849 | 70108727 | 879 | * | 8,611319 | 4,398903517 | 1,455325 | 3,022626 | 0,002506 | 1 | 0 | Exon (ENSMUST00000( | 12 | 70093682 | 70098058 | 4377 | 2 | 18080 | ENSMUST00000223316.2 | -9791 | ENSMUSG00000021068 | Nin | ninein |
| chrX | 1,57E+08 | 1,57E+08 | 892 | * | 8,371778 | 4,399429784 | 1,674808 | 2,626826 | 0,008619 | 1 | 0 | Intron (ENSMUST0000( | 20 | 1,57E+08 | 1,57E+08 | 8294 | 2 | 245684 | ENSMUST00000139664.2 | -57265 | ENSMUSG00000025658 | Cnksr2 | connector enhancer of kinase suppressor of Ras 2 |
| chr2 | 1,27E+08 | 1,27E+08 | 1066 | * | 9,194578 | 4,399640151 | 1,419053 | 3,100405 | 0,001933 | 0,518149475 | 2,237403 | Promoter (<=1kb) | 2 | 1,26E+08 | 1,27E+08 | 37702 | 2 | 14391 | ENSMUST00000039978.13 | 0 | ENSMUSG00000027361 | Gabpb1 | GA repeat binding protein, beta 1 |
| chr13 | 64310027 | 64311105 | 1079 | * | 12,69217 | 4,401402841 | 1,489517 | 2,95492 | 0,003128 | 0,75938015 | 0,918255 | Promoter (<=1kb) | 13 | 64310060 | 64311432 | 1373 | 1 | 56541 | ENSMUST00000221264.2 | 0 | ENSMUSG00000021476 | Habp4 | hyaluronic acid binding protein 4 |
| chr13 | 36461495 | 36462443 | 949 | * | 8,401073 | 4,404574577 | 1,46251 | 3,011653 | 0,002598 | 0,658200216 | 1,425929 | Intron (ENSMUST0000( | 13 | 36416253 | 36597724 | 181472 | 1 | 69955 | ENSMUST00000224619.2 | 45242 | ENSMUSG00000021420 | Fars2 | phenylalanine-tRNA synthetase 2 (mitochondrial) |
| chr1 | 1,66E+08 | 1,66E+08 | 2150 | * | 14,67048 | 4,408744066 | 1,086131 | 4,059128 | 4,93E-05 | 0,115291293 | 1,214904 | Promoter (<=1kb) | 1 | 1,66E+08 | 1,66E+08 | 62685 | 1 | 1E+08 | ENSMUST00000111416.7 | -271 | ENSMUSG00000040612 | Ildr2 | immunoglobulin-like domain containing receptor 2 |
| chr1 | 1,55E+08 | 1,55E+08 | 1022 | * | 9,11835 | 4,409276497 | 1,32698 | 3,322791 | 0,000891 | 0,470960971 | 1,360622 | Distal Intergenic | 1 | 1,55E+08 | 1,55E+08 | 11027 | 2 | 15064 | ENSMUST00000195579.2 | -4223 | ENSMUSG00000026471 | Mr1 | major histocompatibility complex, class I-related |
| chr10 | 1,28E+08 | 1,28E+08 | 841 | * | 7,588939 | 4,410341028 | 1,409439 | 3,129146 | 0,001753 | 0,594676756 | 1,360622 | Distal Intergenic | 10 | 1,28E+08 | 1,28E+08 | 7119 | 1 | 11947 | ENSMUST00000026459.6 | -3746 | ENSMUSG00000025393 | Atp5b | ATP synthase, H+ transporting mitochondrial F1 complex, beta subunit |
| chr13 | 61610650 | 61611385 | 736 | * | 6,146281 | 4,410424783 | 1,595482 | 2,764321 | 0,005704 | 1 | 0 | Distal Intergenic | 13 | 61684076 | 61686282 | 2207 | 2 | 64139 | ENSMUST00000225268.2 | 74897 | ENSMUSG00000074871 | Ctsm | cathepsin M |
| chr18 | 60883345 | 60884184 | 840 | * | 8,82806 | 4,411894546 | 1,631464 | 2,704255 | 0,006846 | 0,830992446 | 1,360622 | Promoter (1-2kb) | 18 | 60819035 | 60881722 | 62688 | 2 | 15531 | ENSMUST00000237783.2 | -1623 | ENSMUSG00000054008 | Ndst1 | N-deacetylase/N-sulfotransferase (heparan glucosaminyl) 1 |
| chr4 | 1,33E+08 | 1,33E+08 | 1470 | * | 11,56646 | 4,411933702 | 1,202573 | 3,668745 | 0,000244 | 0,352029189 | 0,738441 | Promoter (2-3kb) | 4 | 1,33E+08 | 1,33E+08 | 16241 | 2 | 100336 | ENSMUST00000030702.14 | -2209 | ENSMUSG00000028882 | Ppp1r8 | protein phosphatase 1, regulatory subunit 8 |
| chrX | 17013104 | 17013739 | 636 | * | 5,57705 | 4,413780886 | 1,659864 | 2,659121 | 0,007834 | 1 | 0 | Intron (ENSMUST0000( | 20 | 16998288 | 17185607 | 187320 | 2 | 74405 | ENSMUST00000026014.8 | 171868 | ENSMUSG00000025038 | Efhc2 | EF-hand domain (C-terminal) containing 2 |
| chr14 | 1,08E+08 | 1,08E+08 | 740 | * | 5,932322 | 4,417202439 | 1,597653 | 2,764807 | 0,005696 | 1 | 0 | Distal Intergenic | 14 | 1,07E+08 | 1,07E+08 | 127 | 1 | 1,15E+08 | ENSMUST00000104036.3 | 613420 | ENSMUSG00000077224 | Gm23437 | predicted gene, 23437 |
| chr12 | 75869371 | 75869930 | 560 | * | 7,406889 | 4,419363775 | 1,477944 | 2,990211 | 0,002788 | 1 | 0 | Intron (ENSMUST0000( | 12 | 75865124 | 75952677 | 87554 | 1 | 319565 | ENSMUST00000157033.2 | 4247 | ENSMUSG00000063450 | Syne2 | spectrin repeat containing, nuclear envelope 2 |
| chr1 | 99927137 | 99927748 | 612 | * | 6,429665 | 4,419465596 | 1,572147 | 2,811101 | 0,004937 | 1 | 0 | Intron (ENSMUST0000( | 1 | 1E+08 | 1E+08 | 388182 | 1 | 241175 | ENSMUST00000188735.2 | -97738 | ENSMUSG00000067028 | Cntnap5b | contactin associated protein-like 5B |
| chr1 | 94801201 | 94802042 | 842 | * | 6,970026 | 4,420418591 | 1,561749 | 2,830428 | 0,004649 | 1 | 0 | Distal Intergenic | 1 | 94725167 | 94730197 | 5031 | 1 | 74689 | ENSMUST00000185490.2 | 76034 | ENSMUSG00000100518 | 4930440C22Rik | RIKEN cDNA 4930440C22 gene |
| chr5 | 65548353 | 65549927 | 1575 | * | 11,5493 | 4,420891392 | 1,222623 | 3,615907 | 0,000299 | 0,392614537 | 0,738441 | Promoter (<=1kb) | 5 | 65545707 | 65548742 | 3036 | 2 | 20005 | ENSMUST00000127874.6 | 0 | ENSMUSG00000047215 | Rpl9 | ribosomal protein L9 |
| chr15 | 12032501 | 12033344 | 844 | * | 5,588974 | 4,421050197 | 1,623855 | 2,722564 | 0,006478 | 1 | 0 | Distal Intergenic | 15 | 11984127 | 11997069 | 12943 | 2 | 20024 | ENSMUST00000110504.2 | -35432 | ENSMUSG00000022205 | Sub1 | SUB1 homolog, transcriptional regulator |
| chr3 | 1,08E+08 | 1,08E+08 | 890 | * | 7,995527 | 4,422300363 | 1,624413 | 2,722399 | 0,006481 | 1 | 0 | Promoter (<=1kb) | 3 | 1,08E+08 | 1,08E+08 | 19189 | 2 | 12977 | ENSMUST00000118593.8 | 0 | ENSMUSG00000014599 | Csf1 | colony stimulating factor 1 (macrophage) |
| chr13 | 1,13E+08 | 1,13E+08 | 616 | * | 5,662226 | 4,423990757 | 1,674189 | 2,642467 | 0,00823 | 1 | 0 | Intron (ENSMUST0000( | 13 | 1,13E+08 | 1,13E+08 | 66780 | 1 | 19012 | ENSMUST00000070951.8 | 30513 | ENSMUSG00000021759 | Plpp1 | phospholipid phosphatase 1 |
| chr3 | 33642977 | 33644060 | 1084 | * | 8,819426 | 4,42472704 | 1,516871 | 2,917009 | 0,003534 | 0,631820259 | 2,237403 | Distal Intergenic | 3 | 33447206 | 33447348 | 143 | 2 | 1,15E+08 | ENSMUST00000157635.3 | -195629 | ENSMUSG00000088260 | Gm24780 | predicted gene, 24780 |
| chr5 | 32188336 | 32189469 | 1134 | * | 8,645584 | 4,424735682 | 1,443527 | 3,065224 | 0,002175 | 0,63491343 | 1,360622 | Intron (ENSMUST0000( | 5 | 32203322 | 32242097 | 38776 | 1 | 107976 | ENSMUST00000202080.2 | -13853 | ENSMUSG00000052139 | Babam2 | BRISC and BRCA1 A complex member 2 |
| chr4 | 130588143 | 130589841 | 1699 | * | 26,303589 | 4,42478777 | 0,8465402 | 5,2269081 | 1,724E-07 | 0,001047051 | 2,8800171 | Promoter (2-3kb) | 4 | 130585062 | 130585191 | 130 | 2 | 115486250 | ENSMUST00000104193.3 | -2952 | ENSMUSG00000077381 | Gm22660 | predicted gene, 22660 |
| chr9 | 44164154 | 44165874 | 1721 | * | 12,20204 | 4,425732165 | 1,701355 | 2,601298 | 0,009287 | 0,893554647 | 1,2763 | Promoter (1-2kb) | 9 | 44158604 | 44162768 | 4165 | 2 | 170761 | ENSMUST00000034618.6 | -1386 | ENSMUSG00000032105 | Pdzd3 | PDZ domain containing 3 |
| chr18 | 63531268 | 63531853 | 586 | * | 5,242045 | 4,426064162 | 1,628753 | 2,717455 | 0,006579 | 1 | 0 | Promoter (<=1kb) | 18 | 63532285 | 63554268 | 21984 | 1 | 1,05E+08 | ENSMUST00000183218.3 | -432 | | Gm26972 | predicted gene, 26972 |
| chr9 | 64778279 | 64779144 | 866 | * | 5,727252 | 4,427233 | 1,622423 | 2,728779 | 0,006357 | 1 | 0 | Exon (ENSMUST00000( | 9 | 64796242 | 64808257 | 12016 | 1 | 102442 | ENSMUST00000216098.2 | -17098 | ENSMUSG00000053641 | Dennd4a | DENN/MADD domain containing 4A |
| chr11 | 66488269 | 66489081 | 813 | * | 7,465195 | 4,42728277 | 1,459101 | 3,034254 | 0,002411 | 1 | 0 | Distal Intergenic | 11 | 66108298 | 66416790 | 308493 | 2 | 380702 | ENSMUST00000066679.7 | -71479 | ENSMUSG00000053930 | Shisa6 | shisa family member 6 |
| chr10 | 80007634 | 80008382 | 749 | * | 6,043215 | 4,429029397 | 1,616189 | 2,740415 | 0,006136 | 1 | 0 | Promoter (<=1kb) | 10 | 80007816 | 80008532 | 717 | 1 | 12696 | ENSMUST00000147190.2 | 0 | ENSMUSG00000045193 | Cirbp | cold inducible RNA binding protein |
| chr3 | 1,33E+08 | 1,33E+08 | 708 | * | 8,850385 | 4,429837539 | 1,468044 | 3,017511 | 0,002549 | 0,681421811 | 1,211315 | Intron (ENSMUST0000( | 3 | 1,33E+08 | 1,33E+08 | 5525 | 2 | 214133 | ENSMUST00000195913.2 | 17352 | ENSMUSG00000040943 | Tet2 | tet methylcytosine dioxygenase 2 |
| chr17 | 47237309 | 47237977 | 669 | * | 6,222489 | 4,432426986 | 1,661564 | 2,667624 | 0,007639 | 1 | 0 | Distal Intergenic | 17 | 47241169 | 47244001 | 2833 | 2 | 224826 | ENSMUST00000224442.2 | 6024 | ENSMUSG00000023977 | Ubr2 | ubiquitin protein ligase E3 component n-recognin 2 |
| chr4 | 1,33E+08 | 1,33E+08 | 708 | * | 5,623023 | 4,432720094 | 1,649896 | 2,686667 | 0,007217 | 1 | 0 | Intron (ENSMUST0000( | 4 | 1,33E+08 | 1,33E+08 | 28171 | 2 | 503610 | ENSMUST00000084238.5 | 8127 | ENSMUSG00000037553 | Zdhhc18 | zinc finger, DHHC domain containing 18 |
| chr9 | 53533156 | 53533793 | 638 | * | 6,427527 | 4,432775413 | 1,571749 | 2,820282 | 0,004798 | 1 | 0 | Promoter (<=1kb) | 9 | 53527900 | 53533334 | 5435 | 2 | 75717 | ENSMUST00000141803.2 | 0 | ENSMUSG00000032030 | Cul5 | cullin 5 |
| chr5 | 1,22E+08 | 1,22E+08 | 1212 | * | 8,241449 | 4,433056931 | 1,305141 | 3,396612 | 0,000682 | 0,426043749 | 1,360622 | Promoter (<=1kb) | 5 | 1,22E+08 | 1,22E+08 | 37847 | 1 | 20239 | ENSMUST00000162611.8 | 0 | ENSMUSG00000042605 | Atxn2 | ataxin 2 |
| chr11 | 100404140 | 100405350 | 1211 | * | 13,580293 | 4,436786624 | 0,9964511 | 4,4525886 | 8,484E-06 | 0,04517794 | 0,9386821 | Exon (ENSMUST000001( | 11 | 100367184 | 100414424 | 47241 | 2 | 104112 | ENSMUST00000165111.8 | 9074 | ENSMUSG00000020917 | Acly | ATP citrate lyase |
| chr1 | 1,19E+08 | 1,19E+08 | 1439 | * | 11,74075 | 4,437486564 | 1,214441 | 3,653932 | 0,000258 | 0,284932697 | 1,2763 | Distal Intergenic | 1 | 1,19E+08 | 1,19E+08 | 219488 | 2 | 14633 | ENSMUST00000062483.15 | -83911 | ENSMUSG00000048402 | Gli2 | GLI-Kruppel family member GLI2 |
| chr8 | 23216144 | 23217218 | 1075 | * | 9,227901 | 4,437972192 | 1,419075 | 3,12737 | 0,001764 | 0,582425528 | 1,449706 | Distal Intergenic | 8 | 23149784 | 23196605 | 46822 | 2 | 16150 | ENSMUST00000063401.10 | -19539 | ENSMUSG00000031537 | Ikbkb | inhibitor of kappaB kinase beta |
| chr2 | 1,52E+08 | 1,52E+08 | 780 | * | 5,329409 | 4,438013524 | 1,673552 | 2,651853 | 0,008005 | 1 | 0 | Exon (ENSMUST00000: | 2 | 1,52E+08 | 1,52E+08 | 25930 | 1 | 12995 | ENSMUST00000141597.2 | 24286 | ENSMUSG00000074698 | Csnk2a1 | casein kinase 2, alpha 1 polypeptide |
| chr7 | 1,1E+08 | 1,1E+08 | 1137 | * | 12,24 | 4,438036162 | 1,079547 | 4,111018 | 3,94E-05 | 0,137154233 | 0,738441 | Intron (ENSMUST0000( | 7 | 1,1E+08 | 1,1E+08 | 13625 | 1 | 11717 | ENSMUST00000148292.2 | 5183 | ENSMUSG00000005686 | Ampd3 | adenosine monophosphate deaminase 3 |
| chr12 | 13323616 | 13324084 | 469 | * | 4,733055 | 4,438755934 | 1,695318 | 2,618243 | 0,008838 | 1 | 0 | Exon (ENSMUST00000( | 12 | 13319134 | 13633812 | 314679 | 1 | 71169 | ENSMUST00000042953.10 | 4482 | ENSMUSG00000020576 | Nbas | neuroblastoma amplified sequence |
| chr5 | 83537864 | 83538831 | 968 | * | 6,547332 | 4,43959054 | 1,592424 | 2,787944 | 0,005304 | 1 | 0 | Distal Intergenic | 5 | 83425992 | 83503042 | 77051 | 2 | 243078 | ENSMUST00000053543.11 | -34822 | ENSMUSG00000049537 | Tecrl | trans-2,3-enoyl-CoA reductase-like |
| chrX | 16948271 | 16949490 | 1220 | * | 7,267547 | 4,43981263 | 1,533415 | 2,895376 | 0,003787 | 1 | 0 | Distal Intergenic | 20 | 16751760 | 16778013 | 26254 | 2 | 17986 | ENSMUST00000040134.8 | -170258 | ENSMUSG00000040138 | Ndp | Norrie disease (pseudoglioma) (human) |
| chr3 | 1,56E+08 | 1,56E+08 | 593 | * | 7,33458 | 4,440590169 | 1,607409 | 2,762576 | 0,005735 | 1 | 0 | Intron (ENSMUST0000( | 3 | 1,56E+08 | 1,57E+08 | 454095 | 1 | 320840 | ENSMUST00000175773.2 | 76266 | ENSMUSG00000040037 | Negr1 | neuronal growth regulator 1 |
| chr3 | 37365495 | 37367207 | 1713 | * | 15,833971 | 4,441190085 | 1,0394255 | 4,2727352 | 1,931E-05 | 0,037516355 | 2,6636191 | Promoter (<=1kb) | 3 | 37366668 | 37459015 | 92348 | 1 | 14173 | ENSMUST00000138710.3 | 0 | ENSMUSG00000037225 | Fgf2 | fibroblast growth factor 2 |
| chr1 | 45403594 | 45404475 | 882 | * | 7,02856 | 4,441308847 | 1,480456 | 2,999961 | 0,0027 | 1 | 0 | Distal Intergenic | 1 | 45379118 | 45386926 | 7809 | 1 | 12825 | ENSMUST00000186021.2 | 24476 | ENSMUSG00000026043 | Col3a1 | collagen, type III, alpha 1 |
| chr2 | 126849721 | 126852390 | 2670 | * | 19,022105 | 4,445400106 | 1,0106348 | 4,3986218 | 1,089E-05 | 0,026722594 | 2,3860078 | Promoter (<=1kb) | 2 | 126850637 | 126879264 | 28628 | 1 | 108011 | ENSMUST00000149254.9 | 0 | ENSMUSG00000001998 | Ap4e1 | adaptor-related protein complex AP-4, epsilon 1 |
| chr12 | 32146849 | 32148611 | 1763 | * | 10,28743 | 4,445481521 | 1,419118 | 3,132566 | 0,001733 | 0,578908869 | 1,449706 | Exon (ENSMUST00000: | 12 | 32123285 | 32130849 | 7565 | 2 | 1,03E+08 | ENSMUST00000219954.3 | -16000 | ENSMUSG00000112569 | Gm33111 | predicted gene, 33111 |
| chr18 | 56614687 | 56615250 | 564 | * | 5,734632 | 4,44617943 | 1,65642 | 2,68421 | 0,00727 | 1 | 0 | Intron (ENSMUST0000( | 18 | 56602339 | 56625402 | 23064 | 1 | 107022 | ENSMUST00000237716.2 | 12348 | ENSMUSG00000001700 | Gramd3 | GRAM domain containing 3 |
| chr10 | 15361917 | 15362658 | 742 | * | 6,393945 | 4,4474448 | 1,554237 | 2,861498 | 0,004216 | 1 | 0 | Distal Intergenic | 10 | 15610660 | 15614570 | 3911 | 2 | 1,03E+08 | ENSMUST00000218861.2 | 251912 | | Gm32283 | predicted gene, 32283 |
| chr16 | 13921234 | 13922044 | 811 | * | 6,134695 | 4,449210308 | 1,523131 | 2,921096 | 0,003488 | 1 | 0 | Distal Intergenic | 16 | 13956641 | 13960404 | 3764 | 2 | 223989 | ENSMUST00000231032.2 | 38360 | ENSMUSG00000060657 | Marf1 | meiosis regulator and mRNA stability 1 |
| chr18 | 46623038 | 46623972 | 935 | * | 8,247644 | 4,450759951 | 1,457139 | 3,054451 | 0,002255 | 0,542967057 | 2,237403 | Distal Intergenic | 18 | 46598282 | 46598857 | 576 | 2 | 72076 | ENSMUST00000180618.2 | -24181 | | Mospd4 | motile sperm domain containing 4 |
| chr17 | 74406831 | 74407365 | 535 | * | 4,989561 | 4,453442694 | 1,721012 | 2,587688 | 0,009662 | 1 | 0 | Distal Intergenic | 17 | 74385744 | 74387288 | 1545 | 2 | 1,03E+08 | ENSMUST00000233356.2 | -19543 | ENSMUSG00000117226 | Gm31235 | predicted gene, 31235 |
| chr11 | 7801651 | 7802462 | 812 | * | 7,286311 | 4,454218816 | 1,549038 | 2,875474 | 0,004034 | 1 | 0 | Distal Intergenic | 11 | 7836444 | 7836505 | 62 | 2 | 1,15E+08 | ENSMUST00000157263.4 | 34043 | ENSMUSG00000098853 | Gm27393 | predicted gene, 27393 |
| chr5 | 1,38E+08 | 1,38E+08 | 1245 | * | 9,984113 | 4,455377109 | 1,384543 | 3,217941 | 0,001291 | 0,563032042 | 1,173325 | Distal Intergenic | 5 | 1,38E+08 | 1,38E+08 | 9688 | 1 | 83766 | ENSMUST00000136088.8 | -7842 | ENSMUSG00000029712 | Actl6b | actin-like 6B |
| chr12 | 41122588 | 41123381 | 794 | * | 7,704803 | 4,455692286 | 1,465421 | 3,040555 | 0,002361 | 1 | 0 | Intron (ENSMUST0000( | 12 | 41118311 | 41160942 | 42632 | 1 | 93757 | ENSMUST00000080160.4 | 4277 | ENSMUSG00000056899 | Immp2l | IMP2 inner mitochondrial membrane peptidase-like (S. cerevisiae) |
| chr1 | 21488904 | 21489647 | 744 | * | 8,09067 | 4,455779973 | 1,341448 | 3,321618 | 0,000895 | 0,471637333 | 1,360622 | Intron (ENSMUST0000( | 1 | 21453780 | 21456597 | 2818 | 1 | 98582 | ENSMUST00000147615.2 | 35124 | ENSMUSG00000085079 | Khdc1b | KH domain containing 1B |
| chr1 | 1,83E+08 | 1,83E+08 | 743 | * | 7,096229 | 4,458787593 | 1,499012 | 2,974485 | 0,002935 | 1 | 0 | Distal Intergenic | 1 | 1,83E+08 | 1,83E+08 | 61196 | 2 | 68897 | ENSMUST00000171366.7 | 67724 | ENSMUSG00000030768 | Disp1 | dispatched RND transporter family member 1 |
| chr3 | 1,37E+08 | 1,37E+08 | 828 | * | 6,966403 | 4,460569231 | 1,577587 | 2,827463 | 0,004692 | 1 | 0 | Intron (ENSMUST0000( | 3 | 1,37E+08 | 1,37E+08 | 86151 | 1 | 59308 | ENSMUST00000197511.2 | 83702 | ENSMUSG00000054690 | Emcn | endomucin |
| chr3 | 1,27E+08 | 1,27E+08 | 837 | * | 6,722054 | 4,462197849 | 1,533743 | 2,909352 | 0,003622 | 1 | 0 | Intron (ENSMUST0000( | 3 | 1,27E+08 | 1,27E+08 | 329010 | 2 | 109676 | ENSMUST00000183095.3 | 28110 | ENSMUSG00000032826 | Ank2 | ankyrin 2, brain |
| chr1 | 1,58E+08 | 1,58E+08 | 581 | * | 5,903118 | 4,462298042 | 1,620848 | 2,753063 | 0,005904 | 1 | 0 | Distal Intergenic | 1 | 1,58E+08 | 1,58E+08 | 761 | 2 | 240843 | ENSMUST00000194578.2 | 357158 | ENSMUSG00000004031 | Brinp2 | bone morphogenic protein/retinoic acid inducible neural-specific 2 |
| chr18 | 18024566 | 18025142 | 577 | * | 6,675187 | 4,463347581 | 1,608463 | 2,774914 | 0,005522 | 1 | 0 | Distal Intergenic | 18 | 17701803 | 17714047 | 12245 | 2 | 75162 | ENSMUST00000234987.2 | -310519 | ENSMUSG00000117615 | 4930545E07Rik | RIKEN cDNA 4930545E07 gene |
| chr16 | 21512603 | 21513110 | 508 | * | 5,421327 | 4,463377652 | 1,689176 | 2,642339 | 0,008234 | 1 | 0 | Promoter (<=1kb) | 16 | 21495475 | 21513330 | 17856 | 2 | 72190 | ENSMUST00000232634.2 | 220 | ENSMUSG00000043391 | 2510009E07Rik | RIKEN cDNA 2510009E07 gene |
| chrX | 72652745 | 72654260 | 1516 | * | 11,03462 | 4,464209223 | 1,32079 | 3,379953 | 0,000725 | 0,446344042 | 1,2763 | Distal Intergenic | 20 | 72683025 | 72687120 | 4096 | 1 | 75590 | ENSMUST00000019701.9 | -28765 | ENSMUSG00000031383 | Dusp9 | dual specificity phosphatase 9 |
| chr13 | 42976686 | 42977386 | 701 | * | 7,53423 | 4,466491394 | 1,444938 | 3,09113 | 0,001994 | 1 | 0 | Intron (ENSMUST0000( | 13 | 43019723 | 43288780 | 269058 | 1 | 218194 | ENSMUST00000131942.2 | -42337 | ENSMUSG00000054728 | Phactr1 | phosphatase and actin regulator 1 |
| chr13 | 32911603 | 32912537 | 935 | * | 6,249354 | 4,468960952 | 1,589835 | 2,81096 | 0,004939 | 1 | 0 | Exon (ENSMUST00000: | 13 | 32901558 | 32960396 | 58839 | 2 | 238564 | ENSMUST00000230119.2 | 47859 | ENSMUSG00000044951 | Mylk4 | myosin light chain kinase family, member 4 |
| chr1 | 75559564 | 75560389 | 826 | * | 6,969059 | 4,470002404 | 1,535333 | 2,911421 | 0,003598 | 1 | 0 | Distal Intergenic | 1 | 75528353 | 75530376 | 2024 | 1 | 20536 | ENSMUST00000132110.2 | 31211 | ENSMUSG00000006576 | Slc4a3 | solute carrier family 4 (anion exchanger), member 3 |
| chr13 | 111065124 | 111067048 | 1925 | * | 15,278 | 4,471188025 | 1,0190536 | 4,3875886 | 1,146E-05 | 0,046577192 | 1,214904 | Distal Intergenic | 13 | 111019386 | 111040380 | 20995 | 2 | 102636142 | ENSMUST00000225424.2 | -24744 | | 4930526H09Rik | RIKEN cDNA 4930526H09 gene |
| chr2 | 24414670 | 24415340 | 671 | * | 8,618489 | 4,472919101 | 1,419279 | 3,151543 | 0,001624 | 0,568412386 | 1,425929 | Distal Intergenic | 2 | 24310572 | 24365611 | 55040 | 2 | 18510 | ENSMUST00000028355.11 | -49059 | ENSMUSG00000026976 | Pax8 | paired box 8 |
| chr13 | 1,07E+08 | 1,07E+08 | 808 | * | 7,4026 | 4,473970683 | 1,495772 | 2,991078 | 0,00278 | 1 | 0 | Intron (ENSMUST0000( | 13 | 1,07E+08 | 1,07E+08 | 13047 | 1 | 66254 | ENSMUST00000223870.2 | 10248 | ENSMUSG00000021692 | Dimt1 | DIM1 dimethyladenosine transferase 1-like (S. cerevisiae) |
| chr13 | 23769101 | 23770129 | 1029 | * | 12,85893 | 4,477025143 | 1,339883 | 3,341356 | 0,000834 | 0,470829715 | 1,2763 | 5' UTR | 13 | 23765581 | 23766718 | 1138 | 1 | 319156 | ENSMUST00000102968.3 | 3520 | ENSMUSG00000061482 | H4c4 | H4 clustered histone 4 |
| chr4 | 45976395 | 45977168 | 774 | * | 7,498819 | 4,477884771 | 1,574867 | 2,843342 | 0,004464 | 0,752744767 | 1,360622 | Intron (ENSMUST0000( | 4 | 45972834 | 45987622 | 14789 | 1 | 100121 | ENSMUST00000151422.2 | 3561 | ENSMUSG00000035517 | Tdrd7 | tudor domain containing 7 |
| chr18 | 78374458 | 78375171 | 714 | * | 7,380121 | 4,477949308 | 1,561451 | 2,867813 | 0,004133 | 1 | 0 | Intron (ENSMUST0000( | 18 | 78190155 | 78252309 | 62155 | 2 | 27411 | ENSMUST00000025434.14 | -122149 | ENSMUSG00000024552 | Slc14a2 | solute carrier family 14 (urea transporter), member 2 |
| chr3 | 13066561 | 13067158 | 598 | * | 6,432315 | 4,478312443 | 1,665429 | 2,688985 | 0,007167 | 1 | 0 | Distal Intergenic | 3 | 13536715 | 13842163 | 305449 | 1 | 76897 | ENSMUST00000191806.3 | -469557 | ENSMUSG00000039717 | Ralyl | RALY RNA binding protein-like |
| chr9 | 5980464 | 5981414 | 951 | * | 7,843154 | 4,479641175 | 1,467585 | 3,05239 | 0,00227 | 0,642870233 | 1,360622 | Distal Intergenic | 9 | 6168584 | 6378845 | 210262 | 1 | 71785 | ENSMUST00000058692.9 | -187170 | ENSMUSG00000032006 | Pdgfd | platelet-derived growth factor, D polypeptide |
| chrX | 99642667 | 99643629 | 963 | * | 5,817559 | 4,480714155 | 1,638365 | 2,734869 | 0,006241 | 1 | 0 | Distal Intergenic | 20 | 99633760 | 99638475 | 4716 | 2 | 57385 | ENSMUST00000053373.2 | -4192 | ENSMUSG00000044359 | P2ry4 | pyrimidinergic receptor P2Y, G-protein coupled, 4 |
| chrX | 1,47E+08 | 1,47E+08 | 596 | * | 6,495481 | 4,482866768 | 1,709159 | 2,622849 | 0,00872 | 1 | 0 | Promoter (<=1kb) | 20 | 1,47E+08 | 1,47E+08 | 40246 | 1 | 434866 | ENSMUST00000101190.11 | 116 | ENSMUSG00000096834 | Gm15127 | predicted gene 15127 |
| chr11 | 77488439 | 77489590 | 1152 | * | 9,628855 | 4,485006688 | 1,399506 | 3,204708 | 0,001352 | 0,587540639 | 1,086118 | Intron (ENSMUST0000( | 11 | 77419988 | 77498641 | 78654 | 2 | 216965 | ENSMUST00000058496.8 | 9051 | ENSMUSG00000017291 | Taok1 | TAO kinase 1 |
| chrX | 96218081 | 96219058 | 978 | * | 6,688496 | 4,485183408 | 1,588956 | 2,822724 | 0,004762 | 1 | 0 | Distal Intergenic | 20 | 96116202 | 96125710 | 9509 | 1 | 245526 | ENSMUST00000033556.4 | 101879 | ENSMUSG00000031212 | Pgr15l | G protein-coupled receptor 15-like |
| chr2 | 1,31E+08 | 1,31E+08 | 830 | * | 6,878362 | 4,486025375 | 1,580111 | 2,839058 | 0,004525 | 1 | 0 | Distal Intergenic | 2 | 1,31E+08 | 1,31E+08 | 16434 | 2 | 11550 | ENSMUST00000103184.4 | -41571 | ENSMUSG00000027335 | Adra1d | adrenergic receptor, alpha 1d |
| chr18 | 10614216 | 10615430 | 1215 | * | 8,712989 | 4,486117505 | 1,447439 | 3,099347 | 0,001939 | 0,604577026 | 1,425929 | Promoter (2-3kb) | 18 | 10617775 | 10628230 | 10456 | 1 | 20641 | ENSMUST00000002551.5 | -2345 | ENSMUSG00000002477 | Snrpd1 | small nuclear ribonucleoprotein D1 |
| chr15 | 79107089 | 79107788 | 700 | * | 6,919749 | 4,4882035 | 1,621587 | 2,767784 | 0,005644 | 1 | 0 | Intron (ENSMUST0000( | 15 | 79113373 | 79133684 | 20312 | 1 | 18693 | ENSMUST00000163571.8 | -5585 | ENSMUSG00000068206 | Pick1 | protein interacting with C kinase 1 |
| chr9 | 44309803 | 44310480 | 678 | * | 6,933665 | 4,48853283 | 1,565086 | 2,867915 | 0,004132 | 1 | 0 | Promoter (<=1kb) | 9 | 44309771 | 44313765 | 3995 | 1 | 14385 | ENSMUST00000213268.2 | 32 | ENSMUSG00000032114 | Slc37a4 | solute carrier family 37 (glucose-6-phosphate transporter), member 4 |
| chr16 | 24513976 | 24514710 | 735 | * | 7,834926 | 4,488867121 | 1,495331 | 3,001922 | 0,002683 | 1 | 0 | Intron (ENSMUST0000( | 16 | 24540605 | 24798280 | 257676 | 1 | 210126 | ENSMUST00000115314.4 | -25895 | ENSMUSG00000033306 | Lpp | LIM domain containing preferred translocation partner in lipoma |
| chr15 | 94874648 | 94875380 | 733 | * | 10,1634 | 4,488978232 | 1,472296 | 3,048964 | 0,002296 | 0,663630601 | 1,211315 | Intron (ENSMUST0000( | 15 | 94884888 | 94885019 | 132 | 2 | 1,15E+08 | ENSMUST00000104259.3 | 9639 | ENSMUSG00000077447 | Gm23129 | predicted gene, 23129 |
| chr4 | 44376564 | 44377842 | 1279 | * | 9,811109 | 4,489091427 | 1,506363 | 2,980086 | 0,002882 | 0,705098751 | 1,173325 | Distal Intergenic | 4 | 44412469 | 44412595 | 127 | 2 | 1,15E+08 | ENSMUST00000104399.2 | 34753 | ENSMUSG00000077587 | Gm24376 | predicted gene, 24376 |
| chr9 | 44652570 | 44653191 | 622 | * | 5,231317 | 4,489959624 | 1,683348 | 2,66728 | 0,007647 | 1 | 0 | 3' UTR | 9 | 44648110 | 44648244 | 135 | 2 | 1,15E+08 | ENSMUST00000082724.3 | -4326 | ENSMUSG00000064658 | Gm24166 | predicted gene, 24166 |
| chr12 | 70049885 | 70050863 | 979 | * | 5,917694 | 4,493836056 | 1,575075 | 2,853093 | 0,00433 | 1 | 0 | Distal Intergenic | 12 | 70041564 | 70041666 | 103 | 2 | 1,15E+08 | ENSMUST00000158528.4 | -8219 | ENSMUSG00002075853 | Gm23579 | predicted gene, 23579 |
| chr4 | 1,41E+08 | 1,41E+08 | 711 | * | 5,141101 | 4,496767049 | 1,706668 | 2,634823 | 0,008418 | 1 | 0 | Distal Intergenic | 4 | 1,41E+08 | 1,41E+08 | 1430 | 1 | 1,01E+08 | ENSMUST00000142429.3 | 6051 | ENSMUSG00000085395 | Gm13056 | predicted gene 13056 |
| chr7 | 1,23E+08 | 1,23E+08 | 936 | * | 6,483336 | 4,499511435 | 1,547236 | 2,908097 | 0,003636 | 1 | 0 | Intron (ENSMUST0000( | 7 | 1,23E+08 | 1,23E+08 | 53437 | 1 | 233833 | ENSMUST00000206888.2 | 3525 | ENSMUSG00000052707 | Tnrc6a | trinucleotide repeat containing 6a |
| chrX | 1,33E+08 | 1,33E+08 | 1337 | * | 12,0734 | 4,499883093 | 1,150603 | 3,910892 | 9,2E-05 | 0,223981956 | 0,744375 | Promoter (<=1kb) | 20 | 1,33E+08 | 1,33E+08 | 14497 | 2 | 215201 | ENSMUST00000087540.4 | -533 | ENSMUSG00000067369 | Trmt2b | TRM2 tRNA methyltransferase 2B |
| chr5 | 28638294 | 28639357 | 1064 | * | 7,224329 | 4,500071669 | 1,539742 | 2,922614 | 0,003471 | 1 | 0 | Distal Intergenic | 5 | 28615659 | 28619027 | 3369 | 1 | 381626 | ENSMUST00000133313.2 | 22635 | ENSMUSG00000048271 | Rbm33 | RNA binding motif protein 33 |

| | | | | | | | | | | | | | | | | | | | | | | | |
|---|---|---|---|---|---|---|---|---|---|---|---|---|---|---|---|---|---|---|---|---|---|---|---|
| chr1 | 15558281 | 15559073 | 793 | * | 6,051132 | 4,502002464 | 1,703137 | 2,643359 | 0,008209 | 1 | 0 | Intron (ENSMUST0000( | 1 | 15382676 | 15793974 | 411299 | 1 | 98741 | ENSMUST00000170146.3 | 175605 | ENSMUSG00000092083 | Kcnb2 | potassium voltage gated channel, Shab-related subfamily, member 2 |
| chr11 | 1,19E+08 | 1,19E+08 | 2535 | * | 17,3415 | 4,502770646 | 1,306304 | 3,446956 | 0,000567 | 0,306095127 | 2,386008 | Promoter (<=1kb) | 11 | 1,19E+08 | 1,19E+08 | 18401 | 1 | 268512 | ENSMUST00000151612.2 | 0 | ENSMUSG00000039908 | Slc26a11 | solute carrier family 26, member 11 |
| chr9 | 53737906 | 53738915 | 1010 | * | 8,717553 | 4,502806238 | 1,538132 | 2,927452 | 0,003418 | 0,698318058 | 1,425929 | Distal Intergenic | 9 | 53757448 | 53761844 | 4397 | 1 | 66402 | ENSMUST00000048485.7 | -18533 | ENSMUSG00000042045 | Sln | sarcolipin |
| chr1 | 65039434 | 65040225 | 792 | * | 5,180463 | 4,504560002 | 1,660935 | 2,712063 | 0,006687 | 1 | 0 | Promoter (<=1kb) | 1 | 65040466 | 65040621 | 156 | 2 | 1,15E+08 | ENSMUST00000104283.4 | 396 | ENSMUSG00000077471 | Gm24788 | predicted gene, 24788 |
| chr4 | 1,48E+08 | 1,48E+08 | 833 | * | 6,436162 | 4,504876281 | 1,657409 | 2,718023 | 0,006567 | 1 | 0 | Promoter (1-2kb) | 4 | 1,48E+08 | 1,48E+08 | 1418 | 1 | 18158 | ENSMUST00000103231.5 | 1482 | ENSMUSG00000029019 | Nppb | natriuretic peptide type B |
| chr16 | 9622364 | 9623542 | 1179 | * | 7,788007 | 4,505650056 | 1,472637 | 3,05958 | 0,002216 | 1 | 0 | Intron (ENSMUST0000( | 16 | 9579363 | 9615197 | 35835 | 2 | 14811 | ENSMUST00000199267.2 | -7167 | ENSMUSG00000059003 | Grin2a | glutamate receptor, ionotropic, NMDA2A (epsilon 1) |
| chr1 | 1,92E+08 | 1,92E+08 | 1368 | * | 11,83008 | 4,506704653 | 1,325317 | 3,400473 | 0,000673 | 0,434848027 | 1,2763 | Distal Intergenic | 1 | 1,92E+08 | 1,92E+08 | 10914 | 1 | 74023 | ENSMUST00000180463.3 | -3400 | ENSMUSG00000049353 | Rd3 | retinal degeneration 3 |
| chr13 | 38534819 | 38535643 | 825 | * | 5,487987 | 4,509601023 | 1,677448 | 2,68837 | 0,00718 | 1 | 0 | Intron (ENSMUST0000( | 13 | 38530129 | 38541945 | 11817 | 1 | 12161 | ENSMUST00000224452.2 | 4690 | ENSMUSG00000039004 | Bmp6 | bone morphogenetic protein 6 |
| chr11 | 54760030 | 54761278 | 1249 | * | 9,683921 | 4,51192444 | 1,28459 | 3,512347 | 0,000444 | 0,339611164 | 1,425929 | Promoter (2-3kb) | 11 | 54757307 | 54761133 | 3827 | 1 | 15254 | ENSMUST00000117710.2 | 2723 | ENSMUSG00000020267 | Hint1 | histidine triad nucleotide binding protein 1 |
| chr4 | 1,36E+08 | 1,36E+08 | 1180 | * | 5,390418 | 4,512796741 | 1,749307 | 2,579762 | 0,009887 | 1 | 0 | Promoter (1-2kb) | 4 | 1,36E+08 | 1,36E+08 | 3100 | 1 | 14105 | ENSMUST00000129718.2 | 1524 | ENSMUSG00000028676 | Srsf10 | serine and arginine-rich splicing factor 10 |
| chr16 | 8608350 | 8609397 | 1048 | * | 10,13766 | 4,513250988 | 1,408574 | 3,204127 | 0,001355 | 0,534851486 | 1,425929 | Promoter (<=1kb) | 16 | 8507459 | 8610172 | 102714 | 2 | 252870 | ENSMUST00000160405.8 | 775 | ENSMUSG00000022710 | Usp7 | ubiquitin specific peptidase 7 |
| chr9 | 64817821 | 64818992 | 1172 | * | 8,577184 | 4,514482802 | 1,465915 | 3,079635 | 0,002073 | 0,627505501 | 1,360622 | Promoter (<=1kb) | 9 | 64819518 | 64826686 | 7169 | 1 | 102442 | ENSMUST00000213744.2 | -526 | ENSMUSG00000053641 | Dennd4a | DENN/MADD domain containing 4A |
| chrX | 73296432 | 73297513 | 1082 | * | 5,925024 | 4,514690944 | 1,537238 | 2,936885 | 0,003315 | 1 | 0 | Promoter (<=1kb) | 20 | 73298293 | 73301493 | 3201 | 1 | 13726 | ENSMUST00000002029.13 | -780 | ENSMUSG00000001964 | Emd | emerin |
| chr5 | 1,24E+08 | 1,24E+08 | 1532 | * | 11,13471 | 4,51626642 | 1,272261 | 3,549795 | 0,000386 | 0,35147835 | 1,173325 | Distal Intergenic | 5 | 1,24E+08 | 1,24E+08 | 21661 | 1 | 68184 | ENSMUST00000023869.15 | -4532 | ENSMUSG00000023106 | Denr | density-regulated protein |
| chrX | 1,2E+08 | 1,2E+08 | 885 | * | 6,860707 | 4,517397846 | 1,660406 | 2,720658 | 0,006515 | 1 | 0 | Distal Intergenic | 20 | 1,2E+08 | 1,2E+08 | 128 | 1 | 1,15E+08 | ENSMUST00000104247.3 | 82029 | ENSMUSG00000077435 | Gm22590 | predicted gene, 22590 |
| chr15 | 1,01E+08 | 1,01E+08 | 737 | * | 8,058065 | 4,518416619 | 1,580015 | 2,85973 | 0,00424 | 1 | 0 | Intron (ENSMUST0000( | 15 | 1,01E+08 | 1,01E+08 | 63 | 1 | 1,02E+08 | ENSMUST00000199512.3 | 5842 | ENSMUSG00000106563 | Mir6962 | microRNA 6962 |
| chrX | 88854667 | 88855476 | 810 | * | 5,33569 | 4,518579922 | 1,725235 | 2,61911 | 0,008816 | 1 | 0 | Distal Intergenic | 20 | 88795874 | 88799097 | 3224 | 1 | 245509 | ENSMUST00000113958.2 | 58793 | ENSMUSG00000079513 | 4932429P05Rik | RIKEN cDNA 4932429P05 gene |
| chr17 | 50124453 | 50124869 | 417 | * | 5,239672 | 4,51948632 | 1,726808 | 2,617249 | 0,008864 | 1 | 0 | Exon (ENSMUST000001 | 17 | 50124654 | 50128853 | 4200 | 2 | 1,03E+08 | ENSMUST00000159971.2 | 3984 | | Gm16555 | predicted gene 16555 |
| chr4 | 70727422 | 70727988 | 567 | * | 6,108475 | 4,520322928 | 1,623311 | 2,784632 | 0,005359 | 1 | 0 | Distal Intergenic | 4 | 70380063 | 70453232 | 73170 | 2 | 230316 | ENSMUST00000037881.3 | -274190 | ENSMUSG00000039270 | Megf9 | multiple EGF-like-domains 9 |
| chr2 | 1,57E+08 | 1,57E+08 | 1295 | * | 8,330542 | 4,521324551 | 1,551473 | 2,914214 | 0,003566 | 0,705187732 | 1,449706 | Promoter (<=1kb) | 2 | 1,57E+08 | 1,57E+08 | 58620 | 2 | 19650 | ENSMUST00000029170.8 | 0 | ENSMUSG00000027641 | Rbl1 | RB transcriptional corepressor like 1 |
| chr1 | 3215041 | 3216141 | 1101 | * | 8,624466 | 4,521614588 | 1,522453 | 2,969954 | 0,002978 | 0,676922067 | 1,449706 | Distal Intergenic | 1 | 3172239 | 3172348 | 110 | 1 | 1,15E+08 | ENSMUST00000082908.3 | 42802 | ENSMUSG00000064842 | Gm26206 | predicted gene, 26206 |
| chr10 | 34920513 | 34921710 | 1198 | * | 8,026168 | 4,521661051 | 1,615805 | 2,798395 | 0,005136 | 1 | 0 | Intron (ENSMUST0000( | 10 | 34988194 | 34988327 | 134 | 1 | 1,15E+08 | ENSMUST00000104561.3 | -66484 | ENSMUSG00000077749 | Gm26341 | predicted gene, 26341 |
| chr2 | 81429171 | 81430010 | 840 | * | 5,749291 | 4,522941119 | 1,672923 | 2,703615 | 0,006859 | 1 | 0 | Distal Intergenic | 2 | 81883566 | 82090223 | 206658 | 1 | 241514 | ENSMUST00000047527.8 | -453556 | ENSMUSG00000070866 | Zfp804a | zinc finger protein 804A |
| chr18 | 70661520 | 70662833 | 1314 | * | 9,293304 | 4,523972561 | 1,741779 | 2,597329 | 0,009395 | 0,797848249 | 2,237403 | Promoter (<=1kb) | 18 | 70641751 | 70663209 | 21459 | 2 | 26447 | ENSMUST00000161542.8 | 376 | ENSMUSG00000038425 | Poli | polymerase (DNA directed), iota |
| chr16 | 94190815 | 94192713 | 1899 | * | 11,13129 | 4,524826822 | 1,203094 | 3,760993 | 0,000169 | 0,275349208 | 0,918255 | 3' UTR | 16 | 94184731 | 94269098 | 84368 | 1 | 22129 | ENSMUST00000231915.2 | 6084 | ENSMUSG00000040785 | Ttc3 | tetratricopeptide repeat domain 3 |
| chr6 | 98029554 | 98030256 | 703 | * | 7,055913 | 4,525439461 | 1,601287 | 2,826126 | 0,004711 | 1 | 0 | Distal Intergenic | 6 | 97968882 | 97995136 | 26255 | 1 | 17342 | ENSMUST00000203938.2 | 60672 | ENSMUSG00000035158 | Mitf | melanogenesis associated transcription factor |
| chr2 | 12658951 | 12660291 | 1341 | * | 12,35432 | 4,527131008 | 1,211591 | 3,736519 | 0,000187 | 0,312833142 | 0,738441 | Distal Intergenic | 2 | 12352074 | 12424281 | 72208 | 2 | 66960 | ENSMUST00000028105.13 | -234670 | ENSMUSG00000026767 | Mindy3 | MINDY lysine 48 deubiquitinase 3 |
| chr19 | 27390491 | 27391409 | 919 | * | 6,492011 | 4,530103388 | 1,664851 | 2,721026 | 0,006508 | 1 | 0 | Promoter (<=1kb) | 19 | 27367461 | 27389664 | 22204 | 2 | 52874 | ENSMUST00000236585.2 | -827 | ENSMUSG00000041360 | Pum3 | pumilio RNA-binding family member 3 |
| chr4 | 1,37E+08 | 1,37E+08 | 1869 | * | 14,62149 | 4,530160764 | 1,169949 | 3,872102 | 0,000108 | 0,170809454 | 1,41696 | Promoter (<=1kb) | 4 | 1,37E+08 | 1,37E+08 | 34324 | 2 | 230848 | ENSMUST00000145505.2 | 0 | ENSMUSG00000060862 | Zbtb40 | zinc finger and BTB domain containing 40 |
| chr2 | 1,73E+08 | 1,73E+08 | 596 | * | 5,2583 | 4,534340299 | 1,697634 | 2,670976 | 0,007563 | 1 | 0 | Intron (ENSMUST0000( | 2 | 1,73E+08 | 1,73E+08 | 11651 | 1 | 56190 | ENSMUST00000130430.2 | 3071 | ENSMUSG00000027510 | Rbm38 | RNA binding motif protein 38 |
| chr2 | 15073684 | 15074257 | 574 | * | 5,221561 | 4,534814897 | 1,65155 | 2,745793 | 0,006036 | 1 | 0 | Promoter (<=1kb) | 2 | 15072950 | 15081416 | 8467 | 1 | 75869 | ENSMUST00000129509.3 | 734 | ENSMUSG00000017418 | Arl5b | ADP-ribosylation factor-like 5B |
| chr1 | 72272639 | 72273979 | 1341 | * | 10,8572 | 4,535680211 | 1,367711 | 3,316256 | 0,000912 | 0,586304784 | 0,738441 | Promoter (1-2kb) | 1 | 72275642 | 72305891 | 30250 | 1 | 320933 | ENSMUST00000247711.1 | -1663 | | D230017M19Rik | RIKEN cDNA D230017M19 gene |
| chr1 | 17926414 | 17927118 | 705 | * | 6,532873 | 4,537431986 | 1,687095 | 2,689494 | 0,007156 | 1 | 0 | Distal Intergenic | 1 | 17763128 | 17797870 | 34743 | 2 | 1,03E+08 | ENSMUST00000127328.2 | -128544 | ENSMUSG00000085125 | Gm16070 | predicted gene 16070 |
| chr18 | 35023360 | 35023935 | 576 | * | 6,713559 | 4,537811929 | 1,560925 | 2,90713 | 0,003648 | 1 | 0 | Distal Intergenic | 18 | 34994260 | 34998009 | 3750 | 1 | 13653 | ENSMUST00000165033.2 | 29100 | ENSMUSG00000038418 | Egr1 | early growth response 1 |
| chr2 | 1,28E+08 | 1,28E+08 | 1300 | * | 12,79262 | 4,539637237 | 1,224039 | 3,708737 | 0,000208 | 0,259309257 | 1,2763 | Promoter (<=1kb) | 2 | 1,28E+08 | 1,28E+08 | 607 | 1 | 12125 | ENSMUST00000132201.2 | 0 | ENSMUSG00000027381 | Bcl2l11 | BCL2-like 11 (apoptosis facilitator) |
| chr3 | 1,06E+08 | 1,06E+08 | 1042 | * | 8,979278 | 4,539638699 | 1,287738 | 3,525283 | 0,000423 | 0,339611164 | 1,360622 | Promoter (2-3kb) | 3 | 1,06E+08 | 1,06E+08 | 9305 | 2 | 53975 | ENSMUST00000090680.11 | -2955 | ENSMUSG00000027905 | Ddx20 | DEAD box helicase 20 |
| chr7 | 1,1E+08 | 1,1E+08 | 605 | * | 6,803383 | 4,539715325 | 1,728303 | 2,62669 | 0,008622 | 1 | 0 | Distal Intergenic | 7 | 1,1E+08 | 1,1E+08 | 152791 | 2 | 69781 | ENSMUST00000185931.7 | -24193 | ENSMUSG00000101585 | 1600010M07Rik | RIKEN cDNA 1600010M07 gene |
| chr12 | 79550628 | 79551341 | 714 | * | 5,10842 | 4,540421506 | 1,703724 | 2,664998 | 0,007699 | 1 | 0 | Intron (ENSMUST0000( | 12 | 79374127 | 79748630 | 374504 | 1 | 19363 | ENSMUST00000221257.2 | 176501 | ENSMUSG00000059060 | Rad51b | RAD51 paralog B |
| chr3 | 1,41E+08 | 1,41E+08 | 474 | * | 6,49459 | 4,541977352 | 1,626605 | 2,792305 | 0,005233 | 1 | 0 | Distal Intergenic | 3 | 1,41E+08 | 1,41E+08 | 2352 | 2 | 18598 | ENSMUST00000057860.3 | 40357 | ENSMUSG00000047674 | Pdha2 | pyruvate dehydrogenase E1 alpha 2 |
| chr8 | 1,27E+08 | 1,27E+08 | 652 | * | 5,186409 | 4,54407484 | 1,72303 | 2,637257 | 0,008358 | 1 | 0 | Intron (ENSMUST0000( | 8 | 1,27E+08 | 1,27E+08 | 5691 | 2 | 270110 | ENSMUST00000054960.9 | 16653 | ENSMUSG00000051495 | Irf2bp2 | interferon regulatory factor 2 binding protein 2 |
| chr11 | 86833564 | 86834370 | 807 | * | 6,928806 | 4,546847328 | 1,542631 | 2,947464 | 0,003204 | 1 | 0 | Intron (ENSMUST0000( | 11 | 86831425 | 86862850 | 31426 | 2 | 77864 | ENSMUST00000238921.2 | 28480 | ENSMUSG00000018427 | Ypel2 | yippee like 2 |
| chr2 | 86068133 | 86068771 | 639 | * | 5,157487 | 4,547517604 | 1,737848 | 2,616752 | 0,008877 | 1 | 0 | Promoter (<=1kb) | 2 | 86066215 | 86067177 | 963 | 2 | 259016 | ENSMUST00000099900.2 | -956 | ENSMUSG00000075195 | Olfr1048 | olfactory receptor 1048 |
| chr17 | 44037548 | 44038319 | 772 | * | 7,432633 | 4,547864106 | 1,53246 | 2,967688 | 0,003 | 0,684274806 | 1,360622 | Intron (ENSMUST0000( | 17 | 44042834 | 44062322 | 19489 | 1 | 56050 | ENSMUST00000233165.2 | -4515 | ENSMUSG00000023963 | Cyp39a1 | cytochrome P450, family 39, subfamily a, polypeptide 1 |
| chr11 | 53990945 | 53992397 | 1453 | * | 23,989466 | 4,550548835 | 0,8814461 | 5,1625947 | 2,436E-07 | 0,001237282 | 2,9510621 | Promoter (<=1kb) | 11 | 53991750 | 54022478 | 30729 | 1 | 18452 | ENSMUST00000093107.12 | 0 | ENSMUSG00000018906 | P4ha2 | procollagen-proline, 2-oxoglutarate 4-dioxygenase (proline 4-hydroxylase), alpha II polypeptide |
| chr16 | 48768592 | 48769798 | 1207 | * | 7,894471 | 4,551158013 | 1,462237 | 3,112463 | 0,001855 | 1 | 0 | Exon (ENSMUST000001 | 16 | 48765905 | 48774319 | 8415 | 2 | 224170 | ENSMUST00000133377.2 | 4521 | ENSMUSG00000064061 | Dzip3 | DAZ interacting protein 3, zinc finger |
| chr18 | 65714906 | 65716221 | 1316 | * | 11,19491 | 4,551499168 | 1,234904 | 3,685712 | 0,000228 | 0,272208984 | 1,2763 | Promoter (<=1kb) | 18 | 65715156 | 65820938 | 105783 | 1 | 328977 | ENSMUST00000237553.2 | 0 | ENSMUSG00000042439 | Zfp532 | zinc finger protein 532 |
| chr13 | 18031069 | 18031511 | 443 | * | 6,679818 | 4,552896426 | 1,610412 | 2,827162 | 0,004696 | 1 | 0 | Distal Intergenic | 13 | 17984176 | 17984282 | 107 | 2 | 1,15E+08 | ENSMUST00000083232.3 | -46787 | ENSMUSG00000118773 | Gm24018 | predicted gene, 24018 |
| chr9 | 7184073 | 7184915 | 843 | * | 6,237969 | 4,553472442 | 1,676971 | 2,715296 | 0,006622 | 1 | 0 | Promoter (<=1kb) | 9 | 6928503 | 7184446 | 255944 | 2 | 110350 | ENSMUST00000140466.8 | 0 | ENSMUSG00000047193 | Dync2h1 | dynein cytoplasmic 2 heavy chain 1 |
| chr6 | 52691242 | 52692056 | 815 | * | 7,962738 | 4,553949137 | 1,478601 | 3,079905 | 0,002071 | 1 | 0 | Promoter (<=1kb) | 6 | 52691335 | 52704284 | 12950 | 1 | 52440 | ENSMUST00000129660.2 | 0 | ENSMUSG00000004535 | Tax1bp1 | Tax1 (human T cell leukemia virus type I) binding protein 1 |
| chr5 | 79017460 | 79018019 | 560 | * | 6,149774 | 4,554185855 | 1,678878 | 2,712637 | 0,006675 | 1 | 0 | Distal Intergenic | 5 | 78927706 | 78927841 | 136 | 1 | 1,15E+08 | ENSMUST00000103990.3 | 89754 | ENSMUSG00000077178 | Gm25756 | predicted gene, 25756 |
| chr15 | 43750221 | 43750671 | 451 | * | 6,329201 | 4,555228861 | 1,640849 | 2,776142 | 0,005501 | 1 | 0 | Distal Intergenic | 15 | 43728042 | 43733432 | 5391 | 2 | 239408 | ENSMUST00000067469.6 | -16789 | ENSMUSG00000054409 | Tmem74 | transmembrane protein 74 |
| chr14 | 69894014 | 69895347 | 1334 | * | 9,105929 | 4,556306535 | 1,271647 | 3,582995 | 0,00034 | 0,311240408 | 1,360622 | Intron (ENSMUST0000( | 14 | 69875118 | 69882201 | 7084 | 2 | 1,03E+08 | ENSMUST00000184491.2 | -11813 | | Gm27177 | predicted gene 27177 |
| chrX | 1,26E+08 | 1,26E+08 | 846 | * | 5,245228 | 4,557838811 | 1,622726 | 2,808755 | 0,004973 | 1 | 0 | Distal Intergenic | 20 | 1,26E+08 | 1,26E+08 | 2037 | 2 | 1,19E+08 | ENSMUST00000117443.2 | 108749 | ENSMUSG00000082772 | LOC118568449 | vigilin-like |
| chr5 | 66388884 | 66389996 | 1113 | * | 8,228709 | 4,564118863 | 1,452897 | 3,141392 | 0,001681 | 0,57149888 | 1,449706 | Distal Intergenic | 5 | 66417240 | 66454405 | 37166 | 1 | 70918 | ENSMUST00000202994.4 | -27244 | ENSMUSG00000029206 | Nsun7 | NOL1/NOP2/Sun domain family, member 7 |
| chr14 | 73680607 | 73682168 | 1562 | * | 6,748258 | 4,566667777 | 1,59275 | 2,867159 | 0,004142 | 1 | 0 | Distal Intergenic | 14 | 73599666 | 73622729 | 23064 | 2 | 16432 | ENSMUST00000022704.9 | -57878 | ENSMUSG00000022108 | Itm2b | integral membrane protein 2B |
| chr9 | 59393934 | 59395409 | 1476 | * | 23,562459 | 4,56695703 | 0,8768677 | 5,2082622 | 1,906E-07 | 0,001087015 | 2,8800171 | Promoter (<=1kb) | 9 | 59300396 | 59393901 | 93506 | 2 | 23806 | ENSMUST00000026264.12 | -33 | ENSMUSG00000025234 | Arih1 | ariadne RBR E3 ubiquitin protein ligase 1 |
| chr13 | 80381969 | 80383037 | 1069 | * | 5,099579 | 4,567096248 | 1,752264 | 2,606397 | 0,00915 | 1 | 0 | Distal Intergenic | 13 | 80306782 | 80306909 | 128 | 1 | 1,15E+08 | ENSMUST00000122712.4 | 75187 | ENSMUSG00000084661 | Gm24635 | predicted gene, 24635 |
| chr4 | 50822540 | 50823940 | 1401 | * | 11,71052 | 4,569892753 | 1,360778 | 3,358293 | 0,000784 | 0,551069862 | 0,744375 | Distal Intergenic | 4 | 51216678 | 51229928 | 13251 | 1 | 74914 | ENSMUST00000166749.2 | -392738 | ENSMUSG00000039555 | Cylc2 | cylicin, basic protein of sperm head cytoskeleton 2 |
| chr6 | 76919738 | 76920165 | 428 | * | 6,586571 | 4,570804761 | 1,6137 | 2,8325 | 0,004619 | 1 | 0 | Intron (ENSMUST0000( | 6 | 76859739 | 76879871 | 20133 | 2 | 12386 | ENSMUST00000162991.2 | -39867 | ENSMUSG00000063063 | Ctnna2 | catenin (cadherin associated protein), alpha 2 |
| chr16 | 31272428 | 31274166 | 1739 | * | 14,26443 | 4,572463273 | 1,151048 | 3,972436 | 7,11E-05 | 0,17036682 | 0,938682 | 3' UTR | 16 | 31254909 | 31275568 | 20660 | 1 | 71911 | ENSMUST00000149039.2 | 17519 | ENSMUSG00000046598 | Bdh1 | 3-hydroxybutyrate dehydrogenase, type 1 |
| chr17 | 44962537 | 44963375 | 839 | * | 7,278177 | 4,574687818 | 1,562051 | 2,928641 | 0,003404 | 1 | 0 | Intron (ENSMUST0000( | 17 | 44950568 | 45045580 | 95013 | 2 | 12393 | ENSMUST00000160199.2 | 82205 | ENSMUSG00000039153 | Runx2 | runt related transcription factor 2 |
| chr5 | 122467796 | 122471313 | 3518 | * | 27,870756 | 4,57741198 | 0,8426258 | 5,432319 | 5,563E-08 | 0,000412118 | 2,8585108 | Exon (ENSMUST0000011 | 5 | 122463569 | 122489357 | 25789 | 2 | 231724 | ENSMUST00000149600.8 | 18044 | ENSMUSG00000038569 | Rad9b | RAD9 checkpoint clamp component B |
| chr9 | 5430616 | 5431109 | 494 | * | 4,684213 | 4,578060535 | 1,775041 | 2,57913 | 0,009905 | 1 | 0 | Distal Intergenic | 9 | 5345504 | 5358823 | 13320 | 1 | 12364 | ENSMUST00000151332.2 | 85112 | ENSMUSG00000025887 | Casp12 | caspase 12 |
| chrX | 160696618 | 160699263 | 2646 | * | 18,421587 | 4,57835415 | 1,0203751 | 4,4869323 | 7,226E-06 | 0,021335371 | 2,1545044 | Intron (ENSMUST0000008 | 20 | 160597686 | 160615172 | 17487 | 2 | 195726 | ENSMUST00000136860.3 | -81446 | ENSMUSG00000086540 | Scml1 | Scm polycomb group protein like 1 |
| chr16 | 33086460 | 33087893 | 1434 | * | 13,58911 | 4,579466997 | 1,347917 | 3,397441 | 0,00068 | 0,487376501 | 0,938682 | Promoter (<=1kb) | 16 | 33088128 | 33090215 | 2088 | 1 | 69150 | ENSMUST00000231242.2 | -235 | ENSMUSG00000022808 | Snx4 | sorting nexin 4 |
| chr17 | 39738006 | 39738737 | 732 | * | 5,019203 | 4,57977696 | 1,730069 | 2,647165 | 0,008117 | 1 | 0 | Distal Intergenic | 17 | 39717250 | 39721883 | 4634 | 1 | 1E+08 | ENSMUST00000179848.2 | 20756 | ENSMUSG00000094024 | Esp18 | exocrine gland secreted peptide 18 |
| chr5 | 33209516 | 33210964 | 1449 | * | 11,78535 | 4,580225597 | 1,494125 | 3,065491 | 0,002173 | 0,732373293 | 0,744375 | Distal Intergenic | 5 | 33179135 | 33184325 | 5191 | 1 | 22629 | ENSMUST00000136667.2 | 30381 | ENSMUSG00000018965 | Ywhah | tyrosine 3-monooxygenase/tryptophan 5-monooxygenase activation protein, eta polypeptide |
| chr4 | 1,39E+08 | 1,39E+08 | 1463 | * | 11,59831 | 4,581854608 | 1,349708 | 3,394702 | 0,000687 | 0,531203841 | 0,744375 | Promoter (1-2kb) | 4 | 1,39E+08 | 1,39E+08 | 98920 | 1 | 12345 | ENSMUST00000102508.10 | -1282 | ENSMUSG00000028745 | Capzb | capping protein (actin filament) muscle Z-line, beta |
| chr2 | 1,54E+08 | 1,54E+08 | 875 | * | 9,836014 | 4,582438138 | 1,468721 | 3,12002 | 0,001808 | 0,627785088 | 1,173325 | Distal Intergenic | 2 | 1,54E+08 | 1,54E+08 | 8021 | 1 | 73388 | ENSMUST00000028984.13 | -21129 | ENSMUSG00000027482 | Bpifa3 | BPI fold containing family A, member 3 |
| chr1 | 75164915 | 75165784 | 870 | * | 6,327834 | 4,58297254 | 1,688097 | 2,714875 | 0,00663 | 1 | 0 | Promoter (1-2kb) | 1 | 75157509 | 75167026 | 9518 | 2 | 245860 | ENSMUST00000189820.7 | 1242 | ENSMUSG00000033124 | Atg9a | autophagy related 9A |
| chrX | 80149903 | 80150809 | 907 | * | 7,002401 | 4,58299464 | 1,544267 | 2,967747 | 0,003 | 1 | 0 | Distal Intergenic | 20 | 80114322 | 80141478 | 27157 | 1 | 192216 | ENSMUST00000026760.3 | 35581 | ENSMUSG00000025666 | Tmem47 | transmembrane protein 47 |
| chr17 | 36054523 | 36055501 | 979 | * | 7,789863 | 4,583608582 | 1,537483 | 2,981242 | 0,002871 | 0,681459438 | 1,360622 | Intron (ENSMUST0000( | 17 | 36048626 | 36050698 | 2073 | 2 | 1,19E+08 | ENSMUST00000173840.2 | -3825 | | Gm20443 | predicted gene 20443 |
| chr9 | 1,22E+08 | 1,22E+08 | 795 | * | 7,077461 | 4,583850031 | 1,534747 | 2,986713 | 0,00282 | 1 | 0 | Distal Intergenic | 9 | 1,22E+08 | 1,22E+08 | 10441 | 1 | 1,19E+08 | ENSMUST00000213591.2 | -3233 | ENSMUSG00000111465 | Gm47113 | predicted gene, 47113 |
| chr10 | 95368681 | 95369529 | 849 | * | 7,464892 | 4,584344819 | 1,511045 | 3,03389 | 0,002414 | 1 | 0 | Intron (ENSMUST0000( | 10 | 95377254 | 95378832 | 1579 | 1 | 93765 | ENSMUST00000219741.2 | -7725 | ENSMUSG00000074781 | Ube2n | ubiquitin-conjugating enzyme E2N |
| chr14 | 35758636 | 35759252 | 617 | * | 5,763076 | 4,586546092 | 1,644441 | 2,789121 | 0,005285 | 1 | 0 | Distal Intergenic | 14 | 35816874 | 35820238 | 3365 | 1 | 218921 | ENSMUST00000226305.2 | -57622 | ENSMUSG00000096405 | 4930474N05Rik | RIKEN cDNA 4930474N05 gene |
| chrX | 12254233 | 12255441 | 1209 | * | 8,70562 | 4,587343116 | 1,746631 | 2,626395 | 0,008629 | 0,78560377 | 2,237403 | Distal Intergenic | 20 | 12206021 | 12221235 | 15215 | 2 | 1E+08 | ENSMUST00000146535.8 | -32998 | ENSMUSG00000087591 | Gm14635 | predicted gene 14635 |
| chr12 | 80475139 | 80476656 | 1518 | * | 10,31423 | 4,58905658 | 1,377216 | 3,332125 | 0,000862 | 0,483145429 | 1,211315 | Intron (ENSMUST0000( | 12 | 80445609 | 80483219 | 37611 | 2 | 320808 | ENSMUST00000218498.2 | 6563 | ENSMUSG00000049106 | Dcaf5 | DDB1 and CUL4 associated factor 5 |
| chr3 | 1,08E+08 | 1,08E+08 | 811 | * | 7,58524 | 4,589300907 | 1,470471 | 3,120974 | 0,001803 | 1 | 0 | Distal Intergenic | 3 | 1,08E+08 | 1,08E+08 | 1644 | 2 | 229709 | ENSMUST00000153530.2 | 10397 | ENSMUSG00000027893 | Ahcyl1 | S-adenosylhomocysteine hydrolase-like 1 |
| chr15 | 81977397 | 81978965 | 1569 | * | 15,69541 | 4,590191132 | 1,703004 | 2,69535 | 0,007031 | 0,847259603 | 1,2763 | Exon (ENSMUST000002 | 15 | 81967364 | 82010937 | 43574 | 1 | 74369 | ENSMUST00000186125.2 | 10033 | ENSMUSG00000068117 | Mei1 | meiotic double-stranded break formation protein 1 |
| chr1 | 1,65E+08 | 1,65E+08 | 807 | * | 5,876732 | 4,590851905 | 1,688324 | 2,719178 | 0,006544 | 1 | 0 | Intron (ENSMUST0000( | 1 | 1,65E+08 | 1,65E+08 | 3884 | 2 | 16963 | ENSMUST00000027860.8 | 11439 | ENSMUSG00000026573 | Xcl1 | chemokine (C motif) ligand 1 |
| chr13 | 34143735 | 34145102 | 1368 | * | 8,50823 | 4,59089829 | 1,422047 | 3,228374 | 0,001245 | 0,516516173 | 1,449706 | Exon (ENSMUST000001 | 13 | 34148670 | 34169629 | 20960 | 1 | 18105 | ENSMUST00000220844.2 | -3568 | ENSMUSG00000046949 | Nqo2 | N-ribosyldihydronicotinamide quinone reductase 2 |
| chr16 | 6223144 | 6224130 | 987 | * | 5,563369 | 4,594868569 | 1,620334 | 2,835754 | 0,004572 | 1 | 0 | Intron (ENSMUST0000( | 16 | 6167002 | 7229390 | 1062389 | 1 | 268859 | ENSMUST00000056416.9 | 56142 | ENSMUSG00000008658 | Rbfox1 | RNA binding protein, fox-1 homolog (C. elegans) 1 |
| chr19 | 10153180 | 10154391 | 1212 | * | 7,97055 | 4,594958897 | 1,518221 | 3,026542 | 0,002474 | 1 | 0 | Exon (ENSMUST000002 | 19 | 10160252 | 10174238 | 13987 | 1 | 76267 | ENSMUST00000010807.6 | -5861 | ENSMUSG00000010663 | Fads1 | fatty acid desaturase 1 |
| chr16 | 20444197 | 20445023 | 827 | * | 8,83489 | 4,595397274 | 1,774889 | 2,589117 | 0,009622 | 0,883511731 | 1,425929 | Intron (ENSMUST0000( | 16 | 20448578 | 20464665 | 16088 | 1 | 107522 | ENSMUST00000003898.12 | -3555 | ENSMUSG00000022842 | Ece2 | endothelin converting enzyme 2 |
| chr8 | 1,12E+08 | 1,12E+08 | 391 | * | 5,24021 | 4,59548671 | 1,695262 | 2,710782 | 0,006712 | 1 | 0 | Intron (ENSMUST0000( | 8 | 1,12E+08 | 1,12E+08 | 27146 | 1 | 22640 | ENSMUST00000211926.2 | 9860 | ENSMUSG00000055835 | Zfp1 | zinc finger protein 1 |
| chr10 | 4969216 | 4970219 | 1004 | * | 7,041907 | 4,59582808 | 1,609071 | 2,856199 | 0,004287 | 1 | 0 | Distal Intergenic | 10 | 4971579 | 5019063 | 47485 | 2 | 64009 | ENSMUST00000095899.5 | 48844 | ENSMUSG00000096054 | Syne1 | spectrin repeat containing, nuclear envelope 1 |
| chr8 | 13797017 | 13797559 | 543 | * | 6,629474 | 4,596320271 | 1,673343 | 2,746789 | 0,006018 | 1 | 0 | Exon (ENSMUST000002 | 8 | 13807676 | 13816497 | 8822 | 1 | 69957 | ENSMUST00000130173.9 | -10117 | ENSMUSG00000038416 | Cdc16 | CDC16 cell division cycle 16 |
| chrX | 1,09E+08 | 1,09E+08 | 1524 | * | 7,87832 | 4,596441367 | 1,658669 | 2,771163 | 0,005586 | 1 | 0 | Distal Intergenic | 20 | 1,08E+08 | 1,08E+08 | 3682 | 1 | 622976 | ENSMUST00000060013.4 | 479681 | ENSMUSG00000048621 | Gm6377 | predicted gene 6377 |
| chr4 | 94870960 | 94871763 | 804 | * | 6,884688 | 4,596555212 | 1,525822 | 3,01251 | 0,002591 | 1 | 0 | Intron (ENSMUST0000( | 4 | 94867440 | 94916781 | 49342 | 1 | 1,05E+08 | ENSMUST00000149367.2 | 3520 | | Gm12694 | predicted gene 12694 |
| chr14 | 122064721 | 122066607 | 1887 | * | 15,217005 | 4,59672502 | 1,0550161 | 4,357019 | 1,318E-05 | 0,033029217 | 2,1416125 | Distal Intergenic | 14 | 121780838 | 122035249 | 254412 | 2 | 105445 | ENSMUST00000100299.11 | -29472 | ENSMUSG00000025558 | Dock9 | dedicator of cytokinesis 9 |
| chr18 | 58723523 | 58725028 | 1506 | * | 12,24162 | 4,598078772 | 1,312154 | 3,504221 | 0,000458 | 0,40037769 | 1,075735 | Intron (ENSMUST0000( | 18 | 58748431 | 58748561 | 131 | 2 | 1,15E+08 | ENSMUST00000104055.3 | 23533 | ENSMUSG00000077243 | Gm23270 | predicted gene, 23270 |
| chrX | 37705441 | 37706154 | 714 | * | 6,84042 | 4,598287217 | 1,645774 | 2,793996 | 0,005206 | 1 | 0 | Distal Intergenic | 20 | 37698680 | 37702277 | 3598 | 1 | 68995 | ENSMUST00000148938.2 | 6761 | ENSMUSG00000000355 | Mcts1 | malignant T cell amplified sequence 1 |
| chr3 | 86527033 | 86528866 | 1834 | * | 9,778633 | 4,60040219 | 1,257494 | 3,658388 | 0,000254 | 0,271805507 | 1,425929 | Exon (ENSMUST000001 | 3 | 86453357 | 86455590 | 2234 | 2 | 23937 | ENSMUST00000191695.3 | -71443 | ENSMUSG00000057777 | Mab21l2 | mab-21-like 2 |
| chr18 | 44542226 | 44543201 | 976 | * | 8,017452 | 4,601478672 | 1,472825 | 3,124253 | 0,001783 | 1 | 0 | Intron (ENSMUST0000( | 18 | 44513594 | 44558036 | 44443 | 1 | 70640 | ENSMUST00000025350.10 | 28632 | ENSMUSG00000024472 | Dcp2 | decapping mRNA 2 |
| chr16 | 10185876 | 10186300 | 425 | * | 6,864485 | 4,603184189 | 1,444344 | 3,187043 | 0,001437 | 1 | 0 | Intron (ENSMUST0000( | 16 | 10146207 | 10165455 | 19249 | 2 | 381033 | ENSMUST00000229862.2 | -20421 | ENSMUSG00000116270 | Gm1600 | predicted gene 1600 |
| chr14 | 39056276 | 39057514 | 1239 | * | 7,645573 | 4,605736244 | 1,503536 | 3,06327 | 0,002189 | 1 | 0 | Intron (ENSMUST0000( | 14 | 38942028 | 38972387 | 30360 | 1 | 1,03E+08 | ENSMUST00000177185.2 | 114248 | | Gm20642 | predicted gene 20642 |
| chr16 | 90484672 | 90485572 | 901 | * | 8,433509 | 4,606523625 | 1,423006 | 3,237177 | 0,001207 | 1 | 0 | Distal Intergenic | 16 | 90464327 | 90473888 | 9562 | 2 | 1,03E+08 | ENSMUST00000232154.2 | -10784 | | Gm36169 | predicted gene, 36169 |
| chr6 | 1,21E+08 | 1,21E+08 | 582 | * | 6,428217 | 4,60817714 | 1,528053 | 3,015718 | 0,002564 | 1 | 0 | Distal Intergenic | 6 | 1,21E+08 | 1,21E+08 | 37507 | 1 | 14412 | ENSMUST00000064580.14 | -11484 | ENSMUSG00000030108 | Slc6a13 | solute carrier family 6 (neurotransmitter transporter, GABA), member 13 |
| chr4 | 1,2E+08 | 1,2E+08 | 1244 | * | 9,004744 | 4,608245956 | 1,746121 | 2,639133 | 0,008312 | 0,85599412 | 1,425929 | Intron (ENSMUST0000( | 4 | 1,2E+08 | 1,2E+08 | 20928 | 1 | 667063 | ENSMUST00000238451.2 | 7472 | ENSMUSG00000091297 | Gm8439 | predicted gene 8439 |
| chr8 | 50899049 | 50899708 | 660 | * | 7,547852 | 4,6114447 | 1,551305 | 2,972623 | 0,002953 | 1 | 0 | Intron (ENSMUST0000( | 8 | 50845745 | 50970677 | 124933 | 2 | 23965 | ENSMUST00000211156.2 | 70969 | ENSMUSG00000031561 | Tenm3 | teneurin transmembrane protein 3 |
| chr1 | 26698046 | 26698856 | 811 | * | 6,867309 | 4,61167801 | 1,635693 | 2,819403 | 0,004811 | 1 | 0 | Distal Intergenic | 1 | 26720895 | 26726541 | 5647 | 2 | 210940 | ENSMUST00000097801.4 | 27685 | ENSMUSG00000073722 | 4931408C20Rik | RIKEN cDNA 4931408C20 gene |

| | | | | | | | | | | | | | | | | | | | | | | | |
|---|---|---|---|---|---|---|---|---|---|---|---|---|---|---|---|---|---|---|---|---|---|---|---|
| chr1 | 1,32E+08 | 1,32E+08 | 929 | * | 8,948867 | 4,614774456 | 1,396749 | 3,303939 | 0,000953 | 0,471637333 | 1,449706 | 3' UTR | 1 | 1,32E+08 | 1,32E+08 | 8732 | 1 | 213417 | ENSMUST00000046071.5 | 7488 | ENSMUSG00000042115 | Klhdc8a | kelch domain containing 8A |
| chr4 | 1,17E+08 | 1,17E+08 | 1556 | * | 9,360452 | 4,614820699 | 1,521814 | 3,032448 | 0,002426 | 0,642870233 | 1,449706 | Intron (ENSMUST00000 | 4 | 1,17E+08 | 1,17E+08 | 107 | 2 | 1,15E+08 | ENSMUST00000240534.1 | -31422 | ENSMUSG00000119735 | Gm22092 | predicted gene, 22092 |
| chr5 | 3274845 | 3275740 | 896 | * | 9,96245 | 4,616964602 | 1,404833 | 3,286487 | 0,001014 | 0,532146359 | 1,086118 | Distal Intergenic | 5 | 3286431 | 3286868 | 438 | 1 | 1E+08 | ENSMUST00000118533.2 | -10691 | ENSMUSG00000062353 | Gm15772 | ribosomal protein L26 pseudogene |
| chr10 | 33520661 | 33521423 | 763 | * | 7,411709 | 4,617902317 | 1,53083 | 3,0166 | 0,002556 | 0,66269276 | 1,360622 | Intron (ENSMUST00000 | 10 | 33388282 | 33500765 | 112484 | 2 | 215890 | ENSMUST00000019920.13 | -19896 | ENSMUSG00000019785 | Clvs2 | clavesin 2 |
| chr13 | 95857508 | 95858676 | 1169 | * | 9,520287 | 4,618407224 | 1,326758 | 3,480971 | 0,0005 | 0,396791886 | 1,211315 | Exon (ENSMUST000000 | 13 | 95844002 | 95862055 | 18054 | 1 | 1,15E+08 | ENSMUST00000238396.2 | 13506 | | Gm52062 | predicted gene, 52062 |
| chr16 | 32986315 | 32987552 | 1238 | * | 8,061727 | 4,618814986 | 1,452723 | 3,17942 | 0,001476 | 1 | 0 | Distal Intergenic | 16 | 33005441 | 33063636 | 58196 | 1 | 106326 | ENSMUST00000232100.2 | -17889 | ENSMUSG00000022807 | Osbpl11 | oxysterol binding protein-like 11 |
| chr11 | 1,2E+08 | 1,2E+08 | 1119 | * | 9,112471 | 4,620023529 | 1,417289 | 3,25976 | 0,001115 | 0,507398906 | 1,360622 | Distal Intergenic | 11 | 1,2E+08 | 1,2E+08 | 596 | 2 | 11465 | ENSMUST00000089616.6 | 13125 | ENSMUSG00000062825 | Actg1 | actin, gamma, cytoplasmic 1 |
| chr5 | 53024925 | 53025932 | 1008 | * | 7,703782 | 4,620977927 | 1,491395 | 3,098427 | 0,001946 | 1 | 0 | Promoter (1-2kb) | 5 | 53022955 | 53024076 | 1122 | 1 | 52206 | ENSMUST00000138629.2 | 1970 | ENSMUSG00000029176 | Anapc4 | anaphase promoting complex subunit 4 |
| chr16 | 49675309 | 49675922 | 614 | * | 7,393344 | 4,621514259 | 1,523935 | 3,032619 | 0,002424 | 1 | 0 | Promoter (<=1kb) | 16 | 49675729 | 49731454 | 55726 | 1 | 16423 | ENSMUST00000114496.3 | 0 | ENSMUSG00000055447 | Cd47 | CD47 antigen (Rh-related antigen, integrin-associated signal transducer) |
| chr2 | 1,25E+08 | 1,25E+08 | 992 | * | 8,905922 | 4,626501597 | 1,419056 | 3,260267 | 0,001113 | 0,507398906 | 1,360622 | Promoter (2-3kb) | 2 | 1,25E+08 | 1,25E+08 | 11127 | 1 | 110074 | ENSMUST00000051605.9 | 2536 | ENSMUSG00000027203 | Dut | deoxyuridine triphosphatase |
| chr4 | 1,06E+08 | 1,06E+08 | 960 | * | 9,032186 | 4,626800963 | 1,416488 | 3,266388 | 0,001089 | 0,525184597 | 1,211315 | Promoter (1-2kb) | 4 | 1,06E+08 | 1,06E+08 | 1652 | 1 | 230576 | ENSMUST00000136505.2 | -1551 | ENSMUSG00000034919 | Ttc22 | tetratricopeptide repeat domain 22 |
| chr7 | 1E+08 | 1E+08 | 915 | * | 8,104025 | 4,626822033 | 1,526366 | 3,031267 | 0,002435 | 0,646274978 | 1,425929 | Intron (ENSMUST00000 | 7 | 1E+08 | 1E+08 | 25278 | 2 | 52443 | ENSMUST00000107053.11 | 6568 | ENSMUSG00000030706 | Mrpl48 | mitochondrial ribosomal protein L48 |
| chr9 | 1,05E+08 | 1,05E+08 | 959 | * | 7,296244 | 4,627774816 | 1,513351 | 3,057966 | 0,002228 | 0,64068479 | 1,360622 | Distal Intergenic | 9 | 1,05E+08 | 1,05E+08 | 67660 | 2 | 235574 | ENSMUST00000190802.2 | -26340 | ENSMUSG00000032570 | Atp2c1 | ATPase, Ca++-sequestering |
| chrX | 78061913 | 78062820 | 908 | * | 5,167067 | 4,629880664 | 1,696226 | 2,729519 | 0,006343 | 1 | 0 | Distal Intergenic | 20 | 77781369 | 77783007 | 1639 | 1 | 70864 | ENSMUST00000105110.4 | 280544 | ENSMUSG00000078315 | Fam47c | family with sequence similarity 47, member C |
| chr7 | 18743471 | 18746328 | 2858 | * | 47,328253 | 4,63287013 | 0,6377778 | 7,2640821 | 3,756E-13 | 2,1688E-08 | 2,7872553 | Distal Intergenic | 7 | 18737969 | 18740702 | 2734 | 1 | 272359 | ENSMUST00000053713.5 | 5502 | ENSMUSG00000044030 | Irf2bp1 | interferon regulatory factor 2 binding protein 1 |
| chr4 | 1,07E+08 | 1,07E+08 | 867 | * | 8,55977 | 4,634076815 | 1,764805 | 2,62583 | 0,008644 | 1 | 0 | Intron (ENSMUST00000 | 4 | 1,07E+08 | 1,07E+08 | 8831 | 1 | 72787 | ENSMUST00000132950.2 | 21964 | ENSMUSG00000028614 | Ndc1 | NDC1 transmembrane nucleoporin |
| chr1 | 4928257 | 4929170 | 914 | * | 6,082343 | 4,635616487 | 1,558315 | 2,974761 | 0,002932 | 1 | 0 | Promoter (<=1kb) | 1 | 4928261 | 4968132 | 39872 | 1 | 21399 | ENSMUST00000165720.3 | 0 | ENSMUSG00000033813 | Tcea1 | transcription elongation factor A (SII) 1 |
| chr10 | 85879214 | 85879988 | 775 | * | 6,769053 | 4,636975948 | 1,521584 | 3,047467 | 0,002308 | 1 | 0 | Promoter (<=1kb) | 10 | 85878316 | 85882673 | 4358 | 1 | 69754 | ENSMUST00000130383.2 | 898 | ENSMUSG00000001786 | Fbxo7 | F-box protein 7 |
| chr3 | 56376166 | 56377119 | 954 | * | 8,172021 | 4,639896549 | 1,487971 | 3,118271 | 0,001819 | 0,600307614 | 1,360622 | Distal Intergenic | 3 | 56352703 | 56352833 | 131 | 2 | 1,15E+08 | ENSMUST00000104522.3 | -23333 | ENSMUSG00000119391 | Gm25727 | predicted gene, 25727 |
| chr6 | 30379742 | 30380663 | 922 | * | 13,02938 | 4,640590223 | 1,315469 | 3,527709 | 0,000419 | 0,365691948 | 1,173325 | Promoter (2-3kb) | 6 | 30374822 | 30382667 | 7846 | 2 | 232679 | ENSMUST00000126945.8 | 2004 | ENSMUSG00000039130 | Zc3hc1 | zinc finger, C3HC type 1 |
| chr4 | 14714965 | 14715547 | 583 | * | 6,088555 | 4,641689171 | 1,775888 | 2,613728 | 0,008956 | 1 | 0 | Intron (ENSMUST00000 | 4 | 14665754 | 14756845 | 91092 | 2 | 73314 | ENSMUST00000143565.2 | 41298 | ENSMUSG00000023151 | Lrrc69 | leucine rich repeat containing 69 |
| chr3 | 95211600 | 95213234 | 1635 | * | 12,91652 | 4,642489086 | 1,223978 | 3,792951 | 0,000149 | 0,284660899 | 0,738441 | Promoter (<=1kb) | 3 | 95203407 | 95214115 | 10709 | 2 | 71790 | ENSMUST00000107183.8 | 881 | ENSMUSG00000015702 | Anxa9 | annexin A9 |
| chr10 | 80415964 | 80416797 | 834 | * | 6,55703 | 4,642568292 | 1,617773 | 2,869728 | 0,004108 | 1 | 0 | Promoter (2-3kb) | 10 | 80402958 | 80413155 | 10198 | 2 | 118445 | ENSMUST00000038558.9 | -2809 | ENSMUSG00000035397 | Klf16 | Kruppel-like factor 16 |
| chr1 | 72751063 | 72751705 | 643 | * | 5,523735 | 4,644440157 | 1,760586 | 2,638008 | 0,008339 | 1 | 0 | Promoter (<=1kb) | 1 | 72750940 | 72751729 | 790 | 1 | 19981 | ENSMUST00000211963.2 | 123 | ENSMUSG00000046330 | Rpl37a | ribosomal protein L37a |
| chrX | 81808266 | 81808735 | 470 | * | 5,582675 | 4,646458974 | 1,642017 | 2,829726 | 0,004659 | 1 | 0 | Distal Intergenic | 20 | 81735845 | 81735976 | 132 | 2 | 1,15E+08 | ENSMUST00000104486.3 | -72290 | ENSMUSG00000077674 | Gm24470 | predicted gene, 24470 |
| chr16 | 33724475 | 33725141 | 667 | * | 6,86237 | 4,646674589 | 1,573519 | 2,953047 | 0,003147 | 1 | 0 | Intron (ENSMUST00000 | 16 | 33712317 | 33719772 | 7456 | 1 | 16419 | ENSMUST00000148462.2 | 12158 | ENSMUSG00000022817 | Itgb5 | integrin beta 5 |
| chr19 | 10674485 | 10675120 | 636 | * | 7,873139 | 4,647417909 | 1,474761 | 3,151302 | 0,001625 | 1 | 0 | Intron (ENSMUST00000 | 19 | 10666129 | 10691991 | 25863 | 1 | 107305 | ENSMUST00000235921.2 | 8356 | ENSMUSG00000048832 | Vps37c | vacuolar protein sorting 37C |
| chr19 | 52971962 | 52972465 | 504 | * | 6,763913 | 4,651686589 | 1,506154 | 3,088453 | 0,002012 | 1 | 0 | Intron (ENSMUST00000 | 19 | 52983795 | 52985064 | 1270 | 2 | 170750 | ENSMUST00000182728.2 | 12599 | ENSMUSG00000025027 | Xpnpep1 | X-prolyl aminopeptidase (aminopeptidase P) 1, soluble |
| chr1 | 1,05E+08 | 1,05E+08 | 711 | * | 5,819272 | 4,652879834 | 1,635694 | 2,844591 | 0,004447 | 1 | 0 | Intron (ENSMUST00000 | 1 | 1,05E+08 | 1,05E+08 | 226953 | 1 | 23836 | ENSMUST00000062528.9 | 106066 | ENSMUSG00000050840 | Cdh20 | cadherin 20 |
| chr3 | 1,48E+08 | 1,48E+08 | 678 | * | 5,808692 | 4,654103709 | 1,767083 | 2,633778 | 0,008444 | 1 | 0 | Distal Intergenic | 3 | 1,49E+08 | 1,49E+08 | 7124 | 2 | 99633 | ENSMUST00000168352.2 | 162650 | ENSMUSG00000028184 | Adgrl2 | adhesion G protein-coupled receptor L2 |
| chr4 | 1,35E+08 | 1,35E+08 | 1871 | * | 15,32078 | 4,654479635 | 1,092752 | 4,259411 | 2,05E-05 | 0,065437585 | 1,2763 | Distal Intergenic | 4 | 1,35E+08 | 1,35E+08 | 23969 | 1 | 230828 | ENSMUST00000102546.4 | -7934 | ENSMUSG00000037157 | Il22ra1 | interleukin 22 receptor, alpha 1 |
| chrX | 60537557 | 60538572 | 1016 | * | 6,798859 | 4,654778609 | 1,610019 | 2,891132 | 0,003839 | 1 | 0 | Distal Intergenic | 20 | 60331794 | 60336090 | 4297 | 1 | 245440 | ENSMUST00000232598.2 | 205763 | ENSMUSG00000118608 | Gm4988 | predicted gene 4988 |
| chr19 | 46147198 | 46148295 | 1098 | * | 10,86676 | 4,655323609 | 1,726523 | 2,696358 | 0,00701 | 0,847020973 | 1,2763 | Intron (ENSMUST00000 | 19 | 46141019 | 46160955 | 19937 | 1 | 107338 | ENSMUST00000236167.2 | 6179 | ENSMUSG00000025224 | Gbf1 | golgi-specific brefeldin A-resistance factor 1 |
| chr1 | 90901576 | 90903298 | 1723 | * | 35,8273 | 4,656984718 | 1,312205 | 3,548975 | 0,000387 | 0,258941749 | 2,377674 | Exon (ENSMUST000001 | 1 | 90885855 | 90897383 | 11529 | 2 | 19329 | ENSMUST00000027529.12 | -4193 | ENSMUSG00000026304 | Rab17 | RAB17, member RAS oncogene family |
| chr4 | 1,38E+08 | 1,38E+08 | 1292 | * | 7,149203 | 4,659043225 | 1,538145 | 3,029001 | 0,002454 | 1 | 0 | Promoter (1-2kb) | 4 | 1,38E+08 | 1,38E+08 | 6257 | 1 | 15441 | ENSMUST00000130071.2 | 1933 | ENSMUSG00000028759 | Hp1bp3 | heterochromatin protein 1, binding protein 3 |
| chr12 | 13279598 | 13280472 | 875 | * | 7,447093 | 4,659443247 | 1,668882 | 2,791955 | 0,005239 | 1 | 0 | Promoter (<=1kb) | 12 | 13275162 | 13279380 | 4219 | 2 | 104721 | ENSMUST00000221028.2 | -218 | ENSMUSG00000037149 | Ddx1 | DEAD box helicase 1 |
| chr1 | 48931770 | 48932610 | 841 | * | 5,709945 | 4,664621561 | 1,773847 | 2,629664 | 0,008547 | 1 | 0 | Distal Intergenic | 1 | 49283775 | 49377405 | 93631 | 1 | 442837 | ENSMUST00000056879.9 | -351165 | ENSMUSG00000051616 | C230029F24Rik | RIKEN cDNA C230029F24 gene |
| chr11 | 79658646 | 79660417 | 1772 | * | 11,36426 | 4,667069672 | 1,354428 | 3,445787 | 0,000569 | 0,421090004 | 1,173325 | Distal Intergenic | 11 | 79672805 | 79673713 | 909 | 2 | 1,05E+08 | ENSMUST00000150312.2 | 13296 | | 9130204K15Rik | RIKEN cDNA 9130204K15 gene |
| chr5 | 25114787 | 25115457 | 671 | * | 5,819307 | 4,668084099 | 1,716583 | 2,719405 | 0,00654 | 1 | 0 | Promoter (<=1kb) | 5 | 25094159 | 25113975 | 19817 | 2 | 108099 | ENSMUST00000123749.2 | -812 | ENSMUSG00000028944 | Prkag2 | protein kinase, AMP-activated, gamma 2 non-catalytic subunit |
| chr10 | 93983998 | 93984881 | 884 | * | 8,640437 | 4,668333896 | 1,498159 | 3,116047 | 0,001833 | 0,507398906 | 2,237403 | Promoter (<=1kb) | 10 | 93984281 | 94031543 | 47263 | 1 | 22025 | ENSMUST00000092213.11 | 0 | ENSMUSG00000005897 | Nr2c1 | nuclear receptor subfamily 2, group C, member 1 |
| chr11 | 1,16E+08 | 1,16E+08 | 1053 | * | 11,42505 | 4,669260388 | 1,239173 | 3,768047 | 0,000165 | 0,239082971 | 1,173325 | Distal Intergenic | 11 | 1,16E+08 | 1,16E+08 | 4696 | 2 | 15223 | ENSMUST00000036215.8 | 12973 | ENSMUSG00000034227 | Foxj1 | forkhead box J1 |
| chr6 | 86197726 | 86198714 | 989 | * | 10,1151 | 4,670739419 | 1,465931 | 3,186194 | 0,001442 | 0,59839358 | 1,086118 | Intron (ENSMUST00000 | 6 | 86188576 | 86249201 | 60626 | 1 | 21802 | ENSMUST00000194449.3 | 9150 | ENSMUSG00000029999 | Tgfa | transforming growth factor alpha |
| chr10 | 80082177 | 80083369 | 1193 | * | 10,96025 | 4,672028184 | 1,384354 | 3,374879 | 0,000738 | 0,54190677 | 0,738441 | Promoter (1-2kb) | 10 | 80084955 | 80092627 | 7673 | 1 | 75406 | ENSMUST00000105364.8 | -1586 | ENSMUSG00000020153 | Ndufs7 | NADH:ubiquinone oxidoreductase core subunit S7 |
| chrX | 49421434 | 49422594 | 1161 | * | 6,289615 | 4,674355205 | 1,709353 | 2,734576 | 0,006246 | 1 | 0 | Distal Intergenic | 20 | 49494690 | 49494796 | 107 | 1 | 1,15E+08 | ENSMUST00000240068.1 | -72096 | ENSMUSG00000119917 | Gm22288 | predicted gene, 22288 |
| chr3 | 1,31E+08 | 1,31E+08 | 1837 | * | 21,53465 | 4,674616971 | 1,550255 | 3,015386 | 0,002567 | 0,492273754 | 3,471678 | Promoter (<=1kb) | 3 | 1,3E+08 | 1,31E+08 | 13533 | 2 | 66357 | ENSMUST00000043937.9 | 0 | ENSMUSG00000041084 | Ostc | oligosaccharyltransferase complex subunit (non-catalytic) |
| chr6 | 8659136 | 8659530 | 395 | * | 8,013197 | 4,674776642 | 1,788684 | 2,613529 | 0,008961 | 1 | 0 | Intron (ENSMUST00000 | 6 | 8630699 | 8647819 | 17121 | 2 | 15893 | ENSMUST00000145870.2 | -11317 | ENSMUSG00000062995 | Ica1 | islet cell autoantigen 1 |
| chr2 | 14783999 | 14784392 | 394 | * | 4,943063 | 4,676201283 | 1,750592 | 2,671211 | 0,007558 | 1 | 0 | Intron (ENSMUST00000 | 2 | 14745183 | 14771835 | 26653 | 2 | 1,03E+08 | ENSMUST00000147019.4 | -12164 | | Gm13269 | predicted gene 13269 |
| chr6 | 91448775 | 91449648 | 874 | * | 9,799137 | 4,677695665 | 1,377854 | 3,394913 | 0,000687 | 0,451585113 | 1,173325 | Promoter (<=1kb) | 6 | 91439154 | 91450528 | 11375 | 2 | 72170 | ENSMUST00000040835.9 | 880 | ENSMUSG00000034203 | Chchd4 | coiled-coil-helix-coiled-coil-helix domain containing 4 |
| chr13 | 36735982 | 36736953 | 972 | * | 10,01353 | 4,67776116 | 1,773612 | 2,637421 | 0,008354 | 0,890608507 | 1,173325 | Intron (ENSMUST00000 | 13 | 36796790 | 36804413 | 7624 | 1 | 1,08E+08 | ENSMUST00000225239.2 | -59837 | | Gm46410 | predicted gene, 46410 |
| chr11 | 32405680 | 32406516 | 837 | * | 7,874565 | 4,678439331 | 1,764582 | 2,651302 | 0,008018 | 1 | 0 | Promoter (<=1kb) | 11 | 32405370 | 32466687 | 61318 | 1 | 327900 | ENSMUST00000051053.5 | 310 | ENSMUSG00000044949 | Ubtd2 | ubiquitin domain containing 2 |
| chr2 | 1,02E+08 | 1,02E+08 | 1269 | * | 11,72423 | 4,679433937 | 1,389923 | 3,366686 | 0,000761 | 0,469451136 | 1,173325 | Promoter (2-3kb) | 2 | 1,02E+08 | 1,02E+08 | 50386 | 2 | 80985 | ENSMUST00000144110.2 | 2508 | ENSMUSG00000027189 | Trim44 | tripartite motif-containing 44 |
| chr8 | 46452354 | 46454400 | 2047 | * | 86,917212 | 4,681660972 | 0,4372747 | 10,706452 | 9,493E-27 | 4,44225E-21 | 3,439501 | Promoter (<=1kb) | 8 | 46449939 | 46452904 | 2966 | 2 | 654824 | ENSMUST00000053558.10 | 0 | ENSMUSG00000050914 | Ankrd37 | ankyrin repeat domain 37 |
| chrX | 1,35E+08 | 1,35E+08 | 985 | * | 6,418895 | 4,685739642 | 1,581143 | 2,963514 | 0,003041 | 1 | 0 | Promoter (2-3kb) | 20 | 1,35E+08 | 1,35E+08 | 1457 | 1 | 22381 | ENSMUST00000113115.2 | 2795 | ENSMUSG00000042712 | Tceal9 | transcription elongation factor A like 9 |
| chr15 | 11614198 | 11615201 | 1004 | * | 5,003078 | 4,686681419 | 1,727847 | 2,712439 | 0,006679 | 1 | 0 | Distal Intergenic | 15 | 11696896 | 11703789 | 6894 | 2 | 1,05E+08 | ENSMUST00000228108.2 | 88588 | ENSMUSG00000115585 | Gm41271 | predicted gene, 41271 |
| chr7 | 1,05E+08 | 1,05E+08 | 922 | * | 9,525002 | 4,686890327 | 1,414892 | 3,312542 | 0,000925 | 0,396928896 | 2,237403 | Distal Intergenic | 7 | 1,05E+08 | 1,05E+08 | 954 | 1 | 259045 | ENSMUST00000098164.2 | 8738 | ENSMUSG00000073916 | Olfr669 | olfactory receptor 669 |
| chr14 | 31665656 | 31666149 | 494 | * | 5,838443 | 4,692714892 | 1,638669 | 2,863736 | 0,004187 | 1 | 0 | Distal Intergenic | 14 | 31651545 | 31653675 | 2131 | 1 | 1,03E+08 | ENSMUST00000242479.1 | 14111 | | Gm53172 | predicted gene, 53172 |
| chr12 | 31693533 | 31694334 | 802 | * | 7,34937 | 4,693645336 | 1,557538 | 3,013502 | 0,002583 | 1 | 0 | Intron (ENSMUST00000 | 12 | 31684074 | 31684657 | 584 | 2 | 12033 | ENSMUST00000218543.2 | -8876 | ENSMUSG00000020650 | Bcap29 | B cell receptor associated protein 29 |
| chr8 | 46426925 | 46429254 | 2330 | * | 55,435688 | 4,693850857 | 0,9360867 | 5,0143335 | 5,322E-07 | 0,002656105 | 2,3776742 | Promoter (<=1kb) | 8 | 46428565 | 46449995 | 21431 | 1 | 192169 | ENSMUST00000034051.7 | 0 | ENSMUSG00000031634 | Ufsp2 | UFM1-specific peptidase 2 |
| chr3 | 23089641 | 23090778 | 1138 | * | 8,292046 | 4,693940789 | 1,323898 | 3,545545 | 0,000392 | 0,323988916 | 1,425929 | Distal Intergenic | 3 | 23515214 | 23575562 | 60349 | 1 | 1E+08 | ENSMUST00000194267.2 | -424436 | ENSMUSG00000102203 | 4930412E21Rik | RIKEN cDNA 4930412E21 gene |
| chr4 | 90310387 | 90311385 | 999 | * | 6,417662 | 4,693971256 | 1,693779 | 2,771301 | 0,005583 | 1 | 0 | Distal Intergenic | 4 | 90247859 | 90249563 | 1705 | 2 | 619842 | ENSMUST00000117742.2 | -60824 | ENSMUSG00000080756 | Gm12633 | zinc finger protein 352 pseudogene |
| chr4 | 1,34E+08 | 1,34E+08 | 528 | * | 6,333327 | 4,694038224 | 1,616688 | 2,90349 | 0,00369 | 1 | 0 | Promoter (2-3kb) | 4 | 1,34E+08 | 1,34E+08 | 12929 | 1 | 69885 | ENSMUST00000105866.3 | 2819 | ENSMUSG00000078521 | Aunip | aurora kinase A and ninein interacting protein |
| chr1 | 31079512 | 31080497 | 986 | * | 8,292398 | 4,695203098 | 1,763633 | 2,662234 | 0,007762 | 0,764024179 | 2,237403 | Promoter (<=1kb) | 1 | 31081000 | 31082756 | 1757 | 1 | 1,03E+08 | ENSMUST00000191364.2 | -503 | | Gm28644 | predicted gene 28644 |
| chr13 | 42923890 | 42925035 | 1146 | * | 8,998673 | 4,695222093 | 1,767072 | 2,657063 | 0,007882 | 0,767061106 | 2,237403 | Intron (ENSMUST00000 | 13 | 42863057 | 43292002 | 428946 | 1 | 218194 | ENSMUST00000148891.8 | 60833 | ENSMUSG00000054728 | Phactr1 | phosphatase and actin regulator 1 |
| chr11 | 37023109 | 37024129 | 1021 | * | 9,732636 | 4,695275314 | 1,499297 | 3,131651 | 0,001738 | 0,57954252 | 1,449706 | Intron (ENSMUST00000 | 11 | 35897483 | 37126709 | 1229227 | 2 | 23964 | ENSMUST00000102801.8 | 102580 | ENSMUSG00000049336 | Tenm2 | teneurin transmembrane protein 2 |
| chr5 | 1,18E+08 | 1,18E+08 | 1288 | * | 9,380112 | 4,695896023 | 1,543493 | 3,042382 | 0,002347 | 0,649014805 | 1,360622 | Intron (ENSMUST00000 | 5 | 1,18E+08 | 1,18E+08 | 361004 | 1 | 333050 | ENSMUST00000180430.2 | 16379 | ENSMUSG00000061578 | Ksr2 | kinase suppressor of ras 2 |
| chr16 | 23779090 | 23780006 | 917 | * | 7,996645 | 4,697450364 | 1,517184 | 3,096164 | 0,00196 | 1 | 0 | Distal Intergenic | 16 | 23786444 | 23791130 | 4687 | 2 | 12053 | ENSMUST00000135352.2 | 11124 | ENSMUSG00000022508 | Bcl6 | B cell leukemia/lymphoma 6 |
| chr10 | 80476376 | 80477274 | 899 | * | 7,817095 | 4,698314682 | 1,465433 | 3,206093 | 0,001346 | 0,54122637 | 1,360622 | Promoter (<=1kb) | 10 | 80475473 | 80476092 | 620 | 1 | 103236 | ENSMUST00000220431.2 | 903 | ENSMUSG00000003345 | Csnk1g2 | casein kinase 1, gamma 2 |
| chrX | 1,14E+08 | 1,14E+08 | 497 | * | 5,536025 | 4,700016101 | 1,680527 | 2,796751 | 0,005162 | 1 | 0 | Promoter (<=1kb) | 20 | 1,14E+08 | 1,14E+08 | 348 | 2 | 664969 | ENSMUST00000118664.2 | 868 | ENSMUSG00000082260 | Gm7429 | predicted pseudogene 7429 |
| chr7 | 1,04E+08 | 1,04E+08 | 740 | * | 5,210496 | 4,700919146 | 1,686369 | 2,787599 | 0,00531 | 1 | 0 | Promoter (<=1kb) | 7 | 1,04E+08 | 1,04E+08 | 1244 | 2 | 625321 | ENSMUST00000151294.2 | 0 | ENSMUSG00000073929 | Trim30e-ps1 | tripartite motif-containing 30E, pseudogene 1 |
| chr5 | 1,02E+08 | 1,02E+08 | 654 | * | 8,080239 | 4,703034688 | 1,577986 | 2,980403 | 0,002879 | 1 | 0 | Distal Intergenic | 5 | 1,02E+08 | 1,02E+08 | 140616 | 2 | 72145 | ENSMUST00000174698.2 | -104394 | ENSMUSG00000043940 | Wdfy3 | WD repeat and FYVE domain containing 3 |
| chr10 | 82753255 | 82754189 | 935 | * | 7,061332 | 4,703774335 | 1,634251 | 2,878245 | 0,003999 | 1 | 0 | Distal Intergenic | 10 | 82722680 | 82732146 | 9467 | 1 | 50493 | ENSMUST00000218724.2 | 30575 | ENSMUSG00000020250 | Txnrd1 | thioredoxin reductase 1 |
| chr3 | 1,41E+08 | 1,41E+08 | 946 | * | 8,353564 | 4,706857238 | 1,435075 | 3,279868 | 0,001039 | 0,484674158 | 1,449706 | Distal Intergenic | 3 | 1,41E+08 | 1,41E+08 | 2352 | 2 | 18598 | ENSMUST00000057860.3 | 78235 | ENSMUSG00000047674 | Pdha2 | pyruvate dehydrogenase E1 alpha 2 |
| chr13 | 49548137 | 49548947 | 811 | * | 8,704771 | 4,709345214 | 1,439047 | 3,272545 | 0,001066 | 0,490966492 | 1,449706 | Distal Intergenic | 13 | 49536565 | 49538624 | 2060 | 1 | 76895 | ENSMUST00000220723.2 | 11572 | ENSMUSG00000037933 | Bicd2 | BICD cargo adaptor 2 |
| chr8 | 71043478 | 71044260 | 783 | * | 10,12597 | 4,709708943 | 1,443059 | 3,263699 | 0,0011 | 0,527310742 | 1,211315 | Promoter (2-3kb) | 8 | 71047023 | 71047813 | 791 | 1 | 71780 | ENSMUST00000211117.2 | -2763 | ENSMUSG00000019139 | Isyna1 | myo-inositol 1-phosphate synthase A1 |
| chr6 | 40301661 | 40303002 | 1342 | * | 12,95802 | 4,710261201 | 1,342617 | 3,508269 | 0,000451 | 0,380551476 | 1,173325 | Promoter (<=1kb) | 6 | 40302106 | 40373696 | 71591 | 1 | 69923 | ENSMUST00000031977.12 | 0 | ENSMUSG00000029916 | Agk | acylglycerol kinase |
| chr11 | 1,18E+08 | 1,18E+08 | 933 | * | 12,39215 | 4,711834259 | 1,327616 | 3,549095 | 0,000387 | 0,399754181 | 0,918255 | Distal Intergenic | 11 | 1,18E+08 | 1,18E+08 | 5530 | 2 | 71776 | ENSMUST00000153989.2 | -28537 | ENSMUSG00000017713 | Tha1 | threonine aldolase 1 |
| chr11 | 1,05E+08 | 1,05E+08 | 876 | * | 8,101751 | 4,713747087 | 1,464128 | 3,219491 | 0,001284 | 1 | 0 | Intron (ENSMUST00000 | 11 | 1,05E+08 | 1,05E+08 | 25332 | 1 | 70894 | ENSMUST00000021029.6 | -3679 | ENSMUSG00000020690 | Efcab3 | EF-hand calcium binding domain 3 |
| chr11 | 3154499 | 3155166 | 668 | * | 6,345135 | 4,714890982 | 1,56938 | 3,004302 | 0,002662 | 1 | 0 | Promoter (<=1kb) | 11 | 3153630 | 3194586 | 40957 | 1 | 74203 | ENSMUST00000120721.9 | 869 | ENSMUSG00000020454 | Eif4enif1 | eukaryotic translation initiation factor 4E nuclear import factor 1 |
| chr2 | 1,67E+08 | 1,67E+08 | 991 | * | 7,750413 | 4,71523109 | 1,829099 | 2,577898 | 0,00994 | 1 | 0 | 3' UTR | 2 | 1,67E+08 | 1,67E+08 | 1917 | 1 | 76367 | ENSMUST00000051923.7 | 4100 | ENSMUSG00000042854 | Trp53rkb | transformation related protein 53 regulating kinase B |
| chr11 | 60721010 | 60722172 | 1163 | * | 8,703178 | 4,715852657 | 1,798859 | 2,62158 | 0,008752 | 1 | 0 | Promoter (<=1kb) | 11 | 60721457 | 60749249 | 27793 | 1 | 216820 | ENSMUST00000042281.14 | 0 | ENSMUSG00000042569 | Dhrs7b | dehydrogenase/reductase (SDR family) member 7B |
| chr3 | 1,51E+08 | 1,51E+08 | 785 | * | 7,08249 | 4,716093645 | 1,827334 | 2,580861 | 0,009855 | 1 | 0 | Distal Intergenic | 3 | 1,51E+08 | 1,51E+08 | 3243 | 1 | 1,01E+08 | ENSMUST00000196524.3 | -83374 | | 4930555A03Rik | RIKEN cDNA 4930555A03 gene |
| chr5 | 34091372 | 34092525 | 1154 | * | 7,107323 | 4,71858042 | 1,527126 | 3,089844 | 0,002003 | 1 | 0 | Promoter (1-2kb) | 5 | 34076469 | 34093594 | 17126 | 2 | 24116 | ENSMUST00000132942.2 | 1069 | ENSMUSG00000029111 | Nelfa | negative elongation factor complex member A, Whsc2 |
| chr16 | 88852007 | 88852493 | 487 | * | 4,438292 | 4,722088495 | 1,828909 | 2,581915 | 0,009825 | 1 | 0 | Distal Intergenic | 16 | 88844177 | 88844771 | 595 | 2 | 68484 | ENSMUST00000074637.4 | -7236 | ENSMUSG00000062400 | Krtap6-5 | keratin associated protein 6-5 |
| chr12 | 69244292 | 69245451 | 1160 | * | 11,66692 | 4,722727267 | 1,282285 | 3,683057 | 0,00023 | 0,282588188 | 1,173325 | Promoter (<=1kb) | 12 | 69235861 | 69245203 | 9343 | 2 | 109065 | ENSMUST00000021356.6 | 0 | ENSMUSG00000020973 | Dnaaf2 | dynein, axonemal assembly factor 2 |
| chr19 | 25264098 | 25265505 | 1408 | * | 9,438098 | 4,72560513 | 1,424999 | 3,316217 | 0,000912 | 0,46426665 | 1,449706 | Intron (ENSMUST00000 | 19 | 25214339 | 25411860 | 197522 | 1 | 107351 | ENSMUST00000049400.15 | 49759 | ENSMUSG00000032702 | Kank1 | KN motif and ankyrin repeat domains 1 |
| chrX | 15725659 | 15726640 | 982 | * | 6,249493 | 4,726145587 | 1,633587 | 2,893109 | 0,003814 | 1 | 0 | Distal Intergenic | 20 | 15762902 | 15763008 | 107 | 2 | 1,15E+08 | ENSMUST00000239993.1 | 36368 | ENSMUSG00001118666 | Gm23516 | predicted gene, 23516 |
| chr7 | 98180064 | 98181008 | 945 | * | 9,310127 | 4,727467543 | 1,498879 | 3,154002 | 0,00161 | 0,59912059 | 1,211315 | Distal Intergenic | 7 | 98156161 | 98156323 | 163 | 1 | 1,15E+08 | ENSMUST00000083869.4 | 23903 | ENSMUSG00000065803 | Gm23040 | predicted gene, 23040 |
| chr7 | 65779558 | 65780701 | 1144 | * | 9,246463 | 4,730304948 | 1,409746 | 3,355431 | 0,000792 | 0,470960971 | 1,211315 | Intron (ENSMUST00000 | 7 | 65759263 | 65823546 | 64284 | 1 | 269941 | ENSMUST00000036372.8 | 20295 | ENSMUSG00000032640 | Chsy1 | chondroitin sulfate synthase 1 |
| chr9 | 57067468 | 57068751 | 1284 | * | 9,49451 | 4,730990078 | 1,403593 | 3,370629 | 0,00075 | 0,353771182 | 2,237403 | Promoter (1-2kb) | 9 | 57062323 | 57065615 | 3293 | 2 | 66199 | ENSMUST00000065358.9 | -1853 | ENSMUSG00000032299 | Commd4 | COMM domain containing 4 |
| chr1 | 1,16E+08 | 1,16E+08 | 719 | * | 7,441496 | 4,735834004 | 1,45897 | 3,246013 | 0,00117 | 0,516959323 | 1,360622 | Intron (ENSMUST00000 | 1 | 1,16E+08 | 1,17E+08 | 902568 | 1 | 636808 | ENSMUST00000043725.9 | 579062 | ENSMUSG00000070695 | Cntnap5a | contactin associated protein-like 5A |
| chr12 | 1,01E+08 | 1,01E+08 | 813 | * | 8,350536 | 4,735917793 | 1,787629 | 2,649273 | 0,008067 | 0,771131827 | 2,237403 | Promoter (<=1kb) | 12 | 1,01E+08 | 1,01E+08 | 59566 | 2 | 68339 | ENSMUST00000223097.2 | 489 | ENSMUSG00000021182 | Ccdc88c | coiled-coil domain containing 88C |
| chr12 | 1,14E+08 | 1,14E+08 | 713 | * | 6,924997 | 4,736373718 | 1,80693 | 2,621227 | 0,008761 | 1 | 0 | Distal Intergenic | 12 | 1,14E+08 | 1,14E+08 | 50281 | 1 | 1,19E+08 | ENSMUST00000247233.1 | -62676 | | Gm53771 | predicted gene, 53771 |
| chr11 | 79113893 | 79114977 | 1085 | * | 6,839895 | 4,736515093 | 1,802311 | 2,628023 | 0,008588 | 1 | 0 | Distal Intergenic | 11 | 79131133 | 79133775 | 2643 | 2 | 78889 | ENSMUST00000153585.2 | 18798 | ENSMUSG00000017677 | Wsb1 | WD repeat and SOCS box-containing 1 |
| chr18 | 69804232 | 69805456 | 1225 | * | 9,451085 | 4,738270045 | 1,390138 | 3,408489 | 0,000653 | 0,410544751 | 1,425929 | Intron (ENSMUST00000 | 18 | 69812770 | 69817599 | 4830 | 1 | 21413 | ENSMUST00000146908.2 | -7314 | ENSMUSG00000053477 | Tcf4 | transcription factor 4 |
| chr11 | 1,17E+08 | 1,17E+08 | 1303 | * | 9,306259 | 4,738442873 | 1,738352 | 2,725825 | 0,006414 | 0,811407117 | 1,425929 | Intron (ENSMUST00000 | 11 | 1,17E+08 | 1,17E+08 | 96068 | 1 | 53860 | ENSMUST00000019038.15 | 19836 | ENSMUSG00000059248 | Septin9 | septin 9 |
| chr5 | 23245511 | 23245870 | 360 | * | 6,238717 | 4,739119224 | 1,673005 | 2,832699 | 0,004616 | 1 | 0 | Intron (ENSMUST00000 | 5 | 23305550 | 23480586 | 175037 | 1 | 269629 | ENSMUST00000196406.2 | -59680 | ENSMUSG00000106379 | Lhfpl3 | lipoma HMGIC fusion partner-like 3 |
| chrX | 77907591 | 77908964 | 1374 | * | 10,37196 | 4,741780223 | 1,257184 | 3,771747 | 0,000162 | 0,226355781 | 1,2763 | Distal Intergenic | 20 | 77781369 | 77783007 | 1639 | 1 | 70864 | ENSMUST00000105110.4 | 126222 | ENSMUSG00000078315 | Fam47c | family with sequence similarity 47, member C |
| chr16 | 55753865 | 55754549 | 685 | * | 6,516154 | 4,742625545 | 1,689659 | 2,806853 | 0,005003 | 1 | 0 | Promoter (<=1kb) | 16 | 55725341 | 55754596 | 29256 | 2 | 74201 | ENSMUST00000117468.8 | 47 | ENSMUSG00000022604 | Cep97 | centrosomal protein 97 |
| chr7 | 81477880 | 81478686 | 807 | * | 8,241086 | 4,742935762 | 1,512574 | 3,135672 | 0,001715 | 1 | 0 | Promoter (<=1kb) | 7 | 81450845 | 81478803 | 27959 | 2 | 83962 | ENSMUST00000208566.2 | 117 | ENSMUSG00000025103 | Btbd1 | BTB (POZ) domain containing 1 |
| chr5 | 79672719 | 79673547 | 829 | * | 8,282201 | 4,744025808 | 1,322507 | 3,587145 | 0,000334 | 0,30944653 | 1,360622 | Distal Intergenic | 5 | 80246755 | 80246861 | 107 | 1 | 1,15E+08 | ENSMUST00000239632.1 | -573208 | ENSMUSG00000119601 | Gm24304 | predicted gene, 24304 |
| chr1 | 6203192 | 6204336 | 1145 | * | 10,6212 | 4,745748961 | 1,365618 | 3,475167 | 0,000511 | 0,478766182 | 0,738441 | Distal Intergenic | 1 | 6276421 | 6304535 | 28115 | 1 | 12421 | ENSMUST00000159906.8 | -72085 | ENSMUSG00000025907 | Rb1cc1 | RB1-inducible coiled-coil 1 |

| | | | | | | | | | | | | | | | | | | | | | | | |
|---|---|---|---|---|---|---|---|---|---|---|---|---|---|---|---|---|---|---|---|---|---|---|---|
| chr18 | 38570961 | 38571814 | 854 | * | 7,625959 | 4,746623888 | 1,509954 | 3,143556 | 0,001669 | 1 | 0 | Intron (ENSMUST0000 | 18 | 38580766 | 38598031 | 17266 | 1 | 65113 | ENSMUST00000236171.2 | -8952 | ENSMUSG00000024425 | Ndfip1 | Nedd4 family interacting protein 1 |
| chr16 | 14424100 | 14425194 | 1095 | * | 6,225295 | 4,746813502 | 1,651615 | 2,874044 | 0,004053 | 1 | 0 | Intron (ENSMUST0000 | 16 | 14398085 | 14439164 | 41080 | 1 | 545280 | ENSMUST00000147024.8 | 26015 | | A630010A05Rik | RIKEN cDNA A630010A05 gene |
| chr13 | 18596594 | 18596945 | 352 | * | 5,125705 | 4,74693313 | 1,817986 | 2,611094 | 0,009025 | 1 | 0 | Distal Intergenic | 13 | 18520810 | 18572271 | 51462 | 2 | 218030 | ENSMUST00000223058.2 | -24323 | ENSMUSG00000009734 | Pou6f2 | POU domain, class 6, transcription factor 2 |
| chr11 | 83873925 | 83875499 | 1575 | * | 11,58914 | 4,747205478 | 1,221886 | 3,885147 | 0,000102 | 0,236247134 | 0,744375 | Promoter (<=1kb) | 11 | 83873122 | 83881697 | 8576 | 1 | 217030 | ENSMUST00000138676.2 | 803 | ENSMUSG00000034940 | Synrg | synergin, gamma |
| chr2 | 28412369 | 28412902 | 534 | * | 7,588207 | 4,748542892 | 1,589211 | 2,987987 | 0,002808 | 1 | 0 | Intron (ENSMUST0000 | 2 | 28417862 | 28442279 | 24418 | 1 | 19730 | ENSMUST00000238699.2 | -4960 | ENSMUSG00000026821 | Ralgds | ral guanine nucleotide dissociation stimulator |
| chr17 | 10146647 | 10147800 | 1154 | * | 9,587875 | 4,748728578 | 1,404286 | 3,381595 | 0,000721 | 0,454149353 | 1,211315 | Promoter (<=1kb) | 17 | 10145522 | 10148696 | 3175 | 2 | 1,19E+08 | ENSMUST00000232911.2 | 896 | | Gm34799 | predicted gene, 34799 |
| chr3 | 1,31E+08 | 1,31E+08 | 1089 | * | 10,4713 | 4,752623162 | 1,39734 | 3,401193 | 0,000671 | 0,434788971 | 1,2763 | Distal Intergenic | 3 | 1,31E+08 | 1,32E+08 | 270939 | 1 | 1,03E+08 | ENSMUST00000197726.2 | -31740 | | Gm29865 | predicted gene, 29865 |
| chr13 | 51476489 | 51477388 | 900 | * | 7,621556 | 4,754749583 | 1,826396 | 2,60335 | 0,009232 | 1 | 0 | Distal Intergenic | 13 | 51562675 | 51576833 | 14159 | 1 | 13610 | ENSMUST00000087978.5 | -85287 | ENSMUSG00000067586 | S1pr3 | sphingosine-1-phosphate receptor 3 |
| chr16 | 6696982 | 6698370 | 1389 | * | 13,45634 | 4,75571899 | 1,172906 | 4,054647 | 5,02E-05 | 0,113460495 | 1,2763 | Intron (ENSMUST0000 | 16 | 6627086 | 7227677 | 600592 | 1 | 268859 | ENSMUST00000115841.10 | 69896 | ENSMUSG00000008658 | Rbfox1 | RNA binding protein, fox-1 homolog (C. elegans) 1 |
| chr4 | 1,27E+08 | 1,27E+08 | 1516 | * | 8,417109 | 4,756616731 | 1,444311 | 3,293348 | 0,00099 | 0,477253929 | 1,449706 | Promoter (<=1kb) | 4 | 1,27E+08 | 1,27E+08 | 1564 | 1 | 71514 | ENSMUST00000130659.8 | 0 | ENSMUSG00000028820 | Sfpq | splicing factor proline/glutamine rich (polypyrimidine tract binding protein associated) |
| chr19 | 42573551 | 42574867 | 1317 | * | 9,983136 | 4,757841188 | 1,699923 | 2,798857 | 0,005128 | 0,799316832 | 1,211315 | Intron (ENSMUST0000 | 19 | 42569600 | 42573210 | 3611 | 1 | 52013 | ENSMUST00000162238.2 | 3951 | ENSMUSG00000025184 | R3hcc1l | R3H domain and coiled-coil containing 1 like |
| chr3 | 1,01E+08 | 1,01E+08 | 1233 | * | 8,145624 | 4,758485173 | 1,774863 | 2,681044 | 0,007339 | 1 | 0 | Distal Intergenic | 3 | 1,01E+08 | 1,01E+08 | 7379 | 1 | 1,15E+08 | ENSMUST00000198003.2 | 21342 | | Gm43464 | predicted gene 43464 |
| chr6 | 90609051 | 90609742 | 692 | * | 9,09211 | 4,758843172 | 1,464867 | 3,248651 | 0,00116 | 0,537150047 | 1,211315 | Intron (ENSMUST0000 | 6 | 90596129 | 90623394 | 27266 | 1 | 71699 | ENSMUST00000032177.10 | 12922 | ENSMUSG00000030089 | Slc41a3 | solute carrier family 41, member 3 |
| chr9 | 62531810 | 62533203 | 1394 | * | 24,086331 | 4,760946287 | 0,8090386 | 5,884696 | 3,988E-09 | 4,90153E-05 | 2,6724296 | Distal Intergenic | 9 | 62568156 | 62573363 | 5208 | 2 | 118567713 | ENSMUST00000241674.1 | 40160 | | Gm53533 | predicted gene, 53533 |
| chr9 | 59694792 | 59695761 | 970 | * | 8,444172 | 4,763789024 | 1,457069 | 3,269434 | 0,001078 | 0,502454322 | 1,360622 | Exon (ENSMUST00000 | 9 | 59658179 | 59836149 | 177971 | 1 | 270163 | ENSMUST00000136740.8 | 36613 | ENSMUSG00000039585 | Myo9a | myosin IXa |
| chr4 | 80797756 | 80798584 | 829 | * | 9,295869 | 4,764233058 | 1,771462 | 2,689435 | 0,007157 | 0,749550311 | 2,237403 | Distal Intergenic | 4 | 80828883 | 80873865 | 44983 | 1 | 52829 | ENSMUST00000055922.4 | -30299 | ENSMUSG00000048706 | Lurap1l | leucine rich adaptor protein 1-like |
| chr19 | 15156067 | 15157059 | 993 | * | 8,930532 | 4,76595614 | 1,399049 | 3,406568 | 0,000658 | 0,421090004 | 1,360622 | Distal Intergenic | 19 | 15657368 | 15657474 | 107 | 1 | 1,15E+08 | ENSMUST00000083389.3 | -500309 | ENSMUSG00000118757 | Gm24319 | predicted gene, 24319 |
| chr1 | 35426900 | 35428102 | 1203 | * | 11,28882 | 4,766006314 | 1,686589 | 2,825825 | 0,004716 | 0,872576097 | 0,738441 | Distal Intergenic | 1 | 35218037 | 35218165 | 129 | 2 | 1,15E+08 | ENSMUST00000104043.3 | -208735 | ENSMUSG00000077231 | Gm25096 | predicted gene, 25096 |
| chr4 | 64886407 | 64887162 | 756 | * | 8,01121 | 4,767284359 | 1,521401 | 3,133484 | 0,001727 | 1 | 0 | Distal Intergenic | 4 | 65042411 | 65275746 | 233336 | 1 | 18491 | ENSMUST00000084501.4 | -155249 | ENSMUSG00000028370 | Pappa | pregnancy-associated plasma protein A |
| chr2 | 1,43E+08 | 1,43E+08 | 944 | * | 6,140128 | 4,768062012 | 1,588684 | 3,001265 | 0,002689 | 1 | 0 | Intron (ENSMUST0000 | 2 | 1,43E+08 | 1,43E+08 | 3454 | 2 | 16558 | ENSMUST00000212106.2 | -60776 | ENSMUSG00000038844 | Kif16b | kinesin family member 16B |
| chr1 | 1,14E+08 | 1,14E+08 | 971 | * | 9,085419 | 4,768400438 | 1,459115 | 3,26801 | 0,001083 | 0,524454869 | 1,211315 | Distal Intergenic | 1 | 1,12E+08 | 1,12E+08 | 136 | 1 | 1,15E+08 | ENSMUST00000158066.3 | 1515655 | ENSMUSG00000088691 | Gm22331 | predicted gene, 22331 |
| chr14 | 30693205 | 30694468 | 1264 | * | 10,7421 | 4,770001234 | 1,447782 | 3,294695 | 0,000985 | 0,60003658 | 0,738441 | Intron (ENSMUST0000 | 14 | 30678872 | 30709401 | 30530 | 1 | 23955 | ENSMUST00000226146.2 | 14333 | ENSMUSG00000021918 | Nek4 | NIMA (never in mitosis gene a)-related expressed kinase 4 |
| chr13 | 65091771 | 65093499 | 1729 | * | 14,33522 | 4,771230446 | 1,346216 | 3,544178 | 0,000394 | 0,399583375 | 0,938682 | Distal Intergenic | 13 | 65065268 | 65068711 | 3444 | 1 | 78124 | ENSMUST00000221202.2 | 26503 | ENSMUSG00000056223 | Spata31 | spermatogenesis associated 31 |
| chr1 | 1,65E+08 | 1,65E+08 | 605 | * | 5,818978 | 4,773156031 | 1,850706 | 2,5791 | 0,009906 | 1 | 0 | Distal Intergenic | 1 | 1,65E+08 | 1,65E+08 | 26009 | 1 | 240888 | ENSMUST00000178700.8 | -14143 | ENSMUSG00000040836 | Gpr161 | G protein-coupled receptor 161 |
| chr9 | 1,01E+08 | 1,01E+08 | 823 | * | 6,215336 | 4,773958359 | 1,836647 | 2,599279 | 0,009342 | 1 | 0 | Intron (ENSMUST0000 | 9 | 1,01E+08 | 1,01E+08 | 153344 | 1 | 20842 | ENSMUST00000146312.8 | -7310 | ENSMUSG00000037286 | Stag1 | stromal antigen 1 |
| chr10 | 76733910 | 76735285 | 1376 | * | 13,87125 | 4,774015466 | 1,316281 | 3,626896 | 0,000287 | 0,337811311 | 0,938682 | Intron (ENSMUST0000 | 10 | 76625637 | 76741696 | 116060 | 2 | 59093 | ENSMUST00000171437.8 | 6411 | ENSMUSG00000001120 | Pcbp3 | poly(rC) binding protein 3 |
| chr17 | 56185651 | 56186993 | 1343 | * | 11,62576 | 4,77612146 | 1,794033 | 2,662226 | 0,007763 | 0,861411338 | 1,2763 | Promoter (<=1kb) | 17 | 56171892 | 56185930 | 14039 | 2 | 104349 | ENSMUST00000079642.8 | 0 | ENSMUSG00000057835 | Zfp119a | zinc finger protein 119a |
| chr6 | 88828980 | 88829894 | 915 | * | 10,05888 | 4,778321112 | 1,443086 | 3,311184 | 0,000929 | 0,515735451 | 1,086118 | Promoter (2-3kb) | 6 | 88821158 | 88826359 | 5202 | 2 | 319655 | ENSMUST00000141348.3 | -2621 | ENSMUSG00000033152 | Podxl2 | podocalyxin-like 2 |
| chr7 | 1,27E+08 | 1,27E+08 | 903 | * | 7,042802 | 4,781973619 | 1,616078 | 2,958998 | 0,003086 | 1 | 0 | Intron (ENSMUST0000 | 7 | 1,27E+08 | 1,27E+08 | 9713 | 1 | 1,05E+08 | ENSMUST00000246564.1 | -3972 | | Gm39090 | predicted gene, 39090 |
| chr11 | 96143564 | 96144426 | 863 | * | 7,49346 | 4,78337826 | 1,788951 | 2,673846 | 0,007499 | 1 | 0 | Promoter (<=1kb) | 11 | 96143358 | 96152519 | 9162 | 1 | 193022 | ENSMUST00000139381.2 | 206 | | Gm53 | predicted gene 53 |
| chr1 | 33534036 | 33535004 | 969 | * | 9,384371 | 4,783413637 | 1,487308 | 3,216154 | 0,001299 | 0,554723541 | 1,211315 | Intron (ENSMUST0000 | 1 | 33461814 | 33461919 | 106 | 1 | 1,15E+08 | ENSMUST00000083326.3 | 72222 | ENSMUSG00000065260 | Gm23453 | predicted gene, 23453 |
| chr1 | 1,13E+08 | 1,13E+08 | 760 | * | 5,189199 | 4,786045428 | 1,714542 | 2,791442 | 0,005247 | 1 | 0 | Distal Intergenic | 1 | 1,12E+08 | 1,12E+08 | 136 | 1 | 1,15E+08 | ENSMUST00000158066.3 | 657466 | ENSMUSG00000088691 | Gm22331 | predicted gene, 22331 |
| chr4 | 118962246 | 118964114 | 1869 | * | 39,035865 | 4,786563259 | 0,6800634 | 7,0384076 | 1,944E-12 | 1,02953E-07 | 2,3383871 | Promoter (1-2kb) | 4 | 118965908 | 118995180 | 29273 | 1 | 20525 | ENSMUST00000030398.10 | -1794 | ENSMUSG00000028645 | Slc2a1 | solute carrier family 2 (facilitated glucose transporter), member 1 |
| chr11 | 41074000 | 41074668 | 669 | * | 7,529795 | 4,7877564 | 1,48845 | 3,216606 | 0,001297 | 1 | 0 | Distal Intergenic | 11 | 40639379 | 40646138 | 6760 | 2 | 12450 | ENSMUST00000020576.8 | -427862 | ENSMUSG00000020326 | Ccng1 | cyclin G1 |
| chr10 | 59545406 | 59546647 | 1242 | * | 9,949011 | 4,788468848 | 1,37379 | 3,485591 | 0,000491 | 0,39905285 | 1,173325 | Intron (ENSMUST0000 | 10 | 59538480 | 59587527 | 49048 | 1 | 216001 | ENSMUST00000165405.2 | 6926 | ENSMUSG00000020111 | Micu1 | mitochondrial calcium uptake 1 |
| chr9 | 40778438 | 40779184 | 747 | * | 7,487405 | 4,789464152 | 1,52067 | 3,149574 | 0,001635 | 1 | 0 | Distal Intergenic | 9 | 40785423 | 40791353 | 5931 | 1 | 244813 | ENSMUST00000067375.5 | -6239 | ENSMUSG00000054360 | Bsx | brain specific homeobox |
| chr16 | 55795257 | 55796227 | 971 | * | 7,172216 | 4,793057109 | 1,59432 | 3,006334 | 0,002644 | 1 | 0 | Promoter (<=1kb) | 16 | 55793631 | 55794980 | 1350 | 2 | 66391 | ENSMUST00000186480.2 | -277 | ENSMUSG00000102101 | Zbtb11os1 | zinc finger and BTB domain containing 11, opposite strand 1 |
| chr7 | 1,1E+08 | 1,1E+08 | 754 | * | 6,118496 | 4,793508 | 1,840924 | 2,60386 | 0,009218 | 1 | 0 | Promoter (<=1kb) | 7 | 1,1E+08 | 1,1E+08 | 306910 | 2 | 319934 | ENSMUST00000164759.8 | -447 | ENSMUSG00000038371 | Sbf2 | SET binding factor 2 |
| chr4 | 1,38E+08 | 1,38E+08 | 1066 | * | 8,538441 | 4,79568007 | 1,766781 | 2,71436 | 0,00664 | 0,817552804 | 1,425929 | Distal Intergenic | 4 | 1,38E+08 | 1,38E+08 | 5810 | 1 | 66259 | ENSMUST00000050918.4 | 43014 | ENSMUSG00000046447 | Camk2n1 | calcium/calmodulin-dependent protein kinase II inhibitor 1 |
| chr8 | 56121270 | 56121937 | 668 | * | 9,667828 | 4,795836211 | 1,799309 | 2,665376 | 0,00769 | 0,877892722 | 1,173325 | Distal Intergenic | 8 | 56323947 | 56359983 | 36037 | 2 | 244486 | ENSMUST00000053441.5 | 238046 | ENSMUSG00000046258 | Adam29 | a disintegrin and metallopeptidase domain 29 |
| chr7 | 1,02E+08 | 1,02E+08 | 480 | * | 7,758025 | 4,797111409 | 1,767187 | 2,714547 | 0,006637 | 1 | 0 | Intron (ENSMUST0000 | 7 | 1,02E+08 | 1,02E+08 | 1752 | 1 | 233575 | ENSMUST00000134621.2 | 6843 | ENSMUSG00000030990 | Pgap2 | post-GPI attachment to proteins 2 |
| chr16 | 19495175 | 19496313 | 1139 | * | 9,548025 | 4,798284057 | 1,743703 | 2,751778 | 0,005927 | 0,792545355 | 1,449706 | Intron (ENSMUST0000 | 16 | 19472128 | 19525125 | 52998 | 2 | 239739 | ENSMUST00000081880.7 | 28812 | ENSMUSG00000041247 | Lamp3 | lysosomal-associated membrane protein 3 |
| chr16 | 10079511 | 10080727 | 1217 | * | 8,284255 | 4,798322143 | 1,47798 | 3,246542 | 0,001168 | 0,507398906 | 1,425929 | Distal Intergenic | 16 | 10055782 | 10055884 | 103 | 1 | 1,15E+08 | ENSMUST00000158938.3 | 23729 | ENSMUSG00000089563 | Gm24667 | predicted gene, 24667 |
| chr9 | 49357927 | 49358939 | 1013 | * | 10,59573 | 4,799295276 | 1,723929 | 2,783929 | 0,00537 | 0,798559658 | 1,2763 | Intron (ENSMUST0000 | 9 | 49326494 | 49338321 | 11828 | 2 | 244859 | ENSMUST00000034792.7 | -19606 | ENSMUSG00000032257 | Ankk1 | ankyrin repeat and kinase domain containing 1 |
| chr7 | 39515834 | 39517230 | 1397 | * | 10,78838 | 4,801661081 | 1,420142 | 3,381113 | 0,000722 | 0,540220995 | 0,738441 | Intron (ENSMUST0000 | 7 | 39279816 | 39279926 | 111 | 1 | 1,15E+08 | ENSMUST00000093724.4 | 236018 | ENSMUSG00000070236 | Gm26327 | predicted gene, 26327 |
| chr16 | 18105767 | 18106838 | 1072 | * | 10,72434 | 4,805118524 | 1,39246 | 3,450813 | 0,000559 | 0,406085428 | 1,2763 | Promoter (<=1kb) | 16 | 18105447 | 18107034 | 1588 | 2 | 94223 | ENSMUST00000134431.2 | 196 | ENSMUSG00000022718 | Dgcr8 | DGCR8, microprocessor complex subunit |
| chr13 | 88463531 | 88464539 | 1009 | * | 9,648848 | 4,805595524 | 1,454548 | 3,303842 | 0,000954 | 0,471637333 | 1,449706 | Distal Intergenic | 13 | 88669309 | 88726287 | 56979 | 1 | 1,03E+08 | ENSMUST00000224910.2 | -204770 | ENSMUSG00000114273 | Gm34585 | predicted gene, 34585 |
| chr12 | 81531374 | 81533376 | 2003 | * | 15,55179 | 4,808743854 | 1,166698 | 4,121671 | 3,76E-05 | 0,069348015 | 2,141612 | Promoter (<=1kb) | 12 | 81405800 | 81531884 | 126085 | 2 | 66272 | ENSMUST00000110340.9 | 0 | ENSMUSG00000091803 | Cox16 | cytochrome c oxidase assembly protein 16 |
| chr19 | 22024669 | 22025385 | 717 | * | 8,403123 | 4,809139619 | 1,536261 | 3,130417 | 0,001746 | 1 | 0 | Distal Intergenic | 19 | 22116410 | 22972774 | 856365 | 1 | 226025 | ENSMUST00000237357.2 | -91025 | ENSMUSG00000052387 | Trpm3 | transient receptor potential cation channel, subfamily M, member 3 |
| chr1 | 1,21E+08 | 1,21E+08 | 757 | * | 6,244724 | 4,8134235 | 1,832883 | 2,626148 | 0,008636 | 1 | 0 | Promoter (<=1kb) | 1 | 1,21E+08 | 1,21E+08 | 14155 | 2 | 66942 | ENSMUST00000001724.12 | 0 | ENSMUSG00000001674 | Ddx18 | DEAD box helicase 18 |
| chr6 | 31803618 | 31804315 | 698 | * | 7,661517 | 4,813521169 | 1,830415 | 2,629744 | 0,008545 | 0,868235543 | 1,360622 | Intron (ENSMUST0000 | 6 | 31696316 | 31720029 | 23714 | 1 | 1,05E+08 | ENSMUST00000210985.2 | 107302 | | Gm13846 | predicted gene 13846 |
| chr18 | 69699580 | 69701030 | 1451 | * | 13,54623 | 4,813832722 | 1,292933 | 3,72319 | 0,000197 | 0,252371261 | 1,2763 | Intron (ENSMUST0000 | 18 | 69696406 | 69770035 | 73630 | 1 | 21413 | ENSMUST00000136151.8 | 3174 | ENSMUSG00000053477 | Tcf4 | transcription factor 4 |
| chr12 | 1,08E+08 | 1,08E+08 | 601 | * | 7,172813 | 4,813839411 | 1,822103 | 2,641914 | 0,008244 | 0,860883941 | 1,360622 | Distal Intergenic | 12 | 1,08E+08 | 1,08E+08 | 93200 | 2 | 58208 | ENSMUST00000066060.11 | -37022 | ENSMUSG00000048251 | Bcl11b | B cell leukemia/lymphoma 11B |
| chr3 | 19058891 | 19059449 | 559 | * | 5,839668 | 4,814123346 | 1,690541 | 2,847682 | 0,004404 | 1 | 0 | Distal Intergenic | 3 | 19185566 | 19217229 | 31664 | 2 | 74252 | ENSMUST00000029125.10 | 157780 | ENSMUSG00000027599 | Armc1 | armadillo repeat containing 1 |
| chr19 | 46593400 | 46594332 | 933 | * | 8,427194 | 4,81528927 | 1,801554 | 2,672853 | 0,007521 | 1 | 0 | Intron (ENSMUST0000 | 19 | 46587563 | 46632994 | 45432 | 1 | 226178 | ENSMUST00000175937.2 | 5837 | ENSMUSG00000047731 | Wbp1l | WW domain binding protein 1 like |
| chr2 | 1,18E+08 | 1,18E+08 | 701 | * | 6,384103 | 4,815544404 | 1,848984 | 2,604427 | 0,009203 | 1 | 0 | Distal Intergenic | 2 | 1,18E+08 | 1,18E+08 | 6136 | 2 | 1,01E+08 | ENSMUST00000154409.2 | 69751 | | Gm13985 | predicted gene 13985 |
| chr5 | 98123945 | 98125134 | 1190 | * | 8,33747 | 4,816957561 | 1,768147 | 2,724297 | 0,006444 | 1 | 0 | Intron (ENSMUST0000 | 5 | 98097029 | 98143954 | 46926 | 2 | 71914 | ENSMUST00000202736.2 | 18820 | ENSMUSG00000029338 | Antxr2 | anthrax toxin receptor 2 |
| chr4 | 79275901 | 79276830 | 930 | * | 8,528751 | 4,819018908 | 1,418579 | 3,397074 | 0,000681 | 0,426043749 | 1,360622 | Distal Intergenic | 4 | 78130313 | 78133846 | 3534 | 1 | 1,03E+08 | ENSMUST00000246656.1 | 1145588 | | B230208B08Rik | RIKEN cDNA B230208B08 gene |
| chrX | 1,14E+08 | 1,14E+08 | 726 | * | 6,040776 | 4,824576681 | 1,718136 | 2,80803 | 0,004985 | 1 | 0 | Distal Intergenic | 20 | 1,14E+08 | 1,14E+08 | 930 | 1 | 237009 | ENSMUST00000059509.3 | -117059 | ENSMUSG00000050435 | Ube2dnl1 | ubiquitin-conjugating enzyme E2D N-terminal like 1 |
| chr2 | 1,19E+08 | 1,19E+08 | 934 | * | 8,104367 | 4,824629815 | 1,534956 | 3,143171 | 0,001671 | 1 | 0 | Distal Intergenic | 2 | 1,19E+08 | 1,19E+08 | 25003 | 2 | 241624 | ENSMUST00000171024.8 | 25176 | ENSMUSG00000048647 | Exd1 | exonuclease 3'-5' domain containing 1 |
| chr2 | 1,54E+08 | 1,54E+08 | 1122 | * | 9,197959 | 4,824868844 | 1,866052 | 2,585602 | 0,009721 | 0,890735468 | 1,360622 | Distal Intergenic | 2 | 1,54E+08 | 1,54E+08 | 31787 | 2 | 20648 | ENSMUST00000109728.8 | -8251 | ENSMUSG00000027488 | Snta1 | syntrophin, acidic 1 |
| chr16 | 27379870 | 27380780 | 911 | * | 7,7641 | 4,830023598 | 1,840584 | 2,62418 | 0,008686 | 0,872576097 | 1,360622 | Distal Intergenic | 16 | 27208718 | 27209841 | 1124 | 1 | 67501 | ENSMUST00000149077.2 | 171152 | ENSMUSG00000038127 | Ccdc50 | coiled-coil domain containing 50 |
| chr6 | 1,19E+08 | 1,19E+08 | 714 | * | 7,072681 | 4,831201826 | 1,82386 | 2,648888 | 0,008076 | 1 | 0 | Promoter (2-3kb) | 6 | 1,19E+08 | 1,19E+08 | 16856 | 2 | 22419 | ENSMUST00000117171.8 | -2916 | ENSMUSG00000030170 | Wnt5b | wingless-type MMTV integration site family, member 5B |
| chr9 | 1,01E+08 | 1,01E+08 | 1529 | * | 11,22487 | 4,83185783 | 1,360062 | 3,552675 | 0,000381 | 0,431099484 | 0,738441 | Promoter (2-3kb) | 9 | 1,01E+08 | 1,01E+08 | 153344 | 1 | 20842 | ENSMUST00000146312.8 | -2644 | ENSMUSG00000037286 | Stag1 | stromal antigen 1 |
| chr5 | 32236955 | 32238242 | 1288 | * | 9,942694 | 4,833719825 | 1,768743 | 2,732856 | 0,006279 | 0,854052567 | 1,086118 | Promoter (<=1kb) | 5 | 32237041 | 32242090 | 5050 | 1 | 107976 | ENSMUST00000202087.2 | 0 | ENSMUSG00000052139 | Babam2 | BRISC and BRCA1 A complex member 2 |
| chr11 | 1,02E+08 | 1,02E+08 | 1112 | * | 6,729777 | 4,833831593 | 1,796634 | 2,690493 | 0,007135 | 1 | 0 | Distal Intergenic | 11 | 1,02E+08 | 1,02E+08 | 8153 | 1 | 73887 | ENSMUST00000123249.2 | -21883 | ENSMUSG00000071818 | 4930417O22Rik | RIKEN cDNA 4930417O22 gene |
| chr11 | 1,05E+08 | 1,05E+08 | 1264 | * | 8,315206 | 4,834469506 | 1,779933 | 2,716096 | 0,006606 | 0,816751204 | 1,425929 | Promoter (<=1kb) | 11 | 1,05E+08 | 1,05E+08 | 18365 | 1 | 24086 | ENSMUST00000146910.2 | 959 | ENSMUSG00000020694 | Tlk2 | tousled-like kinase 2 (Arabidopsis) |
| chr11 | 66940651 | 66942265 | 1615 | * | 12,11756 | 4,835809346 | 1,654499 | 2,922824 | 0,003469 | 0,779936001 | 0,918255 | Promoter (1-2kb) | 11 | 66928867 | 66943396 | 14530 | 2 | 66358 | ENSMUST00000116363.2 | 1131 | ENSMUSG00000020910 | Adprm | ADP-ribose/CDP-alcohol diphosphatase, manganese dependent |
| chr14 | 5284941 | 5285597 | 657 | * | 5,003073 | 4,840633507 | 1,786501 | 2,709561 | 0,006737 | 1 | 0 | Distal Intergenic | 14 | 5363820 | 5364135 | 316 | 2 | 1E+08 | ENSMUST00000225472.2 | 78538 | | Pou5f1-rs6 | POU domain, class 5, transcription factor 1, related sequence 6 |
| chr9 | 41949857 | 41950648 | 792 | * | 6,996072 | 4,840712254 | 1,817786 | 2,662971 | 0,007745 | 1 | 0 | Intron (ENSMUST0000 | 9 | 41933911 | 41945409 | 11499 | 1 | 1,03E+08 | ENSMUST00000143710.2 | 15946 | | Gm16214 | predicted gene 16214 |
| chr9 | 55594776 | 55595527 | 752 | * | 7,524098 | 4,842945656 | 1,878105 | 2,578635 | 0,009919 | 1 | 0 | Intron (ENSMUST0000 | 9 | 55650913 | 55699594 | 48682 | 2 | 244891 | ENSMUST00000215323.2 | 104067 | ENSMUSG00000034007 | Scaper | S phase cyclin A-associated protein in the ER |
| chr7 | 110379447 | 110381164 | 1718 | * | 28,919108 | 4,843057591 | 0,7575196 | 6,3933099 | 1,623E-10 | 3,49169E-06 | 2,6724296 | Promoter (2-3kb) | 7 | 110376897 | 110390521 | 13625 | 1 | 11717 | ENSMUST00000148292.2 | 2550 | ENSMUSG00000005686 | Ampd3 | adenosine monophosphate deaminase 3 |
| chrX | 1,25E+08 | 1,25E+08 | 1031 | * | 5,874719 | 4,844277783 | 1,618473 | 2,993116 | 0,002761 | 1 | 0 | Distal Intergenic | 20 | 1,26E+08 | 1,26E+08 | 2037 | 2 | 1,19E+08 | ENSMUST00000117443.2 | 409151 | ENSMUSG00000082772 | LOC118568449 | vigilin-like |
| chr1 | 1,14E+08 | 1,14E+08 | 814 | * | 6,369532 | 4,84881754 | 1,616431 | 2,999707 | 0,002702 | 1 | 0 | Distal Intergenic | 1 | 1,16E+08 | 1,16E+08 | 15632 | 2 | 1,08E+08 | ENSMUST00000186137.7 | 1159083 | | Gm50605 | predicted gene, 50605 |
| chr13 | 95983985 | 95984927 | 943 | * | 8,742324 | 4,848879654 | 1,446468 | 3,352221 | 0,000802 | 0,446187869 | 1,425929 | Intron (ENSMUST0000 | 13 | 95882820 | 96028265 | 145446 | 2 | 544963 | ENSMUST00000238261.2 | 43338 | ENSMUSG00000021676 | Iqgap2 | IQ motif containing GTPase activating protein 2 |
| chr18 | 61032708 | 61033976 | 1269 | * | 10,34809 | 4,85079415 | 1,860928 | 2,606654 | 0,009143 | 0,915421656 | 1,086118 | Distal Intergenic | 18 | 61044852 | 61051633 | 6782 | 1 | 545260 | ENSMUST00000040359.6 | -10876 | ENSMUSG00000036412 | Arsi | arylsulfatase i |
| chr12 | 36093840 | 36094360 | 521 | * | 4,84301 | 4,851260706 | 1,767918 | 2,744052 | 0,006069 | 1 | 0 | Promoter (1-2kb) | 12 | 36071800 | 36092499 | 20700 | 2 | 66109 | ENSMUST00000220641.2 | -1341 | ENSMUSG00000020577 | Tspan13 | tetraspanin 13 |
| chr11 | 84040672 | 84041647 | 976 | * | 6,316496 | 4,851884306 | 1,838281 | 2,639359 | 0,008306 | 1 | 0 | Intron (ENSMUST0000 | 11 | 84039177 | 84058302 | 19126 | 2 | 628813 | ENSMUST00000050771.8 | 16655 | ENSMUSG00000051452 | Gm11437 | predicted gene 11437 |
| chr2 | 1,18E+08 | 1,18E+08 | 417 | * | 5,109134 | 4,852031698 | 1,846532 | 2,627646 | 0,008598 | 1 | 0 | Promoter (<=1kb) | 2 | 1,18E+08 | 1,18E+08 | 86121 | 1 | 27103 | ENSMUST00000110872.8 | 372 | ENSMUSG00000005102 | Eif2ak4 | eukaryotic translation initiation factor 2 alpha kinase 4 |
| chr12 | 45742500 | 45743304 | 805 | * | 7,000375 | 4,852258021 | 1,790335 | 2,710252 | 0,006723 | 1 | 0 | Distal Intergenic | 12 | 45513079 | 45542564 | 29486 | 1 | 1,03E+08 | ENSMUST00000219610.2 | 229421 | ENSMUSG00000112054 | Gm33680 | predicted gene, 33680 |
| chr3 | 1,09E+08 | 1,09E+08 | 848 | * | 7,654823 | 4,852701442 | 1,79589 | 2,702115 | 0,00689 | 0,83238039 | 1,360622 | Intron (ENSMUST0000 | 3 | 1,09E+08 | 1,09E+08 | 14789 | 1 | 329738 | ENSMUST00000147916.2 | 13313 | ENSMUSG00000049565 | Aknad1 | AKNA domain containing 1 |
| chr15 | 54643500 | 54644333 | 834 | * | 7,780201 | 4,852832189 | 1,787597 | 2,714723 | 0,006633 | 1 | 0 | Distal Intergenic | 15 | 54609098 | 54617435 | 8338 | 1 | 18133 | ENSMUST00000050027.9 | 34402 | ENSMUSG00000037362 | Ccn3 | cellular communication network factor 3 |
| chr15 | 63215944 | 63216758 | 815 | * | 6,211125 | 4,853509864 | 1,56339 | 3,104478 | 0,001906 | 1 | 0 | Intron (ENSMUST0000 | 15 | 63224834 | 63228868 | 4035 | 2 | 1,05E+08 | ENSMUST00000227983.2 | 12110 | | Gm41335 | predicted gene, 41335 |
| chr7 | 7301530 | 7302615 | 1086 | * | 11,32261 | 4,85899595 | 1,380243 | 3,520392 | 0,000431 | 0,354474598 | 1,2763 | Promoter (<=1kb) | 7 | 7301890 | 7301945 | 56 | 1 | 1,01E+08 | ENSMUST00000176903.3 | 0 | ENSMUSG00000093599 | Mir5620 | microRNA 5620 |
| chr1 | 80165877 | 80166640 | 764 | * | 5,410151 | 4,859576193 | 1,743435 | 2,787358 | 0,005314 | 1 | 0 | Intron (ENSMUST0000 | 1 | 80176423 | 80191661 | 15239 | 2 | 241128 | ENSMUST00000058748.2 | 25021 | ENSMUSG00000043230 | Fam124b | family with sequence similarity 124, member B |
| chr4 | 1,3E+08 | 1,3E+08 | 462 | * | 6,211515 | 4,859635982 | 1,862787 | 2,608799 | 0,009086 | 1 | 0 | Promoter (<=1kb) | 4 | 1,3E+08 | 1,3E+08 | 12979 | 1 | 107581 | ENSMUST00000146606.8 | 0 | ENSMUSG00000040690 | Col16a1 | collagen, type XVI, alpha 1 |
| chr16 | 50359237 | 50360188 | 952 | * | 6,046906 | 4,860034107 | 1,615316 | 3,00872 | 0,002624 | 1 | 0 | Intron (ENSMUST0000 | 16 | 50348865 | 50379044 | 30180 | 1 | 619312 | ENSMUST00000181884.2 | 10372 | ENSMUSG00000097694 | G730013B05Rik | RIKEN cDNA G730013B05 gene |
| chr6 | 97556723 | 97557328 | 606 | * | 8,759393 | 4,863128518 | 1,44633 | 3,362393 | 0,000773 | 0,439829468 | 1,425929 | Exon (ENSMUST00000 | 6 | 97549037 | 97549199 | 163 | 2 | 1,15E+08 | ENSMUST00000083836.4 | -7524 | ENSMUSG00002076989 | Gm23510 | predicted gene, 23510 |
| chr16 | 31767164 | 31768956 | 1793 | * | 13,83947 | 4,864387934 | 1,367909 | 3,556076 | 0,000376 | 0,36185416 | 1,075735 | Promoter (<=1kb) | 16 | 31765868 | 31767312 | 1445 | 2 | 106264 | ENSMUST00000202722.2 | 0 | ENSMUSG00000107002 | 0610012G03Rik | RIKEN cDNA 0610012G03 gene |
| chr3 | 5897751 | 5898492 | 742 | * | 10,47372 | 4,866740567 | 1,446782 | 3,363839 | 0,000769 | 0,547497234 | 0,744375 | Distal Intergenic | 3 | 5625560 | 5641299 | 15740 | 2 | 19302 | ENSMUST00000059021.10 | -256452 | ENSMUSG00000040374 | Pex2 | peroxisomal biogenesis factor 2 |
| chr15 | 35683510 | 35684196 | 687 | * | 6,93053 | 4,868658163 | 1,829428 | 2,661301 | 0,007784 | 1 | 0 | Intron (ENSMUST0000 | 15 | 35661116 | 35661193 | 78 | 2 | 1E+08 | ENSMUST00000104698.3 | -22317 | ENSMUSG00000077890 | Mir875 | microRNA 875 |
| chr7 | 98814179 | 98814854 | 676 | * | 7,402605 | 4,870327576 | 1,504987 | 3,236127 | 0,001212 | 1 | 0 | 3' UTR | 7 | 98808191 | 98819313 | 11123 | 2 | 67800 | ENSMUST00000207894.2 | 4459 | ENSMUSG00000030747 | Dgat2 | diacylglycerol O-acyltransferase 2 |
| chr1 | 1,95E+08 | 1,95E+08 | 1073 | * | 5,579543 | 4,870374549 | 1,877189 | 2,594504 | 0,009473 | 1 | 0 | Distal Intergenic | 1 | 1,95E+08 | 1,95E+08 | 39906 | 2 | 12902 | ENSMUST00000195120.6 | -32722 | ENSMUSG00000026616 | Cr2 | complement receptor 2 |
| chr8 | 85551864 | 85552485 | 622 | * | 9,823545 | 4,870781481 | 1,522532 | 3,199132 | 0,001378 | 0,593612587 | 1,086118 | Promoter (2-3kb) | 8 | 85542033 | 85549452 | 7420 | 2 | 23863 | ENSMUST00000065539.6 | -2412 | ENSMUSG00000053226 | Dand5 | DAN domain family member 5, BMP antagonist |
| chr6 | 1,19E+08 | 1,19E+08 | 666 | * | 7,383654 | 4,871657661 | 1,77264 | 2,748249 | 0,005991 | 1 | 0 | Promoter (2-3kb) | 6 | 1,18E+08 | 1,19E+08 | 60919 | 2 | 232339 | ENSMUST00000112830.3 | 2967 | ENSMUSG00000007827 | Ankrd26 | ankyrin repeat domain 26 |
| chr7 | 46207458 | 46208483 | 1026 | * | 8,630101 | 4,873212684 | 1,744358 | 2,7937 | 0,005211 | 0,693003605 | 2,237403 | Intron (ENSMUST0000 | 7 | 46092645 | 46252666 | 160022 | 2 | 27414 | ENSMUST00000156147.8 | 44183 | ENSMUSG00000030839 | Sergef | secretion regulating guanine nucleotide exchange factor |
| chr12 | 43487209 | 43487875 | 667 | * | 6,132738 | 4,874245819 | 1,759893 | 2,769626 | 0,005612 | 1 | 0 | Distal Intergenic | 12 | 43789820 | 43789948 | 129 | 1 | 1,15E+08 | ENSMUST00000083070.3 | -301945 | ENSMUSG00000065004 | Gm26326 | predicted gene, 26326 |
| chr11 | 1,12E+08 | 1,12E+08 | 728 | * | 6,998452 | 4,876046001 | 1,684364 | 2,894889 | 0,003793 | 0,72659841 | 1,360622 | Distal Intergenic | 11 | 1,11E+08 | 1,11E+08 | 2269 | 2 | 1,03E+08 | ENSMUST00000247083.1 | -561915 | | Gm30158 | predicted gene, 30158 |
| chr5 | 1,3E+08 | 1,3E+08 | 874 | * | 7,938978 | 4,878808582 | 1,85064 | 2,636282 | 0,008382 | 1 | 0 | Promoter (<=1kb) | 5 | 1,3E+08 | 1,3E+08 | 17753 | 2 | 109900 | ENSMUST00000159619.8 | 0 | ENSMUSG00000025533 | Asl | argininosuccinate lyase |
| chr13 | 82386141 | 82386916 | 776 | * | 8,468227 | 4,880596301 | 1,445043 | 3,377476 | 0,000732 | 0,426343337 | 1,449706 | Intron (ENSMUST0000 | 13 | 82198870 | 82199000 | 131 | 1 | 1,15E+08 | ENSMUST00000158298.3 | 187271 | ENSMUSG00000088923 | Gm24295 | predicted gene, 24295 |
| chr15 | 57989381 | 57989992 | 612 | * | 5,89114 | 4,880902183 | 1,879679 | 2,596668 | 0,009413 | 1 | 0 | Exon (ENSMUST0000 | 15 | 57958800 | 57982711 | 23912 | 2 | 70472 | ENSMUST00000228783.2 | -6670 | ENSMUSG00000022360 | Atad2 | ATPase family, AAA domain containing 2 |

| | | | | | | | | | | | | | | | | | | | | | | | |
|---|---|---|---|---|---|---|---|---|---|---|---|---|---|---|---|---|---|---|---|---|---|---|---|
| chr5 | 1,15E+08 | 1,15E+08 | 973 | * | 6,686062 | 4,882771296 | 1,854843 | 2,632445 | 0,008477 | 1 | 0 | Promoter (<=1kb) | 5 | 1,15E+08 | 1,15E+08 | 46089 | 2 | 26431 | ENSMUST00000112185.9 | 0 | ENSMUSG00000041890 | Git2 | GIT ArfGAP 2 |
| chr2 | 1,55E+08 | 1,55E+08 | 1823 | * | 14,94626 | 4,883222988 | 1,15698 | 4,220662 | 2,44E-05 | 0,052288507 | 2,141612 | Intron (ENSMUST00000 | 2 | 1,55E+08 | 1,55E+08 | 638 | 1 | 60525 | ENSMUST00000143169.2 | -9864 | ENSMUSG00000027605 | Acss2 | acyl-CoA synthetase short-chain family member 2 |
| chr4 | 1,32E+08 | 1,32E+08 | 625 | * | 5,405465 | 4,884736694 | 1,890171 | 2,584283 | 0,009758 | 1 | 0 | Distal Intergenic | 4 | 1,32E+08 | 1,32E+08 | 24287 | 2 | 100340 | ENSMUST00000030709.9 | -5227 | ENSMUSG00000028885 | Smpdl3b | sphingomyelin phosphodiesterase, acid-like 3B |
| chrX | 43990907 | 43991855 | 949 | * | 5,590931 | 4,885054888 | 1,882976 | 2,594327 | 0,009478 | 1 | 0 | Distal Intergenic | 20 | 43875447 | 43879074 | 3628 | 2 | 245404 | ENSMUST00000060481.9 | -111833 | ENSMUSG00000045284 | Dcaf12l1 | DDB1 and CUL4 associated factor 12-like 1 |
| chr10 | 1,17E+08 | 1,17E+08 | 933 | * | 8,324397 | 4,885262174 | 1,490667 | 3,277233 | 0,001048 | 1 | 0 | Intron (ENSMUST00000 | 10 | 1,17E+08 | 1,17E+08 | 65691 | 2 | 327826 | ENSMUST00000219483.2 | 3613 | ENSMUSG00000020170 | Frs2 | fibroblast growth factor receptor substrate 2 |
| chr10 | 16143689 | 16144373 | 685 | * | 5,801527 | 4,886421044 | 1,850643 | 2,640391 | 0,008281 | 1 | 0 | Distal Intergenic | 10 | 15667525 | 15682813 | 15289 | 1 | 77538 | ENSMUST00000218833.2 | 476164 | ENSMUSG00000112227 | C330004P14Rik | RIKEN cDNA C330004P14 gene |
| chr9 | 17638219 | 17639011 | 793 | * | 8,601906 | 4,88691855 | 1,838908 | 2,657511 | 0,007872 | 0,848080889 | 1,425929 | Distal Intergenic | 9 | 18203421 | 18228738 | 25318 | 1 | 66917 | ENSMUST00000001825.9 | -564410 | ENSMUSG00000001774 | Chordc1 | cysteine and histidine-rich domain (CHORD)-containing, zinc-binding protein 1 |
| chr12 | 66929459 | 66930373 | 915 | * | 6,947798 | 4,889186813 | 1,487526 | 3,28679 | 0,001013 | 1 | 0 | Intron (ENSMUST00000 | 12 | 66736270 | 66872503 | 136234 | 2 | 320772 | ENSMUST00000177690.2 | -56956 | ENSMUSG00000034912 | Mdga2 | MAM domain containing glycosylphosphatidylinositol anchor 2 |
| chr5 | 1,29E+08 | 1,29E+08 | 1498 | * | 10,23946 | 4,891215364 | 1,449519 | 3,374371 | 0,00074 | 0,457838671 | 1,211315 | Distal Intergenic | 5 | 1,29E+08 | 1,29E+08 | 4052 | 1 | 19384 | ENSMUST00000111343.2 | 6647 | ENSMUSG00000029430 | Ran | RAN, member RAS oncogene family |
| chr4 | 1,4E+08 | 1,4E+08 | 1379 | * | 12,04996 | 4,891339616 | 1,711125 | 2,858552 | 0,004256 | 0,853738252 | 0,738441 | 3' UTR | 4 | 1,4E+08 | 1,4E+08 | 7807 | 2 | 242721 | ENSMUST00000105031.4 | -57539 | ENSMUSG00000078234 | Klhdc7a | kelch domain containing 7A |
| chr8 | 1,21E+08 | 1,21E+08 | 788 | * | 11,41215 | 4,892158591 | 1,819024 | 2,689441 | 0,007157 | 0,86195649 | 1,173325 | Intron (ENSMUST00000 | 8 | 1,21E+08 | 1,21E+08 | 92944 | 1 | 382034 | ENSMUST00000034279.16 | -126378 | ENSMUSG00000031822 | Gse1 | genetic suppressor element 1, coiled-coil protein |
| chr5 | 87398553 | 87399430 | 878 | * | 6,870778 | 4,893677129 | 1,616887 | 3,026605 | 0,002473 | 1 | 0 | Exon (ENSMUST00000 | 5 | 87388352 | 87402663 | 14312 | 2 | 112417 | ENSMUST00000075858.4 | 3233 | ENSMUSG00000057425 | Ugt2b37 | UDP glucuronosyltransferase 2 family, polypeptide B37 |
| chr1 | 1,15E+08 | 1,15E+08 | 805 | * | 7,354092 | 4,894414104 | 1,889384 | 2,590481 | 0,009584 | 1 | 0 | Distal Intergenic | 1 | 1,16E+08 | 1,16E+08 | 15632 | 2 | 1,08E+08 | ENSMUST00000186137.7 | 726261 | | Gm50605 | predicted gene, 50605 |
| chr1 | 34882204 | 34882921 | 718 | * | 6,96097 | 4,89474953 | 1,509698 | 3,242204 | 0,001186 | 1 | 0 | Promoter (<=1kb) | 1 | 34867723 | 34882146 | 14424 | 2 | 214469 | ENSMUST00000140534.2 | -58 | ENSMUSG00000037503 | Fam168b | family with sequence similarity 168, member B |
| chr5 | 35420435 | 35421517 | 1083 | * | 8,751794 | 4,897296133 | 1,763033 | 2,777768 | 0,005473 | 0,791416391 | 1,360622 | Distal Intergenic | 5 | 35435663 | 35439107 | 3445 | 1 | 11553 | ENSMUST00000049545.7 | -14146 | ENSMUSG00000045318 | Adra2c | adrenergic receptor, alpha 2c |
| chr10 | 81184144 | 81184810 | 667 | * | 7,153566 | 4,897465076 | 1,828804 | 2,677961 | 0,007407 | 0,845014042 | 1,360622 | Promoter (<=1kb) | 10 | 81182593 | 81184919 | 2327 | 2 | 15353 | ENSMUST00000141171.8 | 109 | ENSMUSG00000020232 | Hmg20b | high mobility group 20B |
| chr17 | 32536190 | 32537091 | 902 | * | 6,399572 | 4,897522709 | 1,577136 | 3,105326 | 0,001901 | 1 | 0 | Promoter (<=1kb) | 17 | 32535192 | 32537778 | 2587 | 2 | 56399 | ENSMUST00000236785.2 | 687 | ENSMUSG00000024045 | Akap8 | A kinase (PRKA) anchor protein 8 |
| chr2 | 1,58E+08 | 1,58E+08 | 636 | * | 6,317339 | 4,897665524 | 1,840848 | 2,660549 | 0,007801 | 1 | 0 | Intron (ENSMUST00000 | 2 | 1,58E+08 | 1,58E+08 | 30067 | 1 | 277432 | ENSMUST00000109523.2 | -52195 | ENSMUSG00000037843 | Vstm2l | V-set and transmembrane domain containing 2-like |
| chr9 | 59217612 | 59219122 | 1511 | * | 15,25153 | 4,899037279 | 1,0828178 | 4,5243415 | 6,058E-06 | 0,02642625 | 1,3768303 | Promoter (<=1kb) | 9 | 59218931 | 59231335 | 12405 | 1 | 72141 | ENSMUST00000215994.2 | 0 | ENSMUSG00000025236 | Adpgk | ADP-dependent glucokinase |
| chr9 | 65631738 | 65633434 | 1697 | * | 16,82441 | 4,899777885 | 1,251008 | 3,916663 | 8,98E-05 | 0,117758808 | 2,141612 | Intron (ENSMUST00000 | 9 | 65607152 | 65608426 | 1275 | 2 | 214812 | ENSMUST00000159878.2 | -23312 | ENSMUSG00000040524 | Zfp609 | zinc finger protein 609 |
| chr19 | 8897274 | 8898006 | 733 | * | 8,000307 | 4,902385595 | 1,840217 | 2,664025 | 0,007721 | 1 | 0 | Promoter (<=1kb) | 19 | 8897738 | 8904600 | 6863 | 1 | 72727 | ENSMUST00000096243.7 | 0 | ENSMUSG00000071649 | B3gat3 | beta-1,3-glucuronyltransferase 3 (glucuronosyltransferase I) |
| chr1 | 83450119 | 83451016 | 898 | * | 6,467413 | 4,903591288 | 1,823402 | 2,689254 | 0,007161 | 1 | 0 | Distal Intergenic | 1 | 83231860 | 83385921 | 154062 | 2 | 77629 | ENSMUST00000160953.8 | -64198 | ENSMUSG00000026163 | Sphkap | SPHK1 interactor, AKAP domain containing |
| chr11 | 30128944 | 30129532 | 589 | * | 6,655955 | 4,905537086 | 1,838647 | 2,668015 | 0,00763 | 1 | 0 | Intron (ENSMUST00000 | 11 | 30056785 | 30148257 | 91473 | 2 | 20742 | ENSMUST00000102838.10 | 18725 | ENSMUSG00000020315 | Sptbn1 | spectrin beta, non-erythrocytic 1 |
| chr8 | 13600930 | 13601905 | 976 | * | 7,693548 | 4,905668387 | 1,834647 | 2,673904 | 0,007497 | 1 | 0 | 3' UTR | 8 | 13601011 | 13612405 | 11395 | 2 | 66501 | ENSMUST00000033830.15 | 10500 | ENSMUSG00000031452 | 1700029H14Rik | RIKEN cDNA 1700029H14 gene |
| chrX | 1,68E+08 | 1,68E+08 | 651 | * | 8,705376 | 4,906435977 | 1,461156 | 3,357913 | 0,000785 | 0,439829468 | 1,449706 | Intron (ENSMUST00000 | 20 | 1,68E+08 | 1,68E+08 | 191389 | 1 | 11856 | ENSMUST00000112127.2 | 18708 | ENSMUSG00000031355 | Arhgap6 | Rho GTPase activating protein 6 |
| chr11 | 70512898 | 70513643 | 746 | * | 6,951145 | 4,906806385 | 1,801174 | 2,724227 | 0,006445 | 1 | 0 | Promoter (2-3kb) | 11 | 70513853 | 70516387 | 2535 | 2 | 1,03E+08 | ENSMUST00000151599.2 | 2744 | ENSMUSG00000085952 | Gm12315 | predicted gene 12315 |
| chr10 | 79680343 | 79681652 | 1310 | * | 10,59087 | 4,906828275 | 1,287123 | 3,812245 | 0,000138 | 0,27544967 | 0,744375 | Distal Intergenic | 10 | 79690261 | 79700266 | 10006 | 1 | 19205 | ENSMUST00000172282.8 | -8609 | ENSMUSG00000006498 | Ptbp1 | polypyrimidine tract binding protein 1 |
| chr7 | 39194518 | 39195316 | 799 | * | 6,250756 | 4,908607004 | 1,867458 | 2,628496 | 0,008576 | 1 | 0 | Promoter (<=1kb) | 7 | 39193826 | 39202756 | 8931 | 1 | 1,03E+08 | ENSMUST00000190789.3 | 692 | ENSMUSG00000108621 | Gm33989 | predicted gene, 33989 |
| chr3 | 44380430 | 44381104 | 675 | * | 6,308478 | 4,911263353 | 1,571745 | 3,124719 | 0,00178 | 1 | 0 | Distal Intergenic | 3 | 45281480 | 45332640 | 51161 | 2 | 72511 | ENSMUST00000191843.2 | 951536 | ENSMUSG00000097040 | 2610316D01Rik | RIKEN cDNA 2610316D01 gene |
| chr11 | 1,16E+08 | 1,16E+08 | 1287 | * | 9,496697 | 4,912926877 | 1,714478 | 2,865552 | 0,004163 | 0,761673329 | 1,211315 | Distal Intergenic | 11 | 1,16E+08 | 1,16E+08 | 135 | 1 | 1,15E+08 | ENSMUST00000158835.3 | 10805 | ENSMUSG00000089460 | Gm23699 | predicted gene, 23699 |
| chr4 | 1,34E+08 | 1,34E+08 | 766 | * | 7,535044 | 4,913268416 | 1,637299 | 3,000837 | 0,002692 | 1 | 0 | Intron (ENSMUST00000 | 4 | 1,34E+08 | 1,34E+08 | 129 | 2 | 1,02E+08 | ENSMUST00000185007.3 | -20647 | ENSMUSG00000099289 | Mir6403 | microRNA 6403 |
| chrX | 1,36E+08 | 1,36E+08 | 1061 | * | 8,463202 | 4,913489063 | 1,856036 | 2,647302 | 0,008114 | 0,852340527 | 1,425929 | Distal Intergenic | 20 | 1,36E+08 | 1,36E+08 | 10609 | 2 | 319642 | ENSMUST00000058814.7 | -3211 | ENSMUSG00000043463 | Rab9b | RAB9B, member RAS oncogene family |
| chrX | 1,06E+08 | 1,06E+08 | 1160 | * | 8,53606 | 4,913982715 | 1,475644 | 3,330059 | 0,000868 | 0,382960329 | 2,237403 | Exon (ENSMUST00000 | 20 | 1,06E+08 | 1,06E+08 | 3563 | 2 | 213436 | ENSMUST00000062010.10 | 17915 | ENSMUSG00000047686 | Rtl3 | retrotransposon Gag like 3 |
| chr14 | 30818665 | 30819706 | 1042 | * | 5,286834 | 4,917601841 | 1,891852 | 2,599359 | 0,00934 | 1 | 0 | Intron (ENSMUST00000 | 14 | 30806161 | 30809997 | 3837 | 1 | 66923 | ENSMUST00000156407.2 | 12504 | ENSMUSG00000042323 | Pbrm1 | polybromo 1 |
| chr3 | 98124189 | 98124871 | 683 | * | 7,90185 | 4,917888955 | 1,819965 | 2,702188 | 0,006888 | 1 | 0 | Distal Intergenic | 3 | 98129472 | 98144064 | 14593 | 1 | 67709 | ENSMUST00000029469.5 | -4601 | ENSMUSG00000027876 | Reg4 | regenerating islet-derived family, member 4 |
| chr2 | 1,12E+08 | 1,12E+08 | 1099 | * | 5,826118 | 4,918000521 | 1,885003 | 2,609015 | 0,00908 | 1 | 0 | Promoter (<=1kb) | 2 | 1,12E+08 | 1,12E+08 | 17166 | 1 | 73024 | ENSMUST00000069747.6 | -694 | ENSMUSG00000055943 | Emc7 | ER membrane protein complex subunit 7 |
| chr4 | 1,36E+08 | 1,36E+08 | 1427 | * | 12,43748 | 4,919055731 | 1,320114 | 3,726236 | 0,000194 | 0,293348204 | 0,918255 | Promoter (<=1kb) | 4 | 1,36E+08 | 1,36E+08 | 6345 | 1 | 71665 | ENSMUST00000142033.8 | 0 | ENSMUSG00000028673 | Fuca1 | fucosidase, alpha-L- 1, tissue |
| chr6 | 15852931 | 15853395 | 465 | * | 6,200946 | 4,920369693 | 1,845573 | 2,66604 | 0,007675 | 1 | 0 | Distal Intergenic | 6 | 15727798 | 15770348 | 42551 | 1 | 16543 | ENSMUST00000128849.3 | 125133 | ENSMUSG00000041390 | Mdfic | MyoD family inhibitor domain containing |
| chr4 | 1,39E+08 | 1,39E+08 | 2357 | * | 16,7734 | 4,92043611 | 1,286828 | 3,823693 | 0,000131 | 0,203673596 | 1,2763 | Distal Intergenic | 4 | 1,39E+08 | 1,39E+08 | 1249 | 1 | 66825 | ENSMUST00000116094.5 | -18994 | ENSMUSG00000070661 | Rnf186 | ring finger protein 186 |
| chr4 | 1,56E+08 | 1,56E+08 | 621 | * | 6,910257 | 4,922858595 | 1,806436 | 2,725177 | 0,006427 | 1 | 0 | Promoter (1-2kb) | 4 | 1,56E+08 | 1,56E+08 | 17284 | 1 | 230996 | ENSMUST00000072554.13 | -1518 | ENSMUSG00000059939 | 9430015G10Rik | RIKEN cDNA 9430015G10 gene |
| chr16 | 40986212 | 40986696 | 485 | * | 4,041181 | 4,923080888 | 1,891045 | 2,603365 | 0,009231 | 1 | 0 | Intron (ENSMUST00000 | 16 | 41353220 | 41966576 | 613357 | 1 | 268890 | ENSMUST00000078873.11 | -366524 | ENSMUSG00000061080 | Lsamp | limbic system-associated membrane protein |
| chr9 | 58563319 | 58564320 | 1002 | * | 6,992755 | 4,924866323 | 1,803979 | 2,730003 | 0,006333 | 1 | 0 | Promoter (2-3kb) | 9 | 58560133 | 58566575 | 6443 | 2 | 73673 | ENSMUST00000175792.8 | 2255 | ENSMUSG00000074269 | Rec114 | REC114 meiotic recombination protein |
| chr16 | 31602134 | 31603469 | 1336 | * | 10,03046 | 4,925090527 | 1,413103 | 3,485302 | 0,000492 | 0,392772643 | 1,211315 | Intron (ENSMUST00000 | 16 | 31610589 | 31661636 | 51048 | 1 | 13383 | ENSMUST00000131136.2 | -7120 | ENSMUSG00000022770 | Dlg1 | discs large MAGUK scaffold protein 1 |
| chr16 | 31304949 | 31306173 | 1225 | * | 10,74318 | 4,925771644 | 1,763891 | 2,792561 | 0,005229 | 0,890233764 | 0,738441 | Distal Intergenic | 16 | 31327205 | 31330595 | 3391 | 1 | 1,08E+08 | ENSMUST00000231805.2 | -21032 | | Gm46560 | predicted gene, 46560 |
| chr10 | 1,28E+08 | 1,28E+08 | 778 | * | 11,96397 | 4,931437016 | 1,208031 | 4,082211 | 4,46E-05 | 0,148851477 | 0,744375 | Promoter (<=1kb) | 10 | 1,28E+08 | 1,28E+08 | 15483 | 1 | 216456 | ENSMUST00000044776.13 | 0 | ENSMUSG00000044005 | Gls2 | glutaminase 2 (liver, mitochondrial) |
| chr11 | 75843529 | 75844409 | 881 | * | 7,336419 | 4,932202141 | 1,818242 | 2,712621 | 0,006675 | 1 | 0 | Distal Intergenic | 11 | 75797398 | 75829255 | 31858 | 2 | 380714 | ENSMUST00000152035.2 | -14274 | ENSMUSG00000020847 | Rph3al | rabphilin 3A-like (without C2 domains) |
| chr6 | 1,25E+08 | 1,25E+08 | 750 | * | 9,168609 | 4,93279305 | 1,753139 | 2,813692 | 0,004898 | 0,761559581 | 1,425929 | Intron (ENSMUST00000 | 6 | 1,25E+08 | 1,25E+08 | 27246 | 1 | 269800 | ENSMUST00000112424.2 | 3060 | ENSMUSG00000038346 | Zfp384 | zinc finger protein 384 |
| chr13 | 37724116 | 37724927 | 812 | * | 6,426182 | 4,933171863 | 1,867969 | 2,640928 | 0,008268 | 1 | 0 | Promoter (<=1kb) | 13 | 37725333 | 37732419 | 7087 | 1 | 1,03E+08 | ENSMUST00000224508.3 | -406 | | Lncbate6 | brown adipose tissue enriched long noncoding RNA 6 |
| chr1 | 1,53E+08 | 1,53E+08 | 951 | * | 6,452989 | 4,934125184 | 1,856646 | 2,657548 | 0,007871 | 1 | 0 | Intron (ENSMUST00000 | 1 | 1,53E+08 | 1,53E+08 | 113865 | 2 | 226519 | ENSMUST00000027752.15 | 20573 | ENSMUSG00000026478 | Lamc1 | laminin, gamma 1 |
| chr5 | 1,22E+08 | 1,22E+08 | 1054 | * | 9,103869 | 4,937242342 | 1,511025 | 3,26748 | 0,001085 | 0,504157668 | 1,360622 | Intron (ENSMUST00000 | 5 | 1,22E+08 | 1,22E+08 | 11788 | 2 | 11669 | ENSMUST00000199369.5 | -5274 | ENSMUSG00000029455 | Aldh2 | aldehyde dehydrogenase 2, mitochondrial |
| chrX | 1,09E+08 | 1,09E+08 | 619 | * | 5,553788 | 4,937322442 | 1,86919 | 2,641423 | 0,008256 | 1 | 0 | Distal Intergenic | 20 | 1,09E+08 | 1,09E+08 | 2406 | 1 | 634340 | ENSMUST00000122296.2 | -239896 | ENSMUSG00000082853 | Gm7134 | kinesin family member 18B pseudogene |
| chr5 | 88869180 | 88870142 | 963 | * | 7,252058 | 4,938600606 | 1,904787 | 2,592731 | 0,009522 | 1 | 0 | Promoter (<=1kb) | 5 | 88868836 | 88897868 | 29033 | 1 | 68473 | ENSMUST00000125171.2 | 344 | ENSMUSG00000006262 | Mob1b | MOB kinase activator 1B |
| chr5 | 1,17E+08 | 1,17E+08 | 1356 | * | 12,27737 | 4,939557614 | 1,362655 | 3,62495 | 0,000289 | 0,322225015 | 1,075735 | Intron (ENSMUST00000 | 5 | 1,17E+08 | 1,17E+08 | 5096 | 1 | 231668 | ENSMUST00000139026.2 | -4239 | ENSMUSG00000066894 | Vsig10 | V-set and immunoglobulin domain containing 10 |
| chr2 | 78473327 | 78474070 | 744 | * | 6,377078 | 4,940141541 | 1,817033 | 2,718795 | 0,006552 | 1 | 0 | Distal Intergenic | 2 | 78698468 | 78700951 | 2484 | 1 | 22193 | ENSMUST00000124120.2 | -224398 | ENSMUSG00000027011 | Ube2e3 | ubiquitin-conjugating enzyme E2E 3 |
| chr5 | 66292155 | 66292735 | 581 | * | 5,932381 | 4,940728225 | 1,907153 | 2,590631 | 0,00958 | 1 | 0 | Intron (ENSMUST00000 | 5 | 66182885 | 66308299 | 125415 | 2 | 245945 | ENSMUST00000202794.4 | 15564 | ENSMUSG00000070780 | Rbm47 | RNA binding motif protein 47 |
| chr2 | 69709693 | 69710728 | 1036 | * | 6,946701 | 4,941418267 | 1,878275 | 2,630828 | 0,008518 | 1 | 0 | Intron (ENSMUST00000 | 2 | 69701558 | 69715941 | 14384 | 2 | 75422 | ENSMUST00000142127.8 | 5213 | ENSMUSG00000051730 | Mettl5 | methyltransferase like 5 |
| chr14 | 20474747 | 20475665 | 919 | * | 10,60136 | 4,942131509 | 1,402953 | 3,522664 | 0,000427 | 0,369518512 | 1,173325 | Exon (ENSMUST00000 | 14 | 20475477 | 20480056 | 4580 | 2 | 76670 | ENSMUST00000148692.2 | 4391 | ENSMUSG00000039543 | Cfap70 | cilia and flagella associated protein 70 |
| chr7 | 1,11E+08 | 1,12E+08 | 1684 | * | 12,92133 | 4,943798311 | 1,664647 | 2,969878 | 0,002979 | 0,792151706 | 0,738441 | Distal Intergenic | 7 | 1,11E+08 | 1,11E+08 | 308317 | 2 | 233733 | ENSMUST00000106663.2 | -120692 | ENSMUSG00000038296 | Galnt18 | polypeptide N-acetylgalactosaminyltransferase 18 |
| chr3 | 27456407 | 27457099 | 693 | * | 6,752828 | 4,946028728 | 1,54839 | 3,194304 | 0,001402 | 1 | 0 | Distal Intergenic | 3 | 27425500 | 27432159 | 6660 | 1 | 208188 | ENSMUST00000057186.2 | 30907 | ENSMUSG00000051136 | Ghsr | growth hormone secretagogue receptor |
| chr10 | 1,2E+08 | 1,2E+08 | 1309 | * | 10,14101 | 4,946620048 | 1,707327 | 2,897289 | 0,003764 | 0,764024179 | 1,086118 | Distal Intergenic | 10 | 1,2E+08 | 1,2E+08 | 2894 | 2 | 74912 | ENSMUST00000219022.2 | 24867 | ENSMUSG00000112716 | 4930471E19Rik | RIKEN cDNA 4930471E19 gene |
| chr16 | 86285513 | 86286075 | 563 | * | 4,503676 | 4,947869925 | 1,875942 | 2,637539 | 0,008351 | 1 | 0 | Distal Intergenic | 16 | 86033059 | 86066038 | 32980 | 1 | 1,03E+08 | ENSMUST00000231827.2 | 252454 | | Gm32357 | predicted gene, 32357 |
| chr4 | 92007536 | 92008309 | 774 | * | 5,967862 | 4,948167019 | 1,910744 | 2,589655 | 0,009607 | 1 | 0 | Distal Intergenic | 4 | 92032566 | 92035458 | 2893 | 2 | 69314 | ENSMUST00000107108.8 | 27149 | ENSMUSG00000028533 | Izumo3 | IZUMO family member 3 |
| chr16 | 33524273 | 33525320 | 1048 | * | 9,114167 | 4,949262305 | 1,424559 | 3,474241 | 0,000512 | 0,363898568 | 1,449706 | Intron (ENSMUST00000 | 16 | 33504915 | 33589813 | 84899 | 1 | 77446 | ENSMUST00000126532.2 | 19358 | ENSMUSG00000075254 | Heg1 | heart development protein with EGF-like domains 1 |
| chr3 | 1,46E+08 | 1,46E+08 | 1877 | * | 13,25844 | 4,949758505 | 1,318523 | 3,754018 | 0,000174 | 0,247743734 | 1,173325 | 3' UTR | 3 | 1,46E+08 | 1,46E+08 | 6816 | 1 | 66421 | ENSMUST00000039571.14 | -3677 | ENSMUSG00000036873 | 2410004B18Rik | RIKEN cDNA 2410004B18 gene |
| chr11 | 71070762 | 71071548 | 787 | * | 6,764955 | 4,950351941 | 1,830848 | 2,703858 | 0,006854 | 1 | 0 | Intron (ENSMUST00000 | 11 | 71046819 | 71092124 | 45306 | 2 | 637515 | ENSMUST00000108514.10 | 20576 | ENSMUSG00000070390 | Nlrp1b | NLR family, pyrin domain containing 1B |
| chr15 | 41061904 | 41062588 | 685 | * | 7,155966 | 4,950363302 | 1,831931 | 2,702265 | 0,006887 | 1 | 0 | Distal Intergenic | 15 | 41036883 | 41037264 | 382 | 1 | 1E+08 | ENSMUST00000060066.6 | 25021 | | 4930555K19Rik | leucine rich repeat and sterile alpha motif containing 1 pseudogene |
| chr14 | 31506520 | 31507134 | 615 | * | 5,986493 | 4,950671578 | 1,876133 | 2,638764 | 0,008321 | 1 | 0 | Promoter (2-3kb) | 14 | 31423803 | 31503872 | 80070 | 2 | 105522 | ENSMUST00000227089.2 | -2648 | ENSMUSG00000014496 | Ankrd28 | ankyrin repeat domain 28 |
| chr13 | 49126576 | 49127484 | 909 | * | 7,190243 | 4,950678289 | 1,885329 | 2,625896 | 0,008642 | 1 | 0 | Distal Intergenic | 13 | 49121810 | 49123386 | 1577 | 1 | 68128 | ENSMUST00000241437.1 | 4766 | | Fam120aos | family with sequence similarity 120A, opposite strand |
| chr11 | 1,01E+08 | 1,01E+08 | 804 | * | 6,550257 | 4,951942909 | 1,859344 | 2,663274 | 0,007738 | 1 | 0 | 5' UTR | 11 | 1,01E+08 | 1,01E+08 | 3614 | 2 | 19183 | ENSMUST00000019447.15 | -5950 | ENSMUSG00000019303 | Psmc3ip | proteasome (prosome, macropain) 26S subunit, ATPase 3, interacting protein |
| chr2 | 1,57E+08 | 1,57E+08 | 1753 | * | 13,57948 | 4,952017381 | 1,168976 | 4,2362 | 2,27E-05 | 0,07892476 | 1,075735 | Promoter (<=1kb) | 2 | 1,57E+08 | 1,57E+08 | 8650 | 1 | 20014 | ENSMUST00000156007.8 | 0 | ENSMUSG00000027642 | Rpn2 | ribophorin II |
| chrX | 1,4E+08 | 1,4E+08 | 878 | * | 6,65212 | 4,953780938 | 1,835878 | 2,698317 | 0,006969 | 1 | 0 | Intron (ENSMUST00000 | 20 | 1,4E+08 | 1,4E+08 | 21412 | 2 | 94216 | ENSMUST00000137051.8 | -115412 | ENSMUSG00000031273 | Col4a6 | collagen, type IV, alpha 6 |
| chr4 | 1,3E+08 | 1,3E+08 | 796 | * | 7,136171 | 4,955048695 | 1,820444 | 2,72189 | 0,006491 | 1 | 0 | Intron (ENSMUST00000 | 4 | 1,3E+08 | 1,3E+08 | 13355 | 1 | 319894 | ENSMUST00000139190.2 | 8981 | ENSMUSG00000086369 | E330017L17Rik | RIKEN cDNA E330017L17 gene |
| chr14 | 31599577 | 31600706 | 1130 | * | 8,474399 | 4,960122946 | 1,763266 | 2,813031 | 0,004908 | 1 | 0 | Distal Intergenic | 14 | 31420725 | 31552608 | 131884 | 2 | 105522 | ENSMUST00000014640.9 | -46969 | ENSMUSG00000014496 | Ankrd28 | ankyrin repeat domain 28 |
| chr4 | 1,35E+08 | 1,35E+08 | 1761 | * | 11,71075 | 4,963177436 | 1,549483 | 3,203118 | 0,001359 | 0,658200216 | 0,744375 | Exon (ENSMUST00000 | 4 | 1,35E+08 | 1,35E+08 | 41817 | 1 | 78806 | ENSMUST00000063707.3 | 10852 | ENSMUSG00000028801 | Stpg1 | sperm tail PG rich repeat containing 1 |
| chr2 | 1,28E+08 | 1,28E+08 | 943 | * | 7,039637 | 4,964846869 | 1,917302 | 2,589497 | 0,009612 | 0,890005189 | 1,360622 | Distal Intergenic | 2 | 1,28E+08 | 1,28E+08 | 130651 | 2 | 1E+08 | ENSMUST00000223654.2 | -11862 | ENSMUSG00000074813 | Morrbid | myeloid RNA regulator of BCL2L11 induced cell death |
| chr5 | 25212366 | 25213397 | 1032 | * | 8,77755 | 4,967584599 | 1,764039 | 2,816029 | 0,004862 | 0,769699963 | 1,360622 | Intron (ENSMUST00000 | 5 | 25067746 | 25200751 | 133006 | 2 | 108099 | ENSMUST00000076306.12 | -11615 | ENSMUSG00000028944 | Prkag2 | protein kinase, AMP-activated, gamma 2 non-catalytic subunit |
| chr18 | 5969365 | 5970111 | 747 | * | 6,545513 | 4,968161346 | 1,811372 | 2,742761 | 0,006093 | 1 | 0 | Distal Intergenic | 18 | 5945183 | 5965965 | 20783 | 1 | 1,03E+08 | ENSMUST00000234953.2 | 24182 | | Gm34804 | predicted gene, 34804 |
| chr8 | 94904578 | 94905961 | 1384 | * | 45,162507 | 4,970198203 | 0,6635252 | 7,4905947 | 6,856E-14 | 5,80145E-09 | 2,3776742 | Promoter (<=1kb) | 8 | 94905710 | 94906955 | 1246 | 1 | 17748 | ENSMUST00000034215.8 | 0 | ENSMUSG00000031765 | Mt1 | metallothionein 1 |
| chr19 | 46549619 | 46550500 | 882 | * | 6,597402 | 4,971903896 | 1,830171 | 2,716634 | 0,006595 | 1 | 0 | Promoter (2-3kb) | 19 | 46530534 | 46546831 | 16298 | 2 | 56350 | ENSMUST00000238076.2 | -2788 | ENSMUSG00000025035 | Arl3 | ADP-ribosylation factor-like 3 |
| chr11 | 1,18E+08 | 1,18E+08 | 598 | * | 6,606669 | 4,972478348 | 1,822446 | 2,728465 | 0,006363 | 1 | 0 | Intron (ENSMUST00000 | 11 | 1,18E+08 | 1,18E+08 | 42008 | 2 | 19157 | ENSMUST00000141243.8 | -9649 | ENSMUSG00000017132 | Cyth1 | cytohesin 1 |
| chr7 | 24678116 | 24679094 | 979 | * | 7,784549 | 4,972549489 | 1,89587 | 2,622833 | 0,00872 | 1 | 0 | 3' UTR | 7 | 24669177 | 24672179 | 3003 | 2 | 14470 | ENSMUST00000076961.9 | -5937 | ENSMUSG00000003380 | Rabac1 | Rab acceptor 1 (prenylated) |
| chr3 | 97068820 | 97069732 | 913 | * | 6,417273 | 4,972974923 | 1,88543 | 2,637582 | 0,00835 | 1 | 0 | Promoter (2-3kb) | 3 | 97066419 | 97075626 | 9208 | 1 | 66659 | ENSMUST00000126438.2 | 2401 | ENSMUSG00000028093 | Acp6 | acid phosphatase 6, lysophosphatidic |
| chr7 | 1E+08 | 1E+08 | 864 | * | 5,784495 | 4,973144176 | 1,880956 | 2,643946 | 0,008195 | 1 | 0 | Promoter (2-3kb) | 7 | 1E+08 | 1E+08 | 5016 | 1 | 19346 | ENSMUST00000123960.2 | 2323 | ENSMUSG00000030704 | Rab6a | RAB6A, member RAS oncogene family |
| chr7 | 25035578 | 25036666 | 1089 | * | 9,483635 | 4,97335494 | 1,810434 | 2,747051 | 0,006013 | 0,807330913 | 1,360622 | Intron (ENSMUST00000 | 7 | 25045406 | 25065339 | 19934 | 1 | 269878 | ENSMUST00000153077.2 | -8740 | ENSMUSG00000045039 | Megf8 | multiple EGF-like-domains 8 |
| chr4 | 1,35E+08 | 1,35E+08 | 900 | * | 6,787363 | 4,974372792 | 1,899582 | 2,618667 | 0,008827 | 1 | 0 | Distal Intergenic | 4 | 1,35E+08 | 1,35E+08 | 32838 | 2 | 51796 | ENSMUST00000105861.8 | -7160 | ENSMUSG00000028809 | Srrm1 | serine/arginine repetitive matrix 1 |
| chr13 | 67630052 | 67631440 | 1389 | * | 9,007111 | 4,977154551 | 1,827237 | 2,723869 | 0,006452 | 0,821398149 | 1,360622 | Promoter (<=1kb) | 13 | 67622292 | 67632331 | 10040 | 2 | 408067 | ENSMUST00000225027.2 | 891 | ENSMUSG00000059839 | Zfp874b | zinc finger protein 874b |
| chr4 | 1,35E+08 | 1,35E+08 | 763 | * | 8,390089 | 4,977340346 | 1,77167 | 2,809407 | 0,004963 | 0,774763123 | 1,360622 | Intron (ENSMUST00000 | 4 | 1,35E+08 | 1,35E+08 | 907 | 1 | 12399 | ENSMUST00000127109.2 | 18920 | ENSMUSG00000070691 | Runx3 | runt related transcription factor 3 |
| chr7 | 51243574 | 51244773 | 1200 | * | 8,269945 | 4,977980256 | 1,880962 | 2,646508 | 0,008133 | 0,773936378 | 2,237403 | 3' UTR | 7 | 51271754 | 51320867 | 49114 | 1 | 140919 | ENSMUST00000032710.7 | -26981 | ENSMUSG00000030500 | Slc17a6 | solute carrier family 17 (sodium-dependent inorganic phosphate cotransporter), member 6 |
| chr19 | 36044833 | 36045944 | 1112 | * | 7,408337 | 4,978386452 | 1,834296 | 2,714058 | 0,006646 | 1 | 0 | Distal Intergenic | 19 | 35937220 | 36034907 | 97688 | 2 | 15566 | ENSMUST00000170360.2 | -9926 | ENSMUSG00000024798 | Htr7 | 5-hydroxytryptamine (serotonin) receptor 7 |
| chr4 | 1,33E+08 | 1,33E+08 | 790 | * | 6,51582 | 4,980571807 | 1,637697 | 3,041205 | 0,002356 | 1 | 0 | Promoter (2-3kb) | 4 | 1,33E+08 | 1,33E+08 | 4008 | 2 | 100336 | ENSMUST00000156777.2 | 2400 | ENSMUSG00000028882 | Ppp1r8 | protein phosphatase 1, regulatory subunit 8 |
| chr11 | 70676828 | 70678075 | 1248 | * | 11,56811 | 4,98129545 | 1,656408 | 3,007287 | 0,002636 | 0,730469267 | 0,918255 | Distal Intergenic | 11 | 70655035 | 70663754 | 8720 | 1 | 193043 | ENSMUST00000060444.6 | 21793 | ENSMUSG00000043602 | Zfp3 | zinc finger protein 3 |
| chr9 | 56788121 | 56788657 | 537 | * | 5,444698 | 4,982591983 | 1,923429 | 2,590473 | 0,009584 | 1 | 0 | Intron (ENSMUST00000 | 9 | 56795260 | 56803491 | 8232 | 1 | 121021 | ENSMUST00000215666.2 | -6603 | ENSMUSG00000032911 | Cspg4 | chondroitin sulfate proteoglycan 4 |
| chr18 | 57330846 | 57331446 | 601 | * | 5,087541 | 4,983160992 | 1,933787 | 2,576892 | 0,009969 | 1 | 0 | Intron (ENSMUST00000 | 18 | 57275858 | 57425800 | 149943 | 1 | 70417 | ENSMUST00000139892.2 | 54988 | ENSMUSG00000024593 | Megf10 | multiple EGF-like-domains 10 |
| chrX | 63517181 | 63518185 | 1005 | * | 7,128089 | 4,983291365 | 1,860107 | 2,679035 | 0,007383 | 0,844443332 | 1,360622 | Distal Intergenic | 20 | 63540190 | 63540263 | 74 | 1 | 1,15E+08 | ENSMUST00000122510.3 | -22005 | ENSMUSG00000084459 | Gm24598 | predicted gene, 24598 |
| chr5 | 99140132 | 99140954 | 823 | * | 5,93101 | 4,98489629 | 1,903636 | 2,618619 | 0,008829 | 1 | 0 | Exon (ENSMUST00000 | 5 | 99095252 | 99114615 | 19364 | 2 | 19092 | ENSMUST00000160765.2 | -25517 | ENSMUSG00000029334 | Prkg2 | protein kinase, cGMP-dependent, type II |
| chr8 | 4304917 | 4305501 | 585 | * | 7,23443 | 4,986136296 | 1,882621 | 2,648508 | 0,008085 | 1 | 0 | Promoter (<=1kb) | 8 | 4305071 | 4305825 | 755 | 1 | 102209 | ENSMUST00000208110.2 | 0 | ENSMUSG00000011837 | Snapc2 | small nuclear RNA activating complex, polypeptide 2 |
| chr9 | 8156521 | 8156982 | 462 | * | 5,999 | 4,987918136 | 1,853915 | 2,690478 | 0,007135 | 1 | 0 | Intron (ENSMUST00000 | 9 | 8076462 | 8134295 | 57834 | 2 | 234915 | ENSMUST00000238782.2 | -22226 | ENSMUSG00000040729 | Cep126 | centrosomal protein 126 |
| chr19 | 31204312 | 31205203 | 892 | * | 7,830278 | 4,994110878 | 1,784864 | 2,798034 | 0,005141 | 1 | 0 | Intron (ENSMUST00000 | 19 | 31060237 | 31064469 | 4233 | 1 | 83410 | ENSMUST00000066039.8 | 144075 | ENSMUSG00000053536 | Cstf2t | cleavage stimulation factor, 3' pre-RNA subunit 2, tau |
| chr13 | 1,06E+08 | 1,06E+08 | 1091 | * | 8,244177 | 4,994436171 | 1,773419 | 2,816275 | 0,004858 | 0,683966322 | 2,237403 | Distal Intergenic | 13 | 1,06E+08 | 1,06E+08 | 4484 | 1 | 15550 | ENSMUST00000022235.6 | -67545 | ENSMUSG00000021721 | Htr1a | 5-hydroxytryptamine (serotonin) receptor 1A |
| chr5 | 28947762 | 28948875 | 1114 | * | 10,07651 | 4,995141627 | 1,747727 | 2,85808 | 0,004262 | 0,734161328 | 1,449706 | Intron (ENSMUST00000 | 5 | 28661813 | 28672254 | 10442 | 2 | 20423 | ENSMUST00000002708.5 | -275508 | ENSMUSG00000002633 | Shh | sonic hedgehog |
| chr16 | 84633833 | 84634646 | 814 | * | 6,328413 | 4,997979326 | 1,894723 | 2,637842 | 0,008344 | 1 | 0 | Promoter (1-2kb) | 16 | 84624759 | 84632513 | 7755 | 2 | 11957 | ENSMUST00000023608.14 | -1320 | ENSMUSG00000022890 | Atp5j | ATP synthase, H+ transporting, mitochondrial F0 complex, subunit F |

| | | | | | | | | | | | | | | | | | | | | | | | |
|---|---|---|---|---|---|---|---|---|---|---|---|---|---|---|---|---|---|---|---|---|---|---|---|
| chr8 | 94793719 | 94794565 | 847 | * | 8,479201 | 5,000367753 | 1,865033 | 2,681115 | 0,007338 | 0,842458878 | 1,360622 | Promoter (2-3kb) | 8 | 94794772 | 94796841 | 2070 | 2 | 67378 | ENSMUST00000170208.2 | 2276 | ENSMUSG00000031755 | Bbs2 | Bardet-Biedl syndrome 2 (human) |
| chr2 | 1,3E+08 | 1,3E+08 | 643 | * | 5,040566 | 5,001008513 | 1,937839 | 2,580714 | 0,00986 | 1 | 0 | Distal Intergenic | 2 | 1,3E+08 | 1,3E+08 | 41817 | 1 | 241636 | ENSMUST00000140947.8 | -3522 | ENSMUSG00000027403 | Tgm6 | transglutaminase 6 |
| chr9 | 1,2E+08 | 1,2E+08 | 709 | * | 6,184307 | 5,003004851 | 1,899127 | 2,634371 | 0,008429 | 1 | 0 | Distal Intergenic | 9 | 1,2E+08 | 1,2E+08 | 727 | 1 | 76100 | ENSMUST00000064646.5 | 8028 | ENSMUSG00000052658 | 5830454E08Rik | RIKEN cDNA 5830454E08 gene |
| chr3 | 1,11E+08 | 1,11E+08 | 771 | * | 6,41771 | 5,004132789 | 1,550372 | 3,227698 | 0,001248 | 1 | 0 | Distal Intergenic | 3 | 1,1E+08 | 1,1E+08 | 105 | 2 | 1,15E+08 | ENSMUST00000104065.3 | -572316 | ENSMUSG00000119642 | Gm26076 | predicted gene, 26076 |
| chr6 | 85915436 | 85916096 | 661 | * | 7,148547 | 5,004543298 | 1,817414 | 2,753661 | 0,005893 | 0,802490409 | 1,360622 | Exon (ENSMUST000002 | 6 | 85926692 | 85927865 | 1174 | 2 | 72102 | ENSMUST00000137527.8 | 11769 | ENSMUSG00000030002 | Dusp11 | dual specificity phosphatase 11 (RNA/RNP complex 1-interacting) |
| chr7 | 4504734 | 4505246 | 513 | * | 6,202164 | 5,009131887 | 1,862069 | 2,69009 | 0,007143 | 1 | 0 | Promoter (<=1kb) | 7 | 4484519 | 4504679 | 20161 | 2 | 232807 | ENSMUST00000013886.9 | -55 | ENSMUSG00000019254 | Ppp1r12c | protein phosphatase 1, regulatory subunit 12C |
| chr10 | 18873685 | 18875012 | 1328 | * | 8,643207 | 5,009887619 | 1,442831 | 3,472263 | 0,000516 | 0,365335255 | 1,449706 | Distal Intergenic | 10 | 18883015 | 18884593 | 1579 | 2 | 21929 | ENSMUST00000154749.2 | 9581 | ENSMUSG00000019850 | Tnfaip3 | tumor necrosis factor, alpha-induced protein 3 |
| chr4 | 14005325 | 14006092 | 768 | * | 7,047635 | 5,009976586 | 1,776121 | 2,820741 | 0,004791 | 0,767061106 | 1,360622 | Distal Intergenic | 4 | 13919954 | 13995193 | 75240 | 2 | 329797 | ENSMUST00000130582.2 | -10132 | ENSMUSG00000086927 | Gm11823 | predicted gene 11823 |
| chr2 | 96821249 | 96822137 | 889 | * | 6,176165 | 5,010385612 | 1,824781 | 2,745746 | 0,006037 | 1 | 0 | Intron (ENSMUST00000 | 2 | 97068163 | 97068291 | 129 | 1 | 1,15E+08 | ENSMUST00000158008.3 | -246026 | ENSMUSG00000088633 | Gm22313 | predicted gene, 22313 |
| chr4 | 1,36E+08 | 1,36E+08 | 944 | * | 8,565566 | 5,012077006 | 1,731754 | 2,894219 | 0,003801 | 0,715249603 | 1,449706 | Promoter (<=1kb) | 4 | 1,36E+08 | 1,36E+08 | 534 | 2 | 1,15E+08 | ENSMUST00000153223.8 | 548 | ENSMUSG00000059291 | LOC115489947 | uncharacterized LOC115489947 |
| chrX | 57423551 | 57424343 | 793 | * | 5,806449 | 5,012080527 | 1,945086 | 2,576791 | 0,009972 | 1 | 0 | Distal Intergenic | 20 | 57403312 | 57403418 | 107 | 2 | 1,15E+08 | ENSMUST00000240146.1 | -20133 | ENSMUSG00000119633 | Gm25851 | predicted gene, 25851 |
| chr7 | 81219115 | 81220337 | 1223 | * | 9,361785 | 5,012232954 | 1,739951 | 2,880676 | 0,003968 | 0,73231239 | 1,360622 | Promoter (<=1kb) | 7 | 81221014 | 81241596 | 20583 | 1 | 434204 | ENSMUST00000207123.2 | -677 | ENSMUSG00000045795 | Whamm | WAS protein homolog associated with actin, golgi membranes and microtubules |
| chr15 | 55011789 | 55012641 | 853 | * | 8,784186 | 5,012969205 | 1,78089 | 2,814867 | 0,00488 | 0,683988405 | 2,237403 | Intron (ENSMUST00000 | 15 | 54996849 | 55072281 | 75433 | 1 | 97998 | ENSMUST00000226687.2 | 14940 | ENSMUSG00000022419 | Deptor | DEP domain containing MTOR-interacting protein |
| chr18 | 67630396 | 67631656 | 1261 | * | 6,905866 | 5,01494063 | 1,893795 | 2,648091 | 0,008095 | 0,858257847 | 1,360622 | Promoter (<=1kb) | 18 | 67627197 | 67630618 | 3422 | 2 | 68166 | ENSMUST00000224659.2 | 0 | ENSMUSG00000024533 | Spire1 | spire type actin nucleation factor 1 |
| chr8 | 1,21E+08 | 1,21E+08 | 507 | * | 6,227054 | 5,016553149 | 1,940498 | 2,585188 | 0,009733 | 1 | 0 | Intron (ENSMUST00000 | 8 | 1,21E+08 | 1,21E+08 | 92944 | 1 | 382034 | ENSMUST00000034279.16 | 7288 | ENSMUSG00000031822 | Gse1 | genetic suppressor element 1, coiled-coil protein |
| chr1 | 1,33E+08 | 1,33E+08 | 794 | * | 9,111374 | 5,018481054 | 1,796798 | 2,793013 | 0,005222 | 0,775744126 | 1,425929 | Exon (ENSMUST00000 | 1 | 1,33E+08 | 1,33E+08 | 3526 | 2 | 269116 | ENSMUST00000186539.2 | 8530 | ENSMUSG00000026442 | Nfasc | neurofascin |
| chr1 | 65220810 | 65221519 | 710 | * | 7,651602 | 5,018542215 | 1,826048 | 2,748308 | 0,00599 | 1 | 0 | Promoter (2-3kb) | 1 | 65197775 | 65218340 | 20566 | 2 | 15926 | ENSMUST00000097709.11 | -2470 | ENSMUSG00000025950 | Idh1 | isocitrate dehydrogenase 1 (NADP+), soluble |
| chr4 | 1,08E+08 | 1,08E+08 | 623 | * | 7,443609 | 5,018555454 | 1,834426 | 2,735763 | 0,006224 | 1 | 0 | Intron (ENSMUST00000 | 4 | 1,08E+08 | 1,08E+08 | 573 | 2 | 414872 | ENSMUST00000130508.2 | -13436 | ENSMUSG00000034636 | Zyg11b | zyg-II family member B, cell cycle regulator |
| chr19 | 20539567 | 20540908 | 1342 | * | 8,552473 | 5,0186055 | 1,739892 | 2,884435 | 0,003921 | 1 | 0 | Intron (ENSMUST00000 | 19 | 20511891 | 20529694 | 17804 | 2 | 320155 | ENSMUST00000225910.2 | -9873 | ENSMUSG00000114469 | C730002L08Rik | RIKEN cDNA C730002L08 gene |
| chr13 | 62298166 | 62299031 | 866 | * | 6,614291 | 5,019703414 | 1,840145 | 2,727885 | 0,006374 | 1 | 0 | Exon (ENSMUST000002 | 13 | 62284744 | 62284876 | 133 | 1 | 1,15E+08 | ENSMUST00000083063.4 | 13422 | ENSMUSG00000119163 | Gm24539 | predicted gene, 24539 |
| chr11 | 1,16E+08 | 1,16E+08 | 1404 | * | 7,441696 | 5,02155393 | 1,591494 | 3,155245 | 0,001604 | 1 | 0 | Distal Intergenic | 11 | 1,16E+08 | 1,16E+08 | 483 | 2 | 1,08E+08 | ENSMUST00000118352.3 | 6555 | ENSMUSG00000083394 | Gm11703 | predicted gene 11703 |
| chr11 | 69769172 | 69769869 | 698 | * | 8,492362 | 5,021929957 | 1,541741 | 3,25731 | 0,001125 | 0,426043749 | 2,237403 | Promoter (<=1kb) | 11 | 69766883 | 69768879 | 1997 | 2 | 1,01E+08 | ENSMUST00000178597.3 | -293 | ENSMUSG00000094845 | Tmem95 | transmembrane protein 95 |
| chrX | 35414573 | 35415846 | 1274 | * | 6,361763 | 5,024509179 | 1,820483 | 2,759988 | 0,00578 | 1 | 0 | Exon (ENSMUST00000 | 20 | 35375763 | 35434912 | 59150 | 1 | 16164 | ENSMUST00000033418.8 | 38810 | ENSMUSG00000017057 | Il13ra1 | interleukin 13 receptor, alpha 1 |
| chr19 | 5909458 | 5910247 | 790 | * | 5,406034 | 5,025724595 | 1,919181 | 2,618682 | 0,008827 | 1 | 0 | Intron (ENSMUST00000 | 19 | 5912144 | 5913710 | 1567 | 2 | 67457 | ENSMUST00000133729.2 | 3463 | ENSMUSG00000024816 | Frmd8 | FERM domain containing 8 |
| chr1 | 1,13E+08 | 1,13E+08 | 1449 | * | 11,79994 | 5,028244319 | 1,686798 | 2,980941 | 0,002874 | 0,786955198 | 0,738441 | Distal Intergenic | 1 | 1,12E+08 | 1,12E+08 | 136 | 1 | 1,15E+08 | ENSMUST00000158066.3 | 313092 | ENSMUSG00000088691 | Gm22331 | predicted gene, 22331 |
| chr1 | 61704754 | 61705791 | 1038 | * | 8,718778 | 5,028414136 | 1,764373 | 2,849972 | 0,004372 | 0,749550311 | 1,360622 | Intron (ENSMUST00000 | 1 | 61678402 | 62681443 | 1003042 | 1 | 72823 | ENSMUST00000094906.4 | 26352 | ENSMUSG00000052062 | Pard3b | par-3 family cell polarity regulator beta |
| chrX | 91942085 | 91942777 | 693 | * | 6,7225 | 5,028462507 | 1,570483 | 3,201857 | 0,001365 | 1 | 0 | Distal Intergenic | 20 | 91926057 | 91926161 | 105 | 1 | 1,15E+08 | ENSMUST00000083380.3 | 16028 | ENSMUSG00000119580 | Gm25332 | predicted gene, 25332 |
| chr1 | 71453625 | 71454498 | 874 | * | 7,414128 | 5,029035476 | 1,655383 | 3,037988 | 0,002382 | 0,650529916 | 1,360622 | Promoter (<=1kb) | 1 | 71281435 | 71454069 | 172635 | 2 | 74591 | ENSMUST00000087268.7 | 0 | ENSMUSG00000050296 | Abca12 | ATP-binding cassette, sub-family A (ABC1), member 12 |
| chr13 | 69704728 | 69705372 | 645 | * | 5,280515 | 5,029613171 | 1,947936 | 2,582021 | 0,009822 | 1 | 0 | Intron (ENSMUST00000 | 13 | 69691165 | 69692142 | 978 | 2 | 210373 | ENSMUST00000070942.2 | -12586 | | A530095I07Rik | RIKEN cDNA A530095I07 gene |
| chr3 | 44901102 | 44902362 | 1261 | * | 10,46861 | 5,031068041 | 1,45352 | 3,461298 | 0,000538 | 0,399754181 | 1,2763 | Distal Intergenic | 3 | 45281480 | 45332640 | 51161 | 2 | 72511 | ENSMUST00000191843.2 | 430278 | ENSMUSG00000097040 | 2610316D01Rik | RIKEN cDNA 2610316D01 gene |
| chr17 | 73723210 | 73724119 | 910 | * | 7,969278 | 5,031388964 | 1,843758 | 2,728877 | 0,006355 | 1 | 0 | Distal Intergenic | 17 | 73651917 | 73707861 | 55945 | 2 | 381122 | ENSMUST00000233432.2 | -15349 | ENSMUSG00000043705 | Capn13 | calpain 13 |
| chr11 | 78646014 | 78647375 | 1362 | * | 11,24802 | 5,034819391 | 1,6816 | 2,994064 | 0,002753 | 0,738140609 | 0,918255 | Distal Intergenic | 11 | 78641332 | 78642555 | 1224 | 2 | 69109 | ENSMUST00000059468.6 | -3459 | ENSMUSG00000049489 | Ccnq | cyclin Q |
| chr10 | 1,11E+08 | 1,11E+08 | 788 | * | 10,27155 | 5,036418287 | 1,7621 | 2,858191 | 0,004261 | 0,773870972 | 1,173325 | Distal Intergenic | 10 | 1,11E+08 | 1,11E+08 | 871 | 1 | 432502 | ENSMUST00000171120.3 | -22089 | ENSMUSG00000091086 | Rpl6l | ribosomal protein L6-like |
| chr8 | 71599153 | 71599405 | 253 | * | 5,385345 | 5,036964978 | 1,919762 | 2,623745 | 0,008697 | 1 | 0 | Distal Intergenic | 8 | 71459372 | 71538402 | 79031 | 2 | 69419 | ENSMUST00000212266.2 | -60751 | ENSMUSG00000101854 | 1700026F02Rik | RIKEN cDNA 1700026F02 gene |
| chr12 | 70759462 | 70760291 | 830 | * | 5,962878 | 5,037312058 | 1,92099 | 2,622248 | 0,008735 | 1 | 0 | Intron (ENSMUST00000 | 12 | 70722621 | 70728624 | 6004 | 2 | 1,05E+08 | ENSMUST00000222630.2 | -30838 | | Gm32369 | predicted gene, 32369 |
| chr11 | 53945150 | 53945984 | 835 | * | 5,991073 | 5,039089594 | 1,878428 | 2,68261 | 0,007305 | 1 | 0 | 3' UTR | 11 | 53954549 | 53955394 | 846 | 2 | 30794 | ENSMUST00000151948.2 | 9410 | ENSMUSG00000020388 | Pdlim4 | PDZ and LIM domain 4 |
| chr12 | 80924301 | 80925451 | 1151 | * | 6,470446 | 5,041076864 | 1,831396 | 2,752587 | 0,005913 | 1 | 0 | 3' UTR | 12 | 80958251 | 80958357 | 107 | 1 | 1,15E+08 | ENSMUST00001239512.1 | -32800 | ENSMUSG00000119334 | Gm23298 | predicted gene, 23298 |
| chr5 | 1,43E+08 | 1,43E+08 | 663 | * | 6,674447 | 5,041836605 | 1,81317 | 2,780676 | 0,005425 | 1 | 0 | Exon (ENSMUST00000 | 5 | 1,44E+08 | 1,44E+08 | 2940 | 2 | 19353 | ENSMUST00000131513.2 | 9265 | ENSMUSG00000001847 | Rac1 | Rac family small GTPase 1 |
| chr13 | 58792388 | 58793054 | 667 | * | 6,893465 | 5,044790266 | 1,83019 | 2,75643 | 0,005844 | 1 | 0 | Distal Intergenic | 13 | 58729504 | 58758691 | 29188 | 2 | 114304 | ENSMUST00000148396.2 | -33697 | ENSMUSG00000021553 | Slc28a3 | solute carrier family 28 (sodium-coupled nucleoside transporter), member 3 |
| chr4 | 1,38E+08 | 1,38E+08 | 1186 | * | 6,947727 | 5,04485478 | 1,824053 | 2,765739 | 0,005679 | 1 | 0 | Intron (ENSMUST00000 | 4 | 1,38E+08 | 1,38E+08 | 5226 | 2 | 72269 | ENSMUST00000144986.2 | -5018 | ENSMUSG00000028755 | Cda | cytidine deaminase |
| chr10 | 16840452 | 16841794 | 1343 | * | 8,913044 | 5,045902308 | 1,715828 | 2,940797 | 0,003274 | 0,68680243 | 1,449706 | Promoter (1-2kb) | 10 | 16827043 | 16838928 | 11886 | 2 | 1,01E+08 | ENSMUST00000208191.2 | -1524 | ENSMUSG00000101969 | Gm20125 | predicted gene, 20125 |
| chr3 | 39380181 | 39380941 | 761 | * | 5,721561 | 5,052319946 | 1,861851 | 2,7136 | 0,006656 | 1 | 0 | Distal Intergenic | 3 | 39690113 | 39690231 | 119 | 2 | 1,15E+08 | ENSMUST00000083302.3 | 309290 | ENSMUSG00000118708 | n-R5s195 | nuclear encoded rRNA 5S 195 |
| chr16 | 67831447 | 67832492 | 1046 | * | 8,85429 | 5,053165166 | 1,648422 | 3,065457 | 0,002173 | 0,539604518 | 2,237403 | Distal Intergenic | 16 | 66459076 | 67417796 | 958721 | 2 | 239857 | ENSMUST00000120898.8 | -413651 | ENSMUSG00000064115 | Cadm2 | cell adhesion molecule 2 |
| chr17 | 59986718 | 59987270 | 553 | * | 4,829365 | 5,054087326 | 1,939779 | 2,605496 | 0,009174 | 1 | 0 | Distal Intergenic | 17 | 59876554 | 59876660 | 107 | 2 | 1,15E+08 | ENSMUST00000104374.3 | -110058 | ENSMUSG00000077562 | Gm23203 | predicted gene, 23203 |
| chr7 | 46408984 | 46409736 | 753 | * | 6,705161 | 5,054167789 | 1,67244 | 3,022032 | 0,002511 | 1 | 0 | Distal Intergenic | 7 | 46401257 | 46403737 | 2481 | 1 | 20209 | ENSMUST00000075982.4 | 7727 | ENSMUSG00000057465 | Saa2 | serum amyloid A 2 |
| chr8 | 26669214 | 26670342 | 1129 | * | 8,19979 | 5,054218261 | 1,788545 | 2,825883 | 0,004715 | 0,764024179 | 1,360622 | Distal Intergenic | 8 | 26648821 | 26650840 | 2020 | 1 | 73754 | ENSMUST00000209926.2 | 20393 | ENSMUSG00000037214 | Thap1 | THAP domain containing, apoptosis associated protein 1 |
| chr17 | 71109755 | 71110211 | 457 | * | 5,171799 | 5,05461443 | 1,925631 | 2,624913 | 0,008667 | 1 | 0 | Promoter (2-3kb) | 17 | 71072834 | 71112992 | 40159 | 2 | 1,05E+08 | ENSMUST00000233559.2 | 2781 | ENSMUSG00000117231 | Gm41609 | predicted gene, 41609 |
| chr2 | 57662873 | 57663213 | 341 | * | 5,405039 | 5,055136797 | 1,941947 | 2,603128 | 0,009238 | 1 | 0 | Distal Intergenic | 2 | 57543684 | 57543834 | 151 | 2 | 1,15E+08 | ENSMUST00000157339.4 | -119039 | ENSMUSG00000087964 | Gm25388 | predicted gene, 25388 |
| chr4 | 5795606 | 5796971 | 1366 | * | 8,022529 | 5,055219787 | 1,806123 | 2,798934 | 0,005127 | 1 | 0 | Promoter (1-2kb) | 4 | 5798084 | 5798531 | 448 | 1 | 242297 | ENSMUST00000145365.2 | -1113 | ENSMUSG00000049119 | Fam110b | family with sequence similarity 110, member B |
| chrX | 1,49E+08 | 1,49E+08 | 1024 | * | 6,186103 | 5,05556736 | 1,899727 | 2,661207 | 0,007786 | 1 | 0 | Promoter (<=1kb) | 20 | 1,49E+08 | 1,49E+08 | 53958 | 1 | 18639 | ENSMUST00000080884.11 | 149 | ENSMUSG00000025271 | Pfkfb1 | 6-phosphofructo-2-kinase/fructose-2,6-biphosphatase 1 |
| chr3 | 1,08E+08 | 1,08E+08 | 811 | * | 7,01398 | 5,055676239 | 1,896735 | 2,665462 | 0,007688 | 0,852104582 | 1,360622 | Promoter (2-3kb) | 3 | 1,08E+08 | 1,08E+08 | 13319 | 2 | 1,19E+08 | ENSMUST00000198171.2 | -2785 | | LOC118568636 | uncharacterized LOC118568636 |
| chr10 | 45750596 | 45751954 | 1359 | * | 11,59368 | 5,055941701 | 1,3361 | 3,784105 | 0,000154 | 0,291110136 | 0,738441 | Exon (ENSMUST000002 | 10 | 45713976 | 45728379 | 14404 | 1 | 320363 | ENSMUST00000215085.2 | 36620 | ENSMUSG00000110874 | D030045P18Rik | RIKEN cDNA D030045P18 gene |
| chr5 | 34842078 | 34843011 | 934 | * | 6,346176 | 5,056137115 | 1,946429 | 2,597648 | 0,009386 | 1 | 0 | Intron (ENSMUST00000 | 5 | 34818319 | 34832059 | 13741 | 1 | 14772 | ENSMUST00000148588.2 | 23759 | ENSMUSG00000052783 | Grk4 | G protein-coupled receptor kinase 4 |
| chr5 | 1,22E+08 | 1,22E+08 | 913 | * | 6,985715 | 5,056271725 | 1,801956 | 2,80599 | 0,005016 | 1 | 0 | Distal Intergenic | 5 | 1,22E+08 | 1,22E+08 | 20868 | 2 | 17165 | ENSMUST00000031410.14 | -6749 | ENSMUSG00000029454 | Mapkapk5 | MAP kinase-activated protein kinase 5 |
| chr5 | 38001455 | 38002205 | 751 | * | 6,050219 | 5,056664917 | 1,95547 | 2,585907 | 0,009712 | 1 | 0 | Distal Intergenic | 5 | 37977829 | 37981927 | 4099 | 2 | 17701 | ENSMUST00000063116.10 | -19528 | ENSMUSG00000048450 | Msx1 | msh homeobox 1 |
| chr15 | 88858298 | 88858634 | 337 | * | 6,440925 | 5,058184733 | 1,827119 | 2,768394 | 0,005633 | 1 | 0 | Intron (ENSMUST00000 | 15 | 88840087 | 88863210 | 23124 | 2 | 170790 | ENSMUST00000042594.13 | 4576 | ENSMUSG00000035805 | Mlc1 | megalencephalic leukoencephalopathy with subcortical cysts 1 homolog (human) |
| chr5 | 31858851 | 31859811 | 961 | * | 7,805758 | 5,058935097 | 1,764768 | 2,86663 | 0,004149 | 0,741428016 | 1,360622 | Promoter (<=1kb) | 5 | 31859189 | 32215015 | 355827 | 1 | 107976 | ENSMUST00000200705.2 | 0 | ENSMUSG00000052139 | Babam2 | BRISC and BRCA1 A complex member 2 |
| chr9 | 5429499 | 5430405 | 907 | * | 5,647588 | 5,059165578 | 1,931724 | 2,61899 | 0,008819 | 1 | 0 | Distal Intergenic | 9 | 5345504 | 5358823 | 13320 | 1 | 12364 | ENSMUST00000151332.2 | 83995 | ENSMUSG00000025887 | Casp12 | caspase 12 |
| chr6 | 93506505 | 93507118 | 614 | * | 5,623963 | 5,06032708 | 1,953626 | 2,590223 | 0,009591 | 1 | 0 | Distal Intergenic | 6 | 93486554 | 93486683 | 130 | 1 | 1,15E+08 | ENSMUST00000158005.4 | 19951 | ENSMUSG00000088630 | Gm22312 | predicted gene, 22312 |
| chr15 | 99258251 | 99259030 | 780 | * | 9,725512 | 5,060736709 | 1,404137 | 3,604162 | 0,000313 | 0,295900293 | 1,449706 | Intron (ENSMUST00000 | 15 | 99265910 | 99266016 | 107 | 2 | 1,15E+08 | ENSMUST00000179471.3 | 6986 | ENSMUSG00000119078 | Gm22951 | predicted gene, 22951 |
| chr2 | 31736140 | 31736704 | 565 | * | 9,063018 | 5,061581074 | 1,730604 | 2,924748 | 0,003447 | 0,730469267 | 1,211315 | Promoter (<=1kb) | 2 | 31703302 | 31736017 | 32716 | 2 | 98970 | ENSMUST00000028188.8 | -123 | ENSMUSG00000026841 | Fibcd1 | fibrinogen C domain containing 1 |
| chr4 | 95590402 | 95591148 | 747 | * | 6,251064 | 5,06254546 | 1,88712 | 2,682683 | 0,007303 | 1 | 0 | Intron (ENSMUST00000 | 4 | 95626645 | 95657682 | 31038 | 1 | 75578 | ENSMUST00000153119.2 | -35497 | ENSMUSG00000028573 | Fggy | FGGY carbohydrate kinase domain containing |
| chr4 | 1,18E+08 | 1,18E+08 | 504 | * | 6,536114 | 5,065052953 | 1,856328 | 2,728534 | 0,006362 | 1 | 0 | Intron (ENSMUST00000 | 4 | 1,18E+08 | 1,18E+08 | 27441 | 2 | 19268 | ENSMUST00000150096.8 | -21823 | ENSMUSG00000033295 | Ptprf | protein tyrosine phosphatase, receptor type, F |
| chr18 | 52612708 | 52613431 | 724 | * | 5,100418 | 5,066412968 | 1,913484 | 2,647742 | 0,008103 | 1 | 0 | Exon (ENSMUST000002 | 18 | 52608451 | 52621671 | 13221 | 1 | 67222 | ENSMUST00000237092.2 | 4257 | ENSMUSG00000024528 | Srfbp1 | serum response factor binding protein 1 |
| chr11 | 54594825 | 54595437 | 613 | * | 6,100843 | 5,066469481 | 1,912285 | 2,649432 | 0,008063 | 1 | 0 | Distal Intergenic | 11 | 54582258 | 54587312 | 5055 | 1 | 192786 | ENSMUST00000136494.2 | 12567 | ENSMUSG00000037533 | Rapgef6 | Rap guanine nucleotide exchange factor (GEF) 6 |
| chr17 | 79140207 | 79140959 | 753 | * | 7,771657 | 5,066619188 | 1,522156 | 3,328581 | 0,000873 | 1 | 0 | Promoter (1-2kb) | 17 | 79060327 | 79142753 | 82427 | 2 | 320473 | ENSMUST00000097281.4 | 1794 | ENSMUSG00000039414 | Heatr5b | HEAT repeat containing 5B |
| chr5 | 25056618 | 25057794 | 1177 | * | 7,715999 | 5,0671148 | 1,944463 | 2,60592 | 0,009163 | 1 | 0 | Distal Intergenic | 5 | 25007821 | 25047622 | 39802 | 2 | 19744 | ENSMUST00000030787.9 | -8996 | ENSMUSG00000028945 | Rheb | Ras homolog enriched in brain |
| chr3 | 1,34E+08 | 1,34E+08 | 779 | * | 5,698729 | 5,067639284 | 1,890802 | 2,680153 | 0,007359 | 1 | 0 | Exon (ENSMUST000001 | 3 | 1,34E+08 | 1,34E+08 | 13153 | 1 | 1,03E+08 | ENSMUST00000200648.2 | 11951 | ENSMUSG00000106139 | Gm30648 | predicted gene, 30648 |
| chr2 | 1,33E+08 | 1,33E+08 | 1214 | * | 9,765959 | 5,068013995 | 1,730882 | 2,927995 | 0,003412 | 0,695401133 | 1,449706 | Promoter (1-2kb) | 2 | 1,33E+08 | 1,33E+08 | 64125 | 1 | 73747 | ENSMUST00000061891.11 | -1498 | ENSMUSG00000044991 | Shld1 | shieldin complex subunit 1 |
| chr7 | 1,27E+08 | 1,27E+08 | 994 | * | 8,845827 | 5,068055987 | 1,821791 | 2,781908 | 0,005404 | 0,780222008 | 1,425929 | Promoter (<=1kb) | 7 | 1,27E+08 | 1,27E+08 | 8185 | 2 | 320683 | ENSMUST00000122066.8 | 0 | ENSMUSG00000045639 | Zfp629 | zinc finger protein 629 |
| chr1 | 97979999 | 97980511 | 513 | * | 5,947699 | 5,068285094 | 1,879112 | 2,69717 | 0,006993 | 1 | 0 | Promoter (2-3kb) | 1 | 97977547 | 97977653 | 107 | 1 | 1,15E+08 | ENSMUST00000240225.1 | 2452 | ENSMUSG00000119063 | Gm24465 | predicted gene, 24465 |
| chr4 | 65569440 | 65569884 | 445 | * | 5,547557 | 5,068536872 | 1,87468 | 2,703681 | 0,006858 | 1 | 0 | Intron (ENSMUST00000 | 4 | 65523486 | 65531993 | 8508 | 1 | 69807 | ENSMUST00000156922.2 | 45954 | ENSMUSG00000051675 | Trim32 | tripartite motif-containing 32 |
| chr12 | 80286025 | 80287059 | 1035 | * | 10,2684 | 5,069007789 | 1,482214 | 3,419889 | 0,000626 | 0,431115599 | 1,211315 | Intron (ENSMUST00000 | 12 | 80243723 | 80305558 | 61836 | 2 | 109711 | ENSMUST00000220351.2 | 18499 | ENSMUSG00000015143 | Actn1 | actinin, alpha 1 |
| chr1 | 20964546 | 20965224 | 679 | * | 6,03894 | 5,069469838 | 1,858404 | 2,727862 | 0,006375 | 1 | 0 | Intron (ENSMUST00000 | 1 | 20960830 | 21009874 | 49045 | 1 | 74229 | ENSMUST00000189400.7 | 3716 | ENSMUSG00000025931 | Paqr8 | progestin and adipoQ receptor family member VIII |
| chrX | 67046008 | 67046843 | 836 | * | 5,45235 | 5,07074555 | 1,961914 | 2,584591 | 0,009749 | 1 | 0 | Distal Intergenic | 20 | 67053711 | 67053785 | 75 | 2 | 1E+08 | ENSMUST00000196089.3 | 6942 | ENSMUSG00000106233 | Mir509 | microRNA 509 |
| chr1 | 1,67E+08 | 1,67E+08 | 667 | * | 5,652087 | 5,070957594 | 1,955722 | 2,592882 | 0,009518 | 1 | 0 | Intron (ENSMUST00000 | 1 | 1,67E+08 | 1,67E+08 | 3482 | 1 | 56752 | ENSMUST00000191715.2 | 12096 | ENSMUSG00000026687 | Aldh9a1 | aldehyde dehydrogenase 9, subfamily A1 |
| chr8 | 60494045 | 60494956 | 912 | * | 9,772266 | 5,071985503 | 1,815187 | 2,794194 | 0,005203 | 0,801688106 | 1,211315 | Distal Intergenic | 8 | 60532624 | 60532727 | 104 | 1 | 1,15E+08 | ENSMUST00000083793.3 | -37668 | ENSMUSG00000065727 | Gm26164 | predicted gene, 26164 |
| chr17 | 33511241 | 33512082 | 842 | * | 6,287787 | 5,073840325 | 1,866042 | 2,719039 | 0,006547 | 1 | 0 | Promoter (2-3kb) | 17 | 33508518 | 33526215 | 17698 | 1 | 1E+08 | ENSMUST00000099414.5 | 2723 | ENSMUSG00000096910 | Zfp955b | zinc finger protein 955B |
| chr19 | 8670706 | 8671682 | 977 | * | 7,347698 | 5,074726963 | 1,770668 | 2,865996 | 0,004157 | 0,742014238 | 1,360622 | Distal Intergenic | 19 | 8684951 | 8690349 | 5399 | 2 | 17254 | ENSMUST00000206598.2 | 18667 | ENSMUSG00000010095 | Slc3a2 | solute carrier family 3 (activators of dibasic and neutral amino acid transport), member 2 |
| chr12 | 96109832 | 96110508 | 677 | * | 6,139699 | 5,075589521 | 1,942468 | 2,61296 | 0,008976 | 1 | 0 | Distal Intergenic | 12 | 96013541 | 96013894 | 354 | 2 | 69423 | ENSMUST00000073560.5 | -95938 | | 1700019M22Rik | RIKEN cDNA 1700019M22 gene |
| chrX | 47893850 | 47895108 | 1259 | * | 6,202303 | 5,077854566 | 1,95348 | 2,599388 | 0,009339 | 1 | 0 | Distal Intergenic | 20 | 47930343 | 47966590 | 36248 | 2 | 209005 | ENSMUST00000114928.2 | 71482 | ENSMUSG00000079606 | Fsip2l | fibrous sheath-interacting protein 2-like |
| chr11 | 1,17E+08 | 1,17E+08 | 1203 | * | 9,759073 | 5,078421736 | 1,715739 | 2,959903 | 0,003077 | 0,712074475 | 1,211315 | Promoter (<=1kb) | 11 | 1,17E+08 | 1,17E+08 | 1002 | 2 | 20446 | ENSMUST00000129236.2 | -405 | ENSMUSG00000057286 | St6galnac2 | ST6 (alpha-N-acetyl-neuraminyl-2,3-beta-galactosyl-1,3)-N-acetylgalactosaminide alpha-2,6-sialyltransferase 2 |
| chr16 | 95738789 | 95739327 | 539 | * | 5,569619 | 5,079122341 | 1,891391 | 2,68539 | 0,007245 | 1 | 0 | Distal Intergenic | 16 | 95643086 | 95730277 | 87192 | 2 | 69776 | ENSMUST00000233375.2 | -8512 | ENSMUSG00000117232 | 1600002D24Rik | RIKEN cDNA 1600002D24 gene |
| chr1 | 1,83E+08 | 1,83E+08 | 739 | * | 6,45818 | 5,079337124 | 1,956305 | 2,596393 | 0,009421 | 1 | 0 | Distal Intergenic | 1 | 1,83E+08 | 1,83E+08 | 15197 | 1 | 53791 | ENSMUST00000195614.2 | 45670 | ENSMUSG00000079164 | Tlr5 | toll-like receptor 5 |
| chr1 | 53815898 | 53816867 | 970 | * | 6,708907 | 5,080150576 | 1,944823 | 2,612141 | 0,008998 | 1 | 0 | Intron (ENSMUST00000 | 1 | 53800308 | 53824316 | 24009 | 2 | 98267 | ENSMUST00000185920.2 | 7449 | ENSMUSG00000026094 | Stk17b | serine/threonine kinase 17b (apoptosis-inducing) |
| chr9 | 42175735 | 42176648 | 914 | * | 7,971079 | 5,080714703 | 1,936824 | 2,623219 | 0,00871 | 1 | 0 | Promoter (<=1kb) | 9 | 42162891 | 42175552 | 12662 | 2 | 235293 | ENSMUST00000052725.15 | -183 | ENSMUSG00000032018 | Sc5d | sterol-C5-desaturase |
| chr5 | 1,35E+08 | 1,35E+08 | 1635 | * | 11,80087 | 5,082263313 | 1,664887 | 3,052617 | 0,002269 | 0,668772562 | 1,173325 | Promoter (<=1kb) | 5 | 1,35E+08 | 1,35E+08 | 2293 | 2 | 14371 | ENSMUST00000062572.3 | -48 | ENSMUSG00000049551 | Fzd9 | frizzled class receptor 9 |
| chr3 | 58324406 | 58325070 | 665 | * | 6,587673 | 5,084135017 | 1,953807 | 2,602169 | 0,009264 | 1 | 0 | Promoter (<=1kb) | 3 | 58324905 | 58367730 | 42826 | 1 | 72033 | ENSMUST00000196049.2 | 0 | ENSMUSG00000027806 | Tsc22d2 | TSC22 domain family, member 2 |
| chr7 | 1,44E+08 | 1,44E+08 | 1335 | * | 11,88296 | 5,084644508 | 1,677944 | 3,030282 | 0,002443 | 0,680239471 | 1,173325 | Promoter (1-2kb) | 7 | 1,44E+08 | 1,44E+08 | 12579 | 2 | 233977 | ENSMUST00000182982.8 | -1839 | ENSMUSG00000037519 | Ppfia1 | protein tyrosine phosphatase, receptor type, f polypeptide (PTPRF), interacting protein (liprin), alpha 1 |
| chr11 | 52293244 | 52293975 | 732 | * | 5,713715 | 5,085825285 | 1,866375 | 2,724975 | 0,006431 | 1 | 0 | Exon (ENSMUST000001 | 11 | 52287473 | 52294101 | 6629 | 1 | 213673 | ENSMUST00000136303.2 | 5771 | ENSMUSG00000036275 | 9530068E07Rik | RIKEN cDNA 9530068E07 gene |
| chr12 | 66445943 | 66446491 | 549 | * | 6,581342 | 5,086656995 | 1,815739 | 2,801425 | 0,005088 | 1 | 0 | Distal Intergenic | 12 | 66330153 | 66331175 | 1023 | 2 | 238217 | ENSMUST00000081908.8 | -114768 | ENSMUSG00000060499 | Rpl10l | ribosomal protein L10-like |
| chr10 | 30538376 | 30539352 | 977 | * | 6,122616 | 5,088601018 | 1,876458 | 2,711812 | 0,006692 | 1 | 0 | Exon (ENSMUST000002 | 10 | 30522308 | 30531863 | 9556 | 2 | 211329 | ENSMUST00000216172.2 | -6513 | ENSMUSG00000039697 | Ncoa7 | nuclear receptor coactivator 7 |
| chr2 | 1,64E+08 | 1,64E+08 | 1469 | * | 7,748618 | 5,091801819 | 1,753287 | 2,904146 | 0,003683 | 1 | 0 | Promoter (<=1kb) | 2 | 1,64E+08 | 1,64E+08 | 25089 | 1 | 381404 | ENSMUST00000067715.5 | 0 | ENSMUSG00000054582 | Pabpc1l | poly(A) binding protein, cytoplasmic 1-like |
| chr8 | 41830070 | 41831031 | 962 | * | 8,569376 | 5,091987179 | 1,810879 | 2,811887 | 0,004925 | 0,772898261 | 1,360622 | Promoter (2-3kb) | 8 | 41807064 | 41827810 | 20747 | 2 | 11886 | ENSMUST00000143057.2 | -2260 | ENSMUSG00000031591 | Asah1 | N-acylsphingosine amidohydrolase 1 |
| chr17 | 25637134 | 25637780 | 647 | * | 8,026971 | 5,092853229 | 1,442215 | 3,531272 | 0,000414 | 1 | 0 | Intron (ENSMUST00000 | 17 | 25616599 | 25652252 | 35654 | 2 | 58226 | ENSMUST00000159623.2 | 14472 | ENSMUSG00000024112 | Cacna1h | calcium channel, voltage-dependent, T type, alpha 1H subunit |
| chr11 | 95110552 | 95111201 | 650 | * | 5,140774 | 5,094262222 | 1,91624 | 2,658467 | 0,00785 | 1 | 0 | Distal Intergenic | 11 | 95114116 | 95116458 | 2343 | 2 | 1,03E+08 | ENSMUST00000150818.2 | 5257 | | Gm11515 | predicted gene 11515 |
| chr16 | 33996922 | 33997954 | 1033 | * | 6,939423 | 5,094493428 | 1,939213 | 2,627093 | 0,008612 | 1 | 0 | Intron (ENSMUST00000 | 16 | 33833502 | 33983649 | 150148 | 2 | 545156 | ENSMUST00000114963.8 | -13273 | ENSMUSG00000061751 | Kalrn | kalirin, RhoGEF kinase |
| chr1 | 89377153 | 89378287 | 1135 | * | 9,160019 | 5,094495802 | 1,706414 | 2,985497 | 0,002831 | 0,680148888 | 1,360622 | Distal Intergenic | 1 | 89382528 | 89825339 | 442812 | 1 | 347722 | ENSMUST00000027521.15 | -4241 | ENSMUSG00000055013 | Agap1 | ArfGAP with GTPase domain, ankyrin repeat and PH domain 1 |
| chr16 | 34953888 | 34954548 | 661 | * | 7,098706 | 5,094631875 | 1,940759 | 2,625072 | 0,008663 | 1 | 0 | Distal Intergenic | 16 | 34975247 | 35126108 | 150862 | 1 | 224129 | ENSMUST00000114913.3 | -20699 | ENSMUSG00000022840 | Adcy5 | adenylate cyclase 5 |
| chr5 | 1,21E+08 | 1,21E+08 | 508 | * | 5,552221 | 5,094831871 | 1,907316 | 2,671204 | 0,007558 | 1 | 0 | Intron (ENSMUST00000 | 5 | 1,21E+08 | 1,21E+08 | 11526 | 1 | 23961 | ENSMUST00000183291.2 | 3629 | ENSMUSG00000029605 | Oas1b | 2'-5' oligoadenylate synthetase 1B |
| chr3 | 50557735 | 50558270 | 536 | * | 5,947271 | 5,095410454 | 1,898848 | 2,683422 | 0,007287 | 1 | 0 | Intron (ENSMUST00000 | 3 | 50470848 | 50561672 | 90825 | 2 | 621304 | ENSMUST00000194629.2 | 3402 | | Gm6209 | predicted gene 6209 |

| | | | | | | | | | | | | | | | | | | | | | | | |
|---|---|---|---|---|---|---|---|---|---|---|---|---|---|---|---|---|---|---|---|---|---|---|---|
| chr9 | 15222934 | 15223941 | 1008 | * | 7,189113 | 5,09563065 | 1,845648 | 2,760889 | 0,005764 | 1 | 0 | Promoter (<=1kb) | 9 | 15223848 | 15223970 | 123 | 1 | 1,02E+08 | ENSMUST00020183305.1 | 0 | ENSMUSG00000105847 | Mir7650 | microRNA 7650 |
| chr9 | 44456586 | 44457498 | 913 | * | 5,357755 | 5,095899956 | 1,92223 | 2,651036 | 0,008025 | 1 | 0 | Intron (ENSMUST0000( | 9 | 44466542 | 44467583 | 1042 | 2 | 1,08E+08 | ENSMUST00000213357.2 | 10085 | | Gm50603 | predicted gene, 50603 |
| chr9 | 53121749 | 53122638 | 890 | * | 6,051151 | 5,096385983 | 1,883997 | 2,705092 | 0,006829 | 1 | 0 | Intron (ENSMUST0000( | 9 | 53009935 | 53159353 | 149419 | 2 | 77591 | ENSMUST00000065630.8 | 36715 | ENSMUSG00000053289 | Ddx10 | DEAD box helicase 10 |
| chrX | 35686387 | 35687115 | 729 | * | 4,972506 | 5,099211382 | 1,941034 | 2,62706 | 0,008613 | 1 | 0 | Distal Intergenic | 20 | 35687254 | 35690333 | 3080 | 2 | 1,01E+08 | ENSMUST00000140106.8 | 3218 | ENSMUSG00000091556 | Gm14569 | predicted gene 14569 |
| chr19 | 28657498 | 28658949 | 1452 | * | 12,35351 | 5,099641707 | 1,687241 | 3,022475 | 0,002507 | 0,722798324 | 0,918255 | Promoter (<=1kb) | 19 | 28326973 | 28657477 | 330505 | 2 | 226075 | ENSMUST00000112612.9 | -21 | ENSMUSG00000052942 | Glis3 | GLIS family zinc finger 3 |
| chr15 | 76066115 | 76066967 | 853 | * | 7,288834 | 5,101077903 | 1,910287 | 2,67032 | 0,007578 | 1 | 0 | Intron (ENSMUST0000( | 15 | 76059580 | 76072243 | 12664 | 2 | 18810 | ENSMUST00000165210.8 | 5276 | ENSMUSG00000022565 | Plec | plectin |
| chr19 | 36872311 | 36873176 | 866 | * | 6,540574 | 5,10295347 | 1,856327 | 2,748952 | 0,005979 | 1 | 0 | Distal Intergenic | 19 | 36857973 | 36864773 | 6801 | 1 | 74493 | ENSMUST00000168654.2 | 14338 | ENSMUSG00000024811 | Tnks2 | tankyrase, TRF1-interacting ankyrin-related ADP-ribose polymerase 2 |
| chr5 | 25569937 | 25570824 | 888 | * | 6,322259 | 5,103165515 | 1,852027 | 2,755448 | 0,005861 | 1 | 0 | Intron (ENSMUST0000( | 5 | 25477424 | 25513548 | 36125 | 2 | 231051 | ENSMUST00000172556.8 | -56389 | ENSMUSG00000038056 | Kmt2c | lysine (K)-specific methyltransferase 2C |
| chr13 | 54593441 | 54594231 | 791 | * | 6,336235 | 5,103686783 | 1,840002 | 2,77374 | 0,005542 | 1 | 0 | Distal Intergenic | 13 | 54611495 | 54616601 | 5107 | 2 | 73666 | ENSMUST00000151158.2 | 22370 | ENSMUSG00000025872 | Thoc3 | THO complex 3 |
| chr7 | 66131772 | 66132349 | 578 | * | 6,181688 | 5,105278298 | 1,865545 | 2,736615 | 0,006207 | 1 | 0 | Distal Intergenic | 7 | 66040638 | 66077265 | 36628 | 2 | 56847 | ENSMUST00000015278.15 | -54507 | ENSMUSG00000015134 | Aldh1a3 | aldehyde dehydrogenase family 1, subfamily A3 |
| chr2 | 85266289 | 85266761 | 473 | * | 6,315897 | 5,106791408 | 1,899246 | 2,688853 | 0,00717 | 1 | 0 | Promoter (<=1kb) | 2 | 85259783 | 85266670 | 6888 | 2 | 258425 | ENSMUST00000214679.2 | 0 | ENSMUSG00000075219 | Olfr994 | olfactory receptor 994 |
| chr9 | 1,03E+08 | 1,03E+08 | 2188 | * | 16,54191 | 5,106902462 | 1,272875 | 4,012101 | 6,02E-05 | 0,125710024 | 1,2763 | Distal Intergenic | 9 | 1,03E+08 | 1,03E+08 | 7564 | 1 | 69010 | ENSMUST00000186693.2 | 29706 | ENSMUSG00000035048 | Anapc13 | anaphase promoting complex subunit 13 |
| chr2 | 1,55E+08 | 1,55E+08 | 1260 | * | 8,905671 | 5,108378376 | 1,737125 | 2,940709 | 0,003275 | 0,699278451 | 1,360622 | 3' UTR | 2 | 1,55E+08 | 1,55E+08 | 21365 | 2 | 67204 | ENSMUST00000166171.8 | 20945 | ENSMUSG00000074656 | Eif2s2 | eukaryotic translation initiation factor 2, subunit 2 (beta) |
| chr9 | 1,09E+08 | 1,09E+08 | 891 | * | 6,485362 | 5,112556774 | 1,887004 | 2,709352 | 0,006741 | 1 | 0 | Intron (ENSMUST0000( | 9 | 1,08E+08 | 1,09E+08 | 34121 | 2 | 23807 | ENSMUST00000193552.6 | 3478 | ENSMUSG00000064145 | Arih2 | ariadne RBR E3 ubiquitin protein ligase 2 |
| chr5 | 1,36E+08 | 1,36E+08 | 528 | * | 5,480787 | 5,115668583 | 1,985035 | 2,577118 | 0,009963 | 1 | 0 | Intron (ENSMUST0000( | 5 | 1,36E+08 | 1,36E+08 | 88 | 2 | 723975 | ENSMUST00000200230.3 | -93247 | ENSMUSG00000106479 | Mir721 | microRNA 721 |
| chr19 | 18554061 | 18554845 | 785 | * | 4,910333 | 5,115691095 | 1,952003 | 2,620739 | 0,008774 | 1 | 0 | Intron (ENSMUST0000( | 19 | 18570965 | 18609082 | 38118 | 2 | 20409 | ENSMUST00000138860.2 | 54237 | ENSMUSG00000024725 | Ostf1 | osteoclast stimulating factor 1 |
| chr11 | 57693583 | 57694471 | 889 | * | 6,521468 | 5,116257431 | 1,969032 | 2,598362 | 0,009367 | 1 | 0 | Promoter (1-2kb) | 11 | 57692463 | 57701043 | 8581 | 1 | 50724 | ENSMUST00000020826.6 | 1120 | ENSMUSG00000020519 | Sap30l | SAP30-like |
| chr17 | 36244946 | 36245844 | 899 | * | 6,835314 | 5,116454938 | 1,963889 | 2,605267 | 0,00918 | 1 | 0 | Distal Intergenic | 17 | 36237386 | 36239221 | 1836 | 1 | 52040 | ENSMUST00000151375.2 | 7560 | ENSMUSG00000039220 | Ppp1r10 | protein phosphatase 1, regulatory subunit 10 |
| chr1 | 1,36E+08 | 1,36E+08 | 701 | * | 7,586416 | 5,117117442 | 1,946834 | 2,62843 | 0,008578 | 1 | 0 | Intron (ENSMUST0000( | 1 | 1,36E+08 | 1,36E+08 | 3576 | 1 | 381293 | ENSMUST00000195390.6 | 21599 | ENSMUSG00000041498 | Kif14 | kinesin family member 14 |
| chr2 | 38085116 | 38085931 | 816 | * | 5,890044 | 5,117568045 | 1,904423 | 2,687201 | 0,007205 | 1 | 0 | Intron (ENSMUST0000( | 2 | 37911278 | 38056093 | 144816 | 2 | 227801 | ENSMUST00000126889.8 | -29023 | ENSMUSG00000035392 | Dennd1a | DENN/MADD domain containing 1A |
| chr14 | 21671347 | 21672100 | 754 | * | 6,32114 | 5,117733437 | 1,969429 | 2,598588 | 0,009361 | 1 | 0 | Promoter (<=1kb) | 14 | 21672610 | 21684657 | 12048 | 1 | 54169 | ENSMUST00000182732.2 | -510 | ENSMUSG00000021767 | Kat6b | K(lysine) acetyltransferase 6B |
| chr16 | 31764685 | 31766020 | 1336 | * | 15,26945 | 5,117805478 | 1,374801 | 3,722578 | 0,000197 | 0,238040606 | 1,41696 | Promoter (1-2kb) | 16 | 31765868 | 31767312 | 1445 | 2 | 106264 | ENSMUST00000202722.2 | 1292 | ENSMUSG00000107002 | 0610012G03Rik | RIKEN cDNA 0610012G03 gene |
| chr9 | 42005354 | 42005734 | 381 | * | 4,943632 | 5,118540181 | 1,935411 | 2,644679 | 0,008177 | 1 | 0 | Intron (ENSMUST0000( | 9 | 41876016 | 42035593 | 159578 | 2 | 20660 | ENSMUST00000060989.9 | 29859 | ENSMUSG00000049313 | Sorl1 | sortilin-related receptor, LDLR class A repeats-containing |
| chr13 | 1,15E+08 | 1,15E+08 | 636 | * | 4,7395 | 5,119337647 | 1,987377 | 2,575927 | 0,009997 | 1 | 0 | Distal Intergenic | 13 | 1,15E+08 | 1,15E+08 | 90586 | 1 | 75161 | ENSMUST00000181553.2 | 93548 | ENSMUSG00000051758 | 4930544M13Rik | RIKEN cDNA 4930544M13 gene |
| chr7 | 51606634 | 51607731 | 1098 | * | 8,708791 | 5,121247693 | 1,412119 | 3,62664 | 0,000287 | 0,29306015 | 1,360622 | Intron (ENSMUST0000( | 7 | 51593432 | 51606182 | 12751 | 1 | 14453 | ENSMUST00000145723.2 | 13202 | ENSMUSG00000030498 | Gas2 | growth arrest specific 2 |
| chr4 | 44930781 | 44931870 | 1090 | * | 7,598684 | 5,124838742 | 1,873523 | 2,735401 | 0,00623 | 0,814467366 | 1,360622 | 3' UTR | 4 | 44925995 | 44931362 | 5368 | 1 | 319885 | ENSMUST00000126968.2 | 4786 | ENSMUSG00000035649 | Zcchc7 | zinc finger, CCHC domain containing 7 |
| chr11 | 88235483 | 88236028 | 546 | * | 6,642108 | 5,126088487 | 1,858273 | 2,758523 | 0,005806 | 1 | 0 | 3' UTR | 11 | 88243599 | 88243690 | 92 | 1 | 1,01E+08 | ENSMUST00000175244.3 | -7571 | ENSMUSG00000092985 | Mir378b | microRNA 378b |
| chr9 | 69340187 | 69342185 | 1999 | * | 11,39011 | 5,126170542 | 1,671942 | 3,065999 | 0,002169 | 0,660341484 | 1,173325 | 3' UTR | 9 | 69324155 | 69334306 | 10152 | 1 | 93697 | ENSMUST00000147558.2 | 16032 | ENSMUSG00000032235 | Ice2 | interactor of little elongation complex ELL subunit 2 |
| chr9 | 94418738 | 94419518 | 781 | * | 6,069182 | 5,128911844 | 1,893145 | 2,709202 | 0,006745 | 1 | 0 | Promoter (<=1kb) | 9 | 94399917 | 94420134 | 20218 | 2 | 68861 | ENSMUST00000113028.2 | 616 | ENSMUSG00000045414 | Dipk2a | divergent protein kinase domain 2A |
| chr10 | 21797319 | 21798072 | 754 | * | 6,184489 | 5,128965334 | 1,891791 | 2,711169 | 0,006705 | 1 | 0 | Intron (ENSMUST0000( | 10 | 21804369 | 21866301 | 61933 | 1 | 20393 | ENSMUST00000124203.2 | -6297 | ENSMUSG00000019970 | Sgk1 | serum/glucocorticoid regulated kinase 1 |
| chr4 | 1,36E+08 | 1,36E+08 | 858 | * | 6,980232 | 5,129645878 | 1,813387 | 2,828765 | 0,004673 | 1 | 0 | Intron (ENSMUST0000( | 4 | 1,36E+08 | 1,36E+08 | 534 | 2 | 1,15E+08 | ENSMUST00000153223.8 | -9278 | ENSMUSG00000059291 | LOC115489947 | uncharacterized LOC115489947 |
| chr7 | 91751912 | 91753504 | 1593 | * | 11,44372 | 5,130238948 | 1,705019 | 3,008904 | 0,002622 | 0,685078899 | 1,173325 | Exon (ENSMUST00002 | 7 | 91741414 | 91768719 | 27306 | 2 | 75845 | ENSMUST00000207593.2 | 15215 | ENSMUSG00000109382 | 4930567K12Rik | RIKEN cDNA 4930567K12 gene |
| chr14 | 78651690 | 78653207 | 1518 | * | 7,203714 | 5,130561373 | 1,863825 | 2,752705 | 0,005911 | 1 | 0 | Distal Intergenic | 14 | 78732265 | 78741411 | 9147 | 2 | 219181 | ENSMUST00000227808.2 | 88204 | ENSMUSG00000022016 | Akap11 | A kinase (PRKA) anchor protein 11 |
| chr18 | 54686067 | 54687002 | 936 | * | 10,2336 | 5,133115417 | 1,38489 | 3,706514 | 0,00021 | 0,281843592 | 1,086118 | Distal Intergenic | 18 | 54627765 | 54629656 | 1892 | 2 | 1,05E+08 | ENSMUST00000237202.2 | -56411 | | 4933434P08Rik | RIKEN cDNA 4933434P08 gene |
| chr16 | 54240149 | 54240788 | 640 | * | 6,006468 | 5,133436629 | 1,924878 | 2,666889 | 0,007656 | 1 | 0 | Distal Intergenic | 16 | 54174825 | 54174956 | 132 | 1 | 1,15E+08 | ENSMUST00000082793.3 | 65324 | ENSMUSG00000064727 | Gm25853 | predicted gene, 25853 |
| chr14 | 1,13E+08 | 1,13E+08 | 473 | * | 4,828588 | 5,133884029 | 1,977743 | 2,595829 | 0,009436 | 1 | 0 | Distal Intergenic | 14 | 1,12E+08 | 1,12E+08 | 3494 | 1 | 75409 | ENSMUST00000227891.2 | 998894 | ENSMUSG00000033214 | Slitrk5 | SLIT and NTRK-like family, member 5 |
| chr10 | 17932100 | 17933215 | 1116 | * | 7,962582 | 5,133893203 | 1,947556 | 2,636069 | 0,008387 | 1 | 0 | Promoter (<=1kb) | 10 | 17931752 | 18000892 | 69141 | 1 | 19707 | ENSMUST00000150029.8 | 348 | ENSMUSG00000019854 | Reps1 | RalBP1 associated Eps domain containing protein |
| chr16 | 22183939 | 22184726 | 788 | * | 8,243395 | 5,13452257 | 1,798425 | 2,855011 | 0,004304 | 0,748971518 | 1,360622 | Distal Intergenic | 16 | 22202329 | 22220579 | 18251 | 2 | 104156 | ENSMUST00000170803.2 | 35853 | ENSMUSG00000013089 | Etv5 | ets variant 5 |
| chr6 | 1,42E+08 | 1,42E+08 | 785 | * | 5,867723 | 5,136378612 | 1,977577 | 2,597308 | 0,009396 | 1 | 0 | Distal Intergenic | 6 | 1,42E+08 | 1,42E+08 | 47474 | 2 | 28254 | ENSMUST00000174455.2 | -12582 | ENSMUSG00000079262 | Slco1a6 | solute carrier organic anion transporter family, member 1a6 |
| chr7 | 93820051 | 93820649 | 599 | * | 6,480105 | 5,136383385 | 1,838575 | 2,793676 | 0,005211 | 1 | 0 | Intron (ENSMUST0000( | 7 | 93846417 | 93846686 | 270 | 1 | 545982 | ENSMUST00000071770.4 | -25768 | | Gm5899 | predicted pseudogene 5899 |
| chr5 | 97522356 | 97523744 | 1389 | * | 7,436402 | 5,137048869 | 1,977022 | 2,598377 | 0,009367 | 1 | 0 | Distal Intergenic | 5 | 97530057 | 97540238 | 10182 | 2 | 97243 | ENSMUST00000060265.6 | 16494 | ENSMUSG00000046000 | Naa11 | N(alpha)-acetyltransferase 11, NatA catalytic subunit |
| chr4 | 1,25E+08 | 1,25E+08 | 958 | * | 7,046291 | 5,137177475 | 1,830744 | 2,80606 | 0,005015 | 1 | 0 | Promoter (<=1kb) | 4 | 1,25E+08 | 1,25E+08 | 23270 | 1 | 70088 | ENSMUST00000184118.2 | 210 | ENSMUSG00000028863 | Meaf6 | MYST/Esa1-associated factor 6 |
| chr9 | 77662321 | 77663237 | 917 | * | 9,039537 | 5,138158411 | 1,743344 | 2,9473 | 0,003206 | 0,695401133 | 1,360622 | Promoter (<=1kb) | 9 | 77661817 | 77701767 | 39951 | 1 | 14629 | ENSMUST00000034905.9 | 504 | ENSMUSG00000032350 | Gclc | glutamate-cysteine ligase, catalytic subunit |
| chr5 | 73856961 | 73857668 | 708 | * | 5,781597 | 5,138777265 | 1,904688 | 2,697963 | 0,006977 | 1 | 0 | Distal Intergenic | 5 | 73816604 | 73836827 | 20224 | 1 | 73472 | ENSMUST00000178631.6 | 40357 | ENSMUSG00000029155 | Spata18 | spermatogenesis associated 18 |
| chr12 | 32687510 | 32688730 | 1221 | * | 9,237586 | 5,140374409 | 1,880662 | 2,733279 | 0,006271 | 0,81623383 | 1,360622 | Distal Intergenic | 12 | 32869544 | 32889206 | 19663 | 1 | 59027 | ENSMUST00000220200.2 | -180814 | ENSMUSG00000020572 | Nampt | nicotinamide phosphoribosyltransferase |
| chr2 | 33048251 | 33048929 | 679 | * | 6,36032 | 5,140386154 | 1,898535 | 2,707554 | 0,006778 | 1 | 0 | Promoter (<=1kb) | 2 | 33033319 | 33048182 | 14864 | 2 | 241308 | ENSMUST00000138704.2 | -69 | ENSMUSG00000038831 | Ralgps1 | Ral GEF with PH domain and SH3 binding motif 1 |
| chr4 | 28190595 | 28191159 | 565 | * | 6,239762 | 5,140511038 | 1,880864 | 2,733059 | 0,006275 | 1 | 0 | Distal Intergenic | 4 | 28365316 | 28365444 | 129 | 1 | 1,15E+08 | ENSMUST00000104333.2 | -174157 | ENSMUSG00000077521 | Gm25378 | predicted gene, 25378 |
| chr5 | 36714718 | 36715657 | 940 | * | 7,355345 | 5,14400577 | 1,835135 | 2,803066 | 0,005062 | 0,77749164 | 1,360622 | Intron (ENSMUST0000( | 5 | 36700524 | 36721400 | 20877 | 2 | 100855 | ENSMUST00000146252.2 | 5743 | ENSMUSG00000029192 | Tbc1d14 | TBC1 domain family, member 14 |
| chr9 | 64484671 | 64485208 | 538 | * | 6,308473 | 5,144867704 | 1,900607 | 2,70696 | 0,00679 | 1 | 0 | Intron (ENSMUST0000( | 9 | 64454181 | 64545748 | 91568 | 1 | 214058 | ENSMUST00000147388.2 | 30490 | ENSMUSG00000036466 | Megf11 | multiple EGF-like-domains 11 |
| chrX | 35327035 | 35327828 | 794 | * | 4,879089 | 5,146249016 | 1,962129 | 2,622788 | 0,008721 | 1 | 0 | Intron (ENSMUST0000( | 20 | 35304891 | 35339880 | 34990 | 1 | 75974 | ENSMUST00000115264.2 | 22144 | ENSMUSG00000031093 | Dock11 | dedicator of cytokinesis 11 |
| chr3 | 19842398 | 19843140 | 743 | * | 5,519533 | 5,146401519 | 1,98761 | 2,589241 | 0,009619 | 1 | 0 | Distal Intergenic | 3 | 19747565 | 19749560 | 1996 | 2 | 12918 | ENSMUST00000061294.5 | -92838 | ENSMUSG00000049796 | Crh | corticotropin releasing hormone |
| chr18 | 86107567 | 86108828 | 1262 | * | 12,27881 | 5,14675289 | 1,255893 | 4,098083 | 4,17E-05 | 0,123146349 | 0,918255 | Distal Intergenic | 18 | 86367711 | 86367817 | 107 | 1 | 1,15E+08 | ENSMUST00000082733.3 | -258883 | ENSMUSG00000119275 | Gm25453 | predicted gene, 25453 |
| chr3 | 1,46E+08 | 1,46E+08 | 754 | * | 6,077927 | 5,150724695 | 1,900271 | 2,710521 | 0,006718 | 1 | 0 | 3' UTR | 3 | 1,46E+08 | 1,46E+08 | 55456 | 1 | 22262 | ENSMUST00000199489.5 | 10730 | ENSMUSG00000028186 | Uox | urate oxidase |
| chr3 | 1,46E+08 | 1,46E+08 | 558 | * | 7,172026 | 5,150880183 | 1,787923 | 2,88093 | 0,003965 | 1 | 0 | Promoter (1-2kb) | 3 | 1,46E+08 | 1,46E+08 | 3575 | 1 | 12042 | ENSMUST00000198122.2 | 1958 | ENSMUSG00000028191 | Bcl10 | B cell leukemia/lymphoma 10 |
| chr10 | 1,23E+08 | 1,23E+08 | 544 | * | 6,855656 | 5,151339945 | 1,962821 | 2,624457 | 0,008679 | 1 | 0 | Intron (ENSMUST0000( | 10 | 1,23E+08 | 1,23E+08 | 54274 | 1 | 319468 | ENSMUST00000160315.2 | -19855 | ENSMUSG00000034613 | Ppm1h | protein phosphatase 1H (PP2C domain containing) |
| chr11 | 88554166 | 88555533 | 1368 | * | 12,68426 | 5,151524549 | 1,750497 | 2,942893 | 0,003252 | 0,806557336 | 0,738441 | Intron (ENSMUST0000( | 11 | 88234715 | 88608927 | 374213 | 2 | 76626 | ENSMUST00000092794.12 | 53394 | ENSMUSG00000069769 | Msi2 | musashi RNA-binding protein 2 |
| chr9 | 55636150 | 55636488 | 339 | * | 5,452508 | 5,153397857 | 1,99652 | 2,58119 | 0,009846 | 1 | 0 | Intron (ENSMUST0000( | 9 | 55650913 | 55699594 | 48682 | 2 | 244891 | ENSMUST00000215323.2 | 63106 | ENSMUSG00000034007 | Scaper | S phase cyclin A-associated protein in the ER |
| chr7 | 40317908 | 40318463 | 556 | * | 5,075005 | 5,153408003 | 1,994413 | 2,583923 | 0,009768 | 1 | 0 | Distal Intergenic | 7 | 40324161 | 40332685 | 8525 | 1 | 71029 | ENSMUST00000206289.2 | -5698 | ENSMUSG00000102004 | 4933402C06Rik | RIKEN cDNA 4933402C06 gene |
| chr1 | 1,57E+08 | 1,57E+08 | 445 | * | 5,284529 | 5,153424662 | 1,993468 | 2,585155 | 0,009734 | 1 | 0 | Intron (ENSMUST0000( | 1 | 1,57E+08 | 1,57E+08 | 126546 | 2 | 78255 | ENSMUST00000185198.7 | -70940 | ENSMUSG00000026594 | Ralgps2 | Ral GEF with PH domain and SH3 binding motif 2 |
| chr13 | 54300864 | 54301666 | 803 | * | 6,478472 | 5,1542452 | 1,92447 | 2,678267 | 0,0074 | 1 | 0 | Distal Intergenic | 13 | 54346148 | 54376451 | 30304 | 1 | 15466 | ENSMUST00000038101.4 | -44482 | ENSMUSG00000034987 | Hrh2 | histamine receptor H2 |
| chr2 | 72048479 | 72049587 | 1109 | * | 6,807886 | 5,154338063 | 1,916851 | 2,688961 | 0,007167 | 1 | 0 | Intron (ENSMUST0000( | 2 | 72029143 | 72038781 | 9639 | 1 | 56508 | ENSMUST00000156031.2 | 19336 | ENSMUSG00000049044 | Rapgef4 | Rap guanine nucleotide exchange factor (GEF) 4 |
| chr7 | 1,11E+08 | 1,11E+08 | 626 | * | 8,663181 | 5,154569087 | 1,724993 | 2,988168 | 0,002807 | 1 | 0 | Exon (ENSMUST00000( | 7 | 1,11E+08 | 1,11E+08 | 308317 | 2 | 233733 | ENSMUST00000049430.15 | 163104 | ENSMUSG00000038296 | Galnt18 | polypeptide N-acetylgalactosaminyltransferase 18 |
| chr5 | 1,24E+08 | 1,24E+08 | 1140 | * | 6,812615 | 5,154608663 | 1,897477 | 2,71656 | 0,006596 | 1 | 0 | Promoter (2-3kb) | 5 | 1,24E+08 | 1,24E+08 | 106469 | 2 | 56430 | ENSMUST00000111561.8 | 2606 | ENSMUSG00000049550 | Clip1 | CAP-GLY domain containing linker protein 1 |
| chr5 | 1,23E+08 | 1,23E+08 | 995 | * | 8,961395 | 5,155980519 | 1,817283 | 2,837192 | 0,004551 | 0,778555725 | 1,211315 | Distal Intergenic | 5 | 1,23E+08 | 1,23E+08 | 64315 | 2 | 12589 | ENSMUST00000031426.14 | -8277 | ENSMUSG00000029469 | Ift81 | intraflagellar transport 81 |
| chr2 | 1,56E+08 | 1,56E+08 | 876 | * | 6,943298 | 5,156734417 | 1,885886 | 2,734383 | 0,00625 | 1 | 0 | Intron (ENSMUST0000( | 2 | 1,56E+08 | 1,56E+08 | 67679 | 1 | 13821 | ENSMUST00000029155.16 | -16455 | ENSMUSG00000027624 | Epb41l1 | erythrocyte membrane protein band 4.1 like 1 |
| chr13 | 30826694 | 30827719 | 1026 | * | 7,196048 | 5,156939002 | 1,848546 | 2,789726 | 0,005275 | 1 | 0 | Distal Intergenic | 13 | 30843982 | 30895214 | 51233 | 1 | 105352 | ENSMUST00000091672.13 | -16263 | ENSMUSG00000069255 | Dusp22 | dual specificity phosphatase 22 |
| chr2 | 71804310 | 71805327 | 1018 | * | 10,7514 | 5,15733276 | 1,681664 | 3,066804 | 0,002164 | 0,644704154 | 1,2763 | Distal Intergenic | 2 | 71811584 | 72087818 | 276235 | 1 | 56508 | ENSMUST00000090826.12 | -6257 | ENSMUSG00000049044 | Rapgef4 | Rap guanine nucleotide exchange factor (GEF) 4 |
| chr7 | 65006739 | 65007566 | 828 | * | 6,046857 | 5,158183938 | 1,858361 | 2,775664 | 0,005509 | 1 | 0 | Intron (ENSMUST0000( | 7 | 64946456 | 65020763 | 74308 | 2 | 21872 | ENSMUST00000032729.8 | 13197 | ENSMUSG00000030516 | Tjp1 | tight junction protein 1 |
| chr1 | 1,52E+08 | 1,52E+08 | 630 | * | 5,406285 | 5,158657712 | 1,954666 | 2,63915 | 0,008311 | 1 | 0 | Distal Intergenic | 1 | 1,52E+08 | 1,52E+08 | 7655 | 2 | 1,03E+08 | ENSMUST00000244902.1 | 20537 | | Gm36920 | predicted gene, 36920 |
| chr7 | 1,16E+08 | 1,16E+08 | 798 | * | 4,979421 | 5,159097604 | 1,94529 | 2,652097 | 0,007999 | 1 | 0 | Intron (ENSMUST0000( | 7 | 1,16E+08 | 1,16E+08 | 11209 | 2 | 69350 | ENSMUST00000150518.2 | 15885 | ENSMUSG00000087621 | 1700003G18Rik | RIKEN cDNA 1700003G18 gene |
| chr14 | 33463442 | 33464039 | 598 | * | 5,765091 | 5,160636284 | 1,964318 | 2,627189 | 0,008609 | 1 | 0 | Distal Intergenic | 14 | 33530032 | 33533240 | 3209 | 2 | 218902 | ENSMUST00000228030.2 | 69201 | ENSMUSG00000115751 | 4930503F20Rik | RIKEN cDNA 4930503F20 gene |
| chr8 | 95692351 | 95692881 | 531 | * | 6,934126 | 5,162095424 | 1,805521 | 2,859062 | 0,004249 | 1 | 0 | Exon (ENSMUST00002 | 8 | 95701379 | 95732432 | 31054 | 1 | 14766 | ENSMUST00000211850.2 | -8498 | ENSMUSG00000031785 | Adgrg1 | adhesion G protein-coupled receptor G1 |
| chr5 | 1,5E+08 | 1,5E+08 | 538 | * | 5,690917 | 5,163446692 | 1,926821 | 2,679775 | 0,007367 | 1 | 0 | Intron (ENSMUST0000( | 5 | 1,5E+08 | 1,5E+08 | 143955 | 1 | 320365 | ENSMUST00000202600.4 | 24802 | ENSMUSG00000056602 | Fry | FRY microtubule binding protein |
| chr19 | 14661678 | 14662467 | 790 | * | 6,996795 | 5,165863699 | 1,80018 | 2,869637 | 0,004109 | 1 | 0 | Distal Intergenic | 19 | 14425514 | 14575415 | 149902 | 2 | 21888 | ENSMUST00000167776.3 | -86263 | ENSMUSG00000024642 | Tle4 | transducin-like enhancer of split 4 |
| chr1 | 91009332 | 91010061 | 730 | * | 5,070028 | 5,166487636 | 1,920428 | 2,690279 | 0,007139 | 1 | 0 | Exon (ENSMUST00001 | 1 | 90981308 | 91056009 | 74702 | 1 | 16978 | ENSMUST00000189617.3 | 28024 | ENSMUSG00000026305 | Lrrfip1 | leucine rich repeat (in FLII) interacting protein 1 |
| chr12 | 79668781 | 79669817 | 1037 | * | 7,058511 | 5,167077483 | 1,775275 | 2,910579 | 0,003608 | 0,716908183 | 1,360622 | Intron (ENSMUST0000( | 12 | 79938062 | 79938677 | 616 | 1 | 1,08E+08 | ENSMUST00000243367.1 | -268245 | | Gm46362 | predicted gene, 46362 |
| chr1 | 1,11E+08 | 1,11E+08 | 875 | * | 6,030389 | 5,167200982 | 1,871244 | 2,761372 | 0,005756 | 1 | 0 | Intron (ENSMUST0000( | 1 | 1,12E+08 | 1,12E+08 | 6067 | 2 | 319901 | ENSMUST00000186365.2 | 435838 | ENSMUSG00000038702 | Dsel | dermatan sulfate epimerase-like |
| chr14 | 31085452 | 31087026 | 1575 | * | 9,215946 | 5,167210969 | 1,744633 | 2,961775 | 0,003059 | 0,683210259 | 1,425929 | Promoter (<=1kb) | 14 | 31087513 | 31091970 | 4458 | 1 | 12339 | ENSMUST00000228237.2 | -487 | ENSMUSG00000021893 | Capn7 | calpain 7 |
| chr13 | 73753758 | 73754351 | 594 | * | 7,829011 | 5,167289744 | 1,770878 | 2,917925 | 0,003524 | 1 | 0 | Promoter (1-2kb) | 13 | 73752125 | 73768724 | 16600 | 1 | 218335 | ENSMUST00000022102.9 | 1633 | ENSMUSG00000021610 | Clptm1l | CLPTM1-like |
| chr1 | 91418052 | 91418628 | 577 | * | 6,65479 | 5,167688915 | 1,873942 | 2,757657 | 0,005822 | 1 | 0 | Distal Intergenic | 1 | 91422369 | 91455678 | 33310 | 1 | 74019 | ENSMUST00000189341.7 | -3741 | ENSMUSG00000034292 | Traf3ip1 | TRAF3 interacting protein 1 |
| chr3 | 87824256 | 87825284 | 1029 | * | 6,742575 | 5,167999652 | 1,870891 | 2,76232 | 0,005739 | 1 | 0 | Promoter (1-2kb) | 3 | 87826813 | 87830969 | 4157 | 1 | 67707 | ENSMUST00000119968.8 | -1529 | ENSMUSG00000019710 | Mrpl24 | mitochondrial ribosomal protein L24 |
| chr12 | 16875157 | 16875931 | 775 | * | 7,043435 | 5,168426949 | 1,841461 | 2,806699 | 0,005005 | 1 | 0 | Promoter (1-2kb) | 12 | 16874152 | 16875510 | 1359 | 1 | 50496 | ENSMUST00000221600.2 | 1005 | ENSMUSG00000057469 | E2f6 | E2F transcription factor 6 |
| chr6 | 52577764 | 52578637 | 874 | * | 8,885589 | 5,169717457 | 1,722689 | 3,000959 | 0,002691 | 0,661081489 | 1,449706 | Exon (ENSMUST00002 | 6 | 52523681 | 52607243 | 83563 | 2 | 58875 | ENSMUST00000155948.2 | 28606 | ENSMUSG00000029776 | Hibadh | 3-hydroxyisobutyrate dehydrogenase |
| chr16 | 32448771 | 32449412 | 642 | * | 6,235678 | 5,170509688 | 1,914743 | 2,700368 | 0,006926 | 1 | 0 | 3' UTR | 16 | 32443260 | 32451608 | 8349 | 1 | 22042 | ENSMUST00000128285.2 | 5511 | ENSMUSG00000022797 | Tfrc | transferrin receptor |
| chr16 | 57618659 | 57619727 | 1069 | * | 5,933028 | 5,172688169 | 1,931342 | 2,678287 | 0,0074 | 1 | 0 | Distal Intergenic | 16 | 57444621 | 57575100 | 130480 | 2 | 12837 | ENSMUST00000089332.5 | -43559 | ENSMUSG00000068196 | Col8a1 | collagen, type VIII, alpha 1 |
| chr5 | 1,44E+08 | 1,44E+08 | 737 | * | 7,571244 | 5,172901366 | 1,903965 | 2,71691 | 0,006589 | 1 | 0 | Intron (ENSMUST0000( | 5 | 1,44E+08 | 1,44E+08 | 6918 | 2 | 66898 | ENSMUST00000129287.2 | -21475 | ENSMUSG00000038859 | Baiap2l1 | BAI1-associated protein 2-like 1 |
| chr11 | 53740003 | 53741943 | 1941 | * | 13,47464 | 5,174345324 | 1,63495 | 3,164833 | 0,001552 | 0,644704154 | 0,918255 | Promoter (<=1kb) | 11 | 53676315 | 53740874 | 64560 | 2 | 622459 | ENSMUST00000220191.2 | 0 | ENSMUSG00000081769 | Gm12216 | predicted gene 12216 |
| chr11 | 50017531 | 50018829 | 1299 | * | 9,307965 | 5,175128357 | 1,794952 | 2,883157 | 0,003937 | 0,722439117 | 1,449706 | Exon (ENSMUST00001 | 11 | 50022223 | 50062857 | 40635 | 1 | 76795 | ENSMUST00000093138.13 | -3394 | ENSMUSG00000036644 | Tbc1d9b | TBC1 domain family, member 9B |
| chr9 | 44214871 | 44216519 | 1649 | * | 11,13916 | 5,177178251 | 1,703792 | 3,038621 | 0,002377 | 0,661775851 | 1,2763 | Promoter (<=1kb) | 9 | 44211587 | 44216942 | 5356 | 2 | 102423 | ENSMUST00000217605.2 | 423 | ENSMUSG00000032119 | Hinfp | histone H4 transcription factor |
| chr16 | 20137992 | 20138872 | 881 | * | 5,808056 | 5,178451164 | 1,871536 | 2,766952 | 0,005658 | 1 | 0 | Promoter (2-3kb) | 16 | 20128526 | 20135500 | 6975 | 2 | 224044 | ENSMUST00000182741.2 | -2492 | ENSMUSG00000022818 | Cyp2ab1 | cytochrome P450, family 2, subfamily ab, polypeptide 1 |
| chr5 | 65771988 | 65772806 | 819 | * | 5,97935 | 5,179817422 | 1,838596 | 2,817268 | 0,004843 | 1 | 0 | 3' UTR | 5 | 65765113 | 65777943 | 12831 | 2 | 71521 | ENSMUST00000202648.4 | 5137 | ENSMUSG00000029202 | Pds5a | PDS5 cohesin associated factor A |
| chr1 | 1,71E+08 | 1,71E+08 | 474 | * | 4,898267 | 5,180219573 | 1,999872 | 2,590276 | 0,00959 | 1 | 0 | Distal Intergenic | 1 | 1,71E+08 | 1,71E+08 | 12912 | 1 | 75472 | ENSMUST00000129651.8 | -11488 | ENSMUSG00000026649 | Cfap126 | cilia and flagella associated protein 126 |
| chr14 | 1,03E+08 | 1,03E+08 | 1219 | * | 7,840324 | 5,181603491 | 1,823899 | 2,840949 | 0,004498 | 1 | 0 | Promoter (<=1kb) | 14 | 1,03E+08 | 1,03E+08 | 7413 | 1 | 211286 | ENSMUST00000022721.8 | -21 | ENSMUSG00000022125 | Cln5 | ceroid-lipofuscinosis, neuronal 5 |
| chrX | 41607683 | 41608654 | 972 | * | 6,843041 | 5,181684793 | 1,825904 | 2,837873 | 0,004542 | 1 | 0 | Intron (ENSMUST0000( | 20 | 41603190 | 41610847 | 7658 | 1 | 20400 | ENSMUST00000128393.2 | 4493 | ENSMUSG00000005696 | Sh2d1a | SH2 domain containing 1A |
| chr10 | 35266341 | 35267220 | 880 | * | 6,971559 | 5,181686688 | 1,793922 | 2,888469 | 0,003871 | 1 | 0 | Distal Intergenic | 10 | 34988194 | 34988327 | 134 | 1 | 1,15E+08 | ENSMUST00000104561.3 | 278147 | ENSMUSG00000077749 | Gm26341 | predicted gene, 26341 |
| chrX | 1,53E+08 | 1,53E+08 | 855 | * | 5,051117 | 5,184352484 | 1,954079 | 2,653092 | 0,007976 | 1 | 0 | Distal Intergenic | 20 | 1,53E+08 | 1,53E+08 | 2681 | 1 | 75185 | ENSMUST00000026029.2 | 41597 | ENSMUSG00000025051 | Samt4 | spermatogenesis associated multipass transmembrane protein 4 |
| chr2 | 1,54E+08 | 1,54E+08 | 1431 | * | 12,25964 | 5,185069973 | 1,712322 | 3,028093 | 0,002461 | 0,718585444 | 0,918255 | Promoter (<=1kb) | 2 | 1,54E+08 | 1,54E+08 | 17161 | 1 | 228802 | ENSMUST00000045959.8 | 0 | ENSMUSG00000038572 | Bpifb5 | BPI fold containing family B, member 5 |
| chr10 | 21702737 | 21703725 | 989 | * | 9,452264 | 5,185591652 | 1,697944 | 3,054041 | 0,002258 | 0,64236196 | 1,360622 | Distal Intergenic | 10 | 21706398 | 21720750 | 14353 | 2 | 628133 | ENSMUST00000180731.2 | 17025 | | Platr6 | pluripotency associated transcript 6 |
| chr1 | 1,93E+08 | 1,93E+08 | 673 | * | 5,164874 | 5,185718001 | 1,93549 | 2,679279 | 0,007378 | 1 | 0 | Distal Intergenic | 1 | 1,93E+08 | 1,93E+08 | 23953 | 2 | 215303 | ENSMUST00000016323.11 | -50359 | ENSMUSG00000016179 | Camk1g | calcium/calmodulin-dependent protein kinase I gamma |
| chr7 | 97171044 | 97171583 | 540 | * | 7,647944 | 5,186610055 | 1,836398 | 2,824339 | 0,004738 | 0,764803394 | 1,360622 | Exon (ENSMUST00000( | 7 | 97179361 | 97184450 | 5090 | 1 | 101861 | ENSMUST00000205387.2 | -7778 | ENSMUSG00000025133 | Ints4 | integrator complex subunit 4 |
| chr3 | 1,04E+08 | 1,04E+08 | 759 | * | 5,996681 | 5,186974723 | 1,906198 | 2,721111 | 0,006506 | 1 | 0 | Intron (ENSMUST0000( | 3 | 1,04E+08 | 1,04E+08 | 6048 | 1 | 229672 | ENSMUST00000062945.12 | -3773 | ENSMUSG00000044165 | Bcl2l15 | BCLl2-like 15 |

| | | | | | | | | | | | | | | | | | | | | | | | |
|---|---|---|---|---|---|---|---|---|---|---|---|---|---|---|---|---|---|---|---|---|---|---|---|
| chr19 | 38064485 | 38065144 | 660 | * | 7,457401 | 5,188228791 | 1,884031 | 2,753792 | 0,005891 | 1 | 0 | Distal Intergenic | 19 | 38060377 | 38060501 | 125 | 1 | 1,15E+08 | ENSMUST00000082712.3 | 4108 | ENSMUSG00000064646 | Gm23300 | predicted gene, 23300 |
| chr3 | 54310700 | 54311748 | 1049 | * | 7,695257 | 5,189117336 | 1,475083 | 3,517847 | 0,000435 | 1 | 0 | Distal Intergenic | 3 | 54296310 | 54297814 | 1505 | 1 | 50706 | ENSMUST00000150868.2 | 14390 | ENSMUSG00000027750 | Postn | periostin, osteoblast specific factor |
| chr5 | 1,37E+08 | 1,37E+08 | 855 | * | 6,07327 | 5,189345138 | 1,893575 | 2,740501 | 0,006135 | 1 | 0 | Promoter (<=1kb) | 5 | 1,37E+08 | 1,37E+08 | 1795 | 2 | 140709 | ENSMUST00000141744.2 | -408 | ENSMUSG00000004415 | Col26a1 | collagen, type XXVI, alpha 1 |
| chr12 | 83447796 | 83448852 | 1057 | * | 6,887718 | 5,189348771 | 1,878146 | 2,763016 | 0,005727 | 1 | 0 | Intron (ENSMUST0000( | 12 | 83316089 | 83486694 | 170606 | 2 | 70127 | ENSMUST00000177801.2 | 37842 | ENSMUSG00000021221 | Dpf3 | double PHD fingers 3 |
| chr7 | 1,23E+08 | 1,23E+08 | 1416 | * | 7,55485 | 5,190130393 | 1,925577 | 2,695364 | 0,007031 | 1 | 0 | Intron (ENSMUST0000( | 7 | 1,23E+08 | 1,23E+08 | 23348 | 1 | 233833 | ENSMUST00000205760.2 | -9485 | ENSMUSG00000052707 | Tnrc6a | trinucleotide repeat containing 6a |
| chr7 | 1,18E+08 | 1,18E+08 | 1102 | * | 7,104089 | 5,190190593 | 1,929598 | 2,689778 | 0,00715 | 1 | 0 | Promoter (<=1kb) | 7 | 1,18E+08 | 1,18E+08 | 1687 | 1 | 67745 | ENSMUST00000138180.2 | 727 | ENSMUSG00000055159 | 4930583K01Rik | RIKEN cDNA 4930583K01 gene |
| chr11 | 1,2E+08 | 1,2E+08 | 808 | * | 5,936971 | 5,190209062 | 1,930212 | 2,688932 | 0,007168 | 1 | 0 | Intron (ENSMUST0000( | 11 | 1,2E+08 | 1,2E+08 | 48183 | 1 | 74370 | ENSMUST00000139728.8 | -50839 | ENSMUSG00000025583 | Rptor | regulatory associated protein of MTOR, complex 1 |
| chr8 | 11548324 | 11549050 | 727 | * | 7,211702 | 5,190291184 | 1,936394 | 2,68039 | 0,007354 | 1 | 0 | Promoter (<=1kb) | 8 | 11547633 | 11560236 | 12604 | 1 | 69225 | ENSMUST00000180269.2 | 691 | ENSMUSG00000031505 | Naxd | NAD(P)HX dehydratase |
| chr11 | 1,06E+08 | 1,06E+08 | 959 | * | 6,359228 | 5,1903884 | 1,942928 | 2,671426 | 0,007553 | 1 | 0 | Distal Intergenic | 11 | 1,06E+08 | 1,06E+08 | 90402 | 2 | 78943 | ENSMUST00000131895.2 | -5507 | ENSMUSG00000020715 | Ern1 | endoplasmic reticulum (ER) to nucleus signalling 1 |
| chr9 | 56934763 | 56935915 | 1153 | * | 7,326673 | 5,190611479 | 1,958246 | 2,650643 | 0,008034 | 0,857326444 | 1,360622 | Exon (ENSMUST00000( | 9 | 56929516 | 56943903 | 14388 | 1 | 56294 | ENSMUST00000216034.2 | 5247 | ENSMUSG00000032290 | Ptpn9 | protein tyrosine phosphatase, non-receptor type 9 |
| chr10 | 1,11E+08 | 1,11E+08 | 636 | * | 5,726699 | 5,19063267 | 1,960346 | 2,647814 | 0,008101 | 1 | 0 | Distal Intergenic | 10 | 1,11E+08 | 1,11E+08 | 2919 | 1 | 1,03E+08 | ENSMUST00000245392.1 | 16535 | | Gm30262 | predicted gene, 30262 |
| chr2 | 1,32E+08 | 1,32E+08 | 720 | * | 5,590428 | 5,190959464 | 1,99465 | 2,602441 | 0,009256 | 1 | 0 | Intron (ENSMUST0000( | 2 | 1,31E+08 | 1,32E+08 | 68942 | 2 | 1,19E+08 | ENSMUST00000138080.2 | 35853 | | Gm14280 | predicted gene 14280 |
| chr5 | 76991417 | 76992108 | 692 | * | 5,571946 | 5,191023874 | 1,991494 | 2,606598 | 0,009145 | 1 | 0 | Promoter (2-3kb) | 5 | 76988466 | 76990438 | 1973 | 1 | 320827 | ENSMUST00000121851.2 | 2951 | ENSMUSG00000036377 | Cracd | capping protein inhibiting regulator of actin |
| chr9 | 26913941 | 26914573 | 633 | * | 6,139935 | 5,191201492 | 2,00711 | 2,586406 | 0,009698 | 1 | 0 | Promoter (<=1kb) | 9 | 26914356 | 26917743 | 3388 | 1 | 77862 | ENSMUST00000217143.2 | 0 | ENSMUSG00000035443 | Thyn1 | thymocyte nuclear protein 1 |
| chr4 | 1,35E+08 | 1,35E+08 | 1123 | * | 6,883942 | 5,192434456 | 1,920923 | 2,703093 | 0,00687 | 1 | 0 | Intron (ENSMUST0000( | 4 | 1,35E+08 | 1,35E+08 | 41956 | 2 | 66146 | ENSMUST00000137707.8 | 4608 | ENSMUSG00000028826 | Maco1 | macoilin 1 |
| chr12 | 1,04E+08 | 1,04E+08 | 517 | * | 5,210908 | 5,193647471 | 1,889015 | 2,749394 | 0,005971 | 1 | 0 | Distal Intergenic | 12 | 1,04E+08 | 1,04E+08 | 2729 | 2 | 12401 | ENSMUST00000152517.2 | 11640 | ENSMUSG00000060807 | Serpina6 | serine (or cysteine) peptidase inhibitor, clade A, member 6 |
| chr7 | 1,45E+08 | 1,45E+08 | 857 | * | 6,901321 | 5,194934503 | 1,853965 | 2,802068 | 0,005078 | 0,777919825 | 1,360622 | Distal Intergenic | 7 | 1,45E+08 | 1,45E+08 | 794 | 2 | 1E+08 | ENSMUST00000207703.2 | 107942 | ENSMUSG00000109305 | Smim38 | small integral membrane protein 38 |
| chr6 | 55426413 | 55426800 | 388 | * | 5,914572 | 5,195089088 | 1,902283 | 2,730976 | 0,006315 | 1 | 0 | Promoter (2-3kb) | 6 | 55428963 | 55475890 | 46928 | 1 | 11517 | ENSMUST00000070736.12 | -2163 | ENSMUSG00000029778 | Adcyap1r1 | adenylate cyclase activating polypeptide 1 receptor 1 |
| chr3 | 1,01E+08 | 1,01E+08 | 804 | * | 8,864764 | 5,195593377 | 1,76623 | 2,941628 | 0,003265 | 0,698702076 | 1,360622 | Distal Intergenic | 3 | 1,01E+08 | 1,01E+08 | 7379 | 1 | 1,15E+08 | ENSMUST00000198003.2 | 40242 | | Gm43464 | predicted gene 43464 |
| chr7 | 1,1E+08 | 1,1E+08 | 621 | * | 4,696171 | 5,196835321 | 2,006867 | 2,589527 | 0,009611 | 1 | 0 | Intron (ENSMUST0000( | 7 | 1,1E+08 | 1,1E+08 | 305364 | 2 | 319934 | ENSMUST00000166020.8 | 25502 | ENSMUSG00000038371 | Sbf2 | SET binding factor 2 |
| chr16 | 50298483 | 50299000 | 518 | * | 4,988679 | 5,198165724 | 1,999065 | 2,600299 | 0,009314 | 1 | 0 | Distal Intergenic | 16 | 50012207 | 50252753 | 240547 | 2 | 70508 | ENSMUST00000138166.8 | -45730 | ENSMUSG00000022641 | Bbx | bobby sox HMG box containing |
| chr1 | 1,67E+08 | 1,67E+08 | 1173 | * | 7,301222 | 5,199533674 | 1,909487 | 2,723 | 0,006469 | 0,821417341 | 1,360622 | Promoter (<=1kb) | 1 | 1,67E+08 | 1,67E+08 | 10609 | 2 | 80914 | ENSMUST00000195443.6 | -160 | ENSMUSG00000026558 | Uck2 | uridine-cytidine kinase 2 |
| chr2 | 12236561 | 12237476 | 916 | * | 6,213196 | 5,199628762 | 1,872675 | 2,776578 | 0,005493 | 1 | 0 | Intron (ENSMUST0000( | 2 | 12224428 | 12247275 | 22848 | 2 | 241226 | ENSMUST00000172716.3 | 9799 | ENSMUSG00000026768 | Itga8 | integrin alpha 8 |
| chr11 | 84070791 | 84071983 | 1193 | * | 9,179484 | 5,199728659 | 1,709465 | 3,041729 | 0,002352 | 0,639383182 | 1,449706 | Promoter (<=1kb) | 11 | 84070684 | 84105010 | 34327 | 1 | 107476 | ENSMUST00000136463.9 | 107 | ENSMUSG00000020532 | Acaca | acetyl-Coenzyme A carboxylase alpha |
| chr3 | 79789670 | 79791002 | 1333 | * | 10,49318 | 5,200214651 | 1,374393 | 3,783644 | 0,000155 | 0,290582894 | 0,744375 | Promoter (<=1kb) | 3 | 79791840 | 79853587 | 61748 | 1 | 68659 | ENSMUST00000118853.8 | -838 | ENSMUSG00000027955 | Gask1b | golgi associated kinase 1B |
| chr18 | 7116681 | 7117941 | 1261 | * | 10,32019 | 5,200656706 | 1,697681 | 3,063389 | 0,002188 | 0,625646014 | 1,449706 | Intron (ENSMUST0000( | 18 | 7129389 | 7184073 | 54685 | 2 | 74934 | ENSMUST00000235068.2 | 66132 | ENSMUSG00000061802 | Odad2 | outer dynein arm docking complex subunit 2 |
| chr3 | 66887307 | 66888515 | 1209 | * | 7,722884 | 5,205077634 | 1,79982 | 2,891999 | 0,003828 | 0,728477043 | 1,360622 | Promoter (<=1kb) | 3 | 66880592 | 66888474 | 7883 | 2 | 20429 | ENSMUST00000162439.8 | 0 | ENSMUSG00000027833 | Shox2 | short stature homeobox 2 |
| chr10 | 73584686 | 73585424 | 739 | * | 6,859246 | 5,205314543 | 1,96962 | 2,642802 | 0,008222 | 1 | 0 | Intron (ENSMUST0000( | 10 | 73657699 | 74470597 | 812899 | 1 | 11994 | ENSMUST00000064562.14 | -72275 | ENSMUSG00000052613 | Pcdh15 | protocadherin 15 |
| chr3 | 58157923 | 58158759 | 837 | * | 5,04632 | 5,207439691 | 1,987273 | 2,620395 | 0,008783 | 1 | 0 | Distal Intergenic | 3 | 58197672 | 58197778 | 107 | 2 | 1,15E+08 | ENSMUST00000083788.3 | 39019 | ENSMUSG00000065722 | Gm26166 | predicted gene, 26166 |
| chr3 | 30360655 | 30361109 | 455 | * | 6,210634 | 5,208930234 | 1,903926 | 2,735889 | 0,006221 | 1 | 0 | Intron (ENSMUST0000( | 3 | 30326471 | 30328534 | 2064 | 2 | 545510 | ENSMUST00000192999.2 | -32121 | | Gm10258 | predicted gene 10258 |
| chr3 | 1,07E+08 | 1,07E+08 | 648 | * | 4,943417 | 5,209619855 | 1,951057 | 2,670152 | 0,007582 | 1 | 0 | Distal Intergenic | 3 | 1,07E+08 | 1,07E+08 | 914 | 1 | 321000 | ENSMUST00000150888.2 | 6352 | ENSMUSG00000056260 | Lrif1 | ligand dependent nuclear receptor interacting factor 1 |
| chr17 | 40836895 | 40837599 | 705 | * | 6,193523 | 5,211201533 | 1,833196 | 2,842686 | 0,004474 | 1 | 0 | Intron (ENSMUST0000( | 17 | 40830913 | 40841312 | 10400 | 1 | 1E+08 | ENSMUST00000171813.4 | 5982 | ENSMUSG00000090747 | Esp8 | exocrine gland secreted peptide 8 |
| chr13 | 46208026 | 46208641 | 616 | * | 5,826309 | 5,214058805 | 1,882684 | 2,769482 | 0,005615 | 1 | 0 | Distal Intergenic | 13 | 46214073 | 46222238 | 8166 | 1 | 1,08E+08 | ENSMUST00000221550.2 | -5432 | | Gm45949 | predicted gene, 45949 |
| chr7 | 30669220 | 30670172 | 953 | * | 6,829669 | 5,214429041 | 1,87719 | 2,777784 | 0,005473 | 1 | 0 | Promoter (2-3kb) | 7 | 30657204 | 30672747 | 15544 | 2 | 54135 | ENSMUST00000205961.2 | 2575 | ENSMUSG00000001247 | Lsr | lipolysis stimulated lipoprotein receptor |
| chr2 | 78704118 | 78704876 | 759 | * | 4,674272 | 5,215366014 | 1,973934 | 2,642118 | 0,008239 | 1 | 0 | Intron (ENSMUST0000( | 2 | 78700022 | 78750641 | 50620 | 1 | 22193 | ENSMUST00000121433.2 | 4096 | ENSMUSG00000027011 | Ube2e3 | ubiquitin-conjugating enzyme E2E 3 |
| chr3 | 88497799 | 88498062 | 264 | * | 5,063987 | 5,215928681 | 1,961353 | 2,659352 | 0,007829 | 1 | 0 | Distal Intergenic | 3 | 88486978 | 88495726 | 8749 | 1 | 66256 | ENSMUST00000035785.9 | 10821 | ENSMUSG00000041355 | Ssr2 | signal sequence receptor, beta |
| chr4 | 1,41E+08 | 1,41E+08 | 1123 | * | 9,233948 | 5,219544615 | 1,80811 | 2,886741 | 0,003893 | 0,749550311 | 1,211315 | Intron (ENSMUST0000( | 4 | 1,41E+08 | 1,41E+08 | 12895 | 1 | 67680 | ENSMUST00000125780.2 | 6438 | ENSMUSG00000009863 | Sdhb | succinate dehydrogenase complex, subunit B, iron sulfur (Ip) |
| chr4 | 66038693 | 66039503 | 811 | * | 7,743031 | 5,220264233 | 1,79823 | 2,903001 | 0,003696 | 1 | 0 | Intron (ENSMUST0000( | 4 | 65299040 | 66322720 | 1023681 | 2 | 56079 | ENSMUST00000084496.3 | 283217 | ENSMUSG00000028373 | Astn2 | astrotactin 2 |
| chr7 | 19580703 | 19581807 | 1105 | * | 6,666149 | 5,220739813 | 2,001773 | 2,608057 | 0,009106 | 1 | 0 | Exon (ENSMUST000002 | 7 | 19585928 | 19592275 | 6348 | 1 | 1,01E+08 | ENSMUST00000172815.4 | -4121 | ENSMUSG00000092216 | Gm19345 | predicted gene, 19345 |
| chr15 | 82886115 | 82886610 | 496 | * | 6,366215 | 5,222130293 | 1,831796 | 2,850825 | 0,004361 | 1 | 0 | Intron (ENSMUST0000( | 15 | 82741331 | 82872073 | 130743 | 2 | 21411 | ENSMUST00000229439.2 | -14042 | ENSMUSG00000041852 | Tcf20 | transcription factor 20 |
| chr10 | 21690566 | 21690885 | 320 | * | 4,801901 | 5,222380849 | 1,978648 | 2,639368 | 0,008306 | 1 | 0 | Distal Intergenic | 10 | 21706398 | 21720750 | 14353 | 2 | 628133 | ENSMUST00000180731.2 | 29865 | | Platr6 | pluripotency associated transcript 6 |
| chr8 | 1,19E+08 | 1,19E+08 | 188 | * | 5,023562 | 5,223624239 | 1,981399 | 2,636331 | 0,008381 | 1 | 0 | Distal Intergenic | 8 | 1,19E+08 | 1,19E+08 | 10299 | 2 | 68533 | ENSMUST00000034303.3 | -114111 | ENSMUSG00000031843 | Mphosph6 | M phase phosphoprotein 6 |
| chr19 | 12728221 | 12729070 | 850 | * | 7,093781 | 5,22423666 | 1,891684 | 2,761685 | 0,00575 | 1 | 0 | Distal Intergenic | 19 | 12741024 | 12742996 | 1973 | 2 | 12803 | ENSMUST00000112933.2 | 13926 | ENSMUSG00000079415 | Cntf | ciliary neurotrophic factor |
| chr11 | 61084880 | 61085937 | 1058 | * | 7,100225 | 5,224282682 | 1,89262 | 2,760344 | 0,005774 | 0,799782439 | 1,360622 | Distal Intergenic | 11 | 61098363 | 61103159 | 4797 | 1 | 11670 | ENSMUST00000147792.2 | -12426 | ENSMUSG00000019102 | Aldh3a1 | aldehyde dehydrogenase family 3, subfamily A1 |
| chr4 | 1,32E+08 | 1,32E+08 | 1023 | * | 6,031785 | 5,224285278 | 1,969391 | 2,652742 | 0,007984 | 1 | 0 | Intron (ENSMUST0000( | 4 | 1,32E+08 | 1,32E+08 | 18617 | 1 | 19204 | ENSMUST00000070690.8 | 4928 | ENSMUSG00000056529 | Ptafr | platelet-activating factor receptor |
| chr3 | 57194007 | 57194950 | 944 | * | 7,148191 | 5,225094704 | 1,86856 | 2,796321 | 0,005169 | 1 | 0 | 3' UTR | 3 | 57200240 | 57201124 | 885 | 2 | 17112 | ENSMUST00000196704.2 | 6174 | ENSMUSG00000027800 | Tm4sf1 | transmembrane 4 superfamily member 1 |
| chr13 | 59955707 | 59956689 | 983 | * | 8,825812 | 5,225344482 | 1,768682 | 2,954372 | 0,003133 | 0,690933518 | 1,360622 | Promoter (<=1kb) | 13 | 59948150 | 59957292 | 9143 | 2 | 214290 | ENSMUST00000225241.2 | 603 | ENSMUSG00000035248 | Tut7 | terminal uridylyl transferase 7 |
| chr7 | 35353970 | 35354380 | 411 | * | 6,081631 | 5,226530001 | 1,945318 | 2,686723 | 0,007216 | 1 | 0 | Distal Intergenic | 7 | 35341407 | 35346946 | 5540 | 2 | 56330 | ENSMUST00000118501.8 | -7024 | ENSMUSG00000030417 | Pdcd5 | programmed cell death 5 |
| chr1 | 59512621 | 59513353 | 733 | * | 7,107817 | 5,226850681 | 1,953848 | 2,675157 | 0,007469 | 1 | 0 | Distal Intergenic | 1 | 59521583 | 59526114 | 4532 | 1 | 14369 | ENSMUST00000114246.4 | -8230 | ENSMUSG00000041075 | Fzd7 | frizzled class receptor 7 |
| chr12 | 4813182 | 4814079 | 898 | * | 6,482471 | 5,227105465 | 1,960202 | 2,666616 | 0,007662 | 1 | 0 | Promoter (<=1kb) | 12 | 4789888 | 4812409 | 22522 | 2 | 207921 | ENSMUST00000218905.2 | -773 | ENSMUSG00000050545 | Fam228b | family with sequence similarity 228, member B |
| chr5 | 1,51E+08 | 1,51E+08 | 667 | * | 5,903579 | 5,227425256 | 1,969071 | 2,654767 | 0,007936 | 1 | 0 | Intron (ENSMUST0000( | 5 | 1,51E+08 | 1,51E+08 | 13509 | 1 | 625175 | ENSMUST00000181555.4 | -21463 | ENSMUSG00000097321 | 1700028E10Rik | RIKEN cDNA 1700028E10 gene |
| chr1 | 1,56E+08 | 1,56E+08 | 667 | * | 6,291762 | 5,227946959 | 1,907428 | 2,740836 | 0,006128 | 1 | 0 | Exon (ENSMUST000001 | 1 | 1,56E+08 | 1,56E+08 | 7365 | 2 | 214575 | ENSMUST00000148277.2 | -3088 | ENSMUSG00000060985 | Tdrd5 | tudor domain containing 5 |
| chr19 | 53401707 | 53402639 | 933 | * | 5,412011 | 5,228040512 | 1,987141 | 2,630936 | 0,008515 | 1 | 0 | Distal Intergenic | 19 | 53367827 | 53379009 | 11183 | 2 | 76479 | ENSMUST00000025997.7 | -22698 | ENSMUSG00000025024 | Smndc1 | survival motor neuron domain containing 1 |
| chr11 | 45655317 | 45655977 | 661 | * | 6,596861 | 5,228309326 | 1,994945 | 2,620778 | 0,008773 | 1 | 0 | Distal Intergenic | 11 | 45511198 | 45644439 | 133242 | 2 | 1,05E+08 | ENSMUST00000245312.1 | -10878 | | Gm39816 | predicted gene, 39816 |
| chr5 | 1,48E+08 | 1,48E+08 | 864 | * | 6,017905 | 5,228484726 | 2,000609 | 2,613447 | 0,008963 | 1 | 0 | Intron (ENSMUST0000( | 5 | 1,48E+08 | 1,48E+08 | 31613 | 2 | 24109 | ENSMUST00000201688.4 | -5466 | ENSMUSG00000001687 | Ubl3 | ubiquitin-like 3 |
| chr13 | 70045906 | 70046817 | 912 | * | 4,403692 | 5,22946542 | 2,014472 | 2,595949 | 0,009433 | 1 | 0 | Distal Intergenic | 13 | 70107952 | 70111556 | 3605 | 1 | 67550 | ENSMUST00000220972.2 | -61135 | ENSMUSG00000113802 | 4933416O17Rik | RIKEN cDNA 4933416O17 gene |
| chr18 | 56962787 | 56963993 | 1207 | * | 10,13152 | 5,230081048 | 1,73699 | 3,011003 | 0,002604 | 0,654833721 | 1,449706 | Intron (ENSMUST0000( | 18 | 56908924 | 56945091 | 36168 | 2 | 320253 | ENSMUST00000035278.7 | -17696 | ENSMUSG00000032656 | Marchf3 | membrane associated ring-CH-type finger 3 |
| chr6 | 47435606 | 47436077 | 472 | * | 6,078564 | 5,230933032 | 1,884544 | 2,775702 | 0,005508 | 1 | 0 | Intron (ENSMUST0000( | 6 | 47431322 | 47462111 | 30790 | 1 | 26965 | ENSMUST00000143941.3 | 4284 | ENSMUSG00000029686 | Cul1 | cullin 1 |
| chr5 | 35271197 | 35271529 | 333 | * | 6,14358 | 5,232453308 | 1,837421 | 2,847715 | 0,004403 | 1 | 0 | Distal Intergenic | 5 | 35248845 | 35263110 | 14266 | 2 | 16976 | ENSMUST00000030986.15 | -8087 | ENSMUSG00000029103 | Lrpap1 | low density lipoprotein receptor-related protein associated protein 1 |
| chr19 | 11751967 | 11752469 | 503 | * | 5,302798 | 5,234391742 | 1,899418 | 2,755787 | 0,005855 | 1 | 0 | 3' UTR | 19 | 11747776 | 11752157 | 4382 | 1 | 94063 | ENSMUST00000237728.2 | 4191 | ENSMUSG00000024683 | Mrpl16 | mitochondrial ribosomal protein L16 |
| chr1 | 1,28E+08 | 1,28E+08 | 1113 | * | 7,570784 | 5,235556749 | 1,766416 | 2,963943 | 0,003037 | 1 | 0 | Exon (ENSMUST000001 | 1 | 1,28E+08 | 1,28E+08 | 134413 | 1 | 226412 | ENSMUST00000036288.11 | 7249 | ENSMUSG00000056211 | R3hdm1 | R3H domain containing 1 |
| chr3 | 30947981 | 30948574 | 594 | * | 6,516728 | 5,236178648 | 1,82293 | 2,872397 | 0,004074 | 1 | 0 | 5' UTR | 3 | 30910292 | 30949970 | 39679 | 1 | 71862 | ENSMUST00000147697.2 | 37689 | ENSMUSG00000037661 | Gpr160 | G protein-coupled receptor 160 |
| chr2 | 1,47E+08 | 1,47E+08 | 523 | * | 6,09195 | 5,236834858 | 1,843783 | 2,840266 | 0,004508 | 1 | 0 | Intron (ENSMUST0000( | 2 | 1,47E+08 | 1,47E+08 | 25014 | 1 | 18503 | ENSMUST00000156584.2 | 23068 | ENSMUSG00000037034 | Pax1 | paired box 1 |
| chr5 | 32771977 | 32773437 | 1461 | * | 9,285279 | 5,23747186 | 1,828776 | 2,863922 | 0,004184 | 0,743536224 | 1,360622 | Intron (ENSMUST0000( | 5 | 32768679 | 32844401 | 75723 | 1 | 22612 | ENSMUST00000168707.6 | 3298 | ENSMUSG00000014932 | Yes1 | YES proto-oncogene 1, Src family tyrosine kinase |
| chr16 | 24172968 | 24173696 | 729 | * | 5,036383 | 5,237558911 | 1,933834 | 2,70838 | 0,006761 | 1 | 0 | Intron (ENSMUST0000( | 16 | 24089588 | 24198874 | 109287 | 2 | 1,03E+08 | ENSMUST00000231672.2 | 25178 | | Gm38456 | predicted gene, 38456 |
| chr15 | 83006716 | 83007432 | 717 | * | 7,159246 | 5,237888158 | 1,899497 | 2,757513 | 0,005824 | 1 | 0 | Promoter (<=1kb) | 15 | 82997634 | 83007002 | 9369 | 2 | 74778 | ENSMUST00000018184.10 | 0 | ENSMUSG00000018040 | Rrp7a | ribosomal RNA processing 7 homolog A |
| chr2 | 74838691 | 74839381 | 691 | * | 5,740351 | 5,237975198 | 2,013703 | 2,601166 | 0,009291 | 1 | 0 | Distal Intergenic | 2 | 74955215 | 74968142 | 12928 | 2 | 1,03E+08 | ENSMUST00000143898.2 | 128761 | ENSMUSG00000087333 | Gm13652 | predicted gene 13652 |
| chr10 | 48277246 | 48277780 | 535 | * | 5,211871 | 5,238815975 | 1,990836 | 2,631465 | 0,008502 | 1 | 0 | Distal Intergenic | 10 | 48495408 | 48495491 | 84 | 2 | 1,15E+08 | ENSMUST00000083696.4 | 217711 | | Gm24256 | predicted gene, 24256 |
| chr2 | 1,44E+08 | 1,44E+08 | 506 | * | 6,212621 | 5,239097629 | 1,884076 | 2,780726 | 0,005424 | 1 | 0 | Promoter (1-2kb) | 2 | 1,44E+08 | 1,44E+08 | 1313 | 1 | 228715 | ENSMUST00000136628.2 | 1519 | ENSMUSG00000074754 | Smim26 | small integral membrane protein 26 |
| chr7 | 1,34E+08 | 1,34E+08 | 563 | * | 5,855285 | 5,239233458 | 1,991845 | 2,630342 | 0,00853 | 1 | 0 | Intron (ENSMUST0000( | 7 | 1,34E+08 | 1,35E+08 | 56380 | 2 | 627214 | ENSMUST00000210055.2 | 70422 | ENSMUSG00000073805 | Insyn2a | inhibitory synaptic factor 2A |
| chr11 | 95080721 | 95081483 | 763 | * | 6,132915 | 5,240520963 | 1,970692 | 2,659229 | 0,007832 | 1 | 0 | Distal Intergenic | 11 | 95114116 | 95116458 | 2343 | 2 | 1,03E+08 | ENSMUST00000150818.2 | 34975 | | Gm11515 | predicted gene 11515 |
| chr9 | 63994230 | 63995383 | 1154 | * | 9,302861 | 5,240631355 | 1,441976 | 3,634339 | 0,000279 | 0,282588188 | 1,425929 | Exon (ENSMUST000001 | 9 | 63988043 | 63994243 | 6201 | 1 | 76814 | ENSMUST00000176950.2 | 6187 | | Scarletltr | Scarletltr, erythroid developmental long intergenic non-protein coding transcript |
| chr18 | 56073944 | 56075088 | 1145 | * | 9,484478 | 5,242469425 | 1,716972 | 3,053322 | 0,002263 | 0,642482126 | 1,360622 | Intron (ENSMUST0000( | 18 | 56326071 | 56326205 | 135 | 1 | 1,15E+08 | ENSMUST00000157815.3 | -250983 | ENSMUSG00000088440 | Gm23555 | predicted gene, 23555 |
| chr11 | 70423955 | 70425102 | 1148 | * | 6,82758 | 5,243198915 | 1,931218 | 2,714971 | 0,006628 | 1 | 0 | Promoter (<=1kb) | 11 | 70423058 | 70423765 | 708 | 2 | 1,05E+08 | ENSMUST00000209971.2 | -190 | ENSMUSG00000109833 | Gm40193 | predicted gene, 40193 |
| chr15 | 89316163 | 89316734 | 572 | * | 6,723263 | 5,243411177 | 1,928233 | 2,719283 | 0,006542 | 1 | 0 | Promoter (2-3kb) | 15 | 89311318 | 89314111 | 2794 | 2 | 12651 | ENSMUST00000165623.2 | -2052 | ENSMUSG00000022617 | Chkb | choline kinase beta |
| chr3 | 34763677 | 34764663 | 987 | * | 7,928419 | 5,244380946 | 1,915196 | 2,738301 | 0,006176 | 1 | 0 | Exon (ENSMUST000001 | 3 | 34753789 | 34770809 | 17021 | 2 | 1E+08 | ENSMUST00000196308.5 | 6146 | ENSMUSG00000105153 | Gm3143 | predicted gene 3143 |
| chr10 | 95777756 | 95779173 | 1418 | * | 12,3443 | 5,245320567 | 1,752941 | 2,992296 | 0,002769 | 0,713797407 | 1,075735 | Promoter (1-2kb) | 10 | 95776619 | 95838815 | 62197 | 1 | 216238 | ENSMUST00000218517.2 | 1137 | ENSMUSG00000036499 | Eea1 | early endosome antigen 1 |
| chr4 | 1,23E+08 | 1,23E+08 | 603 | * | 4,50753 | 5,245683286 | 2,013712 | 2,604982 | 0,009188 | 1 | 0 | Intron (ENSMUST0000( | 4 | 1,23E+08 | 1,23E+08 | 14828 | 2 | 12331 | ENSMUST00000140783.2 | 7253 | ENSMUSG00000028656 | Cap1 | CAP, adenylate cyclase-associated protein 1 (yeast) |
| chr18 | 40463468 | 40464026 | 559 | * | 5,241464 | 5,246447701 | 2,001647 | 2,621066 | 0,008766 | 1 | 0 | Intron (ENSMUST0000( | 18 | 40391856 | 40664059 | 272204 | 1 | 383348 | ENSMUST00000235241.2 | 71612 | ENSMUSG00000051401 | Kctd16 | potassium channel tetramerisation domain containing 16 |
| chr9 | 92423609 | 92424604 | 996 | * | 15,0982 | 5,247066577 | 1,683156 | 3,117398 | 0,001825 | 0,594676756 | 1,41696 | Promoter (<=1kb) | 9 | 92424276 | 92490481 | 66206 | 1 | 26432 | ENSMUST00000070522.14 | 0 | ENSMUSG00000032374 | Plod2 | procollagen lysine, 2-oxoglutarate 5-dioxygenase 2 |
| chr9 | 51901913 | 51902584 | 672 | * | 4,819993 | 5,247133795 | 2,019133 | 2,598706 | 0,009358 | 1 | 0 | Distal Intergenic | 9 | 51856280 | 51874856 | 18577 | 2 | 14148 | ENSMUST00000214013.2 | -27057 | ENSMUSG00000032051 | Fdx1 | ferredoxin 1 |
| chr14 | 14202784 | 14203930 | 1147 | * | 7,827994 | 5,247403818 | 1,87818 | 2,793876 | 0,005208 | 0,783008377 | 1,360622 | Distal Intergenic | 14 | 14210420 | 14244262 | 33843 | 1 | 93732 | ENSMUST00000164598.8 | -6490 | ENSMUSG00000021751 | Acox2 | acyl-Coenzyme A oxidase 2, branched chain |
| chr15 | 57575162 | 57575488 | 327 | * | 5,628313 | 5,248504227 | 1,968417 | 2,666358 | 0,007668 | 1 | 0 | Intron (ENSMUST0000( | 15 | 57559125 | 57684506 | 125382 | 1 | 387609 | ENSMUST00000160990.2 | 16037 | ENSMUSG00000071757 | Zhx2 | zinc fingers and homeoboxes 2 |
| chr14 | 64160663 | 64161602 | 940 | * | 5,174197 | 5,248504407 | 1,932367 | 2,716101 | 0,006606 | 1 | 0 | Distal Intergenic | 14 | 64181122 | 64188181 | 7060 | 1 | 20680 | ENSMUST00000079652.7 | -19520 | ENSMUSG00000063060 | Sox7 | SRY (sex determining region Y)-box 7 |
| chrX | 1,4E+08 | 1,4E+08 | 956 | * | 7,035789 | 5,249656289 | 1,950667 | 2,691211 | 0,007119 | 1 | 0 | Intron (ENSMUST0000( | 20 | 1,4E+08 | 1,4E+08 | 308674 | 2 | 94216 | ENSMUST00000101205.3 | 63845 | ENSMUSG00000031273 | Col4a6 | collagen, type IV, alpha 6 |
| chr5 | 31907584 | 31909306 | 1723 | * | 15,282257 | 5,251637491 | 1,1662005 | 4,5032028 | 6,694E-06 | 0,030265668 | 1,214904 | Intron (ENSMUST000002 | 5 | 31942727 | 31977987 | 35261 | 1 | 107976 | ENSMUST00000202270.2 | -33421 | ENSMUSG00000052139 | Babam2 | BRISC and BRCA1 A complex member 2 |
| chrX | 97764575 | 97765764 | 1190 | * | 5,643508 | 5,25191692 | 1,917123 | 2,739479 | 0,006154 | 1 | 0 | Intron (ENSMUST0000( | 20 | 97858381 | 97858487 | 107 | 1 | 1,15E+08 | ENSMUST00000157991.3 | -92617 | ENSMUSG00000088616 | Gm26000 | predicted gene, 26000 |
| chr13 | 23846937 | 23848497 | 1561 | * | 11,89356 | 5,252064701 | 1,363153 | 3,85288 | 0,000117 | 0,253171421 | 0,744375 | Promoter (<=1kb) | 13 | 23847004 | 23847107 | 104 | 1 | 1,15E+08 | ENSMUST00000122804.3 | 0 | ENSMUSG00000084753 | Gm22452 | predicted gene, 22452 |
| chr1 | 92142797 | 92143536 | 740 | * | 6,960745 | 5,252114979 | 1,84035 | 2,853867 | 0,004319 | 1 | 0 | Distal Intergenic | 1 | 91957866 | 92123421 | 165556 | 2 | 208727 | ENSMUST00000212408.2 | -19376 | ENSMUSG00000026313 | Hdac4 | histone deacetylase 4 |
| chr19 | 52811836 | 52812526 | 691 | * | 5,137783 | 5,253453182 | 1,995274 | 2,632948 | 0,008465 | 1 | 0 | Distal Intergenic | 19 | 52911612 | 52915362 | 3751 | 1 | 1E+08 | ENSMUST00000236393.2 | -99086 | ENSMUSG00000117896 | AA387883 | expressed sequence AA387883 |
| chr7 | 99929240 | 99930879 | 1640 | * | 13,30319 | 5,254320749 | 1,321885 | 3,97487 | 7,04E-05 | 0,169105736 | 0,938682 | Distal Intergenic | 7 | 99934727 | 99968906 | 34180 | 1 | 320452 | ENSMUST00000057023.12 | -3848 | ENSMUSG00000051048 | P4ha3 | procollagen-proline, 2-oxoglutarate 4-dioxygenase (proline 4-hydroxylase), alpha polypeptide III |
| chr10 | 26495382 | 26496092 | 711 | * | 5,599708 | 5,255286038 | 1,973474 | 2,662961 | 0,007746 | 1 | 0 | Intron (ENSMUST0000( | 10 | 26549597 | 26550608 | 1012 | 2 | 667572 | ENSMUST00000183230.2 | 54516 | ENSMUSG00000098179 | Gm8709 | glyceraldehyde-3-phosphate dehydrogenase pseudogene |
| chrX | 37527405 | 37528425 | 1021 | * | 5,523751 | 5,256049077 | 1,947569 | 2,698774 | 0,00696 | 1 | 0 | Intron (ENSMUST0000( | 20 | 37518759 | 37532027 | 13269 | 2 | 16784 | ENSMUST00000136817.2 | 3602 | ENSMUSG00000016534 | Lamp2 | lysosomal-associated membrane protein 2 |
| chr18 | 44001783 | 44002906 | 1124 | * | 9,546502 | 5,256840477 | 1,845184 | 2,848952 | 0,004386 | 0,777938745 | 1,173325 | Intron (ENSMUST0000( | 18 | 43973163 | 44003716 | 30554 | 1 | 1,05E+08 | ENSMUST00000237459.2 | 28620 | | Gm41715 | predicted gene, 41715 |
| chr10 | 66018602 | 66019528 | 927 | * | 6,102295 | 5,257085513 | 1,965539 | 2,674628 | 0,007481 | 1 | 0 | Intron (ENSMUST0000( | 10 | 66006180 | 66006285 | 106 | 2 | 1,15E+08 | ENSMUST00000083771.3 | -12317 | ENSMUSG00000065705 | Gm22829 | predicted gene, 22829 |
| chr14 | 27078029 | 27079099 | 1071 | * | 7,842468 | 5,257788708 | 1,880851 | 2,795431 | 0,005183 | 1 | 0 | Intron (ENSMUST0000( | 14 | 27083346 | 27101604 | 18259 | 1 | 71704 | ENSMUST00000225494.2 | -4247 | ENSMUSG00000021895 | Arhgef3 | Rho guanine nucleotide exchange factor (GEF) 3 |
| chr2 | 38704986 | 38705612 | 627 | * | 5,451349 | 5,261253698 | 2,013205 | 2,613372 | 0,008965 | 1 | 0 | Intron (ENSMUST0000( | 2 | 38713269 | 38728612 | 15344 | 1 | 1,03E+08 | ENSMUST00000129089.2 | -7657 | | Nr6a1os | nuclear receptor subfamily 6, group A, member 1, opposite strand |
| chr1 | 13537069 | 13538010 | 942 | * | 8,1346 | 5,262034083 | 1,845419 | 2,851404 | 0,004353 | 0,669491826 | 2,237403 | Distal Intergenic | 1 | 13234725 | 13444307 | 209583 | 2 | 17978 | ENSMUST00000147927.8 | -92762 | ENSMUSG00000005886 | Ncoa2 | nuclear receptor coactivator 2 |
| chr6 | 84831720 | 84832298 | 579 | * | 5,974097 | 5,262547796 | 1,949722 | 2,699127 | 0,006952 | 1 | 0 | 3' UTR | 6 | 84811188 | 85046439 | 235252 | 2 | 75914 | ENSMUST00000046334.15 | 214141 | ENSMUSG00000033769 | Exoc6b | exocyst complex component 6B |
| chr19 | 8769312 | 8770260 | 949 | * | 8,661528 | 5,262561553 | 1,785575 | 2,947264 | 0,003206 | 0,684274806 | 1,449706 | Distal Intergenic | 19 | 8751860 | 8763781 | 11922 | 2 | 225895 | ENSMUST00000176496.8 | -5531 | ENSMUSG00000003680 | Taf6l | TATA-box binding protein associated factor 6 like |
| chr5 | 1,32E+08 | 1,32E+08 | 382 | * | 5,390196 | 5,262884348 | 1,960393 | 2,684606 | 0,007262 | 1 | 0 | Intron (ENSMUST0000( | 5 | 1,31E+08 | 1,32E+08 | 611259 | 2 | 319974 | ENSMUST00000160071.10 | -79247 | ENSMUSG00000029673 | Auts2 | autism susceptibility candidate 2 |

| | | | | | | | | | | | | | | | | | | | | | | | |
|---|---|---|---|---|---|---|---|---|---|---|---|---|---|---|---|---|---|---|---|---|---|---|---|
| chr10 | 93981189 | 93982140 | 952 | * | 8,688757 | 5,264160291 | 1,763825 | 2,984514 | 0,00284 | 1 | 0 | Promoter (1-2kb) | 10 | 93983885 | 94033073 | 49189 | 1 | 22025 | ENSMUST00000105290.9 | -1745 | ENSMUSG00000005897 | Nr2c1 | nuclear receptor subfamily 2, group C, member 1 |
| chr8 | 1,02E+08 | 1,02E+08 | 812 | * | 5,435962 | 5,26500272 | 1,993458 | 2,641141 | 0,008263 | 1 | 0 | Distal Intergenic | 8 | 1,01E+08 | 1,01E+08 | 64447 | 2 | 1,05E+08 | ENSMUST00000210760.2 | -906141 | | Gm39232 | predicted gene, 39232 |
| chr4 | 32544119 | 32544776 | 658 | * | 5,738124 | 5,265786157 | 2,008497 | 2,621754 | 0,008748 | 1 | 0 | Intron (ENSMUST0000( | 4 | 32563097 | 32571617 | 8521 | 2 | 414068 | ENSMUST00000136149.2 | 26841 | ENSMUSG00000086150 | Bach2os | BTB and CNC homology 2, opposite strand |
| chr3 | 1,45E+08 | 1,45E+08 | 454 | * | 5,802155 | 5,266236812 | 2,018177 | 2,609403 | 0,00907 | 1 | 0 | Promoter (1-2kb) | 3 | 1,45E+08 | 1,45E+08 | 9503 | 1 | 52184 | ENSMUST00000196426.5 | -1983 | ENSMUSG00000028256 | Odf2l | outer dense fiber of sperm tails 2-like |
| chr10 | 48594216 | 48595000 | 785 | * | 7,24289 | 5,266422709 | 1,86135 | 2,829356 | 0,004664 | 1 | 0 | Distal Intergenic | 10 | 48495408 | 48495491 | 84 | 2 | 1,15E+08 | ENSMUST00000083696.4 | -98725 | | Gm24256 | predicted gene, 24256 |
| chr7 | 74163583 | 74164440 | 858 | * | 5,809197 | 5,266839262 | 2,030808 | 2,59347 | 0,009501 | 1 | 0 | Intron (ENSMUST0000( | 7 | 74009402 | 74154083 | 144682 | 2 | 108116 | ENSMUST00000136389.2 | -9500 | ENSMUSG00000025790 | Slco3a1 | solute carrier organic anion transporter family, member 3a1 |
| chr1 | 49604207 | 49604778 | 572 | * | 4,854816 | 5,267835648 | 2,042874 | 2,57864 | 0,009919 | 1 | 0 | Distal Intergenic | 1 | 49283775 | 49379602 | 95828 | 1 | 442837 | ENSMUST00000188673.7 | 320432 | ENSMUSG00000051616 | C230029F24Rik | RIKEN cDNA C230029F24 gene |
| chr9 | 63553484 | 63554180 | 697 | * | 6,664494 | 5,268203092 | 1,900483 | 2,772034 | 0,005571 | 1 | 0 | 3' UTR | 9 | 63545935 | 63547610 | 1676 | 1 | 66939 | ENSMUST00000213702.2 | 7549 | ENSMUSG00000037257 | Aagab | alpha- and gamma-adaptin binding protein |
| chr4 | 33248246 | 33248794 | 549 | * | 7,959673 | 5,268445216 | 1,489402 | 3,537288 | 0,000404 | 1 | 0 | Promoter (<=1kb) | 4 | 33245423 | 33248510 | 3088 | 2 | 108767 | ENSMUST00000049357.10 | 0 | ENSMUSG00000040128 | Pnrc1 | proline-rich nuclear receptor coactivator 1 |
| chr7 | 80507517 | 80508505 | 989 | * | 7,48246 | 5,2686986 | 1,793206 | 2,938144 | 0,003302 | 1 | 0 | Promoter (2-3kb) | 7 | 80510668 | 80524717 | 14050 | 1 | 22691 | ENSMUST00000132163.8 | -2163 | ENSMUSG00000038797 | Zscan2 | zinc finger and SCAN domain containing 2 |
| chr12 | 41575736 | 41576503 | 768 | * | 6,677397 | 5,270491653 | 1,931046 | 2,729345 | 0,006346 | 1 | 0 | Intron (ENSMUST0000( | 12 | 41501667 | 41536430 | 34764 | 2 | 16981 | ENSMUST00000043884.6 | -39306 | ENSMUSG00000036295 | Lrrn3 | leucine rich repeat protein 3, neuronal |
| chr2 | 1,04E+08 | 1,04E+08 | 669 | * | 5,703258 | 5,270958629 | 1,907916 | 2,762678 | 0,005733 | 1 | 0 | Distal Intergenic | 2 | 1,04E+08 | 1,04E+08 | 6171 | 2 | 78031 | ENSMUST00000143559.2 | -9397 | ENSMUSG00000086687 | 4930547E08Rik | RIKEN cDNA 4930547E08 gene |
| chr3 | 52629418 | 52630112 | 695 | * | 5,814224 | 5,271391064 | 1,984732 | 2,655971 | 0,007908 | 1 | 0 | Promoter (<=1kb) | 3 | 52630070 | 52630176 | 107 | 2 | 1,15E+08 | ENSMUST00000239592.1 | 64 | ENSMUSG00000119552 | Gm22798 | predicted gene, 22798 |
| chr11 | 1,02E+08 | 1,02E+08 | 617 | * | 5,633911 | 5,272960731 | 1,956719 | 2,694797 | 0,007043 | 1 | 0 | Distal Intergenic | 11 | 1,02E+08 | 1,02E+08 | 5209 | 1 | 75437 | ENSMUST00000010985.8 | 8908 | ENSMUSG00000010841 | Cfap97d1 | CFAP97 domain containing 1 |
| chr13 | 58036097 | 58036611 | 515 | * | 5,894992 | 5,273130283 | 2,036167 | 2,589734 | 0,009605 | 1 | 0 | Intron (ENSMUST0000( | 13 | 57577108 | 58055401 | 478294 | 2 | 20745 | ENSMUST00000172326.3 | 18790 | ENSMUSG00000056222 | Spock1 | sparc/osteonectin, cwcv and kazal-like domains proteoglycan 1 |
| chr14 | 79627437 | 79628649 | 1213 | * | 9,24113 | 5,273738743 | 1,457491 | 3,618369 | 0,000296 | 0,308866996 | 1,211315 | Promoter (<=1kb) | 14 | 79562709 | 79628218 | 65510 | 2 | 66897 | ENSMUST00000022597.15 | 0 | ENSMUSG00000022020 | Naa16 | N(alpha)-acetyltransferase 16, NatA auxiliary subunit |
| chr15 | 7428231 | 7428860 | 630 | * | 5,510603 | 5,27430176 | 2,00219 | 2,634266 | 0,008432 | 1 | 0 | Promoter (<=1kb) | 15 | 7236064 | 7427876 | 191813 | 2 | 268780 | ENSMUST00000160207.8 | -355 | ENSMUSG00000042961 | Egflam | EGF-like, fibronectin type III and laminin G domains |
| chr4 | 1,33E+08 | 1,33E+08 | 641 | * | 5,330599 | 5,274346673 | 2,00127 | 2,635499 | 0,008401 | 1 | 0 | Distal Intergenic | 4 | 1,33E+08 | 1,33E+08 | 69252 | 1 | 242687 | ENSMUST00000084241.12 | -15123 | ENSMUSG00000028868 | Wasf2 | WASP family, member 2 |
| chr12 | 70055531 | 70056807 | 1277 | * | 11,32995 | 5,274347734 | 1,451092 | 3,634744 | 0,000278 | 0,337696055 | 0,918255 | Distal Intergenic | 12 | 70041564 | 70041666 | 103 | 2 | 1,15E+08 | ENSMUST00000158528.4 | -13865 | ENSMUSG00002075853 | Gm23579 | predicted gene, 23579 |
| chr13 | 52487255 | 52488295 | 1041 | * | 8,104361 | 5,274563562 | 1,81132 | 2,912 | 0,003591 | 0,707056607 | 1,449706 | Distal Intergenic | 13 | 52334447 | 52335673 | 1227 | 1 | 1,19E+08 | ENSMUST00000222876.2 | 152808 | | Gm48199 | predicted gene, 48199 |
| chr10 | 7468420 | 7469322 | 903 | * | 6,183929 | 5,275399208 | 1,973196 | 2,673531 | 0,007506 | 1 | 0 | Promoter (2-3kb) | 10 | 7465891 | 7499786 | 33896 | 1 | 237253 | ENSMUST00000135907.8 | 2529 | ENSMUSG00000019796 | Lrp11 | low density lipoprotein receptor-related protein 11 |
| chr8 | 1,06E+08 | 1,06E+08 | 636 | * | 7,213847 | 5,276019914 | 1,793055 | 2,942475 | 0,003256 | 0,698133821 | 1,360622 | Promoter (2-3kb) | 8 | 1,06E+08 | 1,06E+08 | 1355 | 2 | 234686 | ENSMUST00000136439.2 | -2634 | ENSMUSG00000014778 | Fhod1 | formin homology 2 domain containing 1 |
| chr5 | 1,37E+08 | 1,37E+08 | 893 | * | 6,071563 | 5,276144117 | 1,954927 | 2,698896 | 0,006957 | 1 | 0 | Intron (ENSMUST0000( | 5 | 1,37E+08 | 1,37E+08 | 2001 | 2 | 22635 | ENSMUST00000150470.2 | 26815 | ENSMUSG00000079173 | Zan | zonadhesin |
| chr10 | 40765295 | 40766251 | 957 | * | 5,986933 | 5,276686611 | 1,942439 | 2,716526 | 0,006597 | 1 | 0 | Intron (ENSMUST0000( | 10 | 40759823 | 40814566 | 54744 | 1 | 83767 | ENSMUST00000105509.2 | 5472 | ENSMUSG00000019831 | Wasf1 | WASP family, member 1 |
| chr2 | 1,31E+08 | 1,31E+08 | 788 | * | 6,381406 | 5,278447467 | 1,904967 | 2,770886 | 0,00559 | 1 | 0 | Distal Intergenic | 2 | 1,31E+08 | 1,31E+08 | 40336 | 2 | 51902 | ENSMUST00000142982.2 | -19939 | ENSMUSG00000048911 | Rnf24 | ring finger protein 24 |
| chrX | 52363194 | 52363942 | 749 | * | 4,555747 | 5,278706959 | 2,000057 | 2,639278 | 0,008308 | 1 | 0 | Promoter (<=1kb) | 20 | 52362301 | 52420062 | 57762 | 1 | 73866 | ENSMUST00000124137.8 | 893 | ENSMUSG00000036013 | Fam122c | family with sequence similarity 122, member C |
| chr3 | 88914746 | 88915982 | 1237 | * | 7,933417 | 5,279161971 | 1,745804 | 3,023914 | 0,002495 | 1 | 0 | Exon (ENSMUST000001 | 3 | 88930490 | 88935575 | 5086 | 1 | 192195 | ENSMUST00000198846.2 | -14508 | ENSMUSG00000028053 | Ash1l | ASH1 like histone lysine methyltransferase |
| chr4 | 1,08E+08 | 1,08E+08 | 757 | * | 11,07901 | 5,280120468 | 1,77453 | 2,975503 | 0,002925 | 0,7893429 | 0,738441 | Promoter (<=1kb) | 4 | 1,08E+08 | 1,08E+08 | 13405 | 1 | 69893 | ENSMUST00000131656.2 | -555 | ENSMUSG00000048351 | Coa7 | cytochrome c oxidase assembly factor 7 |
| chr4 | 41805113 | 41805995 | 883 | * | 7,575987 | 5,280731131 | 1,862854 | 2,834752 | 0,004586 | 1 | 0 | Distal Intergenic | 4 | 41791989 | 41793601 | 1613 | 2 | 1E+08 | ENSMUST00000180219.2 | -11512 | | Gm12407 | predicted gene 12407 |
| chr17 | 42062675 | 42063336 | 662 | * | 4,956529 | 5,28219687 | 2,047775 | 2,579481 | 0,009895 | 1 | 0 | Distal Intergenic | 17 | 42626838 | 42815968 | 189131 | 1 | 627626 | ENSMUST00000048691.6 | -563502 | ENSMUSG00000042256 | Ptchd4 | patched domain containing 4 |
| chr14 | 1,16E+08 | 1,16E+08 | 768 | * | 4,995829 | 5,28394667 | 2,019409 | 2,61658 | 0,008882 | 1 | 0 | Intron (ENSMUST0000( | 14 | 1,16E+08 | 1,16E+08 | 13002 | 2 | 75088 | ENSMUST00000176582.2 | -351650 | ENSMUSG00000093551 | 4930505G20Rik | RIKEN cDNA 4930505G20 gene |
| chr7 | 26929601 | 26930672 | 1072 | * | 8,741582 | 5,284439862 | 1,74681 | 3,025194 | 0,002485 | 0,658200216 | 1,360622 | Promoter (1-2kb) | 7 | 26906597 | 26928053 | 21457 | 2 | 233011 | ENSMUST00000003850.8 | -1548 | ENSMUSG00000003752 | Itpkc | inositol 1,4,5-trisphosphate 3-kinase C |
| chr7 | 27538037 | 27539027 | 991 | * | 9,634945 | 5,284694944 | 1,703551 | 3,102163 | 0,001921 | 0,59811801 | 1,449706 | Promoter (<=1kb) | 7 | 27538032 | 27555863 | 17832 | 1 | 22717 | ENSMUST00000205701.2 | 5 | ENSMUSG00000078779 | Zfp59 | zinc finger protein 59 |
| chr10 | 80947397 | 80948404 | 1008 | * | 5,879363 | 5,28677996 | 1,954532 | 2,704883 | 0,006833 | 1 | 0 | Intron (ENSMUST0000( | 10 | 80942179 | 80954959 | 12781 | 1 | 26396 | ENSMUST00000136743.2 | 5218 | ENSMUSG00000035027 | Map2k2 | mitogen-activated protein kinase kinase 2 |
| chr9 | 1,09E+08 | 1,09E+08 | 445 | * | 5,025433 | 5,2881973 | 1,922899 | 2,750117 | 0,005957 | 1 | 0 | 3' UTR | 9 | 1,09E+08 | 1,09E+08 | 13995 | 2 | 211305 | ENSMUST00000199102.2 | 11985 | ENSMUSG00000049314 | Fbxw13 | F-box and WD-40 domain protein 13 |
| chr5 | 1,07E+08 | 1,07E+08 | 750 | * | 5,385197 | 5,290930975 | 2,040465 | 2,593003 | 0,009514 | 1 | 0 | Promoter (<=1kb) | 5 | 1,07E+08 | 1,07E+08 | 183060 | 2 | 21814 | ENSMUST00000031224.15 | -255 | ENSMUSG00000029287 | Tgfbr3 | transforming growth factor, beta receptor III |
| chr5 | 1,23E+08 | 1,23E+08 | 1219 | * | 8,295114 | 5,291632377 | 1,747894 | 3,027433 | 0,002466 | 0,649217039 | 1,425929 | Intron (ENSMUST0000( | 5 | 1,23E+08 | 1,23E+08 | 2454 | 1 | 208043 | ENSMUST00000173221.2 | -7280 | ENSMUSG00000038384 | Setd1b | SET domain containing 1B |
| chr7 | 74613100 | 74614106 | 1007 | * | 6,759418 | 5,292515929 | 1,82812 | 2,895059 | 0,003791 | 1 | 0 | Distal Intergenic | 7 | 74679942 | 74707910 | 27969 | 1 | 75205 | ENSMUST00000206433.2 | -65836 | ENSMUSG00000108803 | 4930533N22Rik | RIKEN cDNA 4930533N22 gene |
| chr7 | 45320945 | 45321768 | 824 | * | 7,582823 | 5,292828461 | 1,821252 | 2,906149 | 0,003659 | 1 | 0 | Distal Intergenic | 7 | 45298015 | 45315818 | 17804 | 2 | 14344 | ENSMUST00000069800.6 | -5127 | ENSMUSG00000055978 | Fut2 | fucosyltransferase 2 |
| chr1 | 1,34E+08 | 1,34E+08 | 961 | * | 5,738794 | 5,293374918 | 1,921772 | 2,754423 | 0,00588 | 1 | 0 | Promoter (2-3kb) | 1 | 1,34E+08 | 1,34E+08 | 3951 | 2 | 12227 | ENSMUST00000020692.7 | -2131 | ENSMUSG00000020423 | Btg2 | BTG anti-proliferation factor 2 |
| chr4 | 1,29E+08 | 1,29E+08 | 708 | * | 5,343337 | 5,29385035 | 2,009953 | 2,633818 | 0,008443 | 1 | 0 | Promoter (1-2kb) | 4 | 1,29E+08 | 1,29E+08 | 7897 | 2 | 67349 | ENSMUST00000149472.4 | 1106 | ENSMUSG00000085085 | 1700086P04Rik | RIKEN cDNA 1700086P04 gene |
| chr6 | 1,37E+08 | 1,37E+08 | 848 | * | 8,800361 | 5,297744825 | 1,847585 | 2,867389 | 0,004139 | 0,660341484 | 2,237403 | Exon (ENSMUST00000( | 6 | 1,37E+08 | 1,37E+08 | 2609 | 2 | 13860 | ENSMUST00000152046.2 | 29080 | ENSMUSG00000015766 | Eps8 | epidermal growth factor receptor pathway substrate 8 |
| chr9 | 55306017 | 55306668 | 652 | * | 7,276622 | 5,298149125 | 1,951564 | 2,714821 | 0,006631 | 1 | 0 | Intron (ENSMUST0000( | 9 | 55240485 | 55344642 | 104158 | 1 | 244886 | ENSMUST00000130465.2 | 65532 | ENSMUSG00000032313 | Tmem266 | transmembrane protein 266 |
| chr15 | 76188657 | 76189378 | 722 | * | 7,11686 | 5,298653502 | 1,957956 | 2,706217 | 0,006805 | 1 | 0 | Promoter (1-2kb) | 15 | 76186264 | 76187400 | 1137 | 2 | 75475 | ENSMUST00000163977.2 | -1257 | ENSMUSG00000022562 | Oplah | 5-oxoprolinase (ATP-hydrolysing) |
| chr3 | 1,04E+08 | 1,04E+08 | 522 | * | 4,720904 | 5,302327919 | 2,026006 | 2,617133 | 0,008867 | 1 | 0 | Promoter (1-2kb) | 3 | 1,04E+08 | 1,04E+08 | 21848 | 1 | 229672 | ENSMUST00000106822.2 | 1092 | ENSMUSG00000044165 | Bcl2l15 | BCLI2-like 15 |
| chr14 | 56028500 | 56029411 | 912 | * | 5,807328 | 5,302649561 | 2,013529 | 2,633511 | 0,008451 | 1 | 0 | Promoter (2-3kb) | 14 | 56022452 | 56026322 | 3871 | 2 | 56532 | ENSMUST00000022830.14 | -2178 | ENSMUSG00000022221 | Ripk3 | receptor-interacting serine-threonine kinase 3 |
| chr13 | 41759824 | 41760593 | 770 | * | 5,684675 | 5,304033364 | 1,902266 | 2,788272 | 0,005299 | 1 | 0 | Promoter (<=1kb) | 13 | 41759692 | 41794831 | 35140 | 1 | 621976 | ENSMUST00000129449.3 | 132 | ENSMUSG00000087370 | Tmem170b | transmembrane protein 170B |
| chr1 | 1,9E+08 | 1,9E+08 | 864 | * | 5,439251 | 5,3041689 | 2,037603 | 2,603141 | 0,009237 | 1 | 0 | Intron (ENSMUST0000( | 1 | 1,9E+08 | 1,9E+08 | 52680 | 2 | 19130 | ENSMUST00000175916.8 | -27216 | ENSMUSG00000010175 | Prox1 | prospero homeobox 1 |
| chr14 | 30306730 | 30307559 | 830 | * | 9,459126 | 5,304934016 | 1,753895 | 3,024658 | 0,002489 | 0,658408085 | 1,360622 | Distal Intergenic | 14 | 30318611 | 30319971 | 1361 | 1 | 1,15E+08 | ENSMUST00000246455.1 | -11052 | | Gm52147 | predicted gene, 52147 |
| chr15 | 79840193 | 79840902 | 710 | * | 6,025401 | 5,305215621 | 1,923651 | 2,757888 | 0,005818 | 1 | 0 | Intron (ENSMUST0000( | 15 | 79802650 | 79855319 | 52670 | 2 | 52609 | ENSMUST00000132821.8 | 14417 | ENSMUSG00000053411 | Cbx7 | chromobox 7 |
| chr11 | 39234998 | 39235697 | 700 | * | 5,4182 | 5,305310727 | 1,871542 | 2,834727 | 0,004586 | 1 | 0 | Distal Intergenic | 11 | 39721483 | 39721627 | 145 | 1 | 1,15E+08 | ENSMUST00000157978.4 | -485786 | ENSMUSG00000088603 | Gm24192 | predicted gene, 24192 |
| chr16 | 50252519 | 50253374 | 856 | * | 6,933559 | 5,306291238 | 1,907855 | 2,781286 | 0,005414 | 1 | 0 | Promoter (<=1kb) | 16 | 50012207 | 50252753 | 240547 | 2 | 70508 | ENSMUST00000138166.8 | 0 | ENSMUSG00000022641 | Bbx | bobby sox HMG box containing |
| chr17 | 74962223 | 74963286 | 1064 | * | 6,267803 | 5,307742359 | 1,867244 | 2,842555 | 0,004475 | 1 | 0 | Promoter (<=1kb) | 17 | 74961384 | 75010342 | 48959 | 1 | 12211 | ENSMUST00000183249.2 | 839 | ENSMUSG00000024073 | Birc6 | baculoviral IAP repeat-containing 6 |
| chr14 | 22031621 | 22033267 | 1647 | * | 15,65693 | 5,308516999 | 1,299858 | 4,083922 | 4,43E-05 | 0,096916044 | 1,41696 | Distal Intergenic | 14 | 22034027 | 22039669 | 5643 | 2 | 218820 | ENSMUST00000043409.9 | 6402 | ENSMUSG00000039081 | Zfp503 | zinc finger protein 503 |
| chr7 | 1,17E+08 | 1,17E+08 | 375 | * | 4,58636 | 5,308819373 | 2,052107 | 2,58701 | 0,009681 | 1 | 0 | Distal Intergenic | 7 | 1,17E+08 | 1,17E+08 | 107 | 2 | 1,15E+08 | ENSMUST00000240319.1 | -46748 | ENSMUSG00000119801 | Gm26054 | predicted gene, 26054 |
| chr14 | 1,21E+08 | 1,21E+08 | 559 | * | 5,715225 | 5,308819889 | 2,05177 | 2,587434 | 0,009669 | 1 | 0 | Intron (ENSMUST0000( | 14 | 1,21E+08 | 1,22E+08 | 13014 | 1 | 223254 | ENSMUST00000137971.2 | -27501 | ENSMUSG00000025555 | Farp1 | FERM, RhoGEF (Arhgef) and pleckstrin domain protein 1 (chondrocyte-derived) |
| chr12 | 1,15E+08 | 1,15E+08 | 505 | * | 4,7292 | 5,30887572 | 2,047771 | 2,592514 | 0,009528 | 1 | 0 | Distal Intergenic | 12 | 1,15E+08 | 1,15E+08 | 25225 | 1 | 1,03E+08 | ENSMUST00000195683.2 | -308634 | ENSMUSG00000103168 | Gm30948 | predicted gene, 30948 |
| chrX | 13964170 | 13964888 | 719 | * | 4,914405 | 5,3090078 | 2,039734 | 2,602795 | 0,009247 | 1 | 0 | Distal Intergenic | 20 | 14077387 | 14077900 | 514 | 1 | 385328 | ENSMUST00000105137.4 | -112499 | ENSMUSG00000078346 | H2al1n | H2A histone family member L1N |
| chr11 | 76740123 | 76740592 | 470 | * | 4,992162 | 5,309018967 | 2,037708 | 2,605387 | 0,009177 | 1 | 0 | Promoter (2-3kb) | 11 | 76669250 | 76737844 | 68595 | 2 | 12874 | ENSMUST00000021201.6 | -2279 | ENSMUSG00000020841 | Cpd | carboxypeptidase D |
| chr13 | 67655772 | 67656336 | 565 | * | 6,005414 | 5,309266221 | 2,021026 | 2,627015 | 0,008614 | 1 | 0 | Intron (ENSMUST0000( | 13 | 67638287 | 67648674 | 10388 | 2 | 238693 | ENSMUST00000076123.12 | -7098 | ENSMUSG00000071291 | Zfp58 | zinc finger protein 58 |
| chr7 | 1,15E+08 | 1,15E+08 | 657 | * | 5,089465 | 5,309369245 | 2,015169 | 2,634702 | 0,008421 | 1 | 0 | Distal Intergenic | 7 | 1,15E+08 | 1,15E+08 | 35311 | 1 | 101786 | ENSMUST00000205913.2 | -37021 | ENSMUSG00000108379 | A730082K24Rik | RIKEN cDNA A730082K24 gene |
| chr8 | 1,29E+08 | 1,29E+08 | 695 | * | 5,446718 | 5,309552038 | 2,003459 | 2,650193 | 0,008045 | 1 | 0 | Distal Intergenic | 8 | 1,29E+08 | 1,29E+08 | 78 | 2 | 1,02E+08 | ENSMUST00000184910.3 | 166373 | ENSMUSG00000099326 | Mir21c | microRNA 21c |
| chr1 | 1,65E+08 | 1,65E+08 | 562 | * | 5,510377 | 5,309902506 | 1,98293 | 2,677806 | 0,007411 | 1 | 0 | Exon (ENSMUST000002 | 1 | 1,65E+08 | 1,65E+08 | 8306 | 1 | 433374 | ENSMUST00000191735.6 | -4260 | ENSMUSG00000104117 | Gm20743 | predicted gene, 20743 |
| chr16 | 24227516 | 24228453 | 938 | * | 5,003456 | 5,310067173 | 1,952892 | 2,719078 | 0,006546 | 1 | 0 | Promoter (2-3kb) | 16 | 24225105 | 24225167 | 63 | 2 | 1,15E+08 | ENSMUST00000157236.3 | -2349 | ENSMUSG00000087861 | Gm24440 | predicted gene, 24440 |
| chrX | 1,67E+08 | 1,67E+08 | 946 | * | 5,3919 | 5,310365815 | 2,013117 | 2,637882 | 0,008343 | 1 | 0 | Intron (ENSMUST0000( | 20 | 1,66E+08 | 1,67E+08 | 426246 | 2 | 333605 | ENSMUST00000112145.3 | -100299 | ENSMUSG00000049176 | Frmpd4 | FERM and PDZ domain containing 4 |
| chr11 | 53939321 | 53940894 | 1574 | * | 12,22584 | 5,311111669 | 1,759679 | 3,018227 | 0,002543 | 0,671251031 | 1,2763 | Distal Intergenic | 11 | 53954549 | 53955394 | 846 | 2 | 30794 | ENSMUST00000151948.2 | 14500 | ENSMUSG00000020388 | Pdlim4 | PDZ and LIM domain 4 |
| chr3 | 1,38E+08 | 1,38E+08 | 1001 | * | 6,427187 | 5,311486192 | 1,903944 | 2,789728 | 0,005275 | 1 | 0 | Distal Intergenic | 3 | 1,38E+08 | 1,38E+08 | 12407 | 1 | 11532 | ENSMUST00000005964.7 | -4988 | ENSMUSG00000028138 | Adh5 | alcohol dehydrogenase 5 (class III), chi polypeptide |
| chr9 | 61608630 | 61609617 | 988 | * | 6,908503 | 5,312027291 | 1,881264 | 2,823649 | 0,004748 | 0,765231686 | 1,360622 | Distal Intergenic | 9 | 61417005 | 61441529 | 24525 | 1 | 1,03E+08 | ENSMUST00000214104.2 | 191625 | ENSMUSG00000111381 | Gm34424 | predicted gene, 34424 |
| chr15 | 82318263 | 82319159 | 897 | * | 7,491181 | 5,312328875 | 1,869598 | 2,841428 | 0,004491 | 1 | 0 | Promoter (<=1kb) | 15 | 82317638 | 82338583 | 20946 | 1 | 13105 | ENSMUST00000229793.2 | 625 | ENSMUSG00000068086 | Cyp2d9 | cytochrome P450, family 2, subfamily d, polypeptide 9 |
| chr11 | 1,03E+08 | 1,03E+08 | 1313 | * | 9,490814 | 5,31292688 | 1,718839 | 3,090997 | 0,001995 | 0,641205124 | 1,211315 | Distal Intergenic | 11 | 1,03E+08 | 1,03E+08 | 53889 | 2 | 237954 | ENSMUST00000153273.3 | -10647 | ENSMUSG00000078632 | Lrrc37a | leucine rich repeat containing 37A |
| chrX | 1,51E+08 | 1,51E+08 | 996 | * | 9,67939 | 5,315963437 | 1,778004 | 2,98985 | 0,002791 | 0,691026033 | 1,211315 | Distal Intergenic | 20 | 1,51E+08 | 1,51E+08 | 81040 | 1 | 245666 | ENSMUST00000168786.9 | -26898 | ENSMUSG00000041115 | Iqsec2 | IQ motif and Sec7 domain 2 |
| chr11 | 1,02E+08 | 1,02E+08 | 935 | * | 5,663941 | 5,316001312 | 1,958618 | 2,714159 | 0,006644 | 1 | 0 | Intron (ENSMUST0000( | 11 | 1,02E+08 | 1,02E+08 | 14112 | 1 | 52685 | ENSMUST00000107164.3 | 3620 | ENSMUSG00000017309 | Cd300lg | CD300 molecule like family member G |
| chr4 | 1,35E+08 | 1,35E+08 | 926 | * | 6,400991 | 5,318200031 | 1,950931 | 2,725981 | 0,006411 | 1 | 0 | Intron (ENSMUST0000( | 4 | 1,35E+08 | 1,35E+08 | 10497 | 1 | 19746 | ENSMUST00000136163.2 | -5211 | ENSMUSG00000028825 | Rhd | Rh blood group, D antigen |
| chrX | 1,5E+08 | 1,5E+08 | 643 | * | 5,627531 | 5,318560556 | 1,909131 | 2,785855 | 0,005339 | 1 | 0 | Promoter (2-3kb) | 20 | 1,5E+08 | 1,5E+08 | 2286 | 1 | 14163 | ENSMUST00000124882.2 | -2583 | ENSMUSG00000025265 | Fgd1 | FYVE, RhoGEF and PH domain containing 1 |
| chr4 | 56801437 | 56803040 | 1604 | * | 15,150472 | 5,322944795 | 1,1713996 | 4,5440895 | 5,517E-06 | 0,026722594 | 1,214904 | Promoter (<=1kb) | 4 | 56749680 | 56802331 | 52652 | 2 | 230233 | ENSMUST00000030140.3 | 0 | ENSMUSG00000028431 | Elp1 | elongator complex protein 1 |
| chrX | 1,46E+08 | 1,46E+08 | 650 | * | 4,865243 | 5,323176475 | 1,988781 | 2,676603 | 0,007437 | 1 | 0 | Distal Intergenic | 20 | 1,46E+08 | 1,46E+08 | 129 | 2 | 1,15E+08 | ENSMUST00000122472.3 | 26703 | ENSMUSG00000084421 | Gm25107 | predicted gene, 25107 |
| chr18 | 8656113 | 8656428 | 316 | * | 4,678372 | 5,323653092 | 1,979142 | 2,68988 | 0,007148 | 1 | 0 | Distal Intergenic | 18 | 9212163 | 9218136 | 5974 | 1 | 14370 | ENSMUST00000041080.7 | -555735 | ENSMUSG00000036904 | Fzd8 | frizzled class receptor 8 |
| chr4 | 1,36E+08 | 1,36E+08 | 524 | * | 5,460101 | 5,326097566 | 1,928267 | 2,762117 | 0,005743 | 1 | 0 | Distal Intergenic | 4 | 1,36E+08 | 1,36E+08 | 10069 | 1 | 14105 | ENSMUST00000097844.9 | -16104 | ENSMUSG00000028676 | Srsf10 | serine and arginine-rich splicing factor 10 |
| chr6 | 5381496 | 5382283 | 788 | * | 8,974756 | 5,328465355 | 1,907694 | 2,793145 | 0,00522 | 0,775744126 | 1,425929 | Promoter (1-2kb) | 6 | 5383386 | 5433022 | 49637 | 1 | 65255 | ENSMUST00000043294.12 | -1103 | ENSMUSG00000042607 | Asb4 | ankyrin repeat and SOCS box-containing 4 |
| chr7 | 1,27E+08 | 1,27E+08 | 1029 | * | 8,021094 | 5,328837841 | 1,7986 | 2,96277 | 0,003049 | 1 | 0 | Distal Intergenic | 7 | 1,27E+08 | 1,27E+08 | 7042 | 2 | 320683 | ENSMUST00000132524.8 | -27625 | ENSMUSG00000045639 | Zfp629 | zinc finger protein 629 |
| chr18 | 55397944 | 55398348 | 405 | * | 4,77905 | 5,32889294 | 2,022087 | 2,635342 | 0,008405 | 1 | 0 | Distal Intergenic | 18 | 55190657 | 55212486 | 21830 | 1 | 1E+08 | ENSMUST00000181765.2 | 207287 | | Gm4221 | predicted gene 4221 |
| chr15 | 6277703 | 6279111 | 1409 | * | 8,244375 | 5,330460692 | 1,778764 | 2,996722 | 0,002729 | 0,663408464 | 1,449706 | Distal Intergenic | 15 | 6329269 | 6465001 | 135733 | 1 | 13132 | ENSMUST00000159552.2 | -50158 | ENSMUSG00000022150 | Dab2 | disabled 2, mitogen-responsive phosphoprotein |
| chr1 | 1,66E+08 | 1,66E+08 | 843 | * | 8,121523 | 5,331050348 | 1,930603 | 2,761339 | 0,005756 | 0,713797407 | 2,237403 | Distal Intergenic | 1 | 1,66E+08 | 1,66E+08 | 137525 | 2 | 18986 | ENSMUST00000160260.9 | -59105 | ENSMUSG00000026565 | Pou2f1 | POU domain, class 2, transcription factor 1 |
| chr18 | 16950630 | 16951640 | 1011 | * | 8,425175 | 5,33115927 | 1,875735 | 2,84217 | 0,004481 | 1 | 0 | Promoter (1-2kb) | 18 | 16949464 | 16955840 | 6377 | 1 | 328906 | ENSMUST00000144006.2 | 1166 | ENSMUSG00000086095 | Gm15328 | predicted gene 15328 |
| chr7 | 1,28E+08 | 1,28E+08 | 747 | * | 6,648323 | 5,331293844 | 1,826802 | 2,918376 | 0,003519 | 1 | 0 | Intron (ENSMUST0000( | 7 | 1,28E+08 | 1,28E+08 | 18470 | 1 | 101490 | ENSMUST00000129085.8 | -23084 | ENSMUSG00000042105 | Inpp5f | inositol polyphosphate-5-phosphatase F |
| chr13 | 65172375 | 65173122 | 748 | * | 7,328176 | 5,332511543 | 1,804304 | 2,955439 | 0,003122 | 1 | 0 | Distal Intergenic | 13 | 65180872 | 65185818 | 4947 | 1 | 238683 | ENSMUST00000099427.3 | -7750 | ENSMUSG00000074849 | Spata31d1c | spermatogenesis associated 31 subfamily D, member 1C |
| chr5 | 1,32E+08 | 1,32E+08 | 644 | * | 4,344604 | 5,334094534 | 2,055702 | 2,594779 | 0,009465 | 1 | 0 | Intron (ENSMUST0000( | 5 | 1,31E+08 | 1,32E+08 | 178819 | 2 | 319974 | ENSMUST00000161804.9 | -39232 | ENSMUSG00000029673 | Auts2 | autism susceptibility candidate 2 |
| chr3 | 87600486 | 87601139 | 654 | * | 7,527312 | 5,334411335 | 1,770274 | 3,013326 | 0,002584 | 1 | 0 | Intron (ENSMUST0000( | 3 | 87646287 | 87648355 | 2069 | 2 | 74485 | ENSMUST00000174581.2 | 47216 | ENSMUSG00000023084 | Lrrc71 | leucine rich repeat containing 71 |
| chr8 | 1,07E+08 | 1,07E+08 | 929 | * | 6,589166 | 5,334550564 | 2,052945 | 2,598487 | 0,009364 | 1 | 0 | Distal Intergenic | 8 | 1,07E+08 | 1,07E+08 | 10331 | 1 | 22751 | ENSMUST00000212874.2 | 11850 | ENSMUSG00000031907 | Zfp90 | zinc finger protein 90 |
| chr9 | 71574578 | 71575925 | 1348 | * | 8,190956 | 5,335522827 | 1,975645 | 2,700649 | 0,00692 | 0,744477896 | 2,237403 | Intron (ENSMUST0000( | 9 | 71539915 | 71555632 | 15718 | 2 | 68178 | ENSMUST00000146567.2 | -18946 | ENSMUSG00000032232 | Cgnl1 | cingulin-like 1 |
| chr9 | 23619190 | 23619928 | 739 | * | 8,001039 | 5,336039481 | 1,742355 | 3,062544 | 0,002195 | 1 | 0 | Distal Intergenic | 9 | 23285228 | 23318127 | 32900 | 1 | 73230 | ENSMUST00000214050.2 | 333962 | ENSMUSG00000031963 | Bmper | BMP-binding endothelial regulator |
| chr5 | 1,09E+08 | 1,09E+08 | 806 | * | 7,028416 | 5,336791562 | 1,989768 | 2,682118 | 0,007316 | 1 | 0 | Intron (ENSMUST0000( | 5 | 1,09E+08 | 1,09E+08 | 17316 | 2 | 435864 | ENSMUST00000233835.2 | -17645 | ENSMUSG00000091624 | Vmn2r9 | vomeronasal 2, receptor 9 |
| chr4 | 1,32E+08 | 1,32E+08 | 1134 | * | 7,056651 | 5,337326324 | 1,995873 | 2,674181 | 0,007491 | 0,847125405 | 1,360622 | Promoter (1-2kb) | 4 | 1,32E+08 | 1,32E+08 | 8589 | 1 | 68598 | ENSMUST00000105938.4 | 1804 | ENSMUSG00000054405 | Dnajc8 | DnaJ heat shock protein family (Hsp40) member C8 |
| chr2 | 47307772 | 47308169 | 398 | * | 5,655015 | 5,3373751 | 2,001996 | 2,666027 | 0,007675 | 1 | 0 | Distal Intergenic | 2 | 48274231 | 48274350 | 120 | 2 | 1,15E+08 | ENSMUST00000116849.3 | 966181 | ENSMUSG00000080499 | Gm25338 | predicted gene, 25338 |
| chr7 | 1,1E+08 | 1,1E+08 | 760 | * | 6,73441 | 5,337925078 | 2,003029 | 2,664927 | 0,007701 | 1 | 0 | Promoter (<=1kb) | 7 | 1,1E+08 | 1,1E+08 | 13078 | 2 | 1,15E+08 | ENSMUST00000186888.2 | 563 | | Gm28863 | predicted gene 28863 |
| chr1 | 1,23E+08 | 1,23E+08 | 469 | * | 6,944339 | 5,338370488 | 2,008366 | 2,658067 | 0,007859 | 1 | 0 | Distal Intergenic | 1 | 1,23E+08 | 1,23E+08 | 131 | 2 | 1,15E+08 | ENSMUST00000104284.3 | 312883 | ENSMUSG00000077472 | Gm24791 | predicted gene, 24791 |
| chr2 | 1,55E+08 | 1,55E+08 | 1133 | * | 7,464836 | 5,340842894 | 1,808517 | 2,953161 | 0,003145 | 1 | 0 | Exon (ENSMUST000001 | 2 | 1,55E+08 | 1,55E+08 | 21365 | 2 | 67204 | ENSMUST00000166171.8 | 44411 | ENSMUSG00000074656 | Eif2s2 | eukaryotic translation initiation factor 2, subunit 2 (beta) |
| chr8 | 72617857 | 72618943 | 1087 | * | 8,63655 | 5,340938545 | 1,763299 | 3,028946 | 0,002454 | 0,648067121 | 1,425929 | Distal Intergenic | 8 | 72585746 | 72611125 | 25380 | 2 | 385024 | ENSMUST00000212113.2 | -6732 | | Gm5373 | predicted gene 5373 |
| chr5 | 85470550 | 85471200 | 651 | * | 6,918465 | 5,341557605 | 1,816484 | 2,940603 | 0,003276 | 1 | 0 | Distal Intergenic | 5 | 85977486 | 85977646 | 161 | 1 | 1,15E+08 | ENSMUST00000082621.3 | -506286 | ENSMUSG00000064555 | Gm24626 | predicted gene, 24626 |
| chr6 | 48796861 | 48797343 | 483 | * | 6,744374 | 5,341623648 | 1,815489 | 2,94225 | 0,003258 | 1 | 0 | Exon (ENSMUST000002 | 6 | 48811441 | 48817389 | 5949 | 2 | 65963 | ENSMUST00000204783.3 | 20046 | ENSMUSG00000029810 | Tmem176b | transmembrane protein 176B |
| chr12 | 1,02E+08 | 1,02E+08 | 465 | * | 4,246526 | 5,341730282 | 2,060481 | 2,592468 | 0,009529 | 1 | 0 | 3' UTR | 12 | 1,02E+08 | 1,02E+08 | 79225 | 2 | 109181 | ENSMUST00000021605.14 | -9487 | ENSMUSG00000021188 | Trip11 | thyroid hormone receptor interactor 11 |

| | | | | | | | | | | | | | | | | | | | | | | | |
|---|---|---|---|---|---|---|---|---|---|---|---|---|---|---|---|---|---|---|---|---|---|---|---|
| chr4 | 1,56E+08 | 1,56E+08 | 1011 | * | 9,897103 | 5,344086102 | 1,864812 | 2,865751 | 0,00416 | 0,732350188 | 1,425929 | Promoter (1-2kb) | 4 | 1,56E+08 | 1,56E+08 | 14940 | 1 | 140500 | ENSMUST00000137726.8 | 1907 | ENSMUSG00000029033 | Acap3 | ArfGAP with coiled-coil, ankyrin repeat and PH domains 3 |
| chr9 | 34027067 | 34028116 | 1050 | * | 6,712255 | 5,344121752 | 1,872838 | 2,853489 | 0,004324 | 1 | 0 | Distal Intergenic | 9 | 33953665 | 33970315 | 16651 | 1 | 77119 | ENSMUST00000184229.2 | 73402 | ENSMUSG00000098758 | 7630403G23Rik | RIKEN cDNA 7630403G23 gene |
| chr18 | 38063070 | 38063708 | 639 | * | 6,719235 | 5,344170621 | 1,890165 | 2,827356 | 0,004693 | 1 | 0 | Intron (ENSMUST0000 | 18 | 37976656 | 38068434 | 91779 | 2 | 13367 | ENSMUST00000025337.14 | 4726 | ENSMUSG00000024456 | Diaph1 | diaphanous related formin 1 |
| chr11 | 84299602 | 84300230 | 629 | * | 6,720457 | 5,344221807 | 1,909342 | 2,798986 | 0,005126 | 1 | 0 | Distal Intergenic | 11 | 84333335 | 84340359 | 7025 | 2 | 56321 | ENSMUST00000141093.2 | 40129 | ENSMUSG00000018697 | Aatf | apoptosis antagonizing transcription factor |
| chr14 | 1,06E+08 | 1,06E+08 | 809 | * | 6,235312 | 5,3442509 | 1,917189 | 2,787545 | 0,005311 | 1 | 0 | Distal Intergenic | 14 | 1,06E+08 | 1,06E+08 | 3075 | 1 | 212085 | ENSMUST00000022708.7 | -51594 | ENSMUSG00000022113 | Trim52 | tripartite motif-containing 52 |
| chr13 | 5436754 | 5437475 | 722 | * | 6,170896 | 5,344318139 | 1,9475 | 2,744194 | 0,006066 | 1 | 0 | Distal Intergenic | 13 | 5504908 | 5549701 | 44794 | 1 | 1,05E+08 | ENSMUST00000220659.2 | -67433 | | Gm40660 | predicted gene, 40660 |
| chr5 | 1,15E+08 | 1,15E+08 | 804 | * | 5,665621 | 5,344404945 | 1,981056 | 2,697755 | 0,006981 | 1 | 0 | Promoter (<=1kb) | 5 | 1,15E+08 | 1,15E+08 | 3895 | 2 | 63873 | ENSMUST00000141828.2 | -808 | ENSMUSG00000014158 | Trpv4 | transient receptor potential cation channel, subfamily V, member 4 |
| chr18 | 10280172 | 10280727 | 556 | * | 5,755587 | 5,34440626 | 1,988937 | 2,687066 | 0,007208 | 1 | 0 | Distal Intergenic | 18 | 10325177 | 10502412 | 177236 | 1 | 381157 | ENSMUST00000173356.8 | -44450 | ENSMUSG00000042942 | Greb1l | growth regulation by estrogen in breast cancer-like |
| chrX | 1,33E+08 | 1,33E+08 | 950 | * | 6,473382 | 5,344411688 | 1,987432 | 2,689104 | 0,007164 | 1 | 0 | Promoter (2-3kb) | 20 | 1,33E+08 | 1,33E+08 | 26860 | 1 | 108062 | ENSMUST00000113287.8 | -2362 | ENSMUSG00000031256 | Cstf2 | cleavage stimulation factor, 3' pre-RNA subunit 2 |
| chr11 | 1,15E+08 | 1,15E+08 | 671 | * | 6,051928 | 5,344435709 | 1,992971 | 2,681642 | 0,007326 | 1 | 0 | Promoter (<=1kb) | 11 | 1,15E+08 | 1,15E+08 | 12716 | 2 | 170930 | ENSMUST00000117589.8 | 935 | ENSMUSG00000020738 | Sumo2 | small ubiquitin-like modifier 2 |
| chr17 | 25053234 | 25053877 | 644 | * | 5,004358 | 5,344440126 | 2,000442 | 2,671629 | 0,007548 | 1 | 0 | Intron (ENSMUST0000 | 17 | 25059117 | 25082698 | 23582 | 1 | 14651 | ENSMUST00000164251.8 | -5240 | ENSMUSG00000024158 | Hagh | hydroxyacyl glutathione hydrolase |
| chr1 | 1,91E+08 | 1,91E+08 | 763 | * | 5,233223 | 5,344455314 | 2,001553 | 2,670154 | 0,007582 | 1 | 0 | Exon (ENSMUST00000 | 1 | 1,91E+08 | 1,91E+08 | 15124 | 2 | 76843 | ENSMUST00000194064.2 | 14611 | ENSMUSG00000037474 | Dtl | denticleless E3 ubiquitin protein ligase |
| chr9 | 9649912 | 9650301 | 390 | * | 5,097297 | 5,344456255 | 2,011482 | 2,656975 | 0,007885 | 1 | 0 | Distal Intergenic | 9 | 9894436 | 9894526 | 91 | 1 | 1,02E+08 | ENSMUST00000183808.3 | -244135 | ENSMUSG00000098408 | Mir6237 | microRNA 6237 |
| chr11 | 77494812 | 77495410 | 599 | * | 4,920069 | 5,344513183 | 2,038145 | 2,622243 | 0,008735 | 1 | 0 | Intron (ENSMUST0000 | 11 | 77419988 | 77498641 | 78654 | 2 | 216965 | ENSMUST00000058496.8 | 3231 | ENSMUSG00000017291 | Taok1 | TAO kinase 1 |
| chr9 | 17303702 | 17304221 | 520 | * | 5,083568 | 5,344522561 | 2,048333 | 2,609206 | 0,009075 | 1 | 0 | Distal Intergenic | 9 | 15821485 | 16412581 | 591097 | 2 | 270120 | ENSMUST00000217308.3 | -891121 | ENSMUSG00000074505 | Fat3 | FAT atypical cadherin 3 |
| chr15 | 71071879 | 71072510 | 632 | * | 4,941631 | 5,344526659 | 2,041096 | 2,61846 | 0,008833 | 1 | 0 | Distal Intergenic | 15 | 71317525 | 71334249 | 16725 | 2 | 70363 | ENSMUST00000229634.2 | 261739 | ENSMUSG00000036800 | Fam135b | family with sequence similarity 135, member B |
| chr7 | 29275468 | 29276128 | 661 | * | 4,56107 | 5,344533378 | 2,054095 | 2,601893 | 0,009271 | 1 | 0 | Exon (ENSMUST00000 | 7 | 29259940 | 29274903 | 14964 | 1 | 243900 | ENSMUST00000053635.5 | 15528 | | 4930432E11Rik | RIKEN cDNA 4930432E11 gene |
| chr8 | 36253262 | 36253986 | 725 | * | 5,127793 | 5,344553143 | 2,073459 | 2,577602 | 0,009949 | 1 | 0 | Intron (ENSMUST0000 | 8 | 36232944 | 36285201 | 52258 | 1 | 1,01E+08 | ENSMUST00000207505.2 | 20318 | ENSMUSG00000109372 | Gm19410 | predicted gene, 19410 |
| chrX | 70960782 | 70962056 | 1275 | * | 8,069922 | 5,346563623 | 1,750948 | 3,053526 | 0,002262 | 1 | 0 | Promoter (2-3kb) | 20 | 70964553 | 70984476 | 19924 | 1 | 382221 | ENSMUST00000154780.10 | -2497 | ENSMUSG00000073130 | Gm1141 | predicted gene 1141 |
| chr15 | 85358364 | 85359138 | 775 | * | 10,20316 | 5,348290119 | 1,754486 | 3,048352 | 0,002301 | 0,637823949 | 1,425929 | Distal Intergenic | 15 | 85376937 | 85387428 | 10492 | 2 | 328577 | ENSMUST00000178942.2 | 28290 | ENSMUSG00000096607 | 7530416G11Rik | RIKEN cDNA 7530416G11 gene |
| chr5 | 1,24E+08 | 1,24E+08 | 594 | * | 6,33904 | 5,348855428 | 2,027219 | 2,638519 | 0,008327 | 1 | 0 | Promoter (<=1kb) | 5 | 1,24E+08 | 1,24E+08 | 1423 | 1 | 231727 | ENSMUST00000031384.6 | -719 | ENSMUSG00000029431 | B3gnt4 | UDP-GlcNAc:betaGal beta-1,3-N-acetylglucosaminyltransferase 4 |
| chr6 | 1,29E+08 | 1,29E+08 | 724 | * | 6,834152 | 5,34946733 | 1,917765 | 2,789428 | 0,00528 | 1 | 0 | Distal Intergenic | 6 | 1,29E+08 | 1,29E+08 | 4407 | 2 | 17057 | ENSMUST00000203150.2 | -4510 | ENSMUSG00000030361 | Klrb1a | killer cell lectin-like receptor subfamily B member 1A |
| chr14 | 75784103 | 75785502 | 1400 | * | 7,450229 | 5,350674472 | 2,001869 | 2,67284 | 0,007521 | 1 | 0 | Distal Intergenic | 14 | 75820323 | 75829389 | 9067 | 2 | 67926 | ENSMUST00000165569.3 | 43887 | ENSMUSG00000034913 | Cby2 | chibby family member 2 |
| chr7 | 35452396 | 35453635 | 1240 | * | 9,615798 | 5,351109991 | 1,71524 | 3,119744 | 0,00181 | 0,593612587 | 1,425929 | Promoter (<=1kb) | 7 | 35450744 | 35453780 | 3037 | 2 | 233115 | ENSMUST00000124116.2 | 145 | ENSMUSG00000043671 | Dpy19l3 | dpy-19-like 3 (C. elegans) |
| chr1 | 89602885 | 89603737 | 853 | * | 7,425186 | 5,351153903 | 1,790625 | 2,988429 | 0,002804 | 0,677651646 | 1,360622 | Intron (ENSMUST0000 | 1 | 89608807 | 89611610 | 2804 | 2 | 74403 | ENSMUST00000187011.2 | 7873 | | 4933400F21Rik | RIKEN cDNA 4933400F21 gene |
| chr9 | 38487888 | 38488520 | 633 | * | 5,071688 | 5,354440089 | 1,947077 | 2,749989 | 0,00596 | 1 | 0 | Promoter (2-3kb) | 9 | 38491474 | 38498172 | 6699 | 1 | 258806 | ENSMUST00000217160.3 | -2954 | ENSMUSG00000111448 | Olfr912 | olfactory receptor 912 |
| chr5 | 22347570 | 22348286 | 717 | * | 4,876159 | 5,355705793 | 1,98155 | 2,702785 | 0,006876 | 1 | 0 | Intron (ENSMUST0000 | 5 | 22265311 | 22339667 | 74357 | 2 | 19699 | ENSMUST00000162876.2 | -7903 | ENSMUSG00000042453 | Reln | reelin |
| chr19 | 56482021 | 56482818 | 798 | * | 7,735399 | 5,3562524 | 1,803887 | 2,969284 | 0,002985 | 1 | 0 | Distal Intergenic | 19 | 56468119 | 56475184 | 7066 | 1 | 226245 | ENSMUST00000225008.2 | 13902 | ENSMUSG00000035818 | Plekhs1 | pleckstrin homology domain containing, family S member 1 |
| chr1 | 26094243 | 26095050 | 808 | * | 6,145511 | 5,358520881 | 1,901799 | 2,817607 | 0,004838 | 1 | 0 | Distal Intergenic | 1 | 25106631 | 25868788 | 762158 | 2 | 210933 | ENSMUST00000151309.8 | -225455 | ENSMUSG00000033569 | Adgrb3 | adhesion G protein-coupled receptor B3 |
| chr11 | 67535440 | 67536105 | 666 | * | 6,534439 | 5,359532001 | 1,99653 | 2,684423 | 0,007266 | 1 | 0 | Intron (ENSMUST0000 | 11 | 67543403 | 67574514 | 31112 | 1 | 14457 | ENSMUST00000140863.2 | -7298 | ENSMUSG00000033066 | Gas7 | growth arrest specific 7 |
| chr10 | 61469556 | 61471022 | 1467 | * | 9,08923 | 5,360212061 | 1,940502 | 2,762282 | 0,00574 | 0,713633527 | 2,237403 | Distal Intergenic | 10 | 61484331 | 61509947 | 25617 | 1 | 67895 | ENSMUST00000020286.7 | -13309 | ENSMUSG00000020089 | Ppa1 | pyrophosphatase (inorganic) 1 |
| chr2 | 30372725 | 30373270 | 546 | * | 8,173445 | 5,360987335 | 1,79527 | 2,986173 | 0,002825 | 1 | 0 | Distal Intergenic | 2 | 30361549 | 30364231 | 2683 | 2 | 72500 | ENSMUST00000065134.4 | -8494 | ENSMUSG00000089762 | Ier5l | immediate early response 5-like |
| chr1 | 51923853 | 51924482 | 630 | * | 5,301089 | 5,363052059 | 2,058974 | 2,604721 | 0,009195 | 1 | 0 | Intron (ENSMUST0000 | 1 | 51836562 | 51935460 | 98899 | 2 | 17912 | ENSMUST00000144694.2 | 10978 | ENSMUSG00000018417 | Myo1b | myosin IB |
| chr11 | 75902221 | 75902698 | 478 | * | 6,158244 | 5,363673602 | 1,918443 | 2,795847 | 0,005176 | 1 | 0 | Distal Intergenic | 11 | 75890738 | 75894395 | 3658 | 1 | 74230 | ENSMUST00000066408.6 | 11483 | ENSMUSG00000053783 | 1700016K19Rik | RIKEN cDNA 1700016K19 gene |
| chr10 | 1,21E+08 | 1,21E+08 | 651 | * | 5,609593 | 5,365975595 | 1,876865 | 2,859009 | 0,00425 | 1 | 0 | Intron (ENSMUST0000 | 10 | 1,21E+08 | 1,21E+08 | 47827 | 2 | 192678 | ENSMUST00000219500.2 | 27935 | ENSMUSG00000025795 | Rassf3 | Ras association (RalGDS/AF-6) domain family member 3 |
| chrX | 1,55E+08 | 1,55E+08 | 925 | * | 7,050893 | 5,366612968 | 1,865378 | 2,876957 | 0,004015 | 1 | 0 | Intron (ENSMUST0000 | 20 | 1,54E+08 | 1,55E+08 | 721113 | 1 | 1,03E+08 | ENSMUST00000112542.2 | 362392 | | Gm15155 | predicted gene 15155 |
| chr1 | 1,8E+08 | 1,8E+08 | 341 | * | 4,725014 | 5,366776457 | 1,997452 | 2,686811 | 0,007214 | 1 | 0 | Intron (ENSMUST0000 | 1 | 1,8E+08 | 1,8E+08 | 17801 | 2 | 19165 | ENSMUST00000133340.2 | -6628 | ENSMUSG00000010609 | Psen2 | presenilin 2 |
| chr12 | 50011321 | 50012346 | 1026 | * | 6,203609 | 5,368199299 | 1,838162 | 2,920416 | 0,003496 | 1 | 0 | Distal Intergenic | 12 | 50167092 | 50192019 | 24928 | 1 | 1,05E+08 | ENSMUST00000218806.2 | -154746 | ENSMUSG00000112353 | Gm40418 | predicted gene, 40418 |
| chr2 | 64695132 | 64695445 | 314 | * | 6,052247 | 5,370468656 | 1,978487 | 2,714432 | 0,006639 | 1 | 0 | Distal Intergenic | 2 | 64742820 | 64745363 | 2544 | 2 | 50915 | ENSMUST00000134020.2 | 49918 | ENSMUSG00000026888 | Grb14 | growth factor receptor bound protein 14 |
| chrX | 42465153 | 42466130 | 978 | * | 7,861451 | 5,370578722 | 1,934657 | 2,775984 | 0,005503 | 0,792545355 | 1,360622 | Intron (ENSMUST0000 | 20 | 41913235 | 42517880 | 604646 | 2 | 23963 | ENSMUST00000154433.8 | 51750 | ENSMUSG00000016150 | Tenm1 | teneurin transmembrane protein 1 |
| chr13 | 89587784 | 89588309 | 526 | * | 5,081576 | 5,371440643 | 2,052945 | 2,616456 | 0,008885 | 1 | 0 | Distal Intergenic | 13 | 89687915 | 89759771 | 71857 | 1 | 12950 | ENSMUST00000022108.9 | -99606 | ENSMUSG00000021613 | Hapln1 | hyaluronan and proteoglycan link protein 1 |
| chr5 | 1,19E+08 | 1,19E+08 | 1493 | * | 9,054057 | 5,372781263 | 1,79069 | 3,000397 | 0,002696 | 0,685026746 | 1,211315 | Promoter (<=1kb) | 5 | 1,19E+08 | 1,19E+08 | 31965 | 1 | 76199 | ENSMUST00000201696.2 | 49 | ENSMUSG00000018076 | Med13l | mediator complex subunit 13-like |
| chr4 | 34507991 | 34508805 | 815 | * | 5,745686 | 5,374181951 | 1,879905 | 2,858751 | 0,004253 | 1 | 0 | Distal Intergenic | 4 | 34550937 | 34566908 | 15972 | 1 | 433693 | ENSMUST00000084299.6 | -42132 | ENSMUSG00000028291 | Akirin2 | akirin 2 |
| chr8 | 21397350 | 21397983 | 634 | * | 7,404962 | 5,374587504 | 1,81719 | 2,957636 | 0,0031 | 0,688021707 | 1,360622 | Distal Intergenic | 8 | 21390622 | 21392643 | 2022 | 2 | 1,01E+08 | ENSMUST00000122025.9 | -4707 | ENSMUSG00000082976 | Gm15056 | predicted gene 15056 |
| chr9 | 1,05E+08 | 1,05E+08 | 618 | * | 6,098845 | 5,374742506 | 1,825557 | 2,944165 | 0,003238 | 1 | 0 | Exon (ENSMUST00000 | 9 | 1,05E+08 | 1,05E+08 | 8184 | 2 | 235574 | ENSMUST00000177000.2 | -5214 | ENSMUSG00000032570 | Atp2c1 | ATPase, Ca++-sequestering |
| chr9 | 1,08E+08 | 1,08E+08 | 1206 | * | 7,247 | 5,376118206 | 1,884739 | 2,852447 | 0,004338 | 0,749550311 | 1,360622 | 3' UTR | 9 | 1,08E+08 | 1,08E+08 | 3351 | 1 | 69232 | ENSMUST00000194139.2 | -9951 | ENSMUSG00000006673 | Qrich1 | glutamine-rich 1 |
| chr19 | 16732256 | 16733237 | 982 | * | 6,03057 | 5,377140038 | 1,922624 | 2,796771 | 0,005162 | 1 | 0 | Intron (ENSMUST0000 | 19 | 16699702 | 16728302 | 28601 | 2 | 271564 | ENSMUST00000223846.2 | -3954 | ENSMUSG00000046230 | Vps13a | vacuolar protein sorting 13A |
| chr7 | 1,27E+08 | 1,27E+08 | 944 | * | 5,801286 | 5,377191371 | 1,924235 | 2,794457 | 0,005199 | 1 | 0 | Distal Intergenic | 7 | 1,27E+08 | 1,27E+08 | 15058 | 1 | 1E+08 | ENSMUST00000185401.7 | -3960 | ENSMUSG00000053877 | Srcap | Snf2-related CREBBP activator protein |
| chr10 | 65413099 | 65413840 | 742 | * | 6,668266 | 5,378357067 | 1,909769 | 2,816234 | 0,004859 | 1 | 0 | Distal Intergenic | 10 | 65370433 | 65371056 | 624 | 1 | 216036 | ENSMUST00000219802.2 | 42666 | ENSMUSG00000112341 | Gm4796 | predicted gene 4796 |
| chr19 | 5123489 | 5124348 | 860 | * | 5,457038 | 5,37836924 | 1,976684 | 2,720905 | 0,00651 | 1 | 0 | Exon (ENSMUST00002 | 19 | 5118106 | 5120710 | 2605 | 1 | 70445 | ENSMUST00000070630.8 | 5383 | ENSMUSG00000056481 | Cd248 | CD248 antigen, endosialin |
| chr13 | 96688356 | 96689096 | 741 | * | 5,267449 | 5,378605235 | 1,905715 | 2,822356 | 0,004767 | 1 | 0 | Intron (ENSMUST0000 | 13 | 96679263 | 96748247 | 68985 | 1 | 68018 | ENSMUST00000222434.2 | 9093 | ENSMUSG00000021669 | Cert1 | ceramide transporter 1 |
| chr15 | 4401766 | 4402525 | 760 | * | 5,713552 | 5,37876651 | 1,995596 | 2,695319 | 0,007032 | 1 | 0 | Promoter (2-3kb) | 15 | 4404973 | 4549896 | 144924 | 1 | 239318 | ENSMUST00000130332.2 | -2448 | ENSMUSG00000049148 | Plcxd3 | phosphatidylinositol-specific phospholipase C, X domain containing 3 |
| chr1 | 16728356 | 16729554 | 1199 | * | 7,540509 | 5,378921933 | 1,814735 | 2,964027 | 0,003036 | 1 | 0 | Promoter (<=1kb) | 1 | 16727776 | 16732502 | 4727 | 1 | 329101 | ENSMUST00000181371.2 | 580 | ENSMUSG00000097744 | D030040B21Rik | RIKEN cDNA D030040B21 gene |
| chr11 | 97857399 | 97857985 | 587 | * | 5,50491 | 5,37951312 | 2,037886 | 2,639752 | 0,008297 | 1 | 0 | Promoter (1-2kb) | 11 | 97859618 | 97860061 | 444 | 1 | 1,03E+08 | ENSMUST00000118176.2 | -1633 | | Gm11633 | predicted gene 11633 |
| chr9 | 23480733 | 23481534 | 802 | * | 5,078291 | 5,379850523 | 2,058323 | 2,613706 | 0,008957 | 1 | 0 | Distal Intergenic | 9 | 23285228 | 23318127 | 32900 | 1 | 73230 | ENSMUST00000214050.2 | 195505 | ENSMUSG00000031963 | Bmper | BMP-binding endothelial regulator |
| chr11 | 6615236 | 6615657 | 422 | * | 4,515699 | 5,379907007 | 2,06168 | 2,609478 | 0,009068 | 1 | 0 | Intron (ENSMUST0000 | 11 | 6600148 | 6610233 | 10086 | 2 | 1E+08 | ENSMUST00000127510.9 | -5003 | ENSMUSG00000086373 | Gm11978 | predicted gene 11978 |
| chr2 | 77178838 | 77179332 | 495 | * | 4,779515 | 5,380019061 | 2,068436 | 2,601008 | 0,009295 | 1 | 0 | Distal Intergenic | 2 | 77016899 | 77110936 | 94038 | 2 | 228071 | ENSMUST00000102659.2 | -67902 | ENSMUSG00000042272 | Sestd1 | SEC14 and spectrin domains 1 |
| chr17 | 27185972 | 27186523 | 552 | * | 6,367208 | 5,380405245 | 1,852964 | 2,903676 | 0,003688 | 1 | 0 | Intron (ENSMUST0000 | 17 | 27182000 | 27189561 | 7562 | 1 | 240057 | ENSMUST00000202208.2 | 3972 | ENSMUSG00000067629 | Syngap1 | synaptic Ras GTPase activating protein 1 homolog (rat) |
| chr9 | 64059872 | 64060701 | 830 | * | 5,338724 | 5,380698218 | 2,070974 | 2,598148 | 0,009373 | 1 | 0 | Promoter (2-3kb) | 9 | 64051469 | 64062784 | 11316 | 2 | 68014 | ENSMUST00000138075.9 | 2083 | ENSMUSG00000032400 | Zwilch | zwilch kinetochore protein |
| chr18 | 66043075 | 66043514 | 440 | * | 4,854532 | 5,381051245 | 2,06345 | 2,607793 | 0,009113 | 1 | 0 | Distal Intergenic | 18 | 65996404 | 66027826 | 31423 | 2 | 1,03E+08 | ENSMUST00000237028.2 | -15249 | ENSMUSG00000118170 | Gm30094 | predicted gene, 30094 |
| chr18 | 24256067 | 24256731 | 665 | * | 5,356758 | 5,3833801 | 2,015966 | 2,670373 | 0,007577 | 1 | 0 | Promoter (1-2kb) | 18 | 24237814 | 24255010 | 17197 | 2 | 225280 | ENSMUST00000153360.8 | -1057 | ENSMUSG00000047989 | Ino80c | INO80 complex subunit C |
| chr11 | 1,08E+08 | 1,08E+08 | 1097 | * | 11,3481 | 5,384132935 | 1,751693 | 3,073674 | 0,002114 | 0,655368941 | 1,173325 | Promoter (<=1kb) | 11 | 1,08E+08 | 1,08E+08 | 52552 | 1 | 11818 | ENSMUST00000133383.8 | 0 | ENSMUSG00000000049 | Apoh | apolipoprotein H |
| chr18 | 73945706 | 73946571 | 866 | * | 6,95266 | 5,384496657 | 1,795321 | 2,999183 | 0,002707 | 1 | 0 | Promoter (1-2kb) | 18 | 73902974 | 73948509 | 45536 | 2 | 107029 | ENSMUST00000025439.5 | 1938 | ENSMUSG00000024556 | Me2 | malic enzyme 2, NAD(+)-dependent, mitochondrial |
| chr1 | 1,78E+08 | 1,78E+08 | 701 | * | 6,197871 | 5,384856503 | 1,933331 | 2,785274 | 0,005348 | 1 | 0 | Intron (ENSMUST0000 | 1 | 1,78E+08 | 1,78E+08 | 61492 | 1 | 78825 | ENSMUST00000159284.2 | 20705 | ENSMUSG00000026502 | Desi2 | desumoylating isopeptidase 2 |
| chr11 | 83466490 | 83467412 | 923 | * | 9,481725 | 5,386235599 | 1,729156 | 3,114951 | 0,00184 | 0,508068302 | 2,237403 | Promoter (<=1kb) | 11 | 83466144 | 83467968 | 1825 | 2 | 20308 | ENSMUST00000136131.2 | 556 | ENSMUSG00000019122 | Ccl9 | chemokine (C-C motif) ligand 9 |
| chr6 | 50494525 | 50494873 | 349 | * | 5,029578 | 5,387311528 | 2,0219 | 2,66448 | 0,007711 | 1 | 0 | Distal Intergenic | 6 | 50476529 | 50482710 | 6182 | 2 | 1,05E+08 | ENSMUST00000247433.1 | -11815 | | Gm40372 | predicted gene, 40372 |
| chr1 | 1,57E+08 | 1,57E+08 | 531 | * | 6,883561 | 5,390002671 | 1,85183 | 2,910636 | 0,003607 | 1 | 0 | Distal Intergenic | 1 | 1,57E+08 | 1,57E+08 | 40516 | 2 | 215015 | ENSMUST00000086153.8 | -46063 | ENSMUSG00000033557 | Fam20b | FAM20B, glycosaminoglycan xylosylkinase |
| chr6 | 83443566 | 83444333 | 768 | * | 11,42999 | 5,393419591 | 1,654994 | 3,258875 | 0,001119 | 0,626412475 | 0,738441 | Distal Intergenic | 6 | 83381473 | 83436066 | 54594 | 2 | 194388 | ENSMUST00000190295.2 | -7500 | ENSMUSG00000034832 | Tet3 | tet methylcytosine dioxygenase 3 |
| chr1 | 36160702 | 36161568 | 867 | * | 6,77082 | 5,395321155 | 1,892808 | 2,850432 | 0,004366 | 1 | 0 | Distal Intergenic | 1 | 36171099 | 36173635 | 2537 | 1 | 1,03E+08 | ENSMUST00000245804.1 | -9531 | | Gm33280 | predicted gene, 33280 |
| chr4 | 24492040 | 24493411 | 1372 | * | 7,917596 | 5,397837891 | 1,765825 | 3,056836 | 0,002237 | 1 | 0 | Distal Intergenic | 4 | 24496451 | 24581338 | 84888 | 1 | 212377 | ENSMUST00000173079.8 | -3040 | ENSMUSG00000045751 | Mms22l | MMS22-like, DNA repair protein |
| chr4 | 55316282 | 55317520 | 1239 | * | 7,000888 | 5,398381182 | 1,790104 | 3,01568 | 0,002564 | 1 | 0 | Distal Intergenic | 4 | 55350043 | 55392237 | 42195 | 1 | 19359 | ENSMUST00000030134.9 | -32523 | ENSMUSG00000028426 | Rad23b | RAD23 homolog B, nucleotide excision repair protein |
| chr3 | 23771541 | 23772408 | 868 | * | 6,80777 | 5,398766662 | 1,782659 | 3,02849 | 0,002458 | 1 | 0 | Distal Intergenic | 3 | 23858499 | 23993641 | 135143 | 2 | 635702 | ENSMUST00000192246.4 | 221233 | ENSMUSG00000102758 | Naaladl2 | N-acetylated alpha-linked acidic dipeptidase-like 2 |
| chr13 | 45447518 | 45448330 | 813 | * | 6,347648 | 5,398816493 | 1,934723 | 2,790485 | 0,005263 | 1 | 0 | Exon (ENSMUST00020 | 13 | 45400108 | 45421187 | 21080 | 2 | 1,03E+08 | ENSMUST00000222218.2 | -26331 | ENSMUSG00000113919 | Gm34466 | predicted gene, 34466 |
| chr9 | 65793545 | 65794251 | 707 | * | 5,613422 | 5,399384493 | 1,92491 | 2,805006 | 0,005032 | 1 | 0 | Promoter (1-2kb) | 9 | 65740438 | 65792338 | 51901 | 2 | 56404 | ENSMUST00000117083.2 | -1207 | ENSMUSG00000032386 | Trip4 | thyroid hormone receptor interactor 4 |
| chr2 | 92670371 | 92671375 | 1005 | * | 9,633946 | 5,399471029 | 1,721954 | 3,135666 | 0,001715 | 0,619070779 | 1,173325 | Distal Intergenic | 2 | 92686512 | 92744008 | 57497 | 2 | 1,03E+08 | ENSMUST00000123263.2 | 72633 | | Gm13791 | predicted gene 13791 |
| chr1 | 72683291 | 72684181 | 891 | * | 9,403965 | 5,399535026 | 1,767799 | 3,054383 | 0,002255 | 0,642322948 | 1,360622 | Promoter (2-3kb) | 1 | 72680402 | 72682122 | 1721 | 1 | 1,05E+08 | ENSMUST00000211909.2 | 2889 | | Gm39662 | predicted gene, 39662 |
| chr17 | 70147009 | 70147971 | 963 | * | 10,4559 | 5,399999146 | 1,736739 | 3,109275 | 0,001875 | 0,63491343 | 1,173325 | Distal Intergenic | 17 | 70200755 | 70230990 | 30236 | 2 | 1,03E+08 | ENSMUST00000232745.2 | 83019 | | Gm36594 | predicted gene, 36594 |
| chr19 | 43099971 | 43101136 | 1166 | * | 10,14601 | 5,400903901 | 1,711833 | 3,155042 | 0,001605 | 0,566484927 | 1,425929 | Intron (ENSMUST0000 | 19 | 42897023 | 43212128 | 315106 | 2 | 545291 | ENSMUST00000238153.2 | 110992 | ENSMUSG00000074852 | Hpse2 | heparanase 2 |
| chr1 | 81290325 | 81290850 | 526 | * | 4,655375 | 5,403172442 | 1,9903 | 2,714752 | 0,006633 | 1 | 0 | Intron (ENSMUST0000 | 1 | 81055300 | 81319479 | 264180 | 1 | 241134 | ENSMUST00000123285.2 | 235025 | ENSMUSG00000054976 | Nyap2 | neuronal tyrosine-phophorylated phosphoinositide 3-kinase adaptor 2 |
| chr14 | 96700403 | 96701054 | 652 | * | 5,060751 | 5,403394561 | 1,979707 | 2,729391 | 0,006345 | 1 | 0 | Intron (ENSMUST0000 | 14 | 96682391 | 96682506 | 116 | 1 | 1,15E+08 | ENSMUST00000083108.3 | 18012 | ENSMUSG00000065042 | Gm24043 | predicted gene, 24043 |
| chr12 | 79345314 | 79346956 | 1643 | * | 10,61782 | 5,406421963 | 1,757128 | 3,076851 | 0,002092 | 0,7299084 | 0,738441 | Promoter (<=1kb) | 12 | 79344337 | 79861152 | 516816 | 1 | 19363 | ENSMUST00000218039.3 | 977 | ENSMUSG00000059060 | Rad51b | RAD51 paralog B |
| chr16 | 45732071 | 45733372 | 1302 | * | 13,52054 | 5,406670532 | 1,655489 | 3,265906 | 0,001091 | 0,571756101 | 0,938682 | Intron (ENSMUST0000 | 16 | 45765926 | 45777142 | 11217 | 1 | 1,01E+08 | ENSMUST00000241345.1 | -32554 | ENSMUSG00000085826 | Gm15638 | predicted gene 15638 |
| chr14 | 1,21E+08 | 1,21E+08 | 1159 | * | 6,855212 | 5,407145577 | 1,921444 | 2,814105 | 0,004891 | 1 | 0 | Exon (ENSMUST00000 | 14 | 1,21E+08 | 1,22E+08 | 13014 | 1 | 223254 | ENSMUST00000137971.2 | -4334 | ENSMUSG00000025555 | Farp1 | FERM, RhoGEF (Arhgef) and pleckstrin domain protein 1 (chondrocyte-derived) |
| chr5 | 32238518 | 32239176 | 659 | * | 7,275701 | 5,407933691 | 1,886864 | 2,866096 | 0,004156 | 1 | 0 | Promoter (1-2kb) | 5 | 32237041 | 32242090 | 5050 | 1 | 107976 | ENSMUST00000202087.2 | 1477 | ENSMUSG00000052139 | Babam2 | BRISC and BRCA1 A complex member 2 |
| chr5 | 31594001 | 31595232 | 1232 | * | 9,1209 | 5,407941031 | 1,780619 | 3,037114 | 0,002389 | 0,650529916 | 1,360622 | Distal Intergenic | 5 | 31609775 | 31624001 | 14227 | 1 | 269639 | ENSMUST00000202061.4 | -14543 | ENSMUSG00000062761 | Zfp512 | zinc finger protein 512 |
| chr16 | 13767696 | 13768341 | 646 | * | 6,654681 | 5,408177193 | 1,891424 | 2,859315 | 0,004246 | 1 | 0 | Downstream (<=300bp | 16 | 13758958 | 13764709 | 5752 | 1 | 93734 | ENSMUST00000156143.2 | 8738 | ENSMUSG00000022679 | Mpv17l | Mpv17 transgene, kidney disease mutant-like |
| chr1 | 63420682 | 63421473 | 792 | * | 6,009363 | 5,410013375 | 1,880613 | 2,876729 | 0,004018 | 1 | 0 | Distal Intergenic | 1 | 63485050 | 63612376 | 127327 | 1 | 23792 | ENSMUST00000182642.8 | -63577 | ENSMUSG00000025964 | Adam23 | a disintegrin and metallopeptidase domain 23 |
| chr7 | 1,22E+08 | 1,22E+08 | 802 | * | 5,881021 | 5,410590992 | 1,942637 | 2,785179 | 0,00535 | 1 | 0 | Intron (ENSMUST0000 | 7 | 1,22E+08 | 1,22E+08 | 52785 | 2 | 233824 | ENSMUST00000148634.2 | 5626 | ENSMUSG00000034951 | Cog7 | component of oligomeric golgi complex 7 |
| chr15 | 77338325 | 77338912 | 588 | * | 5,874827 | 5,410986875 | 1,952111 | 2,771864 | 0,005574 | 1 | 0 | Intron (ENSMUST0000 | 15 | 77309498 | 77331692 | 22195 | 2 | 278679 | ENSMUST00000149824.2 | -6633 | ENSMUSG00000068252 | Apol7b | apolipoprotein L 7b |
| chr11 | 52324788 | 52325457 | 670 | * | 7,325559 | 5,411260818 | 1,966884 | 2,751184 | 0,005938 | 1 | 0 | Distal Intergenic | 11 | 52287473 | 52294101 | 6629 | 1 | 213673 | ENSMUST00000136303.2 | 37315 | ENSMUSG00000036275 | 9530068E07Rik | RIKEN cDNA 9530068E07 gene |
| chr3 | 81239995 | 81240785 | 791 | * | 7,620643 | 5,411323893 | 1,763175 | 3,06908 | 0,002147 | 1 | 0 | Distal Intergenic | 3 | 80944731 | 81118077 | 173347 | 1 | 54635 | ENSMUST00000143721.2 | 295264 | ENSMUSG00000028019 | Pdgfc | platelet-derived growth factor, C polypeptide |
| chr1 | 50390912 | 50391656 | 745 | * | 4,975953 | 5,413111797 | 2,008585 | 2,694988 | 0,007039 | 1 | 0 | Distal Intergenic | 1 | 50939806 | 50954589 | 14784 | 1 | 56363 | ENSMUST00000139157.8 | -548150 | ENSMUSG00000026109 | Tmeff2 | transmembrane protein with EGF-like and two follistatin-like domains 2 |
| chr19 | 23824984 | 23825500 | 517 | * | 4,844045 | 5,414308281 | 2,04557 | 2,646845 | 0,008125 | 1 | 0 | Intron (ENSMUST0000 | 19 | 23881821 | 23924740 | 42920 | 1 | 319924 | ENSMUST00000237688.2 | -56321 | ENSMUSG00000024897 | Apba1 | amyloid beta (A4) precursor protein binding, family A, member 1 |
| chr6 | 64566320 | 64566732 | 413 | * | 5,083589 | 5,414481318 | 2,051301 | 2,639536 | 0,008302 | 1 | 0 | Intron (ENSMUST0000 | 6 | 64640172 | 64681307 | 41136 | 1 | 14804 | ENSMUST00000210324.2 | -73440 | ENSMUSG00000071424 | Grid2 | glutamate receptor, ionotropic, delta 2 |
| chr11 | 53765858 | 53766406 | 549 | * | 4,929163 | 5,41493186 | 2,067008 | 2,619695 | 0,008801 | 1 | 0 | 3' UTR | 11 | 53756291 | 53762503 | 6213 | 2 | 20520 | ENSMUST00000152084.2 | -3355 | ENSMUSG00000018900 | Slc22a5 | solute carrier family 22 (organic cation transporter), member 5 |
| chr17 | 30591573 | 30592419 | 847 | * | 5,762596 | 5,415002099 | 2,069977 | 2,615972 | 0,008897 | 1 | 0 | Intron (ENSMUST0000 | 17 | 30614594 | 30615007 | 414 | 2 | 623169 | ENSMUST00000235315.2 | 22588 | | Gm6402 | ribosomal protein S17 pseudogene |
| chr13 | 90240160 | 90240939 | 780 | * | 4,439472 | 5,415072079 | 2,07157 | 2,613995 | 0,008949 | 1 | 0 | Promoter (2-3kb) | 13 | 90237881 | 90252780 | 14900 | 1 | 66074 | ENSMUST00000012566.9 | 2279 | ENSMUSG00000012422 | Tmem167 | transmembrane protein 167 |
| chr16 | 11969501 | 11970091 | 591 | * | 4,938138 | 5,415121514 | 2,018432 | 2,682835 | 0,0073 | 1 | 0 | Intron (ENSMUST0000 | 16 | 11802584 | 11813260 | 10677 | 1 | 72555 | ENSMUST00000229804.2 | 166917 | ENSMUSG00000022494 | Shisa9 | shisa family member 9 |
| chr11 | 20404064 | 20404671 | 608 | * | 5,648114 | 5,415136003 | 2,063832 | 2,623826 | 0,008695 | 1 | 0 | Promoter (<=1kb) | 11 | 20403102 | 20404101 | 1000 | 1 | 1,05E+08 | ENSMUST00000247465.1 | 962 | | Gm39606 | predicted gene, 39606 |
| chr11 | 59739117 | 59740148 | 1032 | * | 8,338783 | 5,415163785 | 1,901476 | 2,847875 | 0,004401 | 1 | 0 | Promoter (<=1kb) | 11 | 59738930 | 59767359 | 28430 | 1 | 103850 | ENSMUST00000137695.2 | 187 | ENSMUSG00000032615 | Nt5m | 5',3'-nucleotidase, mitochondrial |
| chr11 | 7286182 | 7286519 | 338 | * | 4,72001 | 5,415222723 | 2,061566 | 2,626752 | 0,00862 | 1 | 0 | Distal Intergenic | 11 | 7157686 | 7163923 | 6238 | 2 | 16009 | ENSMUST00000135887.3 | -122259 | ENSMUSG00000020427 | Igfbp3 | insulin-like growth factor binding protein 3 |

| | | | | | | | | | | | | | | | | | | | | | | | |
|---|---|---|---|---|---|---|---|---|---|---|---|---|---|---|---|---|---|---|---|---|---|---|---|
| chr1 | 26118071 | 26118716 | 646 | * | 4,368794 | 5,415795009 | 2,098803 | 2,58042 | 0,009868 | 1 | 0 | Distal Intergenic | 1 | 25106631 | 25868788 | 762158 | 2 | 210933 | ENSMUST00000151309.8 | -249283 | ENSMUSG00000033569 | Adgrb3 | adhesion G protein-coupled receptor B3 |
| chr13 | 51137243 | 51137641 | 399 | * | 5,033462 | 5,416407087 | 2,025253 | 2,674434 | 0,007486 | 1 | 0 | Distal Intergenic | 13 | 51254916 | 51306582 | 51667 | 1 | 20729 | ENSMUST00000095797.6 | -117275 | ENSMUSG00000021395 | Spin1 | spindlin 1 |
| chr1 | 18433572 | 18434348 | 777 | * | 6,632226 | 5,418388028 | 1,972533 | 2,746919 | 0,006016 | 1 | 0 | Distal Intergenic | 1 | 18321204 | 18335362 | 14159 | 2 | 77673 | ENSMUST00000088463.2 | -98210 | ENSMUSG00000067773 | Defb41 | defensin beta 41 |
| chr5 | 1,15E+08 | 1,15E+08 | 990 | * | 6,661829 | 5,418548242 | 1,968687 | 2,752367 | 0,005917 | 1 | 0 | Intron (ENSMUST0000( | 5 | 1,15E+08 | 1,15E+08 | 73389 | 1 | 74585 | ENSMUST00000128590.3 | 7855 | ENSMUSG00000029550 | Sppl3 | signal peptide peptidase 3 |
| chr5 | 33145684 | 33147286 | 1603 | * | 13,85428 | 5,419285183 | 1,63392 | 3,316738 | 0,000911 | 0,54122637 | 0,918255 | Intron (ENSMUST0000( | 5 | 33125063 | 33131520 | 6458 | 1 | 277854 | ENSMUST00000127560.2 | 20621 | ENSMUSG00000037426 | Depdc5 | DEP domain containing 5 |
| chr5 | 74091035 | 74091807 | 773 | * | 6,551965 | 5,420265503 | 1,930032 | 2,808382 | 0,004979 | 1 | 0 | Distal Intergenic | 5 | 74070239 | 74077764 | 7526 | 1 | 69402 | ENSMUST00000202487.2 | 20796 | ENSMUSG00000107027 | 1700019F05Rik | RIKEN cDNA 1700019F05 gene |
| chr15 | 62196054 | 62196436 | 383 | * | 6,864419 | 5,42087919 | 1,917615 | 2,826886 | 0,0047 | 1 | 0 | Intron (ENSMUST0000( | 15 | 62094452 | 62114354 | 19903 | 1 | 19296 | ENSMUST00000182141.2 | 101602 | ENSMUSG00000097039 | Pvt1 | Pvt1 oncogene |
| chr11 | 34523239 | 34524649 | 1411 | * | 7,34477 | 5,421240874 | 1,807784 | 2,998833 | 0,00271 | 1 | 0 | Intron (ENSMUST0000( | 11 | 34311959 | 34579306 | 267348 | 2 | 94176 | ENSMUST00000101364.3 | 54657 | ENSMUSG00000020143 | Dock2 | dedicator of cyto-kinesis 2 |
| chr8 | 90870835 | 90872052 | 1218 | * | 9,260972 | 5,422356104 | 1,772462 | 3,059222 | 0,002219 | 0,628564376 | 1,425929 | Distal Intergenic | 8 | 90984574 | 91074754 | 90181 | 2 | 244579 | ENSMUST00000176034.2 | 202702 | ENSMUSG00000043668 | Tox3 | TOX high mobility group box family member 3 |
| chr4 | 29145077 | 29145703 | 627 | * | 4,218575 | 5,425370269 | 2,088348 | 2,597925 | 0,009379 | 1 | 0 | Distal Intergenic | 4 | 29068079 | 29088811 | 20733 | 1 | 1,03E+08 | ENSMUST00000154813.2 | 76998 | | Gm11917 | predicted gene 11917 |
| chr18 | 56020251 | 56021057 | 807 | * | 7,185837 | 5,42564408 | 1,781073 | 3,046278 | 0,002317 | 1 | 0 | Distal Intergenic | 18 | 56326071 | 56326205 | 135 | 1 | 1,15E+08 | ENSMUST00000157815.3 | -305014 | ENSMUSG00000088440 | Gm23555 | predicted gene, 23555 |
| chr11 | 48726555 | 48728643 | 2089 | * | 18,77116 | 5,426115869 | 1,293365 | 4,195349 | 2,72E-05 | 0,054226803 | 2,221611 | Promoter (<=1kb) | 11 | 48728292 | 48741382 | 13091 | 1 | 94089 | ENSMUST00000046903.6 | 0 | ENSMUSG00000040350 | Trim7 | tripartite motif-containing 7 |
| chr17 | 5476810 | 5477451 | 642 | * | 6,29385 | 5,426331886 | 1,855506 | 2,924448 | 0,003451 | 1 | 0 | Intron (ENSMUST0000( | 17 | 5467598 | 5469042 | 1445 | 2 | 106557 | ENSMUST00000189788.2 | -7768 | ENSMUSG00000101959 | Ldhal6b | lactate dehydrogenase A-like 6B |
| chr3 | 1,04E+08 | 1,04E+08 | 962 | * | 5,257863 | 5,427679996 | 1,934899 | 2,805149 | 0,005029 | 1 | 0 | Intron (ENSMUST0000( | 3 | 1,04E+08 | 1,04E+08 | 47514 | 2 | 15257 | ENSMUST00000106845.9 | 3583 | ENSMUSG00000008730 | Hipk1 | homeodomain interacting protein kinase 1 |
| chr17 | 42599373 | 42600049 | 677 | * | 5,285242 | 5,428031198 | 1,955646 | 2,77557 | 0,005511 | 1 | 0 | Distal Intergenic | 17 | 42626838 | 42815968 | 189131 | 1 | 627626 | ENSMUST00000048691.6 | -26789 | ENSMUSG00000042256 | Ptchd4 | patched domain containing 4 |
| chr12 | 1,08E+08 | 1,08E+08 | 1320 | * | 10,81968 | 5,430871836 | 1,784245 | 3,043793 | 0,002336 | 0,71238788 | 0,918255 | Distal Intergenic | 12 | 1,08E+08 | 1,08E+08 | 24600 | 1 | 214305 | ENSMUST00000021685.8 | -7557 | ENSMUSG00000021260 | Hhipl1 | hedgehog interacting protein-like 1 |
| chr9 | 15272085 | 15272441 | 357 | * | 5,302263 | 5,432433635 | 1,95958 | 2,772244 | 0,005567 | 1 | 0 | Distal Intergenic | 9 | 15228393 | 15269084 | 40692 | 2 | 319675 | ENSMUST00000098979.12 | -3001 | ENSMUSG00000046111 | Cep295 | centrosomal protein 295 |
| chrX | 1,69E+08 | 1,69E+08 | 1179 | * | 6,479514 | 5,434837986 | 1,846208 | 2,943784 | 0,003242 | 1 | 0 | Promoter (<=1kb) | 20 | 1,69E+08 | 1,69E+08 | 111177 | 1 | 17318 | ENSMUST00000112105.8 | 0 | ENSMUSG00000035299 | Mid1 | midline 1 |
| chr5 | 72009206 | 72009930 | 725 | * | 7,109089 | 5,436365534 | 1,850423 | 2,937904 | 0,003304 | 1 | 0 | Intron (ENSMUST0000( | 5 | 71857275 | 72306380 | 449106 | 1 | 14400 | ENSMUST00000031122.9 | 151931 | ENSMUSG00000029212 | Gabrb1 | gamma-aminobutyric acid (GABA) A receptor, subunit beta 1 |
| chr2 | 1,49E+08 | 1,49E+08 | 729 | * | 5,358502 | 5,442216097 | 1,897251 | 2,868474 | 0,004125 | 1 | 0 | Distal Intergenic | 2 | 1,49E+08 | 1,49E+08 | 3929 | 1 | 78609 | ENSMUST00000028933.3 | -3818 | ENSMUSG00000027444 | Cstdc1 | cystatin domain containing 1 |
| chr12 | 6993839 | 6994445 | 607 | * | 5,989532 | 5,443668582 | 1,867672 | 2,914682 | 0,003561 | 1 | 0 | Distal Intergenic | 12 | 7422039 | 7430330 | 8292 | 2 | 73575 | ENSMUST00000187396.2 | 435885 | ENSMUSG00000102096 | 1700101O22Rik | RIKEN cDNA 1700101O22 gene |
| chr9 | 65633437 | 65634637 | 1201 | * | 7,141709 | 5,443906015 | 1,954748 | 2,784966 | 0,005353 | 1 | 0 | Intron (ENSMUST0000( | 9 | 65607152 | 65608426 | 1275 | 2 | 214812 | ENSMUST00000159878.2 | -25011 | ENSMUSG00000040524 | Zfp609 | zinc finger protein 609 |
| chr4 | 1,33E+08 | 1,33E+08 | 904 | * | 6,408894 | 5,444468885 | 1,964525 | 2,771392 | 0,005582 | 1 | 0 | Intron (ENSMUST0000( | 4 | 1,33E+08 | 1,33E+08 | 612 | 1 | 242687 | ENSMUST00000123578.2 | -16552 | ENSMUSG00000028868 | Wasf2 | WASP family, member 2 |
| chr13 | 58294937 | 58295872 | 936 | * | 6,734167 | 5,445085481 | 1,9756 | 2,756168 | 0,005848 | 1 | 0 | Promoter (<=1kb) | 13 | 58294763 | 58296789 | 2027 | 1 | 67339 | ENSMUST00000224280.2 | 174 | ENSMUSG00000097717 | 1700066J03Rik | RIKEN cDNA 1700066J03 gene |
| chr7 | 50985795 | 50986621 | 827 | * | 6,346533 | 5,446018453 | 1,821615 | 2,989665 | 0,002793 | 1 | 0 | Distal Intergenic | 7 | 51160777 | 51248457 | 87681 | 1 | 233246 | ENSMUST00000043944.6 | -174156 | ENSMUSG00000055489 | Ano5 | anoctamin 5 |
| chr10 | 81056372 | 81057116 | 745 | * | 5,439335 | 5,446677537 | 1,949544 | 2,793822 | 0,005209 | 1 | 0 | Intron (ENSMUST0000( | 10 | 81045646 | 81049110 | 3465 | 2 | 16467 | ENSMUST00000133555.2 | -7262 | ENSMUSG00000034958 | Atcay | ataxia, cerebellar, Cayman type |
| chr7 | 75268120 | 75268626 | 507 | * | 5,920378 | 5,446873333 | 2,010401 | 2,709347 | 0,006742 | 1 | 0 | Intron (ENSMUST0000( | 7 | 75264436 | 75326894 | 62459 | 1 | 75547 | ENSMUST00000208053.2 | 3684 | ENSMUSG00000066406 | Akap13 | A kinase (PRKA) anchor protein 13 |
| chr4 | 1,36E+08 | 1,36E+08 | 655 | * | 5,537123 | 5,447207491 | 2,017518 | 2,699955 | 0,006935 | 1 | 0 | Distal Intergenic | 4 | 1,36E+08 | 1,36E+08 | 24814 | 1 | 12802 | ENSMUST00000097843.9 | -8163 | ENSMUSG00000062585 | Cnr2 | cannabinoid receptor 2 (macrophage) |
| chrX | 64338602 | 64339194 | 593 | * | 5,792561 | 5,447355867 | 1,922349 | 2,833698 | 0,004601 | 1 | 0 | Distal Intergenic | 20 | 64091250 | 64092620 | 1371 | 1 | 70062 | ENSMUST00000033524.3 | 247352 | ENSMUSG00000031181 | Ctag2 | cancer/testis antigen 2 |
| chr5 | 24655172 | 24655883 | 712 | * | 5,35872 | 5,447488441 | 2,022965 | 2,692824 | 0,007085 | 1 | 0 | Promoter (1-2kb) | 5 | 24657175 | 24707045 | 49871 | 1 | 213990 | ENSMUST00000024123.9 | -1292 | ENSMUSG00000023353 | Agap3 | ArfGAP with GTPase domain, ankyrin repeat and PH domain 3 |
| chrX | 1,68E+08 | 1,68E+08 | 864 | * | 5,130607 | 5,447639135 | 2,026548 | 2,688138 | 0,007185 | 1 | 0 | Promoter (<=1kb) | 20 | 1,68E+08 | 1,68E+08 | 8751 | 2 | 15159 | ENSMUST00000033717.9 | 210 | ENSMUSG00000031352 | Hccs | holocytochrome c synthetase |
| chr12 | 1,16E+08 | 1,16E+08 | 772 | * | 4,968584 | 5,44801511 | 2,034856 | 2,677347 | 0,007421 | 1 | 0 | Intron (ENSMUST0000( | 12 | 1,16E+08 | 1,16E+08 | 90971 | 1 | 52635 | ENSMUST00000220720.2 | 13249 | ENSMUSG00000021171 | Esyt2 | extended synaptotagmin-like protein 2 |
| chrX | 36072794 | 36073502 | 709 | * | 4,519974 | 5,448147961 | 2,104833 | 2,588399 | 0,009642 | 1 | 0 | Intron (ENSMUST0000( | 20 | 36064828 | 36147875 | 83048 | 1 | 22209 | ENSMUST00000240546.1 | 7966 | ENSMUSG00000016308 | Ube2a | ubiquitin-conjugating enzyme E2A |
| chr7 | 1,22E+08 | 1,22E+08 | 516 | * | 5,057303 | 5,448392648 | 2,088426 | 2,608851 | 0,009085 | 1 | 0 | Promoter (<=1kb) | 7 | 1,22E+08 | 1,22E+08 | 1533 | 1 | 28018 | ENSMUST00000124733.2 | 280 | ENSMUSG00000030870 | Ubfd1 | ubiquitin family domain containing 1 |
| chr9 | 36796520 | 36797049 | 530 | * | 5,041334 | 5,448441027 | 2,086043 | 2,611854 | 0,009005 | 1 | 0 | Distal Intergenic | 9 | 36781799 | 36790216 | 8418 | 1 | 235180 | ENSMUST00000161978.2 | 14721 | ENSMUSG00000032118 | Fez1 | fasciculation and elongation protein zeta 1 (zygin I) |
| chr10 | 57651237 | 57651779 | 543 | * | 4,664182 | 5,448689873 | 2,071801 | 2,629929 | 0,00854 | 1 | 0 | Distal Intergenic | 10 | 57660977 | 57664546 | 3570 | 1 | 12140 | ENSMUST00000020024.12 | -9198 | ENSMUSG00000019874 | Fabp7 | fatty acid binding protein 7, brain |
| chr18 | 50050011 | 50050987 | 977 | * | 4,635376 | 5,448968168 | 2,054724 | 2,651923 | 0,008003 | 1 | 0 | Exon (ENSMUST000001 | 18 | 50055780 | 50064981 | 9202 | 1 | 240283 | ENSMUST00000224938.2 | -4793 | ENSMUSG00000037416 | Dmxl1 | Dmx-like 1 |
| chr6 | 98867304 | 98867792 | 489 | * | 5,920537 | 5,449069169 | 2,060022 | 2,645151 | 0,008165 | 1 | 0 | Distal Intergenic | 6 | 98907132 | 98912506 | 5375 | 2 | 108655 | ENSMUST00000175880.2 | 44714 | ENSMUSG00000030067 | Foxp1 | forkhead box P1 |
| chrX | 1,64E+08 | 1,64E+08 | 705 | * | 5,252997 | 5,449072969 | 2,099664 | 2,595212 | 0,009453 | 1 | 0 | Distal Intergenic | 20 | 1,64E+08 | 1,64E+08 | 198377 | 2 | 237213 | ENSMUST00000058787.9 | -52577 | ENSMUSG00000018589 | Glra2 | glycine receptor, alpha 2 subunit |
| chr14 | 1,2E+08 | 1,2E+08 | 1072 | * | 5,372458 | 5,449418119 | 2,029411 | 2,685221 | 0,007248 | 1 | 0 | Intron (ENSMUST0000( | 14 | 1,2E+08 | 1,2E+08 | 13417 | 2 | 1E+08 | ENSMUST00000226598.2 | 6081 | | Gm4681 | predicted gene 4681 |
| chr5 | 1,38E+08 | 1,38E+08 | 829 | * | 5,1443 | 5,449436569 | 2,068951 | 2,633913 | 0,008441 | 1 | 0 | Intron (ENSMUST0000( | 5 | 1,38E+08 | 1,38E+08 | 657 | 2 | 545812 | ENSMUST00000199773.2 | -13490 | ENSMUSG00000066682 | Pilrb2 | paired immunoglobin-like type 2 receptor beta 2 |
| chr19 | 7028098 | 7028625 | 528 | * | 4,550564 | 5,450096672 | 2,0855 | 2,613328 | 0,008967 | 1 | 0 | Distal Intergenic | 19 | 7034178 | 7175422 | 141245 | 1 | 107227 | ENSMUST00000040261.7 | -5553 | ENSMUSG00000036278 | Macrod1 | mono-ADP ribosylhydrolase 1 |
| chrX | 1,11E+08 | 1,11E+08 | 876 | * | 5,14137 | 5,450215398 | 2,088959 | 2,609058 | 0,009079 | 1 | 0 | Distal Intergenic | 20 | 1,11E+08 | 1,11E+08 | 349202 | 1 | 71030 | ENSMUST00000207962.2 | -14390 | ENSMUSG00000079460 | 4933403O08Rik | RIKEN cDNA 4933403O08 gene |
| chr10 | 86077514 | 86077961 | 448 | * | 5,278708 | 5,450295885 | 1,986303 | 2,74394 | 0,006071 | 1 | 0 | Intron (ENSMUST0000( | 10 | 86136236 | 86185370 | 49135 | 1 | 21859 | ENSMUST00000020234.14 | -58275 | ENSMUSG00000020044 | Timp3 | tissue inhibitor of metalloproteinase 3 |
| chr15 | 76784912 | 76785438 | 527 | * | 5,420302 | 5,450380326 | 1,982403 | 2,74938 | 0,005971 | 1 | 0 | Promoter (<=1kb) | 15 | 76784136 | 76785391 | 1256 | 1 | 66398 | ENSMUST00000109793.3 | 776 | ENSMUSG00000055041 | Commd5 | COMM domain containing 5 |
| chr7 | 65908238 | 65908826 | 589 | * | 5,792097 | 5,450391573 | 1,981518 | 2,750614 | 0,005948 | 1 | 0 | 3' UTR | 7 | 65876660 | 65912218 | 35559 | 2 | 233328 | ENSMUST00000167705.8 | 3392 | ENSMUSG00000015133 | Lrrk1 | leucine-rich repeat kinase 1 |
| chr12 | 72479935 | 72480465 | 531 | * | 5,588859 | 5,450450759 | 2,095283 | 2,601296 | 0,009287 | 1 | 0 | Distal Intergenic | 12 | 72488640 | 72544013 | 55374 | 1 | 78257 | ENSMUST00000161284.3 | -8175 | ENSMUSG00000021090 | Lrrc9 | leucine rich repeat containing 9 |
| chr10 | 59706796 | 59707308 | 513 | * | 5,668199 | 5,450492907 | 1,976941 | 2,757034 | 0,005833 | 1 | 0 | Exon (ENSMUST000002 | 10 | 59715378 | 59735124 | 19747 | 1 | 56709 | ENSMUST00000147914.8 | -8070 | ENSMUSG00000020109 | Dnajb12 | DnaJ heat shock protein family (Hsp40) member B12 |
| chr4 | 1,34E+08 | 1,34E+08 | 710 | * | 5,703716 | 5,450499413 | 1,976631 | 2,757469 | 0,005825 | 1 | 0 | Intron (ENSMUST0000( | 4 | 1,34E+08 | 1,34E+08 | 135879 | 2 | 230815 | ENSMUST00000176606.2 | 39145 | ENSMUSG00000037306 | Man1c1 | mannosidase, alpha, class 1C, member 1 |
| chrX | 41377283 | 41377961 | 679 | * | 5,304647 | 5,450688414 | 1,996086 | 2,730688 | 0,00632 | 1 | 0 | Distal Intergenic | 20 | 41241057 | 41309906 | 68850 | 1 | 20843 | ENSMUST00000128799.3 | 136226 | ENSMUSG00000025862 | Stag2 | stromal antigen 2 |
| chr14 | 63716809 | 63717313 | 505 | * | 5,64947 | 5,450770538 | 1,964791 | 2,774224 | 0,005533 | 1 | 0 | Distal Intergenic | 14 | 63731148 | 63737321 | 6174 | 1 | 1,05E+08 | ENSMUST00000164515.2 | -13835 | | Gm17232 | predicted gene 17232 |
| chr7 | 99047652 | 99048263 | 612 | * | 6,232475 | 5,451745468 | 1,925499 | 2,831341 | 0,004635 | 1 | 0 | Intron (ENSMUST0000( | 7 | 99036217 | 99073366 | 37150 | 1 | 233552 | ENSMUST00000207530.2 | 11435 | ENSMUSG00000035314 | Gdpd5 | glycerophosphodiester phosphodiesterase domain containing 5 |
| chr12 | 84355510 | 84357446 | 1937 | * | 13,94489 | 5,451966873 | 1,748298 | 3,118442 | 0,001818 | 0,614021623 | 1,2763 | 3' UTR | 12 | 84363626 | 84390497 | 26872 | 1 | 52708 | ENSMUST00000045931.12 | -6180 | ENSMUSG00000042472 | Zfp410 | zinc finger protein 410 |
| chr14 | 69677482 | 69678875 | 1394 | * | 5,413079 | 5,452775582 | 2,063746 | 2,642174 | 0,008238 | 1 | 0 | Intron (ENSMUST0000( | 14 | 69656843 | 69663926 | 7084 | 2 | 1,01E+08 | ENSMUST00000184039.2 | -13556 | | Gm16867 | predicted gene, 16867 |
| chr9 | 90041701 | 90042598 | 898 | * | 6,982941 | 5,453322817 | 1,871926 | 2,913215 | 0,003577 | 1 | 0 | Promoter (2-3kb) | 9 | 90045122 | 90082153 | 37032 | 1 | 108153 | ENSMUST00000113059.8 | -2524 | ENSMUSG00000032363 | Adamts7 | a disintegrin-like and metallopeptidase (reprolysin type) with thrombospondin type 1 motif, 7 |
| chr12 | 1,08E+08 | 1,08E+08 | 879 | * | 6,939242 | 5,453570297 | 1,864392 | 2,925119 | 0,003443 | 0,711199341 | 1,360622 | Promoter (1-2kb) | 12 | 1,08E+08 | 1,08E+08 | 23622 | 1 | 12454 | ENSMUST00000101055.5 | 1065 | ENSMUSG00000021258 | Ccnk | cyclin K |
| chr5 | 78836844 | 78838135 | 1292 | * | 8,252782 | 5,454766946 | 1,817011 | 3,002054 | 0,002682 | 0,579477808 | 2,237403 | Distal Intergenic | 5 | 78927706 | 78927841 | 136 | 1 | 1,15E+08 | ENSMUST00000103990.3 | -89571 | ENSMUSG00000077178 | Gm25756 | predicted gene, 25756 |
| chr15 | 5171056 | 5171982 | 927 | * | 7,97896 | 5,454767815 | 1,831023 | 2,979081 | 0,002891 | 1 | 0 | Promoter (1-2kb) | 15 | 5173343 | 5211380 | 38038 | 1 | 105787 | ENSMUST00000051186.9 | -1361 | ENSMUSG00000050697 | Prkaa1 | protein kinase, AMP-activated, alpha 1 catalytic subunit |
| chr7 | 68038237 | 68038657 | 421 | * | 5,248369 | 5,456642884 | 1,936844 | 2,817286 | 0,004843 | 1 | 0 | Intron (ENSMUST0000( | 7 | 68067150 | 68075881 | 8732 | 1 | 71244 | ENSMUST00000207304.2 | -28493 | ENSMUSG00000108924 | 4933436H12Rik | RIKEN cDNA 4933436H12 gene |
| chr14 | 5611919 | 5612640 | 722 | * | 4,241587 | 5,458716127 | 2,096552 | 2,603664 | 0,009223 | 1 | 0 | Distal Intergenic | 14 | 5650540 | 5960698 | 310159 | 1 | 218772 | ENSMUST00000223976.2 | -37900 | ENSMUSG00000017491 | Rarb | retinoic acid receptor, beta |
| chr4 | 1,37E+08 | 1,37E+08 | 558 | * | 4,262507 | 5,460802663 | 2,097683 | 2,603254 | 0,009234 | 1 | 0 | Distal Intergenic | 4 | 1,36E+08 | 1,37E+08 | 188450 | 2 | 13844 | ENSMUST00000105845.9 | -8818 | ENSMUSG00000028664 | Ephb2 | Eph receptor B2 |
| chr1 | 1,35E+08 | 1,35E+08 | 1500 | * | 11,49656 | 5,462181286 | 1,687125 | 3,237568 | 0,001206 | 0,639271276 | 0,744375 | Promoter (<=1kb) | 1 | 1,35E+08 | 1,35E+08 | 1758 | 2 | 329251 | ENSMUST00000146639.2 | 0 | ENSMUSG00000073557 | Ppp1r12b | protein phosphatase 1, regulatory subunit 12B |
| chr12 | 1,1E+08 | 1,1E+08 | 492 | * | 4,426463 | 5,463124658 | 2,041141 | 2,676505 | 0,007439 | 1 | 0 | Promoter (1-2kb) | 12 | 1,1E+08 | 1,1E+08 | 20327 | 1 | 75745 | ENSMUST00000183180.8 | -1148 | | Rian | RNA imprinted and accumulated in nucleus |
| chr19 | 37402294 | 37403225 | 932 | * | 8,012873 | 5,463343938 | 1,737172 | 3,144965 | 0,001661 | 1 | 0 | Exon (ENSMUST000002 | 19 | 37390629 | 37394219 | 3591 | 1 | 16551 | ENSMUST00000236027.2 | 11665 | ENSMUSG00000012443 | Kif11 | kinesin family member 11 |
| chr5 | 10806814 | 10807578 | 765 | * | 6,227497 | 5,468189134 | 2,078548 | 2,630773 | 0,008519 | 1 | 0 | Distal Intergenic | 5 | 10914996 | 10920775 | 5780 | 1 | 623849 | ENSMUST00000095015.11 | -107418 | ENSMUSG00000091422 | Gm6455 | predicted gene 6455 |
| chr10 | 69708543 | 69709343 | 801 | * | 7,377406 | 5,468949714 | 1,799782 | 3,038673 | 0,002376 | 1 | 0 | Intron (ENSMUST0000( | 10 | 69733961 | 69859751 | 125791 | 1 | 11735 | ENSMUST00000182972.8 | -24618 | ENSMUSG00000069601 | Ank3 | ankyrin 3, epithelial |
| chr10 | 12802012 | 12802658 | 647 | * | 8,339942 | 5,471508886 | 1,521454 | 3,596238 | 0,000323 | 1 | 0 | Distal Intergenic | 10 | 12792387 | 12798860 | 6474 | 2 | 320273 | ENSMUST00000181073.10 | -3152 | ENSMUSG00000097378 | B230208H11Rik | RIKEN cDNA B230208H11 gene |
| chr15 | 81154629 | 81155661 | 1033 | * | 5,955702 | 5,472692553 | 2,006516 | 2,72746 | 0,006382 | 1 | 0 | Intron (ENSMUST0000( | 15 | 81126858 | 81136063 | 9206 | 2 | 78102 | ENSMUST00000230860.2 | -18566 | ENSMUSG00000115970 | 8430426J06Rik | RIKEN cDNA 8430426J06 gene |
| chr5 | 33426116 | 33427561 | 1446 | * | 11,4874 | 5,474581323 | 1,770615 | 3,091909 | 0,001989 | 0,644704154 | 1,173325 | Promoter (<=1kb) | 5 | 33407727 | 33426832 | 19106 | 2 | 13016 | ENSMUST00000201275.4 | 0 | ENSMUSG00000037373 | Ctbp1 | C-terminal binding protein 1 |
| chr5 | 33960648 | 33961538 | 891 | * | 5,320397 | 5,474899008 | 1,907842 | 2,869682 | 0,004109 | 1 | 0 | Distal Intergenic | 5 | 33978069 | 34055310 | 77242 | 1 | 107823 | ENSMUST00000075812.11 | -16531 | ENSMUSG00000057406 | Nsd2 | nuclear receptor binding SET domain protein 2 |
| chr2 | 1,65E+08 | 1,65E+08 | 1253 | * | 12,69224 | 5,476229879 | 1,385866 | 3,951485 | 7,77E-05 | 0,205693058 | 0,738441 | Promoter (<=1kb) | 2 | 1,65E+08 | 1,65E+08 | 7062 | 1 | 101113 | ENSMUST00000172577.8 | 0 | ENSMUSG00000050373 | Snx21 | sorting nexin family member 21 |
| chr18 | 67334054 | 67334719 | 666 | * | 5,261394 | 5,476961685 | 2,011067 | 2,72341 | 0,006461 | 1 | 0 | Intron (ENSMUST0000( | 18 | 67338437 | 67340960 | 2524 | 1 | 67064 | ENSMUST00000210564.3 | -3718 | ENSMUSG00000109901 | Chmp1b | charged multivesicular body protein 1B |
| chr12 | 1,11E+08 | 1,11E+08 | 893 | * | 6,704014 | 5,477426761 | 1,900493 | 2,882108 | 0,00395 | 1 | 0 | Intron (ENSMUST0000( | 12 | 1,11E+08 | 1,11E+08 | 73587 | 1 | 217864 | ENSMUST00000116388.4 | 21828 | ENSMUSG00000037896 | Rcor1 | REST corepressor 1 |
| chrX | 72815037 | 72815672 | 636 | * | 4,899102 | 5,477447869 | 2,002346 | 2,735515 | 0,006228 | 1 | 0 | Promoter (2-3kb) | 20 | 72818011 | 72822531 | 4521 | 1 | 56504 | ENSMUST00000002081.6 | -2339 | ENSMUSG00000002007 | Srpk3 | serine/arginine-rich protein specific kinase 3 |
| chr13 | 5031340 | 5031783 | 444 | * | 5,250755 | 5,478185524 | 2,115305 | 2,589785 | 0,009604 | 1 | 0 | Distal Intergenic | 13 | 5239802 | 5239933 | 132 | 2 | 1,15E+08 | ENSMUST00000104285.3 | 208150 | ENSMUSG00000077473 | Gm24790 | predicted gene, 24790 |
| chr14 | 25223151 | 25223794 | 644 | * | 5,810449 | 5,478795176 | 1,978352 | 2,769373 | 0,005616 | 1 | 0 | Distal Intergenic | 14 | 25140219 | 25143665 | 3447 | 2 | 75862 | ENSMUST00000225429.2 | -79486 | ENSMUSG00000114916 | 4930572O13Rik | RIKEN cDNA 4930572O13 gene |
| chr9 | 1,1E+08 | 1,1E+08 | 334 | * | 4,825177 | 5,479577462 | 2,071721 | 2,64494 | 0,008171 | 1 | 0 | Intron (ENSMUST0000( | 9 | 1,1E+08 | 1,1E+08 | 25967 | 2 | 546161 | ENSMUST00000098359.4 | 5271 | ENSMUSG00000074059 | Fbxw18 | F-box and WD-40 domain protein 18 |
| chr18 | 36501054 | 36501909 | 856 | * | 6,46874 | 5,480235615 | 2,012474 | 2,723134 | 0,006467 | 1 | 0 | Promoter (2-3kb) | 18 | 36498834 | 36535593 | 36760 | 1 | 66060 | ENSMUST00000144158.2 | 2220 | ENSMUSG00000046727 | Cystm1 | cysteine-rich transmembrane module containing 1 |
| chr15 | 31491163 | 31491973 | 811 | * | 5,818007 | 5,480524182 | 2,01704 | 2,717112 | 0,006585 | 1 | 0 | Intron (ENSMUST0000( | 15 | 31465397 | 31472137 | 6741 | 2 | 223455 | ENSMUST00000226868.2 | -19026 | ENSMUSG00000039100 | Marchf6 | membrane associated ring-CH-type finger 6 |
| chr7 | 1,4E+08 | 1,4E+08 | 1004 | * | 9,242479 | 5,480836259 | 1,85734 | 2,950906 | 0,003168 | 0,685026746 | 1,425929 | 3' UTR | 7 | 1,4E+08 | 1,4E+08 | 8709 | 2 | 11501 | ENSMUST00000148670.8 | -3540 | ENSMUSG00000025473 | Adam8 | a disintegrin and metallopeptidase domain 8 |
| chr13 | 35257366 | 35258103 | 738 | * | 5,565462 | 5,481215396 | 2,028946 | 2,701509 | 0,006903 | 1 | 0 | Intron (ENSMUST0000( | 13 | 35262472 | 35267243 | 4772 | 1 | 1,05E+08 | ENSMUST00000223077.2 | -4369 | ENSMUSG00000113738 | Gm40910 | predicted gene, 40910 |
| chr11 | 86892029 | 86892770 | 742 | * | 4,539601 | 5,481791488 | 2,126997 | 2,577245 | 0,009959 | 1 | 0 | Distal Intergenic | 11 | 86887959 | 86888089 | 131 | 2 | 1,15E+08 | ENSMUST00000158191.3 | -3940 | ENSMUSG00000088816 | Gm22387 | predicted gene, 22387 |
| chr19 | 6913196 | 6913621 | 426 | * | 4,336853 | 5,481816447 | 2,120596 | 2,585036 | 0,009737 | 1 | 0 | Promoter (1-2kb) | 19 | 6903030 | 6911883 | 8854 | 2 | 16528 | ENSMUST00000025908.8 | -1313 | ENSMUSG00000024957 | Kcnk4 | potassium channel, subfamily K, member 4 |
| chr7 | 1E+08 | 1E+08 | 801 | * | 4,270307 | 5,481872362 | 2,108041 | 2,600458 | 0,00931 | 1 | 0 | Intron (ENSMUST0000( | 7 | 1E+08 | 1E+08 | 843 | 2 | 1,05E+08 | ENSMUST00000209160.2 | -14391 | | Gm39059 | predicted gene, 39059 |
| chr10 | 1,08E+08 | 1,08E+08 | 520 | * | 4,175201 | 5,481876721 | 2,102067 | 2,607851 | 0,009111 | 1 | 0 | Distal Intergenic | 10 | 1,08E+08 | 1,08E+08 | 115159 | 1 | 17931 | ENSMUST00000219263.2 | -102202 | ENSMUSG00000019907 | Ppp1r12a | protein phosphatase 1, regulatory subunit 12A |
| chr11 | 95336573 | 95337225 | 653 | * | 4,611385 | 5,481885553 | 2,098931 | 2,611751 | 0,009008 | 1 | 0 | Intron (ENSMUST0000( | 11 | 95365145 | 95381141 | 15997 | 1 | 20747 | ENSMUST00000150922.2 | -27920 | ENSMUSG00000057522 | Spop | speckle-type BTB/POZ protein |
| chr6 | 51951367 | 51951802 | 436 | * | 4,459336 | 5,481915866 | 2,094184 | 2,617686 | 0,008853 | 1 | 0 | Intron (ENSMUST0000( | 6 | 51836146 | 51989495 | 153350 | 2 | 54353 | ENSMUST00000078214.8 | 37693 | ENSMUSG00000059182 | Skap2 | src family associated phosphoprotein 2 |
| chr1 | 1,87E+08 | 1,87E+08 | 1174 | * | 5,029274 | 5,482059615 | 2,053418 | 2,669724 | 0,007591 | 1 | 0 | Intron (ENSMUST0000( | 1 | 1,87E+08 | 1,87E+08 | 125248 | 1 | 67769 | ENSMUST00000160570.2 | 76123 | ENSMUSG00000039210 | Gpatch2 | G patch domain containing 2 |
| chr5 | 22748848 | 22749594 | 747 | * | 4,512599 | 5,482067616 | 2,051999 | 2,671574 | 0,00755 | 1 | 0 | Intron (ENSMUST0000( | 5 | 22728514 | 22754516 | 26003 | 2 | 26429 | ENSMUST00000141263.3 | 4922 | ENSMUSG00000029012 | Orc5 | origin recognition complex, subunit 5 |
| chr4 | 53967392 | 53968018 | 627 | * | 4,761443 | 5,482164798 | 2,029704 | 2,700967 | 0,006914 | 1 | 0 | Distal Intergenic | 4 | 54004057 | 54004187 | 131 | 1 | 1,15E+08 | ENSMUST00000158927.2 | -36039 | ENSMUSG00000089552 | Gm24177 | predicted gene, 24177 |
| chr19 | 57465916 | 57466445 | 530 | * | 4,724434 | 5,482169182 | 2,029203 | 2,701636 | 0,0069 | 1 | 0 | Intron (ENSMUST0000( | 19 | 57485822 | 57501216 | 15395 | 1 | 78073 | ENSMUST00000234319.2 | -19377 | ENSMUSG00000117655 | 6720468P15Rik | RIKEN cDNA 6720468P15 gene |
| chr5 | 33300282 | 33300887 | 606 | * | 4,951539 | 5,48218813 | 2,025475 | 2,706619 | 0,006797 | 1 | 0 | Intron (ENSMUST0000( | 5 | 33261563 | 33320214 | 58652 | 1 | 20537 | ENSMUST00000011178.5 | 38719 | ENSMUSG00000011034 | Slc5a1 | solute carrier family 5 (sodium/glucose cotransporter), member 1 |
| chr15 | 1E+08 | 1E+08 | 539 | * | 4,860089 | 5,482207147 | 2,019983 | 2,713987 | 0,006648 | 1 | 0 | Promoter (2-3kb) | 15 | 1E+08 | 1E+08 | 49262 | 2 | 21422 | ENSMUST00000229696.2 | -2066 | ENSMUSG00000009733 | Tfcp2 | transcription factor CP2 |
| chr4 | 41125313 | 41126174 | 862 | * | 5,219237 | 5,482211619 | 2,018828 | 2,715541 | 0,006617 | 1 | 0 | Promoter (<=1kb) | 4 | 41114427 | 41124455 | 10029 | 2 | 230082 | ENSMUST00000030138.9 | -858 | ENSMUSG00000028430 | Nol6 | nucleolar protein family 6 (RNA-associated) |
| chr4 | 98812560 | 98813012 | 453 | * | 5,116501 | 5,482219185 | 2,021289 | 2,712239 | 0,006683 | 1 | 0 | Promoter (<=1kb) | 4 | 98812201 | 98816507 | 4307 | 1 | 230484 | ENSMUST00000169053.2 | 359 | ENSMUSG00000028560 | Usp1 | ubiquitin specific peptidase 1 |
| chr3 | 1,01E+08 | 1,01E+08 | 659 | * | 6,163565 | 5,482259578 | 2,01017 | 2,727261 | 0,006386 | 1 | 0 | Intron (ENSMUST0000( | 3 | 1E+08 | 1,01E+08 | 64683 | 2 | 17156 | ENSMUST00000133642.2 | 36102 | ENSMUSG00000008763 | Man1a2 | mannosidase, alpha, class 1A, member 2 |
| chr14 | 26554725 | 26555356 | 632 | * | 5,011623 | 5,482301823 | 2,000708 | 2,740181 | 0,006141 | 1 | 0 | Intron (ENSMUST0000( | 14 | 26616514 | 26637214 | 20701 | 1 | 142687 | ENSMUST00000238987.2 | -61158 | ENSMUSG00000021898 | Asb14 | ankyrin repeat and SOCS box-containing 14 |
| chr12 | 15438577 | 15439126 | 550 | * | 5,630636 | 5,482303894 | 2,000172 | 2,740916 | 0,006127 | 1 | 0 | Distal Intergenic | 12 | 15656325 | 15779692 | 123368 | 1 | 1,05E+08 | ENSMUST00000222542.3 | -217199 | | Gm40287 | predicted gene, 40287 |
| chrX | 37055520 | 37056431 | 912 | * | 5,134347 | 5,482374332 | 1,985268 | 2,761529 | 0,005753 | 1 | 0 | Distal Intergenic | 20 | 37029693 | 37034479 | 4787 | 2 | 1,03E+08 | ENSMUST00000117962.3 | -21041 | | Rhox11-ps2 | reproductive homeobox 11, pseudogene 2 |
| chr15 | 4688004 | 4688902 | 899 | * | 5,425994 | 5,482385949 | 1,984177 | 2,763052 | 0,005726 | 1 | 0 | Distal Intergenic | 15 | 4756657 | 4833527 | 76871 | 1 | 12274 | ENSMUST00000162585.8 | -67755 | ENSMUSG00000022181 | C6 | complement component 6 |
| chr4 | 1,21E+08 | 1,21E+08 | 660 | * | 5,536798 | 5,482467138 | 1,96766 | 2,786288 | 0,005332 | 1 | 0 | Promoter (<=1kb) | 4 | 1,21E+08 | 1,21E+08 | 1821 | 1 | 12840 | ENSMUST00000140119.2 | 555 | ENSMUSG00000028626 | Col9a2 | collagen, type IX, alpha 2 |

| | | | | | | | | | | | | | | | | | | | | | | | |
|---|---|---|---|---|---|---|---|---|---|---|---|---|---|---|---|---|---|---|---|---|---|---|---|
| chr5 | 1,44E+08 | 1,44E+08 | 721 | * | 6,186905 | 5,482705647 | 1,926734 | 2,845595 | 0,004433 | 1 | 0 | Distal Intergenic | 5 | 1,44E+08 | 1,44E+08 | 87769 | 1 | 231876 | ENSMUST00000041804.8 | -29804 | ENSMUSG00000038970 | Lmtk2 | lemur tyrosine kinase 2 |
| chr19 | 37186510 | 37187329 | 820 | * | 6,205937 | 5,482778089 | 1,915185 | 2,862793 | 0,004199 | 1 | 0 | Promoter (<=1kb) | 19 | 37152023 | 37186000 | 33978 | 2 | 208922 | ENSMUST00000131917.2 | -510 | ENSMUSG00000039652 | Cpeb3 | cytoplasmic polyadenylation element binding protein 3 |
| chr12 | 71858204 | 71859205 | 1002 | * | 6,506874 | 5,482782943 | 1,914711 | 2,863504 | 0,00419 | 1 | 0 | Distal Intergenic | 12 | 71877852 | 72039107 | 161256 | 1 | 208846 | ENSMUST00000223272.2 | -18647 | ENSMUSG00000034574 | Daam1 | dishevelled associated activator of morphogenesis 1 |
| chr8 | 1,12E+08 | 1,12E+08 | 761 | * | 7,538513 | 5,482890073 | 1,898314 | 2,888294 | 0,003873 | 1 | 0 | Distal Intergenic | 8 | 1,12E+08 | 1,12E+08 | 29279 | 2 | 234736 | ENSMUST00000038739.5 | -7556 | ENSMUSG00000033596 | Rfwd3 | ring finger and WD repeat domain 3 |
| chr10 | 19105442 | 19106192 | 751 | * | 5,259312 | 5,483094407 | 2,063292 | 2,65745 | 0,007873 | 1 | 0 | Distal Intergenic | 10 | 19042986 | 19082450 | 39465 | 1 | 1,01E+08 | ENSMUST00000216683.2 | 62456 | | Gm20139 | predicted gene, 20139 |
| chr9 | 55407925 | 55409183 | 1259 | * | 8,719203 | 5,483095744 | 1,870605 | 2,931188 | 0,003377 | 0,693003605 | 1,449706 | Exon (ENSMUST00000 | 9 | 55396110 | 55419092 | 22983 | 2 | 110842 | ENSMUST00000142587.2 | 9909 | ENSMUSG00000032314 | Etfa | electron transferring flavoprotein, alpha polypeptide |
| chr10 | 36262644 | 36263390 | 747 | * | 8,251872 | 5,48383486 | 1,822614 | 3,008774 | 0,002623 | 0,668446397 | 1,360622 | Distal Intergenic | 10 | 36382810 | 36709769 | 326960 | 1 | 319415 | ENSMUST00000167191.8 | -119420 | ENSMUSG00000044499 | Hs3st5 | heparan sulfate (glucosamine) 3-O-sulfotransferase 5 |
| chr9 | 60472199 | 60472749 | 551 | * | 5,62461 | 5,484278772 | 1,993738 | 2,750753 | 0,005946 | 1 | 0 | Distal Intergenic | 9 | 60432993 | 60453185 | 20193 | 2 | 1,01E+08 | ENSMUST00000242602.1 | -19014 | ENSMUSG00000111761 | 9230112J17Rik | RIKEN cDNA 9230112J17 gene |
| chr9 | 8214997 | 8215673 | 677 | * | 4,754785 | 5,484403086 | 1,990271 | 2,755606 | 0,005858 | 1 | 0 | Distal Intergenic | 9 | 8221890 | 8241988 | 20099 | 1 | 76669 | ENSMUST00000034483.5 | -6217 | ENSMUSG00000031998 | 1700128F08Rik | U1-snRNP binding protein homolog pseudogene |
| chr17 | 79746073 | 79747092 | 1020 | * | 6,67097 | 5,48508806 | 1,998885 | 2,744074 | 0,006068 | 1 | 0 | Exon (ENSMUST00000 | 17 | 79746334 | 79759191 | 12858 | 2 | 1,03E+08 | ENSMUST00000242898.1 | 12099 | | Gm32954 | predicted gene, 32954 |
| chr5 | 22877463 | 22877997 | 535 | * | 4,95078 | 5,485294933 | 2,108628 | 2,601357 | 0,009286 | 1 | 0 | Intron (ENSMUST00000 | 5 | 22850117 | 22850181 | 65 | 2 | 1,15E+08 | ENSMUST00000174963.3 | -27282 | ENSMUSG00000092704 | Gm24512 | predicted gene, 24512 |
| chr18 | 57621976 | 57622480 | 505 | * | 6,436746 | 5,48757216 | 2,037537 | 2,693237 | 0,007076 | 1 | 0 | Intron (ENSMUST00000 | 18 | 57605525 | 57611204 | 5680 | 1 | 629147 | ENSMUST00000209786.2 | 16451 | ENSMUSG00000069372 | Ctxn3 | cortexin 3 |
| chr7 | 1,44E+08 | 1,44E+08 | 600 | * | 7,006158 | 5,489025999 | 1,878161 | 2,922554 | 0,003472 | 1 | 0 | Promoter (<=1kb) | 7 | 1,44E+08 | 1,44E+08 | 76972 | 2 | 233977 | ENSMUST00000168134.10 | -307 | ENSMUSG00000037519 | Ppfia1 | protein tyrosine phosphatase, receptor type, f polypeptide (PTPRF), interacting protein (liprin), alpha 1 |
| chr1 | 87919683 | 87920536 | 854 | * | 7,788474 | 5,489178567 | 1,945375 | 2,821656 | 0,004778 | 0,766939286 | 1,360622 | Intron (ENSMUST00000 | 1 | 87913477 | 87936250 | 22774 | 2 | 227334 | ENSMUST00000186315.2 | 15714 | ENSMUSG00000005501 | Usp40 | ubiquitin specific peptidase 40 |
| chr3 | 24378738 | 24379433 | 696 | * | 5,84221 | 5,491859421 | 2,014633 | 2,725985 | 0,006411 | 1 | 0 | Intron (ENSMUST00000 | 3 | 24616238 | 24616344 | 107 | 1 | 1,15E+08 | ENSMUST00000093715.3 | -236805 | ENSMUSG00000070227 | Gm24704 | predicted gene, 24704 |
| chr17 | 29572520 | 29573335 | 816 | * | 7,04799 | 5,493102432 | 1,866714 | 2,942658 | 0,003254 | 1 | 0 | Distal Intergenic | 17 | 29551050 | 29566908 | 15859 | 2 | 56462 | ENSMUST00000234948.2 | -5612 | ENSMUSG00000024012 | Mtch1 | mitochondrial carrier 1 |
| chr10 | 12840061 | 12840655 | 595 | * | 7,881174 | 5,495989611 | 1,496433 | 3,672726 | 0,00024 | 1 | 0 | Promoter (<=1kb) | 10 | 12817471 | 12840042 | 22572 | 2 | 74732 | ENSMUST00000218685.2 | -19 | ENSMUSG00000039232 | Stx11 | syntaxin 11 |
| chr11 | 1,16E+08 | 1,16E+08 | 770 | * | 6,127114 | 5,497028783 | 1,872982 | 2,934907 | 0,003336 | 1 | 0 | Promoter (2-3kb) | 11 | 1,16E+08 | 1,16E+08 | 4855 | 2 | 217335 | ENSMUST00000126861.8 | 2011 | ENSMUSG00000020776 | Fbf1 | Fas (TNFRSF6) binding factor 1 |
| chr12 | 95520224 | 95520614 | 391 | * | 4,122974 | 5,497754294 | 2,134127 | 2,576114 | 0,009992 | 1 | 0 | Distal Intergenic | 12 | 95586592 | 95586698 | 107 | 1 | 1,15E+08 | ENSMUST00000240474.1 | -65978 | ENSMUSG00000118868 | Gm22246 | predicted gene, 22246 |
| chr11 | 57409061 | 57410451 | 1391 | * | 12,76692 | 5,498744518 | 1,362042 | 4,037133 | 5,41E-05 | 0,166885401 | 0,738441 | Promoter (<=1kb) | 11 | 57409490 | 57424641 | 15152 | 1 | 216760 | ENSMUST00000108849.8 | 0 | ENSMUSG00000020522 | Mfap3 | microfibrillar-associated protein 3 |
| chr17 | 50777338 | 50778100 | 763 | * | 10,64433 | 5,499767973 | 1,7873 | 3,077137 | 0,00209 | 0,64068479 | 1,2763 | Distal Intergenic | 17 | 50816431 | 50995512 | 179082 | 1 | 224860 | ENSMUST00000043938.8 | -38331 | ENSMUSG00000038910 | Plcl2 | phospholipase C-like 2 |
| chr5 | 54699825 | 54700632 | 808 | * | 6,183752 | 5,503342598 | 1,886275 | 2,917572 | 0,003528 | 1 | 0 | Distal Intergenic | 5 | 54507333 | 54515885 | 8553 | 1 | 330086 | ENSMUST00000197424.2 | 192492 | ENSMUSG00000105912 | Gm10440 | predicted gene 10440 |
| chrX | 73313549 | 73314564 | 1016 | * | 7,974616 | 5,504039637 | 1,785858 | 3,082014 | 0,002056 | 1 | 0 | Promoter (<=1kb) | 20 | 73314418 | 73316741 | 2324 | 1 | 110954 | ENSMUST00000008826.14 | 0 | ENSMUSG00000008682 | Rpl10 | ribosomal protein L10 |
| chr8 | 65143309 | 65144327 | 1019 | * | 9,183064 | 5,504430131 | 1,913658 | 2,876392 | 0,004023 | 0,728793055 | 1,425929 | Promoter (1-2kb) | 8 | 65045576 | 65146088 | 100513 | 2 | 12876 | ENSMUST00000048967.9 | 1761 | ENSMUSG00000037852 | Cpe | carboxypeptidase E |
| chr13 | 23884565 | 23885642 | 1078 | * | 8,789214 | 5,50509205 | 1,849137 | 2,977114 | 0,00291 | 0,675797279 | 1,425929 | Promoter (2-3kb) | 13 | 23881618 | 23882437 | 820 | 2 | 319155 | ENSMUST00000102967.3 | -2128 | ENSMUSG00000060678 | H4c3 | H4 clustered histone 3 |
| chr18 | 63598617 | 63599571 | 955 | * | 6,38203 | 5,507388367 | 1,941198 | 2,837108 | 0,004552 | 1 | 0 | Distal Intergenic | 18 | 63641240 | 63641302 | 63 | 1 | 1,15E+08 | ENSMUST00000158531.3 | -41669 | ENSMUSG00000089156 | Gm23581 | predicted gene, 23581 |
| chr11 | 80140402 | 80141649 | 1248 | * | 8,134368 | 5,507949904 | 1,949848 | 2,824811 | 0,004731 | 1 | 0 | Promoter (<=1kb) | 11 | 80140500 | 80157564 | 17065 | 1 | 59040 | ENSMUST00000135947.2 | 0 | ENSMUSG00000017686 | Rhot1 | ras homolog family member T1 |
| chr10 | 81405900 | 81407303 | 1404 | * | 9,550784 | 5,509049564 | 1,740968 | 3,164361 | 0,001554 | 0,615591422 | 1,086118 | Distal Intergenic | 10 | 81410395 | 81426300 | 15906 | 1 | 21886 | ENSMUST00000140433.8 | -3092 | ENSMUSG00000034771 | Tle2 | transducin-like enhancer of split 2 |
| chr7 | 92486209 | 92487219 | 1011 | * | 7,402099 | 5,510000384 | 1,973311 | 2,792262 | 0,005234 | 1 | 0 | 3' UTR | 7 | 92469128 | 92485852 | 16725 | 1 | 75985 | ENSMUST00000107179.2 | 17081 | ENSMUSG00000030643 | Rab30 | RAB30, member RAS oncogene family |
| chr14 | 18384137 | 18384435 | 299 | * | 4,424596 | 5,510316427 | 2,13367 | 2,582553 | 0,009807 | 1 | 0 | Promoter (1-2kb) | 14 | 18370467 | 18385653 | 15187 | 2 | 1E+08 | ENSMUST00000166928.8 | 1218 | ENSMUSG00000091022 | Gm3115 | predicted gene 3115 |
| chr1 | 58505430 | 58506768 | 1339 | * | 7,635339 | 5,510340343 | 1,977461 | 2,786573 | 0,005327 | 1 | 0 | Promoter (<=1kb) | 1 | 58505674 | 58505780 | 107 | 2 | 1,15E+08 | ENSMUST00000179823.3 | 0 | ENSMUSG00000119442 | Gm23966 | predicted gene, 23966 |
| chrX | 12729779 | 12731054 | 1276 | * | 5,55563 | 5,510397039 | 2,131206 | 2,585577 | 0,009722 | 1 | 0 | Distal Intergenic | 20 | 12754466 | 12862153 | 107688 | 1 | 75037 | ENSMUST00000245073.1 | -23412 | | 4930500G05Rik | RIKEN cDNA 4930500G05 gene |
| chr6 | 1,18E+08 | 1,18E+08 | 650 | * | 7,62308 | 5,510406683 | 1,978267 | 2,785472 | 0,005345 | 1 | 0 | Promoter (2-3kb) | 6 | 1,18E+08 | 1,18E+08 | 7651 | 1 | 22685 | ENSMUST00000112857.3 | 2033 | ENSMUSG00000042097 | Zfp239 | zinc finger protein 239 |
| chr4 | 6034685 | 6035690 | 1006 | * | 7,201982 | 5,510587305 | 1,980447 | 2,782497 | 0,005394 | 1 | 0 | Intron (ENSMUST00000 | 4 | 6191098 | 6221688 | 30591 | 1 | 68053 | ENSMUST00000029907.6 | -155408 | ENSMUSG00000028243 | Ubxn2b | UBX domain protein 2B |
| chr2 | 1,71E+08 | 1,71E+08 | 335 | * | 4,485172 | 5,510982427 | 2,115803 | 2,604677 | 0,009196 | 1 | 0 | Intron (ENSMUST00000 | 2 | 1,71E+08 | 1,71E+08 | 2894 | 2 | 1,05E+08 | ENSMUST00000242846.1 | 14061 | | Gm40003 | predicted gene, 40003 |
| chr18 | 5338849 | 5340043 | 1195 | * | 4,634321 | 5,511206114 | 2,110027 | 2,611912 | 0,009004 | 1 | 0 | Distal Intergenic | 18 | 5210124 | 5334807 | 124684 | 2 | 240186 | ENSMUST00000234241.2 | -4042 | ENSMUSG00000050945 | Zfp438 | zinc finger protein 438 |
| chr9 | 71161299 | 71162406 | 1108 | * | 8,397343 | 5,51125432 | 1,798 | 3,065213 | 0,002175 | 0,624579927 | 1,449706 | Exon (ENSMUST00000 | 9 | 71123071 | 71203525 | 80455 | 1 | 19378 | ENSMUST00000034723.6 | 38228 | ENSMUSG00000013584 | Aldh1a2 | aldehyde dehydrogenase family 1, subfamily A2 |
| chr9 | 45788552 | 45789339 | 788 | * | 8,350795 | 5,511257639 | 1,988774 | 2,771184 | 0,005585 | 0,792963293 | 1,360622 | Intron (ENSMUST00000 | 9 | 45782879 | 45783559 | 681 | 2 | 235315 | ENSMUST00000160868.2 | -4993 | ENSMUSG00000042790 | Rnf214 | ring finger protein 214 |
| chr15 | 76029703 | 76030398 | 696 | * | 5,229926 | 5,511578736 | 2,100292 | 2,624196 | 0,008685 | 1 | 0 | Distal Intergenic | 15 | 76003717 | 76010796 | 7080 | 2 | 271278 | ENSMUST00000146157.8 | -18907 | ENSMUSG00000044361 | BC024139 | cDNA sequence BC024139 |
| chr2 | 1,32E+08 | 1,32E+08 | 1309 | * | 8,132993 | 5,511661977 | 1,818032 | 3,031663 | 0,002432 | 0,654754826 | 1,360622 | Distal Intergenic | 2 | 1,31E+08 | 1,32E+08 | 68942 | 2 | 1,19E+08 | ENSMUST00000138080.2 | -50582 | | Gm14280 | predicted gene 14280 |
| chr18 | 32014310 | 32015004 | 695 | * | 4,62119 | 5,511801861 | 2,094878 | 2,631085 | 0,008511 | 1 | 0 | Promoter (<=1kb) | 18 | 32015195 | 32019423 | 4229 | 1 | 74320 | ENSMUST00000234137.2 | -191 | ENSMUSG00000024400 | Wdr33 | WD repeat domain 33 |
| chr4 | 1,41E+08 | 1,41E+08 | 1282 | * | 10,02072 | 5,51183729 | 1,827416 | 3,016192 | 0,00256 | 0,683988405 | 1,173325 | Promoter (2-3kb) | 4 | 1,41E+08 | 1,41E+08 | 3324 | 1 | 18600 | ENSMUST00000130958.2 | 2926 | ENSMUSG00000028927 | Padi2 | peptidyl arginine deiminase, type II |
| chr17 | 64810584 | 64810978 | 395 | * | 5,494702 | 5,512762382 | 1,960451 | 2,811986 | 0,004924 | 1 | 0 | Distal Intergenic | 17 | 64821134 | 64862498 | 41365 | 2 | 1E+08 | ENSMUST00000168379.3 | 51520 | ENSMUSG00000073377 | AU016765 | expressed sequence AU016765 |
| chr4 | 1,05E+08 | 1,05E+08 | 736 | * | 6,565809 | 5,513061678 | 1,902179 | 2,898287 | 0,003752 | 1 | 0 | Distal Intergenic | 4 | 1,05E+08 | 1,05E+08 | 493 | 2 | 433742 | ENSMUST00000118507.2 | 52931 | ENSMUSG00000081819 | Gm12722 | predicted gene 12722 |
| chr11 | 86886568 | 86887154 | 587 | * | 5,417141 | 5,513082833 | 2,064419 | 2,670525 | 0,007573 | 1 | 0 | Promoter (<=1kb) | 11 | 86887959 | 86888089 | 131 | 2 | 1,15E+08 | ENSMUST00000158191.3 | 935 | ENSMUSG00000088816 | Gm22387 | predicted gene, 22387 |
| chr4 | 1,4E+08 | 1,4E+08 | 586 | * | 6,00774 | 5,51327251 | 1,917635 | 2,875038 | 0,00404 | 1 | 0 | Intron (ENSMUST00000 | 4 | 1,4E+08 | 1,4E+08 | 219939 | 2 | 230868 | ENSMUST00000039331.9 | 4933 | ENSMUSG00000040972 | Igsf21 | immunoglobulin superfamily, member 21 |
| chr2 | 92525949 | 92526706 | 758 | * | 7,006497 | 5,513321412 | 2,016118 | 2,734623 | 0,006245 | 0,814686254 | 1,360622 | Distal Intergenic | 2 | 92572575 | 92589553 | 16979 | 1 | 1,05E+08 | ENSMUST00000152230.2 | -45869 | | Gm13816 | predicted gene 13816 |
| chr1 | 1,71E+08 | 1,71E+08 | 1190 | * | 5,846763 | 5,513602784 | 1,943666 | 2,836702 | 0,004558 | 1 | 0 | Distal Intergenic | 1 | 1,71E+08 | 1,71E+08 | 6881 | 2 | 14131 | ENSMUST00000168817.2 | 14008 | ENSMUSG00000059498 | Fcgr3 | Fc receptor, IgG, low affinity III |
| chr18 | 59013147 | 59013755 | 609 | * | 5,678088 | 5,513631799 | 1,946349 | 2,832807 | 0,004614 | 1 | 0 | Intron (ENSMUST00000 | 18 | 58970389 | 59037486 | 67098 | 1 | 240322 | ENSMUST00000236901.2 | 42758 | ENSMUSG00000053441 | Adamts19 | a disintegrin-like and metallopeptidase (reprolysin type) with thrombospondin type 1 motif, 19 |
| chr6 | 25029003 | 25029581 | 579 | * | 5,959243 | 5,513765226 | 1,956534 | 2,818129 | 0,00483 | 1 | 0 | Distal Intergenic | 6 | 25078771 | 25078902 | 132 | 2 | 1,15E+08 | ENSMUST00000157242.3 | 49321 | ENSMUSG00000087867 | Gm24443 | predicted gene, 24443 |
| chr13 | 1,17E+08 | 1,17E+08 | 689 | * | 5,050228 | 5,5141628 | 1,993725 | 2,765759 | 0,005679 | 1 | 0 | Distal Intergenic | 13 | 1,17E+08 | 1,17E+08 | 36084 | 2 | 74680 | ENSMUST00000224552.2 | 123383 | ENSMUSG00000114290 | 4930435F18Rik | RIKEN cDNA 4930435F18 gene |
| chr6 | 1,02E+08 | 1,02E+08 | 628 | * | 5,205884 | 5,514366107 | 1,934715 | 2,850222 | 0,004369 | 1 | 0 | Intron (ENSMUST00000 | 6 | 1,02E+08 | 1,02E+08 | 301362 | 2 | 18488 | ENSMUST00000032159.7 | -7461 | ENSMUSG00000030075 | Cntn3 | contactin 3 |
| chrX | 13761379 | 13761773 | 395 | * | 4,69722 | 5,514748274 | 2,059679 | 2,677479 | 0,007418 | 1 | 0 | Distal Intergenic | 20 | 13580970 | 13717606 | 136637 | 2 | 12361 | ENSMUST00000132240.2 | -43773 | ENSMUSG00000031012 | Cask | calcium/calmodulin-dependent serine protein kinase (MAGUK family) |
| chr4 | 1,07E+08 | 1,07E+08 | 654 | * | 4,441079 | 5,514796724 | 2,0661 | 2,669182 | 0,007604 | 1 | 0 | Promoter (<=1kb) | 4 | 1,07E+08 | 1,07E+08 | 39420 | 2 | 66526 | ENSMUST00000141165.2 | -679 | ENSMUSG00000028619 | Tceanc2 | transcription elongation factor A (SII) N-terminal and central domain containing 2 |
| chr6 | 1,08E+08 | 1,08E+08 | 436 | * | 4,495956 | 5,514823951 | 2,069593 | 2,66469 | 0,007706 | 1 | 0 | Promoter (<=1kb) | 6 | 1,08E+08 | 1,08E+08 | 12078 | 1 | 74729 | ENSMUST00000049246.7 | -870 | ENSMUSG00000034639 | Setmar | SET domain without mariner transposase fusion |
| chr5 | 1,21E+08 | 1,21E+08 | 376 | * | 4,778048 | 5,514862038 | 2,073132 | 2,660159 | 0,00781 | 1 | 0 | Intron (ENSMUST00000 | 5 | 1,21E+08 | 1,21E+08 | 9534 | 2 | 100764 | ENSMUST00000156356.2 | 3079 | ENSMUSG00000029600 | Rita1 | RBPJ interacting and tubulin associated 1 |
| chr13 | 1,01E+08 | 1,01E+08 | 704 | * | 4,683332 | 5,514941949 | 2,083308 | 2,647204 | 0,008116 | 1 | 0 | Promoter (2-3kb) | 13 | 1,01E+08 | 1,01E+08 | 130483 | 1 | 1,05E+08 | ENSMUST00000185979.2 | -2705 | | Gm41042 | predicted gene, 41042 |
| chr16 | 20559723 | 20560097 | 375 | * | 4,192476 | 5,51510857 | 2,106938 | 2,617594 | 0,008855 | 1 | 0 | 3' UTR | 16 | 20564058 | 20564222 | 165 | 1 | 1,15E+08 | ENSMUST00000083438.3 | -3961 | ENSMUSG00000065372 | Gm22741 | predicted gene, 22741 |
| chr12 | 73443808 | 73444173 | 366 | * | 4,418883 | 5,515189255 | 2,116052 | 2,606358 | 0,009151 | 1 | 0 | Intron (ENSMUST00000 | 12 | 73441279 | 73456327 | 15049 | 2 | 380765 | ENSMUST00000243711.1 | 12154 | ENSMUSG00000056359 | D830013O20Rik | RIKEN cDNA D830013O20 gene |
| chr8 | 85467293 | 85467603 | 311 | * | 3,957307 | 5,515301891 | 2,134973 | 2,583312 | 0,009786 | 1 | 0 | Intron (ENSMUST00000 | 8 | 85449636 | 85453473 | 3838 | 1 | 619292 | ENSMUST00000098571.5 | 17657 | ENSMUSG00000074203 | G430095P16Rik | RIKEN cDNA G430095P16 gene |
| chr12 | 94450148 | 94450723 | 576 | * | 4,372854 | 5,515310177 | 2,135645 | 2,582503 | 0,009809 | 1 | 0 | Distal Intergenic | 12 | 93894112 | 93894238 | 127 | 1 | 1,02E+08 | ENSMUST00000183568.3 | 556036 | ENSMUSG00000098931 | Mir8099-2 | microRNA 8099-2 |
| chr9 | 17864200 | 17864736 | 537 | * | 4,227902 | 5,515312388 | 2,137707 | 2,580014 | 0,00988 | 1 | 0 | Distal Intergenic | 9 | 18203421 | 18228738 | 25318 | 1 | 66917 | ENSMUST00000001825.9 | -338685 | ENSMUSG00000001774 | Chordc1 | cysteine and histidine-rich domain (CHORD)-containing, zinc-binding protein 1 |
| chr8 | 27773226 | 27774077 | 852 | * | 6,52747 | 5,515471196 | 2,047443 | 2,693833 | 0,007064 | 1 | 0 | Distal Intergenic | 8 | 27786083 | 27786829 | 747 | 1 | 732482 | ENSMUST00000209480.2 | -12006 | | Gm9731 | Fam58a pseudogene |
| chr7 | 67169645 | 67170207 | 563 | * | 5,910599 | 5,517126574 | 1,983145 | 2,782008 | 0,005402 | 1 | 0 | Intron (ENSMUST00000 | 7 | 67179459 | 67185145 | 5687 | 2 | 67867 | ENSMUST00000185818.2 | 14938 | ENSMUSG00000030556 | Lrrc28 | leucine rich repeat containing 28 |
| chr9 | 21228790 | 21229198 | 409 | * | 4,43775 | 5,517335471 | 2,034501 | 2,711886 | 0,00669 | 1 | 0 | Promoter (2-3kb) | 9 | 21229366 | 21231913 | 2548 | 2 | 1,05E+08 | ENSMUST00000245945.1 | 2715 | ENSMUSG00000111172 | Gm39302 | predicted gene, 39302 |
| chr5 | 23691778 | 23692524 | 747 | * | 6,112185 | 5,517338573 | 2,077429 | 2,655849 | 0,007911 | 1 | 0 | Promoter (1-2kb) | 5 | 23690547 | 23707860 | 17314 | 1 | 69188 | ENSMUST00000196260.2 | 1231 | ENSMUSG00000029004 | Kmt2e | lysine (K)-specific methyltransferase 2E |
| chr19 | 58198616 | 58199396 | 781 | * | 6,314234 | 5,517680284 | 2,083082 | 2,648806 | 0,008078 | 1 | 0 | Intron (ENSMUST00000 | 19 | 58080246 | 58084463 | 4218 | 2 | 1,15E+08 | ENSMUST00000235250.2 | -114153 | | Gm50287 | predicted gene, 50287 |
| chr4 | 82730949 | 82731819 | 871 | * | 6,284119 | 5,518323074 | 1,962718 | 2,811572 | 0,00493 | 1 | 0 | Intron (ENSMUST00000 | 4 | 82756549 | 82769051 | 12503 | 2 | 68268 | ENSMUST00000139401.2 | 37232 | ENSMUSG00000028403 | Zdhhc21 | zinc finger, DHHC domain containing 21 |
| chr15 | 79385076 | 79385656 | 581 | * | 6,093515 | 5,518370088 | 1,961764 | 2,812963 | 0,004909 | 1 | 0 | Intron (ENSMUST00000 | 15 | 79367915 | 79389442 | 21528 | 2 | 16520 | ENSMUST00000057801.8 | 3786 | ENSMUSG00000044216 | Kcnj4 | potassium inwardly-rectifying channel, subfamily J, member 4 |
| chr8 | 1,07E+08 | 1,07E+08 | 889 | * | 5,529401 | 5,5187784 | 2,102457 | 2,624919 | 0,008667 | 1 | 0 | Promoter (2-3kb) | 8 | 1,07E+08 | 1,07E+08 | 46407 | 1 | 12560 | ENSMUST00000080797.8 | 2846 | ENSMUSG00000061048 | Cdh3 | cadherin 3 |
| chr1 | 1,42E+08 | 1,42E+08 | 416 | * | 4,264279 | 5,51907678 | 2,083117 | 2,649432 | 0,008063 | 1 | 0 | Distal Intergenic | 1 | 1,41E+08 | 1,41E+08 | 107 | 1 | 1,15E+08 | ENSMUST00000239951.1 | 556730 | ENSMUSG00000118759 | Gm25457 | predicted gene, 25457 |
| chr3 | 1,22E+08 | 1,22E+08 | 650 | * | 5,606958 | 5,519247037 | 2,111284 | 2,614166 | 0,008945 | 1 | 0 | Exon (ENSMUST00000 | 3 | 1,22E+08 | 1,22E+08 | 25163 | 2 | 214459 | ENSMUST00000197259.5 | -10966 | ENSMUSG00000039735 | Fnbp1l | formin binding protein 1-like |
| chrX | 64566587 | 64567369 | 783 | * | 5,144088 | 5,520003874 | 1,950093 | 2,830637 | 0,004646 | 1 | 0 | Distal Intergenic | 20 | 64091250 | 64092620 | 1371 | 1 | 70062 | ENSMUST00000033524.3 | 475337 | ENSMUSG00000031181 | Ctag2 | cancer/testis antigen 2 |
| chr16 | 24423187 | 24423888 | 702 | * | 6,759895 | 5,520069035 | 1,934952 | 2,85282 | 0,004333 | 1 | 0 | Intron (ENSMUST00000 | 16 | 24348144 | 24349112 | 969 | 1 | 624582 | ENSMUST00000231737.2 | 75043 | | Morf4l1-ps1 | mortality factor 4 like 1, pseudogene 1 |
| chr5 | 1,4E+08 | 1,4E+08 | 929 | * | 7,041904 | 5,520200285 | 1,799426 | 3,067756 | 0,002157 | 1 | 0 | Promoter (1-2kb) | 5 | 1,4E+08 | 1,4E+08 | 24033 | 1 | 27979 | ENSMUST00000100507.8 | -1420 | ENSMUSG00000056076 | Eif3b | eukaryotic translation initiation factor 3, subunit B |
| chr8 | 1,12E+08 | 1,12E+08 | 543 | * | 5,346076 | 5,520793916 | 2,141311 | 2,578231 | 0,009931 | 1 | 0 | Promoter (<=1kb) | 8 | 1,12E+08 | 1,12E+08 | 132 | 1 | 1,15E+08 | ENSMUST00000104289.3 | -631 | ENSMUSG00000077477 | Gm24793 | predicted gene, 24793 |
| chr17 | 79808057 | 79809132 | 1076 | * | 10,52021 | 5,523947845 | 1,717073 | 3,217072 | 0,001295 | 0,611687401 | 0,918255 | Distal Intergenic | 17 | 79746334 | 79759191 | 12858 | 2 | 1,03E+08 | ENSMUST00000242898.1 | -48866 | | Gm32954 | predicted gene, 32954 |
| chr5 | 1,22E+08 | 1,22E+08 | 542 | * | 5,701984 | 5,524508553 | 2,017659 | 2,738078 | 0,00618 | 1 | 0 | Distal Intergenic | 5 | 1,22E+08 | 1,22E+08 | 23174 | 2 | 654470 | ENSMUST00000141281.4 | -5877 | ENSMUSG00000038593 | Tctn1 | tectonic family member 1 |
| chr8 | 72116169 | 72116539 | 371 | * | 6,218808 | 5,524610215 | 1,968409 | 2,806637 | 0,005006 | 1 | 0 | Intron (ENSMUST00000 | 8 | 72112990 | 72120726 | 7737 | 2 | 382018 | ENSMUST00000175780.2 | 4187 | ENSMUSG00000034799 | Unc13a | unc-13 homolog A |
| chr11 | 97355930 | 97356746 | 817 | * | 6,145096 | 5,525059116 | 1,8556 | 2,977505 | 0,002906 | 1 | 0 | Exon (ENSMUST00000 | 11 | 97342956 | 97393228 | 50273 | 1 | 58996 | ENSMUST00000142465.8 | 12974 | ENSMUSG00000049807 | Arhgap23 | Rho GTPase activating protein 23 |
| chr9 | 62029859 | 62031135 | 1277 | * | 12,84509 | 5,525369494 | 1,686308 | 3,276608 | 0,001051 | 0,539997763 | 1,075735 | Promoter (<=1kb) | 9 | 62029800 | 62029876 | 77 | 2 | 1,01E+08 | ENSMUST00000174955.3 | 0 | ENSMUSG00000092696 | Mir5133 | microRNA 5133 |
| chr16 | 74091285 | 74091980 | 696 | * | 6,082177 | 5,525769266 | 1,843067 | 2,998138 | 0,002716 | 1 | 0 | Intron (ENSMUST00000 | 16 | 74138878 | 74138955 | 78 | 2 | 751561 | ENSMUST00000199719.3 | 46975 | ENSMUSG00000104758 | Mir691 | microRNA 691 |
| chr18 | 77407316 | 77408092 | 777 | * | 7,905059 | 5,526443894 | 1,831003 | 3,01826 | 0,002542 | 0,661775851 | 1,360622 | Intron (ENSMUST00000 | 18 | 77395446 | 77470024 | 74579 | 1 | 240411 | ENSMUST00000035501.11 | 11870 | ENSMUSG00000032818 | Loxhd1 | lipoxygenase homology domains 1 |
| chr5 | 23680592 | 23681719 | 1128 | * | 7,253032 | 5,52748017 | 2,015706 | 2,742205 | 0,006103 | 1 | 0 | Promoter (1-2kb) | 5 | 23683553 | 23690449 | 6897 | 1 | 69188 | ENSMUST00000200330.2 | -1834 | ENSMUSG00000029004 | Kmt2e | lysine (K)-specific methyltransferase 2E |
| chr4 | 40716601 | 40717524 | 924 | * | 8,617833 | 5,527846158 | 1,837556 | 3,008259 | 0,002627 | 0,659642963 | 1,425929 | Exon (ENSMUST00000 | 4 | 40722150 | 40723795 | 1646 | 1 | 15502 | ENSMUST00000139406.2 | -4626 | ENSMUSG00000028410 | Dnaja1 | DnaJ heat shock protein family (Hsp40) member A1 |
| chr17 | 51981518 | 51981932 | 415 | * | 5,248835 | 5,528071511 | 2,023914 | 2,731377 | 0,006307 | 1 | 0 | Distal Intergenic | 17 | 52047016 | 52049294 | 2279 | 2 | 20230 | ENSMUST00000137697.2 | 67362 | ENSMUSG00000023927 | Satb1 | special AT-rich sequence binding protein 1 |
| chr8 | 25028726 | 25029599 | 874 | * | 5,431395 | 5,528796507 | 2,012048 | 2,747845 | 0,005999 | 1 | 0 | Promoter (1-2kb) | 8 | 25023472 | 25030812 | 7341 | 2 | 209176 | ENSMUST00000138335.2 | 1213 | ENSMUSG00000031549 | Ido2 | indoleamine 2,3-dioxygenase 2 |
| chr12 | 85410579 | 85411518 | 940 | * | 7,046596 | 5,528998352 | 1,824836 | 3,029861 | 0,002447 | 1 | 0 | Intron (ENSMUST00000 | 12 | 85390139 | 85420729 | 30591 | 2 | 68581 | ENSMUST00000222585.2 | 9211 | ENSMUSG00000021248 | Tmed10 | transmembrane p24 trafficking protein 10 |
| chr6 | 72041674 | 72042332 | 659 | * | 6,924898 | 5,530905216 | 1,887445 | 2,930366 | 0,003386 | 1 | 0 | Distal Intergenic | 6 | 71990880 | 72019512 | 28633 | 2 | 330350 | ENSMUST00000206867.2 | -22162 | ENSMUSG00000108356 | 9130221F21Rik | RIKEN cDNA 9130221F21 gene |
| chrX | 22879513 | 22880595 | 1083 | * | 6,523159 | 5,531451749 | 2,060852 | 2,68406 | 0,007273 | 1 | 0 | Distal Intergenic | 20 | 23085525 | 23133041 | 47517 | 2 | 67455 | ENSMUST00000115316.9 | 252446 | ENSMUSG00000036782 | Klhl13 | kelch-like 13 |
| chr5 | 1,3E+08 | 1,3E+08 | 685 | * | 6,48811 | 5,533003541 | 1,845913 | 2,997435 | 0,002723 | 1 | 0 | Promoter (2-3kb) | 5 | 1,3E+08 | 1,3E+08 | 38619 | 1 | 56715 | ENSMUST00000148264.8 | 2518 | ENSMUSG00000025340 | Rabgef1 | RAB guanine nucleotide exchange factor (GEF) 1 |
| chr16 | 62598461 | 62599205 | 745 | * | 7,568039 | 5,534626851 | 1,75418 | 3,155107 | 0,001604 | 0,575786437 | 1,360622 | Intron (ENSMUST00000 | 16 | 62599614 | 62607076 | 7463 | 2 | 106338 | ENSMUST00000152842.2 | 7871 | ENSMUSG00000050312 | Nsun3 | NOL1/NOP2/Sun domain family member 3 |
| chr9 | 84456895 | 84457331 | 437 | * | 4,853292 | 5,53635447 | 2,107755 | 2,626659 | 0,008623 | 1 | 0 | Promoter (2-3kb) | 9 | 84460013 | 84495737 | 35725 | 1 | 1,05E+08 | ENSMUST00000214502.2 | -2682 | | Gm39388 | predicted gene, 39388 |
| chr5 | 1,19E+08 | 1,19E+08 | 1144 | * | 7,310793 | 5,53738248 | 1,862774 | 2,972654 | 0,002952 | 1 | 0 | Distal Intergenic | 5 | 1,19E+08 | 1,19E+08 | 5535 | 2 | 665194 | ENSMUST00000200918.2 | -25983 | ENSMUSG00000107334 | Gm7538 | predicted gene 7538 |
| chr5 | 1,43E+08 | 1,43E+08 | 914 | * | 9,084985 | 5,537652867 | 1,780657 | 3,109894 | 0,001872 | 0,594676756 | 1,449706 | Exon (ENSMUST00000 | 5 | 1,43E+08 | 1,43E+08 | 13019 | 2 | 1,05E+08 | ENSMUST00000176473.2 | 9109 | | Gm20635 | predicted gene 20635 |
| chr18 | 3214393 | 3214934 | 542 | * | 5,229946 | 5,537864145 | 1,953096 | 2,835429 | 0,004576 | 1 | 0 | Distal Intergenic | 18 | 3267546 | 3281078 | 13533 | 2 | 12916 | ENSMUST00000049942.13 | 66144 | ENSMUSG00000063889 | Crem | cAMP responsive element modulator |
| chr2 | 79855174 | 79856256 | 1083 | * | 8,419307 | 5,539189315 | 1,809291 | 3,061524 | 0,002202 | 0,627785088 | 1,425929 | Intron (ENSMUST00000 | 2 | 79736334 | 79869373 | 133040 | 2 | 18573 | ENSMUST00000146076.8 | 13117 | ENSMUSG00000059173 | Pde1a | phosphodiesterase 1A, calmodulin-dependent |
| chr11 | 72685802 | 72687057 | 1256 | * | 9,826681 | 5,539483982 | 1,776037 | 3,119014 | 0,001815 | 0,624307941 | 1,211315 | Promoter (<=1kb) | 11 | 72668058 | 72686682 | 18625 | 2 | 192986 | ENSMUST00000079681.6 | 0 | ENSMUSG00000057778 | Cyb5d2 | cytochrome b5 domain containing 2 |
| chr15 | 80092366 | 80092811 | 446 | * | 4,139952 | 5,542616685 | 2,143743 | 2,585485 | 0,009724 | 1 | 0 | Intron (ENSMUST00000 | 15 | 80118219 | 80137572 | 19354 | 1 | 239555 | ENSMUST00000023048.12 | -25408 | ENSMUSG00000022412 | Mief1 | mitochondrial elongation factor 1 |
| chr10 | 47992103 | 47992981 | 879 | * | 6,529544 | 5,542703015 | 1,892117 | 2,929365 | 0,003397 | 1 | 0 | Distal Intergenic | 10 | 48495408 | 48495491 | 84 | 2 | 1,15E+08 | ENSMUST00000083696.4 | 502510 | | Gm24256 | predicted gene, 24256 |
| chr10 | 76899754 | 76900559 | 806 | * | 7,0796 | 5,542728777 | 1,895006 | 2,924914 | 0,003446 | 0,711218709 | 1,360622 | Exon (ENSMUST00000 | 10 | 76891441 | 76895831 | 4391 | 2 | 12822 | ENSMUST00000218407.2 | -3923 | ENSMUSG00000001435 | Col18a1 | collagen, type XVIII, alpha 1 |

| | | | | | | | | | | | | | | | | | | | | | | | |
|---|---|---|---|---|---|---|---|---|---|---|---|---|---|---|---|---|---|---|---|---|---|---|---|
| chr6 | 85388804 | 85389497 | 694 | * | 7,157747 | 5,542907402 | 1,900168 | 2,917061 | 0,003533 | 1 | 0 | Distal Intergenic | 6 | 85400868 | 85405859 | 4992 | 1 | 384452 | ENSMUST00000089578.9 | -11371 | ENSMUSG00000068302 | Noto | notochord homeobox |
| chr12 | 72564121 | 72565634 | 1514 | * | 6,282453 | 5,543007069 | 1,902748 | 2,913158 | 0,003578 | 1 | 0 | Intron (ENSMUST0000( | 12 | 72583157 | 72626893 | 43737 | 1 | 67708 | ENSMUST00000044352.8 | -17523 | ENSMUSG00000034501 | Pcnx4 | pecanex homolog 4 |
| chr13 | 1,01E+08 | 1,01E+08 | 1416 | * | 7,701857 | 5,543092685 | 1,759145 | 3,151016 | 0,001627 | 1 | 0 | Distal Intergenic | 13 | 1,01E+08 | 1,01E+08 | 15425 | 1 | 328324 | ENSMUST00000223648.2 | -74782 | ENSMUSG00000114590 | 5930438M14Rik | RIKEN cDNA 5930438M14 gene |
| chr13 | 48677716 | 48678211 | 496 | * | 4,410069 | 5,543399282 | 2,113605 | 2,622723 | 0,008723 | 1 | 0 | Distal Intergenic | 13 | 48667046 | 48668335 | 1290 | 1 | 442803 | ENSMUST00000065465.2 | 10670 | ENSMUSG00000053181 | A830005F24Rik | RIKEN cDNA A830005F24 gene |
| chr16 | 21270480 | 21271093 | 614 | * | 4,51328 | 5,543690125 | 2,102885 | 2,636231 | 0,008383 | 1 | 0 | Intron (ENSMUST0000( | 16 | 21288937 | 21460322 | 171386 | 1 | 209018 | ENSMUST00000125487.8 | -17844 | ENSMUSG00000033653 | Vps8 | VPS8 CORVET complex subunit |
| chr9 | 64805866 | 64807883 | 2018 | * | 15,17377 | 5,543777849 | 1,306273 | 4,243966 | 2,2E-05 | 0,068345002 | 1,2763 | Exon (ENSMUST000002 | 9 | 64814479 | 64819393 | 4915 | 1 | 102442 | ENSMUST00000214029.2 | -6596 | ENSMUSG00000053641 | Dennd4a | DENN/MADD domain containing 4A |
| chr13 | 47835200 | 47836004 | 805 | * | 6,28991 | 5,543871828 | 1,930488 | 2,871747 | 0,004082 | 1 | 0 | Distal Intergenic | 13 | 47821026 | 47821868 | 843 | 1 | 74939 | ENSMUST00000223996.2 | 14174 | ENSMUSG00000114509 | 4930471G24Rik | RIKEN cDNA 4930471G24 gene |
| chr11 | 85015831 | 85016585 | 755 | * | 6,174459 | 5,543975048 | 1,835554 | 3,020329 | 0,002525 | 1 | 0 | Intron (ENSMUST0000( | 11 | 84994506 | 85030755 | 36250 | 2 | 237898 | ENSMUST00000174809.2 | 14170 | ENSMUSG00000000804 | Usp32 | ubiquitin specific peptidase 32 |
| chr10 | 95447589 | 95448173 | 585 | * | 5,764016 | 5,544384386 | 1,947518 | 2,846897 | 0,004415 | 1 | 0 | Distal Intergenic | 10 | 95466099 | 95468101 | 2003 | 2 | 1,03E+08 | ENSMUST00000219656.2 | 19928 | | Gm33336 | predicted gene, 33336 |
| chr13 | 36364676 | 36365621 | 946 | * | 6,121196 | 5,544399176 | 1,824107 | 3,039515 | 0,00237 | 1 | 0 | Intron (ENSMUST0000( | 13 | 36416105 | 36430592 | 14488 | 1 | 69955 | ENSMUST00000224540.2 | -50484 | ENSMUSG00000021420 | Fars2 | phenylalanine-tRNA synthetase 2 (mitochondrial) |
| chr5 | 43994133 | 43994877 | 745 | * | 6,585652 | 5,54505713 | 1,972853 | 2,81068 | 0,004944 | 1 | 0 | Promoter (<=1kb) | 5 | 43994456 | 43999752 | 5297 | 1 | 12182 | ENSMUST00000125838.3 | 0 | ENSMUSG00000029082 | Bst1 | bone marrow stromal cell antigen 1 |
| chr2 | 1,21E+08 | 1,21E+08 | 863 | * | 5,751525 | 5,545222898 | 1,979373 | 2,801505 | 0,005086 | 1 | 0 | 3' UTR | 2 | 1,21E+08 | 1,21E+08 | 39731 | 2 | 74176 | ENSMUST00000028721.8 | -7604 | ENSMUSG00000053675 | Tgm5 | transglutaminase 5 |
| chr17 | 29523651 | 29524607 | 957 | * | 7,954583 | 5,545434728 | 1,795763 | 3,088066 | 0,002015 | 0,621585803 | 1,360622 | 3' UTR | 17 | 29517258 | 29520499 | 3242 | 1 | 80748 | ENSMUST00000150874.2 | 6393 | ENSMUSG00000052712 | BC004004 | cDNA sequence BC004004 |
| chr17 | 46559790 | 46560801 | 1012 | * | 6,548198 | 5,54543817 | 1,98824 | 2,789119 | 0,005285 | 1 | 0 | Promoter (<=1kb) | 17 | 46559955 | 46562830 | 2876 | 1 | 28064 | ENSMUST00000127378.2 | 0 | ENSMUSG00000071074 | Yipf3 | Yip1 domain family, member 3 |
| chr10 | 97643751 | 97644511 | 761 | * | 5,452382 | 5,545572211 | 1,993351 | 2,782036 | 0,005402 | 1 | 0 | Intron (ENSMUST0000( | 10 | 97580619 | 97580746 | 128 | 2 | 1,15E+08 | ENSMUST00000158472.3 | -63005 | ENSMUSG00000089097 | Gm25658 | predicted gene, 25658 |
| chr13 | 41478482 | 41479232 | 751 | * | 5,014978 | 5,545582608 | 1,994 | 2,781135 | 0,005417 | 1 | 0 | Intron (ENSMUST0000( | 13 | 41463396 | 41512723 | 49328 | 2 | 18003 | ENSMUST00000163623.3 | 33491 | ENSMUSG00000021365 | Nedd9 | neural precursor cell expressed, developmentally down-regulated gene 9 |
| chr5 | 1,04E+08 | 1,04E+08 | 919 | * | 5,183611 | 5,545784786 | 2,035002 | 2,725198 | 0,006426 | 1 | 0 | Promoter (<=1kb) | 5 | 1,04E+08 | 1,04E+08 | 124055 | 1 | 17355 | ENSMUST00000126335.2 | -609 | ENSMUSG00000029313 | Aff1 | AF4/FMR2 family, member 1 |
| chr4 | 1,54E+08 | 1,54E+08 | 401 | * | 4,960959 | 5,546028603 | 2,028036 | 2,734679 | 0,006244 | 1 | 0 | Distal Intergenic | 4 | 1,53E+08 | 1,54E+08 | 109591 | 2 | 230959 | ENSMUST00000105646.3 | -111644 | ENSMUSG00000039546 | Ajap1 | adherens junction associated protein 1 |
| chr11 | 82120804 | 82121310 | 507 | * | 6,672647 | 5,546058355 | 1,838453 | 3,016698 | 0,002555 | 1 | 0 | Intron (ENSMUST0000( | 11 | 82067485 | 82070638 | 3154 | 2 | 20290 | ENSMUST00000021043.5 | -50166 | ENSMUSG00000020702 | Ccl1 | chemokine (C-C motif) ligand 1 |
| chr7 | 79837869 | 79838446 | 578 | * | 5,040318 | 5,546118683 | 2,016808 | 2,749949 | 0,00596 | 1 | 0 | Promoter (1-2kb) | 7 | 79836626 | 79871580 | 34955 | 1 | 20352 | ENSMUST00000107383.8 | 1243 | ENSMUSG00000030539 | Sema4b | sema domain, immunoglobulin domain (Ig), transmembrane domain (TM) and short cytoplasmic domain, (semaphorin) 4B |
| chr13 | 31827297 | 31827729 | 433 | * | 4,362543 | 5,547251596 | 2,072454 | 2,676659 | 0,007436 | 1 | 0 | Distal Intergenic | 13 | 31809799 | 31815386 | 5588 | 1 | 14238 | ENSMUST00000042054.3 | 17498 | ENSMUSG00000038402 | Foxf2 | forkhead box F2 |
| chr10 | 17519688 | 17520491 | 804 | * | 5,401578 | 5,547301235 | 2,07537 | 2,672922 | 0,007519 | 1 | 0 | Intron (ENSMUST0000( | 10 | 17438491 | 17534261 | 95771 | 2 | 1,19E+08 | ENSMUST00000219966.2 | 13770 | | Gm53271 | predicted gene, 53271 |
| chr14 | 62032402 | 62033568 | 1167 | * | 6,140623 | 5,547480144 | 1,915894 | 2,895505 | 0,003785 | 1 | 0 | Distal Intergenic | 14 | 61840265 | 61919822 | 79558 | 2 | 668253 | ENSMUST00000182768.9 | -112580 | ENSMUSG00000097589 | Dleu2 | deleted in lymphocytic leukemia, 2 |
| chr1 | 81220398 | 81221180 | 783 | * | 8,377963 | 5,547880204 | 1,726148 | 3,214023 | 0,001309 | 0,537550416 | 1,360622 | Intron (ENSMUST0000( | 1 | 81055300 | 81319479 | 264180 | 1 | 241134 | ENSMUST00000123285.2 | 165098 | ENSMUSG00000054976 | Nyap2 | neuronal tyrosine-phophorylated phosphoinositide 3-kinase adaptor 2 |
| chr18 | 34867892 | 34868545 | 654 | * | 4,544547 | 5,547890677 | 2,108663 | 2,630999 | 0,008513 | 1 | 0 | 3' UTR | 18 | 34868030 | 34877116 | 9087 | 2 | 12532 | ENSMUST00000236366.2 | 8571 | ENSMUSG00000044201 | Cdc25c | cell division cycle 25C |
| chr3 | 80904269 | 80904850 | 582 | * | 5,076039 | 5,548362406 | 2,138004 | 2,595113 | 0,009456 | 1 | 0 | Distal Intergenic | 3 | 80943723 | 81121347 | 177625 | 1 | 54635 | ENSMUST00000029652.4 | -38873 | ENSMUSG00000028019 | Pdgfc | platelet-derived growth factor, C polypeptide |
| chr13 | 91642600 | 91643555 | 956 | * | 5,584938 | 5,548544865 | 1,893725 | 2,929964 | 0,00339 | 1 | 0 | Intron (ENSMUST0000( | 13 | 91609300 | 91672985 | 63686 | 1 | 66970 | ENSMUST00000141381.2 | 33300 | ENSMUSG00000003992 | Ssbp2 | single-stranded DNA binding protein 2 |
| chr17 | 41298535 | 41299122 | 588 | * | 5,983095 | 5,549831347 | 1,886511 | 2,94185 | 0,003263 | 1 | 0 | Distal Intergenic | 17 | 41266079 | 41269872 | 3794 | 1 | 17850 | ENSMUST00000233222.2 | 32456 | ENSMUSG00000023921 | Mmut | methylmalonyl-Coenzyme A mutase |
| chr12 | 39173402 | 39173959 | 558 | * | 4,766688 | 5,550249575 | 1,985482 | 2,795417 | 0,005183 | 1 | 0 | Intron (ENSMUST0000( | 12 | 39163815 | 39204718 | 40904 | 2 | 1,03E+08 | ENSMUST00000221271.2 | 30759 | ENSMUSG00000113272 | Gm34809 | predicted gene, 34809 |
| chr1 | 88124826 | 88125837 | 1012 | * | 6,531655 | 5,551122225 | 1,907996 | 2,9094 | 0,003621 | 1 | 0 | Promoter (2-3kb) | 1 | 88128323 | 88146719 | 18397 | 1 | 22236 | ENSMUST00000049289.9 | -2486 | ENSMUSG00000090171 | Ugt1a2 | UDP glucuronosyltransferase 1 family, polypeptide A2 |
| chr19 | 13915603 | 13916370 | 768 | * | 7,35705 | 5,551269154 | 1,905091 | 2,913912 | 0,003569 | 0,715249603 | 1,360622 | Distal Intergenic | 19 | 13896306 | 13897408 | 1103 | 1 | 258151 | ENSMUST00000081520.3 | 19297 | ENSMUSG00000062314 | Olfr1505 | olfactory receptor 1505 |
| chr1 | 88797519 | 88797958 | 440 | * | 6,650058 | 5,552389945 | 1,815698 | 3,057993 | 0,002228 | 1 | 0 | Intron (ENSMUST0000( | 1 | 88682652 | 88683458 | 807 | 1 | 78650 | ENSMUST00000180596.3 | 114867 | ENSMUSG00000097398 | Platr5 | pluripotency associated transcript 5 |
| chr2 | 1,3E+08 | 1,3E+08 | 1052 | * | 7,711695 | 5,552434627 | 1,883925 | 2,94727 | 0,003206 | 1 | 0 | Distal Intergenic | 2 | 1,3E+08 | 1,3E+08 | 28458 | 1 | 67333 | ENSMUST00000165413.9 | -41393 | ENSMUSG00000037885 | Stk35 | serine/threonine kinase 35 |
| chr12 | 26853974 | 26854838 | 865 | * | 8,207358 | 5,556344575 | 1,823293 | 3,047423 | 0,002308 | 0,63917784 | 1,425929 | Distal Intergenic | 12 | 26864370 | 26884858 | 20489 | 2 | 74668 | ENSMUST00000221910.2 | 30020 | ENSMUSG00000113218 | 4933409F18Rik | RIKEN cDNA 4933409F18 gene |
| chrX | 1,35E+08 | 1,35E+08 | 486 | * | 4,897059 | 5,556648872 | 1,977279 | 2,81025 | 0,00495 | 1 | 0 | Promoter (2-3kb) | 20 | 1,35E+08 | 1,35E+08 | 2060 | 1 | 1E+08 | ENSMUST00000113116.3 | 2508 | ENSMUSG00000079428 | Tceal7 | transcription elongation factor A (SII)-like 7 |
| chr5 | 36682423 | 36683508 | 1086 | * | 6,585527 | 5,55880153 | 1,960081 | 2,836006 | 0,004568 | 1 | 0 | Intron (ENSMUST0000( | 5 | 36682285 | 36687851 | 5567 | 2 | 100855 | ENSMUST00000141626.2 | 4343 | ENSMUSG00000029192 | Tbc1d14 | TBC1 domain family, member 14 |
| chr4 | 5253991 | 5254859 | 869 | * | 9,140744 | 5,562299087 | 1,779802 | 3,125236 | 0,001777 | 0,620083984 | 1,211315 | Distal Intergenic | 4 | 5632328 | 5642284 | 9957 | 1 | 73872 | ENSMUST00000156903.8 | -377469 | ENSMUSG00000086066 | 4930423M02Rik | RIKEN cDNA 4930423M02 gene |
| chr14 | 20495496 | 20496207 | 712 | * | 4,331401 | 5,563561774 | 2,083738 | 2,66999 | 0,007585 | 1 | 0 | Promoter (2-3kb) | 14 | 20497428 | 20498769 | 1342 | 2 | 76670 | ENSMUST00000143940.2 | 2562 | ENSMUSG00000039543 | Cfap70 | cilia and flagella associated protein 70 |
| chr14 | 30668373 | 30669310 | 938 | * | 7,583435 | 5,564452656 | 1,856283 | 2,997631 | 0,002721 | 1 | 0 | Distal Intergenic | 14 | 30651137 | 30665246 | 14110 | 2 | 16424 | ENSMUST00000163118.2 | -3127 | ENSMUSG00000006529 | Itih1 | inter-alpha trypsin inhibitor, heavy chain 1 |
| chr5 | 1,48E+08 | 1,48E+08 | 636 | * | 5,917722 | 5,566073525 | 1,925498 | 2,890719 | 0,003844 | 1 | 0 | Distal Intergenic | 5 | 1,48E+08 | 1,48E+08 | 935 | 2 | 71706 | ENSMUST00000138244.2 | -23334 | ENSMUSG00000029650 | Slc46a3 | solute carrier family 46, member 3 |
| chr5 | 5470006 | 5470866 | 861 | * | 7,889583 | 5,568528711 | 1,834082 | 3,03614 | 0,002396 | 1 | 0 | Promoter (<=1kb) | 5 | 5277215 | 5470312 | 193098 | 2 | 18647 | ENSMUST00000167567.2 | 0 | ENSMUSG00000028926 | Cdk14 | cyclin-dependent kinase 14 |
| chr13 | 43739794 | 43740694 | 901 | * | 7,962159 | 5,569513668 | 1,848894 | 3,012348 | 0,002592 | 0,665162479 | 1,360622 | Distal Intergenic | 13 | 43699799 | 43713667 | 13869 | 2 | 76137 | ENSMUST00000223353.2 | -26127 | ENSMUSG00000021371 | Mcur1 | mitochondrial calcium uniporter regulator 1 |
| chr3 | 87073818 | 87074590 | 773 | * | 5,586166 | 5,57019491 | 2,069941 | 2,690992 | 0,007124 | 1 | 0 | Intron (ENSMUST0000( | 3 | 86991935 | 87081857 | 89923 | 2 | 170643 | ENSMUST00000041732.9 | 7267 | ENSMUSG00000041734 | Kirrel | kirre like nephrin family adhesion molecule 1 |
| chr6 | 48525858 | 48526769 | 912 | * | 7,138243 | 5,57175424 | 1,886345 | 2,953729 | 0,00314 | 1 | 0 | Distal Intergenic | 6 | 48531730 | 48547656 | 15927 | 1 | 243371 | ENSMUST00000114545.8 | -4961 | ENSMUSG00000073096 | Lrrc61 | leucine rich repeat containing 61 |
| chr10 | 93952480 | 93953672 | 1193 | * | 6,870253 | 5,572324963 | 1,896673 | 2,937947 | 0,003304 | 1 | 0 | Intron (ENSMUST0000( | 10 | 93969025 | 93975733 | 6709 | 1 | 13998 | ENSMUST00000125535.2 | -15353 | ENSMUSG00000020021 | Fgd6 | FYVE, RhoGEF and PH domain containing 6 |
| chr7 | 80439024 | 80439724 | 701 | * | 6,049202 | 5,57396658 | 1,888955 | 2,95082 | 0,003169 | 1 | 0 | Intron (ENSMUST0000( | 7 | 80399125 | 80453011 | 53887 | 2 | 29875 | ENSMUST00000206149.2 | 13287 | ENSMUSG00000030536 | Iqgap1 | IQ motif containing GTPase activating protein 1 |
| chrX | 1,68E+08 | 1,68E+08 | 825 | * | 4,160667 | 5,574564664 | 2,159807 | 2,581047 | 0,00985 | 1 | 0 | 3' UTR | 20 | 1,68E+08 | 1,68E+08 | 8751 | 2 | 15159 | ENSMUST00000033717.9 | 18347 | ENSMUSG00000031352 | Hccs | holocytochrome c synthetase |
| chrX | 10400547 | 10401098 | 552 | * | 4,113221 | 5,574589434 | 2,15809 | 2,583112 | 0,009791 | 1 | 0 | Intron (ENSMUST0000( | 20 | 10351397 | 10462844 | 111448 | 1 | 21912 | ENSMUST00000149982.2 | 49150 | ENSMUSG00000058254 | Tspan7 | tetraspanin 7 |
| chrX | 1E+08 | 1E+08 | 1158 | * | 4,671862 | 5,574799016 | 2,144531 | 2,599542 | 0,009335 | 1 | 0 | Distal Intergenic | 20 | 1E+08 | 1E+08 | 10605 | 2 | 55988 | ENSMUST00000077876.4 | -7861 | ENSMUSG00000046032 | Snx12 | sorting nexin 12 |
| chr8 | 1,04E+08 | 1,04E+08 | 370 | * | 6,62757 | 5,574875568 | 1,949758 | 2,859265 | 0,004246 | 1 | 0 | Intron (ENSMUST0000( | 8 | 1,04E+08 | 1,04E+08 | 68740 | 1 | 1,03E+08 | ENSMUST00000209324.2 | 5088 | | Gm32531 | predicted gene, 32531 |
| chr3 | 95128143 | 95128603 | 461 | * | 5,069454 | 5,57489278 | 2,138146 | 2,607348 | 0,009125 | 1 | 0 | Promoter (2-3kb) | 3 | 95106881 | 95125227 | 18347 | 2 | 213054 | ENSMUST00000138862.8 | -2916 | ENSMUSG00000038766 | Gabpb2 | GA repeat binding protein, beta 2 |
| chr19 | 29481051 | 29481637 | 587 | * | 4,943444 | 5,574956287 | 2,133147 | 2,613489 | 0,008962 | 1 | 0 | Exon (ENSMUST000001 | 19 | 29442150 | 29498370 | 56221 | 2 | 77779 | ENSMUST00000236439.2 | 16733 | ENSMUSG00000097855 | A930007I19Rik | RIKEN cDNA A930007I19 gene |
| chr1 | 37328743 | 37329464 | 722 | * | 4,041239 | 5,575001862 | 2,131047 | 2,616086 | 0,008894 | 1 | 0 | Distal Intergenic | 1 | 37338946 | 37387593 | 48648 | 1 | 269180 | ENSMUST00000146676.2 | -9482 | ENSMUSG00000026113 | Inpp4a | inositol polyphosphate-4-phosphatase, type I |
| chr14 | 22828862 | 22829605 | 744 | * | 5,747243 | 5,575203285 | 1,957999 | 2,847399 | 0,004408 | 1 | 0 | Intron (ENSMUST0000( | 14 | 22646581 | 22649336 | 2756 | 1 | 76633 | ENSMUST00000226003.2 | 182281 | ENSMUSG00000063458 | Lrmda | leucine rich melanocyte differentiation associated |
| chr1 | 1,32E+08 | 1,32E+08 | 822 | * | 5,541473 | 5,575253068 | 1,959161 | 2,845734 | 0,004431 | 1 | 0 | Promoter (2-3kb) | 1 | 1,32E+08 | 1,32E+08 | 12944 | 1 | 213464 | ENSMUST00000189786.3 | 2337 | ENSMUSG00000026439 | Rbbp5 | retinoblastoma binding protein 5, histone lysine methyltransferase complex subunit |
| chr1 | 1,81E+08 | 1,81E+08 | 1071 | * | 5,563522 | 5,575387676 | 2,106619 | 2,646604 | 0,00813 | 1 | 0 | Promoter (2-3kb) | 1 | 1,81E+08 | 1,81E+08 | 57067 | 2 | 67459 | ENSMUST00000027797.9 | 2948 | ENSMUSG00000026516 | Nvl | nuclear VCP-like |
| chr8 | 86564625 | 86565295 | 671 | * | 6,775409 | 5,575671597 | 1,969284 | 2,83132 | 0,004636 | 1 | 0 | Promoter (2-3kb) | 8 | 86444207 | 86567550 | 123344 | 2 | 71927 | ENSMUST00000034140.9 | 2255 | ENSMUSG00000031703 | Itfg1 | integrin alpha FG-GAP repeat containing 1 |
| chr9 | 70017595 | 70018134 | 540 | * | 4,722855 | 5,575939217 | 2,076884 | 2,684762 | 0,007258 | 1 | 0 | Exon (ENSMUST00000( | 9 | 70009832 | 70010358 | 527 | 2 | 76886 | ENSMUST00000145427.2 | -7237 | ENSMUSG00000032224 | Fam81a | family with sequence similarity 81, member A |
| chr17 | 27751563 | 27752017 | 455 | * | 4,716261 | 5,576010707 | 2,072651 | 2,69028 | 0,007139 | 1 | 0 | Distal Intergenic | 17 | 27644881 | 27740377 | 95497 | 2 | 268934 | ENSMUST00000231416.2 | -11186 | ENSMUSG00000063239 | Grm4 | glutamate receptor, metabotropic 4 |
| chr7 | 92317471 | 92318790 | 1320 | * | 5,340204 | 5,576337159 | 2,054959 | 2,7136 | 0,006656 | 1 | 0 | Promoter (<=1kb) | 7 | 92292751 | 92319142 | 26392 | 2 | 74737 | ENSMUST00000119954.9 | 352 | ENSMUSG00000041328 | Pcf11 | PCF11 cleavage and polyadenylation factor subunit |
| chr4 | 1,31E+08 | 1,31E+08 | 495 | * | 4,616928 | 5,576346995 | 2,054615 | 2,714059 | 0,006646 | 1 | 0 | Distal Intergenic | 4 | 1,31E+08 | 1,31E+08 | 15902 | 2 | 1,05E+08 | ENSMUST00000152128.2 | -8359 | | Gm42321 | predicted gene, 42321 |
| chr7 | 1,09E+08 | 1,09E+08 | 697 | * | 4,565809 | 5,576361079 | 2,054166 | 2,714659 | 0,006634 | 1 | 0 | Promoter (<=1kb) | 7 | 1,09E+08 | 1,09E+08 | 8535 | 1 | 66085 | ENSMUST00000033342.7 | 394 | ENSMUSG00000031029 | Eif3f | eukaryotic translation initiation factor 3, subunit F |
| chr1 | 59064902 | 59065645 | 744 | * | 4,887217 | 5,576366905 | 2,053546 | 2,715482 | 0,006618 | 1 | 0 | Intron (ENSMUST0000( | 1 | 59075214 | 59077065 | 1852 | 1 | 1,05E+08 | ENSMUST00000191356.2 | -9569 | | Gm29018 | predicted gene 29018 |
| chr3 | 28833519 | 28834188 | 670 | * | 4,646797 | 5,576591014 | 2,042822 | 2,729847 | 0,006336 | 1 | 0 | Promoter (1-2kb) | 3 | 28792931 | 28835244 | 42314 | 2 | 73611 | ENSMUST00000241072.1 | 1056 | | 1700112D23Rik | RIKEN cDNA 1700112D23 gene |
| chrX | 35325721 | 35326792 | 1072 | * | 4,979565 | 5,576675407 | 2,038669 | 2,735449 | 0,00623 | 1 | 0 | Exon (ENSMUST00000( | 20 | 35304891 | 35339880 | 34990 | 1 | 75974 | ENSMUST00000115264.2 | 20830 | ENSMUSG00000031093 | Dock11 | dedicator of cytokinesis 11 |
| chrX | 1,64E+08 | 1,64E+08 | 956 | * | 5,828847 | 5,576802051 | 1,938972 | 2,876165 | 0,004025 | 1 | 0 | Intron (ENSMUST0000( | 20 | 1,64E+08 | 1,64E+08 | 198377 | 2 | 237213 | ENSMUST00000058787.9 | 33101 | ENSMUSG00000018589 | Glra2 | glycine receptor, alpha 2 subunit |
| chr12 | 28841237 | 28841565 | 329 | * | 5,395218 | 5,577059068 | 2,020379 | 2,760402 | 0,005773 | 1 | 0 | Exon (ENSMUST000002 | 12 | 28860137 | 28861190 | 1054 | 1 | 380752 | ENSMUST00000222829.2 | -18572 | ENSMUSG00000036613 | Eipr1 | EARP complex and GARP complex interacting protein 1 |
| chr4 | 1,29E+08 | 1,29E+08 | 309 | * | 5,208304 | 5,577147363 | 2,016602 | 2,765616 | 0,005682 | 1 | 0 | Distal Intergenic | 4 | 1,29E+08 | 1,29E+08 | 6916 | 1 | 75234 | ENSMUST00000130803.3 | 10412 | ENSMUSG00000028793 | Rnf19b | ring finger protein 19B |
| chrX | 50163456 | 50164348 | 893 | * | 5,236021 | 5,577341551 | 2,014112 | 2,769132 | 0,005621 | 1 | 0 | Distal Intergenic | 20 | 50209329 | 50216247 | 6919 | 2 | 171170 | ENSMUST00000150014.2 | 51899 | ENSMUSG00000036109 | Mbnl3 | muscleblind like splicing factor 3 |
| chr7 | 1,06E+08 | 1,06E+08 | 543 | * | 7,015962 | 5,577553126 | 1,805032 | 3,090003 | 0,002002 | 1 | 0 | Exon (ENSMUST000002 | 7 | 1,06E+08 | 1,06E+08 | 136 | 1 | 1,15E+08 | ENSMUST00000104589.3 | 80236 | ENSMUSG00000077777 | Gm22372 | predicted gene, 22372 |
| chr10 | 59660802 | 59662104 | 1303 | * | 35,919376 | 5,577840681 | 0,7524218 | 7,4131835 | 1,233E-13 | 9,27413E-09 | 2,3776742 | Promoter (<=1kb) | 10 | 59663043 | 59676189 | 13147 | 1 | 216001 | ENSMUST00000169842.2 | -939 | ENSMUSG00000020111 | Micu1 | mitochondrial calcium uptake 1 |
| chr12 | 85503348 | 85503925 | 578 | * | 5,072959 | 5,577890792 | 2,029912 | 2,747848 | 0,005999 | 1 | 0 | Distal Intergenic | 12 | 85520664 | 85524047 | 3384 | 1 | 14281 | ENSMUST00000021674.7 | -16739 | ENSMUSG00000021250 | Fos | FBJ osteosarcoma oncogene |
| chr13 | 99818388 | 99819411 | 1024 | * | 5,921115 | 5,578049668 | 2,035345 | 2,740592 | 0,006133 | 1 | 0 | Distal Intergenic | 13 | 99828907 | 99829037 | 131 | 2 | 1,15E+08 | ENSMUST00000104485.3 | 9626 | ENSMUSG00000077673 | Gm24471 | predicted gene, 24471 |
| chr19 | 36621201 | 36622563 | 1363 | * | 11,5654 | 5,578105497 | 1,468122 | 3,799483 | 0,000145 | 0,223981956 | 1,173325 | Intron (ENSMUST0000( | 19 | 36591448 | 36603501 | 12054 | 2 | 668215 | ENSMUST00000246331.1 | -17700 | ENSMUSG00000087579 | Hectd2os | Hectd2, opposite strand |
| chr8 | 34148240 | 34149264 | 1025 | * | 10,58172 | 5,578770915 | 1,714794 | 3,253318 | 0,001141 | 0,586831305 | 0,918255 | Intron (ENSMUST0000( | 8 | 34143266 | 34188191 | 44926 | 1 | 14782 | ENSMUST00000033992.9 | 4974 | ENSMUSG00000031584 | Gsr | glutathione reductase |
| chr15 | 98968588 | 98969064 | 477 | * | 6,955778 | 5,579287286 | 1,820613 | 3,064511 | 0,00218 | 0,635792383 | 1,360622 | Intron (ENSMUST0000( | 15 | 98972456 | 98981290 | 8835 | 1 | 78733 | ENSMUST00000230054.2 | -3392 | ENSMUSG00000032783 | Troap | trophinin associated protein |
| chrX | 1,11E+08 | 1,11E+08 | 678 | * | 5,097637 | 5,579291236 | 2,076086 | 2,687409 | 0,007201 | 1 | 0 | Distal Intergenic | 20 | 1,11E+08 | 1,11E+08 | 63070 | 1 | 67715 | ENSMUST00000026602.9 | -18700 | ENSMUSG00000025528 | 2010106E10Rik | RIKEN cDNA 2010106E10 gene |
| chr1 | 1,93E+08 | 1,93E+08 | 881 | * | 6,083355 | 5,57947381 | 1,926263 | 2,896527 | 0,003773 | 1 | 0 | Distal Intergenic | 1 | 1,93E+08 | 1,93E+08 | 18913 | 1 | 54139 | ENSMUST00000076521.7 | -4396 | ENSMUSG00000026638 | Irf6 | interferon regulatory factor 6 |
| chr15 | 12082641 | 12083567 | 927 | * | 5,301187 | 5,579483117 | 2,082909 | 2,678698 | 0,007391 | 1 | 0 | Promoter (1-2kb) | 15 | 12068334 | 12085283 | 16950 | 2 | 1,03E+08 | ENSMUST00000227908.2 | 1716 | | Gm34759 | predicted gene, 34759 |
| chrX | 20502224 | 20502618 | 395 | * | 4,519589 | 5,579779736 | 2,093525 | 2,665256 | 0,007693 | 1 | 0 | Intron (ENSMUST0000( | 20 | 20486337 | 20516923 | 30587 | 1 | 236732 | ENSMUST00000177738.8 | 15887 | ENSMUSG00000031060 | Rbm10 | RNA binding motif protein 10 |
| chr18 | 49069135 | 49069887 | 753 | * | 7,613958 | 5,580403476 | 1,775226 | 3,143489 | 0,001669 | 1 | 0 | Distal Intergenic | 18 | 49656356 | 49657507 | 1152 | 1 | 67321 | ENSMUST00000190201.2 | -586469 | ENSMUSG00000100297 | 1700044K03Rik | RIKEN cDNA 1700044K03 gene |
| chr1 | 1,9E+08 | 1,9E+08 | 526 | * | 6,415351 | 5,580425443 | 2,118754 | 2,633825 | 0,008443 | 1 | 0 | Promoter (<=1kb) | 1 | 1,9E+08 | 1,9E+08 | 37208 | 2 | 226830 | ENSMUST00000144452.2 | 287 | ENSMUSG00000026603 | Smyd2 | SET and MYND domain containing 2 |
| chr5 | 98407707 | 98408419 | 713 | * | 4,689823 | 5,580505579 | 2,120351 | 2,631878 | 0,008491 | 1 | 0 | Intron (ENSMUST0000( | 5 | 98402271 | 98423320 | 21050 | 1 | 14176 | ENSMUST00000200059.2 | 5436 | ENSMUSG00000029337 | Fgf5 | fibroblast growth factor 5 |
| chr4 | 1,05E+08 | 1,05E+08 | 377 | * | 4,667409 | 5,580848243 | 2,135685 | 2,613142 | 0,008971 | 1 | 0 | Distal Intergenic | 4 | 1,05E+08 | 1,05E+08 | 493 | 2 | 433742 | ENSMUST00000118507.2 | -34072 | ENSMUSG00000081819 | Gm12722 | predicted gene 12722 |
| chr12 | 1,08E+08 | 1,08E+08 | 551 | * | 4,54296 | 5,580918051 | 2,138273 | 2,610012 | 0,009054 | 1 | 0 | Exon (ENSMUST000002 | 12 | 1,08E+08 | 1,08E+08 | 27854 | 1 | 13116 | ENSMUST00000021684.6 | -7722 | ENSMUSG00000021259 | Cyp46a1 | cytochrome P450, family 46, subfamily a, polypeptide 1 |
| chr1 | 61828170 | 61828626 | 457 | * | 6,872024 | 5,580919 | 1,805049 | 3,091838 | 0,001989 | 0,619070779 | 1,360622 | Intron (ENSMUST0000( | 1 | 61806574 | 61890621 | 84048 | 2 | 74661 | ENSMUST00000138535.8 | 61995 | | Pard3bos1 | par-3 family cell polarity regulator beta, opposite strand 1 |
| chr10 | 75741765 | 75742463 | 699 | * | 8,250886 | 5,58103815 | 1,891121 | 2,951179 | 0,003166 | 0,684171718 | 1,449706 | 3' UTR | 10 | 75732607 | 75749996 | 17390 | 2 | 20587 | ENSMUST00000133189.8 | 7533 | ENSMUSG00000000902 | Smarcb1 | SWI/SNF related, matrix associated, actin dependent regulator of chromatin, subfamily b, member 1 |
| chr1 | 86104450 | 86105329 | 880 | * | 7,488051 | 5,58256812 | 1,839384 | 3,035021 | 0,002405 | 1 | 0 | Exon (ENSMUST000001 | 1 | 86124112 | 86180114 | 56003 | 1 | 78795 | ENSMUST00000150059.8 | -18783 | ENSMUSG00000062590 | Armc9 | armadillo repeat containing 9 |
| chr4 | 82302815 | 82303902 | 1088 | * | 7,324502 | 5,582569679 | 1,839678 | 3,034536 | 0,002409 | 1 | 0 | Intron (ENSMUST0000( | 4 | 82416781 | 82417553 | 773 | 2 | 18028 | ENSMUST00000152588.2 | 113651 | ENSMUSG00000008575 | Nfib | nuclear factor I/B |
| chr13 | 83147177 | 83148392 | 1216 | * | 7,923737 | 5,583188453 | 1,762594 | 3,167598 | 0,001537 | 1 | 0 | Distal Intergenic | 13 | 82846394 | 82846452 | 59 | 2 | 1,01E+08 | ENSMUST00000198795.3 | -300725 | ENSMUSG00000106655 | Mir3961 | microRNA 3961 |
| chr5 | 7217519 | 7218364 | 846 | * | 5,719497 | 5,584850629 | 2,043506 | 2,732975 | 0,006277 | 1 | 0 | Intron (ENSMUST0000( | 5 | 7227706 | 7227856 | 151 | 2 | 1,15E+08 | ENSMUST00000082917.3 | 9492 | ENSMUSG00000064851 | Gm24525 | predicted gene, 24525 |
| chr7 | 45030873 | 45032110 | 1238 | * | 11,62991 | 5,586095812 | 1,768467 | 3,158722 | 0,001585 | 0,683067851 | 0,744375 | Promoter (<=1kb) | 7 | 45030026 | 45030741 | 716 | 2 | 20637 | ENSMUST00000210661.2 | -132 | ENSMUSG00000063511 | Snrnp70 | small nuclear ribonucleoprotein 70 (U1) |
| chr1 | 56932306 | 56933352 | 1047 | * | 9,844578 | 5,588024777 | 1,712767 | 3,262571 | 0,001104 | 0,533665288 | 1,173325 | Intron (ENSMUST0000( | 1 | 56987293 | 57008999 | 21707 | 2 | 212712 | ENSMUST00000176759.2 | 75647 | ENSMUSG00000038331 | Satb2 | special AT-rich sequence binding protein 2 |
| chr7 | 1,1E+08 | 1,1E+08 | 753 | * | 6,783237 | 5,59038731 | 1,836232 | 3,044488 | 0,002331 | 1 | 0 | Intron (ENSMUST0000( | 7 | 1,1E+08 | 1,1E+08 | 21459 | 2 | 319934 | ENSMUST00000166885.2 | 4076 | ENSMUSG00000038371 | Sbf2 | SET binding factor 2 |
| chr19 | 7185537 | 7186793 | 1257 | * | 10,49086 | 5,591772716 | 1,738193 | 3,217004 | 0,001295 | 0,563566358 | 1,173325 | Promoter (1-2kb) | 19 | 7175567 | 7183681 | 8115 | 2 | 107260 | ENSMUST00000025679.11 | -1856 | ENSMUSG00000024767 | Otub1 | OTU domain, ubiquitin aldehyde binding 1 |
| chr14 | 14294919 | 14296863 | 1945 | * | 18,725297 | 5,595735493 | 1,2716669 | 4,4003153 | 1,081E-05 | 0,026722594 | 2,3860078 | Promoter (<=1kb) | 14 | 14296748 | 14298918 | 2171 | 1 | 68263 | ENSMUST00000224864.2 | 0 | ENSMUSG00000021748 | Pdhb | pyruvate dehydrogenase (lipoamide) beta |
| chr8 | 10028264 | 10029191 | 928 | * | 8,278179 | 5,596959235 | 1,839207 | 3,043137 | 0,002341 | 0,54964001 | 2,237403 | Promoter (<=1kb) | 8 | 10027717 | 10042155 | 14439 | 1 | 68904 | ENSMUST00000048216.6 | 547 | ENSMUSG00000040396 | Abhd13 | abhydrolase domain containing 13 |
| chr10 | 1,01E+08 | 1,01E+08 | 704 | * | 4,686778 | 5,59759951 | 2,113778 | 2,648149 | 0,008093 | 1 | 0 | Distal Intergenic | 10 | 1,01E+08 | 1,01E+08 | 132 | 1 | 1,15E+08 | ENSMUST00000157098.3 | -159190 | ENSMUSG00000087723 | Gm26278 | predicted gene, 26278 |
| chrX | 56480266 | 56481550 | 1285 | * | 8,611563 | 5,598126878 | 1,628698 | 3,43718 | 0,000588 | 1 | 0 | Intron (ENSMUST0000( | 20 | 56454514 | 56534127 | 79614 | 1 | 245423 | ENSMUST00000114725.3 | 25752 | ENSMUSG00000079584 | Tm9sf5 | transmembrane 9 superfamily member 5 |
| chr10 | 21677499 | 21678577 | 1079 | * | 8,523866 | 5,599903016 | 1,753702 | 3,19319 | 0,001407 | 0,54122637 | 1,425929 | Distal Intergenic | 10 | 21706398 | 21720750 | 14353 | 2 | 628133 | ENSMUST00000180731.2 | 42173 | | Platr6 | pluripotency associated transcript 6 |
| chr11 | 86046660 | 86048107 | 1448 | * | 7,629355 | 5,600924005 | 1,888064 | 2,966491 | 0,003012 | 1 | 0 | Intron (ENSMUST0000( | 11 | 86085859 | 86091990 | 6132 | 2 | 237911 | ENSMUST00000149748.8 | 43883 | ENSMUSG00000034329 | Brip1 | BRCA1 interacting protein C-terminal helicase 1 |

| | | | | | | | | | | | | | | | | | | | | | | | |
|---|---|---|---|---|---|---|---|---|---|---|---|---|---|---|---|---|---|---|---|---|---|---|---|
| chr1 | 71138631 | 71139977 | 1347 | * | 8,695913 | 5,601661738 | 1,722358 | 3,252321 | 0,001145 | 1 | 0 | Promoter (2-3kb) | 1 | 71066657 | 71142305 | 75649 | 2 | 12021 | ENSMUST00000027393.8 | 2328 | ENSMUSG00000026196 | Bard1 | BRCA1 associated RING domain 1 |
| chr17 | 66088778 | 66089932 | 1155 | * | 6,711112 | 5,601723603 | 1,888788 | 2,965777 | 0,003019 | 1 | 0 | Promoter (<=1kb) | 17 | 66089568 | 66110907 | 21340 | 1 | 70351 | ENSMUST00000161324.8 | 0 | ENSMUSG00000061950 | Ppp4r1 | protein phosphatase 4, regulatory subunit 1 |
| chr18 | 5934983 | 5936641 | 1659 | * | 11,18365 | 5,601845779 | 1,687668 | 3,319281 | 0,000902 | 0,497788447 | 1,173325 | Distal Intergenic | 18 | 5945183 | 5965965 | 20783 | 1 | 1,03E+08 | ENSMUST00000234953.2 | -8542 | | Gm34804 | predicted gene, 34804 |
| chr6 | 31293558 | 31294149 | 592 | * | 6,85113 | 5,602916659 | 1,917055 | 2,922669 | 0,00347 | 1 | 0 | Intron (ENSMUST00000 | 6 | 31298906 | 31314320 | 15415 | 1 | 73652 | ENSMUST00000124811.2 | -4757 | ENSMUSG00000087380 | 2210408F21Rik | RIKEN cDNA 2210408F21 gene |
| chr18 | 70815682 | 70816532 | 851 | * | 7,599625 | 5,603043192 | 1,856434 | 3,018176 | 0,002543 | 0,661775851 | 1,360622 | Distal Intergenic | 18 | 70750844 | 70758848 | 8005 | 1 | 17191 | ENSMUST00000128555.2 | 64838 | ENSMUSG00000024513 | Mbd2 | methyl-CpG binding domain protein 2 |
| chr4 | 1,08E+08 | 1,08E+08 | 882 | * | 6,928884 | 5,6040095 | 1,934156 | 2,897393 | 0,003763 | 1 | 0 | Distal Intergenic | 4 | 1,08E+08 | 1,08E+08 | 15573 | 2 | 545667 | ENSMUST00000053157.7 | -6212 | ENSMUSG00000059816 | Shisal2a | shisa like 2A |
| chr4 | 1,33E+08 | 1,33E+08 | 887 | * | 7,929781 | 5,604645637 | 1,772018 | 3,16286 | 0,001562 | 1 | 0 | Promoter (1-2kb) | 4 | 1,33E+08 | 1,33E+08 | 868 | 1 | 53608 | ENSMUST00000123612.2 | -1685 | ENSMUSG00000028862 | Map3k6 | mitogen-activated protein kinase kinase kinase 6 |
| chr11 | 8669980 | 8670740 | 761 | * | 6,772368 | 5,605685487 | 2,03251 | 2,758011 | 0,005815 | 1 | 0 | Distal Intergenic | 11 | 8381652 | 8614681 | 233030 | 2 | 319939 | ENSMUST00000239111.2 | -55299 | ENSMUSG00000020422 | Tns3 | tensin 3 |
| chr7 | 70784620 | 70785541 | 922 | * | 8,252131 | 5,606054281 | 1,781632 | 3,146583 | 0,001652 | 0,582232389 | 1,360622 | Distal Intergenic | 7 | 70738175 | 70766287 | 28113 | 2 | 1,03E+08 | ENSMUST00000208671.2 | -18333 | ENSMUSG00000109003 | Gm36011 | predicted gene, 36011 |
| chr4 | 53711446 | 53712236 | 791 | * | 6,278436 | 5,606326209 | 1,975462 | 2,837982 | 0,00454 | 1 | 0 | Promoter (1-2kb) | 4 | 53713998 | 53765785 | 51788 | 1 | 246179 | ENSMUST00000128667.8 | -1762 | ENSMUSG00000028414 | Fktn | fukutin |
| chr14 | 45628926 | 45629609 | 684 | * | 3,904787 | 5,606488763 | 2,14881 | 2,609113 | 0,009078 | 1 | 0 | Promoter (2-3kb) | 14 | 45613791 | 45626418 | 12628 | 2 | 54342 | ENSMUST00000046191.9 | -2508 | ENSMUSG00000037722 | Gnpnat1 | glucosamine-phosphate N-acetyltransferase 1 |
| chr12 | 77451229 | 77451714 | 486 | * | 4,688722 | 5,606492857 | 2,146003 | 2,612528 | 0,008988 | 1 | 0 | Intron (ENSMUST00000 | 12 | 77456915 | 77522763 | 65849 | 1 | 53618 | ENSMUST00000218851.2 | -5201 | ENSMUSG00000021065 | Fut8 | fucosyltransferase 8 |
| chr15 | 70558573 | 70559164 | 592 | * | 5,456271 | 5,606632901 | 2,127383 | 2,63546 | 0,008402 | 1 | 0 | Distal Intergenic | 15 | 69883472 | 69949690 | 66219 | 1 | 1,01E+08 | ENSMUST00000230802.2 | 675101 | ENSMUSG00000116483 | Gm19782 | predicted gene, 19782 |
| chr2 | 30404105 | 30404585 | 481 | * | 4,522223 | 5,606636143 | 2,126461 | 2,636605 | 0,008374 | 1 | 0 | Distal Intergenic | 2 | 30424741 | 30429034 | 4294 | 1 | 1,03E+08 | ENSMUST00000145413.2 | -20156 | ENSMUSG00000085275 | Gm14487 | predicted gene 14487 |
| chrX | 71470095 | 71470579 | 485 | * | 4,262394 | 5,60664707 | 2,123802 | 2,639911 | 0,008293 | 1 | 0 | Distal Intergenic | 20 | 71425476 | 71430464 | 4989 | 2 | 236852 | ENSMUST00000088481.5 | -39631 | ENSMUSG00000043453 | Magea10 | MAGE family member A10 |
| chr11 | 1,02E+08 | 1,02E+08 | 420 | * | 4,331622 | 5,606652301 | 2,124634 | 2,638879 | 0,008318 | 1 | 0 | Distal Intergenic | 11 | 1,02E+08 | 1,02E+08 | 16865 | 2 | 17285 | ENSMUST00000057054.8 | -3593 | ENSMUSG00000001493 | Meox1 | mesenchyme homeobox 1 |
| chr6 | 1,27E+08 | 1,27E+08 | 588 | * | 4,512189 | 5,60677356 | 2,106505 | 2,661648 | 0,007776 | 1 | 0 | Promoter (2-3kb) | 6 | 1,27E+08 | 1,27E+08 | 47401 | 1 | 101187 | ENSMUST00000112191.8 | -2063 | ENSMUSG00000037997 | Parp11 | poly (ADP-ribose) polymerase family, member 11 |
| chr17 | 35294882 | 35295393 | 512 | * | 4,555124 | 5,606820717 | 2,100211 | 2,669646 | 0,007593 | 1 | 0 | Promoter (<=1kb) | 17 | 35295878 | 35297270 | 1393 | 1 | 70274 | ENSMUST00000172494.8 | -485 | ENSMUSG00000013766 | Ly6g6e | lymphocyte antigen 6 complex, locus G6E |
| chr14 | 5791117 | 5792024 | 908 | * | 5,360478 | 5,606913513 | 2,086468 | 2,687275 | 0,007204 | 1 | 0 | Intron (ENSMUST00000 | 14 | 5893936 | 6038426 | 144491 | 1 | 218772 | ENSMUST00000225921.2 | -101912 | ENSMUSG00000017491 | Rarb | retinoic acid receptor, beta |
| chr14 | 1,22E+08 | 1,22E+08 | 484 | * | 4,801079 | 5,606922465 | 2,084304 | 2,690069 | 0,007144 | 1 | 0 | Distal Intergenic | 14 | 1,22E+08 | 1,22E+08 | 4033 | 2 | 1,03E+08 | ENSMUST00000227451.2 | 14170 | | Gm33299 | predicted gene, 33299 |
| chr17 | 24878830 | 24879357 | 528 | * | 4,96572 | 5,607169087 | 2,053432 | 2,730634 | 0,006321 | 1 | 0 | Promoter (1-2kb) | 17 | 24876304 | 24877431 | 1128 | 2 | 381073 | ENSMUST00000095544.6 | -1399 | ENSMUSG00000071230 | Npw | neuropeptide W |
| chr10 | 35890173 | 35891125 | 953 | * | 5,862883 | 5,607276403 | 1,994025 | 2,812039 | 0,004923 | 1 | 0 | Distal Intergenic | 10 | 35714486 | 35730471 | 15986 | 1 | 1,05E+08 | ENSMUST00000213846.2 | 175687 | | Gm40623 | predicted gene, 40623 |
| chr5 | 1,37E+08 | 1,37E+08 | 786 | * | 5,576479 | 5,607333554 | 2,032818 | 2,758404 | 0,005808 | 1 | 0 | 3' UTR | 5 | 1,37E+08 | 1,37E+08 | 3263 | 1 | 59310 | ENSMUST00000197849.2 | 3108 | ENSMUSG00000005474 | Myl10 | myosin, light chain 10, regulatory |
| chr1 | 1,81E+08 | 1,81E+08 | 699 | * | 5,627338 | 5,607431433 | 2,022033 | 2,773165 | 0,005551 | 1 | 0 | Distal Intergenic | 1 | 1,81E+08 | 1,82E+08 | 238183 | 1 | 240960 | ENSMUST00000208001.2 | -28112 | ENSMUSG00000047369 | Dnah14 | dynein, axonemal, heavy chain 14 |
| chr12 | 93633480 | 93634124 | 645 | * | 4,895009 | 5,607457112 | 2,019788 | 2,77626 | 0,005499 | 1 | 0 | Distal Intergenic | 12 | 93713276 | 93713436 | 161 | 1 | 1,15E+08 | ENSMUST00000083369.4 | -79152 | ENSMUSG00000065303 | Gm23249 | predicted gene, 23249 |
| chr11 | 79352929 | 79353770 | 842 | * | 5,348616 | 5,60752194 | 2,013228 | 2,785338 | 0,005347 | 1 | 0 | Intron (ENSMUST00000 | 11 | 79338979 | 79364238 | 25260 | 1 | 18015 | ENSMUST00000130979.2 | 13950 | ENSMUSG00000020716 | Nf1 | neurofibromin 1 |
| chr12 | 84055704 | 84056359 | 656 | * | 5,051372 | 5,60764805 | 1,999343 | 2,804745 | 0,005036 | 1 | 0 | Promoter (<=1kb) | 12 | 84056264 | 84065145 | 8882 | 1 | 26897 | ENSMUST00000168120.3 | 0 | ENSMUSG00000072949 | Acot1 | acyl-CoA thioesterase 1 |
| chr3 | 69254740 | 69255080 | 341 | * | 5,26908 | 5,607734348 | 2,003479 | 2,798999 | 0,005126 | 1 | 0 | Intron (ENSMUST00000 | 3 | 69224194 | 69468135 | 243942 | 1 | 242083 | ENSMUST00000029355.9 | 30546 | ENSMUSG00000027784 | Ppm1l | protein phosphatase 1 (formerly 2C)-like |
| chr19 | 7970223 | 7970682 | 460 | * | 6,287638 | 5,607959258 | 1,830476 | 3,063663 | 0,002186 | 1 | 0 | Distal Intergenic | 19 | 7841753 | 7943392 | 101640 | 2 | 171405 | ENSMUST00000075619.5 | -26831 | ENSMUSG00000067656 | Slc22a27 | solute carrier family 22, member 27 |
| chr15 | 79863780 | 79864559 | 780 | * | 6,592411 | 5,607969882 | 1,967609 | 2,850144 | 0,00437 | 1 | 0 | Distal Intergenic | 15 | 79802618 | 79855320 | 52703 | 2 | 52609 | ENSMUST00000128931.8 | -8460 | ENSMUSG00000053411 | Cbx7 | chromobox 7 |
| chr14 | 6003067 | 6003903 | 837 | * | 5,550029 | 5,608038792 | 1,961683 | 2,85879 | 0,004253 | 1 | 0 | Intron (ENSMUST00000 | 14 | 6041258 | 6046475 | 5218 | 2 | 21974 | ENSMUST00000163032.2 | 42572 | ENSMUSG00000017485 | Top2b | topoisomerase (DNA) II beta |
| chr3 | 84379759 | 84380426 | 668 | * | 5,624529 | 5,608053398 | 1,960334 | 2,860764 | 0,004226 | 1 | 0 | Intron (ENSMUST00000 | 3 | 84381830 | 84386289 | 4460 | 2 | 229474 | ENSMUST00000131065.2 | 5863 | ENSMUSG00000041842 | Fhdc1 | FH2 domain containing 1 |
| chr12 | 87467346 | 87468371 | 1026 | * | 5,960845 | 5,608172321 | 1,949818 | 2,876254 | 0,004024 | 1 | 0 | Exon (ENSMUST000002 | 12 | 87484449 | 87484527 | 79 | 2 | 1,01E+08 | ENSMUST00000197755.3 | 16156 | ENSMUSG00000105428 | Mir3068 | microRNA 3068 |
| chrX | 12039599 | 12040579 | 981 | * | 5,847161 | 5,608312149 | 1,938137 | 2,893662 | 0,003808 | 1 | 0 | Distal Intergenic | 20 | 11902982 | 12026594 | 123613 | 2 | 71458 | ENSMUST00000043441.13 | -13005 | ENSMUSG00000040363 | Bcor | BCL6 interacting corepressor |
| chr10 | 28448968 | 28449652 | 685 | * | 7,470132 | 5,60840409 | 1,852632 | 3,027263 | 0,002468 | 1 | 0 | Exon (ENSMUST000001 | 10 | 28456455 | 28467901 | 11447 | 1 | 19272 | ENSMUST00000220357.2 | -6803 | ENSMUSG00000019889 | Ptprk | protein tyrosine phosphatase, receptor type, K |
| chr7 | 67078780 | 67079622 | 843 | * | 6,649022 | 5,608460073 | 1,932442 | 2,902265 | 0,003705 | 1 | 0 | Exon (ENSMUST000002 | 7 | 67078821 | 67093681 | 14861 | 2 | 1,03E+08 | ENSMUST00000208755.2 | 14059 | | Gm33926 | predicted gene, 33926 |
| chr2 | 36024602 | 36025309 | 708 | * | 6,408862 | 5,608470781 | 1,925252 | 2,91311 | 0,003578 | 1 | 0 | Promoter (1-2kb) | 2 | 36026401 | 36080659 | 54259 | 1 | 67871 | ENSMUST00000028250.9 | -1092 | ENSMUSG00000026887 | Mrrf | mitochondrial ribosome recycling factor |
| chr19 | 4038349 | 4038812 | 464 | * | 6,211703 | 5,608617544 | 1,914001 | 2,93031 | 0,003386 | 1 | 0 | Promoter (1-2kb) | 19 | 4037203 | 4038651 | 1449 | 1 | 71670 | ENSMUST00000235592.2 | 1146 | ENSMUSG00000024866 | Acy3 | aspartoacylase (aminoacylase) 3 |
| chr12 | 3478098 | 3479205 | 1108 | * | 7,244338 | 5,608689853 | 1,908806 | 2,938324 | 0,0033 | 1 | 0 | Promoter (<=1kb) | 12 | 3456227 | 3477528 | 21302 | 2 | 1E+08 | ENSMUST00000180149.3 | -570 | ENSMUSG00000071456 | 1110002L01Rik | RIKEN cDNA 1110002L01 gene |
| chr9 | 1,08E+08 | 1,08E+08 | 751 | * | 6,334279 | 5,608858249 | 1,896038 | 2,9582 | 0,003094 | 1 | 0 | Intron (ENSMUST00000 | 9 | 1,08E+08 | 1,08E+08 | 45494 | 2 | 13138 | ENSMUST00000194698.2 | 32040 | ENSMUSG00000039952 | Dag1 | dystroglycan 1 |
| chr11 | 78954383 | 78955155 | 773 | * | 6,598849 | 5,609123585 | 1,87781 | 2,987056 | 0,002817 | 1 | 0 | Intron (ENSMUST00000 | 11 | 78935746 | 78950701 | 14956 | 2 | 16706 | ENSMUST00000141409.8 | -3682 | ENSMUSG00000018334 | Ksr1 | kinase suppressor of ras 1 |
| chr4 | 1,49E+08 | 1,49E+08 | 846 | * | 5,476599 | 5,60919541 | 2,035719 | 2,755387 | 0,005862 | 1 | 0 | Promoter (<=1kb) | 4 | 1,49E+08 | 1,49E+08 | 108 | 1 | 1,15E+08 | ENSMUST00000158000.3 | 931 | ENSMUSG00000088625 | Gm23318 | predicted gene, 23318 |
| chr9 | 23016889 | 23018302 | 1414 | * | 8,987232 | 5,609366376 | 1,735218 | 3,232657 | 0,001226 | 0,543024297 | 1,211315 | Distal Intergenic | 9 | 23134372 | 23396496 | 262125 | 1 | 73230 | ENSMUST00000071982.7 | -116070 | ENSMUSG00000031963 | Bmper | BMP-binding endothelial regulator |
| chr11 | 37581084 | 37581537 | 454 | * | 4,694089 | 5,609786795 | 2,120149 | 2,645939 | 0,008146 | 1 | 0 | Distal Intergenic | 11 | 37549816 | 37563993 | 14178 | 1 | 1E+08 | ENSMUST00000131013.2 | 31268 | ENSMUSG00000087388 | Gm12128 | predicted gene 12128 |
| chr11 | 87913126 | 87914075 | 950 | * | 9,592536 | 5,611488929 | 1,758323 | 3,191387 | 0,001416 | 0,54122637 | 1,425929 | Intron (ENSMUST00000 | 11 | 87938199 | 87944581 | 6383 | 1 | 110809 | ENSMUST00000139129.9 | -24124 | ENSMUSG00000018379 | Srsf1 | serine and arginine-rich splicing factor 1 |
| chr16 | 25079070 | 25079401 | 332 | * | 4,505645 | 5,611666404 | 2,099537 | 2,672812 | 0,007522 | 1 | 0 | Distal Intergenic | 16 | 25105567 | 25241094 | 135528 | 1 | 71338 | ENSMUST00000056087.4 | -26166 | ENSMUSG00000048399 | Tprg | transformation related protein 63 regulated |
| chr14 | 55022764 | 55024331 | 1568 | * | 12,34128 | 5,611994256 | 1,651326 | 3,398477 | 0,000678 | 0,436654797 | 1,2763 | Distal Intergenic | 14 | 54974092 | 55019403 | 45312 | 2 | 50934 | ENSMUST00000226646.2 | -3361 | ENSMUSG00000022180 | Slc7a8 | solute carrier family 7 (cationic amino acid transporter, y+ system), member 8 |
| chr7 | 46337077 | 46337499 | 423 | * | 4,954123 | 5,612433058 | 2,121463 | 2,645549 | 0,008156 | 1 | 0 | 3' UTR | 7 | 46294065 | 46321961 | 27897 | 2 | 21990 | ENSMUST00000049298.15 | -15116 | ENSMUSG00000040046 | Tph1 | tryptophan hydroxylase 1 |
| chr14 | 67495997 | 67496726 | 730 | * | 5,052448 | 5,612486517 | 2,123803 | 2,642659 | 0,008226 | 1 | 0 | Intron (ENSMUST00000 | 14 | 67472093 | 67666136 | 194044 | 1 | 13592 | ENSMUST00000176029.2 | 23904 | ENSMUSG00000022053 | Ebf2 | early B cell factor 2 |
| chr3 | 1,08E+08 | 1,08E+08 | 535 | * | 4,901125 | 5,613237636 | 2,147319 | 2,614068 | 0,008947 | 1 | 0 | Distal Intergenic | 3 | 1,08E+08 | 1,08E+08 | 4752 | 2 | 14863 | ENSMUST00000012348.9 | -8758 | ENSMUSG00000040562 | Gstm2 | glutathione S-transferase, mu 2 |
| chr4 | 41271972 | 41272459 | 488 | * | 4,475228 | 5,613332168 | 2,149663 | 2,611261 | 0,009021 | 1 | 0 | Promoter (2-3kb) | 4 | 41221592 | 41275116 | 53525 | 2 | 68926 | ENSMUST00000132499.2 | 2657 | ENSMUSG00000028433 | Ubap2 | ubiquitin-associated protein 2 |
| chr1 | 1,71E+08 | 1,71E+08 | 532 | * | 4,427488 | 5,613970417 | 2,171914 | 2,584803 | 0,009743 | 1 | 0 | Promoter (2-3kb) | 1 | 1,71E+08 | 1,71E+08 | 17931 | 1 | 71740 | ENSMUST00000094325.5 | 2685 | ENSMUSG00000006411 | Nectin4 | nectin cell adhesion molecule 4 |
| chr11 | 80299781 | 80300465 | 685 | * | 4,784686 | 5,614078094 | 2,17558 | 2,580498 | 0,009866 | 1 | 0 | Distal Intergenic | 11 | 80319441 | 80325364 | 5924 | 1 | 69077 | ENSMUST00000172641.2 | -18976 | ENSMUSG00000017428 | Psmd11 | proteasome (prosome, macropain) 26S subunit, non-ATPase, 11 |
| chr14 | 77188670 | 77189262 | 593 | * | 5,166149 | 5,615086716 | 1,956608 | 2,869806 | 0,004107 | 1 | 0 | Distal Intergenic | 14 | 77272390 | 77273521 | 1132 | 2 | 210808 | ENSMUST00000142300.2 | 84259 | ENSMUSG00000044350 | Lacc1 | laccase domain containing 1 |
| chr9 | 63869216 | 63870300 | 1085 | * | 7,532924 | 5,620677763 | 1,787816 | 3,143879 | 0,001667 | 1 | 0 | Intron (ENSMUST00000 | 9 | 63843190 | 63849084 | 5895 | 2 | 1,03E+08 | ENSMUST00000244845.1 | -20132 | | Gm36033 | predicted gene, 36033 |
| chr5 | 19471604 | 19472512 | 909 | * | 4,681868 | 5,62276486 | 2,014973 | 2,790491 | 0,005263 | 1 | 0 | Intron (ENSMUST00000 | 5 | 19432053 | 20909771 | 1477719 | 1 | 50791 | ENSMUST00000197443.5 | 39551 | ENSMUSG00000040003 | Magi2 | membrane associated guanylate kinase, WW and PDZ domain containing 2 |
| chr18 | 69136697 | 69137297 | 601 | * | 7,55182 | 5,626514624 | 1,779638 | 3,161606 | 0,001569 | 0,570876293 | 1,360622 | Distal Intergenic | 18 | 69155270 | 69155344 | 75 | 2 | 1,02E+08 | ENSMUST00000184434.3 | 18047 | ENSMUSG00000098366 | Mir145b | microRNA 145b |
| chr12 | 1,13E+08 | 1,13E+08 | 673 | * | 5,557306 | 5,628276799 | 1,929829 | 2,916464 | 0,00354 | 1 | 0 | Promoter (2-3kb) | 12 | 1,13E+08 | 1,13E+08 | 35322 | 1 | 217893 | ENSMUST00000222406.2 | 2102 | ENSMUSG00000021143 | Pacs2 | phosphofurin acidic cluster sorting protein 2 |
| chr13 | 52999928 | 53001129 | 1202 | * | 10,36209 | 5,629005917 | 1,880521 | 2,993323 | 0,00276 | 0,777938745 | 0,738441 | Intron (ENSMUST00000 | 13 | 53064359 | 53066982 | 2624 | 2 | 11992 | ENSMUST00000230563.2 | 65853 | ENSMUSG00000021460 | Auh | AU RNA binding protein/enoyl-coenzyme A hydratase |
| chr2 | 34766177 | 34766580 | 404 | * | 4,653126 | 5,629953103 | 2,174461 | 2,589125 | 0,009622 | 1 | 0 | Promoter (1-2kb) | 2 | 34756951 | 34764980 | 8030 | 2 | 66998 | ENSMUST00000189715.2 | -1197 | ENSMUSG00000026869 | Psmd5 | proteasome (prosome, macropain) 26S subunit, non-ATPase, 5 |
| chr19 | 46802221 | 46802841 | 621 | * | 4,438178 | 5,630371366 | 2,161809 | 2,604473 | 0,009202 | 1 | 0 | Intron (ENSMUST00000 | 19 | 46854099 | 46867018 | 12920 | 1 | 94219 | ENSMUST00000235671.2 | -51258 | ENSMUSG00000064105 | Cnnm2 | cyclin M2 |
| chr7 | 79662852 | 79664158 | 1307 | * | 7,788611 | 5,630783479 | 1,900151 | 2,963335 | 0,003043 | 1 | 0 | Distal Intergenic | 7 | 79674562 | 79743921 | 69360 | 1 | 209225 | ENSMUST00000164056.9 | -10404 | ENSMUSG00000048897 | Zfp710 | zinc finger protein 710 |
| chr12 | 17272447 | 17273186 | 740 | * | 4,414181 | 5,631111758 | 2,139261 | 2,63227 | 0,008482 | 1 | 0 | Distal Intergenic | 12 | 17316546 | 17331811 | 15266 | 1 | 71853 | ENSMUST00000162936.8 | -43360 | ENSMUSG00000020571 | Pdia6 | protein disulfide isomerase associated 6 |
| chr4 | 1,26E+08 | 1,26E+08 | 697 | * | 5,698965 | 5,631431202 | 1,864549 | 3,020264 | 0,002526 | 1 | 0 | Distal Intergenic | 4 | 1,26E+08 | 1,26E+08 | 38679 | 2 | 230753 | ENSMUST00000080919.12 | -5937 | ENSMUSG00000043962 | Thrap3 | thyroid hormone receptor associated protein 3 |
| chr4 | 1,45E+08 | 1,45E+08 | 323 | * | 4,621742 | 5,631774362 | 2,120207 | 2,656237 | 0,007902 | 1 | 0 | Distal Intergenic | 4 | 1,45E+08 | 1,45E+08 | 22321 | 1 | 1,05E+08 | ENSMUST00000143127.8 | -11706 | | Gm42346 | predicted gene, 42346 |
| chrX | 20683627 | 20684822 | 1196 | * | 4,929943 | 5,632622622 | 2,097515 | 2,685379 | 0,007245 | 1 | 0 | Intron (ENSMUST00000 | 20 | 20666571 | 20666675 | 105 | 2 | 1,15E+08 | ENSMUST00000158038.3 | -16952 | ENSMUSG00000118865 | Gm26065 | predicted gene, 26065 |
| chr1 | 88455287 | 88456042 | 756 | * | 4,995877 | 5,632887869 | 2,090889 | 2,694016 | 0,00706 | 1 | 0 | Distal Intergenic | 1 | 88427593 | 88437788 | 10196 | 2 | 14659 | ENSMUST00000067625.9 | -17499 | ENSMUSG00000062310 | Glrp1 | glutamine repeat protein 1 |
| chr3 | 86234896 | 86235748 | 853 | * | 7,638813 | 5,633043904 | 1,949051 | 2,890147 | 0,003851 | 1 | 0 | Exon (ENSMUST000001 | 3 | 86271764 | 86271932 | 169 | 1 | 1,15E+08 | ENSMUST00000083830.3 | -36016 | ENSMUSG00000065764 | Gm23850 | predicted gene, 23850 |
| chr5 | 76955545 | 76956260 | 716 | * | 6,345693 | 5,633174958 | 2,0831 | 2,704227 | 0,006846 | 1 | 0 | Promoter (1-2kb) | 5 | 76957835 | 76990436 | 32602 | 1 | 320827 | ENSMUST00000120818.8 | -1575 | ENSMUSG00000036377 | Cracd | capping protein inhibiting regulator of actin |
| chr1 | 5446254 | 5446883 | 630 | * | 4,948542 | 5,633289971 | 2,081127 | 2,706846 | 0,006793 | 1 | 0 | Distal Intergenic | 1 | 5658716 | 5676354 | 17639 | 1 | 18387 | ENSMUST00000160777.8 | -211833 | ENSMUSG00000025905 | Oprk1 | opioid receptor, kappa 1 |
| chr1 | 87057205 | 87057835 | 631 | * | 5,582453 | 5,633334678 | 2,079431 | 2,709076 | 0,006747 | 1 | 0 | Distal Intergenic | 1 | 87052730 | 87055634 | 2905 | 1 | 11648 | ENSMUST00000044878.4 | 4475 | ENSMUSG00000036500 | Akp3 | alkaline phosphatase 3, intestine, not Mn requiring |
| chr5 | 1,16E+08 | 1,16E+08 | 893 | * | 6,796888 | 5,633447623 | 1,933386 | 2,913773 | 0,003571 | 1 | 0 | Distal Intergenic | 5 | 1,16E+08 | 1,16E+08 | 164345 | 2 | 269693 | ENSMUST00000050178.13 | -8808 | ENSMUSG00000043913 | Ccdc60 | coiled-coil domain containing 60 |
| chr3 | 85303129 | 85303832 | 704 | * | 5,829254 | 5,633564996 | 2,073495 | 2,716942 | 0,006589 | 1 | 0 | Intron (ENSMUST00000 | 3 | 85304831 | 85335435 | 30605 | 2 | 1,03E+08 | ENSMUST00000200057.2 | 31603 | | Gm31359 | predicted gene, 31359 |
| chr15 | 34238803 | 34239347 | 545 | * | 5,018723 | 5,633660136 | 2,071241 | 2,719945 | 0,006529 | 1 | 0 | Promoter (<=1kb) | 15 | 34238313 | 34276314 | 38002 | 1 | 114128 | ENSMUST00000226627.2 | 490 | ENSMUSG00000022257 | Laptm4b | lysosomal-associated protein transmembrane 4B |
| chr12 | 25777571 | 25778338 | 768 | * | 5,248559 | 5,633785852 | 2,06835 | 2,723807 | 0,006453 | 1 | 0 | Distal Intergenic | 12 | 25444754 | 25521978 | 77225 | 1 | 328066 | ENSMUST00000221270.2 | 332817 | | Gm36723 | predicted gene, 36723 |
| chr16 | 74314299 | 74315178 | 880 | * | 6,660078 | 5,634477985 | 1,912308 | 2,946428 | 0,003215 | 1 | 0 | Distal Intergenic | 16 | 74404645 | 74404770 | 126 | 1 | 1,15E+08 | ENSMUST00000157767.3 | -89467 | ENSMUSG00000088392 | Gm26307 | predicted gene, 26307 |
| chrX | 1,04E+08 | 1,04E+08 | 847 | * | 5,401042 | 5,634491507 | 2,051773 | 2,746158 | 0,00603 | 1 | 0 | Intron (ENSMUST00000 | 20 | 1,04E+08 | 1,04E+08 | 24673 | 1 | 331487 | ENSMUST00000087867.6 | 4802 | ENSMUSG00000073016 | Uprt | uracil phosphoribosyltransferase |
| chr6 | 37279561 | 37280105 | 545 | * | 6,378391 | 5,637152258 | 1,925546 | 2,927561 | 0,003416 | 1 | 0 | Promoter (2-3kb) | 6 | 36823916 | 37277119 | 453204 | 2 | 320127 | ENSMUST00000150300.9 | -2442 | ENSMUSG00000038665 | Dgki | diacylglycerol kinase, iota |
| chr10 | 61649886 | 61650564 | 679 | * | 6,190385 | 5,63718225 | 1,935127 | 2,913081 | 0,003579 | 1 | 0 | Distal Intergenic | 10 | 61574444 | 61619926 | 45483 | 2 | 404634 | ENSMUST00000020283.5 | -29960 | ENSMUSG00000020086 | Macroh2a2 | macroH2A.2 histone |
| chr8 | 52779154 | 52779830 | 677 | * | 6,931759 | 5,637247062 | 1,95393 | 2,885081 | 0,003913 | 0,730607562 | 1,360622 | Distal Intergenic | 8 | 52789950 | 52790140 | 191 | 2 | 1,15E+08 | ENSMUST00000082748.3 | 10310 | ENSMUSG00000064682 | Gm25813 | predicted gene, 25813 |
| chr9 | 56878387 | 56879064 | 678 | * | 5,770146 | 5,637264991 | 1,954097 | 2,884845 | 0,003916 | 1 | 0 | Intron (ENSMUST00000 | 9 | 56858162 | 56890490 | 32329 | 1 | 66069 | ENSMUST00000068856.5 | 20225 | ENSMUSG00000055334 | Snupn | snurportin 1 |
| chr4 | 53451337 | 53451812 | 476 | * | 5,130011 | 5,637332981 | 1,973199 | 2,856951 | 0,004277 | 1 | 0 | Intron (ENSMUST00000 | 4 | 53440684 | 53563450 | 122767 | 1 | 100434 | ENSMUST00000107647.8 | 10653 | ENSMUSG00000028412 | Slc44a1 | solute carrier family 44, member 1 |
| chr12 | 76252269 | 76253113 | 845 | * | 6,432232 | 5,637380906 | 1,995007 | 2,825744 | 0,004717 | 1 | 0 | Intron (ENSMUST00000 | 12 | 76261260 | 76276238 | 14979 | 2 | 80384 | ENSMUST00000220068.2 | 23125 | ENSMUSG00000021056 | Tex21 | testis expressed gene 21 |
| chr1 | 48918937 | 48920105 | 1169 | * | 5,77972 | 5,637405537 | 1,999021 | 2,820083 | 0,004801 | 1 | 0 | Distal Intergenic | 1 | 49283775 | 49377405 | 93631 | 1 | 442837 | ENSMUST00000056879.9 | -363670 | ENSMUSG00000051616 | C230029F24Rik | RIKEN cDNA C230029F24 gene |
| chrX | 36445527 | 36446413 | 887 | * | 4,911716 | 5,637457793 | 2,018124 | 2,793415 | 0,005215 | 1 | 0 | Distal Intergenic | 20 | 36451241 | 36454816 | 3576 | 2 | 54405 | ENSMUST00000016571.8 | 8403 | ENSMUSG00000016427 | Ndufa1 | NADH:ubiquinone oxidoreductase subunit A1 |
| chr7 | 58328163 | 58328792 | 630 | * | 5,106689 | 5,637460878 | 2,019205 | 2,79192 | 0,00524 | 1 | 0 | Intron (ENSMUST00000 | 7 | 58307994 | 58479168 | 171175 | 1 | 11982 | ENSMUST00000168747.3 | 20169 | ENSMUSG00000025324 | Atp10a | ATPase, class V, type 10A |
| chrX | 1,14E+08 | 1,14E+08 | 507 | * | 4,996701 | 5,637478341 | 2,02466 | 2,784408 | 0,005363 | 1 | 0 | Distal Intergenic | 20 | 1,14E+08 | 1,14E+08 | 381 | 1 | 1,15E+08 | ENSMUST00000185033.3 | -178267 | ENSMUSG00000098405 | Gm27414 | predicted gene, 27414 |
| chr2 | 5540323 | 5540935 | 613 | * | 5,451611 | 5,637478634 | 2,02629 | 2,782168 | 0,0054 | 1 | 0 | Intron (ENSMUST00000 | 2 | 5303698 | 5680857 | 377160 | 2 | 227541 | ENSMUST00000114987.4 | 139922 | ENSMUSG00000039145 | Camk1d | calcium/calmodulin-dependent protein kinase ID |
| chr4 | 1,34E+08 | 1,34E+08 | 479 | * | 4,844573 | 5,637480899 | 2,026682 | 2,781631 | 0,005409 | 1 | 0 | Intron (ENSMUST00000 | 4 | 1,34E+08 | 1,34E+08 | 4704 | 1 | 1,03E+08 | ENSMUST00000133742.2 | -3302 | | Gm12977 | predicted gene 12977 |
| chr18 | 74913481 | 74914188 | 708 | * | 5,517467 | 5,637515898 | 2,03785 | 2,766404 | 0,005668 | 1 | 0 | Promoter (1-2kb) | 18 | 74912268 | 74939279 | 27012 | 1 | 52538 | ENSMUST00000041053.11 | 1213 | ENSMUSG00000036880 | Acaa2 | acetyl-Coenzyme A acyltransferase 2 (mitochondrial 3-oxoacyl-Coenzyme A thiolase) |
| chr7 | 41803279 | 41803980 | 702 | * | 4,513073 | 5,637561596 | 2,055358 | 2,742861 | 0,006091 | 1 | 0 | Intron (ENSMUST00000 | 7 | 41774828 | 41845632 | 70805 | 1 | 637898 | ENSMUST00000233266.2 | 28451 | ENSMUSG00000090619 | Vmn2r60 | vomeronasal 2, receptor 60 |
| chr5 | 21258317 | 21258777 | 461 | * | 4,827018 | 5,637605999 | 2,068461 | 2,725507 | 0,00642 | 1 | 0 | Promoter (1-2kb) | 5 | 21214442 | 21260723 | 46282 | 2 | 19248 | ENSMUST00000140057.2 | 1946 | ENSMUSG00000028771 | Ptpn12 | protein tyrosine phosphatase, non-receptor type 12 |
| chr11 | 70755009 | 70755716 | 708 | * | 5,191035 | 5,637634569 | 2,090765 | 2,696446 | 0,007008 | 1 | 0 | Exon (ENSMUST000001 | 11 | 70735801 | 70831309 | 95509 | 1 | 54189 | ENSMUST00000178245.2 | 19208 | ENSMUSG00000020817 | Rabep1 | rabaptin, RAB GTPase binding effector protein 1 |
| chr1 | 58641661 | 58642110 | 450 | * | 4,825299 | 5,637644434 | 2,091394 | 2,69564 | 0,007025 | 1 | 0 | Distal Intergenic | 1 | 58625543 | 58635123 | 9581 | 1 | 66495 | ENSMUST00000027193.9 | 16118 | ENSMUSG00000026032 | Ndufb3 | NADH:ubiquinone oxidoreductase subunit B3 |
| chr11 | 1,12E+08 | 1,12E+08 | 488 | * | 4,544421 | 5,637756366 | 2,136019 | 2,639375 | 0,008306 | 1 | 0 | Distal Intergenic | 11 | 1,13E+08 | 1,13E+08 | 4560 | 2 | 217294 | ENSMUST00000141399.2 | 502714 | ENSMUSG00000041674 | BC006965 | cDNA sequence BC006965 |
| chr4 | 1,3E+08 | 1,3E+08 | 393 | * | 3,923821 | 5,637824102 | 2,174052 | 2,593233 | 0,009508 | 1 | 0 | Promoter (2-3kb) | 4 | 1,3E+08 | 1,3E+08 | 27867 | 1 | 622480 | ENSMUST00000084263.6 | 2370 | ENSMUSG00000028784 | Spocd1 | SPOC domain containing 1 |
| chr9 | 98848539 | 98849318 | 780 | * | 5,767414 | 5,637888356 | 1,981919 | 2,844662 | 0,004446 | 1 | 0 | Distal Intergenic | 9 | 98831834 | 98837352 | 5519 | 2 | 768252 | ENSMUST00000181706.8 | -11187 | ENSMUSG00000097072 | Foxl2os | forkhead box L2, opposite strand |
| chr19 | 11598597 | 11599412 | 816 | * | 6,491138 | 5,639480943 | 1,885046 | 2,991693 | 0,002774 | 1 | 0 | Promoter (1-2kb) | 19 | 11594557 | 11601007 | 6451 | 2 | 14126 | ENSMUST00000164792.8 | 1595 | ENSMUSG00000024680 | Ms4a2 | membrane-spanning 4-domains, subfamily A, member 2 |
| chr15 | 9453033 | 9453687 | 655 | * | 5,561844 | 5,641916697 | 2,076919 | 2,716484 | 0,006598 | 1 | 0 | Intron (ENSMUST00000 | 15 | 9436114 | 9466124 | 30011 | 1 | 75568 | ENSMUST00000226688.2 | 16919 | ENSMUSG00000039676 | Capsl | calcyphosine-like |
| chr5 | 31324995 | 31325737 | 743 | * | 5,814314 | 5,643154797 | 2,105196 | 2,680583 | 0,007349 | 1 | 0 | Promoter (1-2kb) | 5 | 31316839 | 31323476 | 6638 | 2 | 71752 | ENSMUST00000201087.2 | -1519 | ENSMUSG00000106864 | Gtf3c2 | general transcription factor IIIC, polypeptide 2, beta |
| chr8 | 80439395 | 80439784 | 390 | * | 4,457953 | 5,644740572 | 2,144166 | 2,632604 | 0,008473 | 1 | 0 | Promoter (<=1kb) | 8 | 80438480 | 80505688 | 67209 | 1 | 68999 | ENSMUST00000210812.2 | 915 | ENSMUSG00000036977 | Anapc10 | anaphase promoting complex subunit 10 |

| | | | | | | | | | | | | | | | | | | | | | | | |
|---|---|---|---|---|---|---|---|---|---|---|---|---|---|---|---|---|---|---|---|---|---|---|---|
| chr2 | 91055036 | 91056036 | 1001 | * | 9,351743 | 5,644996921 | 1,716947 | 3,287812 | 0,00101 | 0,510701052 | 1,211315 | Intron (ENSMUST00000 | 2 | 91041917 | 91065676 | 23760 | 2 | 107986 | ENSMUST00000152277.8 | 9640 | ENSMUSG00000002109 | Ddb2 | damage specific DNA binding protein 2 |
| chr1 | 89191400 | 89191720 | 321 | * | 4,394913 | 5,645346517 | 2,160416 | 2,613083 | 0,008973 | 1 | 0 | Distal Intergenic | 1 | 89288064 | 89333855 | 45792 | 2 | 77331 | ENSMUST00000186258.2 | 142135 | ENSMUSG00000099938 | C030007H22Rik | RIKEN cDNA C030007H22 gene |
| chr16 | 46394170 | 46394879 | 710 | * | 4,312977 | 5,645597942 | 2,167377 | 2,604807 | 0,009193 | 1 | 0 | Distal Intergenic | 16 | 46279215 | 46318888 | 39674 | 2 | 58998 | ENSMUST00000133935.2 | -75282 | ENSMUSG00000022656 | Nectin3 | nectin cell adhesion molecule 3 |
| chr19 | 16759657 | 16760287 | 631 | * | 4,778764 | 5,64596753 | 2,178078 | 2,592178 | 0,009537 | 1 | 0 | Promoter (1-2kb) | 19 | 16729684 | 16758297 | 28614 | 2 | 271564 | ENSMUST00000225764.2 | -1360 | ENSMUSG00000046230 | Vps13a | vacuolar protein sorting 13A |
| chr11 | 72661440 | 72661940 | 501 | * | 5,582089 | 5,646430539 | 2,192075 | 2,575839 | 0,01 | 1 | 0 | 3' UTR | 11 | 72655293 | 72658579 | 3287 | 1 | 11736 | ENSMUST00000132889.2 | 6147 | ENSMUSG00000020790 | Ankfy1 | ankyrin repeat and FYVE domain containing 1 |
| chr11 | 1E+08 | 1E+08 | 1368 | * | 8,945551 | 5,651319417 | 1,743658 | 3,241071 | 0,001191 | 0,541133622 | 1,211315 | Promoter (<=1kb) | 11 | 1E+08 | 1E+08 | 15839 | 1 | 12799 | ENSMUST00000147403.2 | 287 | ENSMUSG00000006782 | Cnp | 2',3'-cyclic nucleotide 3' phosphodiesterase |
| chr17 | 29652807 | 29654031 | 1225 | * | 18,965504 | 5,651752223 | 1,0007371 | 5,6475891 | 1,627E-08 | 0,000164635 | 2,6636191 | Promoter (<=1kb) | 17 | 29653791 | 29685056 | 31266 | 1 | 102640029 | ENSMUST00000234282.2 | 0 | ENSMUSG00000117427 | Gm36199 | predicted gene, 36199 |
| chr14 | 3167976 | 3168757 | 782 | * | 6,560008 | 5,655217292 | 1,84179 | 3,0705 | 0,002137 | 1 | 0 | Distal Intergenic | 14 | 3575429 | 3839361 | 263933 | 1 | 218793 | ENSMUST00000124353.2 | -406672 | ENSMUSG00000058317 | Ube2e2 | ubiquitin-conjugating enzyme E2E 2 |
| chr13 | 74504326 | 74504929 | 604 | * | 5,565909 | 5,655631166 | 2,04646 | 2,763617 | 0,005716 | 1 | 0 | Exon (ENSMUST00000 | 13 | 74498484 | 74505167 | 6684 | 1 | 67433 | ENSMUST00000162376.2 | 5842 | ENSMUSG00000021578 | Ccdc127 | coiled-coil domain containing 127 |
| chr19 | 19150368 | 19151324 | 957 | * | 6,20664 | 5,655821673 | 1,852491 | 3,053091 | 0,002265 | 1 | 0 | Distal Intergenic | 19 | 18907969 | 19088560 | 180592 | 2 | 225998 | ENSMUST00000112832.8 | -61808 | ENSMUSG00000036192 | Rorb | RAR-related orphan receptor beta |
| chr2 | 1,31E+08 | 1,31E+08 | 599 | * | 5,129536 | 5,656503398 | 1,958575 | 2,888071 | 0,003876 | 1 | 0 | Distal Intergenic | 2 | 1,31E+08 | 1,31E+08 | 40336 | 2 | 51902 | ENSMUST00000142982.2 | -9736 | ENSMUSG00000048911 | Rnf24 | ring finger protein 24 |
| chr4 | 4502985 | 4503526 | 542 | * | 5,563848 | 5,656808194 | 2,025829 | 2,792342 | 0,005233 | 1 | 0 | Distal Intergenic | 4 | 4528204 | 4528719 | 516 | 2 | 622019 | ENSMUST00000136880.2 | 25193 | | Gm11780 | predicted gene 11780 |
| chr8 | 1,21E+08 | 1,21E+08 | 1117 | * | 9,543107 | 5,657908545 | 1,686932 | 3,353963 | 0,000797 | 0,441780915 | 1,449706 | Intron (ENSMUST00000 | 8 | 1,21E+08 | 1,21E+08 | 51152 | 1 | 234797 | ENSMUST00000108951.8 | 7398 | ENSMUSG00000031824 | 6430548M08Rik | RIKEN cDNA 6430548M08 gene |
| chr18 | 63108054 | 63109098 | 1045 | * | 9,677411 | 5,659388495 | 1,899437 | 2,979509 | 0,002887 | 0,717133806 | 1,086118 | Promoter (1-2kb) | 18 | 63110902 | 63132521 | 21620 | 1 | 108123 | ENSMUST00000025474.14 | -1804 | ENSMUSG00000024581 | Napg | N-ethylmaleimide sensitive fusion protein attachment protein gamma |
| chr18 | 57495280 | 57496067 | 788 | * | 5,558985 | 5,659721575 | 1,885809 | 3,001217 | 0,002689 | 1 | 0 | 5' UTR | 18 | 57487962 | 57498388 | 10427 | 1 | 73137 | ENSMUST00000238069.2 | 7318 | ENSMUSG00000024594 | Prrc1 | proline-rich coiled-coil 1 |
| chr13 | 96304059 | 96304726 | 668 | * | 4,216078 | 5,660594156 | 2,178598 | 2,598274 | 0,009369 | 1 | 0 | Distal Intergenic | 13 | 96091102 | 96269085 | 177984 | 2 | 75209 | ENSMUST00000161263.8 | -34974 | ENSMUSG00000051111 | Sv2c | synaptic vesicle glycoprotein 2c |
| chr17 | 34579603 | 34580111 | 509 | * | 4,554422 | 5,660703583 | 2,173665 | 2,60422 | 0,009208 | 1 | 0 | Exon (ENSMUST00001 | 17 | 34573796 | 34588469 | 14674 | 1 | 547431 | ENSMUST00000025198.15 | 5807 | ENSMUSG00000024340 | Btnl2 | butyrophilin-like 2 |
| chr18 | 8591243 | 8591879 | 637 | * | 4,525319 | 5,660834504 | 2,1681 | 2,610966 | 0,009029 | 1 | 0 | Distal Intergenic | 18 | 9212163 | 9218136 | 5974 | 1 | 14370 | ENSMUST00000041080.7 | -620284 | ENSMUSG00000036904 | Fzd8 | frizzled class receptor 8 |
| chr19 | 28787877 | 28788702 | 826 | * | 4,880218 | 5,661007488 | 2,161541 | 2,618969 | 0,00882 | 1 | 0 | Distal Intergenic | 19 | 28812449 | 28856701 | 44253 | 1 | 20510 | ENSMUST00000161340.2 | -23747 | ENSMUSG00000024935 | Slc1a1 | solute carrier family 1 (neuronal/epithelial high affinity glutamate transporter, system Xag), member 1 |
| chr4 | 55655247 | 55655933 | 687 | * | 6,894337 | 5,661498896 | 1,862204 | 3,040214 | 0,002364 | 0,64945399 | 1,360622 | Distal Intergenic | 4 | 55605149 | 55636901 | 31753 | 1 | 1,03E+08 | ENSMUST00000123620.2 | 50098 | | Gm12506 | predicted gene 12506 |
| chr1 | 1,64E+08 | 1,64E+08 | 498 | * | 4,391008 | 5,661611679 | 2,136579 | 2,649849 | 0,008053 | 1 | 0 | Promoter (<=1kb) | 1 | 1,64E+08 | 1,64E+08 | 137527 | 1 | 16579 | ENSMUST00000077642.12 | 331 | ENSMUSG00000026585 | Kifap3 | kinesin-associated protein 3 |
| chr4 | 88038169 | 88038720 | 552 | * | 4,302759 | 5,66190535 | 2,124838 | 2,66463 | 0,007707 | 1 | 0 | Promoter (<=1kb) | 4 | 88039536 | 88329241 | 289706 | 1 | 230393 | ENSMUST00000159342.8 | -816 | ENSMUSG00000038368 | Focad | focadhesin |
| chr2 | 1,53E+08 | 1,53E+08 | 559 | * | 4,855977 | 5,662382085 | 2,106758 | 2,687723 | 0,007194 | 1 | 0 | Intron (ENSMUST00000 | 2 | 1,53E+08 | 1,53E+08 | 1052 | 1 | 78151 | ENSMUST00000109783.2 | 5688 | | 4930404H24Rik | RIKEN cDNA 4930404H24 gene |
| chr7 | 1,18E+08 | 1,18E+08 | 569 | * | 4,3256 | 5,662644756 | 2,09768 | 2,69948 | 0,006945 | 1 | 0 | Distal Intergenic | 7 | 1,18E+08 | 1,18E+08 | 1687 | 1 | 67745 | ENSMUST00000138180.2 | 12166 | ENSMUSG00000055159 | 4930583K01Rik | RIKEN cDNA 4930583K01 gene |
| chr10 | 82075686 | 82076562 | 877 | * | 7,86217 | 5,662721751 | 1,812019 | 3,125089 | 0,001778 | 1 | 0 | Promoter (<=1kb) | 10 | 82060684 | 82077107 | 16424 | 2 | 237411 | ENSMUST00000041264.15 | 545 | ENSMUSG00000062931 | Zfp938 | zinc finger protein 938 |
| chr17 | 48575103 | 48575721 | 619 | * | 5,428384 | 5,663114793 | 1,936286 | 2,92473 | 0,003448 | 1 | 0 | Intron (ENSMUST00000 | 17 | 48571401 | 48581528 | 10128 | 1 | 224840 | ENSMUST00000153420.2 | 3702 | ENSMUSG00000051682 | Treml4 | triggering receptor expressed on myeloid cells-like 4 |
| chr3 | 1,42E+08 | 1,42E+08 | 861 | * | 4,524402 | 5,663364412 | 2,08216 | 2,719947 | 0,006529 | 1 | 0 | Intron (ENSMUST00000 | 3 | 1,42E+08 | 1,42E+08 | 55016 | 2 | 56376 | ENSMUST00000197808.5 | -8836 | ENSMUSG00000028273 | Pdlim5 | PDZ and LIM domain 5 |
| chr4 | 21729207 | 21729919 | 713 | * | 4,728906 | 5,663503454 | 2,068815 | 2,737559 | 0,00619 | 1 | 0 | Promoter (1-2kb) | 4 | 21727767 | 21748554 | 20788 | 1 | 51813 | ENSMUST00000133712.2 | 1440 | ENSMUSG00000028252 | Ccnc | cyclin C |
| chr10 | 90691498 | 90692146 | 649 | * | 4,755419 | 5,663508489 | 2,068798 | 2,737584 | 0,006189 | 1 | 0 | Intron (ENSMUST00000 | 10 | 90665705 | 90757200 | 91496 | 1 | 77531 | ENSMUST00000183145.2 | 25793 | ENSMUSG00000058589 | Anks1b | ankyrin repeat and sterile alpha motif domain containing 1B |
| chrX | 1,11E+08 | 1,11E+08 | 791 | * | 5,05727 | 5,663522505 | 2,067908 | 2,738769 | 0,006167 | 1 | 0 | Intron (ENSMUST00000 | 20 | 1,1E+08 | 1,1E+08 | 149768 | 2 | 67071 | ENSMUST00000123951.9 | -52392 | ENSMUSG00000025665 | Rps6ka6 | ribosomal protein S6 kinase polypeptide 6 |
| chr17 | 7578997 | 7579900 | 904 | * | 7,129667 | 5,663677327 | 1,844839 | 3,070011 | 0,002141 | 1 | 0 | Distal Intergenic | 17 | 7592045 | 7613373 | 21329 | 1 | 21461 | ENSMUST00000095726.11 | -12145 | ENSMUSG00000071322 | Tcp10a | t-complex protein 10a |
| chr15 | 97678442 | 97678868 | 427 | * | 5,425665 | 5,663733829 | 2,060612 | 2,748569 | 0,005986 | 1 | 0 | Promoter (2-3kb) | 15 | 97676401 | 97688940 | 12540 | 1 | 67739 | ENSMUST00000230531.2 | 2041 | ENSMUSG00000081534 | Slc48a1 | solute carrier family 48 (heme transporter), member 1 |
| chr10 | 88433811 | 88434655 | 845 | * | 4,953784 | 5,664165914 | 2,047316 | 2,76663 | 0,005664 | 1 | 0 | Intron (ENSMUST00000 | 10 | 88354201 | 88440887 | 86687 | 2 | 109272 | ENSMUST00000119185.8 | 6232 | ENSMUSG00000020061 | Mybpc1 | myosin binding protein C, slow-type |
| chr12 | 84530644 | 84531526 | 883 | * | 6,560556 | 5,664306292 | 1,869042 | 3,030593 | 0,002441 | 1 | 0 | Intron (ENSMUST00000 | 12 | 84504170 | 84576483 | 72314 | 1 | 217708 | ENSMUST00000221172.2 | 26474 | ENSMUSG00000085793 | Lin52 | lin-52 DREAM MuvB core complex component |
| chr16 | 75992726 | 75993460 | 735 | * | 6,313231 | 5,664672094 | 1,884762 | 3,005511 | 0,002651 | 1 | 0 | Distal Intergenic | 16 | 75929429 | 75929559 | 131 | 2 | 1,15E+08 | ENSMUST00000104009.3 | -63167 | ENSMUSG00000077197 | Gm22268 | predicted gene, 22268 |
| chr14 | 70538115 | 70539011 | 897 | * | 5,605069 | 5,664751049 | 2,02976 | 2,790848 | 0,005257 | 1 | 0 | Exon (ENSMUST00002 | 14 | 70546221 | 70548689 | 2469 | 2 | 213053 | ENSMUST00000155825.2 | 9678 | ENSMUSG00000022094 | Slc39a14 | solute carrier family 39 (zinc transporter), member 14 |
| chr3 | 1,17E+08 | 1,17E+08 | 706 | * | 4,951113 | 5,664883379 | 2,026146 | 2,79589 | 0,005176 | 1 | 0 | Promoter (<=1kb) | 3 | 1,17E+08 | 1,17E+08 | 50730 | 2 | 114301 | ENSMUST00000040097.14 | -961 | ENSMUSG00000033377 | Palmd | palmdelphin |
| chr1 | 91806520 | 91807532 | 1013 | * | 7,748077 | 5,665009319 | 1,962165 | 2,887121 | 0,003888 | 1 | 0 | Distal Intergenic | 1 | 91860925 | 91862586 | 1662 | 2 | 208727 | ENSMUST00000191327.2 | 55054 | ENSMUSG00000026313 | Hdac4 | histone deacetylase 4 |
| chr1 | 1,83E+08 | 1,83E+08 | 740 | * | 6,038418 | 5,665298805 | 1,913616 | 2,96052 | 0,003071 | 1 | 0 | Distal Intergenic | 1 | 1,83E+08 | 1,83E+08 | 15197 | 1 | 53791 | ENSMUST00000195614.2 | 54908 | ENSMUSG00000079164 | Tlr5 | toll-like receptor 5 |
| chr1 | 37336733 | 37337570 | 838 | * | 6,119771 | 5,665392326 | 1,918402 | 2,953184 | 0,003145 | 1 | 0 | Promoter (1-2kb) | 1 | 37338946 | 37387593 | 48648 | 1 | 269180 | ENSMUST00000146676.2 | -1376 | ENSMUSG00000026113 | Inpp4a | inositol polyphosphate-4-phosphatase, type I |
| chr1 | 1,86E+08 | 1,86E+08 | 1249 | * | 5,498029 | 5,665412739 | 2,011153 | 2,816997 | 0,004847 | 1 | 0 | Distal Intergenic | 1 | 1,86E+08 | 1,86E+08 | 46635 | 2 | 320449 | ENSMUST00000193084.2 | 57670 | ENSMUSG00000103690 | A730004F24Rik | RIKEN cDNA A730004F24 gene |
| chr14 | 20879463 | 20880686 | 1224 | * | 6,710451 | 5,665502323 | 2,008635 | 2,820574 | 0,004794 | 1 | 0 | Distal Intergenic | 14 | 20886728 | 20893453 | 6726 | 1 | 18792 | ENSMUST00000022368.4 | -6042 | ENSMUSG00000021822 | Plau | plasminogen activator, urokinase |
| chr16 | 32170620 | 32171254 | 635 | * | 5,216666 | 5,665745693 | 2,086576 | 2,715332 | 0,006621 | 1 | 0 | Intron (ENSMUST00000 | 16 | 32151086 | 32206713 | 55628 | 1 | 224111 | ENSMUST00000232137.2 | 19534 | ENSMUSG00000053774 | Ubxn7 | UBX domain protein 7 |
| chr1 | 1,56E+08 | 1,56E+08 | 729 | * | 5,867065 | 5,665797451 | 1,939751 | 2,920889 | 0,00349 | 1 | 0 | Promoter (1-2kb) | 1 | 1,56E+08 | 1,56E+08 | 34732 | 2 | 104009 | ENSMUST00000035325.15 | -1407 | ENSMUSG00000033684 | Qsox1 | quiescin Q6 sulfhydryl oxidase 1 |
| chr11 | 69674961 | 69676156 | 1196 | * | 6,637728 | 5,66585021 | 1,942921 | 2,916151 | 0,003544 | 1 | 0 | 3' UTR | 11 | 69683605 | 69684646 | 1042 | 2 | 11443 | ENSMUST00000129381.2 | 8490 | ENSMUSG00000041189 | Chrnb1 | cholinergic receptor, nicotinic, beta polypeptide 1 (muscle) |
| chr5 | 32675951 | 32677079 | 1129 | * | 5,625445 | 5,666064761 | 1,954637 | 2,898781 | 0,003746 | 1 | 0 | Intron (ENSMUST00000 | 5 | 32643314 | 32650100 | 6787 | 1 | 19046 | ENSMUST00000201880.2 | 32637 | ENSMUSG00000014956 | Ppp1cb | protein phosphatase 1 catalytic subunit beta |
| chr12 | 21458374 | 21459205 | 832 | * | 6,458006 | 5,666250996 | 1,965742 | 2,8825 | 0,003945 | 1 | 0 | Intron (ENSMUST00000 | 12 | 21445571 | 21467131 | 21561 | 2 | 22630 | ENSMUST00000140710.2 | 7926 | ENSMUSG00000076432 | Ywhaq | tyrosine 3-monooxygenase/tryptophan 5-monooxygenase activation protein theta |
| chr17 | 25582262 | 25583048 | 787 | * | 6,569538 | 5,666794519 | 1,999235 | 2,834482 | 0,00459 | 1 | 0 | Promoter (2-3kb) | 17 | 25585279 | 25587068 | 1790 | 1 | 17229 | ENSMUST00000234477.2 | -2231 | ENSMUSG00000033825 | Tpsb2 | tryptase beta 2 |
| chr11 | 96864325 | 96865011 | 687 | * | 6,31622 | 5,667047863 | 2,017564 | 2,808857 | 0,004972 | 1 | 0 | Intron (ENSMUST00000 | 11 | 96844167 | 96868537 | 24371 | 2 | 78912 | ENSMUST00000107624.8 | 3526 | ENSMUSG00000018678 | Sp2 | Sp2 transcription factor |
| chr7 | 27028776 | 27029411 | 636 | * | 5,562365 | 5,667225757 | 1,961922 | 2,888609 | 0,003869 | 1 | 0 | Exon (ENSMUST00000 | 7 | 27005423 | 27033038 | 27616 | 2 | 108075 | ENSMUST00000118583.8 | 3627 | ENSMUSG00000040488 | Ltbp4 | latent transforming growth factor beta binding protein 4 |
| chr2 | 27541966 | 27542382 | 417 | * | 5,784714 | 5,667288846 | 1,963673 | 2,886066 | 0,003901 | 1 | 0 | Distal Intergenic | 2 | 27566452 | 27631910 | 65459 | 1 | 20181 | ENSMUST00000129514.8 | -24070 | ENSMUSG00000015846 | Rxra | retinoid X receptor alpha |
| chr2 | 1,27E+08 | 1,27E+08 | 706 | * | 4,683634 | 5,667313585 | 2,035514 | 2,784217 | 0,005366 | 1 | 0 | Distal Intergenic | 2 | 1,27E+08 | 1,27E+08 | 84666 | 2 | 58800 | ENSMUST00000103224.10 | -3456 | ENSMUSG00000027365 | Trpm7 | transient receptor potential cation channel, subfamily M, member 7 |
| chr12 | 40881202 | 40881597 | 396 | * | 4,526391 | 5,667446155 | 2,04602 | 2,769985 | 0,005606 | 1 | 0 | Intron (ENSMUST00000 | 12 | 40844233 | 40848081 | 3849 | 1 | 238130 | ENSMUST00000220850.2 | 36969 | ENSMUSG00000035954 | Dock4 | dedicator of cytokinesis 4 |
| chr5 | 1,21E+08 | 1,21E+08 | 754 | * | 5,043891 | 5,667470039 | 2,048165 | 2,767097 | 0,005656 | 1 | 0 | Intron (ENSMUST00000 | 5 | 1,21E+08 | 1,21E+08 | 60789 | 2 | 19247 | ENSMUST00000054547.9 | 12694 | ENSMUSG00000043733 | Ptpn11 | protein tyrosine phosphatase, non-receptor type 11 |
| chr11 | 1,03E+08 | 1,03E+08 | 802 | * | 8,03235 | 5,66773881 | 1,861299 | 3,045044 | 0,002326 | 1 | 0 | Distal Intergenic | 11 | 1,03E+08 | 1,03E+08 | 9273 | 1 | 1,05E+08 | ENSMUST00000212331.3 | 36649 | | Gm39397 | predicted gene, 39397 |
| chr2 | 28779107 | 28779603 | 497 | * | 5,266517 | 5,66785639 | 2,079218 | 2,725956 | 0,006412 | 1 | 0 | Intron (ENSMUST00000 | 2 | 28797696 | 28802507 | 4812 | 2 | 54422 | ENSMUST00000124059.2 | 22904 | ENSMUSG00000026805 | Barhl1 | BarH like homeobox 1 |
| chr8 | 50328083 | 50328478 | 396 | * | 5,415783 | 5,667932645 | 2,084106 | 2,7196 | 0,006536 | 1 | 0 | Intron (ENSMUST00000 | 8 | 49938516 | 49974130 | 35615 | 1 | 75359 | ENSMUST00000126285.2 | 389567 | ENSMUSG00000031559 | 4930555F03Rik | RIKEN cDNA 4930555F03 gene |
| chr19 | 12833890 | 12834668 | 779 | * | 4,790593 | 5,66809034 | 2,100605 | 2,698313 | 0,006969 | 1 | 0 | 5' UTR | 19 | 12839106 | 12840234 | 1129 | 1 | 258697 | ENSMUST00000059675.4 | -4438 | ENSMUSG00000046272 | Olfr1444 | olfactory receptor 1444 |
| chr15 | 71798193 | 71798575 | 383 | * | 4,305835 | 5,668124502 | 2,102391 | 2,696038 | 0,007017 | 1 | 0 | Intron (ENSMUST00000 | 15 | 71776577 | 71826223 | 49647 | 2 | 69700 | ENSMUST00000162081.2 | 27648 | ENSMUSG00000079022 | Col22a1 | collagen, type XXII, alpha 1 |
| chr12 | 85643105 | 85643708 | 604 | * | 4,834655 | 5,668379873 | 2,127023 | 2,664936 | 0,0077 | 1 | 0 | Promoter (2-3kb) | 12 | 85645801 | 85686652 | 40852 | 1 | 81703 | ENSMUST00000177587.9 | -2093 | ENSMUSG00000034271 | Jdp2 | Jun dimerization protein 2 |
| chr4 | 89972131 | 89972612 | 482 | * | 4,190632 | 5,668431796 | 2,132603 | 2,657987 | 0,007861 | 1 | 0 | Distal Intergenic | 4 | 90107057 | 90113939 | 6883 | 1 | 236537 | ENSMUST00000107129.2 | -134445 | ENSMUSG00000070902 | Zfp352 | zinc finger protein 352 |
| chr6 | 43643221 | 43643698 | 478 | * | 4,099411 | 5,668471636 | 2,136957 | 2,652591 | 0,007988 | 1 | 0 | Promoter (<=1kb) | 6 | 43321935 | 43643212 | 321278 | 2 | 29807 | ENSMUST00000067888.14 | -9 | ENSMUSG00000029735 | Tpk1 | thiamine pyrophosphokinase |
| chr18 | 80115321 | 80115796 | 476 | * | 4,597997 | 5,668623494 | 2,153097 | 2,632777 | 0,008469 | 1 | 0 | Intron (ENSMUST00000 | 18 | 80090105 | 80162854 | 72750 | 1 | 93737 | ENSMUST00000070219.9 | 25216 | ENSMUSG00000056214 | Pard6g | par-6 family cell polarity regulator gamma |
| chrX | 70329957 | 70330424 | 468 | * | 4,564538 | 5,668626356 | 2,153517 | 2,632264 | 0,008482 | 1 | 0 | Intron (ENSMUST00000 | 20 | 70344975 | 70359297 | 14323 | 1 | 17772 | ENSMUST00000129722.2 | -14551 | ENSMUSG00000031337 | Mtm1 | X-linked myotubular myopathy gene 1 |
| chr3 | 1,25E+08 | 1,25E+08 | 667 | * | 4,149315 | 5,66870352 | 2,159976 | 2,624429 | 0,008679 | 1 | 0 | Distal Intergenic | 3 | 1,24E+08 | 1,24E+08 | 7885 | 2 | 384775 | ENSMUST00000180033.8 | -205092 | ENSMUSG00000039174 | 1700003H04Rik | RIKEN cDNA 1700003H04 gene |
| chr3 | 1,31E+08 | 1,31E+08 | 629 | * | 4,628267 | 5,668718716 | 2,163167 | 2,620564 | 0,008778 | 1 | 0 | Promoter (2-3kb) | 3 | 1,31E+08 | 1,31E+08 | 9913 | 2 | 71519 | ENSMUST00000200236.2 | 2270 | ENSMUSG00000027983 | Cyp2u1 | cytochrome P450, family 2, subfamily u, polypeptide 1 |
| chr4 | 11203122 | 11203590 | 469 | * | 4,210085 | 5,668891127 | 2,1815 | 2,598621 | 0,00936 | 1 | 0 | 3' UTR | 4 | 11204110 | 11213185 | 9076 | 2 | 72656 | ENSMUST00000136545.8 | 9595 | ENSMUSG00000040738 | Ints8 | integrator complex subunit 8 |
| chr4 | 1,16E+08 | 1,16E+08 | 726 | * | 3,700192 | 5,668968724 | 2,190906 | 2,5875 | 0,009668 | 1 | 0 | Distal Intergenic | 4 | 1,16E+08 | 1,16E+08 | 30687 | 1 | 16351 | ENSMUST00000106479.8 | -11751 | ENSMUSG00000028696 | Ipp | IAP promoted placental gene |
| chr5 | 97149143 | 97150052 | 910 | * | 6,365345 | 5,670276237 | 2,037735 | 2,782637 | 0,005392 | 1 | 0 | Intron (ENSMUST00000 | 5 | 97145548 | 97239726 | 94179 | 1 | 140780 | ENSMUST00000035635.10 | 3595 | ENSMUSG00000034663 | Bmp2k | BMP2 inducible kinase |
| chr4 | 1,49E+08 | 1,49E+08 | 1100 | * | 7,170023 | 5,670597864 | 1,894819 | 2,992686 | 0,002765 | 1 | 0 | Promoter (1-2kb) | 4 | 1,49E+08 | 1,49E+08 | 3117 | 1 | 20810 | ENSMUST00000006611.9 | -1535 | ENSMUSG00000006442 | Srm | spermidine synthase |
| chr12 | 85515835 | 85516473 | 639 | * | 5,067494 | 5,670763966 | 1,979302 | 2,865032 | 0,00417 | 1 | 0 | Distal Intergenic | 12 | 85520664 | 85524047 | 3384 | 1 | 14281 | ENSMUST00000021674.7 | -4191 | ENSMUSG00000021250 | Fos | FBJ osteosarcoma oncogene |
| chr9 | 62342619 | 62344057 | 1439 | * | 19,736853 | 5,670891902 | 1,1426102 | 4,9631029 | 6,938E-07 | 0,003457727 | 2,2216115 | Intron (ENSMUST00000 | 9 | 62335545 | 62336532 | 988 | 2 | 235431 | ENSMUST00000151604.2 | -6087 | ENSMUSG00000041729 | Coro2b | coronin, actin binding protein, 2B |
| chr9 | 45209717 | 45210491 | 775 | * | 6,157193 | 5,671017115 | 2,049976 | 2,766382 | 0,005668 | 1 | 0 | Distal Intergenic | 9 | 45230398 | 45258879 | 28482 | 1 | 214531 | ENSMUST00000034597.8 | -19907 | ENSMUSG00000037129 | Tmprss13 | transmembrane protease, serine 13 |
| chr4 | 90774974 | 90775904 | 931 | * | 6,845228 | 5,672926209 | 2,084261 | 2,721793 | 0,006493 | 1 | 0 | Distal Intergenic | 4 | 90886099 | 90886230 | 132 | 2 | 1,15E+08 | ENSMUST00000178089.3 | 110326 | ENSMUSG00000096896 | Gm25244 | predicted gene, 25244 |
| chr3 | 57371433 | 57373356 | 1924 | * | 11,6764 | 5,673270146 | 1,693042 | 3,350933 | 0,000805 | 0,523637575 | 0,918255 | Intron (ENSMUST00000 | 3 | 57332735 | 57349098 | 16364 | 1 | 229302 | ENSMUST00000029377.8 | 38698 | ENSMUSG00000027801 | Tm4sf4 | transmembrane 4 superfamily member 4 |
| chr4 | 39491724 | 39492353 | 630 | * | 5,739305 | 5,673497642 | 2,095071 | 2,708022 | 0,006769 | 1 | 0 | Distal Intergenic | 4 | 39450295 | 39451776 | 1482 | 1 | 75471 | ENSMUST00000029955.5 | 41429 | ENSMUSG00000028287 | 1700009N14Rik | RIKEN cDNA 1700009N14 gene |
| chr5 | 1,43E+08 | 1,43E+08 | 647 | * | 7,563531 | 5,673577308 | 1,85548 | 3,05774 | 0,00223 | 1 | 0 | Exon (ENSMUST00001 | 5 | 1,43E+08 | 1,43E+08 | 13870 | 2 | 231868 | ENSMUST00000159406.8 | 7435 | ENSMUSG00000039244 | Ents15 | integrator complex subunit 15 |
| chr4 | 1,25E+08 | 1,25E+08 | 841 | * | 5,706426 | 5,673773799 | 2,100635 | 2,70098 | 0,006914 | 1 | 0 | Intron (ENSMUST00000 | 4 | 1,25E+08 | 1,25E+08 | 10366 | 2 | 75563 | ENSMUST00000036383.4 | 3007 | ENSMUSG00000042707 | Dnali1 | dynein, axonemal, light intermediate polypeptide 1 |
| chr12 | 14816611 | 14817185 | 575 | * | 5,938047 | 5,673926377 | 2,103562 | 2,697295 | 0,006991 | 1 | 0 | Distal Intergenic | 12 | 14556588 | 14556694 | 107 | 1 | 1,15E+08 | ENSMUST00000240467.1 | 260023 | ENSMUSG00000119624 | Gm25414 | predicted gene, 25414 |
| chr15 | 1,02E+08 | 1,02E+08 | 555 | * | 5,608935 | 5,674919462 | 2,124023 | 2,671778 | 0,007545 | 1 | 0 | Distal Intergenic | 15 | 1,02E+08 | 1,02E+08 | 8570 | 2 | 77055 | ENSMUST00000100179.2 | -10445 | ENSMUSG00000075402 | Krt76 | keratin 76 |
| chr9 | 70768614 | 70769120 | 507 | * | 5,542113 | 5,674926919 | 2,124318 | 2,671411 | 0,007553 | 1 | 0 | Intron (ENSMUST00000 | 9 | 70837770 | 70841712 | 3943 | 1 | 1,03E+08 | ENSMUST00000215190.2 | -68650 | | Gm32017 | predicted gene, 32017 |
| chr7 | 35041612 | 35042184 | 573 | * | 5,572673 | 5,675047403 | 2,126982 | 2,668122 | 0,007628 | 1 | 0 | Intron (ENSMUST00000 | 7 | 35033755 | 35091710 | 57956 | 1 | 52428 | ENSMUST00000085556.4 | 7857 | ENSMUSG00000030494 | Rhpn2 | rhophilin, Rho GTPase binding protein 2 |
| chr12 | 79576275 | 79577094 | 820 | * | 5,931194 | 5,675950677 | 2,146867 | 2,643829 | 0,008197 | 1 | 0 | Intron (ENSMUST00000 | 12 | 79374127 | 79748630 | 374504 | 1 | 19363 | ENSMUST00000221257.2 | 202148 | ENSMUSG00000059060 | Rad51b | RAD51 paralog B |
| chr17 | 8089324 | 8090244 | 921 | * | 5,428145 | 5,676400128 | 2,157182 | 2,631396 | 0,008503 | 1 | 0 | Distal Intergenic | 17 | 8001276 | 8046134 | 44859 | 2 | 68655 | ENSMUST00000167580.8 | -43190 | ENSMUSG00000071984 | Fndc1 | fibronectin type III domain containing 1 |
| chr1 | 63165138 | 63166060 | 923 | * | 5,684764 | 5,676918405 | 2,169407 | 2,616806 | 0,008876 | 1 | 0 | Exon (ENSMUST00001 | 1 | 63153899 | 63196404 | 42506 | 1 | 1E+08 | ENSMUST00000188602.2 | 11239 | ENSMUSG00000084799 | Ino80dos | INO80 complex subunit D, opposite strand |
| chr9 | 94777044 | 94777692 | 649 | * | 5,063409 | 5,67731739 | 2,179207 | 2,605222 | 0,009181 | 1 | 0 | Intron (ENSMUST00000 | 9 | 94633882 | 94633943 | 62 | 1 | 1,02E+08 | ENSMUST00000183864.3 | 143162 | ENSMUSG00000098251 | Mir7656 | microRNA 7656 |
| chr4 | 1,06E+08 | 1,06E+08 | 354 | * | 5,065701 | 5,677482209 | 2,183239 | 2,600486 | 0,009309 | 1 | 0 | Distal Intergenic | 4 | 1,06E+08 | 1,06E+08 | 3229 | 2 | 72563 | ENSMUST00000137882.2 | 5343 | | 2700068H02Rik | RIKEN cDNA 2700068H02 gene |
| chrX | 49966832 | 49967517 | 686 | * | 4,813362 | 5,677609815 | 2,186229 | 2,596988 | 0,009405 | 1 | 0 | Promoter (<=1kb) | 20 | 49967265 | 49980725 | 13461 | 1 | 70415 | ENSMUST00000146956.2 | 0 | ENSMUSG00000031112 | Stk26 | serine/threonine kinase 26 |
| chr1 | 81264950 | 81265709 | 760 | * | 4,9827 | 5,678191274 | 2,201162 | 2,579633 | 0,009891 | 1 | 0 | Intron (ENSMUST00000 | 1 | 81055300 | 81319479 | 264180 | 1 | 241134 | ENSMUST00000123285.2 | 209650 | ENSMUSG00000054976 | Nyap2 | neuronal tyrosine-phophorylated phosphoinositide 3-kinase adaptor 2 |
| chr12 | 82274973 | 82276578 | 1606 | * | 13,77496 | 5,682031149 | 1,63191 | 3,48183 | 0,000498 | 0,475863747 | 0,738441 | Intron (ENSMUST00000 | 12 | 82303359 | 82497045 | 193687 | 1 | 217692 | ENSMUST00000222714.2 | -26781 | ENSMUSG00000042700 | Sipa1l1 | signal-induced proliferation-associated 1 like 1 |
| chr7 | 98313415 | 98314280 | 866 | * | 8,19736 | 5,687118299 | 1,847172 | 3,078825 | 0,002078 | 0,531203841 | 2,237403 | Distal Intergenic | 7 | 98292264 | 98305990 | 13727 | 2 | 233545 | ENSMUST00000206565.2 | -7425 | ENSMUSG00000035401 | Emsy | EMSY, BRCA2-interacting transcriptional repressor |
| chr15 | 79478406 | 79479124 | 719 | * | 4,556754 | 5,688482522 | 1,989213 | 2,859665 | 0,004241 | 1 | 0 | Exon (ENSMUST00020 | 15 | 79446513 | 79470097 | 23585 | 2 | 13404 | ENSMUST00000229472.2 | -8309 | ENSMUSG00000022429 | Dmc1 | DNA meiotic recombinase 1 |
| chr10 | 41164850 | 41165671 | 822 | * | 6,275084 | 5,689624337 | 1,879313 | 3,027502 | 0,002466 | 1 | 0 | Intron (ENSMUST00000 | 10 | 41064168 | 41179256 | 115089 | 2 | 103199 | ENSMUST00000043814.5 | 13585 | ENSMUSG00000038417 | Fig4 | FIG4 phosphoinositide 5-phosphatase |
| chrX | 46137311 | 46137863 | 553 | * | 4,180779 | 5,690503717 | 2,208336 | 2,576829 | 0,009971 | 1 | 0 | Distal Intergenic | 20 | 45417884 | 45419222 | 1339 | 1 | 73360 | ENSMUST00000059466.3 | 719427 | ENSMUSG00000046615 | Actrt1 | actin-related protein T1 |
| chr15 | 1,01E+08 | 1,01E+08 | 695 | * | 8,242209 | 5,690654071 | 1,851064 | 3,07426 | 0,00211 | 1 | 0 | Distal Intergenic | 15 | 1,01E+08 | 1,01E+08 | 66470 | 1 | 20273 | ENSMUST00000201484.4 | -13818 | ENSMUSG00000023033 | Scn8a | sodium channel, voltage-gated, type VIII, alpha |
| chr12 | 48567887 | 48568470 | 584 | * | 4,094414 | 5,690668014 | 2,196233 | 2,591104 | 0,009567 | 1 | 0 | Distal Intergenic | 12 | 49056666 | 49056772 | 107 | 1 | 1,15E+08 | ENSMUST00000239848.1 | -488196 | ENSMUSG00001118661 | Gm26454 | predicted gene, 26454 |
| chr3 | 56694424 | 56694653 | 230 | * | 4,153208 | 5,690719389 | 2,192769 | 2,59522 | 0,009453 | 1 | 0 | Distal Intergenic | 3 | 56961571 | 56961700 | 130 | 1 | 1,15E+08 | ENSMUST00000239689.1 | -266918 | ENSMUSG00000118960 | Gm22269 | predicted gene, 22269 |
| chr8 | 26277474 | 26278194 | 721 | * | 7,222731 | 5,690775258 | 1,85352 | 3,070252 | 0,002139 | 1 | 0 | Promoter (1-2kb) | 8 | 26275615 | 26292137 | 16523 | 1 | 67207 | ENSMUST00000211670.2 | 1859 | ENSMUSG00000037296 | Lsm1 | LSM1 homolog, mRNA degradation associated |
| chr5 | 84230787 | 84231611 | 825 | * | 6,390633 | 5,690780535 | 1,96253 | 2,899717 | 0,003735 | 1 | 0 | Intron (ENSMUST00000 | 5 | 84205650 | 84564665 | 359016 | 2 | 13839 | ENSMUST00000113403.8 | 333054 | ENSMUSG00000029245 | Epha5 | Eph receptor A5 |
| chr13 | 97573521 | 97574108 | 588 | * | 4,040323 | 5,690879246 | 2,180752 | 2,609595 | 0,009065 | 1 | 0 | Promoter (2-3kb) | 13 | 97571478 | 97633515 | 62038 | 1 | 1E+08 | ENSMUST00000109431.10 | 2043 | ENSMUSG00000078952 | Lncenc1 | long non-coding RNA, embryonic stem cells expressed 1 |
| chr2 | 1,1E+08 | 1,1E+08 | 504 | * | 3,993422 | 5,690911466 | 2,179849 | 2,61069 | 0,009036 | 1 | 0 | Intron (ENSMUST00000 | 2 | 1,1E+08 | 1,1E+08 | 22111 | 1 | 107515 | ENSMUST00000152584.2 | -16089 | ENSMUSG00000050199 | Lgr4 | leucine-rich repeat-containing G protein-coupled receptor 4 |

| | | | | | | | | | | | | | | | | | | | | | | | |
|---|---|---|---|---|---|---|---|---|---|---|---|---|---|---|---|---|---|---|---|---|---|---|---|
| chr5 | 68901976 | 68902564 | 589 | * | 4,669831 | 5,690927669 | 2,177889 | 2,613047 | 0,008974 | 1 | 0 | Distal Intergenic | 5 | 68723513 | 68723639 | 127 | 2 | 1,15E+08 | ENSMUST00000157255.3 | -178337 | ENSMUSG00000087880 | Gm23605 | predicted gene, 23605 |
| chr11 | 3536790 | 3537341 | 552 | * | 3,881942 | 5,691052141 | 2,169607 | 2,62308 | 0,008714 | 1 | 0 | Distal Intergenic | 11 | 3482460 | 3490612 | 8153 | 2 | 29856 | ENSMUST00000156201.8 | -46178 | ENSMUSG00000020439 | Smtn | smoothelin |
| chr7 | 1,29E+08 | 1,29E+08 | 368 | * | 4,336173 | 5,691107634 | 2,166576 | 2,626775 | 0,00862 | 1 | 0 | Distal Intergenic | 7 | 1,29E+08 | 1,29E+08 | 20157 | 2 | 1,03E+08 | ENSMUST00000246223.1 | -70604 | | Gm33027 | predicted gene, 33027 |
| chr4 | 1,35E+08 | 1,35E+08 | 521 | * | 3,869012 | 5,691128017 | 2,164558 | 2,629233 | 0,008558 | 1 | 0 | Promoter (2-3kb) | 4 | 1,35E+08 | 1,35E+08 | 3544 | 1 | 73937 | ENSMUST00000153347.2 | -2614 | ENSMUSG00000086788 | 1700029M20Rik | RIKEN cDNA 1700029M20 gene |
| chr5 | 13681672 | 13682106 | 435 | * | 4,309289 | 5,691183758 | 2,162235 | 2,632083 | 0,008486 | 1 | 0 | Distal Intergenic | 5 | 13818676 | 13821901 | 3226 | 1 | 626053 | ENSMUST00000170514.2 | -136570 | | Gm6647 | predicted gene 6647 |
| chr8 | 9705126 | 9705458 | 333 | * | 4,60203 | 5,691203472 | 2,159549 | 2,635367 | 0,008405 | 1 | 0 | Intron (ENSMUST0000( | 8 | 9479733 | 9820104 | 340372 | 2 | 270028 | ENSMUST00000207869.2 | 114646 | ENSMUSG00000079157 | Nalf1 | NALCN channel auxiliary factor 1 |
| chr4 | 1,07E+08 | 1,07E+08 | 408 | * | 4,660219 | 5,691273341 | 2,155386 | 2,640489 | 0,008279 | 1 | 0 | Promoter (<=1kb) | 4 | 1,07E+08 | 1,07E+08 | 2471 | 1 | 230577 | ENSMUST00000106782.2 | 0 | ENSMUSG00000043572 | Pars2 | prolyl-tRNA synthetase (mitochondrial)(putative) |
| chr14 | 71063627 | 71064401 | 775 | * | 4,413263 | 5,6913812 | 2,149282 | 2,648039 | 0,008096 | 1 | 0 | Distal Intergenic | 14 | 71090373 | 71111394 | 21022 | 1 | 73648 | ENSMUST00000190896.2 | -25972 | ENSMUSG00000099528 | 2410012E07Rik | RIKEN cDNA 2410012E07 gene |
| chr3 | 15258178 | 15258955 | 778 | * | 4,426981 | 5,691551903 | 2,138168 | 2,661882 | 0,007771 | 1 | 0 | Distal Intergenic | 3 | 15218125 | 15236502 | 18378 | 2 | 1,03E+08 | ENSMUST00000194008.2 | -21676 | | Gm33819 | predicted gene, 33819 |
| chr11 | 1,21E+08 | 1,21E+08 | 473 | * | 4,272722 | 5,691572635 | 2,136972 | 2,663382 | 0,007736 | 1 | 0 | Intron (ENSMUST0000( | 11 | 1,21E+08 | 1,21E+08 | 22751 | 1 | 68938 | ENSMUST00000168510.2 | -3648 | ENSMUSG00000025142 | Aspscr1 | alveolar soft part sarcoma chromosome region, candidate 1 (human) |
| chr12 | 87436668 | 87437310 | 643 | * | 5,116613 | 5,691596142 | 2,135022 | 2,665826 | 0,00768 | 1 | 0 | Promoter (1-2kb) | 12 | 87351832 | 87435129 | 83298 | 2 | 20773 | ENSMUST00000021424.5 | -1539 | ENSMUSG00000021036 | Sptlc2 | serine palmitoyltransferase, long chain base subunit 2 |
| chr1 | 1,83E+08 | 1,83E+08 | 675 | * | 5,124095 | 5,691696266 | 2,129231 | 2,673123 | 0,007515 | 1 | 0 | Intron (ENSMUST0000( | 1 | 1,83E+08 | 1,83E+08 | 61196 | 2 | 68897 | ENSMUST00000171366.7 | 53649 | ENSMUSG00000030768 | Disp1 | dispatched RND transporter family member 1 |
| chr10 | 59973960 | 59974279 | 320 | * | 4,619186 | 5,691700441 | 2,129094 | 2,673296 | 0,007511 | 1 | 0 | Distal Intergenic | 10 | 59962598 | 59962655 | 58 | 1 | 1,02E+08 | ENSMUST00000198998.3 | 11362 | ENSMUSG00000105468 | Mir6906 | microRNA 6906 |
| chr8 | 41909113 | 41909681 | 569 | * | 6,815033 | 5,692018883 | 1,882151 | 3,02421 | 0,002493 | 1 | 0 | Distal Intergenic | 8 | 41850496 | 41870111 | 19616 | 2 | 14300 | ENSMUST00000033999.8 | -39002 | ENSMUSG00000031590 | Frg1 | FSHD region gene 1 |
| chr2 | 3362127 | 3362585 | 459 | * | 4,472308 | 5,692037656 | 2,111251 | 2,69605 | 0,007017 | 1 | 0 | Intron (ENSMUST0000( | 2 | 3343054 | 3367606 | 24553 | 2 | 99035 | ENSMUST00000115087.4 | 5021 | ENSMUSG00000026645 | Olah | oleoyl-ACP hydrolase |
| chr5 | 1E+08 | 1E+08 | 578 | * | 5,071845 | 5,692259747 | 2,097767 | 2,713485 | 0,006658 | 1 | 0 | Promoter (<=1kb) | 5 | 1E+08 | 1E+08 | 15549 | 2 | 11991 | ENSMUST00000171106.8 | 479 | ENSMUSG00000000568 | Hnrnpd | heterogeneous nuclear ribonucleoprotein D |
| chr19 | 59626994 | 59627403 | 410 | * | 4,284661 | 5,692547439 | 2,083256 | 2,732525 | 0,006285 | 1 | 0 | Distal Intergenic | 19 | 59649755 | 59669638 | 19884 | 1 | 1,01E+08 | ENSMUST00000236333.2 | -22352 | | Gm19956 | predicted gene, 19956 |
| chr19 | 42562814 | 42563210 | 397 | * | 4,767585 | 5,692877936 | 2,066435 | 2,754927 | 0,005871 | 1 | 0 | Intron (ENSMUST0000( | 19 | 42569600 | 42573210 | 3611 | 1 | 52013 | ENSMUST00000162238.2 | -6390 | ENSMUSG00000025184 | R3hcc1l | R3H domain and coiled-coil containing 1 like |
| chr7 | 24402644 | 24403134 | 491 | * | 5,880236 | 5,693017483 | 2,059371 | 2,764445 | 0,005702 | 1 | 0 | Distal Intergenic | 7 | 24408667 | 24412948 | 4282 | 1 | 232972 | ENSMUST00000206826.2 | -5533 | ENSMUSG00000045587 | Lypd10 | Ly6/PLAUR domain containing 10 |
| chr15 | 3641044 | 3641367 | 324 | * | 4,847287 | 5,693135846 | 2,05418 | 2,771488 | 0,00558 | 1 | 0 | Distal Intergenic | 15 | 3632754 | 3632879 | 126 | 2 | 1,15E+08 | ENSMUST00000116945.4 | -8165 | ENSMUSG00000080595 | Gm22031 | predicted gene, 22031 |
| chr14 | 79720387 | 79721203 | 817 | * | 5,113965 | 5,693257115 | 2,048152 | 2,779705 | 0,005441 | 1 | 0 | Promoter (1-2kb) | 14 | 79697377 | 79718960 | 21584 | 2 | 22380 | ENSMUST00000022601.7 | -1427 | ENSMUSG00000022023 | Wbp4 | WW domain binding protein 4 |
| chrX | 35144900 | 35145733 | 834 | * | 4,871801 | 5,693311481 | 2,046057 | 2,782577 | 0,005393 | 1 | 0 | Distal Intergenic | 20 | 35152485 | 35340215 | 187731 | 1 | 75974 | ENSMUST00000033419.13 | -6752 | ENSMUSG00000031093 | Dock11 | dedicator of cytokinesis 11 |
| chr3 | 96389934 | 96390618 | 685 | * | 6,028245 | 5,693314006 | 1,91694 | 2,970001 | 0,002978 | 1 | 0 | Distal Intergenic | 3 | 96393871 | 96394035 | 165 | 2 | 1,15E+08 | ENSMUST00000239775.1 | 3417 | ENSMUSG00000118698 | Gm25890 | predicted gene, 25890 |
| chr4 | 45199442 | 45200070 | 629 | * | 4,715204 | 5,693477575 | 2,038465 | 2,793021 | 0,005222 | 1 | 0 | Intron (ENSMUST0000( | 4 | 45203925 | 45285936 | 82012 | 1 | 666060 | ENSMUST00000107804.2 | -3855 | ENSMUSG00000035615 | Frmpd1 | FERM and PDZ domain containing 1 |
| chr17 | 43659203 | 43659972 | 770 | * | 5,776529 | 5,693609078 | 2,033108 | 2,800446 | 0,005103 | 1 | 0 | Distal Intergenic | 17 | 43671342 | 43766963 | 95622 | 1 | 224792 | ENSMUST00000226087.2 | -11370 | ENSMUSG00000056492 | Adgrf5 | adhesion G protein-coupled receptor F5 |
| chr1 | 59803515 | 59804303 | 789 | * | 7,226758 | 5,693640576 | 1,820825 | 3,126956 | 0,001766 | 0,595550945 | 1,360622 | Promoter (<=1kb) | 1 | 59802559 | 59918173 | 115615 | 1 | 12168 | ENSMUST00000087435.7 | 956 | ENSMUSG00000067336 | Bmpr2 | bone morphogenetic protein receptor, type II (serine/threonine kinase) |
| chr17 | 34611893 | 34612482 | 590 | * | 4,99844 | 5,693714293 | 2,028041 | 2,807494 | 0,004993 | 1 | 0 | Intron (ENSMUST0000( | 17 | 34617794 | 34679708 | 61915 | 1 | 407788 | ENSMUST00000114175.8 | -5312 | ENSMUSG00000057246 | BC051142 | cDNA sequence BC051142 |
| chr1 | 1,35E+08 | 1,35E+08 | 752 | * | 5,771641 | 5,693767107 | 2,025465 | 2,811092 | 0,004937 | 1 | 0 | Promoter (<=1kb) | 1 | 1,35E+08 | 1,35E+08 | 4608 | 2 | 215615 | ENSMUST00000130069.2 | 0 | ENSMUSG00000041926 | Rnpep | arginyl aminopeptidase (aminopeptidase B) |
| chr1 | 29380899 | 29381606 | 708 | * | 5,796398 | 5,693831458 | 1,985481 | 2,867734 | 0,004134 | 1 | 0 | Distal Intergenic | 1 | 29460569 | 29460693 | 125 | 1 | 1,02E+08 | ENSMUST00000183870.3 | -78963 | ENSMUSG00000099257 | Mir6342 | microRNA 6342 |
| chr14 | 92756932 | 92757837 | 906 | * | 5,284448 | 5,693935831 | 2,018134 | 2,821387 | 0,004782 | 1 | 0 | Distal Intergenic | 14 | 92605936 | 92606065 | 130 | 2 | 1,15E+08 | ENSMUST00000104110.3 | -150867 | ENSMUSG00000077298 | Gm24127 | predicted gene, 24127 |
| chrX | 72100827 | 72101538 | 712 | * | 4,921514 | 5,69404813 | 2,01356 | 2,827851 | 0,004686 | 1 | 0 | Distal Intergenic | 20 | 72108393 | 72111797 | 3405 | 1 | 245468 | ENSMUST00000060418.8 | -6855 | ENSMUSG00000046287 | Pnma3 | paraneoplastic antigen MA3 |
| chr13 | 99266004 | 99266852 | 849 | * | 5,795266 | 5,694061119 | 2,012739 | 2,829011 | 0,004669 | 1 | 0 | Distal Intergenic | 13 | 99237269 | 99242680 | 5412 | 1 | 328320 | ENSMUST00000224929.2 | 28735 | ENSMUSG00000097848 | Gm807 | predicted gene 807 |
| chr10 | 1,1E+08 | 1,1E+08 | 509 | * | 5,33125 | 5,694099006 | 2,011976 | 2,830103 | 0,004653 | 1 | 0 | Intron (ENSMUST0000( | 10 | 1,1E+08 | 1,1E+08 | 9709 | 2 | 260315 | ENSMUST00000163071.2 | -26793 | ENSMUSG00000020181 | Nav3 | neuron navigator 3 |
| chr11 | 83580380 | 83580975 | 596 | * | 5,810363 | 5,694173207 | 1,942345 | 2,931598 | 0,003372 | 1 | 0 | Distal Intergenic | 11 | 83586101 | 83587011 | 911 | 1 | 1,03E+08 | ENSMUST00000121317.2 | -5126 | | Gm11430 | predicted gene 11430 |
| chr5 | 35958569 | 35959532 | 964 | * | 8,601442 | 5,694263692 | 1,945766 | 2,926489 | 0,003428 | 1 | 0 | Intron (ENSMUST0000( | 5 | 35971703 | 36000548 | 28846 | 1 | 231148 | ENSMUST00000130233.8 | -12171 | ENSMUSG00000029095 | Ablim2 | actin-binding LIM protein 2 |
| chr14 | 1,17E+08 | 1,17E+08 | 709 | * | 5,242943 | 5,69552676 | 1,988394 | 2,864385 | 0,004178 | 1 | 0 | Intron (ENSMUST0000( | 14 | 1,17E+08 | 1,18E+08 | 1049151 | 1 | 23888 | ENSMUST00000078849.12 | 199868 | ENSMUSG00000058571 | Gpc6 | glypican 6 |
| chr1 | 1,92E+08 | 1,92E+08 | 1179 | * | 5,58862 | 5,695758232 | 1,949861 | 2,921109 | 0,003488 | 1 | 0 | Intron (ENSMUST0000( | 1 | 1,92E+08 | 1,92E+08 | 75420 | 1 | 16510 | ENSMUST00000151152.3 | 15020 | ENSMUSG00000058248 | Kcnh1 | potassium voltage-gated channel, subfamily H (eag-related), member 1 |
| chr9 | 54908858 | 54909504 | 647 | * | 5,686181 | 5,695767036 | 1,949812 | 2,921188 | 0,003487 | 1 | 0 | Intron (ENSMUST0000( | 9 | 54888817 | 54914109 | 25293 | 1 | 110835 | ENSMUST00000213960.2 | 20041 | ENSMUSG00000035594 | Chrna5 | cholinergic receptor, nicotinic, alpha polypeptide 5 |
| chr17 | 48068950 | 48069629 | 680 | * | 5,817396 | 5,695804589 | 1,948108 | 2,923762 | 0,003458 | 1 | 0 | Promoter (<=1kb) | 17 | 48070100 | 48097121 | 27022 | 1 | 21425 | ENSMUST00000125177.2 | -471 | ENSMUSG00000023990 | Tfeb | transcription factor EB |
| chr11 | 1,01E+08 | 1,01E+08 | 898 | * | 5,732776 | 5,696224415 | 1,934584 | 2,944418 | 0,003236 | 1 | 0 | Promoter (<=1kb) | 11 | 1,01E+08 | 1,01E+08 | 2266 | 1 | 19192 | ENSMUST00000131170.2 | 0 | ENSMUSG00000078652 | Psme3 | proteaseome (prosome, macropain) activator subunit 3 (PA28 gamma, Ki) |
| chr8 | 1,07E+08 | 1,07E+08 | 1025 | * | 7,703307 | 5,696376222 | 1,92937 | 2,952454 | 0,003153 | 1 | 0 | Promoter (<=1kb) | 8 | 1,07E+08 | 1,07E+08 | 14314 | 1 | 192654 | ENSMUST00000212963.2 | -561 | ENSMUSG00000031903 | Pla2g15 | phospholipase A2, group XV |
| chr2 | 1,2E+08 | 1,2E+08 | 755 | * | 5,562196 | 5,696892502 | 2,043664 | 2,787587 | 0,00531 | 1 | 0 | Promoter (1-2kb) | 2 | 1,2E+08 | 1,2E+08 | 20372 | 1 | 22174 | ENSMUST00000110783.8 | -1349 | ENSMUSG00000027298 | Tyro3 | TYRO3 protein tyrosine kinase 3 |
| chr11 | 1,06E+08 | 1,06E+08 | 617 | * | 4,82181 | 5,696919612 | 2,043861 | 2,787332 | 0,005314 | 1 | 0 | Intron (ENSMUST0000( | 11 | 1,06E+08 | 1,06E+08 | 93147 | 2 | 78943 | ENSMUST00000001059.9 | 8015 | ENSMUSG00000020715 | Ern1 | endoplasmic reticulum (ER) to nucleus signalling 1 |
| chr6 | 1,18E+08 | 1,18E+08 | 406 | * | 4,467766 | 5,697545798 | 2,073584 | 2,74768 | 0,006002 | 1 | 0 | Distal Intergenic | 6 | 1,18E+08 | 1,18E+08 | 132 | 2 | 1,15E+08 | ENSMUST00000104599.3 | 5705 | ENSMUSG00000077787 | Gm23908 | predicted gene, 23908 |
| chr1 | 1,08E+08 | 1,08E+08 | 570 | * | 5,356654 | 5,697626279 | 2,076621 | 2,743701 | 0,006075 | 1 | 0 | Intron (ENSMUST0000( | 1 | 1,08E+08 | 1,08E+08 | 91629 | 1 | 319596 | ENSMUST00000189497.2 | 67029 | ENSMUSG00000100457 | D830032E09Rik | RIKEN cDNA D830032E09 gene |
| chr5 | 25502501 | 25503189 | 689 | * | 4,450771 | 5,697824713 | 2,086376 | 2,730967 | 0,006315 | 1 | 0 | Promoter (1-2kb) | 5 | 25476796 | 25505073 | 28278 | 2 | 231051 | ENSMUST00000174734.8 | 1884 | ENSMUSG00000038056 | Kmt2c | lysine (K)-specific methyltransferase 2C |
| chr16 | 15543358 | 15543912 | 555 | * | 4,542337 | 5,69796074 | 2,093525 | 2,721706 | 0,006495 | 1 | 0 | Intron (ENSMUST0000( | 16 | 15455753 | 15660099 | 204347 | 1 | 19090 | ENSMUST00000023352.9 | 87605 | ENSMUSG00000022672 | Prkdc | protein kinase, DNA activated, catalytic polypeptide |
| chr17 | 5889362 | 5890519 | 1158 | * | 7,382558 | 5,698610572 | 1,867502 | 3,051462 | 0,002277 | 1 | 0 | Promoter (1-2kb) | 17 | 5891604 | 5982229 | 90626 | 1 | 66616 | ENSMUST00000002436.11 | -1085 | ENSMUSG00000002365 | Snx9 | sorting nexin 9 |
| chr1 | 59206100 | 59206660 | 561 | * | 4,815449 | 5,698764344 | 2,138584 | 2,664737 | 0,007705 | 1 | 0 | Exon (ENSMUST00001 | 1 | 59160094 | 59202548 | 42455 | 2 | 227157 | ENSMUST00000114275.9 | -3552 | ENSMUSG00000079550 | Mpp4 | membrane protein, palmitoylated 4 (MAGUK p55 subfamily member 4) |
| chr4 | 1,51E+08 | 1,51E+08 | 539 | * | 4,896549 | 5,698849545 | 2,143726 | 2,658386 | 0,007852 | 1 | 0 | Distal Intergenic | 4 | 1,51E+08 | 1,51E+08 | 23603 | 2 | 242773 | ENSMUST00000037827.10 | -3411 | ENSMUSG00000039838 | Slc45a1 | solute carrier family 45, member 1 |
| chr15 | 40894047 | 40894532 | 486 | * | 3,950197 | 5,699413774 | 2,177079 | 2,617918 | 0,008847 | 1 | 0 | Intron (ENSMUST0000( | 15 | 41036883 | 41037264 | 382 | 1 | 1E+08 | ENSMUST00000060066.6 | -142351 | | 4930555K19Rik | leucine rich repeat and sterile alpha motif containing 1 pseudogene |
| chr5 | 1,08E+08 | 1,08E+08 | 1336 | * | 10,6259 | 5,699647681 | 1,687483 | 3,377604 | 0,000731 | 0,541111532 | 0,744375 | Distal Intergenic | 5 | 1,08E+08 | 1,08E+08 | 130313 | 2 | 14020 | ENSMUST00000112642.8 | -9988 | ENSMUSG00000011831 | Evi5 | ecotropic viral integration site 5 |
| chr1 | 1,33E+08 | 1,33E+08 | 447 | * | 4,00687 | 5,699708118 | 2,197962 | 2,593179 | 0,009509 | 1 | 0 | Intron (ENSMUST0000( | 1 | 1,32E+08 | 1,33E+08 | 66669 | 2 | 269116 | ENSMUST00000186389.7 | 4138 | ENSMUSG00000026442 | Nfasc | neurofascin |
| chr10 | 19278220 | 19278703 | 484 | * | 5,530457 | 5,701706221 | 1,954768 | 2,91682 | 0,003536 | 1 | 0 | Distal Intergenic | 10 | 19232281 | 19234352 | 2072 | 1 | 94222 | ENSMUST00000053225.7 | 45939 | ENSMUSG00000045591 | Olig3 | oligodendrocyte transcription factor 3 |
| chr5 | 29369816 | 29370767 | 952 | * | 9,269567 | 5,702558474 | 1,786394 | 3,192217 | 0,001412 | 0,539620856 | 1,449706 | Distal Intergenic | 5 | 29400990 | 29408133 | 7144 | 1 | 56874 | ENSMUST00000160888.8 | -30223 | ENSMUSG00000029130 | Rnf32 | ring finger protein 32 |
| chrX | 1,17E+08 | 1,17E+08 | 1572 | * | 7,259019 | 5,704435837 | 1,829612 | 3,117839 | 0,001822 | 1 | 0 | Exon (ENSMUST00001 | 20 | 1,17E+08 | 1,17E+08 | 1030 | 1 | 1E+08 | ENSMUST00000072518.8 | -536455 | ENSMUSG00000100194 | Tgif2lx2 | TGFB-induced factor homeobox 2-like, X-linked 2 |
| chr19 | 7156127 | 7157258 | 1132 | * | 10,67231 | 5,706427192 | 1,762535 | 3,237624 | 0,001205 | 0,640427178 | 0,738441 | Intron (ENSMUST0000( | 19 | 7176905 | 7177829 | 925 | 2 | 107260 | ENSMUST00000127241.2 | 20571 | ENSMUSG00000024767 | Otub1 | OTU domain, ubiquitin aldehyde binding 1 |
| chr15 | 40916445 | 40917058 | 614 | * | 5,595949 | 5,706777865 | 1,970343 | 2,896337 | 0,003775 | 1 | 0 | Intron (ENSMUST0000( | 15 | 41036883 | 41037264 | 382 | 1 | 1E+08 | ENSMUST00000060066.6 | -119825 | | 4930555K19Rik | leucine rich repeat and sterile alpha motif containing 1 pseudogene |
| chr3 | 95484870 | 95485659 | 790 | * | 4,238593 | 5,707171704 | 2,169955 | 2,630088 | 0,008536 | 1 | 0 | Exon (ENSMUST0000( | 3 | 95496245 | 95526553 | 30309 | 1 | 229593 | ENSMUST00000177390.8 | -10586 | ENSMUSG00000046519 | Golph3l | golgi phosphoprotein 3-like |
| chrX | 72435425 | 72436477 | 1053 | * | 5,117796 | 5,708808087 | 1,997304 | 2,858257 | 0,00426 | 1 | 0 | Distal Intergenic | 20 | 72454702 | 72470604 | 15903 | 1 | 22754 | ENSMUST00000033740.12 | -18225 | ENSMUSG00000031374 | Zfp92 | zinc finger protein 92 |
| chr12 | 26625533 | 26626105 | 573 | * | 4,734763 | 5,710849217 | 2,21673 | 2,576249 | 0,009988 | 1 | 0 | Intron (ENSMUST0000( | 12 | 26622107 | 26652308 | 30202 | 2 | 69437 | ENSMUST00000221140.2 | 26203 | | 1700020D12Rik | RIKEN cDNA 1700020D12 gene |
| chr11 | 60809801 | 60810371 | 571 | * | 6,062947 | 5,711812059 | 1,900954 | 3,004708 | 0,002658 | 1 | 0 | Distal Intergenic | 11 | 60796780 | 60805576 | 8797 | 2 | 24083 | ENSMUST00000143800.2 | -4225 | ENSMUSG00000018931 | Natd1 | N-acetyltransferase domain containing 1 |
| chr13 | 87563566 | 87564265 | 700 | * | 5,313106 | 5,713502314 | 2,14513 | 2,663477 | 0,007734 | 1 | 0 | Distal Intergenic | 13 | 88669309 | 88726287 | 56979 | 1 | 1,03E+08 | ENSMUST00000224910.2 | -1105044 | ENSMUSG00000114273 | Gm34585 | predicted gene, 34585 |
| chr12 | 93836710 | 93837620 | 911 | * | 8,37448 | 5,714724381 | 1,744521 | 3,275813 | 0,001054 | 1 | 0 | Distal Intergenic | 12 | 93894112 | 93894238 | 127 | 1 | 1,02E+08 | ENSMUST00000183568.3 | -56492 | ENSMUSG00000098931 | Mir8099-2 | microRNA 8099-2 |
| chr6 | 85585487 | 85586530 | 1044 | * | 6,871009 | 5,719254849 | 1,884193 | 3,035388 | 0,002402 | 1 | 0 | Exon (ENSMUST0000( | 6 | 85564628 | 85679620 | 114993 | 1 | 236266 | ENSMUST00000213058.2 | 20859 | ENSMUSG00000063810 | Alms1 | ALMS1, centrosome and basal body associated |
| chr9 | 64616726 | 64617421 | 696 | * | 6,507737 | 5,720437545 | 1,906093 | 3,001133 | 0,00269 | 1 | 0 | Downstream (<=300bp | 9 | 64623887 | 64633002 | 9116 | 2 | 53869 | ENSMUST00000172444.2 | 15581 | ENSMUSG00000004771 | Rab11a | RAB11A, member RAS oncogene family |
| chr17 | 36098978 | 36099498 | 521 | * | 4,196274 | 5,72059501 | 2,202373 | 2,597469 | 0,009391 | 1 | 0 | Distal Intergenic | 17 | 36083342 | 36091579 | 8238 | 1 | 74601 | ENSMUST00000172874.2 | 15636 | ENSMUSG00000092171 | 4833427F10Rik | RIKEN cDNA 4833427F10 gene |
| chr9 | 64168917 | 64169334 | 418 | * | 3,714341 | 5,720707251 | 2,189073 | 2,613302 | 0,008967 | 1 | 0 | Distal Intergenic | 9 | 64093052 | 64160913 | 67862 | 2 | 26395 | ENSMUST00000005066.9 | -8004 | ENSMUSG00000004936 | Map2k1 | mitogen-activated protein kinase kinase 1 |
| chr2 | 96954295 | 96954738 | 444 | * | 4,623827 | 5,72090167 | 2,164762 | 2,642739 | 0,008224 | 1 | 0 | Intron (ENSMUST0000( | 2 | 97068163 | 97068291 | 129 | 1 | 1,15E+08 | ENSMUST00000158008.3 | -113425 | ENSMUSG00000088633 | Gm22313 | predicted gene, 22313 |
| chr17 | 51256635 | 51256883 | 249 | * | 4,243432 | 5,720932321 | 2,159476 | 2,649222 | 0,008068 | 1 | 0 | Intron (ENSMUST0000( | 17 | 51261021 | 51291654 | 30634 | 2 | 72238 | ENSMUST00000224123.2 | 34771 | ENSMUSG00000023923 | Tbc1d5 | TBC1 domain family, member 5 |
| chr1 | 1,57E+08 | 1,57E+08 | 466 | * | 4,781683 | 5,720971621 | 2,154556 | 2,65529 | 0,007924 | 1 | 0 | Distal Intergenic | 1 | 1,57E+08 | 1,57E+08 | 9076 | 2 | 73435 | ENSMUST00000119526.2 | 33167 | ENSMUSG00000026592 | Tex35 | testis expressed 35 |
| chr2 | 22015290 | 22015741 | 452 | * | 3,884159 | 5,720974467 | 2,155655 | 2,653938 | 0,007956 | 1 | 0 | Distal Intergenic | 2 | 22232314 | 22508264 | 275951 | 1 | 667663 | ENSMUST00000044749.14 | -216573 | ENSMUSG00000025716 | Myo3a | myosin IIIA |
| chr6 | 1,27E+08 | 1,27E+08 | 405 | * | 4,168274 | 5,721246083 | 2,125854 | 2,69127 | 0,007118 | 1 | 0 | 3' UTR | 6 | 1,27E+08 | 1,27E+08 | 8644 | 2 | 12444 | ENSMUST00000201564.2 | 9558 | ENSMUSG00000000184 | Ccnd2 | cyclin D2 |
| chr17 | 7196042 | 7196403 | 362 | * | 4,901755 | 5,721309112 | 2,116251 | 2,703512 | 0,006861 | 1 | 0 | Intron (ENSMUST0000( | 17 | 7188268 | 7215680 | 27413 | 2 | 1E+08 | ENSMUST00000232602.2 | 19277 | ENSMUSG00000023806 | Rsph3b | radial spoke 3B homolog (Chlamydomonas) |
| chr15 | 34363817 | 34364741 | 925 | * | 4,703579 | 5,721536959 | 2,093367 | 2,733175 | 0,006273 | 1 | 0 | Intron (ENSMUST0000( | 15 | 34349197 | 34358018 | 8822 | 2 | 77358 | ENSMUST00000227772.2 | -5799 | ENSMUSG00000094447 | 9430069I07Rik | RIKEN cDNA 9430069I07 gene |
| chrX | 36470002 | 36470355 | 354 | * | 4,229163 | 5,721570135 | 2,08995 | 2,737659 | 0,006188 | 1 | 0 | Distal Intergenic | 20 | 36473001 | 36474694 | 1694 | 2 | 194854 | ENSMUST00000101636.3 | 4339 | ENSMUSG00000073243 | Gm9 | predicted gene 9 |
| chr3 | 1,47E+08 | 1,47E+08 | 641 | * | 4,566476 | 5,721623015 | 2,085685 | 2,743283 | 0,006083 | 1 | 0 | Distal Intergenic | 3 | 1,47E+08 | 1,47E+08 | 4401 | 1 | 1,05E+08 | ENSMUST00000195873.2 | 345939 | | Gm37041 | predicted gene, 37041 |
| chr4 | 21795575 | 21796057 | 483 | * | 5,194748 | 5,7216831 | 2,079602 | 2,751335 | 0,005935 | 1 | 0 | Intron (ENSMUST0000( | 4 | 21784724 | 21816893 | 32170 | 1 | 77593 | ENSMUST00000148647.3 | 10851 | ENSMUSG00000040455 | Usp45 | ubiquitin specific petidase 45 |
| chr14 | 1,05E+08 | 1,05E+08 | 1043 | * | 5,273953 | 5,721873662 | 2,060385 | 2,77709 | 0,005485 | 1 | 0 | Promoter (2-3kb) | 14 | 1,05E+08 | 1,05E+08 | 16340 | 2 | 72486 | ENSMUST00000228210.2 | 2995 | ENSMUSG00000022120 | Obi1 | ORC ubiquitin ligase 1 |
| chr15 | 1E+08 | 1E+08 | 409 | * | 5,211012 | 5,722029158 | 2,046509 | 2,795996 | 0,005174 | 1 | 0 | Intron (ENSMUST0000( | 15 | 1E+08 | 1E+08 | 2494 | 1 | 239667 | ENSMUST00000135658.3 | 10964 | ENSMUSG00000023026 | Dip2b | disco interacting protein 2 homolog B |
| chr9 | 70548115 | 70548845 | 731 | * | 4,7582 | 5,722157865 | 2,037278 | 2,808728 | 0,004974 | 1 | 0 | Intron (ENSMUST0000( | 9 | 70533518 | 70564429 | 30912 | 2 | 235461 | ENSMUST00000214293.2 | 15584 | ENSMUSG00000042444 | Mindy2 | MINDY lysine 48 deubiquitinase 2 |
| chr13 | 97169039 | 97170042 | 1004 | * | 5,372595 | 5,722640187 | 1,99834 | 2,863697 | 0,004187 | 1 | 0 | Promoter (<=1kb) | 13 | 97158643 | 97170841 | 12199 | 2 | 66479 | ENSMUST00000221277.2 | 799 | ENSMUSG00000052075 | 1700029F12Rik | RIKEN cDNA 1700029F12 gene |
| chr1 | 56437587 | 56438192 | 606 | * | 5,196181 | 5,72273589 | 1,990422 | 2,875137 | 0,004039 | 1 | 0 | Distal Intergenic | 1 | 56675203 | 56676594 | 1392 | 2 | 71066 | ENSMUST00000062085.6 | 238402 | ENSMUSG00000045336 | Hsfy2 | heat shock transcription factor, Y-linked 2 |
| chr9 | 66512498 | 66513160 | 663 | * | 5,689695 | 5,722776189 | 1,987445 | 2,879464 | 0,003984 | 1 | 0 | Distal Intergenic | 9 | 66421919 | 66500424 | 78506 | 2 | 235441 | ENSMUST00000127569.8 | -12074 | ENSMUSG00000032376 | Usp3 | ubiquitin specific peptidase 3 |
| chr1 | 8981183 | 8982158 | 976 | * | 5,742402 | 5,723068773 | 1,968062 | 2,907972 | 0,003638 | 1 | 0 | Intron (ENSMUST0000( | 1 | 8433699 | 8874167 | 440469 | 2 | 71096 | ENSMUST00000115488.3 | -107016 | ENSMUSG00000025909 | Sntg1 | syntrophin, gamma 1 |
| chr7 | 1,15E+08 | 1,15E+08 | 583 | * | 5,491014 | 5,723254775 | 1,954576 | 2,92813 | 0,00341 | 1 | 0 | Intron (ENSMUST0000( | 7 | 1,15E+08 | 1,15E+08 | 85153 | 2 | 20679 | ENSMUST00000206775.2 | -15287 | ENSMUSG00000051910 | Sox6 | SRY (sex determining region Y)-box 6 |
| chr14 | 1,23E+08 | 1,23E+08 | 553 | * | 6,09191 | 5,723304674 | 1,952218 | 2,931694 | 0,003371 | 1 | 0 | Intron (ENSMUST0000( | 14 | 1,23E+08 | 1,23E+08 | 2926 | 1 | 110821 | ENSMUST00000138836.2 | 10551 | ENSMUSG00000041650 | Pcca | propionyl-Coenzyme A carboxylase, alpha polypeptide |
| chr3 | 41516311 | 41516927 | 617 | * | 5,493697 | 5,723316084 | 1,950591 | 2,934145 | 0,003345 | 1 | 0 | Promoter (<=1kb) | 3 | 41517804 | 41571293 | 53490 | 1 | 269424 | ENSMUST00000163764.8 | -877 | ENSMUSG00000025764 | Jade1 | jade family PHD finger 1 |
| chr7 | 1,33E+08 | 1,33E+08 | 722 | * | 6,367 | 5,723340899 | 1,949708 | 2,935486 | 0,00333 | 1 | 0 | Promoter (<=1kb) | 7 | 1,33E+08 | 1,33E+08 | 1908 | 1 | 360216 | ENSMUST00000153335.2 | 423 | ENSMUSG00000030967 | Zranb1 | zinc finger, RAN-binding domain containing 1 |
| chr7 | 1,27E+08 | 1,27E+08 | 778 | * | 6,332974 | 5,72345977 | 1,941064 | 2,94862 | 0,003192 | 1 | 0 | Promoter (1-2kb) | 7 | 1,27E+08 | 1,27E+08 | 2310 | 1 | 233900 | ENSMUST00000206699.2 | -1283 | ENSMUSG00000030816 | Rnf40 | ring finger protein 40 |
| chr13 | 43610461 | 43611181 | 721 | * | 5,476213 | 5,723555112 | 1,935714 | 2,956818 | 0,003108 | 1 | 0 | Intron (ENSMUST0000( | 13 | 43573055 | 43615207 | 42153 | 2 | 56705 | ENSMUST00000222239.2 | 4026 | ENSMUSG00000038546 | Ranbp9 | RAN binding protein 9 |
| chr2 | 1,55E+08 | 1,55E+08 | 810 | * | 7,336707 | 5,724221024 | 1,897625 | 3,016519 | 0,002557 | 1 | 0 | Promoter (<=1kb) | 2 | 1,55E+08 | 1,55E+08 | 1325 | 1 | 668940 | ENSMUST00000124415.2 | 0 | ENSMUSG00000074652 | Myh7b | myosin, heavy chain 7B, cardiac muscle, beta |
| chr3 | 69102861 | 69103730 | 870 | * | 6,893515 | 5,724776537 | 1,870145 | 3,061141 | 0,002205 | 1 | 0 | Distal Intergenic | 3 | 69129752 | 69130951 | 1200 | 1 | 71619 | ENSMUST00000183126.2 | -26022 | ENSMUSG00000098207 | Arl14 | ADP-ribosylation factor-like 14 |
| chr11 | 1,1E+08 | 1,1E+08 | 914 | * | 8,276964 | 5,724824152 | 1,862691 | 3,073416 | 0,002116 | 0,622205297 | 1,425929 | Distal Intergenic | 11 | 1,1E+08 | 1,1E+08 | 15324 | 2 | 76184 | ENSMUST00000150988.2 | -13788 | ENSMUSG00000044749 | Abca6 | ATP-binding cassette, sub-family A (ABC1), member 6 |
| chr7 | 27976724 | 27977543 | 820 | * | 4,998006 | 5,725385367 | 2,024902 | 2,827488 | 0,004691 | 1 | 0 | Promoter (<=1kb) | 7 | 27977164 | 27978914 | 1751 | 1 | 434156 | ENSMUST00000094651.4 | 0 | ENSMUSG00000070705 | Eid2b | EP300 interacting inhibitor of differentiation 2B |
| chr2 | 1,71E+08 | 1,71E+08 | 681 | * | 7,132648 | 5,725618367 | 1,849849 | 3,095181 | 0,001967 | 0,617289329 | 1,360622 | Distal Intergenic | 2 | 1,71E+08 | 1,71E+08 | 264 | 2 | 1,15E+08 | ENSMUST00000157880.3 | -197544 | ENSMUSG00000088505 | Gm26489 | predicted gene, 26489 |
| chrX | 1,4E+08 | 1,4E+08 | 580 | * | 4,596577 | 5,726751971 | 2,068321 | 2,768793 | 0,005626 | 1 | 0 | Intron (ENSMUST0000( | 20 | 1,4E+08 | 1,4E+08 | 21412 | 2 | 94216 | ENSMUST00000137051.8 | -58378 | ENSMUSG00000031273 | Col4a6 | collagen, type IV, alpha 6 |
| chr10 | 58312966 | 58313484 | 519 | * | 4,721039 | 5,727238151 | 2,085691 | 2,745967 | 0,006033 | 1 | 0 | Exon (ENSMUST0000( | 10 | 58333770 | 58412066 | 78297 | 1 | 76138 | ENSMUST00000036576.10 | -20286 | ENSMUSG00000038010 | Ccdc138 | coiled-coil domain containing 138 |
| chr1 | 37936298 | 37936912 | 615 | * | 5,046731 | 5,727606406 | 2,099078 | 2,72863 | 0,00636 | 1 | 0 | Promoter (<=1kb) | 1 | 37936337 | 37937206 | 870 | 1 | 107734 | ENSMUST00000162513.2 | 0 | ENSMUSG00000026087 | Mrpl30 | mitochondrial ribosomal protein L30 |
| chr1 | 1,66E+08 | 1,66E+08 | 753 | * | 4,533381 | 5,728245303 | 2,123111 | 2,698044 | 0,006975 | 1 | 0 | Exon (ENSMUST00001 | 1 | 1,66E+08 | 1,66E+08 | 3340 | 1 | 12503 | ENSMUST00000159070.3 | -13651 | ENSMUSG00000005763 | Cd247 | CD247 antigen |
| chr17 | 26966526 | 26967544 | 1019 | * | 5,470336 | 5,728249881 | 1,884026 | 3,04043 | 0,002362 | 1 | 0 | Intron (ENSMUST0000( | 17 | 26961958 | 26977161 | 15204 | 1 | 77128 | ENSMUST00000132972.2 | 4568 | ENSMUSG00000048249 | Crebrf | CREB3 regulatory factor |

| | | | | | | | | | | | | | | | | | | | | | | | |
|---|---|---|---|---|---|---|---|---|---|---|---|---|---|---|---|---|---|---|---|---|---|---|---|
| chrX | 1,41E+08 | 1,41E+08 | 381 | * | 4,181223 | 5,728574015 | 2,137191 | 2,680422 | 0,007353 | 1 | 0 | Promoter (<=1kb) | 20 | 1,41E+08 | 1,41E+08 | 72352 | 2 | 50790 | ENSMUST00000112907.8 | 135 | ENSMUSG00000031278 | Acsl4 | acyl-CoA synthetase long-chain family member 4 |
| chr15 | 88770863 | 88771177 | 315 | * | 4,31121 | 5,728668017 | 2,141332 | 2,675282 | 0,007467 | 1 | 0 | Distal Intergenic | 15 | 88774716 | 88775686 | 971 | 2 | 239591 | ENSMUST00000176338.2 | 4509 | ENSMUSG00000022388 | Ttll8 | tubulin tyrosine ligase-like family, member 8 |
| chr17 | 43456199 | 43456501 | 303 | * | 4,075257 | 5,728777213 | 2,145642 | 2,66996 | 0,007586 | 1 | 0 | Intron (ENSMUST0000( | 17 | 43450543 | 43468823 | 18281 | 2 | 545207 | ENSMUST00000246147.1 | 12322 | ENSMUSG00000117011 | E130008D07Rik | RIKEN cDNA E130008D07 gene |
| chr2 | 1,26E+08 | 1,26E+08 | 567 | * | 3,71311 | 5,730319771 | 2,217967 | 2,583591 | 0,009778 | 1 | 0 | Intron (ENSMUST0000( | 2 | 1,26E+08 | 1,26E+08 | 55549 | 1 | 14178 | ENSMUST00000110442.2 | 11250 | ENSMUSG00000027208 | Fgf7 | fibroblast growth factor 7 |
| chr7 | 66681918 | 66682769 | 852 | * | 11,21596 | 5,7339996 | 1,673292 | 3,426777 | 0,000611 | 0,419066905 | 1,2763 | Exon (ENSMUST00000: | 7 | 66697219 | 66728052 | 30834 | 1 | 233332 | ENSMUST00000154103.2 | -14450 | ENSMUSG00000058145 | Adamts17 | a disintegrin-like and metallopeptidase (reprolysin type) with thrombospondin type 1 motif, 17 |
| chr5 | 1,38E+08 | 1,38E+08 | 653 | * | 6,248092 | 5,734168018 | 1,896239 | 3,023969 | 0,002495 | 1 | 0 | Intron (ENSMUST0000( | 5 | 1,38E+08 | 1,38E+08 | 34196 | 2 | 231801 | ENSMUST00000100544.11 | 5988 | ENSMUSG00000029722 | Agfg2 | ArfGAP with FG repeats 2 |
| chr3 | 1,26E+08 | 1,26E+08 | 285 | * | 5,24726 | 5,736422649 | 1,948587 | 2,943889 | 0,003241 | 1 | 0 | Distal Intergenic | 3 | 1,26E+08 | 1,26E+08 | 14617 | 1 | 64580 | ENSMUST00000198101.2 | 46810 | ENSMUSG00000027971 | Ndst4 | N-deacetylase/N-sulfotransferase (heparin glucosaminyl) 4 |
| chr1 | 1,66E+08 | 1,66E+08 | 684 | * | 7,033435 | 5,739641123 | 1,803323 | 3,182813 | 0,001459 | 1 | 0 | Intron (ENSMUST0000( | 1 | 1,66E+08 | 1,66E+08 | 107 | 2 | 1,15E+08 | ENSMUST00000239609.1 | 9307 | ENSMUSG00000118799 | Gm23402 | predicted gene, 23402 |
| chr10 | 26707677 | 26708291 | 615 | * | 4,758673 | 5,740189268 | 2,222673 | 2,582562 | 0,009807 | 1 | 0 | Intron (ENSMUST0000( | 10 | 26698598 | 26725997 | 27400 | 1 | 73910 | ENSMUST00000135866.2 | 9079 | ENSMUSG00000039031 | Arhgap18 | Rho GTPase activating protein 18 |
| chr19 | 8846813 | 8847300 | 488 | * | 5,612011 | 5,740343076 | 1,920103 | 2,989602 | 0,002793 | 1 | 0 | Promoter (1-2kb) | 19 | 8848860 | 8853027 | 4168 | 1 | 225896 | ENSMUST00000237220.2 | -1560 | ENSMUSG00000071655 | Ubxn1 | UBX domain protein 1 |
| chr4 | 1,37E+08 | 1,37E+08 | 496 | * | 4,661009 | 5,740890075 | 2,197227 | 2,612789 | 0,008981 | 1 | 0 | Promoter (2-3kb) | 4 | 1,37E+08 | 1,37E+08 | 947 | 1 | 15530 | ENSMUST00000155648.2 | -2350 | ENSMUSG00000028763 | Hspg2 | perlecan (heparan sulfate proteoglycan 2) |
| chr5 | 49957700 | 49958357 | 658 | * | 4,442656 | 5,741159211 | 2,188463 | 2,623375 | 0,008706 | 1 | 0 | Distal Intergenic | 5 | 50123452 | 50129417 | 5966 | 2 | 70693 | ENSMUST00000196177.2 | 171060 | ENSMUSG00000029090 | Adgra3 | adhesion G protein-coupled receptor A3 |
| chr5 | 9017379 | 9018066 | 688 | * | 4,154595 | 5,741659076 | 2,170963 | 2,644752 | 0,008175 | 1 | 0 | Intron (ENSMUST0000( | 5 | 9023603 | 9032144 | 8542 | 2 | 74114 | ENSMUST00000146115.2 | 14078 | ENSMUSG00000003623 | Crot | carnitine O-octanoyltransferase |
| chr14 | 7836905 | 7837962 | 1058 | * | 4,715912 | 5,741931359 | 2,162326 | 2,655442 | 0,00792 | 1 | 0 | Distal Intergenic | 14 | 7669823 | 7767484 | 97662 | 2 | 218756 | ENSMUST00000057015.8 | -69421 | ENSMUSG00000021733 | Slc4a7 | solute carrier family 4, sodium bicarbonate cotransporter, member 7 |
| chrX | 23551039 | 23552203 | 1165 | * | 4,441231 | 5,742286755 | 2,151 | 2,66959 | 0,007594 | 1 | 0 | Distal Intergenic | 20 | 23559290 | 23672229 | 112940 | 1 | 72404 | ENSMUST00000035766.13 | -7087 | ENSMUSG00000036769 | Wdr44 | WD repeat domain 44 |
| chr4 | 1,52E+08 | 1,52E+08 | 410 | * | 5,304633 | 5,743168315 | 2,123638 | 2,704401 | 0,006843 | 1 | 0 | Intron (ENSMUST0000( | 4 | 1,52E+08 | 1,52E+08 | 391475 | 2 | 1,03E+08 | ENSMUST00000155685.8 | 88768 | | Gm38442 | predicted gene, 38442 |
| chr2 | 90747109 | 90747660 | 552 | * | 5,794977 | 5,74386613 | 2,103557 | 2,730549 | 0,006323 | 1 | 0 | Promoter (<=1kb) | 2 | 90741338 | 90747940 | 6603 | 2 | 66461 | ENSMUST00000155621.4 | 280 | ENSMUSG00000063235 | Ptpmt1 | protein tyrosine phosphatase, mitochondrial 1 |
| chr4 | 90940051 | 90940611 | 561 | * | 5,5033 | 5,745153055 | 2,069742 | 2,775782 | 0,005507 | 1 | 0 | Distal Intergenic | 4 | 90926137 | 90926510 | 374 | 1 | 1,08E+08 | ENSMUST00000116078.3 | 13914 | | Gm12643 | predicted gene 12643 |
| chr9 | 74859588 | 74860230 | 643 | * | 5,407108 | 5,746209333 | 2,044303 | 2,810841 | 0,004941 | 1 | 0 | Promoter (<=1kb) | 9 | 74860166 | 74862966 | 2801 | 1 | 235493 | ENSMUST00000216809.2 | 0 | ENSMUSG00000034858 | Fam214a | family with sequence similarity 214, member A |
| chr6 | 48061222 | 48061630 | 409 | * | 5,772652 | 5,747006329 | 2,026002 | 2,836625 | 0,004559 | 1 | 0 | Promoter (1-2kb) | 6 | 48039329 | 48063527 | 24199 | 2 | 69228 | ENSMUST00000073124.9 | 1897 | ENSMUSG00000057691 | Zfp746 | zinc finger protein 746 |
| chr17 | 25150934 | 25151533 | 600 | * | 6,733387 | 5,749335256 | 1,978345 | 2,906134 | 0,003659 | 1 | 0 | Intron (ENSMUST0000( | 17 | 25116507 | 25155868 | 39362 | 2 | 30957 | ENSMUST00000115228.9 | 4335 | ENSMUSG00000024163 | Mapk8ip3 | mitogen-activated protein kinase 8 interacting protein 3 |
| chr7 | 97419671 | 97420499 | 829 | * | 6,190922 | 5,749411428 | 1,976857 | 2,90836 | 0,003633 | 1 | 0 | Distal Intergenic | 7 | 97437748 | 97504119 | 66372 | 1 | 18479 | ENSMUST00000205757.2 | -17249 | ENSMUSG00000030774 | Pak1 | p21 (RAC1) activated kinase 1 |
| chr11 | 1,04E+08 | 1,04E+08 | 1037 | * | 6,387492 | 5,74945184 | 1,935327 | 2,970791 | 0,00297 | 1 | 0 | Intron (ENSMUST0000( | 11 | 1,04E+08 | 1,04E+08 | 108791 | 2 | 76719 | ENSMUST00000106972.8 | -3596 | ENSMUSG00000018412 | Kansl1 | KAT8 regulatory NSL complex subunit 1 |
| chr8 | 87028386 | 87028948 | 563 | * | 6,058588 | 5,749825077 | 1,969387 | 2,919602 | 0,003505 | 1 | 0 | Distal Intergenic | 8 | 87231212 | 87261376 | 30165 | 2 | 244562 | ENSMUST00000156066.8 | 232428 | ENSMUSG00000036872 | Abcc12 | ATP-binding cassette, sub-family C (CFTR/MRP), member 12 |
| chr3 | 1,4E+08 | 1,4E+08 | 829 | * | 7,113611 | 5,750127284 | 1,947186 | 2,953044 | 0,003147 | 1 | 0 | Distal Intergenic | 3 | 1,4E+08 | 1,4E+08 | 122 | 1 | 1,02E+08 | ENSMUST00000184427.3 | 55039 | ENSMUSG00000099169 | Mir7j | microRNA 7j |
| chrX | 22629644 | 22630268 | 625 | * | 3,921656 | 5,750191494 | 2,204467 | 2,608427 | 0,009096 | 1 | 0 | Distal Intergenic | 20 | 22327968 | 22328576 | 609 | 1 | 76705 | ENSMUST00000140102.2 | 301676 | | Ppp1r2-ps7 | protein phosphatase 1, regulatory (inhibitor) subunit 2, pseudogene 7 |
| chr17 | 31625429 | 31625836 | 408 | * | 3,713884 | 5,750204655 | 2,184329 | 2,632481 | 0,008476 | 1 | 0 | Intron (ENSMUST0000( | 17 | 31633130 | 31695282 | 62153 | 1 | 18585 | ENSMUST00000235639.2 | -7294 | ENSMUSG00000041119 | Pde9a | phosphodiesterase 9A |
| chrX | 1,45E+08 | 1,45E+08 | 317 | * | 4,180323 | 5,75020604 | 2,207646 | 2,604678 | 0,009196 | 1 | 0 | Distal Intergenic | 20 | 1,44E+08 | 1,44E+08 | 29462 | 2 | 27494 | ENSMUST00000143610.3 | -627580 | ENSMUSG00000041688 | Amot | angiomotin |
| chr5 | 23656491 | 23656811 | 321 | * | 3,967983 | 5,75021625 | 2,171516 | 2,648019 | 0,008097 | 1 | 0 | Intron (ENSMUST0000( | 5 | 23639447 | 23709233 | 69787 | 1 | 69188 | ENSMUST00000115128.8 | 17044 | ENSMUSG00000029004 | Kmt2e | lysine (K)-specific methyltransferase 2E |
| chr17 | 47425479 | 47425932 | 454 | * | 3,76787 | 5,750227449 | 2,168477 | 2,651735 | 0,008008 | 1 | 0 | Distal Intergenic | 17 | 47451801 | 47640462 | 188662 | 1 | 224829 | ENSMUST00000188947.7 | -25869 | ENSMUSG00000064043 | Trerf1 | transcriptional regulating factor 1 |
| chr9 | 65701002 | 65701526 | 525 | * | 4,49561 | 5,750265657 | 2,134466 | 2,694007 | 0,00706 | 1 | 0 | Intron (ENSMUST0000( | 9 | 65702512 | 65705194 | 2683 | 2 | 214812 | ENSMUST00000160747.2 | 3668 | ENSMUSG00000040524 | Zfp609 | zinc finger protein 609 |
| chr11 | 86958618 | 86959219 | 602 | * | 4,341026 | 5,750267957 | 2,14016 | 2,68684 | 0,007213 | 1 | 0 | Intron (ENSMUST0000( | 11 | 86924693 | 86964888 | 40196 | 2 | 66569 | ENSMUST00000020804.8 | 5669 | ENSMUSG00000061666 | Gdpd1 | glycerophosphodiester phosphodiesterase domain containing 1 |
| chr1 | 39471097 | 39471589 | 493 | * | 4,268856 | 5,750274648 | 2,12914 | 2,70075 | 0,006918 | 1 | 0 | Intron (ENSMUST0000( | 1 | 39430279 | 39431712 | 1434 | 2 | 54610 | ENSMUST00000195854.2 | -39385 | ENSMUSG00000003134 | Tbc1d8 | TBC1 domain family, member 8 |
| chr11 | 51672978 | 51673637 | 660 | * | 4,102249 | 5,750319477 | 2,102341 | 2,735198 | 0,006234 | 1 | 0 | Exon (ENSMUST00000( | 11 | 51654521 | 51673628 | 19108 | 1 | 66397 | ENSMUST00000136363.2 | 18457 | ENSMUSG00000020386 | Sar1b | secretion associated Ras related GTPase 1B |
| chr7 | 81076125 | 81076554 | 430 | * | 5,400606 | 5,750332207 | 2,092893 | 2,747552 | 0,006004 | 1 | 0 | Intron (ENSMUST0000( | 7 | 80996776 | 81086125 | 89350 | 2 | 12877 | ENSMUST00000130310.8 | 9571 | ENSMUSG00000025586 | Cpeb1 | cytoplasmic polyadenylation element binding protein 1 |
| chr11 | 84852569 | 84853230 | 662 | * | 5,332955 | 5,750337901 | 2,086435 | 2,756059 | 0,00585 | 1 | 0 | Promoter (1-2kb) | 11 | 84855173 | 84856870 | 1698 | 1 | 12351 | ENSMUST00000108076.3 | -1943 | ENSMUSG00000000805 | Car4 | carbonic anhydrase 4 |
| chr10 | 62435780 | 62436318 | 539 | * | 4,792517 | 5,75039629 | 2,053227 | 2,800662 | 0,0051 | 1 | 0 | Promoter (1-2kb) | 10 | 62416030 | 62438060 | 22031 | 2 | 56200 | ENSMUST00000045866.9 | 1742 | ENSMUSG00000020075 | Ddx21 | DExD box helicase 21 |
| chr3 | 5570909 | 5571535 | 627 | * | 4,703191 | 5,750467276 | 2,019069 | 2,848078 | 0,004398 | 1 | 0 | Distal Intergenic | 3 | 5578055 | 5580393 | 2339 | 2 | 78039 | ENSMUST00000176666.2 | 8858 | ENSMUSG00000093557 | 4930555M17Rik | RIKEN cDNA 4930555M17 gene |
| chr3 | 86933586 | 86934263 | 678 | * | 4,821261 | 5,75049193 | 2,005625 | 2,867182 | 0,004141 | 1 | 0 | Distal Intergenic | 3 | 86903141 | 86906748 | 3608 | 2 | 12479 | ENSMUST00000029717.4 | -26838 | ENSMUSG00000028076 | Cd1d1 | CD1d1 antigen |
| chr12 | 42306265 | 42306718 | 454 | * | 5,159878 | 5,750542505 | 1,981861 | 2,901587 | 0,003713 | 1 | 0 | Distal Intergenic | 12 | 41965491 | 42063539 | 98049 | 2 | 1,08E+08 | ENSMUST00000222917.2 | -242726 | | Gm46348 | predicted gene, 46348 |
| chr1 | 59975091 | 59975959 | 869 | * | 6,132989 | 5,750544117 | 1,974094 | 2,913004 | 0,00358 | 1 | 0 | Intron (ENSMUST0000( | 1 | 59952165 | 60024505 | 72341 | 1 | 72750 | ENSMUST00000036540.12 | 22926 | ENSMUSG00000041040 | Fam117b | family with sequence similarity 117, member B |
| chr2 | 75244767 | 75245494 | 728 | * | 5,914151 | 5,750550803 | 1,971437 | 2,916934 | 0,003535 | 1 | 0 | Distal Intergenic | 2 | 75134918 | 75143195 | 8278 | 1 | 1,03E+08 | ENSMUST00000229575.2 | 109849 | | 2600014E21Rik | RIKEN cDNA 2600014E21 gene |
| chr16 | 92972752 | 92973477 | 726 | * | 5,951889 | 5,75063029 | 1,941914 | 2,96132 | 0,003063 | 1 | 0 | Distal Intergenic | 16 | 93140612 | 93150077 | 9466 | 2 | 69857 | ENSMUST00000185971.7 | 176600 | ENSMUSG00000100277 | 1810053B23Rik | RIKEN cDNA 1810053B23 gene |
| chr9 | 38675275 | 38675751 | 477 | * | 5,97757 | 5,750683278 | 1,918398 | 2,997648 | 0,002721 | 1 | 0 | Intron (ENSMUST0000( | 9 | 38684364 | 38690317 | 5954 | 1 | 258778 | ENSMUST00000217114.3 | -8613 | ENSMUSG00000049926 | Olfr921 | olfactory receptor 921 |
| chr4 | 45941307 | 45941973 | 667 | * | 6,217361 | 5,750690857 | 1,916308 | 3,000923 | 0,002692 | 1 | 0 | Exon (ENSMUST00000: | 4 | 45965334 | 46034761 | 69428 | 1 | 100121 | ENSMUST00000107777.9 | -23361 | ENSMUSG00000035517 | Tdrd7 | tudor domain containing 7 |
| chr7 | 83916818 | 83917401 | 584 | * | 6,977093 | 5,750702465 | 1,914858 | 3,003201 | 0,002672 | 1 | 0 | Exon (ENSMUST00000( | 7 | 83899284 | 83912983 | 13700 | 2 | 11864 | ENSMUST00000208564.2 | -3835 | ENSMUSG00000015709 | Arnt2 | aryl hydrocarbon receptor nuclear translocator 2 |
| chr19 | 20464985 | 20465550 | 566 | * | 6,286022 | 5,75079351 | 1,880157 | 3,058678 | 0,002223 | 1 | 0 | Intron (ENSMUST0000( | 19 | 20470079 | 20620829 | 150751 | 1 | 11668 | ENSMUST00000225337.3 | -4529 | ENSMUSG00000053279 | Aldh1a1 | aldehyde dehydrogenase family 1, subfamily A1 |
| chr14 | 72426476 | 72427381 | 906 | * | 7,530556 | 5,750850622 | 1,86251 | 3,087689 | 0,002017 | 1 | 0 | Distal Intergenic | 14 | 72150531 | 72157878 | 7348 | 1 | 74667 | ENSMUST00000228197.2 | 275945 | ENSMUSG00000115304 | 4930434J06Rik | RIKEN cDNA 4930434J06 gene |
| chr11 | 1,14E+08 | 1,14E+08 | 826 | * | 7,66947 | 5,750977595 | 1,82699 | 3,147789 | 0,001645 | 1 | 0 | Intron (ENSMUST0000( | 11 | 1,14E+08 | 1,14E+08 | 217414 | 2 | 237979 | ENSMUST00000141943.2 | 116601 | ENSMUSG00000041592 | Sdk2 | sidekick cell adhesion molecule 2 |
| chrX | 1,26E+08 | 1,26E+08 | 773 | * | 5,870451 | 5,751259729 | 1,976298 | 2,910118 | 0,003613 | 1 | 0 | Distal Intergenic | 20 | 1,26E+08 | 1,26E+08 | 2031 | 1 | 245598 | ENSMUST00000249240.1 | 87476 | ENSMUSG00000049815 | 4921511C20Rik | high density lipoprotein (HDL) binding protein pseudogene |
| chr10 | 61333247 | 61334285 | 1039 | * | 6,992944 | 5,751544892 | 1,973918 | 2,91377 | 0,003571 | 1 | 0 | Promoter (<=1kb) | 10 | 61332379 | 61333876 | 1498 | 2 | 1,15E+08 | ENSMUST00000220094.2 | 0 | | Gm47593 | predicted gene, 47593 |
| chr2 | 1,54E+08 | 1,54E+08 | 815 | * | 5,864453 | 5,753966344 | 2,025882 | 2,840228 | 0,004508 | 1 | 0 | Distal Intergenic | 2 | 1,54E+08 | 1,54E+08 | 102876 | 1 | 12396 | ENSMUST00000109725.8 | -6087 | ENSMUSG00000038533 | Cbfa2t2 | CBFA2/RUNX1 translocation partner 2 |
| chr2 | 1,57E+08 | 1,57E+08 | 606 | * | 6,041595 | 5,754560501 | 2,04048 | 2,820199 | 0,004799 | 1 | 0 | Exon (ENSMUST00000: | 2 | 1,57E+08 | 1,57E+08 | 3854 | 2 | 29812 | ENSMUST00000149599.2 | 22258 | ENSMUSG00000027634 | Ndrg3 | N-myc downstream regulated gene 3 |
| chr2 | 51011648 | 51012397 | 750 | * | 5,979156 | 5,755554518 | 2,065944 | 2,78592 | 0,005338 | 1 | 0 | Distal Intergenic | 2 | 51027081 | 51039106 | 12026 | 2 | 74194 | ENSMUST00000140864.2 | 26709 | ENSMUSG00000017144 | Rnd3 | Rho family GTPase 3 |
| chr8 | 72775825 | 72776942 | 1118 | * | 7,364953 | 5,755555637 | 1,798559 | 3,200093 | 0,001374 | 0,542967057 | 1,360622 | Distal Intergenic | 8 | 72742326 | 72763470 | 21145 | 2 | 72054 | ENSMUST00000003574.5 | -12355 | ENSMUSG00000003484 | Cyp4f18 | cytochrome P450, family 4, subfamily f, polypeptide 18 |
| chr17 | 45818141 | 45818802 | 662 | * | 5,002682 | 5,755876017 | 2,074517 | 2,774562 | 0,005528 | 1 | 0 | Promoter (<=1kb) | 17 | 45817767 | 45831769 | 14003 | 1 | 224805 | ENSMUST00000024733.9 | 374 | ENSMUSG00000023938 | Aars2 | alanyl-tRNA synthetase 2, mitochondrial |
| chr3 | 1,15E+08 | 1,15E+08 | 548 | * | 4,608877 | 5,756209373 | 2,083627 | 2,762592 | 0,005734 | 1 | 0 | Distal Intergenic | 3 | 1,15E+08 | 1,15E+08 | 44292 | 1 | 229759 | ENSMUST00000051309.9 | 83156 | ENSMUSG00000027965 | Olfm3 | olfactomedin 3 |
| chr5 | 77441008 | 77441826 | 819 | * | 9,178426 | 5,756725241 | 1,778447 | 3,23694 | 0,001208 | 0,439573569 | 2,237403 | Distal Intergenic | 5 | 77451226 | 77452162 | 937 | 2 | 56412 | ENSMUST00000150451.2 | 10336 | ENSMUSG00000036285 | Noa1 | nitric oxide associated 1 |
| chr3 | 37691833 | 37692373 | 541 | * | 4,948755 | 5,756856478 | 2,101719 | 2,739119 | 0,00616 | 1 | 0 | Promoter (1-2kb) | 3 | 37694096 | 37698747 | 4652 | 1 | 24063 | ENSMUST00000108109.8 | -1723 | ENSMUSG00000037211 | Spry1 | sprouty RTK signaling antagonist 1 |
| chr4 | 8536455 | 8537391 | 937 | * | 5,313326 | 5,757123324 | 2,110798 | 2,727464 | 0,006382 | 1 | 0 | Promoter (<=1kb) | 4 | 8535644 | 8607778 | 72135 | 1 | 59021 | ENSMUST00000060232.8 | 811 | ENSMUSG00000047187 | Rab2a | RAB2A, member RAS oncogene family |
| chr1 | 1,42E+08 | 1,42E+08 | 616 | * | 5,794312 | 5,759632397 | 2,196583 | 2,622088 | 0,008739 | 1 | 0 | Distal Intergenic | 1 | 1,41E+08 | 1,41E+08 | 107 | 1 | 1,15E+08 | ENSMUST00000239951.1 | 579410 | ENSMUSG00000118759 | Gm25457 | predicted gene, 25457 |
| chrX | 1,62E+08 | 1,62E+08 | 672 | * | 4,370035 | 5,759912604 | 2,208106 | 2,60853 | 0,009093 | 1 | 0 | Distal Intergenic | 20 | 1,62E+08 | 1,62E+08 | 120 | 2 | 1,15E+08 | ENSMUST00000158798.3 | -25476 | ENSMUSG00000089423 | Gm26007 | predicted gene, 26007 |
| chr19 | 13304897 | 13305335 | 439 | * | 4,352632 | 5,760264237 | 2,222749 | 2,591505 | 0,009556 | 1 | 0 | Distal Intergenic | 19 | 13290724 | 13291647 | 924 | 2 | 258121 | ENSMUST00000080142.2 | -13250 | ENSMUSG00000062199 | Olfr1465 | olfactory receptor 1465 |
| chr6 | 93198873 | 93199272 | 400 | * | 3,80656 | 5,760395825 | 2,227575 | 2,585949 | 0,009711 | 1 | 0 | Intron (ENSMUST0000( | 6 | 93312439 | 93312569 | 131 | 2 | 1,15E+08 | ENSMUST00000103992.3 | 113297 | ENSMUSG00000077180 | Gm25094 | predicted gene, 25094 |
| chr5 | 36831843 | 36833157 | 1315 | * | 8,638875 | 5,762283641 | 1,726163 | 3,338204 | 0,000843 | 1 | 0 | Intron (ENSMUST0000( | 5 | 36823753 | 36823856 | 104 | 2 | 1,15E+08 | ENSMUST00000158782.3 | -7987 | ENSMUSG00000089407 | Gm24878 | predicted gene, 24878 |
| chr12 | 73589346 | 73590392 | 1047 | * | 9,23151 | 5,76393492 | 1,746648 | 3,299998 | 0,000967 | 0,474532331 | 1,449706 | Promoter (2-3kb) | 12 | 73590176 | 73593166 | 2991 | 2 | 238257 | ENSMUST00000042975.7 | 2774 | ENSMUSG00000034435 | Tmem30b | transmembrane protein 30B |
| chr3 | 88182311 | 88183316 | 1006 | * | 10,69501 | 5,764085873 | 1,800711 | 3,201006 | 0,001369 | 0,659642963 | 0,744375 | Intron (ENSMUST0000( | 3 | 88150687 | 88194517 | 43831 | 2 | 58176 | ENSMUST00000165196.8 | 11201 | ENSMUSG00000104445 | Rhbg | Rhesus blood group-associated B glycoprotein |
| chr9 | 20874518 | 20875624 | 1107 | * | 8,000339 | 5,764303859 | 1,831027 | 3,148126 | 0,001643 | 1 | 0 | 3' UTR | 9 | 20818505 | 20871184 | 52680 | 2 | 13433 | ENSMUST00000177754.9 | -3334 | ENSMUSG00000004099 | Dnmt1 | DNA methyltransferase (cytosine-5) 1 |
| chr7 | 60994397 | 60995234 | 838 | * | 7,310183 | 5,764673796 | 1,841105 | 3,131094 | 0,001742 | 1 | 0 | Distal Intergenic | 7 | 60932487 | 60961469 | 28983 | 2 | 27493 | ENSMUST00000154958.2 | -32928 | | A230006K03Rik | RIKEN cDNA A230006K03 gene |
| chr13 | 24945556 | 24946572 | 1017 | * | 4,233158 | 5,768994903 | 2,239044 | 2,576544 | 0,009979 | 1 | 0 | Promoter (<=1kb) | 13 | 24935828 | 24945844 | 10017 | 2 | 57441 | ENSMUST00000006898.10 | 0 | ENSMUSG00000006715 | Gmnn | geminin |
| chr14 | 64827864 | 64828525 | 662 | * | 4,394034 | 5,76911004 | 2,232915 | 2,583668 | 0,009776 | 1 | 0 | Promoter (<=1kb) | 14 | 64828106 | 64828190 | 85 | 1 | 387233 | ENSMUST00000083663.3 | 0 | ENSMUSG00000065597 | Mir124a-1 | microRNA 124a-1 |
| chr11 | 84207455 | 84208202 | 748 | * | 4,10888 | 5,769760655 | 2,201061 | 2,621355 | 0,008758 | 1 | 0 | Intron (ENSMUST0000( | 11 | 84171281 | 84202227 | 30947 | 1 | 107476 | ENSMUST00000183887.2 | 36174 | ENSMUSG00000020532 | Acaca | acetyl-Coenzyme A carboxylase alpha |
| chr10 | 30987556 | 30987849 | 294 | * | 3,980271 | 5,769865189 | 2,195887 | 2,627578 | 0,0086 | 1 | 0 | Distal Intergenic | 10 | 30882408 | 30942240 | 59833 | 2 | 1,03E+08 | ENSMUST00000214464.2 | -45316 | ENSMUSG00000111778 | Gm30676 | predicted gene, 30676 |
| chr4 | 1,51E+08 | 1,51E+08 | 662 | * | 4,118434 | 5,770071685 | 2,186466 | 2,638995 | 0,008315 | 1 | 0 | Distal Intergenic | 4 | 1,51E+08 | 1,51E+08 | 14974 | 1 | 74155 | ENSMUST00000073600.9 | -52632 | ENSMUSG00000028967 | Errfi1 | ERBB receptor feedback inhibitor 1 |
| chr14 | 5039890 | 5040396 | 507 | * | 4,504226 | 5,770191426 | 2,180506 | 2,646263 | 0,008139 | 1 | 0 | Distal Intergenic | 14 | 4506571 | 4809435 | 302865 | 2 | 21834 | ENSMUST00000224470.2 | -230455 | ENSMUSG00000021779 | Thrb | thyroid hormone receptor beta |
| chr8 | 90595294 | 90595649 | 356 | * | 4,184518 | 5,770284992 | 2,176476 | 2,651206 | 0,00802 | 1 | 0 | Distal Intergenic | 8 | 90393801 | 90393932 | 132 | 2 | 1,15E+08 | ENSMUST00000158258.3 | -201362 | ENSMUSG00000088883 | Gm24212 | predicted gene, 24212 |
| chr4 | 1,32E+08 | 1,32E+08 | 541 | * | 4,574379 | 5,770743553 | 2,156414 | 2,676083 | 0,007449 | 1 | 0 | Distal Intergenic | 4 | 1,32E+08 | 1,32E+08 | 18617 | 1 | 19204 | ENSMUST00000070690.8 | 30764 | ENSMUSG00000056529 | Ptafr | platelet-activating factor receptor |
| chr9 | 31130865 | 31131582 | 718 | * | 4,747511 | 5,771209393 | 2,137242 | 2,700307 | 0,006928 | 1 | 0 | Distal Intergenic | 9 | 31067486 | 31123111 | 55626 | 2 | 11804 | ENSMUST00000213254.2 | -7754 | ENSMUSG00000031996 | Aplp2 | amyloid beta (A4) precursor-like protein 2 |
| chr2 | 1,04E+08 | 1,04E+08 | 601 | * | 4,144664 | 5,77126437 | 2,135546 | 2,702478 | 0,006882 | 1 | 0 | Promoter (<=1kb) | 2 | 1,04E+08 | 1,04E+08 | 139 | 1 | 1,15E+08 | ENSMUST00000157114.3 | 960 | ENSMUSG00000087739 | Gm24644 | predicted gene, 24644 |
| chr3 | 1,08E+08 | 1,08E+08 | 447 | * | 4,626591 | 5,771822091 | 2,114094 | 2,730164 | 0,00633 | 1 | 0 | Distal Intergenic | 3 | 1,08E+08 | 1,08E+08 | 2200 | 2 | 12977 | ENSMUST00000156820.2 | 24969 | ENSMUSG00000014599 | Csf1 | colony stimulating factor 1 (macrophage) |
| chr16 | 65232992 | 65233569 | 578 | * | 4,716661 | 5,772232926 | 2,09842 | 2,750752 | 0,005946 | 1 | 0 | Distal Intergenic | 16 | 65317397 | 65331183 | 13787 | 1 | 18736 | ENSMUST00000176330.8 | -83828 | ENSMUSG00000004842 | Pou1f1 | POU domain, class 1, transcription factor 1 |
| chr10 | 36844885 | 36845540 | 656 | * | 4,696718 | 5,772293315 | 2,096366 | 2,753476 | 0,005897 | 1 | 0 | Distal Intergenic | 10 | 36850540 | 36877885 | 27346 | 1 | 15182 | ENSMUST00000019911.14 | -5000 | ENSMUSG00000019777 | Hdac2 | histone deacetylase 2 |
| chr17 | 32566023 | 32566756 | 734 | * | 4,573312 | 5,772379944 | 2,093392 | 2,757429 | 0,005826 | 1 | 0 | Promoter (2-3kb) | 17 | 32555447 | 32569480 | 14034 | 2 | 54194 | ENSMUST00000235265.2 | 2724 | ENSMUSG00000002625 | Akap8l | A kinase (PRKA) anchor protein 8-like |
| chr8 | 1,24E+08 | 1,24E+08 | 340 | * | 4,6473 | 5,772493487 | 2,089227 | 2,762981 | 0,005728 | 1 | 0 | Promoter (<=1kb) | 8 | 1,24E+08 | 1,24E+08 | 8039 | 2 | 77087 | ENSMUST00000212337.2 | -853 | ENSMUSG00000035569 | Ankrd11 | ankyrin repeat domain 11 |
| chr5 | 1,39E+08 | 1,39E+08 | 1087 | * | 11,01563 | 5,772862576 | 1,696663 | 3,40248 | 0,000668 | 0,524856133 | 0,744375 | Promoter (<=1kb) | 5 | 1,39E+08 | 1,39E+08 | 15835 | 1 | 80290 | ENSMUST00000051293.8 | 0 | ENSMUSG00000044197 | Gpr146 | G protein-coupled receptor 146 |
| chr5 | 96563477 | 96563972 | 496 | * | 4,907604 | 5,773752295 | 2,048087 | 2,819095 | 0,004816 | 1 | 0 | Intron (ENSMUST0000( | 5 | 96522294 | 96535493 | 13200 | 1 | 231470 | ENSMUST00000196751.2 | 41183 | ENSMUSG00000034687 | Fras1 | Fraser extracellular matrix complex subunit 1 |
| chr13 | 36747075 | 36747912 | 838 | * | 5,267309 | 5,77498315 | 2,01181 | 2,87054 | 0,004098 | 1 | 0 | Intron (ENSMUST0000( | 13 | 36796790 | 36804413 | 7624 | 1 | 1,08E+08 | ENSMUST00000225239.2 | -48878 | | Gm46410 | predicted gene, 46410 |
| chr9 | 55394967 | 55395600 | 634 | * | 5,25248 | 5,775069998 | 2,008958 | 2,87466 | 0,004045 | 1 | 0 | Promoter (<=1kb) | 9 | 55381821 | 55394797 | 12977 | 2 | 110842 | ENSMUST00000139342.2 | -170 | ENSMUSG00000032314 | Etfa | electron transferring flavoprotein, alpha polypeptide |
| chr5 | 31577419 | 31578543 | 1125 | * | 7,539732 | 5,77591892 | 1,822031 | 3,170044 | 0,001524 | 1 | 0 | Distal Intergenic | 5 | 31609775 | 31624001 | 14227 | 1 | 269639 | ENSMUST00000202061.4 | -31232 | ENSMUSG00000062761 | Zfp512 | zinc finger protein 512 |
| chr2 | 1,74E+08 | 1,74E+08 | 387 | * | 6,316568 | 5,776112962 | 2,008044 | 2,876487 | 0,004021 | 1 | 0 | Exon (ENSMUST00000( | 2 | 1,74E+08 | 1,74E+08 | 7188 | 1 | 545486 | ENSMUST00000016399.6 | 5005 | ENSMUSG00000016255 | Tubb1 | tubulin, beta 1 class VI |
| chr10 | 36647235 | 36647878 | 644 | * | 5,85075 | 5,776498193 | 1,971848 | 2,929485 | 0,003395 | 1 | 0 | Intron (ENSMUST0000( | 10 | 36651781 | 36651927 | 147 | 2 | 1,15E+08 | ENSMUST00000157798.3 | 4049 | ENSMUSG00000088423 | Gm22554 | predicted gene, 22554 |
| chr19 | 21069284 | 21070251 | 968 | * | 6,456295 | 5,77664763 | 1,968234 | 2,934939 | 0,003336 | 1 | 0 | Distal Intergenic | 19 | 21081866 | 21084521 | 2656 | 1 | 75354 | ENSMUST00000236278.2 | -11615 | ENSMUSG00000118047 | 4930554I06Rik | RIKEN cDNA 4930554I06 gene |
| chr7 | 39910329 | 39911126 | 798 | * | 7,084149 | 5,776730465 | 1,876277 | 3,078826 | 0,002078 | 0,627785088 | 1,360622 | Distal Intergenic | 7 | 40324161 | 40332685 | 8525 | 1 | 71029 | ENSMUST00000206289.2 | -413035 | ENSMUSG00000102004 | 4933402C06Rik | RIKEN cDNA 4933402C06 gene |
| chrX | 1,66E+08 | 1,66E+08 | 1078 | * | 6,54686 | 5,776746899 | 1,877531 | 3,076778 | 0,002093 | 1 | 0 | Distal Intergenic | 20 | 1,66E+08 | 1,66E+08 | 1536 | 2 | 19241 | ENSMUST00000112175.2 | 25004 | ENSMUSG00000049775 | Tmsb4x | thymosin, beta 4, X chromosome |
| chr5 | 1,19E+08 | 1,19E+08 | 746 | * | 6,212032 | 5,776770645 | 1,879336 | 3,073836 | 0,002113 | 1 | 0 | Distal Intergenic | 5 | 1,19E+08 | 1,19E+08 | 5535 | 2 | 665194 | ENSMUST00000200918.2 | 6350 | ENSMUSG00000107334 | Gm7538 | predicted gene 7538 |
| chr6 | 1,01E+08 | 1,01E+08 | 603 | * | 5,969388 | 5,777418576 | 1,934215 | 2,986957 | 0,002818 | 1 | 0 | Exon (ENSMUST00000( | 6 | 1,01E+08 | 1,01E+08 | 36313 | 1 | 232314 | ENSMUST00000063854.7 | -35107 | ENSMUSG00000052144 | Ppp4r2 | protein phosphatase 4, regulatory subunit 2 |
| chr17 | 39295456 | 39296297 | 842 | * | 6,221445 | 5,777425681 | 1,934674 | 2,986254 | 0,002824 | 1 | 0 | Distal Intergenic | 17 | 39347586 | 39351118 | 3533 | 1 | 1E+08 | ENSMUST00000232955.2 | -51289 | ENSMUSG00000095625 | Esp24 | exocrine gland secreted peptide 24 |
| chr5 | 61062424 | 61062994 | 571 | * | 5,83687 | 5,777741073 | 1,966219 | 2,938504 | 0,003298 | 1 | 0 | Distal Intergenic | 5 | 60658176 | 60658283 | 108 | 2 | 1,15E+08 | ENSMUST00000157423.3 | -404141 | ENSMUSG00000088048 | Gm23331 | predicted gene, 23331 |
| chrX | 91576024 | 91576787 | 764 | * | 5,249985 | 5,777798551 | 1,971802 | 2,930212 | 0,003387 | 1 | 0 | Intron (ENSMUST0000( | 20 | 91533555 | 91534287 | 733 | 1 | 546347 | ENSMUST00000096371.3 | 42469 | ENSMUSG00000071726 | Gm5941 | predicted gene 5941 |
| chr15 | 62419906 | 62420486 | 581 | * | 6,233683 | 5,777967349 | 1,938163 | 2,981157 | 0,002872 | 1 | 0 | Distal Intergenic | 15 | 62439824 | 62654636 | 214813 | 1 | 1,05E+08 | ENSMUST00000230029.2 | -19338 | | Gm41333 | predicted gene, 41333 |
| chr9 | 22598107 | 22598659 | 553 | * | 4,914625 | 5,778007129 | 1,994555 | 2,89689 | 0,003769 | 1 | 0 | Intron (ENSMUST0000( | 9 | 22590046 | 22799570 | 209525 | 1 | 319845 | ENSMUST00000124076.2 | 8061 | ENSMUSG00000035919 | Bbs9 | Bardet-Biedl syndrome 9 (human) |

| | | | | | | | | | | | | | | | | | | | | | | | |
|---|---|---|---|---|---|---|---|---|---|---|---|---|---|---|---|---|---|---|---|---|---|---|---|
| chr15 | 93233005 | 93233491 | 487 | * | 5,341809 | 5,778136024 | 2,009104 | 2,875977 | 0,004028 | 1 | 0 | Promoter (1-2kb) | 15 | 93182159 | 93234689 | 52531 | 2 | 67057 | ENSMUST00000080299.7 | 1198 | ENSMUSG00000022634 | Yaf2 | YY1 associated factor 2 |
| chr3 | 1,59E+08 | 1,59E+08 | 751 | * | 4,830048 | 5,778309568 | 2,032513 | 2,842938 | 0,00447 | 1 | 0 | Intron (ENSMUST0000( | 3 | 1,59E+08 | 1,59E+08 | 34002 | 1 | 76131 | ENSMUST00000106041.3 | 14078 | ENSMUSG00000028175 | Depdc1a | DEP domain containing 1a |
| chr5 | 36904083 | 36904720 | 638 | * | 4,629774 | 5,778497136 | 2,056056 | 2,810476 | 0,004947 | 1 | 0 | Promoter (1-2kb) | 5 | 36904722 | 36905994 | 1273 | 2 | 117197 | ENSMUST00000071949.5 | 1274 | ENSMUSG00000060708 | Bloc1s4 | biogenesis of lysosomal organelles complex-1, subunit 4, cappuccino |
| chr18 | 90608990 | 90609484 | 495 | * | 5,072352 | 5,778517902 | 2,058156 | 2,807619 | 0,004991 | 1 | 0 | 3' UTR | 18 | 90603531 | 90609199 | 5669 | 1 | 1,08E+08 | ENSMUST00000210766.2 | 5459 | ENSMUSG00000110277 | Gm45871 | predicted gene 45871 |
| chr17 | 9817181 | 9817940 | 760 | * | 4,600124 | 5,778535771 | 2,061404 | 2,803203 | 0,00506 | 1 | 0 | Distal Intergenic | 17 | 9885424 | 9888646 | 3223 | 2 | 67543 | ENSMUST00000057190.4 | 70706 | ENSMUSG00000046173 | Pabpc6 | poly(A) binding protein, cytoplasmic 6 |
| chr13 | 24813181 | 24814224 | 1044 | * | 5,822052 | 5,778542839 | 2,066774 | 2,795924 | 0,005175 | 1 | 0 | Intron (ENSMUST0000( | 13 | 24822631 | 24917799 | 95169 | 1 | 193385 | ENSMUST00000110384.9 | -8407 | ENSMUSG00000036006 | Ripor2 | RHO family interacting cell polarization regulator 2 |
| chr4 | 1,38E+08 | 1,38E+08 | 528 | * | 4,562126 | 5,778628 | 2,075426 | 2,78431 | 0,005364 | 1 | 0 | Distal Intergenic | 4 | 1,38E+08 | 1,38E+08 | 29465 | 2 | 72269 | ENSMUST00000030535.4 | -8344 | ENSMUSG00000028755 | Cda | cytidine deaminase |
| chr7 | 1,36E+08 | 1,36E+08 | 389 | * | 4,584308 | 5,778670389 | 2,082778 | 2,774501 | 0,005529 | 1 | 0 | Distal Intergenic | 7 | 1,36E+08 | 1,36E+08 | 50199 | 1 | 77383 | ENSMUST00000210438.3 | -36674 | ENSMUSG00000110027 | C030029H02Rik | RIKEN cDNA C030029H02 gene |
| chr10 | 83581961 | 83582463 | 503 | * | 4,652538 | 5,778672816 | 2,084606 | 2,77207 | 0,00557 | 1 | 0 | Intron (ENSMUST0000( | 10 | 83573773 | 83598027 | 24255 | 1 | 69784 | ENSMUST00000218623.2 | 8188 | ENSMUSG00000087651 | 1500009L16Rik | RIKEN cDNA 1500009L16 gene |
| chr1 | 1,7E+08 | 1,7E+08 | 403 | * | 4,688643 | 5,778695708 | 2,086632 | 2,769389 | 0,005616 | 1 | 0 | Intron (ENSMUST0000( | 1 | 1,7E+08 | 1,7E+08 | 2988 | 1 | 67303 | ENSMUST00000242643.1 | 8959 | ENSMUSG00000097503 | 3110045C21Rik | RIKEN cDNA 3110045C21 gene |
| chr10 | 1,3E+08 | 1,3E+08 | 494 | * | 4,341971 | 5,778732982 | 2,091734 | 2,762652 | 0,005733 | 1 | 0 | 3' UTR | 10 | 1,3E+08 | 1,3E+08 | 15640 | 2 | 623734 | ENSMUST00000171811.4 | 14298 | ENSMUSG00000092048 | Vmn2r85 | vomeronasal 2, receptor 85 |
| chr5 | 36875754 | 36876386 | 633 | * | 4,627535 | 5,778882509 | 2,116601 | 2,730266 | 0,006328 | 1 | 0 | Distal Intergenic | 5 | 36828743 | 36853368 | 24626 | 2 | 320661 | ENSMUST00000140653.2 | -22386 | ENSMUSG00000029190 | D5Ertd579e | DNA segment, Chr 5, ERATO Doi 579, expressed |
| chr11 | 59041848 | 59042299 | 452 | * | 4,199918 | 5,778913796 | 2,123215 | 2,721775 | 0,006493 | 1 | 0 | Distal Intergenic | 11 | 59051949 | 59054143 | 2195 | 2 | 216792 | ENSMUST00000069631.3 | 11844 | ENSMUSG00000049287 | Iba57 | IBA57 homolog, iron-sulfur cluster assembly |
| chr3 | 30671767 | 30672274 | 508 | * | 4,478448 | 5,778932321 | 2,126811 | 2,717181 | 0,006584 | 1 | 0 | 3' UTR | 3 | 30667660 | 30670944 | 3285 | 1 | 80732 | ENSMUST00000193736.2 | 4107 | ENSMUSG00000037730 | Mynn | myoneurin |
| chr12 | 9814576 | 9814957 | 382 | * | 4,043976 | 5,778944138 | 2,12924 | 2,714088 | 0,006646 | 1 | 0 | Distal Intergenic | 12 | 9915347 | 9915450 | 104 | 2 | 1,15E+08 | ENSMUST00000157712.3 | 100493 | ENSMUSG00000088337 | Gm22845 | predicted gene, 22845 |
| chr13 | 99192234 | 99192822 | 589 | * | 5,066531 | 5,778978845 | 2,134719 | 2,707138 | 0,006787 | 1 | 0 | Distal Intergenic | 13 | 99219536 | 99224412 | 4877 | 2 | 69603 | ENSMUST00000243956.1 | 31590 | ENSMUSG00000100410 | 2310020H05Rik | RIKEN cDNA 2310020H05 gene |
| chr19 | 41838239 | 41838625 | 387 | * | 3,786585 | 5,779117885 | 2,160224 | 2,67524 | 0,007468 | 1 | 0 | Promoter (1-2kb) | 19 | 41834411 | 41836571 | 2161 | 2 | 212398 | ENSMUST00000059231.4 | -1668 | ENSMUSG00000047604 | Frat2 | frequently rearranged in advanced T cell lymphomas 2 |
| chr14 | 10686683 | 10687226 | 544 | * | 3,884795 | 5,779177177 | 2,171922 | 2,660858 | 0,007794 | 1 | 0 | Intron (ENSMUST0000( | 14 | 10505632 | 10916192 | 410561 | 2 | 19270 | ENSMUST00000248718.1 | 228966 | | Ptprg | protein tyrosine phosphatase, receptor type, G |
| chr12 | 40699214 | 40700013 | 800 | * | 6,682258 | 5,780392319 | 1,993115 | 2,900181 | 0,003729 | 1 | 0 | Exon (ENSMUST00000( | 12 | 40837591 | 40861340 | 23750 | 1 | 238130 | ENSMUST00000220783.2 | -137578 | ENSMUSG00000035954 | Dock4 | dedicator of cytokinesis 4 |
| chr1 | 1,88E+08 | 1,88E+08 | 887 | * | 7,350834 | 5,781678567 | 1,833116 | 3,154017 | 0,00161 | 1 | 0 | Intron (ENSMUST0000( | 1 | 1,88E+08 | 1,89E+08 | 702008 | 1 | 22283 | ENSMUST00000060479.14 | 52934 | ENSMUSG00000026609 | Ush2a | usherin |
| chr11 | 87981760 | 87982565 | 806 | * | 5,711619 | 5,782714681 | 2,037214 | 2,838541 | 0,004532 | 1 | 0 | Distal Intergenic | 11 | 87988884 | 87990934 | 2051 | 1 | 70393 | ENSMUST00000131964.2 | -6319 | ENSMUSG00000018378 | 2210416O15Rik | RIKEN cDNA 2210416O15 gene |
| chr4 | 1,3E+08 | 1,3E+08 | 895 | * | 7,043638 | 5,783465032 | 2,053497 | 2,816398 | 0,004857 | 0,769692849 | 1,360622 | Promoter (<=1kb) | 4 | 1,3E+08 | 1,3E+08 | 1641 | 2 | 66264 | ENSMUST00000141361.8 | 0 | ENSMUSG00000028795 | Ccdc28b | coiled coil domain containing 28B |
| chrX | 78218615 | 78219603 | 989 | * | 5,716902 | 5,786713306 | 1,924429 | 3,006976 | 0,002639 | 1 | 0 | Distal Intergenic | 20 | 78374119 | 78393916 | 19798 | 2 | 636104 | ENSMUST00000199433.2 | 174313 | ENSMUSG00000073077 | Cfap47 | cilia and flagella associated protein 47 |
| chr5 | 1,36E+08 | 1,36E+08 | 704 | * | 5,54002 | 5,787556497 | 2,155907 | 2,684511 | 0,007264 | 1 | 0 | Intron (ENSMUST0000( | 5 | 1,36E+08 | 1,36E+08 | 8964 | 2 | 215210 | ENSMUST00000043378.9 | -7919 | ENSMUSG00000039886 | Tmem120a | transmembrane protein 120A |
| chr6 | 74471244 | 74471922 | 679 | * | 4,26867 | 5,788053128 | 2,169845 | 2,667496 | 0,007642 | 1 | 0 | Distal Intergenic | 6 | 74428780 | 74428887 | 108 | 2 | 1,15E+08 | ENSMUST00000082474.3 | -42357 | ENSMUSG00000064408 | Gm23924 | predicted gene, 23924 |
| chr1 | 34499162 | 34499750 | 589 | * | 4,653044 | 5,788206705 | 2,175055 | 2,661177 | 0,007787 | 1 | 0 | Promoter (<=1kb) | 1 | 34498873 | 34510997 | 12125 | 1 | 19253 | ENSMUST00000188884.3 | 289 | ENSMUSG00000026126 | Ptpn18 | protein tyrosine phosphatase, non-receptor type 18 |
| chr15 | 36887474 | 36887850 | 377 | * | 4,430686 | 5,789474312 | 2,216268 | 2,612263 | 0,008994 | 1 | 0 | Distal Intergenic | 15 | 36870744 | 36880121 | 9378 | 2 | 1E+08 | ENSMUST00000100713.4 | -7353 | ENSMUSG00000072618 | Gm10384 | predicted gene 10384 |
| chr16 | 82447589 | 82448107 | 519 | * | 5,683114 | 5,789476554 | 1,96925 | 2,93994 | 0,003283 | 1 | 0 | Distal Intergenic | 16 | 82340117 | 82340223 | 107 | 1 | 1,15E+08 | ENSMUST00000240414.1 | 107472 | ENSMUSG00000119122 | Gm23355 | predicted gene, 23355 |
| chr11 | 54550139 | 54550846 | 708 | * | 5,090273 | 5,789678368 | 2,223533 | 2,603819 | 0,009219 | 1 | 0 | Intron (ENSMUST0000( | 11 | 54582258 | 54587312 | 5055 | 1 | 192786 | ENSMUST00000136494.2 | -31412 | ENSMUSG00000037533 | Rapgef6 | Rap guanine nucleotide exchange factor (GEF) 6 |
| chr18 | 32422398 | 32423919 | 1522 | * | 11,385 | 5,791699956 | 1,643778 | 3,523408 | 0,000426 | 0,44905823 | 0,738441 | Distal Intergenic | 18 | 32390322 | 32403144 | 12823 | 1 | 13872 | ENSMUST00000129023.2 | 32076 | ENSMUSG00000024382 | Ercc3 | excision repair cross-complementing rodent repair deficiency, complementation group 3 |
| chrX | 1,4E+08 | 1,4E+08 | 1061 | * | 7,208359 | 5,79243207 | 1,81179 | 3,197077 | 0,001388 | 1 | 0 | Intron (ENSMUST0000( | 20 | 1,4E+08 | 1,4E+08 | 21412 | 2 | 94216 | ENSMUST00000137051.8 | -131351 | ENSMUSG00000031273 | Col4a6 | collagen, type IV, alpha 6 |
| chr11 | 1,05E+08 | 1,05E+08 | 672 | * | 7,075468 | 5,795172789 | 1,775112 | 3,264681 | 0,001096 | 1 | 0 | Distal Intergenic | 11 | 1,05E+08 | 1,05E+08 | 105153 | 1 | 24086 | ENSMUST00000106939.9 | -3108 | ENSMUSG00000020694 | Tlk2 | tousled-like kinase 2 (Arabidopsis) |
| chr10 | 39944385 | 39945024 | 640 | * | 6,360758 | 5,796065828 | 1,870922 | 3,097973 | 0,001948 | 1 | 0 | Intron (ENSMUST0000( | 10 | 39913295 | 39952601 | 39307 | 2 | 72472 | ENSMUST00000213827.2 | 7577 | ENSMUSG00000019838 | Slc16a10 | solute carrier family 16 (monocarboxylic acid transporters), member 10 |
| chr17 | 45935095 | 45935714 | 620 | * | 4,235082 | 5,797669715 | 2,246163 | 2,581144 | 0,009847 | 1 | 0 | Distal Intergenic | 17 | 45912262 | 45913028 | 767 | 2 | 653016 | ENSMUST00000113529.4 | -22067 | ENSMUSG00000079471 | Mymx | myomixer, myoblast fusion factor |
| chr7 | 67236074 | 67236596 | 523 | * | 4,038447 | 5,797725589 | 2,241592 | 2,586432 | 0,009698 | 1 | 0 | Intron (ENSMUST0000( | 7 | 67164154 | 67229498 | 65345 | 2 | 67867 | ENSMUST00000190115.7 | -6576 | ENSMUSG00000030556 | Lrrc28 | leucine rich repeat containing 28 |
| chrX | 23714291 | 23715227 | 937 | * | 4,888342 | 5,797908038 | 2,229426 | 2,600628 | 0,009305 | 1 | 0 | Distal Intergenic | 20 | 23748792 | 23773846 | 25055 | 1 | 236749 | ENSMUST00000116614.3 | -33565 | ENSMUSG00000068113 | Tesl1 | testin LIM domain protein like 1 |
| chr13 | 74330859 | 74331481 | 623 | * | 3,90177 | 5,798051712 | 2,219755 | 2,612023 | 0,009001 | 1 | 0 | Intron (ENSMUST0000( | 13 | 74320118 | 74324598 | 4481 | 2 | 211446 | ENSMUST00000220679.2 | -6261 | ENSMUSG00000034152 | Exoc3 | exocyst complex component 3 |
| chr4 | 1,33E+08 | 1,33E+08 | 616 | * | 4,393937 | 5,798677986 | 2,176757 | 2,663906 | 0,007724 | 1 | 0 | Promoter (1-2kb) | 4 | 1,33E+08 | 1,33E+08 | 12112 | 1 | 53608 | ENSMUST00000030677.7 | -1029 | ENSMUSG00000028862 | Map3k6 | mitogen-activated protein kinase kinase kinase 6 |
| chr3 | 1,26E+08 | 1,26E+08 | 571 | * | 3,877696 | 5,798892894 | 2,16335 | 2,680515 | 0,007351 | 1 | 0 | Intron (ENSMUST0000( | 3 | 1,26E+08 | 1,26E+08 | 1716 | 2 | 1,03E+08 | ENSMUST00000132762.3 | -25419 | ENSMUSG00000086679 | Gm15551 | predicted gene 15551 |
| chr12 | 14515591 | 14515947 | 357 | * | 3,949737 | 5,798981448 | 2,157406 | 2,687942 | 0,007189 | 1 | 0 | Distal Intergenic | 12 | 14556588 | 14556694 | 107 | 1 | 1,15E+08 | ENSMUST00000240467.1 | -40641 | ENSMUSG00000119624 | Gm25414 | predicted gene, 25414 |
| chr10 | 1,1E+08 | 1,1E+08 | 475 | * | 4,371245 | 5,799035336 | 2,155195 | 2,690724 | 0,00713 | 1 | 0 | Intron (ENSMUST0000( | 10 | 1,1E+08 | 1,1E+08 | 150077 | 2 | 260315 | ENSMUST00000238286.2 | -20553 | ENSMUSG00000020181 | Nav3 | neuron navigator 3 |
| chr2 | 34936423 | 34936872 | 450 | * | 5,150754 | 5,799121243 | 2,149455 | 2,69795 | 0,006977 | 1 | 0 | Intron (ENSMUST0000( | 2 | 34873343 | 34951450 | 78108 | 2 | 15139 | ENSMUST00000028233.7 | 14578 | ENSMUSG00000026874 | Hc | hemolytic complement |
| chr4 | 46131447 | 46131895 | 449 | * | 3,905759 | 5,799173851 | 2,146611 | 2,701549 | 0,006902 | 1 | 0 | Promoter (<=1kb) | 4 | 46128966 | 46131923 | 2958 | 2 | 272027 | ENSMUST00000137399.2 | 28 | ENSMUSG00000035495 | Tstd2 | thiosulfate sulfurtransferase (rhodanese)-like domain containing 2 |
| chr3 | 87879304 | 87879771 | 468 | * | 4,05377 | 5,79934105 | 2,136206 | 2,714786 | 0,006632 | 1 | 0 | Promoter (<=1kb) | 3 | 87878436 | 87887758 | 9323 | 1 | 18008 | ENSMUST00000160694.2 | 868 | ENSMUSG00000004891 | Nes | nestin |
| chr13 | 8935255 | 8936453 | 1199 | * | 9,298279 | 5,799728838 | 1,799002 | 3,223859 | 0,001265 | 0,524368688 | 1,425929 | Promoter (<=1kb) | 13 | 8935537 | 8942487 | 6951 | 1 | 319554 | ENSMUST00000169314.9 | 0 | ENSMUSG00000058258 | Idi1 | isopentenyl-diphosphate delta isomerase |
| chr16 | 91959001 | 91959461 | 461 | * | 4,199638 | 5,799982286 | 2,101386 | 2,760075 | 0,005779 | 1 | 0 | Intron (ENSMUST0000( | 16 | 91951135 | 91953179 | 2045 | 1 | 1,05E+08 | ENSMUST00000232211.2 | 7866 | | Gm41495 | predicted gene, 41495 |
| chr9 | 57431401 | 57432152 | 752 | * | 4,551196 | 5,800093667 | 2,095782 | 2,767508 | 0,005649 | 1 | 0 | Promoter (2-3kb) | 9 | 57428616 | 57437928 | 9313 | 1 | 12858 | ENSMUST00000214154.2 | 2785 | ENSMUSG00000000088 | Cox5a | cytochrome c oxidase subunit 5A |
| chr15 | 52154639 | 52154937 | 299 | * | 4,483559 | 5,800142283 | 2,093294 | 2,770821 | 0,005592 | 1 | 0 | Distal Intergenic | 15 | 52158949 | 52199194 | 40246 | 1 | 239436 | ENSMUST00000037240.3 | -4012 | ENSMUSG00000022315 | Slc30a8 | solute carrier family 30 (zinc transporter), member 8 |
| chr10 | 94952348 | 94952740 | 393 | * | 4,806068 | 5,800156338 | 2,092287 | 2,772161 | 0,005569 | 1 | 0 | Exon (ENSMUST000002 | 10 | 94956444 | 94981687 | 25244 | 2 | 628062 | ENSMUST00000220248.2 | 28947 | ENSMUSG00000112481 | Gm29684 | predicted gene, 29684 |
| chr2 | 76452725 | 76453649 | 925 | * | 5,602952 | 5,800235564 | 2,088686 | 2,776978 | 0,005487 | 1 | 0 | Distal Intergenic | 2 | 76467139 | 76471077 | 3939 | 2 | 23992 | ENSMUST00000140493.2 | 17428 | ENSMUSG00000002731 | Prkra | protein kinase, interferon inducible double stranded RNA dependent activator |
| chr16 | 59588651 | 59589557 | 907 | * | 4,715096 | 5,800301758 | 2,085226 | 2,781618 | 0,005409 | 1 | 0 | Intron (ENSMUST0000( | 16 | 59433691 | 59459754 | 26064 | 2 | 56297 | ENSMUST00000099646.10 | -128897 | ENSMUSG00000022722 | Arl6 | ADP-ribosylation factor-like 6 |
| chr3 | 1,47E+08 | 1,47E+08 | 485 | * | 5,580714 | 5,800445499 | 2,078629 | 2,790515 | 0,005262 | 1 | 0 | Distal Intergenic | 3 | 1,47E+08 | 1,47E+08 | 4401 | 1 | 1,05E+08 | ENSMUST00000195873.2 | 307957 | | Gm37041 | predicted gene, 37041 |
| chr14 | 30438790 | 30439379 | 590 | * | 5,047425 | 5,80047638 | 2,077444 | 2,792122 | 0,005236 | 1 | 0 | Promoter (<=1kb) | 14 | 30438448 | 30540303 | 101856 | 1 | 54650 | ENSMUST00000228006.2 | 342 | ENSMUSG00000006527 | Sfmbt1 | Scm-like with four mbt domains 1 |
| chr1 | 1,23E+08 | 1,23E+08 | 341 | * | 4,680354 | 5,800731704 | 2,064871 | 2,809246 | 0,004966 | 1 | 0 | Distal Intergenic | 1 | 1,23E+08 | 1,23E+08 | 131 | 2 | 1,15E+08 | ENSMUST00000104284.3 | 179925 | ENSMUSG00000077472 | Gm24791 | predicted gene, 24791 |
| chrX | 61609397 | 61610276 | 880 | * | 4,642158 | 5,800780547 | 2,062305 | 2,812766 | 0,004912 | 1 | 0 | Distal Intergenic | 20 | 61554145 | 61570617 | 16473 | 2 | 75545 | ENSMUST00000129479.2 | -38780 | ENSMUSG00000085757 | Tslrn1 | testis specific long noncoding RNA 1 |
| chr7 | 1,1E+08 | 1,1E+08 | 622 | * | 4,669026 | 5,801078243 | 2,048758 | 2,83151 | 0,004633 | 1 | 0 | Promoter (1-2kb) | 7 | 1,1E+08 | 1,1E+08 | 1168 | 2 | 67150 | ENSMUST00000148367.2 | 1046 | ENSMUSG00000030788 | Rnf141 | ring finger protein 141 |
| chr7 | 45337918 | 45338416 | 499 | * | 5,094772 | 5,80125498 | 2,041513 | 2,841645 | 0,004488 | 1 | 0 | Intron (ENSMUST0000( | 7 | 45333853 | 45343851 | 9999 | 1 | 243967 | ENSMUST00000183120.4 | 4065 | ENSMUSG00000070564 | Ntn5 | netrin 5 |
| chr10 | 66901163 | 66901655 | 493 | * | 6,6526 | 5,801355152 | 2,03644 | 2,848772 | 0,004389 | 1 | 0 | Intron (ENSMUST0000( | 10 | 66931904 | 67027527 | 95624 | 1 | 108829 | ENSMUST00000174062.8 | -30249 | ENSMUSG00000037876 | Jmjd1c | jumonji domain containing 1C |
| chr11 | 77487138 | 77487948 | 811 | * | 5,100498 | 5,801547118 | 2,028524 | 2,859984 | 0,004237 | 1 | 0 | Intron (ENSMUST0000( | 11 | 77419988 | 77498641 | 78654 | 2 | 216965 | ENSMUST00000058496.8 | 10693 | ENSMUSG00000017291 | Taok1 | TAO kinase 1 |
| chr5 | 1,43E+08 | 1,43E+08 | 1029 | * | 9,241343 | 5,801789368 | 1,912659 | 3,033364 | 0,002418 | 0,644704154 | 1,425929 | Promoter (1-2kb) | 5 | 1,43E+08 | 1,43E+08 | 34131 | 1 | 170935 | ENSMUST00000110733.9 | -1209 | ENSMUSG00000010825 | Grid2ip | glutamate receptor, ionotropic, delta 2 (Grid2) interacting protein 1 |
| chr6 | 1,18E+08 | 1,18E+08 | 528 | * | 4,976659 | 5,801898786 | 2,014358 | 2,880272 | 0,003973 | 1 | 0 | Promoter (<=1kb) | 6 | 1,18E+08 | 1,18E+08 | 4594 | 1 | 1,05E+08 | ENSMUST00000247298.1 | 778 | | Gm38882 | predicted gene, 38882 |
| chr8 | 86299793 | 86300265 | 473 | * | 5,372765 | 5,802034049 | 2,008922 | 2,888133 | 0,003875 | 1 | 0 | Distal Intergenic | 8 | 86264262 | 86281973 | 17712 | 2 | 56445 | ENSMUST00000034138.7 | -17820 | ENSMUSG00000031701 | Dnaja2 | DnaJ heat shock protein family (Hsp40) member A2 |
| chr15 | 38232622 | 38233264 | 643 | * | 7,468915 | 5,802091004 | 1,865585 | 3,110065 | 0,00187 | 1 | 0 | Distal Intergenic | 15 | 38219447 | 38226979 | 7533 | 1 | 18285 | ENSMUST00000081966.5 | 13175 | ENSMUSG00000061923 | Odf1 | outer dense fiber of sperm tails 1 |
| chr4 | 1,37E+08 | 1,37E+08 | 495 | * | 5,60818 | 5,802408152 | 1,930972 | 3,004915 | 0,002657 | 1 | 0 | Intron (ENSMUST0000( | 4 | 1,37E+08 | 1,37E+08 | 27398 | 2 | 13842 | ENSMUST00000030420.9 | 13709 | ENSMUSG00000028661 | Epha8 | Eph receptor A8 |
| chrX | 97569424 | 97570202 | 779 | * | 5,236947 | 5,802428405 | 1,993665 | 2,910433 | 0,003609 | 1 | 0 | Distal Intergenic | 20 | 97607175 | 97622041 | 14867 | 2 | 94190 | ENSMUST00000154920.2 | 51839 | ENSMUSG00000031214 | Ophn1 | oligophrenin 1 |
| chr11 | 1,17E+08 | 1,17E+08 | 896 | * | 6,539408 | 5,802610171 | 1,986603 | 2,92087 | 0,003491 | 1 | 0 | Distal Intergenic | 11 | 1,17E+08 | 1,17E+08 | 1362 | 1 | 1,03E+08 | ENSMUST00000246631.1 | -12436 | | Gm11734 | predicted gene 11734 |
| chr6 | 17144315 | 17145080 | 766 | * | 5,838904 | 5,802623471 | 1,9867 | 2,920734 | 0,003492 | 1 | 0 | Intron (ENSMUST0000( | 6 | 17097940 | 17171464 | 73525 | 2 | 1,19E+08 | ENSMUST00000136503.2 | 26384 | | Gm53251 | predicted gene, 53251 |
| chr7 | 1,26E+08 | 1,26E+08 | 727 | * | 5,387308 | 5,802695883 | 1,983785 | 2,925063 | 0,003444 | 1 | 0 | Promoter (2-3kb) | 7 | 1,26E+08 | 1,26E+08 | 13884 | 2 | 18020 | ENSMUST00000075671.5 | -2205 | ENSMUSG00000030722 | Nfatc2ip | nuclear factor of activated T cells, cytoplasmic, calcineurin dependent 2 interacting protein |
| chr6 | 43055360 | 43055734 | 375 | * | 6,619978 | 5,803098302 | 1,900871 | 3,052862 | 0,002267 | 1 | 0 | Intron (ENSMUST0000( | 6 | 43092678 | 43093610 | 933 | 1 | 258649 | ENSMUST00000095954.4 | -36944 | ENSMUSG00000094669 | Olfr441 | olfactory receptor 441 |
| chr3 | 138448352 | 138449886 | 1535 | * | 18,665948 | 5,803248053 | 1,1007736 | 5,2719726 | 1,35E-07 | 0,001000466 | 2,3860078 | Promoter (<=1kb) | 3 | 138448530 | 138600245 | 151716 | 1 | 56224 | ENSMUST00000153336.6 | 0 | ENSMUSG00000028152 | Tspan5 | tetraspanin 5 |
| chr15 | 11158512 | 11159359 | 848 | * | 7,065086 | 5,803574434 | 1,920174 | 3,022421 | 0,002508 | 1 | 0 | Intron (ENSMUST0000( | 15 | 11064904 | 11217207 | 152304 | 1 | 239337 | ENSMUST00000228940.2 | 93608 | ENSMUSG00000047497 | Adamts12 | a disintegrin-like and metallopeptidase (reprolysin type) with thrombospondin type 1 motif, 12 |
| chr11 | 1,17E+08 | 1,17E+08 | 1124 | * | 8,398282 | 5,803602166 | 1,934051 | 3,00075 | 0,002693 | 1 | 0 | Promoter (2-3kb) | 11 | 1,17E+08 | 1,17E+08 | 2529 | 1 | 69900 | ENSMUST00000147404.2 | 2354 | ENSMUSG00000020818 | Mfsd11 | major facilitator superfamily domain containing 11 |
| chr15 | 37004626 | 37005346 | 721 | * | 5,283577 | 5,804398245 | 1,955858 | 2,967699 | 0,003 | 1 | 0 | Promoter (<=1kb) | 15 | 37002083 | 37004302 | 2220 | 2 | 68036 | ENSMUST00000228275.2 | -324 | ENSMUSG00000062397 | Zfp706 | zinc finger protein 706 |
| chr15 | 41581193 | 41581825 | 633 | * | 5,578095 | 5,804740985 | 1,971951 | 2,943654 | 0,003244 | 1 | 0 | Intron (ENSMUST0000( | 15 | 41574000 | 41683776 | 109777 | 1 | 170719 | ENSMUST00000229769.2 | 7193 | ENSMUSG00000022307 | Oxr1 | oxidation resistance 1 |
| chr17 | 29392257 | 29393129 | 873 | * | 5,09524 | 5,805133161 | 1,992408 | 2,913627 | 0,003573 | 1 | 0 | Promoter (2-3kb) | 17 | 29381241 | 29395630 | 14390 | 2 | 240058 | ENSMUST00000124917.3 | 2501 | ENSMUSG00000024008 | Cpne5 | copine V |
| chr3 | 88593937 | 88594847 | 911 | * | 7,286673 | 5,805174348 | 1,805512 | 3,215251 | 0,001303 | 0,537150047 | 1,360622 | Promoter (<=1kb) | 3 | 88593144 | 88619310 | 26167 | 1 | 74200 | ENSMUST00000198078.2 | 793 | ENSMUSG00000028060 | Khdc4 | KH domain containing 4, pre-mRNA splicing factor |
| chr7 | 1,45E+08 | 1,45E+08 | 762 | * | 4,834758 | 5,805695394 | 2,024032 | 2,868381 | 0,004126 | 1 | 0 | Distal Intergenic | 7 | 1,44E+08 | 1,44E+08 | 6728 | 2 | 12443 | ENSMUST00000208193.2 | -112416 | ENSMUSG00000070348 | Ccnd1 | cyclin D1 |
| chr17 | 7218088 | 7218853 | 766 | * | 4,907368 | 5,805895474 | 2,036306 | 2,85119 | 0,004356 | 1 | 0 | Promoter (2-3kb) | 17 | 7171812 | 7216081 | 44270 | 2 | 1E+08 | ENSMUST00000231794.2 | -2007 | ENSMUSG00000023806 | Rsph3b | radial spoke 3B homolog (Chlamydomonas) |
| chr2 | 1,53E+08 | 1,53E+08 | 575 | * | 4,541242 | 5,806095348 | 2,049014 | 2,833605 | 0,004603 | 1 | 0 | Distal Intergenic | 2 | 1,53E+08 | 1,53E+08 | 13898 | 2 | 99311 | ENSMUST00000147822.8 | 41076 | ENSMUSG00000056941 | Commd7 | COMM domain containing 7 |
| chr11 | 77506873 | 77507344 | 472 | * | 4,806624 | 5,806302813 | 2,061505 | 2,816536 | 0,004854 | 1 | 0 | Distal Intergenic | 11 | 77419988 | 77498641 | 78654 | 2 | 216965 | ENSMUST00000058496.8 | -8232 | ENSMUSG00000017291 | Taok1 | TAO kinase 1 |
| chr17 | 24109322 | 24109907 | 586 | * | 4,766035 | 5,806452363 | 2,072382 | 2,801826 | 0,005081 | 1 | 0 | Intron (ENSMUST0000( | 17 | 24117695 | 24119738 | 2044 | 1 | 630537 | ENSMUST00000088598.7 | -7788 | ENSMUSG00000096278 | Dcpp2 | demilune cell and parotid protein 2 |
| chr7 | 1,41E+08 | 1,41E+08 | 781 | * | 5,723429 | 5,806456853 | 2,072179 | 2,802102 | 0,005077 | 1 | 0 | Distal Intergenic | 7 | 1,41E+08 | 1,41E+08 | 12530 | 2 | 107702 | ENSMUST00000167493.9 | -6605 | ENSMUSG00000038650 | Rnh1 | ribonuclease/angiogenin inhibitor 1 |
| chr6 | 31797235 | 31797713 | 479 | * | 6,391307 | 5,806619174 | 1,919186 | 3,025563 | 0,002482 | 1 | 0 | Intron (ENSMUST0000( | 6 | 31696316 | 31720029 | 23714 | 1 | 1,05E+08 | ENSMUST00000210985.2 | 100919 | | Gm13846 | predicted gene 13846 |
| chr10 | 75403410 | 75403834 | 425 | * | 5,215586 | 5,806734638 | 2,092525 | 2,774989 | 0,00552 | 1 | 0 | Promoter (<=1kb) | 10 | 75404468 | 75415116 | 10649 | 1 | 14598 | ENSMUST00000145928.8 | -634 | ENSMUSG00000006345 | Ggt1 | gamma-glutamyltransferase 1 |
| chr9 | 1,08E+08 | 1,08E+08 | 338 | * | 4,329966 | 5,806946717 | 2,108187 | 2,754474 | 0,005879 | 1 | 0 | Intron (ENSMUST0000( | 9 | 1,08E+08 | 1,08E+08 | 5440 | 2 | 20350 | ENSMUST00000193665.6 | -6070 | ENSMUSG00000034684 | Sema3f | sema domain, immunoglobulin domain (Ig), short basic domain, secreted, (semaphorin) 3F |
| chr2 | 91607919 | 91608609 | 691 | * | 4,301623 | 5,806969235 | 2,110107 | 2,751978 | 0,005924 | 1 | 0 | Intron (ENSMUST0000( | 2 | 91603577 | 91654864 | 51288 | 1 | 228361 | ENSMUST00000133490.2 | 4342 | ENSMUSG00000040506 | Ambra1 | autophagy/beclin 1 regulator 1 |
| chr11 | 1,14E+08 | 1,14E+08 | 581 | * | 4,434703 | 5,807172946 | 2,125406 | 2,732266 | 0,00629 | 1 | 0 | Exon (ENSMUST00000( | 11 | 1,14E+08 | 1,14E+08 | 1531 | 1 | 1,05E+08 | ENSMUST00000246991.1 | 7320 | | Gm39468 | predicted gene, 39468 |
| chr12 | 1,02E+08 | 1,02E+08 | 720 | * | 4,469661 | 5,807192716 | 2,127506 | 2,729578 | 0,006342 | 1 | 0 | Promoter (<=1kb) | 12 | 1,02E+08 | 1,02E+08 | 25200 | 2 | 110616 | ENSMUST00000159883.2 | 310 | ENSMUSG00000021189 | Atxn3 | ataxin 3 |
| chr10 | 35543189 | 35544005 | 817 | * | 6,298859 | 5,807239952 | 1,901877 | 3,053426 | 0,002262 | 1 | 0 | Distal Intergenic | 10 | 35584671 | 35587888 | 3218 | 2 | 1E+08 | ENSMUST00000080898.4 | 43883 | ENSMUSG00000063953 | Amd2 | S-adenosylmethionine decarboxylase 2 |
| chr13 | 76219925 | 76220375 | 451 | * | 3,973111 | 5,807260803 | 2,133112 | 2,722436 | 0,00648 | 1 | 0 | Exon (ENSMUST00000: | 13 | 76220274 | 76225499 | 5226 | 2 | 77041 | ENSMUST00000224262.2 | 5124 | ENSMUSG00000021592 | Arsk | arylsulfatase K |
| chr7 | 64897097 | 64897493 | 397 | * | 4,135908 | 5,8072941 | 2,135922 | 2,71887 | 0,006551 | 1 | 0 | Distal Intergenic | 7 | 64862565 | 64863177 | 613 | 2 | 75601 | ENSMUST00000205499.2 | -33920 | ENSMUSG00000108587 | 1810049I09Rik | RIKEN cDNA 1810049I09 gene |
| chr5 | 1,16E+08 | 1,16E+08 | 570 | * | 3,930163 | 5,807469587 | 2,15071 | 2,700257 | 0,006929 | 1 | 0 | Exon (ENSMUST00000: | 5 | 1,16E+08 | 1,16E+08 | 5022 | 1 | 231659 | ENSMUST00000125299.2 | 4675 | ENSMUSG00000041638 | Gcn1 | GCN1 activator of EIF2AK4 |
| chr12 | 81019356 | 81020238 | 883 | * | 7,556574 | 5,80749584 | 1,986867 | 2,922942 | 0,003467 | 1 | 0 | Distal Intergenic | 12 | 80999957 | 81015479 | 15523 | 2 | 20493 | ENSMUST00000218162.2 | -3877 | ENSMUSG00000021135 | Slc10a1 | solute carrier family 10 (sodium/bile acid cotransporter family), member 1 |
| chr10 | 85105368 | 85105918 | 551 | * | 3,853714 | 5,807695759 | 2,17044 | 2,675815 | 0,007455 | 1 | 0 | Distal Intergenic | 10 | 84967564 | 85020928 | 53365 | 2 | 12952 | ENSMUST00000020227.11 | -84440 | ENSMUSG00000020038 | Cry1 | cryptochrome 1 (photolyase-like) |
| chr19 | 40329967 | 40330888 | 922 | * | 7,249792 | 5,807808905 | 1,991612 | 2,916135 | 0,003544 | 1 | 0 | Promoter (<=1kb) | 19 | 40322476 | 40329038 | 6563 | 2 | 20411 | ENSMUST00000226028.2 | -929 | ENSMUSG00000025006 | Sorbs1 | sorbin and SH3 domain containing 1 |
| chr3 | 79067477 | 79068165 | 689 | * | 4,169496 | 5,807841974 | 2,1832 | 2,660243 | 0,007808 | 1 | 0 | Intron (ENSMUST0000( | 3 | 79016373 | 79080505 | 64133 | 2 | 76089 | ENSMUST00000152275.8 | 12340 | ENSMUSG00000062232 | Rapgef2 | Rap guanine nucleotide exchange factor (GEF) 2 |
| chr4 | 1,3E+08 | 1,3E+08 | 440 | * | 3,775609 | 5,807866226 | 2,186509 | 2,656228 | 0,007902 | 1 | 0 | Distal Intergenic | 4 | 1,3E+08 | 1,3E+08 | 28014 | 1 | 19244 | ENSMUST00000132386.2 | -22902 | ENSMUSG00000028788 | Ptp4a2 | protein tyrosine phosphatase 4a2 |
| chr9 | 90822373 | 90822966 | 594 | * | 4,093581 | 5,80818181 | 2,218034 | 2,618617 | 0,008829 | 1 | 0 | Distal Intergenic | 9 | 90707355 | 90710913 | 3559 | 2 | 1,03E+08 | ENSMUST00000185312.2 | -111460 | | Gm28703 | predicted gene 28703 |
| chr6 | 33683141 | 33683576 | 436 | * | 4,106735 | 5,808194723 | 2,218316 | 2,61829 | 0,008837 | 1 | 0 | Intron (ENSMUST0000( | 6 | 33678369 | 33839095 | 160727 | 1 | 20336 | ENSMUST00000143190.2 | 4772 | ENSMUSG00000029763 | Exoc4 | exocyst complex component 4 |
| chr2 | 8118592 | 8119266 | 675 | * | 3,719614 | 5,808385174 | 2,237842 | 2,59553 | 0,009445 | 1 | 0 | Distal Intergenic | 2 | 7398887 | 7514374 | 115488 | 2 | 14007 | ENSMUST00000182037.2 | -604218 | ENSMUSG00000002107 | Celf2 | CUGBP, Elav-like family member 2 |
| chr4 | 1,27E+08 | 1,27E+08 | 262 | * | 3,815015 | 5,808442046 | 2,24262 | 2,590025 | 0,009597 | 1 | 0 | Promoter (1-2kb) | 4 | 1,27E+08 | 1,27E+08 | 133 | 2 | 1,15E+08 | ENSMUST00000158359.3 | -1151 | ENSMUSG00000088984 | Gm25604 | predicted gene, 25604 |

| | | | | | | | | | | | | | | | | | | | | | | | |
|---|---|---|---|---|---|---|---|---|---|---|---|---|---|---|---|---|---|---|---|---|---|---|---|
| chr5 | 1,16E+08 | 1,16E+08 | 1060 | * | 7,082225 | 5,809654098 | 2,021102 | 2,874499 | 0,004047 | 1 | 0 | Promoter (<=1kb) | 5 | 1,16E+08 | 1,16E+08 | 1452 | 1 | 11837 | ENSMUST00000155584.2 | 232 | ENSMUSG00000067274 | Rplp0 | ribosomal protein, large, P0 |
| chr14 | 57897026 | 57897780 | 755 | * | 6,549237 | 5,810434837 | 2,034616 | 2,855789 | 0,004293 | 1 | 0 | Intron (ENSMUST0000( | 14 | 57819728 | 57902413 | 82686 | 2 | 57258 | ENSMUST00000089482.12 | 4633 | ENSMUSG00000021952 | Xpo4 | exportin 4 |
| chr6 | 87854885 | 87855639 | 755 | * | 6,189821 | 5,81205655 | 2,064436 | 2,815324 | 0,004873 | 1 | 0 | Distal Intergenic | 6 | 87864796 | 87890577 | 25782 | 1 | 54161 | ENSMUST00000113607.10 | -9157 | ENSMUSG00000030058 | Copg1 | coatomer protein complex, subunit gamma 1 |
| chr4 | 1,35E+08 | 1,35E+08 | 782 | * | 7,087808 | 5,813926505 | 2,103101 | 2,764454 | 0,005702 | 1 | 0 | Distal Intergenic | 4 | 1,35E+08 | 1,35E+08 | 2887 | 2 | 74552 | ENSMUST00000183397.2 | 10013 | ENSMUSG00000028803 | Nipal3 | NIPA-like domain containing 3 |
| chr6 | 37358224 | 37358763 | 540 | * | 5,640179 | 5,814353572 | 2,11236 | 2,752539 | 0,005914 | 1 | 0 | Intron (ENSMUST0000( | 6 | 37311227 | 37313470 | 2244 | 2 | 208647 | ENSMUST00000128534.2 | -44754 | ENSMUSG00000038648 | Creb3l2 | cAMP responsive element binding protein 3-like 2 |
| chr9 | 14075250 | 14075753 | 504 | * | 5,576138 | 5,814526658 | 2,116256 | 2,747553 | 0,006004 | 1 | 0 | Distal Intergenic | 9 | 14014888 | 14020767 | 5880 | 1 | 76412 | ENSMUST00000214881.2 | 60362 | ENSMUSG00000111489 | 1700019J19Rik | RIKEN cDNA 1700019J19 gene |
| chr4 | 1,35E+08 | 1,35E+08 | 579 | * | 5,556637 | 5,815060121 | 2,128503 | 2,731995 | 0,006295 | 1 | 0 | Promoter (<=1kb) | 4 | 1,35E+08 | 1,35E+08 | 31743 | 2 | 230824 | ENSMUST00000105855.2 | 300 | ENSMUSG00000037188 | Grhl3 | grainyhead like transcription factor 3 |
| chr11 | 48709642 | 48710504 | 863 | * | 6,293027 | 5,815708075 | 2,144031 | 2,712512 | 0,006678 | 1 | 0 | Promoter (1-2kb) | 11 | 48697231 | 48708180 | 10950 | 2 | 211007 | ENSMUST00000047145.14 | -1462 | ENSMUSG00000040365 | Trim41 | tripartite motif-containing 41 |
| chr19 | 40685095 | 40685657 | 563 | * | 5,468353 | 5,815752609 | 2,145055 | 2,711237 | 0,006703 | 1 | 0 | Intron (ENSMUST0000( | 19 | 40670111 | 40711637 | 41527 | 1 | 12495 | ENSMUST00000137508.2 | 14984 | ENSMUSG00000048120 | Entpd1 | ectonucleoside triphosphate diphosphohydrolase 1 |
| chr7 | 83359141 | 83359772 | 632 | * | 4,489364 | 5,815777898 | 2,231112 | 2,606672 | 0,009143 | 1 | 0 | Intron (ENSMUST0000( | 7 | 83371718 | 83381615 | 9898 | 2 | 16170 | ENSMUST00000131916.2 | 21843 | ENSMUSG00000001741 | Il16 | interleukin 16 |
| chr7 | 96273161 | 96273873 | 713 | * | 5,150344 | 5,815916647 | 2,149169 | 2,706124 | 0,006807 | 1 | 0 | Intron (ENSMUST0000( | 7 | 96293379 | 96381487 | 88109 | 1 | 23966 | ENSMUST00000131541.2 | -19506 | ENSMUSG00000048078 | Tenm4 | teneurin transmembrane protein 4 |
| chr1 | 96923826 | 96924383 | 558 | * | 5,281882 | 5,816604913 | 2,166641 | 2,684619 | 0,007261 | 1 | 0 | Promoter (<=1kb) | 1 | 96849643 | 96924962 | 75320 | 2 | 67854 | ENSMUST00000178359.3 | 579 | ENSMUSG00000045463 | Slco6b1 | solute carrier organic anion transporter family, member 6b1 |
| chr10 | 59905650 | 59906341 | 692 | * | 4,906703 | 5,816807237 | 2,198635 | 2,645645 | 0,008154 | 1 | 0 | Intron (ENSMUST0000( | 10 | 59942041 | 59963160 | 21120 | 1 | 94214 | ENSMUST00000138976.3 | -35700 | ENSMUSG00000058297 | Spock2 | sparc/osteonectin, cwcv and kazal-like domains proteoglycan 2 |
| chr16 | 37203176 | 37204164 | 989 | * | 5,695739 | 5,816859424 | 2,173514 | 2,676247 | 0,007445 | 1 | 0 | Promoter (1-2kb) | 16 | 36935304 | 37205285 | 269982 | 2 | 207227 | ENSMUST00000114787.8 | 1121 | ENSMUSG00000022829 | Stxbp5l | syntaxin binding protein 5-like |
| chr7 | 1,34E+08 | 1,34E+08 | 679 | * | 5,776613 | 5,816937317 | 2,175254 | 2,674142 | 0,007492 | 1 | 0 | Distal Intergenic | 7 | 1,34E+08 | 1,34E+08 | 2970 | 2 | 71007 | ENSMUST00000205334.2 | -69935 | ENSMUSG00000030994 | D7Ertd443e | DNA segment, Chr 7, ERATO Doi 443, expressed |
| chr5 | 1,24E+08 | 1,24E+08 | 747 | * | 5,73842 | 5,818109444 | 2,208453 | 2,634473 | 0,008427 | 1 | 0 | Promoter (1-2kb) | 5 | 1,24E+08 | 1,24E+08 | 18556 | 2 | 56325 | ENSMUST00000126856.8 | 1085 | ENSMUSG00000029408 | Abcb9 | ATP-binding cassette, sub-family B (MDR/TAP), member 9 |
| chr3 | 87773873 | 87774376 | 504 | * | 4,649501 | 5,818537166 | 2,220873 | 2,619932 | 0,008795 | 1 | 0 | Intron (ENSMUST0000( | 3 | 87766210 | 87792915 | 26706 | 2 | 94315 | ENSMUST00000005015.10 | 18539 | ENSMUSG00000004895 | Prcc | papillary renal cell carcinoma (translocation-associated) |
| chr13 | 91510950 | 91511715 | 766 | * | 4,825422 | 5,819153956 | 2,239597 | 2,598304 | 0,009369 | 1 | 0 | Distal Intergenic | 13 | 91518937 | 91536204 | 17268 | 2 | 320297 | ENSMUST00000244414.1 | 24489 | ENSMUSG00000097182 | A830009L08Rik | RIKEN cDNA A830009L08 gene |
| chr9 | 24988885 | 24989523 | 639 | * | 4,612226 | 5,819675732 | 2,256914 | 2,578599 | 0,00992 | 1 | 0 | Distal Intergenic | 9 | 25019428 | 25063116 | 43689 | 2 | 80517 | ENSMUST00000008573.9 | 73593 | ENSMUSG00000008429 | Herpud2 | HERPUD family member 2 |
| chr1 | 42337689 | 42338397 | 709 | * | 4,556634 | 5,819732172 | 2,25868 | 2,576607 | 0,009978 | 1 | 0 | Exon (ENSMUST000002 | 1 | 42301088 | 42343568 | 42481 | 1 | 1,03E+08 | ENSMUST00000244990.1 | 36601 | | Gm28175 | predicted gene 28175 |
| chr10 | 1,15E+08 | 1,15E+08 | 822 | * | 6,250523 | 5,820093287 | 2,110396 | 2,75782 | 0,005819 | 1 | 0 | Exon (ENSMUST000002 | 10 | 1,15E+08 | 1,15E+08 | 47814 | 1 | 216345 | ENSMUST00000036044.9 | 10121 | ENSMUSG00000034163 | Zfc3h1 | zinc finger, C3H1-type containing |
| chr16 | 10852222 | 10852948 | 727 | * | 6,535514 | 5,820399602 | 2,103018 | 2,767642 | 0,005646 | 1 | 0 | Promoter (2-3kb) | 16 | 10849820 | 10854284 | 4465 | 1 | 1E+08 | ENSMUST00000229764.2 | 2402 | | Gm10832 | predicted gene 10832 |
| chr13 | 97100873 | 97102491 | 1619 | * | 9,56873 | 5,822025778 | 1,772451 | 3,284732 | 0,001021 | 0,484531896 | 1,425929 | Distal Intergenic | 13 | 97061197 | 97087414 | 26218 | 1 | 218476 | ENSMUST00000171324.3 | 39676 | ENSMUSG00000091387 | Gcnt4 | glucosaminyl (N-acetyl) transferase 4, core 2 (beta-1,6-N-acetylglucosaminyltransferase) |
| chr9 | 59659351 | 59659916 | 566 | * | 4,521471 | 5,825958948 | 2,259281 | 2,578678 | 0,009918 | 1 | 0 | Promoter (1-2kb) | 9 | 59658179 | 59836149 | 177971 | 1 | 270163 | ENSMUST00000136740.8 | 1172 | ENSMUSG00000039585 | Myo9a | myosin IXa |
| chr11 | 1,15E+08 | 1,15E+08 | 249 | * | 3,800774 | 5,826093016 | 2,24231 | 2,598255 | 0,00937 | 1 | 0 | Promoter (2-3kb) | 11 | 1,15E+08 | 1,15E+08 | 17893 | 2 | 328035 | ENSMUST00000056153.8 | 2122 | ENSMUSG00000044788 | Fads6 | fatty acid desaturase domain family, member 6 |
| chrX | 40467139 | 40467747 | 609 | * | 3,845294 | 5,826118973 | 2,23877 | 2,602375 | 0,009258 | 1 | 0 | Distal Intergenic | 20 | 40461136 | 40461391 | 256 | 1 | 1,08E+08 | ENSMUST00000121134.2 | 6003 | ENSMUSG00000083515 | Gm14577 | predicted gene 14577 |
| chr15 | 14698704 | 14699273 | 570 | * | 4,445084 | 5,826140861 | 2,098734 | 2,776026 | 0,005503 | 1 | 0 | Distal Intergenic | 15 | 15780080 | 15823510 | 43431 | 1 | 1,03E+08 | ENSMUST00000226912.2 | -1080807 | | 4930557F08Rik | RIKEN cDNA 4930557F08 gene |
| chr1 | 1,71E+08 | 1,71E+08 | 724 | * | 4,244318 | 5,826375837 | 2,208669 | 2,637958 | 0,008341 | 1 | 0 | Distal Intergenic | 1 | 1,71E+08 | 1,71E+08 | 796 | 1 | 16456 | ENSMUST00000155913.2 | 5906 | ENSMUSG00000038235 | F11r | F11 receptor |
| chr4 | 81511526 | 81512058 | 533 | * | 4,244211 | 5,826410496 | 2,203515 | 2,644144 | 0,00819 | 1 | 0 | Distal Intergenic | 4 | 81661383 | 81664938 | 3556 | 1 | 1,03E+08 | ENSMUST00000144875.2 | -149325 | | Gm11412 | predicted gene 11412 |
| chr11 | 63781419 | 63782000 | 582 | * | 7,06501 | 5,826457744 | 1,984642 | 2,935773 | 0,003327 | 1 | 0 | Intron (ENSMUST0000( | 11 | 63776618 | 63813116 | 36499 | 2 | 54710 | ENSMUST00000094103.4 | 31116 | ENSMUSG00000070407 | Hs3st3b1 | heparan sulfate (glucosamine) 3-O-sulfotransferase 3B1 |
| chr16 | 20058966 | 20059571 | 606 | * | 4,377747 | 5,826494002 | 2,194329 | 2,655251 | 0,007925 | 1 | 0 | Promoter (<=1kb) | 16 | 20059777 | 20060864 | 1088 | 1 | 1,15E+08 | ENSMUST00000231237.2 | -206 | | Gm49566 | predicted gene, 49566 |
| chr2 | 95737761 | 95738196 | 436 | * | 4,089549 | 5,826594117 | 2,185123 | 2,666483 | 0,007665 | 1 | 0 | Distal Intergenic | 2 | 95668433 | 95670939 | 2507 | 1 | 1,03E+08 | ENSMUST00000138883.2 | 69328 | ENSMUSG00000087537 | Gm13799 | predicted gene 13799 |
| chr18 | 34752035 | 34752870 | 836 | * | 4,199898 | 5,826706852 | 2,172102 | 2,68252 | 0,007307 | 1 | 0 | Intron (ENSMUST0000( | 18 | 34732883 | 34757606 | 24724 | 2 | 78656 | ENSMUST00000152612.8 | 4736 | ENSMUSG00000003778 | Brd8 | bromodomain containing 8 |
| chrX | 1,29E+08 | 1,29E+08 | 500 | * | 4,025738 | 5,826806696 | 2,163213 | 2,693589 | 0,007069 | 1 | 0 | Distal Intergenic | 20 | 1,3E+08 | 1,3E+08 | 129 | 2 | 1,15E+08 | ENSMUST00000104191.3 | 318528 | ENSMUSG00000077379 | Gm25170 | predicted gene, 25170 |
| chr11 | 99198233 | 99198888 | 656 | * | 3,867061 | 5,826883667 | 2,153577 | 2,705678 | 0,006817 | 1 | 0 | Distal Intergenic | 11 | 99206342 | 99213777 | 7436 | 2 | 70810 | ENSMUST00000038004.3 | 14889 | ENSMUSG00000035831 | Krt25 | keratin 25 |
| chrX | 85610194 | 85611081 | 888 | * | 4,884159 | 5,827137093 | 2,130194 | 2,735496 | 0,006229 | 1 | 0 | Distal Intergenic | 20 | 85293862 | 85348699 | 54838 | 2 | 434903 | ENSMUST00000096378.3 | -261495 | ENSMUSG00000035427 | Mageb4 | MAGE family member B4 |
| chr2 | 1,19E+08 | 1,19E+08 | 574 | * | 4,926685 | 5,827297162 | 2,113602 | 2,757046 | 0,005833 | 1 | 0 | Intron (ENSMUST0000( | 2 | 1,19E+08 | 1,19E+08 | 11726 | 2 | 72112 | ENSMUST00000110820.3 | 3330 | ENSMUSG00000027317 | Ppp1r14d | protein phosphatase 1, regulatory inhibitor subunit 14D |
| chr1 | 59290464 | 59291042 | 579 | * | 4,219343 | 5,827341724 | 2,110741 | 2,760804 | 0,005766 | 1 | 0 | Distal Intergenic | 1 | 59296065 | 59392152 | 96088 | 1 | 271697 | ENSMUST00000160662.8 | -5023 | ENSMUSG00000026023 | Cdk15 | cyclin-dependent kinase 15 |
| chr3 | 44353817 | 44354459 | 643 | * | 5,684034 | 5,827511922 | 2,096707 | 2,779364 | 0,005447 | 1 | 0 | Distal Intergenic | 3 | 45281480 | 45332640 | 51161 | 2 | 72511 | ENSMUST00000191843.2 | 978181 | ENSMUSG00000097040 | 2610316D01Rik | RIKEN cDNA 2610316D01 gene |
| chr4 | 61430275 | 61430635 | 361 | * | 4,14296 | 5,82754968 | 2,093269 | 2,783947 | 0,00537 | 1 | 0 | Distal Intergenic | 4 | 61434757 | 61437704 | 2948 | 2 | 1E+08 | ENSMUST00000095051.6 | 7069 | ENSMUSG00000078675 | Mup16 | major urinary protein 16 |
| chr14 | 73846704 | 73847793 | 1090 | * | 4,974439 | 5,827579044 | 2,089769 | 2,788624 | 0,005293 | 1 | 0 | Distal Intergenic | 14 | 73827948 | 73828473 | 526 | 1 | 20916 | ENSMUST00000161082.2 | 18756 | ENSMUSG00000022110 | Sucla2 | succinate-Coenzyme A ligase, ADP-forming, beta subunit |
| chr10 | 44873488 | 44874147 | 660 | * | 4,567732 | 5,828065911 | 2,051766 | 2,840512 | 0,004504 | 1 | 0 | Distal Intergenic | 10 | 44900398 | 44904248 | 3851 | 1 | 1,05E+08 | ENSMUST00000214642.2 | -26251 | | Gm40639 | predicted gene, 40639 |
| chr4 | 1,24E+08 | 1,24E+08 | 671 | * | 7,124249 | 5,828140434 | 1,882811 | 3,095446 | 0,001965 | 1 | 0 | Promoter (<=1kb) | 4 | 1,23E+08 | 1,24E+08 | 101938 | 2 | 11426 | ENSMUST00000238866.2 | 0 | ENSMUSG00000028649 | Macf1 | microtubule-actin crosslinking factor 1 |
| chr11 | 96682242 | 96683192 | 951 | * | 5,583842 | 5,828468536 | 2,023569 | 2,880292 | 0,003973 | 1 | 0 | Promoter (2-3kb) | 11 | 96680075 | 96699366 | 19292 | 1 | 12412 | ENSMUST00000079702.4 | 2167 | ENSMUSG00000018666 | Cbx1 | chromobox 1 |
| chr5 | 24408221 | 24408825 | 605 | * | 5,088136 | 5,828677807 | 2,009405 | 2,900699 | 0,003723 | 1 | 0 | Distal Intergenic | 5 | 24382902 | 24386430 | 3529 | 1 | 231042 | ENSMUST00000124150.2 | 25319 | ENSMUSG00000048439 | Nupl2 | nucleoporin like 2 |
| chr11 | 84779672 | 84780464 | 793 | * | 6,037745 | 5,82896864 | 1,903051 | 3,062961 | 0,002192 | 1 | 0 | Intron (ENSMUST0000( | 11 | 84773412 | 84801948 | 28537 | 1 | 66196 | ENSMUST00000141173.2 | 6260 | ENSMUSG00000020527 | Myo19 | myosin XIX |
| chr1 | 1,79E+08 | 1,79E+08 | 478 | * | 5,220631 | 5,829155529 | 1,978759 | 2,945865 | 0,003221 | 1 | 0 | Intron (ENSMUST0000( | 1 | 1,79E+08 | 1,79E+08 | 32096 | 2 | 69726 | ENSMUST00000129393.2 | -84511 | ENSMUSG00000055067 | Smyd3 | SET and MYND domain containing 3 |
| chr14 | 52266121 | 52266921 | 801 | * | 6,144957 | 5,829250975 | 1,973226 | 2,954172 | 0,003135 | 1 | 0 | Distal Intergenic | 14 | 52272539 | 52273600 | 1062 | 2 | 258420 | ENSMUST00000214342.2 | 6679 | ENSMUSG00000044286 | Olfr221 | olfactory receptor 221 |
| chr18 | 69733053 | 69734149 | 1097 | * | 5,38591 | 5,829461672 | 1,961497 | 2,971946 | 0,002959 | 1 | 0 | Promoter (<=1kb) | 18 | 69732607 | 69785485 | 52879 | 1 | 21413 | ENSMUST00000145918.5 | 446 | ENSMUSG00000053477 | Tcf4 | transcription factor 4 |
| chr16 | 33474247 | 33475049 | 803 | * | 5,556356 | 5,829568464 | 1,956114 | 2,980178 | 0,002881 | 1 | 0 | Intron (ENSMUST0000( | 16 | 33504740 | 33587183 | 82444 | 1 | 77446 | ENSMUST00000232568.2 | -29691 | ENSMUSG00000075254 | Heg1 | heart development protein with EGF-like domains 1 |
| chr9 | 1,14E+08 | 1,14E+08 | 943 | * | 6,420258 | 5,829795877 | 1,924722 | 3,028902 | 0,002454 | 1 | 0 | Promoter (<=1kb) | 9 | 1,13E+08 | 1,14E+08 | 56516 | 2 | 18571 | ENSMUST00000035086.13 | -183 | ENSMUSG00000032504 | Pdcd6ip | programmed cell death 6 interacting protein |
| chr5 | 37664903 | 37665626 | 724 | * | 5,865833 | 5,830140431 | 1,925801 | 3,027385 | 0,002467 | 1 | 0 | Intron (ENSMUST0000( | 5 | 37573203 | 37575259 | 2057 | 1 | 68525 | ENSMUST00000159854.2 | 91700 | ENSMUSG00000050248 | Evc2 | EvC ciliary complex subunit 2 |
| chr4 | 56761034 | 56761667 | 634 | * | 6,057407 | 5,830195064 | 1,923524 | 3,030996 | 0,002437 | 1 | 0 | Promoter (1-2kb) | 4 | 56751015 | 56763330 | 12316 | 2 | 230233 | ENSMUST00000150002.8 | 1663 | ENSMUSG00000028431 | Elp1 | elongator complex protein 1 |
| chr16 | 32897246 | 32898069 | 824 | * | 5,68982 | 5,830503475 | 1,945342 | 2,997161 | 0,002725 | 1 | 0 | Intron (ENSMUST0000( | 16 | 32911656 | 32925252 | 13597 | 1 | 239833 | ENSMUST00000153549.2 | -13587 | ENSMUSG00000022802 | Lmln | leishmanolysin-like (metallopeptidase M8 family) |
| chr15 | 74878849 | 74879632 | 784 | * | 5,614398 | 5,830531416 | 1,946091 | 2,996022 | 0,002735 | 1 | 0 | Promoter (<=1kb) | 15 | 74879152 | 74896259 | 17108 | 1 | 1,03E+08 | ENSMUST00000190861.2 | 0 | | Gm28068 | predicted gene 28068 |
| chr9 | 51387570 | 51388401 | 832 | * | 5,869633 | 5,831912152 | 1,990937 | 2,92923 | 0,003398 | 1 | 0 | Distal Intergenic | 9 | 51200986 | 51240224 | 39239 | 2 | 69809 | ENSMUST00000238450.3 | -147346 | ENSMUSG00000036027 | 1810046K07Rik | RIKEN cDNA 1810046K07 gene |
| chr11 | 1,02E+08 | 1,02E+08 | 598 | * | 5,044258 | 5,834356667 | 2,092546 | 2,788162 | 0,005301 | 1 | 0 | Intron (ENSMUST0000( | 11 | 1,02E+08 | 1,02E+08 | 8220 | 2 | 72349 | ENSMUST00000125794.2 | 3599 | ENSMUSG00000003518 | Dusp3 | dual specificity phosphatase 3 (vaccinia virus phosphatase VH1-related) |
| chr4 | 47140566 | 47141397 | 832 | * | 5,443045 | 5,835043063 | 2,12696 | 2,743372 | 0,006081 | 1 | 0 | Distal Intergenic | 4 | 47091923 | 47123043 | 31121 | 1 | 230145 | ENSMUST00000107744.2 | 48643 | ENSMUSG00000039774 | Galnt12 | polypeptide N-acetylgalactosaminyltransferase 12 |
| chr3 | 94730945 | 94731687 | 743 | * | 4,622456 | 5,835385369 | 2,145782 | 2,719468 | 0,006539 | 1 | 0 | Distal Intergenic | 3 | 94744878 | 94789637 | 44760 | 1 | 229584 | ENSMUST00000107270.9 | -13191 | ENSMUSG00000038902 | Pogz | pogo transposable element with ZNF domain |
| chr3 | 86217658 | 86218422 | 765 | * | 4,551515 | 5,835898231 | 2,175719 | 2,682285 | 0,007312 | 1 | 0 | Exon (ENSMUST000001 | 3 | 86182539 | 86182669 | 131 | 2 | 1,15E+08 | ENSMUST00000104556.3 | -34989 | ENSMUSG00000077744 | Gm26343 | predicted gene, 26343 |
| chr5 | 70876221 | 70876790 | 570 | * | 5,384141 | 5,838547762 | 1,947129 | 2,998542 | 0,002713 | 1 | 0 | Distal Intergenic | 5 | 70909118 | 70999547 | 90430 | 2 | 14405 | ENSMUST00000199705.2 | 122757 | ENSMUSG00000001260 | Gabrg1 | gamma-aminobutyric acid (GABA) A receptor, subunit gamma 1 |
| chr4 | 1,32E+08 | 1,32E+08 | 990 | * | 8,221173 | 5,839847627 | 1,846358 | 3,162902 | 0,001562 | 1 | 0 | 3' UTR | 4 | 1,32E+08 | 1,32E+08 | 18470 | 2 | 230784 | ENSMUST00000030724.9 | 16382 | ENSMUSG00000028893 | Sesn2 | sestrin 2 |
| chr16 | 58862383 | 58862716 | 334 | * | 3,895725 | 5,843247268 | 2,244262 | 2,603639 | 0,009224 | 1 | 0 | Promoter (2-3kb) | 16 | 58855926 | 58860130 | 4205 | 2 | 258319 | ENSMUST00000207673.4 | -2253 | ENSMUSG00000043357 | Olfr187 | olfactory receptor 187 |
| chr5 | 73973930 | 73974426 | 497 | * | 4,714325 | 5,843929491 | 2,216442 | 2,636626 | 0,008374 | 1 | 0 | Distal Intergenic | 5 | 74070239 | 74077764 | 7526 | 1 | 69402 | ENSMUST00000202487.2 | -95813 | ENSMUSG00000107027 | 1700019F05Rik | RIKEN cDNA 1700019F05 gene |
| chrX | 1,03E+08 | 1,03E+08 | 571 | * | 4,218686 | 5,844383814 | 2,199646 | 2,656966 | 0,007885 | 1 | 0 | Exon (ENSMUST000002 | 20 | 1,03E+08 | 1,03E+08 | 2895 | 1 | 70252 | ENSMUST00000235019.2 | 24349 | ENSMUSG00000097571 | Jpx | Jpx transcript, Xist activator (non-protein coding) |
| chr17 | 63945289 | 63946018 | 730 | * | 4,49056 | 5,844601114 | 2,19207 | 2,666247 | 0,00767 | 1 | 0 | Distal Intergenic | 17 | 63364447 | 63807012 | 442566 | 2 | 50758 | ENSMUST00000024761.13 | -138277 | ENSMUSG00000023965 | Fbxl17 | F-box and leucine-rich repeat protein 17 |
| chr16 | 75097621 | 75098146 | 526 | * | 4,354727 | 5,845115974 | 2,172964 | 2,689928 | 0,007147 | 1 | 0 | Distal Intergenic | 16 | 75179597 | 75244082 | 64486 | 2 | 268902 | ENSMUST00000243278.1 | 145936 | ENSMUSG00000052516 | Robo2 | roundabout guidance receptor 2 |
| chr3 | 86132294 | 86132959 | 666 | * | 4,799636 | 5,845605091 | 2,156535 | 2,710647 | 0,006715 | 1 | 0 | Promoter (<=1kb) | 3 | 86132596 | 86450018 | 317423 | 1 | 80877 | ENSMUST00000212390.2 | 0 | ENSMUSG00000028080 | Lrba | LPS-responsive beige-like anchor |
| chr11 | 1,04E+08 | 1,04E+08 | 467 | * | 4,957907 | 5,846070716 | 2,140832 | 2,730747 | 0,006319 | 1 | 0 | Intron (ENSMUST0000( | 11 | 1,04E+08 | 1,04E+08 | 43808 | 1 | 22415 | ENSMUST00000000127.6 | 13866 | ENSMUSG00000000125 | Wnt3 | wingless-type MMTV integration site family, member 3 |
| chr2 | 92859666 | 92860171 | 506 | * | 5,138757 | 5,847920203 | 2,086455 | 2,802802 | 0,005066 | 1 | 0 | Promoter (<=1kb) | 2 | 92859693 | 92863784 | 4092 | 1 | 433456 | ENSMUST00000099699.8 | 0 | ENSMUSG00000075027 | 4631405J19Rik | RIKEN cDNA 4631405J19 gene |
| chr2 | 73664634 | 73665185 | 552 | * | 5,343012 | 5,848518151 | 2,07016 | 2,825153 | 0,004726 | 1 | 0 | Intron (ENSMUST0000( | 2 | 73646853 | 73675849 | 28997 | 2 | 11909 | ENSMUST00000124737.8 | 10664 | ENSMUSG00000027104 | Atf2 | activating transcription factor 2 |
| chr6 | 1,11E+08 | 1,11E+08 | 455 | * | 5,72943 | 5,848955484 | 2,05866 | 2,841147 | 0,004495 | 1 | 0 | Distal Intergenic | 6 | 1,11E+08 | 1,11E+08 | 851510 | 1 | 108073 | ENSMUST00000174018.8 | -34215 | ENSMUSG00000056755 | Grm7 | glutamate receptor, metabotropic 7 |
| chr2 | 72992165 | 72992898 | 734 | * | 6,996555 | 5,85302426 | 1,875677 | 3,120486 | 0,001806 | 1 | 0 | Intron (ENSMUST0000( | 2 | 72923546 | 73043304 | 119759 | 2 | 67059 | ENSMUST00000112055.8 | 50406 | ENSMUSG00000027108 | Ola1 | Obg-like ATPase 1 |
| chr18 | 65735347 | 65736299 | 953 | * | 6,847562 | 5,853606593 | 1,885874 | 3,103923 | 0,00191 | 1 | 0 | Intron (ENSMUST0000( | 18 | 65717008 | 65756264 | 39257 | 1 | 328977 | ENSMUST00000182973.2 | 18339 | ENSMUSG00000042439 | Zfp532 | zinc finger protein 532 |
| chr2 | 1,65E+08 | 1,65E+08 | 591 | * | 3,689983 | 5,853988605 | 2,262546 | 2,587346 | 0,009672 | 1 | 0 | Promoter (<=1kb) | 2 | 1,65E+08 | 1,65E+08 | 479 | 1 | 19025 | ENSMUST00000140213.2 | -499 | ENSMUSG00000017760 | Ctsa | cathepsin A |
| chr13 | 58781008 | 58781498 | 491 | * | 4,020706 | 5,854001709 | 2,257552 | 2,593075 | 0,009512 | 1 | 0 | Distal Intergenic | 13 | 58729504 | 58758691 | 29188 | 2 | 114304 | ENSMUST00000148396.2 | -22317 | ENSMUSG00000021553 | Slc28a3 | solute carrier family 28 (sodium-coupled nucleoside transporter), member 3 |
| chrX | 1,51E+08 | 1,51E+08 | 517 | * | 3,732299 | 5,8540408 | 2,248295 | 2,603769 | 0,00922 | 1 | 0 | Distal Intergenic | 20 | 1,51E+08 | 1,51E+08 | 1484 | 1 | 20591 | ENSMUST00000155793.2 | 7401 | ENSMUSG00000025332 | Kdm5c | lysine (K)-specific demethylase 5C |
| chr8 | 1,2E+08 | 1,2E+08 | 297 | * | 3,936275 | 5,85414884 | 2,214987 | 2,642973 | 0,008218 | 1 | 0 | Intron (ENSMUST0000( | 8 | 1,2E+08 | 1,2E+08 | 34299 | 1 | 69047 | ENSMUST00000212454.2 | 16305 | ENSMUSG00000034112 | Atp2c2 | ATPase, Ca++ transporting, type 2C, member 2 |
| chr2 | 1,27E+08 | 1,27E+08 | 484 | * | 4,129145 | 5,85415071 | 2,209407 | 2,649648 | 0,008058 | 1 | 0 | Intron (ENSMUST0000( | 2 | 1,27E+08 | 1,27E+08 | 84547 | 2 | 58800 | ENSMUST00000028843.12 | 12157 | ENSMUSG00000027365 | Trpm7 | transient receptor potential cation channel, subfamily M, member 7 |
| chr15 | 74427186 | 74427591 | 406 | * | 4,146783 | 5,854165272 | 2,200798 | 2,660019 | 0,007814 | 1 | 0 | Intron (ENSMUST0000( | 15 | 74433133 | 74445968 | 12836 | 1 | 107831 | ENSMUST00000189353.7 | -5542 | ENSMUSG00000034730 | Adgrb1 | adhesion G protein-coupled receptor B1 |
| chr18 | 33961694 | 33961966 | 273 | * | 3,634584 | 5,854176523 | 2,202791 | 2,657618 | 0,00787 | 1 | 0 | Exon (ENSMUST00000( | 18 | 33935593 | 33965847 | 30255 | 2 | 13824 | ENSMUST00000236514.2 | 3881 | ENSMUSG00000024376 | Epb41l4a | erythrocyte membrane protein band 4.1 like 4a |
| chrX | 1,03E+08 | 1,03E+08 | 710 | * | 4,286913 | 5,854196414 | 2,191794 | 2,670961 | 0,007563 | 1 | 0 | Intron (ENSMUST0000( | 20 | 1,03E+08 | 1,03E+08 | 133190 | 2 | 11306 | ENSMUST00000033695.6 | 92138 | ENSMUSG00000031333 | Abcb7 | ATP-binding cassette, sub-family B (MDR/TAP), member 7 |
| chr4 | 1,09E+08 | 1,09E+08 | 459 | * | 4,404237 | 5,854301964 | 2,163433 | 2,706024 | 0,006809 | 1 | 0 | Promoter (<=1kb) | 4 | 1,09E+08 | 1,09E+08 | 4864 | 2 | 70533 | ENSMUST00000139464.2 | 941 | ENSMUSG00000028568 | Btf3l4 | basic transcription factor 3-like 4 |
| chr9 | 63343921 | 63344530 | 610 | * | 4,377791 | 5,854357456 | 2,14682 | 2,72699 | 0,006391 | 1 | 0 | Intron (ENSMUST0000( | 9 | 63349601 | 63349742 | 142 | 1 | 1,15E+08 | ENSMUST00000175396.3 | -5071 | ENSMUSG00000093137 | Gm25995 | predicted gene, 25995 |
| chr18 | 55539187 | 55539418 | 232 | * | 3,900491 | 5,854384606 | 2,142034 | 2,733096 | 0,006274 | 1 | 0 | Distal Intergenic | 18 | 55190657 | 55212486 | 21830 | 1 | 1E+08 | ENSMUST00000181765.2 | 348530 | | Gm4221 | predicted gene 4221 |
| chr16 | 79673934 | 79674441 | 508 | * | 3,961527 | 5,85443156 | 2,131906 | 2,746102 | 0,006031 | 1 | 0 | Distal Intergenic | 16 | 79796264 | 79797434 | 1171 | 1 | 78654 | ENSMUST00000191029.2 | -121823 | ENSMUSG00000101810 | 1700066C05Rik | RIKEN cDNA 1700066C05 gene |
| chr13 | 9528869 | 9529459 | 591 | * | 4,201956 | 5,854486726 | 2,114966 | 2,768123 | 0,005638 | 1 | 0 | Intron (ENSMUST0000( | 13 | 9660842 | 9716897 | 56056 | 1 | 208440 | ENSMUST00000222280.2 | -131383 | ENSMUSG00000048264 | Dip2c | disco interacting protein 2 homolog C |
| chr12 | 81192627 | 81193197 | 571 | * | 4,972138 | 5,854489867 | 2,11343 | 2,770137 | 0,005603 | 1 | 0 | Intron (ENSMUST0000( | 12 | 81212925 | 81213008 | 84 | 1 | 1,01E+08 | ENSMUST00000174932.3 | -19728 | ENSMUSG00000092673 | Mir3067 | microRNA 3067 |
| chr1 | 1,92E+08 | 1,92E+08 | 403 | * | 4,677478 | 5,854516263 | 2,108555 | 2,776554 | 0,005494 | 1 | 0 | Intron (ENSMUST0000( | 1 | 1,92E+08 | 1,92E+08 | 75420 | 1 | 16510 | ENSMUST00000151152.3 | -40357 | ENSMUSG00000058248 | Kcnh1 | potassium voltage-gated channel, subfamily H (eag-related), member 1 |
| chr9 | 65509966 | 65510569 | 604 | * | 4,23323 | 5,854560944 | 2,099132 | 2,789039 | 0,005286 | 1 | 0 | Distal Intergenic | 9 | 65494508 | 65495562 | 1055 | 1 | 208084 | ENSMUST00000141046.3 | 15458 | ENSMUSG00000041064 | Pif1 | PIF1 5'-to-3' DNA helicase |
| chrX | 46868767 | 46869473 | 707 | * | 4,860995 | 5,854577503 | 2,096147 | 2,793018 | 0,005222 | 1 | 0 | Distal Intergenic | 20 | 46930807 | 46981273 | 50467 | 2 | 93761 | ENSMUST00000153548.9 | 111800 | ENSMUSG00000031099 | Smarca1 | SWI/SNF related, matrix associated, actin dependent regulator of chromatin, subfamily a, member 1 |
| chr10 | 42950081 | 42950738 | 658 | * | 4,446323 | 5,854638294 | 2,083038 | 2,810625 | 0,004945 | 1 | 0 | Exon (ENSMUST00000( | 10 | 42911069 | 42924186 | 13118 | 1 | 1,15E+08 | ENSMUST00000213469.2 | 39012 | | Gm47815 | predicted gene, 47815 |
| chr9 | 1,23E+08 | 1,23E+08 | 590 | * | 5,160907 | 5,854644113 | 2,079094 | 2,815959 | 0,004863 | 1 | 0 | Intron (ENSMUST0000( | 9 | 1,23E+08 | 1,23E+08 | 62717 | 1 | 83493 | ENSMUST00000238984.2 | 8355 | ENSMUSG00000025240 | Sacm1l | SAC1 suppressor of actin mutations 1-like (yeast) |
| chr4 | 6222634 | 6223689 | 1056 | * | 5,131384 | 5,854667627 | 2,072429 | 2,825026 | 0,004728 | 1 | 0 | Distal Intergenic | 4 | 6191098 | 6221688 | 30591 | 1 | 68053 | ENSMUST00000029907.6 | 31536 | ENSMUSG00000028243 | Ubxn2b | UBX domain protein 2B |
| chrX | 1,34E+08 | 1,34E+08 | 669 | * | 5,64602 | 5,854830229 | 1,968673 | 2,973999 | 0,002939 | 1 | 0 | Distal Intergenic | 20 | 1,34E+08 | 1,34E+08 | 11825 | 2 | 83454 | ENSMUST00000143059.2 | 38819 | ENSMUSG00000009941 | Nxf2 | nuclear RNA export factor 2 |
| chrX | 1,66E+08 | 1,66E+08 | 702 | * | 4,868786 | 5,854892043 | 2,028657 | 2,886092 | 0,003901 | 1 | 0 | Promoter (<=1kb) | 20 | 1,66E+08 | 1,66E+08 | 36428 | 2 | 110639 | ENSMUST00000026839.5 | 0 | ENSMUSG00000025742 | Prps2 | phosphoribosyl pyrophosphate synthetase 2 |
| chr3 | 62053198 | 62053685 | 488 | * | 5,081747 | 5,854904984 | 2,028097 | 2,886895 | 0,003891 | 1 | 0 | Distal Intergenic | 3 | 62245765 | 62369642 | 123878 | 1 | 622434 | ENSMUST00000079300.13 | -192080 | ENSMUSG00000036885 | Arhgef26 | Rho guanine nucleotide exchange factor (GEF) 26 |
| chr17 | 30948979 | 30949485 | 507 | * | 6,784758 | 5,854983526 | 1,93353 | 3,028131 | 0,002461 | 1 | 0 | Intron (ENSMUST0000( | 17 | 30940015 | 30948858 | 8844 | 1 | 13417 | ENSMUST00000236948.2 | 8964 | ENSMUSG00000033826 | Dnah8 | dynein, axonemal, heavy chain 8 |
| chr3 | 66455921 | 66456428 | 508 | * | 4,973474 | 5,855179931 | 1,979186 | 2,958377 | 0,003093 | 1 | 0 | Distal Intergenic | 3 | 65960979 | 66204258 | 243280 | 2 | 72789 | ENSMUST00000029419.8 | -251663 | ENSMUSG00000027831 | Veph1 | ventricular zone expressed PH domain-containing 1 |
| chr14 | 20743438 | 20744215 | 778 | * | 5,725808 | 5,855223542 | 1,971576 | 2,96982 | 0,00298 | 1 | 0 | Promoter (<=1kb) | 14 | 20745036 | 20751084 | 6049 | 1 | 73068 | ENSMUST00000048016.3 | -821 | ENSMUSG00000039357 | Fut11 | fucosyltransferase 11 |
| chr12 | 88825174 | 88825877 | 704 | * | 5,179644 | 5,855262151 | 1,966171 | 2,978003 | 0,002901 | 1 | 0 | Intron (ENSMUST0000( | 12 | 88762209 | 88820013 | 57805 | 1 | 18191 | ENSMUST00000164072.2 | 62965 | ENSMUSG00000066392 | Nrxn3 | neurexin III |

| | | | | | | | | | | | | | | | | | | | | | | | |
|---|---|---|---|---|---|---|---|---|---|---|---|---|---|---|---|---|---|---|---|---|---|---|---|
| chr2 | 1,53E+08 | 1,53E+08 | 1014 | * | 9,226388 | 5,855508659 | 1,721099 | 3,402192 | 0,000668 | 0,416183238 | 1,425929 | Intron (ENSMUST0000 | 2 | 1,53E+08 | 1,53E+08 | 16182 | 2 | 329540 | ENSMUST00000156786.2 | 12621 | ENSMUSG00000061411 | Nol4l | nucleolar protein 4-like |
| chr10 | 84749521 | 84750131 | 611 | * | 6,435118 | 5,855863631 | 1,887326 | 3,10273 | 0,001917 | 1 | 0 | Distal Intergenic | 10 | 84753480 | 84852311 | 98832 | 1 | 237422 | ENSMUST00000038523.15 | -3349 | ENSMUSG00000035620 | Ric8b | RIC8 guanine nucleotide exchange factor B |
| chr10 | 21875822 | 21876706 | 885 | * | 6,451418 | 5,855869936 | 1,886513 | 3,104071 | 0,001909 | 1 | 0 | Downstream (<=300bp | 10 | 21872490 | 21874139 | 1650 | 1 | 20393 | ENSMUST00000135195.2 | 3332 | ENSMUSG00000019970 | Sgk1 | serum/glucocorticoid regulated kinase 1 |
| chr9 | 1,2E+08 | 1,2E+08 | 1259 | * | 6,845005 | 5,856047238 | 1,868171 | 3,134642 | 0,001721 | 1 | 0 | Promoter (<=1kb) | 9 | 1,2E+08 | 1,2E+08 | 847 | 1 | 67115 | ENSMUST00000215063.2 | 0 | ENSMUSG00000025794 | Rpl14 | ribosomal protein L14 |
| chr13 | 93604799 | 93605494 | 696 | * | 7,052526 | 5,856278026 | 1,845615 | 3,173077 | 0,001508 | 1 | 0 | Intron (ENSMUST0000 | 13 | 93598354 | 93598692 | 339 | 1 | 1,03E+08 | ENSMUST00000223458.2 | 6445 | ENSMUSG00000113247 | Gm30155 | predicted gene, 30155 |
| chr8 | 87229385 | 87230152 | 768 | * | 7,200085 | 5,856331003 | 1,839251 | 3,184085 | 0,001452 | 1 | 0 | Intron (ENSMUST0000 | 8 | 87231212 | 87261376 | 30165 | 2 | 244562 | ENSMUST00000156066.8 | 31224 | ENSMUSG00000036872 | Abcc12 | ATP-binding cassette, sub-family C (CFTR/MRP), member 12 |
| chr10 | 86542944 | 86543759 | 816 | * | 6,007835 | 5,856726727 | 1,949442 | 3,004309 | 0,002662 | 1 | 0 | Promoter (1-2kb) | 10 | 86541708 | 86560431 | 18724 | 1 | 103220 | ENSMUST00000061458.9 | 1236 | ENSMUSG00000044937 | Ttc41 | tetratricopeptide repeat domain 41 |
| chr4 | 11088366 | 11089646 | 1281 | * | 8,68471 | 5,856795188 | 1,799468 | 3,254736 | 0,001135 | 0,500829828 | 1,449706 | Distal Intergenic | 4 | 11070254 | 11076205 | 5952 | 2 | 76947 | ENSMUST00000148127.2 | -12161 | ENSMUSG00000050323 | Ndufaf6 | NADH:ubiquinone oxidoreductase complex assembly factor 6 |
| chr18 | 69817756 | 69819174 | 1419 | * | 8,83955 | 5,857015783 | 1,783369 | 3,284241 | 0,001023 | 0,410415129 | 2,237403 | Promoter (2-3kb) | 18 | 69815451 | 69816704 | 1254 | 1 | 21413 | ENSMUST00000136019.3 | 2305 | ENSMUSG00000053477 | Tcf4 | transcription factor 4 |
| chr16 | 31276263 | 31276868 | 606 | * | 6,145385 | 5,857317827 | 1,963593 | 2,982959 | 0,002855 | 1 | 0 | 3' UTR | 16 | 31254909 | 31275568 | 20660 | 1 | 71911 | ENSMUST00000149039.2 | 21354 | ENSMUSG00000046598 | Bdh1 | 3-hydroxybutyrate dehydrogenase, type 1 |
| chr6 | 28546899 | 28547276 | 378 | * | 5,491856 | 5,857827892 | 1,975784 | 2,964812 | 0,003029 | 1 | 0 | Intron (ENSMUST0000 | 6 | 28561816 | 28724970 | 163155 | 1 | 56463 | ENSMUST00000171532.8 | -14540 | ENSMUSG00000001424 | Snd1 | staphylococcal nuclease and tudor domain containing 1 |
| chr19 | 12741338 | 12742198 | 861 | * | 5,429566 | 5,858473723 | 1,993142 | 2,939316 | 0,003289 | 1 | 0 | Promoter (<=1kb) | 19 | 12741024 | 12742996 | 1973 | 2 | 12803 | ENSMUST00000112933.2 | 798 | ENSMUSG00000079415 | Cntf | ciliary neurotrophic factor |
| chr5 | 3865350 | 3866079 | 730 | * | 6,151776 | 5,860262408 | 1,850234 | 3,167308 | 0,001539 | 1 | 0 | Promoter (2-3kb) | 5 | 3862778 | 3868569 | 5792 | 1 | 79264 | ENSMUST00000197742.2 | 2572 | ENSMUSG00000000600 | Krit1 | KRIT1, ankyrin repeat containing |
| chr14 | 75457043 | 75457904 | 862 | * | 5,140993 | 5,861602152 | 2,092387 | 2,801395 | 0,005088 | 1 | 0 | Intron (ENSMUST0000 | 14 | 75435340 | 75436806 | 1467 | 1 | 18826 | ENSMUST00000130510.2 | 21703 | ENSMUSG00000021998 | Lcp1 | lymphocyte cytosolic protein 1 |
| chr5 | 92370682 | 92371580 | 899 | * | 4,688035 | 5,862543806 | 2,128403 | 2,754433 | 0,005879 | 1 | 0 | Distal Intergenic | 5 | 92374548 | 92376880 | 2333 | 2 | 19023 | ENSMUST00000201236.2 | 5300 | ENSMUSG00000029410 | Ppef2 | protein phosphatase, EF hand calcium-binding domain 2 |
| chr14 | 8887220 | 8887891 | 672 | * | 4,378491 | 5,862642269 | 2,132877 | 2,748702 | 0,005983 | 1 | 0 | Intron (ENSMUST0000 | 14 | 8786615 | 8798887 | 12273 | 2 | 218739 | ENSMUST00000061045.3 | -88333 | ENSMUSG00000044772 | Sntn | sentan, cilia apical structure protein |
| chr10 | 68629341 | 68629822 | 482 | * | 4,296519 | 5,863116726 | 2,153459 | 2,72265 | 0,006476 | 1 | 0 | Distal Intergenic | 10 | 68559654 | 68615031 | 55378 | 1 | 327766 | ENSMUST00000218918.2 | 69687 | ENSMUSG00000060044 | Tmem26 | transmembrane protein 26 |
| chr7 | 45189580 | 45190112 | 533 | * | 4,813998 | 5,863959687 | 2,192482 | 2,674576 | 0,007482 | 1 | 0 | Exon (ENSMUST000002 | 7 | 45199259 | 45203653 | 4395 | 1 | 69217 | ENSMUST00000210797.2 | -9147 | ENSMUSG00000040428 | Plekha4 | pleckstrin homology domain containing, family A (phosphoinositide binding specific) member 4 |
| chr15 | 99352093 | 99352649 | 557 | * | 4,064603 | 5,864101328 | 2,199801 | 2,665742 | 0,007682 | 1 | 0 | Promoter (2-3kb) | 15 | 99327302 | 99355593 | 28292 | 2 | 380969 | ENSMUST00000161948.2 | 2944 | ENSMUSG00000023009 | Nckap5l | NCK-associated protein 5-like |
| chr16 | 33783107 | 33783540 | 434 | * | 4,179144 | 5,864386493 | 2,213154 | 2,649787 | 0,008054 | 1 | 0 | Intron (ENSMUST0000 | 16 | 33775164 | 33787373 | 12210 | 2 | 22247 | ENSMUST00000131990.2 | 3833 | ENSMUSG00000022814 | Umps | uridine monophosphate synthetase |
| chr19 | 16271609 | 16271972 | 364 | * | 3,843987 | 5,865147368 | 2,254969 | 2,600988 | 0,009296 | 1 | 0 | Intron (ENSMUST0000 | 19 | 16292256 | 16292368 | 113 | 2 | 1,02E+08 | ENSMUST00000184935.3 | 20396 | ENSMUSG00000098718 | Mir496b | microRNA 496b |
| chr3 | 1,16E+08 | 1,16E+08 | 485 | * | 3,772344 | 5,865192159 | 2,257875 | 2,59766 | 0,009386 | 1 | 0 | 3' UTR | 3 | 1,16E+08 | 1,16E+08 | 12566 | 1 | 1,03E+08 | ENSMUST00000198269.2 | -4314 | | 5330425B07Rik | RIKEN cDNA 5330425B07 gene |
| chr7 | 19530553 | 19531054 | 502 | * | 4,418253 | 5,870123417 | 2,27766 | 2,57726 | 0,009959 | 1 | 0 | Promoter (<=1kb) | 7 | 19513643 | 19530718 | 17076 | 2 | 80794 | ENSMUST00000108449.9 | 0 | ENSMUSG00000040525 | Cblc | Casitas B-lineage lymphoma c |
| chr12 | 83236114 | 83236651 | 538 | * | 5,001418 | 5,870183593 | 2,274888 | 2,580427 | 0,009868 | 1 | 0 | Distal Intergenic | 12 | 83162999 | 83184397 | 21399 | 1 | 50779 | ENSMUST00000201512.2 | 73115 | ENSMUSG00000021219 | Rgs6 | regulator of G-protein signaling 6 |
| chr1 | 1,65E+08 | 1,65E+08 | 412 | * | 3,52071 | 5,870190847 | 2,273621 | 2,581869 | 0,009827 | 1 | 0 | Exon (ENSMUST000001 | 1 | 1,65E+08 | 1,65E+08 | 19744 | 1 | 70456 | ENSMUST00000193575.2 | 6562 | ENSMUSG00000026568 | Mpc2 | mitochondrial pyruvate carrier 2 |
| chr1 | 1,42E+08 | 1,42E+08 | 659 | * | 4,109588 | 5,870260882 | 2,271316 | 2,584519 | 0,009751 | 1 | 0 | Distal Intergenic | 1 | 1,41E+08 | 1,41E+08 | 107 | 1 | 1,15E+08 | ENSMUST00000239951.1 | 835711 | ENSMUSG00000118759 | Gm25457 | predicted gene, 25457 |
| chr11 | 29996197 | 29996718 | 522 | * | 5,337404 | 5,870466703 | 2,258117 | 2,599717 | 0,00933 | 1 | 0 | Distal Intergenic | 11 | 29693048 | 29976033 | 282986 | 2 | 237711 | ENSMUST00000058902.6 | -20164 | ENSMUSG00000044072 | Eml6 | echinoderm microtubule associated protein like 6 |
| chrX | 91441966 | 91442571 | 606 | * | 3,947142 | 5,870513076 | 2,25678 | 2,601278 | 0,009288 | 1 | 0 | Intron (ENSMUST0000 | 20 | 91533555 | 91534287 | 733 | 1 | 546347 | ENSMUST00000096371.3 | -90984 | ENSMUSG00000071726 | Gm5941 | predicted gene 5941 |
| chr13 | 5920496 | 5921038 | 543 | * | 4,012847 | 5,870713601 | 2,243951 | 2,61624 | 0,00889 | 1 | 0 | Downstream (<=300bp | 13 | 5914677 | 5916769 | 2093 | 1 | 23849 | ENSMUST00000221734.2 | 5819 | ENSMUSG00000000078 | Klf6 | Kruppel-like factor 6 |
| chr9 | 62670388 | 62670761 | 374 | * | 4,265998 | 5,87080855 | 2,239242 | 2,621784 | 0,008747 | 1 | 0 | Intron (ENSMUST0000 | 9 | 62699103 | 62719216 | 20114 | 2 | 14155 | ENSMUST00000034775.10 | 48455 | ENSMUSG00000032244 | Fem1b | fem 1 homolog b |
| chr8 | 22416982 | 22417306 | 325 | * | 4,775111 | 5,870906251 | 2,234148 | 2,627806 | 0,008594 | 1 | 0 | Intron (ENSMUST0000 | 8 | 22419815 | 22422726 | 2912 | 2 | 246082 | ENSMUST00000062586.3 | 5420 | ENSMUSG00000048500 | Defb15 | defensin beta 15 |
| chr6 | 1,16E+08 | 1,16E+08 | 596 | * | 5,5562 | 5,872372264 | 2,164107 | 2,713532 | 0,006657 | 1 | 0 | Promoter (<=1kb) | 6 | 1,16E+08 | 1,16E+08 | 40195 | 2 | 67784 | ENSMUST00000015511.15 | 741 | ENSMUSG00000030123 | Plxnd1 | plexin D1 |
| chr9 | 55658927 | 55659487 | 561 | * | 4,239796 | 5,872425332 | 2,161346 | 2,717022 | 0,006587 | 1 | 0 | Intron (ENSMUST0000 | 9 | 55650913 | 55699594 | 48682 | 2 | 244891 | ENSMUST00000215323.2 | 40107 | ENSMUSG00000034007 | Scaper | S phase cyclin A-associated protein in the ER |
| chr6 | 1,12E+08 | 1,12E+08 | 297 | * | 4,495445 | 5,873154482 | 2,131431 | 2,755498 | 0,00586 | 1 | 0 | Distal Intergenic | 6 | 1,12E+08 | 1,12E+08 | 1812 | 1 | 67336 | ENSMUST00000186102.2 | -8526 | ENSMUSG00000100219 | 1700054K19Rik | RIKEN cDNA 1700054K19 gene |
| chr17 | 28805853 | 28806376 | 524 | * | 5,197264 | 5,873307371 | 2,124828 | 2,764133 | 0,005707 | 1 | 0 | Distal Intergenic | 17 | 28808564 | 28811537 | 2974 | 2 | 20815 | ENSMUST00000136504.3 | 5161 | ENSMUSG00000004865 | Srpk1 | serine/arginine-rich protein specific kinase 1 |
| chr19 | 28898358 | 28898846 | 489 | * | 4,519435 | 5,873822352 | 2,104831 | 2,790638 | 0,00526 | 1 | 0 | Intron (ENSMUST0000 | 19 | 28900933 | 28906627 | 5695 | 2 | 381218 | ENSMUST00000159655.2 | 7781 | ENSMUSG00000064202 | Spata6l | spermatogenesis associated 6 like |
| chr1 | 1,67E+08 | 1,67E+08 | 865 | * | 4,835578 | 5,874437973 | 2,08302 | 2,820154 | 0,0048 | 1 | 0 | Distal Intergenic | 1 | 1,67E+08 | 1,67E+08 | 89421 | 1 | 226610 | ENSMUST00000128373.2 | 90325 | ENSMUSG00000060568 | Fam78b | family with sequence similarity 78, member B |
| chr1 | 1,82E+08 | 1,82E+08 | 675 | * | 5,584279 | 5,87611922 | 2,02827 | 2,897109 | 0,003766 | 1 | 0 | Promoter (<=1kb) | 1 | 1,82E+08 | 1,82E+08 | 7033 | 2 | 13244 | ENSMUST00000035295.6 | 0 | ENSMUSG00000038633 | Degs1 | delta(4)-desaturase, sphingolipid 1 |
| chr19 | 53380843 | 53381479 | 637 | * | 5,195991 | 5,876142739 | 2,027453 | 2,898288 | 0,003752 | 1 | 0 | Promoter (1-2kb) | 19 | 53367827 | 53379009 | 11183 | 2 | 76479 | ENSMUST00000025997.7 | -1834 | ENSMUSG00000025024 | Smndc1 | survival motor neuron domain containing 1 |
| chr17 | 29455941 | 29456841 | 901 | * | 6,106975 | 5,876212867 | 2,025435 | 2,90121 | 0,003717 | 1 | 0 | Promoter (<=1kb) | 17 | 29436836 | 29456686 | 19851 | 2 | 240058 | ENSMUST00000126755.2 | 0 | ENSMUSG00000024008 | Cpne5 | copine V |
| chr2 | 1,67E+08 | 1,67E+08 | 767 | * | 10,67909 | 5,876377055 | 1,692125 | 3,472779 | 0,000515 | 0,406085428 | 1,173325 | Distal Intergenic | 2 | 1,67E+08 | 1,67E+08 | 55289 | 1 | 77031 | ENSMUST00000047815.13 | -18209 | ENSMUSG00000039463 | Slc9a8 | solute carrier family 9 (sodium/hydrogen exchanger), member 8 |
| chr15 | 78085185 | 78085685 | 501 | * | 5,898692 | 5,877436541 | 1,990996 | 2,952009 | 0,003157 | 1 | 0 | Promoter (2-3kb) | 15 | 78075314 | 78088437 | 13124 | 2 | 19293 | ENSMUST00000005860.16 | 2752 | ENSMUSG00000005716 | Pvalb | parvalbumin |
| chr15 | 8271313 | 8272246 | 934 | * | 6,835386 | 5,880153897 | 1,908629 | 3,080826 | 0,002064 | 1 | 0 | Exon (ENSMUST000001 | 15 | 8276587 | 8282460 | 5874 | 1 | 73692 | ENSMUST00000128772.2 | -4341 | ENSMUSG00000039801 | Cplane1 | ciliogenesis and planar polarity effector 1 |
| chr12 | 11366471 | 11367451 | 981 | * | 7,069016 | 5,880216508 | 1,909645 | 3,07922 | 0,002075 | 1 | 0 | Promoter (1-2kb) | 12 | 11364705 | 11368044 | 3340 | 1 | 67241 | ENSMUST00000218815.2 | 1766 | ENSMUSG00000020608 | Smc6 | structural maintenance of chromosomes 6 |
| chr1 | 1,84E+08 | 1,84E+08 | 1193 | * | 8,657471 | 5,880228291 | 1,840671 | 3,194611 | 0,0014 | 0,541111532 | 1,425929 | Distal Intergenic | 1 | 1,84E+08 | 1,84E+08 | 41256 | 1 | 63953 | ENSMUST00000048655.8 | 75188 | ENSMUSG00000039384 | Dusp10 | dual specificity phosphatase 10 |
| chr4 | 57943594 | 57944658 | 1065 | * | 6,245495 | 5,880447188 | 1,875779 | 3,134936 | 0,001719 | 1 | 0 | 3' UTR | 4 | 57943373 | 57956411 | 13039 | 2 | 22166 | ENSMUST00000030051.6 | 11753 | ENSMUSG00000028367 | Txn1 | thioredoxin 1 |
| chr11 | 29475636 | 29476458 | 823 | * | 6,497129 | 5,880574155 | 1,899954 | 3,095114 | 0,001967 | 1 | 0 | Promoter (<=1kb) | 11 | 29476408 | 29483397 | 6990 | 1 | 76784 | ENSMUST00000133452.8 | 0 | ENSMUSG00000020459 | Mtif2 | mitochondrial translational initiation factor 2 |
| chr13 | 21345573 | 21346104 | 532 | * | 5,887469 | 5,880584168 | 1,901664 | 3,092336 | 0,001986 | 1 | 0 | Distal Intergenic | 13 | 21363615 | 21378894 | 15280 | 1 | 19720 | ENSMUST00000222544.2 | -17511 | ENSMUSG00000021326 | Trim27 | tripartite motif-containing 27 |
| chr11 | 1,19E+08 | 1,19E+08 | 1249 | * | 7,641418 | 5,880613055 | 1,908805 | 3,080782 | 0,002065 | 1 | 0 | Promoter (1-2kb) | 11 | 1,19E+08 | 1,19E+08 | 11043 | 1 | 338371 | ENSMUST00000134873.9 | 1135 | ENSMUSG00000039850 | Endov | endonuclease V |
| chr11 | 1E+08 | 1E+08 | 817 | * | 6,588877 | 5,880726758 | 1,934846 | 3,039377 | 0,002371 | 1 | 0 | Intron (ENSMUST0000 | 11 | 1E+08 | 1E+08 | 6825 | 1 | 12799 | ENSMUST00000103120.5 | -10129 | ENSMUSG00000006782 | Cnp | 2',3'-cyclic nucleotide 3' phosphodiesterase |
| chr5 | 36045084 | 36045844 | 761 | * | 5,216623 | 5,880817988 | 1,954824 | 3,008362 | 0,002627 | 1 | 0 | Distal Intergenic | 5 | 36050663 | 36161267 | 110605 | 1 | 70292 | ENSMUST00000064571.11 | -4819 | ENSMUSG00000029094 | Afap1 | actin filament associated protein 1 |
| chr11 | 26796190 | 26796941 | 752 | * | 5,626895 | 5,880907875 | 1,978447 | 2,972486 | 0,002954 | 1 | 0 | Distal Intergenic | 11 | 26497848 | 26543999 | 46152 | 2 | 69922 | ENSMUST00000156264.2 | -252191 | ENSMUSG00000064090 | Vrk2 | vaccinia related kinase 2 |
| chr12 | 83662892 | 83663712 | 821 | * | 6,184774 | 5,880961895 | 1,995442 | 2,947197 | 0,003207 | 1 | 0 | Distal Intergenic | 12 | 83678010 | 83721191 | 43182 | 1 | 67039 | ENSMUST00000181983.8 | -14298 | ENSMUSG00000010608 | Rbm25 | RNA binding motif protein 25 |
| chr1 | 1,8E+08 | 1,8E+08 | 942 | * | 4,890503 | 5,880979414 | 1,996412 | 2,945774 | 0,003221 | 1 | 0 | Intron (ENSMUST0000 | 1 | 1,8E+08 | 1,81E+08 | 12617 | 1 | 72568 | ENSMUST00000194638.2 | -6102 | ENSMUSG00000058729 | Lin9 | lin-9 DREAM MuvB core complex component |
| chr5 | 57229588 | 57230212 | 625 | * | 4,885706 | 5,880988568 | 1,996424 | 2,945761 | 0,003222 | 1 | 0 | Distal Intergenic | 5 | 57285987 | 57286114 | 128 | 2 | 1,15E+08 | ENSMUST00000104145.3 | 55902 | ENSMUSG00000077333 | Gm23015 | predicted gene, 23015 |
| chr10 | 50394750 | 50395314 | 565 | * | 4,696368 | 5,881125992 | 2,041409 | 2,880914 | 0,003965 | 1 | 0 | Distal Intergenic | 10 | 50468765 | 50727581 | 258817 | 1 | 77987 | ENSMUST00000035606.10 | -73451 | ENSMUSG00000038774 | Ascc3 | activating signal cointegrator 1 complex subunit 3 |
| chr14 | 78263771 | 78264202 | 432 | * | 4,407551 | 5,881205865 | 2,068402 | 2,843357 | 0,004464 | 1 | 0 | Intron (ENSMUST0000 | 14 | 78320183 | 78326421 | 6239 | 2 | 219170 | ENSMUST00000173826.2 | 62219 | ENSMUSG00000045655 | Fam216b | family with sequence similarity 216, member B |
| chr14 | 66794798 | 66795391 | 594 | * | 4,688931 | 5,88124065 | 2,083886 | 2,822247 | 0,004769 | 1 | 0 | Distal Intergenic | 14 | 66872700 | 66970911 | 98212 | 1 | 11549 | ENSMUST00000159365.8 | -77309 | ENSMUSG00000045875 | Adra1a | adrenergic receptor, alpha 1a |
| chr12 | 1,11E+08 | 1,11E+08 | 334 | * | 4,115475 | 5,881407363 | 2,153439 | 2,73117 | 0,006311 | 1 | 0 | Distal Intergenic | 12 | 1,11E+08 | 1,11E+08 | 6175 | 2 | 1,05E+08 | ENSMUST00000246597.1 | -15067 | | Gm40578 | predicted gene, 40578 |
| chr11 | 44219891 | 44220253 | 363 | * | 4,186442 | 5,881407936 | 2,14771 | 2,738455 | 0,006173 | 1 | 0 | Distal Intergenic | 11 | 44290890 | 44304837 | 13948 | 1 | 16160 | ENSMUST00000170513.3 | -70637 | ENSMUSG00000004296 | Il12b | interleukin 12b |
| chr9 | 53483676 | 53484332 | 657 | * | 4,155163 | 5,881410173 | 2,154541 | 2,729774 | 0,006338 | 1 | 0 | 3' UTR | 9 | 53474834 | 53478510 | 3677 | 1 | 244879 | ENSMUST00000148336.2 | 8842 | ENSMUSG00000033054 | Npat | nuclear protein in the AT region |
| chr7 | 99613940 | 99614295 | 356 | * | 3,809107 | 5,881433226 | 2,163828 | 2,718069 | 0,006566 | 1 | 0 | Promoter (<=1kb) | 7 | 99584485 | 99614941 | 30457 | 2 | 108937 | ENSMUST00000174264.2 | 646 | ENSMUSG00000058761 | Rnf169 | ring finger protein 169 |
| chr12 | 9992659 | 9993172 | 514 | * | 4,355383 | 5,881439198 | 2,164349 | 2,717418 | 0,006579 | 1 | 0 | Distal Intergenic | 12 | 9915347 | 9915450 | 104 | 2 | 1,15E+08 | ENSMUST00000157712.3 | -77209 | ENSMUSG00000088337 | Gm22845 | predicted gene, 22845 |
| chr12 | 79140003 | 79140569 | 567 | * | 3,67358 | 5,881463296 | 2,177243 | 2,701335 | 0,006906 | 1 | 0 | Intron (ENSMUST0000 | 12 | 79127438 | 79136425 | 8988 | 2 | 110417 | ENSMUST00000072154.9 | -3578 | ENSMUSG00000021120 | Pigh | phosphatidylinositol glycan anchor biosynthesis, class H |
| chr2 | 1,37E+08 | 1,37E+08 | 592 | * | 4,08685 | 5,881494081 | 2,193235 | 2,681653 | 0,007326 | 1 | 0 | Distal Intergenic | 2 | 1,37E+08 | 1,37E+08 | 40624 | 1 | 1,03E+08 | ENSMUST00000130262.2 | 186843 | | Gm14055 | predicted gene 14055 |
| chr3 | 69001859 | 69002215 | 357 | * | 3,789419 | 5,881533245 | 2,211133 | 2,659963 | 0,007815 | 1 | 0 | Exon (ENSMUST00000 | 3 | 68992438 | 68992780 | 343 | 2 | 1,15E+08 | ENSMUST00000158792.3 | -9079 | ENSMUSG00000089417 | Gm22009 | predicted gene, 22009 |
| chr10 | 13266011 | 13266320 | 310 | * | 3,481895 | 5,881550548 | 2,221099 | 2,648036 | 0,008096 | 1 | 0 | Promoter (1-2kb) | 10 | 13090002 | 13264714 | 174713 | 2 | 215789 | ENSMUST00000079698.7 | -1297 | ENSMUSG00000062866 | Phactr2 | phosphatase and actin regulator 2 |
| chrX | 63357538 | 63358138 | 601 | * | 3,940616 | 5,881579067 | 2,228623 | 2,639109 | 0,008312 | 1 | 0 | Distal Intergenic | 20 | 63308491 | 63320602 | 12112 | 2 | 245446 | ENSMUST00000069926.14 | -36936 | ENSMUSG00000046699 | Slitrk4 | SLIT and NTRK-like family, member 4 |
| chr14 | 39490700 | 39491254 | 555 | * | 3,691912 | 5,881621418 | 2,26907 | 2,592085 | 0,00954 | 1 | 0 | Distal Intergenic | 14 | 39377620 | 39407250 | 29631 | 1 | 73591 | ENSMUST00000185680.2 | 113080 | ENSMUSG00000101261 | 1700109I08Rik | RIKEN cDNA 1700109I08 gene |
| chr1 | 1,49E+08 | 1,49E+08 | 449 | * | 3,679465 | 5,881638026 | 2,265814 | 2,595817 | 0,009437 | 1 | 0 | Distal Intergenic | 1 | 1,49E+08 | 1,49E+08 | 36698 | 2 | 1,05E+08 | ENSMUST00000191253.7 | -92064 | | Gm29398 | predicted gene 29398 |
| chr12 | 78964941 | 78965313 | 373 | * | 3,739222 | 5,881644955 | 2,277994 | 2,581941 | 0,009825 | 1 | 0 | Distal Intergenic | 12 | 78935465 | 78953738 | 18274 | 2 | 27260 | ENSMUST00000021544.8 | -11203 | ENSMUSG00000021118 | Plek2 | pleckstrin 2 |
| chr14 | 26962821 | 26963573 | 753 | * | 6,591891 | 5,881685682 | 1,93451 | 3,040401 | 0,002363 | 1 | 0 | Promoter (2-3kb) | 14 | 26959996 | 27125861 | 165866 | 1 | 71704 | ENSMUST00000049206.6 | 2825 | ENSMUSG00000021895 | Arhgef3 | Rho guanine nucleotide exchange factor (GEF) 3 |
| chr9 | 71540006 | 71540639 | 634 | * | 6,329675 | 5,881730121 | 1,935261 | 3,039244 | 0,002372 | 1 | 0 | Promoter (1-2kb) | 9 | 71537620 | 71538612 | 993 | 2 | 68178 | ENSMUST00000136044.2 | -1394 | ENSMUSG00000032232 | Cgnl1 | cingulin-like 1 |
| chr11 | 5208776 | 5210029 | 1254 | * | 6,846732 | 5,883581165 | 1,970678 | 2,985562 | 0,002831 | 1 | 0 | Promoter (1-2kb) | 11 | 5141552 | 5211558 | 70007 | 2 | 84035 | ENSMUST00000020662.15 | 1529 | ENSMUSG00000020393 | Kremen1 | kringle containing transmembrane protein 1 |
| chr4 | 16418954 | 16419557 | 604 | * | 5,716818 | 5,883893887 | 1,976865 | 2,976376 | 0,002917 | 1 | 0 | Distal Intergenic | 4 | 16164112 | 16219193 | 55082 | 1 | 1E+08 | ENSMUST00000244567.1 | 254842 | ENSMUSG00000085112 | A530072M11Rik | RIKEN cDNA gene A530072M11 |
| chr4 | 22132276 | 22132914 | 639 | * | 6,067724 | 5,884678282 | 1,993899 | 2,951342 | 0,003164 | 1 | 0 | Distal Intergenic | 4 | 21958456 | 21960807 | 2352 | 1 | 76132 | ENSMUST00000155220.2 | 173820 | ENSMUSG00000028246 | Faxc | failed axon connections homolog |
| chr17 | 36346241 | 36347134 | 894 | * | 5,522191 | 5,886093171 | 2,026655 | 2,904339 | 0,00368 | 1 | 0 | Promoter (2-3kb) | 17 | 36340868 | 36343747 | 2880 | 2 | 15040 | ENSMUST00000102678.5 | -2494 | ENSMUSG00000067212 | H2-T23 | histocompatibility 2, T region locus 23 |
| chr2 | 11839232 | 11839744 | 513 | * | 5,492392 | 5,886548344 | 2,038291 | 2,887982 | 0,003877 | 1 | 0 | Distal Intergenic | 2 | 11790969 | 11794933 | 3965 | 1 | 320816 | ENSMUST00000142622.2 | 48263 | ENSMUSG00000047909 | Ankrd16 | ankyrin repeat domain 16 |
| chr1 | 37832914 | 37834112 | 1199 | * | 8,544552 | 5,887241978 | 1,736783 | 3,389739 | 0,0007 | 0,430115816 | 1,360622 | Intron (ENSMUST0000 | 1 | 37879143 | 37882802 | 3660 | 2 | 211484 | ENSMUST00000134024.2 | 48690 | ENSMUSG00000060771 | Tsga10 | testis specific 10 |
| chr9 | 66503787 | 66504349 | 563 | * | 4,47593 | 5,887587438 | 2,159638 | 2,726192 | 0,006407 | 1 | 0 | Distal Intergenic | 9 | 66421919 | 66500424 | 78506 | 2 | 235441 | ENSMUST00000127569.8 | -3363 | ENSMUSG00000032376 | Usp3 | ubiquitin specific peptidase 3 |
| chr18 | 37722007 | 37722383 | 377 | * | 4,657665 | 5,888955498 | 2,106778 | 2,795242 | 0,005186 | 1 | 0 | Intron (ENSMUST0000 | 18 | 37770730 | 37771527 | 798 | 2 | 83885 | ENSMUST00000239189.2 | 49144 | ENSMUSG00000050304 | Slc25a2 | solute carrier family 25 (mitochondrial carrier, ornithine transporter) member 2 |
| chr16 | 20470441 | 20471148 | 708 | * | 5,140901 | 5,889203445 | 2,115126 | 2,784327 | 0,005364 | 1 | 0 | Promoter (<=1kb) | 16 | 20470454 | 20473738 | 3285 | 1 | 21762 | ENSMUST00000232629.2 | 0 | ENSMUSG00000006998 | Psmd2 | proteasome (prosome, macropain) 26S subunit, non-ATPase, 2 |
| chr16 | 4347725 | 4348499 | 775 | * | 4,431406 | 5,889802619 | 2,134452 | 2,759398 | 0,005791 | 1 | 0 | Intron (ENSMUST0000 | 16 | 4298092 | 4340939 | 42848 | 2 | 106393 | ENSMUST00000023161.8 | -6786 | ENSMUSG00000022519 | Srl | sarcalumenin |
| chr5 | 1,08E+08 | 1,08E+08 | 504 | * | 4,343901 | 5,890751846 | 2,168844 | 2,716079 | 0,006606 | 1 | 0 | Intron (ENSMUST0000 | 5 | 1,08E+08 | 1,08E+08 | 9007 | 1 | 231571 | ENSMUST00000129483.3 | 12919 | ENSMUSG00000033773 | Rpap2 | RNA polymerase II associated protein 2 |
| chr15 | 10763430 | 10763814 | 385 | * | 4,202042 | 5,890802645 | 2,171002 | 2,713403 | 0,00666 | 1 | 0 | Intron (ENSMUST0000 | 15 | 10717937 | 10722022 | 4086 | 1 | 75259 | ENSMUST00000181739.3 | 45493 | ENSMUSG00000096971 | 4930556M19Rik | RIKEN cDNA 4930556M19 gene |
| chr4 | 9137115 | 9137751 | 637 | * | 4,886857 | 5,891518704 | 2,19954 | 2,678524 | 0,007395 | 1 | 0 | Distal Intergenic | 4 | 9210003 | 9210108 | 106 | 1 | 1,15E+08 | ENSMUST00000082782.2 | -72252 | ENSMUSG00000075958 | Gm23423 | predicted gene, 23423 |
| chrX | 1,03E+08 | 1,03E+08 | 541 | * | 3,890528 | 5,891817981 | 2,211454 | 2,664228 | 0,007717 | 1 | 0 | Distal Intergenic | 20 | 1,03E+08 | 1,03E+08 | 107 | 2 | 1,15E+08 | ENSMUST00000082942.3 | -12851 | ENSMUSG00000064876 | Gm23378 | predicted gene, 23378 |
| chr1 | 1,78E+08 | 1,78E+08 | 744 | * | 5,939582 | 5,893160784 | 1,900577 | 3,100722 | 0,00193 | 1 | 0 | Exon (ENSMUST00000 | 1 | 1,78E+08 | 1,78E+08 | 23511 | 1 | 69428 | ENSMUST00000016106.6 | 10944 | ENSMUSG00000015962 | 1700016C15Rik | RIKEN cDNA 1700016C15 gene |
| chr15 | 1,03E+08 | 1,03E+08 | 472 | * | 3,934928 | 5,893252459 | 2,277634 | 2,587445 | 0,009669 | 1 | 0 | Intron (ENSMUST0000 | 15 | 1,03E+08 | 1,03E+08 | 48273 | 2 | 12419 | ENSMUST00000118152.8 | 8284 | ENSMUSG00000009575 | Cbx5 | chromobox 5 |
| chr1 | 1,48E+08 | 1,48E+08 | 672 | * | 5,138073 | 5,893568127 | 1,973056 | 2,987025 | 0,002817 | 1 | 0 | Distal Intergenic | 1 | 1,48E+08 | 1,48E+08 | 129 | 2 | 1,15E+08 | ENSMUST00000157957.3 | 161575 | ENSMUSG00000088582 | Gm22966 | predicted gene, 22966 |
| chr4 | 93854609 | 93855257 | 649 | * | 3,741829 | 5,897357956 | 2,278384 | 2,588395 | 0,009642 | 1 | 0 | Distal Intergenic | 4 | 93341495 | 93559307 | 217813 | 2 | 1,03E+08 | ENSMUST00000137509.2 | -295302 | | Gm31828 | predicted gene, 31828 |
| chrX | 1,16E+08 | 1,16E+08 | 688 | * | 4,118396 | 5,897393411 | 2,27493 | 2,59234 | 0,009533 | 1 | 0 | Distal Intergenic | 20 | 1,16E+08 | 1,16E+08 | 348 | 1 | 1E+08 | ENSMUST00000122446.2 | 335322 | ENSMUSG00000067441 | H2ab1 | H2A.B variant histone 1 |
| chrX | 99865871 | 99866170 | 300 | * | 3,698868 | 5,897506849 | 2,265953 | 2,602661 | 0,00925 | 1 | 0 | Distal Intergenic | 20 | 99851606 | 99858436 | 6831 | 1 | 53310 | ENSMUST00000146772.2 | 14265 | ENSMUSG00000000881 | Dlg3 | discs large MAGUK scaffold protein 3 |
| chr15 | 63639794 | 63640161 | 368 | * | 3,831916 | 5,897822255 | 2,242148 | 2,630433 | 0,008528 | 1 | 0 | Distal Intergenic | 15 | 63612533 | 63617216 | 4684 | 1 | 1,05E+08 | ENSMUST00000228483.2 | 27261 | ENSMUSG00000115711 | Gm41336 | predicted gene, 41336 |
| chr5 | 32611306 | 32612162 | 857 | * | 3,885614 | 5,898230369 | 2,214038 | 2,664015 | 0,007721 | 1 | 0 | Distal Intergenic | 5 | 32616187 | 32652867 | 36681 | 1 | 19046 | ENSMUST00000015100.15 | -4025 | ENSMUSG00000014956 | Ppp1cb | protein phosphatase 1 catalytic subunit beta |
| chr14 | 71066619 | 71067142 | 524 | * | 3,779751 | 5,898526842 | 2,193672 | 2,688882 | 0,007169 | 1 | 0 | Distal Intergenic | 14 | 71090373 | 71111394 | 21022 | 1 | 73648 | ENSMUST00000190896.2 | -23231 | ENSMUSG00000099528 | 2410012E07Rik | RIKEN cDNA 2410012E07 gene |
| chr5 | 1,26E+08 | 1,26E+08 | 665 | * | 4,662757 | 5,89885583 | 2,173399 | 2,714115 | 0,006645 | 1 | 0 | Distal Intergenic | 5 | 1,26E+08 | 1,26E+08 | 23861 | 1 | 73396 | ENSMUST00000199057.2 | 201800 | ENSMUSG00000105274 | 1700048F04Rik | RIKEN cDNA 1700048F04 gene |
| chr7 | 34105363 | 34106173 | 811 | * | 4,636304 | 5,898913592 | 2,169119 | 2,719498 | 0,006538 | 1 | 0 | Distal Intergenic | 7 | 34045140 | 34092740 | 47601 | 2 | 67070 | ENSMUST00000206388.2 | -12623 | ENSMUSG00000066568 | Lsm14a | LSM14A mRNA processing body assembly factor |
| chr4 | 1,36E+08 | 1,36E+08 | 660 | * | 4,722232 | 5,898980222 | 2,164517 | 2,72531 | 0,006424 | 1 | 0 | Promoter (<=1kb) | 4 | 1,36E+08 | 1,36E+08 | 9516 | 1 | 269593 | ENSMUST00000168936.2 | 203 | ENSMUSG00000001089 | Luzp1 | leucine zipper protein 1 |
| chr11 | 96861433 | 96861857 | 425 | * | 4,232081 | 5,899105183 | 2,157664 | 2,734024 | 0,006257 | 1 | 0 | Promoter (1-2kb) | 11 | 96844299 | 96859491 | 15193 | 2 | 78912 | ENSMUST00000107623.8 | -1942 | ENSMUSG00000018678 | Sp2 | Sp2 transcription factor |
| chr14 | 72075451 | 72076132 | 682 | * | 4,10267 | 5,899110915 | 2,157191 | 2,734627 | 0,006245 | 1 | 0 | Distal Intergenic | 14 | 72094142 | 72157879 | 63738 | 1 | 74667 | ENSMUST00000228500.2 | -18010 | ENSMUSG00000115304 | 4930434J06Rik | RIKEN cDNA 4930434J06 gene |

| | | | | | | | | | | | | | | | | | | | | | | | |
|---|---|---|---|---|---|---|---|---|---|---|---|---|---|---|---|---|---|---|---|---|---|---|---|
| chr13 | 1,04E+08 | 1,04E+08 | 555 | * | 5,273147 | 5,899326168 | 2,144122 | 2,751395 | 0,005934 | 1 | 0 | Promoter (2-3kb) | 13 | 1,04E+08 | 1,04E+08 | 22951 | 1 | 81003 | ENSMUST00000069174.6 | 2267 | ENSMUSG00000021712 | Trim23 | tripartite motif-containing 23 |
| chr8 | 40749645 | 40750735 | 1091 | * | 12,07035 | 5,899383444 | 1,702099 | 3,465946 | 0,000528 | 0,409763611 | 1,173325 | Intron (ENSMUST0000( | 8 | 40732207 | 40739994 | 7788 | 2 | 80857 | ENSMUST00000034014.14 | -9651 | ENSMUSG00000031603 | Fgf20 | fibroblast growth factor 20 |
| chr14 | 69895798 | 69896474 | 677 | * | 4,404865 | 5,899532605 | 2,133157 | 2,765635 | 0,005681 | 1 | 0 | Intron (ENSMUST0000( | 14 | 69875118 | 69882201 | 7084 | 2 | 1,03E+08 | ENSMUST00000184491.2 | -13597 | | Gm27177 | predicted gene 27177 |
| chr4 | 63958491 | 63958794 | 304 | * | 4,307327 | 5,899675306 | 2,125279 | 2,775954 | 0,005504 | 1 | 0 | Intron (ENSMUST0000( | 4 | 63878022 | 63965252 | 87231 | 2 | 21923 | ENSMUST00000107377.10 | 6458 | ENSMUSG00000028364 | Tnc | tenascin C |
| chr11 | 84030670 | 84031204 | 535 | * | 4,425177 | 5,899935513 | 2,112002 | 2,793527 | 0,005214 | 1 | 0 | Intron (ENSMUST0000( | 11 | 84020498 | 84105116 | 84619 | 1 | 107476 | ENSMUST00000133811.3 | 10172 | ENSMUSG00000020532 | Acaca | acetyl-Coenzyme A carboxylase alpha |
| chr9 | 72437184 | 72437812 | 629 | * | 4,633666 | 5,899982092 | 2,109605 | 2,796724 | 0,005162 | 1 | 0 | Promoter (1-2kb) | 9 | 72425532 | 72439140 | 13609 | 2 | 75115 | ENSMUST00000181405.3 | 1328 | ENSMUSG00000096986 | 4930509E16Rik | RIKEN cDNA 4930509E16 gene |
| chr11 | 77918655 | 77919297 | 643 | * | 5,073288 | 5,900271789 | 2,094705 | 2,816756 | 0,004851 | 1 | 0 | Promoter (<=1kb) | 11 | 77918936 | 77920403 | 1468 | 1 | 268448 | ENSMUST00000125079.2 | 0 | ENSMUSG00000037791 | Phf12 | PHD finger protein 12 |
| chr17 | 6090362 | 6090927 | 566 | * | 4,820306 | 5,900580389 | 2,079489 | 2,837514 | 0,004547 | 1 | 0 | Intron (ENSMUST0000( | 17 | 6094179 | 6100330 | 6152 | 2 | 321007 | ENSMUST00000153325.2 | 9403 | ENSMUSG00000015659 | Serac1 | serine active site containing 1 |
| chr12 | 20160499 | 20160789 | 291 | * | 4,477633 | 5,900723559 | 2,072402 | 2,847287 | 0,004409 | 1 | 0 | Promoter (<=1kb) | 12 | 20161165 | 20194846 | 33682 | 1 | 1,03E+08 | ENSMUST00000222981.2 | -376 | | Gm3993 | predicted gene 3993 |
| chr2 | 1,18E+08 | 1,18E+08 | 628 | * | 4,568112 | 5,900818537 | 2,06816 | 2,853173 | 0,004328 | 1 | 0 | Intron (ENSMUST0000( | 2 | 1,18E+08 | 1,18E+08 | 12614 | 2 | 171543 | ENSMUST00000143583.2 | 6811 | ENSMUSG00000040093 | Bmf | BCL2 modifying factor |
| chr8 | 47988017 | 47988491 | 475 | * | 4,64137 | 5,900921931 | 2,0643 | 2,858559 | 0,004256 | 1 | 0 | Promoter (1-2kb) | 8 | 47986721 | 47997178 | 10458 | 1 | 192174 | ENSMUST00000176379.2 | 1296 | ENSMUSG00000031568 | Rwdd4a | RWD domain containing 4A |
| chr7 | 1,11E+08 | 1,11E+08 | 645 | * | 5,279285 | 5,901183684 | 2,051519 | 2,876494 | 0,004021 | 1 | 0 | Distal Intergenic | 7 | 1,11E+08 | 1,11E+08 | 308317 | 2 | 233733 | ENSMUST00000106663.2 | -62247 | ENSMUSG00000038296 | Galnt18 | polypeptide N-acetylgalactosaminyltransferase 18 |
| chr4 | 26925653 | 26926743 | 1091 | * | 5,540037 | 5,901249679 | 2,049192 | 2,879793 | 0,003979 | 1 | 0 | Distal Intergenic | 4 | 26635820 | 26705449 | 69630 | 2 | 75360 | ENSMUST00000154459.2 | -220204 | ENSMUSG00000086257 | 4930548K13Rik | RIKEN cDNA 4930548K13 gene |
| chr12 | 3335418 | 3336250 | 833 | * | 5,419562 | 5,901776931 | 2,026701 | 2,912012 | 0,003591 | 1 | 0 | Intron (ENSMUST0000( | 12 | 3297428 | 3359969 | 62542 | 2 | 19325 | ENSMUST00000021001.10 | 23719 | ENSMUSG00000020671 | Rab10 | RAB10, member RAS oncogene family |
| chrX | 6353184 | 6354349 | 1166 | * | 5,167547 | 5,902005097 | 2,018085 | 2,924557 | 0,003449 | 1 | 0 | Intron (ENSMUST0000( | 20 | 6327796 | 6495594 | 167799 | 1 | 208431 | ENSMUST00000143641.4 | 25388 | ENSMUSG00000068270 | Shroom4 | shroom family member 4 |
| chr11 | 50138837 | 50139560 | 724 | * | 5,50458 | 5,902355129 | 2,004243 | 2,94493 | 0,00323 | 1 | 0 | Intron (ENSMUST0000( | 11 | 50127299 | 50129443 | 2145 | 2 | 17001 | ENSMUST00000125555.2 | -9394 | ENSMUSG00000020377 | Ltc4s | leukotriene C4 synthase |
| chr6 | 38821962 | 38822834 | 873 | * | 8,048762 | 5,903099153 | 1,805903 | 3,26878 | 0,00108 | 1 | 0 | Intron (ENSMUST0000( | 6 | 38794842 | 38814246 | 19405 | 2 | 15258 | ENSMUST00000114855.3 | -7716 | ENSMUSG00000061436 | Hipk2 | homeodomain interacting protein kinase 2 |
| chr4 | 45971734 | 45972376 | 643 | * | 5,675324 | 5,903596911 | 1,960487 | 3,01129 | 0,002601 | 1 | 0 | Promoter (<=1kb) | 4 | 45972233 | 46034761 | 62529 | 1 | 100121 | ENSMUST00000102929.2 | 0 | ENSMUSG00000035517 | Tdrd7 | tudor domain containing 7 |
| chr12 | 77248131 | 77248964 | 834 | * | 6,186909 | 5,904974728 | 1,890409 | 3,12365 | 0,001786 | 1 | 0 | Distal Intergenic | 12 | 77284899 | 77522759 | 237861 | 1 | 53618 | ENSMUST00000177595.9 | -35935 | ENSMUSG00000021065 | Fut8 | fucosyltransferase 8 |
| chr7 | 1,1E+08 | 1,1E+08 | 943 | * | 6,359649 | 5,90509624 | 1,897349 | 3,112288 | 0,001856 | 1 | 0 | Intron (ENSMUST0000( | 7 | 1,1E+08 | 1,1E+08 | 21241 | 1 | 22390 | ENSMUST00000033326.10 | -3671 | ENSMUSG00000031016 | Wee1 | WEE 1 homolog 1 (S. pombe) |
| chr15 | 5836127 | 5837136 | 1010 | * | 6,900617 | 5,905421402 | 1,916141 | 3,081935 | 0,002057 | 1 | 0 | Distal Intergenic | 15 | 5525799 | 5624464 | 98666 | 1 | 71432 | ENSMUST00000228069.2 | 310328 | ENSMUSG00000110018 | 5430437J10Rik | RIKEN cDNA 5430437J10 gene |
| chr5 | 1,44E+08 | 1,44E+08 | 657 | * | 6,205466 | 5,905566297 | 1,924637 | 3,068406 | 0,002152 | 1 | 0 | Promoter (1-2kb) | 5 | 1,44E+08 | 1,44E+08 | 56204 | 1 | 19159 | ENSMUST00000177281.8 | 1287 | ENSMUSG00000018001 | Cyth3 | cytohesin 3 |
| chr5 | 85114165 | 85114880 | 716 | * | 7,157808 | 5,905770014 | 1,938336 | 3,046825 | 0,002313 | 0,645382905 | 1,360622 | Distal Intergenic | 5 | 84478634 | 84565241 | 86608 | 2 | 13839 | ENSMUST00000136775.2 | -548924 | ENSMUSG00000029245 | Epha5 | Eph receptor A5 |
| chr8 | 91637335 | 91638162 | 828 | * | 6,238304 | 5,905963408 | 1,951314 | 3,026659 | 0,002473 | 1 | 0 | Promoter (2-3kb) | 8 | 91635242 | 91703578 | 68337 | 1 | 109151 | ENSMUST00000211403.2 | 2093 | ENSMUSG00000056608 | Chd9 | chromodomain helicase DNA binding protein 9 |
| chr1 | 1,69E+08 | 1,69E+08 | 768 | * | 6,169886 | 5,906648011 | 2,002833 | 2,949146 | 0,003187 | 1 | 0 | Promoter (<=1kb) | 1 | 1,69E+08 | 1,69E+08 | 33531 | 2 | 66977 | ENSMUST00000111368.8 | 0 | ENSMUSG00000026683 | Nuf2 | NUF2, NDC80 kinetochore complex component |
| chr19 | 9612843 | 9613355 | 513 | * | 4,690311 | 5,906911819 | 2,025212 | 2,916689 | 0,003538 | 1 | 0 | Distal Intergenic | 19 | 9535156 | 9536547 | 1392 | 2 | 619371 | ENSMUST00000090527.5 | -76296 | | Stxbp3-ps | syntaxin-binding protein 3, pseudogene |
| chr1 | 1,75E+08 | 1,75E+08 | 400 | * | 4,771078 | 5,90693771 | 2,027528 | 2,91337 | 0,003576 | 1 | 0 | Distal Intergenic | 1 | 1,75E+08 | 1,75E+08 | 19689 | 2 | 24012 | ENSMUST00000192203.2 | 43716 | ENSMUSG00000026527 | Rgs7 | regulator of G protein signaling 7 |
| chr11 | 69370269 | 69370759 | 491 | * | 4,793483 | 5,90697567 | 2,031174 | 2,908158 | 0,003636 | 1 | 0 | Exon (ENSMUST00000( | 11 | 69355436 | 69356520 | 1085 | 2 | 327954 | ENSMUST00000151811.2 | -13749 | ENSMUSG00000005237 | Dnah2 | dynein, axonemal, heavy chain 2 |
| chr11 | 86875684 | 86876323 | 640 | * | 4,892493 | 5,907176525 | 2,048432 | 2,883755 | 0,00393 | 1 | 0 | Intron (ENSMUST0000( | 11 | 86827251 | 86884533 | 57283 | 2 | 77864 | ENSMUST00000018571.5 | 8210 | ENSMUSG00000018427 | Ypel2 | yippee like 2 |
| chr1 | 1,5E+08 | 1,5E+08 | 885 | * | 5,83002 | 5,907180736 | 2,04923 | 2,882634 | 0,003944 | 1 | 0 | Distal Intergenic | 1 | 1,5E+08 | 1,5E+08 | 7974 | 2 | 96875 | ENSMUST00000159035.2 | -34319 | ENSMUSG00000006014 | Prg4 | proteoglycan 4 (megakaryocyte stimulating factor, articular superficial zone protein) |
| chr7 | 97006859 | 97008143 | 1285 | * | 9,281376 | 5,907188273 | 1,86358 | 3,169806 | 0,001525 | 0,550933624 | 1,449706 | Distal Intergenic | 7 | 97020813 | 97041392 | 20580 | 1 | 381903 | ENSMUST00000098300.6 | -12670 | ENSMUSG00000035704 | Alg8 | asparagine-linked glycosylation 8 (alpha-1,3-glucosyltransferase) |
| chr11 | 60343449 | 60343990 | 542 | * | 5,012133 | 5,907576051 | 2,090688 | 2,825661 | 0,004718 | 1 | 0 | Promoter (1-2kb) | 11 | 60345417 | 60352679 | 7263 | 1 | 13495 | ENSMUST00000138713.8 | -1427 | ENSMUSG00000020537 | Drg2 | developmentally regulated GTP binding protein 2 |
| chr4 | 1,08E+08 | 1,08E+08 | 620 | * | 4,74031 | 5,907718834 | 2,105125 | 2,806351 | 0,005011 | 1 | 0 | Distal Intergenic | 4 | 1,08E+08 | 1,08E+08 | 4455 | 1 | 69893 | ENSMUST00000147054.2 | 23841 | ENSMUSG00000048351 | Coa7 | cytochrome c oxidase assembly factor 7 |
| chr19 | 47158837 | 47159289 | 453 | * | 4,371519 | 5,907789931 | 2,114284 | 2,794227 | 0,005202 | 1 | 0 | Exon (ENSMUST00000 | 19 | 47167259 | 47247880 | 80622 | 1 | 18011 | ENSMUST00000111808.11 | -7970 | ENSMUSG00000006435 | Neurl1a | neuralized E3 ubiquitin protein ligase 1A |
| chrX | 5834007 | 5834758 | 752 | * | 4,321374 | 5,907819523 | 2,118115 | 2,789187 | 0,005284 | 1 | 0 | Distal Intergenic | 20 | 5959507 | 5967164 | 7658 | 1 | 58242 | ENSMUST00000103007.4 | -124749 | ENSMUSG00000073295 | Nudt11 | nudix (nucleoside diphosphate linked moiety X)-type motif 11 |
| chr9 | 1,09E+08 | 1,09E+08 | 605 | * | 4,07406 | 5,907853398 | 2,122514 | 2,783422 | 0,005379 | 1 | 0 | Distal Intergenic | 9 | 1,09E+08 | 1,09E+08 | 107 | 2 | 1,15E+08 | ENSMUST00000083740.3 | -21158 | ENSMUSG00000065674 | Gm24259 | predicted gene, 24259 |
| chr3 | 35985270 | 35986637 | 1368 | * | 7,82323 | 5,907968326 | 1,846464 | 3,199611 | 0,001376 | 1 | 0 | Promoter (<=1kb) | 3 | 35946254 | 35986483 | 40230 | 2 | 114893 | ENSMUST00000178098.8 | 0 | ENSMUSG00000027708 | Dcun1d1 | DCN1, defective in cullin neddylation 1, domain containing 1 (S. cerevisiae) |
| chr3 | 1,05E+08 | 1,05E+08 | 704 | * | 4,499969 | 5,908062673 | 2,149297 | 2,748835 | 0,005981 | 1 | 0 | Exon (ENSMUST00000( | 3 | 1,05E+08 | 1,05E+08 | 19795 | 1 | 20501 | ENSMUST00000046212.2 | 14482 | ENSMUSG00000032902 | Slc16a1 | solute carrier family 16 (monocarboxylic acid transporters), member 1 |
| chr9 | 43501174 | 43501564 | 391 | * | 4,037171 | 5,908664866 | 2,235784 | 2,642771 | 0,008223 | 1 | 0 | Distal Intergenic | 9 | 43519481 | 43531715 | 12235 | 1 | 1,03E+08 | ENSMUST00000215154.2 | -17917 | ENSMUSG00000110988 | Gm29909 | predicted gene, 29909 |
| chr7 | 1,12E+08 | 1,12E+08 | 1118 | * | 8,096114 | 5,909521256 | 1,975418 | 2,991529 | 0,002776 | 0,675778938 | 1,360622 | Intron (ENSMUST0000( | 7 | 1,12E+08 | 1,12E+08 | 69270 | 1 | 57342 | ENSMUST00000106640.2 | -37134 | ENSMUSG00000030770 | Parva | parvin, alpha |
| chr11 | 82284451 | 82284944 | 494 | * | 7,070945 | 5,911556702 | 1,799642 | 3,284852 | 0,00102 | 1 | 0 | Intron (ENSMUST0000( | 11 | 82279949 | 82325549 | 45601 | 1 | 270893 | ENSMUST00000132477.2 | 4502 | ENSMUSG00000020701 | Tmem132e | transmembrane protein 132E |
| chr9 | 84124412 | 84124937 | 526 | * | 4,636145 | 5,912643848 | 2,279876 | 2,593405 | 0,009503 | 1 | 0 | Distal Intergenic | 9 | 84070408 | 84070592 | 185 | 1 | 1,15E+08 | ENSMUST00000104478.4 | 54004 | ENSMUSG00000077666 | Gm26146 | predicted gene, 26146 |
| chr4 | 1,3E+08 | 1,3E+08 | 1002 | * | 5,772196 | 5,913452415 | 2,05468 | 2,87804 | 0,004002 | 1 | 0 | Promoter (<=1kb) | 4 | 1,3E+08 | 1,3E+08 | 7351 | 1 | 19244 | ENSMUST00000141754.2 | 231 | ENSMUSG00000028788 | Ptp4a2 | protein tyrosine phosphatase 4a2 |
| chr15 | 12398346 | 12399233 | 888 | * | 6,250377 | 5,91520043 | 2,097878 | 2,819611 | 0,004808 | 1 | 0 | Intron (ENSMUST0000( | 15 | 12405018 | 12412466 | 7449 | 2 | 68070 | ENSMUST00000191339.2 | 13233 | ENSMUSG00000022197 | Pdzd2 | PDZ domain containing 2 |
| chr9 | 1,21E+08 | 1,21E+08 | 867 | * | 5,80128 | 5,915218414 | 2,189871 | 2,701173 | 0,00691 | 1 | 0 | Intron (ENSMUST0000( | 9 | 1,21E+08 | 1,21E+08 | 958 | 1 | 12387 | ENSMUST00000139138.2 | 14023 | ENSMUSG00000006932 | Ctnnb1 | catenin (cadherin associated protein), beta 1 |
| chr19 | 7216098 | 7216748 | 651 | * | 4,921449 | 5,915643184 | 2,109498 | 2,80429 | 0,005043 | 1 | 0 | Promoter (1-2kb) | 19 | 7203033 | 7218399 | 15367 | 2 | 70999 | ENSMUST00000236628.2 | 1651 | ENSMUSG00000024764 | Naa40 | N(alpha)-acetyltransferase 40, NatD catalytic subunit |
| chr19 | 5594037 | 5594961 | 925 | * | 8,476062 | 5,916645056 | 1,763843 | 3,354406 | 0,000795 | 1 | 0 | Intron (ENSMUST0000( | 19 | 5603588 | 5609431 | 5844 | 2 | 18426 | ENSMUST00000238060.2 | 14470 | ENSMUSG00000024922 | Ovol1 | ovo like zinc finger 1 |
| chr17 | 13160213 | 13160726 | 514 | * | 4,189014 | 5,918543362 | 2,199039 | 2,691423 | 0,007115 | 1 | 0 | Promoter (<=1kb) | 17 | 13142700 | 13159487 | 16788 | 2 | 224530 | ENSMUST00000160378.3 | -726 | ENSMUSG00000062480 | Acat3 | acetyl-Coenzyme A acetyltransferase 3 |
| chr14 | 32165236 | 32165586 | 351 | * | 4,569991 | 5,919435429 | 2,232052 | 2,652015 | 0,008001 | 1 | 0 | Intron (ENSMUST0000( | 14 | 32172825 | 32182868 | 10044 | 2 | 12647 | ENSMUST00000227675.2 | 17282 | ENSMUSG00000021919 | Chat | choline acetyltransferase |
| chr13 | 38706914 | 38707731 | 818 | * | 4,259866 | 5,920697492 | 2,283273 | 2,593075 | 0,009512 | 1 | 0 | 5' UTR | 13 | 38711379 | 38716896 | 5518 | 1 | 1,03E+08 | ENSMUST00000224471.2 | -3648 | | Gm31834 | predicted gene, 31834 |
| chr11 | 79166713 | 79167801 | 1089 | * | 9,171443 | 5,922923803 | 1,810962 | 3,270595 | 0,001073 | 0,492478878 | 1,449706 | Distal Intergenic | 11 | 79130198 | 79145497 | 15300 | 2 | 78889 | ENSMUST00000017821.12 | -21216 | ENSMUSG00000017677 | Wsb1 | WD repeat and SOCS box-containing 1 |
| chr6 | 1,31E+08 | 1,31E+08 | 733 | * | 7,26262 | 5,923452609 | 1,882537 | 3,146527 | 0,001652 | 0,582232389 | 1,360622 | Intron (ENSMUST0000( | 6 | 1,31E+08 | 1,31E+08 | 20880 | 2 | 1,19E+08 | ENSMUST00000205067.2 | 9173 | | Gm53051 | predicted gene, 53051 |
| chrX | 24301460 | 24302029 | 570 | * | 4,076482 | 5,92422392 | 2,273475 | 2,605801 | 0,009166 | 1 | 0 | Distal Intergenic | 20 | 24340547 | 24366999 | 26453 | 2 | 546272 | ENSMUST00000120734.2 | 64970 | ENSMUSG00000084063 | Gm5934 | predicted gene 5934 |
| chr8 | 1,12E+08 | 1,12E+08 | 306 | * | 3,479387 | 5,924251081 | 2,272134 | 2,607351 | 0,009125 | 1 | 0 | 3' UTR | 8 | 1,12E+08 | 1,12E+08 | 16405 | 2 | 434350 | ENSMUST00000143719.2 | -15183 | | 9430091E24Rik | RIKEN cDNA 9430091E24 gene |
| chr10 | 99891226 | 99891639 | 414 | * | 3,878837 | 5,924254151 | 2,27385 | 2,605385 | 0,009177 | 1 | 0 | Exon (ENSMUST00000 | 10 | 99923918 | 99933043 | 9126 | 1 | 17311 | ENSMUST00000219881.2 | -32279 | ENSMUSG00000019966 | Kitl | kit ligand |
| chr1 | 1,51E+08 | 1,51E+08 | 725 | * | 3,836875 | 5,924290195 | 2,268104 | 2,612001 | 0,009001 | 1 | 0 | Promoter (<=1kb) | 1 | 1,51E+08 | 1,51E+08 | 18788 | 1 | 117198 | ENSMUST00000186745.2 | 0 | ENSMUSG00000023150 | Ivns1abp | influenza virus NS1A binding protein |
| chrX | 35764943 | 35765549 | 607 | * | 4,206925 | 5,924385703 | 2,256899 | 2,625011 | 0,008665 | 1 | 0 | Distal Intergenic | 20 | 35687254 | 35755483 | 68230 | 2 | 1,01E+08 | ENSMUST00000169499.2 | -9460 | ENSMUSG00000091556 | Gm14569 | predicted gene 14569 |
| chr4 | 3297387 | 3297978 | 592 | * | 4,207199 | 5,924916861 | 2,198246 | 2,695293 | 0,007033 | 1 | 0 | Distal Intergenic | 4 | 3247518 | 3248527 | 1010 | 1 | 1,08E+08 | ENSMUST00000144933.2 | 49869 | | Gm46840 | predicted gene, 46840 |
| chrX | 1,34E+08 | 1,34E+08 | 485 | * | 3,872212 | 5,925111929 | 2,179614 | 2,718423 | 0,006559 | 1 | 0 | Distal Intergenic | 20 | 1,34E+08 | 1,34E+08 | 5075 | 2 | 67416 | ENSMUST00000168264.2 | -70014 | ENSMUSG00000033436 | Armcx2 | armadillo repeat containing, X-linked 2 |
| chr5 | 1,34E+08 | 1,34E+08 | 401 | * | 3,946317 | 5,92511662 | 2,179862 | 2,718115 | 0,006565 | 1 | 0 | Distal Intergenic | 5 | 1,34E+08 | 1,34E+08 | 797 | 2 | 1,03E+08 | ENSMUST00000140421.2 | 78472 | | Gm15627 | predicted gene 15627 |
| chr12 | 1,16E+08 | 1,16E+08 | 444 | * | 3,891076 | 5,925225144 | 2,16768 | 2,733441 | 0,006268 | 1 | 0 | Intron (ENSMUST0000( | 12 | 1,16E+08 | 1,16E+08 | 90971 | 1 | 52635 | ENSMUST00000220720.2 | 17771 | ENSMUSG00000021171 | Esyt2 | extended synaptotagmin-like protein 2 |
| chr15 | 73460477 | 73460728 | 252 | * | 4,305367 | 5,925238617 | 2,166442 | 2,735009 | 0,006238 | 1 | 0 | Promoter (<=1kb) | 15 | 73458743 | 73460811 | 2069 | 2 | 106068 | ENSMUST00000152558.2 | 83 | ENSMUSG00000079020 | Slc45a4 | solute carrier family 45, member 4 |
| chr3 | 83571064 | 83571518 | 455 | * | 4,344184 | 5,925272459 | 2,164557 | 2,737407 | 0,006193 | 1 | 0 | Distal Intergenic | 3 | 83609559 | 83609665 | 107 | 1 | 1,15E+08 | ENSMUST00000240217.1 | -38041 | ENSMUSG00000119833 | Gm24584 | predicted gene, 24584 |
| chr2 | 1,31E+08 | 1,31E+08 | 377 | * | 4,201953 | 5,925276387 | 2,165222 | 2,736567 | 0,006208 | 1 | 0 | Intron (ENSMUST0000( | 2 | 1,31E+08 | 1,31E+08 | 15250 | 2 | 51902 | ENSMUST00000165420.8 | 5204 | ENSMUSG00000048911 | Rnf24 | ring finger protein 24 |
| chrX | 7212438 | 7212882 | 445 | * | 4,167299 | 5,925298168 | 2,161374 | 2,741449 | 0,006117 | 1 | 0 | Distal Intergenic | 20 | 7037438 | 7185597 | 148160 | 2 | 12728 | ENSMUST00000128319.2 | -26841 | ENSMUSG00000004317 | Clcn5 | chloride channel, voltage-sensitive 5 |
| chr14 | 21388155 | 21388675 | 521 | * | 4,710028 | 5,925323943 | 2,159446 | 2,74391 | 0,006071 | 1 | 0 | Intron (ENSMUST0000( | 14 | 21368193 | 21376861 | 8669 | 1 | 11534 | ENSMUST00000223645.2 | 19962 | ENSMUSG00000039197 | Adk | adenosine kinase |
| chr4 | 1,56E+08 | 1,56E+08 | 442 | * | 3,983766 | 5,925352818 | 2,156543 | 2,747617 | 0,006003 | 1 | 0 | Promoter (<=1kb) | 4 | 1,56E+08 | 1,56E+08 | 5036 | 1 | 66448 | ENSMUST00000143503.2 | 309 | ENSMUSG00000029066 | Mrpl20 | mitochondrial ribosomal protein L20 |
| chr11 | 11755273 | 11756103 | 831 | * | 4,034487 | 5,925432085 | 2,149641 | 2,756475 | 0,005843 | 1 | 0 | Promoter (2-3kb) | 11 | 11737431 | 11758925 | 21495 | 2 | 60530 | ENSMUST00000152345.8 | 2822 | ENSMUSG00000035455 | Fignl1 | fidgetin-like 1 |
| chr11 | 48745280 | 48745918 | 639 | * | 4,844616 | 5,925590193 | 2,135133 | 2,775279 | 0,005515 | 1 | 0 | Distal Intergenic | 11 | 48738729 | 48743036 | 4308 | 1 | 94089 | ENSMUST00000131270.2 | 6551 | ENSMUSG00000040350 | Trim7 | tripartite motif-containing 7 |
| chr14 | 92406857 | 92407750 | 894 | * | 5,136916 | 5,925620194 | 2,132261 | 2,779031 | 0,005452 | 1 | 0 | Distal Intergenic | 14 | 92605936 | 92606065 | 130 | 2 | 1,15E+08 | ENSMUST00000104110.3 | 198315 | ENSMUSG00000077298 | Gm24127 | predicted gene, 24127 |
| chr4 | 97038356 | 97038937 | 582 | * | 6,663255 | 5,925907412 | 1,85145 | 3,200685 | 0,001371 | 1 | 0 | Distal Intergenic | 4 | 96805257 | 96805320 | 64 | 1 | 1,15E+08 | ENSMUST00000184022.3 | 233099 | ENSMUSG00000098261 | Gm27521 | predicted gene, 27521 |
| chr11 | 85204048 | 85205396 | 1349 | * | 8,967193 | 5,925928445 | 1,803556 | 3,285692 | 0,001017 | 0,410415129 | 2,237403 | Promoter (1-2kb) | 11 | 85202990 | 85228029 | 25040 | 1 | 53892 | ENSMUST00000127717.2 | 1058 | ENSMUSG00000020525 | Ppm1d | protein phosphatase 1D magnesium-dependent, delta isoform |
| chr14 | 1,05E+08 | 1,05E+08 | 665 | * | 4,360589 | 5,926592934 | 2,05918 | 2,878133 | 0,004 | 1 | 0 | Promoter (2-3kb) | 14 | 1,05E+08 | 1,05E+08 | 107 | 1 | 1,15E+08 | ENSMUST00000239836.1 | 2254 | ENSMUSG00000119509 | Gm22290 | predicted gene, 22290 |
| chr11 | 1,19E+08 | 1,19E+08 | 467 | * | 4,803341 | 5,92674078 | 2,048868 | 2,892691 | 0,00382 | 1 | 0 | Promoter (1-2kb) | 11 | 1,19E+08 | 1,19E+08 | 18401 | 1 | 268512 | ENSMUST00000151612.2 | 1687 | ENSMUSG00000039908 | Slc26a11 | solute carrier family 26, member 11 |
| chr6 | 39208658 | 39208888 | 231 | * | 5,01009 | 5,926820631 | 2,043234 | 2,900706 | 0,003723 | 1 | 0 | Distal Intergenic | 6 | 39217048 | 39219475 | 2428 | 1 | 1,05E+08 | ENSMUST00000200801.2 | -8160 | | Gm38794 | predicted gene, 38794 |
| chr11 | 77676143 | 77676547 | 405 | * | 4,722238 | 5,927058582 | 2,028194 | 2,922333 | 0,003474 | 1 | 0 | Promoter (2-3kb) | 11 | 77676279 | 77678573 | 2295 | 2 | 791303 | ENSMUST00000092883.3 | 2026 | ENSMUSG00000069804 | Gm10277 | predicted gene 10277 |
| chr8 | 1,23E+08 | 1,23E+08 | 614 | * | 5,251957 | 5,927408196 | 2,007727 | 2,952298 | 0,003154 | 1 | 0 | Promoter (1-2kb) | 8 | 1,23E+08 | 1,23E+08 | 69632 | 2 | 234839 | ENSMUST00000156333.9 | -1485 | ENSMUSG00000014444 | Piezo1 | piezo-type mechanosensitive ion channel component 1 |
| chrX | 76417841 | 76418789 | 949 | * | 6,164766 | 5,927479829 | 2,002998 | 2,959303 | 0,003083 | 1 | 0 | Distal Intergenic | 20 | 76554619 | 76706589 | 151971 | 1 | 21372 | ENSMUST00000088217.12 | -135830 | ENSMUSG00000025246 | Tbl1x | transducin (beta)-like 1 X-linked |
| chr1 | 1,87E+08 | 1,87E+08 | 940 | * | 5,201957 | 5,927561745 | 1,99855 | 2,965931 | 0,003018 | 1 | 0 | Intron (ENSMUST0000( | 1 | 1,87E+08 | 1,87E+08 | 14788 | 1 | 329316 | ENSMUST00000195611.2 | -109252 | ENSMUSG00000103923 | Gm37896 | predicted gene, 37896 |
| chr3 | 83528978 | 83529747 | 770 | * | 5,234325 | 5,927680317 | 1,992435 | 2,975093 | 0,002929 | 1 | 0 | Distal Intergenic | 3 | 83462670 | 83481429 | 18760 | 2 | 69460 | ENSMUST00000185938.2 | -47549 | ENSMUSG00000101334 | 1700028M03Rik | RIKEN cDNA 1700028M03 gene |
| chr17 | 10765097 | 10765877 | 781 | * | 8,417673 | 5,928812123 | 1,878585 | 3,155999 | 0,001599 | 1 | 0 | Intron (ENSMUST0000( | 17 | 10715382 | 10772098 | 56717 | 1 | 1,03E+08 | ENSMUST00000161091.2 | 49715 | | Gm16168 | predicted gene 16168 |
| chr7 | 1,27E+08 | 1,27E+08 | 586 | * | 5,875047 | 5,929128263 | 1,921077 | 3,086357 | 0,002026 | 1 | 0 | Distal Intergenic | 7 | 1,27E+08 | 1,27E+08 | 7042 | 2 | 320683 | ENSMUST00000132524.8 | -30215 | ENSMUSG00000045639 | Zfp629 | zinc finger protein 629 |
| chr7 | 91103613 | 91104412 | 800 | * | 6,148993 | 5,929167382 | 1,9196 | 3,088752 | 0,00201 | 1 | 0 | Intron (ENSMUST0000( | 7 | 91192018 | 91521636 | 329619 | 1 | 23859 | ENSMUST00000239136.2 | -87606 | ENSMUSG00000052572 | Dlg2 | discs large MAGUK scaffold protein 2 |
| chr5 | 1,2E+08 | 1,2E+08 | 625 | * | 5,784439 | 5,93040093 | 1,933078 | 3,067854 | 0,002156 | 1 | 0 | Distal Intergenic | 5 | 1,2E+08 | 1,2E+08 | 8956 | 1 | 21388 | ENSMUST00000202504.3 | 50347 | ENSMUSG00000018263 | Tbx5 | T-box 5 |
| chr10 | 1,17E+08 | 1,17E+08 | 750 | * | 5,648431 | 5,930535738 | 1,938398 | 3,059504 | 0,002217 | 1 | 0 | Distal Intergenic | 10 | 1,17E+08 | 1,17E+08 | 531 | 2 | 432508 | ENSMUST00000175888.2 | 16656 | ENSMUSG00000055531 | Cpsf6 | cleavage and polyadenylation specific factor 6 |
| chr12 | 70419404 | 70420077 | 674 | * | 6,672629 | 5,930538952 | 1,866447 | 3,177448 | 0,001486 | 1 | 0 | Distal Intergenic | 12 | 70312923 | 70394388 | 81466 | 2 | 94090 | ENSMUST00000221370.2 | -25016 | ENSMUSG00000021071 | Trim9 | tripartite motif-containing 9 |
| chr12 | 4816708 | 4817379 | 672 | * | 5,936948 | 5,930723267 | 1,945933 | 3,047753 | 0,002306 | 1 | 0 | Promoter (1-2kb) | 12 | 4789888 | 4818521 | 28634 | 2 | 207921 | ENSMUST00000219898.2 | 1142 | ENSMUSG00000050545 | Fam228b | family with sequence similarity 228, member B |
| chr2 | 1,21E+08 | 1,21E+08 | 970 | * | 7,639142 | 5,930913761 | 1,853743 | 3,199426 | 0,001377 | 1 | 0 | Promoter (<=1kb) | 2 | 1,21E+08 | 1,21E+08 | 9269 | 1 | 96957 | ENSMUST00000130010.8 | -285 | ENSMUSG00000054484 | Tmem62 | transmembrane protein 62 |
| chr2 | 32258841 | 32259268 | 428 | * | 4,865874 | 5,933513006 | 2,087897 | 2,841861 | 0,004485 | 1 | 0 | Promoter (<=1kb) | 2 | 32260074 | 32261762 | 1689 | 1 | 68379 | ENSMUST00000132290.2 | -806 | ENSMUSG00000039205 | Ciz1 | CDKN1A interacting zinc finger protein 1 |
| chr11 | 1,22E+08 | 1,22E+08 | 639 | * | 4,959357 | 5,933698417 | 2,100656 | 2,824688 | 0,004733 | 1 | 0 | Promoter (2-3kb) | 11 | 1,22E+08 | 1,22E+08 | 13177 | 1 | 74675 | ENSMUST00000036690.7 | 2381 | ENSMUSG00000039198 | Ptchd3 | patched domain containing 3 |
| chr1 | 51872931 | 51873448 | 518 | * | 4,574172 | 5,934089228 | 2,127895 | 2,788714 | 0,005292 | 1 | 0 | Intron (ENSMUST0000( | 1 | 51838695 | 51859705 | 21011 | 2 | 17912 | ENSMUST00000135125.8 | -13226 | ENSMUSG00000018417 | Myo1b | myosin IB |
| chr3 | 22730726 | 22731187 | 462 | * | 4,584248 | 5,934327182 | 2,144693 | 2,766982 | 0,005658 | 1 | 0 | Distal Intergenic | 3 | 22464087 | 22464215 | 129 | 2 | 1,15E+08 | ENSMUST00000158221.3 | -266511 | ENSMUSG00000088846 | Gm26354 | predicted gene, 26354 |
| chr18 | 24439341 | 24439718 | 378 | * | 4,05375 | 5,934332659 | 2,145857 | 2,765484 | 0,005684 | 1 | 0 | Distal Intergenic | 18 | 24413090 | 24416248 | 3159 | 1 | 14423 | ENSMUST00000170191.2 | 26251 | ENSMUSG00000000420 | Galnt1 | polypeptide N-acetylgalactosaminyltransferase 1 |
| chr14 | 63519034 | 63519729 | 696 | * | 5,115599 | 5,934334352 | 2,146975 | 2,764044 | 0,005709 | 1 | 0 | Distal Intergenic | 14 | 63478551 | 63509141 | 30591 | 2 | 14463 | ENSMUST00000132122.2 | -9893 | ENSMUSG00000021944 | Gata4 | GATA binding protein 4 |
| chr12 | 53504341 | 53504755 | 415 | * | 4,074417 | 5,934603858 | 2,166101 | 2,739764 | 0,006148 | 1 | 0 | Intron (ENSMUST0000( | 12 | 53350135 | 53356629 | 6495 | 2 | 75576 | ENSMUST00000222223.2 | -147712 | | 1700030L22Rik | RIKEN cDNA 1700030L22 gene |
| chr18 | 34897518 | 34898338 | 821 | * | 4,977608 | 5,934951185 | 2,197433 | 2,700856 | 0,006916 | 1 | 0 | Intron (ENSMUST0000( | 18 | 34892626 | 34906813 | 14188 | 1 | 66306 | ENSMUST00000235929.2 | 4892 | ENSMUSG00000034300 | Fam53c | family with sequence similarity 53, member C |
| chr2 | 60378723 | 60379016 | 294 | * | 3,825201 | 5,93501214 | 2,201408 | 2,696007 | 0,007018 | 1 | 0 | Promoter (2-3kb) | 2 | 60381101 | 60384267 | 3167 | 1 | 1,01E+08 | ENSMUST00000151931.2 | -2085 | | Gm13583 | predicted gene 13583 |
| chr11 | 1,07E+08 | 1,07E+08 | 583 | * | 3,77446 | 5,935816362 | 2,280753 | 2,602568 | 0,009253 | 1 | 0 | Intron (ENSMUST0000( | 11 | 1,07E+08 | 1,07E+08 | 8711 | 2 | 66313 | ENSMUST00000144027.2 | 4631 | ENSMUSG00000018363 | Smurf2 | SMAD specific E3 ubiquitin protein ligase 2 |
| chr12 | 14157368 | 14158005 | 638 | * | 4,031753 | 5,935935728 | 2,292721 | 2,589036 | 0,009625 | 1 | 0 | Distal Intergenic | 12 | 14200535 | 14201350 | 816 | 2 | 105005 | ENSMUST00000221405.2 | 43345 | ENSMUSG00000020607 | Lratd1 | LRAT domain containing 1 |
| chr8 | 79445783 | 79446446 | 664 | * | 7,266438 | 5,936643842 | 2,003438 | 2,963229 | 0,003044 | 1 | 0 | Intron (ENSMUST0000( | 8 | 79423803 | 79455205 | 31403 | 1 | 76775 | ENSMUST00000210132.2 | 21980 | ENSMUSG00000031684 | Slc10a7 | solute carrier family 10 (sodium/bile acid cotransporter family), member 7 |
| chr3 | 1,46E+08 | 1,46E+08 | 807 | * | 7,238039 | 5,936730041 | 2,004903 | 2,961105 | 0,003065 | 1 | 0 | Intron (ENSMUST0000( | 3 | 1,46E+08 | 1,46E+08 | 15628 | 2 | 56629 | ENSMUST00000029836.9 | 10751 | ENSMUSG00000028185 | Dnase2b | deoxyribonuclease II beta |

| | | | | | | | | | | | | | | | | | | | | | | | |
|---|---|---|---|---|---|---|---|---|---|---|---|---|---|---|---|---|---|---|---|---|---|---|---|
| chr11 | 1,02E+08 | 1,02E+08 | 906 | * | 6,930739 | 5,938607124 | 2,038993 | 2,91252 | 0,003585 | 1 | 0 | Distal Intergenic | 11 | 1,02E+08 | 1,02E+08 | 2292 | 1 | 80981 | ENSMUST00000039388.3 | 30933 | ENSMUSG00000034936 | Arl4d | ADP-ribosylation factor-like 4D |
| chr14 | 76245910 | 76246461 | 552 | * | 4,067209 | 5,938642067 | 2,300646 | 2,581293 | 0,009843 | 1 | 0 | Promoter (1-2kb) | 14 | 76134377 | 76248305 | 113929 | 2 | 68705 | ENSMUST00000088922.5 | 1844 | ENSMUSG00000067995 | Gtf2f2 | general transcription factor IIF, polypeptide 2 |
| chr15 | 11856456 | 11857134 | 679 | * | 4,144006 | 5,938993419 | 2,28351 | 2,600818 | 0,0093 | 1 | 0 | Intron (ENSMUST0000 | 15 | 11842346 | 11891074 | 48729 | 2 | 18162 | ENSMUST00000226139.2 | 33940 | ENSMUSG00000022206 | Npr3 | natriuretic peptide receptor 3 |
| chr7 | 53171059 | 53171305 | 247 | * | 3,676046 | 5,939071204 | 2,280281 | 2,604535 | 0,0092 | 1 | 0 | Distal Intergenic | 7 | 53541509 | 53541637 | 129 | 2 | 1,02E+08 | ENSMUST00000184140.3 | 370332 | ENSMUSG00000099176 | Mir6238 | microRNA 6238 |
| chrX | 1,5E+08 | 1,5E+08 | 445 | * | 4,432344 | 5,939140843 | 2,277534 | 2,607707 | 0,009115 | 1 | 0 | Intron (ENSMUST0000 | 20 | 1,5E+08 | 1,5E+08 | 52873 | 2 | 331547 | ENSMUST00000175942.8 | -81602 | ENSMUSG00000084771 | A230072E10Rik | RIKEN cDNA A230072E10 gene |
| chr18 | 29386512 | 29386932 | 421 | * | 4,080168 | 5,939944611 | 2,240792 | 2,650824 | 0,00803 | 1 | 0 | Distal Intergenic | 18 | 29575253 | 29599160 | 23908 | 1 | 78199 | ENSMUST00000234621.2 | -188321 | ENSMUSG00000117554 | 4930527G23Rik | RIKEN cDNA 4930527G23 gene |
| chr10 | 1,04E+08 | 1,04E+08 | 640 | * | 3,79001 | 5,940006939 | 2,296857 | 2,586145 | 0,009706 | 1 | 0 | Distal Intergenic | 10 | 1,04E+08 | 1,04E+08 | 157 | 2 | 1,15E+08 | ENSMUST00000158557.3 | 23730 | ENSMUSG00000089182 | Gm22945 | predicted gene, 22945 |
| chr4 | 1,15E+08 | 1,15E+08 | 850 | * | 6,270077 | 5,940072258 | 2,06869 | 2,871417 | 0,004086 | 1 | 0 | Promoter (<=1kb) | 4 | 1,15E+08 | 1,15E+08 | 27906 | 2 | 66588 | ENSMUST00000030491.9 | -16 | ENSMUSG00000028719 | Cmpk1 | cytidine monophosphate (UMP-CMP) kinase 1 |
| chr16 | 16019652 | 16020324 | 673 | * | 6,513218 | 5,940822592 | 2,085105 | 2,849172 | 0,004383 | 1 | 0 | Distal Intergenic | 16 | 16031182 | 16045151 | 13970 | 1 | 67451 | ENSMUST00000162150.8 | -10858 | ENSMUSG00000041957 | Pkp2 | plakophilin 2 |
| chr9 | 63681345 | 63681823 | 479 | * | 4,521044 | 5,940828949 | 2,204286 | 2,695126 | 0,007036 | 1 | 0 | Distal Intergenic | 9 | 63554049 | 63665276 | 111228 | 2 | 17127 | ENSMUST00000034973.10 | -16069 | ENSMUSG00000032402 | Smad3 | SMAD family member 3 |
| chr12 | 79089028 | 79089614 | 587 | * | 4,560865 | 5,941083549 | 2,194215 | 2,707612 | 0,006777 | 1 | 0 | Intron (ENSMUST0000 | 12 | 79082410 | 79127403 | 44994 | 1 | 211945 | ENSMUST00000219956.2 | 6618 | ENSMUSG00000060716 | Plekhh1 | pleckstrin homology domain containing, family H (with MyTH4 domain) member 1 |
| chr4 | 58377977 | 58378635 | 659 | * | 5,693638 | 5,941507023 | 2,100676 | 2,828379 | 0,004678 | 1 | 0 | Distal Intergenic | 4 | 58441012 | 58441098 | 87 | 2 | 1,01E+08 | ENSMUST00000175552.2 | 62463 | ENSMUSG00000093293 | Mir3095 | microRNA 3095 |
| chr4 | 1,38E+08 | 1,38E+08 | 476 | * | 5,018395 | 5,941625852 | 2,173653 | 2,733475 | 0,006267 | 1 | 0 | Distal Intergenic | 4 | 1,38E+08 | 1,38E+08 | 10038 | 2 | 75718 | ENSMUST00000137206.2 | 57840 | ENSMUSG00000028753 | Vwa5b1 | von Willebrand factor A domain containing 5B1 |
| chr8 | 1,15E+08 | 1,15E+08 | 516 | * | 4,373627 | 5,941716191 | 2,171399 | 2,736354 | 0,006212 | 1 | 0 | Promoter (<=1kb) | 8 | 1,15E+08 | 1,16E+08 | 913053 | 1 | 80707 | ENSMUST00000004756.14 | -601 | ENSMUSG00000004637 | Wwox | WW domain-containing oxidoreductase |
| chr5 | 66630126 | 66630582 | 457 | * | 4,532957 | 5,941883559 | 2,165141 | 2,74434 | 0,006063 | 1 | 0 | Intron (ENSMUST0000 | 5 | 66459548 | 66649514 | 189967 | 2 | 11787 | ENSMUST00000160063.5 | 18932 | ENSMUSG00000029207 | Apbb2 | amyloid beta (A4) precursor protein-binding, family B, member 2 |
| chrX | 71504398 | 71505034 | 637 | * | 4,916999 | 5,941931017 | 2,163316 | 2,746677 | 0,00602 | 1 | 0 | Intron (ENSMUST0000 | 20 | 71425476 | 71430464 | 4989 | 2 | 236852 | ENSMUST00000088481.5 | -73934 | ENSMUSG00000043453 | Magea10 | MAGE family member A10 |
| chr8 | 46759358 | 46759767 | 410 | * | 4,298748 | 5,942233828 | 2,152513 | 2,760603 | 0,005769 | 1 | 0 | Exon (ENSMUST00000 | 8 | 46752168 | 46778711 | 26544 | 1 | 1,05E+08 | ENSMUST00000210181.2 | 7190 | ENSMUSG00000110179 | Gm39168 | predicted gene, 39168 |
| chr15 | 11127647 | 11128440 | 794 | * | 5,947345 | 5,942244023 | 2,118564 | 2,804846 | 0,005034 | 1 | 0 | Intron (ENSMUST0000 | 15 | 11064904 | 11217207 | 152304 | 1 | 239337 | ENSMUST00000228940.2 | 62743 | ENSMUSG00000047497 | Adamts12 | a disintegrin-like and metallopeptidase (reprolysin type) with thrombospondin type 1 motif, 12 |
| chr14 | 9543702 | 9544378 | 677 | * | 5,319562 | 5,942382238 | 2,147162 | 2,767552 | 0,005648 | 1 | 0 | Distal Intergenic | 14 | 9646684 | 10097198 | 450515 | 1 | 27062 | ENSMUST00000112657.9 | -102306 | ENSMUSG00000054423 | Cadps | Ca2+-dependent secretion activator |
| chr7 | 25225810 | 25226343 | 534 | * | 5,301019 | 5,943045961 | 2,125317 | 2,796311 | 0,005169 | 1 | 0 | Intron (ENSMUST0000 | 7 | 25217489 | 25236518 | 19030 | 2 | 26367 | ENSMUST00000145681.2 | 10175 | ENSMUSG00000054385 | Ceacam2 | carcinoembryonic antigen-related cell adhesion molecule 2 |
| chr1 | 42468119 | 42468719 | 601 | * | 5,462261 | 5,943321565 | 2,11623 | 2,808448 | 0,004978 | 1 | 0 | Distal Intergenic | 1 | 42530973 | 42534823 | 3851 | 1 | 1,19E+08 | ENSMUST00000191992.2 | -62254 | | Gm37047 | predicted gene, 37047 |
| chr11 | 88334332 | 88334770 | 439 | * | 4,562403 | 5,943339184 | 2,116054 | 2,808689 | 0,004974 | 1 | 0 | Intron (ENSMUST0000 | 11 | 88243599 | 88243690 | 92 | 1 | 1,01E+08 | ENSMUST00000175244.3 | 90733 | ENSMUSG00000092985 | Mir378b | microRNA 378b |
| chr11 | 1,02E+08 | 1,02E+08 | 688 | * | 6,168692 | 5,943466634 | 2,150319 | 2,763993 | 0,00571 | 1 | 0 | Distal Intergenic | 11 | 1,02E+08 | 1,02E+08 | 18616 | 2 | 237943 | ENSMUST00000125754.2 | -9633 | ENSMUSG00000034621 | Gpatch8 | G patch domain containing 8 |
| chr3 | 97561881 | 97562716 | 836 | * | 5,291263 | 5,943527091 | 2,151655 | 2,762305 | 0,005739 | 1 | 0 | Promoter (2-3kb) | 3 | 97565509 | 97578115 | 12607 | 1 | 108097 | ENSMUST00000143927.8 | -2793 | ENSMUSG00000038205 | Prkab2 | protein kinase, AMP-activated, beta 2 non-catalytic subunit |
| chr11 | 1,06E+08 | 1,06E+08 | 747 | * | 5,049133 | 5,944282981 | 2,172702 | 2,735894 | 0,006221 | 1 | 0 | Promoter (1-2kb) | 11 | 1,06E+08 | 1,06E+08 | 7654 | 2 | 83796 | ENSMUST00000155243.2 | 1475 | ENSMUSG00000078619 | Smarcd2 | SWI/SNF related, matrix associated, actin dependent regulator of chromatin, subfamily d, member 2 |
| chr3 | 1,08E+08 | 1,08E+08 | 1181 | * | 6,023947 | 5,944794785 | 2,072774 | 2,868038 | 0,00413 | 1 | 0 | Distal Intergenic | 3 | 1,08E+08 | 1,08E+08 | 13319 | 2 | 1,19E+08 | ENSMUST00000198171.2 | 33700 | | LOC118568636 | uncharacterized LOC118568636 |
| chr2 | 76351212 | 76352181 | 970 | * | 5,218059 | 5,944906259 | 2,191211 | 2,713069 | 0,006666 | 1 | 0 | Intron (ENSMUST0000 | 2 | 76329441 | 76371874 | 42434 | 1 | 99031 | ENSMUST00000128034.2 | 21771 | ENSMUSG00000042359 | Osbpl6 | oxysterol binding protein-like 6 |
| chr11 | 1,19E+08 | 1,19E+08 | 569 | * | 4,853334 | 5,945693974 | 2,215857 | 2,683248 | 0,007291 | 1 | 0 | Intron (ENSMUST0000 | 11 | 1,18E+08 | 1,19E+08 | 271278 | 2 | 52897 | ENSMUST00000103023.8 | 31761 | ENSMUSG00000025576 | Rbfox3 | RNA binding protein, fox-1 homolog (C. elegans) 3 |
| chr18 | 24083147 | 24083930 | 784 | * | 5,277544 | 5,946162747 | 2,230855 | 2,665419 | 0,007689 | 1 | 0 | Intron (ENSMUST0000 | 18 | 24087762 | 24089633 | 1872 | 1 | 69256 | ENSMUST00000225682.2 | -3832 | ENSMUSG00000024276 | Zfp397 | zinc finger protein 397 |
| chr12 | 4393267 | 4393989 | 723 | * | 4,877998 | 5,946298609 | 2,235269 | 2,660216 | 0,007809 | 1 | 0 | Promoter (<=1kb) | 12 | 4388894 | 4394470 | 5577 | 2 | 17977 | ENSMUST00000217822.2 | 481 | ENSMUSG00000020647 | Ncoa1 | nuclear receptor coactivator 1 |
| chr15 | 11729106 | 11730038 | 933 | * | 4,823975 | 5,946712552 | 2,249842 | 2,643169 | 0,008213 | 1 | 0 | Distal Intergenic | 15 | 11696896 | 11703789 | 6894 | 2 | 1,05E+08 | ENSMUST00000228108.2 | -25317 | ENSMUSG00000115585 | Gm41271 | predicted gene, 41271 |
| chr1 | 1,91E+08 | 1,91E+08 | 350 | * | 4,761286 | 5,947866324 | 2,293058 | 2,593858 | 0,009491 | 1 | 0 | Intron (ENSMUST0000 | 1 | 1,91E+08 | 1,91E+08 | 15124 | 2 | 76843 | ENSMUST00000194064.2 | 10826 | ENSMUSG00000037474 | Dtl | denticleless E3 ubiquitin protein ligase |
| chr9 | 57229076 | 57229518 | 443 | * | 4,756063 | 5,947951913 | 2,295997 | 2,590575 | 0,009582 | 1 | 0 | Distal Intergenic | 9 | 57160400 | 57169895 | 9496 | 2 | 74211 | ENSMUST00000034846.7 | -59181 | ENSMUSG00000032300 | 1700017B05Rik | RIKEN cDNA 1700017B05 gene |
| chr8 | 7255101 | 7256071 | 971 | * | 7,278265 | 5,947970528 | 1,80713 | 3,29139 | 0,000997 | 1 | 0 | Distal Intergenic | 8 | 7431015 | 7478569 | 47555 | 1 | 1,03E+08 | ENSMUST00000170074.2 | -174944 | | Gm17215 | predicted gene 17215 |
| chr10 | 18109193 | 18109723 | 531 | * | 4,748351 | 5,947999521 | 2,297658 | 2,588723 | 0,009633 | 1 | 0 | Promoter (1-2kb) | 10 | 18092532 | 18110726 | 18195 | 2 | 215814 | ENSMUST00000052648.9 | 1003 | ENSMUSG00000059554 | Ccdc28a | coiled-coil domain containing 28A |
| chr16 | 49866077 | 49866304 | 228 | * | 3,589619 | 5,95063354 | 2,283272 | 2,606187 | 0,009156 | 1 | 0 | Promoter (<=1kb) | 16 | 49862852 | 49866652 | 3801 | 2 | 224180 | ENSMUST00000241286.1 | 348 | | Gm4827 | predicted gene 4827 |
| chr13 | 1,13E+08 | 1,13E+08 | 602 | * | 3,862431 | 5,950633542 | 2,285652 | 2,603473 | 0,009228 | 1 | 0 | Intron (ENSMUST0000 | 13 | 1,13E+08 | 1,13E+08 | 6937 | 2 | 72198 | ENSMUST00000224639.2 | -11847 | ENSMUSG00000016018 | Mtrex | Mtr4 exosome RNA helicase |
| chr6 | 27380492 | 27380935 | 444 | * | 3,845728 | 5,95063642 | 2,286481 | 2,602531 | 0,009254 | 1 | 0 | Intron (ENSMUST0000 | 6 | 27936654 | 27936749 | 96 | 2 | 735266 | ENSMUST00000102113.3 | 555814 | ENSMUSG00000076067 | Mir592 | microRNA 592 |
| chr2 | 86656760 | 86657581 | 822 | * | 4,109052 | 5,950692369 | 2,271891 | 2,619269 | 0,008812 | 1 | 0 | Promoter (1-2kb) | 2 | 86659020 | 86660166 | 1147 | 1 | 258362 | ENSMUST00000105211.4 | -1439 | ENSMUSG00000044213 | Olfr1094 | olfactory receptor 1094 |
| chr7 | 99073420 | 99073724 | 305 | * | 4,436188 | 5,950710311 | 2,26006 | 2,632988 | 0,008464 | 1 | 0 | Promoter (<=1kb) | 7 | 99073320 | 99110192 | 36873 | 1 | 233552 | ENSMUST00000213887.2 | 100 | ENSMUSG00000035314 | Gdpd5 | glycerophosphodiester phosphodiesterase domain containing 5 |
| chr5 | 4266979 | 4268033 | 1055 | * | 9,058144 | 5,950730717 | 1,839406 | 3,235137 | 0,001216 | 0,514468591 | 1,425929 | Distal Intergenic | 5 | 4242367 | 4247651 | 5285 | 1 | 208595 | ENSMUST00000177258.2 | 24612 | ENSMUSG00000053178 | Mterf1b | mitochondrial transcription termination factor 1b |
| chr8 | 35250496 | 35250769 | 274 | * | 3,649221 | 5,95077093 | 2,251979 | 2,642463 | 0,008231 | 1 | 0 | Distal Intergenic | 8 | 35274451 | 35287048 | 12598 | 1 | 319520 | ENSMUST00000033930.5 | -23682 | ENSMUSG00000031530 | Dusp4 | dual specificity phosphatase 4 |
| chr11 | 51837644 | 51837951 | 308 | * | 4,332953 | 5,950785508 | 2,247566 | 2,647658 | 0,008105 | 1 | 0 | Distal Intergenic | 11 | 51855209 | 51855316 | 108 | 1 | 1,15E+08 | ENSMUST00000175584.3 | -17258 | ENSMUSG00000093325 | Gm26061 | predicted gene, 26061 |
| chr9 | 7761531 | 7762134 | 604 | * | 4,05557 | 5,95092661 | 2,214258 | 2,687548 | 0,007198 | 1 | 0 | Promoter (1-2kb) | 9 | 7764042 | 7794334 | 30293 | 1 | 71929 | ENSMUST00000052865.16 | -1908 | ENSMUSG00000050912 | Tmem123 | transmembrane protein 123 |
| chr15 | 89056107 | 89056417 | 311 | * | 4,217838 | 5,950988723 | 2,200841 | 2,703961 | 0,006852 | 1 | 0 | Promoter (1-2kb) | 15 | 89039752 | 89054891 | 15140 | 2 | 140570 | ENSMUST00000060808.10 | -1216 | ENSMUSG00000036606 | Plxnb2 | plexin B2 |
| chr16 | 14995566 | 14995983 | 418 | * | 4,109516 | 5,951021683 | 2,194092 | 2,712293 | 0,006682 | 1 | 0 | Distal Intergenic | 16 | 14710192 | 14809747 | 99556 | 2 | 1,03E+08 | ENSMUST00000229379.2 | -185819 | | Gm33696 | predicted gene, 33696 |
| chr2 | 1,27E+08 | 1,27E+08 | 439 | * | 3,970818 | 5,951070515 | 2,189382 | 2,718151 | 0,006565 | 1 | 0 | Intron (ENSMUST0000 | 2 | 1,27E+08 | 1,27E+08 | 16727 | 1 | 109778 | ENSMUST00000110389.10 | 3153 | ENSMUSG00000001999 | Blvra | biliverdin reductase A |
| chr14 | 56700356 | 56701069 | 714 | * | 4,322377 | 5,951076419 | 2,187532 | 2,720452 | 0,006519 | 1 | 0 | 3' UTR | 14 | 56688030 | 56724523 | 36494 | 1 | 30054 | ENSMUST00000225621.2 | 12326 | ENSMUSG00000000365 | Rnf17 | ring finger protein 17 |
| chr13 | 93258482 | 93258992 | 511 | * | 3,703155 | 5,951111447 | 2,180082 | 2,729765 | 0,006338 | 1 | 0 | Intron (ENSMUST0000 | 13 | 93177221 | 93281232 | 104012 | 2 | 76469 | ENSMUST00000062122.4 | 22240 | ENSMUSG00000047419 | Cmya5 | cardiomyopathy associated 5 |
| chrX | 1,66E+08 | 1,66E+08 | 307 | * | 4,517544 | 5,951117486 | 2,183403 | 2,725616 | 0,006418 | 1 | 0 | Exon (ENSMUST00000 | 20 | 1,66E+08 | 1,66E+08 | 12764 | 2 | 170743 | ENSMUST00000149507.2 | 22160 | ENSMUSG00000044583 | Tlr7 | toll-like receptor 7 |
| chr3 | 39493542 | 39493941 | 400 | * | 3,881363 | 5,951159899 | 2,170941 | 2,741281 | 0,00612 | 1 | 0 | Distal Intergenic | 3 | 39690113 | 39690231 | 119 | 2 | 1,15E+08 | ENSMUST00000083302.3 | 196290 | ENSMUSG00000118708 | n-R5s195 | nuclear encoded rRNA 5S 195 |
| chr15 | 75820895 | 75821216 | 322 | * | 3,983825 | 5,951204696 | 2,163212 | 2,751097 | 0,00594 | 1 | 0 | 3' UTR | 15 | 75812801 | 75821249 | 8449 | 1 | 78834 | ENSMUST00000037260.8 | 8094 | ENSMUSG00000050846 | Zfp623 | zinc finger protein 623 |
| chr11 | 6661236 | 6661609 | 374 | * | 4,361717 | 5,95121671 | 2,157317 | 2,758619 | 0,005805 | 1 | 0 | Distal Intergenic | 11 | 6666001 | 6670037 | 4037 | 2 | 626419 | ENSMUST00000139769.2 | 8428 | ENSMUSG00000086808 | Gm11981 | predicted gene 11981 |
| chr8 | 56860275 | 56860975 | 701 | * | 7,142065 | 5,951319081 | 1,850577 | 3,215926 | 0,0013 | 1 | 0 | Distal Intergenic | 8 | 56918657 | 56938003 | 19347 | 2 | 75123 | ENSMUST00000209557.2 | 77028 | ENSMUSG00000110310 | 4930518J21Rik | RIKEN cDNA 4930518J21 gene |
| chr13 | 69681273 | 69681614 | 342 | * | 4,063662 | 5,951319325 | 2,143832 | 2,776019 | 0,005503 | 1 | 0 | Promoter (<=1kb) | 13 | 69681865 | 69763587 | 81723 | 1 | 28114 | ENSMUST00000143716.8 | -251 | ENSMUSG00000021595 | Nsun2 | NOL1/NOP2/Sun domain family member 2 |
| chrX | 1,68E+08 | 1,68E+08 | 431 | * | 4,676342 | 5,951343604 | 2,135607 | 2,786722 | 0,005324 | 1 | 0 | Intron (ENSMUST0000 | 20 | 1,68E+08 | 1,68E+08 | 214917 | 1 | 11856 | ENSMUST00000143048.2 | -92683 | ENSMUSG00000031355 | Arhgap6 | Rho GTPase activating protein 6 |
| chr17 | 84152592 | 84152841 | 250 | * | 4,035779 | 5,951358647 | 2,133403 | 2,789608 | 0,005277 | 1 | 0 | Promoter (1-2kb) | 17 | 84138585 | 84154178 | 15594 | 2 | 107766 | ENSMUST00000000687.9 | 1337 | ENSMUSG00000000673 | Haao | 3-hydroxyanthranilate 3,4-dioxygenase |
| chr9 | 71393159 | 71393819 | 661 | * | 4,417051 | 5,95150683 | 2,109095 | 2,821829 | 0,004775 | 1 | 0 | Promoter (<=1kb) | 9 | 71385719 | 71393217 | 7499 | 2 | 28015 | ENSMUST00000034720.12 | 0 | ENSMUSG00000032199 | Polr2m | polymerase (RNA) II (DNA directed) polypeptide M |
| chr10 | 99279314 | 99279767 | 454 | * | 4,271549 | 5,951590896 | 2,095127 | 2,840683 | 0,004502 | 1 | 0 | Distal Intergenic | 10 | 99295551 | 99296958 | 1408 | 1 | 1,03E+08 | ENSMUST00000220432.2 | -15784 | | Gm34983 | predicted gene, 34983 |
| chr14 | 65625614 | 65626565 | 952 | * | 5,286452 | 5,951768255 | 2,068156 | 2,877814 | 0,004004 | 1 | 0 | Exon (ENSMUST00000 | 14 | 65633129 | 65634199 | 1071 | 1 | 380912 | ENSMUST00000225512.2 | -6564 | ENSMUSG00000034522 | Zfp395 | zinc finger protein 395 |
| chr16 | 63892639 | 63893374 | 736 | * | 5,099332 | 5,951897145 | 2,05173 | 2,900917 | 0,003721 | 1 | 0 | Distal Intergenic | 16 | 63662281 | 63684538 | 22258 | 2 | 13837 | ENSMUST00000232370.2 | -208101 | ENSMUSG00000052504 | Epha3 | Eph receptor A3 |
| chr5 | 19614788 | 19615812 | 1025 | * | 8,288763 | 5,951903745 | 1,75159 | 3,398001 | 0,000679 | 0,337696055 | 2,237403 | Intron (ENSMUST0000 | 5 | 19720403 | 19720505 | 103 | 2 | 1,15E+08 | ENSMUST00000158043.3 | 104693 | ENSMUSG00000088668 | Gm25335 | predicted gene, 25335 |
| chr2 | 1,54E+08 | 1,54E+08 | 532 | * | 4,503085 | 5,951946342 | 2,046228 | 2,90874 | 0,003629 | 1 | 0 | Intron (ENSMUST0000 | 2 | 1,54E+08 | 1,54E+08 | 67936 | 1 | 12396 | ENSMUST00000155202.2 | 20911 | ENSMUSG00000038533 | Cbfa2t2 | CBFA2/RUNX1 translocation partner 2 |
| chr9 | 43025971 | 43026989 | 1019 | * | 5,96361 | 5,952464361 | 1,982655 | 3,00227 | 0,00268 | 1 | 0 | Promoter (<=1kb) | 9 | 43020130 | 43027828 | 7699 | 2 | 235300 | ENSMUST00000213648.2 | 839 | ENSMUSG00000048503 | Tlcd5 | TLC domain containing 5 |
| chr18 | 10032188 | 10033016 | 829 | * | 8,026348 | 5,952608924 | 1,970054 | 3,021546 | 0,002515 | 1 | 0 | Promoter (2-3kb) | 18 | 9993066 | 10030141 | 37076 | 2 | 59025 | ENSMUST00000092096.14 | -2047 | ENSMUSG00000047879 | Usp14 | ubiquitin specific peptidase 14 |
| chr12 | 1,11E+08 | 1,11E+08 | 393 | * | 5,30263 | 5,952785161 | 1,950295 | 3,052248 | 0,002271 | 1 | 0 | Intron (ENSMUST0000 | 12 | 1,11E+08 | 1,11E+08 | 18608 | 1 | 72805 | ENSMUST00000222460.2 | 3422 | ENSMUSG00000021271 | Zfp839 | zinc finger protein 839 |
| chr2 | 1,56E+08 | 1,56E+08 | 703 | * | 5,436383 | 5,952850858 | 1,94414 | 3,061946 | 0,002199 | 1 | 0 | Intron (ENSMUST0000 | 2 | 1,56E+08 | 1,56E+08 | 28931 | 1 | 228829 | ENSMUST00000153822.2 | -4066 | ENSMUSG00000038116 | Phf20 | PHD finger protein 20 |
| chr5 | 65751727 | 65752699 | 973 | * | 5,825439 | 5,952850866 | 1,945591 | 3,059662 | 0,002216 | 1 | 0 | Promoter (2-3kb) | 5 | 65748939 | 65752488 | 3550 | 1 | 53323 | ENSMUST00000202572.2 | 2788 | ENSMUSG00000029203 | Ube2k | ubiquitin-conjugating enzyme E2K |
| chr11 | 23473369 | 23473895 | 527 | * | 6,316141 | 5,952985503 | 1,931879 | 3,081448 | 0,00206 | 1 | 0 | Promoter (2-3kb) | 11 | 23468817 | 23471155 | 2339 | 2 | 67358 | ENSMUST00000156629.3 | -2214 | ENSMUSG00000020286 | 1700093K21Rik | RIKEN cDNA 1700093K21 gene |
| chr13 | 23685816 | 23686585 | 770 | * | 7,352379 | 5,95353008 | 1,887482 | 3,154218 | 0,001609 | 1 | 0 | Distal Intergenic | 13 | 23661846 | 23673027 | 11182 | 2 | 238555 | ENSMUST00000110433.10 | -12789 | ENSMUSG00000053216 | Btn2a2 | butyrophilin, subfamily 2, member A2 |
| chr11 | 1,19E+08 | 1,19E+08 | 947 | * | 6,851241 | 5,954002199 | 1,853366 | 3,212535 | 0,001316 | 1 | 0 | Promoter (<=1kb) | 11 | 1,19E+08 | 1,19E+08 | 9361 | 1 | 672511 | ENSMUST00000172235.2 | 828 | ENSMUSG00000070327 | Rnf213 | ring finger protein 213 |
| chr13 | 93581567 | 93582519 | 953 | * | 6,695355 | 5,954253457 | 1,917217 | 3,105675 | 0,001898 | 1 | 0 | Promoter (2-3kb) | 13 | 93584996 | 93585343 | 348 | 1 | 619883 | ENSMUST00000223297.2 | -2477 | | Gm6109 | predicted gene 6109 |
| chr2 | 37342607 | 37343190 | 584 | * | 7,375011 | 5,954436338 | 1,826574 | 3,259893 | 0,001115 | 0,507398906 | 1,360622 | Promoter (<=1kb) | 2 | 37342279 | 37456466 | 114188 | 1 | 227800 | ENSMUST00000112920.2 | 328 | ENSMUSG00000035437 | Rabgap1 | RAB GTPase activating protein 1 |
| chr19 | 41848436 | 41849126 | 691 | * | 6,011584 | 5,954512636 | 1,924116 | 3,094675 | 0,00197 | 1 | 0 | Distal Intergenic | 19 | 41834411 | 41836571 | 2161 | 2 | 212398 | ENSMUST00000059231.4 | -11865 | ENSMUSG00000047604 | Frat2 | frequently rearranged in advanced T cell lymphomas 2 |
| chr2 | 1,12E+08 | 1,12E+08 | 864 | * | 8,015847 | 5,954709357 | 1,811255 | 3,287615 | 0,00101 | 0,492832383 | 1,360622 | Distal Intergenic | 2 | 1,12E+08 | 1,12E+08 | 17166 | 1 | 73024 | ENSMUST00000069747.6 | -10836 | ENSMUSG00000055943 | Emc7 | ER membrane protein complex subunit 7 |
| chr19 | 8951330 | 8952463 | 1134 | * | 10,87839 | 5,955020381 | 1,794961 | 3,317632 | 0,000908 | 0,54122637 | 0,918255 | Promoter (1-2kb) | 19 | 8949768 | 8955843 | 6076 | 1 | 67160 | ENSMUST00000135025.2 | 1562 | ENSMUSG00000071644 | Eef1g | eukaryotic translation elongation factor 1 gamma |
| chr4 | 1,49E+08 | 1,49E+08 | 656 | * | 5,531775 | 5,956345318 | 1,978731 | 3,010185 | 0,002611 | 1 | 0 | Exon (ENSMUST00000 | 4 | 1,49E+08 | 1,49E+08 | 9432 | 2 | 69928 | ENSMUST00000030813.10 | -5721 | ENSMUSG00000073705 | Cenps | centromere protein S |
| chr4 | 1,18E+08 | 1,18E+08 | 681 | * | 6,198045 | 5,956778312 | 1,993455 | 2,988168 | 0,002807 | 1 | 0 | Promoter (1-2kb) | 4 | 1,18E+08 | 1,18E+08 | 21734 | 1 | 14664 | ENSMUST00000136476.2 | 1709 | ENSMUSG00000028542 | Slc6a9 | solute carrier family 6 (neurotransmitter transporter, glycine), member 9 |
| chr7 | 83129624 | 83130522 | 899 | * | 4,937516 | 5,957850713 | 2,033163 | 2,930336 | 0,003386 | 1 | 0 | Intron (ENSMUST0000 | 7 | 83192078 | 83199464 | 7387 | 2 | 1E+08 | ENSMUST00000208877.2 | 68942 | | Gm10610 | predicted gene 10610 |
| chr15 | 96269403 | 96270029 | 627 | * | 4,5621 | 5,958573881 | 2,062743 | 2,888665 | 0,003869 | 1 | 0 | Exon (ENSMUST00000 | 15 | 96276695 | 96301482 | 24788 | 1 | 77044 | ENSMUST00000176739.3 | -6666 | ENSMUSG00000033237 | Arid2 | AT rich interactive domain 2 (ARID, RFX-like) |
| chr16 | 91730935 | 91731835 | 901 | * | 5,315685 | 5,959165552 | 2,089603 | 2,851817 | 0,004347 | 1 | 0 | Promoter (2-3kb) | 16 | 91722102 | 91728575 | 6474 | 2 | 28080 | ENSMUST00000023677.10 | -2360 | ENSMUSG00000022956 | Atp5o | ATP synthase, H+ transporting, mitochondrial F1 complex, O subunit |
| chr2 | 1,2E+08 | 1,2E+08 | 482 | * | 4,64949 | 5,959395937 | 2,100059 | 2,837728 | 0,004544 | 1 | 0 | Intron (ENSMUST0000 | 2 | 1,2E+08 | 1,2E+08 | 30247 | 2 | 20402 | ENSMUST00000171215.8 | -3960 | ENSMUSG00000027288 | Zfp106 | zinc finger protein 106 |
| chr10 | 1,28E+08 | 1,28E+08 | 671 | * | 3,934891 | 5,960414639 | 2,154341 | 2,766699 | 0,005663 | 1 | 0 | Promoter (1-2kb) | 10 | 1,28E+08 | 1,28E+08 | 5010 | 1 | 216456 | ENSMUST00000152928.2 | 1106 | ENSMUSG00000044005 | Gls2 | glutaminase 2 (liver, mitochondrial) |
| chrX | 93267285 | 93267868 | 584 | * | 4,053405 | 5,961555287 | 2,22583 | 2,678352 | 0,007399 | 1 | 0 | Distal Intergenic | 20 | 93278205 | 93281763 | 3559 | 1 | 236904 | ENSMUST00000150999.8 | -10337 | ENSMUSG00000043929 | Klhl15 | kelch-like 15 |
| chr19 | 14007353 | 14007909 | 557 | * | 3,91922 | 5,961634876 | 2,231553 | 2,671518 | 0,007551 | 1 | 0 | Distal Intergenic | 19 | 13896306 | 13897408 | 1103 | 1 | 258151 | ENSMUST00000081520.3 | 111047 | ENSMUSG00000062314 | Olfr1505 | olfactory receptor 1505 |
| chr14 | 66624395 | 66625211 | 817 | * | 6,40295 | 5,962040485 | 1,842813 | 3,235293 | 0,001215 | 1 | 0 | Distal Intergenic | 14 | 66581830 | 66595551 | 13722 | 1 | 56471 | ENSMUST00000147477.2 | 42565 | ENSMUSG00000022044 | Stmn4 | stathmin-like 4 |
| chrX | 81840859 | 81841669 | 811 | * | 3,68139 | 5,962069396 | 2,261192 | 2,636694 | 0,008372 | 1 | 0 | Distal Intergenic | 20 | 81735845 | 81735976 | 132 | 2 | 1,15E+08 | ENSMUST00000104486.3 | -104883 | ENSMUSG00000077674 | Gm24470 | predicted gene, 24470 |
| chr13 | 36994961 | 36995558 | 598 | * | 4,117381 | 5,962196764 | 2,270514 | 2,625923 | 0,008641 | 1 | 0 | Distal Intergenic | 13 | 37020340 | 37069570 | 49231 | 1 | 1,03E+08 | ENSMUST00000225846.2 | -24782 | | Gm30489 | predicted gene, 30489 |
| chr5 | 1,12E+08 | 1,12E+08 | 335 | * | 3,439516 | 5,962522637 | 2,298073 | 2,594575 | 0,009471 | 1 | 0 | Distal Intergenic | 5 | 1,12E+08 | 1,12E+08 | 11716 | 1 | 54723 | ENSMUST00000031288.14 | -5448 | ENSMUSG00000029345 | Tfip11 | tuftelin interacting protein 11 |
| chr1 | 1,46E+08 | 1,46E+08 | 578 | * | 3,982899 | 5,964675542 | 2,310152 | 2,581941 | 0,009825 | 1 | 0 | Distal Intergenic | 1 | 1,46E+08 | 1,47E+08 | 407712 | 1 | 215378 | ENSMUST00000074622.11 | -83937 | ENSMUSG00000035131 | Brinp3 | bone morphogenetic protein/retinoic acid inducible neural specific 3 |
| chr1 | 61075976 | 61076400 | 425 | * | 3,907258 | 5,964750027 | 2,305628 | 2,587039 | 0,00968 | 1 | 0 | Distal Intergenic | 1 | 61017086 | 61039479 | 22394 | 1 | 54167 | ENSMUST00000102827.4 | 58890 | ENSMUSG00000026009 | Icos | inducible T cell co-stimulator |
| chr4 | 16375199 | 16375771 | 573 | * | 4,060922 | 5,96482864 | 2,299481 | 2,593989 | 0,009487 | 1 | 0 | Distal Intergenic | 4 | 16164112 | 16219193 | 55082 | 1 | 1E+08 | ENSMUST00000244567.1 | 211087 | ENSMUSG00000085112 | A530072M11Rik | RIKEN cDNA gene A530072M11 |
| chr2 | 92051623 | 92052120 | 498 | * | 3,968765 | 5,964978905 | 2,289272 | 2,605623 | 0,009171 | 1 | 0 | Promoter (<=1kb) | 2 | 92051907 | 92195011 | 143105 | 1 | 192285 | ENSMUST00000090586.12 | 0 | ENSMUSG00000058318 | Phf21a | PHD finger protein 21A |
| chr17 | 17966626 | 17967221 | 596 | * | 3,688397 | 5,965191193 | 2,277584 | 2,619087 | 0,008817 | 1 | 0 | Distal Intergenic | 17 | 17924009 | 17955418 | 31410 | 1 | 626942 | ENSMUST00000232113.2 | 42617 | ENSMUSG00000092120 | Vmn2r90 | vomeronasal 2, receptor 90 |
| chr11 | 15767194 | 15767641 | 448 | * | 3,873025 | 5,965396983 | 2,266026 | 2,632537 | 0,008475 | 1 | 0 | Distal Intergenic | 11 | 15402422 | 15402551 | 130 | 1 | 1,15E+08 | ENSMUST00000158734.3 | 364772 | ENSMUSG00000089359 | Gm25490 | predicted gene, 25490 |
| chr3 | 67099442 | 67100202 | 761 | * | 4,150388 | 5,965451431 | 2,262283 | 2,636916 | 0,008366 | 1 | 0 | Intron (ENSMUST0000 | 3 | 66963291 | 66963397 | 107 | 1 | 1,15E+08 | ENSMUST00000240229.1 | 136151 | ENSMUSG00000118721 | Gm22295 | predicted gene, 22295 |
| chr5 | 3852113 | 3852628 | 516 | * | 4,784642 | 5,965706731 | 2,246609 | 2,655428 | 0,007921 | 1 | 0 | Promoter (<=1kb) | 5 | 3783680 | 3852787 | 69108 | 2 | 70797 | ENSMUST00000199763.2 | 159 | ENSMUSG00000040351 | Ankib1 | ankyrin repeat and IBR domain containing 1 |
| chr1 | 1,89E+08 | 1,89E+08 | 509 | * | 3,994341 | 5,965875551 | 2,238032 | 2,665679 | 0,007683 | 1 | 0 | Distal Intergenic | 1 | 1,89E+08 | 1,89E+08 | 6161 | 2 | 108000 | ENSMUST00000165798.2 | 106060 | ENSMUSG00000026605 | Cenpf | centromere protein F |
| chr10 | 84715287 | 84716087 | 801 | * | 6,149645 | 5,965919285 | 1,902079 | 3,136525 | 0,00171 | 1 | 0 | Exon (ENSMUST00000 | 10 | 84753480 | 84852311 | 98832 | 1 | 237422 | ENSMUST00000038523.15 | -37393 | ENSMUSG00000035620 | Ric8b | RIC8 guanine nucleotide exchange factor B |
| chr10 | 1,18E+08 | 1,18E+08 | 469 | * | 3,879941 | 5,965939671 | 2,235189 | 2,669098 | 0,007606 | 1 | 0 | Distal Intergenic | 10 | 1,18E+08 | 1,18E+08 | 22834 | 1 | 71067 | ENSMUST00000217950.2 | 107783 | ENSMUSG00000112500 | 4933411E08Rik | RIKEN cDNA 4933411E08 gene |
| chr5 | 1,2E+08 | 1,2E+08 | 554 | * | 4,741993 | 5,966984855 | 2,182362 | 2,734187 | 0,006253 | 1 | 0 | Distal Intergenic | 5 | 1,2E+08 | 1,2E+08 | 9160 | 2 | 1,03E+08 | ENSMUST00000201435.2 | 139544 | ENSMUSG00000106938 | Gm31314 | predicted gene, 31314 |

| | | | | | | | | | | | | | | | | | | | | | | | |
|---|---|---|---|---|---|---|---|---|---|---|---|---|---|---|---|---|---|---|---|---|---|---|---|
| chr1 | 1,85E+08 | 1,85E+08 | 938 | * | 4,548979 | 5,96703436 | 2,180098 | 2,737049 | 0,006199 | 1 | 0 | Intron (ENSMUST00000 | 1 | 1,85E+08 | 1,85E+08 | 11293 | 2 | 226777 | ENSMUST00000048308.6 | -27166 | ENSMUSG00000039349 | C130074G19Rik | RIKEN cDNA C130074G19 gene |
| chr14 | 79754099 | 79754947 | 849 | * | 5,376066 | 5,967115968 | 2,175416 | 2,742977 | 0,006088 | 1 | 0 | Promoter (<=1kb) | 14 | 79753138 | 79776267 | 23130 | 1 | 13709 | ENSMUST00000227104.2 | 961 | ENSMUSG00000036461 | Elf1 | E74 like ETS transcription factor 1 |
| chr4 | 81947015 | 81947723 | 709 | * | 4,166177 | 5,967236037 | 2,170276 | 2,749529 | 0,005968 | 1 | 0 | Distal Intergenic | 4 | 81983668 | 82021046 | 37379 | 1 | 545641 | ENSMUST00000137810.4 | -35945 | ENSMUSG00000085022 | Gm5860 | predicted gene 5860 |
| chr11 | 5814269 | 5814872 | 604 | * | 4,254907 | 5,967517417 | 2,157781 | 2,76558 | 0,005682 | 1 | 0 | Promoter (2-3kb) | 11 | 5811947 | 5822088 | 10142 | 1 | 11568 | ENSMUST00000102923.10 | 2322 | ENSMUSG00000020473 | Aebp1 | AE binding protein 1 |
| chr16 | 72921026 | 72921406 | 381 | * | 4,264735 | 5,967658396 | 2,152175 | 2,77285 | 0,005557 | 1 | 0 | Intron (ENSMUST00000 | 16 | 72895876 | 73220464 | 324589 | 1 | 74991 | ENSMUST00000231486.2 | 25150 | ENSMUSG00000116976 | 4930500H12Rik | RIKEN cDNA 4930500H12 gene |
| chr14 | 1,07E+08 | 1,07E+08 | 980 | * | 4,697636 | 5,967758774 | 2,147369 | 2,779102 | 0,005451 | 1 | 0 | Promoter (1-2kb) | 14 | 1,07E+08 | 1,07E+08 | 5957 | 1 | 1,03E+08 | ENSMUST00000227226.2 | 1824 | | Gm36107 | predicted gene, 36107 |
| chr15 | 40711666 | 40712063 | 398 | * | 4,372677 | 5,967857646 | 2,143733 | 2,783862 | 0,005372 | 1 | 0 | Intron (ENSMUST00000 | 15 | 40528386 | 40967982 | 439597 | 1 | 22762 | ENSMUST00000230319.2 | 183280 | ENSMUSG00000022306 | Zfpm2 | zinc finger protein, multitype 2 |
| chr17 | 78099854 | 78100341 | 488 | * | 4,800504 | 5,968081564 | 2,134065 | 2,796579 | 0,005165 | 1 | 0 | Distal Intergenic | 17 | 78506685 | 78507454 | 770 | 2 | 1,03E+08 | ENSMUST00000234450.2 | 407113 | ENSMUSG00000117648 | Gm32282 | predicted gene, 32282 |
| chr9 | 13910034 | 13910454 | 421 | * | 4,672333 | 5,968102809 | 2,132906 | 2,798108 | 0,00514 | 1 | 0 | Distal Intergenic | 9 | 14000916 | 14029597 | 28682 | 1 | 76412 | ENSMUST00000216531.2 | -90462 | ENSMUSG00000111489 | 1700019J19Rik | RIKEN cDNA 1700019J19 gene |
| chr18 | 4040783 | 4041109 | 327 | * | 4,333467 | 5,968564488 | 2,114132 | 2,823175 | 0,004755 | 1 | 0 | Distal Intergenic | 18 | 4165832 | 4182232 | 16401 | 1 | 67328 | ENSMUST00000025076.10 | -124723 | ENSMUSG00000024233 | Lyzl1 | lysozyme-like 1 |
| chr18 | 15203929 | 15204318 | 390 | * | 4,530321 | 5,968708203 | 2,108883 | 2,83027 | 0,004651 | 1 | 0 | Intron (ENSMUST00000 | 18 | 15107112 | 15197155 | 90044 | 2 | 106931 | ENSMUST00000234864.2 | -6774 | ENSMUSG00000036225 | Kctd1 | potassium channel tetramerisation domain containing 1 |
| chrX | 83872413 | 83873386 | 974 | * | 7,550861 | 5,96888577 | 1,854073 | 3,219338 | 0,001285 | 1 | 0 | Intron (ENSMUST00000 | 20 | 83816317 | 84019313 | 202997 | 1 | 13405 | ENSMUST00000156107.2 | 56096 | ENSMUSG00000045103 | Dmd | dystrophin, muscular dystrophy |
| chr7 | 34599916 | 34600594 | 679 | * | 5,877004 | 5,968924498 | 2,100398 | 2,841806 | 0,004486 | 1 | 0 | Distal Intergenic | 7 | 34611804 | 34744133 | 132330 | 1 | 18624 | ENSMUST00000075068.14 | -11210 | ENSMUSG00000063931 | Pepd | peptidase D |
| chr16 | 90524185 | 90524577 | 393 | * | 4,799073 | 5,970051695 | 2,060194 | 2,897811 | 0,003758 | 1 | 0 | Promoter (<=1kb) | 16 | 90516200 | 90524292 | 8093 | 2 | 66578 | ENSMUST00000099554.5 | 0 | ENSMUSG00000022978 | Mis18a | MIS18 kinetochore protein A |
| chr1 | 1,93E+08 | 1,93E+08 | 581 | * | 5,149877 | 5,9701524 | 2,056681 | 2,902809 | 0,003698 | 1 | 0 | Distal Intergenic | 1 | 1,93E+08 | 1,93E+08 | 1551 | 2 | 320952 | ENSMUST00000057543.3 | 12429 | ENSMUSG00000047384 | A730013G03Rik | RIKEN cDNA A730013G03 gene |
| chr4 | 38891440 | 38891945 | 506 | * | 5,641647 | 5,970161937 | 2,057047 | 2,902297 | 0,003704 | 1 | 0 | Distal Intergenic | 4 | 38571667 | 38855483 | 283817 | 2 | 1,03E+08 | ENSMUST00000126314.2 | -35957 | ENSMUSG00000085632 | Gm12381 | predicted gene 12381 |
| chr3 | 53430612 | 53431094 | 483 | * | 4,899908 | 5,970293076 | 2,052198 | 2,909219 | 0,003623 | 1 | 0 | Promoter (<=1kb) | 3 | 53423377 | 53430441 | 7065 | 2 | 242022 | ENSMUST00000199543.2 | -171 | ENSMUSG00000037016 | Frem2 | Fras1 related extracellular matrix protein 2 |
| chr17 | 35450907 | 35451613 | 707 | * | 5,538054 | 5,971020646 | 2,029521 | 2,942084 | 0,00326 | 1 | 0 | Promoter (1-2kb) | 17 | 35452636 | 35457743 | 5108 | 1 | 66237 | ENSMUST00000146711.2 | -1023 | ENSMUSG00000024403 | Atp6v1g2 | ATPase, H+ transporting, lysosomal V1 subunit G2 |
| chr13 | 18590830 | 18591650 | 821 | * | 5,25425 | 5,97167102 | 2,0099 | 2,971129 | 0,002967 | 1 | 0 | Distal Intergenic | 13 | 18520810 | 18572271 | 51462 | 2 | 218030 | ENSMUST00000223058.2 | -18559 | ENSMUSG00000009734 | Pou6f2 | POU domain, class 6, transcription factor 2 |
| chr7 | 1,17E+08 | 1,17E+08 | 752 | * | 7,021915 | 5,973752329 | 1,95531 | 3,055143 | 0,00225 | 1 | 0 | Intron (ENSMUST00000 | 7 | 1,17E+08 | 1,17E+08 | 73868 | 1 | 233781 | ENSMUST00000161889.2 | 127031 | ENSMUSG00000030657 | Xylt1 | xylosyltransferase 1 |
| chr19 | 21490082 | 21490955 | 874 | * | 6,640907 | 5,975567004 | 1,915177 | 3,120112 | 0,001808 | 1 | 0 | Distal Intergenic | 19 | 21529568 | 21534105 | 4538 | 1 | 1,19E+08 | ENSMUST00000236172.2 | -38613 | ENSMUSG00000079382 | C19H9orf57 | chromosome 19 C9orf57 homolog |
| chr7 | 44892002 | 44892773 | 772 | * | 7,691801 | 5,976351777 | 1,880405 | 3,178226 | 0,001482 | 0,560372961 | 1,360622 | Promoter (1-2kb) | 7 | 44890509 | 44922791 | 32283 | 1 | 381884 | ENSMUST00000213347.2 | 1493 | ENSMUSG00000094152 | Slc6a16 | solute carrier family 6, member 16 |
| chr18 | 32679281 | 32679553 | 273 | * | 3,561444 | 5,976714776 | 2,296046 | 2,603047 | 0,00924 | 1 | 0 | Intron (ENSMUST00000 | 18 | 32662845 | 32668471 | 5627 | 2 | 71683 | ENSMUST00000172434.3 | -10810 | ENSMUSG00000090523 | Gypc | glycophorin C |
| chrX | 12736571 | 12737023 | 453 | * | 3,52708 | 5,976722382 | 2,288722 | 2,61138 | 0,009018 | 1 | 0 | Exon (ENSMUST00020 | 20 | 12754466 | 12862153 | 107688 | 1 | 75037 | ENSMUST00000245073.1 | -17443 | | 4930500G05Rik | RIKEN cDNA 4930500G05 gene |
| chr14 | 22230040 | 22230249 | 210 | * | 3,521153 | 5,976722476 | 2,310705 | 2,586536 | 0,009695 | 1 | 0 | Intron (ENSMUST00000 | 14 | 22360233 | 22634507 | 274275 | 1 | 76633 | ENSMUST00000159107.2 | -129984 | ENSMUSG00000063458 | Lrmda | leucine rich melanocyte differentiation associated |
| chr18 | 87930057 | 87930547 | 491 | * | 4,507874 | 5,976723265 | 2,241185 | 2,666769 | 0,007658 | 1 | 0 | Distal Intergenic | 18 | 87943555 | 87943688 | 134 | 2 | 1,15E+08 | ENSMUST00000104453.3 | 13141 | ENSMUSG00000077641 | Gm24987 | predicted gene, 24987 |
| chr15 | 84038028 | 84038458 | 431 | * | 3,434609 | 5,976723441 | 2,292281 | 2,607326 | 0,009125 | 1 | 0 | Distal Intergenic | 15 | 84052038 | 84070407 | 18370 | 1 | 116939 | ENSMUST00000045289.6 | -13580 | ENSMUSG00000041653 | Pnpla3 | patatin-like phospholipase domain containing 3 |
| chr3 | 36541907 | 36542498 | 592 | * | 3,648216 | 5,976734849 | 2,301097 | 2,597342 | 0,009395 | 1 | 0 | Distal Intergenic | 3 | 36530086 | 36536448 | 6363 | 1 | 72282 | ENSMUST00000029268.7 | 11821 | ENSMUSG00000027713 | 1810062G17Rik | RIKEN cDNA 1810062G17 gene |
| chrX | 59370372 | 59370850 | 479 | * | 4,090975 | 5,976735339 | 2,195563 | 2,722188 | 0,006485 | 1 | 0 | Intron (ENSMUST00000 | 20 | 59270983 | 59312190 | 41208 | 2 | 320940 | ENSMUST00000133295.8 | -58182 | ENSMUSG00000062949 | Atp11c | ATPase, class VI, type 11C |
| chr10 | 1,12E+08 | 1,12E+08 | 438 | * | 4,019513 | 5,976744414 | 2,196225 | 2,721372 | 0,006501 | 1 | 0 | Promoter (<=1kb) | 10 | 1,12E+08 | 1,12E+08 | 17167 | 2 | 73690 | ENSMUST00000162508.9 | 683 | ENSMUSG00000056888 | Glipr1 | GLI pathogenesis-related 1 (glioma) |
| chr1 | 1,06E+08 | 1,06E+08 | 666 | * | 4,834046 | 5,976761383 | 2,148816 | 2,78142 | 0,005412 | 1 | 0 | Intron (ENSMUST00000 | 1 | 1,06E+08 | 1,06E+08 | 222499 | 1 | 98432 | ENSMUST00000061047.7 | 10572 | ENSMUSG00000044340 | Phlpp1 | PH domain and leucine rich repeat protein phosphatase 1 |
| chr3 | 65294961 | 65295635 | 675 | * | 5,50057 | 5,976780221 | 2,065804 | 2,893198 | 0,003813 | 1 | 0 | Exon (ENSMUST000000 | 3 | 65289101 | 65290802 | 1702 | 2 | 67437 | ENSMUST00000139785.2 | -4159 | ENSMUSG00000027828 | Ssr3 | signal sequence receptor, gamma |
| chr14 | 63114608 | 63115889 | 1282 | * | 4,448177 | 5,976782408 | 2,117402 | 2,822696 | 0,004762 | 1 | 0 | Intron (ENSMUST00000 | 14 | 63102991 | 63103100 | 110 | 1 | 1,02E+08 | ENSMUST00000183649.3 | 11617 | ENSMUSG00000098858 | Mir6541 | microRNA 6541 |
| chr4 | 78659363 | 78659939 | 577 | * | 4,765668 | 5,976794472 | 2,057729 | 2,904559 | 0,003678 | 1 | 0 | Distal Intergenic | 4 | 78130313 | 78133846 | 3534 | 1 | 1,03E+08 | ENSMUST00000246656.1 | 529050 | | B230208B08Rik | RIKEN cDNA B230208B08 gene |
| chr16 | 92230900 | 92231319 | 420 | * | 5,129108 | 5,976796233 | 2,034564 | 2,93763 | 0,003307 | 1 | 0 | Intron (ENSMUST00000 | 16 | 92254365 | 92259378 | 5014 | 1 | 1,08E+08 | ENSMUST00000231968.2 | -23046 | | Gm46555 | predicted gene, 46555 |
| chr5 | 70676116 | 70676460 | 345 | * | 4,736422 | 5,976811819 | 2,008597 | 2,975615 | 0,002924 | 1 | 0 | Distal Intergenic | 5 | 70708184 | 70708290 | 107 | 2 | 1,15E+08 | ENSMUST00000240250.2 | 31830 | ENSMUSG00000118789 | Gm26072 | predicted gene, 26072 |
| chr10 | 10930997 | 10931585 | 589 | * | 5,181156 | 5,976813436 | 1,989115 | 3,00476 | 0,002658 | 1 | 0 | Intron (ENSMUST00000 | 10 | 10564545 | 10956700 | 392156 | 2 | 14816 | ENSMUST00000105560.2 | 25115 | ENSMUSG00000019828 | Grm1 | glutamate receptor, metabotropic 1 |
| chr9 | 13659759 | 13660481 | 723 | * | 5,716321 | 5,976821336 | 2,005265 | 2,980565 | 0,002877 | 1 | 0 | Promoter (<=1kb) | 9 | 13660477 | 13717777 | 57301 | 1 | 77116 | ENSMUST00000155679.8 | 0 | ENSMUSG00000031918 | Mtmr2 | myotubularin related protein 2 |
| chr18 | 10997421 | 10998274 | 854 | * | 5,260824 | 5,976827746 | 1,992304 | 2,999958 | 0,0027 | 1 | 0 | Intron (ENSMUST00000 | 18 | 10987944 | 11047637 | 59694 | 2 | 68434 | ENSMUST00000232314.2 | 49363 | ENSMUSG00000097222 | Gata6os | GATA binding protein 6, opposite strand |
| chr1 | 1,16E+08 | 1,16E+08 | 468 | * | 5,97504 | 5,976835633 | 1,941082 | 3,079126 | 0,002076 | 1 | 0 | Intron (ENSMUST00000 | 1 | 1,16E+08 | 1,17E+08 | 902568 | 1 | 636808 | ENSMUST00000043725.9 | 767828 | ENSMUSG00000070695 | Cntnap5a | contactin associated protein-like 5A |
| chr7 | 1,16E+08 | 1,16E+08 | 909 | * | 5,440101 | 5,976836299 | 1,949256 | 3,066215 | 0,002168 | 1 | 0 | Distal Intergenic | 7 | 1,16E+08 | 1,16E+08 | 7222 | 1 | 53322 | ENSMUST00000183335.2 | 46812 | ENSMUSG00000030659 | Nucb2 | nucleobindin 2 |
| chr11 | 76147215 | 76147976 | 762 | * | 6,268865 | 5,97684426 | 1,927154 | 3,101384 | 0,001926 | 1 | 0 | Distal Intergenic | 11 | 76140038 | 76141334 | 1297 | 1 | 67390 | ENSMUST00000155526.3 | 7177 | ENSMUSG00000038046 | Mrm3 | mitochondrial rRNA methyltransferase 3 |
| chr10 | 52257164 | 52257936 | 773 | * | 5,975918 | 5,976850725 | 1,914648 | 3,121645 | 0,001798 | 1 | 0 | Promoter (<=1kb) | 10 | 52213156 | 52257814 | 44659 | 2 | 94221 | ENSMUST00000217753.2 | 0 | ENSMUSG00000019861 | Gopc | golgi associated PDZ and coiled-coil motif containing |
| chr13 | 36814465 | 36815339 | 875 | * | 6,832129 | 5,976878813 | 1,86954 | 3,196979 | 0,001389 | 1 | 0 | Intron (ENSMUST00000 | 13 | 36796946 | 36837047 | 40102 | 1 | 1,08E+08 | ENSMUST00000226032.2 | 17519 | | Gm46410 | predicted gene, 46410 |
| chr14 | 1,06E+08 | 1,06E+08 | 1054 | * | 6,964387 | 5,976880648 | 1,870411 | 3,19549 | 0,001396 | 1 | 0 | Distal Intergenic | 14 | 1,06E+08 | 1,06E+08 | 4873 | 2 | 24064 | ENSMUST00000022709.6 | -40705 | ENSMUSG00000022114 | Spry2 | sprouty RTK signaling antagonist 2 |
| chrX | 75725848 | 75727332 | 1485 | * | 7,708732 | 5,976906067 | 1,852777 | 3,225918 | 0,001256 | 1 | 0 | Distal Intergenic | 20 | 75626214 | 75646530 | 20317 | 2 | 74987 | ENSMUST00000114054.3 | -79318 | ENSMUSG00000079525 | Cldn34d | claudin 34D |
| chr8 | 1,22E+08 | 1,22E+08 | 869 | * | 7,635357 | 5,977907224 | 1,870927 | 3,195159 | 0,001398 | 1 | 0 | Promoter (1-2kb) | 8 | 1,22E+08 | 1,22E+08 | 29367 | 2 | 76454 | ENSMUST00000059018.14 | -1264 | ENSMUSG00000052934 | Fbxo31 | F-box protein 31 |
| chr4 | 1,29E+08 | 1,29E+08 | 723 | * | 7,902897 | 5,980211816 | 1,826555 | 3,274039 | 0,00106 | 1 | 0 | Promoter (<=1kb) | 4 | 1,29E+08 | 1,29E+08 | 36045 | 2 | 97130 | ENSMUST00000145261.8 | -133 | ENSMUSG00000050390 | C77080 | expressed sequence C77080 |
| chr7 | 99647727 | 99648330 | 604 | * | 6,296222 | 5,980543347 | 1,969352 | 3,036808 | 0,002391 | 1 | 0 | Distal Intergenic | 7 | 99655611 | 99683933 | 28323 | 1 | 69121 | ENSMUST00000032977.11 | -7281 | ENSMUSG00000030732 | Chrdl2 | chordin-like 2 |
| chr9 | 43553899 | 43554767 | 869 | * | 5,369805 | 5,981646766 | 1,998598 | 2,992922 | 0,002763 | 1 | 0 | Promoter (<=1kb) | 9 | 43554016 | 43555375 | 1360 | 2 | 1,03E+08 | ENSMUST00000214692.2 | 608 | ENSMUSG00000111400 | Gm29961 | predicted gene, 29961 |
| chr12 | 9990440 | 9990877 | 438 | * | 4,993861 | 5,982257168 | 2,016034 | 2,967339 | 0,003004 | 1 | 0 | Distal Intergenic | 12 | 9915347 | 9915450 | 104 | 2 | 1,15E+08 | ENSMUST00000157712.3 | -74990 | ENSMUSG00000088337 | Gm22845 | predicted gene, 22845 |
| chr2 | 1,67E+08 | 1,67E+08 | 767 | * | 4,977165 | 5,982748778 | 2,031113 | 2,945552 | 0,003224 | 1 | 0 | Intron (ENSMUST00000 | 2 | 1,67E+08 | 1,67E+08 | 21315 | 1 | 99371 | ENSMUST00000144847.3 | 13862 | ENSMUSG00000074582 | Arfgef2 | ADP-ribosylation factor guanine nucleotide-exchange factor 2 (brefeldin A-inhibited) |
| chr3 | 95642665 | 95643884 | 1220 | * | 8,590606 | 5,983880158 | 1,756874 | 3,405982 | 0,000659 | 0,410415129 | 1,449706 | Promoter (<=1kb) | 3 | 95641461 | 95642330 | 870 | 2 | 13601 | ENSMUST00000144958.3 | -335 | ENSMUSG00000028108 | Ecm1 | extracellular matrix protein 1 |
| chr1 | 1,49E+08 | 1,49E+08 | 934 | * | 4,642255 | 5,984053197 | 2,074417 | 2,884692 | 0,003918 | 1 | 0 | Distal Intergenic | 1 | 1,49E+08 | 1,49E+08 | 36562 | 2 | 1,05E+08 | ENSMUST00000245751.1 | 437291 | | Gm29398 | predicted gene 29398 |
| chr4 | 1,5E+08 | 1,5E+08 | 662 | * | 5,13602 | 5,984516295 | 2,092187 | 2,860412 | 0,004231 | 1 | 0 | Distal Intergenic | 4 | 1,5E+08 | 1,51E+08 | 340321 | 1 | 68703 | ENSMUST00000105682.9 | -9392 | ENSMUSG00000039852 | Rere | arginine glutamic acid dipeptide (RE) repeats |
| chr11 | 6044587 | 6045515 | 929 | * | 9,334653 | 5,985847086 | 1,812456 | 3,302616 | 0,000958 | 0,502272318 | 1,211315 | Distal Intergenic | 11 | 6055986 | 6069409 | 13424 | 2 | 209586 | ENSMUST00000109805.8 | 23894 | ENSMUSG00000053838 | Nudcd3 | NudC domain containing 3 |
| chr11 | 70781701 | 70782914 | 1214 | * | 8,227434 | 5,986133468 | 1,805935 | 3,314701 | 0,000917 | 0,395182737 | 2,237403 | Intron (ENSMUST00000 | 11 | 70735801 | 70831309 | 95509 | 1 | 54189 | ENSMUST00000178245.2 | 45900 | ENSMUSG00000020817 | Rabep1 | rabaptin, RAB GTPase binding effector protein 1 |
| chr14 | 6267988 | 6268422 | 435 | * | 4,304433 | 5,986706474 | 2,186119 | 2,738509 | 0,006172 | 1 | 0 | Distal Intergenic | 14 | 6226476 | 6227493 | 1018 | 1 | 71147 | ENSMUST00000148121.3 | 41512 | ENSMUSG00000021786 | Oxsm | 3-oxoacyl-ACP synthase, mitochondrial |
| chr6 | 70822140 | 70822489 | 350 | * | 3,608977 | 5,988547669 | 2,288657 | 2,616621 | 0,00888 | 1 | 0 | Promoter (<=1kb) | 6 | 70821537 | 70841809 | 20273 | 1 | 13666 | ENSMUST00000162950.2 | 603 | ENSMUSG00000031668 | Eif2ak3 | eukaryotic translation initiation factor 2 alpha kinase 3 |
| chr6 | 86445871 | 86446307 | 437 | * | 3,856861 | 5,988761896 | 2,301673 | 2,601917 | 0,00927 | 1 | 0 | Intron (ENSMUST00000 | 6 | 86441022 | 86446929 | 5908 | 1 | 232196 | ENSMUST00000148768.2 | 4849 | ENSMUSG00000046679 | C87436 | expressed sequence C87436 |
| chr2 | 38652907 | 38653291 | 385 | * | 3,817605 | 5,98879439 | 2,305732 | 2,59735 | 0,009395 | 1 | 0 | Promoter (<=1kb) | 2 | 38632969 | 38653777 | 20809 | 2 | 14536 | ENSMUST00000203202.2 | 486 | ENSMUSG00000063972 | Nr6a1 | nuclear receptor subfamily 6, group A, member 1 |
| chr16 | 30257177 | 30257714 | 538 | * | 3,646506 | 5,990349085 | 2,320477 | 2,581516 | 0,009837 | 1 | 0 | Distal Intergenic | 16 | 30181397 | 30224793 | 43397 | 2 | 224088 | ENSMUST00000229616.2 | -32384 | ENSMUSG00000022533 | Atp13a3 | ATPase type 13A3 |
| chr14 | 1,07E+08 | 1,07E+08 | 359 | * | 3,682394 | 5,990432161 | 2,31523 | 2,587402 | 0,00967 | 1 | 0 | Intron (ENSMUST00000 | 14 | 1,07E+08 | 1,07E+08 | 43282 | 2 | 73631 | ENSMUST00000186643.2 | 53740 | ENSMUSG00000099794 | 1700128A07Rik | RIKEN cDNA 1700128A07 gene |
| chr3 | 34292688 | 34293033 | 346 | * | 3,489737 | 5,990456491 | 2,311398 | 2,591703 | 0,00955 | 1 | 0 | Intron (ENSMUST00000 | 3 | 34250900 | 34692632 | 441733 | 1 | 320478 | ENSMUST00000196058.5 | 41788 | ENSMUSG00000105265 | Sox2ot | SOX2 overlapping transcript (non-protein coding) |
| chr2 | 1,32E+08 | 1,32E+08 | 540 | * | 3,489226 | 5,990613082 | 2,300228 | 2,604357 | 0,009205 | 1 | 0 | Intron (ENSMUST00000 | 2 | 1,32E+08 | 1,32E+08 | 34031 | 1 | 110911 | ENSMUST00000147456.2 | 3364 | ENSMUSG00000058793 | Cds2 | CDP-diacylglycerol synthase (phosphatidate cytidylyltransferase) 2 |
| chrX | 91496785 | 91497476 | 692 | * | 4,273154 | 5,990621874 | 2,297104 | 2,607902 | 0,00911 | 1 | 0 | Intron (ENSMUST00000 | 20 | 91533555 | 91534287 | 733 | 1 | 546347 | ENSMUST00000096371.3 | -36079 | ENSMUSG00000071726 | Gm5941 | predicted gene 5941 |
| chr18 | 34786081 | 34786635 | 555 | * | 4,208347 | 5,99085668 | 2,27859 | 2,629194 | 0,008559 | 1 | 0 | Promoter (1-2kb) | 18 | 34764736 | 34784788 | 20053 | 2 | 52563 | ENSMUST00000133181.2 | -1293 | ENSMUSG00000024370 | Cdc23 | CDC23 cell division cycle 23 |
| chr7 | 1,44E+08 | 1,44E+08 | 397 | * | 4,608697 | 5,990872577 | 2,275901 | 2,632308 | 0,008481 | 1 | 0 | Promoter (1-2kb) | 7 | 1,44E+08 | 1,44E+08 | 4768 | 2 | 1,15E+08 | ENSMUST00000207398.2 | 1859 | | Gm34964 | predicted gene, 34964 |
| chr3 | 72955737 | 72956206 | 470 | * | 4,163701 | 5,990956273 | 2,270938 | 2,638098 | 0,008337 | 1 | 0 | 3' UTR | 3 | 72954598 | 72964276 | 9679 | 2 | 386750 | ENSMUST00000192477.2 | 8070 | ENSMUSG00000048304 | Slitrk3 | SLIT and NTRK-like family, member 3 |
| chr14 | 75435987 | 75436774 | 788 | * | 3,835483 | 5,991016978 | 2,265443 | 2,644524 | 0,008181 | 1 | 0 | Promoter (<=1kb) | 14 | 75435340 | 75436806 | 1467 | 1 | 18826 | ENSMUST00000130510.2 | 647 | ENSMUSG00000021998 | Lcp1 | lymphocyte cytosolic protein 1 |
| chr11 | 11863272 | 11863838 | 567 | * | 3,904747 | 5,991161384 | 2,255489 | 2,656258 | 0,007901 | 1 | 0 | Promoter (<=1kb) | 11 | 11863621 | 11863748 | 128 | 2 | 1,15E+08 | ENSMUST00000157365.4 | 0 | ENSMUSG00000087990 | Gm25058 | predicted gene, 25058 |
| chrX | 7536031 | 7536500 | 470 | * | 3,789867 | 5,991163406 | 2,255208 | 2,65659 | 0,007894 | 1 | 0 | Promoter (1-2kb) | 20 | 7535569 | 7537552 | 1984 | 2 | 18824 | ENSMUST00000143854.2 | 1052 | ENSMUSG00000031146 | Plp2 | proteolipid protein 2 |
| chr2 | 1,3E+08 | 1,3E+08 | 500 | * | 3,449446 | 5,991194722 | 2,252693 | 2,65957 | 0,007824 | 1 | 0 | Distal Intergenic | 2 | 1,3E+08 | 1,3E+08 | 28458 | 1 | 67333 | ENSMUST00000165413.9 | -30676 | ENSMUSG00000037885 | Stk35 | serine/threonine kinase 35 |
| chrX | 1,01E+08 | 1,01E+08 | 749 | * | 4,257682 | 5,991406495 | 2,237889 | 2,677258 | 0,007423 | 1 | 0 | Intron (ENSMUST00000 | 20 | 1,01E+08 | 1,01E+08 | 21116 | 1 | 1E+08 | ENSMUST00000144397.2 | 6013 | ENSMUSG00000079481 | Nhsl2 | NHS-like 2 |
| chr7 | 1,34E+08 | 1,34E+08 | 457 | * | 4,04364 | 5,991613621 | 2,221918 | 2,696595 | 0,007005 | 1 | 0 | Exon (ENSMUST000000 | 7 | 1,33E+08 | 1,34E+08 | 75070 | 2 | 11489 | ENSMUST00000138363.2 | 31352 | ENSMUSG00000054555 | Adam12 | a disintegrin and metallopeptidase domain 12 (meltrin alpha) |
| chrX | 1,62E+08 | 1,62E+08 | 530 | * | 4,265893 | 5,991776183 | 2,211724 | 2,709098 | 0,006747 | 1 | 0 | Promoter (2-3kb) | 20 | 1,62E+08 | 1,62E+08 | 107 | 2 | 1,15E+08 | ENSMUST00000082494.3 | -2029 | ENSMUSG00000118912 | Gm22746 | predicted gene, 22746 |
| chr2 | 71063377 | 71064010 | 634 | * | 4,200619 | 5,992068992 | 2,193099 | 2,732239 | 0,006291 | 1 | 0 | Promoter (1-2kb) | 2 | 71065143 | 71090693 | 25551 | 1 | 13427 | ENSMUST00000137683.2 | -1133 | ENSMUSG00000027012 | Dync1i2 | dynein cytoplasmic 1 intermediate chain 2 |
| chr11 | 94608073 | 94608592 | 520 | * | 4,054373 | 5,992262952 | 2,180336 | 2,748321 | 0,00599 | 1 | 0 | Distal Intergenic | 11 | 94585324 | 94595325 | 10002 | 2 | 432589 | ENSMUST00000069852.2 | -12748 | ENSMUSG00000056008 | Gm11541 | predicted gene 11541 |
| chr17 | 56404148 | 56404417 | 270 | * | 3,940574 | 5,992470927 | 2,167065 | 2,765247 | 0,005688 | 1 | 0 | Promoter (<=1kb) | 17 | 56403858 | 56404891 | 1034 | 1 | 15193 | ENSMUST00000224655.2 | 290 | ENSMUSG00000002833 | Hdgfl2 | HDGF like 2 |
| chr3 | 49925813 | 49926128 | 316 | * | 4,218203 | 5,992606166 | 2,159239 | 2,775333 | 0,005515 | 1 | 0 | Intron (ENSMUST00000 | 3 | 49699720 | 49711774 | 12055 | 2 | 73173 | ENSMUST00000193600.2 | -214039 | ENSMUSG00000037892 | Pcdh18 | protocadherin 18 |
| chr6 | 1,36E+08 | 1,36E+08 | 399 | * | 4,452983 | 5,992628277 | 2,158198 | 2,776682 | 0,005492 | 1 | 0 | Distal Intergenic | 6 | 1,36E+08 | 1,36E+08 | 115 | 1 | 1,15E+08 | ENSMUST00000083253.3 | -46530 | ENSMUSG00000065187 | Gm25136 | predicted gene, 25136 |
| chr4 | 26627988 | 26628616 | 629 | * | 4,314074 | 5,992740945 | 2,151434 | 2,785464 | 0,005345 | 1 | 0 | Distal Intergenic | 4 | 26635820 | 26705449 | 69630 | 2 | 75360 | ENSMUST00000154459.2 | 76833 | ENSMUSG00000086257 | 4930548K13Rik | RIKEN cDNA 4930548K13 gene |
| chr6 | 33958011 | 33958906 | 896 | * | 13,72125 | 5,992966106 | 1,662541 | 3,604703 | 0,000313 | 0,400053238 | 0,738441 | Distal Intergenic | 6 | 33961924 | 33969346 | 7423 | 2 | 1,03E+08 | ENSMUST00000155496.2 | 10440 | | Gm13853 | predicted gene 13853 |
| chrX | 1,13E+08 | 1,13E+08 | 834 | * | 4,79523 | 5,993127336 | 2,070286 | 2,894831 | 0,003794 | 1 | 0 | Intron (ENSMUST00000 | 20 | 1,13E+08 | 1,13E+08 | 109908 | 1 | 93837 | ENSMUST00000123974.2 | -41456 | ENSMUSG00000025592 | Dach2 | dachshund family transcription factor 2 |
| chr3 | 1,41E+08 | 1,41E+08 | 788 | * | 5,570629 | 5,993177448 | 2,127951 | 2,816407 | 0,004856 | 1 | 0 | Distal Intergenic | 3 | 1,41E+08 | 1,41E+08 | 2352 | 2 | 18598 | ENSMUST00000057860.3 | -78626 | ENSMUSG00000047674 | Pdha2 | pyruvate dehydrogenase E1 alpha 2 |
| chr13 | 1,04E+08 | 1,04E+08 | 466 | * | 4,676378 | 5,993906395 | 2,090094 | 2,867769 | 0,004134 | 1 | 0 | Intron (ENSMUST00000 | 13 | 1,04E+08 | 1,05E+08 | 68399 | 1 | 108154 | ENSMUST00000224504.2 | 17995 | ENSMUSG00000046169 | Adamts6 | a disintegrin-like and metallopeptidase (reprolysin type) with thrombospondin type 1 motif, 6 |
| chr2 | 1,21E+08 | 1,21E+08 | 746 | * | 6,998312 | 5,994443137 | 2,064022 | 2,904253 | 0,003681 | 1 | 0 | Intron (ENSMUST00000 | 2 | 1,21E+08 | 1,21E+08 | 8908 | 2 | 22222 | ENSMUST00000154023.2 | -18084 | ENSMUSG00000027272 | Ubr1 | ubiquitin protein ligase E3 component n-recognin 1 |
| chr2 | 91065496 | 91066114 | 619 | * | 5,333702 | 5,994476761 | 2,062832 | 2,905946 | 0,003661 | 1 | 0 | Promoter (<=1kb) | 2 | 91041917 | 91065676 | 23760 | 2 | 107986 | ENSMUST00000152277.8 | 0 | ENSMUSG00000002109 | Ddb2 | damage specific DNA binding protein 2 |
| chr2 | 1,25E+08 | 1,25E+08 | 728 | * | 5,457725 | 5,995252663 | 2,030194 | 2,953044 | 0,003147 | 1 | 0 | Intron (ENSMUST00000 | 2 | 1,25E+08 | 1,25E+08 | 116317 | 2 | 14118 | ENSMUST00000148272.2 | 106163 | ENSMUSG00000027204 | Fbn1 | fibrillin 1 |
| chr8 | 88431061 | 88432090 | 1030 | * | 12,41318 | 5,995509603 | 1,709659 | 3,506844 | 0,000453 | 0,424243345 | 0,918255 | Intron (ENSMUST00000 | 8 | 88509775 | 88530451 | 20677 | 2 | 94187 | ENSMUST00000174249.2 | 98361 | ENSMUSG00000045333 | Zfp423 | zinc finger protein 423 |
| chr4 | 1,36E+08 | 1,36E+08 | 839 | * | 5,613596 | 5,995521686 | 2,019257 | 2,969172 | 0,002986 | 1 | 0 | Promoter (<=1kb) | 4 | 1,36E+08 | 1,36E+08 | 2269 | 1 | 74326 | ENSMUST00000180855.2 | -51 | ENSMUSG00000066037 | Hnrnpr | heterogeneous nuclear ribonucleoprotein R |
| chr18 | 83991515 | 83991916 | 402 | * | 5,200297 | 5,995992128 | 2,001211 | 2,996182 | 0,002734 | 1 | 0 | Distal Intergenic | 18 | 84033943 | 84103200 | 69258 | 2 | 110796 | ENSMUST00000235188.2 | 111284 | ENSMUSG00000046982 | Tshz1 | teashirt zinc finger family member 1 |
| chr16 | 59977242 | 59977895 | 654 | * | 5,653398 | 5,99601168 | 2,001044 | 2,996442 | 0,002732 | 1 | 0 | Intron (ENSMUST00000 | 16 | 59580524 | 59826084 | 245561 | 2 | 13840 | ENSMUST00000161358.2 | -151158 | ENSMUSG00000055540 | Epha6 | Eph receptor A6 |
| chr9 | 59493991 | 59494688 | 698 | * | 5,306086 | 5,997082653 | 1,963936 | 3,053604 | 0,002261 | 1 | 0 | Promoter (1-2kb) | 9 | 59496571 | 59514574 | 18004 | 1 | 76183 | ENSMUST00000121266.8 | -1883 | ENSMUSG00000032297 | Celf6 | CUGBP, Elav-like family member 6 |
| chr12 | 77437441 | 77438214 | 774 | * | 6,025152 | 5,997656131 | 1,945863 | 3,08226 | 0,002054 | 1 | 0 | Promoter (1-2kb) | 12 | 77435638 | 77522763 | 87126 | 1 | 53618 | ENSMUST00000219299.2 | 1803 | ENSMUSG00000021065 | Fut8 | fucosyltransferase 8 |
| chr7 | 43274888 | 43275705 | 818 | * | 9,058174 | 5,998197176 | 1,767537 | 3,393535 | 0,00069 | 0,418252864 | 1,449706 | Distal Intergenic | 7 | 43284131 | 43298719 | 14589 | 1 | 72431 | ENSMUST00000032663.10 | -8426 | ENSMUSG00000030472 | Ceacam18 | carcinoembryonic antigen-related cell adhesion molecule 18 |
| chr14 | 57161917 | 57163066 | 1150 | * | 8,78614 | 5,998500371 | 1,920962 | 3,122654 | 0,001792 | 1 | 0 | Intron (ENSMUST00000 | 14 | 57158222 | 57197809 | 39588 | 1 | 76007 | ENSMUST00000224922.2 | 3695 | ENSMUSG00000021945 | Zmym2 | zinc finger, MYM-type 2 |
| chr13 | 95814107 | 95814822 | 716 | * | 6,008916 | 5,998690967 | 1,91572 | 3,131298 | 0,00174 | 1 | 0 | Intron (ENSMUST00000 | 13 | 95811801 | 95818832 | 7032 | 2 | 544963 | ENSMUST00000238263.2 | 4010 | ENSMUSG00000021676 | Iqgap2 | IQ motif containing GTPase activating protein 2 |
| chr1 | 1,82E+08 | 1,82E+08 | 870 | * | 7,751999 | 5,999308432 | 1,833581 | 3,271908 | 0,001068 | 1 | 0 | Promoter (2-3kb) | 1 | 1,82E+08 | 1,82E+08 | 10706 | 1 | 1,05E+08 | ENSMUST00000246968.1 | -2850 | | Gm33887 | predicted gene, 33887 |
| chr4 | 1,25E+08 | 1,25E+08 | 850 | * | 7,003302 | 5,99961761 | 1,856213 | 3,232182 | 0,001228 | 1 | 0 | Promoter (<=1kb) | 4 | 1,25E+08 | 1,25E+08 | 1963 | 1 | 230737 | ENSMUST00000125078.2 | 0 | ENSMUSG00000028869 | Gnl2 | guanine nucleotide binding protein-like 2 (nucleolar) |
| chr7 | 99359180 | 99360043 | 864 | * | 6,706469 | 6,000035702 | 1,891068 | 3,172829 | 0,00151 | 1 | 0 | Promoter (<=1kb) | 7 | 99307011 | 99360547 | 53537 | 2 | 101488 | ENSMUST00000032985.11 | 504 | ENSMUSG00000030737 | Slco2b1 | solute carrier organic anion transporter family, member 2b1 |

| | | | | | | | | | | | | | | | | | | | | | | |
|---|---|---|---|---|---|---|---|---|---|---|---|---|---|---|---|---|---|---|---|---|---|---|
| chr1 | 60857696 | 60858442 | 747 | * | 5,913754 | 6,000102962 | 1,896883 | 3,163138 | 0,001561 | 1 | 0 | Distal Intergenic | 1 | 60817905 | 60842949 | 25045 | 1 | 76443 | ENSMUST00000130082.3 | 39791 | | 2310016D23Rik | RIKEN cDNA 2310016D23 gene |
| chr17 | 6040307 | 6040863 | 557 | * | 5,82554 | 6,000320192 | 1,917833 | 3,128698 | 0,001756 | 1 | 0 | Promoter (<=1kb) | 17 | 6040338 | 6094565 | 54228 | 1 | 20975 | ENSMUST00000115788.8 | 0 | ENSMUSG00000023805 | Synj2 | synaptojanin 2 |
| chr13 | 8229955 | 8230698 | 744 | * | 5,67468 | 6,00042465 | 1,927863 | 3,112474 | 0,001855 | 1 | 0 | Distal Intergenic | 13 | 8252902 | 8811566 | 558665 | 1 | 94191 | ENSMUST00000064473.13 | -22204 | ENSMUSG00000052551 | Adarb2 | adenosine deaminase, RNA-specific, B2 |
| chr8 | 71930829 | 71931417 | 589 | * | 5,43857 | 6,000509697 | 1,937266 | 3,097411 | 0,001952 | 1 | 0 | Exon (ENSMUST00000( | 8 | 71929487 | 71936382 | 6896 | 2 | 382014 | ENSMUST00000212768.2 | 4965 | ENSMUSG00000034863 | Ano8 | anoctamin 8 |
| chr17 | 74942115 | 74942607 | 493 | * | 6,060139 | 6,000614924 | 1,949331 | 3,078294 | 0,002082 | 1 | 0 | Promoter (<=1kb) | 17 | 74942717 | 74947234 | 4518 | 1 | 12211 | ENSMUST00000234521.2 | -110 | ENSMUSG00000024073 | Birc6 | baculoviral IAP repeat-containing 6 |
| chr18 | 14273171 | 14273492 | 322 | * | 5,36712 | 6,00092333 | 1,985884 | 3,021789 | 0,002513 | 1 | 0 | Distal Intergenic | 18 | 14365366 | 14411599 | 46234 | 1 | 74529 | ENSMUST00000234218.2 | -91874 | ENSMUSG00000117633 | 8430422H06Rik | RIKEN cDNA 8430422H06 gene |
| chr9 | 86573993 | 86574973 | 981 | * | 4,998254 | 6,001102446 | 2,007917 | 2,98872 | 0,002801 | 1 | 0 | Promoter (2-3kb) | 9 | 86577513 | 86583306 | 5794 | 1 | 319556 | ENSMUST00000150367.4 | -2540 | ENSMUSG00000087466 | A330041J22Rik | RIKEN cDNA A330041J22 gene |
| chr3 | 87876556 | 87877249 | 694 | * | 4,983942 | 6,001166757 | 2,018082 | 2,973698 | 0,002942 | 1 | 0 | Promoter (1-2kb) | 3 | 87878385 | 87880698 | 2314 | 1 | 18008 | ENSMUST00000159830.2 | -1136 | ENSMUSG00000004891 | Nes | nestin |
| chr18 | 63519820 | 63520615 | 796 | * | 5,924518 | 6,001225891 | 2,029186 | 2,957455 | 0,003102 | 1 | 0 | Promoter (<=1kb) | 18 | 63143284 | 63520254 | 376971 | 2 | 667742 | ENSMUST00000047480.13 | 0 | ENSMUSG00000041482 | Piezo2 | piezo-type mechanosensitive ion channel component 2 |
| chr5 | 80759002 | 80760059 | 1058 | * | 7,103669 | 6,001226353 | 1,85478 | 3,235546 | 0,001214 | 1 | 0 | Distal Intergenic | 5 | 81041501 | 81041603 | 103 | 1 | 1,15E+08 | ENSMUST00000083301.3 | -281442 | ENSMUSG00000065235 | Gm22974 | predicted gene, 22974 |
| chr9 | 1,1E+08 | 1,1E+08 | 557 | * | 4,520023 | 6,001267934 | 2,032943 | 2,952009 | 0,003157 | 1 | 0 | Promoter (2-3kb) | 9 | 1,1E+08 | 1,1E+08 | 18793 | 1 | 29873 | ENSMUST00000035058.10 | -2076 | ENSMUSG00000032482 | Cspg5 | chondroitin sulfate proteoglycan 5 |
| chr5 | 1,24E+08 | 1,24E+08 | 719 | * | 5,021516 | 6,001371195 | 2,050808 | 2,926345 | 0,00343 | 1 | 0 | Intron (ENSMUST0000( | 5 | 1,24E+08 | 1,24E+08 | 63133 | 1 | 208104 | ENSMUST00000068237.12 | 5399 | ENSMUSG00000038342 | Mlxip | MLX interacting protein |
| chr1 | 42167050 | 42167972 | 923 | * | 6,683105 | 6,00143532 | 2,06158 | 2,911086 | 0,003602 | 1 | 0 | Distal Intergenic | 1 | 42298929 | 42343568 | 44640 | 1 | 1,03E+08 | ENSMUST00000240955.1 | -130957 | | Gm28175 | predicted gene 28175 |
| chr18 | 65507311 | 65507804 | 494 | * | 4,471483 | 6,001514322 | 2,072354 | 2,895989 | 0,00378 | 1 | 0 | Intron (ENSMUST0000( | 18 | 65496120 | 65526965 | 30846 | 2 | 225638 | ENSMUST00000182629.3 | 19161 | ENSMUSG00000032845 | Alpk2 | alpha-kinase 2 |
| chr3 | 1,44E+08 | 1,44E+08 | 1156 | * | 6,049403 | 6,001752043 | 2,116378 | 2,83586 | 0,00457 | 1 | 0 | Promoter (2-3kb) | 3 | 1,44E+08 | 1,44E+08 | 115919 | 2 | 23908 | ENSMUST00000160690.2 | 2481 | ENSMUSG00000040151 | Hs2st1 | heparan sulfate 2-O-sulfotransferase 1 |
| chr7 | 29354068 | 29354644 | 577 | * | 4,492295 | 6,001847575 | 2,137524 | 2,807851 | 0,004987 | 1 | 0 | Distal Intergenic | 7 | 29397116 | 29424675 | 27560 | 2 | 76718 | ENSMUST00000208371.2 | 70031 | ENSMUSG00000049123 | Catsperg2 | cation channel sperm associated auxiliary subunit gamma 2 |
| chr14 | 8621624 | 8622479 | 856 | * | 4,54156 | 6,001871696 | 2,138515 | 2,806561 | 0,005007 | 1 | 0 | Intron (ENSMUST0000( | 14 | 8555285 | 8655145 | 99861 | 1 | 238939 | ENSMUST00000144914.3 | 66339 | ENSMUSG00000084902 | Gm281 | predicted gene 281 |
| chr10 | 61483509 | 61483807 | 299 | * | 4,024432 | 6,001984151 | 2,164534 | 2,772875 | 0,005556 | 1 | 0 | Promoter (<=1kb) | 10 | 61484331 | 61509947 | 25617 | 1 | 67895 | ENSMUST00000020286.7 | -524 | ENSMUSG00000020089 | Ppa1 | pyrophosphatase (inorganic) 1 |
| chr13 | 49160753 | 49161260 | 508 | * | 4,277343 | 6,002053201 | 2,182601 | 2,749954 | 0,00596 | 1 | 0 | Distal Intergenic | 13 | 49121810 | 49123386 | 1577 | 1 | 68128 | ENSMUST00000241437.1 | 38943 | | Fam120aos | family with sequence similarity 120A, opposite strand |
| chr9 | 54804188 | 54804619 | 432 | * | 4,148251 | 6,002093111 | 2,190406 | 2,740173 | 0,006141 | 1 | 0 | Promoter (<=1kb) | 9 | 54804757 | 54811271 | 6515 | 1 | 64602 | ENSMUST00000214023.2 | -138 | ENSMUSG00000032293 | Ireb2 | iron responsive element binding protein 2 |
| chr18 | 31194169 | 31194485 | 317 | * | 3,625697 | 6,002330756 | 2,249307 | 2,668525 | 0,007619 | 1 | 0 | Intron (ENSMUST0000( | 18 | 31286324 | 31345785 | 59462 | 2 | 19762 | ENSMUST00000153196.2 | 151300 | ENSMUSG00000057455 | Rit2 | Ras-like without CAAX 2 |
| chr14 | 61772290 | 61772789 | 500 | * | 3,821246 | 6,002331883 | 2,248812 | 2,669113 | 0,007605 | 1 | 0 | 3' UTR | 14 | 61771754 | 61794207 | 22454 | 2 | 66674 | ENSMUST00000100496.5 | 21418 | ENSMUSG00000021930 | Spryd7 | SPRY domain containing 7 |
| chr4 | 28930078 | 28930700 | 623 | * | 4,077418 | 6,002396875 | 2,269201 | 2,645159 | 0,008165 | 1 | 0 | Intron (ENSMUST0000( | 4 | 28871924 | 28946334 | 74411 | 1 | 13841 | ENSMUST00000129912.2 | 58154 | ENSMUSG00000028289 | Epha7 | Eph receptor A7 |
| chr11 | 97075906 | 97076448 | 543 | * | 3,597156 | 6,002489735 | 2,293575 | 2,617088 | 0,008868 | 1 | 0 | Promoter (2-3kb) | 11 | 97050540 | 97078707 | 28168 | 2 | 16211 | ENSMUST00000001479.5 | 2259 | ENSMUSG00000001440 | Kpnb1 | karyopherin (importin) beta 1 |
| chr12 | 71857767 | 71858199 | 433 | * | 3,523364 | 6,002501599 | 2,300984 | 2,608667 | 0,00909 | 1 | 0 | Distal Intergenic | 12 | 71877852 | 72039107 | 161256 | 1 | 208846 | ENSMUST00000223272.2 | -19653 | ENSMUSG00000034574 | Daam1 | dishevelled associated activator of morphogenesis 1 |
| chr15 | 81472564 | 81473248 | 685 | * | 3,655727 | 6,002521788 | 2,301771 | 2,607784 | 0,009113 | 1 | 0 | Promoter (2-3kb) | 15 | 81470421 | 81508419 | 37999 | 1 | 328572 | ENSMUST00000189338.2 | 2143 | ENSMUSG00000055024 | Ep300 | E1A binding protein p300 |
| chr10 | 6876520 | 6876806 | 287 | * | 4,045731 | 6,003977522 | 2,308288 | 2,601052 | 0,009294 | 1 | 0 | Intron (ENSMUST0000( | 10 | 6840439 | 6929711 | 89273 | 2 | 320495 | ENSMUST00000078070.10 | 52905 | ENSMUSG00000064065 | Ipcef1 | interaction protein for cytohesin exchange factors 1 |
| chr1 | 1,56E+08 | 1,56E+08 | 780 | * | 4,802467 | 6,004332221 | 2,29379 | 2,617647 | 0,008854 | 1 | 0 | Exon (ENSMUST00000 | 1 | 1,56E+08 | 1,56E+08 | 37021 | 2 | 74081 | ENSMUST00000124495.3 | 32025 | ENSMUSG00000033671 | Cep350 | centrosomal protein 350 |
| chr16 | 17561051 | 17561901 | 851 | * | 6,762064 | 6,004739557 | 1,969064 | 3,049539 | 0,002292 | 1 | 0 | Distal Intergenic | 16 | 17498554 | 17550755 | 52202 | 2 | 94112 | ENSMUST00000231413.2 | -10296 | ENSMUSG00000012114 | Med15 | mediator complex subunit 15 |
| chr17 | 45087610 | 45087977 | 368 | * | 4,398212 | 6,005235929 | 2,256664 | 2,661112 | 0,007788 | 1 | 0 | Promoter (<=1kb) | 17 | 45088039 | 45430177 | 342139 | 1 | 109115 | ENSMUST00000050630.14 | -62 | ENSMUSG00000038954 | Supt3 | SPT3, SAGA and STAGA complex component |
| chr16 | 58850417 | 58850868 | 452 | * | 4,856189 | 6,005365385 | 2,25141 | 2,66738 | 0,007645 | 1 | 0 | Promoter (2-3kb) | 16 | 58847286 | 58848290 | 1005 | 2 | 258318 | ENSMUST00000062380.3 | -2127 | ENSMUSG00000047960 | Olfr186 | olfactory receptor 186 |
| chr2 | 1,13E+08 | 1,13E+08 | 696 | * | 5,624797 | 6,007323061 | 2,029177 | 2,960473 | 0,003072 | 1 | 0 | Intron (ENSMUST0000( | 2 | 1,12E+08 | 1,13E+08 | 398952 | 2 | 20192 | ENSMUST00000091818.6 | -14066 | ENSMUSG00000057378 | Ryr3 | ryanodine receptor 3 |
| chr7 | 1,06E+08 | 1,06E+08 | 806 | * | 5,057837 | 6,007488134 | 2,033704 | 2,953964 | 0,003137 | 1 | 0 | Distal Intergenic | 7 | 1,06E+08 | 1,06E+08 | 936 | 2 | 258595 | ENSMUST00000074981.2 | -8057 | ENSMUSG00000059087 | Olfr698 | olfactory receptor 698 |
| chr1 | 1,66E+08 | 1,66E+08 | 718 | * | 5,355654 | 6,007882656 | 2,165218 | 2,774725 | 0,005525 | 1 | 0 | Distal Intergenic | 1 | 1,66E+08 | 1,66E+08 | 13976 | 2 | 226601 | ENSMUST00000143922.2 | -11922 | ENSMUSG00000086056 | Gm4846 | predicted gene 4846 |
| chr7 | 1,04E+08 | 1,04E+08 | 558 | * | 5,547946 | 6,008043841 | 2,159597 | 2,78202 | 0,005402 | 1 | 0 | Promoter (2-3kb) | 7 | 1,04E+08 | 1,04E+08 | 1061 | 1 | 259050 | ENSMUST00000098175.3 | -2760 | ENSMUSG00000073927 | Olfr652 | olfactory receptor 652 |
| chr19 | 53515184 | 53515839 | 656 | * | 4,777071 | 6,008686047 | 2,067282 | 2,906563 | 0,003654 | 1 | 0 | Promoter (1-2kb) | 19 | 53517540 | 53530862 | 13323 | 1 | 240672 | ENSMUST00000038287.7 | -1701 | ENSMUSG00000034765 | Dusp5 | dual specificity phosphatase 5 |
| chr15 | 78891221 | 78891839 | 619 | * | 5,534671 | 6,009469504 | 2,118669 | 2,836436 | 0,004562 | 1 | 0 | Intron (ENSMUST0000( | 15 | 78879155 | 78895241 | 16087 | 2 | 1E+08 | ENSMUST00000230425.2 | 3402 | ENSMUSG00000116505 | Gm10865 | predicted gene 10865 |
| chrX | 35301394 | 35302185 | 792 | * | 5,159051 | 6,009597328 | 2,095608 | 2,86771 | 0,004135 | 1 | 0 | Promoter (2-3kb) | 20 | 35304891 | 35339880 | 34990 | 1 | 75974 | ENSMUST00000115264.2 | -2706 | ENSMUSG00000031093 | Dock11 | dedicator of cytokinesis 11 |
| chrX | 48884846 | 48885459 | 614 | * | 5,361201 | 6,009653366 | 2,113672 | 2,843228 | 0,004466 | 1 | 0 | Promoter (<=1kb) | 20 | 48875224 | 48884241 | 9018 | 2 | 209268 | ENSMUST00000072037.13 | -605 | ENSMUSG00000031111 | Igsf1 | immunoglobulin superfamily, member 1 |
| chr4 | 15644093 | 15645021 | 929 | * | 10,66276 | 6,010950689 | 1,745904 | 3,442887 | 0,000576 | 0,422869079 | 1,173325 | Distal Intergenic | 4 | 15881264 | 15908064 | 26801 | 1 | 12307 | ENSMUST00000029876.2 | -236243 | ENSMUSG00000028222 | Calb1 | calbindin 1 |
| chr7 | 1,14E+08 | 1,14E+08 | 824 | * | 7,277632 | 6,01128991 | 1,779641 | 3,37781 | 0,000731 | 1 | 0 | Intron (ENSMUST0000( | 7 | 1,14E+08 | 1,14E+08 | 132 | 2 | 1,15E+08 | ENSMUST00000104168.3 | 13239 | ENSMUSG00000118946 | Gm24982 | predicted gene, 24982 |
| chr13 | 1,14E+08 | 1,14E+08 | 607 | * | 4,924789 | 6,012076004 | 2,185992 | 2,750274 | 0,005955 | 1 | 0 | Intron (ENSMUST0000( | 13 | 1,14E+08 | 1,14E+08 | 27245 | 2 | 319803 | ENSMUST00000225044.2 | 52918 | ENSMUSG00000114213 | A430090L17Rik | RIKEN cDNA A430090L17 gene |
| chrX | 9139698 | 9140223 | 526 | * | 3,943798 | 6,012966524 | 2,224146 | 2,703495 | 0,006861 | 1 | 0 | Promoter (<=1kb) | 20 | 9138995 | 9179489 | 40495 | 1 | 22439 | ENSMUST00000015486.7 | 703 | ENSMUSG00000015342 | Xk | X-linked Kx blood group |
| chr9 | 41606104 | 41607157 | 1054 | * | 7,360332 | 6,013532115 | 1,906451 | 3,154308 | 0,001609 | 1 | 0 | Intron (ENSMUST0000( | 9 | 41570460 | 41586350 | 15891 | 2 | 1,03E+08 | ENSMUST00000214294.2 | -19754 | | Gm35835 | predicted gene, 35835 |
| chr12 | 32838542 | 32839475 | 934 | * | 10,95193 | 6,014224014 | 1,701897 | 3,533835 | 0,00041 | 0,408596045 | 0,918255 | Distal Intergenic | 12 | 32869544 | 32889206 | 19663 | 1 | 59027 | ENSMUST00000220200.2 | -30069 | ENSMUSG00000020572 | Nampt | nicotinamide phosphoribosyltransferase |
| chr1 | 1,36E+08 | 1,36E+08 | 807 | * | 7,108144 | 6,014896605 | 1,995714 | 3,013907 | 0,002579 | 0,663630601 | 1,360622 | Distal Intergenic | 1 | 1,36E+08 | 1,36E+08 | 47813 | 2 | 18772 | ENSMUST00000027667.13 | -20102 | ENSMUSG00000026413 | Pkp1 | plakophilin 1 |
| chr14 | 95781632 | 95782412 | 781 | * | 4,213681 | 6,016061986 | 2,295618 | 2,620672 | 0,008776 | 1 | 0 | Distal Intergenic | 14 | 96118690 | 96120216 | 1527 | 1 | 66732 | ENSMUST00000228913.2 | -336278 | ENSMUSG00000034689 | Spertl | spermatid associated like |
| chr14 | 4965091 | 4965555 | 465 | * | 3,884853 | 6,016107085 | 2,289074 | 2,628183 | 0,008584 | 1 | 0 | Distal Intergenic | 14 | 4506571 | 4809435 | 302865 | 2 | 21834 | ENSMUST00000224470.2 | -155656 | ENSMUSG00000021779 | Thrb | thyroid hormone receptor beta |
| chr12 | 1,14E+08 | 1,14E+08 | 238 | * | 3,487082 | 6,016113032 | 2,288557 | 2,628781 | 0,008569 | 1 | 0 | Distal Intergenic | 12 | 1,14E+08 | 1,14E+08 | 50281 | 1 | 1,19E+08 | ENSMUST00000247233.1 | -280194 | | Gm53771 | predicted gene, 53771 |
| chr14 | 66449459 | 66450051 | 593 | * | 3,939954 | 6,016193323 | 2,276946 | 2,642221 | 0,008236 | 1 | 0 | Promoter (<=1kb) | 14 | 66390707 | 66451001 | 60295 | 2 | 19229 | ENSMUST00000178730.8 | 950 | ENSMUSG00000059456 | Ptk2b | PTK2 protein tyrosine kinase 2 beta |
| chrX | 83935289 | 83935849 | 561 | * | 4,021318 | 6,016568765 | 2,237554 | 2,688905 | 0,007169 | 1 | 0 | Intron (ENSMUST0000( | 20 | 83816317 | 84019313 | 202997 | 1 | 13405 | ENSMUST00000156107.2 | 118972 | ENSMUSG00000045103 | Dmd | dystrophin, muscular dystrophy |
| chr1 | 1,42E+08 | 1,42E+08 | 326 | * | 4,068121 | 6,016695629 | 2,223417 | 2,706058 | 0,006809 | 1 | 0 | Distal Intergenic | 1 | 1,41E+08 | 1,41E+08 | 107 | 1 | 1,15E+08 | ENSMUST00000239951.1 | 337722 | ENSMUSG00000118759 | Gm25457 | predicted gene, 25457 |
| chr9 | 66530512 | 66530910 | 399 | * | 3,816744 | 6,017177852 | 2,178441 | 2,762149 | 0,005742 | 1 | 0 | Distal Intergenic | 9 | 66421919 | 66500424 | 78506 | 2 | 235441 | ENSMUST00000127569.8 | -30088 | ENSMUSG00000032376 | Usp3 | ubiquitin specific peptidase 3 |
| chr16 | 26707791 | 26708373 | 583 | * | 4,612394 | 6,017199255 | 2,176727 | 2,764333 | 0,005704 | 1 | 0 | Distal Intergenic | 16 | 26732303 | 26784356 | 52054 | 2 | 239789 | ENSMUST00000231417.2 | 75983 | ENSMUSG00000068428 | Gmnc | geminin coiled-coil domain containing |
| chr10 | 51669396 | 51669800 | 405 | * | 4,379786 | 6,017213859 | 2,174311 | 2,767412 | 0,00565 | 1 | 0 | Distal Intergenic | 10 | 51632270 | 51633290 | 1021 | 2 | 71090 | ENSMUST00000219766.2 | -36106 | ENSMUSG00000062224 | 4933411G06Rik | Sumo1 pseudogene |
| chr5 | 98640520 | 98640988 | 469 | * | 4,494797 | 6,017218426 | 2,174108 | 2,767672 | 0,005646 | 1 | 0 | Intron (ENSMUST0000( | 5 | 98623415 | 98623523 | 109 | 2 | 735265 | ENSMUST00000197105.3 | -16997 | ENSMUSG00000105814 | Mir703 | microRNA 703 |
| chr1 | 1,71E+08 | 1,71E+08 | 485 | * | 4,273521 | 6,017325978 | 2,164546 | 2,779948 | 0,005437 | 1 | 0 | Distal Intergenic | 1 | 1,71E+08 | 1,71E+08 | 6462 | 2 | 66155 | ENSMUST00000080001.9 | -5332 | ENSMUSG00000062963 | Ufc1 | ubiquitin-fold modifier conjugating enzyme 1 |
| chr3 | 40557451 | 40557889 | 439 | * | 4,219289 | 6,017351253 | 2,162918 | 2,782052 | 0,005402 | 1 | 0 | Distal Intergenic | 3 | 40562579 | 40563923 | 1345 | 2 | 1E+08 | ENSMUST00000202778.2 | 6034 | ENSMUSG00000037884 | 1700017G19Rik | RIKEN cDNA 1700017G19 gene |
| chrX | 14181271 | 14182094 | 824 | * | 4,0804 | 6,017488272 | 2,153745 | 2,793965 | 0,005207 | 1 | 0 | Promoter (<=1kb) | 20 | 14181640 | 14181830 | 191 | 2 | 1,15E+08 | ENSMUST00000082838.3 | 0 | ENSMUSG00000064772 | Gm25202 | predicted gene, 25202 |
| chr12 | 79949520 | 79950475 | 956 | * | 5,589482 | 6,017547844 | 2,145144 | 2,805195 | 0,005029 | 1 | 0 | Distal Intergenic | 12 | 79938062 | 79938677 | 616 | 1 | 1,08E+08 | ENSMUST00000243367.1 | 11458 | | Gm46362 | predicted gene, 46362 |
| chr17 | 51892387 | 51893287 | 901 | * | 4,565602 | 6,017583543 | 2,142987 | 2,808036 | 0,004984 | 1 | 0 | Distal Intergenic | 17 | 51841305 | 51841886 | 582 | 2 | 1,03E+08 | ENSMUST00000233527.2 | -50501 | ENSMUSG00000117181 | Gm31143 | predicted gene, 31143 |
| chr10 | 99113822 | 99114323 | 502 | * | 4,773902 | 6,017715315 | 2,132247 | 2,822241 | 0,004769 | 1 | 0 | Intron (ENSMUST0000( | 10 | 99110658 | 99114935 | 4278 | 1 | 1,19E+08 | ENSMUST00000218105.2 | 3164 | | Gm48089 | predicted gene, 48089 |
| chr8 | 22674572 | 22674946 | 375 | * | 5,144816 | 6,018269829 | 2,092272 | 2,876428 | 0,004022 | 1 | 0 | Promoter (<=1kb) | 8 | 22658176 | 22675835 | 17660 | 2 | 80986 | ENSMUST00000046916.9 | 889 | ENSMUSG00000037725 | Ckap2 | cytoskeleton associated protein 2 |
| chr5 | 35845729 | 35846463 | 735 | * | 4,992952 | 6,018509914 | 2,075546 | 2,899724 | 0,003735 | 1 | 0 | Distal Intergenic | 5 | 35809367 | 35837126 | 27760 | 2 | 78558 | ENSMUST00000087629.10 | -8603 | ENSMUSG00000029096 | Htra3 | HtrA serine peptidase 3 |
| chr4 | 42927362 | 42927904 | 543 | * | 4,965821 | 6,01859889 | 2,068964 | 2,908992 | 0,003626 | 1 | 0 | Intron (ENSMUST0000( | 4 | 42922268 | 42933797 | 11530 | 1 | 230085 | ENSMUST00000139100.2 | 5094 | ENSMUSG00000036062 | Phf24 | PHD finger protein 24 |
| chr16 | 4360825 | 4361339 | 515 | * | 4,893479 | 6,018970493 | 2,045738 | 2,9422 | 0,003259 | 1 | 0 | Promoter (1-2kb) | 16 | 4305443 | 4359680 | 54238 | 2 | 106393 | ENSMUST00000133440.2 | -1145 | ENSMUSG00000022519 | Srl | sarcalumenin |
| chr10 | 18896727 | 18897225 | 499 | * | 5,190417 | 6,019103035 | 2,037767 | 2,953774 | 0,003139 | 1 | 0 | Distal Intergenic | 10 | 18884014 | 18891405 | 7392 | 2 | 21929 | ENSMUST00000122863.2 | -5322 | ENSMUSG00000019850 | Tnfaip3 | tumor necrosis factor, alpha-induced protein 3 |
| chr4 | 1,37E+08 | 1,37E+08 | 765 | * | 4,947576 | 6,019910664 | 1,992791 | 3,020844 | 0,002521 | 1 | 0 | Distal Intergenic | 4 | 1,37E+08 | 1,37E+08 | 5985 | 2 | 1,01E+08 | ENSMUST00000151040.3 | 11409 | ENSMUSG00000085833 | Gm13003 | predicted gene 13003 |
| chr7 | 1,18E+08 | 1,18E+08 | 587 | * | 5,985633 | 6,020214048 | 1,977052 | 3,045046 | 0,002326 | 1 | 0 | Promoter (<=1kb) | 7 | 1,18E+08 | 1,18E+08 | 41351 | 1 | 74424 | ENSMUST00000057320.8 | 789 | ENSMUSG00000030650 | Tmc5 | transmembrane channel-like gene family 5 |
| chr4 | 6040812 | 6041460 | 649 | * | 5,161516 | 6,020277228 | 1,974398 | 3,049171 | 0,002295 | 1 | 0 | Intron (ENSMUST0000( | 4 | 6191098 | 6221688 | 30591 | 1 | 68053 | ENSMUST00000029907.6 | -149638 | ENSMUSG00000028243 | Ubxn2b | UBX domain protein 2B |
| chr17 | 25943494 | 25944446 | 953 | * | 6,894619 | 6,020716724 | 1,953171 | 3,082535 | 0,002052 | 1 | 0 | Promoter (1-2kb) | 17 | 25943755 | 25946330 | 2576 | 2 | 214901 | ENSMUST00000237158.2 | 1884 | ENSMUSG00000019214 | Chtf18 | CTF18, chromosome transmission fidelity factor 18 |
| chr11 | 1,07E+08 | 1,07E+08 | 780 | * | 5,823386 | 6,02073033 | 1,952871 | 3,083015 | 0,002049 | 1 | 0 | Intron (ENSMUST0000( | 11 | 1,07E+08 | 1,07E+08 | 2565 | 2 | 18613 | ENSMUST00000135481.2 | 14070 | ENSMUSG00000020717 | Pecam1 | platelet/endothelial cell adhesion molecule 1 |
| chr11 | 8201191 | 8201627 | 437 | * | 5,970247 | 6,020814513 | 1,949029 | 3,089135 | 0,002007 | 1 | 0 | Distal Intergenic | 11 | 8401070 | 8418675 | 17606 | 2 | 319939 | ENSMUST00000129074.2 | 217048 | ENSMUSG00000020422 | Tns3 | tensin 3 |
| chr9 | 56567750 | 56568244 | 495 | * | 6,007476 | 6,021363889 | 1,925678 | 3,12688 | 0,001767 | 1 | 0 | Intron (ENSMUST0000( | 9 | 56526389 | 56554944 | 28556 | 2 | 235402 | ENSMUST00000114247.2 | -12806 | ENSMUSG00000049556 | Lingo1 | leucine rich repeat and Ig domain containing 1 |
| chr19 | 19920159 | 19921182 | 1024 | * | 6,049238 | 6,022093545 | 1,89788 | 3,173064 | 0,001508 | 1 | 0 | Distal Intergenic | 19 | 19983304 | 19987508 | 4205 | 2 | 1E+08 | ENSMUST00000243433.1 | 66326 | | Gm3728 | predicted gene 3728 |
| chr18 | 55033279 | 55034062 | 784 | * | 7,496921 | 6,022462374 | 1,86865 | 3,222895 | 0,001269 | 1 | 0 | 3' UTR | 18 | 55025553 | 55028634 | 3082 | 2 | 269023 | ENSMUST00000237009.2 | -4645 | ENSMUSG00000052713 | Zfp608 | zinc finger protein 608 |
| chr9 | 22064400 | 22065201 | 802 | * | 7,073875 | 6,022843423 | 1,871836 | 3,217613 | 0,001293 | 1 | 0 | Promoter (2-3kb) | 9 | 22068142 | 22069654 | 1513 | 1 | 66268 | ENSMUST00000123680.2 | -2941 | ENSMUSG00000010607 | Pigyl | phosphatidylinositol glycan anchor biosynthesis, class Y-like |
| chr5 | 1,43E+08 | 1,43E+08 | 980 | * | 7,69433 | 6,023074287 | 1,863945 | 3,231358 | 0,001232 | 1 | 0 | Promoter (<=1kb) | 5 | 1,43E+08 | 1,43E+08 | 105889 | 2 | 108086 | ENSMUST00000197296.5 | 0 | ENSMUSG00000045078 | Rnf216 | ring finger protein 216 |
| chr7 | 1,04E+08 | 1,04E+08 | 672 | * | 5,80461 | 6,024046958 | 1,943802 | 3,099105 | 0,001941 | 1 | 0 | Distal Intergenic | 7 | 1,04E+08 | 1,04E+08 | 936 | 2 | 18367 | ENSMUST00000079117.2 | 9275 | ENSMUSG00000058200 | Olfr66 | olfactory receptor 66 |
| chr1 | 1,82E+08 | 1,82E+08 | 847 | * | 5,816285 | 6,024068847 | 1,944838 | 3,097466 | 0,001952 | 1 | 0 | Distal Intergenic | 1 | 1,82E+08 | 1,82E+08 | 115591 | 2 | 13800 | ENSMUST00000111024.10 | -5914 | ENSMUSG00000022995 | Enah | ENAH actin regulator |
| chr10 | 10252209 | 10252795 | 587 | * | 5,953325 | 6,024441807 | 1,966483 | 3,063562 | 0,002187 | 1 | 0 | Intron (ENSMUST0000( | 10 | 10211453 | 10287488 | 76036 | 2 | 215772 | ENSMUST00000148816.8 | 34693 | ENSMUSG00000050994 | Adgb | androglobin |
| chr8 | 1,09E+08 | 1,09E+08 | 697 | * | 5,172995 | 6,02449377 | 1,969188 | 3,059379 | 0,002218 | 1 | 0 | Intron (ENSMUST0000( | 8 | 1,09E+08 | 1,09E+08 | 9318 | 2 | 1,05E+08 | ENSMUST00000210837.2 | 148886 | | Gm39244 | predicted gene, 39244 |
| chr11 | 77716887 | 77717478 | 592 | * | 4,789287 | 6,02510373 | 2,009306 | 2,998599 | 0,002712 | 1 | 0 | Promoter (1-2kb) | 11 | 77715380 | 77718467 | 3088 | 1 | 360013 | ENSMUST00000154892.2 | 1507 | ENSMUSG00000000631 | Myo18a | myosin XVIIIA |
| chr17 | 31476219 | 31476804 | 586 | * | 5,382891 | 6,025169995 | 2,013706 | 2,99208 | 0,002771 | 1 | 0 | Exon (ENSMUST00000 | 17 | 31466638 | 31472395 | 5758 | 1 | 328795 | ENSMUST00000237156.2 | 9581 | ENSMUSG00000042345 | Ubash3a | ubiquitin associated and SH3 domain containing, A |
| chr15 | 78298150 | 78298781 | 632 | * | 4,606103 | 6,026184279 | 2,09531 | 2,876035 | 0,004027 | 1 | 0 | 3' UTR | 15 | 78294182 | 78298206 | 4025 | 1 | 246221 | ENSMUST00000229739.2 | 3968 | ENSMUSG00000071711 | Mpst | mercaptopyruvate sulfurtransferase |
| chr1 | 19487018 | 19487359 | 342 | * | 4,478059 | 6,026628617 | 2,138955 | 2,817557 | 0,004839 | 1 | 0 | Distal Intergenic | 1 | 19284103 | 19289477 | 5375 | 1 | 21419 | ENSMUST00000186972.2 | 202915 | ENSMUSG00000025927 | Tfap2b | transcription factor AP-2 beta |
| chr11 | 24832379 | 24832992 | 614 | * | 5,155924 | 6,02667329 | 2,143344 | 2,811809 | 0,004926 | 1 | 0 | Distal Intergenic | 11 | 24853983 | 24876050 | 22068 | 2 | 1,03E+08 | ENSMUST00000130092.2 | 43058 | ENSMUSG00000087145 | Gm12665 | predicted gene 12665 |
| chr18 | 71476082 | 71476626 | 545 | * | 4,789682 | 6,026740627 | 2,14898 | 2,804465 | 0,00504 | 1 | 0 | Intron (ENSMUST0000( | 18 | 70750844 | 70758848 | 8005 | 1 | 17191 | ENSMUST00000128555.2 | 725238 | ENSMUSG00000024513 | Mbd2 | methyl-CpG binding domain protein 2 |
| chr9 | 55990864 | 55991267 | 404 | * | 4,835563 | 6,026764297 | 2,151406 | 2,801314 | 0,005089 | 1 | 0 | Distal Intergenic | 9 | 55997246 | 56036172 | 38927 | 1 | 19200 | ENSMUST00000059206.8 | -5979 | ENSMUSG00000032322 | Pstpip1 | proline-serine-threonine phosphatase-interacting protein 1 |
| chr16 | 31587985 | 31588536 | 552 | * | 4,458929 | 6,027039576 | 2,182434 | 2,761614 | 0,005752 | 1 | 0 | Intron (ENSMUST0000( | 16 | 31573168 | 31691079 | 117912 | 1 | 13383 | ENSMUST00000115196.8 | 14817 | ENSMUSG00000022770 | Dlg1 | discs large MAGUK scaffold protein 1 |
| chr1 | 1,65E+08 | 1,65E+08 | 484 | * | 4,065929 | 6,027112841 | 2,192736 | 2,748672 | 0,005984 | 1 | 0 | Intron (ENSMUST0000( | 1 | 1,65E+08 | 1,65E+08 | 73418 | 1 | 433374 | ENSMUST00000194322.2 | 5199 | ENSMUSG00000104117 | Gm20743 | predicted gene, 20743 |
| chr1 | 1,56E+08 | 1,56E+08 | 294 | * | 3,990879 | 6,02737991 | 2,223966 | 2,710194 | 0,006724 | 1 | 0 | Promoter (2-3kb) | 1 | 1,56E+08 | 1,56E+08 | 7365 | 2 | 214575 | ENSMUST00000148277.2 | 2929 | ENSMUSG00000060985 | Tdrd5 | tudor domain containing 5 |
| chr3 | 57543779 | 57544105 | 327 | * | 3,845119 | 6,027508138 | 2,243219 | 2,686991 | 0,00721 | 1 | 0 | Distal Intergenic | 3 | 57530245 | 57533245 | 3001 | 2 | 1,03E+08 | ENSMUST00000199419.2 | -10534 | ENSMUSG00000097339 | Gm26671 | predicted gene, 26671 |
| chr12 | 25745699 | 25746173 | 475 | * | 4,025772 | 6,027587102 | 2,253009 | 2,675349 | 0,007465 | 1 | 0 | Distal Intergenic | 12 | 25444754 | 25521978 | 77225 | 1 | 328066 | ENSMUST00000221270.2 | 300945 | | Gm36723 | predicted gene, 36723 |
| chr7 | 31626060 | 31626648 | 589 | * | 3,978373 | 6,027633803 | 2,25675 | 2,670935 | 0,007564 | 1 | 0 | Distal Intergenic | 7 | 31671530 | 31671992 | 463 | 2 | 1,03E+08 | ENSMUST00000186898.2 | 45344 | ENSMUSG00000100926 | Gm5329 | predicted gene 5329 |
| chr7 | 65406143 | 65406708 | 566 | * | 3,896258 | 6,027824833 | 2,289247 | 2,633104 | 0,008461 | 1 | 0 | Distal Intergenic | 7 | 65452758 | 65455998 | 3241 | 1 | 1,01E+08 | ENSMUST00000243649.1 | -46050 | ENSMUSG00000100094 | 1810008I18Rik | RIKEN cDNA 1810008I18 gene |
| chr5 | 1,36E+08 | 1,36E+08 | 534 | * | 3,848298 | 6,027967174 | 2,312393 | 2,606809 | 0,009139 | 1 | 0 | Distal Intergenic | 5 | 1,36E+08 | 1,36E+08 | 35064 | 1 | 58212 | ENSMUST00000146038.2 | -10487 | ENSMUSG00000039860 | Srrm3 | serine/arginine repetitive matrix 3 |
| chr5 | 1,16E+08 | 1,16E+08 | 329 | * | 3,790162 | 6,028005663 | 2,317703 | 2,600854 | 0,009299 | 1 | 0 | Promoter (2-3kb) | 5 | 1,16E+08 | 1,16E+08 | 13979 | 1 | 77407 | ENSMUST00000138885.2 | 2465 | ENSMUSG00000029518 | Rab35 | RAB35, member RAS oncogene family |
| chr1 | 1,71E+08 | 1,71E+08 | 356 | * | 3,730147 | 6,028022788 | 2,318612 | 2,599841 | 0,009327 | 1 | 0 | Promoter (<=1kb) | 1 | 1,71E+08 | 1,71E+08 | 2911 | 2 | 226646 | ENSMUST00000131286.3 | 358 | ENSMUSG00000013593 | Ndufs2 | NADH:ubiquinone oxidoreductase core subunit S2 |
| chr7 | 89651519 | 89651938 | 420 | * | 3,761964 | 6,028068073 | 2,325356 | 2,59232 | 0,009533 | 1 | 0 | Intron (ENSMUST0000( | 7 | 89603862 | 89630191 | 26330 | 2 | 13626 | ENSMUST00000107234.3 | -21328 | ENSMUSG00000030619 | Eed | embryonic ectoderm development |
| chr7 | 1,27E+08 | 1,27E+08 | 461 | * | 3,988787 | 6,028855948 | 2,320839 | 2,597705 | 0,009385 | 1 | 0 | Distal Intergenic | 7 | 1,27E+08 | 1,27E+08 | 1522 | 2 | 269997 | ENSMUST00000205832.2 | 9061 | ENSMUSG00000054381 | Zfp747 | zinc finger protein 747 |

| | | | | | | | | | | | | | | | | | | | | | | | |
|---|---|---|---|---|---|---|---|---|---|---|---|---|---|---|---|---|---|---|---|---|---|---|---|
| chr7 | 98104989 | 98105452 | 464 | * | 4,246234 | 6,028900075 | 2,317172 | 2,601835 | 0,009273 | 1 | 0 | Exon (ENSMUST000002 | 7 | 98116901 | 98124386 | 7486 | 1 | 14918 | ENSMUST00000205350.2 | -11449 | ENSMUSG00000074003 | Gucy2d | guanylate cyclase 2d |
| chr6 | 86305268 | 86305686 | 419 | * | 4,350203 | 6,029139252 | 2,305147 | 2,615512 | 0,008909 | 1 | 0 | Distal Intergenic | 6 | 86342628 | 86347040 | 4413 | 1 | 66488 | ENSMUST00000071492.9 | -36942 | ENSMUSG00000057497 | Fam136a | family with sequence similarity 136, member A |
| chr2 | 78633254 | 78634019 | 766 | * | 6,573023 | 6,029217873 | 1,979055 | 3,046514 | 0,002315 | 1 | 0 | Distal Intergenic | 2 | 78698468 | 78700951 | 2484 | 1 | 22193 | ENSMUST00000124120.2 | -64449 | ENSMUSG00000027011 | Ube2e3 | ubiquitin-conjugating enzyme E2E 3 |
| chr5 | 34756585 | 34757172 | 588 | * | 4,169445 | 6,029515098 | 2,285491 | 2,63817 | 0,008335 | 1 | 0 | Promoter (1-2kb) | 5 | 34758686 | 34788160 | 29475 | 1 | 11518 | ENSMUST00000052836.14 | -1514 | ENSMUSG00000029106 | Add1 | adducin 1 (alpha) |
| chr15 | 78632979 | 78633526 | 548 | * | 6,696633 | 6,029705335 | 1,988875 | 3,031717 | 0,002432 | 1 | 0 | Distal Intergenic | 15 | 78648084 | 78651364 | 3281 | 2 | 17305 | ENSMUST00000136795.2 | 17838 | ENSMUSG00000018169 | Mfng | MFNG O-fucosylpeptide 3-beta-N-acetylglucosaminyltransferase |
| chr4 | 1,32E+08 | 1,32E+08 | 454 | * | 4,011482 | 6,029851083 | 2,269219 | 2,657236 | 0,007878 | 1 | 0 | Promoter (<=1kb) | 4 | 1,32E+08 | 1,32E+08 | 21392 | 1 | 66464 | ENSMUST00000030731.11 | -964 | ENSMUSG00000028899 | Taf12 | TATA-box binding protein associated factor 12 |
| chr14 | 75670617 | 75671106 | 490 | * | 4,298344 | 6,030146741 | 2,256203 | 2,672697 | 0,007524 | 1 | 0 | Distal Intergenic | 14 | 75693422 | 75767148 | 73727 | 1 | 380918 | ENSMUST00000164848.3 | -22316 | ENSMUSG00000091722 | Siah3 | siah E3 ubiquitin protein ligase family member 3 |
| chr1 | 1,84E+08 | 1,84E+08 | 401 | * | 3,781847 | 6,030359376 | 2,2453 | 2,685769 | 0,007236 | 1 | 0 | Intron (ENSMUST00000 | 1 | 1,83E+08 | 1,84E+08 | 60770 | 1 | 1,03E+08 | ENSMUST00000193310.2 | 49602 | ENSMUSG00000103305 | Gm34068 | predicted gene, 34068 |
| chrX | 80055258 | 80055799 | 542 | * | 4,127504 | 6,030789913 | 2,226968 | 2,708072 | 0,006768 | 1 | 0 | Distal Intergenic | 20 | 80114304 | 80138382 | 24079 | 1 | 192216 | ENSMUST00000150052.2 | -58505 | ENSMUSG00000025666 | Tmem47 | transmembrane protein 47 |
| chr5 | 1,22E+08 | 1,22E+08 | 403 | * | 3,92227 | 6,031020487 | 2,216522 | 2,720939 | 0,00651 | 1 | 0 | Promoter (2-3kb) | 5 | 1,22E+08 | 1,22E+08 | 25789 | 2 | 231724 | ENSMUST00000149600.8 | 2491 | ENSMUSG00000038569 | Rad9b | RAD9 checkpoint clamp component B |
| chr4 | 37847306 | 37847969 | 664 | * | 4,945311 | 6,032585377 | 2,155295 | 2,79896 | 0,005127 | 1 | 0 | Distal Intergenic | 4 | 37249403 | 37332515 | 83113 | 1 | 1,03E+08 | ENSMUST00000147108.2 | 597903 | ENSMUSG00000086026 | Gm12374 | predicted gene 12374 |
| chr12 | 15452742 | 15453252 | 511 | * | 4,786272 | 6,03339104 | 2,127144 | 2,83638 | 0,004563 | 1 | 0 | Distal Intergenic | 12 | 15656325 | 15779692 | 123368 | 1 | 1,05E+08 | ENSMUST00000222542.3 | -203073 | | Gm40287 | predicted gene, 40287 |
| chr10 | 80517832 | 80518634 | 803 | * | 4,837432 | 6,034920949 | 2,114356 | 2,854259 | 0,004314 | 1 | 0 | Distal Intergenic | 10 | 80507780 | 80513946 | 6167 | 2 | 17347 | ENSMUST00000199949.2 | -3886 | ENSMUSG00000020190 | Mknk2 | MAP kinase-interacting serine/threonine kinase 2 |
| chr1 | 1,69E+08 | 1,69E+08 | 723 | * | 4,696785 | 6,036037001 | 2,148792 | 2,809038 | 0,004969 | 1 | 0 | Intron (ENSMUST00000 | 1 | 1,69E+08 | 1,69E+08 | 16286 | 2 | 66977 | ENSMUST00000195342.6 | 183159 | ENSMUSG00000026683 | Nuf2 | NUF2, NDC80 kinetochore complex component |
| chr7 | 29794836 | 29795726 | 891 | * | 5,714866 | 6,036251846 | 2,042181 | 2,955786 | 0,003119 | 1 | 0 | Promoter (<=1kb) | 7 | 29794610 | 29807039 | 12430 | 1 | 26466 | ENSMUST00000108200.2 | 226 | ENSMUSG00000049421 | Zfp260 | zinc finger protein 260 |
| chr11 | 31432966 | 31433538 | 573 | * | 5,448344 | 6,037174991 | 2,018418 | 2,991043 | 0,00278 | 1 | 0 | Distal Intergenic | 11 | 31307307 | 31320074 | 12768 | 2 | 20856 | ENSMUST00000020546.3 | -112892 | ENSMUSG00000020303 | Stc2 | stanniocalcin 2 |
| chr7 | 65733281 | 65734003 | 723 | * | 6,234806 | 6,037266974 | 2,01643 | 2,994038 | 0,002753 | 1 | 0 | Intron (ENSMUST00000 | 7 | 65729709 | 65738935 | 9227 | 1 | 109815 | ENSMUST00000206044.2 | 3572 | ENSMUSG00000075701 | Selenos | selenoprotein S |
| chr17 | 9324926 | 9325350 | 425 | * | 4,215435 | 6,039256562 | 2,271965 | 2,658165 | 0,007857 | 1 | 0 | Distal Intergenic | 17 | 9366551 | 9386668 | 20118 | 2 | 67791 | ENSMUST00000181618.2 | 61318 | ENSMUSG00000097049 | 6530411M01Rik | RIKEN cDNA 6530411M01 gene |
| chr11 | 13160725 | 13160988 | 264 | * | 3,734992 | 6,039561228 | 2,285511 | 2,642543 | 0,008229 | 1 | 0 | Distal Intergenic | 11 | 12642813 | 12642942 | 130 | 1 | 1,15E+08 | ENSMUST00000157884.3 | 517912 | ENSMUSG00000088509 | Gm22789 | predicted gene, 22789 |
| chr7 | 18519767 | 18520251 | 485 | * | 3,638745 | 6,040902629 | 2,334526 | 2,587635 | 0,009664 | 1 | 0 | Promoter (<=1kb) | 7 | 18520257 | 18523024 | 2768 | 1 | 20958 | ENSMUST00000207882.2 | -6 | ENSMUSG00000109116 | Sycp1-ps1 | synaptonemal complex protein 1, pseudogene 1 |
| chr5 | 1,05E+08 | 1,05E+08 | 238 | * | 3,365723 | 6,040982733 | 2,316694 | 2,607588 | 0,009118 | 1 | 0 | Distal Intergenic | 5 | 1,05E+08 | 1,05E+08 | 14615 | 2 | 626391 | ENSMUST00000089906.7 | 102936 | ENSMUSG00000072774 | Zfp951 | zinc finger protein 951 |
| chr2 | 11672080 | 11672677 | 598 | * | 3,649076 | 6,041079633 | 2,300842 | 2,625596 | 0,00865 | 1 | 0 | Intron (ENSMUST00000 | 2 | 11647660 | 11653579 | 5920 | 1 | 16184 | ENSMUST00000150890.3 | 24420 | ENSMUSG00000026770 | Il2ra | interleukin 2 receptor, alpha chain |
| chr3 | 57586082 | 57586395 | 314 | * | 3,832367 | 6,041288813 | 2,257325 | 2,676304 | 0,007444 | 1 | 0 | Intron (ENSMUST00000 | 3 | 57564814 | 57599956 | 35143 | 2 | 242037 | ENSMUST00000238789.2 | 13561 | ENSMUSG00000074591 | Ankub1 | ankyrin repeat and ubiquitin domain containing 1 |
| chr9 | 69343664 | 69344240 | 577 | * | 4,091738 | 6,041413334 | 2,237243 | 2,700384 | 0,006926 | 1 | 0 | Distal Intergenic | 9 | 69360902 | 69399077 | 38176 | 1 | 12306 | ENSMUST00000034756.15 | -16662 | ENSMUSG00000032231 | Anxa2 | annexin A2 |
| chr13 | 33217687 | 33217930 | 244 | * | 3,834816 | 6,041689286 | 2,186024 | 2,76378 | 0,005714 | 1 | 0 | Promoter (<=1kb) | 13 | 33215185 | 33218543 | 3359 | 2 | 1,03E+08 | ENSMUST00000221946.2 | 613 | | Gm36099 | predicted gene, 36099 |
| chr14 | 1,01E+08 | 1,01E+08 | 688 | * | 3,853072 | 6,04170801 | 2,184852 | 2,765271 | 0,005688 | 1 | 0 | Distal Intergenic | 14 | 1,01E+08 | 1,01E+08 | 101 | 2 | 1,15E+08 | ENSMUST00000157535.4 | -44320 | ENSMUSG00000088160 | Gm23538 | predicted gene, 23538 |
| chr2 | 1,74E+08 | 1,74E+08 | 403 | * | 4,010826 | 6,04178884 | 2,171915 | 2,78178 | 0,005406 | 1 | 0 | Distal Intergenic | 2 | 1,74E+08 | 1,74E+08 | 16126 | 1 | 228960 | ENSMUST00000155000.8 | -20259 | ENSMUSG00000027522 | Stx16 | syntaxin 16 |
| chr15 | 23699570 | 23700155 | 586 | * | 5,225967 | 6,042381124 | 2,095227 | 2,883879 | 0,003928 | 1 | 0 | Distal Intergenic | 15 | 23646506 | 23646663 | 158 | 2 | 1,15E+08 | ENSMUST00000157135.4 | -52907 | ENSMUSG00000087760 | Gm23980 | predicted gene, 23980 |
| chr12 | 80188032 | 80189420 | 1389 | * | 9,396232 | 6,04239407 | 1,737033 | 3,478572 | 0,000504 | 0,300935339 | 2,237403 | Distal Intergenic | 12 | 80170745 | 80179640 | 8896 | 2 | 69548 | ENSMUST00000180643.4 | -8392 | | 2310015A10Rik | RIKEN cDNA 2310015A10 gene |
| chr15 | 77193129 | 77193520 | 392 | * | 5,024065 | 6,042464329 | 2,082575 | 2,901439 | 0,003715 | 1 | 0 | Promoter (1-2kb) | 15 | 76997806 | 77191204 | 193399 | 2 | 93686 | ENSMUST00000228582.2 | -1925 | ENSMUSG00000033565 | Rbfox2 | RNA binding protein, fox-1 homolog (C. elegans) 2 |
| chr6 | 66546823 | 66547179 | 357 | * | 4,980818 | 6,042587072 | 2,068897 | 2,920681 | 0,003493 | 1 | 0 | Distal Intergenic | 6 | 66529164 | 66537080 | 7917 | 2 | 171188 | ENSMUST00000227014.2 | -9743 | ENSMUSG00000062905 | Vmn1r32 | vomeronasal 1 receptor 32 |
| chr13 | 52496078 | 52496611 | 534 | * | 4,770337 | 6,042875448 | 2,035483 | 2,968767 | 0,00299 | 1 | 0 | Distal Intergenic | 13 | 52334447 | 52335673 | 1227 | 1 | 1,19E+08 | ENSMUST00000222876.2 | 161631 | | Gm48199 | predicted gene, 48199 |
| chr7 | 24881282 | 24881780 | 499 | * | 5,42499 | 6,042959578 | 2,026809 | 2,981515 | 0,002868 | 1 | 0 | Promoter (2-3kb) | 7 | 24820043 | 24879151 | 59109 | 2 | 18987 | ENSMUST00000175754.4 | -2131 | ENSMUSG00000008496 | Pou2f2 | POU domain, class 2, transcription factor 2 |
| chr15 | 71717180 | 71717674 | 495 | * | 4,75552 | 6,042968506 | 2,026803 | 2,981527 | 0,002868 | 1 | 0 | Intron (ENSMUST00000 | 15 | 71666037 | 71666214 | 178 | 1 | 1,15E+08 | ENSMUST00000158270.3 | 51143 | ENSMUSG00000088895 | Gm22519 | predicted gene, 22519 |
| chr15 | 40598832 | 40599657 | 826 | * | 5,010411 | 6,042979781 | 2,025864 | 2,982915 | 0,002855 | 1 | 0 | Intron (ENSMUST00000 | 15 | 40528386 | 40967982 | 439597 | 1 | 22762 | ENSMUST00000230319.2 | 70446 | ENSMUSG00000022306 | Zfpm2 | zinc finger protein, multitype 2 |
| chr3 | 30474827 | 30475591 | 765 | * | 4,960294 | 6,043166665 | 2,007679 | 3,010026 | 0,002612 | 1 | 0 | Intron (ENSMUST00000 | 3 | 30290718 | 30563636 | 272919 | 2 | 14013 | ENSMUST00000173022.2 | 88045 | ENSMUSG00000027684 | Mecom | MDS1 and EVI1 complex locus |
| chr6 | 48103445 | 48103929 | 485 | * | 7,381605 | 6,043276692 | 1,826815 | 3,308094 | 0,000939 | 1 | 0 | Distal Intergenic | 6 | 48107840 | 48136214 | 28375 | 2 | 1,01E+08 | ENSMUST00000180741.3 | 32285 | ENSMUSG00000107875 | Gm16630 | predicted gene, 16630 |
| chr19 | 4375916 | 4376983 | 1068 | * | 5,527681 | 6,043369086 | 1,989011 | 3,038379 | 0,002379 | 1 | 0 | Promoter (1-2kb) | 19 | 4367734 | 4374893 | 7160 | 2 | 225876 | ENSMUST00000235335.2 | -1023 | ENSMUSG00000054611 | Kdm2a | lysine (K)-specific demethylase 2A |
| chr4 | 41216882 | 41217927 | 1046 | * | 10,60358 | 6,043650332 | 1,835708 | 3,292272 | 0,000994 | 0,499240204 | 1,2763 | Intron (ENSMUST00000 | 4 | 41213776 | 41213853 | 78 | 2 | 1,15E+08 | ENSMUST00000177331.2 | -3029 | ENSMUSG00000093562 | Gm23090 | predicted gene, 23090 |
| chrX | 1,62E+08 | 1,62E+08 | 959 | * | 5,609348 | 6,043698617 | 1,961523 | 3,081125 | 0,002062 | 1 | 0 | Distal Intergenic | 20 | 1,62E+08 | 1,62E+08 | 1536 | 2 | 1,03E+08 | ENSMUST00000136472.2 | 18389 | ENSMUSG00000086034 | Gm15201 | predicted gene 15201 |
| chr19 | 37190146 | 37191151 | 1006 | * | 5,718621 | 6,043844655 | 1,950768 | 3,098188 | 0,001947 | 1 | 0 | Intron (ENSMUST00000 | 19 | 37194589 | 37199376 | 4788 | 1 | 69104 | ENSMUST00000128530.2 | -3438 | ENSMUSG00000023307 | Marchf5 | membrane associated ring-CH-type finger 5 |
| chr17 | 44325834 | 44326620 | 787 | * | 5,980585 | 6,044094103 | 1,932885 | 3,126981 | 0,001766 | 1 | 0 | Intron (ENSMUST00000 | 17 | 44337601 | 44348283 | 10683 | 1 | 53901 | ENSMUST00000229939.2 | -10981 | ENSMUSG00000039601 | Rcan2 | regulator of calcineurin 2 |
| chr19 | 40293171 | 40293943 | 773 | * | 7,464428 | 6,044278253 | 1,851173 | 3,265107 | 0,001094 | 1 | 0 | Intron (ENSMUST00000 | 19 | 40283463 | 40288270 | 4808 | 2 | 20411 | ENSMUST00000225306.2 | -4901 | ENSMUSG00000025006 | Sorbs1 | sorbin and SH3 domain containing 1 |
| chr18 | 68904141 | 68904838 | 698 | * | 7,040094 | 6,045169823 | 1,864004 | 3,243111 | 0,001182 | 1 | 0 | Distal Intergenic | 18 | 68872471 | 68872572 | 102 | 1 | 1,02E+08 | ENSMUST00000183541.3 | 31670 | ENSMUSG00000099300 | Mir6356 | microRNA 6356 |
| chrX | 1,68E+08 | 1,68E+08 | 1086 | * | 6,842377 | 6,045202897 | 1,86271 | 3,245381 | 0,001173 | 1 | 0 | Intron (ENSMUST00000 | 20 | 1,68E+08 | 1,68E+08 | 191389 | 1 | 11856 | ENSMUST00000112127.2 | 29651 | ENSMUSG00000031355 | Arhgap6 | Rho GTPase activating protein 6 |
| chr11 | 97489252 | 97489852 | 601 | * | 7,456401 | 6,045963089 | 1,824589 | 3,313602 | 0,000921 | 1 | 0 | Promoter (<=1kb) | 11 | 97489306 | 97514726 | 25421 | 1 | 69729 | ENSMUST00000131687.3 | 0 | ENSMUSG00000085860 | 2410003L11Rik | RIKEN cDNA 2410003L11 gene |
| chr4 | 6790702 | 6791547 | 846 | * | 5,920491 | 6,046654309 | 1,921533 | 3,146786 | 0,001651 | 1 | 0 | Intron (ENSMUST00000 | 4 | 6688046 | 6689351 | 1306 | 2 | 252838 | ENSMUST00000125974.2 | -101351 | ENSMUSG00000041272 | Tox | thymocyte selection-associated high mobility group box |
| chr1 | 1,52E+08 | 1,52E+08 | 891 | * | 6,008906 | 6,047329099 | 1,94591 | 3,107713 | 0,001885 | 1 | 0 | Intron (ENSMUST00000 | 1 | 1,52E+08 | 1,52E+08 | 42176 | 1 | 63913 | ENSMUST00000086267.6 | 7857 | ENSMUSG00000026483 | Niban1 | niban apoptosis regulator 1 |
| chr1 | 1,14E+08 | 1,14E+08 | 704 | * | 5,890806 | 6,047433217 | 1,949356 | 3,102272 | 0,00192 | 1 | 0 | Distal Intergenic | 1 | 1,16E+08 | 1,16E+08 | 15632 | 2 | 1,08E+08 | ENSMUST00000186137.7 | 1403468 | | Gm50605 | predicted gene, 50605 |
| chr1 | 63208378 | 63208798 | 421 | * | 5,391511 | 6,047591083 | 1,955853 | 3,092048 | 0,001988 | 1 | 0 | Exon (ENSMUST00000 | 1 | 63189225 | 63215822 | 26598 | 2 | 227197 | ENSMUST00000185847.2 | 7024 | ENSMUSG00000025968 | Ndufs1 | NADH:ubiquinone oxidoreductase core subunit S1 |
| chr9 | 1,03E+08 | 1,03E+08 | 675 | * | 5,419568 | 6,048405399 | 1,988868 | 3,041129 | 0,002357 | 1 | 0 | Intron (ENSMUST00000 | 9 | 1,03E+08 | 1,03E+08 | 2982 | 2 | 22041 | ENSMUST00000190051.2 | 4050 | ENSMUSG00000032554 | Trf | transferrin |
| chr9 | 83375176 | 83375603 | 428 | * | 4,17073 | 6,05072389 | 2,1141 | 2,86208 | 0,004209 | 1 | 0 | Distal Intergenic | 9 | 83273417 | 83323180 | 49764 | 2 | 75782 | ENSMUST00000034793.15 | -51996 | ENSMUSG00000032258 | Lca5 | Leber congenital amaurosis 5 (human) |
| chr4 | 1,27E+08 | 1,27E+08 | 609 | * | 4,758433 | 6,051223211 | 2,148539 | 2,816436 | 0,004856 | 1 | 0 | Distal Intergenic | 4 | 1,27E+08 | 1,27E+08 | 1993 | 1 | 73488 | ENSMUST00000129084.8 | -38350 | | 1700080G11Rik | RIKEN cDNA 1700080G11 gene |
| chr5 | 92272868 | 92273259 | 392 | * | 4,023498 | 6,051232906 | 2,149523 | 2,815151 | 0,004875 | 1 | 0 | Promoter (1-2kb) | 5 | 92275107 | 92277972 | 2866 | 1 | 1,05E+08 | ENSMUST00000243223.1 | -1848 | | LOC105246914 | uncharacterized LOC105246914 |
| chr15 | 48202310 | 48202689 | 380 | * | 4,031351 | 6,051400593 | 2,160874 | 2,800441 | 0,005103 | 1 | 0 | Intron (ENSMUST00000 | 15 | 47445119 | 47865017 | 419899 | 2 | 239420 | ENSMUST00000161653.2 | -337293 | ENSMUSG00000022311 | Csmd3 | CUB and Sushi multiple domains 3 |
| chr15 | 81615005 | 81615581 | 577 | * | 3,86605 | 6,052037537 | 2,212325 | 2,735601 | 0,006227 | 1 | 0 | Promoter (<=1kb) | 15 | 81588454 | 81614204 | 25751 | 2 | 19387 | ENSMUST00000052374.13 | -801 | ENSMUSG00000022391 | Rangap1 | RAN GTPase activating protein 1 |
| chr11 | 58209153 | 58209844 | 692 | * | 3,716744 | 6,052199304 | 2,227636 | 2,716871 | 0,00659 | 1 | 0 | Promoter (<=1kb) | 11 | 58208153 | 58210780 | 2628 | 2 | 319475 | ENSMUST00000155662.2 | 936 | ENSMUSG00000049755 | Zfp672 | zinc finger protein 672 |
| chr19 | 4183065 | 4183539 | 475 | * | 4,321558 | 6,052525207 | 2,256441 | 2,682333 | 0,007311 | 1 | 0 | Distal Intergenic | 19 | 4177571 | 4181256 | 3686 | 1 | 66990 | ENSMUST00000127555.2 | 5494 | ENSMUSG00000024845 | Tmem134 | transmembrane protein 134 |
| chrX | 45440968 | 45441550 | 583 | * | 3,760921 | 6,052838635 | 2,286172 | 2,647587 | 0,008107 | 1 | 0 | Distal Intergenic | 20 | 45417884 | 45419222 | 1339 | 1 | 73360 | ENSMUST00000059466.3 | 23084 | ENSMUSG00000046615 | Actrt1 | actin-related protein T1 |
| chr18 | 19521590 | 19522027 | 438 | * | 4,267668 | 6,052949761 | 2,299397 | 2,632407 | 0,008478 | 1 | 0 | Distal Intergenic | 18 | 19266274 | 19334773 | 68500 | 1 | 1,03E+08 | ENSMUST00000234128.2 | 255316 | | Gm31086 | predicted gene, 31086 |
| chr8 | 1,02E+08 | 1,02E+08 | 409 | * | 3,900024 | 6,0531342 | 2,317054 | 2,612426 | 0,00899 | 1 | 0 | Distal Intergenic | 8 | 1,01E+08 | 1,01E+08 | 64447 | 2 | 1,05E+08 | ENSMUST00000210760.2 | -944809 | | Gm39232 | predicted gene, 39232 |
| chr14 | 1,16E+08 | 1,16E+08 | 394 | * | 3,400068 | 6,053226031 | 2,329101 | 2,598954 | 0,009351 | 1 | 0 | Intron (ENSMUST00000 | 14 | 1,16E+08 | 1,16E+08 | 13002 | 2 | 75088 | ENSMUST00000176582.2 | -394376 | ENSMUSG00000093551 | 4930505G20Rik | RIKEN cDNA 4930505G20 gene |
| chr4 | 1,26E+08 | 1,26E+08 | 881 | * | 7,379861 | 6,053852298 | 1,995389 | 3,03392 | 0,002414 | 0,652809396 | 1,360622 | Distal Intergenic | 4 | 1,26E+08 | 1,26E+08 | 11132 | 2 | 72395 | ENSMUST00000123368.8 | 179622 | ENSMUSG00000085562 | 2610028E06Rik | RIKEN cDNA 2610028E06 gene |
| chr13 | 91511733 | 91512254 | 522 | * | 3,850401 | 6,053876755 | 2,3065 | 2,624703 | 0,008672 | 1 | 0 | Distal Intergenic | 13 | 91518937 | 91536204 | 17268 | 2 | 320297 | ENSMUST00000244414.1 | 23950 | ENSMUSG00000097182 | A830009L08Rik | RIKEN cDNA A830009L08 gene |
| chr10 | 77362997 | 77363231 | 235 | * | 3,952962 | 6,054005385 | 2,297686 | 2,634827 | 0,008418 | 1 | 0 | Promoter (2-3kb) | 10 | 77366086 | 77401520 | 35435 | 1 | 16414 | ENSMUST00000000299.14 | -2855 | ENSMUSG00000000290 | Itgb2 | integrin beta 2 |
| chr1 | 1,41E+08 | 1,41E+08 | 453 | * | 4,012065 | 6,054113792 | 2,290136 | 2,643561 | 0,008204 | 1 | 0 | Distal Intergenic | 1 | 1,41E+08 | 1,41E+08 | 107 | 1 | 1,15E+08 | ENSMUST00000239951.1 | 241787 | ENSMUSG00000118759 | Gm25457 | predicted gene, 25457 |
| chrX | 93660763 | 93661356 | 594 | * | 4,45133 | 6,054369217 | 2,273851 | 2,662606 | 0,007754 | 1 | 0 | Promoter (1-2kb) | 20 | 93659406 | 93659731 | 326 | 1 | 1,08E+08 | ENSMUST00000117321.3 | 1357 | ENSMUSG00000080974 | Gm9009 | predicted pseudogene 9009 |
| chr11 | 4254066 | 4254405 | 340 | * | 3,756542 | 6,054523554 | 2,265481 | 2,672511 | 0,007529 | 1 | 0 | Distal Intergenic | 11 | 4235460 | 4236939 | 1480 | 1 | 1,03E+08 | ENSMUST00000127224.2 | 18606 | | Gm11958 | predicted gene 11958 |
| chr5 | 1,32E+08 | 1,32E+08 | 711 | * | 3,820464 | 6,054561741 | 2,263594 | 2,674756 | 0,007478 | 1 | 0 | Intron (ENSMUST00000 | 5 | 1,32E+08 | 1,32E+08 | 3500 | 1 | 75369 | ENSMUST00000198640.2 | -39294 | ENSMUSG00000105773 | 4930563F08Rik | RIKEN cDNA 4930563F08 gene |
| chr7 | 27053402 | 27053886 | 485 | * | 3,745407 | 6,054852205 | 2,245924 | 2,69593 | 0,007019 | 1 | 0 | Promoter (1-2kb) | 7 | 27041562 | 27055168 | 13607 | 2 | 192192 | ENSMUST00000132684.8 | 1282 | ENSMUSG00000089832 | Shkbp1 | Sh3kbp1 binding protein 1 |
| chr7 | 80295156 | 80296070 | 915 | * | 6,889175 | 6,055195973 | 2,020443 | 2,996965 | 0,002727 | 1 | 0 | Intron (ENSMUST00000 | 7 | 80285765 | 80294132 | 8368 | 1 | 1,01E+08 | ENSMUST00000142678.2 | 9391 | ENSMUSG00000084821 | Gm15880 | predicted gene 15880 |
| chr18 | 46410370 | 46410950 | 581 | * | 3,904977 | 6,055838601 | 2,193786 | 2,760451 | 0,005772 | 1 | 0 | Promoter (2-3kb) | 18 | 46374587 | 46413886 | 39300 | 2 | 225467 | ENSMUST00000236999.2 | 2936 | ENSMUSG00000024477 | Pggt1b | protein geranylgeranyltransferase type I, beta subunit |
| chrX | 73206514 | 73207302 | 789 | * | 4,458829 | 6,056197773 | 2,17602 | 2,783153 | 0,005383 | 1 | 0 | Intron (ENSMUST00000 | 20 | 73194550 | 73211444 | 16895 | 2 | 385380 | ENSMUST00000078060.4 | 4142 | ENSMUSG00000062564 | Tex28 | testis expressed 28 |
| chr6 | 29199204 | 29199545 | 342 | * | 4,601978 | 6,057701963 | 2,111055 | 2,869514 | 0,004111 | 1 | 0 | Distal Intergenic | 6 | 29200444 | 29203419 | 2976 | 2 | 23917 | ENSMUST00000162242.2 | 3874 | ENSMUSG00000003500 | Impdh1 | inosine monophosphate dehydrogenase 1 |
| chr10 | 47477177 | 47477978 | 802 | * | 4,77882 | 6,057794928 | 2,107571 | 2,874301 | 0,004049 | 1 | 0 | Distal Intergenic | 10 | 46892675 | 46978928 | 86254 | 2 | 1,03E+08 | ENSMUST00000214355.3 | -498249 | | Gm35552 | predicted gene, 35552 |
| chrX | 11726503 | 11727230 | 728 | * | 5,072919 | 6,057845007 | 2,104663 | 2,878297 | 0,003998 | 1 | 0 | Distal Intergenic | 20 | 11716723 | 11716829 | 107 | 1 | 1,15E+08 | ENSMUST00000083891.3 | 9780 | ENSMUSG00000065825 | Gm26314 | predicted gene, 26314 |
| chr5 | 1,23E+08 | 1,23E+08 | 505 | * | 4,532174 | 6,058091666 | 2,095634 | 2,890816 | 0,003842 | 1 | 0 | Distal Intergenic | 5 | 1,23E+08 | 1,23E+08 | 10634 | 2 | 12589 | ENSMUST00000135956.3 | 28389 | ENSMUSG00000029469 | Ift81 | intraflagellar transport 81 |
| chr4 | 1,24E+08 | 1,24E+08 | 679 | * | 4,663183 | 6,05867667 | 2,073972 | 2,921291 | 0,003486 | 1 | 0 | Promoter (2-3kb) | 4 | 1,24E+08 | 1,24E+08 | 3617 | 2 | 595136 | ENSMUST00000106207.8 | 2464 | ENSMUSG00000028648 | Ndufs5 | NADH:ubiquinone oxidoreductase core subunit S5 |
| chr2 | 1,35E+08 | 1,35E+08 | 348 | * | 4,858536 | 6,059460074 | 2,046913 | 2,960291 | 0,003073 | 1 | 0 | Distal Intergenic | 2 | 1,34E+08 | 1,34E+08 | 49961 | 2 | 52837 | ENSMUST00000038228.11 | -36609 | ENSMUSG00000034723 | Tmx4 | thioredoxin-related transmembrane protein 4 |
| chr13 | 78711533 | 78712134 | 602 | * | 4,911859 | 6,059752802 | 2,03746 | 2,97417 | 0,002938 | 1 | 0 | Distal Intergenic | 13 | 78631473 | 78710689 | 79217 | 1 | 1,03E+08 | ENSMUST00000224561.2 | 80060 | | Gm32067 | predicted gene, 32067 |
| chr11 | 1,02E+08 | 1,02E+08 | 734 | * | 5,373954 | 6,06050603 | 2,146249 | 2,823766 | 0,004746 | 1 | 0 | Distal Intergenic | 11 | 1,02E+08 | 1,02E+08 | 8153 | 1 | 73887 | ENSMUST00000123249.2 | -15384 | ENSMUSG00000071818 | 4930417O22Rik | RIKEN cDNA 4930417O22 gene |
| chr1 | 1,76E+08 | 1,76E+08 | 490 | * | 5,455299 | 6,061195844 | 2,166426 | 2,797786 | 0,005145 | 1 | 0 | Promoter (1-2kb) | 1 | 1,76E+08 | 1,76E+08 | 6546 | 2 | 13603 | ENSMUST00000210367.2 | -1123 | ENSMUSG00000026525 | Opn3 | opsin 3 |
| chr9 | 65792167 | 65792762 | 596 | * | 5,444631 | 6,063078905 | 2,229372 | 2,719636 | 0,006535 | 1 | 0 | Promoter (<=1kb) | 9 | 65738418 | 65792337 | 53920 | 2 | 56404 | ENSMUST00000122410.8 | 0 | ENSMUSG00000032386 | Trip4 | thyroid hormone receptor interactor 4 |
| chr2 | 1,68E+08 | 1,68E+08 | 489 | * | 6,674692 | 6,063907339 | 1,927208 | 3,146473 | 0,001653 | 1 | 0 | Distal Intergenic | 2 | 1,68E+08 | 1,68E+08 | 5103 | 1 | 1E+08 | ENSMUST00000139927.2 | -15860 | | Gm14321 | predicted gene 14321 |
| chrX | 81237016 | 81238135 | 1120 | * | 7,270585 | 6,064653502 | 1,801813 | 3,365863 | 0,000763 | 1 | 0 | Distal Intergenic | 20 | 81444697 | 81444833 | 137 | 2 | 1,15E+08 | ENSMUST00000083047.3 | 206698 | ENSMUSG00000119611 | Gm24706 | predicted gene, 24706 |
| chr4 | 1,54E+08 | 1,54E+08 | 711 | * | 6,689756 | 6,064722151 | 1,909744 | 3,175672 | 0,001495 | 1 | 0 | Promoter (1-2kb) | 4 | 1,54E+08 | 1,54E+08 | 2527 | 2 | 67808 | ENSMUST00000134492.2 | -1938 | ENSMUSG00000029030 | Tprgl | transformation related protein 63 regulated like |
| chr1 | 1,32E+08 | 1,32E+08 | 659 | * | 4,696463 | 6,065211154 | 2,314642 | 2,620366 | 0,008784 | 1 | 0 | Distal Intergenic | 1 | 1,32E+08 | 1,32E+08 | 9600 | 2 | 21367 | ENSMUST00000186530.3 | -7840 | ENSMUSG00000053024 | Cntn2 | contactin 2 |
| chr17 | 63567394 | 63568123 | 730 | * | 4,929571 | 6,065413683 | 2,322765 | 2,61129 | 0,00902 | 1 | 0 | Intron (ENSMUST00000 | 17 | 63612441 | 63619966 | 7526 | 2 | 414079 | ENSMUST00000164605.8 | 51843 | ENSMUSG00000090565 | 4930405O22Rik | RIKEN cDNA 4930405O22 gene |
| chr16 | 12133982 | 12134528 | 547 | * | 3,40557 | 6,065704091 | 2,349338 | 2,581878 | 0,009826 | 1 | 0 | Distal Intergenic | 16 | 11802584 | 11813260 | 10677 | 1 | 72555 | ENSMUST00000229804.2 | 331398 | ENSMUSG00000022494 | Shisa9 | shisa family member 9 |
| chr14 | 96011174 | 96011667 | 494 | * | 3,379119 | 6,065743788 | 2,323072 | 2,611088 | 0,009025 | 1 | 0 | Distal Intergenic | 14 | 96118690 | 96120216 | 1527 | 1 | 66732 | ENSMUST00000228913.2 | -107023 | ENSMUSG00000034689 | Spertl | spermatid associated like |
| chr16 | 92941419 | 92941829 | 411 | * | 3,834525 | 6,065773193 | 2,289099 | 2,649852 | 0,008053 | 1 | 0 | Distal Intergenic | 16 | 93140612 | 93150077 | 9466 | 2 | 69857 | ENSMUST00000185971.7 | 208248 | ENSMUSG00000100277 | 1810053B23Rik | RIKEN cDNA 1810053B23 gene |
| chr9 | 1,07E+08 | 1,07E+08 | 565 | * | 4,019632 | 6,065814186 | 2,264855 | 2,678235 | 0,007401 | 1 | 0 | Promoter (<=1kb) | 9 | 1,07E+08 | 1,07E+08 | 34813 | 2 | 102626 | ENSMUST00000143487.8 | 0 | ENSMUSG00000032577 | Mapkapk3 | mitogen-activated protein kinase-activated protein kinase 3 |
| chr15 | 8478690 | 8479242 | 553 | * | 3,734932 | 6,065861491 | 2,248542 | 2,697687 | 0,006982 | 1 | 0 | Distal Intergenic | 15 | 8320101 | 8473947 | 153847 | 2 | 71175 | ENSMUST00000052965.8 | -4743 | ENSMUSG00000022141 | Nipbl | NIPBL cohesin loading factor |
| chr6 | 35099713 | 35100002 | 290 | * | 3,959403 | 6,065899278 | 2,220986 | 2,731174 | 0,006311 | 1 | 0 | Intron (ENSMUST00000 | 6 | 34999000 | 35110560 | 111561 | 2 | 53621 | ENSMUST00000202143.4 | 10558 | ENSMUSG00000038784 | Cnot4 | CCR4-NOT transcription complex, subunit 4 |
| chr1 | 1,62E+08 | 1,62E+08 | 353 | * | 4,029326 | 6,065908245 | 2,220532 | 2,731737 | 0,0063 | 1 | 0 | Intron (ENSMUST00000 | 1 | 1,62E+08 | 1,62E+08 | 27058 | 2 | 75368 | ENSMUST00000141397.2 | 6761 | ENSMUSG00000086277 | 4930558K02Rik | RIKEN cDNA 4930558K02 gene |
| chr12 | 83258077 | 83258603 | 527 | * | 4,972426 | 6,065942315 | 2,190833 | 2,768783 | 0,005627 | 1 | 0 | Distal Intergenic | 12 | 83307267 | 83341202 | 33936 | 2 | 70127 | ENSMUST00000140327.8 | 82599 | ENSMUSG00000021221 | Dpf3 | double PHD fingers 3 |
| chr14 | 1,19E+08 | 1,19E+08 | 487 | * | 4,05061 | 6,065943914 | 2,196307 | 2,761883 | 0,005747 | 1 | 0 | 3' UTR | 14 | 1,19E+08 | 1,19E+08 | 767 | 2 | 66435 | ENSMUST00000227218.2 | 3185 | ENSMUSG00000042104 | Uggt2 | UDP-glucose glycoprotein glucosyltransferase 2 |
| chr12 | 1,11E+08 | 1,11E+08 | 511 | * | 4,033961 | 6,06601535 | 2,169892 | 2,795537 | 0,005181 | 1 | 0 | Promoter (1-2kb) | 12 | 1,11E+08 | 1,11E+08 | 7859 | 1 | 74190 | ENSMUST00000222126.2 | -1809 | ENSMUSG00000021280 | Exoc3l4 | exocyst complex component 3-like 4 |
| chr15 | 56829636 | 56829962 | 327 | * | 4,708636 | 6,066066189 | 2,144186 | 2,829076 | 0,004668 | 1 | 0 | Intron (ENSMUST00000 | 15 | 56628856 | 56641474 | 12619 | 1 | 1,03E+08 | ENSMUST00000165880.2 | 200780 | | Gm38516 | predicted gene, 38516 |
| chr15 | 8454526 | 8454975 | 450 | * | 4,014835 | 6,066109984 | 2,125636 | 2,853786 | 0,00432 | 1 | 0 | Intron (ENSMUST00000 | 15 | 8320101 | 8473947 | 153847 | 2 | 71175 | ENSMUST00000052965.8 | 18972 | ENSMUSG00000022141 | Nipbl | NIPBL cohesin loading factor |
| chr3 | 1,15E+08 | 1,15E+08 | 661 | * | 5,503087 | 6,066112794 | 2,118129 | 2,863901 | 0,004185 | 1 | 0 | 3' UTR | 3 | 1,15E+08 | 1,15E+08 | 44292 | 1 | 229759 | ENSMUST00000051309.9 | 43682 | ENSMUSG00000027965 | Olfm3 | olfactomedin 3 |

| | | | | | | | | | | | | | | | | | | | | | | | |
|---|---|---|---|---|---|---|---|---|---|---|---|---|---|---|---|---|---|---|---|---|---|---|---|
| chr9 | 31000277 | 31000800 | 524 | * | 4,580189 | 6,066116107 | 2,115646 | 2,867264 | 0,00414 | 1 | 0 | 3' UTR | 9 | 30991913 | 30993771 | 1859 | 2 | 1,05E+08 | ENSMUST00000181215.2 | -6506 | | Gm17508 | predicted gene, 17508 |
| chrX | 91833648 | 91834238 | 591 | * | 4,33465 | 6,066204178 | 2,091437 | 2,900496 | 0,003726 | 1 | 0 | Distal Intergenic | 20 | 91926057 | 91926161 | 105 | 1 | 1,15E+08 | ENSMUST00000083380.3 | -91819 | ENSMUSG00000119580 | Gm25332 | predicted gene, 25332 |
| chr9 | 15189606 | 15190657 | 1052 | * | 8,431727 | 6,066210532 | 1,848492 | 3,281708 | 0,001032 | 0,412357885 | 2,237403 | Promoter (<=1kb) | 9 | 15183024 | 15191199 | 8176 | 2 | 234959 | ENSMUST00000216406.2 | 542 | ENSMUSG00000031935 | Med17 | mediator complex subunit 17 |
| chr11 | 83371539 | 83372267 | 729 | * | 4,761587 | 6,066212297 | 2,082101 | 2,913506 | 0,003574 | 1 | 0 | Exon (ENSMUST00000 | 11 | 83379853 | 83397569 | 17717 | 1 | 70439 | ENSMUST00000133170.2 | -7586 | ENSMUSG00000020680 | Taf15 | TATA-box binding protein associated factor 15 |
| chr7 | 28053861 | 28054458 | 598 | * | 5,303306 | 6,066214033 | 2,079185 | 2,917592 | 0,003527 | 1 | 0 | Promoter (2-3kb) | 7 | 28050933 | 28051057 | 125 | 1 | 1E+08 | ENSMUST00000104523.3 | 2928 | | AF357399 | snoRNA AF357399 |
| chr9 | 29453943 | 29454619 | 677 | * | 4,165783 | 6,06622109 | 2,079381 | 2,91732 | 0,003531 | 1 | 0 | Intron (ENSMUST0000 | 9 | 29501780 | 29503806 | 2027 | 2 | 1,05E+08 | ENSMUST00000141752.2 | 49187 | | Gm15521 | predicted gene 15521 |
| chrX | 1,04E+08 | 1,04E+08 | 673 | * | 4,168613 | 6,066221828 | 2,079143 | 2,917655 | 0,003527 | 1 | 0 | Intron (ENSMUST0000 | 20 | 1,04E+08 | 1,04E+08 | 88 | 2 | 723861 | ENSMUST00000083565.3 | -8696 | ENSMUSG00000065499 | Mir384 | microRNA 384 |
| chr19 | 8734783 | 8735364 | 582 | * | 4,426676 | 6,066246264 | 2,069732 | 2,930933 | 0,003379 | 1 | 0 | Promoter (<=1kb) | 19 | 8734514 | 8742655 | 8142 | 1 | 53319 | ENSMUST00000183939.2 | 269 | ENSMUSG00000010097 | Nxf1 | nuclear RNA export factor 1 |
| chrX | 8205601 | 8206222 | 622 | * | 4,652325 | 6,06625507 | 2,068094 | 2,933259 | 0,003354 | 1 | 0 | Distal Intergenic | 20 | 8193663 | 8202476 | 8814 | 1 | 385312 | ENSMUST00000103003.4 | 11938 | ENSMUSG00000068219 | Ssxb10 | synovial sarcoma, X member B10 |
| chr2 | 1,66E+08 | 1,66E+08 | 591 | * | 6,079956 | 6,066359737 | 2,028076 | 2,991189 | 0,002779 | 1 | 0 | Distal Intergenic | 2 | 1,66E+08 | 1,66E+08 | 17330 | 2 | 1,05E+08 | ENSMUST00000151070.2 | 35495 | | Gm14268 | predicted gene 14268 |
| chr11 | 1,21E+08 | 1,21E+08 | 1142 | * | 7,550068 | 6,066362346 | 1,851434 | 3,276576 | 0,001051 | 0,497895643 | 1,360622 | Promoter (1-2kb) | 11 | 1,21E+08 | 1,21E+08 | 4726 | 2 | 217370 | ENSMUST00000147490.6 | -1247 | ENSMUSG00000039294 | Cybc1 | cytochrome b 245 chaperone 1 |
| chr1 | 1,9E+08 | 1,9E+08 | 796 | * | 5,587122 | 6,066501358 | 1,988378 | 3,05098 | 0,002281 | 1 | 0 | Distal Intergenic | 1 | 1,9E+08 | 1,9E+08 | 41872 | 2 | 226830 | ENSMUST00000027897.8 | -82719 | ENSMUSG00000026603 | Smyd2 | SET and MYND domain containing 2 |
| chr5 | 64502086 | 64502670 | 585 | * | 5,581957 | 6,066626932 | 1,956409 | 3,100899 | 0,001929 | 1 | 0 | Exon (ENSMUST00000 | 5 | 64540797 | 64543548 | 2752 | 1 | 1,08E+08 | ENSMUST00000200306.2 | -38127 | | Gm43837 | predicted gene 43837 |
| chr9 | 34167580 | 34168664 | 1085 | * | 6,500867 | 6,066709669 | 1,937599 | 3,131045 | 0,001742 | 1 | 0 | Distal Intergenic | 9 | 33953665 | 33970315 | 16651 | 1 | 77119 | ENSMUST00000184229.2 | 213915 | ENSMUSG00000098758 | 7630403G23Rik | RIKEN cDNA 7630403G23 gene |
| chr7 | 1,25E+08 | 1,25E+08 | 753 | * | 7,445803 | 6,06671928 | 1,871735 | 3,241228 | 0,00119 | 1 | 0 | Exon (ENSMUST00000 | 7 | 1,25E+08 | 1,25E+08 | 53779 | 1 | 233865 | ENSMUST00000148701.4 | 5995 | ENSMUSG00000032743 | Katnip | katanin interacting protein |
| chr3 | 1,31E+08 | 1,31E+08 | 779 | * | 5,901148 | 6,066852767 | 1,909858 | 3,176599 | 0,00149 | 1 | 0 | Promoter (1-2kb) | 3 | 1,31E+08 | 1,31E+08 | 26924 | 2 | 15107 | ENSMUST00000152427.2 | 1208 | ENSMUSG00000027984 | Hadh | hydroxyacyl-Coenzyme A dehydrogenase |
| chr6 | 1,02E+08 | 1,02E+08 | 581 | * | 6,428153 | 6,067009159 | 1,875288 | 3,235242 | 0,001215 | 1 | 0 | Distal Intergenic | 6 | 1,02E+08 | 1,02E+08 | 5386 | 1 | 207157 | ENSMUST00000064002.6 | 199490 | ENSMUSG00000052236 | Gm9871 | predicted gene 9871 |
| chr7 | 19117819 | 19118815 | 997 | * | 8,757805 | 6,067157161 | 1,849127 | 3,281092 | 0,001034 | 1 | 0 | Promoter (1-2kb) | 7 | 19116194 | 19128386 | 12193 | 1 | 13871 | ENSMUST00000129249.2 | 1625 | ENSMUSG00000030400 | Ercc2 | excision repair cross-complementing rodent repair deficiency, complementation group 2 |
| chr13 | 92274709 | 92275538 | 830 | * | 7,253365 | 6,067188581 | 1,845942 | 3,286771 | 0,001013 | 1 | 0 | Distal Intergenic | 13 | 92158557 | 92268164 | 109608 | 2 | 19418 | ENSMUST00000216219.2 | -6545 | ENSMUSG00000021708 | Rasgrf2 | RAS protein-specific guanine nucleotide-releasing factor 2 |
| chr14 | 21831966 | 21832692 | 727 | * | 7,40232 | 6,067269802 | 1,832905 | 3,310194 | 0,000932 | 1 | 0 | Intron (ENSMUST0000 | 14 | 21841940 | 21843043 | 1104 | 1 | 67630 | ENSMUST00000142023.2 | -9248 | ENSMUSG00000021770 | Samd8 | sterile alpha motif domain containing 8 |
| chr12 | 59178528 | 59179476 | 949 | * | 6,717426 | 6,067640775 | 1,877923 | 3,231038 | 0,001233 | 1 | 0 | Promoter (<=1kb) | 12 | 59178288 | 59237008 | 58721 | 1 | 338320 | ENSMUST00000177162.8 | 240 | ENSMUSG00000021000 | Mia2 | MIA SH3 domain ER export factor 2 |
| chr1 | 21341424 | 21341921 | 498 | * | 6,61262 | 6,068432847 | 1,896057 | 3,200554 | 0,001372 | 1 | 0 | Promoter (<=1kb) | 1 | 21320583 | 21341817 | 21235 | 2 | 1,03E+08 | ENSMUST00000185842.2 | 0 | | Gm28836 | predicted gene 28836 |
| chr3 | 1,27E+08 | 1,27E+08 | 1042 | * | 10,20344 | 6,07187673 | 1,762823 | 3,444405 | 0,000572 | 0,434848027 | 1,086118 | Distal Intergenic | 3 | 1,27E+08 | 1,27E+08 | 540344 | 2 | 109676 | ENSMUST00000182711.8 | -19764 | ENSMUSG00000032826 | Ank2 | ankyrin 2, brain |
| chr10 | 67869744 | 67870431 | 688 | * | 5,101461 | 6,074477125 | 2,101649 | 2,890338 | 0,003848 | 1 | 0 | Intron (ENSMUST0000 | 10 | 67879798 | 67886876 | 7079 | 1 | 170799 | ENSMUST00000123147.2 | -9367 | ENSMUSG00000037846 | Rtkn2 | rhotekin 2 |
| chr3 | 1,49E+08 | 1,49E+08 | 702 | * | 4,749468 | 6,075935466 | 2,180349 | 2,78668 | 0,005325 | 1 | 0 | Distal Intergenic | 3 | 1,49E+08 | 1,49E+08 | 97515 | 2 | 1,03E+08 | ENSMUST00000197948.2 | -41543 | ENSMUSG00000104785 | Gm31121 | predicted gene, 31121 |
| chr14 | 98906933 | 98907584 | 652 | * | 4,142381 | 6,076270623 | 2,200668 | 2,761102 | 0,005761 | 1 | 0 | Distal Intergenic | 14 | 98861626 | 98861732 | 107 | 1 | 1,15E+08 | ENSMUST00000180098.3 | 45307 | ENSMUSG00000119462 | Gm26139 | predicted gene, 26139 |
| chr3 | 16023715 | 16024185 | 471 | * | 4,20946 | 6,077465375 | 2,287509 | 2,656805 | 0,007889 | 1 | 0 | Intron (ENSMUST0000 | 3 | 16002234 | 16060496 | 58263 | 2 | 381484 | ENSMUST00000108347.3 | 36311 | ENSMUSG00000078780 | Gm5150 | predicted gene 5150 |
| chr9 | 32626214 | 32626448 | 235 | * | 3,753945 | 6,077661756 | 2,302226 | 2,639907 | 0,008293 | 1 | 0 | Intron (ENSMUST0000 | 9 | 32607680 | 32665724 | 58045 | 1 | 23871 | ENSMUST00000184364.2 | 18534 | ENSMUSG00000032035 | Ets1 | E26 avian leukemia oncogene 1, 5' domain |
| chr2 | 71438446 | 71438739 | 294 | * | 3,314051 | 6,077770249 | 2,350658 | 2,585561 | 0,009722 | 1 | 0 | Distal Intergenic | 2 | 71373752 | 71377098 | 3347 | 2 | 13392 | ENSMUST00000024159.8 | -61348 | ENSMUSG00000023391 | Dlx2 | distal-less homeobox 2 |
| chr13 | 41760911 | 41761279 | 369 | * | 3,732888 | 6,077866354 | 2,320205 | 2,619539 | 0,008805 | 1 | 0 | Promoter (1-2kb) | 13 | 41759692 | 41794831 | 35140 | 1 | 621976 | ENSMUST00000129449.3 | 1219 | ENSMUSG00000087370 | Tmem170b | transmembrane protein 170B |
| chr6 | 20078619 | 20079025 | 407 | * | 5,247448 | 6,077919491 | 2,336596 | 2,601185 | 0,00929 | 1 | 0 | Distal Intergenic | 6 | 19624608 | 19624714 | 107 | 1 | 1,15E+08 | ENSMUST00000083768.3 | 454011 | ENSMUSG00000065702 | Gm22828 | predicted gene, 22828 |
| chr3 | 1,26E+08 | 1,26E+08 | 609 | * | 4,198128 | 6,077924173 | 2,32591 | 2,613138 | 0,008972 | 1 | 0 | Distal Intergenic | 3 | 1,26E+08 | 1,26E+08 | 88387 | 1 | 108058 | ENSMUST00000143383.8 | -5517 | ENSMUSG00000053819 | Camk2d | calcium/calmodulin-dependent protein kinase II, delta |
| chr13 | 42060489 | 42060951 | 463 | * | 3,642561 | 6,078065669 | 2,322614 | 2,616907 | 0,008873 | 1 | 0 | Intron (ENSMUST0000 | 13 | 42055127 | 42055230 | 104 | 1 | 1,15E+08 | ENSMUST00000158410.3 | 5362 | ENSMUSG00000089035 | Gm22157 | predicted gene, 22157 |
| chr19 | 14452309 | 14452824 | 516 | * | 3,948271 | 6,078070064 | 2,322799 | 2,616701 | 0,008878 | 1 | 0 | Intron (ENSMUST0000 | 19 | 14521090 | 14572903 | 51814 | 2 | 21888 | ENSMUST00000235975.2 | 120079 | ENSMUSG00000024642 | Tle4 | transducin-like enhancer of split 4 |
| chr14 | 96776157 | 96776365 | 209 | * | 3,533287 | 6,078149995 | 2,317013 | 2,623269 | 0,008709 | 1 | 0 | Distal Intergenic | 14 | 96340172 | 96756538 | 416367 | 2 | 93688 | ENSMUST00000022666.9 | -19619 | ENSMUSG00000022076 | Klhl1 | kelch-like 1 |
| chr12 | 33711144 | 33711494 | 351 | * | 3,496122 | 6,07832923 | 2,302257 | 2,640161 | 0,008287 | 1 | 0 | Distal Intergenic | 12 | 33502711 | 33502842 | 132 | 2 | 1,15E+08 | ENSMUST00000239981.1 | -208302 | ENSMUSG00000119573 | Gm23529 | predicted gene, 23529 |
| chr14 | 1,2E+08 | 1,2E+08 | 564 | * | 3,471593 | 6,078334207 | 2,301349 | 2,641205 | 0,008261 | 1 | 0 | Intron (ENSMUST0000 | 14 | 1,2E+08 | 1,2E+08 | 2340 | 1 | 75432 | ENSMUST00000188582.2 | -113882 | ENSMUSG00000100274 | 1700006F04Rik | RIKEN cDNA 1700006F04 gene |
| chr7 | 1,02E+08 | 1,02E+08 | 1403 | * | 11,01827 | 6,078674673 | 1,722928 | 3,528107 | 0,000419 | 0,351174296 | 1,2763 | Promoter (<=1kb) | 7 | 1,02E+08 | 1,02E+08 | 553 | 1 | 75430 | ENSMUST00000129641.2 | 0 | ENSMUSG00000030649 | Anapc15 | anaphase promoting complex C subunit 15 |
| chr5 | 71151246 | 71151627 | 382 | * | 3,491945 | 6,079020457 | 2,247103 | 2,70527 | 0,006825 | 1 | 0 | Intron (ENSMUST0000 | 5 | 71118298 | 71171027 | 52730 | 2 | 14395 | ENSMUST00000197124.5 | 19400 | ENSMUSG00000000560 | Gabra2 | gamma-aminobutyric acid (GABA) A receptor, subunit alpha 2 |
| chr2 | 1,44E+08 | 1,44E+08 | 291 | * | 4,060522 | 6,079117443 | 2,241626 | 2,711923 | 0,006689 | 1 | 0 | Intron (ENSMUST0000 | 2 | 1,44E+08 | 1,44E+08 | 41604 | 2 | 378775 | ENSMUST00000153878.8 | 18426 | ENSMUSG00000087115 | Pcsk2os2 | proprotein convertase subtilisin/kexin type 2, opposite strand 2 |
| chr4 | 66124280 | 66124705 | 426 | * | 3,805562 | 6,079241829 | 2,230943 | 2,724965 | 0,006431 | 1 | 0 | Intron (ENSMUST0000 | 4 | 65299040 | 66322720 | 1023681 | 2 | 56079 | ENSMUST00000084496.3 | 198015 | ENSMUSG00000028373 | Astn2 | astrotactin 2 |
| chr1 | 1,84E+08 | 1,84E+08 | 627 | * | 3,790598 | 6,079369655 | 2,222886 | 2,734899 | 0,00624 | 1 | 0 | Distal Intergenic | 1 | 1,84E+08 | 1,84E+08 | 98 | 2 | 1,15E+08 | ENSMUST00000158432.4 | -20805 | ENSMUSG00000089057 | Gm23349 | predicted gene, 23349 |
| chr11 | 96440944 | 96441476 | 533 | * | 4,420776 | 6,079790298 | 2,194218 | 2,770823 | 0,005591 | 1 | 0 | Intron (ENSMUST0000 | 11 | 96380687 | 96595257 | 214571 | 1 | 78473 | ENSMUST00000127639.2 | 60257 | ENSMUSG00000057058 | Skap1 | src family associated phosphoprotein 1 |
| chr10 | 65681547 | 65682388 | 842 | * | 4,904195 | 6,079842713 | 2,191214 | 2,774646 | 0,005526 | 1 | 0 | Distal Intergenic | 10 | 65711567 | 65744018 | 32452 | 2 | 1,03E+08 | ENSMUST00000218367.2 | 61630 | ENSMUSG00000112913 | Gm30853 | predicted gene, 30853 |
| chr7 | 1,1E+08 | 1,1E+08 | 580 | * | 4,18087 | 6,079921341 | 2,185339 | 2,782141 | 0,0054 | 1 | 0 | Distal Intergenic | 7 | 1,1E+08 | 1,1E+08 | 43714 | 1 | 11717 | ENSMUST00000170374.8 | -17205 | ENSMUSG00000005686 | Ampd3 | adenosine monophosphate deaminase 3 |
| chr14 | 72438641 | 72439197 | 557 | * | 4,34429 | 6,079943856 | 2,183818 | 2,784089 | 0,005368 | 1 | 0 | Distal Intergenic | 14 | 72150531 | 72157878 | 7348 | 1 | 74667 | ENSMUST00000228197.2 | 288110 | ENSMUSG00000115304 | 4930434J06Rik | RIKEN cDNA 4930434J06 gene |
| chr10 | 1,27E+08 | 1,27E+08 | 503 | * | 4,651162 | 6,080161829 | 2,170734 | 2,80097 | 0,005095 | 1 | 0 | Promoter (<=1kb) | 10 | 1,27E+08 | 1,27E+08 | 31677 | 1 | 71750 | ENSMUST00000171434.8 | 0 | ENSMUSG00000025404 | R3hdm2 | R3H domain containing 2 |
| chr5 | 1,22E+08 | 1,22E+08 | 517 | * | 5,267268 | 6,080590541 | 2,145407 | 2,834237 | 0,004594 | 1 | 0 | Promoter (<=1kb) | 5 | 1,22E+08 | 1,22E+08 | 9156 | 1 | 20239 | ENSMUST00000159928.2 | -414 | ENSMUSG00000042605 | Atxn2 | ataxin 2 |
| chr2 | 1,8E+08 | 1,8E+08 | 731 | * | 4,365177 | 6,080968001 | 2,125752 | 2,86062 | 0,004228 | 1 | 0 | Promoter (2-3kb) | 2 | 1,8E+08 | 1,8E+08 | 107 | 1 | 1,15E+08 | ENSMUST00000082488.3 | -2584 | ENSMUSG00000119719 | Gm22502 | predicted gene, 22502 |
| chr2 | 90484842 | 90485392 | 551 | * | 4,42824 | 6,081575745 | 2,093989 | 2,904302 | 0,003681 | 1 | 0 | Distal Intergenic | 2 | 90507559 | 90566672 | 59114 | 1 | 59015 | ENSMUST00000057481.7 | -22167 | ENSMUSG00000051329 | Nup160 | nucleoporin 160 |
| chr16 | 18192280 | 18192774 | 495 | * | 4,802042 | 6,081583954 | 2,093169 | 2,905444 | 0,003667 | 1 | 0 | Intron (ENSMUST0000 | 16 | 18198648 | 18225497 | 26850 | 1 | 11877 | ENSMUST00000239535.1 | -5874 | ENSMUSG00000118669 | Arvcf | armadillo repeat gene deleted in velocardiofacial syndrome |
| chr3 | 60757039 | 60757407 | 369 | * | 4,663003 | 6,081675011 | 2,088973 | 2,911324 | 0,003599 | 1 | 0 | Distal Intergenic | 3 | 60695782 | 60702151 | 6370 | 1 | 1,03E+08 | ENSMUST00000194030.2 | 61257 | ENSMUSG00000103587 | Gm38326 | predicted gene, 38326 |
| chr11 | 23108031 | 23108628 | 598 | * | 4,61786 | 6,08197235 | 2,075746 | 2,930018 | 0,003389 | 1 | 0 | Intron (ENSMUST0000 | 11 | 23102140 | 23116375 | 14236 | 1 | 76993 | ENSMUST00000141678.2 | 5891 | ENSMUSG00000086366 | 1700061J23Rik | RIKEN cDNA 1700061J23 gene |
| chr11 | 1,17E+08 | 1,17E+08 | 527 | * | 6,065002 | 6,082480674 | 2,051906 | 2,964307 | 0,003034 | 1 | 0 | Intron (ENSMUST0000 | 11 | 1,17E+08 | 1,17E+08 | 130029 | 1 | 53860 | ENSMUST00000106354.9 | 5119 | ENSMUSG00000059248 | Septin9 | septin 9 |
| chr7 | 37804030 | 37804531 | 502 | * | 4,7769 | 6,082595943 | 2,047634 | 2,970548 | 0,002973 | 1 | 0 | Promoter (1-2kb) | 7 | 37800083 | 37806333 | 6251 | 2 | 12447 | ENSMUST00000130329.2 | 1802 | ENSMUSG00000002068 | Ccne1 | cyclin E1 |
| chr1 | 36124394 | 36125171 | 778 | * | 5,515394 | 6,08310026 | 2,026186 | 3,002242 | 0,00268 | 1 | 0 | Intron (ENSMUST0000 | 1 | 36107481 | 36145527 | 38047 | 1 | 50785 | ENSMUST00000088174.4 | 16913 | ENSMUSG00000045216 | Hs6st1 | heparan sulfate 6-O-sulfotransferase 1 |
| chr7 | 30168662 | 30169126 | 465 | * | 5,428813 | 6,083400382 | 2,014519 | 3,019778 | 0,00253 | 1 | 0 | Intron (ENSMUST0000 | 7 | 30161589 | 30165016 | 3428 | 2 | 445267 | ENSMUST00000152625.2 | -3646 | ENSMUSG00000087009 | Nphs1os | nephrosis 1 homolog, nephrin, opposite strand |
| chr4 | 72126215 | 72126802 | 588 | * | 4,911365 | 6,083463487 | 2,012471 | 3,022883 | 0,002504 | 1 | 0 | Intron (ENSMUST0000 | 4 | 72119481 | 72122138 | 2658 | 1 | 68285 | ENSMUST00000246336.1 | 6734 | | C630043F03Rik | RIKEN cDNA C630043F03 gene |
| chr9 | 41072224 | 41073398 | 1175 | * | 8,366979 | 6,085758718 | 1,935206 | 3,14476 | 0,001662 | 0,569763832 | 1,449706 | Promoter (<=1kb) | 9 | 40961822 | 41072816 | 110995 | 2 | 72828 | ENSMUST00000124819.2 | 0 | ENSMUSG00000032020 | Ubash3b | ubiquitin associated and SH3 domain containing, B |
| chr1 | 1,56E+08 | 1,56E+08 | 629 | * | 6,77057 | 6,087829447 | 1,881261 | 3,236037 | 0,001212 | 1 | 0 | Promoter (2-3kb) | 1 | 1,56E+08 | 1,56E+08 | 27787 | 2 | 20652 | ENSMUST00000187073.2 | 2549 | ENSMUSG00000026600 | Soat1 | sterol O-acyltransferase 1 |
| chr12 | 4243465 | 4243937 | 473 | * | 5,922269 | 6,08848915 | 1,940272 | 3,137956 | 0,001701 | 1 | 0 | Promoter (<=1kb) | 12 | 4244821 | 4246904 | 2084 | 1 | 104111 | ENSMUST00000150944.2 | -884 | ENSMUSG00000020654 | Adcy3 | adenylate cyclase 3 |
| chr8 | 85624210 | 85624784 | 575 | * | 7,136187 | 6,088805559 | 1,867255 | 3,260832 | 0,001111 | 1 | 0 | Intron (ENSMUST0000 | 8 | 85620627 | 85626780 | 6154 | 1 | 1,05E+08 | ENSMUST00000240952.1 | 3583 | | Gm39213 | predicted gene, 39213 |
| chr16 | 20618773 | 20619769 | 997 | * | 6,29654 | 6,088893877 | 1,881785 | 3,235702 | 0,001213 | 1 | 0 | Exon (ENSMUST00000 | 16 | 20564058 | 20564222 | 165 | 1 | 1,15E+08 | ENSMUST00000083438.3 | 54715 | ENSMUSG00000065372 | Gm22741 | predicted gene, 22741 |
| chr10 | 19209864 | 19210783 | 920 | * | 6,255675 | 6,089141705 | 1,929015 | 3,156606 | 0,001596 | 1 | 0 | Distal Intergenic | 10 | 19232281 | 19234352 | 2072 | 1 | 94222 | ENSMUST00000053225.7 | -21498 | ENSMUSG00000045591 | Olig3 | oligodendrocyte transcription factor 3 |
| chr2 | 1,66E+08 | 1,66E+08 | 700 | * | 6,801863 | 6,089244706 | 1,956908 | 3,111667 | 0,00186 | 1 | 0 | Exon (ENSMUST00000 | 2 | 1,66E+08 | 1,66E+08 | 6379 | 2 | 228880 | ENSMUST00000139202.2 | 36647 | ENSMUSG00000039671 | Zmynd8 | zinc finger, MYND-type containing 8 |
| chr10 | 40289431 | 40290215 | 785 | * | 5,408297 | 6,089373986 | 1,987726 | 3,063488 | 0,002188 | 1 | 0 | Intron (ENSMUST0000 | 10 | 40225522 | 40358097 | 132576 | 1 | 78334 | ENSMUST00000214659.2 | 63909 | ENSMUSG00000038481 | Cdk19 | cyclin-dependent kinase 19 |
| chr15 | 50068402 | 50068929 | 528 | * | 4,334958 | 6,089397914 | 2,362167 | 2,577886 | 0,009941 | 1 | 0 | Distal Intergenic | 15 | 50682895 | 50690917 | 8023 | 2 | 83925 | ENSMUST00000183421.2 | 621988 | ENSMUSG00000038679 | Trps1 | transcriptional repressor GATA binding 1 |
| chr5 | 66415200 | 66415651 | 452 | * | 4,809273 | 6,089416211 | 1,998632 | 3,046793 | 0,002313 | 1 | 0 | Promoter (1-2kb) | 5 | 66417240 | 66454405 | 37166 | 1 | 70918 | ENSMUST00000202994.4 | -1589 | ENSMUSG00000029206 | Nsun7 | NOL1/NOP2/Sun domain family, member 7 |
| chr13 | 31946783 | 31947509 | 727 | * | 5,406881 | 6,089431227 | 2,003905 | 3,038782 | 0,002375 | 1 | 0 | Distal Intergenic | 13 | 31921616 | 31921723 | 108 | 2 | 1,15E+08 | ENSMUST00000083909.3 | -25060 | ENSMUSG00000065843 | Gm25186 | predicted gene, 25186 |
| chr8 | 70610381 | 70610811 | 431 | * | 4,4277 | 6,08949976 | 2,355899 | 2,584788 | 0,009744 | 1 | 0 | Intron (ENSMUST0000 | 8 | 70605431 | 70620137 | 14707 | 1 | 17259 | ENSMUST00000110141.9 | 4950 | ENSMUSG00000079033 | Mef2b | myocyte enhancer factor 2B |
| chr10 | 83839350 | 83839879 | 530 | * | 4,904601 | 6,089547332 | 2,037676 | 2,988476 | 0,002804 | 1 | 0 | Intron (ENSMUST0000 | 10 | 83928130 | 83932113 | 3984 | 1 | 1,03E+08 | ENSMUST00000213796.2 | -88251 | ENSMUSG00000111524 | Gm34113 | predicted gene, 34113 |
| chr1 | 54706440 | 54706767 | 328 | * | 4,388165 | 6,089579792 | 2,046157 | 2,976106 | 0,002919 | 1 | 0 | Exon (ENSMUST00000 | 1 | 54688256 | 54696846 | 8591 | 2 | 329154 | ENSMUST00000178156.8 | -9594 | ENSMUSG00000052331 | Ankrd44 | ankyrin repeat domain 44 |
| chr16 | 93023221 | 93023587 | 367 | * | 4,982612 | 6,089681088 | 2,084171 | 2,921872 | 0,003479 | 1 | 0 | Distal Intergenic | 16 | 93140612 | 93150077 | 9466 | 2 | 69857 | ENSMUST00000185971.7 | 126490 | ENSMUSG00000100277 | 1810053B23Rik | RIKEN cDNA 1810053B23 gene |
| chr15 | 80395841 | 80396645 | 805 | * | 4,965969 | 6,08973755 | 2,104811 | 2,893246 | 0,003813 | 1 | 0 | Intron (ENSMUST0000 | 15 | 80336441 | 80444671 | 108231 | 2 | 383075 | ENSMUST00000096347.4 | 48026 | ENSMUSG00000050439 | Enthd1 | ENTH domain containing 1 |
| chr9 | 65350979 | 65351607 | 629 | * | 5,028175 | 6,089738598 | 2,107417 | 2,88967 | 0,003856 | 1 | 0 | Promoter (<=1kb) | 9 | 65350823 | 65359407 | 8585 | 1 | 69606 | ENSMUST00000130057.2 | 156 | ENSMUSG00000059183 | Mtfmt | mitochondrial methionyl-tRNA formyltransferase |
| chr5 | 1,2E+08 | 1,2E+08 | 422 | * | 4,283449 | 6,089793486 | 2,119634 | 2,873041 | 0,004065 | 1 | 0 | Promoter (<=1kb) | 5 | 1,2E+08 | 1,2E+08 | 584 | 1 | 21386 | ENSMUST00000154680.2 | -76 | ENSMUSG00000018604 | Tbx3 | T-box 3 |
| chr16 | 32748840 | 32749451 | 612 | * | 4,07932 | 6,089891534 | 2,16609 | 2,811468 | 0,004932 | 1 | 0 | Intron (ENSMUST0000 | 16 | 32734512 | 32833069 | 98558 | 1 | 70144 | ENSMUST00000135193.9 | 14328 | ENSMUSG00000022801 | Lrch3 | leucine-rich repeats and calponin homology (CH) domain containing 3 |
| chr3 | 97906738 | 97907357 | 620 | * | 4,541157 | 6,089911434 | 2,16286 | 2,815676 | 0,004867 | 1 | 0 | Distal Intergenic | 3 | 97920843 | 98007975 | 87133 | 1 | 18129 | ENSMUST00000198324.2 | -13486 | ENSMUSG00000027878 | Notch2 | notch 2 |
| chr10 | 65304874 | 65305268 | 395 | * | 3,838083 | 6,08993383 | 2,181325 | 2,791851 | 0,005241 | 1 | 0 | Distal Intergenic | 10 | 65370433 | 65371056 | 624 | 1 | 216036 | ENSMUST00000219802.2 | -65165 | ENSMUSG00000112341 | Gm4796 | predicted gene 4796 |
| chr4 | 16841966 | 16842261 | 296 | * | 4,54186 | 6,089960353 | 2,208835 | 2,757091 | 0,005832 | 1 | 0 | Distal Intergenic | 4 | 16861689 | 16861794 | 106 | 2 | 1,15E+08 | ENSMUST00000082685.2 | 19533 | ENSMUSG00000064619 | Gm23975 | predicted gene, 23975 |
| chrX | 1,43E+08 | 1,43E+08 | 462 | * | 4,821336 | 6,089987485 | 2,218683 | 2,744866 | 0,006054 | 1 | 0 | Promoter (1-2kb) | 20 | 1,43E+08 | 1,43E+08 | 11271 | 2 | 12338 | ENSMUST00000151154.2 | 1195 | ENSMUSG00000067276 | Capn6 | calpain 6 |
| chr13 | 1,02E+08 | 1,02E+08 | 449 | * | 3,734679 | 6,090013216 | 2,224431 | 2,737784 | 0,006185 | 1 | 0 | Distal Intergenic | 13 | 1,02E+08 | 1,02E+08 | 4376 | 2 | 1,18E+08 | ENSMUST00000225534.2 | -6541 | ENSMUSG00000114652 | Gm41043 | predicted gene, 41043 |
| chr10 | 87408065 | 87408336 | 272 | * | 3,607213 | 6,09003849 | 2,239555 | 2,719307 | 0,006542 | 1 | 0 | Intron (ENSMUST0000 | 10 | 87382535 | 87406194 | 23660 | 1 | 18478 | ENSMUST00000217864.2 | 25530 | ENSMUSG00000020051 | Pah | phenylalanine hydroxylase |
| chr7 | 82008194 | 82008560 | 367 | * | 3,34059 | 6,090070635 | 2,260323 | 2,694337 | 0,007053 | 1 | 0 | Intron (ENSMUST0000 | 7 | 81986426 | 82261638 | 275213 | 1 | 269959 | ENSMUST00000173828.3 | 21768 | ENSMUSG00000070469 | Adamtsl3 | ADAMTS-like 3 |
| chr8 | 75364092 | 75364427 | 336 | * | 3,520554 | 6,090078227 | 2,261889 | 2,692474 | 0,007092 | 1 | 0 | Intron (ENSMUST0000 | 8 | 75599801 | 75620134 | 20334 | 1 | 71597 | ENSMUST00000034034.10 | -235374 | ENSMUSG00000031621 | Isx | intestine specific homeobox |
| chr5 | 31962692 | 31963081 | 390 | * | 3,737571 | 6,090083778 | 2,265053 | 2,688716 | 0,007173 | 1 | 0 | Intron (ENSMUST0000 | 5 | 31942727 | 31977987 | 35261 | 1 | 107976 | ENSMUST00000202270.2 | 19965 | ENSMUSG00000052139 | Babam2 | BRISC and BRCA1 A complex member 2 |
| chr14 | 82021254 | 82021714 | 461 | * | 3,483456 | 6,090092483 | 2,266702 | 2,686763 | 0,007215 | 1 | 0 | Distal Intergenic | 14 | 81923250 | 81958568 | 35319 | 1 | 1,05E+08 | ENSMUST00000227374.2 | 98004 | | Gm41219 | predicted gene, 41219 |
| chr19 | 22306697 | 22307181 | 485 | * | 3,744862 | 6,090165193 | 2,330656 | 2,613069 | 0,008973 | 1 | 0 | Intron (ENSMUST0000 | 19 | 22415884 | 22425506 | 9623 | 2 | 1,03E+08 | ENSMUST00000183473.2 | 118325 | | Gm27151 | predicted gene 27151 |
| chr4 | 1,25E+08 | 1,25E+08 | 438 | * | 3,562341 | 6,090193879 | 2,343589 | 2,598661 | 0,009359 | 1 | 0 | Promoter (2-3kb) | 4 | 1,25E+08 | 1,25E+08 | 2230 | 1 | 230737 | ENSMUST00000147550.2 | -2459 | ENSMUSG00000028869 | Gnl2 | guanine nucleotide binding protein-like 2 (nucleolar) |
| chr16 | 49713540 | 49714100 | 561 | * | 4,353116 | 6,09027917 | 2,318816 | 2,626461 | 0,008628 | 1 | 0 | Intron (ENSMUST0000 | 16 | 49687196 | 49731454 | 44259 | 1 | 16423 | ENSMUST00000229104.2 | 26344 | ENSMUSG00000055447 | Cd47 | CD47 antigen (Rh-related antigen, integrin-associated signal transducer) |
| chr9 | 51395700 | 51396410 | 711 | * | 6,915807 | 6,090356042 | 1,888287 | 3,225335 | 0,001258 | 0,531203841 | 1,360622 | Distal Intergenic | 9 | 51200986 | 51240224 | 39239 | 2 | 69809 | ENSMUST00000238450.3 | -155476 | ENSMUSG00000036027 | 1810046K07Rik | RIKEN cDNA 1810046K07 gene |
| chr3 | 1,14E+08 | 1,14E+08 | 793 | * | 7,164001 | 6,090448533 | 1,92108 | 3,170326 | 0,001523 | 1 | 0 | Intron (ENSMUST0000 | 3 | 1,14E+08 | 1,14E+08 | 54040 | 1 | 12814 | ENSMUST00000184978.2 | 33524 | ENSMUSG00000027966 | Col11a1 | collagen, type XI, alpha 1 |
| chr15 | 78037890 | 78038772 | 883 | * | 9,055742 | 6,090986932 | 1,900777 | 3,204471 | 0,001353 | 0,456947326 | 2,237403 | Distal Intergenic | 15 | 78048012 | 78051136 | 3125 | 2 | 67042 | ENSMUST00000231069.2 | 12364 | ENSMUSG00000016637 | Ift27 | intraflagellar transport 27 |
| chr18 | 77988150 | 77988857 | 708 | * | 5,98477 | 6,092239337 | 1,92654 | 3,162269 | 0,001565 | 1 | 0 | Intron (ENSMUST0000 | 18 | 77981680 | 78078228 | 96549 | 1 | 1,01E+08 | ENSMUST00000044622.7 | 6470 | ENSMUSG00000039840 | Epg5 | ectopic P-granules autophagy protein 5 homolog (C. elegans) |
| chr15 | 29727061 | 29727616 | 556 | * | 4,79968 | 6,09287868 | 2,210032 | 2,756919 | 0,005835 | 1 | 0 | Distal Intergenic | 15 | 29713477 | 29713550 | 74 | 2 | 1,01E+08 | ENSMUST00000174898.3 | -13511 | ENSMUSG00000092639 | Mir3964 | microRNA 3964 |
| chr15 | 39210195 | 39211119 | 925 | * | 6,052864 | 6,092897725 | 1,941619 | 3,138049 | 0,001701 | 1 | 0 | Intron (ENSMUST0000 | 15 | 39255196 | 39315628 | 60433 | 1 | 116838 | ENSMUST00000228839.2 | -44077 | ENSMUSG00000037386 | Rims2 | regulating synaptic membrane exocytosis 2 |
| chr2 | 1,19E+08 | 1,19E+08 | 642 | * | 5,273851 | 6,094605768 | 2,151436 | 2,832808 | 0,004614 | 1 | 0 | Promoter (2-3kb) | 2 | 1,19E+08 | 1,19E+08 | 2072 | 2 | 228543 | ENSMUST00000037360.8 | 2742 | ENSMUSG00000034226 | Rhov | ras homolog family member V |
| chr11 | 74919102 | 74919697 | 596 | * | 6,073595 | 6,095492851 | 2,010328 | 3,032088 | 0,002429 | 1 | 0 | Intron (ENSMUST0000 | 11 | 74884679 | 75030195 | 145517 | 1 | 103677 | ENSMUST00000140197.3 | 34423 | ENSMUSG00000038290 | Smg6 | SMG6 nonsense mediated mRNA decay factor |
| chrX | 98333725 | 98334634 | 910 | * | 5,09753 | 6,095769034 | 2,018419 | 3,020071 | 0,002527 | 1 | 0 | Distal Intergenic | 20 | 98181737 | 98190836 | 9100 | 1 | 13641 | ENSMUST00000151750.2 | 151988 | ENSMUSG00000031217 | Efnb1 | ephrin B1 |
| chr17 | 79758413 | 79758947 | 535 | * | 5,581656 | 6,0959438 | 2,11177 | 2,886652 | 0,003894 | 1 | 0 | Promoter (<=1kb) | 17 | 79746334 | 79759191 | 12858 | 2 | 1,03E+08 | ENSMUST00000242898.1 | 244 | | Gm32954 | predicted gene, 32954 |
| chr7 | 29978030 | 29978639 | 610 | * | 4,623212 | 6,09869476 | 2,124792 | 2,870255 | 0,004101 | 1 | 0 | Promoter (<=1kb) | 7 | 29979519 | 29989662 | 10144 | 1 | 1,08E+08 | ENSMUST00000014072.6 | -880 | ENSMUSG00000013928 | 1700020L13Rik | RIKEN cDNA 1700020L13 gene |

| | | | | | | | | | | | | | | | | | | | | | | | |
|---|---|---|---|---|---|---|---|---|---|---|---|---|---|---|---|---|---|---|---|---|---|---|---|
| chr2 | 26557845 | 26558214 | 370 | * | 4,675398 | 6,09879861 | 2,129222 | 2,864332 | 0,004179 | 1 | 0 | 3' UTR | 2 | 26557686 | 26561294 | 3609 | 2 | 16821 | ENSMUST00000028283.2 | 3080 | ENSMUSG00000026919 | Lcn4 | lipocalin 4 |
| chr3 | 52005228 | 52005753 | 526 | * | 4,896977 | 6,098818551 | 2,128821 | 2,864881 | 0,004172 | 1 | 0 | Intron (ENSMUST0000( | 3 | 51593328 | 52012497 | 419170 | 2 | 433586 | ENSMUST00000121440.4 | 6744 | ENSMUSG00000061143 | Maml3 | mastermind like transcriptional coactivator 3 |
| chr7 | 89752051 | 89752757 | 707 | * | 4,880809 | 6,09883655 | 2,12921 | 2,864366 | 0,004178 | 1 | 0 | Distal Intergenic | 7 | 89774030 | 89778685 | 4656 | 2 | 78329 | ENSMUST00000181189.3 | 25928 | ENSMUSG00000097162 | 2310010J17Rik | RIKEN cDNA 2310010J17 gene |
| chr7 | 1,09E+08 | 1,09E+08 | 700 | * | 4,510226 | 6,099562382 | 2,164079 | 2,818549 | 0,004824 | 1 | 0 | Promoter (2-3kb) | 7 | 1,09E+08 | 1,09E+08 | 4920 | 2 | 57355 | ENSMUST00000145211.3 | 2255 | ENSMUSG00000031022 | BC051019 | cDNA sequence BC051019 |
| chr4 | 82017341 | 82017778 | 438 | * | 4,006544 | 6,100006272 | 2,185423 | 2,791224 | 0,005251 | 1 | 0 | Intron (ENSMUST0000( | 4 | 81952907 | 81984130 | 31224 | 2 | 1,03E+08 | ENSMUST00000243987.1 | -33211 | ENSMUSG00000087110 | Gm11264 | predicted gene 11264 |
| chr4 | 25608621 | 25609167 | 547 | * | 4,092689 | 6,101074 | 2,243648 | 2,719266 | 0,006543 | 1 | 0 | Distal Intergenic | 4 | 25566660 | 25566822 | 163 | 2 | 1,15E+08 | ENSMUST00000083106.2 | -41799 | ENSMUSG00000065040 | Gm24977 | predicted gene, 24977 |
| chr19 | 37116215 | 37116774 | 560 | * | 4,410045 | 6,101686097 | 2,2816 | 2,674302 | 0,007489 | 1 | 0 | Exon (ENSMUST00000( | 19 | 37151243 | 37173940 | 22698 | 1 | 319931 | ENSMUST00000180609.2 | -34469 | ENSMUSG00000097591 | A330032B11Rik | RIKEN cDNA A330032B11 gene |
| chr2 | 1,56E+08 | 1,56E+08 | 349 | * | 3,524177 | 6,10182571 | 2,290199 | 2,664321 | 0,007714 | 1 | 0 | Intron (ENSMUST0000( | 2 | 1,56E+08 | 1,57E+08 | 150659 | 1 | 228836 | ENSMUST00000169464.9 | 34495 | ENSMUSG00000061689 | Dlgap4 | DLG associated protein 4 |
| chr17 | 30708661 | 30709461 | 801 | * | 7,308437 | 6,101845697 | 1,976448 | 3,087279 | 0,00202 | 1 | 0 | Promoter (1-2kb) | 17 | 30439527 | 30710684 | 271158 | 2 | 224671 | ENSMUST00000236153.2 | 1223 | ENSMUSG00000062202 | Btbd9 | BTB (POZ) domain containing 9 |
| chrX | 1,67E+08 | 1,67E+08 | 659 | * | 3,732885 | 6,102028173 | 2,347441 | 2,599439 | 0,009338 | 1 | 0 | Promoter (2-3kb) | 20 | 1,66E+08 | 1,67E+08 | 426246 | 2 | 333605 | ENSMUST00000112145.3 | -2880 | ENSMUSG00000049176 | Frmpd4 | FERM and PDZ domain containing 4 |
| chr13 | 93470422 | 93470904 | 483 | * | 4,236062 | 6,102054944 | 2,306335 | 2,64578 | 0,00815 | 1 | 0 | Intron (ENSMUST0000( | 13 | 93444542 | 93541637 | 97096 | 1 | 26556 | ENSMUST00000079086.8 | 25880 | ENSMUSG00000007617 | Homer1 | homer scaffolding protein 1 |
| chr2 | 29899119 | 29899412 | 294 | * | 3,187998 | 6,102090424 | 2,335936 | 2,612268 | 0,008994 | 1 | 0 | Intron (ENSMUST0000( | 2 | 29894100 | 29896658 | 2559 | 1 | 20740 | ENSMUST00000202844.2 | 5019 | ENSMUSG00000057738 | Sptan1 | spectrin alpha, non-erythrocytic 1 |
| chr8 | 41645188 | 41645398 | 211 | * | 3,205827 | 6,102204993 | 2,322385 | 2,62756 | 0,0086 | 1 | 0 | Intron (ENSMUST0000( | 8 | 41650613 | 41668190 | 17578 | 2 | 234199 | ENSMUST00000134510.2 | 22792 | ENSMUSG00000031594 | Fgl1 | fibrinogen-like protein 1 |
| chr14 | 26125011 | 26125442 | 432 | * | 3,815667 | 6,102233894 | 2,317374 | 2,633254 | 0,008457 | 1 | 0 | Distal Intergenic | 14 | 26148480 | 26151256 | 2777 | 2 | 83997 | ENSMUST00000136625.2 | 25814 | ENSMUSG00000021870 | Slmap | sarcolemma associated protein |
| chr10 | 24535191 | 24535759 | 569 | * | 3,808578 | 6,102239142 | 2,317391 | 2,633237 | 0,008458 | 1 | 0 | Intron (ENSMUST0000( | 10 | 24544384 | 24550156 | 5773 | 2 | 18605 | ENSMUST00000142974.2 | 14397 | ENSMUSG00000037370 | Enpp1 | ectonucleotide pyrophosphatase/phosphodiesterase 1 |
| chr4 | 56546652 | 56547167 | 516 | * | 3,645308 | 6,102270506 | 2,314281 | 2,636789 | 0,008369 | 1 | 0 | Distal Intergenic | 4 | 56678904 | 56679035 | 132 | 2 | 1,15E+08 | ENSMUST00000104475.2 | 131868 | ENSMUSG00000077663 | Gm26144 | predicted gene, 26144 |
| chr16 | 32125598 | 32125949 | 352 | * | 3,716053 | 6,102334971 | 2,306146 | 2,646118 | 0,008142 | 1 | 0 | Distal Intergenic | 16 | 32151075 | 32171881 | 20807 | 1 | 224111 | ENSMUST00000232016.2 | -25126 | ENSMUSG00000053774 | Ubxn7 | UBX domain protein 7 |
| chr5 | 1,22E+08 | 1,22E+08 | 348 | * | 3,836985 | 6,102527431 | 2,283928 | 2,671944 | 0,007541 | 1 | 0 | Distal Intergenic | 5 | 1,22E+08 | 1,22E+08 | 26737 | 1 | 74096 | ENSMUST00000145854.8 | -13945 | ENSMUSG00000064267 | Hvcn1 | hydrogen voltage-gated channel 1 |
| chrX | 98317640 | 98318314 | 675 | * | 3,838776 | 6,102785351 | 2,257774 | 2,70301 | 0,006871 | 1 | 0 | Distal Intergenic | 20 | 98181737 | 98190836 | 9100 | 1 | 13641 | ENSMUST00000151750.2 | 135903 | ENSMUSG00000031217 | Efnb1 | ephrin B1 |
| chr5 | 65799365 | 65799884 | 520 | * | 3,778084 | 6,102879012 | 2,247018 | 2,71599 | 0,006608 | 1 | 0 | Exon (ENSMUST000002 | 5 | 65795085 | 65796259 | 1175 | 2 | 71521 | ENSMUST00000201046.2 | -3106 | ENSMUSG00000029202 | Pds5a | PDS5 cohesin associated factor A |
| chr13 | 55821537 | 55821956 | 420 | * | 3,748958 | 6,102950114 | 2,242044 | 2,722048 | 0,006488 | 1 | 0 | Exon (ENSMUST00000( | 13 | 55817577 | 55823730 | 6154 | 1 | 212880 | ENSMUST00000231097.2 | 3960 | ENSMUSG00000021500 | Ddx46 | DEAD box helicase 46 |
| chr11 | 70436533 | 70436969 | 437 | * | 4,244051 | 6,102958304 | 2,241169 | 2,723114 | 0,006467 | 1 | 0 | 3' UTR | 11 | 70431182 | 70437224 | 6043 | 1 | 18806 | ENSMUST00000108556.2 | 5351 | ENSMUSG00000020828 | Pld2 | phospholipase D2 |
| chr15 | 1,03E+08 | 1,03E+08 | 286 | * | 4,044095 | 6,103001264 | 2,234473 | 2,731293 | 0,006309 | 1 | 0 | Distal Intergenic | 15 | 1,03E+08 | 1,03E+08 | 23759 | 2 | 432989 | ENSMUST00000181306.2 | -3074 | ENSMUSG00000097003 | D930007P13Rik | Riken cDNA D930007P13 gene |
| chr14 | 24618012 | 24618430 | 419 | * | 3,710654 | 6,103044478 | 2,230535 | 2,736134 | 0,006217 | 1 | 0 | Distal Intergenic | 14 | 24667625 | 24788944 | 121320 | 1 | 1,03E+08 | ENSMUST00000225654.2 | -49195 | | Gm31775 | predicted gene, 31775 |
| chr1 | 51271420 | 51271795 | 376 | * | 4,754185 | 6,103284352 | 2,211414 | 2,759901 | 0,005782 | 1 | 0 | Intron (ENSMUST0000( | 1 | 51328285 | 51342119 | 13835 | 1 | 20324 | ENSMUST00000051572.8 | -56490 | ENSMUSG00000045954 | Cavin2 | caveolae associated 2 |
| chr4 | 33982551 | 33983061 | 511 | * | 3,694189 | 6,10347136 | 2,191796 | 2,784689 | 0,005358 | 1 | 0 | Distal Intergenic | 4 | 33926202 | 33948819 | 22618 | 1 | 12801 | ENSMUST00000084736.3 | 56349 | ENSMUSG00000044288 | Cnr1 | cannabinoid receptor 1 (brain) |
| chr4 | 94389143 | 94389413 | 271 | * | 3,975351 | 6,103617909 | 2,178768 | 2,801408 | 0,005088 | 1 | 0 | 3' UTR | 4 | 94348510 | 94365475 | 16966 | 2 | 1,15E+08 | ENSMUST00000133884.9 | -23668 | | Gm12655 | predicted gene 12655 |
| chr10 | 26567586 | 26567880 | 295 | * | 3,814201 | 6,103669966 | 2,17451 | 2,806918 | 0,005002 | 1 | 0 | Distal Intergenic | 10 | 26571570 | 26618856 | 47287 | 1 | 1,15E+08 | ENSMUST00000218733.2 | -3690 | | Gm48893 | predicted gene, 48893 |
| chr11 | 51923463 | 51924211 | 749 | * | 4,091885 | 6,104124344 | 2,137632 | 2,855555 | 0,004296 | 1 | 0 | Promoter (<=1kb) | 11 | 51924354 | 51980603 | 56250 | 1 | 213084 | ENSMUST00000136021.2 | -143 | ENSMUSG00000020389 | Cdkl3 | cyclin-dependent kinase-like 3 |
| chr11 | 21007305 | 21007815 | 511 | * | 5,164186 | 6,104442276 | 2,113596 | 2,888178 | 0,003875 | 1 | 0 | Distal Intergenic | 11 | 21041291 | 21093005 | 51715 | 1 | 67245 | ENSMUST00000156122.8 | -33476 | ENSMUSG00000020134 | Peli1 | pellino 1 |
| chr5 | 1,44E+08 | 1,44E+08 | 444 | * | 5,018078 | 6,104584459 | 2,103404 | 2,902241 | 0,003705 | 1 | 0 | Distal Intergenic | 5 | 1,44E+08 | 1,44E+08 | 87804 | 1 | 19159 | ENSMUST00000116456.10 | -20138 | ENSMUSG00000018001 | Cyth3 | cytohesin 3 |
| chr3 | 52307117 | 52307755 | 639 | * | 4,506628 | 6,10465824 | 2,099053 | 2,908291 | 0,003634 | 1 | 0 | Distal Intergenic | 3 | 52332829 | 52333069 | 241 | 1 | 1,03E+08 | ENSMUST00000194115.2 | -25074 | | Gm34303 | predicted gene, 34303 |
| chrX | 60258674 | 60259711 | 1038 | * | 5,138426 | 6,104724562 | 2,093699 | 2,91576 | 0,003548 | 1 | 0 | Intron (ENSMUST0000( | 20 | 60227611 | 60229164 | 1554 | 2 | 631990 | ENSMUST00000166381.3 | -29510 | ENSMUSG00000090546 | Cdr1 | cerebellar degeneration related antigen 1 |
| chr14 | 76107255 | 76108304 | 1050 | * | 5,645919 | 6,105066415 | 2,072617 | 2,945584 | 0,003223 | 1 | 0 | Distal Intergenic | 14 | 76085345 | 76085474 | 130 | 1 | 1E+08 | ENSMUST00000083213.3 | 21910 | ENSMUSG00000065147 | Snora31 | small nucleolar RNA, H/ACA box 31 |
| chr9 | 1,22E+08 | 1,22E+08 | 605 | * | 5,386104 | 6,105158912 | 2,065489 | 2,955793 | 0,003119 | 1 | 0 | Exon (ENSMUST000002 | 9 | 1,22E+08 | 1,22E+08 | 41835 | 2 | 1,01E+08 | ENSMUST00000135986.9 | -6811 | ENSMUSG00000079235 | Ccdc13 | coiled-coil domain containing 13 |
| chr2 | 40506807 | 40507211 | 405 | * | 5,2629 | 6,105343785 | 2,054854 | 2,971182 | 0,002967 | 1 | 0 | Promoter (1-2kb) | 2 | 40485258 | 40505263 | 20006 | 2 | 94217 | ENSMUST00000129974.3 | -1544 | ENSMUSG00000049252 | Lrp1b | low density lipoprotein-related protein 1B |
| chr12 | 80894309 | 80894934 | 626 | * | 5,311641 | 6,105435711 | 2,049123 | 2,979535 | 0,002887 | 1 | 0 | Intron (ENSMUST0000( | 12 | 80837333 | 80927609 | 90277 | 1 | 217684 | ENSMUST00000068519.7 | 56976 | ENSMUSG00000021133 | Susd6 | sushi domain containing 6 |
| chr17 | 57558563 | 57558718 | 156 | * | 4,855761 | 6,105549369 | 2,04262 | 2,989077 | 0,002798 | 1 | 0 | Promoter (1-2kb) | 17 | 57560111 | 57562328 | 2218 | 1 | 106628 | ENSMUST00000224586.2 | -1393 | ENSMUSG00000019487 | Trip10 | thyroid hormone receptor interactor 10 |
| chr8 | 84168232 | 84168931 | 700 | * | 4,857835 | 6,105849213 | 2,026718 | 3,012678 | 0,00259 | 1 | 0 | Promoter (<=1kb) | 8 | 84157139 | 84169079 | 11941 | 2 | 56367 | ENSMUST00000212449.2 | 148 | ENSMUSG00000063253 | Scoc | short coiled-coil protein |
| chr1 | 13416191 | 13416733 | 543 | * | 5,252049 | 6,105917022 | 2,021933 | 3,019842 | 0,002529 | 1 | 0 | Exon (ENSMUST000001 | 1 | 13404722 | 13404833 | 112 | 2 | 1,15E+08 | ENSMUST00000157157.3 | -11358 | ENSMUSG00000087782 | Gm23169 | predicted gene, 23169 |
| chr5 | 46878368 | 46878922 | 555 | * | 4,727568 | 6,105937795 | 2,020989 | 3,021263 | 0,002517 | 1 | 0 | Distal Intergenic | 5 | 46084160 | 46089751 | 5592 | 2 | 78106 | ENSMUST00000198088.2 | -788617 | ENSMUSG00000106602 | 4930405L22Rik | RIKEN cDNA 4930405L22 gene |
| chr1 | 1,86E+08 | 1,86E+08 | 806 | * | 5,0461 | 6,105958945 | 2,020266 | 3,022354 | 0,002508 | 1 | 0 | Intron (ENSMUST0000( | 1 | 1,86E+08 | 1,86E+08 | 24411 | 2 | 21808 | ENSMUST00000191766.6 | 6216 | ENSMUSG00000039239 | Tgfb2 | transforming growth factor, beta 2 |
| chr17 | 15163223 | 15164282 | 1060 | * | 9,460156 | 6,108014344 | 1,701878 | 3,588985 | 0,000332 | 0,302204121 | 1,449706 | Promoter (<=1kb) | 17 | 15163446 | 15166198 | 2753 | 1 | 67912 | ENSMUST00000052691.9 | 0 | ENSMUSG00000050088 | 1600012H06Rik | RIKEN cDNA 1600012H06 gene |
| chr3 | 1,23E+08 | 1,23E+08 | 866 | * | 5,733007 | 6,108034909 | 1,925238 | 3,172612 | 0,001511 | 1 | 0 | Intron (ENSMUST0000( | 3 | 1,23E+08 | 1,23E+08 | 36292 | 2 | 118449 | ENSMUST00000139160.8 | 9099 | ENSMUSG00000050315 | Synpo2 | synaptopodin 2 |
| chr17 | 28755543 | 28756646 | 1104 | * | 9,049395 | 6,109215338 | 1,809211 | 3,376729 | 0,000734 | 0,430115816 | 1,425929 | 3' UTR | 17 | 28749841 | 28758084 | 8244 | 1 | 67645 | ENSMUST00000233114.2 | 5702 | ENSMUSG00000024223 | Armc12 | armadillo repeat containing 12 |
| chr1 | 92118094 | 92118977 | 884 | * | 6,986718 | 6,109863703 | 1,862601 | 3,280285 | 0,001037 | 1 | 0 | Intron (ENSMUST0000( | 1 | 91957866 | 92123416 | 165551 | 2 | 208727 | ENSMUST00000212867.2 | 4439 | ENSMUSG00000026313 | Hdac4 | histone deacetylase 4 |
| chr11 | 62771962 | 62772803 | 842 | * | 6,865178 | 6,110031562 | 1,848983 | 3,304537 | 0,000951 | 1 | 0 | Promoter (1-2kb) | 11 | 62770440 | 62776172 | 5733 | 1 | 67510 | ENSMUST00000154176.2 | 1522 | ENSMUSG00000014177 | Tvp23b | trans-golgi network vesicle protein 23B |
| chr6 | 1,16E+08 | 1,16E+08 | 384 | * | 5,580634 | 6,11192219 | 1,981404 | 3,084643 | 0,002038 | 1 | 0 | Promoter (<=1kb) | 6 | 1,16E+08 | 1,16E+08 | 20723 | 1 | 67027 | ENSMUST00000000449.9 | -139 | ENSMUSG00000000439 | Mkrn2 | makorin, ring finger protein, 2 |
| chr13 | 53462439 | 53462870 | 432 | * | 4,995406 | 6,111942062 | 1,982673 | 3,082678 | 0,002051 | 1 | 0 | Distal Intergenic | 13 | 53263353 | 53440160 | 176808 | 2 | 26564 | ENSMUST00000021918.10 | -22279 | ENSMUSG00000021464 | Ror2 | receptor tyrosine kinase-like orphan receptor 2 |
| chr13 | 78189654 | 78190462 | 809 | * | 5,52167 | 6,11206681 | 1,994772 | 3,064043 | 0,002184 | 1 | 0 | Intron (ENSMUST0000( | 13 | 78173021 | 78174414 | 1394 | 1 | 75507 | ENSMUST00000175955.4 | 16633 | ENSMUSG00000093668 | Pou5f2 | POU domain class 5, transcription factor 2 |
| chr12 | 1,12E+08 | 1,12E+08 | 822 | * | 8,468966 | 6,112272112 | 1,799844 | 3,396002 | 0,000684 | 0,419066905 | 1,425929 | Promoter (1-2kb) | 12 | 1,12E+08 | 1,12E+08 | 1208 | 1 | 16593 | ENSMUST00000147203.2 | 1498 | ENSMUSG00000021288 | Klc1 | kinesin light chain 1 |
| chr19 | 9544931 | 9545501 | 571 | * | 4,884788 | 6,112698288 | 2,061757 | 2,964801 | 0,003029 | 1 | 0 | Distal Intergenic | 19 | 9535156 | 9536547 | 1392 | 2 | 619371 | ENSMUST00000090527.5 | -8384 | | Stxbp3-ps | syntaxin-binding protein 3, pseudogene |
| chr11 | 80611924 | 80612438 | 515 | * | 4,374129 | 6,112711873 | 2,06287 | 2,963207 | 0,003045 | 1 | 0 | Intron (ENSMUST0000( | 11 | 80564493 | 80670829 | 106337 | 2 | 338367 | ENSMUST00000125944.2 | 58391 | ENSMUSG00000035441 | Myo1d | myosin ID |
| chr17 | 30658572 | 30659083 | 512 | * | 4,788831 | 6,112874127 | 2,083499 | 2,933947 | 0,003347 | 1 | 0 | Intron (ENSMUST0000( | 17 | 30515928 | 30665114 | 149187 | 2 | 224671 | ENSMUST00000235555.2 | 6031 | ENSMUSG00000062202 | Btbd9 | BTB (POZ) domain containing 9 |
| chr8 | 50089351 | 50090020 | 670 | * | 4,691876 | 6,112917622 | 2,087874 | 2,927819 | 0,003413 | 1 | 0 | Distal Intergenic | 8 | 49938516 | 49974130 | 35615 | 1 | 75359 | ENSMUST00000126285.2 | 150835 | ENSMUSG00000031559 | 4930555F03Rik | RIKEN cDNA 4930555F03 gene |
| chrX | 39879985 | 39880790 | 806 | * | 4,190774 | 6,112958695 | 2,093771 | 2,919593 | 0,003505 | 1 | 0 | Distal Intergenic | 20 | 40461136 | 40461391 | 256 | 1 | 1,08E+08 | ENSMUST00000121134.2 | -580346 | ENSMUSG00000083515 | Gm14577 | predicted gene 14577 |
| chr3 | 40520720 | 40521146 | 427 | * | 4,320112 | 6,113414432 | 2,358096 | 2,592522 | 0,009528 | 1 | 0 | Distal Intergenic | 3 | 40562579 | 40563923 | 1345 | 2 | 1E+08 | ENSMUST00000202778.2 | 42777 | ENSMUSG00000037884 | 1700017G19Rik | RIKEN cDNA 1700017G19 gene |
| chr10 | 68745983 | 68746365 | 383 | * | 3,898751 | 6,113690655 | 2,206773 | 2,770421 | 0,005598 | 1 | 0 | Distal Intergenic | 10 | 68856570 | 68869718 | 13149 | 2 | 1,03E+08 | ENSMUST00000219854.2 | 123353 | ENSMUSG00000112331 | Gm33035 | predicted gene, 33035 |
| chr18 | 61151720 | 61152173 | 454 | * | 4,625259 | 6,113846324 | 2,238409 | 2,731336 | 0,006308 | 1 | 0 | 3' UTR | 18 | 61128452 | 61147294 | 18843 | 2 | 240332 | ENSMUST00000025520.10 | -4426 | ENSMUSG00000052026 | Slc6a7 | solute carrier family 6 (neurotransmitter transporter, L-proline), member 7 |
| chr9 | 93086766 | 93087252 | 487 | * | 4,278625 | 6,113930877 | 2,253646 | 2,712907 | 0,00667 | 1 | 0 | Distal Intergenic | 9 | 92862048 | 92887792 | 25745 | 2 | 78349 | ENSMUST00000190562.7 | -198974 | ENSMUSG00000100294 | 1700034K08Rik | RIKEN cDNA 1700034K08 gene |
| chr10 | 45363464 | 45363773 | 310 | * | 3,91288 | 6,114011255 | 2,267992 | 2,695782 | 0,007022 | 1 | 0 | Promoter (<=1kb) | 10 | 45294459 | 45362491 | 68033 | 2 | 380669 | ENSMUST00000215708.2 | -973 | ENSMUSG00000063804 | Lin28b | lin-28 homolog B |
| chr14 | 83413972 | 83414541 | 570 | * | 4,260415 | 6,114039558 | 2,321813 | 2,633304 | 0,008456 | 1 | 0 | Distal Intergenic | 14 | 83643071 | 83643204 | 134 | 1 | 1,15E+08 | ENSMUST00000157384.3 | -228530 | ENSMUSG00000088009 | Gm24774 | predicted gene, 24774 |
| chr7 | 91342981 | 91343584 | 604 | * | 3,862308 | 6,114341887 | 2,344122 | 2,608372 | 0,009097 | 1 | 0 | Intron (ENSMUST0000( | 7 | 91360628 | 91589349 | 228722 | 1 | 23859 | ENSMUST00000128592.3 | -17044 | ENSMUSG00000052572 | Dlg2 | discs large MAGUK scaffold protein 2 |
| chr11 | 57992196 | 57992556 | 361 | * | 4,140439 | 6,115253776 | 2,257748 | 2,708563 | 0,006758 | 1 | 0 | Promoter (2-3kb) | 11 | 57994979 | 58009420 | 14442 | 1 | 69125 | ENSMUST00000020822.12 | -2423 | ENSMUSG00000020515 | Cnot8 | CCR4-NOT transcription complex, subunit 8 |
| chr1 | 1,52E+08 | 1,52E+08 | 887 | * | 8,021713 | 6,115376005 | 1,808072 | 3,382263 | 0,000719 | 1 | 0 | Promoter (2-3kb) | 1 | 1,52E+08 | 1,52E+08 | 6073 | 2 | 1,03E+08 | ENSMUST00000243781.1 | 2629 | ENSMUSG00000100594 | 2810414N06Rik | RIKEN cDNA 2810414N06 gene |
| chr2 | 25854469 | 25854878 | 410 | * | 4,071532 | 6,115528779 | 2,24479 | 2,724321 | 0,006443 | 1 | 0 | Promoter (<=1kb) | 2 | 25835044 | 25855762 | 20719 | 2 | 227634 | ENSMUST00000151593.8 | 884 | ENSMUSG00000026933 | Camsap1 | calmodulin regulated spectrin-associated protein 1 |
| chrX | 1,02E+08 | 1,02E+08 | 719 | * | 4,291335 | 6,115617687 | 2,24068 | 2,729357 | 0,006346 | 1 | 0 | 3' UTR | 20 | 1,02E+08 | 1,02E+08 | 10114 | 1 | 22127 | ENSMUST00000033691.5 | -24196 | ENSMUSG00000031329 | Tsx | testis specific X-linked gene |
| chr13 | 36994115 | 36994858 | 744 | * | 5,269172 | 6,116222941 | 2,212867 | 2,763936 | 0,005711 | 1 | 0 | Distal Intergenic | 13 | 37020340 | 37069570 | 49231 | 1 | 1,03E+08 | ENSMUST00000225846.2 | -25482 | | Gm30489 | predicted gene, 30489 |
| chr17 | 34052166 | 34052733 | 568 | * | 4,220545 | 6,116278157 | 2,210478 | 2,766948 | 0,005658 | 1 | 0 | Intron (ENSMUST0000( | 17 | 34048367 | 34057288 | 8922 | 1 | 66416 | ENSMUST00000173946.2 | 3799 | ENSMUSG00000041881 | Ndufa7 | NADH:ubiquinone oxidoreductase subunit A7 |
| chr12 | 51735378 | 51736691 | 1314 | * | 6,778001 | 6,116817686 | 1,972245 | 3,101449 | 0,001926 | 1 | 0 | Promoter (1-2kb) | 12 | 51737816 | 51791569 | 53754 | 1 | 11782 | ENSMUST00000218820.2 | -1125 | ENSMUSG00000020955 | Ap4s1 | adaptor-related protein complex AP-4, sigma 1 |
| chr4 | 72542278 | 72542993 | 716 | * | 6,063699 | 6,117458658 | 1,986722 | 3,079172 | 0,002076 | 1 | 0 | Distal Intergenic | 4 | 72768820 | 72770869 | 2050 | 2 | 353204 | ENSMUST00000179234.2 | 227876 | ENSMUSG00000059343 | Aldoart1 | aldolase 1 A, retrogene 1 |
| chr7 | 1,26E+08 | 1,26E+08 | 481 | * | 6,067426 | 6,117874879 | 2,146299 | 2,850429 | 0,004366 | 1 | 0 | Promoter (1-2kb) | 7 | 1,26E+08 | 1,26E+08 | 22398 | 1 | 104175 | ENSMUST00000056028.11 | -1497 | ENSMUSG00000042978 | Sbk1 | SH3-binding kinase 1 |
| chr15 | 79770645 | 79771258 | 614 | * | 4,90472 | 6,118262651 | 2,132126 | 2,86956 | 0,00411 | 1 | 0 | Distal Intergenic | 15 | 79775860 | 79789748 | 13889 | 1 | 80287 | ENSMUST00000177350.8 | -4602 | ENSMUSG00000009585 | Apobec3 | apolipoprotein B mRNA editing enzyme, catalytic polypeptide 3 |
| chr2 | 1,07E+08 | 1,07E+08 | 552 | * | 4,893412 | 6,118725014 | 2,115932 | 2,89174 | 0,003831 | 1 | 0 | Distal Intergenic | 2 | 1,07E+08 | 1,07E+08 | 7864 | 1 | 16492 | ENSMUST00000037012.3 | -78729 | ENSMUSG00000042604 | Kcna4 | potassium voltage-gated channel, shaker-related subfamily, member 4 |
| chr4 | 1,5E+08 | 1,5E+08 | 611 | * | 5,822204 | 6,122246836 | 2,013833 | 3,040097 | 0,002365 | 1 | 0 | Distal Intergenic | 4 | 1,5E+08 | 1,5E+08 | 3738 | 2 | 63954 | ENSMUST00000148596.8 | 9433 | ENSMUSG00000028996 | Rbp7 | retinol binding protein 7, cellular |
| chr3 | 1,08E+08 | 1,08E+08 | 763 | * | 5,789869 | 6,123188784 | 1,990854 | 3,075659 | 0,0021 | 1 | 0 | Promoter (<=1kb) | 3 | 1,08E+08 | 1,08E+08 | 77421 | 1 | 20661 | ENSMUST00000102632.7 | 748 | ENSMUSG00000068747 | Sort1 | sortilin 1 |
| chr11 | 97694630 | 97694901 | 272 | * | 3,349005 | 6,125993899 | 2,325903 | 2,633813 | 0,008443 | 1 | 0 | Promoter (1-2kb) | 11 | 97696018 | 97726945 | 30928 | 1 | 16796 | ENSMUST00000148280.3 | -1117 | ENSMUSG00000038366 | Lasp1 | LIM and SH3 protein 1 |
| chr11 | 78180605 | 78181002 | 398 | * | 3,402669 | 6,126106146 | 2,305311 | 2,657388 | 0,007875 | 1 | 0 | Promoter (<=1kb) | 11 | 78181667 | 78184571 | 2905 | 1 | 216976 | ENSMUST00000046361.5 | -665 | ENSMUSG00000037593 | Rskr | ribosomal protein S6 kinase related |
| chr17 | 5705619 | 5706080 | 462 | * | 3,886526 | 6,126119865 | 2,300712 | 2,662706 | 0,007752 | 1 | 0 | Intron (ENSMUST0000( | 17 | 5848932 | 5853168 | 4237 | 1 | 78512 | ENSMUST00000180924.8 | -142852 | ENSMUSG00000096965 | 3300005D01Rik | RIKEN cDNA 3300005D01 gene |
| chrX | 90318835 | 90319321 | 487 | * | 4,040471 | 6,126174775 | 2,290674 | 2,674399 | 0,007486 | 1 | 0 | Distal Intergenic | 20 | 90375361 | 90376503 | 1143 | 1 | 17146 | ENSMUST00000101397.2 | -56040 | ENSMUSG00000073069 | Mageb2 | MAGE family member B2 |
| chr8 | 14871578 | 14871949 | 372 | * | 3,76374 | 6,12621993 | 2,281176 | 2,685554 | 0,007241 | 1 | 0 | Intron (ENSMUST0000( | 8 | 14827773 | 14897398 | 69626 | 1 | 244310 | ENSMUST00000129119.2 | 43805 | ENSMUSG00000047495 | Dlgap2 | DLG associated protein 2 |
| chr13 | 30790177 | 30791116 | 940 | * | 4,685789 | 6,126493396 | 2,23525 | 2,740854 | 0,006128 | 1 | 0 | Distal Intergenic | 13 | 30799026 | 30808242 | 9217 | 1 | 75090 | ENSMUST00000151471.2 | -7910 | ENSMUSG00000086263 | 4930519D14Rik | RIKEN cDNA 4930519D14 gene |
| chr5 | 1,08E+08 | 1,08E+08 | 338 | * | 3,901774 | 6,126760187 | 2,199352 | 2,785712 | 0,005341 | 1 | 0 | Intron (ENSMUST0000( | 5 | 1,08E+08 | 1,08E+08 | 14266 | 1 | 76421 | ENSMUST00000162591.2 | 5151 | ENSMUSG00000089798 | 1700028K03Rik | RIKEN cDNA 1700028K03 gene |
| chr16 | 77644773 | 77645184 | 412 | * | 3,977418 | 6,126893586 | 2,181785 | 2,808202 | 0,004982 | 1 | 0 | Intron (ENSMUST0000( | 16 | 77443228 | 77446314 | 3087 | 1 | 723952 | ENSMUST00000193537.2 | 201545 | ENSMUSG00000065472 | Mir125b-2 | microRNA 125b-2 |
| chr13 | 9383303 | 9383609 | 307 | * | 3,793411 | 6,126922073 | 2,176426 | 2,81513 | 0,004876 | 1 | 0 | Intron (ENSMUST0000( | 13 | 9327018 | 9716214 | 389197 | 1 | 208440 | ENSMUST00000166299.2 | 56285 | ENSMUSG00000048264 | Dip2c | disco interacting protein 2 homolog C |
| chr5 | 1,25E+08 | 1,25E+08 | 414 | * | 3,745748 | 6,126931777 | 2,17512 | 2,816825 | 0,00485 | 1 | 0 | Intron (ENSMUST0000( | 5 | 1,25E+08 | 1,25E+08 | 7472 | 2 | 20602 | ENSMUST00000199561.2 | -3540 | ENSMUSG00000029478 | Ncor2 | nuclear receptor co-repressor 2 |
| chr10 | 26712614 | 26713067 | 454 | * | 4,54111 | 6,127311712 | 2,130317 | 2,876245 | 0,004024 | 1 | 0 | Intron (ENSMUST0000( | 10 | 26698598 | 26725997 | 27400 | 1 | 73910 | ENSMUST00000135866.2 | 14016 | ENSMUSG00000039031 | Arhgap18 | Rho GTPase activating protein 18 |
| chr11 | 98660476 | 98660912 | 437 | * | 5,069033 | 6,127314947 | 2,126759 | 2,881058 | 0,003963 | 1 | 0 | Promoter (<=1kb) | 11 | 98660173 | 98661252 | 1080 | 2 | 217166 | ENSMUST00000139220.2 | 340 | ENSMUSG00000020889 | Nr1d1 | nuclear receptor subfamily 1, group D, member 1 |
| chr4 | 1,35E+08 | 1,35E+08 | 506 | * | 4,976101 | 6,127423129 | 2,113975 | 2,898532 | 0,003749 | 1 | 0 | 3' UTR | 4 | 1,35E+08 | 1,35E+08 | 2887 | 2 | 74552 | ENSMUST00000183397.2 | 5284 | ENSMUSG00000028803 | Nipal3 | NIPA-like domain containing 3 |
| chr9 | 62714903 | 62715350 | 448 | * | 5,012989 | 6,127502831 | 2,107206 | 2,907881 | 0,003639 | 1 | 0 | Intron (ENSMUST0000( | 9 | 62699103 | 62719216 | 20114 | 2 | 14155 | ENSMUST00000034775.10 | 3866 | ENSMUSG00000032244 | Fem1b | fem 1 homolog b |
| chrX | 71277702 | 71278186 | 485 | * | 4,820141 | 6,127578811 | 2,098047 | 2,92061 | 0,003493 | 1 | 0 | Distal Intergenic | 20 | 71265623 | 71266570 | 948 | 1 | 17140 | ENSMUST00000048675.2 | 12079 | ENSMUSG00000033343 | Magea4 | MAGE family member A4 |
| chr9 | 66810846 | 66811312 | 467 | * | 4,468034 | 6,127615226 | 2,093628 | 2,926793 | 0,003425 | 1 | 0 | Intron (ENSMUST0000( | 9 | 66750946 | 66826969 | 76024 | 2 | 235442 | ENSMUST00000041139.9 | 15657 | ENSMUSG00000036943 | Rab8b | RAB8B, member RAS oncogene family |
| chr1 | 1,35E+08 | 1,35E+08 | 799 | * | 4,66 | 6,127662186 | 2,089461 | 2,932652 | 0,003361 | 1 | 0 | Intron (ENSMUST0000( | 1 | 1,35E+08 | 1,35E+08 | 43755 | 2 | 329251 | ENSMUST00000238280.2 | 4705 | ENSMUSG00000073557 | Ppp1r12b | protein phosphatase 1, regulatory subunit 12B |
| chr1 | 1,91E+08 | 1,91E+08 | 605 | * | 4,900443 | 6,128147497 | 2,041685 | 3,001515 | 0,002686 | 1 | 0 | Intron (ENSMUST0000( | 1 | 1,91E+08 | 1,91E+08 | 11406 | 2 | 66208 | ENSMUST00000046770.10 | 38699 | ENSMUSG00000037499 | Nenf | neuron derived neurotrophic factor |
| chr9 | 67515038 | 67515701 | 664 | * | 4,756848 | 6,128416587 | 2,01935 | 3,034847 | 0,002407 | 1 | 0 | Distal Intergenic | 9 | 67124369 | 67466985 | 342617 | 2 | 70549 | ENSMUST00000040025.14 | -48053 | ENSMUSG00000052698 | Tln2 | talin 2 |
| chr19 | 37209177 | 37209871 | 695 | * | 4,8139 | 6,128747861 | 1,99321 | 3,074813 | 0,002106 | 1 | 0 | Promoter (1-2kb) | 19 | 37210781 | 37210883 | 103 | 2 | 1,15E+08 | ENSMUST00000158518.4 | 1012 | ENSMUSG00000089143 | Gm25268 | predicted gene, 25268 |
| chr2 | 1,57E+08 | 1,57E+08 | 671 | * | 5,165232 | 6,128751135 | 1,991402 | 3,077607 | 0,002087 | 1 | 0 | Intron (ENSMUST0000( | 2 | 1,57E+08 | 1,57E+08 | 43667 | 2 | 29812 | ENSMUST00000156651.2 | 11452 | ENSMUSG00000027634 | Ndrg3 | N-myc downstream regulated gene 3 |
| chr17 | 49566484 | 49566946 | 463 | * | 5,670909 | 6,12878045 | 1,990345 | 3,079256 | 0,002075 | 1 | 0 | Distal Intergenic | 17 | 49599770 | 49599929 | 160 | 1 | 1,15E+08 | ENSMUST00000082607.3 | -32824 | ENSMUSG00000064541 | Gm22978 | predicted gene, 22978 |
| chr13 | 34292363 | 34292780 | 418 | * | 5,099906 | 6,1291181 | 1,965062 | 3,119046 | 0,001814 | 1 | 0 | Intron (ENSMUST0000( | 13 | 34268138 | 34297255 | 29118 | 2 | 432745 | ENSMUST00000222745.2 | 4475 | ENSMUSG00000113591 | 4930447K03Rik | RIKEN cDNA 4930447K03 gene |
| chr4 | 16646677 | 16647462 | 786 | * | 9,389901 | 6,129345667 | 1,749779 | 3,502925 | 0,00046 | 0,344155199 | 1,449706 | Distal Intergenic | 4 | 16675923 | 16676029 | 107 | 1 | 1,15E+08 | ENSMUST00000179782.2 | -28461 | ENSMUSG00000096423 | Gm24978 | predicted gene, 24978 |
| chr3 | 88415786 | 88416318 | 533 | * | 5,411909 | 6,129364217 | 1,948074 | 3,146371 | 0,001653 | 1 | 0 | Promoter (<=1kb) | 3 | 88410060 | 88417263 | 7204 | 2 | 16905 | ENSMUST00000149068.2 | 945 | ENSMUSG00000028063 | Lmna | lamin A |

| | | | | | | | | | | | | | | | | | | | | | | | |
|---|---|---|---|---|---|---|---|---|---|---|---|---|---|---|---|---|---|---|---|---|---|---|---|
| chr3 | 18691431 | 18692008 | 578 | * | 5,708384 | 6,129369281 | 1,948498 | 3,145689 | 0,001657 | 1 | 0 | Distal Intergenic | 3 | 18522451 | 18534641 | 12191 | 2 | 73959 | ENSMUST00000192010.2 | -156790 | ENSMUSG00000102699 | 4930433B08Rik | RIKEN cDNA 4930433B08 gene |
| chr11 | 50567365 | 50567890 | 526 | * | 6,371773 | 6,130207547 | 1,897677 | 3,230375 | 0,001236 | 1 | 0 | Intron (ENSMUST0000( | 11 | 50608497 | 50609742 | 1246 | 2 | 1,19E+08 | ENSMUST00000137567.2 | 41852 | | Gm12199 | predicted gene 12199 |
| chr15 | 69593438 | 69594053 | 616 | * | 6,917799 | 6,131307277 | 1,845171 | 3,322894 | 0,000891 | 1 | 0 | Distal Intergenic | 15 | 69557427 | 69557555 | 129 | 2 | 1,15E+08 | ENSMUST00000117039.3 | -35883 | ENSMUSG00000080689 | Gm23987 | predicted gene, 23987 |
| chr15 | 75644323 | 75645283 | 961 | * | 8,088441 | 6,132539156 | 1,898582 | 3,230062 | 0,001238 | 0,527676616 | 1,360622 | Intron (ENSMUST0000( | 15 | 75631708 | 75631838 | 131 | 1 | 1,15E+08 | ENSMUST00000104228.3 | 12615 | ENSMUSG00000077416 | Gm23747 | predicted gene, 23747 |
| chr15 | 97954654 | 97955147 | 494 | * | 6,245792 | 6,132730007 | 1,907002 | 3,215902 | 0,0013 | 1 | 0 | 3' UTR | 15 | 97939113 | 97962118 | 23006 | 2 | 223870 | ENSMUST00000181855.8 | 6971 | ENSMUSG00000033075 | Senp1 | SUMO1/sentrin specific peptidase 1 |
| chr14 | 76053342 | 76054163 | 822 | * | 7,090543 | 6,132755966 | 1,908144 | 3,213991 | 0,001309 | 1 | 0 | Distal Intergenic | 14 | 76080819 | 76082404 | 1586 | 2 | 1E+08 | ENSMUST00000131222.2 | 28241 | | Gm4285 | predicted gene 4285 |
| chr18 | 71303513 | 71304006 | 494 | * | 5,808043 | 6,133117816 | 1,923354 | 3,188762 | 0,001429 | 1 | 0 | Distal Intergenic | 18 | 70750844 | 70758848 | 8005 | 1 | 17191 | ENSMUST00000128555.2 | 552669 | ENSMUSG00000024513 | Mbd2 | methyl-CpG binding domain protein 2 |
| chr3 | 1,09E+08 | 1,09E+08 | 602 | * | 5,604757 | 6,133550598 | 1,944506 | 3,154297 | 0,001609 | 1 | 0 | Promoter (1-2kb) | 3 | 1,09E+08 | 1,09E+08 | 30496 | 2 | 20912 | ENSMUST00000196679.5 | 1903 | ENSMUSG00000027882 | Stxbp3 | syntaxin binding protein 3 |
| chr3 | 26201235 | 26201850 | 616 | * | 5,417249 | 6,133564489 | 1,944842 | 3,153759 | 0,001612 | 1 | 0 | Intron (ENSMUST0000( | 3 | 25487595 | 26207456 | 719862 | 2 | 192167 | ENSMUST00000191835.2 | 5606 | ENSMUSG00000063887 | Nlgn1 | neuroligin 1 |
| chr5 | 1,24E+08 | 1,24E+08 | 987 | * | 7,274698 | 6,13364344 | 1,949563 | 3,146164 | 0,001654 | 0,582425528 | 1,360622 | Distal Intergenic | 5 | 1,24E+08 | 1,24E+08 | 3285 | 2 | 243270 | ENSMUST00000164267.3 | 4241 | ENSMUSG00000049241 | Hcar1 | hydrocarboxylic acid receptor 1 |
| chr17 | 74497011 | 74498157 | 1147 | * | 6,024089 | 6,133659278 | 1,949922 | 3,145593 | 0,001658 | 1 | 0 | Exon (ENSMUST000002 | 17 | 74508111 | 74537496 | 29386 | 2 | 76890 | ENSMUST00000233495.2 | 39339 | ENSMUSG00000058704 | Memo1 | mediator of cell motility 1 |
| chr11 | 4148924 | 4149603 | 680 | * | 5,327213 | 6,134032793 | 1,969339 | 3,114768 | 0,001841 | 1 | 0 | Intron (ENSMUST0000( | 11 | 4155423 | 4159990 | 4568 | 1 | 103724 | ENSMUST00000151672.2 | -5820 | ENSMUSG00000034412 | Tbc1d10a | TBC1 domain family, member 10a |
| chr14 | 25186680 | 25187294 | 615 | * | 6,426789 | 6,134036124 | 1,969055 | 3,115218 | 0,001838 | 1 | 0 | Distal Intergenic | 14 | 25140219 | 25143665 | 3447 | 2 | 75862 | ENSMUST00000225429.2 | -43015 | ENSMUSG00000114916 | 4930572O13Rik | RIKEN cDNA 4930572O13 gene |
| chr9 | 58858718 | 58859505 | 788 | * | 5,580281 | 6,134097238 | 1,972054 | 3,110512 | 0,001868 | 1 | 0 | Intron (ENSMUST0000( | 9 | 58820119 | 58889391 | 69273 | 2 | 18007 | ENSMUST00000215026.2 | 29886 | ENSMUSG00000032340 | Neo1 | neogenin |
| chr3 | 46008568 | 46009152 | 585 | * | 4,775515 | 6,134891951 | 2,019133 | 3,038379 | 0,002379 | 1 | 0 | Distal Intergenic | 3 | 45933013 | 45933130 | 118 | 1 | 1,15E+08 | ENSMUST00000158959.4 | 75555 | ENSMUSG00000089584 | Gm23337 | predicted gene, 23337 |
| chr9 | 57534087 | 57534846 | 760 | * | 5,551818 | 6,135271092 | 2,045496 | 2,999404 | 0,002705 | 1 | 0 | Promoter (1-2kb) | 9 | 57534601 | 57536044 | 1444 | 2 | 12988 | ENSMUST00000213660.2 | 1198 | ENSMUSG00000032312 | Csk | c-src tyrosine kinase |
| chr11 | 1E+08 | 1E+08 | 1111 | * | 10,27253 | 6,135379352 | 1,798872 | 3,410681 | 0,000648 | 0,410415129 | 1,425929 | Promoter (2-3kb) | 11 | 1E+08 | 1E+08 | 4090 | 1 | 66720 | ENSMUST00000107395.3 | 2498 | ENSMUSG00000001558 | Klhl10 | kelch-like 10 |
| chr15 | 1,03E+08 | 1,03E+08 | 664 | * | 4,76784 | 6,135394995 | 2,053676 | 2,987518 | 0,002813 | 1 | 0 | Intron (ENSMUST0000( | 15 | 1,03E+08 | 1,03E+08 | 48273 | 2 | 12419 | ENSMUST00000118152.8 | 9895 | ENSMUSG00000009575 | Cbx5 | chromobox 5 |
| chr2 | 34251408 | 34252064 | 657 | * | 4,530318 | 6,135909785 | 2,092392 | 2,932485 | 0,003363 | 1 | 0 | Intron (ENSMUST0000( | 2 | 34066871 | 34260858 | 193988 | 2 | 18516 | ENSMUST00000175855.3 | 8794 | ENSMUSG00000038718 | Pbx3 | pre B cell leukemia homeobox 3 |
| chr13 | 45708864 | 45709359 | 496 | * | 4,485605 | 6,135925327 | 2,093439 | 2,931026 | 0,003378 | 1 | 0 | 3' UTR | 13 | 45692285 | 45696026 | 3742 | 1 | 66355 | ENSMUST00000129176.2 | 16579 | ENSMUSG00000000253 | Gmpr | guanosine monophosphate reductase |
| chr5 | 31371985 | 31372734 | 750 | * | 4,798238 | 6,136083748 | 2,107615 | 2,911387 | 0,003598 | 1 | 0 | Intron (ENSMUST0000( | 5 | 31362434 | 31377663 | 15230 | 2 | 14208 | ENSMUST00000200778.5 | 4929 | ENSMUSG00000029147 | Ppm1g | protein phosphatase 1G (formerly 2C), magnesium-dependent, gamma isoform |
| chr11 | 68855108 | 68855581 | 474 | * | 4,310882 | 6,136126231 | 2,111451 | 2,906118 | 0,003659 | 1 | 0 | Distal Intergenic | 11 | 68858957 | 68863342 | 4386 | 1 | 71998 | ENSMUST00000102606.10 | -3376 | ENSMUSG00000018740 | Slc25a35 | solute carrier family 25, member 35 |
| chr4 | 1,3E+08 | 1,3E+08 | 834 | * | 4,872287 | 6,136488165 | 2,143233 | 2,863192 | 0,004194 | 1 | 0 | Exon (ENSMUST000001 | 4 | 1,3E+08 | 1,3E+08 | 28478 | 2 | 20218 | ENSMUST00000066257.6 | 12167 | ENSMUSG00000028790 | Khdrbs1 | KH domain containing, RNA binding, signal transduction associated 1 |
| chr13 | 67742458 | 67742934 | 477 | * | 5,339841 | 6,13666519 | 2,160734 | 2,840083 | 0,00451 | 1 | 0 | Intron (ENSMUST0000( | 13 | 67738231 | 67757764 | 19534 | 2 | 1E+08 | ENSMUST00000223599.2 | 14830 | ENSMUSG00000058093 | Zfp729b | zinc finger protein 729b |
| chr2 | 53791792 | 53792317 | 526 | * | 4,376537 | 6,13680447 | 2,377395 | 2,581314 | 0,009842 | 1 | 0 | Distal Intergenic | 2 | 53974105 | 53975564 | 1460 | 2 | 67874 | ENSMUST00000100089.3 | 183247 | ENSMUSG00000075334 | Rprm | reprimo, TP53 dependent G2 arrest mediator candidate |
| chr1 | 23364025 | 23364544 | 520 | * | 4,281903 | 6,136895347 | 2,368418 | 2,591137 | 0,009566 | 1 | 0 | Distal Intergenic | 1 | 23330782 | 23330865 | 84 | 1 | 723964 | ENSMUST00000083633.3 | 33243 | ENSMUSG00000065567 | Mir30c-2 | microRNA 30c-2 |
| chrX | 70240329 | 70240861 | 533 | * | 3,894904 | 6,137042797 | 2,199072 | 2,790742 | 0,005259 | 1 | 0 | Exon (ENSMUST000001 | 20 | 70254373 | 70358901 | 104529 | 1 | 17772 | ENSMUST00000171933.8 | -13512 | ENSMUSG00000031337 | Mtm1 | X-linked myotubular myopathy gene 1 |
| chr5 | 9559220 | 9559501 | 282 | * | 3,490171 | 6,137281079 | 2,226537 | 2,756425 | 0,005844 | 1 | 0 | Intron (ENSMUST0000( | 5 | 9467615 | 9472330 | 4716 | 1 | 231014 | ENSMUST00000142202.2 | 91605 | ENSMUSG00000056004 | Elapor2 | endosome-lysosome associated apoptosis and autophagy regulator family member 2 |
| chr14 | 57904291 | 57904649 | 359 | * | 3,938992 | 6,137339382 | 2,230911 | 2,751046 | 0,005941 | 1 | 0 | Promoter (1-2kb) | 14 | 57879486 | 57902887 | 23402 | 2 | 57258 | ENSMUST00000172524.2 | -1404 | ENSMUSG00000021952 | Xpo4 | exportin 4 |
| chr6 | 1,42E+08 | 1,42E+08 | 560 | * | 4,420324 | 6,137365101 | 2,234241 | 2,746958 | 0,006015 | 1 | 0 | Promoter (2-3kb) | 6 | 1,42E+08 | 1,42E+08 | 17604 | 1 | 232491 | ENSMUST00000041852.8 | -2293 | ENSMUSG00000041671 | Pyroxd1 | pyridine nucleotide-disulphide oxidoreductase domain 1 |
| chr11 | 1,17E+08 | 1,17E+08 | 2121 | * | 14,47125 | 6,13740612 | 1,611863 | 3,807649 | 0,00014 | 0,209871787 | 1,2763 | Promoter (<=1kb) | 11 | 1,17E+08 | 1,17E+08 | 2529 | 1 | 69900 | ENSMUST00000147404.2 | -364 | ENSMUSG00000020818 | Mfsd11 | major facilitator superfamily domain containing 11 |
| chr3 | 1,11E+08 | 1,11E+08 | 419 | * | 3,702729 | 6,137421362 | 2,330873 | 2,6331 | 0,008461 | 1 | 0 | Distal Intergenic | 3 | 1,1E+08 | 1,1E+08 | 105 | 2 | 1,15E+08 | ENSMUST00000104065.3 | -150061 | ENSMUSG00000119642 | Gm26076 | predicted gene, 26076 |
| chr13 | 52630966 | 52631321 | 356 | * | 3,577146 | 6,137437355 | 2,330261 | 2,633798 | 0,008444 | 1 | 0 | Distal Intergenic | 13 | 52658416 | 52685315 | 26900 | 2 | 68203 | ENSMUST00000057442.8 | 53994 | ENSMUSG00000047842 | Diras2 | DIRAS family, GTP-binding RAS-like 2 |
| chr5 | 1,39E+08 | 1,39E+08 | 294 | * | 3,588077 | 6,13753292 | 2,323142 | 2,64191 | 0,008244 | 1 | 0 | Distal Intergenic | 5 | 1,39E+08 | 1,39E+08 | 53747 | 2 | 231821 | ENSMUST00000110865.2 | -5696 | ENSMUSG00000056413 | Adap1 | ArfGAP with dual PH domains 1 |
| chr11 | 14872199 | 14872822 | 624 | * | 4,093041 | 6,137539913 | 2,322661 | 2,64246 | 0,008231 | 1 | 0 | Distal Intergenic | 11 | 14549276 | 14550314 | 1039 | 1 | 432536 | ENSMUST00000117584.3 | 322923 | ENSMUSG00000084135 | Pom121l12 | POM121 membrane glycoprotein-like 12 |
| chr19 | 55950553 | 55950987 | 435 | * | 3,899385 | 6,137593098 | 2,261999 | 2,713349 | 0,006661 | 1 | 0 | Distal Intergenic | 19 | 55886741 | 55908036 | 21296 | 1 | 21416 | ENSMUST00000148666.3 | 63812 | ENSMUSG00000024985 | Tcf7l2 | transcription factor 7 like 2, T cell specific, HMG box |
| chr11 | 1,04E+08 | 1,04E+08 | 642 | * | 4,045225 | 6,137647362 | 2,315385 | 2,65081 | 0,00803 | 1 | 0 | Promoter (<=1kb) | 11 | 1,04E+08 | 1,05E+08 | 62477 | 1 | 16416 | ENSMUST00000021028.5 | 149 | ENSMUSG00000020689 | Itgb3 | integrin beta 3 |
| chr14 | 1,01E+08 | 1,01E+08 | 441 | * | 4,125034 | 6,137904448 | 2,298398 | 2,670514 | 0,007574 | 1 | 0 | Intron (ENSMUST0000( | 14 | 1E+08 | 1,01E+08 | 261653 | 2 | 16597 | ENSMUST00000226774.2 | 12694 | ENSMUSG00000072294 | Klf12 | Kruppel-like factor 12 |
| chr11 | 69152184 | 69152663 | 480 | * | 3,560169 | 6,137979833 | 2,318954 | 2,646874 | 0,008124 | 1 | 0 | Distal Intergenic | 11 | 69131425 | 69131529 | 105 | 1 | 1,15E+08 | ENSMUST00000158813.3 | 20759 | ENSMUSG00000089438 | Gm23194 | predicted gene, 23194 |
| chr7 | 75912138 | 75912744 | 607 | * | 3,544553 | 6,137987339 | 2,319334 | 2,646444 | 0,008134 | 1 | 0 | Intron (ENSMUST0000( | 7 | 75879635 | 76774446 | 894812 | 1 | 244071 | ENSMUST00000156166.8 | 32503 | ENSMUSG00000025754 | Agbl1 | ATP/GTP binding protein-like 1 |
| chr12 | 1,09E+08 | 1,09E+08 | 709 | * | 4,604512 | 6,138023944 | 2,326159 | 2,638695 | 0,008323 | 1 | 0 | Distal Intergenic | 12 | 1,09E+08 | 1,09E+08 | 1970 | 2 | 1,05E+08 | ENSMUST00000243496.1 | 3447 | | Gm40572 | predicted gene, 40572 |
| chr14 | 56934612 | 56934959 | 348 | * | 3,938485 | 6,138282716 | 2,273227 | 2,700251 | 0,006929 | 1 | 0 | Promoter (1-2kb) | 14 | 56933737 | 56936818 | 3082 | 2 | 1,01E+08 | ENSMUST00000180752.2 | 1859 | ENSMUSG00000097743 | Gm16973 | predicted gene, 16973 |
| chrX | 10618280 | 10619022 | 743 | * | 3,595914 | 6,13830214 | 2,272552 | 2,701062 | 0,006912 | 1 | 0 | Distal Intergenic | 20 | 10583707 | 10584623 | 917 | 1 | 68041 | ENSMUST00000156321.2 | 34573 | ENSMUSG00000008035 | Mid1ip1 | Mid1 interacting protein 1 (gastrulation specific G12-like (zebrafish)) |
| chr17 | 43716582 | 43716937 | 356 | * | 4,013993 | 6,138350178 | 2,270283 | 2,703782 | 0,006856 | 1 | 0 | Intron (ENSMUST0000( | 17 | 43700379 | 43733478 | 33100 | 1 | 224792 | ENSMUST00000225466.2 | 16203 | ENSMUSG00000056492 | Adgrf5 | adhesion G protein-coupled receptor F5 |
| chr14 | 1,22E+08 | 1,22E+08 | 629 | * | 4,069923 | 6,138407581 | 2,266218 | 2,708657 | 0,006756 | 1 | 0 | Intron (ENSMUST0000( | 14 | 1,22E+08 | 1,22E+08 | 2599 | 2 | 319727 | ENSMUST00000131416.3 | 30772 | | A330035P11Rik | RIKEN cDNA A330035P11 gene |
| chrX | 1,65E+08 | 1,65E+08 | 360 | * | 3,476843 | 6,138414373 | 2,381485 | 2,577557 | 0,00995 | 1 | 0 | Promoter (1-2kb) | 20 | 1,65E+08 | 1,65E+08 | 39309 | 1 | 14758 | ENSMUST00000123408.8 | -1935 | ENSMUSG00000031342 | Gpm6b | glycoprotein m6b |
| chr8 | 93385904 | 93386075 | 172 | * | 3,76366 | 6,138657969 | 2,251146 | 2,726903 | 0,006393 | 1 | 0 | Distal Intergenic | 8 | 93400916 | 93407584 | 6669 | 1 | 64379 | ENSMUST00000034185.13 | -14841 | ENSMUSG00000031738 | Irx6 | Iroquois homeobox 6 |
| chrX | 94948264 | 94948977 | 714 | * | 4,035663 | 6,139108889 | 2,22515 | 2,758963 | 0,005799 | 1 | 0 | Intron (ENSMUST0000( | 20 | 94942726 | 94976243 | 33518 | 1 | 547176 | ENSMUST00000145362.4 | 5538 | ENSMUSG00000035045 | Zc3h12b | zinc finger CCCH-type containing 12B |
| chr16 | 10186620 | 10187050 | 431 | * | 4,581528 | 6,139341061 | 2,213747 | 2,773281 | 0,005549 | 1 | 0 | Intron (ENSMUST0000( | 16 | 10146207 | 10165455 | 19249 | 2 | 381033 | ENSMUST00000229862.2 | -21165 | ENSMUSG00000116270 | Gm1600 | predicted gene 1600 |
| chr2 | 75450603 | 75450983 | 381 | * | 4,354417 | 6,139432514 | 2,207714 | 2,7809 | 0,005421 | 1 | 0 | Distal Intergenic | 2 | 75464373 | 75465119 | 747 | 1 | 1,05E+08 | ENSMUST00000247606.1 | -13390 | | Gm39858 | predicted gene, 39858 |
| chr13 | 28977636 | 28978068 | 433 | * | 4,659131 | 6,140286503 | 2,165718 | 2,835219 | 0,004579 | 1 | 0 | Intron (ENSMUST0000( | 13 | 28644785 | 28941618 | 296834 | 2 | 72518 | ENSMUST00000244648.1 | -36018 | | 2610307P16Rik | RIKEN cDNA 2610307P16 gene |
| chr18 | 54070132 | 54070873 | 742 | * | 4,527247 | 6,141275916 | 2,120957 | 2,895521 | 0,003785 | 1 | 0 | Intron (ENSMUST0000( | 18 | 54065316 | 54081743 | 16428 | 1 | 70425 | ENSMUST00000236399.2 | 4816 | ENSMUSG00000073563 | Csnk1g3 | casein kinase 1, gamma 3 |
| chr4 | 1,39E+08 | 1,39E+08 | 455 | * | 4,471662 | 6,141666862 | 2,104717 | 2,918049 | 0,003522 | 1 | 0 | Distal Intergenic | 4 | 1,39E+08 | 1,39E+08 | 18567 | 2 | 73162 | ENSMUST00000097830.4 | -18928 | ENSMUSG00000041161 | Otud3 | OTU domain containing 3 |
| chr11 | 1,22E+08 | 1,22E+08 | 695 | * | 4,835076 | 6,143081265 | 2,052426 | 2,993082 | 0,002762 | 1 | 0 | Exon (ENSMUST000001 | 11 | 1,22E+08 | 1,22E+08 | 20748 | 2 | 1,19E+08 | ENSMUST00000243379.1 | -13790 | | Gm53660 | predicted gene, 53660 |
| chr3 | 30067027 | 30068455 | 1429 | * | 12,0071 | 6,144154213 | 1,680771 | 3,655558 | 0,000257 | 0,358680655 | 0,744375 | Promoter (<=1kb) | 3 | 30005445 | 30067305 | 61861 | 2 | 14013 | ENSMUST00000172694.8 | 0 | ENSMUSG00000027684 | Mecom | MDS1 and EVI1 complex locus |
| chr14 | 74581698 | 74582713 | 1016 | * | 5,154419 | 6,14431187 | 2,012713 | 3,052752 | 0,002268 | 1 | 0 | Intron (ENSMUST0000( | 14 | 74592967 | 74607181 | 14215 | 1 | 74425 | ENSMUST00000226927.2 | -10254 | ENSMUSG00000115845 | 4933402J15Rik | RIKEN cDNA 4933402J15 gene |
| chr18 | 88314343 | 88314803 | 461 | * | 5,498685 | 6,145406394 | 1,98121 | 3,101845 | 0,001923 | 1 | 0 | Distal Intergenic | 18 | 88627526 | 88676931 | 49406 | 1 | 1,19E+08 | ENSMUST00000237266.2 | -312723 | | Gm50401 | predicted gene, 50401 |
| chr1 | 1,51E+08 | 1,51E+08 | 640 | * | 5,812418 | 6,146118936 | 1,962208 | 3,132247 | 0,001735 | 1 | 0 | Intron (ENSMUST0000( | 1 | 1,51E+08 | 1,52E+08 | 118687 | 1 | 63913 | ENSMUST00000111875.2 | 51629 | ENSMUSG00000026483 | Niban1 | niban apoptosis regulator 1 |
| chr18 | 75634887 | 75635454 | 568 | * | 5,996269 | 6,146412578 | 1,95477 | 3,144314 | 0,001665 | 1 | 0 | Intron (ENSMUST0000( | 18 | 75647716 | 75655877 | 8162 | 1 | 1E+08 | ENSMUST00000181913.2 | -12262 | ENSMUSG00000097712 | Gm10532 | predicted gene 10532 |
| chr11 | 78046728 | 78047423 | 696 | * | 6,339392 | 6,146490495 | 1,952878 | 3,147402 | 0,001647 | 1 | 0 | 3' UTR | 11 | 78051619 | 78056342 | 4724 | 2 | 22032 | ENSMUST00000148218.2 | 8919 | ENSMUSG00000017386 | Traf4 | TNF receptor associated factor 4 |
| chr9 | 54048756 | 54049871 | 1116 | * | 7,322288 | 6,146734875 | 1,946964 | 3,157087 | 0,001594 | 1 | 0 | Distal Intergenic | 9 | 54073221 | 54100726 | 27506 | 2 | 13075 | ENSMUST00000034811.8 | 50855 | ENSMUSG00000032274 | Cyp19a1 | cytochrome P450, family 19, subfamily a, polypeptide 1 |
| chr4 | 1,36E+08 | 1,36E+08 | 1309 | * | 7,741257 | 6,146950519 | 1,941695 | 3,165765 | 0,001547 | 1 | 0 | Exon (ENSMUST00000( | 4 | 1,36E+08 | 1,36E+08 | 18396 | 2 | 27224 | ENSMUST00000030427.6 | 12215 | ENSMUSG00000028668 | Eloa | elongin A |
| chr12 | 68471750 | 68472340 | 591 | * | 6,480393 | 6,14837444 | 1,91002 | 3,21901 | 0,001286 | 1 | 0 | Intron (ENSMUST0000( | 12 | 69176610 | 69191850 | 15241 | 1 | 1,03E+08 | ENSMUST00000222345.2 | -704270 | ENSMUSG00000113158 | Gm31447 | predicted gene, 31447 |
| chr9 | 96974205 | 96974571 | 367 | * | 3,001354 | 6,149344574 | 2,377701 | 2,586257 | 0,009702 | 1 | 0 | Promoter (2-3kb) | 9 | 96962412 | 96972197 | 9786 | 2 | 192287 | ENSMUST00000137372.2 | -2008 | ENSMUSG00000032449 | Slc25a36 | solute carrier family 25, member 36 |
| chr15 | 66691243 | 66691502 | 260 | * | 3,619846 | 6,149363295 | 2,376977 | 2,587053 | 0,00968 | 1 | 0 | Intron (ENSMUST0000( | 15 | 66652713 | 66684442 | 31730 | 2 | 20491 | ENSMUST00000100572.10 | -6801 | ENSMUSG00000022372 | Sla | src-like adaptor |
| chr13 | 97288265 | 97288614 | 350 | * | 3,686301 | 6,14937723 | 2,361692 | 2,603802 | 0,00922 | 1 | 0 | Promoter (1-2kb) | 13 | 97286858 | 97299410 | 12553 | 1 | 320806 | ENSMUST00000160981.2 | 1407 | ENSMUSG00000021666 | Gfm2 | G elongation factor, mitochondrial 2 |
| chr3 | 64581403 | 64581889 | 487 | * | 3,525014 | 6,149463326 | 2,325514 | 2,644346 | 0,008185 | 1 | 0 | Intron (ENSMUST0000( | 3 | 64539995 | 64569682 | 29688 | 2 | 319210 | ENSMUST00000176568.2 | -11721 | ENSMUSG00000097538 | Vmn2r-ps11 | vomeronasal 2, receptor, pseudogene 11 |
| chrX | 69921935 | 69922635 | 701 | * | 3,782191 | 6,149519908 | 2,305749 | 2,667037 | 0,007652 | 1 | 0 | Distal Intergenic | 20 | 70093846 | 70198553 | 104708 | 1 | 333639 | ENSMUST00000239162.2 | -171211 | ENSMUSG00000059401 | Mamld1 | mastermind-like domain containing 1 |
| chr1 | 1,78E+08 | 1,78E+08 | 523 | * | 3,819615 | 6,149534567 | 2,301505 | 2,671962 | 0,007541 | 1 | 0 | Distal Intergenic | 1 | 1,78E+08 | 1,78E+08 | 23511 | 1 | 69428 | ENSMUST00000016106.6 | -21804 | ENSMUSG00000015962 | 1700016C15Rik | RIKEN cDNA 1700016C15 gene |
| chr11 | 98127801 | 98128427 | 627 | * | 3,761995 | 6,149647325 | 2,261474 | 2,71931 | 0,006542 | 1 | 0 | Intron (ENSMUST0000( | 11 | 98132563 | 98169330 | 36768 | 1 | 69131 | ENSMUST00000147441.8 | -4136 | ENSMUSG00000003119 | Cdk12 | cyclin-dependent kinase 12 |
| chrX | 1,38E+08 | 1,38E+08 | 566 | * | 4,874962 | 6,149823721 | 2,198631 | 2,797115 | 0,005156 | 1 | 0 | Distal Intergenic | 20 | 1,38E+08 | 1,38E+08 | 21170 | 1 | 245631 | ENSMUST00000113042.3 | 21882 | ENSMUSG00000042515 | Pwwp3b | PWWP domain containing 3B |
| chr11 | 75932908 | 75933518 | 611 | * | 7,596502 | 6,149910435 | 1,879776 | 3,271618 | 0,001069 | 1 | 0 | Distal Intergenic | 11 | 75910020 | 75918608 | 8589 | 2 | 76566 | ENSMUST00000021207.7 | -14300 | ENSMUSG00000020846 | Rflnb | refilin B |
| chr1 | 86658363 | 86658788 | 426 | * | 3,861476 | 6,149915134 | 2,173268 | 2,8298 | 0,004658 | 1 | 0 | Intron (ENSMUST0000( | 1 | 86631565 | 86977814 | 346250 | 1 | 208718 | ENSMUST00000065694.8 | 26798 | ENSMUSG00000053333 | Dis3l2 | DIS3 like 3'-5' exoribonuclease 2 |
| chr17 | 27947987 | 27948463 | 477 | * | 4,167726 | 6,149915221 | 2,171639 | 2,831924 | 0,004627 | 1 | 0 | Promoter (<=1kb) | 17 | 27933633 | 27947929 | 14297 | 2 | 30051 | ENSMUST00000114870.9 | -58 | ENSMUSG00000024215 | Spdef | SAM pointed domain containing ets transcription factor |
| chr3 | 27839482 | 27839824 | 343 | * | 3,7959 | 6,149972812 | 2,157435 | 2,850595 | 0,004364 | 1 | 0 | Distal Intergenic | 3 | 27548581 | 27765456 | 216876 | 2 | 72007 | ENSMUST00000191684.2 | -74026 | ENSMUSG00000039286 | Fndc3b | fibronectin type III domain containing 3B |
| chr10 | 61131099 | 61131552 | 454 | * | 4,539823 | 6,149973399 | 2,155873 | 2,852661 | 0,004335 | 1 | 0 | Promoter (2-3kb) | 10 | 61133612 | 61140459 | 6848 | 1 | 18646 | ENSMUST00000219375.2 | -2060 | ENSMUSG00000037202 | Prf1 | perforin 1 (pore forming protein) |
| chr12 | 42356764 | 42357470 | 707 | * | 4,055904 | 6,150047496 | 2,140545 | 2,873123 | 0,004064 | 1 | 0 | Distal Intergenic | 12 | 41965491 | 42063539 | 98049 | 2 | 1,08E+08 | ENSMUST00000222917.2 | -293225 | | Gm46348 | predicted gene, 46348 |
| chr19 | 48262599 | 48263050 | 452 | * | 4,755714 | 6,150137573 | 2,118698 | 2,902791 | 0,003699 | 1 | 0 | Intron (ENSMUST0000( | 19 | 48194464 | 48793944 | 599481 | 1 | 66673 | ENSMUST00000078880.6 | 68135 | ENSMUSG00000063434 | Sorcs3 | sortilin-related VPS10 domain containing receptor 3 |
| chr18 | 83859483 | 83859909 | 427 | * | 4,235454 | 6,150334403 | 2,072474 | 2,96763 | 0,003001 | 1 | 0 | Distal Intergenic | 18 | 83652026 | 83656210 | 4185 | 2 | 1,03E+08 | ENSMUST00000237865.2 | -203273 | | Gm31621 | predicted gene, 31621 |
| chr19 | 20744560 | 20744938 | 379 | * | 4,992728 | 6,150368959 | 2,063293 | 2,980851 | 0,002874 | 1 | 0 | Distal Intergenic | 19 | 20670317 | 20704926 | 34610 | 2 | 26358 | ENSMUST00000025656.4 | -39634 | ENSMUSG00000024747 | Aldh1a7 | aldehyde dehydrogenase family 1, subfamily A7 |
| chr6 | 36043409 | 36043737 | 329 | * | 4,622538 | 6,15042201 | 2,057009 | 2,989983 | 0,00279 | 1 | 0 | Distal Intergenic | 6 | 35952265 | 35952399 | 135 | 1 | 1,15E+08 | ENSMUST00000104058.3 | 91144 | ENSMUSG00000077246 | Gm23273 | predicted gene, 23273 |
| chr1 | 54945139 | 54945672 | 534 | * | 5,417673 | 6,150738129 | 1,994626 | 3,083655 | 0,002045 | 1 | 0 | Intron (ENSMUST0000( | 1 | 54774461 | 54965513 | 191053 | 2 | 329154 | ENSMUST00000178935.8 | 19841 | ENSMUSG00000052331 | Ankrd44 | ankyrin repeat domain 44 |
| chr11 | 78986690 | 78987510 | 821 | * | 6,494235 | 6,150771342 | 1,988615 | 3,092992 | 0,001981 | 1 | 0 | Intron (ENSMUST0000( | 11 | 78921451 | 78965312 | 43862 | 2 | 16706 | ENSMUST00000208969.2 | -21378 | ENSMUSG00000018334 | Ksr1 | kinase suppressor of ras 1 |
| chr15 | 13290945 | 13291725 | 781 | * | 5,024124 | 6,150796099 | 1,987584 | 3,094609 | 0,001971 | 1 | 0 | Distal Intergenic | 15 | 13059337 | 13173761 | 114425 | 2 | 12563 | ENSMUST00000226917.2 | -117184 | ENSMUSG00000039385 | Cdh6 | cadherin 6 |
| chr18 | 55309778 | 55310424 | 647 | * | 5,287421 | 6,150817526 | 1,982068 | 3,103233 | 0,001914 | 1 | 0 | Distal Intergenic | 18 | 55190657 | 55212486 | 21830 | 1 | 1E+08 | ENSMUST00000181765.2 | 119121 | | Gm4221 | predicted gene 4221 |
| chr13 | 1,04E+08 | 1,04E+08 | 698 | * | 5,583701 | 6,150892898 | 1,972462 | 3,118384 | 0,001818 | 1 | 0 | Distal Intergenic | 13 | 1,03E+08 | 1,03E+08 | 1453 | 1 | 432800 | ENSMUST00000052030.6 | 52715 | | Gm5454 | predicted gene 5454 |
| chr13 | 96675609 | 96676310 | 702 | * | 5,741061 | 6,150901557 | 1,970425 | 3,121612 | 0,001799 | 1 | 0 | Promoter (2-3kb) | 13 | 96641047 | 96678848 | 37802 | 2 | 27015 | ENSMUST00000223475.2 | 2538 | ENSMUSG00000021668 | Polk | polymerase (DNA directed), kappa |
| chr11 | 95973010 | 95973874 | 865 | * | 6,364727 | 6,150936412 | 1,962498 | 3,134239 | 0,001723 | 1 | 0 | Promoter (<=1kb) | 11 | 95974499 | 95974682 | 184 | 2 | 1,15E+08 | ENSMUST00000082861.3 | 808 | ENSMUSG00000064795 | Gm22423 | predicted gene, 22423 |
| chr13 | 96251991 | 96252762 | 772 | * | 6,019596 | 6,151384665 | 1,903891 | 3,230954 | 0,001234 | 1 | 0 | Intron (ENSMUST0000( | 13 | 96091102 | 96269085 | 177984 | 2 | 75209 | ENSMUST00000161263.8 | 16323 | ENSMUSG00000051111 | Sv2c | synaptic vesicle glycoprotein 2c |
| chr11 | 77583627 | 77584182 | 556 | * | 6,018371 | 6,151442504 | 1,897308 | 3,242196 | 0,001186 | 1 | 0 | Exon (ENSMUST000001 | 11 | 77578427 | 77582544 | 4118 | 1 | 68564 | ENSMUST00000155238.2 | 5200 | ENSMUSG00000037857 | Nufip2 | nuclear fragile X mental retardation protein interacting protein 2 |
| chr8 | 45391286 | 45391932 | 647 | * | 6,35166 | 6,151535543 | 1,886465 | 3,26088 | 0,001111 | 1 | 0 | Promoter (2-3kb) | 8 | 45388484 | 45505294 | 116811 | 1 | 14107 | ENSMUST00000191428.7 | 2802 | ENSMUSG00000070047 | Fat1 | FAT atypical cadherin 1 |
| chr5 | 28335108 | 28335734 | 627 | * | 6,004324 | 6,15156953 | 1,883107 | 3,266713 | 0,001088 | 1 | 0 | Distal Intergenic | 5 | 28370692 | 28377164 | 6473 | 1 | 13799 | ENSMUST00000036177.9 | -34958 | ENSMUSG00000039095 | En2 | engrailed 2 |
| chr11 | 97892149 | 97892974 | 826 | * | 7,196482 | 6,151786914 | 1,860121 | 3,307198 | 0,000942 | 1 | 0 | 3' UTR | 11 | 97880404 | 97887007 | 6604 | 2 | 217151 | ENSMUST00000042971.16 | -5142 | ENSMUSG00000038352 | Arl5c | ADP-ribosylation factor-like 5C |
| chr1 | 1,8E+08 | 1,8E+08 | 1046 | * | 6,567377 | 6,154175061 | 1,904215 | 3,231869 | 0,00123 | 1 | 0 | Intron (ENSMUST0000( | 1 | 1,8E+08 | 1,8E+08 | 10364 | 1 | 226751 | ENSMUST00000139002.2 | 7710 | ENSMUSG00000026490 | Cdc42bpa | CDC42 binding protein kinase alpha |
| chr4 | 1,56E+08 | 1,56E+08 | 691 | * | 6,085888 | 6,154883551 | 1,927011 | 3,194005 | 0,001403 | 1 | 0 | Promoter (<=1kb) | 4 | 1,56E+08 | 1,56E+08 | 8460 | 2 | 1,01E+08 | ENSMUST00000154931.2 | 310 | ENSMUSG00000086682 | Gm16023 | predicted gene 16023 |
| chr5 | 93274730 | 93275508 | 779 | * | 5,818662 | 6,155882715 | 1,963312 | 3,135458 | 0,001716 | 1 | 0 | Intron (ENSMUST0000( | 5 | 93296362 | 93305561 | 9200 | 1 | 52398 | ENSMUST00000201695.2 | -20854 | ENSMUSG00000058013 | Septin11 | septin 11 |
| chr9 | 22229080 | 22229990 | 911 | * | 5,38881 | 6,156356731 | 1,981974 | 3,106174 | 0,001895 | 1 | 0 | Distal Intergenic | 9 | 22190363 | 22218944 | 28582 | 2 | 235050 | ENSMUST00000215202.2 | -10136 | ENSMUSG00000066829 | Zfp810 | zinc finger protein 810 |
| chr2 | 1,26E+08 | 1,26E+08 | 759 | * | 5,516296 | 6,157315972 | 2,024563 | 3,041306 | 0,002356 | 1 | 0 | Intron (ENSMUST0000( | 2 | 1,26E+08 | 1,26E+08 | 81083 | 1 | 69976 | ENSMUST00000110454.2 | 75880 | ENSMUSG00000027207 | Galk2 | galactokinase 2 |
| chr1 | 1,5E+08 | 1,5E+08 | 495 | * | 4,576911 | 6,157739466 | 2,045861 | 3,009852 | 0,002614 | 1 | 0 | Intron (ENSMUST0000( | 1 | 1,5E+08 | 1,5E+08 | 16016 | 2 | 18783 | ENSMUST00000155438.2 | 65084 | ENSMUSG00000056220 | Pla2g4a | phospholipase A2, group IVA (cytosolic, calcium-dependent) |
| chr9 | 29851900 | 29852690 | 791 | * | 4,936948 | 6,158532498 | 2,090312 | 2,946226 | 0,003217 | 1 | 0 | Intron (ENSMUST0000( | 9 | 29322884 | 29874415 | 551532 | 2 | 235106 | ENSMUST00000146651.2 | 21725 | ENSMUSG00000059974 | Ntm | neurotrimin |
| chr2 | 8784698 | 8785105 | 408 | * | 3,379867 | 6,160104328 | 2,381898 | 2,586217 | 0,009704 | 1 | 0 | Distal Intergenic | 2 | 9194326 | 9202288 | 7963 | 1 | 73438 | ENSMUST00000132414.2 | -409221 | ENSMUSG00000085580 | 1700061F12Rik | RIKEN cDNA 1700061F12 gene |

| | | | | | | | | | | | | | | | | | | | | | | | |
|---|---|---|---|---|---|---|---|---|---|---|---|---|---|---|---|---|---|---|---|---|---|---|---|
| chrX | 1,06E+08 | 1,06E+08 | 805 | * | 4,078151 | 6,160124917 | 2,38155 | 2,586603 | 0,009693 | 1 | 0 | Promoter (<=1kb) | 20 | 1,06E+08 | 1,06E+08 | 13276 | 1 | 78134 | ENSMUST00000060576.8 | 414 | ENSMUSG00000049929 | Lpar4 | lysophosphatidic acid receptor 4 |
| chr14 | 1,18E+08 | 1,18E+08 | 713 | * | 4,023079 | 6,160302917 | 2,212739 | 2,784017 | 0,005369 | 1 | 0 | Intron (ENSMUST0000( | 14 | 1,18E+08 | 1,18E+08 | 105 | 2 | 1,02E+08 | ENSMUST00000183564.3 | 158569 | ENSMUSG00000098648 | Mir6239 | microRNA 6239 |
| chrX | 1,45E+08 | 1,45E+08 | 576 | * | 3,720578 | 6,160307187 | 2,362107 | 2,607971 | 0,009108 | 1 | 0 | Distal Intergenic | 20 | 1,46E+08 | 1,46E+08 | 26816 | 1 | 15560 | ENSMUST00000149260.8 | -309009 | ENSMUSG00000041380 | Htr2c | 5-hydroxytryptamine (serotonin) receptor 2C |
| chr13 | 64311933 | 64312167 | 235 | * | 3,341086 | 6,160430182 | 2,351625 | 2,619648 | 0,008802 | 1 | 0 | Promoter (1-2kb) | 13 | 64310060 | 64320610 | 10551 | 1 | 56541 | ENSMUST00000223328.2 | 1873 | ENSMUSG00000021476 | Habp4 | hyaluronic acid binding protein 4 |
| chr9 | 1,08E+08 | 1,08E+08 | 680 | * | 4,229463 | 6,160788422 | 2,257241 | 2,729345 | 0,006346 | 1 | 0 | Intron (ENSMUST0000( | 9 | 1,08E+08 | 1,08E+08 | 38246 | 1 | 27399 | ENSMUST00000175874.2 | 6883 | ENSMUSG00000032594 | Ip6k1 | inositol hexaphosphate kinase 1 |
| chrX | 36255420 | 36256154 | 735 | * | 3,837709 | 6,160936185 | 2,270088 | 2,713964 | 0,006648 | 1 | 0 | Promoter (<=1kb) | 20 | 36230249 | 36255447 | 25199 | 2 | 56526 | ENSMUST00000152291.3 | 0 | ENSMUSG00000050379 | Septin6 | septin 6 |
| chrX | 59282413 | 59283104 | 692 | * | 3,41714 | 6,161096006 | 2,293002 | 2,686913 | 0,007212 | 1 | 0 | Intron (ENSMUST0000( | 20 | 59302912 | 59303018 | 107 | 1 | 1,15E+08 | ENSMUST00000239780.1 | -19808 | ENSMUSG00000119039 | Gm22287 | predicted gene, 22287 |
| chr14 | 1,21E+08 | 1,21E+08 | 536 | * | 4,043939 | 6,161398053 | 2,268664 | 2,715871 | 0,00661 | 1 | 0 | Intron (ENSMUST0000( | 14 | 1,21E+08 | 1,21E+08 | 133025 | 1 | 105559 | ENSMUST00000226800.2 | -6115 | ENSMUSG00000022139 | Mbnl2 | muscleblind like splicing factor 2 |
| chr1 | 1,36E+08 | 1,36E+08 | 482 | * | 4,427322 | 6,161566773 | 2,335264 | 2,638488 | 0,008328 | 1 | 0 | Exon (ENSMUST00000( | 1 | 1,36E+08 | 1,36E+08 | 13499 | 2 | 67886 | ENSMUST00000194808.6 | -22200 | ENSMUSG00000041570 | Camsap2 | calmodulin regulated spectrin-associated protein family, member 2 |
| chr9 | 20956203 | 20956461 | 259 | * | 3,753726 | 6,161569443 | 2,254995 | 2,732409 | 0,006287 | 1 | 0 | Distal Intergenic | 9 | 20943547 | 20943624 | 78 | 2 | 1E+08 | ENSMUST00000122714.3 | -12579 | ENSMUSG00000084663 | Mir1900 | microRNA 1900 |
| chrX | 73276341 | 73277375 | 1035 | * | 4,15156 | 6,161699003 | 2,245199 | 2,744389 | 0,006062 | 1 | 0 | Promoter (2-3kb) | 20 | 73271986 | 73274176 | 2191 | 2 | 192176 | ENSMUST00000127463.2 | -2165 | ENSMUSG00000031328 | Flna | filamin, alpha |
| chr18 | 34566993 | 34567332 | 340 | * | 3,614673 | 6,161946727 | 2,3822 | 2,586662 | 0,009691 | 1 | 0 | Intron (ENSMUST0000( | 18 | 34575809 | 34584490 | 8682 | 2 | 225358 | ENSMUST00000235364.2 | 17158 | ENSMUSG00000036501 | Fam13b | family with sequence similarity 13, member B |
| chr11 | 40046056 | 40046659 | 604 | * | 4,827453 | 6,162009446 | 2,223026 | 2,771902 | 0,005573 | 1 | 0 | Intron (ENSMUST0000( | 11 | 39899021 | 40228658 | 329638 | 1 | 1,03E+08 | ENSMUST00000130945.2 | 147035 | ENSMUSG00000085301 | Gm12132 | predicted gene 12132 |
| chr9 | 45646667 | 45646978 | 312 | * | 4,231857 | 6,162021305 | 2,222631 | 2,7724 | 0,005564 | 1 | 0 | Intron (ENSMUST0000( | 9 | 45653707 | 45656546 | 2840 | 1 | 114873 | ENSMUST00000215027.2 | -6729 | ENSMUSG00000032087 | Dscaml1 | DS cell adhesion molecule like 1 |
| chr7 | 19241148 | 19241658 | 511 | * | 4,020056 | 6,162249222 | 2,207173 | 2,791919 | 0,00524 | 1 | 0 | Promoter (<=1kb) | 7 | 19238411 | 19242292 | 3882 | 2 | 232946 | ENSMUST00000077408.8 | 634 | ENSMUSG00000057667 | Bloc1s3 | biogenesis of lysosomal organelles complex-1, subunit 3 |
| chr19 | 56460308 | 56461056 | 749 | * | 4,079083 | 6,162274573 | 2,205035 | 2,794638 | 0,005196 | 1 | 0 | Intron (ENSMUST0000( | 19 | 56467333 | 56468296 | 964 | 1 | 226245 | ENSMUST00000225391.2 | -6277 | ENSMUSG00000035818 | Plekhs1 | pleckstrin homology domain containing, family S member 1 |
| chr15 | 41505472 | 41506082 | 611 | * | 4,51218 | 6,162650594 | 2,182392 | 2,823805 | 0,004746 | 1 | 0 | Intron (ENSMUST0000( | 15 | 41574000 | 41683776 | 109777 | 1 | 170719 | ENSMUST00000229769.2 | -67918 | ENSMUSG00000022307 | Oxr1 | oxidation resistance 1 |
| chr8 | 1,27E+08 | 1,27E+08 | 448 | * | 3,965792 | 6,162651344 | 2,181003 | 2,825604 | 0,004719 | 1 | 0 | Distal Intergenic | 8 | 1,27E+08 | 1,27E+08 | 2764 | 2 | 1,15E+08 | ENSMUST00000212695.2 | 32003 | | Gm45805 | predicted gene 45805 |
| chr6 | 83045434 | 83045670 | 237 | * | 4,255096 | 6,16389521 | 2,110267 | 2,920908 | 0,00349 | 1 | 0 | Promoter (1-2kb) | 6 | 83045305 | 83047206 | 1902 | 2 | 21909 | ENSMUST00000174674.3 | 1536 | ENSMUSG00000068327 | Tlx2 | T cell leukemia, homeobox 2 |
| chr5 | 1,19E+08 | 1,19E+08 | 586 | * | 4,742492 | 6,164502314 | 2,080445 | 2,963069 | 0,003046 | 1 | 0 | Promoter (1-2kb) | 5 | 1,19E+08 | 1,19E+08 | 13530 | 1 | 76199 | ENSMUST00000202787.2 | 1664 | ENSMUSG00000018076 | Med13l | mediator complex subunit 13-like |
| chr6 | 65628493 | 65630269 | 1777 | * | 14,46417 | 6,165618064 | 1,741781 | 3,539836 | 0,0004 | 0,327375387 | 1,41696 | Distal Intergenic | 6 | 65648574 | 65689310 | 40737 | 1 | 68169 | ENSMUST00000054351.6 | -18305 | ENSMUSG00000049001 | Ndnf | neuron-derived neurotrophic factor |
| chr3 | 1,28E+08 | 1,28E+08 | 533 | * | 6,093889 | 6,167559078 | 2,107702 | 2,9262 | 0,003431 | 1 | 0 | Promoter (<=1kb) | 3 | 1,28E+08 | 1,28E+08 | 30517 | 2 | 211556 | ENSMUST00000200409.5 | -42 | ENSMUSG00000074238 | Ap1ar | adaptor-related protein complex 1 associated regulatory protein |
| chr5 | 80748051 | 80749036 | 986 | * | 5,563893 | 6,169642211 | 2,170291 | 2,842772 | 0,004472 | 1 | 0 | Distal Intergenic | 5 | 81041501 | 81041603 | 103 | 1 | 1,15E+08 | ENSMUST00000083301.3 | -292465 | ENSMUSG00000065235 | Gm22974 | predicted gene, 22974 |
| chr15 | 36170606 | 36171291 | 686 | * | 4,943884 | 6,170292107 | 2,192263 | 2,814576 | 0,004884 | 1 | 0 | Promoter (<=1kb) | 15 | 36169942 | 36170100 | 159 | 1 | 1,15E+08 | ENSMUST00000082609.3 | 664 | ENSMUSG00000064543 | Gm22979 | predicted gene, 22979 |
| chr2 | 88790463 | 88790927 | 465 | * | 3,862635 | 6,170478419 | 2,394458 | 2,576983 | 0,009967 | 1 | 0 | Promoter (1-2kb) | 2 | 88788812 | 88789747 | 936 | 1 | 258241 | ENSMUST00000055895.3 | 1651 | ENSMUSG00000048226 | Olfr1212 | olfactory receptor 1212 |
| chr13 | 38663070 | 38663432 | 363 | * | 3,742037 | 6,170604784 | 2,387346 | 2,584713 | 0,009746 | 1 | 0 | Intron (ENSMUST0000( | 13 | 38644232 | 38645781 | 1550 | 2 | 75854 | ENSMUST00000225893.2 | -17289 | ENSMUSG00000114895 | 4930579J19Rik | RIKEN cDNA 4930579J19 gene |
| chr19 | 28620922 | 28621731 | 810 | * | 5,632168 | 6,170796275 | 2,210577 | 2,791486 | 0,005247 | 1 | 0 | Intron (ENSMUST0000( | 19 | 28509247 | 28644085 | 134839 | 2 | 226075 | ENSMUST00000159178.2 | 22354 | ENSMUSG00000052942 | Glis3 | GLIS family zinc finger 3 |
| chr19 | 43197992 | 43198719 | 728 | * | 5,12571 | 6,171088356 | 2,221622 | 2,77774 | 0,005474 | 1 | 0 | Intron (ENSMUST0000( | 19 | 42897023 | 43212128 | 315106 | 2 | 545291 | ENSMUST00000238153.2 | 13409 | ENSMUSG00000074852 | Hpse2 | heparanase 2 |
| chr15 | 34081875 | 34082484 | 610 | * | 5,517525 | 6,171703405 | 2,245098 | 2,748968 | 0,005978 | 1 | 0 | Promoter (<=1kb) | 15 | 34081906 | 34082779 | 874 | 2 | 1,05E+08 | ENSMUST00000247181.1 | 295 | | Gm41291 | predicted gene, 41291 |
| chr5 | 98116505 | 98117294 | 790 | * | 4,821382 | 6,171796672 | 2,24916 | 2,744045 | 0,006069 | 1 | 0 | Intron (ENSMUST0000( | 5 | 98097029 | 98143954 | 46926 | 2 | 71914 | ENSMUST00000202736.2 | 26660 | ENSMUSG00000029338 | Antxr2 | anthrax toxin receptor 2 |
| chr18 | 66088703 | 66089338 | 636 | * | 7,121912 | 6,171854056 | 1,850554 | 3,335139 | 0,000853 | 0,463890057 | 1,360622 | 3' UTR | 18 | 66089844 | 66100753 | 10910 | 2 | 225644 | ENSMUST00000236894.2 | 11415 | ENSMUSG00000024519 | Cplx4 | complexin 4 |
| chr11 | 79076353 | 79076883 | 531 | * | 5,51752 | 6,171920273 | 2,254544 | 2,737547 | 0,00619 | 1 | 0 | Distal Intergenic | 11 | 78917533 | 79037233 | 119701 | 2 | 16706 | ENSMUST00000108264.8 | -39120 | ENSMUSG00000018334 | Ksr1 | kinase suppressor of ras 1 |
| chr3 | 73442099 | 73442910 | 812 | * | 5,065406 | 6,172348474 | 2,272306 | 2,716336 | 0,006601 | 1 | 0 | Intron (ENSMUST0000( | 3 | 73258539 | 73388984 | 130446 | 2 | 74693 | ENSMUST00000194514.2 | -53115 | ENSMUSG00000102362 | 4930509J09Rik | RIKEN cDNA 4930509J09 gene |
| chr9 | 42072023 | 42072872 | 850 | * | 6,123727 | 6,17245578 | 1,873106 | 3,295305 | 0,000983 | 1 | 0 | Distal Intergenic | 9 | 42053657 | 42055560 | 1904 | 1 | 1,05E+08 | ENSMUST00000217090.2 | 18366 | | Gm39321 | predicted gene, 39321 |
| chr13 | 53252566 | 53253313 | 748 | * | 7,274727 | 6,172471662 | 1,908046 | 3,23497 | 0,001217 | 0,524454869 | 1,360622 | Distal Intergenic | 13 | 53155899 | 53172529 | 16631 | 1 | 1,03E+08 | ENSMUST00000222317.3 | 96667 | | Gm33424 | predicted gene, 33424 |
| chr19 | 12835997 | 12836641 | 645 | * | 6,034368 | 6,172494765 | 1,929346 | 3,199267 | 0,001378 | 1 | 0 | Promoter (2-3kb) | 19 | 12839106 | 12840234 | 1129 | 1 | 258697 | ENSMUST00000059675.4 | -2465 | ENSMUSG00000046272 | Olfr1444 | olfactory receptor 1444 |
| chr3 | 86923485 | 86924502 | 1018 | * | 6,341274 | 6,172507678 | 1,959593 | 3,149893 | 0,001633 | 1 | 0 | Distal Intergenic | 3 | 86903141 | 86906748 | 3608 | 2 | 12479 | ENSMUST00000029717.4 | -16737 | ENSMUSG00000028076 | Cd1d1 | CD1d1 antigen |
| chr2 | 45857144 | 45857912 | 769 | * | 6,0933 | 6,172512969 | 1,961936 | 3,146134 | 0,001654 | 1 | 0 | Distal Intergenic | 2 | 45586812 | 45588079 | 1268 | 1 | 69405 | ENSMUST00000134308.2 | 270332 | ENSMUSG00000086202 | 1700019E08Rik | RIKEN cDNA 1700019E08 gene |
| chr5 | 43935882 | 43936399 | 518 | * | 4,545052 | 6,172552271 | 2,039305 | 3,026792 | 0,002472 | 1 | 0 | Intron (ENSMUST0000( | 5 | 43925451 | 43939442 | 13992 | 2 | 242960 | ENSMUST00000199055.2 | 3043 | ENSMUSG00000039753 | Fbxl5 | F-box and leucine-rich repeat protein 5 |
| chr17 | 36186465 | 36187150 | 686 | * | 4,715121 | 6,172563524 | 2,057791 | 2,999606 | 0,002703 | 1 | 0 | 3' UTR | 17 | 36190711 | 36203562 | 12852 | 1 | 69192 | ENSMUST00000025292.15 | -3561 | ENSMUSG00000024422 | Dhx16 | DEAH (Asp-Glu-Ala-His) box polypeptide 16 |
| chrX | 51602775 | 51603575 | 801 | * | 4,728456 | 6,17257071 | 2,041238 | 3,023935 | 0,002495 | 1 | 0 | Intron (ENSMUST0000( | 20 | 51664720 | 51707924 | 43205 | 1 | 1,01E+08 | ENSMUST00000068507.2 | -61145 | ENSMUSG00000055110 | A630012P03Rik | RIKEN cDNA A630012P03 gene |
| chrX | 1,13E+08 | 1,13E+08 | 676 | * | 5,042012 | 6,172573754 | 2,092852 | 2,949359 | 0,003184 | 1 | 0 | Intron (ENSMUST0000( | 20 | 1,13E+08 | 1,13E+08 | 109908 | 1 | 93837 | ENSMUST00000123974.2 | -8186 | ENSMUSG00000025592 | Dach2 | dachshund family transcription factor 2 |
| chr14 | 77554739 | 77555275 | 537 | * | 4,487031 | 6,172574875 | 2,282161 | 2,704706 | 0,006836 | 1 | 0 | Intron (ENSMUST0000( | 14 | 77549516 | 77815137 | 265622 | 1 | 239188 | ENSMUST00000228148.2 | 5223 | ENSMUSG00000022012 | Enox1 | ecto-NOX disulfide-thiol exchanger 1 |
| chr17 | 34025921 | 34026716 | 796 | * | 4,625838 | 6,172577861 | 2,108827 | 2,92702 | 0,003422 | 1 | 0 | Promoter (2-3kb) | 17 | 34029493 | 34041155 | 11663 | 1 | 80880 | ENSMUST00000234426.2 | -2777 | ENSMUSG00000042099 | Kank3 | KN motif and ankyrin repeat domains 3 |
| chr3 | 1,46E+08 | 1,46E+08 | 484 | * | 4,652504 | 6,172579138 | 2,097722 | 2,942516 | 0,003256 | 1 | 0 | Intron (ENSMUST0000( | 3 | 1,46E+08 | 1,46E+08 | 78817 | 1 | 69219 | ENSMUST00000127193.2 | 47809 | ENSMUSG00000028194 | Ddah1 | dimethylarginine dimethylaminohydrolase 1 |
| chr13 | 24986435 | 24987157 | 723 | * | 4,556178 | 6,172590398 | 2,139797 | 2,884662 | 0,003918 | 1 | 0 | Promoter (<=1kb) | 13 | 24986003 | 24996643 | 10641 | 1 | 79555 | ENSMUST00000155575.2 | 432 | ENSMUSG00000019132 | BC005537 | cDNA sequence BC005537 |
| chr12 | 1,03E+08 | 1,03E+08 | 253 | * | 4,736856 | 6,172596342 | 2,144998 | 2,87767 | 0,004006 | 1 | 0 | Distal Intergenic | 12 | 1,03E+08 | 1,04E+08 | 81549 | 1 | 74521 | ENSMUST00000021631.12 | -19908 | ENSMUSG00000021209 | Ppp4r4 | protein phosphatase 4, regulatory subunit 4 |
| chr4 | 1,45E+08 | 1,45E+08 | 634 | * | 4,375175 | 6,172598174 | 2,104864 | 2,93254 | 0,003362 | 1 | 0 | Distal Intergenic | 4 | 1,45E+08 | 1,45E+08 | 1094 | 1 | 20148 | ENSMUST00000133265.2 | 13105 | ENSMUSG00000066026 | Dhrs3 | dehydrogenase/reductase (SDR family) member 3 |
| chr9 | 45914791 | 45915346 | 556 | * | 4,301021 | 6,172602797 | 2,170559 | 2,843784 | 0,004458 | 1 | 0 | Intron (ENSMUST0000( | 9 | 45886402 | 45923988 | 37587 | 2 | 18475 | ENSMUST00000217514.2 | 8642 | ENSMUSG00000003131 | Pafah1b2 | platelet-activating factor acetylhydrolase, isoform 1b, subunit 2 |
| chr1 | 62499694 | 62499940 | 247 | * | 4,049082 | 6,172605674 | 2,181972 | 2,828912 | 0,004671 | 1 | 0 | Intron (ENSMUST0000( | 1 | 62742444 | 62779091 | 36648 | 1 | 18187 | ENSMUST00000126344.2 | -242504 | ENSMUSG00000025969 | Nrp2 | neuropilin 2 |
| chr12 | 1,09E+08 | 1,09E+08 | 434 | * | 3,630131 | 6,172612796 | 2,221453 | 2,778638 | 0,005459 | 1 | 0 | Promoter (<=1kb) | 12 | 1,09E+08 | 1,09E+08 | 36031 | 2 | 380785 | ENSMUST00000190647.3 | -716 | ENSMUSG00000040867 | Begain | brain-enriched guanylate kinase-associated |
| chr11 | 1,08E+08 | 1,08E+08 | 509 | * | 3,705349 | 6,17261595 | 2,241355 | 2,753966 | 0,005888 | 1 | 0 | Distal Intergenic | 11 | 1,08E+08 | 1,08E+08 | 62201 | 2 | 54377 | ENSMUST00000021066.4 | -21375 | ENSMUSG00000020723 | Cacng4 | calcium channel, voltage-dependent, gamma subunit 4 |
| chr4 | 1,34E+08 | 1,34E+08 | 262 | * | 3,804903 | 6,172626498 | 2,311865 | 2,669977 | 0,007586 | 1 | 0 | Intron (ENSMUST0000( | 4 | 1,34E+08 | 1,34E+08 | 129 | 2 | 1,02E+08 | ENSMUST00000185007.3 | -24635 | ENSMUSG00000099289 | Mir6403 | microRNA 6403 |
| chr11 | 60072863 | 60073232 | 370 | * | 3,575935 | 6,172627179 | 2,261869 | 2,728994 | 0,006353 | 1 | 0 | Intron (ENSMUST0000( | 11 | 60066705 | 60090023 | 23319 | 1 | 19377 | ENSMUST00000102688.2 | 6158 | ENSMUSG00000062115 | Rai1 | retinoic acid induced 1 |
| chr16 | 19916156 | 19916489 | 334 | * | 3,366439 | 6,172629526 | 2,324236 | 2,655767 | 0,007913 | 1 | 0 | Promoter (<=1kb) | 16 | 19916292 | 19947971 | 31680 | 1 | 75785 | ENSMUST00000023509.5 | 0 | ENSMUSG00000062901 | Klhl24 | kelch-like 24 |
| chr13 | 35126012 | 35126302 | 291 | * | 3,244864 | 6,172656495 | 2,382951 | 2,590341 | 0,009588 | 1 | 0 | Intron (ENSMUST0000( | 13 | 35108392 | 35139858 | 31467 | 1 | 66761 | ENSMUST00000160905.2 | 17620 | ENSMUSG00000021415 | 4933417A18Rik | RIKEN cDNA 4933417A18 gene |
| chr9 | 59553448 | 59554487 | 1040 | * | 4,995098 | 6,17274156 | 2,289299 | 2,696346 | 0,00701 | 1 | 0 | Promoter (<=1kb) | 9 | 59555480 | 59556815 | 1336 | 1 | 67287 | ENSMUST00000217284.2 | -993 | ENSMUSG00000025237 | Parp6 | poly (ADP-ribose) polymerase family, member 6 |
| chr15 | 79084908 | 79085426 | 519 | * | 4,932905 | 6,172823842 | 2,293879 | 2,690998 | 0,007124 | 1 | 0 | Exon (ENSMUST00000: | 15 | 79093371 | 79095211 | 1841 | 1 | 1E+08 | ENSMUST00000242492.1 | -7945 | ENSMUSG00000075555 | Gm10863 | predicted gene 10863 |
| chr11 | 51781464 | 51781902 | 439 | * | 5,21621 | 6,172916595 | 2,297346 | 2,686978 | 0,00721 | 1 | 0 | Distal Intergenic | 11 | 51789793 | 51800451 | 10659 | 1 | 1,05E+08 | ENSMUST00000212253.2 | -7891 | ENSMUSG00000110618 | Gm39822 | predicted gene, 39822 |
| chr16 | 91853953 | 91855501 | 1549 | * | 9,679108 | 6,173043457 | 1,826963 | 3,378855 | 0,000728 | 0,47475665 | 1,086118 | Promoter (<=1kb) | 16 | 91855204 | 91984229 | 129026 | 1 | 1,05E+08 | ENSMUST00000232677.2 | 0 | | Gm41496 | predicted gene, 41496 |
| chr9 | 82824324 | 82825000 | 677 | * | 4,704213 | 6,173048393 | 2,30358 | 2,679763 | 0,007367 | 1 | 0 | Intron (ENSMUST0000( | 9 | 82812019 | 82815085 | 3067 | 2 | 83946 | ENSMUST00000190405.2 | -9239 | ENSMUSG00000032253 | Phip | pleckstrin homology domain interacting protein |
| chrX | 71505878 | 71506747 | 870 | * | 4,637336 | 6,174030409 | 2,351707 | 2,62534 | 0,008656 | 1 | 0 | Intron (ENSMUST0000( | 20 | 71425476 | 71430464 | 4989 | 2 | 236852 | ENSMUST00000088481.5 | -75414 | ENSMUSG00000043453 | Magea10 | MAGE family member A10 |
| chr16 | 93581429 | 93581825 | 397 | * | 4,815926 | 6,174449407 | 2,373874 | 2,601001 | 0,009295 | 1 | 0 | Intron (ENSMUST0000( | 16 | 93574289 | 93595808 | 21520 | 1 | 70028 | ENSMUST00000228261.2 | 7140 | ENSMUSG00000022946 | Dop1b | DOP1 leucine zipper like protein B |
| chr8 | 22254440 | 22254947 | 508 | * | 5,401789 | 6,175691431 | 2,162669 | 2,855588 | 0,004296 | 1 | 0 | Distal Intergenic | 8 | 22266615 | 22285201 | 18587 | 1 | 13214 | ENSMUST00000051017.9 | -11668 | ENSMUSG00000044748 | Defb1 | defensin beta 1 |
| chr4 | 1,4E+08 | 1,4E+08 | 849 | * | 5,422597 | 6,175956302 | 2,153552 | 2,8678 | 0,004133 | 1 | 0 | Distal Intergenic | 4 | 1,4E+08 | 1,4E+08 | 7807 | 2 | 242721 | ENSMUST00000105031.4 | -15289 | ENSMUSG00000078234 | Klhdc7a | kelch domain containing 7A |
| chr11 | 32453292 | 32453977 | 686 | * | 5,250552 | 6,175958657 | 2,153571 | 2,867776 | 0,004134 | 1 | 0 | Intron (ENSMUST0000( | 11 | 32472751 | 32477579 | 4829 | 2 | 69306 | ENSMUST00000054327.3 | 23602 | ENSMUSG00000044056 | Efcab9 | EF-hand calcium binding domain 9 |
| chr15 | 62197449 | 62198001 | 553 | * | 5,759949 | 6,176676134 | 1,935498 | 3,19126 | 0,001417 | 1 | 0 | Intron (ENSMUST0000( | 15 | 62094452 | 62114354 | 19903 | 1 | 19296 | ENSMUST00000182141.2 | 102997 | ENSMUSG00000097039 | Pvt1 | Pvt1 oncogene |
| chr15 | 21791902 | 21792555 | 654 | * | 5,779072 | 6,176954921 | 1,943682 | 3,177967 | 0,001483 | 1 | 0 | Distal Intergenic | 15 | 21114455 | 21114561 | 107 | 1 | 1,15E+08 | ENSMUST00000240163.1 | 677447 | ENSMUSG00000119035 | Gm25101 | predicted gene, 25101 |
| chrX | 1,34E+08 | 1,34E+08 | 686 | * | 5,683272 | 6,177318552 | 1,95441 | 3,160708 | 0,001574 | 1 | 0 | Distal Intergenic | 20 | 1,34E+08 | 1,34E+08 | 5769 | 1 | 56258 | ENSMUST00000059297.6 | 14182 | ENSMUSG00000045427 | Hnrnph2 | heterogeneous nuclear ribonucleoprotein H2 |
| chr10 | 26658307 | 26658926 | 620 | * | 5,149099 | 6,178513157 | 1,992657 | 3,10064 | 0,001931 | 1 | 0 | Intron (ENSMUST0000( | 10 | 26648473 | 26794644 | 146172 | 1 | 73910 | ENSMUST00000039557.9 | 9834 | ENSMUSG00000039031 | Arhgap18 | Rho GTPase activating protein 18 |
| chr12 | 16857499 | 16858566 | 1068 | * | 5,892575 | 6,179133156 | 2,061396 | 2,997548 | 0,002722 | 1 | 0 | Promoter (2-3kb) | 12 | 16860964 | 16889743 | 28780 | 1 | 50496 | ENSMUST00000220794.2 | -2398 | ENSMUSG00000057469 | E2f6 | E2F transcription factor 6 |
| chr5 | 40358776 | 40359434 | 659 | * | 4,854454 | 6,179288829 | 2,020665 | 3,058047 | 0,002228 | 1 | 0 | Distal Intergenic | 5 | 40133014 | 40142971 | 9958 | 2 | 1,03E+08 | ENSMUST00000200925.2 | -215805 | ENSMUSG00000107070 | Gm35191 | predicted gene, 35191 |
| chrX | 1,11E+08 | 1,11E+08 | 495 | * | 5,007198 | 6,179588272 | 2,032016 | 3,041112 | 0,002357 | 1 | 0 | Intron (ENSMUST0000( | 20 | 1,1E+08 | 1,11E+08 | 127149 | 2 | 245596 | ENSMUST00000113422.9 | 69583 | ENSMUSG00000034551 | Hdx | highly divergent homeobox |
| chr19 | 10463134 | 10464209 | 1076 | * | 7,497487 | 6,182583328 | 1,983174 | 3,117519 | 0,001824 | 1 | 0 | Promoter (2-3kb) | 19 | 10451603 | 10460292 | 8690 | 2 | 67752 | ENSMUST00000038842.5 | -2842 | ENSMUSG00000035179 | Ppp1r32 | protein phosphatase 1, regulatory subunit 32 |
| chr2 | 59350110 | 59350424 | 315 | * | 3,21493 | 6,183050544 | 2,397287 | 2,579187 | 0,009903 | 1 | 0 | Distal Intergenic | 2 | 59315013 | 59335364 | 20352 | 1 | 76747 | ENSMUST00000144415.2 | 35097 | ENSMUSG00000026989 | Dapl1 | death associated protein-like 1 |
| chr7 | 91475367 | 91475970 | 604 | * | 3,595474 | 6,183246921 | 2,374852 | 2,603634 | 0,009224 | 1 | 0 | Intron (ENSMUST0000( | 7 | 91360628 | 92096211 | 735584 | 1 | 23859 | ENSMUST00000107193.8 | 114739 | ENSMUSG00000052572 | Dlg2 | discs large MAGUK scaffold protein 2 |
| chr11 | 1,2E+08 | 1,2E+08 | 443 | * | 3,661434 | 6,183327202 | 2,360013 | 2,620039 | 0,008792 | 1 | 0 | Intron (ENSMUST0000( | 11 | 1,2E+08 | 1,2E+08 | 48183 | 1 | 74370 | ENSMUST00000139728.8 | -15674 | ENSMUSG00000025583 | Rptor | regulatory associated protein of MTOR, complex 1 |
| chr11 | 23396099 | 23396682 | 584 | * | 3,705675 | 6,183950365 | 2,287507 | 2,703357 | 0,006864 | 1 | 0 | Exon (ENSMUST00000: | 11 | 23407938 | 23410897 | 2960 | 1 | 17847 | ENSMUST00000129521.8 | -11256 | ENSMUSG00000056342 | Usp34 | ubiquitin specific peptidase 34 |
| chr9 | 44440758 | 44441532 | 775 | * | 3,692878 | 6,183955218 | 2,272164 | 2,721615 | 0,006496 | 1 | 0 | Promoter (2-3kb) | 9 | 44425502 | 44437879 | 12378 | 2 | 12145 | ENSMUST00000215661.2 | -2879 | ENSMUSG00000047880 | Cxcr5 | chemokine (C-X-C motif) receptor 5 |
| chr19 | 41401742 | 41402094 | 353 | * | 3,835344 | 6,184184168 | 2,263874 | 2,731682 | 0,006301 | 1 | 0 | Distal Intergenic | 19 | 41295791 | 41373541 | 77751 | 2 | 83490 | ENSMUST00000235978.2 | -28201 | ENSMUSG00000025017 | Pik3ap1 | phosphoinositide-3-kinase adaptor protein 1 |
| chr2 | 1,61E+08 | 1,61E+08 | 390 | * | 4,28873 | 6,184219736 | 2,259588 | 2,73688 | 0,006202 | 1 | 0 | Distal Intergenic | 2 | 1,62E+08 | 1,62E+08 | 102147 | 2 | 19281 | ENSMUST00000153770.2 | 294697 | ENSMUSG00000053141 | Ptprt | protein tyrosine phosphatase, receptor type, T |
| chr7 | 1E+08 | 1E+08 | 650 | * | 4,430002 | 6,184369481 | 2,243288 | 2,756833 | 0,005836 | 1 | 0 | Intron (ENSMUST0000( | 7 | 1,01E+08 | 1,01E+08 | 4318 | 2 | 320100 | ENSMUST00000156855.8 | 25136 | ENSMUSG00000008318 | Relt | RELT tumor necrosis factor receptor |
| chr11 | 1,21E+08 | 1,21E+08 | 481 | * | 3,918716 | 6,18486219 | 2,199073 | 2,812486 | 0,004916 | 1 | 0 | Promoter (<=1kb) | 11 | 1,21E+08 | 1,21E+08 | 1549 | 2 | 68671 | ENSMUST00000145356.2 | -319 | ENSMUSG00000025137 | Pcyt2 | phosphate cytidylyltransferase 2, ethanolamine |
| chr4 | 45250716 | 45250965 | 250 | * | 4,233365 | 6,184891544 | 2,19669 | 2,81555 | 0,004869 | 1 | 0 | Intron (ENSMUST0000( | 4 | 45234723 | 45243990 | 9268 | 2 | 78508 | ENSMUST00000134640.2 | -6726 | ENSMUSG00000086284 | Frmpd1os | FERM and PDZ domain containing 1, opposite strand |
| chr6 | 86858573 | 86858973 | 401 | * | 4,178961 | 6,185036805 | 2,183658 | 2,83242 | 0,00462 | 1 | 0 | Intron (ENSMUST0000( | 6 | 86828167 | 86912489 | 84323 | 1 | 269774 | ENSMUST00000156885.2 | 30406 | ENSMUSG00000057230 | Aak1 | AP2 associated kinase 1 |
| chr11 | 99116625 | 99117375 | 751 | * | 4,247165 | 6,18526919 | 2,164531 | 2,857556 | 0,004269 | 1 | 0 | Exon (ENSMUST00000: | 11 | 99110535 | 99121239 | 10705 | 2 | 57376 | ENSMUST00000128707.8 | 3864 | ENSMUSG00000037935 | Smarce1 | SWI/SNF related, matrix associated, actin dependent regulator of chromatin, subfamily e, member 1 |
| chr5 | 23741743 | 23742015 | 273 | * | 4,155471 | 6,185427594 | 2,152742 | 2,873279 | 0,004062 | 1 | 0 | Intron (ENSMUST0000( | 5 | 23730729 | 23747116 | 16388 | 2 | 20817 | ENSMUST00000200173.2 | 5101 | ENSMUSG00000062604 | Srpk2 | serine/arginine-rich protein specific kinase 2 |
| chr12 | 38396226 | 38396861 | 636 | * | 4,285992 | 6,185428812 | 2,151917 | 2,87438 | 0,004048 | 1 | 0 | Intron (ENSMUST0000( | 12 | 38602807 | 38664520 | 61714 | 2 | 73867 | ENSMUST00000222132.2 | 267659 | ENSMUSG00000112967 | 4930428E07Rik | RIKEN cDNA 4930428E07 gene |
| chr18 | 88751347 | 88751766 | 420 | * | 4,419914 | 6,18550257 | 2,146131 | 2,882164 | 0,00395 | 1 | 0 | Intron (ENSMUST0000( | 18 | 88683348 | 88776615 | 93268 | 2 | 1,19E+08 | ENSMUST00000155169.2 | 24849 | | LOC118568389 | uncharacterized LOC118568389 |
| chr9 | 66791048 | 66791505 | 458 | * | 4,017309 | 6,185659219 | 2,135309 | 2,896845 | 0,003769 | 1 | 0 | Exon (ENSMUST00000: | 9 | 66779871 | 66802988 | 23118 | 1 | 1,05E+08 | ENSMUST00000247274.1 | 11177 | | Gm40528 | predicted gene, 40528 |
| chr14 | 31536167 | 31537051 | 885 | * | 4,352027 | 6,185665206 | 2,135354 | 2,896787 | 0,00377 | 1 | 0 | Intron (ENSMUST0000( | 14 | 31467232 | 31552335 | 85104 | 2 | 105522 | ENSMUST00000228037.2 | 15284 | ENSMUSG00000014496 | Ankrd28 | ankyrin repeat domain 28 |
| chr9 | 99571809 | 99572351 | 543 | * | 4,111648 | 6,185696949 | 2,132209 | 2,901074 | 0,003719 | 1 | 0 | 3' UTR | 9 | 99574235 | 99592062 | 17828 | 2 | 56492 | ENSMUST00000136429.8 | 19711 | ENSMUSG00000032473 | Cldn18 | claudin 18 |
| chr4 | 15270961 | 15271528 | 568 | * | 4,50968 | 6,185814394 | 2,123145 | 2,913514 | 0,003574 | 1 | 0 | Intron (ENSMUST0000( | 4 | 15265831 | 15286753 | 20923 | 1 | 100201 | ENSMUST00000062684.9 | 5130 | ENSMUSG00000043252 | Tmem64 | transmembrane protein 64 |
| chr14 | 73166486 | 73167299 | 814 | * | 4,830833 | 6,185935963 | 2,114906 | 2,924922 | 0,003445 | 1 | 0 | Distal Intergenic | 14 | 73078081 | 73100995 | 22915 | 2 | 1E+08 | ENSMUST00000226844.2 | -65491 | ENSMUSG00000115611 | Gm4606 | predicted gene 4606 |
| chr12 | 57351089 | 57351518 | 430 | * | 5,544895 | 6,186803072 | 2,061129 | 3,001657 | 0,002685 | 1 | 0 | Promoter (1-2kb) | 12 | 57352869 | 57354462 | 1594 | 1 | 73490 | ENSMUST00000136006.2 | -1351 | ENSMUSG00000047022 | Mipol1 | mirror-image polydactyly 1 |
| chrX | 1,34E+08 | 1,34E+08 | 620 | * | 5,239839 | 6,188481033 | 1,973352 | 3,136025 | 0,001713 | 1 | 0 | Exon (ENSMUST00000: | 20 | 1,34E+08 | 1,34E+08 | 13415 | 2 | 320606 | ENSMUST00000153610.8 | 12432 | ENSMUSG00000089759 | 3632454L22Rik | RIKEN cDNA 3632454L22 gene |
| chr2 | 1,65E+08 | 1,65E+08 | 669 | * | 6,030741 | 6,188485384 | 1,972716 | 3,137039 | 0,001707 | 1 | 0 | Distal Intergenic | 2 | 1,65E+08 | 1,65E+08 | 2764 | 1 | 641368 | ENSMUST00000180193.2 | 11968 | ENSMUSG00000074593 | Spint5 | serine protease inhibitor, Kunitz type 5 |
| chr5 | 30741054 | 30741723 | 670 | * | 6,495248 | 6,19151654 | 1,863944 | 3,321728 | 0,000895 | 1 | 0 | Intron (ENSMUST0000( | 5 | 30745514 | 30782615 | 37102 | 1 | 16527 | ENSMUST00000066295.5 | -3791 | ENSMUSG00000049265 | Kcnk3 | potassium channel, subfamily K, member 3 |
| chr19 | 44345413 | 44346476 | 1064 | * | 8,131901 | 6,192028457 | 1,79396 | 3,451597 | 0,000557 | 0,384274727 | 1,425929 | Distal Intergenic | 19 | 44321531 | 44335182 | 13652 | 1 | 329065 | ENSMUST00000058856.9 | 23882 | ENSMUSG00000050195 | Scd4 | stearoyl-coenzyme A desaturase 4 |
| chr19 | 43659053 | 43659730 | 678 | * | 8,55532 | 6,192781768 | 1,849393 | 3,348549 | 0,000812 | 0,369395048 | 2,237403 | Intron (ENSMUST0000( | 19 | 43652240 | 43655582 | 3343 | 2 | 246696 | ENSMUST00000237364.2 | -3471 | ENSMUSG00000040414 | Slc25a28 | solute carrier family 25, member 28 |
| chr3 | 95919642 | 95920441 | 800 | * | 6,007363 | 6,193134449 | 1,880328 | 3,293646 | 0,000989 | 1 | 0 | Intron (ENSMUST0000( | 3 | 95895741 | 95903313 | 7573 | 2 | 67220 | ENSMUST00000015889.10 | -16329 | ENSMUSG00000015745 | Plekho1 | pleckstrin homology domain containing, family O member 1 |

| | | | | | | | | | | | | | | | | | | | | | | | |
|---|---|---|---|---|---|---|---|---|---|---|---|---|---|---|---|---|---|---|---|---|---|---|---|
| chr9 | 55113498 | 55114535 | 1038 | * | 7,738897 | 6,193206837 | 1,886122 | 3,283566 | 0,001025 | 1 | 0 | Promoter (1-2kb) | 9 | 55116209 | 55131717 | 15509 | 1 | 71999 | ENSMUST00000034859.15 | -1674 | ENSMUSG00000032309 | Fbxo22 | F-box protein 22 |
| chr16 | 12926110 | 12927114 | 1005 | * | 6,37104 | 6,193367916 | 1,903907 | 3,252978 | 0,001142 | 1 | 0 | Promoter (<=1kb) | 16 | 12927548 | 12966598 | 39051 | 1 | 50505 | ENSMUST00000023206.14 | -434 | ENSMUSG00000022545 | Ercc4 | excision repair cross-complementing rodent repair deficiency, complementation group 4 |
| chr4 | 1,09E+08 | 1,09E+08 | 611 | * | 5,992325 | 6,193419518 | 1,909724 | 3,243096 | 0,001182 | 1 | 0 | Promoter (<=1kb) | 4 | 1,09E+08 | 1,09E+08 | 7408 | 2 | 70533 | ENSMUST00000125707.2 | -400 | ENSMUSG00000028568 | Btf3l4 | basic transcription factor 3-like 4 |
| chr3 | 1,18E+08 | 1,18E+08 | 549 | * | 5,715491 | 6,193450473 | 1,912915 | 3,237704 | 0,001205 | 1 | 0 | Distal Intergenic | 3 | 1,18E+08 | 1,18E+08 | 87290 | 2 | 76561 | ENSMUST00000029639.12 | -259780 | ENSMUSG00000028007 | Snx7 | sorting nexin 7 |
| chrX | 23829262 | 23829741 | 480 | * | 3,570008 | 6,19363839 | 2,374189 | 2,608738 | 0,009088 | 1 | 0 | Distal Intergenic | 20 | 23759023 | 23773736 | 14714 | 1 | 236749 | ENSMUST00000163718.3 | 70239 | ENSMUSG00000068113 | Tesl1 | testin LIM domain protein like 1 |
| chr5 | 12982354 | 12982947 | 594 | * | 5,870378 | 6,193910921 | 1,969128 | 3,14551 | 0,001658 | 1 | 0 | Distal Intergenic | 5 | 13175381 | 13360721 | 185341 | 1 | 20346 | ENSMUST00000196514.5 | -192434 | ENSMUSG00000028883 | Sema3a | sema domain, immunoglobulin domain (Ig), short basic domain, secreted, (semaphorin) 3A |
| chrX | 92246897 | 92247495 | 599 | * | 3,564083 | 6,193969353 | 2,353781 | 2,631498 | 0,008501 | 1 | 0 | Distal Intergenic | 20 | 92226833 | 92227552 | 720 | 1 | 69357 | ENSMUST00000088133.4 | 20064 | ENSMUSG00000096153 | BC061195 | cDNA sequence BC061195 |
| chr7 | 1,07E+08 | 1,07E+08 | 425 | * | 3,522486 | 6,19418028 | 2,339519 | 2,64763 | 0,008106 | 1 | 0 | Intron (ENSMUST0000( | 7 | 1,07E+08 | 1,07E+08 | 44592 | 1 | 60510 | ENSMUST00000144003.8 | -4982 | ENSMUSG00000062542 | Syt9 | synaptotagmin IX |
| chr14 | 79422286 | 79423326 | 1041 | * | 4,986978 | 6,194283435 | 2,025213 | 3,058583 | 0,002224 | 1 | 0 | Intron (ENSMUST0000( | 14 | 79416334 | 79436630 | 20297 | 2 | 1E+08 | ENSMUST00000228380.2 | 13304 | ENSMUSG00000115111 | Gm4632 | predicted gene 4632 |
| chr13 | 96911776 | 96912399 | 624 | * | 4,808198 | 6,194310637 | 2,029769 | 3,051731 | 0,002275 | 1 | 0 | Intron (ENSMUST0000( | 13 | 96923104 | 96957929 | 34826 | 1 | 625662 | ENSMUST00000208004.2 | -10705 | ENSMUSG00000109561 | Ankrd31 | ankyrin repeat domain 31 |
| chr17 | 30324528 | 30325135 | 608 | * | 4,621528 | 6,194362805 | 2,037205 | 3,040619 | 0,002361 | 1 | 0 | Intron (ENSMUST0000( | 17 | 30355164 | 30428994 | 73831 | 1 | 21769 | ENSMUST00000234902.2 | -30029 | ENSMUSG00000044477 | Zfand3 | zinc finger, AN1-type domain 3 |
| chr3 | 1,46E+08 | 1,46E+08 | 522 | * | 5,348232 | 6,194467852 | 2,057791 | 3,010252 | 0,00261 | 1 | 0 | Distal Intergenic | 3 | 1,46E+08 | 1,46E+08 | 10095 | 1 | 12042 | ENSMUST00000029842.9 | -3859 | ENSMUSG00000028191 | Bcl10 | B cell leukemia/lymphoma 10 |
| chr1 | 1,71E+08 | 1,71E+08 | 592 | * | 4,521358 | 6,194855103 | 2,1343 | 2,902523 | 0,003702 | 1 | 0 | Promoter (1-2kb) | 1 | 1,71E+08 | 1,71E+08 | 5669 | 2 | 435653 | ENSMUST00000094337.2 | 1628 | ENSMUSG00000070524 | Fcrlb | Fc receptor-like B |
| chr19 | 42765974 | 42766404 | 431 | * | 4,353023 | 6,194912099 | 2,150687 | 2,880434 | 0,003971 | 1 | 0 | Promoter (<=1kb) | 19 | 42746919 | 42766702 | 19784 | 2 | 192236 | ENSMUST00000162061.2 | 298 | ENSMUSG00000025188 | Hps1 | HPS1, biogenesis of lysosomal organelles complex 3 subunit 1 |
| chr12 | 1E+08 | 1E+08 | 353 | * | 4,380698 | 6,194927723 | 2,155737 | 2,873694 | 0,004057 | 1 | 0 | Intron (ENSMUST0000( | 12 | 1E+08 | 1E+08 | 5340 | 1 | 19179 | ENSMUST00000223078.2 | 3137 | ENSMUSG00000021178 | Psmc1 | protease (prosome, macropain) 26S subunit, ATPase 1 |
| chr2 | 32148362 | 32148898 | 537 | * | 4,871059 | 6,194959768 | 2,164637 | 2,861893 | 0,004211 | 1 | 0 | Promoter (<=1kb) | 2 | 32147859 | 32149645 | 1787 | 2 | 22245 | ENSMUST00000142797.2 | 747 | ENSMUSG00000002550 | Uck1 | uridine-cytidine kinase 1 |
| chr7 | 29052920 | 29053191 | 272 | * | 3,769838 | 6,195192433 | 2,226056 | 2,783035 | 0,005385 | 1 | 0 | Promoter (<=1kb) | 7 | 29020759 | 29053644 | 32886 | 2 | 74206 | ENSMUST00000182009.8 | 453 | ENSMUSG00000030583 | Sipa1l3 | signal-induced proliferation-associated 1 like 3 |
| chr14 | 1,18E+08 | 1,18E+08 | 561 | * | 3,842047 | 6,195283648 | 2,252993 | 2,749802 | 0,005963 | 1 | 0 | Intron (ENSMUST0000( | 14 | 1,18E+08 | 1,18E+08 | 105 | 2 | 1,02E+08 | ENSMUST00000183564.3 | 293865 | ENSMUSG00000098648 | Mir6239 | microRNA 6239 |
| chr5 | 35348692 | 35349305 | 614 | * | 4,046092 | 6,195338616 | 2,273596 | 2,724908 | 0,006432 | 1 | 0 | Distal Intergenic | 5 | 35248845 | 35263110 | 14266 | 2 | 16976 | ENSMUST00000030986.15 | -85582 | ENSMUSG00000029103 | Lrpap1 | low density lipoprotein receptor-related protein associated protein 1 |
| chr18 | 71340330 | 71340794 | 465 | * | 3,620409 | 6,195382899 | 2,281214 | 2,715827 | 0,006611 | 1 | 0 | Distal Intergenic | 18 | 70750844 | 70758848 | 8005 | 1 | 17191 | ENSMUST00000128555.2 | 589486 | ENSMUSG00000024513 | Mbd2 | methyl-CpG binding domain protein 2 |
| chr7 | 12646751 | 12647321 | 571 | * | 4,116683 | 6,195397723 | 2,287602 | 2,70825 | 0,006764 | 1 | 0 | Distal Intergenic | 7 | 12656217 | 12660612 | 4396 | 1 | 20103 | ENSMUST00000108539.8 | -8896 | ENSMUSG00000012848 | Rps5 | ribosomal protein S5 |
| chr2 | 43976072 | 43976501 | 430 | * | 3,91398 | 6,195659173 | 2,392132 | 2,590016 | 0,009597 | 1 | 0 | Intron (ENSMUST0000( | 2 | 43949250 | 43955336 | 6087 | 2 | 320872 | ENSMUST00000139948.2 | -20736 | ENSMUSG00000085089 | Arhgap15os | Rho GTPase activating protein 15, opposite strand |
| chr1 | 33941079 | 33941806 | 728 | * | 6,841747 | 6,196275229 | 1,898158 | 3,264363 | 0,001097 | 1 | 0 | Intron (ENSMUST0000( | 1 | 33891133 | 33946849 | 55717 | 2 | 320705 | ENSMUST00000115161.8 | 5043 | ENSMUSG00000042182 | Bend6 | BEN domain containing 6 |
| chrX | 99687825 | 99688409 | 585 | * | 4,510484 | 6,196426131 | 2,219721 | 2,791534 | 0,005246 | 1 | 0 | Intron (ENSMUST0000( | 20 | 99677566 | 99679913 | 2348 | 1 | 16571 | ENSMUST00000156386.2 | 10259 | ENSMUSG00000034311 | Kif4 | kinesin family member 4 |
| chr10 | 24818324 | 24819059 | 736 | * | 4,903062 | 6,196876578 | 2,199061 | 2,817965 | 0,004833 | 1 | 0 | Distal Intergenic | 10 | 24803468 | 24804659 | 1192 | 1 | 1,05E+08 | ENSMUST00000244341.1 | 14856 | | Gm40616 | predicted gene, 40616 |
| chr16 | 32683971 | 32684531 | 561 | * | 4,18889 | 6,197214953 | 2,184605 | 2,836766 | 0,004557 | 1 | 0 | Intron (ENSMUST0000( | 16 | 32681830 | 32688708 | 6879 | 2 | 1,01E+08 | ENSMUST00000149621.2 | 4177 | ENSMUSG00000035629 | Rubcn | RUN domain and cysteine-rich domain containing, Beclin 1-interacting protein |
| chr11 | 79932075 | 79932985 | 911 | * | 6,945523 | 6,197709821 | 1,929433 | 3,212192 | 0,001317 | 0,538977324 | 1,360622 | Distal Intergenic | 11 | 79947043 | 79948725 | 1683 | 2 | 54394 | ENSMUST00000147164.2 | 15740 | ENSMUSG00000017561 | Crlf3 | cytokine receptor-like factor 3 |
| chr4 | 35387760 | 35388239 | 480 | * | 5,275308 | 6,197886006 | 2,157788 | 2,872333 | 0,004075 | 1 | 0 | Distal Intergenic | 4 | 35212423 | 35226175 | 13753 | 2 | 73205 | ENSMUST00000156472.2 | -161585 | ENSMUSG00000028300 | C9orf72 | C9orf72, member of C9orf72-SMCR8 complex |
| chr8 | 24458138 | 24458859 | 722 | * | 6,087732 | 6,198404044 | 2,13677 | 2,900828 | 0,003722 | 1 | 0 | Intron (ENSMUST0000( | 8 | 24419069 | 24505208 | 86140 | 1 | 320158 | ENSMUST00000146774.2 | 39069 | ENSMUSG00000037492 | Zmat4 | zinc finger, matrin type 4 |
| chr17 | 72642941 | 72643450 | 510 | * | 5,491869 | 6,198818327 | 2,121745 | 2,921566 | 0,003483 | 1 | 0 | Intron (ENSMUST0000( | 17 | 72175967 | 72910822 | 734856 | 2 | 11682 | ENSMUST00000086639.6 | 267372 | ENSMUSG00000055471 | Alk | anaplastic lymphoma kinase |
| chr6 | 1,1E+08 | 1,1E+08 | 316 | * | 4,817682 | 6,199327805 | 2,104011 | 2,946434 | 0,003215 | 1 | 0 | Distal Intergenic | 6 | 1,11E+08 | 1,11E+08 | 851510 | 1 | 108073 | ENSMUST00000174018.8 | -716378 | ENSMUSG00000056755 | Grm7 | glutamate receptor, metabotropic 7 |
| chrX | 31269524 | 31270006 | 483 | * | 6,054619 | 6,199353916 | 2,103478 | 2,947192 | 0,003207 | 1 | 0 | Distal Intergenic | 20 | 31276303 | 31278014 | 1712 | 2 | 1,01E+08 | ENSMUST00000238728.2 | 8008 | ENSMUSG00000096645 | Spin2e | spindlin family, member 2E |
| chr8 | 25650942 | 25651517 | 576 | * | 5,905755 | 6,200934448 | 2,015681 | 3,076348 | 0,002096 | 1 | 0 | Promoter (1-2kb) | 8 | 25647745 | 25652587 | 4843 | 2 | 320165 | ENSMUST00000209441.2 | 1070 | ENSMUSG00000065954 | Tacc1 | transforming, acidic coiled-coil containing protein 1 |
| chr13 | 1,04E+08 | 1,04E+08 | 655 | * | 3,386975 | 6,205910254 | 2,383041 | 2,604198 | 0,009209 | 1 | 0 | Promoter (1-2kb) | 13 | 1,04E+08 | 1,04E+08 | 70502 | 2 | 59079 | ENSMUST00000053927.12 | -1532 | ENSMUSG00000021709 | Erbin | Erbb2 interacting protein |
| chr13 | 1,16E+08 | 1,16E+08 | 417 | * | 3,619701 | 6,206009666 | 2,366875 | 2,622027 | 0,008741 | 1 | 0 | Distal Intergenic | 13 | 1,16E+08 | 1,16E+08 | 3957 | 2 | 16392 | ENSMUST00000176444.2 | 502247 | ENSMUSG00000042258 | Isl1 | ISL1 transcription factor, LIM/homeodomain |
| chr7 | 1,2E+08 | 1,2E+08 | 295 | * | 3,924568 | 6,20620559 | 2,328847 | 2,664927 | 0,007701 | 1 | 0 | Intron (ENSMUST0000( | 7 | 1,2E+08 | 1,2E+08 | 7854 | 1 | 233813 | ENSMUST00000165055.2 | -16843 | ENSMUSG00000030889 | Vwa3a | von Willebrand factor A domain containing 3A |
| chr14 | 97117972 | 97118579 | 608 | * | 3,870001 | 6,206308283 | 2,310144 | 2,686546 | 0,007219 | 1 | 0 | Distal Intergenic | 14 | 97280037 | 97290404 | 10368 | 2 | 1,03E+08 | ENSMUST00000228946.2 | 171825 | | Gm33525 | predicted gene, 33525 |
| chr2 | 1,63E+08 | 1,63E+08 | 472 | * | 3,672865 | 6,206329282 | 2,305982 | 2,691404 | 0,007115 | 1 | 0 | Distal Intergenic | 2 | 1,63E+08 | 1,63E+08 | 21354 | 1 | 1,01E+08 | ENSMUST00000144092.2 | -19623 | | Gm11454 | predicted gene 11454 |
| chr8 | 11433439 | 11433701 | 263 | * | 3,674453 | 6,206337221 | 2,307739 | 2,689358 | 0,007159 | 1 | 0 | Intron (ENSMUST0000( | 8 | 11454908 | 11471254 | 16347 | 1 | 12827 | ENSMUST00000145295.2 | -21207 | ENSMUSG00000031503 | Col4a2 | collagen, type IV, alpha 2 |
| chr10 | 1,27E+08 | 1,27E+08 | 458 | * | 3,601322 | 6,206686701 | 2,255092 | 2,752299 | 0,005918 | 1 | 0 | Promoter (<=1kb) | 10 | 1,27E+08 | 1,27E+08 | 938 | 2 | 216443 | ENSMUST00000128369.2 | -551 | ENSMUSG00000040354 | Mars1 | methionine-tRNA synthetase 1 |
| chr17 | 25219113 | 25219656 | 544 | * | 3,868629 | 6,206717669 | 2,246362 | 2,763008 | 0,005727 | 1 | 0 | Intron (ENSMUST0000( | 17 | 25180200 | 25234204 | 54005 | 2 | 57354 | ENSMUST00000073337.8 | 14548 | ENSMUSG00000038002 | Cramp1 | cramped chromatin regulator 1 |
| chr5 | 41923178 | 41923594 | 417 | * | 3,877509 | 6,206734577 | 2,246389 | 2,762983 | 0,005728 | 1 | 0 | Promoter (1-2kb) | 5 | 41918826 | 41921844 | 3019 | 2 | 12020 | ENSMUST00000060820.8 | -1334 | ENSMUSG00000049691 | Nkx3-2 | NK3 homeobox 2 |
| chrX | 17395858 | 17396203 | 346 | * | 3,950397 | 6,206814889 | 2,232331 | 2,780419 | 0,005429 | 1 | 0 | Distal Intergenic | 20 | 17424137 | 17437863 | 13727 | 2 | 72018 | ENSMUST00000127690.2 | 41660 | ENSMUSG00000025040 | Fundc1 | FUN14 domain containing 1 |
| chr10 | 16857796 | 16858625 | 830 | * | 4,139927 | 6,206822223 | 2,232565 | 2,78013 | 0,005434 | 1 | 0 | Intron (ENSMUST0000( | 10 | 16827041 | 16852896 | 25856 | 2 | 1,01E+08 | ENSMUST00000209129.2 | -4900 | ENSMUSG00000101969 | Gm20125 | predicted gene, 20125 |
| chr4 | 1,39E+08 | 1,39E+08 | 597 | * | 4,134279 | 6,206927528 | 2,217597 | 2,798943 | 0,005127 | 1 | 0 | Intron (ENSMUST0000( | 4 | 1,39E+08 | 1,39E+08 | 4385 | 2 | 18784 | ENSMUST00000140100.2 | 13704 | ENSMUSG00000041193 | Pla2g5 | phospholipase A2, group V |
| chr4 | 41677541 | 41677846 | 306 | * | 4,239448 | 6,206943292 | 2,215692 | 2,801356 | 0,005089 | 1 | 0 | Intron (ENSMUST0000( | 4 | 41663268 | 41664034 | 767 | 2 | 12804 | ENSMUST00000151181.2 | -13507 | ENSMUSG00000028444 | Cntfr | ciliary neurotrophic factor receptor |
| chr15 | 72153576 | 72153910 | 335 | * | 4,257001 | 6,207142722 | 2,190901 | 2,833146 | 0,004609 | 1 | 0 | Intron (ENSMUST0000( | 15 | 72127199 | 72127305 | 107 | 2 | 1,15E+08 | ENSMUST00001239540.1 | -26271 | ENSMUSG00000119376 | Gm23217 | predicted gene, 23217 |
| chr11 | 1,05E+08 | 1,05E+08 | 571 | * | 3,612624 | 6,207144601 | 2,305719 | 2,692065 | 0,007101 | 1 | 0 | Distal Intergenic | 11 | 1,05E+08 | 1,05E+08 | 431688 | 1 | 1,05E+08 | ENSMUST00000212287.2 | -3173 | | Gm11639 | predicted gene 11639 |
| chr1 | 14833415 | 14833931 | 517 | * | 4,70928 | 6,207453258 | 2,15254 | 2,883781 | 0,003929 | 1 | 0 | Intron (ENSMUST0000( | 1 | 14823570 | 14826214 | 2645 | 2 | 17681 | ENSMUST00000027062.7 | -7201 | ENSMUSG00000025930 | Msc | musculin |
| chr15 | 36876946 | 36877363 | 418 | * | 4,865627 | 6,207533527 | 2,142969 | 2,896698 | 0,003771 | 1 | 0 | Promoter (2-3kb) | 15 | 36870744 | 36880121 | 9378 | 2 | 1E+08 | ENSMUST00000100713.4 | 2758 | ENSMUSG00000072618 | Gm10384 | predicted gene 10384 |
| chr3 | 73036182 | 73036837 | 656 | * | 5,142765 | 6,207744222 | 2,121034 | 2,926754 | 0,003425 | 1 | 0 | Intron (ENSMUST0000( | 3 | 72973714 | 73205464 | 231751 | 1 | 626082 | ENSMUST00000244216.1 | 62468 | ENSMUSG00000103428 | Gm20754 | predicted gene, 20754 |
| chr5 | 24265311 | 24265796 | 486 | * | 4,462228 | 6,207812687 | 2,115577 | 2,934336 | 0,003343 | 1 | 0 | Distal Intergenic | 5 | 24165929 | 24235688 | 69760 | 2 | 84652 | ENSMUST00000030849.13 | -29623 | ENSMUSG00000028995 | Fam126a | family with sequence similarity 126, member A |
| chr7 | 1,26E+08 | 1,26E+08 | 484 | * | 4,901947 | 6,207820526 | 2,11279 | 2,938209 | 0,003301 | 1 | 0 | Promoter (<=1kb) | 7 | 1,26E+08 | 1,26E+08 | 5914 | 2 | 56312 | ENSMUST00000191413.2 | 0 | ENSMUSG00000030717 | Nupr1 | nuclear protein transcription regulator 1 |
| chr11 | 5498624 | 5499107 | 484 | * | 4,116004 | 6,207935445 | 2,102489 | 2,952661 | 0,00315 | 1 | 0 | Distal Intergenic | 11 | 5478887 | 5492187 | 13301 | 2 | 104479 | ENSMUST00000020776.5 | -6437 | ENSMUSG00000020482 | Ccdc117 | coiled-coil domain containing 117 |
| chr2 | 1,58E+08 | 1,58E+08 | 1156 | * | 11,33596 | 6,208236623 | 1,706593 | 3,637795 | 0,000275 | 0,372510154 | 0,744375 | Exon (ENSMUST00000: | 2 | 1,58E+08 | 1,58E+08 | 26438 | 1 | 329547 | ENSMUST00000065039.3 | 13382 | ENSMUSG00000052922 | Bpi | bactericidal permeablility increasing protein |
| chr9 | 84937204 | 84937902 | 699 | * | 4,625771 | 6,208237107 | 2,073991 | 2,993377 | 0,002759 | 1 | 0 | Distal Intergenic | 9 | 85202492 | 85209177 | 6686 | 2 | 212943 | ENSMUST00000034802.15 | 271275 | ENSMUSG00000032265 | Tent5a | terminal nucleotidyltransferase 5A |
| chr8 | 32783224 | 32783687 | 464 | * | 4,538415 | 6,208602644 | 2,042855 | 3,039179 | 0,002372 | 1 | 0 | Promoter (<=1kb) | 8 | 32339339 | 32782941 | 443603 | 2 | 211323 | ENSMUST00000238824.2 | -283 | ENSMUSG00000062991 | Nrg1 | neuregulin 1 |
| chr14 | 66285838 | 66286513 | 676 | * | 8,2374 | 6,208766079 | 1,882849 | 3,297538 | 0,000975 | 0,475365233 | 1,449706 | Intron (ENSMUST0000( | 14 | 66283438 | 66315146 | 31709 | 2 | 11495 | ENSMUST00000225667.2 | 28633 | ENSMUSG00000022039 | Adam2 | a disintegrin and metallopeptidase domain 2 |
| chr2 | 1,63E+08 | 1,63E+08 | 801 | * | 5,130221 | 6,209021241 | 2,011763 | 3,086358 | 0,002026 | 1 | 0 | Promoter (<=1kb) | 2 | 1,63E+08 | 1,63E+08 | 951 | 1 | 68386 | ENSMUST00000076667.2 | -172 | ENSMUSG00000058812 | 0610039K10Rik | RIKEN cDNA 0610039K10 gene |
| chr18 | 34756096 | 34756689 | 594 | * | 5,700623 | 6,20939835 | 1,983478 | 3,13056 | 0,001745 | 1 | 0 | Promoter (<=1kb) | 18 | 34732883 | 34757606 | 24724 | 2 | 78656 | ENSMUST00000152612.8 | 917 | ENSMUSG00000003778 | Brd8 | bromodomain containing 8 |
| chr18 | 77065957 | 77066707 | 751 | * | 5,438069 | 6,209612544 | 1,968749 | 3,154091 | 0,00161 | 1 | 0 | 3' UTR | 18 | 77066157 | 77094568 | 28412 | 2 | 71206 | ENSMUST00000154053.8 | 27861 | ENSMUSG00000025420 | Katnal2 | katanin p60 subunit A-like 2 |
| chr12 | 61782540 | 61783546 | 1007 | * | 7,506306 | 6,209661036 | 1,965452 | 3,159406 | 0,001581 | 1 | 0 | Intron (ENSMUST0000( | 12 | 61570734 | 61904714 | 333981 | 1 | 238205 | ENSMUST00000119481.2 | 211806 | ENSMUSG00000035653 | Lrfn5 | leucine rich repeat and fibronectin type III domain containing 5 |
| chr4 | 1,25E+08 | 1,25E+08 | 600 | * | 5,445601 | 6,209836184 | 1,954937 | 3,176489 | 0,001491 | 1 | 0 | Promoter (<=1kb) | 4 | 1,25E+08 | 1,25E+08 | 2610 | 1 | 16330 | ENSMUST00000135971.2 | 971 | ENSMUSG00000028894 | Inpp5b | inositol polyphosphate-5-phosphatase B |
| chr3 | 1,05E+08 | 1,05E+08 | 848 | * | 5,901195 | 6,210472908 | 1,918684 | 3,23684 | 0,001209 | 1 | 0 | 5' UTR | 3 | 1,05E+08 | 1,05E+08 | 19795 | 1 | 20501 | ENSMUST00000046212.2 | 10058 | ENSMUSG00000032902 | Slc16a1 | solute carrier family 16 (monocarboxylic acid transporters), member 1 |
| chr10 | 84269306 | 84269840 | 535 | * | 6,623516 | 6,210616968 | 1,911053 | 3,24984 | 0,001155 | 1 | 0 | Intron (ENSMUST0000( | 10 | 84206769 | 84276461 | 69693 | 2 | 77976 | ENSMUST00000020220.15 | 6621 | ENSMUSG00000020032 | Nuak1 | NUAK family, SNF1-like kinase, 1 |
| chr11 | 79603589 | 79604772 | 1184 | * | 8,969048 | 6,21077397 | 1,769367 | 3,510167 | 0,000448 | 0,284660899 | 2,237403 | Promoter (<=1kb) | 11 | 79602795 | 79602860 | 66 | 1 | 387188 | ENSMUST00000083461.3 | 794 | ENSMUSG00000065395 | Mir193a | microRNA 193a |
| chr18 | 55911505 | 55912515 | 1011 | * | 6,352362 | 6,210844395 | 1,900188 | 3,268542 | 0,001081 | 1 | 0 | Distal Intergenic | 18 | 56326071 | 56326205 | 135 | 1 | 1,15E+08 | ENSMUST00000157815.3 | -413556 | ENSMUSG00000088440 | Gm23555 | predicted gene, 23555 |
| chr5 | 24327742 | 24328671 | 930 | * | 8,258795 | 6,211445345 | 1,790358 | 3,469387 | 0,000522 | 0,304096805 | 2,237403 | Intron (ENSMUST0000( | 5 | 24305646 | 24341001 | 35356 | 1 | 52323 | ENSMUST00000199530.2 | 22096 | ENSMUSG00000028986 | Klhl7 | kelch-like 7 |
| chr2 | 1,74E+08 | 1,74E+08 | 854 | * | 6,613509 | 6,211814916 | 1,855794 | 3,347255 | 0,000816 | 1 | 0 | Intron (ENSMUST0000( | 2 | 1,74E+08 | 1,74E+08 | 17672 | 1 | 14683 | ENSMUST00000124935.8 | -13038 | ENSMUSG00000027523 | Gnas | GNAS (guanine nucleotide binding protein, alpha stimulating) complex locus |
| chr8 | 48887230 | 48888494 | 1265 | * | 9,270722 | 6,211853629 | 1,804903 | 3,441655 | 0,000578 | 0,40037769 | 1,360622 | Intron (ENSMUST0000( | 8 | 48788228 | 49008348 | 220121 | 2 | 23965 | ENSMUST00000110346.9 | 119854 | ENSMUSG00000031561 | Tenm3 | teneurin transmembrane protein 3 |
| chr11 | 1,06E+08 | 1,06E+08 | 923 | * | 7,154691 | 6,2136052 | 1,879204 | 3,306509 | 0,000945 | 0,479393375 | 1,360622 | Promoter (<=1kb) | 11 | 1,06E+08 | 1,06E+08 | 1229 | 2 | 1,15E+08 | ENSMUST00000140736.2 | -408 | | Taco1os | translational activator of mitochondrially encoded cytochrome c oxidase I, opposite strand |
| chr11 | 6402800 | 6403560 | 761 | * | 6,56076 | 6,214186495 | 1,910817 | 3,252109 | 0,001146 | 1 | 0 | Promoter (<=1kb) | 11 | 6403151 | 6403438 | 288 | 1 | 1,03E+08 | ENSMUST00000117715.2 | 0 | | Gm11971 | predicted gene 11971 |
| chrX | 1,35E+08 | 1,35E+08 | 884 | * | 6,474129 | 6,214504275 | 1,930834 | 3,218559 | 0,001288 | 1 | 0 | 5' UTR | 20 | 1,35E+08 | 1,35E+08 | 54 | 1 | 1,02E+08 | ENSMUST00000196502.3 | 4539 | ENSMUSG00000106264 | Mir7646 | microRNA 7646 |
| chr12 | 77549019 | 77549697 | 679 | * | 8,417962 | 6,214608401 | 1,858417 | 3,344033 | 0,000826 | 0,449809164 | 1,425929 | Distal Intergenic | 12 | 77600121 | 77616483 | 16363 | 1 | 1,03E+08 | ENSMUST00000218335.2 | -50424 | | Gm35240 | predicted gene, 35240 |
| chr19 | 14424861 | 14425733 | 873 | * | 5,497738 | 6,215048583 | 1,96592 | 3,161395 | 0,00157 | 1 | 0 | 3' UTR | 19 | 14521090 | 14572903 | 51814 | 2 | 21888 | ENSMUST00000235975.2 | 147170 | ENSMUSG00000024642 | Tle4 | transducin-like enhancer of split 4 |
| chr13 | 34415754 | 34416223 | 470 | * | 4,921203 | 6,215402992 | 1,993685 | 3,117545 | 0,001824 | 1 | 0 | Intron (ENSMUST0000( | 13 | 34366850 | 34377514 | 10665 | 2 | 73102 | ENSMUST00000128392.2 | -38240 | ENSMUSG00000038267 | Slc22a23 | solute carrier family 22, member 23 |
| chr2 | 46817572 | 46817845 | 274 | * | 4,195566 | 6,21568387 | 2,412879 | 2,576045 | 0,009994 | 1 | 0 | Distal Intergenic | 2 | 46309545 | 46332892 | 23348 | 2 | 1,03E+08 | ENSMUST00000243386.1 | -484680 | | Gm13470 | predicted gene 13470 |
| chr6 | 91643987 | 91644311 | 325 | * | 3,635071 | 6,215709138 | 2,409928 | 2,579209 | 0,009903 | 1 | 0 | Distal Intergenic | 6 | 91661034 | 91736047 | 75014 | 1 | 21366 | ENSMUST00000032185.9 | -16723 | ENSMUSG00000030096 | Slc6a6 | solute carrier family 6 (neurotransmitter transporter, taurine), member 6 |
| chr9 | 1,13E+08 | 1,13E+08 | 317 | * | 3,253765 | 6,215811758 | 2,402425 | 2,587308 | 0,009673 | 1 | 0 | Distal Intergenic | 9 | 1,13E+08 | 1,13E+08 | 13892 | 2 | 1,03E+08 | ENSMUST00000214286.2 | -3388 | | Gm36485 | predicted gene, 36485 |
| chr5 | 1,05E+08 | 1,05E+08 | 625 | * | 3,113327 | 6,215850185 | 2,39784 | 2,59227 | 0,009534 | 1 | 0 | Distal Intergenic | 5 | 1,05E+08 | 1,05E+08 | 23820 | 2 | 17472 | ENSMUST00000196204.5 | -16957 | ENSMUSG00000079363 | Gbp4 | guanylate binding protein 4 |
| chr4 | 1,37E+08 | 1,37E+08 | 468 | * | 4,847963 | 6,215916014 | 2,037334 | 3,051004 | 0,002281 | 1 | 0 | Intron (ENSMUST0000( | 4 | 1,37E+08 | 1,37E+08 | 101827 | 1 | 15530 | ENSMUST00000171332.2 | 24321 | ENSMUSG00000028763 | Hspg2 | perlecan (heparan sulfate proteoglycan 2) |
| chr5 | 1,49E+08 | 1,49E+08 | 550 | * | 4,596657 | 6,215932081 | 2,038453 | 3,049337 | 0,002293 | 1 | 0 | Intron (ENSMUST0000( | 5 | 1,49E+08 | 1,5E+08 | 83216 | 1 | 381693 | ENSMUST00000110502.7 | -13941 | ENSMUSG00000029658 | Wdr95 | WD40 repeat domain 95 |
| chrX | 1E+08 | 1E+08 | 969 | * | 3,492247 | 6,216204909 | 2,36915 | 2,623813 | 0,008695 | 1 | 0 | Exon (ENSMUST00000( | 20 | 99882254 | 1E+08 | 220992 | 2 | 83558 | ENSMUST00000009814.10 | 15505 | ENSMUSG00000009670 | Tex11 | testis expressed gene 11 |
| chrX | 1E+08 | 1E+08 | 478 | * | 3,909585 | 6,216293155 | 2,360906 | 2,633012 | 0,008463 | 1 | 0 | Promoter (1-2kb) | 20 | 1E+08 | 1E+08 | 1894 | 1 | 14618 | ENSMUST00000119190.2 | 1027 | ENSMUSG00000047797 | Gjb1 | gap junction protein, beta 1 |
| chr1 | 68792076 | 68792515 | 440 | * | 3,833077 | 6,216421273 | 2,351915 | 2,643132 | 0,008214 | 1 | 0 | Intron (ENSMUST0000( | 1 | 68071345 | 69146915 | 1075571 | 2 | 13869 | ENSMUST00000119142.8 | 354400 | ENSMUSG00000062209 | Erbb4 | erb-b2 receptor tyrosine kinase 4 |
| chr2 | 1,29E+08 | 1,29E+08 | 426 | * | 3,782124 | 6,216514791 | 2,344955 | 2,651017 | 0,008025 | 1 | 0 | Distal Intergenic | 2 | 1,29E+08 | 1,29E+08 | 151 | 2 | 1,15E+08 | ENSMUST00000083123.3 | -16957 | ENSMUSG00000065057 | Gm25703 | predicted gene, 25703 |
| chr2 | 81101982 | 81102434 | 453 | * | 3,916528 | 6,216637002 | 2,337935 | 2,659029 | 0,007837 | 1 | 0 | Distal Intergenic | 2 | 80814427 | 80814533 | 107 | 2 | 1,15E+08 | ENSMUST00000240007.1 | -287449 | ENSMUSG00000119241 | Gm23452 | predicted gene, 23452 |
| chr12 | 73969187 | 73969804 | 618 | * | 3,695606 | 6,216769826 | 2,327869 | 2,670584 | 0,007572 | 1 | 0 | Intron (ENSMUST0000( | 12 | 73973376 | 73975306 | 1931 | 1 | 15251 | ENSMUST00000149740.2 | -3572 | ENSMUSG00000021109 | Hif1a | hypoxia inducible factor 1, alpha subunit |
| chr5 | 22708815 | 22709655 | 841 | * | 4,14901 | 6,216923413 | 2,315424 | 2,685004 | 0,007253 | 1 | 0 | Intron (ENSMUST0000( | 5 | 22733319 | 22733717 | 399 | 2 | 1E+08 | ENSMUST00000118224.2 | 24062 | | Gm15421 | ribosomal protein L22 like 1 pseudogene |
| chr8 | 9211374 | 9211862 | 489 | * | 4,656563 | 6,217008768 | 2,309252 | 2,692218 | 0,007098 | 1 | 0 | Distal Intergenic | 8 | 9202600 | 9207221 | 4622 | 2 | 1,03E+08 | ENSMUST00000208794.2 | -4153 | | Gm31726 | predicted gene, 31726 |
| chr15 | 22567728 | 22568453 | 726 | * | 4,62166 | 6,217046398 | 2,160291 | 2,877875 | 0,004004 | 1 | 0 | Intron (ENSMUST0000( | 15 | 22549108 | 23400653 | 851546 | 1 | 320865 | ENSMUST00000163361.8 | 18620 | ENSMUSG00000040420 | Cdh18 | cadherin 18 |
| chr2 | 1,32E+08 | 1,32E+08 | 527 | * | 4,499381 | 6,217125039 | 2,172265 | 2,862047 | 0,004209 | 1 | 0 | Intron (ENSMUST0000( | 2 | 1,32E+08 | 1,32E+08 | 46174 | 1 | 111368 | ENSMUST00000136783.2 | 3769 | ENSMUSG00000098754 | Prn | prion protein readthrough transcript |
| chr7 | 1,26E+08 | 1,26E+08 | 379 | * | 3,928032 | 6,217127975 | 2,300895 | 2,702048 | 0,006891 | 1 | 0 | Distal Intergenic | 7 | 1,26E+08 | 1,26E+08 | 11730 | 2 | 233871 | ENSMUST00000040202.15 | -20026 | ENSMUSG00000032637 | Atxn2l | ataxin 2-like |
| chr7 | 29992030 | 29992417 | 388 | * | 3,451798 | 6,217169049 | 2,298256 | 2,705168 | 0,006827 | 1 | 0 | Promoter (<=1kb) | 7 | 29991383 | 29996359 | 4977 | 1 | 76686 | ENSMUST00000144508.2 | 647 | ENSMUSG00000013921 | Clip3 | CAP-GLY domain containing linker protein 3 |
| chr9 | 18206111 | 18206552 | 442 | * | 4,045783 | 6,217207374 | 2,182999 | 2,848012 | 0,004399 | 1 | 0 | Promoter (2-3kb) | 9 | 18203684 | 18221140 | 17457 | 1 | 66917 | ENSMUST00000217083.2 | 2427 | ENSMUSG00000001774 | Chordc1 | cysteine and histidine-rich domain (CHORD)-containing, zinc-binding protein 1 |
| chr2 | 75593766 | 75594258 | 493 | * | 3,930751 | 6,217247854 | 2,188251 | 2,841194 | 0,004494 | 1 | 0 | Distal Intergenic | 2 | 75607495 | 75609446 | 1952 | 1 | 1,03E+08 | ENSMUST00000242477.1 | -13237 | | Gm13657 | predicted gene 13657 |
| chr4 | 32856033 | 32856402 | 370 | * | 4,416237 | 6,217373925 | 2,207531 | 2,816437 | 0,004856 | 1 | 0 | Intron (ENSMUST0000( | 4 | 32805740 | 32860442 | 54703 | 2 | 140577 | ENSMUST00000108166.2 | 4040 | ENSMUSG00000040183 | Ankrd6 | ankyrin repeat domain 6 |
| chr13 | 1,17E+08 | 1,17E+08 | 414 | * | 3,744128 | 6,217612606 | 2,269647 | 2,739462 | 0,006154 | 1 | 0 | Distal Intergenic | 13 | 1,17E+08 | 1,17E+08 | 4469 | 2 | 52552 | ENSMUST00000223833.2 | -51824 | ENSMUSG00000021725 | Parp8 | poly (ADP-ribose) polymerase family, member 8 |
| chr5 | 1,24E+08 | 1,24E+08 | 472 | * | 3,584249 | 6,217634365 | 2,268173 | 2,741253 | 0,006121 | 1 | 0 | Promoter (<=1kb) | 5 | 1,24E+08 | 1,24E+08 | 2438 | 2 | 76399 | ENSMUST00000198901.2 | -311 | ENSMUSG00000029437 | Il31 | interleukin 31 |
| chr8 | 86753853 | 86754122 | 270 | * | 4,056408 | 6,217794278 | 2,274039 | 2,734251 | 0,006252 | 1 | 0 | Intron (ENSMUST0000( | 8 | 86762275 | 86787270 | 24996 | 1 | 102093 | ENSMUST00000162305.2 | -8153 | ENSMUSG00000036879 | Phkb | phosphorylase kinase beta |
| chr13 | 1,06E+08 | 1,06E+08 | 465 | * | 3,831649 | 6,218016916 | 2,24518 | 2,769497 | 0,005614 | 1 | 0 | Distal Intergenic | 13 | 1,07E+08 | 1,07E+08 | 14510 | 2 | 1,03E+08 | ENSMUST00000225713.2 | 542197 | | Gm30411 | predicted gene, 30411 |

| | | | | | | | | | | | | | | | | | | | | | | | |
|---|---|---|---|---|---|---|---|---|---|---|---|---|---|---|---|---|---|---|---|---|---|---|---|
| chr18 | 51692958 | 51693462 | 505 | * | 3,853093 | 6,218162579 | 2,343008 | 2,653923 | 0,007956 | 1 | 0 | Distal Intergenic | 18 | 51740616 | 51740740 | 125 | 2 | 1,15E+08 | ENSMUST00000104129.3 | 47278 | ENSMUSG00000077317 | Gm25785 | predicted gene, 25785 |
| chr11 | 98829456 | 98829732 | 277 | * | 3,696846 | 6,218216806 | 2,360034 | 2,634799 | 0,008419 | 1 | 0 | Promoter (<=1kb) | 11 | 98829154 | 98841187 | 12034 | 1 | 19401 | ENSMUST00000139399.2 | 302 | ENSMUSG00000037992 | Rara | retinoic acid receptor, alpha |
| chr5 | 1,2E+08 | 1,2E+08 | 470 | * | 4,358803 | 6,218263125 | 2,229853 | 2,788643 | 0,005293 | 1 | 0 | Distal Intergenic | 5 | 1,2E+08 | 1,2E+08 | 9160 | 2 | 1,03E+08 | ENSMUST00000201435.2 | 88230 | ENSMUSG00000106938 | Gm31314 | predicted gene, 31314 |
| chr15 | 50476338 | 50476711 | 374 | * | 3,169819 | 6,218290887 | 2,374207 | 2,619103 | 0,008816 | 1 | 0 | Distal Intergenic | 15 | 50682895 | 50690917 | 8023 | 2 | 83925 | ENSMUST00000183421.2 | 214206 | ENSMUSG00000038679 | Trps1 | transcriptional repressor GATA binding 1 |
| chr7 | 9199362 | 9199628 | 267 | * | 4,0427 | 6,218374443 | 2,224216 | 2,795761 | 0,005178 | 1 | 0 | Distal Intergenic | 7 | 9208545 | 9226652 | 18108 | 2 | 22305 | ENSMUST00000072787.5 | 27024 | ENSMUSG00000066828 | Vmn2r37 | vomeronasal 2, receptor 37 |
| chr10 | 89048241 | 89048627 | 387 | * | 4,723238 | 6,218392542 | 2,223204 | 2,797041 | 0,005157 | 1 | 0 | Intron (ENSMUST0000 | 10 | 88903594 | 89092998 | 189405 | 2 | 320091 | ENSMUST00000181976.8 | 44371 | ENSMUSG00000035189 | Ano4 | anoctamin 4 |
| chr4 | 1,4E+08 | 1,4E+08 | 704 | * | 4,873122 | 6,218428229 | 2,220358 | 2,800642 | 0,0051 | 1 | 0 | Intron (ENSMUST0000 | 4 | 1,4E+08 | 1,4E+08 | 1761 | 1 | 1,03E+08 | ENSMUST00000137283.2 | -73039 | | Gm13029 | predicted gene 13029 |
| chr7 | 39838472 | 39838886 | 415 | * | 3,79644 | 6,218516176 | 2,216174 | 2,805969 | 0,005017 | 1 | 0 | Distal Intergenic | 7 | 40324161 | 40332685 | 8525 | 1 | 71029 | ENSMUST00000206289.2 | -485275 | ENSMUSG00000102004 | 4933402C06Rik | RIKEN cDNA 4933402C06 gene |
| chr5 | 66454869 | 66455436 | 568 | * | 4,305981 | 6,218607005 | 2,211473 | 2,811975 | 0,004924 | 1 | 0 | 3' UTR | 5 | 66456046 | 66471071 | 15026 | 2 | 11787 | ENSMUST00000162955.8 | 15635 | ENSMUSG00000029207 | Apbb2 | amyloid beta (A4) precursor protein-binding, family B, member 2 |
| chr11 | 29920356 | 29920949 | 594 | * | 4,90757 | 6,219079363 | 2,185777 | 2,845249 | 0,004438 | 1 | 0 | Intron (ENSMUST0000 | 11 | 29693048 | 29976033 | 282986 | 2 | 237711 | ENSMUST00000058902.6 | 55084 | ENSMUSG00000044072 | Eml6 | echinoderm microtubule associated protein like 6 |
| chr19 | 48515862 | 48516286 | 425 | * | 4,298818 | 6,21913585 | 2,182836 | 2,849108 | 0,004384 | 1 | 0 | Intron (ENSMUST0000 | 19 | 48418399 | 48643749 | 225351 | 1 | 66673 | ENSMUST00000236094.2 | 97463 | ENSMUSG00000063434 | Sorcs3 | sortilin-related VPS10 domain containing receptor 3 |
| chr16 | 31633919 | 31634685 | 767 | * | 5,032999 | 6,219851952 | 2,147927 | 2,895747 | 0,003783 | 1 | 0 | Intron (ENSMUST0000 | 16 | 31648117 | 31655309 | 7193 | 1 | 13383 | ENSMUST00000137322.2 | -13432 | ENSMUSG00000022770 | Dlg1 | discs large MAGUK scaffold protein 1 |
| chr8 | 37900506 | 37900836 | 331 | * | 4,368796 | 6,220608482 | 2,114405 | 2,942013 | 0,003261 | 1 | 0 | Distal Intergenic | 8 | 37784631 | 37784732 | 102 | 2 | 1,15E+08 | ENSMUST00000083343.3 | -115774 | ENSMUSG00000065277 | Gm25126 | predicted gene, 25126 |
| chr13 | 53541800 | 53542181 | 382 | * | 4,588009 | 6,220798128 | 2,106356 | 2,953346 | 0,003143 | 1 | 0 | Distal Intergenic | 13 | 53547654 | 53594528 | 46875 | 1 | 1E+08 | ENSMUST00000246406.1 | -5473 | ENSMUSG00000107008 | Gm2762 | predicted gene 2762 |
| chr10 | 58106513 | 58107211 | 699 | * | 4,686536 | 6,220949757 | 2,100959 | 2,961005 | 0,003066 | 1 | 0 | Promoter (2-3kb) | 10 | 58109268 | 58112116 | 2849 | 1 | 70297 | ENSMUST00000160677.2 | -2057 | ENSMUSG00000038039 | Gcc2 | GRIP and coiled-coil domain containing 2 |
| chr10 | 1,28E+08 | 1,28E+08 | 685 | * | 5,010192 | 6,221751862 | 2,069146 | 3,006917 | 0,002639 | 1 | 0 | Distal Intergenic | 10 | 1,28E+08 | 1,28E+08 | 131 | 2 | 1,15E+08 | ENSMUST00000158062.3 | 10312 | ENSMUSG00000088687 | Gm23996 | predicted gene, 23996 |
| chr14 | 1,19E+08 | 1,19E+08 | 571 | * | 5,839351 | 6,223723715 | 2,003176 | 3,106928 | 0,00189 | 1 | 0 | Intron (ENSMUST0000 | 14 | 1,19E+08 | 1,19E+08 | 193121 | 2 | 239273 | ENSMUST00000226703.2 | 5066 | ENSMUSG00000032849 | Abcc4 | ATP-binding cassette, sub-family C (CFTR/MRP), member 4 |
| chr17 | 30647267 | 30647783 | 517 | * | 5,676999 | 6,224146801 | 1,990587 | 3,126789 | 0,001767 | 1 | 0 | Intron (ENSMUST0000 | 17 | 30439320 | 30633749 | 194430 | 2 | 224671 | ENSMUST00000235342.2 | -13518 | ENSMUSG00000062202 | Btbd9 | BTB (POZ) domain containing 9 |
| chr8 | 74398661 | 74400083 | 1423 | * | 5,662648 | 6,224402768 | 1,983568 | 3,137983 | 0,001701 | 1 | 0 | Distal Intergenic | 8 | 73541227 | 74080168 | 538942 | 2 | 16795 | ENSMUST00000212459.2 | -318493 | ENSMUSG00000004383 | Large1 | LARGE xylosyl- and glucuronyltransferase 1 |
| chrX | 38003337 | 38003749 | 413 | * | 4,940628 | 6,225633364 | 2,115812 | 2,942433 | 0,003256 | 1 | 0 | Intron (ENSMUST0000 | 20 | 38046961 | 38048432 | 1472 | 1 | 77883 | ENSMUST00000168144.3 | -43212 | ENSMUSG00000051361 | 6030498E09Rik | RIKEN cDNA 6030498E09 gene |
| chr14 | 1,07E+08 | 1,07E+08 | 996 | * | 6,327202 | 6,226618582 | 1,927045 | 3,231174 | 0,001233 | 1 | 0 | Distal Intergenic | 14 | 1,07E+08 | 1,07E+08 | 69334 | 2 | 73631 | ENSMUST00000186311.2 | -14815 | ENSMUSG00000099794 | 1700128A07Rik | RIKEN cDNA 1700128A07 gene |
| chr2 | 36076942 | 36077471 | 530 | * | 3,16982 | 6,228317308 | 2,404189 | 2,59061 | 0,009581 | 1 | 0 | Intron (ENSMUST0000 | 2 | 36106733 | 36113030 | 6298 | 1 | 19224 | ENSMUST00000147012.3 | -29262 | ENSMUSG00000047250 | Ptgs1 | prostaglandin-endoperoxide synthase 1 |
| chr3 | 1,13E+08 | 1,13E+08 | 379 | * | 4,187459 | 6,228519631 | 2,328896 | 2,674452 | 0,007485 | 1 | 0 | Distal Intergenic | 3 | 1,13E+08 | 1,13E+08 | 872 | 2 | 109959 | ENSMUST00000144352.2 | 34375 | ENSMUSG00000074268 | Amy2a5 | amylase 2a5 |
| chr10 | 5424755 | 5425253 | 499 | * | 3,368983 | 6,228525874 | 2,329519 | 2,673739 | 0,007501 | 1 | 0 | Intron (ENSMUST0000 | 10 | 4970917 | 5500678 | 529762 | 2 | 64009 | ENSMUST00000215295.2 | 75425 | ENSMUSG00000096054 | Syne1 | spectrin repeat containing, nuclear envelope 1 |
| chr11 | 3601703 | 3602052 | 350 | * | 3,697169 | 6,228580144 | 2,313592 | 2,692169 | 0,007099 | 1 | 0 | Promoter (1-2kb) | 11 | 3599759 | 3619410 | 19652 | 1 | 74522 | ENSMUST00000140242.8 | 1944 | ENSMUSG00000034543 | Morc2a | microrchidia 2A |
| chr5 | 1,25E+08 | 1,25E+08 | 569 | * | 3,752701 | 6,22859012 | 2,248117 | 2,770581 | 0,005596 | 1 | 0 | Promoter (1-2kb) | 5 | 1,25E+08 | 1,25E+08 | 7650 | 1 | 76167 | ENSMUST00000111453.2 | 1023 | ENSMUSG00000029402 | Snrnp35 | small nuclear ribonucleoprotein 35 (U11/U12) |
| chr10 | 4334438 | 4334767 | 330 | * | 3,509378 | 6,228775409 | 2,260381 | 2,75563 | 0,005858 | 1 | 0 | Promoter (2-3kb) | 10 | 4317075 | 4337475 | 20401 | 2 | 381990 | ENSMUST00000100077.5 | 2708 | ENSMUSG00000075327 | Zbtb2 | zinc finger and BTB domain containing 2 |
| chr10 | 40749135 | 40749763 | 629 | * | 4,327186 | 6,22878297 | 2,259152 | 2,757134 | 0,005831 | 1 | 0 | Intron (ENSMUST0000 | 10 | 40721160 | 40759088 | 37929 | 2 | 71713 | ENSMUST00000215051.2 | 9325 | ENSMUSG00000038446 | Cdc40 | cell division cycle 40 |
| chr11 | 73078988 | 73079365 | 378 | * | 4,763539 | 6,228854359 | 2,232942 | 2,789528 | 0,005278 | 1 | 0 | Promoter (<=1kb) | 11 | 73077178 | 73079632 | 2455 | 2 | 83429 | ENSMUST00000130101.2 | 267 | ENSMUSG00000005949 | Ctns | cystinosis, nephropathic |
| chrX | 15416406 | 15416994 | 589 | * | 3,591501 | 6,229011782 | 2,201378 | 2,829597 | 0,004661 | 1 | 0 | Distal Intergenic | 20 | 15551524 | 15551630 | 107 | 2 | 1,15E+08 | ENSMUST00000239881.1 | 134636 | ENSMUSG00000118898 | Gm25207 | predicted gene, 25207 |
| chrX | 1,62E+08 | 1,62E+08 | 430 | * | 4,118724 | 6,229012331 | 2,198564 | 2,833219 | 0,004608 | 1 | 0 | Distal Intergenic | 20 | 1,62E+08 | 1,62E+08 | 62215 | 1 | 55936 | ENSMUST00000112300.2 | 68836 | ENSMUSG00000031360 | Ctps2 | cytidine 5'-triphosphate synthase 2 |
| chr12 | 4560788 | 4561208 | 421 | * | 4,38433 | 6,229153933 | 2,168114 | 2,873075 | 0,004065 | 1 | 0 | Promoter (1-2kb) | 12 | 4559317 | 4572370 | 13054 | 1 | 1,05E+08 | ENSMUST00000218374.2 | 1471 | | LOC105244649 | uncharacterized LOC105244649 |
| chr18 | 10786974 | 10787868 | 895 | * | 4,83927 | 6,229239639 | 2,14844 | 2,899424 | 0,003738 | 1 | 0 | Promoter (<=1kb) | 18 | 10781718 | 10787714 | 5997 | 2 | 1,05E+08 | ENSMUST00000240973.1 | 0 | | Mir133a-1hg | Mir133a-1, Mir1b and Mir1a-2 host gene |
| chr11 | 75290686 | 75291200 | 515 | * | 3,904059 | 6,229314604 | 2,289387 | 2,720953 | 0,006509 | 1 | 0 | Promoter (<=1kb) | 11 | 75290435 | 75291459 | 1025 | 1 | 319822 | ENSMUST00000135774.2 | 251 | ENSMUSG00000018809 | Smyd4 | SET and MYND domain containing 4 |
| chr3 | 1,56E+08 | 1,56E+08 | 449 | * | 4,1586 | 6,229374517 | 2,124356 | 2,932359 | 0,003364 | 1 | 0 | Distal Intergenic | 3 | 1,56E+08 | 1,56E+08 | 32239 | 1 | 329780 | ENSMUST00000196033.2 | 46909 | ENSMUSG00000104553 | 9330178D15Rik | RIKEN cDNA 9330178D15 gene |
| chr7 | 60581187 | 60582081 | 895 | * | 7,489948 | 6,229399668 | 1,869904 | 3,331402 | 0,000864 | 0,465630924 | 1,360622 | Intron (ENSMUST0000 | 7 | 60567505 | 60572658 | 5154 | 2 | 1,01E+08 | ENSMUST00000207994.2 | -8529 | | Gm29729 | predicted gene, 29729 |
| chrX | 1,63E+08 | 1,63E+08 | 701 | * | 4,809397 | 6,229561593 | 2,094156 | 2,974737 | 0,002932 | 1 | 0 | Distal Intergenic | 20 | 1,63E+08 | 1,63E+08 | 1191 | 2 | 74264 | ENSMUST00000059320.4 | 34796 | ENSMUSG00000083695 | Rnf138rt1 | ring finger protein 138, retrogene 1 |
| chr3 | 88267457 | 88267869 | 413 | * | 4,126346 | 6,229572119 | 2,089263 | 2,981707 | 0,002866 | 1 | 0 | Promoter (<=1kb) | 3 | 88267843 | 88268398 | 556 | 1 | 229512 | ENSMUST00000194359.2 | 0 | ENSMUSG00000001415 | Smg5 | SMG5 nonsense mediated mRNA decay factor |
| chr17 | 28705370 | 28706222 | 853 | * | 5,470448 | 6,229759427 | 2,058271 | 3,026696 | 0,002472 | 1 | 0 | Promoter (<=1kb) | 17 | 28618068 | 28705123 | 87056 | 2 | 14229 | ENSMUST00000079413.11 | -247 | ENSMUSG00000024222 | Fkbp5 | FK506 binding protein 5 |
| chr9 | 43864592 | 43865186 | 595 | * | 5,040226 | 6,229857563 | 2,045739 | 3,045285 | 0,002325 | 1 | 0 | Intron (ENSMUST0000 | 9 | 43873156 | 43881468 | 8313 | 2 | 1,03E+08 | ENSMUST00000217507.2 | 16282 | | Gm30373 | predicted gene, 30373 |
| chr7 | 1,1E+08 | 1,1E+08 | 552 | * | 4,615838 | 6,229887685 | 2,04191 | 3,051009 | 0,002281 | 1 | 0 | Promoter (<=1kb) | 7 | 1,1E+08 | 1,1E+08 | 9572 | 2 | 1,01E+08 | ENSMUST00000210822.2 | -997 | ENSMUSG00000109700 | Gm21123 | predicted gene, 21123 |
| chr2 | 12578049 | 12578589 | 541 | * | 4,875106 | 6,230020688 | 2,020624 | 3,083216 | 0,002048 | 1 | 0 | Distal Intergenic | 2 | 12352074 | 12424281 | 72208 | 2 | 66960 | ENSMUST00000028105.13 | -153768 | ENSMUSG00000026767 | Mindy3 | MINDY lysine 48 deubiquitinase 3 |
| chr6 | 44506356 | 44506911 | 556 | * | 4,864505 | 6,230028319 | 2,019512 | 3,084917 | 0,002036 | 1 | 0 | Distal Intergenic | 6 | 44057422 | 44060289 | 2868 | 2 | 1,03E+08 | ENSMUST00000203777.2 | -446067 | ENSMUSG00000107598 | Gm35216 | predicted gene, 35216 |
| chrX | 73316391 | 73317451 | 1061 | * | 4,936822 | 6,230033018 | 2,020569 | 3,083306 | 0,002047 | 1 | 0 | Promoter (<=1kb) | 20 | 73317633 | 73317711 | 79 | 2 | 1,02E+08 | ENSMUST00000184075.3 | 260 | ENSMUSG00000099226 | Mir7091 | microRNA 7091 |
| chr9 | 59564988 | 59565328 | 341 | * | 4,785038 | 6,230113129 | 2,008295 | 3,102191 | 0,001921 | 1 | 0 | Promoter (<=1kb) | 9 | 59565526 | 59577881 | 12356 | 1 | 18746 | ENSMUST00000213930.2 | -198 | ENSMUSG00000032294 | Pkm | pyruvate kinase, muscle |
| chr9 | 74942564 | 74943398 | 835 | * | 5,069808 | 6,230403634 | 1,973268 | 3,157404 | 0,001592 | 1 | 0 | Promoter (1-2kb) | 9 | 74944896 | 74967576 | 22681 | 1 | 59046 | ENSMUST00000168166.8 | -1498 | ENSMUSG00000007656 | Arpp19 | cAMP-regulated phosphoprotein 19 |
| chr10 | 1,23E+08 | 1,23E+08 | 1023 | * | 6,754832 | 6,230541243 | 1,957667 | 3,182636 | 0,001459 | 1 | 0 | Intron (ENSMUST0000 | 10 | 1,23E+08 | 1,23E+08 | 4628 | 1 | 319468 | ENSMUST00000159885.2 | -6094 | ENSMUSG00000034613 | Ppm1h | protein phosphatase 1H (PP2C domain containing) |
| chr14 | 22043169 | 22044012 | 844 | * | 6,811135 | 6,230959588 | 1,914319 | 3,254922 | 0,001134 | 1 | 0 | Exon (ENSMUST00000 | 14 | 22034027 | 22039669 | 5643 | 2 | 218820 | ENSMUST00000043409.9 | -3500 | ENSMUSG00000039081 | Zfp503 | zinc finger protein 503 |
| chrX | 1,06E+08 | 1,06E+08 | 1046 | * | 7,501023 | 6,23110934 | 1,902597 | 3,275054 | 0,001056 | 1 | 0 | Promoter (<=1kb) | 20 | 1,06E+08 | 1,06E+08 | 8108 | 1 | 213439 | ENSMUST00000156709.8 | 0 | ENSMUSG00000073008 | Gpr174 | G protein-coupled receptor 174 |
| chr14 | 44479065 | 44479952 | 888 | * | 6,421754 | 6,231444759 | 1,872911 | 3,327145 | 0,000877 | 1 | 0 | Intron (ENSMUST0000 | 14 | 44463766 | 44470269 | 6504 | 2 | 1,03E+08 | ENSMUST00000227517.2 | -8796 | ENSMUSG00000115536 | Gm32857 | predicted gene, 32857 |
| chr12 | 84076104 | 84076929 | 826 | * | 6,665376 | 6,231681566 | 1,855186 | 3,359061 | 0,000782 | 1 | 0 | Distal Intergenic | 12 | 84085153 | 84095375 | 10223 | 1 | 171282 | ENSMUST00000021652.5 | -8224 | ENSMUSG00000052392 | Acot4 | acyl-CoA thioesterase 4 |
| chr3 | 1,45E+08 | 1,45E+08 | 897 | * | 6,261656 | 6,233972616 | 1,882278 | 3,31193 | 0,000927 | 1 | 0 | Intron (ENSMUST0000 | 3 | 1,45E+08 | 1,45E+08 | 23256 | 1 | 71355 | ENSMUST00000127379.2 | -38319 | ENSMUSG00000028197 | Col24a1 | collagen, type XXIV, alpha 1 |
| chr6 | 85926940 | 85927457 | 518 | * | 6,805905 | 6,234027293 | 1,884313 | 3,308382 | 0,000938 | 1 | 0 | Promoter (<=1kb) | 6 | 85926692 | 85927865 | 1174 | 2 | 72102 | ENSMUST00000137527.8 | 408 | ENSMUSG00000030002 | Dusp11 | dual specificity phosphatase 11 (RNA/RNP complex 1-interacting) |
| chr13 | 95432306 | 95433718 | 1413 | * | 10,32658 | 6,234103184 | 1,727599 | 3,608536 | 0,000308 | 0,295900293 | 1,425929 | Distal Intergenic | 13 | 95460120 | 95474349 | 14230 | 1 | 72114 | ENSMUST00000221807.2 | -26402 | ENSMUSG00000041995 | Zbed3 | zinc finger, BED type containing 3 |
| chrX | 66013619 | 66014212 | 594 | * | 4,5656 | 6,237840791 | 2,078322 | 3,001383 | 0,002688 | 1 | 0 | Exon (ENSMUST00000 | 20 | 65882658 | 65882731 | 74 | 2 | 723888 | ENSMUST00000093563.3 | -130888 | ENSMUSG00000070071 | Mir465 | microRNA 465 |
| chrX | 91105728 | 91106026 | 299 | * | 3,066526 | 6,237952529 | 2,414261 | 2,583794 | 0,009772 | 1 | 0 | Distal Intergenic | 20 | 91058797 | 91059939 | 1143 | 2 | 17145 | ENSMUST00000072269.2 | -45789 | ENSMUSG00000062162 | Mageb1 | MAGE family member B1 |
| chr10 | 77626700 | 77627095 | 396 | * | 4,516124 | 6,237980525 | 2,08835 | 2,987038 | 0,002817 | 1 | 0 | Promoter (2-3kb) | 10 | 77623324 | 77624034 | 711 | 2 | 634504 | ENSMUST00000180161.2 | -2666 | ENSMUSG00000095721 | Gm7137 | predicted gene 7137 |
| chr19 | 44216423 | 44216842 | 420 | * | 3,579763 | 6,238028833 | 2,405295 | 2,593457 | 0,009502 | 1 | 0 | Intron (ENSMUST0000 | 19 | 44226254 | 44230457 | 4204 | 1 | 30049 | ENSMUST00000237324.2 | -9412 | ENSMUSG00000025202 | Scd3 | stearoyl-coenzyme A desaturase 3 |
| chr3 | 34939196 | 34939364 | 169 | * | 4,184586 | 6,238119734 | 2,0986 | 2,972515 | 0,002954 | 1 | 0 | Distal Intergenic | 3 | 34976694 | 34976800 | 107 | 2 | 1,02E+08 | ENSMUST00000183357.3 | 37436 | ENSMUSG00000098862 | Mir6378 | microRNA 6378 |
| chr18 | 62071471 | 62071751 | 281 | * | 3,245695 | 6,238132299 | 2,396472 | 2,603048 | 0,00924 | 1 | 0 | Distal Intergenic | 18 | 62086146 | 62148786 | 62641 | 1 | 225608 | ENSMUST00000051720.6 | -14395 | ENSMUSG00000045629 | Sh3tc2 | SH3 domain and tetratricopeptide repeats 2 |
| chr14 | 81381226 | 81381602 | 377 | * | 3,225661 | 6,238186238 | 2,390079 | 2,610034 | 0,009053 | 1 | 0 | Distal Intergenic | 14 | 81923250 | 81958568 | 35319 | 1 | 1,05E+08 | ENSMUST00000227374.2 | -541648 | | Gm41219 | predicted gene, 41219 |
| chr17 | 19685718 | 19686041 | 324 | * | 3,762297 | 6,238201742 | 2,390583 | 2,609489 | 0,009068 | 1 | 0 | Intron (ENSMUST0000 | 17 | 19724994 | 19755493 | 30500 | 1 | 627537 | ENSMUST00000166081.3 | -38953 | ENSMUSG00000091859 | Vmn2r100 | vomeronasal 2, receptor 100 |
| chrX | 1,27E+08 | 1,27E+08 | 885 | * | 3,807302 | 6,238415826 | 2,367957 | 2,634514 | 0,008426 | 1 | 0 | Distal Intergenic | 20 | 1,27E+08 | 1,27E+08 | 107 | 2 | 1,15E+08 | ENSMUST00000240076.1 | -161205 | ENSMUSG00000119321 | Gm25061 | predicted gene, 25061 |
| chrX | 31411718 | 31412712 | 995 | * | 5,265019 | 6,238438191 | 2,125306 | 2,935313 | 0,003332 | 1 | 0 | Distal Intergenic | 20 | 31373674 | 31375374 | 1701 | 2 | 1,01E+08 | ENSMUST00000238772.2 | -36344 | ENSMUSG00000118431 | Btbd35f25 | BTB domain containing 35, family member 25 |
| chr3 | 87456607 | 87456920 | 314 | * | 3,371293 | 6,238673117 | 2,342416 | 2,66335 | 0,007737 | 1 | 0 | Promoter (<=1kb) | 3 | 87457406 | 87468416 | 11011 | 1 | 546801 | ENSMUST00000238894.2 | -486 | ENSMUSG00000118505 | Etv3l | ets variant 3-like |
| chr1 | 1,71E+08 | 1,71E+08 | 412 | * | 4,40874 | 6,238813176 | 2,155316 | 2,894617 | 0,003796 | 1 | 0 | 3' UTR | 1 | 1,71E+08 | 1,71E+08 | 4335 | 2 | 80915 | ENSMUST00000172042.2 | 4185 | ENSMUSG00000026659 | Dusp12 | dual specificity phosphatase 12 |
| chrX | 17743887 | 17744645 | 759 | * | 3,883399 | 6,23887962 | 2,322654 | 2,686099 | 0,007229 | 1 | 0 | Distal Intergenic | 20 | 18012096 | 18012940 | 845 | 1 | 73547 | ENSMUST00000026018.4 | -267451 | ENSMUSG00000025043 | Dusp21 | dual specificity phosphatase 21 |
| chr6 | 35975273 | 35975571 | 299 | * | 4,186746 | 6,239033095 | 2,309484 | 2,701484 | 0,006903 | 1 | 0 | Distal Intergenic | 6 | 35952265 | 35952399 | 135 | 1 | 1,15E+08 | ENSMUST00000104058.3 | 23008 | ENSMUSG00000077246 | Gm23273 | predicted gene, 23273 |
| chr13 | 51107863 | 51108329 | 467 | * | 3,683551 | 6,239146346 | 2,299707 | 2,713017 | 0,006667 | 1 | 0 | Distal Intergenic | 13 | 51254916 | 51306582 | 51667 | 1 | 20729 | ENSMUST00000095797.6 | -146587 | ENSMUSG00000021395 | Spin1 | spindlin 1 |
| chr14 | 29341681 | 29342017 | 337 | * | 3,709303 | 6,239204193 | 2,297323 | 2,715858 | 0,00661 | 1 | 0 | Intron (ENSMUST0000 | 14 | 29439424 | 29443752 | 4329 | 2 | 12294 | ENSMUST00000224128.2 | 101735 | ENSMUSG00000021991 | Cacna2d3 | calcium channel, voltage-dependent, alpha2/delta subunit 3 |
| chr12 | 81502969 | 81503628 | 660 | * | 4,063312 | 6,239303426 | 2,287447 | 2,727628 | 0,006379 | 1 | 0 | Intron (ENSMUST0000 | 12 | 81518867 | 81531603 | 12737 | 2 | 66272 | ENSMUST00000171237.8 | 27975 | ENSMUSG00000091803 | Cox16 | cytochrome c oxidase assembly protein 16 |
| chr19 | 14340423 | 14340728 | 306 | * | 3,881731 | 6,239568528 | 2,26546 | 2,754217 | 0,005883 | 1 | 0 | Distal Intergenic | 19 | 14521090 | 14572903 | 51814 | 2 | 21888 | ENSMUST00000235975.2 | 232175 | ENSMUSG00000024642 | Tle4 | transducin-like enhancer of split 4 |
| chr4 | 10597749 | 10598145 | 397 | * | 4,241686 | 6,239617989 | 2,262117 | 2,758309 | 0,00581 | 1 | 0 | Intron (ENSMUST0000 | 4 | 10479962 | 10742904 | 262943 | 2 | 73624 | ENSMUST00000145856.2 | 144759 | ENSMUSG00000086080 | 1700123O12Rik | RIKEN cDNA 1700123O12 gene |
| chr13 | 47215113 | 47215606 | 494 | * | 3,717284 | 6,239778617 | 2,256424 | 2,765339 | 0,005686 | 1 | 0 | Promoter (<=1kb) | 13 | 47216482 | 47238072 | 21591 | 1 | 218214 | ENSMUST00000143518.3 | -876 | ENSMUSG00000038080 | Kdm1b | lysine (K)-specific demethylase 1B |
| chr18 | 46658098 | 46658559 | 462 | * | 3,993666 | 6,23985447 | 2,244495 | 2,78007 | 0,005435 | 1 | 0 | Promoter (<=1kb) | 18 | 46634813 | 46659038 | 24226 | 2 | 240263 | ENSMUST00000036226.6 | 479 | ENSMUSG00000033319 | Fem1c | fem 1 homolog c |
| chrX | 1,04E+08 | 1,04E+08 | 366 | * | 3,961089 | 6,239861319 | 2,265612 | 2,754162 | 0,005884 | 1 | 0 | Distal Intergenic | 20 | 1,04E+08 | 1,04E+08 | 88 | 2 | 723861 | ENSMUST00000083565.3 | 80550 | ENSMUSG00000065499 | Mir384 | microRNA 384 |
| chr3 | 1,04E+08 | 1,04E+08 | 379 | * | 3,645439 | 6,239876488 | 2,266693 | 2,752855 | 0,005908 | 1 | 0 | Intron (ENSMUST0000 | 3 | 1,04E+08 | 1,04E+08 | 52494 | 1 | 229675 | ENSMUST00000051139.13 | -4918 | ENSMUSG00000044098 | Rsbn1 | rosbin, round spermatid basic protein 1 |
| chr12 | 47263722 | 47264256 | 535 | * | 4,257713 | 6,239949925 | 2,237898 | 2,788309 | 0,005298 | 1 | 0 | Distal Intergenic | 12 | 46741300 | 46865712 | 124413 | 2 | 664883 | ENSMUST00000021438.8 | -398010 | ENSMUSG00000021047 | Nova1 | NOVA alternative splicing regulator 1 |
| chr3 | 51356136 | 51356679 | 544 | * | 3,515157 | 6,239960844 | 2,277585 | 2,739727 | 0,006149 | 1 | 0 | Intron (ENSMUST0000 | 3 | 51349327 | 51355980 | 6654 | 1 | 74838 | ENSMUST00000192197.2 | 6809 | ENSMUSG00000063273 | Naa15 | N(alpha)-acetyltransferase 15, NatA auxiliary subunit |
| chr12 | 1,15E+08 | 1,15E+08 | 255 | * | 3,792257 | 6,239970091 | 2,276657 | 2,740848 | 0,006128 | 1 | 0 | Distal Intergenic | 12 | 1,15E+08 | 1,15E+08 | 25225 | 1 | 1,03E+08 | ENSMUST00000195683.2 | -236259 | ENSMUSG00000103168 | Gm30948 | predicted gene, 30948 |
| chr14 | 10069932 | 10070557 | 626 | * | 3,850213 | 6,239997149 | 2,283036 | 2,733201 | 0,006272 | 1 | 0 | Intron (ENSMUST0000 | 14 | 10063475 | 10096615 | 33141 | 1 | 27062 | ENSMUST00000225807.2 | 6457 | ENSMUSG00000054423 | Cadps | Ca2+-dependent secretion activator |
| chr17 | 24683990 | 24684465 | 476 | * | 3,772586 | 6,24020024 | 2,220047 | 2,810842 | 0,004941 | 1 | 0 | Exon (ENSMUST00000 | 17 | 24684432 | 24689165 | 4734 | 2 | 74844 | ENSMUST00000226743.2 | 4700 | | 4833447I15Rik | RIKEN cDNA 4833447I15 gene |
| chr11 | 1,06E+08 | 1,06E+08 | 394 | * | 3,46536 | 6,240283358 | 2,322087 | 2,68736 | 0,007202 | 1 | 0 | Intron (ENSMUST0000 | 11 | 1,06E+08 | 1,06E+08 | 133 | 1 | 1,15E+08 | ENSMUST00000116890.4 | 6520 | ENSMUSG00000080540 | Gm22711 | predicted gene, 22711 |
| chr6 | 86823328 | 86823616 | 289 | * | 4,247835 | 6,24071918 | 2,185113 | 2,856016 | 0,00429 | 1 | 0 | Promoter (2-3kb) | 6 | 86825381 | 86826422 | 1042 | 2 | 67170 | ENSMUST00000185414.2 | 2806 | ENSMUSG00000100164 | 2610306M01Rik | RIKEN cDNA 2610306M01 gene |
| chr2 | 76632829 | 76633293 | 465 | * | 3,32484 | 6,24093798 | 2,421836 | 2,576945 | 0,009968 | 1 | 0 | Exon (ENSMUST00000 | 2 | 76651475 | 76656886 | 5412 | 2 | 22138 | ENSMUST00000142251.8 | 23593 | ENSMUSG00000051747 | Ttn | titin |
| chr1 | 62584767 | 62585318 | 552 | * | 4,805715 | 6,241312461 | 2,14848 | 2,904989 | 0,003673 | 1 | 0 | Intron (ENSMUST0000 | 1 | 62742444 | 62779091 | 36648 | 1 | 18187 | ENSMUST00000126344.2 | -157126 | ENSMUSG00000025969 | Nrp2 | neuropilin 2 |
| chrX | 55537739 | 55538657 | 919 | * | 5,408433 | 6,241431183 | 2,142593 | 2,913027 | 0,003579 | 1 | 0 | Exon (ENSMUST00000 | 20 | 55529178 | 55553198 | 24021 | 1 | 236790 | ENSMUST00000143142.8 | 8561 | ENSMUSG00000035967 | Ints6l | integrator complex subunit 6 like |
| chr12 | 10047473 | 10048249 | 777 | * | 4,474774 | 6,242190175 | 2,102111 | 2,969487 | 0,002983 | 1 | 0 | Distal Intergenic | 12 | 9915347 | 9915450 | 104 | 2 | 1,15E+08 | ENSMUST00000157712.3 | -132023 | ENSMUSG00000088337 | Gm22845 | predicted gene, 22845 |
| chr13 | 91995811 | 91996286 | 476 | * | 5,135834 | 6,242452746 | 2,088195 | 2,989401 | 0,002795 | 1 | 0 | Distal Intergenic | 13 | 92001506 | 92025004 | 23499 | 2 | 76722 | ENSMUST00000022122.4 | 28718 | ENSMUSG00000021622 | Ckmt2 | creatine kinase, mitochondrial 2 |
| chr17 | 77184410 | 77184837 | 428 | * | 5,239778 | 6,242506552 | 2,085755 | 2,992924 | 0,002763 | 1 | 0 | Distal Intergenic | 17 | 76381541 | 76381646 | 106 | 1 | 1,15E+08 | ENSMUST00000104111.3 | 802869 | ENSMUSG00000077299 | Gm24126 | predicted gene, 24126 |
| chr2 | 22989863 | 22990308 | 446 | * | 4,817432 | 6,242682374 | 2,078402 | 3,003597 | 0,002668 | 1 | 0 | Promoter (<=1kb) | 2 | 22989002 | 23003376 | 14375 | 1 | 74159 | ENSMUST00000227612.2 | 861 | ENSMUSG00000026781 | Acbd5 | acyl-Coenzyme A binding domain containing 5 |
| chr8 | 85952818 | 85953401 | 584 | * | 5,083399 | 6,242783269 | 2,072406 | 3,012335 | 0,002592 | 1 | 0 | Intron (ENSMUST0000 | 8 | 85957126 | 85958064 | 939 | 1 | 258858 | ENSMUST00000070849.6 | -3725 | ENSMUSG00000051952 | Olfr371 | olfactory receptor 371 |
| chr3 | 97205402 | 97206023 | 622 | * | 6,167673 | 6,242984996 | 2,063259 | 3,025788 | 0,00248 | 1 | 0 | Promoter (<=1kb) | 3 | 97123869 | 97205233 | 81365 | 2 | 77578 | ENSMUST00000196586.2 | -169 | ENSMUSG00000038256 | Bcl9 | B cell CLL/lymphoma 9 |
| chr16 | 37689197 | 37689861 | 665 | * | 5,216109 | 6,243144858 | 2,056266 | 3,036157 | 0,002396 | 1 | 0 | Promoter (<=1kb) | 16 | 37688751 | 37709220 | 20470 | 1 | 320184 | ENSMUST00000078717.7 | 446 | ENSMUSG00000034158 | Lrrc58 | leucine rich repeat containing 58 |
| chr8 | 73057961 | 73058615 | 655 | * | 5,397528 | 6,244353947 | 2,007004 | 3,111281 | 0,001863 | 1 | 0 | 3' UTR | 8 | 73072877 | 73075500 | 2624 | 1 | 16598 | ENSMUST00000067912.8 | -14262 | ENSMUSG00000055148 | Klf2 | Kruppel-like factor 2 (lung) |
| chr19 | 5316873 | 5317695 | 823 | * | 6,280738 | 6,244454322 | 2,002738 | 3,117959 | 0,001821 | 1 | 0 | Intron (ENSMUST0000 | 19 | 5210428 | 5322735 | 112308 | 2 | 107975 | ENSMUST00000236470.2 | 5040 | ENSMUSG00000024855 | Pacs1 | phosphofurin acidic cluster sorting protein 1 |
| chr18 | 18870406 | 18871242 | 837 | * | 6,223875 | 6,244862495 | 1,988033 | 3,141226 | 0,001682 | 1 | 0 | Distal Intergenic | 18 | 19266274 | 19334773 | 68500 | 1 | 1,03E+08 | ENSMUST00000234128.2 | -395032 | | Gm31086 | predicted gene, 31086 |
| chr5 | 1,14E+08 | 1,14E+08 | 642 | * | 6,207379 | 6,245126642 | 1,978695 | 3,156184 | 0,001598 | 1 | 0 | Intron (ENSMUST0000 | 5 | 1,14E+08 | 1,14E+08 | 21053 | 2 | 381707 | ENSMUST00000047891.6 | 19490 | ENSMUSG00000042184 | 1700069L16Rik | RIKEN cDNA 1700069L16 gene |
| chrX | 1,03E+08 | 1,03E+08 | 866 | * | 6,984021 | 6,245662257 | 1,961644 | 3,183891 | 0,001453 | 1 | 0 | Distal Intergenic | 20 | 1,03E+08 | 1,03E+08 | 124570 | 2 | 20502 | ENSMUST00000042664.10 | -22480 | ENSMUSG00000033965 | Slc16a2 | solute carrier family 16 (monocarboxylic acid transporters), member 2 |
| chr13 | 1,17E+08 | 1,17E+08 | 549 | * | 6,188794 | 6,2457327 | 1,959287 | 3,187759 | 0,001434 | 1 | 0 | Intron (ENSMUST0000 | 13 | 1,17E+08 | 1,17E+08 | 36084 | 2 | 74680 | ENSMUST00000224552.2 | 239627 | ENSMUSG00000114290 | 4930435F18Rik | RIKEN cDNA 4930435F18 gene |
| chr1 | 92483593 | 92483920 | 328 | * | 4,352065 | 6,24909231 | 2,321603 | 2,691715 | 0,007109 | 1 | 0 | Distal Intergenic | 1 | 92500847 | 92501928 | 1082 | 1 | 259039 | ENSMUST00000074859.3 | -16927 | ENSMUSG00000058904 | Olfr1413 | olfactory receptor 1413 |

| | | | | | | | | | | | | | | | | | | | | | | | |
|---|---|---|---|---|---|---|---|---|---|---|---|---|---|---|---|---|---|---|---|---|---|---|---|
| chr1 | 43972351 | 43973586 | 1236 | * | 18,2088 | 6,249877319 | 1,2593234 | 4,9628853 | 6,945E-07 | 0,003162373 | 2,5070128 | Promoter (<=1kb) | 1 | 43972807 | 43988565 | 15759 | 1 | 22019 | ENSMUST00000185579.7 | 0 | ENSMUSG00000041763 | Tpp2 | tripeptidyl peptidase II |
| chr11 | 61222394 | 61222941 | 548 | * | 3,430302 | 6,250535423 | 2,369821 | 2,637556 | 0,008351 | 1 | 0 | Promoter (2-3kb) | 11 | 61225587 | 61225693 | 107 | 1 | 1,15E+08 | ENSMUST00000240018.1 | -2646 | ENSMUSG00000119377 | Gm25929 | predicted gene, 25929 |
| chr14 | 1,06E+08 | 1,06E+08 | 377 | * | 3,576959 | 6,250551422 | 2,345784 | 2,66459 | 0,007708 | 1 | 0 | Distal Intergenic | 14 | 1,06E+08 | 1,06E+08 | 14324 | 1 | 76273 | ENSMUST00000149358.2 | 31842 | ENSMUSG00000053253 | Ndfip2 | Nedd4 family interacting protein 2 |
| chr7 | 31451608 | 31452018 | 411 | * | 3,177706 | 6,250570942 | 2,317067 | 2,697622 | 0,006984 | 1 | 0 | Distal Intergenic | 7 | 31466377 | 31466839 | 463 | 1 | 1,03E+08 | ENSMUST00000187984.2 | -14359 | ENSMUSG00000100517 | Gm29627 | predicted gene 29627 |
| chr19 | 17017355 | 17017738 | 384 | * | 3,864831 | 6,250600923 | 2,292911 | 2,726055 | 0,00641 | 1 | 0 | Intron (ENSMUST0000( | 19 | 16933482 | 17201296 | 267815 | 1 | 353211 | ENSMUST00000087689.5 | 83873 | ENSMUSG00000039126 | Prune2 | prune homolog 2 |
| chr15 | 46707969 | 46708425 | 457 | * | 4,827558 | 6,250627215 | 2,245284 | 2,783892 | 0,005371 | 1 | 0 | Distal Intergenic | 15 | 46807783 | 46807844 | 62 | 2 | 1,15E+08 | ENSMUST00000158054.3 | 99419 | ENSMUSG00000088679 | Gm23665 | predicted gene, 23665 |
| chr11 | 1,02E+08 | 1,02E+08 | 646 | * | 3,960148 | 6,250647093 | 2,241403 | 2,788721 | 0,005292 | 1 | 0 | Promoter (<=1kb) | 11 | 1,02E+08 | 1,02E+08 | 1398 | 1 | 217207 | ENSMUST00000141461.2 | 0 | ENSMUSG00000034931 | Dhx8 | DEAH (Asp-Glu-Ala-His) box polypeptide 8 |
| chr8 | 25636880 | 25637274 | 395 | * | 3,94257 | 6,250654854 | 2,232852 | 2,799404 | 0,00512 | 1 | 0 | Distal Intergenic | 8 | 25649141 | 25649249 | 109 | 2 | 1,02E+08 | ENSMUST00000184998.3 | 11975 | ENSMUSG00000099142 | Mir8108 | microRNA 8108 |
| chr17 | 6156452 | 6156862 | 411 | * | 3,941774 | 6,250703136 | 2,18074 | 2,866322 | 0,004153 | 1 | 0 | Promoter (<=1kb) | 17 | 6156712 | 6189067 | 32356 | 1 | 68842 | ENSMUST00000149838.8 | 0 | ENSMUSG00000034377 | Tulp4 | tubby like protein 4 |
| chr13 | 38410854 | 38411754 | 901 | * | 4,896027 | 6,250710703 | 2,184326 | 2,86162 | 0,004215 | 1 | 0 | 3' UTR | 13 | 38405095 | 38410956 | 5862 | 1 | 67797 | ENSMUST00000223893.2 | 5759 | ENSMUSG00000021431 | Snrnp48 | small nuclear ribonucleoprotein 48 (U11/U12) |
| chr13 | 95511949 | 95512296 | 348 | * | 3,872637 | 6,25073186 | 2,159601 | 2,894392 | 0,003799 | 1 | 0 | Promoter (<=1kb) | 13 | 95487191 | 95511860 | 24670 | 2 | 66549 | ENSMUST00000022189.9 | -89 | ENSMUSG00000021681 | Aggf1 | angiogenic factor with G patch and FHA domains 1 |
| chr10 | 72625065 | 72625445 | 381 | * | 4,234831 | 6,250826855 | 2,100072 | 2,976482 | 0,002916 | 1 | 0 | Distal Intergenic | 10 | 72492076 | 72492927 | 852 | 1 | 52696 | ENSMUST00000219352.2 | 132989 | ENSMUSG00000019923 | Zwint | ZW10 interactor |
| chr1 | 1,56E+08 | 1,56E+08 | 455 | * | 4,576158 | 6,250833329 | 2,093347 | 2,986047 | 0,002826 | 1 | 0 | Intron (ENSMUST0000( | 1 | 1,56E+08 | 1,56E+08 | 10531 | 2 | 77352 | ENSMUST00000212747.2 | -16892 | ENSMUSG00000026601 | Axdnd1 | axonemal dynein light chain domain containing 1 |
| chr1 | 1,57E+08 | 1,57E+08 | 663 | * | 5,205955 | 6,25086455 | 2,062488 | 3,030741 | 0,00244 | 1 | 0 | Intron (ENSMUST0000( | 1 | 1,57E+08 | 1,57E+08 | 126546 | 2 | 78255 | ENSMUST00000185198.7 | -61114 | ENSMUSG00000026594 | Ralgps2 | Ral GEF with PH domain and SH3 binding motif 2 |
| chr14 | 27477584 | 27478332 | 749 | * | 4,859068 | 6,250966947 | 2,010263 | 3,109527 | 0,001874 | 1 | 0 | Intron (ENSMUST0000( | 14 | 27446593 | 27733724 | 287132 | 1 | 238988 | ENSMUST00000211627.2 | 30991 | ENSMUSG00000040640 | Erc2 | ELKS/RAB6-interacting/CAST family member 2 |
| chr1 | 90935781 | 90936192 | 412 | * | 5,221238 | 6,251044872 | 1,969435 | 3,17403 | 0,001503 | 1 | 0 | Intron (ENSMUST0000( | 1 | 90926501 | 91056662 | 130162 | 1 | 16978 | ENSMUST00000068116.13 | 9280 | ENSMUSG00000026305 | Lrrfip1 | leucine rich repeat (in FLII) interacting protein 1 |
| chr7 | 1,11E+08 | 1,11E+08 | 759 | * | 5,865566 | 6,251060538 | 1,969298 | 3,174258 | 0,001502 | 1 | 0 | Intron (ENSMUST0000( | 7 | 1,11E+08 | 1,11E+08 | 1357 | 1 | 22083 | ENSMUST00000140220.2 | 3436 | ENSMUSG00000005609 | Ctr9 | CTR9 homolog, Paf1/RNA polymerase II complex component |
| chrX | 42637571 | 42638480 | 910 | * | 6,83551 | 6,251077669 | 1,959733 | 3,18976 | 0,001424 | 1 | 0 | Distal Intergenic | 20 | 42680037 | 42682409 | 2373 | 1 | 637093 | ENSMUST00000105113.4 | -41557 | ENSMUSG00000078320 | Tex13c1 | TEX13 family member C1 |
| chr11 | 1,06E+08 | 1,06E+08 | 1090 | * | 5,512711 | 6,251089968 | 1,949857 | 3,205921 | 0,001346 | 1 | 0 | Intron (ENSMUST0000( | 11 | 1,06E+08 | 1,06E+08 | 1290 | 1 | 77097 | ENSMUST00000167695.2 | 75786 | ENSMUSG00000053580 | Tanc2 | tetratricopeptide repeat, ankyrin repeat and coiled-coil containing 2 |
| chr9 | 52113047 | 52113744 | 698 | * | 6,078295 | 6,251226048 | 1,898944 | 3,291948 | 0,000995 | 1 | 0 | Distal Intergenic | 9 | 52022644 | 52079872 | 57229 | 2 | 244871 | ENSMUST00000213645.2 | -33175 | ENSMUSG00000035164 | Zc3h12c | zinc finger CCCH type containing 12C |
| chr12 | 84904336 | 84905252 | 917 | * | 6,58046 | 6,251250442 | 1,888706 | 3,309806 | 0,000934 | 1 | 0 | Intron (ENSMUST0000( | 12 | 84922543 | 84924906 | 2364 | 1 | 1E+08 | ENSMUST00000171853.2 | -17291 | ENSMUSG00000092130 | D030025P21Rik | RIKEN cDNA D030025P21 gene |
| chr4 | 1,33E+08 | 1,33E+08 | 864 | * | 6,999857 | 6,251437058 | 1,835424 | 3,405992 | 0,000659 | 1 | 0 | Promoter (<=1kb) | 4 | 1,33E+08 | 1,33E+08 | 3307 | 1 | 242691 | ENSMUST00000154648.2 | -107 | ENSMUSG00000028850 | Gpatch3 | G patch domain containing 3 |
| chr1 | 1,34E+08 | 1,34E+08 | 1381 | * | 7,358865 | 6,251437427 | 1,837693 | 3,401786 | 0,000669 | 1 | 0 | Promoter (<=1kb) | 1 | 1,34E+08 | 1,34E+08 | 6717 | 2 | 70579 | ENSMUST00000192775.2 | -21 | ENSMUSG00000102976 | Zc3h11a | zinc finger CCCH type containing 11A |
| chr14 | 49195063 | 49195914 | 852 | * | 7,525535 | 6,251552114 | 1,805574 | 3,462364 | 0,000535 | 1 | 0 | Distal Intergenic | 14 | 49239174 | 49239293 | 120 | 2 | 1,15E+08 | ENSMUST00000157129.4 | 43379 | | Gm23470 | predicted gene, 23470 |
| chr5 | 75527199 | 75527869 | 671 | * | 5,713763 | 6,252200117 | 2,183467 | 2,863428 | 0,004191 | 1 | 0 | Distal Intergenic | 5 | 75337492 | 75337565 | 74 | 1 | 1,02E+08 | ENSMUST00000183797.3 | 189707 | ENSMUSG00000099060 | Mir7025 | microRNA 7025 |
| chr8 | 1,06E+08 | 1,06E+08 | 647 | * | 9,70106 | 6,252240269 | 1,836444 | 3,404536 | 0,000663 | 0,410415129 | 1,449706 | Promoter (<=1kb) | 8 | 1,06E+08 | 1,06E+08 | 6861 | 1 | 102075 | ENSMUST00000160191.9 | 0 | ENSMUSG00000014782 | Plekhg4 | pleckstrin homology domain containing, family G (with RhoGef domain) member 4 |
| chr11 | 23721266 | 23721953 | 688 | * | 9,101603 | 6,252282906 | 1,823679 | 3,42839 | 0,000607 | 0,40037769 | 1,425929 | Promoter (<=1kb) | 11 | 23686847 | 23720970 | 34124 | 2 | 19696 | ENSMUST00000102864.5 | -296 | ENSMUSG00000020275 | Rel | reticuloendotheliosis oncogene |
| chr9 | 62568003 | 62568646 | 644 | * | 5,715289 | 6,252331946 | 2,178167 | 2,870456 | 0,004099 | 1 | 0 | Exon (ENSMUST000002 | 9 | 62568156 | 62573363 | 5208 | 2 | 1,19E+08 | ENSMUST00000241674.1 | 4717 | | Gm53533 | predicted gene, 53533 |
| chr12 | 84456444 | 84457368 | 925 | * | 6,862206 | 6,254635721 | 1,897813 | 3,295708 | 0,000982 | 1 | 0 | Promoter (<=1kb) | 12 | 84455845 | 84480554 | 24710 | 1 | 72873 | ENSMUST00000081828.13 | 599 | ENSMUSG00000057265 | Bbof1 | basal body orientation factor 1 |
| chr12 | 76812779 | 76813665 | 887 | * | 6,550241 | 6,25637739 | 1,955251 | 3,199783 | 0,001375 | 1 | 0 | Promoter (<=1kb) | 12 | 76812372 | 76967005 | 154634 | 1 | 110606 | ENSMUST00000137826.8 | 407 | ENSMUSG00000033373 | Fntb | farnesyltransferase, CAAX box, beta |
| chr19 | 3546901 | 3547592 | 692 | * | 5,064102 | 6,257201733 | 1,987241 | 3,148688 | 0,00164 | 1 | 0 | Promoter (<=1kb) | 19 | 3533528 | 3546530 | 13003 | 2 | 52036 | ENSMUST00000225705.2 | -371 | ENSMUSG00000024908 | Ppp6r3 | protein phosphatase 6, regulatory subunit 3 |
| chrX | 6486447 | 6487487 | 1041 | * | 7,309498 | 6,258039203 | 2,011882 | 3,11054 | 0,001867 | 1 | 0 | Promoter (1-2kb) | 20 | 6489340 | 6549502 | 60163 | 1 | 208431 | ENSMUST00000089520.3 | -1853 | ENSMUSG00000068270 | Shroom4 | shroom family member 4 |
| chr11 | 5017773 | 5018628 | 856 | * | 5,036378 | 6,258246681 | 2,033918 | 3,076942 | 0,002091 | 1 | 0 | Promoter (2-3kb) | 11 | 5019974 | 5020891 | 918 | 2 | 14030 | ENSMUST00000130118.2 | 2263 | ENSMUSG00000009079 | Ewsr1 | Ewing sarcoma breakpoint region 1 |
| chr10 | 8239251 | 8240044 | 794 | * | 4,523234 | 6,259466255 | 2,09904 | 2,982061 | 0,002863 | 1 | 0 | Intron (ENSMUST0000( | 10 | 8080520 | 8178006 | 97487 | 2 | 338362 | ENSMUST00000219116.2 | -61245 | ENSMUSG00000047712 | Ust | uronyl-2-sulfotransferase |
| chr5 | 30031337 | 30031973 | 637 | * | 4,818089 | 6,259664078 | 2,112129 | 2,963676 | 0,00304 | 1 | 0 | Distal Intergenic | 5 | 29970188 | 29971685 | 1498 | 1 | 23950 | ENSMUST00000139126.2 | 61149 | ENSMUSG00000029131 | Dnajb6 | DnaJ heat shock protein family (Hsp40) member B6 |
| chr2 | 35079482 | 35079935 | 454 | * | 4,38048 | 6,259817078 | 2,120507 | 2,952037 | 0,003157 | 1 | 0 | Promoter (<=1kb) | 2 | 35072413 | 35079972 | 7560 | 2 | 68365 | ENSMUST00000230657.2 | 37 | ENSMUSG00000026878 | Rab14 | RAB14, member RAS oncogene family |
| chr18 | 13071479 | 13072027 | 549 | * | 4,479561 | 6,259867958 | 2,124852 | 2,946026 | 0,003219 | 1 | 0 | Promoter (2-3kb) | 18 | 13031174 | 13074834 | 43661 | 2 | 64291 | ENSMUST00000122175.8 | 2807 | ENSMUSG00000044252 | Osbpl1a | oxysterol binding protein-like 1A |
| chr12 | 1,11E+08 | 1,11E+08 | 556 | * | 4,936168 | 6,259928019 | 2,128028 | 2,941657 | 0,003265 | 1 | 0 | Promoter (2-3kb) | 12 | 1,11E+08 | 1,11E+08 | 2884 | 2 | 217866 | ENSMUST00000220619.2 | 2168 | ENSMUSG00000021279 | Cdc42bpb | CDC42 binding protein kinase beta |
| chr7 | 49507818 | 49508317 | 500 | * | 4,677935 | 6,260199894 | 2,14624 | 2,916822 | 0,003536 | 1 | 0 | 3' UTR | 7 | 49559894 | 49613604 | 53711 | 1 | 104245 | ENSMUST00000056442.12 | -51577 | ENSMUSG00000039728 | Slc6a5 | solute carrier family 6 (neurotransmitter transporter, glycine), member 5 |
| chr1 | 1,06E+08 | 1,06E+08 | 510 | * | 3,767293 | 6,260211561 | 2,382853 | 2,627192 | 0,008609 | 1 | 0 | Intron (ENSMUST0000( | 1 | 1,06E+08 | 1,06E+08 | 28327 | 1 | 227449 | ENSMUST00000186983.7 | -5047 | ENSMUSG00000038866 | Zcchc2 | zinc finger, CCHC domain containing 2 |
| chr5 | 39771125 | 39771861 | 737 | * | 4,086156 | 6,260888251 | 2,300711 | 2,721284 | 0,006503 | 1 | 0 | 3' UTR | 5 | 39619908 | 39760917 | 141010 | 2 | 75095 | ENSMUST00000202742.4 | -10208 | ENSMUSG00000106741 | 4930513D17Rik | RIKEN cDNA 4930513D17 gene |
| chr14 | 66110713 | 66111442 | 730 | * | 3,546933 | 6,260902008 | 2,297432 | 2,725174 | 0,006427 | 1 | 0 | Distal Intergenic | 14 | 66074809 | 66078178 | 3370 | 1 | 67179 | ENSMUST00000226027.2 | 35904 | ENSMUSG00000022035 | Ccdc25 | coiled-coil domain containing 25 |
| chr14 | 79611609 | 79612336 | 728 | * | 3,894448 | 6,260920584 | 2,198179 | 2,848231 | 0,004396 | 1 | 0 | 3' UTR | 14 | 79578417 | 79624546 | 46130 | 2 | 66897 | ENSMUST00000163486.2 | 12210 | ENSMUSG00000022020 | Naa16 | N(alpha)-acetyltransferase 16, NatA auxiliary subunit |
| chr10 | 86621834 | 86622331 | 498 | * | 4,281237 | 6,261057482 | 2,279846 | 2,746264 | 0,006028 | 1 | 0 | Intron (ENSMUST0000( | 10 | 86614869 | 86674253 | 59385 | 1 | 103466 | ENSMUST00000099396.3 | 6965 | ENSMUSG00000054027 | Nt5dc3 | 5'-nucleotidase domain containing 3 |
| chr12 | 51876723 | 51876973 | 251 | * | 3,614634 | 6,261131746 | 2,274394 | 2,752879 | 0,005907 | 1 | 0 | Promoter (<=1kb) | 12 | 51790505 | 51876319 | 85815 | 2 | 207304 | ENSMUST00000179265.8 | -404 | ENSMUSG00000035247 | Hectd1 | HECT domain E3 ubiquitin protein ligase 1 |
| chr11 | 1,19E+08 | 1,19E+08 | 476 | * | 4,156959 | 6,261254967 | 2,260775 | 2,769517 | 0,005614 | 1 | 0 | Distal Intergenic | 11 | 1,19E+08 | 1,19E+08 | 5098 | 2 | 1,05E+08 | ENSMUST00000156576.2 | 7386 | | Gm11754 | predicted gene 11754 |
| chr4 | 18421252 | 18421582 | 331 | * | 3,993941 | 6,261569766 | 2,230516 | 2,80723 | 0,004997 | 1 | 0 | Intron (ENSMUST0000( | 4 | 18051757 | 18093719 | 41963 | 1 | 17389 | ENSMUST00000139418.2 | 369495 | ENSMUSG00000028226 | Mmp16 | matrix metallopeptidase 16 |
| chr17 | 13981792 | 13982407 | 616 | * | 4,393739 | 6,261582666 | 2,256858 | 2,774469 | 0,005529 | 1 | 0 | Promoter (<=1kb) | 17 | 13981867 | 14074487 | 92621 | 1 | 17356 | ENSMUST00000137495.8 | 0 | ENSMUSG00000068036 | Afdn | afadin, adherens junction formation factor |
| chr9 | 56642947 | 56643397 | 451 | * | 4,562532 | 6,262394144 | 2,160931 | 2,898008 | 0,003755 | 1 | 0 | Intron (ENSMUST0000( | 9 | 56592474 | 56676407 | 83934 | 2 | 235402 | ENSMUST00000210995.2 | 33010 | ENSMUSG00000049556 | Lingo1 | leucine rich repeat and Ig domain containing 1 |
| chr17 | 27405437 | 27405725 | 289 | * | 3,89437 | 6,262540697 | 2,355141 | 2,659094 | 0,007835 | 1 | 0 | Distal Intergenic | 17 | 27409086 | 27412454 | 3369 | 2 | 224640 | ENSMUST00000237625.2 | 6729 | ENSMUSG00000044857 | Lemd2 | LEM domain containing 2 |
| chr1 | 93735546 | 93736262 | 717 | * | 5,152477 | 6,263391025 | 2,090129 | 2,996653 | 0,00273 | 1 | 0 | Promoter (<=1kb) | 1 | 93736292 | 93736393 | 102 | 2 | 1,15E+08 | ENSMUST00000158336.3 | 131 | ENSMUSG00000088961 | Gm22089 | predicted gene, 22089 |
| chr11 | 75632297 | 75632869 | 573 | * | 4,313855 | 6,263404006 | 2,09002 | 2,996815 | 0,002728 | 1 | 0 | Intron (ENSMUST0000( | 11 | 75623974 | 75647698 | 23725 | 1 | 22627 | ENSMUST00000131398.2 | 8323 | ENSMUSG00000020849 | Ywhae | tyrosine 3-monooxygenase/tryptophan 5-monooxygenase activation protein, epsilon polypeptide |
| chr12 | 26931669 | 26932377 | 709 | * | 4,229139 | 6,263493442 | 2,084438 | 3,004884 | 0,002657 | 1 | 0 | Distal Intergenic | 12 | 26869784 | 26884872 | 15089 | 2 | 74668 | ENSMUST00000222026.2 | -46797 | ENSMUSG00000113218 | 4933409F18Rik | RIKEN cDNA 4933409F18 gene |
| chr4 | 1,33E+08 | 1,33E+08 | 562 | * | 5,205925 | 6,263970564 | 2,055027 | 3,04812 | 0,002303 | 1 | 0 | Promoter (<=1kb) | 4 | 1,33E+08 | 1,33E+08 | 2035 | 1 | 19891 | ENSMUST00000156968.2 | 84 | ENSMUSG00000028884 | Rpa2 | replication protein A2 |
| chr13 | 43533644 | 43534467 | 824 | * | 5,288916 | 6,264095455 | 2,048065 | 3,058543 | 0,002224 | 1 | 0 | Promoter (<=1kb) | 13 | 43533149 | 43541132 | 7984 | 1 | 68346 | ENSMUST00000222106.2 | 495 | ENSMUSG00000054021 | Sirt5 | sirtuin 5 |
| chr6 | 1,18E+08 | 1,18E+08 | 412 | * | 4,76239 | 6,264289982 | 2,038488 | 3,073009 | 0,002119 | 1 | 0 | Promoter (1-2kb) | 6 | 1,18E+08 | 1,18E+08 | 31689 | 2 | 78752 | ENSMUST00000049344.15 | -1222 | ENSMUSG00000042042 | Csgalnact2 | chondroitin sulfate N-acetylgalactosaminyltransferase 2 |
| chr2 | 30028346 | 30028900 | 555 | * | 4,786129 | 6,264573863 | 2,022749 | 3,097059 | 0,001955 | 1 | 0 | Promoter (1-2kb) | 2 | 30027042 | 30032536 | 5495 | 1 | 70296 | ENSMUST00000129789.2 | 1304 | ENSMUSG00000039678 | Tbc1d13 | TBC1 domain family, member 13 |
| chr8 | 40294904 | 40295937 | 1034 | * | 8,703442 | 6,266077503 | 1,716334 | 3,650851 | 0,000261 | 0,275159027 | 1,425929 | Distal Intergenic | 8 | 40056495 | 40095714 | 39220 | 2 | 20288 | ENSMUST00000170091.3 | -199190 | ENSMUSG00000025044 | Msr1 | macrophage scavenger receptor 1 |
| chrX | 1,49E+08 | 1,49E+08 | 635 | * | 6,137556 | 6,267034745 | 1,917466 | 3,268394 | 0,001082 | 1 | 0 | Distal Intergenic | 20 | 1,49E+08 | 1,49E+08 | 12280 | 2 | 56191 | ENSMUST00000087258.10 | -13771 | ENSMUSG00000025272 | Tro | trophinin |
| chr9 | 1,03E+08 | 1,03E+08 | 796 | * | 6,069571 | 6,267148133 | 1,913605 | 3,275048 | 0,001056 | 1 | 0 | Intron (ENSMUST0000( | 9 | 1,03E+08 | 1,03E+08 | 17505 | 2 | 71775 | ENSMUST00000150524.2 | -6820 | ENSMUSG00000033688 | Inhca | inhibitor of carbonic anhydrase |
| chr16 | 12977205 | 12977752 | 548 | * | 5,951056 | 6,26767398 | 2,085381 | 3,00553 | 0,002651 | 1 | 0 | Distal Intergenic | 16 | 12950255 | 12962355 | 12101 | 1 | 50505 | ENSMUST00000156393.2 | 26950 | ENSMUSG00000022545 | Ercc4 | excision repair cross-complementing rodent repair deficiency, complementation group 4 |
| chr16 | 24827974 | 24829334 | 1361 | * | 8,21928 | 6,2685618 | 1,868087 | 3,355605 | 0,000792 | 0,443906448 | 1,425929 | Distal Intergenic | 16 | 24878389 | 24887808 | 9420 | 1 | 319487 | ENSMUST00000181655.2 | -49055 | ENSMUSG00000096960 | A230028O05Rik | RIKEN cDNA A230028O05 gene |
| chr11 | 87418085 | 87418558 | 474 | * | 3,813137 | 6,269401625 | 2,407886 | 2,603695 | 0,009222 | 1 | 0 | Intron (ENSMUST0000( | 11 | 87427550 | 87445783 | 18234 | 1 | 83560 | ENSMUST00000131973.2 | -8992 | ENSMUSG00000010342 | Tex14 | testis expressed gene 14 |
| chr5 | 1,17E+08 | 1,17E+08 | 302 | * | 3,395073 | 6,269656755 | 2,387676 | 2,62584 | 0,008644 | 1 | 0 | Intron (ENSMUST0000( | 5 | 1,17E+08 | 1,17E+08 | 2598 | 1 | 1,03E+08 | ENSMUST00000201443.2 | -21130 | | Gm29926 | predicted gene, 29926 |
| chr1 | 58040034 | 58040455 | 422 | * | 4,177455 | 6,270276746 | 2,346072 | 2,67267 | 0,007525 | 1 | 0 | Exon (ENSMUST0000( | 1 | 58035130 | 58065058 | 29929 | 1 | 68549 | ENSMUST00000027202.9 | 4904 | ENSMUSG00000026039 | Sgo2a | shugoshin 2A |
| chr11 | 3622763 | 3623764 | 1002 | * | 6,800432 | 6,270773394 | 1,839113 | 3,409673 | 0,00065 | 1 | 0 | Intron (ENSMUST0000( | 11 | 3629907 | 3633907 | 4001 | 1 | 74522 | ENSMUST00000126035.8 | -6143 | ENSMUSG00000034543 | Morc2a | microrchidia 2A |
| chr11 | 67979285 | 67979878 | 594 | * | 5,780773 | 6,271059399 | 2,194281 | 2,857911 | 0,004264 | 1 | 0 | Intron (ENSMUST0000( | 11 | 67857332 | 67864467 | 7136 | 1 | 55943 | ENSMUST00000142649.2 | 121953 | ENSMUSG00000020903 | Stx8 | syntaxin 8 |
| chr9 | 43233356 | 43233770 | 415 | * | 4,274528 | 6,271129288 | 2,292973 | 2,734934 | 0,006239 | 1 | 0 | Exon (ENSMUST0000( | 9 | 43230256 | 43231668 | 1413 | 1 | 72169 | ENSMUST00000217418.2 | 3100 | ENSMUSG00000032013 | Trim29 | tripartite motif-containing 29 |
| chr19 | 18914561 | 18915234 | 674 | * | 4,318997 | 6,271295335 | 2,285224 | 2,74428 | 0,006064 | 1 | 0 | 3' UTR | 19 | 18953326 | 18978464 | 25139 | 2 | 225998 | ENSMUST00000149635.8 | 63230 | ENSMUSG00000036192 | Rorb | RAR-related orphan receptor beta |
| chr9 | 65167175 | 65167951 | 777 | * | 6,151528 | 6,27143379 | 1,967654 | 3,187264 | 0,001436 | 1 | 0 | Distal Intergenic | 9 | 65172462 | 65187887 | 15426 | 1 | 214425 | ENSMUST00000048762.8 | -4511 | ENSMUSG00000042254 | Cilp | cartilage intermediate layer protein, nucleotide pyrophosphohydrolase |
| chr10 | 33788665 | 33789329 | 665 | * | 5,463745 | 6,271502444 | 1,981837 | 3,164489 | 0,001554 | 1 | 0 | Exon (ENSMUST00000 | 10 | 33781107 | 33792017 | 10911 | 1 | 212892 | ENSMUST00000169670.8 | 7558 | ENSMUSG00000039552 | Rsph4a | radial spoke head 4 homolog A (Chlamydomonas) |
| chr4 | 1,19E+08 | 1,19E+08 | 599 | * | 4,676552 | 6,2716104 | 2,01181 | 3,117397 | 0,001825 | 1 | 0 | Intron (ENSMUST0000( | 4 | 1,19E+08 | 1,19E+08 | 14322 | 2 | 433751 | ENSMUST00000128998.2 | 4559 | ENSMUSG00000066060 | Gm12866 | predicted gene 12866 |
| chr7 | 82069587 | 82070072 | 486 | * | 4,90293 | 6,271685768 | 2,035698 | 3,080852 | 0,002064 | 1 | 0 | Intron (ENSMUST0000( | 7 | 82057682 | 82115052 | 57371 | 1 | 269959 | ENSMUST00000174495.2 | 11905 | ENSMUSG00000070469 | Adamtsl3 | ADAMTS-like 3 |
| chr2 | 1,64E+08 | 1,64E+08 | 562 | * | 4,699532 | 6,271782633 | 2,069686 | 3,030306 | 0,002443 | 1 | 0 | Distal Intergenic | 2 | 1,64E+08 | 1,64E+08 | 23629 | 1 | 54401 | ENSMUST00000018470.10 | -6317 | ENSMUSG00000018326 | Ywhab | tyrosine 3-monooxygenase/tryptophan 5-monooxygenase activation protein, beta polypeptide |
| chr7 | 1,28E+08 | 1,28E+08 | 606 | * | 4,893043 | 6,27179156 | 2,066647 | 3,034767 | 0,002407 | 1 | 0 | Intron (ENSMUST0000( | 7 | 1,28E+08 | 1,28E+08 | 4959 | 1 | 207352 | ENSMUST00000206986.2 | 6133 | ENSMUSG00000055319 | Sec23ip | Sec23 interacting protein |
| chrX | 36424693 | 36425621 | 929 | * | 5,089898 | 6,271889385 | 2,109145 | 2,973664 | 0,002943 | 1 | 0 | Intron (ENSMUST0000( | 20 | 36414351 | 36432495 | 18145 | 2 | 434756 | ENSMUST00000046557.6 | 6874 | ENSMUSG00000036551 | Akap14 | A kinase (PRKA) anchor protein 14 |
| chr5 | 1,07E+08 | 1,07E+08 | 998 | * | 5,751924 | 6,271981775 | 2,154334 | 2,911332 | 0,003599 | 1 | 0 | Distal Intergenic | 5 | 1,07E+08 | 1,07E+08 | 14727 | 2 | 21814 | ENSMUST00000136882.2 | 36161 | ENSMUSG00000029287 | Tgfbr3 | transforming growth factor, beta receptor III |
| chr13 | 1,11E+08 | 1,11E+08 | 712 | * | 5,449678 | 6,271982316 | 2,150341 | 2,916738 | 0,003537 | 1 | 0 | Distal Intergenic | 13 | 1,11E+08 | 1,11E+08 | 1924 | 1 | 1,03E+08 | ENSMUST00000209229.2 | -60612 | ENSMUSG00000109853 | Gm33045 | predicted gene, 33045 |
| chr16 | 25889509 | 25889852 | 344 | * | 4,709012 | 6,272038434 | 2,160581 | 2,90294 | 0,003697 | 1 | 0 | Intron (ENSMUST0000( | 16 | 25778038 | 25924534 | 146497 | 2 | 210530 | ENSMUST00000039990.6 | 34682 | ENSMUSG00000038168 | P3h2 | prolyl 3-hydroxylase 2 |
| chr4 | 57749119 | 57749563 | 445 | * | 4,03698 | 6,272103376 | 2,205629 | 2,84368 | 0,00446 | 1 | 0 | Intron (ENSMUST0000( | 4 | 57742095 | 57896982 | 154888 | 1 | 677884 | ENSMUST00000107600.8 | 7024 | ENSMUSG00000038729 | Pakap | paralemmin A kinase anchor protein |
| chr10 | 66930930 | 66931561 | 632 | * | 3,694019 | 6,272128689 | 2,211619 | 2,83599 | 0,004568 | 1 | 0 | Promoter (<=1kb) | 10 | 66931904 | 67027527 | 95624 | 1 | 108829 | ENSMUST00000174062.8 | -343 | ENSMUSG00000037876 | Jmjd1c | jumonji domain containing 1C |
| chr13 | 35657566 | 35657905 | 340 | * | 3,777194 | 6,272255514 | 2,273903 | 2,758365 | 0,005809 | 1 | 0 | Distal Intergenic | 13 | 35477130 | 35477260 | 131 | 2 | 1,15E+08 | ENSMUST00000104413.3 | -180306 | ENSMUSG00000077601 | Gm22674 | predicted gene, 22674 |
| chrX | 15826410 | 15826702 | 293 | * | 3,855777 | 6,272256301 | 2,298068 | 2,729361 | 0,006346 | 1 | 0 | Distal Intergenic | 20 | 15762902 | 15763008 | 107 | 2 | 1,15E+08 | ENSMUST00000239993.1 | -63402 | ENSMUSG00001118666 | Gm23516 | predicted gene, 23516 |
| chr13 | 18015563 | 18016181 | 619 | * | 3,806071 | 6,272266563 | 2,298433 | 2,728932 | 0,006354 | 1 | 0 | Distal Intergenic | 13 | 17984176 | 17984282 | 107 | 2 | 1,15E+08 | ENSMUST00000083232.3 | -31281 | ENSMUSG00000118773 | Gm24018 | predicted gene, 24018 |
| chr8 | 53284582 | 53285312 | 731 | * | 5,160762 | 6,272279554 | 2,3009 | 2,726011 | 0,00641 | 1 | 0 | Distal Intergenic | 8 | 52809646 | 52856374 | 46729 | 1 | 330767 | ENSMUST00000210977.2 | 474936 | ENSMUSG00000110013 | 1190028D05Rik | RIKEN cDNA 1190028D05 gene |
| chr5 | 12838104 | 12838538 | 435 | * | 3,88674 | 6,272371532 | 2,368505 | 2,648241 | 0,008091 | 1 | 0 | Distal Intergenic | 5 | 12625761 | 12638719 | 12959 | 1 | 108151 | ENSMUST00000195923.2 | 212343 | ENSMUSG00000040254 | Sema3d | sema domain, immunoglobulin domain (Ig), short basic domain, secreted, (semaphorin) 3D |
| chr5 | 1,44E+08 | 1,44E+08 | 351 | * | 3,788747 | 6,272375146 | 2,385473 | 2,629406 | 0,008553 | 1 | 0 | Promoter (<=1kb) | 5 | 1,44E+08 | 1,44E+08 | 4327 | 2 | 66729 | ENSMUST00000110717.9 | -884 | ENSMUSG00000029607 | Ankrd61 | ankyrin repeat domain 61 |
| chr14 | 67295733 | 67296292 | 560 | * | 3,546862 | 6,272432515 | 2,424279 | 2,587339 | 0,009672 | 1 | 0 | Intron (ENSMUST0000( | 14 | 67253123 | 67309631 | 56509 | 2 | 71978 | ENSMUST00000225251.2 | 13339 | ENSMUSG00000022052 | Ppp2r2a | protein phosphatase 2, regulatory subunit B, alpha |
| chr9 | 51936837 | 51937954 | 1118 | * | 9,861997 | 6,272980625 | 1,767355 | 3,549361 | 0,000386 | 0,351510656 | 1,173325 | Distal Intergenic | 9 | 51958473 | 52000035 | 41563 | 1 | 19684 | ENSMUST00000163153.9 | -20519 | ENSMUSG00000032050 | Rdx | radixin |
| chrX | 83269043 | 83269800 | 758 | * | 4,43671 | 6,273135606 | 2,192193 | 2,86158 | 0,004215 | 1 | 0 | Intron (ENSMUST0000( | 20 | 83120255 | 83353500 | 233246 | 1 | 13405 | ENSMUST00000139998.2 | 148788 | ENSMUSG00000045103 | Dmd | dystrophin, muscular dystrophy |
| chr18 | 25838150 | 25838660 | 511 | * | 5,114894 | 6,273489454 | 2,175868 | 2,883212 | 0,003936 | 1 | 0 | Intron (ENSMUST0000( | 18 | 25637277 | 25885749 | 248473 | 2 | 108013 | ENSMUST00000225927.2 | 47089 | ENSMUSG00000024268 | Celf4 | CUGBP, Elav-like family member 4 |
| chr14 | 14110722 | 14111296 | 575 | * | 4,957495 | 6,273601558 | 2,170678 | 2,890157 | 0,00385 | 1 | 0 | Intron (ENSMUST0000( | 14 | 14098763 | 14102333 | 3571 | 2 | 1,03E+08 | ENSMUST00000225127.2 | -8389 | | 2610318M16Rik | RIKEN cDNA 2610318M16 gene |
| chr10 | 1,15E+08 | 1,15E+08 | 602 | * | 4,051377 | 6,274362273 | 2,342885 | 2,678049 | 0,007405 | 1 | 0 | Promoter (2-3kb) | 10 | 1,15E+08 | 1,15E+08 | 53596 | 2 | 66687 | ENSMUST00000020339.10 | -2345 | ENSMUSG00000020130 | Tbc1d15 | TBC1 domain family, member 15 |
| chr15 | 81108155 | 81108721 | 567 | * | 4,22255 | 6,274930401 | 2,375385 | 2,641648 | 0,00825 | 1 | 0 | Distal Intergenic | 15 | 81119700 | 81123165 | 3466 | 1 | 207911 | ENSMUST00000166855.3 | -10979 | ENSMUSG00000050164 | Mchr1 | melanin-concentrating hormone receptor 1 |
| chr6 | 39514313 | 39515287 | 975 | * | 12,76812 | 6,275043095 | 1,642794 | 3,819737 | 0,000134 | 0,273126507 | 0,738441 | Intron (ENSMUST0000( | 6 | 39504456 | 39521420 | 16965 | 1 | 77209 | ENSMUST00000135363.2 | 9857 | ENSMUSG00000085058 | 8030453O22Rik | RIKEN cDNA 8030453O22 gene |
| chr13 | 8960255 | 8961016 | 762 | * | 5,23153 | 6,276527556 | 2,064007 | 3,040943 | 0,002358 | 1 | 0 | Distal Intergenic | 13 | 8937105 | 8940481 | 3377 | 1 | 319554 | ENSMUST00000176851.2 | 23150 | ENSMUSG00000058258 | Idi1 | isopentenyl-diphosphate delta isomerase |
| chr19 | 55660264 | 55660758 | 495 | * | 6,22005 | 6,278403989 | 2,009427 | 3,124474 | 0,001781 | 1 | 0 | Distal Intergenic | 19 | 55730252 | 55921661 | 191410 | 1 | 21416 | ENSMUST00000111662.11 | -69494 | ENSMUSG00000024985 | Tcf7l2 | transcription factor 7 like 2, T cell specific, HMG box |
| chr4 | 1,38E+08 | 1,38E+08 | 1015 | * | 5,815451 | 6,278772471 | 2,022435 | 3,10456 | 0,001906 | 1 | 0 | 3' UTR | 4 | 1,38E+08 | 1,38E+08 | 4289 | 1 | 68350 | ENSMUST00000105815.2 | 6443 | ENSMUSG00000041241 | Mul1 | mitochondrial ubiquitin ligase activator of NFKB 1 |
| chr2 | 1,09E+08 | 1,09E+08 | 428 | * | 4,586478 | 6,280364874 | 2,089567 | 3,005582 | 0,002651 | 1 | 0 | Distal Intergenic | 2 | 1,09E+08 | 1,09E+08 | 2758 | 1 | 1,03E+08 | ENSMUST00000153283.2 | -45850 | | Gm13923 | predicted gene 13923 |
| chr17 | 66359084 | 66359493 | 410 | * | 3,492845 | 6,281619426 | 2,42371 | 2,591737 | 0,009549 | 1 | 0 | Intron (ENSMUST0000( | 17 | 66344609 | 66383459 | 38851 | 2 | 106585 | ENSMUST00000233045.2 | 23966 | ENSMUSG00000034647 | Ankrd12 | ankyrin repeat domain 12 |
| chr8 | 25693107 | 25693433 | 327 | * | 3,57759 | 6,281680924 | 2,413701 | 2,60251 | 0,009254 | 1 | 0 | Promoter (1-2kb) | 8 | 25644568 | 25691558 | 46991 | 2 | 320165 | ENSMUST00000084030.6 | -1549 | ENSMUSG00000065954 | Tacc1 | transforming, acidic coiled-coil containing protein 1 |

| | | | | | | | | | | | | | | | | | | | | | | | |
|---|---|---|---|---|---|---|---|---|---|---|---|---|---|---|---|---|---|---|---|---|---|---|---|
| chr10 | 82827128 | 82827668 | 541 | * | 4,695299 | 6,281689384 | 2,160548 | 2,907452 | 0,003644 | 1 | 0 | Intron (ENSMUST0000 | 10 | 82821332 | 83031734 | 210403 | 1 | 58250 | ENSMUST00000040110.8 | 5796 | ENSMUSG00000034612 | Chst11 | carbohydrate sulfotransferase 11 |
| chr4 | 25934715 | 25935197 | 483 | * | 4,668705 | 6,281702305 | 2,160932 | 2,906941 | 0,00365 | 1 | 0 | Distal Intergenic | 4 | 25940365 | 25940470 | 106 | 1 | 1,15E+08 | ENSMUST00000157922.4 | -5168 | ENSMUSG00000088547 | Gm24965 | predicted gene, 24965 |
| chr4 | 46750781 | 46751114 | 334 | * | 3,349521 | 6,281784599 | 2,392348 | 2,625783 | 0,008645 | 1 | 0 | Intron (ENSMUST0000 | 4 | 46677559 | 46724391 | 46833 | 2 | 242425 | ENSMUST00000205643.2 | -26390 | ENSMUSG00000039809 | Gabbr2 | gamma-aminobutyric acid (GABA) B receptor, 2 |
| chr9 | 96944748 | 96945425 | 678 | * | 4,841174 | 6,281935297 | 2,175109 | 2,888102 | 0,003876 | 1 | 0 | Intron (ENSMUST0000 | 9 | 96929497 | 96949360 | 19864 | 2 | 1,01E+08 | ENSMUST00000186238.2 | 3935 | | Gm19325 | predicted gene, 19325 |
| chrX | 23480861 | 23481542 | 682 | * | 4,000187 | 6,282249349 | 2,318913 | 2,709136 | 0,006746 | 1 | 0 | Distal Intergenic | 20 | 23559290 | 23672229 | 112940 | 1 | 72404 | ENSMUST00000035766.13 | -77748 | ENSMUSG00000036769 | Wdr44 | WD repeat domain 44 |
| chr9 | 95919398 | 95920292 | 895 | * | 7,138395 | 6,282324144 | 1,920729 | 3,270802 | 0,001072 | 1 | 0 | Intron (ENSMUST0000 | 9 | 95855570 | 95860386 | 4817 | 1 | 24127 | ENSMUST00000189806.2 | 63828 | ENSMUSG00000032410 | Xrn1 | 5'-3' exoribonuclease 1 |
| chrX | 76531471 | 76531864 | 394 | * | 3,545231 | 6,28233269 | 2,30039 | 2,730986 | 0,006315 | 1 | 0 | Distal Intergenic | 20 | 76554619 | 76706589 | 151971 | 1 | 21372 | ENSMUST00000088217.12 | -22755 | ENSMUSG00000025246 | Tbl1x | transducin (beta)-like 1 X-linked |
| chr15 | 8462635 | 8462963 | 329 | * | 3,774291 | 6,282594405 | 2,261646 | 2,777886 | 0,005471 | 1 | 0 | Intron (ENSMUST0000 | 15 | 8320101 | 8473947 | 153847 | 2 | 71175 | ENSMUST00000052965.8 | 10984 | ENSMUSG00000022141 | Nipbl | NIPBL cohesin loading factor |
| chr11 | 75139402 | 75139788 | 387 | * | 3,952949 | 6,282768212 | 2,238501 | 2,806686 | 0,005005 | 1 | 0 | Intron (ENSMUST0000 | 11 | 75186192 | 75188651 | 2460 | 2 | 1,03E+08 | ENSMUST00000241351.1 | 48863 | | Gm35117 | predicted gene, 35117 |
| chr15 | 1,01E+08 | 1,01E+08 | 164 | * | 3,888627 | 6,28277024 | 2,239726 | 2,805151 | 0,005029 | 1 | 0 | 3' UTR | 15 | 1,01E+08 | 1,01E+08 | 5025 | 1 | 11482 | ENSMUST00000128525.2 | 7171 | ENSMUSG00000000530 | Acvrl1 | activin A receptor, type II-like 1 |
| chr11 | 79337035 | 79337690 | 656 | * | 3,716628 | 6,282796226 | 2,236124 | 2,809682 | 0,004959 | 1 | 0 | Promoter (1-2kb) | 11 | 79338979 | 79364238 | 25260 | 1 | 18015 | ENSMUST00000122917.8 | -1289 | ENSMUSG00000020716 | Nf1 | neurofibromin 1 |
| chr18 | 3331850 | 3332133 | 284 | * | 3,640924 | 6,282796279 | 2,237395 | 2,808085 | 0,004984 | 1 | 0 | Intron (ENSMUST0000 | 18 | 3266048 | 3327591 | 61544 | 2 | 12916 | ENSMUST00000151311.8 | -4259 | ENSMUSG00000063889 | Crem | cAMP responsive element modulator |
| chr3 | 43250592 | 43250942 | 351 | * | 4,144903 | 6,282851106 | 2,229618 | 2,817904 | 0,004834 | 1 | 0 | Distal Intergenic | 3 | 41705631 | 41710927 | 5297 | 1 | 73852 | ENSMUST00000195030.2 | 1544961 | ENSMUSG00000025766 | D3Ertd751e | DNA segment, Chr 3, ERATO Doi 751, expressed |
| chrX | 17254557 | 17255415 | 859 | * | 3,589451 | 6,283009034 | 2,247134 | 2,79601 | 0,005174 | 1 | 0 | Distal Intergenic | 20 | 17191554 | 17191657 | 104 | 1 | 1,15E+08 | ENSMUST00000083308.3 | 63003 | ENSMUSG00000119745 | Gm22294 | predicted gene, 22294 |
| chr6 | 92869730 | 92869982 | 253 | * | 4,103591 | 6,283293144 | 2,174649 | 2,889336 | 0,003861 | 1 | 0 | Intron (ENSMUST0000 | 6 | 92749683 | 92878422 | 128740 | 2 | 101401 | ENSMUST00000167391.2 | 8440 | ENSMUSG00000030022 | Adamts9 | a disintegrin-like and metallopeptidase (reprolysin type) with thrombospondin type 1 motif, 9 |
| chrX | 8094135 | 8094784 | 650 | * | 4,563012 | 6,28347146 | 2,154656 | 2,91623 | 0,003543 | 1 | 0 | Distal Intergenic | 20 | 8111186 | 8112045 | 860 | 2 | 54632 | ENSMUST00000126924.2 | 17261 | ENSMUSG00000031171 | Ftsj1 | FtsJ RNA methyltransferase homolog 1 (E. coli) |
| chr7 | 1,21E+08 | 1,21E+08 | 426 | * | 3,922488 | 6,283563266 | 2,146067 | 2,927944 | 0,003412 | 1 | 0 | Promoter (1-2kb) | 7 | 1,21E+08 | 1,21E+08 | 32201 | 1 | 74466 | ENSMUST00000216241.2 | 1552 | ENSMUSG00000030877 | Mfsd13b | major facilitator superfamily domain containing 13B |
| chr19 | 20567087 | 20567882 | 796 | * | 3,634588 | 6,283785404 | 2,309729 | 2,720572 | 0,006517 | 1 | 0 | Intron (ENSMUST0000 | 19 | 20579325 | 20620826 | 41502 | 1 | 11668 | ENSMUST00000087638.4 | -11443 | ENSMUSG00000053279 | Aldh1a1 | aldehyde dehydrogenase family 1, subfamily A1 |
| chr15 | 79963182 | 79963892 | 711 | * | 4,507901 | 6,284433676 | 2,066357 | 3,041311 | 0,002356 | 1 | 0 | Promoter (<=1kb) | 15 | 79962699 | 79962792 | 94 | 2 | 1E+08 | ENSMUST00000104546.3 | -390 | ENSMUSG00000077734 | Snord83b | small nucleolar RNA, C/D box 83B |
| chr4 | 1,54E+08 | 1,54E+08 | 301 | * | 4,877309 | 6,284602122 | 2,051982 | 3,062698 | 0,002194 | 1 | 0 | Intron (ENSMUST0000 | 4 | 1,54E+08 | 1,54E+08 | 34215 | 1 | 230971 | ENSMUST00000152159.8 | 4230 | ENSMUSG00000057751 | Megf6 | multiple EGF-like-domains 6 |
| chr8 | 68240429 | 68240853 | 425 | * | 5,491424 | 6,284703159 | 2,044811 | 3,073488 | 0,002116 | 1 | 0 | Intron (ENSMUST0000 | 8 | 68164178 | 68270885 | 106708 | 2 | 234353 | ENSMUST00000059374.5 | 30032 | ENSMUSG00000030465 | Psd3 | pleckstrin and Sec7 domain containing 3 |
| chr15 | 53069440 | 53070012 | 573 | * | 5,239009 | 6,284710899 | 2,042997 | 3,076221 | 0,002096 | 1 | 0 | Intron (ENSMUST0000 | 15 | 52946542 | 53059651 | 113110 | 2 | 14042 | ENSMUST00000110244.2 | -9789 | ENSMUSG00000061731 | Ext1 | exostosin glycosyltransferase 1 |
| chr16 | 31304314 | 31304779 | 466 | * | 4,651099 | 6,284729179 | 2,042265 | 3,077333 | 0,002089 | 1 | 0 | Distal Intergenic | 16 | 31327205 | 31330595 | 3391 | 1 | 1,08E+08 | ENSMUST00000231805.2 | -22426 | | Gm46560 | predicted gene, 46560 |
| chr14 | 5733351 | 5734224 | 874 | * | 4,828724 | 6,284732771 | 2,041652 | 3,078258 | 0,002082 | 1 | 0 | Intron (ENSMUST0000 | 14 | 5650691 | 5921244 | 270554 | 1 | 218772 | ENSMUST00000225245.2 | 82660 | ENSMUSG00000017491 | Rarb | retinoic acid receptor, beta |
| chrX | 10342075 | 10342907 | 833 | * | 5,45336 | 6,285012172 | 2,021894 | 3,108478 | 0,001881 | 1 | 0 | Distal Intergenic | 20 | 10351397 | 10462844 | 111448 | 1 | 21912 | ENSMUST00000076354.13 | -8490 | ENSMUSG00000058254 | Tspan7 | tetraspanin 7 |
| chr15 | 8001219 | 8001899 | 681 | * | 3,692426 | 6,285019743 | 2,429321 | 2,587151 | 0,009677 | 1 | 0 | Intron (ENSMUST0000 | 15 | 7902536 | 7915858 | 13323 | 2 | 545085 | ENSMUST00000226928.2 | -85361 | ENSMUSG00000039828 | Wdr70 | WD repeat domain 70 |
| chr5 | 1,2E+08 | 1,2E+08 | 567 | * | 4,880343 | 6,285228863 | 2,00613 | 3,133011 | 0,00173 | 1 | 0 | Distal Intergenic | 5 | 1,2E+08 | 1,2E+08 | 22245 | 1 | 74111 | ENSMUST00000202788.2 | 112204 | ENSMUSG00000029594 | Rbm19 | RNA binding motif protein 19 |
| chr2 | 1,33E+08 | 1,33E+08 | 724 | * | 6,819734 | 6,286185235 | 1,947041 | 3,228584 | 0,001244 | 1 | 0 | Distal Intergenic | 2 | 1,33E+08 | 1,33E+08 | 13802 | 1 | 12653 | ENSMUST00000028826.4 | -3751 | ENSMUSG00000027350 | Chgb | chromogranin B |
| chr11 | 87325412 | 87326268 | 857 | * | 6,243893 | 6,287197898 | 1,896353 | 3,315415 | 0,000915 | 1 | 0 | Intron (ENSMUST0000 | 11 | 87318417 | 87375592 | 57176 | 1 | 83560 | ENSMUST00000156483.2 | 6995 | ENSMUSG00000010342 | Tex14 | testis expressed gene 14 |
| chr7 | 19371750 | 19372274 | 525 | * | 6,356781 | 6,287332406 | 1,890516 | 3,325724 | 0,000882 | 1 | 0 | Promoter (<=1kb) | 7 | 19368881 | 19371671 | 2791 | 2 | 56457 | ENSMUST00000208868.2 | -79 | ENSMUSG00000002981 | Clptm1 | cleft lip and palate associated transmembrane protein 1 |
| chr7 | 80523363 | 80524138 | 776 | * | 7,284045 | 6,287480644 | 1,883633 | 3,337954 | 0,000844 | 0,46163452 | 1,360622 | Intron (ENSMUST0000 | 7 | 80512644 | 80524846 | 12203 | 1 | 22691 | ENSMUST00000155128.3 | 10719 | ENSMUSG00000038797 | Zscan2 | zinc finger and SCAN domain containing 2 |
| chr19 | 38577251 | 38577512 | 262 | * | 3,54321 | 6,290836926 | 2,412824 | 2,60725 | 0,009127 | 1 | 0 | Intron (ENSMUST0000 | 19 | 38571812 | 38640451 | 68640 | 1 | 74055 | ENSMUST00000182999.2 | 5439 | ENSMUSG00000024998 | Plce1 | phospholipase C, epsilon 1 |
| chr19 | 59831482 | 59832109 | 628 | * | 3,930805 | 6,290902627 | 2,407411 | 2,61314 | 0,008971 | 1 | 0 | Distal Intergenic | 19 | 59895348 | 59931432 | 36085 | 2 | 74998 | ENSMUST00000170819.2 | 99323 | ENSMUSG00000040022 | Rab11fip2 | RAB11 family interacting protein 2 (class I) |
| chr7 | 1,39E+08 | 1,39E+08 | 552 | * | 6,07336 | 6,291036004 | 1,91387 | 3,287076 | 0,001012 | 1 | 0 | Promoter (2-3kb) | 7 | 1,39E+08 | 1,39E+08 | 16325 | 1 | 212111 | ENSMUST00000131781.2 | 2745 | ENSMUSG00000025477 | Inpp5a | inositol polyphosphate-5-phosphatase A |
| chr4 | 1,37E+08 | 1,37E+08 | 1478 | * | 14,00175 | 6,291080882 | 1,765901 | 3,562533 | 0,000367 | 0,425046984 | 0,738441 | 3' UTR | 4 | 1,37E+08 | 1,37E+08 | 22238 | 1 | 22417 | ENSMUST00000045747.5 | 18034 | ENSMUSG00000036856 | Wnt4 | wingless-type MMTV integration site family, member 4 |
| chr5 | 1,49E+08 | 1,49E+08 | 276 | * | 3,349441 | 6,291138511 | 2,386974 | 2,635612 | 0,008399 | 1 | 0 | Distal Intergenic | 5 | 1,49E+08 | 1,49E+08 | 1312 | 1 | 70138 | ENSMUST00000244021.1 | -77543 | ENSMUSG00000086494 | 2210417A02Rik | RIKEN cDNA 2210417A02 gene |
| chr10 | 30073771 | 30074425 | 655 | * | 4,00689 | 6,291636091 | 2,34767 | 2,679949 | 0,007363 | 1 | 0 | Promoter (2-3kb) | 10 | 30070565 | 30076488 | 5924 | 2 | 66311 | ENSMUST00000216853.2 | 2063 | ENSMUSG00000075266 | Cenpw | centromere protein W |
| chr9 | 43906080 | 43906944 | 865 | * | 5,529644 | 6,291743343 | 1,979176 | 3,178971 | 0,001478 | 1 | 0 | Exon (ENSMUST00002 | 9 | 43836048 | 43881550 | 45503 | 2 | 1,03E+08 | ENSMUST00000215774.2 | -24530 | | Gm30373 | predicted gene, 30373 |
| chr8 | 1,13E+08 | 1,13E+08 | 247 | * | 4,097335 | 6,291773535 | 2,337515 | 2,69165 | 0,00711 | 1 | 0 | Distal Intergenic | 8 | 1,13E+08 | 1,13E+08 | 5909 | 2 | 672498 | ENSMUST00000247388.1 | -11867 | | Gm9564 | predicted gene 9564 |
| chr14 | 20531594 | 20532085 | 492 | * | 3,536821 | 6,291815972 | 2,332077 | 2,697945 | 0,006977 | 1 | 0 | Promoter (1-2kb) | 14 | 20505328 | 20530201 | 24874 | 2 | 11750 | ENSMUST00000065504.17 | -1393 | ENSMUSG00000021814 | Anxa7 | annexin A7 |
| chr5 | 12631717 | 12632693 | 977 | * | 6,069141 | 6,291873965 | 1,993259 | 3,156576 | 0,001596 | 1 | 0 | Intron (ENSMUST0000 | 5 | 12625761 | 12638719 | 12959 | 1 | 108151 | ENSMUST00000195923.2 | 5956 | ENSMUSG00000040254 | Sema3d | sema domain, immunoglobulin domain (Ig), short basic domain, secreted, (semaphorin) 3D |
| chr11 | 1,18E+08 | 1,18E+08 | 667 | * | 4,695979 | 6,292011032 | 2,009024 | 3,131875 | 0,001737 | 1 | 0 | Intron (ENSMUST0000 | 11 | 1,18E+08 | 1,18E+08 | 37720 | 1 | 74451 | ENSMUST00000132676.8 | -7079 | ENSMUSG00000017715 | Pgs1 | phosphatidylglycerophosphate synthase 1 |
| chr11 | 3640828 | 3641348 | 521 | * | 3,915152 | 6,292083653 | 2,313515 | 2,719708 | 0,006534 | 1 | 0 | Distal Intergenic | 11 | 3630362 | 3633689 | 3328 | 1 | 74522 | ENSMUST00000128666.2 | 10466 | ENSMUSG00000034543 | Morc2a | microrchidia 2A |
| chr3 | 90375373 | 90375898 | 526 | * | 4,223169 | 6,292553696 | 2,2805 | 2,759287 | 0,005793 | 1 | 0 | Promoter (2-3kb) | 3 | 90357898 | 90373173 | 15276 | 2 | 18160 | ENSMUST00000029540.13 | -2200 | ENSMUSG00000027931 | Npr1 | natriuretic peptide receptor 1 |
| chr2 | 18769625 | 18770019 | 395 | * | 4,990199 | 6,292621296 | 2,087648 | 3,014216 | 0,002576 | 1 | 0 | Distal Intergenic | 2 | 18747526 | 18765673 | 18148 | 2 | 1E+08 | ENSMUST00000174811.2 | -3952 | ENSMUSG00000092353 | Gm3363 | predicted gene 3363 |
| chr12 | 81286338 | 81286993 | 656 | * | 4,081639 | 6,292963991 | 2,143202 | 2,936244 | 0,003322 | 1 | 0 | Intron (ENSMUST0000 | 12 | 81275058 | 81290032 | 14975 | 1 | 1E+08 | ENSMUST00000182077.2 | 11280 | ENSMUSG00000098128 | Gm3693 | predicted gene 3693 |
| chr2 | 18972588 | 18972923 | 336 | * | 4,026834 | 6,293155247 | 2,243224 | 2,805407 | 0,005025 | 1 | 0 | Intron (ENSMUST0000 | 2 | 18847066 | 19002937 | 155872 | 2 | 18718 | ENSMUST00000006912.12 | 30014 | ENSMUSG00000026737 | Pip4k2a | phosphatidylinositol-5-phosphate 4-kinase, type II, alpha |
| chr11 | 97822190 | 97822694 | 505 | * | 4,014233 | 6,293320214 | 2,231567 | 2,820135 | 0,0048 | 1 | 0 | Intron (ENSMUST0000 | 11 | 97859618 | 97860061 | 444 | 1 | 1,03E+08 | ENSMUST00000118176.2 | -36924 | | Gm11633 | predicted gene 11633 |
| chr3 | 5348637 | 5349318 | 682 | * | 4,538343 | 6,293434393 | 2,225839 | 2,827444 | 0,004692 | 1 | 0 | Intron (ENSMUST0000 | 3 | 5306732 | 5480917 | 174186 | 1 | 80892 | ENSMUST00000176383.3 | 41905 | ENSMUSG00000025255 | Zfhx4 | zinc finger homeodomain 4 |
| chr11 | 60860782 | 60861287 | 506 | * | 3,800266 | 6,293501291 | 2,256252 | 2,789361 | 0,005281 | 1 | 0 | Distal Intergenic | 11 | 60841488 | 60842647 | 1160 | 1 | 26397 | ENSMUST00000137609.2 | 19294 | ENSMUSG00000018932 | Map2k3 | mitogen-activated protein kinase kinase 3 |
| chr14 | 96843159 | 96843862 | 704 | * | 3,379049 | 6,293516579 | 2,257854 | 2,787389 | 0,005313 | 1 | 0 | Distal Intergenic | 14 | 96826263 | 96826392 | 130 | 1 | 1,15E+08 | ENSMUST00000082647.3 | 16896 | ENSMUSG00000064581 | Gm25133 | predicted gene, 25133 |
| chr10 | 18309320 | 18309634 | 315 | * | 3,505774 | 6,293775649 | 2,319871 | 2,712985 | 0,006668 | 1 | 0 | Intron (ENSMUST0000 | 10 | 18283423 | 18409640 | 126218 | 1 | 215819 | ENSMUST00000037341.14 | 25897 | ENSMUSG00000039835 | Nhsl1 | NHS-like 1 |
| chr11 | 12896277 | 12896763 | 487 | * | 5,433624 | 6,293799473 | 2,204206 | 2,855359 | 0,004299 | 1 | 0 | Distal Intergenic | 11 | 12642813 | 12642942 | 130 | 1 | 1,15E+08 | ENSMUST00000157884.3 | 253464 | ENSMUSG00000088509 | Gm22789 | predicted gene, 22789 |
| chr12 | 1,2E+08 | 1,2E+08 | 543 | * | 4,820585 | 6,293898521 | 2,200321 | 2,860445 | 0,00423 | 1 | 0 | Distal Intergenic | 12 | 1,19E+08 | 1,19E+08 | 23525 | 1 | 238455 | ENSMUST00000048880.7 | 238792 | ENSMUSG00000041886 | Macc1 | metastasis associated in colon cancer 1 |
| chr13 | 20269429 | 20269857 | 429 | * | 4,776708 | 6,294731705 | 2,155809 | 2,919894 | 0,003502 | 1 | 0 | Distal Intergenic | 13 | 20274766 | 20448177 | 173412 | 1 | 140580 | ENSMUST00000181429.8 | -4909 | ENSMUSG00000041112 | Elmo1 | engulfment and cell motility 1 |
| chr12 | 44337244 | 44337836 | 593 | * | 5,30508 | 6,29478816 | 2,153894 | 2,922515 | 0,003472 | 1 | 0 | Promoter (<=1kb) | 12 | 44337393 | 44358408 | 21016 | 1 | 67452 | ENSMUST00000218803.2 | 0 | ENSMUSG00000036257 | Pnpla8 | patatin-like phospholipase domain containing 8 |
| chr7 | 1,1E+08 | 1,1E+08 | 648 | * | 4,684522 | 6,296665602 | 2,072256 | 3,038556 | 0,002377 | 1 | 0 | Intron (ENSMUST0000 | 7 | 1,1E+08 | 1,1E+08 | 305364 | 2 | 319934 | ENSMUST00000166020.8 | 54687 | ENSMUSG00000038371 | Sbf2 | SET binding factor 2 |
| chr19 | 25995535 | 25996400 | 866 | * | 5,533113 | 6,297825253 | 2,030729 | 3,101263 | 0,001927 | 1 | 0 | Distal Intergenic | 19 | 26015816 | 26023831 | 8016 | 2 | 73433 | ENSMUST00000237184.2 | 27431 | ENSMUSG00000118016 | 1700048O14Rik | RIKEN cDNA 1700048O14 gene |
| chr11 | 53263811 | 53264625 | 815 | * | 5,348908 | 6,300043735 | 2,047086 | 3,077566 | 0,002087 | 1 | 0 | Promoter (<=1kb) | 11 | 53263801 | 53290452 | 26652 | 1 | 93736 | ENSMUST00000130152.2 | 10 | ENSMUSG00000049470 | Aff4 | AF4/FMR2 family, member 4 |
| chr4 | 1,17E+08 | 1,17E+08 | 680 | * | 4,896077 | 6,300071792 | 2,048007 | 3,076197 | 0,002097 | 1 | 0 | Promoter (1-2kb) | 4 | 1,17E+08 | 1,17E+08 | 3212 | 1 | 68777 | ENSMUST00000145836.2 | 1435 | ENSMUSG00000048772 | Tmem53 | transmembrane protein 53 |
| chr2 | 6159689 | 6160635 | 947 | * | 4,708525 | 6,300592361 | 2,067244 | 3,047822 | 0,002305 | 1 | 0 | Distal Intergenic | 2 | 6137694 | 6145324 | 7631 | 1 | 1E+08 | ENSMUST00000195480.2 | 21995 | ENSMUSG00000102887 | Gm10857 | predicted gene 10857 |
| chr2 | 92394176 | 92394800 | 625 | * | 6,184715 | 6,300801339 | 1,943835 | 3,241428 | 0,001189 | 1 | 0 | Distal Intergenic | 2 | 92360039 | 92371325 | 11287 | 2 | 1,03E+08 | ENSMUST00000142249.3 | -22851 | | Gm13817 | predicted gene 13817 |
| chr9 | 58867339 | 58868455 | 1117 | * | 7,591174 | 6,301136414 | 1,935605 | 3,255384 | 0,001132 | 1 | 0 | Intron (ENSMUST0000 | 9 | 58820119 | 58889391 | 69273 | 2 | 18007 | ENSMUST00000215026.2 | 20936 | ENSMUSG00000032340 | Neo1 | neogenin |
| chr16 | 21498803 | 21499165 | 363 | * | 3,053606 | 6,303091417 | 2,423157 | 2,60119 | 0,00929 | 1 | 0 | Intron (ENSMUST0000 | 16 | 21472068 | 21511981 | 39914 | 2 | 72190 | ENSMUST00000232164.2 | 12816 | ENSMUSG00000043391 | 2510009E07Rik | RIKEN cDNA 2510009E07 gene |
| chr11 | 1,15E+08 | 1,15E+08 | 312 | * | 3,707302 | 6,303146274 | 2,411725 | 2,613542 | 0,008961 | 1 | 0 | Promoter (1-2kb) | 11 | 1,15E+08 | 1,15E+08 | 14217 | 1 | 237988 | ENSMUST00000053288.6 | -1337 | ENSMUSG00000050910 | Cdr2l | cerebellar degeneration-related protein 2-like |
| chrX | 12033420 | 12034225 | 806 | * | 3,406016 | 6,303180114 | 2,394431 | 2,632433 | 0,008478 | 1 | 0 | Distal Intergenic | 20 | 11902982 | 12026594 | 123613 | 2 | 71458 | ENSMUST00000043441.13 | -6826 | ENSMUSG00000040363 | Bcor | BCL6 interacting corepressor |
| chr11 | 24871927 | 24872375 | 449 | * | 3,419826 | 6,303287661 | 2,36629 | 2,663785 | 0,007727 | 1 | 0 | Intron (ENSMUST0000 | 11 | 24853983 | 24876050 | 22068 | 2 | 1,03E+08 | ENSMUST00000130092.2 | 3675 | ENSMUSG00000087145 | Gm12665 | predicted gene 12665 |
| chr12 | 68541526 | 68542097 | 572 | * | 4,507826 | 6,303963411 | 2,204246 | 2,859918 | 0,004238 | 1 | 0 | Distal Intergenic | 12 | 69176610 | 69191850 | 15241 | 1 | 1,03E+08 | ENSMUST00000222345.2 | -634513 | ENSMUSG00000113158 | Gm31447 | predicted gene, 31447 |
| chr4 | 1,13E+08 | 1,13E+08 | 523 | * | 3,978092 | 6,304026656 | 2,193714 | 2,873677 | 0,004057 | 1 | 0 | Intron (ENSMUST0000 | 4 | 1,13E+08 | 1,13E+08 | 1212 | 2 | 433745 | ENSMUST00000118069.2 | -48236 | ENSMUSG00000083011 | Gm12816 | predicted gene 12816 |
| chr4 | 1,33E+08 | 1,33E+08 | 321 | * | 4,787985 | 6,304094588 | 2,236478 | 2,81876 | 0,004821 | 1 | 0 | Intron (ENSMUST0000 | 4 | 1,33E+08 | 1,33E+08 | 5815 | 1 | 230793 | ENSMUST00000148518.2 | -6138 | ENSMUSG00000037692 | Ahdc1 | AT hook, DNA binding motif, containing 1 |
| chr13 | 46852362 | 46853046 | 685 | * | 4,055033 | 6,304104126 | 2,183699 | 2,886892 | 0,003891 | 1 | 0 | Promoter (2-3kb) | 13 | 46847243 | 46850324 | 3082 | 2 | 218210 | ENSMUST00000224186.2 | -2038 | ENSMUSG00000021374 | Nup153 | nucleoporin 153 |
| chr2 | 23826788 | 23827359 | 572 | * | 5,297902 | 6,304524912 | 2,110712 | 2,986918 | 0,002818 | 1 | 0 | Distal Intergenic | 2 | 23893541 | 23938878 | 45338 | 2 | 140483 | ENSMUST00000114497.2 | 111519 | ENSMUSG00000026986 | Hnmt | histamine N-methyltransferase |
| chrX | 67160698 | 67161137 | 440 | * | 4,777379 | 6,304546923 | 2,107877 | 2,990947 | 0,002781 | 1 | 0 | Distal Intergenic | 20 | 67053711 | 67053785 | 75 | 2 | 1E+08 | ENSMUST00000196089.3 | -106913 | ENSMUSG00000106233 | Mir509 | microRNA 509 |
| chr17 | 74799540 | 74800032 | 493 | * | 4,444517 | 6,304561156 | 2,106136 | 2,993425 | 0,002759 | 1 | 0 | Promoter (<=1kb) | 17 | 74798839 | 74801073 | 2235 | 1 | 67864 | ENSMUST00000235064.2 | 701 | ENSMUSG00000024072 | Yipf4 | Yip1 domain family, member 4 |
| chr12 | 78023711 | 78024328 | 618 | * | 4,122186 | 6,304634561 | 2,093984 | 3,010832 | 0,002605 | 1 | 0 | Distal Intergenic | 12 | 77861454 | 77861583 | 130 | 2 | 1,15E+08 | ENSMUST00000104519.3 | -162128 | ENSMUSG00000077707 | Gm24070 | predicted gene, 24070 |
| chr9 | 30303103 | 30303557 | 455 | * | 4,94451 | 6,304643551 | 2,092918 | 3,012369 | 0,002592 | 1 | 0 | Intron (ENSMUST0000 | 9 | 30297276 | 30338272 | 40997 | 2 | 1,03E+08 | ENSMUST00000217174.2 | 34715 | ENSMUSG00000111109 | Gm30933 | predicted gene, 30933 |
| chr13 | 28403808 | 28404462 | 655 | * | 4,858539 | 6,304787226 | 2,074164 | 3,039676 | 0,002368 | 1 | 0 | Distal Intergenic | 13 | 28326467 | 28335594 | 9128 | 1 | 28078 | ENSMUST00000017208.5 | 77341 | ENSMUSG00000017064 | Prl5a1 | prolactin family 5, subfamily a, member 1 |
| chr9 | 59008347 | 59009474 | 1128 | * | 4,7896 | 6,305170092 | 2,026867 | 3,110796 | 0,001866 | 1 | 0 | Distal Intergenic | 9 | 58781970 | 58943724 | 161755 | 2 | 18007 | ENSMUST00000068664.7 | -64623 | ENSMUSG00000032340 | Neo1 | neogenin |
| chr11 | 39471352 | 39472178 | 827 | * | 5,266368 | 6,30524964 | 2,017955 | 3,124574 | 0,001781 | 1 | 0 | Distal Intergenic | 11 | 39721483 | 39721627 | 145 | 1 | 1,15E+08 | ENSMUST00000157978.4 | -249305 | ENSMUSG00000088603 | Gm24192 | predicted gene, 24192 |
| chr6 | 26261523 | 26262013 | 491 | * | 4,939675 | 6,305299952 | 2,012287 | 3,1334 | 0,001728 | 1 | 0 | Distal Intergenic | 6 | 25954484 | 25958158 | 3675 | 1 | 626904 | ENSMUST00000201080.2 | 307039 | ENSMUSG00000106799 | Gm20756 | predicted gene, 20756 |
| chr15 | 83337662 | 83338338 | 677 | * | 4,904343 | 6,305375427 | 2,003393 | 3,147349 | 0,001648 | 1 | 0 | Intron (ENSMUST0000 | 15 | 83274703 | 83348753 | 74051 | 2 | 23970 | ENSMUST00000229337.2 | 10415 | ENSMUSG00000016664 | Pacsin2 | protein kinase C and casein kinase substrate in neurons 2 |
| chr14 | 73831357 | 73831894 | 538 | * | 3,768541 | 6,305565987 | 2,339463 | 2,695305 | 0,007032 | 1 | 0 | Intron (ENSMUST0000 | 14 | 73827948 | 73828473 | 526 | 1 | 20916 | ENSMUST00000161082.2 | 3409 | ENSMUSG00000022110 | Sucla2 | succinate-Coenzyme A ligase, ADP-forming, beta subunit |
| chr4 | 33986974 | 33987636 | 663 | * | 5,352697 | 6,305985642 | 1,943271 | 3,245036 | 0,001174 | 1 | 0 | Distal Intergenic | 4 | 34024876 | 34041040 | 16165 | 2 | 67652 | ENSMUST00000108148.8 | 53404 | ENSMUSG00000028264 | Spaca1 | sperm acrosome associated 1 |
| chr5 | 1,26E+08 | 1,26E+08 | 463 | * | 5,477743 | 6,306167284 | 1,927833 | 3,271117 | 0,001071 | 1 | 0 | Distal Intergenic | 5 | 1,26E+08 | 1,26E+08 | 107 | 2 | 1,15E+08 | ENSMUST00000239978.1 | -14003 | ENSMUSG00000119346 | Gm23151 | predicted gene, 23151 |
| chr13 | 34104421 | 34105406 | 986 | * | 8,230876 | 6,306518228 | 1,83027 | 3,445676 | 0,00057 | 0,3894426 | 1,425929 | Promoter (<=1kb) | 13 | 34102215 | 34106042 | 3828 | 2 | 20719 | ENSMUST00000171951.2 | 636 | ENSMUSG00000060147 | Serpinb6a | serine (or cysteine) peptidase inhibitor, clade B, member 6a |
| chr3 | 1,1E+08 | 1,1E+08 | 656 | * | 6,460643 | 6,306583682 | 1,89551 | 3,327118 | 0,000877 | 1 | 0 | Intron (ENSMUST0000 | 3 | 1,1E+08 | 1,1E+08 | 304511 | 2 | 80883 | ENSMUST00000189335.2 | 133307 | ENSMUSG00000059857 | Ntng1 | netrin G1 |
| chr6 | 1,13E+08 | 1,13E+08 | 708 | * | 7,989394 | 6,309063913 | 1,813296 | 3,479334 | 0,000503 | 1 | 0 | Exon (ENSMUST00000 | 6 | 1,13E+08 | 1,13E+08 | 24108 | 1 | 72895 | ENSMUST00000150107.2 | -6287 | ENSMUSG00000034269 | Setd5 | SET domain containing 5 |
| chr18 | 62625757 | 62626429 | 673 | * | 7,260922 | 6,309287779 | 1,850479 | 3,409542 | 0,000651 | 1 | 0 | Intron (ENSMUST0000 | 18 | 62652448 | 62653800 | 1353 | 2 | 107035 | ENSMUST00000237378.2 | 27371 | ENSMUSG00000042211 | Fbxo38 | F-box protein 38 |
| chr7 | 1,27E+08 | 1,27E+08 | 1124 | * | 8,507076 | 6,310685624 | 1,914329 | 3,296552 | 0,000979 | 0,406085428 | 2,237403 | Distal Intergenic | 7 | 1,27E+08 | 1,27E+08 | 1570 | 1 | 101602 | ENSMUST00000206102.2 | 4716 | ENSMUSG00000045165 | AI467606 | expressed sequence AI467606 |
| chr2 | 25129962 | 25130444 | 483 | * | 5,466309 | 6,311340506 | 1,950765 | 3,235315 | 0,001215 | 1 | 0 | Promoter (<=1kb) | 2 | 25128830 | 25130000 | 1171 | 2 | 67859 | ENSMUST00000043379.5 | 0 | ENSMUSG00000036731 | Cysrt1 | cysteine rich tail 1 |
| chr1 | 1,9E+08 | 1,9E+08 | 440 | * | 3,331081 | 6,31178153 | 2,445754 | 2,580709 | 0,00986 | 1 | 0 | Distal Intergenic | 1 | 1,9E+08 | 1,9E+08 | 10223 | 1 | 1,03E+08 | ENSMUST00000191893.2 | 102961 | | Gm30932 | predicted gene, 30932 |
| chrX | 1,32E+08 | 1,32E+08 | 175 | * | 3,093867 | 6,311964735 | 2,427153 | 2,600564 | 0,009307 | 1 | 0 | Distal Intergenic | 20 | 1,33E+08 | 1,33E+08 | 3672 | 2 | 279653 | ENSMUST00000193485.2 | 272474 | ENSMUSG00000051323 | Pcdh19 | protocadherin 19 |
| chr3 | 13927306 | 13927704 | 399 | * | 3,305542 | 6,312033412 | 2,419593 | 2,608717 | 0,009088 | 1 | 0 | Intron (ENSMUST0000 | 3 | 14011453 | 14246184 | 234732 | 1 | 76897 | ENSMUST00000192209.6 | -83749 | ENSMUSG00000039717 | Ralyl | RALY RNA binding protein-like |
| chr11 | 59091686 | 59092207 | 522 | * | 4,91722 | 6,312101349 | 2,001236 | 3,154101 | 0,00161 | 1 | 0 | Promoter (1-2kb) | 11 | 59090655 | 59093211 | 2557 | 2 | 72203 | ENSMUST00000243015.1 | 1004 | | 2610507I01Rik | RIKEN cDNA 2610507I01 gene |
| chr11 | 20682948 | 20683568 | 621 | * | 4,938475 | 6,312583605 | 2,03701 | 3,098945 | 0,001942 | 1 | 0 | Intron (ENSMUST0000 | 11 | 20635084 | 20677607 | 42524 | 2 | 216549 | ENSMUST00000146722.9 | -5341 | ENSMUSG00000049659 | Aftph | aftiphilin |
| chr1 | 1,15E+08 | 1,15E+08 | 428 | * | 4,135616 | 6,312603873 | 2,359915 | 2,674928 | 0,007475 | 1 | 0 | Distal Intergenic | 1 | 1,16E+08 | 1,16E+08 | 15632 | 2 | 1,08E+08 | ENSMUST00000186137.7 | 601223 | | Gm50605 | predicted gene, 50605 |
| chr4 | 1,26E+08 | 1,26E+08 | 393 | * | 4,73208 | 6,312627374 | 2,040585 | 3,093538 | 0,001978 | 1 | 0 | Intron (ENSMUST0000 | 4 | 1,26E+08 | 1,26E+08 | 28617 | 2 | 214150 | ENSMUST00000151179.8 | 11961 | ENSMUSG00000028842 | Ago3 | argonaute RISC catalytic subunit 3 |
| chr3 | 30309050 | 30309702 | 653 | * | 4,68108 | 6,312764304 | 2,05276 | 3,075257 | 0,002103 | 1 | 0 | Intron (ENSMUST0000 | 3 | 30321974 | 30324096 | 2123 | 2 | 545510 | ENSMUST00000091270.3 | 14394 | | Gm10258 | predicted gene 10258 |
| chr14 | 50025354 | 50026069 | 716 | * | 4,435428 | 6,313590264 | 2,132992 | 2,959969 | 0,003077 | 1 | 0 | Exon (ENSMUST00020 | 14 | 49913409 | 50020840 | 107432 | 2 | 67419 | ENSMUST00000036972.14 | -4514 | ENSMUSG00000036242 | Armh4 | armadillo-like helical domain containing 4 |
| chr1 | 67026097 | 67026610 | 514 | * | 3,751287 | 6,313843356 | 2,253008 | 2,802406 | 0,005072 | 1 | 0 | Distal Intergenic | 1 | 67046177 | 67060224 | 14048 | 2 | 14768 | ENSMUST00000133508.2 | 33614 | ENSMUSG00000026000 | Lancl1 | LanC (bacterial lantibiotic synthetase component C)-like 1 |
| chr14 | 1,17E+08 | 1,17E+08 | 796 | * | 3,851247 | 6,314166564 | 2,228983 | 2,832757 | 0,004615 | 1 | 0 | Intron (ENSMUST0000 | 14 | 1,17E+08 | 1,18E+08 | 1049151 | 1 | 23888 | ENSMUST00000078849.12 | 140421 | ENSMUSG00000058571 | Gpc6 | glypican 6 |

| | | | | | | | | | | | | | | | | | | | | | | | |
|---|---|---|---|---|---|---|---|---|---|---|---|---|---|---|---|---|---|---|---|---|---|---|---|
| chr12 | 18864564 | 18864999 | 436 | * | 4,105887 | 6,31422506 | 2,22326 | 2,840075 | 0,00451 | 1 | 0 | Distal Intergenic | 12 | 18903841 | 18956504 | 52664 | 1 | 1,05E+08 | ENSMUST00000222888.2 | -38842 | | Gm40847 | predicted gene, 40847 |
| chr5 | 1,4E+08 | 1,4E+08 | 412 | * | 3,406004 | 6,314545892 | 2,254872 | 2,800401 | 0,005104 | 1 | 0 | Intron (ENSMUST0000( | 5 | 1,4E+08 | 1,4E+08 | 24033 | 1 | 27979 | ENSMUST00000100507.8 | 3532 | ENSMUSG00000056076 | Eif3b | eukaryotic translation initiation factor 3, subunit B |
| chr6 | 41272096 | 41272423 | 328 | * | 4,063822 | 6,31461828 | 2,264134 | 2,788977 | 0,005287 | 1 | 0 | Distal Intergenic | 6 | 41279203 | 41282466 | 3264 | 1 | 22074 | ENSMUST00000031913.5 | -6780 | ENSMUSG00000054106 | Try4 | trypsin 4 |
| chr10 | 99065161 | 99065715 | 555 | * | 4,468488 | 6,314662112 | 2,194325 | 2,877725 | 0,004006 | 1 | 0 | Promoter (<=1kb) | 10 | 99052304 | 99064768 | 12465 | 2 | 1,03E+08 | ENSMUST00000219984.2 | -393 | ENSMUSG00000112805 | Gm34574 | predicted gene, 34574 |
| chr14 | 67502900 | 67503365 | 466 | * | 3,479813 | 6,314913715 | 2,314879 | 2,727968 | 0,006373 | 1 | 0 | Intron (ENSMUST0000( | 14 | 67472093 | 67666136 | 194044 | 1 | 13592 | ENSMUST00000176029.2 | 30807 | ENSMUSG00000022053 | Ebf2 | early B cell factor 2 |
| chr11 | 1,16E+08 | 1,16E+08 | 552 | * | 3,707 | 6,314963188 | 2,32263 | 2,718884 | 0,00655 | 1 | 0 | Intron (ENSMUST0000( | 11 | 1,16E+08 | 1,16E+08 | 55263 | 2 | 14784 | ENSMUST00000106499.8 | 11822 | ENSMUSG00000059923 | Grb2 | growth factor receptor bound protein 2 |
| chr13 | 1,03E+08 | 1,03E+08 | 584 | * | 3,447079 | 6,314991539 | 2,33 | 2,710297 | 0,006722 | 1 | 0 | Distal Intergenic | 13 | 1,03E+08 | 1,03E+08 | 27099 | 2 | 1,03E+08 | ENSMUST00000225128.2 | 86594 | | Gm29927 | predicted gene, 29927 |
| chr16 | 33394900 | 33395116 | 217 | * | 3,502679 | 6,315057481 | 2,342133 | 2,696285 | 0,007012 | 1 | 0 | Intron (ENSMUST0000( | 16 | 33416932 | 33484505 | 67574 | 1 | 171286 | ENSMUST00000122314.8 | -21816 | ENSMUSG00000035506 | Slc12a8 | solute carrier family 12 (potassium/chloride transporters), member 8 |
| chr14 | 82460187 | 82460838 | 652 | * | 4,618819 | 6,315186174 | 2,160912 | 2,922463 | 0,003473 | 1 | 0 | Distal Intergenic | 14 | 81923250 | 81958568 | 35319 | 1 | 1,05E+08 | ENSMUST00000227374.2 | 536937 | | Gm41219 | predicted gene, 41219 |
| chr11 | 1,05E+08 | 1,05E+08 | 601 | * | 3,66532 | 6,315226789 | 2,372067 | 2,662331 | 0,00776 | 1 | 0 | Distal Intergenic | 11 | 1,05E+08 | 1,05E+08 | 431688 | 1 | 1,05E+08 | ENSMUST00000212287.2 | -12727 | | Gm11639 | predicted gene 11639 |
| chr3 | 1,05E+08 | 1,05E+08 | 421 | * | 3,548343 | 6,315330216 | 2,396686 | 2,635026 | 0,008413 | 1 | 0 | Promoter (<=1kb) | 3 | 1,05E+08 | 1,05E+08 | 3074 | 2 | 12340 | ENSMUST00000196924.2 | 394 | ENSMUSG00000070372 | Capza1 | capping protein (actin filament) muscle Z-line, alpha 1 |
| chr3 | 40608163 | 40608480 | 318 | * | 3,517547 | 6,315398802 | 2,411053 | 2,619353 | 0,00881 | 1 | 0 | Exon (ENSMUST00000( | 3 | 40594916 | 40659204 | 64289 | 1 | 380614 | ENSMUST00000091186.7 | 13247 | ENSMUSG00000060798 | Intu | inturned planar cell polarity protein |
| chr10 | 82675032 | 82675291 | 260 | * | 3,673449 | 6,315507862 | 2,433818 | 2,594897 | 0,009462 | 1 | 0 | Intron (ENSMUST0000( | 10 | 82669785 | 82733546 | 63762 | 1 | 50493 | ENSMUST00000219368.3 | 5247 | ENSMUSG00000020250 | Txnrd1 | thioredoxin reductase 1 |
| chr1 | 1,86E+08 | 1,86E+08 | 876 | * | 5,041411 | 6,315631241 | 2,134958 | 2,958199 | 0,003094 | 1 | 0 | Promoter (<=1kb) | 1 | 1,86E+08 | 1,86E+08 | 83198 | 2 | 21808 | ENSMUST00000045288.14 | 0 | ENSMUSG00000039239 | Tgfb2 | transforming growth factor, beta 2 |
| chr19 | 46954582 | 46955245 | 664 | * | 4,412511 | 6,315735241 | 2,128809 | 2,966793 | 0,003009 | 1 | 0 | Promoter (2-3kb) | 19 | 46875270 | 46957913 | 82644 | 2 | 76952 | ENSMUST00000086961.15 | 2668 | ENSMUSG00000025041 | Nt5c2 | 5'-nucleotidase, cytosolic II |
| chrX | 47165167 | 47165834 | 668 | * | 4,77921 | 6,315794289 | 2,12587 | 2,970923 | 0,002969 | 1 | 0 | Distal Intergenic | 20 | 47197602 | 47225858 | 28257 | 1 | 170745 | ENSMUST00000077775.11 | -31768 | ENSMUSG00000037005 | Xpnpep2 | X-prolyl aminopeptidase (aminopeptidase P) 2, membrane-bound |
| chr14 | 1,14E+08 | 1,14E+08 | 931 | * | 4,493857 | 6,316299185 | 2,099061 | 3,009107 | 0,00262 | 1 | 0 | Distal Intergenic | 14 | 1,15E+08 | 1,15E+08 | 29363 | 1 | 75822 | ENSMUST00000226121.2 | -889765 | ENSMUSG00000115320 | 4930524C18Rik | RIKEN cDNA 4930524C18 gene |
| chr1 | 59120634 | 59121534 | 901 | * | 5,106835 | 6,317055775 | 2,06275 | 3,062444 | 0,002195 | 1 | 0 | Promoter (2-3kb) | 1 | 59108979 | 59123682 | 14704 | 2 | 73463 | ENSMUST00000149784.2 | 2148 | ENSMUSG00000072295 | C2cd6 | C2 calcium dependent domain containing 6 |
| chr8 | 88589049 | 88589802 | 754 | * | 7,645146 | 6,317212136 | 1,959196 | 3,22439 | 0,001262 | 1 | 0 | Intron (ENSMUST0000( | 8 | 88388440 | 88622363 | 233924 | 2 | 94187 | ENSMUST00000052250.15 | 32561 | ENSMUSG00000045333 | Zfp423 | zinc finger protein 423 |
| chrX | 62778053 | 62778548 | 496 | * | 4,634324 | 6,317424191 | 2,046247 | 3,087323 | 0,00202 | 1 | 0 | Distal Intergenic | 20 | 63195011 | 63195804 | 794 | 2 | 627470 | ENSMUST00000114682.4 | 417256 | ENSMUSG00000079579 | Gm6760 | predicted gene 6760 |
| chr19 | 4352342 | 4352906 | 565 | * | 5,253887 | 6,318064386 | 2,019991 | 3,127768 | 0,001761 | 1 | 0 | Promoter (2-3kb) | 19 | 4342595 | 4355236 | 12642 | 2 | 110355 | ENSMUST00000167511.3 | 2330 | ENSMUSG00000024858 | Grk2 | G protein-coupled receptor kinase 2 |
| chr12 | 32337171 | 32338068 | 898 | * | 6,025428 | 6,319255173 | 2,011218 | 3,142004 | 0,001678 | 1 | 0 | Distal Intergenic | 12 | 32296329 | 32298620 | 2292 | 1 | 1,03E+08 | ENSMUST00000218566.2 | 40842 | ENSMUSG00000112578 | Gm33308 | predicted gene, 33308 |
| chr13 | 95617569 | 95618119 | 551 | * | 3,99045 | 6,320571032 | 2,43619 | 2,594449 | 0,009474 | 1 | 0 | Promoter (2-3kb) | 13 | 95613776 | 95615229 | 1454 | 2 | 268686 | ENSMUST00000022186.5 | -2340 | ENSMUSG00000021679 | S100z | S100 calcium binding protein, zeta |
| chr13 | 35564090 | 35564351 | 262 | * | 3,979343 | 6,321055392 | 2,402895 | 2,6306 | 0,008523 | 1 | 0 | Distal Intergenic | 13 | 35477130 | 35477260 | 131 | 2 | 1,15E+08 | ENSMUST00000104413.3 | -86830 | ENSMUSG00000077601 | Gm22674 | predicted gene, 22674 |
| chr14 | 64968098 | 64968910 | 813 | * | 6,271823 | 6,321139148 | 1,917539 | 3,296485 | 0,000979 | 1 | 0 | Intron (ENSMUST0000( | 14 | 64991367 | 64991501 | 135 | 1 | 1,15E+08 | ENSMUST00000082508.3 | -22457 | ENSMUSG00000064442 | Gm26225 | predicted gene, 26225 |
| chr18 | 77155426 | 77156353 | 928 | * | 6,656435 | 6,321660799 | 2,087729 | 3,028008 | 0,002462 | 1 | 0 | Promoter (1-2kb) | 18 | 77153515 | 77240813 | 87299 | 1 | 17344 | ENSMUST00000168882.9 | 1911 | ENSMUSG00000025423 | Pias2 | protein inhibitor of activated STAT 2 |
| chr16 | 31944095 | 31944657 | 563 | * | 4,976551 | 6,322273276 | 2,109378 | 2,997222 | 0,002725 | 1 | 0 | Distal Intergenic | 16 | 31922696 | 31926873 | 4178 | 1 | 66994 | ENSMUST00000169186.3 | 21399 | ENSMUSG00000035790 | Cep19 | centrosomal protein 19 |
| chr15 | 11405119 | 11405759 | 641 | * | 4,111193 | 6,322326444 | 2,326744 | 2,717242 | 0,006583 | 1 | 0 | Distal Intergenic | 15 | 11382387 | 11399751 | 17365 | 2 | 110960 | ENSMUST00000022849.7 | -5368 | ENSMUSG00000022241 | Tars | threonyl-tRNA synthetase |
| chr8 | 70081960 | 70082807 | 848 | * | 8,557321 | 6,322543879 | 1,881143 | 3,361012 | 0,000777 | 0,447238105 | 1,360622 | Distal Intergenic | 8 | 70063508 | 70078199 | 14692 | 2 | 234362 | ENSMUST00000121886.8 | -3761 | ENSMUSG00000060427 | Zfp868 | zinc finger protein 868 |
| chrX | 43910107 | 43910481 | 375 | * | 3,083204 | 6,324292167 | 2,415824 | 2,617861 | 0,008848 | 1 | 0 | Distal Intergenic | 20 | 43875447 | 43879074 | 3628 | 2 | 245404 | ENSMUST00000060481.9 | -31033 | ENSMUSG00000045284 | Dcaf12l1 | DDB1 and CUL4 associated factor 12-like 1 |
| chrX | 5504226 | 5504738 | 513 | * | 3,894651 | 6,32442349 | 2,323103 | 2,722404 | 0,006481 | 1 | 0 | Distal Intergenic | 20 | 5581121 | 5583080 | 1960 | 1 | 434725 | ENSMUST00000105007.3 | -76383 | ENSMUSG00000095814 | Btbd35f7 | BTB domain containing 35, family member 7 |
| chr4 | 98306307 | 98306502 | 196 | * | 3,351242 | 6,324448572 | 2,310262 | 2,737546 | 0,00619 | 1 | 0 | Promoter (1-2kb) | 4 | 98304836 | 98320111 | 15276 | 1 | 12695 | ENSMUST00000136675.2 | 1471 | ENSMUSG00000061859 | Patj | PATJ, crumbs cell polarity complex component |
| chr4 | 28146784 | 28147363 | 580 | * | 4,275169 | 6,324653386 | 2,21819 | 2,851267 | 0,004355 | 1 | 0 | Distal Intergenic | 4 | 28365316 | 28365444 | 129 | 1 | 1,15E+08 | ENSMUST00000104333.2 | -217953 | ENSMUSG00000077521 | Gm25378 | predicted gene, 25378 |
| chr8 | 89715766 | 89716117 | 352 | * | 4,964603 | 6,324686155 | 2,212093 | 2,859141 | 0,004248 | 1 | 0 | Distal Intergenic | 8 | 89751363 | 89751459 | 97 | 1 | 1,02E+08 | ENSMUST00000183771.3 | -35246 | ENSMUSG00000099078 | Mir8110 | microRNA 8110 |
| chr19 | 16920854 | 16921565 | 712 | * | 3,899309 | 6,324776661 | 2,159388 | 2,928967 | 0,003401 | 1 | 0 | Distal Intergenic | 19 | 16933482 | 17033146 | 99665 | 1 | 353211 | ENSMUST00000224145.2 | -11917 | ENSMUSG00000039126 | Prune2 | prune homolog 2 |
| chr6 | 72882800 | 72883738 | 939 | * | 4,668195 | 6,324815389 | 2,136306 | 2,960632 | 0,00307 | 1 | 0 | Distal Intergenic | 6 | 72818097 | 72876962 | 58866 | 2 | 74287 | ENSMUST00000068697.11 | -5838 | ENSMUSG00000055239 | Kcmf1 | potassium channel modulatory factor 1 |
| chr1 | 66817154 | 66817927 | 774 | * | 4,539894 | 6,324864412 | 2,117727 | 2,986628 | 0,002821 | 1 | 0 | Exon (ENSMUST00001 | 1 | 66758407 | 66840369 | 81963 | 2 | 68691 | ENSMUST00000129190.8 | 22442 | ENSMUSG00000026004 | Kansl1l | KAT8 regulatory NSL complex subunit 1-like |
| chr1 | 60194010 | 60194457 | 448 | * | 4,341799 | 6,324901589 | 2,1094 | 2,998437 | 0,002714 | 1 | 0 | Distal Intergenic | 1 | 60180607 | 60187186 | 6580 | 1 | 241066 | ENSMUST00000132949.3 | 13403 | ENSMUSG00000026017 | Carf | calcium response factor |
| chr7 | 93569164 | 93569597 | 434 | * | 4,299511 | 6,324907598 | 2,108367 | 2,999908 | 0,002701 | 1 | 0 | Distal Intergenic | 7 | 93691898 | 93998607 | 306710 | 1 | 1,03E+08 | ENSMUST00000206116.2 | -122301 | | Gm32647 | predicted gene, 32647 |
| chr11 | 1,18E+08 | 1,18E+08 | 576 | * | 4,69198 | 6,324955438 | 2,087325 | 3,030173 | 0,002444 | 1 | 0 | Intron (ENSMUST0000( | 11 | 1,18E+08 | 1,18E+08 | 11669 | 1 | 74451 | ENSMUST00000155507.2 | -4334 | ENSMUSG00000017715 | Pgs1 | phosphatidylglycerophosphate synthase 1 |
| chr11 | 76398357 | 76398865 | 509 | * | 5,016849 | 6,32506726 | 2,052856 | 3,081106 | 0,002062 | 1 | 0 | Promoter (<=1kb) | 11 | 76307561 | 76399821 | 92261 | 2 | 109934 | ENSMUST00000108407.10 | 956 | ENSMUSG00000017631 | Abr | active BCR-related gene |
| chr14 | 9205950 | 9206780 | 831 | * | 4,676189 | 6,325102691 | 2,04983 | 3,085672 | 0,002031 | 1 | 0 | Exon (ENSMUST00001 | 14 | 9185133 | 9206630 | 21498 | 1 | 328354 | ENSMUST00000140967.2 | 20817 | ENSMUSG00000051729 | Gm5087 | predicted gene 5087 |
| chr15 | 6655888 | 6656861 | 974 | * | 4,713308 | 6,32513991 | 2,034171 | 3,109443 | 0,001874 | 1 | 0 | Exon (ENSMUST00001 | 15 | 6649075 | 6661873 | 12799 | 1 | 23880 | ENSMUST00000162430.2 | 6813 | ENSMUSG00000022148 | Fyb | FYN binding protein |
| chrX | 10975078 | 10976023 | 946 | * | 5,273297 | 6,325189225 | 2,02067 | 3,130243 | 0,001747 | 1 | 0 | Distal Intergenic | 20 | 11165496 | 11165996 | 501 | 1 | 1E+08 | ENSMUST00000188439.2 | -189473 | ENSMUSG00000100626 | H2al1a | H2A histone family member L1A |
| chr17 | 46157747 | 46158130 | 384 | * | 5,185356 | 6,325216079 | 2,012096 | 3,143596 | 0,001669 | 1 | 0 | Distal Intergenic | 17 | 46117130 | 46133249 | 16120 | 2 | 1,03E+08 | ENSMUST00000233237.2 | -24498 | | Gm35692 | predicted gene, 35692 |
| chr16 | 5026540 | 5027101 | 562 | * | 5,024347 | 6,325274019 | 1,998849 | 3,164458 | 0,001554 | 1 | 0 | Distal Intergenic | 16 | 5013153 | 5021876 | 8724 | 2 | 27426 | ENSMUST00000023911.11 | -4664 | ENSMUSG00000023143 | Nagpa | N-acetylglucosamine-1-phosphodiester alpha-N-acetylglucosaminidase |
| chrX | 14231428 | 14232065 | 638 | * | 5,046828 | 6,325356449 | 1,979511 | 3,195413 | 0,001396 | 1 | 0 | Distal Intergenic | 20 | 14181640 | 14181830 | 191 | 2 | 1,15E+08 | ENSMUST00000082838.3 | -49598 | ENSMUSG00000064772 | Gm25202 | predicted gene, 25202 |
| chr11 | 87007686 | 87008655 | 970 | * | 6,521144 | 6,325641787 | 1,918898 | 3,296497 | 0,000979 | 1 | 0 | Promoter (<=1kb) | 11 | 87006695 | 87012933 | 6239 | 1 | 66140 | ENSMUST00000131202.2 | 991 | ENSMUSG00000020492 | Ska2 | spindle and kinetochore associated complex subunit 2 |
| chr5 | 32652891 | 32653734 | 844 | * | 5,90043 | 6,325666911 | 1,917896 | 3,298233 | 0,000973 | 1 | 0 | Intron (ENSMUST0000( | 5 | 32643314 | 32650100 | 6787 | 1 | 19046 | ENSMUST00000201880.2 | 9577 | ENSMUSG00000014956 | Ppp1cb | protein phosphatase 1 catalytic subunit beta |
| chr9 | 70646917 | 70647793 | 877 | * | 6,030801 | 6,325797195 | 1,89335 | 3,34106 | 0,000835 | 1 | 0 | Exon (ENSMUST00001 | 9 | 70683335 | 70685660 | 2326 | 1 | 11487 | ENSMUST00000145377.2 | -35542 | ENSMUSG00000054693 | Adam10 | a disintegrin and metallopeptidase domain 10 |
| chr2 | 29984304 | 29984756 | 453 | * | 5,828596 | 6,325834312 | 1,887338 | 3,351723 | 0,000803 | 1 | 0 | Promoter (<=1kb) | 2 | 29980956 | 29983660 | 2705 | 2 | 66220 | ENSMUST00000102865.11 | -644 | ENSMUSG00000015335 | Zdhhc12 | zinc finger, DHHC domain containing 12 |
| chr9 | 54750511 | 54751226 | 716 | * | 7,237244 | 6,32623114 | 1,829281 | 3,458316 | 0,000544 | 1 | 0 | Distal Intergenic | 9 | 54771073 | 54819814 | 48742 | 1 | 64602 | ENSMUST00000034843.9 | -19847 | ENSMUSG00000032293 | Ireb2 | iron responsive element binding protein 2 |
| chr11 | 45815061 | 45815828 | 768 | * | 7,118476 | 6,326265096 | 1,825589 | 3,465329 | 0,00053 | 1 | 0 | Distal Intergenic | 11 | 45819096 | 45835762 | 16667 | 2 | 72290 | ENSMUST00000129820.8 | 19934 | ENSMUSG00000044847 | Lsm11 | U7 snRNP-specific Sm-like protein LSM11 |
| chr8 | 1,21E+08 | 1,21E+08 | 809 | * | 7,612335 | 6,326405093 | 1,808886 | 3,497403 | 0,00047 | 1 | 0 | Distal Intergenic | 8 | 1,21E+08 | 1,21E+08 | 60356 | 1 | 78892 | ENSMUST00000132583.8 | -7270 | ENSMUSG00000031825 | Crispld2 | cysteine-rich secretory protein LCCL domain containing 2 |
| chr9 | 35952715 | 35953503 | 789 | * | 5,265039 | 6,32698165 | 2,358082 | 2,683105 | 0,007294 | 1 | 0 | Distal Intergenic | 9 | 36031116 | 36040075 | 8960 | 2 | 546123 | ENSMUST00000168452.3 | 86572 | ENSMUSG00000091411 | Gm5916 | predicted gene 5916 |
| chr8 | 1,27E+08 | 1,27E+08 | 439 | * | 5,956784 | 6,331041879 | 1,951484 | 3,244219 | 0,001178 | 1 | 0 | Distal Intergenic | 8 | 1,27E+08 | 1,27E+08 | 49737 | 2 | 212728 | ENSMUST00000170518.3 | -41228 | ENSMUSG00000090290 | Tarbp1 | TAR RNA binding protein 1 |
| chr8 | 1,24E+08 | 1,24E+08 | 657 | * | 5,949296 | 6,331262304 | 1,961566 | 3,227657 | 0,001248 | 1 | 0 | Distal Intergenic | 8 | 1,24E+08 | 1,24E+08 | 9956 | 1 | 69581 | ENSMUST00000045487.4 | 22923 | ENSMUSG00000039960 | Rhou | ras homolog family member U |
| chr7 | 1,32E+08 | 1,32E+08 | 658 | * | 4,959672 | 6,331897326 | 1,992833 | 3,177334 | 0,001486 | 1 | 0 | Promoter (2-3kb) | 7 | 1,32E+08 | 1,32E+08 | 25217 | 2 | 72096 | ENSMUST00000033257.15 | -2238 | ENSMUSG00000030960 | Eef1akmt2 | EEF1A lysine methyltransferase 2 |
| chr7 | 80137417 | 80138027 | 611 | * | 5,029936 | 6,33253767 | 2,02845 | 3,121861 | 0,001797 | 1 | 0 | Intron (ENSMUST0000( | 7 | 80123886 | 80131652 | 7767 | 2 | 12144 | ENSMUST00000205584.2 | -5765 | ENSMUSG00000030528 | Blm | Bloom syndrome, RecQ like helicase |
| chr9 | 44285957 | 44286686 | 730 | * | 5,068858 | 6,332832604 | 2,046399 | 3,094622 | 0,001971 | 1 | 0 | Exon (ENSMUST00000( | 9 | 44290787 | 44303666 | 12880 | 1 | 12282 | ENSMUST00000066601.13 | -4101 | ENSMUSG00000032115 | Hyou1 | hypoxia up-regulated 1 |
| chr2 | 1,35E+08 | 1,35E+08 | 338 | * | 3,487734 | 6,333051831 | 2,414746 | 2,622658 | 0,008725 | 1 | 0 | Intron (ENSMUST0000( | 2 | 1,35E+08 | 1,35E+08 | 46044 | 2 | 78805 | ENSMUST00000138303.2 | 123736 | ENSMUSG00000085247 | 4930545L23Rik | RIKEN cDNA 4930545L23 gene |
| chr5 | 54118038 | 54118543 | 506 | * | 4,789884 | 6,333094484 | 2,062333 | 3,07084 | 0,002135 | 1 | 0 | Distal Intergenic | 5 | 54155841 | 54278399 | 122559 | 1 | 116873 | ENSMUST00000117661.9 | -37298 | ENSMUSG00000039156 | Stim2 | stromal interaction molecule 2 |
| chr8 | 71434998 | 71435463 | 466 | * | 3,850332 | 6,333364772 | 2,373917 | 2,667897 | 0,007633 | 1 | 0 | Intron (ENSMUST0000( | 8 | 71479240 | 71578962 | 99723 | 1 | 665858 | ENSMUST00000245061.1 | -43777 | | Gm7827 | predicted gene 7827 |
| chr8 | 66621130 | 66621579 | 450 | * | 4,062635 | 6,333592054 | 2,346935 | 2,698665 | 0,006962 | 1 | 0 | Intron (ENSMUST0000( | 8 | 66574394 | 66924277 | 349884 | 1 | 72925 | ENSMUST00000178982.8 | 46736 | ENSMUSG00000036469 | Marchf1 | membrane associated ring-CH-type finger 1 |
| chr1 | 92570510 | 92571259 | 750 | * | 5,255044 | 6,333760303 | 2,110604 | 3,000924 | 0,002692 | 1 | 0 | Promoter (<=1kb) | 1 | 92567092 | 92569707 | 2616 | 2 | 66915 | ENSMUST00000129730.2 | -803 | ENSMUSG00000073616 | Cops9 | COP9 signalosome subunit 9 |
| chr17 | 31529822 | 31530883 | 1062 | * | 8,982831 | 6,333967146 | 1,947584 | 3,252217 | 0,001145 | 0,504427013 | 1,425929 | Intron (ENSMUST0000( | 17 | 31524849 | 31569669 | 44821 | 1 | 224674 | ENSMUST00000235564.2 | 4973 | ENSMUSG00000024036 | Slc37a1 | solute carrier family 37 (glycerol-3-phosphate transporter), member 1 |
| chr2 | 30592546 | 30592780 | 235 | * | 4,252876 | 6,334012928 | 2,296059 | 2,758645 | 0,005804 | 1 | 0 | Distal Intergenic | 2 | 30603162 | 30607055 | 3894 | 2 | 1E+08 | ENSMUST00000131207.2 | 14275 | | Gm14488 | predicted gene 14488 |
| chr6 | 1,41E+08 | 1,41E+08 | 572 | * | 3,955563 | 6,334052477 | 2,292425 | 2,763035 | 0,005727 | 1 | 0 | Promoter (1-2kb) | 6 | 1,41E+08 | 1,41E+08 | 2185 | 1 | 109135 | ENSMUST00000204528.2 | -1867 | ENSMUSG00000030231 | Plekha5 | pleckstrin homology domain containing, family A member 5 |
| chr17 | 21668822 | 21669467 | 646 | * | 3,964309 | 6,33455572 | 2,240884 | 2,826811 | 0,004701 | 1 | 0 | Promoter (1-2kb) | 17 | 21670636 | 21685849 | 15214 | 1 | 22709 | ENSMUST00000039577.8 | -1169 | ENSMUSG00000023892 | Zfp51 | zinc finger protein 51 |
| chr4 | 1,49E+08 | 1,49E+08 | 385 | * | 3,864962 | 6,334637312 | 2,234624 | 2,834767 | 0,004586 | 1 | 0 | Exon (ENSMUST00000( | 4 | 1,49E+08 | 1,49E+08 | 2624 | 2 | 16561 | ENSMUST00000140749.2 | -14019 | ENSMUSG00000063077 | Kif1b | kinesin family member 1B |
| chr11 | 1,2E+08 | 1,2E+08 | 579 | * | 3,946994 | 6,334794257 | 2,219 | 2,854797 | 0,004306 | 1 | 0 | Intron (ENSMUST0000( | 11 | 1,2E+08 | 1,2E+08 | 367 | 1 | 108100 | ENSMUST00000146960.2 | 12641 | ENSMUSG00000025372 | Baiap2 | brain-specific angiogenesis inhibitor 1-associated protein 2 |
| chr11 | 94153187 | 94153640 | 454 | * | 4,185603 | 6,335278572 | 2,250645 | 2,814873 | 0,00488 | 1 | 0 | Distal Intergenic | 11 | 94133273 | 94136831 | 3559 | 2 | 278507 | ENSMUST00000131352.2 | -16356 | ENSMUSG00000044177 | Wfikkn2 | WAP, follistatin/kazal, immunoglobulin, kunitz and netrin domain containing 2 |
| chr10 | 77027207 | 77027900 | 694 | * | 5,041229 | 6,335417281 | 2,166306 | 2,924526 | 0,00345 | 1 | 0 | Distal Intergenic | 10 | 76888012 | 77002382 | 114371 | 2 | 12822 | ENSMUST00000081654.13 | -24825 | ENSMUSG00000001435 | Col18a1 | collagen, type XVIII, alpha 1 |
| chrX | 57076481 | 57077358 | 878 | * | 4,692028 | 6,335507607 | 2,159803 | 2,933373 | 0,003353 | 1 | 0 | Promoter (<=1kb) | 20 | 57076370 | 57087096 | 10727 | 1 | 22773 | ENSMUST00000088629.4 | 111 | ENSMUSG00000067860 | Zic3 | zinc finger protein of the cerebellum 3 |
| chr11 | 38324687 | 38325238 | 552 | * | 4,327352 | 6,3358875 | 2,133196 | 2,970139 | 0,002977 | 1 | 0 | Distal Intergenic | 11 | 38384157 | 38410840 | 26684 | 2 | 1,01E+08 | ENSMUST00000133599.2 | 85602 | | Gm12130 | predicted gene 12130 |
| chr7 | 44205397 | 44205693 | 297 | * | 3,534375 | 6,335960031 | 2,33359 | 2,715113 | 0,006625 | 1 | 0 | Promoter (2-3kb) | 7 | 44200481 | 44203375 | 2895 | 2 | 22260 | ENSMUST00000132769.8 | -2022 | ENSMUSG00000060601 | Nr1h2 | nuclear receptor subfamily 1, group H, member 2 |
| chr11 | 83634519 | 83634960 | 442 | * | 4,834982 | 6,335966633 | 2,126374 | 2,979705 | 0,002885 | 1 | 0 | Promoter (2-3kb) | 11 | 83637766 | 83643468 | 5703 | 1 | 66107 | ENSMUST00000070832.3 | -2806 | ENSMUSG00000051748 | Wfdc21 | WAP four-disulfide core domain 21 |
| chrX | 62892956 | 62893759 | 804 | * | 4,457642 | 6,336322487 | 2,103612 | 3,012115 | 0,002594 | 1 | 0 | Distal Intergenic | 20 | 63195011 | 63195804 | 794 | 2 | 627470 | ENSMUST00000114682.4 | 302045 | ENSMUSG00000079579 | Gm6760 | predicted gene 6760 |
| chr11 | 44656593 | 44657300 | 708 | * | 4,803488 | 6,336595365 | 2,085459 | 3,038465 | 0,002378 | 1 | 0 | Intron (ENSMUST0000( | 11 | 44512588 | 44534286 | 21699 | 1 | 13591 | ENSMUST00000138452.2 | 144005 | ENSMUSG00000057098 | Ebf1 | early B cell factor 1 |
| chrX | 92888741 | 92889407 | 667 | * | 4,987119 | 6,337642123 | 2,027227 | 3,126262 | 0,00177 | 1 | 0 | Distal Intergenic | 20 | 92808213 | 92875807 | 67595 | 2 | 236900 | ENSMUST00000045748.7 | -12934 | ENSMUSG00000035232 | Pdk3 | pyruvate dehydrogenase kinase, isoenzyme 3 |
| chr1 | 1,94E+08 | 1,94E+08 | 644 | * | 4,78142 | 6,337643901 | 2,026586 | 3,127251 | 0,001764 | 1 | 0 | Distal Intergenic | 1 | 1,94E+08 | 1,94E+08 | 10573 | 2 | 74942 | ENSMUST00000193395.2 | 55239 | ENSMUSG00000103692 | 4930503O07Rik | RIKEN cDNA 4930503O07 gene |
| chr14 | 57520053 | 57520640 | 588 | * | 5,475795 | 6,33780427 | 2,018075 | 3,140519 | 0,001686 | 1 | 0 | Intron (ENSMUST0000( | 14 | 57570823 | 57570942 | 120 | 2 | 1,15E+08 | ENSMUST00000104309.3 | 50302 | ENSMUSG00000077497 | Gm23593 | predicted gene, 23593 |
| chr1 | 1,42E+08 | 1,42E+08 | 576 | * | 5,13864 | 6,338138773 | 2,002549 | 3,165035 | 0,001551 | 1 | 0 | Distal Intergenic | 1 | 1,41E+08 | 1,41E+08 | 107 | 1 | 1,15E+08 | ENSMUST00000239951.1 | 306360 | ENSMUSG00000118759 | Gm25457 | predicted gene, 25457 |
| chr3 | 52878575 | 52879139 | 565 | * | 5,067933 | 6,338317641 | 1,99359 | 3,179349 | 0,001476 | 1 | 0 | Distal Intergenic | 3 | 52889543 | 52902106 | 12564 | 2 | 67542 | ENSMUST00000194779.2 | 22967 | ENSMUSG00000027742 | Cog6 | component of oligomeric golgi complex 6 |
| chr3 | 88463549 | 88464190 | 642 | * | 5,657724 | 6,341446168 | 2,070084 | 3,063376 | 0,002189 | 1 | 0 | Promoter (<=1kb) | 3 | 88464010 | 88471821 | 7812 | 1 | 94232 | ENSMUST00000192687.2 | 0 | ENSMUSG00000008604 | Ubqln4 | ubiquilin 4 |
| chr18 | 61625075 | 61625629 | 555 | * | 4,168087 | 6,341824515 | 2,403989 | 2,638042 | 0,008339 | 1 | 0 | 3' UTR | 18 | 61617837 | 61622291 | 4455 | 1 | 1,08E+08 | ENSMUST00000237683.2 | 7238 | ENSMUSG00000117713 | Gm46637 | predicted gene, 46637 |
| chr2 | 50240869 | 50241325 | 457 | * | 3,787554 | 6,341915147 | 2,396627 | 2,646183 | 0,008141 | 1 | 0 | Intron (ENSMUST0000( | 2 | 50270412 | 50302622 | 32211 | 1 | 1,03E+08 | ENSMUST00000227437.2 | -29087 | | Gm48908 | predicted gene 48908 |
| chr14 | 75179329 | 75180314 | 986 | * | 5,972641 | 6,34243805 | 2,102299 | 3,016906 | 0,002554 | 1 | 0 | Intron (ENSMUST0000( | 14 | 74992115 | 75185306 | 193192 | 2 | 380916 | ENSMUST00000088970.7 | 4992 | ENSMUSG00000068015 | Lrch1 | leucine-rich repeats and calponin homology (CH) domain containing 1 |
| chr6 | 41216651 | 41216875 | 225 | * | 3,821171 | 6,342551096 | 2,351995 | 2,696669 | 0,007004 | 1 | 0 | Distal Intergenic | 6 | 41279203 | 41282466 | 3264 | 1 | 22074 | ENSMUST00000031913.5 | -62328 | ENSMUSG00000054106 | Try4 | trypsin 4 |
| chr1 | 19671961 | 19672757 | 797 | * | 3,638203 | 6,343131563 | 2,31381 | 2,741422 | 0,006117 | 1 | 0 | Distal Intergenic | 1 | 19284103 | 19289477 | 5375 | 1 | 21419 | ENSMUST00000186972.2 | 387858 | ENSMUSG00000025927 | Tfap2b | transcription factor AP-2 beta |
| chr4 | 1,5E+08 | 1,5E+08 | 764 | * | 7,694332 | 6,343320343 | 1,827691 | 3,470674 | 0,000519 | 1 | 0 | Distal Intergenic | 4 | 1,5E+08 | 1,5E+08 | 671 | 1 | 69151 | ENSMUST00000146427.2 | 14332 | ENSMUSG00000028990 | Lzic | leucine zipper and CTNNBIP1 domain containing |
| chr13 | 1,07E+08 | 1,07E+08 | 543 | * | 4,419806 | 6,343631015 | 2,283481 | 2,778053 | 0,005469 | 1 | 0 | Promoter (<=1kb) | 13 | 1,07E+08 | 1,07E+08 | 946 | 1 | 66254 | ENSMUST00000223851.2 | 0 | ENSMUSG00000021692 | Dimt1 | DIM1 dimethyladenosine transferase 1-like (S. cerevisiae) |
| chr10 | 83854811 | 83855463 | 653 | * | 4,168516 | 6,344100508 | 2,256339 | 2,811679 | 0,004928 | 1 | 0 | Distal Intergenic | 10 | 83928130 | 83932113 | 3984 | 1 | 1,03E+08 | ENSMUST00000213796.2 | -72667 | ENSMUSG00000111524 | Gm34113 | predicted gene, 34113 |
| chr6 | 92580231 | 92580973 | 743 | * | 11,19849 | 6,344269573 | 1,700451 | 3,730933 | 0,000191 | 0,315367183 | 0,738441 | Intron (ENSMUST0000( | 6 | 92347889 | 92544347 | 196459 | 2 | 243548 | ENSMUST00000113447.8 | -35884 | ENSMUSG00000030020 | Prickle2 | prickle planar cell polarity protein 2 |
| chrX | 1,62E+08 | 1,62E+08 | 993 | * | 7,321002 | 6,344636698 | 1,812594 | 3,500308 | 0,000465 | 1 | 0 | Distal Intergenic | 20 | 1,62E+08 | 1,62E+08 | 1536 | 2 | 1,03E+08 | ENSMUST00000136472.2 | 16760 | ENSMUSG00000086034 | Gm15201 | predicted gene 15201 |
| chrX | 1,03E+08 | 1,03E+08 | 1353 | * | 7,015285 | 6,344697985 | 1,840739 | 3,446821 | 0,000567 | 1 | 0 | Distal Intergenic | 20 | 1,03E+08 | 1,03E+08 | 123752 | 2 | 245555 | ENSMUST00000118314.8 | -61927 | ENSMUSG00000046449 | Nexmif | neurite extension and migration factor |
| chr10 | 12662381 | 12663312 | 932 | * | 6,485644 | 6,344757598 | 1,869063 | 3,39462 | 0,000687 | 1 | 0 | Intron (ENSMUST0000( | 10 | 12623671 | 12689318 | 65648 | 2 | 22288 | ENSMUST00000219163.2 | 26006 | ENSMUSG00000019820 | Utrn | utrophin |
| chr1 | 38171232 | 38172164 | 933 | * | 7,268657 | 6,344760632 | 1,874831 | 3,384177 | 0,000714 | 0,434643701 | 1,360622 | Promoter (<=1kb) | 1 | 38171164 | 38198973 | 27810 | 1 | 329123 | ENSMUST00000194892.2 | 68 | ENSMUSG00000102716 | Gm5099 | predicted gene 5099 |
| chr5 | 13280749 | 13281402 | 654 | * | 6,165601 | 6,34478591 | 1,886284 | 3,363644 | 0,000769 | 1 | 0 | Intron (ENSMUST0000( | 5 | 13175560 | 13574674 | 399115 | 1 | 20346 | ENSMUST00000195907.5 | 105189 | ENSMUSG00000028883 | Sema3a | sema domain, immunoglobulin domain (Ig), short basic domain, secreted, (semaphorin) 3A |
| chr6 | 83189670 | 83190205 | 536 | * | 7,092023 | 6,344818909 | 1,911505 | 3,319279 | 0,000903 | 1 | 0 | Distal Intergenic | 6 | 83196810 | 83280458 | 83649 | 1 | 232156 | ENSMUST00000113900.8 | -6605 | ENSMUSG00000068323 | Slc4a5 | solute carrier family 4, sodium bicarbonate cotransporter, member 5 |
| chr11 | 1,03E+08 | 1,03E+08 | 854 | * | 5,952067 | 6,344881454 | 1,957007 | 3,242136 | 0,001186 | 1 | 0 | Promoter (<=1kb) | 11 | 1,03E+08 | 1,03E+08 | 8502 | 1 | 268491 | ENSMUST00000155813.2 | 98 | ENSMUSG00000051455 | Meioc | meiosis specific with coiled-coil domain |

| | | | | | | | | | | | | | | | | | | | | | | | |
|---|---|---|---|---|---|---|---|---|---|---|---|---|---|---|---|---|---|---|---|---|---|---|---|
| chr17 | 75564372 | 75564927 | 556 | * | 5,366776 | 6,34488211 | 1,947922 | 3,257257 | 0,001125 | 1 | 0 | Intron (ENSMUST00000 | 17 | 75596971 | 75601364 | 4394 | 1 | 268977 | ENSMUST00000127458.2 | -32044 | ENSMUSG00000001870 | Ltbp1 | latent transforming growth factor beta binding protein 1 |
| chr2 | 1,5E+08 | 1,5E+08 | 610 | * | 4,776612 | 6,344933553 | 1,993301 | 3,183129 | 0,001457 | 1 | 0 | Intron (ENSMUST00000 | 2 | 1,5E+08 | 1,5E+08 | 14101 | 2 | 545471 | ENSMUST00000109914.2 | 4800 | ENSMUSG00000074731 | Zfp345 | zinc finger protein 345 |
| chr3 | 98135710 | 98136225 | 516 | * | 5,221533 | 6,344936111 | 2,00383 | 3,166404 | 0,001543 | 1 | 0 | Intron (ENSMUST00000 | 3 | 98129472 | 98144064 | 14593 | 1 | 67709 | ENSMUST00000029469.5 | 6238 | ENSMUSG00000027876 | Reg4 | regenerating islet-derived family, member 4 |
| chr4 | 1,29E+08 | 1,29E+08 | 618 | * | 5,042762 | 6,344988357 | 2,053609 | 3,089677 | 0,002004 | 1 | 0 | Distal Intergenic | 4 | 1,29E+08 | 1,29E+08 | 15054 | 2 | 215627 | ENSMUST00000053042.6 | 22970 | ENSMUSG00000048485 | Zbtb8b | zinc finger and BTB domain containing 8b |
| chr5 | 1E+08 | 1E+08 | 629 | * | 4,98545 | 6,34499443 | 2,062257 | 3,076723 | 0,002093 | 1 | 0 | Promoter (1-2kb) | 5 | 1E+08 | 1E+08 | 21289 | 1 | 67870 | ENSMUST00000199518.2 | 1412 | ENSMUSG00000029326 | Enoph1 | enolase-phosphatase 1 |
| chr11 | 1,04E+08 | 1,04E+08 | 562 | * | 5,019478 | 6,345004277 | 2,079804 | 3,05077 | 0,002283 | 1 | 0 | Intron (ENSMUST00000 | 11 | 1,04E+08 | 1,04E+08 | 96636 | 1 | 17762 | ENSMUST00000100347.11 | 5506 | ENSMUSG00000018411 | Mapt | microtubule-associated protein tau |
| chr6 | 51299798 | 51300222 | 425 | * | 4,39053 | 6,345011006 | 2,071641 | 3,062794 | 0,002193 | 1 | 0 | Distal Intergenic | 6 | 51260512 | 51264835 | 4324 | 2 | 1,03E+08 | ENSMUST00000204226.2 | -34963 | | Gm32479 | predicted gene, 32479 |
| chr4 | 97698575 | 97698957 | 383 | * | 4,252091 | 6,345012368 | 2,073217 | 3,060468 | 0,00221 | 1 | 0 | Intron (ENSMUST00000 | 4 | 97456370 | 97666285 | 209916 | 2 | 319865 | ENSMUST00000146447.8 | -32290 | | E130114P18Rik | RIKEN cDNA E130114P18 gene |
| chr18 | 35326767 | 35327611 | 845 | * | 4,00899 | 6,345065823 | 2,143852 | 2,959657 | 0,00308 | 1 | 0 | Intron (ENSMUST00000 | 18 | 35347983 | 35372523 | 24541 | 1 | 12385 | ENSMUST00000235449.2 | -20372 | ENSMUSG00000037815 | Ctnna1 | catenin (cadherin associated protein), alpha 1 |
| chr14 | 21330685 | 21331110 | 426 | * | 4,146339 | 6,345074419 | 2,123563 | 2,987937 | 0,002809 | 1 | 0 | Intron (ENSMUST00000 | 14 | 21368193 | 21376861 | 8669 | 1 | 11534 | ENSMUST00000223645.2 | -37083 | ENSMUSG00000039197 | Adk | adenosine kinase |
| chr7 | 63651983 | 63652445 | 463 | * | 4,35777 | 6,345091757 | 2,175931 | 2,916036 | 0,003545 | 1 | 0 | Promoter (<=1kb) | 7 | 63651603 | 63671126 | 19524 | 1 | 1,03E+08 | ENSMUST00000205726.3 | 380 | | Gm32633 | predicted gene, 32633 |
| chr14 | 27746246 | 27747074 | 829 | * | 3,759546 | 6,345167869 | 2,35839 | 2,690466 | 0,007135 | 1 | 0 | Exon (ENSMUST00002 | 14 | 27768385 | 28200281 | 431897 | 1 | 238988 | ENSMUST00000209800.2 | -21311 | ENSMUSG00000040640 | Erc2 | ELKS/RAB6-interacting/CAST family member 2 |
| chr11 | 5423925 | 5424469 | 545 | * | 3,575935 | 6,345173031 | 2,356995 | 2,69206 | 0,007101 | 1 | 0 | Distal Intergenic | 11 | 5435868 | 5445931 | 10064 | 1 | 1,03E+08 | ENSMUST00000241583.1 | -11399 | | Gm11963 | predicted gene 11963 |
| chr11 | 70045947 | 70046507 | 561 | * | 4,723544 | 6,34549464 | 2,186342 | 2,902335 | 0,003704 | 1 | 0 | Promoter (<=1kb) | 11 | 70047023 | 70061089 | 14067 | 1 | 17312 | ENSMUST00000144935.9 | -516 | ENSMUSG00000000318 | Clec10a | C-type lectin domain family 10, member A |
| chr12 | 53344692 | 53345242 | 551 | * | 5,185739 | 6,346651362 | 2,136719 | 2,970279 | 0,002975 | 1 | 0 | Intron (ENSMUST00000 | 12 | 53350427 | 53356623 | 6197 | 2 | 75576 | ENSMUST00000221214.2 | 11381 | | 1700030L22Rik | RIKEN cDNA 1700030L22 gene |
| chr7 | 88844924 | 88845501 | 578 | * | 4,595259 | 6,346925595 | 2,125933 | 2,985478 | 0,002831 | 1 | 0 | Exon (ENSMUST000000 | 7 | 88803335 | 88808825 | 5491 | 2 | 72759 | ENSMUST00000136267.8 | -36099 | ENSMUSG00000039428 | Tmem135 | transmembrane protein 135 |
| chr3 | 5282479 | 5284156 | 1678 | * | 11,47016 | 6,347426215 | 1,747768 | 3,631733 | 0,000282 | 0,338031787 | 0,918255 | Promoter (<=1kb) | 3 | 5283586 | 5478225 | 194640 | 1 | 80892 | ENSMUST00000175866.8 | 0 | ENSMUSG00000025255 | Zfhx4 | zinc finger homeodomain 4 |
| chr1 | 58807870 | 58808980 | 1111 | * | 7,476505 | 6,347439596 | 1,859414 | 3,413677 | 0,000641 | 1 | 0 | Distal Intergenic | 1 | 58834533 | 58886662 | 52130 | 1 | 12370 | ENSMUST00000027189.15 | -25553 | ENSMUSG00000026029 | Casp8 | caspase 8 |
| chr6 | 1,28E+08 | 1,28E+08 | 472 | * | 4,456444 | 6,347534534 | 2,335089 | 2,718326 | 0,006561 | 1 | 0 | Distal Intergenic | 6 | 1,28E+08 | 1,28E+08 | 1561 | 2 | 109246 | ENSMUST00000202141.2 | 9272 | ENSMUSG00000030352 | Tspan9 | tetraspanin 9 |
| chr8 | 26602879 | 26603488 | 610 | * | 6,534391 | 6,348133576 | 1,876877 | 3,382285 | 0,000719 | 1 | 0 | Intron (ENSMUST00000 | 8 | 26511449 | 26609252 | 97804 | 2 | 320191 | ENSMUST00000037182.14 | 5764 | ENSMUSG00000037234 | Hook3 | hook microtubule tethering protein 3 |
| chr18 | 67303824 | 67304552 | 729 | * | 5,325827 | 6,348623461 | 2,065144 | 3,07418 | 0,002111 | 1 | 0 | Intron (ENSMUST00000 | 18 | 67307956 | 67308062 | 107 | 2 | 1,15E+08 | ENSMUST00000240361.1 | 3510 | ENSMUSG00000119766 | Gm23512 | predicted gene, 23512 |
| chr4 | 1,4E+08 | 1,4E+08 | 612 | * | 6,432766 | 6,350736353 | 2,002044 | 3,172126 | 0,001513 | 1 | 0 | Intron (ENSMUST00000 | 4 | 1,39E+08 | 1,4E+08 | 92332 | 2 | 18509 | ENSMUST00000174681.2 | 52433 | ENSMUSG00000028736 | Pax7 | paired box 7 |
| chr11 | 75568501 | 75568938 | 438 | * | 4,782312 | 6,352323621 | 2,024943 | 3,137038 | 0,001707 | 1 | 0 | Promoter (1-2kb) | 11 | 75570085 | 75596891 | 26807 | 1 | 12928 | ENSMUST00000017920.14 | -1147 | ENSMUSG00000017776 | Crk | v-crk avian sarcoma virus CT10 oncogene homolog |
| chr18 | 11424047 | 11424574 | 528 | * | 6,905815 | 6,352407084 | 1,960541 | 3,24013 | 0,001195 | 0,523637575 | 1,360622 | Distal Intergenic | 18 | 11766333 | 11876264 | 109932 | 1 | 225182 | ENSMUST00000115861.9 | -341759 | ENSMUSG00000041238 | Rbbp8 | retinoblastoma binding protein 8, endonuclease |
| chr4 | 1,29E+08 | 1,29E+08 | 400 | * | 3,423433 | 6,353660631 | 2,433876 | 2,610511 | 0,009041 | 1 | 0 | Promoter (<=1kb) | 4 | 1,29E+08 | 1,29E+08 | 7074 | 2 | 54709 | ENSMUST00000135055.8 | 861 | ENSMUSG00000028798 | Eif3i | eukaryotic translation initiation factor 3, subunit I |
| chr18 | 30745838 | 30746283 | 446 | * | 3,189307 | 6,35390846 | 2,391083 | 2,657335 | 0,007876 | 1 | 0 | Distal Intergenic | 18 | 30405980 | 30481173 | 75194 | 1 | 225326 | ENSMUST00000131405.2 | 339858 | ENSMUSG00000033628 | Pik3c3 | phosphatidylinositol 3-kinase catalytic subunit type 3 |
| chr14 | 95200532 | 95201327 | 796 | * | 3,702001 | 6,35412915 | 2,351085 | 2,702637 | 0,006879 | 1 | 0 | Distal Intergenic | 14 | 95167137 | 95167262 | 126 | 1 | 1,15E+08 | ENSMUST00000158759.3 | 33395 | ENSMUSG00000089384 | Gm24680 | predicted gene, 24680 |
| chr11 | 1E+08 | 1E+08 | 699 | * | 4,663017 | 6,354662258 | 2,158035 | 2,944652 | 0,003233 | 1 | 0 | Distal Intergenic | 11 | 1E+08 | 1E+08 | 4387 | 2 | 16664 | ENSMUST00000007272.8 | -17757 | ENSMUSG00000045545 | Krt14 | keratin 14 |
| chr1 | 1,67E+08 | 1,67E+08 | 372 | * | 3,922733 | 6,35508821 | 2,216486 | 2,86719 | 0,004141 | 1 | 0 | 3' UTR | 1 | 1,67E+08 | 1,67E+08 | 107 | 1 | 1,15E+08 | ENSMUST00000240139.1 | -15857 | ENSMUSG00000119799 | Gm24182 | predicted gene, 24182 |
| chrX | 73330263 | 73331350 | 1088 | * | 5,10531 | 6,355191212 | 2,199736 | 2,88907 | 0,003864 | 1 | 0 | Promoter (<=1kb) | 20 | 73331271 | 73332215 | 945 | 1 | 66826 | ENSMUST00000153459.2 | 0 | ENSMUSG00000009995 | Tafazzin | tafazzin, phospholipid-lysophospholipid transacylase |
| chr4 | 1,39E+08 | 1,39E+08 | 656 | * | 4,830845 | 6,355241633 | 2,198108 | 2,891233 | 0,003837 | 1 | 0 | Intron (ENSMUST00000 | 4 | 1,39E+08 | 1,39E+08 | 88691 | 1 | 212632 | ENSMUST00000040023.16 | 18185 | ENSMUSG00000041025 | Iffo2 | intermediate filament family orphan 2 |
| chr15 | 99707235 | 99708104 | 870 | * | 9,205437 | 6,355328377 | 1,935069 | 3,28429 | 0,001022 | 0,410415129 | 2,237403 | Intron (ENSMUST00000 | 15 | 99700901 | 99701007 | 107 | 2 | 1,15E+08 | ENSMUST00000083709.3 | -6228 | ENSMUSG00000065643 | Gm25897 | predicted gene, 25897 |
| chrX | 59889234 | 59889641 | 408 | * | 3,932828 | 6,355387757 | 2,182659 | 2,911764 | 0,003594 | 1 | 0 | Distal Intergenic | 20 | 59934972 | 59937036 | 2065 | 2 | 20675 | ENSMUST00000135107.4 | 47395 | ENSMUSG00000045179 | Sox3 | SRY (sex determining region Y)-box 3 |
| chr16 | 24812598 | 24813017 | 420 | * | 4,289437 | 6,355413332 | 2,182724 | 2,911689 | 0,003595 | 1 | 0 | Distal Intergenic | 16 | 24878389 | 24887808 | 9420 | 1 | 319487 | ENSMUST00000181655.2 | -65372 | ENSMUSG00000096960 | A230028O05Rik | RIKEN cDNA A230028O05 gene |
| chr3 | 43192071 | 43192610 | 540 | * | 4,188957 | 6,355575399 | 2,165128 | 2,935427 | 0,003331 | 1 | 0 | Distal Intergenic | 3 | 41705631 | 41710927 | 5297 | 1 | 73852 | ENSMUST00000195030.2 | 1486440 | ENSMUSG00000025766 | D3Ertd751e | DNA segment, Chr 3, ERATO Doi 751, expressed |
| chr19 | 29038129 | 29038650 | 522 | * | 3,960527 | 6,355773314 | 2,142566 | 2,966431 | 0,003013 | 1 | 0 | Distal Intergenic | 19 | 29044636 | 29046918 | 2283 | 1 | 1,01E+08 | ENSMUST00000235636.3 | -5986 | ENSMUSG00000118038 | Gm9895 | predicted gene 9895 |
| chr8 | 91595082 | 91595647 | 566 | * | 3,852717 | 6,355859998 | 2,253228 | 2,82078 | 0,004791 | 1 | 0 | Intron (ENSMUST00000 | 8 | 91576295 | 91598718 | 22424 | 1 | 109151 | ENSMUST00000211795.2 | 18787 | ENSMUSG00000056608 | Chd9 | chromodomain helicase DNA binding protein 9 |
| chr2 | 1,81E+08 | 1,81E+08 | 537 | * | 4,165535 | 6,355905891 | 2,260116 | 2,812204 | 0,00492 | 1 | 0 | Promoter (<=1kb) | 2 | 1,81E+08 | 1,81E+08 | 17974 | 1 | 66314 | ENSMUST00000184588.8 | -345 | ENSMUSG00000000827 | Tpd52l2 | tumor protein D52-like 2 |
| chr17 | 60902621 | 60903070 | 450 | * | 4,869462 | 6,356598766 | 2,07102 | 3,069308 | 0,002146 | 1 | 0 | Distal Intergenic | 17 | 60793002 | 60793164 | 163 | 1 | 1,15E+08 | ENSMUST00000082798.3 | 109619 | ENSMUSG00000064732 | Gm23447 | predicted gene, 23447 |
| chr14 | 69433878 | 69434332 | 455 | * | 4,388009 | 6,356632763 | 2,332211 | 2,725582 | 0,006419 | 1 | 0 | Distal Intergenic | 14 | 69428087 | 69432111 | 4025 | 1 | 18095 | ENSMUST00000022646.9 | 5791 | ENSMUSG00000022061 | Nkx3-1 | NK3 homeobox 1 |
| chr3 | 35778126 | 35778563 | 438 | * | 4,766405 | 6,356776668 | 2,056288 | 3,091385 | 0,001992 | 1 | 0 | Distal Intergenic | 3 | 35789522 | 35789626 | 105 | 1 | 1,15E+08 | ENSMUST00000082900.3 | -10959 | ENSMUSG00000119567 | Gm25696 | predicted gene, 25696 |
| chr6 | 47810059 | 47810657 | 599 | * | 4,488766 | 6,356999492 | 2,040123 | 3,115989 | 0,001833 | 1 | 0 | Promoter (1-2kb) | 6 | 47812595 | 47820581 | 7987 | 1 | 272347 | ENSMUST00000146202.2 | -1938 | ENSMUSG00000062519 | Zfp398 | zinc finger protein 398 |
| chr14 | 92267280 | 92267967 | 688 | * | 3,871843 | 6,35713668 | 2,3887 | 2,661338 | 0,007783 | 1 | 0 | Distal Intergenic | 14 | 92605936 | 92606065 | 130 | 2 | 1,15E+08 | ENSMUST00000104110.3 | 338098 | ENSMUSG00000077298 | Gm24127 | predicted gene, 24127 |
| chr1 | 87090700 | 87091206 | 507 | * | 5,440849 | 6,35760827 | 1,998161 | 3,18173 | 0,001464 | 1 | 0 | Distal Intergenic | 1 | 87075377 | 87084243 | 8867 | 2 | 13599 | ENSMUST00000161002.8 | -6457 | ENSMUSG00000026247 | Ecel1 | endothelin converting enzyme-like 1 |
| chr12 | 10319260 | 10319911 | 652 | * | 5,354513 | 6,358115613 | 1,968232 | 3,23037 | 0,001236 | 1 | 0 | Distal Intergenic | 12 | 10419973 | 10425935 | 5963 | 1 | 70881 | ENSMUST00000143739.9 | -100062 | ENSMUSG00000020622 | Nt5c1b | 5'-nucleotidase, cytosolic IB |
| chr10 | 18379531 | 18380533 | 1003 | * | 6,787336 | 6,358780067 | 1,932667 | 3,290159 | 0,001001 | 1 | 0 | Intron (ENSMUST00000 | 10 | 18345729 | 18409637 | 63909 | 1 | 215819 | ENSMUST00000100054.4 | 33802 | ENSMUSG00000039835 | Nhsl1 | NHS-like 1 |
| chr12 | 1,1E+08 | 1,1E+08 | 667 | * | 6,509493 | 6,359455368 | 1,900934 | 3,345437 | 0,000822 | 1 | 0 | Promoter (1-2kb) | 12 | 1,1E+08 | 1,1E+08 | 6424 | 2 | 67280 | ENSMUST00000223217.2 | 1268 | ENSMUSG00000113331 | 3110009F21Rik | RIKEN cDNA 3110009F21 gene |
| chr3 | 1,46E+08 | 1,46E+08 | 835 | * | 6,731967 | 6,359988534 | 2,031008 | 3,131444 | 0,001739 | 1 | 0 | Intron (ENSMUST00000 | 3 | 1,46E+08 | 1,46E+08 | 78817 | 1 | 69219 | ENSMUST00000127193.2 | 31313 | ENSMUSG00000028194 | Ddah1 | dimethylarginine dimethylaminohydrolase 1 |
| chr10 | 26627871 | 26628618 | 748 | * | 6,959063 | 6,360030293 | 1,87694 | 3,388511 | 0,000703 | 1 | 0 | Promoter (<=1kb) | 10 | 26629417 | 26736728 | 107312 | 1 | 73910 | ENSMUST00000176060.8 | -799 | ENSMUSG00000039031 | Arhgap18 | Rho GTPase activating protein 18 |
| chr6 | 97730910 | 97731429 | 520 | * | 6,617788 | 6,360437894 | 1,86197 | 3,415971 | 0,000636 | 1 | 0 | Distal Intergenic | 6 | 97707400 | 97707506 | 107 | 1 | 1,15E+08 | ENSMUST00000239830.1 | 23510 | ENSMUSG00000118937 | Gm24565 | predicted gene, 24565 |
| chr10 | 59607946 | 59609031 | 1086 | * | 6,905677 | 6,36060261 | 1,855182 | 3,428559 | 0,000607 | 1 | 0 | Intron (ENSMUST00000 | 10 | 59598235 | 59624858 | 26624 | 1 | 216001 | ENSMUST00000172034.2 | 9711 | ENSMUSG00000020111 | Micu1 | mitochondrial calcium uptake 1 |
| chr2 | 1,49E+08 | 1,49E+08 | 1274 | * | 8,66796 | 6,360826596 | 1,847445 | 3,44304 | 0,000575 | 0,400053238 | 1,360622 | Distal Intergenic | 2 | 1,49E+08 | 1,49E+08 | 2845 | 2 | 13010 | ENSMUST00000144845.2 | -57479 | ENSMUSG00000027447 | Cst3 | cystatin C |
| chr3 | 1,05E+08 | 1,05E+08 | 451 | * | 3,403204 | 6,361928043 | 2,445413 | 2,601577 | 0,00928 | 1 | 0 | Promoter (1-2kb) | 3 | 1,05E+08 | 1,05E+08 | 19795 | 1 | 20501 | ENSMUST00000046212.2 | -1142 | ENSMUSG00000032902 | Slc16a1 | solute carrier family 16 (monocarboxylic acid transporters), member 1 |
| chr5 | 30055767 | 30056448 | 682 | * | 3,928244 | 6,362244678 | 2,413678 | 2,635912 | 0,008391 | 1 | 0 | Distal Intergenic | 5 | 29970188 | 29971685 | 1498 | 1 | 23950 | ENSMUST00000139126.2 | 85579 | ENSMUSG00000029131 | Dnajb6 | DnaJ heat shock protein family (Hsp40) member B6 |
| chrX | 1,41E+08 | 1,41E+08 | 563 | * | 5,5887 | 6,363374719 | 2,127756 | 2,99065 | 0,002784 | 1 | 0 | Distal Intergenic | 20 | 1,4E+08 | 1,41E+08 | 14266 | 2 | 16370 | ENSMUST00000067841.8 | -212031 | ENSMUSG00000054667 | Irs4 | insulin receptor substrate 4 |
| chr3 | 4831080 | 4831858 | 779 | * | 6,131365 | 6,363399235 | 1,868224 | 3,406122 | 0,000659 | 1 | 0 | Exon (ENSMUST00001 | 3 | 4862388 | 4879967 | 17580 | 1 | 1,01E+08 | ENSMUST00000245577.1 | -30530 | ENSMUSG00000098659 | 1110015O18Rik | RIKEN cDNA 1110015O18 gene |
| chr6 | 71418809 | 71419405 | 597 | * | 7,04586 | 6,363473595 | 1,875847 | 3,39232 | 0,000693 | 1 | 0 | Promoter (1-2kb) | 6 | 71365618 | 71417621 | 52004 | 2 | 68477 | ENSMUST00000002292.15 | -1188 | ENSMUSG00000002222 | Rmnd5a | required for meiotic nuclear division 5 homolog A |
| chr1 | 58594086 | 58594478 | 393 | * | 3,47691 | 6,363515235 | 2,309703 | 2,755123 | 0,005867 | 1 | 0 | Intron (ENSMUST00000 | 1 | 58566902 | 58605113 | 38212 | 2 | 213056 | ENSMUST00000187717.7 | 10635 | ENSMUSG00000038174 | Fam126b | family with sequence similarity 126, member B |
| chr13 | 78774718 | 78775184 | 467 | * | 3,49618 | 6,363568033 | 2,305175 | 2,760558 | 0,00577 | 1 | 0 | Distal Intergenic | 13 | 78631473 | 78710689 | 79217 | 1 | 1,03E+08 | ENSMUST00000224561.2 | 143245 | | Gm32067 | predicted gene, 32067 |
| chr8 | 85053709 | 85054216 | 508 | * | 6,435523 | 6,363735223 | 1,905782 | 3,339173 | 0,00084 | 1 | 0 | Distal Intergenic | 8 | 85065268 | 85366875 | 301608 | 1 | 12286 | ENSMUST00000238701.2 | -11052 | ENSMUSG00000034656 | Cacna1a | calcium channel, voltage-dependent, P/Q type, alpha 1A subunit |
| chr8 | 22981341 | 22981791 | 451 | * | 5,935124 | 6,363926443 | 1,928471 | 3,299986 | 0,000967 | 1 | 0 | Intron (ENSMUST00000 | 8 | 22996141 | 23025237 | 29097 | 1 | 20516 | ENSMUST00000209347.2 | -14350 | ENSMUSG00000037656 | Slc20a2 | solute carrier family 20, member 2 |
| chr3 | 23348155 | 23348991 | 837 | * | 7,337397 | 6,364105914 | 1,951691 | 3,260817 | 0,001111 | 1 | 0 | Distal Intergenic | 3 | 23515214 | 23575562 | 60349 | 1 | 1E+08 | ENSMUST00000194267.2 | -166223 | ENSMUSG00000102203 | 4930412E21Rik | RIKEN cDNA 4930412E21 gene |
| chr15 | 38420039 | 38420482 | 444 | * | 5,10469 | 6,364314666 | 1,984364 | 3,207231 | 0,00134 | 1 | 0 | Distal Intergenic | 15 | 38385878 | 38404398 | 18521 | 2 | 1,05E+08 | ENSMUST00000227422.2 | -15641 | | Gm41307 | predicted gene, 41307 |
| chr13 | 14218382 | 14218830 | 449 | * | 4,464763 | 6,364746895 | 2,226995 | 2,857998 | 0,004263 | 1 | 0 | Distal Intergenic | 13 | 14172538 | 14214223 | 41686 | 2 | 70430 | ENSMUST00000162326.2 | -4159 | ENSMUSG00000039233 | Tbce | tubulin-specific chaperone E |
| chr11 | 1,2E+08 | 1,2E+08 | 432 | * | 4,082503 | 6,365050383 | 2,13024 | 2,98795 | 0,002809 | 1 | 0 | Promoter (2-3kb) | 11 | 1,2E+08 | 1,2E+08 | 12956 | 2 | 18453 | ENSMUST00000026122.11 | -2316 | ENSMUSG00000025130 | P4hb | prolyl 4-hydroxylase, beta polypeptide |
| chr14 | 1,07E+08 | 1,07E+08 | 798 | * | 4,534022 | 6,365552035 | 2,181276 | 2,918269 | 0,00352 | 1 | 0 | Distal Intergenic | 14 | 1,07E+08 | 1,07E+08 | 69334 | 2 | 73631 | ENSMUST00000186311.2 | -79246 | ENSMUSG00000099794 | 1700128A07Rik | RIKEN cDNA 1700128A07 gene |
| chr9 | 1,1E+08 | 1,1E+08 | 363 | * | 3,445895 | 6,365702519 | 2,348807 | 2,710186 | 0,006725 | 1 | 0 | Intron (ENSMUST00000 | 9 | 1,1E+08 | 1,1E+08 | 4448 | 1 | 72341 | ENSMUST00000196855.2 | 3377 | ENSMUSG00000054836 | Elp6 | elongator acetyltransferase complex subunit 6 |
| chr3 | 1,1E+08 | 1,1E+08 | 361 | * | 3,760179 | 6,365725654 | 2,359712 | 2,69767 | 0,006983 | 1 | 0 | Intron (ENSMUST00000 | 3 | 1,1E+08 | 1,1E+08 | 106 | 1 | 1,15E+08 | ENSMUST00000083938.3 | 11816 | ENSMUSG00000065872 | Gm25681 | predicted gene, 25681 |
| chr7 | 1,05E+08 | 1,05E+08 | 279 | * | 2,866247 | 6,365888698 | 2,441874 | 2,606968 | 0,009135 | 1 | 0 | Promoter (<=1kb) | 7 | 1,05E+08 | 1,05E+08 | 4982 | 2 | 101867 | ENSMUST00000098148.6 | -907 | ENSMUSG00000030888 | Rrp8 | ribosomal RNA processing 8 |
| chr18 | 52634529 | 52635180 | 652 | * | 4,863927 | 6,366401914 | 2,244724 | 2,836163 | 0,004566 | 1 | 0 | Distal Intergenic | 18 | 52620522 | 52621728 | 1207 | 1 | 67222 | ENSMUST00000237434.2 | 14007 | ENSMUSG00000024528 | Srfbp1 | serum response factor binding protein 1 |
| chr3 | 81564789 | 81565705 | 917 | * | 4,952079 | 6,368130622 | 2,064214 | 3,085015 | 0,002035 | 1 | 0 | Distal Intergenic | 3 | 81709332 | 81712468 | 3137 | 1 | 320566 | ENSMUST00000193672.2 | -143627 | ENSMUSG00000103695 | A830029E22Rik | RIKEN cDNA A830029E22 gene |
| chr3 | 8363425 | 8364029 | 605 | * | 5,289011 | 6,368143762 | 2,063648 | 3,085867 | 0,00203 | 1 | 0 | Distal Intergenic | 3 | 8494218 | 8494324 | 107 | 1 | 1,15E+08 | ENSMUST00000083269.3 | -130189 | ENSMUSG00000065203 | Gm24614 | predicted gene, 24614 |
| chr14 | 70613285 | 70614222 | 938 | * | 6,864554 | 6,368208823 | 1,906249 | 3,340702 | 0,000836 | 1 | 0 | Exon (ENSMUST000000 | 14 | 70568526 | 70588874 | 20349 | 2 | 213053 | ENSMUST00000142598.2 | -24411 | ENSMUSG00000022094 | Slc39a14 | solute carrier family 39 (zinc transporter), member 14 |
| chr6 | 62313521 | 62314026 | 506 | * | 5,320253 | 6,368220158 | 2,060958 | 3,089931 | 0,002002 | 1 | 0 | Intron (ENSMUST00000 | 6 | 61933673 | 62359849 | 426177 | 1 | 232035 | ENSMUST00000232648.2 | 379848 | ENSMUSG00000039578 | Ccser1 | coiled-coil serine rich 1 |
| chr2 | 1,37E+08 | 1,37E+08 | 513 | * | 4,109897 | 6,368495851 | 2,354124 | 2,705251 | 0,006825 | 1 | 0 | Intron (ENSMUST00000 | 2 | 1,37E+08 | 1,37E+08 | 40624 | 1 | 1,03E+08 | ENSMUST00000130262.2 | 11766 | | Gm14055 | predicted gene 14055 |
| chr4 | 53781553 | 53782401 | 849 | * | 7,897441 | 6,36871157 | 1,919136 | 3,318531 | 0,000905 | 1 | 0 | Promoter (1-2kb) | 4 | 53779705 | 53788712 | 9008 | 1 | 21350 | ENSMUST00000030124.4 | 1848 | ENSMUSG00000028417 | Tal2 | T cell acute lymphocytic leukemia 2 |
| chr5 | 1,24E+08 | 1,24E+08 | 636 | * | 5,083049 | 6,368722695 | 2,042454 | 3,118173 | 0,00182 | 1 | 0 | Promoter (1-2kb) | 5 | 1,24E+08 | 1,24E+08 | 56943 | 2 | 19679 | ENSMUST00000159677.8 | -1576 | ENSMUSG00000029406 | Pitpnm2 | phosphatidylinositol transfer protein, membrane-associated 2 |
| chr13 | 6599624 | 6600258 | 635 | * | 5,493514 | 6,369094628 | 2,02909 | 3,138891 | 0,001696 | 1 | 0 | Promoter (1-2kb) | 13 | 6598271 | 6606740 | 8470 | 1 | 69617 | ENSMUST00000220454.2 | 1353 | ENSMUSG00000021193 | Pitrm1 | pitrilysin metallepetidase 1 |
| chr1 | 90263768 | 90264267 | 500 | * | 5,960274 | 6,369828058 | 2,004843 | 3,177221 | 0,001487 | 1 | 0 | Distal Intergenic | 1 | 90208180 | 90211634 | 3455 | 1 | 1,03E+08 | ENSMUST00000188640.2 | 55588 | | 2700003A03Rik | RIKEN cDNA 2700003A03 gene |
| chr19 | 4885751 | 4886501 | 751 | * | 6,416362 | 6,371593347 | 1,953503 | 3,261625 | 0,001108 | 1 | 0 | Promoter (2-3kb) | 19 | 4875575 | 4889247 | 13673 | 2 | 12460 | ENSMUST00000238136.2 | 2746 | ENSMUSG00000034108 | Ccs | copper chaperone for superoxide dismutase |
| chr1 | 1,9E+08 | 1,9E+08 | 1566 | * | 7,588687 | 6,373915096 | 1,897693 | 3,358771 | 0,000783 | 1 | 0 | Intron (ENSMUST00000 | 1 | 1,9E+08 | 1,9E+08 | 46947 | 2 | 19130 | ENSMUST00000010319.14 | 43290 | ENSMUSG00000010175 | Prox1 | prospero homeobox 1 |
| chr4 | 76405015 | 76405343 | 329 | * | 3,293716 | 6,374102154 | 2,447961 | 2,603842 | 0,009219 | 1 | 0 | Intron (ENSMUST00000 | 4 | 76369129 | 76408926 | 39798 | 1 | 1,05E+08 | ENSMUST00000209960.3 | 35886 | | Gm42303 | predicted gene, 42303 |
| chr12 | 39869616 | 39870085 | 470 | * | 3,347903 | 6,374186317 | 2,429171 | 2,624017 | 0,00869 | 1 | 0 | Distal Intergenic | 12 | 39998628 | 40000631 | 2004 | 2 | 1,05E+08 | ENSMUST00000222820.2 | 130546 | | Gm40399 | predicted gene, 40399 |
| chr3 | 1,1E+08 | 1,1E+08 | 510 | * | 3,395581 | 6,374354183 | 2,379174 | 2,67923 | 0,007379 | 1 | 0 | Distal Intergenic | 3 | 1,1E+08 | 1,1E+08 | 337 | 2 | 1,15E+08 | ENSMUST00000198642.2 | -66584 | ENSMUSG00000104944 | Gm43405 | predicted gene 43405 |
| chr18 | 17581356 | 17582133 | 778 | * | 4,631654 | 6,374564743 | 2,149978 | 2,964945 | 0,003027 | 1 | 0 | Distal Intergenic | 18 | 17367381 | 17452787 | 85407 | 2 | 1,03E+08 | ENSMUST00000234727.2 | -128569 | | Gm30960 | predicted gene, 30960 |
| chr4 | 1,38E+08 | 1,38E+08 | 717 | * | 4,01713 | 6,374665974 | 2,303158 | 2,767793 | 0,005644 | 1 | 0 | Distal Intergenic | 4 | 1,38E+08 | 1,38E+08 | 5810 | 1 | 66259 | ENSMUST00000050918.4 | 44584 | ENSMUSG00000046447 | Camk2n1 | calcium/calmodulin-dependent protein kinase II inhibitor 1 |
| chrX | 69320857 | 69321269 | 413 | * | 3,740375 | 6,374771376 | 2,283141 | 2,792106 | 0,005237 | 1 | 0 | Distal Intergenic | 20 | 69394676 | 69404697 | 10022 | 2 | 15931 | ENSMUST00000130519.2 | 83428 | ENSMUSG00000035847 | Ids | iduronate 2-sulfatase |
| chr15 | 10725940 | 10726354 | 415 | * | 4,291778 | 6,375311274 | 2,184636 | 2,918249 | 0,00352 | 1 | 0 | Intron (ENSMUST00000 | 15 | 10717937 | 10722022 | 4086 | 1 | 75259 | ENSMUST00000181739.3 | 8003 | ENSMUSG00000096971 | 4930556M19Rik | RIKEN cDNA 4930556M19 gene |
| chr4 | 8223360 | 8223818 | 459 | * | 4,163478 | 6,375314664 | 2,185519 | 2,917071 | 0,003533 | 1 | 0 | Intron (ENSMUST00000 | 4 | 8143362 | 8239041 | 95680 | 2 | 12319 | ENSMUST00000066674.8 | 15223 | ENSMUSG00000041261 | Car8 | carbonic anhydrase 8 |
| chr1 | 1,32E+08 | 1,32E+08 | 573 | * | 4,179347 | 6,375586592 | 2,142073 | 2,976363 | 0,002917 | 1 | 0 | Promoter (<=1kb) | 1 | 1,32E+08 | 1,32E+08 | 10613 | 2 | 1,15E+08 | ENSMUST00000180835.2 | 0 | ENSMUSG00000097760 | 6030442K20Rik | RIKEN cDNA 6030442K20 gene |
| chr11 | 94648617 | 94649084 | 468 | * | 5,42494 | 6,375913049 | 2,095778 | 3,042266 | 0,002348 | 1 | 0 | Distal Intergenic | 11 | 94668590 | 94673326 | 4737 | 2 | 544806 | ENSMUST00000143620.9 | 24242 | ENSMUSG00000075610 | Tmem92 | transmembrane protein 92 |
| chr7 | 98613976 | 98614415 | 440 | * | 4,732625 | 6,376053743 | 2,079708 | 3,065841 | 0,002171 | 1 | 0 | Intron (ENSMUST00000 | 7 | 98537716 | 98593112 | 55397 | 2 | 78610 | ENSMUST00000207919.2 | -20864 | ENSMUSG00000035354 | Uvrag | UV radiation resistance associated gene |
| chr17 | 11820666 | 11821270 | 605 | * | 4,651527 | 6,376156232 | 2,068753 | 3,082126 | 0,002055 | 1 | 0 | Intron (ENSMUST00000 | 17 | 12057489 | 12086448 | 28960 | 1 | 50873 | ENSMUST00000179624.8 | -236219 | ENSMUSG00000023826 | Prkn | parkin RBR E3 ubiquitin protein ligase |
| chr11 | 76713514 | 76714008 | 495 | * | 4,547177 | 6,376311154 | 2,051529 | 3,108078 | 0,001883 | 1 | 0 | Intron (ENSMUST00000 | 11 | 76696542 | 76699652 | 3111 | 2 | 12874 | ENSMUST00000132796.2 | -13862 | ENSMUSG00000020841 | Cpd | carboxypeptidase D |
| chr2 | 31285579 | 31285883 | 305 | * | 5,777903 | 6,377144441 | 1,970034 | 3,237073 | 0,001208 | 1 | 0 | Intron (ENSMUST00000 | 2 | 31290289 | 31295826 | 5538 | 1 | 665700 | ENSMUST00000130375.2 | -4406 | ENSMUSG00000055632 | Hmcn2 | hemicentin 2 |
| chr11 | 72378987 | 72379928 | 942 | * | 8,523798 | 6,377497621 | 1,830884 | 3,483288 | 0,000495 | 0,369518512 | 1,360622 | Promoter (<=1kb) | 11 | 72342464 | 72380730 | 38267 | 2 | 216892 | ENSMUST00000045303.10 | 802 | ENSMUSG00000040447 | Spns2 | spinster homolog 2 |
| chr14 | 23472700 | 23473130 | 431 | * | 5,329425 | 6,377516461 | 1,939896 | 3,287555 | 0,001011 | 1 | 0 | Intron (ENSMUST00000 | 14 | 23574930 | 23593354 | 18425 | 2 | 16531 | ENSMUST00000223847.2 | 120224 | ENSMUSG00000063142 | Kcnma1 | potassium large conductance calcium-activated channel, subfamily M, alpha member 1 |
| chr4 | 1,54E+08 | 1,54E+08 | 871 | * | 8,000074 | 6,379063055 | 1,842983 | 3,46127 | 0,000538 | 1 | 0 | Promoter (<=1kb) | 4 | 1,54E+08 | 1,54E+08 | 3406 | 1 | 97159 | ENSMUST00000133524.2 | 178 | ENSMUSG00000047613 | A430005L14Rik | RIKEN cDNA A430005L14 gene |
| chr10 | 42120973 | 42121616 | 644 | * | 6,645794 | 6,379066274 | 1,843093 | 3,461067 | 0,000538 | 1 | 0 | Intron (ENSMUST00000 | 10 | 42073550 | 42134362 | 60813 | 2 | 56484 | ENSMUST00000177542.2 | 12746 | ENSMUSG00000048756 | Foxo3 | forkhead box O3 |
| chr8 | 26465816 | 26466400 | 585 | * | 7,598437 | 6,381425994 | 1,844229 | 3,460214 | 0,00054 | 1 | 0 | Promoter (<=1kb) | 8 | 26434481 | 26466781 | 32301 | 2 | 52120 | ENSMUST00000037609.8 | 381 | ENSMUSG00000037260 | Hgsnat | heparan-alpha-glucosaminide N-acetyltransferase |
| chr8 | 10038959 | 10039716 | 758 | * | 7,11236 | 6,381435978 | 1,843926 | 3,460787 | 0,000539 | 1 | 0 | 3' UTR | 8 | 10027717 | 10042155 | 14439 | 1 | 68904 | ENSMUST00000048216.6 | 11242 | ENSMUSG00000040396 | Abhd13 | abhydrolase domain containing 13 |

| | | | | | | | | | | | | | | | | | | | | | | | |
|---|---|---|---|---|---|---|---|---|---|---|---|---|---|---|---|---|---|---|---|---|---|---|---|
| chr11 | 53153613 | 53154711 | 1099 | * | 6,519522 | 6,382093637 | 1,88079 | 3,393305 | 0,000691 | 1 | 0 | Exon (ENSMUST00000 | 11 | 53153707 | 53158068 | 4362 | 2 | 15525 | ENSMUST00000139322.8 | 3357 | ENSMUSG00000020361 | Hspa4 | heat shock protein 4 |
| chr9 | 62355699 | 62356081 | 383 | * | 3,252181 | 6,382167345 | 2,45796 | 2,59653 | 0,009417 | 1 | 0 | Promoter (<=1kb) | 9 | 62336616 | 62356430 | 19815 | 2 | 235431 | ENSMUST00000174439.2 | 349 | ENSMUSG00000041729 | Coro2b | coronin, actin binding protein, 2B |
| chr9 | 21461342 | 21461769 | 428 | * | 3,521901 | 6,382515939 | 2,417243 | 2,640411 | 0,008281 | 1 | 0 | Promoter (2-3kb) | 9 | 21458454 | 21490763 | 32310 | 1 | 59035 | ENSMUST00000147749.2 | 2888 | ENSMUSG00000032185 | Carm1 | coactivator-associated arginine methyltransferase 1 |
| chr7 | 1,23E+08 | 1,23E+08 | 293 | * | 3,464065 | 6,382639559 | 2,404751 | 2,654179 | 0,00795 | 1 | 0 | Intron (ENSMUST0000 | 7 | 1,23E+08 | 1,23E+08 | 38779 | 1 | 233833 | ENSMUST00000206458.2 | -27022 | ENSMUSG00000052707 | Tnrc6a | trinucleotide repeat containing 6a |
| chrX | 40430901 | 40431640 | 740 | * | 4,560708 | 6,383486676 | 2,319104 | 2,752565 | 0,005913 | 1 | 0 | Distal Intergenic | 20 | 40461136 | 40461391 | 256 | 1 | 1,08E+08 | ENSMUST00000121134.2 | -29496 | ENSMUSG00000083515 | Gm14577 | predicted gene 14577 |
| chr2 | 1,59E+08 | 1,59E+08 | 547 | * | 4,540072 | 6,383889427 | 2,282194 | 2,79726 | 0,005154 | 1 | 0 | Distal Intergenic | 2 | 1,59E+08 | 1,59E+08 | 132 | 1 | 1,15E+08 | ENSMUST00000239735.1 | 141785 | ENSMUSG00000119093 | Gm25090 | predicted gene, 25090 |
| chr10 | 85847866 | 85848392 | 527 | * | 4,243983 | 6,384080334 | 2,268085 | 2,814744 | 0,004882 | 1 | 0 | Promoter (<=1kb) | 10 | 85795204 | 85847759 | 52556 | 2 | 270757 | ENSMUST00000061699.12 | -107 | ENSMUSG00000050108 | Bpifc | BPI fold containing family C |
| chr18 | 10289656 | 10290165 | 510 | * | 4,383346 | 6,384374453 | 2,074036 | 3,078237 | 0,002082 | 1 | 0 | Distal Intergenic | 18 | 10325177 | 10502412 | 177236 | 1 | 381157 | ENSMUST00000173356.8 | -35012 | ENSMUSG00000042942 | Greb1l | growth regulation by estrogen in breast cancer-like |
| chr9 | 56292462 | 56292946 | 485 | * | 3,748214 | 6,384952739 | 2,150925 | 2,968468 | 0,002993 | 1 | 0 | Intron (ENSMUST0000 | 9 | 56167841 | 56325326 | 157486 | 2 | 244895 | ENSMUST00000186735.2 | 32380 | ENSMUSG00000074305 | Peak1 | pseudopodium-enriched atypical kinase 1 |
| chr11 | 3875470 | 3875771 | 302 | * | 3,938512 | 6,385227506 | 2,197636 | 2,905498 | 0,003667 | 1 | 0 | Intron (ENSMUST0000 | 11 | 3867192 | 3881789 | 14598 | 2 | 21452 | ENSMUST00000109992.8 | 6018 | ENSMUSG00000020432 | Tcn2 | transcobalamin 2 |
| chr7 | 92404070 | 92404688 | 619 | * | 4,294076 | 6,385427268 | 2,173141 | 2,93834 | 0,0033 | 1 | 0 | Intron (ENSMUST0000 | 7 | 92390875 | 92478824 | 87950 | 1 | 75985 | ENSMUST00000208470.2 | 13195 | ENSMUSG00000030643 | Rab30 | RAB30, member RAS oncogene family |
| chr14 | 32342901 | 32343461 | 561 | * | 4,526843 | 6,385546534 | 2,16477 | 2,949757 | 0,00318 | 1 | 0 | Intron (ENSMUST0000 | 14 | 32321915 | 32370559 | 48645 | 1 | 107751 | ENSMUST00000228878.2 | 20986 | ENSMUSG00000041730 | Prrxl1 | paired related homeobox protein-like 1 |
| chrX | 1,06E+08 | 1,06E+08 | 695 | * | 3,863617 | 6,385782147 | 2,307787 | 2,767059 | 0,005656 | 1 | 0 | Distal Intergenic | 20 | 1,06E+08 | 1,06E+08 | 107 | 2 | 1,15E+08 | ENSMUST00000083799.3 | -92958 | ENSMUSG00000065733 | Gm24491 | predicted gene, 24491 |
| chr3 | 88932475 | 88933069 | 595 | * | 3,973888 | 6,386033744 | 2,360183 | 2,705737 | 0,006815 | 1 | 0 | Promoter (1-2kb) | 3 | 88930490 | 88935575 | 5086 | 1 | 192195 | ENSMUST00000198846.2 | 1985 | ENSMUSG00000028053 | Ash1l | ASH1 like histone lysine methyltransferase |
| chr2 | 1,26E+08 | 1,26E+08 | 307 | * | 3,117349 | 6,38636422 | 2,460123 | 2,595953 | 0,009433 | 1 | 0 | Intron (ENSMUST0000 | 2 | 1,26E+08 | 1,26E+08 | 10732 | 1 | 26458 | ENSMUST00000150947.2 | 6695 | ENSMUSG00000027359 | Slc27a2 | solute carrier family 27 (fatty acid transporter), member 2 |
| chr14 | 53954098 | 53954699 | 602 | * | 4,596649 | 6,386726472 | 2,097913 | 3,044323 | 0,002332 | 1 | 0 | 5' UTR | 14 | 54155784 | 54198972 | 43189 | 2 | 1,03E+08 | ENSMUST00000198142.2 | 244273 | | Gm30392 | predicted gene, 30392 |
| chr4 | 1,49E+08 | 1,49E+08 | 409 | * | 5,129847 | 6,387238978 | 2,072439 | 3,081991 | 0,002056 | 1 | 0 | Distal Intergenic | 4 | 1,49E+08 | 1,49E+08 | 8273 | 2 | 71707 | ENSMUST00000156686.2 | 13161 | ENSMUSG00000047719 | Ubiad1 | UbiA prenyltransferase domain containing 1 |
| chr17 | 34017523 | 34018294 | 772 | * | 5,656573 | 6,389331168 | 1,986423 | 3,216501 | 0,001298 | 1 | 0 | Distal Intergenic | 17 | 34029493 | 34041155 | 11663 | 1 | 80880 | ENSMUST00000234426.2 | -11199 | ENSMUSG00000042099 | Kank3 | KN motif and ankyrin repeat domains 3 |
| chr4 | 1,36E+08 | 1,36E+08 | 282 | * | 3,4971 | 6,390105855 | 2,459805 | 2,59781 | 0,009382 | 1 | 0 | Distal Intergenic | 4 | 1,36E+08 | 1,36E+08 | 1896 | 1 | 15903 | ENSMUST00000008016.3 | -21859 | ENSMUSG00000007872 | Id3 | inhibitor of DNA binding 3 |
| chr8 | 1,05E+08 | 1,05E+08 | 354 | * | 4,089769 | 6,390127508 | 2,459549 | 2,598089 | 0,009374 | 1 | 0 | Promoter (1-2kb) | 8 | 1,05E+08 | 1,05E+08 | 10379 | 2 | 234664 | ENSMUST00000160913.8 | -1460 | ENSMUSG00000031878 | Nae1 | NEDD8 activating enzyme E1 subunit 1 |
| chrX | 86026092 | 86026564 | 473 | * | 5,753752 | 6,390290275 | 1,953615 | 3,271009 | 0,001072 | 1 | 0 | Intron (ENSMUST0000 | 20 | 85293862 | 85348699 | 54838 | 2 | 434903 | ENSMUST00000096378.3 | -677393 | ENSMUSG00000035427 | Mageb4 | MAGE family member B4 |
| chr18 | 77796706 | 77797669 | 964 | * | 6,630591 | 6,392534155 | 1,889862 | 3,382541 | 0,000718 | 1 | 0 | Intron (ENSMUST0000 | 18 | 77719193 | 77801633 | 82441 | 2 | 212163 | ENSMUST00000074653.6 | 3964 | ENSMUSG00000047466 | 8030462N17Rik | RIKEN cDNA 8030462N17 gene |
| chr2 | 1,17E+08 | 1,17E+08 | 465 | * | 4,47927 | 6,39346021 | 2,10635 | 3,035326 | 0,002403 | 1 | 0 | Distal Intergenic | 2 | 1,17E+08 | 1,17E+08 | 100 | 1 | 732489 | ENSMUST00000102421.3 | 24345 | ENSMUSG00000076376 | Mir674 | microRNA 674 |
| chr11 | 1,03E+08 | 1,03E+08 | 544 | * | 5,469663 | 6,394034963 | 2,226173 | 2,87221 | 0,004076 | 1 | 0 | Distal Intergenic | 11 | 1,03E+08 | 1,03E+08 | 4621 | 2 | 77132 | ENSMUST00000154798.8 | -3836 | | 2810433D01Rik | RIKEN cDNA 2810433D01 gene |
| chr8 | 1,06E+08 | 1,06E+08 | 442 | * | 3,853436 | 6,394494632 | 2,396416 | 2,668357 | 0,007622 | 1 | 0 | Promoter (1-2kb) | 8 | 1,06E+08 | 1,06E+08 | 13136 | 2 | 546100 | ENSMUST00000182046.2 | -1289 | ENSMUSG00000098120 | Gm5914 | predicted gene 5914 |
| chr14 | 1,05E+08 | 1,05E+08 | 964 | * | 7,518957 | 6,39465343 | 1,841823 | 3,471915 | 0,000517 | 1 | 0 | Intron (ENSMUST0000 | 14 | 1,05E+08 | 1,05E+08 | 327640 | 1 | 320249 | ENSMUST00000228036.2 | 98483 | ENSMUSG00000115432 | D130009I18Rik | RIKEN cDNA D130009I18 gene |
| chrX | 77427902 | 77428614 | 713 | * | 3,707022 | 6,39478805 | 2,266076 | 2,821966 | 0,004773 | 1 | 0 | Distal Intergenic | 20 | 77413249 | 77414734 | 1486 | 1 | 74910 | ENSMUST00000035626.8 | 14653 | ENSMUSG00000035651 | 4930480E11Rik | RIKEN cDNA 4930480E11 gene |
| chrX | 41135798 | 41136860 | 1063 | * | 4,555443 | 6,394898446 | 2,221035 | 2,879243 | 0,003986 | 1 | 0 | Distal Intergenic | 20 | 41148556 | 41161514 | 12959 | 1 | 11798 | ENSMUST00000141316.8 | -11696 | ENSMUSG00000025860 | Xiap | X-linked inhibitor of apoptosis |
| chrX | 1,07E+08 | 1,07E+08 | 517 | * | 3,880779 | 6,394915587 | 2,217853 | 2,883381 | 0,003934 | 1 | 0 | Promoter (<=1kb) | 20 | 1,07E+08 | 1,07E+08 | 80320 | 1 | 213449 | ENSMUST00000101292.9 | 149 | ENSMUSG00000073007 | Tent5d | terminal nucleotidyltransferase 5D |
| chr4 | 34000819 | 34001203 | 385 | * | 3,874154 | 6,394973626 | 2,204028 | 2,901493 | 0,003714 | 1 | 0 | Distal Intergenic | 4 | 34024876 | 34041040 | 16165 | 2 | 67652 | ENSMUST00000108148.8 | 39837 | ENSMUSG00000028264 | Spaca1 | sperm acrosome associated 1 |
| chr2 | 13057840 | 13058429 | 590 | * | 4,058775 | 6,39508943 | 2,168655 | 2,948874 | 0,003189 | 1 | 0 | Distal Intergenic | 2 | 13008268 | 13016617 | 8350 | 2 | 227580 | ENSMUST00000061545.7 | -41223 | ENSMUSG00000049630 | C1ql3 | C1q-like 3 |
| chr12 | 10211870 | 10212384 | 515 | * | 4,359856 | 6,395258756 | 2,125763 | 3,008453 | 0,002626 | 1 | 0 | Distal Intergenic | 12 | 10419973 | 10425935 | 5963 | 1 | 70881 | ENSMUST00000143739.9 | -207589 | ENSMUSG00000020622 | Nt5c1b | 5'-nucleotidase, cytosolic IB |
| chr16 | 23872772 | 23873113 | 342 | * | 4,826167 | 6,395541472 | 2,053088 | 3,115085 | 0,001839 | 1 | 0 | Intron (ENSMUST0000 | 16 | 23895138 | 23901440 | 6303 | 2 | 1,08E+08 | ENSMUST00000230580.2 | 28327 | ENSMUSG00000116437 | Gm46545 | predicted gene, 46545 |
| chr5 | 1,39E+08 | 1,39E+08 | 1110 | * | 8,734011 | 6,395557726 | 1,82404 | 3,506259 | 0,000454 | 0,285767582 | 2,237403 | Intron (ENSMUST0000 | 5 | 1,39E+08 | 1,39E+08 | 15819 | 1 | 67604 | ENSMUST00000110878.2 | 5374 | ENSMUSG00000025858 | Get4 | golgi to ER traffic protein 4 |
| chr1 | 41212456 | 41212889 | 434 | * | 4,41454 | 6,395594738 | 2,043349 | 3,129957 | 0,001748 | 1 | 0 | Intron (ENSMUST0000 | 1 | 41206747 | 41220412 | 13666 | 2 | 74897 | ENSMUST00000194002.2 | 7523 | ENSMUSG00000104072 | 4930448I06Rik | RIKEN cDNA 4930448I06 gene |
| chr9 | 1,15E+08 | 1,15E+08 | 722 | * | 5,533727 | 6,395642999 | 2,03536 | 3,142267 | 0,001676 | 1 | 0 | Distal Intergenic | 9 | 1,15E+08 | 1,15E+08 | 9017 | 1 | 67213 | ENSMUST00000217637.2 | -4441 | ENSMUSG00000032434 | Cmtm6 | CKLF-like MARVEL transmembrane domain containing 6 |
| chr15 | 81701793 | 81702252 | 460 | * | 6,712968 | 6,395895985 | 2,148907 | 2,976348 | 0,002917 | 1 | 0 | Promoter (2-3kb) | 15 | 81704794 | 81705584 | 791 | 1 | 21685 | ENSMUST00000167138.2 | -2542 | ENSMUSG00000022389 | Tef | thyrotroph embryonic factor |
| chrX | 1,62E+08 | 1,62E+08 | 709 | * | 4,141715 | 6,39633122 | 2,263071 | 2,826394 | 0,004708 | 1 | 0 | 3' UTR | 20 | 1,62E+08 | 1,62E+08 | 5564 | 2 | 67043 | ENSMUST00000134629.2 | 5460 | ENSMUSG00000031357 | Syap1 | synapse associated protein 1 |
| chr14 | 9547543 | 9548511 | 969 | * | 6,248355 | 6,396820379 | 1,86522 | 3,429527 | 0,000605 | 1 | 0 | Distal Intergenic | 14 | 9646684 | 10097198 | 450515 | 1 | 27062 | ENSMUST00000112657.9 | -98173 | ENSMUSG00000054423 | Cadps | Ca2+-dependent secretion activator |
| chr5 | 1,15E+08 | 1,15E+08 | 453 | * | 3,750381 | 6,396964745 | 2,308789 | 2,770702 | 0,005594 | 1 | 0 | Promoter (<=1kb) | 5 | 1,15E+08 | 1,15E+08 | 3500 | 1 | 74585 | ENSMUST00000128678.2 | 0 | ENSMUSG00000029550 | Sppl3 | signal peptide peptidase 3 |
| chr3 | 90123336 | 90124637 | 1302 | * | 7,816529 | 6,39743344 | 1,808277 | 3,537861 | 0,000403 | 1 | 0 | Promoter (2-3kb) | 3 | 90121002 | 90133692 | 12691 | 1 | 68328 | ENSMUST00000107373.8 | 2334 | ENSMUSG00000027935 | Rab13 | RAB13, member RAS oncogene family |
| chr11 | 98909841 | 98911654 | 1814 | * | 10,01227 | 6,398224862 | 1,753382 | 3,649076 | 0,000263 | 0,275349208 | 1,425929 | Promoter (1-2kb) | 11 | 98907419 | 98913036 | 5618 | 2 | 21973 | ENSMUST00000139730.2 | 1382 | ENSMUSG00000020914 | Top2a | topoisomerase (DNA) II alpha |
| chr1 | 1,54E+08 | 1,54E+08 | 1060 | * | 10,71163 | 6,398962076 | 1,714097 | 3,73314 | 0,000189 | 0,313797801 | 0,744375 | Distal Intergenic | 1 | 1,54E+08 | 1,54E+08 | 1074 | 1 | 14645 | ENSMUST00000154576.2 | 44390 | ENSMUSG00000026473 | Glul | glutamate-ammonia ligase (glutamine synthetase) |
| chr4 | 1,01E+08 | 1,01E+08 | 759 | * | 6,51943 | 6,400520928 | 1,881823 | 3,401234 | 0,000671 | 1 | 0 | Distal Intergenic | 4 | 1,01E+08 | 1,01E+08 | 227870 | 1 | 320508 | ENSMUST00000030257.15 | -8807 | ENSMUSG00000028532 | Cachd1 | cache domain containing 1 |
| chr15 | 8223581 | 8224365 | 785 | * | 5,935587 | 6,400889348 | 1,897685 | 3,372998 | 0,000744 | 1 | 0 | Promoter (<=1kb) | 15 | 8223785 | 8232090 | 8306 | 1 | 73692 | ENSMUST00000130748.2 | 0 | ENSMUSG00000039801 | Cplane1 | ciliogenesis and planar polarity effector 1 |
| chr1 | 1,35E+08 | 1,35E+08 | 788 | * | 5,733646 | 6,402221644 | 1,966785 | 3,255171 | 0,001133 | 1 | 0 | Distal Intergenic | 1 | 1,35E+08 | 1,35E+08 | 7448 | 2 | 171469 | ENSMUST00000027682.9 | -8630 | ENSMUSG00000026424 | Gpr37l1 | G protein-coupled receptor 37-like 1 |
| chr19 | 47432320 | 47432666 | 347 | * | 3,212227 | 6,402228538 | 2,466543 | 2,595628 | 0,009442 | 1 | 0 | Intron (ENSMUST0000 | 19 | 47255392 | 47452686 | 197295 | 2 | 14218 | ENSMUST00000111800.4 | 20020 | ENSMUSG00000053617 | Sh3pxd2a | SH3 and PX domains 2A |
| chr10 | 93756916 | 93757167 | 252 | * | 3,293551 | 6,402336353 | 2,450076 | 2,613117 | 0,008972 | 1 | 0 | Intron (ENSMUST0000 | 10 | 93725903 | 93732955 | 7053 | 2 | 56307 | ENSMUST00000181434.2 | -23961 | ENSMUSG00000036112 | Metap2 | methionine aminopeptidase 2 |
| chrX | 11357292 | 11357875 | 584 | * | 3,686381 | 6,402467477 | 2,428315 | 2,636589 | 0,008374 | 1 | 0 | Distal Intergenic | 20 | 11194061 | 11194390 | 330 | 1 | 236663 | ENSMUST00000091180.3 | 163231 | ENSMUSG00000069038 | H2al1j | H2A histone family member L1J |
| chr9 | 95898758 | 95899048 | 291 | * | 3,159517 | 6,40266038 | 2,405507 | 2,661667 | 0,007775 | 1 | 0 | Intron (ENSMUST0000 | 9 | 95855570 | 95860386 | 4817 | 1 | 24127 | ENSMUST00000189806.2 | 43188 | ENSMUSG00000032410 | Xrn1 | 5'-3' exoribonuclease 1 |
| chr17 | 38526877 | 38527231 | 355 | * | 3,501902 | 6,403077978 | 2,351113 | 2,723425 | 0,006461 | 1 | 0 | Distal Intergenic | 17 | 38519060 | 38520138 | 1079 | 1 | 258329 | ENSMUST00000076245.3 | 7817 | ENSMUSG00000057801 | Olfr135 | olfactory receptor 135 |
| chr18 | 61834317 | 61834702 | 386 | * | 3,699539 | 6,403190723 | 2,335164 | 2,742073 | 0,006105 | 1 | 0 | Intron (ENSMUST0000 | 18 | 61832546 | 61840684 | 8139 | 2 | 240334 | ENSMUST00000195229.2 | 5982 | ENSMUSG00000024579 | Pcyox1l | prenylcysteine oxidase 1 like |
| chrX | 51435813 | 51436611 | 799 | * | 4,421119 | 6,403536051 | 2,056859 | 3,11326 | 0,00185 | 1 | 0 | Intron (ENSMUST0000 | 20 | 51466998 | 51467058 | 61 | 2 | 1,02E+08 | ENSMUST00000196833.3 | 30447 | ENSMUSG00000105738 | Mir6384 | microRNA 6384 |
| chr10 | 76099708 | 76100245 | 538 | * | 3,968653 | 6,403838966 | 2,265389 | 2,826816 | 0,004701 | 1 | 0 | Promoter (1-2kb) | 10 | 76095263 | 76101342 | 6080 | 2 | 64451 | ENSMUST00000162524.2 | 1097 | ENSMUSG00000020231 | Dip2a | disco interacting protein 2 homolog A |
| chrX | 1,68E+08 | 1,68E+08 | 732 | * | 4,621786 | 6,403998349 | 2,097628 | 3,052972 | 0,002266 | 1 | 0 | Distal Intergenic | 20 | 1,68E+08 | 1,68E+08 | 47569 | 1 | 1,01E+08 | ENSMUST00000149873.2 | 58329 | ENSMUSG00000087159 | Gm15246 | predicted gene 15246 |
| chr10 | 62850101 | 62850594 | 494 | * | 3,711935 | 6,40420384 | 2,231386 | 2,870057 | 0,004104 | 1 | 0 | Promoter (1-2kb) | 10 | 62851210 | 62851947 | 738 | 2 | 432467 | ENSMUST00000141616.2 | 1353 | ENSMUSG00000020069 | Hnrnph3 | heterogeneous nuclear ribonucleoprotein H3 |
| chr11 | 83938377 | 83938843 | 467 | * | 4,114497 | 6,404473183 | 2,207658 | 2,901026 | 0,003719 | 1 | 0 | Distal Intergenic | 11 | 83939240 | 83957906 | 18667 | 2 | 56405 | ENSMUST00000108101.8 | 19063 | ENSMUSG00000018648 | Dusp14 | dual specificity phosphatase 14 |
| chr4 | 1,13E+08 | 1,13E+08 | 424 | * | 3,980182 | 6,404515329 | 2,203989 | 2,905874 | 0,003662 | 1 | 0 | Exon (ENSMUST00000 | 4 | 1,13E+08 | 1,13E+08 | 482358 | 2 | 230622 | ENSMUST00000138966.9 | -342035 | ENSMUSG00000087194 | Skint6 | selection and upkeep of intraepithelial T cells 6 |
| chr2 | 1,12E+08 | 1,12E+08 | 302 | * | 4,20318 | 6,404570619 | 2,198952 | 2,912556 | 0,003585 | 1 | 0 | Promoter (<=1kb) | 2 | 1,12E+08 | 1,12E+08 | 2392 | 1 | 99010 | ENSMUST00000129503.2 | -329 | ENSMUSG00000027134 | Lpcat4 | lysophosphatidylcholine acyltransferase 4 |
| chr12 | 27227000 | 27227483 | 484 | * | 5,101985 | 6,404572805 | 2,15457 | 2,972553 | 0,002953 | 1 | 0 | Distal Intergenic | 12 | 27184079 | 27210632 | 26554 | 2 | 636791 | ENSMUST00000182592.3 | -16368 | ENSMUSG00000094002 | Gm9866 | predicted gene 9866 |
| chr3 | 1,09E+08 | 1,09E+08 | 479 | * | 4,102563 | 6,404749385 | 2,185308 | 2,930822 | 0,003381 | 1 | 0 | Intron (ENSMUST0000 | 3 | 1,09E+08 | 1,09E+08 | 7835 | 2 | 320181 | ENSMUST00000147002.3 | -3271 | ENSMUSG00000045326 | Fndc7 | fibronectin type III domain containing 7 |
| chr19 | 35378530 | 35379062 | 533 | * | 3,690534 | 6,405678495 | 2,294396 | 2,79188 | 0,00524 | 1 | 0 | Distal Intergenic | 19 | 35180635 | 35180766 | 132 | 1 | 1,15E+08 | ENSMUST00000158462.3 | 197895 | ENSMUSG00000089087 | Gm24006 | predicted gene, 24006 |
| chr3 | 34601376 | 34601903 | 528 | * | 4,268799 | 6,405746373 | 2,111332 | 3,033984 | 0,002413 | 1 | 0 | Intron (ENSMUST0000 | 3 | 34614324 | 34732144 | 117821 | 1 | 320478 | ENSMUST00000246089.1 | -12421 | ENSMUSG00000105265 | Sox2ot | SOX2 overlapping transcript (non-protein coding) |
| chr12 | 84467327 | 84468182 | 856 | * | 4,361407 | 6,405881769 | 2,327343 | 2,752444 | 0,005915 | 1 | 0 | Promoter (<=1kb) | 12 | 84466748 | 84477154 | 10407 | 1 | 72873 | ENSMUST00000153540.2 | 579 | ENSMUSG00000057265 | Bbof1 | basal body orientation factor 1 |
| chr11 | 1,02E+08 | 1,02E+08 | 677 | * | 4,167132 | 6,405968388 | 2,097077 | 3,054713 | 0,002253 | 1 | 0 | Exon (ENSMUST00000 | 11 | 1,02E+08 | 1,02E+08 | 4307 | 1 | 72053 | ENSMUST00000036376.13 | -6917 | ENSMUSG00000034757 | Tmub2 | transmembrane and ubiquitin-like domain containing 2 |
| chr12 | 60561759 | 60563030 | 1272 | * | 13,99809 | 6,407910816 | 1,620016 | 3,955461 | 7,64E-05 | 0,178046761 | 0,918255 | Distal Intergenic | 12 | 60364674 | 60364850 | 177 | 1 | 1,03E+08 | ENSMUST00000219590.2 | 197085 | ENSMUSG00000112092 | Gm32936 | predicted gene, 32936 |
| chr4 | 26570949 | 26571555 | 607 | * | 5,919621 | 6,408290281 | 1,974999 | 3,244706 | 0,001176 | 1 | 0 | Distal Intergenic | 4 | 26635820 | 26705449 | 69630 | 2 | 75360 | ENSMUST00000154459.2 | 133894 | ENSMUSG00000086257 | 4930548K13Rik | RIKEN cDNA 4930548K13 gene |
| chr2 | 1,31E+08 | 1,31E+08 | 752 | * | 5,406376 | 6,40902796 | 1,945292 | 3,294636 | 0,000985 | 1 | 0 | Distal Intergenic | 2 | 1,31E+08 | 1,31E+08 | 16434 | 2 | 11550 | ENSMUST00000103184.4 | -76344 | ENSMUSG00000027335 | Adra1d | adrenergic receptor, alpha 1d |
| chr19 | 6382977 | 6383753 | 777 | * | 6,505026 | 6,409044811 | 1,944363 | 3,296219 | 0,00098 | 1 | 0 | Promoter (1-2kb) | 19 | 6385009 | 6390910 | 5902 | 1 | 17283 | ENSMUST00000113502.10 | -1256 | ENSMUSG00000024947 | Men1 | multiple endocrine neoplasia 1 |
| chr9 | 54743045 | 54743908 | 864 | * | 7,15985 | 6,411088871 | 1,876068 | 3,417302 | 0,000632 | 1 | 0 | Distal Intergenic | 9 | 54771073 | 54819814 | 48742 | 1 | 64602 | ENSMUST00000034843.9 | -27165 | ENSMUSG00000032293 | Ireb2 | iron responsive element binding protein 2 |
| chr1 | 72254415 | 72254924 | 510 | * | 3,704294 | 6,411104776 | 2,376275 | 2,697965 | 0,006976 | 1 | 0 | Promoter (2-3kb) | 1 | 72198601 | 72251466 | 52866 | 2 | 381269 | ENSMUST00000048860.9 | -2949 | ENSMUSG00000039395 | Mreg | melanoregulin |
| chr12 | 78924384 | 78925276 | 893 | * | 6,6453 | 6,411203543 | 1,872355 | 3,424139 | 0,000617 | 1 | 0 | Intron (ENSMUST0000 | 12 | 78908904 | 78927948 | 19045 | 1 | 13665 | ENSMUST00000220382.2 | 15480 | ENSMUSG00000021116 | Eif2s1 | eukaryotic translation initiation factor 2, subunit 1 alpha |
| chr7 | 71882674 | 71882915 | 242 | * | 4,128776 | 6,412787003 | 2,266571 | 2,82929 | 0,004665 | 1 | 0 | Intron (ENSMUST0000 | 7 | 71875116 | 71879108 | 3993 | 2 | 244049 | ENSMUST00000206976.2 | -3566 | ENSMUSG00000032776 | Mctp2 | multiple C2 domains, transmembrane 2 |
| chr14 | 76064984 | 76066407 | 1424 | * | 9,752119 | 6,413515647 | 1,81365 | 3,536247 | 0,000406 | 0,359181559 | 1,173325 | Distal Intergenic | 14 | 76080819 | 76082404 | 1586 | 2 | 1E+08 | ENSMUST00000131222.2 | 15997 | | Gm4285 | predicted gene 4285 |
| chr11 | 1,15E+08 | 1,15E+08 | 269 | * | 4,119284 | 6,41379089 | 2,21094 | 2,900933 | 0,003721 | 1 | 0 | Distal Intergenic | 11 | 1,15E+08 | 1,15E+08 | 57217 | 1 | 58222 | ENSMUST00000131046.2 | -9084 | ENSMUSG00000020732 | Rab37 | RAB37, member RAS oncogene family |
| chr9 | 58470836 | 58472069 | 1234 | * | 7,975088 | 6,413850578 | 1,806626 | 3,550181 | 0,000385 | 1 | 0 | Distal Intergenic | 9 | 58431581 | 58462720 | 31140 | 2 | 102657 | ENSMUST00000165365.3 | -8116 | ENSMUSG00000035914 | Cd276 | CD276 antigen |
| chr5 | 69379208 | 69379800 | 593 | * | 3,356462 | 6,414399064 | 2,420996 | 2,649488 | 0,008061 | 1 | 0 | Exon (ENSMUST00000 | 5 | 69453552 | 69499020 | 45469 | 2 | 243043 | ENSMUST00000087231.4 | 119220 | ENSMUSG00000037653 | Kctd8 | potassium channel tetramerisation domain containing 8 |
| chrX | 48766978 | 48767508 | 531 | * | 3,25773 | 6,41441998 | 2,386223 | 2,688106 | 0,007186 | 1 | 0 | Intron (ENSMUST0000 | 20 | 48772381 | 48773340 | 960 | 1 | 236784 | ENSMUST00000096449.2 | -4873 | ENSMUSG00000071764 | Olfr1320 | olfactory receptor 1320 |
| chr14 | 3888159 | 3888956 | 798 | * | 4,09921 | 6,414474382 | 2,276011 | 2,818297 | 0,004828 | 1 | 0 | Intron (ENSMUST0000 | 14 | 3872455 | 3895333 | 22879 | 1 | 218793 | ENSMUST00000176672.2 | 15704 | ENSMUSG00000058317 | Ube2e2 | ubiquitin-conjugating enzyme E2E 2 |
| chrX | 97304640 | 97305244 | 605 | * | 3,713071 | 6,414479446 | 2,274022 | 2,820765 | 0,004791 | 1 | 0 | Intron (ENSMUST0000 | 20 | 97192375 | 97366821 | 174447 | 1 | 11835 | ENSMUST00000052837.9 | 112265 | ENSMUSG00000046532 | Ar | androgen receptor |
| chr11 | 82673754 | 82674467 | 714 | * | 3,955001 | 6,414495238 | 2,213494 | 2,897906 | 0,003757 | 1 | 0 | Promoter (1-2kb) | 11 | 82672029 | 82695024 | 22996 | 1 | 16882 | ENSMUST00000173722.8 | 1725 | ENSMUSG00000020697 | Lig3 | ligase III, DNA, ATP-dependent |
| chr1 | 98374739 | 98375157 | 419 | * | 4,18536 | 6,414550047 | 2,168531 | 2,958017 | 0,003096 | 1 | 0 | Intron (ENSMUST0000 | 1 | 98348879 | 98436972 | 88094 | 1 | 70866 | ENSMUST00000160796.2 | 25860 | ENSMUSG00000026336 | Slco6d1 | solute carrier organic anion transporter family, member 6d1 |
| chr7 | 1,04E+08 | 1,04E+08 | 532 | * | 4,55037 | 6,414583384 | 2,092801 | 3,06507 | 0,002176 | 1 | 0 | Distal Intergenic | 7 | 1,04E+08 | 1,04E+08 | 1141 | 2 | 76681 | ENSMUST00000142641.2 | 5641 | ENSMUSG00000066258 | Trim12a | tripartite motif-containing 12A |
| chr4 | 1,25E+08 | 1,25E+08 | 527 | * | 5,272911 | 6,414623882 | 2,032983 | 3,155277 | 0,001603 | 1 | 0 | Promoter (<=1kb) | 4 | 1,25E+08 | 1,25E+08 | 2230 | 1 | 230737 | ENSMUST00000147550.2 | -406 | ENSMUSG00000028869 | Gnl2 | guanine nucleotide binding protein-like 2 (nucleolar) |
| chr11 | 79492803 | 79493263 | 461 | * | 4,450637 | 6,414625422 | 2,033952 | 3,153774 | 0,001612 | 1 | 0 | Intron (ENSMUST0000 | 11 | 79485991 | 79486094 | 104 | 1 | 1,15E+08 | ENSMUST00000082552.4 | 6812 | ENSMUSG00000064486 | Gm23293 | predicted gene, 23293 |
| chr14 | 1,2E+08 | 1,2E+08 | 779 | * | 5,542788 | 6,414655896 | 1,998101 | 3,210376 | 0,001326 | 1 | 0 | Intron (ENSMUST0000 | 14 | 1,2E+08 | 1,2E+08 | 13417 | 2 | 1E+08 | ENSMUST00000226598.2 | -15388 | | Gm4681 | predicted gene 4681 |
| chr15 | 84038831 | 84039343 | 513 | * | 5,339968 | 6,414716288 | 1,961042 | 3,271076 | 0,001071 | 1 | 0 | Distal Intergenic | 15 | 84052038 | 84070407 | 18370 | 1 | 116939 | ENSMUST00000045289.6 | -12695 | ENSMUSG00000041653 | Pnpla3 | patatin-like phospholipase domain containing 3 |
| chr10 | 21086067 | 21087026 | 960 | * | 6,369815 | 6,41479749 | 1,892468 | 3,389646 | 0,0007 | 1 | 0 | Intron (ENSMUST0000 | 10 | 21053526 | 21065081 | 11556 | 2 | 1,05E+08 | ENSMUST00000213692.2 | -20986 | | Gm40608 | predicted gene, 40608 |
| chr9 | 62865901 | 62866898 | 998 | * | 6,345455 | 6,414811573 | 1,877022 | 3,417548 | 0,000632 | 1 | 0 | Intron (ENSMUST0000 | 9 | 62830189 | 62859278 | 29090 | 2 | 56469 | ENSMUST00000215455.2 | -6623 | ENSMUSG00000032405 | Pias1 | protein inhibitor of activated STAT 1 |
| chr13 | 21396525 | 21397426 | 902 | * | 6,878821 | 6,414815833 | 1,87279 | 3,425273 | 0,000614 | 1 | 0 | Distal Intergenic | 13 | 21374031 | 21374590 | 560 | 1 | 19720 | ENSMUST00000124794.2 | 22494 | ENSMUSG00000021326 | Trim27 | tripartite motif-containing 27 |
| chrX | 8876522 | 8877730 | 1209 | * | 6,691397 | 6,414839734 | 1,857126 | 3,454176 | 0,000552 | 1 | 0 | Distal Intergenic | 20 | 8899732 | 8900572 | 841 | 1 | 83457 | ENSMUST00000096509.7 | -22002 | ENSMUSG00000071815 | Fthl17e | ferritin, heavy polypeptide-like 17, member E |
| chr8 | 1,21E+08 | 1,21E+08 | 441 | * | 5,257657 | 6,415397133 | 2,136692 | 3,002491 | 0,002678 | 1 | 0 | Intron (ENSMUST0000 | 8 | 1,21E+08 | 1,21E+08 | 325179 | 1 | 382034 | ENSMUST00000180448.3 | 123673 | ENSMUSG00000031822 | Gse1 | genetic suppressor element 1, coiled-coil protein |
| chr1 | 1,83E+08 | 1,83E+08 | 442 | * | 4,831931 | 6,415927286 | 2,115372 | 3,033002 | 0,002421 | 1 | 0 | Intron (ENSMUST0000 | 1 | 1,83E+08 | 1,83E+08 | 131697 | 1 | 96935 | ENSMUST00000085724.5 | 34616 | ENSMUSG00000038576 | Susd4 | sushi domain containing 4 |
| chr19 | 26660642 | 26661481 | 840 | * | 6,179984 | 6,416242885 | 2,102763 | 3,051339 | 0,002278 | 1 | 0 | Promoter (2-3kb) | 19 | 26658410 | 26729334 | 70925 | 1 | 67155 | ENSMUST00000175935.8 | 2232 | ENSMUSG00000024921 | Smarca2 | SWI/SNF related, matrix associated, actin dependent regulator of chromatin, subfamily a, member 2 |
| chr15 | 66743479 | 66744244 | 766 | * | 5,544671 | 6,41764631 | 2,053967 | 3,124513 | 0,001781 | 1 | 0 | Distal Intergenic | 15 | 66763169 | 66795050 | 31882 | 1 | 22402 | ENSMUST00000005255.9 | -18925 | ENSMUSG00000005124 | Ccn4 | cellular communication network factor 4 |
| chr13 | 74971926 | 74972576 | 651 | * | 5,704179 | 6,418003512 | 2,042204 | 3,142684 | 0,001674 | 1 | 0 | Distal Intergenic | 13 | 74840487 | 74956929 | 116443 | 2 | 12380 | ENSMUST00000223206.2 | -14997 | ENSMUSG00000021585 | Cast | calpastatin |
| chr19 | 5382263 | 5383517 | 1255 | * | 10,86084 | 6,418328023 | 1,728964 | 3,712239 | 0,000205 | 0,256659223 | 1,2763 | Promoter (2-3kb) | 19 | 5385769 | 5394308 | 8540 | 1 | 225865 | ENSMUST00000043380.5 | -2252 | ENSMUSG00000038498 | Catsper1 | cation channel, sperm associated 1 |
| chr4 | 64762088 | 64763023 | 936 | * | 5,436313 | 6,418586026 | 2,024257 | 3,170836 | 0,00152 | 1 | 0 | Distal Intergenic | 4 | 65042411 | 65275746 | 233336 | 1 | 18491 | ENSMUST00000084501.4 | -279388 | ENSMUSG00000028370 | Pappa | pregnancy-associated plasma protein A |
| chr13 | 74221574 | 74222309 | 736 | * | 5,888894 | 6,419357853 | 1,901458 | 3,376019 | 0,000735 | 1 | 0 | Distal Intergenic | 13 | 74203077 | 74210418 | 7342 | 2 | 74470 | ENSMUST00000220483.2 | -11156 | ENSMUSG00000021572 | Cep72 | centrosomal protein 72 |
| chr16 | 57065145 | 57065750 | 606 | * | 5,624475 | 6,419927919 | 1,922964 | 3,338558 | 0,000842 | 1 | 0 | Distal Intergenic | 16 | 56989225 | 57051867 | 62643 | 2 | 67581 | ENSMUST00000023431.8 | -13278 | ENSMUSG00000022749 | Tbc1d23 | TBC1 domain family, member 23 |
| chr1 | 1,82E+08 | 1,82E+08 | 823 | * | 7,304496 | 6,420096743 | 1,929458 | 3,327409 | 0,000877 | 0,467714818 | 1,360622 | Promoter (1-2kb) | 1 | 1,82E+08 | 1,82E+08 | 53261 | 1 | 209456 | ENSMUST00000117245.2 | -1358 | ENSMUSG00000026510 | Trp53bp2 | transformation related protein 53 binding protein 2 |

| | | | | | | | | | | | | | | | | | | | | | | | |
|---|---|---|---|---|---|---|---|---|---|---|---|---|---|---|---|---|---|---|---|---|---|---|---|
| chr10 | 59784726 | 59785373 | 648 | * | 6,503788 | 6,420517189 | 1,947358 | 3,29704 | 0,000977 | 1 | 0 | Promoter (2-3kb) | 10 | 59785491 | 59787656 | 2166 | 2 | 74747 | ENSMUST00000020308.5 | 2283 | ENSMUSG00000020108 | Ddit4 | DNA-damage-inducible transcript 4 |
| chrX | 5667049 | 5668084 | 1036 | * | 5,618462 | 6,420535038 | 1,947894 | 3,296142 | 0,00098 | 1 | 0 | Distal Intergenic | 20 | 5581121 | 5583080 | 1960 | 1 | 434725 | ENSMUST00000105007.3 | 85928 | ENSMUSG00000095814 | Btbd35f7 | BTB domain containing 35, family member 7 |
| chr14 | 74358831 | 74359305 | 475 | * | 3,262399 | 6,422214872 | 2,446609 | 2,624945 | 0,008666 | 1 | 0 | Intron (ENSMUST0000 | 14 | 74592967 | 74607181 | 14215 | 1 | 74425 | ENSMUST00000226927.2 | -233662 | ENSMUSG00000115845 | 4933402J15Rik | RIKEN cDNA 4933402J15 gene |
| chr16 | 90018061 | 90018771 | 711 | * | 3,956125 | 6,422986013 | 2,315351 | 2,774087 | 0,005536 | 1 | 0 | Promoter (<=1kb) | 16 | 90017642 | 90023217 | 5576 | 1 | 20655 | ENSMUST00000232505.2 | 419 | ENSMUSG00000022982 | Sod1 | superoxide dismutase 1, soluble |
| chr18 | 54019062 | 54020102 | 1041 | * | 8,050496 | 6,423009373 | 1,915331 | 3,353473 | 0,000798 | 0,451739959 | 1,360622 | Intron (ENSMUST0000 | 18 | 53995260 | 54086545 | 91286 | 1 | 70425 | ENSMUST00000237880.2 | 23802 | ENSMUSG00000073563 | Csnk1g3 | casein kinase 1, gamma 3 |
| chr4 | 61451649 | 61451935 | 287 | * | 3,55991 | 6,423220329 | 2,28492 | 2,811136 | 0,004937 | 1 | 0 | Promoter (<=1kb) | 4 | 61452467 | 61455210 | 2744 | 1 | 1E+08 | ENSMUST00000120662.2 | -532 | | Mup-ps12 | major urinary protein, pseudogene 12 |
| chr1 | 1,91E+08 | 1,91E+08 | 520 | * | 4,025067 | 6,423397958 | 2,261152 | 2,840763 | 0,004501 | 1 | 0 | Distal Intergenic | 1 | 1,91E+08 | 1,91E+08 | 1185 | 1 | 1,01E+08 | ENSMUST00000242907.1 | 26966 | | D730003I15Rik | RIKEN cDNA D730003I15 gene |
| chr14 | 67219454 | 67219858 | 405 | * | 3,942895 | 6,423917804 | 2,198251 | 2,922286 | 0,003475 | 1 | 0 | Distal Intergenic | 14 | 67224722 | 67227290 | 2569 | 2 | 12177 | ENSMUST00000138160.8 | 7432 | ENSMUSG00000022051 | Bnip3l | BCL2/adenovirus E1B interacting protein 3-like |
| chr8 | 72035913 | 72036429 | 517 | * | 4,7608 | 6,424674113 | 2,123231 | 3,025895 | 0,002479 | 1 | 0 | Promoter (2-3kb) | 8 | 72033160 | 72039946 | 6787 | 1 | 26457 | ENSMUST00000211811.2 | 2753 | ENSMUSG00000031808 | Slc27a1 | solute carrier family 27 (fatty acid transporter), member 1 |
| chr3 | 97298172 | 97298693 | 522 | * | 4,744056 | 6,424759314 | 2,115714 | 3,036686 | 0,002392 | 1 | 0 | Distal Intergenic | 3 | 97317533 | 97318495 | 963 | 2 | 258272 | ENSMUST00000060912.4 | 19802 | ENSMUSG00000051392 | Olfr1402 | olfactory receptor 1402 |
| chr4 | 33304530 | 33305057 | 528 | * | 4,791625 | 6,425785446 | 2,035924 | 3,156201 | 0,001598 | 1 | 0 | Exon (ENSMUST00000 | 4 | 33310311 | 33502614 | 192304 | 1 | 24018 | ENSMUST00000108153.9 | -5254 | ENSMUSG00000028274 | Rngtt | RNA guanylyltransferase and 5'-phosphatase |
| chr4 | 8539976 | 8540769 | 794 | * | 5,295636 | 6,42634159 | 1,999703 | 3,213649 | 0,001311 | 1 | 0 | Intron (ENSMUST0000 | 4 | 8535644 | 8607778 | 72135 | 1 | 59021 | ENSMUST00000060232.8 | 4332 | ENSMUSG00000047187 | Rab2a | RAB2A, member RAS oncogene family |
| chr2 | 84181112 | 84181582 | 471 | * | 4,92502 | 6,42646627 | 1,993109 | 3,224342 | 0,001263 | 1 | 0 | Exon (ENSMUST00000 | 2 | 84161003 | 84205702 | 44700 | 2 | 54598 | ENSMUST00000099944.4 | 24120 | ENSMUSG00000059588 | Calcrl | calcitonin receptor-like |
| chrX | 5446098 | 5446647 | 550 | * | 3,79376 | 6,42981928 | 2,457962 | 2,615915 | 0,008899 | 1 | 0 | Distal Intergenic | 20 | 5378952 | 5381319 | 2368 | 2 | 17870 | ENSMUST00000058404.5 | -64779 | ENSMUSG00000044597 | Mycs | myc-like oncogene, s-myc protein |
| chr19 | 30533072 | 30533994 | 923 | * | 7,963979 | 6,430196679 | 1,831198 | 3,511469 | 0,000446 | 1 | 0 | Distal Intergenic | 19 | 30523263 | 30527065 | 3803 | 2 | 13380 | ENSMUST00000025803.9 | -6007 | ENSMUSG00000024868 | Dkk1 | dickkopf WNT signaling pathway inhibitor 1 |
| chr13 | 13690938 | 13692170 | 1233 | * | 6,938498 | 6,430278054 | 2,044382 | 3,145341 | 0,001659 | 1 | 0 | Distal Intergenic | 13 | 13683297 | 13684933 | 1637 | 1 | 18073 | ENSMUST00000222436.2 | 7641 | ENSMUSG00000005397 | Nid1 | nidogen 1 |
| chr3 | 1,4E+08 | 1,4E+08 | 209 | * | 3,596629 | 6,430533864 | 2,391185 | 2,689267 | 0,007161 | 1 | 0 | Intron (ENSMUST0000 | 3 | 1,41E+08 | 1,41E+08 | 3470 | 2 | 1,03E+08 | ENSMUST00000199504.2 | 187234 | | Gm32754 | predicted gene, 32754 |
| chr9 | 96602877 | 96604270 | 1394 | * | 10,25689 | 6,431082061 | 1,764251 | 3,645219 | 0,000267 | 0,275349208 | 1,449706 | Promoter (<=1kb) | 9 | 96569451 | 96602353 | 32903 | 2 | 245007 | ENSMUST00000140121.8 | -524 | ENSMUSG00000040433 | Zbtb38 | zinc finger and BTB domain containing 38 |
| chr14 | 73056067 | 73056670 | 604 | * | 3,805535 | 6,431735889 | 2,292567 | 2,805473 | 0,005024 | 1 | 0 | Intron (ENSMUST0000 | 14 | 73026793 | 73097708 | 70916 | 1 | 1,05E+08 | ENSMUST00000227565.2 | 29274 | | Gm41206 | predicted gene, 41206 |
| chrX | 95419507 | 95420210 | 704 | * | 3,864107 | 6,431804315 | 2,288098 | 2,810983 | 0,004939 | 1 | 0 | Intron (ENSMUST0000 | 20 | 95351129 | 95411525 | 60397 | 2 | 245525 | ENSMUST00000059003.5 | -7982 | ENSMUSG00000045802 | Hsf3 | heat shock transcription factor 3 |
| chr1 | 44966315 | 44966865 | 551 | * | 4,05105 | 6,432034075 | 2,27148 | 2,831649 | 0,004631 | 1 | 0 | Distal Intergenic | 1 | 44778836 | 44784109 | 5274 | 2 | 75066 | ENSMUST00000186732.2 | -182206 | ENSMUSG00000101701 | 4930521E06Rik | RIKEN cDNA 4930521E06 gene |
| chr5 | 98521074 | 98521557 | 484 | * | 5,99682 | 6,432903333 | 1,901656 | 3,382791 | 0,000718 | 1 | 0 | Intron (ENSMUST0000 | 5 | 98477229 | 98949906 | 472678 | 1 | 75784 | ENSMUST00000196339.2 | 43845 | ENSMUSG00000057816 | Cfap299 | cilia and flagella associated protein 299 |
| chr1 | 90558092 | 90558662 | 571 | * | 5,970745 | 6,432916387 | 1,90625 | 3,374644 | 0,000739 | 1 | 0 | Distal Intergenic | 1 | 90539302 | 90540344 | 1043 | 1 | 108679 | ENSMUST00000186882.2 | 18790 | ENSMUSG00000034432 | Cops8 | COP9 signalosome subunit 8 |
| chr5 | 1,11E+08 | 1,11E+08 | 608 | * | 5,999448 | 6,432947661 | 1,913178 | 3,362441 | 0,000773 | 1 | 0 | Distal Intergenic | 5 | 1,11E+08 | 1,11E+08 | 11114 | 1 | 231603 | ENSMUST00000243989.1 | -21550 | ENSMUSG00000048215 | A630023P12Rik | RIKEN cDNA A630023P12 gene |
| chr10 | 80700374 | 80700922 | 549 | * | 6,669111 | 6,433068911 | 1,940735 | 3,314759 | 0,000917 | 1 | 0 | Promoter (1-2kb) | 10 | 80699315 | 80701580 | 2266 | 1 | 73218 | ENSMUST00000219136.2 | 1059 | ENSMUSG00000035206 | Sppl2b | signal peptide peptidase like 2B |
| chr9 | 51838489 | 51839139 | 651 | * | 5,145958 | 6,433279598 | 1,993235 | 3,227556 | 0,001249 | 1 | 0 | Intron (ENSMUST0000 | 9 | 51824777 | 51842961 | 18185 | 2 | 629554 | ENSMUST00000214327.2 | 3822 | | Gm6980 | predicted gene 6980 |
| chrX | 37863321 | 37864435 | 1115 | * | 5,133231 | 6,433336171 | 2,004295 | 3,209775 | 0,001328 | 1 | 0 | Promoter (1-2kb) | 20 | 37861548 | 38051563 | 190016 | 1 | 77883 | ENSMUST00000050744.6 | 1773 | ENSMUSG00000051361 | 6030498E09Rik | RIKEN cDNA 6030498E09 gene |
| chr5 | 16906500 | 16906889 | 390 | * | 5,417578 | 6,433725284 | 2,152276 | 2,989267 | 0,002796 | 1 | 0 | Distal Intergenic | 5 | 16996808 | 17087050 | 90243 | 1 | 1,03E+08 | ENSMUST00000211738.2 | -89919 | ENSMUSG00000109903 | Gm28710 | predicted gene 28710 |
| chr9 | 39380020 | 39380569 | 550 | * | 4,62516 | 6,43385946 | 2,203968 | 2,919217 | 0,003509 | 1 | 0 | Promoter (1-2kb) | 9 | 39381076 | 39382020 | 945 | 2 | 258242 | ENSMUST00000073946.2 | 1451 | ENSMUSG00000063176 | Olfr955 | olfactory receptor 955 |
| chr14 | 1,04E+08 | 1,04E+08 | 435 | * | 3,67623 | 6,433946956 | 2,273949 | 2,829416 | 0,004663 | 1 | 0 | Intron (ENSMUST0000 | 14 | 1,04E+08 | 1,04E+08 | 99377 | 1 | 64929 | ENSMUST00000095576.5 | 56125 | ENSMUSG00000022123 | Scel | sciellin |
| chr10 | 48874264 | 48874721 | 458 | * | 4,219095 | 6,433973095 | 2,292703 | 2,806283 | 0,005012 | 1 | 0 | Distal Intergenic | 10 | 48977224 | 48980027 | 2804 | 2 | 14806 | ENSMUST00000220330.2 | 105306 | ENSMUSG00000056073 | Grik2 | glutamate receptor, ionotropic, kainate 2 (beta 2) |
| chr1 | 58657740 | 58658132 | 393 | * | 3,62651 | 6,434036932 | 2,326815 | 2,765169 | 0,005689 | 1 | 0 | Distal Intergenic | 1 | 58625543 | 58635123 | 9581 | 1 | 66495 | ENSMUST00000027193.9 | 32197 | ENSMUSG00000026032 | Ndufb3 | NADH:ubiquinone oxidoreductase subunit B3 |
| chrX | 47368333 | 47368929 | 597 | * | 4,013223 | 6,43403703 | 2,342278 | 2,746914 | 0,006016 | 1 | 0 | Intron (ENSMUST0000 | 20 | 47345795 | 47361961 | 16167 | 1 | 72554 | ENSMUST00000152624.2 | 22538 | ENSMUSG00000063785 | Utp14a | UTP14A small subunit processome component |
| chr1 | 1,66E+08 | 1,66E+08 | 510 | * | 3,30879 | 6,434141832 | 2,413891 | 2,665464 | 0,007688 | 1 | 0 | Intron (ENSMUST0000 | 1 | 1,66E+08 | 1,66E+08 | 137525 | 2 | 18986 | ENSMUST00000160260.9 | -6529 | ENSMUSG00000026565 | Pou2f1 | POU domain, class 2, transcription factor 1 |
| chr5 | 1,24E+08 | 1,24E+08 | 623 | * | 4,395767 | 6,434290346 | 2,140439 | 3,006061 | 0,002647 | 1 | 0 | Intron (ENSMUST0000 | 5 | 1,24E+08 | 1,24E+08 | 38252 | 2 | 56430 | ENSMUST00000144121.2 | 3132 | ENSMUSG00000049550 | Clip1 | CAP-GLY domain containing linker protein 1 |
| chr8 | 1,11E+08 | 1,11E+08 | 569 | * | 4,739107 | 6,434492568 | 2,131003 | 3,019467 | 0,002532 | 1 | 0 | Intron (ENSMUST0000 | 8 | 1,11E+08 | 1,11E+08 | 82453 | 1 | 11765 | ENSMUST00000034171.9 | 7466 | ENSMUSG00000031731 | Ap1g1 | adaptor protein complex AP-1, gamma 1 subunit |
| chr4 | 1,29E+08 | 1,29E+08 | 604 | * | 5,271425 | 6,434553577 | 2,127339 | 3,024696 | 0,002489 | 1 | 0 | Intron (ENSMUST0000 | 4 | 1,29E+08 | 1,29E+08 | 25293 | 2 | 16818 | ENSMUST00000102596.8 | 3272 | ENSMUSG00000000409 | Lck | lymphocyte protein tyrosine kinase |
| chr19 | 39499116 | 39499757 | 642 | * | 5,388561 | 6,435608566 | 2,080566 | 3,093201 | 0,00198 | 1 | 0 | Promoter (<=1kb) | 19 | 39499306 | 39556973 | 57668 | 1 | 13098 | ENSMUST00000025968.5 | 0 | ENSMUSG00000025003 | Cyp2c39 | cytochrome P450, family 2, subfamily c, polypeptide 39 |
| chr13 | 45630386 | 45631057 | 672 | * | 5,911158 | 6,435780945 | 2,073061 | 3,104482 | 0,001906 | 1 | 0 | Distal Intergenic | 13 | 45660920 | 45699858 | 38939 | 1 | 66355 | ENSMUST00000000260.13 | -29863 | ENSMUSG00000000253 | Gmpr | guanosine monophosphate reductase |
| chrX | 1,41E+08 | 1,41E+08 | 866 | * | 4,988911 | 6,436312305 | 2,052426 | 3,135954 | 0,001713 | 1 | 0 | Distal Intergenic | 20 | 1,41E+08 | 1,41E+08 | 432 | 2 | 1,05E+08 | ENSMUST00000117247.2 | 5738 | ENSMUSG00000081750 | Gm4995 | predicted pseudogene 4995 |
| chr5 | 1,44E+08 | 1,44E+08 | 690 | * | 6,563693 | 6,436790215 | 2,034293 | 3,164142 | 0,001555 | 1 | 0 | Promoter (<=1kb) | 5 | 1,44E+08 | 1,44E+08 | 14410 | 2 | 67277 | ENSMUST00000153130.2 | -755 | | 2900089D17Rik | RIKEN cDNA 2900089D17 gene |
| chr11 | 20172405 | 20173267 | 863 | * | 6,485214 | 6,437691178 | 1,905199 | 3,379012 | 0,000727 | 1 | 0 | Exon (ENSMUST00000 | 11 | 20151609 | 20173572 | 21964 | 1 | 19324 | ENSMUST00000152728.2 | 20796 | ENSMUSG00000020149 | Rab1a | RAB1A, member RAS oncogene family |
| chr3 | 1,04E+08 | 1,04E+08 | 696 | * | 5,438061 | 6,437780625 | 2,000943 | 3,217373 | 0,001294 | 1 | 0 | Distal Intergenic | 3 | 1,04E+08 | 1,04E+08 | 15979 | 2 | 433634 | ENSMUST00000056145.3 | 17804 | | Gm5546 | predicted gene 5546 |
| chr6 | 21846234 | 21846677 | 444 | * | 6,244549 | 6,438515539 | 1,978149 | 3,254818 | 0,001135 | 1 | 0 | Intron (ENSMUST0000 | 6 | 21799949 | 21851913 | 51965 | 2 | 269831 | ENSMUST00000134635.2 | 5236 | ENSMUSG00000029669 | Tspan12 | tetraspanin 12 |
| chr3 | 38781342 | 38781998 | 657 | * | 6,118783 | 6,43928082 | 1,956363 | 3,291455 | 0,000997 | 1 | 0 | Distal Intergenic | 3 | 38941089 | 39066134 | 125046 | 1 | 329628 | ENSMUST00000061260.8 | -159091 | ENSMUSG00000046743 | Fat4 | FAT atypical cadherin 4 |
| chrX | 1,42E+08 | 1,42E+08 | 907 | * | 6,645991 | 6,439801814 | 1,942125 | 3,315853 | 0,000914 | 1 | 0 | Distal Intergenic | 20 | 1,42E+08 | 1,42E+08 | 4704 | 1 | 209540 | ENSMUST00000165829.3 | 20343 | ENSMUSG00000085584 | Rtl9 | retrotransposon Gag like 9 |
| chr4 | 45977657 | 45978214 | 558 | * | 5,476851 | 6,43997779 | 1,985345 | 3,243758 | 0,00118 | 1 | 0 | Intron (ENSMUST0000 | 4 | 45972834 | 45987622 | 14789 | 1 | 100121 | ENSMUST00000151422.2 | 4823 | ENSMUSG00000035517 | Tdrd7 | tudor domain containing 7 |
| chr11 | 80271654 | 80272387 | 734 | * | 5,444498 | 6,440037373 | 1,987961 | 3,239519 | 0,001197 | 1 | 0 | Promoter (1-2kb) | 11 | 80274105 | 80287071 | 12967 | 1 | 22680 | ENSMUST00000165565.8 | -1718 | ENSMUSG00000017421 | Zfp207 | zinc finger protein 207 |
| chr6 | 1,37E+08 | 1,37E+08 | 513 | * | 5,957875 | 6,440211458 | 1,995267 | 3,227744 | 0,001248 | 1 | 0 | Exon (ENSMUST00000 | 6 | 1,37E+08 | 1,37E+08 | 49859 | 2 | 66857 | ENSMUST00000032336.7 | -53638 | ENSMUSG00000030214 | Plbd1 | phospholipase B domain containing 1 |
| chr11 | 4462457 | 4463513 | 1057 | * | 6,489076 | 6,440276568 | 1,930057 | 3,336832 | 0,000847 | 1 | 0 | Intron (ENSMUST0000 | 11 | 4430890 | 4478406 | 47517 | 2 | 74302 | ENSMUST00000123506.8 | 14893 | ENSMUSG00000034354 | Mtmr3 | myotubularin related protein 3 |
| chr1 | 1,86E+08 | 1,86E+08 | 626 | * | 3,066274 | 6,441733168 | 2,480005 | 2,597467 | 0,009391 | 1 | 0 | Distal Intergenic | 1 | 1,86E+08 | 1,86E+08 | 46800 | 2 | 320449 | ENSMUST00000195454.2 | -9301 | ENSMUSG00000103690 | A730004F24Rik | RIKEN cDNA A730004F24 gene |
| chr10 | 46568866 | 46569016 | 151 | * | 2,913127 | 6,441772515 | 2,467129 | 2,61104 | 0,009027 | 1 | 0 | Distal Intergenic | 10 | 46534604 | 46534734 | 131 | 2 | 1,15E+08 | ENSMUST00000104173.3 | -34132 | ENSMUSG00000077361 | Gm23481 | predicted gene, 23481 |
| chr14 | 45496932 | 45497299 | 368 | * | 3,381914 | 6,441840796 | 2,450279 | 2,629024 | 0,008563 | 1 | 0 | Intron (ENSMUST0000 | 14 | 45502783 | 45516710 | 13928 | 1 | 70713 | ENSMUST00000151749.2 | -5484 | ENSMUSG00000049092 | Gpr137c | G protein-coupled receptor 137C |
| chr2 | 93170819 | 93171202 | 384 | * | 3,352187 | 6,442006707 | 2,403381 | 2,680393 | 0,007354 | 1 | 0 | Distal Intergenic | 2 | 93141608 | 93164850 | 23243 | 2 | 241556 | ENSMUST00000152141.2 | -5969 | ENSMUSG00000027217 | Tspan18 | tetraspanin 18 |
| chr16 | 40077386 | 40077794 | 409 | * | 3,314965 | 6,442052537 | 2,392898 | 2,692155 | 0,007099 | 1 | 0 | Intron (ENSMUST0000 | 16 | 39804957 | 40478248 | 673292 | 1 | 268890 | ENSMUST00000189829.2 | 272429 | ENSMUSG00000061080 | Lsamp | limbic system-associated membrane protein |
| chr12 | 81000675 | 81001102 | 428 | * | 4,334714 | 6,442503858 | 2,291759 | 2,811161 | 0,004936 | 1 | 0 | Intron (ENSMUST0000 | 12 | 80993150 | 80997271 | 4122 | 1 | 20384 | ENSMUST00000151036.2 | 7525 | ENSMUSG00000021134 | Srsf5 | serine and arginine-rich splicing factor 5 |
| chrX | 70944077 | 70944789 | 713 | * | 4,248979 | 6,442547294 | 2,281745 | 2,823518 | 0,00475 | 1 | 0 | Distal Intergenic | 20 | 70964553 | 70984476 | 19924 | 1 | 382221 | ENSMUST00000154780.10 | -19764 | ENSMUSG00000073130 | Gm1141 | predicted gene 1141 |
| chrX | 20810347 | 20810991 | 645 | * | 4,431723 | 6,442785456 | 2,242941 | 2,872472 | 0,004073 | 1 | 0 | Intron (ENSMUST0000 | 20 | 20799634 | 20816847 | 17214 | 2 | 13712 | ENSMUST00000009550.14 | 5856 | ENSMUSG00000009406 | Elk1 | ELK1, member of ETS oncogene family |
| chr13 | 54934277 | 54934809 | 533 | * | 4,456593 | 6,442982641 | 2,143501 | 3,005823 | 0,002649 | 1 | 0 | Promoter (2-3kb) | 13 | 54937190 | 54944589 | 7400 | 1 | 74257 | ENSMUST00000026993.14 | -2381 | ENSMUSG00000025875 | Tspan17 | tetraspanin 17 |
| chr11 | 21864308 | 21864647 | 340 | * | 4,212282 | 6,442983494 | 2,211457 | 2,913456 | 0,003575 | 1 | 0 | Distal Intergenic | 11 | 21944764 | 21951615 | 6852 | 2 | 18423 | ENSMUST00000006071.14 | 86968 | ENSMUSG00000005917 | Otx1 | orthodenticle homeobox 1 |
| chr3 | 95785092 | 95785442 | 351 | * | 3,634978 | 6,443072627 | 2,194161 | 2,936463 | 0,00332 | 1 | 0 | Intron (ENSMUST0000 | 3 | 95786822 | 95788558 | 1737 | 2 | 229599 | ENSMUST00000161867.8 | 3116 | ENSMUSG00000038550 | Ciart | circadian associated repressor of transcription |
| chr6 | 3953180 | 3953687 | 508 | * | 7,317544 | 6,443104554 | 1,867815 | 3,44954 | 0,000562 | 1 | 0 | Distal Intergenic | 6 | 3962595 | 3968376 | 5782 | 2 | 21789 | ENSMUST00000031674.11 | 14689 | ENSMUSG00000029664 | Tfpi2 | tissue factor pathway inhibitor 2 |
| chr2 | 74703197 | 74703896 | 700 | * | 4,317326 | 6,443699822 | 2,195416 | 2,93507 | 0,003335 | 1 | 0 | Intron (ENSMUST0000 | 2 | 74699325 | 74706848 | 7524 | 1 | 53375 | ENSMUST00000129284.2 | 3872 | ENSMUSG00000027099 | Mtx2 | metaxin 2 |
| chr3 | 40842050 | 40842927 | 878 | * | 4,403606 | 6,44397545 | 2,078912 | 3,099687 | 0,001937 | 1 | 0 | Intron (ENSMUST0000 | 3 | 40840332 | 40848558 | 8227 | 2 | 619719 | ENSMUST00000181813.2 | 5631 | ENSMUSG00000097583 | 6430590A07Rik | RIKEN cDNA 6430590A07 gene |
| chr1 | 1,93E+08 | 1,93E+08 | 663 | * | 4,645098 | 6,443986202 | 2,075663 | 3,104543 | 0,001906 | 1 | 0 | Distal Intergenic | 1 | 1,93E+08 | 1,93E+08 | 129274 | 1 | 1,03E+08 | ENSMUST00000246715.1 | 142178 | | Gm31870 | predicted gene, 31870 |
| chr2 | 1,12E+08 | 1,12E+08 | 468 | * | 4,486506 | 6,444042561 | 2,069922 | 3,113181 | 0,001851 | 1 | 0 | Distal Intergenic | 2 | 1,12E+08 | 1,12E+08 | 17166 | 1 | 73024 | ENSMUST00000069747.6 | -20762 | ENSMUSG00000055943 | Emc7 | ER membrane protein complex subunit 7 |
| chr5 | 1,24E+08 | 1,24E+08 | 468 | * | 4,828802 | 6,444327767 | 2,040251 | 3,158596 | 0,001585 | 1 | 0 | Intron (ENSMUST0000 | 5 | 1,24E+08 | 1,24E+08 | 10971 | 1 | 208908 | ENSMUST00000249271.1 | -6268 | ENSMUSG00000061882 | Ccdc62 | coiled-coil domain containing 62 |
| chr14 | 60480672 | 60481128 | 457 | * | 4,558308 | 6,444374247 | 2,036144 | 3,16499 | 0,001551 | 1 | 0 | Promoter (2-3kb) | 14 | 60476047 | 60478413 | 2367 | 2 | 71844 | ENSMUST00000224575.2 | -2259 | ENSMUSG00000114797 | Nupl1 | nucleoporin like 1 |
| chr8 | 61954644 | 61955267 | 624 | * | 4,876004 | 6,44482887 | 1,995612 | 3,2295 | 0,00124 | 1 | 0 | Intron (ENSMUST0000 | 8 | 61947863 | 61956423 | 8561 | 1 | 234309 | ENSMUST00000150932.2 | 6781 | ENSMUSG00000031641 | Cbr4 | carbonyl reductase 4 |
| chr1 | 1,64E+08 | 1,64E+08 | 818 | * | 5,613069 | 6,44499915 | 1,980349 | 3,254476 | 0,001136 | 1 | 0 | Exon (ENSMUST00000 | 1 | 1,64E+08 | 1,64E+08 | 16340 | 1 | 116914 | ENSMUST00000044021.12 | -41726 | ENSMUSG00000040918 | Slc19a2 | solute carrier family 19 (thiamine transporter), member 2 |
| chr5 | 1,38E+08 | 1,38E+08 | 440 | * | 5,220476 | 6,445335342 | 1,955177 | 3,296549 | 0,000979 | 1 | 0 | Promoter (<=1kb) | 5 | 1,38E+08 | 1,38E+08 | 935 | 2 | 17220 | ENSMUST00000139223.2 | 0 | ENSMUSG00000029730 | Mcm7 | minichromosome maintenance complex component 7 |
| chr5 | 1,43E+08 | 1,43E+08 | 288 | * | 3,583655 | 6,446019727 | 2,432975 | 2,649439 | 0,008063 | 1 | 0 | Intron (ENSMUST0000 | 5 | 1,42E+08 | 1,43E+08 | 60187 | 2 | 231858 | ENSMUST00000085758.10 | 4124 | ENSMUSG00000029576 | Radil | Ras association and DIL domains |
| chr1 | 97993006 | 97994034 | 1029 | * | 7,049242 | 6,44766161 | 1,824566 | 3,533805 | 0,00041 | 0,337054778 | 1,360622 | Intron (ENSMUST0000 | 1 | 97959593 | 97985405 | 25813 | 2 | 18484 | ENSMUST00000159353.2 | -7601 | ENSMUSG00000026335 | Pam | peptidylglycine alpha-amidating monooxygenase |
| chr12 | 66720089 | 66720402 | 314 | * | 3,445655 | 6,44978173 | 2,406455 | 2,6802 | 0,007358 | 1 | 0 | Intron (ENSMUST0000 | 12 | 66553317 | 66702117 | 148801 | 2 | 320772 | ENSMUST00000223289.2 | -17972 | ENSMUSG00000034912 | Mdga2 | MAM domain containing glycosylphosphatidylinositol anchor 2 |
| chr14 | 1E+08 | 1E+08 | 572 | * | 3,759191 | 6,44983054 | 2,401495 | 2,685757 | 0,007237 | 1 | 0 | Intron (ENSMUST0000 | 14 | 1E+08 | 1E+08 | 275279 | 2 | 16597 | ENSMUST00000097079.5 | -30902 | ENSMUSG00000072294 | Klf12 | Kruppel-like factor 12 |
| chr7 | 66997386 | 66997730 | 345 | * | 3,665398 | 6,450009218 | 2,380649 | 2,709349 | 0,006742 | 1 | 0 | Exon (ENSMUST00000 | 7 | 66995340 | 67022575 | 27236 | 2 | 17258 | ENSMUST00000144057.3 | 24845 | ENSMUSG00000030557 | Mef2a | myocyte enhancer factor 2A |
| chr12 | 94518081 | 94518313 | 233 | * | 3,53121 | 6,450067611 | 2,373835 | 2,717151 | 0,006585 | 1 | 0 | Distal Intergenic | 12 | 93894112 | 93894238 | 127 | 1 | 1,02E+08 | ENSMUST00000183568.3 | 623969 | ENSMUSG00000098931 | Mir8099-2 | microRNA 8099-2 |
| chr15 | 34837828 | 34838060 | 233 | * | 3,543347 | 6,450112714 | 2,368473 | 2,723322 | 0,006463 | 1 | 0 | Promoter (<=1kb) | 15 | 34838195 | 34843371 | 5177 | 1 | 16539 | ENSMUST00000228725.2 | -135 | ENSMUSG00000050963 | Kcns2 | K+ voltage-gated channel, subfamily S, 2 |
| chr13 | 53626740 | 53627019 | 280 | * | 3,658335 | 6,450525777 | 2,327934 | 2,770923 | 0,00559 | 1 | 0 | Promoter (<=1kb) | 13 | 53620920 | 53627110 | 6191 | 2 | 17702 | ENSMUST00000021922.10 | 91 | ENSMUSG00000021469 | Msx2 | msh homeobox 2 |
| chr14 | 20976940 | 20977341 | 402 | * | 3,849858 | 6,450824554 | 2,301232 | 2,803205 | 0,00506 | 1 | 0 | Promoter (2-3kb) | 14 | 20979466 | 21048849 | 69384 | 1 | 22330 | ENSMUST00000226113.2 | -2125 | ENSMUSG00000021823 | Vcl | vinculin |
| chr4 | 1,33E+08 | 1,33E+08 | 796 | * | 5,982467 | 6,451288478 | 1,944189 | 3,318241 | 0,000906 | 1 | 0 | Promoter (2-3kb) | 4 | 1,33E+08 | 1,33E+08 | 31009 | 2 | 100226 | ENSMUST00000030698.5 | -2526 | ENSMUSG00000028879 | Stx12 | syntaxin 12 |
| chr11 | 1,18E+08 | 1,18E+08 | 589 | * | 3,908019 | 6,451367182 | 2,255953 | 2,859708 | 0,00424 | 1 | 0 | Promoter (2-3kb) | 11 | 1,18E+08 | 1,18E+08 | 30592 | 2 | 72344 | ENSMUST00000092382.10 | -2213 | ENSMUSG00000033909 | Usp36 | ubiquitin specific peptidase 36 |
| chr4 | 1,36E+08 | 1,36E+08 | 964 | * | 5,831746 | 6,45143927 | 1,959559 | 3,292292 | 0,000994 | 1 | 0 | Intron (ENSMUST0000 | 4 | 1,36E+08 | 1,36E+08 | 70818 | 1 | 269593 | ENSMUST00000129230.3 | 11538 | ENSMUSG00000001089 | Luzp1 | leucine zipper protein 1 |
| chr4 | 1,03E+08 | 1,03E+08 | 708 | * | 4,963603 | 6,451730878 | 1,994282 | 3,235115 | 0,001216 | 1 | 0 | Intron (ENSMUST0000 | 4 | 1,03E+08 | 1,03E+08 | 42961 | 1 | 67013 | ENSMUST00000133178.2 | 16932 | ENSMUSG00000035069 | Oma1 | OMA1 zinc metallopeptidase |
| chr14 | 55766795 | 55767529 | 735 | * | 5,682753 | 6,451874932 | 2,128156 | 3,031674 | 0,002432 | 1 | 0 | Exon (ENSMUST00000 | 14 | 55756435 | 55762438 | 6004 | 2 | 18185 | ENSMUST00000062232.15 | -4357 | ENSMUSG00000040632 | Nrl | neural retina leucine zipper gene |
| chr10 | 97706494 | 97706856 | 363 | * | 4,073948 | 6,452434492 | 2,179155 | 2,96098 | 0,003067 | 1 | 0 | Intron (ENSMUST0000 | 10 | 97517276 | 97802952 | 285677 | 2 | 1E+08 | ENSMUST00000220233.2 | 96096 | ENSMUSG00000074776 | Gm10754 | predicted gene 10754 |
| chr9 | 86690285 | 86690846 | 562 | * | 4,295154 | 6,452530204 | 2,117226 | 3,047633 | 0,002307 | 1 | 0 | Promoter (<=1kb) | 9 | 86647986 | 86690794 | 42809 | 2 | 20616 | ENSMUST00000190388.7 | 0 | ENSMUSG00000033419 | Snap91 | synaptosomal-associated protein 91 |
| chr3 | 1,05E+08 | 1,05E+08 | 861 | * | 4,394193 | 6,452531586 | 2,118089 | 3,046393 | 0,002316 | 1 | 0 | Intron (ENSMUST0000 | 3 | 1,05E+08 | 1,05E+08 | 12805 | 2 | 17454 | ENSMUST00000196161.2 | 5500 | ENSMUSG00000002227 | Mov10 | Mov10 RISC complex RNA helicase |
| chr12 | 19284221 | 19284712 | 492 | * | 3,975578 | 6,452586462 | 2,127396 | 3,033092 | 0,002421 | 1 | 0 | Distal Intergenic | 12 | 19342814 | 19351265 | 8452 | 1 | 546524 | ENSMUST00000101568.5 | -58102 | | Gm5953 | predicted gene 5953 |
| chr2 | 4942142 | 4942907 | 766 | * | 8,324531 | 6,45280968 | 1,839718 | 3,507499 | 0,000452 | 1 | 0 | Promoter (<=1kb) | 2 | 4942294 | 4943154 | 861 | 1 | 16922 | ENSMUST00000128738.2 | 0 | ENSMUSG00000026664 | Phyh | phytanoyl-CoA hydroxylase |
| chr15 | 10665770 | 10666324 | 555 | * | 4,515865 | 6,452842134 | 2,15295 | 2,997209 | 0,002725 | 1 | 0 | Intron (ENSMUST0000 | 15 | 10569055 | 10713626 | 144572 | 2 | 75646 | ENSMUST00000090339.11 | 47302 | ENSMUSG00000022246 | Rai14 | retinoic acid induced 14 |
| chr1 | 87743751 | 87744175 | 425 | * | 4,547073 | 6,452982837 | 2,145033 | 3,008337 | 0,002627 | 1 | 0 | Intron (ENSMUST0000 | 1 | 87739886 | 87752181 | 12296 | 1 | 20215 | ENSMUST00000144334.8 | 3865 | ENSMUSG00000056055 | Sag | S-antigen, retina and pineal gland (arrestin) |
| chr2 | 1,13E+08 | 1,13E+08 | 408 | * | 3,80592 | 6,453047354 | 2,229197 | 2,894786 | 0,003794 | 1 | 0 | Distal Intergenic | 2 | 1,13E+08 | 1,13E+08 | 166 | 1 | 1,15E+08 | ENSMUST00000176208.3 | 13036 | ENSMUSG00000093660 | Gm26505 | predicted gene, 26505 |
| chrX | 1,3E+08 | 1,3E+08 | 655 | * | 3,901899 | 6,453052353 | 2,231716 | 2,891521 | 0,003834 | 1 | 0 | Distal Intergenic | 20 | 1,3E+08 | 1,3E+08 | 129 | 2 | 1,15E+08 | ENSMUST00000104191.3 | 244683 | ENSMUSG00000077379 | Gm25170 | predicted gene, 25170 |
| chr11 | 79091628 | 79092526 | 899 | * | 7,873903 | 6,45326008 | 1,848527 | 3,491028 | 0,000481 | 1 | 0 | Distal Intergenic | 11 | 79131133 | 79133775 | 2643 | 2 | 78889 | ENSMUST00000153585.2 | 41249 | ENSMUSG00000017677 | Wsb1 | WD repeat and SOCS box-containing 1 |
| chrX | 1,5E+08 | 1,5E+08 | 465 | * | 5,021354 | 6,453285163 | 2,127836 | 3,032793 | 0,002423 | 1 | 0 | Distal Intergenic | 20 | 1,5E+08 | 1,5E+08 | 7063 | 2 | 69499 | ENSMUST00000166010.2 | -19151 | ENSMUSG00000025264 | Tsr2 | TSR2 20S rRNA accumulation |
| chr2 | 98181367 | 98182001 | 635 | * | 3,702511 | 6,453313272 | 2,297508 | 2,808832 | 0,004972 | 1 | 0 | Distal Intergenic | 2 | 97298434 | 97459408 | 160975 | 1 | 241568 | ENSMUST00000170144.2 | 882933 | ENSMUSG00000050587 | Lrrc4c | leucine rich repeat containing 4C |
| chr10 | 79621779 | 79622052 | 274 | * | 3,610518 | 6,453342867 | 2,315575 | 2,786929 | 0,005321 | 1 | 0 | Promoter (1-2kb) | 10 | 79618930 | 79620417 | 1488 | 2 | 73106 | ENSMUST00000171681.2 | -1362 | ENSMUSG00000020323 | Prss57 | protease, serine 57 |
| chr3 | 41726650 | 41727052 | 403 | * | 3,794089 | 6,453496983 | 2,356765 | 2,738286 | 0,006176 | 1 | 0 | Intron (ENSMUST0000 | 3 | 41705631 | 41710927 | 5297 | 1 | 73852 | ENSMUST00000195030.2 | 21019 | ENSMUSG00000025766 | D3Ertd751e | DNA segment, Chr 3, ERATO Doi 751, expressed |
| chr5 | 28050093 | 28050385 | 293 | * | 4,280297 | 6,453497862 | 2,115577 | 3,050467 | 0,002285 | 1 | 0 | Promoter (2-3kb) | 5 | 28047147 | 28060086 | 12940 | 1 | 15563 | ENSMUST00000036227.7 | 2946 | ENSMUSG00000039106 | Htr5a | 5-hydroxytryptamine (serotonin) receptor 5A |
| chr8 | 28766372 | 28766926 | 555 | * | 3,195464 | 6,453517938 | 2,366172 | 2,727409 | 0,006383 | 1 | 0 | Distal Intergenic | 8 | 28573358 | 28574948 | 1591 | 2 | 666422 | ENSMUST00000209921.2 | -191424 | ENSMUSG00000110372 | Gm8096 | 3-phosphoglycerate dehydrogenase pseudogene |

| | | | | | | | | | | | | | | | | | | | | | | | |
|---|---|---|---|---|---|---|---|---|---|---|---|---|---|---|---|---|---|---|---|---|---|---|---|
| chr5 | 34789778 | 34790214 | 437 | * | 4,763134 | 6,454086077 | 2,084256 | 3,096591 | 0,001958 | 1 | 0 | Promoter (1-2kb) | 5 | 34790992 | 34791973 | 982 | 2 | 68294 | ENSMUST00000128670.2 | 1759 | ENSMUSG00000001082 | Mfsd10 | major facilitator superfamily domain containing 10 |
| chr11 | 96065990 | 96066594 | 605 | * | 4,736463 | 6,454393911 | 2,069485 | 3,11884 | 0,001816 | 1 | 0 | Distal Intergenic | 11 | 96085142 | 96088273 | 3132 | 1 | 15408 | ENSMUST00000062709.4 | -18548 | ENSMUSG00000049604 | Hoxb13 | homeobox B13 |
| chr11 | 1,01E+08 | 1,01E+08 | 599 | * | 5,495821 | 6,455579365 | 2,017392 | 3,199962 | 0,001374 | 1 | 0 | Promoter (<=1kb) | 11 | 1,01E+08 | 1,01E+08 | 10343 | 1 | 14377 | ENSMUST00000019469.3 | -297 | ENSMUSG00000078650 | G6pc | glucose-6-phosphatase, catalytic |
| chr11 | 1,17E+08 | 1,17E+08 | 822 | * | 5,186094 | 6,455948683 | 2,002509 | 3,22393 | 0,001264 | 1 | 0 | Distal Intergenic | 11 | 1,17E+08 | 1,17E+08 | 68086 | 1 | 268510 | ENSMUST00000103027.10 | -7133 | ENSMUSG00000043857 | Mgat5b | mannoside acetylglucosaminyltransferase 5, isoenzyme B |
| chr5 | 1,47E+08 | 1,47E+08 | 919 | * | 5,613264 | 6,456189301 | 1,993518 | 3,238591 | 0,001201 | 1 | 0 | Promoter (<=1kb) | 5 | 1,47E+08 | 1,47E+08 | 781 | 1 | 19933 | ENSMUST00000135345.2 | 310 | ENSMUSG00000041453 | Rpl21 | ribosomal protein L21 |
| chrX | 1,32E+08 | 1,32E+08 | 883 | * | 5,626715 | 6,456436444 | 1,984961 | 3,252677 | 0,001143 | 1 | 0 | Distal Intergenic | 20 | 1,31E+08 | 1,31E+08 | 131 | 2 | 1,15E+08 | ENSMUST00000104440.3 | -404910 | ENSMUSG00000077628 | Gm23768 | predicted gene, 23768 |
| chr2 | 23822410 | 23822696 | 287 | * | 4,208693 | 6,457222063 | 2,387911 | 2,704131 | 0,006848 | 1 | 0 | Distal Intergenic | 2 | 23893541 | 23938878 | 45338 | 2 | 140483 | ENSMUST00000114497.2 | 116182 | ENSMUSG00000026986 | Hnmt | histamine N-methyltransferase |
| chr4 | 1,36E+08 | 1,36E+08 | 400 | * | 4,495579 | 6,458643938 | 2,326892 | 2,775653 | 0,005509 | 1 | 0 | Exon (ENSMUST00000 | 4 | 1,35E+08 | 1,36E+08 | 55850 | 1 | 242702 | ENSMUST00000105854.2 | 18307 | ENSMUSG00000037139 | Myom3 | myomesin family, member 3 |
| chr10 | 23023811 | 23024616 | 806 | * | 7,567942 | 6,46003108 | 1,877866 | 3,440092 | 0,000582 | 1 | 0 | Intron (ENSMUST00000 | 10 | 23042763 | 23042869 | 107 | 2 | 1,15E+08 | ENSMUST00000239688.1 | 18253 | ENSMUSG00000119293 | Gm24655 | predicted gene, 24655 |
| chr14 | 27321075 | 27321750 | 676 | * | 5,509112 | 6,460878765 | 2,209244 | 2,924474 | 0,00345 | 1 | 0 | Distal Intergenic | 14 | 27344385 | 28200492 | 856108 | 1 | 238988 | ENSMUST00000210135.2 | -22635 | ENSMUSG00000040640 | Erc2 | ELKS/RAB6-interacting/CAST family member 2 |
| chr9 | 51896970 | 51897413 | 444 | * | 3,569907 | 6,461211622 | 2,450562 | 2,636624 | 0,008374 | 1 | 0 | Distal Intergenic | 9 | 51856280 | 51874856 | 18577 | 2 | 14148 | ENSMUST00000214013.2 | -22114 | ENSMUSG00000032051 | Fdx1 | ferredoxin 1 |
| chrX | 54701197 | 54701840 | 644 | * | 3,60527 | 6,461350139 | 2,396665 | 2,695976 | 0,007018 | 1 | 0 | Distal Intergenic | 20 | 54598507 | 54613718 | 15212 | 1 | 1E+08 | ENSMUST00000177739.2 | 102690 | ENSMUSG00000095398 | Gm16404 | predicted gene 16404 |
| chr16 | 24712165 | 24712516 | 352 | * | 4,078222 | 6,46137591 | 2,393868 | 2,699136 | 0,006952 | 1 | 0 | Intron (ENSMUST00000 | 16 | 24702923 | 24703056 | 134 | 1 | 1,15E+08 | ENSMUST00000104414.3 | 9242 | ENSMUSG00000077602 | Gm22672 | predicted gene, 22672 |
| chr2 | 70573924 | 70574309 | 386 | * | 3,399931 | 6,461408613 | 2,374172 | 2,721542 | 0,006498 | 1 | 0 | Intron (ENSMUST00000 | 2 | 70544485 | 70548568 | 4084 | 2 | 228012 | ENSMUST00000141707.2 | -25356 | ENSMUSG00000041997 | Tlk1 | tousled-like kinase 1 |
| chr3 | 20396449 | 20396929 | 481 | * | 3,434591 | 6,461504078 | 2,339308 | 2,762143 | 0,005742 | 1 | 0 | Intron (ENSMUST00000 | 3 | 20369061 | 20421288 | 52228 | 2 | 11608 | ENSMUST00000163776.3 | 24359 | ENSMUSG00000054988 | Agtr1b | angiotensin II receptor, type 1b |
| chrX | 21132681 | 21133324 | 644 | * | 4,392641 | 6,461864754 | 2,223956 | 2,905572 | 0,003666 | 1 | 0 | Distal Intergenic | 20 | 21072819 | 21087904 | 15086 | 2 | 1,01E+08 | ENSMUST00000125569.8 | -44777 | ENSMUSG00000087358 | 4930453H23Rik | RIKEN cDNA 4930453H23 gene |
| chr16 | 4288562 | 4289157 | 596 | * | 4,30968 | 6,462143044 | 2,154749 | 2,999023 | 0,002708 | 1 | 0 | Distal Intergenic | 16 | 4105393 | 4238362 | 132970 | 2 | 11515 | ENSMUST00000120080.8 | -50200 | ENSMUSG00000005580 | Adcy9 | adenylate cyclase 9 |
| chrX | 95004820 | 95005322 | 503 | * | 5,754913 | 6,462211335 | 2,152437 | 3,002276 | 0,00268 | 1 | 0 | Distal Intergenic | 20 | 94993041 | 95000568 | 7528 | 2 | 76130 | ENSMUST00000139717.2 | -4252 | ENSMUSG00000057421 | Las1l | LAS1-like (S. cerevisiae) |
| chr2 | 1,65E+08 | 1,65E+08 | 619 | * | 5,803094 | 6,462826359 | 2,02615 | 3,189707 | 0,001424 | 1 | 0 | Promoter (1-2kb) | 2 | 1,65E+08 | 1,65E+08 | 542 | 2 | 228869 | ENSMUST00000126685.2 | 1607 | ENSMUSG00000039804 | Ncoa5 | nuclear receptor coactivator 5 |
| chr11 | 85083350 | 85083898 | 549 | * | 5,817464 | 6,463025884 | 1,996737 | 3,236793 | 0,001209 | 1 | 0 | Intron (ENSMUST00000 | 11 | 85061922 | 85071971 | 10050 | 1 | 73634 | ENSMUST00000018623.4 | 21428 | ENSMUSG00000018479 | 1700125H20Rik | RIKEN cDNA 1700125H20 gene |
| chr4 | 1,27E+08 | 1,27E+08 | 987 | * | 6,803485 | 6,463835613 | 2,09375 | 3,087205 | 0,00202 | 1 | 0 | Promoter (1-2kb) | 4 | 1,27E+08 | 1,27E+08 | 42340 | 2 | 67785 | ENSMUST00000154821.2 | 1478 | ENSMUSG00000042446 | Zmym4 | zinc finger, MYM-type 4 |
| chr17 | 51723016 | 51723732 | 717 | * | 6,278075 | 6,463912284 | 1,894688 | 3,411597 | 0,000646 | 1 | 0 | Distal Intergenic | 17 | 51732237 | 51742866 | 10630 | 2 | 328837 | ENSMUST00000233466.2 | 19134 | ENSMUSG00000117204 | C330011F03Rik | RIKEN cDNA C330011F03 gene |
| chrX | 50880917 | 50881759 | 843 | * | 3,800292 | 6,463958535 | 2,285229 | 2,828582 | 0,004675 | 1 | 0 | Intron (ENSMUST00000 | 20 | 50842836 | 50890021 | 47186 | 2 | 83563 | ENSMUST00000069509.4 | 8262 | ENSMUSG00000055780 | Usp26 | ubiquitin specific peptidase 26 |
| chr2 | 1,63E+08 | 1,63E+08 | 848 | * | 10,76617 | 6,465008525 | 1,701671 | 3,799211 | 0,000145 | 0,223981956 | 1,173325 | Promoter (<=1kb) | 2 | 1,63E+08 | 1,63E+08 | 602 | 1 | 1,08E+08 | ENSMUST00000190930.3 | 0 | ENSMUSG00000099632 | 2900093K20Rik | RIKEN cDNA 2900093K20 gene |
| chr12 | 3315438 | 3316195 | 758 | * | 8,501868 | 6,465697656 | 1,768531 | 3,655971 | 0,000256 | 0,278016366 | 1,360622 | Intron (ENSMUST00000 | 12 | 3286611 | 3299999 | 13389 | 1 | 74173 | ENSMUST00000049684.7 | 28827 | ENSMUSG00000079179 | Rab10os | RAB10, member RAS oncogene family, opposite strand |
| chr2 | 97242568 | 97243013 | 446 | * | 3,9475 | 6,468445404 | 2,482792 | 2,605311 | 0,009179 | 1 | 0 | Intron (ENSMUST00000 | 2 | 97298002 | 97462009 | 164008 | 1 | 241568 | ENSMUST00000059049.8 | -54989 | ENSMUSG00000050587 | Lrrc4c | leucine rich repeat containing 4C |
| chr13 | 98343757 | 98344186 | 430 | * | 3,285868 | 6,468635421 | 2,451153 | 2,639017 | 0,008315 | 1 | 0 | Promoter (<=1kb) | 13 | 98035977 | 98342947 | 306971 | 2 | 110596 | ENSMUST00000225884.2 | -810 | ENSMUSG00000021662 | Arhgef28 | Rho guanine nucleotide exchange factor (GEF) 28 |
| chr3 | 88423221 | 88423609 | 389 | * | 3,462217 | 6,468843636 | 2,420836 | 2,672153 | 0,007537 | 1 | 0 | Distal Intergenic | 3 | 88410060 | 88417263 | 7204 | 2 | 16905 | ENSMUST00000149068.2 | -5958 | ENSMUSG00000028063 | Lmna | lamin A |
| chrX | 16931377 | 16931675 | 299 | * | 3,197461 | 6,469065495 | 2,390549 | 2,7061 | 0,006808 | 1 | 0 | Distal Intergenic | 20 | 16751760 | 16778013 | 26254 | 2 | 17986 | ENSMUST00000040134.8 | -153364 | ENSMUSG00000040138 | Ndp | Norrie disease (pseudoglioma) (human) |
| chr9 | 62408426 | 62408718 | 293 | * | 3,401584 | 6,469357035 | 2,354118 | 2,748102 | 0,005994 | 1 | 0 | Intron (ENSMUST00000 | 9 | 62335218 | 62417780 | 82563 | 2 | 235431 | ENSMUST00000164246.9 | 9062 | ENSMUSG00000041729 | Coro2b | coronin, actin binding protein, 2B |
| chr1 | 97589849 | 97590474 | 626 | * | 3,777067 | 6,469916904 | 2,289287 | 2,826171 | 0,004711 | 1 | 0 | Promoter (<=1kb) | 1 | 97571628 | 97589799 | 18172 | 2 | 52392 | ENSMUST00000149927.2 | -50 | ENSMUSG00000044768 | Macir | macrophage immunometabolism regulator |
| chr18 | 42680897 | 42681521 | 625 | * | 5,216052 | 6,470173188 | 2,010419 | 3,218321 | 0,001289 | 1 | 0 | Exon (ENSMUST00000 | 18 | 42685573 | 42701098 | 15526 | 1 | 56070 | ENSMUST00000237646.2 | -4052 | ENSMUSG00000024498 | Tcerg1 | transcription elongation regulator 1 (CA150) |
| chr14 | 1,04E+08 | 1,04E+08 | 558 | * | 5,231312 | 6,47021346 | 2,012875 | 3,214414 | 0,001307 | 1 | 0 | Distal Intergenic | 14 | 1,04E+08 | 1,04E+08 | 49515 | 2 | 73877 | ENSMUST00000228770.2 | -16652 | ENSMUSG00000115727 | 4930432J09Rik | RIKEN cDNA 4930432J09 gene |
| chr2 | 1,02E+08 | 1,02E+08 | 630 | * | 4,454956 | 6,470226739 | 2,257903 | 2,865592 | 0,004162 | 1 | 0 | Promoter (<=1kb) | 2 | 1,02E+08 | 1,02E+08 | 100782 | 2 | 80985 | ENSMUST00000102573.8 | -937 | ENSMUSG00000027189 | Trim44 | tripartite motif-containing 44 |
| chr13 | 52919794 | 52920209 | 416 | * | 3,876381 | 6,470459251 | 2,236128 | 2,8936 | 0,003809 | 1 | 0 | Distal Intergenic | 13 | 52883958 | 52911241 | 27284 | 2 | 105404 | ENSMUST00000223209.2 | -8553 | | BB123696 | expressed sequence BB123696 |
| chr4 | 1,32E+08 | 1,32E+08 | 523 | * | 3,898795 | 6,470632544 | 2,220229 | 2,914398 | 0,003564 | 1 | 0 | Distal Intergenic | 4 | 1,32E+08 | 1,32E+08 | 2934 | 2 | 11983 | ENSMUST00000152993.8 | 3933 | ENSMUSG00000054428 | Atpif1 | ATPase inhibitory factor 1 |
| chr18 | 7807442 | 7808194 | 753 | * | 4,128751 | 6,47082384 | 2,204929 | 2,934709 | 0,003339 | 1 | 0 | Distal Intergenic | 18 | 7835177 | 7835283 | 107 | 1 | 1,15E+08 | ENSMUST00000082888.2 | -26983 | ENSMUSG00000064822 | Gm24292 | predicted gene, 24292 |
| chr2 | 16831429 | 16832085 | 657 | * | 4,590949 | 6,470824169 | 2,072801 | 3,121777 | 0,001798 | 1 | 0 | Distal Intergenic | 2 | 16643507 | 16756501 | 112995 | 1 | 67448 | ENSMUST00000126173.2 | 187922 | ENSMUSG00000026748 | Plxdc2 | plexin domain containing 2 |
| chr13 | 5028956 | 5029541 | 586 | * | 4,346806 | 6,471004475 | 2,189582 | 2,955361 | 0,003123 | 1 | 0 | Distal Intergenic | 13 | 5239802 | 5239933 | 132 | 2 | 1,15E+08 | ENSMUST00000104285.3 | 210392 | ENSMUSG00000077473 | Gm24790 | predicted gene, 24790 |
| chr4 | 99784993 | 99785201 | 209 | * | 4,235086 | 6,471354126 | 2,161286 | 2,994215 | 0,002752 | 1 | 0 | Promoter (1-2kb) | 4 | 99786611 | 99844491 | 57881 | 1 | 72157 | ENSMUST00000058351.16 | -1410 | ENSMUSG00000025791 | Pgm1 | phosphoglucomutase 1 |
| chr4 | 1,37E+08 | 1,37E+08 | 601 | * | 5,22124 | 6,471374906 | 2,14015 | 3,023795 | 0,002496 | 1 | 0 | Intron (ENSMUST00000 | 4 | 1,37E+08 | 1,37E+08 | 9542 | 2 | 67868 | ENSMUST00000102522.5 | 3394 | ENSMUSG00000023433 | Cela3b | chymotrypsin-like elastase family, member 3B |
| chr5 | 1,16E+08 | 1,16E+08 | 607 | * | 4,712314 | 6,471407165 | 2,157529 | 2,999454 | 0,002705 | 1 | 0 | Intron (ENSMUST00000 | 5 | 1,16E+08 | 1,16E+08 | 53202 | 1 | 12704 | ENSMUST00000122877.8 | -4803 | ENSMUSG00000029516 | Cit | citron |
| chrX | 38195070 | 38195612 | 543 | * | 4,315009 | 6,471650095 | 2,139701 | 3,024558 | 0,00249 | 1 | 0 | Distal Intergenic | 20 | 38046961 | 38048432 | 1472 | 1 | 77883 | ENSMUST00000168144.3 | 148109 | ENSMUSG00000051361 | 6030498E09Rik | RIKEN cDNA 6030498E09 gene |
| chr18 | 6831869 | 6832259 | 391 | * | 4,293437 | 6,471721162 | 2,1359 | 3,029974 | 0,002446 | 1 | 0 | Distal Intergenic | 18 | 6765220 | 6789240 | 24021 | 1 | 19330 | ENSMUST00000097680.7 | 66649 | ENSMUSG00000073639 | Rab18 | RAB18, member RAS oncogene family |
| chr13 | 36223738 | 36224348 | 611 | * | 4,830589 | 6,472470631 | 2,084656 | 3,104814 | 0,001904 | 1 | 0 | Intron (ENSMUST00000 | 13 | 36174844 | 36175503 | 660 | 2 | 1E+08 | ENSMUST00000224867.2 | -48235 | | Tmed10-ps | transmembrane p24 trafficking protein 10, pseudogene |
| chr1 | 1,25E+08 | 1,25E+08 | 430 | * | 3,875038 | 6,472607071 | 2,344218 | 2,761094 | 0,005761 | 1 | 0 | Distal Intergenic | 1 | 1,24E+08 | 1,25E+08 | 340283 | 2 | 269109 | ENSMUST00000187286.2 | -150434 | ENSMUSG00000036815 | Dpp10 | dipeptidylpeptidase 10 |
| chrX | 1,12E+08 | 1,12E+08 | 509 | * | 3,5784 | 6,472798 | 2,385148 | 2,713793 | 0,006652 | 1 | 0 | Distal Intergenic | 20 | 1,12E+08 | 1,12E+08 | 60221 | 2 | 69693 | ENSMUST00000039887.4 | -119525 | ENSMUSG00000034607 | Pof1b | premature ovarian failure 1B |
| chr7 | 1,42E+08 | 1,42E+08 | 467 | * | 4,621852 | 6,472943001 | 2,056982 | 3,146816 | 0,001651 | 1 | 0 | Intron (ENSMUST00000 | 7 | 1,42E+08 | 1,42E+08 | 758 | 2 | 1,15E+08 | ENSMUST00000209533.2 | 4027 | | Gm45417 | predicted gene 45417 |
| chr14 | 25985968 | 25986349 | 382 | * | 4,954296 | 6,472945463 | 2,057637 | 3,145815 | 0,001656 | 1 | 0 | Promoter (2-3kb) | 14 | 25983429 | 25990512 | 7084 | 1 | 278672 | ENSMUST00000049793.9 | 2539 | ENSMUSG00000048502 | Duxbl1 | double homeobox B-like 1 |
| chr19 | 8231430 | 8232028 | 599 | * | 5,786356 | 6,47310857 | 2,048385 | 3,160104 | 0,001577 | 1 | 0 | Promoter (2-3kb) | 19 | 8229332 | 8294543 | 65212 | 1 | 1,19E+08 | ENSMUST00000235607.2 | 2098 | | LOC118568404 | uncharacterized LOC118568404 |
| chr7 | 80396036 | 80396850 | 815 | * | 5,24123 | 6,473372842 | 2,034014 | 3,18256 | 0,00146 | 1 | 0 | Intron (ENSMUST00000 | 7 | 80376527 | 80379996 | 3470 | 2 | 29875 | ENSMUST00000205606.2 | -16040 | ENSMUSG00000030536 | Iqgap1 | IQ motif containing GTPase activating protein 1 |
| chr1 | 93386335 | 93387129 | 795 | * | 4,794082 | 6,473701733 | 2,016924 | 3,20969 | 0,001329 | 1 | 0 | Intron (ENSMUST00000 | 1 | 93335484 | 93373408 | 37925 | 2 | 110611 | ENSMUST00000042498.14 | -12927 | ENSMUSG00000034088 | Hdlbp | high density lipoprotein (HDL) binding protein |
| chr9 | 20413421 | 20414120 | 700 | * | 5,041765 | 6,474028798 | 2,001117 | 3,235208 | 0,001216 | 1 | 0 | Intron (ENSMUST00000 | 9 | 20379845 | 20404042 | 24198 | 2 | 235028 | ENSMUST00000167457.8 | -9379 | ENSMUSG00000059475 | Zfp426 | zinc finger protein 426 |
| chr17 | 37103136 | 37104019 | 884 | * | 6,427515 | 6,47504003 | 1,938716 | 3,339861 | 0,000838 | 1 | 0 | Distal Intergenic | 17 | 37123063 | 37126456 | 3394 | 1 | 399549 | ENSMUST00000041398.5 | -19044 | ENSMUSG00000037130 | H2-M10.6 | histocompatibility 2, M region locus 10.6 |
| chr7 | 1,28E+08 | 1,28E+08 | 525 | * | 3,273282 | 6,475364865 | 2,510187 | 2,579634 | 0,009891 | 1 | 0 | Intron (ENSMUST00000 | 7 | 1,28E+08 | 1,28E+08 | 10781 | 1 | 16409 | ENSMUST00000134694.2 | -18344 | ENSMUSG00000030786 | Itgam | integrin alpha M |
| chrX | 99513299 | 99514744 | 1446 | * | 5,800362 | 6,475980929 | 1,921633 | 3,37004 | 0,000752 | 1 | 0 | Distal Intergenic | 20 | 99537897 | 99559731 | 21835 | 1 | 18518 | ENSMUST00000033570.6 | -23153 | ENSMUSG00000031221 | Igbp1 | immunoglobulin (CD79A) binding protein 1 |
| chr6 | 83401769 | 83402403 | 635 | * | 8,458658 | 6,476251204 | 1,911921 | 3,3873 | 0,000706 | 0,421090004 | 1,449706 | Intron (ENSMUST00000 | 6 | 83339355 | 83418657 | 79303 | 2 | 194388 | ENSMUST00000089622.11 | 16254 | ENSMUSG00000034832 | Tet3 | tet methylcytosine dioxygenase 3 |
| chr11 | 95680931 | 95681744 | 814 | * | 6,709164 | 6,476527811 | 1,903335 | 3,402725 | 0,000667 | 1 | 0 | Intron (ENSMUST00000 | 11 | 95715325 | 95721527 | 6203 | 1 | 237928 | ENSMUST00000150134.2 | -33581 | ENSMUSG00000050860 | Phospho1 | phosphatase, orphan 1 |
| chr11 | 1,03E+08 | 1,03E+08 | 655 | * | 3,756187 | 6,477197836 | 2,351415 | 2,754596 | 0,005876 | 1 | 0 | Intron (ENSMUST00000 | 11 | 1,03E+08 | 1,03E+08 | 11197 | 2 | 52715 | ENSMUST00000124869.8 | 10213 | ENSMUSG00000020925 | Ccdc43 | coiled-coil domain containing 43 |
| chr18 | 58036683 | 58037136 | 454 | * | 3,713506 | 6,477336392 | 2,342126 | 2,76558 | 0,005682 | 1 | 0 | Intron (ENSMUST00000 | 18 | 58011750 | 58079893 | 68144 | 1 | 20496 | ENSMUST00000115366.3 | 24933 | ENSMUSG00000024597 | Slc12a2 | solute carrier family 12, member 2 |
| chr5 | 34726812 | 34727156 | 345 | * | 5,018088 | 6,478973208 | 2,066535 | 3,135187 | 0,001717 | 1 | 0 | Distal Intergenic | 5 | 34731008 | 34789652 | 58645 | 1 | 11518 | ENSMUST00000114338.9 | -3852 | ENSMUSG00000029106 | Add1 | adducin 1 (alpha) |
| chr1 | 52556407 | 52556818 | 412 | * | 5,504371 | 6,479090354 | 2,07111 | 3,128318 | 0,001758 | 1 | 0 | Distal Intergenic | 1 | 52562061 | 52621838 | 59778 | 1 | 1,18E+08 | ENSMUST00000189527.2 | -5243 | ENSMUSG00000100120 | Gm53065 | predicted gene 53065 |
| chr18 | 32767468 | 32768114 | 647 | * | 7,832475 | 6,479293395 | 1,826443 | 3,547492 | 0,000389 | 1 | 0 | Distal Intergenic | 18 | 32754973 | 32755109 | 137 | 2 | 1,15E+08 | ENSMUST00000157090.3 | -12359 | ENSMUSG00000087715 | Gm25826 | predicted gene, 25826 |
| chr11 | 85946383 | 85947173 | 791 | * | 5,321144 | 6,480314535 | 2,168432 | 2,988479 | 0,002804 | 1 | 0 | Distal Intergenic | 11 | 86085859 | 86091990 | 6132 | 2 | 237911 | ENSMUST00000149748.8 | 144817 | ENSMUSG00000034329 | Brip1 | BRCA1 interacting protein C-terminal helicase 1 |
| chr1 | 1,89E+08 | 1,89E+08 | 466 | * | 3,173951 | 6,480353404 | 2,500461 | 2,591664 | 0,009551 | 1 | 0 | Distal Intergenic | 1 | 1,89E+08 | 1,89E+08 | 194337 | 2 | 16526 | ENSMUST00000180044.8 | -93991 | ENSMUSG00000037624 | Kcnk2 | potassium channel, subfamily K, member 2 |
| chr7 | 61333476 | 61334063 | 588 | * | 3,285698 | 6,480375029 | 2,446735 | 2,648581 | 0,008083 | 1 | 0 | Promoter (<=1kb) | 7 | 61332872 | 61332940 | 69 | 2 | 1,01E+08 | ENSMUST00000175106.3 | -536 | ENSMUSG00000092847 | Mir344d-1 | microRNA 344d-1 |
| chr11 | 61732514 | 61733128 | 615 | * | 4,449646 | 6,480776348 | 2,143652 | 3,023241 | 0,002501 | 1 | 0 | Intron (ENSMUST00000 | 11 | 61668328 | 61745328 | 77001 | 2 | 29869 | ENSMUST00000129025.8 | 12200 | ENSMUSG00000004798 | Ulk2 | unc-51 like kinase 2 |
| chrX | 1,21E+08 | 1,21E+08 | 654 | * | 4,846721 | 6,480882986 | 2,084231 | 3,109484 | 0,001874 | 1 | 0 | Distal Intergenic | 20 | 1,21E+08 | 1,21E+08 | 136 | 1 | 1,15E+08 | ENSMUST00000157914.3 | -53367 | ENSMUSG00000088539 | Gm23277 | predicted gene, 23277 |
| chr2 | 69570617 | 69571353 | 737 | * | 5,355863 | 6,480936513 | 2,066168 | 3,136694 | 0,001709 | 1 | 0 | Promoter (<=1kb) | 2 | 69572013 | 69579892 | 7880 | 1 | 228005 | ENSMUST00000143954.2 | -660 | ENSMUSG00000042133 | Ppig | peptidyl-prolyl isomerase G (cyclophilin G) |
| chr10 | 26859466 | 26859857 | 392 | * | 5,148978 | 6,480939102 | 2,061862 | 3,143246 | 0,001671 | 1 | 0 | Exon (ENSMUST00000 | 10 | 26856032 | 26864510 | 8479 | 2 | 16773 | ENSMUST00000186279.2 | 4653 | ENSMUSG00000019899 | Lama2 | laminin, alpha 2 |
| chr8 | 23839027 | 23839712 | 686 | * | 5,49282 | 6,480962476 | 2,159933 | 3,000539 | 0,002695 | 1 | 0 | Distal Intergenic | 8 | 23780049 | 23780180 | 132 | 2 | 1,15E+08 | ENSMUST00000104211.3 | -58847 | ENSMUSG00000077399 | Gm24335 | predicted gene, 24335 |
| chr16 | 39653353 | 39654055 | 703 | * | 5,407804 | 6,481002535 | 2,029969 | 3,192661 | 0,00141 | 1 | 0 | Distal Intergenic | 16 | 39804723 | 40800071 | 995349 | 1 | 268890 | ENSMUST00000189229.7 | -150668 | ENSMUSG00000061080 | Lsamp | limbic system-associated membrane protein |
| chr19 | 6146914 | 6147771 | 858 | * | 5,394817 | 6,4812722 | 1,937742 | 3,344754 | 0,000824 | 1 | 0 | Promoter (1-2kb) | 19 | 6145867 | 6148264 | 2398 | 1 | 225852 | ENSMUST00000237778.2 | 1047 | ENSMUSG00000117666 | Gm550 | predicted gene 550 |
| chr14 | 23096952 | 23097683 | 732 | * | 7,195231 | 6,48153419 | 1,868327 | 3,469165 | 0,000522 | 1 | 0 | Intron (ENSMUST00000 | 14 | 23088262 | 23144639 | 56378 | 2 | 791307 | ENSMUST00000090409.2 | 46956 | ENSMUSG00000068680 | Gm10248 | predicted gene 10248 |
| chr19 | 40673499 | 40674176 | 678 | * | 5,027178 | 6,481660358 | 2,109561 | 3,072516 | 0,002123 | 1 | 0 | Intron (ENSMUST00000 | 19 | 40670111 | 40711637 | 41527 | 1 | 12495 | ENSMUST00000137508.2 | 3388 | ENSMUSG00000048120 | Entpd1 | ectonucleoside triphosphate diphosphohydrolase 1 |
| chr12 | 28317355 | 28318263 | 909 | * | 9,342983 | 6,481807479 | 1,815482 | 3,570296 | 0,000357 | 0,311294276 | 1,425929 | Intron (ENSMUST00000 | 12 | 28400262 | 28400393 | 132 | 1 | 1,15E+08 | ENSMUST00000240083.1 | -81999 | ENSMUSG00000118901 | Gm25564 | predicted gene, 25564 |
| chr9 | 51639809 | 51641048 | 1240 | * | 9,39155 | 6,483736998 | 1,811664 | 3,578885 | 0,000345 | 0,313746033 | 1,360622 | Distal Intergenic | 9 | 51676637 | 51701549 | 24913 | 1 | 244867 | ENSMUST00000124907.2 | -35589 | ENSMUSG00000053199 | Arhgap20 | Rho GTPase activating protein 20 |
| chr5 | 24499206 | 24499842 | 637 | * | 7,640023 | 6,484002152 | 1,819193 | 3,56422 | 0,000365 | 0,322225015 | 1,360622 | Distal Intergenic | 5 | 24525081 | 24527252 | 2172 | 2 | 16511 | ENSMUST00000129246.2 | 27410 | ENSMUSG00000038319 | Kcnh2 | potassium voltage-gated channel, subfamily H (eag-related), member 2 |
| chr1 | 1,91E+08 | 1,91E+08 | 360 | * | 3,537554 | 6,48405262 | 2,374852 | 2,730297 | 0,006328 | 1 | 0 | 3' UTR | 1 | 1,91E+08 | 1,91E+08 | 15124 | 2 | 76843 | ENSMUST00000194064.2 | 22078 | ENSMUSG00000037474 | Dtl | denticleless E3 ubiquitin protein ligase |
| chr2 | 69728406 | 69729183 | 778 | * | 7,186849 | 6,484708279 | 2,005357 | 3,233693 | 0,001222 | 1 | 0 | Promoter (<=1kb) | 2 | 69727647 | 69769003 | 41357 | 1 | 68795 | ENSMUST00000131553.2 | 759 | ENSMUSG00000044308 | Ubr3 | ubiquitin protein ligase E3 component n-recognin 3 |
| chr11 | 95187898 | 95188687 | 790 | * | 6,125096 | 6,486963866 | 1,933195 | 3,355567 | 0,000792 | 1 | 0 | Intron (ENSMUST00000 | 11 | 95174967 | 95181863 | 6897 | 2 | 217127 | ENSMUST00000149356.2 | -6035 | ENSMUSG00000038909 | Kat7 | K(lysine) acetyltransferase 7 |
| chr11 | 1,13E+08 | 1,13E+08 | 424 | * | 4,072576 | 6,487668355 | 2,448441 | 2,649714 | 0,008056 | 1 | 0 | Distal Intergenic | 11 | 1,13E+08 | 1,13E+08 | 6278 | 2 | 1E+08 | ENSMUST00000127337.2 | -125300 | ENSMUSG00000075437 | Gm11681 | predicted gene 11681 |
| chr10 | 67589426 | 67589809 | 384 | * | 3,314469 | 6,487751455 | 2,435395 | 2,663942 | 0,007723 | 1 | 0 | Distal Intergenic | 10 | 67621579 | 67672231 | 50653 | 1 | 1,03E+08 | ENSMUST00000232007.2 | -31770 | | 4930543I11Rik | RIKEN cDNA 4930543I11 gene |
| chr3 | 1,13E+08 | 1,13E+08 | 419 | * | 4,281978 | 6,487847539 | 2,419904 | 2,681035 | 0,007339 | 1 | 0 | Promoter (<=1kb) | 3 | 1,13E+08 | 1,13E+08 | 41713 | 2 | 11722 | ENSMUST00000174147.2 | -766 | ENSMUSG00000074264 | Amy1 | amylase 1, salivary |
| chr9 | 5466378 | 5466782 | 405 | * | 3,166226 | 6,487922125 | 2,404095 | 2,698697 | 0,006961 | 1 | 0 | Distal Intergenic | 9 | 5345504 | 5358823 | 13320 | 1 | 12364 | ENSMUST00000151332.2 | 120874 | ENSMUSG00000025887 | Casp12 | caspase 12 |
| chr12 | 38547182 | 38547428 | 247 | * | 3,282617 | 6,488167077 | 2,363838 | 2,74476 | 0,006056 | 1 | 0 | Intron (ENSMUST00000 | 12 | 38602807 | 38664520 | 61714 | 2 | 73867 | ENSMUST00000222132.2 | 117092 | ENSMUSG00000112967 | 4930428E07Rik | RIKEN cDNA 4930428E07 gene |
| chr19 | 20449368 | 20450019 | 652 | * | 4,111843 | 6,488293858 | 2,341291 | 2,771246 | 0,005584 | 1 | 0 | Intron (ENSMUST00000 | 19 | 20470079 | 20620829 | 150751 | 1 | 11668 | ENSMUST00000225337.3 | -20060 | ENSMUSG00000053279 | Aldh1a1 | aldehyde dehydrogenase family 1, subfamily A1 |
| chr8 | 37421177 | 37421532 | 356 | * | 4,330304 | 6,488847797 | 2,26724 | 2,862003 | 0,00421 | 1 | 0 | Promoter (<=1kb) | 8 | 37405709 | 37420297 | 14589 | 2 | 50768 | ENSMUST00000179501.2 | -880 | ENSMUSG00000031523 | Dlc1 | deleted in liver cancer 1 |
| chr8 | 24712814 | 24713299 | 486 | * | 4,706078 | 6,489097725 | 2,065818 | 3,141175 | 0,001683 | 1 | 0 | Intron (ENSMUST00000 | 8 | 24902195 | 24906953 | 4759 | 1 | 1,03E+08 | ENSMUST00000208709.2 | -188896 | | Gm30692 | predicted gene, 30692 |
| chr3 | 99961572 | 99961897 | 326 | * | 3,598443 | 6,489343353 | 2,208997 | 2,937688 | 0,003307 | 1 | 0 | Intron (ENSMUST00000 | 3 | 99934924 | 99946267 | 11344 | 1 | 74362 | ENSMUST00000143050.2 | 26648 | ENSMUSG00000027867 | Spag17 | sperm associated antigen 17 |
| chr15 | 80105469 | 80105876 | 408 | * | 4,891416 | 6,489571146 | 2,185782 | 2,968993 | 0,002988 | 1 | 0 | Distal Intergenic | 15 | 80118219 | 80137572 | 19354 | 1 | 239555 | ENSMUST00000023048.12 | -12343 | ENSMUSG00000022412 | Mief1 | mitochondrial elongation factor 1 |
| chr11 | 1,12E+08 | 1,12E+08 | 438 | * | 4,29831 | 6,490159684 | 2,127871 | 3,050072 | 0,002288 | 1 | 0 | Distal Intergenic | 11 | 1,13E+08 | 1,13E+08 | 4560 | 2 | 217294 | ENSMUST00000141399.2 | 350184 | ENSMUSG00000041674 | BC006965 | cDNA sequence BC006965 |
| chr9 | 20791232 | 20791677 | 446 | * | 4,612322 | 6,490214774 | 2,122037 | 3,058483 | 0,002225 | 1 | 0 | Promoter (<=1kb) | 9 | 20785104 | 20791023 | 5920 | 2 | 70726 | ENSMUST00000043726.8 | -209 | ENSMUSG00000038742 | Angptl6 | angiopoietin-like 6 |
| chr19 | 47298386 | 47298839 | 454 | * | 5,00366 | 6,490519229 | 2,096534 | 3,095833 | 0,001963 | 1 | 0 | Intron (ENSMUST00000 | 19 | 47255352 | 47303190 | 47839 | 2 | 14218 | ENSMUST00000237796.2 | 4351 | ENSMUSG00000053617 | Sh3pxd2a | SH3 and PX domains 2A |
| chr18 | 40713255 | 40714157 | 903 | * | 7,169665 | 6,491348638 | 1,904154 | 3,409047 | 0,000652 | 1 | 0 | Distal Intergenic | 18 | 40656152 | 40664232 | 8081 | 1 | 383348 | ENSMUST00000236525.2 | 57103 | ENSMUSG00000051401 | Kctd16 | potassium channel tetramerisation domain containing 16 |
| chr15 | 27024542 | 27025039 | 498 | * | 6,682751 | 6,491502439 | 2,024988 | 3,205699 | 0,001347 | 1 | 0 | Distal Intergenic | 15 | 27061822 | 27061952 | 131 | 2 | 1,15E+08 | ENSMUST00000157566.3 | 36913 | ENSMUSG00000088191 | Gm23903 | predicted gene, 23903 |
| chr5 | 1,36E+08 | 1,36E+08 | 775 | * | 6,001295 | 6,492049071 | 1,99193 | 3,259176 | 0,001117 | 1 | 0 | 3' UTR | 5 | 1,36E+08 | 1,36E+08 | 8964 | 2 | 215210 | ENSMUST00000043378.9 | -10362 | ENSMUSG00000039886 | Tmem120a | transmembrane protein 120A |
| chr1 | 1,87E+08 | 1,87E+08 | 360 | * | 3,287305 | 6,492051891 | 2,440092 | 2,660577 | 0,007801 | 1 | 0 | Intron (ENSMUST00000 | 1 | 1,87E+08 | 1,87E+08 | 125248 | 1 | 67769 | ENSMUST00000160570.2 | 90388 | ENSMUSG00000039210 | Gpatch2 | G patch domain containing 2 |
| chr5 | 1,35E+08 | 1,35E+08 | 489 | * | 5,505328 | 6,492650877 | 1,959246 | 3,313852 | 0,00092 | 1 | 0 | Distal Intergenic | 5 | 1,35E+08 | 1,35E+08 | 2293 | 2 | 14371 | ENSMUST00000062572.3 | -19161 | ENSMUSG00000049551 | Fzd9 | frizzled class receptor 9 |
| chr12 | 1,05E+08 | 1,05E+08 | 649 | * | 5,466869 | 6,492753785 | 1,954419 | 3,322089 | 0,000893 | 1 | 0 | Distal Intergenic | 12 | 1,05E+08 | 1,05E+08 | 79877 | 2 | 212073 | ENSMUST00000067005.11 | -5319 | ENSMUSG00000054150 | Syne3 | spectrin repeat containing, nuclear envelope family member 3 |
| chr6 | 67150234 | 67151266 | 1033 | * | 6,021677 | 6,492994174 | 1,943044 | 3,341661 | 0,000833 | 1 | 0 | Intron (ENSMUST00000 | 6 | 67102993 | 67163578 | 60586 | 2 | 1,05E+08 | ENSMUST00000246733.1 | 12312 | | Gm38825 | predicted gene, 38825 |

| | | | | | | | | | | | | | | | | | | | | | | | |
|---|---|---|---|---|---|---|---|---|---|---|---|---|---|---|---|---|---|---|---|---|---|---|---|
| chr10 | 16083613 | 16084310 | 698 | * | 3,561129 | 6,494532976 | 2,486123 | 2,612313 | 0,008993 | 1 | 0 | Distal Intergenic | 10 | 15667525 | 15682813 | 15289 | 1 | 77538 | ENSMUST00000218833.2 | 416088 | ENSMUSG00000112227 | C330004P14Rik | RIKEN cDNA C330004P14 gene |
| chr11 | 26589636 | 26590123 | 488 | * | 3,736509 | 6,494655469 | 2,47301 | 2,626215 | 0,008634 | 1 | 0 | Distal Intergenic | 11 | 26497848 | 26543999 | 46152 | 2 | 69922 | ENSMUST00000156264.2 | -45637 | ENSMUSG00000064090 | Vrk2 | vaccinia related kinase 2 |
| chr17 | 19290303 | 19290581 | 279 | * | 4,327866 | 6,494909403 | 2,446249 | 2,655048 | 0,00793 | 1 | 0 | Intron (ENSMUST00000 | 17 | 19273722 | 19302673 | 28952 | 1 | 224552 | ENSMUST00000170424.3 | 16581 | ENSMUSG00000096717 | Vmn2r98 | vomeronasal 2, receptor 98 |
| chr7 | 10328634 | 10329126 | 493 | * | 3,420342 | 6,495139597 | 2,425351 | 2,678021 | 0,007406 | 1 | 0 | Intron (ENSMUST00000 | 7 | 10312857 | 10315414 | 2558 | 2 | 252904 | ENSMUST00000163658.3 | -13220 | ENSMUSG00000091662 | Vmn1r69 | vomeronasal 1 receptor 69 |
| chr12 | 49920072 | 49920948 | 877 | * | 7,177975 | 6,49525551 | 1,851082 | 3,508896 | 0,00045 | 0,352029189 | 1,360622 | Distal Intergenic | 12 | 50167092 | 50192019 | 24928 | 1 | 1,05E+08 | ENSMUST00000218806.2 | -246144 | ENSMUSG00000112353 | Gm40418 | predicted gene, 40418 |
| chr14 | 1,18E+08 | 1,18E+08 | 481 | * | 3,660636 | 6,495419437 | 2,395296 | 2,71174 | 0,006693 | 1 | 0 | Distal Intergenic | 14 | 1,18E+08 | 1,18E+08 | 105 | 2 | 1,02E+08 | ENSMUST00000183564.3 | -36754 | ENSMUSG00000098648 | Mir6239 | microRNA 6239 |
| chr12 | 1,07E+08 | 1,07E+08 | 315 | * | 3,567099 | 6,495643186 | 2,375228 | 2,734745 | 0,006243 | 1 | 0 | Distal Intergenic | 12 | 1,07E+08 | 1,07E+08 | 2117 | 2 | 1,05E+08 | ENSMUST00000124395.2 | 36712 | ENSMUSG00000086319 | Gm16087 | predicted gene 16087 |
| chr9 | 62865329 | 62865674 | 346 | * | 5,160984 | 6,498021954 | 2,200952 | 2,952369 | 0,003153 | 1 | 0 | Intron (ENSMUST00000 | 9 | 62830189 | 62859278 | 29090 | 2 | 56469 | ENSMUST00000215455.2 | -6051 | ENSMUSG00000032405 | Pias1 | protein inhibitor of activated STAT 1 |
| chr14 | 21400704 | 21401358 | 655 | * | 5,287744 | 6,498132492 | 2,098131 | 3,097105 | 0,001954 | 1 | 0 | Intron (ENSMUST00000 | 14 | 21419138 | 21420402 | 1265 | 1 | 11534 | ENSMUST00000224974.2 | -17780 | ENSMUSG00000039197 | Adk | adenosine kinase |
| chr5 | 32320201 | 32322007 | 1807 | * | 13,35475 | 6,498380085 | 1,762786 | 3,686427 | 0,000227 | 0,338590591 | 0,738441 | Distal Intergenic | 5 | 32332443 | 32332549 | 107 | 1 | 1,15E+08 | ENSMUST00000240394.1 | -10436 | ENSMUSG00000118998 | Gm26335 | predicted gene, 26335 |
| chr10 | 16997068 | 16998177 | 1110 | * | 6,64571 | 6,498810545 | 1,90288 | 3,415249 | 0,000637 | 1 | 0 | Distal Intergenic | 10 | 16857052 | 16971555 | 114504 | 2 | 1,01E+08 | ENSMUST00000208870.2 | -25513 | ENSMUSG00000101969 | Gm20125 | predicted gene, 20125 |
| chr13 | 12594616 | 12595390 | 775 | * | 6,277998 | 6,498919338 | 1,967094 | 3,303817 | 0,000954 | 1 | 0 | Exon (ENSMUST000002 | 13 | 12581339 | 12621990 | 40652 | 1 | 67475 | ENSMUST00000221739.2 | 13277 | ENSMUSG00000057069 | Ero1b | endoplasmic reticulum oxidoreductase 1 beta |
| chr1 | 96267226 | 96267979 | 754 | * | 5,906327 | 6,498962814 | 1,985721 | 3,272847 | 0,001065 | 1 | 0 | Distal Intergenic | 1 | 96544824 | 96544930 | 107 | 1 | 1,15E+08 | ENSMUST00001239562.1 | -276845 | ENSMUSG00000119190 | Gm26245 | predicted gene, 26245 |
| chr17 | 18648930 | 18649567 | 638 | * | 5,552183 | 6,49898012 | 1,991178 | 3,263888 | 0,001099 | 1 | 0 | Intron (ENSMUST00000 | 17 | 18644340 | 18681167 | 36828 | 1 | 328759 | ENSMUST00000232090.2 | 4590 | ENSMUSG00000091631 | Vmn2r95 | vomeronasal 2, receptor 95 |
| chrX | 77512865 | 77513326 | 462 | * | 5,034087 | 6,499043302 | 2,044546 | 3,178722 | 0,001479 | 1 | 0 | Intron (ENSMUST00000 | 20 | 77413249 | 77414734 | 1486 | 1 | 74910 | ENSMUST00000035626.8 | 99616 | ENSMUSG00000035651 | 4930480E11Rik | RIKEN cDNA 4930480E11 gene |
| chr17 | 74612903 | 74613502 | 600 | * | 4,236631 | 6,499100836 | 2,089706 | 3,110055 | 0,001871 | 1 | 0 | Intron (ENSMUST00000 | 17 | 74606477 | 74623198 | 16722 | 2 | 66310 | ENSMUST00000232687.2 | 9696 | ENSMUSG00000024067 | Dpy30 | dpy-30, histone methyltransferase complex regulatory subunit |
| chr14 | 88630665 | 88631016 | 352 | * | 5,013727 | 6,49911281 | 2,09868 | 3,096762 | 0,001956 | 1 | 0 | Distal Intergenic | 14 | 88520693 | 88574980 | 54288 | 2 | 1,03E+08 | ENSMUST00000226380.2 | -55685 | | Gm33203 | predicted gene, 33203 |
| chr1 | 64761040 | 64761525 | 486 | * | 4,160231 | 6,499135015 | 2,127434 | 3,054918 | 0,002251 | 1 | 0 | Intron (ENSMUST00000 | 1 | 64773628 | 64776908 | 3281 | 2 | 14367 | ENSMUST00000116133.4 | 15383 | ENSMUSG00000045005 | Fzd5 | frizzled class receptor 5 |
| chr4 | 9249593 | 9250125 | 533 | * | 4,107252 | 6,499223363 | 2,268588 | 2,864877 | 0,004172 | 1 | 0 | Distal Intergenic | 4 | 9269293 | 9282272 | 12980 | 1 | 74438 | ENSMUST00000147285.2 | -19168 | ENSMUSG00000041216 | Clvs1 | clavesin 1 |
| chr5 | 68662267 | 68662691 | 425 | * | 3,722925 | 6,499348387 | 2,418638 | 2,687194 | 0,007206 | 1 | 0 | Distal Intergenic | 5 | 68723513 | 68723639 | 127 | 2 | 1,15E+08 | ENSMUST00000157255.3 | 60948 | ENSMUSG00000087880 | Gm23605 | predicted gene, 23605 |
| chr7 | 79300302 | 79301053 | 752 | * | 5,334727 | 6,500491062 | 2,077573 | 3,128886 | 0,001755 | 1 | 0 | Distal Intergenic | 7 | 79309944 | 79347896 | 37953 | 1 | 77011 | ENSMUST00000035977.9 | -8891 | ENSMUSG00000046591 | Ticrr | TOPBP1-interacting checkpoint and replication regulator |
| chr16 | 58926689 | 58927285 | 597 | * | 4,410408 | 6,50107165 | 2,242341 | 2,899235 | 0,003741 | 1 | 0 | Distal Intergenic | 16 | 58929988 | 58930996 | 1009 | 2 | 257972 | ENSMUST00000076262.4 | 3711 | ENSMUSG00000060057 | Olfr193 | olfactory receptor 193 |
| chrX | 1,49E+08 | 1,49E+08 | 780 | * | 6,890218 | 6,502715381 | 1,867308 | 3,4824 | 0,000497 | 1 | 0 | Distal Intergenic | 20 | 1,49E+08 | 1,49E+08 | 42193 | 2 | 382245 | ENSMUST00000155740.2 | -17194 | ENSMUSG00000041353 | Tmem29 | transmembrane protein 29 |
| chr2 | 1,18E+08 | 1,18E+08 | 647 | * | 5,742015 | 6,503541476 | 1,969581 | 3,301992 | 0,00096 | 1 | 0 | Intron (ENSMUST00000 | 2 | 1,18E+08 | 1,18E+08 | 96310 | 2 | 381413 | ENSMUST00000039160.3 | 12149 | ENSMUSG00000040133 | Gpr176 | G protein-coupled receptor 176 |
| chr12 | 59117458 | 59118525 | 1068 | * | 7,777603 | 6,504657673 | 1,935702 | 3,36036 | 0,000778 | 1 | 0 | Exon (ENSMUST00000 | 12 | 59113750 | 59115477 | 1728 | 1 | 18949 | ENSMUST00000217676.2 | 3708 | ENSMUSG00000020994 | Pnn | pinin |
| chrX | 12979166 | 12980160 | 995 | * | 5,29842 | 6,505948256 | 1,992161 | 3,265774 | 0,001092 | 1 | 0 | Promoter (<=1kb) | 20 | 12979923 | 12989472 | 9550 | 1 | 22284 | ENSMUST00000149021.3 | 0 | ENSMUSG00000031010 | Usp9x | ubiquitin specific peptidase 9, X chromosome |
| chrX | 72972056 | 72972531 | 476 | * | 3,101879 | 6,506285961 | 2,505509 | 2,596792 | 0,00941 | 1 | 0 | Promoter (<=1kb) | 20 | 72968869 | 72972735 | 3867 | 2 | 19703 | ENSMUST00000144771.2 | 204 | ENSMUSG00000031387 | Renbp | renin binding protein |
| chr14 | 54871000 | 54871501 | 502 | * | 4,743435 | 6,507415119 | 2,077783 | 3,131903 | 0,001737 | 1 | 0 | Exon (ENSMUST00000 | 14 | 54870010 | 54874998 | 4989 | 2 | 239096 | ENSMUST00000227124.2 | 3497 | ENSMUSG00000059674 | Cdh24 | cadherin-like 24 |
| chr5 | 31878117 | 31878689 | 573 | * | 4,077118 | 6,507441658 | 2,25374 | 2,887396 | 0,003884 | 1 | 0 | Intron (ENSMUST00000 | 5 | 31859189 | 32215015 | 355827 | 1 | 107976 | ENSMUST00000200705.2 | 18928 | ENSMUSG00000052139 | Babam2 | BRISC and BRCA1 A complex member 2 |
| chr16 | 80091863 | 80092455 | 593 | * | 3,944274 | 6,507604191 | 2,2278 | 2,92109 | 0,003488 | 1 | 0 | Distal Intergenic | 16 | 79796264 | 79797434 | 1171 | 1 | 78654 | ENSMUST00000191029.2 | 295599 | ENSMUSG00000101810 | 1700066C05Rik | RIKEN cDNA 1700066C05 gene |
| chr14 | 73261157 | 73261881 | 725 | * | 4,794265 | 6,507846786 | 2,107554 | 3,087867 | 0,002016 | 1 | 0 | Distal Intergenic | 14 | 73266568 | 73286512 | 19945 | 2 | 70086 | ENSMUST00000044664.12 | 24631 | ENSMUSG00000033470 | Cysltr2 | cysteinyl leukotriene receptor 2 |
| chr16 | 76023801 | 76024432 | 632 | * | 5,342121 | 6,507974503 | 2,169074 | 3,000346 | 0,002697 | 1 | 0 | Distal Intergenic | 16 | 75929429 | 75929559 | 131 | 2 | 1,15E+08 | ENSMUST00000104009.3 | -94242 | ENSMUSG00000077197 | Gm22268 | predicted gene, 22268 |
| chr14 | 9284287 | 9285677 | 1391 | * | 8,805991 | 6,508225983 | 1,858087 | 3,502648 | 0,000461 | 0,377927525 | 1,211315 | Intron (ENSMUST00000 | 14 | 9298861 | 9311189 | 12329 | 1 | 328354 | ENSMUST00000161835.3 | -13184 | ENSMUSG00000051729 | Gm5087 | predicted gene 5087 |
| chr19 | 46708955 | 46709350 | 396 | * | 4,385804 | 6,508809037 | 2,073893 | 3,13845 | 0,001698 | 1 | 0 | Intron (ENSMUST00000 | 19 | 46700411 | 46708860 | 8450 | 1 | 57344 | ENSMUST00000235266.2 | 8544 | ENSMUSG00000003559 | As3mt | arsenite methyltransferase |
| chrX | 87346828 | 87347613 | 786 | * | 5,104575 | 6,508858305 | 2,067496 | 3,148185 | 0,001643 | 1 | 0 | Distal Intergenic | 20 | 87236223 | 87236329 | 107 | 1 | 1,15E+08 | ENSMUST00000083813.3 | 110605 | ENSMUSG00000065747 | Gm25006 | predicted gene, 25006 |
| chr12 | 1,06E+08 | 1,06E+08 | 668 | * | 4,945413 | 6,509002557 | 2,053204 | 3,170168 | 0,001524 | 1 | 0 | Intron (ENSMUST00000 | 12 | 1,06E+08 | 1,06E+08 | 52402 | 1 | 18789 | ENSMUST00000170002.8 | 3661 | ENSMUSG00000021111 | Papola | poly (A) polymerase alpha |
| chr1 | 87859082 | 87859774 | 693 | * | 4,686274 | 6,50900677 | 2,054356 | 3,168393 | 0,001533 | 1 | 0 | Promoter (<=1kb) | 1 | 87859704 | 87865975 | 6272 | 1 | 227333 | ENSMUST00000185260.7 | 0 | ENSMUSG00000070738 | Dgkd | diacylglycerol kinase, delta |
| chr2 | 38508326 | 38508879 | 554 | * | 5,097172 | 6,50990104 | 1,977568 | 3,291872 | 0,000995 | 1 | 0 | Intron (ENSMUST00000 | 2 | 38501183 | 38533932 | 32750 | 2 | 19177 | ENSMUST00000151981.3 | 25053 | ENSMUSG00000026750 | Psmb7 | proteasome (prosome, macropain) subunit, beta type 7 |
| chr5 | 31081321 | 31082138 | 818 | * | 6,733934 | 6,51129834 | 1,889722 | 3,445639 | 0,00057 | 1 | 0 | Promoter (2-3kb) | 5 | 31079202 | 31088137 | 8936 | 1 | 16548 | ENSMUST00000202752.2 | 2119 | ENSMUSG00000029162 | Khk | ketohexokinase |
| chr11 | 69263437 | 69263904 | 468 | * | 3,692783 | 6,513702142 | 2,430848 | 2,679601 | 0,007371 | 1 | 0 | Distal Intergenic | 11 | 69234099 | 69260232 | 26134 | 2 | 216848 | ENSMUST00000108661.8 | -3205 | ENSMUSG00000018474 | Chd3 | chromodomain helicase DNA binding protein 3 |
| chr16 | 97763392 | 97764552 | 1161 | * | 9,290158 | 6,514163382 | 1,739989 | 3,743795 | 0,000181 | 0,17530799 | 2,237403 | Promoter (<=1kb) | 16 | 97744557 | 97763815 | 19259 | 2 | 114565 | ENSMUST00000232010.2 | 0 | ENSMUSG00000046962 | Zbtb21 | zinc finger and BTB domain containing 21 |
| chr14 | 1,17E+08 | 1,17E+08 | 560 | * | 4,040777 | 6,514226462 | 2,371907 | 2,746409 | 0,006025 | 1 | 0 | Intron (ENSMUST00000 | 14 | 1,17E+08 | 1,18E+08 | 701884 | 1 | 23888 | ENSMUST00000123239.2 | -573072 | ENSMUSG00000058571 | Gpc6 | glypican 6 |
| chr3 | 84572625 | 84573187 | 563 | * | 3,884322 | 6,514463313 | 2,349572 | 2,772617 | 0,005561 | 1 | 0 | Promoter (<=1kb) | 3 | 84573499 | 84611882 | 38384 | 1 | 320782 | ENSMUST00000107682.2 | -312 | ENSMUSG00000056498 | Tmem154 | transmembrane protein 154 |
| chr1 | 1,93E+08 | 1,93E+08 | 730 | * | 3,59854 | 6,514608635 | 2,334434 | 2,790659 | 0,00526 | 1 | 0 | Intron (ENSMUST00000 | 1 | 1,93E+08 | 1,93E+08 | 23572 | 2 | 215193 | ENSMUST00000194412.3 | 12149 | ENSMUSG00000016181 | Utp25 | UTP25 small subunit processome component |
| chr1 | 88630085 | 88630596 | 512 | * | 4,242395 | 6,51461094 | 2,334095 | 2,791065 | 0,005253 | 1 | 0 | Promoter (<=1kb) | 1 | 88625947 | 88629943 | 3997 | 2 | 320982 | ENSMUST00000159814.2 | -142 | ENSMUSG00000049866 | Arl4c | ADP-ribosylation factor-like 4C |
| chr9 | 96661835 | 96662535 | 701 | * | 7,7641 | 6,515215165 | 1,81387 | 3,591887 | 0,000328 | 1 | 0 | Exon (ENSMUST000001 | 9 | 96647654 | 96655343 | 7690 | 1 | 319707 | ENSMUST00000190054.2 | 14181 | ENSMUSG00000101581 | C430002N11Rik | RIKEN cDNA C430002N11 gene |
| chr6 | 29731380 | 29731877 | 498 | * | 3,830353 | 6,515882625 | 2,227448 | 2,925268 | 0,003442 | 1 | 0 | Distal Intergenic | 6 | 29735693 | 29761364 | 25672 | 1 | 319757 | ENSMUST00000001812.5 | -3816 | ENSMUSG00000001761 | Smo | smoothened, frizzled class receptor |
| chr11 | 1,03E+08 | 1,03E+08 | 775 | * | 6,214184 | 6,515882835 | 1,880347 | 3,465256 | 0,00053 | 1 | 0 | Distal Intergenic | 11 | 1,03E+08 | 1,03E+08 | 11593 | 1 | 268491 | ENSMUST00000156590.8 | -9118 | ENSMUSG00000051455 | Meioc | meiosis specific with coiled-coil domain |
| chr10 | 1,15E+08 | 1,15E+08 | 752 | * | 7,677873 | 6,516085711 | 1,90566 | 3,419332 | 0,000628 | 1 | 0 | Promoter (<=1kb) | 10 | 1,15E+08 | 1,15E+08 | 53596 | 2 | 66687 | ENSMUST00000020339.10 | 0 | ENSMUSG00000020130 | Tbc1d15 | TBC1 domain family, member 15 |
| chr3 | 4333531 | 4334099 | 569 | * | 6,620064 | 6,516372681 | 1,945696 | 3,349121 | 0,000811 | 1 | 0 | Distal Intergenic | 3 | 4404058 | 4404176 | 119 | 1 | 1,15E+08 | ENSMUST00000239917.1 | -69959 | ENSMUSG00000119546 | n-R5s194 | nuclear encoded rRNA 5S 194 |
| chr18 | 66147220 | 66147732 | 513 | * | 5,27498 | 6,51660823 | 2,17448 | 2,996859 | 0,002728 | 1 | 0 | Intron (ENSMUST00000 | 18 | 66126199 | 66152849 | 26651 | 2 | 70361 | ENSMUST00000150133.3 | 5117 | ENSMUSG00000041891 | Lman1 | lectin, mannose-binding, 1 |
| chr13 | 51794211 | 51794635 | 425 | * | 4,72925 | 6,516722807 | 2,004497 | 3,251051 | 0,00115 | 1 | 0 | Distal Intergenic | 13 | 51799268 | 51804696 | 5429 | 1 | 66197 | ENSMUST00000075853.6 | -4633 | ENSMUSG00000062248 | Cks2 | CDC28 protein kinase regulatory subunit 2 |
| chrX | 86543752 | 86544309 | 558 | * | 4,906735 | 6,516913073 | 2,155285 | 3,02369 | 0,002497 | 1 | 0 | Intron (ENSMUST00000 | 20 | 85790848 | 86933841 | 1142994 | 2 | 331461 | ENSMUST00000078875.8 | 389532 | ENSMUSG00000052372 | Il1rapl1 | interleukin 1 receptor accessory protein-like 1 |
| chrX | 85237108 | 85237996 | 889 | * | 3,994433 | 6,516925365 | 2,15522 | 3,023786 | 0,002496 | 1 | 0 | Promoter (1-2kb) | 20 | 85235370 | 85239553 | 4184 | 1 | 11614 | ENSMUST00000026036.5 | 1738 | ENSMUSG00000025056 | Nr0b1 | nuclear receptor subfamily 0, group B, member 1 |
| chrX | 1,01E+08 | 1,01E+08 | 806 | * | 4,635075 | 6,517158832 | 2,101596 | 3,101053 | 0,001928 | 1 | 0 | Intron (ENSMUST00000 | 20 | 1,01E+08 | 1,01E+08 | 95 | 2 | 1,15E+08 | ENSMUST00000157087.4 | -8031 | ENSMUSG00000087712 | Gm25828 | predicted gene, 25828 |
| chr1 | 23532826 | 23533243 | 418 | * | 4,599147 | 6,517335574 | 2,143306 | 3,040786 | 0,00236 | 1 | 0 | Distal Intergenic | 1 | 23613668 | 23635745 | 22078 | 1 | 1,05E+08 | ENSMUST00000191140.2 | -80425 | ENSMUSG00000099503 | Gm28822 | predicted gene 28822 |
| chr11 | 80291637 | 80292155 | 519 | * | 3,937495 | 6,517576409 | 2,229442 | 2,923412 | 0,003462 | 1 | 0 | Intron (ENSMUST00000 | 11 | 80279957 | 80295647 | 15691 | 1 | 22680 | ENSMUST00000123726.2 | 11680 | ENSMUSG00000017421 | Zfp207 | zinc finger protein 207 |
| chr9 | 55790878 | 55791225 | 348 | * | 3,632711 | 6,517639685 | 2,247552 | 2,899884 | 0,003733 | 1 | 0 | Exon (ENSMUST000002 | 9 | 55457167 | 55826889 | 369723 | 2 | 244891 | ENSMUST00000037408.9 | 35664 | ENSMUSG00000034007 | Scaper | S phase cyclin A-associated protein in the ER |
| chr1 | 1,85E+08 | 1,85E+08 | 523 | * | 3,806054 | 6,517707754 | 2,278808 | 2,860139 | 0,004235 | 1 | 0 | Distal Intergenic | 1 | 1,85E+08 | 1,85E+08 | 630 | 1 | 114668 | ENSMUST00000192616.2 | 242968 | ENSMUSG00000102185 | 5033404E19Rik | NSA2 ribosome biogenesis homolog pseudogene |
| chr7 | 81568958 | 81569362 | 405 | * | 4,266767 | 6,517791583 | 2,311842 | 2,819306 | 0,004813 | 1 | 0 | Exon (ENSMUST000002 | 7 | 81547559 | 81583771 | 36213 | 2 | 29877 | ENSMUST00000149288.2 | 14409 | ENSMUSG00000025104 | Hdgfl3 | HDGF like 3 |
| chr1 | 1,83E+08 | 1,83E+08 | 370 | * | 3,200072 | 6,518035262 | 2,431792 | 2,680342 | 0,007355 | 1 | 0 | Distal Intergenic | 1 | 1,83E+08 | 1,83E+08 | 90181 | 1 | 96935 | ENSMUST00000155229.7 | -7130 | ENSMUSG00000038576 | Susd4 | sushi domain containing 4 |
| chr4 | 29775358 | 29775810 | 453 | * | 2,866452 | 6,518112105 | 2,479377 | 2,628932 | 0,008565 | 1 | 0 | Distal Intergenic | 4 | 29068079 | 29088811 | 20733 | 1 | 1,03E+08 | ENSMUST00000154813.2 | 707279 | | Gm11917 | predicted gene 11917 |
| chr7 | 89317664 | 89318306 | 643 | * | 6,006349 | 6,519261119 | 2,033208 | 3,206392 | 0,001344 | 1 | 0 | Intron (ENSMUST00000 | 7 | 89282094 | 89498360 | 216267 | 1 | 109264 | ENSMUST00000159167.8 | 35570 | ENSMUSG00000030621 | Me3 | malic enzyme 3, NADP(+)-dependent, mitochondrial |
| chr15 | 26240817 | 26241482 | 666 | * | 5,931639 | 6,519390929 | 2,027774 | 3,215048 | 0,001304 | 1 | 0 | Distal Intergenic | 15 | 26309134 | 26312457 | 3324 | 1 | 211147 | ENSMUST00000126304.2 | -67652 | ENSMUSG00000022269 | Marchf11 | membrane associated ring-CH-type finger 11 |
| chr11 | 84841765 | 84842504 | 740 | * | 6,249622 | 6,520168417 | 1,996123 | 3,266416 | 0,001089 | 1 | 0 | Distal Intergenic | 11 | 84848612 | 84853381 | 4770 | 1 | 12351 | ENSMUST00000139416.2 | -6108 | ENSMUSG00000000805 | Car4 | carbonic anhydrase 4 |
| chr6 | 43580216 | 43580607 | 392 | * | 4,123489 | 6,521172009 | 2,403058 | 2,713697 | 0,006654 | 1 | 0 | Intron (ENSMUST00000 | 6 | 43322007 | 43602900 | 280894 | 2 | 29807 | ENSMUST00000203649.3 | 22293 | ENSMUSG00000029735 | Tpk1 | thiamine pyrophosphokinase |
| chr2 | 1,32E+08 | 1,32E+08 | 558 | * | 5,846596 | 6,522283655 | 1,939759 | 3,362419 | 0,000773 | 1 | 0 | Distal Intergenic | 2 | 1,32E+08 | 1,32E+08 | 8163 | 2 | 70612 | ENSMUST00000110164.8 | 24153 | ENSMUSG00000027341 | Tmem230 | transmembrane protein 230 |
| chr7 | 18605463 | 18606120 | 658 | * | 6,309636 | 6,522389852 | 1,92225 | 3,393101 | 0,000691 | 1 | 0 | Promoter (<=1kb) | 7 | 18605295 | 18624384 | 19090 | 1 | 21946 | ENSMUST00000206569.2 | 168 | ENSMUSG00000030413 | Pglyrp1 | peptidoglycan recognition protein 1 |
| chr1 | 1,24E+08 | 1,24E+08 | 423 | * | 4,704917 | 6,523126972 | 2,275763 | 2,866348 | 0,004152 | 1 | 0 | Intron (ENSMUST00000 | 1 | 1,24E+08 | 1,24E+08 | 104 | 1 | 1,15E+08 | ENSMUST00000083406.3 | -72645 | ENSMUSG00000065340 | Gm25578 | predicted gene, 25578 |
| chr12 | 73245928 | 73246345 | 418 | * | 4,964403 | 6,524277414 | 2,215899 | 2,944302 | 0,003237 | 1 | 0 | Intron (ENSMUST00000 | 12 | 73192548 | 73192679 | 132 | 2 | 1,15E+08 | ENSMUST00000104562.3 | -53249 | ENSMUSG00000119928 | Gm23910 | predicted gene, 23910 |
| chr12 | 26715834 | 26716675 | 842 | * | 8,172731 | 6,524895025 | 1,857721 | 3,512312 | 0,000444 | 0,284054467 | 2,237403 | Distal Intergenic | 12 | 26622107 | 26652308 | 30202 | 2 | 69437 | ENSMUST00000221140.2 | -63526 | | 1700020D12Rik | RIKEN cDNA 1700020D12 gene |
| chr4 | 1,27E+08 | 1,27E+08 | 833 | * | 4,286434 | 6,525118026 | 2,427526 | 2,68797 | 0,007189 | 1 | 0 | Distal Intergenic | 4 | 1,27E+08 | 1,27E+08 | 106197 | 2 | 67785 | ENSMUST00000106108.9 | -18906 | ENSMUSG00000042446 | Zmym4 | zinc finger, MYM-type 4 |
| chr9 | 52056201 | 52056987 | 787 | * | 5,859771 | 6,525315796 | 2,16791 | 3,009957 | 0,002613 | 1 | 0 | Intron (ENSMUST00000 | 9 | 52023285 | 52079411 | 56127 | 2 | 244871 | ENSMUST00000165519.2 | 22424 | ENSMUSG00000035164 | Zc3h12c | zinc finger CCCH type containing 12C |
| chr7 | 45088102 | 45088536 | 435 | * | 4,144599 | 6,525789307 | 2,227915 | 2,929101 | 0,003399 | 1 | 0 | Promoter (<=1kb) | 7 | 45071184 | 45087520 | 16337 | 2 | 20174 | ENSMUST00000211214.2 | -582 | ENSMUSG00000003868 | Ruvbl2 | RuvB-like protein 2 |
| chr3 | 77400976 | 77401492 | 517 | * | 3,562878 | 6,525885946 | 2,208522 | 2,954866 | 0,003128 | 1 | 0 | Distal Intergenic | 3 | 77012516 | 77012592 | 77 | 2 | 1,15E+08 | ENSMUST00000157856.4 | -388384 | | Gm23644 | predicted gene, 23644 |
| chr5 | 9092457 | 9092813 | 357 | * | 4,007946 | 6,526327855 | 2,120427 | 3,077837 | 0,002085 | 1 | 0 | Intron (ENSMUST00000 | 5 | 9085124 | 9111316 | 26193 | 1 | 1,05E+08 | ENSMUST00000224996.2 | 7333 | | Gm40264 | predicted gene, 40264 |
| chr8 | 1,12E+08 | 1,12E+08 | 485 | * | 4,21835 | 6,526495457 | 2,092543 | 3,11893 | 0,001815 | 1 | 0 | Distal Intergenic | 8 | 1,12E+08 | 1,12E+08 | 29279 | 2 | 234736 | ENSMUST00000038739.5 | -5906 | ENSMUSG00000033596 | Rfwd3 | ring finger and WD repeat domain 3 |
| chr10 | 73653907 | 73654432 | 526 | * | 5,125917 | 6,526586886 | 2,075263 | 3,144945 | 0,001661 | 1 | 0 | Intron (ENSMUST00000 | 10 | 73657699 | 74470597 | 812899 | 1 | 11994 | ENSMUST00000064562.14 | -3267 | ENSMUSG00000052613 | Pcdh15 | protocadherin 15 |
| chr12 | 51797913 | 51798391 | 479 | * | 4,944278 | 6,527095513 | 2,005993 | 3,253798 | 0,001139 | 1 | 0 | Intron (ENSMUST00000 | 12 | 51804821 | 51812014 | 7194 | 2 | 207304 | ENSMUST00000218626.2 | 13623 | ENSMUSG00000035247 | Hectd1 | HECT domain E3 ubiquitin protein ligase 1 |
| chr10 | 81211068 | 81211344 | 277 | * | 4,607186 | 6,527395887 | 2,21311 | 2,949423 | 0,003184 | 1 | 0 | Promoter (2-3kb) | 10 | 81211197 | 81214214 | 3018 | 2 | 56371 | ENSMUST00000127203.2 | 2870 | ENSMUSG00000020235 | Fzr1 | fizzy and cell division cycle 20 related 1 |
| chr1 | 64646475 | 64647104 | 630 | * | 6,293743 | 6,527656378 | 1,942636 | 3,360205 | 0,000779 | 1 | 0 | 3' UTR | 1 | 64647115 | 64654790 | 7676 | 2 | 67099 | ENSMUST00000162806.8 | 7686 | ENSMUSG00000025956 | Mettl21a | methyltransferase like 21A |
| chr17 | 32160985 | 32161854 | 870 | * | 9,047875 | 6,528090991 | 1,795717 | 3,635368 | 0,000278 | 0,282235674 | 1,425929 | Distal Intergenic | 17 | 32126853 | 32128003 | 1151 | 2 | 69561 | ENSMUST00000235350.2 | -32982 | | 2310015A16Rik | RIKEN cDNA 2310015A16 gene |
| chr2 | 1,74E+08 | 1,74E+08 | 730 | * | 6,098108 | 6,528099396 | 1,902274 | 3,431735 | 0,0006 | 1 | 0 | Distal Intergenic | 2 | 1,74E+08 | 1,74E+08 | 46829 | 1 | 56491 | ENSMUST00000067530.6 | -5910 | ENSMUSG00000054455 | Vapb | vesicle-associated membrane protein, associated protein B and C |
| chr5 | 36566592 | 36567214 | 623 | * | 7,237118 | 6,528603755 | 1,862859 | 3,504616 | 0,000457 | 0,353999513 | 1,360622 | Distal Intergenic | 5 | 36307547 | 36555483 | 247937 | 2 | 81840 | ENSMUST00000070720.8 | -11109 | ENSMUSG00000029093 | Sorcs2 | sortilin-related VPS10 domain containing receptor 2 |
| chr4 | 86531860 | 86532940 | 1081 | * | 8,28038 | 6,529507031 | 1,806199 | 3,615054 | 0,0003 | 0,233277523 | 2,237403 | Promoter (1-2kb) | 4 | 86503264 | 86530292 | 27029 | 2 | 230376 | ENSMUST00000128381.8 | -1568 | ENSMUSG00000038047 | Haus6 | HAUS augmin-like complex, subunit 6 |
| chr17 | 33800063 | 33801607 | 1545 | * | 10,47176 | 6,529640834 | 1,731327 | 3,771467 | 0,000162 | 0,270529135 | 0,918255 | Promoter (<=1kb) | 17 | 33801577 | 33826738 | 25162 | 1 | 17916 | ENSMUST00000173426.2 | 0 | ENSMUSG00000024300 | Myo1f | myosin IF |
| chr11 | 62738030 | 62739044 | 1015 | * | 9,022863 | 6,530403039 | 1,757133 | 3,71651 | 0,000202 | 0,188932071 | 2,237403 | Promoter (<=1kb) | 11 | 62737958 | 62768269 | 30312 | 1 | 213980 | ENSMUST00000176577.2 | 72 | ENSMUSG00000090173 | Fbxw10 | F-box and WD-40 domain protein 10 |
| chr1 | 84834568 | 84835353 | 786 | * | 11,03872 | 6,530510518 | 1,758994 | 3,712639 | 0,000205 | 0,300106667 | 0,918255 | Intron (ENSMUST00000 | 1 | 84844157 | 84878208 | 34052 | 1 | 66153 | ENSMUST00000139562.8 | -8804 | ENSMUSG00000073633 | Fbxo36 | F-box protein 36 |
| chr9 | 75723688 | 75724441 | 754 | * | 7,47579 | 6,531310615 | 1,97659 | 3,304332 | 0,000952 | 1 | 0 | Intron (ENSMUST00000 | 9 | 75735304 | 75750882 | 15579 | 1 | 12160 | ENSMUST00000137213.2 | -10863 | ENSMUSG00000032179 | Bmp5 | bone morphogenetic protein 5 |
| chr4 | 1,01E+08 | 1,01E+08 | 1410 | * | 7,944311 | 6,531704676 | 1,967051 | 3,320557 | 0,000898 | 1 | 0 | Promoter (<=1kb) | 4 | 1,01E+08 | 1,01E+08 | 83312 | 2 | 16451 | ENSMUST00000132292.2 | 56 | ENSMUSG00000028530 | Jak1 | Janus kinase 1 |
| chr2 | 1,67E+08 | 1,67E+08 | 1092 | * | 7,941511 | 6,53204668 | 1,830968 | 3,567538 | 0,00036 | 1 | 0 | Promoter (1-2kb) | 2 | 1,67E+08 | 1,67E+08 | 6291 | 2 | 56336 | ENSMUST00000133557.2 | -1605 | ENSMUSG00000017929 | B4galt5 | UDP-Gal:betaGlcNAc beta 1,4-galactosyltransferase, polypeptide 5 |
| chr19 | 58905375 | 58906093 | 719 | * | 8,91782 | 6,532136409 | 1,957235 | 3,337431 | 0,000846 | 0,453798789 | 1,425929 | Intron (ENSMUST00000 | 19 | 58900105 | 58900191 | 87 | 2 | 1,01E+08 | ENSMUST00000175072.3 | -5184 | ENSMUSG00000092813 | Mir3086 | microRNA 3086 |
| chr17 | 64261734 | 64262269 | 536 | * | 3,685933 | 6,532288036 | 2,409583 | 2,710962 | 0,006709 | 1 | 0 | Intron (ENSMUST00000 | 17 | 64244967 | 64446192 | 201226 | 1 | 14158 | ENSMUST00000038080.7 | 16767 | ENSMUSG00000000127 | Fer | fer (fms/fps related) protein kinase |
| chrX | 64494440 | 64494999 | 560 | * | 3,616611 | 6,532714271 | 2,356941 | 2,771692 | 0,005577 | 1 | 0 | Distal Intergenic | 20 | 64091250 | 64092620 | 1371 | 1 | 70062 | ENSMUST00000033524.3 | 403190 | ENSMUSG00000031181 | Ctag2 | cancer/testis antigen 2 |
| chrX | 1,09E+08 | 1,09E+08 | 983 | * | 4,083214 | 6,532840414 | 2,340621 | 2,791072 | 0,005253 | 1 | 0 | Distal Intergenic | 20 | 1,09E+08 | 1,09E+08 | 2406 | 1 | 634340 | ENSMUST00000122296.2 | -69500 | ENSMUSG00000082853 | Gm7134 | kinesin family member 18B pseudogene |
| chr18 | 66081611 | 66082416 | 806 | * | 6,157024 | 6,532887404 | 1,881793 | 3,471629 | 0,000517 | 1 | 0 | Distal Intergenic | 18 | 66071098 | 66072858 | 1761 | 2 | 19434 | ENSMUST00000237506.2 | -8753 | ENSMUSG00000024518 | Rax | retina and anterior neural fold homeobox |
| chr4 | 74686363 | 74686732 | 370 | * | 3,537419 | 6,532949745 | 2,330448 | 2,803302 | 0,005058 | 1 | 0 | Distal Intergenic | 4 | 74553456 | 74553587 | 132 | 1 | 1,15E+08 | ENSMUST00000200225.3 | 132907 | ENSMUSG00000104700 | Gm26379 | predicted gene, 26379 |
| chr4 | 58879732 | 58880599 | 868 | * | 5,707458 | 6,533529594 | 1,930247 | 3,384816 | 0,000712 | 1 | 0 | Intron (ENSMUST00000 | 4 | 58869539 | 58911902 | 42364 | 2 | 230249 | ENSMUST00000134848.2 | 31303 | ENSMUSG00000050812 | Ecpas | Ecm29 proteasome adaptor and scaffold |
| chr1 | 38510755 | 38511369 | 615 | * | 5,682414 | 6,53394054 | 1,9683 | 3,319585 | 0,000902 | 1 | 0 | Intron (ENSMUST00000 | 1 | 38427910 | 38429325 | 1416 | 1 | 1,03E+08 | ENSMUST00000131663.2 | 82845 | | Gm16152 | predicted gene 16152 |
| chr3 | 1,04E+08 | 1,04E+08 | 693 | * | 4,956715 | 6,534796385 | 2,155351 | 3,031894 | 0,00243 | 1 | 0 | Intron (ENSMUST00000 | 3 | 1,04E+08 | 1,04E+08 | 16201 | 1 | 19260 | ENSMUST00000198530.2 | 10521 | ENSMUSG00000027843 | Ptpn22 | protein tyrosine phosphatase, non-receptor type 22 (lymphoid) |
| chr4 | 44778330 | 44778984 | 655 | * | 4,169175 | 6,534857978 | 2,072557 | 3,153042 | 0,001616 | 1 | 0 | Intron (ENSMUST00000 | 4 | 44760444 | 44912059 | 151616 | 1 | 319885 | ENSMUST00000067521.8 | 17886 | ENSMUSG00000035649 | Zcchc7 | zinc finger, CCHC domain containing 7 |

| | | | | | | | | | | | | | | | | | | | | | | | |
|---|---|---|---|---|---|---|---|---|---|---|---|---|---|---|---|---|---|---|---|---|---|---|---|
| chr14 | 1,24E+08 | 1,24E+08 | 706 | * | 4,351297 | 6,535761482 | 2,089966 | 3,127209 | 0,001765 | 1 | 0 | Intron (ENSMUST0000 | 14 | 1,24E+08 | 1,24E+08 | 214612 | 2 | 14169 | ENSMUST00000026631.6 | -22978 | ENSMUSG00000025551 | Fgf14 | fibroblast growth factor 14 |
| chr3 | 51139495 | 51140610 | 1116 | * | 7,697722 | 6,536285937 | 1,855446 | 3,522758 | 0,000427 | 1 | 0 | Exon (ENSMUST00000 | 3 | 51147627 | 51159045 | 11419 | 1 | 12457 | ENSMUST00000144826.2 | -7017 | ENSMUSG00000023087 | Noct | nocturnin |
| chr4 | 57148573 | 57149322 | 750 | * | 5,813963 | 6,53642118 | 2,051869 | 3,185595 | 0,001445 | 1 | 0 | Distal Intergenic | 4 | 57061726 | 57143437 | 81712 | 2 | 54357 | ENSMUST00000030142.4 | -5136 | ENSMUSG00000028434 | Epb41l4b | erythrocyte membrane protein band 4.1 like 4b |
| chr7 | 66580904 | 66581552 | 649 | * | 5,688755 | 6,537233453 | 2,008307 | 3,255097 | 0,001134 | 1 | 0 | Intron (ENSMUST0000 | 7 | 66559216 | 66619303 | 60088 | 1 | 233332 | ENSMUST00000138009.2 | 21688 | ENSMUSG00000058145 | Adamts17 | a disintegrin-like and metallopeptidase (reprolysin type) with thrombospondin type 1 motif, 17 |
| chr2 | 1,8E+08 | 1,8E+08 | 565 | * | 5,48036 | 6,537391363 | 2,218779 | 2,946391 | 0,003215 | 1 | 0 | Intron (ENSMUST0000 | 2 | 1,8E+08 | 1,8E+08 | 39889 | 1 | 228983 | ENSMUST00000142376.2 | 4185 | ENSMUSG00000039050 | Osbpl2 | oxysterol binding protein-like 2 |
| chr19 | 6106257 | 6106854 | 598 | * | 5,813722 | 6,537544122 | 1,994289 | 3,278133 | 0,001045 | 1 | 0 | Promoter (<=1kb) | 19 | 6104961 | 6107109 | 2149 | 2 | 18120 | ENSMUST00000236412.2 | 255 | ENSMUSG00000007338 | Mrpl49 | mitochondrial ribosomal protein L49 |
| chr1 | 18475642 | 18476178 | 537 | * | 5,252215 | 6,53763421 | 1,990485 | 3,284443 | 0,001022 | 1 | 0 | Distal Intergenic | 1 | 18321204 | 18335362 | 14159 | 2 | 77673 | ENSMUST00000088463.2 | -140280 | ENSMUSG00000067773 | Defb41 | defensin beta 41 |
| chr18 | 24077873 | 24078171 | 299 | * | 3,557283 | 6,538076807 | 2,533677 | 2,58047 | 0,009867 | 1 | 0 | Exon (ENSMUST00002 | 18 | 24087762 | 24089633 | 1872 | 1 | 69256 | ENSMUST00000225682.2 | -9591 | ENSMUSG00000024276 | Zfp397 | zinc finger protein 397 |
| chr7 | 95962345 | 95962594 | 250 | * | 3,65954 | 6,539053678 | 2,435807 | 2,684554 | 0,007263 | 1 | 0 | Intron (ENSMUST0000 | 7 | 95954874 | 95954972 | 99 | 1 | 1,02E+08 | ENSMUST00000184738.3 | 7471 | ENSMUSG00000099030 | Mir6394 | microRNA 6394 |
| chr4 | 1,26E+08 | 1,26E+08 | 788 | * | 6,680252 | 6,539280123 | 1,923638 | 3,399433 | 0,000675 | 1 | 0 | Exon (ENSMUST00000 | 4 | 1,26E+08 | 1,26E+08 | 1496 | 1 | 230751 | ENSMUST00000153138.2 | -9020 | ENSMUSG00000042616 | Oscp1 | organic solute carrier partner 1 |
| chr5 | 1,08E+08 | 1,08E+08 | 881 | * | 6,03283 | 6,539781344 | 1,906615 | 3,430048 | 0,000603 | 1 | 0 | Promoter (<=1kb) | 5 | 1,08E+08 | 1,08E+08 | 13829 | 1 | 17765 | ENSMUST00000137996.8 | -874 | ENSMUSG00000029267 | Mtf2 | metal response element binding transcription factor 2 |
| chr13 | 51511885 | 51512610 | 726 | * | 6,276661 | 6,540362895 | 1,888211 | 3,463788 | 0,000533 | 1 | 0 | Distal Intergenic | 13 | 51562675 | 51576833 | 14159 | 1 | 13610 | ENSMUST00000087978.5 | -50065 | ENSMUSG00000067586 | S1pr3 | sphingosine-1-phosphate receptor 3 |
| chr5 | 1,15E+08 | 1,15E+08 | 690 | * | 6,823121 | 6,540577469 | 1,881366 | 3,476505 | 0,000508 | 1 | 0 | Intron (ENSMUST0000 | 5 | 1,15E+08 | 1,15E+08 | 5904 | 1 | 108014 | ENSMUST00000031513.14 | -10748 | ENSMUSG00000029538 | Srsf9 | serine and arginine-rich splicing factor 9 |
| chr10 | 24227802 | 24228695 | 894 | * | 6,713394 | 6,540598507 | 1,881238 | 3,476753 | 0,000508 | 1 | 0 | Distal Intergenic | 10 | 24099443 | 24107097 | 7655 | 1 | 59012 | ENSMUST00000154916.2 | 128359 | ENSMUSG00000020000 | Moxd1 | monooxygenase, DBH-like 1 |
| chr1 | 1,67E+08 | 1,67E+08 | 685 | * | 4,304954 | 6,540676624 | 2,415628 | 2,707651 | 0,006776 | 1 | 0 | Promoter (1-2kb) | 1 | 1,67E+08 | 1,67E+08 | 16464 | 1 | 68944 | ENSMUST00000193101.2 | -1370 | ENSMUSG00000052428 | Tmco1 | transmembrane and coiled-coil domains 1 |
| chr15 | 6572818 | 6573468 | 651 | * | 4,553629 | 6,541935451 | 2,228584 | 2,935467 | 0,00333 | 1 | 0 | Intron (ENSMUST0000 | 15 | 6552334 | 6609774 | 57441 | 1 | 23880 | ENSMUST00000226412.2 | 20484 | ENSMUSG00000022148 | Fyb | FYN binding protein |
| chr2 | 1,08E+08 | 1,08E+08 | 268 | * | 4,270823 | 6,542675757 | 2,188685 | 2,989319 | 0,002796 | 1 | 0 | Distal Intergenic | 2 | 1,08E+08 | 1,08E+08 | 131 | 2 | 1,15E+08 | ENSMUST00000104223.3 | 85147 | ENSMUSG00000077411 | Gm23749 | predicted gene, 23749 |
| chr18 | 40405839 | 40406439 | 601 | * | 5,200256 | 6,542733431 | 2,185413 | 2,99382 | 0,002755 | 1 | 0 | Intron (ENSMUST0000 | 18 | 40391856 | 40664059 | 272204 | 1 | 383348 | ENSMUST00000235241.2 | 13983 | ENSMUSG00000051401 | Kctd16 | potassium channel tetramerisation domain containing 16 |
| chr7 | 45049591 | 45050201 | 611 | * | 4,685693 | 6,542974847 | 2,173303 | 3,010613 | 0,002607 | 1 | 0 | Distal Intergenic | 7 | 45025877 | 45045166 | 19290 | 2 | 20637 | ENSMUST00000074575.11 | -4425 | ENSMUSG00000063511 | Snrnp70 | small nuclear ribonucleoprotein 70 (U1) |
| chr2 | 1,65E+08 | 1,65E+08 | 280 | * | 2,870179 | 6,543402555 | 2,486784 | 2,631271 | 0,008507 | 1 | 0 | Promoter (<=1kb) | 2 | 1,65E+08 | 1,65E+08 | 4209 | 1 | 57138 | ENSMUST00000135918.2 | -889 | ENSMUSG00000017740 | Slc12a5 | solute carrier family 12, member 5 |
| chrX | 82045669 | 82046084 | 416 | * | 3,308784 | 6,543468016 | 2,439452 | 2,682352 | 0,007311 | 1 | 0 | Intron (ENSMUST0000 | 20 | 82087008 | 82773937 | 686930 | 1 | 13405 | ENSMUST00000147740.2 | -40924 | ENSMUSG00000045103 | Dmd | dystrophin, muscular dystrophy |
| chr1 | 70694508 | 70695033 | 526 | * | 3,772924 | 6,543565765 | 2,378523 | 2,751105 | 0,005939 | 1 | 0 | Intron (ENSMUST0000 | 1 | 70635974 | 70636080 | 107 | 1 | 1,15E+08 | ENSMUST00000102000.3 | 58534 | ENSMUSG00000075952 | Gm23422 | predicted gene, 23422 |
| chr9 | 1,08E+08 | 1,08E+08 | 828 | * | 4,180386 | 6,543816111 | 2,226613 | 2,93891 | 0,003294 | 1 | 0 | Distal Intergenic | 9 | 1,08E+08 | 1,08E+08 | 59325 | 2 | 13138 | ENSMUST00000191899.6 | -3332 | ENSMUSG00000039952 | Dag1 | dystroglycan 1 |
| chr19 | 35960508 | 35961061 | 554 | * | 4,102038 | 6,544005386 | 2,155776 | 3,035568 | 0,002401 | 1 | 0 | Intron (ENSMUST0000 | 19 | 35812621 | 35901978 | 89358 | 2 | 329056 | ENSMUST00000178904.2 | -58530 | ENSMUSG00000094707 | A830019P07Rik | RIKEN cDNA A830019P07 gene |
| chr4 | 13672881 | 13673407 | 527 | * | 4,083113 | 6,544035174 | 2,143402 | 3,053107 | 0,002265 | 1 | 0 | Distal Intergenic | 4 | 13743436 | 13893649 | 150214 | 1 | 12395 | ENSMUST00000006761.10 | -70029 | ENSMUSG00000006586 | Runx1t1 | RUNX1 translocation partner 1 |
| chr14 | 1,03E+08 | 1,03E+08 | 672 | * | 4,393782 | 6,544095248 | 2,118273 | 3,089354 | 0,002006 | 1 | 0 | Intron (ENSMUST0000 | 14 | 1,03E+08 | 1,03E+08 | 1380 | 1 | 1,03E+08 | ENSMUST00000244208.1 | -4995 | ENSMUSG00000115855 | Gm34643 | predicted gene, 34643 |
| chr19 | 30745030 | 30745265 | 236 | * | 4,363301 | 6,544120789 | 2,113557 | 3,096259 | 0,00196 | 1 | 0 | Intron (ENSMUST0000 | 19 | 30686285 | 30758852 | 72568 | 2 | 19091 | ENSMUST00000182401.2 | 13587 | ENSMUSG00000052920 | Prkg1 | protein kinase, cGMP-dependent, type I |
| chr13 | 21897231 | 21897968 | 738 | * | 6,281586 | 6,544871684 | 1,922887 | 3,40367 | 0,000665 | 1 | 0 | Promoter (2-3kb) | 13 | 21899933 | 21900313 | 381 | 2 | 319185 | ENSMUST00000091756.2 | 2345 | ENSMUSG00000094338 | H2bc13 | H2B clustered histone 13 |
| chr4 | 86445972 | 86446796 | 825 | * | 7,285394 | 6,545054888 | 1,892093 | 3,459162 | 0,000542 | 0,388942349 | 1,360622 | Intron (ENSMUST0000 | 4 | 86362878 | 86457354 | 94477 | 2 | 75811 | ENSMUST00000151481.8 | 10558 | ENSMUSG00000028492 | Saxo1 | stabilizer of axonemal microtubules 1 |
| chr4 | 52438938 | 52439900 | 963 | * | 8,089698 | 6,545786624 | 1,797878 | 3,64084 | 0,000272 | 1 | 0 | Promoter (<=1kb) | 4 | 52439243 | 52488260 | 49018 | 1 | 14211 | ENSMUST00000102915.10 | 0 | ENSMUSG00000028312 | Smc2 | structural maintenance of chromosomes 2 |
| chr3 | 44774489 | 44775293 | 805 | * | 6,636167 | 6,546873867 | 2,020274 | 3,240587 | 0,001193 | 1 | 0 | Distal Intergenic | 3 | 45281480 | 45332640 | 51161 | 2 | 72511 | ENSMUST00000191843.2 | 557347 | ENSMUSG00000097040 | 2610316D01Rik | RIKEN cDNA 2610316D01 gene |
| chr2 | 1,68E+08 | 1,68E+08 | 307 | * | 3,248297 | 6,546963816 | 2,334029 | 2,805005 | 0,005032 | 1 | 0 | Exon (ENSMUST00020 | 2 | 1,68E+08 | 1,68E+08 | 1911 | 2 | 77017 | ENSMUST00000129713.2 | -23651 | ENSMUSG00000087648 | E130018N17Rik | RIKEN cDNA E130018N17 gene |
| chr1 | 97833821 | 97834645 | 825 | * | 5,825392 | 6,549896322 | 1,889836 | 3,465854 | 0,000529 | 1 | 0 | Intron (ENSMUST0000 | 1 | 97782359 | 97822121 | 39763 | 2 | 18484 | ENSMUST00000159841.2 | -11700 | ENSMUSG00000026335 | Pam | peptidylglycine alpha-amidating monooxygenase |
| chr17 | 57838224 | 57838469 | 246 | * | 3,165348 | 6,549964369 | 2,51271 | 2,606733 | 0,009141 | 1 | 0 | Intron (ENSMUST0000 | 17 | 57813453 | 57852505 | 39053 | 2 | 224916 | ENSMUST00000165781.3 | 14036 | ENSMUSG00000090655 | Vmn2r120 | vomeronasal 2, receptor 120 |
| chr3 | 13601860 | 13602465 | 606 | * | 4,316576 | 6,550827691 | 2,358109 | 2,778 | 0,005469 | 1 | 0 | Intron (ENSMUST0000 | 3 | 13538280 | 14180787 | 642508 | 1 | 76897 | ENSMUST00000211860.2 | 63580 | ENSMUSG00000039717 | Ralyl | RALY RNA binding protein-like |
| chr11 | 1,02E+08 | 1,02E+08 | 554 | * | 4,419447 | 6,551546588 | 2,25812 | 2,901329 | 0,003716 | 1 | 0 | Distal Intergenic | 11 | 1,02E+08 | 1,02E+08 | 3965 | 1 | 217214 | ENSMUST00000129799.8 | -6627 | ENSMUSG00000048217 | Nags | N-acetylglutamate synthase |
| chr17 | 88099753 | 88100115 | 363 | * | 4,708888 | 6,551889907 | 2,019186 | 3,244818 | 0,001175 | 1 | 0 | Intron (ENSMUST0000 | 17 | 88053229 | 88105422 | 52194 | 2 | 210741 | ENSMUST00000055221.9 | 5307 | ENSMUSG00000050138 | Kcnk12 | potassium channel, subfamily K, member 12 |
| chr7 | 1E+08 | 1E+08 | 679 | * | 4,109649 | 6,551978069 | 2,20733 | 2,968282 | 0,002995 | 1 | 0 | Intron (ENSMUST0000 | 7 | 1E+08 | 1E+08 | 19327 | 1 | 277939 | ENSMUST00000208753.2 | 7573 | ENSMUSG00000047248 | C2cd3 | C2 calcium-dependent domain containing 3 |
| chr3 | 1,46E+08 | 1,46E+08 | 692 | * | 4,408086 | 6,55273162 | 2,134136 | 3,070438 | 0,002137 | 1 | 0 | Intron (ENSMUST0000 | 3 | 1,46E+08 | 1,46E+08 | 36319 | 2 | 75015 | ENSMUST00000124931.8 | 3477 | ENSMUSG00000048652 | Samd13 | sterile alpha motif domain containing 13 |
| chrX | 88151167 | 88151949 | 783 | * | 4,334822 | 6,553125237 | 2,144486 | 3,055802 | 0,002245 | 1 | 0 | Distal Intergenic | 20 | 88447454 | 88453295 | 5842 | 2 | 18630 | ENSMUST00000113959.8 | 301346 | ENSMUSG00000035395 | Dcaf8l | DDB1 and CUL4 associated factor 8 like |
| chr8 | 4799763 | 4800214 | 452 | * | 3,883866 | 6,553530779 | 2,196613 | 2,983471 | 0,00285 | 1 | 0 | Promoter (<=1kb) | 8 | 4789595 | 4799675 | 10081 | 2 | 20419 | ENSMUST00000207876.2 | -88 | ENSMUSG00000022322 | Shcbp1 | Shc SH2-domain binding protein 1 |
| chr14 | 86061711 | 86062634 | 924 | * | 5,473453 | 6,554301911 | 2,017795 | 3,248249 | 0,001161 | 1 | 0 | Distal Intergenic | 14 | 86410363 | 86417021 | 6659 | 1 | 1,03E+08 | ENSMUST00000227116.2 | -347729 | | Gm32729 | predicted gene, 32729 |
| chr11 | 5675596 | 5676143 | 548 | * | 4,875911 | 6,55451143 | 2,005325 | 3,268554 | 0,001081 | 1 | 0 | Intron (ENSMUST0000 | 11 | 5654523 | 5665680 | 11158 | 2 | 64660 | ENSMUST00000149980.2 | -9916 | ENSMUSG00000020477 | Mrps24 | mitochondrial ribosomal protein S24 |
| chr3 | 10173893 | 10174368 | 476 | * | 3,651622 | 6,554527481 | 2,369108 | 2,766664 | 0,005663 | 1 | 0 | Distal Intergenic | 3 | 10153170 | 10157625 | 4456 | 1 | 1E+08 | ENSMUST00000061419.9 | 20723 | ENSMUSG00000049230 | Myef2l | myelin expression factor 2 like |
| chr1 | 88466534 | 88467199 | 666 | * | 5,434197 | 6,55463411 | 1,997491 | 3,281434 | 0,001033 | 1 | 0 | Distal Intergenic | 1 | 88479239 | 88489889 | 10651 | 1 | 1,01E+08 | ENSMUST00000189394.2 | -12040 | ENSMUSG00000100798 | Gm19589 | predicted gene, 19589 |
| chr14 | 71991187 | 71991661 | 475 | * | 3,242362 | 6,554779598 | 2,427926 | 2,699745 | 0,006939 | 1 | 0 | Distal Intergenic | 14 | 72094142 | 72157879 | 63738 | 1 | 74667 | ENSMUST00000228500.2 | -102481 | ENSMUSG00000115304 | 4930434J06Rik | RIKEN cDNA 4930434J06 gene |
| chr3 | 1,02E+08 | 1,02E+08 | 595 | * | 5,25383 | 6,554792855 | 1,989537 | 3,294633 | 0,000986 | 1 | 0 | Distal Intergenic | 3 | 1,02E+08 | 1,02E+08 | 23148 | 2 | 242125 | ENSMUST00000061831.11 | 52916 | ENSMUSG00000044313 | Mab21l3 | mab-21-like 3 |
| chr11 | 29881159 | 29881550 | 392 | * | 2,945426 | 6,555176184 | 2,534023 | 2,586865 | 0,009685 | 1 | 0 | Intron (ENSMUST0000 | 11 | 29799461 | 29799600 | 140 | 1 | 1,15E+08 | ENSMUST00000157510.3 | 81698 | ENSMUSG00000088135 | Gm22588 | predicted gene, 22588 |
| chr4 | 93045505 | 93046417 | 913 | * | 5,796009 | 6,55573653 | 1,940107 | 3,37906 | 0,000727 | 1 | 0 | Distal Intergenic | 4 | 93222375 | 93223748 | 1374 | 2 | 69136 | ENSMUST00000066774.6 | 177331 | ENSMUSG00000054000 | Tusc1 | tumor suppressor candidate 1 |
| chr9 | 57007500 | 57007956 | 457 | * | 3,642989 | 6,556728337 | 2,483157 | 2,640481 | 0,008279 | 1 | 0 | Exon (ENSMUST00000 | 9 | 57002532 | 57005164 | 2633 | 1 | 20466 | ENSMUST00000165927.2 | 4968 | ENSMUSG00000042557 | Sin3a | transcriptional regulator, SIN3A (yeast) |
| chr14 | 1,19E+08 | 1,19E+08 | 365 | * | 3,470383 | 6,556765893 | 2,479073 | 2,644846 | 0,008173 | 1 | 0 | Promoter (<=1kb) | 14 | 1,19E+08 | 1,19E+08 | 20758 | 1 | 58187 | ENSMUST00000100314.4 | 451 | ENSMUSG00000022132 | Cldn10 | claudin 10 |
| chr2 | 1,24E+08 | 1,24E+08 | 589 | * | 3,925651 | 6,558287271 | 2,33358 | 2,810397 | 0,004948 | 1 | 0 | Intron (ENSMUST0000 | 2 | 1,24E+08 | 1,25E+08 | 52780 | 1 | 214968 | ENSMUST00000132088.2 | -48398 | ENSMUSG00000027200 | Sema6d | sema domain, transmembrane domain (TM), and cytoplasmic domain, (semaphorin) 6D |
| chr11 | 75409419 | 75410223 | 805 | * | 8,73112 | 6,558988478 | 1,82097 | 3,601921 | 0,000316 | 0,303890196 | 1,360622 | Intron (ENSMUST0000 | 11 | 75404401 | 75417404 | 13004 | 1 | 380713 | ENSMUST00000123819.2 | 5018 | ENSMUSG00000038188 | Scarf1 | scavenger receptor class F, member 1 |
| chr7 | 88826750 | 88827308 | 559 | * | 4,519218 | 6,560375311 | 2,185477 | 3,001805 | 0,002684 | 1 | 0 | Intron (ENSMUST0000 | 7 | 88803335 | 88808825 | 5491 | 2 | 72759 | ENSMUST00000136267.8 | -17925 | ENSMUSG00000039428 | Tmem135 | transmembrane protein 135 |
| chr14 | 92764660 | 92765147 | 488 | * | 4,599715 | 6,561326129 | 2,133962 | 3,074715 | 0,002107 | 1 | 0 | Distal Intergenic | 14 | 92605936 | 92606065 | 130 | 2 | 1,15E+08 | ENSMUST00000104110.3 | -158595 | ENSMUSG00000077298 | Gm24127 | predicted gene, 24127 |
| chr13 | 25212331 | 25212563 | 233 | * | 3,293742 | 6,561628233 | 2,403965 | 2,729503 | 0,006343 | 1 | 0 | Distal Intergenic | 13 | 25171466 | 25204362 | 32897 | 2 | 380836 | ENSMUST00000021772.4 | -7969 | ENSMUSG00000021339 | Mrs2 | MRS2 magnesium transporter |
| chr10 | 36251098 | 36251495 | 398 | * | 3,887257 | 6,561628829 | 2,349022 | 2,793344 | 0,005217 | 1 | 0 | Distal Intergenic | 10 | 36382810 | 36709769 | 326960 | 1 | 319415 | ENSMUST00000167191.8 | -131315 | ENSMUSG00000044499 | Hs3st5 | heparan sulfate (glucosamine) 3-O-sulfotransferase 5 |
| chrX | 57854443 | 57855204 | 762 | * | 4,23925 | 6,561670516 | 2,135271 | 3,072992 | 0,002119 | 1 | 0 | Distal Intergenic | 20 | 57954720 | 57965664 | 10945 | 1 | 75352 | ENSMUST00000118305.8 | -99516 | ENSMUSG00000046180 | Magea13 | MAGE family member A13 |
| chr13 | 5998452 | 5999001 | 550 | * | 4,39835 | 6,561672881 | 2,131948 | 3,077782 | 0,002085 | 1 | 0 | Intron (ENSMUST0000 | 13 | 6027784 | 6064284 | 36501 | 1 | 1,08E+08 | ENSMUST00000221291.2 | -28783 | | Gm46401 | predicted gene, 46401 |
| chr13 | 12150900 | 12151453 | 554 | * | 4,231857 | 6,561689922 | 2,154965 | 3,044917 | 0,002327 | 1 | 0 | Distal Intergenic | 13 | 11785926 | 12121831 | 335906 | 2 | 20191 | ENSMUST00000220597.2 | -29069 | ENSMUSG00000021313 | Ryr2 | ryanodine receptor 2, cardiac |
| chr19 | 41463255 | 41464078 | 824 | * | 6,282458 | 6,561695481 | 2,020966 | 3,246812 | 0,001167 | 1 | 0 | Exon (ENSMUST00002 | 19 | 41471076 | 41550700 | 79625 | 1 | 212391 | ENSMUST00000067795.13 | -6998 | ENSMUSG00000025019 | Lcor | ligand dependent nuclear receptor corepressor |
| chr9 | 63577205 | 63577882 | 678 | * | 4,672045 | 6,561696323 | 2,097731 | 3,127997 | 0,00176 | 1 | 0 | Intron (ENSMUST0000 | 9 | 63562053 | 63573829 | 11777 | 2 | 17127 | ENSMUST00000137713.8 | -3376 | ENSMUSG00000032402 | Smad3 | SMAD family member 3 |
| chr3 | 76785999 | 76786625 | 627 | * | 4,942512 | 6,561699244 | 2,043633 | 3,210801 | 0,001324 | 1 | 0 | Distal Intergenic | 3 | 77012516 | 77012592 | 77 | 2 | 1,15E+08 | ENSMUST00000157856.4 | 225967 | | Gm23644 | predicted gene, 23644 |
| chr12 | 1,02E+08 | 1,02E+08 | 823 | * | 5,936461 | 6,561719961 | 1,959254 | 3,34909 | 0,000811 | 1 | 0 | Intron (ENSMUST0000 | 12 | 1,02E+08 | 1,02E+08 | 29686 | 2 | 19141 | ENSMUST00000110020.8 | -5968 | ENSMUSG00000021190 | Lgmn | legumain |
| chr3 | 1,28E+08 | 1,28E+08 | 386 | * | 5,228966 | 6,561729679 | 1,942465 | 3,378043 | 0,00073 | 1 | 0 | Distal Intergenic | 3 | 1,28E+08 | 1,28E+08 | 2173 | 1 | 1,03E+08 | ENSMUST00000199122.2 | 73320 | | Gm35667 | predicted gene, 35667 |
| chr13 | 1,03E+08 | 1,03E+08 | 473 | * | 5,710401 | 6,561756193 | 1,892866 | 3,466571 | 0,000527 | 1 | 0 | Distal Intergenic | 13 | 1,02E+08 | 1,02E+08 | 8467 | 1 | 1,03E+08 | ENSMUST00000244780.1 | 97020 | | Gm36703 | predicted gene, 36703 |
| chr10 | 77056884 | 77057798 | 915 | * | 8,506144 | 6,561761636 | 1,846436 | 3,553745 | 0,00038 | 0,326832856 | 1,360622 | Distal Intergenic | 10 | 77095052 | 77105409 | 10358 | 1 | 80294 | ENSMUST00000020493.9 | -37254 | ENSMUSG00000020260 | Pofut2 | protein O-fucosyltransferase 2 |
| chr13 | 95727834 | 95729036 | 1203 | * | 8,594579 | 6,561761715 | 1,844534 | 3,557409 | 0,000375 | 0,262086394 | 2,237403 | Distal Intergenic | 13 | 95738311 | 95754995 | 16685 | 2 | 14062 | ENSMUST00000059193.7 | 25959 | ENSMUSG00000048376 | F2r | coagulation factor II (thrombin) receptor |
| chr11 | 53848643 | 53849521 | 879 | * | 7,377017 | 6,561795766 | 1,820758 | 3,603882 | 0,000313 | 1 | 0 | 3' UTR | 11 | 53841389 | 53843972 | 2584 | 2 | 56517 | ENSMUST00000129301.2 | -4671 | ENSMUSG00000063652 | Slc22a21 | solute carrier family 22 (organic cation transporter), member 21 |
| chrX | 97615747 | 97616497 | 751 | * | 5,145223 | 6,561861948 | 2,125245 | 3,087579 | 0,002018 | 1 | 0 | Intron (ENSMUST0000 | 20 | 97607175 | 97622041 | 14867 | 2 | 94190 | ENSMUST00000154920.2 | 5544 | ENSMUSG00000031214 | Ophn1 | oligophrenin 1 |
| chr3 | 89073652 | 89074353 | 702 | * | 5,30332 | 6,564011605 | 2,018192 | 3,252422 | 0,001144 | 1 | 0 | Promoter (<=1kb) | 3 | 89074471 | 89077716 | 3246 | 1 | 1,06E+08 | ENSMUST00000133323.2 | -118 | | Gm45927 | predicted gene, 45927 |
| chr19 | 36340936 | 36341702 | 767 | * | 6,707299 | 6,564233986 | 2,009852 | 3,266028 | 0,001091 | 1 | 0 | Intron (ENSMUST0000 | 19 | 36356468 | 36433604 | 77137 | 1 | 76073 | ENSMUST00000071267.14 | -14766 | ENSMUSG00000024805 | Pcgf5 | polycomb group ring finger 5 |
| chr1 | 67362231 | 67362964 | 734 | * | 7,683246 | 6,564629187 | 1,821244 | 3,604475 | 0,000313 | 1 | 0 | Exon (ENSMUST00000 | 1 | 67296383 | 67296511 | 129 | 1 | 1,15E+08 | ENSMUST00000104557.3 | 65848 | ENSMUSG00000077745 | Gm26342 | predicted gene, 26342 |
| chr11 | 10570807 | 10571641 | 835 | * | 6,714766 | 6,56575123 | 1,961418 | 3,347452 | 0,000816 | 1 | 0 | Distal Intergenic | 11 | 11064223 | 11213526 | 149304 | 1 | 319922 | ENSMUST00000109681.8 | -492582 | ENSMUSG00000050830 | Vwc2 | von Willebrand factor C domain containing 2 |
| chr10 | 11486119 | 11486865 | 747 | * | 8,247588 | 6,566350797 | 1,878795 | 3,494979 | 0,000474 | 0,352527836 | 1,425929 | Exon (ENSMUST00000 | 10 | 11523749 | 11523878 | 130 | 1 | 1,15E+08 | ENSMUST00000104351.3 | -36884 | ENSMUSG00000077539 | Gm22546 | predicted gene, 22546 |
| chr11 | 30979566 | 30980698 | 1133 | * | 7,064223 | 6,567735877 | 1,90848 | 3,441344 | 0,000579 | 1 | 0 | Exon (ENSMUST00000 | 11 | 31008595 | 31011626 | 3032 | 1 | 65257 | ENSMUST00000148808.2 | -27897 | ENSMUSG00000020305 | Asb3 | ankyrin repeat and SOCS box-containing 3 |
| chr1 | 23442834 | 23443318 | 485 | * | 2,874628 | 6,568171934 | 2,490903 | 2,636863 | 0,008368 | 1 | 0 | Distal Intergenic | 1 | 23430830 | 23436853 | 6024 | 2 | 1,01E+08 | ENSMUST00000189537.2 | -5981 | | Gm19673 | predicted gene, 19673 |
| chr1 | 85586242 | 85586632 | 391 | * | 3,262651 | 6,568762986 | 2,354355 | 2,790048 | 0,00527 | 1 | 0 | Intron (ENSMUST0000 | 1 | 85577771 | 85637719 | 59949 | 1 | 20684 | ENSMUST00000066427.11 | 8471 | ENSMUSG00000026222 | Sp100 | nuclear antigen Sp100 |
| chr11 | 59363557 | 59364245 | 689 | * | 6,125095 | 6,568796091 | 1,996165 | 3,290709 | 0,000999 | 1 | 0 | Promoter (<=1kb) | 11 | 59352023 | 59363300 | 11278 | 2 | 237775 | ENSMUST00000057799.8 | -257 | ENSMUSG00000054519 | Zfp867 | zinc finger protein 867 |
| chr14 | 1,1E+08 | 1,1E+08 | 828 | * | 3,492509 | 6,568975631 | 2,316171 | 2,836135 | 0,004566 | 1 | 0 | Distal Intergenic | 14 | 1,1E+08 | 1,1E+08 | 130 | 2 | 1,15E+08 | ENSMUST00000158494.3 | 282766 | ENSMUSG00000089119 | Gm25901 | predicted gene, 25901 |
| chr8 | 10173176 | 10173625 | 450 | * | 4,090497 | 6,569589351 | 2,214914 | 2,96607 | 0,003016 | 1 | 0 | Distal Intergenic | 8 | 10203911 | 10476702 | 272792 | 1 | 244281 | ENSMUST00000208309.2 | -30286 | ENSMUSG00000039057 | Myo16 | myosin XVI |
| chr13 | 47105166 | 47105827 | 662 | * | 4,588935 | 6,569598478 | 2,210037 | 2,972619 | 0,002953 | 1 | 0 | Distal Intergenic | 13 | 47054574 | 47083343 | 28770 | 2 | 16553 | ENSMUST00000224923.2 | -21823 | ENSMUSG00000021375 | Kif13a | kinesin family member 13A |
| chrX | 84662985 | 84663720 | 736 | * | 3,820304 | 6,570102695 | 2,146743 | 3,060498 | 0,00221 | 1 | 0 | Intron (ENSMUST0000 | 20 | 84617756 | 84657589 | 39834 | 1 | 66724 | ENSMUST00000146063.2 | 45229 | ENSMUSG00000035476 | Tab3 | TGF-beta activated kinase 1/MAP3K7 binding protein 3 |
| chr19 | 17625086 | 17625440 | 355 | * | 4,110153 | 6,570212829 | 2,13511 | 3,077224 | 0,002089 | 1 | 0 | Intron (ENSMUST0000 | 19 | 17650816 | 17814981 | 164166 | 2 | 18552 | ENSMUST00000235505.2 | 189541 | ENSMUSG00000024713 | Pcsk5 | proprotein convertase subtilisin/kexin type 5 |
| chr7 | 34765462 | 34766047 | 586 | * | 4,80339 | 6,570247209 | 2,128299 | 3,087089 | 0,002021 | 1 | 0 | Exon (ENSMUST00002 | 7 | 34745847 | 34755998 | 10152 | 2 | 12611 | ENSMUST00000130491.3 | -9464 | ENSMUSG00000056216 | Cebpg | CCAAT/enhancer binding protein (C/EBP), gamma |
| chr12 | 1,05E+08 | 1,05E+08 | 567 | * | 4,792928 | 6,570637719 | 2,134931 | 3,077682 | 0,002086 | 1 | 0 | Intron (ENSMUST0000 | 12 | 1,05E+08 | 1,05E+08 | 39240 | 1 | 268595 | ENSMUST00000178224.2 | 11892 | ENSMUSG00000094910 | D430019H16Rik | RIKEN cDNA D430019H16 gene |
| chr1 | 91386909 | 91387452 | 544 | * | 5,378322 | 6,571187435 | 2,03518 | 3,2288 | 0,001243 | 1 | 0 | Promoter (<=1kb) | 1 | 91343704 | 91387046 | 43343 | 2 | 18627 | ENSMUST00000069620.10 | 0 | ENSMUSG00000055866 | Per2 | period circadian clock 2 |
| chr6 | 1,14E+08 | 1,14E+08 | 269 | * | 4,057539 | 6,571200046 | 2,19108 | 2,999068 | 0,002708 | 1 | 0 | Intron (ENSMUST0000 | 6 | 1,14E+08 | 1,14E+08 | 118712 | 1 | 243616 | ENSMUST00000032451.9 | 55640 | ENSMUSG00000030307 | Slc6a11 | solute carrier family 6 (neurotransmitter transporter, GABA), member 11 |
| chr18 | 54596697 | 54597709 | 1013 | * | 6,420574 | 6,571552098 | 2,005725 | 3,276397 | 0,001051 | 1 | 0 | Distal Intergenic | 18 | 54627765 | 54629656 | 1892 | 2 | 1,05E+08 | ENSMUST00000237202.2 | 31947 | | 4933434P08Rik | RIKEN cDNA 4933434P08 gene |
| chr5 | 1,34E+08 | 1,34E+08 | 418 | * | 3,841633 | 6,572044735 | 2,298286 | 2,859541 | 0,004243 | 1 | 0 | Promoter (<=1kb) | 5 | 1,34E+08 | 1,34E+08 | 28721 | 2 | 94254 | ENSMUST00000076228.3 | 887 | ENSMUSG00000061979 | Rcc1l | reculator of chromosome condensation 1 like |
| chr13 | 19721849 | 19723199 | 1351 | * | 7,870585 | 6,572648324 | 1,912065 | 3,437461 | 0,000587 | 1 | 0 | Distal Intergenic | 13 | 19647017 | 19688537 | 41521 | 2 | 319466 | ENSMUST00000221607.2 | -33312 | ENSMUSG00000114189 | 9330199G10Rik | RIKEN cDNA 9330199G10 gene |
| chr7 | 1,3E+08 | 1,3E+08 | 419 | * | 5,944131 | 6,572935447 | 1,913906 | 3,434304 | 0,000594 | 1 | 0 | Promoter (1-2kb) | 7 | 1,3E+08 | 1,3E+08 | 115207 | 1 | 57752 | ENSMUST00000209108.2 | 1121 | ENSMUSG00000030852 | Tacc2 | transforming, acidic coiled-coil containing protein 2 |
| chrX | 72865576 | 72866985 | 1410 | * | 6,324232 | 6,57350228 | 1,884276 | 3,488609 | 0,000486 | 1 | 0 | Promoter (1-2kb) | 20 | 72836965 | 72868575 | 31611 | 2 | 245469 | ENSMUST00000002080.12 | 1590 | ENSMUSG00000002006 | Pdzd4 | PDZ domain containing 4 |
| chr12 | 1,13E+08 | 1,13E+08 | 607 | * | 6,803031 | 6,573612203 | 1,878619 | 3,499172 | 0,000467 | 1 | 0 | Distal Intergenic | 12 | 1,13E+08 | 1,13E+08 | 26174 | 1 | 70435 | ENSMUST00000101029.4 | -29127 | ENSMUSG00000037679 | Inf2 | inverted formin, FH2 and WH2 domain containing |
| chr1 | 1,84E+08 | 1,84E+08 | 551 | * | 3,152321 | 6,574557695 | 2,5059 | 2,623631 | 0,0087 | 1 | 0 | Distal Intergenic | 1 | 1,84E+08 | 1,84E+08 | 5480 | 2 | 15284 | ENSMUST00000048572.7 | -24369 | ENSMUSG00000039377 | Hlx | H2.0-like homeobox |
| chr4 | 93974034 | 93974597 | 564 | * | 3,40134 | 6,574906704 | 2,45942 | 2,673356 | 0,00751 | 1 | 0 | Distal Intergenic | 4 | 93977813 | 94313825 | 336013 | 2 | 1,08E+08 | ENSMUST00000126270.2 | 339228 | | Gm12648 | predicted gene 12648 |
| chr3 | 56544747 | 56545345 | 599 | * | 4,244904 | 6,575076198 | 2,438091 | 2,696814 | 0,007001 | 1 | 0 | Distal Intergenic | 3 | 56352703 | 56352833 | 131 | 2 | 1,15E+08 | ENSMUST00000104522.3 | -191914 | ENSMUSG00000119391 | Gm25727 | predicted gene, 25727 |
| chr1 | 1,15E+08 | 1,15E+08 | 409 | * | 3,839224 | 6,575143427 | 2,430741 | 2,704996 | 0,006831 | 1 | 0 | Distal Intergenic | 1 | 1,16E+08 | 1,16E+08 | 15632 | 2 | 1,08E+08 | ENSMUST00000186137.7 | 794899 | | Gm50605 | predicted gene, 50605 |
| chr14 | 1,01E+08 | 1,01E+08 | 629 | * | 3,961855 | 6,575148933 | 2,430528 | 2,705235 | 0,006826 | 1 | 0 | Distal Intergenic | 14 | 1,01E+08 | 1,01E+08 | 68647 | 2 | 16597 | ENSMUST00000241711.1 | -35608 | ENSMUSG00000072294 | Klf12 | Kruppel-like factor 12 |
| chrX | 73302740 | 73303109 | 370 | * | 3,487103 | 6,575226497 | 2,417958 | 2,719331 | 0,006541 | 1 | 0 | Intron (ENSMUST0000 | 20 | 73298436 | 73305154 | 6719 | 1 | 13726 | ENSMUST00000088313.5 | 4304 | ENSMUSG00000001964 | Emd | emerin |
| chr1 | 1,93E+08 | 1,93E+08 | 464 | * | 3,9221 | 6,575743626 | 2,361993 | 2,783982 | 0,00537 | 1 | 0 | Exon (ENSMUST00002 | 1 | 1,93E+08 | 1,93E+08 | 129274 | 1 | 1,03E+08 | ENSMUST00000246715.1 | 47136 | | Gm31870 | predicted gene, 31870 |

| | | | | | | | | | | | | | | | | | | | | | | |
|---|---|---|---|---|---|---|---|---|---|---|---|---|---|---|---|---|---|---|---|---|---|---|
| chr19 | 10081558 | 10081801 | 244 | * | 3,927406 | 6,575912515 | 2,342996 | 2,806626 | 0,005006 | 1 | 0 | Promoter (2-3kb) | 19 | 10040129 | 10079110 | 38982 | 2 | 56473 | ENSMUST00000025567.9 | -2448 | ENSMUSG00000024665 | Fads2 | fatty acid desaturase 2 |
| chr14 | 86088212 | 86088597 | 386 | * | 3,614441 | 6,576429499 | 2,295891 | 2,864435 | 0,004178 | 1 | 0 | Distal Intergenic | 14 | 86410363 | 86417021 | 6659 | 1 | 1,03E+08 | ENSMUST00000227116.2 | -321766 | | Gm32729 | predicted gene, 32729 |
| chrX | 44186574 | 44187637 | 1064 | * | 3,822316 | 6,577332002 | 2,221249 | 2,961096 | 0,003065 | 1 | 0 | Distal Intergenic | 20 | 44179781 | 44181668 | 1888 | 1 | 68800 | ENSMUST00000040002.4 | 6793 | ENSMUSG00000037086 | Prr32 | proline rich 32 |
| chr2 | 1,28E+08 | 1,28E+08 | 374 | * | 4,339705 | 6,57746426 | 2,211475 | 2,974244 | 0,002937 | 1 | 0 | Intron (ENSMUST0000( | 2 | 1,28E+08 | 1,29E+08 | 5457 | 2 | 17222 | ENSMUST00000123503.2 | 15529 | ENSMUSG00000014355 | Anapc1 | anaphase promoting complex subunit 1 |
| chr9 | 59674701 | 59676513 | 1813 | * | 9,468758 | 6,577878061 | 1,822016 | 3,61022 | 0,000306 | 0,294898446 | 1,425929 | Intron (ENSMUST0000( | 9 | 59658179 | 59836149 | 177971 | 1 | 270163 | ENSMUST00000136740.8 | 16522 | ENSMUSG00000039585 | Myo9a | myosin IXa |
| chr14 | 76867116 | 76868424 | 1309 | * | 7,545212 | 6,578088856 | 1,854567 | 3,546967 | 0,00039 | 1 | 0 | Distal Intergenic | 14 | 76915006 | 76924532 | 9527 | 1 | 73272 | ENSMUST00000190818.7 | -46582 | ENSMUSG00000101009 | 1700108F19Rik | RIKEN cDNA 1700108F19 gene |
| chr2 | 1,68E+08 | 1,68E+08 | 838 | * | 6,368364 | 6,578288099 | 1,889695 | 3,481138 | 0,000499 | 1 | 0 | Distal Intergenic | 2 | 1,68E+08 | 1,68E+08 | 5103 | 1 | 1E+08 | ENSMUST00000139927.2 | -9137 | | Gm14321 | predicted gene 14321 |
| chr10 | 77846312 | 77846996 | 685 | * | 6,720493 | 6,578364617 | 1,905824 | 3,451717 | 0,000557 | 1 | 0 | Promoter (<=1kb) | 10 | 77836514 | 77845917 | 9404 | 2 | 18641 | ENSMUST00000145716.2 | -395 | ENSMUSG00000020277 | Pfkl | phosphofructokinase, liver, B-type |
| chr11 | 81578997 | 81580594 | 1598 | * | 11,79064 | 6,578402062 | 1,723392 | 3,817124 | 0,000135 | 0,205744841 | 1,2763 | Intron (ENSMUST0000( | 11 | 81478187 | 81484304 | 6118 | 1 | 1,03E+08 | ENSMUST00000156193.2 | 100810 | | Gm11418 | predicted gene 11418 |
| chr4 | 99096858 | 99097268 | 411 | * | 5,539047 | 6,578740325 | 2,095095 | 3,140068 | 0,001689 | 1 | 0 | Intron (ENSMUST0000( | 4 | 99106328 | 99109932 | 3605 | 1 | 242557 | ENSMUST00000152121.2 | -9060 | ENSMUSG00000028550 | Atg4c | autophagy related 4C, cysteine peptidase |
| chr10 | 1,16E+08 | 1,16E+08 | 881 | * | 4,973176 | 6,578822902 | 2,017962 | 3,260133 | 0,001114 | 1 | 0 | Intron (ENSMUST0000( | 10 | 1,16E+08 | 1,16E+08 | 40866 | 1 | 19279 | ENSMUST00000124369.2 | 13537 | ENSMUSG00000020151 | Ptprr | protein tyrosine phosphatase, receptor type, R |
| chr14 | 5360130 | 5360886 | 757 | * | 5,847729 | 6,578867313 | 2,035614 | 3,231884 | 0,00123 | 1 | 0 | Distal Intergenic | 14 | 5363820 | 5364135 | 316 | 2 | 1E+08 | ENSMUST00000225472.2 | 3249 | | Pou5f1-rs6 | POU domain, class 5, transcription factor 1, related sequence 6 |
| chr17 | 8286782 | 8287934 | 1153 | * | 8,690323 | 6,578958865 | 1,780719 | 3,694552 | 0,00022 | 0,250313148 | 1,449706 | Intron (ENSMUST0000( | 17 | 8302907 | 8302994 | 88 | 1 | 1,15E+08 | ENSMUST00000158815.3 | -14973 | ENSMUSG00000118778 | Gm22416 | predicted gene, 22416 |
| chr13 | 41309173 | 41309526 | 354 | * | 4,613642 | 6,579077449 | 2,107976 | 3,12104 | 0,001802 | 1 | 0 | Promoter (2-3kb) | 13 | 41306864 | 41325160 | 18297 | 1 | 637277 | ENSMUST00000145810.2 | 2309 | ENSMUSG00000038651 | Sycp2l | synaptonemal complex protein 2-like |
| chr13 | 38141947 | 38142869 | 923 | * | 5,083659 | 6,579202967 | 2,101277 | 3,13105 | 0,001742 | 1 | 0 | Distal Intergenic | 13 | 38130992 | 38133182 | 2191 | 1 | 68750 | ENSMUST00000162849.2 | 10955 | ENSMUSG00000039087 | Rreb1 | ras responsive element binding protein 1 |
| chrX | 1,01E+08 | 1,01E+08 | 512 | * | 3,73246 | 6,579453193 | 2,314294 | 2,842963 | 0,00447 | 1 | 0 | Distal Intergenic | 20 | 1,01E+08 | 1,01E+08 | 4454 | 2 | 20102 | ENSMUST00000033683.8 | -24624 | ENSMUSG00000031320 | Rps4x | ribosomal protein S4, X-linked |
| chr1 | 1,84E+08 | 1,84E+08 | 623 | * | 3,276856 | 6,57945931 | 2,328614 | 2,825483 | 0,004721 | 1 | 0 | Distal Intergenic | 1 | 1,84E+08 | 1,84E+08 | 41256 | 1 | 63953 | ENSMUST00000048655.8 | 129452 | ENSMUSG00000039384 | Dusp10 | dual specificity phosphatase 10 |
| chrX | 1,4E+08 | 1,4E+08 | 566 | * | 3,72717 | 6,579514021 | 2,371695 | 2,774183 | 0,005534 | 1 | 0 | Intron (ENSMUST0000( | 20 | 1,4E+08 | 1,4E+08 | 59111 | 1 | 12830 | ENSMUST00000130732.2 | 3272 | ENSMUSG00000031274 | Col4a5 | collagen, type IV, alpha 5 |
| chr7 | 27221174 | 27221432 | 259 | * | 3,432464 | 6,579517975 | 2,372012 | 2,773813 | 0,00554 | 1 | 0 | Promoter (1-2kb) | 7 | 27222692 | 27230600 | 7909 | 1 | 233020 | ENSMUST00000108353.9 | -1260 | ENSMUSG00000040424 | Hipk4 | homeodomain interacting protein kinase 4 |
| chr19 | 10505619 | 10505903 | 285 | * | 3,36212 | 6,579546782 | 2,392838 | 2,749683 | 0,005965 | 1 | 0 | Promoter (2-3kb) | 19 | 10480181 | 10503506 | 23326 | 2 | 66072 | ENSMUST00000235222.2 | -2113 | ENSMUSG00000024668 | Sdhaf2 | succinate dehydrogenase complex assembly factor 2 |
| chr9 | 1,15E+08 | 1,15E+08 | 252 | * | 3,932644 | 6,581000816 | 2,49931 | 2,633127 | 0,00846 | 1 | 0 | 3' UTR | 9 | 1,15E+08 | 1,15E+08 | 96274 | 1 | 74486 | ENSMUST00000182413.2 | -10803 | ENSMUSG00000040875 | Osbpl10 | oxysterol binding protein-like 10 |
| chr2 | 1,55E+08 | 1,55E+08 | 782 | * | 5,774486 | 6,581325074 | 2,002069 | 3,287263 | 0,001012 | 1 | 0 | Intron (ENSMUST0000( | 2 | 1,55E+08 | 1,55E+08 | 20509 | 2 | 56406 | ENSMUST00000109669.8 | -3847 | ENSMUSG00000038369 | Ncoa6 | nuclear receptor coactivator 6 |
| chrX | 9065805 | 9066555 | 751 | * | 5,698022 | 6,58251019 | 1,958502 | 3,360993 | 0,000777 | 1 | 0 | Promoter (<=1kb) | 20 | 9066141 | 9134324 | 68184 | 1 | 236285 | ENSMUST00000069763.3 | 0 | ENSMUSG00000047344 | Lancl3 | LanC lantibiotic synthetase component C-like 3 (bacterial) |
| chr16 | 23188127 | 23188738 | 612 | * | 6,13463 | 6,582543646 | 1,957297 | 3,363078 | 0,000771 | 1 | 0 | Distal Intergenic | 16 | 23247883 | 23252710 | 4828 | 1 | 239766 | ENSMUST00000038730.7 | -59145 | ENSMUSG00000033383 | Rtp1 | receptor transporter protein 1 |
| chr3 | 95614347 | 95614971 | 625 | * | 6,082976 | 6,583183585 | 1,936455 | 3,399606 | 0,000675 | 1 | 0 | Distal Intergenic | 3 | 95583511 | 95595228 | 11718 | 2 | 229595 | ENSMUST00000117782.8 | -19119 | ENSMUSG00000015850 | Adamtsl4 | ADAMTS-like 4 |
| chr4 | 19079430 | 19079879 | 450 | * | 3,86111 | 6,585905467 | 2,554835 | 2,57782 | 0,009943 | 1 | 0 | Intron (ENSMUST0000( | 4 | 18767980 | 19122565 | 354586 | 2 | 435772 | ENSMUST00000137780.3 | 42686 | ENSMUSG00000073991 | Cnbd1 | cyclic nucleotide binding domain containing 1 |
| chr10 | 30683577 | 30684030 | 454 | * | 2,960396 | 6,585932011 | 2,553823 | 2,578852 | 0,009913 | 1 | 0 | Distal Intergenic | 10 | 30571887 | 30679322 | 107436 | 2 | 211329 | ENSMUST00000215725.2 | -4255 | ENSMUSG00000039697 | Ncoa7 | nuclear receptor coactivator 7 |
| chr17 | 32982272 | 32982641 | 370 | * | 3,215734 | 6,586141476 | 2,467553 | 2,669099 | 0,007606 | 1 | 0 | Distal Intergenic | 17 | 32992226 | 33006521 | 14296 | 2 | 208292 | ENSMUST00000057501.9 | 23880 | ENSMUSG00000024298 | Zfp871 | zinc finger protein 871 |
| chr2 | 96743861 | 96744404 | 544 | * | 3,330763 | 6,586279069 | 2,422185 | 2,719148 | 0,006545 | 1 | 0 | Intron (ENSMUST0000( | 2 | 97068163 | 97068291 | 129 | 1 | 1,15E+08 | ENSMUST00000158008.3 | -323759 | ENSMUSG00000088633 | Gm22313 | predicted gene, 22313 |
| chr2 | 1,55E+08 | 1,55E+08 | 517 | * | 3,887168 | 6,586483801 | 2,354395 | 2,797527 | 0,00515 | 1 | 0 | Promoter (1-2kb) | 2 | 1,55E+08 | 1,55E+08 | 16082 | 2 | 1,19E+08 | ENSMUST00000155334.2 | -1125 | | LOC118568528 | uncharacterized LOC118568528 |
| chrX | 88132924 | 88133619 | 696 | * | 3,546417 | 6,586645877 | 2,309406 | 2,852096 | 0,004343 | 1 | 0 | Distal Intergenic | 20 | 88447454 | 88453295 | 5842 | 2 | 18630 | ENSMUST00000113959.8 | 319676 | ENSMUSG00000035395 | Dcaf8l | DDB1 and CUL4 associated factor 8 like |
| chrX | 73443654 | 73444130 | 477 | * | 3,718538 | 6,586852582 | 2,259489 | 2,915195 | 0,003555 | 1 | 0 | Intron (ENSMUST0000( | 20 | 73436933 | 73483494 | 46562 | 1 | 16151 | ENSMUST00000130802.8 | 6721 | ENSMUSG00000004221 | Ikbkg | inhibitor of kappaB kinase gamma |
| chr9 | 1,07E+08 | 1,07E+08 | 600 | * | 6,09869 | 6,586975249 | 2,131987 | 3,089595 | 0,002004 | 1 | 0 | Distal Intergenic | 9 | 1,07E+08 | 1,07E+08 | 2637 | 1 | 1,03E+08 | ENSMUST00000187786.2 | -6823 | ENSMUSG00000101411 | Gm28111 | predicted gene 28111 |
| chrX | 72722445 | 72723339 | 895 | * | 4,765077 | 6,587060923 | 2,063826 | 3,191674 | 0,001415 | 1 | 0 | Promoter (<=1kb) | 20 | 72722484 | 72723605 | 1122 | 1 | 102857 | ENSMUST00000166200.8 | 0 | ENSMUSG00000019558 | Slc6a8 | solute carrier family 6 (neurotransmitter transporter, creatine), member 8 |
| chr11 | 21979552 | 21980369 | 818 | * | 4,631253 | 6,587219775 | 2,181547 | 3,019518 | 0,002532 | 1 | 0 | Intron (ENSMUST0000( | 11 | 21946858 | 21952897 | 6040 | 2 | 18423 | ENSMUST00000147486.3 | -26655 | ENSMUSG00000005917 | Otx1 | orthodenticle homeobox 1 |
| chr10 | 69357983 | 69358721 | 739 | * | 5,048655 | 6,587989109 | 2,060302 | 3,197584 | 0,001386 | 1 | 0 | Intron (ENSMUST0000( | 10 | 69369602 | 69556690 | 187089 | 1 | 11735 | ENSMUST00000182379.7 | -10881 | ENSMUSG00000069601 | Ank3 | ankyrin 3, epithelial |
| chr9 | 65095934 | 65096716 | 783 | * | 5,294614 | 6,588058581 | 2,052325 | 3,210047 | 0,001327 | 1 | 0 | Distal Intergenic | 9 | 65102635 | 65145712 | 43078 | 1 | 214424 | ENSMUST00000216702.2 | -5919 | ENSMUSG00000032392 | Parp16 | poly (ADP-ribose) polymerase family, member 16 |
| chr18 | 45499944 | 45500437 | 494 | * | 5,326411 | 6,588270479 | 2,151526 | 3,062138 | 0,002198 | 1 | 0 | Intron (ENSMUST0000( | 18 | 45402072 | 45819091 | 417020 | 1 | 140492 | ENSMUST00000066890.14 | 97872 | ENSMUSG00000054477 | Kcnn2 | potassium intermediate/small conductance calcium-activated channel, subfamily N, member 2 |
| chrX | 1,51E+08 | 1,51E+08 | 774 | * | 5,012348 | 6,588303182 | 2,021412 | 3,259258 | 0,001117 | 1 | 0 | Intron (ENSMUST0000( | 20 | 1,51E+08 | 1,51E+08 | 4924 | 1 | 59026 | ENSMUST00000150426.8 | -9066 | ENSMUSG00000025261 | Huwe1 | HECT, UBA and WWE domain containing 1 |
| chr13 | 60449500 | 60449999 | 500 | * | 3,861322 | 6,588947283 | 2,211259 | 2,979726 | 0,002885 | 1 | 0 | Distal Intergenic | 13 | 60353285 | 60363837 | 10553 | 1 | 1,01E+08 | ENSMUST00000224064.2 | 96215 | ENSMUSG00000050765 | Gm5084 | predicted gene 5084 |
| chr4 | 1,51E+08 | 1,51E+08 | 745 | * | 6,857376 | 6,589290055 | 1,924326 | 3,424207 | 0,000617 | 1 | 0 | Promoter (1-2kb) | 4 | 1,51E+08 | 1,51E+08 | 160 | 1 | 1,15E+08 | ENSMUST00000082542.4 | 1291 | ENSMUSG00000064476 | Gm23405 | predicted gene, 23405 |
| chr2 | 70949313 | 70949739 | 427 | * | 3,720691 | 6,590082347 | 2,338593 | 2,817969 | 0,004833 | 1 | 0 | Promoter (1-2kb) | 2 | 70948267 | 70973270 | 25004 | 1 | 73649 | ENSMUST00000028403.3 | 1046 | ENSMUSG00000027015 | Cybrd1 | cytochrome b reductase 1 |
| chr6 | 25662409 | 25662827 | 419 | * | 3,354368 | 6,5905152 | 2,397877 | 2,748479 | 0,005987 | 1 | 0 | Distal Intergenic | 6 | 25665877 | 25689791 | 23915 | 2 | 14763 | ENSMUST00000200812.2 | 26964 | ENSMUSG00000039904 | Gpr37 | G protein-coupled receptor 37 |
| chr11 | 71323394 | 71324275 | 882 | * | 7,110756 | 6,590691951 | 1,829258 | 3,602932 | 0,000315 | 1 | 0 | Promoter (2-3kb) | 11 | 71320654 | 71327006 | 6353 | 2 | 1,03E+08 | ENSMUST00000152249.2 | 2731 | | Gm16013 | predicted gene 16013 |
| chr16 | 5752668 | 5753317 | 650 | * | 7,507301 | 6,590978082 | 1,813766 | 3,633863 | 0,000279 | 1 | 0 | Intron (ENSMUST0000( | 16 | 5793449 | 5816360 | 22912 | 1 | 68238 | ENSMUST00000190515.2 | -40132 | ENSMUSG00000101481 | 1700123O21Rik | RIKEN cDNA 1700123O21 gene |
| chr1 | 50235863 | 50236382 | 520 | * | 2,887744 | 6,592094811 | 2,554768 | 2,58031 | 0,009871 | 1 | 0 | Distal Intergenic | 1 | 50939806 | 50954589 | 14784 | 1 | 56363 | ENSMUST00000139157.8 | -703424 | ENSMUSG00000026109 | Tmeff2 | transmembrane protein with EGF-like and two follistatin-like domains 2 |
| chr4 | 1,27E+08 | 1,27E+08 | 354 | * | 3,521101 | 6,592543265 | 2,475516 | 2,663098 | 0,007742 | 1 | 0 | Promoter (<=1kb) | 4 | 1,27E+08 | 1,27E+08 | 32085 | 1 | 26445 | ENSMUST00000030642.3 | -829 | ENSMUSG00000028837 | Psmb2 | proteasome (prosome, macropain) subunit, beta type 2 |
| chr19 | 36175925 | 36176184 | 260 | * | 3,658807 | 6,592585129 | 2,472707 | 2,666141 | 0,007673 | 1 | 0 | Intron (ENSMUST0000( | 19 | 36159980 | 36199018 | 39039 | 1 | 1,03E+08 | ENSMUST00000236660.2 | 15945 | ENSMUSG00000117998 | Gm32081 | predicted gene, 32081 |
| chr7 | 1,17E+08 | 1,17E+08 | 439 | * | 3,717782 | 6,594007008 | 2,286655 | 2,883691 | 0,00393 | 1 | 0 | Intron (ENSMUST0000( | 7 | 1,17E+08 | 1,17E+08 | 292590 | 1 | 233781 | ENSMUST00000032892.7 | 22453 | ENSMUSG00000030657 | Xylt1 | xylosyltransferase 1 |
| chr7 | 42968019 | 42969021 | 1003 | * | 7,412431 | 6,594238926 | 1,850835 | 3,562846 | 0,000367 | 1 | 0 | Distal Intergenic | 7 | 42950900 | 42962718 | 11819 | 2 | 69930 | ENSMUST00000151659.2 | -5301 | ENSMUSG00000012640 | Zfp715 | zinc finger protein 715 |
| chrX | 11403436 | 11404104 | 669 | * | 3,992533 | 6,594456217 | 2,242774 | 2,940312 | 0,003279 | 1 | 0 | Distal Intergenic | 20 | 11194061 | 11194390 | 330 | 1 | 236663 | ENSMUST00000091180.3 | 209375 | ENSMUSG00000069038 | H2al1j | H2A histone family member L1J |
| chr8 | 61900821 | 61901136 | 316 | * | 4,051019 | 6,594648091 | 2,225303 | 2,963483 | 0,003042 | 1 | 0 | Distal Intergenic | 8 | 61931704 | 61933555 | 1852 | 2 | 1,19E+08 | ENSMUST00000240963.1 | 32419 | | Gm53435 | predicted gene, 53435 |
| chr9 | 42988764 | 42989091 | 328 | * | 4,280907 | 6,595150574 | 2,181057 | 3,023833 | 0,002496 | 1 | 0 | Intron (ENSMUST0000( | 9 | 42875139 | 43016781 | 141643 | 2 | 69632 | ENSMUST00000072767.7 | 27690 | ENSMUSG00000059495 | Arhgef12 | Rho guanine nucleotide exchange factor (GEF) 12 |
| chr5 | 97297032 | 97297827 | 796 | * | 4,547868 | 6,595249993 | 2,175317 | 3,031857 | 0,002431 | 1 | 0 | Exon (ENSMUST000001 | 5 | 97290885 | 97297707 | 6823 | 1 | 1,03E+08 | ENSMUST00000198559.2 | 6147 | | Gm33609 | predicted gene, 33609 |
| chr18 | 69183903 | 69184791 | 889 | * | 4,883423 | 6,596076483 | 2,072046 | 3,183364 | 0,001456 | 1 | 0 | Distal Intergenic | 18 | 69155270 | 69155344 | 75 | 2 | 1,02E+08 | ENSMUST00000184434.3 | -28559 | ENSMUSG00000098366 | Mir145b | microRNA 145b |
| chr14 | 4270159 | 4270724 | 566 | * | 4,490767 | 6,596538963 | 2,166027 | 3,045456 | 0,002323 | 1 | 0 | Distal Intergenic | 14 | 4257324 | 4261659 | 4336 | 1 | 353187 | ENSMUST00000225308.2 | 12835 | ENSMUSG00000021775 | Nr1d2 | nuclear receptor subfamily 1, group D, member 2 |
| chr9 | 1,07E+08 | 1,07E+08 | 262 | * | 4,097223 | 6,59678588 | 2,234913 | 2,951697 | 0,00316 | 1 | 0 | Promoter (<=1kb) | 9 | 1,07E+08 | 1,07E+08 | 1910 | 1 | 109685 | ENSMUST00000149487.2 | -488 | ENSMUSG00000036091 | Hyal3 | hyaluronoglucosaminidase 3 |
| chrX | 51720743 | 51721092 | 350 | * | 3,808208 | 6,597217325 | 2,383407 | 2,767977 | 0,005641 | 1 | 0 | Distal Intergenic | 20 | 51476541 | 51702827 | 226287 | 2 | 14734 | ENSMUST00000114857.2 | -17916 | ENSMUSG00000055653 | Gpc3 | glypican 3 |
| chr13 | 40641511 | 40642105 | 595 | * | 5,375841 | 6,597583742 | 2,024266 | 3,259247 | 0,001117 | 1 | 0 | Distal Intergenic | 13 | 40433365 | 40514926 | 81562 | 2 | 218165 | ENSMUST00000224813.2 | -126585 | ENSMUSG00000047094 | Ofcc1 | orofacial cleft 1 candidate 1 |
| chrX | 92504892 | 92505356 | 465 | * | 3,253435 | 6,598515296 | 2,520351 | 2,618094 | 0,008842 | 1 | 0 | Exon (ENSMUST00000( | 20 | 92524096 | 92534552 | 10457 | 2 | 18968 | ENSMUST00000142479.2 | 29196 | ENSMUSG00000006678 | Pola1 | polymerase (DNA directed), alpha 1 |
| chr16 | 20467483 | 20467955 | 473 | * | 3,151544 | 6,598562624 | 2,513454 | 2,625297 | 0,008657 | 1 | 0 | Promoter (2-3kb) | 16 | 20470402 | 20482164 | 11763 | 1 | 21762 | ENSMUST00000007212.9 | -2447 | ENSMUSG00000006998 | Psmd2 | proteasome (prosome, macropain) 26S subunit, non-ATPase, 2 |
| chrX | 86198501 | 86199122 | 622 | * | 3,877121 | 6,59926981 | 2,440955 | 2,703561 | 0,00686 | 1 | 0 | Intron (ENSMUST0000( | 20 | 85790848 | 86933841 | 1142994 | 2 | 331461 | ENSMUST00000078875.8 | 734719 | ENSMUSG00000052372 | Il1rapl1 | interleukin 1 receptor accessory protein-like 1 |
| chr13 | 95214979 | 95215688 | 710 | * | 6,553157 | 6,599925844 | 1,924095 | 3,430145 | 0,000603 | 1 | 0 | Intron (ENSMUST0000( | 13 | 95179976 | 95189526 | 9551 | 2 | 218461 | ENSMUST00000160412.2 | -25453 | ENSMUSG00000021684 | Pde8b | phosphodiesterase 8B |
| chr8 | 1,1E+08 | 1,1E+08 | 355 | * | 4,409627 | 6,60175733 | 2,250276 | 2,933755 | 0,003349 | 1 | 0 | Intron (ENSMUST0000( | 8 | 1,1E+08 | 1,1E+08 | 79 | 1 | 1,01E+08 | ENSMUST00000175500.3 | -46639 | ENSMUSG00000093241 | Mir3108 | microRNA 3108 |
| chr19 | 26300124 | 26300846 | 723 | * | 4,320416 | 6,601957595 | 2,238742 | 2,948959 | 0,003188 | 1 | 0 | Distal Intergenic | 19 | 26460957 | 26461850 | 894 | 2 | 1,03E+08 | ENSMUST00000237584.2 | 161004 | | Gm34808 | predicted gene, 34808 |
| chr11 | 1,02E+08 | 1,02E+08 | 485 | * | 4,737236 | 6,602226011 | 2,2227 | 2,970363 | 0,002974 | 1 | 0 | Promoter (<=1kb) | 11 | 1,02E+08 | 1,02E+08 | 928 | 1 | 51799 | ENSMUST00000132876.2 | 894 | ENSMUSG00000006575 | Rundc3a | RUN domain containing 3A |
| chr17 | 25498267 | 25499277 | 1011 | * | 5,872341 | 6,60245096 | 1,984577 | 3,326881 | 0,000878 | 1 | 0 | Distal Intergenic | 17 | 25481367 | 25493596 | 12230 | 2 | 22196 | ENSMUST00000174001.8 | -4671 | ENSMUSG00000015120 | Ube2i | ubiquitin-conjugating enzyme E2I |
| chr12 | 41121846 | 41122445 | 600 | * | 3,573001 | 6,603806688 | 2,482437 | 2,660211 | 0,007809 | 1 | 0 | Intron (ENSMUST0000( | 12 | 41118311 | 41160942 | 42632 | 1 | 93757 | ENSMUST00000080160.4 | 3535 | ENSMUSG00000056899 | Immp2l | IMP2 inner mitochondrial membrane peptidase-like (S. cerevisiae) |
| chr1 | 1,28E+08 | 1,28E+08 | 387 | * | 4,22415 | 6,604534656 | 2,161348 | 3,055748 | 0,002245 | 1 | 0 | 3' UTR | 1 | 1,28E+08 | 1,28E+08 | 20439 | 2 | 1,01E+08 | ENSMUST00000188163.2 | 6700 | ENSMUSG00000102014 | 2900009J06Rik | RIKEN cDNA 2900009J06 gene |
| chr11 | 29380868 | 29381501 | 634 | * | 4,367416 | 6,604574028 | 2,149871 | 3,072079 | 0,002126 | 1 | 0 | Intron (ENSMUST0000( | 11 | 29406496 | 29411228 | 4733 | 1 | 108686 | ENSMUST00000137227.2 | -24995 | ENSMUSG00000032740 | Ccdc88a | coiled coil domain containing 88A |
| chr1 | 1,12E+08 | 1,12E+08 | 676 | * | 5,16358 | 6,604789186 | 2,09167 | 3,157664 | 0,00159 | 1 | 0 | Distal Intergenic | 1 | 1,12E+08 | 1,12E+08 | 136 | 1 | 1,15E+08 | ENSMUST00000158066.3 | -257651 | ENSMUSG00000088691 | Gm22331 | predicted gene, 22331 |
| chr16 | 53694565 | 53695185 | 621 | * | 5,433661 | 6,605002036 | 2,040434 | 3,237057 | 0,001208 | 1 | 0 | Distal Intergenic | 16 | 54174825 | 54174956 | 132 | 1 | 1,15E+08 | ENSMUST00000082793.3 | -479640 | ENSMUSG00000064727 | Gm25853 | predicted gene, 25853 |
| chr4 | 1,09E+08 | 1,09E+08 | 681 | * | 5,736615 | 6,605348832 | 1,973393 | 3,347204 | 0,000816 | 1 | 0 | Intron (ENSMUST0000( | 4 | 1,09E+08 | 1,09E+08 | 4061 | 1 | 13858 | ENSMUST00000141751.2 | 8161 | ENSMUSG00000028552 | Eps15 | epidermal growth factor receptor pathway substrate 15 |
| chr3 | 5481496 | 5482117 | 622 | * | 5,110116 | 6,605394687 | 1,967514 | 3,357229 | 0,000787 | 1 | 0 | Distal Intergenic | 3 | 5578055 | 5580393 | 2339 | 2 | 78039 | ENSMUST00000176666.2 | 98276 | ENSMUSG00000093557 | 4930555M17Rik | RIKEN cDNA 4930555M17 gene |
| chr3 | 42434999 | 42435627 | 629 | * | 5,709207 | 6,605904238 | 1,894039 | 3,487734 | 0,000487 | 1 | 0 | Distal Intergenic | 3 | 41705631 | 41710927 | 5297 | 1 | 73852 | ENSMUST00000195030.2 | 729368 | ENSMUSG00000025766 | D3Ertd751e | DNA segment, Chr 3, ERATO Doi 751, expressed |
| chr3 | 90529237 | 90530255 | 1019 | * | 7,507061 | 6,606134626 | 1,865825 | 3,540596 | 0,000399 | 1 | 0 | Intron (ENSMUST0000( | 3 | 90521067 | 90531488 | 10422 | 1 | 20200 | ENSMUST00000200289.2 | 8170 | ENSMUSG00000001025 | S100a6 | S100 calcium binding protein A6 (calcyclin) |
| chr4 | 1,08E+08 | 1,08E+08 | 1185 | * | 7,642083 | 6,606565722 | 1,824062 | 3,621898 | 0,000292 | 1 | 0 | Promoter (<=1kb) | 4 | 1,08E+08 | 1,08E+08 | 107 | 2 | 1,15E+08 | ENSMUST00000122644.3 | 478 | ENSMUSG00000084593 | Gm22345 | predicted gene, 22345 |
| chr12 | 55033964 | 55034576 | 613 | * | 6,220151 | 6,607768555 | 1,992968 | 3,315541 | 0,000915 | 1 | 0 | Promoter (<=1kb) | 12 | 54940336 | 55033292 | 92957 | 2 | 217578 | ENSMUST00000173433.8 | -672 | ENSMUSG00000035021 | Baz1a | bromodomain adjacent to zinc finger domain 1A |
| chr11 | 48692643 | 48693809 | 1167 | * | 8,233963 | 6,609832967 | 1,828394 | 3,615103 | 0,0003 | 0,29289663 | 1,425929 | Promoter (<=1kb) | 11 | 48692859 | 48692939 | 81 | 1 | 1E+08 | ENSMUST00000082791.3 | 0 | ENSMUSG00000064725 | Snord96a | small nucleolar RNA, C/D box 96A |
| chr14 | 65379583 | 65380142 | 560 | * | 3,301355 | 6,610328757 | 2,43798 | 2,711396 | 0,0067 | 1 | 0 | Intron (ENSMUST0000( | 14 | 65315199 | 65387304 | 72106 | 2 | 54616 | ENSMUST00000223989.2 | 7162 | ENSMUSG00000021978 | Extl3 | exostosin-like glycosyltransferase 3 |
| chr14 | 45464156 | 45464543 | 388 | * | 3,415714 | 6,610354163 | 2,430462 | 2,719793 | 0,006532 | 1 | 0 | Intron (ENSMUST0000( | 14 | 45372364 | 45457785 | 85422 | 2 | 70561 | ENSMUST00000123879.8 | -6371 | ENSMUSG00000021830 | Txndc16 | thioredoxin domain containing 16 |
| chr3 | 1,48E+08 | 1,48E+08 | 543 | * | 3,567943 | 6,611016053 | 2,328129 | 2,839626 | 0,004517 | 1 | 0 | Distal Intergenic | 3 | 1,48E+08 | 1,48E+08 | 132 | 2 | 1,15E+08 | ENSMUST00000104253.3 | 340697 | ENSMUSG00000077441 | Gm23131 | predicted gene, 23131 |
| chr7 | 69564085 | 69564412 | 328 | * | 3,94111 | 6,611018419 | 2,327801 | 2,840027 | 0,004511 | 1 | 0 | Distal Intergenic | 7 | 69607524 | 69607652 | 129 | 2 | 1,15E+08 | ENSMUST00000103978.3 | 43240 | ENSMUSG00000077166 | Gm24120 | predicted gene, 24120 |
| chr19 | 41254130 | 41254461 | 332 | * | 3,657357 | 6,61115067 | 2,308482 | 2,863852 | 0,004185 | 1 | 0 | Promoter (1-2kb) | 19 | 41199283 | 41252436 | 53154 | 2 | 107358 | ENSMUST00000025989.10 | -1694 | ENSMUSG00000025016 | Tm9sf3 | transmembrane 9 superfamily member 3 |
| chrX | 71850624 | 71851286 | 663 | * | 4,30646 | 6,611258234 | 2,295233 | 2,88043 | 0,003971 | 1 | 0 | Distal Intergenic | 20 | 71868784 | 71886208 | 17425 | 1 | 57249 | ENSMUST00000114553.8 | -17498 | ENSMUSG00000031344 | Gabrq | gamma-aminobutyric acid (GABA) A receptor, subunit theta |
| chr9 | 34888097 | 34888694 | 598 | * | 4,636312 | 6,612165495 | 2,189241 | 3,0203 | 0,002525 | 1 | 0 | Intron (ENSMUST0000( | 9 | 34927748 | 34942172 | 14425 | 1 | 67703 | ENSMUST00000188658.2 | -39054 | ENSMUSG00000032036 | Kirrel3 | kirre like nephrin family adhesion molecule 3 |
| chr7 | 92377150 | 92377541 | 392 | * | 3,877947 | 6,612444289 | 2,162164 | 3,058253 | 0,002226 | 1 | 0 | Distal Intergenic | 7 | 92388086 | 92390642 | 2557 | 2 | 666737 | ENSMUST00000245044.1 | 13101 | ENSMUSG00000074024 | 4632427E13Rik | RIKEN cDNA 4632427E13 gene |
| chr2 | 1,57E+08 | 1,57E+08 | 417 | * | 4,264167 | 6,61281637 | 2,128525 | 3,106761 | 0,001891 | 1 | 0 | Intron (ENSMUST0000( | 2 | 1,57E+08 | 1,57E+08 | 16168 | 1 | 20779 | ENSMUST00000109531.8 | -8794 | ENSMUSG00000027646 | Src | Rous sarcoma oncogene |
| chr5 | 1,3E+08 | 1,3E+08 | 506 | * | 5,549956 | 6,613336907 | 2,085318 | 3,171381 | 0,001517 | 1 | 0 | Promoter (<=1kb) | 5 | 1,3E+08 | 1,3E+08 | 42545 | 1 | 56715 | ENSMUST00000119797.8 | 0 | ENSMUSG00000025340 | Rabgef1 | RAB guanine nucleotide exchange factor (GEF) 1 |
| chr15 | 62614334 | 62614841 | 508 | * | 4,20164 | 6,613552271 | 2,143522 | 3,085368 | 0,002033 | 1 | 0 | Intron (ENSMUST0000( | 15 | 62665050 | 62665183 | 134 | 1 | 1,15E+08 | ENSMUST00000158272.3 | -50209 | ENSMUSG00000088897 | Gm22521 | predicted gene, 22521 |
| chr11 | 72632671 | 72633343 | 673 | * | 4,457451 | 6,613714721 | 2,168993 | 3,04921 | 0,002294 | 1 | 0 | Intron (ENSMUST0000( | 11 | 72650742 | 72651716 | 975 | 1 | 11736 | ENSMUST00000147195.2 | -17399 | ENSMUSG00000020790 | Ankfy1 | ankyrin repeat and FYVE domain containing 1 |
| chr19 | 16580618 | 16581080 | 463 | * | 3,415047 | 6,614484492 | 2,323699 | 2,846533 | 0,00442 | 1 | 0 | Exon (ENSMUST00000( | 19 | 16592730 | 16608711 | 15982 | 2 | 271564 | ENSMUST00000223667.2 | 27631 | ENSMUSG00000046230 | Vps13a | vacuolar protein sorting 13A |
| chr19 | 5028966 | 5030167 | 1202 | * | 8,284311 | 6,614711243 | 1,85912 | 3,55798 | 0,000374 | 0,323988916 | 1,360622 | Distal Intergenic | 19 | 5035030 | 5039932 | 4903 | 2 | 225872 | ENSMUST00000237881.2 | 9765 | ENSMUSG00000045903 | Npas4 | neuronal PAS domain protein 4 |
| chr1 | 24726127 | 24726512 | 386 | * | 3,203237 | 6,615049991 | 2,479545 | 2,667848 | 0,007634 | 1 | 0 | Intron (ENSMUST0000( | 1 | 24718007 | 24805382 | 87376 | 1 | 68421 | ENSMUST00000095062.10 | 8120 | ENSMUSG00000073725 | Lmbrd1 | LMBR1 domain containing 1 |
| chr12 | 99779637 | 99779917 | 281 | * | 3,493884 | 6,616059182 | 2,517938 | 2,627571 | 0,0086 | 1 | 0 | Intron (ENSMUST0000( | 12 | 99684981 | 99849116 | 164136 | 2 | 78767 | ENSMUST00000220875.2 | 69199 | ENSMUSG00000021176 | Efcab11 | EF-hand calcium binding domain 11 |
| chr19 | 43043717 | 43044046 | 330 | * | 3,535441 | 6,616337857 | 2,483783 | 2,663815 | 0,007726 | 1 | 0 | Intron (ENSMUST0000( | 19 | 42897023 | 43212128 | 315106 | 2 | 545291 | ENSMUST00000238153.2 | 168082 | ENSMUSG00000074852 | Hpse2 | heparanase 2 |
| chr11 | 18278908 | 18279259 | 352 | * | 4,561236 | 6,619291415 | 2,421425 | 2,733634 | 0,006264 | 1 | 0 | Distal Intergenic | 11 | 18417479 | 18443255 | 25777 | 2 | 1,03E+08 | ENSMUST00000143311.2 | 163996 | ENSMUSG00000085675 | Gm12022 | predicted gene 12022 |
| chr10 | 77280164 | 77280616 | 453 | * | 4,271882 | 6,619732245 | 2,194348 | 3,01672 | 0,002555 | 1 | 0 | Distal Intergenic | 10 | 77293751 | 77307463 | 13713 | 1 | 1,03E+08 | ENSMUST00000218522.2 | -13135 | | Gm35920 | predicted gene, 35920 |
| chr9 | 49147934 | 49148601 | 668 | * | 5,470993 | 6,620505146 | 2,039092 | 3,246791 | 0,001167 | 1 | 0 | Distal Intergenic | 9 | 49161831 | 49191894 | 30064 | 1 | 235327 | ENSMUST00000216428.2 | -13230 | ENSMUSG00000054156 | Gm4894 | predicted gene 4894 |
| chr7 | 30418243 | 30418753 | 511 | * | 4,912225 | 6,621229484 | 2,111731 | 3,135452 | 0,001716 | 1 | 0 | Promoter (<=1kb) | 7 | 30417257 | 30420045 | 2789 | 1 | 11944 | ENSMUST00000171014.2 | 986 | ENSMUSG00000005553 | Atp4a | ATPase, H+/K+ exchanging, gastric, alpha polypeptide |
| chr14 | 66519210 | 66519866 | 657 | * | 3,929649 | 6,621329296 | 2,411858 | 2,745323 | 0,006045 | 1 | 0 | Promoter (<=1kb) | 14 | 66390711 | 66518501 | 127791 | 2 | 19229 | ENSMUST00000089250.9 | -709 | ENSMUSG00000059456 | Ptk2b | PTK2 protein tyrosine kinase 2 beta |

| | | | | | | | | | | | | | | | | | | | | | | | |
|---|---|---|---|---|---|---|---|---|---|---|---|---|---|---|---|---|---|---|---|---|---|---|---|
| chr5 | 72905450 | 72905918 | 469 | * | 3,585596 | 6,621400577 | 2,336346 | 2,834084 | 0,004596 | 1 | 0 | Exon (ENSMUST00000 | 5 | 72853321 | 72910116 | 56796 | 2 | 22165 | ENSMUST00000197313.5 | 4198 | ENSMUSG00000054892 | Txk | TXK tyrosine kinase |
| chr12 | 1,1E+08 | 1,1E+08 | 270 | * | 3,682619 | 6,621406457 | 2,308257 | 2,868574 | 0,004123 | 1 | 0 | Distal Intergenic | 12 | 1,1E+08 | 1,1E+08 | 26388 | 2 | 1,03E+08 | ENSMUST00000244432.1 | -9498 | | Gm34667 | predicted gene, 34667 |
| chr12 | 83907606 | 83907983 | 378 | * | 3,644869 | 6,621425104 | 2,27813 | 2,906517 | 0,003655 | 1 | 0 | Intron (ENSMUST00000 | 12 | 83878602 | 83923572 | 44971 | 2 | 18222 | ENSMUST00000125089.2 | 15589 | ENSMUSG00000021224 | Numb | NUMB endocytic adaptor protein |
| chr16 | 32158870 | 32159427 | 558 | * | 4,550872 | 6,62166847 | 2,058147 | 3,217297 | 0,001294 | 1 | 0 | Intron (ENSMUST00000 | 16 | 32151086 | 32206713 | 55628 | 1 | 224111 | ENSMUST00000232137.2 | 7784 | ENSMUSG00000053774 | Ubxn7 | UBX domain protein 7 |
| chr19 | 24429748 | 24430400 | 653 | * | 4,906311 | 6,621762649 | 2,003609 | 3,304917 | 0,00095 | 1 | 0 | Intron (ENSMUST00000 | 19 | 24453866 | 24454720 | 855 | 2 | 68034 | ENSMUST00000099556.2 | 24320 | ENSMUSG00000074922 | Fam122a | family with sequence similarity 122, member A |
| chr15 | 31794513 | 31795146 | 634 | * | 6,287644 | 6,621866569 | 1,953236 | 3,390203 | 0,000698 | 1 | 0 | Distal Intergenic | 15 | 31602264 | 31618755 | 16492 | 1 | 68073 | ENSMUST00000161061.3 | 192249 | ENSMUSG00000039065 | Atpsckmt | ATP synthase C subunit lysine N-methyltransferase |
| chr4 | 43700939 | 43702267 | 1329 | * | 8,08068 | 6,622378786 | 1,782798 | 3,714598 | 0,000204 | 0,247743734 | 1,360622 | Promoter (<=1kb) | 4 | 43694784 | 43700807 | 6024 | 2 | 56014 | ENSMUST00000055545.5 | -132 | ENSMUSG00000050215 | Olfr70 | olfactory receptor 70 |
| chr19 | 5155467 | 5156061 | 595 | * | 4,204369 | 6,624757552 | 2,313601 | 2,863396 | 0,004191 | 1 | 0 | Promoter (<=1kb) | 19 | 5150437 | 5156273 | 5837 | 2 | 76308 | ENSMUST00000237201.2 | 212 | ENSMUSG00000024870 | Rab1b | RAB1B, member RAS oncogene family |
| chr4 | 1,25E+08 | 1,25E+08 | 725 | * | 6,036481 | 6,624893048 | 1,968123 | 3,366098 | 0,000762 | 1 | 0 | Intron (ENSMUST00000 | 4 | 1,25E+08 | 1,25E+08 | 155288 | 1 | 1,08E+08 | ENSMUST00000245356.1 | 34747 | | Gm46869 | predicted gene, 46869 |
| chr15 | 3142890 | 3143273 | 384 | * | 3,320464 | 6,627169219 | 2,567932 | 2,580742 | 0,009859 | 1 | 0 | Distal Intergenic | 15 | 3298029 | 3306755 | 8727 | 1 | 20363 | ENSMUST00000160787.10 | -154756 | ENSMUSG00000064373 | Selenop | selenoprotein P |
| chr10 | 1,18E+08 | 1,18E+08 | 425 | * | 3,346743 | 6,627681905 | 2,432986 | 2,724094 | 0,006448 | 1 | 0 | 3' UTR | 10 | 1,18E+08 | 1,18E+08 | 1278 | 2 | 17246 | ENSMUST00000131627.2 | 3907 | ENSMUSG00000020184 | Mdm2 | transformed mouse 3T3 cell double minute 2 |
| chr7 | 58236928 | 58237486 | 559 | * | 3,986592 | 6,628180082 | 2,331265 | 2,843169 | 0,004467 | 1 | 0 | Distal Intergenic | 7 | 58305914 | 58308626 | 2713 | 1 | 11982 | ENSMUST00000207668.2 | -68428 | ENSMUSG00000025324 | Atp10a | ATPase, class V, type 10A |
| chr5 | 47945197 | 47945822 | 626 | * | 4,211724 | 6,628344556 | 2,303702 | 2,877257 | 0,004011 | 1 | 0 | Distal Intergenic | 5 | 48140480 | 48465075 | 324596 | 1 | 20563 | ENSMUST00000173107.8 | -194658 | ENSMUSG00000031558 | Slit2 | slit guidance ligand 2 |
| chr10 | 49446438 | 49446749 | 312 | * | 3,831906 | 6,628445738 | 2,287617 | 2,897533 | 0,003761 | 1 | 0 | Intron (ENSMUST00000 | 10 | 49232129 | 49411449 | 179321 | 2 | 14806 | ENSMUST00000218669.2 | -34989 | ENSMUSG00000056073 | Grik2 | glutamate receptor, ionotropic, kainate 2 (beta 2) |
| chr19 | 38748229 | 38749195 | 967 | * | 8,738264 | 6,630490802 | 1,835571 | 3,612223 | 0,000304 | 0,291564571 | 1,449706 | Exon (ENSMUST00000 | 19 | 38723358 | 38739567 | 16210 | 1 | 74055 | ENSMUST00000183131.2 | 24871 | ENSMUSG00000024998 | Plce1 | phospholipase C, epsilon 1 |
| chr5 | 77555907 | 77556298 | 392 | * | 4,90205 | 6,630688931 | 2,026859 | 3,271411 | 0,00107 | 1 | 0 | Promoter (<=1kb) | 5 | 77497087 | 77555888 | 58802 | 2 | 29817 | ENSMUST00000046746.10 | -19 | ENSMUSG00000036256 | Igfbp7 | insulin-like growth factor binding protein 7 |
| chr16 | 33939438 | 33940325 | 888 | * | 6,749114 | 6,630823058 | 2,014789 | 3,291075 | 0,000998 | 1 | 0 | Intron (ENSMUST00000 | 16 | 33833507 | 33917460 | 83954 | 2 | 545156 | ENSMUST00000114966.8 | -21978 | ENSMUSG00000061751 | Kalrn | kalirin, RhoGEF kinase |
| chr11 | 61787442 | 61788485 | 1044 | * | 8,632425 | 6,630951146 | 1,863228 | 3,558852 | 0,000372 | 0,316831371 | 1,425929 | Exon (ENSMUST00000 | 11 | 61762498 | 61769276 | 6779 | 2 | 56697 | ENSMUST00000137323.2 | -18166 | ENSMUSG00000047804 | Akap10 | A kinase (PRKA) anchor protein 10 |
| chr15 | 75565588 | 75566144 | 557 | * | 5,752684 | 6,632262439 | 1,921227 | 3,452097 | 0,000556 | 1 | 0 | Distal Intergenic | 15 | 75576129 | 75587334 | 11206 | 1 | 14787 | ENSMUST00000121137.9 | -9985 | ENSMUSG00000022580 | Rhpn1 | rhophilin, Rho GTPase binding protein 1 |
| chr5 | 92109980 | 92110875 | 896 | * | 6,239537 | 6,632388446 | 1,915071 | 3,463259 | 0,000534 | 1 | 0 | Promoter (<=1kb) | 5 | 92110248 | 92114457 | 4210 | 1 | 381650 | ENSMUST00000193925.6 | 0 | ENSMUSG00000102644 | Thap6 | THAP domain containing 6 |
| chr18 | 57564664 | 57565205 | 542 | * | 6,28538 | 6,632526042 | 1,907621 | 3,476857 | 0,000507 | 1 | 0 | Intron (ENSMUST00000 | 18 | 57538457 | 57547308 | 8852 | 1 | 1,01E+08 | ENSMUST00000242320.1 | 26207 | ENSMUSG00000117864 | Gm19500 | predicted gene, 19500 |
| chr17 | 13987105 | 13987715 | 611 | * | 3,466281 | 6,633200149 | 2,54337 | 2,608035 | 0,009106 | 1 | 0 | Intron (ENSMUST00000 | 17 | 13936698 | 13982087 | 45390 | 2 | 21646 | ENSMUST00000148430.2 | -5018 | ENSMUSG00000038347 | Tcte2 | t-complex-associated testis expressed 2 |
| chrX | 1,61E+08 | 1,61E+08 | 255 | * | 3,235377 | 6,633385167 | 2,519774 | 2,632532 | 0,008475 | 1 | 0 | Distal Intergenic | 20 | 1,61E+08 | 1,61E+08 | 326435 | 2 | 195727 | ENSMUST00000087085.10 | -32628 | ENSMUSG00000059493 | Nhs | NHS actin remodeling regulator |
| chr5 | 98326353 | 98326710 | 358 | * | 3,393189 | 6,634482559 | 2,378695 | 2,789127 | 0,005285 | 1 | 0 | Promoter (2-3kb) | 5 | 98328837 | 98336850 | 8014 | 1 | 77630 | ENSMUST00000112959.4 | -2127 | ENSMUSG00000035456 | Prdm8 | PR domain containing 8 |
| chrX | 55709349 | 55709800 | 452 | * | 3,937802 | 6,634519356 | 2,37752 | 2,79052 | 0,005262 | 1 | 0 | 3' UTR | 20 | 55703605 | 55709189 | 5585 | 1 | 236794 | ENSMUST00000202670.2 | 5744 | ENSMUSG00000060681 | Slc9a6 | solute carrier family 9 (sodium/hydrogen exchanger), member 6 |
| chr13 | 50155888 | 50156451 | 564 | * | 3,724589 | 6,634566451 | 2,368978 | 2,800603 | 0,005101 | 1 | 0 | Exon (ENSMUST00000 | 13 | 50291105 | 50300706 | 9602 | 1 | 71252 | ENSMUST00000222134.2 | -134654 | | 4933433N18Rik | RIKEN cDNA 4933433N18 gene |
| chr5 | 1,17E+08 | 1,17E+08 | 1155 | * | 7,945538 | 6,634567012 | 1,819291 | 3,646787 | 0,000266 | 1 | 0 | Promoter (<=1kb) | 5 | 1,17E+08 | 1,17E+08 | 14639 | 2 | 68955 | ENSMUST00000150127.2 | 0 | ENSMUSG00000063919 | Srrm4 | serine/arginine repetitive matrix 4 |
| chr14 | 54766639 | 54767507 | 869 | * | 5,008111 | 6,637589495 | 2,055124 | 3,229776 | 0,001239 | 1 | 0 | Distal Intergenic | 14 | 54744644 | 54754982 | 10339 | 2 | 27374 | ENSMUST00000023873.12 | -11657 | ENSMUSG00000023110 | Prmt5 | protein arginine N-methyltransferase 5 |
| chr17 | 26368676 | 26369520 | 845 | * | 5,873475 | 6,638014786 | 1,917245 | 3,462267 | 0,000536 | 1 | 0 | Intron (ENSMUST00000 | 17 | 26361592 | 26409205 | 47614 | 1 | 12005 | ENSMUST00000163421.9 | 7084 | ENSMUSG00000024182 | Axin1 | axin 1 |
| chr3 | 86046228 | 86047032 | 805 | * | 5,010678 | 6,63809187 | 2,098422 | 3,163373 | 0,00156 | 1 | 0 | Promoter (<=1kb) | 3 | 86046098 | 86046165 | 68 | 2 | 19870 | ENSMUST00000083050.3 | -63 | ENSMUSG00000064984 | Snord73a | small nucleolar RNA, C/D box U73A |
| chr2 | 1,28E+08 | 1,28E+08 | 595 | * | 5,000135 | 6,638180466 | 2,001256 | 3,317007 | 0,00091 | 1 | 0 | Intron (ENSMUST00000 | 2 | 1,28E+08 | 1,28E+08 | 15009 | 2 | 1E+08 | ENSMUST00000239358.2 | 50599 | ENSMUSG00000074813 | Morrbid | myeloid RNA regulator of BCL2L11 induced cell death |
| chr2 | 1,7E+08 | 1,7E+08 | 749 | * | 5,666706 | 6,638195861 | 2,008562 | 3,304949 | 0,00095 | 1 | 0 | Distal Intergenic | 2 | 1,7E+08 | 1,7E+08 | 894 | 2 | 1,01E+08 | ENSMUST00000241007.1 | 16266 | | Gm17619 | predicted gene, 17619 |
| chr12 | 5977163 | 5977591 | 429 | * | 5,356037 | 6,638224424 | 2,090299 | 3,17573 | 0,001495 | 1 | 0 | Intron (ENSMUST00000 | 12 | 5858234 | 5906142 | 47909 | 1 | 1,03E+08 | ENSMUST00000219472.2 | 118929 | | Gm32442 | predicted gene, 32442 |
| chr5 | 50285319 | 50285754 | 436 | * | 5,113438 | 6,638241152 | 2,042069 | 3,250743 | 0,001151 | 1 | 0 | Distal Intergenic | 5 | 50308675 | 50313437 | 4763 | 1 | 73965 | ENSMUST00000197958.2 | -22921 | ENSMUSG00000104870 | 4930448I18Rik | RIKEN cDNA 4930448I18 gene |
| chr9 | 28116115 | 28116650 | 536 | * | 4,643429 | 6,638241891 | 2,090116 | 3,176017 | 0,001493 | 1 | 0 | Intron (ENSMUST00000 | 9 | 28314445 | 28317247 | 2803 | 1 | 330908 | ENSMUST00000133142.2 | -197795 | ENSMUSG00000062257 | Opcml | opioid binding protein/cell adhesion molecule-like |
| chr10 | 32442547 | 32443051 | 505 | * | 5,0767 | 6,63827059 | 2,056198 | 3,228421 | 0,001245 | 1 | 0 | Intron (ENSMUST00000 | 10 | 32205761 | 32286331 | 80571 | 2 | 432450 | ENSMUST00000092603.5 | -156216 | ENSMUSG00000069670 | Nkain2 | Na+/K+ transporting ATPase interacting 2 |
| chr13 | 1,13E+08 | 1,13E+08 | 740 | * | 4,137381 | 6,638315436 | 2,094563 | 3,169308 | 0,001528 | 1 | 0 | Distal Intergenic | 13 | 1,13E+08 | 1,13E+08 | 46017 | 1 | 16195 | ENSMUST00000183663.8 | -19033 | ENSMUSG00000021756 | Il6st | interleukin 6 signal transducer |
| chr10 | 18376980 | 18377832 | 853 | * | 5,158286 | 6,638340618 | 2,128905 | 3,118195 | 0,00182 | 1 | 0 | Intron (ENSMUST00000 | 10 | 18345729 | 18409637 | 63909 | 1 | 215819 | ENSMUST00000100054.4 | 31251 | ENSMUSG00000039835 | Nhsl1 | NHS-like 1 |
| chr12 | 83941783 | 83942346 | 564 | * | 5,346256 | 6,638342826 | 2,105035 | 3,153555 | 0,001613 | 1 | 0 | Intron (ENSMUST00000 | 12 | 83840810 | 83923584 | 82775 | 2 | 18222 | ENSMUST00000110298.9 | -18199 | ENSMUSG00000021224 | Numb | NUMB endocytic adaptor protein |
| chr7 | 99217890 | 99218224 | 335 | * | 3,773995 | 6,638488834 | 2,290421 | 2,898371 | 0,003751 | 1 | 0 | Exon (ENSMUST00000 | 7 | 99222271 | 99237335 | 15065 | 1 | 109689 | ENSMUST00000161268.2 | -4047 | ENSMUSG00000018909 | Arrb1 | arrestin, beta 1 |
| chr5 | 1,25E+08 | 1,25E+08 | 883 | * | 4,646806 | 6,638671289 | 2,104041 | 3,1552 | 0,001604 | 1 | 0 | Exon (ENSMUST00000 | 5 | 1,25E+08 | 1,25E+08 | 5750 | 2 | 243272 | ENSMUST00000200674.2 | -3580 | ENSMUSG00000038095 | Sbno1 | strawberry notch 1 |
| chr19 | 12991652 | 12991974 | 323 | * | 3,534033 | 6,638866968 | 2,568464 | 2,584762 | 0,009745 | 1 | 0 | Distal Intergenic | 19 | 13004768 | 13005691 | 924 | 2 | 258695 | ENSMUST00000087773.2 | 13717 | ENSMUSG00000094755 | Olfr1453 | olfactory receptor 1453 |
| chr16 | 48838157 | 48838950 | 794 | * | 5,24892 | 6,639033059 | 2,050179 | 3,23827 | 0,001203 | 1 | 0 | Promoter (1-2kb) | 16 | 48836200 | 48840072 | 3873 | 1 | 224171 | ENSMUST00000149004.2 | 1957 | ENSMUSG00000033031 | Cip2a | cell proliferation regulating inhibitor of protein phosphatase 2A |
| chr1 | 66439581 | 66440086 | 506 | * | 4,120685 | 6,641231117 | 2,257121 | 2,942347 | 0,003257 | 1 | 0 | Intron (ENSMUST00000 | 1 | 66426170 | 66440734 | 14565 | 1 | 17756 | ENSMUST00000174590.2 | 13411 | ENSMUSG00000015222 | Map2 | microtubule-associated protein 2 |
| chr7 | 1,1E+08 | 1,1E+08 | 465 | * | 6,791959 | 6,641478966 | 1,951387 | 3,403465 | 0,000665 | 1 | 0 | Distal Intergenic | 7 | 1,1E+08 | 1,1E+08 | 19545 | 2 | 319934 | ENSMUST00000165992.2 | 38712 | ENSMUSG00000038371 | Sbf2 | SET binding factor 2 |
| chr15 | 59613015 | 59613773 | 759 | * | 7,828292 | 6,641559571 | 1,836902 | 3,61563 | 0,0003 | 1 | 0 | Distal Intergenic | 15 | 59660266 | 59666808 | 6543 | 2 | 1,01E+08 | ENSMUST00000228865.2 | 53035 | ENSMUSG00000115426 | Gm19510 | predicted gene, 19510 |
| chr5 | 1,47E+08 | 1,47E+08 | 606 | * | 5,929529 | 6,641901851 | 1,93792 | 3,427335 | 0,00061 | 1 | 0 | Intron (ENSMUST00000 | 5 | 1,47E+08 | 1,47E+08 | 60198 | 2 | 22217 | ENSMUST00000085614.6 | 8669 | ENSMUSG00000029640 | Usp12 | ubiquitin specific peptidase 12 |
| chr7 | 45025156 | 45026189 | 1034 | * | 7,209055 | 6,642739308 | 1,877064 | 3,538898 | 0,000402 | 1 | 0 | 3' UTR | 7 | 45030026 | 45030741 | 716 | 2 | 20637 | ENSMUST00000210661.2 | 4552 | ENSMUSG00000063511 | Snrnp70 | small nuclear ribonucleoprotein 70 (U1) |
| chr17 | 29020669 | 29021619 | 951 | * | 7,338466 | 6,644084316 | 1,875756 | 3,542084 | 0,000397 | 1 | 0 | Promoter (<=1kb) | 17 | 29020758 | 29025282 | 4525 | 1 | 268936 | ENSMUST00000136233.2 | 0 | ENSMUSG00000063952 | Brpf3 | bromodomain and PHD finger containing, 3 |
| chrX | 43737883 | 43738385 | 503 | * | 3,544316 | 6,64507992 | 2,369655 | 2,804239 | 0,005044 | 1 | 0 | Distal Intergenic | 20 | 43875447 | 43879038 | 3592 | 2 | 245404 | ENSMUST00000115056.2 | 140653 | ENSMUSG00000045284 | Dcaf12l1 | DDB1 and CUL4 associated factor 12-like 1 |
| chr12 | 56845372 | 56845752 | 381 | * | 3,823879 | 6,645405028 | 2,285948 | 2,907068 | 0,003648 | 1 | 0 | Intron (ENSMUST00000 | 12 | 56773912 | 56822145 | 48234 | 2 | 217593 | ENSMUST00000160709.2 | -23227 | ENSMUSG00000035472 | Slc25a21 | solute carrier family 25 (mitochondrial oxodicarboxylate carrier), member 21 |
| chr17 | 42248940 | 42249464 | 525 | * | 4,916564 | 6,646812574 | 2,052293 | 3,238726 | 0,001201 | 1 | 0 | Distal Intergenic | 17 | 42626838 | 42815968 | 189131 | 1 | 627626 | ENSMUST00000048691.6 | -377374 | ENSMUSG00000042256 | Ptchd4 | patched domain containing 4 |
| chr12 | 38992611 | 38993308 | 698 | * | 4,424733 | 6,646846865 | 2,048769 | 3,244312 | 0,001177 | 1 | 0 | Distal Intergenic | 12 | 38956054 | 38956952 | 899 | 2 | 1,03E+08 | ENSMUST00000223252.2 | -35659 | | Gm34611 | predicted gene, 34611 |
| chr5 | 83604330 | 83604873 | 544 | * | 3,782653 | 6,648470208 | 2,277081 | 2,919733 | 0,003503 | 1 | 0 | Distal Intergenic | 5 | 83425992 | 83503042 | 77051 | 2 | 243078 | ENSMUST00000053543.11 | -101288 | ENSMUSG00000049537 | Tecrl | trans-2,3-enoyl-CoA reductase-like |
| chrX | 74302607 | 74303609 | 1003 | * | 5,988077 | 6,648543466 | 1,896829 | 3,505084 | 0,000456 | 1 | 0 | Exon (ENSMUST00000 | 20 | 74192727 | 74207376 | 14650 | 2 | 434800 | ENSMUST00000141041.2 | -95231 | ENSMUSG00000073094 | Smim9 | small integral membrane protein 9 |
| chr3 | 82002446 | 82002894 | 449 | * | 3,546793 | 6,648937362 | 2,345998 | 2,834162 | 0,004595 | 1 | 0 | Intron (ENSMUST00000 | 3 | 82001859 | 82009418 | 7560 | 2 | 60596 | ENSMUST00000192289.2 | 6524 | ENSMUSG00000033910 | Gucy1a1 | guanylate cyclase 1, soluble, alpha 1 |
| chr1 | 1,65E+08 | 1,65E+08 | 1619 | * | 10,09092 | 6,64900557 | 1,778329 | 3,738906 | 0,000185 | 0,253278065 | 1,173325 | Intron (ENSMUST00000 | 1 | 1,65E+08 | 1,65E+08 | 22494 | 2 | 74106 | ENSMUST00000193353.2 | 6017 | ENSMUSG00000026571 | Dcaf6 | DDB1 and CUL4 associated factor 6 |
| chr10 | 1,17E+08 | 1,17E+08 | 1710 | * | 13,93265 | 6,649874985 | 1,642587 | 4,048416 | 5,16E-05 | 0,14233608 | 0,918255 | Distal Intergenic | 10 | 1,17E+08 | 1,17E+08 | 1356 | 1 | 1E+08 | ENSMUST00000217799.2 | -12476 | | Gm10747 | predicted gene 10747 |
| chrX | 95169963 | 95170487 | 525 | * | 3,06814 | 6,649907526 | 2,528345 | 2,630142 | 0,008535 | 1 | 0 | Intron (ENSMUST00000 | 20 | 95194443 | 95194578 | 136 | 2 | 1,15E+08 | ENSMUST00000157521.3 | 24091 | ENSMUSG00000088146 | Gm24718 | predicted gene, 24718 |
| chr9 | 1,1E+08 | 1,1E+08 | 710 | * | 8,038236 | 6,650380101 | 1,796182 | 3,702508 | 0,000213 | 1 | 0 | Intron (ENSMUST00000 | 9 | 1,1E+08 | 1,1E+08 | 10819 | 1 | 235626 | ENSMUST00000198697.2 | -7318 | ENSMUSG00000044791 | Setd2 | SET domain containing 2 |
| chr14 | 67948480 | 67949023 | 544 | * | 2,990417 | 6,650631754 | 2,499792 | 2,660474 | 0,007803 | 1 | 0 | Intron (ENSMUST00000 | 14 | 67936385 | 67952824 | 16440 | 2 | 108912 | ENSMUST00000155312.2 | 3801 | ENSMUSG00000048922 | Cdca2 | cell division cycle associated 2 |
| chr5 | 1,07E+08 | 1,07E+08 | 528 | * | 3,312224 | 6,650639785 | 2,502848 | 2,657229 | 0,007879 | 1 | 0 | Promoter (1-2kb) | 5 | 1,07E+08 | 1,07E+08 | 55136 | 2 | 330149 | ENSMUST00000148495.2 | 1749 | ENSMUSG00000043410 | Hfm1 | HFM1, ATP-dependent DNA helicase homolog |
| chr19 | 39367793 | 39368121 | 329 | * | 3,429142 | 6,650738899 | 2,480708 | 2,680984 | 0,007341 | 1 | 0 | Distal Intergenic | 19 | 39378000 | 39451519 | 73520 | 2 | 13097 | ENSMUST00000035488.3 | 83398 | ENSMUSG00000032808 | Cyp2c38 | cytochrome P450, family 2, subfamily c, polypeptide 38 |
| chr3 | 1,3E+08 | 1,3E+08 | 1437 | * | 11,69239 | 6,651084823 | 1,748201 | 3,804532 | 0,000142 | 0,280031433 | 0,744375 | Distal Intergenic | 3 | 1,3E+08 | 1,31E+08 | 13533 | 2 | 66357 | ENSMUST00000043937.9 | 18177 | ENSMUSG00000041084 | Ostc | oligosaccharyltransferase complex subunit (non-catalytic) |
| chr16 | 3642856 | 3643129 | 274 | * | 3,739651 | 6,651208472 | 2,410347 | 2,75944 | 0,00579 | 1 | 0 | Distal Intergenic | 16 | 3656839 | 3657777 | 939 | 1 | 18312 | ENSMUST00000080917.2 | -13710 | ENSMUSG00000059043 | Olfr15 | olfactory receptor 15 |
| chr14 | 9991114 | 9991513 | 400 | * | 3,205232 | 6,651324927 | 2,397363 | 2,774433 | 0,00553 | 1 | 0 | Intron (ENSMUST00000 | 14 | 9980246 | 10093085 | 112840 | 1 | 27062 | ENSMUST00000224581.2 | 10868 | ENSMUSG00000054423 | Cadps | Ca2+-dependent secretion activator |
| chr3 | 10674540 | 10674868 | 329 | * | 3,753978 | 6,651450021 | 2,380367 | 2,794296 | 0,005201 | 1 | 0 | Distal Intergenic | 3 | 10482877 | 10505162 | 22286 | 2 | 74718 | ENSMUST00000099223.11 | -169378 | ENSMUSG00000027534 | Snx16 | sorting nexin 16 |
| chr15 | 91582424 | 91582726 | 303 | * | 3,588727 | 6,651589608 | 2,361572 | 2,816594 | 0,004854 | 1 | 0 | Intron (ENSMUST00000 | 15 | 91557378 | 91700323 | 142946 | 1 | 66725 | ENSMUST00000060642.7 | 25046 | ENSMUSG00000036273 | Lrrk2 | leucine-rich repeat kinase 2 |
| chr14 | 1,02E+08 | 1,02E+08 | 573 | * | 3,972194 | 6,651918253 | 2,323709 | 2,86263 | 0,004201 | 1 | 0 | Distal Intergenic | 14 | 1,02E+08 | 1,02E+08 | 3606 | 1 | 380928 | ENSMUST00000160038.2 | 12471 | ENSMUSG00000033060 | Lmo7 | LIM domain only 7 |
| chrX | 21390870 | 21391520 | 651 | * | 3,638812 | 6,652399351 | 2,270772 | 2,929576 | 0,003394 | 1 | 0 | Distal Intergenic | 20 | 21366605 | 21366735 | 131 | 2 | 1,15E+08 | ENSMUST00000104242.3 | -24135 | ENSMUSG00000077430 | Gm22592 | predicted gene, 22592 |
| chr2 | 71810983 | 71811498 | 516 | * | 3,755687 | 6,652563172 | 2,253787 | 2,951727 | 0,00316 | 1 | 0 | Promoter (<=1kb) | 2 | 71811584 | 72087818 | 276235 | 1 | 56508 | ENSMUST00000090826.12 | -86 | ENSMUSG00000049044 | Rapgef4 | Rap guanine nucleotide exchange factor (GEF) 4 |
| chrX | 1,59E+08 | 1,59E+08 | 534 | * | 4,124847 | 6,652600318 | 2,249602 | 2,957235 | 0,003104 | 1 | 0 | Intron (ENSMUST00000 | 20 | 1,59E+08 | 1,59E+08 | 1675 | 2 | 18597 | ENSMUST00000156531.2 | 24343 | ENSMUSG00000031299 | Pdha1 | pyruvate dehydrogenase E1 alpha 1 |
| chr15 | 81929163 | 81929532 | 370 | * | 3,848134 | 6,652641163 | 2,244559 | 2,963897 | 0,003038 | 1 | 0 | Promoter (2-3kb) | 15 | 81925935 | 81931720 | 5786 | 2 | 20826 | ENSMUST00000230953.2 | 2188 | ENSMUSG00000063480 | Snu13 | SNU13 homolog, small nuclear ribonucleoprotein (U4/U6.U5) |
| chr11 | 51157097 | 51158141 | 1045 | * | 7,718529 | 6,653215269 | 1,827306 | 3,640997 | 0,000272 | 1 | 0 | 5' UTR | 11 | 51154013 | 51172593 | 18581 | 1 | 12750 | ENSMUST00000109113.8 | 3084 | ENSMUSG00000020385 | Clk4 | CDC like kinase 4 |
| chr11 | 70114438 | 70115077 | 640 | * | 5,001675 | 6,653688642 | 2,155729 | 3,086515 | 0,002025 | 1 | 0 | Promoter (2-3kb) | 11 | 70110002 | 70111890 | 1889 | 2 | 69309 | ENSMUST00000190533.2 | -2548 | ENSMUSG00000044367 | Slc16a13 | solute carrier family 16 (monocarboxylic acid transporters), member 13 |
| chr15 | 7582374 | 7582918 | 545 | * | 4,607769 | 6,654208377 | 2,117523 | 3,142449 | 0,001675 | 1 | 0 | Distal Intergenic | 15 | 7236064 | 7427876 | 191813 | 2 | 268780 | ENSMUST00000160207.8 | -154498 | ENSMUSG00000042961 | Egflam | EGF-like, fibronectin type III and laminin G domains |
| chr8 | 53668131 | 53668541 | 411 | * | 6,450095 | 6,655177924 | 2,05626 | 3,236544 | 0,00121 | 1 | 0 | Distal Intergenic | 8 | 53889135 | 53933533 | 44399 | 1 | 1,05E+08 | ENSMUST00000210964.2 | -220594 | | Gm39180 | predicted gene, 39180 |
| chr9 | 1,08E+08 | 1,08E+08 | 1277 | * | 11,04696 | 6,655446915 | 2,337208 | 2,847606 | 0,004405 | 0,763005183 | 1,2763 | Distal Intergenic | 9 | 1,08E+08 | 1,08E+08 | 94363 | 2 | 12217 | ENSMUST00000035208.14 | -5417 | ENSMUSG00000032589 | Bsn | bassoon |
| chrX | 1,09E+08 | 1,09E+08 | 485 | * | 4,047298 | 6,655579361 | 2,385994 | 2,789437 | 0,00528 | 1 | 0 | Distal Intergenic | 20 | 1,09E+08 | 1,09E+08 | 2406 | 1 | 634340 | ENSMUST00000122296.2 | -247259 | ENSMUSG00000082853 | Gm7134 | kinesin family member 18B pseudogene |
| chr6 | 86327545 | 86327900 | 356 | * | 3,652077 | 6,655664737 | 2,444094 | 2,723162 | 0,006466 | 1 | 0 | Distal Intergenic | 6 | 86342628 | 86347040 | 4413 | 1 | 66488 | ENSMUST00000071492.9 | -14728 | ENSMUSG00000057497 | Fam136a | family with sequence similarity 136, member A |
| chr4 | 1,06E+08 | 1,06E+08 | 327 | * | 3,707583 | 6,656214839 | 2,542574 | 2,617904 | 0,008847 | 1 | 0 | Distal Intergenic | 4 | 1,06E+08 | 1,06E+08 | 125105 | 1 | 329908 | ENSMUST00000165709.8 | -12132 | ENSMUSG00000028514 | Usp24 | ubiquitin specific peptidase 24 |
| chr7 | 33895889 | 33897006 | 1118 | * | 8,637052 | 6,657146099 | 1,837304 | 3,623322 | 0,000291 | 0,228721288 | 2,237403 | Promoter (<=1kb) | 7 | 33883924 | 33896086 | 12163 | 2 | 68079 | ENSMUST00000002710.10 | 0 | ENSMUSG00000002635 | Pdcd2l | programmed cell death 2-like |
| chr1 | 1,67E+08 | 1,67E+08 | 367 | * | 3,817537 | 6,657526306 | 2,40447 | 2,768812 | 0,005626 | 1 | 0 | Intron (ENSMUST00000 | 1 | 1,67E+08 | 1,67E+08 | 89421 | 1 | 226610 | ENSMUST00000128373.2 | 45376 | ENSMUSG00000060568 | Fam78b | family with sequence similarity 78, member B |
| chr4 | 99893780 | 99894438 | 659 | * | 6,298326 | 6,657734948 | 1,938198 | 3,435013 | 0,000593 | 1 | 0 | Distal Intergenic | 4 | 99952988 | 1E+08 | 348975 | 1 | 26563 | ENSMUST00000039630.6 | -58550 | ENSMUSG00000035305 | Ror1 | receptor tyrosine kinase-like orphan receptor 1 |
| chrX | 1,02E+08 | 1,02E+08 | 762 | * | 4,266762 | 6,658236629 | 2,344281 | 2,840204 | 0,004508 | 1 | 0 | Intron (ENSMUST00000 | 20 | 1,01E+08 | 1,02E+08 | 220721 | 2 | 70315 | ENSMUST00000087916.11 | -12500 | ENSMUSG00000067567 | Hdac8 | histone deacetylase 8 |
| chr2 | 90703386 | 90703904 | 519 | * | 5,408127 | 6,660980838 | 2,166969 | 3,073871 | 0,002113 | 1 | 0 | Distal Intergenic | 2 | 90693803 | 90695732 | 1930 | 1 | 56428 | ENSMUST00000142350.2 | 9583 | ENSMUSG00000027282 | Mtch2 | mitochondrial carrier 2 |
| chr11 | 47374261 | 47374738 | 478 | * | 4,742793 | 6,661621014 | 2,135355 | 3,119678 | 0,00181 | 1 | 0 | Intron (ENSMUST00000 | 11 | 46787080 | 47469792 | 682713 | 2 | 24052 | ENSMUST00000109220.8 | 95054 | ENSMUSG00000020354 | Sgcd | sarcoglycan, delta (dystrophin-associated glycoprotein) |
| chr6 | 71581537 | 71582526 | 990 | * | 7,99684 | 6,661821936 | 1,818232 | 3,663902 | 0,000248 | 1 | 0 | Promoter (<=1kb) | 6 | 71580116 | 71580964 | 849 | 2 | 104263 | ENSMUST00000206050.2 | -573 | ENSMUSG00000053470 | Kdm3a | lysine (K)-specific demethylase 3A |
| chr19 | 3740240 | 3740862 | 623 | * | 5,220122 | 6,66217945 | 2,029586 | 3,282531 | 0,001029 | 1 | 0 | Intron (ENSMUST00000 | 19 | 3671671 | 3736564 | 64894 | 2 | 16973 | ENSMUST00000176867.2 | -3676 | ENSMUSG00000024913 | Lrp5 | low density lipoprotein receptor-related protein 5 |
| chr2 | 55843830 | 55844211 | 382 | * | 3,895539 | 6,662347764 | 2,231626 | 2,985424 | 0,002832 | 1 | 0 | Distal Intergenic | 2 | 55327177 | 55488152 | 160976 | 1 | 16519 | ENSMUST00000112632.2 | 516653 | ENSMUSG00000026824 | Kcnj3 | potassium inwardly-rectifying channel, subfamily J, member 3 |
| chr9 | 1,01E+08 | 1,01E+08 | 454 | * | 4,916623 | 6,663005368 | 2,069396 | 3,219783 | 0,001283 | 1 | 0 | Promoter (<=1kb) | 9 | 1,01E+08 | 1,01E+08 | 5645 | 2 | 235542 | ENSMUST00000191226.2 | -4 | ENSMUSG00000043154 | Ppp2r3a | protein phosphatase 2, regulatory subunit B'', alpha |
| chr2 | 73740348 | 73741013 | 666 | * | 5,209259 | 6,663245414 | 2,024968 | 3,290544 | 0,001 | 1 | 0 | Promoter (<=1kb) | 2 | 73739527 | 73740803 | 1277 | 2 | 228033 | ENSMUST00000131045.2 | 0 | ENSMUSG00000018770 | Atp5g3 | ATP synthase, H+ transporting, mitochondrial F0 complex, subunit C3 (subunit 9) |
| chr10 | 59536072 | 59536876 | 805 | * | 5,278172 | 6,663375693 | 2,003856 | 3,325277 | 0,000883 | 1 | 0 | Promoter (1-2kb) | 10 | 59538299 | 59699954 | 161656 | 1 | 216001 | ENSMUST00000165563.8 | -1423 | ENSMUSG00000020111 | Micu1 | mitochondrial calcium uptake 1 |
| chr5 | 1,35E+08 | 1,35E+08 | 740 | * | 5,855448 | 6,663937261 | 1,926598 | 3,458914 | 0,000542 | 1 | 0 | Promoter (<=1kb) | 5 | 1,35E+08 | 1,35E+08 | 1145 | 2 | 69812 | ENSMUST00000136022.9 | -921 | ENSMUSG00000085042 | Abhd11os | abhydrolase domain containing 11, opposite strand |
| chr10 | 61559678 | 61560953 | 1276 | * | 7,6007 | 6,664393374 | 1,876636 | 3,551245 | 0,000383 | 1 | 0 | Promoter (2-3kb) | 10 | 61563927 | 61570349 | 6423 | 1 | 71361 | ENSMUST00000140664.2 | -2974 | ENSMUSG00000020085 | Aifm2 | apoptosis-inducing factor, mitochondrion-associated 2 |
| chr13 | 97036186 | 97037538 | 1353 | * | 7,095295 | 6,664532742 | 1,863875 | 3,575633 | 0,000349 | 1 | 0 | Intron (ENSMUST00000 | 13 | 97061197 | 97087414 | 26218 | 1 | 218476 | ENSMUST00000171324.3 | -23659 | ENSMUSG00000091387 | Gcnt4 | glucosaminyl (N-acetyl) transferase 4, core 2 (beta-1,6-N-acetylglucosaminyltransferase) |
| chr2 | 70079841 | 70080419 | 579 | * | 6,878938 | 6,664791536 | 1,841377 | 3,619461 | 0,000295 | 1 | 0 | Intron (ENSMUST00000 | 2 | 69926642 | 70013611 | 86970 | 1 | 329421 | ENSMUST00000112241.2 | 153199 | ENSMUSG00000042064 | Myo3b | myosin IIIB |
| chr12 | 26018493 | 26019795 | 1303 | * | 10,28888 | 6,66640953 | 1,729248 | 3,855092 | 0,000116 | 0,17530799 | 1,425929 | Distal Intergenic | 12 | 26141961 | 26260504 | 118544 | 2 | 629481 | ENSMUST00000223022.2 | 240709 | ENSMUSG00000113550 | Gm29687 | predicted gene, 29687 |
| chr18 | 27575969 | 27576471 | 503 | * | 3,325247 | 6,667670665 | 2,479917 | 2,688667 | 0,007174 | 1 | 0 | Distal Intergenic | 18 | 27942785 | 27960738 | 17954 | 1 | 1,03E+08 | ENSMUST00000234286.2 | -366314 | | Gm33674 | predicted gene, 33674 |
| chr4 | 38823358 | 38824379 | 1022 | * | 6,800068 | 6,668343226 | 1,985837 | 3,357952 | 0,000785 | 1 | 0 | Intron (ENSMUST00000 | 4 | 38645427 | 38855463 | 210037 | 2 | 1,03E+08 | ENSMUST00000244574.1 | 31084 | ENSMUSG00000085632 | Gm12381 | predicted gene 12381 |
| chr2 | 99333538 | 99334495 | 958 | * | 4,742622 | 6,668390031 | 2,354168 | 2,832588 | 0,004617 | 1 | 0 | Distal Intergenic | 2 | 97298434 | 97459408 | 160975 | 1 | 241568 | ENSMUST00000170144.2 | 2035104 | ENSMUSG00000050587 | Lrrc4c | leucine rich repeat containing 4C |

| | | | | | | | | | | | | | | | | | | | | | | | |
|---|---|---|---|---|---|---|---|---|---|---|---|---|---|---|---|---|---|---|---|---|---|---|---|
| chr13 | 35598588 | 35599446 | 859 | * | 6,847417 | 6,668489035 | 1,823186 | 3,657601 | 0,000255 | 1 | 0 | Intron (ENSMUST0000( | 13 | 35477130 | 35477260 | 131 | 2 | 1,15E+08 | ENSMUST00000104413.3 | -121328 | ENSMUSG00000077601 | Gm22674 | predicted gene, 22674 |
| chr1 | 68861831 | 68862929 | 1099 | * | 7,070696 | 6,668725785 | 1,838221 | 3,627815 | 0,000286 | 1 | 0 | Intron (ENSMUST0000( | 1 | 68071345 | 69146915 | 1075571 | 2 | 13869 | ENSMUST00000119142.8 | 283986 | ENSMUSG00000062209 | Erbb4 | erb-b2 receptor tyrosine kinase 4 |
| chr10 | 12567886 | 12568315 | 430 | * | 4,047427 | 6,670056866 | 2,160292 | 3,087572 | 0,002018 | 1 | 0 | Intron (ENSMUST0000( | 10 | 12542167 | 12564606 | 22440 | 2 | 22288 | ENSMUST00000219724.2 | -3280 | ENSMUSG00000019820 | Utrn | utrophin |
| chr1 | 1,84E+08 | 1,84E+08 | 742 | * | 4,813374 | 6,670306713 | 2,137017 | 3,121316 | 0,0018 | 1 | 0 | Distal Intergenic | 1 | 1,84E+08 | 1,84E+08 | 23973 | 1 | 63953 | ENSMUST00000139839.2 | -102909 | ENSMUSG00000039384 | Dusp10 | dual specificity phosphatase 10 |
| chr16 | 46321711 | 46322240 | 530 | * | 5,16461 | 6,670394188 | 1,980531 | 3,367983 | 0,000757 | 1 | 0 | Promoter (2-3kb) | 16 | 46279215 | 46318888 | 39674 | 2 | 58998 | ENSMUST00000133935.2 | -2823 | ENSMUSG00000022656 | Nectin3 | nectin cell adhesion molecule 3 |
| chr12 | 84143659 | 84144265 | 607 | * | 5,771238 | 6,670558239 | 2,113965 | 3,155473 | 0,001602 | 1 | 0 | Distal Intergenic | 12 | 84147428 | 84158123 | 10696 | 1 | 217700 | ENSMUST00000056822.4 | -3163 | ENSMUSG00000043487 | Acot6 | acyl-CoA thioesterase 6 |
| chr11 | 10868828 | 10869255 | 428 | * | 5,707834 | 6,670804236 | 2,031179 | 3,284203 | 0,001023 | 1 | 0 | Distal Intergenic | 11 | 11064223 | 11213526 | 149304 | 1 | 319922 | ENSMUST00000109681.8 | -194968 | ENSMUSG00000050830 | Vwc2 | von Willebrand factor C domain containing 2 |
| chr9 | 13332224 | 13332743 | 520 | * | 4,512084 | 6,671206077 | 2,066242 | 3,228667 | 0,001244 | 1 | 0 | Intron (ENSMUST0000( | 9 | 13342657 | 13344032 | 1376 | 1 | 18689 | ENSMUST00000071254.7 | -9914 | ENSMUSG00000031802 | Phxr4 | per-hexamer repeat gene 4 |
| chr19 | 13126484 | 13126957 | 474 | * | 4,759871 | 6,672045244 | 2,011195 | 3,317453 | 0,000908 | 1 | 0 | Promoter (2-3kb) | 19 | 13123006 | 13124107 | 1102 | 2 | 258684 | ENSMUST00000078299.5 | -2377 | ENSMUSG00000057503 | Olfr1459 | olfactory receptor 1459 |
| chr8 | 1,12E+08 | 1,12E+08 | 336 | * | 3,255891 | 6,673377644 | 2,505471 | 2,663522 | 0,007733 | 1 | 0 | Intron (ENSMUST0000( | 8 | 1,12E+08 | 1,12E+08 | 71780 | 2 | 319481 | ENSMUST00000211981.2 | 4310 | ENSMUSG00000031959 | Wdr59 | WD repeat domain 59 |
| chr11 | 1,2E+08 | 1,2E+08 | 231 | * | 3,426061 | 6,673470857 | 2,493811 | 2,676013 | 0,00745 | 1 | 0 | Distal Intergenic | 11 | 1,2E+08 | 1,2E+08 | 2375 | 1 | 268515 | ENSMUST00000143667.2 | 24405 | ENSMUSG00000039741 | Bahcc1 | BAH domain and coiled-coil containing 1 |
| chr13 | 1,03E+08 | 1,03E+08 | 1175 | * | 7,630059 | 6,676106164 | 1,839344 | 3,629612 | 0,000284 | 0,290961988 | 1,360622 | Promoter (1-2kb) | 13 | 1,03E+08 | 1,03E+08 | 204219 | 2 | 328329 | ENSMUST00000172264.8 | 1477 | ENSMUSG00000034751 | Mast4 | microtubule associated serine/threonine kinase family member 4 |
| chr19 | 26135262 | 26136138 | 877 | * | 4,766671 | 6,676956404 | 2,188708 | 3,050639 | 0,002284 | 1 | 0 | Distal Intergenic | 19 | 26015816 | 26023831 | 8016 | 2 | 73433 | ENSMUST00000237184.2 | -111431 | ENSMUSG00000118016 | 1700048O14Rik | RIKEN cDNA 1700048O14 gene |
| chr14 | 1,19E+08 | 1,19E+08 | 579 | * | 4,504287 | 6,677513186 | 2,155287 | 3,098202 | 0,001947 | 1 | 0 | Exon (ENSMUST00000( | 14 | 1,19E+08 | 1,19E+08 | 767 | 2 | 66435 | ENSMUST00000227218.2 | 5623 | ENSMUSG00000042104 | Uggt2 | UDP-glucose glycoprotein glucosyltransferase 2 |
| chr11 | 8615580 | 8616174 | 595 | * | 4,737313 | 6,678461634 | 2,03287 | 3,285238 | 0,001019 | 1 | 0 | Promoter (<=1kb) | 11 | 8381652 | 8614681 | 233030 | 2 | 319939 | ENSMUST00000239111.2 | -899 | ENSMUSG00000020422 | Tns3 | tensin 3 |
| chr14 | 70065176 | 70065503 | 328 | * | 3,185813 | 6,678608549 | 2,441018 | 2,735993 | 0,006219 | 1 | 0 | Distal Intergenic | 14 | 70077869 | 70297367 | 219499 | 1 | 73523 | ENSMUST00000022678.5 | -12366 | ENSMUSG00000022085 | Pebp4 | phosphatidylethanolamine binding protein 4 |
| chrX | 69804528 | 69804927 | 400 | * | 3,800113 | 6,678746336 | 2,310433 | 2,890691 | 0,003844 | 1 | 0 | Distal Intergenic | 20 | 69737082 | 69737213 | 132 | 1 | 1,15E+08 | ENSMUST00000104087.3 | 67446 | ENSMUSG00000077275 | Gm22785 | predicted gene, 22785 |
| chr10 | 39959313 | 39959754 | 442 | * | 4,934587 | 6,678761361 | 2,090231 | 3,195226 | 0,001397 | 1 | 0 | Intron (ENSMUST0000( | 10 | 39913295 | 39952601 | 39307 | 2 | 72472 | ENSMUST00000213827.2 | -6712 | ENSMUSG00000019838 | Slc16a10 | solute carrier family 16 (monocarboxylic acid transporters), member 10 |
| chr11 | 89718517 | 89718901 | 385 | * | 3,687867 | 6,678762769 | 2,298527 | 2,905671 | 0,003665 | 1 | 0 | Intron (ENSMUST0000( | 11 | 89282064 | 89732092 | 450029 | 2 | 382543 | ENSMUST00000238273.3 | 13191 | ENSMUSG00000047773 | Ankfn1 | ankyrin-repeat and fibronectin type III domain containing 1 |
| chr14 | 94938625 | 94939447 | 823 | * | 3,735377 | 6,67888199 | 2,218786 | 3,010152 | 0,002611 | 1 | 0 | Distal Intergenic | 14 | 94953131 | 94972901 | 19771 | 2 | 1,03E+08 | ENSMUST00000227716.2 | 33454 | ENSMUSG00000115822 | Gm33450 | predicted gene, 33450 |
| chrX | 35273974 | 35274461 | 488 | * | 4,561785 | 6,679005973 | 2,158252 | 3,094637 | 0,001971 | 1 | 0 | Promoter (<=1kb) | 20 | 35275080 | 35280492 | 5413 | 1 | 75974 | ENSMUST00000140175.2 | -619 | ENSMUSG00000031093 | Dock11 | dedicator of cytokinesis 11 |
| chr11 | 1,01E+08 | 1,01E+08 | 914 | * | 5,143758 | 6,679496894 | 1,961783 | 3,40481 | 0,000662 | 1 | 0 | Promoter (2-3kb) | 11 | 1,01E+08 | 1,01E+08 | 10263 | 1 | 68107 | ENSMUST00000153438.2 | 2009 | ENSMUSG00000078653 | Cntd1 | cyclin N-terminal domain containing 1 |
| chr16 | 73299838 | 73300396 | 559 | * | 5,944771 | 6,679635011 | 1,923835 | 3,472041 | 0,000517 | 1 | 0 | Distal Intergenic | 16 | 73219681 | 73219847 | 167 | 2 | 1,15E+08 | ENSMUST00000082896.3 | -79991 | ENSMUSG00000064830 | Gm25699 | predicted gene, 25699 |
| chr13 | 51359245 | 51360069 | 825 | * | 7,45858 | 6,679738537 | 1,897364 | 3,520536 | 0,000431 | 1 | 0 | Promoter (2-3kb) | 13 | 51356763 | 51357101 | 339 | 2 | 667728 | ENSMUST00000091708.6 | -2144 | | H2af-ps2 | H2A histone family, pseudogene 2 |
| chr11 | 74827502 | 74828432 | 931 | * | 8,037426 | 6,680022574 | 1,842237 | 3,626038 | 0,000288 | 1 | 0 | Intron (ENSMUST0000( | 11 | 74821346 | 75055274 | 233929 | 1 | 103677 | ENSMUST00000130145.8 | 6156 | ENSMUSG00000038290 | Smg6 | SMG6 nonsense mediated mRNA decay factor |
| chr1 | 71452165 | 71452924 | 760 | * | 8,063686 | 6,68427347 | 1,850671 | 3,611811 | 0,000304 | 1 | 0 | Promoter (1-2kb) | 1 | 71400902 | 71454055 | 53154 | 2 | 74591 | ENSMUST00000189102.2 | 1131 | ENSMUSG00000050296 | Abca12 | ATP-binding cassette, sub-family A (ABC1), member 12 |
| chr19 | 4374286 | 4375215 | 930 | * | 7,490742 | 6,684442727 | 1,859406 | 3,594934 | 0,000324 | 1 | 0 | Promoter (<=1kb) | 19 | 4367734 | 4374893 | 7160 | 2 | 225876 | ENSMUST00000235335.2 | 0 | ENSMUSG00000054611 | Kdm2a | lysine (K)-specific demethylase 2A |
| chr1 | 75455107 | 75455502 | 396 | * | 3,147196 | 6,684444883 | 2,476534 | 2,699113 | 0,006952 | 1 | 0 | Promoter (<=1kb) | 1 | 75451220 | 75455390 | 4171 | 2 | 74241 | ENSMUST00000143496.8 | 0 | ENSMUSG00000032997 | Chpf | chondroitin polymerizing factor |
| chr14 | 1,14E+08 | 1,14E+08 | 472 | * | 3,548691 | 6,6851855 | 2,322094 | 2,878948 | 0,00399 | 1 | 0 | Distal Intergenic | 14 | 1,15E+08 | 1,15E+08 | 29363 | 1 | 75822 | ENSMUST00000226121.2 | -834532 | ENSMUSG00000115320 | 4930524C18Rik | RIKEN cDNA 4930524C18 gene |
| chr5 | 1,08E+08 | 1,08E+08 | 610 | * | 4,141197 | 6,686092912 | 2,18664 | 3,057701 | 0,00223 | 1 | 0 | Promoter (1-2kb) | 5 | 1,08E+08 | 1,08E+08 | 12110 | 1 | 73254 | ENSMUST00000195973.2 | -1913 | ENSMUSG00000056531 | Ccdc18 | coiled-coil domain containing 18 |
| chr4 | 86605763 | 86606723 | 961 | * | 6,026079 | 6,686259582 | 1,977027 | 3,381977 | 0,00072 | 1 | 0 | Distal Intergenic | 4 | 86574802 | 86588297 | 13496 | 2 | 11520 | ENSMUST00000000466.13 | -17466 | ENSMUSG00000028494 | Plin2 | perilipin 2 |
| chr11 | 22104043 | 22104642 | 600 | * | 4,175974 | 6,686277843 | 2,166386 | 3,086375 | 0,002026 | 1 | 0 | Intron (ENSMUST0000( | 11 | 22119037 | 22121410 | 2374 | 2 | 216565 | ENSMUST00000136762.2 | 16768 | ENSMUSG00000042302 | Ehbp1 | EH domain binding protein 1 |
| chr11 | 1,09E+08 | 1,09E+08 | 698 | * | 4,773849 | 6,686456562 | 2,144957 | 3,117291 | 0,001825 | 1 | 0 | Promoter (<=1kb) | 11 | 1,09E+08 | 1,09E+08 | 177737 | 1 | 76380 | ENSMUST00000106724.10 | 817 | ENSMUSG00000020728 | Cep112 | centrosomal protein 112 |
| chr7 | 44517665 | 44518295 | 631 | * | 6,094251 | 6,687931523 | 2,008843 | 3,329245 | 0,000871 | 1 | 0 | Promoter (<=1kb) | 7 | 44514392 | 44518481 | 4090 | 2 | 84113 | ENSMUST00000209139.2 | 186 | ENSMUSG00000038502 | Ptov1 | prostate tumor over expressed gene 1 |
| chr7 | 34431500 | 34432104 | 605 | * | 3,833775 | 6,688517984 | 2,259184 | 2,96059 | 0,003071 | 1 | 0 | Intron (ENSMUST0000( | 7 | 34373893 | 34453237 | 79345 | 2 | 68947 | ENSMUST00000238947.2 | 21133 | ENSMUSG00000060402 | Chst8 | carbohydrate sulfotransferase 8 |
| chr5 | 1,07E+08 | 1,07E+08 | 648 | * | 6,01258 | 6,688718768 | 2,124777 | 3,147963 | 0,001644 | 1 | 0 | Intron (ENSMUST0000( | 5 | 1,07E+08 | 1,07E+08 | 4086 | 1 | 12545 | ENSMUST00000123546.2 | 6881 | ENSMUSG00000029283 | Cdc7 | cell division cycle 7 (S. cerevisiae) |
| chr6 | 29532657 | 29533004 | 348 | * | 3,468093 | 6,68932017 | 2,449101 | 2,731338 | 0,006308 | 1 | 0 | Promoter (1-2kb) | 6 | 29531247 | 29535345 | 4099 | 1 | 27056 | ENSMUST00000164922.3 | 1410 | ENSMUSG00000029771 | Irf5 | interferon regulatory factor 5 |
| chr1 | 11090124 | 11090432 | 309 | * | 2,902715 | 6,6896539 | 2,567969 | 2,605037 | 0,009186 | 1 | 0 | Intron (ENSMUST0000( | 1 | 11063689 | 11373905 | 310217 | 1 | 109294 | ENSMUST00000027056.12 | 26435 | ENSMUSG00000048960 | Prex2 | phosphatidylinositol-3,4,5-trisphosphate-dependent Rac exchange factor 2 |
| chr1 | 1,82E+08 | 1,82E+08 | 708 | * | 7,032178 | 6,690382209 | 1,869951 | 3,577839 | 0,000346 | 0,313797801 | 1,360622 | Promoter (<=1kb) | 1 | 1,82E+08 | 1,82E+08 | 33704 | 2 | 12334 | ENSMUST00000192483.2 | -752 | ENSMUSG00000026509 | Capn2 | calpain 2 |
| chr18 | 16878750 | 16879297 | 548 | * | 3,438377 | 6,690514579 | 2,443304 | 2,738307 | 0,006176 | 1 | 0 | Intron (ENSMUST0000( | 18 | 16904454 | 16941519 | 37066 | 2 | 12558 | ENSMUST00000152779.2 | 62222 | ENSMUSG00000024304 | Cdh2 | cadherin 2 |
| chrX | 17911666 | 17912050 | 385 | * | 3,228104 | 6,690697516 | 2,419488 | 2,765337 | 0,005686 | 1 | 0 | Distal Intergenic | 20 | 18012096 | 18012940 | 845 | 1 | 73547 | ENSMUST00000026018.4 | -100046 | ENSMUSG00000025043 | Dusp21 | dual specificity phosphatase 21 |
| chr7 | 77510916 | 77511438 | 523 | * | 3,546841 | 6,691069263 | 2,37774 | 2,814046 | 0,004892 | 1 | 0 | Distal Intergenic | 7 | 77830631 | 77842491 | 11861 | 2 | 18213 | ENSMUST00000205354.2 | 331053 | ENSMUSG00000059146 | Ntrk3 | neurotrophic tyrosine kinase, receptor, type 3 |
| chr9 | 70416614 | 70417355 | 742 | * | 7,059189 | 6,691379905 | 1,830369 | 3,655754 | 0,000256 | 1 | 0 | Distal Intergenic | 9 | 70332711 | 70411007 | 78297 | 2 | 93836 | ENSMUST00000034739.12 | -5607 | ENSMUSG00000032217 | Rnf111 | ring finger 111 |
| chr6 | 14827779 | 14828128 | 350 | * | 3,94764 | 6,692245982 | 2,260391 | 2,960659 | 0,00307 | 1 | 0 | Distal Intergenic | 6 | 14713976 | 14755273 | 41298 | 2 | 140491 | ENSMUST00000045096.6 | -72506 | ENSMUSG00000042717 | Ppp1r3a | protein phosphatase 1, regulatory subunit 3A |
| chr1 | 1,82E+08 | 1,82E+08 | 572 | * | 4,663857 | 6,693134463 | 2,189543 | 3,056864 | 0,002237 | 1 | 0 | Promoter (2-3kb) | 1 | 1,82E+08 | 1,82E+08 | 4308 | 2 | 13244 | ENSMUST00000132551.2 | -2283 | ENSMUSG00000038633 | Degs1 | delta(4)-desaturase, sphingolipid 1 |
| chr1 | 1,45E+08 | 1,45E+08 | 450 | * | 4,203672 | 6,693364092 | 2,174135 | 3,078633 | 0,00208 | 1 | 0 | Distal Intergenic | 1 | 1,45E+08 | 1,45E+08 | 127 | 2 | 1,15E+08 | ENSMUST00000157954.3 | -149513 | ENSMUSG00000088579 | Gm25460 | predicted gene, 25460 |
| chr14 | 72051718 | 72052577 | 860 | * | 4,505778 | 6,693662432 | 2,153396 | 3,108421 | 0,001881 | 1 | 0 | Distal Intergenic | 14 | 72094142 | 72157879 | 63738 | 1 | 74667 | ENSMUST00000228500.2 | -41565 | ENSMUSG00000115304 | 4930434J06Rik | RIKEN cDNA 4930434J06 gene |
| chr1 | 30906279 | 30906897 | 619 | * | 3,806202 | 6,693736677 | 2,417199 | 2,769212 | 0,005619 | 1 | 0 | Exon (ENSMUST00000: | 1 | 30841423 | 30902337 | 60915 | 2 | 213109 | ENSMUST00000088310.10 | -3942 | ENSMUSG00000048874 | Phf3 | PHD finger protein 3 |
| chr11 | 61454867 | 61455533 | 667 | * | 5,258212 | 6,694963151 | 2,077977 | 3,221865 | 0,001274 | 1 | 0 | Intron (ENSMUST0000( | 11 | 61437332 | 61470484 | 33153 | 2 | 13855 | ENSMUST00000147501.8 | 14951 | ENSMUSG00000001036 | Epn2 | epsin 2 |
| chr3 | 69144657 | 69145637 | 981 | * | 8,797248 | 6,69514031 | 1,813829 | 3,691164 | 0,000223 | 0,252371261 | 1,449706 | Distal Intergenic | 3 | 69129752 | 69130951 | 1200 | 1 | 71619 | ENSMUST00000183126.2 | 14905 | ENSMUSG00000098207 | Arl14 | ADP-ribosylation factor-like 14 |
| chr6 | 83817220 | 83817649 | 430 | * | 6,181684 | 6,695176949 | 1,930538 | 3,468036 | 0,000524 | 1 | 0 | Distal Intergenic | 6 | 83813919 | 83814048 | 130 | 2 | 1,15E+08 | ENSMUST00000104144.3 | -3172 | ENSMUSG00000077332 | Gm23014 | predicted gene, 23014 |
| chr7 | 86240415 | 86241078 | 664 | * | 5,712334 | 6,69518256 | 1,8999 | 3,523966 | 0,000425 | 1 | 0 | Distal Intergenic | 7 | 86234485 | 86235414 | 930 | 1 | 258850 | ENSMUST00000078447.2 | 5930 | ENSMUSG00000059319 | Olfr295 | olfactory receptor 295 |
| chr9 | 62088784 | 62089223 | 440 | * | 4,580275 | 6,695205058 | 2,12601 | 3,149188 | 0,001637 | 1 | 0 | Distal Intergenic | 9 | 61977750 | 62029937 | 52188 | 2 | 93683 | ENSMUST00000185873.2 | -58847 | ENSMUSG00000032252 | Glce | glucuronyl C5-epimerase |
| chrX | 48770645 | 48771592 | 948 | * | 5,971389 | 6,695207175 | 2,028813 | 3,300061 | 0,000967 | 1 | 0 | Promoter (<=1kb) | 20 | 48772381 | 48773340 | 960 | 1 | 236784 | ENSMUST00000096449.2 | -789 | ENSMUSG00000071764 | Olfr1320 | olfactory receptor 1320 |
| chr5 | 38670783 | 38671194 | 412 | * | 4,441356 | 6,695207933 | 2,06064 | 3,249092 | 0,001158 | 1 | 0 | Distal Intergenic | 5 | 38506982 | 38660486 | 153505 | 2 | 117591 | ENSMUST00000155634.8 | -10297 | ENSMUSG00000005107 | Slc2a9 | solute carrier family 2 (facilitated glucose transporter), member 9 |
| chr5 | 1,35E+08 | 1,35E+08 | 595 | * | 3,96648 | 6,695216416 | 2,363015 | 2,833337 | 0,004606 | 1 | 0 | Intron (ENSMUST0000( | 5 | 1,35E+08 | 1,35E+08 | 12510 | 2 | 94244 | ENSMUST00000201784.4 | 16812 | ENSMUSG00000040013 | Fkbp6 | FK506 binding protein 6 |
| chr14 | 24111152 | 24112291 | 1140 | * | 10,77299 | 6,695701888 | 1,716122 | 3,901647 | 9,55E-05 | 0,22808412 | 0,738441 | Intron (ENSMUST0000( | 14 | 24119802 | 24120705 | 904 | 1 | 620499 | ENSMUST00000109467.4 | -7511 | | Gm6158 | predicted gene 6158 |
| chr3 | 20989173 | 20989511 | 339 | * | 3,896776 | 6,69657091 | 2,427306 | 2,758849 | 0,005801 | 1 | 0 | Distal Intergenic | 3 | 20557262 | 20560372 | 3111 | 2 | 1,08E+08 | ENSMUST00000195457.2 | -428801 | | Gm31466 | predicted gene, 31466 |
| chr3 | 78153282 | 78153636 | 355 | * | 3,892717 | 6,696849343 | 2,403456 | 2,786341 | 0,005331 | 1 | 0 | Distal Intergenic | 3 | 78887229 | 78887358 | 130 | 1 | 1,15E+08 | ENSMUST00000240248.1 | -733593 | ENSMUSG00000119445 | Gm23440 | predicted gene, 23440 |
| chr14 | 1,19E+08 | 1,19E+08 | 676 | * | 3,874299 | 6,699057029 | 2,337872 | 2,865451 | 0,004164 | 1 | 0 | Intron (ENSMUST0000( | 14 | 1,19E+08 | 1,2E+08 | 731475 | 1 | 50787 | ENSMUST00000065904.5 | 45881 | ENSMUSG00000053465 | Hs6st3 | heparan sulfate 6-O-sulfotransferase 3 |
| chr18 | 56860738 | 56861658 | 921 | * | 6,717797 | 6,6998143 | 1,893498 | 3,538327 | 0,000403 | 1 | 0 | Promoter (<=1kb) | 18 | 56861513 | 56866528 | 5016 | 1 | 16906 | ENSMUST00000128651.2 | 0 | ENSMUSG00000024590 | Lmnb1 | lamin B1 |
| chr9 | 19483036 | 19483560 | 525 | * | 3,365125 | 6,700957577 | 2,490467 | 2,690643 | 0,007131 | 1 | 0 | Distal Intergenic | 9 | 19477731 | 19478678 | 948 | 2 | 258515 | ENSMUST00000072419.3 | -4358 | ENSMUSG00000095667 | Olfr854 | olfactory receptor 854 |
| chr8 | 4755408 | 4756015 | 608 | * | 5,896292 | 6,701074613 | 1,921089 | 3,488165 | 0,000486 | 1 | 0 | Distal Intergenic | 8 | 4728431 | 4738497 | 10067 | 1 | 623223 | ENSMUST00000181337.3 | 26977 | | Gm6410 | predicted gene 6410 |
| chr18 | 89850700 | 89851045 | 346 | * | 4,234628 | 6,70130906 | 2,390111 | 2,803764 | 0,005051 | 1 | 0 | Distal Intergenic | 18 | 89310548 | 89787652 | 477105 | 2 | 623279 | ENSMUST00000097495.5 | -63048 | ENSMUSG00000073514 | Dok6 | docking protein 6 |
| chrX | 1,28E+08 | 1,28E+08 | 718 | * | 3,518472 | 6,701461273 | 2,3502 | 2,851443 | 0,004352 | 1 | 0 | Distal Intergenic | 20 | 1,29E+08 | 1,29E+08 | 430730 | 1 | 54004 | ENSMUST00000127577.8 | -270074 | ENSMUSG00000034480 | Diaph2 | diaphanous related formin 2 |
| chr2 | 1,31E+08 | 1,31E+08 | 655 | * | 5,924488 | 6,701863517 | 2,110502 | 3,175483 | 0,001496 | 1 | 0 | 3' UTR | 2 | 1,31E+08 | 1,31E+08 | 13407 | 2 | 269356 | ENSMUST00000099362.11 | -10351 | ENSMUSG00000074796 | Slc4a11 | solute carrier family 4, sodium bicarbonate transporter-like, member 11 |
| chr2 | 14890916 | 14891217 | 302 | * | 3,858521 | 6,701876387 | 2,255058 | 2,971931 | 0,002959 | 1 | 0 | Intron (ENSMUST0000( | 2 | 14879563 | 14991907 | 112345 | 1 | 12296 | ENSMUST00000193452.2 | 11353 | ENSMUSG00000057914 | Cacnb2 | calcium channel, voltage-dependent, beta 2 subunit |
| chr9 | 1,2E+08 | 1,2E+08 | 625 | * | 4,434015 | 6,702851966 | 2,099131 | 3,193156 | 0,001407 | 1 | 0 | Promoter (1-2kb) | 9 | 1,2E+08 | 1,2E+08 | 4501 | 2 | 74498 | ENSMUST00000213409.2 | 1151 | ENSMUSG00000032513 | Gorasp1 | golgi reassembly stacking protein 1 |
| chr15 | 43247061 | 43247445 | 385 | * | 5,696691 | 6,704178967 | 1,962212 | 3,416643 | 0,000634 | 1 | 0 | Distal Intergenic | 15 | 43294339 | 43340432 | 46094 | 2 | 1E+08 | ENSMUST00000100683.3 | 92987 | ENSMUSG00000072592 | Gm10373 | predicted gene 10373 |
| chr8 | 65136187 | 65136649 | 463 | * | 5,278616 | 6,704265609 | 1,954786 | 3,429668 | 0,000604 | 1 | 0 | Intron (ENSMUST0000( | 8 | 65045576 | 65146088 | 100513 | 2 | 12876 | ENSMUST00000048967.9 | 9439 | ENSMUSG00000037852 | Cpe | carboxypeptidase E |
| chr17 | 56446133 | 56446719 | 587 | * | 6,723354 | 6,704509272 | 1,934666 | 3,465461 | 0,000529 | 1 | 0 | Promoter (<=1kb) | 17 | 56430085 | 56447343 | 17259 | 2 | 20359 | ENSMUST00000001256.11 | 624 | ENSMUSG00000001227 | Sema6b | sema domain, transmembrane domain (TM), and cytoplasmic domain, (semaphorin) 6B |
| chr14 | 47335728 | 47336335 | 608 | * | 6,635668 | 6,705544823 | 1,867556 | 3,590545 | 0,00033 | 1 | 0 | Exon (ENSMUST00000: | 14 | 47324592 | 47326558 | 1967 | 1 | 74480 | ENSMUST00000130046.2 | 11136 | ENSMUSG00000021838 | Samd4 | sterile alpha motif domain containing 4 |
| chr19 | 6884259 | 6885083 | 825 | * | 7,432656 | 6,706182377 | 1,834162 | 3,656265 | 0,000256 | 1 | 0 | Promoter (<=1kb) | 19 | 6884118 | 6885669 | 1552 | 2 | 54683 | ENSMUST00000149261.8 | 586 | ENSMUSG00000024953 | Prdx5 | peroxiredoxin 5 |
| chr10 | 39641110 | 39641381 | 272 | * | 3,540154 | 6,70624834 | 2,563169 | 2,61639 | 0,008887 | 1 | 0 | Intron (ENSMUST0000( | 10 | 39608360 | 39751202 | 142843 | 1 | 19714 | ENSMUST00000164763.8 | 32750 | ENSMUSG00000019841 | Rev3l | REV3 like, DNA directed polymerase zeta catalytic subunit |
| chr14 | 1,16E+08 | 1,16E+08 | 711 | * | 3,518711 | 6,70745573 | 2,375365 | 2,823758 | 0,004746 | 1 | 0 | Intron (ENSMUST0000( | 14 | 1,16E+08 | 1,16E+08 | 13002 | 2 | 75088 | ENSMUST00000176582.2 | -655900 | ENSMUSG00000093551 | 4930505G20Rik | RIKEN cDNA 4930505G20 gene |
| chrX | 5391008 | 5391539 | 532 | * | 3,7507 | 6,708297826 | 2,273818 | 2,950236 | 0,003175 | 1 | 0 | Distal Intergenic | 20 | 5378952 | 5381319 | 2368 | 2 | 17870 | ENSMUST00000058404.5 | -9689 | ENSMUSG00000044597 | Mycs | myc-like oncogene, s-myc protein |
| chr14 | 1,13E+08 | 1,13E+08 | 905 | * | 4,659537 | 6,708796483 | 2,224423 | 3,015972 | 0,002562 | 1 | 0 | Distal Intergenic | 14 | 1,12E+08 | 1,12E+08 | 3494 | 1 | 75409 | ENSMUST00000227891.2 | 603187 | ENSMUSG00000033214 | Slitrk5 | SLIT and NTRK-like family, member 5 |
| chr18 | 24848328 | 24848852 | 525 | * | 4,267241 | 6,708862829 | 2,220587 | 3,021211 | 0,002518 | 1 | 0 | Intron (ENSMUST0000( | 18 | 24842502 | 25266558 | 424057 | 1 | 225288 | ENSMUST00000234834.2 | 5826 | ENSMUSG00000034295 | Fhod3 | formin homology 2 domain containing 3 |
| chr12 | 99531660 | 99532064 | 405 | * | 4,275316 | 6,709139047 | 2,195622 | 3,05569 | 0,002245 | 1 | 0 | Promoter (1-2kb) | 12 | 99530100 | 99586297 | 56198 | 1 | 1,03E+08 | ENSMUST00000180686.2 | 1560 | ENSMUSG00000097917 | Gm26839 | predicted gene, 26839 |
| chr1 | 1,78E+08 | 1,78E+08 | 436 | * | 4,453161 | 6,709203404 | 2,188378 | 3,065834 | 0,002171 | 1 | 0 | Promoter (1-2kb) | 1 | 1,78E+08 | 1,79E+08 | 187376 | 1 | 269152 | ENSMUST00000193931.2 | -1288 | ENSMUSG00000026494 | Kif26b | kinesin family member 26B |
| chr3 | 96155186 | 96155779 | 594 | * | 4,750203 | 6,709618108 | 2,1565 | 3,111346 | 0,001862 | 1 | 0 | Promoter (1-2kb) | 3 | 96154001 | 96155825 | 1825 | 1 | 15077 | ENSMUST00000177796.2 | 1185 | ENSMUSG00000093769 | H3c14 | H3 clustered histone 14 |
| chr5 | 1,14E+08 | 1,14E+08 | 481 | * | 4,782958 | 6,709871778 | 2,135963 | 3,14138 | 0,001682 | 1 | 0 | Intron (ENSMUST0000( | 5 | 1,14E+08 | 1,14E+08 | 21206 | 1 | 1,03E+08 | ENSMUST00000160164.2 | 9931 | ENSMUSG00000029592 | Gm16108 | predicted gene 16108 |
| chr4 | 1,09E+08 | 1,09E+08 | 1540 | * | 11,30439 | 6,710080652 | 1,741199 | 3,853712 | 0,000116 | 0,198990239 | 1,173325 | Distal Intergenic | 4 | 1,09E+08 | 1,09E+08 | 107550 | 1 | 13858 | ENSMUST00000102729.10 | -14559 | ENSMUSG00000028552 | Eps15 | epidermal growth factor receptor pathway substrate 15 |
| chr1 | 1,14E+08 | 1,14E+08 | 511 | * | 4,604723 | 6,710367174 | 2,103197 | 3,190556 | 0,00142 | 1 | 0 | Distal Intergenic | 1 | 1,16E+08 | 1,16E+08 | 15632 | 2 | 1,08E+08 | ENSMUST00000186137.7 | 1534943 | | Gm50605 | predicted gene, 50605 |
| chr13 | 38287534 | 38288307 | 774 | * | 6,160085 | 6,710432662 | 1,889473 | 3,551484 | 0,000383 | 1 | 0 | Distal Intergenic | 13 | 38241191 | 38245289 | 4099 | 1 | 71340 | ENSMUST00000225174.2 | 46343 | ENSMUSG00000021428 | Riok1 | RIO kinase 1 |
| chr4 | 18757696 | 18758299 | 604 | * | 5,742672 | 6,710567914 | 1,917428 | 3,499776 | 0,000466 | 1 | 0 | Intron (ENSMUST0000( | 4 | 18767980 | 19122565 | 354586 | 2 | 435772 | ENSMUST00000137780.3 | 364266 | ENSMUSG00000073991 | Cnbd1 | cyclic nucleotide binding domain containing 1 |
| chr9 | 9221284 | 9221783 | 500 | * | 4,540964 | 6,711116646 | 2,074332 | 3,235315 | 0,001215 | 1 | 0 | Intron (ENSMUST0000( | 9 | 9236293 | 9260888 | 24596 | 1 | 1,01E+08 | ENSMUST00000189204.2 | -14510 | ENSMUSG00000085253 | Gm16833 | predicted gene, 16833 |
| chr12 | 37700412 | 37700768 | 357 | * | 4,193286 | 6,711161834 | 2,099128 | 3,197119 | 0,001388 | 1 | 0 | Distal Intergenic | 12 | 37867725 | 38032009 | 164285 | 1 | 217480 | ENSMUST00000222337.2 | -166957 | ENSMUSG00000036095 | Dgkb | diacylglycerol kinase, beta |
| chr13 | 13878015 | 13878324 | 310 | * | 3,44702 | 6,711583244 | 2,33043 | 2,879977 | 0,003977 | 1 | 0 | Intron (ENSMUST0000( | 13 | 13883936 | 13922114 | 38179 | 1 | 17101 | ENSMUST00000221717.2 | -5612 | ENSMUSG00000019726 | Lyst | lysosomal trafficking regulator |
| chr3 | 79356398 | 79356796 | 399 | * | 3,643171 | 6,711598533 | 2,371187 | 2,83048 | 0,004648 | 1 | 0 | Exon (ENSMUST00000: | 3 | 79364194 | 79373512 | 9319 | 2 | 329679 | ENSMUST00000149475.8 | 16716 | ENSMUSG00000061175 | Fnip2 | folliculin interacting protein 2 |
| chr14 | 31919160 | 31920040 | 881 | * | 5,89562 | 6,71166607 | 2,025455 | 3,313659 | 0,000921 | 1 | 0 | Intron (ENSMUST0000( | 14 | 31913244 | 31923785 | 10542 | 2 | 53600 | ENSMUST00000172052.8 | 3745 | ENSMUSG00000013701 | Timm23 | translocase of inner mitochondrial membrane 23 |
| chr5 | 57729207 | 57729684 | 478 | * | 3,808594 | 6,712865342 | 2,430489 | 2,76194 | 0,005746 | 1 | 0 | Intron (ENSMUST0000( | 5 | 57727424 | 57875262 | 147839 | 2 | 319216 | ENSMUST00000180708.3 | 145578 | ENSMUSG00000097216 | 4932441J04Rik | RIKEN cDNA 4932441J04 gene |
| chr7 | 1,13E+08 | 1,13E+08 | 951 | * | 7,01752 | 6,71313581 | 1,957416 | 3,429591 | 0,000604 | 1 | 0 | Intron (ENSMUST0000( | 7 | 1,13E+08 | 1,13E+08 | 41782 | 2 | 75239 | ENSMUST00000210408.2 | 3966 | ENSMUSG00000110009 | 4930543E12Rik | RIKEN cDNA 4930543E12 gene |
| chr7 | 67041769 | 67042227 | 459 | * | 3,648365 | 6,713399605 | 2,379571 | 2,821265 | 0,004783 | 1 | 0 | Distal Intergenic | 7 | 66901557 | 67022606 | 121050 | 2 | 17258 | ENSMUST00000133074.2 | -19163 | ENSMUSG00000030557 | Mef2a | myocyte enhancer factor 2A |
| chrX | 47296906 | 47297900 | 995 | * | 7,629617 | 6,713549451 | 1,940721 | 3,459307 | 0,000542 | 1 | 0 | Promoter (<=1kb) | 20 | 47260846 | 47297564 | 36719 | 2 | 208884 | ENSMUST00000037960.11 | 0 | ENSMUSG00000036985 | Zdhhc9 | zinc finger, DHHC domain containing 9 |
| chr6 | 47811742 | 47813342 | 1601 | * | 17,368922 | 6,714245386 | 1,5953476 | 4,208641 | 2,569E-05 | 0,047042565 | 2,6636191 | Promoter (<=1kb) | 6 | 47812595 | 47820581 | 7987 | 1 | 272347 | ENSMUST00000146202.2 | 0 | ENSMUSG00000062519 | Zfp398 | zinc finger protein 398 |
| chr1 | 36108056 | 36108754 | 699 | * | 7,410342 | 6,714374048 | 1,910867 | 3,513784 | 0,000442 | 1 | 0 | Promoter (<=1kb) | 1 | 36107481 | 36145527 | 38047 | 1 | 50785 | ENSMUST00000088174.4 | 575 | ENSMUSG00000045216 | Hs6st1 | heparan sulfate 6-O-sulfotransferase 1 |
| chr6 | 39423905 | 39424305 | 401 | * | 4,155936 | 6,714636491 | 2,281053 | 2,943656 | 0,003244 | 1 | 0 | Distal Intergenic | 6 | 39445138 | 39447258 | 2121 | 2 | 209773 | ENSMUST00000145541.2 | 22953 | ENSMUSG00000038456 | Dennd2a | DENN/MADD domain containing 2A |
| chr5 | 1,09E+08 | 1,09E+08 | 787 | * | 4,727493 | 6,714865079 | 2,264889 | 2,964766 | 0,003029 | 1 | 0 | Distal Intergenic | 5 | 1,09E+08 | 1,09E+08 | 11975 | 2 | 211223 | ENSMUST00000167133.3 | 26374 | ENSMUSG00000091375 | Vmn2r15 | vomeronasal 2, receptor 15 |
| chr17 | 30308129 | 30308958 | 830 | * | 6,40229 | 6,71539631 | 1,876392 | 3,578887 | 0,000345 | 1 | 0 | Intron (ENSMUST0000( | 17 | 30279356 | 30427127 | 147772 | 1 | 21769 | ENSMUST00000234790.2 | 28773 | ENSMUSG00000044477 | Zfand3 | zinc finger, AN1-type domain 3 |
| chr9 | 25233933 | 25234339 | 407 | * | 4,804063 | 6,716305127 | 2,177195 | 3,084843 | 0,002037 | 1 | 0 | Distal Intergenic | 9 | 25210632 | 25215363 | 4732 | 1 | 235072 | ENSMUST00000213435.2 | 23301 | ENSMUSG00000001833 | Septin7 | septin 7 |

| | | | | | | | | | | | | | | | | | | | | | | | |
|---|---|---|---|---|---|---|---|---|---|---|---|---|---|---|---|---|---|---|---|---|---|---|---|
| chr4 | 6299177 | 6300228 | 1052 | * | 6,307137 | 6,716432 | 1,918075 | 3,501653 | 0,000462 | 1 | 0 | Distal Intergenic | 4 | 6272805 | 6275633 | 2829 | 2 | 13122 | ENSMUST00000147346.2 | -23544 | ENSMUSG00000028240 | Cyp7a1 | cytochrome P450, family 7, subfamily a, polypeptide 1 |
| chr14 | 1,06E+08 | 1,06E+08 | 578 | * | 4,03295 | 6,718247735 | 2,19118 | 3,06604 | 0,002169 | 1 | 0 | Distal Intergenic | 14 | 1,06E+08 | 1,06E+08 | 14324 | 1 | 76273 | ENSMUST00000149358.2 | 30665 | ENSMUSG00000053253 | Ndfip2 | Nedd4 family interacting protein 2 |
| chr9 | 69914636 | 69915327 | 692 | * | 6,517359 | 6,719583268 | 1,965094 | 3,419472 | 0,000627 | 1 | 0 | 3' UTR | 9 | 69910661 | 69912045 | 1385 | 1 | 12175 | ENSMUST00000137472.2 | 3975 | ENSMUSG00000011958 | Bnip2 | BCL2/adenovirus E1B interacting protein 2 |
| chr19 | 59275763 | 59276535 | 773 | * | 6,430819 | 6,719802917 | 1,940366 | 3,463163 | 0,000534 | 1 | 0 | Exon (ENSMUST00000( | 19 | 59249391 | 59283675 | 34285 | 1 | 214084 | ENSMUST00000236270.2 | 26372 | ENSMUSG00000025094 | Slc18a2 | solute carrier family 18 (vesicular monoamine), member 2 |
| chr19 | 51361123 | 51361733 | 611 | * | 4,263776 | 6,721424704 | 2,308617 | 2,91145 | 0,003598 | 1 | 0 | Distal Intergenic | 19 | 51228128 | 51228234 | 107 | 2 | 1,15E+08 | ENSMUST00000083209.3 | -132889 | ENSMUSG00000065143 | Gm25644 | predicted gene, 25644 |
| chr18 | 10612722 | 10614135 | 1414 | * | 10,74573 | 6,722162397 | 1,764101 | 3,810532 | 0,000139 | 0,276598083 | 0,744375 | Promoter (2-3kb) | 18 | 10566602 | 10610352 | 43751 | 2 | 77805 | ENSMUST00000127099.9 | -2370 | ENSMUSG00000024293 | Esco1 | establishment of sister chromatid cohesion N-acetyltransferase 1 |
| chr17 | 19725251 | 19725633 | 383 | * | 3,18534 | 6,722581543 | 2,57632 | 2,609374 | 0,009071 | 1 | 0 | Promoter (<=1kb) | 17 | 19725062 | 19755493 | 30432 | 1 | 627537 | ENSMUST00000231465.2 | 189 | ENSMUSG00000091859 | Vmn2r100 | vomeronasal 2, receptor 100 |
| chr3 | 97240461 | 97240721 | 261 | * | 2,969512 | 6,722710668 | 2,546626 | 2,63985 | 0,008294 | 1 | 0 | Distal Intergenic | 3 | 97123869 | 97205233 | 81365 | 2 | 77578 | ENSMUST00000196586.2 | -35228 | ENSMUSG00000038256 | Bcl9 | B cell CLL/lymphoma 9 |
| chr9 | 7150222 | 7150530 | 309 | * | 3,055849 | 6,722847296 | 2,517686 | 2,670249 | 0,00758 | 1 | 0 | Intron (ENSMUST0000( | 9 | 6929304 | 7176957 | 247654 | 2 | 110350 | ENSMUST00000147193.8 | 26427 | ENSMUSG00000047193 | Dync2h1 | dynein cytoplasmic 2 heavy chain 1 |
| chr16 | 18630140 | 18631751 | 1612 | * | 11,38815 | 6,723407993 | 1,712211 | 3,92674 | 8,61E-05 | 0,166885401 | 1,173325 | Promoter (<=1kb) | 16 | 18599197 | 18630722 | 31526 | 2 | 12544 | ENSMUST00000000028.14 | 0 | ENSMUSG00000000028 | Cdc45 | cell division cycle 45 |
| chr2 | 1,27E+08 | 1,27E+08 | 524 | * | 3,637607 | 6,723797523 | 2,35015 | 2,861008 | 0,004223 | 1 | 0 | Promoter (<=1kb) | 2 | 1,27E+08 | 1,27E+08 | 11039 | 1 | 108011 | ENSMUST00000142740.2 | -454 | ENSMUSG00000001998 | Ap4e1 | adaptor-related protein complex AP-4, epsilon 1 |
| chr4 | 1,36E+08 | 1,36E+08 | 378 | * | 3,991811 | 6,724533752 | 2,251467 | 2,986734 | 0,00282 | 1 | 0 | Distal Intergenic | 4 | 1,36E+08 | 1,36E+08 | 20875 | 1 | 15552 | ENSMUST00000088677.5 | -18142 | ENSMUSG00000070687 | Htr1d | 5-hydroxytryptamine (serotonin) receptor 1D |
| chr16 | 4286458 | 4287524 | 1067 | * | 6,574444 | 6,724970634 | 1,832942 | 3,668949 | 0,000244 | 1 | 0 | Distal Intergenic | 16 | 4105393 | 4238362 | 132970 | 2 | 11515 | ENSMUST00000120080.8 | -48096 | ENSMUSG00000005580 | Adcy9 | adenylate cyclase 9 |
| chr7 | 1,45E+08 | 1,45E+08 | 1127 | * | 7,642149 | 6,725980725 | 1,93737 | 3,471707 | 0,000517 | 0,377927525 | 1,360622 | 3' UTR | 7 | 1,45E+08 | 1,45E+08 | 1364 | 2 | 1E+08 | ENSMUST00000207841.2 | -6834 | ENSMUSG00000109305 | Smim38 | small integral membrane protein 38 |
| chr17 | 5737148 | 5737763 | 616 | * | 4,187466 | 6,727425271 | 2,223372 | 3,025775 | 0,00248 | 1 | 0 | Intron (ENSMUST0000( | 17 | 5848932 | 5853168 | 4237 | 1 | 78512 | ENSMUST00000180924.8 | -111169 | ENSMUSG00000096965 | 3300005D01Rik | RIKEN cDNA 3300005D01 gene |
| chr5 | 83801456 | 83801870 | 415 | * | 5,391748 | 6,728054353 | 1,975113 | 3,406415 | 0,000658 | 1 | 0 | Distal Intergenic | 5 | 83796670 | 83796801 | 132 | 2 | 1,15E+08 | ENSMUST00000104100.3 | -4655 | ENSMUSG00000077288 | Gm25765 | predicted gene, 25765 |
| chr12 | 15610026 | 15611304 | 1279 | * | 11,60353 | 6,72830045 | 1,823767 | 3,689232 | 0,000225 | 0,337512625 | 0,744375 | Distal Intergenic | 12 | 15656325 | 15779692 | 123368 | 1 | 1,05E+08 | ENSMUST00000222542.3 | -45021 | | Gm40287 | predicted gene, 40287 |
| chr10 | 98750203 | 98750438 | 236 | * | 3,932124 | 6,728392343 | 2,507986 | 2,682787 | 0,007301 | 1 | 0 | Promoter (<=1kb) | 10 | 98750268 | 98822984 | 72717 | 1 | 67972 | ENSMUST00000219557.2 | 0 | ENSMUSG00000019943 | Atp2b1 | ATPase, Ca++ transporting, plasma membrane 1 |
| chr13 | 5925097 | 5925426 | 330 | * | 3,782483 | 6,729054287 | 2,431368 | 2,767601 | 0,005647 | 1 | 0 | Distal Intergenic | 13 | 5914677 | 5916769 | 2093 | 1 | 23849 | ENSMUST00000221734.2 | 10420 | ENSMUSG00000000078 | Klf6 | Kruppel-like factor 6 |
| chr4 | 1,34E+08 | 1,34E+08 | 1034 | * | 7,576652 | 6,729178531 | 1,921367 | 3,502288 | 0,000461 | 1 | 0 | Distal Intergenic | 4 | 1,34E+08 | 1,34E+08 | 9497 | 1 | 230810 | ENSMUST00000105873.8 | -8297 | ENSMUSG00000028836 | Slc30a2 | solute carrier family 30 (zinc transporter), member 2 |
| chrX | 39885928 | 39886381 | 454 | * | 3,588436 | 6,729240451 | 2,409966 | 2,792255 | 0,005234 | 1 | 0 | Distal Intergenic | 20 | 40461136 | 40461391 | 256 | 1 | 1,08E+08 | ENSMUST00000121134.2 | -574755 | ENSMUSG00000083515 | Gm14577 | predicted gene 14577 |
| chr14 | 62286941 | 62287508 | 568 | * | 4,275913 | 6,729590181 | 2,372268 | 2,836775 | 0,004557 | 1 | 0 | Distal Intergenic | 14 | 62350569 | 62408575 | 58007 | 2 | 1,03E+08 | ENSMUST00000226702.2 | 121067 | | Gm35419 | predicted gene, 35419 |
| chr11 | 62254159 | 62255045 | 887 | * | 4,914197 | 6,732690244 | 2,14024 | 3,145765 | 0,001657 | 1 | 0 | Intron (ENSMUST0000( | 11 | 62208623 | 62250617 | 41995 | 2 | 20185 | ENSMUST00000037575.15 | -3542 | ENSMUSG00000018501 | Ncor1 | nuclear receptor co-repressor 1 |
| chr13 | 29679930 | 29680479 | 550 | * | 6,198737 | 6,733467848 | 2,099281 | 3,207512 | 0,001339 | 1 | 0 | Intron (ENSMUST0000( | 13 | 29627759 | 29701437 | 73679 | 2 | 68916 | ENSMUST00000154209.2 | 20958 | ENSMUSG00000006191 | Cdkal1 | CDK5 regulatory subunit associated protein 1-like 1 |
| chr5 | 1,47E+08 | 1,47E+08 | 562 | * | 3,083898 | 6,733731057 | 2,404248 | 2,800763 | 0,005098 | 1 | 0 | Promoter (2-3kb) | 5 | 1,47E+08 | 1,47E+08 | 96131 | 1 | 72587 | ENSMUST00000176466.8 | 2914 | ENSMUSG00000029647 | Pan3 | PAN3 poly(A) specific ribonuclease subunit |
| chr13 | 1,11E+08 | 1,11E+08 | 434 | * | 3,650228 | 6,733746718 | 2,401178 | 2,804351 | 0,005042 | 1 | 0 | Distal Intergenic | 13 | 1,11E+08 | 1,11E+08 | 1239 | 1 | 20620 | ENSMUST00000225156.2 | 20074 | ENSMUSG00000021701 | Plk2 | polo like kinase 2 |
| chr8 | 48142739 | 48143201 | 463 | * | 5,57239 | 6,735143573 | 1,911531 | 3,523428 | 0,000426 | 1 | 0 | Distal Intergenic | 8 | 48121790 | 48128591 | 6802 | 2 | 69260 | ENSMUST00000125536.2 | -14148 | ENSMUSG00000063049 | Ing2 | inhibitor of growth family, member 2 |
| chr4 | 91696837 | 91697435 | 599 | * | 4,21683 | 6,736637132 | 2,212874 | 3,044294 | 0,002332 | 1 | 0 | Promoter (2-3kb) | 4 | 91693784 | 91694792 | 1009 | 2 | 620016 | ENSMUST00000122162.2 | -2045 | ENSMUSG00000100863 | Gm12669 | glutaredoxin 3 pseudogene |
| chr17 | 67617278 | 67617680 | 403 | * | 3,499589 | 6,737127571 | 2,263668 | 2,976199 | 0,002918 | 1 | 0 | Intron (ENSMUST0000( | 17 | 66973942 | 67661068 | 687127 | 2 | 19274 | ENSMUST00000037974.10 | 43388 | ENSMUSG00000033278 | Ptprm | protein tyrosine phosphatase, receptor type, M |
| chr1 | 1,53E+08 | 1,53E+08 | 2085 | * | 12,22429 | 6,73771193 | 1,675893 | 4,02037 | 5,81E-05 | 0,153389938 | 0,918255 | Intron (ENSMUST0000( | 1 | 1,53E+08 | 1,53E+08 | 113865 | 2 | 226519 | ENSMUST00000027752.15 | 3552 | ENSMUSG00000026478 | Lamc1 | laminin, gamma 1 |
| chr12 | 17886189 | 17886499 | 311 | * | 3,121233 | 6,738752548 | 2,590011 | 2,601824 | 0,009273 | 1 | 0 | Distal Intergenic | 12 | 17950735 | 18112465 | 161731 | 1 | 668525 | ENSMUST00000221149.2 | -64236 | | Gm9222 | predicted gene 9222 |
| chr18 | 41228182 | 41228474 | 293 | * | 3,335801 | 6,738960362 | 2,54065 | 2,652455 | 0,007991 | 1 | 0 | Distal Intergenic | 18 | 40854557 | 40866926 | 12370 | 1 | 1,03E+08 | ENSMUST00000237626.2 | 373625 | ENSMUSG00000117847 | Gm31205 | predicted gene, 31205 |
| chrX | 61471715 | 61472237 | 523 | * | 3,559355 | 6,739775133 | 2,354648 | 2,862328 | 0,004205 | 1 | 0 | Distal Intergenic | 20 | 61511597 | 61512418 | 822 | 1 | 70936 | ENSMUST00000033537.2 | -39360 | ENSMUSG00000031194 | 4931400O07Rik | RIKEN cDNA 4931400O07 gene |
| chrX | 1,38E+08 | 1,38E+08 | 488 | * | 3,515986 | 6,739850277 | 2,340731 | 2,879379 | 0,003985 | 1 | 0 | Distal Intergenic | 20 | 1,38E+08 | 1,38E+08 | 74216 | 1 | 102871 | ENSMUST00000046763.13 | -55469 | ENSMUSG00000042498 | Radx | RPA1 related single stranded DNA binding protein, X-linked |
| chrX | 59019875 | 59020169 | 295 | * | 3,733915 | 6,739941388 | 2,322156 | 2,902449 | 0,003703 | 1 | 0 | Distal Intergenic | 20 | 59044824 | 59076119 | 31296 | 1 | 14071 | ENSMUST00000033477.5 | -24655 | ENSMUSG00000031138 | F9 | coagulation factor IX |
| chr15 | 20922212 | 20922945 | 734 | * | 4,715837 | 6,742071354 | 2,065236 | 3,264553 | 0,001096 | 1 | 0 | Intron (ENSMUST0000( | 15 | 20908958 | 20909085 | 128 | 1 | 1,15E+08 | ENSMUST00000158633.3 | 13254 | ENSMUSG00000089258 | Gm25711 | predicted gene, 25711 |
| chr19 | 46422308 | 46422659 | 352 | * | 6,23917 | 6,743264869 | 1,974824 | 3,414616 | 0,000639 | 1 | 0 | Intron (ENSMUST0000( | 19 | 46446345 | 46464073 | 17729 | 1 | 24069 | ENSMUST00000123252.2 | -23686 | ENSMUSG00000025231 | Sufu | SUFU negative regulator of hedgehog signaling |
| chr11 | 23577703 | 23578429 | 727 | * | 6,057245 | 6,743758181 | 1,945204 | 3,466864 | 0,000527 | 1 | 0 | Promoter (<=1kb) | 11 | 23523251 | 23579027 | 55777 | 2 | 71675 | ENSMUST00000043356.13 | 598 | ENSMUSG00000042208 | Sanbr | SANT and BTB domain regulator of CSR |
| chr9 | 66903547 | 66903941 | 395 | * | 3,171249 | 6,744883822 | 2,452769 | 2,749906 | 0,005961 | 1 | 0 | Distal Intergenic | 9 | 66920349 | 66929969 | 9621 | 1 | 1,03E+08 | ENSMUST00000240821.1 | -16408 | | Gm30639 | predicted gene, 30639 |
| chr12 | 22812219 | 22812613 | 395 | * | 3,540444 | 6,745152089 | 2,415626 | 2,792299 | 0,005233 | 1 | 0 | Distal Intergenic | 12 | 22635890 | 22673346 | 37457 | 2 | 668662 | ENSMUST00000223403.2 | -138873 | | Gm9292 | predicted gene 9292 |
| chr11 | 4483298 | 4484208 | 911 | * | 3,915118 | 6,746343067 | 2,289177 | 2,94706 | 0,003208 | 1 | 0 | Intron (ENSMUST0000( | 11 | 4432752 | 4478406 | 45655 | 2 | 74302 | ENSMUST00000128256.2 | -4892 | ENSMUSG00000034354 | Mtmr3 | myotubularin related protein 3 |
| chr4 | 17640366 | 17641481 | 1116 | * | 9,602266 | 6,74700666 | 1,806639 | 3,734563 | 0,000188 | 0,252389304 | 1,211315 | Distal Intergenic | 4 | 17546447 | 17636191 | 89745 | 2 | 1,03E+08 | ENSMUST00000245354.1 | -4175 | | Gm30092 | predicted gene, 30092 |
| chr11 | 87883661 | 87884332 | 672 | * | 4,364296 | 6,747943391 | 2,161028 | 3,122562 | 0,001793 | 1 | 0 | Distal Intergenic | 11 | 87870351 | 87878359 | 8009 | 2 | 68097 | ENSMUST00000020775.9 | -5302 | ENSMUSG00000020483 | Dynll2 | dynein light chain LC8-type 2 |
| chr17 | 64837856 | 64838101 | 246 | * | 3,40373 | 6,749676333 | 2,563387 | 2,633109 | 0,008461 | 1 | 0 | Intron (ENSMUST0000( | 17 | 64821134 | 64862498 | 41365 | 2 | 1E+08 | ENSMUST00000168379.3 | 24397 | ENSMUSG00000073377 | AU016765 | expressed sequence AU016765 |
| chr16 | 11831070 | 11831462 | 393 | * | 3,382701 | 6,749772012 | 2,306696 | 2,926165 | 0,003432 | 1 | 0 | Intron (ENSMUST0000( | 16 | 11802584 | 11813260 | 10677 | 1 | 72555 | ENSMUST00000229804.2 | 28486 | ENSMUSG00000022494 | Shisa9 | shisa family member 9 |
| chr15 | 53175948 | 53176469 | 522 | * | 4,816121 | 6,749904925 | 2,107274 | 3,203145 | 0,001359 | 1 | 0 | Intron (ENSMUST0000( | 15 | 52932060 | 53209055 | 276996 | 2 | 14042 | ENSMUST00000133362.8 | 32586 | ENSMUSG00000061731 | Ext1 | exostosin glycosyltransferase 1 |
| chr1 | 1,9E+08 | 1,9E+08 | 791 | * | 4,950896 | 6,74994041 | 2,050085 | 3,292517 | 0,000993 | 1 | 0 | Distal Intergenic | 1 | 1,9E+08 | 1,9E+08 | 52680 | 2 | 19130 | ENSMUST00000175916.8 | -50486 | ENSMUSG00000010175 | Prox1 | prospero homeobox 1 |
| chr10 | 81098601 | 81099239 | 639 | * | 6,503546 | 6,750020665 | 1,971534 | 3,42374 | 0,000618 | 1 | 0 | Promoter (1-2kb) | 10 | 81097004 | 81098501 | 1498 | 1 | 17179 | ENSMUST00000148735.2 | 1597 | ENSMUSG00000004933 | Matk | megakaryocyte-associated tyrosine kinase |
| chr1 | 1,75E+08 | 1,75E+08 | 609 | * | 7,193855 | 6,750099008 | 1,91208 | 3,530238 | 0,000415 | 1 | 0 | 3' UTR | 1 | 1,75E+08 | 1,75E+08 | 2910 | 2 | 14194 | ENSMUST00000130996.2 | 3068 | ENSMUSG00000026526 | Fh1 | fumarate hydratase 1 |
| chr12 | 84275725 | 84276531 | 807 | * | 6,809471 | 6,750105937 | 1,910767 | 3,532668 | 0,000411 | 1 | 0 | Distal Intergenic | 12 | 84199233 | 84265655 | 66423 | 2 | 238317 | ENSMUST00000220974.2 | -10070 | ENSMUSG00000042507 | Mideas | mitotic deacetylase associated SANT domain protein |
| chr7 | 1,14E+08 | 1,14E+08 | 1186 | * | 7,271343 | 6,750301971 | 1,81395 | 3,721327 | 0,000198 | 1 | 0 | Distal Intergenic | 7 | 1,14E+08 | 1,14E+08 | 106690 | 1 | 233752 | ENSMUST00000161800.3 | -10199 | ENSMUSG00000048782 | Insc | INSC spindle orientation adaptor protein |
| chr4 | 44754461 | 44755799 | 1339 | * | 10,49219 | 6,7505372 | 1,732557 | 3,896285 | 9,77E-05 | 0,170809454 | 1,2763 | Promoter (<=1kb) | 4 | 44755877 | 44761971 | 6095 | 1 | 319885 | ENSMUST00000154121.2 | -78 | ENSMUSG00000035649 | Zcchc7 | zinc finger, CCHC domain containing 7 |
| chr19 | 45233792 | 45234734 | 943 | * | 10,42768 | 6,752716104 | 1,774288 | 3,805874 | 0,000141 | 0,221323359 | 1,173325 | Distal Intergenic | 19 | 45221123 | 45224251 | 3129 | 2 | 16814 | ENSMUST00000099401.6 | -9541 | ENSMUSG00000025216 | Lbx1 | ladybird homeobox 1 |
| chr2 | 5320273 | 5320564 | 292 | * | 4,762862 | 6,75276653 | 2,268847 | 2,976298 | 0,002918 | 1 | 0 | Intron (ENSMUST0000( | 2 | 5264740 | 5264926 | 187 | 2 | 1,15E+08 | ENSMUST00000122657.3 | -55347 | ENSMUSG00000084606 | Gm23118 | predicted gene, 23118 |
| chr8 | 11598558 | 11599148 | 591 | * | 7,196759 | 6,754536166 | 1,84165 | 3,667655 | 0,000245 | 1 | 0 | Promoter (1-2kb) | 8 | 11579612 | 11600391 | 20780 | 2 | 71941 | ENSMUST00000209236.2 | 1243 | ENSMUSG00000056228 | Cars2 | cysteinyl-tRNA synthetase 2 (mitochondrial)(putative) |
| chr12 | 45121213 | 45122180 | 968 | * | 7,85232 | 6,755411842 | 1,859039 | 3,633819 | 0,000279 | 1 | 0 | Promoter (<=1kb) | 12 | 44899267 | 45121492 | 222226 | 2 | 217517 | ENSMUST00000053768.14 | 0 | ENSMUSG00000046314 | Stxbp6 | syntaxin binding protein 6 (amisyn) |
| chr12 | 21926776 | 21927657 | 882 | * | 7,591309 | 6,75554285 | 1,855857 | 3,64012 | 0,000273 | 1 | 0 | Distal Intergenic | 12 | 21826160 | 21826708 | 549 | 2 | 434693 | ENSMUST00000164169.2 | -100068 | | Mrto4-ps2 | mRNA turnover 4, pseudogene 2 |
| chrX | 60876194 | 60876836 | 643 | * | 4,046583 | 6,756749254 | 2,147562 | 3,146241 | 0,001654 | 1 | 0 | Distal Intergenic | 20 | 60982497 | 60982603 | 107 | 1 | 1,15E+08 | ENSMUST00000240328.1 | -105661 | ENSMUSG00000118848 | Gm25412 | predicted gene, 25412 |
| chr14 | 74415650 | 74416401 | 752 | * | 4,811337 | 6,757030194 | 2,108745 | 3,204289 | 0,001354 | 1 | 0 | Intron (ENSMUST0000( | 14 | 74592967 | 74607181 | 14215 | 1 | 74425 | ENSMUST00000226927.2 | -176566 | ENSMUSG00000115845 | 4933402J15Rik | RIKEN cDNA 4933402J15 gene |
| chr5 | 1,47E+08 | 1,47E+08 | 673 | * | 4,926167 | 6,758090191 | 1,995428 | 3,386788 | 0,000707 | 1 | 0 | Promoter (2-3kb) | 5 | 1,47E+08 | 1,47E+08 | 4143 | 1 | 19933 | ENSMUST00000035983.12 | -2807 | ENSMUSG00000041453 | Rpl21 | ribosomal protein L21 |
| chr12 | 54143911 | 54144546 | 636 | * | 5,395459 | 6,758208057 | 1,985195 | 3,404304 | 0,000663 | 1 | 0 | Intron (ENSMUST0000( | 12 | 54138213 | 54153668 | 15456 | 2 | 1,03E+08 | ENSMUST00000219480.2 | 9122 | | Gm35818 | predicted gene, 35818 |
| chr3 | 1,22E+08 | 1,22E+08 | 1069 | * | 6,986407 | 6,75839937 | 2,070627 | 3,263939 | 0,001099 | 1 | 0 | Promoter (<=1kb) | 3 | 1,22E+08 | 1,22E+08 | 56396 | 2 | 19299 | ENSMUST00000029770.8 | -46 | ENSMUSG00000028127 | Abcd3 | ATP-binding cassette, sub-family D (ALD), member 3 |
| chr4 | 1,33E+08 | 1,33E+08 | 812 | * | 6,181688 | 6,758641412 | 1,94978 | 3,466362 | 0,000528 | 1 | 0 | Intron (ENSMUST0000( | 4 | 1,33E+08 | 1,33E+08 | 45932 | 1 | 242687 | ENSMUST00000136132.2 | 9783 | ENSMUSG00000028868 | Wasf2 | WASP family, member 2 |
| chr19 | 27291791 | 27292496 | 706 | * | 6,43712 | 6,758772175 | 1,940803 | 3,482461 | 0,000497 | 1 | 0 | Promoter (1-2kb) | 19 | 27290299 | 27299593 | 9295 | 1 | 1,03E+08 | ENSMUST00000243868.1 | 1492 | | Gm35268 | predicted gene, 35268 |
| chr8 | 3261410 | 3262058 | 649 | * | 6,889981 | 6,759170354 | 1,912545 | 3,534123 | 0,000409 | 0,337054778 | 1,360622 | Exon (ENSMUST00000( | 8 | 3224795 | 3242717 | 17923 | 2 | 16337 | ENSMUST00000207100.2 | -18693 | ENSMUSG00000005534 | Insr | insulin receptor |
| chr5 | 93115395 | 93115974 | 580 | * | 4,043835 | 6,759243472 | 2,263045 | 2,986791 | 0,002819 | 1 | 0 | Distal Intergenic | 5 | 93109898 | 93112649 | 2752 | 1 | 27428 | ENSMUST00000172752.2 | 5497 | ENSMUSG00000029381 | Shroom3 | shroom family member 3 |
| chr10 | 88636903 | 88637533 | 631 | * | 6,828002 | 6,760133138 | 1,857443 | 3,639484 | 0,000273 | 1 | 0 | Exon (ENSMUST00000( | 10 | 88612041 | 88618528 | 6488 | 2 | 70683 | ENSMUST00000219662.2 | -18375 | ENSMUSG00000004356 | Utp20 | UTP20 small subunit processome component |
| chr4 | 72193733 | 72194702 | 970 | * | 7,257958 | 6,760658169 | 1,883159 | 3,590062 | 0,000331 | 1 | 0 | Distal Intergenic | 4 | 72119481 | 72122138 | 2658 | 1 | 68285 | ENSMUST00000246336.1 | 74252 | | C630043F03Rik | RIKEN cDNA C630043F03 gene |
| chrX | 37010720 | 37011994 | 1275 | * | 7,597853 | 6,760982329 | 1,817099 | 3,720756 | 0,000199 | 1 | 0 | Distal Intergenic | 20 | 36987974 | 36990647 | 2674 | 2 | 104384 | ENSMUST00000089062.8 | -20073 | ENSMUSG00000068048 | Rhox9 | reproductive homeobox 9 |
| chr1 | 1,4E+08 | 1,4E+08 | 925 | * | 8,732394 | 6,761121801 | 1,811985 | 3,731334 | 0,00019 | 0,238928695 | 1,360622 | Distal Intergenic | 1 | 1,4E+08 | 1,4E+08 | 142 | 1 | 1,15E+08 | ENSMUST00000157305.3 | 46554 | ENSMUSG00000087930 | Gm26048 | predicted gene, 26048 |
| chr4 | 43487558 | 43487851 | 294 | * | 3,510813 | 6,764888812 | 2,618192 | 2,583802 | 0,009772 | 1 | 0 | Distal Intergenic | 4 | 43482081 | 43483734 | 1654 | 2 | 54390 | ENSMUST00000030180.7 | -3824 | ENSMUSG00000028460 | Sit1 | suppression inducing transmembrane adaptor 1 |
| chr7 | 1,13E+08 | 1,13E+08 | 429 | * | 5,078682 | 6,764929671 | 2,055688 | 3,290834 | 0,000999 | 1 | 0 | Exon (ENSMUST00000( | 7 | 1,13E+08 | 1,13E+08 | 7419 | 1 | 11865 | ENSMUST00000211547.2 | -6073 | ENSMUSG00000055116 | Arntl | aryl hydrocarbon receptor nuclear translocator-like |
| chr16 | 34524480 | 34525263 | 784 | * | 6,222903 | 6,764946894 | 2,028 | 3,335772 | 0,000851 | 1 | 0 | Promoter (<=1kb) | 16 | 34524413 | 34528625 | 4213 | 1 | 239839 | ENSMUST00000232341.2 | 67 | ENSMUSG00000022833 | Ccdc14 | coiled-coil domain containing 14 |
| chr4 | 1,02E+08 | 1,02E+08 | 402 | * | 5,676066 | 6,765133733 | 2,044172 | 3,309474 | 0,000935 | 1 | 0 | Intron (ENSMUST0000( | 4 | 1,02E+08 | 1,02E+08 | 469857 | 1 | 18578 | ENSMUST00000173119.8 | 73095 | ENSMUSG00000028525 | Pde4b | phosphodiesterase 4B, cAMP specific |
| chr14 | 1,06E+08 | 1,06E+08 | 563 | * | 3,729502 | 6,76554813 | 2,527578 | 2,676692 | 0,007435 | 1 | 0 | Distal Intergenic | 14 | 1,06E+08 | 1,06E+08 | 3075 | 1 | 212085 | ENSMUST00000022708.7 | -55854 | ENSMUSG00000022113 | Trim52 | tripartite motif-containing 52 |
| chr14 | 51543528 | 51543835 | 308 | * | 3,561706 | 6,765568071 | 2,428856 | 2,785495 | 0,005345 | 1 | 0 | Distal Intergenic | 14 | 51554792 | 51582946 | 28155 | 2 | 75377 | ENSMUST00000228344.2 | 39111 | ENSMUSG00000115540 | 4930597G03Rik | RIKEN cDNA 4930597G03 gene |
| chr4 | 58323480 | 58324297 | 818 | * | 6,060919 | 6,765616197 | 2,018171 | 3,352351 | 0,000801 | 1 | 0 | Intron (ENSMUST0000( | 4 | 58286175 | 58304324 | 18150 | 1 | 18198 | ENSMUST00000135879.2 | 37305 | ENSMUSG00000057280 | Musk | muscle, skeletal, receptor tyrosine kinase |
| chr11 | 6418873 | 6419644 | 772 | * | 4,974063 | 6,765727572 | 2,012762 | 3,361415 | 0,000775 | 1 | 0 | Promoter (2-3kb) | 11 | 6422225 | 6422277 | 53 | 2 | 19291 | ENSMUST00000175144.4 | 2633 | ENSMUSG00000094483 | Purb | purine rich element binding protein B |
| chr17 | 59994793 | 59995305 | 513 | * | 3,492791 | 6,766639045 | 2,408103 | 2,809946 | 0,004955 | 1 | 0 | Distal Intergenic | 17 | 59876554 | 59876660 | 107 | 2 | 1,15E+08 | ENSMUST00000104374.3 | -118133 | ENSMUSG00000077562 | Gm23203 | predicted gene, 23203 |
| chr4 | 56846560 | 56847062 | 503 | * | 4,166984 | 6,766943639 | 2,37973 | 2,843577 | 0,004461 | 1 | 0 | Promoter (<=1kb) | 4 | 56844467 | 56846092 | 1626 | 2 | 54366 | ENSMUST00000132764.2 | -468 | ENSMUSG00000038816 | Ctnnal1 | catenin (cadherin associated protein), alpha-like 1 |
| chr8 | 12918062 | 12918621 | 560 | * | 4,130096 | 6,768680712 | 2,243441 | 3,017098 | 0,002552 | 1 | 0 | 3' UTR | 8 | 12923806 | 12930146 | 6341 | 1 | 17207 | ENSMUST00000156560.2 | -5185 | ENSMUSG00000031442 | Mcf2l | mcf.2 transforming sequence-like |
| chr7 | 1,41E+08 | 1,41E+08 | 656 | * | 7,255452 | 6,769254634 | 1,87565 | 3,609018 | 0,000307 | 0,301441602 | 1,360622 | Promoter (<=1kb) | 7 | 1,41E+08 | 1,41E+08 | 91 | 1 | 1,02E+08 | ENSMUST00000184158.3 | -808 | ENSMUSG00000098759 | Mir7063 | microRNA 7063 |
| chr1 | 57449249 | 57449775 | 527 | * | 4,658857 | 6,77188961 | 2,197075 | 3,08223 | 0,002055 | 1 | 0 | Promoter (2-3kb) | 1 | 57427396 | 57446260 | 18865 | 2 | 68736 | ENSMUST00000162686.8 | -2989 | ENSMUSG00000048495 | Tyw5 | tRNA-yW synthesizing protein 5 |
| chr10 | 88457259 | 88457925 | 667 | * | 4,374902 | 6,772014403 | 2,172958 | 3,116496 | 0,00183 | 1 | 0 | Distal Intergenic | 10 | 88354141 | 88441014 | 86874 | 2 | 109272 | ENSMUST00000121629.8 | -16245 | ENSMUSG00000020061 | Mybpc1 | myosin binding protein C, slow-type |
| chr11 | 97322916 | 97323384 | 469 | * | 4,820577 | 6,772295685 | 2,114952 | 3,202103 | 0,001364 | 1 | 0 | Intron (ENSMUST0000( | 11 | 97327111 | 97335054 | 7944 | 1 | 58996 | ENSMUST00000152933.2 | -3727 | ENSMUSG00000049807 | Arhgap23 | Rho GTPase activating protein 23 |
| chr9 | 44495685 | 44496109 | 425 | * | 5,172424 | 6,772402371 | 2,008653 | 3,371613 | 0,000747 | 1 | 0 | Exon (ENSMUST00000 2 | 9 | 44516189 | 44552028 | 35840 | 1 | 13209 | ENSMUST00000170489.2 | -20080 | ENSMUSG00000032097 | Ddx6 | DEAD (Asp-Glu-Ala-Asp) box polypeptide 6 |
| chrX | 63558394 | 63559196 | 803 | * | 4,554998 | 6,772699809 | 2,05176 | 3,300922 | 0,000964 | 1 | 0 | Distal Intergenic | 20 | 63540190 | 63540263 | 74 | 1 | 1,15E+08 | ENSMUST00000122510.3 | 18204 | ENSMUSG00000084459 | Gm24598 | predicted gene, 24598 |
| chrX | 16006087 | 16006619 | 533 | * | 5,594554 | 6,773608269 | 1,939729 | 3,492038 | 0,000479 | 1 | 0 | Distal Intergenic | 20 | 15762902 | 15763008 | 107 | 2 | 1,15E+08 | ENSMUST00000239993.1 | -243079 | ENSMUSG00001118666 | Gm23516 | predicted gene, 23516 |
| chr18 | 33464782 | 33465596 | 815 | * | 6,079378 | 6,774140348 | 1,89117 | 3,581984 | 0,000341 | 1 | 0 | Distal Intergenic | 18 | 33332408 | 33346915 | 14508 | 2 | 170459 | ENSMUST00000025236.9 | -117867 | ENSMUSG00000024378 | Stard4 | StAR-related lipid transfer (START) domain containing 4 |
| chr5 | 32744869 | 32746039 | 1171 | * | 8,188538 | 6,774954497 | 1,832295 | 3,697524 | 0,000218 | 1 | 0 | Intron (ENSMUST0000( | 5 | 32768515 | 32842921 | 74407 | 1 | 22612 | ENSMUST00000202543.4 | -22476 | ENSMUSG00000014932 | Yes1 | YES proto-oncogene 1, Src family tyrosine kinase |
| chr3 | 1,04E+08 | 1,04E+08 | 435 | * | 3,86428 | 6,775665582 | 2,40181 | 2,821066 | 0,004786 | 1 | 0 | Intron (ENSMUST0000( | 3 | 1,04E+08 | 1,04E+08 | 206479 | 2 | 99470 | ENSMUST00000121198.8 | 40860 | ENSMUSG00000052539 | Magi3 | membrane associated guanylate kinase, WW and PDZ domain containing 3 |
| chrX | 53198982 | 53199384 | 403 | * | 4,029898 | 6,778513804 | 2,178343 | 3,111776 | 0,00186 | 1 | 0 | Intron (ENSMUST0000( | 20 | 53120962 | 53121068 | 107 | 1 | 1,15E+08 | ENSMUST00000240118.1 | 78020 | ENSMUSG00000118772 | Gm22550 | predicted gene, 22550 |
| chr5 | 1,38E+08 | 1,38E+08 | 691 | * | 4,404597 | 6,778784006 | 2,151809 | 3,150273 | 0,001631 | 1 | 0 | Promoter (<=1kb) | 5 | 1,38E+08 | 1,38E+08 | 9638 | 2 | 118446 | ENSMUST00000077119.8 | 879 | ENSMUSG00000056966 | Gjc3 | gap junction protein, gamma 3 |
| chr11 | 87169720 | 87170711 | 992 | * | 7,439126 | 6,779694222 | 1,913292 | 3,54347 | 0,000395 | 1 | 0 | Intron (ENSMUST0000( | 11 | 87209520 | 87209625 | 106 | 2 | 1,15E+08 | ENSMUST00000157376.3 | 38914 | ENSMUSG00000088001 | Gm22883 | predicted gene, 22883 |
| chr16 | 17419412 | 17420263 | 852 | * | 5,285217 | 6,779841428 | 2,067236 | 3,279665 | 0,001039 | 1 | 0 | Distal Intergenic | 16 | 17437218 | 17462692 | 25475 | 1 | 77626 | ENSMUST00000006053.13 | -16955 | ENSMUSG00000005899 | Smpd4 | sphingomyelin phosphodiesterase 4 |
| chr2 | 1,73E+08 | 1,73E+08 | 317 | * | 5,566739 | 6,780260063 | 2,021357 | 3,354311 | 0,000796 | 1 | 0 | Distal Intergenic | 2 | 1,73E+08 | 1,73E+08 | 23975 | 2 | 12162 | ENSMUST00000137247.2 | 84335 | ENSMUSG00000008999 | Bmp7 | bone morphogenetic protein 7 |
| chr15 | 84551268 | 84551730 | 463 | * | 5,072294 | 6,780321762 | 2,035603 | 3,330867 | 0,000866 | 1 | 0 | Promoter (2-3kb) | 15 | 84553821 | 84587874 | 34054 | 1 | 109270 | ENSMUST00000171460.8 | -2091 | ENSMUSG00000036106 | Prr5 | proline rich 5 (renal) |
| chr3 | 50277461 | 50277957 | 497 | * | 3,891139 | 6,780937727 | 2,232877 | 3,036861 | 0,002391 | 1 | 0 | Intron (ENSMUST0000( | 3 | 50319387 | 50398060 | 78674 | 2 | 26570 | ENSMUST00000029297.6 | 120103 | ENSMUSG00000027737 | Slc7a11 | solute carrier family 7 (cationic amino acid transporter, y+ system), member 11 |
| chr1 | 63484298 | 63484800 | 503 | * | 5,363559 | 6,78106335 | 1,990342 | 3,406984 | 0,000657 | 1 | 0 | Promoter (<=1kb) | 1 | 63485050 | 63612376 | 127327 | 1 | 23792 | ENSMUST00000182642.8 | -250 | ENSMUSG00000025964 | Adam23 | a disintegrin and metallopeptidase domain 23 |
| chr19 | 10070863 | 10071287 | 425 | * | 3,625989 | 6,782878153 | 2,342355 | 2,895752 | 0,003783 | 1 | 0 | Intron (ENSMUST0000( | 19 | 10043937 | 10066080 | 22144 | 2 | 56473 | ENSMUST00000235937.2 | -4783 | ENSMUSG00000024665 | Fads2 | fatty acid desaturase 2 |
| chr2 | 32859472 | 32859789 | 318 | * | 4,269364 | 6,783005744 | 2,328902 | 2,912534 | 0,003585 | 1 | 0 | Distal Intergenic | 2 | 32853303 | 32853432 | 130 | 1 | 104367 | ENSMUST00000083190.3 | 6169 | ENSMUSG00000065124 | Snora65 | small nucleolar RNA, H/ACA box 65 |

| | | | | | | | | | | | | | | | | | | | | | | | |
|---|---|---|---|---|---|---|---|---|---|---|---|---|---|---|---|---|---|---|---|---|---|---|---|
| chr3 | 15214487 | 15214831 | 345 | * | 4,253156 | 6,784172204 | 2,234426 | 3,036204 | 0,002396 | 1 | 0 | Distal Intergenic | 3 | 15218125 | 15236502 | 18378 | 2 | 1,03E+08 | ENSMUST00000194008.2 | 21671 | | Gm33819 | predicted gene, 33819 |
| chr14 | 65617898 | 65618568 | 671 | * | 4,933748 | 6,785501665 | 2,145521 | 3,162636 | 0,001563 | 1 | 0 | Intron (ENSMUST0000( | 14 | 65612842 | 65623812 | 10971 | 1 | 380912 | ENSMUST00000224687.2 | 5056 | ENSMUSG00000034522 | Zfp395 | zinc finger protein 395 |
| chr17 | 46143508 | 46144222 | 715 | * | 5,122314 | 6,785631427 | 2,13782 | 3,174089 | 0,001503 | 1 | 0 | Distal Intergenic | 17 | 46117130 | 46133249 | 16120 | 2 | 1,03E+08 | ENSMUST00000233237.2 | -10259 | | Gm35692 | predicted gene, 35692 |
| chr10 | 20548143 | 20548561 | 419 | * | 4,780143 | 6,786679188 | 2,083098 | 3,257973 | 0,001122 | 1 | 0 | Intron (ENSMUST0000( | 10 | 20276164 | 20600442 | 324279 | 2 | 29863 | ENSMUST00000170265.8 | 51881 | ENSMUSG00000019990 | Pde7b | phosphodiesterase 7B |
| chrX | 98352696 | 98353550 | 855 | * | 3,435038 | 6,786711384 | 2,443652 | 2,777282 | 0,005482 | 1 | 0 | Distal Intergenic | 20 | 98448937 | 98487403 | 38467 | 2 | 1E+08 | ENSMUST00000156494.2 | 133853 | ENSMUSG00000073051 | Gm14812 | predicted gene 14812 |
| chr10 | 1,23E+08 | 1,23E+08 | 650 | * | 4,598425 | 6,787387121 | 2,144528 | 3,16498 | 0,001551 | 1 | 0 | Distal Intergenic | 10 | 1,23E+08 | 1,23E+08 | 758 | 2 | 387251 | ENSMUST00000220378.2 | -4708 | ENSMUSG00000065406 | Mirlet7i | microRNA let7i |
| chr1 | 1,14E+08 | 1,14E+08 | 496 | * | 5,736793 | 6,787513332 | 2,10487 | 3,224672 | 0,001261 | 1 | 0 | Distal Intergenic | 1 | 1,12E+08 | 1,12E+08 | 136 | 1 | 1,15E+08 | ENSMUST00000158066.3 | 1502479 | ENSMUSG00000088691 | Gm22331 | predicted gene, 22331 |
| chr3 | 1,19E+08 | 1,19E+08 | 823 | * | 5,958506 | 6,788776886 | 1,875618 | 3,619487 | 0,000295 | 1 | 0 | Intron (ENSMUST0000( | 3 | 1,19E+08 | 1,19E+08 | 88610 | 1 | 99586 | ENSMUST00000129148.2 | 213239 | ENSMUSG00000033308 | Dpyd | dihydropyrimidine dehydrogenase |
| chr14 | 21743487 | 21744482 | 996 | * | 10,62555 | 6,79018636 | 1,749556 | 3,881092 | 0,000104 | 0,239591648 | 0,738441 | Intron (ENSMUST0000( | 14 | 21735014 | 21753170 | 18157 | 2 | 435391 | ENSMUST00000224735.2 | 8688 | ENSMUSG00000063821 | Dusp29 | dual specificity phosphatase 29 |
| chr14 | 1,21E+08 | 1,21E+08 | 2024 | * | 11,92562 | 6,790769073 | 1,715934 | 3,957477 | 7,57E-05 | 0,203319961 | 0,738441 | Promoter (1-2kb) | 14 | 1,21E+08 | 1,21E+08 | 52677 | 1 | 105559 | ENSMUST00000227508.2 | 1260 | ENSMUSG00000022139 | Mbnl2 | muscleblind like splicing factor 2 |
| chr9 | 90014419 | 90014870 | 452 | * | 3,336536 | 6,791430861 | 2,604989 | 2,607086 | 0,009132 | 1 | 0 | Distal Intergenic | 9 | 89973718 | 89996827 | 23110 | 2 | 21761 | ENSMUST00000169860.8 | -17592 | ENSMUSG00000062270 | Morf4l1 | mortality factor 4 like 1 |
| chr3 | 1,56E+08 | 1,56E+08 | 384 | * | 3,325589 | 6,791448483 | 2,430504 | 2,794256 | 0,005202 | 1 | 0 | Distal Intergenic | 3 | 1,55E+08 | 1,55E+08 | 104 | 2 | 1,15E+08 | ENSMUST00000083873.3 | -69317 | ENSMUSG00000065807 | Gm23038 | predicted gene, 23038 |
| chr11 | 1,18E+08 | 1,18E+08 | 1034 | * | 9,577765 | 6,791896948 | 1,751639 | 3,877452 | 0,000106 | 0,184528934 | 1,211315 | Promoter (<=1kb) | 11 | 1,18E+08 | 1,18E+08 | 23131 | 1 | 74451 | ENSMUST00000100185.10 | 0 | ENSMUSG00000017715 | Pgs1 | phosphatidylglycerophosphate synthase 1 |
| chr11 | 41974804 | 41975134 | 331 | * | 3,728278 | 6,792864131 | 2,294766 | 2,960155 | 0,003075 | 1 | 0 | Distal Intergenic | 11 | 41801030 | 41891684 | 90655 | 2 | 14406 | ENSMUST00000070725.11 | -83120 | ENSMUSG00000020436 | Gabrg2 | gamma-aminobutyric acid (GABA) A receptor, subunit gamma 2 |
| chr14 | 23260930 | 23261736 | 807 | * | 6,695193 | 6,79298658 | 1,954243 | 3,476019 | 0,000509 | 1 | 0 | Distal Intergenic | 14 | 23088262 | 23144639 | 56378 | 2 | 791307 | ENSMUST00000090409.2 | -116291 | ENSMUSG00000068680 | Gm10248 | predicted gene 10248 |
| chr3 | 93428104 | 93428561 | 458 | * | 4,663327 | 6,794172805 | 2,126292 | 3,195314 | 0,001397 | 1 | 0 | Promoter (<=1kb) | 3 | 93427795 | 93433594 | 5800 | 1 | 20195 | ENSMUST00000029515.5 | 309 | ENSMUSG00000027907 | S100a11 | S100 calcium binding protein A11 |
| chrX | 1,55E+08 | 1,55E+08 | 639 | * | 6,052532 | 6,796020317 | 1,976712 | 3,438042 | 0,000586 | 1 | 0 | Distal Intergenic | 20 | 1,54E+08 | 1,55E+08 | 721113 | 1 | 1,03E+08 | ENSMUST00000112542.2 | 729904 | | Gm15155 | predicted gene 15155 |
| chr4 | 72919565 | 72920194 | 630 | * | 4,133356 | 6,798103724 | 2,354981 | 2,886692 | 0,003893 | 1 | 0 | Distal Intergenic | 4 | 72824438 | 72824575 | 138 | 2 | 1,15E+08 | ENSMUST00000158473.3 | -94990 | ENSMUSG00000089098 | Gm25653 | predicted gene, 25653 |
| chr16 | 25124353 | 25125341 | 989 | * | 6,841378 | 6,798757715 | 1,843931 | 3,687101 | 0,000227 | 1 | 0 | Intron (ENSMUST0000( | 16 | 25105571 | 25136985 | 31415 | 1 | 71338 | ENSMUST00000128595.2 | 18782 | ENSMUSG00000048399 | Tprg | transformation related protein 63 regulated |
| chr10 | 28143421 | 28144157 | 737 | * | 8,383752 | 6,799037414 | 1,83403 | 3,707157 | 0,00021 | 0,252371261 | 1,360622 | Intron (ENSMUST0000( | 10 | 27951006 | 28140669 | 189664 | 1 | 19272 | ENSMUST00000219761.2 | 192415 | ENSMUSG00000019889 | Ptprk | protein tyrosine phosphatase, receptor type, K |
| chr1 | 1,77E+08 | 1,77E+08 | 853 | * | 4,316016 | 6,799816693 | 2,271501 | 2,993535 | 0,002758 | 1 | 0 | Promoter (2-3kb) | 1 | 1,77E+08 | 1,77E+08 | 18741 | 2 | 545389 | ENSMUST00000192643.2 | -2284 | ENSMUSG00000057335 | Cep170 | centrosomal protein 170 |
| chr4 | 1,49E+08 | 1,49E+08 | 608 | * | 4,321045 | 6,8024112 | 2,210055 | 3,077938 | 0,002084 | 1 | 0 | Promoter (2-3kb) | 4 | 1,49E+08 | 1,49E+08 | 867 | 2 | 63958 | ENSMUST00000131986.2 | -2070 | ENSMUSG00000028960 | Ube4b | ubiquitination factor E4B |
| chr1 | 3590511 | 3591059 | 549 | * | 4,962448 | 6,802629978 | 2,063202 | 3,297123 | 0,000977 | 1 | 0 | Intron (ENSMUST0000( | 1 | 3284705 | 3741721 | 457017 | 2 | 497097 | ENSMUST00000070533.5 | 150662 | ENSMUSG00000051951 | Xkr4 | X-linked Kx blood group related 4 |
| chr14 | 25244052 | 25244696 | 645 | * | 5,036549 | 6,802666542 | 2,037598 | 3,338571 | 0,000842 | 1 | 0 | Distal Intergenic | 14 | 25140219 | 25143665 | 3447 | 2 | 75862 | ENSMUST00000225429.2 | -100387 | ENSMUSG00000114916 | 4930572O13Rik | RIKEN cDNA 4930572O13 gene |
| chr13 | 43611933 | 43612633 | 701 | * | 5,691536 | 6,802838885 | 1,963247 | 3,465096 | 0,00053 | 1 | 0 | Promoter (2-3kb) | 13 | 43573055 | 43615207 | 42153 | 2 | 56705 | ENSMUST00000222239.2 | 2574 | ENSMUSG00000038546 | Ranbp9 | RAN binding protein 9 |
| chr3 | 1,02E+08 | 1,02E+08 | 705 | * | 6,278748 | 6,802887271 | 1,941371 | 3,504166 | 0,000458 | 1 | 0 | Promoter (1-2kb) | 3 | 1,02E+08 | 1,02E+08 | 7051 | 2 | 229658 | ENSMUST00000161722.2 | 1426 | ENSMUSG00000027860 | Vangl1 | VANGL planar cell polarity 1 |
| chr2 | 1,19E+08 | 1,19E+08 | 630 | * | 6,625685 | 6,803010549 | 1,899488 | 3,581498 | 0,000342 | 1 | 0 | Intron (ENSMUST0000( | 2 | 1,19E+08 | 1,19E+08 | 3153 | 1 | 69065 | ENSMUST00000028780.4 | 39805 | ENSMUSG00000027313 | Chac1 | ChaC, cation transport regulator 1 |
| chr1 | 89193240 | 89193879 | 640 | * | 7,259296 | 6,803327831 | 1,818917 | 3,740317 | 0,000184 | 1 | 0 | Distal Intergenic | 1 | 89288064 | 89333855 | 45792 | 2 | 77331 | ENSMUST00000186258.2 | 139976 | ENSMUSG00000099938 | C030007H22Rik | RIKEN cDNA C030007H22 gene |
| chr11 | 35660772 | 35661430 | 659 | * | 5,909792 | 6,804184755 | 1,964502 | 3,463568 | 0,000533 | 1 | 0 | Promoter (<=1kb) | 11 | 35660311 | 35682112 | 21802 | 1 | 211347 | ENSMUST00000018990.8 | 461 | ENSMUSG00000018846 | Pank3 | pantothenate kinase 3 |
| chr8 | 5231685 | 5232235 | 551 | * | 6,136959 | 6,804360241 | 1,957824 | 3,475471 | 0,00051 | 1 | 0 | Distal Intergenic | 8 | 5133219 | 5155351 | 22133 | 2 | 20494 | ENSMUST00000023835.3 | -76334 | ENSMUSG00000023073 | Slc10a2 | solute carrier family 10, member 2 |
| chr15 | 99467909 | 99468685 | 777 | * | 8,447671 | 6,808292828 | 1,844244 | 3,691644 | 0,000223 | 0,25250467 | 1,425929 | Distal Intergenic | 15 | 99476936 | 99482428 | 5493 | 1 | 11827 | ENSMUST00000023752.6 | -8251 | ENSMUSG00000023013 | Aqp2 | aquaporin 2 |
| chr12 | 83914013 | 83914588 | 576 | * | 4,418812 | 6,809709115 | 2,060364 | 3,305101 | 0,000949 | 1 | 0 | Intron (ENSMUST0000( | 12 | 83878602 | 83923572 | 44971 | 2 | 18222 | ENSMUST00000125089.2 | 8984 | ENSMUSG00000021224 | Numb | NUMB endocytic adaptor protein |
| chr14 | 1,2E+08 | 1,2E+08 | 591 | * | 5,203018 | 6,810147066 | 2,01653 | 3,377161 | 0,000732 | 1 | 0 | Intron (ENSMUST0000( | 14 | 1,2E+08 | 1,2E+08 | 2340 | 1 | 75432 | ENSMUST00000188582.2 | 13774 | ENSMUSG00000100274 | 1700006F04Rik | RIKEN cDNA 1700006F04 gene |
| chr10 | 42154820 | 42155650 | 831 | * | 5,180224 | 6,810238667 | 2,008295 | 3,391054 | 0,000696 | 1 | 0 | Promoter (2-3kb) | 10 | 42123427 | 42152751 | 29325 | 2 | 56484 | ENSMUST00000105501.2 | -2069 | ENSMUSG00000048756 | Foxo3 | forkhead box O3 |
| chrX | 12181178 | 12181548 | 371 | * | 3,685881 | 6,811802659 | 2,310063 | 2,948751 | 0,003191 | 1 | 0 | Distal Intergenic | 20 | 12206876 | 12211461 | 4586 | 2 | 1E+08 | ENSMUST00000137281.2 | 29913 | ENSMUSG00000087591 | Gm14635 | predicted gene 14635 |
| chr9 | 64808706 | 64809295 | 590 | * | 6,799338 | 6,812090322 | 1,879102 | 3,625184 | 0,000289 | 1 | 0 | Intron (ENSMUST0000( | 9 | 64814479 | 64819393 | 4915 | 1 | 102442 | ENSMUST00000214029.2 | -5184 | ENSMUSG00000053641 | Dennd4a | DENN/MADD domain containing 4A |
| chr18 | 42652846 | 42653432 | 587 | * | 3,556798 | 6,812227433 | 2,422425 | 2,812153 | 0,004921 | 1 | 0 | Intron (ENSMUST0000( | 18 | 42657004 | 42669354 | 12351 | 1 | 56070 | ENSMUST00000237390.2 | -3572 | ENSMUSG00000024498 | Tcerg1 | transcription elongation regulator 1 (CA150) |
| chr9 | 65265125 | 65265794 | 670 | * | 6,836675 | 6,812309343 | 1,867519 | 3,647786 | 0,000265 | 1 | 0 | Promoter (2-3kb) | 9 | 65268323 | 65273930 | 5608 | 1 | 208080 | ENSMUST00000217103.2 | -2529 | ENSMUSG00000086228 | Ubap1l | ubiquitin-associated protein 1-like |
| chr10 | 49108413 | 49109066 | 654 | * | 3,785671 | 6,812448998 | 2,496907 | 2,728355 | 0,006365 | 1 | 0 | Intron (ENSMUST0000( | 10 | 49095518 | 49101749 | 6232 | 2 | 319306 | ENSMUST00000180729.2 | -6664 | | A730099G02Rik | RIKEN cDNA A730099G02 gene |
| chr11 | 75477163 | 75477738 | 576 | * | 4,116278 | 6,813963342 | 2,283531 | 2,983959 | 0,002845 | 1 | 0 | Promoter (1-2kb) | 11 | 75478923 | 75519630 | 40708 | 1 | 18738 | ENSMUST00000143219.8 | -1185 | ENSMUSG00000017781 | Pitpna | phosphatidylinositol transfer protein, alpha |
| chr3 | 82038218 | 82038571 | 354 | * | 5,121262 | 6,815654996 | 2,130967 | 3,198385 | 0,001382 | 1 | 0 | Intron (ENSMUST0000( | 3 | 81999734 | 82052380 | 52647 | 2 | 60596 | ENSMUST00000193924.6 | 13809 | ENSMUSG00000033910 | Gucy1a1 | guanylate cyclase 1, soluble, alpha 1 |
| chrX | 52510915 | 52511304 | 390 | * | 3,38319 | 6,817975856 | 2,439136 | 2,795242 | 0,005186 | 1 | 0 | Distal Intergenic | 20 | 52523795 | 52532453 | 8659 | 1 | 69501 | ENSMUST00000074232.7 | -12491 | ENSMUSG00000060967 | Etd | embryonic testis differentiation |
| chr12 | 37782661 | 37783367 | 707 | * | 3,746057 | 6,818997883 | 2,341419 | 2,912336 | 0,003587 | 1 | 0 | Distal Intergenic | 12 | 37867725 | 38032009 | 164285 | 1 | 217480 | ENSMUST00000222337.2 | -84358 | ENSMUSG00000036095 | Dgkb | diacylglycerol kinase, beta |
| chr9 | 21712519 | 21712909 | 391 | * | 4,270968 | 6,820205312 | 2,246475 | 3,035959 | 0,002398 | 1 | 0 | Promoter (2-3kb) | 9 | 21713021 | 21715297 | 2277 | 2 | 319899 | ENSMUST00000217310.2 | 2388 | ENSMUSG00000032198 | Dock6 | dedicator of cytokinesis 6 |
| chr10 | 32133246 | 32133875 | 630 | * | 6,328258 | 6,820309508 | 1,900107 | 3,589434 | 0,000331 | 1 | 0 | Intron (ENSMUST0000( | 10 | 32205761 | 32286331 | 80571 | 2 | 432450 | ENSMUST00000092603.5 | 152456 | ENSMUSG00000069670 | Nkain2 | Na+/K+ transporting ATPase interacting 2 |
| chr18 | 39073366 | 39073939 | 574 | * | 6,410967 | 6,822560066 | 1,877466 | 3,633919 | 0,000279 | 1 | 0 | Intron (ENSMUST0000( | 18 | 38979630 | 39062525 | 82896 | 2 | 14164 | ENSMUST00000235922.2 | -10841 | ENSMUSG00000036585 | Fgf1 | fibroblast growth factor 1 |
| chrX | 14477423 | 14478198 | 776 | * | 6,301065 | 6,822817556 | 2,096961 | 3,253669 | 0,001139 | 1 | 0 | Distal Intergenic | 20 | 14181640 | 14181830 | 191 | 2 | 1,15E+08 | ENSMUST00000082838.3 | -295593 | ENSMUSG00000064772 | Gm25202 | predicted gene, 25202 |
| chr15 | 6930305 | 6930718 | 414 | * | 5,288747 | 6,823015481 | 2,088155 | 3,267485 | 0,001085 | 1 | 0 | Distal Intergenic | 15 | 6844518 | 6904450 | 59933 | 2 | 18414 | ENSMUST00000176826.2 | -25855 | ENSMUSG00000022146 | Osmr | oncostatin M receptor |
| chr12 | 14967008 | 14967268 | 261 | * | 3,396162 | 6,823139239 | 2,575092 | 2,649668 | 0,008057 | 1 | 0 | Distal Intergenic | 12 | 15046390 | 15100390 | 54001 | 2 | 73972 | ENSMUST00000221781.2 | 133122 | ENSMUSG00000113644 | 4930448C13Rik | RIKEN cDNA 4930448C13 gene |
| chr7 | 1,43E+08 | 1,43E+08 | 371 | * | 4,108237 | 6,823256699 | 2,278076 | 2,995184 | 0,002743 | 1 | 0 | Intron (ENSMUST0000( | 7 | 1,43E+08 | 1,43E+08 | 5578 | 2 | 1,15E+08 | ENSMUST00000185637.2 | -21364 | ENSMUSG00000101062 | Gm28821 | predicted gene 28821 |
| chr19 | 37314245 | 37314739 | 495 | * | 4,322966 | 6,823547777 | 2,213058 | 3,083311 | 0,002047 | 1 | 0 | Promoter (<=1kb) | 19 | 37281049 | 37315251 | 34203 | 2 | 15925 | ENSMUST00000154308.2 | 512 | ENSMUSG00000056999 | Ide | insulin degrading enzyme |
| chr16 | 27208073 | 27208633 | 561 | * | 4,776245 | 6,823942548 | 2,134847 | 3,196456 | 0,001391 | 1 | 0 | Promoter (<=1kb) | 16 | 27208686 | 27257099 | 48414 | 1 | 67501 | ENSMUST00000143823.2 | -53 | ENSMUSG00000038127 | Ccdc50 | coiled-coil domain containing 50 |
| chr8 | 86470591 | 86471189 | 599 | * | 4,968393 | 6,824410928 | 2,060547 | 3,311941 | 0,000927 | 1 | 0 | Promoter (1-2kb) | 8 | 86457873 | 86469147 | 11275 | 2 | 71927 | ENSMUST00000210572.2 | -1444 | ENSMUSG00000031703 | Itfg1 | integrin alpha FG-GAP repeat containing 1 |
| chr9 | 57214622 | 57215408 | 787 | * | 5,600382 | 6,82515806 | 1,968945 | 3,466403 | 0,000527 | 1 | 0 | Distal Intergenic | 9 | 57160400 | 57169895 | 9496 | 2 | 74211 | ENSMUST00000034846.7 | -44727 | ENSMUSG00000032300 | 1700017B05Rik | RIKEN cDNA 1700017B05 gene |
| chr9 | 61604290 | 61604892 | 603 | * | 5,288892 | 6,825197609 | 1,966283 | 3,471117 | 0,000518 | 1 | 0 | Distal Intergenic | 9 | 61417005 | 61441529 | 24525 | 1 | 1,03E+08 | ENSMUST00000214104.2 | 187285 | ENSMUSG00000111381 | Gm34424 | predicted gene, 34424 |
| chr18 | 38652840 | 38653573 | 734 | * | 6,44406 | 6,825529174 | 1,990572 | 3,428929 | 0,000606 | 1 | 0 | Distal Intergenic | 18 | 38584363 | 38597436 | 13074 | 1 | 65113 | ENSMUST00000238135.2 | 68477 | ENSMUSG00000024425 | Ndfip1 | Nedd4 family interacting protein 1 |
| chr7 | 1,02E+08 | 1,02E+08 | 545 | * | 7,448934 | 6,826027001 | 1,890649 | 3,610416 | 0,000306 | 1 | 0 | Promoter (1-2kb) | 7 | 1,02E+08 | 1,02E+08 | 10966 | 2 | 56212 | ENSMUST00000106923.2 | 1015 | ENSMUSG00000073982 | Rhog | ras homolog family member G |
| chr15 | 3975686 | 3976144 | 459 | * | 3,158532 | 6,827106733 | 2,611353 | 2,614394 | 0,008939 | 1 | 0 | Distal Intergenic | 15 | 4000766 | 4001464 | 699 | 2 | 106052 | ENSMUST00000148169.2 | 25320 | ENSMUSG00000022184 | Fbxo4 | F-box protein 4 |
| chr13 | 78353147 | 78353632 | 486 | * | 3,467795 | 6,82869058 | 2,319449 | 2,9441 | 0,003239 | 1 | 0 | Promoter (2-3kb) | 13 | 78350672 | 78383779 | 33108 | 1 | 320174 | ENSMUST00000134705.3 | 2475 | ENSMUSG00000087143 | A830082K12Rik | RIKEN cDNA A830082K12 gene |
| chr14 | 57047590 | 57048681 | 1092 | * | 6,301787 | 6,83156318 | 1,902289 | 3,591233 | 0,000329 | 1 | 0 | Promoter (<=1kb) | 14 | 57028045 | 57048921 | 20877 | 2 | 219105 | ENSMUST00000173954.3 | 240 | ENSMUSG00000040123 | Zmym5 | zinc finger, MYM-type 5 |
| chr10 | 56710466 | 56711400 | 935 | * | 5,994199 | 6,831795857 | 1,959729 | 3,486093 | 0,00049 | 1 | 0 | Distal Intergenic | 10 | 56846178 | 56858295 | 12118 | 2 | 1,03E+08 | ENSMUST00000241975.1 | 146895 | | Gm36827 | predicted gene, 36827 |
| chr1 | 1,64E+08 | 1,64E+08 | 836 | * | 5,894277 | 6,831884313 | 1,986407 | 3,439317 | 0,000583 | 1 | 0 | Distal Intergenic | 1 | 1,64E+08 | 1,64E+08 | 11574 | 1 | 1,03E+08 | ENSMUST00000181831.3 | 51900 | ENSMUSG00000097281 | Gm26685 | predicted gene, 26685 |
| chr7 | 1,01E+08 | 1,01E+08 | 662 | * | 6,286881 | 6,832130941 | 2,008115 | 3,402262 | 0,000668 | 1 | 0 | Intron (ENSMUST0000( | 7 | 1,01E+08 | 1,01E+08 | 12955 | 2 | 1,03E+08 | ENSMUST00000209330.2 | -3150 | | Gm35363 | predicted gene, 35363 |
| chrX | 23293574 | 23294266 | 693 | * | 3,952077 | 6,832750381 | 2,479645 | 2,755536 | 0,00586 | 1 | 0 | Distal Intergenic | 20 | 23085510 | 23231321 | 145812 | 2 | 67455 | ENSMUST00000115313.8 | -62253 | ENSMUSG00000036782 | Klhl13 | kelch-like 13 |
| chr5 | 1,24E+08 | 1,24E+08 | 638 | * | 5,616612 | 6,833031645 | 1,959983 | 3,486272 | 0,00049 | 1 | 0 | Intron (ENSMUST0000( | 5 | 1,24E+08 | 1,24E+08 | 16709 | 2 | 56325 | ENSMUST00000153433.8 | 7522 | ENSMUSG00000029408 | Abcb9 | ATP-binding cassette, sub-family B (MDR/TAP), member 9 |
| chr7 | 24807268 | 24807763 | 496 | * | 6,636156 | 6,833306587 | 1,94576 | 3,511896 | 0,000445 | 1 | 0 | Intron (ENSMUST0000( | 7 | 24796491 | 24796600 | 110 | 2 | 1,02E+08 | ENSMUST00000184150.3 | -10668 | ENSMUSG00000098441 | Mir6537 | microRNA 6537 |
| chr9 | 1,12E+08 | 1,12E+08 | 622 | * | 7,146654 | 6,834091332 | 1,911416 | 3,575407 | 0,00035 | 1 | 0 | Distal Intergenic | 9 | 1,12E+08 | 1,12E+08 | 6356 | 2 | 78028 | ENSMUST00000215688.2 | 87837 | ENSMUSG00000111054 | 4930545L08Rik | RIKEN cDNA 4930545L08 gene |
| chr4 | 1,41E+08 | 1,41E+08 | 751 | * | 7,894925 | 6,834741872 | 1,885988 | 3,623959 | 0,00029 | 1 | 0 | Distal Intergenic | 4 | 1,41E+08 | 1,41E+08 | 6420 | 2 | 74202 | ENSMUST00000137491.2 | 21123 | ENSMUSG00000006219 | Fblim1 | filamin binding LIM protein 1 |
| chr4 | 9299476 | 9299879 | 404 | * | 4,297906 | 6,834771397 | 2,26954 | 3,011522 | 0,002599 | 1 | 0 | Intron (ENSMUST0000( | 4 | 9311756 | 9429803 | 118048 | 1 | 74438 | ENSMUST00000141085.8 | -11877 | ENSMUSG00000041216 | Clvs1 | clavesin 1 |
| chr1 | 1,56E+08 | 1,56E+08 | 639 | * | 4,477633 | 6,836542209 | 2,145307 | 3,186744 | 0,001439 | 1 | 0 | Intron (ENSMUST0000( | 1 | 1,56E+08 | 1,56E+08 | 1630 | 1 | 240832 | ENSMUST00000125371.2 | 23252 | ENSMUSG00000050565 | Tor1aip2 | torsin A interacting protein 2 |
| chrX | 89250354 | 89251022 | 669 | * | 5,220346 | 6,837673915 | 2,083781 | 3,281379 | 0,001033 | 1 | 0 | Distal Intergenic | 20 | 88973704 | 88976458 | 2755 | 2 | 245511 | ENSMUST00000088146.3 | -273896 | ENSMUSG00000035387 | 4930415L06Rik | RIKEN cDNA 4930415L06 gene |
| chr2 | 1,09E+08 | 1,09E+08 | 302 | * | 4,718594 | 6,838855396 | 2,088005 | 3,275306 | 0,001055 | 1 | 0 | Distal Intergenic | 2 | 1,09E+08 | 1,09E+08 | 2758 | 1 | 1,03E+08 | ENSMUST00000153283.2 | -179025 | | Gm13923 | predicted gene 13923 |
| chr1 | 63939459 | 63940335 | 877 | * | 5,370079 | 6,839065666 | 2,02027 | 3,385224 | 0,000711 | 1 | 0 | Distal Intergenic | 1 | 64003886 | 64008158 | 4273 | 1 | 433315 | ENSMUST00000133244.2 | -63551 | ENSMUSG00000044689 | Gm13749 | predicted gene 13749 |
| chr11 | 77532387 | 77532728 | 342 | * | 3,763213 | 6,843416686 | 2,346597 | 2,916316 | 0,003542 | 1 | 0 | Distal Intergenic | 11 | 77549402 | 77549533 | 132 | 2 | 1,15E+08 | ENSMUST00000104254.3 | 16805 | ENSMUSG00000118748 | Gm23132 | predicted gene, 23132 |
| chr4 | 1,24E+08 | 1,24E+08 | 501 | * | 4,363118 | 6,844610972 | 2,176389 | 3,144939 | 0,001661 | 1 | 0 | Promoter (<=1kb) | 4 | 1,24E+08 | 1,24E+08 | 1905 | 1 | 54170 | ENSMUST00000143635.2 | -246 | ENSMUSG00000028646 | Rragc | Ras-related GTP binding C |
| chr1 | 1,35E+08 | 1,35E+08 | 1017 | * | 6,583034 | 6,845285704 | 1,858411 | 3,683408 | 0,00023 | 1 | 0 | Distal Intergenic | 1 | 1,35E+08 | 1,35E+08 | 95153 | 1 | 20980 | ENSMUST00000187725.7 | -6381 | ENSMUSG00000026452 | Syt2 | synaptotagmin II |
| chr3 | 65790238 | 65790817 | 580 | * | 5,267444 | 6,845947309 | 1,936878 | 3,534526 | 0,000409 | 1 | 0 | Distal Intergenic | 3 | 65854203 | 65855419 | 1217 | 2 | 56706 | ENSMUST00000149729.2 | 64602 | ENSMUSG00000027829 | Ccnl1 | cyclin L1 |
| chr10 | 30459306 | 30460383 | 1078 | * | 5,606486 | 6,846553094 | 2,002035 | 3,419797 | 0,000627 | 1 | 0 | Intron (ENSMUST0000( | 10 | 30435117 | 30466105 | 30989 | 2 | 73681 | ENSMUST00000215041.2 | 5722 | ENSMUSG00000019792 | Trmt11 | tRNA methyltransferase 11 |
| chr5 | 1,05E+08 | 1,05E+08 | 390 | * | 3,647095 | 6,847936554 | 2,453296 | 2,791321 | 0,005249 | 1 | 0 | Intron (ENSMUST0000( | 5 | 1,05E+08 | 1,05E+08 | 14615 | 2 | 626391 | ENSMUST00000089906.7 | -15107 | ENSMUSG00000072774 | Zfp951 | zinc finger protein 951 |
| chr18 | 21133302 | 21134364 | 1063 | * | 6,349973 | 6,848425166 | 1,896441 | 3,611198 | 0,000305 | 1 | 0 | Promoter (<=1kb) | 18 | 21134398 | 21160592 | 26195 | 1 | 56515 | ENSMUST00000234107.2 | -34 | ENSMUSG00000024317 | Rnf138 | ring finger protein 138 |
| chr7 | 1,27E+08 | 1,27E+08 | 509 | * | 6,412453 | 6,84870354 | 1,883778 | 3,635621 | 0,000277 | 1 | 0 | Promoter (<=1kb) | 7 | 1,27E+08 | 1,27E+08 | 2632 | 1 | 101502 | ENSMUST00000136823.8 | 106 | ENSMUSG00000042289 | Hsd3b7 | hydroxy-delta-5-steroid dehydrogenase, 3 beta- and steroid delta-isomerase 7 |
| chrX | 1,08E+08 | 1,08E+08 | 690 | * | 4,637986 | 6,848959545 | 2,134313 | 3,208976 | 0,001332 | 1 | 0 | Intron (ENSMUST0000( | 20 | 1,08E+08 | 1,08E+08 | 92730 | 1 | 69964 | ENSMUST00000207966.2 | -7303 | ENSMUSG00000086629 | 2810403D21Rik | RIKEN cDNA 2810403D21 gene |
| chr4 | 78539533 | 78540108 | 576 | * | 3,793678 | 6,852864836 | 2,415001 | 2,837624 | 0,004545 | 1 | 0 | Distal Intergenic | 4 | 78130313 | 78133846 | 3534 | 1 | 1,03E+08 | ENSMUST00000246656.1 | 409220 | | B230208B08Rik | RIKEN cDNA B230208B08 gene |
| chr8 | 18840695 | 18841276 | 582 | * | 4,616795 | 6,853455063 | 2,049914 | 3,343289 | 0,000828 | 1 | 0 | Intron (ENSMUST0000( | 8 | 18740279 | 18791578 | 51300 | 2 | 11601 | ENSMUST00000033846.7 | -49117 | ENSMUSG00000031465 | Angpt2 | angiopoietin 2 |
| chr7 | 23896481 | 23897345 | 865 | * | 7,127695 | 6,854036834 | 1,888511 | 3,629334 | 0,000284 | 1 | 0 | 3' UTR | 7 | 23896302 | 23903008 | 6707 | 2 | 56707 | ENSMUST00000056683.3 | 5663 | ENSMUSG00000087598 | Zfp111 | zinc finger protein 111 |
| chr9 | 88317411 | 88318065 | 655 | * | 5,001932 | 6,85430212 | 2,056218 | 3,333451 | 0,000858 | 1 | 0 | Promoter (2-3kb) | 9 | 88258805 | 88320951 | 62147 | 2 | 244962 | ENSMUST00000173011.9 | 2886 | ENSMUSG00000032422 | Snx14 | sorting nexin 14 |
| chr8 | 7828342 | 7828929 | 588 | * | 5,931444 | 6,854456598 | 1,972202 | 3,475535 | 0,00051 | 1 | 0 | Distal Intergenic | 8 | 7688764 | 7688892 | 129 | 1 | 1,15E+08 | ENSMUST00000104190.3 | 139578 | ENSMUSG00000077378 | Gm25169 | predicted gene, 25169 |
| chr7 | 1,43E+08 | 1,43E+08 | 681 | * | 5,897892 | 6,854725404 | 1,961901 | 3,493919 | 0,000476 | 1 | 0 | Promoter (1-2kb) | 7 | 1,43E+08 | 1,43E+08 | 12817 | 2 | 17955 | ENSMUST00000208093.2 | 1431 | ENSMUSG00000059119 | Nap1l4 | nucleosome assembly protein 1-like 4 |
| chr10 | 1E+08 | 1E+08 | 1591 | * | 9,532309 | 6,854937853 | 1,755783 | 3,904207 | 9,45E-05 | 0,161408969 | 1,360622 | Intron (ENSMUST0000( | 10 | 1E+08 | 1E+08 | 10628 | 2 | 237500 | ENSMUST00000128009.2 | 17640 | ENSMUSG00000036676 | Tmtc3 | transmembrane and tetratricopeptide repeat containing 3 |
| chr9 | 1,03E+08 | 1,03E+08 | 991 | * | 9,447111 | 6,855116583 | 1,738016 | 3,944219 | 8,01E-05 | 0,14233608 | 1,425929 | Distal Intergenic | 9 | 1,03E+08 | 1,03E+08 | 72626 | 1 | 20187 | ENSMUST00000175883.8 | -6100 | ENSMUSG00000032547 | Ryk | receptor-like tyrosine kinase |
| chr9 | 75311266 | 75311937 | 672 | * | 5,923689 | 6,8565998 | 2,179056 | 3,146591 | 0,001652 | 1 | 0 | Intron (ENSMUST0000( | 9 | 75294184 | 75316641 | 22458 | 2 | 50772 | ENSMUST00000168937.8 | 4704 | ENSMUSG00000042688 | Mapk6 | mitogen-activated protein kinase 6 |
| chr5 | 1,23E+08 | 1,23E+08 | 698 | * | 5,042851 | 6,860232143 | 2,049493 | 3,347282 | 0,000816 | 1 | 0 | Promoter (<=1kb) | 5 | 1,23E+08 | 1,23E+08 | 33881 | 2 | 59008 | ENSMUST00000086216.9 | -963 | ENSMUSG00000029472 | Anapc5 | anaphase-promoting complex subunit 5 |
| chr11 | 78644662 | 78646007 | 1346 | * | 9,905685 | 6,860694885 | 1,77822 | 3,858181 | 0,000114 | 0,173497184 | 1,449706 | Promoter (2-3kb) | 11 | 78641332 | 78642555 | 1224 | 2 | 69109 | ENSMUST00000059468.6 | -2107 | ENSMUSG00000049489 | Ccnq | cyclin Q |
| chr3 | 1,06E+08 | 1,06E+08 | 962 | * | 6,103134 | 6,861233189 | 1,964306 | 3,492955 | 0,000478 | 1 | 0 | Intron (ENSMUST0000( | 3 | 1,06E+08 | 1,06E+08 | 72643 | 2 | 109905 | ENSMUST00000197094.5 | 23393 | ENSMUSG00000068798 | Rap1a | RAS-related protein 1a |
| chr7 | 28055061 | 28055692 | 632 | * | 5,670369 | 6,86192726 | 1,91694 | 3,579625 | 0,000344 | 1 | 0 | Distal Intergenic | 7 | 28050933 | 28051057 | 125 | 1 | 1E+08 | ENSMUST00000104523.3 | 4128 | | AF357399 | snoRNA AF357399 |
| chr3 | 83580095 | 83580903 | 809 | * | 6,010466 | 6,862413157 | 1,888492 | 3,633805 | 0,000279 | 1 | 0 | Distal Intergenic | 3 | 83609559 | 83609665 | 107 | 1 | 1,15E+08 | ENSMUST00000240217.1 | -28656 | ENSMUSG00000119833 | Gm24584 | predicted gene, 24584 |
| chr18 | 40479603 | 40480029 | 427 | * | 4,350249 | 6,864000405 | 2,287631 | 3,000484 | 0,002696 | 1 | 0 | Exon (ENSMUST000002 | 18 | 40541138 | 40664683 | 123546 | 1 | 383348 | ENSMUST00000236301.2 | -61109 | ENSMUSG00000051401 | Kctd16 | potassium channel tetramerisation domain containing 16 |
| chr19 | 42016531 | 42017620 | 1090 | * | 9,36768 | 6,870190998 | 1,750693 | 3,924269 | 8,7E-05 | 0,149914039 | 1,425929 | Intron (ENSMUST0000( | 19 | 42024439 | 42033549 | 9111 | 1 | 56642 | ENSMUST00000238137.2 | -6819 | ENSMUSG00000025172 | Ankrd2 | ankyrin repeat domain 2 (stretch responsive muscle) |
| chrX | 1,67E+08 | 1,67E+08 | 553 | * | 5,354813 | 6,870658718 | 2,20222 | 3,119878 | 0,001809 | 1 | 0 | Intron (ENSMUST0000( | 20 | 1,66E+08 | 1,67E+08 | 426246 | 2 | 333605 | ENSMUST00000112145.3 | -96475 | ENSMUSG00000049176 | Frmpd4 | FERM and PDZ domain containing 4 |
| chr11 | 88619638 | 88620369 | 732 | * | 4,356446 | 6,870796937 | 2,192096 | 3,134351 | 0,001722 | 1 | 0 | Exon (ENSMUST000001 | 11 | 88611315 | 88614999 | 3685 | 1 | 193280 | ENSMUST00000125967.2 | 8323 | | C030037D09Rik | RIKEN cDNA C030037D09 gene |
| chr16 | 7216837 | 7217644 | 808 | * | 7,341406 | 6,87209313 | 1,820824 | 3,774167 | 0,000161 | 1 | 0 | Intron (ENSMUST0000( | 16 | 7170834 | 7227833 | 57000 | 1 | 268859 | ENSMUST00000231031.2 | 46003 | ENSMUSG00000008658 | Rbfox1 | RNA binding protein, fox-1 homolog (C. elegans) 1 |

| | | | | | | | | | | | | | | | | | | | | | | | |
|---|---|---|---|---|---|---|---|---|---|---|---|---|---|---|---|---|---|---|---|---|---|---|---|
| chr5 | 44494609 | 44494973 | 365 | * | 3,772667 | 6,873483971 | 2,205227 | 3,116905 | 0,001828 | 1 | 0 | Distal Intergenic | 5 | 44570135 | 44584934 | 14800 | 2 | 1,03E+08 | ENSMUST00000199005.2 | 89961 | ENSMUSG00000104800 | Gm36017 | predicted gene, 36017 |
| chr15 | 79539298 | 79540081 | 784 | * | 6,033241 | 6,873561516 | 2,050937 | 3,351425 | 0,000804 | 1 | 0 | Promoter (2-3kb) | 15 | 79496572 | 79542950 | 46379 | 2 | 75729 | ENSMUST00000109646.9 | 2869 | ENSMUSG00000042564 | Fam227a | family with sequence similarity 227, member A |
| chrX | 69382748 | 69383502 | 755 | * | 4,425003 | 6,873895957 | 2,128968 | 3,228746 | 0,001243 | 1 | 0 | Distal Intergenic | 20 | 69394676 | 69404697 | 10022 | 2 | 15931 | ENSMUST00000130519.2 | 21195 | ENSMUSG00000035847 | Ids | iduronate 2-sulfatase |
| chr5 | 31006236 | 31006962 | 727 | * | 6,190938 | 6,876485693 | 1,864029 | 3,689045 | 0,000225 | 1 | 0 | Intron (ENSMUST0000( | 5 | 31011143 | 31020702 | 9560 | 1 | 100732 | ENSMUST00000202501.2 | -4181 | ENSMUSG00000029166 | Mapre3 | microtubule-associated protein, RP/EB family, member 3 |
| chr17 | 73392421 | 73392765 | 345 | * | 4,011968 | 6,876520096 | 2,2265 | 3,088489 | 0,002012 | 1 | 0 | Distal Intergenic | 17 | 73414980 | 73468986 | 54007 | 1 | 225010 | ENSMUST00000130574.4 | -22215 | ENSMUSG00000054469 | Lclat1 | lysocardiolipin acyltransferase 1 |
| chrX | 49772032 | 49772642 | 611 | * | 3,787472 | 6,878000287 | 2,362089 | 2,911829 | 0,003593 | 1 | 0 | Distal Intergenic | 20 | 49647012 | 49724198 | 77187 | 2 | 103012 | ENSMUST00000143999.9 | -47834 | ENSMUSG00000085396 | Firre | functional intergenic repeating RNA element |
| chr19 | 37583139 | 37583564 | 426 | * | 5,85266 | 6,879978373 | 2,132162 | 3,226762 | 0,001252 | 1 | 0 | Intron (ENSMUST0000( | 19 | 37538907 | 37671703 | 132797 | 1 | 107371 | ENSMUST00000238817.2 | 44232 | ENSMUSG00000053799 | Exoc6 | exocyst complex component 6 |
| chr9 | 1,08E+08 | 1,08E+08 | 986 | * | 7,806235 | 6,880176345 | 1,930758 | 3,56346 | 0,000366 | 0,322374461 | 1,360622 | Intron (ENSMUST0000( | 9 | 1,08E+08 | 1,08E+08 | 24088 | 1 | 11848 | ENSMUST00000193490.2 | 5476 | ENSMUSG00000007815 | Rhoa | ras homolog family member A |
| chr7 | 44648080 | 44648440 | 361 | * | 5,099148 | 6,880291128 | 2,106984 | 3,26547 | 0,001093 | 1 | 0 | Promoter (<=1kb) | 7 | 44640646 | 44648136 | 7491 | 2 | 75736 | ENSMUST00000208128.2 | 0 | ENSMUSG00000003190 | Bcl2l12 | BCL2-like 12 (proline rich) |
| chrX | 97285028 | 97285856 | 829 | * | 5,46621 | 6,880733046 | 2,072231 | 3,320447 | 0,000899 | 1 | 0 | Intron (ENSMUST0000( | 20 | 97192375 | 97366821 | 174447 | 1 | 11835 | ENSMUST00000052837.9 | 92653 | ENSMUSG00000046532 | Ar | androgen receptor |
| chr11 | 25255974 | 25256538 | 565 | * | 5,426333 | 6,881742117 | 1,942012 | 3,543614 | 0,000395 | 1 | 0 | Intron (ENSMUST0000( | 11 | 25251263 | 25272786 | 21524 | 1 | 1,03E+08 | ENSMUST00000145127.2 | 4711 | | Gm12069 | predicted gene 12069 |
| chrX | 95937552 | 95938178 | 627 | * | 5,286 | 6,881921581 | 2,03884 | 3,37541 | 0,000737 | 1 | 0 | Distal Intergenic | 20 | 96116202 | 96125710 | 9509 | 1 | 245526 | ENSMUST00000033556.4 | -178024 | ENSMUSG00000031212 | Pgr15l | G protein-coupled receptor 15-like |
| chr1 | 37276494 | 37276970 | 477 | * | 4,575549 | 6,882168013 | 2,236698 | 3,076932 | 0,002091 | 1 | 0 | Intron (ENSMUST0000( | 1 | 37258286 | 37302465 | 44180 | 1 | 12790 | ENSMUST00000027288.10 | 18208 | ENSMUSG00000026114 | Cnga3 | cyclic nucleotide gated channel alpha 3 |
| chr3 | 1,23E+08 | 1,23E+08 | 583 | * | 4,140922 | 6,882904409 | 2,385169 | 2,885709 | 0,003905 | 1 | 0 | Exon (ENSMUST00000 | 3 | 1,23E+08 | 1,23E+08 | 27342 | 2 | 99526 | ENSMUST00000199923.5 | -10988 | ENSMUSG00000039701 | Usp53 | ubiquitin specific peptidase 53 |
| chr19 | 40821957 | 40822705 | 749 | * | 6,249567 | 6,88330045 | 1,927517 | 3,571072 | 0,000356 | 1 | 0 | Promoter (2-3kb) | 19 | 40819752 | 40836453 | 16702 | 1 | 240665 | ENSMUST00000119316.2 | 2205 | ENSMUSG00000025010 | Ccnj | cyclin J |
| chr13 | 1,08E+08 | 1,08E+08 | 591 | * | 6,365372 | 6,88431241 | 1,886669 | 3,648924 | 0,000263 | 1 | 0 | Distal Intergenic | 13 | 1,08E+08 | 1,08E+08 | 71280 | 1 | 74559 | ENSMUST00000022207.10 | -10584 | ENSMUSG00000021696 | Elovl7 | ELOVL family member 7, elongation of long chain fatty acids (yeast) |
| chr2 | 90335065 | 90336017 | 953 | * | 8,809207 | 6,887903109 | 1,782608 | 3,863948 | 0,000112 | 0,131142082 | 2,237403 | Intron (ENSMUST0000( | 2 | 90260100 | 90309514 | 49415 | 2 | 19271 | ENSMUST00000111495.9 | -25551 | ENSMUSG00000025314 | Ptprj | protein tyrosine phosphatase, receptor type, J |
| chr3 | 1,27E+08 | 1,27E+08 | 609 | * | 6,215441 | 6,888782529 | 2,006556 | 3,433138 | 0,000597 | 1 | 0 | 3' UTR | 3 | 1,27E+08 | 1,27E+08 | 6821 | 2 | 71481 | ENSMUST00000159720.2 | 15110 | ENSMUSG00000028028 | Alpk1 | alpha-kinase 1 |
| chr5 | 59257060 | 59257923 | 864 | * | 5,147589 | 6,889210541 | 1,948325 | 3,535966 | 0,000406 | 1 | 0 | Distal Intergenic | 5 | 59372289 | 59372395 | 107 | 2 | 1,15E+08 | ENSMUST00000239835.1 | 114472 | ENSMUSG00000119859 | Gm22308 | predicted gene, 22308 |
| chr5 | 1,32E+08 | 1,32E+08 | 918 | * | 8,115573 | 6,891010226 | 1,788122 | 3,853769 | 0,000116 | 0,174628563 | 1,449706 | Intron (ENSMUST0000( | 5 | 1,31E+08 | 1,32E+08 | 178819 | 2 | 319974 | ENSMUST00000161804.9 | -26926 | ENSMUSG00000029673 | Auts2 | autism susceptibility candidate 2 |
| chr3 | 46621960 | 46622266 | 307 | * | 3,626561 | 6,892314548 | 2,392149 | 2,881223 | 0,003961 | 1 | 0 | Distal Intergenic | 3 | 46396632 | 46402654 | 6023 | 2 | 241989 | ENSMUST00000192793.6 | -219306 | ENSMUSG00000090919 | Pabpc4l | poly(A) binding protein, cytoplasmic 4-like |
| chr11 | 30816060 | 30816485 | 426 | * | 4,032039 | 6,900773302 | 2,543641 | 2,712951 | 0,006669 | 1 | 0 | Promoter (<=1kb) | 11 | 30815469 | 30824204 | 8736 | 1 | 103554 | ENSMUST00000155719.2 | 591 | ENSMUSG00000040850 | Psme4 | proteasome (prosome, macropain) activator subunit 4 |
| chr11 | 77238439 | 77239040 | 602 | * | 5,775236 | 6,90212104 | 2,019593 | 3,41758 | 0,000632 | 1 | 0 | Promoter (<=1kb) | 11 | 77239106 | 77304461 | 65356 | 1 | 237860 | ENSMUST00000127890.9 | -66 | ENSMUSG00000037926 | Ssh2 | slingshot protein phosphatase 2 |
| chr3 | 97673123 | 97673515 | 393 | * | 4,658265 | 6,902451677 | 2,068014 | 3,337721 | 0,000845 | 1 | 0 | Promoter (1-2kb) | 3 | 97648133 | 97675247 | 27115 | 2 | 83679 | ENSMUST00000175751.8 | 1732 | ENSMUSG00000038170 | Pde4dip | phosphodiesterase 4D interacting protein (myomegalin) |
| chr12 | 85013466 | 85015720 | 2255 | * | 14,03942 | 6,905866678 | 1,664675 | 4,148477 | 3,35E-05 | 0,088115747 | 1,2763 | Promoter (<=1kb) | 12 | 84997856 | 85012804 | 14949 | 2 | 68497 | ENSMUST00000168713.2 | -662 | ENSMUSG00000042350 | Arel1 | apoptosis resistant E3 ubiquitin protein ligase 1 |
| chr6 | 1,13E+08 | 1,13E+08 | 377 | * | 3,176022 | 6,905999095 | 2,58758 | 2,668903 | 0,00761 | 1 | 0 | Promoter (1-2kb) | 6 | 1,13E+08 | 1,13E+08 | 4227 | 2 | 101206 | ENSMUST00000125414.2 | 1237 | ENSMUSG00000048930 | Tada3 | transcriptional adaptor 3 |
| chr1 | 1,16E+08 | 1,16E+08 | 447 | * | 4,261569 | 6,907136402 | 2,298387 | 3,00521 | 0,002654 | 1 | 0 | Intron (ENSMUST0000( | 1 | 1,16E+08 | 1,17E+08 | 902568 | 1 | 636808 | ENSMUST00000043725.9 | 22957 | ENSMUSG00000070695 | Cntnap5a | contactin associated protein-like 5A |
| chr7 | 1,27E+08 | 1,27E+08 | 969 | * | 8,478836 | 6,907291358 | 1,778414 | 3,883961 | 0,000103 | 1 | 0 | Promoter (<=1kb) | 7 | 1,27E+08 | 1,27E+08 | 2555 | 2 | 27973 | ENSMUST00000033074.8 | -955 | ENSMUSG00000096145 | Vkorc1 | vitamin K epoxide reductase complex, subunit 1 |
| chr3 | 13854557 | 13855733 | 1177 | * | 8,630386 | 6,907795775 | 1,810088 | 3,816277 | 0,000135 | 1 | 0 | Intron (ENSMUST0000( | 3 | 13841908 | 13842367 | 460 | 1 | 76897 | ENSMUST00000108373.4 | 12649 | ENSMUSG00000039717 | Ralyl | RALY RNA binding protein-like |
| chr11 | 86460466 | 86460925 | 460 | * | 4,705228 | 6,9085245 | 2,092638 | 3,301348 | 0,000962 | 1 | 0 | Distal Intergenic | 11 | 86474893 | 86474984 | 92 | 2 | 387140 | ENSMUST00000083521.3 | 14059 | ENSMUSG00000065455 | Mir21a | microRNA 21a |
| chr1 | 83886423 | 83886792 | 370 | * | 3,754611 | 6,912580644 | 2,268786 | 3,046819 | 0,002313 | 1 | 0 | Intron (ENSMUST0000( | 1 | 84015647 | 84030974 | 15328 | 2 | 98496 | ENSMUST00000175948.2 | 144182 | ENSMUSG00000045658 | Pid1 | phosphotyrosine interaction domain containing 1 |
| chr8 | 39122374 | 39122981 | 608 | * | 4,195392 | 6,913009383 | 2,22253 | 3,110424 | 0,001868 | 1 | 0 | Intron (ENSMUST0000( | 8 | 39126917 | 39138220 | 11304 | 1 | 1,05E+08 | ENSMUST00000210223.2 | -3936 | | Gm40493 | predicted gene, 40493 |
| chr9 | 60562667 | 60563062 | 396 | * | 4,559272 | 6,914172654 | 2,126142 | 3,25198 | 0,001146 | 1 | 0 | Intron (ENSMUST0000( | 9 | 60476455 | 60557084 | 80630 | 2 | 102747 | ENSMUST00000053171.14 | -5583 | ENSMUSG00000047766 | Lrrc49 | leucine rich repeat containing 49 |
| chr2 | 60684845 | 60685608 | 764 | * | 3,930804 | 6,914744935 | 2,603932 | 2,655501 | 0,007919 | 1 | 0 | Intron (ENSMUST0000( | 2 | 60582297 | 60711782 | 129486 | 2 | 56878 | ENSMUST00000164147.8 | 26174 | ENSMUSG00000026970 | Rbms1 | RNA binding motif, single stranded interacting protein 1 |
| chr5 | 45420249 | 45420690 | 442 | * | 4,478876 | 6,914865597 | 2,077138 | 3,329036 | 0,000871 | 1 | 0 | Intron (ENSMUST0000( | 5 | 45267971 | 45333035 | 65065 | 2 | 52401 | ENSMUST00000198493.2 | -87214 | ENSMUSG00000029092 | D5Ertd615e | DNA segment, Chr 5, ERATO Doi 615, expressed |
| chrX | 1,07E+08 | 1,07E+08 | 857 | * | 5,415323 | 6,91526964 | 2,052709 | 3,368851 | 0,000755 | 1 | 0 | Distal Intergenic | 20 | 1,07E+08 | 1,07E+08 | 1155 | 1 | 71011 | ENSMUST00000139758.2 | 19482 | ENSMUSG00000055700 | 4933401B06Rik | GLE1 RNA export mediator pseudogene |
| chr5 | 13753877 | 13754216 | 340 | * | 3,333238 | 6,915322434 | 2,524085 | 2,739734 | 0,006149 | 1 | 0 | Distal Intergenic | 5 | 13818676 | 13821901 | 3226 | 1 | 626053 | ENSMUST00000170514.2 | -64460 | | Gm6647 | predicted gene 6647 |
| chr7 | 67298099 | 67298742 | 644 | * | 5,463395 | 6,915972358 | 1,961671 | 3,525552 | 0,000423 | 1 | 0 | Promoter (<=1kb) | 7 | 67298898 | 67328820 | 29923 | 1 | 67009 | ENSMUST00000130681.8 | -156 | ENSMUSG00000030555 | Ttc23 | tetratricopeptide repeat domain 23 |
| chr9 | 8134557 | 8135076 | 520 | * | 5,30688 | 6,916314213 | 1,995844 | 3,465358 | 0,00053 | 1 | 0 | Promoter (<=1kb) | 9 | 8076462 | 8134295 | 57834 | 2 | 234915 | ENSMUST00000238782.2 | -262 | ENSMUSG00000040729 | Cep126 | centrosomal protein 126 |
| chr17 | 24973504 | 24974326 | 823 | * | 8,28922 | 6,916672607 | 1,879353 | 3,680347 | 0,000233 | 0,256539258 | 1,449706 | Promoter (1-2kb) | 17 | 24971990 | 24977622 | 5633 | 1 | 328779 | ENSMUST00000235111.2 | 1514 | ENSMUSG00000039628 | Hs3st6 | heparan sulfate (glucosamine) 3-O-sulfotransferase 6 |
| chr12 | 93976506 | 93977171 | 666 | * | 6,969774 | 6,916768181 | 1,850557 | 3,737668 | 0,000186 | 1 | 0 | Distal Intergenic | 12 | 93894112 | 93894238 | 127 | 1 | 1,02E+08 | ENSMUST00000183568.3 | 82394 | ENSMUSG00000098931 | Mir8099-2 | microRNA 8099-2 |
| chr15 | 5240643 | 5241221 | 579 | * | 6,261958 | 6,917559191 | 1,940531 | 3,564776 | 0,000364 | 1 | 0 | Intron (ENSMUST0000( | 15 | 5216835 | 5241524 | 24690 | 1 | 67515 | ENSMUST00000141167.2 | 23808 | ENSMUSG00000022151 | Ttc33 | tetratricopeptide repeat domain 33 |
| chr4 | 79082541 | 79083105 | 565 | * | 4,503281 | 6,918067458 | 2,249856 | 3,074893 | 0,002106 | 1 | 0 | Distal Intergenic | 4 | 78130313 | 78133846 | 3534 | 1 | 1,03E+08 | ENSMUST00000246656.1 | 952228 | | B230208B08Rik | RIKEN cDNA B230208B08 gene |
| chr12 | 15781782 | 15782235 | 454 | * | 4,65663 | 6,918938273 | 2,189481 | 3,160081 | 0,001577 | 1 | 0 | Distal Intergenic | 12 | 15767601 | 15778403 | 10803 | 1 | 1,05E+08 | ENSMUST00000221185.2 | 14181 | | Gm40287 | predicted gene, 40287 |
| chr11 | 1,17E+08 | 1,17E+08 | 674 | * | 6,053699 | 6,919539683 | 2,151655 | 3,215915 | 0,0013 | 1 | 0 | Promoter (<=1kb) | 11 | 1,17E+08 | 1,17E+08 | 110 | 1 | 1,02E+08 | ENSMUST00000196236.3 | 41 | ENSMUSG00000104740 | Mir6516 | microRNA 6516 |
| chr7 | 28656042 | 28656334 | 293 | * | 3,542292 | 6,920963839 | 2,388253 | 2,897919 | 0,003756 | 1 | 0 | Intron (ENSMUST0000( | 7 | 28598045 | 28661648 | 63604 | 2 | 60595 | ENSMUST00000127210.8 | 5314 | ENSMUSG00000054808 | Actn4 | actinin alpha 4 |
| chr16 | 69391149 | 69392264 | 1116 | * | 6,880332 | 6,921233471 | 1,827147 | 3,788 | 0,000152 | 1 | 0 | Distal Intergenic | 16 | 69593023 | 69597269 | 4247 | 2 | 78110 | ENSMUST00000201886.2 | 205005 | ENSMUSG00000107073 | 4930428D20Rik | RIKEN cDNA 4930428D20 gene |
| chr10 | 81439820 | 81440597 | 778 | * | 7,721077 | 6,92153619 | 1,84533 | 3,750839 | 0,000176 | 1 | 0 | Promoter (1-2kb) | 10 | 81442104 | 81456537 | 14434 | 1 | 268319 | ENSMUST00000126940.2 | -1507 | ENSMUSG00000074862 | BC025920 | zinc finger protein pseudogene |
| chr14 | 1,09E+08 | 1,09E+08 | 543 | * | 3,400646 | 6,924881317 | 2,56447 | 2,700317 | 0,006927 | 1 | 0 | Distal Intergenic | 14 | 1,09E+08 | 1,09E+08 | 5359 | 2 | 76965 | ENSMUST00000100322.4 | 7250 | ENSMUSG00000075478 | Slitrk1 | SLIT and NTRK-like family, member 1 |
| chr14 | 1,1E+08 | 1,1E+08 | 454 | * | 3,121808 | 6,928619104 | 2,680868 | 2,584469 | 0,009753 | 1 | 0 | Distal Intergenic | 14 | 1,1E+08 | 1,1E+08 | 130 | 2 | 1,15E+08 | ENSMUST00000158494.3 | 340064 | ENSMUSG00000089119 | Gm25901 | predicted gene, 25901 |
| chr3 | 12543669 | 12544402 | 734 | * | 6,863363 | 6,929907888 | 1,825365 | 3,796451 | 0,000147 | 1 | 0 | Distal Intergenic | 3 | 11888289 | 11909419 | 21131 | 2 | 1E+08 | ENSMUST00000192022.2 | -634250 | ENSMUSG00000104015 | Gm10745 | predicted gene 10745 |
| chr15 | 81799500 | 81800015 | 516 | * | 5,659728 | 6,930097423 | 1,982345 | 3,495909 | 0,000472 | 1 | 0 | 3' UTR | 15 | 81794490 | 81797494 | 3005 | 1 | 11429 | ENSMUST00000144324.2 | 5010 | ENSMUSG00000022477 | Aco2 | aconitase 2, mitochondrial |
| chrX | 75542154 | 75542857 | 704 | * | 3,793221 | 6,930359868 | 2,412726 | 2,872419 | 0,004073 | 1 | 0 | Distal Intergenic | 20 | 75626214 | 75646530 | 20317 | 2 | 74987 | ENSMUST00000114054.3 | 103673 | ENSMUSG00000079525 | Cldn34d | claudin 34D |
| chr3 | 1,15E+08 | 1,15E+08 | 684 | * | 5,779779 | 6,931124691 | 1,932217 | 3,587136 | 0,000334 | 1 | 0 | Distal Intergenic | 3 | 1,15E+08 | 1,15E+08 | 132 | 1 | 1,15E+08 | ENSMUST00000157721.3 | -111550 | ENSMUSG00000088346 | Gm22337 | predicted gene, 22337 |
| chr1 | 77573092 | 77573859 | 768 | * | 7,188894 | 6,93238092 | 1,881265 | 3,684957 | 0,000229 | 1 | 0 | Distal Intergenic | 1 | 77498961 | 77533210 | 34250 | 1 | 1,03E+08 | ENSMUST00000246015.1 | 74131 | | Gm32793 | predicted gene, 32793 |
| chr12 | 51594053 | 51594431 | 379 | * | 4,66365 | 6,934158705 | 2,103905 | 3,295851 | 0,000981 | 1 | 0 | Distal Intergenic | 12 | 51640124 | 51652554 | 12431 | 1 | 12810 | ENSMUST00000164782.10 | -45693 | ENSMUSG00000020953 | Coch | cochlin |
| chr2 | 1,65E+08 | 1,65E+08 | 994 | * | 9,280878 | 6,93548826 | 1,790123 | 3,874307 | 0,000107 | 0,167380886 | 1,449706 | Distal Intergenic | 2 | 1,65E+08 | 1,65E+08 | 5470 | 1 | 629777 | ENSMUST00000142338.2 | 7079 | ENSMUSG00000085436 | Zfp335os | zinc finger protein 335, opposite strand |
| chr16 | 72894636 | 72895341 | 706 | * | 5,046416 | 6,936483117 | 2,033281 | 3,411474 | 0,000646 | 1 | 0 | Promoter (<=1kb) | 16 | 72895795 | 73000900 | 105106 | 1 | 74991 | ENSMUST00000231434.2 | -454 | ENSMUSG00000116976 | 4930500H12Rik | RIKEN cDNA 4930500H12 gene |
| chr16 | 51851295 | 51852576 | 1282 | * | 16,87397 | 6,936939347 | 1,653614 | 4,195016 | 2,73E-05 | 0,052288507 | 2,386008 | Promoter (<=1kb) | 16 | 51851588 | 52028411 | 176824 | 1 | 208650 | ENSMUST00000114471.3 | 0 | ENSMUSG00000022637 | Cblb | Casitas B-lineage lymphoma b |
| chr2 | 1,23E+08 | 1,23E+08 | 469 | * | 4,405608 | 6,943911794 | 2,119239 | 3,276607 | 0,001051 | 1 | 0 | Promoter (1-2kb) | 2 | 1,23E+08 | 1,23E+08 | 5425 | 2 | 75915 | ENSMUST00000150154.3 | 1916 | | Bloc1s6os | biogenesis of lysosomal organelles complex-1, subunit 6, pallidin, opposite strand |
| chr9 | 96677076 | 96677834 | 759 | * | 4,671289 | 6,943922725 | 2,128398 | 3,262511 | 0,001104 | 1 | 0 | Distal Intergenic | 9 | 96647654 | 96655343 | 7690 | 1 | 319707 | ENSMUST00000190054.2 | 29422 | ENSMUSG00000101581 | C430002N11Rik | RIKEN cDNA C430002N11 gene |
| chr15 | 69239751 | 69240275 | 525 | * | 4,269432 | 6,945652343 | 2,269527 | 3,060397 | 0,00221 | 1 | 0 | Distal Intergenic | 15 | 69322864 | 69322995 | 132 | 2 | 1,15E+08 | ENSMUST00000104314.3 | 82720 | ENSMUSG00000077502 | Gm24232 | predicted gene, 24232 |
| chr16 | 25140580 | 25141445 | 866 | * | 6,908654 | 6,948445514 | 1,873294 | 3,709212 | 0,000208 | 0,25121113 | 1,360622 | Intron (ENSMUST0000( | 16 | 25105571 | 25136985 | 31415 | 1 | 71338 | ENSMUST00000128595.2 | 35009 | ENSMUSG00000048399 | Tprg | transformation related protein 63 regulated |
| chr10 | 1,12E+08 | 1,12E+08 | 251 | * | 3,20032 | 6,948926293 | 2,470107 | 2,813208 | 0,004905 | 1 | 0 | Distal Intergenic | 10 | 1,12E+08 | 1,12E+08 | 19856 | 2 | 629206 | ENSMUST00000220339.2 | -12357 | ENSMUSG00000112533 | Gm20758 | predicted gene, 20758 |
| chr7 | 1,21E+08 | 1,21E+08 | 1127 | * | 8,598004 | 6,952029636 | 1,7641 | 3,940837 | 8,12E-05 | 1 | 0 | Intron (ENSMUST0000( | 7 | 1,21E+08 | 1,21E+08 | 65233 | 2 | 76179 | ENSMUST00000046929.7 | 10749 | ENSMUSG00000063317 | Usp31 | ubiquitin specific peptidase 31 |
| chr12 | 1,11E+08 | 1,11E+08 | 1150 | * | 11,5882 | 6,952909067 | 1,708669 | 4,069197 | 4,72E-05 | 0,153389938 | 0,744375 | Promoter (<=1kb) | 12 | 1,11E+08 | 1,11E+08 | 83081 | 1 | 104859 | ENSMUST00000169597.9 | 261 | ENSMUSG00000021275 | Tecpr2 | tectonin beta-propeller repeat containing 2 |
| chr11 | 1,16E+08 | 1,16E+08 | 1402 | * | 10,46079 | 6,952994231 | 1,704516 | 4,079161 | 4,52E-05 | 0,105241527 | 1,2763 | Promoter (<=1kb) | 11 | 1,16E+08 | 1,16E+08 | 40883 | 2 | 170472 | ENSMUST00000021097.10 | 0 | ENSMUSG00000020752 | Recql5 | RecQ protein-like 5 |
| chr10 | 77984429 | 77985188 | 760 | * | 6,723258 | 6,956218507 | 1,949335 | 3,568509 | 0,000359 | 1 | 0 | Distal Intergenic | 10 | 77957267 | 77973931 | 16665 | 2 | 1,15E+08 | ENSMUST00000218616.2 | -10498 | | Gm47922 | predicted gene, 47922 |
| chr5 | 1,23E+08 | 1,23E+08 | 559 | * | 4,462997 | 6,959909347 | 2,21385 | 3,143803 | 0,001668 | 1 | 0 | Exon (ENSMUST00000 | 5 | 1,23E+08 | 1,23E+08 | 5604 | 2 | 30841 | ENSMUST00000139674.2 | 4776 | ENSMUSG00000029475 | Kdm2b | lysine (K)-specific demethylase 2B |
| chrX | 11119263 | 11119904 | 642 | * | 4,743035 | 6,961355998 | 2,096748 | 3,320072 | 0,0009 | 1 | 0 | Distal Intergenic | 20 | 11165496 | 11165996 | 501 | 1 | 1E+08 | ENSMUST00000188439.2 | -45592 | ENSMUSG00000100626 | H2al1a | H2A histone family member L1A |
| chr15 | 1,01E+08 | 1,01E+08 | 616 | * | 6,515642 | 6,961864377 | 1,887099 | 3,689189 | 0,000225 | 1 | 0 | Intron (ENSMUST0000( | 15 | 1,01E+08 | 1,01E+08 | 34682 | 1 | 59033 | ENSMUST00000162049.2 | -8181 | ENSMUSG00000023032 | Slc4a8 | solute carrier family 4 (anion exchanger), member 8 |
| chrX | 47518526 | 47519104 | 579 | * | 3,88799 | 6,961999655 | 2,516951 | 2,766045 | 0,005674 | 1 | 0 | Intron (ENSMUST0000( | 20 | 47499923 | 47543046 | 43124 | 2 | 56501 | ENSMUST00000114958.8 | 23942 | ENSMUSG00000031103 | Elf4 | E74-like factor 4 (ets domain transcription factor) |
| chr4 | 1,38E+08 | 1,38E+08 | 878 | * | 5,312665 | 6,962048622 | 2,05188 | 3,39301 | 0,000691 | 1 | 0 | Promoter (2-3kb) | 4 | 1,38E+08 | 1,38E+08 | 7094 | 2 | 26971 | ENSMUST00000030526.7 | -2197 | ENSMUSG00000028749 | Pla2g2f | phospholipase A2, group IIF |
| chr14 | 1,17E+08 | 1,17E+08 | 746 | * | 3,400468 | 6,962964123 | 2,334497 | 2,98264 | 0,002858 | 1 | 0 | Intron (ENSMUST0000( | 14 | 1,17E+08 | 1,18E+08 | 1049151 | 1 | 23888 | ENSMUST00000078849.12 | 51570 | ENSMUSG00000058571 | Gpc6 | glypican 6 |
| chr18 | 37302343 | 37302835 | 493 | * | 4,095009 | 6,962979453 | 2,308877 | 3,015743 | 0,002564 | 1 | 0 | Intron (ENSMUST0000( | 18 | 37276556 | 37320710 | 44155 | 1 | 353237 | ENSMUST00000047479.3 | 25787 | ENSMUSG00000102697 | Pcdhac2 | protocadherin alpha subfamily C, 2 |
| chr10 | 79874728 | 79875417 | 690 | * | 6,11198 | 6,963543018 | 1,975714 | 3,524569 | 0,000424 | 1 | 0 | Promoter (<=1kb) | 10 | 79871785 | 79875484 | 3700 | 2 | 66420 | ENSMUST00000131743.8 | 67 | ENSMUSG00000004667 | Polr2e | polymerase (RNA) II (DNA directed) polypeptide E |
| chrX | 1,3E+08 | 1,3E+08 | 512 | * | 4,325128 | 6,963950963 | 2,304963 | 3,021285 | 0,002517 | 1 | 0 | Distal Intergenic | 20 | 1,3E+08 | 1,3E+08 | 129 | 2 | 1,15E+08 | ENSMUST00000104191.3 | 311642 | ENSMUSG00000077379 | Gm25170 | predicted gene, 25170 |
| chr9 | 71057900 | 71058958 | 1059 | * | 10,16388 | 6,970372432 | 1,772319 | 3,932912 | 8,39E-05 | 0,161376593 | 1,211315 | Intron (ENSMUST0000( | 9 | 71017944 | 71069915 | 51972 | 2 | 64008 | ENSMUST00000060917.12 | 10957 | ENSMUSG00000032204 | Aqp9 | aquaporin 9 |
| chrX | 17246007 | 17247224 | 1218 | * | 7,658584 | 6,972301401 | 1,82759 | 3,815026 | 0,000136 | 1 | 0 | Distal Intergenic | 20 | 17191554 | 17191657 | 104 | 1 | 1,15E+08 | ENSMUST00000083308.3 | 54453 | ENSMUSG00000119745 | Gm22294 | predicted gene, 22294 |
| chr9 | 42902008 | 42902374 | 367 | * | 3,760736 | 6,973061496 | 2,261539 | 3,083326 | 0,002047 | 1 | 0 | Intron (ENSMUST0000( | 9 | 42937300 | 42939114 | 1815 | 2 | 69632 | ENSMUST00000214175.2 | 36740 | ENSMUSG00000059495 | Arhgef12 | Rho guanine nucleotide exchange factor (GEF) 12 |
| chrX | 68946392 | 68946865 | 474 | * | 4,549436 | 6,974551233 | 2,111464 | 3,303183 | 0,000956 | 1 | 0 | Distal Intergenic | 20 | 68988906 | 68989584 | 679 | 1 | 75509 | ENSMUST00000075654.2 | -42041 | ENSMUSG00000059690 | 1700020N15Rik | RIKEN cDNA 1700020N15 gene |
| chr2 | 80384936 | 80385494 | 559 | * | 5,743302 | 6,975261512 | 2,058116 | 3,389149 | 0,000701 | 1 | 0 | Promoter (<=1kb) | 2 | 80375457 | 80385414 | 9958 | 2 | 50884 | ENSMUST00000131872.8 | 0 | ENSMUSG00000027002 | Nckap1 | NCK-associated protein 1 |
| chr19 | 21549933 | 21550614 | 682 | * | 4,978858 | 6,975488643 | 2,041689 | 3,416528 | 0,000634 | 1 | 0 | Distal Intergenic | 19 | 21530305 | 21535021 | 4717 | 1 | 1,19E+08 | ENSMUST00000112772.4 | 19628 | ENSMUSG00000079382 | C19H9orf57 | chromosome 19 C9orf57 homolog |
| chr8 | 1,16E+08 | 1,16E+08 | 573 | * | 6,585979 | 6,97568877 | 1,919043 | 3,634983 | 0,000278 | 1 | 0 | Intron (ENSMUST0000( | 8 | 1,15E+08 | 1,16E+08 | 639976 | 1 | 80707 | ENSMUST00000162052.8 | 170707 | ENSMUSG00000004637 | Wwox | WW domain-containing oxidoreductase |
| chr1 | 92620973 | 92621713 | 741 | * | 5,887369 | 6,976380675 | 1,987223 | 3,510619 | 0,000447 | 1 | 0 | Distal Intergenic | 1 | 92641956 | 92644924 | 2969 | 2 | 1,03E+08 | ENSMUST00000188188.2 | 23211 | | Gm29483 | predicted gene 29483 |
| chr18 | 35258606 | 35259143 | 538 | * | 5,133788 | 6,976734528 | 1,959194 | 3,561023 | 0,000369 | 1 | 0 | Intron (ENSMUST0000( | 18 | 35252495 | 35307472 | 54978 | 1 | 12385 | ENSMUST00000237154.2 | 6111 | ENSMUSG00000037815 | Ctnna1 | catenin (cadherin associated protein), alpha 1 |
| chr3 | 12052055 | 12052997 | 943 | * | 6,442889 | 6,976739996 | 1,908875 | 3,654897 | 0,000257 | 1 | 0 | Distal Intergenic | 3 | 11888289 | 11909419 | 21131 | 2 | 1E+08 | ENSMUST00000192022.2 | -142636 | ENSMUSG00000104015 | Gm10745 | predicted gene 10745 |
| chr2 | 10084570 | 10085796 | 1227 | * | 9,529631 | 6,978565122 | 1,887115 | 3,698008 | 0,000217 | 0,247946394 | 1,449706 | Promoter (<=1kb) | 2 | 10060827 | 10085238 | 24412 | 2 | 11949 | ENSMUST00000026887.14 | 0 | ENSMUSG00000025781 | Atp5c1 | ATP synthase, H+ transporting, mitochondrial F1 complex, gamma polypeptide 1 |
| chr4 | 34242524 | 34243153 | 630 | * | 3,464918 | 6,981142313 | 2,430975 | 2,871746 | 0,004082 | 1 | 0 | Distal Intergenic | 4 | 34086126 | 34086233 | 108 | 1 | 1,15E+08 | ENSMUST00000157216.2 | 156398 | ENSMUSG00000087841 | Gm25623 | predicted gene, 25623 |
| chr4 | 89795634 | 89796658 | 1025 | * | 6,026988 | 6,983045726 | 1,975411 | 3,534984 | 0,000408 | 1 | 0 | Distal Intergenic | 4 | 89576435 | 89583009 | 6575 | 1 | 242523 | ENSMUST00000052478.3 | 219199 | ENSMUSG00000043753 | Dmrta1 | doublesex and mab-3 related transcription factor like family A1 |
| chr7 | 1,16E+08 | 1,16E+08 | 1266 | * | 10,67342 | 6,984472977 | 1,736441 | 4,022292 | 5,76E-05 | 0,172125866 | 0,744375 | Distal Intergenic | 7 | 1,16E+08 | 1,16E+08 | 7222 | 1 | 53322 | ENSMUST00000183335.2 | 45516 | ENSMUSG00000030659 | Nucb2 | nucleobindin 2 |
| chrX | 16724078 | 16724568 | 491 | * | 3,33726 | 6,986210145 | 2,569222 | 2,719193 | 0,006544 | 1 | 0 | Distal Intergenic | 20 | 16575521 | 16683605 | 108085 | 2 | 109731 | ENSMUST00000040820.13 | -40473 | ENSMUSG00000040147 | Maob | monoamine oxidase B |
| chrX | 61354615 | 61355303 | 689 | * | 4,474767 | 6,98727716 | 2,171429 | 3,217825 | 0,001292 | 1 | 0 | Distal Intergenic | 20 | 61330748 | 61335684 | 4937 | 1 | 74437 | ENSMUST00000051414.2 | 23867 | ENSMUSG00000045330 | Magec2 | MAGE family member C2 |
| chr13 | 99685255 | 99685777 | 523 | * | 5,4946 | 6,989009812 | 2,014616 | 3,469152 | 0,000522 | 1 | 0 | Distal Intergenic | 13 | 99653137 | 99658268 | 5132 | 1 | 1,03E+08 | ENSMUST00000181939.3 | 32118 | | Gm26559 | predicted gene, 26559 |
| chr5 | 1,49E+08 | 1,49E+08 | 615 | * | 6,650528 | 6,989533377 | 1,871352 | 3,735018 | 0,000188 | 1 | 0 | Distal Intergenic | 5 | 1,49E+08 | 1,49E+08 | 5118 | 1 | 78015 | ENSMUST00000155504.2 | 28617 | ENSMUSG00000085740 | 4930505K14Rik | RIKEN cDNA 4930505K14 gene |
| chr4 | 65869838 | 65870379 | 542 | * | 4,822474 | 6,9899002 | 2,028932 | 3,445112 | 0,000571 | 1 | 0 | Intron (ENSMUST0000( | 4 | 65523486 | 65531993 | 8508 | 1 | 69807 | ENSMUST00000156922.2 | 346352 | ENSMUSG00000051675 | Trim32 | tripartite motif-containing 32 |
| chr7 | 78954756 | 78955317 | 562 | * | 6,045154 | 6,989923894 | 1,972025 | 3,544542 | 0,000393 | 1 | 0 | Intron (ENSMUST0000( | 7 | 78923002 | 78995119 | 72118 | 1 | 54608 | ENSMUST00000135333.3 | 31754 | ENSMUSG00000039202 | Abhd2 | abhydrolase domain containing 2 |
| chr1 | 1,9E+08 | 1,9E+08 | 610 | * | 5,429218 | 6,994140367 | 2,063159 | 3,390015 | 0,000699 | 1 | 0 | Distal Intergenic | 1 | 1,9E+08 | 1,9E+08 | 10223 | 1 | 1,03E+08 | ENSMUST00000191893.2 | 87658 | | Gm30932 | predicted gene, 30932 |
| chr9 | 1,01E+08 | 1,01E+08 | 1189 | * | 8,326748 | 6,997899123 | 1,784504 | 3,921482 | 8,8E-05 | 0,151196729 | 1,425929 | Distal Intergenic | 9 | 1,01E+08 | 1,01E+08 | 853 | 2 | 382099 | ENSMUST00000075243.8 | -54850 | ENSMUSG00000061486 | Gm5161 | predicted pseudogene 5161 |
| chr19 | 56429417 | 56429812 | 396 | * | 3,263364 | 6,998348298 | 2,677457 | 2,613804 | 0,008954 | 1 | 0 | 3' UTR | 19 | 56425128 | 56430747 | 5620 | 1 | 12369 | ENSMUST00000237765.2 | 4289 | ENSMUSG00000025076 | Casp7 | caspase 7 |
| chr14 | 1,21E+08 | 1,21E+08 | 1379 | * | 10,02871 | 7,000833662 | 1,728218 | 4,050896 | 5,1E-05 | 0,120486853 | 1,173325 | Distal Intergenic | 14 | 1,21E+08 | 1,21E+08 | 84 | 1 | 1,15E+08 | ENSMUST00000122527.4 | -136321 | | Gm25781 | predicted gene, 25781 |

| | | | | | | | | | | | | | | | | | | | | | | | |
|---|---|---|---|---|---|---|---|---|---|---|---|---|---|---|---|---|---|---|---|---|---|---|---|
| chr17 | 46581307 | 46581811 | 505 | * | 5,496457 | 7,001139928 | 2,097865 | 3,337268 | 0,000846 | 1 | 0 | Promoter (<=1kb) | 17 | 46572451 | 46581697 | 9247 | 2 | 74094 | ENSMUST00000224055.2 | 0 | ENSMUSG00000012296 | Tjap1 | tight junction associated protein 1 |
| chr5 | 1,38E+08 | 1,38E+08 | 497 | * | 4,795859 | 7,002860131 | 1,992723 | 3,514216 | 0,000441 | 1 | 0 | Distal Intergenic | 5 | 1,38E+08 | 1,38E+08 | 8037 | 2 | 243300 | ENSMUST00000212152.2 | 26984 | ENSMUSG00000045348 | Nyap1 | neuronal tyrosine-phosphorylated phosphoinositide 3-kinase adaptor 1 |
| chr1 | 1,84E+08 | 1,84E+08 | 998 | * | 5,524809 | 7,004518679 | 2,048579 | 3,419208 | 0,000628 | 1 | 0 | Distal Intergenic | 1 | 1,84E+08 | 1,84E+08 | 41256 | 1 | 63953 | ENSMUST00000048655.8 | 48209 | ENSMUSG00000039384 | Dusp10 | dual specificity phosphatase 10 |
| chr15 | 12875919 | 12876521 | 603 | * | 7,241251 | 7,005154264 | 1,832536 | 3,822655 | 0,000132 | 1 | 0 | Promoter (2-3kb) | 15 | 12878794 | 12885316 | 6523 | 1 | 14000 | ENSMUST00000141010.2 | -2273 | ENSMUSG00000022191 | Drosha | drosha, ribonuclease type III |
| chr9 | 66003446 | 66003899 | 454 | * | 4,755293 | 7,005857738 | 2,159369 | 3,2444 | 0,001177 | 1 | 0 | Promoter (<=1kb) | 9 | 66001469 | 66003873 | 2405 | 2 | 56440 | ENSMUST00000143148.2 | 0 | ENSMUSG00000032382 | Snx1 | sorting nexin 1 |
| chr11 | 52249459 | 52249811 | 353 | * | 7,210004 | 7,007571422 | 1,872986 | 3,741389 | 0,000183 | 1 | 0 | Promoter (1-2kb) | 11 | 52251687 | 52280224 | 28538 | 1 | 22333 | ENSMUST00000102758.8 | -1876 | ENSMUSG00000020402 | Vdac1 | voltage-dependent anion channel 1 |
| chr12 | 69432490 | 69433337 | 848 | * | 5,506116 | 7,007797313 | 2,025736 | 3,459382 | 0,000541 | 1 | 0 | Distal Intergenic | 12 | 69418924 | 69420557 | 1634 | 1 | 11845 | ENSMUST00000050063.9 | 13566 | ENSMUSG00000044147 | Arf6 | ADP-ribosylation factor 6 |
| chrX | 21206237 | 21206883 | 647 | * | 4,075708 | 7,008332964 | 2,32326 | 3,016594 | 0,002556 | 1 | 0 | Distal Intergenic | 20 | 21178449 | 21200966 | 22518 | 2 | 629024 | ENSMUST00000123014.2 | -5271 | ENSMUSG00000087586 | Gm6938 | predicted gene 6938 |
| chr1 | 50082110 | 50082766 | 657 | * | 4,152869 | 7,008653179 | 2,149263 | 3,260957 | 0,00111 | 1 | 0 | Distal Intergenic | 1 | 49283775 | 49379602 | 95828 | 1 | 442837 | ENSMUST00000188673.7 | 798335 | ENSMUSG00000051616 | C230029F24Rik | RIKEN cDNA C230029F24 gene |
| chrX | 1,11E+08 | 1,11E+08 | 585 | * | 4,835224 | 7,008972471 | 2,012227 | 3,483192 | 0,000495 | 1 | 0 | Intron (ENSMUST0000( | 20 | 1,11E+08 | 1,11E+08 | 61065 | 1 | 68117 | ENSMUST00000113415.2 | 41056 | ENSMUSG00000025525 | Apool | apolipoprotein O-like |
| chr5 | 54597998 | 54598742 | 745 | * | 5,342196 | 7,010077976 | 2,028324 | 3,456093 | 0,000548 | 1 | 0 | Distal Intergenic | 5 | 54507333 | 54515885 | 8553 | 1 | 330086 | ENSMUST00000197424.2 | 90665 | ENSMUSG00000105912 | Gm10440 | predicted gene 10440 |
| chr5 | 1,16E+08 | 1,16E+08 | 595 | * | 4,934558 | 7,010988039 | 2,162373 | 3,242266 | 0,001186 | 1 | 0 | Promoter (<=1kb) | 5 | 1,16E+08 | 1,16E+08 | 6716 | 2 | 75387 | ENSMUST00000112066.2 | -995 | ENSMUSG00000029524 | Sirt4 | sirtuin 4 |
| chr11 | 1,2E+08 | 1,2E+08 | 681 | * | 5,728805 | 7,014768644 | 2,029901 | 3,45572 | 0,000549 | 1 | 0 | Intron (ENSMUST0000( | 11 | 1,2E+08 | 1,2E+08 | 27524 | 1 | 74370 | ENSMUST00000131217.2 | 10252 | ENSMUSG00000025583 | Rptor | regulatory associated protein of MTOR, complex 1 |
| chr12 | 16941262 | 16941989 | 728 | * | 5,744016 | 7,015036786 | 2,002337 | 3,503425 | 0,000459 | 1 | 0 | Promoter (2-3kb) | 12 | 16944896 | 17037824 | 92929 | 1 | 19878 | ENSMUST00000020904.8 | -2907 | ENSMUSG00000020580 | Rock2 | Rho-associated coiled-coil containing protein kinase 2 |
| chr13 | 31236317 | 31236928 | 612 | * | 6,595996 | 7,01695182 | 1,860477 | 3,771588 | 0,000162 | 1 | 0 | Distal Intergenic | 13 | 31107639 | 31158076 | 50438 | 2 | 66482 | ENSMUST00000222133.2 | -78241 | ENSMUSG00000021357 | Exoc2 | exocyst complex component 2 |
| chr7 | 1,1E+08 | 1,1E+08 | 785 | * | 5,035048 | 7,018477824 | 2,100116 | 3,341948 | 0,000832 | 1 | 0 | Intron (ENSMUST0000( | 7 | 1,1E+08 | 1,1E+08 | 305364 | 2 | 319934 | ENSMUST00000166020.8 | 19773 | ENSMUSG00000038371 | Sbf2 | SET binding factor 2 |
| chr2 | 1,67E+08 | 1,67E+08 | 1869 | * | 15,97733 | 7,018780537 | 1,720035 | 4,080604 | 4,49E-05 | 0,072320047 | 2,386008 | Promoter (<=1kb) | 2 | 1,67E+08 | 1,67E+08 | 92465 | 1 | 99371 | ENSMUST00000099078.10 | 126 | ENSMUSG00000074582 | Arfgef2 | ADP-ribosylation factor guanine nucleotide-exchange factor 2 (brefeldin A-inhibited) |
| chrX | 37734259 | 37734878 | 620 | * | 4,172837 | 7,021408734 | 2,399688 | 2,925968 | 0,003434 | 1 | 0 | Distal Intergenic | 20 | 37719662 | 37723964 | 4303 | 2 | 59048 | ENSMUST00000058265.8 | -10295 | ENSMUSG00000048970 | C1galt1c1 | C1GALT1-specific chaperone 1 |
| chr4 | 1,49E+08 | 1,49E+08 | 1061 | * | 6,166694 | 7,025946173 | 2,048768 | 3,429352 | 0,000605 | 1 | 0 | Promoter (1-2kb) | 4 | 1,49E+08 | 1,49E+08 | 19632 | 2 | 16561 | ENSMUST00000139123.8 | 1873 | ENSMUSG00000063077 | Kif1b | kinesin family member 1B |
| chr12 | 57324518 | 57324914 | 397 | * | 4,56863 | 7,026595129 | 2,211801 | 3,176865 | 0,001489 | 1 | 0 | Intron (ENSMUST0000( | 12 | 57328959 | 57513827 | 184869 | 1 | 73490 | ENSMUST00000153137.6 | -4045 | ENSMUSG00000047022 | Mipol1 | mirror-image polydactyly 1 |
| chr18 | 57003457 | 57004474 | 1018 | * | 6,707398 | 7,02802933 | 1,960742 | 3,584373 | 0,000338 | 1 | 0 | Intron (ENSMUST0000( | 18 | 56936741 | 57058587 | 121847 | 2 | 320253 | ENSMUST00000153044.2 | 54113 | ENSMUSG00000032656 | Marchf3 | membrane associated ring-CH-type finger 3 |
| chr14 | 83346709 | 83347637 | 929 | * | 4,975828 | 7,028678049 | 1,999707 | 3,514855 | 0,00044 | 1 | 0 | Distal Intergenic | 14 | 83643071 | 83643204 | 134 | 1 | 1,15E+08 | ENSMUST00000157384.3 | -295434 | ENSMUSG00000088009 | Gm24774 | predicted gene, 24774 |
| chr3 | 58482577 | 58483194 | 618 | * | 4,819453 | 7,028997582 | 2,046689 | 3,434326 | 0,000594 | 1 | 0 | Promoter (<=1kb) | 3 | 58484057 | 58500554 | 16498 | 1 | 69227 | ENSMUST00000107924.3 | -863 | ENSMUSG00000075700 | Selenot | selenoprotein T |
| chr3 | 1,34E+08 | 1,34E+08 | 645 | * | 7,436017 | 7,02928296 | 1,863294 | 3,772504 | 0,000162 | 1 | 0 | Distal Intergenic | 3 | 1,34E+08 | 1,34E+08 | 19493 | 2 | 75208 | ENSMUST00000198231.2 | 53801 | ENSMUSG00000105765 | 4930539C22Rik | RIKEN cDNA 4930539C22 gene |
| chr10 | 11584822 | 11585502 | 681 | * | 6,874285 | 7,029604173 | 1,841821 | 3,81666 | 0,000135 | 1 | 0 | Distal Intergenic | 10 | 11523749 | 11523878 | 130 | 1 | 1,15E+08 | ENSMUST00000104351.3 | 61073 | ENSMUSG00000077539 | Gm22546 | predicted gene, 22546 |
| chr3 | 15067898 | 15068221 | 324 | * | 4,03085 | 7,030481597 | 2,362939 | 2,975312 | 0,002927 | 1 | 0 | Downstream (<=300bp | 3 | 15036261 | 15067787 | 31527 | 1 | 77127 | ENSMUST00000182586.2 | 31637 | ENSMUSG00000098008 | A930001A20Rik | RIKEN cDNA A930001A20 gene |
| chr4 | 12943886 | 12944411 | 526 | * | 5,749721 | 7,032068061 | 2,141926 | 3,283059 | 0,001027 | 1 | 0 | Intron (ENSMUST0000( | 4 | 12917967 | 12981070 | 63104 | 1 | 208820 | ENSMUST00000156536.3 | 25919 | ENSMUSG00000055963 | Triqk | triple QxxK/R motif containing |
| chr19 | 7492983 | 7493325 | 343 | * | 3,395153 | 7,033254924 | 2,626378 | 2,677929 | 0,007408 | 1 | 0 | Intron (ENSMUST0000( | 19 | 7471900 | 7511565 | 39666 | 1 | 109168 | ENSMUST00000236382.2 | 21083 | ENSMUSG00000024759 | Atl3 | atlastin GTPase 3 |
| chr3 | 44323400 | 44323987 | 588 | * | 5,059434 | 7,033803755 | 1,999119 | 3,518452 | 0,000434 | 1 | 0 | Distal Intergenic | 3 | 45281480 | 45332640 | 51161 | 2 | 72511 | ENSMUST00000191843.2 | 1008653 | ENSMUSG00000097040 | 2610316D01Rik | RIKEN cDNA 2610316D01 gene |
| chr17 | 56971599 | 56972202 | 604 | * | 7,160068 | 7,036307728 | 1,868343 | 3,766068 | 0,000166 | 1 | 0 | Downstream (<=300bp | 17 | 56980294 | 57014879 | 34586 | 1 | 71810 | ENSMUST00000232975.2 | -8092 | ENSMUSG00000002372 | Ranbp3 | RAN binding protein 3 |
| chr2 | 1,27E+08 | 1,27E+08 | 1294 | * | 12,4239 | 7,036518855 | 1,83963 | 3,824965 | 0,000131 | 0,270724242 | 0,738441 | Intron (ENSMUST0000( | 2 | 1,26E+08 | 1,27E+08 | 9599 | 2 | 14391 | ENSMUST00000137335.2 | -4537 | ENSMUSG00000027361 | Gabpb1 | GA repeat binding protein, beta 1 |
| chr14 | 64321797 | 64322476 | 680 | * | 5,556155 | 7,038464026 | 2,061554 | 3,414154 | 0,00064 | 1 | 0 | Promoter (<=1kb) | 14 | 64312894 | 64322857 | 9964 | 2 | 71037 | ENSMUST00000089338.6 | 381 | ENSMUSG00000034623 | Prss55 | protease, serine 55 |
| chr14 | 65348921 | 65349481 | 561 | * | 5,310333 | 7,040500162 | 2,003668 | 3,513806 | 0,000442 | 1 | 0 | Intron (ENSMUST0000( | 14 | 65289509 | 65335555 | 46047 | 2 | 54616 | ENSMUST00000022550.8 | -13366 | ENSMUSG00000021978 | Extl3 | exostosin-like glycosyltransferase 3 |
| chr10 | 62063065 | 62063475 | 411 | * | 6,824709 | 7,041436478 | 1,927478 | 3,653187 | 0,000259 | 1 | 0 | Intron (ENSMUST0000( | 10 | 62021175 | 62067030 | 45856 | 2 | 70423 | ENSMUST00000047883.11 | 3555 | ENSMUSG00000037031 | Tspan15 | tetraspanin 15 |
| chr12 | 80424495 | 80424833 | 339 | * | 3,030875 | 7,042623875 | 2,629322 | 2,678495 | 0,007395 | 1 | 0 | Promoter (1-2kb) | 12 | 80413366 | 80423466 | 10101 | 2 | 320808 | ENSMUST00000218728.2 | -1029 | ENSMUSG00000049106 | Dcaf5 | DDB1 and CUL4 associated factor 5 |
| chr12 | 99527094 | 99527974 | 881 | * | 9,241064 | 7,042781034 | 1,770619 | 3,977581 | 6,96E-05 | 0,096870739 | 2,237403 | Promoter (1-2kb) | 12 | 99354604 | 99529767 | 175164 | 2 | 71375 | ENSMUST00000176928.3 | 1793 | ENSMUSG00000033713 | Foxn3 | forkhead box N3 |
| chr10 | 65866461 | 65866814 | 354 | * | 4,487257 | 7,045069863 | 2,126198 | 3,313459 | 0,000921 | 1 | 0 | Distal Intergenic | 10 | 65956455 | 65958855 | 2401 | 2 | 69433 | ENSMUST00000188279.2 | 92041 | | 1700023F02Rik | RIKEN cDNA 1700023F02 gene |
| chr12 | 1,09E+08 | 1,09E+08 | 325 | * | 5,428514 | 7,046160127 | 2,018472 | 3,490839 | 0,000482 | 1 | 0 | Promoter (<=1kb) | 12 | 1,09E+08 | 1,09E+08 | 27176 | 1 | 22632 | ENSMUST00000021692.9 | 0 | ENSMUSG00000021264 | Yy1 | YY1 transcription factor |
| chr16 | 31034567 | 31035538 | 972 | * | 7,834463 | 7,046243408 | 1,780053 | 3,958446 | 7,54E-05 | 1 | 0 | Distal Intergenic | 16 | 30911230 | 31020063 | 108834 | 2 | 78618 | ENSMUST00000230698.2 | -14504 | ENSMUSG00000049076 | Acap2 | ArfGAP with coiled-coil, ankyrin repeat and PH domains 2 |
| chr16 | 17984421 | 17985251 | 831 | * | 5,480922 | 7,047130006 | 1,946195 | 3,620978 | 0,000293 | 1 | 0 | Promoter (1-2kb) | 16 | 17982953 | 17983034 | 82 | 2 | 1,02E+08 | ENSMUST00000184472.3 | -1387 | ENSMUSG00000099293 | Mir6366 | microRNA 6366 |
| chr5 | 1,39E+08 | 1,39E+08 | 779 | * | 6,368264 | 7,047347032 | 1,932244 | 3,647235 | 0,000265 | 1 | 0 | Promoter (2-3kb) | 5 | 1,39E+08 | 1,39E+08 | 5352 | 2 | 100494 | ENSMUST00000143889.2 | 2115 | ENSMUSG00000053581 | Zfand2a | zinc finger, AN1-type domain 2A |
| chr9 | 10764621 | 10765131 | 511 | * | 6,051213 | 7,047580094 | 1,918496 | 3,673493 | 0,000239 | 1 | 0 | Intron (ENSMUST0000( | 9 | 10904350 | 10907823 | 3474 | 1 | 1,03E+08 | ENSMUST00000210677.2 | -139219 | | Gm32710 | predicted gene, 32710 |
| chrX | 61185793 | 61186502 | 710 | * | 4,622373 | 7,052991191 | 2,034162 | 3,467271 | 0,000526 | 1 | 0 | Distal Intergenic | 20 | 61327105 | 61335684 | 8580 | 1 | 74437 | ENSMUST00000178085.8 | -140603 | ENSMUSG00000045330 | Magec2 | MAGE family member C2 |
| chr8 | 79784491 | 79785102 | 612 | * | 5,858816 | 7,053309066 | 1,953075 | 3,611387 | 0,000305 | 1 | 0 | Intron (ENSMUST0000( | 8 | 79755216 | 79917485 | 162270 | 1 | 622675 | ENSMUST00000238669.2 | 29275 | ENSMUSG00000071064 | Zfp827 | zinc finger protein 827 |
| chr8 | 88905210 | 88905936 | 727 | * | 5,791842 | 7,053628803 | 1,892166 | 3,727807 | 0,000193 | 1 | 0 | Distal Intergenic | 8 | 88891110 | 88893673 | 2564 | 1 | 234549 | ENSMUST00000137452.2 | 14100 | ENSMUSG00000031657 | Heatr3 | HEAT repeat containing 3 |
| chr6 | 22356124 | 22356698 | 575 | * | 7,335394 | 7,053905259 | 1,850497 | 3,811899 | 0,000138 | 1 | 0 | Promoter (<=1kb) | 6 | 22306519 | 22356165 | 49647 | 2 | 27999 | ENSMUST00000165576.8 | 0 | ENSMUSG00000029672 | Fam3c | FAM3 metabolism regulating signaling molecule C |
| chr19 | 19191419 | 19192000 | 582 | * | 3,874227 | 7,057420177 | 2,188659 | 3,224541 | 0,001262 | 1 | 0 | Distal Intergenic | 19 | 19253263 | 19253394 | 132 | 1 | 1,15E+08 | ENSMUST00000158282.3 | -61263 | ENSMUSG00000088907 | Gm25437 | predicted gene, 25437 |
| chr2 | 1,2E+08 | 1,2E+08 | 622 | * | 5,439117 | 7,059540303 | 1,944442 | 3,630625 | 0,000283 | 1 | 0 | Promoter (1-2kb) | 2 | 1,2E+08 | 1,2E+08 | 2527 | 2 | 20402 | ENSMUST00000130303.2 | -1639 | ENSMUSG00000027288 | Zfp106 | zinc finger protein 106 |
| chr2 | 1,14E+08 | 1,14E+08 | 712 | * | 7,015704 | 7,064107469 | 2,031296 | 3,477635 | 0,000506 | 0,373994686 | 1,360622 | Distal Intergenic | 2 | 1,14E+08 | 1,14E+08 | 27747 | 2 | 66632 | ENSMUST00000138106.8 | 92121 | ENSMUSG00000057147 | Dph6 | diphthamine biosynthesis 6 |
| chr16 | 93586656 | 93587599 | 944 | * | 8,720598 | 7,066446767 | 1,747668 | 4,043358 | 5,27E-05 | 1 | 0 | Promoter (2-3kb) | 16 | 93589677 | 93596903 | 7227 | 1 | 70028 | ENSMUST00000232210.2 | -2078 | ENSMUSG00000022946 | Dop1b | DOP1 leucine zipper like protein B |
| chr10 | 84989685 | 84990472 | 788 | * | 4,995082 | 7,067103266 | 2,040837 | 3,462846 | 0,000534 | 1 | 0 | Promoter (2-3kb) | 10 | 84984301 | 84987220 | 2920 | 2 | 12952 | ENSMUST00000214805.2 | -2465 | ENSMUSG00000020038 | Cry1 | cryptochrome 1 (photolyase-like) |
| chr6 | 49978939 | 49979863 | 925 | * | 8,723141 | 7,067990655 | 1,852218 | 3,815961 | 0,000136 | 0,14942522 | 2,237403 | Distal Intergenic | 6 | 50087221 | 50175578 | 88358 | 1 | 56524 | ENSMUST00000166318.8 | -107358 | ENSMUSG00000038388 | Pals2 | protein associated with LIN7 2, MAGUK family member |
| chr9 | 70462257 | 70463272 | 1016 | * | 8,742539 | 7,0697364 | 1,768199 | 3,99827 | 6,38E-05 | 0,120888649 | 1,449706 | Intron (ENSMUST0000( | 9 | 70450180 | 70469088 | 18909 | 1 | 66660 | ENSMUST00000213362.2 | 12077 | ENSMUSG00000032212 | Sltm | SAFB-like, transcription modulator |
| chr15 | 34105886 | 34106831 | 946 | * | 8,065388 | 7,07052434 | 1,816 | 3,89346 | 9,88E-05 | 1 | 0 | Intron (ENSMUST0000( | 15 | 34114233 | 34115348 | 1116 | 1 | 67154 | ENSMUST00000226331.2 | -7402 | ENSMUSG00000022255 | Mtdh | metadherin |
| chrX | 1,19E+08 | 1,19E+08 | 878 | * | 6,099364 | 7,071742204 | 1,953573 | 3,619902 | 0,000295 | 1 | 0 | Distal Intergenic | 20 | 1,19E+08 | 1,19E+08 | 2970 | 1 | 93728 | ENSMUST00000040961.3 | -65825 | ENSMUSG00000034732 | Pabpc5 | poly(A) binding protein, cytoplasmic 5 |
| chrX | 1,02E+08 | 1,02E+08 | 1525 | * | 7,374576 | 7,073441301 | 1,824738 | 3,876416 | 0,000106 | 1 | 0 | Intron (ENSMUST0000( | 20 | 1,02E+08 | 1,02E+08 | 130272 | 2 | 18679 | ENSMUST00000043596.12 | 31659 | ENSMUSG00000034055 | Phka1 | phosphorylase kinase alpha 1 |
| chr16 | 29999266 | 30000452 | 1187 | * | 11,9862 | 7,075132034 | 1,745723 | 4,052837 | 5,06E-05 | 0,159721691 | 0,744375 | Distal Intergenic | 16 | 30013889 | 30019117 | 5229 | 1 | 75531 | ENSMUST00000187479.2 | -13437 | | 1700025H01Rik | RIKEN cDNA 1700025H01 gene |
| chr7 | 75304503 | 75305750 | 1248 | * | 10,23242 | 7,075894343 | 1,71975 | 4,11449 | 3,88E-05 | 0,089194938 | 1,449706 | Intron (ENSMUST0000( | 7 | 75293933 | 75327078 | 33146 | 1 | 75547 | ENSMUST00000209040.2 | 10570 | ENSMUSG00000066406 | Akap13 | A kinase (PRKA) anchor protein 13 |
| chr18 | 43372430 | 43373027 | 598 | * | 4,103268 | 7,078832114 | 2,260114 | 3,132068 | 0,001736 | 1 | 0 | Intron (ENSMUST0000( | 18 | 43376105 | 43396737 | 20633 | 1 | 269019 | ENSMUST00000236899.2 | -3078 | ENSMUSG00000039954 | Stk32a | serine/threonine kinase 32A |
| chr2 | 44149793 | 44150230 | 438 | * | 4,355855 | 7,081717988 | 2,400611 | 2,949965 | 0,003178 | 1 | 0 | Intron (ENSMUST0000( | 2 | 43949250 | 43955336 | 6087 | 2 | 320872 | ENSMUST00000139948.2 | -194457 | ENSMUSG00000085089 | Arhgap15os | Rho GTPase activating protein 15, opposite strand |
| chr11 | 94958438 | 94959110 | 673 | * | 5,576617 | 7,083435682 | 2,033853 | 3,482767 | 0,000496 | 1 | 0 | Promoter (<=1kb) | 11 | 94935300 | 94960024 | 24725 | 2 | 16400 | ENSMUST00000107739.8 | 914 | ENSMUSG00000001507 | Itga3 | integrin alpha 3 |
| chrX | 70349260 | 70349938 | 679 | * | 5,1156 | 7,083548508 | 2,019335 | 3,507862 | 0,000452 | 1 | 0 | Intron (ENSMUST0000( | 20 | 70344975 | 70359297 | 14323 | 1 | 17772 | ENSMUST00000129722.2 | 4285 | ENSMUSG00000031337 | Mtm1 | X-linked myotubular myopathy gene 1 |
| chr3 | 7659935 | 7660567 | 633 | * | 7,089302 | 7,08458194 | 1,883342 | 3,761708 | 0,000169 | 1 | 0 | Intron (ENSMUST0000( | 3 | 7637088 | 7670924 | 33837 | 2 | 16196 | ENSMUST00000168269.8 | 10357 | ENSMUSG00000040329 | Il7 | interleukin 7 |
| chr7 | 1,01E+08 | 1,01E+08 | 858 | * | 7,875447 | 7,085220145 | 1,897323 | 3,734324 | 0,000188 | 1 | 0 | Promoter (<=1kb) | 7 | 1,01E+08 | 1,01E+08 | 68 | 1 | 387157 | ENSMUST00000083512.3 | 962 | ENSMUSG00000065446 | Mir139 | microRNA 139 |
| chr10 | 33507883 | 33508562 | 680 | * | 5,704999 | 7,089147382 | 2,006583 | 3,532944 | 0,000411 | 1 | 0 | Intron (ENSMUST0000( | 10 | 33388282 | 33500765 | 112484 | 2 | 215890 | ENSMUST00000019920.13 | -7118 | ENSMUSG00000019785 | Clvs2 | clavesin 2 |
| chr14 | 12360155 | 12360671 | 517 | * | 4,437312 | 7,095116237 | 2,252327 | 3,150127 | 0,001632 | 1 | 0 | Intron (ENSMUST0000( | 14 | 12583144 | 12583747 | 604 | 2 | 1E+08 | ENSMUST00000162462.2 | 223076 | ENSMUSG00000089686 | Mnd1-ps | Mnd1 retrotransposed pseudogene |
| chr3 | 75421358 | 75421836 | 479 | * | 5,007995 | 7,09568984 | 2,04977 | 3,4617 | 0,000537 | 1 | 0 | Distal Intergenic | 3 | 75424583 | 75428337 | 3755 | 2 | 56426 | ENSMUST00000159519.2 | 6501 | ENSMUSG00000027835 | Pdcd10 | programmed cell death 10 |
| chr10 | 9936097 | 9936665 | 569 | * | 5,430893 | 7,096826872 | 1,982375 | 3,579962 | 0,000344 | 1 | 0 | Intron (ENSMUST0000( | 10 | 9976697 | 9976860 | 164 | 1 | 1,15E+08 | ENSMUST00000083942.3 | -40032 | ENSMUSG00000065876 | Gm25682 | predicted gene, 25682 |
| chr19 | 16679585 | 16680735 | 1151 | * | 7,010324 | 7,096911602 | 1,835928 | 3,865572 | 0,000111 | 1 | 0 | Intron (ENSMUST0000( | 19 | 16662532 | 16663937 | 1406 | 2 | 271564 | ENSMUST00000225349.2 | -15648 | ENSMUSG00000046230 | Vps13a | vacuolar protein sorting 13A |
| chr18 | 24822882 | 24823379 | 498 | * | 5,630447 | 7,098654274 | 2,072686 | 3,424858 | 0,000615 | 1 | 0 | Exon (ENSMUST000002 | 18 | 24841680 | 24904593 | 62914 | 1 | 225288 | ENSMUST00000234462.2 | -18301 | ENSMUSG00000034295 | Fhod3 | formin homology 2 domain containing 3 |
| chrX | 76884252 | 76884819 | 568 | * | 3,818004 | 7,099049569 | 2,362886 | 3,004397 | 0,002661 | 1 | 0 | Intron (ENSMUST0000( | 20 | 76881508 | 76897136 | 15629 | 2 | 54192 | ENSMUST00000114041.3 | 12317 | ENSMUSG00000000003 | Pbsn | probasin |
| chr3 | 29015983 | 29016271 | 289 | * | 3,69094 | 7,09920019 | 2,323203 | 3,055782 | 0,002245 | 1 | 0 | Distal Intergenic | 3 | 28946766 | 28980873 | 34108 | 1 | 385263 | ENSMUST00000099170.2 | 69217 | ENSMUSG00000074655 | Gm1527 | predicted gene 1527 |
| chr19 | 10154943 | 10155430 | 488 | * | 5,346189 | 7,105630043 | 2,08585 | 3,406588 | 0,000658 | 1 | 0 | Distal Intergenic | 19 | 10160252 | 10174238 | 13987 | 1 | 76267 | ENSMUST00000010807.6 | -4822 | ENSMUSG00000010663 | Fads1 | fatty acid desaturase 1 |
| chr4 | 1,07E+08 | 1,07E+08 | 714 | * | 5,411556 | 7,106014983 | 2,057464 | 3,453774 | 0,000553 | 1 | 0 | Intron (ENSMUST0000( | 4 | 1,07E+08 | 1,07E+08 | 24166 | 2 | 329910 | ENSMUST00000140541.8 | -5592 | ENSMUSG00000034853 | Acot11 | acyl-CoA thioesterase 11 |
| chr13 | 93878028 | 93878862 | 835 | * | 8,908916 | 7,115592095 | 1,830523 | 3,88719 | 0,000101 | 0,168300715 | 1,360622 | Intron (ENSMUST0000( | 13 | 93842690 | 93855683 | 12994 | 2 | 105364 | ENSMUST00000149839.2 | -22345 | ENSMUSG00000087334 | AW495222 | expressed sequence AW495222 |
| chr14 | 99169341 | 99169916 | 576 | * | 4,899539 | 7,115646391 | 2,257525 | 3,151968 | 0,001622 | 1 | 0 | Distal Intergenic | 14 | 99125410 | 99158381 | 32972 | 2 | 1,19E+08 | ENSMUST00000226462.2 | -10960 | | Gm49018 | predicted gene, 49018 |
| chr7 | 1,1E+08 | 1,1E+08 | 967 | * | 8,414665 | 7,116985411 | 1,794023 | 3,967053 | 7,28E-05 | 1 | 0 | Exon (ENSMUST00000( | 7 | 1,1E+08 | 1,1E+08 | 21241 | 1 | 22390 | ENSMUST00000033326.10 | 4151 | ENSMUSG00000031016 | Wee1 | WEE 1 homolog 1 (S. pombe) |
| chr11 | 12939541 | 12940356 | 816 | * | 5,605936 | 7,119624508 | 1,955233 | 3,641318 | 0,000271 | 1 | 0 | Distal Intergenic | 11 | 12642813 | 12642942 | 130 | 1 | 1,15E+08 | ENSMUST00000157884.3 | 296728 | ENSMUSG00000088509 | Gm22789 | predicted gene, 22789 |
| chr1 | 33795927 | 33796588 | 662 | * | 4,827462 | 7,120041684 | 2,135387 | 3,334309 | 0,000855 | 1 | 0 | Promoter (<=1kb) | 1 | 33785883 | 33796822 | 10940 | 2 | 213539 | ENSMUST00000187602.2 | 234 | ENSMUSG00000042215 | Bag2 | BCL2-associated athanogene 2 |
| chr9 | 1,14E+08 | 1,14E+08 | 1381 | * | 10,7279 | 7,120583149 | 1,694452 | 4,202293 | 2,64E-05 | 0,092441431 | 0,918255 | Promoter (<=1kb) | 9 | 1,14E+08 | 1,14E+08 | 6172 | 1 | 1,01E+08 | ENSMUST00000111820.4 | 0 | ENSMUSG00000079260 | Tmppe | transmembrane protein with metallophosphoesterase domain |
| chr14 | 68070728 | 68071257 | 530 | * | 4,27772 | 7,122756862 | 2,129398 | 3,344963 | 0,000823 | 1 | 0 | Intron (ENSMUST0000( | 14 | 68097085 | 68097194 | 110 | 2 | 1,02E+08 | ENSMUST00000183732.3 | 25937 | ENSMUSG00000098866 | Mir6539 | microRNA 6539 |
| chr17 | 10428802 | 10429663 | 862 | * | 6,883351 | 7,1228082 | 2,080588 | 3,423459 | 0,000618 | 1 | 0 | Promoter (2-3kb) | 17 | 10428022 | 10432503 | 4482 | 2 | 19317 | ENSMUST00000232721.2 | 2840 | ENSMUSG00000062078 | Qk | quaking, KH domain containing RNA binding |
| chr17 | 34625623 | 34626030 | 408 | * | 5,555463 | 7,124493906 | 1,989801 | 3,580507 | 0,000343 | 1 | 0 | Intron (ENSMUST0000( | 17 | 34617796 | 34640123 | 22328 | 1 | 407788 | ENSMUST00000142208.8 | 7827 | ENSMUSG00000057246 | BC051142 | cDNA sequence BC051142 |
| chr16 | 9123083 | 9123764 | 682 | * | 6,047305 | 7,124889405 | 1,972309 | 3,61246 | 0,000303 | 1 | 0 | Distal Intergenic | 16 | 8867836 | 8873262 | 5427 | 1 | 1,03E+08 | ENSMUST00000229201.2 | 255247 | | Gm30567 | predicted gene, 30567 |
| chr16 | 14071807 | 14072381 | 575 | * | 6,144385 | 7,125346395 | 1,952631 | 3,649101 | 0,000263 | 1 | 0 | Intron (ENSMUST0000( | 16 | 14012764 | 14109227 | 96464 | 2 | 17880 | ENSMUST00000230397.2 | 36846 | ENSMUSG00000018830 | Myh11 | myosin, heavy polypeptide 11, smooth muscle |
| chr19 | 56809158 | 56810086 | 929 | * | 7,075876 | 7,12571622 | 1,886715 | 3,776784 | 0,000159 | 1 | 0 | Promoter (<=1kb) | 19 | 56775913 | 56810622 | 34710 | 2 | 213993 | ENSMUST00000118592.8 | 536 | ENSMUSG00000035173 | Ccdc186 | coiled-coil domain containing 186 |
| chr16 | 35460280 | 35461132 | 853 | * | 5,730507 | 7,126024451 | 1,933897 | 3,684801 | 0,000229 | 1 | 0 | Intron (ENSMUST0000( | 16 | 35478465 | 35478857 | 393 | 1 | 20357 | ENSMUST00000139805.2 | -17333 | ENSMUSG00000052133 | Sema5b | sema domain, seven thrombospondin repeats (type 1 and type 1-like), transmembrane domain (TM) and short cytoplasmic domain, (semaphorin) 5B |
| chr16 | 49795061 | 49796060 | 1000 | * | 10,34753 | 7,128732458 | 1,740001 | 4,09697 | 4,19E-05 | 0,103768001 | 1,211315 | Distal Intergenic | 16 | 49862852 | 49866652 | 3801 | 2 | 224180 | ENSMUST00000241286.1 | 70592 | | Gm4827 | predicted gene 4827 |
| chr17 | 18564643 | 18565188 | 546 | * | 4,633638 | 7,130005561 | 2,074459 | 3,437044 | 0,000588 | 1 | 0 | Distal Intergenic | 17 | 18518361 | 18552176 | 33816 | 1 | 627132 | ENSMUST00000231879.3 | 46282 | ENSMUSG00000079698 | Vmn2r93 | vomeronasal 2, receptor 93 |
| chr17 | 20885431 | 20886133 | 703 | * | 4,93083 | 7,133493189 | 2,189789 | 3,257616 | 0,001124 | 1 | 0 | Distal Intergenic | 17 | 20907770 | 20908666 | 897 | 1 | 171225 | ENSMUST00000061756.4 | -21637 | ENSMUSG00000042848 | Vmn1r226 | vomeronasal 1 receptor 226 |
| chr16 | 32236406 | 32236864 | 459 | * | 5,218253 | 7,134632953 | 2,094211 | 3,406835 | 0,000657 | 1 | 0 | Promoter (1-2kb) | 16 | 32238520 | 32247917 | 9398 | 1 | 66061 | ENSMUST00000014220.15 | -1656 | ENSMUSG00000014075 | Dynlt2b | dynein light chain Tctex-type 2B |
| chr11 | 56435194 | 56435867 | 674 | * | 8,969906 | 7,134637536 | 2,53053 | 2,819424 | 0,004811 | 0,683210259 | 2,237403 | Distal Intergenic | 11 | 56153991 | 56154066 | 76 | 2 | 1,15E+08 | ENSMUST00000083701.3 | -281128 | ENSMUSG00000065635 | Gm24253 | predicted gene, 24253 |
| chrX | 23715305 | 23716155 | 851 | * | 4,344179 | 7,136059554 | 2,26734 | 3,147326 | 0,001648 | 1 | 0 | Distal Intergenic | 20 | 23748792 | 23773846 | 25055 | 1 | 236749 | ENSMUST00000116614.3 | -32637 | ENSMUSG00000068113 | Tesl1 | testin LIM domain protein like 1 |
| chr11 | 28944485 | 28945014 | 530 | * | 6,305605 | 7,137579126 | 1,895354 | 3,765829 | 0,000166 | 1 | 0 | Distal Intergenic | 11 | 28865667 | 28876743 | 11077 | 1 | 216616 | ENSMUST00000139713.2 | 78818 | ENSMUSG00000020467 | Efemp1 | epidermal growth factor-containing fibulin-like extracellular matrix protein 1 |
| chr10 | 30591189 | 30592380 | 1192 | * | 7,484799 | 7,13856701 | 1,894072 | 3,768899 | 0,000164 | 0,221323359 | 1,360622 | Intron (ENSMUST0000( | 10 | 30570161 | 30598701 | 28541 | 2 | 211329 | ENSMUST00000217398.2 | 6321 | ENSMUSG00000039697 | Ncoa7 | nuclear receptor coactivator 7 |
| chr12 | 21201368 | 21201775 | 408 | * | 7,662743 | 7,139928454 | 1,850485 | 3,85841 | 0,000114 | 1 | 0 | Intron (ENSMUST0000( | 12 | 21161955 | 21318180 | 156226 | 1 | 211914 | ENSMUST00000050990.10 | 39413 | ENSMUSG00000052632 | Asap2 | ArfGAP with SH3 domain, ankyrin repeat and PH domain 2 |
| chr11 | 86532143 | 86532728 | 586 | * | 6,42376 | 7,143670729 | 1,900753 | 3,758336 | 0,000171 | 1 | 0 | Intron (ENSMUST0000( | 11 | 86521254 | 86521852 | 599 | 2 | 1,01E+08 | ENSMUST00000117851.2 | -10291 | ENSMUSG00000083992 | Gm11478 | predicted gene 11478 |
| chr14 | 26254142 | 26255114 | 973 | * | 8,644301 | 7,146256433 | 1,76041 | 4,059427 | 4,92E-05 | 0,080427024 | 2,237403 | Promoter (<=1kb) | 14 | 26134330 | 26254895 | 120566 | 2 | 83997 | ENSMUST00000038522.10 | 0 | ENSMUSG00000021870 | Slmap | sarcolemma associated protein |
| chr11 | 79356456 | 79357570 | 1115 | * | 6,89982 | 7,148589057 | 1,857152 | 3,849221 | 0,000118 | 1 | 0 | Intron (ENSMUST0000( | 11 | 79372545 | 79374857 | 2313 | 2 | 104522 | ENSMUST00000165610.2 | 17287 | ENSMUSG00000091523 | AU040972 | expressed sequence AU040972 |
| chrX | 1,6E+08 | 1,6E+08 | 327 | * | 4,464274 | 7,153287898 | 2,169753 | 3,296821 | 0,000978 | 1 | 0 | Promoter (<=1kb) | 20 | 1,6E+08 | 1,6E+08 | 210397 | 2 | 382253 | ENSMUST00000087104.11 | -38 | ENSMUSG00000031292 | Cdkl5 | cyclin-dependent kinase-like 5 |
| chr1 | 59954098 | 59954790 | 693 | * | 4,81577 | 7,156020564 | 2,04846 | 3,493365 | 0,000477 | 1 | 0 | Promoter (1-2kb) | 1 | 59952165 | 60024505 | 72341 | 1 | 72750 | ENSMUST00000036540.12 | 1933 | ENSMUSG00000041040 | Fam117b | family with sequence similarity 117, member B |

| | | | | | | | | | | | | | | | | | | | | | | | |
|---|---|---|---|---|---|---|---|---|---|---|---|---|---|---|---|---|---|---|---|---|---|---|---|
| chrX | 96704404 | 96705078 | 675 | * | 5,178753 | 7,158504629 | 2,08897 | 3,42681 | 0,000611 | 1 | 0 | Distal Intergenic | 20 | 96381397 | 96420822 | 39426 | 2 | 245527 | ENSMUST00000126283.8 | -283582 | ENSMUSG00000034457 | Eda2r | ectodysplasin A2 receptor |
| chr9 | 1,06E+08 | 1,06E+08 | 1352 | * | 11,69447 | 7,16028972 | 1,695221 | 4,22381 | 2,4E-05 | 0,072320047 | 1,2763 | Distal Intergenic | 9 | 1,06E+08 | 1,06E+08 | 69461 | 1 | 70235 | ENSMUST00000072206.14 | -8578 | ENSMUSG00000023345 | Poc1a | POC1 centriolar protein A |
| chr9 | 63228444 | 63229469 | 1026 | * | 11,04969 | 7,161843459 | 1,707371 | 4,194662 | 2,73E-05 | 0,108693506 | 0,738441 | Intron (ENSMUST0000 | 9 | 63123553 | 63210425 | 86873 | 2 | 23938 | ENSMUST00000213604.2 | -18019 | ENSMUSG00000058444 | Map2k5 | mitogen-activated protein kinase kinase 5 |
| chr14 | 79893687 | 79894160 | 474 | * | 4,116205 | 7,163265556 | 2,462743 | 2,908653 | 0,00363 | 1 | 0 | Exon (ENSMUST00000 | 14 | 79875130 | 79899524 | 24395 | 2 | 16840 | ENSMUST00000165835.8 | 5364 | ENSMUSG00000022025 | Cnmd | chondromodulin |
| chr12 | 79582476 | 79583740 | 1265 | * | 9,197929 | 7,164488384 | 1,790846 | 4,000616 | 6,32E-05 | 0,120486853 | 1,449706 | Intron (ENSMUST0000 | 12 | 79374127 | 79748630 | 374504 | 1 | 19363 | ENSMUST00000221257.2 | 208349 | ENSMUSG00000059060 | Rad51b | RAD51 paralog B |
| chr1 | 1,9E+08 | 1,9E+08 | 1345 | * | 6,456666 | 7,166550835 | 1,873296 | 3,825638 | 0,00013 | 1 | 0 | Promoter (2-3kb) | 1 | 1,9E+08 | 1,9E+08 | 775 | 1 | 19250 | ENSMUST00000148129.2 | 2856 | ENSMUSG00000026604 | Ptpn14 | protein tyrosine phosphatase, non-receptor type 14 |
| chr11 | 1,16E+08 | 1,16E+08 | 604 | * | 5,347088 | 7,167146407 | 1,940487 | 3,693479 | 0,000221 | 1 | 0 | Promoter (<=1kb) | 11 | 1,16E+08 | 1,16E+08 | 27158 | 2 | 11430 | ENSMUST00000066587.12 | -771 | ENSMUSG00000020777 | Acox1 | acyl-Coenzyme A oxidase 1, palmitoyl |
| chr4 | 32799975 | 32800915 | 941 | * | 5,576554 | 7,170826977 | 2,037931 | 3,518679 | 0,000434 | 1 | 0 | Promoter (<=1kb) | 4 | 32800253 | 32801383 | 1131 | 1 | 108755 | ENSMUST00000062802.5 | 0 | ENSMUSG00000045854 | Lyrm2 | LYR motif containing 2 |
| chr17 | 26472409 | 26473279 | 871 | * | 5,812368 | 7,171844269 | 1,95892 | 3,661121 | 0,000251 | 1 | 0 | Promoter (<=1kb) | 17 | 26472196 | 26485367 | 13172 | 1 | 66978 | ENSMUST00000155151.3 | 213 | ENSMUSG00000024188 | Luc7l | Luc7-like |
| chr5 | 32393149 | 32393888 | 740 | * | 6,213795 | 7,172025346 | 1,946921 | 3,683779 | 0,00023 | 1 | 0 | Promoter (2-3kb) | 5 | 32396428 | 32479588 | 83161 | 1 | 665270 | ENSMUST00000202201.4 | -2540 | ENSMUSG00000029134 | Plb1 | phospholipase B1 |
| chr15 | 1,03E+08 | 1,03E+08 | 614 | * | 6,670606 | 7,172606429 | 1,826818 | 3,926285 | 8,63E-05 | 1 | 0 | Promoter (2-3kb) | 15 | 1,03E+08 | 1,03E+08 | 4462 | 1 | 15382 | ENSMUST00000231141.2 | 2351 | ENSMUSG00000046434 | Hnrnpa1 | heterogeneous nuclear ribonucleoprotein A1 |
| chr1 | 33027278 | 33027928 | 651 | * | 4,823702 | 7,17452986 | 2,119778 | 3,384566 | 0,000713 | 1 | 0 | Distal Intergenic | 1 | 33231074 | 33231183 | 110 | 1 | 1,15E+08 | ENSMUST00000158125.3 | -203146 | ENSMUSG00000088750 | Gm24901 | predicted gene, 24901 |
| chr1 | 1,84E+08 | 1,84E+08 | 900 | * | 4,977832 | 7,177369898 | 2,086484 | 3,439935 | 0,000582 | 1 | 0 | Intron (ENSMUST0000 | 1 | 1,84E+08 | 1,84E+08 | 41256 | 1 | 63953 | ENSMUST00000048655.8 | 24545 | ENSMUSG00000039384 | Dusp10 | dual specificity phosphatase 10 |
| chrX | 57700871 | 57701390 | 520 | * | 5,576438 | 7,178245595 | 1,92572 | 3,727564 | 0,000193 | 1 | 0 | Distal Intergenic | 20 | 57954720 | 57965664 | 10945 | 1 | 75352 | ENSMUST00000118305.8 | -253330 | ENSMUSG00000046180 | Magea13 | MAGE family member A13 |
| chrX | 12422746 | 12423739 | 994 | * | 5,767862 | 7,178650082 | 1,937498 | 3,705113 | 0,000211 | 1 | 0 | Distal Intergenic | 20 | 12454040 | 12462820 | 8781 | 1 | 70495 | ENSMUST00000137277.2 | -30301 | ENSMUSG00000031007 | Atp6ap2 | ATPase, H+ transporting, lysosomal accessory protein 2 |
| chrX | 57097739 | 57098592 | 854 | * | 6,364558 | 7,184036924 | 1,904292 | 3,772551 | 0,000162 | 1 | 0 | Distal Intergenic | 20 | 57076370 | 57087096 | 10727 | 1 | 22773 | ENSMUST00000088629.4 | 21369 | ENSMUSG00000067860 | Zic3 | zinc finger protein of the cerebellum 3 |
| chr7 | 93377514 | 93378297 | 784 | * | 4,941443 | 7,189408763 | 1,972671 | 3,644504 | 0,000268 | 1 | 0 | Distal Intergenic | 7 | 93691898 | 93998607 | 306710 | 1 | 1,03E+08 | ENSMUST00000206116.2 | -313601 | | Gm32647 | predicted gene, 32647 |
| chr4 | 1,56E+08 | 1,56E+08 | 1223 | * | 10,49139 | 7,191520666 | 1,706301 | 4,214684 | 2,5E-05 | 0,074746651 | 1,2763 | Promoter (1-2kb) | 4 | 1,56E+08 | 1,56E+08 | 13881 | 1 | 140499 | ENSMUST00000103175.8 | -1509 | ENSMUSG00000023286 | Ube2j2 | ubiquitin-conjugating enzyme E2J 2 |
| chr14 | 31386137 | 31386943 | 807 | * | 8,721388 | 7,197590593 | 1,779985 | 4,043625 | 5,26E-05 | 1 | 0 | Intron (ENSMUST0000 | 14 | 31357605 | 31363243 | 5639 | 2 | 56794 | ENSMUST00000135751.2 | -22894 | ENSMUSG00000021884 | Hacl1 | 2-hydroxyacyl-CoA lyase 1 |
| chr2 | 97933596 | 97934232 | 637 | * | 5,443446 | 7,200939072 | 1,923102 | 3,74444 | 0,000181 | 1 | 0 | Distal Intergenic | 2 | 97298434 | 97459408 | 160975 | 1 | 241568 | ENSMUST00000170144.2 | 635162 | ENSMUSG00000050587 | Lrrc4c | leucine rich repeat containing 4C |
| chr11 | 1,21E+08 | 1,21E+08 | 556 | * | 5,69498 | 7,20292818 | 2,070053 | 3,479586 | 0,000502 | 1 | 0 | Intron (ENSMUST0000 | 11 | 1,21E+08 | 1,21E+08 | 106344 | 2 | 71276 | ENSMUST00000056781.5 | 4302 | ENSMUSG00000048445 | Ccdc57 | coiled-coil domain containing 57 |
| chr11 | 86707251 | 86708083 | 833 | * | 6,591157 | 7,2065075 | 1,910148 | 3,772748 | 0,000161 | 1 | 0 | Distal Intergenic | 11 | 86659672 | 86698572 | 38901 | 2 | 67487 | ENSMUST00000018569.14 | -8679 | ENSMUSG00000018425 | Dhx40 | DEAH (Asp-Glu-Ala-His) box polypeptide 40 |
| chr3 | 1,08E+08 | 1,08E+08 | 846 | * | 7,659863 | 7,206784068 | 1,894349 | 3,80436 | 0,000142 | 0,204595855 | 1,360622 | Promoter (<=1kb) | 3 | 1,08E+08 | 1,08E+08 | 414 | 2 | 1E+08 | ENSMUST00000157560.3 | -45 | | Scarna2 | small Cajal body-specific RNA 2 |
| chr4 | 74030882 | 74031720 | 839 | * | 5,247563 | 7,213929194 | 2,100219 | 3,434847 | 0,000593 | 1 | 0 | Intron (ENSMUST0000 | 4 | 74087174 | 74107238 | 20065 | 1 | 242506 | ENSMUST00000154134.2 | -55454 | ENSMUSG00000049122 | Frmd3 | FERM domain containing 3 |
| chr2 | 1,66E+08 | 1,66E+08 | 461 | * | 3,667643 | 7,215673778 | 2,385433 | 3,02489 | 0,002487 | 1 | 0 | Intron (ENSMUST0000 | 2 | 1,66E+08 | 1,66E+08 | 72009 | 2 | 277360 | ENSMUST00000099080.9 | -8129 | ENSMUSG00000039621 | Prex1 | phosphatidylinositol-3,4,5-trisphosphate-dependent Rac exchange factor 1 |
| chrX | 1,33E+08 | 1,33E+08 | 686 | * | 6,009281 | 7,216109994 | 1,967198 | 3,668217 | 0,000244 | 1 | 0 | Distal Intergenic | 20 | 1,32E+08 | 1,33E+08 | 106128 | 2 | 279653 | ENSMUST00000149154.8 | -6441 | ENSMUSG00000051323 | Pcdh19 | protocadherin 19 |
| chr1 | 1,8E+08 | 1,8E+08 | 1018 | * | 7,564503 | 7,217768261 | 1,902493 | 3,793848 | 0,000148 | 1 | 0 | Promoter (2-3kb) | 1 | 1,8E+08 | 1,8E+08 | 14412 | 2 | 67426 | ENSMUST00000160482.8 | 2617 | ENSMUSG00000026489 | Coq8a | coenzyme Q8A |
| chr12 | 79667632 | 79668583 | 952 | * | 8,310701 | 7,219944304 | 1,827407 | 3,950923 | 7,79E-05 | 0,143892051 | 1,360622 | Intron (ENSMUST0000 | 12 | 79938062 | 79938677 | 616 | 1 | 1,08E+08 | ENSMUST00000243367.1 | -269479 | | Gm46362 | predicted gene, 46362 |
| chr14 | 47482060 | 47482998 | 939 | * | 9,050386 | 7,222901976 | 1,748842 | 4,130106 | 3,63E-05 | 0,065868596 | 2,237403 | Promoter (<=1kb) | 14 | 47478989 | 47482727 | 3739 | 2 | 218973 | ENSMUST00000228810.2 | 0 | ENSMUSG00000037572 | Wdhd1 | WD repeat and HMG-box DNA binding protein 1 |
| chr4 | 1,3E+08 | 1,3E+08 | 1177 | * | 8,650494 | 7,223009968 | 1,780613 | 4,056473 | 4,98E-05 | 0,103578649 | 1,449706 | Intron (ENSMUST0000 | 4 | 1,3E+08 | 1,3E+08 | 18646 | 1 | 19244 | ENSMUST00000133662.2 | 7747 | ENSMUSG00000028788 | Ptp4a2 | protein tyrosine phosphatase 4a2 |
| chr13 | 1,04E+08 | 1,04E+08 | 539 | * | 7,534817 | 7,223369312 | 1,890344 | 3,821194 | 0,000133 | 0,197827068 | 1,360622 | Intron (ENSMUST0000 | 13 | 1,04E+08 | 1,04E+08 | 38633 | 2 | 75805 | ENSMUST00000225324.2 | -3865 | ENSMUSG00000021710 | Nln | neurolysin (metallopeptidase M3 family) |
| chr3 | 67132970 | 67134299 | 1330 | * | 12,5397 | 7,225244126 | 1,715096 | 4,212734 | 2,52E-05 | 0,079560837 | 1,173325 | Intron (ENSMUST0000 | 3 | 67281430 | 67307331 | 25902 | 1 | 17349 | ENSMUST00000077916.12 | -147131 | ENSMUSG00000048416 | Mlf1 | myeloid leukemia factor 1 |
| chr1 | 1,67E+08 | 1,67E+08 | 980 | * | 6,946138 | 7,228066299 | 1,925534 | 3,753798 | 0,000174 | 1 | 0 | Distal Intergenic | 1 | 1,68E+08 | 1,68E+08 | 159505 | 1 | 110648 | ENSMUST00000111377.8 | -26940 | ENSMUSG00000026686 | Lmx1a | LIM homeobox transcription factor 1 alpha |
| chr2 | 1,2E+08 | 1,2E+08 | 1356 | * | 9,599698 | 7,229863015 | 1,770026 | 4,084609 | 4,42E-05 | 0,115300307 | 1,086118 | Promoter (2-3kb) | 2 | 1,2E+08 | 1,2E+08 | 2410 | 1 | 29808 | ENSMUST00000150742.2 | 2362 | ENSMUSG00000033943 | Mga | MAX gene associated |
| chrX | 1,69E+08 | 1,69E+08 | 636 | * | 5,327842 | 7,233045211 | 1,999915 | 3,616677 | 0,000298 | 1 | 0 | Intron (ENSMUST0000 | 20 | 1,69E+08 | 1,69E+08 | 3258 | 2 | 66747 | ENSMUST00000068874.8 | -9842 | ENSMUSG00000055357 | 4933400A11Rik | capping protein (actin filament) muscle Z-line, alpha 1 pseudogene |
| chr14 | 31610795 | 31611814 | 1020 | * | 7,21097 | 7,239581401 | 1,899319 | 3,811672 | 0,000138 | 1 | 0 | Distal Intergenic | 14 | 31651545 | 31653675 | 2131 | 1 | 1,03E+08 | ENSMUST00000242479.1 | -39731 | | Gm53172 | predicted gene, 53172 |
| chr16 | 94009839 | 94011549 | 1711 | * | 13,35602 | 7,249234556 | 1,725565 | 4,201078 | 2,66E-05 | 0,09162632 | 0,938682 | Intron (ENSMUST0000 | 16 | 93932798 | 94006493 | 73696 | 2 | 110948 | ENSMUST00000227447.2 | -3346 | ENSMUSG00000040820 | Hlcs | holocarboxylase synthetase (biotin- [propriony-Coenzyme A-carboxylase (ATP-hydrolysing)] ligase) |
| chr16 | 92679471 | 92680234 | 764 | * | 7,009036 | 7,250783518 | 1,820272 | 3,983352 | 6,79E-05 | 1 | 0 | Distal Intergenic | 16 | 92398354 | 92623037 | 224684 | 2 | 12394 | ENSMUST00000023673.14 | -56434 | ENSMUSG00000022952 | Runx1 | runt related transcription factor 1 |
| chr9 | 19371920 | 19372487 | 568 | * | 3,963021 | 7,252107105 | 2,503423 | 2,896877 | 0,003769 | 1 | 0 | Distal Intergenic | 9 | 19388597 | 19389543 | 947 | 2 | 258516 | ENSMUST00000211832.2 | 17056 | ENSMUSG00000094535 | Olfr850 | olfactory receptor 850 |
| chr13 | 52859616 | 52860713 | 1098 | * | 9,041375 | 7,252872618 | 1,798025 | 4,033801 | 5,49E-05 | 0,112224124 | 1,425929 | Distal Intergenic | 13 | 52883651 | 52889644 | 5994 | 2 | 105404 | ENSMUST00000222580.2 | 28931 | | BB123696 | expressed sequence BB123696 |
| chr11 | 78066179 | 78066578 | 400 | * | 4,083703 | 7,254003622 | 2,292056 | 3,164845 | 0,001552 | 1 | 0 | Promoter (<=1kb) | 11 | 78056932 | 78067437 | 10506 | 2 | 140859 | ENSMUST00000152611.8 | 859 | ENSMUSG00000017405 | Nek8 | NIMA (never in mitosis gene a)-related expressed kinase 8 |
| chrX | 62962784 | 62963594 | 811 | * | 5,531599 | 7,259497049 | 2,034113 | 3,568877 | 0,000359 | 1 | 0 | Distal Intergenic | 20 | 63195011 | 63195804 | 794 | 2 | 627470 | ENSMUST00000114682.4 | 232210 | ENSMUSG00000079579 | Gm6760 | predicted gene 6760 |
| chr5 | 1,44E+08 | 1,44E+08 | 982 | * | 7,148858 | 7,261434559 | 1,829103 | 3,969943 | 7,19E-05 | 1 | 0 | Intron (ENSMUST0000 | 5 | 1,44E+08 | 1,44E+08 | 4156 | 1 | 17341 | ENSMUST00000060747.8 | -35197 | ENSMUSG00000052271 | Bhlha15 | basic helix-loop-helix family, member a15 |
| chr3 | 89665148 | 89665795 | 648 | * | 5,530179 | 7,263675877 | 2,054384 | 3,535696 | 0,000407 | 1 | 0 | Intron (ENSMUST0000 | 3 | 89659740 | 89670708 | 10969 | 2 | 11444 | ENSMUST00000199372.2 | 4913 | ENSMUSG00000027950 | Chrnb2 | cholinergic receptor, nicotinic, beta polypeptide 2 (neuronal) |
| chr19 | 5930835 | 5931681 | 847 | * | 7,529135 | 7,267526741 | 1,804506 | 4,027433 | 5,64E-05 | 1 | 0 | Promoter (2-3kb) | 19 | 5928684 | 5934513 | 5830 | 1 | 107375 | ENSMUST00000141362.2 | 2151 | ENSMUSG00000024818 | Slc25a45 | solute carrier family 25, member 45 |
| chr5 | 37981801 | 37982661 | 861 | * | 9,506643 | 7,267527144 | 1,803974 | 4,028621 | 5,61E-05 | 0,113460495 | 1,425929 | Promoter (<=1kb) | 5 | 37977829 | 37981927 | 4099 | 2 | 17701 | ENSMUST00000063116.10 | 0 | ENSMUSG00000048450 | Msx1 | msh homeobox 1 |
| chr4 | 48043266 | 48043884 | 619 | * | 8,839718 | 7,272402649 | 1,86822 | 3,89269 | 9,91E-05 | 0,121267917 | 2,237403 | Promoter (1-2kb) | 4 | 48045153 | 48086445 | 41293 | 1 | 18124 | ENSMUST00000030025.10 | -1269 | ENSMUSG00000028341 | Nr4a3 | nuclear receptor subfamily 4, group A, member 3 |
| chr11 | 3867829 | 3868323 | 495 | * | 6,643506 | 7,276783144 | 1,915698 | 3,798502 | 0,000146 | 1 | 0 | Intron (ENSMUST0000 | 11 | 3857022 | 3864664 | 7643 | 2 | 103710 | ENSMUST00000051207.2 | -3165 | ENSMUSG00000048807 | Slc35e4 | solute carrier family 35, member E4 |
| chr9 | 3342809 | 3343735 | 927 | * | 7,363763 | 7,28176747 | 1,980193 | 3,677302 | 0,000236 | 1 | 0 | 5' UTR | 9 | 3338445 | 3385842 | 47398 | 1 | 67667 | ENSMUST00000165105.2 | 4364 | ENSMUSG00000025899 | Alkbh8 | alkB homolog 8, tRNA methyltransferase |
| chr18 | 8651208 | 8651659 | 452 | * | 4,868793 | 7,284930789 | 2,111815 | 3,449606 | 0,000561 | 1 | 0 | Distal Intergenic | 18 | 9212163 | 9218136 | 5974 | 1 | 14370 | ENSMUST00000041080.7 | -560504 | ENSMUSG00000036904 | Fzd8 | frizzled class receptor 8 |
| chr17 | 91298143 | 91298942 | 800 | * | 7,383168 | 7,287039893 | 1,818653 | 4,006834 | 6,15E-05 | 1 | 0 | Exon (ENSMUST00000 | 17 | 90842634 | 91395911 | 553278 | 2 | 18189 | ENSMUST00000161637.8 | 96969 | ENSMUSG00000024109 | Nrxn1 | neurexin I |
| chrX | 59600161 | 59600918 | 758 | * | 5,468181 | 7,292016743 | 2,181761 | 3,342262 | 0,000831 | 1 | 0 | Promoter (<=1kb) | 20 | 59601334 | 59851599 | 250266 | 1 | 1,05E+08 | ENSMUST00000152669.2 | -416 | | Gm14661 | predicted gene 14661 |
| chr19 | 12576626 | 12577013 | 388 | * | 4,304296 | 7,298323669 | 2,279395 | 3,201869 | 0,001365 | 1 | 0 | Promoter (<=1kb) | 19 | 12577338 | 12604237 | 26900 | 1 | 240549 | ENSMUST00000181868.8 | -325 | ENSMUSG00000071633 | Gm4952 | predicted gene 4952 |
| chr17 | 74079695 | 74080049 | 355 | * | 5,884899 | 7,30141319 | 1,994152 | 3,661412 | 0,000251 | 1 | 0 | Distal Intergenic | 17 | 74111836 | 74139089 | 27254 | 1 | 57440 | ENSMUST00000024860.9 | -31787 | ENSMUSG00000024065 | Ehd3 | EH-domain containing 3 |
| chr13 | 51144209 | 51145352 | 1144 | * | 8,354563 | 7,30376932 | 1,878577 | 3,887926 | 0,000101 | 0,168212038 | 1,360622 | Distal Intergenic | 13 | 51254916 | 51306582 | 51667 | 1 | 20729 | ENSMUST00000095797.6 | -109564 | ENSMUSG00000021395 | Spin1 | spindlin 1 |
| chr7 | 65566529 | 65567638 | 1110 | * | 11,59584 | 7,308513083 | 1,747384 | 4,182544 | 2,88E-05 | 0,113207887 | 0,738441 | Intron (ENSMUST0000 | 7 | 65602760 | 65612941 | 10182 | 1 | 18553 | ENSMUST00000177369.2 | -35122 | ENSMUSG00000030513 | Pcsk6 | proprotein convertase subtilisin/kexin type 6 |
| chr12 | 79560394 | 79561082 | 689 | * | 5,054393 | 7,346390987 | 2,099111 | 3,499763 | 0,000466 | 1 | 0 | Intron (ENSMUST0000 | 12 | 79374127 | 79748630 | 374504 | 1 | 19363 | ENSMUST00000221257.2 | 186267 | ENSMUSG00000059060 | Rad51b | RAD51 paralog B |
| chr19 | 7006015 | 7006889 | 875 | * | 6,213668 | 7,35318833 | 2,064557 | 3,561631 | 0,000369 | 1 | 0 | Exon (ENSMUST00000 | 19 | 6976326 | 6996837 | 20512 | 2 | 108101 | ENSMUST00000040772.9 | -9178 | ENSMUSG00000024965 | Fermt3 | fermitin family member 3 |
| chrX | 1,16E+08 | 1,16E+08 | 1288 | * | 7,061073 | 7,355131293 | 1,830087 | 4,019007 | 5,84E-05 | 1 | 0 | Distal Intergenic | 20 | 1,16E+08 | 1,16E+08 | 348 | 1 | 624153 | ENSMUST00000118616.3 | 6537 | ENSMUSG00000082482 | H2ab2 | H2A.B variant histone 2 |
| chr12 | 73024480 | 73025371 | 892 | * | 7,101648 | 7,355464355 | 1,871925 | 3,929358 | 8,52E-05 | 1 | 0 | Distal Intergenic | 12 | 72958433 | 72987548 | 29116 | 2 | 75801 | ENSMUST00000136075.8 | -36932 | ENSMUSG00000021098 | 4930447C04Rik | RIKEN cDNA 4930447C04 gene |
| chr5 | 1,24E+08 | 1,24E+08 | 1344 | * | 8,021697 | 7,356044723 | 1,812877 | 4,057662 | 4,96E-05 | 1 | 0 | Promoter (1-2kb) | 5 | 1,24E+08 | 1,24E+08 | 4496 | 2 | 330192 | ENSMUST00000199741.2 | 1553 | ENSMUSG00000066278 | Vps37b | vacuolar protein sorting 37B |
| chr14 | 58037347 | 58038561 | 1215 | * | 9,877951 | 7,356347579 | 1,796046 | 4,095858 | 4,21E-05 | 0,10388 | 1,211315 | Promoter (1-2kb) | 14 | 58036095 | 58039792 | 3698 | 1 | 20220 | ENSMUST00000111267.2 | 1252 | ENSMUSG00000021963 | Sap18 | Sin3-associated polypeptide 18 |
| chr2 | 1,13E+08 | 1,13E+08 | 895 | * | 6,467254 | 7,364246393 | 1,963747 | 3,750098 | 0,000177 | 1 | 0 | Intron (ENSMUST0000 | 2 | 1,13E+08 | 1,14E+08 | 345767 | 1 | 14260 | ENSMUST00000161731.5 | 11954 | ENSMUSG00000044042 | Fmn1 | formin 1 |
| chr1 | 51518563 | 51519288 | 726 | * | 6,252447 | 7,370290562 | 1,858392 | 3,965951 | 7,31E-05 | 1 | 0 | Promoter (<=1kb) | 1 | 51510288 | 51517584 | 7297 | 2 | 109019 | ENSMUST00000188051.7 | -979 | ENSMUSG00000026107 | Nabp1 | nucleic acid binding protein 1 |
| chr9 | 63299734 | 63300797 | 1064 | * | 9,051168 | 7,370466495 | 1,793157 | 4,110329 | 3,95E-05 | 0,099805478 | 1,211315 | Distal Intergenic | 9 | 63305883 | 63400817 | 94935 | 1 | 1,03E+08 | ENSMUST00000191455.7 | -5086 | | Gm16759 | predicted gene, 16759 |
| chrX | 73466677 | 73467559 | 883 | * | 5,694616 | 7,375158741 | 2,067368 | 3,567414 | 0,000361 | 1 | 0 | Promoter (<=1kb) | 20 | 73468140 | 73483556 | 15417 | 1 | 16151 | ENSMUST00000135165.8 | -581 | ENSMUSG00000004221 | Ikbkg | inhibitor of kappaB kinase gamma |
| chr2 | 25993831 | 25994535 | 705 | * | 10,1341 | 7,378705968 | 1,760132 | 4,192131 | 2,76E-05 | 0,072876072 | 1,449706 | Intron (ENSMUST0000 | 2 | 25945547 | 25982189 | 36643 | 2 | 67991 | ENSMUST00000114159.9 | -11642 | ENSMUSG00000026932 | Nacc2 | nucleus accumbens associated 2, BEN and BTB (POZ) domain containing |
| chr11 | 86636915 | 86637726 | 812 | * | 8,416067 | 7,3789763 | 1,794695 | 4,11155 | 3,93E-05 | 1 | 0 | Intron (ENSMUST0000 | 11 | 86585481 | 86648327 | 62847 | 2 | 67300 | ENSMUST00000060766.16 | 10601 | ENSMUSG00000047126 | Cltc | clathrin, heavy polypeptide (Hc) |
| chr2 | 57010138 | 57010904 | 767 | * | 6,317251 | 7,38024236 | 1,968258 | 3,749631 | 0,000177 | 1 | 0 | Intron (ENSMUST0000 | 2 | 56996842 | 57014015 | 17174 | 2 | 18227 | ENSMUST00000112629.8 | 3111 | ENSMUSG00000026826 | Nr4a2 | nuclear receptor subfamily 4, group A, member 2 |
| chr1 | 1,67E+08 | 1,67E+08 | 686 | * | 5,552181 | 7,380244355 | 1,987987 | 3,712421 | 0,000205 | 1 | 0 | Distal Intergenic | 1 | 1,66E+08 | 1,67E+08 | 19791 | 2 | 240894 | ENSMUST00000027843.11 | -113683 | ENSMUSG00000026560 | Fmo9 | flavin containing monooxygenase 9 |
| chr16 | 38245547 | 38246289 | 743 | * | 6,54476 | 7,380256396 | 1,870644 | 3,945302 | 7,97E-05 | 1 | 0 | Intron (ENSMUST0000 | 16 | 38216479 | 38253507 | 37029 | 2 | 85031 | ENSMUST00000002926.8 | 7218 | ENSMUSG00000002847 | Pla1a | phospholipase A1 member A |
| chr13 | 53428338 | 53429397 | 1060 | * | 8,956432 | 7,391224348 | 1,791196 | 4,126418 | 3,68E-05 | 0,066764601 | 2,237403 | Promoter (<=1kb) | 13 | 53285996 | 53429694 | 143699 | 2 | 26564 | ENSMUST00000149299.2 | 297 | ENSMUSG00000021464 | Ror2 | receptor tyrosine kinase-like orphan receptor 2 |
| chr7 | 1,01E+08 | 1,01E+08 | 874 | * | 7,763796 | 7,395327339 | 1,875994 | 3,942084 | 8,08E-05 | 1 | 0 | Distal Intergenic | 7 | 1,01E+08 | 1,01E+08 | 104041 | 1 | 207278 | ENSMUST00000208439.2 | -27014 | ENSMUSG00000030691 | Fchsd2 | FCH and double SH3 domains 2 |
| chr4 | 44616246 | 44617716 | 1471 | * | 11,57837 | 7,403390195 | 1,761165 | 4,20369 | 2,63E-05 | 0,080985792 | 1,173325 | Intron (ENSMUST0000 | 4 | 44607491 | 44607568 | 78 | 2 | 1,01E+08 | ENSMUST00000175020.2 | -8678 | ENSMUSG00000092761 | Mir5120 | microRNA 5120 |
| chrX | 16349883 | 16350650 | 768 | * | 6,118016 | 7,409633841 | 2,042742 | 3,627298 | 0,000286 | 1 | 0 | Distal Intergenic | 20 | 16389108 | 16389928 | 821 | 1 | 66742 | ENSMUST00000115422.2 | -38458 | ENSMUSG00000023257 | Cypt1 | cysteine-rich perinuclear theca 1 |
| chr14 | 54667951 | 54669112 | 1162 | * | 11,46113 | 7,41573921 | 2,441139 | 3,03782 | 0,002383 | 0,749931768 | 0,738441 | Promoter (<=1kb) | 14 | 54669069 | 54678722 | 9654 | 1 | 17387 | ENSMUST00000089688.6 | 0 | ENSMUSG00000000957 | Mmp14 | matrix metallopeptidase 14 (membrane-inserted) |
| chr9 | 1,05E+08 | 1,05E+08 | 599 | * | 7,25081 | 7,423294141 | 1,905482 | 3,895758 | 9,79E-05 | 1 | 0 | Intron (ENSMUST0000 | 9 | 1,05E+08 | 1,05E+08 | 149743 | 1 | 74020 | ENSMUST00000213452.2 | -10290 | ENSMUSG00000032564 | Cpne4 | copine IV |
| chr16 | 86714684 | 86715033 | 350 | * | 4,273785 | 7,424080639 | 2,241106 | 3,312687 | 0,000924 | 1 | 0 | Distal Intergenic | 16 | 86613780 | 86674259 | 60480 | 1 | 1,03E+08 | ENSMUST00000242045.1 | 100904 | | Gm32624 | predicted gene, 32624 |
| chr9 | 34187626 | 34188634 | 1009 | * | 9,360566 | 7,428429863 | 1,866409 | 3,980065 | 6,89E-05 | 0,14371218 | 1,211315 | Distal Intergenic | 9 | 34397190 | 34400454 | 3265 | 1 | 67703 | ENSMUST00000184203.8 | -208556 | ENSMUSG00000032036 | Kirrel3 | kirre like nephrin family adhesion molecule 3 |
| chr18 | 32304857 | 32306074 | 1218 | * | 9,653414 | 7,428522773 | 1,806069 | 4,113089 | 3,9E-05 | 0,106347561 | 1,086118 | Intron (ENSMUST0000 | 18 | 32296249 | 32369804 | 73556 | 1 | 26405 | ENSMUST00000096575.5 | 8608 | ENSMUSG00000024383 | Map3k2 | mitogen-activated protein kinase kinase kinase 2 |
| chr11 | 6421451 | 6422594 | 1144 | * | 8,286461 | 7,429668893 | 1,758185 | 4,225762 | 2,38E-05 | 0,050236155 | 2,237403 | Promoter (<=1kb) | 11 | 6422225 | 6422277 | 53 | 2 | 19291 | ENSMUST00000175144.4 | 0 | ENSMUSG00000094483 | Purb | purine rich element binding protein B |
| chr10 | 77589997 | 77590364 | 368 | * | 4,928371 | 7,431379403 | 2,161573 | 3,437951 | 0,000586 | 1 | 0 | Promoter (2-3kb) | 10 | 77586355 | 77587188 | 834 | 2 | 77763 | ENSMUST00000219969.2 | -2809 | ENSMUSG00000112380 | Gm9736 | predicted gene 9736 |
| chr2 | 1,49E+08 | 1,49E+08 | 371 | * | 5,669406 | 7,431736066 | 2,133055 | 3,484081 | 0,000494 | 1 | 0 | Distal Intergenic | 2 | 1,49E+08 | 1,49E+08 | 4961 | 1 | 58214 | ENSMUST00000109939.3 | 138910 | ENSMUSG00000033156 | Cst10 | cystatin 10 (chondrocytes) |
| chr14 | 1,24E+08 | 1,24E+08 | 684 | * | 5,39955 | 7,432784603 | 1,925994 | 3,859194 | 0,000114 | 1 | 0 | Intron (ENSMUST0000 | 14 | 1,24E+08 | 1,24E+08 | 28050 | 1 | 69393 | ENSMUST00000190088.2 | 5146 | ENSMUSG00000099994 | 1700024B18Rik | RIKEN cDNA 1700024B18 gene |
| chrX | 1,33E+08 | 1,33E+08 | 384 | * | 5,379338 | 7,459201693 | 2,000473 | 3,728719 | 0,000192 | 1 | 0 | Distal Intergenic | 20 | 1,33E+08 | 1,33E+08 | 14598 | 1 | 64103 | ENSMUST00000033602.9 | -52668 | ENSMUSG00000031250 | Tnmd | tenomodulin |
| chr12 | 26930735 | 26931610 | 876 | * | 6,058673 | 7,461942508 | 1,865789 | 3,999349 | 6,35E-05 | 1 | 0 | Distal Intergenic | 12 | 26869784 | 26884872 | 15089 | 2 | 74668 | ENSMUST00000222026.2 | -45863 | ENSMUSG00000113218 | 4933409F18Rik | RIKEN cDNA 4933409F18 gene |
| chr15 | 79794680 | 79796783 | 2104 | * | 15,92389 | 7,464646864 | 1,649543 | 4,5252818 | 6,032E-06 | 0,027558645 | 1,2763004 | 3' UTR | 15 | 79790324 | 79791292 | 969 | 1 | 80287 | ENSMUST00000175827.2 | 4356 | ENSMUSG00000009585 | Apobec3 | apolipoprotein B mRNA editing enzyme, catalytic polypeptide 3 |
| chr1 | 1,35E+08 | 1,35E+08 | 1001 | * | 10,0543 | 7,464811521 | 1,74994 | 4,265752 | 1,99E-05 | 0,059300632 | 1,449706 | 3' UTR | 1 | 1,35E+08 | 1,35E+08 | 902 | 1 | 67196 | ENSMUST00000139617.2 | 10859 | ENSMUSG00000026429 | Ube2t | ubiquitin-conjugating enzyme E2T |
| chr5 | 1,22E+08 | 1,22E+08 | 804 | * | 6,679108 | 7,471351135 | 1,86162 | 4,01336 | 5,99E-05 | 1 | 0 | 3' UTR | 5 | 1,22E+08 | 1,22E+08 | 26569 | 1 | 320717 | ENSMUST00000119015.2 | 26535 | ENSMUSG00000038582 | Pptc7 | PTC7 protein phosphatase homolog |
| chr7 | 77587904 | 77588689 | 786 | * | 6,948962 | 7,475152774 | 2,06113 | 3,626725 | 0,000287 | 0,29306015 | 1,360622 | Distal Intergenic | 7 | 77830631 | 77842491 | 11861 | 2 | 18213 | ENSMUST00000205354.2 | 253802 | ENSMUSG00000059146 | Ntrk3 | neurotrophic tyrosine kinase, receptor, type 3 |
| chr5 | 1,35E+08 | 1,35E+08 | 953 | * | 9,531002 | 7,477325514 | 1,755588 | 4,259158 | 2,05E-05 | 0,067993447 | 1,211315 | Promoter (1-2kb) | 5 | 1,35E+08 | 1,35E+08 | 4476 | 2 | 16885 | ENSMUST00000137897.2 | -1743 | ENSMUSG00000029674 | Limk1 | LIM-domain containing, protein kinase |
| chr8 | 122317825 | 122319115 | 1291 | * | 16,203832 | 7,477361772 | 1,6739962 | 4,4667736 | 7,941E-06 | 0,021335371 | 2,3860078 | Promoter (<=1kb) | 8 | 122317451 | 122323734 | 6284 | 1 | 67443 | ENSMUST00000181521.2 | 374 | ENSMUSG00000031812 | Map1lc3b | microtubule-associated protein 1 light chain 3 beta |
| chr4 | 1,33E+08 | 1,33E+08 | 850 | * | 8,184072 | 7,48062574 | 1,819778 | 4,110735 | 3,94E-05 | 0,090530822 | 1,425929 | Promoter (<=1kb) | 4 | 1,33E+08 | 1,33E+08 | 7169 | 2 | 230789 | ENSMUST00000148667.2 | -204 | ENSMUSG00000028878 | Fam76a | family with sequence similarity 76, member A |
| chr11 | 95273675 | 95274964 | 1290 | * | 10,48149 | 7,492552542 | 1,7235 | 4,347289 | 1,38E-05 | 0,063243151 | 0,918255 | Promoter (<=1kb) | 11 | 95275518 | 95282602 | 7085 | 1 | 110172 | ENSMUST00000021243.16 | -554 | ENSMUSG00000020873 | Slc35b1 | solute carrier family 35, member B1 |
| chr9 | 1,14E+08 | 1,14E+08 | 807 | * | 7,447834 | 7,494556401 | 1,824638 | 4,107421 | 4E-05 | 1 | 0 | Promoter (2-3kb) | 9 | 1,14E+08 | 1,14E+08 | 177325 | 1 | 76499 | ENSMUST00000213663.2 | -2837 | ENSMUSG00000033392 | Clasp2 | CLIP associating protein 2 |
| chr6 | 34756432 | 34758583 | 2152 | * | 18,192705 | 7,498395095 | 1,6113653 | 4,6534422 | 3,264E-06 | 0,011612601 | 2,1545044 | Promoter (<=1kb) | 6 | 34757367 | 34824049 | 66683 | 1 | 76223 | ENSMUST00000202017.4 | 0 | ENSMUSG00000038836 | Agbl3 | ATP/GTP binding protein-like 3 |
| chr9 | 64934466 | 64935056 | 591 | * | 6,38961 | 7,502561438 | 1,879347 | 3,992111 | 6,55E-05 | 1 | 0 | Distal Intergenic | 9 | 64939696 | 64989933 | 50238 | 1 | 74388 | ENSMUST00000034960.14 | -4640 | ENSMUSG00000032393 | Dpp8 | dipeptidylpeptidase 8 |
| chr3 | 94554013 | 94555764 | 1752 | * | 12,74549 | 7,520248067 | 1,74348 | 4,313355 | 1,61E-05 | 0,080010047 | 0,738441 | Intron (ENSMUST0000 | 3 | 94520303 | 94566107 | 45805 | 2 | 22156 | ENSMUST00000196655.5 | 10343 | ENSMUSG00000005968 | Tuft1 | tuftelin 1 |
| chr19 | 44268947 | 44270124 | 1178 | * | 11,744278 | 7,533129482 | 1,6864871 | 4,4667579 | 7,941E-06 | 0,033161239 | 1,2763004 | Distal Intergenic | 19 | 44260710 | 44266660 | 5951 | 1 | 102638677 | ENSMUST00000237187.2 | 8237 | | Gm35183 | predicted gene, 35183 |
| chr4 | 1,5E+08 | 1,5E+08 | 792 | * | 8,908873 | 7,544400912 | 1,762114 | 4,281448 | 1,86E-05 | 0,063587778 | 1,211315 | Promoter (1-2kb) | 4 | 1,5E+08 | 1,5E+08 | 4443 | 1 | 13806 | ENSMUST00000141931.2 | 1859 | ENSMUSG00000063524 | Eno1 | enolase 1, alpha non-neuron |
| chrX | 1,4E+08 | 1,4E+08 | 985 | * | 7,491081 | 7,549362505 | 1,855168 | 4,069369 | 4,71E-05 | 1 | 0 | Exon (ENSMUST00000 | 20 | 1,4E+08 | 1,4E+08 | 308674 | 2 | 94216 | ENSMUST00000101205.3 | 117467 | ENSMUSG00000031273 | Col4a6 | collagen, type IV, alpha 6 |

| | | | | | | | | | | | | | | | | | | | | | | | |
|---|---|---|---|---|---|---|---|---|---|---|---|---|---|---|---|---|---|---|---|---|---|---|---|
| chr15 | 83060004 | 83060941 | 938 | * | 10,26659 | 7,566948711 | 1,777215 | 4,257756 | 2,06E-05 | 0,061899193 | 1,425929 | Promoter (<=1kb) | 15 | 83057397 | 83060646 | 3250 | 2 | 109754 | ENSMUST00000229317.2 | 0 | ENSMUSG00000018042 | Cyb5r3 | cytochrome b5 reductase 3 |
| chr14 | 55958473 | 55958909 | 437 | * | 6,10201 | 7,568029127 | 2,100431 | 3,603084 | 0,000314 | 1 | 0 | Promoter (<=1kb) | 14 | 55953334 | 55959633 | 6300 | 2 | 56187 | ENSMUST00000062861.15 | 724 | ENSMUSG00000040472 | Rabggta | Rab geranylgeranyl transferase, a subunit |
| chr2 | 1,27E+08 | 1,27E+08 | 543 | * | 7,310221 | 7,568798974 | 1,864498 | 4,059431 | 4,92E-05 | 1 | 0 | Promoter (1-2kb) | 2 | 1,27E+08 | 1,27E+08 | 8815 | 2 | 66552 | ENSMUST00000125592.2 | 1035 | ENSMUSG00000027366 | Sppl2a | signal peptide peptidase like 2A |
| chr1 | 34159053 | 34160566 | 1514 | * | 11,8465 | 7,577329969 | 1,721311 | 4,40207 | 1,07E-05 | 0,05285139 | 0,918255 | Promoter (<=1kb) | 1 | 34160331 | 34193602 | 33272 | 1 | 13518 | ENSMUST00000183006.5 | 0 | ENSMUSG00000026131 | Dst | dystonin |
| chr11 | 75535925 | 75537036 | 1112 | * | 11,44724 | 7,595701301 | 1,749449 | 4,341768 | 1,41E-05 | 0,051605231 | 1,2763 | Promoter (<=1kb) | 11 | 75537612 | 75538557 | 946 | 1 | 19062 | ENSMUST00000149134.2 | -576 | ENSMUSG00000006127 | Inpp5k | inositol polyphosphate 5-phosphatase K |
| chr11 | 4655985 | 4656660 | 676 | * | 6,044618 | 7,609274146 | 1,907483 | 3,989169 | 6,63E-05 | 1 | 0 | Promoter (1-2kb) | 11 | 4654708 | 4687395 | 32688 | 1 | 67178 | ENSMUST00000154341.8 | 1277 | ENSMUSG00000009076 | Zmat5 | zinc finger, matrin type 5 |
| chr11 | 90143131 | 90144270 | 1140 | * | 12,66429 | 7,631669431 | 1,752438 | 4,354886 | 1,33E-05 | 0,061899193 | 0,918255 | Promoter (2-3kb) | 11 | 90140302 | 90169415 | 29114 | 1 | 67468 | ENSMUST00000004050.7 | 2829 | ENSMUSG00000003948 | Mmd | monocyte to macrophage differentiation-associated |
| chr18 | 67839109 | 67839795 | 687 | * | 7,548166 | 7,646551236 | 1,79401 | 4,262268 | 2,02E-05 | 0,062829128 | 1,360622 | Intron (ENSMUST00000 | 18 | 67808888 | 67823294 | 14407 | 2 | 19255 | ENSMUST00000128169.2 | -15815 | ENSMUSG00000024539 | Ptpn2 | protein tyrosine phosphatase, non-receptor type 2 |
| chr17 | 47320208 | 47321291 | 1084 | * | 8,262428 | 7,659185999 | 1,773685 | 4,318233 | 1,57E-05 | 0,052557724 | 1,360622 | Promoter (<=1kb) | 17 | 47241174 | 47321405 | 80232 | 2 | 224826 | ENSMUST00000224759.2 | 114 | ENSMUSG00000023977 | Ubr2 | ubiquitin protein ligase E3 component n-recognin 2 |
| chr4 | 79291640 | 79292608 | 969 | * | 9,376125 | 7,692361452 | 1,890311 | 4,069363 | 4,71E-05 | 0,099733893 | 1,449706 | Distal Intergenic | 4 | 78130313 | 78133846 | 3534 | 1 | 1,03E+08 | ENSMUST00000246656.1 | 1161327 | | B230208B08Rik | RIKEN cDNA B230208B08 gene |
| chr14 | 31122426 | 31123453 | 1028 | * | 10,872138 | 7,719566133 | 1,724334 | 4,4768393 | 7,576E-06 | 0,048727933 | 0,7443745 | Intron (ENSMUST000001 | 14 | 31099552 | 31129342 | 29791 | 2 | 24056 | ENSMUST00000147586.2 | 5889 | ENSMUSG00000021892 | Sh3bp5 | SH3-domain binding protein 5 (BTK-associated) |
| chr17 | 24637241 | 24638250 | 1010 | * | 9,9306411 | 7,735401154 | 1,7388446 | 4,4485868 | 8,644E-06 | 0,037168825 | 1,2113146 | Promoter (<=1kb) | 17 | 24637466 | 24639559 | 2094 | 1 | 19826 | ENSMUST00000234154.2 | 0 | ENSMUSG00000034681 | Rnps1 | RNA binding protein with serine rich domain 1 |
| chr7 | 110385727 | 110386884 | 1158 | * | 10,498311 | 7,739521459 | 1,7050696 | 4,5391235 | 5,649E-06 | 0,027873477 | 1,1733253 | Intron (ENSMUST000001 | 7 | 110376897 | 110390521 | 13625 | 1 | 11717 | ENSMUST00000148292.2 | 8830 | ENSMUSG00000005686 | Ampd3 | adenosine monophosphate deaminase 3 |
| chr15 | 81726247 | 81727899 | 1653 | * | 15,153451 | 7,777182159 | 1,6234957 | 4,7903927 | 1,665E-06 | 0,007032136 | 2,1416125 | Exon (ENSMUST0000009 | 15 | 81732473 | 81742686 | 10214 | 2 | 57259 | ENSMUST00000050467.9 | 14787 | ENSMUSG00000048546 | Tob2 | transducer of ERBB2, 2 |
| chr19 | 43488388 | 43489496 | 1109 | * | 9,2992374 | 7,824346639 | 1,7673979 | 4,427043 | 9,553E-06 | 0,034889687 | 1,4497059 | 3' UTR | 19 | 43488754 | 43493235 | 4482 | 2 | 14718 | ENSMUST00000132504.2 | 3739 | ENSMUSG00000025190 | Got1 | glutamic-oxaloacetic transaminase 1, soluble |
| chr19 | 24001038 | 24001830 | 793 | * | 9,6138139 | 7,828738603 | 1,7374914 | 4,5057712 | 6,613E-06 | 0,026722594 | 1,4497059 | Intron (ENSMUST000000 | 19 | 23950114 | 24008383 | 58270 | 2 | 381217 | ENSMUST00000096164.6 | 6553 | ENSMUSG00000071604 | Fam189a2 | family with sequence similarity 189, member A2 |
| chr13 | 38533180 | 38534336 | 1157 | * | 11,230535 | 7,842818209 | 1,7378492 | 4,5129452 | 6,393E-06 | 0,030165351 | 1,1733253 | Intron (ENSMUST000001 | 13 | 38530129 | 38541945 | 11817 | 1 | 12161 | ENSMUST00000224452.2 | 3051 | ENSMUSG00000039004 | Bmp6 | bone morphogenetic protein 6 |
| chr1 | 30988845 | 30989619 | 775 | * | 8,009555 | 7,863821976 | 1,794413 | 4,382393 | 1,17E-05 | 1 | 0 | Promoter (<=1kb) | 1 | 30979383 | 30988851 | 9469 | 2 | 19243 | ENSMUST00000233506.2 | 0 | ENSMUSG00000026064 | Ptp4a1 | protein tyrosine phosphatase 4a1 |
| chr17 | 58098260 | 58099693 | 1434 | * | 10,271565 | 7,943507839 | 1,7341033 | 4,580758 | 4,633E-06 | 0,023585918 | 1,2113146 | Intron (ENSMUST000000 | 17 | 58076565 | 58717350 | 640786 | 1 | 620292 | ENSMUST00000076038.7 | 21695 | ENSMUSG00000038048 | Cntnap5c | contactin associated protein-like 5C |
| chr2 | 101864498 | 101865552 | 1055 | * | 10,412106 | 7,950124047 | 1,725719 | 4,6068473 | 4,088E-06 | 0,026638542 | 0,9182553 | Intron (ENSMUST000001 | 2 | 101869974 | 101897070 | 27097 | 1 | 102643083 | ENSMUST00000140238.2 | -4422 | | Gm13919 | predicted gene 13919 |
| chr4 | 135116431 | 135117646 | 1216 | * | 10,232419 | 8,10614452 | 1,7714753 | 4,5759286 | 4,741E-06 | 0,024635007 | 1,1733253 | Intron (ENSMUST000001 | 4 | 135097963 | 135112983 | 15021 | 2 | 230822 | ENSMUST00000123632.2 | -3448 | ENSMUSG00000043924 | Ncmap | noncompact myelin associated protein |
| chr5 | 33159635 | 33161531 | 1897 | * | 21,926357 | 8,275079464 | 1,5882084 | 5,2103236 | 1,885E-07 | 0,001081418 | 2,9519335 | Distal Intergenic | 5 | 33176160 | 33185310 | 9151 | 1 | 22629 | ENSMUST00000019109.8 | -14629 | ENSMUSG00000018965 | Ywhah | tyrosine 3-monooxygenase/tryptophan 5-monooxygenase activation protein, eta polypeptide |
| chr2 | 154734263 | 154735700 | 1438 | * | 15,120896 | 8,340777433 | 1,6288392 | 5,1206881 | 3,044E-07 | 0,002655742 | 1,3768303 | Promoter (<=1kb) | 2 | 154713330 | 154734855 | 21526 | 2 | 67204 | ENSMUST00000099173.11 | 0 | ENSMUSG00000074656 | Eif2s2 | eukaryotic translation initiation factor 2, subunit 2 (beta) |
| chr15 | 84920146 | 84921297 | 1152 | * | 19,907131 | 8,389048667 | 1,6087926 | 5,2144999 | 1,843E-07 | 0,001120595 | 2,6636191 | Promoter (<=1kb) | 15 | 84921290 | 84947031 | 25742 | 1 | 73225 | ENSMUST00000023069.9 | 0 | ENSMUSG00000022434 | Fam118a | family with sequence similarity 118, member A |
| chr5 | 115629741 | 115630838 | 1098 | * | 15,627472 | 8,535145193 | 1,6462359 | 5,1846429 | 2,164E-07 | 0,001425027 | 2,1416125 | Promoter (1-2kb) | 5 | 115628313 | 115628453 | 141 | 1 | 115490316 | ENSMUST00000240114.1 | 1428 | ENSMUSG00000119132 | Gm24407 | predicted gene, 24407 |
| chr19 | 44371082 | 44371885 | 804 | * | 14,014164 | 8,811109276 | 1,648309 | 5,3455445 | 9,015E-08 | 0,001125549 | 1,2763004 | Distal Intergenic | 19 | 44386432 | 44395129 | 8698 | 2 | 20249 | ENSMUST00000236824.2 | 23244 | ENSMUSG00000037071 | Scd1 | stearoyl-Coenzyme A desaturase 1 |
| chr14 | 55112972 | 55114314 | 1343 | * | 17,786654 | 8,900911355 | 1,6145505 | 5,5129346 | 3,529E-08 | 0,00031467 | 2,5070128 | Promoter (<=1kb) | 14 | 55113054 | 55114995 | 1942 | 2 | 105651 | ENSMUST00000168622.2 | 681 | ENSMUSG00000072494 | Ppp1r3e | protein phosphatase 1, regulatory subunit 3E |

| regions | DMOG UP | DMOG UP % | DMOG DOWN | DMOG DOWN % |
|---|---|---|---|---|
| 3' UTR | 260 | 2,62546703 | 53 | 1,534008683 |
| 5' UTR | 26 | 0,262546703 | 6 | 0,17366136 |
| distal intergenic | 3341 | 33,73725134 | 1715 | 49,6382055 |
| downstream (<= 300 bp) | 15 | 0,151469252 | 6 | 0,17366136 |
| exon | 626 | 6,321316773 | 200 | 5,788712012 |
| intron | 2782 | 28,09249722 | 1106 | 32,01157742 |
| promoter (<= 1kb) | 1762 | 17,7925881 | 167 | 4,83357453 |
| promoter (1-2 kb) | 590 | 5,957790569 | 113 | 3,270622287 |
| promoter (2-3 kb) | 501 | 5,059073008 | 89 | 2,575976845 |
| promoter total | 2853 | 28,80945168 | 369 | 10,68017366 |
| TOTAL | 9903 | 100 | 3455 | 100 |

### Add. Table 3

| geneSet | description | link | size | overlap | expect | enrichmentRatio | pValue | FDR | overlapId | userId |
|---|---|---|---|---|---|---|---|---|---|---|
| mmu00534 | Glycosaminoglycan biosynthesis | http://www.kegg.jp/kegg-bin/show_pathway?mmu00534+14042+14043+15478+15531+217119+218271+233781+23908+319415+50785+50787+54616+54710+64580+72727+93683 | 24 | 16 | 5,805257 | 2,756122599 | 1,25E-05 | 3,52E-04 | 14042;14043;15478;15531;217119;218271;233781;23908;319415;50785;50787;54616;54710;64580;72727;93683 | B4galt7;Hs3st5;Xylt2;Glce;Ndst4;Ext1;Ndst1;Hs3st3a1;Ext2;Extl3;B3gat3;Hs3st3b1;Xylt1;Hs2st1;Hs6st1;Hs6st3 |
| mmu04710 | Circadian rhythm | http://www.kegg.jp/kegg-bin/show_pathway?mmu04710+105787+108097+108099+12753+12912+12952+12953+18030+18626+18627+18628+20893+213311+217166+225998+26965+27373+353187+56438 | 34 | 19 | 8,22411408 | 2,310279237 | 7,14E-05 | 0,00138421 | 105787;108097;108099;12753;12912;12952;12953;18030;18626;18627;18628;20893;213311;217166;225998;26965;27373;353187;56438 | Bhlhe40;Per1;Creb1;Rbx1;Nfil3;Nr1d1;Csnk1e;Fbxl21;Cry2;Per3;Prkag2;Clock;Cry1;Cul1;Prkaa1;Rorb;Prkab2;Per2;Nr1d2 |
| mmu00532 | Glycosaminoglycan biosynthesis | http://www.kegg.jp/kegg-bin/show_pathway?mmu00532+217119+218271+233781+269941+319901+338362+53374+58250+72727+74241+78752 | 21 | 11 | 5,07959987 | 2,165524899 | 0,00485948 | 0,02886363 | 217119;218271;233781;269941;319901;338362;53374;58250;72727;74241;78752 | Chsy1;Chst3;Chpf;B4galt7;Xylt2;Chst11;Dsel;Ust;B3gat3;Xylt1;Csgalnact2 |
| mmu04520 | Adherens junction | http://www.kegg.jp/kegg-bin/show_pathway?mmu04520+108100+109711+109905+11465+11848+12385+12386+12387+12388+12995+14158+16001+16337+17127+17295+17356+17906+19268+19271+19274+19353+19878+20411+20779+21414+21416+215449+21812+21872+22330+22612+231858+242687+26413+29875+328572+380928+56426+58998+59310+60595+66313+71740+75788+77446+79264+83767+93742 | 92 | 48 | 22,2534852 | 2,156965512 | 5,73E-09 | 2,00E-06 | 108100;109711;109905;11465;11848;12385;12386;12387;12388;12995;14158;16001;16337;17127;17295;17356;17906;19268;19271;19274;19353;19878;20411;20779;21414;21416;215449;21812;21872;22330;22612;231858;242687;26413;29875;328572;380928;56426;58998;59310;60595;66313;71740;75788;77446;79264;83767;93742 | Smad3;Mapk1;Iqgap1;Ptprj;Rac1;Met;Rhoa;Insr;Heg1;Actn1;Pard3;Actg1;Smurf2;Sorbs1;Myl2;Vcl;Tcf7;Actn4;Rap1b;Yes1;Smurf1;Tgfbr1;Fer;Igf1r;Wasf2;Myl10;Nectin3;Ptprm;Ctnnd1;Csnk2a1;Ctnna2;Ptprf;Tjp1;Wasf1;Nectin4;Krit1;Ctnnb1;Ep300;Tcf7l2;Afdn;Baiap2;Ctnna1;Radil;Src;Lmo7;Rap1a;Rock2;Pdcd10 |
| mmu05211 | Renal cell carcinoma | http://www.kegg.jp/kegg-bin/show_pathway?mmu05211+107746+109905+110157+112405+112406+11863+11864+12575+12928+14194+14784+15251+17295+18479+18706+19247+19353+20525+209446+215449+21802+21808+22339+224105+23797+23871+241656+26395+26396+26413+328572+56438+71745+94315 | 68 | 34 | 16,4482282 | 2,067091949 | 3,36E-06 | 1,30E-04 | 107746;109905;110157;112405;112406;11863;11864;12575;12928;14194;14784;15251;17295;18479;18706;19247;19353;20525;209446;215449;21802;21808;22339;224105;23797;23871;241656;26395;26396;26413;328572;56438;71745;94315 | Slc2a1;Vegfa;Tgfa;Mapk1;Rac1;Map2k1;Egln1;Met;Egln2;Rbx1;Ptpn11;Pik3ca;Pak5;Cdkn1a;Rapgef1;Rap1b;Arnt;Raf1;Tfe3;Akt3;Pak1;Pak2;Cul2;Map2k2;Arnt2;Prcc;Ep300;Ets1;Tgfb2;Hif1a;Grb2;Crk;Fh1;Rap1a |

| | | | | | | | | | | |
|---|---|---|---|---|---|---|---|---|---|---|
| mmu00620 | Pyruvate metabolism | http://www.kegg.jp/kegg-bin/show_pathway?mmu00620+106557+107029+107476+109264+11532+11669+11671+14194+14651+16828+16833+17448+17449+18563+18597+18598+18746+224530+56752+60525+68263 | 44 | 21 | 10,6429712 | 1,973133224 | 5,61E-04 | 0,00674546 | 106557;107029;107476;109264;11532;11669;11671;14194;14651;16828;16833;17448;17449;18563;18597;18598;18746;224530;56752;60525;68263 | Acss2;Pdhb;Pkm;Pcx;Pdha2;Aldh2;Me2;Ldhc;Aldh3a2;Acaca;Ldha;Mdh1;Me3;Adh5;Mdh2;Hagh;Aldh9a1;Ldhal6b;Acat3;Pdha1;Fh1 |
| mmu04216 | Ferroptosis | http://www.kegg.jp/kegg-bin/show_pathway?mmu04216+14629+14630+15368+17254+213053+22041+22042+22334+23983+26570+27057+50790+53945+66734+67443+67634+68428+71775+74205 | 40 | 19 | 9,67542833 | 1,963737352 | 0,00109416 | 0,0097913 | 14629;14630;15368;17254;213053;22041;22042;22334;23983;26570;27057;50790;53945;66734;67443;67634;68428;71775;74205 | Hmox1;Map1lc3b;Slc7a11;Gclm;Pcbp1;Ncoa4;Map1lc3a;Slc40a1;Acsl3;Steap3;Gclc;Vdac2;Slc3a2;Tfrc;Ftmt;Slc39a14;Acsl4;Trf;Inhca |
| mmu05220 | Chronic myeloid leukemia | http://www.kegg.jp/kegg-bin/show_pathway?mmu05220+107986+110157+110279+11350+12394+12402+12443+12571+12575+12928+13016+13555+13557+14013+14389+14784+15182+16150+16151+17127+17246+17873+18035+18706+19247+20418+21808+21812+23797+23882+242705+26395+26396+26413+27015+433759 | 76 | 36 | 18,3833138 | 1,958297636 | 8,55E-06 | 2,71E-04 | 107986;110157;110279;11350;12394;12402;12443;12571;12575;12928;13016;13555;13557;14013;14389;14784;15182;16150;16151;17127;17246;17873;18035;18706;19247;20418;21808;21812;23797;23882;242705;26395;26396;26413;27015;433759 | E2f1;Gadd45b;Smad3;Mapk1;Mdm2;Gab2;Map2k1;Ctbp1;Mecom;Shc3;Abl1;Ddb2;Ptpn11;Pik3ca;Ikbkb;E2f2;Nfkbia;Cdkn1a;Hdac1;Raf1;Cbl;Tgfbr1;Akt3;E2f3;Ikbkg;Gadd45g;Bcr;Cdk6;Runx1;Map2k2;Hdac2;Ccnd1;Tgfb2;Polk;Grb2;Crk |
| mmu01040 | Biosynthesis of unsaturated fatty acids | http://www.kegg.jp/kegg-bin/show_pathway?mmu01040+11430+171210+171282+20249+20250+20280+230161+26897+30049+30963+329065+56473+57874+74559+76267+80911 | 34 | 16 | 8,22411408 | 1,945498305 | 0,00300435 | 0,01978339 | 11430;171210;171282;20249;20250;20280;230161;26897;30049;30963;329065;56473;57874;74559;76267;80911 | Scd1;Scd2;Scp2;Acox3;Scd4;Hacd1;Fads2;Hacd3;Acnat1;Acot2;Scd3;Fads1;Acot1;Acot4;Elovl7;Acox1 |
| mmu04066 | HIF-1 signaling pathway | http://www.kegg.jp/kegg-bin/show_pathway?mmu04066+106557+108058+112405+112406+11600+11601+11863+12325+12575+13614+13685+13806+13856+14433+15251+15275+15368+16001+16333+16337+16828+16833+17347+18127+18597+18598+18641+18655+18706+18751+18787+20525+21898+22041+22042+22339+226265+228026+234779+23797+26395+26396+26413+328572+353204+433182+56421+56438+68263+71745+71775 | 114 | 51 | 27,5749707 | 1,849503323 | 1,10E-06 | 9,36E-05 | 106557;108058;112405;112406;11600;11601;11863;12325;12575;13614;13685;13806;13856;14433;15251;15275;15368;16001;16333;16337;16828;16833;17347;18127;18597;18598;18641;18655;18706;18751;18787;20525;21898;22041;22042;22339;226265;228026;234779;23797;26395;26396;26413;328572;353204;433182;56421;56438;68263;71745;71775 | Slc2a1;Pgk1;Eno1;Hmox1;Pdk1;Eif4ebp1;Gapdh;Vegfa;Epo;Hk1;Pdhb;Mapk1;Pfkl;Map2k1;Egln1;Camk2g;Plcg2;Insr;Egln2;Eno4;Pdha2;Rbx1;Ldhc;Pik3ca;Pfkp;Cdkn1a;Ldha;Ins1;Mknk2;Aldoart1;Tlr4;Arnt;Eno1b;Angpt1;Nos3;Akt3;Prkcb;Tfrc;Igf1r;Serpine1;Edn1;Cul2;Map2k2;Ldhal6b;Ep300;Trf;Camk2d;Hif1a;Inhca;Pdha1;Angpt2 |

| | | | | | | | | | | |
|---|---|---|---|---|---|---|---|---|---|---|
| mmu04927 | Cortisol synthesis and secretion | http://www.kegg.jp/kegg-bin/show_pathway?mmu04927+104111+109305+11512+11515+11607+11608+11614+11909+12289+12912+13070+13074+14672+14683+16440+16492+16526+16527+16835+17200+18514+18584+18795+18796+20778+208647+218461+224129+239556+58226+77037+78284 | 72 | 32 | 17,415771 | 1,837415066 | 1,25E-04 | 0,00208105 | 104111;109305;11512;11515;11607;11608;11614;11909;12289;12912;13070;13074;14672;14683;16440;16492;16526;16527;16835;17200;18514;18584;18795;18796;20778;208647;218461;224129;239556;58226;77037;78284 | Adcy6;Scarb1;Pbx1;Cacna1h;Plcb2;Mc2r;Pde8a;Gna11;Itpr3;Orai1;Creb3l4;Creb1;Mrap;Agtr1a;Kcnk2;Cacna1d;Pde8b;Cyp17a1;Nr0b1;Cyp11a1;Ldlr;Plcb1;Kcna4;Cacna1i;Adcy5;Creb3l2;Atf2;Adcy3;Kcnk3;Gnas;Agtr1b;Adcy9 |
| mmu05216 | Thyroid cancer | http://www.kegg.jp/kegg-bin/show_pathway?mmu05216+107986+108989+12387+12443+12575+17873+18510+20181+21414+21416+23882+26395+26396+26413+27015+27057 | 37 | 16 | 8,9497712 | 1,787755199 | 0,00829491 | 0,03965647 | 107986;108989;12387;12443;12575;17873;18510;20181;21414;21416;23882;26395;26396;26413;27015;27057 | Gadd45b;Mapk1;Tpr;Map2k1;Ddb2;Pax8;Ncoa4;Cdkn1a;Tcf7;Gadd45g;Map2k2;Rxra;Ccnd1;Ctnnb1;Tcf7l2;Polk |
| mmu04934 | Cushing syndrome | http://www.kegg.jp/kegg-bin/show_pathway?mmu04934+104111+108058+109305+109905+11512+11515+11607+11608+11622+11789+11863+11909+12005+12289+12325+12387+12443+12447+12571+12575+12912+12918+13070+13074+13555+13557+14194+14362+14367+14369+14370+14371+14672+14679+14683+16440+16492+16526+16527+16835+17200+17283+18514+18584+18795+18796+20778+208647+213464+21414+21416+215449+218461+22408+224129+22414+22415+22417+22419+23805+239556+241489+242705+26395+26396+26413+58226+77037+78284 | 162 | 69 | 39,1854847 | 1,760856105 | 1,58E-07 | 1,83E-05 | 104111;108058;109305;109905;11512;11515;11607;11608;11622;11789;11863;11909;12005;12289;12325;12387;12443;12447;12571;12575;12912;12918;13070;13074;13555;13557;14194;14362;14367;14369;14370;14371;14672;14679;14683;16440;16492;16526;16527;16835;17200;17283;18514;18584;18795;18796;20778;208647;213464;21414;21416;215449;218461;22408;224129;22414;22415;22417;22419;23805;239556;241489;242705;26395;26396;26413;58226;77037;78284 | E2f1;Adcy6;Scarb1;Pbx1;Mapk1;Cacna1h;Plcb2;Map2k1;Wnt2b;Mc2r;Camk2g;Pde8a;Gna11;Itpr3;Orai1;Creb3l4;Men1;Wnt4;Gnai3;Creb1;E2f2;Mrap;Cdkn1a;Tcf7;Fzd8;Fzd9;Agtr1a;Fzd1;Rap1b;Ahr;Arnt;Axin1;E2f3;Fzd7;Kcnk2;Cacna1d;Apc;Pde8b;Cyp17a1;Wnt1;Cyp11a1;Cdk6;Pde11a;Ldlr;Plcb1;Kcna4;Wnt5b;Apc2;Cacna1i;Adcy5;Crh;Map2k2;Rbbp5;Ccnd1;Creb3l2;Wnt3;Atf2;Ctnnb1;Camk2d;Ccne1;Adcy3;Tcf7l2;Kcnk3;Gnas;Agtr1b;Adcy9;Fzd5;Fh1;Rap1a |
| mmu04137 | Mitophagy | http://www.kegg.jp/kegg-bin/show_pathway?mmu04137+12176+12177+12995+13555+13665+13666+14232+14479+15251+17342+19345+19349+20779+209446+213948+21425+22030+22186+22190+228361+23806+245860+26420+271457+50873+52440+53375+56480+56484+56715+59026+59040+66313+66687+66734+67443+68350+68943+69104+72018+75788+94044 | 99 | 42 | 23,9466851 | 1,753896199 | 4,57E-05 | 9,39E-04 | 12176;12177;12995;13555;13665;13666;14232;14479;15251;17342;19345;19349;20779;209446;213948;21425;22030;22186;22190;228361;23806;245860;26420;271457;50873;52440;53375;56480;56484;56715;59026;59040;66313;66687;66734;67443;68350;68943;69104;72018;75788;94044 | E2f1;Bnip3;Atg9b;Arih1;Traf2;Map1lc3b;Atg9a;Uba52;Mapk9;Rab5a;Bcl2l13;Fkbp8;Rab5c;Foxo3;Bnip3l;Smurf2;Map1lc3a;Huwe1;Pink1;Ubc;Smurf1;Tfe3;Rab7;Usp15;Prkn;Tbk1;Rhot1;Csnk2a1;Mitf;Tax1bp1;Rabgef1;Tfeb;Ambra1;Eif2ak3;Marchf5;Fundc1;Hif1a;Tbc1d15;Mul1;Eif2s1;Mtx2;Src |

| | | | | | | | | | | |
|---|---|---|---|---|---|---|---|---|---|---|
| mmu05412 | Arrhythmogenic right ventricular cardiomyopathy | http://www.kegg.jp/kegg-bin/show_pathway?mmu05412+11465+11938+12289+12293+12294+12296+12385+12386+12387+12558+13138+13405+13527+13726+14609+16400+16410+16416+16419+16421+16772+16773+16905+20191+20648+20649+20650+21414+21416+24052+241226+319734+320910+54377+67451 | 84 | 35 | 20,3183995 | 1,722576624 | 2,92E-04 | 0,00407477 | 11465;11938;12289;12293;12294;12296;12385;12386;12387;12558;13138;13405;13527;13726;14609;16400;16410;16416;16419;16421;16772;16773;16905;20191;20648;20649;20650;21414;21416;24052;241226;319734;320910;54377;67451 | Dag1;Ryr2;Atp2a2;Sntb2;Dtna;Actg1;Itgb8;Cacna2d4;Dmd;Tcf7;Lmna;Lama1;Cdh2;Cacnb2;Sntb1;Cacna2d1;Cacna1d;Sgcd;Itgav;Snta1;Lama2;Itgb7;Itgb5;Gja1;Emd;Ctnna2;Itga8;Ctnnb1;Pkp2;Tcf7l2;Itgb3;Cacng4;Cacna2d3;Ctnna1;Itga3 |
| mmu05217 | Basal cell carcinoma | http://www.kegg.jp/kegg-bin/show_pathway?mmu05217+107986+11789+12005+12387+12575+14362+14367+14369+14370+14371+14633+17873+19206+20423+21414+21416+22408+22414+22415+22417+22419+23805+23882+24069+27015+319757 | 63 | 26 | 15,2387996 | 1,706171132 | 0,00198918 | 0,0147707 | 107986;11789;12005;12387;12575;14362;14367;14369;14370;14371;14633;17873;19206;20423;21414;21416;22408;22414;22415;22417;22419;23805;23882;24069;27015;319757 | Ptch1;Gadd45b;Wnt2b;Gli2;Wnt4;Ddb2;Cdkn1a;Tcf7;Fzd8;Fzd9;Fzd1;Shh;Smo;Axin1;Fzd7;Gadd45g;Apc;Wnt1;Wnt5b;Apc2;Wnt3;Ctnnb1;Tcf7l2;Polk;Fzd5;Sufu |
| mmu04120 | Ubiquitin mediated proteolysis | http://www.kegg.jp/kegg-bin/show_pathway?mmu04120+106052+107986+109161+11797+11798+12189+12211+12402+140499+14897+17222+17246+17318+17344+17999+208650+216150+217232+217342+218793+22186+22190+22193+22196+22209+22214+230904+237009+26401+26965+50754+50868+50873+52206+52563+53323+56371+56438+56469+56550+59004+59008+59026+63958+66053+66313+67030+67345+67455+68999+69010+69807+69957+70790+71745+75430+75717+75788+80794+83814+93765 | 148 | 61 | 35,7990848 | 1,703954174 | 3,05E-06 | 1,30E-04 | 106052;107986;109161;11797;11798;12189;12211;12402;140499;14897;17222;17246;17318;17344;17999;208650;216150;217232;217342;218793;22186;22190;22193;22196;22209;22214;230904;237009;26401;26965;50754;50868;50873;52206;52563;53323;56371;56438;56469;56550;59004;59008;59026;63958;66053;66313;67030;67345;67455;68999;69010;69807;69957;70790;71745;75430;75717;75788;80794;83814;93765 | Keap1;Map3k1;Cblb;Anapc4;Ube2j2;Trip12;Mdm2;Anapc13;Ube4b;Uba52;Anapc15;Ube2d2a;Pias4;Birc6;Rbx1;Ddb2;Ube2q2;Smurf2;Ube2h;Trim32;Huwe1;Ppil2;Brca1;Cdc34;Herc4;Fzr1;Nedd4;Nedd4l;Birc2;Ubc;Smurf1;Cdc23;Cbl;Ube2i;Cdc16;Fbxw7;Ube2o;Cdc27;Prkn;Ubr5;Ube2e2;Fancl;Fbxo2;Mid1;Anapc1;Cul2;Cul5;Ube2n;Ube2dnl1;Ube2e3;Cul1;Ube2a;Klhl13;Anapc10;Cblc;Ube2k;Pias2;Xiap;Pias1;Fbxo4;Anapc5 |

| | | | | | | | | | | |
|---|---|---|---|---|---|---|---|---|---|---|
| mmu04916 | Melanogenesis | http://www.kegg.jp/kegg-bin/show_pathway?mmu04916+104111+108058+110157+11512+11515+12313+12315+12325+12387+12912+13614+14362+14367+14369+14370+14371+14679+14683+16590+17311+17342+18751+18795+18796+208647+21414+21416+22178+22408+224129+22414+22415+22417+22419+26395+26396+26413+328572+50518+78284+80796 | 100 | 41 | 24,1885708 | 1,695015398 | 1,41E-04 | 0,00213493 | 104111;108058;110157;11512;11515;12313;12315;12325;12387;12912;13614;14362;14367;14369;14370;14371;14679;14683;16590;17311;17342;18751;18795;18796;208647;21414;21416;22178;22408;224129;22414;22415;22417;22419;26395;26396;26413;328572;50518;78284;80796 | Adcy6;a;Mapk1;Plcb2;Map2k1;Wnt2b;Calm3;Camk2g;Creb3l4;Wnt4;Tyrp1;Gnai3;Creb1;Tcf7;Fzd8;Fzd9;Fzd1;Raf1;Prkcb;Fzd7;Kit;Edn1;Wnt1;Plcb1;Calm1;Calm4;Wnt5b;Mitf;Adcy5;Map2k2;Creb3l2;Wnt3;Ctnnb1;Kitl;Ep300;Camk2d;Adcy3;Tcf7l2;Gnas;Adcy9;Fzd5 |
| mmu04962 | Vasopressin-regulated water reabsorption | http://www.kegg.jp/kegg-bin/show_pathway?mmu04962+104111+110350+11512+11515+11827+12912+13191+13427+14683+18195+19345+208647+22318+234663+271457+53869+68097+78284 | 44 | 18 | 10,6429712 | 1,691257049 | 0,01036514 | 0,04637734 | 104111;110350;11512;11515;11827;12912;13191;13427;14683;18195;19345;208647;22318;234663;271457;53869;68097;78284 | Adcy6;Aqp2;Creb3l4;Vamp2;Rab5a;Creb1;Rab5c;Dctn1;Nsf;Dync1li2;Dync2h1;Rab11a;Creb3l2;Dync1i2;Adcy3;Gnas;Adcy9;Dynll2 |
| mmu05230 | Central carbon metabolism in cancer | http://www.kegg.jp/kegg-bin/show_pathway?mmu05230+106557+110157+14183+14380+15251+15275+15926+16590+16828+16833+17295+18213+18597+18598+18641+18648+18706+18746+20514+20525+216456+228026+23797+26395+26396+26413+56421+68263 | 69 | 28 | 16,6901139 | 1,677639843 | 0,00184293 | 0,01429292 | 106557;110157;14183;14380;15251;15275;15926;16590;16828;16833;17295;18213;18597;18598;18641;18648;18706;18746;20514;20525;216456;228026;23797;26395;26396;26413;56421;68263 | Slc2a1;Pdk1;Hk1;Pdhb;Mapk1;Pfkl;Gls2;Map2k1;Met;Ntrk3;Fgfr2;Pkm;Pdha2;Ldhc;Pik3ca;Pfkp;Pgam1;Ldha;Raf1;Akt3;Kit;Slc1a5;G6pd2;Idh1;Map2k2;Ldhal6b;Hif1a;Pdha1 |
| mmu03018 | RNA degradation | http://www.kegg.jp/kegg-bin/show_pathway?mmu03018+104625+109075+12227+13209+13806+15526+18459+18641+18983+210106+213550+22057+225929+226265+227715+24127+241989+319885+381404+433182+50912+53621+56421+57259+57316+67207+67543+68533+69125+70640+72198+72587+75901+93728 | 84 | 34 | 20,3183995 | 1,673360149 | 6,68E-04 | 0,00728847 | 104625;109075;12227;13209;13806;15526;18459;18641;18983;210106;213550;22057;225929;226265;227715;24127;241989;319885;381404;433182;50912;53621;56421;57259;57316;67207;67543;68533;69125;70640;72198;72587;75901;93728 | Eno1;Tob2;Hspa9;Btg2;Pfkl;Tent4a;Zcchc7;Exosc2;Exosc10;Dcp2;Tob1;Xrn1;Eno4;Pfkp;Pabpc2;C1d;Dcp1a;Patl1;Dis3l;Eno1b;Mtrex;Exosc4;Pabpc4l;Cnot6;Cnot7;Pabpc1l;Mphosph6;Lsm1;Pabpc6;Cnot4;Cnot8;Pan3;Ddx6;Pabpc5 |

| | | | | | | | | | | |
|---|---|---|---|---|---|---|---|---|---|---|
| mmu05225 | Hepatocellular carcinoma | http://www.kegg.jp/kegg-bin/show_pathway?mmu05225+107986+110157+11465+11789+12005+12387+12443+12571+12575+13555+13557+13712+14362+14367+14369+14370+14371+14784+14858+14862+14863+14872+15368+16001+16973+17126+17127+17295+17873+18024+18706+18751+20418+20587+211666+212398+21414+21416+21802+21808+21812+22408+22414+22415+22417+22419+234779+23797+23805+23882+242705+26395+26396+26413+26992+27015+50493+50868+57376+66923+67155+68094+68214+70127+77044+83766+83796+83797+93687+93760 | 174 | 70 | 42,0881132 | 1,66317743 | 1,69E-06 | 9,83E-05 | 107986;110157;11465;11789;12005;12387;12443;12571;12575;13555;13557;13712;14362;14367;14369;14370;14371;14784;14858;14862;14863;14872;15368;16001;16973;17126;17127;17295;17873;18024;18706;18751;20418;20587;211666;212398;21414;21416;21802;21808;21812;22408;22414;22415;22417;22419;234779;23797;23805;23882;242705;26395;26396;26413;26992;27015;50493;50868;57376;66923;67155;68094;68214;70127;77044;83766;83796;83797;93687;93760 | Keap1;E2f1;Hmox1;Gadd45b;Smad3;Tgfa;Mapk1;Lrp5;Map2k1;Wnt2b;Smarcc2;Met;Plcg2;Shc3;Wnt4;Arid2;Actg1;Ddb2;Gstt2;Gsta2;Actl6b;Csnk1a1;Pik3ca;Mgst2;E2f2;Gsto2;Cdkn1a;Tcf7;Fzd8;Fzd9;Smarcd1;Fzd1;Smarcb1;Smad2;Axin1;Nfe2l2;Gstm1;Raf1;Tgfbr1;Akt3;Prkcb;E2f3;Fzd7;Arid1a;Igf1r;Gadd45g;Apc;Brd7;Wnt1;Cdk6;Wnt5b;Apc2;Txnrd1;Pbrm1;Dpf3;Map2k2;Gstm2;Frat2;Ccnd1;Wnt3;Ctnnb1;Smarcd2;Tgfb2;Tcf7l2;Polk;Smarce1;Grb2;Smarca2;Elk1;Fzd5 |
| mmu05210 | Colorectal cancer | http://www.kegg.jp/kegg-bin/show_pathway?mmu05210+107986+110157+11789+11848+12005+12125+12387+12443+12575+13063+13176+14281+14784+17126+17127+17688+17873+18706+19353+19730+21414+21416+21802+21808+21812+23797+23805+23882+26395+26396+26413+26420+27015+58801+72993 | 88 | 35 | 21,2859423 | 1,644277687 | 8,20E-04 | 0,0081803 | 107986;110157;11789;11848;12005;12125;12387;12443;12575;13063;13176;14281;14784;17126;17127;17688;17873;18706;19353;19730;21414;21416;21802;21808;21812;23797;23805;23882;26395;26396;26413;26420;27015;58801;72993 | Gadd45b;Smad3;Tgfa;Mapk1;Fos;Cycs;Rac1;Map2k1;Msh6;Appl1;Bcl2l11;Rhoa;Mapk9;Ddb2;Pik3ca;Cdkn1a;Tcf7;Smad2;Axin1;Raf1;Tgfbr1;Akt3;Dcc;Gadd45g;Apc;Pmaip1;Apc2;Ralgds;Map2k2;Ccnd1;Ctnnb1;Tgfb2;Tcf7l2;Polk;Grb2 |
| mmu05213 | Endometrial cancer | http://www.kegg.jp/kegg-bin/show_pathway?mmu05213+107986+110157+11789+12005+12385+12386+12387+12443+12575+13712+14784+17873+18706+21414+21416+23797+23805+23882+26395+26396+26413+27015+56484 | 58 | 23 | 14,0293711 | 1,639417753 | 0,00634417 | 0,03354721 | 107986;110157;11789;12005;12385;12386;12387;12443;12575;13712;14784;17873;18706;21414;21416;23797;23805;23882;26395;26396;26413;27015;56484 | Gadd45b;Mapk1;Map2k1;Ddb2;Pik3ca;Foxo3;Cdkn1a;Tcf7;Axin1;Raf1;Akt3;Gadd45g;Apc;Apc2;Ctnna2;Map2k2;Ccnd1;Ctnnb1;Tcf7l2;Polk;Grb2;Ctnna1;Elk1 |

| | | | | | | | | | |
|---|---|---|---|---|---|---|---|---|---|
| mmu04390 | Hippo signaling pathway | http://www.kegg.jp/kegg-bin/show_pathway?mmu04390+11465+11789+11797+12005+12160+12161+12162+12166+12168+12385+12386+12387+12443+12444+12445+12695+13383+14164+14362+14367+14369+14370+14371+14633+15902+16414+17126+17127+17131+18016+18787+19046+19052+209456+211652+21414+21416+21676+21808+21812+22408+22414+22415+22417+22419+22627+22629+22630+22631+23805+23859+269643+27373+27494+50523+53310+54401+68473+71978+73246+93737+93742 | 157 | 62 | 37,9760562 | 1,632607654 | 1,31E-05 | 3,52E-04 | 11465;11789;11797;12005;12160;12161;12162;12166;12168;12385;12386;12387;12443;12444;12445;12695;13383;14164;14362;14367;14369;14370;14371;14633;15902;16414;17126;17127;17131;18016;18787;19046;19052;209456;211652;21414;21416;21676;21808;21812;22408;22414;22415;22417;22419;22627;22629;22630;22631;23805;23859;269643;27373;27494;50523;53310;54401;68473;71978;73246;93737;93742 | Ywhah;Ppp2r2a;Smad3;Bmp6;Wwc1;Ppp1cb;Id2;Wnt2b;Gli2;Smad7;Dlg2;Dlg1;Wnt4;Trp53bp2;Pard3;Actg1;Ywhae;Bmpr2;Csnk1e;Ywhaz;Tcf7;Nf2;Fzd8;Fzd9;Fzd1;Smad2;Birc2;Ppp2ca;Axin1;Tead1;Tgfbr1;Fzd7;Itgb2;Serpine1;Ppp2r2c;Apc;Lats2;Ccnd3;Wnt1;Amot;Wnt5b;Bmpr1a;Rassf6;Apc2;Ctnna2;Mob1b;Ywhaq;Pard6g;Ccnd2;Ccnd1;Wnt3;Dlg3;Ctnnb1;Tgfb2;Tcf7l2;Ywhab;Patj;Ctnna1;Fzd5;Bmp5;Bmp7;Fgf1 |
| mmu05214 | Glioma | http://www.kegg.jp/kegg-bin/show_pathway?mmu05214+107986+108058+110157+12313+12315+12325+12443+12571+12575+13555+13557+14784+16001+17246+17873+18706+18751+20418+215303+21802+234779+23797+23882+242705+26395+26396+26413+27015+80796 | 74 | 29 | 17,8995424 | 1,620153149 | 0,00287832 | 0,01931798 | 107986;108058;110157;12313;12315;12325;12443;12571;12575;13555;13557;14784;16001;17246;17873;18706;18751;20418;215303;21802;234779;23797;23882;242705;26395;26396;26413;27015;80796 | E2f1;Gadd45b;Tgfa;Mapk1;Mdm2;Map2k1;Calm3;Camk2g;Plcg2;Shc3;Ddb2;Pik3ca;E2f2;Cdkn1a;Raf1;Akt3;Prkcb;E2f3;Igf1r;Gadd45g;Cdk6;Calm1;Calm4;Camk1g;Map2k2;Ccnd1;Camk2d;Polk;Grb2 |
| mmu05205 | Proteoglycans in cancer | http://www.kegg.jp/kegg-bin/show_pathway?mmu05205+108058+109676+110157+11465+11735+11848+12325+12387+12402+12443+12575+12842+13712+13869+14000+14083+14173+14362+14367+14369+14370+14371+14733+14734+14784+15200+15251+15530+16001+16160+16410+16416+16419+16440+17126+17246+17295+17931+18479+18706+18751+18792+19046+19206+192176+19247+19353+19684+19878+20423+20544+20779+21808+21844+21859+21898+22160+22339+22408+22414+22415+22417+22419+232807+234779+23797+24088+26395+26396+26413+29875+319757+327826+329251+387140+545291+57257+68794+69632+74055 | 205 | 80 | 49,5865702 | 1,613340058 | 1,34E-06 | 9,36E-05 | 108058;109676;110157;11465;11735;11848;12325;12387;12402;12443;12575;12842;13712;13869;14000;14083;14173;14362;14367;14369;14370;14371;14733;14734;14784;15200;15251;15530;16001;16160;16410;16416;16419;16440;17126;17246;17295;17931;18479;18706;18751;18792;19046;19206;192176;19247;19353;19684;19878;20423;20544;20779;21808;21844;21859;21898;22160;22339;22408;22414;22415;22417;22419;232807;234779;23797;24088;26395;26396;26413;29875;319757;327826;329251;387140;545291;57257;68794;69632;74055 | Ptch1;Vegfa;Fgf2;Ppp1cb;Mapk1;Slc9a1;Mdm2;Iqgap1;Rac1;Map2k1;Wnt2b;Met;Ppp1r12b;Hspg2;Camk2g;Plcg2;Flnc;Plce1;Itpr3;Hbegf;Rhoa;Twist1;Rdx;Wnt4;Ank2;Ptk2;Arhgef12;Actg1;Ptpn11;Hpse2;Pik3ca;Cdkn1a;Tiam1;Fzd8;Tlr2;Fzd9;Fzd1;Vav3;Tlr4;Smad2;Shh;Smo;Raf1;Cbl;Akt3;Prkcb;Gpc1;Fzd7;Igf1r;Pak1;Col1a1;Itgav;Wnt1;Itgb5;Wnt5b;Frs2;Ppp1r12c;Map2k2;Timp3;Ank3;Ppp1r12a;Plau;Ccnd1;Wnt3;Il12b;Ctnnb1;Camk2d;Tgfb2;Itgb3;Flna;Erbb4;Hif1a;Grb2;Elk1;Fzd5;Gpc3;Src;Mir21a;Drosha;Rock2 |

| | | | | | | | | | | |
|---|---|---|---|---|---|---|---|---|---|---|
| mmu05218 | Melanoma | http://www.kegg.jp/kegg-bin/show_pathway?mmu05218+107986+110157+12443+12571+12575+13555+13557+14164+14173+14176+14178+14180+16001+17246+17295+17342+17873+18706+23797+23882+242705+26395+26396+26413+27015+54635+71785+80857 | 72 | 28 | 17,415771 | 1,607738183 | 0,00383571 | 0,0243393 | 107986;110157;12443;12571;12575;13555;13557;14164;14173;14176;14178;14180;16001;17246;17295;17342;17873;18706;23797;23882;242705;26395;26396;26413;27015;54635;71785;80857 | E2f1;Gadd45b;Fgf2;Mapk1;Mdm2;Map2k1;Met;Fgf20;Ddb2;Pik3ca;E2f2;Cdkn1a;Pdgfd;Raf1;Akt3;E2f3;Igf1r;Gadd45g;Fgf9;Cdk6;Mitf;Map2k2;Pdgfc;Fgf5;Fgf7;Ccnd1;Polk;Fgf1 |
| mmu01200 | Carbon metabolism | http://www.kegg.jp/kegg-bin/show_pathway?mmu01200+100198+100678+107029+107272+109264+110639+110821+11429+11430+11532+12359+13806+14194+14380+14433+14718+14719+14751+15275+15926+17448+17449+17850+18563+18597+18598+18641+18648+18655+18746+20425+20916+21881+224530+226265+235582+353204+433182+56421+60525+67680+68263+71832+72141+73724+76615+80911 | 121 | 47 | 29,2681707 | 1,605840026 | 0,00022279 | 0,00323977 | 100198;100678;107029;107272;109264;110639;110821;11429;11430;11532;12359;13806;14194;14380;14433;14718;14719;14751;15275;15926;17448;17449;17850;18563;18597;18598;18641;18648;18655;18746;20425;20916;21881;224530;226265;235582;353204;433182;56421;60525;67680;68263;71832;72141;73724;76615;80911 | Pgk1;Eno1;Acss2;Gapdh;Hk1;Adpgk;Pdhb;Got1;Pfkl;Got2;H6pd;Acox3;Pkm;Gpi1;Pcx;Eno4;Pdha2;Me2;Pfkp;Mcee;Pgam1;Glyctk;Aldoart1;Mdh1;Me3;Got1l1;Csl;Eno1b;Sdhb;Adh5;Psat1;Mdh2;Psph;G6pd2;Tkt;Cat;Shmt1;Idh1;Mmut;Pcca;Sucla2;Prps2;Acat3;Pdha1;Fh1;Aco2;Acox1 |
| mmu00010 | Glycolysis / Gluconeogenesis | http://www.kegg.jp/kegg-bin/show_pathway?mmu00010+106557+11532+11669+11670+11671+13806+14433+14751+15275+16828+16833+18597+18598+18641+18648+18655+18746+226265+353204+433182+56421+56752+60525+68263+72141+72157 | 67 | 26 | 16,2063424 | 1,604310169 | 0,00540045 | 0,03039931 | 106557;11532;11669;11670;11671;13806;14433;14751;15275;16828;16833;18597;18598;18641;18648;18655;18746;226265;353204;433182;56421;56752;60525;68263;72141;72157 | Pgk1;Eno1;Acss2;Gapdh;Hk1;Adpgk;Pdhb;Pfkl;Pkm;Gpi1;Eno4;Pdha2;Aldh2;Ldhc;Pfkp;Aldh3a2;Pgam1;Ldha;Aldoart1;Eno1b;Adh5;Aldh3a1;Aldh9a1;Ldhal6b;Pgm1;Pdha1 |

| | | | | | | | | | | |
|---|---|---|---|---|---|---|---|---|---|---|
| mmu04110 | Cell cycle | http://www.kegg.jp/kegg-bin/show_pathway?mmu04110+11350+11920+12443+12444+12445+12447+12532+12544+12545+12571+12575+13555+13557+15182+17120+17126+17127+17220+17222+17246+17873+19052+19090+19650+20842+20843+217232+21808+218294+22390+22627+22629+22630+22631+229776+23882+242705+26429+26931+26932+26965+320209+328572+433759+52206+52563+54401+56371+56438+59008+68999+69010+69957+70315+71175+71521+72415+75430+77011+77805 | 155 | 60 | 37,4922848 | 1,600329251 | 3,63E-05 | 7,93E-04 | 11350;11920;12443;12444;12445;12447;12532;12544;12545;12571;12575;13555;13557;15182;17120;17126;17127;17220;17222;17246;17873;19052;19090;19650;20842;20843;217232;21808;218294;22390;22627;22629;22630;22631;229776;23882;242705;26429;26931;26932;26965;320209;328572;433759;52206;52563;54401;56371;56438;59008;68999;69010;69957;70315;71175;71521;72415;75430;77011;77805 | E2f1;Ywhah;Gadd45b;Smad3;Anapc4;Mdm2;Anapc13;Sgo1;Cdc45;Esco1;Ddx11;Anapc15;Orc5;Nipbl;Stag1;Mad1l1;Cdc7;Rbx1;Abl1;Ywhae;E2f2;Cdkn1a;Ppp2r5c;Hdac1;Ywhaz;Rbl1;Cdc14a;Smad2;Fzr1;Ppp2ca;Cdc23;Wee1;Cdc14b;Atm;E2f3;Cdc16;Ppp2r5e;Cdc27;Gadd45g;Ccnd3;Cdk6;Anapc1;Pds5a;Cul1;Stag2;Cdc25c;Anapc10;Ywhaq;Prkdc;Ccnd2;Hdac2;Ccnd1;Ep300;Ccne1;Tgfb2;Ywhab;Mcm7;Ticrr;Hdac8;Anapc5 |
| mmu04015 | Rap1 signaling pathway | http://www.kegg.jp/kegg-bin/show_pathway?mmu04015+101540+104111+107746+109905+110157+11465+11512+11515+11600+11601+11848+12313+12315+12387+12388+12801+12927+12928+12977+13800+14062+14164+14173+14176+14178+14180+14183+14679+14683+14811+16001+16333+16337+16409+16414+16416+16590+16797+17295+17311+17356+18053+18645+18654+18706+18751+18795+18796+192786+19353+19730+20779+215449+217692+21844+22339+224129+23797+238130+244668+26395+26396+26397+26413+50791+54354+54635+56508+57257+70549+71785+74055+74206+76089+78134+78473+79264+80796+80857+93737+93742+99470 | 214 | 82 | 51,7635416 | 1,58412654 | 2,34E-06 | 1,17E-04 | 101540;104111;107746;109905;110157;11465;11512;11515;11600;11601;11848;12313;12315;12387;12388;12801;12927;12928;12977;13800;14062;14164;14173;14176;14178;14180;14183;14679;14683;14811;16001;16333;16337;16409;16414;16416;16590;16797;17295;17311;17356;18053;18645;18654;18706;18751;18795;18796;192786;19353;19730;20779;215449;217692;21844;22339;224129;23797;238130;244668;26395;26396;26397;26413;50791;54354;54635;56508;57257;70549;71785;74055;74206;76089;78134;78473;79264;80796;80857;93737;93742;99470 | Sipa1l3;Adcy6;Vegfa;Fgf2;Mapk1;Pgf;Sipa1l2;Plcb2;Magi3;F2r;Rac1;Map2k1;Met;Bcar1;Calm3;Plce1;Rhoa;Insr;Fgfr2;Fgf20;Gnai3;Sipa1l1;Pard3;Actg1;Lat;Pik3ca;Magi2;Pdgfd;Tiam1;Rapgef4;Ins1;Dock4;Enah;Vav3;Rapgef1;Rap1b;Prkd2;Angpt1;Raf1;Akt3;Ngfr;Prkcb;Pfn2;Itgb2;Igf1r;Kit;Itgam;Grin2a;Fgf9;Skap1;Rapgef2;Rapgef6;Plcb1;Ctnnd1;Calm1;Rassf5;Calm4;Csf1;Ralgds;Adcy5;Map2k2;Pdgfc;Fgf5;Pard6g;Fgf7;Krit1;Ctnnb1;Kitl;Adcy3;Cnr1;Tln2;Itgb3;Lpar4;Gnas;Afdn;Map2k3;Crk;Adcy9;Src;Fgf1;Angpt2;Rap1a |

| | | | | | | | | | | |
|---|---|---|---|---|---|---|---|---|---|---|
| mmu04350 | TGF-beta signaling pathway | http://www.kegg.jp/kegg-bin/show_pathway?mmu04350+11481+11848+12160+12161+12162+12166+12168+14118+14313+15182+15902+15903+15904+17126+17127+17131+18007+18119+18741+19052+19650+20185+20466+20481+207596+209268+21808+21812+22041+22042+230597+26413+268977+26965+328572+433759+434215+50765+56438+66313+71775+75788 | 110 | 42 | 26,6074279 | 1,578506579 | 7,13E-04 | 0,00753708 | 11481;11848;12160;12161;12162;12166;12168;14118;14313;15182;15902;15903;15904;17126;17127;17131;18007;18119;18741;19052;19650;20185;20466;20481;207596;209268;21808;21812;22041;22042;230597;26413;268977;26965;328572;433759;434215;50765;56438;66313;71775;75788 | Id3;Smad3;Bmp6;Id2;Mapk1;Thsd4;Smad7;Rhoa;Tfr2;Neo1;Rbx1;Acvr2b;Ski;Ncor1;Fst;Bmpr2;Smurf2;Nodal;Pitx2;Hdac1;Lrrc32;Rbl1;Smad2;Id4;Sin3a;Ppp2ca;Smurf1;Tgfbr1;Tfrc;Zfyve9;Bmpr1a;Cul1;Hdac2;Fbn1;Ep300;Igsf1;Trf;Tgfb2;Inhca;Ltbp1;Bmp5;Bmp7 |
| mmu04928 | Parathyroid hormone synthesis, secretion and action | http://www.kegg.jp/kegg-bin/show_pathway?mmu04928+104111+109689+110157+11512+11515+11848+11909+12393+12575+12912+13653+14281+14672+14679+14683+15200+16440+16973+17258+17260+17261+17387+17389+17391+18227+18578+18751+18795+18796+18806+19228+20181+20531+208647+213498+224129+238871+26395+26413+75547+78284 | 108 | 41 | 26,1236565 | 1,569458702 | 9,37E-04 | 0,00863059 | 104111;109689;110157;11512;11515;11848;11909;12393;12575;12912;13653;14281;14672;14679;14683;15200;16440;16973;17258;17260;17261;17387;17389;17391;18227;18578;18751;18795;18796;18806;19228;20181;20531;208647;213498;224129;238871;26395;26413;75547;78284 | Adcy6;Pde4d;Mapk1;Lrp5;Fos;Akap13;Plcb2;Map2k1;Mmp24;Gna11;Itpr3;Nr4a2;Hbegf;Rhoa;Creb3l4;Gnai3;Creb1;Runx2;Arrb1;Cdkn1a;Slc34a2;Pth1r;Mef2d;Mef2c;Mmp14;Raf1;Prkcb;Arhgef11;Plcb1;Egr1;Adcy5;Rxra;Creb3l2;Atf2;Adcy3;Pld2;Gnas;Mmp16;Mef2a;Adcy9;Pde4b |
| mmu04510 | Focal adhesion | http://www.kegg.jp/kegg-bin/show_pathway?mmu04510+107746+109711+109905+110157+11465+11797+11798+11848+12334+12387+12443+12444+12445+12826+12827+12830+12839+12840+12842+12927+12928+13367+13712+13731+14083+14784+16001+16400+16410+16416+16419+16421+16772+16773+170736+17295+17906+17931+18479+18654+18706+18720+18751+19046+192176+19353+19417+19699+19878+20418+20779+215449+21923+22330+22339+224105+226519+232807+23797+238564+241226+241656+26395+26413+26420+320910+329251+54635+57257+57342+59310+60595+68553+68794+70549+71785+94216 | 203 | 77 | 49,1027988 | 1,56813872 | 7,33E-06 | 2,56E-04 | 107746;109711;109905;110157;11465;11797;11798;11848;12334;12387;12443;12444;12445;12826;12827;12830;12839;12840;12842;12927;12928;13367;13712;13731;14083;14784;16001;16400;16410;16416;16419;16421;16772;16773;170736;17295;17906;17931;18479;18654;18706;18720;18751;19046;192176;19353;19417;19699;19878;20418;20779;215449;21923;22330;22339;224105;226519;232807;23797;238564;241226;241656;26395;26413;26420;320910;329251;54635;57257;57342;59310;60595;68553;68794;70549;71785;94216 | Vegfa;Parvb;Ppp1cb;Mapk1;Pgf;Rac1;Lamc1;Map2k1;Emp2;Met;Ppp1r12b;Bcar1;Flnc;Capn2;Rhoa;Mapk9;Shc3;Actn1;Ptk2;Actg1;Rasgrf1;Pip5k1a;Pik3ca;Itgb8;Pak5;Myl2;Pdgfd;Vcl;Col6a4;Parva;Vav3;Actn4;Lama1;Rapgef1;Rap1b;Birc2;Raf1;Akt3;Reln;Prkcb;Col4a1;Igf1r;Pak1;Col1a1;Itgav;Myl10;Diaph1;Ccnd3;Col9a1;Lama2;Itgb7;Itgb5;Col4a6;Pak2;Mylk4;Ppp1r12c;Itga8;Pdgfc;Ppp1r12a;Col9a2;Ccnd2;Ccnd1;Tnc;Ctnnb1;Tln2;Itgb3;Flna;Col4a2;Grb2;Crk;Xiap;Elk1;Col4a5;Src;Rap1a;Rock2;Itga3 |

| | | | | | | | | | | |
|---|---|---|---|---|---|---|---|---|---|---|
| mmu04912 | GnRH signaling pathway | http://www.kegg.jp/kegg-bin/show_pathway?mmu04912+104111+108058+110157+11512+11515+12289+12313+12315+12325+13653+13712+14672+14683+14784+15200+16440+17387+18751+18753+18783+18795+18796+18806+19229+20779+224129+26395+26396+26397+26401+26405+26413+26420+80796 | 90 | 34 | 21,7697137 | 1,561802806 | 0,0027012 | 0,01860065 | 104111;108058;110157;11512;11515;12289;12313;12315;12325;13653;13712;14672;14683;14784;15200;16440;17387;18751;18753;18783;18795;18796;18806;19229;20779;224129;26395;26396;26397;26401;26405;26413;26420;80796 | Adcy6;Map3k1;Mapk1;Plcb2;Map3k2;Map2k1;Calm3;Camk2g;Gna11;Itpr3;Hbegf;Mapk9;Mmp14;Raf1;Prkcb;Cacna1d;Pla2g4a;Prkcd;Plcb1;Calm1;Calm4;Egr1;Adcy5;Map2k2;Ptk2b;Camk2d;Adcy3;Pld2;Gnas;Map2k3;Grb2;Elk1;Adcy9;Src |
| mmu05222 | Small cell lung cancer | http://www.kegg.jp/kegg-bin/show_pathway?mmu05222+107986+11797+11798+12443+12447+12571+12575+12826+12827+12830+13063+13555+13557+14083+16150+16151+16400+16410+16772+16773+17873+18035+18706+20181+218772+22030+22031+22032+226519+23797+23882+242705+27015+66197+94216 | 93 | 35 | 22,4953709 | 1,555875661 | 0,00254005 | 0,01809135 | 107986;11797;11798;12443;12447;12571;12575;12826;12827;12830;13063;13555;13557;14083;16150;16151;16400;16410;16772;16773;17873;18035;18706;20181;218772;22030;22031;22032;226519;23797;23882;242705;27015;66197;94216 | E2f1;Traf2;Gadd45b;Cycs;Lamc1;Ptk2;Ddb2;Pik3ca;Ikbkb;E2f2;Nfkbia;Cdkn1a;Traf3;Lama1;Birc2;Akt3;E2f3;Ikbkg;Col4a1;Gadd45g;Traf4;Itgav;Lama2;Cdk6;Col4a6;Rarb;Rxra;Ccnd1;Ccne1;Polk;Col4a2;Xiap;Cks2;Col4a5;Itga3 |
| mmu04360 | Axon guidance | http://www.kegg.jp/kegg-bin/show_pathway?mmu04360+107449+108058+108151+110157+11350+117600+11848+12162+12168+12325+12632+13176+13641+13800+13837+13839+13840+13841+13842+13844+140570+14083+14270+14679+16885+16886+17295+18007+18186+18479+18706+19206+19247+19353+19878+20187+20346+20349+20350+20352+20356+20357+20358+20359+20361+20423+20563+20779+214968+224105+22417+22419+226251+231148+234779+237860+241568+241656+26413+268902+319757+53972+54712+69632+73181+80883+93737+93742 | 181 | 68 | 43,7813132 | 1,553174061 | 3,53E-05 | 7,93E-04 | 107449;108058;108151;110157;11350;117600;11848;12162;12168;12325;12632;13176;13641;13800;13837;13839;13840;13841;13842;13844;140570;14083;14270;14679;16885;16886;17295;18007;18186;18479;18706;19206;19247;19353;19878;20187;20346;20349;20350;20352;20356;20357;20358;20359;20361;20423;20563;20779;214968;224105;22417;22419;226251;231148;234779;237860;241568;241656;26413;268902;319757;53972;54712;69632;73181;80883;93737;93742 | Ptch1;Limk2;Mapk1;Limk1;Ryk;Rac1;Ablim2;Ngef;Met;Camk2g;Plcg2;Rhoa;Robo2;Sema6a;Wnt4;Gnai3;Neo1;Ptk2;Ablim1;Abl1;Pard3;Ephb2;Arhgef12;Ptpn11;Epha8;Pik3ca;Plxnc1;Nfatc4;Pak5;Bmpr2;Srgap2;Sema7a;Epha5;Sema3e;Sema5a;Enah;Unc5b;Sema3d;Efnb1;Shh;Lrrc4c;Smo;Cfl2;Raf1;Dcc;Pak1;Srgap1;Pak2;Nrp1;Wnt5b;Sema4b;Pard6g;Sema3f;Plxnb2;Epha3;Epha6;Epha7;Camk2d;Sema3a;Ntng1;Sema6d;Src;Slit2;Sema6b;Bmp7;Ssh2;Rock2;Sema5b |

| | | | | | | | | | | |
|---|---|---|---|---|---|---|---|---|---|---|
| mmu04919 | Thyroid hormone signaling pathway | http://www.kegg.jp/kegg-bin/show_pathway?mmu04919+110157+11465+11938+12387+12443+14463+15182+15251+16410+16416+17246+17977+17978+18129+18132+18640+18641+18706+18751+18795+18796+19744+20181+20185+20466+20502+20525+20544+20779+21834+22417+234779+234959+23797+26395+26396+26413+328572+433759+53901+56421+68975+72472+74055+76199 | 120 | 45 | 29,026285 | 1,550318962 | 7,36E-04 | 0,00755446 | 110157;11465;11938;12387;12443;14463;15182;15251;16410;16416;17246;17977;17978;18129;18132;18640;18641;18706;18751;18795;18796;19744;20181;20185;20466;20502;20525;20544;20779;21834;22417;234779;234959;23797;26395;26396;26413;328572;433759;53901;56421;68975;72472;74055;76199 | Slc2a1;Mapk1;Atp2a2;Slc9a1;Mdm2;Pfkl;Plcb2;Map2k1;Plcg2;Plce1;Med17;Wnt4;Actg1;Ncor1;Pik3ca;Pfkp;Hdac1;Ncoa2;Med13l;Pfkfb2;Sin3a;Med27;Raf1;Akt3;Prkcb;Notch4;Itgav;Plcb1;Rheb;Map2k2;Rxra;Thrb;Hdac2;Slc16a10;Ccnd1;Ctnnb1;Gata4;Ncoa1;Ep300;Rcan2;Notch2;Itgb3;Hif1a;Slc16a2;Src |
| mmu04070 | Phosphatidylinositol signaling system | http://www.kegg.jp/kegg-bin/show_pathway?mmu04070+101490+110197+110911+12313+12315+16330+16440+17772+18706+18711+18718+18720+18751+18795+18796+20975+212111+217480+217837+225326+227333+233011+234515+234779+269180+27399+320127+64436+69718+74055+74302+74596+77116+80796+83493+84095 | 96 | 36 | 23,221028 | 1,550318962 | 0,00238695 | 0,01735512 | 101490;110197;110911;12313;12315;16330;16440;17772;18706;18711;18718;18720;18751;18795;18796;20975;212111;217480;217837;225326;227333;233011;234515;234779;269180;27399;320127;64436;69718;74055;74302;74596;77116;80796;83493;84095 | Itpk1;Pik3c3;Plcb2;Calm3;Plcg2;Ipmk;Plce1;Itpr3;Pip5k1a;Dgki;Pik3ca;Inpp5e;Itpkc;Dgkb;Prkcb;Pi4k2a;Dgkg;Inpp5a;Cds1;Pikfyve;Plcb1;Calm1;Calm4;Inpp4b;Inpp5f;Inpp4a;Mtm1;Sacm1l;Mtmr2;Cds2;Synj2;Ip6k1;Inpp5b;Pip4k2a;Mtmr3;Dgkd |
| mmu00562 | Inositol phosphate metabolism | http://www.kegg.jp/kegg-bin/show_pathway?mmu00562+101490+103199+16330+17772+18706+18711+18718+18720+18795+18796+19062+20975+212111+217837+225326+233011+234515+234779+269180+64436+69718+71780+74055+74302+77116+83493+84095 | 72 | 27 | 17,415771 | 1,550318962 | 0,00789417 | 0,03935809 | 101490;103199;16330;17772;18706;18711;18718;18720;18795;18796;19062;20975;212111;217837;225326;233011;234515;234779;269180;64436;69718;71780;74055;74302;77116;83493;84095 | Itpk1;Pik3c3;Inpp5k;Plcb2;Plcg2;Ipmk;Plce1;Isyna1;Pip5k1a;Pik3ca;Inpp5e;Itpkc;Pi4k2a;Inpp5a;Pikfyve;Plcb1;Inpp4b;Inpp5f;Inpp4a;Mtm1;Fig4;Sacm1l;Mtmr2;Synj2;Inpp5b;Pip4k2a;Mtmr3 |
| mmu05223 | Non-small cell lung cancer | http://www.kegg.jp/kegg-bin/show_pathway?mmu05223+107986+110157+11682+12443+12571+12575+13555+13557+14784+16572+17295+17873+18706+18751+20181+21802+218772+234779+23797+23882+242705+26395+26396+26413+27015+54354+56484 | 72 | 27 | 17,415771 | 1,550318962 | 0,00789417 | 0,03935809 | 107986;110157;11682;12443;12571;12575;13555;13557;14784;16572;17295;17873;18706;18751;20181;21802;218772;234779;23797;23882;242705;26395;26396;26413;27015;54354;56484 | E2f1;Gadd45b;Tgfa;Mapk1;Map2k1;Met;Plcg2;Kif5a;Ddb2;Pik3ca;Foxo3;E2f2;Cdkn1a;Raf1;Akt3;Prkcb;E2f3;Gadd45g;Cdk6;Rassf5;Map2k2;Rarb;Rxra;Ccnd1;Polk;Alk;Grb2 |

| | | | | | | | | | | |
|---|---|---|---|---|---|---|---|---|---|---|
| mmu04540 | Gap junction | http://www.kegg.jp/kegg-bin/show_pathway?mmu04540+104111+108071+110157+11512+11515+14609+14672+14679+14683+14784+14816+15560+16440+18751+18795+18796+19091+19092+20779+21872+22143+224129+227613+23938+26395+26396+26405+26413+545486+54635+60596+71785 | 86 | 32 | 20,8021709 | 1,538300985 | 0,00464552 | 0,02844363 | 104111;108071;110157;11512;11515;14609;14672;14679;14683;14784;14816;15560;16440;18751;18795;18796;19091;19092;20779;21872;22143;224129;227613;23938;26395;26396;26405;26413;545486;54635;60596;71785 | Adcy6;Tubb4b;Mapk1;Plcb2;Map3k2;Map2k5;Map2k1;Gna11;Itpr3;Gnai3;Tuba1b;Pdgfd;Raf1;Prkcb;Grm5;Prkg1;Gja1;Plcb1;Prkg2;Adcy5;Tjp1;Map2k2;Pdgfc;Tubb1;Grm1;Adcy3;Htr2c;Gnas;Grb2;Adcy9;Src;Gucy1a1 |
| mmu04530 | Tight junction | http://www.kegg.jp/kegg-bin/show_pathway?mmu04530+104027+105787+108097+108099+109711+109905+11465+11848+12289+12394+12443+12479+12695+12741+13383+14463+15525+16456+17356+17880+17886+17906+17999+18016+19052+192786+19353+19684+19878+20779+20869+21844+21872+22143+227157+231830+235442+23859+26401+26420+269643+27375+27494+434204+53310+54357+56378+56492+58187+60595+67771+68178+68328+71978+73608+74094+74987+76089+77579+83814+93737+93742 | 168 | 62 | 40,636799 | 1,525710724 | 1,39E-04 | 0,00213493 | 104027;105787;108097;108099;109711;109905;11465;11848;12289;12394;12443;12479;12695;12741;13383;14463;15525;16456;17356;17880;17886;17906;17999;18016;19052;192786;19353;19684;19878;20779;20869;21844;21872;22143;227157;231830;235442;23859;26401;26420;269643;27375;27494;434204;53310;54357;56378;56492;58187;60595;67771;68178;68328;71978;73608;74094;74987;76089;77579;83814;93737;93742 | Ppp2r2a;Map3k1;Stk11;Cgnl1;Synpo;Rac1;Rhoa;Dlg2;Mapk9;Rdx;Dlg1;Myh9;Actn1;Mpp4;Pard3;Actg1;Tuba1b;Myl2;Tiam1;Nf2;Cldn5;Micall2;Actn4;Nedd4;Nedd4l;Whamm;Marveld3;Ppp2ca;Arpc5;Prkag2;Arpc3;Cacna1d;Ppp2r2c;Tjp3;Myh10;Rapgef2;Tjap1;Rapgef6;Amot;Runx1;Tjp1;Prkaa1;Pard6g;Cd1d1;Ccnd1;F11r;Dlg3;Gata4;Prkab2;Cldn34d;Rab8b;Cldn18;Afdn;Patj;Hspa4;Rab13;Epb41l4b;Cldn10;Src;Rap1a;Rock2;Myh11 |
| mmu05212 | Pancreatic cancer | http://www.kegg.jp/kegg-bin/show_pathway?mmu05212+107986+110157+12443+12571+12575+13555+13557+16150+16151+16451+17126+17127+17873+18706+18806+19353+19730+21802+21808+21812+22339+23797+23882+242705+26395+26413+26420+27015 | 76 | 28 | 18,3833138 | 1,523120383 | 0,00906876 | 0,04110387 | 107986;110157;12443;12571;12575;13555;13557;16150;16151;16451;17126;17127;17873;18706;18806;19353;19730;21802;21808;21812;22339;23797;23882;242705;26395;26413;26420;27015 | E2f1;Gadd45b;Vegfa;Smad3;Tgfa;Mapk1;Rac1;Map2k1;Mapk9;Ddb2;Pik3ca;Ikbkb;E2f2;Cdkn1a;Smad2;Raf1;Tgfbr1;Akt3;E2f3;Ikbkg;Gadd45g;Cdk6;Ralgds;Ccnd1;Pld2;Tgfb2;Polk;Jak1 |

| | | | | | | | | | | |
|---|---|---|---|---|---|---|---|---|---|---|
| mmu03082 | ATP-dependent chromatin remodeling | http://www.kegg.jp/kegg-bin/show_pathway?mmu03082+100039319+100042922+100043597+100683+11465+116870+12054+15182+17191+20174+20587+210982+216848+217578+21761+225280+22632+229542+233532+235134+236663+243842+26992+319164+319165+319168+319170+319171+319173+385328+404634+407823+433759+57376+624153+665433+66923+67155+68094+70088+70127+77044+78656+81601+83766+83796+83797+93696+93760+93761 | 136 | 50 | 32,8964563 | 1,519920551 | 6,55E-04 | 0,00728847 | 100039319;100042922;100043597;100683;11465;116870;12054;15182;17191;20174;20587;210982;216848;217578;21761;225280;22632;229542;233532;235134;236663;243842;26992;319164;319165;319168;319170;319171;319173;385328;404634;407823;433759;57376;624153;665433;66923;67155;68094;70088;70127;77044;78656;81601;83766;83796;83797;93696;93760;93761 | H2ac12;H2ac10;H2ac24;H2ac7;Bcl7b;Smarcc2;Mta1;H2ac22;H2ac6;Chrac1;Trrap;Arid2;Kat5;Actg1;H2ac23;Actl6b;Bicra;Rsf1;Hdac1;Baz2b;Mbd2;Smarcd1;Smarcb1;Srcap;Bicral;Gatad2b;Nfrkb;Arid1a;Brd7;Smarca1;Baz1a;Pbrm1;Meaf6;Dpf3;H2al1n;Ino80c;Macroh2a2;Hdac2;Brd8;H2ab1;Smarcd2;Smarce1;H2al1a;H2al1j;Smarca2;Chd3;Ruvbl2;Morf4l1;Yy1;H2ab2 |
| mmu01230 | Biosynthesis of amino acids | http://www.kegg.jp/kegg-bin/show_pathway?mmu01230+100678+107272+109900+110639+11429+11898+12036+13806+14433+14645+14718+14719+15926+18478+18563+18641+18648+18655+18746+20425+217214+21881+226265+353204+433182+56421+71776+71832+76615 | 79 | 29 | 19,1089709 | 1,517611811 | 0,00847289 | 0,03995999 | 100678;107272;109900;110639;11429;11898;12036;13806;14433;14645;14718;14719;15926;18478;18563;18641;18648;18655;18746;20425;217214;21881;226265;353204;433182;56421;71776;71832;76615 | Pgk1;Eno1;Ass1;Gapdh;Got1;Pfkl;Got2;Glul;Tha1;Pkm;Bcat2;Pcx;Eno4;Pfkp;Pgam1;Aldoart1;Got1l1;Csl;Eno1b;Psat1;Nags;Psph;Tkt;Shmt1;Asl;Idh1;Prps2;Pah;Aco2 |
| mmu05226 | Gastric cancer | http://www.kegg.jp/kegg-bin/show_pathway?mmu05226+107986+110157+11789+12005+12385+12386+12387+12443+12447+12575+13555+13557+14164+14173+14176+14178+14180+14183+14362+14367+14369+14370+14371+14784+16973+17126+17127+17295+17873+18706+20181+20418+20423+21239 8+21414+21416+21808+21812+218772+22408+22414+22415+22417+22419+23797+23805+23882+242705+26395+26396+26413+27015+67709+80857+93687 | 150 | 55 | 36,2828562 | 1,515867429 | 3,90E-04 | 0,00504134 | 107986;110157;11789;12005;12385;12386;12387;12443;12447;12575;13555;13557;14164;14173;14176;14178;14180;14183;14362;14367;14369;14370;14371;14784;16973;17126;17127;17295;17873;18706;20181;20418;20423;21239 8;21414;21416;21808;21812;218772;22408;22414;22415;22417;22419;23797;23805;23882;242705;26395;26396;26413;27015;67709;80857;93687 | E2f1;Gadd45b;Smad3;Fgf2;Mapk1;Lrp5;Map2k1;Wnt2b;Met;Fgfr2;Shc3;Fgf20;Wnt4;Ddb2;Csnk1a1;Pik3ca;E2f2;Cdkn1a;Tcf7;Fzd8;Fzd9;Fzd1;Smad2;Shh;Axin1;Raf1;Tgfbr1;Akt3;E2f3;Fzd7;Gadd45g;Apc;Fgf9;Wnt1;Wnt5b;Apc2;Ctnna2;Reg4;Map2k2;Rarb;Fgf5;Rxra;Fgf7;Frat2;Ccnd1;Wnt3;Ctnnb1;Ccne1;Tgfb2;Tcf7l2;Polk;Grb2;Ctnna1;Fzd5;Fgf1 |

| | | | | | | | | | |
|---|---|---|---|---|---|---|---|---|---|
| mmu04666 | Fc gamma R-mediated phagocytosis | http://www.kegg.jp/kegg-bin/show_pathway?mmu04666+110157+11845+12632+12928+13196+14130+14131+14389+16797+16885+16886+17909+18479+18706+18720+18751+18753+18783+18806+19012+19353+20963+211914+227753+234779+23797+242687+26395+26413+56378+57257+67771+67916+83767 | 93 | 34 | 22,4953709 | 1,51142207 | 0,00496223 | 0,02886363 | 110157;11845;12632;12928;13196;14130;14131;14389;16797;16885;16886;17909;18479;18706;18720;18751;18753;18783;18806;19012;19353;20963;211914;227753;234779;23797;242687;26395;26413;56378;57257;67771;67916;83767 | Limk2;Mapk1;Limk1;Gab2;Rac1;Map2k1;Gsn;Plpp3;Plcg2;Arf6;Syk;Myo10;Pip5k1a;Lat;Pik3ca;Vav3;Cfl2;Arpc5;Raf1;Akt3;Prkcb;Arpc3;Wasf2;Asap1;Pla2g4a;Pak1;Prkcd;Fcgr2b;Plpp1;Wasf1;Fcgr3;Pld2;Crk;Asap2 |
| mmu04922 | Glucagon signaling pathway | http://www.kegg.jp/kegg-bin/show_pathway?mmu04922+102093+105787+106557+107476+108058+108097+108099+11909+12313+12315+12325+12894+12912+14527+14683+16440+16828+16833+17691+18597+18598+18639+18641+18648+18679+18746+18795+18796+19017+20525+208647+23797+328572+56420+56421+68263+78284+80796 | 104 | 38 | 25,1561137 | 1,510567193 | 0,00312456 | 0,02019389 | 102093;105787;106557;107476;108058;108097;108099;11909;12313;12315;12325;12894;12912;14527;14683;16440;16828;16833;17691;18597;18598;18639;18641;18648;18679;18746;18795;18796;19017;20525;208647;23797;328572;56420;56421;68263;78284;80796 | Slc2a1;Sik1;Pdhb;Pfkl;Plcb2;Calm3;Camk2g;Itpr3;Cpt1a;Creb3l4;Pkm;Pdha2;Creb1;Ppp4c;Gcgr;Ldhc;Pfkp;Acaca;Pgam1;Ldha;Prkag2;Akt3;Ppargc1a;Phkb;Plcb1;Calm1;Calm4;Pfkfb1;Ldhal6b;Prkaa1;Creb3l2;Atf2;Prkab2;Ep300;Camk2d;Gnas;Pdha1;Phka1 |
| mmu04810 | Regulation of actin cytoskeleton | http://www.kegg.jp/kegg-bin/show_pathway?mmu04810+108100+109711+110157+11465+11789+11848+12274+12632+12927+12928+13367+13800+14062+14083+14163+14164+14173+14176+14178+14180+14183+15139+16333+16400+16409+16410+16414+16416+16419+16421+16885+16886+17880+17886+17906+17931+18479+18645+18706+18711+18718+18720+19046+19241+19353+19684+19878+20544+20779+21844+219140+22330+224105+227753+232807+237860+23797+23805+238564+241226+241656+242687+26395+26396+26413+29875+320910+329165+329251+50884+54004+544963+54635+56378+57257+59310+60595+67771+69632+71785+77579+78134+80857+83767 | 231 | 84 | 55,8755986 | 1,503339599 | 1,88E-05 | 4,69E-04 | 108100;109711;110157;11465;11789;11848;12274;12632;12927;12928;13367;13800;14062;14083;14163;14164;14173;14176;14178;14180;14183;15139;16333;16400;16409;16410;16414;16416;16419;16421;16885;16886;17880;17886;17906;17931;18479;18645;18706;18711;18718;18720;19046;19241;19353;19684;19878;20544;20779;21844;219140;22330;224105;227753;232807;237860;23797;23805;238564;241226;241656;242687;26395;26396;26413;29875;320910;329165;329251;50884;54004;544963;54635;56378;57257;59310;60595;67771;69632;71785;77579;78134;80857;83767 | Limk2;Fgf2;Ppp1cb;Mapk1;Limk1;Slc9a1;Iqgap2;Iqgap1;F2r;Rac1;Map2k1;Gsn;Ppp1r12b;Bcar1;Rhoa;Fgfr2;Rdx;Fgf20;Myh9;Actn1;Spata13;Ptk2;Arhgef12;Actg1;Pip5k1a;Pik3ca;Itgb8;Pak5;Myl2;Pdgfd;Tiam1;Vcl;Ins1;Enah;Vav3;Actn4;Cfl2;Arpc5;Raf1;Akt3;Pfn2;Itgb2;Arpc3;Wasf2;Itgam;Apc;Pak1;Fgf9;Itgav;Myl10;Diaph1;Myh10;Pikfyve;Itgb7;Itgb5;Pak2;Abi2;Apc2;Mylk4;Ppp1r12c;Itga8;Wasf1;Map2k2;Fgd1;Pdgfc;Ppp1r12a;C6;Fgf5;Fgf7;Tmsb4x;Hc;Itgb3;Lpar4;Pip4k2a;Baiap2;Crk;Src;Diaph2;Fgf1;Ssh2;Nckap1;Rock2;Itga3;Myh11 |

| | | | | | | | | | | |
|---|---|---|---|---|---|---|---|---|---|---|
| mmu04114 | Oocyte meiosis | http://www.kegg.jp/kegg-bin/show_pathway?mmu04114+104111+108058+110651+11512+11515+11835+12313+12315+12325+12447+12532+12877+16001+16333+16440+17120+17222+19046+19052+208922+217232+224129+22627+22629+22630+22631+231207+26395+26413+26931+26932+26965+52206+52563+54401+56438+59008+67071+68999+69010+69957+72415+75430+80796 | 121 | 44 | 29,2681707 | 1,503339599 | 0,00172626 | 0,01369234 | 104111;108058;110651;11512;11515;11835;12313;12315;12325;12447;12532;12877;16001;16333;16440;17120;17222;19046;19052;208922;217232;224129;22627;22629;22630;22631;231207;26395;26413;26931;26932;26965;52206;52563;54401;56438;59008;67071;68999;69010;69957;72415;75430;80796 | Ywhah;Adcy6;Ppp1cb;Mapk1;Anapc4;Anapc13;Sgo1;Map2k1;Calm3;Camk2g;Itpr3;Anapc15;Mad1l1;Rbx1;Ywhae;Rps6ka6;Ppp2r5c;Ywhaz;Ins1;Ppp2ca;Cdc23;Cdc16;Rps6ka3;Ppp2r5e;Cdc27;Igf1r;Calm1;Cpeb2;Calm4;Anapc1;Adcy5;Cul1;Cpeb3;Cdc25c;Anapc10;Ywhaq;Cpeb1;Camk2d;Ccne1;Adcy3;Ywhab;Ar;Adcy9;Anapc5 |
| mmu04072 | Phospholipase D signaling pathway | http://www.kegg.jp/kegg-bin/show_pathway?mmu04072+103967+104111+108071+108073+110157+110197+11512+11515+11607+11608+11841+11845+11848+14062+14126+14389+14683+14784+14816+16333+16337+16590+17311+18706+18720+18783+18795+18796+18806+19012+19157+19159+19229+19247+19730+19744+20418+20963+217480+224129+227333+234779+23797+26395+26396+26413+268934+320127+54635+56508+67916+71785+72318+78134 | 149 | 54 | 36,0409705 | 1,498294835 | 6,07E-04 | 0,00706634 | 103967;104111;108071;108073;110157;110197;11512;11515;11607;11608;11841;11845;11848;14062;14126;14389;14683;14784;14816;16333;16337;16590;17311;18706;18720;18783;18795;18796;18806;19012;19157;19159;19229;19247;19730;19744;20418;20963;217480;224129;227333;234779;23797;26395;26396;26413;268934;320127;54635;56508;67916;71785;72318;78134 | Adcy6;Mapk1;Plcb2;F2r;Gab2;Map2k1;Plpp3;Plcg2;Rhoa;Insr;Arf6;Shc3;Ptpn11;Cyth4;Syk;Pip5k1a;Dgki;Pik3ca;Pdgfd;Rapgef4;Agtr1a;Ins1;Dnm3;Dgkb;Arf2;Raf1;Akt3;Dgkg;Kit;Pla2g4a;Grm5;Plcb1;Plpp1;Ralgds;Cyth1;Rheb;Adcy5;Map2k2;Pdgfc;Grm4;Ms4a2;Grm7;Cyth3;Kitl;Grm1;Ptk2b;Adcy3;Pld2;Lpar4;Gnas;Grb2;Agtr1b;Adcy9;Dgkd |
| mmu04140 | Autophagy | http://www.kegg.jp/kegg-bin/show_pathway?mmu04140+100502698+100502841+105787+110157+12176+12211+12421+13143+13665+13666+15251+15289+16001+16333+16370+16784+18518+18706+18753+19052+19324+19349+19744+20869+213948+217695+218035+218460+22186+22190+225326+228361+23797+242557+245860+26395+26396+26413+26420+27103+29869+329015+52440+54170+56480+66734+66840+67443+67727+68441+72145+73205+74302+74370+74498+74747+75669+78610+78943+97998 | 166 | 60 | 40,1530276 | 1,494283337 | 3,38E-04 | 0,00453398 | 100502698;100502841;105787;110157;12176;12211;12421;13143;13665;13666;15251;15289;16001;16333;16370;16784;18518;18706;18753;19052;19324;19349;19744;20869;213948;217695;218035;218460;22186;22190;225326;228361;23797;242557;245860;26395;26396;26413;26420;27103;29869;329015;52440;54170;56480;66734;66840;67443;67727;68441;72145;73205;74302;74370;74498;74747;75669;78610;78943;97998 | Bnip3;Atg9b;Pik3c3;Map1lc3b;Stk11;Mapk1;Rubcn;Map2k1;Atg9a;Uba52;Ddit4;Dapk2;Mapk9;Rb1cc1;Wdr45b;Birc6;Deptor;Pik3ca;Rraga;Wdr41;Map1lc3a;Hmgb1;Stx17;Ins1;Vps41;Atg2a;Pik3r4;Ppp2ca;Rragc;Ubc;Zfyve1;Raf1;Akt3;Uvrag;Rab7;Igf1r;Tbk1;Rab1a;Prkcd;Irs4;Tax1bp1;Wdfy3;Eif2ak4;Rheb;Rptor;Ern1;Lamp2;Map2k2;Prkaa1;Ambra1;Eif2ak3;Epg5;C9orf72;Hif1a;Eif2s1;Mtmr3;Igbp1;Ulk2;Atg4c;Gorasp1 |

| | | | | | | | | | | |
|---|---|---|---|---|---|---|---|---|---|---|
| mmu04713 | Circadian entrainment | http://www.kegg.jp/kegg-bin/show_pathway?mmu04713+104111+108058+11512+11515+11517+12289+12313+12315+12325+12912+14281+14679+14683+14700+14800+14802+14811+16440+16519+18626+18627+18628+18751+18795+18796+19091+19092+20191+20192+224129+239556+26413+58226+60596+80796 | 98 | 35 | 23,7047994 | 1,476494249 | 0,00672864 | 0,03453378 | 104111;108058;11512;11515;11517;12289;12313;12315;12325;12912;14281;14679;14683;14700;14800;14802;14811;16440;16519;18626;18627;18628;18751;18795;18796;19091;19092;20191;20192;224129;239556;26413;58226;60596;80796 | Adcy6;Mapk1;Cacna1h;Fos;Ryr2;Plcb2;Calm3;Camk2g;Per1;Itpr3;Gnai3;Creb1;Gria2;Kcnj3;Per3;Gng10;Prkcb;Cacna1d;Grin2a;Prkg1;Plcb1;Gria4;Calm1;Calm4;Cacna1i;Prkg2;Adcy5;Adcyap1r1;Ryr3;Camk2d;Adcy3;Gnas;Adcy9;Per2;Gucy1a1 |
| mmu05410 | Hypertrophic cardiomyopathy | http://www.kegg.jp/kegg-bin/show_pathway?mmu05410+105787+108097+108099+11465+11938+12289+12293+12294+12296+13138+13405+13527+13614+13726+16400+16410+16416+16419+16421+16772+16773+16905+17906+20191+20648+20649+20650+21808+22003+22138+24052+241226+319734+320910+54377 | 98 | 35 | 23,7047994 | 1,476494249 | 0,00672864 | 0,03453378 | 105787;108097;108099;11465;11938;12289;12293;12294;12296;13138;13405;13527;13614;13726;16400;16410;16416;16419;16421;16772;16773;16905;17906;20191;20648;20649;20650;21808;22003;22138;24052;241226;319734;320910;54377 | Dag1;Ryr2;Atp2a2;Sntb2;Tpm1;Dtna;Actg1;Itgb8;Cacna2d4;Myl2;Dmd;Lmna;Lama1;Cacnb2;Prkag2;Sntb1;Cacna2d1;Cacna1d;Sgcd;Edn1;Itgav;Snta1;Lama2;Itgb7;Itgb5;Emd;Itga8;Prkaa1;Prkab2;Tgfb2;Itgb3;Cacng4;Cacna2d3;Ttn;Itga3 |
| mmu03083 | Polycomb repressive complex | http://www.kegg.jp/kegg-bin/show_pathway?mmu03083+13626+15182+17765+211961+212391+214669+225876+22632+237009+252870+29808+29871+30841+30951+319974+320858+328572+433759+494448+50496+52609+52615+54650+56550+67057+67933+71458+75302+76073+77593 | 84 | 30 | 20,3183995 | 1,476494249 | 0,01154156 | 0,04972844 | 13626;15182;17765;211961;212391;214669;225876;22632;237009;252870;29808;29871;30841;30951;319974;320858;328572;433759;494448;50496;52609;52615;54650;56550;67057;67933;71458;75302;76073;77593 | Mga;Scmh1;Auts2;L3mbtl2;Hcfc2;Ube2d2a;Kdm2a;Usp7;Hdac1;Asxl3;Suz12;L3mbtl4;Cbx8;Asxl2;Lcor;Cbx6;Ube2dnl1;E2f6;Cbx7;Bcor;Usp45;Hdac2;Yaf2;Sfmbt1;Ep300;Eed;Mtf2;Pcgf5;Kdm2b;Yy1 |
| mmu04012 | ErbB signaling pathway | http://www.kegg.jp/kegg-bin/show_pathway?mmu04012+108058+110157+11350+12325+12402+12575+12928+13685+13712+13869+14083+14784+15200+18183+18479+18706+18751+20418+20779+208650+211323+21802+224105+234779+23797+241656+26395+26396+26413+26420 | 84 | 30 | 20,3183995 | 1,476494249 | 0,01154156 | 0,04972844 | 108058;110157;11350;12325;12402;12575;12928;13685;13712;13869;14083;14784;15200;18183;18479;18706;18751;20418;20779;208650;211323;21802;224105;234779;23797;241656;26395;26396;26413;26420 | Eif4ebp1;Tgfa;Mapk1;Cblb;Map2k1;Camk2g;Plcg2;Hbegf;Mapk9;Shc3;Ptk2;Abl1;Pik3ca;Pak5;Cdkn1a;Nrg3;Raf1;Cbl;Akt3;Prkcb;Pak1;Pak2;Map2k2;Camk2d;Nrg1;Erbb4;Grb2;Crk;Elk1;Src |

| | | | | | | | | | | |
|---|---|---|---|---|---|---|---|---|---|---|
| mmu04141 | Protein processing in endoplasmic reticulum | http://www.kegg.jp/kegg-bin/show_pathway?mmu04141+105245+108159+108687+109815+110616+116891+12282+12334+12364+12954+13002+13665+13666+140499+15502+17156+18024+18453+19359+193740+20014+20335+213539+216440+22030+223455+22393+225896+230815+230904+235416+237009+26420+26965+320011+50527+50873+54609+56438+56445+56550+56709+56812+57377+63958+66256+66357+66397+66435+66753+67437+67475+67819+69162+70361+71853+78943+80286+81500+94232+99683 | 171 | 61 | 41,3624561 | 1,474767355 | 4,52E-04 | 0,00563261 | 105245;108159;108687;109815;110616;116891;12282;12334;12364;12954;13002;13665;13666;140499;15502;17156;18024;18453;19359;193740;20014;20335;213539;216440;22030;223455;22393;225896;230815;230904;235416;237009;26420;26965;320011;50527;50873;54609;56438;56445;56550;56709;56812;57377;63958;66256;66357;66397;66435;66753;67437;67475;67819;69162;70361;71853;78943;80286;81500;94232;99683 | Ero1a;Sec24b;Traf2;Cryaa;Os9;Dnajc5;Ube2j2;Rpn2;Ubxn8;Ube4b;Tusc3;Lman1l;Sil1;Hspa1a;Ostc;Capn2;Mapk9;Ube2d2a;Derl2;Rbx1;Mogs;Selenos;P4hb;Dnaja1;Sec31a;Sec61g;Erlec1;Dnajb2;Txndc5;Bag2;Nfe2l2;Wfs1;Man1c1;Uggt1;Edem2;Prkn;Derl1;Ubqln2;Atxn3;Fbxo2;Casp12;Ube2dnl1;Ern1;Ssr2;Cul1;Rad23b;Dnajb12;Marchf6;Man1a2;Pdia6;Ubxn1;Sar1b;Dnaja2;Ssr3;Eif2ak3;Uggt2;Hyou1;Ubqln4;Eif2s1;Ero1b;Lman1 |
| mmu05414 | Dilated cardiomyopathy | http://www.kegg.jp/kegg-bin/show_pathway?mmu05414+104111+11465+11512+11515+11938+12289+12293+12294+12296+13138+13405+13527+13726+14683+16400+16410+16416+16419+16421+16772+16773+16905+17906+20191+20648+20649+20650+21808+22003+22138+224129+24052+241226+319734+320910+54377 | 101 | 36 | 24,4304565 | 1,473570498 | 0,00626671 | 0,03354721 | 104111;11465;11512;11515;11938;12289;12293;12294;12296;13138;13405;13527;13726;14683;16400;16410;16416;16419;16421;16772;16773;16905;17906;20191;20648;20649;20650;21808;22003;22138;224129;24052;241226;319734;320910;54377 | Adcy6;Dag1;Ryr2;Atp2a2;Sntb2;Tpm1;Dtna;Actg1;Itgb8;Cacna2d4;Myl2;Dmd;Lmna;Lama1;Cacnb2;Sntb1;Cacna2d1;Cacna1d;Sgcd;Itgav;Snta1;Lama2;Itgb7;Itgb5;Emd;Adcy5;Itga8;Adcy3;Tgfb2;Itgb3;Cacng4;Gnas;Cacna2d3;Ttn;Adcy9;Itga3 |
| mmu04148 | Efferocytosis | http://www.kegg.jp/kegg-bin/show_pathway?mmu04148+107831+108058+11487+11491+11502+11863+11938+12215+12325+12369+12608+12894+12912+12927+12928+13435+13856+140580+14083+15251+16410+16416+16419+16423+18514+18613+19247+192654+19329+19345+19349+19353+20181+20393+20501+20525+209018+212555+214058+218035+22174+22260+233246+233571+240672+26395+26396+26413+271457+319520+320091+320940+380713+497097+68365+70417+73181 | 161 | 57 | 38,943599 | 1,463655169 | 8,45E-04 | 0,00819165 | 107831;108058;11487;11491;11502;11863;11938;12215;12325;12369;12608;12894;12912;12927;12928;13435;13856;140580;14083;15251;16410;16416;16419;16423;18514;18613;19247;192654;19329;19345;19349;19353;20181;20393;20501;20525;209018;212555;214058;218035;22174;22260;233246;233571;240672;26395;26396;26413;271457;319520;320091;320940;380713;497097;68365;70417;73181 | Slc2a1;Epo;Bsg;Pbx1;Dnmt3a;Mapk1;Dusp4;Atp2a2;Rac1;Map2k1;Bcar1;Rab17;Camk2g;Vps8;Scarf1;Adam9;Cpt1a;Cd47;P2ry6;Rab5a;Cebpb;Ptk2;Creb1;Ptpn11;Rab5c;Nfatc4;Vps41;Adam17;Megf10;Arnt;Slc66a1;Rab7;Xkr4;Itgav;Itgb5;Sgk1;Megf11;Map2k2;Ano5;Rxra;Pla2g15;Tyro3;Adgrb1;Slc16a1;Atp11c;Dusp5;Pecam1;Camk2d;Itgb3;Hif1a;Ano4;Rab14;Elmo1;Adam10;Nr1h2;Crk;Casp7 |

| | | | | | | | | | | |
|---|---|---|---|---|---|---|---|---|---|---|
| mmu05224 | Breast cancer | http://www.kegg.jp/kegg-bin/show_pathway?mmu05224+107986+110157+11789+12005+12189+12387+12443+12571+12575+13388+13555+13557+14164+14173+14176+14178+14180+14281+14362+14367+14369+14370+14371+14784+15205+16001+16590+16973+17873+17977+18129+18132+18706+20418+212398+21414+21416+22408+22414+22415+22417+22419+23797+23805+23882+242705+26395+26396+26413+27015+80857+93687 | 147 | 52 | 35,5571991 | 1,462432399 | 0,00142512 | 0,01192985 | 107986;110157;11789;12005;12189;12387;12443;12571;12575;13388;13555;13557;14164;14173;14176;14178;14180;14281;14362;14367;14369;14370;14371;14784;15205;16001;16590;16973;17873;17977;18129;18132;18706;20418;212398;21414;21416;22408;22414;22415;22417;22419;23797;23805;23882;242705;26395;26396;26413;27015;80857;93687 | E2f1;Hes1;Gadd45b;Fgf2;Mapk1;Lrp5;Fos;Map2k1;Wnt2b;Shc3;Fgf20;Wnt4;Dll1;Ddb2;Csnk1a1;Pik3ca;E2f2;Cdkn1a;Tcf7;Fzd8;Fzd9;Fzd1;Brca1;Axin1;Raf1;Akt3;E2f3;Notch4;Fzd7;Igf1r;Gadd45g;Kit;Apc;Fgf9;Wnt1;Cdk6;Wnt5b;Apc2;Map2k2;Fgf5;Fgf7;Frat2;Ccnd1;Wnt3;Ctnnb1;Ncoa1;Notch2;Tcf7l2;Polk;Grb2;Fzd5;Fgf1 |
| mmu05215 | Prostate cancer | http://www.kegg.jp/kegg-bin/show_pathway?mmu05215+104156+110157+11835+12387+12443+12447+12575+12912+13555+13557+14183+14784+16001+16150+16151+16333+17246+18035+18095+18706+18792+208647+21414+21416+21417+21802+23797+242705+26395+26396+26413+328572+54635+71785+78284 | 99 | 35 | 23,9466851 | 1,461580166 | 0,00803968 | 0,03951901 | 104156;110157;11835;12387;12443;12447;12575;12912;13555;13557;14183;14784;16001;16150;16151;16333;17246;18035;18095;18706;18792;208647;21414;21416;21417;21802;23797;242705;26395;26396;26413;328572;54635;71785;78284 | E2f1;Tgfa;Mapk1;Mdm2;Map2k1;Zeb1;Fgfr2;Creb3l4;Creb1;Pik3ca;Ikbkb;E2f2;Nfkbia;Cdkn1a;Pdgfd;Tcf7;Ins1;Etv5;Raf1;Akt3;E2f3;Ikbkg;Igf1r;Map2k2;Pdgfc;Plau;Ccnd1;Creb3l2;Ctnnb1;Ep300;Ccne1;Tcf7l2;Grb2;Nkx3-1;Ar |
| mmu04921 | Oxytocin signaling pathway | http://www.kegg.jp/kegg-bin/show_pathway?mmu04921+104111+105787+108058+108097+108099+110157+11465+11512+11515+11848+12289+12293+12294+12296+12313+12315+12325+12443+12575+13712+14281+14679+14683+16440+16519+16520+17260+17931+18127+18160+18751+18783+18795+18796+19046+19878+20191+20192+20779+215303+224129+230103+232807+238564+23938+26395+26396+26413+319734+329251+54377+60596+73181+80796 | 153 | 54 | 37,0085134 | 1,459123729 | 0,00123814 | 0,01080276 | 104111;105787;108058;108097;108099;110157;11465;11512;11515;11848;12289;12293;12294;12296;12313;12315;12325;12443;12575;13712;14281;14679;14683;16440;16519;16520;17260;17931;18127;18160;18751;18783;18795;18796;19046;19878;20191;20192;20779;215303;224129;230103;232807;238564;23938;26395;26396;26413;319734;329251;54377;60596;73181;80796 | Adcy6;Ppp1cb;Mapk1;Fos;Ryr2;Plcb2;Map2k5;Map2k1;Npr2;Ppp1r12b;Calm3;Camk2g;Itpr3;Rhoa;Gnai3;Actg1;Nfatc4;Cacna2d4;Cdkn1a;Kcnj3;Mef2c;Cacnb2;Nos3;Prkag2;Raf1;Cacna2d1;Prkcb;Cacna1d;Pla2g4a;Plcb1;Calm1;Calm4;Mylk4;Ppp1r12c;Adcy5;Camk1g;Map2k2;Prkaa1;Ppp1r12a;Kcnj4;Ccnd1;Prkab2;Ryr3;Camk2d;Adcy3;Cacng4;Gnas;Cacna2d3;Npr1;Elk1;Adcy9;Src;Gucy1a1;Rock2 |

| | | | | | | | | | |
|---|---|---|---|---|---|---|---|---|---|
| mmu04935 | Growth hormone synthesis, secretion and action | http://www.kegg.jp/kegg-bin/show_pathway?mmu04935+104111+110157+11512+11515+11909+12289+12912+12927+12928+14083+14281+14600+14672+14679+14683+14784+16009+16370+16440+18706+18736+18751+18795+18796+20418+20604+20606+20609+208188+208647+224129+234779+23797+26395+26396+26397+26401+26413+26420+328572+78284 | 117 | 41 | 28,3006279 | 1,44873111 | 0,00520852 | 0,02979957 | 104111;110157;11512;11515;11909;12289;12912;12927;12928;14083;14281;14600;14672;14679;14683;14784;16009;16370;16440;18706;18736;18751;18795;18796;20418;20604;20606;20609;208188;208647;224129;234779;23797;26395;26396;26397;26401;26413;26420;328572;78284 | Adcy6;Map3k1;Mapk1;Fos;Plcb2;Map2k1;Sstr5;Bcar1;Plcg2;Gna11;Itpr3;Mapk9;Shc3;Creb3l4;Gnai3;Ptk2;Creb1;Pik3ca;Ghr;Sst;Raf1;Akt3;Prkcb;Sstr2;Cacna1d;Plcb1;Irs4;Ghsr;Adcy5;Map2k2;Igfbp3;Pou1f1;Creb3l2;Atf2;Ep300;Adcy3;Gnas;Map2k3;Grb2;Crk;Adcy9 |
| mmu03015 | mRNA surveillance pathway | http://www.kegg.jp/kegg-bin/show_pathway?mmu03015+103677+108062+17690+18459+18789+18949+19046+19052+19826+20220+226517+229512+230908+233908+235542+24018+241989+26931+26932+269643+326622+381404+432508+433702+51796+53319+54196+67543+70248+71978+74320+74737+76626+83410+83454+93728 | 103 | 36 | 24,9142279 | 1,444957479 | 0,00887932 | 0,04077479 | 103677;108062;17690;18459;18789;18949;19046;19052;19826;20220;226517;229512;230908;233908;235542;24018;241989;26931;26932;269643;326622;381404;432508;433702;51796;53319;54196;67543;70248;71978;74320;74737;76626;83410;83454;93728 | Sap18;Ppp2r2a;Rnps1;Ppp1cb;Msi2;Ppp2r3a;Wdr33;Dazap1;Cstf2t;Pabpc2;Ncbp1;Ppp2r5c;Ppp2ca;Papola;Upf2;Pabpc4l;Ppp2r5e;Pabpn1;Ppp2r2c;Nxf1;Msi1;Smg7;Tardbp;Fus;Srrm1;Pabpc1l;Cstf2;Pcf11;Pabpc6;Nxf2;Cpsf6;Smg6;Smg5;Rngtt;Pnn;Pabpc5 |
| mmu04014 | Ras signaling pathway | http://www.kegg.jp/kegg-bin/show_pathway?mmu04014+109905+110157+11350+11600+11601+11845+11848+12313+12315+12977+13712+14164+14173+14176+14178+14180+14183+14389+14700+14784+14811+15566+16001+16150+16151+16333+16337+16590+16706+16797+17295+17311+17356+18015+18053+18479+18654+18706+18751+18783+18784+18806+19247+19345+19353+19417+19418+19696+19730+19731+20418+215449+21802+21844+22339+224105+226525+234779+23797+23871+23872+240057+241656+241694+26395+26396+26413+26420+26970+26971+271457+333050+54354+54635+56480+56784+66482+71785+74055+80796+80857+85031 | 235 | 82 | 56,8431414 | 1,442566296 | 1,23E-04 | 0,00208105 | 109905;110157;11350;11600;11601;11845;11848;12313;12315;12977;13712;14164;14173;14176;14178;14180;14183;14389;14700;14784;14811;15566;16001;16150;16151;16333;16337;16590;16706;16797;17295;17311;17356;18015;18053;18479;18654;18706;18751;18783;18784;18806;19247;19345;19353;19417;19418;19696;19730;19731;20418;215449;21802;21844;22339;224105;226525;234779;23797;23871;23872;240057;241656;241694;26395;26396;26413;26420;26970;26971;271457;333050;54354;54635;56480;56784;66482;71785;74055;80796;80857;85031 | Vegfa;Tgfa;Fgf2;Mapk1;Pgf;Ksr1;Gab2;Rac1;Map2k1;Met;Ralgapa1;Calm3;Plcg2;Plce1;Rhoa;Insr;Arf6;Fgfr2;Mapk9;Shc3;Rel;Ets2;Fgf20;Rab5a;Abl1;Ptpn11;Rasgrf1;Exoc2;Rgl1;Rab5c;Lat;Pik3ca;Ikbkb;Pak5;Pdgfd;Tiam1;Ksr2;Ins1;Syngap1;Rap1b;Angpt1;Ralgapa2;Gng10;Raf1;Akt3;Ngfr;Prkcb;Ikbkg;Pla2g2e;Pla1a;Igf1r;Kit;Pla2g4a;Grin2a;Pak1;Fgf9;Pla2g2f;Tbk1;Rasal2;Pak2;Calm1;Rassf5;Calm4;Csf1;Ralgds;Htr7;Map2k2;Pdgfc;Fgf5;Nf1;Fgf7;Kitl;Rasgrf2;Ets1;Pld2;Pla2g5;Afdn;Grb2;Elk1;Fgf1;Angpt2;Rap1a |

| | | | | | | | | | | |
|---|---|---|---|---|---|---|---|---|---|---|
| mmu04270 | Vascular smooth muscle contraction | http://www.kegg.jp/kegg-bin/show_pathway?mmu04270+104111+109624+110157+11512+11515+11535+11549+11550+11607+11608+11848+12289+12313+12315+13614+13615+14672+14683+16440+16531+17880+17886+17931+18158+18160+18751+18753+18783+18784+18795+18796+19046+19091+19878+213498+224129+230103+232807+238564+26395+26396+26413+26970+26971+329251+54598+60596+69632+77579+80796 | 144 | 50 | 34,831542 | 1,43548052 | 0,00271815 | 0,01860065 | 104111;109624;110157;11512;11515;11535;11549;11550;11607;11608;11848;12289;12313;12315;13614;13615;14672;14683;16440;16531;17880;17886;17931;18158;18160;18751;18753;18783;18784;18795;18796;19046;19091;19878;213498;224129;230103;232807;238564;26395;26396;26413;26970;26971;329251;54598;60596;69632;77579;80796 | Adm;Adcy6;Ppp1cb;Mapk1;Plcb2;Map2k1;Npr2;Ppp1r12b;Calm3;Gna11;Itpr3;Rhoa;Myh9;Arhgef12;Cald1;Kcnma1;Edn2;Agtr1a;Raf1;Prkcb;Pla2g2e;Cacna1d;Pla2g4a;Edn1;Pla2g2f;Prkcd;Myh10;Prkg1;Arhgef11;Plcb1;Calm1;Calm4;Mylk4;Adra1d;Nppb;Ppp1r12c;Adcy5;Map2k2;Ppp1r12a;Adra1a;Adcy3;Pla2g5;Gnas;Npr1;Calcrl;Agtr1b;Adcy9;Gucy1a1;Rock2;Myh11 |
| mmu04152 | AMPK signaling pathway | http://www.kegg.jp/kegg-bin/show_pathway?mmu04152+105787+107476+108097+108099+11549+12443+12894+12912+13685+14104+15357+16001+16333+16337+16370+18639+18640+18641+18706+19017+19052+19325+19744+20249+20250+208647+20869+235542+23797+26931+26932+269643+30049+329065+56421+56484+57915+59021+68365+68465+69008+71978+74370+78284 | 127 | 44 | 30,7194849 | 1,432315681 | 0,00494447 | 0,02886363 | 105787;107476;108097;108099;11549;12443;12894;12912;13685;14104;15357;16001;16333;16337;16370;18639;18640;18641;18706;19017;19052;19325;19744;20249;20250;208647;20869;235542;23797;26931;26932;269643;30049;329065;56421;56484;57915;59021;68365;68465;69008;71978;74370;78284 | Scd1;Scd2;Eif4ebp1;Adipor2;Ppp2r2a;Stk11;Pfkl;Ppp2r3a;Insr;Cpt1a;Scd4;Creb3l4;Fasn;Creb1;Pik3ca;Foxo3;Pfkp;Ppp2r5c;Acaca;Rab2a;Ins1;Pfkfb2;Tbc1d1;Ppp2ca;Prkag2;Akt3;Hmgcr;Ppargc1a;Ppp2r5e;Igf1r;Ppp2r2c;Cab39l;Scd3;Irs4;Pfkfb1;Rheb;Rptor;Prkaa1;Ccnd1;Creb3l2;Adra1a;Rab10;Prkab2;Rab14 |

| | | | | | | | | | | |
|---|---|---|---|---|---|---|---|---|---|---|
| mmu05200 | Pathways in cancer | bin/show_pathway?mmu05200+104111+107986+108058+108989+110157+110279+112405+112406+11350+11512+11515+11607+11608+11682+11789+11797+11798+11835+11848+11863+11864+12005+12125+12313+12315+12325+12369+12370+12385+12386+12387+12394+12395+12402+12443+12444+12445+12447+12571+12575+12826+12827+12830+12928+13016+13063+13143+13176+13388+13555+13557+13614+13712+13856+14013+14062+14082+14083+14164+14173+14176+14178+14180+14183+14194+14281+14362+14367+14369+14370+14371+14633+14672+14679+14683+14700+14784+14858+14862+14863+14872+15182+15205+15251+15368+15975+16001+16150+16151+16160+16164+16169+16184+16185+ | 543 | 187 | 131,34394 | 1,423742889 | 1,98E-08 | 3,45E-06 | 89;110157;110279;112405;112406;11350;11512;11515;11607;11608;11682;11789;11797;11798;11835;11848;11863;11864;12005;12125;12313;12315;12325;12369;12370;12385;12386;12387;12394;12395;12402;12443;12444;12445;12447;12571;12575;12826;12827;12830;12928;13016;13063;13143;13176;13388;13555;13557;13614;13712;13856;14013;14062;14082;14083;14164;14173;14176;14178;14180;14183;14194;14281;14362;14367;14369;14370;14371;14633;14672;14679;14683;14700;14784;14858;14862;14863;14872;15182;15205;15251;15368;15975;16001;16150;16151;16160;16164;16169;16184;16185;16187;16190;16195;16196;16400;16410;16451;16590;16772;16773;16973;17126;17127;17246;17295;17311;17342;17688; | Slc2a1;Keap1;E2f1;Hes1;Hmox1;Ptch1;Adcy6;Traf2;Gadd45b;Vegfa;Smad3;Epo;Tgfa;Fgf2;Mapk1;Lrp5;Tpr;Fos;Cycs;Pgf;Mdm2;Plcb2;Rara;F2r;Rac1;Lamc1;Map2k1;Wnt2b;Msh6;Egln1;Met;Appl1;Ctbp1;Dapk2;Bcl2l11;Calm3;Camk2g;Plcg2;Gna11;Gli2;Rhoa;Fgfr2;Mapk9;Mecom;Il3;Egln2;Fgf20;Wnt4;Stat6;Il2rb;Gnai3;Ptk2;Dll1;Rbx1;Abl1;Arhgef12;Ddb2;Gstt2;Gsta2;Pik3ca;Pax8;Mgst2;Ikbkb;Pim1;E2f2;Ncoa4;Gsto2;Nfkbia;Cdkn1a;Traf3;Hdac1;Tcf7;Fzd8;Fzd9;Agtr1a;Fzd1;Lama1;Smad2;Arnt;Ifnar1;Il4ra;Fadd;Shh;Birc2;Smo;Gng10;Axin1;Nfe2l2;Gstm1;Raf1;Cbl;Tgfbr1;Akt3;Il15ra;Prkcb;E2f3;Notch4;Ikbkg;Fzd7;Col4a1;Dcc;Igf1r;Gadd45g;Kit;Traf4;Apc;Fgf9;Edn1;Itgav;Bcr;Ccnd3;Wnt1;Lama2;Cdk6;Arhgef11;Plcb1;Col4a6;Calm1;Pmaip1;Runx1;Rassf5;Calm4;Cul2;Wnt5b;Apc2;Mitf;Ctnna2;Txnrd1;Ralgds;Il13ra1;Adcy5;Cul1;Map2k2;Rarb;Fgf5;Gstm2;Rxra;Ccnd2;Fgf7;Arnt2;Hdac2;Frat2;Ccnd1;Wnt3;Il12b;Ctnnb1;Kitl;Ncoa1;Ep300;Il2ra;Ets1;Camk2d;Ccne1;Adcy3;Notch2;Pld2;Tgfb2;Tcf7l2;Polk;Lpar4;Alk;Col4a2;Gnas;Hif1a;Grb2;Ctnna1;Casp8;Crk;Nkx3-1;Xiap;Ar;Elk1;Agtr1b;Adcy9;Fzd5;Cks2;Jak1;Runx1t1;Col4a5;Il6st;Sufu;Fh1;Fgf1;Casp7;Rock2;Itga3;Il7 |
| mmu04144 | Endocytosis | bin/show_pathway?mmu04144+103967+107305+107338+109689+110355+11841+11843+11845+11848+12340+12345+12402+13132+13196+13660+13855+13858+140500+14183+14772+15040+16001+16184+16185+16572+16835+17126+17127+17246+17999+18571+18720+18806+19157+19159+19325+19345+19349+193740+20405+20779+208092+208650+211673+211914+213990+216238+216724+21812+22042+22088+225861+230597+232227+234353+245666+26431+271457+27965+30930+330192+330319+347722+399549+53869+54189+54198+55988+56378+56440+57440+58194+66251+66313+67064+67300+67771+69150+69710+70432+72318+74998+75608+75788+77407+78618+80794+83814+93737+93742+98878+99 | 268 | 92 | 64,8253698 | 1,419197457 | 9,70E-05 | 0,00178118 | 103967;107305;107338;109689;110355;11841;11843;11845;11848;12340;12345;12402;13132;13196;13660;13855;13858;140500;14183;14772;15040;16001;16184;16185;16572;16835;17126;17127;17246;17999;18571;18720;18806;19157;19159;19325;19345;19349;193740;20405;20779;208092;208650;211673;211914;213990;216238;216724;21812;22042;22088;225861;230597;232227;234353;245666;26431;271457;27965;30930;330192;330319;347722;399549;53869;54189;54198;55988;56378;56440;57440;58194;66251;66313;67064;67300;67771;69150;69710;70432;72318;74998;75608;75788;77407;78618;80794;83814;93737;93742;98878;99371 | Snx32;Vps37b;Sh3gl1;Smad3;Cblb;Arfgef2;Capzb;Mdm2;Snx3;Eps15;Arfgap3;Vps26a;Ehd1;Hspa1a;Rhoa;Arf6;Fgfr2;Arap1;Rabep1;Kif5a;Il2rb;Rab11fip2;Rab5a;Snx4;Pard3;Cyth4;Chmp4b;Pip5k1a;Rab5c;Epn2;Smurf2;Arrb1;Agap1;Arf4;Spg21;Dab2;Wipf3;Sh3kbp1;Iqsec1;Iqsec2;Smad2;Eea1;Dnm3;Nedd4;Rufy1;Nedd4l;Acap3;Arf2;Arpc5;Ehd4;Smurf1;Cbl;Grk2;Tgfbr1;Gbf1;Chmp6;Rab7;Tfrc;Arpc3;Igf1r;Asap1;Tsg101;Pdcd6ip;Zfyve9;Ldlr;Arfgef1;Rufy2;Vps37c;Git2;Cyth1;Grk4;Agap3;Chmp1b;Snx12;Pard6g;Rab11a;Cblc;H2-T23;Rab10;Cyth3;Rab35;Il2ra;Pld2;Psd3;Capza1;H2-M10.6;Src;Snx1;Acap2;Asap2;Ehd3;Cltc |

| | | | | | | | | | | |
|---|---|---|---|---|---|---|---|---|---|---|
| mmu04310 | Wnt signaling pathway | http://www.kegg.jp/kegg-bin/show_pathway?mmu04310+106042+107515+108058+11789+11848+12005+12325+12387+12443+12444+12445+12995+13016+13380+14283+14362+14367+14369+14370+14371+16973+17127+18751+18795+18796+19353+19878+20187+20319+20379+207742+208846+212398+21372+21414+21416+21886+21888+22402+22408+22414+22415+22417+22419+229658+23805+243548+26420+26563+26564+26965+27373+328572+329252+407821+56438+56811+68339+73181+93687 | 175 | 60 | 42,3299989 | 1,417434479 | 0,00154567 | 0,01254505 | 106042;107515;108058;11789;11848;12005;12325;12387;12443;12444;12445;12995;13016;13380;14283;14362;14367;14369;14370;14371;16973;17127;18751;18795;18796;19353;19878;20187;20319;20379;207742;208846;212398;21372;21414;21416;21886;21888;22402;22408;22414;22415;22417;22419;229658;23805;243548;26420;26563;26564;26965;27373;328572;329252;407821;56438;56811;68339;73181;93687 | Smad3;Lrp5;Ror2;Plcb2;Ryk;Rac1;Wnt2b;Prickle1;Ctbp1;Camk2g;Prickle2;Rhoa;Mapk9;Wnt4;Rbx1;Lgr6;Csnk1a1;Lgr4;Nfatc4;Tle4;Znrf3;Tle2;Csnk1e;Tcf7;Fzd8;Fzd9;Fzd1;Axin1;Ccdc88c;Dkk2;Sfrp2;Prkcb;Fzd7;Rnf43;Fosl1;Apc;Ccnd3;Wnt1;Plcb1;Sfrp4;Wnt5b;Apc2;Csnk2a1;Cul1;Daam1;Ccnd2;Frat2;Ccnd1;Wnt3;Ctnnb1;Tbl1x;Ep300;Camk2d;Tcf7l2;Ccn4;Dkk1;Fzd5;Ror1;Vangl1;Rock2 |
| mmu04022 | cGMP-PKG signaling pathway | http://www.kegg.jp/kegg-bin/show_pathway?mmu04022+104111+110157+11512+11515+11539+11549+11550+11552+11553+11555+11607+11608+11739+11848+11909+11938+12289+12313+12315+12912+14463+14672+14679+16333+16337+16370+16440+16531+17258+17259+17260+17261+17931+18127+18158+18160+18795+18796+19046+19091+19092+19878+208647+22333+22334+224129+230103+23797+238564+242202+26395+26396+26413+333329+60596+67972+73181+78284+80796 | 173 | 59 | 41,8462275 | 1,409923988 | 0,00193901 | 0,0147112 | 104111;110157;11512;11515;11539;11549;11550;11552;11553;11555;11607;11608;11739;11848;11909;11938;12289;12313;12315;12912;14463;14672;14679;16333;16337;16370;16440;16531;17258;17259;17260;17261;17931;18127;18158;18160;18795;18796;19046;19091;19092;19878;208647;22333;22334;224129;230103;23797;238564;242202;26395;26396;26413;333329;60596;67972;73181;78284;80796 | Adcy6;Ppp1cb;Mapk1;Atp2a2;Plcb2;Map2k1;Npr2;Calm3;Gna11;Itpr3;Cngb1;Rhoa;Vdac1;Insr;Creb3l4;Adora1;Gnai3;Creb1;Adrb2;Kcnma1;Nfatc4;Agtr1a;Ins1;Mef2d;Vdac2;Mef2c;Nos3;Slc25a4;Raf1;Akt3;Adra2c;Adra2b;Cacna1d;Pde5a;Prkg1;Plcb1;Irs4;Calm1;Calm4;Atp2b1;Mylk4;Adra1d;Nppb;Prkg2;Adcy5;Map2k2;Ppp1r12a;Creb3l2;Atf2;Adra1a;Gata4;Adcy3;Mef2b;Npr1;Mef2a;Agtr1b;Adcy9;Gucy1a1;Rock2 |
| mmu05135 | Yersinia infection | http://www.kegg.jp/kegg-bin/show_pathway?mmu05135+106759+108100+109333+110596+110651+11465+11845+11848+12927+12928+140580+14083+14131+14281+16150+16151+16797+16818+16885+18035+18706+18720+19229+19353+19878+20779+21898+22030+23797+242687+26395+26396+26397+26413+26420+26431+330319+54353+56212+56378+56480+57257+67071+67771+68652+69632 | 135 | 46 | 32,6545706 | 1,408684884 | 0,00586024 | 0,03234306 | 106759;108100;109333;110596;110651;11465;11845;11848;12927;12928;140580;14083;14131;14281;16150;16151;16797;16818;16885;18035;18706;18720;19229;19353;19878;20779;21898;22030;23797;242687;26395;26396;26397;26413;26420;26431;330319;54353;56212;56378;56480;57257;67071;67771;68652;69632 | Traf2;Mapk1;Limk1;Fos;Arhgef28;Rac1;Map2k1;Bcar1;Rhoa;Arf6;Mapk9;Skap2;Ptk2;Arhgef12;Actg1;Ticam1;Pip5k1a;Lat;Pik3ca;Ikbkb;Rps6ka6;Nfkbia;Wipf3;Vav3;Tlr4;Arpc5;Akt3;Ikbkg;Rps6ka3;Pkn2;Arpc3;Wasf2;Tbk1;Tab2;Git2;Map2k2;Fcgr3;Ptk2b;Map2k3;Elmo1;Baiap2;Crk;Lck;Src;Rhog;Rock2 |

| | | | | | | | | | | |
|---|---|---|---|---|---|---|---|---|---|---|
| mmu04024 | cAMP signaling pathway | http://www.kegg.jp/kegg-bin/show_pathway?mmu04024+104111+108058+109305+109905+110157+11430+11512+11515+11517+11539+11555+11848+11938+12289+12313+12315+12325+12790+12912+12918+13614+13615+14062+14281+14679+14683+14800+14802+14811+15166+15550+15552+16526+17200+17356+17931+18035+18160+18479+18578+18706+18806+19046+19206+19353+19878+20191+20544+20604+20606+20609+208188+208647+215449+21844+224129+23797+238871+239273+23984+242425+243270+26395+26396+26413+26420+328572+333329+56508+57257+67972+74055+78284+80796+80911 | 224 | 75 | 54,1823986 | 1,384213359 | 9,40E-04 | 0,00863059 | 104111;108058;109305;109905;110157;11430;11512;11515;11517;11539;11555;11848;11938;12289;12313;12315;12325;12790;12912;12918;13614;13615;14062;14281;14679;14683;14800;14802;14811;15166;15550;15552;16526;17200;17356;17931;18035;18160;18479;18578;18706;18806;19046;19206;19353;19878;20191;20544;20604;20606;20609;208188;208647;215449;21844;224129;23797;238871;239273;23984;242425;243270;26395;26396;26413;26420;328572;333329;56508;57257;67972;74055;78284;80796;80911 | Ptch1;Adcy6;Pde4d;Ppp1cb;Mapk1;Fos;Ryr2;Atp2a2;Slc9a1;Gabbr2;F2r;Rac1;Map2k1;Sstr5;Mc2r;Calm3;Camk2g;Plce1;Acox3;Cngb1;Rhoa;Cnga3;Mapk9;Orai1;Creb3l4;Adora1;Gnai3;Creb1;Adrb2;Hcn2;Pik3ca;Hcar1;Gria2;Nfkbia;Tiam1;Rapgef4;Edn2;Sst;Htr1a;Vav3;Rap1b;Raf1;Akt3;Pde10a;Sstr2;Kcnk2;Cacna1d;Grin2a;Pak1;Edn1;Gria4;Calm1;Calm4;Atp2b1;Ghsr;Adcy5;Crh;Adcyap1r1;Map2k2;Ppp1r12a;Creb3l2;Ep300;Camk2d;Adcy3;Pld2;Gnas;Abcc4;Afdn;Npr1;Adcy9;Htr1d;Pde4b;Rap1a;Rock2;Acox1 |
| mmu04150 | mTOR signaling pathway | http://www.kegg.jp/kegg-bin/show_pathway?mmu04150+105787+109270+110157+110651+11848+13685+14362+14367+14369+14370+14371+14784+16001+16150+16333+16337+16973+17254+18706+18751+19744+20393+20869+216742+22408+22414+22415+22417+22419+227743+230784+23797+26395+26396+26413+277854+29869+319481+329679+338375+54170+56430+66237+67071+68441+69008+72124+72446+74370+74747+97998 | 157 | 51 | 37,9760562 | 1,342951457 | 0,01089589 | 0,04813503 | 105787;109270;110157;110651;11848;13685;14362;14367;14369;14370;14371;14784;16001;16150;16333;16337;16973;17254;18706;18751;19744;20393;20869;216742;22408;22414;22415;22417;22419;227743;230784;23797;26395;26396;26413;277854;29869;319481;329679;338375;54170;56430;66237;67071;68441;69008;72124;72446;74370;74747;97998 | Eif4ebp1;Stk11;Seh1l;Mapk1;Lrp5;Map2k1;Wnt2b;Ddit4;Rhoa;Insr;Wnt4;Atp6v1g3;Deptor;Depdc5;Pik3ca;Rraga;Ikbkb;Rps6ka6;Fzd8;Fzd9;Ins1;Fzd1;Slc3a2;Rragc;Raf1;Fnip1;Akt3;Prkcb;Rps6ka3;Fzd7;Igf1r;Clip1;Cab39l;Mapkap1;Wnt1;Prr5l;Wnt5b;Rheb;Sgk1;Rptor;Map2k2;Prkaa1;Sesn2;Wnt3;Atp6v1g2;Grb2;Ulk2;Fzd5;Wdr59;Fnip2;Prr5 |

| | | | | | | | | | | |
|---|---|---|---|---|---|---|---|---|---|---|
| mmu05132 | Salmonella infection | http://www.kegg.jp/kegg-bin/show_pathway?mmu05132+110157+110350+11465+11797+11841+11845+11848+12306+12369+12370+12387+13063+13191+13427+140580+14082+14281+14433+16150+16151+16572+16593+17281+17906+18035+18479+18645+18706+19157+19159+192176+19345+19349+19353+19878+20194+20336+21414+21416+218035+21898+22030+22143+22166+225326+227613+234663+234686+23797+24088+26395+26397+26413+26420+271457+319642+50884+53413+53791+545486+56212+56378+56532+56551+59310+622434+66395+66482+66616+66724+67771+68097+68652+68794+70297+71999+72318+80837 | 253 | 79 | 61,1970842 | 1,290911178 | 0,0059311 | 0,03234306 | 110157;110350;11465;11797;11841;11845;11848;12306;12369;12370;12387;13063;13191;13427;140580;14082;14281;14433;16150;16151;16572;16593;17281;17906;18035;18479;18645;18706;19157;19159;192176;19345;19349;19353;19878;20194;20336;21414;21416;218035;21898;22030;22143;22166;225326;227613;234663;234686;23797;24088;26395;26396;26397;26413;26420;271457;319642;50884;53413;53791;545486;56212;56378;56532;56551;59310;622434;66395;66482;66616;66724;67771;68097;68652;68794;70297;71999;72318;80837 | Traf2;Tubb4b;Gapdh;Fyco1;Pik3c3;Mapk1;Fos;Cycs;Exoc7;Rac1;Map2k1;Flnc;Rhoa;Arf6;Mapk9;Anxa2;Klc1;Kif5a;Rab5a;Txn2;Actg1;Tuba1b;Cyth4;Exoc2;Rab5c;Pik3ca;Ikbkb;Dctn1;Nfkbia;Myl2;Tcf7;S100a10;Tlr2;Vps41;Fhod1;Tlr4;Ahnak;Fadd;Birc2;Exoc4;Arf2;Rhoj;Arpc5;Raf1;Akt3;Pfn2;Ikbkg;Dync1li2;Rab7;Arpc3;Rab9b;Pak1;Myl10;Tab2;Arhgef26;Dync2h1;Cyth1;Tlr5;Map2k2;Ripk3;Snx9;Tubb1;Txn1;Cyth3;Ctnnb1;Dync1i2;Tcf7l2;Flna;Fbxo22;Gcc2;Map2k3;Elmo1;Casp8;Tab3;Dynll2;Rhog;Nckap1;Casp7;Rock2 |
| mmu04151 | PI3K-Akt signaling pathway | [illegible]bin/show_pathway?mmu04151+105787+109333+110157+11600+11601+11909+12125+12189+12443+12444+12445+12447+12571+12575+12826+12827+12830+12839+12840+12842+12912+12977+13685+13856+13869+14062+14083+14164+14173+14176+14178+14180+14183+14573+14600+14700+14784+15975+16001+16150+16151+16184+16185+16187+16190+16196+16333+16337+16400+16410+16416+16419+16421+16451+16590+16772+16773+17210+17246+17295+17311+18053+18127+18414+18654+18706+19052+19116+19353+19699+19744+20181+20393+208647+20869+20963+21802+21898+21923+22339+22627+22629+22630+22631+226519+235542+23797+24088+241226+263803+26395+26396+26413+26931+ | 364 | 113 | 88,0463978 | 1,283414232 | 0,00143568 | 0,01192985 | [illegible]0;11601;11909;12125;12189;12443;12444;12445;12447;12571;12575;12826;12827;12830;12839;12840;12842;12912;12977;13685;13856;13869;14062;14083;14164;14173;14176;14178;14180;14183;14573;14600;14700;14784;15975;16001;16150;16151;16184;16185;16187;16190;16196;16333;16337;16400;16410;16416;16419;16421;16451;16590;16772;16773;17210;17246;17295;17311;18053;18127;18414;18654;18706;19052;19116;19353;19699;19744;20181;20393;208647;20869;20963;21802;21898;21923;22339;22627;22629;22630;22631;226519;235542;23797;24088;241226;263803;26395;26396;26413;26931;26932;269643;28078;320910;50791;54401;54635;56484;68553;71785;71978;74370;74747;78134;78284;80857;83490;942 | Mcl1;Ywhah;Eif4ebp1;Ppp2r2a;Vegfa;Epo;Tgfa;Stk11;Fgf2;Mapk1;Pgf;Mdm2;F2r;Rac1;Lamc1;Map2k1;Met;Ddit4;Bcl2l11;Ppp2r3a;Insr;Fgfr2;Creb3l4;Il3;Fgf20;Il2rb;Ptk2;Creb1;Syk;Pik3ca;Foxo3;Ywhae;Ghr;Itgb8;Ikbkb;Magi2;Cdkn1a;Ppp2r5c;Pdgfd;Ywhaz;Col6a4;Osmr;Tlr2;Pkn3;Ins1;Brca1;Lama1;Tlr4;Gdnf;Ifnar1;Il4ra;Pik3ap1;Angpt1;Ppp2ca;Gng10;Nos3;Raf1;Akt3;Reln;Ngfr;Ikbkg;Ppp2r5e;Pkn2;Col4a1;Prl5a1;Igf1r;Kit;Ppp2r2c;Fgf9;Col1a1;Prlr;Itgav;Ccnd3;Col9a1;Lama2;Cdk6;Itgb7;Itgb5;Col4a6;Csf1;Rheb;Sgk1;Rptor;Itga8;Map2k2;Pdgfc;Prkaa1;Col9a2;Fgf5;Ywhaq;Rxra;Ccnd2;Fgf7;Ccnd1;Creb3l2;Atf2;Tnc;Kitl;Phlpp1;Il2ra;Ccne1;Itgb3;Lpar4;Col4a2;Erbb4;Ywhab;Grb2;Jak1;Col4a5;Fgf1;Angpt2;Itga3;Il7 |

| | | | | | | | | | |
|---|---|---|---|---|---|---|---|---|---|
| mmu04010 | MAPK signaling pathway | [illegible]bin/show_pathway?mmu04010+102626+109689+109905+110157+110651+11600+11601+11909+12286+12289+12293+12294+12296+12928+12977+13712+13869+14013+14164+14173+14176+14178+14180+14183+14281+14573+14784+16001+16150+16151+16333+16337+16590+17165+17260+17295+17311+17347+17762+17873+18015+18053+18479+18654+18751+18783+19042+192176+19252+19279+19353+193740+19417+19418+215449+216965+21802+21808+21812+22030+22339+224105+225028+23797+23882+23938+239556+240672+26395+26396+26397+26401+26405+26413+26420+30957+319520+319734+338372+53608+54377+54635+58226+63953+65964+67071+68652+68794+71785+72349+75590+76089+8085 | 301 | 93 | 72,8075982 | 1,277339211 | 0,00421824 | 0,02628867 | 102626;109689;109905;110157;110651;11600;11601;11909;12286;12289;12293;12294;12296;12928;12977;13712;13869;14013;14164;14173;14176;14178;14180;14183;14281;14573;14784;16001;16150;16151;16333;16337;16590;17165;17260;17295;17311;17347;17762;17873;18015;18053;18479;18654;18751;18783;19042;192176;19252;19279;19353;193740;19417;19418;215449;216965;21802;21808;21812;22030;22339;224105;225028;23797;23882;23938;239556;240672;26395;26396;26397;26401;26405;26413;26420;30957;319520;319734;338372;53608;54377;54635;58226;63953;65964;67071;68652;68794;71785;72349;75590;76089;80857 | Traf2;Gadd45b;Vegfa;Map3k1;Tgfa;Fgf2;Mapk1;Dusp4;Cacna1h;Fos;Pgf;Map3k2;Map2k5;Rac1;Map2k1;Map4k3;Met;Hspa1a;Flnc;Insr;Fgfr2;Mapk9;Mecom;Fgf20;Cacna1a;Dusp9;Rasgrf1;Dusp10;Ikbkb;Taok1;Rps6ka6;Cacna2d4;Arrb1;Pdgfd;Ins1;Map3k20;Mknk2;Gdnf;Rap1b;Dusp1;Angpt1;Mef2c;Map3k9;Cacnb2;Mapk8ip3;Raf1;Tgfbr1;Akt3;Ngfr;Cacna2d1;Prkcb;Ikbkg;Rps6ka3;Igf1r;Gadd45g;Kit;Cacna1d;Pla2g4a;Pak1;Fgf9;Rapgef2;Tab2;Pak2;Ppm1a;Cacna1i;Csf1;Mapkapk5;Map2k2;Pdgfc;Fgf5;Map3k6;Nf1;Fgf7;Dusp3;Atf2;Kitl;Dusp5;Mapkapk3;Rasgrf2;Tgfb2;Flna;Cacng4;Erbb4;Cacna2d3;Map2k3;Grb2;Mapt;Crk;Elk1;Ptprr;Fgf1;Angpt2;Rap1a |
| mmu04020 | Calcium signaling pathway | http://www.kegg.jp/kegg-bin/show_pathway?mmu04020+102093+104111+108058+108071+109305+11515+11549+11550+11555+11607+11608+116873+11739+11938+12286+12289+12313+12315+12325+13869+14062+14164+14173+14176+14178+14180+14183+14573+14672+14680+14683+14811+14816+15466+15560+15563+15566+16440+170732+17295+18127+18213+18573+18679+18751+18795+18796+19204+19229+20191+20192+21338+21425+215303+215999+21802+22333+22334+22339+233011+233979+234779+238564+239556+252972+269717+54635+58226+65973+66402+67972+68528+70086+71785+73181+74055+80796+80857 | 253 | 78 | 61,1970842 | 1,27457053 | 0,00877836 | 0,04077479 | 102093;104111;108058;108071;109305;11515;11549;11550;11555;11607;11608;116873;11739;11938;12286;12289;12313;12315;12325;13869;14062;14164;14173;14176;14178;14180;14183;14573;14672;14680;14683;14811;14816;15466;15560;15563;15566;16440;170732;17295;18127;18213;18573;18679;18751;18795;18796;19204;19229;20191;20192;21338;21425;215303;215999;21802;22333;22334;22339;233011;233979;234779;238564;239556;252972;269717;54635;58226;65973;66402;67972;68528;70086;71785;73181;74055;80796;80857 | Asph;Vegfa;Tgfa;Fgf2;Cacna1h;Ryr2;Atp2a2;Plcb2;F2r;Met;Trhr2;Calm3;Camk2g;Plcg2;Gna11;Plce1;Ntrk3;Itpr3;Vdac1;Fgfr2;Orai1;Fgf20;Cacna1a;Adrb2;Nfatc4;Smim6;Pde1a;Mcu;Pdgfd;Itpkc;Agtr1a;Tpcn2;Orai2;Sln;Gdnf;Vdac2;Nos3;Slc25a4;Tacr3;Tpcn1;Prkcb;Cacna1d;Grin2a;Fgf9;Grm5;Phkb;Plcb1;Gnal;Calm1;Calm4;Atp2b1;Cacna1i;Mylk4;Adra1d;Htr7;Hrh2;Camk1g;Ptafr;Pdgfc;Fgf5;Tfeb;Fgf7;Adra1a;Grm1;Ryr3;Ptk2b;Camk2d;Adcy3;Htr2c;Gnas;Erbb4;Stim2;Htr5a;Agtr1b;Adcy9;Cysltr2;Fgf1;Phka1 |

| | | | | | | | | | | |
|---|---|---|---|---|---|---|---|---|---|---|
| mmu05022 | Pathways of neurodegeneration | [illegible]bin/show_pathway?mmu05022+103199+108058+108071+110082+110157+110616+11739+11789+11820+11938+12005+12289+12313+12315+12325+12334+12359+12364+12369+12370+12387+12421+12460+12858+12977+12995+13063+13191+13380+13417+13665+13666+140499+14082+14362+14367+14369+14370+14371+14763+14800+14802+14811+14816+15194+16440+16572+16593+16973+17762+17992+18751+18795+18796+19165+19177+19179+19324+19353+20191+20192+20238+20239+20655+20907+212398+215114+215999+21762+217695+218460+22030+22143+22186+22190+22333+22334+22408+22414+22415+22417+22419+225326+226646+227197+227613+228361+230908+233871+233908+23805+240960 | 473 | 137 | 114,41194 | 1,197427471 | 0,00829145 | 0,03965647 | [illegible]82;110157;110616;11739;11789;11820;11938;12005;12289;12313;12315;12325;12334;12359;12364;12369;12370;12387;12421;12460;12858;12977;12995;13063;13191;13380;13417;13665;13666;140499;14082;14362;14367;14369;14370;14371;14763;14800;14802;14811;14816;15194;16440;16572;16593;16973;17762;17992;18751;18795;18796;19165;19177;19179;19324;19353;20191;20192;20238;20239;20655;20907;212398;215114;215999;21762;217695;218460;22030;22143;22186;22190;22333;22334;22408;22414;22415;22417;22419;225326;226646;227197;227613;228361;230908;233871;233908;23805;240960;26395;26396;26397;26413;26420;26442;26443;26445;271457;27373;29869;327954;329015;432611;50873;54405;54548 | Traf2;Psma6;Tubb4b;Ndufa12;Pik3c3;Map1lc3b;Psmc6;Mapk1;Lrp5;Atxn1;Ube2j2;Ryr2;Cycs;Atp2a2;Plcb2;Rac1;Ccs;Map2k1;Wnt2b;Atxn2;Uba52;Calm3;Camk2g;Psmb7;Itpr3;Capn2;Vdac1;Mapk9;Klc1;Wnt4;Lrrk2;Kif5a;Dnah14;Cox7b2;Rab5a;Rb1cc1;Dnali1;Gpx7;Tuba1b;Ndufs7;Psma5;Csnk1a1;Dctn1;Trap1;Gria2;Wdr41;Map1lc3a;Mcu;Psmd11;Csnk1e;Ndufs5;Fzd8;Dnah5;Fzd9;Fzd1;Atg2a;Htt;Pink1;Stx1a;Vdac2;App;Pik3r4;Dnah2;Fadd;Psmd2;Dnah7a;Hip1;Sdhb;Axin1;Ubc;Zfyve1;Slc25a4;Raf1;Dkk2;Atxn2l;Uqcr11;Prkcb;Fzd7;Dnah10;Cacna1d;Prkn;Grin2a;Apc;Derl1;Dnai2;Ndufs2;Tbk1;Wnt1;Rab1a;Grm5;Atxn3;Cat;Plcb1;Gria4;Calm1;Calm4;Tardbp;Fus;Ndufa4;Wnt5b;Apc2;Csf1;Csnk2a1;Casp12;Ern1;Map2k2;Psen2;Ndufa1;Ndufb3;Fig4;Tubb1;Frat2;Cox5a;Ambra1;Wnt3;Dnah8;Ctnnb1;Grm1;Eif2ak3;Ryr3;Ndufs1;Camk2d;Ndufa7;Psmc1;C9orf72;Map2k3;Mapt;Casp8;Eif2s1;Sod1;Dkk1;Ulk2;Fzd5;Vapb;Gpr37;Psmb2;Casp7 |

### Add. Table 4

| motif Name | group | dataset | method | Consensus | P-value | Log P-value | q-value (Benjamini) | # of Target Sequences v | % of Target Sequences | # of Background Sequ | % of Background Sequences with Motif |
|---|---|---|---|---|---|---|---|---|---|---|---|
| JunB | bZIP | DendriticCells-Junb-ChIP-Seq(GSE36099) | Homer | RATGASTCAT | 1,00E-11 | -2,56E+01 | 0 | 937 | 5,91% | 3984,8 | 4,73% |
| BATF | bZIP | Th17-BATF-ChIP-Seq(GSE39756) | Homer | DATGASTCAT | 1,00E-09 | -2,28E+01 | 0 | 1141 | 7,20% | 5028,2 | 5,97% |
| Fos | bZIP | TSC-Fos-ChIP-Seq(GSE110950) | Homer | NDATGASTCAYN | 1,00E-09 | -2,24E+01 | 0 | 988 | 6,23% | 4295,9 | 5,10% |
| Fra2 | bZIP | Striatum-Fra2-ChIP-Seq(GSE43429) | Homer | GGATGACTCATC | 1,00E-09 | -2,24E+01 | 0 | 783 | 4,94% | 3313,6 | 3,93% |
| Fra1 | bZIP | BT549-Fra1-ChIP-Seq(GSE46166) | Homer | NNATGASTCATH | 1,00E-09 | -2,23E+01 | 0 | 940 | 5,93% | 4067,6 | 4,83% |
| Atf3 | bZIP | GBM-ATF3-ChIP-Seq(GSE33912) | Homer | DATGASTCATHN | 1,00E-08 | -1,85E+01 | 0 | 1092 | 6,89% | 4894,1 | 5,81% |
| Fosl2 | bZIP | 3T3L1-Fosl2-ChIP-Seq(GSE56872) | Homer | NATGASTCABNN | 1,00E-07 | -1,80E+01 | 0 | 527 | 3,32% | 2185,5 | 2,60% |
| AP-1 | bZIP | ThioMac-PU.1-ChIP-Seq(GSE21512) | Homer | VTGACTCATC | 1,00E-07 | -1,72E+01 | 0 | 1220 | 7,70% | 5563,9 | 6,61% |
| Jun-AP1 | bZIP | K562-cJun-ChIP-Seq(GSE31477) | Homer | GATGASTCATCN | 1,00E-07 | -1,64E+01 | 0 | 381 | 2,40% | 1531,5 | 1,82% |
| THRb | NR | Liver-NR1A2-ChIP-Seq(GSE52613) | Homer | TRAGGTCA | 1,00E-06 | -1,43E+01 | 0 | 7021 | 44,30% | 35696,6 | 42,39% |
| RARa | NR | K562-RARa-ChIP-Seq(Encode) | Homer | TTGAMCTTTG | 1,00E-05 | -1,21E+01 | 0,0002 | 5173 | 32,64% | 26117,2 | 31,01% |
| NR5A2 | NR | Pancreas-LRH1-ChIP-Seq(GSE34295) | Homer | BTCAAGGTCA | 1,00E-04 | -9,57E+00 | 0,0027 | 1324 | 8,35% | 6349,9 | 7,54% |
| EAR2/NR2F6 | NR | K562-NR2F6-ChIP-Seq(Encode) | Homer | NRBCARRGGTCA | 1,00E-04 | -9,29E+00 | 0,0033 | 2089 | 13,18% | 10269,8 | 12,20% |
| ERb,IR3 | NR | Ovary-ERb-ChIP-Seq(GSE203391) | Homer | RGGTCAGGGTGACCT | 1,00E-03 | -8,26E+00 | 0,0087 | 672 | 4,24% | 3120,5 | 3,71% |
| ERRA/ESRRA/NR3B1 | NR | HepG2-Erra-ChIP-Seq(GSE31477) | Homer | CAAAGGTCAG | 1,00E-03 | -7,98E+00 | 0,0108 | 3576 | 22,56% | 18059,5 | 21,45% |
| SF1/NR5A1 | NR | H295R-Nr5a1-ChIP-Seq(GSE44220) | Homer | CAAGGHCANV | 1,00E-03 | -7,70E+00 | 0,0133 | 889 | 5,61% | 4228,4 | 5,02% |
| COUP-TFII | NR | K562-NR2F1-ChIP-Seq(Encode) | Homer | GKBCARAGGTCA | 1,00E-03 | -7,48E+00 | 0,0156 | 2265 | 14,29% | 11283,3 | 13,40% |
| HOXB4 | Homeobox | ES-Hoxb4-ChIP-Seq(GSE34014) | Homer | TGATTRATGGCY | 1,00E-03 | -7,12E+00 | 0,0212 | 364 | 2,30% | 1633,9 | 1,94% |
| COUP-TFII | NR | Artia-Nr2f2-ChIP-Seq(GSE46497) | Homer | AGRGGTCA | 1,00E-02 | -6,84E+00 | 0,0267 | 2723 | 17,18% | 13702,7 | 16,27% |
| HOXA2 | Homeobox | mES-Hoxa2-ChIP-Seq(Donaldson_et_al.) | Homer | GYCATCMATCAT | 1,00E-02 | -6,68E+00 | 0,0295 | 212 | 1,34% | 908 | 1,08% |
| Bach2 | bZIP | OCILy7-Bach2-ChIP-Seq(GSE44420) | Homer | TGCTGAGTCA | 1,00E-02 | -6,44E+00 | 0,036 | 294 | 1,85% | 1309,5 | 1,56% |
| GFY | ? | Promoter | Homer | ACTACAATTCCC | 1,00E-02 | -6,29E+00 | 0,0396 | 101 | 0,64% | 395,4 | 0,47% |
| TR4,DR1 | NR | Hela-TR4-ChIP-Seq(GSE24685) | Homer | GAGGTCAAAGGTCA | 1,00E-02 | -6,12E+00 | 0,0451 | 183 | 1,15% | 781,2 | 0,93% |
| GLI3 | Zf | Limb-GLI3-ChIP-Chip(GSE11077) | Homer | CGTGGGTGGTCC | 1,00E-02 | -6,09E+00 | 0,0451 | 173 | 1,09% | 734,4 | 0,87% |
| BORIS | Zf | K562-CTCFL-ChIP-Seq(GSE32465) | Homer | CNNBRGCGCCCCCTGSTGGC | 1,00E-02 | -6,01E+00 | 0,0462 | 292 | 1,84% | 1310,6 | 1,56% |
| ERRg | NR | Kidney-ESRRG-ChIP-Seq(GSE104905) | Homer | GTGACCTTGRVN | 1,00E-02 | -5,90E+00 | 0,0497 | 1272 | 8,03% | 6263,4 | 7,44% |
| RXR,DR1 | NR | 3T3L1-RXR-ChIP-Seq(GSE13511) | Homer | TAGGGCAAAGGTCA | 1,00E-02 | -5,81E+00 | 0,0525 | 1613 | 10,18% | 8023,7 | 9,53% |
| c-Myc | bHLH | mES-cMyc-ChIP-Seq(GSE11431) | Homer | VVCCACGTGG | 1,00E-02 | -5,78E+00 | 0,0525 | 829 | 5,23% | 4007,6 | 4,76% |
| HIF-2a | bHLH | 785_O-HIF2a-ChIP-Seq(GSE34871) | Homer | GCACGTACCC | 1,00E-02 | -5,58E+00 | 0,0614 | 404 | 2,55% | 1875,6 | 2,23% |
| HIF-1a | bHLH | MCF7-HIF1a-ChIP-Seq(GSE28352) | Homer | TACGTGCV | 1,00E-02 | -5,33E+00 | 0,0764 | 285 | 1,80% | 1294,8 | 1,54% |
| NPAS4 | bHLH | Neuron-Npas4-ChIP-Seq(GSE127793) | Homer | NHRTCACGACDN | 1,00E-02 | -5,24E+00 | 0,0811 | 863 | 5,44% | 4207 | 5,00% |
| Zscan4c | Zf | ESC-Zscan4c-ChIP-seq(GSE140619) | Homer | ATRTGTGCAC | 1,00E-02 | -5,22E+00 | 0,0811 | 800 | 5,05% | 3886 | 4,61% |

| target | 5' primer sequence | 3' primer sequence | reference |
|---|---|---|---|
| miR-29b-1-5p | GCTGGTTTCATATGGTGGTTTA | mRQ 3’ Primer (Takara Bio USA) | https://mirbase.org/mature/MIMAT0004523 |
| Drosha | ATGCAAGGCAATACGTGTCAT | TTTTGGGGTCTGAAAGCTGGT | Spandidos et al., 2010 (Nucleic Acids Research); PrimerBank ID: 29123082a1 |
| Dgcr8 (set 1) | GCAGGAGAAGCGATGATGGAG | CCGTAGAAGTTGAATGGGTCG | Spandidos et al., 2010 (Nucleic Acids Research); PrimerBank ID: 15277314a1 |
| Dgcr8 (set 2) | GATGGTGTGACTTACGGATCTGG | TAGGCTTCTCCTCAGAGGTCTG | Origene, MP203668 |
| Ago1 | CTGCCTTCTACAAAGCACAGCC | TCTGTCCACAGTGGGTCACTTC | Origene, MP204395 |
| Ago2 | TGCCATGGTACGAGAGTTGCTCAT | ACGATGAACGTGATTCCTGGCTGA | Martinez and Gregory, 2013 (RNA) |
| Ago3 | CTTCTGTGTTCCAGCAACCAGTG | GGCACAGTATCTGCTTGGATGG | Origene, MP204397 |
| Ago4 | CACACGCATCATCTACTACCGC | GCCGATAGTCTTCCTCCAAGCT | Origene, MP204398 |
| Mtdh | TGCTCTTCCTTCTAGGTTACGG | CCATTTGGTTTGGGCTTTTCAG | Spandidos et al., 2010 (Nucleic Acids Research); PrimerBank ID: 31982233a1 |
| Snd1 | TCTGGGTGCGCCATAATTGTC | TCAGCTTCTTGCGAAGGAACT | Spandidos et al., 2010 (Nucleic Acids Research); PrimerBank ID: 9790067a1 |
| Tarbp1 | CCAAAGAAGGCGTGATCCACT | CGGGCCAGTAATAAACTCTGAGA | Spandidos et al., 2010 (Nucleic Acids Research); PrimerBank ID: 229577449c3 |
| Tnrc6a | ATGCTCCTGAAAGCAAACCAG | CCTTTTAGGGCAAGTCCATTGT | Spandidos et al., 2010 (Nucleic Acids Research); PrimerBank ID: 26350805a1 |
| Tnrc6b | CCTCTTGCCAAACAGTTTTCAAA | CCCAGGTCGATCCTTCCTCA | Spandidos et al., 2010 (Nucleic Acids Research); PrimerBank ID: 26342470a1 |
| Hif1a | GGCGAGAACGAGAAGAAAAA | AAGTGGCAACTGATGAGCAA | Watts, Stein et al., 2021 |

| ID | Accession | Target | TargetID | Experiment | Literature | Tissue |
|---|---|---|---|---|---|---|
| mmu-let-7i | MI0000138 | Apc2 | 23805 | CLIP-Seq dataset analysis | 23142080\|25083871\|21258322 | Unspecified |
| mmu-let-7i | MI0000138 | Armc2 | 213402 | CLIP-Seq dataset analysis | 21258322\|23597149 | Unspecified |
| mmu-let-7i | MI0000138 | Atp2b2 | 11941 | CLIP-Seq dataset analysis | 21258322 | Unspecified |
| mmu-let-7i | MI0000138 | Cts8 | 56094 | CLIP-Seq dataset analysis | 25083871\|21258322 | Unspecified |
| mmu-let-7i | MI0000138 | Fbxl14 | 101358 | CLIP-Seq dataset analysis | 25083871 | Unspecified |
| mmu-let-7i | MI0000138 | Gnl3l | 237107 | CLIP-Seq dataset analysis | 21258322\|25083871 | Unspecified |
| mmu-let-7i | MI0000138 | Ifnar1 | 15975 | CLIP-Seq dataset analysis | 21258322 | Unspecified |
| mmu-let-7i | MI0000138 | Lins | 72635 | CLIP-Seq dataset analysis | 25083871 | Unspecified |
| mmu-let-7i | MI0000138 | Meis2 | 17536 | CLIP-Seq dataset analysis | 23142080 | Unspecified |
| mmu-let-7i | MI0000138 | Msi2 | 76626 | CLIP-Seq dataset analysis | 25083871 | Unspecified |
| mmu-let-7i | MI0000138 | Mup11 | 100039028 | CLIP-Seq dataset analysis | 23597149 | Unspecified |
| mmu-let-7i | MI0000138 | Mup17 | 100039206 | CLIP-Seq dataset analysis | 23597149 | Unspecified |
| mmu-let-7i | MI0000138 | Nf2 | 18016 | CLIP-Seq dataset analysis | 25083871\|23597149 | Unspecified |
| mmu-let-7i | MI0000138 | Nup214 | 227720 | CLIP-Seq dataset analysis | 21258322 | Unspecified |
| mmu-let-7i | MI0000138 | Tnfrsf26 | 244237 | CLIP-Seq dataset analysis | 21258322 | Unspecified |
| mmu-let-7i | MI0000138 | Zfp444 | 72667 | CLIP-Seq dataset analysis | 21258322 | Unspecified |
| mmu-miR-24-3p | MIMAT0000219 | 0610010F05Rik | 71675 | Sequencing | 37986224\|19536157 | Unspecified |
| mmu-miR-24-3p | MIMAT0000219 | 1200011I18Rik | 67467 | HITS-CLIP | 21258322 | Unspecified |
| mmu-miR-24-3p | MIMAT0000219 | 1700025G04Rik | 69399 | Sequencing | 19536157\|37986224 | Unspecified |
| mmu-miR-24-3p | MIMAT0000219 | 1700052N19Rik | 73419 | Sequencing | 19536157\|37986224 | Unspecified |
| mmu-miR-24-3p | MIMAT0000219 | 2510039O18Rik | 77034 | Sequencing | 19536157\|37986224 | Unspecified |
| mmu-miR-24-3p | MIMAT0000219 | 2610008E11Rik | 72128 | Sequencing | 19536157\|37986224 | Unspecified |
| mmu-miR-24-3p | MIMAT0000219 | 2700089E24Rik | 381820 | Sequencing | 19536157\|37986224 | Unspecified |
| mmu-miR-24-3p | MIMAT0000219 | 4930444A02Rik | 74653 | HITS-CLIP | 23142080 | Unspecified |
| mmu-miR-24-3p | MIMAT0000219 | 4931428F04Rik | 74356 | HITS-CLIP | 21258322 | Unspecified |
| mmu-miR-24-3p | MIMAT0000219 | 4933426M11Rik | 217684 | Sequencing | 19536157\|37986224 | Unspecified |
| mmu-miR-24-3p | MIMAT0000219 | 5730559C18Rik | 67313 | Sequencing | 37986224\|19536157 | Unspecified |
| mmu-miR-24-3p | MIMAT0000219 | 9430020K01Rik | 240185 | Sequencing | 19536157\|37986224 | Unspecified |
| mmu-miR-24-3p | MIMAT0000219 | AU019823 | 270156 | Sequencing | 37986224\|19536157 | Unspecified |
| mmu-miR-24-3p | MIMAT0000219 | AW549877 | 106064 | Sequencing//HITS-CLIP | 19536157\|23142080\|37986224 | Unspecified |
| mmu-miR-24-3p | MIMAT0000219 | Abcd1 | 11666 | Sequencing | 37986224\|19536157 | Unspecified |
| mmu-miR-24-3p | MIMAT0000219 | Abcg1 | 11307 | Sequencing | 37986224\|19536157 | Unspecified |
| mmu-miR-24-3p | MIMAT0000219 | Abi2 | 329165 | Sequencing//HITS-CLIP | 37986224\|25753418\|19536157 | Unspecified |
| mmu-miR-24-3p | MIMAT0000219 | Abl1 | 11350 | Sequencing | 19536157\|37986224 | Unspecified |
| mmu-miR-24-3p | MIMAT0000219 | Abr | 109934 | Sequencing | 19536157\|37986224 | Unspecified |
| mmu-miR-24-3p | MIMAT0000219 | Acsl4 | 50790 | Sequencing | 37986224\|19536157 | Unspecified |
| mmu-miR-24-3p | MIMAT0000219 | Actr10 | 56444 | HITS-CLIP | 21258322 | Unspecified |
| mmu-miR-24-3p | MIMAT0000219 | Acvr1b | 11479 | Sequencing//HITS-CLIP | 19536157\|23142080\|37986224 | Unspecified |

| | | | | | | |
|---|---|---|---|---|---|---|
| mmu-miR-24-3p | MIMAT0000219 | Adam19 | 11492 | Sequencing//HITS-CLIP | 19536157\|25753418\|37986224 | Unspecified |
| mmu-miR-24-3p | MIMAT0000219 | Adcy1 | 432530 | Sequencing | 37986224\|19536157 | Unspecified |
| mmu-miR-24-3p | MIMAT0000219 | Adcyap1r1 | 11517 | Sequencing | 19536157\|37986224 | Unspecified |
| mmu-miR-24-3p | MIMAT0000219 | Add3 | 27360 | Sequencing | 37986224\|19536157 | Unspecified |
| mmu-miR-24-3p | MIMAT0000219 | Adpgk | 72141 | Sequencing//HITS-CLIP | 37986224\|23142080\|19536157 | Unspecified |
| mmu-miR-24-3p | MIMAT0000219 | Agpat3 | 28169 | Sequencing | 19536157\|37986224 | Unspecified |
| mmu-miR-24-3p | MIMAT0000219 | Als2 | 74018 | Sequencing | 19536157\|37986224 | Unspecified |
| mmu-miR-24-3p | MIMAT0000219 | Ank3 | 11735 | Sequencing | 37986224\|19536157 | Unspecified |
| mmu-miR-24-3p | MIMAT0000219 | Ankrd52 | 237615 | Sequencing//HITS-CLIP | 18353861\|23142080\|37986224 | Unspecified |
| mmu-miR-24-3p | MIMAT0000219 | Apba1 | 319924 | Western blot//qRT-PCR//Western blot;qR | 18353861\|37986224 | Unspecified |
| mmu-miR-24-3p | MIMAT0000219 | Arhgap26 | 71302 | Sequencing | 19536157\|37986224 | Unspecified |
| mmu-miR-24-3p | MIMAT0000219 | Armc2 | 213402 | HITS-CLIP | 21258322 | Unspecified |
| mmu-miR-24-3p | MIMAT0000219 | Arxes2 | 76976 | Sequencing | 19536157 | Unspecified |
| mmu-miR-24-3p | MIMAT0000219 | Astn1 | 11899 | Sequencing | 37986224\|19536157 | Unspecified |
| mmu-miR-24-3p | MIMAT0000219 | Atcay | 16467 | Sequencing | 19536157\|37986224 | Unspecified |
| mmu-miR-24-3p | MIMAT0000219 | Atp1a2 | 98660 | Sequencing//HITS-CLIP | 37986224\|25753418\|19536157 | Unspecified |
| mmu-miR-24-3p | MIMAT0000219 | Atp2a2 | 11938 | Sequencing//HITS-CLIP | 37986224\|23142080\|19536157 | Unspecified |
| mmu-miR-24-3p | MIMAT0000219 | Avl9 | 78937 | Sequencing//HITS-CLIP | 19536157\|23142080\|37986224 | Unspecified |
| mmu-miR-24-3p | MIMAT0000219 | B2m | 12010 | Luciferase reporter assay//Western blottin | 31729180 | Unspecified |
| mmu-miR-24-3p | MIMAT0000219 | B3gnt5 | 108105 | HITS-CLIP | 23142080 | Unspecified |
| mmu-miR-24-3p | MIMAT0000219 | B4galnt2 | 14422 | HITS-CLIP | 21258322 | Unspecified |
| mmu-miR-24-3p | MIMAT0000219 | Bahcc1 | 268515 | Sequencing | 37986224\|19536157 | Unspecified |
| mmu-miR-24-3p | MIMAT0000219 | Bbc3 | 170770 | HITS-CLIP//Luciferase reporter assay | 23142080\|27797453 | Unspecified |
| mmu-miR-24-3p | MIMAT0000219 | Bcl2l11 | 12125 | Immunohistochemistry//Luciferase assay/ | 21383058\|23142080 | Unspecified |
| mmu-miR-24-3p | MIMAT0000219 | Birc5 | 11799 | HITS-CLIP | 21258322 | Unspecified |
| mmu-miR-24-3p | MIMAT0000219 | Bpnt1 | 23827 | Sequencing | 37986224\|19536157 | Unspecified |
| mmu-miR-24-3p | MIMAT0000219 | Bsn | 12217 | Sequencing | 19536157\|37986224 | Unspecified |
| mmu-miR-24-3p | MIMAT0000219 | Calu | 12321 | Sequencing | 19536157\|37986224 | Unspecified |
| mmu-miR-24-3p | MIMAT0000219 | Camk2d | 108058 | Sequencing//HITS-CLIP | 37986224\|23142080\|19536157 | Unspecified |
| mmu-miR-24-3p | MIMAT0000219 | Camkk1 | 55984 | Sequencing | 19536157\|37986224 | Unspecified |
| mmu-miR-24-3p | MIMAT0000219 | Casc4 | 319996 | Sequencing | 37986224\|19536157 | Unspecified |
| mmu-miR-24-3p | MIMAT0000219 | Casp9 | 12371 | Sequencing | 37986224\|19536157 | Unspecified |
| mmu-miR-24-3p | MIMAT0000219 | Cbfa2t2 | 12396 | Sequencing | 37986224\|19536157 | Unspecified |
| mmu-miR-24-3p | MIMAT0000219 | Cbx1 | 12412 | Sequencing | 37986224\|19536157 | Unspecified |
| mmu-miR-24-3p | MIMAT0000219 | Ccdc55 | 237859 | HITS-CLIP | 21258322 | Unspecified |
| mmu-miR-24-3p | MIMAT0000219 | Ccdc85b | 240514 | HITS-CLIP | 21258322\|25753418 | Unspecified |
| mmu-miR-24-3p | MIMAT0000219 | Ccnt2 | 72949 | Sequencing//HITS-CLIP | 19536157\|23142080\|37986224 | Unspecified |
| mmu-miR-24-3p | MIMAT0000219 | Ccser2 | 72972 | Sequencing//HITS-CLIP | 19536157\|23142080 | Unspecified |
| mmu-miR-24-3p | MIMAT0000219 | Cd300a | 217303 | HITS-CLIP | 21258322\|23597149 | Unspecified |

| | | | | | | |
|---|---|---|---|---|---|---|
| mmu-miR-24-3p | MIMAT0000219 | Cd93 | 17064 | Sequencing | 37986224\|19536157 | Unspecified |
| mmu-miR-24-3p | MIMAT0000219 | Cds2 | 110911 | Sequencing//HITS-CLIP | 37986224\|23142080\|19536157 | Unspecified |
| mmu-miR-24-3p | MIMAT0000219 | Celf4 | 108013 | Sequencing | 19536157 | Unspecified |
| mmu-miR-24-3p | MIMAT0000219 | Celf5 | 319586 | Sequencing | 19536157 | Unspecified |
| mmu-miR-24-3p | MIMAT0000219 | Cep120 | 225523 | Sequencing | 19536157 | Unspecified |
| mmu-miR-24-3p | MIMAT0000219 | Cep170 | 545389 | Sequencing//HITS-CLIP | 19536157\|23142080\|37986224 | Unspecified |
| mmu-miR-24-3p | MIMAT0000219 | Cep95 | 320162 | Sequencing//HITS-CLIP | 19536157\|23142080 | Unspecified |
| mmu-miR-24-3p | MIMAT0000219 | Chl1 | 12661 | Sequencing | 37986224\|19536157 | Unspecified |
| mmu-miR-24-3p | MIMAT0000219 | Chrna1 | 11435 | HITS-CLIP | 21258322 | Unspecified |
| mmu-miR-24-3p | MIMAT0000219 | Clcn3 | 12725 | Sequencing//HITS-CLIP | 18353861\|23142080\|37986224 | Unspecified |
| mmu-miR-24-3p | MIMAT0000219 | Clmn | 94040 | Sequencing | 37986224\|19536157 | Unspecified |
| mmu-miR-24-3p | MIMAT0000219 | Cmtm6 | 67213 | Sequencing//HITS-CLIP | 19536157\|25753418\|23142080\|3798 | Unspecified |
| mmu-miR-24-3p | MIMAT0000219 | Cnbd2 | 70873 | Sequencing | 19536157 | Unspecified |
| mmu-miR-24-3p | MIMAT0000219 | Cnnm3 | 94218 | Sequencing | 19536157\|37986224 | Unspecified |
| mmu-miR-24-3p | MIMAT0000219 | Cnot6 | 104625 | Sequencing//HITS-CLIP | 19536157\|23142080\|37986224 | Unspecified |
| mmu-miR-24-3p | MIMAT0000219 | Cnst | 226744 | Sequencing//HITS-CLIP | 19536157\|23142080 | Unspecified |
| mmu-miR-24-3p | MIMAT0000219 | Coa3 | 52469 | Sequencing | 19536157 | Unspecified |
| mmu-miR-24-3p | MIMAT0000219 | Cog3 | 338337 | Sequencing//HITS-CLIP | 19536157\|23142080\|37986224 | Unspecified |
| mmu-miR-24-3p | MIMAT0000219 | Col4a1 | 12826 | Sequencing//HITS-CLIP | 19536157\|25753418\|37986224 | Unspecified |
| mmu-miR-24-3p | MIMAT0000219 | Commd7 | 99311 | HITS-CLIP | 21258322 | Unspecified |
| mmu-miR-24-3p | MIMAT0000219 | Cplx2 | 12890 | Sequencing | 19536157\|37986224 | Unspecified |
| mmu-miR-24-3p | MIMAT0000219 | Crhr1 | 12921 | Sequencing | 37986224\|19536157 | Unspecified |
| mmu-miR-24-3p | MIMAT0000219 | Crtc1 | 382056 | Sequencing | 19536157\|37986224 | Unspecified |
| mmu-miR-24-3p | MIMAT0000219 | Csnk1a1 | 93687 | Sequencing//HITS-CLIP | 37986224\|23142080\|19536157 | Unspecified |
| mmu-miR-24-3p | MIMAT0000219 | Csnk1g1 | 214897 | Sequencing | 19536157\|37986224 | Unspecified |
| mmu-miR-24-3p | MIMAT0000219 | Ctnnbip1 | 67087 | Sequencing | 37986224\|19536157 | Unspecified |
| mmu-miR-24-3p | MIMAT0000219 | Cyld | 74256 | HITS-CLIP | 21258322 | Unspecified |
| mmu-miR-24-3p | MIMAT0000219 | D14Abb1e | 218850 | Sequencing | 19536157\|37986224 | Unspecified |
| mmu-miR-24-3p | MIMAT0000219 | Dcaf8 | 98193 | Sequencing//HITS-CLIP | 19536157\|23142080 | Unspecified |
| mmu-miR-24-3p | MIMAT0000219 | Dpagt1 | 13478 | Sequencing | 37986224\|19536157 | Unspecified |
| mmu-miR-24-3p | MIMAT0000219 | Dph7 | 67228 | HITS-CLIP | 25083871\|21258322 | Unspecified |
| mmu-miR-24-3p | MIMAT0000219 | Dpp6 | 13483 | Sequencing | 19536157\|37986224 | Unspecified |
| mmu-miR-24-3p | MIMAT0000219 | Drd1a | 13488 | Sequencing | 19536157\|37986224 | Unspecified |
| mmu-miR-24-3p | MIMAT0000219 | Dtna | 13527 | Sequencing | 37986224\|19536157 | Unspecified |
| mmu-miR-24-3p | MIMAT0000219 | Ece1 | 230857 | Sequencing//HITS-CLIP | 37986224\|23142080\|19536157 | Unspecified |
| mmu-miR-24-3p | MIMAT0000219 | Eif4g2 | 13690 | HITS-CLIP | 21258322\|31433990\|25753418 | Unspecified |
| mmu-miR-24-3p | MIMAT0000219 | Emc10 | 69683 | Sequencing | 19536157 | Unspecified |
| mmu-miR-24-3p | MIMAT0000219 | Eno2 | 13807 | Sequencing | 37986224\|19536157 | Unspecified |
| mmu-miR-24-3p | MIMAT0000219 | Entpd6 | 12497 | Sequencing//HITS-CLIP | 37986224\|23142080\|19536157 | Unspecified |

| | | | | | | |
|---|---|---|---|---|---|---|
| mmu-miR-24-3p | MIMAT0000219 | Epb4.1l1 | 13821 | Sequencing | 19536157\|37986224 | Unspecified |
| mmu-miR-24-3p | MIMAT0000219 | Epn1 | 13854 | Sequencing//HITS-CLIP | 19536157\|25753418\|23142080\|3798[illegible] | Unspecified |
| mmu-miR-24-3p | MIMAT0000219 | Ept1 | 28042 | Sequencing | 19536157 | Unspecified |
| mmu-miR-24-3p | MIMAT0000219 | Esyt2 | 52635 | Sequencing//HITS-CLIP | 19536157\|23142080 | Unspecified |
| mmu-miR-24-3p | MIMAT0000219 | Exoc6b | 75914 | Sequencing//HITS-CLIP | 37986224\|25753418\|19536157 | Unspecified |
| mmu-miR-24-3p | MIMAT0000219 | Extl2 | 58193 | Sequencing | 37986224\|19536157 | Unspecified |
| mmu-miR-24-3p | MIMAT0000219 | Fam115a | 77574 | Sequencing | 19536157\|37986224 | Unspecified |
| mmu-miR-24-3p | MIMAT0000219 | Fam117b | 72750 | Sequencing | 37986224\|19536157 | Unspecified |
| mmu-miR-24-3p | MIMAT0000219 | Fam131a | 78408 | Sequencing | 19536157\|37986224 | Unspecified |
| mmu-miR-24-3p | MIMAT0000219 | Fam134c | 67998 | Sequencing | 19536157\|37986224 | Unspecified |
| mmu-miR-24-3p | MIMAT0000219 | Fam161b | 217705 | HITS-CLIP | 25083871 | Unspecified |
| mmu-miR-24-3p | MIMAT0000219 | Fam168b | 214469 | Sequencing//HITS-CLIP | 37986224\|23142080\|19536157 | Unspecified |
| mmu-miR-24-3p | MIMAT0000219 | Fam76b | 72826 | Sequencing | 37986224\|19536157 | Unspecified |
| mmu-miR-24-3p | MIMAT0000219 | Fat1 | 14107 | Sequencing | 37986224\|19536157 | Unspecified |
| mmu-miR-24-3p | MIMAT0000219 | Fbxl20 | 72194 | Sequencing//HITS-CLIP | 19536157\|23142080\|37986224 | Unspecified |
| mmu-miR-24-3p | MIMAT0000219 | Fgfr1 | 14182 | Sequencing | 19536157\|37986224 | Unspecified |
| mmu-miR-24-3p | MIMAT0000219 | Fkbp1b | 14226 | Sequencing | 37986224\|19536157 | Unspecified |
| mmu-miR-24-3p | MIMAT0000219 | Flna | 192176 | Sequencing | 19536157\|37986224 | Unspecified |
| mmu-miR-24-3p | MIMAT0000219 | Foxn3 | 71375 | Sequencing//HITS-CLIP | 19536157\|23142080\|37986224 | Unspecified |
| mmu-miR-24-3p | MIMAT0000219 | Frem3 | 333315 | HITS-CLIP | 23142080 | Unspecified |
| mmu-miR-24-3p | MIMAT0000219 | Fut9 | 14348 | Sequencing | 19536157\|37986224 | Unspecified |
| mmu-miR-24-3p | MIMAT0000219 | Gabpb2 | 213054 | Sequencing//HITS-CLIP | 37986224\|23142080\|19536157 | Unspecified |
| mmu-miR-24-3p | MIMAT0000219 | Gabrb2 | 14401 | Sequencing | 37986224\|19536157 | Unspecified |
| mmu-miR-24-3p | MIMAT0000219 | Gal3st3 | 545276 | Sequencing | 19536157\|37986224 | Unspecified |
| mmu-miR-24-3p | MIMAT0000219 | Gap43 | 14432 | Sequencing | 37986224\|19536157 | Unspecified |
| mmu-miR-24-3p | MIMAT0000219 | Gas2l3 | 237436 | HITS-CLIP | 21258322 | Unspecified |
| mmu-miR-24-3p | MIMAT0000219 | Gatsl2 | 80909 | Sequencing | 19536157 | Unspecified |
| mmu-miR-24-3p | MIMAT0000219 | Gba2 | 230101 | Sequencing//HITS-CLIP | 19536157\|25753418\|23142080\|3798[illegible] | Unspecified |
| mmu-miR-24-3p | MIMAT0000219 | Ggps1 | 14593 | HITS-CLIP | 21258322 | Unspecified |
| mmu-miR-24-3p | MIMAT0000219 | Ggta1 | 14594 | Sequencing//HITS-CLIP | 37986224\|23142080\|19536157 | Unspecified |
| mmu-miR-24-3p | MIMAT0000219 | Gm16532 | 100042450 | HITS-CLIP | 21258322 | Unspecified |
| mmu-miR-24-3p | MIMAT0000219 | Gm5113 | 330503 | HITS-CLIP | 21258322 | Unspecified |
| mmu-miR-24-3p | MIMAT0000219 | Gna14 | 14675 | Sequencing | 19536157\|37986224 | Unspecified |
| mmu-miR-24-3p | MIMAT0000219 | Gnal | 14680 | Sequencing | 19536157\|37986224 | Unspecified |
| mmu-miR-24-3p | MIMAT0000219 | Gnaq | 14682 | Sequencing//HITS-CLIP | 37986224\|23142080\|19536157 | Unspecified |
| mmu-miR-24-3p | MIMAT0000219 | Gnb5 | 14697 | Sequencing | 37986224\|19536157 | Unspecified |
| mmu-miR-24-3p | MIMAT0000219 | Gpalpp1 | 67467 | HITS-CLIP | 21258322 | Unspecified |
| mmu-miR-24-3p | MIMAT0000219 | Gpr37l1 | 171469 | Sequencing | 19536157\|37986224 | Unspecified |
| mmu-miR-24-3p | MIMAT0000219 | Gpr88 | 64378 | Sequencing | 19536157\|37986224 | Unspecified |

| | | | | | | |
|---|---|---|---|---|---|---|
| mmu-miR-24-3p | MIMAT0000219 | Gprasp1 | 67298 | Sequencing//HITS-CLIP | 37986224\|23142080\|19536157 | Unspecified |
| mmu-miR-24-3p | MIMAT0000219 | Gpt2 | 108682 | Sequencing | 37986224\|19536157 | Unspecified |
| mmu-miR-24-3p | MIMAT0000219 | Gria3 | 53623 | Sequencing | 19536157\|37986224 | Unspecified |
| mmu-miR-24-3p | MIMAT0000219 | Grin1 | 14810 | Sequencing | 19536157\|37986224 | Unspecified |
| mmu-miR-24-3p | MIMAT0000219 | Grina | 66168 | Sequencing | 37986224\|19536157 | Unspecified |
| mmu-miR-24-3p | MIMAT0000219 | H2-T24 | 15042 | HITS-CLIP | 25083871\|23142080\|21258322 | Unspecified |
| mmu-miR-24-3p | MIMAT0000219 | H2afx | 15270 | Sequencing//HITS-CLIP | 37986224\|23142080\|19536157 | Unspecified |
| mmu-miR-24-3p | MIMAT0000219 | Has3 | 15118 | Sequencing | 37986224\|19536157 | Unspecified |
| mmu-miR-24-3p | MIMAT0000219 | Hbegf | 15200 | Sequencing//HITS-CLIP | 19536157\|25753418\|37986224 | Unspecified |
| mmu-miR-24-3p | MIMAT0000219 | Hdgfrp3 | 29877 | Sequencing | 19536157\|37986224 | Unspecified |
| mmu-miR-24-3p | MIMAT0000219 | Hiat1 | 15247 | Sequencing | 19536157\|37986224 | Unspecified |
| mmu-miR-24-3p | MIMAT0000219 | Hic2 | 58180 | Sequencing//HITS-CLIP | 37986224\|25753418\|19536157 | Unspecified |
| mmu-miR-24-3p | MIMAT0000219 | Hmox1 | 15368 | Sequencing//qCLASH//HITS-CLIP | 19536157\|31393953\|25753418\|3798 | Unspecified |
| mmu-miR-24-3p | MIMAT0000219 | Hnf1a | 21405 | Luciferase assay//qRT-PCR//Western blot/ | 23761103 | Unspecified |
| mmu-miR-24-3p | MIMAT0000219 | Homez | 239099 | HITS-CLIP | 21258322\|25753418 | Unspecified |
| mmu-miR-24-3p | MIMAT0000219 | Hprt | 15452 | Sequencing | 19536157 | Unspecified |
| mmu-miR-24-3p | MIMAT0000219 | Htr5a | 15563 | Sequencing | 37986224\|19536157 | Unspecified |
| mmu-miR-24-3p | MIMAT0000219 | Ifnar1 | 15975 | Sequencing//HITS-CLIP | 37986224\|23142080\|19536157 | Unspecified |
| mmu-miR-24-3p | MIMAT0000219 | Igf2 | 16002 | Sequencing | 37986224\|19536157 | Unspecified |
| mmu-miR-24-3p | MIMAT0000219 | Igfbp5 | 16011 | Sequencing | 37986224\|19536157 | Unspecified |
| mmu-miR-24-3p | MIMAT0000219 | Ikbkg | 16151 | HITS-CLIP | 25083871\|21258322 | Unspecified |
| mmu-miR-24-3p | MIMAT0000219 | Il6st | 16195 | Sequencing//HITS-CLIP | 37986224\|23142080\|19536157 | Unspecified |
| mmu-miR-24-3p | MIMAT0000219 | Itsn1 | 16443 | Sequencing//HITS-CLIP | 19536157\|21258322\|37986224 | Unspecified |
| mmu-miR-24-3p | MIMAT0000219 | Jph2 | 59091 | Luciferase assay//Luciferase reporter assay | 22891046 | Unspecified |
| mmu-miR-24-3p | MIMAT0000219 | Kansl3 | 226976 | Sequencing//HITS-CLIP | 19536157\|25753418 | Unspecified |
| mmu-miR-24-3p | MIMAT0000219 | Kcnc3 | 16504 | Sequencing | 37986224\|19536157 | Unspecified |
| mmu-miR-24-3p | MIMAT0000219 | Kcnj2 | 16518 | Sequencing | 19536157\|37986224 | Unspecified |
| mmu-miR-24-3p | MIMAT0000219 | Kcnj3 | 16519 | Sequencing | 37986224\|19536157 | Unspecified |
| mmu-miR-24-3p | MIMAT0000219 | Kcnk6 | 52150 | Sequencing | 19536157\|37986224 | Unspecified |
| mmu-miR-24-3p | MIMAT0000219 | Kdm4c | 76804 | Sequencing//HITS-CLIP | 19536157\|23142080 | Unspecified |
| mmu-miR-24-3p | MIMAT0000219 | Kdm5b | 75605 | Sequencing//HITS-CLIP | 19536157\|23142080 | Unspecified |
| mmu-miR-24-3p | MIMAT0000219 | Kif21b | 16565 | Sequencing//HITS-CLIP | 37986224\|23142080\|19536157 | Unspecified |
| mmu-miR-24-3p | MIMAT0000219 | Kif3b | 16569 | Sequencing//HITS-CLIP | 19536157\|23142080\|37986224 | Unspecified |
| mmu-miR-24-3p | MIMAT0000219 | Kifap3 | 16579 | Sequencing | 37986224\|19536157 | Unspecified |
| mmu-miR-24-3p | MIMAT0000219 | Kitl | 17311 | Sequencing | 37986224\|19536157 | Unspecified |
| mmu-miR-24-3p | MIMAT0000219 | Klc2 | 16594 | Sequencing | 19536157\|37986224 | Unspecified |
| mmu-miR-24-3p | MIMAT0000219 | Klf6 | 23849 | Sequencing//HITS-CLIP | 19536157\|23142080\|25753418\|3798 | Unspecified |
| mmu-miR-24-3p | MIMAT0000219 | Klhdc9 | 68874 | HITS-CLIP | 21258322 | Unspecified |
| mmu-miR-24-3p | MIMAT0000219 | Klhl24 | 75785 | Sequencing//HITS-CLIP | 37986224\|23142080\|25753418\|1953 | Unspecified |

| | | | | | | |
|---|---|---|---|---|---|---|
| mmu-miR-24-3p | MIMAT0000219 | L1cam | 16728 | Sequencing | 19536157\|37986224 | Unspecified |
| mmu-miR-24-3p | MIMAT0000219 | Lbh | 77889 | Sequencing | 19536157\|37986224 | Unspecified |
| mmu-miR-24-3p | MIMAT0000219 | Ldb2 | 16826 | Sequencing | 37986224\|19536157 | Unspecified |
| mmu-miR-24-3p | MIMAT0000219 | Ldha | 16828 | Sequencing//HITS-CLIP | 37986224\|23142080\|19536157 | Unspecified |
| mmu-miR-24-3p | MIMAT0000219 | Letmd1 | 68614 | Sequencing | 37986224\|19536157 | Unspecified |
| mmu-miR-24-3p | MIMAT0000219 | Lims1 | 110829 | Sequencing//HITS-CLIP | 19536157\|23142080\|37986224 | Unspecified |
| mmu-miR-24-3p | MIMAT0000219 | Lipa | 16889 | Sequencing | 37986224\|19536157 | Unspecified |
| mmu-miR-24-3p | MIMAT0000219 | Lmbr1 | 56873 | Sequencing | 19536157\|37986224 | Unspecified |
| mmu-miR-24-3p | MIMAT0000219 | Lmtk2 | 231876 | Sequencing//HITS-CLIP | 37986224\|23142080\|19536157 | Unspecified |
| mmu-miR-24-3p | MIMAT0000219 | Lrch1 | 380916 | Sequencing//HITS-CLIP | 19536157\|23142080\|37986224 | Unspecified |
| mmu-miR-24-3p | MIMAT0000219 | Lrrc3 | 237387 | Sequencing | 37986224\|19536157 | Unspecified |
| mmu-miR-24-3p | MIMAT0000219 | Lrrc39 | 109245 | HITS-CLIP | 21258322 | Unspecified |
| mmu-miR-24-3p | MIMAT0000219 | Lrrc58 | 320184 | Sequencing//HITS-CLIP | 19536157\|23142080\|37986224 | Unspecified |
| mmu-miR-24-3p | MIMAT0000219 | Lrtm2 | 211187 | Sequencing | 19536157\|37986224 | Unspecified |
| mmu-miR-24-3p | MIMAT0000219 | Lsamp | 268890 | Sequencing | 19536157\|37986224 | Unspecified |
| mmu-miR-24-3p | MIMAT0000219 | Ly6g6d | 114654 | HITS-CLIP | 21258322 | Unspecified |
| mmu-miR-24-3p | MIMAT0000219 | MAPK7 | 5598 | HITS-CLIP//Luciferase reporter assay | 23142080\|27103442 | Unspecified |
| mmu-miR-24-3p | MIMAT0000219 | Macf1 | 11426 | Sequencing//HITS-CLIP | 37986224\|23142080\|25753418\|1953€ | Unspecified |
| mmu-miR-24-3p | MIMAT0000219 | Mak16 | 67920 | Sequencing | 19536157\|37986224 | Unspecified |
| mmu-miR-24-3p | MIMAT0000219 | Man1b1 | 227619 | HITS-CLIP | 23597149 | Unspecified |
| mmu-miR-24-3p | MIMAT0000219 | Marchf5 | 69104 | HITS-CLIP | 21258322 | Unspecified |
| mmu-miR-24-3p | MIMAT0000219 | Mbd6 | 110962 | Sequencing//HITS-CLIP | 19536157\|23142080\|37986224 | Unspecified |
| mmu-miR-24-3p | MIMAT0000219 | Mcm2 | 17216 | Sequencing | 19536157\|37986224 | Unspecified |
| mmu-miR-24-3p | MIMAT0000219 | Mdh1 | 17449 | Sequencing//HITS-CLIP | 37986224\|23142080\|19536157 | Unspecified |
| mmu-miR-24-3p | MIMAT0000219 | Mdk | 17242 | Sequencing | 19536157\|37986224 | Unspecified |
| mmu-miR-24-3p | MIMAT0000219 | Med10 | 28077 | Sequencing | 19536157\|37986224 | Unspecified |
| mmu-miR-24-3p | MIMAT0000219 | Mef2c | 17260 | Western blot//qRT-PCR//Western blot;qR | 18353861 | Unspecified |
| mmu-miR-24-3p | MIMAT0000219 | Mllt11 | 56772 | Sequencing | 37986224\|19536157 | Unspecified |
| mmu-miR-24-3p | MIMAT0000219 | Mmp24 | 17391 | Sequencing | 37986224\|19536157 | Unspecified |
| mmu-miR-24-3p | MIMAT0000219 | Mrpl35 | 66223 | HITS-CLIP | 21258322 | Unspecified |
| mmu-miR-24-3p | MIMAT0000219 | Myh10 | 77579 | Sequencing | 37986224\|19536157 | Unspecified |
| mmu-miR-24-3p | MIMAT0000219 | Mylip | 218203 | HITS-CLIP | 23597149\|21258322 | Unspecified |
| mmu-miR-24-3p | MIMAT0000219 | Myog | 17928 | Western blot//qRT-PCR//Western blot;qR | 18353861 | Unspecified |
| mmu-miR-24-3p | MIMAT0000219 | N28178 | 230085 | Sequencing | 19536157\|37986224 | Unspecified |
| mmu-miR-24-3p | MIMAT0000219 | Naa40 | 70999 | Sequencing//HITS-CLIP | 19536157\|23142080 | Unspecified |
| mmu-miR-24-3p | MIMAT0000219 | Nacc2 | 67991 | Sequencing | 37986224\|19536157 | Unspecified |
| mmu-miR-24-3p | MIMAT0000219 | Nat8l | 269642 | Sequencing | 19536157\|37986224 | Unspecified |
| mmu-miR-24-3p | MIMAT0000219 | Ncl | 17975 | HITS-CLIP | 21258322\|23142080 | Unspecified |
| mmu-miR-24-3p | MIMAT0000219 | Ndfip1 | 65113 | Sequencing//HITS-CLIP | 37986224\|23142080\|25753418\|1953€ | Unspecified |

| | | | | | | |
|---|---|---|---|---|---|---|
| mmu-miR-24-3p | MIMAT0000219 | Ndrg3 | 29812 | Sequencing//HITS-CLIP | 37986224\|23142080\|19536157 | Unspecified |
| mmu-miR-24-3p | MIMAT0000219 | Nedd4 | 17999 | Sequencing | 37986224\|19536157 | Unspecified |
| mmu-miR-24-3p | MIMAT0000219 | Negr1 | 320840 | Sequencing | 19536157\|37986224 | Unspecified |
| mmu-miR-24-3p | MIMAT0000219 | Neu3 | 50877 | HITS-CLIP | 21258322\|23142080 | Unspecified |
| mmu-miR-24-3p | MIMAT0000219 | Neurod1 | 18012 | Luciferase assay//qRT-PCR//Western blot/ | 23761103 | Unspecified |
| mmu-miR-24-3p | MIMAT0000219 | Nf1 | 18015 | Sequencing | 37986224\|19536157 | Unspecified |
| mmu-miR-24-3p | MIMAT0000219 | Nfat5 | 54446 | Sequencing//HITS-CLIP | 19536157\|23142080\|37986224 | Unspecified |
| mmu-miR-24-3p | MIMAT0000219 | Nfia | 18027 | Sequencing | 19536157\|37986224 | Unspecified |
| mmu-miR-24-3p | MIMAT0000219 | Nfix | 18032 | Sequencing | 19536157\|37986224 | Unspecified |
| mmu-miR-24-3p | MIMAT0000219 | Nme7 | 171567 | HITS-CLIP | 21258322 | Unspecified |
| mmu-miR-24-3p | MIMAT0000219 | Nr2c2 | 22026 | Sequencing | 37986224\|19536157 | Unspecified |
| mmu-miR-24-3p | MIMAT0000219 | Nrgn | 64011 | Sequencing | 37986224\|19536157 | Unspecified |
| mmu-miR-24-3p | MIMAT0000219 | Nrp2 | 18187 | Sequencing | 37986224\|19536157 | Unspecified |
| mmu-miR-24-3p | MIMAT0000219 | Nrxn3 | 18191 | Sequencing | 19536157\|37986224 | Unspecified |
| mmu-miR-24-3p | MIMAT0000219 | Nsd1 | 18193 | Sequencing//HITS-CLIP | 19536157\|23142080\|25753418\|3798€ | Unspecified |
| mmu-miR-24-3p | MIMAT0000219 | Nsrp1 | 237859 | HITS-CLIP | 21258322 | Unspecified |
| mmu-miR-24-3p | MIMAT0000219 | Nucks1 | 98415 | Sequencing//HITS-CLIP | 19536157\|23142080\|37986224 | Unspecified |
| mmu-miR-24-3p | MIMAT0000219 | Nup210 | 54563 | Sequencing//HITS-CLIP | 19536157\|23142080\|37986224 | Unspecified |
| mmu-miR-24-3p | MIMAT0000219 | Ogt | 108155 | Sequencing//HITS-CLIP | 19536157\|23142080\|37986224 | Unspecified |
| mmu-miR-24-3p | MIMAT0000219 | Os9 | 216440 | Sequencing | 19536157\|37986224 | Unspecified |
| mmu-miR-24-3p | MIMAT0000219 | Oxt | 18429 | ELISA//Luciferase assay//qRT-PCR//Lucifer | 23682839 | Unspecified |
| mmu-miR-24-3p | MIMAT0000219 | Padi2 | 18600 | HITS-CLIP | 21258322 | Unspecified |
| mmu-miR-24-3p | MIMAT0000219 | Pak7 | 241656 | Sequencing | 19536157\|37986224 | Unspecified |
| mmu-miR-24-3p | MIMAT0000219 | Pclo | 26875 | Sequencing | 37986224\|19536157 | Unspecified |
| mmu-miR-24-3p | MIMAT0000219 | Pcsk1n | 30052 | Sequencing | 37986224\|19536157 | Unspecified |
| mmu-miR-24-3p | MIMAT0000219 | Pde1a | 18573 | Sequencing | 19536157\|37986224 | Unspecified |
| mmu-miR-24-3p | MIMAT0000219 | Pgrmc2 | 70804 | Sequencing//HITS-CLIP | 19536157\|23142080\|37986224 | Unspecified |
| mmu-miR-24-3p | MIMAT0000219 | Pianp | 319352 | Sequencing | 19536157 | Unspecified |
| mmu-miR-24-3p | MIMAT0000219 | Pign | 27392 | Sequencing | 37986224\|19536157 | Unspecified |
| mmu-miR-24-3p | MIMAT0000219 | Pigv | 230801 | HITS-CLIP | 21258322 | Unspecified |
| mmu-miR-24-3p | MIMAT0000219 | Pknox2 | 208076 | Sequencing | 19536157\|37986224 | Unspecified |
| mmu-miR-24-3p | MIMAT0000219 | Plch1 | 269437 | Sequencing | 19536157\|37986224 | Unspecified |
| mmu-miR-24-3p | MIMAT0000219 | Plec | 18810 | Sequencing//HITS-CLIP | 19536157\|23142080\|25753418 | Unspecified |
| mmu-miR-24-3p | MIMAT0000219 | Plod2 | 26432 | Sequencing//HITS-CLIP | 19536157\|23142080\|37986224 | Unspecified |
| mmu-miR-24-3p | MIMAT0000219 | Plxna2 | 18845 | Sequencing | 19536157\|37986224 | Unspecified |
| mmu-miR-24-3p | MIMAT0000219 | Pml | 18854 | Sequencing//HITS-CLIP | 37986224\|23142080\|19536157 | Unspecified |
| mmu-miR-24-3p | MIMAT0000219 | Pnoc | 18155 | HITS-CLIP | 21258322 | Unspecified |
| mmu-miR-24-3p | MIMAT0000219 | Pogz | 229584 | Sequencing | 37986224\|19536157 | Unspecified |
| mmu-miR-24-3p | MIMAT0000219 | Polr1e | 64424 | HITS-CLIP | 21258322 | Unspecified |

| | | | | | | |
|---|---|---|---|---|---|---|
| mmu-miR-24-3p | MIMAT0000219 | Pomk | 74653 | HITS-CLIP | 23142080 | Unspecified |
| mmu-miR-24-3p | MIMAT0000219 | Pou5f1 | 18999 | HITS-CLIP//qRT-PCR//Luciferase reporter a | 37986224\|25519956 | Unspecified |
| mmu-miR-24-3p | MIMAT0000219 | Ppm1k | 243382 | Sequencing | 19536157\|37986224 | Unspecified |
| mmu-miR-24-3p | MIMAT0000219 | Preb | 50907 | Sequencing | 19536157\|37986224 | Unspecified |
| mmu-miR-24-3p | MIMAT0000219 | Psmb11 | 73902 | HITS-CLIP | 21258322 | Unspecified |
| mmu-miR-24-3p | MIMAT0000219 | Rab31 | 106572 | Sequencing | 19536157\|37986224 | Unspecified |
| mmu-miR-24-3p | MIMAT0000219 | Rab3d | 19340 | Sequencing | 37986224\|19536157 | Unspecified |
| mmu-miR-24-3p | MIMAT0000219 | Rab3gap1 | 226407 | Sequencing//HITS-CLIP | 19536157\|25753418\|37986224 | Unspecified |
| mmu-miR-24-3p | MIMAT0000219 | Rab43 | 69834 | Sequencing | 37986224\|19536157 | Unspecified |
| mmu-miR-24-3p | MIMAT0000219 | Rasal1 | 19415 | Sequencing | 19536157\|37986224 | Unspecified |
| mmu-miR-24-3p | MIMAT0000219 | Reep1 | 52250 | Sequencing | 19536157\|37986224 | Unspecified |
| mmu-miR-24-3p | MIMAT0000219 | Reep2 | 225362 | Sequencing | 37986224\|19536157 | Unspecified |
| mmu-miR-24-3p | MIMAT0000219 | Rhbdl1 | 214951 | Sequencing//HITS-CLIP | 19536157\|23142080\|37986224 | Unspecified |
| mmu-miR-24-3p | MIMAT0000219 | Rictor | 78757 | Sequencing//HITS-CLIP | 19536157\|23142080 | Unspecified |
| mmu-miR-24-3p | MIMAT0000219 | Rnf138 | 56515 | Sequencing//HITS-CLIP | 37986224\|23142080\|19536157 | Unspecified |
| mmu-miR-24-3p | MIMAT0000219 | Rnf170 | 77733 | Sequencing | 37986224\|19536157 | Unspecified |
| mmu-miR-24-3p | MIMAT0000219 | Rnf24 | 51902 | Sequencing | 19536157\|37986224 | Unspecified |
| mmu-miR-24-3p | MIMAT0000219 | Rpl12 | 269261 | Sequencing | 19536157\|37986224 | Unspecified |
| mmu-miR-24-3p | MIMAT0000219 | Rps6kl1 | 238323 | Sequencing | 37986224\|19536157 | Unspecified |
| mmu-miR-24-3p | MIMAT0000219 | Rragc | 54170 | Sequencing | 37986224\|19536157 | Unspecified |
| mmu-miR-24-3p | MIMAT0000219 | Rtn1 | 104001 | Sequencing | 37986224\|19536157 | Unspecified |
| mmu-miR-24-3p | MIMAT0000219 | Sacs | 50720 | Sequencing//HITS-CLIP | 37986224\|23142080\|19536157 | Unspecified |
| mmu-miR-24-3p | MIMAT0000219 | Saysd1 | 67509 | HITS-CLIP | 21258322 | Unspecified |
| mmu-miR-24-3p | MIMAT0000219 | Scd2 | 20250 | Sequencing//HITS-CLIP | 19536157\|23142080\|37986224 | Unspecified |
| mmu-miR-24-3p | MIMAT0000219 | Scn2a1 | 110876 | Sequencing | 37986224\|19536157 | Unspecified |
| mmu-miR-24-3p | MIMAT0000219 | Scrt2 | 545474 | Sequencing | 19536157\|37986224 | Unspecified |
| mmu-miR-24-3p | MIMAT0000219 | Sec23ip | 207352 | HITS-CLIP | 21258322 | Unspecified |
| mmu-miR-24-3p | MIMAT0000219 | Sema4a | 20351 | Sequencing | 19536157\|37986224 | Unspecified |
| mmu-miR-24-3p | MIMAT0000219 | Sema4g | 26456 | Sequencing | 19536157\|37986224 | Unspecified |
| mmu-miR-24-3p | MIMAT0000219 | Setd7 | 73251 | Sequencing//HITS-CLIP | 19536157\|25753418\|23142080\|3798( | Unspecified |
| mmu-miR-24-3p | MIMAT0000219 | Sez6 | 20370 | Sequencing | 19536157\|37986224 | Unspecified |
| mmu-miR-24-3p | MIMAT0000219 | Sfxn4 | 94281 | HITS-CLIP | 21258322 | Unspecified |
| mmu-miR-24-3p | MIMAT0000219 | Sfxn5 | 94282 | Sequencing//HITS-CLIP | 19536157\|25753418\|37986224 | Unspecified |
| mmu-miR-24-3p | MIMAT0000219 | Sgo2a | 68549 | HITS-CLIP | 21258322 | Unspecified |
| mmu-miR-24-3p | MIMAT0000219 | Sgol2 | 68549 | HITS-CLIP | 21258322 | Unspecified |
| mmu-miR-24-3p | MIMAT0000219 | Sh3d19 | 27059 | Sequencing | 19536157\|37986224 | Unspecified |
| mmu-miR-24-3p | MIMAT0000219 | Shisa6 | 380702 | Sequencing | 19536157 | Unspecified |
| mmu-miR-24-3p | MIMAT0000219 | Ska3 | 219114 | Sequencing | 19536157 | Unspecified |
| mmu-miR-24-3p | MIMAT0000219 | Skida1 | 72668 | Sequencing//HITS-CLIP | 19536157\|23142080 | Unspecified |

| | | | | | | |
|---|---|---|---|---|---|---|
| mmu-miR-24-3p | MIMAT0000219 | Slc12a6 | 107723 | Sequencing//HITS-CLIP | 37986224\|23142080\|19536157 | Unspecified |
| mmu-miR-24-3p | MIMAT0000219 | Slc15a4 | 100561 | Sequencing | 19536157\|37986224 | Unspecified |
| mmu-miR-24-3p | MIMAT0000219 | Slc18a2 | 214084 | Sequencing | 37986224\|19536157 | Unspecified |
| mmu-miR-24-3p | MIMAT0000219 | Slc22a23 | 73102 | Sequencing | 19536157\|37986224 | Unspecified |
| mmu-miR-24-3p | MIMAT0000219 | Slc25a12 | 78830 | Sequencing//HITS-CLIP | 19536157\|23142080\|37986224 | Unspecified |
| mmu-miR-24-3p | MIMAT0000219 | Slc25a25 | 227731 | Sequencing//HITS-CLIP | 37986224\|31433990\|25753418\|2620 | Unspecified |
| mmu-miR-24-3p | MIMAT0000219 | Slc30a10 | 226781 | Sequencing | 37986224\|19536157 | Unspecified |
| mmu-miR-24-3p | MIMAT0000219 | Slc35d2 | 70484 | HITS-CLIP | 21258322 | Unspecified |
| mmu-miR-24-3p | MIMAT0000219 | Slc35e2 | 320541 | Sequencing//HITS-CLIP | 19536157\|23142080 | Unspecified |
| mmu-miR-24-3p | MIMAT0000219 | Slc4a8 | 59033 | Sequencing | 37986224\|19536157 | Unspecified |
| mmu-miR-24-3p | MIMAT0000219 | Slc7a1 | 11987 | Sequencing//HITS-CLIP | 37986224\|23142080\|19536157 | Unspecified |
| mmu-miR-24-3p | MIMAT0000219 | Slitrk5 | 75409 | Sequencing | 19536157\|37986224 | Unspecified |
| mmu-miR-24-3p | MIMAT0000219 | Smad3 | 17127 | Sequencing//HITS-CLIP//In situ hybridizati | 19536157\|23142080\|19582816 | Unspecified |
| mmu-miR-24-3p | MIMAT0000219 | Smad4 | 17128 | In situ hybridization//Luciferase assay//Mi | 19582816\|23142080\|25519956 | Unspecified |
| mmu-miR-24-3p | MIMAT0000219 | Smad5 | 17129 | In situ hybridization//Luciferase assay//Mi | 19582816 | Unspecified |
| mmu-miR-24-3p | MIMAT0000219 | Smim17 | 100042450 | HITS-CLIP | 21258322 | Unspecified |
| mmu-miR-24-3p | MIMAT0000219 | Snap47 | 67826 | Sequencing | 37986224\|19536157 | Unspecified |
| mmu-miR-24-3p | MIMAT0000219 | Snap91 | 20616 | Sequencing | 19536157\|37986224 | Unspecified |
| mmu-miR-24-3p | MIMAT0000219 | Snhg11 | 319317 | HITS-CLIP | 21258322 | Unspecified |
| mmu-miR-24-3p | MIMAT0000219 | Snx27 | 76742 | Sequencing//HITS-CLIP | 19536157\|23142080\|37986224 | Unspecified |
| mmu-miR-24-3p | MIMAT0000219 | Soat1 | 20652 | Sequencing//HITS-CLIP | 37986224\|23142080\|19536157 | Unspecified |
| mmu-miR-24-3p | MIMAT0000219 | Sp1 | 20683 | Sequencing//HITS-CLIP | 37986224\|23142080\|25753418\|1953 | Unspecified |
| mmu-miR-24-3p | MIMAT0000219 | Sp8 | 320145 | Sequencing | 19536157\|37986224 | Unspecified |
| mmu-miR-24-3p | MIMAT0000219 | Sparc | 20692 | Sequencing | 37986224\|19536157 | Unspecified |
| mmu-miR-24-3p | MIMAT0000219 | Spsb4 | 211949 | Sequencing | 19536157\|37986224 | Unspecified |
| mmu-miR-24-3p | MIMAT0000219 | Sptbn2 | 20743 | Sequencing | 19536157 | Unspecified |
| mmu-miR-24-3p | MIMAT0000219 | Srgap2 | 14270 | Sequencing//qCLASH//HITS-CLIP | 37986224\|31393953\|23142080\|1953 | Unspecified |
| mmu-miR-24-3p | MIMAT0000219 | Sri | 109552 | Sequencing//HITS-CLIP | 37986224\|25753418\|23142080\|1953 | Unspecified |
| mmu-miR-24-3p | MIMAT0000219 | Srsf1 | 110809 | Sequencing | 19536157 | Unspecified |
| mmu-miR-24-3p | MIMAT0000219 | Srsf11 | 69207 | Sequencing//HITS-CLIP | 19536157\|23142080 | Unspecified |
| mmu-miR-24-3p | MIMAT0000219 | Srxn1 | 76650 | Sequencing | 37986224\|19536157 | Unspecified |
| mmu-miR-24-3p | MIMAT0000219 | Ssfa2 | 70599 | Sequencing | 37986224\|19536157 | Unspecified |
| mmu-miR-24-3p | MIMAT0000219 | St8sia2 | 20450 | Sequencing | 37986224\|19536157 | Unspecified |
| mmu-miR-24-3p | MIMAT0000219 | Stk38 | 106504 | Sequencing//HITS-CLIP | 37986224\|23142080\|19536157 | Unspecified |
| mmu-miR-24-3p | MIMAT0000219 | Sumf2 | 67902 | HITS-CLIP | 21258322 | Unspecified |
| mmu-miR-24-3p | MIMAT0000219 | Syp | 20977 | Sequencing | 19536157\|37986224 | Unspecified |
| mmu-miR-24-3p | MIMAT0000219 | Syt1 | 20979 | Sequencing | 37986224\|19536157 | Unspecified |
| mmu-miR-24-3p | MIMAT0000219 | Syt2 | 20980 | Sequencing | 37986224\|19536157 | Unspecified |
| mmu-miR-24-3p | MIMAT0000219 | T2 | 21331 | HITS-CLIP | 21258322 | Unspecified |

| | | | | | | |
|---|---|---|---|---|---|---|
| mmu-miR-24-3p | MIMAT0000219 | Tcf7 | 21414 | HITS-CLIP//Luciferase reporter assay | 37986224\|23142080\|26006243 | Unspecified |
| mmu-miR-24-3p | MIMAT0000219 | Tmco3 | 234076 | HITS-CLIP | 25083871 | Unspecified |
| mmu-miR-24-3p | MIMAT0000219 | Tmed7 | 66676 | Sequencing | 19536157 | Unspecified |
| mmu-miR-24-3p | MIMAT0000219 | Tmem135 | 72759 | Sequencing//HITS-CLIP | 19536157\|23142080\|37986224 | Unspecified |
| mmu-miR-24-3p | MIMAT0000219 | Tmem206 | 66950 | HITS-CLIP | 21258322 | Unspecified |
| mmu-miR-24-3p | MIMAT0000219 | Tmem69 | 230657 | HITS-CLIP | 21258322 | Unspecified |
| mmu-miR-24-3p | MIMAT0000219 | Tmem87b | 72477 | Sequencing | 19536157\|37986224 | Unspecified |
| mmu-miR-24-3p | MIMAT0000219 | Tmem88b | 320587 | HITS-CLIP | 21258322 | Unspecified |
| mmu-miR-24-3p | MIMAT0000219 | Tnfsf10 | 22035 | HITS-CLIP | 21258322 | Unspecified |
| mmu-miR-24-3p | MIMAT0000219 | Tnks | 21951 | Sequencing//HITS-CLIP | 19536157\|25753418\|23142080\|3798€ | Unspecified |
| mmu-miR-24-3p | MIMAT0000219 | Tpmt | 22017 | HITS-CLIP | 25083871 | Unspecified |
| mmu-miR-24-3p | MIMAT0000219 | Tppp | 72948 | Sequencing | 19536157\|37986224 | Unspecified |
| mmu-miR-24-3p | MIMAT0000219 | Tpst1 | 22021 | Sequencing | 19536157\|37986224 | Unspecified |
| mmu-miR-24-3p | MIMAT0000219 | Trappc4 | 60409 | Sequencing//HITS-CLIP | 37986224\|25753418\|19536157 | Unspecified |
| mmu-miR-24-3p | MIMAT0000219 | Trib3 | 228775 | qRT-PCR | 25634354 | Unspecified |
| mmu-miR-24-3p | MIMAT0000219 | Trip10 | 106628 | Sequencing | 37986224\|19536157 | Unspecified |
| mmu-miR-24-3p | MIMAT0000219 | Trmt10a | 108943 | HITS-CLIP | 21258322\|23142080 | Unspecified |
| mmu-miR-24-3p | MIMAT0000219 | Tspyl4 | 72480 | Sequencing | 19536157\|37986224 | Unspecified |
| mmu-miR-24-3p | MIMAT0000219 | Ttc3 | 22129 | Sequencing//HITS-CLIP | 19536157\|23142080\|37986224 | Unspecified |
| mmu-miR-24-3p | MIMAT0000219 | Ttyh3 | 78339 | Sequencing//HITS-CLIP | 19536157\|25753418\|37986224 | Unspecified |
| mmu-miR-24-3p | MIMAT0000219 | Txnl1 | 53382 | Sequencing | 19536157\|37986224 | Unspecified |
| mmu-miR-24-3p | MIMAT0000219 | Tyr | 22173 | HITS-CLIP | 21258322 | Unspecified |
| mmu-miR-24-3p | MIMAT0000219 | Ubn1 | 170644 | Sequencing | 37986224\|19536157 | Unspecified |
| mmu-miR-24-3p | MIMAT0000219 | Ubr4 | 69116 | Sequencing//HITS-CLIP | 37986224\|25753418\|19536157 | Unspecified |
| mmu-miR-24-3p | MIMAT0000219 | Ugcg | 22234 | Sequencing//HITS-CLIP | 37986224\|31433990\|25753418\|23142 | Unspecified |
| mmu-miR-24-3p | MIMAT0000219 | Unc5b | 107449 | Sequencing | 19536157\|37986224 | Unspecified |
| mmu-miR-24-3p | MIMAT0000219 | Usp40 | 227334 | Sequencing | 19536157\|37986224 | Unspecified |
| mmu-miR-24-3p | MIMAT0000219 | VSIVgp5 | 1489835 | Luciferase reporter assay | 17613256 | Unspecified |
| mmu-miR-24-3p | MIMAT0000219 | Vav1 | 22324 | HITS-CLIP | 21258322\|23142080 | Unspecified |
| mmu-miR-24-3p | MIMAT0000219 | Vhl | 22346 | Sequencing//HITS-CLIP | 37986224\|23142080\|19536157 | Unspecified |
| mmu-miR-24-3p | MIMAT0000219 | Wdr13 | 73447 | Sequencing//HITS-CLIP | 37986224\|23142080\|19536157 | Unspecified |
| mmu-miR-24-3p | MIMAT0000219 | Wdr59 | 319481 | Sequencing | 19536157\|37986224 | Unspecified |
| mmu-miR-24-3p | MIMAT0000219 | Wdr85 | 67228 | HITS-CLIP | 21258322\|25083871 | Unspecified |
| mmu-miR-24-3p | MIMAT0000219 | Xpo7 | 65246 | Sequencing//HITS-CLIP | 37986224\|23142080\|19536157\|2125€ | Unspecified |
| mmu-miR-24-3p | MIMAT0000219 | Xpr1 | 19775 | Sequencing//HITS-CLIP | 19536157\|23142080\|37986224 | Unspecified |
| mmu-miR-24-3p | MIMAT0000219 | Xxylt1 | 268880 | Sequencing | 19536157 | Unspecified |
| mmu-miR-24-3p | MIMAT0000219 | Zcchc24 | 71918 | Sequencing | 19536157 | Unspecified |
| mmu-miR-24-3p | MIMAT0000219 | Zfp113 | 56314 | HITS-CLIP | 25083871 | Unspecified |
| mmu-miR-24-3p | MIMAT0000219 | Zfp395 | 380912 | Sequencing | 19536157\|37986224 | Unspecified |

| | | | | | | |
|---|---|---|---|---|---|---|
| mmu-miR-24-3p | MIMAT0000219 | Zfp397 | 69256 | Sequencing//HITS-CLIP | 37986224\|25753418\|19536157 | Unspecified |
| mmu-miR-24-3p | MIMAT0000219 | Zfp605 | 675812 | Sequencing | 19536157 | Unspecified |
| mmu-miR-24-3p | MIMAT0000219 | Zfp664 | 269704 | Sequencing | 19536157\|37986224 | Unspecified |
| mmu-miR-24-3p | MIMAT0000219 | Zfp697 | 242109 | Sequencing | 37986224\|19536157 | Unspecified |
| mmu-miR-24-3p | MIMAT0000219 | Zfp704 | 170753 | Sequencing | 19536157\|37986224 | Unspecified |
| mmu-miR-24-3p | MIMAT0000219 | Zfp799 | 240064 | HITS-CLIP | 21258322 | Unspecified |
| mmu-miR-24-3p | MIMAT0000219 | Zfp933 | 242747 | HITS-CLIP | 21258322 | Unspecified |
| mmu-miR-24-3p | MIMAT0000219 | Zhx3 | 320799 | Sequencing | 37986224\|19536157 | Unspecified |
| mmu-miR-24-3p | MIMAT0000219 | Zmat3 | 22401 | Sequencing | 19536157\|37986224 | Unspecified |
| mmu-miR-24-3p | MIMAT0000219 | Zmym3 | 56364 | Sequencing//HITS-CLIP | 37986224\|23142080\|19536157 | Unspecified |
| mmu-miR-24-3p | MIMAT0000219 | Zxdc | 80292 | Sequencing | 37986224\|19536157 | Unspecified |
| mmu-miR-9-5p | MIMAT0000142 | 08-Mar | 71779 | Sequencing | 19536157 | Unspecified |
| mmu-miR-9-5p | MIMAT0000142 | 2700089E24Rik | 381820 | Sequencing | 19536157\|37986224 | Unspecified |
| mmu-miR-9-5p | MIMAT0000142 | 4930402H24Rik | 228602 | Sequencing | 19536157\|37986224 | Unspecified |
| mmu-miR-9-5p | MIMAT0000142 | 4931406P16Rik | 233103 | Sequencing | 19536157\|37986224 | Unspecified |
| mmu-miR-9-5p | MIMAT0000142 | AI504432 | 229694 | Sequencing | 19536157\|37986224 | Unspecified |
| mmu-miR-9-5p | MIMAT0000142 | AK010878 | 100233175 | Sequencing | 19536157 | Unspecified |
| mmu-miR-9-5p | MIMAT0000142 | AW549877 | 106064 | Sequencing | 37986224\|19536157 | Unspecified |
| mmu-miR-9-5p | MIMAT0000142 | AW554918 | 225289 | Sequencing | 37986224\|19536157 | Unspecified |
| mmu-miR-9-5p | MIMAT0000142 | Aatk | 11302 | Sequencing | 19536157\|37986224 | Unspecified |
| mmu-miR-9-5p | MIMAT0000142 | Abca1 | 11303 | Sequencing//HITS-CLIP | 37986224\|19536157 | Unspecified |
| mmu-miR-9-5p | MIMAT0000142 | Abcd1 | 11666 | Sequencing | 19536157\|37986224 | Unspecified |
| mmu-miR-9-5p | MIMAT0000142 | Abcf1 | 224742 | Sequencing | 37986224\|19536157 | Unspecified |
| mmu-miR-9-5p | MIMAT0000142 | Abhd10 | 213012 | Sequencing | 37986224\|19536157 | Unspecified |
| mmu-miR-9-5p | MIMAT0000142 | Abr | 109934 | Sequencing | 37986224\|19536157 | Unspecified |
| mmu-miR-9-5p | MIMAT0000142 | Acer1 | 171168 | HITS-CLIP | 21258322 | Unspecified |
| mmu-miR-9-5p | MIMAT0000142 | Acer3 | 66190 | Sequencing | 37986224\|19536157 | Unspecified |
| mmu-miR-9-5p | MIMAT0000142 | Acly | 104112 | Sequencing | 37986224\|19536157 | Unspecified |
| mmu-miR-9-5p | MIMAT0000142 | Actn4 | 60595 | Sequencing | 19536157\|37986224 | Unspecified |
| mmu-miR-9-5p | MIMAT0000142 | Actr1a | 54130 | Sequencing | 37986224\|19536157 | Unspecified |
| mmu-miR-9-5p | MIMAT0000142 | Adam10 | 11487 | Sequencing | 37986224\|19536157 | Unspecified |
| mmu-miR-9-5p | MIMAT0000142 | Adam11 | 11488 | Sequencing | 37986224\|19536157 | Unspecified |
| mmu-miR-9-5p | MIMAT0000142 | Adamts3 | 330119 | Sequencing | 19536157\|37986224 | Unspecified |
| mmu-miR-9-5p | MIMAT0000142 | Adamts9 | 101401 | Sequencing | 37986224\|19536157 | Unspecified |
| mmu-miR-9-5p | MIMAT0000142 | Adcy1 | 432530 | Sequencing | 37986224\|19536157 | Unspecified |
| mmu-miR-9-5p | MIMAT0000142 | Adcy6 | 11512 | Sequencing | 19536157\|37986224 | Unspecified |
| mmu-miR-9-5p | MIMAT0000142 | Adcyap1r1 | 11517 | Sequencing | 37986224\|19536157 | Unspecified |
| mmu-miR-9-5p | MIMAT0000142 | Agfg1 | 15463 | Sequencing//HITS-CLIP | 37986224\|19536157 | Unspecified |
| mmu-miR-9-5p | MIMAT0000142 | Ak4 | 11639 | Sequencing | 19536157 | Unspecified |

| | | | | | | |
|---|---|---|---|---|---|---|
| mmu-miR-9-5p | MIMAT0000142 | Akap9 | 100986 | Sequencing | 37986224\|19536157 | Unspecified |
| mmu-miR-9-5p | MIMAT0000142 | Akt3 | 23797 | Sequencing | 37986224\|19536157 | Unspecified |
| mmu-miR-9-5p | MIMAT0000142 | Aldh3a2 | 11671 | Sequencing | 37986224\|19536157 | Unspecified |
| mmu-miR-9-5p | MIMAT0000142 | Ambra1 | 228361 | Sequencing | 37986224\|19536157 | Unspecified |
| mmu-miR-9-5p | MIMAT0000142 | Amer1 | 72345 | Sequencing | 19536157 | Unspecified |
| mmu-miR-9-5p | MIMAT0000142 | Amer2 | 72125 | Sequencing | 19536157 | Unspecified |
| mmu-miR-9-5p | MIMAT0000142 | Ank2 | 109676 | Sequencing | 19536157\|37986224 | Unspecified |
| mmu-miR-9-5p | MIMAT0000142 | Ank3 | 11735 | Sequencing | 19536157\|37986224 | Unspecified |
| mmu-miR-9-5p | MIMAT0000142 | Ankrd12 | 106585 | Sequencing | 37986224\|19536157 | Unspecified |
| mmu-miR-9-5p | MIMAT0000142 | Ankrd40 | 71452 | Sequencing | 19536157\|37986224 | Unspecified |
| mmu-miR-9-5p | MIMAT0000142 | Ankrd63 | 383787 | Sequencing | 19536157 | Unspecified |
| mmu-miR-9-5p | MIMAT0000142 | Antxr2 | 71914 | Sequencing | 37986224\|19536157 | Unspecified |
| mmu-miR-9-5p | MIMAT0000142 | Ap1s2 | 108012 | Sequencing | 19536157\|37986224 | Unspecified |
| mmu-miR-9-5p | MIMAT0000142 | Aqp4 | 11829 | Sequencing | 37986224\|19536157 | Unspecified |
| mmu-miR-9-5p | MIMAT0000142 | Arf2 | 11841 | Sequencing | 37986224\|19536157 | Unspecified |
| mmu-miR-9-5p | MIMAT0000142 | Arhgap6 | 11856 | Sequencing | 19536157\|37986224 | Unspecified |
| mmu-miR-9-5p | MIMAT0000142 | Arhgdia | 192662 | Sequencing | 37986224\|19536157 | Unspecified |
| mmu-miR-9-5p | MIMAT0000142 | Arid1a | 93760 | Sequencing | 19536157\|37986224 | Unspecified |
| mmu-miR-9-5p | MIMAT0000142 | Arid4a | 238247 | Sequencing | 19536157\|37986224 | Unspecified |
| mmu-miR-9-5p | MIMAT0000142 | Armcx2 | 67416 | Sequencing | 19536157\|37986224 | Unspecified |
| mmu-miR-9-5p | MIMAT0000142 | Arpc1b | 11867 | Sequencing | 19536157\|37986224 | Unspecified |
| mmu-miR-9-5p | MIMAT0000142 | Arpp21 | 74100 | Sequencing | 37986224\|19536157 | Unspecified |
| mmu-miR-9-5p | MIMAT0000142 | Asb7 | 117589 | Sequencing | 19536157\|37986224 | Unspecified |
| mmu-miR-9-5p | MIMAT0000142 | Asxl3 | 211961 | Sequencing | 19536157\|37986224 | Unspecified |
| mmu-miR-9-5p | MIMAT0000142 | Atf1 | 11908 | Sequencing | 37986224\|19536157 | Unspecified |
| mmu-miR-9-5p | MIMAT0000142 | Atp1a2 | 98660 | Sequencing | 37986224\|19536157 | Unspecified |
| mmu-miR-9-5p | MIMAT0000142 | Atp1b1 | 11931 | Sequencing//HITS-CLIP | 19536157\|37986224 | Unspecified |
| mmu-miR-9-5p | MIMAT0000142 | Atp2b3 | 320707 | Sequencing | 37986224\|19536157 | Unspecified |
| mmu-miR-9-5p | MIMAT0000142 | Atp6v0a1 | 11975 | Sequencing | 19536157\|37986224 | Unspecified |
| mmu-miR-9-5p | MIMAT0000142 | Atp8a1 | 11980 | Sequencing | 19536157\|37986224 | Unspecified |
| mmu-miR-9-5p | MIMAT0000142 | B3galt1 | 26877 | Sequencing | 37986224\|19536157 | Unspecified |
| mmu-miR-9-5p | MIMAT0000142 | B4galnt1 | 14421 | Sequencing | 37986224\|19536157 | Unspecified |
| mmu-miR-9-5p | MIMAT0000142 | B4galt1 | 14595 | Sequencing | 37986224\|19536157 | Unspecified |
| mmu-miR-9-5p | MIMAT0000142 | B4galt5 | 56336 | Sequencing | 37986224\|19536157 | Unspecified |
| mmu-miR-9-5p | MIMAT0000142 | Bbs1 | 52028 | Sequencing | 19536157\|37986224 | Unspecified |
| mmu-miR-9-5p | MIMAT0000142 | Bche | 12038 | Sequencing | 19536157\|37986224 | Unspecified |
| mmu-miR-9-5p | MIMAT0000142 | Bcl2l11 | 12125 | HITS-CLIP | 23597149 | Unspecified |
| mmu-miR-9-5p | MIMAT0000142 | Bcl6 | 12053 | Sequencing | 37986224\|19536157 | Unspecified |
| mmu-miR-9-5p | MIMAT0000142 | Bclaf1 | 72567 | Sequencing | 37986224\|19536157 | Unspecified |

| | | | | | | | |
|---|---|---|---|---|---|---|---|
| mmu-miR-9-5p | MIMAT0000142 | Bend6 | 320705 | Sequencing | 37986224\|19536157 | | Unspecified |
| mmu-miR-9-5p | MIMAT0000142 | Bmp2 | 12156 | Sequencing | 37986224\|19536157 | | Unspecified |
| mmu-miR-9-5p | MIMAT0000142 | Bptf | 207165 | Sequencing | 19536157\|37986224 | | Unspecified |
| mmu-miR-9-5p | MIMAT0000142 | Bsn | 12217 | Sequencing | 37986224\|19536157 | | Unspecified |
| mmu-miR-9-5p | MIMAT0000142 | C1ql3 | 227580 | Sequencing | 19536157\|37986224 | | Unspecified |
| mmu-miR-9-5p | MIMAT0000142 | Caly | 68566 | Sequencing | 37986224\|19536157 | | Unspecified |
| mmu-miR-9-5p | MIMAT0000142 | Camkk2 | 207565 | Luciferase assay//Western blot//Luciferase | | 24603903 | Unspecified |
| mmu-miR-9-5p | MIMAT0000142 | Camta1 | 100072 | Sequencing | 37986224\|19536157 | | Unspecified |
| mmu-miR-9-5p | MIMAT0000142 | Car7 | 12354 | Sequencing | 37986224\|19536157 | | Unspecified |
| mmu-miR-9-5p | MIMAT0000142 | Carm1 | 59035 | Sequencing | 37986224\|19536157 | | Unspecified |
| mmu-miR-9-5p | MIMAT0000142 | Cbln4 | 228942 | Sequencing | 19536157\|37986224 | | Unspecified |
| mmu-miR-9-5p | MIMAT0000142 | Ccdc50 | 67501 | Sequencing | 37986224\|19536157 | | Unspecified |
| mmu-miR-9-5p | MIMAT0000142 | Ccdc6 | 76551 | Sequencing | | 19536157 | Unspecified |
| mmu-miR-9-5p | MIMAT0000142 | Ccdc91 | 67015 | Sequencing | 37986224\|19536157 | | Unspecified |
| mmu-miR-9-5p | MIMAT0000142 | Ccnf | 12449 | Sequencing | 19536157\|37986224 | | Unspecified |
| mmu-miR-9-5p | MIMAT0000142 | Ccser2 | 72972 | Sequencing//HITS-CLIP | | 19536157 | Unspecified |
| mmu-miR-9-5p | MIMAT0000142 | Cd151 | 12476 | Sequencing | 19536157\|37986224 | | Unspecified |
| mmu-miR-9-5p | MIMAT0000142 | Cdan1 | 68968 | Sequencing | 37986224\|19536157 | | Unspecified |
| mmu-miR-9-5p | MIMAT0000142 | Cdc14b | 218294 | HITS-CLIP | | 21258322 | Unspecified |
| mmu-miR-9-5p | MIMAT0000142 | Cdc73 | 214498 | Sequencing | 37986224\|19536157 | | Unspecified |
| mmu-miR-9-5p | MIMAT0000142 | Cdh10 | 320873 | Sequencing | 19536157\|37986224 | | Unspecified |
| mmu-miR-9-5p | MIMAT0000142 | Cdk13 | 69562 | Sequencing | | 19536157 | Unspecified |
| mmu-miR-9-5p | MIMAT0000142 | Cds2 | 110911 | Sequencing | 37986224\|19536157 | | Unspecified |
| mmu-miR-9-5p | MIMAT0000142 | Cecr2 | 330409 | Sequencing | 37986224\|19536157 | | Unspecified |
| mmu-miR-9-5p | MIMAT0000142 | Celf6 | 76183 | Sequencing | | 19536157 | Unspecified |
| mmu-miR-9-5p | MIMAT0000142 | Celsr2 | 53883 | Sequencing | 19536157\|37986224 | | Unspecified |
| mmu-miR-9-5p | MIMAT0000142 | Chl1 | 12661 | Sequencing | 37986224\|19536157 | | Unspecified |
| mmu-miR-9-5p | MIMAT0000142 | Chpt1 | 212862 | Sequencing | 19536157\|37986224 | | Unspecified |
| mmu-miR-9-5p | MIMAT0000142 | Chst11 | 58250 | Sequencing | 37986224\|19536157 | | Unspecified |
| mmu-miR-9-5p | MIMAT0000142 | Chst15 | 77590 | Sequencing | | 19536157 | Unspecified |
| mmu-miR-9-5p | MIMAT0000142 | Cib2 | 56506 | Sequencing | 19536157\|37986224 | | Unspecified |
| mmu-miR-9-5p | MIMAT0000142 | Clasp1 | 76707 | Sequencing | 19536157\|37986224 | | Unspecified |
| mmu-miR-9-5p | MIMAT0000142 | Clip1 | 56430 | Sequencing//HITS-CLIP | 37986224\|19536157 | | Unspecified |
| mmu-miR-9-5p | MIMAT0000142 | Clock | 12753 | Sequencing | 19536157\|37986224 | | Unspecified |
| mmu-miR-9-5p | MIMAT0000142 | Clp1 | 98985 | Sequencing | 37986224\|19536157 | | Unspecified |
| mmu-miR-9-5p | MIMAT0000142 | Clstn1 | 65945 | Sequencing | 19536157\|37986224 | | Unspecified |
| mmu-miR-9-5p | MIMAT0000142 | Cnnm2 | 94219 | Sequencing//HITS-CLIP | 37986224\|19536157 | | Unspecified |
| mmu-miR-9-5p | MIMAT0000142 | Cnrip1 | 380686 | Sequencing | 37986224\|19536157 | | Unspecified |
| mmu-miR-9-5p | MIMAT0000142 | Cntn3 | 18488 | Sequencing//HITS-CLIP | 37986224\|19536157 | | Unspecified |

| | | | | | | |
|---|---|---|---|---|---|---|
| mmu-miR-9-5p | MIMAT0000142 | Col4a1 | 12826 | Sequencing | 37986224\|19536157 | Unspecified |
| mmu-miR-9-5p | MIMAT0000142 | Col4a2 | 12827 | Sequencing | 37986224\|19536157 | Unspecified |
| mmu-miR-9-5p | MIMAT0000142 | Colec12 | 140792 | Sequencing//HITS-CLIP | 19536157\|37986224 | Unspecified |
| mmu-miR-9-5p | MIMAT0000142 | Cox6c | 12864 | Sequencing | 19536157\|37986224 | Unspecified |
| mmu-miR-9-5p | MIMAT0000142 | Cpeb2 | 231207 | Sequencing | 19536157\|37986224 | Unspecified |
| mmu-miR-9-5p | MIMAT0000142 | Cplx2 | 12890 | Sequencing | 19536157\|37986224 | Unspecified |
| mmu-miR-9-5p | MIMAT0000142 | Creb1 | 12912 | Sequencing | 19536157\|37986224 | Unspecified |
| mmu-miR-9-5p | MIMAT0000142 | Creb3l2 | 208647 | Sequencing | 19536157\|37986224 | Unspecified |
| mmu-miR-9-5p | MIMAT0000142 | Crebrf | 77128 | Sequencing//HITS-CLIP | 19536157 | Unspecified |
| mmu-miR-9-5p | MIMAT0000142 | Cspp1 | 211660 | Sequencing | 37986224\|19536157 | Unspecified |
| mmu-miR-9-5p | MIMAT0000142 | Csrnp2 | 207785 | Sequencing | 37986224\|19536157 | Unspecified |
| mmu-miR-9-5p | MIMAT0000142 | Ctdsp1 | 227292 | Sequencing | 37986224\|19536157 | Unspecified |
| mmu-miR-9-5p | MIMAT0000142 | Ctdsp2 | 52468 | Sequencing | 37986224\|19536157 | Unspecified |
| mmu-miR-9-5p | MIMAT0000142 | Ctgf | 14219 | Sequencing | 37986224\|19536157 | Unspecified |
| mmu-miR-9-5p | MIMAT0000142 | Ctnnb1 | 12387 | Sequencing | 37986224\|19536157 | Unspecified |
| mmu-miR-9-5p | MIMAT0000142 | Cxcl12 | 20315 | Sequencing | 19536157\|37986224 | Unspecified |
| mmu-miR-9-5p | MIMAT0000142 | Cxcr4 | 12767 | Sequencing | 37986224\|19536157 | Unspecified |
| mmu-miR-9-5p | MIMAT0000142 | Cxxc5 | 67393 | Sequencing | 37986224\|19536157 | Unspecified |
| mmu-miR-9-5p | MIMAT0000142 | D3Bwg0562e | 229791 | Sequencing | 37986224\|19536157 | Unspecified |
| mmu-miR-9-5p | MIMAT0000142 | D430041D05Rik | 241589 | Sequencing | 37986224\|19536157 | Unspecified |
| mmu-miR-9-5p | MIMAT0000142 | Dact3 | 629378 | Sequencing | 19536157\|37986224 | Unspecified |
| mmu-miR-9-5p | MIMAT0000142 | Dbt | 13171 | Sequencing | 19536157\|37986224 | Unspecified |
| mmu-miR-9-5p | MIMAT0000142 | Ddx17 | 67040 | Sequencing | 19536157\|37986224 | Unspecified |
| mmu-miR-9-5p | MIMAT0000142 | Dhcr24 | 74754 | Sequencing | 37986224\|19536157 | Unspecified |
| mmu-miR-9-5p | MIMAT0000142 | Diras2 | 68203 | Sequencing | 19536157\|37986224 | Unspecified |
| mmu-miR-9-5p | MIMAT0000142 | Disp2 | 214240 | Sequencing | 37986224\|19536157 | Unspecified |
| mmu-miR-9-5p | MIMAT0000142 | Dlg4 | 13385 | Sequencing | 37986224\|19536157 | Unspecified |
| mmu-miR-9-5p | MIMAT0000142 | Dmtf1 | 23857 | Sequencing | 19536157\|37986224 | Unspecified |
| mmu-miR-9-5p | MIMAT0000142 | Dnajb4 | 67035 | Sequencing | 19536157\|37986224 | Unspecified |
| mmu-miR-9-5p | MIMAT0000142 | Dnajc14 | 74330 | Sequencing//HITS-CLIP | 37986224\|19536157 | Unspecified |
| mmu-miR-9-5p | MIMAT0000142 | Dock9 | 105445 | Sequencing | 37986224\|19536157 | Unspecified |
| mmu-miR-9-5p | MIMAT0000142 | Dram2 | 67171 | Sequencing | 19536157 | Unspecified |
| mmu-miR-9-5p | MIMAT0000142 | Dtna | 13527 | Sequencing | 37986224\|19536157 | Unspecified |
| mmu-miR-9-5p | MIMAT0000142 | Dusp6 | 67603 | Sequencing | 19536157\|37986224 | Unspecified |
| mmu-miR-9-5p | MIMAT0000142 | Dzank1 | 241688 | Sequencing | 19536157 | Unspecified |
| mmu-miR-9-5p | MIMAT0000142 | E130309D14Rik | 432582 | Sequencing | 37986224\|19536157 | Unspecified |
| mmu-miR-9-5p | MIMAT0000142 | Eaf1 | 74427 | Sequencing | 19536157\|37986224 | Unspecified |
| mmu-miR-9-5p | MIMAT0000142 | Eef1d | 66656 | Sequencing | 37986224\|19536157 | Unspecified |
| mmu-miR-9-5p | MIMAT0000142 | Efnb2 | 13642 | Sequencing | 19536157\|37986224 | Unspecified |

| | | | | | | | |
|---|---|---|---|---|---|---|---|
| mmu-miR-9-5p | MIMAT0000142 | Egr1 | 13653 | Sequencing | 19536157\|37986224 | | Unspecified |
| mmu-miR-9-5p | MIMAT0000142 | Eif1a | 13664 | Sequencing//HITS-CLIP | 19536157\|37986224 | | Unspecified |
| mmu-miR-9-5p | MIMAT0000142 | Elfn2 | 207393 | Sequencing | 19536157\|37986224 | | Unspecified |
| mmu-miR-9-5p | MIMAT0000142 | Elmod2 | 244548 | Sequencing | 19536157\|37986224 | | Unspecified |
| mmu-miR-9-5p | MIMAT0000142 | Elmod3 | 232089 | Sequencing | 37986224\|19536157 | | Unspecified |
| mmu-miR-9-5p | MIMAT0000142 | Elovl2 | 54326 | Sequencing | 37986224\|19536157 | | Unspecified |
| mmu-miR-9-5p | MIMAT0000142 | Elovl5 | 68801 | Sequencing | 37986224\|19536157 | | Unspecified |
| mmu-miR-9-5p | MIMAT0000142 | Elovl6 | 170439 | Sequencing | 19536157\|37986224 | | Unspecified |
| mmu-miR-9-5p | MIMAT0000142 | Eml1 | 68519 | Sequencing | 37986224\|19536157 | | Unspecified |
| mmu-miR-9-5p | MIMAT0000142 | Eml5 | 319670 | Sequencing | 19536157\|37986224 | | Unspecified |
| mmu-miR-9-5p | MIMAT0000142 | En2 | 13799 | Sequencing | 37986224\|19536157 | | Unspecified |
| mmu-miR-9-5p | MIMAT0000142 | Epas1 | 13819 | Sequencing | 19536157\|37986224 | | Unspecified |
| mmu-miR-9-5p | MIMAT0000142 | Epm2aip1 | 77781 | Sequencing | 19536157\|37986224 | | Unspecified |
| mmu-miR-9-5p | MIMAT0000142 | Eps15 | 13858 | Sequencing | 37986224\|19536157 | | Unspecified |
| mmu-miR-9-5p | MIMAT0000142 | Erc2 | 238988 | Sequencing | 19536157\|37986224 | | Unspecified |
| mmu-miR-9-5p | MIMAT0000142 | Erlin1 | 226144 | Sequencing | 37986224\|19536157 | | Unspecified |
| mmu-miR-9-5p | MIMAT0000142 | Esyt2 | 52635 | Sequencing | | 19536157 | Unspecified |
| mmu-miR-9-5p | MIMAT0000142 | Exoc2 | 66482 | Sequencing | 19536157\|37986224 | | Unspecified |
| mmu-miR-9-5p | MIMAT0000142 | Fads2 | 56473 | Sequencing | 37986224\|19536157 | | Unspecified |
| mmu-miR-9-5p | MIMAT0000142 | Fam101b | 76566 | Sequencing | 19536157\|37986224 | | Unspecified |
| mmu-miR-9-5p | MIMAT0000142 | Fam115a | 77574 | Sequencing | 19536157\|37986224 | | Unspecified |
| mmu-miR-9-5p | MIMAT0000142 | Fam219a | 71901 | Sequencing | | 19536157 | Unspecified |
| mmu-miR-9-5p | MIMAT0000142 | Fam43a | 224093 | Sequencing | 37986224\|19536157 | | Unspecified |
| mmu-miR-9-5p | MIMAT0000142 | Fam53c | 66306 | Sequencing | | 19536157 | Unspecified |
| mmu-miR-9-5p | MIMAT0000142 | Fam63b | 235461 | Sequencing | | 19536157 | Unspecified |
| mmu-miR-9-5p | MIMAT0000142 | Fam76a | 230789 | Sequencing | 37986224\|19536157 | | Unspecified |
| mmu-miR-9-5p | MIMAT0000142 | Fbxl16 | 214931 | Sequencing | 19536157\|37986224 | | Unspecified |
| mmu-miR-9-5p | MIMAT0000142 | Fbxl17 | 50758 | HITS-CLIP | | 21258322 | Unspecified |
| mmu-miR-9-5p | MIMAT0000142 | Fbxw11 | 103583 | Sequencing//HITS-CLIP | 19536157\|37986224 | | Unspecified |
| mmu-miR-9-5p | MIMAT0000142 | Fem1c | 240263 | Sequencing | 37986224\|19536157 | | Unspecified |
| mmu-miR-9-5p | MIMAT0000142 | Flnb | 286940 | Sequencing | 37986224\|19536157 | | Unspecified |
| mmu-miR-9-5p | MIMAT0000142 | Flt1 | 14254 | Sequencing | 19536157\|37986224 | | Unspecified |
| mmu-miR-9-5p | MIMAT0000142 | Flt3 | 14255 | Sequencing | 37986224\|19536157 | | Unspecified |
| mmu-miR-9-5p | MIMAT0000142 | Fmn1 | 14260 | Sequencing | 37986224\|19536157 | | Unspecified |
| mmu-miR-9-5p | MIMAT0000142 | Fmnl2 | 71409 | Sequencing | 19536157\|37986224 | | Unspecified |
| mmu-miR-9-5p | MIMAT0000142 | Fnip1 | 216742 | Sequencing | 37986224\|19536157 | | Unspecified |
| mmu-miR-9-5p | MIMAT0000142 | Fnip2 | 329679 | Sequencing | 37986224\|19536157 | | Unspecified |
| mmu-miR-9-5p | MIMAT0000142 | Foxg1 | 15228 | Luciferase assay//Luciferase reporter assay | | 18842901 | Unspecified |
| mmu-miR-9-5p | MIMAT0000142 | Foxk2 | 68837 | Sequencing | 37986224\|19536157 | | Unspecified |

| | | | | | | |
|---|---|---|---|---|---|---|
| mmu-miR-9-5p | MIMAT0000142 | Foxn2 | 14236 | Sequencing | 37986224\|19536157 | Unspecified |
| mmu-miR-9-5p | MIMAT0000142 | Foxo3 | 56484 | HITS-CLIP//Immunoblot//Immunohistoche | 37986224\|19536157\|26470727 | Unspecified |
| mmu-miR-9-5p | MIMAT0000142 | Foxp4 | 74123 | Sequencing//HITS-CLIP | 37986224\|19536157 | Unspecified |
| mmu-miR-9-5p | MIMAT0000142 | Frmd4a | 209630 | Sequencing | 19536157\|37986224 | Unspecified |
| mmu-miR-9-5p | MIMAT0000142 | Fstl1 | 14314 | Sequencing | 37986224\|19536157 | Unspecified |
| mmu-miR-9-5p | MIMAT0000142 | Furin | 18550 | Sequencing | 19536157\|37986224 | Unspecified |
| mmu-miR-9-5p | MIMAT0000142 | Fut9 | 14348 | Sequencing | 37986224\|19536157 | Unspecified |
| mmu-miR-9-5p | MIMAT0000142 | Fxr1 | 14359 | Sequencing | 37986224\|19536157 | Unspecified |
| mmu-miR-9-5p | MIMAT0000142 | Fyttd1 | 69823 | Sequencing//HITS-CLIP | 37986224\|19536157 | Unspecified |
| mmu-miR-9-5p | MIMAT0000142 | Gabra1 | 14394 | Sequencing | 37986224\|19536157 | Unspecified |
| mmu-miR-9-5p | MIMAT0000142 | Gfap | 14580 | Sequencing | 37986224\|19536157 | Unspecified |
| mmu-miR-9-5p | MIMAT0000142 | Ggt7 | 207182 | Sequencing | 19536157\|37986224 | Unspecified |
| mmu-miR-9-5p | MIMAT0000142 | Gjc3 | 118446 | Sequencing | 19536157\|37986224 | Unspecified |
| mmu-miR-9-5p | MIMAT0000142 | Gm12258 | 237769 | Sequencing | 19536157 | Unspecified |
| mmu-miR-9-5p | MIMAT0000142 | Gm2a | 14667 | Sequencing | 37986224\|19536157 | Unspecified |
| mmu-miR-9-5p | MIMAT0000142 | Gnai1 | 14677 | Sequencing | 37986224\|19536157 | Unspecified |
| mmu-miR-9-5p | MIMAT0000142 | Gpr88 | 64378 | Sequencing | 37986224\|19536157 | Unspecified |
| mmu-miR-9-5p | MIMAT0000142 | Gramd1b | 235283 | Sequencing | 37986224\|19536157 | Unspecified |
| mmu-miR-9-5p | MIMAT0000142 | Grhl1 | 195733 | Sequencing | 37986224\|19536157 | Unspecified |
| mmu-miR-9-5p | MIMAT0000142 | Gria2 | 14800 | Sequencing | 19536157\|37986224 | Unspecified |
| mmu-miR-9-5p | MIMAT0000142 | Gria4 | 14802 | Sequencing | 19536157\|37986224 | Unspecified |
| mmu-miR-9-5p | MIMAT0000142 | Grin2b | 14812 | Sequencing//HITS-CLIP | 19536157\|37986224 | Unspecified |
| mmu-miR-9-5p | MIMAT0000142 | Gtdc1 | 227835 | Sequencing | 19536157\|37986224 | Unspecified |
| mmu-miR-9-5p | MIMAT0000142 | Gtf3c4 | 269252 | Sequencing | 19536157\|37986224 | Unspecified |
| mmu-miR-9-5p | MIMAT0000142 | Gucy1b3 | 54195 | Sequencing | 37986224\|19536157 | Unspecified |
| mmu-miR-9-5p | MIMAT0000142 | Guk1 | 14923 | Sequencing | 19536157\|37986224 | Unspecified |
| mmu-miR-9-5p | MIMAT0000142 | Gzf1 | 74533 | Sequencing | 19536157\|37986224 | Unspecified |
| mmu-miR-9-5p | MIMAT0000142 | Hapln1 | 12950 | Sequencing | 37986224\|19536157 | Unspecified |
| mmu-miR-9-5p | MIMAT0000142 | Hdac2 | 15182 | Sequencing | 37986224\|19536157 | Unspecified |
| mmu-miR-9-5p | MIMAT0000142 | Hdgfrp3 | 29877 | Sequencing | 37986224\|19536157 | Unspecified |
| mmu-miR-9-5p | MIMAT0000142 | Heatr5a | 320487 | Sequencing | 19536157\|37986224 | Unspecified |
| mmu-miR-9-5p | MIMAT0000142 | Hecw2 | 329152 | Sequencing | 37986224\|19536157 | Unspecified |
| mmu-miR-9-5p | MIMAT0000142 | Hes1 | 15205 | Luciferase assay//Luciferase reporter assay | 21238922\|22840391 | Unspecified |
| mmu-miR-9-5p | MIMAT0000142 | Hipk1 | 15257 | Sequencing//HITS-CLIP | 19536157\|37986224 | Unspecified |
| mmu-miR-9-5p | MIMAT0000142 | Hist3h2ba | 78303 | Sequencing | 37986224\|19536157 | Unspecified |
| mmu-miR-9-5p | MIMAT0000142 | Hpcal4 | 170638 | Sequencing | 19536157\|37986224 | Unspecified |
| mmu-miR-9-5p | MIMAT0000142 | Hs6st1 | 50785 | Sequencing | 37986224\|19536157 | Unspecified |
| mmu-miR-9-5p | MIMAT0000142 | Igf2 | 16002 | Sequencing | 37986224\|19536157 | Unspecified |
| mmu-miR-9-5p | MIMAT0000142 | Il1rap | 16180 | Sequencing | 37986224\|19536157 | Unspecified |

| | | | | | | | |
|---|---|---|---|---|---|---|---|
| mmu-miR-9-5p | MIMAT0000142 | Ilvbl | 216136 | HITS-CLIP | | 21258322 | Unspecified |
| mmu-miR-9-5p | MIMAT0000142 | Impa1 | 55980 | Sequencing//HITS-CLIP | 37986224\|19536157 | | Unspecified |
| mmu-miR-9-5p | MIMAT0000142 | Ist1 | 71955 | Sequencing | | 19536157 | Unspecified |
| mmu-miR-9-5p | MIMAT0000142 | Isyna1 | 71780 | Sequencing | 37986224\|19536157 | | Unspecified |
| mmu-miR-9-5p | MIMAT0000142 | Itga4 | 16401 | Sequencing | 37986224\|19536157 | | Unspecified |
| mmu-miR-9-5p | MIMAT0000142 | Itm2c | 64294 | Sequencing | 37986224\|19536157 | | Unspecified |
| mmu-miR-9-5p | MIMAT0000142 | Jakmip2 | 76217 | Sequencing | 19536157\|37986224 | | Unspecified |
| mmu-miR-9-5p | MIMAT0000142 | Kalrn | 545156 | Sequencing | 19536157\|37986224 | | Unspecified |
| mmu-miR-9-5p | MIMAT0000142 | Kcnc1 | 16502 | Sequencing//HITS-CLIP | 19536157\|37986224 | | Unspecified |
| mmu-miR-9-5p | MIMAT0000142 | Kcnh5 | 238271 | Sequencing | 37986224\|19536157 | | Unspecified |
| mmu-miR-9-5p | MIMAT0000142 | Kcns2 | 16539 | Sequencing | 19536157\|37986224 | | Unspecified |
| mmu-miR-9-5p | MIMAT0000142 | Kif21a | 16564 | Sequencing | 19536157\|37986224 | | Unspecified |
| mmu-miR-9-5p | MIMAT0000142 | Kif5a | 16572 | Sequencing | 19536157\|37986224 | | Unspecified |
| mmu-miR-9-5p | MIMAT0000142 | Kif5c | 16574 | Sequencing | 19536157\|37986224 | | Unspecified |
| mmu-miR-9-5p | MIMAT0000142 | Klf7 | 93691 | Sequencing | 37986224\|19536157 | | Unspecified |
| mmu-miR-9-5p | MIMAT0000142 | Klf8 | 245671 | Sequencing | 37986224\|19536157 | | Unspecified |
| mmu-miR-9-5p | MIMAT0000142 | Klhl21 | 242785 | Sequencing | 37986224\|19536157 | | Unspecified |
| mmu-miR-9-5p | MIMAT0000142 | Kmt2a | 214162 | Sequencing | | 19536157 | Unspecified |
| mmu-miR-9-5p | MIMAT0000142 | Ky | 16716 | HITS-CLIP | | 21258322 | Unspecified |
| mmu-miR-9-5p | MIMAT0000142 | Lancl3 | 236285 | Sequencing | 37986224\|19536157 | | Unspecified |
| mmu-miR-9-5p | MIMAT0000142 | Larp4 | 207214 | Sequencing | 37986224\|19536157 | | Unspecified |
| mmu-miR-9-5p | MIMAT0000142 | Lfng | 16848 | Sequencing | 19536157\|37986224 | | Unspecified |
| mmu-miR-9-5p | MIMAT0000142 | Lgi1 | 56839 | Sequencing | 19536157\|37986224 | | Unspecified |
| mmu-miR-9-5p | MIMAT0000142 | Lin28b | 380669 | Sequencing | 19536157\|37986224 | | Unspecified |
| mmu-miR-9-5p | MIMAT0000142 | Lmbrd1 | 68421 | Sequencing | 37986224\|19536157 | | Unspecified |
| mmu-miR-9-5p | MIMAT0000142 | Lmbrd2 | 320506 | Sequencing | 19536157\|37986224 | | Unspecified |
| mmu-miR-9-5p | MIMAT0000142 | Lonrf2 | 381338 | Sequencing | 37986224\|19536157 | | Unspecified |
| mmu-miR-9-5p | MIMAT0000142 | Lphn3 | 319387 | Sequencing | 19536157\|37986224 | | Unspecified |
| mmu-miR-9-5p | MIMAT0000142 | Lrig2 | 269473 | Sequencing | 37986224\|19536157 | | Unspecified |
| mmu-miR-9-5p | MIMAT0000142 | Lrrc58 | 320184 | Sequencing | 19536157\|37986224 | | Unspecified |
| mmu-miR-9-5p | MIMAT0000142 | Lsamp | 268890 | Sequencing//HITS-CLIP | 37986224\|19536157 | | Unspecified |
| mmu-miR-9-5p | MIMAT0000142 | Lzts3 | 241638 | Sequencing//HITS-CLIP | | 19536157 | Unspecified |
| mmu-miR-9-5p | MIMAT0000142 | M6pr | 17113 | Sequencing | 37986224\|19536157 | | Unspecified |
| mmu-miR-9-5p | MIMAT0000142 | Maea | 59003 | Sequencing | 37986224\|19536157 | | Unspecified |
| mmu-miR-9-5p | MIMAT0000142 | Magt1 | 67075 | Sequencing | 37986224\|19536157 | | Unspecified |
| mmu-miR-9-5p | MIMAT0000142 | Man1a2 | 17156 | Sequencing | 19536157\|37986224 | | Unspecified |
| mmu-miR-9-5p | MIMAT0000142 | Map6 | 17760 | Sequencing | | 19536157 | Unspecified |
| mmu-miR-9-5p | MIMAT0000142 | Mapk4 | 225724 | Sequencing | 37986224\|19536157 | | Unspecified |
| mmu-miR-9-5p | MIMAT0000142 | Mapt | 17762 | Sequencing | 37986224\|19536157 | | Unspecified |

| | | | | | | |
|---|---|---|---|---|---|---|
| mmu-miR-9-5p | MIMAT0000142 | 08-Mar | 71779 | Sequencing | 37986224\|19536157 | Unspecified |
| mmu-miR-9-5p | MIMAT0000142 | Marchf6 | 223455 | Sequencing//HITS-CLIP | 37986224\|19536157 | Unspecified |
| mmu-miR-9-5p | MIMAT0000142 | Mdga2 | 320772 | Sequencing//HITS-CLIP | 37986224\|19536157 | Unspecified |
| mmu-miR-9-5p | MIMAT0000142 | Mdk | 17242 | Sequencing | 37986224\|19536157 | Unspecified |
| mmu-miR-9-5p | MIMAT0000142 | Megf10 | 70417 | Sequencing | 19536157\|37986224 | Unspecified |
| mmu-miR-9-5p | MIMAT0000142 | Megf9 | 230316 | Sequencing | 19536157\|37986224 | Unspecified |
| mmu-miR-9-5p | MIMAT0000142 | Mkx | 210719 | Sequencing | 19536157\|37986224 | Unspecified |
| mmu-miR-9-5p | MIMAT0000142 | Mmd | 67468 | Sequencing | 19536157\|37986224 | Unspecified |
| mmu-miR-9-5p | MIMAT0000142 | Mmp16 | 17389 | Sequencing | 37986224\|19536157 | Unspecified |
| mmu-miR-9-5p | MIMAT0000142 | Morc2a | 74522 | Sequencing | 19536157\|37986224 | Unspecified |
| mmu-miR-9-5p | MIMAT0000142 | Mtmr4 | 170749 | Sequencing | 19536157\|37986224 | Unspecified |
| mmu-miR-9-5p | MIMAT0000142 | Mycn | 18109 | Immunofluorescence//qRT-PCR//Western | 26238256 | Unspecified |
| mmu-miR-9-5p | MIMAT0000142 | Myo10 | 17909 | Sequencing | 19536157\|37986224 | Unspecified |
| mmu-miR-9-5p | MIMAT0000142 | Myo18a | 360013 | Sequencing | 37986224\|19536157 | Unspecified |
| mmu-miR-9-5p | MIMAT0000142 | Myo5a | 17918 | Sequencing | 19536157\|37986224 | Unspecified |
| mmu-miR-9-5p | MIMAT0000142 | Myo6 | 17920 | Sequencing | 37986224\|19536157 | Unspecified |
| mmu-miR-9-5p | MIMAT0000142 | N28178 | 230085 | Sequencing | 19536157\|37986224 | Unspecified |
| mmu-miR-9-5p | MIMAT0000142 | Narf | 67608 | Sequencing | 19536157\|37986224 | Unspecified |
| mmu-miR-9-5p | MIMAT0000142 | Ncor1 | 20185 | Sequencing | 19536157\|37986224 | Unspecified |
| mmu-miR-9-5p | MIMAT0000142 | Ncor2 | 20602 | Sequencing | 19536157\|37986224 | Unspecified |
| mmu-miR-9-5p | MIMAT0000142 | Ndrg2 | 29811 | Sequencing | 19536157\|37986224 | Unspecified |
| mmu-miR-9-5p | MIMAT0000142 | Nedd4 | 17999 | Sequencing//HITS-CLIP | 37986224\|19536157 | Unspecified |
| mmu-miR-9-5p | MIMAT0000142 | Nek9 | 217718 | Sequencing | 19536157\|37986224 | Unspecified |
| mmu-miR-9-5p | MIMAT0000142 | Nelfb | 58202 | Sequencing | 19536157 | Unspecified |
| mmu-miR-9-5p | MIMAT0000142 | Neto2 | 74513 | Sequencing | 19536157\|37986224 | Unspecified |
| mmu-miR-9-5p | MIMAT0000142 | Nf1 | 18015 | Sequencing//HITS-CLIP | 19536157\|37986224 | Unspecified |
| mmu-miR-9-5p | MIMAT0000142 | Nhsl1 | 215819 | Sequencing | 37986224\|19536157 | Unspecified |
| mmu-miR-9-5p | MIMAT0000142 | Nkrf | 77286 | Sequencing | 37986224\|19536157 | Unspecified |
| mmu-miR-9-5p | MIMAT0000142 | Nmt2 | 18108 | Sequencing | 37986224\|19536157 | Unspecified |
| mmu-miR-9-5p | MIMAT0000142 | Nos1 | 18125 | Sequencing | 19536157\|37986224 | Unspecified |
| mmu-miR-9-5p | MIMAT0000142 | Npc1 | 18145 | Sequencing | 37986224\|19536157 | Unspecified |
| mmu-miR-9-5p | MIMAT0000142 | Npc2 | 67963 | Sequencing | 19536157\|37986224 | Unspecified |
| mmu-miR-9-5p | MIMAT0000142 | Nptx1 | 18164 | Sequencing//HITS-CLIP | 37986224\|19536157 | Unspecified |
| mmu-miR-9-5p | MIMAT0000142 | Nr2c1 | 22025 | Sequencing | 19536157\|37986224 | Unspecified |
| mmu-miR-9-5p | MIMAT0000142 | Nrarp | 67122 | Sequencing | 19536157\|37986224 | Unspecified |
| mmu-miR-9-5p | MIMAT0000142 | Nup50 | 18141 | Sequencing | 19536157\|37986224 | Unspecified |
| mmu-miR-9-5p | MIMAT0000142 | Ocrl | 320634 | Sequencing | 37986224\|19536157 | Unspecified |
| mmu-miR-9-5p | MIMAT0000142 | Opcml | 330908 | Sequencing | 37986224\|19536157 | Unspecified |
| mmu-miR-9-5p | MIMAT0000142 | Opn3 | 13603 | Sequencing | 37986224\|19536157 | Unspecified |

| | | | | | | |
|---|---|---|---|---|---|---|
| mmu-miR-9-5p | MIMAT0000142 | P4ha2 | 18452 | Sequencing | 19536157\|37986224 | Unspecified |
| mmu-miR-9-5p | MIMAT0000142 | Pacs2 | 217893 | Sequencing | 37986224\|19536157 | Unspecified |
| mmu-miR-9-5p | MIMAT0000142 | Padi2 | 18600 | Sequencing | 37986224\|19536157 | Unspecified |
| mmu-miR-9-5p | MIMAT0000142 | Paip2 | 67869 | Sequencing | 19536157\|37986224 | Unspecified |
| mmu-miR-9-5p | MIMAT0000142 | Pank3 | 211347 | Sequencing | 19536157\|37986224 | Unspecified |
| mmu-miR-9-5p | MIMAT0000142 | Paqr8 | 74229 | Sequencing | 19536157\|37986224 | Unspecified |
| mmu-miR-9-5p | MIMAT0000142 | Pard6g | 93737 | Sequencing | 19536157\|37986224 | Unspecified |
| mmu-miR-9-5p | MIMAT0000142 | Pbrm1 | 66923 | Sequencing | 19536157\|37986224 | Unspecified |
| mmu-miR-9-5p | MIMAT0000142 | Pcdh7 | 54216 | Sequencing//HITS-CLIP | 19536157\|37986224 | Unspecified |
| mmu-miR-9-5p | MIMAT0000142 | Pcdh8 | 18530 | Sequencing | 37986224\|19536157 | Unspecified |
| mmu-miR-9-5p | MIMAT0000142 | Pcmtd2 | 245867 | Sequencing | 19536157\|37986224 | Unspecified |
| mmu-miR-9-5p | MIMAT0000142 | Pcnx | 54604 | Sequencing | 19536157\|37986224 | Unspecified |
| mmu-miR-9-5p | MIMAT0000142 | Pdhx | 27402 | Sequencing | 37986224\|19536157 | Unspecified |
| mmu-miR-9-5p | MIMAT0000142 | Pds5b | 100710 | Sequencing | 37986224\|19536157 | Unspecified |
| mmu-miR-9-5p | MIMAT0000142 | Pea15a | 18611 | Sequencing | 19536157\|37986224 | Unspecified |
| mmu-miR-9-5p | MIMAT0000142 | Pgrmc1 | 53328 | Sequencing | 19536157\|37986224 | Unspecified |
| mmu-miR-9-5p | MIMAT0000142 | Pik3cb | 74769 | Sequencing | 37986224\|19536157 | Unspecified |
| mmu-miR-9-5p | MIMAT0000142 | Pik3cd | 18707 | Sequencing | 19536157\|37986224 | Unspecified |
| mmu-miR-9-5p | MIMAT0000142 | Pikfyve | 18711 | Sequencing | 19536157 | Unspecified |
| mmu-miR-9-5p | MIMAT0000142 | Pitpnc1 | 71795 | Sequencing | 19536157\|37986224 | Unspecified |
| mmu-miR-9-5p | MIMAT0000142 | Plat | 18791 | Sequencing | 19536157\|37986224 | Unspecified |
| mmu-miR-9-5p | MIMAT0000142 | Plbd2 | 71772 | Sequencing | 19536157 | Unspecified |
| mmu-miR-9-5p | MIMAT0000142 | Pld3 | 18807 | Sequencing | 19536157\|37986224 | Unspecified |
| mmu-miR-9-5p | MIMAT0000142 | Plscr3 | 70310 | Sequencing//HITS-CLIP | 19536157\|37986224 | Unspecified |
| mmu-miR-9-5p | MIMAT0000142 | Plxnb3 | 140571 | Sequencing | 37986224\|19536157 | Unspecified |
| mmu-miR-9-5p | MIMAT0000142 | Pnma2 | 239157 | Sequencing | 19536157\|37986224 | Unspecified |
| mmu-miR-9-5p | MIMAT0000142 | Polr3k | 67005 | Sequencing | 37986224\|19536157 | Unspecified |
| mmu-miR-9-5p | MIMAT0000142 | Ppm1l | 242083 | Sequencing | 37986224\|19536157 | Unspecified |
| mmu-miR-9-5p | MIMAT0000142 | Ppp1r13b | 21981 | Sequencing | 37986224\|19536157 | Unspecified |
| mmu-miR-9-5p | MIMAT0000142 | Ppp2r5d | 21770 | Sequencing | 19536157\|37986224 | Unspecified |
| mmu-miR-9-5p | MIMAT0000142 | Ppp3r1 | 19058 | Sequencing | 19536157\|37986224 | Unspecified |
| mmu-miR-9-5p | MIMAT0000142 | Prkar2a | 19087 | Sequencing | 37986224\|19536157 | Unspecified |
| mmu-miR-9-5p | MIMAT0000142 | Prkcb | 18751 | Sequencing | 37986224\|19536157 | Unspecified |
| mmu-miR-9-5p | MIMAT0000142 | Prnp | 19122 | Sequencing | 37986224\|19536157 | Unspecified |
| mmu-miR-9-5p | MIMAT0000142 | Prom1 | 19126 | Sequencing | 19536157\|37986224 | Unspecified |
| mmu-miR-9-5p | MIMAT0000142 | Prps2 | 110639 | Sequencing | 37986224\|19536157 | Unspecified |
| mmu-miR-9-5p | MIMAT0000142 | Prrc2b | 227723 | Sequencing | 19536157 | Unspecified |
| mmu-miR-9-5p | MIMAT0000142 | Prrg3 | 208748 | Sequencing | 19536157\|37986224 | Unspecified |
| mmu-miR-9-5p | MIMAT0000142 | Prrt3 | 210673 | Sequencing | 37986224\|19536157 | Unspecified |

| | | | | | | |
|---|---|---|---|---|---|---|
| mmu-miR-9-5p | MIMAT0000142 | Psd3 | 234353 | Sequencing | 37986224\|19536157 | Unspecified |
| mmu-miR-9-5p | MIMAT0000142 | Psen1 | 19164 | Sequencing | 19536157\|37986224 | Unspecified |
| mmu-miR-9-5p | MIMAT0000142 | Psme4 | 103554 | Sequencing | 37986224\|19536157 | Unspecified |
| mmu-miR-9-5p | MIMAT0000142 | Ptbp2 | 56195 | Sequencing | 37986224\|19536157 | Unspecified |
| mmu-miR-9-5p | MIMAT0000142 | Ptcd1 | 71799 | Sequencing | 19536157\|37986224 | Unspecified |
| mmu-miR-9-5p | MIMAT0000142 | Ptch1 | 19206 | Sequencing | 19536157\|37986224 | Unspecified |
| mmu-miR-9-5p | MIMAT0000142 | Ptgds | 19215 | Sequencing | 19536157\|37986224 | Unspecified |
| mmu-miR-9-5p | MIMAT0000142 | Ptprd | 19266 | Sequencing | 19536157\|37986224 | Unspecified |
| mmu-miR-9-5p | MIMAT0000142 | Ptprg | 19270 | Sequencing | 19536157\|37986224 | Unspecified |
| mmu-miR-9-5p | MIMAT0000142 | Pvrl1 | 58235 | Sequencing | 19536157\|37986224 | Unspecified |
| mmu-miR-9-5p | MIMAT0000142 | Pxdn | 69675 | Sequencing | 37986224\|19536157 | Unspecified |
| mmu-miR-9-5p | MIMAT0000142 | R3hdm1 | 226412 | Sequencing | 19536157\|37986224 | Unspecified |
| mmu-miR-9-5p | MIMAT0000142 | Rab9b | 319642 | Sequencing | 19536157\|37986224 | Unspecified |
| mmu-miR-9-5p | MIMAT0000142 | Rad9b | 231724 | Sequencing | 37986224\|19536157 | Unspecified |
| mmu-miR-9-5p | MIMAT0000142 | Rap1b | 215449 | Sequencing | 37986224\|19536157 | Unspecified |
| mmu-miR-9-5p | MIMAT0000142 | Rapgef6 | 192786 | Sequencing | 19536157\|37986224 | Unspecified |
| mmu-miR-9-5p | MIMAT0000142 | Rasa1 | 218397 | Sequencing | 19536157\|37986224 | Unspecified |
| mmu-miR-9-5p | MIMAT0000142 | Rasd2 | 75141 | Sequencing//HITS-CLIP | 37986224\|19536157 | Unspecified |
| mmu-miR-9-5p | MIMAT0000142 | Rasgrp1 | 19419 | Sequencing | 37986224\|19536157 | Unspecified |
| mmu-miR-9-5p | MIMAT0000142 | Rassf3 | 192678 | Sequencing//HITS-CLIP | 19536157\|25083871\|37986224 | Unspecified |
| mmu-miR-9-5p | MIMAT0000142 | Rbfox1 | 268859 | Sequencing | 19536157 | Unspecified |
| mmu-miR-9-5p | MIMAT0000142 | Rbm24 | 666794 | Sequencing | 37986224\|19536157 | Unspecified |
| mmu-miR-9-5p | MIMAT0000142 | Rbm25 | 67039 | Sequencing | 37986224\|19536157 | Unspecified |
| mmu-miR-9-5p | MIMAT0000142 | Rbm27 | 225432 | Sequencing | 37986224\|19536157 | Unspecified |
| mmu-miR-9-5p | MIMAT0000142 | Rcan2 | 53901 | Sequencing | 37986224\|19536157 | Unspecified |
| mmu-miR-9-5p | MIMAT0000142 | Reep1 | 52250 | Sequencing | 37986224\|19536157 | Unspecified |
| mmu-miR-9-5p | MIMAT0000142 | Reep4 | 72549 | Sequencing | 37986224\|19536157 | Unspecified |
| mmu-miR-9-5p | MIMAT0000142 | Reep5 | 13476 | Sequencing | 19536157\|37986224 | Unspecified |
| mmu-miR-9-5p | MIMAT0000142 | Rela | 19697 | Sequencing | 19536157\|37986224 | Unspecified |
| mmu-miR-9-5p | MIMAT0000142 | Rfx7 | 319758 | Sequencing//HITS-CLIP | 37986224\|19536157 | Unspecified |
| mmu-miR-9-5p | MIMAT0000142 | Rgp1 | 242406 | Sequencing | 19536157\|37986224 | Unspecified |
| mmu-miR-9-5p | MIMAT0000142 | Rgs7bp | 52882 | Sequencing | 37986224\|19536157 | Unspecified |
| mmu-miR-9-5p | MIMAT0000142 | Rictor | 78757 | Sequencing//qCLASH | 19536157\|31393953 | Unspecified |
| mmu-miR-9-5p | MIMAT0000142 | Rnf128 | 66889 | Sequencing | 19536157\|37986224 | Unspecified |
| mmu-miR-9-5p | MIMAT0000142 | Rnf144a | 108089 | Sequencing | 37986224\|19536157 | Unspecified |
| mmu-miR-9-5p | MIMAT0000142 | Rnf19a | 30945 | Sequencing | 37986224\|19536157 | Unspecified |
| mmu-miR-9-5p | MIMAT0000142 | Rnf2 | 19821 | Sequencing | 19536157\|37986224 | Unspecified |
| mmu-miR-9-5p | MIMAT0000142 | Rnf4 | 19822 | Sequencing | 19536157\|37986224 | Unspecified |
| mmu-miR-9-5p | MIMAT0000142 | Ror1 | 26563 | Sequencing | 19536157\|37986224 | Unspecified |

| | | | | | | | |
|---|---|---|---|---|---|---|---|
| mmu-miR-9-5p | MIMAT0000142 | Ror2 | 26564 | Sequencing | 19536157\|37986224 | | Unspecified |
| mmu-miR-9-5p | MIMAT0000142 | Rpgrip1l | 244585 | Sequencing | 37986224\|19536157 | | Unspecified |
| mmu-miR-9-5p | MIMAT0000142 | Rptor | 74370 | Sequencing | | 19536157 | Unspecified |
| mmu-miR-9-5p | MIMAT0000142 | Rundc3b | 242819 | Sequencing//HITS-CLIP | 19536157\|37986224 | | Unspecified |
| mmu-miR-9-5p | MIMAT0000142 | Runx1 | 12394 | Luciferase assay//Western blot//Luciferase | 19114653\|26091714 | | Unspecified |
| mmu-miR-9-5p | MIMAT0000142 | Sacm1l | 83493 | Sequencing | 37986224\|19536157 | | Unspecified |
| mmu-miR-9-5p | MIMAT0000142 | Scamp1 | 107767 | Sequencing | 19536157\|37986224 | | Unspecified |
| mmu-miR-9-5p | MIMAT0000142 | Scd1 | 20249 | Sequencing | 37986224\|19536157 | | Unspecified |
| mmu-miR-9-5p | MIMAT0000142 | Scn2b | 72821 | Sequencing | 37986224\|19536157 | | Unspecified |
| mmu-miR-9-5p | MIMAT0000142 | Scn3b | 235281 | Sequencing | 37986224\|19536157 | | Unspecified |
| mmu-miR-9-5p | MIMAT0000142 | Scn8a | 20273 | Sequencing | 37986224\|19536157 | | Unspecified |
| mmu-miR-9-5p | MIMAT0000142 | Scrn1 | 69938 | Sequencing | 19536157\|37986224 | | Unspecified |
| mmu-miR-9-5p | MIMAT0000142 | Sec31a | 69162 | Sequencing | 19536157\|37986224 | | Unspecified |
| mmu-miR-9-5p | MIMAT0000142 | Sel1l | 20338 | Sequencing | 19536157\|37986224 | | Unspecified |
| mmu-miR-9-5p | MIMAT0000142 | Sema6d | 214968 | Sequencing | 19536157\|37986224 | | Unspecified |
| mmu-miR-9-5p | MIMAT0000142 | Senp8 | 71599 | Sequencing | 37986224\|19536157 | | Unspecified |
| mmu-miR-9-5p | MIMAT0000142 | Serpini1 | 20713 | Sequencing | 19536157\|37986224 | | Unspecified |
| mmu-miR-9-5p | MIMAT0000142 | Sertm1 | 329641 | Sequencing | | 19536157 | Unspecified |
| mmu-miR-9-5p | MIMAT0000142 | Sestd1 | 228071 | Sequencing | 19536157\|37986224 | | Unspecified |
| mmu-miR-9-5p | MIMAT0000142 | Sfrp1 | 20377 | Sequencing | 37986224\|19536157 | | Unspecified |
| mmu-miR-9-5p | MIMAT0000142 | Shank2 | 210274 | Sequencing | 37986224\|19536157 | | Unspecified |
| mmu-miR-9-5p | MIMAT0000142 | Shroom4 | 208431 | Sequencing | 19536157\|37986224 | | Unspecified |
| mmu-miR-9-5p | MIMAT0000142 | Sidt2 | 214597 | Sequencing | 19536157\|37986224 | | Unspecified |
| mmu-miR-9-5p | MIMAT0000142 | Sirt1 | 93759 | ELISA//Luciferase assay//qRT-PCR//Wester | | 21288303 | Unspecified |
| mmu-miR-9-5p | MIMAT0000142 | Slc10a2 | 20494 | HITS-CLIP | 23142080\|21258322\|19536157\|23597 | | Unspecified |
| mmu-miR-9-5p | MIMAT0000142 | Slc10a7 | 76775 | Sequencing | 37986224\|19536157 | | Unspecified |
| mmu-miR-9-5p | MIMAT0000142 | Slc12a5 | 57138 | Sequencing | 37986224\|19536157 | | Unspecified |
| mmu-miR-9-5p | MIMAT0000142 | Slc14a2 | 27411 | HITS-CLIP | | 21258322 | Unspecified |
| mmu-miR-9-5p | MIMAT0000142 | Slc16a7 | 20503 | HITS-CLIP | | 21258322 | Unspecified |
| mmu-miR-9-5p | MIMAT0000142 | Slc1a1 | 20510 | Sequencing | 37986224\|19536157 | | Unspecified |
| mmu-miR-9-5p | MIMAT0000142 | Slc24a2 | 76376 | Sequencing | 19536157\|37986224 | | Unspecified |
| mmu-miR-9-5p | MIMAT0000142 | Slc30a10 | 226781 | Sequencing | 19536157\|37986224 | | Unspecified |
| mmu-miR-9-5p | MIMAT0000142 | Slc30a7 | 66500 | HITS-CLIP | | 25083871 | Unspecified |
| mmu-miR-9-5p | MIMAT0000142 | Slc31a2 | 20530 | Sequencing | 37986224\|19536157 | | Unspecified |
| mmu-miR-9-5p | MIMAT0000142 | Slc33a1 | 11416 | Sequencing | 37986224\|19536157 | | Unspecified |
| mmu-miR-9-5p | MIMAT0000142 | Slc35g1 | 240660 | HITS-CLIP | | 25083871 | Unspecified |
| mmu-miR-9-5p | MIMAT0000142 | Slc38a7 | 234595 | Sequencing | 19536157\|37986224 | | Unspecified |
| mmu-miR-9-5p | MIMAT0000142 | Slc6a6 | 21366 | Sequencing | 19536157\|37986224 | | Unspecified |
| mmu-miR-9-5p | MIMAT0000142 | Slc9a1 | 20544 | Sequencing | 19536157\|37986224 | | Unspecified |

| | | | | | | |
|---|---|---|---|---|---|---|
| mmu-miR-9-5p | MIMAT0000142 | Smap2 | 69780 | Sequencing | 37986224\|19536157 | Unspecified |
| mmu-miR-9-5p | MIMAT0000142 | Smarcd2 | 83796 | Sequencing | 37986224\|19536157 | Unspecified |
| mmu-miR-9-5p | MIMAT0000142 | Smc2 | 14211 | Sequencing | 37986224\|19536157 | Unspecified |
| mmu-miR-9-5p | MIMAT0000142 | Smyd1 | 12180 | Sequencing | 37986224\|19536157 | Unspecified |
| mmu-miR-9-5p | MIMAT0000142 | Snap25 | 20614 | Sequencing | 37986224\|19536157 | Unspecified |
| mmu-miR-9-5p | MIMAT0000142 | Snph | 241727 | Sequencing | 37986224\|19536157 | Unspecified |
| mmu-miR-9-5p | MIMAT0000142 | Snx25 | 102141 | Sequencing//HITS-CLIP | 19536157\|37986224 | Unspecified |
| mmu-miR-9-5p | MIMAT0000142 | Snx27 | 76742 | Sequencing | 19536157\|37986224 | Unspecified |
| mmu-miR-9-5p | MIMAT0000142 | Soat1 | 20652 | Sequencing | 19536157\|37986224 | Unspecified |
| mmu-miR-9-5p | MIMAT0000142 | Socs5 | 56468 | Sequencing//HITS-CLIP | 19536157\|37986224 | Unspecified |
| mmu-miR-9-5p | MIMAT0000142 | Sos1 | 20662 | HITS-CLIP | 25083871 | Unspecified |
| mmu-miR-9-5p | MIMAT0000142 | Specc1l | 74392 | Sequencing | 37986224\|19536157 | Unspecified |
| mmu-miR-9-5p | MIMAT0000142 | Sphkap | 77629 | Sequencing | 37986224\|19536157 | Unspecified |
| mmu-miR-9-5p | MIMAT0000142 | Spire1 | 68166 | Sequencing | 19536157\|37986224 | Unspecified |
| mmu-miR-9-5p | MIMAT0000142 | Spred3 | 101809 | Sequencing | 37986224\|19536157 | Unspecified |
| mmu-miR-9-5p | MIMAT0000142 | Sprn | 212518 | Sequencing | 37986224\|19536157 | Unspecified |
| mmu-miR-9-5p | MIMAT0000142 | Sptb | 20741 | Sequencing | 19536157 | Unspecified |
| mmu-miR-9-5p | MIMAT0000142 | Sra1 | 24068 | Sequencing | 19536157\|37986224 | Unspecified |
| mmu-miR-9-5p | MIMAT0000142 | Srcin1 | 56013 | Sequencing | 19536157 | Unspecified |
| mmu-miR-9-5p | MIMAT0000142 | Srek1 | 218543 | Sequencing | 19536157 | Unspecified |
| mmu-miR-9-5p | MIMAT0000142 | Srrm2 | 75956 | Sequencing | 37986224\|19536157 | Unspecified |
| mmu-miR-9-5p | MIMAT0000142 | Ssr1 | 107513 | Sequencing | 19536157\|37986224 | Unspecified |
| mmu-miR-9-5p | MIMAT0000142 | St8sia3 | 20451 | Sequencing | 37986224\|19536157 | Unspecified |
| mmu-miR-9-5p | MIMAT0000142 | Stam2 | 56324 | HITS-CLIP | 25083871 | Unspecified |
| mmu-miR-9-5p | MIMAT0000142 | Stat2 | 20847 | Sequencing | 37986224\|19536157 | Unspecified |
| mmu-miR-9-5p | MIMAT0000142 | Stk11ip | 71728 | Sequencing | 19536157\|37986224 | Unspecified |
| mmu-miR-9-5p | MIMAT0000142 | Stmn1 | 16765 | qRT-PCR//Luciferase assay//Western blot/ | 20362537 | Unspecified |
| mmu-miR-9-5p | MIMAT0000142 | Sufu | 24069 | Sequencing | 19536157\|37986224 | Unspecified |
| mmu-miR-9-5p | MIMAT0000142 | Sumf2 | 67902 | Sequencing | 19536157\|37986224 | Unspecified |
| mmu-miR-9-5p | MIMAT0000142 | Surf6 | 20935 | Sequencing | 19536157\|37986224 | Unspecified |
| mmu-miR-9-5p | MIMAT0000142 | Syap1 | 67043 | Sequencing | 37986224\|19536157 | Unspecified |
| mmu-miR-9-5p | MIMAT0000142 | Sybu | 319613 | Sequencing | 19536157 | Unspecified |
| mmu-miR-9-5p | MIMAT0000142 | Syne1 | 64009 | Sequencing//HITS-CLIP | 37986224\|19536157 | Unspecified |
| mmu-miR-9-5p | MIMAT0000142 | Syt11 | 229521 | Sequencing | 37986224\|19536157 | Unspecified |
| mmu-miR-9-5p | MIMAT0000142 | Tacc1 | 320165 | Sequencing | 19536157\|37986224 | Unspecified |
| mmu-miR-9-5p | MIMAT0000142 | Tanc2 | 77097 | Sequencing | 37986224\|19536157 | Unspecified |
| mmu-miR-9-5p | MIMAT0000142 | Tbc1d22a | 223754 | Sequencing | 19536157\|37986224 | Unspecified |
| mmu-miR-9-5p | MIMAT0000142 | Tbc1d24 | 224617 | Sequencing | 19536157\|37986224 | Unspecified |
| mmu-miR-9-5p | MIMAT0000142 | Tef | 21685 | Sequencing | 37986224\|19536157 | Unspecified |

| | | | | | | | |
|---|---|---|---|---|---|---|---|
| mmu-miR-9-5p | MIMAT0000142 | Tenm4 | 23966 | Sequencing//HITS-CLIP | | 19536157 | Unspecified |
| mmu-miR-9-5p | MIMAT0000142 | Tgfbr2 | 21813 | Sequencing | 19536157\|37986224 | | Unspecified |
| mmu-miR-9-5p | MIMAT0000142 | Tgfbr3 | 21814 | Sequencing | 37986224\|19536157 | | Unspecified |
| mmu-miR-9-5p | MIMAT0000142 | Thrsp | 21835 | Sequencing | 19536157\|37986224 | | Unspecified |
| mmu-miR-9-5p | MIMAT0000142 | Tln2 | 70549 | Sequencing | 19536157\|37986224 | | Unspecified |
| mmu-miR-9-5p | MIMAT0000142 | Tmc7 | 209760 | Sequencing | 19536157\|37986224 | | Unspecified |
| mmu-miR-9-5p | MIMAT0000142 | Tmed8 | 382620 | Sequencing | 37986224\|19536157 | | Unspecified |
| mmu-miR-9-5p | MIMAT0000142 | Tmem109 | 68539 | Sequencing | 37986224\|19536157 | | Unspecified |
| mmu-miR-9-5p | MIMAT0000142 | Tmem136 | 235300 | Sequencing | 37986224\|19536157 | | Unspecified |
| mmu-miR-9-5p | MIMAT0000142 | Tmem184b | 223693 | Sequencing | 37986224\|19536157 | | Unspecified |
| mmu-miR-9-5p | MIMAT0000142 | Tmem248 | 71667 | Sequencing | 37986224\|19536157 | | Unspecified |
| mmu-miR-9-5p | MIMAT0000142 | Tmem40 | 94346 | Sequencing | 37986224\|19536157 | | Unspecified |
| mmu-miR-9-5p | MIMAT0000142 | Tmem65 | 74868 | Sequencing//HITS-CLIP | 19536157\|37986224 | | Unspecified |
| mmu-miR-9-5p | MIMAT0000142 | Tnks | 21951 | Sequencing | 37986224\|19536157 | | Unspecified |
| mmu-miR-9-5p | MIMAT0000142 | Tomm6 | 66119 | Sequencing | 37986224\|19536157 | | Unspecified |
| mmu-miR-9-5p | MIMAT0000142 | Tor1aip2 | 240832 | Sequencing | | 19536157 | Unspecified |
| mmu-miR-9-5p | MIMAT0000142 | Tpm1 | 22003 | Sequencing | 37986224\|19536157 | | Unspecified |
| mmu-miR-9-5p | MIMAT0000142 | Traf3 | 22031 | Sequencing//HITS-CLIP | 37986224\|19536157 | | Unspecified |
| mmu-miR-9-5p | MIMAT0000142 | Tram1 | 72265 | Sequencing | 19536157\|37986224 | | Unspecified |
| mmu-miR-9-5p | MIMAT0000142 | Trim25 | 217069 | Sequencing | 37986224\|19536157 | | Unspecified |
| mmu-miR-9-5p | MIMAT0000142 | Tsc1 | 64930 | Sequencing | 19536157\|37986224 | | Unspecified |
| mmu-miR-9-5p | MIMAT0000142 | Tspan31 | 67125 | Sequencing | 19536157\|37986224 | | Unspecified |
| mmu-miR-9-5p | MIMAT0000142 | Tspan7 | 21912 | Sequencing | 19536157\|37986224 | | Unspecified |
| mmu-miR-9-5p | MIMAT0000142 | Ttc14 | 67120 | Sequencing | 37986224\|19536157 | | Unspecified |
| mmu-miR-9-5p | MIMAT0000142 | Ttr | 22139 | Sequencing | 37986224\|19536157 | | Unspecified |
| mmu-miR-9-5p | MIMAT0000142 | Ttyh2 | 117160 | Sequencing | 37986224\|19536157 | | Unspecified |
| mmu-miR-9-5p | MIMAT0000142 | Ttyh3 | 78339 | Sequencing | 37986224\|19536157 | | Unspecified |
| mmu-miR-9-5p | MIMAT0000142 | Tubgcp4 | 51885 | Sequencing | 19536157\|37986224 | | Unspecified |
| mmu-miR-9-5p | MIMAT0000142 | Ube3c | 100763 | Sequencing | 37986224\|19536157 | | Unspecified |
| mmu-miR-9-5p | MIMAT0000142 | Ufl1 | 67490 | Sequencing | | 19536157 | Unspecified |
| mmu-miR-9-5p | MIMAT0000142 | Unc13c | 208898 | Sequencing | 37986224\|19536157 | | Unspecified |
| mmu-miR-9-5p | MIMAT0000142 | Unc5c | 22253 | Sequencing | 19536157\|37986224 | | Unspecified |
| mmu-miR-9-5p | MIMAT0000142 | Unk | 217331 | Sequencing | 19536157\|37986224 | | Unspecified |
| mmu-miR-9-5p | MIMAT0000142 | Usp46 | 69727 | Sequencing | 19536157\|37986224 | | Unspecified |
| mmu-miR-9-5p | MIMAT0000142 | Utp23 | 78581 | HITS-CLIP | | 21258322 | Unspecified |
| mmu-miR-9-5p | MIMAT0000142 | Utrn | 22288 | Sequencing | 37986224\|19536157 | | Unspecified |
| mmu-miR-9-5p | MIMAT0000142 | Uvssa | 71101 | Sequencing | | 19536157 | Unspecified |
| mmu-miR-9-5p | MIMAT0000142 | Vamp3 | 22319 | Sequencing | 19536157\|37986224 | | Unspecified |
| mmu-miR-9-5p | MIMAT0000142 | Vbp1 | 22327 | Sequencing | 19536157\|37986224 | | Unspecified |

| | | | | | | |
|---|---|---|---|---|---|---|
| mmu-miR-9-5p | MIMAT0000142 | Vipr1 | 22354 | Sequencing | 19536157\|37986224 | Unspecified |
| mmu-miR-9-5p | MIMAT0000142 | Vps37a | 52348 | Sequencing | 19536157\|37986224 | Unspecified |
| mmu-miR-9-5p | MIMAT0000142 | Wasf2 | 242687 | Sequencing | 37986224\|19536157 | Unspecified |
| mmu-miR-9-5p | MIMAT0000142 | Wdr37 | 207615 | Sequencing | 37986224\|19536157 | Unspecified |
| mmu-miR-9-5p | MIMAT0000142 | Wipf1 | 215280 | Sequencing | 19536157\|37986224 | Unspecified |
| mmu-miR-9-5p | MIMAT0000142 | Wnk1 | 232341 | Sequencing//qCLASH//HITS-CLIP | 37986224\|31393953\|19536157\|26203 | Unspecified |
| mmu-miR-9-5p | MIMAT0000142 | Wtip | 101543 | Sequencing | 19536157\|37986224 | Unspecified |
| mmu-miR-9-5p | MIMAT0000142 | Xxylt1 | 268880 | Sequencing | 19536157 | Unspecified |
| mmu-miR-9-5p | MIMAT0000142 | Ybx3 | 56449 | Sequencing//HITS-CLIP | 19536157 | Unspecified |
| mmu-miR-9-5p | MIMAT0000142 | Zbed3 | 72114 | Sequencing | 19536157\|37986224 | Unspecified |
| mmu-miR-9-5p | MIMAT0000142 | Zbtb24 | 268294 | Sequencing | 37986224\|19536157 | Unspecified |
| mmu-miR-9-5p | MIMAT0000142 | Zbtb34 | 241311 | Sequencing//HITS-CLIP | 37986224\|19536157 | Unspecified |
| mmu-miR-9-5p | MIMAT0000142 | Zbtb41 | 226470 | Sequencing | 37986224\|19536157 | Unspecified |
| mmu-miR-9-5p | MIMAT0000142 | Zcchc14 | 142682 | Sequencing | 19536157\|37986224 | Unspecified |
| mmu-miR-9-5p | MIMAT0000142 | Zdhhc20 | 75965 | Sequencing | 19536157\|37986224 | Unspecified |
| mmu-miR-9-5p | MIMAT0000142 | Zdhhc7 | 102193 | Sequencing | 19536157\|37986224 | Unspecified |
| mmu-miR-9-5p | MIMAT0000142 | Zfand4 | 67492 | Sequencing | 19536157 | Unspecified |
| mmu-miR-9-5p | MIMAT0000142 | Zfhx3 | 11906 | Sequencing | 37986224\|19536157 | Unspecified |
| mmu-miR-9-5p | MIMAT0000142 | Zfhx4 | 80892 | Sequencing | 19536157\|37986224 | Unspecified |
| mmu-miR-9-5p | MIMAT0000142 | Zfp275 | 27081 | Sequencing | 37986224\|19536157 | Unspecified |
| mmu-miR-9-5p | MIMAT0000142 | Zfp282 | 101095 | Sequencing//HITS-CLIP | 19536157\|37986224 | Unspecified |
| mmu-miR-9-5p | MIMAT0000142 | Zfp36l2 | 12193 | Sequencing | 37986224\|19536157 | Unspecified |
| mmu-miR-9-5p | MIMAT0000142 | Zfp521 | 225207 | Immunocytochemistry//Luciferase assay//( | 22465325 | Unspecified |
| mmu-miR-9-5p | MIMAT0000142 | Zfp704 | 170753 | Sequencing | 19536157\|37986224 | Unspecified |
| mmu-miR-9-5p | MIMAT0000142 | Zic1 | 22771 | Sequencing | 19536157\|37986224 | Unspecified |
| mmu-miR-9-5p | MIMAT0000142 | Zic3 | 22773 | Sequencing | 19536157\|37986224 | Unspecified |
| mmu-miR-9-5p | MIMAT0000142 | Zkscan1 | 74570 | Sequencing | 19536157\|37986224 | Unspecified |
| mmu-miR-9-5p | MIMAT0000142 | Zmat3 | 22401 | Sequencing | 37986224\|19536157 | Unspecified |
| mmu-miR-9-5p | MIMAT0000142 | Zmynd8 | 228880 | Sequencing | 19536157 | Unspecified |
| mmu-miR-9-5p | MIMAT0000142 | Zzz3 | 108946 | Sequencing | 37986224\|19536157 | Unspecified |
| mmu-miR-703 | MIMAT0003493 | 08-Sept | 20362 | HITS-CLIP | 25083871 | Unspecified |
| mmu-miR-703 | MIMAT0003493 | 4933427D06Rik | 232217 | HITS-CLIP | 25083871 | Unspecified |
| mmu-miR-703 | MIMAT0003493 | Arhgap11a | 228482 | HITS-CLIP | 25083871 | Unspecified |
| mmu-miR-703 | MIMAT0003493 | Asxl2 | 75302 | HITS-CLIP | 25083871\|21258322 | Unspecified |
| mmu-miR-703 | MIMAT0003493 | Braf | 109880 | HITS-CLIP | 25083871 | Unspecified |
| mmu-miR-703 | MIMAT0003493 | Cep97 | 74201 | HITS-CLIP | 19536157\|26203562\|23597149\|25083 | Unspecified |
| mmu-miR-703 | MIMAT0003493 | Cnnm3 | 94218 | HITS-CLIP | 21258322 | Unspecified |
| mmu-miR-703 | MIMAT0003493 | Cnot4 | 53621 | HITS-CLIP | 25083871 | Unspecified |
| mmu-miR-703 | MIMAT0003493 | Commd8 | 27784 | HITS-CLIP | 21258322 | Unspecified |

| | | | | | | |
|---|---|---|---|---|---|---|
| mmu-miR-703 | MIMAT0003493 | Epas1 | 13819 | HITS-CLIP | 25083871 | Unspecified |
| mmu-miR-703 | MIMAT0003493 | Faah | 14073 | HITS-CLIP | 21258322 | Unspecified |
| mmu-miR-703 | MIMAT0003493 | Fam96a | 68250 | HITS-CLIP | 21258322 | Unspecified |
| mmu-miR-703 | MIMAT0003493 | Foxj3 | 230700 | HITS-CLIP | 23597149\|21258322\|23142080\|2508 | Unspecified |
| mmu-miR-703 | MIMAT0003493 | Git2 | 26431 | HITS-CLIP | 21258322 | Unspecified |
| mmu-miR-703 | MIMAT0003493 | Gpr3 | 14748 | HITS-CLIP | 25083871 | Unspecified |
| mmu-miR-703 | MIMAT0003493 | Il1r1 | 16177 | HITS-CLIP | 25083871 | Unspecified |
| mmu-miR-703 | MIMAT0003493 | Ilkap | 67444 | HITS-CLIP | 25083871 | Unspecified |
| mmu-miR-703 | MIMAT0003493 | Kif23 | 71819 | HITS-CLIP | 23142080\|25083871 | Unspecified |
| mmu-miR-703 | MIMAT0003493 | Larp4b | 217980 | HITS-CLIP | 25083871 | Unspecified |
| mmu-miR-703 | MIMAT0003493 | Mars2 | 212679 | HITS-CLIP | 25083871 | Unspecified |
| mmu-miR-703 | MIMAT0003493 | Mau2 | 74549 | HITS-CLIP | 25083871 | Unspecified |
| mmu-miR-703 | MIMAT0003493 | Mcart1 | 230125 | CLIP-Seq dataset analysis | 25083871\|21258322\|23597149 | Unspecified |
| mmu-miR-703 | MIMAT0003493 | Mmab | 77697 | HITS-CLIP | 21258322 | Unspecified |
| mmu-miR-703 | MIMAT0003493 | Mnat1 | 17420 | HITS-CLIP | 25083871 | Unspecified |
| mmu-miR-703 | MIMAT0003493 | Mxra7 | 67622 | HITS-CLIP | 25083871 | Unspecified |
| mmu-miR-703 | MIMAT0003493 | Narf | 67608 | HITS-CLIP | 25083871 | Unspecified |
| mmu-miR-703 | MIMAT0003493 | Pcnp | 76302 | HITS-CLIP | 25083871 | Unspecified |
| mmu-miR-703 | MIMAT0003493 | Pctp | 18559 | HITS-CLIP | 23597149\|25083871\|21258322 | Unspecified |
| mmu-miR-703 | MIMAT0003493 | Pde8b | 218461 | HITS-CLIP | 23142080\|25083871 | Unspecified |
| mmu-miR-703 | MIMAT0003493 | Pdxk | 216134 | HITS-CLIP | 23597149\|21258322 | Unspecified |
| mmu-miR-703 | MIMAT0003493 | Phex | 18675 | HITS-CLIP | 25083871 | Unspecified |
| mmu-miR-703 | MIMAT0003493 | Pitpnm2 | 19679 | HITS-CLIP | 25083871 | Unspecified |
| mmu-miR-703 | MIMAT0003493 | Prex2 | 109294 | HITS-CLIP | 25083871 | Unspecified |
| mmu-miR-703 | MIMAT0003493 | Prr18 | 320111 | HITS-CLIP | 25083871 | Unspecified |
| mmu-miR-703 | MIMAT0003493 | Prtg | 235472 | HITS-CLIP | 25083871 | Unspecified |
| mmu-miR-703 | MIMAT0003493 | Ptcd1 | 71799 | HITS-CLIP | 25083871 | Unspecified |
| mmu-miR-703 | MIMAT0003493 | Rem1 | 19700 | HITS-CLIP | 25083871 | Unspecified |
| mmu-miR-703 | MIMAT0003493 | Ret | 19713 | HITS-CLIP | 25083871 | Unspecified |
| mmu-miR-703 | MIMAT0003493 | Ric3 | 320360 | HITS-CLIP | 21258322 | Unspecified |
| mmu-miR-703 | MIMAT0003493 | Rnf165 | 225743 | HITS-CLIP | 23142080 | Unspecified |
| mmu-miR-703 | MIMAT0003493 | Rpp14 | 67053 | HITS-CLIP | 25083871 | Unspecified |
| mmu-miR-703 | MIMAT0003493 | Rras2 | 66922 | HITS-CLIP | 25083871 | Unspecified |
| mmu-miR-703 | MIMAT0003493 | Rsf1 | 233532 | HITS-CLIP | 25083871 | Unspecified |
| mmu-miR-703 | MIMAT0003493 | Sec62 | 69276 | HITS-CLIP | 28735896 | Unspecified |
| mmu-miR-703 | MIMAT0003493 | 08-Sept | 20362 | HITS-CLIP | 25083871 | Unspecified |
| mmu-miR-703 | MIMAT0003493 | Slc12a2 | 20496 | HITS-CLIP | 25083871 | Unspecified |
| mmu-miR-703 | MIMAT0003493 | Slc25a51 | 230125 | HITS-CLIP | 25083871\|23597149\|21258322 | Unspecified |
| mmu-miR-703 | MIMAT0003493 | Slc30a7 | 66500 | HITS-CLIP | 25083871 | Unspecified |

| | | | | | | | |
|---|---|---|---|---|---|---|---|
| mmu-miR-703 | MIMAT0003493 | Tcf7l2 | 21416 | HITS-CLIP | | 25083871 | Unspecified |
| mmu-miR-703 | MIMAT0003493 | Tctn3 | 67590 | HITS-CLIP | | 25083871 | Unspecified |
| mmu-miR-703 | MIMAT0003493 | Tlx2 | 21909 | HITS-CLIP | | 25083871 | Unspecified |
| mmu-miR-703 | MIMAT0003493 | Tmem170b | 621976 | HITS-CLIP | | 21258322 | Unspecified |
| mmu-miR-703 | MIMAT0003493 | Ttc17 | 74569 | HITS-CLIP | | 25083871 | Unspecified |
| mmu-miR-703 | MIMAT0003493 | Ttc32 | 75516 | HITS-CLIP | | 25083871 | Unspecified |
| mmu-miR-703 | MIMAT0003493 | Usp29 | 57775 | HITS-CLIP | | 25083871 | Unspecified |
| mmu-miR-703 | MIMAT0003493 | Usp43 | 216835 | HITS-CLIP | | 25083871 | Unspecified |
| mmu-miR-703 | MIMAT0003493 | Vps26a | 30930 | HITS-CLIP | | 25083871 | Unspecified |
| mmu-miR-703 | MIMAT0003493 | Xpnpep3 | 321003 | HITS-CLIP | | 21258322 | Unspecified |
| mmu-miR-703 | MIMAT0003493 | Zbed4 | 223773 | HITS-CLIP | | 25083871 | Unspecified |
| mmu-miR-703 | MIMAT0003493 | Zbtb14 | 22666 | HITS-CLIP | | 25083871 | Unspecified |
| mmu-miR-703 | MIMAT0003493 | Zfp106 | 20402 | HITS-CLIP | | 25083871 | Unspecified |
| mmu-miR-703 | MIMAT0003493 | Zfp11 | 22648 | HITS-CLIP | | 25083871 | Unspecified |
| mmu-miR-703 | MIMAT0003493 | Zfp161 | 22666 | CLIP-Seq dataset analysis | | 25083871 | Unspecified |
| mmu-miR-703 | MIMAT0003493 | Zfp53 | 24132 | HITS-CLIP | | 25083871 | Unspecified |
| mmu-miR-703 | MIMAT0003493 | Zfp84 | 74352 | HITS-CLIP | | 25083871 | Unspecified |
| mmu-miR-466q | MIMAT0020631 | 4932443I19Rik | 403185 | HITS-CLIP | | 21258322 | Unspecified |
| mmu-miR-466q | MIMAT0020631 | 6330578E17Rik | 76178 | CLIP-Seq dataset analysis | 23142080\|25083871 | | Unspecified |
| mmu-miR-466q | MIMAT0020631 | AI429214 | 621080 | HITS-CLIP | | 28735896 | Unspecified |
| mmu-miR-466q | MIMAT0020631 | AI597468 | 103266 | HITS-CLIP | | 21258322 | Unspecified |
| mmu-miR-466q | MIMAT0020631 | Aak1 | 269774 | HITS-CLIP | | 25083871 | Unspecified |
| mmu-miR-466q | MIMAT0020631 | Acsf3 | 257633 | HITS-CLIP | | 28735896 | Unspecified |
| mmu-miR-466q | MIMAT0020631 | Adat1 | 30947 | HITS-CLIP | | 28735896 | Unspecified |
| mmu-miR-466q | MIMAT0020631 | Adig | 246747 | HITS-CLIP | | 28735896 | Unspecified |
| mmu-miR-466q | MIMAT0020631 | Agrn | 11603 | HITS-CLIP | | 28735896 | Unspecified |
| mmu-miR-466q | MIMAT0020631 | Akap5 | 238276 | HITS-CLIP | | 28735896 | Unspecified |
| mmu-miR-466q | MIMAT0020631 | Aldh1l1 | 107747 | HITS-CLIP | | 28735896 | Unspecified |
| mmu-miR-466q | MIMAT0020631 | Alkbh1 | 211064 | HITS-CLIP | | 28735896 | Unspecified |
| mmu-miR-466q | MIMAT0020631 | Ankrd28 | 105522 | HITS-CLIP | | 21258322 | Unspecified |
| mmu-miR-466q | MIMAT0020631 | Arhgef9 | 236915 | HITS-CLIP | | 21258322 | Unspecified |
| mmu-miR-466q | MIMAT0020631 | Arl14epl | 381142 | HITS-CLIP | 23597149\|25083871 | | Unspecified |
| mmu-miR-466q | MIMAT0020631 | Arl4c | 320982 | HITS-CLIP | | 21258322 | Unspecified |
| mmu-miR-466q | MIMAT0020631 | Armc8 | 74125 | HITS-CLIP | 23597149\|28735896 | | Unspecified |
| mmu-miR-466q | MIMAT0020631 | Arsi | 545260 | HITS-CLIP | | 28735896 | Unspecified |
| mmu-miR-466q | MIMAT0020631 | Atf1 | 11908 | HITS-CLIP | | 28735896 | Unspecified |
| mmu-miR-466q | MIMAT0020631 | Atf3 | 11910 | HITS-CLIP | | 28735896 | Unspecified |
| mmu-miR-466q | MIMAT0020631 | Atg10 | 66795 | HITS-CLIP | | 28735896 | Unspecified |
| mmu-miR-466q | MIMAT0020631 | Atm | 11920 | HITS-CLIP | | 28735896 | Unspecified |

| | | | | | | |
|---|---|---|---|---|---|---|
| mmu-miR-466q | MIMAT0020631 | Atpbd4 | 66632 | HITS-CLIP | 21258322 | Unspecified |
| mmu-miR-466q | MIMAT0020631 | Auts2 | 319974 | HITS-CLIP | 25083871\|21258322 | Unspecified |
| mmu-miR-466q | MIMAT0020631 | Bace1 | 23821 | HITS-CLIP | 28735896 | Unspecified |
| mmu-miR-466q | MIMAT0020631 | Bach2 | 12014 | HITS-CLIP | 25083871\|21258322\|23597149\|23142 | Unspecified |
| mmu-miR-466q | MIMAT0020631 | Bcl2l11 | 12125 | HITS-CLIP | 25083871 | Unspecified |
| mmu-miR-466q | MIMAT0020631 | Bcl2l15 | 229672 | HITS-CLIP | 25083871\|21258322 | Unspecified |
| mmu-miR-466q | MIMAT0020631 | Bloc1s6 | 18457 | HITS-CLIP | 21258322\|25083871\|23597149 | Unspecified |
| mmu-miR-466q | MIMAT0020631 | Brf1 | 72308 | HITS-CLIP | 28735896 | Unspecified |
| mmu-miR-466q | MIMAT0020631 | Btg2 | 12227 | HITS-CLIP | 23142080\|23597149 | Unspecified |
| mmu-miR-466q | MIMAT0020631 | Cavin3 | 109042 | HITS-CLIP | 28735896 | Unspecified |
| mmu-miR-466q | MIMAT0020631 | Cd69 | 12515 | HITS-CLIP | 23142080 | Unspecified |
| mmu-miR-466q | MIMAT0020631 | Cfap65 | 241116 | HITS-CLIP | 28735896 | Unspecified |
| mmu-miR-466q | MIMAT0020631 | Chrna7 | 11441 | HITS-CLIP | 28735896 | Unspecified |
| mmu-miR-466q | MIMAT0020631 | Chst15 | 77590 | HITS-CLIP | 28735896 | Unspecified |
| mmu-miR-466q | MIMAT0020631 | Clec1a | 243653 | HITS-CLIP | 28735896 | Unspecified |
| mmu-miR-466q | MIMAT0020631 | Clic5 | 224796 | HITS-CLIP | 28735896 | Unspecified |
| mmu-miR-466q | MIMAT0020631 | Coa5 | 76178 | HITS-CLIP | 23142080\|25083871 | Unspecified |
| mmu-miR-466q | MIMAT0020631 | Col22a1 | 69700 | HITS-CLIP | 28735896 | Unspecified |
| mmu-miR-466q | MIMAT0020631 | Crem | 12916 | HITS-CLIP | 28735896 | Unspecified |
| mmu-miR-466q | MIMAT0020631 | Cst3 | 13010 | HITS-CLIP | 28735896 | Unspecified |
| mmu-miR-466q | MIMAT0020631 | Ctif | 269037 | HITS-CLIP | 23597149\|19536157\|25083871 | Unspecified |
| mmu-miR-466q | MIMAT0020631 | Cyb5r2 | 320635 | HITS-CLIP | 21258322 | Unspecified |
| mmu-miR-466q | MIMAT0020631 | Dagla | 269060 | HITS-CLIP | 21258322 | Unspecified |
| mmu-miR-466q | MIMAT0020631 | Dcaf17 | 75763 | HITS-CLIP | 28735896 | Unspecified |
| mmu-miR-466q | MIMAT0020631 | Dcaf5 | 320808 | HITS-CLIP | 23142080 | Unspecified |
| mmu-miR-466q | MIMAT0020631 | Ddx21 | 56200 | HITS-CLIP | 21258322\|25083871\|23142080 | Unspecified |
| mmu-miR-466q | MIMAT0020631 | Dennd4b | 229541 | HITS-CLIP | 28735896 | Unspecified |
| mmu-miR-466q | MIMAT0020631 | Dhx33 | 216877 | HITS-CLIP | 21258322 | Unspecified |
| mmu-miR-466q | MIMAT0020631 | Dicer1 | 192119 | HITS-CLIP | 21258322 | Unspecified |
| mmu-miR-466q | MIMAT0020631 | Dnajb4 | 67035 | HITS-CLIP | 25083871\|21258322\|23597149 | Unspecified |
| mmu-miR-466q | MIMAT0020631 | Dph6 | 66632 | HITS-CLIP | 21258322 | Unspecified |
| mmu-miR-466q | MIMAT0020631 | Eif1ad | 69860 | HITS-CLIP | 25083871 | Unspecified |
| mmu-miR-466q | MIMAT0020631 | Eif4e | 13684 | HITS-CLIP | 28735896 | Unspecified |
| mmu-miR-466q | MIMAT0020631 | Enah | 13800 | HITS-CLIP | 28735896 | Unspecified |
| mmu-miR-466q | MIMAT0020631 | Ercc6 | 319955 | HITS-CLIP | 23597149 | Unspecified |
| mmu-miR-466q | MIMAT0020631 | Erlec1 | 66753 | HITS-CLIP | 28735896 | Unspecified |
| mmu-miR-466q | MIMAT0020631 | Etv3 | 27049 | HITS-CLIP | 25083871 | Unspecified |
| mmu-miR-466q | MIMAT0020631 | F2rl1 | 14063 | HITS-CLIP | 23597149\|19536157\|26203562\|25083 | Unspecified |
| mmu-miR-466q | MIMAT0020631 | Fam160a1 | 229488 | HITS-CLIP | 21258322 | Unspecified |

| | | | | | | |
|---|---|---|---|---|---|---|
| mmu-miR-466q | MIMAT0020631 | Fam178a | 226151 | HITS-CLIP | 25083871\|23597149 | Unspecified |
| mmu-miR-466q | MIMAT0020631 | Fam189a1 | 70638 | HITS-CLIP | 28735896 | Unspecified |
| mmu-miR-466q | MIMAT0020631 | Fam212b | 109050 | HITS-CLIP | 28735896 | Unspecified |
| mmu-miR-466q | MIMAT0020631 | Fam53c | 66306 | HITS-CLIP | 25083871\|21258322 | Unspecified |
| mmu-miR-466q | MIMAT0020631 | Fam69c | 240479 | HITS-CLIP | 23142080\|23597149\|25083871 | Unspecified |
| mmu-miR-466q | MIMAT0020631 | Fezf2 | 54713 | HITS-CLIP | 28735896 | Unspecified |
| mmu-miR-466q | MIMAT0020631 | Ffar3 | 233080 | HITS-CLIP | 25083871\|21258322 | Unspecified |
| mmu-miR-466q | MIMAT0020631 | Fhod1 | 234686 | HITS-CLIP | 28735896 | Unspecified |
| mmu-miR-466q | MIMAT0020631 | Fkbp6 | 94244 | HITS-CLIP | 28735896 | Unspecified |
| mmu-miR-466q | MIMAT0020631 | Fosl2 | 14284 | HITS-CLIP | 19536157\|23142080 | Unspecified |
| mmu-miR-466q | MIMAT0020631 | Foxf2 | 14238 | HITS-CLIP | 28735896 | Unspecified |
| mmu-miR-466q | MIMAT0020631 | Foxk1 | 17425 | HITS-CLIP | 25083871\|21258322 | Unspecified |
| mmu-miR-466q | MIMAT0020631 | Gabarapl2 | 93739 | HITS-CLIP | 28735896 | Unspecified |
| mmu-miR-466q | MIMAT0020631 | Gcfc1 | 67367 | CLIP-Seq dataset analysis | 25083871\|23597149\|21258322 | Unspecified |
| mmu-miR-466q | MIMAT0020631 | Gja5 | 14613 | HITS-CLIP | 21258322\|25083871\|23597149\|28735 | Unspecified |
| mmu-miR-466q | MIMAT0020631 | Gm13476 | 433424 | HITS-CLIP | 25083871 | Unspecified |
| mmu-miR-466q | MIMAT0020631 | Gm13889 | 620695 | HITS-CLIP | 28735896 | Unspecified |
| mmu-miR-466q | MIMAT0020631 | Gm17174 | 102631744 | HITS-CLIP | 28735896 | Unspecified |
| mmu-miR-466q | MIMAT0020631 | Gm5148 | 381438 | HITS-CLIP | 23142080\|25083871 | Unspecified |
| mmu-miR-466q | MIMAT0020631 | Gm949 | 381142 | CLIP-Seq dataset analysis | 23597149\|25083871 | Unspecified |
| mmu-miR-466q | MIMAT0020631 | Gmeb1 | 56809 | HITS-CLIP | 21258322 | Unspecified |
| mmu-miR-466q | MIMAT0020631 | Grik3 | 14807 | HITS-CLIP | 23597149\|25083871\|19536157 | Unspecified |
| mmu-miR-466q | MIMAT0020631 | Gsk3b | 56637 | HITS-CLIP | 28735896 | Unspecified |
| mmu-miR-466q | MIMAT0020631 | Hist4h4 | 320332 | HITS-CLIP | 21258322 | Unspecified |
| mmu-miR-466q | MIMAT0020631 | Homer2 | 26557 | HITS-CLIP | 25083871\|19536157 | Unspecified |
| mmu-miR-466q | MIMAT0020631 | Hrk | 12123 | HITS-CLIP | 19536157\|23597149 | Unspecified |
| mmu-miR-466q | MIMAT0020631 | Hs3st3b1 | 54710 | HITS-CLIP | 25083871\|21258322 | Unspecified |
| mmu-miR-466q | MIMAT0020631 | Ido2 | 209176 | HITS-CLIP | 28735896 | Unspecified |
| mmu-miR-466q | MIMAT0020631 | Igsf9 | 93842 | HITS-CLIP | 28735896 | Unspecified |
| mmu-miR-466q | MIMAT0020631 | Ilf3 | 16201 | HITS-CLIP | 25083871 | Unspecified |
| mmu-miR-466q | MIMAT0020631 | Iqsec3 | 243621 | HITS-CLIP | 21258322 | Unspecified |
| mmu-miR-466q | MIMAT0020631 | Itpk1 | 217837 | HITS-CLIP | 28735896 | Unspecified |
| mmu-miR-466q | MIMAT0020631 | Itpripl2 | 319622 | HITS-CLIP | 21258322 | Unspecified |
| mmu-miR-466q | MIMAT0020631 | Iws1 | 73473 | HITS-CLIP | 21258322\|23597149 | Unspecified |
| mmu-miR-466q | MIMAT0020631 | Kcnb1 | 16500 | HITS-CLIP | 28735896 | Unspecified |
| mmu-miR-466q | MIMAT0020631 | Kcng3 | 225030 | HITS-CLIP | 28735896 | Unspecified |
| mmu-miR-466q | MIMAT0020631 | Kcnj12 | 16515 | HITS-CLIP | 28735896 | Unspecified |
| mmu-miR-466q | MIMAT0020631 | Klhl5 | 71778 | HITS-CLIP | 28735896 | Unspecified |
| mmu-miR-466q | MIMAT0020631 | Lin54 | 231506 | HITS-CLIP | 25083871 | Unspecified |

| | | | | | | |
|---|---|---|---|---|---|---|
| mmu-miR-466q | MIMAT0020631 | Llgl1 | 16897 | HITS-CLIP | 21258322 | Unspecified |
| mmu-miR-466q | MIMAT0020631 | Lrrc4c | 241568 | HITS-CLIP | 28735896 | Unspecified |
| mmu-miR-466q | MIMAT0020631 | Ltf | 17002 | HITS-CLIP | 25083871 | Unspecified |
| mmu-miR-466q | MIMAT0020631 | Mcph1 | 244329 | HITS-CLIP | 28735896 | Unspecified |
| mmu-miR-466q | MIMAT0020631 | Mdga1 | 74762 | HITS-CLIP | 28735896 | Unspecified |
| mmu-miR-466q | MIMAT0020631 | Med20 | 56771 | HITS-CLIP | 28735896 | Unspecified |
| mmu-miR-466q | MIMAT0020631 | Memo1 | 76890 | HITS-CLIP | 28735896 | Unspecified |
| mmu-miR-466q | MIMAT0020631 | Mfsd7b | 226844 | HITS-CLIP | 23142080 | Unspecified |
| mmu-miR-466q | MIMAT0020631 | Mrc2 | 17534 | HITS-CLIP | 28735896 | Unspecified |
| mmu-miR-466q | MIMAT0020631 | Mrps24 | 64660 | HITS-CLIP | 25083871\|23597149\|21258322 | Unspecified |
| mmu-miR-466q | MIMAT0020631 | Mtpn | 14489 | HITS-CLIP | 28735896 | Unspecified |
| mmu-miR-466q | MIMAT0020631 | Mtx3 | 382793 | HITS-CLIP | 25083871 | Unspecified |
| mmu-miR-466q | MIMAT0020631 | Mxi1 | 17859 | HITS-CLIP | 21258322 | Unspecified |
| mmu-miR-466q | MIMAT0020631 | Mylk | 107589 | HITS-CLIP | 25083871 | Unspecified |
| mmu-miR-466q | MIMAT0020631 | Myo7a | 17921 | HITS-CLIP | 28735896 | Unspecified |
| mmu-miR-466q | MIMAT0020631 | Nacc2 | 67991 | HITS-CLIP | 21258322\|23597149\|25083871 | Unspecified |
| mmu-miR-466q | MIMAT0020631 | Nav2 | 78286 | HITS-CLIP | 28735896 | Unspecified |
| mmu-miR-466q | MIMAT0020631 | Ndufc1 | 66377 | HITS-CLIP | 28735896 | Unspecified |
| mmu-miR-466q | MIMAT0020631 | Nectin1 | 58235 | HITS-CLIP | 23142080 | Unspecified |
| mmu-miR-466q | MIMAT0020631 | Neu3 | 50877 | HITS-CLIP | 21258322 | Unspecified |
| mmu-miR-466q | MIMAT0020631 | Neurl1b | 240055 | HITS-CLIP | 23597149\|25083871\|23142080 | Unspecified |
| mmu-miR-466q | MIMAT0020631 | Nfatc2 | 18019 | HITS-CLIP | 23142080\|21258322\|23597149 | Unspecified |
| mmu-miR-466q | MIMAT0020631 | Nfatc4 | 73181 | HITS-CLIP | 28735896 | Unspecified |
| mmu-miR-466q | MIMAT0020631 | Nmnat3 | 74080 | HITS-CLIP | 23142080 | Unspecified |
| mmu-miR-466q | MIMAT0020631 | Npbwr1 | 226304 | HITS-CLIP | 28735896 | Unspecified |
| mmu-miR-466q | MIMAT0020631 | Nras | 18176 | HITS-CLIP | 19536157\|23597149\|25083871\|26203 | Unspecified |
| mmu-miR-466q | MIMAT0020631 | Nrg3 | 18183 | HITS-CLIP | 19536157\|23597149 | Unspecified |
| mmu-miR-466q | MIMAT0020631 | Ntrk3 | 18213 | HITS-CLIP | 28735896 | Unspecified |
| mmu-miR-466q | MIMAT0020631 | Nudt18 | 213484 | HITS-CLIP | 28735896 | Unspecified |
| mmu-miR-466q | MIMAT0020631 | Nup93 | 71805 | HITS-CLIP | 28735896 | Unspecified |
| mmu-miR-466q | MIMAT0020631 | Nyap2 | 241134 | HITS-CLIP | 28735896 | Unspecified |
| mmu-miR-466q | MIMAT0020631 | Olfr1342 | 258708 | HITS-CLIP | 28735896 | Unspecified |
| mmu-miR-466q | MIMAT0020631 | Olfr1393 | 258463 | HITS-CLIP | 28735896 | Unspecified |
| mmu-miR-466q | MIMAT0020631 | Olfr1512 | 258424 | HITS-CLIP | 28735896 | Unspecified |
| mmu-miR-466q | MIMAT0020631 | Pard3b | 72823 | HITS-CLIP | 25083871\|23597149 | Unspecified |
| mmu-miR-466q | MIMAT0020631 | Parg | 26430 | HITS-CLIP | 28735896 | Unspecified |
| mmu-miR-466q | MIMAT0020631 | Paxbp1 | 67367 | HITS-CLIP | 25083871\|21258322\|23597149 | Unspecified |
| mmu-miR-466q | MIMAT0020631 | Pbx1 | 18514 | HITS-CLIP | 28735896 | Unspecified |
| mmu-miR-466q | MIMAT0020631 | Pck1 | 18534 | HITS-CLIP | 28735896 | Unspecified |

| | | | | | | |
|---|---|---|---|---|---|---|
| mmu-miR-466q | MIMAT0020631 | Pde1a | 18573 | HITS-CLIP | 28735896 | Unspecified |
| mmu-miR-466q | MIMAT0020631 | Pkib | 18768 | HITS-CLIP | 21258322\|23597149 | Unspecified |
| mmu-miR-466q | MIMAT0020631 | Plcl1 | 227120 | HITS-CLIP | 28735896 | Unspecified |
| mmu-miR-466q | MIMAT0020631 | Pldn | 18457 | CLIP-Seq dataset analysis | 25083871\|23597149\|21258322 | Unspecified |
| mmu-miR-466q | MIMAT0020631 | Plekhg3 | 263406 | HITS-CLIP | 28735896 | Unspecified |
| mmu-miR-466q | MIMAT0020631 | Plet1 | 76509 | HITS-CLIP | 28735896 | Unspecified |
| mmu-miR-466q | MIMAT0020631 | Pnrc2 | 52830 | HITS-CLIP | 28735896 | Unspecified |
| mmu-miR-466q | MIMAT0020631 | Pou3f3 | 18993 | HITS-CLIP | 28735896 | Unspecified |
| mmu-miR-466q | MIMAT0020631 | Prkab2 | 108097 | HITS-CLIP | 21258322 | Unspecified |
| mmu-miR-466q | MIMAT0020631 | Prnd | 26434 | HITS-CLIP | 23142080\|23597149\|21258322 | Unspecified |
| mmu-miR-466q | MIMAT0020631 | Prpf38b | 66921 | HITS-CLIP | 28735896 | Unspecified |
| mmu-miR-466q | MIMAT0020631 | Prrg3 | 208748 | HITS-CLIP | 26203562\|21258322\|19536157\|23597 | Unspecified |
| mmu-miR-466q | MIMAT0020631 | Psma4 | 26441 | HITS-CLIP | 28735896 | Unspecified |
| mmu-miR-466q | MIMAT0020631 | Pten | 19211 | HITS-CLIP | 25083871\|23142080 | Unspecified |
| mmu-miR-466q | MIMAT0020631 | Pvrl1 | 58235 | HITS-CLIP | 23142080 | Unspecified |
| mmu-miR-466q | MIMAT0020631 | Rad51c | 114714 | HITS-CLIP | 25083871\|23142080\|28735896 | Unspecified |
| mmu-miR-466q | MIMAT0020631 | Rapgefl1 | 268480 | HITS-CLIP | 28735896 | Unspecified |
| mmu-miR-466q | MIMAT0020631 | Rasal2 | 226525 | HITS-CLIP | 28735896 | Unspecified |
| mmu-miR-466q | MIMAT0020631 | Rbbp8nl | 271887 | HITS-CLIP | 28735896 | Unspecified |
| mmu-miR-466q | MIMAT0020631 | Rflnb | 76566 | HITS-CLIP | 28735896 | Unspecified |
| mmu-miR-466q | MIMAT0020631 | Rhbdl3 | 246104 | HITS-CLIP | 25083871 | Unspecified |
| mmu-miR-466q | MIMAT0020631 | Rhobtb1 | 69288 | HITS-CLIP | 25083871 | Unspecified |
| mmu-miR-466q | MIMAT0020631 | Rmi2 | 223970 | HITS-CLIP | 28735896 | Unspecified |
| mmu-miR-466q | MIMAT0020631 | Rmnd5a | 68477 | HITS-CLIP | 23597149 | Unspecified |
| mmu-miR-466q | MIMAT0020631 | Rnf130 | 59044 | HITS-CLIP | 28735896 | Unspecified |
| mmu-miR-466q | MIMAT0020631 | Rnf41 | 67588 | HITS-CLIP | 23597149\|21258322 | Unspecified |
| mmu-miR-466q | MIMAT0020631 | Rpf2 | 67239 | HITS-CLIP | 28735896 | Unspecified |
| mmu-miR-466q | MIMAT0020631 | Rrn3 | 106298 | HITS-CLIP | 21258322 | Unspecified |
| mmu-miR-466q | MIMAT0020631 | Rsf1 | 233532 | HITS-CLIP | 28735896 | Unspecified |
| mmu-miR-466q | MIMAT0020631 | S100pbp | 74648 | HITS-CLIP | 28735896 | Unspecified |
| mmu-miR-466q | MIMAT0020631 | Sec62 | 69276 | HITS-CLIP | 28735896 | Unspecified |
| mmu-miR-466q | MIMAT0020631 | Serpina1a | 20700 | HITS-CLIP | 28735896 | Unspecified |
| mmu-miR-466q | MIMAT0020631 | Sgms1 | 208449 | HITS-CLIP | 25083871 | Unspecified |
| mmu-miR-466q | MIMAT0020631 | Slamf7 | 75345 | HITS-CLIP | 28735896 | Unspecified |
| mmu-miR-466q | MIMAT0020631 | Slc10a5 | 241877 | HITS-CLIP | 23597149 | Unspecified |
| mmu-miR-466q | MIMAT0020631 | Slc39a14 | 213053 | HITS-CLIP | 25083871 | Unspecified |
| mmu-miR-466q | MIMAT0020631 | Slc7a13 | 74087 | HITS-CLIP | 28735896 | Unspecified |
| mmu-miR-466q | MIMAT0020631 | Slc7a2 | 11988 | HITS-CLIP | 23597149\|28735896 | Unspecified |
| mmu-miR-466q | MIMAT0020631 | Slc8a2 | 110891 | HITS-CLIP | 28735896 | Unspecified |

| | | | | | | |
|---|---|---|---|---|---|---|
| mmu-miR-466q | MIMAT0020631 | Slf2 | 226151 | HITS-CLIP | 23597149\|25083871 | Unspecified |
| mmu-miR-466q | MIMAT0020631 | Smad2 | 17126 | HITS-CLIP | 37986224\|28735896 | Unspecified |
| mmu-miR-466q | MIMAT0020631 | Spc24 | 67629 | HITS-CLIP | 21258322 | Unspecified |
| mmu-miR-466q | MIMAT0020631 | Spg7 | 234847 | HITS-CLIP | 28735896 | Unspecified |
| mmu-miR-466q | MIMAT0020631 | Spock2 | 94214 | HITS-CLIP | 28735896 | Unspecified |
| mmu-miR-466q | MIMAT0020631 | Srek1 | 218543 | HITS-CLIP | 28735896 | Unspecified |
| mmu-miR-466q | MIMAT0020631 | Srfbp1 | 67222 | HITS-CLIP | 28735896 | Unspecified |
| mmu-miR-466q | MIMAT0020631 | St3gal5 | 20454 | HITS-CLIP | 28735896 | Unspecified |
| mmu-miR-466q | MIMAT0020631 | St6galnac5 | 26938 | HITS-CLIP | 28735896 | Unspecified |
| mmu-miR-466q | MIMAT0020631 | Svopl | 320590 | HITS-CLIP | 21258322 | Unspecified |
| mmu-miR-466q | MIMAT0020631 | Synrg | 217030 | HITS-CLIP | 28735896 | Unspecified |
| mmu-miR-466q | MIMAT0020631 | Sytl4 | 27359 | HITS-CLIP | 25083871\|21258322\|23142080 | Unspecified |
| mmu-miR-466q | MIMAT0020631 | Tacr2 | 21337 | HITS-CLIP | 21258322 | Unspecified |
| mmu-miR-466q | MIMAT0020631 | Tagap | 72536 | HITS-CLIP | 23142080 | Unspecified |
| mmu-miR-466q | MIMAT0020631 | Tagap1 | 380608 | HITS-CLIP | 23142080 | Unspecified |
| mmu-miR-466q | MIMAT0020631 | Tcf12 | 21406 | HITS-CLIP | 28735896 | Unspecified |
| mmu-miR-466q | MIMAT0020631 | Them7 | 74088 | HITS-CLIP | 28735896 | Unspecified |
| mmu-miR-466q | MIMAT0020631 | Tll1 | 21892 | HITS-CLIP | 21258322\|23597149 | Unspecified |
| mmu-miR-466q | MIMAT0020631 | Tlr6 | 21899 | HITS-CLIP | 28735896 | Unspecified |
| mmu-miR-466q | MIMAT0020631 | Tmc7 | 209760 | HITS-CLIP | 23142080\|21258322 | Unspecified |
| mmu-miR-466q | MIMAT0020631 | Tmem151b | 210573 | HITS-CLIP | 28735896 | Unspecified |
| mmu-miR-466q | MIMAT0020631 | Tmem196 | 217951 | HITS-CLIP | 28735896 | Unspecified |
| mmu-miR-466q | MIMAT0020631 | Tmem220 | 338369 | HITS-CLIP | 28735896 | Unspecified |
| mmu-miR-466q | MIMAT0020631 | Tmem263 | 103266 | HITS-CLIP | 21258322 | Unspecified |
| mmu-miR-466q | MIMAT0020631 | Tmprss3 | 140765 | HITS-CLIP | 21258322 | Unspecified |
| mmu-miR-466q | MIMAT0020631 | Tnfrsf9 | 21942 | HITS-CLIP | 25083871\|26203562\|23597149\|1953( | Unspecified |
| mmu-miR-466q | MIMAT0020631 | Tnfsf10 | 22035 | HITS-CLIP | 21258322 | Unspecified |
| mmu-miR-466q | MIMAT0020631 | Tns4 | 217169 | HITS-CLIP | 25083871 | Unspecified |
| mmu-miR-466q | MIMAT0020631 | Trim25 | 217069 | HITS-CLIP | 28735896 | Unspecified |
| mmu-miR-466q | MIMAT0020631 | Trim36 | 28105 | HITS-CLIP | 28735896 | Unspecified |
| mmu-miR-466q | MIMAT0020631 | Trim44 | 80985 | HITS-CLIP | 28735896 | Unspecified |
| mmu-miR-466q | MIMAT0020631 | Trim62 | 67525 | HITS-CLIP | 28735896 | Unspecified |
| mmu-miR-466q | MIMAT0020631 | Trim71 | 636931 | HITS-CLIP | 21258322 | Unspecified |
| mmu-miR-466q | MIMAT0020631 | Trmt61a | 328162 | HITS-CLIP | 28735896 | Unspecified |
| mmu-miR-466q | MIMAT0020631 | Trpc6 | 22068 | HITS-CLIP | 28735896 | Unspecified |
| mmu-miR-466q | MIMAT0020631 | Tubal3 | 238463 | HITS-CLIP | 19536157\|23597149\|25083871 | Unspecified |
| mmu-miR-466q | MIMAT0020631 | Ubxn2b | 68053 | HITS-CLIP | 23142080 | Unspecified |
| mmu-miR-466q | MIMAT0020631 | Uggt1 | 320011 | HITS-CLIP | 28735896 | Unspecified |
| mmu-miR-466q | MIMAT0020631 | Uggt2 | 66435 | HITS-CLIP | 28735896 | Unspecified |

| | | | | | | |
|---|---|---|---|---|---|---|
| mmu-miR-466q | MIMAT0020631 | Uhrf1bp1l | 75089 | HITS-CLIP | 25083871 | Unspecified |
| mmu-miR-466q | MIMAT0020631 | Unc5b | 107449 | HITS-CLIP | 25083871\|19536157\|23597149 | Unspecified |
| mmu-miR-466q | MIMAT0020631 | Vangl2 | 93840 | HITS-CLIP | 28735896 | Unspecified |
| mmu-miR-466q | MIMAT0020631 | Vwa5a | 67776 | HITS-CLIP | 28735896 | Unspecified |
| mmu-miR-466q | MIMAT0020631 | Wbp1l | 226178 | HITS-CLIP | 37986224\|23597149\|28735896 | Unspecified |
| mmu-miR-466q | MIMAT0020631 | Wdr12 | 57750 | HITS-CLIP | 28735896 | Unspecified |
| mmu-miR-466q | MIMAT0020631 | Wdr26 | 226757 | HITS-CLIP | 28735896 | Unspecified |
| mmu-miR-466q | MIMAT0020631 | Wwp2 | 66894 | HITS-CLIP | 25083871\|21258322 | Unspecified |
| mmu-miR-466q | MIMAT0020631 | Xkr9 | 381246 | HITS-CLIP | 28735896 | Unspecified |
| mmu-miR-466q | MIMAT0020631 | Xylt1 | 233781 | HITS-CLIP | 28735896 | Unspecified |
| mmu-miR-466q | MIMAT0020631 | Xylt2 | 217119 | HITS-CLIP | 28735896 | Unspecified |
| mmu-miR-466q | MIMAT0020631 | Yod1 | 226418 | HITS-CLIP | 23142080 | Unspecified |
| mmu-miR-466q | MIMAT0020631 | Zbtb16 | 235320 | HITS-CLIP | 28735896 | Unspecified |
| mmu-miR-466q | MIMAT0020631 | Zbtb7b | 22724 | HITS-CLIP | 25083871\|23597149 | Unspecified |
| mmu-miR-466q | MIMAT0020631 | Zc3h12d | 237256 | HITS-CLIP | 21258322 | Unspecified |
| mmu-miR-466q | MIMAT0020631 | Zeb2os | 433424 | HITS-CLIP | 25083871 | Unspecified |
| mmu-miR-466q | MIMAT0020631 | Zfand5 | 22682 | HITS-CLIP | 28735896 | Unspecified |
| mmu-miR-466q | MIMAT0020631 | Zfp157 | 72154 | HITS-CLIP | 21258322 | Unspecified |
| mmu-miR-466q | MIMAT0020631 | Zfp26 | 22688 | HITS-CLIP | 25083871 | Unspecified |
| mmu-miR-466q | MIMAT0020631 | Zfp317 | 244713 | HITS-CLIP | 21258322 | Unspecified |
| mmu-miR-466q | MIMAT0020631 | Zfp326 | 54367 | HITS-CLIP | 28735896 | Unspecified |
| mmu-miR-466q | MIMAT0020631 | Zfp648 | 100503355 | HITS-CLIP | 28735896 | Unspecified |
| mmu-miR-466q | MIMAT0020631 | Zfyve26 | 211978 | HITS-CLIP | 37986224\|28735896 | Unspecified |
| mmu-miR-466q | MIMAT0020631 | Zgrf1 | 71643 | HITS-CLIP | 28735896 | Unspecified |
| mmu-miR-344d-3p | MIMAT0014808 | 1200011I18Rik | 67467 | HITS-CLIP | 25083871 | Unspecified |
| mmu-miR-344d-3p | MIMAT0014808 | 4921524J17Rik | 66714 | HITS-CLIP | 25083871 | Unspecified |
| mmu-miR-344d-3p | MIMAT0014808 | Acbd5 | 74159 | HITS-CLIP | 25083871 | Unspecified |
| mmu-miR-344d-3p | MIMAT0014808 | Akna | 100182 | HITS-CLIP | 25083871 | Unspecified |
| mmu-miR-344d-3p | MIMAT0014808 | Anln | 68743 | HITS-CLIP | 25083871\|23142080\|21258322 | Unspecified |
| mmu-miR-344d-3p | MIMAT0014808 | Arhgap11a | 228482 | HITS-CLIP | 25083871 | Unspecified |
| mmu-miR-344d-3p | MIMAT0014808 | Arl15 | 218639 | HITS-CLIP | 25083871 | Unspecified |
| mmu-miR-344d-3p | MIMAT0014808 | Atp2b2 | 11941 | HITS-CLIP | 25083871 | Unspecified |
| mmu-miR-344d-3p | MIMAT0014808 | Cacnb2 | 12296 | HITS-CLIP | 25083871 | Unspecified |
| mmu-miR-344d-3p | MIMAT0014808 | Calm1 | 12313 | HITS-CLIP | 25083871 | Unspecified |
| mmu-miR-344d-3p | MIMAT0014808 | Casc4 | 319996 | HITS-CLIP | 25083871 | Unspecified |
| mmu-miR-344d-3p | MIMAT0014808 | Cbx3 | 12417 | HITS-CLIP | 25083871 | Unspecified |
| mmu-miR-344d-3p | MIMAT0014808 | Ccdc90b | 66365 | HITS-CLIP | 25083871 | Unspecified |
| mmu-miR-344d-3p | MIMAT0014808 | Cd69 | 12515 | HITS-CLIP | 23142080 | Unspecified |
| mmu-miR-344d-3p | MIMAT0014808 | Ceacam1 | 26365 | HITS-CLIP | 23142080 | Unspecified |

| | | | | | | |
|---|---|---|---|---|---|---|
| mmu-miR-344d-3p | MIMAT0014808 | Cops7b | 26895 | HITS-CLIP | 25083871 | Unspecified |
| mmu-miR-344d-3p | MIMAT0014808 | Crebzf | 233490 | HITS-CLIP | 25083871 | Unspecified |
| mmu-miR-344d-3p | MIMAT0014808 | Crp | 12944 | HITS-CLIP | 25083871 | Unspecified |
| mmu-miR-344d-3p | MIMAT0014808 | Ctxn3 | 629147 | HITS-CLIP | 25083871 | Unspecified |
| mmu-miR-344d-3p | MIMAT0014808 | Cxxc5 | 67393 | HITS-CLIP | 25083871 | Unspecified |
| mmu-miR-344d-3p | MIMAT0014808 | Dimt1 | 66254 | HITS-CLIP | 25083871 | Unspecified |
| mmu-miR-344d-3p | MIMAT0014808 | Dlx3 | 13393 | HITS-CLIP | 25083871 | Unspecified |
| mmu-miR-344d-3p | MIMAT0014808 | Eif2s1 | 13665 | HITS-CLIP | 25083871 | Unspecified |
| mmu-miR-344d-3p | MIMAT0014808 | Elavl4 | 15572 | HITS-CLIP | 25083871 | Unspecified |
| mmu-miR-344d-3p | MIMAT0014808 | Ep300 | 328572 | HITS-CLIP | 25083871 | Unspecified |
| mmu-miR-344d-3p | MIMAT0014808 | Erc1 | 111173 | HITS-CLIP | 25083871\|23597149\|19536157 | Unspecified |
| mmu-miR-344d-3p | MIMAT0014808 | Erlec1 | 66753 | HITS-CLIP | 28735896 | Unspecified |
| mmu-miR-344d-3p | MIMAT0014808 | Etfbkmt | 320204 | HITS-CLIP | 25083871 | Unspecified |
| mmu-miR-344d-3p | MIMAT0014808 | Etv3 | 27049 | HITS-CLIP | 23142080 | Unspecified |
| mmu-miR-344d-3p | MIMAT0014808 | Fam151b | 73942 | HITS-CLIP | 25083871 | Unspecified |
| mmu-miR-344d-3p | MIMAT0014808 | Fam46a | 212943 | HITS-CLIP | 25083871 | Unspecified |
| mmu-miR-344d-3p | MIMAT0014808 | Fbxl20 | 72194 | HITS-CLIP | 25083871 | Unspecified |
| mmu-miR-344d-3p | MIMAT0014808 | Fgf16 | 80903 | HITS-CLIP | 25083871 | Unspecified |
| mmu-miR-344d-3p | MIMAT0014808 | Folh1 | 53320 | HITS-CLIP | 25083871 | Unspecified |
| mmu-miR-344d-3p | MIMAT0014808 | Fsd1l | 319636 | HITS-CLIP | 25083871 | Unspecified |
| mmu-miR-344d-3p | MIMAT0014808 | Fzd1 | 14362 | HITS-CLIP | 25083871 | Unspecified |
| mmu-miR-344d-3p | MIMAT0014808 | Fzd5 | 14367 | HITS-CLIP | 25083871 | Unspecified |
| mmu-miR-344d-3p | MIMAT0014808 | Gm8369 | 666926 | HITS-CLIP | 25083871 | Unspecified |
| mmu-miR-344d-3p | MIMAT0014808 | Gpalpp1 | 67467 | HITS-CLIP | 25083871 | Unspecified |
| mmu-miR-344d-3p | MIMAT0014808 | Gpc6 | 23888 | HITS-CLIP | 25083871 | Unspecified |
| mmu-miR-344d-3p | MIMAT0014808 | Grhl3 | 230824 | HITS-CLIP | 25083871 | Unspecified |
| mmu-miR-344d-3p | MIMAT0014808 | Has2 | 15117 | HITS-CLIP | 25083871 | Unspecified |
| mmu-miR-344d-3p | MIMAT0014808 | Hexim1 | 192231 | HITS-CLIP | 25083871 | Unspecified |
| mmu-miR-344d-3p | MIMAT0014808 | Hiat1 | 15247 | HITS-CLIP | 25083871 | Unspecified |
| mmu-miR-344d-3p | MIMAT0014808 | Hist1h1d | 14957 | HITS-CLIP | 25083871 | Unspecified |
| mmu-miR-344d-3p | MIMAT0014808 | Hopx | 74318 | HITS-CLIP | 25083871 | Unspecified |
| mmu-miR-344d-3p | MIMAT0014808 | Hoxa11 | 15396 | HITS-CLIP | 25083871 | Unspecified |
| mmu-miR-344d-3p | MIMAT0014808 | Id2 | 15902 | HITS-CLIP | 25083871 | Unspecified |
| mmu-miR-344d-3p | MIMAT0014808 | Igfbp3 | 16009 | HITS-CLIP | 25083871 | Unspecified |
| mmu-miR-344d-3p | MIMAT0014808 | Ikzf5 | 67143 | HITS-CLIP | 25083871 | Unspecified |
| mmu-miR-344d-3p | MIMAT0014808 | Ints6 | 18130 | HITS-CLIP | 23142080 | Unspecified |
| mmu-miR-344d-3p | MIMAT0014808 | Ism1 | 319909 | HITS-CLIP | 25083871 | Unspecified |
| mmu-miR-344d-3p | MIMAT0014808 | Jade1 | 269424 | HITS-CLIP | 25083871 | Unspecified |
| mmu-miR-344d-3p | MIMAT0014808 | Klf12 | 16597 | HITS-CLIP | 25083871 | Unspecified |

| | | | | | | |
|---|---|---|---|---|---|---|
| mmu-miR-344d-3p | MIMAT0014808 | L3mbtl4 | 320858 | HITS-CLIP | 25083871\|23142080 | Unspecified |
| mmu-miR-344d-3p | MIMAT0014808 | Larp1 | 73158 | HITS-CLIP | 25083871 | Unspecified |
| mmu-miR-344d-3p | MIMAT0014808 | Lin54 | 231506 | HITS-CLIP | 25083871 | Unspecified |
| mmu-miR-344d-3p | MIMAT0014808 | Lrig2 | 269473 | HITS-CLIP | 23142080 | Unspecified |
| mmu-miR-344d-3p | MIMAT0014808 | Lrrc71 | 74485 | HITS-CLIP | 25083871 | Unspecified |
| mmu-miR-344d-3p | MIMAT0014808 | Lrrtm2 | 107065 | HITS-CLIP | 25083871 | Unspecified |
| mmu-miR-344d-3p | MIMAT0014808 | Lzts3 | 241638 | HITS-CLIP | 25083871 | Unspecified |
| mmu-miR-344d-3p | MIMAT0014808 | Map1a | 17754 | HITS-CLIP | 25083871 | Unspecified |
| mmu-miR-344d-3p | MIMAT0014808 | Mettl20 | 320204 | HITS-CLIP | 25083871 | Unspecified |
| mmu-miR-344d-3p | MIMAT0014808 | Mex3b | 108797 | HITS-CLIP | 23142080 | Unspecified |
| mmu-miR-344d-3p | MIMAT0014808 | Mfsd14a | 15247 | HITS-CLIP | 25083871 | Unspecified |
| mmu-miR-344d-3p | MIMAT0014808 | Mgea5 | 76055 | HITS-CLIP | 25083871 | Unspecified |
| mmu-miR-344d-3p | MIMAT0014808 | Mllt3 | 70122 | HITS-CLIP | 25083871 | Unspecified |
| mmu-miR-344d-3p | MIMAT0014808 | Mpped2 | 77015 | HITS-CLIP | 25083871 | Unspecified |
| mmu-miR-344d-3p | MIMAT0014808 | Nadsyn1 | 78914 | HITS-CLIP | 25083871 | Unspecified |
| mmu-miR-344d-3p | MIMAT0014808 | Nck2 | 17974 | HITS-CLIP | 25083871 | Unspecified |
| mmu-miR-344d-3p | MIMAT0014808 | Nckap5l | 380969 | HITS-CLIP | 25083871 | Unspecified |
| mmu-miR-344d-3p | MIMAT0014808 | Ndnf | 68169 | HITS-CLIP | 25083871 | Unspecified |
| mmu-miR-344d-3p | MIMAT0014808 | Nf2 | 18016 | HITS-CLIP | 23597149 | Unspecified |
| mmu-miR-344d-3p | MIMAT0014808 | Nfib | 18028 | HITS-CLIP | 25083871 | Unspecified |
| mmu-miR-344d-3p | MIMAT0014808 | Nln | 75805 | HITS-CLIP | 25083871 | Unspecified |
| mmu-miR-344d-3p | MIMAT0014808 | Nol7 | 70078 | HITS-CLIP | 25083871 | Unspecified |
| mmu-miR-344d-3p | MIMAT0014808 | Nsd3 | 234135 | HITS-CLIP | 25083871 | Unspecified |
| mmu-miR-344d-3p | MIMAT0014808 | Ntrk3 | 18213 | HITS-CLIP | 25083871 | Unspecified |
| mmu-miR-344d-3p | MIMAT0014808 | Nupl2 | 231042 | HITS-CLIP | 25083871 | Unspecified |
| mmu-miR-344d-3p | MIMAT0014808 | Ocrl | 320634 | HITS-CLIP | 23142080 | Unspecified |
| mmu-miR-344d-3p | MIMAT0014808 | Olr1 | 108078 | HITS-CLIP | 21258322 | Unspecified |
| mmu-miR-344d-3p | MIMAT0014808 | Osbp | 76303 | HITS-CLIP | 25083871 | Unspecified |
| mmu-miR-344d-3p | MIMAT0014808 | Pcdh20 | 219257 | HITS-CLIP | 25083871 | Unspecified |
| mmu-miR-344d-3p | MIMAT0014808 | Pcdhb17 | 93888 | HITS-CLIP | 25083871 | Unspecified |
| mmu-miR-344d-3p | MIMAT0014808 | Pcsk5 | 18552 | HITS-CLIP | 25083871 | Unspecified |
| mmu-miR-344d-3p | MIMAT0014808 | Pfkl | 18641 | HITS-CLIP | 25083871\|21258322 | Unspecified |
| mmu-miR-344d-3p | MIMAT0014808 | Phf17 | 269424 | HITS-CLIP | 25083871 | Unspecified |
| mmu-miR-344d-3p | MIMAT0014808 | Pkib | 18768 | HITS-CLIP | 25083871 | Unspecified |
| mmu-miR-344d-3p | MIMAT0014808 | Plau | 18792 | HITS-CLIP | 25083871 | Unspecified |
| mmu-miR-344d-3p | MIMAT0014808 | Plaur | 18793 | HITS-CLIP | 25083871 | Unspecified |
| mmu-miR-344d-3p | MIMAT0014808 | Plxnc1 | 54712 | HITS-CLIP | 25083871 | Unspecified |
| mmu-miR-344d-3p | MIMAT0014808 | Pm20d2 | 242377 | HITS-CLIP | 25083871 | Unspecified |
| mmu-miR-344d-3p | MIMAT0014808 | Pou4f2 | 18997 | HITS-CLIP | 25083871 | Unspecified |

| | | | | | | |
|---|---|---|---|---|---|---|
| mmu-miR-344d-3p | MIMAT0014808 | Ppp4r2 | 232314 | HITS-CLIP | 25083871 | Unspecified |
| mmu-miR-344d-3p | MIMAT0014808 | Ppp6r1 | 243819 | HITS-CLIP | 25083871 | Unspecified |
| mmu-miR-344d-3p | MIMAT0014808 | Prrx1 | 18933 | HITS-CLIP | 25083871 | Unspecified |
| mmu-miR-344d-3p | MIMAT0014808 | Pten | 19211 | HITS-CLIP | 25083871 | Unspecified |
| mmu-miR-344d-3p | MIMAT0014808 | Ranbp10 | 74334 | HITS-CLIP | 25083871 | Unspecified |
| mmu-miR-344d-3p | MIMAT0014808 | Rasal2 | 226525 | HITS-CLIP | 25083871 | Unspecified |
| mmu-miR-344d-3p | MIMAT0014808 | Rassf5 | 54354 | HITS-CLIP | 25083871 | Unspecified |
| mmu-miR-344d-3p | MIMAT0014808 | Rbms3 | 207181 | HITS-CLIP | 25083871 | Unspecified |
| mmu-miR-344d-3p | MIMAT0014808 | Rccd1 | 269955 | HITS-CLIP | 25083871 | Unspecified |
| mmu-miR-344d-3p | MIMAT0014808 | Reep5 | 13476 | HITS-CLIP | 25083871 | Unspecified |
| mmu-miR-344d-3p | MIMAT0014808 | Ret | 19713 | HITS-CLIP | 25083871 | Unspecified |
| mmu-miR-344d-3p | MIMAT0014808 | Rnf166 | 68718 | HITS-CLIP | 25083871 | Unspecified |
| mmu-miR-344d-3p | MIMAT0014808 | Rnf6 | 74132 | HITS-CLIP | 23142080 | Unspecified |
| mmu-miR-344d-3p | MIMAT0014808 | Rragb | 245670 | HITS-CLIP | 25083871 | Unspecified |
| mmu-miR-344d-3p | MIMAT0014808 | Rufy2 | 70432 | HITS-CLIP | 25083871 | Unspecified |
| mmu-miR-344d-3p | MIMAT0014808 | Runx1t1 | 12395 | HITS-CLIP | 25083871 | Unspecified |
| mmu-miR-344d-3p | MIMAT0014808 | Senp3 | 80886 | HITS-CLIP | 25083871 | Unspecified |
| mmu-miR-344d-3p | MIMAT0014808 | Shh | 20423 | HITS-CLIP | 25083871 | Unspecified |
| mmu-miR-344d-3p | MIMAT0014808 | Slc12a8 | 171286 | HITS-CLIP | 25083871 | Unspecified |
| mmu-miR-344d-3p | MIMAT0014808 | Slc35a1 | 24060 | HITS-CLIP | 25083871 | Unspecified |
| mmu-miR-344d-3p | MIMAT0014808 | Slc7a11 | 26570 | HITS-CLIP | 25083871 | Unspecified |
| mmu-miR-344d-3p | MIMAT0014808 | Slc8a1 | 20541 | HITS-CLIP | 25083871 | Unspecified |
| mmu-miR-344d-3p | MIMAT0014808 | Slitrk2 | 245450 | HITS-CLIP | 23142080 | Unspecified |
| mmu-miR-344d-3p | MIMAT0014808 | Smurf2 | 66313 | HITS-CLIP | 25083871 | Unspecified |
| mmu-miR-344d-3p | MIMAT0014808 | Snx27 | 76742 | HITS-CLIP | 25083871 | Unspecified |
| mmu-miR-344d-3p | MIMAT0014808 | Sort1 | 20661 | HITS-CLIP | 25083871 | Unspecified |
| mmu-miR-344d-3p | MIMAT0014808 | Sox11 | 20666 | HITS-CLIP | 23142080 | Unspecified |
| mmu-miR-344d-3p | MIMAT0014808 | Sri | 109552 | HITS-CLIP | 25083871 | Unspecified |
| mmu-miR-344d-3p | MIMAT0014808 | Strbp | 20744 | HITS-CLIP | 25083871 | Unspecified |
| mmu-miR-344d-3p | MIMAT0014808 | Synpr | 72003 | HITS-CLIP | 25083871 | Unspecified |
| mmu-miR-344d-3p | MIMAT0014808 | Syp | 20977 | HITS-CLIP | 25083871 | Unspecified |
| mmu-miR-344d-3p | MIMAT0014808 | Tbx4 | 21387 | HITS-CLIP | 25083871 | Unspecified |
| mmu-miR-344d-3p | MIMAT0014808 | Tcf21 | 21412 | HITS-CLIP | 25083871 | Unspecified |
| mmu-miR-344d-3p | MIMAT0014808 | Tcf7l2 | 21416 | HITS-CLIP | 25083871 | Unspecified |
| mmu-miR-344d-3p | MIMAT0014808 | Tead3 | 21678 | HITS-CLIP | 25083871 | Unspecified |
| mmu-miR-344d-3p | MIMAT0014808 | Tet2 | 214133 | HITS-CLIP | 25083871 | Unspecified |
| mmu-miR-344d-3p | MIMAT0014808 | Tex12 | 66654 | HITS-CLIP | 23142080 | Unspecified |
| mmu-miR-344d-3p | MIMAT0014808 | Tfam | 21780 | HITS-CLIP | 25083871 | Unspecified |
| mmu-miR-344d-3p | MIMAT0014808 | Timp3 | 21859 | HITS-CLIP | 25083871 | Unspecified |

| | | | | | | |
|---|---|---|---|---|---|---|
| mmu-miR-344d-3p | MIMAT0014808 | Tmem100 | 67888 | HITS-CLIP | 25083871 | Unspecified |
| mmu-miR-344d-3p | MIMAT0014808 | Tmem161b | 72745 | HITS-CLIP | 25083871 | Unspecified |
| mmu-miR-344d-3p | MIMAT0014808 | Tmem56 | 99887 | HITS-CLIP | 23142080 | Unspecified |
| mmu-miR-344d-3p | MIMAT0014808 | Trak2 | 70827 | HITS-CLIP | 25083871 | Unspecified |
| mmu-miR-344d-3p | MIMAT0014808 | Trpc7 | 26946 | HITS-CLIP | 25083871 | Unspecified |
| mmu-miR-344d-3p | MIMAT0014808 | Trps1 | 83925 | HITS-CLIP | 25083871 | Unspecified |
| mmu-miR-344d-3p | MIMAT0014808 | Ttc17 | 74569 | HITS-CLIP | 25083871 | Unspecified |
| mmu-miR-344d-3p | MIMAT0014808 | Ttc39b | 69863 | HITS-CLIP | 25083871 | Unspecified |
| mmu-miR-344d-3p | MIMAT0014808 | Ugt2b35 | 243085 | HITS-CLIP | 25083871 | Unspecified |
| mmu-miR-344d-3p | MIMAT0014808 | Usp6nl | 98910 | HITS-CLIP | 25083871 | Unspecified |
| mmu-miR-344d-3p | MIMAT0014808 | Whsc1l1 | 234135 | HITS-CLIP | 25083871 | Unspecified |
| mmu-miR-344d-3p | MIMAT0014808 | Wnt11 | 22411 | HITS-CLIP | 25083871 | Unspecified |
| mmu-miR-344d-3p | MIMAT0014808 | Wnt3 | 22415 | HITS-CLIP | 25083871 | Unspecified |
| mmu-miR-344d-3p | MIMAT0014808 | Wwtr1 | 97064 | HITS-CLIP | 25083871 | Unspecified |
| mmu-miR-344d-3p | MIMAT0014808 | Zbed6 | 667118 | HITS-CLIP | 25083871 | Unspecified |
| mmu-miR-344d-3p | MIMAT0014808 | Zc3hav1l | 209032 | HITS-CLIP | 25083871 | Unspecified |
| mmu-miR-344d-3p | MIMAT0014808 | Zer1 | 227693 | HITS-CLIP | 25083871 | Unspecified |
| mmu-miR-344d-3p | MIMAT0014808 | Zfand5 | 22682 | HITS-CLIP | 25083871 | Unspecified |
| mmu-miR-344d-3p | MIMAT0014808 | Zfp148 | 22661 | HITS-CLIP | 25083871 | Unspecified |
| mmu-miR-344d-3p | MIMAT0014808 | Zfp384 | 269800 | HITS-CLIP | 25083871 | Unspecified |
| mmu-miR-344d-3p | MIMAT0014808 | Zfp518b | 100515 | HITS-CLIP | 25083871 | Unspecified |
| mmu-miR-344d-3p | MIMAT0014808 | Zfp644 | 52397 | HITS-CLIP | 25083871 | Unspecified |
| mmu-miR-344d-3p | MIMAT0014808 | Zmynd8 | 228880 | HITS-CLIP | 25083871 | Unspecified |
| mmu-miR-344d-3p | MIMAT0014808 | Znrf1 | 170737 | HITS-CLIP | 23142080 | Unspecified |
| mmu-miR-344d-3p | MIMAT0014808 | Zwint | 52696 | HITS-CLIP | 23142080 | Unspecified |
| mmu-miR-3108-5p | MIMAT0014947 | Akap5 | 238276 | HITS-CLIP | 21258322 | Unspecified |
| mmu-miR-3108-5p | MIMAT0014947 | Aph1b | 208117 | HITS-CLIP | 23142080 | Unspecified |
| mmu-miR-3108-5p | MIMAT0014947 | Arid4a | 238247 | HITS-CLIP | 25083871 | Unspecified |
| mmu-miR-3108-5p | MIMAT0014947 | Csf1 | 12977 | HITS-CLIP | 21258322 | Unspecified |
| mmu-miR-3108-5p | MIMAT0014947 | Ctla2a | 13024 | HITS-CLIP | 21258322 | Unspecified |
| mmu-miR-3108-5p | MIMAT0014947 | Dcp2 | 70640 | HITS-CLIP | 25083871 | Unspecified |
| mmu-miR-3108-5p | MIMAT0014947 | Dusp3 | 72349 | HITS-CLIP | 21258322 | Unspecified |
| mmu-miR-3108-5p | MIMAT0014947 | Eif2ak2 | 19106 | HITS-CLIP | 21258322 | Unspecified |
| mmu-miR-3108-5p | MIMAT0014947 | Fam49a | 76820 | HITS-CLIP | 21258322 | Unspecified |
| mmu-miR-3108-5p | MIMAT0014947 | Glt28d2 | 320302 | HITS-CLIP | 25083871 | Unspecified |
| mmu-miR-3108-5p | MIMAT0014947 | Hist1h2ac | 319164 | HITS-CLIP | 21258322 | Unspecified |
| mmu-miR-3108-5p | MIMAT0014947 | Hnrnpd | 11991 | HITS-CLIP | 25083871 | Unspecified |
| mmu-miR-3108-5p | MIMAT0014947 | Itga4 | 16401 | HITS-CLIP | 21258322 | Unspecified |
| mmu-miR-3108-5p | MIMAT0014947 | Lzts3 | 241638 | HITS-CLIP | 26203562\|19536157\|25083871\|23597 | Unspecified |

| | | | | | | |
|---|---|---|---|---|---|---|
| mmu-miR-3108-5p | MIMAT0014947 | Mcmdc2 | 240697 | HITS-CLIP | 21258322 | Unspecified |
| mmu-miR-3108-5p | MIMAT0014947 | Mob3c | 100465 | HITS-CLIP | 25083871 | Unspecified |
| mmu-miR-3108-5p | MIMAT0014947 | Nab1 | 17936 | HITS-CLIP | 25083871\|23597149 | Unspecified |
| mmu-miR-3108-5p | MIMAT0014947 | Nars2 | 244141 | HITS-CLIP | 21258322 | Unspecified |
| mmu-miR-3108-5p | MIMAT0014947 | Neu3 | 50877 | HITS-CLIP | 21258322 | Unspecified |
| mmu-miR-3108-5p | MIMAT0014947 | Osbp | 76303 | HITS-CLIP | 25083871 | Unspecified |
| mmu-miR-3108-5p | MIMAT0014947 | Pak6 | 214230 | HITS-CLIP | 19536157 | Unspecified |
| mmu-miR-3108-5p | MIMAT0014947 | Phf14 | 75725 | HITS-CLIP | 21258322 | Unspecified |
| mmu-miR-3108-5p | MIMAT0014947 | Prrg3 | 208748 | HITS-CLIP | 25083871\|26203562\|28735896\|23597 | Unspecified |
| mmu-miR-3108-5p | MIMAT0014947 | Rc3h1 | 381305 | HITS-CLIP | 23142080 | Unspecified |
| mmu-miR-3108-5p | MIMAT0014947 | Steap2 | 74051 | HITS-CLIP | 25083871 | Unspecified |
| mmu-miR-3108-5p | MIMAT0014947 | Uevld | 54122 | HITS-CLIP | 21258322 | Unspecified |
| mmu-miR-3108-5p | MIMAT0014947 | Ung | 22256 | HITS-CLIP | 25083871 | Unspecified |
| mmu-miR-3108-5p | MIMAT0014947 | Vmn1r58 | 81014 | HITS-CLIP | 25083871\|21258322 | Unspecified |
| mmu-miR-3108-5p | MIMAT0014947 | Zfp263 | 74120 | HITS-CLIP | 23142080 | Unspecified |
| mmu-miR-3108-5p | MIMAT0014947 | Zfp568 | 243905 | HITS-CLIP | 25083871 | Unspecified |
| mmu-miR-3108-5p | MIMAT0014947 | Zfp933 | 242747 | HITS-CLIP | 25083871\|21258322 | Unspecified |
| mmu-miR-1187 | MIMAT0005837 | 1700019G17Rik | 75541 | HITS-CLIP | 25083871 | Unspecified |
| mmu-miR-1187 | MIMAT0005837 | Abcc9 | 20928 | HITS-CLIP | 23597149 | Unspecified |
| mmu-miR-1187 | MIMAT0005837 | Ado | 211488 | HITS-CLIP | 25083871 | Unspecified |
| mmu-miR-1187 | MIMAT0005837 | Adra1b | 11548 | HITS-CLIP | 23597149 | Unspecified |
| mmu-miR-1187 | MIMAT0005837 | Akap10 | 56697 | HITS-CLIP | 25083871 | Unspecified |
| mmu-miR-1187 | MIMAT0005837 | Akap2 | 11641 | HITS-CLIP | 23597149 | Unspecified |
| mmu-miR-1187 | MIMAT0005837 | Ap1ar | 211556 | HITS-CLIP | 23597149 | Unspecified |
| mmu-miR-1187 | MIMAT0005837 | Apba1 | 319924 | HITS-CLIP | 25083871 | Unspecified |
| mmu-miR-1187 | MIMAT0005837 | Aplnr | 23796 | HITS-CLIP | 25083871 | Unspecified |
| mmu-miR-1187 | MIMAT0005837 | Appl1 | 72993 | HITS-CLIP | 25083871 | Unspecified |
| mmu-miR-1187 | MIMAT0005837 | Arhgef9 | 236915 | HITS-CLIP | 21258322 | Unspecified |
| mmu-miR-1187 | MIMAT0005837 | Asah2 | 54447 | HITS-CLIP | 25083871 | Unspecified |
| mmu-miR-1187 | MIMAT0005837 | Asb7 | 117589 | HITS-CLIP | 23597149 | Unspecified |
| mmu-miR-1187 | MIMAT0005837 | B4galt6 | 56386 | HITS-CLIP | 25083871 | Unspecified |
| mmu-miR-1187 | MIMAT0005837 | BC021785 | 215928 | HITS-CLIP | 25083871 | Unspecified |
| mmu-miR-1187 | MIMAT0005837 | Bcl2 | 12043 | HITS-CLIP | 23597149 | Unspecified |
| mmu-miR-1187 | MIMAT0005837 | Calcoco1 | 67488 | HITS-CLIP | 25083871 | Unspecified |
| mmu-miR-1187 | MIMAT0005837 | Calcr | 12311 | HITS-CLIP | 23142080 | Unspecified |
| mmu-miR-1187 | MIMAT0005837 | Casp8 | 12370 | HITS-CLIP | 23597149\|25083871 | Unspecified |
| mmu-miR-1187 | MIMAT0005837 | Ccdc85a | 216613 | HITS-CLIP | 25083871 | Unspecified |
| mmu-miR-1187 | MIMAT0005837 | Ccpg1 | 72278 | HITS-CLIP | 25083871 | Unspecified |
| mmu-miR-1187 | MIMAT0005837 | Cd200r1 | 57781 | HITS-CLIP | 25083871 | Unspecified |

| | | | | | | |
|---|---|---|---|---|---|---|
| mmu-miR-1187 | MIMAT0005837 | Cd28 | 12487 | HITS-CLIP | 25083871 | Unspecified |
| mmu-miR-1187 | MIMAT0005837 | Cdk7 | 12572 | HITS-CLIP | 25083871 | Unspecified |
| mmu-miR-1187 | MIMAT0005837 | Cer1 | 12622 | HITS-CLIP | 23597149 | Unspecified |
| mmu-miR-1187 | MIMAT0005837 | Chrna1 | 11435 | HITS-CLIP | 25083871 | Unspecified |
| mmu-miR-1187 | MIMAT0005837 | Clca2 | 80797 | HITS-CLIP | 25083871 | Unspecified |
| mmu-miR-1187 | MIMAT0005837 | Clca3a2 | 80797 | HITS-CLIP | 25083871 | Unspecified |
| mmu-miR-1187 | MIMAT0005837 | Cml2 | 93673 | HITS-CLIP | 25083871 | Unspecified |
| mmu-miR-1187 | MIMAT0005837 | Cnih3 | 72978 | HITS-CLIP | 25083871\|23597149 | Unspecified |
| mmu-miR-1187 | MIMAT0005837 | Cox15 | 226139 | HITS-CLIP | 23597149\|25083871 | Unspecified |
| mmu-miR-1187 | MIMAT0005837 | D630003M21Rik | 228846 | HITS-CLIP | 25083871 | Unspecified |
| mmu-miR-1187 | MIMAT0005837 | D630045J12Rik | 330286 | HITS-CLIP | 19536157\|21258322\|23597149\|25083 | Unspecified |
| mmu-miR-1187 | MIMAT0005837 | D730040F13Rik | 242474 | CLIP-Seq dataset analysis | 23142080 | Unspecified |
| mmu-miR-1187 | MIMAT0005837 | Dnaaf5 | 433956 | HITS-CLIP | 25083871\|23597149 | Unspecified |
| mmu-miR-1187 | MIMAT0005837 | Drp2 | 13497 | HITS-CLIP | 25083871 | Unspecified |
| mmu-miR-1187 | MIMAT0005837 | Egfl6 | 54156 | HITS-CLIP | 23597149 | Unspecified |
| mmu-miR-1187 | MIMAT0005837 | Eif2s1 | 13665 | HITS-CLIP | 25083871 | Unspecified |
| mmu-miR-1187 | MIMAT0005837 | Ell2 | 192657 | HITS-CLIP | 25083871 | Unspecified |
| mmu-miR-1187 | MIMAT0005837 | Enpp4 | 224794 | HITS-CLIP | 23597149 | Unspecified |
| mmu-miR-1187 | MIMAT0005837 | Entpd1 | 12495 | HITS-CLIP | 25083871 | Unspecified |
| mmu-miR-1187 | MIMAT0005837 | Epas1 | 13819 | HITS-CLIP | 21258322 | Unspecified |
| mmu-miR-1187 | MIMAT0005837 | Fam107a | 268709 | HITS-CLIP | 21258322\|25083871\|23142080 | Unspecified |
| mmu-miR-1187 | MIMAT0005837 | Fam46a | 212943 | HITS-CLIP | 25083871 | Unspecified |
| mmu-miR-1187 | MIMAT0005837 | Fam55c | 385658 | CLIP-Seq dataset analysis | 25083871 | Unspecified |
| mmu-miR-1187 | MIMAT0005837 | Fblim1 | 74202 | HITS-CLIP | 23142080 | Unspecified |
| mmu-miR-1187 | MIMAT0005837 | Fbrs | 14123 | HITS-CLIP | 23597149 | Unspecified |
| mmu-miR-1187 | MIMAT0005837 | Flrt1 | 396184 | HITS-CLIP | 23597149 | Unspecified |
| mmu-miR-1187 | MIMAT0005837 | Fsd1l | 319636 | HITS-CLIP | 21258322 | Unspecified |
| mmu-miR-1187 | MIMAT0005837 | Gabpb2 | 213054 | HITS-CLIP | 25083871 | Unspecified |
| mmu-miR-1187 | MIMAT0005837 | Gatc | 384281 | HITS-CLIP | 23597149 | Unspecified |
| mmu-miR-1187 | MIMAT0005837 | Gcnt4 | 218476 | HITS-CLIP | 25083871 | Unspecified |
| mmu-miR-1187 | MIMAT0005837 | Gfod1 | 328232 | HITS-CLIP | 25083871 | Unspecified |
| mmu-miR-1187 | MIMAT0005837 | Gfpt1 | 14583 | HITS-CLIP | 25083871 | Unspecified |
| mmu-miR-1187 | MIMAT0005837 | Gm14326 | 665211 | HITS-CLIP | 23142080 | Unspecified |
| mmu-miR-1187 | MIMAT0005837 | Gnb4 | 14696 | HITS-CLIP | 23142080\|25083871 | Unspecified |
| mmu-miR-1187 | MIMAT0005837 | Gramd1c | 207798 | HITS-CLIP | 25083871 | Unspecified |
| mmu-miR-1187 | MIMAT0005837 | Gstt3 | 103140 | HITS-CLIP | 25083871 | Unspecified |
| mmu-miR-1187 | MIMAT0005837 | Gtf2h2 | 23894 | HITS-CLIP | 25083871 | Unspecified |
| mmu-miR-1187 | MIMAT0005837 | Has2 | 15117 | HITS-CLIP | 23597149 | Unspecified |
| mmu-miR-1187 | MIMAT0005837 | Heatr2 | 433956 | HITS-CLIP | 23597149\|25083871 | Unspecified |

| | | | | | | |
|---|---|---|---|---|---|---|
| mmu-miR-1187 | MIMAT0005837 | Hecw1 | 94253 | HITS-CLIP | 25083871 | Unspecified |
| mmu-miR-1187 | MIMAT0005837 | Hif1an | 319594 | HITS-CLIP | 25083871 | Unspecified |
| mmu-miR-1187 | MIMAT0005837 | Htr1f | 15557 | HITS-CLIP | 25083871 | Unspecified |
| mmu-miR-1187 | MIMAT0005837 | Htt | 15194 | HITS-CLIP | 25083871 | Unspecified |
| mmu-miR-1187 | MIMAT0005837 | Igf2 | 16002 | HITS-CLIP | 23597149\|25083871 | Unspecified |
| mmu-miR-1187 | MIMAT0005837 | Iglon5 | 210094 | HITS-CLIP | 23597149 | Unspecified |
| mmu-miR-1187 | MIMAT0005837 | Il18r1 | 16182 | HITS-CLIP | 23597149 | Unspecified |
| mmu-miR-1187 | MIMAT0005837 | Insig2 | 72999 | HITS-CLIP | 25083871\|21258322 | Unspecified |
| mmu-miR-1187 | MIMAT0005837 | Ints8 | 72656 | HITS-CLIP | 25083871 | Unspecified |
| mmu-miR-1187 | MIMAT0005837 | Klf8 | 245671 | HITS-CLIP | 25083871 | Unspecified |
| mmu-miR-1187 | MIMAT0005837 | LOC100048884 | 100048884 | HITS-CLIP | 23597149 | Unspecified |
| mmu-miR-1187 | MIMAT0005837 | Lifr | 16880 | HITS-CLIP | 23597149 | Unspecified |
| mmu-miR-1187 | MIMAT0005837 | Lin7a | 108030 | HITS-CLIP | 23597149 | Unspecified |
| mmu-miR-1187 | MIMAT0005837 | Lrrc40 | 67144 | HITS-CLIP | 25083871 | Unspecified |
| mmu-miR-1187 | MIMAT0005837 | Ly96 | 17087 | HITS-CLIP | 23597149 | Unspecified |
| mmu-miR-1187 | MIMAT0005837 | Mavs | 228607 | HITS-CLIP | 25083871 | Unspecified |
| mmu-miR-1187 | MIMAT0005837 | Mfsd4b5 | 215928 | HITS-CLIP | 25083871 | Unspecified |
| mmu-miR-1187 | MIMAT0005837 | Mocs1 | 56738 | HITS-CLIP | 23597149 | Unspecified |
| mmu-miR-1187 | MIMAT0005837 | Mrgpre | 244238 | HITS-CLIP | 25083871 | Unspecified |
| mmu-miR-1187 | MIMAT0005837 | Mup18 | 100048884 | HITS-CLIP | 23597149 | Unspecified |
| mmu-miR-1187 | MIMAT0005837 | Mylk4 | 238564 | HITS-CLIP | 21258322 | Unspecified |
| mmu-miR-1187 | MIMAT0005837 | Nat8f2 | 93673 | HITS-CLIP | 25083871 | Unspecified |
| mmu-miR-1187 | MIMAT0005837 | Nat8f4 | 75541 | HITS-CLIP | 25083871 | Unspecified |
| mmu-miR-1187 | MIMAT0005837 | Nbeal1 | 269198 | HITS-CLIP | 21258322 | Unspecified |
| mmu-miR-1187 | MIMAT0005837 | Neu1 | 18010 | HITS-CLIP | 25083871 | Unspecified |
| mmu-miR-1187 | MIMAT0005837 | Nqo2 | 18105 | HITS-CLIP | 25083871 | Unspecified |
| mmu-miR-1187 | MIMAT0005837 | Nsd3 | 234135 | HITS-CLIP | 25083871\|23597149 | Unspecified |
| mmu-miR-1187 | MIMAT0005837 | Nsun3 | 106338 | HITS-CLIP | 23597149 | Unspecified |
| mmu-miR-1187 | MIMAT0005837 | Nxpe3 | 385658 | HITS-CLIP | 25083871 | Unspecified |
| mmu-miR-1187 | MIMAT0005837 | Onecut2 | 225631 | HITS-CLIP | 25083871 | Unspecified |
| mmu-miR-1187 | MIMAT0005837 | Oxtr | 18430 | HITS-CLIP | 23597149 | Unspecified |
| mmu-miR-1187 | MIMAT0005837 | Pgm3 | 109785 | HITS-CLIP | 25083871 | Unspecified |
| mmu-miR-1187 | MIMAT0005837 | Plag1 | 56711 | HITS-CLIP | 25083871 | Unspecified |
| mmu-miR-1187 | MIMAT0005837 | Prkcd | 18753 | HITS-CLIP | 25083871 | Unspecified |
| mmu-miR-1187 | MIMAT0005837 | Pstpip2 | 19201 | HITS-CLIP | 21258322 | Unspecified |
| mmu-miR-1187 | MIMAT0005837 | Ptprr | 19279 | HITS-CLIP | 25083871 | Unspecified |
| mmu-miR-1187 | MIMAT0005837 | Rab3c | 67295 | HITS-CLIP | 21258322 | Unspecified |
| mmu-miR-1187 | MIMAT0005837 | Rab7 | 19349 | HITS-CLIP//Western blotting//RNA-seq | 23597149\|34014398 | Unspecified |
| mmu-miR-1187 | MIMAT0005837 | Rab9b | 319642 | HITS-CLIP | 21258322\|23597149 | Unspecified |

| | | | | | | |
|---|---|---|---|---|---|---|
| mmu-miR-1187 | MIMAT0005837 | Rabgap1 | 227800 | HITS-CLIP | 25083871 | Unspecified |
| mmu-miR-1187 | MIMAT0005837 | Rasa2 | 114713 | HITS-CLIP | 25083871 | Unspecified |
| mmu-miR-1187 | MIMAT0005837 | Rasal2 | 226525 | HITS-CLIP | 25083871 | Unspecified |
| mmu-miR-1187 | MIMAT0005837 | Rassf2 | 215653 | HITS-CLIP | 23597149 | Unspecified |
| mmu-miR-1187 | MIMAT0005837 | Rassf5 | 54354 | HITS-CLIP | 25083871 | Unspecified |
| mmu-miR-1187 | MIMAT0005837 | Rfx3 | 19726 | HITS-CLIP | 23597149 | Unspecified |
| mmu-miR-1187 | MIMAT0005837 | Rgs8 | 67792 | HITS-CLIP | 25083871 | Unspecified |
| mmu-miR-1187 | MIMAT0005837 | Rorb | 225998 | HITS-CLIP | 23597149 | Unspecified |
| mmu-miR-1187 | MIMAT0005837 | Rpusd2 | 271842 | HITS-CLIP | 23597149 | Unspecified |
| mmu-miR-1187 | MIMAT0005837 | Runx1 | 12394 | HITS-CLIP | 23597149 | Unspecified |
| mmu-miR-1187 | MIMAT0005837 | Scd3 | 30049 | HITS-CLIP | 23597149 | Unspecified |
| mmu-miR-1187 | MIMAT0005837 | Scn2a | 110876 | HITS-CLIP | 25083871 | Unspecified |
| mmu-miR-1187 | MIMAT0005837 | Scn2a1 | 110876 | HITS-CLIP | 25083871 | Unspecified |
| mmu-miR-1187 | MIMAT0005837 | Sema6a | 20358 | HITS-CLIP | 21258322 | Unspecified |
| mmu-miR-1187 | MIMAT0005837 | Septin3 | 24050 | HITS-CLIP | 25083871 | Unspecified |
| mmu-miR-1187 | MIMAT0005837 | Sh2d2a | 27371 | HITS-CLIP | 23597149 | Unspecified |
| mmu-miR-1187 | MIMAT0005837 | Sike1 | 66641 | HITS-CLIP | 23597149\|21258322\|25083871 | Unspecified |
| mmu-miR-1187 | MIMAT0005837 | Six4 | 20474 | HITS-CLIP | 25083871 | Unspecified |
| mmu-miR-1187 | MIMAT0005837 | Ski | 20481 | HITS-CLIP | 23597149 | Unspecified |
| mmu-miR-1187 | MIMAT0005837 | Slc17a5 | 235504 | HITS-CLIP | 25083871 | Unspecified |
| mmu-miR-1187 | MIMAT0005837 | Slc22a8 | 19879 | HITS-CLIP | 25083871\|23597149\|21258322\|1953( | Unspecified |
| mmu-miR-1187 | MIMAT0005837 | Slc39a14 | 213053 | HITS-CLIP | 25083871\|19536157\|23597149 | Unspecified |
| mmu-miR-1187 | MIMAT0005837 | Slc5a8 | 216225 | HITS-CLIP | 21258322 | Unspecified |
| mmu-miR-1187 | MIMAT0005837 | Slc6a6 | 21366 | HITS-CLIP | 23597149 | Unspecified |
| mmu-miR-1187 | MIMAT0005837 | Smarca2 | 67155 | HITS-CLIP | 25083871\|23597149 | Unspecified |
| mmu-miR-1187 | MIMAT0005837 | St18 | 240690 | HITS-CLIP | 21258322 | Unspecified |
| mmu-miR-1187 | MIMAT0005837 | Steap2 | 74051 | HITS-CLIP | 25083871 | Unspecified |
| mmu-miR-1187 | MIMAT0005837 | Supt7l | 72195 | HITS-CLIP | 23597149 | Unspecified |
| mmu-miR-1187 | MIMAT0005837 | Tacr1 | 21336 | HITS-CLIP | 19536157 | Unspecified |
| mmu-miR-1187 | MIMAT0005837 | Tacr2 | 21337 | HITS-CLIP | 23597149 | Unspecified |
| mmu-miR-1187 | MIMAT0005837 | Tbc1d24 | 224617 | HITS-CLIP | 25083871 | Unspecified |
| mmu-miR-1187 | MIMAT0005837 | Tbx22 | 245572 | HITS-CLIP | 25083871 | Unspecified |
| mmu-miR-1187 | MIMAT0005837 | Tmco1 | 68944 | HITS-CLIP | 23597149 | Unspecified |
| mmu-miR-1187 | MIMAT0005837 | Tmem170b | 621976 | HITS-CLIP | 25083871 | Unspecified |
| mmu-miR-1187 | MIMAT0005837 | Tmem245 | 242474 | HITS-CLIP | 23142080 | Unspecified |
| mmu-miR-1187 | MIMAT0005837 | Tmem26 | 327766 | HITS-CLIP | 25083871 | Unspecified |
| mmu-miR-1187 | MIMAT0005837 | Trim12c | 319236 | HITS-CLIP | 25083871 | Unspecified |
| mmu-miR-1187 | MIMAT0005837 | Tsc22d3 | 14605 | HITS-CLIP | 25083871 | Unspecified |
| mmu-miR-1187 | MIMAT0005837 | Txlnb | 378431 | HITS-CLIP | 25083871 | Unspecified |

| | | | | | | | |
|---|---|---|---|---|---|---|---|
| mmu-miR-1187 | MIMAT0005837 | Ubn2 | 320538 | HITS-CLIP | | 25083871 | Unspecified |
| mmu-miR-1187 | MIMAT0005837 | Uhrf1bp1l | 75089 | HITS-CLIP | | 25083871 | Unspecified |
| mmu-miR-1187 | MIMAT0005837 | Vdr | 22337 | HITS-CLIP | | 23142080 | Unspecified |
| mmu-miR-1187 | MIMAT0005837 | Vps33b | 233405 | HITS-CLIP | 21258322\|23597149 | | Unspecified |
| mmu-miR-1187 | MIMAT0005837 | Vps37a | 52348 | HITS-CLIP | | 23597149 | Unspecified |
| mmu-miR-1187 | MIMAT0005837 | Wars2 | 70560 | HITS-CLIP | | 25083871 | Unspecified |
| mmu-miR-1187 | MIMAT0005837 | Wdr46 | 57315 | HITS-CLIP | | 23597149 | Unspecified |
| mmu-miR-1187 | MIMAT0005837 | Whsc1l1 | 234135 | HITS-CLIP | 23597149\|25083871 | | Unspecified |
| mmu-miR-1187 | MIMAT0005837 | Xrcc3 | 74335 | HITS-CLIP | | 23597149 | Unspecified |
| mmu-miR-1187 | MIMAT0005837 | Zeb2 | 24136 | HITS-CLIP | | 25083871 | Unspecified |
| mmu-miR-1187 | MIMAT0005837 | Zfhx3 | 11906 | HITS-CLIP | 23597149\|23142080 | | Unspecified |
| mmu-miR-1187 | MIMAT0005837 | Zfp157 | 72154 | HITS-CLIP | | 25083871 | Unspecified |
| mmu-miR-1187 | MIMAT0005837 | Zfp26 | 22688 | HITS-CLIP | | 25083871 | Unspecified |
| mmu-miR-1187 | MIMAT0005837 | Zfp518b | 100515 | HITS-CLIP | | 25083871 | Unspecified |
| mmu-miR-1187 | MIMAT0005837 | Zfp92 | 22754 | HITS-CLIP | | 21258322 | Unspecified |
| mmu-miR-1187 | MIMAT0005837 | Zfp931 | 353208 | HITS-CLIP | | 23142080 | Unspecified |
| mmu-miR-1187 | MIMAT0005837 | Znrf3 | 407821 | HITS-CLIP | | 23597149 | Unspecified |
| mmu-miR-326-3p | MIMAT0000559 | Ccdc25 | 67179 | HITS-CLIP | | 23597149 | Unspecified |
| mmu-miR-326-3p | MIMAT0000559 | Clgn | 12745 | HITS-CLIP | | 21258322 | Unspecified |
| mmu-miR-326-3p | MIMAT0000559 | Dync1li2 | 234663 | HITS-CLIP | 21258322\|25083871 | | Unspecified |
| mmu-miR-326-3p | MIMAT0000559 | Efr3b | 668212 | HITS-CLIP | | 21258322 | Unspecified |
| mmu-miR-326-3p | MIMAT0000559 | Ephb3 | 13845 | HITS-CLIP | 21258322\|23142080\|23597149\|19536 | | Unspecified |
| mmu-miR-326-3p | MIMAT0000559 | Ept1 | 28042 | HITS-CLIP | | 21258322 | Unspecified |
| mmu-miR-326-3p | MIMAT0000559 | Ets1 | 23871 | ELISA//Flow//GFP reporter assay//Immuno | | 19838199 | Unspecified |
| mmu-miR-326-3p | MIMAT0000559 | Exoc8 | 102058 | HITS-CLIP | | 21258322 | Unspecified |
| mmu-miR-326-3p | MIMAT0000559 | Fbxl16 | 214931 | HITS-CLIP | | 21258322 | Unspecified |
| mmu-miR-326-3p | MIMAT0000559 | Ildr2 | 100039795 | HITS-CLIP | | 25083871 | Unspecified |
| mmu-miR-326-3p | MIMAT0000559 | Maff | 17133 | HITS-CLIP | | 25083871 | Unspecified |
| mmu-miR-326-3p | MIMAT0000559 | Map3k7 | 26409 | HITS-CLIP | | 25083871 | Unspecified |
| mmu-miR-326-3p | MIMAT0000559 | Mylip | 218203 | HITS-CLIP | | 21258322 | Unspecified |
| mmu-miR-326-3p | MIMAT0000559 | Phactr1 | 218194 | HITS-CLIP | | 21258322 | Unspecified |
| mmu-miR-326-3p | MIMAT0000559 | Rnf150 | 330812 | HITS-CLIP | 25083871\|21258322 | | Unspecified |
| mmu-miR-326-3p | MIMAT0000559 | Selenoi | 28042 | HITS-CLIP | | 21258322 | Unspecified |
| mmu-miR-326-3p | MIMAT0000559 | Smo | 319757 | Luciferase assay//Western blot//Luciferase | | 20216554 | Unspecified |
| mmu-miR-326-3p | MIMAT0000559 | Tars2 | 71807 | HITS-CLIP | | 21258322 | Unspecified |
| mmu-miR-326-3p | MIMAT0000559 | Tbx5 | 21388 | HITS-CLIP | | 21258322 | Unspecified |
| mmu-miR-326-3p | MIMAT0000559 | Tmem179 | 104885 | HITS-CLIP | | 21258322 | Unspecified |
| mmu-miR-326-3p | MIMAT0000559 | Usp22 | 216825 | HITS-CLIP | | 21258322 | Unspecified |
| mmu-miR-326-3p | MIMAT0000559 | Vsx1 | 114889 | HITS-CLIP | 23597149\|21258322 | | Unspecified |

| | | | | | | |
|---|---|---|---|---|---|---|
| mmu-miR-875-3p | MIMAT0004938 | 3110082D06Rik | 627626 | CLIP-Seq dataset analysis | 23142080\|25083871 | Unspecified |
| mmu-miR-875-3p | MIMAT0004938 | 4931406B18Rik | 74054 | HITS-CLIP | 23597149\|21258322 | Unspecified |
| mmu-miR-875-3p | MIMAT0004938 | Arl16 | 70317 | HITS-CLIP | 21258322 | Unspecified |
| mmu-miR-875-3p | MIMAT0004938 | Atp2b1 | 67972 | HITS-CLIP | 21258322 | Unspecified |
| mmu-miR-875-3p | MIMAT0004938 | Bnc2 | 242509 | HITS-CLIP | 23142080 | Unspecified |
| mmu-miR-875-3p | MIMAT0004938 | Cd63 | 12512 | HITS-CLIP | 25083871 | Unspecified |
| mmu-miR-875-3p | MIMAT0004938 | Cdh12 | 215654 | HITS-CLIP | 25083871 | Unspecified |
| mmu-miR-875-3p | MIMAT0004938 | Celsr2 | 53883 | HITS-CLIP | 23142080 | Unspecified |
| mmu-miR-875-3p | MIMAT0004938 | Cep68 | 216543 | HITS-CLIP | 25083871 | Unspecified |
| mmu-miR-875-3p | MIMAT0004938 | Commd7 | 99311 | HITS-CLIP | 21258322 | Unspecified |
| mmu-miR-875-3p | MIMAT0004938 | Dennd4a | 102442 | HITS-CLIP | 25083871 | Unspecified |
| mmu-miR-875-3p | MIMAT0004938 | Dnajc1 | 13418 | HITS-CLIP | 21258322 | Unspecified |
| mmu-miR-875-3p | MIMAT0004938 | Fam114a1 | 68303 | HITS-CLIP | 25083871 | Unspecified |
| mmu-miR-875-3p | MIMAT0004938 | Gmip | 78816 | HITS-CLIP | 25083871\|21258322 | Unspecified |
| mmu-miR-875-3p | MIMAT0004938 | Gng12 | 14701 | HITS-CLIP | 25083871 | Unspecified |
| mmu-miR-875-3p | MIMAT0004938 | Gpr82 | 319200 | HITS-CLIP | 25083871 | Unspecified |
| mmu-miR-875-3p | MIMAT0004938 | Inpp5d | 16331 | HITS-CLIP | 25083871 | Unspecified |
| mmu-miR-875-3p | MIMAT0004938 | Lars2 | 102436 | HITS-CLIP | 21258322\|23597149 | Unspecified |
| mmu-miR-875-3p | MIMAT0004938 | Mboat1 | 218121 | HITS-CLIP | 25083871 | Unspecified |
| mmu-miR-875-3p | MIMAT0004938 | Nab1 | 17936 | HITS-CLIP | 23597149 | Unspecified |
| mmu-miR-875-3p | MIMAT0004938 | Nat8l | 269642 | HITS-CLIP | 25083871 | Unspecified |
| mmu-miR-875-3p | MIMAT0004938 | Olig3 | 94222 | HITS-CLIP | 21258322 | Unspecified |
| mmu-miR-875-3p | MIMAT0004938 | Ppp4r2 | 232314 | HITS-CLIP | 25083871 | Unspecified |
| mmu-miR-875-3p | MIMAT0004938 | Ptchd4 | 627626 | HITS-CLIP | 25083871\|23142080 | Unspecified |
| mmu-miR-875-3p | MIMAT0004938 | Rbm47 | 245945 | HITS-CLIP | 25083871 | Unspecified |
| mmu-miR-875-3p | MIMAT0004938 | Ret | 19713 | HITS-CLIP | 25083871 | Unspecified |
| mmu-miR-875-3p | MIMAT0004938 | Serpine1 | 18787 | HITS-CLIP | 23142080 | Unspecified |
| mmu-miR-875-3p | MIMAT0004938 | Slc15a2 | 57738 | HITS-CLIP | 25083871 | Unspecified |
| mmu-miR-875-3p | MIMAT0004938 | Spry4 | 24066 | HITS-CLIP | 25083871 | Unspecified |
| mmu-miR-875-3p | MIMAT0004938 | Sspn | 16651 | HITS-CLIP | 21258322 | Unspecified |
| mmu-miR-875-3p | MIMAT0004938 | Strbp | 20744 | HITS-CLIP | 25083871 | Unspecified |
| mmu-miR-875-3p | MIMAT0004938 | Tbx4 | 21387 | HITS-CLIP | 23142080\|19536157\|21258322\|25083 | Unspecified |
| mmu-miR-875-3p | MIMAT0004938 | Tet2 | 214133 | HITS-CLIP | 25083871 | Unspecified |
| mmu-miR-875-3p | MIMAT0004938 | Tmem101 | 76547 | HITS-CLIP | 21258322\|23142080 | Unspecified |
| mmu-miR-875-3p | MIMAT0004938 | Tmod2 | 50876 | HITS-CLIP | 23142080 | Unspecified |
| mmu-miR-875-3p | MIMAT0004938 | Ttpa | 50500 | HITS-CLIP | 25083871 | Unspecified |
| mmu-miR-875-3p | MIMAT0004938 | Tube1 | 71924 | HITS-CLIP | 25083871 | Unspecified |
| mmu-miR-875-3p | MIMAT0004938 | Wdr25 | 212198 | HITS-CLIP | 21258322 | Unspecified |
| mmu-miR-875-3p | MIMAT0004938 | Wnk1 | 232341 | HITS-CLIP | 21258322 | Unspecified |

| | | | | | | |
|---|---|---|---|---|---|---|
| mmu-miR-875-3p | MIMAT0004938 | Wtap | 60532 | HITS-CLIP | 25083871 | Unspecified |
| mmu-miR-875-3p | MIMAT0004938 | Zfhx3 | 11906 | HITS-CLIP | 26203562 | Unspecified |
| mmu-miR-875-3p | MIMAT0004938 | Zfp192 | 93681 | CLIP-Seq dataset analysis | 25083871 | Unspecified |
| mmu-miR-875-3p | MIMAT0004938 | Zfp882 | 382019 | HITS-CLIP | 25083871 | Unspecified |
| mmu-miR-875-3p | MIMAT0004938 | Zfyve27 | 319740 | HITS-CLIP | 25083871 | Unspecified |
| mmu-miR-875-3p | MIMAT0004938 | Zkscan8 | 93681 | HITS-CLIP | 25083871 | Unspecified |
| mmu-miR-216b-5p | MIMAT0003729 | Acad9 | 229211 | HITS-CLIP | 25083871 | Unspecified |
| mmu-miR-216b-5p | MIMAT0003729 | Adat1 | 30947 | HITS-CLIP | 21258322 | Unspecified |
| mmu-miR-216b-5p | MIMAT0003729 | Ccdc32 | 269336 | HITS-CLIP | 21258322 | Unspecified |
| mmu-miR-216b-5p | MIMAT0003729 | Cep97 | 74201 | HITS-CLIP | 21258322 | Unspecified |
| mmu-miR-216b-5p | MIMAT0003729 | Dcaf12l2 | 245403 | HITS-CLIP | 25083871\|21258322\|23597149 | Unspecified |
| mmu-miR-216b-5p | MIMAT0003729 | Dzank1 | 241688 | HITS-CLIP | 23597149 | Unspecified |
| mmu-miR-216b-5p | MIMAT0003729 | Efr3b | 668212 | HITS-CLIP | 21258322 | Unspecified |
| mmu-miR-216b-5p | MIMAT0003729 | Fbxl17 | 50758 | HITS-CLIP | 21258322 | Unspecified |
| mmu-miR-216b-5p | MIMAT0003729 | Glt28d2 | 320302 | HITS-CLIP | 25083871 | Unspecified |
| mmu-miR-216b-5p | MIMAT0003729 | Gm5464 | 432870 | HITS-CLIP | 25083871\|23597149 | Unspecified |
| mmu-miR-216b-5p | MIMAT0003729 | Gng2 | 14702 | HITS-CLIP | 21258322 | Unspecified |
| mmu-miR-216b-5p | MIMAT0003729 | Golga7b | 71146 | HITS-CLIP | 25083871\|21258322\|23597149 | Unspecified |
| mmu-miR-216b-5p | MIMAT0003729 | Grin2b | 14812 | HITS-CLIP | 21258322 | Unspecified |
| mmu-miR-216b-5p | MIMAT0003729 | Higd1a | 56295 | HITS-CLIP | 23142080 | Unspecified |
| mmu-miR-216b-5p | MIMAT0003729 | Hmgb1 | 15289 | HITS-CLIP | 25083871 | Unspecified |
| mmu-miR-216b-5p | MIMAT0003729 | Ildr2 | 100039795 | HITS-CLIP | 25083871 | Unspecified |
| mmu-miR-216b-5p | MIMAT0003729 | Ilf3 | 16201 | HITS-CLIP | 25083871\|21258322 | Unspecified |
| mmu-miR-216b-5p | MIMAT0003729 | Map3k7 | 26409 | HITS-CLIP | 25083871 | Unspecified |
| mmu-miR-216b-5p | MIMAT0003729 | Nhsl1 | 215819 | HITS-CLIP | 25083871 | Unspecified |
| mmu-miR-216b-5p | MIMAT0003729 | P2rx7 | 18439 | HITS-CLIP | 21258322 | Unspecified |
| mmu-miR-216b-5p | MIMAT0003729 | Pacs2 | 217893 | HITS-CLIP | 21258322 | Unspecified |
| mmu-miR-216b-5p | MIMAT0003729 | Pax3 | 18505 | HITS-CLIP | 23597149\|21258322 | Unspecified |
| mmu-miR-216b-5p | MIMAT0003729 | Pgm5 | 226041 | HITS-CLIP | 25083871 | Unspecified |
| mmu-miR-216b-5p | MIMAT0003729 | Pigv | 230801 | HITS-CLIP | 25083871\|21258322 | Unspecified |
| mmu-miR-216b-5p | MIMAT0003729 | Ranbp10 | 74334 | HITS-CLIP | 25083871 | Unspecified |
| mmu-miR-216b-5p | MIMAT0003729 | Rnf150 | 330812 | HITS-CLIP | 21258322\|25083871 | Unspecified |
| mmu-miR-216b-5p | MIMAT0003729 | Scamp1 | 107767 | HITS-CLIP | 25083871 | Unspecified |
| mmu-miR-216b-5p | MIMAT0003729 | Synb | 239167 | HITS-CLIP | 23597149\|25083871 | Unspecified |
| mmu-miR-216b-5p | MIMAT0003729 | Tmem242 | 70544 | HITS-CLIP | 21258322 | Unspecified |
| mmu-miR-216b-5p | MIMAT0003729 | Ung | 22256 | HITS-CLIP | 25083871 | Unspecified |
| mmu-miR-216b-5p | MIMAT0003729 | Vsx1 | 114889 | HITS-CLIP | 23597149\|21258322 | Unspecified |
| mmu-miR-216b-5p | MIMAT0003729 | Zfp280d | 235469 | HITS-CLIP | 25083871 | Unspecified |
| mmu-miR-216b-5p | MIMAT0003729 | Zfp951 | 626391 | HITS-CLIP | 23142080\|25083871\|21258322 | Unspecified |

| | | | | | | |
|---|---|---|---|---|---|---|
| mmu-miR-139-5p | MIMAT0000656 | 2310008H09Rik | 66356 | CLIP-Seq dataset analysis | 21258322 | Unspecified |
| mmu-miR-139-5p | MIMAT0000656 | 2310010M20Rik | 69576 | CLIP-Seq dataset analysis | 21258322\|25083871 | Unspecified |
| mmu-miR-139-5p | MIMAT0000656 | 2510049J12Rik | 70291 | HITS-CLIP | 21258322 | Unspecified |
| mmu-miR-139-5p | MIMAT0000656 | 2610027L16Rik | 67842 | CLIP-Seq dataset analysis | 21258322 | Unspecified |
| mmu-miR-139-5p | MIMAT0000656 | 2810021J22Rik | 69944 | HITS-CLIP | 21258322\|25083871 | Unspecified |
| mmu-miR-139-5p | MIMAT0000656 | 6030458C11Rik | 77877 | HITS-CLIP | 21258322 | Unspecified |
| mmu-miR-139-5p | MIMAT0000656 | A230051G13Rik | 216792 | CLIP-Seq dataset analysis | 21258322 | Unspecified |
| mmu-miR-139-5p | MIMAT0000656 | Acvr1c | 269275 | HITS-CLIP | 23142080 | Unspecified |
| mmu-miR-139-5p | MIMAT0000656 | Ajuba | 16475 | HITS-CLIP | 21258322\|26203562 | Unspecified |
| mmu-miR-139-5p | MIMAT0000656 | Akap5 | 238276 | HITS-CLIP | 21258322 | Unspecified |
| mmu-miR-139-5p | MIMAT0000656 | Akap8 | 56399 | HITS-CLIP | 21258322 | Unspecified |
| mmu-miR-139-5p | MIMAT0000656 | Akirin2 | 433693 | HITS-CLIP | 25083871 | Unspecified |
| mmu-miR-139-5p | MIMAT0000656 | Akr1d1 | 208665 | HITS-CLIP | 21258322 | Unspecified |
| mmu-miR-139-5p | MIMAT0000656 | Atp5g3 | 228033 | HITS-CLIP | 23597149 | Unspecified |
| mmu-miR-139-5p | MIMAT0000656 | Atrn | 11990 | HITS-CLIP | 23142080\|29167373 | Unspecified |
| mmu-miR-139-5p | MIMAT0000656 | B230120H23Rik | 65964 | HITS-CLIP | 21258322 | Unspecified |
| mmu-miR-139-5p | MIMAT0000656 | Bach2 | 12014 | HITS-CLIP | 21258322 | Unspecified |
| mmu-miR-139-5p | MIMAT0000656 | Bbs1 | 52028 | HITS-CLIP | 21258322 | Unspecified |
| mmu-miR-139-5p | MIMAT0000656 | Bub1 | 12235 | HITS-CLIP | 21258322 | Unspecified |
| mmu-miR-139-5p | MIMAT0000656 | Bysl | 53414 | HITS-CLIP | 21258322 | Unspecified |
| mmu-miR-139-5p | MIMAT0000656 | Ccdc157 | 216516 | HITS-CLIP | 25083871 | Unspecified |
| mmu-miR-139-5p | MIMAT0000656 | Cdk7 | 12572 | HITS-CLIP | 21258322 | Unspecified |
| mmu-miR-139-5p | MIMAT0000656 | Cep68 | 216543 | HITS-CLIP | 21258322 | Unspecified |
| mmu-miR-139-5p | MIMAT0000656 | Cep97 | 74201 | HITS-CLIP | 21258322 | Unspecified |
| mmu-miR-139-5p | MIMAT0000656 | Chrna1 | 11435 | HITS-CLIP | 21258322 | Unspecified |
| mmu-miR-139-5p | MIMAT0000656 | Clns1a | 12729 | HITS-CLIP | 21258322 | Unspecified |
| mmu-miR-139-5p | MIMAT0000656 | Commd8 | 27784 | HITS-CLIP//sRNA-Seq | 21258322\|33184237\|29167373\|25083 | Unspecified |
| mmu-miR-139-5p | MIMAT0000656 | Cracr2b | 213573 | HITS-CLIP | 25083871\|21258322 | Unspecified |
| mmu-miR-139-5p | MIMAT0000656 | Cts8 | 56094 | HITS-CLIP | 21258322 | Unspecified |
| mmu-miR-139-5p | MIMAT0000656 | Ctsa | 19025 | HITS-CLIP | 25083871\|29167373 | Unspecified |
| mmu-miR-139-5p | MIMAT0000656 | Cwc25 | 67480 | HITS-CLIP | 25083871\|21258322 | Unspecified |
| mmu-miR-139-5p | MIMAT0000656 | Cyb5d2 | 192986 | HITS-CLIP | 21258322 | Unspecified |
| mmu-miR-139-5p | MIMAT0000656 | D17H6S53E | 114585 | HITS-CLIP | 21258322 | Unspecified |
| mmu-miR-139-5p | MIMAT0000656 | D330012F22Rik | 269952 | CLIP-Seq dataset analysis | 21258322 | Unspecified |
| mmu-miR-139-5p | MIMAT0000656 | Dars2 | 226539 | HITS-CLIP | 21258322 | Unspecified |
| mmu-miR-139-5p | MIMAT0000656 | Ddhd2 | 72108 | HITS-CLIP | 21258322 | Unspecified |
| mmu-miR-139-5p | MIMAT0000656 | Decr2 | 26378 | HITS-CLIP | 21258322 | Unspecified |
| mmu-miR-139-5p | MIMAT0000656 | Dennd6a | 211922 | HITS-CLIP | 21258322 | Unspecified |
| mmu-miR-139-5p | MIMAT0000656 | Dhx37 | 208144 | HITS-CLIP | 21258322 | Unspecified |

| | | | | | | |
|---|---|---|---|---|---|---|
| mmu-miR-139-5p | MIMAT0000656 | Dll3 | 13389 | HITS-CLIP | 25083871\|21258322 | Unspecified |
| mmu-miR-139-5p | MIMAT0000656 | Dlx3 | 13393 | HITS-CLIP | 25083871 | Unspecified |
| mmu-miR-139-5p | MIMAT0000656 | Dnase2a | 13423 | HITS-CLIP | 21258322\|25083871 | Unspecified |
| mmu-miR-139-5p | MIMAT0000656 | Dtwd2 | 68857 | HITS-CLIP | 25083871 | Unspecified |
| mmu-miR-139-5p | MIMAT0000656 | Dusp3 | 72349 | HITS-CLIP//sRNA-Seq | 21258322\|33184237\|29167373 | Unspecified |
| mmu-miR-139-5p | MIMAT0000656 | Dynll1 | 56455 | HITS-CLIP | 21258322 | Unspecified |
| mmu-miR-139-5p | MIMAT0000656 | Efcab4a | 213573 | HITS-CLIP | 21258322\|25083871 | Unspecified |
| mmu-miR-139-5p | MIMAT0000656 | Entpd5 | 12499 | HITS-CLIP | 25083871 | Unspecified |
| mmu-miR-139-5p | MIMAT0000656 | Exosc4 | 109075 | HITS-CLIP | 21258322 | Unspecified |
| mmu-miR-139-5p | MIMAT0000656 | Fam116a | 211922 | CLIP-Seq dataset analysis | 21258322 | Unspecified |
| mmu-miR-139-5p | MIMAT0000656 | Fam198b | 68659 | HITS-CLIP | 21258322 | Unspecified |
| mmu-miR-139-5p | MIMAT0000656 | Fitm2 | 228859 | HITS-CLIP | 21258322 | Unspecified |
| mmu-miR-139-5p | MIMAT0000656 | FoxO1 | 56458 | Western Blot//RT-PCR//Luciferase assay//I | 19883627\|29167373 | Unspecified |
| mmu-miR-139-5p | MIMAT0000656 | Foxo1 | 56458 | Luciferase assay//qRT-PCR//Western blot/ | 19883627\|29167373 | Unspecified |
| mmu-miR-139-5p | MIMAT0000656 | Fzd7 | 14369 | HITS-CLIP | 21258322 | Unspecified |
| mmu-miR-139-5p | MIMAT0000656 | Gas2l3 | 237436 | HITS-CLIP | 21258322\|29167373\|25083871 | Unspecified |
| mmu-miR-139-5p | MIMAT0000656 | Gcnt3 | 72077 | HITS-CLIP | 21258322 | Unspecified |
| mmu-miR-139-5p | MIMAT0000656 | Gdpgp1 | 269952 | HITS-CLIP | 21258322\|26203562 | Unspecified |
| mmu-miR-139-5p | MIMAT0000656 | Gfpt1 | 14583 | HITS-CLIP | 21258322\|25083871 | Unspecified |
| mmu-miR-139-5p | MIMAT0000656 | Gje1 | 76743 | HITS-CLIP | 21258322 | Unspecified |
| mmu-miR-139-5p | MIMAT0000656 | Gm5531 | 433365 | HITS-CLIP | 25083871 | Unspecified |
| mmu-miR-139-5p | MIMAT0000656 | Gng2 | 14702 | HITS-CLIP | 21258322\|25083871 | Unspecified |
| mmu-miR-139-5p | MIMAT0000656 | Gngt1 | 14699 | HITS-CLIP | 25083871\|21258322 | Unspecified |
| mmu-miR-139-5p | MIMAT0000656 | Gpr68 | 238377 | HITS-CLIP | 21258322 | Unspecified |
| mmu-miR-139-5p | MIMAT0000656 | Gtpbp3 | 70359 | HITS-CLIP | 21258322 | Unspecified |
| mmu-miR-139-5p | MIMAT0000656 | Hacd3 | 57874 | HITS-CLIP | 21258322 | Unspecified |
| mmu-miR-139-5p | MIMAT0000656 | Hist1h4i | 319158 | HITS-CLIP | 21258322 | Unspecified |
| mmu-miR-139-5p | MIMAT0000656 | Hnrnpab | 15384 | HITS-CLIP//sRNA-Seq | 21258322\|33184237\|31433990\|29167 | Unspecified |
| mmu-miR-139-5p | MIMAT0000656 | Iba57 | 216792 | HITS-CLIP | 21258322 | Unspecified |
| mmu-miR-139-5p | MIMAT0000656 | Inpp5e | 64436 | HITS-CLIP | 21258322\|25083871 | Unspecified |
| mmu-miR-139-5p | MIMAT0000656 | Irgm2 | 54396 | HITS-CLIP | 25083871\|31433990 | Unspecified |
| mmu-miR-139-5p | MIMAT0000656 | Irs1 | 16367 | Luciferase reporter assay//qRT-PCR//West | 25536154 | Unspecified |
| mmu-miR-139-5p | MIMAT0000656 | Isg20l2 | 229504 | HITS-CLIP | 21258322 | Unspecified |
| mmu-miR-139-5p | MIMAT0000656 | Itgb1bp3 | 69564 | CLIP-Seq dataset analysis | 21258322 | Unspecified |
| mmu-miR-139-5p | MIMAT0000656 | Jub | 16475 | CLIP-Seq dataset analysis | 21258322 | Unspecified |
| mmu-miR-139-5p | MIMAT0000656 | Knop1 | 66356 | HITS-CLIP | 21258322 | Unspecified |
| mmu-miR-139-5p | MIMAT0000656 | Ldlrad2 | 435811 | HITS-CLIP | 21258322 | Unspecified |
| mmu-miR-139-5p | MIMAT0000656 | Lix1 | 66643 | HITS-CLIP | 25083871\|21258322 | Unspecified |
| mmu-miR-139-5p | MIMAT0000656 | Lrrc58 | 320184 | HITS-CLIP//sRNA-Seq | 25083871\|33184237\|29167373\|2125ε | Unspecified |

| | | | | | | | |
|---|---|---|---|---|---|---|---|
| mmu-miR-139-5p | MIMAT0000656 | Luc7l2 | 192196 | HITS-CLIP | | 21258322 | Unspecified |
| mmu-miR-139-5p | MIMAT0000656 | Man2a2 | 140481 | HITS-CLIP | | 21258322 | Unspecified |
| mmu-miR-139-5p | MIMAT0000656 | Map3k20 | 65964 | HITS-CLIP | | 21258322 | Unspecified |
| mmu-miR-139-5p | MIMAT0000656 | Marchf5 | 69104 | HITS-CLIP | | 21258322 | Unspecified |
| mmu-miR-139-5p | MIMAT0000656 | Mbd4 | 17193 | HITS-CLIP | | 21258322 | Unspecified |
| mmu-miR-139-5p | MIMAT0000656 | Mcart1 | 230125 | CLIP-Seq dataset analysis | | 21258322 | Unspecified |
| mmu-miR-139-5p | MIMAT0000656 | Mga | 29808 | HITS-CLIP | | 21258322 | Unspecified |
| mmu-miR-139-5p | MIMAT0000656 | Mkrn2os | 70291 | HITS-CLIP | | 21258322 | Unspecified |
| mmu-miR-139-5p | MIMAT0000656 | Mmp25 | 240047 | HITS-CLIP | | 21258322 | Unspecified |
| mmu-miR-139-5p | MIMAT0000656 | Mrpl15 | 27395 | HITS-CLIP | | 21258322 | Unspecified |
| mmu-miR-139-5p | MIMAT0000656 | Mrpl40 | 18100 | HITS-CLIP | 25083871\|21258322 | | Unspecified |
| mmu-miR-139-5p | MIMAT0000656 | Msi2 | 76626 | HITS-CLIP | | 25083871 | Unspecified |
| mmu-miR-139-5p | MIMAT0000656 | Mzt1 | 76789 | HITS-CLIP | 21258322\|25083871 | | Unspecified |
| mmu-miR-139-5p | MIMAT0000656 | Nanog | 71950 | HITS-CLIP | 23597149\|21258322 | | Unspecified |
| mmu-miR-139-5p | MIMAT0000656 | Nip7 | 66164 | HITS-CLIP | 21258322\|31433990 | | Unspecified |
| mmu-miR-139-5p | MIMAT0000656 | Nmrk2 | 69564 | HITS-CLIP | | 21258322 | Unspecified |
| mmu-miR-139-5p | MIMAT0000656 | Nop9 | 67842 | HITS-CLIP | 21258322\|29167373 | | Unspecified |
| mmu-miR-139-5p | MIMAT0000656 | Notch1 | 18128 | Luciferase reporter assay//qRT-PCR//West | | 25536154 | Unspecified |
| mmu-miR-139-5p | MIMAT0000656 | Nup214 | 227720 | HITS-CLIP | | 21258322 | Unspecified |
| mmu-miR-139-5p | MIMAT0000656 | Ocln | 18260 | HITS-CLIP | | 21258322 | Unspecified |
| mmu-miR-139-5p | MIMAT0000656 | Onecut3 | 246086 | HITS-CLIP | | 21258322 | Unspecified |
| mmu-miR-139-5p | MIMAT0000656 | Orai3 | 269999 | HITS-CLIP | | 21258322 | Unspecified |
| mmu-miR-139-5p | MIMAT0000656 | Osbpl8 | 237542 | HITS-CLIP//sRNA-Seq | 25083871\|33184237\|29167373\|2125{ | | Unspecified |
| mmu-miR-139-5p | MIMAT0000656 | Ostf1 | 20409 | HITS-CLIP | 21258322\|29167373 | | Unspecified |
| mmu-miR-139-5p | MIMAT0000656 | Parp11 | 101187 | HITS-CLIP | 21258322\|29167373 | | Unspecified |
| mmu-miR-139-5p | MIMAT0000656 | Pcdhb13 | 93884 | HITS-CLIP | | 21258322 | Unspecified |
| mmu-miR-139-5p | MIMAT0000656 | Pfkl | 18641 | HITS-CLIP | 25083871\|29167373\|21258322 | | Unspecified |
| mmu-miR-139-5p | MIMAT0000656 | Pigv | 230801 | HITS-CLIP | | 21258322 | Unspecified |
| mmu-miR-139-5p | MIMAT0000656 | Pmaip1 | 58801 | HITS-CLIP | | 21258322 | Unspecified |
| mmu-miR-139-5p | MIMAT0000656 | Pphln1 | 223828 | HITS-CLIP | 21258322\|29167373 | | Unspecified |
| mmu-miR-139-5p | MIMAT0000656 | Prdx1 | 18477 | HITS-CLIP//sRNA-Seq | 21258322\|33184237\|29167373 | | Unspecified |
| mmu-miR-139-5p | MIMAT0000656 | Prpf4 | 70052 | HITS-CLIP | | 21258322 | Unspecified |
| mmu-miR-139-5p | MIMAT0000656 | Ptp4a3 | 19245 | HITS-CLIP | | 21258322 | Unspecified |
| mmu-miR-139-5p | MIMAT0000656 | Ptplad1 | 57874 | HITS-CLIP | | 21258322 | Unspecified |
| mmu-miR-139-5p | MIMAT0000656 | Pycr1 | 209027 | HITS-CLIP | | 21258322 | Unspecified |
| mmu-miR-139-5p | MIMAT0000656 | Rasal2 | 226525 | HITS-CLIP | | 21258322 | Unspecified |
| mmu-miR-139-5p | MIMAT0000656 | Rbak | 57782 | HITS-CLIP | | 21258322 | Unspecified |
| mmu-miR-139-5p | MIMAT0000656 | Rchy1 | 68098 | HITS-CLIP | | 25083871 | Unspecified |
| mmu-miR-139-5p | MIMAT0000656 | Rg9mtd2 | 108943 | CLIP-Seq dataset analysis | | 21258322 | Unspecified |

| | | | | | | |
|---|---|---|---|---|---|---|
| mmu-miR-139-5p | MIMAT0000656 | Rho | 212541 | Luciferase reporter assay | 25425000 | Unspecified |
| mmu-miR-139-5p | MIMAT0000656 | Rnaseh2a | 69724 | HITS-CLIP | 25083871 | Unspecified |
| mmu-miR-139-5p | MIMAT0000656 | Rnd1 | 223881 | HITS-CLIP | 21258322 | Unspecified |
| mmu-miR-139-5p | MIMAT0000656 | Rnf150 | 330812 | HITS-CLIP | 23142080 | Unspecified |
| mmu-miR-139-5p | MIMAT0000656 | Rprd1a | 225283 | HITS-CLIP | 21258322 | Unspecified |
| mmu-miR-139-5p | MIMAT0000656 | Rrp36 | 224823 | HITS-CLIP | 21258322 | Unspecified |
| mmu-miR-139-5p | MIMAT0000656 | Sc5d | 235293 | HITS-CLIP | 21258322 | Unspecified |
| mmu-miR-139-5p | MIMAT0000656 | Sco1 | 52892 | HITS-CLIP | 21258322 | Unspecified |
| mmu-miR-139-5p | MIMAT0000656 | Sec22c | 215474 | HITS-CLIP | 21258322 | Unspecified |
| mmu-miR-139-5p | MIMAT0000656 | Sec23ip | 207352 | HITS-CLIP | 25083871 | Unspecified |
| mmu-miR-139-5p | MIMAT0000656 | Sema3e | 20349 | HITS-CLIP | 25083871 | Unspecified |
| mmu-miR-139-5p | MIMAT0000656 | Sfxn4 | 94281 | HITS-CLIP | 25083871 | Unspecified |
| mmu-miR-139-5p | MIMAT0000656 | Sgo2a | 68549 | HITS-CLIP | 21258322 | Unspecified |
| mmu-miR-139-5p | MIMAT0000656 | Sgol2 | 68549 | HITS-CLIP | 21258322 | Unspecified |
| mmu-miR-139-5p | MIMAT0000656 | Sirpa | 19261 | HITS-CLIP | 25083871\|29167373 | Unspecified |
| mmu-miR-139-5p | MIMAT0000656 | Slc15a1 | 56643 | HITS-CLIP | 21258322 | Unspecified |
| mmu-miR-139-5p | MIMAT0000656 | Slc1a7 | 242607 | HITS-CLIP | 25083871 | Unspecified |
| mmu-miR-139-5p | MIMAT0000656 | Slc25a51 | 230125 | HITS-CLIP | 21258322 | Unspecified |
| mmu-miR-139-5p | MIMAT0000656 | Slc35e1 | 270066 | HITS-CLIP | 25083871 | Unspecified |
| mmu-miR-139-5p | MIMAT0000656 | Smco1 | 69576 | HITS-CLIP | 25083871\|21258322 | Unspecified |
| mmu-miR-139-5p | MIMAT0000656 | Spib | 272382 | HITS-CLIP | 21258322 | Unspecified |
| mmu-miR-139-5p | MIMAT0000656 | Stx17 | 67727 | HITS-CLIP | 21258322 | Unspecified |
| mmu-miR-139-5p | MIMAT0000656 | Sumf2 | 67902 | HITS-CLIP | 21258322 | Unspecified |
| mmu-miR-139-5p | MIMAT0000656 | T2 | 21331 | HITS-CLIP | 25083871 | Unspecified |
| mmu-miR-139-5p | MIMAT0000656 | Tars2 | 71807 | HITS-CLIP | 21258322 | Unspecified |
| mmu-miR-139-5p | MIMAT0000656 | Tcfcp2 | 21422 | CLIP-Seq dataset analysis | 21258322 | Unspecified |
| mmu-miR-139-5p | MIMAT0000656 | Teddm1b | 433365 | HITS-CLIP | 25083871 | Unspecified |
| mmu-miR-139-5p | MIMAT0000656 | Tex9 | 21778 | HITS-CLIP | 21258322 | Unspecified |
| mmu-miR-139-5p | MIMAT0000656 | Tfcp2 | 21422 | HITS-CLIP | 21258322 | Unspecified |
| mmu-miR-139-5p | MIMAT0000656 | Thap2 | 66816 | HITS-CLIP | 21258322 | Unspecified |
| mmu-miR-139-5p | MIMAT0000656 | Tln2 | 70549 | HITS-CLIP | 21258322 | Unspecified |
| mmu-miR-139-5p | MIMAT0000656 | Tmco3 | 234076 | HITS-CLIP | 25083871 | Unspecified |
| mmu-miR-139-5p | MIMAT0000656 | Tmem206 | 66950 | HITS-CLIP | 21258322\|29167373\|25083871 | Unspecified |
| mmu-miR-139-5p | MIMAT0000656 | Tmem69 | 230657 | HITS-CLIP | 25083871 | Unspecified |
| mmu-miR-139-5p | MIMAT0000656 | Tmem88b | 320587 | HITS-CLIP | 21258322 | Unspecified |
| mmu-miR-139-5p | MIMAT0000656 | Tmx4 | 52837 | HITS-CLIP | 21258322 | Unspecified |
| mmu-miR-139-5p | MIMAT0000656 | Trim36 | 28105 | HITS-CLIP | 21258322 | Unspecified |
| mmu-miR-139-5p | MIMAT0000656 | Trmt10a | 108943 | HITS-CLIP | 21258322 | Unspecified |
| mmu-miR-139-5p | MIMAT0000656 | Tspyl1 | 22110 | HITS-CLIP | 21258322 | Unspecified |

| | | | | | | |
|---|---|---|---|---|---|---|
| mmu-miR-139-5p | MIMAT0000656 | Ttc26 | 264134 | HITS-CLIP | 21258322 | Unspecified |
| mmu-miR-139-5p | MIMAT0000656 | Ttc5 | 219022 | HITS-CLIP | 21258322 | Unspecified |
| mmu-miR-139-5p | MIMAT0000656 | Tulp3 | 22158 | HITS-CLIP | 21258322 | Unspecified |
| mmu-miR-139-5p | MIMAT0000656 | Ubac1 | 98766 | HITS-CLIP | 21258322\|25083871 | Unspecified |
| mmu-miR-139-5p | MIMAT0000656 | Vsig10 | 231668 | HITS-CLIP | 21258322 | Unspecified |
| mmu-miR-139-5p | MIMAT0000656 | Xpo7 | 65246 | HITS-CLIP | 25083871\|29167373 | Unspecified |
| mmu-miR-139-5p | MIMAT0000656 | Xrcc2 | 57434 | HITS-CLIP | 25083871 | Unspecified |
| mmu-miR-139-5p | MIMAT0000656 | Ybey | 216119 | HITS-CLIP | 21258322\|25083871 | Unspecified |
| mmu-miR-139-5p | MIMAT0000656 | Zc3hav1l | 209032 | HITS-CLIP | 25083871 | Unspecified |
| mmu-miR-139-5p | MIMAT0000656 | Zdhhc3 | 69035 | HITS-CLIP | 21258322\|29167373\|23597149 | Unspecified |
| mmu-miR-139-5p | MIMAT0000656 | Zfp11 | 22648 | HITS-CLIP | 25083871\|21258322 | Unspecified |
| mmu-miR-139-5p | MIMAT0000656 | Zfp113 | 56314 | HITS-CLIP | 25083871 | Unspecified |
| mmu-miR-139-5p | MIMAT0000656 | Zfp322a | 218100 | HITS-CLIP | 25083871\|29167373 | Unspecified |
| mmu-miR-139-5p | MIMAT0000656 | Zfp345 | 545471 | HITS-CLIP | 21258322 | Unspecified |
| mmu-miR-139-5p | MIMAT0000656 | Zfp697 | 242109 | HITS-CLIP | 21258322 | Unspecified |
| mmu-miR-139-5p | MIMAT0000656 | Zfp799 | 240064 | HITS-CLIP | 21258322 | Unspecified |
| mmu-miR-139-5p | MIMAT0000656 | Zfp830 | 66983 | HITS-CLIP | 21258322 | Unspecified |
| mmu-miR-139-5p | MIMAT0000656 | Zfp872 | 619310 | HITS-CLIP | 25083871 | Unspecified |
| mmu-miR-3109-3p | MIMAT0014950 | Ap1s3 | 252903 | HITS-CLIP | 21258322 | Unspecified |
| mmu-miR-3109-3p | MIMAT0014950 | Fam96a | 68250 | HITS-CLIP | 21258322 | Unspecified |
| mmu-miR-3109-3p | MIMAT0014950 | Gaa | 14387 | HITS-CLIP | 21258322 | Unspecified |
| mmu-miR-3109-3p | MIMAT0014950 | Gjc3 | 118446 | HITS-CLIP | 21258322\|25083871 | Unspecified |
| mmu-miR-3109-3p | MIMAT0014950 | Isg20l2 | 229504 | HITS-CLIP | 21258322 | Unspecified |
| mmu-miR-3109-3p | MIMAT0014950 | Onecut3 | 246086 | HITS-CLIP | 21258322 | Unspecified |
| mmu-miR-3109-3p | MIMAT0014950 | Rmi2 | 223970 | HITS-CLIP | 21258322 | Unspecified |
| mmu-miR-3109-3p | MIMAT0014950 | Six3 | 20473 | HITS-CLIP | 21258322 | Unspecified |
| mmu-miR-3109-3p | MIMAT0014950 | Slc18b1 | 76306 | HITS-CLIP | 21258322 | Unspecified |
| mmu-miR-3109-3p | MIMAT0014950 | Ston2 | 108800 | HITS-CLIP | 21258322 | Unspecified |
| mmu-miR-3109-3p | MIMAT0014950 | Tnfrsf10b | 21933 | HITS-CLIP | 25083871 | Unspecified |
| mmu-miR-3109-3p | MIMAT0014950 | Zbtb8a | 73680 | HITS-CLIP | 21258322 | Unspecified |
| mmu-miR-3109-3p | MIMAT0014950 | Zfp488 | 382867 | HITS-CLIP | 21258322 | Unspecified |
| mmu-miR-691 | MIMAT0003470 | 1700025G04Rik | 69399 | HITS-CLIP | 21258322 | Unspecified |
| mmu-miR-691 | MIMAT0003470 | Adamtsl1 | 77739 | HITS-CLIP | 23142080 | Unspecified |
| mmu-miR-691 | MIMAT0003470 | Arl6ip1 | 54208 | HITS-CLIP | 23142080 | Unspecified |
| mmu-miR-691 | MIMAT0003470 | Atp2b1 | 67972 | HITS-CLIP | 21258322 | Unspecified |
| mmu-miR-691 | MIMAT0003470 | Cbln2 | 12405 | HITS-CLIP | 23142080 | Unspecified |
| mmu-miR-691 | MIMAT0003470 | Cdk7 | 12572 | HITS-CLIP | 21258322 | Unspecified |
| mmu-miR-691 | MIMAT0003470 | Cnnm1 | 83674 | HITS-CLIP | 23142080 | Unspecified |
| mmu-miR-691 | MIMAT0003470 | D330012F22Rik | 269952 | CLIP-Seq dataset analysis | 25083871\|21258322 | Unspecified |

| | | | | | | |
|---|---|---|---|---|---|---|
| mmu-miR-691 | MIMAT0003470 | D930020B18Rik | 216393 | HITS-CLIP | 23142080 | Unspecified |
| mmu-miR-691 | MIMAT0003470 | Dapl1 | 76747 | HITS-CLIP | 23142080 | Unspecified |
| mmu-miR-691 | MIMAT0003470 | Fam38b | 667742 | CLIP-Seq dataset analysis | 23142080 | Unspecified |
| mmu-miR-691 | MIMAT0003470 | Gdpgp1 | 269952 | HITS-CLIP | 21258322\|25083871 | Unspecified |
| mmu-miR-691 | MIMAT0003470 | Hlcs | 110948 | HITS-CLIP | 21258322 | Unspecified |
| mmu-miR-691 | MIMAT0003470 | Il25 | 140806 | Luciferase reporter assay//RNA-seq//qRT-I | 32779953 | Unspecified |
| mmu-miR-691 | MIMAT0003470 | Inpp5d | 16331 | HITS-CLIP | 25083871 | Unspecified |
| mmu-miR-691 | MIMAT0003470 | Intu | 380614 | HITS-CLIP | 21258322 | Unspecified |
| mmu-miR-691 | MIMAT0003470 | Lhfpl3 | 269629 | HITS-CLIP | 23142080 | Unspecified |
| mmu-miR-691 | MIMAT0003470 | Ndrg1 | 17988 | HITS-CLIP | 21258322 | Unspecified |
| mmu-miR-691 | MIMAT0003470 | Pcgf2 | 22658 | HITS-CLIP | 25083871\|23142080\|21258322\|23597 | Unspecified |
| mmu-miR-691 | MIMAT0003470 | Piezo2 | 667742 | HITS-CLIP | 23142080 | Unspecified |
| mmu-miR-691 | MIMAT0003470 | Scd3 | 30049 | HITS-CLIP | 25083871 | Unspecified |
| mmu-miR-691 | MIMAT0003470 | Sik3 | 70661 | HITS-CLIP | 23142080 | Unspecified |
| mmu-miR-691 | MIMAT0003470 | Slc7a14 | 241919 | HITS-CLIP | 21258322 | Unspecified |
| mmu-miR-691 | MIMAT0003470 | Sp4 | 20688 | HITS-CLIP | 23142080 | Unspecified |
| mmu-miR-691 | MIMAT0003470 | Syndig1l | 627191 | HITS-CLIP | 23142080 | Unspecified |
| mmu-miR-691 | MIMAT0003470 | Tmem90a | 627191 | CLIP-Seq dataset analysis | 23142080 | Unspecified |
| mmu-miR-691 | MIMAT0003470 | Tnfrsf26 | 244237 | HITS-CLIP | 21258322\|25083871 | Unspecified |
| mmu-miR-691 | MIMAT0003470 | Utp23 | 78581 | HITS-CLIP | 21258322 | Unspecified |
| mmu-miR-691 | MIMAT0003470 | Wtap | 60532 | HITS-CLIP | 25083871 | Unspecified |
| mmu-miR-3095-3p | MIMAT0014912 | Ago4 | 76850 | HITS-CLIP | 23142080 | Unspecified |
| mmu-miR-3095-3p | MIMAT0014912 | Ccdc97 | 52132 | HITS-CLIP | 23142080 | Unspecified |
| mmu-miR-3095-3p | MIMAT0014912 | Dusp8 | 18218 | HITS-CLIP | 23142080 | Unspecified |
| mmu-miR-3095-3p | MIMAT0014912 | Eif2c4 | 76850 | CLIP-Seq dataset analysis | 23142080 | Unspecified |
| mmu-miR-3095-3p | MIMAT0014912 | Gcet2 | 14525 | CLIP-Seq dataset analysis | 23142080 | Unspecified |
| mmu-miR-3095-3p | MIMAT0014912 | Gcsam | 14525 | HITS-CLIP | 23142080 | Unspecified |
| mmu-miR-3095-3p | MIMAT0014912 | Gpx5 | 14780 | HITS-CLIP | 26203562 | Unspecified |
| mmu-miR-3095-3p | MIMAT0014912 | Hecw1 | 94253 | HITS-CLIP | 23142080 | Unspecified |
| mmu-miR-3095-3p | MIMAT0014912 | Hiat1 | 15247 | HITS-CLIP | 25083871 | Unspecified |
| mmu-miR-3095-3p | MIMAT0014912 | Mfsd14a | 15247 | HITS-CLIP | 25083871 | Unspecified |
| mmu-miR-3095-3p | MIMAT0014912 | Mkl2 | 239719 | HITS-CLIP | 23142080 | Unspecified |
| mmu-miR-3095-3p | MIMAT0014912 | Nebl | 74103 | HITS-CLIP | 23142080 | Unspecified |
| mmu-miR-3095-3p | MIMAT0014912 | Nfatc4 | 73181 | HITS-CLIP | 23597149\|25083871\|21258322 | Unspecified |
| mmu-miR-3095-3p | MIMAT0014912 | Nr2c1 | 22025 | HITS-CLIP | 23142080 | Unspecified |
| mmu-miR-3095-3p | MIMAT0014912 | Sik3 | 70661 | HITS-CLIP | 23142080 | Unspecified |
| mmu-miR-3095-3p | MIMAT0014912 | Slc25a41 | 103775 | HITS-CLIP | 23142080 | Unspecified |
| mmu-miR-3095-3p | MIMAT0014912 | Slc38a8 | 234788 | HITS-CLIP | 23142080 | Unspecified |
| mmu-miR-3095-3p | MIMAT0014912 | Spn | 20737 | HITS-CLIP | 23142080 | Unspecified |

| | | | | | | |
|---|---|---|---|---|---|---|
| mmu-miR-3095-3p | MIMAT0014912 | Tgfbr3 | 21814 | HITS-CLIP | 23142080 | Unspecified |
| mmu-miR-3095-3p | MIMAT0014912 | Tnfrsf1b | 21938 | HITS-CLIP | 25083871\|21258322\|23597149 | Unspecified |
| mmu-miR-669n | MIMAT0009427 | 1200014J11Rik | 66874 | HITS-CLIP | 23142080 | Unspecified |
| mmu-miR-669n | MIMAT0009427 | 4933432B09Rik | 75729 | CLIP-Seq dataset analysis | 23142080 | Unspecified |
| mmu-miR-669n | MIMAT0009427 | 5730437N04Rik | 70544 | CLIP-Seq dataset analysis | 21258322 | Unspecified |
| mmu-miR-669n | MIMAT0009427 | 5730507C01Rik | 236366 | HITS-CLIP | 23142080 | Unspecified |
| mmu-miR-669n | MIMAT0009427 | Acadm | 11364 | HITS-CLIP | 25083871 | Unspecified |
| mmu-miR-669n | MIMAT0009427 | Ackr4 | 252837 | HITS-CLIP | 25083871\|23597149 | Unspecified |
| mmu-miR-669n | MIMAT0009427 | Asap3 | 230837 | HITS-CLIP | 25083871 | Unspecified |
| mmu-miR-669n | MIMAT0009427 | Atxn1 | 20238 | HITS-CLIP | 25083871 | Unspecified |
| mmu-miR-669n | MIMAT0009427 | Atxn3 | 110616 | HITS-CLIP | 23142080 | Unspecified |
| mmu-miR-669n | MIMAT0009427 | Barhl2 | 104382 | HITS-CLIP | 23597149 | Unspecified |
| mmu-miR-669n | MIMAT0009427 | Bean1 | 65115 | HITS-CLIP | 23142080 | Unspecified |
| mmu-miR-669n | MIMAT0009427 | Ccrl1 | 252837 | HITS-CLIP | 23597149\|25083871 | Unspecified |
| mmu-miR-669n | MIMAT0009427 | Cdc14b | 218294 | HITS-CLIP | 23142080 | Unspecified |
| mmu-miR-669n | MIMAT0009427 | Cdk13 | 69562 | HITS-CLIP | 23142080 | Unspecified |
| mmu-miR-669n | MIMAT0009427 | Creg2 | 263764 | HITS-CLIP | 23597149 | Unspecified |
| mmu-miR-669n | MIMAT0009427 | Edaradd | 171211 | HITS-CLIP | 23142080 | Unspecified |
| mmu-miR-669n | MIMAT0009427 | Faf2 | 76577 | HITS-CLIP | 21258322 | Unspecified |
| mmu-miR-669n | MIMAT0009427 | Fam178a | 226151 | HITS-CLIP | 23142080 | Unspecified |
| mmu-miR-669n | MIMAT0009427 | Fam227a | 75729 | HITS-CLIP | 23142080 | Unspecified |
| mmu-miR-669n | MIMAT0009427 | Fbxl14 | 101358 | HITS-CLIP | 25083871\|23142080 | Unspecified |
| mmu-miR-669n | MIMAT0009427 | Fbxl18 | 231863 | HITS-CLIP | 23142080 | Unspecified |
| mmu-miR-669n | MIMAT0009427 | Frk | 14302 | HITS-CLIP | 25083871 | Unspecified |
| mmu-miR-669n | MIMAT0009427 | Fyn | 14360 | HITS-CLIP | 23142080 | Unspecified |
| mmu-miR-669n | MIMAT0009427 | Gcm1 | 14531 | HITS-CLIP | 23597149 | Unspecified |
| mmu-miR-669n | MIMAT0009427 | Gcnt4 | 218476 | HITS-CLIP | 25083871 | Unspecified |
| mmu-miR-669n | MIMAT0009427 | Gla | 11605 | HITS-CLIP | 25083871 | Unspecified |
| mmu-miR-669n | MIMAT0009427 | Gm4841 | 225594 | HITS-CLIP | 25083871 | Unspecified |
| mmu-miR-669n | MIMAT0009427 | Gpr146 | 80290 | HITS-CLIP | 21258322\|23597149 | Unspecified |
| mmu-miR-669n | MIMAT0009427 | Gpr35 | 64095 | HITS-CLIP | 23142080 | Unspecified |
| mmu-miR-669n | MIMAT0009427 | Gria3 | 53623 | HITS-CLIP | 23597149 | Unspecified |
| mmu-miR-669n | MIMAT0009427 | Haus2 | 66296 | HITS-CLIP | 25083871 | Unspecified |
| mmu-miR-669n | MIMAT0009427 | Hecw1 | 94253 | HITS-CLIP | 23142080 | Unspecified |
| mmu-miR-669n | MIMAT0009427 | Hs3st3b1 | 54710 | HITS-CLIP | 23142080 | Unspecified |
| mmu-miR-669n | MIMAT0009427 | Ints6 | 18130 | HITS-CLIP | 21258322\|23142080 | Unspecified |
| mmu-miR-669n | MIMAT0009427 | Ly6g6c | 68468 | HITS-CLIP | 23597149 | Unspecified |
| mmu-miR-669n | MIMAT0009427 | Man2a2 | 140481 | HITS-CLIP | 23597149 | Unspecified |
| mmu-miR-669n | MIMAT0009427 | Mapkbp1 | 26390 | HITS-CLIP | 23597149 | Unspecified |

| | | | | | | | |
|---|---|---|---|---|---|---|---|
| mmu-miR-669n | MIMAT0009427 | Mctp2 | 244049 | HITS-CLIP | | 23142080 | Unspecified |
| mmu-miR-669n | MIMAT0009427 | Mill1 | 266815 | HITS-CLIP | | 25083871 | Unspecified |
| mmu-miR-669n | MIMAT0009427 | Mlec | 109154 | HITS-CLIP | | 23597149 | Unspecified |
| mmu-miR-669n | MIMAT0009427 | Nab1 | 17936 | HITS-CLIP | | 23597149 | Unspecified |
| mmu-miR-669n | MIMAT0009427 | Ncbp3 | 66874 | HITS-CLIP | | 23142080 | Unspecified |
| mmu-miR-669n | MIMAT0009427 | Npr3 | 18162 | HITS-CLIP | | 25083871 | Unspecified |
| mmu-miR-669n | MIMAT0009427 | Nrxn1 | 18189 | HITS-CLIP | | 23142080 | Unspecified |
| mmu-miR-669n | MIMAT0009427 | Nrxn3 | 18191 | HITS-CLIP | 23142080\|25083871 | | Unspecified |
| mmu-miR-669n | MIMAT0009427 | Nus1 | 52014 | HITS-CLIP | 25083871\|23142080 | | Unspecified |
| mmu-miR-669n | MIMAT0009427 | Ocrl | 320634 | HITS-CLIP | | 23142080 | Unspecified |
| mmu-miR-669n | MIMAT0009427 | Palm2 | 242481 | HITS-CLIP | 25083871\|23597149 | | Unspecified |
| mmu-miR-669n | MIMAT0009427 | Pgm5 | 226041 | HITS-CLIP | | 23597149 | Unspecified |
| mmu-miR-669n | MIMAT0009427 | Ppp4r2 | 232314 | HITS-CLIP | | 25083871 | Unspecified |
| mmu-miR-669n | MIMAT0009427 | Prkab2 | 108097 | HITS-CLIP | | 25083871 | Unspecified |
| mmu-miR-669n | MIMAT0009427 | Prkd3 | 75292 | HITS-CLIP | | 23142080 | Unspecified |
| mmu-miR-669n | MIMAT0009427 | Rnf11 | 29864 | HITS-CLIP | | 23142080 | Unspecified |
| mmu-miR-669n | MIMAT0009427 | Rnf6 | 74132 | HITS-CLIP | | 23142080 | Unspecified |
| mmu-miR-669n | MIMAT0009427 | Rspry1 | 67610 | HITS-CLIP | | 23142080 | Unspecified |
| mmu-miR-669n | MIMAT0009427 | Senp6 | 215351 | Luciferase reporter assay//qRT-PCR//West | | 26807181 | Unspecified |
| mmu-miR-669n | MIMAT0009427 | Slc25a21 | 217593 | HITS-CLIP | | 23597149 | Unspecified |
| mmu-miR-669n | MIMAT0009427 | Slco2a1 | 24059 | HITS-CLIP | | 21258322 | Unspecified |
| mmu-miR-669n | MIMAT0009427 | Slf2 | 226151 | HITS-CLIP | | 23142080 | Unspecified |
| mmu-miR-669n | MIMAT0009427 | Ssr1 | 107513 | HITS-CLIP | | 23597149 | Unspecified |
| mmu-miR-669n | MIMAT0009427 | Tmem151b | 210573 | HITS-CLIP | | 23597149 | Unspecified |
| mmu-miR-669n | MIMAT0009427 | Tmem242 | 70544 | HITS-CLIP | | 21258322 | Unspecified |
| mmu-miR-669n | MIMAT0009427 | Tmem47 | 192216 | HITS-CLIP | | 23597149 | Unspecified |
| mmu-miR-669n | MIMAT0009427 | Tmem88b | 320587 | HITS-CLIP | | 21258322 | Unspecified |
| mmu-miR-669n | MIMAT0009427 | Txndc9 | 98258 | HITS-CLIP | | 21258322 | Unspecified |
| mmu-miR-669n | MIMAT0009427 | Uncx | 22255 | HITS-CLIP | | 23597149 | Unspecified |
| mmu-miR-669n | MIMAT0009427 | Usp2 | 53376 | HITS-CLIP | | 23142080 | Unspecified |
| mmu-miR-669n | MIMAT0009427 | Yipf5 | 67180 | HITS-CLIP | | 23142080 | Unspecified |
| mmu-miR-669n | MIMAT0009427 | Zfhx3 | 11906 | HITS-CLIP | | 23142080 | Unspecified |
| mmu-miR-669n | MIMAT0009427 | Zfp68 | 24135 | HITS-CLIP | | 25083871 | Unspecified |
| mmu-miR-669n | MIMAT0009427 | Zfp92 | 22754 | HITS-CLIP | 25083871\|23597149 | | Unspecified |
| mmu-miR-669n | MIMAT0009427 | Zscan29 | 99334 | HITS-CLIP | | 23597149 | Unspecified |
| mmu-miR-592-5p | MIMAT0003730 | 5730507C01Rik | 236366 | HITS-CLIP | | 23142080 | Unspecified |
| mmu-miR-592-5p | MIMAT0003730 | Ackr4 | 252837 | HITS-CLIP | 23597149\|25083871 | | Unspecified |
| mmu-miR-592-5p | MIMAT0003730 | Atxn1 | 20238 | HITS-CLIP | | 25083871 | Unspecified |
| mmu-miR-592-5p | MIMAT0003730 | Barhl2 | 104382 | HITS-CLIP | | 23597149 | Unspecified |

| | | | | | | |
|---|---|---|---|---|---|---|
| mmu-miR-592-5p | MIMAT0003730 | Ccrl1 | 252837 | HITS-CLIP | 25083871\|23597149 | Unspecified |
| mmu-miR-592-5p | MIMAT0003730 | Cdk13 | 69562 | HITS-CLIP | 23142080 | Unspecified |
| mmu-miR-592-5p | MIMAT0003730 | Creg2 | 263764 | HITS-CLIP | 23597149 | Unspecified |
| mmu-miR-592-5p | MIMAT0003730 | Fam227a | 75729 | HITS-CLIP | 23142080 | Unspecified |
| mmu-miR-592-5p | MIMAT0003730 | Frk | 14302 | HITS-CLIP | 25083871 | Unspecified |
| mmu-miR-592-5p | MIMAT0003730 | Gcnt4 | 218476 | HITS-CLIP | 25083871 | Unspecified |
| mmu-miR-592-5p | MIMAT0003730 | Gla | 11605 | HITS-CLIP | 25083871 | Unspecified |
| mmu-miR-592-5p | MIMAT0003730 | Gm4841 | 225594 | HITS-CLIP | 25083871 | Unspecified |
| mmu-miR-592-5p | MIMAT0003730 | Gpr146 | 80290 | HITS-CLIP | 23597149\|21258322 | Unspecified |
| mmu-miR-592-5p | MIMAT0003730 | Gpr35 | 64095 | HITS-CLIP | 23142080 | Unspecified |
| mmu-miR-592-5p | MIMAT0003730 | Gria3 | 53623 | HITS-CLIP | 23597149 | Unspecified |
| mmu-miR-592-5p | MIMAT0003730 | Haus2 | 66296 | HITS-CLIP | 25083871 | Unspecified |
| mmu-miR-592-5p | MIMAT0003730 | Hs3st3b1 | 54710 | HITS-CLIP | 23142080 | Unspecified |
| mmu-miR-592-5p | MIMAT0003730 | Ly6g6c | 68468 | HITS-CLIP | 23597149 | Unspecified |
| mmu-miR-592-5p | MIMAT0003730 | Man2a2 | 140481 | HITS-CLIP | 23597149 | Unspecified |
| mmu-miR-592-5p | MIMAT0003730 | Mapkbp1 | 26390 | HITS-CLIP | 23597149 | Unspecified |
| mmu-miR-592-5p | MIMAT0003730 | Mctp2 | 244049 | HITS-CLIP | 23142080 | Unspecified |
| mmu-miR-592-5p | MIMAT0003730 | Mill1 | 266815 | HITS-CLIP | 25083871 | Unspecified |
| mmu-miR-592-5p | MIMAT0003730 | Mlec | 109154 | HITS-CLIP | 23597149 | Unspecified |
| mmu-miR-592-5p | MIMAT0003730 | Nab1 | 17936 | HITS-CLIP | 23597149 | Unspecified |
| mmu-miR-592-5p | MIMAT0003730 | Npr3 | 18162 | HITS-CLIP | 25083871 | Unspecified |
| mmu-miR-592-5p | MIMAT0003730 | Nrxn1 | 18189 | HITS-CLIP | 23142080 | Unspecified |
| mmu-miR-592-5p | MIMAT0003730 | Nrxn3 | 18191 | HITS-CLIP | 23142080 | Unspecified |
| mmu-miR-592-5p | MIMAT0003730 | Nus1 | 52014 | HITS-CLIP | 25083871 | Unspecified |
| mmu-miR-592-5p | MIMAT0003730 | Palm2 | 242481 | HITS-CLIP | 25083871\|23597149 | Unspecified |
| mmu-miR-592-5p | MIMAT0003730 | Pgm5 | 226041 | HITS-CLIP | 23597149 | Unspecified |
| mmu-miR-592-5p | MIMAT0003730 | Ppp4r2 | 232314 | HITS-CLIP | 25083871 | Unspecified |
| mmu-miR-592-5p | MIMAT0003730 | Prkab2 | 108097 | HITS-CLIP | 25083871 | Unspecified |
| mmu-miR-592-5p | MIMAT0003730 | Rnf6 | 74132 | HITS-CLIP | 23142080 | Unspecified |
| mmu-miR-592-5p | MIMAT0003730 | Slc24a2 | 76376 | HITS-CLIP | 25083871 | Unspecified |
| mmu-miR-592-5p | MIMAT0003730 | Slc25a21 | 217593 | HITS-CLIP | 23597149 | Unspecified |
| mmu-miR-592-5p | MIMAT0003730 | Ssr1 | 107513 | HITS-CLIP | 23597149 | Unspecified |
| mmu-miR-592-5p | MIMAT0003730 | Tmem151b | 210573 | HITS-CLIP | 23597149 | Unspecified |
| mmu-miR-592-5p | MIMAT0003730 | Tmem47 | 192216 | HITS-CLIP | 23597149 | Unspecified |
| mmu-miR-592-5p | MIMAT0003730 | Txndc9 | 98258 | HITS-CLIP | 21258322 | Unspecified |
| mmu-miR-592-5p | MIMAT0003730 | Uba1y | 22202 | HITS-CLIP | 21258322 | Unspecified |
| mmu-miR-592-5p | MIMAT0003730 | Uncx | 22255 | HITS-CLIP | 23597149 | Unspecified |
| mmu-miR-592-5p | MIMAT0003730 | Zfp385c | 278304 | HITS-CLIP | 21258322 | Unspecified |
| mmu-miR-592-5p | MIMAT0003730 | Zfp68 | 24135 | HITS-CLIP | 25083871 | Unspecified |

| | | | | | | |
|---|---|---|---|---|---|---|
| mmu-miR-592-5p | MIMAT0003730 | Zfp92 | 22754 | HITS-CLIP | 23597149\|25083871 | Unspecified |
| mmu-miR-592-5p | MIMAT0003730 | Zscan29 | 99334 | HITS-CLIP | 23597149 | Unspecified |
| mmu-miR-3086-3p | MIMAT0014881 | Cadm2 | 239857 | HITS-CLIP | 21258322\|23142080\|25083871 | Unspecified |
| mmu-miR-3086-3p | MIMAT0014881 | Ccdc82 | 66396 | HITS-CLIP | 25083871 | Unspecified |
| mmu-miR-3086-3p | MIMAT0014881 | Ccl9 | 20308 | HITS-CLIP | 21258322 | Unspecified |
| mmu-miR-3086-3p | MIMAT0014881 | Cml3 | 93674 | HITS-CLIP | 21258322 | Unspecified |
| mmu-miR-3086-3p | MIMAT0014881 | Glrx2 | 69367 | HITS-CLIP | 21258322 | Unspecified |
| mmu-miR-3086-3p | MIMAT0014881 | Gm11128 | 100504710 | HITS-CLIP | 21258322 | Unspecified |
| mmu-miR-3086-3p | MIMAT0014881 | Isg15 | 100038882 | HITS-CLIP | 21258322\|25083871 | Unspecified |
| mmu-miR-3086-3p | MIMAT0014881 | Maoa | 17161 | HITS-CLIP | 21258322\|25083871 | Unspecified |
| mmu-miR-3086-3p | MIMAT0014881 | N4bp1 | 80750 | HITS-CLIP | 23597149 | Unspecified |
| mmu-miR-3086-3p | MIMAT0014881 | Nat8f3 | 93674 | HITS-CLIP | 21258322 | Unspecified |
| mmu-miR-3086-3p | MIMAT0014881 | Nat8f6 | 100504710 | HITS-CLIP | 21258322 | Unspecified |
| mmu-miR-3086-3p | MIMAT0014881 | Nkain3 | 269513 | HITS-CLIP | 25083871\|21258322 | Unspecified |
| mmu-miR-3086-3p | MIMAT0014881 | Nol12 | 97961 | HITS-CLIP | 25083871 | Unspecified |
| mmu-miR-3086-3p | MIMAT0014881 | Nr2f2 | 11819 | HITS-CLIP | 25083871 | Unspecified |
| mmu-miR-3086-3p | MIMAT0014881 | Oxsm | 71147 | HITS-CLIP | 25083871 | Unspecified |
| mmu-miR-3086-3p | MIMAT0014881 | Phf14 | 75725 | HITS-CLIP | 25083871 | Unspecified |
| mmu-miR-3086-3p | MIMAT0014881 | Plxnc1 | 54712 | HITS-CLIP | 25083871 | Unspecified |
| mmu-miR-3086-3p | MIMAT0014881 | Pramef8 | 242736 | HITS-CLIP | 21258322 | Unspecified |
| mmu-miR-3086-3p | MIMAT0014881 | Prkca | 18750 | HITS-CLIP | 23142080 | Unspecified |
| mmu-miR-3086-3p | MIMAT0014881 | Prrxl1 | 107751 | HITS-CLIP | 21258322 | Unspecified |
| mmu-miR-3086-3p | MIMAT0014881 | Slc5a3 | 53881 | HITS-CLIP | 21258322 | Unspecified |
| mmu-miR-3086-3p | MIMAT0014881 | Ybey | 216119 | HITS-CLIP | 25083871 | Unspecified |
| mmu-miR-3086-3p | MIMAT0014881 | Zfp65 | 235907 | HITS-CLIP | 21258322 | Unspecified |
| mmu-miR-3086-3p | MIMAT0014881 | Zfp71-rs1 | 235907 | HITS-CLIP | 21258322 | Unspecified |
| mmu-miR-5133 | MIMAT0020644 | 2010109I03Rik | 67038 | HITS-CLIP | 21258322 | Unspecified |
| mmu-miR-5133 | MIMAT0020644 | Arhgef39 | 230098 | HITS-CLIP | 21258322 | Unspecified |
| mmu-miR-5133 | MIMAT0020644 | Ccdc78 | 381077 | HITS-CLIP | 25083871\|21258322 | Unspecified |
| mmu-miR-5133 | MIMAT0020644 | Chrna1 | 11435 | HITS-CLIP | 21258322 | Unspecified |
| mmu-miR-5133 | MIMAT0020644 | Clns1a | 12729 | HITS-CLIP | 21258322 | Unspecified |
| mmu-miR-5133 | MIMAT0020644 | Cspg4 | 121021 | HITS-CLIP | 23142080 | Unspecified |
| mmu-miR-5133 | MIMAT0020644 | Dync1li2 | 234663 | HITS-CLIP | 21258322\|25083871 | Unspecified |
| mmu-miR-5133 | MIMAT0020644 | E130306D19Rik | 230098 | CLIP-Seq dataset analysis | 21258322 | Unspecified |
| mmu-miR-5133 | MIMAT0020644 | Efr3b | 668212 | HITS-CLIP | 21258322 | Unspecified |
| mmu-miR-5133 | MIMAT0020644 | Ggps1 | 14593 | HITS-CLIP | 21258322 | Unspecified |
| mmu-miR-5133 | MIMAT0020644 | H2-T24 | 15042 | HITS-CLIP | 25083871\|21258322 | Unspecified |
| mmu-miR-5133 | MIMAT0020644 | Hcls1 | 15163 | HITS-CLIP | 21258322 | Unspecified |
| mmu-miR-5133 | MIMAT0020644 | Ikbkg | 16151 | HITS-CLIP | 21258322\|25083871 | Unspecified |

| | | | | | | |
|---|---|---|---|---|---|---|
| mmu-miR-5133 | MIMAT0020644 | Madd | 228355 | HITS-CLIP | 26203562\|23597149\|21258322\|1953[illegible] | Unspecified |
| mmu-miR-5133 | MIMAT0020644 | Marchf5 | 69104 | HITS-CLIP | 25083871\|21258322 | Unspecified |
| mmu-miR-5133 | MIMAT0020644 | Nanog | 71950 | HITS-CLIP | 21258322\|23597149 | Unspecified |
| mmu-miR-5133 | MIMAT0020644 | Paxip1 | 55982 | HITS-CLIP | 23597149 | Unspecified |
| mmu-miR-5133 | MIMAT0020644 | Ppp1r16b | 228852 | HITS-CLIP | 21258322 | Unspecified |
| mmu-miR-5133 | MIMAT0020644 | Prss43 | 272643 | HITS-CLIP | 21258322 | Unspecified |
| mmu-miR-5133 | MIMAT0020644 | Rmnd5a | 68477 | HITS-CLIP | 23597149 | Unspecified |
| mmu-miR-5133 | MIMAT0020644 | Rnf150 | 330812 | HITS-CLIP | 21258322\|25083871 | Unspecified |
| mmu-miR-5133 | MIMAT0020644 | Sec23ip | 207352 | HITS-CLIP | 25083871 | Unspecified |
| mmu-miR-5133 | MIMAT0020644 | Sgo2a | 68549 | HITS-CLIP | 21258322 | Unspecified |
| mmu-miR-5133 | MIMAT0020644 | Sgol2 | 68549 | HITS-CLIP | 21258322 | Unspecified |
| mmu-miR-5133 | MIMAT0020644 | Slc35d2 | 70484 | HITS-CLIP | 21258322 | Unspecified |
| mmu-miR-5133 | MIMAT0020644 | Slc4a7 | 218756 | HITS-CLIP | 21258322 | Unspecified |
| mmu-miR-5133 | MIMAT0020644 | Slc7a2 | 11988 | HITS-CLIP | 21258322 | Unspecified |
| mmu-miR-5133 | MIMAT0020644 | Slx1b | 75764 | HITS-CLIP | 21258322 | Unspecified |
| mmu-miR-5133 | MIMAT0020644 | Svopl | 320590 | HITS-CLIP | 21258322 | Unspecified |
| mmu-miR-5133 | MIMAT0020644 | Timm17b | 21855 | HITS-CLIP | 25083871\|21258322 | Unspecified |
| mmu-miR-5133 | MIMAT0020644 | Tmem206 | 66950 | HITS-CLIP | 25083871\|21258322 | Unspecified |
| mmu-miR-5133 | MIMAT0020644 | Tmppe | 100504715 | HITS-CLIP | 21258322 | Unspecified |
| mmu-miR-5133 | MIMAT0020644 | Xpo7 | 65246 | HITS-CLIP | 21258322 | Unspecified |
| mmu-miR-5133 | MIMAT0020644 | Zfp113 | 56314 | HITS-CLIP | 25083871 | Unspecified |
| mmu-miR-674-5p | MIMAT0003740 | 1700016D06Rik | 76413 | HITS-CLIP | 21258322 | Unspecified |
| mmu-miR-674-5p | MIMAT0003740 | 1700047I17Rik2 | 100101807 | HITS-CLIP | 21258322 | Unspecified |
| mmu-miR-674-5p | MIMAT0003740 | A230046K03Rik | 319277 | HITS-CLIP | 23597149\|21258322 | Unspecified |
| mmu-miR-674-5p | MIMAT0003740 | AK010878 | 100233175 | HITS-CLIP | 21258322 | Unspecified |
| mmu-miR-674-5p | MIMAT0003740 | Aipl1 | 114230 | HITS-CLIP | 21258322 | Unspecified |
| mmu-miR-674-5p | MIMAT0003740 | Ankfy1 | 11736 | HITS-CLIP | 21258322 | Unspecified |
| mmu-miR-674-5p | MIMAT0003740 | Anks1b | 77531 | HITS-CLIP | 23142080 | Unspecified |
| mmu-miR-674-5p | MIMAT0003740 | Arl16 | 70317 | HITS-CLIP | 21258322 | Unspecified |
| mmu-miR-674-5p | MIMAT0003740 | Asxl2 | 75302 | HITS-CLIP | 21258322 | Unspecified |
| mmu-miR-674-5p | MIMAT0003740 | Atrn | 11990 | HITS-CLIP | 21258322 | Unspecified |
| mmu-miR-674-5p | MIMAT0003740 | Bach2 | 12014 | HITS-CLIP | 25083871 | Unspecified |
| mmu-miR-674-5p | MIMAT0003740 | Banp | 53325 | HITS-CLIP | 21258322 | Unspecified |
| mmu-miR-674-5p | MIMAT0003740 | Brip1 | 237911 | HITS-CLIP | 21258322 | Unspecified |
| mmu-miR-674-5p | MIMAT0003740 | Bzrap1 | 207777 | HITS-CLIP | 25083871 | Unspecified |
| mmu-miR-674-5p | MIMAT0003740 | Casc4 | 319996 | HITS-CLIP | 21258322\|23597149 | Unspecified |
| mmu-miR-674-5p | MIMAT0003740 | Ccdc25 | 67179 | HITS-CLIP | 23597149\|25083871\|23142080\|2125[illegible] | Unspecified |
| mmu-miR-674-5p | MIMAT0003740 | Ccr9 | 12769 | HITS-CLIP | 21258322 | Unspecified |
| mmu-miR-674-5p | MIMAT0003740 | Cenpf | 108000 | HITS-CLIP | 21258322 | Unspecified |

| | | | | | | | |
|---|---|---|---|---|---|---|---|
| mmu-miR-674-5p | MIMAT0003740 | Cflar | 12633 | HITS-CLIP | | 21258322 | Unspecified |
| mmu-miR-674-5p | MIMAT0003740 | Churc1 | 211151 | HITS-CLIP | 23597149\|21258322 | | Unspecified |
| mmu-miR-674-5p | MIMAT0003740 | Clca2 | 80797 | HITS-CLIP | | 21258322 | Unspecified |
| mmu-miR-674-5p | MIMAT0003740 | Clca3a2 | 80797 | HITS-CLIP | | 21258322 | Unspecified |
| mmu-miR-674-5p | MIMAT0003740 | Commd7 | 99311 | HITS-CLIP | | 21258322 | Unspecified |
| mmu-miR-674-5p | MIMAT0003740 | Comt | 12846 | HITS-CLIP | | 21258322 | Unspecified |
| mmu-miR-674-5p | MIMAT0003740 | Cpne9 | 211232 | HITS-CLIP | | 21258322 | Unspecified |
| mmu-miR-674-5p | MIMAT0003740 | Cx3cr1 | 13051 | HITS-CLIP | | 21258322 | Unspecified |
| mmu-miR-674-5p | MIMAT0003740 | Cxcr2 | 12765 | HITS-CLIP | | 21258322 | Unspecified |
| mmu-miR-674-5p | MIMAT0003740 | Cyp2e1 | 13106 | HITS-CLIP | | 21258322 | Unspecified |
| mmu-miR-674-5p | MIMAT0003740 | D3Ertd254e | 241944 | HITS-CLIP | | 21258322 | Unspecified |
| mmu-miR-674-5p | MIMAT0003740 | Dcaf4 | 73828 | HITS-CLIP | | 21258322 | Unspecified |
| mmu-miR-674-5p | MIMAT0003740 | Ddx3x | 13205 | HITS-CLIP | | 23597149 | Unspecified |
| mmu-miR-674-5p | MIMAT0003740 | Dennd5b | 320560 | HITS-CLIP | | 21258322 | Unspecified |
| mmu-miR-674-5p | MIMAT0003740 | Dph5 | 69740 | HITS-CLIP | | 21258322 | Unspecified |
| mmu-miR-674-5p | MIMAT0003740 | Exosc2 | 227715 | HITS-CLIP | | 21258322 | Unspecified |
| mmu-miR-674-5p | MIMAT0003740 | Fam177a | 73385 | HITS-CLIP | | 21258322 | Unspecified |
| mmu-miR-674-5p | MIMAT0003740 | Gm10778 | 100233208 | HITS-CLIP | | 21258322 | Unspecified |
| mmu-miR-674-5p | MIMAT0003740 | Gon7 | 100233175 | HITS-CLIP | | 21258322 | Unspecified |
| mmu-miR-674-5p | MIMAT0003740 | H2-T23 | 15040 | HITS-CLIP | | 21258322 | Unspecified |
| mmu-miR-674-5p | MIMAT0003740 | Il15ra | 16169 | HITS-CLIP | | 25083871 | Unspecified |
| mmu-miR-674-5p | MIMAT0003740 | Irak1 | 16179 | HITS-CLIP | | 21258322 | Unspecified |
| mmu-miR-674-5p | MIMAT0003740 | Lcmt2 | 329504 | HITS-CLIP | | 21258322 | Unspecified |
| mmu-miR-674-5p | MIMAT0003740 | Loxl2 | 94352 | HITS-CLIP | 21258322\|23597149 | | Unspecified |
| mmu-miR-674-5p | MIMAT0003740 | Magt1 | 67075 | HITS-CLIP | | 25083871 | Unspecified |
| mmu-miR-674-5p | MIMAT0003740 | Mbp | 17196 | HITS-CLIP | 25083871\|21258322 | | Unspecified |
| mmu-miR-674-5p | MIMAT0003740 | Med22 | 20933 | HITS-CLIP | | 23142080 | Unspecified |
| mmu-miR-674-5p | MIMAT0003740 | Mfsd11 | 69900 | HITS-CLIP | | 21258322 | Unspecified |
| mmu-miR-674-5p | MIMAT0003740 | Mfsd2b | 432628 | HITS-CLIP | | 21258322 | Unspecified |
| mmu-miR-674-5p | MIMAT0003740 | Mpv17l | 93734 | HITS-CLIP | | 21258322 | Unspecified |
| mmu-miR-674-5p | MIMAT0003740 | Myh10 | 77579 | HITS-CLIP | | 23142080 | Unspecified |
| mmu-miR-674-5p | MIMAT0003740 | Myo1c | 17913 | HITS-CLIP | | 21258322 | Unspecified |
| mmu-miR-674-5p | MIMAT0003740 | Naip5 | 17951 | HITS-CLIP | | 21258322 | Unspecified |
| mmu-miR-674-5p | MIMAT0003740 | Nip7 | 66164 | HITS-CLIP | | 21258322 | Unspecified |
| mmu-miR-674-5p | MIMAT0003740 | Nol3 | 78688 | HITS-CLIP | | 21258322 | Unspecified |
| mmu-miR-674-5p | MIMAT0003740 | Nup133 | 234865 | HITS-CLIP | | 21258322 | Unspecified |
| mmu-miR-674-5p | MIMAT0003740 | Ogdh | 18293 | HITS-CLIP | 21258322\|23597149 | | Unspecified |
| mmu-miR-674-5p | MIMAT0003740 | Pbld1 | 68371 | HITS-CLIP | | 21258322 | Unspecified |
| mmu-miR-674-5p | MIMAT0003740 | Pcsk6 | 18553 | HITS-CLIP | | 23142080 | Unspecified |

| | | | | | | |
|---|---|---|---|---|---|---|
| mmu-miR-674-5p | MIMAT0003740 | Pla2g4b | 211429 | HITS-CLIP | 21258322 | Unspecified |
| mmu-miR-674-5p | MIMAT0003740 | Pola2 | 18969 | HITS-CLIP | 21258322\|23597149 | Unspecified |
| mmu-miR-674-5p | MIMAT0003740 | Polr2l | 66491 | HITS-CLIP | 21258322 | Unspecified |
| mmu-miR-674-5p | MIMAT0003740 | Ppil4 | 67418 | HITS-CLIP | 21258322 | Unspecified |
| mmu-miR-674-5p | MIMAT0003740 | Ppm1k | 243382 | HITS-CLIP | 21258322 | Unspecified |
| mmu-miR-674-5p | MIMAT0003740 | Ppp2r2c | 269643 | HITS-CLIP | 25083871 | Unspecified |
| mmu-miR-674-5p | MIMAT0003740 | Prps2 | 110639 | HITS-CLIP | 25083871\|23142080\|26203562 | Unspecified |
| mmu-miR-674-5p | MIMAT0003740 | Rab33b | 19338 | HITS-CLIP | 21258322 | Unspecified |
| mmu-miR-674-5p | MIMAT0003740 | Rbbp4 | 19646 | HITS-CLIP | 21258322 | Unspecified |
| mmu-miR-674-5p | MIMAT0003740 | Rbm34 | 52202 | HITS-CLIP | 21258322 | Unspecified |
| mmu-miR-674-5p | MIMAT0003740 | Rin1 | 225870 | HITS-CLIP | 21258322 | Unspecified |
| mmu-miR-674-5p | MIMAT0003740 | Rmi2 | 223970 | HITS-CLIP | 21258322 | Unspecified |
| mmu-miR-674-5p | MIMAT0003740 | Rpl7l1 | 66229 | HITS-CLIP | 25083871 | Unspecified |
| mmu-miR-674-5p | MIMAT0003740 | Sap18 | 20220 | HITS-CLIP | 21258322 | Unspecified |
| mmu-miR-674-5p | MIMAT0003740 | Scd3 | 30049 | HITS-CLIP | 21258322 | Unspecified |
| mmu-miR-674-5p | MIMAT0003740 | Sh3rf3 | 237353 | HITS-CLIP | 23142080 | Unspecified |
| mmu-miR-674-5p | MIMAT0003740 | Slc22a15 | 242126 | HITS-CLIP | 21258322 | Unspecified |
| mmu-miR-674-5p | MIMAT0003740 | Slc23a1 | 20522 | HITS-CLIP | 21258322\|23597149 | Unspecified |
| mmu-miR-674-5p | MIMAT0003740 | Slc7a14 | 241919 | HITS-CLIP | 21258322 | Unspecified |
| mmu-miR-674-5p | MIMAT0003740 | Slfn5 | 327978 | HITS-CLIP | 23597149\|21258322 | Unspecified |
| mmu-miR-674-5p | MIMAT0003740 | Spc24 | 67629 | HITS-CLIP | 21258322 | Unspecified |
| mmu-miR-674-5p | MIMAT0003740 | Srp72 | 66661 | HITS-CLIP | 25083871\|23142080 | Unspecified |
| mmu-miR-674-5p | MIMAT0003740 | Ssr1 | 107513 | HITS-CLIP | 21258322 | Unspecified |
| mmu-miR-674-5p | MIMAT0003740 | Synrg | 217030 | HITS-CLIP | 21258322 | Unspecified |
| mmu-miR-674-5p | MIMAT0003740 | T2 | 21331 | HITS-CLIP | 21258322 | Unspecified |
| mmu-miR-674-5p | MIMAT0003740 | Tapbpl | 213233 | HITS-CLIP | 21258322 | Unspecified |
| mmu-miR-674-5p | MIMAT0003740 | Tbc1d24 | 224617 | HITS-CLIP | 21258322 | Unspecified |
| mmu-miR-674-5p | MIMAT0003740 | Tmem69 | 230657 | HITS-CLIP | 23597149\|21258322 | Unspecified |
| mmu-miR-674-5p | MIMAT0003740 | Tspan15 | 70423 | HITS-CLIP | 21258322 | Unspecified |
| mmu-miR-674-5p | MIMAT0003740 | Tspoap1 | 207777 | HITS-CLIP | 25083871 | Unspecified |
| mmu-miR-674-5p | MIMAT0003740 | Tyk2 | 54721 | HITS-CLIP | 21258322 | Unspecified |
| mmu-miR-674-5p | MIMAT0003740 | Ubp1 | 22221 | HITS-CLIP | 23142080 | Unspecified |
| mmu-miR-674-5p | MIMAT0003740 | Vwa5a | 67776 | HITS-CLIP | 28735896 | Unspecified |
| mmu-miR-674-5p | MIMAT0003740 | Washc4 | 319277 | HITS-CLIP | 29167373\|21258322\|23597149 | Unspecified |
| mmu-miR-674-5p | MIMAT0003740 | Wdr89 | 72338 | HITS-CLIP | 21258322 | Unspecified |
| mmu-miR-674-5p | MIMAT0003740 | Ypel2 | 77864 | HITS-CLIP | 21258322 | Unspecified |
| mmu-miR-674-5p | MIMAT0003740 | Ywhag | 22628 | HITS-CLIP | 23142080 | Unspecified |
| mmu-miR-674-5p | MIMAT0003740 | Zfp202 | 80902 | HITS-CLIP | 21258322 | Unspecified |
| mmu-miR-674-5p | MIMAT0003740 | Zfp605 | 675812 | HITS-CLIP | 21258322 | Unspecified |

| | | | | | | |
|---|---|---|---|---|---|---|
| mmu-miR-674-5p | MIMAT0003740 | Zfp607 | 545938 | HITS-CLIP | 21258322 | Unspecified |
| mmu-miR-674-5p | MIMAT0003740 | Zfp607a | 545938 | HITS-CLIP | 21258322 | Unspecified |
| mmu-miR-674-5p | MIMAT0003740 | Zfp935 | 71508 | HITS-CLIP | 21258322 | Unspecified |
| mmu-miR-674-5p | MIMAT0003740 | Zyg11b | 414872 | HITS-CLIP | 21258322 | Unspecified |
| mmu-miR-5114 | MIMAT0020622 | 1200014J11Rik | 66874 | HITS-CLIP | 23142080 | Unspecified |
| mmu-miR-5114 | MIMAT0020622 | 4931428F04Rik | 74356 | HITS-CLIP | 21258322 | Unspecified |
| mmu-miR-5114 | MIMAT0020622 | Arhgef39 | 230098 | HITS-CLIP | 21258322 | Unspecified |
| mmu-miR-5114 | MIMAT0020622 | Armc2 | 213402 | HITS-CLIP | 21258322 | Unspecified |
| mmu-miR-5114 | MIMAT0020622 | Ccdc78 | 381077 | HITS-CLIP | 21258322 | Unspecified |
| mmu-miR-5114 | MIMAT0020622 | Cd63 | 12512 | HITS-CLIP | 25083871 | Unspecified |
| mmu-miR-5114 | MIMAT0020622 | Cdc14b | 218294 | HITS-CLIP | 21258322 | Unspecified |
| mmu-miR-5114 | MIMAT0020622 | Chrna1 | 11435 | HITS-CLIP | 21258322 | Unspecified |
| mmu-miR-5114 | MIMAT0020622 | Cmtm6 | 67213 | HITS-CLIP | 25083871 | Unspecified |
| mmu-miR-5114 | MIMAT0020622 | Cpeb3 | 208922 | HITS-CLIP | 25083871 | Unspecified |
| mmu-miR-5114 | MIMAT0020622 | Cspg4 | 121021 | HITS-CLIP | 23142080 | Unspecified |
| mmu-miR-5114 | MIMAT0020622 | Dync1li2 | 234663 | HITS-CLIP | 25083871\|21258322 | Unspecified |
| mmu-miR-5114 | MIMAT0020622 | E130306D19Rik | 230098 | CLIP-Seq dataset analysis | 21258322 | Unspecified |
| mmu-miR-5114 | MIMAT0020622 | Efr3b | 668212 | HITS-CLIP | 21258322 | Unspecified |
| mmu-miR-5114 | MIMAT0020622 | Fam161b | 217705 | HITS-CLIP | 25083871 | Unspecified |
| mmu-miR-5114 | MIMAT0020622 | Ggps1 | 14593 | HITS-CLIP | 21258322 | Unspecified |
| mmu-miR-5114 | MIMAT0020622 | Glrx2 | 69367 | HITS-CLIP | 21258322 | Unspecified |
| mmu-miR-5114 | MIMAT0020622 | H2-T24 | 15042 | HITS-CLIP | 25083871\|21258322 | Unspecified |
| mmu-miR-5114 | MIMAT0020622 | Ikbkg | 16151 | HITS-CLIP | 25083871\|21258322 | Unspecified |
| mmu-miR-5114 | MIMAT0020622 | Madd | 228355 | HITS-CLIP | 23597149\|26203562\|21258322\|1953[illegible] | Unspecified |
| mmu-miR-5114 | MIMAT0020622 | Marchf5 | 69104 | HITS-CLIP | 21258322\|25083871 | Unspecified |
| mmu-miR-5114 | MIMAT0020622 | Mtf1 | 17764 | HITS-CLIP | 23142080 | Unspecified |
| mmu-miR-5114 | MIMAT0020622 | Nanog | 71950 | HITS-CLIP | 21258322\|23597149 | Unspecified |
| mmu-miR-5114 | MIMAT0020622 | Nav2 | 78286 | HITS-CLIP | 25083871 | Unspecified |
| mmu-miR-5114 | MIMAT0020622 | Ncbp3 | 66874 | HITS-CLIP | 23142080 | Unspecified |
| mmu-miR-5114 | MIMAT0020622 | Padi2 | 18600 | HITS-CLIP | 21258322 | Unspecified |
| mmu-miR-5114 | MIMAT0020622 | Paxip1 | 55982 | HITS-CLIP | 23597149 | Unspecified |
| mmu-miR-5114 | MIMAT0020622 | Ppp1r16b | 228852 | HITS-CLIP | 21258322 | Unspecified |
| mmu-miR-5114 | MIMAT0020622 | Prtg | 235472 | HITS-CLIP | 25083871 | Unspecified |
| mmu-miR-5114 | MIMAT0020622 | Rmnd5a | 68477 | HITS-CLIP | 23597149 | Unspecified |
| mmu-miR-5114 | MIMAT0020622 | Rnf150 | 330812 | HITS-CLIP | 25083871\|21258322 | Unspecified |
| mmu-miR-5114 | MIMAT0020622 | Sec23ip | 207352 | HITS-CLIP | 25083871 | Unspecified |
| mmu-miR-5114 | MIMAT0020622 | Sgo2a | 68549 | HITS-CLIP | 21258322 | Unspecified |
| mmu-miR-5114 | MIMAT0020622 | Sgol2 | 68549 | HITS-CLIP | 21258322 | Unspecified |
| mmu-miR-5114 | MIMAT0020622 | Slc35d2 | 70484 | HITS-CLIP | 21258322 | Unspecified |

| | | | | | | |
|---|---|---|---|---|---|---|
| mmu-miR-5114 | MIMAT0020622 | Slc4a7 | 218756 | HITS-CLIP | 21258322 | Unspecified |
| mmu-miR-5114 | MIMAT0020622 | Srrm4 | 68955 | HITS-CLIP | 25083871 | Unspecified |
| mmu-miR-5114 | MIMAT0020622 | Svopl | 320590 | HITS-CLIP | 21258322 | Unspecified |
| mmu-miR-5114 | MIMAT0020622 | T2 | 21331 | HITS-CLIP | 21258322 | Unspecified |
| mmu-miR-5114 | MIMAT0020622 | Tmem206 | 66950 | HITS-CLIP | 25083871\|21258322 | Unspecified |
| mmu-miR-5114 | MIMAT0020622 | Xpo7 | 65246 | HITS-CLIP | 21258322 | Unspecified |
| mmu-miR-5114 | MIMAT0020622 | Zfp113 | 56314 | HITS-CLIP | 25083871 | Unspecified |
| mmu-miR-3106-5p | MIMAT0014817 | 1200011I18Rik | 67467 | HITS-CLIP | 21258322 | Unspecified |
| mmu-miR-3106-5p | MIMAT0014817 | 4930444A02Rik | 74653 | HITS-CLIP | 23142080 | Unspecified |
| mmu-miR-3106-5p | MIMAT0014817 | 4931428F04Rik | 74356 | HITS-CLIP | 21258322 | Unspecified |
| mmu-miR-3106-5p | MIMAT0014817 | Actr10 | 56444 | HITS-CLIP | 21258322 | Unspecified |
| mmu-miR-3106-5p | MIMAT0014817 | Armc2 | 213402 | HITS-CLIP | 21258322 | Unspecified |
| mmu-miR-3106-5p | MIMAT0014817 | B3gnt5 | 108105 | HITS-CLIP | 23142080 | Unspecified |
| mmu-miR-3106-5p | MIMAT0014817 | B4galnt2 | 14422 | HITS-CLIP | 21258322 | Unspecified |
| mmu-miR-3106-5p | MIMAT0014817 | Birc5 | 11799 | HITS-CLIP | 21258322 | Unspecified |
| mmu-miR-3106-5p | MIMAT0014817 | Ccdc55 | 237859 | HITS-CLIP | 21258322 | Unspecified |
| mmu-miR-3106-5p | MIMAT0014817 | Ccdc85b | 240514 | HITS-CLIP | 21258322 | Unspecified |
| mmu-miR-3106-5p | MIMAT0014817 | Cd300a | 217303 | HITS-CLIP | 23597149\|21258322 | Unspecified |
| mmu-miR-3106-5p | MIMAT0014817 | Chrna1 | 11435 | HITS-CLIP | 21258322 | Unspecified |
| mmu-miR-3106-5p | MIMAT0014817 | Commd7 | 99311 | HITS-CLIP | 21258322 | Unspecified |
| mmu-miR-3106-5p | MIMAT0014817 | Cyld | 74256 | HITS-CLIP | 21258322 | Unspecified |
| mmu-miR-3106-5p | MIMAT0014817 | Dph7 | 67228 | HITS-CLIP | 21258322\|25083871 | Unspecified |
| mmu-miR-3106-5p | MIMAT0014817 | Eif4g2 | 13690 | HITS-CLIP | 21258322 | Unspecified |
| mmu-miR-3106-5p | MIMAT0014817 | Fam161b | 217705 | HITS-CLIP | 25083871 | Unspecified |
| mmu-miR-3106-5p | MIMAT0014817 | Frem3 | 333315 | HITS-CLIP | 23142080 | Unspecified |
| mmu-miR-3106-5p | MIMAT0014817 | Gas2l3 | 237436 | HITS-CLIP | 21258322 | Unspecified |
| mmu-miR-3106-5p | MIMAT0014817 | Ggps1 | 14593 | HITS-CLIP | 21258322 | Unspecified |
| mmu-miR-3106-5p | MIMAT0014817 | Gm16532 | 100042450 | HITS-CLIP | 21258322 | Unspecified |
| mmu-miR-3106-5p | MIMAT0014817 | Gm5113 | 330503 | HITS-CLIP | 21258322 | Unspecified |
| mmu-miR-3106-5p | MIMAT0014817 | Gpalpp1 | 67467 | HITS-CLIP | 21258322 | Unspecified |
| mmu-miR-3106-5p | MIMAT0014817 | H2-T24 | 15042 | HITS-CLIP | 21258322\|25083871 | Unspecified |
| mmu-miR-3106-5p | MIMAT0014817 | Homez | 239099 | HITS-CLIP | 21258322 | Unspecified |
| mmu-miR-3106-5p | MIMAT0014817 | Ikbkg | 16151 | HITS-CLIP | 25083871\|21258322 | Unspecified |
| mmu-miR-3106-5p | MIMAT0014817 | Itsn1 | 16443 | HITS-CLIP | 21258322 | Unspecified |
| mmu-miR-3106-5p | MIMAT0014817 | Klhdc9 | 68874 | HITS-CLIP | 21258322 | Unspecified |
| mmu-miR-3106-5p | MIMAT0014817 | Lrrc39 | 109245 | HITS-CLIP | 21258322 | Unspecified |
| mmu-miR-3106-5p | MIMAT0014817 | Ly6g6d | 114654 | HITS-CLIP | 21258322 | Unspecified |
| mmu-miR-3106-5p | MIMAT0014817 | Marchf5 | 69104 | HITS-CLIP | 21258322 | Unspecified |
| mmu-miR-3106-5p | MIMAT0014817 | Mrpl35 | 66223 | HITS-CLIP | 21258322 | Unspecified |

| | | | | | | |
|---|---|---|---|---|---|---|
| mmu-miR-3106-5p | MIMAT0014817 | Mylip | 218203 | HITS-CLIP | 23597149\|21258322 | Unspecified |
| mmu-miR-3106-5p | MIMAT0014817 | Neu3 | 50877 | HITS-CLIP | 21258322 | Unspecified |
| mmu-miR-3106-5p | MIMAT0014817 | Nsrp1 | 237859 | HITS-CLIP | 21258322 | Unspecified |
| mmu-miR-3106-5p | MIMAT0014817 | Padi2 | 18600 | HITS-CLIP | 21258322 | Unspecified |
| mmu-miR-3106-5p | MIMAT0014817 | Pigv | 230801 | HITS-CLIP | 21258322 | Unspecified |
| mmu-miR-3106-5p | MIMAT0014817 | Pnoc | 18155 | HITS-CLIP | 21258322 | Unspecified |
| mmu-miR-3106-5p | MIMAT0014817 | Pomk | 74653 | HITS-CLIP | 23142080 | Unspecified |
| mmu-miR-3106-5p | MIMAT0014817 | Psmb11 | 73902 | HITS-CLIP | 21258322 | Unspecified |
| mmu-miR-3106-5p | MIMAT0014817 | Saysd1 | 67509 | HITS-CLIP | 21258322 | Unspecified |
| mmu-miR-3106-5p | MIMAT0014817 | Sec23ip | 207352 | HITS-CLIP | 21258322 | Unspecified |
| mmu-miR-3106-5p | MIMAT0014817 | Sfxn4 | 94281 | HITS-CLIP | 21258322 | Unspecified |
| mmu-miR-3106-5p | MIMAT0014817 | Sgo2a | 68549 | HITS-CLIP | 21258322 | Unspecified |
| mmu-miR-3106-5p | MIMAT0014817 | Sgol2 | 68549 | HITS-CLIP | 21258322 | Unspecified |
| mmu-miR-3106-5p | MIMAT0014817 | Slc35d2 | 70484 | HITS-CLIP | 21258322 | Unspecified |
| mmu-miR-3106-5p | MIMAT0014817 | Smim17 | 100042450 | HITS-CLIP | 21258322 | Unspecified |
| mmu-miR-3106-5p | MIMAT0014817 | Snhg11 | 319317 | HITS-CLIP | 21258322 | Unspecified |
| mmu-miR-3106-5p | MIMAT0014817 | Sumf2 | 67902 | HITS-CLIP | 21258322 | Unspecified |
| mmu-miR-3106-5p | MIMAT0014817 | T2 | 21331 | HITS-CLIP | 21258322 | Unspecified |
| mmu-miR-3106-5p | MIMAT0014817 | Tmco3 | 234076 | HITS-CLIP | 25083871 | Unspecified |
| mmu-miR-3106-5p | MIMAT0014817 | Tmem206 | 66950 | HITS-CLIP | 21258322 | Unspecified |
| mmu-miR-3106-5p | MIMAT0014817 | Tmem69 | 230657 | HITS-CLIP | 21258322 | Unspecified |
| mmu-miR-3106-5p | MIMAT0014817 | Tmem88b | 320587 | HITS-CLIP | 21258322 | Unspecified |
| mmu-miR-3106-5p | MIMAT0014817 | Tnfsf10 | 22035 | HITS-CLIP | 21258322 | Unspecified |
| mmu-miR-3106-5p | MIMAT0014817 | Tpmt | 22017 | HITS-CLIP | 25083871 | Unspecified |
| mmu-miR-3106-5p | MIMAT0014817 | Trmt10a | 108943 | HITS-CLIP | 21258322 | Unspecified |
| mmu-miR-3106-5p | MIMAT0014817 | Tyr | 22173 | HITS-CLIP | 21258322 | Unspecified |
| mmu-miR-3106-5p | MIMAT0014817 | Vav1 | 22324 | HITS-CLIP | 21258322 | Unspecified |
| mmu-miR-3106-5p | MIMAT0014817 | Wdr85 | 67228 | HITS-CLIP | 25083871\|21258322 | Unspecified |
| mmu-miR-3106-5p | MIMAT0014817 | Xpo7 | 65246 | HITS-CLIP | 21258322 | Unspecified |
| mmu-miR-3106-5p | MIMAT0014817 | Zfp113 | 56314 | HITS-CLIP | 25083871 | Unspecified |
| mmu-miR-3106-5p | MIMAT0014817 | Zfp799 | 240064 | HITS-CLIP | 21258322 | Unspecified |
| mmu-miR-3106-5p | MIMAT0014817 | Zfp933 | 242747 | HITS-CLIP | 21258322 | Unspecified |
| mmu-miR-29b-3p | MIMAT0000127 | 4921536K21Rik | 67430 | HITS-CLIP | 21258322 | Unspecified |
| mmu-miR-29b-3p | MIMAT0000127 | Acvr2a | 11480 | Luciferase assay//Microarray//Northern bl | 19342382\|25083871\|23597149\|23142 | Unspecified |
| mmu-miR-29b-3p | MIMAT0000127 | Akt3 | 23797 | Luciferase assay//qRT-PCR//Western blot/ | 23764849\|25083871\|33184237\|23142 | Unspecified |
| mmu-miR-29b-3p | MIMAT0000127 | Bace1 | 23821 | Luciferase assay//qRT-PCR//Western blot/ | 22664922\|25083871\|23597149\|33184 | Unspecified |
| mmu-miR-29b-3p | MIMAT0000127 | Bak1 | 12018 | Luciferase assay//Microarray//qRT-PCR//V | 21245165\|25083871\|23597149\|33184 | Unspecified |
| mmu-miR-29b-3p | MIMAT0000127 | Bbc3 | 170770 | Luciferase assay//Microarray//qRT-PCR//V | 21245165\|23597149\|23142080\|25083 | Unspecified |
| mmu-miR-29b-3p | MIMAT0000127 | Bcl2l11 | 12125 | Luciferase assay//Microarray//qRT-PCR//V | 21245165\|25083871\|23142080 | Unspecified |

| | | | | | | | |
|---|---|---|---|---|---|---|---|
| mmu-miR-29b-3p | MIMAT0000127 | Bicd1 | 12121 | HITS-CLIP | 23597149\|19536157 | | Unspecified |
| mmu-miR-29b-3p | MIMAT0000127 | Bmf | 171543 | Luciferase assay//Microarray//qRT-PCR//V | 21245165\|25083871\|23597149\|23142 | | Unspecified |
| mmu-miR-29b-3p | MIMAT0000127 | Cav2 | 12390 | reporter_gene//HITS-CLIP//Reporter assay | 37986224\|25083871\|17652184 | | Unspecified |
| mmu-miR-29b-3p | MIMAT0000127 | Cisd2 | 67006 | HITS-CLIP | 21258322\|25083871\|23597149\|23142 | | Unspecified |
| mmu-miR-29b-3p | MIMAT0000127 | Col1a1 | 12842 | Luciferase assay//Microarray//Northern bl | 19342382\|23597149\|33184237\|25083 | | Unspecified |
| mmu-miR-29b-3p | MIMAT0000127 | Col1a2 | 12843 | Luciferase assay//Microarray//Northern bl | 18723672\|25083871\|23597149\|33184 | | Unspecified |
| mmu-miR-29b-3p | MIMAT0000127 | Col2a1 | 12824 | Luciferase assay//Luciferase reporter assay | | 21665270 | Unspecified |
| mmu-miR-29b-3p | MIMAT0000127 | Col3a1 | 12825 | Luciferase assay//Microarray//Northern bl | 18723672\|25083871\|33184237\|23597 | | Unspecified |
| mmu-miR-29b-3p | MIMAT0000127 | Col4a1 | 12826 | Luciferase assay//qRT-PCR//Western blot/ | 23139829\|23597149\|25083871\|33184 | | Unspecified |
| mmu-miR-29b-3p | MIMAT0000127 | Col4a2 | 12827 | Luciferase assay//qRT-PCR//Western blot/ | 23139829\|25083871\|23597149\|33184 | | Unspecified |
| mmu-miR-29b-3p | MIMAT0000127 | Col4a3 | 12828 | Luciferase assay//qRT-PCR//Western blot/ | | 23139829 | Unspecified |
| mmu-miR-29b-3p | MIMAT0000127 | Col4a4 | 12829 | Luciferase assay//qRT-PCR//Western blot/ | | 23139829 | Unspecified |
| mmu-miR-29b-3p | MIMAT0000127 | Col4a5 | 12830 | Luciferase assay//qRT-PCR//Western blot/ | 23139829\|25083871\|33184237 | | Unspecified |
| mmu-miR-29b-3p | MIMAT0000127 | Col4a6 | 94216 | Luciferase assay//qRT-PCR//Western blot/ | | 23139829 | Unspecified |
| mmu-miR-29b-3p | MIMAT0000127 | Col5a3 | 53867 | Luciferase assay//Microarray//Northern bl | 19342382\|25083871\|33184237 | | Unspecified |
| mmu-miR-29b-3p | MIMAT0000127 | Ctnnbip1 | 67087 | Luciferase assay//Microarray//Northern bl | 19342382\|25083871 | | Unspecified |
| mmu-miR-29b-3p | MIMAT0000127 | Dnmt3a | 13435 | Luciferase assay//mRAP//Northern blot//H | 19509302\|23597149\|23142080\|25083 | | Unspecified |
| mmu-miR-29b-3p | MIMAT0000127 | Dnmt3b | 13436 | qRT-PCR//Western blot//Luciferase report | 19959559\|19509302 | | Unspecified |
| mmu-miR-29b-3p | MIMAT0000127 | Dusp2 | 13537 | Luciferase assay//Microarray//Northern bl | 19342382\|25083871\|31393953\|23142 | | Unspecified |
| mmu-miR-29b-3p | MIMAT0000127 | Eln | 13717 | Luciferase assay//Microarray//Northern bl | | 18723672 | Unspecified |
| mmu-miR-29b-3p | MIMAT0000127 | Eomes | 13813 | Luciferase assay//Microarray//qRT-PCR//H | 21820330\|23142080 | | Unspecified |
| mmu-miR-29b-3p | MIMAT0000127 | Eps15 | 13858 | HITS-CLIP//sRNA-Seq | 21258322\|25083871\|33184237\|23142 | | Unspecified |
| mmu-miR-29b-3p | MIMAT0000127 | Fam126b | 213056 | HITS-CLIP | 21258322\|25083871\|23142080 | | Unspecified |
| mmu-miR-29b-3p | MIMAT0000127 | Fbn1 | 14118 | Luciferase assay//Microarray//Northern bl | 18723672\|25083871\|33184237 | | Unspecified |
| mmu-miR-29b-3p | MIMAT0000127 | Gm14137 | 623781 | HITS-CLIP | | 21258322 | Unspecified |
| mmu-miR-29b-3p | MIMAT0000127 | Hdac4 | 208727 | Luciferase assay//qRT-PCR//Western blot/ | 21324893\|25083871\|23142080\|19342 | | Unspecified |
| mmu-miR-29b-3p | MIMAT0000127 | Hif3a | 53417 | HITS-CLIP | 21258322\|23142080\|25083871\|23597 | | Unspecified |
| mmu-miR-29b-3p | MIMAT0000127 | Hrk | 12123 | Luciferase assay//Microarray//qRT-PCR//V | | 21245165 | Unspecified |
| mmu-miR-29b-3p | MIMAT0000127 | Igf1 | 16000 | ELISA//Luciferase assay//qRT-PCR//HITS-C | 23880139\|25083871\|23597149 | | Unspecified |
| mmu-miR-29b-3p | MIMAT0000127 | Insig1 | 231070 | reporter_gene//sRNA-Seq//HITS-CLIP//Re | 37986224\|33184237\|23597149\|17652 | | Unspecified |
| mmu-miR-29b-3p | MIMAT0000127 | Lactb2 | 212442 | HITS-CLIP | 21258322\|23597149 | | Unspecified |
| mmu-miR-29b-3p | MIMAT0000127 | Mcl1 | 17210 | HITS-CLIP//sRNA-Seq | 23597149\|25083871\|33184237\|23142 | | Unspecified |
| mmu-miR-29b-3p | MIMAT0000127 | Mrpl35 | 66223 | HITS-CLIP | | 21258322 | Unspecified |
| mmu-miR-29b-3p | MIMAT0000127 | Pigt | 78928 | HITS-CLIP//sRNA-Seq | 21258322\|33184237\|23142080 | | Unspecified |
| mmu-miR-29b-3p | MIMAT0000127 | Prr5l | 72446 | HITS-CLIP | | 21258322 | Unspecified |
| mmu-miR-29b-3p | MIMAT0000127 | RARB | 218772 | Immunoprecipitaion (IP)//Luciferase repor | | 31420607 | Unspecified |
| mmu-miR-29b-3p | MIMAT0000127 | Rsad1 | 237926 | HITS-CLIP | | 21258322 | Unspecified |
| mmu-miR-29b-3p | MIMAT0000127 | Scap | 235623 | sRNA-Seq//HITS-CLIP//Luciferase reporter | 33184237\|23597149\|23142080\|27477 | | Unspecified |
| mmu-miR-29b-3p | MIMAT0000127 | Sirt1 | 93759 | Luciferase assay//qRT-PCR//Western blot/ | 24657470\|25083871\|23142080 | | Unspecified |

| | | | | | | |
|---|---|---|---|---|---|---|
| mmu-miR-29b-3p | MIMAT0000127 | Sox2 | 20674 | Luciferase assay//Luciferase reporter assay | 23266889 | Unspecified |
| mmu-miR-29b-3p | MIMAT0000127 | Sparc | 20692 | HITS-CLIP | 23142080 | Unspecified |
| mmu-miR-29b-3p | MIMAT0000127 | Tbx21 | 57765 | Luciferase assay//Microarray//qRT-PCR//H | 21820330\|23142080 | Unspecified |
| mmu-miR-29b-3p | MIMAT0000127 | Tgfb3 | 21809 | Luciferase assay//Microarray//Northern bl | 19342382\|25083871\|33184237 | Unspecified |
| mmu-miR-29b-3p | MIMAT0000127 | Tnfaip3 | 21929 | Immunoblot//Immunocytochemistry//Luci | 24236890\|23142080\|25083871 | Unspecified |
| mmu-miR-29b-3p | MIMAT0000127 | Vegfa | 22339 | HITS-CLIP//ELISA//Luciferase reporter assa | 37986224\|25083871\|23142080\|27589 | Unspecified |
| mmu-miR-29b-3p | MIMAT0000127 | Yy1 | 22632 | HITS-CLIP//Other | 37986224\|23142080\|25083871\|18977 | Unspecified |
| mmu-miR-29b-3p | MIMAT0000127 | Zbtb5 | 230119 | HITS-CLIP | 23142080\|25083871\|23597149\|28735 | Unspecified |
| mmu-miR-29b-3p | MIMAT0000127 | Zfp26 | 22688 | HITS-CLIP | 25083871\|23142080 | Unspecified |
| mmu-miR-5120 | MIMAT0020628 | 1810041L15Rik | 72301 | HITS-CLIP | 23142080 | Unspecified |
| mmu-miR-5120 | MIMAT0020628 | Abca5 | 217265 | HITS-CLIP | 25083871 | Unspecified |
| mmu-miR-5120 | MIMAT0020628 | Cpox | 12892 | HITS-CLIP | 23142080 | Unspecified |
| mmu-miR-5120 | MIMAT0020628 | Ddx27 | 228889 | HITS-CLIP | 21258322 | Unspecified |
| mmu-miR-5120 | MIMAT0020628 | Epas1 | 13819 | HITS-CLIP | 25083871 | Unspecified |
| mmu-miR-5120 | MIMAT0020628 | Fam172a | 68675 | HITS-CLIP | 23142080 | Unspecified |
| mmu-miR-5120 | MIMAT0020628 | Gpr180 | 58245 | HITS-CLIP | 21258322\|23597149 | Unspecified |
| mmu-miR-5120 | MIMAT0020628 | Itga9 | 104099 | HITS-CLIP | 23142080 | Unspecified |
| mmu-miR-5120 | MIMAT0020628 | Ncan | 13004 | HITS-CLIP | 21258322 | Unspecified |
| mmu-miR-5120 | MIMAT0020628 | Ncl | 17975 | HITS-CLIP | 21258322 | Unspecified |
| mmu-miR-5120 | MIMAT0020628 | Nsg2 | 18197 | HITS-CLIP | 25083871 | Unspecified |
| mmu-miR-5120 | MIMAT0020628 | Pm20d2 | 242377 | HITS-CLIP | 25083871\|23142080 | Unspecified |
| mmu-miR-5120 | MIMAT0020628 | Prpf31 | 68988 | HITS-CLIP | 21258322 | Unspecified |
| mmu-miR-5120 | MIMAT0020628 | Zic4 | 22774 | HITS-CLIP | 23142080 | Unspecified |
| mmu-miR-21a-5p | MIMAT0000530 | AK010878 | 100233175 | HITS-CLIP | 21258322 | Unspecified |
| mmu-miR-21a-5p | MIMAT0000530 | Btg2 | 12227 | HITS-CLIP, qPCR//HITS-CLIP//qCLASH//Oth | 37986224\|23597149\|31393953\|29167 | Unspecified |
| mmu-miR-21a-5p | MIMAT0000530 | Cdk6 | 12571 | Microarrays//HITS-CLIP//qCLASH//Lucifera | 37986224\|23142080\|25263593\|23597 | Unspecified |
| mmu-miR-21a-5p | MIMAT0000530 | Cyfip1 | 20430 | HITS-CLIP | 25083871\|23597149\|29167373 | Unspecified |
| mmu-miR-21a-5p | MIMAT0000530 | E2f2 | 242705 | HITS-CLIP//Microarrays | 23142080\|25263593\|23597149 | Unspecified |
| mmu-miR-21a-5p | MIMAT0000530 | Eif4e3 | 66892 | Luciferase assay//qRT-PCR//Luciferase rep | 24891504 | Unspecified |
| mmu-miR-21a-5p | MIMAT0000530 | Elavl4 | 15572 | Microarrays//Other | 37986224\|20167875 | Unspecified |
| mmu-miR-21a-5p | MIMAT0000530 | Fasl | 14103 | Luciferase assay//Western blot//Northern | 20404348\|23142080 | Unspecified |
| mmu-miR-21a-5p | MIMAT0000530 | Gid4 | 66771 | HITS-CLIP | 23142080\|23597149 | Unspecified |
| mmu-miR-21a-5p | MIMAT0000530 | Gon7 | 100233175 | HITS-CLIP | 21258322 | Unspecified |
| mmu-miR-21a-5p | MIMAT0000530 | Gt(ROSA)26Sor | 14910 | Immunohistochemistry//Luciferase assay/ | 23238710 | Unspecified |
| mmu-miR-21a-5p | MIMAT0000530 | Il12a | 16159 | qPCR//RNA-seq | 37986224\|31710308 | Unspecified |
| mmu-miR-21a-5p | MIMAT0000530 | Kcnk6 | 52150 | HITS-CLIP | 25083871 | Unspecified |
| mmu-miR-21a-5p | MIMAT0000530 | MAP2K3 | 5606 | HITS-CLIP//Luciferase reporter assay//Flov | 23597149\|23142080\|26996129\|31585 | Unspecified |
| mmu-miR-21a-5p | MIMAT0000530 | Map3k1 | 26401 | HITS-CLIP | 25083871\|23597149\|19536157\|23142 | Unspecified |
| mmu-miR-21a-5p | MIMAT0000530 | Mmp9 | 17395 | Immunohistochemistry//In situ hybridizati | 23443810\|25263593 | Unspecified |

| | | | | | | |
|---|---|---|---|---|---|---|
| mmu-miR-21a-5p | MIMAT0000530 | Moap1 | 64113 | HITS-CLIP | 21258322 | Unspecified |
| mmu-miR-21a-5p | MIMAT0000530 | Pdcd10 | 56426 | Luciferase assay//qRT-PCR//HITS-CLIP//Lu | 24891504\|23142080 | Unspecified |
| mmu-miR-21a-5p | MIMAT0000530 | Peli1 | 67245 | Luciferase assay//Reporter assay//qRT-PCI | 20167875\|23597149\|23142080\|29167 | Unspecified |
| mmu-miR-21a-5p | MIMAT0000530 | Pias3 | 229615 | Western blot | 22517757 | Unspecified |
| mmu-miR-21a-5p | MIMAT0000530 | Pten | 19211 | Reporter assay//Western Blot//Luciferase | 20167875\|20404348\|18508928\|26744 | Unspecified |
| mmu-miR-21a-5p | MIMAT0000530 | Rc3h1 | 381305 | HITS-CLIP//In situ hybridization//Luciferase | 29167373\|25697406 | Unspecified |
| mmu-miR-21a-5p | MIMAT0000530 | Reck | 53614 | Luciferase assay//HITS-CLIP//Microarrays/ | 18556655\|23597149\|25263593\|31393 | Unspecified |
| mmu-miR-21a-5p | MIMAT0000530 | Rmnd5a | 68477 | HITS-CLIP | 23597149\|29167373 | Unspecified |
| mmu-miR-21a-5p | MIMAT0000530 | Rpp40 | 208366 | HITS-CLIP | 21258322 | Unspecified |
| mmu-miR-21a-5p | MIMAT0000530 | SORBS2 | 8470 | HITS-CLIP//Western blot | 19536157\|23597149\|27033308 | Unspecified |
| mmu-miR-21a-5p | MIMAT0000530 | Smad7 | 17131 | Luciferase assay//Northern blot//QRTPCR/ | 23784029\|23597149\|25263593\|23142 | Unspecified |
| mmu-miR-21a-5p | MIMAT0000530 | Sod2 | 20656 | HITS-CLIP//ChIP-seq//qRT-PCR//Western b | 23597149\|23142080\|26222667 | Unspecified |
| mmu-miR-21a-5p | MIMAT0000530 | Sox2 | 20674 | qRT-PCR | 26209818 | Unspecified |
| mmu-miR-21a-5p | MIMAT0000530 | Sox5 | 20678 | HITS-CLIP//Luciferase reporter assay//qRT- | 37986224\|23597149\|23142080\|27347 | Unspecified |
| mmu-miR-21a-5p | MIMAT0000530 | Spry1 | 24063 | Luciferase assay//Western blot//qPCR//Mi | 20216554\|23416424\|23142080\|18508 | Unspecified |
| mmu-miR-21a-5p | MIMAT0000530 | Spry2 | 24064 | Luciferase assay//Western blot//Reporter | 18508928 | Unspecified |
| mmu-miR-21a-5p | MIMAT0000530 | Spry3 | 236576 | Western blot//Other | 19147652\|20404348\|18508928 | Unspecified |
| mmu-miR-21a-5p | MIMAT0000530 | Spry4 | 24066 | qRT-PCR//HITS-CLIP//Other | 19043405\|23597149\|18508928 | Unspecified |
| mmu-miR-21a-5p | MIMAT0000530 | Tgfbi | 21810 | ELISA//FACS//Luciferase assay//qRT-PCR// | 21494432\|23597149\|29167373 | Unspecified |
| mmu-miR-21a-5p | MIMAT0000530 | Tgfbr3 | 21814 | Luciferase assay//qRT-PCR//Western blot/ | 22960625\|23142080 | Unspecified |
| mmu-miR-21a-5p | MIMAT0000530 | Timp3 | 21859 | qRT-PCR//HITS-CLIP | 23797704\|19536157 | Unspecified |
| mmu-miR-21a-5p | MIMAT0000530 | Tnfaip8l2 | 69769 | ChIP-seq//Flow//Luciferase assay//qRT-PC | 24577093\|23142080 | Unspecified |
| mmu-miR-21a-5p | MIMAT0000530 | Tns1 | 21961 | HITS-CLIP | 23142080\|23597149 | Unspecified |
| mmu-miR-21a-5p | MIMAT0000530 | Yy1 | 22632 | Luciferase assay//qRT-PCR//HITS-CLIP//Lu | 24891504\|23597149\|23142080 | Unspecified |
| mmu-miR-3069-3p | MIMAT0014845 | Cd69 | 12515 | HITS-CLIP | 23142080 | Unspecified |
| mmu-miR-3069-3p | MIMAT0014845 | Glrx2 | 69367 | HITS-CLIP | 21258322 | Unspecified |
| mmu-miR-3069-3p | MIMAT0014845 | Gpx5 | 14780 | HITS-CLIP | 26203562 | Unspecified |
| mmu-miR-3069-3p | MIMAT0014845 | Hecw1 | 94253 | HITS-CLIP | 23142080 | Unspecified |
| mmu-miR-3069-3p | MIMAT0014845 | Hiat1 | 15247 | HITS-CLIP | 25083871 | Unspecified |
| mmu-miR-3069-3p | MIMAT0014845 | Mfsd14a | 15247 | HITS-CLIP | 25083871 | Unspecified |
| mmu-miR-3069-3p | MIMAT0014845 | Nfatc4 | 73181 | HITS-CLIP | 23597149\|25083871\|21258322 | Unspecified |
| mmu-miR-3069-3p | MIMAT0014845 | Nudt18 | 213484 | HITS-CLIP | 23597149 | Unspecified |
| mmu-miR-3069-3p | MIMAT0014845 | Spn | 20737 | HITS-CLIP | 23142080 | Unspecified |
| mmu-miR-3069-3p | MIMAT0014845 | Tgfbr3 | 21814 | HITS-CLIP | 23142080 | Unspecified |
| mmu-miR-3069-3p | MIMAT0014845 | Tnfrsf1b | 21938 | HITS-CLIP | 23597149\|25083871\|21258322 | Unspecified |
| mmu-miR-3069-5p | MIMAT0014844 | 9930013L23Rik | 80982 | HITS-CLIP | 23142080 | Unspecified |
| mmu-miR-3069-5p | MIMAT0014844 | Cemip | 80982 | HITS-CLIP | 23142080 | Unspecified |
| mmu-miR-3069-5p | MIMAT0014844 | Chmp5 | 76959 | HITS-CLIP | 25083871 | Unspecified |
| mmu-miR-3069-5p | MIMAT0014844 | Dcun1d3 | 233805 | HITS-CLIP | 21258322 | Unspecified |

| | | | | | | |
|---|---|---|---|---|---|---|
| mmu-miR-3069-5p | MIMAT0014844 | Dusp11 | 72102 | HITS-CLIP | 21258322 | Unspecified |
| mmu-miR-3069-5p | MIMAT0014844 | Fyco1 | 17281 | HITS-CLIP | 23142080 | Unspecified |
| mmu-miR-3069-5p | MIMAT0014844 | Med22 | 20933 | HITS-CLIP | 23142080 | Unspecified |
| mmu-miR-3069-5p | MIMAT0014844 | Mrm3 | 67390 | HITS-CLIP | 25083871 | Unspecified |
| mmu-miR-3069-5p | MIMAT0014844 | Peg10 | 170676 | HITS-CLIP | 23142080 | Unspecified |
| mmu-miR-3069-5p | MIMAT0014844 | Phc3 | 241915 | HITS-CLIP | 21258322 | Unspecified |
| mmu-miR-3069-5p | MIMAT0014844 | Prex2 | 109294 | HITS-CLIP | 25083871\|19536157 | Unspecified |
| mmu-miR-3069-5p | MIMAT0014844 | Rab11b | 19326 | HITS-CLIP | 21258322 | Unspecified |
| mmu-miR-3069-5p | MIMAT0014844 | Rnmtl1 | 67390 | HITS-CLIP | 25083871 | Unspecified |
| mmu-miR-3069-5p | MIMAT0014844 | Stxbp5 | 78808 | HITS-CLIP | 25083871 | Unspecified |
| mmu-miR-3069-5p | MIMAT0014844 | Ubp1 | 22221 | HITS-CLIP | 23142080 | Unspecified |
| mmu-miR-34a-5p | MIMAT0000542 | 9930013L23Rik | 80982 | HITS-CLIP | 23142080 | Unspecified |
| mmu-miR-34a-5p | MIMAT0000542 | Actb | 11461 | Luciferase assay//HITS-CLIP//Luciferase re | 18835850\|19536157\|23142080 | Unspecified |
| mmu-miR-34a-5p | MIMAT0000542 | Ankrd28 | 105522 | HITS-CLIP | 21258322\|23142080\|29167373 | Unspecified |
| mmu-miR-34a-5p | MIMAT0000542 | Atg9a | 245860 | HITS-CLIP//Immunoblot//Luciferase report | 23142080\|19536157\|26203562\|24973 | Unspecified |
| mmu-miR-34a-5p | MIMAT0000542 | B3galtl | 381694 | HITS-CLIP | 21258322 | Unspecified |
| mmu-miR-34a-5p | MIMAT0000542 | B3glct | 381694 | HITS-CLIP | 21258322 | Unspecified |
| mmu-miR-34a-5p | MIMAT0000542 | Bcl2 | 12043 | qRT-PCR//Western blot//HITS-CLIP//Other | 19683563\|23142080\|31433990\|29167 | Unspecified |
| mmu-miR-34a-5p | MIMAT0000542 | Bcl6 | 12053 | Luciferase assay//Northern blot//qRT-PCR | 23047694\|23142080\|29167373 | Unspecified |
| mmu-miR-34a-5p | MIMAT0000542 | Bloc1s3 | 232946 | HITS-CLIP | 21258322\|23142080 | Unspecified |
| mmu-miR-34a-5p | MIMAT0000542 | Brca1 | 12189 | HITS-CLIP//Next Generation Sequencing (N | 23142080\|26403328 | Unspecified |
| mmu-miR-34a-5p | MIMAT0000542 | Ccnd1 | 12443 | Northern blot//qRT-PCR//Western blot//N | 24587330\|23027903\|29167373 | Unspecified |
| mmu-miR-34a-5p | MIMAT0000542 | Cd93 | 17064 | HITS-CLIP | 21258322\|29167373 | Unspecified |
| mmu-miR-34a-5p | MIMAT0000542 | Cemip | 80982 | HITS-CLIP | 23142080 | Unspecified |
| mmu-miR-34a-5p | MIMAT0000542 | Cenpl | 70454 | HITS-CLIP//Microarrays | 21258322\|23027903\|23142080 | Unspecified |
| mmu-miR-34a-5p | MIMAT0000542 | Cercam | 99151 | HITS-CLIP | 21258322 | Unspecified |
| mmu-miR-34a-5p | MIMAT0000542 | Cnnm3 | 94218 | HITS-CLIP | 21258322\|26203562\|29167373 | Unspecified |
| mmu-miR-34a-5p | MIMAT0000542 | Csf1r | 12978 | HITS-CLIP//qRT-PCR | 29167373\|32612530 | Unspecified |
| mmu-miR-34a-5p | MIMAT0000542 | Ctsa | 19025 | HITS-CLIP | 21258322\|23142080\|29167373 | Unspecified |
| mmu-miR-34a-5p | MIMAT0000542 | Dll1 | 13388 | Luciferase assay//qRT-PCR//Western blot/ | 21931765 | Unspecified |
| mmu-miR-34a-5p | MIMAT0000542 | Dnase2a | 13423 | HITS-CLIP | 25083871 | Unspecified |
| mmu-miR-34a-5p | MIMAT0000542 | Dusp11 | 72102 | HITS-CLIP | 21258322\|29167373\|23142080 | Unspecified |
| mmu-miR-34a-5p | MIMAT0000542 | Epb4.2 | 13828 | HITS-CLIP | 21258322 | Unspecified |
| mmu-miR-34a-5p | MIMAT0000542 | Epb42 | 13828 | HITS-CLIP | 21258322 | Unspecified |
| mmu-miR-34a-5p | MIMAT0000542 | Fam118a | 73225 | HITS-CLIP | 21258322\|23142080 | Unspecified |
| mmu-miR-34a-5p | MIMAT0000542 | Fndc5 | 252995 | Microarrays//Luciferase reporter assay//q | 37986224\|27297797 | Unspecified |
| mmu-miR-34a-5p | MIMAT0000542 | Foxp1 | 108655 | Western blot//Western Blot//HITS-CLIP | 37986224\|20598588\|19536157\|23142 | Unspecified |
| mmu-miR-34a-5p | MIMAT0000542 | Foxp3 | 20371 | qRT-PCR | 32612530 | Unspecified |
| mmu-miR-34a-5p | MIMAT0000542 | Fyco1 | 17281 | HITS-CLIP | 23142080\|29167373 | Unspecified |

| | | | | | | |
|---|---|---|---|---|---|---|
| mmu-miR-34a-5p | MIMAT0000542 | Gas1 | 14451 | Immunohistochemistry//Luciferase assay/, | 24560136 | Unspecified |
| mmu-miR-34a-5p | MIMAT0000542 | MSN | 17698 | HITS-CLIP//qRT-PCR//Western blotting | 31433990\|23142080\|29167373\|32319 | Unspecified |
| mmu-miR-34a-5p | MIMAT0000542 | Mapre1 | 13589 | HITS-CLIP | 21258322\|31433990\|23142080\|29167 | Unspecified |
| mmu-miR-34a-5p | MIMAT0000542 | Mbd4 | 17193 | HITS-CLIP | 21258322\|23142080 | Unspecified |
| mmu-miR-34a-5p | MIMAT0000542 | Med22 | 20933 | HITS-CLIP//Microarrays | 23142080\|23027903 | Unspecified |
| mmu-miR-34a-5p | MIMAT0000542 | Mmp12 | 17381 | HITS-CLIP//Microarray//qRT-PCR | 29167373\|32855961 | Unspecified |
| mmu-miR-34a-5p | MIMAT0000542 | Mrm3 | 67390 | HITS-CLIP | 25083871 | Unspecified |
| mmu-miR-34a-5p | MIMAT0000542 | Nos1 | 18125 | qRT-PCR | 32612530 | Unspecified |
| mmu-miR-34a-5p | MIMAT0000542 | Notch1 | 18128 | ELISA//Immunofluorescence//Luciferase a | 22483937\|23142080\|29167373\|24587 | Unspecified |
| mmu-miR-34a-5p | MIMAT0000542 | Phc3 | 241915 | HITS-CLIP | 21258322\|23142080\|31433990\|29167 | Unspecified |
| mmu-miR-34a-5p | MIMAT0000542 | Pin1 | 23988 | HITS-CLIP | 21258322\|23142080 | Unspecified |
| mmu-miR-34a-5p | MIMAT0000542 | Pofut1 | 140484 | Luciferase assay//Northern blot//qRT-PCR, | 23047694\|23027903\|23142080\|24587 | Unspecified |
| mmu-miR-34a-5p | MIMAT0000542 | Ppp1r10 | 52040 | Northern blot//qRT-PCR//Western blot//H | 24587330\|23142080 | Unspecified |
| mmu-miR-34a-5p | MIMAT0000542 | Prex2 | 109294 | HITS-CLIP | 25083871\|19536157 | Unspecified |
| mmu-miR-34a-5p | MIMAT0000542 | Rab11b | 19326 | HITS-CLIP | 21258322\|23142080 | Unspecified |
| mmu-miR-34a-5p | MIMAT0000542 | Rgs17 | 56533 | HITS-CLIP | 23142080 | Unspecified |
| mmu-miR-34a-5p | MIMAT0000542 | Rheb | 19744 | HITS-CLIP | 21258322\|23142080\|29167373 | Unspecified |
| mmu-miR-34a-5p | MIMAT0000542 | Rnmtl1 | 67390 | HITS-CLIP | 25083871 | Unspecified |
| mmu-miR-34a-5p | MIMAT0000542 | Sco1 | 52892 | HITS-CLIP | 21258322\|23142080 | Unspecified |
| mmu-miR-34a-5p | MIMAT0000542 | Sema4b | 20352 | HITS-CLIP//Microarrays//Luciferase report | 21258322\|23027903\|23142080\|29167 | Unspecified |
| mmu-miR-34a-5p | MIMAT0000542 | Sgk3 | 170755 | HITS-CLIP | 21258322\|29167373 | Unspecified |
| mmu-miR-34a-5p | MIMAT0000542 | Shisa7 | 232813 | HITS-CLIP | 23142080 | Unspecified |
| mmu-miR-34a-5p | MIMAT0000542 | Slc22a12 | 20521 | qRT-PCR//Luciferase reporter assay | 25535087 | Unspecified |
| mmu-miR-34a-5p | MIMAT0000542 | Slc35e2 | 320541 | HITS-CLIP | 21258322\|23630078\|30971389\|23142 | Unspecified |
| mmu-miR-34a-5p | MIMAT0000542 | Smco1 | 69576 | HITS-CLIP | 21258322\|25083871 | Unspecified |
| mmu-miR-34a-5p | MIMAT0000542 | St6galnac3 | 20447 | HITS-CLIP | 23142080\|19536157 | Unspecified |
| mmu-miR-34a-5p | MIMAT0000542 | Syt1 | 20979 | HITS-CLIP//Immunoblot//Luciferase report | 19536157\|24973144 | Unspecified |
| mmu-miR-34a-5p | MIMAT0000542 | TGIF2 | 60436 | Luciferase assay//qRT-PCR////Luciferase re | 25043055 | Unspecified |
| mmu-miR-34a-5p | MIMAT0000542 | Tbc1d2 | 381605 | HITS-CLIP | 23142080 | Unspecified |
| mmu-miR-34a-5p | MIMAT0000542 | Trem2 | 83433 | Western blot | 23778113 | Unspecified |
| mmu-miR-34a-5p | MIMAT0000542 | Trim36 | 28105 | HITS-CLIP | 21258322\|23142080 | Unspecified |
| mmu-miR-34a-5p | MIMAT0000542 | Trmt10a | 108943 | HITS-CLIP | 21258322\|23142080 | Unspecified |
| mmu-miR-34a-5p | MIMAT0000542 | Ubp1 | 22221 | HITS-CLIP | 23142080 | Unspecified |
| mmu-miR-34a-5p | MIMAT0000542 | Vcl | 22330 | Luciferase assay//Northern blot//qRT-PCR, | 23047694\|23630078\|23142080\|29167 | Unspecified |
| mmu-miR-34a-5p | MIMAT0000542 | Vegfb | 22340 | Northern blot//qRT-PCR//Western blot//H | 24587330\|23142080 | Unspecified |
| mmu-miR-34a-5p | MIMAT0000542 | Vps9d1 | 72325 | HITS-CLIP | 23142080 | Unspecified |
| mmu-miR-34a-5p | MIMAT0000542 | Wasf1 | 83767 | HITS-CLIP//Luciferase reporter assay//qRT- | 19536157\|28363996 | Unspecified |
| mmu-miR-34a-5p | MIMAT0000542 | Wipf2 | 68524 | HITS-CLIP | 21258322\|23142080 | Unspecified |
| mmu-miR-34a-5p | MIMAT0000542 | Ywhag | 22628 | HITS-CLIP | 23142080\|19536157\|29167373 | Unspecified |

| | | | | | | |
|---|---|---|---|---|---|---|
| mmu-miR-34a-5p | MIMAT0000542 | Zfp317 | 244713 | HITS-CLIP | 21258322\|23142080 | Unspecified |
| mmu-miR-34a-5p | MIMAT0000542 | Zfp446 | 269870 | HITS-CLIP | 21258322 | Unspecified |
| mmu-miR-509-5p | MIMAT0004890 | A230046K03Rik | 319277 | HITS-CLIP | 21258322 | Unspecified |
| mmu-miR-509-5p | MIMAT0004890 | Abcb7 | 11306 | HITS-CLIP | 21258322 | Unspecified |
| mmu-miR-509-5p | MIMAT0004890 | Arpc4 | 68089 | HITS-CLIP | 23597149 | Unspecified |
| mmu-miR-509-5p | MIMAT0004890 | BC016495 | 225994 | CLIP-Seq dataset analysis | 21258322 | Unspecified |
| mmu-miR-509-5p | MIMAT0004890 | Bbs1 | 52028 | HITS-CLIP | 21258322 | Unspecified |
| mmu-miR-509-5p | MIMAT0004890 | Bcas2 | 68183 | HITS-CLIP | 21258322 | Unspecified |
| mmu-miR-509-5p | MIMAT0004890 | Bri3bp | 76809 | HITS-CLIP | 21258322 | Unspecified |
| mmu-miR-509-5p | MIMAT0004890 | Brip1 | 237911 | HITS-CLIP | 21258322 | Unspecified |
| mmu-miR-509-5p | MIMAT0004890 | Cflar | 12633 | HITS-CLIP | 21258322 | Unspecified |
| mmu-miR-509-5p | MIMAT0004890 | Chrnb4 | 108015 | HITS-CLIP | 21258322 | Unspecified |
| mmu-miR-509-5p | MIMAT0004890 | Churc1 | 211151 | HITS-CLIP | 21258322 | Unspecified |
| mmu-miR-509-5p | MIMAT0004890 | Cxcr2 | 12765 | HITS-CLIP | 21258322 | Unspecified |
| mmu-miR-509-5p | MIMAT0004890 | Cyp2e1 | 13106 | HITS-CLIP | 25083871 | Unspecified |
| mmu-miR-509-5p | MIMAT0004890 | Dusp26 | 66959 | HITS-CLIP | 25083871 | Unspecified |
| mmu-miR-509-5p | MIMAT0004890 | Exosc2 | 227715 | HITS-CLIP | 21258322 | Unspecified |
| mmu-miR-509-5p | MIMAT0004890 | H2-T23 | 15040 | HITS-CLIP | 21258322 | Unspecified |
| mmu-miR-509-5p | MIMAT0004890 | Hist2h3c1 | 15077 | HITS-CLIP | 21258322 | Unspecified |
| mmu-miR-509-5p | MIMAT0004890 | Irak1 | 16179 | HITS-CLIP | 21258322 | Unspecified |
| mmu-miR-509-5p | MIMAT0004890 | Klhl7 | 52323 | HITS-CLIP | 21258322 | Unspecified |
| mmu-miR-509-5p | MIMAT0004890 | Lcmt2 | 329504 | HITS-CLIP | 21258322 | Unspecified |
| mmu-miR-509-5p | MIMAT0004890 | Man1b1 | 227619 | HITS-CLIP | 23597149 | Unspecified |
| mmu-miR-509-5p | MIMAT0004890 | Mpv17 | 17527 | HITS-CLIP | 21258322 | Unspecified |
| mmu-miR-509-5p | MIMAT0004890 | Mrpl11 | 66419 | HITS-CLIP | 21258322 | Unspecified |
| mmu-miR-509-5p | MIMAT0004890 | Mrpl9 | 78523 | HITS-CLIP | 21258322 | Unspecified |
| mmu-miR-509-5p | MIMAT0004890 | Mylk2 | 228785 | HITS-CLIP | 21258322 | Unspecified |
| mmu-miR-509-5p | MIMAT0004890 | Nip7 | 66164 | HITS-CLIP | 21258322 | Unspecified |
| mmu-miR-509-5p | MIMAT0004890 | Ogdh | 18293 | HITS-CLIP | 25083871\|21258322 | Unspecified |
| mmu-miR-509-5p | MIMAT0004890 | Pdzd8 | 107368 | HITS-CLIP | 21258322 | Unspecified |
| mmu-miR-509-5p | MIMAT0004890 | Plxnc1 | 54712 | HITS-CLIP | 25083871 | Unspecified |
| mmu-miR-509-5p | MIMAT0004890 | Pramef8 | 242736 | HITS-CLIP | 21258322 | Unspecified |
| mmu-miR-509-5p | MIMAT0004890 | Rab33b | 19338 | HITS-CLIP | 21258322 | Unspecified |
| mmu-miR-509-5p | MIMAT0004890 | Rs1 | 20147 | HITS-CLIP | 21258322 | Unspecified |
| mmu-miR-509-5p | MIMAT0004890 | S100pbp | 74648 | HITS-CLIP | 28735896 | Unspecified |
| mmu-miR-509-5p | MIMAT0004890 | Sap18 | 20220 | HITS-CLIP | 21258322 | Unspecified |
| mmu-miR-509-5p | MIMAT0004890 | Slc25a34 | 384071 | HITS-CLIP | 21258322 | Unspecified |
| mmu-miR-509-5p | MIMAT0004890 | Slc7a2 | 11988 | HITS-CLIP | 23142080\|23597149\|25083871\|28735 | Unspecified |
| mmu-miR-509-5p | MIMAT0004890 | Tapbpl | 213233 | HITS-CLIP | 21258322 | Unspecified |

| | | | | | | |
|---|---|---|---|---|---|---|
| mmu-miR-509-5p | MIMAT0004890 | Tmem69 | 230657 | HITS-CLIP | 21258322 | Unspecified |
| mmu-miR-509-5p | MIMAT0004890 | Trim44 | 80985 | HITS-CLIP | 19536157\|23597149\|23142080\|21258 | Unspecified |
| mmu-miR-509-5p | MIMAT0004890 | Ttc9c | 70387 | HITS-CLIP | 21258322 | Unspecified |
| mmu-miR-509-5p | MIMAT0004890 | Ttf1 | 22130 | HITS-CLIP | 21258322 | Unspecified |
| mmu-miR-509-5p | MIMAT0004890 | Washc4 | 319277 | HITS-CLIP | 21258322 | Unspecified |
| mmu-miR-509-5p | MIMAT0004890 | Wdr89 | 72338 | HITS-CLIP | 21258322 | Unspecified |
| mmu-miR-509-5p | MIMAT0004890 | Zfp459 | 328274 | HITS-CLIP | 21258322 | Unspecified |
| mmu-miR-384-5p | MIMAT0004745 | 2410017P09Rik | 69709 | CLIP-Seq dataset analysis | 23142080 | Unspecified |
| mmu-miR-384-5p | MIMAT0004745 | 4933432B09Rik | 75729 | CLIP-Seq dataset analysis | 23142080 | Unspecified |
| mmu-miR-384-5p | MIMAT0004745 | 4933433P14Rik | 66787 | CLIP-Seq dataset analysis | 25083871 | Unspecified |
| mmu-miR-384-5p | MIMAT0004745 | AW555464 | 217882 | CLIP-Seq dataset analysis | 23597149\|25083871\|23142080 | Unspecified |
| mmu-miR-384-5p | MIMAT0004745 | Amer1 | 72345 | HITS-CLIP | 25083871 | Unspecified |
| mmu-miR-384-5p | MIMAT0004745 | Camk4 | 12326 | HITS-CLIP | 23142080 | Unspecified |
| mmu-miR-384-5p | MIMAT0004745 | Cd300a | 217303 | HITS-CLIP | 21258322\|25083871 | Unspecified |
| mmu-miR-384-5p | MIMAT0004745 | Cdk13 | 69562 | HITS-CLIP | 23142080 | Unspecified |
| mmu-miR-384-5p | MIMAT0004745 | Cep170b | 217882 | HITS-CLIP | 23142080\|23597149\|25083871 | Unspecified |
| mmu-miR-384-5p | MIMAT0004745 | Csf1 | 12977 | HITS-CLIP | 21258322 | Unspecified |
| mmu-miR-384-5p | MIMAT0004745 | Dpy19l1 | 244745 | HITS-CLIP | 25083871 | Unspecified |
| mmu-miR-384-5p | MIMAT0004745 | Eno2 | 13807 | HITS-CLIP | 25083871 | Unspecified |
| mmu-miR-384-5p | MIMAT0004745 | Fam123b | 72345 | CLIP-Seq dataset analysis | 25083871 | Unspecified |
| mmu-miR-384-5p | MIMAT0004745 | Fam227a | 75729 | HITS-CLIP | 23142080 | Unspecified |
| mmu-miR-384-5p | MIMAT0004745 | Fam40a | 229707 | CLIP-Seq dataset analysis | 25083871 | Unspecified |
| mmu-miR-384-5p | MIMAT0004745 | Gskip | 66787 | HITS-CLIP | 25083871 | Unspecified |
| mmu-miR-384-5p | MIMAT0004745 | Hoxb3 | 15410 | HITS-CLIP | 25083871 | Unspecified |
| mmu-miR-384-5p | MIMAT0004745 | Ice2 | 93697 | HITS-CLIP | 25083871 | Unspecified |
| mmu-miR-384-5p | MIMAT0004745 | Il18r1 | 16182 | HITS-CLIP | 23597149\|25083871 | Unspecified |
| mmu-miR-384-5p | MIMAT0004745 | Ing1 | 26356 | HITS-CLIP | 23142080 | Unspecified |
| mmu-miR-384-5p | MIMAT0004745 | Ints12 | 71793 | HITS-CLIP | 25083871 | Unspecified |
| mmu-miR-384-5p | MIMAT0004745 | Lpar3 | 65086 | HITS-CLIP | 23597149\|21258322 | Unspecified |
| mmu-miR-384-5p | MIMAT0004745 | Mesd | 67943 | HITS-CLIP | 25083871 | Unspecified |
| mmu-miR-384-5p | MIMAT0004745 | Mesdc2 | 67943 | HITS-CLIP | 25083871 | Unspecified |
| mmu-miR-384-5p | MIMAT0004745 | Mgea5 | 76055 | HITS-CLIP | 25083871 | Unspecified |
| mmu-miR-384-5p | MIMAT0004745 | Narg2 | 93697 | HITS-CLIP | 25083871 | Unspecified |
| mmu-miR-384-5p | MIMAT0004745 | Nfatc3 | 18021 | HITS-CLIP | 25083871 | Unspecified |
| mmu-miR-384-5p | MIMAT0004745 | Pgr | 18667 | HITS-CLIP | 25083871 | Unspecified |
| mmu-miR-384-5p | MIMAT0004745 | Pigh | 110417 | HITS-CLIP | 25083871 | Unspecified |
| mmu-miR-384-5p | MIMAT0004745 | Ptrhd1 | 69709 | HITS-CLIP | 23142080 | Unspecified |
| mmu-miR-384-5p | MIMAT0004745 | Reep4 | 72549 | HITS-CLIP | 25083871\|23597149 | Unspecified |
| mmu-miR-384-5p | MIMAT0004745 | Rrad | 56437 | HITS-CLIP | 25083871 | Unspecified |

| | | | | | | |
|---|---|---|---|---|---|---|
| mmu-miR-384-5p | MIMAT0004745 | Serpine1 | 18787 | HITS-CLIP | 23142080 | Unspecified |
| mmu-miR-384-5p | MIMAT0004745 | Six4 | 20474 | HITS-CLIP | 25083871 | Unspecified |
| mmu-miR-384-5p | MIMAT0004745 | Slc16a1 | 20501 | HITS-CLIP | 23142080\|19536157 | Unspecified |
| mmu-miR-384-5p | MIMAT0004745 | Strip1 | 229707 | HITS-CLIP | 25083871 | Unspecified |
| mmu-miR-384-5p | MIMAT0004745 | Tbc1d5 | 72238 | HITS-CLIP | 23142080 | Unspecified |
| mmu-miR-384-5p | MIMAT0004745 | Tnrc6b | 213988 | HITS-CLIP | 19536157\|25083871\|23597149 | Unspecified |
| mmu-miR-384-5p | MIMAT0004745 | Tomm70a | 28185 | HITS-CLIP | 25083871 | Unspecified |
| mmu-miR-384-5p | MIMAT0004745 | Trp53rk | 76367 | HITS-CLIP | 21258322 | Unspecified |
| mmu-miR-384-5p | MIMAT0004745 | Trp53rkb | 76367 | HITS-CLIP | 21258322 | Unspecified |
| mmu-miR-384-5p | MIMAT0004745 | Wdr44 | 72404 | HITS-CLIP | 25083871 | Unspecified |
| mmu-miR-384-5p | MIMAT0004745 | Wdr89 | 72338 | HITS-CLIP | 25083871\|19536157\|23597149 | Unspecified |
| mmu-miR-384-5p | MIMAT0004745 | Zfp446 | 269870 | HITS-CLIP | 25083871 | Unspecified |
| mmu-miR-384-5p | MIMAT0004745 | Zfp711 | 245595 | HITS-CLIP | 25083871 | Unspecified |
| mmu-miR-384-5p | MIMAT0004745 | Zfp9 | 22750 | HITS-CLIP | 21258322 | Unspecified |
| mmu-miR-30c-5p | MIMAT0000514 | Amer1 | 72345 | HITS-CLIP | 25083871\|23142080 | Unspecified |
| mmu-miR-30c-5p | MIMAT0000514 | Camk4 | 12326 | HITS-CLIP | 23142080 | Unspecified |
| mmu-miR-30c-5p | MIMAT0000514 | Ccne2 | 12448 | Luciferase assay//qRT-PCR//HITS-CLIP//Lu | 24555094\|23142080\|23597149\|1953 | Unspecified |
| mmu-miR-30c-5p | MIMAT0000514 | Cd300a | 217303 | HITS-CLIP | 25083871\|21258322 | Unspecified |
| mmu-miR-30c-5p | MIMAT0000514 | Cdk13 | 69562 | HITS-CLIP | 23142080 | Unspecified |
| mmu-miR-30c-5p | MIMAT0000514 | Celsr3 | 107934 | Luciferase assay//qRT-PCR//Luciferase rep | 24555094 | Unspecified |
| mmu-miR-30c-5p | MIMAT0000514 | Cep170b | 217882 | HITS-CLIP | 23142080\|23597149\|25083871 | Unspecified |
| mmu-miR-30c-5p | MIMAT0000514 | Csf1 | 12977 | HITS-CLIP | 21258322 | Unspecified |
| mmu-miR-30c-5p | MIMAT0000514 | Dpy19l1 | 244745 | HITS-CLIP | 25083871\|23142080 | Unspecified |
| mmu-miR-30c-5p | MIMAT0000514 | Egfr | 13649 | Luciferase assay//qRT-PCR//Luciferase rep | 24555094 | Unspecified |
| mmu-miR-30c-5p | MIMAT0000514 | Eno2 | 13807 | HITS-CLIP | 25083871 | Unspecified |
| mmu-miR-30c-5p | MIMAT0000514 | FASN | 2194 | Immunoprecipitaion//qRT-PCR//Western | 28088781 | Unspecified |
| mmu-miR-30c-5p | MIMAT0000514 | Fam227a | 75729 | HITS-CLIP | 23142080 | Unspecified |
| mmu-miR-30c-5p | MIMAT0000514 | Gskip | 66787 | HITS-CLIP | 25083871\|23142080 | Unspecified |
| mmu-miR-30c-5p | MIMAT0000514 | Hoxb3 | 15410 | HITS-CLIP | 25083871 | Unspecified |
| mmu-miR-30c-5p | MIMAT0000514 | Ice2 | 93697 | HITS-CLIP | 25083871 | Unspecified |
| mmu-miR-30c-5p | MIMAT0000514 | Il18r1 | 16182 | HITS-CLIP | 25083871\|23142080\|23597149 | Unspecified |
| mmu-miR-30c-5p | MIMAT0000514 | Ing1 | 26356 | HITS-CLIP | 23142080 | Unspecified |
| mmu-miR-30c-5p | MIMAT0000514 | Ints12 | 71793 | HITS-CLIP | 25083871 | Unspecified |
| mmu-miR-30c-5p | MIMAT0000514 | Lpar3 | 65086 | HITS-CLIP | 21258322\|23597149 | Unspecified |
| mmu-miR-30c-5p | MIMAT0000514 | Mdm2 | 17246 | Luciferase assay//qRT-PCR//Luciferase rep | 24555094 | Unspecified |
| mmu-miR-30c-5p | MIMAT0000514 | Mesd | 67943 | HITS-CLIP | 25083871 | Unspecified |
| mmu-miR-30c-5p | MIMAT0000514 | Mesdc2 | 67943 | HITS-CLIP | 25083871 | Unspecified |
| mmu-miR-30c-5p | MIMAT0000514 | Mgea5 | 76055 | HITS-CLIP | 25083871\|23142080 | Unspecified |
| mmu-miR-30c-5p | MIMAT0000514 | Mtdh | 67154 | Luciferase assay//qRT-PCR//HITS-CLIP//Lu | 24555094\|23597149\|19536157\|2314 | Unspecified |

| | | | | | | |
|---|---|---|---|---|---|---|
| mmu-miR-30c-5p | MIMAT0000514 | Narg2 | 93697 | HITS-CLIP | 25083871 | Unspecified |
| mmu-miR-30c-5p | MIMAT0000514 | Nfatc3 | 18021 | HITS-CLIP | 25083871\|23142080\|25753418 | Unspecified |
| mmu-miR-30c-5p | MIMAT0000514 | Pgr | 18667 | HITS-CLIP | 25083871 | Unspecified |
| mmu-miR-30c-5p | MIMAT0000514 | Pigh | 110417 | HITS-CLIP | 25083871 | Unspecified |
| mmu-miR-30c-5p | MIMAT0000514 | Ptrhd1 | 69709 | HITS-CLIP | 23142080 | Unspecified |
| mmu-miR-30c-5p | MIMAT0000514 | Reep4 | 72549 | HITS-CLIP | 23597149\|23142080\|25083871 | Unspecified |
| mmu-miR-30c-5p | MIMAT0000514 | Rrad | 56437 | HITS-CLIP | 25083871\|23142080 | Unspecified |
| mmu-miR-30c-5p | MIMAT0000514 | Runx1 | 12394 | Luciferase assay//Western blot//Luciferase | 19114653 | Unspecified |
| mmu-miR-30c-5p | MIMAT0000514 | Serpine1 | 18787 | HITS-CLIP | 23142080 | Unspecified |
| mmu-miR-30c-5p | MIMAT0000514 | Six4 | 20474 | HITS-CLIP | 25083871 | Unspecified |
| mmu-miR-30c-5p | MIMAT0000514 | Slc16a1 | 20501 | HITS-CLIP | 23142080 | Unspecified |
| mmu-miR-30c-5p | MIMAT0000514 | Smad1 | 17125 | Luciferase assay//qRT-PCR//Western Blot/ | 24555094\|22253433 | Unspecified |
| mmu-miR-30c-5p | MIMAT0000514 | Snai1 | 20613 | Luciferase assay//Western blot//Luciferase | 21945074 | Unspecified |
| mmu-miR-30c-5p | MIMAT0000514 | Strip1 | 229707 | HITS-CLIP | 25083871\|23597149\|23142080 | Unspecified |
| mmu-miR-30c-5p | MIMAT0000514 | Tbc1d5 | 72238 | HITS-CLIP | 23142080 | Unspecified |
| mmu-miR-30c-5p | MIMAT0000514 | Timp3 | 21859 | Luciferase assay//qRT-PCR//Luciferase rep | 24555094 | Unspecified |
| mmu-miR-30c-5p | MIMAT0000514 | Tnrc6b | 213988 | HITS-CLIP | 25083871\|23597149\|23142080\|1953( | Unspecified |
| mmu-miR-30c-5p | MIMAT0000514 | Tomm70a | 28185 | HITS-CLIP | 25083871\|23142080 | Unspecified |
| mmu-miR-30c-5p | MIMAT0000514 | Trp53rk | 76367 | HITS-CLIP | 21258322 | Unspecified |
| mmu-miR-30c-5p | MIMAT0000514 | Trp53rkb | 76367 | HITS-CLIP | 21258322 | Unspecified |
| mmu-miR-30c-5p | MIMAT0000514 | Twf1 | 19230 | Luciferase assay//qRT-PCR//HITS-CLIP//Lu | 24555094\|23597149\|23142080 | Unspecified |
| mmu-miR-30c-5p | MIMAT0000514 | Wdr44 | 72404 | HITS-CLIP | 25083871 | Unspecified |
| mmu-miR-30c-5p | MIMAT0000514 | Wdr89 | 72338 | HITS-CLIP | 25083871\|23597149\|19536157 | Unspecified |
| mmu-miR-30c-5p | MIMAT0000514 | Zfp446 | 269870 | HITS-CLIP | 25083871 | Unspecified |
| mmu-miR-30c-5p | MIMAT0000514 | Zfp711 | 245595 | HITS-CLIP | 25083871 | Unspecified |
| mmu-miR-30c-5p | MIMAT0000514 | Zfp9 | 22750 | HITS-CLIP | 21258322 | Unspecified |
| mmu-miR-875-5p | MIMAT0004937 | 09-Sept | 53860 | HITS-CLIP | 25083871 | Unspecified |
| mmu-miR-875-5p | MIMAT0004937 | 4833442J19Rik | 320204 | CLIP-Seq dataset analysis | 25083871 | Unspecified |
| mmu-miR-875-5p | MIMAT0004937 | 6030419C18Rik | 319477 | HITS-CLIP | 25083871 | Unspecified |
| mmu-miR-875-5p | MIMAT0004937 | C2cd3 | 277939 | HITS-CLIP | 25083871 | Unspecified |
| mmu-miR-875-5p | MIMAT0004937 | Capza1 | 12340 | HITS-CLIP | 21258322\|25083871 | Unspecified |
| mmu-miR-875-5p | MIMAT0004937 | Ddx17 | 67040 | HITS-CLIP | 25083871 | Unspecified |
| mmu-miR-875-5p | MIMAT0004937 | Etfbkmt | 320204 | HITS-CLIP | 25083871 | Unspecified |
| mmu-miR-875-5p | MIMAT0004937 | Fign | 60344 | HITS-CLIP | 21258322\|23597149\|19536157 | Unspecified |
| mmu-miR-875-5p | MIMAT0004937 | Gfod1 | 328232 | HITS-CLIP | 25083871 | Unspecified |
| mmu-miR-875-5p | MIMAT0004937 | Golt1a | 68338 | HITS-CLIP | 25083871 | Unspecified |
| mmu-miR-875-5p | MIMAT0004937 | Mettl20 | 320204 | HITS-CLIP | 25083871 | Unspecified |
| mmu-miR-875-5p | MIMAT0004937 | Mfsd2b | 432628 | HITS-CLIP | 21258322 | Unspecified |
| mmu-miR-875-5p | MIMAT0004937 | Nhlrc2 | 66866 | HITS-CLIP | 25083871 | Unspecified |

| | | | | | | |
|---|---|---|---|---|---|---|
| mmu-miR-875-5p | MIMAT0004937 | Nufip2 | 68564 | HITS-CLIP | 25083871 | Unspecified |
| mmu-miR-875-5p | MIMAT0004937 | Prdx3 | 11757 | Luciferase assay//qRT-PCR//Western blot/ | 23846856 | Unspecified |
| mmu-miR-875-5p | MIMAT0004937 | Prkaa1 | 105787 | HITS-CLIP | 25083871 | Unspecified |
| mmu-miR-875-5p | MIMAT0004937 | Rbms3 | 207181 | HITS-CLIP | 25083871 | Unspecified |
| mmu-miR-875-5p | MIMAT0004937 | Rwdd4a | 192174 | HITS-CLIP | 21258322 | Unspecified |
| mmu-miR-875-5p | MIMAT0004937 | Satb2 | 212712 | HITS-CLIP | 25083871 | Unspecified |
| mmu-miR-875-5p | MIMAT0004937 | 09-Sept | 53374 | HITS-CLIP | 25083871 | Unspecified |
| mmu-miR-875-5p | MIMAT0004937 | Slc15a2 | 57738 | HITS-CLIP | 25083871 | Unspecified |
| mmu-miR-875-5p | MIMAT0004937 | Stat1 | 20846 | HITS-CLIP | 25083871 | Unspecified |
| mmu-miR-875-5p | MIMAT0004937 | Synj2bp | 24071 | HITS-CLIP | 21258322 | Unspecified |
| mmu-miR-875-5p | MIMAT0004937 | Vmn2r123 | 22300 | HITS-CLIP | 25083871 | Unspecified |
| mmu-miR-875-5p | MIMAT0004937 | Zcchc8 | 70650 | HITS-CLIP | 25083871 | Unspecified |
| mmu-miR-139-3p | MIMAT0004662 | Mtf1 | 17764 | HITS-CLIP | 23142080 | Unspecified |
| mmu-miR-139-3p | MIMAT0004662 | Nav2 | 78286 | HITS-CLIP | 25083871 | Unspecified |
| mmu-miR-139-3p | MIMAT0004662 | Opa3 | 403187 | HITS-CLIP | 21258322 | Unspecified |
| mmu-miR-139-3p | MIMAT0004662 | Plxna4 | 243743 | HITS-CLIP | 23142080 | Unspecified |
| mmu-miR-139-3p | MIMAT0004662 | Sco1 | 52892 | HITS-CLIP | 23142080 | Unspecified |
| mmu-miR-152-3p | MIMAT0000162 | Adam19 | 11492 | HITS-CLIP | 25083871 | Unspecified |
| mmu-miR-152-3p | MIMAT0000162 | Arrdc3 | 105171 | HITS-CLIP | 25083871 | Unspecified |
| mmu-miR-152-3p | MIMAT0000162 | Camk2a | 12322 | Luciferase assay//qRT-PCR//Western blot/ | 21068402 | Unspecified |
| mmu-miR-152-3p | MIMAT0000162 | Ccna2 | 12428 | HITS-CLIP | 25083871 | Unspecified |
| mmu-miR-152-3p | MIMAT0000162 | Chrna7 | 11441 | HITS-CLIP | 28735896 | Unspecified |
| mmu-miR-152-3p | MIMAT0000162 | Clca2 | 80797 | HITS-CLIP | 21258322 | Unspecified |
| mmu-miR-152-3p | MIMAT0000162 | Clca3a2 | 80797 | HITS-CLIP | 21258322 | Unspecified |
| mmu-miR-152-3p | MIMAT0000162 | Cpm | 70574 | HITS-CLIP | 25083871 | Unspecified |
| mmu-miR-152-3p | MIMAT0000162 | Dnmt1 | 13433 | Luciferase assay//Western blot//Luciferas | 22905112\|34006907 | Unspecified |
| mmu-miR-152-3p | MIMAT0000162 | Ets1 | 23871 | HITS-CLIP | 23597149\|25083871 | Unspecified |
| mmu-miR-152-3p | MIMAT0000162 | Gpr157 | 269604 | HITS-CLIP | 21258322 | Unspecified |
| mmu-miR-152-3p | MIMAT0000162 | Grik3 | 14807 | HITS-CLIP | 19536157\|23597149\|25083871 | Unspecified |
| mmu-miR-152-3p | MIMAT0000162 | H2-T23 | 15040 | HITS-CLIP | 21258322 | Unspecified |
| mmu-miR-152-3p | MIMAT0000162 | Hpca | 15444 | HITS-CLIP | 23142080 | Unspecified |
| mmu-miR-152-3p | MIMAT0000162 | Itgb3 | 16416 | HITS-CLIP | 21258322\|25083871\|23597149\|23142 | Unspecified |
| mmu-miR-152-3p | MIMAT0000162 | Lypd1 | 72585 | HITS-CLIP | 28735896 | Unspecified |
| mmu-miR-152-3p | MIMAT0000162 | Mbp | 17196 | HITS-CLIP | 25083871 | Unspecified |
| mmu-miR-152-3p | MIMAT0000162 | Mrpl51 | 66493 | HITS-CLIP | 21258322 | Unspecified |
| mmu-miR-152-3p | MIMAT0000162 | Ms4a15 | 545279 | HITS-CLIP | 23142080 | Unspecified |
| mmu-miR-152-3p | MIMAT0000162 | N6amt1 | 67768 | HITS-CLIP | 21258322 | Unspecified |
| mmu-miR-152-3p | MIMAT0000162 | Naip5 | 17951 | HITS-CLIP | 21258322 | Unspecified |
| mmu-miR-152-3p | MIMAT0000162 | Ociad2 | 433904 | HITS-CLIP | 21258322 | Unspecified |

| | | | | | | |
|---|---|---|---|---|---|---|
| mmu-miR-152-3p | MIMAT0000162 | Psma4 | 26441 | HITS-CLIP | 37986224\|28735896 | Unspecified |
| mmu-miR-152-3p | MIMAT0000162 | Pten | 19211 | HITS-CLIP//Luciferase reporter assay//Wes | 25083871\|26996529 | Unspecified |
| mmu-miR-152-3p | MIMAT0000162 | Rad51c | 114714 | HITS-CLIP | 25083871\|23142080 | Unspecified |
| mmu-miR-152-3p | MIMAT0000162 | Robo1 | 19876 | HITS-CLIP | 25083871 | Unspecified |
| mmu-miR-152-3p | MIMAT0000162 | Scml4 | 268297 | HITS-CLIP | 21258322\|25083871 | Unspecified |
| mmu-miR-152-3p | MIMAT0000162 | Snx27 | 76742 | HITS-CLIP | 25083871\|23597149 | Unspecified |
| mmu-miR-152-3p | MIMAT0000162 | Stxbp2 | 20911 | HITS-CLIP | 23597149 | Unspecified |
| mmu-miR-152-3p | MIMAT0000162 | Tbc1d24 | 224617 | HITS-CLIP | 21258322 | Unspecified |
| mmu-miR-152-3p | MIMAT0000162 | Tmem178b | 434008 | HITS-CLIP | 25083871\|23597149\|21258322\|19536 | Unspecified |
| mmu-miR-152-3p | MIMAT0000162 | Trim43b | 666747 | HITS-CLIP | 21258322 | Unspecified |
| mmu-miR-152-3p | MIMAT0000162 | Ubap2l | 74383 | HITS-CLIP | 25083871 | Unspecified |
| mmu-miR-152-3p | MIMAT0000162 | Zfp607 | 545938 | HITS-CLIP | 21258322 | Unspecified |
| mmu-miR-152-3p | MIMAT0000162 | Zfp607a | 545938 | HITS-CLIP | 21258322 | Unspecified |
| mmu-miR-1946a | MIMAT0009412 | 2310010M20Rik | 69576 | CLIP-Seq dataset analysis | 25083871\|21258322 | Unspecified |
| mmu-miR-1946a | MIMAT0009412 | 9830001H06Rik | 320706 | CLIP-Seq dataset analysis | 21258322 | Unspecified |
| mmu-miR-1946a | MIMAT0009412 | B4galnt2 | 14422 | HITS-CLIP | 21258322 | Unspecified |
| mmu-miR-1946a | MIMAT0009412 | Bach2 | 12014 | HITS-CLIP | 21258322 | Unspecified |
| mmu-miR-1946a | MIMAT0009412 | Cbln3 | 56410 | HITS-CLIP | 21258322 | Unspecified |
| mmu-miR-1946a | MIMAT0009412 | Cdnf | 227526 | HITS-CLIP | 21258322 | Unspecified |
| mmu-miR-1946a | MIMAT0009412 | Cnnm3 | 94218 | HITS-CLIP | 21258322 | Unspecified |
| mmu-miR-1946a | MIMAT0009412 | Cybrd1 | 73649 | HITS-CLIP | 21258322 | Unspecified |
| mmu-miR-1946a | MIMAT0009412 | Ddhd2 | 72108 | HITS-CLIP | 21258322 | Unspecified |
| mmu-miR-1946a | MIMAT0009412 | Dnase2a | 13423 | HITS-CLIP | 25083871 | Unspecified |
| mmu-miR-1946a | MIMAT0009412 | Ercc6 | 319955 | HITS-CLIP | 25083871\|21258322 | Unspecified |
| mmu-miR-1946a | MIMAT0009412 | Fbxo43 | 78803 | HITS-CLIP | 25083871 | Unspecified |
| mmu-miR-1946a | MIMAT0009412 | Gnl3l | 237107 | HITS-CLIP | 21258322\|25083871 | Unspecified |
| mmu-miR-1946a | MIMAT0009412 | Il17ra | 16172 | HITS-CLIP | 25083871\|23597149 | Unspecified |
| mmu-miR-1946a | MIMAT0009412 | Kdelc2 | 68304 | HITS-CLIP | 21258322 | Unspecified |
| mmu-miR-1946a | MIMAT0009412 | Mapre1 | 13589 | HITS-CLIP | 21258322 | Unspecified |
| mmu-miR-1946a | MIMAT0009412 | Mdc1 | 240087 | HITS-CLIP | 21258322 | Unspecified |
| mmu-miR-1946a | MIMAT0009412 | Mga | 29808 | HITS-CLIP | 25083871\|21258322 | Unspecified |
| mmu-miR-1946a | MIMAT0009412 | Mrpl43 | 94067 | HITS-CLIP | 25083871 | Unspecified |
| mmu-miR-1946a | MIMAT0009412 | Nosip | 66394 | HITS-CLIP | 21258322 | Unspecified |
| mmu-miR-1946a | MIMAT0009412 | Pfas | 237823 | HITS-CLIP | 21258322 | Unspecified |
| mmu-miR-1946a | MIMAT0009412 | Pik3c2a | 18704 | HITS-CLIP | 21258322 | Unspecified |
| mmu-miR-1946a | MIMAT0009412 | Rab11b | 19326 | HITS-CLIP | 21258322 | Unspecified |
| mmu-miR-1946a | MIMAT0009412 | Ranbp3l | 223332 | HITS-CLIP | 21258322 | Unspecified |
| mmu-miR-1946a | MIMAT0009412 | Rpl7l1 | 66229 | HITS-CLIP | 21258322 | Unspecified |
| mmu-miR-1946a | MIMAT0009412 | Sema4b | 20352 | HITS-CLIP | 21258322 | Unspecified |

| | | | | | | | |
|---|---|---|---|---|---|---|---|
| mmu-miR-1946a | MIMAT0009412 | Sfxn2 | 94279 | HITS-CLIP | | 21258322 | Unspecified |
| mmu-miR-1946a | MIMAT0009412 | Sgk3 | 170755 | HITS-CLIP | | 21258322 | Unspecified |
| mmu-miR-1946a | MIMAT0009412 | Slc28a3 | 114304 | HITS-CLIP | 21258322\|25083871 | | Unspecified |
| mmu-miR-1946a | MIMAT0009412 | Slc35e2 | 320541 | HITS-CLIP | | 21258322 | Unspecified |
| mmu-miR-1946a | MIMAT0009412 | Smco1 | 69576 | HITS-CLIP | 21258322\|25083871 | | Unspecified |
| mmu-miR-1946a | MIMAT0009412 | Soga1 | 320706 | HITS-CLIP | | 21258322 | Unspecified |
| mmu-miR-1946a | MIMAT0009412 | TDRD7 | 23424 | Microarray | | 33665188 | Unspecified |
| mmu-miR-1946a | MIMAT0009412 | Thap2 | 66816 | HITS-CLIP | | 21258322 | Unspecified |
| mmu-miR-1946a | MIMAT0009412 | Tnfrsf10b | 21933 | HITS-CLIP | | 21258322 | Unspecified |
| mmu-miR-1946a | MIMAT0009412 | Trim27 | 19720 | HITS-CLIP | 25083871\|21258322 | | Unspecified |
| mmu-miR-1946a | MIMAT0009412 | Trim36 | 28105 | HITS-CLIP | | 21258322 | Unspecified |
| mmu-miR-1946a | MIMAT0009412 | Ttc26 | 264134 | HITS-CLIP | | 21258322 | Unspecified |
| mmu-miR-1946a | MIMAT0009412 | Wipf2 | 68524 | HITS-CLIP | | 21258322 | Unspecified |
| mmu-miR-1946a | MIMAT0009412 | Zbtb44 | 235132 | HITS-CLIP | | 21258322 | Unspecified |
| mmu-miR-1946a | MIMAT0009412 | Zfp830 | 66983 | HITS-CLIP | | 21258322 | Unspecified |
| mmu-miR-1249-3p | MIMAT0010560 | Auts2 | 319974 | HITS-CLIP | | 21258322 | Unspecified |
| mmu-miR-1249-3p | MIMAT0010560 | Foxl1 | 14241 | HITS-CLIP | | 25083871 | Unspecified |
| mmu-miR-1249-3p | MIMAT0010560 | Tnfsf10 | 22035 | HITS-CLIP | | 21258322 | Unspecified |
| mmu-miR-1249-3p | MIMAT0010560 | Uggt2 | 66435 | HITS-CLIP | | 28735896 | Unspecified |
| mmu-miR-1249-3p | MIMAT0010560 | Zfp941 | 407812 | HITS-CLIP | | 25083871 | Unspecified |
| mmu-miR-680 | MIMAT0003457 | Cd59a | 12509 | HITS-CLIP | | 21258322 | Unspecified |
| mmu-miR-680 | MIMAT0003457 | Cldn19 | 242653 | HITS-CLIP | | 21258322 | Unspecified |
| mmu-miR-680 | MIMAT0003457 | Ctnnb1 | 12387 | Immunohistochemistry//Luciferase assay/ | | 24594381 | Unspecified |
| mmu-miR-680 | MIMAT0003457 | Hist2h3c1 | 15077 | HITS-CLIP | | 21258322 | Unspecified |
| mmu-miR-680 | MIMAT0003457 | Rnase4 | 58809 | HITS-CLIP | | 21258322 | Unspecified |
| mmu-miR-680 | MIMAT0003457 | Slc17a5 | 235504 | HITS-CLIP | | 21258322 | Unspecified |
| mmu-miR-680 | MIMAT0003457 | Vcan | 13003 | Immunohistochemistry//Luciferase assay/ | | 24594381 | Unspecified |
| mmu-miR-3067-5p | MIMAT0014840 | Adamtsl1 | 77739 | HITS-CLIP | | 23142080 | Unspecified |
| mmu-miR-3067-5p | MIMAT0014840 | Auts2 | 319974 | HITS-CLIP | | 23597149 | Unspecified |
| mmu-miR-3067-5p | MIMAT0014840 | B3galtl | 381694 | HITS-CLIP | | 21258322 | Unspecified |
| mmu-miR-3067-5p | MIMAT0014840 | B3glct | 381694 | HITS-CLIP | | 21258322 | Unspecified |
| mmu-miR-3067-5p | MIMAT0014840 | Cd300lf | 246746 | HITS-CLIP | 21258322\|23597149 | | Unspecified |
| mmu-miR-3067-5p | MIMAT0014840 | Cdc27 | 217232 | HITS-CLIP | | 25083871 | Unspecified |
| mmu-miR-3067-5p | MIMAT0014840 | Chrna2 | 110902 | HITS-CLIP | | 21258322 | Unspecified |
| mmu-miR-3067-5p | MIMAT0014840 | Dsg2 | 13511 | HITS-CLIP | 23597149\|21258322 | | Unspecified |
| mmu-miR-3067-5p | MIMAT0014840 | E4f1 | 13560 | HITS-CLIP | 23597149\|19536157 | | Unspecified |
| mmu-miR-3067-5p | MIMAT0014840 | Edaradd | 171211 | HITS-CLIP | | 23142080 | Unspecified |
| mmu-miR-3067-5p | MIMAT0014840 | Faah | 14073 | HITS-CLIP | | 21258322 | Unspecified |
| mmu-miR-3067-5p | MIMAT0014840 | Fam81a | 76886 | HITS-CLIP | 21258322\|25083871 | | Unspecified |

| | | | | | | |
|---|---|---|---|---|---|---|
| mmu-miR-3067-5p | MIMAT0014840 | Hook3 | 320191 | HITS-CLIP | 21258322\|23142080 | Unspecified |
| mmu-miR-3067-5p | MIMAT0014840 | Opa1 | 74143 | HITS-CLIP | 21258322 | Unspecified |
| mmu-miR-3067-5p | MIMAT0014840 | Pak6 | 214230 | HITS-CLIP | 19536157 | Unspecified |
| mmu-miR-3067-5p | MIMAT0014840 | Pbx1 | 18514 | HITS-CLIP | 25083871 | Unspecified |
| mmu-miR-3067-5p | MIMAT0014840 | Pkn2 | 109333 | HITS-CLIP | 23142080 | Unspecified |
| mmu-miR-3067-5p | MIMAT0014840 | Ptgr2 | 77219 | HITS-CLIP | 21258322 | Unspecified |
| mmu-miR-3067-5p | MIMAT0014840 | Rad54l2 | 81000 | HITS-CLIP | 23597149 | Unspecified |
| mmu-miR-3067-5p | MIMAT0014840 | Stx16 | 228960 | HITS-CLIP | 23597149 | Unspecified |
| mmu-miR-3067-5p | MIMAT0014840 | Tgm3 | 21818 | HITS-CLIP | 21258322 | Unspecified |
| mmu-miR-3067-5p | MIMAT0014840 | Usp40 | 227334 | HITS-CLIP | 25083871\|21258322 | Unspecified |
| mmu-miR-3067-5p | MIMAT0014840 | Vldlr | 22359 | HITS-CLIP | 21258322 | Unspecified |
| mmu-miR-3067-5p | MIMAT0014840 | Zbtb26 | 320633 | HITS-CLIP | 21258322 | Unspecified |
| mmu-miR-465a-5p | MIMAT0002106 | 4930430F08Rik | 68281 | HITS-CLIP | 25083871 | Unspecified |
| mmu-miR-465a-5p | MIMAT0002106 | 4930563E22Rik | 75304 | HITS-CLIP | 25083871 | Unspecified |
| mmu-miR-465a-5p | MIMAT0002106 | 4933439F18Rik | 66771 | CLIP-Seq dataset analysis | 21258322\|25083871 | Unspecified |
| mmu-miR-465a-5p | MIMAT0002106 | Arl4a | 11861 | HITS-CLIP | 23142080 | Unspecified |
| mmu-miR-465a-5p | MIMAT0002106 | Cdc37 | 12539 | HITS-CLIP | 25083871 | Unspecified |
| mmu-miR-465a-5p | MIMAT0002106 | Dapk1 | 69635 | HITS-CLIP | 23142080 | Unspecified |
| mmu-miR-465a-5p | MIMAT0002106 | Gid4 | 66771 | HITS-CLIP | 21258322\|25083871 | Unspecified |
| mmu-miR-465a-5p | MIMAT0002106 | Gm11559 | 100415785 | HITS-CLIP | 25083871 | Unspecified |
| mmu-miR-465a-5p | MIMAT0002106 | Hltf | 20585 | HITS-CLIP | 21258322 | Unspecified |
| mmu-miR-465a-5p | MIMAT0002106 | Map3k1 | 26401 | HITS-CLIP | 25083871 | Unspecified |
| mmu-miR-465a-5p | MIMAT0002106 | Nkain3 | 269513 | HITS-CLIP | 21258322\|25083871 | Unspecified |
| mmu-miR-465a-5p | MIMAT0002106 | PAK1 | 5058 | Luciferase reporter assay//Western blot// | 18922890 | Unspecified |
| mmu-miR-465a-5p | MIMAT0002106 | Ppp1r1c | 75276 | HITS-CLIP | 23597149 | Unspecified |
| mmu-miR-465a-5p | MIMAT0002106 | Setd7 | 73251 | HITS-CLIP | 25083871 | Unspecified |
| mmu-miR-465a-5p | MIMAT0002106 | Sim1 | 20464 | HITS-CLIP | 25083871 | Unspecified |
| mmu-miR-465a-5p | MIMAT0002106 | Tbc1d24 | 224617 | HITS-CLIP | 25083871 | Unspecified |
| mmu-miR-465a-5p | MIMAT0002106 | Tcfap2b | 21419 | CLIP-Seq dataset analysis | 23142080 | Unspecified |
| mmu-miR-465a-5p | MIMAT0002106 | Tfap2b | 21419 | HITS-CLIP | 23142080 | Unspecified |
| mmu-miR-465a-5p | MIMAT0002106 | Thrb | 21834 | HITS-CLIP | 21258322\|23597149 | Unspecified |
| mmu-miR-465a-5p | MIMAT0002106 | Tirap | 117149 | HITS-CLIP | 25083871 | Unspecified |
| mmu-miR-465a-5p | MIMAT0002106 | Veph1 | 72789 | HITS-CLIP | 25083871 | Unspecified |
| mmu-miR-465a-5p | MIMAT0002106 | Ybey | 216119 | HITS-CLIP | 25083871 | Unspecified |
| mmu-miR-465a-5p | MIMAT0002106 | Ybx1 | 22608 | HITS-CLIP | 25083871\|21258322 | Unspecified |
| mmu-miR-193a-3p | MIMAT0000223 | Dph3 | 105638 | HITS-CLIP | 21258322 | Unspecified |
| mmu-miR-193a-3p | MIMAT0000223 | Gla | 11605 | HITS-CLIP | 25083871 | Unspecified |
| mmu-miR-193a-3p | MIMAT0000223 | Ifngr2 | 15980 | HITS-CLIP | 21258322\|31433990 | Unspecified |
| mmu-miR-193a-3p | MIMAT0000223 | Kif15 | 209737 | HITS-CLIP | 21258322 | Unspecified |

| | | | | | | | |
|---|---|---|---|---|---|---|---|
| mmu-miR-193a-3p | MIMAT0000223 | Metap2 | 56307 | HITS-CLIP | | 23142080 | Unspecified |
| mmu-miR-193a-3p | MIMAT0000223 | Mrpl45 | 67036 | HITS-CLIP | | 21258322 | Unspecified |
| mmu-miR-193a-3p | MIMAT0000223 | Tmem245 | 242474 | HITS-CLIP | | 21258322 | Unspecified |
| mmu-miR-193a-3p | MIMAT0000223 | Ubxn2a | 217379 | HITS-CLIP | | 21258322 | Unspecified |
| mmu-miR-378a-3p | MIMAT0003151 | Aoc3 | 11754 | HITS-CLIP | 25083871\|21258322 | | Unspecified |
| mmu-miR-378a-3p | MIMAT0003151 | Arhgef39 | 230098 | HITS-CLIP | | 21258322 | Unspecified |
| mmu-miR-378a-3p | MIMAT0003151 | Casp9 | 12371 | HITS-CLIP//Western blotting//Immunofluo | 37986224\|32913213 | | Unspecified |
| mmu-miR-378a-3p | MIMAT0003151 | Cpox | 12892 | HITS-CLIP | | 25083871 | Unspecified |
| mmu-miR-378a-3p | MIMAT0003151 | Csf2rb | 12983 | HITS-CLIP | | 21258322 | Unspecified |
| mmu-miR-378a-3p | MIMAT0003151 | Dkc1 | 245474 | HITS-CLIP//Microarrays | 21258322\|22855601 | | Unspecified |
| mmu-miR-378a-3p | MIMAT0003151 | Dnajb1 | 81489 | HITS-CLIP | | 25083871 | Unspecified |
| mmu-miR-378a-3p | MIMAT0003151 | Ercc6 | 319955 | HITS-CLIP | | 21258322 | Unspecified |
| mmu-miR-378a-3p | MIMAT0003151 | Gli3 | 2737 | HITS-CLIP//Luciferase reporter assay | 37986224\|27001906 | | Unspecified |
| mmu-miR-378a-3p | MIMAT0003151 | H2-T23 | 15040 | HITS-CLIP | | 21258322 | Unspecified |
| mmu-miR-378a-3p | MIMAT0003151 | Il17rd | 171463 | HITS-CLIP | | 21258322 | Unspecified |
| mmu-miR-378a-3p | MIMAT0003151 | Il17re | 57890 | HITS-CLIP | | 25083871 | Unspecified |
| mmu-miR-378a-3p | MIMAT0003151 | Kif15 | 209737 | HITS-CLIP | 21258322\|23142080 | | Unspecified |
| mmu-miR-378a-3p | MIMAT0003151 | Mapk1 | 26413 | HITS-CLIP//Luciferase reporter assay | 25083871\|23142080\|25529446 | | Unspecified |
| mmu-miR-378a-3p | MIMAT0003151 | Ncoa7 | 211329 | HITS-CLIP | 25083871\|23142080 | | Unspecified |
| mmu-miR-378a-3p | MIMAT0003151 | Nrf1 | 18181 | Luciferase assay//qRT-PCR//Western blot/ | | 23818290 | Unspecified |
| mmu-miR-378a-3p | MIMAT0003151 | Odc1 | 18263 | Other | | 20432256 | Unspecified |
| mmu-miR-378a-3p | MIMAT0003151 | Pdk1 | 228026 | HITS-CLIP//Western blotting//Immunofluo | 37986224\|32913213 | | Unspecified |
| mmu-miR-378a-3p | MIMAT0003151 | Polr2l | 66491 | HITS-CLIP | | 23597149 | Unspecified |
| mmu-miR-378a-3p | MIMAT0003151 | Scfd2 | 212986 | HITS-CLIP | | 21258322 | Unspecified |
| mmu-miR-378a-3p | MIMAT0003151 | Stk32b | 64293 | HITS-CLIP | | 23142080 | Unspecified |
| mmu-miR-378a-3p | MIMAT0003151 | Tgfbr3 | 21814 | HITS-CLIP | | 23142080 | Unspecified |
| mmu-miR-378a-3p | MIMAT0003151 | Tnfrsf1b | 21938 | HITS-CLIP | | 23597149 | Unspecified |
| mmu-miR-378a-3p | MIMAT0003151 | Tsn | 22099 | HITS-CLIP | | 21258322 | Unspecified |
| mmu-miR-378a-3p | MIMAT0003151 | Zfp941 | 407812 | HITS-CLIP | 21258322\|25083871 | | Unspecified |
| mmu-miR-1306-5p | MIMAT0019136 | 4921536K21Rik | 67430 | HITS-CLIP | | 21258322 | Unspecified |
| mmu-miR-1306-5p | MIMAT0019136 | Ap4e1 | 108011 | HITS-CLIP | | 21258322 | Unspecified |
| mmu-miR-1306-5p | MIMAT0019136 | Bbs1 | 52028 | HITS-CLIP | | 21258322 | Unspecified |
| mmu-miR-1306-5p | MIMAT0019136 | Camk2n1 | 66259 | HITS-CLIP | | 21258322 | Unspecified |
| mmu-miR-1306-5p | MIMAT0019136 | Cir1 | 66935 | HITS-CLIP | | 21258322 | Unspecified |
| mmu-miR-1306-5p | MIMAT0019136 | Cyb561d1 | 72023 | HITS-CLIP | | 21258322 | Unspecified |
| mmu-miR-1306-5p | MIMAT0019136 | Elovl1 | 54325 | HITS-CLIP | | 23597149 | Unspecified |
| mmu-miR-1306-5p | MIMAT0019136 | Elovl2 | 54326 | HITS-CLIP | | 21258322 | Unspecified |
| mmu-miR-1306-5p | MIMAT0019136 | Fign | 60344 | HITS-CLIP | | 21258322 | Unspecified |
| mmu-miR-1306-5p | MIMAT0019136 | Frmd3 | 242506 | HITS-CLIP | 21258322\|25083871\|23597149\|23142 | | Unspecified |

| | | | | | | |
|---|---|---|---|---|---|---|
| mmu-miR-1306-5p | MIMAT0019136 | Lhfpl4 | 269788 | HITS-CLIP | 25083871 | Unspecified |
| mmu-miR-1306-5p | MIMAT0019136 | Map3k7 | 26409 | HITS-CLIP | 25083871\|19536157\|23597149 | Unspecified |
| mmu-miR-1306-5p | MIMAT0019136 | Mpg | 268395 | HITS-CLIP | 21258322 | Unspecified |
| mmu-miR-1306-5p | MIMAT0019136 | Ptprb | 19263 | HITS-CLIP | 19536157\|25083871 | Unspecified |
| mmu-miR-1306-5p | MIMAT0019136 | Rab11fip5 | 52055 | HITS-CLIP | 21258322 | Unspecified |
| mmu-miR-1306-5p | MIMAT0019136 | Rinl | 320435 | HITS-CLIP | 25083871 | Unspecified |
| mmu-miR-1306-5p | MIMAT0019136 | S1pr3 | 13610 | HITS-CLIP | 21258322 | Unspecified |
| mmu-miR-1306-5p | MIMAT0019136 | Slc17a7 | 72961 | HITS-CLIP | 23142080 | Unspecified |
| mmu-miR-1306-5p | MIMAT0019136 | Slc1a2 | 20511 | HITS-CLIP | 21258322 | Unspecified |
| mmu-miR-1306-5p | MIMAT0019136 | Slc8a2 | 110891 | HITS-CLIP | 28735896 | Unspecified |
| mmu-miR-1306-5p | MIMAT0019136 | Tpmt | 22017 | HITS-CLIP | 21258322\|25083871 | Unspecified |
| mmu-miR-3961 | MIMAT0019338 | 6030422M02Rik | 240697 | CLIP-Seq dataset analysis | 21258322 | Unspecified |
| mmu-miR-3961 | MIMAT0019338 | A630055G03Rik | 223970 | CLIP-Seq dataset analysis | 21258322 | Unspecified |
| mmu-miR-3961 | MIMAT0019338 | AK010878 | 100233175 | HITS-CLIP | 21258322 | Unspecified |
| mmu-miR-3961 | MIMAT0019338 | Akr1d1 | 208665 | HITS-CLIP | 21258322 | Unspecified |
| mmu-miR-3961 | MIMAT0019338 | Chrna1 | 11435 | HITS-CLIP | 21258322 | Unspecified |
| mmu-miR-3961 | MIMAT0019338 | Dclre1b | 140917 | HITS-CLIP | 21258322 | Unspecified |
| mmu-miR-3961 | MIMAT0019338 | Fam81a | 76886 | HITS-CLIP | 21258322 | Unspecified |
| mmu-miR-3961 | MIMAT0019338 | Fsd1l | 319636 | HITS-CLIP | 21258322 | Unspecified |
| mmu-miR-3961 | MIMAT0019338 | Ggps1 | 14593 | HITS-CLIP | 21258322 | Unspecified |
| mmu-miR-3961 | MIMAT0019338 | Gm5113 | 330503 | HITS-CLIP | 21258322 | Unspecified |
| mmu-miR-3961 | MIMAT0019338 | Gon7 | 100233175 | HITS-CLIP | 21258322 | Unspecified |
| mmu-miR-3961 | MIMAT0019338 | Mcmdc2 | 240697 | HITS-CLIP | 21258322 | Unspecified |
| mmu-miR-3961 | MIMAT0019338 | N4bp2l2 | 381695 | HITS-CLIP | 21258322 | Unspecified |
| mmu-miR-3961 | MIMAT0019338 | Nid1 | 18073 | HITS-CLIP | 25083871 | Unspecified |
| mmu-miR-3961 | MIMAT0019338 | Nufip1 | 27275 | HITS-CLIP | 21258322 | Unspecified |
| mmu-miR-3961 | MIMAT0019338 | Paxip1 | 55982 | HITS-CLIP | 23597149 | Unspecified |
| mmu-miR-3961 | MIMAT0019338 | Pigv | 230801 | HITS-CLIP | 21258322 | Unspecified |
| mmu-miR-3961 | MIMAT0019338 | Prpf31 | 68988 | HITS-CLIP | 21258322 | Unspecified |
| mmu-miR-3961 | MIMAT0019338 | Prrc1 | 73137 | HITS-CLIP | 21258322 | Unspecified |
| mmu-miR-3961 | MIMAT0019338 | Prrxl1 | 107751 | HITS-CLIP | 21258322 | Unspecified |
| mmu-miR-3961 | MIMAT0019338 | Rmi2 | 223970 | HITS-CLIP | 21258322 | Unspecified |
| mmu-miR-3961 | MIMAT0019338 | Rtkn2 | 170799 | HITS-CLIP | 21258322 | Unspecified |
| mmu-miR-3961 | MIMAT0019338 | Sgo2a | 68549 | HITS-CLIP | 21258322 | Unspecified |
| mmu-miR-3961 | MIMAT0019338 | Sgol2 | 68549 | HITS-CLIP | 21258322 | Unspecified |
| mmu-miR-3961 | MIMAT0019338 | Sh3pxd2a | 14218 | HITS-CLIP | 21258322\|23597149\|25083871 | Unspecified |
| mmu-miR-3961 | MIMAT0019338 | Shisa6 | 380702 | HITS-CLIP | 21258322 | Unspecified |
| mmu-miR-3961 | MIMAT0019338 | Slc12a8 | 171286 | HITS-CLIP | 21258322 | Unspecified |
| mmu-miR-3961 | MIMAT0019338 | Slc25a27 | 74011 | HITS-CLIP | 21258322\|23597149 | Unspecified |

| | | | | | | |
|---|---|---|---|---|---|---|
| mmu-miR-3961 | MIMAT0019338 | Sumf2 | 67902 | HITS-CLIP | | 21258322 Unspecified |
| mmu-miR-3961 | MIMAT0019338 | Tmem151a | 381199 | HITS-CLIP | | 21258322 Unspecified |
| mmu-miR-3961 | MIMAT0019338 | Tmem206 | 66950 | HITS-CLIP | 21258322\|25083871 | Unspecified |
| mmu-miR-3961 | MIMAT0019338 | Tmem88b | 320587 | HITS-CLIP | | 21258322 Unspecified |
| mmu-miR-3961 | MIMAT0019338 | Xpo7 | 65246 | HITS-CLIP | | 21258322 Unspecified |
| mmu-miR-3961 | MIMAT0019338 | Zfp1 | 22640 | HITS-CLIP | | 21258322 Unspecified |
| mmu-miR-3961 | MIMAT0019338 | Zfp287 | 170740 | HITS-CLIP | | 21258322 Unspecified |
| mmu-miR-3961 | MIMAT0019338 | Zfp790 | 233056 | HITS-CLIP | | 21258322 Unspecified |
| mmu-miR-3961 | MIMAT0019338 | Zfp941 | 407812 | HITS-CLIP | | 21258322 Unspecified |
| mmu-miR-125b-5p | MIMAT0000136 | 4632428N05Rik | 74048 | Luciferase assay//Western blot//Luciferase | | 22751012 Unspecified |
| mmu-miR-125b-5p | MIMAT0000136 | Abtb1 | 80283 | Microarray//Microarrays//Other | 16495412\|22469780 | Unspecified |
| mmu-miR-125b-5p | MIMAT0000136 | Ajuba | 16475 | B-globin reporter assay//Microarray | | 16495412 Unspecified |
| mmu-miR-125b-5p | MIMAT0000136 | Antxr2 | 71914 | HITS-CLIP | | 25083871 Unspecified |
| mmu-miR-125b-5p | MIMAT0000136 | Aph1c | 68318 | HITS-CLIP | | 21258322 Unspecified |
| mmu-miR-125b-5p | MIMAT0000136 | Apln | 30878 | Microarray//Other | | 16495412 Unspecified |
| mmu-miR-125b-5p | MIMAT0000136 | Arid3a | 13496 | Microarray//Microarrays//Other | 16495412\|22469780 | Unspecified |
| mmu-miR-125b-5p | MIMAT0000136 | Arid3b | 56380 | Microarray//Other | | 16495412 Unspecified |
| mmu-miR-125b-5p | MIMAT0000136 | Bak1 | 12018 | Luciferase assay//qRT-PCR//Western blot/ | 21368288\|21935352\|25083871\|26024 | Unspecified |
| mmu-miR-125b-5p | MIMAT0000136 | Bmf | 171543 | qRT-PCR | | 21118986 Unspecified |
| mmu-miR-125b-5p | MIMAT0000136 | Btg2 | 12227 | HITS-CLIP//Microarray//qRT-PCR | 37986224\|25753418\|33330617 | Unspecified |
| mmu-miR-125b-5p | MIMAT0000136 | Cbfb | 12400 | Luciferase assay//Luciferase reporter assay | | 22689670 Unspecified |
| mmu-miR-125b-5p | MIMAT0000136 | Ddx19b | 234733 | Microarray//Other | | 16495412 Unspecified |
| mmu-miR-125b-5p | MIMAT0000136 | Dqx1 | 93838 | HITS-CLIP | | 21258322 Unspecified |
| mmu-miR-125b-5p | MIMAT0000136 | Dus1l | 68730 | Microarray//HITS-CLIP//qCLASH//Microarr | 16495412\|23597149\|25083871\|3139 | Unspecified |
| mmu-miR-125b-5p | MIMAT0000136 | Entpd4 | 67464 | Microarray//Microarrays | 16495412\|22469780 | Unspecified |
| mmu-miR-125b-5p | MIMAT0000136 | Foxk1 | 17425 | HITS-CLIP | | 25083871 Unspecified |
| mmu-miR-125b-5p | MIMAT0000136 | Gm14137 | 623781 | HITS-CLIP | | 21258322 Unspecified |
| mmu-miR-125b-5p | MIMAT0000136 | Hif1an | 319594 | HITS-CLIP | | 23142080 Unspecified |
| mmu-miR-125b-5p | MIMAT0000136 | Igf2 | 16002 | Northern blot//qRT-PCR//Western blot | | 21200031 Unspecified |
| mmu-miR-125b-5p | MIMAT0000136 | Il1rn | 16181 | HITS-CLIP | 21258322\|23142080 | Unspecified |
| mmu-miR-125b-5p | MIMAT0000136 | Jmy | 57748 | HITS-CLIP | | 21258322 Unspecified |
| mmu-miR-125b-5p | MIMAT0000136 | Jub | 16475 | B-globin reporter assay//Microarray//Othe | | 16495412 Unspecified |
| mmu-miR-125b-5p | MIMAT0000136 | Jun | 16476 | Luciferase assay//qRT-PCR//Western blot/ | | 21368288 Unspecified |
| mmu-miR-125b-5p | MIMAT0000136 | Jund | 16478 | Luciferase assay//qRT-PCR//Western blot/ | | 21368288 Unspecified |
| mmu-miR-125b-5p | MIMAT0000136 | Klf13 | 50794 | qRT-PCR | | 21118986 Unspecified |
| mmu-miR-125b-5p | MIMAT0000136 | Lars2 | 102436 | HITS-CLIP | 23597149\|21258322 | Unspecified |
| mmu-miR-125b-5p | MIMAT0000136 | Lin28a | 83557 | Luciferase assay//qRT-PCR//Western blot/ | 22277001\|16495412\|16227573 | Unspecified |
| mmu-miR-125b-5p | MIMAT0000136 | Madd | 228355 | HITS-CLIP | | 23597149 Unspecified |
| mmu-miR-125b-5p | MIMAT0000136 | Map2k7 | 26400 | B-globin reporter assay//Microarray//HITS | 16495412\|23597149 | Unspecified |

| | | | | | | | |
|---|---|---|---|---|---|---|---|
| mmu-miR-125b-5p | MIMAT0000136 | Mgat4a | 269181 | HITS-CLIP | | 25083871 | Unspecified |
| mmu-miR-125b-5p | MIMAT0000136 | Mtf1 | 17764 | HITS-CLIP | | 25083871 | Unspecified |
| mmu-miR-125b-5p | MIMAT0000136 | Ppp1ca | 19045 | Luciferase assay//qRT-PCR//Western blot/ | | 21935352 | Unspecified |
| mmu-miR-125b-5p | MIMAT0000136 | Ppt2 | 54397 | Microarray//Other | | 16495412 | Unspecified |
| mmu-miR-125b-5p | MIMAT0000136 | Prkra | 23992 | Luciferase assay//qRT-PCR//Western blot/ | | 21935352 | Unspecified |
| mmu-miR-125b-5p | MIMAT0000136 | RYBP | 23429 | Luciferase reporter assay//qRT-PCR//West | | 28111200 | Unspecified |
| mmu-miR-125b-5p | MIMAT0000136 | Rabl6 | 227624 | Microarray//HITS-CLIP//Other | 16495412\|25083871 | | Unspecified |
| mmu-miR-125b-5p | MIMAT0000136 | Rhebl1 | 69159 | Microarray//Microarrays//HITS-CLIP//Othe | 16495412\|22469780\|25083871 | | Unspecified |
| mmu-miR-125b-5p | MIMAT0000136 | Smo | 319757 | Luciferase assay//Western blot//Luciferase | | 20216554 | Unspecified |
| mmu-miR-125b-5p | MIMAT0000136 | Snai1 | 20613 | Luciferase assay//Western blot//Luciferase | | 21945074 | Unspecified |
| mmu-miR-125b-5p | MIMAT0000136 | Snrnp40 | 66585 | HITS-CLIP//Microarrays | 25083871\|22469780 | | Unspecified |
| mmu-miR-125b-5p | MIMAT0000136 | Sox11 | 20666 | HITS-CLIP//Western blotting//qRT-PCR | 37986224\|25083871\|33584192 | | Unspecified |
| mmu-miR-125b-5p | MIMAT0000136 | Stat1 | 20846 | HITS-CLIP | | 21258322 | Unspecified |
| mmu-miR-125b-5p | MIMAT0000136 | Stat3 | 20848 | Luciferase assay//qRT-PCR//Western blot/ | 21368288\|28108732 | | Unspecified |
| mmu-miR-125b-5p | MIMAT0000136 | Suv39h1 | 20937 | Luciferase assay//Western blot//HITS-CLIP | 20699419\|25083871\|22469780 | | Unspecified |
| mmu-miR-125b-5p | MIMAT0000136 | Tnf | 21926 | GFP reporter assay//Luciferase assay//Luci | | 17911593 | Unspecified |
| mmu-miR-125b-5p | MIMAT0000136 | Tor2a | 30933 | Microarray//Microarrays//qCLASH//Other | 16495412\|22469780\|31393953 | | Unspecified |
| mmu-miR-125b-5p | MIMAT0000136 | Triap1 | 69076 | HITS-CLIP//Luciferase reporter assay//Wes | 37986224\|33921710 | | Unspecified |
| mmu-miR-125b-5p | MIMAT0000136 | Trim71 | 636931 | HITS-CLIP//qRT-PCR//Other | 25083871\|19898466 | | Unspecified |
| mmu-miR-125b-5p | MIMAT0000136 | Tspan12 | 269831 | HITS-CLIP | | 25083871 | Unspecified |
| mmu-miR-125b-5p | MIMAT0000136 | Vps4b | 20479 | Luciferase assay//HITS-CLIP//Luciferase re | 25403182\|25083871 | | Unspecified |
| mmu-miR-125b-5p | MIMAT0000136 | Wdr25 | 212198 | HITS-CLIP | | 21258322 | Unspecified |
| mmu-miR-125b-5p | MIMAT0000136 | Zfp385a | 29813 | Microarray//HITS-CLIP//Other | 16495412\|25083871 | | Unspecified |
| mmu-miR-3095-5p | MIMAT0014911 | 4933427D06Rik | 232217 | HITS-CLIP | | 25083871 | Unspecified |
| mmu-miR-3095-5p | MIMAT0014911 | Casp8 | 12370 | HITS-CLIP | | 25083871 | Unspecified |
| mmu-miR-3095-5p | MIMAT0014911 | Cdyl2 | 75796 | HITS-CLIP | | 25083871 | Unspecified |
| mmu-miR-3095-5p | MIMAT0014911 | Csmd1 | 94109 | HITS-CLIP | | 23142080 | Unspecified |
| mmu-miR-3095-5p | MIMAT0014911 | Frmd6 | 319710 | HITS-CLIP | 25083871\|23142080 | | Unspecified |
| mmu-miR-3095-5p | MIMAT0014911 | Gprc5b | 64297 | HITS-CLIP | | 21258322 | Unspecified |
| mmu-miR-3095-5p | MIMAT0014911 | Kcnj15 | 16516 | HITS-CLIP | | 25083871 | Unspecified |
| mmu-miR-3095-5p | MIMAT0014911 | Kitl | 17311 | HITS-CLIP | | 25083871 | Unspecified |
| mmu-miR-3095-5p | MIMAT0014911 | Maml3 | 433586 | HITS-CLIP | | 23142080 | Unspecified |
| mmu-miR-3095-5p | MIMAT0014911 | Neurl1b | 240055 | HITS-CLIP | | 21258322 | Unspecified |
| mmu-miR-3095-5p | MIMAT0014911 | Pik3r1 | 18708 | HITS-CLIP | | 25083871 | Unspecified |
| mmu-miR-3095-5p | MIMAT0014911 | Pknox1 | 18771 | HITS-CLIP | | 23142080 | Unspecified |
| mmu-miR-3095-5p | MIMAT0014911 | Rap2a | 76108 | HITS-CLIP | 23597149\|19536157 | | Unspecified |
| mmu-miR-3095-5p | MIMAT0014911 | Tmem132c | 208213 | HITS-CLIP | | 23142080 | Unspecified |
| mmu-miR-3095-5p | MIMAT0014911 | Tmem65 | 74868 | HITS-CLIP | 26203562\|25083871\|21258322 | | Unspecified |
| mmu-miR-3079-3p | MIMAT0014867 | 2510049J12Rik | 70291 | HITS-CLIP | | 21258322 | Unspecified |

| | | | | | | | |
|---|---|---|---|---|---|---|---|
| mmu-miR-3079-3p | MIMAT0014867 | Armc2 | 213402 | HITS-CLIP | | 21258322 | Unspecified |
| mmu-miR-3079-3p | MIMAT0014867 | Clns1a | 12729 | HITS-CLIP | | 21258322 | Unspecified |
| mmu-miR-3079-3p | MIMAT0014867 | Gm347 | 241289 | CLIP-Seq dataset analysis | 25083871\|23142080\|23597149\|2125E | | Unspecified |
| mmu-miR-3079-3p | MIMAT0014867 | Mkrn2os | 70291 | HITS-CLIP | | 21258322 | Unspecified |
| mmu-miR-3079-3p | MIMAT0014867 | Pkia | 18767 | HITS-CLIP | | 25083871 | Unspecified |
| mmu-miR-3079-3p | MIMAT0014867 | Ppp1r26 | 241289 | HITS-CLIP | 25083871\|21258322\|23142080\|23597 | | Unspecified |
| mmu-miR-3079-3p | MIMAT0014867 | Ptp4a3 | 19245 | HITS-CLIP | | 21258322 | Unspecified |
| mmu-miR-3079-3p | MIMAT0014867 | Slc35e1 | 270066 | HITS-CLIP | | 21258322 | Unspecified |
| mmu-miR-3079-3p | MIMAT0014867 | Tyw1 | 100929 | HITS-CLIP | | 21258322 | Unspecified |
| mmu-miR-3079-3p | MIMAT0014867 | Utp23 | 78581 | HITS-CLIP | | 21258322 | Unspecified |
| mmu-miR-3079-3p | MIMAT0014867 | Zfp11 | 22648 | HITS-CLIP | | 21258322 | Unspecified |
| mmu-miR-509-3p | MIMAT0004891 | Fmo5 | 14263 | HITS-CLIP | | 23597149 | Unspecified |
| mmu-miR-509-3p | MIMAT0004891 | Frmpd4 | 333605 | HITS-CLIP | | 25083871 | Unspecified |
| mmu-miR-509-3p | MIMAT0004891 | MAP3K8 | 1326 | Luciferase reporter assay | | 27001999 | Unspecified |
| mmu-miR-509-3p | MIMAT0004891 | Mrpl41 | 107733 | HITS-CLIP | | 21258322 | Unspecified |
| mmu-miR-509-3p | MIMAT0004891 | Ptpn3 | 545622 | HITS-CLIP | | 21258322 | Unspecified |
| mmu-miR-3068-5p | MIMAT0014842 | 1200014J11Rik | 66874 | HITS-CLIP | | 21258322 | Unspecified |
| mmu-miR-3068-5p | MIMAT0014842 | Arhgef39 | 230098 | HITS-CLIP | | 21258322 | Unspecified |
| mmu-miR-3068-5p | MIMAT0014842 | Ddx19b | 234733 | HITS-CLIP | | 21258322 | Unspecified |
| mmu-miR-3068-5p | MIMAT0014842 | Ggps1 | 14593 | HITS-CLIP | | 21258322 | Unspecified |
| mmu-miR-3068-5p | MIMAT0014842 | Iffo2 | 212632 | HITS-CLIP | | 23597149 | Unspecified |
| mmu-miR-3068-5p | MIMAT0014842 | Ikbkg | 16151 | HITS-CLIP | 25083871\|21258322 | | Unspecified |
| mmu-miR-3068-5p | MIMAT0014842 | Mcc | 328949 | HITS-CLIP | | 25083871 | Unspecified |
| mmu-miR-3068-5p | MIMAT0014842 | Msl1 | 74026 | HITS-CLIP | 25083871\|23597149 | | Unspecified |
| mmu-miR-3068-5p | MIMAT0014842 | Ncbp3 | 66874 | HITS-CLIP | | 21258322 | Unspecified |
| mmu-miR-3068-5p | MIMAT0014842 | Rnf169 | 108937 | HITS-CLIP | | 23142080 | Unspecified |
| mmu-miR-3068-5p | MIMAT0014842 | Samd7 | 75953 | HITS-CLIP | | 21258322 | Unspecified |
| mmu-miR-3092-3p | MIMAT0014906 | 0610030E20Rik | 68364 | HITS-CLIP | | 25083871 | Unspecified |
| mmu-miR-3092-3p | MIMAT0014906 | B4galnt2 | 14422 | HITS-CLIP | | 21258322 | Unspecified |
| mmu-miR-3092-3p | MIMAT0014906 | Cadm2 | 239857 | HITS-CLIP | | 21258322 | Unspecified |
| mmu-miR-3092-3p | MIMAT0014906 | Eif4g2 | 13690 | HITS-CLIP | | 21258322 | Unspecified |
| mmu-miR-3092-3p | MIMAT0014906 | Fubp1 | 51886 | HITS-CLIP | | 25083871 | Unspecified |
| mmu-miR-3092-3p | MIMAT0014906 | Gm10778 | 100233208 | HITS-CLIP | | 21258322 | Unspecified |
| mmu-miR-3092-3p | MIMAT0014906 | Gm5113 | 330503 | HITS-CLIP | | 21258322 | Unspecified |
| mmu-miR-3092-3p | MIMAT0014906 | Klhdc9 | 68874 | HITS-CLIP | | 21258322 | Unspecified |
| mmu-miR-3092-3p | MIMAT0014906 | Lrrc39 | 109245 | HITS-CLIP | | 21258322 | Unspecified |
| mmu-miR-3092-3p | MIMAT0014906 | Onecut2 | 225631 | HITS-CLIP | | 23142080 | Unspecified |
| mmu-miR-3092-3p | MIMAT0014906 | Padi2 | 18600 | HITS-CLIP | | 21258322 | Unspecified |
| mmu-miR-3092-3p | MIMAT0014906 | Psmb11 | 73902 | HITS-CLIP | | 21258322 | Unspecified |

| | | | | | | |
|---|---|---|---|---|---|---|
| mmu-miR-3092-3p | MIMAT0014906 | Sema6d | 214968 | HITS-CLIP | 21258322 | Unspecified |
| mmu-miR-3092-3p | MIMAT0014906 | Slc12a8 | 171286 | HITS-CLIP | 21258322 | Unspecified |
| mmu-miR-3092-3p | MIMAT0014906 | Snhg11 | 319317 | HITS-CLIP | 21258322 | Unspecified |
| mmu-miR-3092-3p | MIMAT0014906 | Syt11 | 229521 | HITS-CLIP | 23142080 | Unspecified |
| mmu-miR-3092-3p | MIMAT0014906 | TDRD7 | 23424 | Microarray | 33665188 | Unspecified |
| mmu-miR-3092-3p | MIMAT0014906 | Zfp287 | 170740 | HITS-CLIP | 21258322 | Unspecified |
| mmu-miR-3092-3p | MIMAT0014906 | Zfp300 | 245368 | HITS-CLIP | 25083871 | Unspecified |
| mmu-miR-3092-3p | MIMAT0014906 | Zfp617 | 170938 | HITS-CLIP | 23142080 | Unspecified |
| mmu-miR-384-3p | MIMAT0001076 | 1600029D21Rik | 76509 | HITS-CLIP | 25083871\|23597149\|23142080 | Unspecified |
| mmu-miR-384-3p | MIMAT0001076 | AK010878 | 100233175 | HITS-CLIP | 21258322 | Unspecified |
| mmu-miR-384-3p | MIMAT0001076 | Atp2a2 | 11938 | HITS-CLIP | 25083871 | Unspecified |
| mmu-miR-384-3p | MIMAT0001076 | Atp2b1 | 67972 | HITS-CLIP | 26203562\|23142080 | Unspecified |
| mmu-miR-384-3p | MIMAT0001076 | Bach2 | 12014 | HITS-CLIP | 23142080 | Unspecified |
| mmu-miR-384-3p | MIMAT0001076 | Crkl | 12929 | HITS-CLIP | 21258322 | Unspecified |
| mmu-miR-384-3p | MIMAT0001076 | Cyld | 74256 | HITS-CLIP | 23142080 | Unspecified |
| mmu-miR-384-3p | MIMAT0001076 | Dlgap1 | 224997 | HITS-CLIP | 23142080 | Unspecified |
| mmu-miR-384-3p | MIMAT0001076 | Fam55b | 78252 | CLIP-Seq dataset analysis | 21258322 | Unspecified |
| mmu-miR-384-3p | MIMAT0001076 | Fbxo32 | 67731 | HITS-CLIP | 25083871\|23142080 | Unspecified |
| mmu-miR-384-3p | MIMAT0001076 | Gm14137 | 623781 | HITS-CLIP | 21258322 | Unspecified |
| mmu-miR-384-3p | MIMAT0001076 | Gon7 | 100233175 | HITS-CLIP | 21258322 | Unspecified |
| mmu-miR-384-3p | MIMAT0001076 | Gpi1 | 14751 | HITS-CLIP | 21258322 | Unspecified |
| mmu-miR-384-3p | MIMAT0001076 | Hdac3 | 15183 | ChIP-seq//Immunofluorescence//Luciferas | 24619412 | Unspecified |
| mmu-miR-384-3p | MIMAT0001076 | Hk2 | 15277 | HITS-CLIP | 21258322 | Unspecified |
| mmu-miR-384-3p | MIMAT0001076 | Hoxb9 | 15417 | HITS-CLIP | 23142080 | Unspecified |
| mmu-miR-384-3p | MIMAT0001076 | Lhfpl4 | 269788 | HITS-CLIP | 23597149 | Unspecified |
| mmu-miR-384-3p | MIMAT0001076 | Mbtps2 | 270669 | HITS-CLIP | 19536157\|23597149\|23142080\|2125 | Unspecified |
| mmu-miR-384-3p | MIMAT0001076 | Moap1 | 64113 | HITS-CLIP | 21258322 | Unspecified |
| mmu-miR-384-3p | MIMAT0001076 | Myo1c | 17913 | HITS-CLIP | 21258322 | Unspecified |
| mmu-miR-384-3p | MIMAT0001076 | Nol3 | 78688 | HITS-CLIP | 21258322 | Unspecified |
| mmu-miR-384-3p | MIMAT0001076 | Nxpe2 | 78252 | HITS-CLIP | 21258322 | Unspecified |
| mmu-miR-384-3p | MIMAT0001076 | Pbld1 | 68371 | HITS-CLIP | 21258322 | Unspecified |
| mmu-miR-384-3p | MIMAT0001076 | Plet1 | 76509 | HITS-CLIP | 23142080\|23597149\|25083871 | Unspecified |
| mmu-miR-384-3p | MIMAT0001076 | Syt13 | 80976 | HITS-CLIP | 21258322 | Unspecified |
| mmu-miR-384-3p | MIMAT0001076 | TDRD7 | 23424 | Microarray | 33665188 | Unspecified |
| mmu-miR-384-3p | MIMAT0001076 | Tacc2 | 57752 | HITS-CLIP | 23142080\|25083871 | Unspecified |
| mmu-miR-384-3p | MIMAT0001076 | Tceal8 | 66684 | HITS-CLIP | 21258322 | Unspecified |
| mmu-miR-384-3p | MIMAT0001076 | Tspan13 | 66109 | HITS-CLIP | 23142080 | Unspecified |
| mmu-miR-28c | MIMAT0019339 | 2010109I03Rik | 67038 | HITS-CLIP | 21258322 | Unspecified |
| mmu-miR-28c | MIMAT0019339 | Ccdc78 | 381077 | HITS-CLIP | 21258322\|25083871 | Unspecified |

| | | | | | | |
|---|---|---|---|---|---|---|
| mmu-miR-28c | MIMAT0019339 | Nr3c1 | 14815 | HITS-CLIP | 23142080 | Unspecified |
| mmu-miR-28c | MIMAT0019339 | Pld5 | 319455 | HITS-CLIP | 23142080 | Unspecified |
| mmu-miR-28c | MIMAT0019339 | Prss43 | 272643 | HITS-CLIP | 21258322 | Unspecified |
| mmu-miR-28c | MIMAT0019339 | Slc7a2 | 11988 | HITS-CLIP | 21258322 | Unspecified |
| mmu-miR-28c | MIMAT0019339 | Slx1b | 75764 | HITS-CLIP | 21258322 | Unspecified |
| mmu-miR-28c | MIMAT0019339 | Tars2 | 71807 | HITS-CLIP | 21258322 | Unspecified |
| mmu-miR-28c | MIMAT0019339 | Timm17b | 21855 | HITS-CLIP | 21258322\|25083871 | Unspecified |
| mmu-miR-28c | MIMAT0019339 | Tmppe | 100504715 | HITS-CLIP | 21258322 | Unspecified |
| mmu-miR-28c | MIMAT0019339 | Tns4 | 217169 | HITS-CLIP | 19536157 | Unspecified |
| mmu-miR-147-3p | MIMAT0004857 | Slco2a1 | 24059 | HITS-CLIP | 21258322 | Unspecified |
| mmu-miR-721 | MIMAT0003515 | Acbd5 | 74159 | HITS-CLIP | 25083871 | Unspecified |
| mmu-miR-721 | MIMAT0003515 | Arrdc3 | 105171 | HITS-CLIP | 25083871 | Unspecified |
| mmu-miR-721 | MIMAT0003515 | Fgfr1op2 | 67529 | HITS-CLIP | 23142080 | Unspecified |
| mmu-miR-721 | MIMAT0003515 | Homez | 239099 | HITS-CLIP | 21258322 | Unspecified |
| mmu-miR-721 | MIMAT0003515 | Inpp5e | 64436 | HITS-CLIP | 21258322 | Unspecified |
| mmu-miR-721 | MIMAT0003515 | Kdm2a | 225876 | HITS-CLIP | 25083871 | Unspecified |
| mmu-miR-721 | MIMAT0003515 | Map3k12 | 26404 | HITS-CLIP | 21258322 | Unspecified |
| mmu-miR-721 | MIMAT0003515 | Meox2 | 17286 | Luciferase assay//qRT-PCR//Western blot/ | 21941297 | Unspecified |
| mmu-miR-721 | MIMAT0003515 | Mup13 | 100039089 | HITS-CLIP | 23597149 | Unspecified |
| mmu-miR-721 | MIMAT0003515 | Mup7 | 100041658 | HITS-CLIP | 23597149 | Unspecified |
| mmu-miR-721 | MIMAT0003515 | Nfia | 18027 | HITS-CLIP | 23142080 | Unspecified |
| mmu-miR-721 | MIMAT0003515 | Snx27 | 76742 | HITS-CLIP | 25083871 | Unspecified |
| mmu-miR-721 | MIMAT0003515 | Timp2 | 21858 | HITS-CLIP | 23142080 | Unspecified |
| mmu-miR-721 | MIMAT0003515 | Tmem25 | 71687 | HITS-CLIP | 25083871 | Unspecified |
| mmu-miR-721 | MIMAT0003515 | Ubap2l | 74383 | HITS-CLIP | 25083871 | Unspecified |
| mmu-miR-383-5p | MIMAT0000748 | 9230110C19Rik | 234912 | HITS-CLIP | 21258322 | Unspecified |
| mmu-miR-383-5p | MIMAT0000748 | Disp2 | 214240 | HITS-CLIP | 23142080 | Unspecified |
| mmu-miR-383-5p | MIMAT0000748 | Kcnj1 | 56379 | HITS-CLIP | 25083871 | Unspecified |
| mmu-miR-383-5p | MIMAT0000748 | Lin28a | 83557 | HITS-CLIP | 21258322 | Unspecified |
| mmu-miR-383-5p | MIMAT0000748 | Rufy2 | 70432 | HITS-CLIP | 21258322 | Unspecified |
| mmu-miR-383-5p | MIMAT0000748 | Sptbn1 | 20742 | HITS-CLIP | 23142080 | Unspecified |
| mmu-miR-3079-5p | MIMAT0014866 | Ddhd2 | 72108 | HITS-CLIP | 21258322 | Unspecified |
| mmu-miR-3079-5p | MIMAT0014866 | Ddx26b | 236790 | HITS-CLIP | 25083871 | Unspecified |
| mmu-miR-3079-5p | MIMAT0014866 | Exoc8 | 102058 | HITS-CLIP | 21258322 | Unspecified |
| mmu-miR-3079-5p | MIMAT0014866 | Fadd | 14082 | HITS-CLIP | 21258322 | Unspecified |
| mmu-miR-3079-5p | MIMAT0014866 | Fam65b | 193385 | HITS-CLIP | 21258322 | Unspecified |
| mmu-miR-3079-5p | MIMAT0014866 | Ints6l | 236790 | HITS-CLIP | 25083871 | Unspecified |
| mmu-miR-3079-5p | MIMAT0014866 | Opcml | 330908 | HITS-CLIP | 23142080 | Unspecified |
| mmu-miR-3079-5p | MIMAT0014866 | Pigy | 66459 | CLIP-Seq dataset analysis | 21258322 | Unspecified |

| | | | | | | | |
|---|---|---|---|---|---|---|---|
| mmu-miR-3079-5p | MIMAT0014866 | Pigyl | 66268 | HITS-CLIP | | 21258322 | Unspecified |
| mmu-miR-3079-5p | MIMAT0014866 | Ppp4r2 | 232314 | HITS-CLIP | | 25083871 | Unspecified |
| mmu-miR-3079-5p | MIMAT0014866 | Ptk6 | 20459 | HITS-CLIP | | 21258322 | Unspecified |
| mmu-miR-3079-5p | MIMAT0014866 | Rg9mtd2 | 108943 | CLIP-Seq dataset analysis | 25083871\|21258322 | | Unspecified |
| mmu-miR-3079-5p | MIMAT0014866 | Rhod | 11854 | HITS-CLIP | | 28735896 | Unspecified |
| mmu-miR-3079-5p | MIMAT0014866 | Ripor2 | 193385 | HITS-CLIP | | 21258322 | Unspecified |
| mmu-miR-3079-5p | MIMAT0014866 | Trmt10a | 108943 | HITS-CLIP | 21258322\|25083871 | | Unspecified |
| mmu-miR-3079-5p | MIMAT0014866 | Vcan | 13003 | HITS-CLIP | | 21258322 | Unspecified |
| mmu-miR-3079-5p | MIMAT0014866 | Zfp830 | 66983 | HITS-CLIP | | 21258322 | Unspecified |
| mmu-miR-3079-5p | MIMAT0014866 | Zfp935 | 71508 | HITS-CLIP | | 25083871 | Unspecified |
| mmu-miR-125b-2-3p | MIMAT0004529 | Ccna2 | 12428 | HITS-CLIP//Luciferase reporter assay | 31433990\|27020049 | | Unspecified |
| mmu-miR-125b-2-3p | MIMAT0004529 | Cercam | 99151 | HITS-CLIP | | 21258322 | Unspecified |
| mmu-miR-125b-2-3p | MIMAT0004529 | Chrnb4 | 108015 | HITS-CLIP | | 21258322 | Unspecified |
| mmu-miR-125b-2-3p | MIMAT0004529 | Ddx3x | 13205 | HITS-CLIP | 23597149\|29167373 | | Unspecified |
| mmu-miR-125b-2-3p | MIMAT0004529 | Dlgap4 | 228836 | HITS-CLIP | | 23142080 | Unspecified |
| mmu-miR-125b-2-3p | MIMAT0004529 | Eef2k | 13631 | Luciferase reporter assay | | 27020049 | Unspecified |
| mmu-miR-125b-2-3p | MIMAT0004529 | Egr2 | 13654 | HITS-CLIP | | 21258322 | Unspecified |
| mmu-miR-125b-2-3p | MIMAT0004529 | Fermt2 | 218952 | HITS-CLIP | | 23597149 | Unspecified |
| mmu-miR-125b-2-3p | MIMAT0004529 | Mrpl45 | 67036 | HITS-CLIP | | 21258322 | Unspecified |
| mmu-miR-125b-2-3p | MIMAT0004529 | Nphp3 | 74025 | HITS-CLIP | | 25083871 | Unspecified |
| mmu-miR-125b-2-3p | MIMAT0004529 | Plekhm3 | 241075 | HITS-CLIP | | 25083871 | Unspecified |
| mmu-miR-125b-2-3p | MIMAT0004529 | Serpine1 | 18787 | HITS-CLIP | 23142080\|23597149 | | Unspecified |
| mmu-miR-125b-2-3p | MIMAT0004529 | Slc35d2 | 70484 | HITS-CLIP | | 21258322 | Unspecified |
| mmu-miR-125b-2-3p | MIMAT0004529 | Sumf2 | 67902 | HITS-CLIP | | 21258322 | Unspecified |
| mmu-miR-125b-2-3p | MIMAT0004529 | Tecpr2 | 104859 | HITS-CLIP | | 25083871 | Unspecified |
| mmu-miR-125b-2-3p | MIMAT0004529 | Utp23 | 78581 | HITS-CLIP | | 21258322 | Unspecified |
| mmu-miR-3964 | MIMAT0019344 | Abce1 | 24015 | HITS-CLIP | | 23142080 | Unspecified |
| mmu-miR-3964 | MIMAT0019344 | Camk4 | 12326 | HITS-CLIP | | 23142080 | Unspecified |
| mmu-miR-3964 | MIMAT0019344 | Csrnp3 | 77771 | HITS-CLIP | | 25083871 | Unspecified |
| mmu-miR-3964 | MIMAT0019344 | Dcun1d1 | 114893 | HITS-CLIP | | 25083871 | Unspecified |
| mmu-miR-3964 | MIMAT0019344 | Fsd1l | 319636 | HITS-CLIP | | 21258322 | Unspecified |
| mmu-miR-3964 | MIMAT0019344 | Glod4 | 67201 | HITS-CLIP | | 23142080 | Unspecified |
| mmu-miR-3964 | MIMAT0019344 | Gm4951 | 240327 | HITS-CLIP | | 21258322 | Unspecified |
| mmu-miR-3964 | MIMAT0019344 | Herpud2 | 80517 | HITS-CLIP | | 23142080 | Unspecified |
| mmu-miR-3964 | MIMAT0019344 | Ptpn11 | 19247 | HITS-CLIP | | 25083871 | Unspecified |
| mmu-miR-3964 | MIMAT0019344 | Rab11fip1 | 75767 | HITS-CLIP | 19536157\|23597149\|25083871 | | Unspecified |
| mmu-miR-3964 | MIMAT0019344 | Rerg | 232441 | HITS-CLIP | | 25083871 | Unspecified |
| mmu-miR-3964 | MIMAT0019344 | Rragb | 245670 | HITS-CLIP | | 25083871 | Unspecified |
| mmu-miR-1900 | MIMAT0007870 | Elovl2 | 54326 | HITS-CLIP | | 21258322 | Unspecified |

| | | | | | | |
|---|---|---|---|---|---|---|
| mmu-miR-1900 | MIMAT0007870 | Ttc38 | 239570 | HITS-CLIP | 21258322 | Unspecified |
| mmu-miR-1900 | MIMAT0007870 | Ubxn2b | 68053 | HITS-CLIP | 25083871\|21258322\|23597149 | Unspecified |
| mmu-miR-100-5p | MIMAT0000655 | Asxl2 | 75302 | HITS-CLIP | 21258322 | Unspecified |
| mmu-miR-100-5p | MIMAT0000655 | Comp | 12845 | HITS-CLIP | 21258322 | Unspecified |
| mmu-miR-100-5p | MIMAT0000655 | Grik3 | 14807 | HITS-CLIP | 23597149 | Unspecified |
| mmu-miR-100-5p | MIMAT0000655 | MTOR | 2475 | Immunohistochemistry//Microarray//qRT- | 21339483 | Unspecified |
| mmu-miR-100-5p | MIMAT0000655 | Slc35d2 | 70484 | HITS-CLIP | 21258322 | Unspecified |
| mmu-miR-3086-5p | MIMAT0014880 | Bach2 | 12014 | HITS-CLIP | 23142080 | Unspecified |
| mmu-miR-3086-5p | MIMAT0014880 | Bri3bp | 76809 | HITS-CLIP | 21258322 | Unspecified |
| mmu-miR-3086-5p | MIMAT0014880 | Fmo5 | 14263 | HITS-CLIP | 23597149\|21258322 | Unspecified |
| mmu-miR-3086-5p | MIMAT0014880 | Gpr35 | 64095 | HITS-CLIP | 23142080 | Unspecified |
| mmu-miR-3086-5p | MIMAT0014880 | Mterf4 | 69821 | HITS-CLIP | 21258322 | Unspecified |
| mmu-miR-3086-5p | MIMAT0014880 | Mterfd2 | 69821 | HITS-CLIP | 21258322 | Unspecified |
| mmu-miR-3086-5p | MIMAT0014880 | Sfmbt2 | 353282 | HITS-CLIP | 25083871 | Unspecified |
| mmu-miR-3086-5p | MIMAT0014880 | Thsd7a | 330267 | HITS-CLIP | 25083871 | Unspecified |
| mmu-miR-3086-5p | MIMAT0014880 | Vezt | 215008 | HITS-CLIP | 23142080 | Unspecified |
| mmu-miR-1306-3p | MIMAT0009411 | Suds3 | 71954 | HITS-CLIP | 21258322 | Unspecified |
| mmu-miR-378b | MIMAT0019348 | Bmp2 | 12156 | HITS-CLIP | 23142080 | Unspecified |
| mmu-miR-378b | MIMAT0019348 | Dcaf6 | 74106 | HITS-CLIP | 23142080 | Unspecified |
| mmu-miR-378b | MIMAT0019348 | LOC100048885 | 100048885 | CLIP-Seq dataset analysis | 23597149 | Unspecified |
| mmu-miR-378b | MIMAT0019348 | Mup10 | 100039008 | HITS-CLIP | 23597149 | Unspecified |
| mmu-miR-378b | MIMAT0019348 | Mup11 | 100039028 | HITS-CLIP | 23597149 | Unspecified |
| mmu-miR-378b | MIMAT0019348 | Mup12 | 100039054 | HITS-CLIP | 23597149 | Unspecified |
| mmu-miR-378b | MIMAT0019348 | Mup14 | 100039116 | HITS-CLIP | 23597149 | Unspecified |
| mmu-miR-378b | MIMAT0019348 | Mup15 | 100039150 | HITS-CLIP | 23597149 | Unspecified |
| mmu-miR-378b | MIMAT0019348 | Mup17 | 100039206 | HITS-CLIP | 23597149 | Unspecified |
| mmu-miR-378b | MIMAT0019348 | Mup19 | 100189605 | HITS-CLIP | 23597149 | Unspecified |
| mmu-miR-378b | MIMAT0019348 | Ncoa7 | 211329 | HITS-CLIP | 25083871 | Unspecified |
| mmu-miR-378b | MIMAT0019348 | Pi15 | 94227 | HITS-CLIP | 21258322 | Unspecified |
| mmu-miR-378b | MIMAT0019348 | TDRD7 | 23424 | Microarray | 33665188 | Unspecified |
| mmu-miR-378b | MIMAT0019348 | Tmcc3 | 319880 | HITS-CLIP | 21258322 | Unspecified |
| mmu-miR-708-5p | MIMAT0004828 | Ccdc50 | 67501 | HITS-CLIP | 23597149\|21258322 | Unspecified |
| mmu-miR-708-5p | MIMAT0004828 | Elovl1 | 54325 | HITS-CLIP | 23597149 | Unspecified |
| mmu-miR-708-5p | MIMAT0004828 | Gjc3 | 118446 | HITS-CLIP | 25083871 | Unspecified |
| mmu-miR-708-5p | MIMAT0004828 | Nav2 | 78286 | HITS-CLIP | 25083871 | Unspecified |
| mmu-miR-708-5p | MIMAT0004828 | Pi15 | 94227 | HITS-CLIP | 21258322 | Unspecified |
| mmu-miR-708-5p | MIMAT0004828 | Prlr | 19116 | HITS-CLIP | 21258322 | Unspecified |
| mmu-miR-708-5p | MIMAT0004828 | Rasal2 | 226525 | HITS-CLIP | 21258322\|23597149 | Unspecified |
| mmu-miR-29b-1-5p | MIMAT0004523 | Acta2 | 11475 | Immunohistochemistry//qRT-PCR//Wester | 26524535 | Unspecified |

| | | | | | | |
|---|---|---|---|---|---|---|
| mmu-miR-29b-1-5p | MIMAT0004523 | Akt1 | 11651 | Immunohistochemistry//qRT-PCR//Wester | 26524535 | Unspecified |
| mmu-miR-29b-1-5p | MIMAT0004523 | Arg1 | 11846 | Microarray//Quantitative proteomic appro | 19878148 | Unspecified |
| mmu-miR-29b-1-5p | MIMAT0004523 | Col1a2 | 12843 | HITS-CLIP//Immunohistochemistry//qRT-P | 25083871\|26524535 | Unspecified |
| mmu-miR-29b-1-5p | MIMAT0004523 | Cxcl12 | 20315 | HITS-CLIP//Luciferase reporter assay//Wes | 37986224\|25083871\|32395570 | Unspecified |
| mmu-miR-29b-1-5p | MIMAT0004523 | Fn1 | 14268 | Immunohistochemistry//qRT-PCR//Wester | 26524535 | Unspecified |
| mmu-miR-29b-1-5p | MIMAT0004523 | Gli1 | 14632 | Immunohistochemistry//qRT-PCR//Wester | 26524535 | Unspecified |
| mmu-miR-29b-1-5p | MIMAT0004523 | Pdgfrb | 18596 | Immunohistochemistry//qRT-PCR//Wester | 26524535 | Unspecified |
| mmu-miR-29b-1-5p | MIMAT0004523 | Sox4 | 20677 | qRT-PCR//Western blot | 27421576 | Unspecified |
| mmu-miR-378a-5p | MIMAT0000742 | VSIVgp2 | 1489832 | Luciferase reporter assay | 17613256 | Unspecified |
| mmu-miR-21a-3p | MIMAT0004628 | GFAP | 2670 | Western blotting//qRT-PCR//Immunocyto | 31419975 | Unspecified |
| mmu-miR-21a-3p | MIMAT0004628 | Ifng | 15978 | Luciferase reporter assay//qRT-PCR//Flow | 33552001 | Unspecified |
| mmu-miR-21a-3p | MIMAT0004628 | RECK | 8434 | HITS-CLIP//Luciferase reporter assay//qRT- | 23142080\|25893734 | Unspecified |
| mmu-miR-21a-3p | MIMAT0004628 | Snca | 20617 | qRT-PCR | 26689812 | Unspecified |
| mmu-miR-24-2-5p | MIMAT0005440 | Atf3 | 11910 | HITS-CLIP | 28735896 | Unspecified |
| mmu-miR-24-2-5p | MIMAT0005440 | PRKCA | 5578 | Luciferase reporter assay//qRT-PCR//West | 22911661 | Unspecified |
| mmu-miR-24-2-5p | MIMAT0005440 | Smad5 | 17129 | qRT-PCR//Western blot | 25476774 | Unspecified |
| mmu-miR-24-2-5p | MIMAT0005440 | Trib3 | 228775 | Western blot | 27084569 | Unspecified |
| mmu-miR-100-3p | MIMAT0017051 | Map3k7 | 26409 | Luciferase reporter assay//Western blottin | 31825165 | Unspecified |
| mmu-miR-100-3p | MIMAT0017051 | Srek1 | 218543 | HITS-CLIP | 28735896 | Unspecified |
| mmu-miR-145b | MIMAT0025105 | Cited2 | 17684 | Luciferase reporter assay//qRT-PCR//West | 27907012 | Unspecified |
| mmu-miR-326-5p | MIMAT0017027 | Lypd1 | 72585 | HITS-CLIP | 28735896 | Unspecified |
| mmu-miR-34a-3p | MIMAT0017022 | Dll1 | 13388 | Immunohistochemistry | 27534418 | Unspecified |
| mmu-miR-34a-3p | MIMAT0017022 | Foxo3 | 56484 | Luciferase reporter assay//qRT-PCR//West | 28321785 | Unspecified |
| mmu-miR-34a-3p | MIMAT0017022 | IRF1 | 3659 | Microarray//qRT-PCR | 33713393 | Unspecified |
| mmu-miR-34a-3p | MIMAT0017022 | Notch1 | 18128 | Immunohistochemistry//Western blotting/ | 27534418\|33942341 | Unspecified |
| mmu-miR-34a-3p | MIMAT0017022 | Rras | 20130 | Western blotting//Immunohistochemistry | 33942341 | Unspecified |
| mmu-miR-383-3p | MIMAT0017082 | Tmem178b | 434008 | HITS-CLIP | 28735896 | Unspecified |
| mmu-miR-5626-3p | MIMAT0022382 | Nfatc4 | 73181 | HITS-CLIP | 28735896 | Unspecified |
| mmu-miR-5626-5p | MIMAT0022381 | Alg12 | 223774 | Western blotting//Microarray//qRT-PCR | 33200806 | Unspecified |
| mmu-miR-6356 | MIMAT0025099 | Dcaf17 | 75763 | HITS-CLIP | 28735896 | Unspecified |
| mmu-miR-6516-3p | MIMAT0027344 | Plcl1 | 227120 | HITS-CLIP | 28735896 | Unspecified |
| mmu-miR-6934-3p | MIMAT0027769 | Xylt2 | 217119 | HITS-CLIP | 28735896 | Unspecified |
| mmu-miR-6934-3p | MIMAT0027769 | Zfp326 | 54367 | HITS-CLIP | 28735896 | Unspecified |
| mmu-miR-7025-3p | MIMAT0027955 | Igf1r | 16001 | Luciferase reporter assay//Western blottin | 31755174 | Unspecified |
| mmu-miR-7025-5p | MIMAT0027954 | Il10 | 16153 | Luciferase reporter assay//Western blottin | 31755174 | Unspecified |
| mmu-miR-7025-5p | MIMAT0027954 | Nfatc4 | 73181 | HITS-CLIP | 28735896 | Unspecified |
| mmu-miR-7650-3p | MIMAT0029803 | Brd7 | 26992 | HITS-CLIP | 28735896 | Unspecified |
| mmu-miR-7650-3p | MIMAT0029803 | Rhod | 11854 | HITS-CLIP | 28735896 | Unspecified |
| mmu-miR-7663-3p | MIMAT0029833 | Olfr1393 | 258463 | HITS-CLIP | 28735896 | Unspecified |

| | | | | | | |
|---|---|---|---|---|---|---|
| mmu-miR-7663-5p | MIMAT0029832 | Fam212b | 109050 | HITS-CLIP | 28735896 | Unspecified |
| mmu-miR-8118 | MIMAT0031424 | Adat1 | 30947 | HITS-CLIP | 28735896 | Unspecified |
| mmu-miR-8118 | MIMAT0031424 | Gja5 | 14613 | HITS-CLIP | 28735896 | Unspecified |
| mmu-miR-9-3p | MIMAT0000143 | Scn2b | 72821 | Luciferase reporter assay | 26259688 | Unspecified |
| mmu-miR-24-3p | MIMAT0000219 | 0610030E20Rik | 68364 | HITS-CLIP | 37986224 | Unspecified |
| mmu-miR-24-3p | MIMAT0000219 | 1110002E22Rik | 102634333 | HITS-CLIP | 25753418 | Unspecified |
| mmu-miR-24-3p | MIMAT0000219 | 1110059E24Rik | 66206 | HITS-CLIP | 23142080 | Unspecified |
| mmu-miR-24-3p | MIMAT0000219 | 1500011B03Rik | 66236 | HITS-CLIP | 37986224\|23142080 | Unspecified |
| mmu-miR-24-3p | MIMAT0000219 | 1700017B05Rik | 74211 | HITS-CLIP | 23142080 | Unspecified |
| mmu-miR-24-3p | MIMAT0000219 | 1700037H04Rik | 67326 | HITS-CLIP | 23142080 | Unspecified |
| mmu-miR-24-3p | MIMAT0000219 | 1700123O20Rik | 58248 | HITS-CLIP | 23142080 | Unspecified |
| mmu-miR-24-3p | MIMAT0000219 | 1810013L24Rik | 69053 | HITS-CLIP | 25753418 | Unspecified |
| mmu-miR-24-3p | MIMAT0000219 | 1810030O07Rik | 69155 | HITS-CLIP | 23142080 | Unspecified |
| mmu-miR-24-3p | MIMAT0000219 | 2010111I01Rik | 72061 | HITS-CLIP | 23142080 | Unspecified |
| mmu-miR-24-3p | MIMAT0000219 | 2310061I04Rik | 69662 | HITS-CLIP | 25753418 | Unspecified |
| mmu-miR-24-3p | MIMAT0000219 | 2510009E07Rik | 72190 | HITS-CLIP | 25753418 | Unspecified |
| mmu-miR-24-3p | MIMAT0000219 | 2610002M06Rik | 67028 | HITS-CLIP | 23142080 | Unspecified |
| mmu-miR-24-3p | MIMAT0000219 | 2610507B11Rik | 72503 | HITS-CLIP | 23142080 | Unspecified |
| mmu-miR-24-3p | MIMAT0000219 | 4833420G17Rik | 67392 | HITS-CLIP | 23142080 | Unspecified |
| mmu-miR-24-3p | MIMAT0000219 | 4921524J17Rik | 66714 | HITS-CLIP | 23142080 | Unspecified |
| mmu-miR-24-3p | MIMAT0000219 | 4930523C07Rik | 67647 | HITS-CLIP | 23142080 | Unspecified |
| mmu-miR-24-3p | MIMAT0000219 | 4932438A13Rik | 229227 | HITS-CLIP | 37986224\|23142080 | Unspecified |
| mmu-miR-24-3p | MIMAT0000219 | 4932438H23Rik | 74387 | HITS-CLIP | 23142080 | Unspecified |
| mmu-miR-24-3p | MIMAT0000219 | 5031439G07Rik | 223739 | HITS-CLIP | 25753418 | Unspecified |
| mmu-miR-24-3p | MIMAT0000219 | 5730455P16Rik | 70591 | HITS-CLIP | 25753418 | Unspecified |
| mmu-miR-24-3p | MIMAT0000219 | 9930021J03Rik | 240613 | HITS-CLIP | 23142080\|25753418 | Unspecified |
| mmu-miR-24-3p | MIMAT0000219 | AB124611 | 382062 | HITS-CLIP | 25753418 | Unspecified |
| mmu-miR-24-3p | MIMAT0000219 | AW549877 | 106064 | Sequencing//HITS-CLIP | 19536157\|23142080\|37986224 | Unspecified |
| mmu-miR-24-3p | MIMAT0000219 | Aacs | 78894 | HITS-CLIP | 23142080 | Unspecified |
| mmu-miR-24-3p | MIMAT0000219 | Aak1 | 269774 | HITS-CLIP | 23142080 | Unspecified |
| mmu-miR-24-3p | MIMAT0000219 | Aars | 234734 | HITS-CLIP//qCLASH | 37986224\|31393953\|25753418\|23142 | Unspecified |
| mmu-miR-24-3p | MIMAT0000219 | Abca1 | 11303 | HITS-CLIP | 25753418\|23142080 | Unspecified |
| mmu-miR-24-3p | MIMAT0000219 | Abca2 | 11305 | HITS-CLIP | 23142080 | Unspecified |
| mmu-miR-24-3p | MIMAT0000219 | Abcb7 | 11306 | HITS-CLIP | 25753418 | Unspecified |
| mmu-miR-24-3p | MIMAT0000219 | Abcb8 | 74610 | HITS-CLIP | 25753418 | Unspecified |
| mmu-miR-24-3p | MIMAT0000219 | Abcb9 | 56325 | HITS-CLIP | 23142080 | Unspecified |
| mmu-miR-24-3p | MIMAT0000219 | Abcc8 | 20927 | HITS-CLIP | 25753418 | Unspecified |
| mmu-miR-24-3p | MIMAT0000219 | Abcc9 | 20928 | HITS-CLIP | 25753418 | Unspecified |
| mmu-miR-24-3p | MIMAT0000219 | Abhd13 | 68904 | HITS-CLIP | 23142080 | Unspecified |

| | | | | | | |
|---|---|---|---|---|---|---|
| mmu-miR-24-3p | MIMAT0000219 | Abhd17a | 216169 | HITS-CLIP | 23142080 | Unspecified |
| mmu-miR-24-3p | MIMAT0000219 | Abhd18 | 269423 | HITS-CLIP | 23142080 | Unspecified |
| mmu-miR-24-3p | MIMAT0000219 | Abhd4 | 105501 | HITS-CLIP | 25753418 | Unspecified |
| mmu-miR-24-3p | MIMAT0000219 | Abi2 | 329165 | Sequencing//HITS-CLIP | 37986224\|25753418\|19536157 | Unspecified |
| mmu-miR-24-3p | MIMAT0000219 | Ablim1 | 226251 | HITS-CLIP | 23142080 | Unspecified |
| mmu-miR-24-3p | MIMAT0000219 | Acaa2 | 52538 | HITS-CLIP | 23142080 | Unspecified |
| mmu-miR-24-3p | MIMAT0000219 | Acacb | 100705 | HITS-CLIP | 25753418 | Unspecified |
| mmu-miR-24-3p | MIMAT0000219 | Acadsb | 66885 | HITS-CLIP | 23142080 | Unspecified |
| mmu-miR-24-3p | MIMAT0000219 | Acadvl | 11370 | HITS-CLIP | 25753418 | Unspecified |
| mmu-miR-24-3p | MIMAT0000219 | Acap2 | 78618 | HITS-CLIP | 23142080 | Unspecified |
| mmu-miR-24-3p | MIMAT0000219 | Acat1 | 110446 | HITS-CLIP | 23142080 | Unspecified |
| mmu-miR-24-3p | MIMAT0000219 | Acbd3 | 170760 | HITS-CLIP | 23142080 | Unspecified |
| mmu-miR-24-3p | MIMAT0000219 | Acbd4 | 67131 | HITS-CLIP | 23142080 | Unspecified |
| mmu-miR-24-3p | MIMAT0000219 | Acin1 | 56215 | HITS-CLIP | 23142080\|25753418 | Unspecified |
| mmu-miR-24-3p | MIMAT0000219 | Acly | 104112 | HITS-CLIP | 23142080 | Unspecified |
| mmu-miR-24-3p | MIMAT0000219 | Acot1 | 26897 | HITS-CLIP | 25753418 | Unspecified |
| mmu-miR-24-3p | MIMAT0000219 | Acot11 | 329910 | HITS-CLIP | 25753418 | Unspecified |
| mmu-miR-24-3p | MIMAT0000219 | Acox1 | 11430 | HITS-CLIP | 25753418 | Unspecified |
| mmu-miR-24-3p | MIMAT0000219 | Actb | 11461 | HITS-CLIP | 23142080 | Unspecified |
| mmu-miR-24-3p | MIMAT0000219 | Actn2 | 11472 | HITS-CLIP | 25753418 | Unspecified |
| mmu-miR-24-3p | MIMAT0000219 | Actn4 | 60595 | HITS-CLIP | 23142080 | Unspecified |
| mmu-miR-24-3p | MIMAT0000219 | Actr2 | 66713 | HITS-CLIP | 23142080 | Unspecified |
| mmu-miR-24-3p | MIMAT0000219 | Acvr1b | 11479 | Sequencing//HITS-CLIP | 19536157\|23142080\|37986224 | Unspecified |
| mmu-miR-24-3p | MIMAT0000219 | Acvrl1 | 11482 | HITS-CLIP | 23142080 | Unspecified |
| mmu-miR-24-3p | MIMAT0000219 | Adam10 | 11487 | HITS-CLIP | 23142080 | Unspecified |
| mmu-miR-24-3p | MIMAT0000219 | Adam19 | 11492 | Sequencing//HITS-CLIP | 19536157\|25753418\|37986224 | Unspecified |
| mmu-miR-24-3p | MIMAT0000219 | Adamts6 | 108154 | HITS-CLIP | 23142080 | Unspecified |
| mmu-miR-24-3p | MIMAT0000219 | Adamtsl4 | 229595 | HITS-CLIP | 25753418\|23142080 | Unspecified |
| mmu-miR-24-3p | MIMAT0000219 | Adar | 56417 | HITS-CLIP | 23142080 | Unspecified |
| mmu-miR-24-3p | MIMAT0000219 | Adck5 | 268822 | HITS-CLIP | 23142080 | Unspecified |
| mmu-miR-24-3p | MIMAT0000219 | Adcy3 | 104111 | HITS-CLIP | 23142080 | Unspecified |
| mmu-miR-24-3p | MIMAT0000219 | Adcy6 | 11512 | HITS-CLIP | 25753418 | Unspecified |
| mmu-miR-24-3p | MIMAT0000219 | Adhfe1 | 76187 | HITS-CLIP | 25753418 | Unspecified |
| mmu-miR-24-3p | MIMAT0000219 | Adipor1 | 72674 | HITS-CLIP | 23142080 | Unspecified |
| mmu-miR-24-3p | MIMAT0000219 | Adipor2 | 68465 | HITS-CLIP | 23142080 | Unspecified |
| mmu-miR-24-3p | MIMAT0000219 | Adnp | 11538 | HITS-CLIP | 37986224 | Unspecified |
| mmu-miR-24-3p | MIMAT0000219 | Adnp2 | 240442 | HITS-CLIP | 23142080 | Unspecified |
| mmu-miR-24-3p | MIMAT0000219 | Adora2a | 11540 | HITS-CLIP | 23142080 | Unspecified |
| mmu-miR-24-3p | MIMAT0000219 | Adpgk | 72141 | Sequencing//HITS-CLIP | 37986224\|23142080\|19536157 | Unspecified |

| | | | | | | |
|---|---|---|---|---|---|---|
| mmu-miR-24-3p | MIMAT0000219 | Adprh | 11544 | HITS-CLIP | 23142080 | Unspecified |
| mmu-miR-24-3p | MIMAT0000219 | Adsl | 11564 | HITS-CLIP | 23142080 | Unspecified |
| mmu-miR-24-3p | MIMAT0000219 | Aebp2 | 11569 | HITS-CLIP | 23142080 | Unspecified |
| mmu-miR-24-3p | MIMAT0000219 | Afdn | 4301 | HITS-CLIP | 23142080 | Unspecified |
| mmu-miR-24-3p | MIMAT0000219 | Aff4 | 93736 | HITS-CLIP | 25753418\|23142080 | Unspecified |
| mmu-miR-24-3p | MIMAT0000219 | Aftph | 216549 | HITS-CLIP | 37986224 | Unspecified |
| mmu-miR-24-3p | MIMAT0000219 | Agfg1 | 15463 | HITS-CLIP | 23142080 | Unspecified |
| mmu-miR-24-3p | MIMAT0000219 | Agfg2 | 231801 | HITS-CLIP | 25753418 | Unspecified |
| mmu-miR-24-3p | MIMAT0000219 | Ago1 | 236511 | HITS-CLIP | 37986224\|23142080 | Unspecified |
| mmu-miR-24-3p | MIMAT0000219 | Ago3 | 214150 | HITS-CLIP | 23142080 | Unspecified |
| mmu-miR-24-3p | MIMAT0000219 | Agpat1 | 55979 | HITS-CLIP | 23142080 | Unspecified |
| mmu-miR-24-3p | MIMAT0000219 | Agpat5 | 52123 | HITS-CLIP | 23142080 | Unspecified |
| mmu-miR-24-3p | MIMAT0000219 | Ahctf1 | 226747 | HITS-CLIP | 23142080 | Unspecified |
| mmu-miR-24-3p | MIMAT0000219 | Ahcyl1 | 229709 | HITS-CLIP | 23142080 | Unspecified |
| mmu-miR-24-3p | MIMAT0000219 | Ahcyl2 | 74340 | HITS-CLIP | 25753418 | Unspecified |
| mmu-miR-24-3p | MIMAT0000219 | Ahdc1 | 230793 | HITS-CLIP | 26203562 | Unspecified |
| mmu-miR-24-3p | MIMAT0000219 | Ahi1 | 52906 | HITS-CLIP | 23142080 | Unspecified |
| mmu-miR-24-3p | MIMAT0000219 | Ahnak | 66395 | HITS-CLIP | 23142080 | Unspecified |
| mmu-miR-24-3p | MIMAT0000219 | Ak2 | 11637 | HITS-CLIP | 37986224\|23142080 | Unspecified |
| mmu-miR-24-3p | MIMAT0000219 | Ak4 | 11639 | HITS-CLIP | 23142080 | Unspecified |
| mmu-miR-24-3p | MIMAT0000219 | Akap10 | 56697 | HITS-CLIP | 23142080 | Unspecified |
| mmu-miR-24-3p | MIMAT0000219 | Akap11 | 219181 | HITS-CLIP | 23142080 | Unspecified |
| mmu-miR-24-3p | MIMAT0000219 | Akap13 | 75547 | HITS-CLIP | 25753418 | Unspecified |
| mmu-miR-24-3p | MIMAT0000219 | Akr1a1 | 58810 | HITS-CLIP | 37986224 | Unspecified |
| mmu-miR-24-3p | MIMAT0000219 | Akt1s1 | 67605 | HITS-CLIP | 23142080 | Unspecified |
| mmu-miR-24-3p | MIMAT0000219 | Aldh3a2 | 11671 | HITS-CLIP | 23142080 | Unspecified |
| mmu-miR-24-3p | MIMAT0000219 | Aldh7a1 | 110695 | HITS-CLIP | 23142080 | Unspecified |
| mmu-miR-24-3p | MIMAT0000219 | Alkbh5 | 268420 | HITS-CLIP | 25753418\|23142080 | Unspecified |
| mmu-miR-24-3p | MIMAT0000219 | Alpk2 | 225638 | HITS-CLIP | 25753418 | Unspecified |
| mmu-miR-24-3p | MIMAT0000219 | Alpk3 | 116904 | HITS-CLIP | 25753418 | Unspecified |
| mmu-miR-24-3p | MIMAT0000219 | Alyref | 21681 | HITS-CLIP | 23142080 | Unspecified |
| mmu-miR-24-3p | MIMAT0000219 | Alyref2 | 56009 | HITS-CLIP | 23142080 | Unspecified |
| mmu-miR-24-3p | MIMAT0000219 | Ambra1 | 228361 | HITS-CLIP | 23142080 | Unspecified |
| mmu-miR-24-3p | MIMAT0000219 | Amigo1 | 229715 | HITS-CLIP | 25753418 | Unspecified |
| mmu-miR-24-3p | MIMAT0000219 | Ammecr1 | 56068 | HITS-CLIP | 23142080 | Unspecified |
| mmu-miR-24-3p | MIMAT0000219 | Ammecr1l | 225339 | HITS-CLIP | 23142080 | Unspecified |
| mmu-miR-24-3p | MIMAT0000219 | Ampd3 | 11717 | HITS-CLIP | 23142080 | Unspecified |
| mmu-miR-24-3p | MIMAT0000219 | Anapc4 | 52206 | HITS-CLIP | 23142080 | Unspecified |
| mmu-miR-24-3p | MIMAT0000219 | Anapc5 | 59008 | HITS-CLIP | 23142080 | Unspecified |

| | | | | | | |
|---|---|---|---|---|---|---|
| mmu-miR-24-3p | MIMAT0000219 | Ankfy1 | 11736 | HITS-CLIP | 23142080 | Unspecified |
| mmu-miR-24-3p | MIMAT0000219 | Ankhd1 | 108857 | HITS-CLIP | 23142080\|25753418 | Unspecified |
| mmu-miR-24-3p | MIMAT0000219 | Ankra2 | 68558 | HITS-CLIP | 25753418 | Unspecified |
| mmu-miR-24-3p | MIMAT0000219 | Ankrd11 | 77087 | HITS-CLIP | 23142080 | Unspecified |
| mmu-miR-24-3p | MIMAT0000219 | Ankrd12 | 106585 | HITS-CLIP | 23142080 | Unspecified |
| mmu-miR-24-3p | MIMAT0000219 | Ankrd17 | 81702 | HITS-CLIP | 23142080 | Unspecified |
| mmu-miR-24-3p | MIMAT0000219 | Ankrd37 | 654824 | HITS-CLIP | 23142080 | Unspecified |
| mmu-miR-24-3p | MIMAT0000219 | Ankrd44 | 329154 | HITS-CLIP | 37986224\|25753418 | Unspecified |
| mmu-miR-24-3p | MIMAT0000219 | Ankrd49 | 56503 | HITS-CLIP | 23142080 | Unspecified |
| mmu-miR-24-3p | MIMAT0000219 | Ankrd52 | 237615 | Sequencing//HITS-CLIP | 18353861\|23142080\|37986224 | Unspecified |
| mmu-miR-24-3p | MIMAT0000219 | Ankrd6 | 140577 | HITS-CLIP | 25753418\|23142080 | Unspecified |
| mmu-miR-24-3p | MIMAT0000219 | Anks1 | 224650 | HITS-CLIP | 23142080\|25753418 | Unspecified |
| mmu-miR-24-3p | MIMAT0000219 | Anln | 68743 | HITS-CLIP | 23142080 | Unspecified |
| mmu-miR-24-3p | MIMAT0000219 | Anp32e | 66471 | HITS-CLIP | 25753418 | Unspecified |
| mmu-miR-24-3p | MIMAT0000219 | Antxr2 | 71914 | HITS-CLIP | 23142080 | Unspecified |
| mmu-miR-24-3p | MIMAT0000219 | Anxa2 | 12306 | HITS-CLIP | 25753418\|23142080 | Unspecified |
| mmu-miR-24-3p | MIMAT0000219 | Anxa5 | 11747 | HITS-CLIP | 25753418 | Unspecified |
| mmu-miR-24-3p | MIMAT0000219 | Ap1ar | 211556 | HITS-CLIP | 23142080 | Unspecified |
| mmu-miR-24-3p | MIMAT0000219 | Ap1b1 | 11764 | HITS-CLIP | 23142080 | Unspecified |
| mmu-miR-24-3p | MIMAT0000219 | Ap1g1 | 11765 | HITS-CLIP | 23142080 | Unspecified |
| mmu-miR-24-3p | MIMAT0000219 | Ap1s1 | 11769 | HITS-CLIP | 23142080 | Unspecified |
| mmu-miR-24-3p | MIMAT0000219 | Ap2a1 | 11771 | HITS-CLIP | 23142080\|25753418 | Unspecified |
| mmu-miR-24-3p | MIMAT0000219 | Ap2a2 | 11772 | qCLASH//HITS-CLIP | 31393953\|25753418 | Unspecified |
| mmu-miR-24-3p | MIMAT0000219 | Ap2b1 | 71770 | HITS-CLIP | 23142080 | Unspecified |
| mmu-miR-24-3p | MIMAT0000219 | Ap4e1 | 108011 | HITS-CLIP | 23142080 | Unspecified |
| mmu-miR-24-3p | MIMAT0000219 | Ap5b1 | 381201 | HITS-CLIP | 23142080 | Unspecified |
| mmu-miR-24-3p | MIMAT0000219 | Ap5m1 | 74385 | HITS-CLIP | 23142080 | Unspecified |
| mmu-miR-24-3p | MIMAT0000219 | Apaf1 | 11783 | HITS-CLIP | 23142080 | Unspecified |
| mmu-miR-24-3p | MIMAT0000219 | Apeh | 235606 | HITS-CLIP | 23142080 | Unspecified |
| mmu-miR-24-3p | MIMAT0000219 | Aplp2 | 11804 | HITS-CLIP | 25753418 | Unspecified |
| mmu-miR-24-3p | MIMAT0000219 | Apobec3 | 80287 | HITS-CLIP | 37986224\|23142080 | Unspecified |
| mmu-miR-24-3p | MIMAT0000219 | Arap1 | 69710 | HITS-CLIP | 23142080 | Unspecified |
| mmu-miR-24-3p | MIMAT0000219 | Arel1 | 68497 | HITS-CLIP | 37986224\|23142080 | Unspecified |
| mmu-miR-24-3p | MIMAT0000219 | Arf1 | 11840 | HITS-CLIP | 25753418 | Unspecified |
| mmu-miR-24-3p | MIMAT0000219 | Arf3 | 11842 | HITS-CLIP | 31433990\|25753418\|23142080 | Unspecified |
| mmu-miR-24-3p | MIMAT0000219 | Arf4 | 11843 | HITS-CLIP | 37986224\|23142080 | Unspecified |
| mmu-miR-24-3p | MIMAT0000219 | Arf6 | 11845 | HITS-CLIP | 37986224\|23142080 | Unspecified |
| mmu-miR-24-3p | MIMAT0000219 | Arfgap1 | 228998 | HITS-CLIP | 23142080 | Unspecified |
| mmu-miR-24-3p | MIMAT0000219 | Arhgap1 | 228359 | HITS-CLIP | 23142080 | Unspecified |

| | | | | | | |
|---|---|---|---|---|---|---|
| mmu-miR-24-3p | MIMAT0000219 | Arhgap17 | 70497 | HITS-CLIP | 23142080 | Unspecified |
| mmu-miR-24-3p | MIMAT0000219 | Arhgap18 | 73910 | HITS-CLIP | 23142080 | Unspecified |
| mmu-miR-24-3p | MIMAT0000219 | Arhgap19 | 71085 | Western Blot, HITS-CLIP//Western Blot//H | 37986224\|23071155\|23142080 | Unspecified |
| mmu-miR-24-3p | MIMAT0000219 | Arhgap21 | 71435 | HITS-CLIP | 25753418 | Unspecified |
| mmu-miR-24-3p | MIMAT0000219 | Arhgap25 | 232201 | HITS-CLIP | 23142080 | Unspecified |
| mmu-miR-24-3p | MIMAT0000219 | Arhgap30 | 226652 | HITS-CLIP | 23142080 | Unspecified |
| mmu-miR-24-3p | MIMAT0000219 | Arhgap35 | 232906 | HITS-CLIP | 23142080 | Unspecified |
| mmu-miR-24-3p | MIMAT0000219 | Arhgap5 | 11855 | HITS-CLIP | 25753418 | Unspecified |
| mmu-miR-24-3p | MIMAT0000219 | Arhgdia | 192662 | HITS-CLIP | 23142080 | Unspecified |
| mmu-miR-24-3p | MIMAT0000219 | Arhgef12 | 69632 | HITS-CLIP | 23142080 | Unspecified |
| mmu-miR-24-3p | MIMAT0000219 | Arhgef18 | 102098 | HITS-CLIP | 23142080 | Unspecified |
| mmu-miR-24-3p | MIMAT0000219 | Arhgef7 | 54126 | HITS-CLIP | 23142080 | Unspecified |
| mmu-miR-24-3p | MIMAT0000219 | Arid1a | 93760 | HITS-CLIP | 25753418 | Unspecified |
| mmu-miR-24-3p | MIMAT0000219 | Arid1b | 239985 | HITS-CLIP | 23142080 | Unspecified |
| mmu-miR-24-3p | MIMAT0000219 | Arid2 | 77044 | HITS-CLIP | 37986224\|23142080\|25753418 | Unspecified |
| mmu-miR-24-3p | MIMAT0000219 | Arid3b | 56380 | HITS-CLIP | 37986224 | Unspecified |
| mmu-miR-24-3p | MIMAT0000219 | Arid5b | 71371 | HITS-CLIP | 23142080 | Unspecified |
| mmu-miR-24-3p | MIMAT0000219 | Arih1 | 23806 | HITS-CLIP | 37986224\|23142080 | Unspecified |
| mmu-miR-24-3p | MIMAT0000219 | Arl13b | 68146 | HITS-CLIP | 23142080 | Unspecified |
| mmu-miR-24-3p | MIMAT0000219 | Arl15 | 218639 | HITS-CLIP | 25753418 | Unspecified |
| mmu-miR-24-3p | MIMAT0000219 | Arl5b | 75869 | HITS-CLIP | 25753418 | Unspecified |
| mmu-miR-24-3p | MIMAT0000219 | Arl6ip5 | 65106 | HITS-CLIP | 23142080 | Unspecified |
| mmu-miR-24-3p | MIMAT0000219 | Armc8 | 74125 | HITS-CLIP | 23142080 | Unspecified |
| mmu-miR-24-3p | MIMAT0000219 | Armcx4 | 100503043 | HITS-CLIP | 31433990 | Unspecified |
| mmu-miR-24-3p | MIMAT0000219 | Armh3 | 79591 | HITS-CLIP | 25753418\|23142080 | Unspecified |
| mmu-miR-24-3p | MIMAT0000219 | Arpc1a | 56443 | HITS-CLIP | 23142080 | Unspecified |
| mmu-miR-24-3p | MIMAT0000219 | Arrdc3 | 105171 | HITS-CLIP | 23142080 | Unspecified |
| mmu-miR-24-3p | MIMAT0000219 | Asb2 | 65256 | HITS-CLIP | 31433990\|25753418 | Unspecified |
| mmu-miR-24-3p | MIMAT0000219 | Asb4 | 65255 | HITS-CLIP | 25753418 | Unspecified |
| mmu-miR-24-3p | MIMAT0000219 | Asb7 | 117589 | HITS-CLIP | 23142080 | Unspecified |
| mmu-miR-24-3p | MIMAT0000219 | Ascc3 | 77987 | HITS-CLIP | 23142080 | Unspecified |
| mmu-miR-24-3p | MIMAT0000219 | Ash1l | 192195 | HITS-CLIP | 37986224\|23142080\|25753418 | Unspecified |
| mmu-miR-24-3p | MIMAT0000219 | Ash2l | 23808 | HITS-CLIP | 25753418 | Unspecified |
| mmu-miR-24-3p | MIMAT0000219 | Asna1 | 56495 | HITS-CLIP | 23142080 | Unspecified |
| mmu-miR-24-3p | MIMAT0000219 | Asph | 65973 | HITS-CLIP | 25753418 | Unspecified |
| mmu-miR-24-3p | MIMAT0000219 | Aspscr1 | 68938 | qCLASH | 31393953 | Unspecified |
| mmu-miR-24-3p | MIMAT0000219 | Asxl1 | 228790 | HITS-CLIP | 23142080 | Unspecified |
| mmu-miR-24-3p | MIMAT0000219 | Asxl2 | 75302 | HITS-CLIP | 23142080 | Unspecified |
| mmu-miR-24-3p | MIMAT0000219 | Atad1 | 67979 | HITS-CLIP | 23142080\|25753418 | Unspecified |

| | | | | | | |
|---|---|---|---|---|---|---|
| mmu-miR-24-3p | MIMAT0000219 | Atad2 | 70472 | HITS-CLIP | 23142080 | Unspecified |
| mmu-miR-24-3p | MIMAT0000219 | Atad2b | 320817 | HITS-CLIP | 23142080 | Unspecified |
| mmu-miR-24-3p | MIMAT0000219 | Atf4 | 11911 | HITS-CLIP | 37986224 | Unspecified |
| mmu-miR-24-3p | MIMAT0000219 | Atf7 | 223922 | HITS-CLIP | 23142080 | Unspecified |
| mmu-miR-24-3p | MIMAT0000219 | Atg12 | 67526 | HITS-CLIP | 23142080 | Unspecified |
| mmu-miR-24-3p | MIMAT0000219 | Atl2 | 56298 | HITS-CLIP | 23142080 | Unspecified |
| mmu-miR-24-3p | MIMAT0000219 | Atm | 11920 | HITS-CLIP | 23142080 | Unspecified |
| mmu-miR-24-3p | MIMAT0000219 | Atmin | 234776 | HITS-CLIP | 23142080 | Unspecified |
| mmu-miR-24-3p | MIMAT0000219 | Atox1 | 11927 | HITS-CLIP | 23142080 | Unspecified |
| mmu-miR-24-3p | MIMAT0000219 | Atp11a | 50770 | HITS-CLIP | 23142080 | Unspecified |
| mmu-miR-24-3p | MIMAT0000219 | Atp11b | 76295 | HITS-CLIP | 23142080 | Unspecified |
| mmu-miR-24-3p | MIMAT0000219 | Atp13a2 | 23400 | HITS-CLIP | 23142080 | Unspecified |
| mmu-miR-24-3p | MIMAT0000219 | Atp13a3 | 224088 | HITS-CLIP | 23142080 | Unspecified |
| mmu-miR-24-3p | MIMAT0000219 | Atp1a1 | 11928 | HITS-CLIP//qCLASH | 37986224\|31393953\|23142080 | Unspecified |
| mmu-miR-24-3p | MIMAT0000219 | Atp1a2 | 98660 | Sequencing//HITS-CLIP | 37986224\|25753418\|19536157 | Unspecified |
| mmu-miR-24-3p | MIMAT0000219 | Atp1b1 | 11931 | HITS-CLIP | 25753418 | Unspecified |
| mmu-miR-24-3p | MIMAT0000219 | Atp1b3 | 11933 | HITS-CLIP | 23142080 | Unspecified |
| mmu-miR-24-3p | MIMAT0000219 | Atp2a2 | 11938 | Sequencing//HITS-CLIP | 37986224\|23142080\|19536157 | Unspecified |
| mmu-miR-24-3p | MIMAT0000219 | Atp2a3 | 53313 | HITS-CLIP | 23142080 | Unspecified |
| mmu-miR-24-3p | MIMAT0000219 | Atp2b1 | 67972 | HITS-CLIP | 37986224\|23142080 | Unspecified |
| mmu-miR-24-3p | MIMAT0000219 | Atp5a1 | 11946 | HITS-CLIP | 23142080 | Unspecified |
| mmu-miR-24-3p | MIMAT0000219 | Atp5b | 11947 | HITS-CLIP | 23142080\|25753418 | Unspecified |
| mmu-miR-24-3p | MIMAT0000219 | Atp5pb | 515 | HITS-CLIP | 25753418 | Unspecified |
| mmu-miR-24-3p | MIMAT0000219 | Atp6v1b2 | 11966 | HITS-CLIP | 23142080 | Unspecified |
| mmu-miR-24-3p | MIMAT0000219 | Atp6v1c1 | 66335 | HITS-CLIP | 23142080 | Unspecified |
| mmu-miR-24-3p | MIMAT0000219 | Atp6v1e1 | 11973 | HITS-CLIP | 25753418 | Unspecified |
| mmu-miR-24-3p | MIMAT0000219 | Atp8b2 | 54667 | HITS-CLIP | 23142080 | Unspecified |
| mmu-miR-24-3p | MIMAT0000219 | Atr | 245000 | HITS-CLIP | 23142080 | Unspecified |
| mmu-miR-24-3p | MIMAT0000219 | Atrx | 22589 | HITS-CLIP | 23142080\|25753418 | Unspecified |
| mmu-miR-24-3p | MIMAT0000219 | Atxn1 | 20238 | HITS-CLIP | 31433990\|23142080 | Unspecified |
| mmu-miR-24-3p | MIMAT0000219 | Atxn1l | 52335 | HITS-CLIP | 37986224\|23142080 | Unspecified |
| mmu-miR-24-3p | MIMAT0000219 | Atxn2l | 233871 | HITS-CLIP//qCLASH | 23142080\|31393953 | Unspecified |
| mmu-miR-24-3p | MIMAT0000219 | Atxn7l3 | 217218 | HITS-CLIP | 23142080 | Unspecified |
| mmu-miR-24-3p | MIMAT0000219 | Atxn7l3b | 382423 | HITS-CLIP | 23142080 | Unspecified |
| mmu-miR-24-3p | MIMAT0000219 | Avl9 | 78937 | Sequencing//HITS-CLIP | 19536157\|23142080\|37986224 | Unspecified |
| mmu-miR-24-3p | MIMAT0000219 | Axin1 | 12005 | HITS-CLIP | 37986224 | Unspecified |
| mmu-miR-24-3p | MIMAT0000219 | Axl | 26362 | HITS-CLIP | 23142080 | Unspecified |
| mmu-miR-24-3p | MIMAT0000219 | Azin1 | 54375 | HITS-CLIP | 23142080 | Unspecified |
| mmu-miR-24-3p | MIMAT0000219 | B230219D22Rik | 78521 | HITS-CLIP | 23142080 | Unspecified |

| | | | | | | |
|---|---|---|---|---|---|---|
| mmu-miR-24-3p | MIMAT0000219 | B3gnt2 | 53625 | HITS-CLIP | 37986224\|23142080 | Unspecified |
| mmu-miR-24-3p | MIMAT0000219 | B4galnt1 | 14421 | HITS-CLIP | 23142080 | Unspecified |
| mmu-miR-24-3p | MIMAT0000219 | B4galt1 | 14595 | HITS-CLIP | 25753418\|23142080 | Unspecified |
| mmu-miR-24-3p | MIMAT0000219 | BC005537 | 79555 | HITS-CLIP | 37986224\|23142080 | Unspecified |
| mmu-miR-24-3p | MIMAT0000219 | BC005561 | 100042165 | HITS-CLIP | 23142080 | Unspecified |
| mmu-miR-24-3p | MIMAT0000219 | BC052040 | 399568 | HITS-CLIP | 23142080 | Unspecified |
| mmu-miR-24-3p | MIMAT0000219 | Bach1 | 12013 | HITS-CLIP | 25753418 | Unspecified |
| mmu-miR-24-3p | MIMAT0000219 | Bad | 12015 | HITS-CLIP | 23142080 | Unspecified |
| mmu-miR-24-3p | MIMAT0000219 | Bag1 | 12017 | HITS-CLIP | 25753418 | Unspecified |
| mmu-miR-24-3p | MIMAT0000219 | Bag3 | 29810 | HITS-CLIP | 25753418 | Unspecified |
| mmu-miR-24-3p | MIMAT0000219 | Bag6 | 224727 | HITS-CLIP | 23142080 | Unspecified |
| mmu-miR-24-3p | MIMAT0000219 | Bak1 | 12018 | HITS-CLIP | 23142080 | Unspecified |
| mmu-miR-24-3p | MIMAT0000219 | Banp | 53325 | HITS-CLIP | 23142080 | Unspecified |
| mmu-miR-24-3p | MIMAT0000219 | Bard1 | 12021 | HITS-CLIP | 23142080 | Unspecified |
| mmu-miR-24-3p | MIMAT0000219 | Baz1a | 217578 | HITS-CLIP | 23142080 | Unspecified |
| mmu-miR-24-3p | MIMAT0000219 | Baz1b | 22385 | HITS-CLIP | 25753418 | Unspecified |
| mmu-miR-24-3p | MIMAT0000219 | Baz2a | 116848 | HITS-CLIP | 37986224\|23142080\|25753418 | Unspecified |
| mmu-miR-24-3p | MIMAT0000219 | Baz2b | 407823 | HITS-CLIP | 23142080 | Unspecified |
| mmu-miR-24-3p | MIMAT0000219 | Bbc3 | 170770 | HITS-CLIP//Luciferase reporter assay | 23142080\|27797453 | Unspecified |
| mmu-miR-24-3p | MIMAT0000219 | Bbx | 70508 | HITS-CLIP | 23142080 | Unspecified |
| mmu-miR-24-3p | MIMAT0000219 | Bcap29 | 12033 | HITS-CLIP | 23142080 | Unspecified |
| mmu-miR-24-3p | MIMAT0000219 | Bcas3 | 192197 | HITS-CLIP | 25753418 | Unspecified |
| mmu-miR-24-3p | MIMAT0000219 | Bcl11b | 58208 | HITS-CLIP | 37986224\|23142080 | Unspecified |
| mmu-miR-24-3p | MIMAT0000219 | Bcl2l1 | 12048 | HITS-CLIP | 37986224\|23142080 | Unspecified |
| mmu-miR-24-3p | MIMAT0000219 | Bcl2l11 | 12125 | Immunohistochemistry//Luciferase assay/ | 21383058\|23142080 | Unspecified |
| mmu-miR-24-3p | MIMAT0000219 | Bcl2l13 | 94044 | HITS-CLIP | 25753418 | Unspecified |
| mmu-miR-24-3p | MIMAT0000219 | Bcl2l2 | 12050 | HITS-CLIP | 37986224\|23142080 | Unspecified |
| mmu-miR-24-3p | MIMAT0000219 | Bcl3 | 12051 | HITS-CLIP | 23142080 | Unspecified |
| mmu-miR-24-3p | MIMAT0000219 | Bcl6 | 12053 | HITS-CLIP | 23142080 | Unspecified |
| mmu-miR-24-3p | MIMAT0000219 | Bcor | 71458 | HITS-CLIP | 23142080\|25753418 | Unspecified |
| mmu-miR-24-3p | MIMAT0000219 | Bcorl1 | 320376 | HITS-CLIP | 23142080 | Unspecified |
| mmu-miR-24-3p | MIMAT0000219 | Bdp1 | 544971 | HITS-CLIP | 23142080 | Unspecified |
| mmu-miR-24-3p | MIMAT0000219 | Bhlhe40 | 20893 | HITS-CLIP | 23142080 | Unspecified |
| mmu-miR-24-3p | MIMAT0000219 | Bik | 12124 | HITS-CLIP | 31433990 | Unspecified |
| mmu-miR-24-3p | MIMAT0000219 | Birc3 | 11796 | HITS-CLIP | 23142080 | Unspecified |
| mmu-miR-24-3p | MIMAT0000219 | Birc6 | 12211 | HITS-CLIP | 23142080\|25753418 | Unspecified |
| mmu-miR-24-3p | MIMAT0000219 | Blmh | 104184 | HITS-CLIP | 23142080 | Unspecified |
| mmu-miR-24-3p | MIMAT0000219 | Bltp2 | 9703 | HITS-CLIP | 25753418 | Unspecified |
| mmu-miR-24-3p | MIMAT0000219 | Bmp2k | 140780 | HITS-CLIP | 23142080 | Unspecified |

| | | | | | | |
|---|---|---|---|---|---|---|
| mmu-miR-24-3p | MIMAT0000219 | Bmpr1a | 12166 | HITS-CLIP | 31433990\|23142080 | Unspecified |
| mmu-miR-24-3p | MIMAT0000219 | Bmpr2 | 12168 | HITS-CLIP | 37986224\|23142080 | Unspecified |
| mmu-miR-24-3p | MIMAT0000219 | Bms1 | 213895 | HITS-CLIP | 23142080 | Unspecified |
| mmu-miR-24-3p | MIMAT0000219 | Bnip2 | 12175 | HITS-CLIP | 23142080 | Unspecified |
| mmu-miR-24-3p | MIMAT0000219 | Bnip3l | 12177 | HITS-CLIP | 23142080 | Unspecified |
| mmu-miR-24-3p | MIMAT0000219 | Bod1l | 665775 | HITS-CLIP | 23142080 | Unspecified |
| mmu-miR-24-3p | MIMAT0000219 | Borcs6 | 54785 | HITS-CLIP | 23142080 | Unspecified |
| mmu-miR-24-3p | MIMAT0000219 | Bptf | 207165 | HITS-CLIP | 25753418 | Unspecified |
| mmu-miR-24-3p | MIMAT0000219 | Braf | 109880 | HITS-CLIP | 23142080 | Unspecified |
| mmu-miR-24-3p | MIMAT0000219 | Brca1 | 12189 | HITS-CLIP | 23142080 | Unspecified |
| mmu-miR-24-3p | MIMAT0000219 | Brd1 | 223770 | HITS-CLIP | 23142080 | Unspecified |
| mmu-miR-24-3p | MIMAT0000219 | Brd4 | 57261 | HITS-CLIP | 23142080 | Unspecified |
| mmu-miR-24-3p | MIMAT0000219 | Brd8 | 78656 | HITS-CLIP | 23142080 | Unspecified |
| mmu-miR-24-3p | MIMAT0000219 | Bri3 | 55950 | HITS-CLIP | 25753418 | Unspecified |
| mmu-miR-24-3p | MIMAT0000219 | Brms1l | 52592 | HITS-CLIP | 25753418 | Unspecified |
| mmu-miR-24-3p | MIMAT0000219 | Bsg | 12215 | HITS-CLIP | 23142080 | Unspecified |
| mmu-miR-24-3p | MIMAT0000219 | Btbd7 | 238386 | HITS-CLIP | 23142080 | Unspecified |
| mmu-miR-24-3p | MIMAT0000219 | Btg2 | 12227 | HITS-CLIP | 37986224\|23142080 | Unspecified |
| mmu-miR-24-3p | MIMAT0000219 | Btla | 208154 | HITS-CLIP | 37986224\|23142080 | Unspecified |
| mmu-miR-24-3p | MIMAT0000219 | Bub1b | 12236 | HITS-CLIP | 23142080 | Unspecified |
| mmu-miR-24-3p | MIMAT0000219 | Bzw2 | 66912 | HITS-CLIP | 23142080 | Unspecified |
| mmu-miR-24-3p | MIMAT0000219 | C1ql3 | 227580 | HITS-CLIP | 23142080 | Unspecified |
| mmu-miR-24-3p | MIMAT0000219 | C1qtnf12 | 67389 | HITS-CLIP | 23142080 | Unspecified |
| mmu-miR-24-3p | MIMAT0000219 | C2cd3 | 277939 | HITS-CLIP | 23142080 | Unspecified |
| mmu-miR-24-3p | MIMAT0000219 | C9orf72 | 203228 | HITS-CLIP | 25753418\|23142080 | Unspecified |
| mmu-miR-24-3p | MIMAT0000219 | Cacfd1 | 381356 | HITS-CLIP | 25753418 | Unspecified |
| mmu-miR-24-3p | MIMAT0000219 | Cacna1c | 12288 | HITS-CLIP | 23142080 | Unspecified |
| mmu-miR-24-3p | MIMAT0000219 | Cacna2d2 | 56808 | HITS-CLIP | 25753418 | Unspecified |
| mmu-miR-24-3p | MIMAT0000219 | Cad | 69719 | HITS-CLIP | 23142080 | Unspecified |
| mmu-miR-24-3p | MIMAT0000219 | Calm1 | 12313 | HITS-CLIP | 23142080 | Unspecified |
| mmu-miR-24-3p | MIMAT0000219 | Camk2b | 12323 | HITS-CLIP | 23142080 | Unspecified |
| mmu-miR-24-3p | MIMAT0000219 | Camk2d | 108058 | Sequencing//HITS-CLIP | 37986224\|23142080\|19536157 | Unspecified |
| mmu-miR-24-3p | MIMAT0000219 | Camk2n1 | 66259 | HITS-CLIP | 23142080 | Unspecified |
| mmu-miR-24-3p | MIMAT0000219 | Camsap2 | 67886 | HITS-CLIP | 23142080 | Unspecified |
| mmu-miR-24-3p | MIMAT0000219 | Cand1 | 71902 | HITS-CLIP | 23142080 | Unspecified |
| mmu-miR-24-3p | MIMAT0000219 | Cand2 | 67088 | HITS-CLIP | 25753418 | Unspecified |
| mmu-miR-24-3p | MIMAT0000219 | Canx | 12330 | HITS-CLIP | 23142080 | Unspecified |
| mmu-miR-24-3p | MIMAT0000219 | Cap1 | 12331 | HITS-CLIP | 23142080 | Unspecified |
| mmu-miR-24-3p | MIMAT0000219 | Capn15 | 50817 | HITS-CLIP | 23142080 | Unspecified |

| | | | | | | | |
|---|---|---|---|---|---|---|---|
| mmu-miR-24-3p | MIMAT0000219 | Caprin1 | 53872 | HITS-CLIP | | 23142080 | Unspecified |
| mmu-miR-24-3p | MIMAT0000219 | Capza2 | 12343 | HITS-CLIP | 23142080\|25753418 | | Unspecified |
| mmu-miR-24-3p | MIMAT0000219 | Capzb | 12345 | HITS-CLIP | 37986224\|23142080 | | Unspecified |
| mmu-miR-24-3p | MIMAT0000219 | Carmil2 | 146206 | HITS-CLIP | | 23142080 | Unspecified |
| mmu-miR-24-3p | MIMAT0000219 | Cars2 | 71941 | HITS-CLIP | | 25753418 | Unspecified |
| mmu-miR-24-3p | MIMAT0000219 | Casc3 | 192160 | HITS-CLIP | | 37986224 | Unspecified |
| mmu-miR-24-3p | MIMAT0000219 | Casd1 | 213819 | HITS-CLIP | | 25753418 | Unspecified |
| mmu-miR-24-3p | MIMAT0000219 | Casp2 | 12366 | HITS-CLIP | | 25753418 | Unspecified |
| mmu-miR-24-3p | MIMAT0000219 | Casp4 | 12363 | HITS-CLIP | | 23142080 | Unspecified |
| mmu-miR-24-3p | MIMAT0000219 | Casp6 | 12368 | HITS-CLIP | 37986224\|23142080 | | Unspecified |
| mmu-miR-24-3p | MIMAT0000219 | Casp8ap2 | 26885 | HITS-CLIP | | 23142080 | Unspecified |
| mmu-miR-24-3p | MIMAT0000219 | Cav1 | 12389 | HITS-CLIP | | 25753418 | Unspecified |
| mmu-miR-24-3p | MIMAT0000219 | Cav2 | 12390 | HITS-CLIP | | 25753418 | Unspecified |
| mmu-miR-24-3p | MIMAT0000219 | Cbfb | 12400 | HITS-CLIP | | 23142080 | Unspecified |
| mmu-miR-24-3p | MIMAT0000219 | Cbl | 12402 | HITS-CLIP | | 23142080 | Unspecified |
| mmu-miR-24-3p | MIMAT0000219 | Cblb | 208650 | HITS-CLIP | 37986224\|23142080 | | Unspecified |
| mmu-miR-24-3p | MIMAT0000219 | Cbwd1 | 226043 | HITS-CLIP | | 23142080 | Unspecified |
| mmu-miR-24-3p | MIMAT0000219 | Cbx2 | 12416 | HITS-CLIP | 37986224\|23142080 | | Unspecified |
| mmu-miR-24-3p | MIMAT0000219 | Cbx4 | 12418 | HITS-CLIP | | 23142080 | Unspecified |
| mmu-miR-24-3p | MIMAT0000219 | Cbx5 | 12419 | HITS-CLIP | | 23142080 | Unspecified |
| mmu-miR-24-3p | MIMAT0000219 | Ccar1 | 67500 | HITS-CLIP | | 23142080 | Unspecified |
| mmu-miR-24-3p | MIMAT0000219 | Ccar2 | 219158 | HITS-CLIP | | 23142080 | Unspecified |
| mmu-miR-24-3p | MIMAT0000219 | Ccdc126 | 57895 | HITS-CLIP | | 23142080 | Unspecified |
| mmu-miR-24-3p | MIMAT0000219 | Ccdc141 | 545428 | HITS-CLIP | 25753418\|23142080 | | Unspecified |
| mmu-miR-24-3p | MIMAT0000219 | Ccdc25 | 67179 | HITS-CLIP | | 23142080 | Unspecified |
| mmu-miR-24-3p | MIMAT0000219 | Ccdc47 | 67163 | HITS-CLIP | | 25753418 | Unspecified |
| mmu-miR-24-3p | MIMAT0000219 | Ccdc50 | 67501 | HITS-CLIP | | 23142080 | Unspecified |
| mmu-miR-24-3p | MIMAT0000219 | Ccdc71l | 72123 | HITS-CLIP | | 23142080 | Unspecified |
| mmu-miR-24-3p | MIMAT0000219 | Ccdc82 | 66396 | HITS-CLIP | | 23142080 | Unspecified |
| mmu-miR-24-3p | MIMAT0000219 | Ccdc85b | 240514 | HITS-CLIP | 21258322\|25753418 | | Unspecified |
| mmu-miR-24-3p | MIMAT0000219 | Ccdc85c | 668158 | HITS-CLIP | | 25753418 | Unspecified |
| mmu-miR-24-3p | MIMAT0000219 | Ccna2 | 12428 | HITS-CLIP | | 23142080 | Unspecified |
| mmu-miR-24-3p | MIMAT0000219 | Ccnb2 | 12442 | HITS-CLIP | | 23142080 | Unspecified |
| mmu-miR-24-3p | MIMAT0000219 | Ccnc | 51813 | HITS-CLIP | 25753418\|23142080 | | Unspecified |
| mmu-miR-24-3p | MIMAT0000219 | Ccnd2 | 12444 | HITS-CLIP | | 23142080 | Unspecified |
| mmu-miR-24-3p | MIMAT0000219 | Ccne1 | 12447 | HITS-CLIP | | 23142080 | Unspecified |
| mmu-miR-24-3p | MIMAT0000219 | Ccne2 | 12448 | HITS-CLIP | | 23142080 | Unspecified |
| mmu-miR-24-3p | MIMAT0000219 | Ccnf | 12449 | HITS-CLIP | | 23142080 | Unspecified |
| mmu-miR-24-3p | MIMAT0000219 | Ccng1 | 12450 | HITS-CLIP | | 23142080 | Unspecified |

| | | | | | | |
|---|---|---|---|---|---|---|
| mmu-miR-24-3p | MIMAT0000219 | Ccnt2 | 72949 | Sequencing//HITS-CLIP | 19536157\|23142080\|37986224 | Unspecified |
| mmu-miR-24-3p | MIMAT0000219 | Ccny | 67974 | HITS-CLIP | 25753418\|23142080 | Unspecified |
| mmu-miR-24-3p | MIMAT0000219 | Ccs | 12460 | HITS-CLIP | 25753418 | Unspecified |
| mmu-miR-24-3p | MIMAT0000219 | Ccsap | 73420 | HITS-CLIP | 23142080 | Unspecified |
| mmu-miR-24-3p | MIMAT0000219 | Ccser2 | 72972 | Sequencing//HITS-CLIP | 19536157\|23142080 | Unspecified |
| mmu-miR-24-3p | MIMAT0000219 | Cct3 | 12462 | HITS-CLIP | 25753418 | Unspecified |
| mmu-miR-24-3p | MIMAT0000219 | Cct4 | 12464 | HITS-CLIP | 23142080 | Unspecified |
| mmu-miR-24-3p | MIMAT0000219 | Cct5 | 12465 | HITS-CLIP | 23142080 | Unspecified |
| mmu-miR-24-3p | MIMAT0000219 | Cct6a | 12466 | HITS-CLIP | 25753418 | Unspecified |
| mmu-miR-24-3p | MIMAT0000219 | Cct7 | 12468 | HITS-CLIP | 23142080 | Unspecified |
| mmu-miR-24-3p | MIMAT0000219 | Cd164 | 53599 | HITS-CLIP | 25753418 | Unspecified |
| mmu-miR-24-3p | MIMAT0000219 | Cd28 | 12487 | HITS-CLIP | 37986224\|23142080 | Unspecified |
| mmu-miR-24-3p | MIMAT0000219 | Cd3g | 12502 | HITS-CLIP | 23142080 | Unspecified |
| mmu-miR-24-3p | MIMAT0000219 | Cd4 | 12504 | HITS-CLIP | 23142080 | Unspecified |
| mmu-miR-24-3p | MIMAT0000219 | Cd44 | 12505 | HITS-CLIP | 23142080 | Unspecified |
| mmu-miR-24-3p | MIMAT0000219 | Cd47 | 16423 | HITS-CLIP | 37986224\|23142080\|25753418 | Unspecified |
| mmu-miR-24-3p | MIMAT0000219 | Cd5 | 12507 | HITS-CLIP | 23142080 | Unspecified |
| mmu-miR-24-3p | MIMAT0000219 | Cd53 | 12508 | HITS-CLIP | 37986224 | Unspecified |
| mmu-miR-24-3p | MIMAT0000219 | Cd69 | 12515 | HITS-CLIP | 37986224 | Unspecified |
| mmu-miR-24-3p | MIMAT0000219 | Cd74 | 16149 | HITS-CLIP | 23142080 | Unspecified |
| mmu-miR-24-3p | MIMAT0000219 | Cd81 | 12520 | qCLASH//HITS-CLIP | 31393953\|23142080 | Unspecified |
| mmu-miR-24-3p | MIMAT0000219 | Cd82 | 12521 | HITS-CLIP | 23142080 | Unspecified |
| mmu-miR-24-3p | MIMAT0000219 | Cdc14b | 218294 | HITS-CLIP | 25753418\|23142080 | Unspecified |
| mmu-miR-24-3p | MIMAT0000219 | Cdc25b | 12531 | HITS-CLIP | 37986224 | Unspecified |
| mmu-miR-24-3p | MIMAT0000219 | Cdc25c | 12532 | HITS-CLIP | 37986224\|23142080 | Unspecified |
| mmu-miR-24-3p | MIMAT0000219 | Cdc27 | 217232 | HITS-CLIP | 37986224\|23142080 | Unspecified |
| mmu-miR-24-3p | MIMAT0000219 | Cdc34 | 216150 | HITS-CLIP | 23142080 | Unspecified |
| mmu-miR-24-3p | MIMAT0000219 | Cdc37l1 | 67072 | HITS-CLIP | 23142080 | Unspecified |
| mmu-miR-24-3p | MIMAT0000219 | Cdc42ep1 | 104445 | HITS-CLIP | 25753418 | Unspecified |
| mmu-miR-24-3p | MIMAT0000219 | Cdc42se1 | 57912 | HITS-CLIP | 31433990 | Unspecified |
| mmu-miR-24-3p | MIMAT0000219 | Cdc5l | 71702 | HITS-CLIP | 23142080\|25753418 | Unspecified |
| mmu-miR-24-3p | MIMAT0000219 | Cdc7 | 12545 | HITS-CLIP | 37986224 | Unspecified |
| mmu-miR-24-3p | MIMAT0000219 | Cdca2 | 108912 | HITS-CLIP | 23142080 | Unspecified |
| mmu-miR-24-3p | MIMAT0000219 | Cdca3 | 14793 | HITS-CLIP//qCLASH | 23142080\|31393953 | Unspecified |
| mmu-miR-24-3p | MIMAT0000219 | Cdh13 | 12554 | HITS-CLIP | 25753418 | Unspecified |
| mmu-miR-24-3p | MIMAT0000219 | Cdip1 | 66626 | HITS-CLIP | 23142080 | Unspecified |
| mmu-miR-24-3p | MIMAT0000219 | Cdk11b | 12537 | HITS-CLIP | 23142080 | Unspecified |
| mmu-miR-24-3p | MIMAT0000219 | Cdk12 | 69131 | HITS-CLIP | 23142080 | Unspecified |
| mmu-miR-24-3p | MIMAT0000219 | Cdk13 | 69562 | HITS-CLIP | 23142080 | Unspecified |

| | | | | | | |
|---|---|---|---|---|---|---|
| mmu-miR-24-3p | MIMAT0000219 | Cdk14 | 18647 | Luciferase Reporter Assay | 37986224 | Unspecified |
| mmu-miR-24-3p | MIMAT0000219 | Cdk16 | 18555 | HITS-CLIP | 25753418 | Unspecified |
| mmu-miR-24-3p | MIMAT0000219 | Cdk17 | 237459 | HITS-CLIP | 37986224 | Unspecified |
| mmu-miR-24-3p | MIMAT0000219 | Cdk2ap1 | 13445 | HITS-CLIP | 37986224 | Unspecified |
| mmu-miR-24-3p | MIMAT0000219 | Cdk5 | 12568 | HITS-CLIP | 23142080 | Unspecified |
| mmu-miR-24-3p | MIMAT0000219 | Cdk6 | 12571 | HITS-CLIP | 23142080\|26203562\|25753418 | Unspecified |
| mmu-miR-24-3p | MIMAT0000219 | Cdkn1b | 12576 | HITS-CLIP | 37986224\|23142080 | Unspecified |
| mmu-miR-24-3p | MIMAT0000219 | Cdkn2aipnl | 52626 | HITS-CLIP | 23142080 | Unspecified |
| mmu-miR-24-3p | MIMAT0000219 | Cdon | 57810 | HITS-CLIP | 23142080 | Unspecified |
| mmu-miR-24-3p | MIMAT0000219 | Cds2 | 110911 | Sequencing//HITS-CLIP | 37986224\|23142080\|19536157 | Unspecified |
| mmu-miR-24-3p | MIMAT0000219 | Cdt1 | 67177 | HITS-CLIP | 23142080 | Unspecified |
| mmu-miR-24-3p | MIMAT0000219 | Cdv3 | 321022 | HITS-CLIP | 23142080 | Unspecified |
| mmu-miR-24-3p | MIMAT0000219 | Cdyl | 12593 | HITS-CLIP | 23142080 | Unspecified |
| mmu-miR-24-3p | MIMAT0000219 | Cdyl2 | 75796 | HITS-CLIP | 37986224 | Unspecified |
| mmu-miR-24-3p | MIMAT0000219 | Celf1 | 13046 | HITS-CLIP | 23142080\|25753418 | Unspecified |
| mmu-miR-24-3p | MIMAT0000219 | Celf2 | 14007 | HITS-CLIP | 23142080\|25753418 | Unspecified |
| mmu-miR-24-3p | MIMAT0000219 | Celsr1 | 12614 | HITS-CLIP | 23142080 | Unspecified |
| mmu-miR-24-3p | MIMAT0000219 | Cenpe | 229841 | HITS-CLIP | 23142080 | Unspecified |
| mmu-miR-24-3p | MIMAT0000219 | Cenpf | 108000 | HITS-CLIP | 23142080 | Unspecified |
| mmu-miR-24-3p | MIMAT0000219 | Cenpk | 60411 | HITS-CLIP | 23142080 | Unspecified |
| mmu-miR-24-3p | MIMAT0000219 | Cenpt | 320394 | HITS-CLIP | 23142080 | Unspecified |
| mmu-miR-24-3p | MIMAT0000219 | Cep135 | 381644 | qCLASH | 31393953 | Unspecified |
| mmu-miR-24-3p | MIMAT0000219 | Cep170 | 545389 | Sequencing//HITS-CLIP | 19536157\|23142080\|37986224 | Unspecified |
| mmu-miR-24-3p | MIMAT0000219 | Cep192 | 70799 | HITS-CLIP | 23142080 | Unspecified |
| mmu-miR-24-3p | MIMAT0000219 | Cep20 | 123811 | HITS-CLIP | 25753418 | Unspecified |
| mmu-miR-24-3p | MIMAT0000219 | Cep295 | 85459 | HITS-CLIP | 23142080 | Unspecified |
| mmu-miR-24-3p | MIMAT0000219 | Cep55 | 74107 | HITS-CLIP | 23142080 | Unspecified |
| mmu-miR-24-3p | MIMAT0000219 | Cep95 | 320162 | Sequencing//HITS-CLIP | 19536157\|23142080 | Unspecified |
| mmu-miR-24-3p | MIMAT0000219 | Cers2 | 76893 | HITS-CLIP | 23142080 | Unspecified |
| mmu-miR-24-3p | MIMAT0000219 | Cfl2 | 12632 | HITS-CLIP | 25753418 | Unspecified |
| mmu-miR-24-3p | MIMAT0000219 | Champ1 | 101994 | HITS-CLIP | 23142080 | Unspecified |
| mmu-miR-24-3p | MIMAT0000219 | Chd2 | 244059 | HITS-CLIP | 23142080 | Unspecified |
| mmu-miR-24-3p | MIMAT0000219 | Chd4 | 107932 | HITS-CLIP | 23142080 | Unspecified |
| mmu-miR-24-3p | MIMAT0000219 | Chd7 | 320790 | HITS-CLIP | 25753418\|23142080 | Unspecified |
| mmu-miR-24-3p | MIMAT0000219 | Chd8 | 67772 | HITS-CLIP | 23142080 | Unspecified |
| mmu-miR-24-3p | MIMAT0000219 | Chd9 | 109151 | HITS-CLIP | 23142080 | Unspecified |
| mmu-miR-24-3p | MIMAT0000219 | Chic2 | 74277 | HITS-CLIP | 25753418 | Unspecified |
| mmu-miR-24-3p | MIMAT0000219 | Chka | 12660 | HITS-CLIP | 25753418 | Unspecified |
| mmu-miR-24-3p | MIMAT0000219 | Chmp1b | 67078 | HITS-CLIP | 25753418 | Unspecified |

| | | | | | | |
|---|---|---|---|---|---|---|
| mmu-miR-24-3p | MIMAT0000219 | Chmp1b2 | 67028 | qCLASH | 31393953 | Unspecified |
| mmu-miR-24-3p | MIMAT0000219 | Chmp4b | 75608 | HITS-CLIP | 23142080 | Unspecified |
| mmu-miR-24-3p | MIMAT0000219 | Chmp5 | 76959 | HITS-CLIP | 25753418\|23142080 | Unspecified |
| mmu-miR-24-3p | MIMAT0000219 | Chmp7 | 105513 | HITS-CLIP | 23142080 | Unspecified |
| mmu-miR-24-3p | MIMAT0000219 | Chordc1 | 66917 | HITS-CLIP | 23142080 | Unspecified |
| mmu-miR-24-3p | MIMAT0000219 | Chp1 | 56398 | HITS-CLIP | 25753418 | Unspecified |
| mmu-miR-24-3p | MIMAT0000219 | Chsy1 | 269941 | HITS-CLIP | 23142080 | Unspecified |
| mmu-miR-24-3p | MIMAT0000219 | Chtop | 66511 | HITS-CLIP | 37986224 | Unspecified |
| mmu-miR-24-3p | MIMAT0000219 | Ciao1 | 26371 | HITS-CLIP | 23142080 | Unspecified |
| mmu-miR-24-3p | MIMAT0000219 | Ciapin1 | 109006 | HITS-CLIP | 25753418 | Unspecified |
| mmu-miR-24-3p | MIMAT0000219 | Cic | 71722 | HITS-CLIP | 23142080 | Unspecified |
| mmu-miR-24-3p | MIMAT0000219 | Cisd2 | 67006 | Luciferase Reporter Assay | 37986224 | Unspecified |
| mmu-miR-24-3p | MIMAT0000219 | Cish | 12700 | HITS-CLIP | 37986224\|23142080 | Unspecified |
| mmu-miR-24-3p | MIMAT0000219 | Cited2 | 17684 | HITS-CLIP | 37986224\|25753418 | Unspecified |
| mmu-miR-24-3p | MIMAT0000219 | Ckap4 | 216197 | HITS-CLIP | 25753418 | Unspecified |
| mmu-miR-24-3p | MIMAT0000219 | Ckap5 | 75786 | HITS-CLIP | 23142080 | Unspecified |
| mmu-miR-24-3p | MIMAT0000219 | Ckb | 12709 | HITS-CLIP | 23142080 | Unspecified |
| mmu-miR-24-3p | MIMAT0000219 | Cks1b | 54124 | HITS-CLIP | 23142080 | Unspecified |
| mmu-miR-24-3p | MIMAT0000219 | Clcf1 | 56708 | HITS-CLIP | 23142080 | Unspecified |
| mmu-miR-24-3p | MIMAT0000219 | Clcn3 | 12725 | Sequencing//HITS-CLIP | 18353861\|23142080\|37986224 | Unspecified |
| mmu-miR-24-3p | MIMAT0000219 | Cldn12 | 64945 | HITS-CLIP | 23142080 | Unspecified |
| mmu-miR-24-3p | MIMAT0000219 | Clint1 | 216705 | HITS-CLIP | 23142080 | Unspecified |
| mmu-miR-24-3p | MIMAT0000219 | Clk4 | 12750 | HITS-CLIP | 23142080 | Unspecified |
| mmu-miR-24-3p | MIMAT0000219 | Clp1 | 98985 | HITS-CLIP | 23142080 | Unspecified |
| mmu-miR-24-3p | MIMAT0000219 | Clpb | 20480 | HITS-CLIP | 25753418 | Unspecified |
| mmu-miR-24-3p | MIMAT0000219 | Clptm1 | 56457 | HITS-CLIP | 25753418 | Unspecified |
| mmu-miR-24-3p | MIMAT0000219 | Clpx | 270166 | HITS-CLIP | 23142080 | Unspecified |
| mmu-miR-24-3p | MIMAT0000219 | Clspn | 269582 | HITS-CLIP | 37986224 | Unspecified |
| mmu-miR-24-3p | MIMAT0000219 | Cltc | 67300 | HITS-CLIP | 37986224\|23142080\|25753418 | Unspecified |
| mmu-miR-24-3p | MIMAT0000219 | Clybl | 69634 | HITS-CLIP | 23142080 | Unspecified |
| mmu-miR-24-3p | MIMAT0000219 | Cmas | 12764 | HITS-CLIP | 23142080 | Unspecified |
| mmu-miR-24-3p | MIMAT0000219 | Cmtm4 | 97487 | HITS-CLIP | 23142080 | Unspecified |
| mmu-miR-24-3p | MIMAT0000219 | Cmtm6 | 67213 | Sequencing//HITS-CLIP | 19536157\|25753418\|23142080\|3798 | Unspecified |
| mmu-miR-24-3p | MIMAT0000219 | Cmtr1 | 74157 | HITS-CLIP | 23142080 | Unspecified |
| mmu-miR-24-3p | MIMAT0000219 | Cnbp | 12785 | HITS-CLIP | 23142080 | Unspecified |
| mmu-miR-24-3p | MIMAT0000219 | Cnep1r1 | 382030 | HITS-CLIP | 23142080 | Unspecified |
| mmu-miR-24-3p | MIMAT0000219 | Cnn2 | 12798 | HITS-CLIP | 37986224 | Unspecified |
| mmu-miR-24-3p | MIMAT0000219 | Cnot1 | 234594 | HITS-CLIP | 23142080\|25753418 | Unspecified |
| mmu-miR-24-3p | MIMAT0000219 | Cnot6 | 104625 | Sequencing//HITS-CLIP | 19536157\|23142080\|37986224 | Unspecified |

| | | | | | | |
|---|---|---|---|---|---|---|
| mmu-miR-24-3p | MIMAT0000219 | Cnst | 226744 | Sequencing//HITS-CLIP | 19536157\|23142080 | Unspecified |
| mmu-miR-24-3p | MIMAT0000219 | Coa5 | 76178 | HITS-CLIP | 25753418 | Unspecified |
| mmu-miR-24-3p | MIMAT0000219 | Cog3 | 338337 | Sequencing//HITS-CLIP | 19536157\|23142080\|37986224 | Unspecified |
| mmu-miR-24-3p | MIMAT0000219 | Cog5 | 238123 | HITS-CLIP | 23142080 | Unspecified |
| mmu-miR-24-3p | MIMAT0000219 | Cog6 | 67542 | HITS-CLIP | 23142080 | Unspecified |
| mmu-miR-24-3p | MIMAT0000219 | Cog7 | 233824 | HITS-CLIP | 23142080\|25753418 | Unspecified |
| mmu-miR-24-3p | MIMAT0000219 | Col15a1 | 12819 | HITS-CLIP | 25753418 | Unspecified |
| mmu-miR-24-3p | MIMAT0000219 | Col4a1 | 12826 | Sequencing//HITS-CLIP | 19536157\|25753418\|37986224 | Unspecified |
| mmu-miR-24-3p | MIMAT0000219 | Col4a2 | 12827 | HITS-CLIP | 25753418 | Unspecified |
| mmu-miR-24-3p | MIMAT0000219 | Colgalt1 | 234407 | HITS-CLIP | 23142080 | Unspecified |
| mmu-miR-24-3p | MIMAT0000219 | Commd9 | 76501 | HITS-CLIP | 23142080 | Unspecified |
| mmu-miR-24-3p | MIMAT0000219 | Copa | 12847 | HITS-CLIP | 23142080 | Unspecified |
| mmu-miR-24-3p | MIMAT0000219 | Copb2 | 50797 | HITS-CLIP | 23142080 | Unspecified |
| mmu-miR-24-3p | MIMAT0000219 | Copg1 | 54161 | HITS-CLIP | 23142080 | Unspecified |
| mmu-miR-24-3p | MIMAT0000219 | Copg2 | 54160 | HITS-CLIP | 23142080 | Unspecified |
| mmu-miR-24-3p | MIMAT0000219 | Cops2 | 12848 | HITS-CLIP | 23142080 | Unspecified |
| mmu-miR-24-3p | MIMAT0000219 | Cops4 | 26891 | HITS-CLIP | 37986224\|23142080 | Unspecified |
| mmu-miR-24-3p | MIMAT0000219 | Copz1 | 56447 | HITS-CLIP | 23142080 | Unspecified |
| mmu-miR-24-3p | MIMAT0000219 | Coq10a | 210582 | HITS-CLIP | 25753418 | Unspecified |
| mmu-miR-24-3p | MIMAT0000219 | Coq3 | 230027 | HITS-CLIP | 25753418\|23142080 | Unspecified |
| mmu-miR-24-3p | MIMAT0000219 | Coro1a | 12721 | HITS-CLIP | 23142080 | Unspecified |
| mmu-miR-24-3p | MIMAT0000219 | Coro1c | 23790 | HITS-CLIP | 25753418 | Unspecified |
| mmu-miR-24-3p | MIMAT0000219 | Cox5a | 12858 | HITS-CLIP | 23142080 | Unspecified |
| mmu-miR-24-3p | MIMAT0000219 | Cox6b2 | 333182 | HITS-CLIP | 25753418 | Unspecified |
| mmu-miR-24-3p | MIMAT0000219 | Cox7a2 | 12866 | HITS-CLIP | 23142080 | Unspecified |
| mmu-miR-24-3p | MIMAT0000219 | Cpd | 12874 | HITS-CLIP | 23142080 | Unspecified |
| mmu-miR-24-3p | MIMAT0000219 | Cpeb3 | 208922 | HITS-CLIP | 25753418 | Unspecified |
| mmu-miR-24-3p | MIMAT0000219 | Cpeb4 | 67579 | HITS-CLIP | 25753418 | Unspecified |
| mmu-miR-24-3p | MIMAT0000219 | Cpm | 70574 | HITS-CLIP | 23142080 | Unspecified |
| mmu-miR-24-3p | MIMAT0000219 | Cpne1 | 266692 | HITS-CLIP | 23142080 | Unspecified |
| mmu-miR-24-3p | MIMAT0000219 | Cpsf4 | 54188 | HITS-CLIP | 23142080 | Unspecified |
| mmu-miR-24-3p | MIMAT0000219 | Cpsf6 | 432508 | HITS-CLIP | 23142080 | Unspecified |
| mmu-miR-24-3p | MIMAT0000219 | Cpt1a | 12894 | HITS-CLIP | 25753418 | Unspecified |
| mmu-miR-24-3p | MIMAT0000219 | Cradd | 12905 | HITS-CLIP | 25753418 | Unspecified |
| mmu-miR-24-3p | MIMAT0000219 | Crcp | 12909 | HITS-CLIP | 23142080 | Unspecified |
| mmu-miR-24-3p | MIMAT0000219 | Creb1 | 12912 | HITS-CLIP | 23142080 | Unspecified |
| mmu-miR-24-3p | MIMAT0000219 | Creb3l2 | 208647 | HITS-CLIP | 25753418 | Unspecified |
| mmu-miR-24-3p | MIMAT0000219 | Crebbp | 12914 | HITS-CLIP | 25753418 | Unspecified |
| mmu-miR-24-3p | MIMAT0000219 | Crebl2 | 232430 | HITS-CLIP | 37986224\|23142080 | Unspecified |

| | | | | | | |
|---|---|---|---|---|---|---|
| mmu-miR-24-3p | MIMAT0000219 | Crebrf | 77128 | HITS-CLIP | 25753418 | Unspecified |
| mmu-miR-24-3p | MIMAT0000219 | Crip1 | 12925 | HITS-CLIP | 23142080 | Unspecified |
| mmu-miR-24-3p | MIMAT0000219 | Crip2 | 68337 | HITS-CLIP | 25753418 | Unspecified |
| mmu-miR-24-3p | MIMAT0000219 | Crk | 12928 | HITS-CLIP | 25753418\|23142080 | Unspecified |
| mmu-miR-24-3p | MIMAT0000219 | Crkl | 12929 | HITS-CLIP | 23142080 | Unspecified |
| mmu-miR-24-3p | MIMAT0000219 | Crls1 | 66586 | HITS-CLIP | 23142080 | Unspecified |
| mmu-miR-24-3p | MIMAT0000219 | Crnkl1 | 66877 | HITS-CLIP | 23142080 | Unspecified |
| mmu-miR-24-3p | MIMAT0000219 | Crtam | 54698 | HITS-CLIP | 23142080 | Unspecified |
| mmu-miR-24-3p | MIMAT0000219 | Cry2 | 12953 | HITS-CLIP | 37986224\|23142080 | Unspecified |
| mmu-miR-24-3p | MIMAT0000219 | Cryab | 12955 | HITS-CLIP | 25753418 | Unspecified |
| mmu-miR-24-3p | MIMAT0000219 | Crybg1 | 202 | HITS-CLIP | 23142080 | Unspecified |
| mmu-miR-24-3p | MIMAT0000219 | Csde1 | 229663 | HITS-CLIP | 23142080 | Unspecified |
| mmu-miR-24-3p | MIMAT0000219 | Csk | 12988 | HITS-CLIP | 25753418\|23142080 | Unspecified |
| mmu-miR-24-3p | MIMAT0000219 | Csnk1a1 | 93687 | Sequencing//HITS-CLIP | 37986224\|23142080\|19536157 | Unspecified |
| mmu-miR-24-3p | MIMAT0000219 | Csnk1d | 104318 | HITS-CLIP | 23142080 | Unspecified |
| mmu-miR-24-3p | MIMAT0000219 | Csnk1g2 | 103236 | HITS-CLIP | 23142080 | Unspecified |
| mmu-miR-24-3p | MIMAT0000219 | Cspp1 | 211660 | HITS-CLIP | 23142080 | Unspecified |
| mmu-miR-24-3p | MIMAT0000219 | Csrnp1 | 215418 | HITS-CLIP | 23142080 | Unspecified |
| mmu-miR-24-3p | MIMAT0000219 | Cst7 | 13011 | HITS-CLIP | 23142080 | Unspecified |
| mmu-miR-24-3p | MIMAT0000219 | Cstf3 | 228410 | HITS-CLIP | 37986224 | Unspecified |
| mmu-miR-24-3p | MIMAT0000219 | Ctbp1 | 13016 | HITS-CLIP | 23142080 | Unspecified |
| mmu-miR-24-3p | MIMAT0000219 | Ctcf | 13018 | HITS-CLIP | 37986224\|25753418 | Unspecified |
| mmu-miR-24-3p | MIMAT0000219 | Ctla4 | 12477 | HITS-CLIP | 23142080 | Unspecified |
| mmu-miR-24-3p | MIMAT0000219 | Ctnna1 | 12385 | HITS-CLIP | 23142080 | Unspecified |
| mmu-miR-24-3p | MIMAT0000219 | Ctnnb1 | 12387 | HITS-CLIP | 37986224\|23142080\|25753418 | Unspecified |
| mmu-miR-24-3p | MIMAT0000219 | Ctps2 | 55936 | HITS-CLIP | 23142080 | Unspecified |
| mmu-miR-24-3p | MIMAT0000219 | Ctr9 | 22083 | HITS-CLIP | 23142080 | Unspecified |
| mmu-miR-24-3p | MIMAT0000219 | Ctsd | 13033 | HITS-CLIP | 23142080 | Unspecified |
| mmu-miR-24-3p | MIMAT0000219 | Cul1 | 26965 | HITS-CLIP | 25753418 | Unspecified |
| mmu-miR-24-3p | MIMAT0000219 | Cul2 | 71745 | HITS-CLIP | 23142080 | Unspecified |
| mmu-miR-24-3p | MIMAT0000219 | Cul3 | 26554 | HITS-CLIP | 23142080\|25753418 | Unspecified |
| mmu-miR-24-3p | MIMAT0000219 | Cul4a | 99375 | HITS-CLIP | 25753418 | Unspecified |
| mmu-miR-24-3p | MIMAT0000219 | Cul5 | 75717 | HITS-CLIP | 23142080 | Unspecified |
| mmu-miR-24-3p | MIMAT0000219 | Cux1 | 13047 | HITS-CLIP | 37986224 | Unspecified |
| mmu-miR-24-3p | MIMAT0000219 | Cwc15 | 66070 | HITS-CLIP | 23142080 | Unspecified |
| mmu-miR-24-3p | MIMAT0000219 | Cxadr | 13052 | HITS-CLIP | 23142080 | Unspecified |
| mmu-miR-24-3p | MIMAT0000219 | Cxcr4 | 12767 | HITS-CLIP | 23142080 | Unspecified |
| mmu-miR-24-3p | MIMAT0000219 | Cxxc1 | 74322 | HITS-CLIP | 23142080 | Unspecified |
| mmu-miR-24-3p | MIMAT0000219 | Cyb561d1 | 72023 | HITS-CLIP | 23142080 | Unspecified |

| | | | | | | |
|---|---|---|---|---|---|---|
| mmu-miR-24-3p | MIMAT0000219 | Cyb5r3 | 109754 | HITS-CLIP | 23142080 | Unspecified |
| mmu-miR-24-3p | MIMAT0000219 | Cyp20a1 | 77951 | HITS-CLIP | 23142080 | Unspecified |
| mmu-miR-24-3p | MIMAT0000219 | Cyp51 | 13121 | HITS-CLIP | 23142080 | Unspecified |
| mmu-miR-24-3p | MIMAT0000219 | Cysltr1 | 58861 | HITS-CLIP | 23142080 | Unspecified |
| mmu-miR-24-3p | MIMAT0000219 | Cyth2 | 19158 | HITS-CLIP | 23142080 | Unspecified |
| mmu-miR-24-3p | MIMAT0000219 | Cyth3 | 19159 | HITS-CLIP | 23142080 | Unspecified |
| mmu-miR-24-3p | MIMAT0000219 | Cytip | 227929 | HITS-CLIP | 23142080 | Unspecified |
| mmu-miR-24-3p | MIMAT0000219 | D030056L22Rik | 225995 | HITS-CLIP | 23142080 | Unspecified |
| mmu-miR-24-3p | MIMAT0000219 | D16Ertd472e | 67102 | HITS-CLIP | 23142080 | Unspecified |
| mmu-miR-24-3p | MIMAT0000219 | Daam1 | 208846 | HITS-CLIP | 25753418\|23142080 | Unspecified |
| mmu-miR-24-3p | MIMAT0000219 | Dab2ip | 69601 | HITS-CLIP | 25753418 | Unspecified |
| mmu-miR-24-3p | MIMAT0000219 | Dag1 | 13138 | HITS-CLIP | 23142080\|25753418 | Unspecified |
| mmu-miR-24-3p | MIMAT0000219 | Dars | 226414 | HITS-CLIP | 25753418 | Unspecified |
| mmu-miR-24-3p | MIMAT0000219 | Dbf4 | 27214 | HITS-CLIP | 23142080 | Unspecified |
| mmu-miR-24-3p | MIMAT0000219 | Dbnl | 13169 | HITS-CLIP | 23142080 | Unspecified |
| mmu-miR-24-3p | MIMAT0000219 | Dbt | 13171 | HITS-CLIP | 23142080 | Unspecified |
| mmu-miR-24-3p | MIMAT0000219 | Dcaf10 | 242418 | HITS-CLIP | 23142080\|25753418 | Unspecified |
| mmu-miR-24-3p | MIMAT0000219 | Dcaf13 | 223499 | HITS-CLIP | 23142080 | Unspecified |
| mmu-miR-24-3p | MIMAT0000219 | Dcaf6 | 74106 | HITS-CLIP | 23142080 | Unspecified |
| mmu-miR-24-3p | MIMAT0000219 | Dcaf7 | 71833 | HITS-CLIP | 37986224\|25753418 | Unspecified |
| mmu-miR-24-3p | MIMAT0000219 | Dcaf8 | 98193 | Sequencing//HITS-CLIP | 19536157\|23142080 | Unspecified |
| mmu-miR-24-3p | MIMAT0000219 | Dcakd | 68087 | HITS-CLIP | 23142080 | Unspecified |
| mmu-miR-24-3p | MIMAT0000219 | Dcp1a | 75901 | HITS-CLIP | 23142080 | Unspecified |
| mmu-miR-24-3p | MIMAT0000219 | Dcp2 | 70640 | HITS-CLIP | 37986224\|23142080 | Unspecified |
| mmu-miR-24-3p | MIMAT0000219 | Dctn1 | 13191 | HITS-CLIP | 23142080 | Unspecified |
| mmu-miR-24-3p | MIMAT0000219 | Dctn4 | 67665 | HITS-CLIP | 23142080 | Unspecified |
| mmu-miR-24-3p | MIMAT0000219 | Dcun1d5 | 76863 | HITS-CLIP | 23142080 | Unspecified |
| mmu-miR-24-3p | MIMAT0000219 | Ddb1 | 13194 | HITS-CLIP | 23142080\|25753418 | Unspecified |
| mmu-miR-24-3p | MIMAT0000219 | Ddhd1 | 114874 | HITS-CLIP | 23142080 | Unspecified |
| mmu-miR-24-3p | MIMAT0000219 | Ddi2 | 68817 | HITS-CLIP | 25753418 | Unspecified |
| mmu-miR-24-3p | MIMAT0000219 | Ddit3 | 13198 | HITS-CLIP | 23142080 | Unspecified |
| mmu-miR-24-3p | MIMAT0000219 | Ddr1 | 12305 | HITS-CLIP | 23142080 | Unspecified |
| mmu-miR-24-3p | MIMAT0000219 | Ddx17 | 67040 | HITS-CLIP | 23142080 | Unspecified |
| mmu-miR-24-3p | MIMAT0000219 | Ddx23 | 74351 | HITS-CLIP | 23142080 | Unspecified |
| mmu-miR-24-3p | MIMAT0000219 | Ddx24 | 27225 | HITS-CLIP | 25753418 | Unspecified |
| mmu-miR-24-3p | MIMAT0000219 | Ddx39 | 68278 | HITS-CLIP | 23142080 | Unspecified |
| mmu-miR-24-3p | MIMAT0000219 | Ddx3x | 13205 | HITS-CLIP | 37986224\|23142080 | Unspecified |
| mmu-miR-24-3p | MIMAT0000219 | Ddx5 | 13207 | HITS-CLIP | 25753418 | Unspecified |
| mmu-miR-24-3p | MIMAT0000219 | Ddx6 | 13209 | HITS-CLIP | 23142080 | Unspecified |

| | | | | | | |
|---|---|---|---|---|---|---|
| mmu-miR-24-3p | MIMAT0000219 | Deaf1 | 54006 | HITS-CLIP | 23142080 | Unspecified |
| mmu-miR-24-3p | MIMAT0000219 | Decr1 | 67460 | HITS-CLIP | 25753418 | Unspecified |
| mmu-miR-24-3p | MIMAT0000219 | Dedd | 21945 | HITS-CLIP | 23142080 | Unspecified |
| mmu-miR-24-3p | MIMAT0000219 | Dennd1b | 329260 | HITS-CLIP | 23142080\|25753418 | Unspecified |
| mmu-miR-24-3p | MIMAT0000219 | Dennd2d | 72121 | HITS-CLIP | 37986224 | Unspecified |
| mmu-miR-24-3p | MIMAT0000219 | Dennd3 | 105841 | HITS-CLIP | 31433990 | Unspecified |
| mmu-miR-24-3p | MIMAT0000219 | Dennd4b | 229541 | HITS-CLIP | 23142080\|31433990\|25753418 | Unspecified |
| mmu-miR-24-3p | MIMAT0000219 | Dennd4c | 329877 | HITS-CLIP | 23142080 | Unspecified |
| mmu-miR-24-3p | MIMAT0000219 | Dennd5a | 19347 | HITS-CLIP | 25753418 | Unspecified |
| mmu-miR-24-3p | MIMAT0000219 | Dennd6a | 211922 | HITS-CLIP | 23142080 | Unspecified |
| mmu-miR-24-3p | MIMAT0000219 | Depdc1b | 218581 | HITS-CLIP | 23142080 | Unspecified |
| mmu-miR-24-3p | MIMAT0000219 | Derl1 | 67819 | HITS-CLIP | 37986224\|23142080 | Unspecified |
| mmu-miR-24-3p | MIMAT0000219 | Desi2 | 78825 | HITS-CLIP | 23142080 | Unspecified |
| mmu-miR-24-3p | MIMAT0000219 | Dgkq | 110524 | HITS-CLIP | 37986224\|23142080 | Unspecified |
| mmu-miR-24-3p | MIMAT0000219 | Dgkz | 104418 | HITS-CLIP | 23142080 | Unspecified |
| mmu-miR-24-3p | MIMAT0000219 | Dguok | 27369 | HITS-CLIP | 31433990 | Unspecified |
| mmu-miR-24-3p | MIMAT0000219 | Dhcr7 | 13360 | HITS-CLIP | 23142080 | Unspecified |
| mmu-miR-24-3p | MIMAT0000219 | Dhrs4 | 28200 | HITS-CLIP | 25753418 | Unspecified |
| mmu-miR-24-3p | MIMAT0000219 | Dhx30 | 72831 | HITS-CLIP | 37986224\|23142080 | Unspecified |
| mmu-miR-24-3p | MIMAT0000219 | Dhx33 | 216877 | HITS-CLIP | 37986224\|23142080 | Unspecified |
| mmu-miR-24-3p | MIMAT0000219 | Dhx36 | 72162 | HITS-CLIP | 37986224\|23142080 | Unspecified |
| mmu-miR-24-3p | MIMAT0000219 | Dhx40 | 67487 | HITS-CLIP | 23142080 | Unspecified |
| mmu-miR-24-3p | MIMAT0000219 | Dhx57 | 106794 | HITS-CLIP | 26203562\|23142080 | Unspecified |
| mmu-miR-24-3p | MIMAT0000219 | Dhx58 | 80861 | HITS-CLIP | 23142080 | Unspecified |
| mmu-miR-24-3p | MIMAT0000219 | Dhx9 | 13211 | HITS-CLIP | 23142080 | Unspecified |
| mmu-miR-24-3p | MIMAT0000219 | Diaph1 | 1729 | HITS-CLIP | 23142080 | Unspecified |
| mmu-miR-24-3p | MIMAT0000219 | Dicer1 | 192119 | HITS-CLIP | 25753418\|23142080 | Unspecified |
| mmu-miR-24-3p | MIMAT0000219 | Dido1 | 23856 | HITS-CLIP | 23142080 | Unspecified |
| mmu-miR-24-3p | MIMAT0000219 | Dip2b | 239667 | HITS-CLIP | 37986224\|23142080 | Unspecified |
| mmu-miR-24-3p | MIMAT0000219 | Disp1 | 68897 | HITS-CLIP | 23142080 | Unspecified |
| mmu-miR-24-3p | MIMAT0000219 | Dlc1 | 50768 | HITS-CLIP | 25753418 | Unspecified |
| mmu-miR-24-3p | MIMAT0000219 | Dlg1 | 13383 | HITS-CLIP | 23142080 | Unspecified |
| mmu-miR-24-3p | MIMAT0000219 | Dlg4 | 13385 | HITS-CLIP | 23142080 | Unspecified |
| mmu-miR-24-3p | MIMAT0000219 | Dlgap5 | 218977 | HITS-CLIP | 23142080 | Unspecified |
| mmu-miR-24-3p | MIMAT0000219 | Dlst | 78920 | HITS-CLIP | 23142080 | Unspecified |
| mmu-miR-24-3p | MIMAT0000219 | Dmtf1 | 23857 | HITS-CLIP | 37986224\|23142080 | Unspecified |
| mmu-miR-24-3p | MIMAT0000219 | Dmxl2 | 235380 | HITS-CLIP | 25753418 | Unspecified |
| mmu-miR-24-3p | MIMAT0000219 | Dna2 | 327762 | HITS-CLIP | 23142080 | Unspecified |
| mmu-miR-24-3p | MIMAT0000219 | Dnaja2 | 56445 | HITS-CLIP | 25753418 | Unspecified |

| | | | | | | | |
|---|---|---|---|---|---|---|---|
| mmu-miR-24-3p | MIMAT0000219 | Dnaja3 | 83945 | HITS-CLIP | | 23142080 | Unspecified |
| mmu-miR-24-3p | MIMAT0000219 | Dnajb12 | 56709 | HITS-CLIP | 37986224\|23142080 | | Unspecified |
| mmu-miR-24-3p | MIMAT0000219 | Dnajb5 | 56323 | HITS-CLIP | 23142080\|25753418 | | Unspecified |
| mmu-miR-24-3p | MIMAT0000219 | Dnajc16 | 214063 | HITS-CLIP | | 23142080 | Unspecified |
| mmu-miR-24-3p | MIMAT0000219 | Dnajc8 | 68598 | HITS-CLIP | | 23142080 | Unspecified |
| mmu-miR-24-3p | MIMAT0000219 | Dnm1l | 74006 | HITS-CLIP | | 23142080 | Unspecified |
| mmu-miR-24-3p | MIMAT0000219 | Dnm2 | 13430 | HITS-CLIP | | 23142080 | Unspecified |
| mmu-miR-24-3p | MIMAT0000219 | Dnmt1 | 13433 | HITS-CLIP | | 23142080 | Unspecified |
| mmu-miR-24-3p | MIMAT0000219 | Dnmt3a | 13435 | HITS-CLIP | | 25753418 | Unspecified |
| mmu-miR-24-3p | MIMAT0000219 | Dock10 | 210293 | HITS-CLIP | | 23142080 | Unspecified |
| mmu-miR-24-3p | MIMAT0000219 | Dock11 | 75974 | HITS-CLIP | | 23142080 | Unspecified |
| mmu-miR-24-3p | MIMAT0000219 | Dock2 | 94176 | HITS-CLIP | 37986224\|23142080 | | Unspecified |
| mmu-miR-24-3p | MIMAT0000219 | Dock8 | 76088 | HITS-CLIP | | 23142080 | Unspecified |
| mmu-miR-24-3p | MIMAT0000219 | Dop1b | 9980 | HITS-CLIP | | 23142080 | Unspecified |
| mmu-miR-24-3p | MIMAT0000219 | Dot1l | 208266 | HITS-CLIP | 37986224\|23142080 | | Unspecified |
| mmu-miR-24-3p | MIMAT0000219 | Dpm3 | 68563 | HITS-CLIP | | 23142080 | Unspecified |
| mmu-miR-24-3p | MIMAT0000219 | Dpp9 | 224897 | HITS-CLIP | | 23142080 | Unspecified |
| mmu-miR-24-3p | MIMAT0000219 | Dpysl2 | 12934 | HITS-CLIP | | 23142080 | Unspecified |
| mmu-miR-24-3p | MIMAT0000219 | Dr1 | 13486 | HITS-CLIP | | 23142080 | Unspecified |
| mmu-miR-24-3p | MIMAT0000219 | Dsp | 109620 | HITS-CLIP | | 25753418 | Unspecified |
| mmu-miR-24-3p | MIMAT0000219 | Dst | 13518 | HITS-CLIP | | 25753418 | Unspecified |
| mmu-miR-24-3p | MIMAT0000219 | Dtnb | 13528 | HITS-CLIP | | 23142080 | Unspecified |
| mmu-miR-24-3p | MIMAT0000219 | Dtx3l | 209200 | HITS-CLIP | | 23142080 | Unspecified |
| mmu-miR-24-3p | MIMAT0000219 | Dusp1 | 19252 | HITS-CLIP | | 23142080 | Unspecified |
| mmu-miR-24-3p | MIMAT0000219 | Dusp16 | 70686 | HITS-CLIP | | 23142080 | Unspecified |
| mmu-miR-24-3p | MIMAT0000219 | Dusp18 | 75219 | HITS-CLIP | | 25753418 | Unspecified |
| mmu-miR-24-3p | MIMAT0000219 | Dusp2 | 13537 | HITS-CLIP | | 23142080 | Unspecified |
| mmu-miR-24-3p | MIMAT0000219 | Dusp22 | 105352 | HITS-CLIP | | 23142080 | Unspecified |
| mmu-miR-24-3p | MIMAT0000219 | Dusp5 | 240672 | HITS-CLIP | | 37986224 | Unspecified |
| mmu-miR-24-3p | MIMAT0000219 | Dusp6 | 67603 | HITS-CLIP | 37986224\|23142080 | | Unspecified |
| mmu-miR-24-3p | MIMAT0000219 | Dym | 69190 | HITS-CLIP | | 23142080 | Unspecified |
| mmu-miR-24-3p | MIMAT0000219 | Dync1h1 | 13424 | HITS-CLIP | 23142080\|25753418 | | Unspecified |
| mmu-miR-24-3p | MIMAT0000219 | Dync1i2 | 13427 | HITS-CLIP | | 25753418 | Unspecified |
| mmu-miR-24-3p | MIMAT0000219 | Dync1li1 | 235661 | HITS-CLIP | | 25753418 | Unspecified |
| mmu-miR-24-3p | MIMAT0000219 | Dync1li2 | 234663 | HITS-CLIP | | 25753418 | Unspecified |
| mmu-miR-24-3p | MIMAT0000219 | Dynll2 | 68097 | HITS-CLIP | | 23142080 | Unspecified |
| mmu-miR-24-3p | MIMAT0000219 | Dynlrb1 | 67068 | HITS-CLIP | | 23142080 | Unspecified |
| mmu-miR-24-3p | MIMAT0000219 | Dyrk1a | 13548 | HITS-CLIP | | 23142080 | Unspecified |
| mmu-miR-24-3p | MIMAT0000219 | Dyrk2 | 69181 | HITS-CLIP | | 23142080 | Unspecified |

| | | | | | | |
|---|---|---|---|---|---|---|
| mmu-miR-24-3p | MIMAT0000219 | Dysf | 26903 | HITS-CLIP | 25753418 | Unspecified |
| mmu-miR-24-3p | MIMAT0000219 | Dzip3 | 224170 | HITS-CLIP | 23142080 | Unspecified |
| mmu-miR-24-3p | MIMAT0000219 | E2f4 | 104394 | HITS-CLIP | 37986224 | Unspecified |
| mmu-miR-24-3p | MIMAT0000219 | E2f7 | 52679 | HITS-CLIP | 23142080 | Unspecified |
| mmu-miR-24-3p | MIMAT0000219 | E2f8 | 108961 | HITS-CLIP | 23142080 | Unspecified |
| mmu-miR-24-3p | MIMAT0000219 | E330009J07Rik | 243780 | HITS-CLIP | 23142080 | Unspecified |
| mmu-miR-24-3p | MIMAT0000219 | E430018J23Rik | 101604 | HITS-CLIP | 25753418 | Unspecified |
| mmu-miR-24-3p | MIMAT0000219 | Eapp | 66266 | HITS-CLIP | 23142080 | Unspecified |
| mmu-miR-24-3p | MIMAT0000219 | Ece1 | 230857 | Sequencing//HITS-CLIP | 37986224\|23142080\|19536157 | Unspecified |
| mmu-miR-24-3p | MIMAT0000219 | Ecpas | 23392 | HITS-CLIP | 25753418\|23142080 | Unspecified |
| mmu-miR-24-3p | MIMAT0000219 | Edc4 | 234699 | HITS-CLIP | 25753418\|23142080 | Unspecified |
| mmu-miR-24-3p | MIMAT0000219 | Edem1 | 192193 | HITS-CLIP | 23142080 | Unspecified |
| mmu-miR-24-3p | MIMAT0000219 | Edem3 | 66967 | HITS-CLIP | 23142080 | Unspecified |
| mmu-miR-24-3p | MIMAT0000219 | Edrf1 | 214764 | HITS-CLIP | 23142080 | Unspecified |
| mmu-miR-24-3p | MIMAT0000219 | Eed | 13626 | HITS-CLIP | 23142080 | Unspecified |
| mmu-miR-24-3p | MIMAT0000219 | Eef1e1 | 66143 | HITS-CLIP | 23142080 | Unspecified |
| mmu-miR-24-3p | MIMAT0000219 | Eef2k | 13631 | HITS-CLIP | 23142080 | Unspecified |
| mmu-miR-24-3p | MIMAT0000219 | Efcab14 | 230648 | HITS-CLIP | 23142080 | Unspecified |
| mmu-miR-24-3p | MIMAT0000219 | Efr3a | 76740 | HITS-CLIP | 23142080 | Unspecified |
| mmu-miR-24-3p | MIMAT0000219 | Egr1 | 13653 | HITS-CLIP | 23142080 | Unspecified |
| mmu-miR-24-3p | MIMAT0000219 | Egr3 | 13655 | HITS-CLIP | 23142080 | Unspecified |
| mmu-miR-24-3p | MIMAT0000219 | Ehbp1l1 | 114601 | HITS-CLIP | 25753418 | Unspecified |
| mmu-miR-24-3p | MIMAT0000219 | Ehhadh | 74147 | HITS-CLIP | 25753418 | Unspecified |
| mmu-miR-24-3p | MIMAT0000219 | Ehmt1 | 77683 | HITS-CLIP | 23142080 | Unspecified |
| mmu-miR-24-3p | MIMAT0000219 | Ehmt2 | 110147 | HITS-CLIP | 23142080 | Unspecified |
| mmu-miR-24-3p | MIMAT0000219 | Ei24 | 13663 | HITS-CLIP | 26203562\|25753418 | Unspecified |
| mmu-miR-24-3p | MIMAT0000219 | Eif2ak1 | 15467 | HITS-CLIP | 23142080\|25753418 | Unspecified |
| mmu-miR-24-3p | MIMAT0000219 | Eif2ak3 | 13666 | HITS-CLIP | 23142080 | Unspecified |
| mmu-miR-24-3p | MIMAT0000219 | Eif2ak4 | 27103 | HITS-CLIP | 23142080 | Unspecified |
| mmu-miR-24-3p | MIMAT0000219 | Eif2s1 | 13665 | HITS-CLIP | 25753418 | Unspecified |
| mmu-miR-24-3p | MIMAT0000219 | Eif3a | 13669 | HITS-CLIP | 37986224\|23142080 | Unspecified |
| mmu-miR-24-3p | MIMAT0000219 | Eif3c | 56347 | HITS-CLIP | 23142080 | Unspecified |
| mmu-miR-24-3p | MIMAT0000219 | Eif3d | 55944 | HITS-CLIP | 25753418 | Unspecified |
| mmu-miR-24-3p | MIMAT0000219 | Eif3f | 66085 | HITS-CLIP | 23142080 | Unspecified |
| mmu-miR-24-3p | MIMAT0000219 | Eif3h | 68135 | HITS-CLIP | 23142080 | Unspecified |
| mmu-miR-24-3p | MIMAT0000219 | Eif4a2 | 13682 | HITS-CLIP | 23142080 | Unspecified |
| mmu-miR-24-3p | MIMAT0000219 | Eif4b | 75705 | HITS-CLIP//qCLASH | 23142080\|31393953 | Unspecified |
| mmu-miR-24-3p | MIMAT0000219 | Eif4e | 13684 | HITS-CLIP | 23142080 | Unspecified |
| mmu-miR-24-3p | MIMAT0000219 | Eif4e2 | 26987 | HITS-CLIP | 23142080\|25753418 | Unspecified |

| | | | | | | |
|---|---|---|---|---|---|---|
| mmu-miR-24-3p | MIMAT0000219 | Eif4ebp2 | 13688 | HITS-CLIP | 23142080 | Unspecified |
| mmu-miR-24-3p | MIMAT0000219 | Eif4enif1 | 74203 | HITS-CLIP | 23142080 | Unspecified |
| mmu-miR-24-3p | MIMAT0000219 | Eif4g1 | 208643 | HITS-CLIP | 25753418\|23142080 | Unspecified |
| mmu-miR-24-3p | MIMAT0000219 | Eif4g2 | 13690 | HITS-CLIP | 21258322\|31433990\|25753418 | Unspecified |
| mmu-miR-24-3p | MIMAT0000219 | Eif4g3 | 230861 | HITS-CLIP | 25753418 | Unspecified |
| mmu-miR-24-3p | MIMAT0000219 | Eif4h | 22384 | HITS-CLIP | 23142080\|25753418 | Unspecified |
| mmu-miR-24-3p | MIMAT0000219 | Eif5b | 226982 | HITS-CLIP | 23142080 | Unspecified |
| mmu-miR-24-3p | MIMAT0000219 | Elavl1 | 15568 | HITS-CLIP | 25753418 | Unspecified |
| mmu-miR-24-3p | MIMAT0000219 | Elf1 | 13709 | HITS-CLIP | 23142080 | Unspecified |
| mmu-miR-24-3p | MIMAT0000219 | Elf2 | 69257 | HITS-CLIP | 23142080 | Unspecified |
| mmu-miR-24-3p | MIMAT0000219 | Elf4 | 56501 | HITS-CLIP | 23142080 | Unspecified |
| mmu-miR-24-3p | MIMAT0000219 | Elk3 | 13713 | HITS-CLIP | 23142080 | Unspecified |
| mmu-miR-24-3p | MIMAT0000219 | Elk4 | 13714 | HITS-CLIP | 23142080 | Unspecified |
| mmu-miR-24-3p | MIMAT0000219 | Ell | 13716 | HITS-CLIP | 37986224\|23142080 | Unspecified |
| mmu-miR-24-3p | MIMAT0000219 | Ell2 | 192657 | HITS-CLIP | 23142080 | Unspecified |
| mmu-miR-24-3p | MIMAT0000219 | Elmo1 | 140580 | HITS-CLIP | 23142080 | Unspecified |
| mmu-miR-24-3p | MIMAT0000219 | Eln | 13717 | HITS-CLIP | 25753418 | Unspecified |
| mmu-miR-24-3p | MIMAT0000219 | Eloa | 6924 | HITS-CLIP | 25753418 | Unspecified |
| mmu-miR-24-3p | MIMAT0000219 | Elp1 | 37455 | HITS-CLIP | 25753418 | Unspecified |
| mmu-miR-24-3p | MIMAT0000219 | Elp3 | 74195 | HITS-CLIP | 23142080 | Unspecified |
| mmu-miR-24-3p | MIMAT0000219 | Elp5 | 54351 | HITS-CLIP | 37986224\|23142080 | Unspecified |
| mmu-miR-24-3p | MIMAT0000219 | Eml3 | 225898 | HITS-CLIP | 23142080 | Unspecified |
| mmu-miR-24-3p | MIMAT0000219 | Eml4 | 78798 | HITS-CLIP | 37986224\|23142080 | Unspecified |
| mmu-miR-24-3p | MIMAT0000219 | Eng | 13805 | HITS-CLIP | 25753418 | Unspecified |
| mmu-miR-24-3p | MIMAT0000219 | Eno3 | 13808 | HITS-CLIP | 25753418\|23142080 | Unspecified |
| mmu-miR-24-3p | MIMAT0000219 | Entpd5 | 12499 | HITS-CLIP | 25753418\|23142080 | Unspecified |
| mmu-miR-24-3p | MIMAT0000219 | Entpd6 | 12497 | Sequencing//HITS-CLIP | 37986224\|23142080\|19536157 | Unspecified |
| mmu-miR-24-3p | MIMAT0000219 | Eomes | 13813 | HITS-CLIP | 23142080 | Unspecified |
| mmu-miR-24-3p | MIMAT0000219 | Ep400 | 75560 | HITS-CLIP | 23142080\|25753418 | Unspecified |
| mmu-miR-24-3p | MIMAT0000219 | Epas1 | 13819 | HITS-CLIP | 23142080 | Unspecified |
| mmu-miR-24-3p | MIMAT0000219 | Epb41 | 269587 | HITS-CLIP | 23142080 | Unspecified |
| mmu-miR-24-3p | MIMAT0000219 | Epc2 | 227867 | HITS-CLIP | 23142080 | Unspecified |
| mmu-miR-24-3p | MIMAT0000219 | Epcam | 17075 | HITS-CLIP | 23142080 | Unspecified |
| mmu-miR-24-3p | MIMAT0000219 | Epg5 | 100502841 | HITS-CLIP | 23142080 | Unspecified |
| mmu-miR-24-3p | MIMAT0000219 | Ephx1 | 13849 | HITS-CLIP | 23142080 | Unspecified |
| mmu-miR-24-3p | MIMAT0000219 | Epm2aip1 | 77781 | HITS-CLIP | 23142080 | Unspecified |
| mmu-miR-24-3p | MIMAT0000219 | Epn1 | 13854 | Sequencing//HITS-CLIP | 19536157\|25753418\|23142080\|3798[illegible] | Unspecified |
| mmu-miR-24-3p | MIMAT0000219 | Eprs | 107508 | HITS-CLIP | 25753418\|23142080 | Unspecified |
| mmu-miR-24-3p | MIMAT0000219 | Eps15 | 13858 | HITS-CLIP | 23142080 | Unspecified |

| | | | | | | |
|---|---|---|---|---|---|---|
| mmu-miR-24-3p | MIMAT0000219 | Erbb2 | 13866 | HITS-CLIP | 25753418 | Unspecified |
| mmu-miR-24-3p | MIMAT0000219 | Erbin | 55914 | HITS-CLIP | 23142080 | Unspecified |
| mmu-miR-24-3p | MIMAT0000219 | Ercc2 | 13871 | HITS-CLIP | 23142080 | Unspecified |
| mmu-miR-24-3p | MIMAT0000219 | Ercc3 | 13872 | HITS-CLIP | 23142080 | Unspecified |
| mmu-miR-24-3p | MIMAT0000219 | Erf | 13875 | HITS-CLIP | 23142080 | Unspecified |
| mmu-miR-24-3p | MIMAT0000219 | Ergic2 | 67456 | HITS-CLIP | 25753418 | Unspecified |
| mmu-miR-24-3p | MIMAT0000219 | Eri1 | 67276 | HITS-CLIP | 23142080 | Unspecified |
| mmu-miR-24-3p | MIMAT0000219 | Eri2 | 71151 | HITS-CLIP | 23142080 | Unspecified |
| mmu-miR-24-3p | MIMAT0000219 | Erlin1 | 226144 | HITS-CLIP | 23142080 | Unspecified |
| mmu-miR-24-3p | MIMAT0000219 | Erlin2 | 244373 | HITS-CLIP | 23142080 | Unspecified |
| mmu-miR-24-3p | MIMAT0000219 | Ern1 | 78943 | HITS-CLIP | 23142080 | Unspecified |
| mmu-miR-24-3p | MIMAT0000219 | Ero1l | 50527 | HITS-CLIP | 23142080 | Unspecified |
| mmu-miR-24-3p | MIMAT0000219 | Erp29 | 67397 | HITS-CLIP | 23142080 | Unspecified |
| mmu-miR-24-3p | MIMAT0000219 | Esco2 | 71988 | HITS-CLIP | 23142080 | Unspecified |
| mmu-miR-24-3p | MIMAT0000219 | Esf1 | 66580 | HITS-CLIP | 23142080 | Unspecified |
| mmu-miR-24-3p | MIMAT0000219 | Espl1 | 105988 | HITS-CLIP | 23142080 | Unspecified |
| mmu-miR-24-3p | MIMAT0000219 | Espn | 56226 | HITS-CLIP | 23142080 | Unspecified |
| mmu-miR-24-3p | MIMAT0000219 | Esyt1 | 23943 | HITS-CLIP | 23142080 | Unspecified |
| mmu-miR-24-3p | MIMAT0000219 | Esyt2 | 52635 | Sequencing//HITS-CLIP | 19536157\|23142080 | Unspecified |
| mmu-miR-24-3p | MIMAT0000219 | Etf1 | 225363 | HITS-CLIP | 25753418 | Unspecified |
| mmu-miR-24-3p | MIMAT0000219 | Etfa | 110842 | HITS-CLIP | 23142080 | Unspecified |
| mmu-miR-24-3p | MIMAT0000219 | Etnk1 | 75320 | HITS-CLIP | 23142080 | Unspecified |
| mmu-miR-24-3p | MIMAT0000219 | Ets1 | 23871 | HITS-CLIP | 37986224\|23142080 | Unspecified |
| mmu-miR-24-3p | MIMAT0000219 | Ets2 | 23872 | HITS-CLIP | 23142080 | Unspecified |
| mmu-miR-24-3p | MIMAT0000219 | Etv5 | 104156 | HITS-CLIP | 23142080 | Unspecified |
| mmu-miR-24-3p | MIMAT0000219 | Etv6 | 14011 | HITS-CLIP | 23142080 | Unspecified |
| mmu-miR-24-3p | MIMAT0000219 | Exoc1 | 69940 | HITS-CLIP | 23142080 | Unspecified |
| mmu-miR-24-3p | MIMAT0000219 | Exoc2 | 66482 | HITS-CLIP | 23142080 | Unspecified |
| mmu-miR-24-3p | MIMAT0000219 | Exoc4 | 20336 | HITS-CLIP | 23142080 | Unspecified |
| mmu-miR-24-3p | MIMAT0000219 | Exoc6 | 107371 | HITS-CLIP | 23142080 | Unspecified |
| mmu-miR-24-3p | MIMAT0000219 | Exoc6b | 75914 | Sequencing//HITS-CLIP | 37986224\|25753418\|19536157 | Unspecified |
| mmu-miR-24-3p | MIMAT0000219 | Ezh1 | 14055 | HITS-CLIP | 23142080 | Unspecified |
| mmu-miR-24-3p | MIMAT0000219 | Ezh2 | 14056 | HITS-CLIP | 23142080 | Unspecified |
| mmu-miR-24-3p | MIMAT0000219 | Fadd | 14082 | HITS-CLIP | 23142080 | Unspecified |
| mmu-miR-24-3p | MIMAT0000219 | Fahd1 | 68636 | HITS-CLIP | 25753418 | Unspecified |
| mmu-miR-24-3p | MIMAT0000219 | Fam102a | 98952 | HITS-CLIP | 23142080 | Unspecified |
| mmu-miR-24-3p | MIMAT0000219 | Fam107b | 66540 | HITS-CLIP | 37986224\|23142080 | Unspecified |
| mmu-miR-24-3p | MIMAT0000219 | Fam118b | 109229 | HITS-CLIP | 23142080 | Unspecified |
| mmu-miR-24-3p | MIMAT0000219 | Fam120a | 218236 | HITS-CLIP | 37986224\|25753418\|23142080 | Unspecified |

| | | | | | | |
|---|---|---|---|---|---|---|
| mmu-miR-24-3p | MIMAT0000219 | Fam120b | 67544 | HITS-CLIP | 23142080 | Unspecified |
| mmu-miR-24-3p | MIMAT0000219 | Fam126b | 213056 | HITS-CLIP | 23142080 | Unspecified |
| mmu-miR-24-3p | MIMAT0000219 | Fam133b | 68152 | HITS-CLIP | 23142080 | Unspecified |
| mmu-miR-24-3p | MIMAT0000219 | Fam160a2 | 74349 | HITS-CLIP | 23142080 | Unspecified |
| mmu-miR-24-3p | MIMAT0000219 | Fam160b1 | 226252 | HITS-CLIP | 23142080 | Unspecified |
| mmu-miR-24-3p | MIMAT0000219 | Fam168b | 214469 | Sequencing//HITS-CLIP | 37986224\|23142080\|19536157 | Unspecified |
| mmu-miR-24-3p | MIMAT0000219 | Fam174b | 100038347 | HITS-CLIP | 25753418 | Unspecified |
| mmu-miR-24-3p | MIMAT0000219 | Fam193b | 212483 | HITS-CLIP | 23142080 | Unspecified |
| mmu-miR-24-3p | MIMAT0000219 | Fam198a | 245050 | HITS-CLIP | 23142080 | Unspecified |
| mmu-miR-24-3p | MIMAT0000219 | Fam199x | 245622 | HITS-CLIP | 23142080 | Unspecified |
| mmu-miR-24-3p | MIMAT0000219 | Fam208a | 218850 | HITS-CLIP | 23142080 | Unspecified |
| mmu-miR-24-3p | MIMAT0000219 | Fam20c | 80752 | HITS-CLIP | 25753418 | Unspecified |
| mmu-miR-24-3p | MIMAT0000219 | Fam219a | 71901 | HITS-CLIP | 23142080 | Unspecified |
| mmu-miR-24-3p | MIMAT0000219 | Fam3c | 27999 | HITS-CLIP | 37986224\|23142080\|25753418 | Unspecified |
| mmu-miR-24-3p | MIMAT0000219 | Fam53b | 77938 | HITS-CLIP | 23142080 | Unspecified |
| mmu-miR-24-3p | MIMAT0000219 | Fam53c | 66306 | HITS-CLIP | 25753418 | Unspecified |
| mmu-miR-24-3p | MIMAT0000219 | Fam69a | 388650 | HITS-CLIP | 23142080 | Unspecified |
| mmu-miR-24-3p | MIMAT0000219 | Fam78a | 241303 | HITS-CLIP | 37986224\|23142080 | Unspecified |
| mmu-miR-24-3p | MIMAT0000219 | Fam83d | 71878 | HITS-CLIP | 23142080 | Unspecified |
| mmu-miR-24-3p | MIMAT0000219 | Fam84a | 105005 | HITS-CLIP | 23142080 | Unspecified |
| mmu-miR-24-3p | MIMAT0000219 | Fam91a1 | 210998 | HITS-CLIP | 23142080 | Unspecified |
| mmu-miR-24-3p | MIMAT0000219 | Farp2 | 227377 | HITS-CLIP | 25753418 | Unspecified |
| mmu-miR-24-3p | MIMAT0000219 | Farsb | 23874 | HITS-CLIP | 25753418 | Unspecified |
| mmu-miR-24-3p | MIMAT0000219 | Fasl | 14103 | HITS-CLIP | 37986224 | Unspecified |
| mmu-miR-24-3p | MIMAT0000219 | Fasn | 14104 | HITS-CLIP | 23142080 | Unspecified |
| mmu-miR-24-3p | MIMAT0000219 | Fastkd3 | 69577 | HITS-CLIP | 23142080 | Unspecified |
| mmu-miR-24-3p | MIMAT0000219 | Fbxl14 | 101358 | HITS-CLIP | 23142080 | Unspecified |
| mmu-miR-24-3p | MIMAT0000219 | Fbxl20 | 72194 | Sequencing//HITS-CLIP | 19536157\|23142080\|37986224 | Unspecified |
| mmu-miR-24-3p | MIMAT0000219 | Fbxl5 | 242960 | HITS-CLIP | 23142080 | Unspecified |
| mmu-miR-24-3p | MIMAT0000219 | Fbxo11 | 225055 | HITS-CLIP | 23142080 | Unspecified |
| mmu-miR-24-3p | MIMAT0000219 | Fbxo28 | 67948 | HITS-CLIP | 23142080 | Unspecified |
| mmu-miR-24-3p | MIMAT0000219 | Fbxo3 | 57443 | HITS-CLIP | 23142080 | Unspecified |
| mmu-miR-24-3p | MIMAT0000219 | Fbxo32 | 67731 | HITS-CLIP | 25753418 | Unspecified |
| mmu-miR-24-3p | MIMAT0000219 | Fbxo33 | 70611 | HITS-CLIP | 23142080 | Unspecified |
| mmu-miR-24-3p | MIMAT0000219 | Fbxo40 | 51725 | HITS-CLIP | 25753418 | Unspecified |
| mmu-miR-24-3p | MIMAT0000219 | Fbxo5 | 67141 | HITS-CLIP | 23142080 | Unspecified |
| mmu-miR-24-3p | MIMAT0000219 | Fchsd2 | 207278 | HITS-CLIP | 25753418 | Unspecified |
| mmu-miR-24-3p | MIMAT0000219 | Fech | 14151 | HITS-CLIP | 25753418 | Unspecified |
| mmu-miR-24-3p | MIMAT0000219 | Fen1 | 14156 | HITS-CLIP | 23142080 | Unspecified |

| | | | | | | |
|---|---|---|---|---|---|---|
| mmu-miR-24-3p | MIMAT0000219 | Fgfr1op2 | 67529 | HITS-CLIP | 23142080 | Unspecified |
| mmu-miR-24-3p | MIMAT0000219 | Fhod1 | 234686 | HITS-CLIP | 25753418 | Unspecified |
| mmu-miR-24-3p | MIMAT0000219 | Fis1 | 66437 | HITS-CLIP | 23142080 | Unspecified |
| mmu-miR-24-3p | MIMAT0000219 | Fitm2 | 228859 | HITS-CLIP | 25753418 | Unspecified |
| mmu-miR-24-3p | MIMAT0000219 | Fkbp11 | 66120 | HITS-CLIP | 23142080 | Unspecified |
| mmu-miR-24-3p | MIMAT0000219 | Fkbp15 | 338355 | HITS-CLIP | 23142080 | Unspecified |
| mmu-miR-24-3p | MIMAT0000219 | Fkbp8 | 14232 | HITS-CLIP | 25753418 | Unspecified |
| mmu-miR-24-3p | MIMAT0000219 | Flcn | 216805 | HITS-CLIP | 37986224\|23142080 | Unspecified |
| mmu-miR-24-3p | MIMAT0000219 | Fli1 | 14247 | HITS-CLIP | 23142080 | Unspecified |
| mmu-miR-24-3p | MIMAT0000219 | Flnb | 286940 | HITS-CLIP | 31433990 | Unspecified |
| mmu-miR-24-3p | MIMAT0000219 | Flnc | 68794 | HITS-CLIP | 25753418 | Unspecified |
| mmu-miR-24-3p | MIMAT0000219 | Flot1 | 14251 | HITS-CLIP | 25753418 | Unspecified |
| mmu-miR-24-3p | MIMAT0000219 | Flt3l | 14256 | HITS-CLIP | 23142080 | Unspecified |
| mmu-miR-24-3p | MIMAT0000219 | Fmnl1 | 57778 | HITS-CLIP | 23142080 | Unspecified |
| mmu-miR-24-3p | MIMAT0000219 | Fnbp1 | 14269 | HITS-CLIP | 23142080 | Unspecified |
| mmu-miR-24-3p | MIMAT0000219 | Fnbp4 | 55935 | HITS-CLIP | 23142080 | Unspecified |
| mmu-miR-24-3p | MIMAT0000219 | Fndc3a | 319448 | HITS-CLIP | 37986224\|23142080 | Unspecified |
| mmu-miR-24-3p | MIMAT0000219 | Fnta | 14272 | HITS-CLIP | 23142080 | Unspecified |
| mmu-miR-24-3p | MIMAT0000219 | Fosl2 | 14284 | HITS-CLIP | 23142080 | Unspecified |
| mmu-miR-24-3p | MIMAT0000219 | Foxm1 | 14235 | HITS-CLIP | 23142080 | Unspecified |
| mmu-miR-24-3p | MIMAT0000219 | Foxn2 | 14236 | HITS-CLIP | 23142080 | Unspecified |
| mmu-miR-24-3p | MIMAT0000219 | Foxn3 | 71375 | Sequencing//HITS-CLIP | 19536157\|23142080\|37986224 | Unspecified |
| mmu-miR-24-3p | MIMAT0000219 | Foxo1 | 56458 | HITS-CLIP | 23142080 | Unspecified |
| mmu-miR-24-3p | MIMAT0000219 | Foxo3 | 56484 | HITS-CLIP | 23142080 | Unspecified |
| mmu-miR-24-3p | MIMAT0000219 | Foxo4 | 54601 | HITS-CLIP | 23142080 | Unspecified |
| mmu-miR-24-3p | MIMAT0000219 | Foxp1 | 108655 | HITS-CLIP | 23142080\|25753418 | Unspecified |
| mmu-miR-24-3p | MIMAT0000219 | Foxred2 | 239554 | HITS-CLIP | 23142080 | Unspecified |
| mmu-miR-24-3p | MIMAT0000219 | Fpgs | 14287 | HITS-CLIP | 25753418 | Unspecified |
| mmu-miR-24-3p | MIMAT0000219 | Fpgt | 75540 | HITS-CLIP | 23142080 | Unspecified |
| mmu-miR-24-3p | MIMAT0000219 | Frmd8 | 67457 | HITS-CLIP | 23142080\|31433990 | Unspecified |
| mmu-miR-24-3p | MIMAT0000219 | Frs2 | 327826 | HITS-CLIP | 31433990 | Unspecified |
| mmu-miR-24-3p | MIMAT0000219 | Fry | 320365 | HITS-CLIP | 25753418 | Unspecified |
| mmu-miR-24-3p | MIMAT0000219 | Fryl | 72313 | HITS-CLIP | 25753418 | Unspecified |
| mmu-miR-24-3p | MIMAT0000219 | Ftl1 | 14325 | HITS-CLIP | 25753418 | Unspecified |
| mmu-miR-24-3p | MIMAT0000219 | Fto | 26383 | HITS-CLIP | 25753418 | Unspecified |
| mmu-miR-24-3p | MIMAT0000219 | Fubp1 | 51886 | HITS-CLIP | 23142080 | Unspecified |
| mmu-miR-24-3p | MIMAT0000219 | Fubp3 | 320267 | HITS-CLIP | 25753418 | Unspecified |
| mmu-miR-24-3p | MIMAT0000219 | Fuca2 | 66848 | HITS-CLIP | 23142080 | Unspecified |
| mmu-miR-24-3p | MIMAT0000219 | Fundc2 | 67391 | HITS-CLIP | 25753418 | Unspecified |

| | | | | | | |
|---|---|---|---|---|---|---|
| mmu-miR-24-3p | MIMAT0000219 | Furin | 18550 | HITS-CLIP | 23142080 | Unspecified |
| mmu-miR-24-3p | MIMAT0000219 | Fus | 233908 | HITS-CLIP | 25753418 | Unspecified |
| mmu-miR-24-3p | MIMAT0000219 | Fut4 | 14345 | HITS-CLIP | 23142080 | Unspecified |
| mmu-miR-24-3p | MIMAT0000219 | Fyco1 | 17281 | HITS-CLIP | 37986224\|25753418 | Unspecified |
| mmu-miR-24-3p | MIMAT0000219 | G2e3 | 217558 | HITS-CLIP | 23142080 | Unspecified |
| mmu-miR-24-3p | MIMAT0000219 | G3bp2 | 23881 | HITS-CLIP | 37986224\|23142080 | Unspecified |
| mmu-miR-24-3p | MIMAT0000219 | G6pdx | 14381 | HITS-CLIP | 23142080 | Unspecified |
| mmu-miR-24-3p | MIMAT0000219 | Gaa | 14387 | HITS-CLIP | 25753418 | Unspecified |
| mmu-miR-24-3p | MIMAT0000219 | Gab2 | 14389 | HITS-CLIP | 25753418 | Unspecified |
| mmu-miR-24-3p | MIMAT0000219 | Gabarap | 56486 | HITS-CLIP | 25753418 | Unspecified |
| mmu-miR-24-3p | MIMAT0000219 | Gabpb2 | 213054 | Sequencing//HITS-CLIP | 37986224\|23142080\|19536157 | Unspecified |
| mmu-miR-24-3p | MIMAT0000219 | Galk1 | 14635 | HITS-CLIP | 31433990 | Unspecified |
| mmu-miR-24-3p | MIMAT0000219 | Galm | 319625 | HITS-CLIP | 25753418 | Unspecified |
| mmu-miR-24-3p | MIMAT0000219 | Galnt7 | 108150 | HITS-CLIP | 23142080 | Unspecified |
| mmu-miR-24-3p | MIMAT0000219 | Ganc | 76051 | HITS-CLIP | 23142080 | Unspecified |
| mmu-miR-24-3p | MIMAT0000219 | Gapvd1 | 66691 | HITS-CLIP | 23142080 | Unspecified |
| mmu-miR-24-3p | MIMAT0000219 | Gars | 353172 | HITS-CLIP | 23142080 | Unspecified |
| mmu-miR-24-3p | MIMAT0000219 | Gata3 | 14462 | HITS-CLIP | 23142080 | Unspecified |
| mmu-miR-24-3p | MIMAT0000219 | Gata4 | 14463 | HITS-CLIP | 25753418 | Unspecified |
| mmu-miR-24-3p | MIMAT0000219 | Gata6 | 14465 | HITS-CLIP | 25753418 | Unspecified |
| mmu-miR-24-3p | MIMAT0000219 | Gatad2a | 234366 | HITS-CLIP | 23142080 | Unspecified |
| mmu-miR-24-3p | MIMAT0000219 | Gatm | 67092 | HITS-CLIP | 23142080 | Unspecified |
| mmu-miR-24-3p | MIMAT0000219 | Gba2 | 230101 | Sequencing//HITS-CLIP | 19536157\|25753418\|23142080\|37986 | Unspecified |
| mmu-miR-24-3p | MIMAT0000219 | Gbf1 | 107338 | HITS-CLIP | 25753418 | Unspecified |
| mmu-miR-24-3p | MIMAT0000219 | Gbp2 | 14469 | HITS-CLIP | 23142080 | Unspecified |
| mmu-miR-24-3p | MIMAT0000219 | Gbp8 | 76074 | HITS-CLIP | 23142080 | Unspecified |
| mmu-miR-24-3p | MIMAT0000219 | Gcc2 | 70297 | HITS-CLIP | 23142080 | Unspecified |
| mmu-miR-24-3p | MIMAT0000219 | Gcfc2 | 330361 | HITS-CLIP | 23142080 | Unspecified |
| mmu-miR-24-3p | MIMAT0000219 | Gch1 | 14528 | HITS-CLIP | 23142080 | Unspecified |
| mmu-miR-24-3p | MIMAT0000219 | Gcn1l1 | 10985 | HITS-CLIP | 23142080 | Unspecified |
| mmu-miR-24-3p | MIMAT0000219 | Gdpgp1 | 269952 | HITS-CLIP | 23142080 | Unspecified |
| mmu-miR-24-3p | MIMAT0000219 | Gemin4 | 276919 | HITS-CLIP | 25753418 | Unspecified |
| mmu-miR-24-3p | MIMAT0000219 | Gfer | 11692 | HITS-CLIP | 37986224\|23142080 | Unspecified |
| mmu-miR-24-3p | MIMAT0000219 | Gfm1 | 28030 | HITS-CLIP | 23142080 | Unspecified |
| mmu-miR-24-3p | MIMAT0000219 | Gfm2 | 320806 | HITS-CLIP | 23142080\|25753418 | Unspecified |
| mmu-miR-24-3p | MIMAT0000219 | Gfpt1 | 14583 | HITS-CLIP | 37986224\|23142080 | Unspecified |
| mmu-miR-24-3p | MIMAT0000219 | Gga2 | 74105 | HITS-CLIP | 25753418 | Unspecified |
| mmu-miR-24-3p | MIMAT0000219 | Ggnbp2 | 217039 | HITS-CLIP | 23142080 | Unspecified |
| mmu-miR-24-3p | MIMAT0000219 | Ggta1 | 14594 | Sequencing//HITS-CLIP | 37986224\|23142080\|19536157 | Unspecified |

| | | | | | | |
|---|---|---|---|---|---|---|
| mmu-miR-24-3p | MIMAT0000219 | Gid8 | 76425 | HITS-CLIP | 25753418 | Unspecified |
| mmu-miR-24-3p | MIMAT0000219 | Gigyf1 | 57330 | HITS-CLIP | 37986224\|25753418\|23142080 | Unspecified |
| mmu-miR-24-3p | MIMAT0000219 | Gigyf2 | 227331 | HITS-CLIP | 25753418 | Unspecified |
| mmu-miR-24-3p | MIMAT0000219 | Gimap1 | 16205 | HITS-CLIP | 23142080 | Unspecified |
| mmu-miR-24-3p | MIMAT0000219 | Gimap7 | 231932 | HITS-CLIP | 37986224\|23142080 | Unspecified |
| mmu-miR-24-3p | MIMAT0000219 | Git2 | 26431 | HITS-CLIP | 23142080 | Unspecified |
| mmu-miR-24-3p | MIMAT0000219 | Gja1 | 14609 | HITS-CLIP | 23142080 | Unspecified |
| mmu-miR-24-3p | MIMAT0000219 | Glg1 | 20340 | HITS-CLIP | 25753418 | Unspecified |
| mmu-miR-24-3p | MIMAT0000219 | Glis1 | 230587 | HITS-CLIP | 23142080 | Unspecified |
| mmu-miR-24-3p | MIMAT0000219 | Gls | 14660 | HITS-CLIP | 37986224\|23142080 | Unspecified |
| mmu-miR-24-3p | MIMAT0000219 | Gltp | 56356 | HITS-CLIP | 31433990\|25753418 | Unspecified |
| mmu-miR-24-3p | MIMAT0000219 | Glyr1 | 74022 | HITS-CLIP | 31433990\|25753418 | Unspecified |
| mmu-miR-24-3p | MIMAT0000219 | Gm17018 | 226162 | HITS-CLIP | 23142080 | Unspecified |
| mmu-miR-24-3p | MIMAT0000219 | Gm20604 | 100859931 | HITS-CLIP | 37986224\|23142080\|25753418 | Unspecified |
| mmu-miR-24-3p | MIMAT0000219 | Gm21970 | 6609959 | HITS-CLIP | 25753418 | Unspecified |
| mmu-miR-24-3p | MIMAT0000219 | Gm2a | 14667 | HITS-CLIP | 23142080 | Unspecified |
| mmu-miR-24-3p | MIMAT0000219 | Gm38394 | 667118 | HITS-CLIP | 23142080 | Unspecified |
| mmu-miR-24-3p | MIMAT0000219 | Gm45713 | 103691134 | HITS-CLIP | 23142080 | Unspecified |
| mmu-miR-24-3p | MIMAT0000219 | Gm49336 | 127230315 | HITS-CLIP | 23142080 | Unspecified |
| mmu-miR-24-3p | MIMAT0000219 | Gm49378 | | HITS-CLIP | 25753418 | Unspecified |
| mmu-miR-24-3p | MIMAT0000219 | Gm9803 | 100042179 | HITS-CLIP | 23142080 | Unspecified |
| mmu-miR-24-3p | MIMAT0000219 | Gmnn | 57441 | HITS-CLIP | 23142080 | Unspecified |
| mmu-miR-24-3p | MIMAT0000219 | Gmppb | 331026 | HITS-CLIP | 23142080 | Unspecified |
| mmu-miR-24-3p | MIMAT0000219 | Gna13 | 14674 | HITS-CLIP | 23142080 | Unspecified |
| mmu-miR-24-3p | MIMAT0000219 | Gnai3 | 14679 | HITS-CLIP | 23142080 | Unspecified |
| mmu-miR-24-3p | MIMAT0000219 | Gnaq | 14682 | Sequencing//HITS-CLIP | 37986224\|23142080\|19536157 | Unspecified |
| mmu-miR-24-3p | MIMAT0000219 | Gnb1 | 14688 | HITS-CLIP | 23142080 | Unspecified |
| mmu-miR-24-3p | MIMAT0000219 | Gnb4 | 14696 | HITS-CLIP | 23142080 | Unspecified |
| mmu-miR-24-3p | MIMAT0000219 | Gne | 50798 | HITS-CLIP | 25753418\|23142080 | Unspecified |
| mmu-miR-24-3p | MIMAT0000219 | Gng12 | 14701 | HITS-CLIP | 23142080 | Unspecified |
| mmu-miR-24-3p | MIMAT0000219 | Gnl1 | 14670 | HITS-CLIP | 23142080 | Unspecified |
| mmu-miR-24-3p | MIMAT0000219 | Gnl3l | 237107 | HITS-CLIP | 23142080 | Unspecified |
| mmu-miR-24-3p | MIMAT0000219 | Gnpat | 14712 | HITS-CLIP | 23142080 | Unspecified |
| mmu-miR-24-3p | MIMAT0000219 | Gnptab | 432486 | HITS-CLIP | 23142080 | Unspecified |
| mmu-miR-24-3p | MIMAT0000219 | Golga1 | 76899 | HITS-CLIP | 23142080 | Unspecified |
| mmu-miR-24-3p | MIMAT0000219 | Golga3 | 269682 | HITS-CLIP | 23142080 | Unspecified |
| mmu-miR-24-3p | MIMAT0000219 | Golga4 | 54214 | HITS-CLIP | 25753418 | Unspecified |
| mmu-miR-24-3p | MIMAT0000219 | Golph3l | 229593 | HITS-CLIP | 23142080 | Unspecified |
| mmu-miR-24-3p | MIMAT0000219 | Gon4l | 76022 | HITS-CLIP | 23142080 | Unspecified |

| | | | | | | |
|---|---|---|---|---|---|---|
| mmu-miR-24-3p | MIMAT0000219 | Gorab | 98376 | HITS-CLIP | 23142080 | Unspecified |
| mmu-miR-24-3p | MIMAT0000219 | Gorasp1 | 74498 | HITS-CLIP | 25753418 | Unspecified |
| mmu-miR-24-3p | MIMAT0000219 | Gosr2 | 56494 | HITS-CLIP | 23142080 | Unspecified |
| mmu-miR-24-3p | MIMAT0000219 | Gpam | 14732 | HITS-CLIP | 25753418 | Unspecified |
| mmu-miR-24-3p | MIMAT0000219 | Gpatch2 | 67769 | HITS-CLIP | 23142080 | Unspecified |
| mmu-miR-24-3p | MIMAT0000219 | Gpd2 | 14571 | HITS-CLIP | 37986224\|23142080 | Unspecified |
| mmu-miR-24-3p | MIMAT0000219 | Gpld1 | 14756 | HITS-CLIP | 23142080 | Unspecified |
| mmu-miR-24-3p | MIMAT0000219 | Gpm6b | 14758 | HITS-CLIP | 23142080 | Unspecified |
| mmu-miR-24-3p | MIMAT0000219 | Gpr107 | 277463 | HITS-CLIP | 25753418 | Unspecified |
| mmu-miR-24-3p | MIMAT0000219 | Gpr132 | 56696 | HITS-CLIP | 37986224\|23142080 | Unspecified |
| mmu-miR-24-3p | MIMAT0000219 | Gpr146 | 80290 | HITS-CLIP | 37986224\|23142080 | Unspecified |
| mmu-miR-24-3p | MIMAT0000219 | Gpr174 | 213439 | HITS-CLIP | 23142080 | Unspecified |
| mmu-miR-24-3p | MIMAT0000219 | Gpr183 | 321019 | HITS-CLIP | 31433990 | Unspecified |
| mmu-miR-24-3p | MIMAT0000219 | Gpr68 | 238377 | HITS-CLIP | 23142080 | Unspecified |
| mmu-miR-24-3p | MIMAT0000219 | Gprasp1 | 67298 | Sequencing//HITS-CLIP | 37986224\|23142080\|19536157 | Unspecified |
| mmu-miR-24-3p | MIMAT0000219 | Gprin3 | 243385 | HITS-CLIP | 31433990\|23142080 | Unspecified |
| mmu-miR-24-3p | MIMAT0000219 | Gramd1b | 235283 | HITS-CLIP | 37986224\|23142080 | Unspecified |
| mmu-miR-24-3p | MIMAT0000219 | Gramd4 | 223752 | HITS-CLIP | 23142080 | Unspecified |
| mmu-miR-24-3p | MIMAT0000219 | Grap | 71520 | HITS-CLIP | 23142080 | Unspecified |
| mmu-miR-24-3p | MIMAT0000219 | Grap2 | 17444 | HITS-CLIP | 23142080 | Unspecified |
| mmu-miR-24-3p | MIMAT0000219 | Grb14 | 50915 | HITS-CLIP | 25753418 | Unspecified |
| mmu-miR-24-3p | MIMAT0000219 | Grb2 | 14784 | HITS-CLIP | 23142080 | Unspecified |
| mmu-miR-24-3p | MIMAT0000219 | Grk2 | 156 | HITS-CLIP | 25753418 | Unspecified |
| mmu-miR-24-3p | MIMAT0000219 | Grk5 | 14773 | HITS-CLIP | 23142080 | Unspecified |
| mmu-miR-24-3p | MIMAT0000219 | Grk6 | 26385 | HITS-CLIP | 23142080 | Unspecified |
| mmu-miR-24-3p | MIMAT0000219 | Grsf1 | 231413 | HITS-CLIP | 23142080 | Unspecified |
| mmu-miR-24-3p | MIMAT0000219 | Gse1 | 382034 | HITS-CLIP | 23142080 | Unspecified |
| mmu-miR-24-3p | MIMAT0000219 | Gskip | 66787 | HITS-CLIP | 25753418 | Unspecified |
| mmu-miR-24-3p | MIMAT0000219 | Gstcd | 67553 | HITS-CLIP | 23142080 | Unspecified |
| mmu-miR-24-3p | MIMAT0000219 | Gtf2e1 | 74197 | HITS-CLIP | 37986224\|23142080 | Unspecified |
| mmu-miR-24-3p | MIMAT0000219 | Gtf2h1 | 14884 | HITS-CLIP | 23142080 | Unspecified |
| mmu-miR-24-3p | MIMAT0000219 | Gtf2h2 | 23894 | HITS-CLIP | 25753418 | Unspecified |
| mmu-miR-24-3p | MIMAT0000219 | Gtf2i | 14886 | HITS-CLIP | 25753418\|23142080 | Unspecified |
| mmu-miR-24-3p | MIMAT0000219 | Gtf3c1 | 233863 | HITS-CLIP | 23142080 | Unspecified |
| mmu-miR-24-3p | MIMAT0000219 | Gtf3c2 | 71752 | HITS-CLIP | 37986224\|23142080 | Unspecified |
| mmu-miR-24-3p | MIMAT0000219 | Gtf3c3 | 98488 | HITS-CLIP | 25753418 | Unspecified |
| mmu-miR-24-3p | MIMAT0000219 | Gtpbp1 | 14904 | HITS-CLIP | 25753418 | Unspecified |
| mmu-miR-24-3p | MIMAT0000219 | Gtpbp2 | 56055 | HITS-CLIP | 23142080 | Unspecified |
| mmu-miR-24-3p | MIMAT0000219 | Gucy1a1 | 60596 | HITS-CLIP | 23142080 | Unspecified |

| | | | | | | | |
|---|---|---|---|---|---|---|---|
| mmu-miR-24-3p | MIMAT0000219 | Gucy2e | 14919 | HITS-CLIP | | 23142080 | Unspecified |
| mmu-miR-24-3p | MIMAT0000219 | Gyg | 27357 | HITS-CLIP | | 23142080 | Unspecified |
| mmu-miR-24-3p | MIMAT0000219 | Gypc | 71683 | HITS-CLIP | | 23142080 | Unspecified |
| mmu-miR-24-3p | MIMAT0000219 | Gzf1 | 74533 | HITS-CLIP | | 23142080 | Unspecified |
| mmu-miR-24-3p | MIMAT0000219 | H1f0 | 14958 | HITS-CLIP | | 25753418 | Unspecified |
| mmu-miR-24-3p | MIMAT0000219 | H1f2 | 3006 | HITS-CLIP | 31433990\|25753418 | | Unspecified |
| mmu-miR-24-3p | MIMAT0000219 | H2-Aa | 14960 | HITS-CLIP | | 23142080 | Unspecified |
| mmu-miR-24-3p | MIMAT0000219 | H2-Eb1 | 14969 | HITS-CLIP | | 23142080 | Unspecified |
| mmu-miR-24-3p | MIMAT0000219 | H2-K1 | 14972 | HITS-CLIP | | 25753418 | Unspecified |
| mmu-miR-24-3p | MIMAT0000219 | H2-Q4 | 15015 | HITS-CLIP | | 23142080 | Unspecified |
| mmu-miR-24-3p | MIMAT0000219 | H2-T24 | 15042 | HITS-CLIP | 25083871\|23142080\|21258322 | | Unspecified |
| mmu-miR-24-3p | MIMAT0000219 | H2afx | 15270 | Sequencing//HITS-CLIP | 37986224\|23142080\|19536157 | | Unspecified |
| mmu-miR-24-3p | MIMAT0000219 | H2afy | 9555 | HITS-CLIP | | 23142080 | Unspecified |
| mmu-miR-24-3p | MIMAT0000219 | H3c14 | 126961 | HITS-CLIP | | 25753418 | Unspecified |
| mmu-miR-24-3p | MIMAT0000219 | H3f3b | 15081 | HITS-CLIP | | 23142080 | Unspecified |
| mmu-miR-24-3p | MIMAT0000219 | Hadha | 97212 | HITS-CLIP | | 23142080 | Unspecified |
| mmu-miR-24-3p | MIMAT0000219 | Hand2 | 15111 | HITS-CLIP | | 25753418 | Unspecified |
| mmu-miR-24-3p | MIMAT0000219 | Hbegf | 15200 | Sequencing//HITS-CLIP | 19536157\|25753418\|37986224 | | Unspecified |
| mmu-miR-24-3p | MIMAT0000219 | Hbp1 | 73389 | HITS-CLIP | 37986224\|23142080 | | Unspecified |
| mmu-miR-24-3p | MIMAT0000219 | Hbs1l | 56422 | HITS-CLIP | | 23142080 | Unspecified |
| mmu-miR-24-3p | MIMAT0000219 | Hcfc1 | 15161 | HITS-CLIP | | 23142080 | Unspecified |
| mmu-miR-24-3p | MIMAT0000219 | Hdac3 | 15183 | HITS-CLIP | | 23142080 | Unspecified |
| mmu-miR-24-3p | MIMAT0000219 | Hdac7 | 56233 | HITS-CLIP | | 23142080 | Unspecified |
| mmu-miR-24-3p | MIMAT0000219 | Hddc2 | 69692 | HITS-CLIP | | 23142080 | Unspecified |
| mmu-miR-24-3p | MIMAT0000219 | Hdgfl2 | 84717 | HITS-CLIP | | 23142080 | Unspecified |
| mmu-miR-24-3p | MIMAT0000219 | Hdgfl3 | 29877 | HITS-CLIP | | 23142080 | Unspecified |
| mmu-miR-24-3p | MIMAT0000219 | Hdlbp | 110611 | HITS-CLIP | | 23142080 | Unspecified |
| mmu-miR-24-3p | MIMAT0000219 | Heatr5b | 320473 | HITS-CLIP | | 23142080 | Unspecified |
| mmu-miR-24-3p | MIMAT0000219 | Heca | 380629 | HITS-CLIP | | 23142080 | Unspecified |
| mmu-miR-24-3p | MIMAT0000219 | Hectd1 | 207304 | HITS-CLIP | | 23142080 | Unspecified |
| mmu-miR-24-3p | MIMAT0000219 | Hectd2 | 226098 | HITS-CLIP | | 23142080 | Unspecified |
| mmu-miR-24-3p | MIMAT0000219 | Hectd4 | 283450 | HITS-CLIP | | 23142080 | Unspecified |
| mmu-miR-24-3p | MIMAT0000219 | Heg1 | 77446 | HITS-CLIP | | 25753418 | Unspecified |
| mmu-miR-24-3p | MIMAT0000219 | Hells | 15201 | HITS-CLIP | | 23142080 | Unspecified |
| mmu-miR-24-3p | MIMAT0000219 | Helz | 78455 | HITS-CLIP | | 25753418 | Unspecified |
| mmu-miR-24-3p | MIMAT0000219 | Helz2 | 229003 | HITS-CLIP | | 23142080 | Unspecified |
| mmu-miR-24-3p | MIMAT0000219 | Herc1 | 235439 | HITS-CLIP | 37986224\|23142080 | | Unspecified |
| mmu-miR-24-3p | MIMAT0000219 | Herc2 | 15204 | HITS-CLIP | | 25753418 | Unspecified |
| mmu-miR-24-3p | MIMAT0000219 | Herpud1 | 64209 | HITS-CLIP | | 25753418 | Unspecified |

| | | | | | | |
|---|---|---|---|---|---|---|
| mmu-miR-24-3p | MIMAT0000219 | Hes1 | 15205 | HITS-CLIP | 25753418 | Unspecified |
| mmu-miR-24-3p | MIMAT0000219 | Hgsnat | 52120 | HITS-CLIP | 23142080 | Unspecified |
| mmu-miR-24-3p | MIMAT0000219 | Hibch | 227095 | HITS-CLIP | 23142080 | Unspecified |
| mmu-miR-24-3p | MIMAT0000219 | Hic2 | 58180 | Sequencing//HITS-CLIP | 37986224\|25753418\|19536157 | Unspecified |
| mmu-miR-24-3p | MIMAT0000219 | Hif1a | 15251 | HITS-CLIP | 23142080 | Unspecified |
| mmu-miR-24-3p | MIMAT0000219 | Hif1an | 319594 | HITS-CLIP | 25753418\|23142080 | Unspecified |
| mmu-miR-24-3p | MIMAT0000219 | Hip1 | 215114 | HITS-CLIP | 23142080 | Unspecified |
| mmu-miR-24-3p | MIMAT0000219 | Hipk1 | 15257 | HITS-CLIP | 25753418\|23142080 | Unspecified |
| mmu-miR-24-3p | MIMAT0000219 | Hipk2 | 15258 | HITS-CLIP | 23142080 | Unspecified |
| mmu-miR-24-3p | MIMAT0000219 | Hipk3 | 15259 | HITS-CLIP | 25753418\|23142080 | Unspecified |
| mmu-miR-24-3p | MIMAT0000219 | Hist1h1b | 3009 | HITS-CLIP | 23142080 | Unspecified |
| mmu-miR-24-3p | MIMAT0000219 | Hist1h2bc | 3017 | HITS-CLIP | 23142080 | Unspecified |
| mmu-miR-24-3p | MIMAT0000219 | Hist1h2bj | 8970 | HITS-CLIP | 23142080 | Unspecified |
| mmu-miR-24-3p | MIMAT0000219 | Hist1h2bl | 8340 | HITS-CLIP | 23142080 | Unspecified |
| mmu-miR-24-3p | MIMAT0000219 | Hist1h2bm | 8342 | HITS-CLIP | 23142080 | Unspecified |
| mmu-miR-24-3p | MIMAT0000219 | Hivep2 | 15273 | HITS-CLIP | 37986224\|23142080 | Unspecified |
| mmu-miR-24-3p | MIMAT0000219 | Hivep3 | 16656 | HITS-CLIP | 23142080 | Unspecified |
| mmu-miR-24-3p | MIMAT0000219 | Hjurp | 381280 | HITS-CLIP | 37986224\|23142080 | Unspecified |
| mmu-miR-24-3p | MIMAT0000219 | Hk1 | 15275 | HITS-CLIP | 37986224\|23142080 | Unspecified |
| mmu-miR-24-3p | MIMAT0000219 | Hk2 | 15277 | HITS-CLIP | 23142080\|25753418 | Unspecified |
| mmu-miR-24-3p | MIMAT0000219 | Hmbox1 | 219150 | HITS-CLIP | 37986224\|23142080 | Unspecified |
| mmu-miR-24-3p | MIMAT0000219 | Hmg20b | 10362 | HITS-CLIP | 25753418 | Unspecified |
| mmu-miR-24-3p | MIMAT0000219 | Hmgcl | 15356 | HITS-CLIP | 23142080 | Unspecified |
| mmu-miR-24-3p | MIMAT0000219 | Hmgcs1 | 208715 | HITS-CLIP | 37986224\|23142080\|25753418 | Unspecified |
| mmu-miR-24-3p | MIMAT0000219 | Hmox1 | 15368 | Sequencing//qCLASH//HITS-CLIP | 19536157\|31393953\|25753418\|3798 | Unspecified |
| mmu-miR-24-3p | MIMAT0000219 | Hnrnpa0 | 77134 | HITS-CLIP | 31433990 | Unspecified |
| mmu-miR-24-3p | MIMAT0000219 | Hnrnpab | 15384 | HITS-CLIP | 23142080 | Unspecified |
| mmu-miR-24-3p | MIMAT0000219 | Hnrnpdl | 50926 | HITS-CLIP | 37986224\|23142080 | Unspecified |
| mmu-miR-24-3p | MIMAT0000219 | Hnrnph2 | 56258 | HITS-CLIP | 23142080 | Unspecified |
| mmu-miR-24-3p | MIMAT0000219 | Hnrnpk | 15387 | HITS-CLIP | 23142080 | Unspecified |
| mmu-miR-24-3p | MIMAT0000219 | Hnrnpll | 72692 | HITS-CLIP | 23142080 | Unspecified |
| mmu-miR-24-3p | MIMAT0000219 | Hnrnpm | 76936 | HITS-CLIP | 23142080 | Unspecified |
| mmu-miR-24-3p | MIMAT0000219 | Hnrnpr | 74326 | HITS-CLIP | 23142080 | Unspecified |
| mmu-miR-24-3p | MIMAT0000219 | Hnrnpul1 | 232989 | HITS-CLIP | 37986224 | Unspecified |
| mmu-miR-24-3p | MIMAT0000219 | Hnrnpul2 | 68693 | HITS-CLIP | 23142080 | Unspecified |
| mmu-miR-24-3p | MIMAT0000219 | Homez | 239099 | HITS-CLIP | 21258322\|25753418 | Unspecified |
| mmu-miR-24-3p | MIMAT0000219 | Hoxc8 | 15426 | qCLASH | 31393953 | Unspecified |
| mmu-miR-24-3p | MIMAT0000219 | Hrc | 15464 | HITS-CLIP | 25753418 | Unspecified |
| mmu-miR-24-3p | MIMAT0000219 | Hs3st2 | 195646 | Luciferase Reporter Assay | 37986224 | Unspecified |

| | | | | | | | |
|---|---|---|---|---|---|---|---|
| mmu-miR-24-3p | MIMAT0000219 | Hsf1 | 15499 | HITS-CLIP | | 25753418 | Unspecified |
| mmu-miR-24-3p | MIMAT0000219 | Hsf2bp | 74377 | HITS-CLIP | | 23142080 | Unspecified |
| mmu-miR-24-3p | MIMAT0000219 | Hsp90ab1 | 15516 | HITS-CLIP | 37986224\|23142080 | | Unspecified |
| mmu-miR-24-3p | MIMAT0000219 | Hspa4l | 18415 | HITS-CLIP | | 23142080 | Unspecified |
| mmu-miR-24-3p | MIMAT0000219 | Hspa5 | 14828 | HITS-CLIP | | 23142080 | Unspecified |
| mmu-miR-24-3p | MIMAT0000219 | Hspa9 | 15526 | HITS-CLIP | | 23142080 | Unspecified |
| mmu-miR-24-3p | MIMAT0000219 | Hspb7 | 29818 | HITS-CLIP | | 25753418 | Unspecified |
| mmu-miR-24-3p | MIMAT0000219 | Hspd1 | 15510 | HITS-CLIP | | 23142080 | Unspecified |
| mmu-miR-24-3p | MIMAT0000219 | Hspg2 | 15530 | HITS-CLIP | | 25753418 | Unspecified |
| mmu-miR-24-3p | MIMAT0000219 | Hsph1 | 15505 | HITS-CLIP | | 23142080 | Unspecified |
| mmu-miR-24-3p | MIMAT0000219 | Htt | 15194 | HITS-CLIP | | 23142080 | Unspecified |
| mmu-miR-24-3p | MIMAT0000219 | Hus1 | 15574 | HITS-CLIP | | 23142080 | Unspecified |
| mmu-miR-24-3p | MIMAT0000219 | Huwe1 | 59026 | HITS-CLIP | | 25753418 | Unspecified |
| mmu-miR-24-3p | MIMAT0000219 | Hyal1 | 15586 | HITS-CLIP | | 26203562 | Unspecified |
| mmu-miR-24-3p | MIMAT0000219 | Iars | 105148 | HITS-CLIP | | 23142080 | Unspecified |
| mmu-miR-24-3p | MIMAT0000219 | Iars2 | 381314 | HITS-CLIP | 31433990\|25753418 | | Unspecified |
| mmu-miR-24-3p | MIMAT0000219 | Ice1 | 822287 | HITS-CLIP | 25753418\|23142080 | | Unspecified |
| mmu-miR-24-3p | MIMAT0000219 | Id2 | 15902 | HITS-CLIP | | 23142080 | Unspecified |
| mmu-miR-24-3p | MIMAT0000219 | Id3 | 15903 | HITS-CLIP | | 23142080 | Unspecified |
| mmu-miR-24-3p | MIMAT0000219 | Ide | 15925 | HITS-CLIP | | 23142080 | Unspecified |
| mmu-miR-24-3p | MIMAT0000219 | Idh1 | 15926 | HITS-CLIP | | 23142080 | Unspecified |
| mmu-miR-24-3p | MIMAT0000219 | Ier5l | 72500 | HITS-CLIP | | 23142080 | Unspecified |
| mmu-miR-24-3p | MIMAT0000219 | Iffo2 | 212632 | HITS-CLIP | | 23142080 | Unspecified |
| mmu-miR-24-3p | MIMAT0000219 | Ifi27l2a | 76933 | HITS-CLIP | | 23142080 | Unspecified |
| mmu-miR-24-3p | MIMAT0000219 | Ifi47 | 15953 | HITS-CLIP | | 23142080 | Unspecified |
| mmu-miR-24-3p | MIMAT0000219 | Ifih1 | 71586 | HITS-CLIP | | 23142080 | Unspecified |
| mmu-miR-24-3p | MIMAT0000219 | Ifitm3 | 66141 | HITS-CLIP | | 25753418 | Unspecified |
| mmu-miR-24-3p | MIMAT0000219 | Ifnar1 | 15975 | Sequencing//HITS-CLIP | 37986224\|23142080\|19536157 | | Unspecified |
| mmu-miR-24-3p | MIMAT0000219 | Ifngr1 | 15979 | HITS-CLIP | | 37986224 | Unspecified |
| mmu-miR-24-3p | MIMAT0000219 | Ifrd1 | 15982 | HITS-CLIP | | 23142080 | Unspecified |
| mmu-miR-24-3p | MIMAT0000219 | Ift20 | 55978 | HITS-CLIP | | 25753418 | Unspecified |
| mmu-miR-24-3p | MIMAT0000219 | Igf1r | 16001 | HITS-CLIP | | 23142080 | Unspecified |
| mmu-miR-24-3p | MIMAT0000219 | Igf2r | 16004 | HITS-CLIP | | 25753418 | Unspecified |
| mmu-miR-24-3p | MIMAT0000219 | Igfbp7 | 29817 | qCLASH | | 31393953 | Unspecified |
| mmu-miR-24-3p | MIMAT0000219 | Ik | 24010 | HITS-CLIP | | 23142080 | Unspecified |
| mmu-miR-24-3p | MIMAT0000219 | Ikbkb | 16150 | HITS-CLIP | | 31433990 | Unspecified |
| mmu-miR-24-3p | MIMAT0000219 | Ikbke | 56489 | HITS-CLIP | | 23142080 | Unspecified |
| mmu-miR-24-3p | MIMAT0000219 | Ikzf3 | 22780 | HITS-CLIP | | 23142080 | Unspecified |
| mmu-miR-24-3p | MIMAT0000219 | Il10rb | 16155 | HITS-CLIP | | 25753418 | Unspecified |

| | | | | | | | |
|---|---|---|---|---|---|---|---|
| mmu-miR-24-3p | MIMAT0000219 | Il12rb2 | 16162 | HITS-CLIP | | 23142080 | Unspecified |
| mmu-miR-24-3p | MIMAT0000219 | Il16 | 16170 | HITS-CLIP | | 23142080 | Unspecified |
| mmu-miR-24-3p | MIMAT0000219 | Il21r | 60504 | HITS-CLIP | | 37986224 | Unspecified |
| mmu-miR-24-3p | MIMAT0000219 | Il24 | 93672 | HITS-CLIP | | 23142080 | Unspecified |
| mmu-miR-24-3p | MIMAT0000219 | Il2ra | 16184 | HITS-CLIP | 37986224\|31433990 | | Unspecified |
| mmu-miR-24-3p | MIMAT0000219 | Il2rb | 16185 | HITS-CLIP | | 23142080 | Unspecified |
| mmu-miR-24-3p | MIMAT0000219 | Il2rg | 16186 | HITS-CLIP | | 23142080 | Unspecified |
| mmu-miR-24-3p | MIMAT0000219 | Il4 | 16189 | HITS-CLIP | | 23142080 | Unspecified |
| mmu-miR-24-3p | MIMAT0000219 | Il4ra | 16190 | HITS-CLIP | | 23142080 | Unspecified |
| mmu-miR-24-3p | MIMAT0000219 | Il6st | 16195 | Sequencing//HITS-CLIP | 37986224\|23142080\|19536157 | | Unspecified |
| mmu-miR-24-3p | MIMAT0000219 | Il7r | 16197 | HITS-CLIP | | 23142080 | Unspecified |
| mmu-miR-24-3p | MIMAT0000219 | Immt | 76614 | HITS-CLIP | | 23142080 | Unspecified |
| mmu-miR-24-3p | MIMAT0000219 | Imp4 | 27993 | HITS-CLIP | | 25753418 | Unspecified |
| mmu-miR-24-3p | MIMAT0000219 | Impa2 | 114663 | HITS-CLIP | | 37986224 | Unspecified |
| mmu-miR-24-3p | MIMAT0000219 | Impad1 | 242291 | HITS-CLIP | | 23142080 | Unspecified |
| mmu-miR-24-3p | MIMAT0000219 | Impdh1 | 23917 | HITS-CLIP | | 25753418 | Unspecified |
| mmu-miR-24-3p | MIMAT0000219 | Inafm2 | 100505573 | HITS-CLIP | | 25753418 | Unspecified |
| mmu-miR-24-3p | MIMAT0000219 | Ing4 | 28019 | HITS-CLIP | | 25753418 | Unspecified |
| mmu-miR-24-3p | MIMAT0000219 | Ino80d | 227195 | HITS-CLIP | | 23142080 | Unspecified |
| mmu-miR-24-3p | MIMAT0000219 | Inpp4b | 234515 | HITS-CLIP | | 23142080 | Unspecified |
| mmu-miR-24-3p | MIMAT0000219 | Inpp5b | 16330 | HITS-CLIP | 23142080\|25753418 | | Unspecified |
| mmu-miR-24-3p | MIMAT0000219 | Inpp5e | 64436 | HITS-CLIP | | 25753418 | Unspecified |
| mmu-miR-24-3p | MIMAT0000219 | Insig1 | 231070 | HITS-CLIP | | 23142080 | Unspecified |
| mmu-miR-24-3p | MIMAT0000219 | Ints10 | 70885 | HITS-CLIP | | 23142080 | Unspecified |
| mmu-miR-24-3p | MIMAT0000219 | Ints3 | 229543 | HITS-CLIP | | 25753418 | Unspecified |
| mmu-miR-24-3p | MIMAT0000219 | Ints5 | 80789 | HITS-CLIP | | 23142080 | Unspecified |
| mmu-miR-24-3p | MIMAT0000219 | Ints6 | 18130 | HITS-CLIP | 37986224\|23142080 | | Unspecified |
| mmu-miR-24-3p | MIMAT0000219 | Ip6k1 | 27399 | HITS-CLIP | | 25753418 | Unspecified |
| mmu-miR-24-3p | MIMAT0000219 | Ipcef1 | 320495 | HITS-CLIP | | 23142080 | Unspecified |
| mmu-miR-24-3p | MIMAT0000219 | Ipo11 | 76582 | HITS-CLIP | | 23142080 | Unspecified |
| mmu-miR-24-3p | MIMAT0000219 | Ipo4 | 75751 | HITS-CLIP | 25753418\|23142080 | | Unspecified |
| mmu-miR-24-3p | MIMAT0000219 | Ipo5 | 70572 | HITS-CLIP | 25753418\|23142080 | | Unspecified |
| mmu-miR-24-3p | MIMAT0000219 | Iqgap1 | 29875 | HITS-CLIP | 37986224\|23142080 | | Unspecified |
| mmu-miR-24-3p | MIMAT0000219 | Iqsec1 | 232227 | HITS-CLIP | | 25753418 | Unspecified |
| mmu-miR-24-3p | MIMAT0000219 | Irak1 | 16179 | HITS-CLIP | | 23142080 | Unspecified |
| mmu-miR-24-3p | MIMAT0000219 | Irf1 | 16362 | HITS-CLIP | | 23142080 | Unspecified |
| mmu-miR-24-3p | MIMAT0000219 | Irf2bp2 | 270110 | HITS-CLIP | | 23142080 | Unspecified |
| mmu-miR-24-3p | MIMAT0000219 | Irf2bpl | 238330 | HITS-CLIP | 25753418\|23142080 | | Unspecified |
| mmu-miR-24-3p | MIMAT0000219 | Irf4 | 16364 | HITS-CLIP | 37986224\|23142080 | | Unspecified |

| | | | | | | |
|---|---|---|---|---|---|---|
| mmu-miR-24-3p | MIMAT0000219 | Irf6 | 54139 | HITS-CLIP | 23142080 | Unspecified |
| mmu-miR-24-3p | MIMAT0000219 | Irf8 | 15900 | HITS-CLIP | 37986224\|23142080 | Unspecified |
| mmu-miR-24-3p | MIMAT0000219 | Irs2 | 384783 | HITS-CLIP | 25753418\|23142080 | Unspecified |
| mmu-miR-24-3p | MIMAT0000219 | Isca1 | 69046 | HITS-CLIP | 25753418 | Unspecified |
| mmu-miR-24-3p | MIMAT0000219 | Isg20l2 | 229504 | HITS-CLIP | 25753418 | Unspecified |
| mmu-miR-24-3p | MIMAT0000219 | Isoc1 | 66307 | HITS-CLIP | 25753418 | Unspecified |
| mmu-miR-24-3p | MIMAT0000219 | Itga3 | 16400 | HITS-CLIP | 23142080 | Unspecified |
| mmu-miR-24-3p | MIMAT0000219 | Itga6 | 16403 | HITS-CLIP | 23142080 | Unspecified |
| mmu-miR-24-3p | MIMAT0000219 | Itga7 | 16404 | HITS-CLIP | 23142080 | Unspecified |
| mmu-miR-24-3p | MIMAT0000219 | Itgal | 16408 | HITS-CLIP | 23142080 | Unspecified |
| mmu-miR-24-3p | MIMAT0000219 | Itgav | 16410 | HITS-CLIP | 23142080 | Unspecified |
| mmu-miR-24-3p | MIMAT0000219 | Itgb1 | 16412 | HITS-CLIP | 23142080 | Unspecified |
| mmu-miR-24-3p | MIMAT0000219 | Itgb7 | 16421 | HITS-CLIP | 23142080 | Unspecified |
| mmu-miR-24-3p | MIMAT0000219 | Itk | 16428 | HITS-CLIP | 37986224\|23142080 | Unspecified |
| mmu-miR-24-3p | MIMAT0000219 | Itm2b | 16432 | HITS-CLIP | 25753418 | Unspecified |
| mmu-miR-24-3p | MIMAT0000219 | Itpk1 | 217837 | HITS-CLIP | 23142080 | Unspecified |
| mmu-miR-24-3p | MIMAT0000219 | Itpr3 | 16440 | HITS-CLIP | 23142080 | Unspecified |
| mmu-miR-24-3p | MIMAT0000219 | Ivd | 56357 | HITS-CLIP | 25753418 | Unspecified |
| mmu-miR-24-3p | MIMAT0000219 | Iws1 | 73473 | HITS-CLIP | 23142080 | Unspecified |
| mmu-miR-24-3p | MIMAT0000219 | Jade1 | 269424 | HITS-CLIP | 23142080 | Unspecified |
| mmu-miR-24-3p | MIMAT0000219 | Jade2 | 76901 | HITS-CLIP | 23142080 | Unspecified |
| mmu-miR-24-3p | MIMAT0000219 | Jagn1 | 67767 | HITS-CLIP | 25753418 | Unspecified |
| mmu-miR-24-3p | MIMAT0000219 | Jak1 | 16451 | HITS-CLIP | 37986224\|23142080\|25753418 | Unspecified |
| mmu-miR-24-3p | MIMAT0000219 | Jak2 | 16452 | HITS-CLIP | 37986224\|23142080 | Unspecified |
| mmu-miR-24-3p | MIMAT0000219 | Jak3 | 16453 | HITS-CLIP | 23142080 | Unspecified |
| mmu-miR-24-3p | MIMAT0000219 | Jarid2 | 16468 | HITS-CLIP | 23142080 | Unspecified |
| mmu-miR-24-3p | MIMAT0000219 | Jcad | 57608 | HITS-CLIP | 25753418\|23142080 | Unspecified |
| mmu-miR-24-3p | MIMAT0000219 | Jmjd1c | 108829 | HITS-CLIP | 23142080 | Unspecified |
| mmu-miR-24-3p | MIMAT0000219 | Jmy | 57748 | HITS-CLIP | 23142080 | Unspecified |
| mmu-miR-24-3p | MIMAT0000219 | Jpt1 | 51155 | HITS-CLIP | 23142080 | Unspecified |
| mmu-miR-24-3p | MIMAT0000219 | Jup | 16480 | HITS-CLIP | 25753418 | Unspecified |
| mmu-miR-24-3p | MIMAT0000219 | Kank2 | 235041 | HITS-CLIP | 25753418 | Unspecified |
| mmu-miR-24-3p | MIMAT0000219 | Kank3 | 80880 | HITS-CLIP | 25753418 | Unspecified |
| mmu-miR-24-3p | MIMAT0000219 | Kansl1 | 76719 | HITS-CLIP | 23142080 | Unspecified |
| mmu-miR-24-3p | MIMAT0000219 | Kansl3 | 226976 | Sequencing//HITS-CLIP | 19536157\|25753418 | Unspecified |
| mmu-miR-24-3p | MIMAT0000219 | Kat6a | 244349 | HITS-CLIP | 23142080 | Unspecified |
| mmu-miR-24-3p | MIMAT0000219 | Kat6b | 54169 | HITS-CLIP | 23142080 | Unspecified |
| mmu-miR-24-3p | MIMAT0000219 | Kat7 | 217127 | HITS-CLIP | 37986224\|23142080\|25753418 | Unspecified |
| mmu-miR-24-3p | MIMAT0000219 | Kbtbd11 | 74901 | HITS-CLIP | 23142080 | Unspecified |

| | | | | | | | |
|---|---|---|---|---|---|---|---|
| mmu-miR-24-3p | MIMAT0000219 | Kbtbd3 | 69149 | HITS-CLIP | | 25753418 | Unspecified |
| mmu-miR-24-3p | MIMAT0000219 | Kbtbd6 | 89890 | HITS-CLIP | | 23142080 | Unspecified |
| mmu-miR-24-3p | MIMAT0000219 | Kcna3 | 16491 | HITS-CLIP | | 23142080 | Unspecified |
| mmu-miR-24-3p | MIMAT0000219 | Kcnb1 | 16500 | HITS-CLIP | | 25753418 | Unspecified |
| mmu-miR-24-3p | MIMAT0000219 | Kcnk5 | 16529 | HITS-CLIP | | 23142080 | Unspecified |
| mmu-miR-24-3p | MIMAT0000219 | Kcnq5 | 226922 | HITS-CLIP | | 23142080 | Unspecified |
| mmu-miR-24-3p | MIMAT0000219 | Kctd5 | 69259 | HITS-CLIP | | 23142080 | Unspecified |
| mmu-miR-24-3p | MIMAT0000219 | Kdelr2 | 66913 | HITS-CLIP | | 23142080 | Unspecified |
| mmu-miR-24-3p | MIMAT0000219 | Kdm2a | 225876 | HITS-CLIP | 26203562\|23142080 | | Unspecified |
| mmu-miR-24-3p | MIMAT0000219 | Kdm3a | 104263 | HITS-CLIP | | 23142080 | Unspecified |
| mmu-miR-24-3p | MIMAT0000219 | Kdm3b | 277250 | HITS-CLIP | | 23142080 | Unspecified |
| mmu-miR-24-3p | MIMAT0000219 | Kdm4c | 76804 | Sequencing//HITS-CLIP | 19536157\|23142080 | | Unspecified |
| mmu-miR-24-3p | MIMAT0000219 | Kdm5a | 214899 | HITS-CLIP | | 23142080 | Unspecified |
| mmu-miR-24-3p | MIMAT0000219 | Kdm5b | 75605 | Sequencing//HITS-CLIP | 19536157\|23142080 | | Unspecified |
| mmu-miR-24-3p | MIMAT0000219 | Kdm5c | 20591 | HITS-CLIP | | 23142080 | Unspecified |
| mmu-miR-24-3p | MIMAT0000219 | Kdm5d | 20592 | HITS-CLIP | | 25753418 | Unspecified |
| mmu-miR-24-3p | MIMAT0000219 | Kdm6b | 216850 | HITS-CLIP | | 23142080 | Unspecified |
| mmu-miR-24-3p | MIMAT0000219 | Kdm7a | 338523 | HITS-CLIP | | 23142080 | Unspecified |
| mmu-miR-24-3p | MIMAT0000219 | Keap1 | 50868 | HITS-CLIP | | 23142080 | Unspecified |
| mmu-miR-24-3p | MIMAT0000219 | Khdc4 | 22889 | HITS-CLIP | | 23142080 | Unspecified |
| mmu-miR-24-3p | MIMAT0000219 | Khdrbs1 | 20218 | HITS-CLIP | | 25753418 | Unspecified |
| mmu-miR-24-3p | MIMAT0000219 | Khsrp | 16549 | HITS-CLIP | | 23142080 | Unspecified |
| mmu-miR-24-3p | MIMAT0000219 | Kidins220 | 77480 | HITS-CLIP | | 37986224 | Unspecified |
| mmu-miR-24-3p | MIMAT0000219 | Kif11 | 16551 | HITS-CLIP | 37986224\|23142080 | | Unspecified |
| mmu-miR-24-3p | MIMAT0000219 | Kif13a | 16553 | HITS-CLIP | | 25753418 | Unspecified |
| mmu-miR-24-3p | MIMAT0000219 | Kif1b | 16561 | HITS-CLIP | 25753418\|23142080 | | Unspecified |
| mmu-miR-24-3p | MIMAT0000219 | Kif20b | 240641 | HITS-CLIP | | 23142080 | Unspecified |
| mmu-miR-24-3p | MIMAT0000219 | Kif21b | 16565 | Sequencing//HITS-CLIP | 37986224\|23142080\|19536157 | | Unspecified |
| mmu-miR-24-3p | MIMAT0000219 | Kif23 | 71819 | HITS-CLIP | 37986224\|23142080 | | Unspecified |
| mmu-miR-24-3p | MIMAT0000219 | Kif24 | 109242 | HITS-CLIP | | 37986224 | Unspecified |
| mmu-miR-24-3p | MIMAT0000219 | Kif2a | 16563 | HITS-CLIP | | 23142080 | Unspecified |
| mmu-miR-24-3p | MIMAT0000219 | Kif3b | 16569 | Sequencing//HITS-CLIP | 19536157\|23142080\|37986224 | | Unspecified |
| mmu-miR-24-3p | MIMAT0000219 | Kif4 | 16571 | HITS-CLIP | | 23142080 | Unspecified |
| mmu-miR-24-3p | MIMAT0000219 | Kif5b | 16573 | HITS-CLIP | | 25753418 | Unspecified |
| mmu-miR-24-3p | MIMAT0000219 | Klc1 | 16593 | HITS-CLIP | | 23142080 | Unspecified |
| mmu-miR-24-3p | MIMAT0000219 | Klf2 | 16598 | HITS-CLIP | 37986224\|23142080 | | Unspecified |
| mmu-miR-24-3p | MIMAT0000219 | Klf3 | 16599 | HITS-CLIP | | 23142080 | Unspecified |
| mmu-miR-24-3p | MIMAT0000219 | Klf6 | 23849 | Sequencing//HITS-CLIP | 19536157\|23142080\|25753418\|3798( | | Unspecified |
| mmu-miR-24-3p | MIMAT0000219 | Klhdc2 | 69554 | HITS-CLIP | | 25753418 | Unspecified |

| | | | | | | |
|---|---|---|---|---|---|---|
| mmu-miR-24-3p | MIMAT0000219 | Klhl15 | 236904 | HITS-CLIP | 23142080 | Unspecified |
| mmu-miR-24-3p | MIMAT0000219 | Klhl23 | 277396 | HITS-CLIP | 25753418 | Unspecified |
| mmu-miR-24-3p | MIMAT0000219 | Klhl24 | 75785 | Sequencing//HITS-CLIP | 37986224\|23142080\|25753418\|19536 | Unspecified |
| mmu-miR-24-3p | MIMAT0000219 | Klhl28 | 66689 | HITS-CLIP | 23142080 | Unspecified |
| mmu-miR-24-3p | MIMAT0000219 | Klhl3 | 100503085 | HITS-CLIP | 25753418 | Unspecified |
| mmu-miR-24-3p | MIMAT0000219 | Klhl30 | 70788 | HITS-CLIP | 25753418 | Unspecified |
| mmu-miR-24-3p | MIMAT0000219 | Klhl31 | 244923 | HITS-CLIP | 25753418 | Unspecified |
| mmu-miR-24-3p | MIMAT0000219 | Klhl34 | 257240 | HITS-CLIP | 25753418 | Unspecified |
| mmu-miR-24-3p | MIMAT0000219 | Klhl42 | 232539 | HITS-CLIP | 37986224\|23142080 | Unspecified |
| mmu-miR-24-3p | MIMAT0000219 | Klhl9 | 242521 | HITS-CLIP | 23142080 | Unspecified |
| mmu-miR-24-3p | MIMAT0000219 | Kmt2a | 214162 | HITS-CLIP | 37986224\|23142080\|25753418 | Unspecified |
| mmu-miR-24-3p | MIMAT0000219 | Kmt2b | 75410 | HITS-CLIP | 23142080\|25753418 | Unspecified |
| mmu-miR-24-3p | MIMAT0000219 | Kmt2c | 231051 | HITS-CLIP | 37986224 | Unspecified |
| mmu-miR-24-3p | MIMAT0000219 | Kmt2d | 381022 | HITS-CLIP | 23142080\|25753418 | Unspecified |
| mmu-miR-24-3p | MIMAT0000219 | Kmt2e | 69188 | HITS-CLIP | 23142080 | Unspecified |
| mmu-miR-24-3p | MIMAT0000219 | Kmt5c | 84787 | HITS-CLIP | 23142080 | Unspecified |
| mmu-miR-24-3p | MIMAT0000219 | Kntc1 | 208628 | HITS-CLIP | 23142080 | Unspecified |
| mmu-miR-24-3p | MIMAT0000219 | Kpna3 | 16648 | HITS-CLIP | 25753418 | Unspecified |
| mmu-miR-24-3p | MIMAT0000219 | Kpna6 | 16650 | HITS-CLIP | 23142080\|25753418 | Unspecified |
| mmu-miR-24-3p | MIMAT0000219 | Kpnb1 | 16211 | HITS-CLIP | 23142080\|25753418 | Unspecified |
| mmu-miR-24-3p | MIMAT0000219 | Kremen2 | 73016 | HITS-CLIP | 23142080 | Unspecified |
| mmu-miR-24-3p | MIMAT0000219 | Krit1 | 79264 | HITS-CLIP | 23142080 | Unspecified |
| mmu-miR-24-3p | MIMAT0000219 | L3mbtl3 | 237339 | HITS-CLIP | 23142080 | Unspecified |
| mmu-miR-24-3p | MIMAT0000219 | Lama4 | 16775 | HITS-CLIP | 25753418 | Unspecified |
| mmu-miR-24-3p | MIMAT0000219 | Lama5 | 16776 | HITS-CLIP | 23142080 | Unspecified |
| mmu-miR-24-3p | MIMAT0000219 | Lamb2 | 16779 | HITS-CLIP | 25753418 | Unspecified |
| mmu-miR-24-3p | MIMAT0000219 | Laptm4a | 17775 | HITS-CLIP | 23142080 | Unspecified |
| mmu-miR-24-3p | MIMAT0000219 | Laptm4b | 114128 | HITS-CLIP | 25753418 | Unspecified |
| mmu-miR-24-3p | MIMAT0000219 | Laptm5 | 16792 | HITS-CLIP | 23142080 | Unspecified |
| mmu-miR-24-3p | MIMAT0000219 | Larp1 | 73158 | HITS-CLIP | 23142080 | Unspecified |
| mmu-miR-24-3p | MIMAT0000219 | Larp4b | 217980 | HITS-CLIP | 23142080\|25753418 | Unspecified |
| mmu-miR-24-3p | MIMAT0000219 | Larp7 | 28036 | HITS-CLIP | 25753418 | Unspecified |
| mmu-miR-24-3p | MIMAT0000219 | Lars | 107045 | HITS-CLIP | 37986224\|23142080 | Unspecified |
| mmu-miR-24-3p | MIMAT0000219 | Lars2 | 102436 | HITS-CLIP | 25753418\|23142080 | Unspecified |
| mmu-miR-24-3p | MIMAT0000219 | Las1l | 76130 | HITS-CLIP | 23142080 | Unspecified |
| mmu-miR-24-3p | MIMAT0000219 | Lats1 | 16798 | HITS-CLIP | 23142080 | Unspecified |
| mmu-miR-24-3p | MIMAT0000219 | Lbr | 98386 | HITS-CLIP | 37986224\|23142080 | Unspecified |
| mmu-miR-24-3p | MIMAT0000219 | Lck | 16818 | HITS-CLIP | 23142080 | Unspecified |
| mmu-miR-24-3p | MIMAT0000219 | Lclat1 | 225010 | HITS-CLIP | 23142080 | Unspecified |

| | | | | | | |
|---|---|---|---|---|---|---|
| mmu-miR-24-3p | MIMAT0000219 | Lcmt1 | 30949 | HITS-CLIP | 23142080 | Unspecified |
| mmu-miR-24-3p | MIMAT0000219 | Lcmt2 | 329504 | HITS-CLIP | 25753418 | Unspecified |
| mmu-miR-24-3p | MIMAT0000219 | Lcor | 212391 | HITS-CLIP | 23142080 | Unspecified |
| mmu-miR-24-3p | MIMAT0000219 | Lcp1 | 18826 | HITS-CLIP | 37986224\|23142080 | Unspecified |
| mmu-miR-24-3p | MIMAT0000219 | Lcp2 | 16822 | HITS-CLIP | 37986224 | Unspecified |
| mmu-miR-24-3p | MIMAT0000219 | Ldha | 16828 | Sequencing//HITS-CLIP | 37986224\|23142080\|19536157 | Unspecified |
| mmu-miR-24-3p | MIMAT0000219 | Ldlrap1 | 100017 | HITS-CLIP | 25753418\|23142080 | Unspecified |
| mmu-miR-24-3p | MIMAT0000219 | Lef1 | 16842 | HITS-CLIP | 23142080 | Unspecified |
| mmu-miR-24-3p | MIMAT0000219 | Lemd2 | 224640 | HITS-CLIP | 23142080 | Unspecified |
| mmu-miR-24-3p | MIMAT0000219 | Lemd3 | 380664 | HITS-CLIP | 23142080 | Unspecified |
| mmu-miR-24-3p | MIMAT0000219 | Leng8 | 232798 | HITS-CLIP | 25753418\|23142080 | Unspecified |
| mmu-miR-24-3p | MIMAT0000219 | Leo1 | 235497 | HITS-CLIP | 23142080 | Unspecified |
| mmu-miR-24-3p | MIMAT0000219 | Lgals3 | 16854 | HITS-CLIP | 23142080 | Unspecified |
| mmu-miR-24-3p | MIMAT0000219 | Lgals8 | 56048 | HITS-CLIP | 37986224 | Unspecified |
| mmu-miR-24-3p | MIMAT0000219 | Lif | 16878 | HITS-CLIP | 23142080 | Unspecified |
| mmu-miR-24-3p | MIMAT0000219 | Limd2 | 67803 | HITS-CLIP | 23142080 | Unspecified |
| mmu-miR-24-3p | MIMAT0000219 | Limk2 | 16886 | HITS-CLIP | 23142080 | Unspecified |
| mmu-miR-24-3p | MIMAT0000219 | Lims1 | 110829 | Sequencing//HITS-CLIP | 19536157\|23142080\|37986224 | Unspecified |
| mmu-miR-24-3p | MIMAT0000219 | Lin9 | 72568 | HITS-CLIP | 23142080 | Unspecified |
| mmu-miR-24-3p | MIMAT0000219 | Lipe | 16890 | HITS-CLIP | 23142080 | Unspecified |
| mmu-miR-24-3p | MIMAT0000219 | Llgl1 | 16897 | HITS-CLIP | 23142080 | Unspecified |
| mmu-miR-24-3p | MIMAT0000219 | Lmbr1l | 74775 | HITS-CLIP | 37986224\|23142080 | Unspecified |
| mmu-miR-24-3p | MIMAT0000219 | Lmnb1 | 16906 | HITS-CLIP | 23142080 | Unspecified |
| mmu-miR-24-3p | MIMAT0000219 | Lmo7 | 380928 | HITS-CLIP | 25753418 | Unspecified |
| mmu-miR-24-3p | MIMAT0000219 | Lmtk2 | 231876 | Sequencing//HITS-CLIP | 37986224\|23142080\|19536157 | Unspecified |
| mmu-miR-24-3p | MIMAT0000219 | Lnpep | 240028 | HITS-CLIP | 23142080 | Unspecified |
| mmu-miR-24-3p | MIMAT0000219 | Lnpk | 69605 | HITS-CLIP | 23142080 | Unspecified |
| mmu-miR-24-3p | MIMAT0000219 | Lonp1 | 74142 | HITS-CLIP | 23142080 | Unspecified |
| mmu-miR-24-3p | MIMAT0000219 | Lox | 16948 | HITS-CLIP | 23142080 | Unspecified |
| mmu-miR-24-3p | MIMAT0000219 | Lpcat3 | 14792 | HITS-CLIP | 37986224\|25753418 | Unspecified |
| mmu-miR-24-3p | MIMAT0000219 | Lpl | 16956 | HITS-CLIP | 25753418 | Unspecified |
| mmu-miR-24-3p | MIMAT0000219 | Lrba | 80877 | HITS-CLIP | 23142080 | Unspecified |
| mmu-miR-24-3p | MIMAT0000219 | Lrch1 | 380916 | Sequencing//HITS-CLIP | 19536157\|23142080\|37986224 | Unspecified |
| mmu-miR-24-3p | MIMAT0000219 | Lrp10 | 65107 | HITS-CLIP | 23142080 | Unspecified |
| mmu-miR-24-3p | MIMAT0000219 | Lrpprc | 72416 | HITS-CLIP | 23142080 | Unspecified |
| mmu-miR-24-3p | MIMAT0000219 | Lrr1 | 69706 | HITS-CLIP | 23142080 | Unspecified |
| mmu-miR-24-3p | MIMAT0000219 | Lrrc2 | 74249 | HITS-CLIP | 25753418 | Unspecified |
| mmu-miR-24-3p | MIMAT0000219 | Lrrc32 | 434215 | HITS-CLIP | 23142080\|25753418 | Unspecified |
| mmu-miR-24-3p | MIMAT0000219 | Lrrc41 | 230654 | HITS-CLIP | 23142080 | Unspecified |

| | | | | | | | |
|---|---|---|---|---|---|---|---|
| mmu-miR-24-3p | MIMAT0000219 | Lrrc42 | 77809 | HITS-CLIP | | 23142080 | Unspecified |
| mmu-miR-24-3p | MIMAT0000219 | Lrrc58 | 320184 | Sequencing//HITS-CLIP | 19536157\|23142080\|37986224 | | Unspecified |
| mmu-miR-24-3p | MIMAT0000219 | Lrrc59 | 98238 | HITS-CLIP | | 23142080 | Unspecified |
| mmu-miR-24-3p | MIMAT0000219 | Lrtm1 | 319476 | HITS-CLIP | | 25753418 | Unspecified |
| mmu-miR-24-3p | MIMAT0000219 | Lrwd1 | 71735 | HITS-CLIP | | 23142080 | Unspecified |
| mmu-miR-24-3p | MIMAT0000219 | Lsm12 | 268490 | HITS-CLIP | | 23142080 | Unspecified |
| mmu-miR-24-3p | MIMAT0000219 | Lsm14a | 67070 | HITS-CLIP | 37986224\|23142080 | | Unspecified |
| mmu-miR-24-3p | MIMAT0000219 | Lsm14b | 241846 | HITS-CLIP | | 31433990 | Unspecified |
| mmu-miR-24-3p | MIMAT0000219 | Lss | 16987 | HITS-CLIP | | 23142080 | Unspecified |
| mmu-miR-24-3p | MIMAT0000219 | Ltb | 16994 | HITS-CLIP | | 23142080 | Unspecified |
| mmu-miR-24-3p | MIMAT0000219 | Ltn1 | 78913 | HITS-CLIP | | 23142080 | Unspecified |
| mmu-miR-24-3p | MIMAT0000219 | Luc7l | 66978 | HITS-CLIP | | 23142080 | Unspecified |
| mmu-miR-24-3p | MIMAT0000219 | Ly6a | 110454 | HITS-CLIP | | 23142080 | Unspecified |
| mmu-miR-24-3p | MIMAT0000219 | Ly6c1 | 17067 | HITS-CLIP | | 25753418 | Unspecified |
| mmu-miR-24-3p | MIMAT0000219 | Ly6e | 17069 | HITS-CLIP | 37986224\|23142080 | | Unspecified |
| mmu-miR-24-3p | MIMAT0000219 | Lyn | 17096 | HITS-CLIP | | 25753418 | Unspecified |
| mmu-miR-24-3p | MIMAT0000219 | Lynx1 | 23936 | HITS-CLIP | | 25753418 | Unspecified |
| mmu-miR-24-3p | MIMAT0000219 | Lypla2 | 26394 | HITS-CLIP | | 23142080 | Unspecified |
| mmu-miR-24-3p | MIMAT0000219 | Lyst | 17101 | HITS-CLIP | 37986224\|23142080 | | Unspecified |
| mmu-miR-24-3p | MIMAT0000219 | Lzic | 69151 | HITS-CLIP | | 25753418 | Unspecified |
| mmu-miR-24-3p | MIMAT0000219 | MAPK7 | 5598 | HITS-CLIP//Luciferase reporter assay | 23142080\|27103442 | | Unspecified |
| mmu-miR-24-3p | MIMAT0000219 | Macf1 | 11426 | Sequencing//HITS-CLIP | 37986224\|23142080\|25753418\|19536 | | Unspecified |
| mmu-miR-24-3p | MIMAT0000219 | Mad1l1 | 17120 | HITS-CLIP | | 23142080 | Unspecified |
| mmu-miR-24-3p | MIMAT0000219 | Maea | 59003 | HITS-CLIP | | 25753418 | Unspecified |
| mmu-miR-24-3p | MIMAT0000219 | Maf | 17132 | HITS-CLIP | | 23142080 | Unspecified |
| mmu-miR-24-3p | MIMAT0000219 | Maf1 | 68877 | HITS-CLIP | | 23142080 | Unspecified |
| mmu-miR-24-3p | MIMAT0000219 | Maff | 17133 | HITS-CLIP | | 23142080 | Unspecified |
| mmu-miR-24-3p | MIMAT0000219 | Mafg | 17134 | HITS-CLIP | | 37986224 | Unspecified |
| mmu-miR-24-3p | MIMAT0000219 | Magee1 | 107528 | HITS-CLIP | | 23142080 | Unspecified |
| mmu-miR-24-3p | MIMAT0000219 | Magi3 | 99470 | HITS-CLIP | 25753418\|23142080 | | Unspecified |
| mmu-miR-24-3p | MIMAT0000219 | Magt1 | 67075 | HITS-CLIP | | 23142080 | Unspecified |
| mmu-miR-24-3p | MIMAT0000219 | Malsu1 | 75593 | HITS-CLIP | | 25753418 | Unspecified |
| mmu-miR-24-3p | MIMAT0000219 | Malt1 | 240354 | HITS-CLIP | | 23142080 | Unspecified |
| mmu-miR-24-3p | MIMAT0000219 | Maml1 | 103806 | HITS-CLIP | | 23142080 | Unspecified |
| mmu-miR-24-3p | MIMAT0000219 | Maml2 | 270118 | HITS-CLIP | | 23142080 | Unspecified |
| mmu-miR-24-3p | MIMAT0000219 | Man1a2 | 17156 | HITS-CLIP | | 23142080 | Unspecified |
| mmu-miR-24-3p | MIMAT0000219 | Man2b1 | 17159 | HITS-CLIP | | 23142080 | Unspecified |
| mmu-miR-24-3p | MIMAT0000219 | Manea | 242362 | HITS-CLIP | | 25753418 | Unspecified |
| mmu-miR-24-3p | MIMAT0000219 | Manf | 74840 | HITS-CLIP | | 23142080 | Unspecified |

| | | | | | | |
|---|---|---|---|---|---|---|
| mmu-miR-24-3p | MIMAT0000219 | Map3k1 | 26401 | HITS-CLIP | 37986224\|23142080 | Unspecified |
| mmu-miR-24-3p | MIMAT0000219 | Map3k11 | 26403 | HITS-CLIP | 23142080 | Unspecified |
| mmu-miR-24-3p | MIMAT0000219 | Map3k12 | 26404 | HITS-CLIP | 23142080 | Unspecified |
| mmu-miR-24-3p | MIMAT0000219 | Map3k2 | 26405 | HITS-CLIP | 23142080 | Unspecified |
| mmu-miR-24-3p | MIMAT0000219 | Map3k5 | 26408 | HITS-CLIP | 23142080 | Unspecified |
| mmu-miR-24-3p | MIMAT0000219 | Map3k8 | 26410 | HITS-CLIP | 37986224 | Unspecified |
| mmu-miR-24-3p | MIMAT0000219 | Map4k2 | 26412 | HITS-CLIP | 23142080 | Unspecified |
| mmu-miR-24-3p | MIMAT0000219 | Mapk1 | 26413 | HITS-CLIP | 23142080 | Unspecified |
| mmu-miR-24-3p | MIMAT0000219 | Mapk14 | 26416 | HITS-CLIP | 23142080 | Unspecified |
| mmu-miR-24-3p | MIMAT0000219 | Mapk6 | 50772 | HITS-CLIP | 23142080\|25753418 | Unspecified |
| mmu-miR-24-3p | MIMAT0000219 | Mapkapk2 | 17164 | HITS-CLIP | 23142080 | Unspecified |
| mmu-miR-24-3p | MIMAT0000219 | Marchf9 | 92979 | HITS-CLIP | 25753418 | Unspecified |
| mmu-miR-24-3p | MIMAT0000219 | Marcks | 17118 | HITS-CLIP | 23142080 | Unspecified |
| mmu-miR-24-3p | MIMAT0000219 | Marf1 | 223989 | HITS-CLIP | 23142080 | Unspecified |
| mmu-miR-24-3p | MIMAT0000219 | Mark3 | 17169 | HITS-CLIP | 23142080 | Unspecified |
| mmu-miR-24-3p | MIMAT0000219 | Mark4 | 232944 | HITS-CLIP | 23142080 | Unspecified |
| mmu-miR-24-3p | MIMAT0000219 | Mars1 | 4141 | HITS-CLIP | 25753418 | Unspecified |
| mmu-miR-24-3p | MIMAT0000219 | Mast2 | 17776 | HITS-CLIP | 25753418 | Unspecified |
| mmu-miR-24-3p | MIMAT0000219 | Mast4 | 328329 | HITS-CLIP | 25753418 | Unspecified |
| mmu-miR-24-3p | MIMAT0000219 | Mat2b | 108645 | HITS-CLIP | 23142080 | Unspecified |
| mmu-miR-24-3p | MIMAT0000219 | Matr3 | 17184 | HITS-CLIP | 37986224\|23142080 | Unspecified |
| mmu-miR-24-3p | MIMAT0000219 | Mau2 | 74549 | HITS-CLIP | 25753418\|23142080 | Unspecified |
| mmu-miR-24-3p | MIMAT0000219 | Maz | 17188 | HITS-CLIP | 37986224\|23142080 | Unspecified |
| mmu-miR-24-3p | MIMAT0000219 | Mbd2 | 17191 | HITS-CLIP | 23142080 | Unspecified |
| mmu-miR-24-3p | MIMAT0000219 | Mbd6 | 110962 | Sequencing//HITS-CLIP | 19536157\|23142080\|37986224 | Unspecified |
| mmu-miR-24-3p | MIMAT0000219 | Mbnl1 | 56758 | HITS-CLIP | 37986224\|23142080 | Unspecified |
| mmu-miR-24-3p | MIMAT0000219 | Mbnl2 | 105559 | HITS-CLIP | 23142080 | Unspecified |
| mmu-miR-24-3p | MIMAT0000219 | Mbnl3 | 171170 | HITS-CLIP | 23142080 | Unspecified |
| mmu-miR-24-3p | MIMAT0000219 | Mbtps2 | 270669 | HITS-CLIP | 23142080 | Unspecified |
| mmu-miR-24-3p | MIMAT0000219 | Mcfd2 | 193813 | HITS-CLIP | 23142080 | Unspecified |
| mmu-miR-24-3p | MIMAT0000219 | Mcl1 | 17210 | HITS-CLIP | 25753418 | Unspecified |
| mmu-miR-24-3p | MIMAT0000219 | Mcm10 | 70024 | HITS-CLIP | 23142080 | Unspecified |
| mmu-miR-24-3p | MIMAT0000219 | Mcm4 | 17217 | HITS-CLIP | 23142080 | Unspecified |
| mmu-miR-24-3p | MIMAT0000219 | Mcm6 | 17219 | HITS-CLIP | 23142080 | Unspecified |
| mmu-miR-24-3p | MIMAT0000219 | Mcmbp | 210711 | HITS-CLIP | 23142080 | Unspecified |
| mmu-miR-24-3p | MIMAT0000219 | Mdh1 | 17449 | Sequencing//HITS-CLIP | 37986224\|23142080\|19536157 | Unspecified |
| mmu-miR-24-3p | MIMAT0000219 | Mdh2 | 17448 | HITS-CLIP | 37986224\|23142080 | Unspecified |
| mmu-miR-24-3p | MIMAT0000219 | Mdm4 | 17248 | HITS-CLIP | 23142080 | Unspecified |
| mmu-miR-24-3p | MIMAT0000219 | Mdn1 | 100019 | HITS-CLIP | 25753418\|23142080 | Unspecified |

| | | | | | | |
|---|---|---|---|---|---|---|
| mmu-miR-24-3p | MIMAT0000219 | Me2 | 107029 | HITS-CLIP | 23142080 | Unspecified |
| mmu-miR-24-3p | MIMAT0000219 | Mecp2 | 17257 | HITS-CLIP | 23142080 | Unspecified |
| mmu-miR-24-3p | MIMAT0000219 | Med1 | 19014 | qCLASH | 31393953 | Unspecified |
| mmu-miR-24-3p | MIMAT0000219 | Med12 | 59024 | HITS-CLIP | 23142080 | Unspecified |
| mmu-miR-24-3p | MIMAT0000219 | Med13 | 327987 | HITS-CLIP | 23142080 | Unspecified |
| mmu-miR-24-3p | MIMAT0000219 | Med13l | 76199 | HITS-CLIP | 37986224\|23142080 | Unspecified |
| mmu-miR-24-3p | MIMAT0000219 | Med14 | 26896 | HITS-CLIP | 23142080 | Unspecified |
| mmu-miR-24-3p | MIMAT0000219 | Med17 | 234959 | HITS-CLIP | 23142080 | Unspecified |
| mmu-miR-24-3p | MIMAT0000219 | Med20 | 56771 | HITS-CLIP | 23142080 | Unspecified |
| mmu-miR-24-3p | MIMAT0000219 | Med25 | 75613 | HITS-CLIP | 23142080 | Unspecified |
| mmu-miR-24-3p | MIMAT0000219 | Med26 | 70625 | HITS-CLIP | 23142080 | Unspecified |
| mmu-miR-24-3p | MIMAT0000219 | Med6 | 69792 | HITS-CLIP | 25753418 | Unspecified |
| mmu-miR-24-3p | MIMAT0000219 | Med7 | 66213 | HITS-CLIP | 23142080 | Unspecified |
| mmu-miR-24-3p | MIMAT0000219 | Med8 | 80509 | HITS-CLIP | 23142080 | Unspecified |
| mmu-miR-24-3p | MIMAT0000219 | Mef2a | 17258 | HITS-CLIP | 37986224\|23142080\|25753418 | Unspecified |
| mmu-miR-24-3p | MIMAT0000219 | Melk | 17279 | HITS-CLIP | 23142080 | Unspecified |
| mmu-miR-24-3p | MIMAT0000219 | Men1 | 17283 | HITS-CLIP | 23142080 | Unspecified |
| mmu-miR-24-3p | MIMAT0000219 | Met | 17295 | HITS-CLIP | 25753418 | Unspecified |
| mmu-miR-24-3p | MIMAT0000219 | Metap1 | 75624 | HITS-CLIP | 23142080 | Unspecified |
| mmu-miR-24-3p | MIMAT0000219 | Mettl9 | 59052 | HITS-CLIP | 25753418 | Unspecified |
| mmu-miR-24-3p | MIMAT0000219 | Mfap3 | 216760 | HITS-CLIP | 25753418 | Unspecified |
| mmu-miR-24-3p | MIMAT0000219 | Mfn1 | 67414 | HITS-CLIP | 23142080 | Unspecified |
| mmu-miR-24-3p | MIMAT0000219 | Mfn2 | 170731 | HITS-CLIP | 25753418 | Unspecified |
| mmu-miR-24-3p | MIMAT0000219 | Mfsd14a | 15247 | HITS-CLIP | 23142080 | Unspecified |
| mmu-miR-24-3p | MIMAT0000219 | Mfsd14b | 84641 | HITS-CLIP | 23142080 | Unspecified |
| mmu-miR-24-3p | MIMAT0000219 | Mfsd2a | 76574 | HITS-CLIP | 37986224 | Unspecified |
| mmu-miR-24-3p | MIMAT0000219 | Mfsd6 | 98682 | HITS-CLIP | 23142080 | Unspecified |
| mmu-miR-24-3p | MIMAT0000219 | Mfsd8 | 72175 | HITS-CLIP | 23142080 | Unspecified |
| mmu-miR-24-3p | MIMAT0000219 | Mga | 29808 | HITS-CLIP | 25753418 | Unspecified |
| mmu-miR-24-3p | MIMAT0000219 | Mgat1 | 17308 | HITS-CLIP | 23142080 | Unspecified |
| mmu-miR-24-3p | MIMAT0000219 | Mgat4b | 103534 | HITS-CLIP | 23142080 | Unspecified |
| mmu-miR-24-3p | MIMAT0000219 | Mgea5 | 76055 | HITS-CLIP | 23142080 | Unspecified |
| mmu-miR-24-3p | MIMAT0000219 | Mia2 | 338320 | HITS-CLIP | 23142080 | Unspecified |
| mmu-miR-24-3p | MIMAT0000219 | Mia3 | 338366 | HITS-CLIP | 37986224 | Unspecified |
| mmu-miR-24-3p | MIMAT0000219 | Micos13 | 125988 | HITS-CLIP | 25753418 | Unspecified |
| mmu-miR-24-3p | MIMAT0000219 | Midn | 59090 | HITS-CLIP | 37986224\|25753418\|23142080 | Unspecified |
| mmu-miR-24-3p | MIMAT0000219 | Mief1 | 239555 | HITS-CLIP | 23142080 | Unspecified |
| mmu-miR-24-3p | MIMAT0000219 | Miga1 | 215708 | HITS-CLIP | 25753418 | Unspecified |
| mmu-miR-24-3p | MIMAT0000219 | Mindy2 | 235461 | HITS-CLIP | 23142080 | Unspecified |

| | | | | | | |
|---|---|---|---|---|---|---|
| mmu-miR-24-3p | MIMAT0000219 | Mindy3 | 80013 | HITS-CLIP | 23142080 | Unspecified |
| mmu-miR-24-3p | MIMAT0000219 | Mlec | 109154 | HITS-CLIP | 23142080 | Unspecified |
| mmu-miR-24-3p | MIMAT0000219 | Mlf2 | 30853 | HITS-CLIP | 23142080 | Unspecified |
| mmu-miR-24-3p | MIMAT0000219 | Mlh3 | 217716 | HITS-CLIP | 23142080 | Unspecified |
| mmu-miR-24-3p | MIMAT0000219 | Mllt3 | 70122 | HITS-CLIP | 23142080 | Unspecified |
| mmu-miR-24-3p | MIMAT0000219 | Mllt6 | 246198 | HITS-CLIP | 23142080 | Unspecified |
| mmu-miR-24-3p | MIMAT0000219 | Mlx | 21428 | HITS-CLIP | 25753418 | Unspecified |
| mmu-miR-24-3p | MIMAT0000219 | Mlxip | 208104 | HITS-CLIP | 23142080 | Unspecified |
| mmu-miR-24-3p | MIMAT0000219 | Mlycd | 56690 | HITS-CLIP | 25753418 | Unspecified |
| mmu-miR-24-3p | MIMAT0000219 | Mmadhc | 109129 | HITS-CLIP | 25753418\|23142080 | Unspecified |
| mmu-miR-24-3p | MIMAT0000219 | Mmgt1 | 236792 | HITS-CLIP | 23142080 | Unspecified |
| mmu-miR-24-3p | MIMAT0000219 | Mmp11 | 17385 | HITS-CLIP | 23142080 | Unspecified |
| mmu-miR-24-3p | MIMAT0000219 | Mms22l | 212377 | HITS-CLIP | 23142080 | Unspecified |
| mmu-miR-24-3p | MIMAT0000219 | Mob1a | 232157 | HITS-CLIP | 25753418\|23142080 | Unspecified |
| mmu-miR-24-3p | MIMAT0000219 | Mogs | 57377 | HITS-CLIP | 23142080 | Unspecified |
| mmu-miR-24-3p | MIMAT0000219 | Mon2 | 67074 | HITS-CLIP | 23142080 | Unspecified |
| mmu-miR-24-3p | MIMAT0000219 | Mov10 | 17454 | HITS-CLIP | 23142080 | Unspecified |
| mmu-miR-24-3p | MIMAT0000219 | Mpdu1 | 24070 | HITS-CLIP | 37986224\|23142080 | Unspecified |
| mmu-miR-24-3p | MIMAT0000219 | Mprip | 26936 | HITS-CLIP | 23142080 | Unspecified |
| mmu-miR-24-3p | MIMAT0000219 | Mpst | 246221 | HITS-CLIP | 25753418 | Unspecified |
| mmu-miR-24-3p | MIMAT0000219 | Mreg | 381269 | HITS-CLIP | 25753418 | Unspecified |
| mmu-miR-24-3p | MIMAT0000219 | Mroh1 | 223658 | HITS-CLIP | 23142080 | Unspecified |
| mmu-miR-24-3p | MIMAT0000219 | Mrpl15 | 27395 | HITS-CLIP | 23142080 | Unspecified |
| mmu-miR-24-3p | MIMAT0000219 | Mrpl17 | 27397 | HITS-CLIP | 23142080 | Unspecified |
| mmu-miR-24-3p | MIMAT0000219 | Mrpl24 | 67707 | HITS-CLIP | 23142080 | Unspecified |
| mmu-miR-24-3p | MIMAT0000219 | Mrpl3 | 94062 | HITS-CLIP | 23142080 | Unspecified |
| mmu-miR-24-3p | MIMAT0000219 | Mrpl33 | 66845 | HITS-CLIP | 23142080 | Unspecified |
| mmu-miR-24-3p | MIMAT0000219 | Mrpl44 | 69163 | HITS-CLIP | 37986224 | Unspecified |
| mmu-miR-24-3p | MIMAT0000219 | Mrps18b | 66973 | HITS-CLIP | 25753418 | Unspecified |
| mmu-miR-24-3p | MIMAT0000219 | Mrps22 | 64655 | HITS-CLIP | 23142080 | Unspecified |
| mmu-miR-24-3p | MIMAT0000219 | Mrps31 | 57312 | HITS-CLIP | 23142080 | Unspecified |
| mmu-miR-24-3p | MIMAT0000219 | Mrtfb | 239719 | HITS-CLIP | 23142080 | Unspecified |
| mmu-miR-24-3p | MIMAT0000219 | Ms4a6d | 68774 | HITS-CLIP | 23142080 | Unspecified |
| mmu-miR-24-3p | MIMAT0000219 | Msh6 | 17688 | HITS-CLIP | 25753418 | Unspecified |
| mmu-miR-24-3p | MIMAT0000219 | Msi2 | 76626 | HITS-CLIP | 25753418 | Unspecified |
| mmu-miR-24-3p | MIMAT0000219 | Msl1 | 74026 | HITS-CLIP | 23142080 | Unspecified |
| mmu-miR-24-3p | MIMAT0000219 | Msn | 17698 | HITS-CLIP | 25753418 | Unspecified |
| mmu-miR-24-3p | MIMAT0000219 | Msrb1 | 27361 | HITS-CLIP | 37986224\|23142080 | Unspecified |
| mmu-miR-24-3p | MIMAT0000219 | Mt1 | 17748 | HITS-CLIP | 25753418 | Unspecified |

| | | | | | | |
|---|---|---|---|---|---|---|
| mmu-miR-24-3p | MIMAT0000219 | Mtbp | 105837 | HITS-CLIP | 23142080 | Unspecified |
| mmu-miR-24-3p | MIMAT0000219 | Mtch1 | 56462 | HITS-CLIP | 23142080 | Unspecified |
| mmu-miR-24-3p | MIMAT0000219 | Mtch2 | 56428 | HITS-CLIP | 25753418 | Unspecified |
| mmu-miR-24-3p | MIMAT0000219 | Mtdh | 67154 | HITS-CLIP | 23142080 | Unspecified |
| mmu-miR-24-3p | MIMAT0000219 | Mtf2 | 17765 | HITS-CLIP | 23142080 | Unspecified |
| mmu-miR-24-3p | MIMAT0000219 | Mthfd2 | 17768 | HITS-CLIP | 23142080 | Unspecified |
| mmu-miR-24-3p | MIMAT0000219 | Mthfr | 17769 | HITS-CLIP | 23142080 | Unspecified |
| mmu-miR-24-3p | MIMAT0000219 | Mthfsl | 100039707 | HITS-CLIP | 23142080 | Unspecified |
| mmu-miR-24-3p | MIMAT0000219 | Mtmr1 | 53332 | HITS-CLIP | 23142080\|25753418 | Unspecified |
| mmu-miR-24-3p | MIMAT0000219 | Mtmr14 | 97287 | HITS-CLIP | 23142080 | Unspecified |
| mmu-miR-24-3p | MIMAT0000219 | Mtmr6 | 219135 | HITS-CLIP | 25753418 | Unspecified |
| mmu-miR-24-3p | MIMAT0000219 | Mtmr7 | 54384 | HITS-CLIP | 23142080 | Unspecified |
| mmu-miR-24-3p | MIMAT0000219 | Mto1 | 68291 | HITS-CLIP | 23142080 | Unspecified |
| mmu-miR-24-3p | MIMAT0000219 | Mtpn | 14489 | HITS-CLIP | 23142080 | Unspecified |
| mmu-miR-24-3p | MIMAT0000219 | Mtr | 238505 | HITS-CLIP | 23142080 | Unspecified |
| mmu-miR-24-3p | MIMAT0000219 | Mtx1 | 17827 | HITS-CLIP | 23142080 | Unspecified |
| mmu-miR-24-3p | MIMAT0000219 | Mxd1 | 17119 | HITS-CLIP | 37986224\|23142080 | Unspecified |
| mmu-miR-24-3p | MIMAT0000219 | Mxd4 | 17122 | HITS-CLIP | 23142080 | Unspecified |
| mmu-miR-24-3p | MIMAT0000219 | Mxi1 | 17859 | HITS-CLIP | 23142080 | Unspecified |
| mmu-miR-24-3p | MIMAT0000219 | Myadm | 50918 | HITS-CLIP | 25753418 | Unspecified |
| mmu-miR-24-3p | MIMAT0000219 | Myb | 17863 | HITS-CLIP | 23142080 | Unspecified |
| mmu-miR-24-3p | MIMAT0000219 | Myc | 17869 | HITS-CLIP | 23142080 | Unspecified |
| mmu-miR-24-3p | MIMAT0000219 | Mycbp2 | 105689 | HITS-CLIP | 23142080 | Unspecified |
| mmu-miR-24-3p | MIMAT0000219 | Mydgf | 56005 | HITS-CLIP | 23142080 | Unspecified |
| mmu-miR-24-3p | MIMAT0000219 | Myh2 | 17882 | Luciferase Reporter Assay | 37986224 | Unspecified |
| mmu-miR-24-3p | MIMAT0000219 | Myh7 | 140781 | HITS-CLIP | 25753418 | Unspecified |
| mmu-miR-24-3p | MIMAT0000219 | Myh9 | 17886 | HITS-CLIP | 23142080\|25753418 | Unspecified |
| mmu-miR-24-3p | MIMAT0000219 | Myl2 | 17906 | HITS-CLIP | 25753418 | Unspecified |
| mmu-miR-24-3p | MIMAT0000219 | Myo18a | 360013 | HITS-CLIP | 25753418 | Unspecified |
| mmu-miR-24-3p | MIMAT0000219 | Myo18b | 74376 | HITS-CLIP | 25753418 | Unspecified |
| mmu-miR-24-3p | MIMAT0000219 | Myo1b | 17912 | HITS-CLIP | 25753418 | Unspecified |
| mmu-miR-24-3p | MIMAT0000219 | Myo1c | 17913 | HITS-CLIP | 31433990\|25753418 | Unspecified |
| mmu-miR-24-3p | MIMAT0000219 | Myo1g | 246177 | HITS-CLIP | 23142080 | Unspecified |
| mmu-miR-24-3p | MIMAT0000219 | Myo6 | 17920 | HITS-CLIP | 23142080 | Unspecified |
| mmu-miR-24-3p | MIMAT0000219 | Myo9b | 17925 | HITS-CLIP | 23142080 | Unspecified |
| mmu-miR-24-3p | MIMAT0000219 | Myom1 | 17929 | HITS-CLIP | 25753418 | Unspecified |
| mmu-miR-24-3p | MIMAT0000219 | Mzt1 | 76789 | HITS-CLIP | 23142080 | Unspecified |
| mmu-miR-24-3p | MIMAT0000219 | N4bp1 | 80750 | HITS-CLIP | 23142080 | Unspecified |
| mmu-miR-24-3p | MIMAT0000219 | N4bp2 | 333789 | HITS-CLIP | 23142080 | Unspecified |

| | | | | | | |
|---|---|---|---|---|---|---|
| mmu-miR-24-3p | MIMAT0000219 | N4bp2l1 | 100637 | HITS-CLIP | 23142080 | Unspecified |
| mmu-miR-24-3p | MIMAT0000219 | N4bp2l2 | 381695 | HITS-CLIP | 37986224\|23142080 | Unspecified |
| mmu-miR-24-3p | MIMAT0000219 | Naa15 | 74838 | HITS-CLIP | 23142080 | Unspecified |
| mmu-miR-24-3p | MIMAT0000219 | Naa35 | 78689 | HITS-CLIP | 23142080\|25753418 | Unspecified |
| mmu-miR-24-3p | MIMAT0000219 | Naa40 | 70999 | Sequencing//HITS-CLIP | 19536157\|23142080 | Unspecified |
| mmu-miR-24-3p | MIMAT0000219 | Naa50 | 72117 | HITS-CLIP | 23142080 | Unspecified |
| mmu-miR-24-3p | MIMAT0000219 | Nab1 | 17936 | HITS-CLIP | 23142080 | Unspecified |
| mmu-miR-24-3p | MIMAT0000219 | Nabp1 | 109019 | HITS-CLIP | 37986224\|23142080 | Unspecified |
| mmu-miR-24-3p | MIMAT0000219 | Nae1 | 234664 | HITS-CLIP | 23142080 | Unspecified |
| mmu-miR-24-3p | MIMAT0000219 | Nagk | 56174 | HITS-CLIP | 23142080 | Unspecified |
| mmu-miR-24-3p | MIMAT0000219 | Naif1 | 71254 | HITS-CLIP | 23142080 | Unspecified |
| mmu-miR-24-3p | MIMAT0000219 | Nampt | 59027 | HITS-CLIP | 23142080 | Unspecified |
| mmu-miR-24-3p | MIMAT0000219 | Nat10 | 98956 | HITS-CLIP | 25753418 | Unspecified |
| mmu-miR-24-3p | MIMAT0000219 | Naxe | 128240 | HITS-CLIP | 25753418 | Unspecified |
| mmu-miR-24-3p | MIMAT0000219 | Nbr1 | 17966 | HITS-CLIP | 23142080\|26203562 | Unspecified |
| mmu-miR-24-3p | MIMAT0000219 | Ncapd2 | 68298 | HITS-CLIP | 25753418\|23142080 | Unspecified |
| mmu-miR-24-3p | MIMAT0000219 | Ncapd3 | 78658 | HITS-CLIP | 23142080 | Unspecified |
| mmu-miR-24-3p | MIMAT0000219 | Ncapg | 54392 | HITS-CLIP | 23142080 | Unspecified |
| mmu-miR-24-3p | MIMAT0000219 | Ncapg2 | 76044 | HITS-CLIP | 23142080 | Unspecified |
| mmu-miR-24-3p | MIMAT0000219 | Ncaph | 215387 | HITS-CLIP | 23142080 | Unspecified |
| mmu-miR-24-3p | MIMAT0000219 | Ncbp3 | 66874 | HITS-CLIP | 25753418 | Unspecified |
| mmu-miR-24-3p | MIMAT0000219 | Nck1 | 17973 | HITS-CLIP | 23142080 | Unspecified |
| mmu-miR-24-3p | MIMAT0000219 | Nckap1 | 50884 | HITS-CLIP | 25753418 | Unspecified |
| mmu-miR-24-3p | MIMAT0000219 | Ncl | 17975 | HITS-CLIP | 21258322\|23142080 | Unspecified |
| mmu-miR-24-3p | MIMAT0000219 | Ncln | 103425 | HITS-CLIP | 37986224 | Unspecified |
| mmu-miR-24-3p | MIMAT0000219 | Ncoa1 | 17977 | HITS-CLIP | 25753418 | Unspecified |
| mmu-miR-24-3p | MIMAT0000219 | Ncoa2 | 17978 | HITS-CLIP | 37986224 | Unspecified |
| mmu-miR-24-3p | MIMAT0000219 | Ncoa3 | 17979 | HITS-CLIP | 23142080 | Unspecified |
| mmu-miR-24-3p | MIMAT0000219 | Ncor2 | 20602 | HITS-CLIP | 23142080 | Unspecified |
| mmu-miR-24-3p | MIMAT0000219 | Ncstn | 59287 | HITS-CLIP | 23142080 | Unspecified |
| mmu-miR-24-3p | MIMAT0000219 | Nde1 | 67203 | HITS-CLIP | 31433990 | Unspecified |
| mmu-miR-24-3p | MIMAT0000219 | Ndfip1 | 65113 | Sequencing//HITS-CLIP | 37986224\|23142080\|25753418\|19536 | Unspecified |
| mmu-miR-24-3p | MIMAT0000219 | Ndrg2 | 29811 | HITS-CLIP | 23142080 | Unspecified |
| mmu-miR-24-3p | MIMAT0000219 | Ndrg3 | 29812 | Sequencing//HITS-CLIP | 37986224\|23142080\|19536157 | Unspecified |
| mmu-miR-24-3p | MIMAT0000219 | Ndst1 | 15531 | HITS-CLIP | 23142080 | Unspecified |
| mmu-miR-24-3p | MIMAT0000219 | Ndst2 | 17423 | HITS-CLIP | 23142080 | Unspecified |
| mmu-miR-24-3p | MIMAT0000219 | Ndufa13 | 67184 | HITS-CLIP | 37986224 | Unspecified |
| mmu-miR-24-3p | MIMAT0000219 | Ndufb2 | 68198 | HITS-CLIP | 25753418 | Unspecified |
| mmu-miR-24-3p | MIMAT0000219 | Ndufb7 | 66916 | HITS-CLIP | 25753418 | Unspecified |

| | | | | | | |
|---|---|---|---|---|---|---|
| mmu-miR-24-3p | MIMAT0000219 | Ndufb8 | 67264 | HITS-CLIP | 23142080 | Unspecified |
| mmu-miR-24-3p | MIMAT0000219 | Ndufc2 | 68197 | HITS-CLIP | 37986224\|23142080 | Unspecified |
| mmu-miR-24-3p | MIMAT0000219 | Ndufs4 | 17993 | HITS-CLIP | 23142080 | Unspecified |
| mmu-miR-24-3p | MIMAT0000219 | Ndufs7 | 75406 | HITS-CLIP | 25753418 | Unspecified |
| mmu-miR-24-3p | MIMAT0000219 | Ndufs8 | 225887 | HITS-CLIP | 23142080 | Unspecified |
| mmu-miR-24-3p | MIMAT0000219 | Ndufv3 | 78330 | HITS-CLIP | 23142080 | Unspecified |
| mmu-miR-24-3p | MIMAT0000219 | Necap2 | 66147 | HITS-CLIP | 23142080 | Unspecified |
| mmu-miR-24-3p | MIMAT0000219 | Nectin2 | 5819 | HITS-CLIP | 25753418 | Unspecified |
| mmu-miR-24-3p | MIMAT0000219 | Nefh | 380684 | HITS-CLIP | 31433990 | Unspecified |
| mmu-miR-24-3p | MIMAT0000219 | Nek4 | 23955 | HITS-CLIP | 23142080 | Unspecified |
| mmu-miR-24-3p | MIMAT0000219 | Nek9 | 217718 | HITS-CLIP | 23142080 | Unspecified |
| mmu-miR-24-3p | MIMAT0000219 | Nelfa | 24116 | HITS-CLIP | 23142080 | Unspecified |
| mmu-miR-24-3p | MIMAT0000219 | Nelfb | 58202 | HITS-CLIP | 23142080 | Unspecified |
| mmu-miR-24-3p | MIMAT0000219 | Nemp1 | 23306 | HITS-CLIP | 23142080 | Unspecified |
| mmu-miR-24-3p | MIMAT0000219 | Nemp2 | 227094 | HITS-CLIP | 23142080 | Unspecified |
| mmu-miR-24-3p | MIMAT0000219 | Neu1 | 18010 | HITS-CLIP | 23142080 | Unspecified |
| mmu-miR-24-3p | MIMAT0000219 | Neu3 | 50877 | HITS-CLIP | 21258322\|23142080 | Unspecified |
| mmu-miR-24-3p | MIMAT0000219 | Neurl3 | 214854 | HITS-CLIP | 23142080 | Unspecified |
| mmu-miR-24-3p | MIMAT0000219 | Nf2 | 18016 | HITS-CLIP | 23142080 | Unspecified |
| mmu-miR-24-3p | MIMAT0000219 | Nfat5 | 54446 | Sequencing//HITS-CLIP | 19536157\|23142080\|37986224 | Unspecified |
| mmu-miR-24-3p | MIMAT0000219 | Nfatc1 | 18018 | HITS-CLIP | 23142080 | Unspecified |
| mmu-miR-24-3p | MIMAT0000219 | Nfatc3 | 18021 | HITS-CLIP | 23142080 | Unspecified |
| mmu-miR-24-3p | MIMAT0000219 | Nfic | 18029 | HITS-CLIP | 25753418 | Unspecified |
| mmu-miR-24-3p | MIMAT0000219 | Nfil3 | 18030 | HITS-CLIP | 23142080 | Unspecified |
| mmu-miR-24-3p | MIMAT0000219 | Nfkb1 | 18033 | HITS-CLIP | 23142080 | Unspecified |
| mmu-miR-24-3p | MIMAT0000219 | Nfrkb | 235134 | HITS-CLIP | 23142080 | Unspecified |
| mmu-miR-24-3p | MIMAT0000219 | Nfs1 | 18041 | HITS-CLIP | 23142080 | Unspecified |
| mmu-miR-24-3p | MIMAT0000219 | Nfx1 | 74164 | HITS-CLIP | 25753418\|23142080 | Unspecified |
| mmu-miR-24-3p | MIMAT0000219 | Nfxl1 | 100978 | HITS-CLIP | 23142080 | Unspecified |
| mmu-miR-24-3p | MIMAT0000219 | Nfya | 18044 | HITS-CLIP | 23142080 | Unspecified |
| mmu-miR-24-3p | MIMAT0000219 | Ngrn | 83485 | HITS-CLIP | 23142080 | Unspecified |
| mmu-miR-24-3p | MIMAT0000219 | Nhsl1 | 215819 | qCLASH | 31393953 | Unspecified |
| mmu-miR-24-3p | MIMAT0000219 | Nifk | 67949 | HITS-CLIP | 23142080 | Unspecified |
| mmu-miR-24-3p | MIMAT0000219 | Nisch | 64652 | HITS-CLIP | 23142080 | Unspecified |
| mmu-miR-24-3p | MIMAT0000219 | Nit2 | 52633 | HITS-CLIP | 23142080 | Unspecified |
| mmu-miR-24-3p | MIMAT0000219 | Nkapd1 | 55216 | HITS-CLIP | 23142080 | Unspecified |
| mmu-miR-24-3p | MIMAT0000219 | Nkrf | 77286 | HITS-CLIP | 23142080 | Unspecified |
| mmu-miR-24-3p | MIMAT0000219 | Nktr | 18087 | HITS-CLIP | 23142080 | Unspecified |
| mmu-miR-24-3p | MIMAT0000219 | Nlk | 18099 | HITS-CLIP | 23142080 | Unspecified |

| | | | | | | |
|---|---|---|---|---|---|---|
| mmu-miR-24-3p | MIMAT0000219 | Nmt1 | 18107 | HITS-CLIP | 23142080 | Unspecified |
| mmu-miR-24-3p | MIMAT0000219 | Noc4l | 100608 | HITS-CLIP | 23142080 | Unspecified |
| mmu-miR-24-3p | MIMAT0000219 | Noct | 12457 | HITS-CLIP | 23142080 | Unspecified |
| mmu-miR-24-3p | MIMAT0000219 | Nod1 | 107607 | HITS-CLIP | 23142080 | Unspecified |
| mmu-miR-24-3p | MIMAT0000219 | Nol6 | 230082 | HITS-CLIP | 25753418 | Unspecified |
| mmu-miR-24-3p | MIMAT0000219 | Nolc1 | 70769 | HITS-CLIP | 25753418 | Unspecified |
| mmu-miR-24-3p | MIMAT0000219 | Nom1 | 433864 | HITS-CLIP | 23142080 | Unspecified |
| mmu-miR-24-3p | MIMAT0000219 | Nono | 53610 | HITS-CLIP | 23142080 | Unspecified |
| mmu-miR-24-3p | MIMAT0000219 | Nop14 | 75416 | HITS-CLIP | 23142080 | Unspecified |
| mmu-miR-24-3p | MIMAT0000219 | Nop58 | 55989 | HITS-CLIP | 37986224 | Unspecified |
| mmu-miR-24-3p | MIMAT0000219 | Nosip | 66394 | HITS-CLIP | 23142080\|25753418 | Unspecified |
| mmu-miR-24-3p | MIMAT0000219 | Notch1 | 18128 | HITS-CLIP | 37986224\|23142080 | Unspecified |
| mmu-miR-24-3p | MIMAT0000219 | Npat | 244879 | HITS-CLIP | 37986224\|23142080 | Unspecified |
| mmu-miR-24-3p | MIMAT0000219 | Npc2 | 67963 | HITS-CLIP | 37986224\|23142080 | Unspecified |
| mmu-miR-24-3p | MIMAT0000219 | Npepps | 19155 | HITS-CLIP | 37986224\|23142080\|25753418 | Unspecified |
| mmu-miR-24-3p | MIMAT0000219 | Nphp3 | 74025 | HITS-CLIP | 23142080 | Unspecified |
| mmu-miR-24-3p | MIMAT0000219 | Nploc4 | 217365 | HITS-CLIP | 25753418\|23142080 | Unspecified |
| mmu-miR-24-3p | MIMAT0000219 | Nr1d1 | 217166 | HITS-CLIP | 25753418 | Unspecified |
| mmu-miR-24-3p | MIMAT0000219 | Nr1h2 | 22260 | HITS-CLIP | 25753418\|23142080 | Unspecified |
| mmu-miR-24-3p | MIMAT0000219 | Nr4a1 | 15370 | HITS-CLIP | 23142080\|25753418 | Unspecified |
| mmu-miR-24-3p | MIMAT0000219 | Nr4a2 | 18227 | HITS-CLIP | 23142080 | Unspecified |
| mmu-miR-24-3p | MIMAT0000219 | Nrap | 18175 | HITS-CLIP | 25753418 | Unspecified |
| mmu-miR-24-3p | MIMAT0000219 | Nrarp | 67122 | HITS-CLIP | 25753418 | Unspecified |
| mmu-miR-24-3p | MIMAT0000219 | Nras | 18176 | HITS-CLIP | 23142080 | Unspecified |
| mmu-miR-24-3p | MIMAT0000219 | Nrbp1 | 192292 | HITS-CLIP | 23142080 | Unspecified |
| mmu-miR-24-3p | MIMAT0000219 | Nrd1 | 230598 | HITS-CLIP | 25753418\|23142080 | Unspecified |
| mmu-miR-24-3p | MIMAT0000219 | Nrf1 | 18181 | HITS-CLIP | 25753418\|23142080 | Unspecified |
| mmu-miR-24-3p | MIMAT0000219 | Nrip1 | 268903 | HITS-CLIP | 23142080 | Unspecified |
| mmu-miR-24-3p | MIMAT0000219 | Nrn1 | 68404 | HITS-CLIP | 23142080 | Unspecified |
| mmu-miR-24-3p | MIMAT0000219 | Nrros | 224109 | HITS-CLIP | 23142080 | Unspecified |
| mmu-miR-24-3p | MIMAT0000219 | Nsd1 | 18193 | Sequencing//HITS-CLIP | 19536157\|23142080\|25753418\|37986 | Unspecified |
| mmu-miR-24-3p | MIMAT0000219 | Nsd3 | 234135 | HITS-CLIP | 23142080\|31433990\|25753418 | Unspecified |
| mmu-miR-24-3p | MIMAT0000219 | Nsmce4a | 67872 | HITS-CLIP | 23142080 | Unspecified |
| mmu-miR-24-3p | MIMAT0000219 | Nsun2 | 28114 | HITS-CLIP | 23142080 | Unspecified |
| mmu-miR-24-3p | MIMAT0000219 | Nsun6 | 74455 | HITS-CLIP | 23142080 | Unspecified |
| mmu-miR-24-3p | MIMAT0000219 | Nt5c2 | 76952 | HITS-CLIP | 23142080 | Unspecified |
| mmu-miR-24-3p | MIMAT0000219 | Nt5dc1 | 319638 | HITS-CLIP | 23142080 | Unspecified |
| mmu-miR-24-3p | MIMAT0000219 | Nucb1 | 18220 | HITS-CLIP | 23142080 | Unspecified |
| mmu-miR-24-3p | MIMAT0000219 | Nucb2 | 53322 | HITS-CLIP | 23142080 | Unspecified |

| | | | | | | |
|---|---|---|---|---|---|---|
| mmu-miR-24-3p | MIMAT0000219 | Nucks1 | 98415 | Sequencing//HITS-CLIP | 19536157\|23142080\|37986224 | Unspecified |
| mmu-miR-24-3p | MIMAT0000219 | Nudt16l1 | 66911 | HITS-CLIP | 25753418 | Unspecified |
| mmu-miR-24-3p | MIMAT0000219 | Nudt22 | 68323 | HITS-CLIP | 23142080 | Unspecified |
| mmu-miR-24-3p | MIMAT0000219 | Nuf2 | 66977 | HITS-CLIP | 23142080 | Unspecified |
| mmu-miR-24-3p | MIMAT0000219 | Nufip2 | 68564 | HITS-CLIP | 37986224\|23142080 | Unspecified |
| mmu-miR-24-3p | MIMAT0000219 | Numa1 | 101706 | HITS-CLIP | 25753418 | Unspecified |
| mmu-miR-24-3p | MIMAT0000219 | Nup107 | 103468 | HITS-CLIP | 23142080 | Unspecified |
| mmu-miR-24-3p | MIMAT0000219 | Nup155 | 170762 | HITS-CLIP | 23142080 | Unspecified |
| mmu-miR-24-3p | MIMAT0000219 | Nup205 | 70699 | HITS-CLIP | 23142080 | Unspecified |
| mmu-miR-24-3p | MIMAT0000219 | Nup210 | 54563 | Sequencing//HITS-CLIP | 19536157\|23142080\|37986224 | Unspecified |
| mmu-miR-24-3p | MIMAT0000219 | Nup50 | 18141 | HITS-CLIP | 26203562 | Unspecified |
| mmu-miR-24-3p | MIMAT0000219 | Nup54 | 269113 | qCLASH | 31393953 | Unspecified |
| mmu-miR-24-3p | MIMAT0000219 | Nup62 | 18226 | HITS-CLIP | 23142080 | Unspecified |
| mmu-miR-24-3p | MIMAT0000219 | Nup88 | 19069 | HITS-CLIP | 23142080 | Unspecified |
| mmu-miR-24-3p | MIMAT0000219 | Nup93 | 71805 | HITS-CLIP | 23142080 | Unspecified |
| mmu-miR-24-3p | MIMAT0000219 | Nup98 | 269966 | HITS-CLIP | 23142080 | Unspecified |
| mmu-miR-24-3p | MIMAT0000219 | Nusap1 | 108907 | HITS-CLIP | 23142080 | Unspecified |
| mmu-miR-24-3p | MIMAT0000219 | Nxn | 18230 | HITS-CLIP | 23142080 | Unspecified |
| mmu-miR-24-3p | MIMAT0000219 | Oas3 | 246727 | HITS-CLIP | 23142080 | Unspecified |
| mmu-miR-24-3p | MIMAT0000219 | Obscn | 380698 | HITS-CLIP | 25753418 | Unspecified |
| mmu-miR-24-3p | MIMAT0000219 | Obsl1 | 98733 | HITS-CLIP | 25753418 | Unspecified |
| mmu-miR-24-3p | MIMAT0000219 | Oga | 10724 | HITS-CLIP | 25753418 | Unspecified |
| mmu-miR-24-3p | MIMAT0000219 | Ogdh | 18293 | HITS-CLIP | 23142080 | Unspecified |
| mmu-miR-24-3p | MIMAT0000219 | Ogdhl | 239017 | HITS-CLIP | 25753418 | Unspecified |
| mmu-miR-24-3p | MIMAT0000219 | Ogfod2 | 66627 | HITS-CLIP | 23142080 | Unspecified |
| mmu-miR-24-3p | MIMAT0000219 | Ogfr | 72075 | HITS-CLIP | 37986224\|25753418 | Unspecified |
| mmu-miR-24-3p | MIMAT0000219 | Ogt | 108155 | Sequencing//HITS-CLIP | 19536157\|23142080\|37986224 | Unspecified |
| mmu-miR-24-3p | MIMAT0000219 | Opa1 | 74143 | HITS-CLIP | 23142080 | Unspecified |
| mmu-miR-24-3p | MIMAT0000219 | Opa3 | 403187 | HITS-CLIP | 25753418 | Unspecified |
| mmu-miR-24-3p | MIMAT0000219 | Oplah | 75475 | HITS-CLIP | 25753418 | Unspecified |
| mmu-miR-24-3p | MIMAT0000219 | Orai2 | 269717 | HITS-CLIP | 23142080 | Unspecified |
| mmu-miR-24-3p | MIMAT0000219 | Osbp | 76303 | HITS-CLIP | 23142080 | Unspecified |
| mmu-miR-24-3p | MIMAT0000219 | Osbpl8 | 237542 | HITS-CLIP | 23142080 | Unspecified |
| mmu-miR-24-3p | MIMAT0000219 | Osbpl9 | 100273 | HITS-CLIP | 23142080 | Unspecified |
| mmu-miR-24-3p | MIMAT0000219 | Otub1 | 107260 | HITS-CLIP | 25753418 | Unspecified |
| mmu-miR-24-3p | MIMAT0000219 | Otud4 | 73945 | HITS-CLIP | 23142080 | Unspecified |
| mmu-miR-24-3p | MIMAT0000219 | Otud5 | 54644 | HITS-CLIP | 23142080 | Unspecified |
| mmu-miR-24-3p | MIMAT0000219 | Otud7b | 229603 | HITS-CLIP | 23142080 | Unspecified |
| mmu-miR-24-3p | MIMAT0000219 | Otx2 | 18424 | Luciferase Reporter Assay | 37986224 | Unspecified |

| | | | | | | |
|---|---|---|---|---|---|---|
| mmu-miR-24-3p | MIMAT0000219 | Oxa1l | 69089 | HITS-CLIP | 25753418 | Unspecified |
| mmu-miR-24-3p | MIMAT0000219 | Oxsm | 71147 | HITS-CLIP | 25753418 | Unspecified |
| mmu-miR-24-3p | MIMAT0000219 | Oxsr1 | 108737 | HITS-CLIP | 23142080 | Unspecified |
| mmu-miR-24-3p | MIMAT0000219 | P2rx4 | 18438 | HITS-CLIP | 25753418 | Unspecified |
| mmu-miR-24-3p | MIMAT0000219 | P2ry1 | 18441 | HITS-CLIP | 23142080 | Unspecified |
| mmu-miR-24-3p | MIMAT0000219 | P4ha1 | 18451 | HITS-CLIP | 23142080 | Unspecified |
| mmu-miR-24-3p | MIMAT0000219 | Pa2g4 | 18813 | HITS-CLIP | 23142080\|25753418 | Unspecified |
| mmu-miR-24-3p | MIMAT0000219 | Pabpc1 | 18458 | HITS-CLIP | 23142080 | Unspecified |
| mmu-miR-24-3p | MIMAT0000219 | Pacsin2 | 23970 | HITS-CLIP | 25753418 | Unspecified |
| mmu-miR-24-3p | MIMAT0000219 | Paf1 | 54624 | HITS-CLIP | 23142080 | Unspecified |
| mmu-miR-24-3p | MIMAT0000219 | Pafah1b1 | 18472 | HITS-CLIP | 23142080 | Unspecified |
| mmu-miR-24-3p | MIMAT0000219 | Pafah1b3 | 18476 | HITS-CLIP | 23142080 | Unspecified |
| mmu-miR-24-3p | MIMAT0000219 | Pagr1a | 67278 | HITS-CLIP | 23142080 | Unspecified |
| mmu-miR-24-3p | MIMAT0000219 | Paics | 67054 | HITS-CLIP | 23142080 | Unspecified |
| mmu-miR-24-3p | MIMAT0000219 | Pak2 | 224105 | HITS-CLIP | 23142080 | Unspecified |
| mmu-miR-24-3p | MIMAT0000219 | Pak4 | 70584 | Western Blot | 37986224\|23071155 | Unspecified |
| mmu-miR-24-3p | MIMAT0000219 | Pakap | 677884 | HITS-CLIP | 25753418 | Unspecified |
| mmu-miR-24-3p | MIMAT0000219 | Palld | 72333 | HITS-CLIP | 25753418 | Unspecified |
| mmu-miR-24-3p | MIMAT0000219 | Pan2 | 103135 | HITS-CLIP | 23142080 | Unspecified |
| mmu-miR-24-3p | MIMAT0000219 | Pank1 | 75735 | HITS-CLIP | 23142080 | Unspecified |
| mmu-miR-24-3p | MIMAT0000219 | Pank3 | 211347 | HITS-CLIP | 23142080 | Unspecified |
| mmu-miR-24-3p | MIMAT0000219 | Panx1 | 55991 | HITS-CLIP | 23142080 | Unspecified |
| mmu-miR-24-3p | MIMAT0000219 | Papss1 | 23971 | HITS-CLIP | 23142080 | Unspecified |
| mmu-miR-24-3p | MIMAT0000219 | Parp14 | 547253 | HITS-CLIP | 25753418 | Unspecified |
| mmu-miR-24-3p | MIMAT0000219 | Parvb | 170736 | HITS-CLIP | 25753418 | Unspecified |
| mmu-miR-24-3p | MIMAT0000219 | Paxip1 | 55982 | HITS-CLIP | 23142080 | Unspecified |
| mmu-miR-24-3p | MIMAT0000219 | Pbk | 52033 | HITS-CLIP | 23142080 | Unspecified |
| mmu-miR-24-3p | MIMAT0000219 | Pbrm1 | 66923 | HITS-CLIP | 37986224\|25753418\|23142080 | Unspecified |
| mmu-miR-24-3p | MIMAT0000219 | Pcbp1 | 23983 | qCLASH//HITS-CLIP | 31393953\|23142080 | Unspecified |
| mmu-miR-24-3p | MIMAT0000219 | Pcca | 110821 | HITS-CLIP | 25753418 | Unspecified |
| mmu-miR-24-3p | MIMAT0000219 | Pcdh7 | 54216 | HITS-CLIP | 25753418 | Unspecified |
| mmu-miR-24-3p | MIMAT0000219 | Pcgf3 | 69587 | HITS-CLIP | 25753418 | Unspecified |
| mmu-miR-24-3p | MIMAT0000219 | Pcgf5 | 76073 | HITS-CLIP | 23142080 | Unspecified |
| mmu-miR-24-3p | MIMAT0000219 | Pcgf6 | 71041 | HITS-CLIP | 25753418 | Unspecified |
| mmu-miR-24-3p | MIMAT0000219 | Pcid2 | 234069 | HITS-CLIP | 23142080 | Unspecified |
| mmu-miR-24-3p | MIMAT0000219 | Pcm1 | 18536 | HITS-CLIP | 23142080 | Unspecified |
| mmu-miR-24-3p | MIMAT0000219 | Pcnt | 18541 | HITS-CLIP | 26203562 | Unspecified |
| mmu-miR-24-3p | MIMAT0000219 | Pcnx4 | 67708 | HITS-CLIP | 23142080 | Unspecified |
| mmu-miR-24-3p | MIMAT0000219 | Pcsk6 | 18553 | HITS-CLIP | 25753418 | Unspecified |

| | | | | | | |
|---|---|---|---|---|---|---|
| mmu-miR-24-3p | MIMAT0000219 | Pdcd1 | 18566 | HITS-CLIP | 23142080 | Unspecified |
| mmu-miR-24-3p | MIMAT0000219 | Pdcd6ip | 18571 | HITS-CLIP | 23142080 | Unspecified |
| mmu-miR-24-3p | MIMAT0000219 | Pde11a | 241489 | HITS-CLIP | 23142080 | Unspecified |
| mmu-miR-24-3p | MIMAT0000219 | Pde2a | 207728 | HITS-CLIP | 23142080 | Unspecified |
| mmu-miR-24-3p | MIMAT0000219 | Pde3b | 18576 | HITS-CLIP | 23142080 | Unspecified |
| mmu-miR-24-3p | MIMAT0000219 | Pde4a | 18577 | HITS-CLIP | 25753418 | Unspecified |
| mmu-miR-24-3p | MIMAT0000219 | Pde4d | 238871 | HITS-CLIP | 23142080 | Unspecified |
| mmu-miR-24-3p | MIMAT0000219 | Pde4dip | 83679 | HITS-CLIP | 23142080 | Unspecified |
| mmu-miR-24-3p | MIMAT0000219 | Pdgfd | 71785 | HITS-CLIP | 25753418 | Unspecified |
| mmu-miR-24-3p | MIMAT0000219 | Pdha1 | 18597 | HITS-CLIP | 23142080 | Unspecified |
| mmu-miR-24-3p | MIMAT0000219 | Pdhx | 27402 | HITS-CLIP | 25753418 | Unspecified |
| mmu-miR-24-3p | MIMAT0000219 | Pdia6 | 71853 | HITS-CLIP | 37986224 | Unspecified |
| mmu-miR-24-3p | MIMAT0000219 | Pdk2 | 18604 | HITS-CLIP | 25753418 | Unspecified |
| mmu-miR-24-3p | MIMAT0000219 | Pdk3 | 236900 | HITS-CLIP | 23142080 | Unspecified |
| mmu-miR-24-3p | MIMAT0000219 | Pdlim1 | 54132 | HITS-CLIP | 25753418 | Unspecified |
| mmu-miR-24-3p | MIMAT0000219 | Pdp1 | 381511 | HITS-CLIP | 25753418 | Unspecified |
| mmu-miR-24-3p | MIMAT0000219 | Pdp2 | 382051 | HITS-CLIP | 25753418 | Unspecified |
| mmu-miR-24-3p | MIMAT0000219 | Pdpk1 | 18607 | HITS-CLIP | 23142080 | Unspecified |
| mmu-miR-24-3p | MIMAT0000219 | Pdpr | 319518 | HITS-CLIP | 25753418 | Unspecified |
| mmu-miR-24-3p | MIMAT0000219 | Pdxdc1 | 94184 | HITS-CLIP | 23142080 | Unspecified |
| mmu-miR-24-3p | MIMAT0000219 | Pdzd8 | 107368 | HITS-CLIP | 37986224\|23142080 | Unspecified |
| mmu-miR-24-3p | MIMAT0000219 | Pea15a | 18611 | HITS-CLIP | 37986224\|23142080 | Unspecified |
| mmu-miR-24-3p | MIMAT0000219 | Peak1 | 244895 | HITS-CLIP | 37986224\|23142080 | Unspecified |
| mmu-miR-24-3p | MIMAT0000219 | Pecam1 | 18613 | HITS-CLIP | 25753418 | Unspecified |
| mmu-miR-24-3p | MIMAT0000219 | Pepd | 18624 | HITS-CLIP | 25753418 | Unspecified |
| mmu-miR-24-3p | MIMAT0000219 | Per1 | 18626 | HITS-CLIP | 23142080\|25753418 | Unspecified |
| mmu-miR-24-3p | MIMAT0000219 | Per3 | 18628 | HITS-CLIP | 25753418 | Unspecified |
| mmu-miR-24-3p | MIMAT0000219 | Pex19 | 19298 | HITS-CLIP | 37986224 | Unspecified |
| mmu-miR-24-3p | MIMAT0000219 | Pfdn5 | 56612 | HITS-CLIP | 25753418 | Unspecified |
| mmu-miR-24-3p | MIMAT0000219 | Pfdn6 | 14976 | HITS-CLIP | 37986224 | Unspecified |
| mmu-miR-24-3p | MIMAT0000219 | Pfkl | 18641 | qCLASH | 31393953 | Unspecified |
| mmu-miR-24-3p | MIMAT0000219 | Pfkp | 56421 | HITS-CLIP | 37986224\|23142080 | Unspecified |
| mmu-miR-24-3p | MIMAT0000219 | Pfn1 | 18643 | HITS-CLIP | 23142080 | Unspecified |
| mmu-miR-24-3p | MIMAT0000219 | Pgap2 | 233575 | HITS-CLIP | 23142080 | Unspecified |
| mmu-miR-24-3p | MIMAT0000219 | Pgap4 | 84302 | HITS-CLIP | 25753418 | Unspecified |
| mmu-miR-24-3p | MIMAT0000219 | Pgrmc2 | 70804 | Sequencing//HITS-CLIP | 19536157\|23142080\|37986224 | Unspecified |
| mmu-miR-24-3p | MIMAT0000219 | Phactr2 | 215789 | HITS-CLIP | 25753418 | Unspecified |
| mmu-miR-24-3p | MIMAT0000219 | Phf2 | 18676 | HITS-CLIP | 25753418 | Unspecified |
| mmu-miR-24-3p | MIMAT0000219 | Phf20 | 228829 | HITS-CLIP | 23142080 | Unspecified |

| | | | | | | | |
|---|---|---|---|---|---|---|---|
| mmu-miR-24-3p | MIMAT0000219 | Phf20l1 | 239510 | HITS-CLIP | | 23142080 | Unspecified |
| mmu-miR-24-3p | MIMAT0000219 | Phf21a | 192285 | HITS-CLIP | | 23142080 | Unspecified |
| mmu-miR-24-3p | MIMAT0000219 | Phf3 | 213109 | HITS-CLIP | 37986224\|23142080 | | Unspecified |
| mmu-miR-24-3p | MIMAT0000219 | Phf6 | 70998 | HITS-CLIP | | 23142080 | Unspecified |
| mmu-miR-24-3p | MIMAT0000219 | Phf8 | 320595 | HITS-CLIP | | 23142080 | Unspecified |
| mmu-miR-24-3p | MIMAT0000219 | Phip | 83946 | HITS-CLIP | | 23142080 | Unspecified |
| mmu-miR-24-3p | MIMAT0000219 | Phlpp2 | 244650 | HITS-CLIP | 37986224\|23142080 | | Unspecified |
| mmu-miR-24-3p | MIMAT0000219 | Pi4ka | 224020 | HITS-CLIP | 23142080\|25753418 | | Unspecified |
| mmu-miR-24-3p | MIMAT0000219 | Pi4kb | 107650 | HITS-CLIP | | 23142080 | Unspecified |
| mmu-miR-24-3p | MIMAT0000219 | Pias2 | 17344 | HITS-CLIP | | 23142080 | Unspecified |
| mmu-miR-24-3p | MIMAT0000219 | Pias4 | 59004 | HITS-CLIP | | 23142080 | Unspecified |
| mmu-miR-24-3p | MIMAT0000219 | Piga | 18700 | HITS-CLIP | | 23142080 | Unspecified |
| mmu-miR-24-3p | MIMAT0000219 | Pigg | 433931 | HITS-CLIP | | 23142080 | Unspecified |
| mmu-miR-24-3p | MIMAT0000219 | Pigm | 67556 | HITS-CLIP | | 23142080 | Unspecified |
| mmu-miR-24-3p | MIMAT0000219 | Pigo | 56703 | HITS-CLIP | | 25753418 | Unspecified |
| mmu-miR-24-3p | MIMAT0000219 | Pik3cd | 18707 | HITS-CLIP | | 23142080 | Unspecified |
| mmu-miR-24-3p | MIMAT0000219 | Pik3ip1 | 216505 | HITS-CLIP | | 23142080 | Unspecified |
| mmu-miR-24-3p | MIMAT0000219 | Pik3r1 | 18708 | HITS-CLIP | | 25753418 | Unspecified |
| mmu-miR-24-3p | MIMAT0000219 | Pik3r5 | 320207 | HITS-CLIP | | 23142080 | Unspecified |
| mmu-miR-24-3p | MIMAT0000219 | Pim1 | 18712 | HITS-CLIP | | 23142080 | Unspecified |
| mmu-miR-24-3p | MIMAT0000219 | Pim2 | 18715 | HITS-CLIP | | 23142080 | Unspecified |
| mmu-miR-24-3p | MIMAT0000219 | Pim3 | 223775 | HITS-CLIP | | 23142080 | Unspecified |
| mmu-miR-24-3p | MIMAT0000219 | Pip5k1a | 18720 | HITS-CLIP | | 23142080 | Unspecified |
| mmu-miR-24-3p | MIMAT0000219 | Pip5k1b | 18719 | HITS-CLIP | | 23142080 | Unspecified |
| mmu-miR-24-3p | MIMAT0000219 | Pisd | 320951 | qCLASH | | 31393953 | Unspecified |
| mmu-miR-24-3p | MIMAT0000219 | Pitpna | 18738 | HITS-CLIP | | 25753418 | Unspecified |
| mmu-miR-24-3p | MIMAT0000219 | Pitpnc1 | 71795 | HITS-CLIP | | 23142080 | Unspecified |
| mmu-miR-24-3p | MIMAT0000219 | Pitpnm1 | 18739 | HITS-CLIP | | 23142080 | Unspecified |
| mmu-miR-24-3p | MIMAT0000219 | Pitpnm2 | 19679 | HITS-CLIP | | 23142080 | Unspecified |
| mmu-miR-24-3p | MIMAT0000219 | Pitrm1 | 69617 | HITS-CLIP | | 37986224 | Unspecified |
| mmu-miR-24-3p | MIMAT0000219 | Piwil2 | 57746 | HITS-CLIP | | 23142080 | Unspecified |
| mmu-miR-24-3p | MIMAT0000219 | Pkdcc | 106522 | HITS-CLIP | | 25753418 | Unspecified |
| mmu-miR-24-3p | MIMAT0000219 | Pkm | 18746 | HITS-CLIP | | 23142080 | Unspecified |
| mmu-miR-24-3p | MIMAT0000219 | Pkmyt1 | 268930 | HITS-CLIP | | 23142080 | Unspecified |
| mmu-miR-24-3p | MIMAT0000219 | Pkn1 | 320795 | HITS-CLIP | | 23142080 | Unspecified |
| mmu-miR-24-3p | MIMAT0000219 | Pkp1 | 18772 | HITS-CLIP | | 26203562 | Unspecified |
| mmu-miR-24-3p | MIMAT0000219 | Pla2g16 | 225845 | HITS-CLIP | | 23142080 | Unspecified |
| mmu-miR-24-3p | MIMAT0000219 | Plagl2 | 54711 | HITS-CLIP | | 23142080 | Unspecified |
| mmu-miR-24-3p | MIMAT0000219 | Plcb3 | 18797 | HITS-CLIP | | 23142080 | Unspecified |

| | | | | | | | |
|---|---|---|---|---|---|---|---|
| mmu-miR-24-3p | MIMAT0000219 | Plcl2 | 224860 | HITS-CLIP | | 23142080 | Unspecified |
| mmu-miR-24-3p | MIMAT0000219 | Plcxd2 | 433022 | HITS-CLIP | 37986224\|23142080 | | Unspecified |
| mmu-miR-24-3p | MIMAT0000219 | Pld2 | 18806 | HITS-CLIP | | 25753418 | Unspecified |
| mmu-miR-24-3p | MIMAT0000219 | Plec | 18810 | Sequencing//HITS-CLIP | 19536157\|23142080\|25753418 | | Unspecified |
| mmu-miR-24-3p | MIMAT0000219 | Plekha2 | 83436 | HITS-CLIP | | 23142080 | Unspecified |
| mmu-miR-24-3p | MIMAT0000219 | Plekhh1 | 211945 | HITS-CLIP | | 25753418 | Unspecified |
| mmu-miR-24-3p | MIMAT0000219 | Plekhm1 | 353047 | HITS-CLIP | 37986224\|23142080 | | Unspecified |
| mmu-miR-24-3p | MIMAT0000219 | Plekho2 | 102595 | HITS-CLIP | | 23142080 | Unspecified |
| mmu-miR-24-3p | MIMAT0000219 | Plin3 | 66905 | HITS-CLIP | | 23142080 | Unspecified |
| mmu-miR-24-3p | MIMAT0000219 | Plk1 | 18817 | HITS-CLIP | | 23142080 | Unspecified |
| mmu-miR-24-3p | MIMAT0000219 | Plk3 | 12795 | HITS-CLIP | 37986224\|23142080 | | Unspecified |
| mmu-miR-24-3p | MIMAT0000219 | Plk4 | 20873 | HITS-CLIP | | 23142080 | Unspecified |
| mmu-miR-24-3p | MIMAT0000219 | Plod2 | 26432 | Sequencing//HITS-CLIP | 19536157\|23142080\|37986224 | | Unspecified |
| mmu-miR-24-3p | MIMAT0000219 | Plrg1 | 53317 | HITS-CLIP | | 23142080 | Unspecified |
| mmu-miR-24-3p | MIMAT0000219 | Plxna1 | 18844 | HITS-CLIP | | 25753418 | Unspecified |
| mmu-miR-24-3p | MIMAT0000219 | Pm20d2 | 242377 | HITS-CLIP | | 25753418 | Unspecified |
| mmu-miR-24-3p | MIMAT0000219 | Pmepa1 | 65112 | HITS-CLIP | | 23142080 | Unspecified |
| mmu-miR-24-3p | MIMAT0000219 | Pml | 18854 | Sequencing//HITS-CLIP | 37986224\|23142080\|19536157 | | Unspecified |
| mmu-miR-24-3p | MIMAT0000219 | Pmm2 | 54128 | HITS-CLIP | | 23142080 | Unspecified |
| mmu-miR-24-3p | MIMAT0000219 | Pms2 | 18861 | HITS-CLIP | | 23142080 | Unspecified |
| mmu-miR-24-3p | MIMAT0000219 | Pnisr | 66625 | HITS-CLIP | | 23142080 | Unspecified |
| mmu-miR-24-3p | MIMAT0000219 | Pnn | 18949 | HITS-CLIP | 37986224\|23142080 | | Unspecified |
| mmu-miR-24-3p | MIMAT0000219 | Pnpla2 | 66853 | HITS-CLIP | | 25753418 | Unspecified |
| mmu-miR-24-3p | MIMAT0000219 | Pnrc1 | 108767 | HITS-CLIP | | 23142080 | Unspecified |
| mmu-miR-24-3p | MIMAT0000219 | Pola1 | 18968 | HITS-CLIP | | 23142080 | Unspecified |
| mmu-miR-24-3p | MIMAT0000219 | Pole | 18973 | HITS-CLIP | | 23142080 | Unspecified |
| mmu-miR-24-3p | MIMAT0000219 | Polm | 54125 | HITS-CLIP | | 23142080 | Unspecified |
| mmu-miR-24-3p | MIMAT0000219 | Polr2a | 20020 | HITS-CLIP | 25753418\|23142080 | | Unspecified |
| mmu-miR-24-3p | MIMAT0000219 | Polr3a | 218832 | HITS-CLIP | | 25753418 | Unspecified |
| mmu-miR-24-3p | MIMAT0000219 | Polr3c | 74414 | HITS-CLIP | | 23142080 | Unspecified |
| mmu-miR-24-3p | MIMAT0000219 | Polr3e | 26939 | HITS-CLIP | | 23142080 | Unspecified |
| mmu-miR-24-3p | MIMAT0000219 | Polr3k | 67005 | HITS-CLIP | | 23142080 | Unspecified |
| mmu-miR-24-3p | MIMAT0000219 | Pom121 | 107939 | HITS-CLIP | | 25753418 | Unspecified |
| mmu-miR-24-3p | MIMAT0000219 | Pon3 | 269823 | HITS-CLIP | | 25753418 | Unspecified |
| mmu-miR-24-3p | MIMAT0000219 | Pou2f2 | 18987 | HITS-CLIP | 37986224\|23142080 | | Unspecified |
| mmu-miR-24-3p | MIMAT0000219 | Ppa2 | 74776 | HITS-CLIP | | 23142080 | Unspecified |
| mmu-miR-24-3p | MIMAT0000219 | Ppan | 235036 | HITS-CLIP | | 23142080 | Unspecified |
| mmu-miR-24-3p | MIMAT0000219 | Ppard | 19015 | HITS-CLIP | | 23142080 | Unspecified |
| mmu-miR-24-3p | MIMAT0000219 | Ppat | 231327 | HITS-CLIP | | 25753418 | Unspecified |

| | | | | | | | |
|---|---|---|---|---|---|---|---|
| mmu-miR-24-3p | MIMAT0000219 | Ppil3 | 53938 | HITS-CLIP | | 23142080 | Unspecified |
| mmu-miR-24-3p | MIMAT0000219 | Ppip5k1 | 327655 | HITS-CLIP | | 25753418 | Unspecified |
| mmu-miR-24-3p | MIMAT0000219 | Ppip5k2 | 227399 | HITS-CLIP | | 25753418 | Unspecified |
| mmu-miR-24-3p | MIMAT0000219 | Ppm1a | 19042 | HITS-CLIP | | 25753418 | Unspecified |
| mmu-miR-24-3p | MIMAT0000219 | Ppm1d | 53892 | HITS-CLIP | | 23142080 | Unspecified |
| mmu-miR-24-3p | MIMAT0000219 | Ppm1g | 14208 | HITS-CLIP | | 23142080 | Unspecified |
| mmu-miR-24-3p | MIMAT0000219 | Ppme1 | 72590 | HITS-CLIP | | 23142080 | Unspecified |
| mmu-miR-24-3p | MIMAT0000219 | Ppp1ca | 19045 | HITS-CLIP | | 23142080 | Unspecified |
| mmu-miR-24-3p | MIMAT0000219 | Ppp1r10 | 52040 | HITS-CLIP | | 23142080 | Unspecified |
| mmu-miR-24-3p | MIMAT0000219 | Ppp1r12b | 329251 | HITS-CLIP | | 25753418 | Unspecified |
| mmu-miR-24-3p | MIMAT0000219 | Ppp1r12c | 232807 | HITS-CLIP | | 23142080 | Unspecified |
| mmu-miR-24-3p | MIMAT0000219 | Ppp1r13b | 21981 | HITS-CLIP | | 23142080 | Unspecified |
| mmu-miR-24-3p | MIMAT0000219 | Ppp1r15b | 108954 | HITS-CLIP | | 23142080 | Unspecified |
| mmu-miR-24-3p | MIMAT0000219 | Ppp1r16b | 228852 | HITS-CLIP | | 23142080 | Unspecified |
| mmu-miR-24-3p | MIMAT0000219 | Ppp1r21 | 73825 | HITS-CLIP | | 23142080 | Unspecified |
| mmu-miR-24-3p | MIMAT0000219 | Ppp1r3a | 140491 | HITS-CLIP | | 25753418 | Unspecified |
| mmu-miR-24-3p | MIMAT0000219 | Ppp1r3b | 244416 | HITS-CLIP | | 23142080 | Unspecified |
| mmu-miR-24-3p | MIMAT0000219 | Ppp1r3e | 90673 | HITS-CLIP | | 23142080 | Unspecified |
| mmu-miR-24-3p | MIMAT0000219 | Ppp2r5a | 226849 | HITS-CLIP | | 23142080 | Unspecified |
| mmu-miR-24-3p | MIMAT0000219 | Ppp2r5d | 21770 | HITS-CLIP | 23142080\|25753418 | | Unspecified |
| mmu-miR-24-3p | MIMAT0000219 | Ppp3cb | 19056 | HITS-CLIP | | 23142080 | Unspecified |
| mmu-miR-24-3p | MIMAT0000219 | Ppp3cc | 19057 | HITS-CLIP | | 23142080 | Unspecified |
| mmu-miR-24-3p | MIMAT0000219 | Ppp4r1 | 70351 | HITS-CLIP | | 23142080 | Unspecified |
| mmu-miR-24-3p | MIMAT0000219 | Ppp4r3b | 57223 | HITS-CLIP | | 23142080 | Unspecified |
| mmu-miR-24-3p | MIMAT0000219 | Ppp5c | 19060 | HITS-CLIP | | 25753418 | Unspecified |
| mmu-miR-24-3p | MIMAT0000219 | Ppp6c | 67857 | HITS-CLIP | | 25753418 | Unspecified |
| mmu-miR-24-3p | MIMAT0000219 | Ppp6r1 | 243819 | HITS-CLIP | | 23142080 | Unspecified |
| mmu-miR-24-3p | MIMAT0000219 | Ppp6r3 | 52036 | HITS-CLIP | | 23142080 | Unspecified |
| mmu-miR-24-3p | MIMAT0000219 | Pprc1 | 226169 | HITS-CLIP | | 23142080 | Unspecified |
| mmu-miR-24-3p | MIMAT0000219 | Pptc7 | 320717 | HITS-CLIP | | 23142080 | Unspecified |
| mmu-miR-24-3p | MIMAT0000219 | Prc1 | 233406 | HITS-CLIP | | 23142080 | Unspecified |
| mmu-miR-24-3p | MIMAT0000219 | Prcc | 94315 | HITS-CLIP | | 23142080 | Unspecified |
| mmu-miR-24-3p | MIMAT0000219 | Prdm1 | 12142 | HITS-CLIP | | 23142080 | Unspecified |
| mmu-miR-24-3p | MIMAT0000219 | Prdm2 | 110593 | HITS-CLIP | | 23142080 | Unspecified |
| mmu-miR-24-3p | MIMAT0000219 | Prdx3 | 11757 | HITS-CLIP | | 23142080 | Unspecified |
| mmu-miR-24-3p | MIMAT0000219 | Prdx6 | 11758 | HITS-CLIP | | 25753418 | Unspecified |
| mmu-miR-24-3p | MIMAT0000219 | Prelid1 | 66494 | HITS-CLIP | | 25753418 | Unspecified |
| mmu-miR-24-3p | MIMAT0000219 | Prkaa2 | 108079 | HITS-CLIP | | 25753418 | Unspecified |
| mmu-miR-24-3p | MIMAT0000219 | Prkacb | 18749 | HITS-CLIP | | 23142080 | Unspecified |

| | | | | | | |
|---|---|---|---|---|---|---|
| mmu-miR-24-3p | MIMAT0000219 | Prkcd | 18753 | HITS-CLIP | 37986224 | Unspecified |
| mmu-miR-24-3p | MIMAT0000219 | Prkcsh | 19089 | HITS-CLIP | 23142080 | Unspecified |
| mmu-miR-24-3p | MIMAT0000219 | Prmt5 | 27374 | HITS-CLIP | 25753418 | Unspecified |
| mmu-miR-24-3p | MIMAT0000219 | Prmt9 | 80204 | HITS-CLIP | 23142080 | Unspecified |
| mmu-miR-24-3p | MIMAT0000219 | Prpf31 | 68988 | HITS-CLIP | 25753418 | Unspecified |
| mmu-miR-24-3p | MIMAT0000219 | Prpf38b | 66921 | HITS-CLIP | 23142080 | Unspecified |
| mmu-miR-24-3p | MIMAT0000219 | Prpf40a | 56194 | HITS-CLIP | 23142080 | Unspecified |
| mmu-miR-24-3p | MIMAT0000219 | Prpf4b | 19134 | HITS-CLIP | 23142080 | Unspecified |
| mmu-miR-24-3p | MIMAT0000219 | Prpf6 | 68879 | HITS-CLIP | 23142080 | Unspecified |
| mmu-miR-24-3p | MIMAT0000219 | Prpf8 | 192159 | HITS-CLIP//qCLASH | 37986224\|23142080\|31393953\|25753 | Unspecified |
| mmu-miR-24-3p | MIMAT0000219 | Prps2 | 110639 | HITS-CLIP | 23142080 | Unspecified |
| mmu-miR-24-3p | MIMAT0000219 | Prr12 | 233210 | HITS-CLIP | 25753418 | Unspecified |
| mmu-miR-24-3p | MIMAT0000219 | Prr14l | 215476 | HITS-CLIP | 23142080 | Unspecified |
| mmu-miR-24-3p | MIMAT0000219 | Prr33 | 102724536 | HITS-CLIP | 25753418 | Unspecified |
| mmu-miR-24-3p | MIMAT0000219 | Prr36 | 80164 | HITS-CLIP | 23142080 | Unspecified |
| mmu-miR-24-3p | MIMAT0000219 | Prr5 | 109270 | HITS-CLIP | 23142080 | Unspecified |
| mmu-miR-24-3p | MIMAT0000219 | Prrc2a | 53761 | HITS-CLIP | 23142080 | Unspecified |
| mmu-miR-24-3p | MIMAT0000219 | Prrc2c | 226562 | HITS-CLIP | 23142080 | Unspecified |
| mmu-miR-24-3p | MIMAT0000219 | Prrt2 | 69017 | HITS-CLIP | 23142080 | Unspecified |
| mmu-miR-24-3p | MIMAT0000219 | Psap | 19156 | Luciferase Reporter Assay//HITS-CLIP | 37986224\|25753418 | Unspecified |
| mmu-miR-24-3p | MIMAT0000219 | Psen1 | 19164 | HITS-CLIP | 23142080 | Unspecified |
| mmu-miR-24-3p | MIMAT0000219 | Psma3 | 19167 | HITS-CLIP | 25753418 | Unspecified |
| mmu-miR-24-3p | MIMAT0000219 | Psmb1 | 19170 | HITS-CLIP | 23142080 | Unspecified |
| mmu-miR-24-3p | MIMAT0000219 | Psmb4 | 19172 | HITS-CLIP | 23142080 | Unspecified |
| mmu-miR-24-3p | MIMAT0000219 | Psmc2 | 19181 | HITS-CLIP | 25753418 | Unspecified |
| mmu-miR-24-3p | MIMAT0000219 | Psmd11 | 69077 | HITS-CLIP | 23142080\|25753418 | Unspecified |
| mmu-miR-24-3p | MIMAT0000219 | Psmd12 | 66997 | HITS-CLIP | 37986224 | Unspecified |
| mmu-miR-24-3p | MIMAT0000219 | Psmd14 | 59029 | HITS-CLIP | 23142080 | Unspecified |
| mmu-miR-24-3p | MIMAT0000219 | Psmd4 | 19185 | HITS-CLIP | 25753418 | Unspecified |
| mmu-miR-24-3p | MIMAT0000219 | Psmd8 | 57296 | HITS-CLIP | 23142080 | Unspecified |
| mmu-miR-24-3p | MIMAT0000219 | Psme2b | 621823 | HITS-CLIP | 23142080 | Unspecified |
| mmu-miR-24-3p | MIMAT0000219 | Psme4 | 103554 | HITS-CLIP | 23142080 | Unspecified |
| mmu-miR-24-3p | MIMAT0000219 | Ptar1 | 72351 | HITS-CLIP | 23142080 | Unspecified |
| mmu-miR-24-3p | MIMAT0000219 | Ptbp1 | 19205 | HITS-CLIP | 23142080\|31433990 | Unspecified |
| mmu-miR-24-3p | MIMAT0000219 | Ptbp3 | 230257 | HITS-CLIP | 23142080 | Unspecified |
| mmu-miR-24-3p | MIMAT0000219 | Ptger4 | 19219 | HITS-CLIP | 37986224\|23142080 | Unspecified |
| mmu-miR-24-3p | MIMAT0000219 | Ptgfrn | 19221 | HITS-CLIP | 23142080 | Unspecified |
| mmu-miR-24-3p | MIMAT0000219 | Ptp4a3 | 19245 | HITS-CLIP | 23142080 | Unspecified |
| mmu-miR-24-3p | MIMAT0000219 | Ptpn11 | 19247 | HITS-CLIP | 25753418\|23142080 | Unspecified |

| | | | | | | |
|---|---|---|---|---|---|---|
| mmu-miR-24-3p | MIMAT0000219 | Ptprc | 19264 | HITS-CLIP | 23142080 | Unspecified |
| mmu-miR-24-3p | MIMAT0000219 | Ptprcap | 19265 | HITS-CLIP | 23142080 | Unspecified |
| mmu-miR-24-3p | MIMAT0000219 | Pum2 | 80913 | HITS-CLIP | 23142080 | Unspecified |
| mmu-miR-24-3p | MIMAT0000219 | Pura | 19290 | HITS-CLIP | 23142080 | Unspecified |
| mmu-miR-24-3p | MIMAT0000219 | Purb | 19291 | HITS-CLIP | 23142080 | Unspecified |
| mmu-miR-24-3p | MIMAT0000219 | Pvr | 52118 | HITS-CLIP | 31433990\|25753418\|23142080 | Unspecified |
| mmu-miR-24-3p | MIMAT0000219 | Pxylp1 | 235534 | HITS-CLIP | 25753418 | Unspecified |
| mmu-miR-24-3p | MIMAT0000219 | Pycard | 66824 | HITS-CLIP | 25753418 | Unspecified |
| mmu-miR-24-3p | MIMAT0000219 | Pycrl | 66194 | HITS-CLIP | 23142080 | Unspecified |
| mmu-miR-24-3p | MIMAT0000219 | Pygb | 110078 | HITS-CLIP | 37986224\|25753418\|23142080 | Unspecified |
| mmu-miR-24-3p | MIMAT0000219 | Pygm | 19309 | HITS-CLIP | 25753418 | Unspecified |
| mmu-miR-24-3p | MIMAT0000219 | Qars | 97541 | HITS-CLIP | 23142080 | Unspecified |
| mmu-miR-24-3p | MIMAT0000219 | Qrich1 | 69232 | HITS-CLIP | 23142080 | Unspecified |
| mmu-miR-24-3p | MIMAT0000219 | Rab10 | 19325 | HITS-CLIP | 23142080 | Unspecified |
| mmu-miR-24-3p | MIMAT0000219 | Rab11fip2 | 74998 | HITS-CLIP | 23142080 | Unspecified |
| mmu-miR-24-3p | MIMAT0000219 | Rab11fip3 | 215445 | HITS-CLIP | 23142080 | Unspecified |
| mmu-miR-24-3p | MIMAT0000219 | Rab21 | 216344 | HITS-CLIP | 23142080 | Unspecified |
| mmu-miR-24-3p | MIMAT0000219 | Rab33b | 19338 | HITS-CLIP | 37986224\|23142080 | Unspecified |
| mmu-miR-24-3p | MIMAT0000219 | Rab35 | 77407 | HITS-CLIP | 23142080 | Unspecified |
| mmu-miR-24-3p | MIMAT0000219 | Rab3gap1 | 226407 | Sequencing//HITS-CLIP | 19536157\|25753418\|37986224 | Unspecified |
| mmu-miR-24-3p | MIMAT0000219 | Rab3gap2 | 98732 | HITS-CLIP | 23142080 | Unspecified |
| mmu-miR-24-3p | MIMAT0000219 | Rab3ip | 216363 | HITS-CLIP | 23142080 | Unspecified |
| mmu-miR-24-3p | MIMAT0000219 | Rab40c | 224624 | HITS-CLIP | 25753418 | Unspecified |
| mmu-miR-24-3p | MIMAT0000219 | Rab5a | 271457 | HITS-CLIP | 23142080 | Unspecified |
| mmu-miR-24-3p | MIMAT0000219 | Rab5c | 19345 | HITS-CLIP | 37986224\|23142080 | Unspecified |
| mmu-miR-24-3p | MIMAT0000219 | Rabac1 | 14470 | HITS-CLIP | 25753418 | Unspecified |
| mmu-miR-24-3p | MIMAT0000219 | Rabgap1 | 227800 | HITS-CLIP | 25753418 | Unspecified |
| mmu-miR-24-3p | MIMAT0000219 | Rabgap1l | 29809 | HITS-CLIP | 23142080 | Unspecified |
| mmu-miR-24-3p | MIMAT0000219 | Rabgef1 | 56715 | HITS-CLIP | 23142080 | Unspecified |
| mmu-miR-24-3p | MIMAT0000219 | Rabggtb | 19352 | HITS-CLIP | 23142080 | Unspecified |
| mmu-miR-24-3p | MIMAT0000219 | Rac1 | 19353 | HITS-CLIP | 31433990\|25753418 | Unspecified |
| mmu-miR-24-3p | MIMAT0000219 | Rac2 | 19354 | HITS-CLIP | 23142080 | Unspecified |
| mmu-miR-24-3p | MIMAT0000219 | Rad21 | 19357 | HITS-CLIP | 23142080 | Unspecified |
| mmu-miR-24-3p | MIMAT0000219 | Rad51b | 5890 | HITS-CLIP | 23142080 | Unspecified |
| mmu-miR-24-3p | MIMAT0000219 | Rad51c | 114714 | HITS-CLIP | 23142080 | Unspecified |
| mmu-miR-24-3p | MIMAT0000219 | Raf1 | 110157 | HITS-CLIP | 23142080 | Unspecified |
| mmu-miR-24-3p | MIMAT0000219 | Rai14 | 75646 | Luciferase Reporter Assay | 37986224 | Unspecified |
| mmu-miR-24-3p | MIMAT0000219 | Rala | 56044 | HITS-CLIP | 23142080 | Unspecified |
| mmu-miR-24-3p | MIMAT0000219 | Ralbp1 | 19765 | HITS-CLIP | 23142080 | Unspecified |

| | | | | | | |
|---|---|---|---|---|---|---|
| mmu-miR-24-3p | MIMAT0000219 | Ralgapa1 | 56784 | HITS-CLIP | 25753418 | Unspecified |
| mmu-miR-24-3p | MIMAT0000219 | Ralgapa2 | 241694 | HITS-CLIP | 25753418 | Unspecified |
| mmu-miR-24-3p | MIMAT0000219 | Ralgapb | 228850 | HITS-CLIP | 25753418 | Unspecified |
| mmu-miR-24-3p | MIMAT0000219 | Ralgps1 | 241308 | HITS-CLIP | 23142080 | Unspecified |
| mmu-miR-24-3p | MIMAT0000219 | Ralgps2 | 78255 | HITS-CLIP | 23142080 | Unspecified |
| mmu-miR-24-3p | MIMAT0000219 | Raly | 19383 | HITS-CLIP | 23142080 | Unspecified |
| mmu-miR-24-3p | MIMAT0000219 | Ramp2 | 54409 | HITS-CLIP | 25753418 | Unspecified |
| mmu-miR-24-3p | MIMAT0000219 | Ranbp2 | 19386 | HITS-CLIP | 23142080\|25753418 | Unspecified |
| mmu-miR-24-3p | MIMAT0000219 | Ranbp6 | 240614 | HITS-CLIP | 37986224\|23142080 | Unspecified |
| mmu-miR-24-3p | MIMAT0000219 | Ranbp9 | 56705 | HITS-CLIP | 25753418 | Unspecified |
| mmu-miR-24-3p | MIMAT0000219 | Rap1b | 215449 | HITS-CLIP | 37986224\|31433990\|25753418\|23142 | Unspecified |
| mmu-miR-24-3p | MIMAT0000219 | Rap1gds1 | 229877 | HITS-CLIP | 23142080 | Unspecified |
| mmu-miR-24-3p | MIMAT0000219 | Rap2c | 72065 | HITS-CLIP | 23142080 | Unspecified |
| mmu-miR-24-3p | MIMAT0000219 | Rapgef2 | 76089 | HITS-CLIP | 23142080 | Unspecified |
| mmu-miR-24-3p | MIMAT0000219 | Rapgef4 | 56508 | HITS-CLIP | 25753418 | Unspecified |
| mmu-miR-24-3p | MIMAT0000219 | Rapgef6 | 192786 | HITS-CLIP | 23142080 | Unspecified |
| mmu-miR-24-3p | MIMAT0000219 | Raph1 | 77300 | HITS-CLIP | 37986224\|23142080\|25753418 | Unspecified |
| mmu-miR-24-3p | MIMAT0000219 | Rarg | 19411 | HITS-CLIP | 23142080 | Unspecified |
| mmu-miR-24-3p | MIMAT0000219 | Rasa1 | 218397 | HITS-CLIP | 37986224\|23142080 | Unspecified |
| mmu-miR-24-3p | MIMAT0000219 | Rasa3 | 19414 | HITS-CLIP | 23142080 | Unspecified |
| mmu-miR-24-3p | MIMAT0000219 | Rasal3 | 320484 | HITS-CLIP | 23142080 | Unspecified |
| mmu-miR-24-3p | MIMAT0000219 | Rasl2-9 | 19428 | HITS-CLIP | 25753418 | Unspecified |
| mmu-miR-24-3p | MIMAT0000219 | Rassf3 | 192678 | HITS-CLIP | 25753418 | Unspecified |
| mmu-miR-24-3p | MIMAT0000219 | Rassf5 | 54354 | HITS-CLIP | 23142080 | Unspecified |
| mmu-miR-24-3p | MIMAT0000219 | Raver1 | 71766 | qCLASH//HITS-CLIP | 31393953\|23142080 | Unspecified |
| mmu-miR-24-3p | MIMAT0000219 | Rb1 | 19645 | HITS-CLIP | 23142080 | Unspecified |
| mmu-miR-24-3p | MIMAT0000219 | Rb1cc1 | 12421 | HITS-CLIP | 23142080 | Unspecified |
| mmu-miR-24-3p | MIMAT0000219 | Rbbp6 | 19647 | HITS-CLIP | 37986224\|23142080 | Unspecified |
| mmu-miR-24-3p | MIMAT0000219 | Rbm15b | 109095 | HITS-CLIP | 25753418 | Unspecified |
| mmu-miR-24-3p | MIMAT0000219 | Rbm18 | 67889 | HITS-CLIP | 23142080 | Unspecified |
| mmu-miR-24-3p | MIMAT0000219 | Rbm20 | 73713 | HITS-CLIP | 25753418 | Unspecified |
| mmu-miR-24-3p | MIMAT0000219 | Rbm25 | 67039 | HITS-CLIP | 25753418\|23142080 | Unspecified |
| mmu-miR-24-3p | MIMAT0000219 | Rbm26 | 74213 | HITS-CLIP | 23142080 | Unspecified |
| mmu-miR-24-3p | MIMAT0000219 | Rbm27 | 225432 | HITS-CLIP | 23142080 | Unspecified |
| mmu-miR-24-3p | MIMAT0000219 | Rbm43 | 71684 | HITS-CLIP | 23142080 | Unspecified |
| mmu-miR-24-3p | MIMAT0000219 | Rbm5 | 83486 | HITS-CLIP | 23142080 | Unspecified |
| mmu-miR-24-3p | MIMAT0000219 | Rbp7 | 33968 | HITS-CLIP | 25753418 | Unspecified |
| mmu-miR-24-3p | MIMAT0000219 | Rbx1 | 56438 | HITS-CLIP | 23142080 | Unspecified |
| mmu-miR-24-3p | MIMAT0000219 | Rc3h1 | 381305 | HITS-CLIP | 23142080 | Unspecified |

| | | | | | | |
|---|---|---|---|---|---|---|
| mmu-miR-24-3p | MIMAT0000219 | Rc3h2 | 319817 | HITS-CLIP | 37986224\|23142080 | Unspecified |
| mmu-miR-24-3p | MIMAT0000219 | Rcc2 | 108911 | HITS-CLIP | 23142080 | Unspecified |
| mmu-miR-24-3p | MIMAT0000219 | Rdx | 19684 | HITS-CLIP | 23142080 | Unspecified |
| mmu-miR-24-3p | MIMAT0000219 | Rela | 19697 | HITS-CLIP | 23142080\|25753418 | Unspecified |
| mmu-miR-24-3p | MIMAT0000219 | Reps1 | 19707 | qCLASH | 31393953 | Unspecified |
| mmu-miR-24-3p | MIMAT0000219 | Rer1 | 67830 | HITS-CLIP | 23142080 | Unspecified |
| mmu-miR-24-3p | MIMAT0000219 | Rev1 | 56210 | HITS-CLIP | 23142080 | Unspecified |
| mmu-miR-24-3p | MIMAT0000219 | Rev3l | 19714 | HITS-CLIP | 23142080 | Unspecified |
| mmu-miR-24-3p | MIMAT0000219 | Rexo4 | 227656 | HITS-CLIP | 23142080 | Unspecified |
| mmu-miR-24-3p | MIMAT0000219 | Rfx1 | 19724 | HITS-CLIP | 23142080 | Unspecified |
| mmu-miR-24-3p | MIMAT0000219 | Rfx3 | 19726 | HITS-CLIP | 23142080 | Unspecified |
| mmu-miR-24-3p | MIMAT0000219 | Rfx5 | 53970 | HITS-CLIP | 23142080 | Unspecified |
| mmu-miR-24-3p | MIMAT0000219 | Rgp1 | 242406 | HITS-CLIP | 25753418 | Unspecified |
| mmu-miR-24-3p | MIMAT0000219 | Rgs16 | 19734 | HITS-CLIP | 23142080 | Unspecified |
| mmu-miR-24-3p | MIMAT0000219 | Rgs2 | 19735 | HITS-CLIP | 25753418 | Unspecified |
| mmu-miR-24-3p | MIMAT0000219 | Rhbdd2 | 215160 | HITS-CLIP | 25753418 | Unspecified |
| mmu-miR-24-3p | MIMAT0000219 | Rhbdl1 | 214951 | Sequencing//HITS-CLIP | 19536157\|23142080\|37986224 | Unspecified |
| mmu-miR-24-3p | MIMAT0000219 | Rhoq | 104215 | HITS-CLIP | 23142080 | Unspecified |
| mmu-miR-24-3p | MIMAT0000219 | Rhot1 | 59040 | HITS-CLIP | 25753418 | Unspecified |
| mmu-miR-24-3p | MIMAT0000219 | Rhpn2 | 52428 | HITS-CLIP | 25753418 | Unspecified |
| mmu-miR-24-3p | MIMAT0000219 | Rictor | 78757 | Sequencing//HITS-CLIP | 19536157\|23142080 | Unspecified |
| mmu-miR-24-3p | MIMAT0000219 | Riok3 | 66878 | HITS-CLIP | 25753418 | Unspecified |
| mmu-miR-24-3p | MIMAT0000219 | Ripk1 | 19766 | HITS-CLIP | 23142080 | Unspecified |
| mmu-miR-24-3p | MIMAT0000219 | Ripk2 | 192656 | HITS-CLIP | 37986224\|23142080 | Unspecified |
| mmu-miR-24-3p | MIMAT0000219 | Rlim | 19820 | HITS-CLIP | 23142080 | Unspecified |
| mmu-miR-24-3p | MIMAT0000219 | Rmi2 | 223970 | HITS-CLIP | 23142080 | Unspecified |
| mmu-miR-24-3p | MIMAT0000219 | Rnaseh2b | 67153 | HITS-CLIP | 23142080 | Unspecified |
| mmu-miR-24-3p | MIMAT0000219 | Rnf10 | 50849 | HITS-CLIP | 25753418\|23142080 | Unspecified |
| mmu-miR-24-3p | MIMAT0000219 | Rnf11 | 29864 | HITS-CLIP | 23142080 | Unspecified |
| mmu-miR-24-3p | MIMAT0000219 | Rnf111 | 93836 | HITS-CLIP | 25753418 | Unspecified |
| mmu-miR-24-3p | MIMAT0000219 | Rnf115 | 67845 | HITS-CLIP | 23142080 | Unspecified |
| mmu-miR-24-3p | MIMAT0000219 | Rnf13 | 24017 | HITS-CLIP | 23142080 | Unspecified |
| mmu-miR-24-3p | MIMAT0000219 | Rnf138 | 56515 | Sequencing//HITS-CLIP | 37986224\|23142080\|19536157 | Unspecified |
| mmu-miR-24-3p | MIMAT0000219 | Rnf14 | 56736 | HITS-CLIP | 23142080 | Unspecified |
| mmu-miR-24-3p | MIMAT0000219 | Rnf144a | 108089 | Luciferase Reporter Assay | 37986224 | Unspecified |
| mmu-miR-24-3p | MIMAT0000219 | Rnf145 | 74315 | HITS-CLIP | 23142080 | Unspecified |
| mmu-miR-24-3p | MIMAT0000219 | Rnf149 | 67702 | HITS-CLIP | 23142080 | Unspecified |
| mmu-miR-24-3p | MIMAT0000219 | Rnf150 | 330812 | HITS-CLIP | 25753418 | Unspecified |
| mmu-miR-24-3p | MIMAT0000219 | Rnf157 | 217340 | HITS-CLIP | 23142080 | Unspecified |

| | | | | | | |
|---|---|---|---|---|---|---|
| mmu-miR-24-3p | MIMAT0000219 | Rnf181 | 66510 | HITS-CLIP | 25753418 | Unspecified |
| mmu-miR-24-3p | MIMAT0000219 | Rnf19a | 30945 | HITS-CLIP | 23142080 | Unspecified |
| mmu-miR-24-3p | MIMAT0000219 | Rnf2 | 19821 | HITS-CLIP | 37986224\|23142080 | Unspecified |
| mmu-miR-24-3p | MIMAT0000219 | Rnf213 | 672511 | HITS-CLIP | 23142080 | Unspecified |
| mmu-miR-24-3p | MIMAT0000219 | Rnf214 | 235315 | HITS-CLIP | 23142080 | Unspecified |
| mmu-miR-24-3p | MIMAT0000219 | Rnf38 | 73469 | HITS-CLIP | 23142080 | Unspecified |
| mmu-miR-24-3p | MIMAT0000219 | Rnf44 | 105239 | HITS-CLIP | 26203562\|25753418 | Unspecified |
| mmu-miR-24-3p | MIMAT0000219 | Rnh1 | 107702 | HITS-CLIP | 23142080 | Unspecified |
| mmu-miR-24-3p | MIMAT0000219 | Rnpep | 215615 | HITS-CLIP | 25753418 | Unspecified |
| mmu-miR-24-3p | MIMAT0000219 | Ro60 | 6738 | HITS-CLIP | 25753418 | Unspecified |
| mmu-miR-24-3p | MIMAT0000219 | Rock2 | 19878 | HITS-CLIP | 23142080 | Unspecified |
| mmu-miR-24-3p | MIMAT0000219 | Ror2 | 26564 | qCLASH | 31393953 | Unspecified |
| mmu-miR-24-3p | MIMAT0000219 | Rora | 19883 | HITS-CLIP | 23142080 | Unspecified |
| mmu-miR-24-3p | MIMAT0000219 | Rpa1 | 68275 | HITS-CLIP | 23142080 | Unspecified |
| mmu-miR-24-3p | MIMAT0000219 | Rpap1 | 68925 | HITS-CLIP | 23142080 | Unspecified |
| mmu-miR-24-3p | MIMAT0000219 | Rpap2 | 231571 | HITS-CLIP | 23142080 | Unspecified |
| mmu-miR-24-3p | MIMAT0000219 | Rpl3l | 66211 | HITS-CLIP | 25753418 | Unspecified |
| mmu-miR-24-3p | MIMAT0000219 | Rpl6 | 19988 | HITS-CLIP | 23142080 | Unspecified |
| mmu-miR-24-3p | MIMAT0000219 | Rpl7 | 19989 | qCLASH | 31393953 | Unspecified |
| mmu-miR-24-3p | MIMAT0000219 | Rpn2 | 20014 | HITS-CLIP | 23142080 | Unspecified |
| mmu-miR-24-3p | MIMAT0000219 | Rprd2 | 75137 | HITS-CLIP | 23142080 | Unspecified |
| mmu-miR-24-3p | MIMAT0000219 | Rps20 | 67427 | HITS-CLIP | 37986224\|23142080\|31433990 | Unspecified |
| mmu-miR-24-3p | MIMAT0000219 | Rps3 | 27050 | HITS-CLIP | 23142080 | Unspecified |
| mmu-miR-24-3p | MIMAT0000219 | Rps6ka1 | 20111 | HITS-CLIP | 31433990\|25753418 | Unspecified |
| mmu-miR-24-3p | MIMAT0000219 | Rps6ka3 | 110651 | HITS-CLIP | 23142080 | Unspecified |
| mmu-miR-24-3p | MIMAT0000219 | Rps6ka5 | 73086 | HITS-CLIP | 23142080 | Unspecified |
| mmu-miR-24-3p | MIMAT0000219 | Rps6kb1 | 72508 | HITS-CLIP | 23142080 | Unspecified |
| mmu-miR-24-3p | MIMAT0000219 | Rpusd2 | 271842 | HITS-CLIP | 25753418 | Unspecified |
| mmu-miR-24-3p | MIMAT0000219 | Rreb1 | 68750 | HITS-CLIP | 23142080 | Unspecified |
| mmu-miR-24-3p | MIMAT0000219 | Rrm1 | 20133 | HITS-CLIP | 23142080 | Unspecified |
| mmu-miR-24-3p | MIMAT0000219 | Rrp1b | 72462 | HITS-CLIP | 23142080 | Unspecified |
| mmu-miR-24-3p | MIMAT0000219 | Rsbn1 | 229675 | HITS-CLIP | 25753418\|23142080 | Unspecified |
| mmu-miR-24-3p | MIMAT0000219 | Rsbn1l | 242860 | HITS-CLIP | 37986224\|23142080 | Unspecified |
| mmu-miR-24-3p | MIMAT0000219 | Rsf1 | 233532 | HITS-CLIP | 31433990 | Unspecified |
| mmu-miR-24-3p | MIMAT0000219 | Rsl1d1 | 66409 | HITS-CLIP | 25753418\|23142080 | Unspecified |
| mmu-miR-24-3p | MIMAT0000219 | Rspry1 | 67610 | HITS-CLIP | 23142080 | Unspecified |
| mmu-miR-24-3p | MIMAT0000219 | Rtcb | 28088 | HITS-CLIP | 23142080 | Unspecified |
| mmu-miR-24-3p | MIMAT0000219 | Rtn4 | 68585 | HITS-CLIP | 37986224\|23142080 | Unspecified |
| mmu-miR-24-3p | MIMAT0000219 | Rubcn | 9711 | HITS-CLIP | 23142080 | Unspecified |

| | | | | | | |
|---|---|---|---|---|---|---|
| mmu-miR-24-3p | MIMAT0000219 | Rundc1 | 217201 | HITS-CLIP | 25753418 | Unspecified |
| mmu-miR-24-3p | MIMAT0000219 | Runx2 | 12393 | HITS-CLIP | 23142080 | Unspecified |
| mmu-miR-24-3p | MIMAT0000219 | Rusc2 | 100213 | HITS-CLIP | 25753418 | Unspecified |
| mmu-miR-24-3p | MIMAT0000219 | Rxrb | 20182 | HITS-CLIP | 23142080 | Unspecified |
| mmu-miR-24-3p | MIMAT0000219 | Ryr2 | 20191 | HITS-CLIP | 25753418 | Unspecified |
| mmu-miR-24-3p | MIMAT0000219 | S100a1 | 20193 | HITS-CLIP | 23142080 | Unspecified |
| mmu-miR-24-3p | MIMAT0000219 | S100pbp | 74648 | HITS-CLIP | 23142080 | Unspecified |
| mmu-miR-24-3p | MIMAT0000219 | S1pr1 | 13609 | HITS-CLIP | 23142080 | Unspecified |
| mmu-miR-24-3p | MIMAT0000219 | S1pr2 | 14739 | HITS-CLIP | 25753418\|23142080 | Unspecified |
| mmu-miR-24-3p | MIMAT0000219 | S1pr4 | 13611 | HITS-CLIP | 23142080 | Unspecified |
| mmu-miR-24-3p | MIMAT0000219 | Sacm1l | 83493 | HITS-CLIP | 23142080\|25753418 | Unspecified |
| mmu-miR-24-3p | MIMAT0000219 | Sacs | 50720 | Sequencing//HITS-CLIP | 37986224\|23142080\|19536157 | Unspecified |
| mmu-miR-24-3p | MIMAT0000219 | Sall2 | 50524 | HITS-CLIP | 23142080 | Unspecified |
| mmu-miR-24-3p | MIMAT0000219 | Samd4b | 233033 | HITS-CLIP | 23142080 | Unspecified |
| mmu-miR-24-3p | MIMAT0000219 | Samd8 | 67630 | HITS-CLIP | 25753418 | Unspecified |
| mmu-miR-24-3p | MIMAT0000219 | Samhd1 | 56045 | HITS-CLIP | 37986224\|23142080 | Unspecified |
| mmu-miR-24-3p | MIMAT0000219 | Samm50 | 68653 | HITS-CLIP | 23142080\|25753418 | Unspecified |
| mmu-miR-24-3p | MIMAT0000219 | Sap130 | 269003 | HITS-CLIP | 23142080 | Unspecified |
| mmu-miR-24-3p | MIMAT0000219 | Satb1 | 20230 | HITS-CLIP | 37986224\|23142080 | Unspecified |
| mmu-miR-24-3p | MIMAT0000219 | Sav1 | 64010 | HITS-CLIP | 23142080 | Unspecified |
| mmu-miR-24-3p | MIMAT0000219 | Sbf1 | 77980 | HITS-CLIP | 25753418 | Unspecified |
| mmu-miR-24-3p | MIMAT0000219 | Sbno1 | 243272 | HITS-CLIP | 23142080\|25753418 | Unspecified |
| mmu-miR-24-3p | MIMAT0000219 | Sc5d | 235293 | HITS-CLIP | 23142080 | Unspecified |
| mmu-miR-24-3p | MIMAT0000219 | Scaf4 | 224432 | HITS-CLIP | 23142080 | Unspecified |
| mmu-miR-24-3p | MIMAT0000219 | Scamp1 | 107767 | HITS-CLIP | 25753418 | Unspecified |
| mmu-miR-24-3p | MIMAT0000219 | Scarb1 | 20778 | HITS-CLIP | 23142080 | Unspecified |
| mmu-miR-24-3p | MIMAT0000219 | Scd2 | 20250 | Sequencing//HITS-CLIP | 19536157\|23142080\|37986224 | Unspecified |
| mmu-miR-24-3p | MIMAT0000219 | Scoc | 56367 | HITS-CLIP | 25753418 | Unspecified |
| mmu-miR-24-3p | MIMAT0000219 | Sdad1 | 231452 | HITS-CLIP | 37986224\|23142080 | Unspecified |
| mmu-miR-24-3p | MIMAT0000219 | Sdf2 | 20316 | HITS-CLIP | 23142080 | Unspecified |
| mmu-miR-24-3p | MIMAT0000219 | Sdf4 | 20318 | HITS-CLIP | 25753418 | Unspecified |
| mmu-miR-24-3p | MIMAT0000219 | Sdhb | 67680 | HITS-CLIP | 23142080 | Unspecified |
| mmu-miR-24-3p | MIMAT0000219 | Sdhc | 66052 | HITS-CLIP | 25753418\|23142080 | Unspecified |
| mmu-miR-24-3p | MIMAT0000219 | Sdr39u1 | 654795 | HITS-CLIP | 23142080 | Unspecified |
| mmu-miR-24-3p | MIMAT0000219 | Sec11a | 56529 | HITS-CLIP | 37986224\|23142080 | Unspecified |
| mmu-miR-24-3p | MIMAT0000219 | Sec14l1 | 74136 | HITS-CLIP | 23142080 | Unspecified |
| mmu-miR-24-3p | MIMAT0000219 | Sec16a | 227648 | HITS-CLIP | 25753418\|23142080 | Unspecified |
| mmu-miR-24-3p | MIMAT0000219 | Sec22b | 20333 | HITS-CLIP | 25753418\|23142080 | Unspecified |
| mmu-miR-24-3p | MIMAT0000219 | Sec23b | 27054 | HITS-CLIP | 23142080 | Unspecified |

| | | | | | | |
|---|---|---|---|---|---|---|
| mmu-miR-24-3p | MIMAT0000219 | Sec24a | 77371 | HITS-CLIP | 37986224\|23142080 | Unspecified |
| mmu-miR-24-3p | MIMAT0000219 | Sec24c | 218811 | HITS-CLIP | 23142080 | Unspecified |
| mmu-miR-24-3p | MIMAT0000219 | Sec62 | 69276 | HITS-CLIP | 23142080 | Unspecified |
| mmu-miR-24-3p | MIMAT0000219 | Sec63 | 140740 | HITS-CLIP | 23142080\|25753418 | Unspecified |
| mmu-miR-24-3p | MIMAT0000219 | Seh1l | 72124 | HITS-CLIP | 23142080 | Unspecified |
| mmu-miR-24-3p | MIMAT0000219 | Selenoh | 280636 | HITS-CLIP | 23142080 | Unspecified |
| mmu-miR-24-3p | MIMAT0000219 | Selenoi | 28042 | HITS-CLIP | 23142080 | Unspecified |
| mmu-miR-24-3p | MIMAT0000219 | Selenop | 6414 | HITS-CLIP | 23142080 | Unspecified |
| mmu-miR-24-3p | MIMAT0000219 | Sema4b | 20352 | HITS-CLIP | 23142080 | Unspecified |
| mmu-miR-24-3p | MIMAT0000219 | Sema4d | 20354 | HITS-CLIP | 37986224\|23142080 | Unspecified |
| mmu-miR-24-3p | MIMAT0000219 | Sema7a | 20361 | HITS-CLIP | 37986224\|25753418\|23142080 | Unspecified |
| mmu-miR-24-3p | MIMAT0000219 | Senp2 | 75826 | HITS-CLIP | 37986224 | Unspecified |
| mmu-miR-24-3p | MIMAT0000219 | Senp5 | 320213 | HITS-CLIP | 23142080 | Unspecified |
| mmu-miR-24-3p | MIMAT0000219 | Sephs2 | 20768 | HITS-CLIP | 23142080 | Unspecified |
| mmu-miR-24-3p | MIMAT0000219 | Sepsecs | 211006 | HITS-CLIP | 23142080 | Unspecified |
| mmu-miR-24-3p | MIMAT0000219 | Septin2 | 18000 | HITS-CLIP | 25753418 | Unspecified |
| mmu-miR-24-3p | MIMAT0000219 | Septin9 | 10801 | HITS-CLIP | 25753418 | Unspecified |
| mmu-miR-24-3p | MIMAT0000219 | Serbp1 | 66870 | HITS-CLIP | 23142080 | Unspecified |
| mmu-miR-24-3p | MIMAT0000219 | Serinc3 | 26943 | HITS-CLIP | 23142080 | Unspecified |
| mmu-miR-24-3p | MIMAT0000219 | Serp1 | 28146 | HITS-CLIP | 37986224 | Unspecified |
| mmu-miR-24-3p | MIMAT0000219 | Serpinf1 | 20317 | HITS-CLIP | 23142080\|25753418 | Unspecified |
| mmu-miR-24-3p | MIMAT0000219 | Sertad2 | 58172 | HITS-CLIP | 25753418 | Unspecified |
| mmu-miR-24-3p | MIMAT0000219 | Sesn1 | 140742 | HITS-CLIP | 37986224\|23142080 | Unspecified |
| mmu-miR-24-3p | MIMAT0000219 | Sesn3 | 75747 | HITS-CLIP | 23142080 | Unspecified |
| mmu-miR-24-3p | MIMAT0000219 | Setd1a | 233904 | HITS-CLIP | 23142080 | Unspecified |
| mmu-miR-24-3p | MIMAT0000219 | Setd7 | 73251 | Sequencing//HITS-CLIP | 19536157\|25753418\|23142080\|3798[illegible] | Unspecified |
| mmu-miR-24-3p | MIMAT0000219 | Setdb2 | 83852 | HITS-CLIP | 23142080 | Unspecified |
| mmu-miR-24-3p | MIMAT0000219 | Setx | 269254 | HITS-CLIP | 23142080 | Unspecified |
| mmu-miR-24-3p | MIMAT0000219 | Sf1 | 22668 | HITS-CLIP | 23142080 | Unspecified |
| mmu-miR-24-3p | MIMAT0000219 | Sf3a2 | 20222 | HITS-CLIP | 23142080 | Unspecified |
| mmu-miR-24-3p | MIMAT0000219 | Sf3b1 | 81898 | HITS-CLIP | 23142080 | Unspecified |
| mmu-miR-24-3p | MIMAT0000219 | Sf3b2 | 319322 | HITS-CLIP | 25753418 | Unspecified |
| mmu-miR-24-3p | MIMAT0000219 | Sf3b3 | 101943 | qCLASH//HITS-CLIP | 31393953\|23142080 | Unspecified |
| mmu-miR-24-3p | MIMAT0000219 | Sfr1 | 67788 | HITS-CLIP | 23142080 | Unspecified |
| mmu-miR-24-3p | MIMAT0000219 | Sfxn2 | 94279 | Luciferase Reporter Assay//HITS-CLIP | 37986224\|23142080 | Unspecified |
| mmu-miR-24-3p | MIMAT0000219 | Sfxn5 | 94282 | Sequencing//HITS-CLIP | 19536157\|25753418\|37986224 | Unspecified |
| mmu-miR-24-3p | MIMAT0000219 | Sgpp1 | 81535 | HITS-CLIP | 23142080 | Unspecified |
| mmu-miR-24-3p | MIMAT0000219 | Sh2b1 | 20399 | HITS-CLIP | 23142080 | Unspecified |
| mmu-miR-24-3p | MIMAT0000219 | Sh3bgrl3 | 73723 | HITS-CLIP | 23142080 | Unspecified |

| | | | | | | | |
|---|---|---|---|---|---|---|---|
| mmu-miR-24-3p | MIMAT0000219 | Sh3gl1 | 20405 | HITS-CLIP | | 23142080 | Unspecified |
| mmu-miR-24-3p | MIMAT0000219 | Sh3glb1 | 54673 | HITS-CLIP | | 23142080 | Unspecified |
| mmu-miR-24-3p | MIMAT0000219 | Sh3pxd2a | 14218 | Western Blot | 37986224\|23071155 | | Unspecified |
| mmu-miR-24-3p | MIMAT0000219 | Siah1a | 20437 | HITS-CLIP | | 23142080 | Unspecified |
| mmu-miR-24-3p | MIMAT0000219 | Siah1b | 20438 | HITS-CLIP | | 23142080 | Unspecified |
| mmu-miR-24-3p | MIMAT0000219 | Siah2 | 20439 | HITS-CLIP | 23142080\|25753418 | | Unspecified |
| mmu-miR-24-3p | MIMAT0000219 | Sigirr | 24058 | HITS-CLIP | | 23142080 | Unspecified |
| mmu-miR-24-3p | MIMAT0000219 | Sik1 | 17691 | HITS-CLIP | | 23142080 | Unspecified |
| mmu-miR-24-3p | MIMAT0000219 | Sik2 | 235344 | HITS-CLIP | | 23142080 | Unspecified |
| mmu-miR-24-3p | MIMAT0000219 | Sik3 | 70661 | HITS-CLIP | | 23142080 | Unspecified |
| mmu-miR-24-3p | MIMAT0000219 | Sin3a | 20466 | HITS-CLIP | | 23142080 | Unspecified |
| mmu-miR-24-3p | MIMAT0000219 | Sirt2 | 64383 | HITS-CLIP | | 25753418 | Unspecified |
| mmu-miR-24-3p | MIMAT0000219 | Skida1 | 72668 | Sequencing//HITS-CLIP | 19536157\|23142080 | | Unspecified |
| mmu-miR-24-3p | MIMAT0000219 | Skil | 20482 | HITS-CLIP | | 23142080 | Unspecified |
| mmu-miR-24-3p | MIMAT0000219 | Skp1a | 21402 | HITS-CLIP | | 23142080 | Unspecified |
| mmu-miR-24-3p | MIMAT0000219 | Skp2 | 27401 | HITS-CLIP | | 23142080 | Unspecified |
| mmu-miR-24-3p | MIMAT0000219 | Sla | 20491 | HITS-CLIP | 37986224\|23142080 | | Unspecified |
| mmu-miR-24-3p | MIMAT0000219 | Slain2 | 75991 | HITS-CLIP | | 25753418 | Unspecified |
| mmu-miR-24-3p | MIMAT0000219 | Slbp | 20492 | HITS-CLIP | | 23142080 | Unspecified |
| mmu-miR-24-3p | MIMAT0000219 | Slc10a3 | 214601 | HITS-CLIP | | 23142080 | Unspecified |
| mmu-miR-24-3p | MIMAT0000219 | Slc12a6 | 107723 | Sequencing//HITS-CLIP | 37986224\|23142080\|19536157 | | Unspecified |
| mmu-miR-24-3p | MIMAT0000219 | Slc12a8 | 171286 | Luciferase Reporter Assay | | 37986224 | Unspecified |
| mmu-miR-24-3p | MIMAT0000219 | Slc12a9 | 83704 | HITS-CLIP | | 23142080 | Unspecified |
| mmu-miR-24-3p | MIMAT0000219 | Slc16a1 | 20501 | HITS-CLIP | | 25753418 | Unspecified |
| mmu-miR-24-3p | MIMAT0000219 | Slc17a7 | 72961 | HITS-CLIP | | 25753418 | Unspecified |
| mmu-miR-24-3p | MIMAT0000219 | Slc1a4 | 55963 | HITS-CLIP | | 23142080 | Unspecified |
| mmu-miR-24-3p | MIMAT0000219 | Slc1a5 | 20514 | HITS-CLIP | | 23142080 | Unspecified |
| mmu-miR-24-3p | MIMAT0000219 | Slc22a5 | 20520 | HITS-CLIP | | 25753418 | Unspecified |
| mmu-miR-24-3p | MIMAT0000219 | Slc25a1 | 13358 | HITS-CLIP | | 37986224 | Unspecified |
| mmu-miR-24-3p | MIMAT0000219 | Slc25a12 | 78830 | Sequencing//HITS-CLIP | 19536157\|23142080\|37986224 | | Unspecified |
| mmu-miR-24-3p | MIMAT0000219 | Slc25a13 | 50799 | HITS-CLIP | | 23142080 | Unspecified |
| mmu-miR-24-3p | MIMAT0000219 | Slc25a15 | 18408 | HITS-CLIP | | 23142080 | Unspecified |
| mmu-miR-24-3p | MIMAT0000219 | Slc25a20 | 57279 | HITS-CLIP | | 23142080 | Unspecified |
| mmu-miR-24-3p | MIMAT0000219 | Slc25a25 | 227731 | Sequencing//HITS-CLIP | 37986224\|31433990\|25753418\|2620 | | Unspecified |
| mmu-miR-24-3p | MIMAT0000219 | Slc25a36 | 192287 | HITS-CLIP | | 23142080 | Unspecified |
| mmu-miR-24-3p | MIMAT0000219 | Slc26a10 | 216441 | HITS-CLIP | | 23142080 | Unspecified |
| mmu-miR-24-3p | MIMAT0000219 | Slc2a1 | 20525 | HITS-CLIP | | 23142080 | Unspecified |
| mmu-miR-24-3p | MIMAT0000219 | Slc2a3 | 20527 | HITS-CLIP | 31433990\|23142080 | | Unspecified |
| mmu-miR-24-3p | MIMAT0000219 | Slc2a6 | 227659 | HITS-CLIP | | 23142080 | Unspecified |

| | | | | | | |
|---|---|---|---|---|---|---|
| mmu-miR-24-3p | MIMAT0000219 | Slc30a1 | 22782 | HITS-CLIP | 25753418 | Unspecified |
| mmu-miR-24-3p | MIMAT0000219 | Slc31a1 | 20529 | HITS-CLIP | 23142080 | Unspecified |
| mmu-miR-24-3p | MIMAT0000219 | Slc33a1 | 11416 | HITS-CLIP | 23142080 | Unspecified |
| mmu-miR-24-3p | MIMAT0000219 | Slc35a3 | 229782 | HITS-CLIP | 25753418 | Unspecified |
| mmu-miR-24-3p | MIMAT0000219 | Slc35b1 | 110172 | HITS-CLIP | 23142080 | Unspecified |
| mmu-miR-24-3p | MIMAT0000219 | Slc35b2 | 73836 | HITS-CLIP | 23142080 | Unspecified |
| mmu-miR-24-3p | MIMAT0000219 | Slc35b3 | 108652 | HITS-CLIP | 23142080 | Unspecified |
| mmu-miR-24-3p | MIMAT0000219 | Slc35e2 | 320541 | Sequencing//HITS-CLIP | 19536157\|23142080 | Unspecified |
| mmu-miR-24-3p | MIMAT0000219 | Slc35f2 | 72022 | HITS-CLIP | 23142080 | Unspecified |
| mmu-miR-24-3p | MIMAT0000219 | Slc35f6 | 74919 | HITS-CLIP | 31433990 | Unspecified |
| mmu-miR-24-3p | MIMAT0000219 | Slc38a1 | 105727 | HITS-CLIP | 23142080 | Unspecified |
| mmu-miR-24-3p | MIMAT0000219 | Slc38a10 | 72055 | HITS-CLIP | 25753418 | Unspecified |
| mmu-miR-24-3p | MIMAT0000219 | Slc39a3 | 106947 | HITS-CLIP | 25753418 | Unspecified |
| mmu-miR-24-3p | MIMAT0000219 | Slc39a7 | 14977 | HITS-CLIP | 23142080 | Unspecified |
| mmu-miR-24-3p | MIMAT0000219 | Slc41a1 | 98396 | HITS-CLIP | 37986224\|23142080 | Unspecified |
| mmu-miR-24-3p | MIMAT0000219 | Slc43a2 | 215113 | HITS-CLIP | 25753418 | Unspecified |
| mmu-miR-24-3p | MIMAT0000219 | Slc44a1 | 100434 | HITS-CLIP | 25753418 | Unspecified |
| mmu-miR-24-3p | MIMAT0000219 | Slc44a2 | 68682 | HITS-CLIP | 25753418 | Unspecified |
| mmu-miR-24-3p | MIMAT0000219 | Slc4a3 | 20536 | HITS-CLIP | 25753418 | Unspecified |
| mmu-miR-24-3p | MIMAT0000219 | Slc4a7 | 218756 | HITS-CLIP | 23142080 | Unspecified |
| mmu-miR-24-3p | MIMAT0000219 | Slc52a2 | 52710 | HITS-CLIP | 23142080 | Unspecified |
| mmu-miR-24-3p | MIMAT0000219 | Slc5a3 | 53881 | HITS-CLIP | 23142080 | Unspecified |
| mmu-miR-24-3p | MIMAT0000219 | Slc6a6 | 21366 | HITS-CLIP | 25753418 | Unspecified |
| mmu-miR-24-3p | MIMAT0000219 | Slc7a1 | 11987 | Sequencing//HITS-CLIP | 37986224\|23142080\|19536157 | Unspecified |
| mmu-miR-24-3p | MIMAT0000219 | Slc9a1 | 20544 | HITS-CLIP | 25753418 | Unspecified |
| mmu-miR-24-3p | MIMAT0000219 | Slc9a3r1 | 26941 | HITS-CLIP | 23142080 | Unspecified |
| mmu-miR-24-3p | MIMAT0000219 | Slc9a6 | 236794 | HITS-CLIP | 23142080 | Unspecified |
| mmu-miR-24-3p | MIMAT0000219 | Slco3a1 | 108116 | HITS-CLIP | 23142080 | Unspecified |
| mmu-miR-24-3p | MIMAT0000219 | Slf2 | 226151 | HITS-CLIP | 23142080 | Unspecified |
| mmu-miR-24-3p | MIMAT0000219 | Slfn5 | 327978 | HITS-CLIP | 25753418 | Unspecified |
| mmu-miR-24-3p | MIMAT0000219 | Slit3 | 20564 | HITS-CLIP | 25753418 | Unspecified |
| mmu-miR-24-3p | MIMAT0000219 | Slk | 20874 | HITS-CLIP | 37986224\|23142080 | Unspecified |
| mmu-miR-24-3p | MIMAT0000219 | Slmap | 83997 | HITS-CLIP | 23142080 | Unspecified |
| mmu-miR-24-3p | MIMAT0000219 | Smad2 | 17126 | HITS-CLIP | 23142080 | Unspecified |
| mmu-miR-24-3p | MIMAT0000219 | Smad3 | 17127 | Sequencing//HITS-CLIP//In situ hybridizati | 19536157\|23142080\|19582816 | Unspecified |
| mmu-miR-24-3p | MIMAT0000219 | Smad4 | 17128 | In situ hybridization//Luciferase assay//Mi | 19582816\|23142080\|25519956 | Unspecified |
| mmu-miR-24-3p | MIMAT0000219 | Smarca5 | 93762 | HITS-CLIP | 23142080 | Unspecified |
| mmu-miR-24-3p | MIMAT0000219 | Smarcad1 | 13990 | HITS-CLIP | 23142080 | Unspecified |
| mmu-miR-24-3p | MIMAT0000219 | Smarcc1 | 20588 | HITS-CLIP | 23142080\|25753418 | Unspecified |

| | | | | | | | |
|---|---|---|---|---|---|---|---|
| mmu-miR-24-3p | MIMAT0000219 | Smarcc2 | 68094 | HITS-CLIP | | 23142080 | Unspecified |
| mmu-miR-24-3p | MIMAT0000219 | Smarcd2 | 83796 | HITS-CLIP | | 37986224 | Unspecified |
| mmu-miR-24-3p | MIMAT0000219 | Smc2 | 14211 | HITS-CLIP | | 23142080 | Unspecified |
| mmu-miR-24-3p | MIMAT0000219 | Smc3 | 13006 | HITS-CLIP | | 23142080 | Unspecified |
| mmu-miR-24-3p | MIMAT0000219 | Smc5 | 226026 | HITS-CLIP | | 23142080 | Unspecified |
| mmu-miR-24-3p | MIMAT0000219 | Smc6 | 67241 | HITS-CLIP | | 31433990 | Unspecified |
| mmu-miR-24-3p | MIMAT0000219 | Smchd1 | 74355 | HITS-CLIP | 37986224\|23142080 | | Unspecified |
| mmu-miR-24-3p | MIMAT0000219 | Smcr8 | 237782 | HITS-CLIP | 37986224\|23142080 | | Unspecified |
| mmu-miR-24-3p | MIMAT0000219 | Smg1 | 233789 | HITS-CLIP | | 23142080 | Unspecified |
| mmu-miR-24-3p | MIMAT0000219 | Smg7 | 226517 | HITS-CLIP | | 23142080 | Unspecified |
| mmu-miR-24-3p | MIMAT0000219 | Smg8 | 74133 | HITS-CLIP | | 23142080 | Unspecified |
| mmu-miR-24-3p | MIMAT0000219 | Smim14 | 68552 | HITS-CLIP | | 23142080 | Unspecified |
| mmu-miR-24-3p | MIMAT0000219 | Smim15 | 75616 | HITS-CLIP | | 25753418 | Unspecified |
| mmu-miR-24-3p | MIMAT0000219 | Smndc1 | 76479 | HITS-CLIP | | 23142080 | Unspecified |
| mmu-miR-24-3p | MIMAT0000219 | Smpd1 | 20597 | HITS-CLIP | 25753418\|23142080 | | Unspecified |
| mmu-miR-24-3p | MIMAT0000219 | Smug1 | 71726 | HITS-CLIP | | 23142080 | Unspecified |
| mmu-miR-24-3p | MIMAT0000219 | Smurf2 | 66313 | HITS-CLIP | 31433990\|23142080 | | Unspecified |
| mmu-miR-24-3p | MIMAT0000219 | Smyd2 | 226830 | HITS-CLIP | | 23142080 | Unspecified |
| mmu-miR-24-3p | MIMAT0000219 | Smyd3 | 69726 | HITS-CLIP | | 23142080 | Unspecified |
| mmu-miR-24-3p | MIMAT0000219 | Smyd5 | 232187 | HITS-CLIP | | 23142080 | Unspecified |
| mmu-miR-24-3p | MIMAT0000219 | Snap29 | 67474 | HITS-CLIP | 37986224\|23142080 | | Unspecified |
| mmu-miR-24-3p | MIMAT0000219 | Snapc5 | 330959 | HITS-CLIP | | 25753418 | Unspecified |
| mmu-miR-24-3p | MIMAT0000219 | Snip1 | 76793 | HITS-CLIP | | 23142080 | Unspecified |
| mmu-miR-24-3p | MIMAT0000219 | Snn | 20621 | HITS-CLIP | | 23142080 | Unspecified |
| mmu-miR-24-3p | MIMAT0000219 | Snrk | 20623 | HITS-CLIP | | 25753418 | Unspecified |
| mmu-miR-24-3p | MIMAT0000219 | Snrnp200 | 320632 | HITS-CLIP | 23142080\|25753418 | | Unspecified |
| mmu-miR-24-3p | MIMAT0000219 | Snrnp40 | 66585 | HITS-CLIP | | 23142080 | Unspecified |
| mmu-miR-24-3p | MIMAT0000219 | Snrpb | 20638 | HITS-CLIP | | 23142080 | Unspecified |
| mmu-miR-24-3p | MIMAT0000219 | Snx12 | 55988 | HITS-CLIP | | 23142080 | Unspecified |
| mmu-miR-24-3p | MIMAT0000219 | Snx13 | 217463 | HITS-CLIP | 37986224\|23142080 | | Unspecified |
| mmu-miR-24-3p | MIMAT0000219 | Snx18 | 170625 | HITS-CLIP | | 23142080 | Unspecified |
| mmu-miR-24-3p | MIMAT0000219 | Snx27 | 76742 | Sequencing//HITS-CLIP | 19536157\|23142080\|37986224 | | Unspecified |
| mmu-miR-24-3p | MIMAT0000219 | Snx30 | 209131 | HITS-CLIP | | 25753418 | Unspecified |
| mmu-miR-24-3p | MIMAT0000219 | Snx5 | 69178 | HITS-CLIP | | 23142080 | Unspecified |
| mmu-miR-24-3p | MIMAT0000219 | Snx6 | 72183 | HITS-CLIP | | 23142080 | Unspecified |
| mmu-miR-24-3p | MIMAT0000219 | Snx7 | 76561 | HITS-CLIP | | 25753418 | Unspecified |
| mmu-miR-24-3p | MIMAT0000219 | Snx9 | 66616 | HITS-CLIP | | 23142080 | Unspecified |
| mmu-miR-24-3p | MIMAT0000219 | Soat1 | 20652 | Sequencing//HITS-CLIP | 37986224\|23142080\|19536157 | | Unspecified |
| mmu-miR-24-3p | MIMAT0000219 | Socs1 | 12703 | HITS-CLIP | | 23142080 | Unspecified |

| | | | | | | | |
|---|---|---|---|---|---|---|---|
| mmu-miR-24-3p | MIMAT0000219 | Socs2 | 216233 | HITS-CLIP | | 23142080 | Unspecified |
| mmu-miR-24-3p | MIMAT0000219 | Socs3 | 12702 | HITS-CLIP | 37986224\|23142080 | | Unspecified |
| mmu-miR-24-3p | MIMAT0000219 | Socs4 | 67296 | HITS-CLIP | | 23142080 | Unspecified |
| mmu-miR-24-3p | MIMAT0000219 | Socs5 | 56468 | HITS-CLIP | | 23142080 | Unspecified |
| mmu-miR-24-3p | MIMAT0000219 | Socs6 | 54607 | HITS-CLIP | | 23142080 | Unspecified |
| mmu-miR-24-3p | MIMAT0000219 | Socs7 | 192157 | HITS-CLIP | | 23142080 | Unspecified |
| mmu-miR-24-3p | MIMAT0000219 | Soga1 | 320706 | HITS-CLIP | | 25753418 | Unspecified |
| mmu-miR-24-3p | MIMAT0000219 | Son | 20658 | HITS-CLIP | 37986224\|23142080\|25753418 | | Unspecified |
| mmu-miR-24-3p | MIMAT0000219 | Sorbs1 | 20411 | HITS-CLIP | | 25753418 | Unspecified |
| mmu-miR-24-3p | MIMAT0000219 | Sorbs2 | 234214 | HITS-CLIP | | 25753418 | Unspecified |
| mmu-miR-24-3p | MIMAT0000219 | Sos1 | 20662 | HITS-CLIP | 23142080\|25753418 | | Unspecified |
| mmu-miR-24-3p | MIMAT0000219 | Sos2 | 20663 | HITS-CLIP | | 23142080 | Unspecified |
| mmu-miR-24-3p | MIMAT0000219 | Sp1 | 20683 | Sequencing//HITS-CLIP | 37986224\|23142080\|25753418\|1953( | | Unspecified |
| mmu-miR-24-3p | MIMAT0000219 | Sp4 | 20688 | HITS-CLIP | 23142080\|25753418 | | Unspecified |
| mmu-miR-24-3p | MIMAT0000219 | Spag9 | 70834 | HITS-CLIP | 25753418\|23142080 | | Unspecified |
| mmu-miR-24-3p | MIMAT0000219 | Spast | 50850 | HITS-CLIP | | 25753418 | Unspecified |
| mmu-miR-24-3p | MIMAT0000219 | Spcs3 | 76687 | HITS-CLIP | | 23142080 | Unspecified |
| mmu-miR-24-3p | MIMAT0000219 | Speg | 11790 | HITS-CLIP | | 25753418 | Unspecified |
| mmu-miR-24-3p | MIMAT0000219 | Spg20 | 229285 | HITS-CLIP | | 23142080 | Unspecified |
| mmu-miR-24-3p | MIMAT0000219 | Spg21 | 27965 | HITS-CLIP | | 23142080 | Unspecified |
| mmu-miR-24-3p | MIMAT0000219 | Spg7 | 234847 | HITS-CLIP | | 25753418 | Unspecified |
| mmu-miR-24-3p | MIMAT0000219 | Spindoc | 144097 | HITS-CLIP | | 25753418 | Unspecified |
| mmu-miR-24-3p | MIMAT0000219 | Spint2 | 20733 | HITS-CLIP | | 23142080 | Unspecified |
| mmu-miR-24-3p | MIMAT0000219 | Spop | 20747 | HITS-CLIP | | 25753418 | Unspecified |
| mmu-miR-24-3p | MIMAT0000219 | Sppl2a | 66552 | HITS-CLIP | 25753418\|23142080 | | Unspecified |
| mmu-miR-24-3p | MIMAT0000219 | Spred1 | 114715 | HITS-CLIP | | 23142080 | Unspecified |
| mmu-miR-24-3p | MIMAT0000219 | Spred2 | 114716 | HITS-CLIP | | 23142080 | Unspecified |
| mmu-miR-24-3p | MIMAT0000219 | Spryd7 | 66674 | HITS-CLIP | | 25753418 | Unspecified |
| mmu-miR-24-3p | MIMAT0000219 | Spsb1 | 74646 | HITS-CLIP | | 25753418 | Unspecified |
| mmu-miR-24-3p | MIMAT0000219 | Sptan1 | 20740 | HITS-CLIP | 37986224\|25753418\|23142080 | | Unspecified |
| mmu-miR-24-3p | MIMAT0000219 | Sptbn1 | 20742 | HITS-CLIP | 23142080\|25753418 | | Unspecified |
| mmu-miR-24-3p | MIMAT0000219 | Sptlc1 | 268656 | HITS-CLIP | | 23142080 | Unspecified |
| mmu-miR-24-3p | MIMAT0000219 | Sptlc2 | 20773 | HITS-CLIP | | 23142080 | Unspecified |
| mmu-miR-24-3p | MIMAT0000219 | Sptssa | 104725 | HITS-CLIP | | 25753418 | Unspecified |
| mmu-miR-24-3p | MIMAT0000219 | Sqle | 20775 | HITS-CLIP | 37986224\|23142080 | | Unspecified |
| mmu-miR-24-3p | MIMAT0000219 | Sra1 | 24068 | HITS-CLIP | | 25753418 | Unspecified |
| mmu-miR-24-3p | MIMAT0000219 | Srd5a1 | 78925 | HITS-CLIP | | 23142080 | Unspecified |
| mmu-miR-24-3p | MIMAT0000219 | Srebf1 | 20787 | HITS-CLIP | | 23142080 | Unspecified |
| mmu-miR-24-3p | MIMAT0000219 | Srebf2 | 20788 | HITS-CLIP | | 23142080 | Unspecified |

| | | | | | | |
|---|---|---|---|---|---|---|
| mmu-miR-24-3p | MIMAT0000219 | Srek1 | 218543 | HITS-CLIP | 23142080 | Unspecified |
| mmu-miR-24-3p | MIMAT0000219 | Srgap2 | 14270 | Sequencing//qCLASH//HITS-CLIP | 37986224\|31393953\|23142080\|19536 | Unspecified |
| mmu-miR-24-3p | MIMAT0000219 | Srgap3 | 259302 | HITS-CLIP | 23142080 | Unspecified |
| mmu-miR-24-3p | MIMAT0000219 | Sri | 109552 | Sequencing//HITS-CLIP | 37986224\|25753418\|23142080\|19536 | Unspecified |
| mmu-miR-24-3p | MIMAT0000219 | Srp14 | 20813 | HITS-CLIP | 25753418 | Unspecified |
| mmu-miR-24-3p | MIMAT0000219 | Srp19 | 66384 | HITS-CLIP | 23142080 | Unspecified |
| mmu-miR-24-3p | MIMAT0000219 | Srpk1 | 20815 | HITS-CLIP | 37986224\|23142080\|25753418 | Unspecified |
| mmu-miR-24-3p | MIMAT0000219 | Srpk2 | 20817 | HITS-CLIP | 23142080 | Unspecified |
| mmu-miR-24-3p | MIMAT0000219 | Srrm2 | 75956 | HITS-CLIP | 37986224\|23142080\|25753418 | Unspecified |
| mmu-miR-24-3p | MIMAT0000219 | Srrt | 83701 | HITS-CLIP | 23142080 | Unspecified |
| mmu-miR-24-3p | MIMAT0000219 | Srsf11 | 69207 | Sequencing//HITS-CLIP | 19536157\|23142080 | Unspecified |
| mmu-miR-24-3p | MIMAT0000219 | Srsf7 | 225027 | HITS-CLIP | 23142080 | Unspecified |
| mmu-miR-24-3p | MIMAT0000219 | Ss18l1 | 269397 | HITS-CLIP | 25753418 | Unspecified |
| mmu-miR-24-3p | MIMAT0000219 | Ssb | 20823 | HITS-CLIP | 23142080 | Unspecified |
| mmu-miR-24-3p | MIMAT0000219 | Ssbp2 | 66970 | HITS-CLIP | 23142080 | Unspecified |
| mmu-miR-24-3p | MIMAT0000219 | Ssh1 | 231637 | HITS-CLIP | 23142080 | Unspecified |
| mmu-miR-24-3p | MIMAT0000219 | Ssh2 | 237860 | HITS-CLIP | 37986224\|23142080 | Unspecified |
| mmu-miR-24-3p | MIMAT0000219 | Sspn | 16651 | HITS-CLIP | 25753418 | Unspecified |
| mmu-miR-24-3p | MIMAT0000219 | Ssr1 | 107513 | HITS-CLIP | 23142080 | Unspecified |
| mmu-miR-24-3p | MIMAT0000219 | Ssr2 | 66256 | HITS-CLIP | 25753418 | Unspecified |
| mmu-miR-24-3p | MIMAT0000219 | Ssr3 | 67437 | HITS-CLIP | 23142080 | Unspecified |
| mmu-miR-24-3p | MIMAT0000219 | Ssu72 | 68991 | HITS-CLIP | 23142080 | Unspecified |
| mmu-miR-24-3p | MIMAT0000219 | St3gal4 | 20443 | HITS-CLIP | 23142080 | Unspecified |
| mmu-miR-24-3p | MIMAT0000219 | St6gal1 | 20440 | HITS-CLIP | 37986224 | Unspecified |
| mmu-miR-24-3p | MIMAT0000219 | St8sia4 | 20452 | HITS-CLIP | 37986224\|23142080 | Unspecified |
| mmu-miR-24-3p | MIMAT0000219 | Stag1 | 20842 | HITS-CLIP | 23142080 | Unspecified |
| mmu-miR-24-3p | MIMAT0000219 | Stag2 | 20843 | HITS-CLIP | 23142080 | Unspecified |
| mmu-miR-24-3p | MIMAT0000219 | Stap1 | 56792 | HITS-CLIP | 23142080 | Unspecified |
| mmu-miR-24-3p | MIMAT0000219 | Stard3 | 59045 | HITS-CLIP | 37986224\|23142080 | Unspecified |
| mmu-miR-24-3p | MIMAT0000219 | Stard3nl | 76205 | HITS-CLIP | 25753418 | Unspecified |
| mmu-miR-24-3p | MIMAT0000219 | Stard4 | 170459 | HITS-CLIP | 23142080 | Unspecified |
| mmu-miR-24-3p | MIMAT0000219 | Stard7 | 99138 | HITS-CLIP | 25753418 | Unspecified |
| mmu-miR-24-3p | MIMAT0000219 | Stat2 | 20847 | HITS-CLIP | 23142080 | Unspecified |
| mmu-miR-24-3p | MIMAT0000219 | Stat3 | 20848 | HITS-CLIP | 23142080 | Unspecified |
| mmu-miR-24-3p | MIMAT0000219 | Stat4 | 20849 | HITS-CLIP | 23142080 | Unspecified |
| mmu-miR-24-3p | MIMAT0000219 | Stat5b | 20851 | HITS-CLIP | 23142080 | Unspecified |
| mmu-miR-24-3p | MIMAT0000219 | Stat6 | 20852 | HITS-CLIP | 23142080 | Unspecified |
| mmu-miR-24-3p | MIMAT0000219 | Stau1 | 20853 | HITS-CLIP | 23142080 | Unspecified |
| mmu-miR-24-3p | MIMAT0000219 | Stc2 | 20856 | HITS-CLIP | 37986224\|23142080 | Unspecified |

| | | | | | | |
|---|---|---|---|---|---|---|
| mmu-miR-24-3p | MIMAT0000219 | Stil | 20460 | HITS-CLIP | 23142080 | Unspecified |
| mmu-miR-24-3p | MIMAT0000219 | Stim2 | 116873 | HITS-CLIP | 23142080 | Unspecified |
| mmu-miR-24-3p | MIMAT0000219 | Stk17b | 98267 | HITS-CLIP | 23142080 | Unspecified |
| mmu-miR-24-3p | MIMAT0000219 | Stk24 | 223255 | HITS-CLIP | 23142080 | Unspecified |
| mmu-miR-24-3p | MIMAT0000219 | Stk26 | 51765 | HITS-CLIP | 23142080 | Unspecified |
| mmu-miR-24-3p | MIMAT0000219 | Stk38 | 106504 | Sequencing//HITS-CLIP | 37986224\|23142080\|19536157 | Unspecified |
| mmu-miR-24-3p | MIMAT0000219 | Stk40 | 74178 | HITS-CLIP | 37986224\|23142080 | Unspecified |
| mmu-miR-24-3p | MIMAT0000219 | Strip2 | 320609 | HITS-CLIP | 25753418 | Unspecified |
| mmu-miR-24-3p | MIMAT0000219 | Strn | 268980 | HITS-CLIP | 23142080\|25753418 | Unspecified |
| mmu-miR-24-3p | MIMAT0000219 | Strn3 | 94186 | qCLASH//HITS-CLIP | 31393953\|23142080 | Unspecified |
| mmu-miR-24-3p | MIMAT0000219 | Stt3a | 16430 | HITS-CLIP | 23142080 | Unspecified |
| mmu-miR-24-3p | MIMAT0000219 | Stt3b | 68292 | HITS-CLIP | 23142080 | Unspecified |
| mmu-miR-24-3p | MIMAT0000219 | Stub1 | 56424 | HITS-CLIP | 25753418 | Unspecified |
| mmu-miR-24-3p | MIMAT0000219 | Stx16 | 228960 | HITS-CLIP | 23142080 | Unspecified |
| mmu-miR-24-3p | MIMAT0000219 | Stx17 | 67727 | HITS-CLIP | 25753418 | Unspecified |
| mmu-miR-24-3p | MIMAT0000219 | Stx18 | 71116 | HITS-CLIP | 23142080 | Unspecified |
| mmu-miR-24-3p | MIMAT0000219 | Stx5a | 56389 | HITS-CLIP | 23142080 | Unspecified |
| mmu-miR-24-3p | MIMAT0000219 | Stxbp3 | 20912 | HITS-CLIP | 23142080 | Unspecified |
| mmu-miR-24-3p | MIMAT0000219 | Sub1 | 20024 | HITS-CLIP | 37986224\|23142080 | Unspecified |
| mmu-miR-24-3p | MIMAT0000219 | Suclg1 | 56451 | HITS-CLIP | 23142080 | Unspecified |
| mmu-miR-24-3p | MIMAT0000219 | Suco | 226551 | HITS-CLIP | 23142080 | Unspecified |
| mmu-miR-24-3p | MIMAT0000219 | Sugp1 | 70616 | HITS-CLIP | 25753418 | Unspecified |
| mmu-miR-24-3p | MIMAT0000219 | Sun1 | 77053 | HITS-CLIP | 23142080 | Unspecified |
| mmu-miR-24-3p | MIMAT0000219 | Supt16 | 114741 | HITS-CLIP | 23142080 | Unspecified |
| mmu-miR-24-3p | MIMAT0000219 | Supv3l1 | 338359 | HITS-CLIP | 25753418 | Unspecified |
| mmu-miR-24-3p | MIMAT0000219 | Surf1 | 20930 | HITS-CLIP | 23142080 | Unspecified |
| mmu-miR-24-3p | MIMAT0000219 | Surf4 | 20932 | HITS-CLIP | 23142080 | Unspecified |
| mmu-miR-24-3p | MIMAT0000219 | Susd1 | 634731 | HITS-CLIP | 23142080 | Unspecified |
| mmu-miR-24-3p | MIMAT0000219 | Susd6 | 9766 | HITS-CLIP | 25753418 | Unspecified |
| mmu-miR-24-3p | MIMAT0000219 | Sv2a | 64051 | HITS-CLIP | 25753418 | Unspecified |
| mmu-miR-24-3p | MIMAT0000219 | Svil | 225115 | HITS-CLIP | 25753418\|23142080 | Unspecified |
| mmu-miR-24-3p | MIMAT0000219 | Swap70 | 20947 | HITS-CLIP | 23142080 | Unspecified |
| mmu-miR-24-3p | MIMAT0000219 | Syap1 | 67043 | HITS-CLIP | 23142080 | Unspecified |
| mmu-miR-24-3p | MIMAT0000219 | Syncrip | 56403 | HITS-CLIP | 23142080 | Unspecified |
| mmu-miR-24-3p | MIMAT0000219 | Synj1 | 104015 | HITS-CLIP | 23142080 | Unspecified |
| mmu-miR-24-3p | MIMAT0000219 | Synj2bp | 24071 | HITS-CLIP | 25753418 | Unspecified |
| mmu-miR-24-3p | MIMAT0000219 | Synrg | 217030 | HITS-CLIP | 23142080 | Unspecified |
| mmu-miR-24-3p | MIMAT0000219 | Sypl | 19027 | HITS-CLIP | 23142080 | Unspecified |
| mmu-miR-24-3p | MIMAT0000219 | Sys1 | 66460 | HITS-CLIP | 23142080 | Unspecified |

| | | | | | | |
|---|---|---|---|---|---|---|
| mmu-miR-24-3p | MIMAT0000219 | Syvn1 | 74126 | HITS-CLIP | 25753418 | Unspecified |
| mmu-miR-24-3p | MIMAT0000219 | Szrd1 | 213491 | HITS-CLIP | 23142080 | Unspecified |
| mmu-miR-24-3p | MIMAT0000219 | Tab2 | 68652 | HITS-CLIP | 23142080\|25753418 | Unspecified |
| mmu-miR-24-3p | MIMAT0000219 | Tacc2 | 57752 | HITS-CLIP | 23142080 | Unspecified |
| mmu-miR-24-3p | MIMAT0000219 | Taf15 | 70439 | HITS-CLIP | 37986224 | Unspecified |
| mmu-miR-24-3p | MIMAT0000219 | Taf1d | 75316 | qCLASH | 31393953 | Unspecified |
| mmu-miR-24-3p | MIMAT0000219 | Taf4 | 6874 | HITS-CLIP | 23142080 | Unspecified |
| mmu-miR-24-3p | MIMAT0000219 | Taf5 | 226182 | HITS-CLIP | 37986224\|23142080 | Unspecified |
| mmu-miR-24-3p | MIMAT0000219 | Taf6 | 21343 | HITS-CLIP | 25753418 | Unspecified |
| mmu-miR-24-3p | MIMAT0000219 | Taf8 | 63856 | HITS-CLIP | 23142080 | Unspecified |
| mmu-miR-24-3p | MIMAT0000219 | Tagln2 | 21346 | HITS-CLIP | 23142080 | Unspecified |
| mmu-miR-24-3p | MIMAT0000219 | Taldo1 | 21351 | HITS-CLIP | 25753418 | Unspecified |
| mmu-miR-24-3p | MIMAT0000219 | Taok1 | 216965 | HITS-CLIP | 37986224\|23142080 | Unspecified |
| mmu-miR-24-3p | MIMAT0000219 | Tap1 | 21354 | HITS-CLIP | 23142080 | Unspecified |
| mmu-miR-24-3p | MIMAT0000219 | Tap2 | 21355 | HITS-CLIP | 23142080 | Unspecified |
| mmu-miR-24-3p | MIMAT0000219 | Tapbp | 21356 | HITS-CLIP | 25753418 | Unspecified |
| mmu-miR-24-3p | MIMAT0000219 | Tapt1 | 231225 | HITS-CLIP | 23142080 | Unspecified |
| mmu-miR-24-3p | MIMAT0000219 | Tarbp1 | 6894 | HITS-CLIP | 23142080 | Unspecified |
| mmu-miR-24-3p | MIMAT0000219 | Tatdn2 | 381801 | HITS-CLIP | 25753418 | Unspecified |
| mmu-miR-24-3p | MIMAT0000219 | Tax1bp1 | 52440 | HITS-CLIP | 23142080\|25753418 | Unspecified |
| mmu-miR-24-3p | MIMAT0000219 | Tbc1d10a | 103724 | HITS-CLIP | 23142080 | Unspecified |
| mmu-miR-24-3p | MIMAT0000219 | Tbc1d14 | 100855 | HITS-CLIP | 23142080 | Unspecified |
| mmu-miR-24-3p | MIMAT0000219 | Tbc1d15 | 66687 | HITS-CLIP | 23142080 | Unspecified |
| mmu-miR-24-3p | MIMAT0000219 | Tbc1d17 | 233204 | HITS-CLIP | 23142080 | Unspecified |
| mmu-miR-24-3p | MIMAT0000219 | Tbc1d30 | 74694 | HITS-CLIP | 23142080 | Unspecified |
| mmu-miR-24-3p | MIMAT0000219 | Tbc1d4 | 210789 | HITS-CLIP | 23142080 | Unspecified |
| mmu-miR-24-3p | MIMAT0000219 | Tbc1d5 | 72238 | HITS-CLIP | 23142080 | Unspecified |
| mmu-miR-24-3p | MIMAT0000219 | Tbcd | 108903 | HITS-CLIP | 25753418 | Unspecified |
| mmu-miR-24-3p | MIMAT0000219 | Tbk1 | 56480 | HITS-CLIP | 23142080 | Unspecified |
| mmu-miR-24-3p | MIMAT0000219 | Tbkbp1 | 73174 | HITS-CLIP | 23142080 | Unspecified |
| mmu-miR-24-3p | MIMAT0000219 | Tbl1xr1 | 81004 | HITS-CLIP | 37986224\|23142080 | Unspecified |
| mmu-miR-24-3p | MIMAT0000219 | Tbpl1 | 237336 | HITS-CLIP | 25753418 | Unspecified |
| mmu-miR-24-3p | MIMAT0000219 | Tbrg4 | 21379 | HITS-CLIP | 25753418 | Unspecified |
| mmu-miR-24-3p | MIMAT0000219 | Tceanc2 | 66526 | HITS-CLIP | 23142080 | Unspecified |
| mmu-miR-24-3p | MIMAT0000219 | Tcerg1 | 56070 | HITS-CLIP | 23142080 | Unspecified |
| mmu-miR-24-3p | MIMAT0000219 | Tcf25 | 66855 | HITS-CLIP | 25753418 | Unspecified |
| mmu-miR-24-3p | MIMAT0000219 | Tcf4 | 21413 | HITS-CLIP | 25753418 | Unspecified |
| mmu-miR-24-3p | MIMAT0000219 | Tcf7 | 21414 | HITS-CLIP//Luciferase reporter assay | 37986224\|23142080\|26006243 | Unspecified |
| mmu-miR-24-3p | MIMAT0000219 | Tcp1 | 21454 | HITS-CLIP | 23142080 | Unspecified |

| | | | | | | |
|---|---|---|---|---|---|---|
| mmu-miR-24-3p | MIMAT0000219 | Tcp11l2 | 216198 | HITS-CLIP | 23142080\|25753418 | Unspecified |
| mmu-miR-24-3p | MIMAT0000219 | Tcta | 102791 | HITS-CLIP | 25753418 | Unspecified |
| mmu-miR-24-3p | MIMAT0000219 | Tdrd7 | 100121 | HITS-CLIP | 25753418 | Unspecified |
| mmu-miR-24-3p | MIMAT0000219 | Tead1 | 21676 | HITS-CLIP | 37986224 | Unspecified |
| mmu-miR-24-3p | MIMAT0000219 | Tecpr2 | 104859 | HITS-CLIP | 23142080 | Unspecified |
| mmu-miR-24-3p | MIMAT0000219 | Tent2 | 167153 | HITS-CLIP | 23142080 | Unspecified |
| mmu-miR-24-3p | MIMAT0000219 | Tent5a | 55603 | HITS-CLIP | 23142080 | Unspecified |
| mmu-miR-24-3p | MIMAT0000219 | Terf2 | 21750 | HITS-CLIP | 23142080 | Unspecified |
| mmu-miR-24-3p | MIMAT0000219 | Tes | 21753 | HITS-CLIP | 37986224 | Unspecified |
| mmu-miR-24-3p | MIMAT0000219 | Tet2 | 214133 | HITS-CLIP | 23142080 | Unspecified |
| mmu-miR-24-3p | MIMAT0000219 | Tet3 | 194388 | HITS-CLIP | 23142080 | Unspecified |
| mmu-miR-24-3p | MIMAT0000219 | Tex2 | 21763 | HITS-CLIP | 25753418 | Unspecified |
| mmu-miR-24-3p | MIMAT0000219 | Tex261 | 21766 | HITS-CLIP | 23142080 | Unspecified |
| mmu-miR-24-3p | MIMAT0000219 | Tex264 | 21767 | HITS-CLIP | 23142080 | Unspecified |
| mmu-miR-24-3p | MIMAT0000219 | Tex30 | 75623 | HITS-CLIP | 23142080 | Unspecified |
| mmu-miR-24-3p | MIMAT0000219 | Tfam | 21780 | HITS-CLIP | 23142080 | Unspecified |
| mmu-miR-24-3p | MIMAT0000219 | Tfdp2 | 211586 | HITS-CLIP | 25753418 | Unspecified |
| mmu-miR-24-3p | MIMAT0000219 | Tfg | 21787 | HITS-CLIP | 23142080 | Unspecified |
| mmu-miR-24-3p | MIMAT0000219 | Tfrc | 22042 | HITS-CLIP | 31433990\|23142080\|25753418 | Unspecified |
| mmu-miR-24-3p | MIMAT0000219 | Tgfbr2 | 21813 | HITS-CLIP | 23142080 | Unspecified |
| mmu-miR-24-3p | MIMAT0000219 | Tgfbr3 | 21814 | HITS-CLIP | 37986224\|23142080 | Unspecified |
| mmu-miR-24-3p | MIMAT0000219 | Tgm2 | 21817 | HITS-CLIP | 23142080 | Unspecified |
| mmu-miR-24-3p | MIMAT0000219 | Tgoln1 | 22134 | HITS-CLIP | 25753418\|23142080 | Unspecified |
| mmu-miR-24-3p | MIMAT0000219 | Tgs1 | 116940 | HITS-CLIP | 25753418\|23142080 | Unspecified |
| mmu-miR-24-3p | MIMAT0000219 | Thada | 240174 | HITS-CLIP | 23142080 | Unspecified |
| mmu-miR-24-3p | MIMAT0000219 | Thoc2 | 331401 | HITS-CLIP | 23142080 | Unspecified |
| mmu-miR-24-3p | MIMAT0000219 | Thoc5 | 8563 | HITS-CLIP | 23142080 | Unspecified |
| mmu-miR-24-3p | MIMAT0000219 | Thoc7 | 66231 | HITS-CLIP | 25753418 | Unspecified |
| mmu-miR-24-3p | MIMAT0000219 | Thsd7a | 330267 | HITS-CLIP | 25753418 | Unspecified |
| mmu-miR-24-3p | MIMAT0000219 | Tial1 | 21843 | HITS-CLIP | 23142080 | Unspecified |
| mmu-miR-24-3p | MIMAT0000219 | Tiam1 | 21844 | HITS-CLIP | 23142080 | Unspecified |
| mmu-miR-24-3p | MIMAT0000219 | Tigar | 319801 | HITS-CLIP | 25753418 | Unspecified |
| mmu-miR-24-3p | MIMAT0000219 | Tigit | 100043314 | HITS-CLIP | 37986224\|23142080 | Unspecified |
| mmu-miR-24-3p | MIMAT0000219 | Timm44 | 21856 | HITS-CLIP | 25753418 | Unspecified |
| mmu-miR-24-3p | MIMAT0000219 | Timp2 | 21858 | HITS-CLIP | 25753418 | Unspecified |
| mmu-miR-24-3p | MIMAT0000219 | Tiparp | 99929 | HITS-CLIP | 37986224\|23142080 | Unspecified |
| mmu-miR-24-3p | MIMAT0000219 | Tk1 | 21877 | HITS-CLIP | 37986224 | Unspecified |
| mmu-miR-24-3p | MIMAT0000219 | Tkt | 21881 | HITS-CLIP | 23142080 | Unspecified |
| mmu-miR-24-3p | MIMAT0000219 | Tle3 | 21887 | HITS-CLIP | 23142080 | Unspecified |

| | | | | | | |
|---|---|---|---|---|---|---|
| mmu-miR-24-3p | MIMAT0000219 | Tlk1 | 228012 | HITS-CLIP | 23142080 | Unspecified |
| mmu-miR-24-3p | MIMAT0000219 | Tlk2 | 24086 | HITS-CLIP | 31433990 | Unspecified |
| mmu-miR-24-3p | MIMAT0000219 | Tln1 | 21894 | HITS-CLIP | 37986224\|25753418\|23142080 | Unspecified |
| mmu-miR-24-3p | MIMAT0000219 | Tln2 | 70549 | HITS-CLIP | 25753418 | Unspecified |
| mmu-miR-24-3p | MIMAT0000219 | Tlr6 | 21899 | HITS-CLIP | 23142080 | Unspecified |
| mmu-miR-24-3p | MIMAT0000219 | Tm2d2 | 69742 | HITS-CLIP | 25753418 | Unspecified |
| mmu-miR-24-3p | MIMAT0000219 | Tm9sf1 | 74140 | HITS-CLIP | 23142080 | Unspecified |
| mmu-miR-24-3p | MIMAT0000219 | Tm9sf2 | 68059 | HITS-CLIP | 25753418 | Unspecified |
| mmu-miR-24-3p | MIMAT0000219 | Tm9sf3 | 107358 | HITS-CLIP | 23142080 | Unspecified |
| mmu-miR-24-3p | MIMAT0000219 | Tmbim6 | 110213 | HITS-CLIP | 23142080 | Unspecified |
| mmu-miR-24-3p | MIMAT0000219 | Tmc6 | 217353 | HITS-CLIP | 23142080 | Unspecified |
| mmu-miR-24-3p | MIMAT0000219 | Tmco1 | 68944 | HITS-CLIP | 37986224\|23142080 | Unspecified |
| mmu-miR-24-3p | MIMAT0000219 | Tmed5 | 73130 | HITS-CLIP | 23142080 | Unspecified |
| mmu-miR-24-3p | MIMAT0000219 | Tmem11 | 216821 | HITS-CLIP | 37986224\|23142080 | Unspecified |
| mmu-miR-24-3p | MIMAT0000219 | Tmem123 | 71929 | HITS-CLIP | 23142080 | Unspecified |
| mmu-miR-24-3p | MIMAT0000219 | Tmem127 | 69470 | HITS-CLIP | 23142080 | Unspecified |
| mmu-miR-24-3p | MIMAT0000219 | Tmem131 | 56030 | HITS-CLIP | 25753418 | Unspecified |
| mmu-miR-24-3p | MIMAT0000219 | Tmem131l | 23240 | HITS-CLIP | 23142080 | Unspecified |
| mmu-miR-24-3p | MIMAT0000219 | Tmem135 | 72759 | Sequencing//HITS-CLIP | 19536157\|23142080\|37986224 | Unspecified |
| mmu-miR-24-3p | MIMAT0000219 | Tmem143 | 70209 | HITS-CLIP | 25753418 | Unspecified |
| mmu-miR-24-3p | MIMAT0000219 | Tmem147 | 69804 | HITS-CLIP | 23142080 | Unspecified |
| mmu-miR-24-3p | MIMAT0000219 | Tmem161b | 72745 | HITS-CLIP | 23142080 | Unspecified |
| mmu-miR-24-3p | MIMAT0000219 | Tmem164 | 209497 | HITS-CLIP | 23142080 | Unspecified |
| mmu-miR-24-3p | MIMAT0000219 | Tmem165 | 21982 | HITS-CLIP | 23142080 | Unspecified |
| mmu-miR-24-3p | MIMAT0000219 | Tmem167 | 66074 | HITS-CLIP | 23142080 | Unspecified |
| mmu-miR-24-3p | MIMAT0000219 | Tmem170b | 621976 | HITS-CLIP | 23142080 | Unspecified |
| mmu-miR-24-3p | MIMAT0000219 | Tmem173 | 72512 | HITS-CLIP | 23142080 | Unspecified |
| mmu-miR-24-3p | MIMAT0000219 | Tmem175 | 72392 | HITS-CLIP | 23142080 | Unspecified |
| mmu-miR-24-3p | MIMAT0000219 | Tmem184b | 223693 | HITS-CLIP | 23142080 | Unspecified |
| mmu-miR-24-3p | MIMAT0000219 | Tmem186 | 66690 | HITS-CLIP | 23142080 | Unspecified |
| mmu-miR-24-3p | MIMAT0000219 | Tmem189 | 407243 | HITS-CLIP | 23142080 | Unspecified |
| mmu-miR-24-3p | MIMAT0000219 | Tmem19 | 67226 | HITS-CLIP | 23142080 | Unspecified |
| mmu-miR-24-3p | MIMAT0000219 | Tmem230 | 70612 | HITS-CLIP | 25753418 | Unspecified |
| mmu-miR-24-3p | MIMAT0000219 | Tmem237 | 381259 | HITS-CLIP | 23142080 | Unspecified |
| mmu-miR-24-3p | MIMAT0000219 | Tmem245 | 242474 | HITS-CLIP | 23142080\|25753418 | Unspecified |
| mmu-miR-24-3p | MIMAT0000219 | Tmem248 | 71667 | HITS-CLIP | 25753418 | Unspecified |
| mmu-miR-24-3p | MIMAT0000219 | Tmem250-ps | 109299 | HITS-CLIP | 25753418 | Unspecified |
| mmu-miR-24-3p | MIMAT0000219 | Tmem268 | 203197 | HITS-CLIP | 23142080 | Unspecified |
| mmu-miR-24-3p | MIMAT0000219 | Tmem30a | 69981 | qCLASH | 31393953 | Unspecified |

| | | | | | | |
|---|---|---|---|---|---|---|
| mmu-miR-24-3p | MIMAT0000219 | Tmem62 | 96957 | HITS-CLIP | 23142080 | Unspecified |
| mmu-miR-24-3p | MIMAT0000219 | Tmem63a | 208795 | HITS-CLIP | 23142080 | Unspecified |
| mmu-miR-24-3p | MIMAT0000219 | Tmem65 | 74868 | HITS-CLIP | 37986224\|25753418\|23142080 | Unspecified |
| mmu-miR-24-3p | MIMAT0000219 | Tmem70 | 70397 | HITS-CLIP | 25753418 | Unspecified |
| mmu-miR-24-3p | MIMAT0000219 | Tmem86a | 67893 | HITS-CLIP | 25753418 | Unspecified |
| mmu-miR-24-3p | MIMAT0000219 | Tmf1 | 232286 | HITS-CLIP | 25753418 | Unspecified |
| mmu-miR-24-3p | MIMAT0000219 | Tmpo | 21917 | HITS-CLIP | 23142080 | Unspecified |
| mmu-miR-24-3p | MIMAT0000219 | Tmsb10 | 19240 | HITS-CLIP | 37986224 | Unspecified |
| mmu-miR-24-3p | MIMAT0000219 | Tmx4 | 52837 | HITS-CLIP | 37986224\|23142080 | Unspecified |
| mmu-miR-24-3p | MIMAT0000219 | Tnfaip1 | 21927 | HITS-CLIP | 23142080 | Unspecified |
| mmu-miR-24-3p | MIMAT0000219 | Tnfrsf1b | 21938 | HITS-CLIP | 37986224\|23142080 | Unspecified |
| mmu-miR-24-3p | MIMAT0000219 | Tnfrsf26 | 244237 | HITS-CLIP | 37986224 | Unspecified |
| mmu-miR-24-3p | MIMAT0000219 | Tnfrsf8 | 21941 | HITS-CLIP | 23142080 | Unspecified |
| mmu-miR-24-3p | MIMAT0000219 | Tnfrsf9 | 21942 | HITS-CLIP | 37986224 | Unspecified |
| mmu-miR-24-3p | MIMAT0000219 | Tnfsf8 | 21949 | HITS-CLIP | 23142080 | Unspecified |
| mmu-miR-24-3p | MIMAT0000219 | Tnik | 665113 | HITS-CLIP | 23142080 | Unspecified |
| mmu-miR-24-3p | MIMAT0000219 | Tnks | 21951 | Sequencing//HITS-CLIP | 19536157\|25753418\|23142080\|3798 | Unspecified |
| mmu-miR-24-3p | MIMAT0000219 | Tnpo1 | 238799 | HITS-CLIP | 23142080 | Unspecified |
| mmu-miR-24-3p | MIMAT0000219 | Tnpo2 | 212999 | HITS-CLIP | 23142080\|25753418 | Unspecified |
| mmu-miR-24-3p | MIMAT0000219 | Tnpo3 | 320938 | HITS-CLIP | 25753418 | Unspecified |
| mmu-miR-24-3p | MIMAT0000219 | Tnrc6a | 233833 | HITS-CLIP | 37986224\|23142080 | Unspecified |
| mmu-miR-24-3p | MIMAT0000219 | Tnrc6b | 213988 | HITS-CLIP | 25753418\|23142080 | Unspecified |
| mmu-miR-24-3p | MIMAT0000219 | Tns1 | 21961 | HITS-CLIP | 25753418 | Unspecified |
| mmu-miR-24-3p | MIMAT0000219 | Tob1 | 22057 | HITS-CLIP | 23142080 | Unspecified |
| mmu-miR-24-3p | MIMAT0000219 | Tob2 | 57259 | HITS-CLIP | 23142080 | Unspecified |
| mmu-miR-24-3p | MIMAT0000219 | Togaram1 | 23116 | HITS-CLIP | 23142080 | Unspecified |
| mmu-miR-24-3p | MIMAT0000219 | Tollip | 54473 | HITS-CLIP | 25753418 | Unspecified |
| mmu-miR-24-3p | MIMAT0000219 | Tomm40 | 53333 | HITS-CLIP | 23142080 | Unspecified |
| mmu-miR-24-3p | MIMAT0000219 | Tomm7 | 66169 | HITS-CLIP | 23142080 | Unspecified |
| mmu-miR-24-3p | MIMAT0000219 | Top1 | 21969 | HITS-CLIP | 23142080 | Unspecified |
| mmu-miR-24-3p | MIMAT0000219 | Top2a | 21973 | HITS-CLIP | 23142080 | Unspecified |
| mmu-miR-24-3p | MIMAT0000219 | Tor1aip2 | 240832 | HITS-CLIP | 23142080 | Unspecified |
| mmu-miR-24-3p | MIMAT0000219 | Tpcn1 | 252972 | HITS-CLIP | 23142080 | Unspecified |
| mmu-miR-24-3p | MIMAT0000219 | Tpi1 | 21991 | HITS-CLIP | 37986224\|23142080 | Unspecified |
| mmu-miR-24-3p | MIMAT0000219 | Tpm3 | 59069 | HITS-CLIP | 23142080 | Unspecified |
| mmu-miR-24-3p | MIMAT0000219 | Tpp1 | 12751 | HITS-CLIP | 23142080 | Unspecified |
| mmu-miR-24-3p | MIMAT0000219 | Tpp2 | 22019 | HITS-CLIP | 25753418\|23142080 | Unspecified |
| mmu-miR-24-3p | MIMAT0000219 | Tpr | 108989 | HITS-CLIP | 23142080 | Unspecified |
| mmu-miR-24-3p | MIMAT0000219 | Tpx2 | 72119 | HITS-CLIP | 23142080 | Unspecified |

| | | | | | | |
|---|---|---|---|---|---|---|
| mmu-miR-24-3p | MIMAT0000219 | Traf1 | 22029 | HITS-CLIP | 23142080 | Unspecified |
| mmu-miR-24-3p | MIMAT0000219 | Trak2 | 70827 | HITS-CLIP | 23142080 | Unspecified |
| mmu-miR-24-3p | MIMAT0000219 | Tram2 | 170829 | HITS-CLIP | 23142080 | Unspecified |
| mmu-miR-24-3p | MIMAT0000219 | Trap1 | 68015 | HITS-CLIP | 25753418 | Unspecified |
| mmu-miR-24-3p | MIMAT0000219 | Trappc11 | 320714 | HITS-CLIP | 23142080 | Unspecified |
| mmu-miR-24-3p | MIMAT0000219 | Trappc13 | 66975 | HITS-CLIP | 23142080 | Unspecified |
| mmu-miR-24-3p | MIMAT0000219 | Trappc3 | 27096 | HITS-CLIP | 23142080 | Unspecified |
| mmu-miR-24-3p | MIMAT0000219 | Trappc4 | 60409 | Sequencing//HITS-CLIP | 37986224\|25753418\|19536157 | Unspecified |
| mmu-miR-24-3p | MIMAT0000219 | Trappc8 | 75964 | HITS-CLIP | 23142080 | Unspecified |
| mmu-miR-24-3p | MIMAT0000219 | Trib1 | 211770 | HITS-CLIP | 23142080 | Unspecified |
| mmu-miR-24-3p | MIMAT0000219 | Trim23 | 81003 | HITS-CLIP | 25753418 | Unspecified |
| mmu-miR-24-3p | MIMAT0000219 | Trim24 | 21848 | HITS-CLIP | 23142080 | Unspecified |
| mmu-miR-24-3p | MIMAT0000219 | Trim25 | 217069 | HITS-CLIP | 23142080 | Unspecified |
| mmu-miR-24-3p | MIMAT0000219 | Trim28 | 21849 | HITS-CLIP | 23142080 | Unspecified |
| mmu-miR-24-3p | MIMAT0000219 | Trim33 | 94093 | qCLASH//HITS-CLIP | 31393953\|23142080 | Unspecified |
| mmu-miR-24-3p | MIMAT0000219 | Trim44 | 80985 | HITS-CLIP | 23142080 | Unspecified |
| mmu-miR-24-3p | MIMAT0000219 | Trim63 | 433766 | HITS-CLIP | 25753418 | Unspecified |
| mmu-miR-24-3p | MIMAT0000219 | Trim71 | 636931 | HITS-CLIP | 37986224 | Unspecified |
| mmu-miR-24-3p | MIMAT0000219 | Trim8 | 93679 | HITS-CLIP | 25753418 | Unspecified |
| mmu-miR-24-3p | MIMAT0000219 | Triobp | 110253 | HITS-CLIP | 23142080 | Unspecified |
| mmu-miR-24-3p | MIMAT0000219 | Trip11 | 109181 | HITS-CLIP | 25753418 | Unspecified |
| mmu-miR-24-3p | MIMAT0000219 | Trip12 | 14897 | HITS-CLIP | 23142080 | Unspecified |
| mmu-miR-24-3p | MIMAT0000219 | Trmt10a | 108943 | HITS-CLIP | 21258322\|23142080 | Unspecified |
| mmu-miR-24-3p | MIMAT0000219 | Trmt11 | 73681 | HITS-CLIP | 23142080 | Unspecified |
| mmu-miR-24-3p | MIMAT0000219 | Trmt44 | 78890 | HITS-CLIP | 37986224 | Unspecified |
| mmu-miR-24-3p | MIMAT0000219 | Trmt5 | 76357 | HITS-CLIP | 23142080 | Unspecified |
| mmu-miR-24-3p | MIMAT0000219 | Trmt6 | 66926 | HITS-CLIP | 23142080 | Unspecified |
| mmu-miR-24-3p | MIMAT0000219 | Trp53 | 22059 | HITS-CLIP | 23142080 | Unspecified |
| mmu-miR-24-3p | MIMAT0000219 | Trp53inp1 | 60599 | HITS-CLIP | 37986224\|31433990\|25753418\|23142 | Unspecified |
| mmu-miR-24-3p | MIMAT0000219 | Trpc4ap | 56407 | HITS-CLIP | 23142080 | Unspecified |
| mmu-miR-24-3p | MIMAT0000219 | Trps1 | 83925 | HITS-CLIP | 23142080 | Unspecified |
| mmu-miR-24-3p | MIMAT0000219 | Trrap | 100683 | HITS-CLIP | 25753418\|23142080 | Unspecified |
| mmu-miR-24-3p | MIMAT0000219 | Tsc22d2 | 72033 | HITS-CLIP | 37986224\|23142080 | Unspecified |
| mmu-miR-24-3p | MIMAT0000219 | Tsc22d3 | 14605 | HITS-CLIP | 25753418\|23142080 | Unspecified |
| mmu-miR-24-3p | MIMAT0000219 | Tsc22d4 | 78829 | HITS-CLIP | 23142080 | Unspecified |
| mmu-miR-24-3p | MIMAT0000219 | Tsg101 | 22088 | HITS-CLIP | 23142080 | Unspecified |
| mmu-miR-24-3p | MIMAT0000219 | Tshz1 | 110796 | HITS-CLIP | 23142080 | Unspecified |
| mmu-miR-24-3p | MIMAT0000219 | Tspan14 | 52588 | HITS-CLIP | 23142080\|31433990 | Unspecified |
| mmu-miR-24-3p | MIMAT0000219 | Tspan5 | 56224 | HITS-CLIP | 23142080 | Unspecified |

| | | | | | | |
|---|---|---|---|---|---|---|
| mmu-miR-24-3p | MIMAT0000219 | Tspyl3 | 241732 | HITS-CLIP | 23142080 | Unspecified |
| mmu-miR-24-3p | MIMAT0000219 | Tsr1 | 104662 | HITS-CLIP | 23142080 | Unspecified |
| mmu-miR-24-3p | MIMAT0000219 | Tsta3 | 22122 | HITS-CLIP | 23142080 | Unspecified |
| mmu-miR-24-3p | MIMAT0000219 | Ttc17 | 74569 | HITS-CLIP | 23142080 | Unspecified |
| mmu-miR-24-3p | MIMAT0000219 | Ttc3 | 22129 | Sequencing//HITS-CLIP | 19536157\|23142080\|37986224 | Unspecified |
| mmu-miR-24-3p | MIMAT0000219 | Ttc30b | 72421 | HITS-CLIP | 25753418 | Unspecified |
| mmu-miR-24-3p | MIMAT0000219 | Ttc39b | 69863 | HITS-CLIP | 37986224\|23142080 | Unspecified |
| mmu-miR-24-3p | MIMAT0000219 | Tti1 | 75425 | HITS-CLIP | 37986224\|23142080 | Unspecified |
| mmu-miR-24-3p | MIMAT0000219 | Ttll4 | 67534 | HITS-CLIP | 23142080 | Unspecified |
| mmu-miR-24-3p | MIMAT0000219 | Ttn | 22138 | HITS-CLIP | 25753418 | Unspecified |
| mmu-miR-24-3p | MIMAT0000219 | Ttpal | 76080 | HITS-CLIP | 23142080 | Unspecified |
| mmu-miR-24-3p | MIMAT0000219 | Ttyh3 | 78339 | Sequencing//HITS-CLIP | 19536157\|25753418\|37986224 | Unspecified |
| mmu-miR-24-3p | MIMAT0000219 | Tuba4a | 22145 | HITS-CLIP | 23142080 | Unspecified |
| mmu-miR-24-3p | MIMAT0000219 | Tubb4b | 227613 | HITS-CLIP | 25753418 | Unspecified |
| mmu-miR-24-3p | MIMAT0000219 | Tubb5 | 22154 | HITS-CLIP | 37986224\|23142080 | Unspecified |
| mmu-miR-24-3p | MIMAT0000219 | Tubgcp3 | 259279 | HITS-CLIP | 23142080 | Unspecified |
| mmu-miR-24-3p | MIMAT0000219 | Tusc1 | 69136 | HITS-CLIP | 23142080 | Unspecified |
| mmu-miR-24-3p | MIMAT0000219 | Twf1 | 19230 | qCLASH | 31393953 | Unspecified |
| mmu-miR-24-3p | MIMAT0000219 | Twf2 | 23999 | HITS-CLIP | 25753418 | Unspecified |
| mmu-miR-24-3p | MIMAT0000219 | Twsg1 | 65960 | HITS-CLIP | 23142080 | Unspecified |
| mmu-miR-24-3p | MIMAT0000219 | Txndc12 | 66073 | HITS-CLIP | 23142080 | Unspecified |
| mmu-miR-24-3p | MIMAT0000219 | Txndc5 | 105245 | HITS-CLIP | 37986224\|23142080 | Unspecified |
| mmu-miR-24-3p | MIMAT0000219 | Txnip | 56338 | HITS-CLIP | 25753418\|23142080 | Unspecified |
| mmu-miR-24-3p | MIMAT0000219 | U2af2 | 22185 | HITS-CLIP | 23142080 | Unspecified |
| mmu-miR-24-3p | MIMAT0000219 | U2surp | 67958 | HITS-CLIP | 23142080 | Unspecified |
| mmu-miR-24-3p | MIMAT0000219 | Uba1 | 22201 | HITS-CLIP | 23142080 | Unspecified |
| mmu-miR-24-3p | MIMAT0000219 | Uba52 | 22186 | HITS-CLIP | 37986224 | Unspecified |
| mmu-miR-24-3p | MIMAT0000219 | Uba6 | 231380 | HITS-CLIP | 23142080 | Unspecified |
| mmu-miR-24-3p | MIMAT0000219 | Ubap2 | 68926 | HITS-CLIP | 23142080\|25753418 | Unspecified |
| mmu-miR-24-3p | MIMAT0000219 | Ubap2l | 74383 | HITS-CLIP | 25753418\|23142080 | Unspecified |
| mmu-miR-24-3p | MIMAT0000219 | Ube2a | 22209 | HITS-CLIP | 25753418 | Unspecified |
| mmu-miR-24-3p | MIMAT0000219 | Ube2b | 22210 | HITS-CLIP | 25753418 | Unspecified |
| mmu-miR-24-3p | MIMAT0000219 | Ube2c | 68612 | HITS-CLIP | 23142080 | Unspecified |
| mmu-miR-24-3p | MIMAT0000219 | Ube2d2a | 56550 | HITS-CLIP | 25753418 | Unspecified |
| mmu-miR-24-3p | MIMAT0000219 | Ube2d3 | 66105 | HITS-CLIP | 23142080 | Unspecified |
| mmu-miR-24-3p | MIMAT0000219 | Ube2j1 | 56228 | HITS-CLIP | 25753418\|23142080 | Unspecified |
| mmu-miR-24-3p | MIMAT0000219 | Ube2k | 53323 | HITS-CLIP | 23142080 | Unspecified |
| mmu-miR-24-3p | MIMAT0000219 | Ube2l3 | 22195 | HITS-CLIP | 23142080 | Unspecified |
| mmu-miR-24-3p | MIMAT0000219 | Ube2q2 | 109161 | HITS-CLIP | 23142080 | Unspecified |

| | | | | | | | |
|---|---|---|---|---|---|---|---|
| mmu-miR-24-3p | MIMAT0000219 | Ube2r2 | 67615 | HITS-CLIP | 25753418\|23142080 | | Unspecified |
| mmu-miR-24-3p | MIMAT0000219 | Ube2v2 | 70620 | HITS-CLIP | | 23142080 | Unspecified |
| mmu-miR-24-3p | MIMAT0000219 | Ube2w | 66799 | HITS-CLIP | | 37986224 | Unspecified |
| mmu-miR-24-3p | MIMAT0000219 | Ube2z | 268470 | HITS-CLIP | 37986224\|23142080 | | Unspecified |
| mmu-miR-24-3p | MIMAT0000219 | Ube3a | 22215 | HITS-CLIP | | 23142080 | Unspecified |
| mmu-miR-24-3p | MIMAT0000219 | Ube3c | 100763 | HITS-CLIP | | 23142080 | Unspecified |
| mmu-miR-24-3p | MIMAT0000219 | Ube4a | 140630 | HITS-CLIP | | 23142080 | Unspecified |
| mmu-miR-24-3p | MIMAT0000219 | Ube4b | 63958 | HITS-CLIP | | 23142080 | Unspecified |
| mmu-miR-24-3p | MIMAT0000219 | Ubfd1 | 28018 | qCLASH//HITS-CLIP | 31393953\|23142080 | | Unspecified |
| mmu-miR-24-3p | MIMAT0000219 | Ubl7 | 69459 | HITS-CLIP | | 23142080 | Unspecified |
| mmu-miR-24-3p | MIMAT0000219 | Ubn2 | 320538 | HITS-CLIP | | 37986224 | Unspecified |
| mmu-miR-24-3p | MIMAT0000219 | Ubp1 | 22221 | HITS-CLIP | | 25753418 | Unspecified |
| mmu-miR-24-3p | MIMAT0000219 | Ubqln1 | 56085 | HITS-CLIP | | 23142080 | Unspecified |
| mmu-miR-24-3p | MIMAT0000219 | Ubqln2 | 54609 | HITS-CLIP | | 23142080 | Unspecified |
| mmu-miR-24-3p | MIMAT0000219 | Ubr1 | 22222 | HITS-CLIP | 37986224\|23142080 | | Unspecified |
| mmu-miR-24-3p | MIMAT0000219 | Ubr2 | 224826 | HITS-CLIP | 26203562\|23142080 | | Unspecified |
| mmu-miR-24-3p | MIMAT0000219 | Ubr3 | 68795 | HITS-CLIP | | 23142080 | Unspecified |
| mmu-miR-24-3p | MIMAT0000219 | Ubr4 | 69116 | Sequencing//HITS-CLIP | 37986224\|25753418\|19536157 | | Unspecified |
| mmu-miR-24-3p | MIMAT0000219 | Ubr5 | 70790 | HITS-CLIP | 25753418\|23142080 | | Unspecified |
| mmu-miR-24-3p | MIMAT0000219 | Ubr7 | 66622 | HITS-CLIP | | 23142080 | Unspecified |
| mmu-miR-24-3p | MIMAT0000219 | Ubtf | 21429 | HITS-CLIP | | 23142080 | Unspecified |
| mmu-miR-24-3p | MIMAT0000219 | Ubxn2b | 68053 | Luciferase Reporter Assay | | 37986224 | Unspecified |
| mmu-miR-24-3p | MIMAT0000219 | Uck2 | 80914 | HITS-CLIP | 37986224\|25753418 | | Unspecified |
| mmu-miR-24-3p | MIMAT0000219 | Ucp2 | 22228 | HITS-CLIP | | 23142080 | Unspecified |
| mmu-miR-24-3p | MIMAT0000219 | Ugcg | 22234 | Sequencing//HITS-CLIP | 37986224\|31433990\|25753418\|2314 | | Unspecified |
| mmu-miR-24-3p | MIMAT0000219 | Uggt1 | 320011 | HITS-CLIP | | 25753418 | Unspecified |
| mmu-miR-24-3p | MIMAT0000219 | Uhmk1 | 16589 | HITS-CLIP//qCLASH | 37986224\|31393953\|23142080 | | Unspecified |
| mmu-miR-24-3p | MIMAT0000219 | Uhrf1bp1 | 224648 | HITS-CLIP | | 25753418 | Unspecified |
| mmu-miR-24-3p | MIMAT0000219 | Unc119 | 22248 | HITS-CLIP | | 25753418 | Unspecified |
| mmu-miR-24-3p | MIMAT0000219 | Unc13a | 382018 | HITS-CLIP | | 23142080 | Unspecified |
| mmu-miR-24-3p | MIMAT0000219 | Unc13b | 22249 | HITS-CLIP | | 23142080 | Unspecified |
| mmu-miR-24-3p | MIMAT0000219 | Unc93b1 | 54445 | HITS-CLIP | | 23142080 | Unspecified |
| mmu-miR-24-3p | MIMAT0000219 | Upf1 | 19704 | HITS-CLIP | 25753418\|23142080 | | Unspecified |
| mmu-miR-24-3p | MIMAT0000219 | Upf2 | 326622 | HITS-CLIP | | 23142080 | Unspecified |
| mmu-miR-24-3p | MIMAT0000219 | Uqcc1 | 56046 | HITS-CLIP | | 23142080 | Unspecified |
| mmu-miR-24-3p | MIMAT0000219 | Uqcr11 | 66594 | HITS-CLIP | | 25753418 | Unspecified |
| mmu-miR-24-3p | MIMAT0000219 | Uqcrc1 | 22273 | HITS-CLIP | | 25753418 | Unspecified |
| mmu-miR-24-3p | MIMAT0000219 | Uqcrc2 | 67003 | HITS-CLIP | | 23142080 | Unspecified |
| mmu-miR-24-3p | MIMAT0000219 | Usf2 | 22282 | HITS-CLIP | | 25753418 | Unspecified |

| | | | | | | |
|---|---|---|---|---|---|---|
| mmu-miR-24-3p | MIMAT0000219 | Usf3 | 205717 | HITS-CLIP | 23142080 | Unspecified |
| mmu-miR-24-3p | MIMAT0000219 | Usp14 | 59025 | HITS-CLIP | 23142080\|25753418 | Unspecified |
| mmu-miR-24-3p | MIMAT0000219 | Usp15 | 14479 | HITS-CLIP | 25753418 | Unspecified |
| mmu-miR-24-3p | MIMAT0000219 | Usp20 | 74270 | HITS-CLIP | 25753418 | Unspecified |
| mmu-miR-24-3p | MIMAT0000219 | Usp22 | 216825 | HITS-CLIP | 25753418 | Unspecified |
| mmu-miR-24-3p | MIMAT0000219 | Usp24 | 329908 | HITS-CLIP | 25753418 | Unspecified |
| mmu-miR-24-3p | MIMAT0000219 | Usp28 | 235323 | HITS-CLIP | 25753418 | Unspecified |
| mmu-miR-24-3p | MIMAT0000219 | Usp3 | 235441 | HITS-CLIP | 23142080 | Unspecified |
| mmu-miR-24-3p | MIMAT0000219 | Usp31 | 76179 | HITS-CLIP | 23142080 | Unspecified |
| mmu-miR-24-3p | MIMAT0000219 | Usp32 | 237898 | HITS-CLIP | 25753418 | Unspecified |
| mmu-miR-24-3p | MIMAT0000219 | Usp34 | 17847 | HITS-CLIP | 37986224\|23142080 | Unspecified |
| mmu-miR-24-3p | MIMAT0000219 | Usp37 | 319651 | HITS-CLIP | 37986224\|23142080 | Unspecified |
| mmu-miR-24-3p | MIMAT0000219 | Usp38 | 74841 | HITS-CLIP | 37986224\|23142080 | Unspecified |
| mmu-miR-24-3p | MIMAT0000219 | Usp47 | 74996 | HITS-CLIP | 23142080 | Unspecified |
| mmu-miR-24-3p | MIMAT0000219 | Usp7 | 252870 | HITS-CLIP | 23142080 | Unspecified |
| mmu-miR-24-3p | MIMAT0000219 | Usp8 | 84092 | HITS-CLIP | 23142080 | Unspecified |
| mmu-miR-24-3p | MIMAT0000219 | Utrn | 22288 | HITS-CLIP | 23142080 | Unspecified |
| mmu-miR-24-3p | MIMAT0000219 | Vamp3 | 22319 | HITS-CLIP | 25753418 | Unspecified |
| mmu-miR-24-3p | MIMAT0000219 | Vav1 | 22324 | HITS-CLIP | 21258322\|23142080 | Unspecified |
| mmu-miR-24-3p | MIMAT0000219 | Vav2 | 22325 | HITS-CLIP | 23142080 | Unspecified |
| mmu-miR-24-3p | MIMAT0000219 | Vcpip1 | 70675 | HITS-CLIP | 37986224\|23142080 | Unspecified |
| mmu-miR-24-3p | MIMAT0000219 | Vdac1 | 22333 | HITS-CLIP | 23142080 | Unspecified |
| mmu-miR-24-3p | MIMAT0000219 | Vegfa | 22339 | HITS-CLIP | 23142080 | Unspecified |
| mmu-miR-24-3p | MIMAT0000219 | Vgll4 | 232334 | HITS-CLIP | 23142080 | Unspecified |
| mmu-miR-24-3p | MIMAT0000219 | Vhl | 22346 | Sequencing//HITS-CLIP | 37986224\|23142080\|19536157 | Unspecified |
| mmu-miR-24-3p | MIMAT0000219 | Vim | 22352 | HITS-CLIP | 23142080 | Unspecified |
| mmu-miR-24-3p | MIMAT0000219 | Virma | 25962 | HITS-CLIP | 25753418\|23142080 | Unspecified |
| mmu-miR-24-3p | MIMAT0000219 | Vmp1 | 75909 | HITS-CLIP | 25753418 | Unspecified |
| mmu-miR-24-3p | MIMAT0000219 | Vps11 | 71732 | HITS-CLIP | 23142080 | Unspecified |
| mmu-miR-24-3p | MIMAT0000219 | Vps13a | 271564 | HITS-CLIP | 23142080 | Unspecified |
| mmu-miR-24-3p | MIMAT0000219 | Vps13d | 230895 | HITS-CLIP | 25753418\|23142080 | Unspecified |
| mmu-miR-24-3p | MIMAT0000219 | Vps25 | 28084 | HITS-CLIP | 25753418 | Unspecified |
| mmu-miR-24-3p | MIMAT0000219 | Vps35l | 57020 | HITS-CLIP | 23142080 | Unspecified |
| mmu-miR-24-3p | MIMAT0000219 | Vps36 | 70160 | HITS-CLIP | 37986224 | Unspecified |
| mmu-miR-24-3p | MIMAT0000219 | Vps52 | 224705 | HITS-CLIP | 25753418 | Unspecified |
| mmu-miR-24-3p | MIMAT0000219 | Vsir | 64115 | HITS-CLIP | 23142080 | Unspecified |
| mmu-miR-24-3p | MIMAT0000219 | Vti1b | 53612 | HITS-CLIP | 23142080 | Unspecified |
| mmu-miR-24-3p | MIMAT0000219 | Vwa5a | 67776 | HITS-CLIP | 23142080 | Unspecified |
| mmu-miR-24-3p | MIMAT0000219 | Vwa8 | 219189 | HITS-CLIP | 25753418 | Unspecified |

| | | | | | | |
|---|---|---|---|---|---|---|
| mmu-miR-24-3p | MIMAT0000219 | Wac | 225131 | HITS-CLIP | 23142080 | Unspecified |
| mmu-miR-24-3p | MIMAT0000219 | Wapl | 23063 | HITS-CLIP | 23142080 | Unspecified |
| mmu-miR-24-3p | MIMAT0000219 | Washc5 | 9897 | HITS-CLIP | 23142080 | Unspecified |
| mmu-miR-24-3p | MIMAT0000219 | Wbp1 | 22377 | HITS-CLIP | 25753418 | Unspecified |
| mmu-miR-24-3p | MIMAT0000219 | Wbp11 | 60321 | HITS-CLIP | 25753418 | Unspecified |
| mmu-miR-24-3p | MIMAT0000219 | Wbp1l | 226178 | HITS-CLIP | 25753418 | Unspecified |
| mmu-miR-24-3p | MIMAT0000219 | Wdfy3 | 72145 | HITS-CLIP | 25753418 | Unspecified |
| mmu-miR-24-3p | MIMAT0000219 | Wdhd1 | 218973 | HITS-CLIP | 23142080 | Unspecified |
| mmu-miR-24-3p | MIMAT0000219 | Wdr11 | 207425 | HITS-CLIP | 23142080 | Unspecified |
| mmu-miR-24-3p | MIMAT0000219 | Wdr13 | 73447 | Sequencing//HITS-CLIP | 37986224\|23142080\|19536157 | Unspecified |
| mmu-miR-24-3p | MIMAT0000219 | Wdr18 | 216156 | HITS-CLIP | 25753418\|23142080 | Unspecified |
| mmu-miR-24-3p | MIMAT0000219 | Wdr26 | 226757 | HITS-CLIP | 37986224\|23142080 | Unspecified |
| mmu-miR-24-3p | MIMAT0000219 | Wdr35 | 74682 | HITS-CLIP | 23142080 | Unspecified |
| mmu-miR-24-3p | MIMAT0000219 | Wdr5 | 140858 | HITS-CLIP | 26203562 | Unspecified |
| mmu-miR-24-3p | MIMAT0000219 | Wdr81 | 192652 | HITS-CLIP | 23142080 | Unspecified |
| mmu-miR-24-3p | MIMAT0000219 | Wdr82 | 77305 | HITS-CLIP | 23142080\|25753418 | Unspecified |
| mmu-miR-24-3p | MIMAT0000219 | Wdr91 | 101240 | HITS-CLIP | 23142080 | Unspecified |
| mmu-miR-24-3p | MIMAT0000219 | Wfs1 | 22393 | HITS-CLIP | 31433990 | Unspecified |
| mmu-miR-24-3p | MIMAT0000219 | Wipf1 | 215280 | HITS-CLIP | 37986224 | Unspecified |
| mmu-miR-24-3p | MIMAT0000219 | Wiz | 22404 | HITS-CLIP | 37986224\|23142080 | Unspecified |
| mmu-miR-24-3p | MIMAT0000219 | Wnk1 | 232341 | HITS-CLIP | 23142080 | Unspecified |
| mmu-miR-24-3p | MIMAT0000219 | Wrnip1 | 78903 | HITS-CLIP//qCLASH | 37986224\|31393953 | Unspecified |
| mmu-miR-24-3p | MIMAT0000219 | Xbp1 | 22433 | HITS-CLIP | 23142080 | Unspecified |
| mmu-miR-24-3p | MIMAT0000219 | Xirp1 | 22437 | HITS-CLIP | 25753418 | Unspecified |
| mmu-miR-24-3p | MIMAT0000219 | Xpnpep1 | 170750 | HITS-CLIP | 37986224 | Unspecified |
| mmu-miR-24-3p | MIMAT0000219 | Xpo1 | 103573 | HITS-CLIP | 23142080 | Unspecified |
| mmu-miR-24-3p | MIMAT0000219 | Xpo4 | 57258 | HITS-CLIP | 23142080 | Unspecified |
| mmu-miR-24-3p | MIMAT0000219 | Xpo6 | 74204 | HITS-CLIP | 23142080 | Unspecified |
| mmu-miR-24-3p | MIMAT0000219 | Xpo7 | 65246 | Sequencing//HITS-CLIP | 37986224\|23142080\|19536157\|2125[illegible] | Unspecified |
| mmu-miR-24-3p | MIMAT0000219 | Xpr1 | 19775 | Sequencing//HITS-CLIP | 19536157\|23142080\|37986224 | Unspecified |
| mmu-miR-24-3p | MIMAT0000219 | Yap1 | 22601 | HITS-CLIP | 25753418 | Unspecified |
| mmu-miR-24-3p | MIMAT0000219 | Ybx1 | 22608 | HITS-CLIP | 23142080 | Unspecified |
| mmu-miR-24-3p | MIMAT0000219 | Yjefn3 | 234365 | HITS-CLIP | 25753418 | Unspecified |
| mmu-miR-24-3p | MIMAT0000219 | Yod1 | 226418 | HITS-CLIP | 23142080 | Unspecified |
| mmu-miR-24-3p | MIMAT0000219 | Ythdc2 | 240255 | HITS-CLIP | 23142080 | Unspecified |
| mmu-miR-24-3p | MIMAT0000219 | Ythdf1 | 228994 | HITS-CLIP | 25753418\|23142080 | Unspecified |
| mmu-miR-24-3p | MIMAT0000219 | Ythdf3 | 229096 | HITS-CLIP | 23142080 | Unspecified |
| mmu-miR-24-3p | MIMAT0000219 | Ywhae | 22627 | HITS-CLIP | 37986224\|23142080 | Unspecified |
| mmu-miR-24-3p | MIMAT0000219 | Ywhag | 22628 | HITS-CLIP | 25753418\|23142080 | Unspecified |

| | | | | | | | |
|---|---|---|---|---|---|---|---|
| mmu-miR-24-3p | MIMAT0000219 | Ywhah | 22629 | HITS-CLIP | | 23142080 | Unspecified |
| mmu-miR-24-3p | MIMAT0000219 | Yy1 | 22632 | HITS-CLIP | | 23142080 | Unspecified |
| mmu-miR-24-3p | MIMAT0000219 | Zap70 | 22637 | HITS-CLIP | | 23142080 | Unspecified |
| mmu-miR-24-3p | MIMAT0000219 | Zbp1 | 58203 | HITS-CLIP | 37986224\|23142080 | | Unspecified |
| mmu-miR-24-3p | MIMAT0000219 | Zbtb11 | 271377 | HITS-CLIP | | 23142080 | Unspecified |
| mmu-miR-24-3p | MIMAT0000219 | Zbtb18 | 30928 | HITS-CLIP | | 23142080 | Unspecified |
| mmu-miR-24-3p | MIMAT0000219 | Zbtb20 | 56490 | HITS-CLIP | | 23142080 | Unspecified |
| mmu-miR-24-3p | MIMAT0000219 | Zbtb37 | 240869 | HITS-CLIP | | 23142080 | Unspecified |
| mmu-miR-24-3p | MIMAT0000219 | Zbtb41 | 226470 | HITS-CLIP | | 23142080 | Unspecified |
| mmu-miR-24-3p | MIMAT0000219 | Zbtb44 | 235132 | HITS-CLIP | 37986224\|23142080 | | Unspecified |
| mmu-miR-24-3p | MIMAT0000219 | Zbtb7b | 22724 | HITS-CLIP | | 23142080 | Unspecified |
| mmu-miR-24-3p | MIMAT0000219 | Zc3h12d | 237256 | HITS-CLIP | | 23142080 | Unspecified |
| mmu-miR-24-3p | MIMAT0000219 | Zc3h18 | 76014 | HITS-CLIP | | 23142080 | Unspecified |
| mmu-miR-24-3p | MIMAT0000219 | Zc3h7a | 106205 | HITS-CLIP | | 23142080 | Unspecified |
| mmu-miR-24-3p | MIMAT0000219 | Zcchc2 | 227449 | HITS-CLIP | | 23142080 | Unspecified |
| mmu-miR-24-3p | MIMAT0000219 | Zcrb1 | 67197 | HITS-CLIP | | 23142080 | Unspecified |
| mmu-miR-24-3p | MIMAT0000219 | Zdhhc17 | 320150 | HITS-CLIP | | 23142080 | Unspecified |
| mmu-miR-24-3p | MIMAT0000219 | Zdhhc18 | 503610 | HITS-CLIP | 37986224\|23142080\|25753418 | | Unspecified |
| mmu-miR-24-3p | MIMAT0000219 | Zdhhc21 | 68268 | HITS-CLIP | | 23142080 | Unspecified |
| mmu-miR-24-3p | MIMAT0000219 | Zdhhc5 | 228136 | HITS-CLIP | | 23142080 | Unspecified |
| mmu-miR-24-3p | MIMAT0000219 | Zdhhc8 | 27801 | HITS-CLIP | | 23142080 | Unspecified |
| mmu-miR-24-3p | MIMAT0000219 | Zeb2 | 24136 | HITS-CLIP | | 23142080 | Unspecified |
| mmu-miR-24-3p | MIMAT0000219 | Zer1 | 227693 | HITS-CLIP | 37986224\|23142080 | | Unspecified |
| mmu-miR-24-3p | MIMAT0000219 | Zfc3h1 | 216345 | HITS-CLIP | | 23142080 | Unspecified |
| mmu-miR-24-3p | MIMAT0000219 | Zfp106 | 20402 | HITS-CLIP | | 23142080 | Unspecified |
| mmu-miR-24-3p | MIMAT0000219 | Zfp148 | 22661 | HITS-CLIP | | 23142080 | Unspecified |
| mmu-miR-24-3p | MIMAT0000219 | Zfp207 | 22680 | HITS-CLIP | | 25753418 | Unspecified |
| mmu-miR-24-3p | MIMAT0000219 | Zfp213 | 449521 | HITS-CLIP | | 23142080 | Unspecified |
| mmu-miR-24-3p | MIMAT0000219 | Zfp217 | 228913 | HITS-CLIP | 37986224\|23142080 | | Unspecified |
| mmu-miR-24-3p | MIMAT0000219 | Zfp219 | 69890 | HITS-CLIP | | 23142080 | Unspecified |
| mmu-miR-24-3p | MIMAT0000219 | Zfp24 | 59057 | HITS-CLIP | | 23142080 | Unspecified |
| mmu-miR-24-3p | MIMAT0000219 | Zfp26 | 22688 | HITS-CLIP | | 23142080 | Unspecified |
| mmu-miR-24-3p | MIMAT0000219 | Zfp266 | 77519 | HITS-CLIP | 23142080\|25753418 | | Unspecified |
| mmu-miR-24-3p | MIMAT0000219 | Zfp280b | 64453 | HITS-CLIP | 37986224\|25753418\|23142080 | | Unspecified |
| mmu-miR-24-3p | MIMAT0000219 | Zfp280d | 235469 | HITS-CLIP | | 31433990 | Unspecified |
| mmu-miR-24-3p | MIMAT0000219 | Zfp281 | 226442 | HITS-CLIP | 37986224\|23142080 | | Unspecified |
| mmu-miR-24-3p | MIMAT0000219 | Zfp292 | 30046 | HITS-CLIP | 25753418\|23142080 | | Unspecified |
| mmu-miR-24-3p | MIMAT0000219 | Zfp296 | 63872 | HITS-CLIP | | 23142080 | Unspecified |
| mmu-miR-24-3p | MIMAT0000219 | Zfp330 | 30932 | HITS-CLIP | | 23142080 | Unspecified |

| | | | | | | |
|---|---|---|---|---|---|---|
| mmu-miR-24-3p | MIMAT0000219 | Zfp35 | 22694 | HITS-CLIP | 25753418 | Unspecified |
| mmu-miR-24-3p | MIMAT0000219 | Zfp397 | 69256 | Sequencing//HITS-CLIP | 37986224\|25753418\|19536157 | Unspecified |
| mmu-miR-24-3p | MIMAT0000219 | Zfp40 | 22700 | HITS-CLIP | 23142080 | Unspecified |
| mmu-miR-24-3p | MIMAT0000219 | Zfp408 | 381410 | HITS-CLIP | 25753418 | Unspecified |
| mmu-miR-24-3p | MIMAT0000219 | Zfp451 | 98403 | HITS-CLIP | 23142080 | Unspecified |
| mmu-miR-24-3p | MIMAT0000219 | Zfp513 | 101023 | HITS-CLIP | 37986224 | Unspecified |
| mmu-miR-24-3p | MIMAT0000219 | Zfp521 | 225207 | HITS-CLIP | 25753418 | Unspecified |
| mmu-miR-24-3p | MIMAT0000219 | Zfp568 | 243905 | HITS-CLIP | 23142080 | Unspecified |
| mmu-miR-24-3p | MIMAT0000219 | Zfp592 | 233410 | HITS-CLIP | 23142080 | Unspecified |
| mmu-miR-24-3p | MIMAT0000219 | Zfp607a | 545938 | HITS-CLIP | 23142080 | Unspecified |
| mmu-miR-24-3p | MIMAT0000219 | Zfp608 | 269023 | HITS-CLIP | 23142080 | Unspecified |
| mmu-miR-24-3p | MIMAT0000219 | Zfp609 | 214812 | HITS-CLIP | 23142080\|25753418 | Unspecified |
| mmu-miR-24-3p | MIMAT0000219 | Zfp654 | 72020 | HITS-CLIP | 37986224\|23142080 | Unspecified |
| mmu-miR-24-3p | MIMAT0000219 | Zfp667 | 384763 | HITS-CLIP | 23142080 | Unspecified |
| mmu-miR-24-3p | MIMAT0000219 | Zfp692 | 103836 | HITS-CLIP | 23142080 | Unspecified |
| mmu-miR-24-3p | MIMAT0000219 | Zfp7 | 223669 | HITS-CLIP | 23142080 | Unspecified |
| mmu-miR-24-3p | MIMAT0000219 | Zfp703 | 353310 | HITS-CLIP | 23142080 | Unspecified |
| mmu-miR-24-3p | MIMAT0000219 | Zfp760 | 240034 | HITS-CLIP | 37986224 | Unspecified |
| mmu-miR-24-3p | MIMAT0000219 | Zfp770 | 228491 | HITS-CLIP | 23142080 | Unspecified |
| mmu-miR-24-3p | MIMAT0000219 | Zfp788 | 67607 | HITS-CLIP | 23142080 | Unspecified |
| mmu-miR-24-3p | MIMAT0000219 | Zfp800 | 627049 | HITS-CLIP | 25753418 | Unspecified |
| mmu-miR-24-3p | MIMAT0000219 | Zfp81 | 224694 | HITS-CLIP | 37986224 | Unspecified |
| mmu-miR-24-3p | MIMAT0000219 | Zfp831 | 100043757 | HITS-CLIP | 37986224\|23142080 | Unspecified |
| mmu-miR-24-3p | MIMAT0000219 | Zfp871 | 208292 | HITS-CLIP | 23142080 | Unspecified |
| mmu-miR-24-3p | MIMAT0000219 | Zfp91 | 109910 | HITS-CLIP | 25753418\|23142080 | Unspecified |
| mmu-miR-24-3p | MIMAT0000219 | Zfp950 | 214368 | HITS-CLIP | 23142080 | Unspecified |
| mmu-miR-24-3p | MIMAT0000219 | Zfp985 | 433804 | HITS-CLIP | 23142080 | Unspecified |
| mmu-miR-24-3p | MIMAT0000219 | Zfpm2 | 22762 | HITS-CLIP | 25753418 | Unspecified |
| mmu-miR-24-3p | MIMAT0000219 | Zfr | 22763 | HITS-CLIP | 23142080 | Unspecified |
| mmu-miR-24-3p | MIMAT0000219 | Zfyve26 | 211978 | HITS-CLIP | 23142080 | Unspecified |
| mmu-miR-24-3p | MIMAT0000219 | Zfyve9 | 230597 | HITS-CLIP | 25753418 | Unspecified |
| mmu-miR-24-3p | MIMAT0000219 | Zik1 | 22775 | HITS-CLIP | 37986224\|23142080 | Unspecified |
| mmu-miR-24-3p | MIMAT0000219 | Zmat5 | 67178 | HITS-CLIP | 25753418 | Unspecified |
| mmu-miR-24-3p | MIMAT0000219 | Zmiz1 | 328365 | HITS-CLIP | 25753418 | Unspecified |
| mmu-miR-24-3p | MIMAT0000219 | Zmpste24 | 230709 | HITS-CLIP | 23142080 | Unspecified |
| mmu-miR-24-3p | MIMAT0000219 | Zmym1 | 68310 | HITS-CLIP | 23142080 | Unspecified |
| mmu-miR-24-3p | MIMAT0000219 | Zmym2 | 76007 | HITS-CLIP | 23142080 | Unspecified |
| mmu-miR-24-3p | MIMAT0000219 | Zmym3 | 56364 | Sequencing//HITS-CLIP | 37986224\|23142080\|19536157 | Unspecified |
| mmu-miR-24-3p | MIMAT0000219 | Zmym4 | 67785 | HITS-CLIP | 23142080 | Unspecified |

| | | | | | | |
|---|---|---|---|---|---|---|
| mmu-miR-24-3p | MIMAT0000219 | Zmym6 | 100177 | HITS-CLIP | 23142080 | Unspecified |
| mmu-miR-24-3p | MIMAT0000219 | Zmynd8 | 228880 | HITS-CLIP | 23142080 | Unspecified |
| mmu-miR-24-3p | MIMAT0000219 | Znfx1 | 98999 | HITS-CLIP | 25753418 | Unspecified |
| mmu-miR-24-3p | MIMAT0000219 | Znrf1 | 170737 | HITS-CLIP | 23142080 | Unspecified |
| mmu-miR-24-3p | MIMAT0000219 | Znrf3 | 407821 | HITS-CLIP | 25753418 | Unspecified |
| mmu-miR-24-3p | MIMAT0000219 | Zswim7 | 69747 | HITS-CLIP | 23142080 | Unspecified |
| mmu-miR-24-3p | MIMAT0000219 | Zwilch | 68014 | HITS-CLIP | 23142080 | Unspecified |
| mmu-miR-24-3p | MIMAT0000219 | Zwint | 52696 | HITS-CLIP | 23142080\|25753418 | Unspecified |
| mmu-miR-24-3p | MIMAT0000219 | Zxdb | 668166 | HITS-CLIP | 23142080 | Unspecified |
| mmu-miR-24-3p | MIMAT0000219 | Zyg11b | 414872 | HITS-CLIP | 23142080 | Unspecified |
| mmu-miR-24-3p | MIMAT0000219 | Zzef1 | 195018 | HITS-CLIP | 23142080 | Unspecified |
| mmu-miR-24-3p | MIMAT0000219 | mt-Nd1 | 4535 | HITS-CLIP | 23142080 | Unspecified |
| mmu-miR-24-3p | MIMAT0000219 | mt-Nd5 | 4540 | HITS-CLIP | 23142080 | Unspecified |
| mmu-miR-9-5p | MIMAT0000142 | 1700030J22Rik | 69528 | HITS-CLIP | 37986224 | Unspecified |
| mmu-miR-9-5p | MIMAT0000142 | 5730507C01Rik | 236366 | qCLASH | 31393953 | Unspecified |
| mmu-miR-9-5p | MIMAT0000142 | A830010M20Rik | 231570 | HITS-CLIP | 37986224 | Unspecified |
| mmu-miR-9-5p | MIMAT0000142 | AI597479 | 98404 | HITS-CLIP | 37986224 | Unspecified |
| mmu-miR-9-5p | MIMAT0000142 | Abca1 | 11303 | Sequencing//HITS-CLIP | 37986224\|19536157 | Unspecified |
| mmu-miR-9-5p | MIMAT0000142 | Abca2 | 11305 | HITS-CLIP | 37986224 | Unspecified |
| mmu-miR-9-5p | MIMAT0000142 | Abcc8 | 20927 | HITS-CLIP | 37986224 | Unspecified |
| mmu-miR-9-5p | MIMAT0000142 | Abhd13 | 68904 | HITS-CLIP | 19536157 | Unspecified |
| mmu-miR-9-5p | MIMAT0000142 | Abhd2 | 54608 | HITS-CLIP | 37986224 | Unspecified |
| mmu-miR-9-5p | MIMAT0000142 | Abhd3 | 106861 | HITS-CLIP | 19536157 | Unspecified |
| mmu-miR-9-5p | MIMAT0000142 | Abl2 | 11352 | HITS-CLIP | 37986224\|19536157 | Unspecified |
| mmu-miR-9-5p | MIMAT0000142 | Ablim2 | 231148 | HITS-CLIP | 37986224 | Unspecified |
| mmu-miR-9-5p | MIMAT0000142 | Acat1 | 110446 | HITS-CLIP | 37986224 | Unspecified |
| mmu-miR-9-5p | MIMAT0000142 | Acin1 | 56215 | qCLASH | 31393953 | Unspecified |
| mmu-miR-9-5p | MIMAT0000142 | Aco2 | 11429 | HITS-CLIP | 37986224 | Unspecified |
| mmu-miR-9-5p | MIMAT0000142 | Acot7 | 70025 | HITS-CLIP | 37986224 | Unspecified |
| mmu-miR-9-5p | MIMAT0000142 | Acsbg1 | 94180 | HITS-CLIP | 37986224 | Unspecified |
| mmu-miR-9-5p | MIMAT0000142 | Acsl1 | 14081 | HITS-CLIP | 37986224 | Unspecified |
| mmu-miR-9-5p | MIMAT0000142 | Acvr2a | 11480 | HITS-CLIP | 37986224 | Unspecified |
| mmu-miR-9-5p | MIMAT0000142 | Add2 | 11519 | HITS-CLIP | 37986224 | Unspecified |
| mmu-miR-9-5p | MIMAT0000142 | Adipor1 | 72674 | HITS-CLIP | 37986224 | Unspecified |
| mmu-miR-9-5p | MIMAT0000142 | Adra2a | 11551 | HITS-CLIP | 37986224 | Unspecified |
| mmu-miR-9-5p | MIMAT0000142 | Afap1 | 70292 | HITS-CLIP | 37986224 | Unspecified |
| mmu-miR-9-5p | MIMAT0000142 | Aff2 | 14266 | HITS-CLIP | 37986224 | Unspecified |
| mmu-miR-9-5p | MIMAT0000142 | Aff4 | 93736 | HITS-CLIP | 37986224 | Unspecified |
| mmu-miR-9-5p | MIMAT0000142 | Afg3l2 | 69597 | HITS-CLIP | 37986224 | Unspecified |

| | | | | | | |
|---|---|---|---|---|---|---|
| mmu-miR-9-5p | MIMAT0000142 | Agap1 | 347722 | HITS-CLIP | 37986224 | Unspecified |
| mmu-miR-9-5p | MIMAT0000142 | Agfg1 | 15463 | Sequencing//HITS-CLIP | 37986224\|19536157 | Unspecified |
| mmu-miR-9-5p | MIMAT0000142 | Ago2 | 239528 | HITS-CLIP | 37986224 | Unspecified |
| mmu-miR-9-5p | MIMAT0000142 | Ago3 | 214150 | HITS-CLIP | 37986224 | Unspecified |
| mmu-miR-9-5p | MIMAT0000142 | Agpat4 | 68262 | HITS-CLIP | 37986224 | Unspecified |
| mmu-miR-9-5p | MIMAT0000142 | Ahctf1 | 226747 | HITS-CLIP | 37986224 | Unspecified |
| mmu-miR-9-5p | MIMAT0000142 | Ahcyl1 | 229709 | HITS-CLIP | 37986224 | Unspecified |
| mmu-miR-9-5p | MIMAT0000142 | Ahi1 | 52906 | HITS-CLIP | 37986224 | Unspecified |
| mmu-miR-9-5p | MIMAT0000142 | Ahnak | 66395 | HITS-CLIP | 37986224 | Unspecified |
| mmu-miR-9-5p | MIMAT0000142 | Ahr | 11622 | HITS-CLIP | 37986224 | Unspecified |
| mmu-miR-9-5p | MIMAT0000142 | Akap12 | 83397 | HITS-CLIP | 37986224 | Unspecified |
| mmu-miR-9-5p | MIMAT0000142 | Aldh2 | 11669 | HITS-CLIP | 37986224 | Unspecified |
| mmu-miR-9-5p | MIMAT0000142 | Aldh6a1 | 104776 | HITS-CLIP | 19536157 | Unspecified |
| mmu-miR-9-5p | MIMAT0000142 | Alg10b | 380959 | HITS-CLIP | 37986224 | Unspecified |
| mmu-miR-9-5p | MIMAT0000142 | Alg5 | 66248 | HITS-CLIP | 37986224 | Unspecified |
| mmu-miR-9-5p | MIMAT0000142 | Amotl2 | 56332 | HITS-CLIP | 37986224 | Unspecified |
| mmu-miR-9-5p | MIMAT0000142 | Ankhd1 | 108857 | HITS-CLIP | 37986224 | Unspecified |
| mmu-miR-9-5p | MIMAT0000142 | Ankib1 | 70797 | HITS-CLIP | 19536157 | Unspecified |
| mmu-miR-9-5p | MIMAT0000142 | Ankrd11 | 77087 | HITS-CLIP | 37986224\|19536157 | Unspecified |
| mmu-miR-9-5p | MIMAT0000142 | Ankrd28 | 105522 | HITS-CLIP | 37986224 | Unspecified |
| mmu-miR-9-5p | MIMAT0000142 | Ankrd52 | 237615 | HITS-CLIP | 37986224 | Unspecified |
| mmu-miR-9-5p | MIMAT0000142 | Ano10 | 102566 | HITS-CLIP | 37986224 | Unspecified |
| mmu-miR-9-5p | MIMAT0000142 | Ano8 | 382014 | HITS-CLIP | 37986224 | Unspecified |
| mmu-miR-9-5p | MIMAT0000142 | Anp32a | 11737 | HITS-CLIP | 37986224 | Unspecified |
| mmu-miR-9-5p | MIMAT0000142 | Anxa5 | 11747 | HITS-CLIP | 37986224 | Unspecified |
| mmu-miR-9-5p | MIMAT0000142 | Apba1 | 319924 | HITS-CLIP | 37986224 | Unspecified |
| mmu-miR-9-5p | MIMAT0000142 | Apc | 11789 | HITS-CLIP | 37986224 | Unspecified |
| mmu-miR-9-5p | MIMAT0000142 | Apln | 30878 | HITS-CLIP | 37986224 | Unspecified |
| mmu-miR-9-5p | MIMAT0000142 | App | 11820 | HITS-CLIP | 37986224\|19536157 | Unspecified |
| mmu-miR-9-5p | MIMAT0000142 | Arfgef3 | 57221 | HITS-CLIP | 19536157 | Unspecified |
| mmu-miR-9-5p | MIMAT0000142 | Arhgap20 | 244867 | HITS-CLIP | 37986224 | Unspecified |
| mmu-miR-9-5p | MIMAT0000142 | Arhgef17 | 207212 | HITS-CLIP | 37986224 | Unspecified |
| mmu-miR-9-5p | MIMAT0000142 | Arl8b | 67166 | HITS-CLIP | 37986224 | Unspecified |
| mmu-miR-9-5p | MIMAT0000142 | Armc1 | 74252 | HITS-CLIP | 37986224 | Unspecified |
| mmu-miR-9-5p | MIMAT0000142 | Arnt2 | 11864 | HITS-CLIP | 37986224 | Unspecified |
| mmu-miR-9-5p | MIMAT0000142 | Arpp19 | 59046 | HITS-CLIP | 37986224 | Unspecified |
| mmu-miR-9-5p | MIMAT0000142 | Arsb | 11881 | HITS-CLIP | 37986224 | Unspecified |
| mmu-miR-9-5p | MIMAT0000142 | Asb8 | 78541 | HITS-CLIP | 19536157 | Unspecified |
| mmu-miR-9-5p | MIMAT0000142 | Ascc3 | 77987 | HITS-CLIP | 37986224 | Unspecified |

| | | | | | | |
|---|---|---|---|---|---|---|
| mmu-miR-9-5p | MIMAT0000142 | Asic1 | 11419 | HITS-CLIP | 37986224 | Unspecified |
| mmu-miR-9-5p | MIMAT0000142 | Astn1 | 11899 | HITS-CLIP | 37986224 | Unspecified |
| mmu-miR-9-5p | MIMAT0000142 | Atad1 | 67979 | HITS-CLIP | 37986224 | Unspecified |
| mmu-miR-9-5p | MIMAT0000142 | Atad2b | 320817 | Sequencing | 37986224 | Unspecified |
| mmu-miR-9-5p | MIMAT0000142 | Atg16l1 | 77040 | HITS-CLIP | 37986224 | Unspecified |
| mmu-miR-9-5p | MIMAT0000142 | Atg2b | 76559 | HITS-CLIP | 37986224 | Unspecified |
| mmu-miR-9-5p | MIMAT0000142 | Atg4b | 66615 | HITS-CLIP | 37986224 | Unspecified |
| mmu-miR-9-5p | MIMAT0000142 | Atl2 | 56298 | HITS-CLIP | 19536157 | Unspecified |
| mmu-miR-9-5p | MIMAT0000142 | Atp1a1 | 11928 | HITS-CLIP | 37986224 | Unspecified |
| mmu-miR-9-5p | MIMAT0000142 | Atp1b1 | 11931 | Sequencing//HITS-CLIP | 19536157\|37986224 | Unspecified |
| mmu-miR-9-5p | MIMAT0000142 | Atp1b2 | 11932 | HITS-CLIP | 37986224 | Unspecified |
| mmu-miR-9-5p | MIMAT0000142 | Atp2b1 | 67972 | HITS-CLIP | 37986224 | Unspecified |
| mmu-miR-9-5p | MIMAT0000142 | Atp5a1 | 11946 | HITS-CLIP | 37986224 | Unspecified |
| mmu-miR-9-5p | MIMAT0000142 | Atp5b | 11947 | HITS-CLIP | 37986224 | Unspecified |
| mmu-miR-9-5p | MIMAT0000142 | Atp6v0e2 | 76252 | HITS-CLIP | 37986224 | Unspecified |
| mmu-miR-9-5p | MIMAT0000142 | Atp6v1b2 | 11966 | HITS-CLIP | 37986224 | Unspecified |
| mmu-miR-9-5p | MIMAT0000142 | Atp6v1h | 108664 | HITS-CLIP | 37986224 | Unspecified |
| mmu-miR-9-5p | MIMAT0000142 | Atrn | 11990 | HITS-CLIP | 37986224 | Unspecified |
| mmu-miR-9-5p | MIMAT0000142 | Atrx | 22589 | HITS-CLIP | 37986224\|19536157 | Unspecified |
| mmu-miR-9-5p | MIMAT0000142 | Atxn1l | 52335 | HITS-CLIP | 37986224 | Unspecified |
| mmu-miR-9-5p | MIMAT0000142 | Atxn7l3b | 382423 | HITS-CLIP | 37986224 | Unspecified |
| mmu-miR-9-5p | MIMAT0000142 | Auts2 | 319974 | HITS-CLIP | 37986224 | Unspecified |
| mmu-miR-9-5p | MIMAT0000142 | B230219D22Rik | 78521 | HITS-CLIP | 37986224 | Unspecified |
| mmu-miR-9-5p | MIMAT0000142 | Bace1 | 23821 | HITS-CLIP | 37986224\|19536157 | Unspecified |
| mmu-miR-9-5p | MIMAT0000142 | Bahd1 | 228536 | HITS-CLIP | 37986224 | Unspecified |
| mmu-miR-9-5p | MIMAT0000142 | Baz2a | 116848 | HITS-CLIP | 19536157 | Unspecified |
| mmu-miR-9-5p | MIMAT0000142 | Bbs4 | 102774 | HITS-CLIP | 37986224 | Unspecified |
| mmu-miR-9-5p | MIMAT0000142 | Bbx | 70508 | HITS-CLIP | 37986224 | Unspecified |
| mmu-miR-9-5p | MIMAT0000142 | Bcap31 | 27061 | HITS-CLIP | 37986224 | Unspecified |
| mmu-miR-9-5p | MIMAT0000142 | Bfar | 67118 | HITS-CLIP | 37986224 | Unspecified |
| mmu-miR-9-5p | MIMAT0000142 | Bicc1 | 83675 | HITS-CLIP | 37986224 | Unspecified |
| mmu-miR-9-5p | MIMAT0000142 | Bicd1 | 12121 | HITS-CLIP | 19536157 | Unspecified |
| mmu-miR-9-5p | MIMAT0000142 | Bmp2k | 140780 | HITS-CLIP | 26203562 | Unspecified |
| mmu-miR-9-5p | MIMAT0000142 | Bmp7 | 12162 | HITS-CLIP | 37986224 | Unspecified |
| mmu-miR-9-5p | MIMAT0000142 | Bmpr2 | 12168 | HITS-CLIP | 37986224 | Unspecified |
| mmu-miR-9-5p | MIMAT0000142 | Bod1l | 665775 | HITS-CLIP | 37986224 | Unspecified |
| mmu-miR-9-5p | MIMAT0000142 | Brwd1 | 93871 | HITS-CLIP | 19536157 | Unspecified |
| mmu-miR-9-5p | MIMAT0000142 | Brwd3 | 382236 | HITS-CLIP | 37986224 | Unspecified |
| mmu-miR-9-5p | MIMAT0000142 | Btbd3 | 228662 | Luciferase Reporter Assay//HITS-CLIP | 37986224\|19536157 | Unspecified |

| | | | | | | | |
|---|---|---|---|---|---|---|---|
| mmu-miR-9-5p | MIMAT0000142 | Btbd8 | 100503185 | HITS-CLIP | | 19536157 | Unspecified |
| mmu-miR-9-5p | MIMAT0000142 | Btg2 | 12227 | HITS-CLIP | | 37986224 | Unspecified |
| mmu-miR-9-5p | MIMAT0000142 | C530008M17Rik | 320827 | HITS-CLIP | | 37986224 | Unspecified |
| mmu-miR-9-5p | MIMAT0000142 | Cacna2d3 | 12294 | HITS-CLIP | | 37986224 | Unspecified |
| mmu-miR-9-5p | MIMAT0000142 | Cadm1 | 54725 | HITS-CLIP | | 37986224 | Unspecified |
| mmu-miR-9-5p | MIMAT0000142 | Camk4 | 12326 | HITS-CLIP | | 37986224 | Unspecified |
| mmu-miR-9-5p | MIMAT0000142 | Cask | 12361 | HITS-CLIP | | 19536157 | Unspecified |
| mmu-miR-9-5p | MIMAT0000142 | Cbx6 | 494448 | HITS-CLIP | | 37986224 | Unspecified |
| mmu-miR-9-5p | MIMAT0000142 | Ccdc112 | 240261 | HITS-CLIP | | 37986224 | Unspecified |
| mmu-miR-9-5p | MIMAT0000142 | Ccdc88a | 108686 | HITS-CLIP | | 37986224 | Unspecified |
| mmu-miR-9-5p | MIMAT0000142 | Ccnd2 | 12444 | HITS-CLIP | | 37986224 | Unspecified |
| mmu-miR-9-5p | MIMAT0000142 | Ccnt2 | 72949 | HITS-CLIP | | 37986224 | Unspecified |
| mmu-miR-9-5p | MIMAT0000142 | Ccpg1os | 546143 | HITS-CLIP | | 37986224 | Unspecified |
| mmu-miR-9-5p | MIMAT0000142 | Ccser2 | 72972 | Sequencing//HITS-CLIP | | 19536157 | Unspecified |
| mmu-miR-9-5p | MIMAT0000142 | Cct2 | 12461 | HITS-CLIP | | 37986224 | Unspecified |
| mmu-miR-9-5p | MIMAT0000142 | Cd93 | 17064 | HITS-CLIP | | 37986224 | Unspecified |
| mmu-miR-9-5p | MIMAT0000142 | Cdc37 | 12539 | qCLASH | | 31393953 | Unspecified |
| mmu-miR-9-5p | MIMAT0000142 | Cdc37l1 | 67072 | HITS-CLIP | | 37986224 | Unspecified |
| mmu-miR-9-5p | MIMAT0000142 | Cdh11 | 12552 | HITS-CLIP | | 37986224 | Unspecified |
| mmu-miR-9-5p | MIMAT0000142 | Cdh12 | 215654 | HITS-CLIP | | 37986224 | Unspecified |
| mmu-miR-9-5p | MIMAT0000142 | Cdh20 | 23836 | HITS-CLIP | | 37986224 | Unspecified |
| mmu-miR-9-5p | MIMAT0000142 | Cdk5r1 | 12569 | HITS-CLIP | | 37986224 | Unspecified |
| mmu-miR-9-5p | MIMAT0000142 | Cdk8 | 264064 | HITS-CLIP | | 37986224 | Unspecified |
| mmu-miR-9-5p | MIMAT0000142 | Cdkl5 | 382253 | HITS-CLIP | | 37986224 | Unspecified |
| mmu-miR-9-5p | MIMAT0000142 | Cdkn2aipnl | 52626 | HITS-CLIP | | 37986224 | Unspecified |
| mmu-miR-9-5p | MIMAT0000142 | Celf2 | 14007 | HITS-CLIP | 37986224\|19536157 | | Unspecified |
| mmu-miR-9-5p | MIMAT0000142 | Cep162 | 382090 | HITS-CLIP | | 37986224 | Unspecified |
| mmu-miR-9-5p | MIMAT0000142 | Cep350 | 74081 | HITS-CLIP | | 37986224 | Unspecified |
| mmu-miR-9-5p | MIMAT0000142 | Chd1 | 12648 | HITS-CLIP | | 37986224 | Unspecified |
| mmu-miR-9-5p | MIMAT0000142 | Chd3 | 216848 | HITS-CLIP | | 19536157 | Unspecified |
| mmu-miR-9-5p | MIMAT0000142 | Chd6 | 71389 | HITS-CLIP | | 37986224 | Unspecified |
| mmu-miR-9-5p | MIMAT0000142 | Chfr | 231600 | qCLASH | | 31393953 | Unspecified |
| mmu-miR-9-5p | MIMAT0000142 | Chgb | 12653 | HITS-CLIP | | 37986224 | Unspecified |
| mmu-miR-9-5p | MIMAT0000142 | Chmp2b | 68942 | HITS-CLIP | | 37986224 | Unspecified |
| mmu-miR-9-5p | MIMAT0000142 | Chn1 | 108699 | HITS-CLIP | | 37986224 | Unspecified |
| mmu-miR-9-5p | MIMAT0000142 | Chsy1 | 269941 | HITS-CLIP | 37986224\|19536157 | | Unspecified |
| mmu-miR-9-5p | MIMAT0000142 | Chsy3 | 78923 | HITS-CLIP | | 37986224 | Unspecified |
| mmu-miR-9-5p | MIMAT0000142 | Cit | 12704 | HITS-CLIP | | 19536157 | Unspecified |
| mmu-miR-9-5p | MIMAT0000142 | Clcn5 | 12728 | HITS-CLIP | | 37986224 | Unspecified |

| | | | | | | | |
|---|---|---|---|---|---|---|---|
| mmu-miR-9-5p | MIMAT0000142 | Cldn11 | 18417 | HITS-CLIP | | 37986224 | Unspecified |
| mmu-miR-9-5p | MIMAT0000142 | Clic4 | 29876 | qCLASH | | 31393953 | Unspecified |
| mmu-miR-9-5p | MIMAT0000142 | Clip1 | 56430 | Sequencing//HITS-CLIP | 37986224\|19536157 | | Unspecified |
| mmu-miR-9-5p | MIMAT0000142 | Clip3 | 76686 | HITS-CLIP | | 37986224 | Unspecified |
| mmu-miR-9-5p | MIMAT0000142 | Clptm1 | 56457 | HITS-CLIP | 37986224\|19536157 | | Unspecified |
| mmu-miR-9-5p | MIMAT0000142 | Cmas | 12764 | HITS-CLIP | | 37986224 | Unspecified |
| mmu-miR-9-5p | MIMAT0000142 | Cmtr2 | 234728 | HITS-CLIP | | 37986224 | Unspecified |
| mmu-miR-9-5p | MIMAT0000142 | Cnnm1 | 83674 | HITS-CLIP | | 37986224 | Unspecified |
| mmu-miR-9-5p | MIMAT0000142 | Cnnm2 | 94219 | Sequencing//HITS-CLIP | 37986224\|19536157 | | Unspecified |
| mmu-miR-9-5p | MIMAT0000142 | Cntn3 | 18488 | Sequencing//HITS-CLIP | 37986224\|19536157 | | Unspecified |
| mmu-miR-9-5p | MIMAT0000142 | Cog5 | 238123 | HITS-CLIP | | 37986224 | Unspecified |
| mmu-miR-9-5p | MIMAT0000142 | Col1a2 | 12843 | HITS-CLIP | | 37986224 | Unspecified |
| mmu-miR-9-5p | MIMAT0000142 | Colec12 | 140792 | Sequencing//HITS-CLIP | 19536157\|37986224 | | Unspecified |
| mmu-miR-9-5p | MIMAT0000142 | Copg1 | 54161 | HITS-CLIP | | 37986224 | Unspecified |
| mmu-miR-9-5p | MIMAT0000142 | Coro1c | 23790 | HITS-CLIP | | 37986224 | Unspecified |
| mmu-miR-9-5p | MIMAT0000142 | Cox10 | 70383 | HITS-CLIP | | 37986224 | Unspecified |
| mmu-miR-9-5p | MIMAT0000142 | Cox7b | 66142 | HITS-CLIP | | 37986224 | Unspecified |
| mmu-miR-9-5p | MIMAT0000142 | Cpd | 12874 | HITS-CLIP | | 37986224 | Unspecified |
| mmu-miR-9-5p | MIMAT0000142 | Cpe | 12876 | HITS-CLIP | | 37986224 | Unspecified |
| mmu-miR-9-5p | MIMAT0000142 | Cplane1 | 65250 | HITS-CLIP | | 19536157 | Unspecified |
| mmu-miR-9-5p | MIMAT0000142 | Cpox | 12892 | HITS-CLIP | | 37986224 | Unspecified |
| mmu-miR-9-5p | MIMAT0000142 | Cpsf6 | 432508 | HITS-CLIP | | 37986224 | Unspecified |
| mmu-miR-9-5p | MIMAT0000142 | Cpsf7 | 269061 | HITS-CLIP | | 37986224 | Unspecified |
| mmu-miR-9-5p | MIMAT0000142 | Crebbp | 12914 | HITS-CLIP | | 37986224 | Unspecified |
| mmu-miR-9-5p | MIMAT0000142 | Crebrf | 77128 | Sequencing//HITS-CLIP | | 19536157 | Unspecified |
| mmu-miR-9-5p | MIMAT0000142 | Crim1 | 50766 | HITS-CLIP | | 37986224 | Unspecified |
| mmu-miR-9-5p | MIMAT0000142 | Csde1 | 229663 | HITS-CLIP | | 37986224 | Unspecified |
| mmu-miR-9-5p | MIMAT0000142 | Cthrc1 | 68588 | HITS-CLIP | | 37986224 | Unspecified |
| mmu-miR-9-5p | MIMAT0000142 | Ctsl | 13039 | HITS-CLIP | 37986224\|19536157 | | Unspecified |
| mmu-miR-9-5p | MIMAT0000142 | Cwf19l2 | 244672 | HITS-CLIP | | 37986224 | Unspecified |
| mmu-miR-9-5p | MIMAT0000142 | Cxadr | 13052 | HITS-CLIP | | 37986224 | Unspecified |
| mmu-miR-9-5p | MIMAT0000142 | Cyb561 | 13056 | HITS-CLIP | | 37986224 | Unspecified |
| mmu-miR-9-5p | MIMAT0000142 | Cyhr1 | 54151 | HITS-CLIP | | 37986224 | Unspecified |
| mmu-miR-9-5p | MIMAT0000142 | D430019H16Rik | 268595 | HITS-CLIP | | 37986224 | Unspecified |
| mmu-miR-9-5p | MIMAT0000142 | Daam1 | 208846 | HITS-CLIP | | 37986224 | Unspecified |
| mmu-miR-9-5p | MIMAT0000142 | Dbx2 | 223843 | HITS-CLIP | | 37986224 | Unspecified |
| mmu-miR-9-5p | MIMAT0000142 | Dcp2 | 70640 | HITS-CLIP | | 37986224 | Unspecified |
| mmu-miR-9-5p | MIMAT0000142 | Dcun1d4 | 100737 | HITS-CLIP | | 37986224 | Unspecified |
| mmu-miR-9-5p | MIMAT0000142 | Ddhd2 | 72108 | HITS-CLIP | | 37986224 | Unspecified |

| | | | | | | |
|---|---|---|---|---|---|---|
| mmu-miR-9-5p | MIMAT0000142 | Ddi2 | 68817 | HITS-CLIP | 37986224 | Unspecified |
| mmu-miR-9-5p | MIMAT0000142 | Ddx10 | 77591 | HITS-CLIP | 37986224 | Unspecified |
| mmu-miR-9-5p | MIMAT0000142 | Ddx20 | 53975 | HITS-CLIP | 37986224 | Unspecified |
| mmu-miR-9-5p | MIMAT0000142 | Ddx6 | 13209 | HITS-CLIP | 37986224 | Unspecified |
| mmu-miR-9-5p | MIMAT0000142 | Derl1 | 67819 | HITS-CLIP | 37986224 | Unspecified |
| mmu-miR-9-5p | MIMAT0000142 | Dgkh | 380921 | HITS-CLIP | 37986224 | Unspecified |
| mmu-miR-9-5p | MIMAT0000142 | Dhdds | 67422 | HITS-CLIP | 37986224 | Unspecified |
| mmu-miR-9-5p | MIMAT0000142 | Dhx29 | 218629 | HITS-CLIP | 37986224 | Unspecified |
| mmu-miR-9-5p | MIMAT0000142 | Dhx36 | 72162 | HITS-CLIP | 37986224 | Unspecified |
| mmu-miR-9-5p | MIMAT0000142 | Diablo | 66593 | HITS-CLIP | 37986224 | Unspecified |
| mmu-miR-9-5p | MIMAT0000142 | Dicer1 | 192119 | HITS-CLIP | 37986224 | Unspecified |
| mmu-miR-9-5p | MIMAT0000142 | Dido1 | 23856 | HITS-CLIP | 37986224 | Unspecified |
| mmu-miR-9-5p | MIMAT0000142 | Dlgap3 | 242667 | HITS-CLIP | 37986224 | Unspecified |
| mmu-miR-9-5p | MIMAT0000142 | Dlgap4 | 228836 | HITS-CLIP | 37986224 | Unspecified |
| mmu-miR-9-5p | MIMAT0000142 | Dnajb9 | 27362 | HITS-CLIP | 37986224 | Unspecified |
| mmu-miR-9-5p | MIMAT0000142 | Dnajc10 | 66861 | HITS-CLIP | 37986224\|19536157 | Unspecified |
| mmu-miR-9-5p | MIMAT0000142 | Dnajc14 | 74330 | Sequencing//HITS-CLIP | 37986224\|19536157 | Unspecified |
| mmu-miR-9-5p | MIMAT0000142 | Dnajc18 | 76594 | HITS-CLIP | 37986224 | Unspecified |
| mmu-miR-9-5p | MIMAT0000142 | Dnajc3 | 100037258 | HITS-CLIP | 37986224 | Unspecified |
| mmu-miR-9-5p | MIMAT0000142 | Dner | 227325 | HITS-CLIP | 37986224 | Unspecified |
| mmu-miR-9-5p | MIMAT0000142 | Dnm1 | 13429 | HITS-CLIP | 37986224 | Unspecified |
| mmu-miR-9-5p | MIMAT0000142 | Dock8 | 76088 | qCLASH | 31393953 | Unspecified |
| mmu-miR-9-5p | MIMAT0000142 | Dok5 | 76829 | HITS-CLIP | 37986224 | Unspecified |
| mmu-miR-9-5p | MIMAT0000142 | Dop1a | 23033 | HITS-CLIP | 19536157 | Unspecified |
| mmu-miR-9-5p | MIMAT0000142 | Dpysl2 | 12934 | HITS-CLIP, Luciferase Reporter Assay | 37986224 | Unspecified |
| mmu-miR-9-5p | MIMAT0000142 | Dusp10 | 63953 | HITS-CLIP | 37986224 | Unspecified |
| mmu-miR-9-5p | MIMAT0000142 | Dusp8 | 18218 | HITS-CLIP | 37986224 | Unspecified |
| mmu-miR-9-5p | MIMAT0000142 | Dync1h1 | 13424 | HITS-CLIP | 37986224\|19536157 | Unspecified |
| mmu-miR-9-5p | MIMAT0000142 | Dyrk1b | 13549 | HITS-CLIP | 19536157 | Unspecified |
| mmu-miR-9-5p | MIMAT0000142 | Dzip1 | 66573 | HITS-CLIP | 19536157 | Unspecified |
| mmu-miR-9-5p | MIMAT0000142 | Dzip1l | 72507 | HITS-CLIP | 37986224 | Unspecified |
| mmu-miR-9-5p | MIMAT0000142 | E2f3 | 13557 | HITS-CLIP | 37986224 | Unspecified |
| mmu-miR-9-5p | MIMAT0000142 | Ecpas | 23392 | HITS-CLIP | 19536157 | Unspecified |
| mmu-miR-9-5p | MIMAT0000142 | Edem3 | 66967 | HITS-CLIP | 37986224 | Unspecified |
| mmu-miR-9-5p | MIMAT0000142 | Edrf1 | 214764 | HITS-CLIP | 19536157 | Unspecified |
| mmu-miR-9-5p | MIMAT0000142 | Efr3a | 76740 | HITS-CLIP | 37986224 | Unspecified |
| mmu-miR-9-5p | MIMAT0000142 | Ehd1 | 13660 | HITS-CLIP | 37986224 | Unspecified |
| mmu-miR-9-5p | MIMAT0000142 | Eif1a | 13664 | Sequencing//HITS-CLIP | 19536157\|37986224 | Unspecified |
| mmu-miR-9-5p | MIMAT0000142 | Eif4e3 | 66892 | HITS-CLIP | 37986224 | Unspecified |

| | | | | | | |
|---|---|---|---|---|---|---|
| mmu-miR-9-5p | MIMAT0000142 | Eif4g2 | 13690 | HITS-CLIP | 37986224 | Unspecified |
| mmu-miR-9-5p | MIMAT0000142 | Eif5 | 217869 | HITS-CLIP | 26203562 | Unspecified |
| mmu-miR-9-5p | MIMAT0000142 | Elavl1 | 15568 | HITS-CLIP | 37986224 | Unspecified |
| mmu-miR-9-5p | MIMAT0000142 | Elovl7 | 74559 | HITS-CLIP | 37986224 | Unspecified |
| mmu-miR-9-5p | MIMAT0000142 | Elp2 | 58523 | HITS-CLIP | 37986224 | Unspecified |
| mmu-miR-9-5p | MIMAT0000142 | Emb | 13723 | HITS-CLIP | 37986224 | Unspecified |
| mmu-miR-9-5p | MIMAT0000142 | Enpp6 | 320981 | HITS-CLIP | 37986224 | Unspecified |
| mmu-miR-9-5p | MIMAT0000142 | Ensa | 56205 | qCLASH | 31393953 | Unspecified |
| mmu-miR-9-5p | MIMAT0000142 | Epha4 | 13838 | HITS-CLIP | 37986224 | Unspecified |
| mmu-miR-9-5p | MIMAT0000142 | Epn1 | 13854 | HITS-CLIP | 37986224 | Unspecified |
| mmu-miR-9-5p | MIMAT0000142 | Ercc6l2 | 76251 | HITS-CLIP | 37986224 | Unspecified |
| mmu-miR-9-5p | MIMAT0000142 | Eri3 | 140546 | HITS-CLIP | 37986224 | Unspecified |
| mmu-miR-9-5p | MIMAT0000142 | Ermp1 | 226090 | HITS-CLIP | 37986224 | Unspecified |
| mmu-miR-9-5p | MIMAT0000142 | Ern1 | 78943 | HITS-CLIP | 37986224 | Unspecified |
| mmu-miR-9-5p | MIMAT0000142 | Esrra | 26379 | HITS-CLIP | 37986224 | Unspecified |
| mmu-miR-9-5p | MIMAT0000142 | Ets2 | 23872 | HITS-CLIP | 19536157 | Unspecified |
| mmu-miR-9-5p | MIMAT0000142 | Exosc9 | 50911 | HITS-CLIP | 19536157 | Unspecified |
| mmu-miR-9-5p | MIMAT0000142 | Ext1 | 14042 | HITS-CLIP | 37986224 | Unspecified |
| mmu-miR-9-5p | MIMAT0000142 | Fam13c | 71721 | HITS-CLIP | 37986224 | Unspecified |
| mmu-miR-9-5p | MIMAT0000142 | Fam199x | 245622 | HITS-CLIP | 37986224 | Unspecified |
| mmu-miR-9-5p | MIMAT0000142 | Fam214b | 230088 | HITS-CLIP | 37986224 | Unspecified |
| mmu-miR-9-5p | MIMAT0000142 | Fam76b | 72826 | HITS-CLIP | 37986224 | Unspecified |
| mmu-miR-9-5p | MIMAT0000142 | Fat3 | 270120 | HITS-CLIP | 37986224 | Unspecified |
| mmu-miR-9-5p | MIMAT0000142 | Fbxl2 | 72179 | HITS-CLIP | 37986224 | Unspecified |
| mmu-miR-9-5p | MIMAT0000142 | Fbxo41 | 330369 | HITS-CLIP | 19536157 | Unspecified |
| mmu-miR-9-5p | MIMAT0000142 | Fbxw11 | 103583 | Sequencing//HITS-CLIP | 19536157\|37986224 | Unspecified |
| mmu-miR-9-5p | MIMAT0000142 | Fh1 | 14194 | HITS-CLIP | 37986224 | Unspecified |
| mmu-miR-9-5p | MIMAT0000142 | Fig4 | 103199 | HITS-CLIP | 37986224 | Unspecified |
| mmu-miR-9-5p | MIMAT0000142 | Fign | 60344 | HITS-CLIP | 37986224 | Unspecified |
| mmu-miR-9-5p | MIMAT0000142 | Fkbp7 | 14231 | HITS-CLIP | 37986224\|19536157 | Unspecified |
| mmu-miR-9-5p | MIMAT0000142 | Fnbp4 | 55935 | HITS-CLIP | 37986224 | Unspecified |
| mmu-miR-9-5p | MIMAT0000142 | Fnta | 14272 | HITS-CLIP | 37986224 | Unspecified |
| mmu-miR-9-5p | MIMAT0000142 | Foxo1 | 56458 | HITS-CLIP | 37986224 | Unspecified |
| mmu-miR-9-5p | MIMAT0000142 | Foxo3 | 56484 | HITS-CLIP//Immunoblot//Immunohistoche | 37986224\|19536157\|26470727 | Unspecified |
| mmu-miR-9-5p | MIMAT0000142 | Foxp2 | 114142 | HITS-CLIP | 37986224 | Unspecified |
| mmu-miR-9-5p | MIMAT0000142 | Foxp4 | 74123 | Sequencing//HITS-CLIP | 37986224\|19536157 | Unspecified |
| mmu-miR-9-5p | MIMAT0000142 | Frem2 | 242022 | HITS-CLIP | 37986224\|19536157 | Unspecified |
| mmu-miR-9-5p | MIMAT0000142 | Fry | 320365 | HITS-CLIP | 37986224 | Unspecified |
| mmu-miR-9-5p | MIMAT0000142 | Fth1 | 14319 | HITS-CLIP | 37986224 | Unspecified |

| | | | | | | | |
|---|---|---|---|---|---|---|---|
| mmu-miR-9-5p | MIMAT0000142 | Fto | 26383 | HITS-CLIP | | 37986224 | Unspecified |
| mmu-miR-9-5p | MIMAT0000142 | Fundc2 | 67391 | qCLASH | | 31393953 | Unspecified |
| mmu-miR-9-5p | MIMAT0000142 | Fut8 | 53618 | HITS-CLIP | | 19536157 | Unspecified |
| mmu-miR-9-5p | MIMAT0000142 | Fyn | 14360 | HITS-CLIP | 37986224\|19536157 | | Unspecified |
| mmu-miR-9-5p | MIMAT0000142 | Fyttd1 | 69823 | Sequencing//HITS-CLIP | 37986224\|19536157 | | Unspecified |
| mmu-miR-9-5p | MIMAT0000142 | Fzd3 | 14365 | HITS-CLIP | | 37986224 | Unspecified |
| mmu-miR-9-5p | MIMAT0000142 | Gabbr2 | 242425 | HITS-CLIP | | 37986224 | Unspecified |
| mmu-miR-9-5p | MIMAT0000142 | Gabra2 | 14395 | HITS-CLIP | | 37986224 | Unspecified |
| mmu-miR-9-5p | MIMAT0000142 | Gabra4 | 14397 | HITS-CLIP | | 37986224 | Unspecified |
| mmu-miR-9-5p | MIMAT0000142 | Gabrb2 | 14401 | HITS-CLIP | | 37986224 | Unspecified |
| mmu-miR-9-5p | MIMAT0000142 | Galnt16 | 108760 | HITS-CLIP | | 37986224 | Unspecified |
| mmu-miR-9-5p | MIMAT0000142 | Gba2 | 230101 | HITS-CLIP | | 37986224 | Unspecified |
| mmu-miR-9-5p | MIMAT0000142 | Gca | 227960 | HITS-CLIP | | 37986224 | Unspecified |
| mmu-miR-9-5p | MIMAT0000142 | Gdpd5 | 233552 | HITS-CLIP | | 37986224 | Unspecified |
| mmu-miR-9-5p | MIMAT0000142 | Gja1 | 14609 | HITS-CLIP | | 37986224 | Unspecified |
| mmu-miR-9-5p | MIMAT0000142 | Glb1l | 74577 | HITS-CLIP | | 37986224 | Unspecified |
| mmu-miR-9-5p | MIMAT0000142 | Gm5283 | 383936 | HITS-CLIP | | 37986224 | Unspecified |
| mmu-miR-9-5p | MIMAT0000142 | Gm9774 | 100043022 | HITS-CLIP | | 37986224 | Unspecified |
| mmu-miR-9-5p | MIMAT0000142 | Gmeb2 | 229004 | Luciferase Reporter Assay | | 37986224 | Unspecified |
| mmu-miR-9-5p | MIMAT0000142 | Gna13 | 14674 | HITS-CLIP | | 37986224 | Unspecified |
| mmu-miR-9-5p | MIMAT0000142 | Gnb4 | 14696 | HITS-CLIP | | 37986224 | Unspecified |
| mmu-miR-9-5p | MIMAT0000142 | Gng12 | 14701 | HITS-CLIP | | 37986224 | Unspecified |
| mmu-miR-9-5p | MIMAT0000142 | Gnptab | 432486 | HITS-CLIP | | 37986224 | Unspecified |
| mmu-miR-9-5p | MIMAT0000142 | Gpatch2 | 67769 | HITS-CLIP | | 37986224 | Unspecified |
| mmu-miR-9-5p | MIMAT0000142 | Gpd1l | 333433 | HITS-CLIP | | 37986224 | Unspecified |
| mmu-miR-9-5p | MIMAT0000142 | Gpm6a | 234267 | HITS-CLIP | | 37986224 | Unspecified |
| mmu-miR-9-5p | MIMAT0000142 | Gpr158 | 241263 | HITS-CLIP | 37986224\|19536157 | | Unspecified |
| mmu-miR-9-5p | MIMAT0000142 | Gpr26 | 233919 | HITS-CLIP | | 37986224 | Unspecified |
| mmu-miR-9-5p | MIMAT0000142 | Gprasp1 | 67298 | HITS-CLIP | | 37986224 | Unspecified |
| mmu-miR-9-5p | MIMAT0000142 | Gprc5b | 64297 | HITS-CLIP | | 37986224 | Unspecified |
| mmu-miR-9-5p | MIMAT0000142 | Gpt2 | 108682 | HITS-CLIP | | 37986224 | Unspecified |
| mmu-miR-9-5p | MIMAT0000142 | Gpx8 | 69590 | HITS-CLIP | | 37986224 | Unspecified |
| mmu-miR-9-5p | MIMAT0000142 | Grin2a | 14811 | HITS-CLIP | | 19536157 | Unspecified |
| mmu-miR-9-5p | MIMAT0000142 | Grin2b | 14812 | Sequencing//HITS-CLIP | 19536157\|37986224 | | Unspecified |
| mmu-miR-9-5p | MIMAT0000142 | Grm4 | 268934 | HITS-CLIP | | 37986224 | Unspecified |
| mmu-miR-9-5p | MIMAT0000142 | H1f0 | 14958 | HITS-CLIP | | 37986224 | Unspecified |
| mmu-miR-9-5p | MIMAT0000142 | Hadha | 97212 | HITS-CLIP | | 37986224 | Unspecified |
| mmu-miR-9-5p | MIMAT0000142 | Has2 | 15117 | HITS-CLIP | | 37986224 | Unspecified |
| mmu-miR-9-5p | MIMAT0000142 | Haus3 | 231123 | HITS-CLIP | | 37986224 | Unspecified |

| | | | | | | |
|---|---|---|---|---|---|---|
| mmu-miR-9-5p | MIMAT0000142 | Hcfc1 | 15161 | HITS-CLIP | 37986224 | Unspecified |
| mmu-miR-9-5p | MIMAT0000142 | Hcfc2 | 67933 | HITS-CLIP | 37986224 | Unspecified |
| mmu-miR-9-5p | MIMAT0000142 | Heatr5b | 320473 | HITS-CLIP | 37986224 | Unspecified |
| mmu-miR-9-5p | MIMAT0000142 | Heatr6 | 217026 | HITS-CLIP | 37986224\|19536157 | Unspecified |
| mmu-miR-9-5p | MIMAT0000142 | Hectd4 | 283450 | HITS-CLIP | 19536157 | Unspecified |
| mmu-miR-9-5p | MIMAT0000142 | Heg1 | 77446 | HITS-CLIP | 37986224 | Unspecified |
| mmu-miR-9-5p | MIMAT0000142 | Herc1 | 235439 | HITS-CLIP | 37986224\|19536157 | Unspecified |
| mmu-miR-9-5p | MIMAT0000142 | Herc6 | 67138 | HITS-CLIP | 37986224 | Unspecified |
| mmu-miR-9-5p | MIMAT0000142 | Herpud2 | 80517 | HITS-CLIP | 19536157 | Unspecified |
| mmu-miR-9-5p | MIMAT0000142 | Hhex | 15242 | HITS-CLIP | 37986224 | Unspecified |
| mmu-miR-9-5p | MIMAT0000142 | Hic1 | 15248 | HITS-CLIP | 19536157 | Unspecified |
| mmu-miR-9-5p | MIMAT0000142 | Hic2 | 58180 | HITS-CLIP | 37986224 | Unspecified |
| mmu-miR-9-5p | MIMAT0000142 | Hint3 | 66847 | HITS-CLIP | 37986224 | Unspecified |
| mmu-miR-9-5p | MIMAT0000142 | Hipk1 | 15257 | Sequencing//HITS-CLIP | 19536157\|37986224 | Unspecified |
| mmu-miR-9-5p | MIMAT0000142 | Hist1h1e | 3008 | HITS-CLIP | 19536157 | Unspecified |
| mmu-miR-9-5p | MIMAT0000142 | Hivep1 | 110521 | HITS-CLIP | 37986224\|19536157 | Unspecified |
| mmu-miR-9-5p | MIMAT0000142 | Hmox2 | 15369 | HITS-CLIP | 37986224 | Unspecified |
| mmu-miR-9-5p | MIMAT0000142 | Hnrnpu | 51810 | HITS-CLIP | 37986224 | Unspecified |
| mmu-miR-9-5p | MIMAT0000142 | Hs2st1 | 23908 | HITS-CLIP | 37986224 | Unspecified |
| mmu-miR-9-5p | MIMAT0000142 | Hs3st2 | 195646 | HITS-CLIP | 37986224 | Unspecified |
| mmu-miR-9-5p | MIMAT0000142 | Hs3st5 | 319415 | HITS-CLIP | 19536157 | Unspecified |
| mmu-miR-9-5p | MIMAT0000142 | Hsf2 | 15500 | HITS-CLIP | 37986224 | Unspecified |
| mmu-miR-9-5p | MIMAT0000142 | Hsp90ab1 | 15516 | HITS-CLIP | 37986224 | Unspecified |
| mmu-miR-9-5p | MIMAT0000142 | Hsp90b1 | 22027 | HITS-CLIP | 37986224 | Unspecified |
| mmu-miR-9-5p | MIMAT0000142 | Hspa13 | 110920 | HITS-CLIP | 37986224 | Unspecified |
| mmu-miR-9-5p | MIMAT0000142 | Hspa4 | 15525 | HITS-CLIP | 37986224 | Unspecified |
| mmu-miR-9-5p | MIMAT0000142 | Hspa9 | 15526 | HITS-CLIP | 37986224\|19536157 | Unspecified |
| mmu-miR-9-5p | MIMAT0000142 | Hunk | 26559 | HITS-CLIP | 37986224 | Unspecified |
| mmu-miR-9-5p | MIMAT0000142 | Id4 | 15904 | HITS-CLIP | 19536157 | Unspecified |
| mmu-miR-9-5p | MIMAT0000142 | Ide | 15925 | HITS-CLIP | 19536157 | Unspecified |
| mmu-miR-9-5p | MIMAT0000142 | Idh1 | 15926 | HITS-CLIP | 19536157 | Unspecified |
| mmu-miR-9-5p | MIMAT0000142 | Idh3a | 67834 | HITS-CLIP | 37986224 | Unspecified |
| mmu-miR-9-5p | MIMAT0000142 | Idh3g | 15929 | HITS-CLIP | 19536157 | Unspecified |
| mmu-miR-9-5p | MIMAT0000142 | Ids | 15931 | HITS-CLIP | 37986224 | Unspecified |
| mmu-miR-9-5p | MIMAT0000142 | Ifngr2 | 15980 | HITS-CLIP | 37986224 | Unspecified |
| mmu-miR-9-5p | MIMAT0000142 | Ift52 | 245866 | HITS-CLIP | 37986224 | Unspecified |
| mmu-miR-9-5p | MIMAT0000142 | Igfbp5 | 16011 | HITS-CLIP | 37986224 | Unspecified |
| mmu-miR-9-5p | MIMAT0000142 | Il6st | 16195 | HITS-CLIP | 37986224 | Unspecified |
| mmu-miR-9-5p | MIMAT0000142 | Ildr2 | 100039795 | HITS-CLIP | 37986224 | Unspecified |

| | | | | | | | |
|---|---|---|---|---|---|---|---|
| mmu-miR-9-5p | MIMAT0000142 | Imp4 | 27993 | HITS-CLIP | | 37986224 | Unspecified |
| mmu-miR-9-5p | MIMAT0000142 | Impa1 | 55980 | Sequencing//HITS-CLIP | 37986224\|19536157 | | Unspecified |
| mmu-miR-9-5p | MIMAT0000142 | Impact | 16210 | HITS-CLIP | | 37986224 | Unspecified |
| mmu-miR-9-5p | MIMAT0000142 | Ints4 | 101861 | HITS-CLIP | | 37986224 | Unspecified |
| mmu-miR-9-5p | MIMAT0000142 | Ipo4 | 75751 | HITS-CLIP | | 37986224 | Unspecified |
| mmu-miR-9-5p | MIMAT0000142 | Iqgap1 | 29875 | HITS-CLIP | | 37986224 | Unspecified |
| mmu-miR-9-5p | MIMAT0000142 | Itgb8 | 320910 | HITS-CLIP | | 37986224 | Unspecified |
| mmu-miR-9-5p | MIMAT0000142 | Itm2b | 16432 | HITS-CLIP | 37986224\|19536157 | | Unspecified |
| mmu-miR-9-5p | MIMAT0000142 | Itpr1 | 16438 | HITS-CLIP | | 37986224 | Unspecified |
| mmu-miR-9-5p | MIMAT0000142 | Itpripl2 | 319622 | HITS-CLIP | | 37986224 | Unspecified |
| mmu-miR-9-5p | MIMAT0000142 | Itsn2 | 20403 | HITS-CLIP | 37986224\|19536157 | | Unspecified |
| mmu-miR-9-5p | MIMAT0000142 | Jag2 | 16450 | HITS-CLIP | | 37986224 | Unspecified |
| mmu-miR-9-5p | MIMAT0000142 | Jak1 | 16451 | HITS-CLIP | | 37986224 | Unspecified |
| mmu-miR-9-5p | MIMAT0000142 | Jmjd1c | 108829 | HITS-CLIP | | 37986224 | Unspecified |
| mmu-miR-9-5p | MIMAT0000142 | Jph1 | 57339 | HITS-CLIP | 37986224\|19536157 | | Unspecified |
| mmu-miR-9-5p | MIMAT0000142 | Jrkl | 77532 | HITS-CLIP | | 37986224 | Unspecified |
| mmu-miR-9-5p | MIMAT0000142 | Kbtbd7 | 211255 | HITS-CLIP | | 37986224 | Unspecified |
| mmu-miR-9-5p | MIMAT0000142 | Kcnc1 | 16502 | Sequencing//HITS-CLIP | 19536157\|37986224 | | Unspecified |
| mmu-miR-9-5p | MIMAT0000142 | Kcnh7 | 170738 | HITS-CLIP | | 37986224 | Unspecified |
| mmu-miR-9-5p | MIMAT0000142 | Kcnj4 | 16520 | HITS-CLIP | | 37986224 | Unspecified |
| mmu-miR-9-5p | MIMAT0000142 | Kcnk10 | 72258 | HITS-CLIP | | 37986224 | Unspecified |
| mmu-miR-9-5p | MIMAT0000142 | Kcnk3 | 16527 | HITS-CLIP | | 37986224 | Unspecified |
| mmu-miR-9-5p | MIMAT0000142 | Kcnk4 | 16528 | HITS-CLIP | | 37986224 | Unspecified |
| mmu-miR-9-5p | MIMAT0000142 | Kcnn2 | 140492 | HITS-CLIP | | 19536157 | Unspecified |
| mmu-miR-9-5p | MIMAT0000142 | Kcnq2 | 16536 | HITS-CLIP | | 37986224 | Unspecified |
| mmu-miR-9-5p | MIMAT0000142 | Kcnq3 | 110862 | HITS-CLIP | | 37986224 | Unspecified |
| mmu-miR-9-5p | MIMAT0000142 | Kcnt2 | 240776 | HITS-CLIP | | 37986224 | Unspecified |
| mmu-miR-9-5p | MIMAT0000142 | Kctd12 | 239217 | HITS-CLIP | | 37986224 | Unspecified |
| mmu-miR-9-5p | MIMAT0000142 | Kdelr2 | 66913 | HITS-CLIP | | 37986224 | Unspecified |
| mmu-miR-9-5p | MIMAT0000142 | Kdm5b | 75605 | qCLASH | | 31393953 | Unspecified |
| mmu-miR-9-5p | MIMAT0000142 | Kdsr | 70750 | HITS-CLIP | | 37986224 | Unspecified |
| mmu-miR-9-5p | MIMAT0000142 | Kif13a | 16553 | HITS-CLIP | 37986224\|19536157 | | Unspecified |
| mmu-miR-9-5p | MIMAT0000142 | Kif5b | 16573 | HITS-CLIP | | 37986224 | Unspecified |
| mmu-miR-9-5p | MIMAT0000142 | Klf13 | 50794 | HITS-CLIP | | 37986224 | Unspecified |
| mmu-miR-9-5p | MIMAT0000142 | Klf3 | 16599 | HITS-CLIP | | 37986224 | Unspecified |
| mmu-miR-9-5p | MIMAT0000142 | Klf9 | 16601 | HITS-CLIP | | 37986224 | Unspecified |
| mmu-miR-9-5p | MIMAT0000142 | Klhl11 | 217194 | HITS-CLIP | | 37986224 | Unspecified |
| mmu-miR-9-5p | MIMAT0000142 | Klhl23 | 277396 | HITS-CLIP | | 37986224 | Unspecified |
| mmu-miR-9-5p | MIMAT0000142 | Klhl42 | 232539 | HITS-CLIP | | 37986224 | Unspecified |

| | | | | | | |
|---|---|---|---|---|---|---|
| mmu-miR-9-5p | MIMAT0000142 | Klhl9 | 242521 | HITS-CLIP | 37986224 | Unspecified |
| mmu-miR-9-5p | MIMAT0000142 | Kmt2c | 231051 | HITS-CLIP | 37986224 | Unspecified |
| mmu-miR-9-5p | MIMAT0000142 | Kpnb1 | 16211 | HITS-CLIP | 37986224 | Unspecified |
| mmu-miR-9-5p | MIMAT0000142 | Ksr2 | 333050 | HITS-CLIP | 37986224 | Unspecified |
| mmu-miR-9-5p | MIMAT0000142 | Lamp2 | 16784 | HITS-CLIP//qCLASH | 37986224\|31393953 | Unspecified |
| mmu-miR-9-5p | MIMAT0000142 | Lca5 | 75782 | HITS-CLIP | 37986224 | Unspecified |
| mmu-miR-9-5p | MIMAT0000142 | Lhx1 | 16869 | HITS-CLIP | 37986224 | Unspecified |
| mmu-miR-9-5p | MIMAT0000142 | Lhx2 | 16870 | HITS-CLIP | 37986224 | Unspecified |
| mmu-miR-9-5p | MIMAT0000142 | Lifr | 16880 | HITS-CLIP | 37986224\|19536157 | Unspecified |
| mmu-miR-9-5p | MIMAT0000142 | Limk2 | 16886 | qCLASH | 31393953 | Unspecified |
| mmu-miR-9-5p | MIMAT0000142 | Lman2 | 66890 | HITS-CLIP | 37986224 | Unspecified |
| mmu-miR-9-5p | MIMAT0000142 | Lnpep | 240028 | HITS-CLIP | 37986224 | Unspecified |
| mmu-miR-9-5p | MIMAT0000142 | Lrch4 | 231798 | HITS-CLIP | 37986224 | Unspecified |
| mmu-miR-9-5p | MIMAT0000142 | Lrp8 | 16975 | HITS-CLIP | 37986224 | Unspecified |
| mmu-miR-9-5p | MIMAT0000142 | Lrrc7 | 242274 | HITS-CLIP | 37986224 | Unspecified |
| mmu-miR-9-5p | MIMAT0000142 | Lrrc8d | 231549 | HITS-CLIP | 37986224 | Unspecified |
| mmu-miR-9-5p | MIMAT0000142 | Lrrn3 | 16981 | HITS-CLIP | 37986224 | Unspecified |
| mmu-miR-9-5p | MIMAT0000142 | Lrrtm4 | 243499 | HITS-CLIP | 37986224 | Unspecified |
| mmu-miR-9-5p | MIMAT0000142 | Lsamp | 268890 | Sequencing//HITS-CLIP | 37986224\|19536157 | Unspecified |
| mmu-miR-9-5p | MIMAT0000142 | Lsm11 | 72290 | HITS-CLIP | 37986224 | Unspecified |
| mmu-miR-9-5p | MIMAT0000142 | Luc7l | 66978 | HITS-CLIP | 37986224 | Unspecified |
| mmu-miR-9-5p | MIMAT0000142 | Luzp1 | 269593 | HITS-CLIP | 37986224 | Unspecified |
| mmu-miR-9-5p | MIMAT0000142 | Lxn | 17035 | HITS-CLIP | 37986224 | Unspecified |
| mmu-miR-9-5p | MIMAT0000142 | Lztfl1 | 93730 | qCLASH | 31393953 | Unspecified |
| mmu-miR-9-5p | MIMAT0000142 | Lzts3 | 241638 | Sequencing//HITS-CLIP | 19536157 | Unspecified |
| mmu-miR-9-5p | MIMAT0000142 | Macf1 | 11426 | HITS-CLIP | 37986224 | Unspecified |
| mmu-miR-9-5p | MIMAT0000142 | Maged1 | 94275 | HITS-CLIP | 19536157 | Unspecified |
| mmu-miR-9-5p | MIMAT0000142 | Magee1 | 107528 | HITS-CLIP | 37986224 | Unspecified |
| mmu-miR-9-5p | MIMAT0000142 | Maml2 | 270118 | HITS-CLIP | 37986224\|19536157 | Unspecified |
| mmu-miR-9-5p | MIMAT0000142 | Man2a1 | 17158 | HITS-CLIP | 37986224 | Unspecified |
| mmu-miR-9-5p | MIMAT0000142 | Man2b1 | 17159 | HITS-CLIP | 37986224 | Unspecified |
| mmu-miR-9-5p | MIMAT0000142 | Maoa | 17161 | HITS-CLIP | 37986224 | Unspecified |
| mmu-miR-9-5p | MIMAT0000142 | Map1a | 17754 | HITS-CLIP | 37986224 | Unspecified |
| mmu-miR-9-5p | MIMAT0000142 | Map1b | 17755 | HITS-CLIP | 37986224 | Unspecified |
| mmu-miR-9-5p | MIMAT0000142 | Map2 | 17756 | HITS-CLIP | 37986224\|19536157 | Unspecified |
| mmu-miR-9-5p | MIMAT0000142 | Map2k4 | 26398 | HITS-CLIP | 37986224 | Unspecified |
| mmu-miR-9-5p | MIMAT0000142 | Map2k7 | 26400 | HITS-CLIP | 37986224 | Unspecified |
| mmu-miR-9-5p | MIMAT0000142 | Map3k1 | 26401 | HITS-CLIP | 37986224 | Unspecified |
| mmu-miR-9-5p | MIMAT0000142 | Map9 | 213582 | HITS-CLIP | 37986224 | Unspecified |

| | | | | | | |
|---|---|---|---|---|---|---|
| mmu-miR-9-5p | MIMAT0000142 | Mapk10 | 26414 | HITS-CLIP | | 37986224 Unspecified |
| mmu-miR-9-5p | MIMAT0000142 | Mapk8ip2 | 60597 | HITS-CLIP | | 37986224 Unspecified |
| mmu-miR-9-5p | MIMAT0000142 | Mapre1 | 13589 | HITS-CLIP | 37986224\|19536157 | Unspecified |
| mmu-miR-9-5p | MIMAT0000142 | Marchf6 | 223455 | Sequencing//HITS-CLIP | 37986224\|19536157 | Unspecified |
| mmu-miR-9-5p | MIMAT0000142 | Marcks | 17118 | HITS-CLIP | | 37986224 Unspecified |
| mmu-miR-9-5p | MIMAT0000142 | Mark3 | 17169 | HITS-CLIP | | 19536157 Unspecified |
| mmu-miR-9-5p | MIMAT0000142 | Mbnl1 | 56758 | HITS-CLIP | | 37986224 Unspecified |
| mmu-miR-9-5p | MIMAT0000142 | Mbnl2 | 105559 | HITS-CLIP | | 37986224 Unspecified |
| mmu-miR-9-5p | MIMAT0000142 | Mbp | 17196 | HITS-CLIP | | 37986224 Unspecified |
| mmu-miR-9-5p | MIMAT0000142 | Mbtd1 | 103537 | HITS-CLIP | | 37986224 Unspecified |
| mmu-miR-9-5p | MIMAT0000142 | Mcf2l | 17207 | HITS-CLIP | | 37986224 Unspecified |
| mmu-miR-9-5p | MIMAT0000142 | Mcm6 | 17219 | qCLASH | | 31393953 Unspecified |
| mmu-miR-9-5p | MIMAT0000142 | Mcm8 | 66634 | HITS-CLIP | | 37986224 Unspecified |
| mmu-miR-9-5p | MIMAT0000142 | Mdga2 | 320772 | Sequencing//HITS-CLIP | 37986224\|19536157 | Unspecified |
| mmu-miR-9-5p | MIMAT0000142 | Med10 | 28077 | HITS-CLIP | 37986224\|19536157 | Unspecified |
| mmu-miR-9-5p | MIMAT0000142 | Med13 | 327987 | HITS-CLIP | | 37986224 Unspecified |
| mmu-miR-9-5p | MIMAT0000142 | Med13l | 76199 | HITS-CLIP | | 37986224 Unspecified |
| mmu-miR-9-5p | MIMAT0000142 | Mef2c | 17260 | HITS-CLIP | | 19536157 Unspecified |
| mmu-miR-9-5p | MIMAT0000142 | Mfap3 | 216760 | HITS-CLIP | | 37986224 Unspecified |
| mmu-miR-9-5p | MIMAT0000142 | Mfap3l | 71306 | HITS-CLIP | | 37986224 Unspecified |
| mmu-miR-9-5p | MIMAT0000142 | Mfsd6 | 98682 | HITS-CLIP | | 19536157 Unspecified |
| mmu-miR-9-5p | MIMAT0000142 | Mical2 | 320878 | HITS-CLIP | | 19536157 Unspecified |
| mmu-miR-9-5p | MIMAT0000142 | Mki67 | 17345 | HITS-CLIP | | 37986224 Unspecified |
| mmu-miR-9-5p | MIMAT0000142 | Mknk2 | 17347 | HITS-CLIP | | 37986224 Unspecified |
| mmu-miR-9-5p | MIMAT0000142 | Mllt3 | 70122 | HITS-CLIP | | 37986224 Unspecified |
| mmu-miR-9-5p | MIMAT0000142 | Mob2 | 101513 | HITS-CLIP | | 37986224 Unspecified |
| mmu-miR-9-5p | MIMAT0000142 | Mpp1 | 17524 | HITS-CLIP | | 37986224 Unspecified |
| mmu-miR-9-5p | MIMAT0000142 | Mtfmt | 69606 | HITS-CLIP | | 37986224 Unspecified |
| mmu-miR-9-5p | MIMAT0000142 | Mtss1 | 211401 | HITS-CLIP | | 37986224 Unspecified |
| mmu-miR-9-5p | MIMAT0000142 | Mtus1 | 102103 | HITS-CLIP | | 19536157 Unspecified |
| mmu-miR-9-5p | MIMAT0000142 | Mtx3 | 382793 | HITS-CLIP | | 37986224 Unspecified |
| mmu-miR-9-5p | MIMAT0000142 | Mul1 | 68350 | HITS-CLIP | | 37986224 Unspecified |
| mmu-miR-9-5p | MIMAT0000142 | Myh9 | 17886 | HITS-CLIP | | 37986224 Unspecified |
| mmu-miR-9-5p | MIMAT0000142 | Myo9a | 270163 | HITS-CLIP | | 19536157 Unspecified |
| mmu-miR-9-5p | MIMAT0000142 | N4bp1 | 80750 | HITS-CLIP | | 37986224 Unspecified |
| mmu-miR-9-5p | MIMAT0000142 | Nacad | 192950 | HITS-CLIP | 37986224\|19536157 | Unspecified |
| mmu-miR-9-5p | MIMAT0000142 | Nampt | 59027 | HITS-CLIP | | 37986224 Unspecified |
| mmu-miR-9-5p | MIMAT0000142 | Nap1l3 | 54561 | HITS-CLIP | | 37986224 Unspecified |
| mmu-miR-9-5p | MIMAT0000142 | Napb | 17957 | HITS-CLIP | | 37986224 Unspecified |

| | | | | | | |
|---|---|---|---|---|---|---|
| mmu-miR-9-5p | MIMAT0000142 | Nasp | 50927 | HITS-CLIP | 37986224 | Unspecified |
| mmu-miR-9-5p | MIMAT0000142 | Nav3 | 260315 | HITS-CLIP | 37986224 | Unspecified |
| mmu-miR-9-5p | MIMAT0000142 | Nbea | 26422 | HITS-CLIP | 37986224 | Unspecified |
| mmu-miR-9-5p | MIMAT0000142 | Nbeal1 | 269198 | HITS-CLIP | 37986224 | Unspecified |
| mmu-miR-9-5p | MIMAT0000142 | Ncapd3 | 78658 | HITS-CLIP | 37986224 | Unspecified |
| mmu-miR-9-5p | MIMAT0000142 | Ncl | 17975 | HITS-CLIP | 37986224\|19536157 | Unspecified |
| mmu-miR-9-5p | MIMAT0000142 | Ncoa1 | 17977 | HITS-CLIP | 37986224 | Unspecified |
| mmu-miR-9-5p | MIMAT0000142 | Ncstn | 59287 | HITS-CLIP | 37986224 | Unspecified |
| mmu-miR-9-5p | MIMAT0000142 | Ndc1 | 72787 | HITS-CLIP | 37986224 | Unspecified |
| mmu-miR-9-5p | MIMAT0000142 | Ndfip1 | 65113 | HITS-CLIP | 37986224 | Unspecified |
| mmu-miR-9-5p | MIMAT0000142 | Ndufb8 | 67264 | HITS-CLIP | 37986224 | Unspecified |
| mmu-miR-9-5p | MIMAT0000142 | Ndufs4 | 17993 | HITS-CLIP | 37986224 | Unspecified |
| mmu-miR-9-5p | MIMAT0000142 | Necap1 | 67602 | HITS-CLIP | 37986224 | Unspecified |
| mmu-miR-9-5p | MIMAT0000142 | Nedd4 | 17999 | Sequencing//HITS-CLIP | 37986224\|19536157 | Unspecified |
| mmu-miR-9-5p | MIMAT0000142 | Nefh | 380684 | HITS-CLIP | 37986224 | Unspecified |
| mmu-miR-9-5p | MIMAT0000142 | Negr1 | 320840 | HITS-CLIP | 19536157 | Unspecified |
| mmu-miR-9-5p | MIMAT0000142 | Neurl4 | 216860 | HITS-CLIP | 37986224 | Unspecified |
| mmu-miR-9-5p | MIMAT0000142 | Nf1 | 18015 | Sequencing//HITS-CLIP | 19536157\|37986224 | Unspecified |
| mmu-miR-9-5p | MIMAT0000142 | Nfasc | 269116 | HITS-CLIP | 37986224 | Unspecified |
| mmu-miR-9-5p | MIMAT0000142 | Nfx1 | 74164 | HITS-CLIP | 19536157 | Unspecified |
| mmu-miR-9-5p | MIMAT0000142 | Ngly1 | 59007 | HITS-CLIP | 37986224 | Unspecified |
| mmu-miR-9-5p | MIMAT0000142 | Nipa1 | 233280 | HITS-CLIP | 37986224 | Unspecified |
| mmu-miR-9-5p | MIMAT0000142 | Nipbl | 71175 | HITS-CLIP | 37986224 | Unspecified |
| mmu-miR-9-5p | MIMAT0000142 | Nisch | 64652 | HITS-CLIP | 37986224 | Unspecified |
| mmu-miR-9-5p | MIMAT0000142 | Nmt1 | 18107 | HITS-CLIP | 37986224 | Unspecified |
| mmu-miR-9-5p | MIMAT0000142 | Nol6 | 230082 | HITS-CLIP | 37986224 | Unspecified |
| mmu-miR-9-5p | MIMAT0000142 | Nop56 | 67134 | qCLASH | 31393953 | Unspecified |
| mmu-miR-9-5p | MIMAT0000142 | Nova1 | 664883 | HITS-CLIP | 37986224 | Unspecified |
| mmu-miR-9-5p | MIMAT0000142 | Nptx1 | 18164 | Sequencing//HITS-CLIP | 37986224\|19536157 | Unspecified |
| mmu-miR-9-5p | MIMAT0000142 | Nr3c1 | 14815 | HITS-CLIP | 37986224 | Unspecified |
| mmu-miR-9-5p | MIMAT0000142 | Nr3c2 | 110784 | HITS-CLIP | 37986224 | Unspecified |
| mmu-miR-9-5p | MIMAT0000142 | Nrgn | 64011 | HITS-CLIP | 37986224 | Unspecified |
| mmu-miR-9-5p | MIMAT0000142 | Nrp1 | 18186 | HITS-CLIP | 37986224 | Unspecified |
| mmu-miR-9-5p | MIMAT0000142 | Nsd1 | 18193 | HITS-CLIP | 37986224 | Unspecified |
| mmu-miR-9-5p | MIMAT0000142 | Nsg2 | 18197 | HITS-CLIP | 37986224 | Unspecified |
| mmu-miR-9-5p | MIMAT0000142 | Nsun5 | 100609 | HITS-CLIP | 37986224 | Unspecified |
| mmu-miR-9-5p | MIMAT0000142 | Ntm | 235106 | HITS-CLIP | 37986224 | Unspecified |
| mmu-miR-9-5p | MIMAT0000142 | Ntn4 | 57764 | HITS-CLIP | 37986224 | Unspecified |
| mmu-miR-9-5p | MIMAT0000142 | Ntrk2 | 18212 | HITS-CLIP | 37986224\|19536157 | Unspecified |

| | | | | | | | |
|---|---|---|---|---|---|---|---|
| mmu-miR-9-5p | MIMAT0000142 | Nub1 | 53312 | HITS-CLIP | | 37986224 | Unspecified |
| mmu-miR-9-5p | MIMAT0000142 | Nudt4 | 71207 | HITS-CLIP | | 37986224 | Unspecified |
| mmu-miR-9-5p | MIMAT0000142 | Nus1 | 52014 | HITS-CLIP | | 37986224 | Unspecified |
| mmu-miR-9-5p | MIMAT0000142 | Onecut2 | 225631 | HITS-CLIP | | 37986224 | Unspecified |
| mmu-miR-9-5p | MIMAT0000142 | Osbpl11 | 106326 | HITS-CLIP | | 37986224 | Unspecified |
| mmu-miR-9-5p | MIMAT0000142 | Otud4 | 73945 | HITS-CLIP | | 19536157 | Unspecified |
| mmu-miR-9-5p | MIMAT0000142 | P4hb | 18453 | HITS-CLIP | | 37986224 | Unspecified |
| mmu-miR-9-5p | MIMAT0000142 | Pacs1 | 107975 | HITS-CLIP | | 37986224 | Unspecified |
| mmu-miR-9-5p | MIMAT0000142 | Pak2 | 224105 | HITS-CLIP | | 37986224 | Unspecified |
| mmu-miR-9-5p | MIMAT0000142 | Papola | 18789 | HITS-CLIP | | 37986224 | Unspecified |
| mmu-miR-9-5p | MIMAT0000142 | Parp1 | 11545 | HITS-CLIP | | 37986224 | Unspecified |
| mmu-miR-9-5p | MIMAT0000142 | Parva | 57342 | HITS-CLIP | | 37986224 | Unspecified |
| mmu-miR-9-5p | MIMAT0000142 | Pcdh10 | 18526 | HITS-CLIP | 37986224\|19536157 | | Unspecified |
| mmu-miR-9-5p | MIMAT0000142 | Pcdh7 | 54216 | Sequencing//HITS-CLIP | 19536157\|37986224 | | Unspecified |
| mmu-miR-9-5p | MIMAT0000142 | Pcdh9 | 211712 | HITS-CLIP | | 37986224 | Unspecified |
| mmu-miR-9-5p | MIMAT0000142 | Pcdhga12 | 93724 | HITS-CLIP | | 37986224 | Unspecified |
| mmu-miR-9-5p | MIMAT0000142 | Pcid2 | 234069 | HITS-CLIP | | 37986224 | Unspecified |
| mmu-miR-9-5p | MIMAT0000142 | Pcm1 | 18536 | HITS-CLIP | 37986224\|19536157 | | Unspecified |
| mmu-miR-9-5p | MIMAT0000142 | Pcsk1 | 18548 | HITS-CLIP | | 37986224 | Unspecified |
| mmu-miR-9-5p | MIMAT0000142 | Pcsk2 | 18549 | HITS-CLIP | | 37986224 | Unspecified |
| mmu-miR-9-5p | MIMAT0000142 | Pde4d | 238871 | HITS-CLIP | | 37986224 | Unspecified |
| mmu-miR-9-5p | MIMAT0000142 | Pde7b | 29863 | HITS-CLIP | | 37986224 | Unspecified |
| mmu-miR-9-5p | MIMAT0000142 | Pdgfrb | 18596 | HITS-CLIP | | 37986224 | Unspecified |
| mmu-miR-9-5p | MIMAT0000142 | Pdia3 | 14827 | HITS-CLIP | | 37986224 | Unspecified |
| mmu-miR-9-5p | MIMAT0000142 | Pds5a | 71521 | HITS-CLIP | 37986224\|19536157 | | Unspecified |
| mmu-miR-9-5p | MIMAT0000142 | Pdxk | 216134 | HITS-CLIP | | 37986224 | Unspecified |
| mmu-miR-9-5p | MIMAT0000142 | Pdzd2 | 68070 | HITS-CLIP | | 37986224 | Unspecified |
| mmu-miR-9-5p | MIMAT0000142 | Peg3 | 18616 | HITS-CLIP | | 37986224 | Unspecified |
| mmu-miR-9-5p | MIMAT0000142 | Per2 | 18627 | HITS-CLIP | | 37986224 | Unspecified |
| mmu-miR-9-5p | MIMAT0000142 | Per3 | 18628 | HITS-CLIP | | 37986224 | Unspecified |
| mmu-miR-9-5p | MIMAT0000142 | Pex1 | 71382 | HITS-CLIP | | 37986224 | Unspecified |
| mmu-miR-9-5p | MIMAT0000142 | Pgm2l1 | 70974 | HITS-CLIP | | 37986224 | Unspecified |
| mmu-miR-9-5p | MIMAT0000142 | Phactr1 | 218194 | HITS-CLIP | | 37986224 | Unspecified |
| mmu-miR-9-5p | MIMAT0000142 | Phf21a | 192285 | HITS-CLIP | | 37986224 | Unspecified |
| mmu-miR-9-5p | MIMAT0000142 | Phip | 83946 | HITS-CLIP | | 37986224 | Unspecified |
| mmu-miR-9-5p | MIMAT0000142 | Phrf1 | 101471 | HITS-CLIP | | 19536157 | Unspecified |
| mmu-miR-9-5p | MIMAT0000142 | Pi4k2a | 84095 | HITS-CLIP | | 37986224 | Unspecified |
| mmu-miR-9-5p | MIMAT0000142 | Picalm | 233489 | HITS-CLIP | | 37986224 | Unspecified |
| mmu-miR-9-5p | MIMAT0000142 | Piga | 18700 | HITS-CLIP | | 37986224 | Unspecified |

| | | | | | | | |
|---|---|---|---|---|---|---|---|
| mmu-miR-9-5p | MIMAT0000142 | Pin1 | 23988 | HITS-CLIP | | 37986224 | Unspecified |
| mmu-miR-9-5p | MIMAT0000142 | Pip4k2a | 18718 | HITS-CLIP | | 37986224 | Unspecified |
| mmu-miR-9-5p | MIMAT0000142 | Pitrm1 | 69617 | HITS-CLIP | | 37986224 | Unspecified |
| mmu-miR-9-5p | MIMAT0000142 | Pkia | 18767 | HITS-CLIP | | 37986224 | Unspecified |
| mmu-miR-9-5p | MIMAT0000142 | Pkp4 | 227937 | HITS-CLIP | 37986224\|19536157 | | Unspecified |
| mmu-miR-9-5p | MIMAT0000142 | Pla2g7 | 27226 | HITS-CLIP | | 37986224 | Unspecified |
| mmu-miR-9-5p | MIMAT0000142 | Plcd4 | 18802 | HITS-CLIP | | 19536157 | Unspecified |
| mmu-miR-9-5p | MIMAT0000142 | Plch2 | 269615 | HITS-CLIP | | 37986224 | Unspecified |
| mmu-miR-9-5p | MIMAT0000142 | Plcl1 | 227120 | HITS-CLIP | | 37986224 | Unspecified |
| mmu-miR-9-5p | MIMAT0000142 | Plcxd3 | 239318 | HITS-CLIP | | 37986224 | Unspecified |
| mmu-miR-9-5p | MIMAT0000142 | Pld1 | 18805 | HITS-CLIP | | 37986224 | Unspecified |
| mmu-miR-9-5p | MIMAT0000142 | Plekha1 | 101476 | HITS-CLIP | | 37986224 | Unspecified |
| mmu-miR-9-5p | MIMAT0000142 | Plp1 | 18823 | HITS-CLIP | | 37986224 | Unspecified |
| mmu-miR-9-5p | MIMAT0000142 | Plpp6 | 403313 | HITS-CLIP | | 19536157 | Unspecified |
| mmu-miR-9-5p | MIMAT0000142 | Plscr3 | 70310 | Sequencing//HITS-CLIP | 19536157\|37986224 | | Unspecified |
| mmu-miR-9-5p | MIMAT0000142 | Plxna2 | 18845 | HITS-CLIP | | 37986224 | Unspecified |
| mmu-miR-9-5p | MIMAT0000142 | Pnmal2 | 434128 | HITS-CLIP | | 37986224 | Unspecified |
| mmu-miR-9-5p | MIMAT0000142 | Pnn | 18949 | HITS-CLIP | | 37986224 | Unspecified |
| mmu-miR-9-5p | MIMAT0000142 | Pnrc1 | 108767 | HITS-CLIP | | 37986224 | Unspecified |
| mmu-miR-9-5p | MIMAT0000142 | Pou2f1 | 18986 | qCLASH//HITS-CLIP | 31393953\|19536157 | | Unspecified |
| mmu-miR-9-5p | MIMAT0000142 | Ppfia2 | 327814 | HITS-CLIP | | 37986224 | Unspecified |
| mmu-miR-9-5p | MIMAT0000142 | Ppif | 105675 | HITS-CLIP | | 37986224 | Unspecified |
| mmu-miR-9-5p | MIMAT0000142 | Ppil4 | 67418 | HITS-CLIP | | 37986224 | Unspecified |
| mmu-miR-9-5p | MIMAT0000142 | Ppm1a | 19042 | HITS-CLIP | 37986224\|19536157 | | Unspecified |
| mmu-miR-9-5p | MIMAT0000142 | Ppm1e | 320472 | HITS-CLIP | | 37986224 | Unspecified |
| mmu-miR-9-5p | MIMAT0000142 | Ppm1k | 243382 | HITS-CLIP | | 37986224 | Unspecified |
| mmu-miR-9-5p | MIMAT0000142 | Ppp1cb | 19046 | HITS-CLIP | | 19536157 | Unspecified |
| mmu-miR-9-5p | MIMAT0000142 | Ppp1r12a | 17931 | HITS-CLIP | | 37986224 | Unspecified |
| mmu-miR-9-5p | MIMAT0000142 | Ppp1r9a | 243725 | HITS-CLIP | | 37986224 | Unspecified |
| mmu-miR-9-5p | MIMAT0000142 | Ppp4r1 | 70351 | HITS-CLIP | | 37986224 | Unspecified |
| mmu-miR-9-5p | MIMAT0000142 | Ppp6r1 | 243819 | HITS-CLIP | | 37986224 | Unspecified |
| mmu-miR-9-5p | MIMAT0000142 | Prdm8 | 77630 | HITS-CLIP | 37986224\|19536157 | | Unspecified |
| mmu-miR-9-5p | MIMAT0000142 | Prdx3 | 11757 | HITS-CLIP | | 37986224 | Unspecified |
| mmu-miR-9-5p | MIMAT0000142 | Prkx | 19108 | HITS-CLIP | | 37986224 | Unspecified |
| mmu-miR-9-5p | MIMAT0000142 | Prpf4b | 19134 | HITS-CLIP | | 37986224 | Unspecified |
| mmu-miR-9-5p | MIMAT0000142 | Prr12 | 233210 | HITS-CLIP | | 37986224 | Unspecified |
| mmu-miR-9-5p | MIMAT0000142 | Prr14l | 215476 | HITS-CLIP | | 37986224 | Unspecified |
| mmu-miR-9-5p | MIMAT0000142 | Prtg | 235472 | HITS-CLIP | | 37986224 | Unspecified |
| mmu-miR-9-5p | MIMAT0000142 | Pskh1 | 244631 | HITS-CLIP | | 37986224 | Unspecified |

| | | | | | | |
|---|---|---|---|---|---|---|
| mmu-miR-9-5p | MIMAT0000142 | Psmc2 | 19181 | HITS-CLIP | 37986224 | Unspecified |
| mmu-miR-9-5p | MIMAT0000142 | Psmd1 | 70247 | HITS-CLIP | 37986224 | Unspecified |
| mmu-miR-9-5p | MIMAT0000142 | Ptchd4 | 627626 | HITS-CLIP | 37986224 | Unspecified |
| mmu-miR-9-5p | MIMAT0000142 | Pten | 19211 | HITS-CLIP | 37986224 | Unspecified |
| mmu-miR-9-5p | MIMAT0000142 | Ptpn13 | 19249 | HITS-CLIP | 37986224 | Unspecified |
| mmu-miR-9-5p | MIMAT0000142 | Ptpn4 | 19258 | HITS-CLIP | 37986224 | Unspecified |
| mmu-miR-9-5p | MIMAT0000142 | Ptprb | 19263 | HITS-CLIP | 37986224\|19536157 | Unspecified |
| mmu-miR-9-5p | MIMAT0000142 | Ptprk | 19272 | HITS-CLIP | 37986224\|19536157 | Unspecified |
| mmu-miR-9-5p | MIMAT0000142 | Ptprn2 | 19276 | HITS-CLIP | 37986224 | Unspecified |
| mmu-miR-9-5p | MIMAT0000142 | Ptprz1 | 19283 | HITS-CLIP | 37986224 | Unspecified |
| mmu-miR-9-5p | MIMAT0000142 | Pura | 19290 | HITS-CLIP | 37986224 | Unspecified |
| mmu-miR-9-5p | MIMAT0000142 | Purb | 19291 | Sequencing | 37986224 | Unspecified |
| mmu-miR-9-5p | MIMAT0000142 | Pwwp2a | 70802 | HITS-CLIP | 37986224 | Unspecified |
| mmu-miR-9-5p | MIMAT0000142 | Pygb | 110078 | HITS-CLIP | 37986224 | Unspecified |
| mmu-miR-9-5p | MIMAT0000142 | Qk | 19317 | HITS-CLIP | 37986224 | Unspecified |
| mmu-miR-9-5p | MIMAT0000142 | Qser1 | 99003 | HITS-CLIP | 37986224 | Unspecified |
| mmu-miR-9-5p | MIMAT0000142 | Rab10 | 19325 | HITS-CLIP | 37986224 | Unspecified |
| mmu-miR-9-5p | MIMAT0000142 | Rab11fip4 | 268451 | HITS-CLIP | 37986224 | Unspecified |
| mmu-miR-9-5p | MIMAT0000142 | Rab24 | 19336 | HITS-CLIP | 37986224 | Unspecified |
| mmu-miR-9-5p | MIMAT0000142 | Rab33a | 19337 | HITS-CLIP | 37986224 | Unspecified |
| mmu-miR-9-5p | MIMAT0000142 | Rabgap1l | 29809 | HITS-CLIP | 37986224 | Unspecified |
| mmu-miR-9-5p | MIMAT0000142 | Rad21 | 19357 | HITS-CLIP | 37986224 | Unspecified |
| mmu-miR-9-5p | MIMAT0000142 | Ralgds | 19730 | HITS-CLIP | 37986224\|19536157 | Unspecified |
| mmu-miR-9-5p | MIMAT0000142 | Ranbp2 | 19386 | HITS-CLIP | 37986224 | Unspecified |
| mmu-miR-9-5p | MIMAT0000142 | Raph1 | 77300 | HITS-CLIP | 37986224 | Unspecified |
| mmu-miR-9-5p | MIMAT0000142 | Rasd2 | 75141 | Sequencing//HITS-CLIP | 37986224\|19536157 | Unspecified |
| mmu-miR-9-5p | MIMAT0000142 | Rbm28 | 68272 | HITS-CLIP | 37986224 | Unspecified |
| mmu-miR-9-5p | MIMAT0000142 | Rbm4 | 19653 | Sequencing | 37986224 | Unspecified |
| mmu-miR-9-5p | MIMAT0000142 | Rbm5 | 83486 | HITS-CLIP | 19536157 | Unspecified |
| mmu-miR-9-5p | MIMAT0000142 | Rbms3 | 207181 | HITS-CLIP | 37986224 | Unspecified |
| mmu-miR-9-5p | MIMAT0000142 | Rbp1 | 19659 | HITS-CLIP | 37986224 | Unspecified |
| mmu-miR-9-5p | MIMAT0000142 | Reep3 | 28193 | HITS-CLIP | 37986224 | Unspecified |
| mmu-miR-9-5p | MIMAT0000142 | Rer1 | 67830 | HITS-CLIP | 37986224 | Unspecified |
| mmu-miR-9-5p | MIMAT0000142 | Rest | 19712 | HITS-CLIP | 37986224 | Unspecified |
| mmu-miR-9-5p | MIMAT0000142 | Rev3l | 19714 | HITS-CLIP | 37986224\|19536157 | Unspecified |
| mmu-miR-9-5p | MIMAT0000142 | Rfx7 | 319758 | Sequencing//HITS-CLIP | 37986224\|19536157 | Unspecified |
| mmu-miR-9-5p | MIMAT0000142 | Rgs19 | 56470 | HITS-CLIP | 37986224 | Unspecified |
| mmu-miR-9-5p | MIMAT0000142 | Rgs4 | 19736 | HITS-CLIP | 37986224 | Unspecified |
| mmu-miR-9-5p | MIMAT0000142 | Rgs5 | 19737 | HITS-CLIP | 37986224 | Unspecified |

| | | | | | | |
|---|---|---|---|---|---|---|
| mmu-miR-9-5p | MIMAT0000142 | Rhoq | 104215 | HITS-CLIP | 37986224 | Unspecified |
| mmu-miR-9-5p | MIMAT0000142 | Rictor | 78757 | Sequencing//qCLASH | 19536157\|31393953 | Unspecified |
| mmu-miR-9-5p | MIMAT0000142 | Rnf111 | 93836 | HITS-CLIP | 19536157 | Unspecified |
| mmu-miR-9-5p | MIMAT0000142 | Rnf150 | 330812 | HITS-CLIP | 37986224 | Unspecified |
| mmu-miR-9-5p | MIMAT0000142 | Rnf167 | 70510 | HITS-CLIP | 37986224 | Unspecified |
| mmu-miR-9-5p | MIMAT0000142 | Rnf216 | 108086 | HITS-CLIP | 37986224 | Unspecified |
| mmu-miR-9-5p | MIMAT0000142 | Rnf219 | 72486 | HITS-CLIP | 37986224 | Unspecified |
| mmu-miR-9-5p | MIMAT0000142 | Rnf6 | 74132 | HITS-CLIP | 19536157 | Unspecified |
| mmu-miR-9-5p | MIMAT0000142 | Rock2 | 19878 | HITS-CLIP | 37986224 | Unspecified |
| mmu-miR-9-5p | MIMAT0000142 | Rph3a | 19894 | HITS-CLIP | 37986224 | Unspecified |
| mmu-miR-9-5p | MIMAT0000142 | Rpl14 | 67115 | HITS-CLIP | 37986224 | Unspecified |
| mmu-miR-9-5p | MIMAT0000142 | Rpl4 | 67891 | HITS-CLIP | 37986224 | Unspecified |
| mmu-miR-9-5p | MIMAT0000142 | Rpn2 | 20014 | HITS-CLIP | 19536157 | Unspecified |
| mmu-miR-9-5p | MIMAT0000142 | Rps16 | 20055 | qCLASH | 31393953 | Unspecified |
| mmu-miR-9-5p | MIMAT0000142 | Rragc | 54170 | HITS-CLIP | 37986224 | Unspecified |
| mmu-miR-9-5p | MIMAT0000142 | Rrm1 | 20133 | qCLASH | 31393953 | Unspecified |
| mmu-miR-9-5p | MIMAT0000142 | Rrp36 | 224823 | HITS-CLIP | 37986224 | Unspecified |
| mmu-miR-9-5p | MIMAT0000142 | Rsbn1l | 242860 | HITS-CLIP | 37986224 | Unspecified |
| mmu-miR-9-5p | MIMAT0000142 | Rsl1d1 | 66409 | HITS-CLIP | 37986224 | Unspecified |
| mmu-miR-9-5p | MIMAT0000142 | Rtf1 | 76246 | HITS-CLIP | 37986224 | Unspecified |
| mmu-miR-9-5p | MIMAT0000142 | Rundc3b | 242819 | Sequencing//HITS-CLIP | 19536157\|37986224 | Unspecified |
| mmu-miR-9-5p | MIMAT0000142 | S100pbp | 74648 | HITS-CLIP | 37986224 | Unspecified |
| mmu-miR-9-5p | MIMAT0000142 | Sacs | 50720 | HITS-CLIP | 37986224 | Unspecified |
| mmu-miR-9-5p | MIMAT0000142 | Sall1 | 58198 | HITS-CLIP | 37986224 | Unspecified |
| mmu-miR-9-5p | MIMAT0000142 | Samd4 | 74480 | HITS-CLIP | 37986224 | Unspecified |
| mmu-miR-9-5p | MIMAT0000142 | Scg2 | 20254 | HITS-CLIP | 37986224 | Unspecified |
| mmu-miR-9-5p | MIMAT0000142 | Scn4b | 399548 | HITS-CLIP | 37986224 | Unspecified |
| mmu-miR-9-5p | MIMAT0000142 | Scrt1 | 170729 | HITS-CLIP | 37986224 | Unspecified |
| mmu-miR-9-5p | MIMAT0000142 | Sdad1 | 231452 | HITS-CLIP | 37986224 | Unspecified |
| mmu-miR-9-5p | MIMAT0000142 | Sdc2 | 15529 | HITS-CLIP | 37986224 | Unspecified |
| mmu-miR-9-5p | MIMAT0000142 | Sdc3 | 20970 | HITS-CLIP | 37986224 | Unspecified |
| mmu-miR-9-5p | MIMAT0000142 | Sec23b | 27054 | HITS-CLIP | 37986224 | Unspecified |
| mmu-miR-9-5p | MIMAT0000142 | Sec63 | 140740 | HITS-CLIP | 37986224 | Unspecified |
| mmu-miR-9-5p | MIMAT0000142 | Secisbp2l | 70354 | HITS-CLIP | 37986224 | Unspecified |
| mmu-miR-9-5p | MIMAT0000142 | Sel1l3 | 231238 | HITS-CLIP | 37986224 | Unspecified |
| mmu-miR-9-5p | MIMAT0000142 | Sema4f | 20355 | HITS-CLIP | 37986224 | Unspecified |
| mmu-miR-9-5p | MIMAT0000142 | Senp5 | 320213 | HITS-CLIP | 37986224 | Unspecified |
| mmu-miR-9-5p | MIMAT0000142 | Sept5 | 18951 | HITS-CLIP | 37986224 | Unspecified |
| mmu-miR-9-5p | MIMAT0000142 | Sept8 | 20362 | HITS-CLIP | 37986224 | Unspecified |

| | | | | | | |
|---|---|---|---|---|---|---|
| mmu-miR-9-5p | MIMAT0000142 | Serinc1 | 56442 | HITS-CLIP | 37986224 | Unspecified |
| mmu-miR-9-5p | MIMAT0000142 | Serinc5 | 218442 | HITS-CLIP | 37986224\|19536157 | Unspecified |
| mmu-miR-9-5p | MIMAT0000142 | Setd2 | 235626 | HITS-CLIP | 37986224\|19536157 | Unspecified |
| mmu-miR-9-5p | MIMAT0000142 | Setx | 269254 | HITS-CLIP | 37986224 | Unspecified |
| mmu-miR-9-5p | MIMAT0000142 | Sez6l2 | 233878 | HITS-CLIP | 37986224 | Unspecified |
| mmu-miR-9-5p | MIMAT0000142 | Sgsm1 | 52850 | HITS-CLIP | 19536157 | Unspecified |
| mmu-miR-9-5p | MIMAT0000142 | Sh2b3 | 16923 | HITS-CLIP | 37986224 | Unspecified |
| mmu-miR-9-5p | MIMAT0000142 | Sh3bgrl3 | 73723 | HITS-CLIP | 37986224 | Unspecified |
| mmu-miR-9-5p | MIMAT0000142 | Sh3d19 | 27059 | HITS-CLIP | 19536157 | Unspecified |
| mmu-miR-9-5p | MIMAT0000142 | Sh3glb1 | 54673 | HITS-CLIP | 37986224 | Unspecified |
| mmu-miR-9-5p | MIMAT0000142 | Sh3pxd2b | 268396 | HITS-CLIP | 37986224 | Unspecified |
| mmu-miR-9-5p | MIMAT0000142 | Sh3rf1 | 59009 | HITS-CLIP | 37986224 | Unspecified |
| mmu-miR-9-5p | MIMAT0000142 | Sh3rf3 | 237353 | HITS-CLIP | 37986224 | Unspecified |
| mmu-miR-9-5p | MIMAT0000142 | Shank1 | 243961 | HITS-CLIP | 19536157 | Unspecified |
| mmu-miR-9-5p | MIMAT0000142 | Skida1 | 72668 | HITS-CLIP | 37986224 | Unspecified |
| mmu-miR-9-5p | MIMAT0000142 | Slc12a6 | 107723 | HITS-CLIP | 37986224 | Unspecified |
| mmu-miR-9-5p | MIMAT0000142 | Slc16a1 | 20501 | HITS-CLIP | 37986224 | Unspecified |
| mmu-miR-9-5p | MIMAT0000142 | Slc16a13 | 69309 | HITS-CLIP | 37986224 | Unspecified |
| mmu-miR-9-5p | MIMAT0000142 | Slc1a2 | 20511 | HITS-CLIP | 37986224 | Unspecified |
| mmu-miR-9-5p | MIMAT0000142 | Slc20a2 | 20516 | HITS-CLIP | 37986224 | Unspecified |
| mmu-miR-9-5p | MIMAT0000142 | Slc22a17 | 59049 | HITS-CLIP | 37986224 | Unspecified |
| mmu-miR-9-5p | MIMAT0000142 | Slc25a12 | 78830 | HITS-CLIP | 37986224 | Unspecified |
| mmu-miR-9-5p | MIMAT0000142 | Slc26a2 | 13521 | HITS-CLIP | 37986224 | Unspecified |
| mmu-miR-9-5p | MIMAT0000142 | Slc2a13 | 239606 | HITS-CLIP | 37986224 | Unspecified |
| mmu-miR-9-5p | MIMAT0000142 | Slc2a8 | 56017 | HITS-CLIP | 37986224\|19536157 | Unspecified |
| mmu-miR-9-5p | MIMAT0000142 | Slc30a1 | 22782 | HITS-CLIP | 37986224 | Unspecified |
| mmu-miR-9-5p | MIMAT0000142 | Slc39a14 | 213053 | HITS-CLIP | 37986224\|19536157 | Unspecified |
| mmu-miR-9-5p | MIMAT0000142 | Slc39a9 | 328133 | HITS-CLIP | 37986224\|19536157 | Unspecified |
| mmu-miR-9-5p | MIMAT0000142 | Slc41a1 | 98396 | HITS-CLIP | 37986224 | Unspecified |
| mmu-miR-9-5p | MIMAT0000142 | Slc44a5 | 242259 | HITS-CLIP | 37986224 | Unspecified |
| mmu-miR-9-5p | MIMAT0000142 | Slc4a7 | 218756 | HITS-CLIP | 37986224 | Unspecified |
| mmu-miR-9-5p | MIMAT0000142 | Slc5a3 | 53881 | HITS-CLIP | 37986224 | Unspecified |
| mmu-miR-9-5p | MIMAT0000142 | Slc7a1 | 11987 | HITS-CLIP | 37986224 | Unspecified |
| mmu-miR-9-5p | MIMAT0000142 | Slc7a11 | 26570 | HITS-CLIP | 37986224 | Unspecified |
| mmu-miR-9-5p | MIMAT0000142 | Slco1c1 | 58807 | HITS-CLIP | 37986224 | Unspecified |
| mmu-miR-9-5p | MIMAT0000142 | Slf1 | 852127 | HITS-CLIP | 19536157 | Unspecified |
| mmu-miR-9-5p | MIMAT0000142 | Slf2 | 226151 | HITS-CLIP | 19536157 | Unspecified |
| mmu-miR-9-5p | MIMAT0000142 | Slmap | 83997 | HITS-CLIP//qCLASH | 37986224\|31393953\|19536157 | Unspecified |
| mmu-miR-9-5p | MIMAT0000142 | Smpd1 | 20597 | HITS-CLIP | 37986224 | Unspecified |

| | | | | | | |
|---|---|---|---|---|---|---|
| mmu-miR-9-5p | MIMAT0000142 | Smpd2 | 20598 | HITS-CLIP | 37986224 | Unspecified |
| mmu-miR-9-5p | MIMAT0000142 | Smpdl3a | 57319 | HITS-CLIP | 37986224 | Unspecified |
| mmu-miR-9-5p | MIMAT0000142 | Smurf2 | 66313 | HITS-CLIP | 37986224 | Unspecified |
| mmu-miR-9-5p | MIMAT0000142 | Snn | 20621 | HITS-CLIP | 37986224 | Unspecified |
| mmu-miR-9-5p | MIMAT0000142 | Snx25 | 102141 | Sequencing//HITS-CLIP | 19536157\|37986224 | Unspecified |
| mmu-miR-9-5p | MIMAT0000142 | Socs5 | 56468 | Sequencing//HITS-CLIP | 19536157\|37986224 | Unspecified |
| mmu-miR-9-5p | MIMAT0000142 | Soga1 | 320706 | HITS-CLIP | 19536157 | Unspecified |
| mmu-miR-9-5p | MIMAT0000142 | Son | 20658 | HITS-CLIP | 37986224 | Unspecified |
| mmu-miR-9-5p | MIMAT0000142 | Sorl1 | 20660 | HITS-CLIP | 37986224 | Unspecified |
| mmu-miR-9-5p | MIMAT0000142 | Sort1 | 20661 | HITS-CLIP | 37986224 | Unspecified |
| mmu-miR-9-5p | MIMAT0000142 | Sos2 | 20663 | qCLASH//HITS-CLIP | 31393953\|19536157 | Unspecified |
| mmu-miR-9-5p | MIMAT0000142 | Sowaha | 237761 | HITS-CLIP | 37986224 | Unspecified |
| mmu-miR-9-5p | MIMAT0000142 | Sparcl1 | 13602 | HITS-CLIP | 19536157 | Unspecified |
| mmu-miR-9-5p | MIMAT0000142 | Spats2 | 72572 | HITS-CLIP | 37986224 | Unspecified |
| mmu-miR-9-5p | MIMAT0000142 | Spin1 | 20729 | HITS-CLIP | 37986224 | Unspecified |
| mmu-miR-9-5p | MIMAT0000142 | Sppl2a | 66552 | HITS-CLIP | 37986224 | Unspecified |
| mmu-miR-9-5p | MIMAT0000142 | Sptlc2 | 20773 | HITS-CLIP | 37986224\|19536157 | Unspecified |
| mmu-miR-9-5p | MIMAT0000142 | Sptssa | 104725 | HITS-CLIP | 37986224 | Unspecified |
| mmu-miR-9-5p | MIMAT0000142 | Spty2d1 | 101685 | HITS-CLIP | 37986224 | Unspecified |
| mmu-miR-9-5p | MIMAT0000142 | Srebf2 | 20788 | HITS-CLIP | 37986224 | Unspecified |
| mmu-miR-9-5p | MIMAT0000142 | Srp54b | 665155 | HITS-CLIP | 37986224 | Unspecified |
| mmu-miR-9-5p | MIMAT0000142 | Srpk1 | 20815 | HITS-CLIP | 37986224 | Unspecified |
| mmu-miR-9-5p | MIMAT0000142 | Srsf10 | 14105 | HITS-CLIP | 37986224 | Unspecified |
| mmu-miR-9-5p | MIMAT0000142 | St6gal1 | 20440 | HITS-CLIP | 37986224 | Unspecified |
| mmu-miR-9-5p | MIMAT0000142 | Stim2 | 116873 | HITS-CLIP | 37986224 | Unspecified |
| mmu-miR-9-5p | MIMAT0000142 | Stk24 | 223255 | HITS-CLIP | 37986224 | Unspecified |
| mmu-miR-9-5p | MIMAT0000142 | Stk38l | 232533 | HITS-CLIP | 37986224 | Unspecified |
| mmu-miR-9-5p | MIMAT0000142 | Stmn2 | 20257 | HITS-CLIP | 37986224 | Unspecified |
| mmu-miR-9-5p | MIMAT0000142 | Strap | 20901 | HITS-CLIP | 37986224 | Unspecified |
| mmu-miR-9-5p | MIMAT0000142 | Stx16 | 228960 | HITS-CLIP | 37986224 | Unspecified |
| mmu-miR-9-5p | MIMAT0000142 | Stx17 | 67727 | HITS-CLIP | 37986224 | Unspecified |
| mmu-miR-9-5p | MIMAT0000142 | Stx3 | 20908 | Luciferase Reporter Assay | 37986224 | Unspecified |
| mmu-miR-9-5p | MIMAT0000142 | Stxbp6 | 217517 | HITS-CLIP | 37986224 | Unspecified |
| mmu-miR-9-5p | MIMAT0000142 | Sucla2 | 20916 | HITS-CLIP | 37986224 | Unspecified |
| mmu-miR-9-5p | MIMAT0000142 | Supt16 | 114741 | HITS-CLIP | 26203562 | Unspecified |
| mmu-miR-9-5p | MIMAT0000142 | Supt6 | 20926 | HITS-CLIP | 37986224\|19536157 | Unspecified |
| mmu-miR-9-5p | MIMAT0000142 | Sv2a | 64051 | HITS-CLIP | 37986224 | Unspecified |
| mmu-miR-9-5p | MIMAT0000142 | Syne1 | 64009 | Sequencing//HITS-CLIP | 37986224\|19536157 | Unspecified |
| mmu-miR-9-5p | MIMAT0000142 | Synj1 | 104015 | Luciferase Reporter Assay | 37986224 | Unspecified |

| | | | | | | |
|---|---|---|---|---|---|---|
| mmu-miR-9-5p | MIMAT0000142 | Synpr | 72003 | Luciferase Reporter Assay | 37986224 | Unspecified |
| mmu-miR-9-5p | MIMAT0000142 | Synrg | 217030 | HITS-CLIP | 19536157 | Unspecified |
| mmu-miR-9-5p | MIMAT0000142 | Syp | 20977 | HITS-CLIP | 37986224 | Unspecified |
| mmu-miR-9-5p | MIMAT0000142 | Syt4 | 20983 | HITS-CLIP | 37986224 | Unspecified |
| mmu-miR-9-5p | MIMAT0000142 | Tab2 | 68652 | HITS-CLIP | 37986224 | Unspecified |
| mmu-miR-9-5p | MIMAT0000142 | Tagln3 | 56370 | HITS-CLIP | 37986224 | Unspecified |
| mmu-miR-9-5p | MIMAT0000142 | Tbc1d13 | 70296 | HITS-CLIP | 37986224 | Unspecified |
| mmu-miR-9-5p | MIMAT0000142 | Tbc1d15 | 66687 | HITS-CLIP | 37986224 | Unspecified |
| mmu-miR-9-5p | MIMAT0000142 | Tbc1d16 | 207592 | HITS-CLIP | 37986224 | Unspecified |
| mmu-miR-9-5p | MIMAT0000142 | Tbc1d8 | 54610 | HITS-CLIP | 37986224 | Unspecified |
| mmu-miR-9-5p | MIMAT0000142 | Tbc1d9b | 76795 | HITS-CLIP | 37986224 | Unspecified |
| mmu-miR-9-5p | MIMAT0000142 | Tbl1x | 21372 | HITS-CLIP | 37986224 | Unspecified |
| mmu-miR-9-5p | MIMAT0000142 | Tbl3 | 213773 | HITS-CLIP | 37986224 | Unspecified |
| mmu-miR-9-5p | MIMAT0000142 | Tbr1 | 21375 | HITS-CLIP | 37986224 | Unspecified |
| mmu-miR-9-5p | MIMAT0000142 | Tceanc2 | 66526 | HITS-CLIP | 19536157 | Unspecified |
| mmu-miR-9-5p | MIMAT0000142 | Tenm1 | 23963 | HITS-CLIP | 37986224 | Unspecified |
| mmu-miR-9-5p | MIMAT0000142 | Tenm4 | 23966 | Sequencing//HITS-CLIP | 19536157 | Unspecified |
| mmu-miR-9-5p | MIMAT0000142 | Tgfbi | 21810 | Luciferase Reporter Assay | 37986224 | Unspecified |
| mmu-miR-9-5p | MIMAT0000142 | Tgfbr1 | 21812 | Luciferase Reporter Assay | 37986224 | Unspecified |
| mmu-miR-9-5p | MIMAT0000142 | Tgoln1 | 22134 | HITS-CLIP | 37986224 | Unspecified |
| mmu-miR-9-5p | MIMAT0000142 | Thap4 | 67026 | HITS-CLIP | 37986224 | Unspecified |
| mmu-miR-9-5p | MIMAT0000142 | Thbs2 | 21826 | HITS-CLIP | 37986224 | Unspecified |
| mmu-miR-9-5p | MIMAT0000142 | Thrb | 21834 | HITS-CLIP | 37986224 | Unspecified |
| mmu-miR-9-5p | MIMAT0000142 | Timm17a | 21854 | HITS-CLIP | 37986224 | Unspecified |
| mmu-miR-9-5p | MIMAT0000142 | Timp2 | 21858 | HITS-CLIP | 37986224 | Unspecified |
| mmu-miR-9-5p | MIMAT0000142 | Timp4 | 110595 | HITS-CLIP | 37986224 | Unspecified |
| mmu-miR-9-5p | MIMAT0000142 | Tln1 | 21894 | HITS-CLIP | 37986224 | Unspecified |
| mmu-miR-9-5p | MIMAT0000142 | Tm2d2 | 69742 | HITS-CLIP | 37986224 | Unspecified |
| mmu-miR-9-5p | MIMAT0000142 | Tm9sf3 | 107358 | HITS-CLIP | 37986224 | Unspecified |
| mmu-miR-9-5p | MIMAT0000142 | Tmcc3 | 319880 | HITS-CLIP | 37986224 | Unspecified |
| mmu-miR-9-5p | MIMAT0000142 | Tmed10 | 68581 | HITS-CLIP | 37986224 | Unspecified |
| mmu-miR-9-5p | MIMAT0000142 | Tmem106b | 71900 | HITS-CLIP | 37986224 | Unspecified |
| mmu-miR-9-5p | MIMAT0000142 | Tmem135 | 72759 | HITS-CLIP | 19536157 | Unspecified |
| mmu-miR-9-5p | MIMAT0000142 | Tmem151b | 210573 | HITS-CLIP | 37986224 | Unspecified |
| mmu-miR-9-5p | MIMAT0000142 | Tmem169 | 271711 | HITS-CLIP | 37986224 | Unspecified |
| mmu-miR-9-5p | MIMAT0000142 | Tmem170b | 621976 | HITS-CLIP | 37986224\|19536157 | Unspecified |
| mmu-miR-9-5p | MIMAT0000142 | Tmem19 | 67226 | HITS-CLIP | 37986224 | Unspecified |
| mmu-miR-9-5p | MIMAT0000142 | Tmem218 | 66279 | HITS-CLIP | 37986224 | Unspecified |
| mmu-miR-9-5p | MIMAT0000142 | Tmem65 | 74868 | Sequencing//HITS-CLIP | 19536157\|37986224 | Unspecified |

| | | | | | | |
|---|---|---|---|---|---|---|
| mmu-miR-9-5p | MIMAT0000142 | Tnrc6a | 233833 | HITS-CLIP | 37986224\|19536157 | Unspecified |
| mmu-miR-9-5p | MIMAT0000142 | Tnrc6b | 213988 | HITS-CLIP | 37986224 | Unspecified |
| mmu-miR-9-5p | MIMAT0000142 | Tob1 | 22057 | HITS-CLIP | 37986224 | Unspecified |
| mmu-miR-9-5p | MIMAT0000142 | Tomm34 | 67145 | qCLASH | 31393953 | Unspecified |
| mmu-miR-9-5p | MIMAT0000142 | Tomm70a | 28185 | HITS-CLIP | 37986224 | Unspecified |
| mmu-miR-9-5p | MIMAT0000142 | Top1 | 21969 | HITS-CLIP | 37986224 | Unspecified |
| mmu-miR-9-5p | MIMAT0000142 | Top2b | 21974 | HITS-CLIP | 37986224 | Unspecified |
| mmu-miR-9-5p | MIMAT0000142 | Traf3 | 22031 | Sequencing//HITS-CLIP | 37986224\|19536157 | Unspecified |
| mmu-miR-9-5p | MIMAT0000142 | Trak2 | 70827 | Luciferase Reporter Assay | 37986224 | Unspecified |
| mmu-miR-9-5p | MIMAT0000142 | Tram1l1 | 229801 | HITS-CLIP | 37986224 | Unspecified |
| mmu-miR-9-5p | MIMAT0000142 | Trib2 | 217410 | HITS-CLIP | 37986224 | Unspecified |
| mmu-miR-9-5p | MIMAT0000142 | Trim2 | 80890 | Luciferase Reporter Assay | 37986224 | Unspecified |
| mmu-miR-9-5p | MIMAT0000142 | Trim28 | 21849 | HITS-CLIP | 37986224 | Unspecified |
| mmu-miR-9-5p | MIMAT0000142 | Trim62 | 67525 | HITS-CLIP | 37986224 | Unspecified |
| mmu-miR-9-5p | MIMAT0000142 | Trim71 | 636931 | HITS-CLIP | 37986224 | Unspecified |
| mmu-miR-9-5p | MIMAT0000142 | Trnp1 | 69539 | HITS-CLIP | 37986224 | Unspecified |
| mmu-miR-9-5p | MIMAT0000142 | Trpm7 | 58800 | qCLASH | 31393953 | Unspecified |
| mmu-miR-9-5p | MIMAT0000142 | Trps1 | 83925 | HITS-CLIP | 37986224\|19536157 | Unspecified |
| mmu-miR-9-5p | MIMAT0000142 | Tsc22d1 | 21807 | HITS-CLIP | 37986224 | Unspecified |
| mmu-miR-9-5p | MIMAT0000142 | Tsc22d2 | 72033 | HITS-CLIP | 37986224 | Unspecified |
| mmu-miR-9-5p | MIMAT0000142 | Tsnax | 53424 | HITS-CLIP | 37986224 | Unspecified |
| mmu-miR-9-5p | MIMAT0000142 | Tspan2 | 70747 | HITS-CLIP | 37986224 | Unspecified |
| mmu-miR-9-5p | MIMAT0000142 | Tspan3 | 56434 | HITS-CLIP | 37986224 | Unspecified |
| mmu-miR-9-5p | MIMAT0000142 | Tspan4 | 64540 | HITS-CLIP | 37986224 | Unspecified |
| mmu-miR-9-5p | MIMAT0000142 | Tspyl5 | 239364 | HITS-CLIP | 37986224 | Unspecified |
| mmu-miR-9-5p | MIMAT0000142 | Ttbk2 | 140810 | HITS-CLIP | 19536157 | Unspecified |
| mmu-miR-9-5p | MIMAT0000142 | Ttc21a | 74052 | HITS-CLIP | 37986224 | Unspecified |
| mmu-miR-9-5p | MIMAT0000142 | Ttc3 | 22129 | HITS-CLIP | 37986224\|19536157 | Unspecified |
| mmu-miR-9-5p | MIMAT0000142 | Ttyh1 | 57776 | HITS-CLIP | 37986224 | Unspecified |
| mmu-miR-9-5p | MIMAT0000142 | Tuba1b | 22143 | HITS-CLIP | 37986224 | Unspecified |
| mmu-miR-9-5p | MIMAT0000142 | Tuba4a | 22145 | HITS-CLIP | 37986224 | Unspecified |
| mmu-miR-9-5p | MIMAT0000142 | Tubb5 | 22154 | HITS-CLIP | 37986224 | Unspecified |
| mmu-miR-9-5p | MIMAT0000142 | Tulp4 | 68842 | HITS-CLIP | 37986224 | Unspecified |
| mmu-miR-9-5p | MIMAT0000142 | Twf1 | 19230 | HITS-CLIP | 37986224 | Unspecified |
| mmu-miR-9-5p | MIMAT0000142 | Twsg1 | 65960 | HITS-CLIP | 37986224 | Unspecified |
| mmu-miR-9-5p | MIMAT0000142 | Txnrd1 | 50493 | HITS-CLIP | 37986224 | Unspecified |
| mmu-miR-9-5p | MIMAT0000142 | Uba5 | 66663 | HITS-CLIP | 37986224 | Unspecified |
| mmu-miR-9-5p | MIMAT0000142 | Ubac1 | 98766 | HITS-CLIP | 37986224 | Unspecified |
| mmu-miR-9-5p | MIMAT0000142 | Ube2k | 53323 | HITS-CLIP | 37986224 | Unspecified |

| | | | | | | | |
|---|---|---|---|---|---|---|---|
| mmu-miR-9-5p | MIMAT0000142 | Ube2o | 217342 | qCLASH | | 31393953 | Unspecified |
| mmu-miR-9-5p | MIMAT0000142 | Ube2z | 268470 | HITS-CLIP | | 37986224 | Unspecified |
| mmu-miR-9-5p | MIMAT0000142 | Ube4b | 63958 | HITS-CLIP | | 37986224 | Unspecified |
| mmu-miR-9-5p | MIMAT0000142 | Ubfd1 | 28018 | HITS-CLIP | | 37986224 | Unspecified |
| mmu-miR-9-5p | MIMAT0000142 | Ubn1 | 170644 | HITS-CLIP | | 37986224 | Unspecified |
| mmu-miR-9-5p | MIMAT0000142 | Ubox5 | 140629 | HITS-CLIP | | 37986224 | Unspecified |
| mmu-miR-9-5p | MIMAT0000142 | Ubqln2 | 54609 | HITS-CLIP | | 37986224 | Unspecified |
| mmu-miR-9-5p | MIMAT0000142 | Ubr5 | 70790 | HITS-CLIP | | 37986224 | Unspecified |
| mmu-miR-9-5p | MIMAT0000142 | Ugcg | 22234 | HITS-CLIP | | 37986224 | Unspecified |
| mmu-miR-9-5p | MIMAT0000142 | Uggt1 | 320011 | HITS-CLIP//qCLASH | 37986224\|31393953 | | Unspecified |
| mmu-miR-9-5p | MIMAT0000142 | Ugt1a6b | 394435 | HITS-CLIP | | 37986224 | Unspecified |
| mmu-miR-9-5p | MIMAT0000142 | Ugt8a | 22239 | HITS-CLIP | | 19536157 | Unspecified |
| mmu-miR-9-5p | MIMAT0000142 | Unc5b | 107449 | HITS-CLIP | | 37986224 | Unspecified |
| mmu-miR-9-5p | MIMAT0000142 | Unc5d | 210801 | HITS-CLIP | | 37986224 | Unspecified |
| mmu-miR-9-5p | MIMAT0000142 | Unc80 | 329178 | HITS-CLIP | 37986224\|19536157 | | Unspecified |
| mmu-miR-9-5p | MIMAT0000142 | Uqcrc2 | 67003 | HITS-CLIP | | 37986224 | Unspecified |
| mmu-miR-9-5p | MIMAT0000142 | Uqcrh | 66576 | HITS-CLIP | | 37986224 | Unspecified |
| mmu-miR-9-5p | MIMAT0000142 | Usp22 | 216825 | HITS-CLIP | | 37986224 | Unspecified |
| mmu-miR-9-5p | MIMAT0000142 | Usp24 | 329908 | HITS-CLIP | | 19536157 | Unspecified |
| mmu-miR-9-5p | MIMAT0000142 | Usp28 | 235323 | HITS-CLIP | | 37986224 | Unspecified |
| mmu-miR-9-5p | MIMAT0000142 | Usp32 | 237898 | HITS-CLIP | | 37986224 | Unspecified |
| mmu-miR-9-5p | MIMAT0000142 | Usp42 | 76800 | HITS-CLIP | | 19536157 | Unspecified |
| mmu-miR-9-5p | MIMAT0000142 | Usp47 | 74996 | HITS-CLIP | | 37986224 | Unspecified |
| mmu-miR-9-5p | MIMAT0000142 | Usp54 | 78787 | HITS-CLIP | | 19536157 | Unspecified |
| mmu-miR-9-5p | MIMAT0000142 | Utp15 | 105372 | HITS-CLIP | | 19536157 | Unspecified |
| mmu-miR-9-5p | MIMAT0000142 | Vamp1 | 22317 | HITS-CLIP | | 37986224 | Unspecified |
| mmu-miR-9-5p | MIMAT0000142 | Vcam1 | 22329 | HITS-CLIP | | 37986224 | Unspecified |
| mmu-miR-9-5p | MIMAT0000142 | Vcl | 22330 | HITS-CLIP | | 37986224 | Unspecified |
| mmu-miR-9-5p | MIMAT0000142 | Vps13a | 271564 | HITS-CLIP | | 19536157 | Unspecified |
| mmu-miR-9-5p | MIMAT0000142 | Vps13c | 320528 | HITS-CLIP | | 37986224 | Unspecified |
| mmu-miR-9-5p | MIMAT0000142 | Vps35 | 65114 | HITS-CLIP//qCLASH | 37986224\|31393953 | | Unspecified |
| mmu-miR-9-5p | MIMAT0000142 | Vsnl1 | 26950 | HITS-CLIP | | 37986224 | Unspecified |
| mmu-miR-9-5p | MIMAT0000142 | Vstm2a | 211739 | HITS-CLIP | | 37986224 | Unspecified |
| mmu-miR-9-5p | MIMAT0000142 | Vti1a | 53611 | HITS-CLIP | | 37986224 | Unspecified |
| mmu-miR-9-5p | MIMAT0000142 | Wapl | 23063 | HITS-CLIP | | 19536157 | Unspecified |
| mmu-miR-9-5p | MIMAT0000142 | Wars2 | 70560 | HITS-CLIP | | 37986224 | Unspecified |
| mmu-miR-9-5p | MIMAT0000142 | Wdr47 | 99512 | HITS-CLIP | | 37986224 | Unspecified |
| mmu-miR-9-5p | MIMAT0000142 | Wdr7 | 104082 | HITS-CLIP | | 37986224 | Unspecified |
| mmu-miR-9-5p | MIMAT0000142 | Wnk1 | 232341 | Sequencing//qCLASH//HITS-CLIP | 37986224\|31393953\|19536157\|26203 | | Unspecified |

| | | | | | | |
|---|---|---|---|---|---|---|
| mmu-miR-9-5p | MIMAT0000142 | Wscd2 | 320916 | HITS-CLIP | 37986224 | Unspecified |
| mmu-miR-9-5p | MIMAT0000142 | Wtap | 60532 | HITS-CLIP | 19536157 | Unspecified |
| mmu-miR-9-5p | MIMAT0000142 | Xpo5 | 72322 | HITS-CLIP | 37986224 | Unspecified |
| mmu-miR-9-5p | MIMAT0000142 | Yars2 | 70120 | HITS-CLIP | 37986224 | Unspecified |
| mmu-miR-9-5p | MIMAT0000142 | Ybx3 | 56449 | Sequencing//HITS-CLIP | 19536157 | Unspecified |
| mmu-miR-9-5p | MIMAT0000142 | Yeats2 | 208146 | HITS-CLIP | 19536157 | Unspecified |
| mmu-miR-9-5p | MIMAT0000142 | Ypel2 | 77864 | HITS-CLIP | 37986224 | Unspecified |
| mmu-miR-9-5p | MIMAT0000142 | Ythdf3 | 229096 | qCLASH | 31393953 | Unspecified |
| mmu-miR-9-5p | MIMAT0000142 | Zbtb25 | 109929 | HITS-CLIP | 37986224 | Unspecified |
| mmu-miR-9-5p | MIMAT0000142 | Zbtb34 | 241311 | Sequencing//HITS-CLIP | 37986224\|19536157 | Unspecified |
| mmu-miR-9-5p | MIMAT0000142 | Zbtb38 | 245007 | HITS-CLIP | 37986224 | Unspecified |
| mmu-miR-9-5p | MIMAT0000142 | Zbtb7a | 16969 | HITS-CLIP | 37986224 | Unspecified |
| mmu-miR-9-5p | MIMAT0000142 | Zc3h13 | 67302 | HITS-CLIP | 37986224 | Unspecified |
| mmu-miR-9-5p | MIMAT0000142 | Zc3hav1l | 209032 | HITS-CLIP | 37986224 | Unspecified |
| mmu-miR-9-5p | MIMAT0000142 | Zcchc8 | 70650 | HITS-CLIP | 37986224 | Unspecified |
| mmu-miR-9-5p | MIMAT0000142 | Zeb1 | 21417 | HITS-CLIP | 37986224 | Unspecified |
| mmu-miR-9-5p | MIMAT0000142 | Zfhx2 | 239102 | HITS-CLIP | 37986224 | Unspecified |
| mmu-miR-9-5p | MIMAT0000142 | Zfp236 | 329002 | HITS-CLIP | 37986224 | Unspecified |
| mmu-miR-9-5p | MIMAT0000142 | Zfp266 | 77519 | HITS-CLIP | 37986224\|19536157 | Unspecified |
| mmu-miR-9-5p | MIMAT0000142 | Zfp273 | 212569 | HITS-CLIP | 37986224\|19536157 | Unspecified |
| mmu-miR-9-5p | MIMAT0000142 | Zfp282 | 101095 | Sequencing//HITS-CLIP | 19536157\|37986224 | Unspecified |
| mmu-miR-9-5p | MIMAT0000142 | Zfp318 | 57908 | HITS-CLIP | 37986224 | Unspecified |
| mmu-miR-9-5p | MIMAT0000142 | Zfp335 | 329559 | HITS-CLIP | 37986224 | Unspecified |
| mmu-miR-9-5p | MIMAT0000142 | Zfp358 | 140482 | HITS-CLIP | 37986224 | Unspecified |
| mmu-miR-9-5p | MIMAT0000142 | Zfp385a | 29813 | HITS-CLIP | 37986224 | Unspecified |
| mmu-miR-9-5p | MIMAT0000142 | Zfp407 | 240476 | HITS-CLIP | 37986224 | Unspecified |
| mmu-miR-9-5p | MIMAT0000142 | Zfp445 | 235682 | HITS-CLIP | 37986224 | Unspecified |
| mmu-miR-9-5p | MIMAT0000142 | Zfp458 | 238690 | HITS-CLIP | 37986224 | Unspecified |
| mmu-miR-9-5p | MIMAT0000142 | Zfp518a | 72672 | HITS-CLIP | 37986224 | Unspecified |
| mmu-miR-9-5p | MIMAT0000142 | Zfp551 | 619331 | HITS-CLIP | 37986224 | Unspecified |
| mmu-miR-9-5p | MIMAT0000142 | Zfp597 | 71063 | HITS-CLIP | 37986224 | Unspecified |
| mmu-miR-9-5p | MIMAT0000142 | Zfp608 | 269023 | HITS-CLIP | 37986224 | Unspecified |
| mmu-miR-9-5p | MIMAT0000142 | Zfp655 | 72611 | HITS-CLIP | 37986224 | Unspecified |
| mmu-miR-9-5p | MIMAT0000142 | Zfp672 | 319475 | HITS-CLIP | 37986224 | Unspecified |
| mmu-miR-9-5p | MIMAT0000142 | Zfp706 | 68036 | HITS-CLIP | 37986224 | Unspecified |
| mmu-miR-9-5p | MIMAT0000142 | Zfp712 | 78251 | HITS-CLIP | 37986224 | Unspecified |
| mmu-miR-9-5p | MIMAT0000142 | Zfp72 | 238722 | HITS-CLIP | 37986224 | Unspecified |
| mmu-miR-9-5p | MIMAT0000142 | Zfp90 | 22751 | HITS-CLIP | 37986224 | Unspecified |
| mmu-miR-9-5p | MIMAT0000142 | Zfp959 | 224893 | HITS-CLIP | 37986224\|19536157 | Unspecified |

| | | | | | | |
|---|---|---|---|---|---|---|
| mmu-miR-9-5p | MIMAT0000142 | Zfyve26 | 211978 | HITS-CLIP | 37986224 | Unspecified |
| mmu-miR-9-5p | MIMAT0000142 | Zhx1 | 22770 | HITS-CLIP | 37986224 | Unspecified |
| mmu-miR-9-5p | MIMAT0000142 | Zmynd11 | 66505 | HITS-CLIP | 37986224 | Unspecified |
| mmu-miR-9-5p | MIMAT0000142 | Zswim8 | 268721 | HITS-CLIP | 37986224 | Unspecified |
| mmu-miR-9-5p | MIMAT0000142 | Zyg11b | 414872 | HITS-CLIP | 37986224 | Unspecified |
| mmu-miR-703 | MIMAT0003493 | Actb | 11461 | HITS-CLIP | 37986224 | Unspecified |
| mmu-miR-703 | MIMAT0003493 | Ago1 | 236511 | HITS-CLIP | 37986224 | Unspecified |
| mmu-miR-703 | MIMAT0003493 | Atxn1l | 52335 | HITS-CLIP | 37986224 | Unspecified |
| mmu-miR-703 | MIMAT0003493 | Brd2 | 14312 | HITS-CLIP | 37986224 | Unspecified |
| mmu-miR-703 | MIMAT0003493 | Brd8 | 78656 | HITS-CLIP | 37986224 | Unspecified |
| mmu-miR-703 | MIMAT0003493 | Casc3 | 192160 | HITS-CLIP | 37986224 | Unspecified |
| mmu-miR-703 | MIMAT0003493 | Cdc42 | 12540 | HITS-CLIP | 37986224 | Unspecified |
| mmu-miR-703 | MIMAT0003493 | Cnot6 | 104625 | HITS-CLIP | 37986224 | Unspecified |
| mmu-miR-703 | MIMAT0003493 | Ddx17 | 67040 | HITS-CLIP | 37986224 | Unspecified |
| mmu-miR-703 | MIMAT0003493 | Eif1a | 13664 | HITS-CLIP | 37986224 | Unspecified |
| mmu-miR-703 | MIMAT0003493 | Epha2 | 13836 | HITS-CLIP | 37986224 | Unspecified |
| mmu-miR-703 | MIMAT0003493 | Erlin2 | 244373 | HITS-CLIP | 37986224 | Unspecified |
| mmu-miR-703 | MIMAT0003493 | Fam169a | 320557 | HITS-CLIP | 37986224 | Unspecified |
| mmu-miR-703 | MIMAT0003493 | Fbxw2 | 30050 | HITS-CLIP | 37986224 | Unspecified |
| mmu-miR-703 | MIMAT0003493 | Iws1 | 73473 | HITS-CLIP | 37986224 | Unspecified |
| mmu-miR-703 | MIMAT0003493 | Lars2 | 102436 | HITS-CLIP | 31433990 | Unspecified |
| mmu-miR-703 | MIMAT0003493 | Lmnb1 | 16906 | HITS-CLIP | 37986224 | Unspecified |
| mmu-miR-703 | MIMAT0003493 | Mcl1 | 17210 | HITS-CLIP | 37986224 | Unspecified |
| mmu-miR-703 | MIMAT0003493 | Mkrn1 | 54484 | HITS-CLIP | 37986224 | Unspecified |
| mmu-miR-703 | MIMAT0003493 | Mprip | 26936 | HITS-CLIP | 31433990 | Unspecified |
| mmu-miR-703 | MIMAT0003493 | Mtx3 | 382793 | HITS-CLIP | 37986224 | Unspecified |
| mmu-miR-703 | MIMAT0003493 | Nacc1 | 66830 | HITS-CLIP | 37986224 | Unspecified |
| mmu-miR-703 | MIMAT0003493 | Nucks1 | 98415 | HITS-CLIP | 37986224 | Unspecified |
| mmu-miR-703 | MIMAT0003493 | Pafah1b2 | 18475 | HITS-CLIP | 21258322 | Unspecified |
| mmu-miR-703 | MIMAT0003493 | Purb | 19291 | HITS-CLIP | 37986224 | Unspecified |
| mmu-miR-703 | MIMAT0003493 | Qser1 | 99003 | HITS-CLIP | 37986224 | Unspecified |
| mmu-miR-703 | MIMAT0003493 | Rai14 | 75646 | HITS-CLIP | 37986224 | Unspecified |
| mmu-miR-703 | MIMAT0003493 | Rnf38 | 73469 | HITS-CLIP | 37986224 | Unspecified |
| mmu-miR-703 | MIMAT0003493 | Sema3e | 20349 | HITS-CLIP | 37986224 | Unspecified |
| mmu-miR-703 | MIMAT0003493 | Slc16a13 | 69309 | HITS-CLIP | 37986224 | Unspecified |
| mmu-miR-703 | MIMAT0003493 | Slc38a2 | 67760 | HITS-CLIP | 37986224 | Unspecified |
| mmu-miR-703 | MIMAT0003493 | Srsf4 | 57317 | HITS-CLIP | 37986224 | Unspecified |
| mmu-miR-703 | MIMAT0003493 | Strbp | 20744 | HITS-CLIP | 37986224 | Unspecified |
| mmu-miR-703 | MIMAT0003493 | Tead1 | 21676 | HITS-CLIP | 37986224 | Unspecified |

| | | | | | | |
|---|---|---|---|---|---|---|
| mmu-miR-703 | MIMAT0003493 | Tle4 | 21888 | HITS-CLIP | 37986224 | Unspecified |
| mmu-miR-703 | MIMAT0003493 | Tnrc6a | 233833 | HITS-CLIP | 31433990 | Unspecified |
| mmu-miR-703 | MIMAT0003493 | Txnip | 56338 | HITS-CLIP | 37986224 | Unspecified |
| mmu-miR-703 | MIMAT0003493 | Wdr3 | 269470 | HITS-CLIP | 21258322 | Unspecified |
| mmu-miR-703 | MIMAT0003493 | Zfp451 | 98403 | HITS-CLIP | 37986224 | Unspecified |
| mmu-miR-703 | MIMAT0003493 | Zfp704 | 170753 | HITS-CLIP | 37986224 | Unspecified |
| mmu-miR-703 | MIMAT0003493 | Zmynd8 | 228880 | HITS-CLIP | 37986224 | Unspecified |
| mmu-miR-466q | MIMAT0020631 | 2610507B11Rik | 72503 | HITS-CLIP | 37986224 | Unspecified |
| mmu-miR-466q | MIMAT0020631 | 4932438A13Rik | 229227 | HITS-CLIP | 37986224 | Unspecified |
| mmu-miR-466q | MIMAT0020631 | Abca2 | 11305 | HITS-CLIP | 37986224 | Unspecified |
| mmu-miR-466q | MIMAT0020631 | Abcb11 | 27413 | HITS-CLIP | 23597149 | Unspecified |
| mmu-miR-466q | MIMAT0020631 | Abcc6 | 27421 | HITS-CLIP | 37986224\|23597149 | Unspecified |
| mmu-miR-466q | MIMAT0020631 | Abcf3 | 27406 | HITS-CLIP | 23597149 | Unspecified |
| mmu-miR-466q | MIMAT0020631 | Acadl | 11363 | HITS-CLIP | 23597149 | Unspecified |
| mmu-miR-466q | MIMAT0020631 | Acly | 104112 | HITS-CLIP | 23597149 | Unspecified |
| mmu-miR-466q | MIMAT0020631 | Actn4 | 60595 | HITS-CLIP | 37986224 | Unspecified |
| mmu-miR-466q | MIMAT0020631 | Adam9 | 11502 | HITS-CLIP | 23597149 | Unspecified |
| mmu-miR-466q | MIMAT0020631 | Adck2 | 57869 | HITS-CLIP | 37986224 | Unspecified |
| mmu-miR-466q | MIMAT0020631 | Adck5 | 268822 | HITS-CLIP | 37986224 | Unspecified |
| mmu-miR-466q | MIMAT0020631 | Adnp | 11538 | HITS-CLIP | 23597149 | Unspecified |
| mmu-miR-466q | MIMAT0020631 | Afdn | 4301 | HITS-CLIP | 23597149 | Unspecified |
| mmu-miR-466q | MIMAT0020631 | Aff4 | 93736 | HITS-CLIP | 23597149 | Unspecified |
| mmu-miR-466q | MIMAT0020631 | Agpat2 | 67512 | HITS-CLIP | 23597149 | Unspecified |
| mmu-miR-466q | MIMAT0020631 | Ahcyl1 | 229709 | HITS-CLIP | 23597149 | Unspecified |
| mmu-miR-466q | MIMAT0020631 | Ahdc1 | 230793 | HITS-CLIP | 23597149 | Unspecified |
| mmu-miR-466q | MIMAT0020631 | Ahr | 11622 | HITS-CLIP | 37986224\|23597149 | Unspecified |
| mmu-miR-466q | MIMAT0020631 | Aifm2 | 71361 | HITS-CLIP | 23597149 | Unspecified |
| mmu-miR-466q | MIMAT0020631 | Akap8l | 54194 | HITS-CLIP | 37986224 | Unspecified |
| mmu-miR-466q | MIMAT0020631 | Akr1a1 | 58810 | HITS-CLIP | 37986224 | Unspecified |
| mmu-miR-466q | MIMAT0020631 | Akr1c20 | 116852 | HITS-CLIP | 23597149 | Unspecified |
| mmu-miR-466q | MIMAT0020631 | Aldh6a1 | 104776 | HITS-CLIP | 37986224 | Unspecified |
| mmu-miR-466q | MIMAT0020631 | Aldh7a1 | 110695 | HITS-CLIP | 37986224 | Unspecified |
| mmu-miR-466q | MIMAT0020631 | Aldh8a1 | 237320 | HITS-CLIP | 37986224 | Unspecified |
| mmu-miR-466q | MIMAT0020631 | Amfr | 23802 | HITS-CLIP | 23597149 | Unspecified |
| mmu-miR-466q | MIMAT0020631 | Amz2 | 13929 | HITS-CLIP | 37986224 | Unspecified |
| mmu-miR-466q | MIMAT0020631 | Anapc1 | 17222 | HITS-CLIP | 37986224 | Unspecified |
| mmu-miR-466q | MIMAT0020631 | Ankrd52 | 237615 | HITS-CLIP | 23597149 | Unspecified |
| mmu-miR-466q | MIMAT0020631 | Arfgef2 | 99371 | HITS-CLIP | 37986224 | Unspecified |
| mmu-miR-466q | MIMAT0020631 | Arhgef19 | 213649 | HITS-CLIP | 37986224 | Unspecified |

| | | | | | | |
|---|---|---|---|---|---|---|
| mmu-miR-466q | MIMAT0020631 | Arid1a | 93760 | HITS-CLIP | 37986224 | Unspecified |
| mmu-miR-466q | MIMAT0020631 | Arid1b | 239985 | HITS-CLIP | 23597149 | Unspecified |
| mmu-miR-466q | MIMAT0020631 | Arrdc3 | 105171 | HITS-CLIP | 23597149 | Unspecified |
| mmu-miR-466q | MIMAT0020631 | Ascc2 | 75452 | HITS-CLIP | 23597149 | Unspecified |
| mmu-miR-466q | MIMAT0020631 | Atp2a2 | 11938 | HITS-CLIP | 37986224 | Unspecified |
| mmu-miR-466q | MIMAT0020631 | Atp6v0e | 11974 | HITS-CLIP | 37986224 | Unspecified |
| mmu-miR-466q | MIMAT0020631 | Avpi1 | 69534 | HITS-CLIP | 23597149 | Unspecified |
| mmu-miR-466q | MIMAT0020631 | BC031181 | 407819 | HITS-CLIP | 37986224\|23597149 | Unspecified |
| mmu-miR-466q | MIMAT0020631 | Bach1 | 12013 | HITS-CLIP | 37986224 | Unspecified |
| mmu-miR-466q | MIMAT0020631 | Bhmt | 12116 | HITS-CLIP | 37986224 | Unspecified |
| mmu-miR-466q | MIMAT0020631 | Btaf1 | 107182 | HITS-CLIP | 37986224 | Unspecified |
| mmu-miR-466q | MIMAT0020631 | Btg2 | 12227 | HITS-CLIP | 23142080\|23597149 | Unspecified |
| mmu-miR-466q | MIMAT0020631 | C4bp | 12269 | HITS-CLIP | 23597149 | Unspecified |
| mmu-miR-466q | MIMAT0020631 | C6 | 12274 | HITS-CLIP | 37986224\|23597149 | Unspecified |
| mmu-miR-466q | MIMAT0020631 | C8b | 110382 | HITS-CLIP | 23597149 | Unspecified |
| mmu-miR-466q | MIMAT0020631 | Cadps2 | 320405 | HITS-CLIP | 37986224 | Unspecified |
| mmu-miR-466q | MIMAT0020631 | Cbfa2t3 | 12398 | HITS-CLIP | 23597149 | Unspecified |
| mmu-miR-466q | MIMAT0020631 | Cbs | 12411 | HITS-CLIP | 37986224 | Unspecified |
| mmu-miR-466q | MIMAT0020631 | Ccdc47 | 67163 | HITS-CLIP | 37986224 | Unspecified |
| mmu-miR-466q | MIMAT0020631 | Ccdc6 | 76551 | HITS-CLIP | 37986224\|23597149 | Unspecified |
| mmu-miR-466q | MIMAT0020631 | Ccnc | 51813 | HITS-CLIP | 37986224 | Unspecified |
| mmu-miR-466q | MIMAT0020631 | Ccs | 12460 | HITS-CLIP | 37986224\|23597149 | Unspecified |
| mmu-miR-466q | MIMAT0020631 | Cd164 | 53599 | HITS-CLIP | 37986224 | Unspecified |
| mmu-miR-466q | MIMAT0020631 | Cd302 | 66205 | HITS-CLIP | 37986224 | Unspecified |
| mmu-miR-466q | MIMAT0020631 | Cdk6 | 12571 | HITS-CLIP | 23597149 | Unspecified |
| mmu-miR-466q | MIMAT0020631 | Cebpa | 12606 | HITS-CLIP | 23597149 | Unspecified |
| mmu-miR-466q | MIMAT0020631 | Cemip2 | 23670 | HITS-CLIP | 23597149 | Unspecified |
| mmu-miR-466q | MIMAT0020631 | Cep350 | 74081 | HITS-CLIP | 23597149 | Unspecified |
| mmu-miR-466q | MIMAT0020631 | Cfh | 12628 | HITS-CLIP | 37986224 | Unspecified |
| mmu-miR-466q | MIMAT0020631 | Cflar | 12633 | HITS-CLIP | 37986224 | Unspecified |
| mmu-miR-466q | MIMAT0020631 | Chd4 | 107932 | HITS-CLIP | 23597149 | Unspecified |
| mmu-miR-466q | MIMAT0020631 | Clic4 | 29876 | HITS-CLIP | 37986224 | Unspecified |
| mmu-miR-466q | MIMAT0020631 | Clu | 12759 | HITS-CLIP | 23597149 | Unspecified |
| mmu-miR-466q | MIMAT0020631 | Cnot1 | 234594 | HITS-CLIP | 37986224 | Unspecified |
| mmu-miR-466q | MIMAT0020631 | Cnot6l | 231464 | HITS-CLIP | 23597149 | Unspecified |
| mmu-miR-466q | MIMAT0020631 | Cobll1 | 319876 | HITS-CLIP | 37986224 | Unspecified |
| mmu-miR-466q | MIMAT0020631 | Col18a1 | 12822 | HITS-CLIP | 37986224 | Unspecified |
| mmu-miR-466q | MIMAT0020631 | Cox19 | 68033 | HITS-CLIP | 37986224 | Unspecified |
| mmu-miR-466q | MIMAT0020631 | Cp | 12870 | HITS-CLIP | 23597149 | Unspecified |

| | | | | | | | |
|---|---|---|---|---|---|---|---|
| mmu-miR-466q | MIMAT0020631 | Cpeb2 | 231207 | HITS-CLIP | | 23597149 | Unspecified |
| mmu-miR-466q | MIMAT0020631 | Cpne2 | 234577 | HITS-CLIP | | 23597149 | Unspecified |
| mmu-miR-466q | MIMAT0020631 | Cps1 | 227231 | HITS-CLIP | | 23597149 | Unspecified |
| mmu-miR-466q | MIMAT0020631 | Creb3l3 | 208677 | HITS-CLIP | | 37986224 | Unspecified |
| mmu-miR-466q | MIMAT0020631 | Ctbp1 | 13016 | HITS-CLIP | | 37986224 | Unspecified |
| mmu-miR-466q | MIMAT0020631 | Cttn | 13043 | HITS-CLIP | | 37986224 | Unspecified |
| mmu-miR-466q | MIMAT0020631 | Cul1 | 26965 | HITS-CLIP | | 23597149 | Unspecified |
| mmu-miR-466q | MIMAT0020631 | Cul3 | 26554 | HITS-CLIP | | 37986224 | Unspecified |
| mmu-miR-466q | MIMAT0020631 | Cxcl12 | 20315 | HITS-CLIP | | 23597149 | Unspecified |
| mmu-miR-466q | MIMAT0020631 | Cyb5b | 66427 | HITS-CLIP | | 37986224 | Unspecified |
| mmu-miR-466q | MIMAT0020631 | Cyp4a14 | 13119 | HITS-CLIP | | 37986224 | Unspecified |
| mmu-miR-466q | MIMAT0020631 | Daam1 | 208846 | HITS-CLIP | | 23597149 | Unspecified |
| mmu-miR-466q | MIMAT0020631 | Dap | 223453 | HITS-CLIP | | 37986224 | Unspecified |
| mmu-miR-466q | MIMAT0020631 | Dazap2 | 23994 | HITS-CLIP | 37986224\|23597149 | | Unspecified |
| mmu-miR-466q | MIMAT0020631 | Dcaf8 | 98193 | HITS-CLIP | | 37986224 | Unspecified |
| mmu-miR-466q | MIMAT0020631 | Dgkd | 227333 | HITS-CLIP | | 37986224 | Unspecified |
| mmu-miR-466q | MIMAT0020631 | Dhrs3 | 20148 | HITS-CLIP | | 37986224 | Unspecified |
| mmu-miR-466q | MIMAT0020631 | Dhrs7 | 66375 | HITS-CLIP | 37986224\|23597149 | | Unspecified |
| mmu-miR-466q | MIMAT0020631 | Dlc1 | 50768 | HITS-CLIP | | 23597149 | Unspecified |
| mmu-miR-466q | MIMAT0020631 | Dlst | 78920 | HITS-CLIP | 37986224\|23597149 | | Unspecified |
| mmu-miR-466q | MIMAT0020631 | Dnm1l | 74006 | HITS-CLIP | | 23597149 | Unspecified |
| mmu-miR-466q | MIMAT0020631 | Dtymk | 21915 | HITS-CLIP | | 37986224 | Unspecified |
| mmu-miR-466q | MIMAT0020631 | Dusp1 | 19252 | HITS-CLIP | | 37986224 | Unspecified |
| mmu-miR-466q | MIMAT0020631 | Dusp16 | 70686 | HITS-CLIP | | 37986224 | Unspecified |
| mmu-miR-466q | MIMAT0020631 | Eaf1 | 74427 | HITS-CLIP | | 23597149 | Unspecified |
| mmu-miR-466q | MIMAT0020631 | Edc4 | 234699 | HITS-CLIP | | 23597149 | Unspecified |
| mmu-miR-466q | MIMAT0020631 | Eea1 | 216238 | HITS-CLIP | | 23597149 | Unspecified |
| mmu-miR-466q | MIMAT0020631 | Eef2 | 13629 | HITS-CLIP | 37986224\|23597149 | | Unspecified |
| mmu-miR-466q | MIMAT0020631 | Ehd1 | 13660 | HITS-CLIP | | 23597149 | Unspecified |
| mmu-miR-466q | MIMAT0020631 | Ehhadh | 74147 | HITS-CLIP | | 37986224 | Unspecified |
| mmu-miR-466q | MIMAT0020631 | Elovl2 | 54326 | HITS-CLIP | | 37986224 | Unspecified |
| mmu-miR-466q | MIMAT0020631 | Enc1 | 13803 | HITS-CLIP | | 37986224 | Unspecified |
| mmu-miR-466q | MIMAT0020631 | Erap1 | 80898 | HITS-CLIP | | 37986224 | Unspecified |
| mmu-miR-466q | MIMAT0020631 | Erlin1 | 226144 | HITS-CLIP | | 23597149 | Unspecified |
| mmu-miR-466q | MIMAT0020631 | Esrp2 | 77411 | HITS-CLIP | | 23597149 | Unspecified |
| mmu-miR-466q | MIMAT0020631 | Esyt2 | 52635 | HITS-CLIP | | 37986224 | Unspecified |
| mmu-miR-466q | MIMAT0020631 | Etfdh | 66841 | HITS-CLIP | | 23597149 | Unspecified |
| mmu-miR-466q | MIMAT0020631 | Ets1 | 23871 | HITS-CLIP | 37986224\|23597149 | | Unspecified |
| mmu-miR-466q | MIMAT0020631 | F2 | 14061 | HITS-CLIP | 37986224\|23597149 | | Unspecified |

| | | | | | | |
|---|---|---|---|---|---|---|
| mmu-miR-466q | MIMAT0020631 | Faah | 14073 | HITS-CLIP | 23597149 | Unspecified |
| mmu-miR-466q | MIMAT0020631 | Fam120a | 218236 | HITS-CLIP | 37986224 | Unspecified |
| mmu-miR-466q | MIMAT0020631 | Fam126b | 213056 | HITS-CLIP | 37986224 | Unspecified |
| mmu-miR-466q | MIMAT0020631 | Fam199x | 245622 | HITS-CLIP | 37986224 | Unspecified |
| mmu-miR-466q | MIMAT0020631 | Fam32a | 67922 | HITS-CLIP | 37986224 | Unspecified |
| mmu-miR-466q | MIMAT0020631 | Farp1 | 223254 | HITS-CLIP | 37986224 | Unspecified |
| mmu-miR-466q | MIMAT0020631 | Fbxl17 | 50758 | HITS-CLIP | 23597149 | Unspecified |
| mmu-miR-466q | MIMAT0020631 | Fbxl20 | 72194 | HITS-CLIP | 37986224 | Unspecified |
| mmu-miR-466q | MIMAT0020631 | Fbxo21 | 231670 | HITS-CLIP | 37986224\|23597149 | Unspecified |
| mmu-miR-466q | MIMAT0020631 | Fibp | 58249 | HITS-CLIP | 23597149 | Unspecified |
| mmu-miR-466q | MIMAT0020631 | Fmo1 | 14261 | HITS-CLIP | 23597149 | Unspecified |
| mmu-miR-466q | MIMAT0020631 | Fn1 | 14268 | HITS-CLIP | 37986224 | Unspecified |
| mmu-miR-466q | MIMAT0020631 | Fnip1 | 216742 | HITS-CLIP | 23597149 | Unspecified |
| mmu-miR-466q | MIMAT0020631 | Fpgs | 14287 | HITS-CLIP | 23597149 | Unspecified |
| mmu-miR-466q | MIMAT0020631 | Fryl | 72313 | HITS-CLIP | 23597149 | Unspecified |
| mmu-miR-466q | MIMAT0020631 | Fth1 | 14319 | HITS-CLIP | 37986224 | Unspecified |
| mmu-miR-466q | MIMAT0020631 | Fto | 26383 | HITS-CLIP | 37986224\|23597149 | Unspecified |
| mmu-miR-466q | MIMAT0020631 | Fyn | 14360 | HITS-CLIP | 23597149 | Unspecified |
| mmu-miR-466q | MIMAT0020631 | Fzd1 | 14362 | HITS-CLIP | 37986224 | Unspecified |
| mmu-miR-466q | MIMAT0020631 | Fzd5 | 14367 | HITS-CLIP | 23597149 | Unspecified |
| mmu-miR-466q | MIMAT0020631 | G0s2 | 14373 | HITS-CLIP | 37986224 | Unspecified |
| mmu-miR-466q | MIMAT0020631 | Galk2 | 69976 | HITS-CLIP | 37986224\|23597149 | Unspecified |
| mmu-miR-466q | MIMAT0020631 | Gc | 14473 | HITS-CLIP | 23597149 | Unspecified |
| mmu-miR-466q | MIMAT0020631 | Gcgr | 14527 | HITS-CLIP | 37986224 | Unspecified |
| mmu-miR-466q | MIMAT0020631 | Gcnt2 | 14538 | HITS-CLIP | 37986224\|23597149 | Unspecified |
| mmu-miR-466q | MIMAT0020631 | Gde1 | 56209 | HITS-CLIP | 37986224 | Unspecified |
| mmu-miR-466q | MIMAT0020631 | Ggact | 223267 | HITS-CLIP | 37986224 | Unspecified |
| mmu-miR-466q | MIMAT0020631 | Ghr | 14600 | HITS-CLIP | 37986224 | Unspecified |
| mmu-miR-466q | MIMAT0020631 | Glg1 | 20340 | HITS-CLIP | 37986224 | Unspecified |
| mmu-miR-466q | MIMAT0020631 | Glyctk | 235582 | HITS-CLIP | 37986224\|23597149 | Unspecified |
| mmu-miR-466q | MIMAT0020631 | Gna12 | 14673 | HITS-CLIP | 37986224 | Unspecified |
| mmu-miR-466q | MIMAT0020631 | Gpalpp1 | 67467 | HITS-CLIP | 37986224 | Unspecified |
| mmu-miR-466q | MIMAT0020631 | Gpihbp1 | 68453 | HITS-CLIP | 37986224 | Unspecified |
| mmu-miR-466q | MIMAT0020631 | Gpr146 | 80290 | HITS-CLIP | 37986224\|23597149 | Unspecified |
| mmu-miR-466q | MIMAT0020631 | Gpr182 | 11536 | HITS-CLIP | 23597149 | Unspecified |
| mmu-miR-466q | MIMAT0020631 | Gprin3 | 243385 | HITS-CLIP | 23597149 | Unspecified |
| mmu-miR-466q | MIMAT0020631 | Gsta3 | 14859 | HITS-CLIP | 37986224\|23597149 | Unspecified |
| mmu-miR-466q | MIMAT0020631 | Gstz1 | 14874 | HITS-CLIP | 23597149 | Unspecified |
| mmu-miR-466q | MIMAT0020631 | Gtf2h4 | 14885 | HITS-CLIP | 37986224 | Unspecified |

| | | | | | | |
|---|---|---|---|---|---|---|
| mmu-miR-466q | MIMAT0020631 | Gtf3c4 | 269252 | HITS-CLIP | 37986224 | Unspecified |
| mmu-miR-466q | MIMAT0020631 | Gucd1 | 68778 | HITS-CLIP | 37986224 | Unspecified |
| mmu-miR-466q | MIMAT0020631 | Gulo | 268756 | HITS-CLIP | 37986224 | Unspecified |
| mmu-miR-466q | MIMAT0020631 | Gzf1 | 74533 | HITS-CLIP | 23597149 | Unspecified |
| mmu-miR-466q | MIMAT0020631 | H1f0 | 14958 | HITS-CLIP | 37986224\|23597149 | Unspecified |
| mmu-miR-466q | MIMAT0020631 | H6pd | 100198 | HITS-CLIP | 37986224\|23597149 | Unspecified |
| mmu-miR-466q | MIMAT0020631 | Hcfc1 | 15161 | HITS-CLIP | 37986224\|23597149 | Unspecified |
| mmu-miR-466q | MIMAT0020631 | Heatr6 | 217026 | HITS-CLIP | 23597149 | Unspecified |
| mmu-miR-466q | MIMAT0020631 | Hectd1 | 207304 | HITS-CLIP | 37986224 | Unspecified |
| mmu-miR-466q | MIMAT0020631 | Herpud1 | 64209 | HITS-CLIP | 23597149 | Unspecified |
| mmu-miR-466q | MIMAT0020631 | Hnrnph3 | 432467 | HITS-CLIP | 37986224 | Unspecified |
| mmu-miR-466q | MIMAT0020631 | Hnrnpu | 51810 | HITS-CLIP | 37986224\|23597149 | Unspecified |
| mmu-miR-466q | MIMAT0020631 | Hp1bp3 | 15441 | HITS-CLIP | 37986224 | Unspecified |
| mmu-miR-466q | MIMAT0020631 | Hprt | 15452 | HITS-CLIP | 23597149 | Unspecified |
| mmu-miR-466q | MIMAT0020631 | Hsd17b10 | 15108 | HITS-CLIP | 37986224 | Unspecified |
| mmu-miR-466q | MIMAT0020631 | Hsd3b7 | 101502 | HITS-CLIP | 23597149 | Unspecified |
| mmu-miR-466q | MIMAT0020631 | Hspa5 | 14828 | HITS-CLIP | 37986224 | Unspecified |
| mmu-miR-466q | MIMAT0020631 | Huwe1 | 59026 | HITS-CLIP | 31433990 | Unspecified |
| mmu-miR-466q | MIMAT0020631 | Hykk | 235386 | HITS-CLIP | 37986224 | Unspecified |
| mmu-miR-466q | MIMAT0020631 | Idh2 | 269951 | HITS-CLIP | 23597149 | Unspecified |
| mmu-miR-466q | MIMAT0020631 | Ifngr1 | 15979 | HITS-CLIP | 37986224 | Unspecified |
| mmu-miR-466q | MIMAT0020631 | Igfbp2 | 16008 | HITS-CLIP | 37986224\|23597149 | Unspecified |
| mmu-miR-466q | MIMAT0020631 | Igfbp4 | 16010 | HITS-CLIP | 23597149 | Unspecified |
| mmu-miR-466q | MIMAT0020631 | Igfbp7 | 29817 | HITS-CLIP | 37986224 | Unspecified |
| mmu-miR-466q | MIMAT0020631 | Insig2 | 72999 | HITS-CLIP | 23597149 | Unspecified |
| mmu-miR-466q | MIMAT0020631 | Ints1 | 68510 | HITS-CLIP | 23597149 | Unspecified |
| mmu-miR-466q | MIMAT0020631 | Ints8 | 72656 | HITS-CLIP | 37986224 | Unspecified |
| mmu-miR-466q | MIMAT0020631 | Ipo5 | 70572 | HITS-CLIP | 37986224 | Unspecified |
| mmu-miR-466q | MIMAT0020631 | Iqgap2 | 544963 | HITS-CLIP | 37986224 | Unspecified |
| mmu-miR-466q | MIMAT0020631 | Itm2b | 16432 | HITS-CLIP | 23597149 | Unspecified |
| mmu-miR-466q | MIMAT0020631 | Itprid2 | 6744 | HITS-CLIP | 23597149 | Unspecified |
| mmu-miR-466q | MIMAT0020631 | Iws1 | 73473 | HITS-CLIP | 21258322\|23597149 | Unspecified |
| mmu-miR-466q | MIMAT0020631 | Jade1 | 269424 | HITS-CLIP | 37986224\|23597149 | Unspecified |
| mmu-miR-466q | MIMAT0020631 | Jak1 | 16451 | HITS-CLIP | 37986224\|23597149 | Unspecified |
| mmu-miR-466q | MIMAT0020631 | Kansl3 | 226976 | HITS-CLIP | 37986224 | Unspecified |
| mmu-miR-466q | MIMAT0020631 | Kat5 | 81601 | HITS-CLIP | 37986224 | Unspecified |
| mmu-miR-466q | MIMAT0020631 | Kdm2a | 225876 | HITS-CLIP | 37986224 | Unspecified |
| mmu-miR-466q | MIMAT0020631 | Kif2a | 16563 | HITS-CLIP | 23597149 | Unspecified |
| mmu-miR-466q | MIMAT0020631 | Klf9 | 16601 | HITS-CLIP | 37986224\|23597149 | Unspecified |

| | | | | | | |
|---|---|---|---|---|---|---|
| mmu-miR-466q | MIMAT0020631 | Klhdc2 | 69554 | HITS-CLIP | 23597149 | Unspecified |
| mmu-miR-466q | MIMAT0020631 | Kmt2a | 214162 | HITS-CLIP | 23597149 | Unspecified |
| mmu-miR-466q | MIMAT0020631 | Kng1 | 16644 | HITS-CLIP | 23597149 | Unspecified |
| mmu-miR-466q | MIMAT0020631 | Kpna4 | 16649 | HITS-CLIP | 23597149 | Unspecified |
| mmu-miR-466q | MIMAT0020631 | L2hgdh | 217666 | HITS-CLIP | 37986224 | Unspecified |
| mmu-miR-466q | MIMAT0020631 | Larp1 | 73158 | HITS-CLIP | 37986224\|23597149 | Unspecified |
| mmu-miR-466q | MIMAT0020631 | Ldha | 16828 | HITS-CLIP | 23597149 | Unspecified |
| mmu-miR-466q | MIMAT0020631 | Ldhd | 52815 | HITS-CLIP | 23597149 | Unspecified |
| mmu-miR-466q | MIMAT0020631 | Letm1 | 56384 | HITS-CLIP | 37986224 | Unspecified |
| mmu-miR-466q | MIMAT0020631 | Letmd1 | 68614 | HITS-CLIP | 23597149 | Unspecified |
| mmu-miR-466q | MIMAT0020631 | Lifr | 16880 | HITS-CLIP | 37986224 | Unspecified |
| mmu-miR-466q | MIMAT0020631 | Lims2 | 225341 | HITS-CLIP | 37986224\|23597149 | Unspecified |
| mmu-miR-466q | MIMAT0020631 | Lipa | 16889 | HITS-CLIP | 37986224 | Unspecified |
| mmu-miR-466q | MIMAT0020631 | Lman1 | 70361 | HITS-CLIP | 37986224 | Unspecified |
| mmu-miR-466q | MIMAT0020631 | Lnpep | 240028 | HITS-CLIP | 37986224 | Unspecified |
| mmu-miR-466q | MIMAT0020631 | Lpin1 | 14245 | HITS-CLIP | 37986224 | Unspecified |
| mmu-miR-466q | MIMAT0020631 | Lpin2 | 64898 | HITS-CLIP | 23597149 | Unspecified |
| mmu-miR-466q | MIMAT0020631 | Lpp | 210126 | HITS-CLIP | 23597149 | Unspecified |
| mmu-miR-466q | MIMAT0020631 | Lrg1 | 76905 | HITS-CLIP | 37986224 | Unspecified |
| mmu-miR-466q | MIMAT0020631 | Lrp1 | 16971 | HITS-CLIP | 37986224 | Unspecified |
| mmu-miR-466q | MIMAT0020631 | Lrrc59 | 98238 | HITS-CLIP | 37986224 | Unspecified |
| mmu-miR-466q | MIMAT0020631 | Lurap1l | 52829 | HITS-CLIP | 23597149 | Unspecified |
| mmu-miR-466q | MIMAT0020631 | Lypla1 | 18777 | HITS-CLIP | 23597149 | Unspecified |
| mmu-miR-466q | MIMAT0020631 | Map3k9 | 338372 | HITS-CLIP | 26203562 | Unspecified |
| mmu-miR-466q | MIMAT0020631 | Mapk9 | 26420 | HITS-CLIP | 37986224 | Unspecified |
| mmu-miR-466q | MIMAT0020631 | Marveld2 | 218518 | HITS-CLIP | 23597149 | Unspecified |
| mmu-miR-466q | MIMAT0020631 | Mat2a | 232087 | HITS-CLIP | 37986224\|23597149 | Unspecified |
| mmu-miR-466q | MIMAT0020631 | Mau2 | 74549 | HITS-CLIP | 37986224 | Unspecified |
| mmu-miR-466q | MIMAT0020631 | Mbnl1 | 56758 | HITS-CLIP | 23597149 | Unspecified |
| mmu-miR-466q | MIMAT0020631 | Mboat7 | 77582 | HITS-CLIP | 37986224 | Unspecified |
| mmu-miR-466q | MIMAT0020631 | Mccc1 | 72039 | HITS-CLIP | 37986224 | Unspecified |
| mmu-miR-466q | MIMAT0020631 | Med24 | 23989 | HITS-CLIP | 37986224 | Unspecified |
| mmu-miR-466q | MIMAT0020631 | Mef2d | 17261 | HITS-CLIP | 37986224\|23597149 | Unspecified |
| mmu-miR-466q | MIMAT0020631 | Melk | 17279 | HITS-CLIP | 23597149 | Unspecified |
| mmu-miR-466q | MIMAT0020631 | Mfsd2a | 76574 | HITS-CLIP | 23597149 | Unspecified |
| mmu-miR-466q | MIMAT0020631 | Mgea5 | 76055 | HITS-CLIP | 23597149 | Unspecified |
| mmu-miR-466q | MIMAT0020631 | Mob2 | 101513 | HITS-CLIP | 37986224 | Unspecified |
| mmu-miR-466q | MIMAT0020631 | Mpdu1 | 24070 | HITS-CLIP | 23597149 | Unspecified |
| mmu-miR-466q | MIMAT0020631 | Myo18a | 360013 | HITS-CLIP | 37986224 | Unspecified |

| | | | | | | |
|---|---|---|---|---|---|---|
| mmu-miR-466q | MIMAT0020631 | Narf | 67608 | HITS-CLIP | 37986224 | Unspecified |
| mmu-miR-466q | MIMAT0020631 | Ndufs3 | 68349 | HITS-CLIP | 37986224 | Unspecified |
| mmu-miR-466q | MIMAT0020631 | Necap1 | 67602 | HITS-CLIP | 23597149 | Unspecified |
| mmu-miR-466q | MIMAT0020631 | Nfil3 | 18030 | HITS-CLIP | 37986224\|23597149 | Unspecified |
| mmu-miR-466q | MIMAT0020631 | Nhlrc3 | 212114 | HITS-CLIP | 37986224 | Unspecified |
| mmu-miR-466q | MIMAT0020631 | Noa1 | 56412 | HITS-CLIP | 37986224 | Unspecified |
| mmu-miR-466q | MIMAT0020631 | Nsd1 | 18193 | HITS-CLIP | 37986224 | Unspecified |
| mmu-miR-466q | MIMAT0020631 | Nudt7 | 67528 | HITS-CLIP | 37986224 | Unspecified |
| mmu-miR-466q | MIMAT0020631 | Ociad1 | 68095 | HITS-CLIP | 37986224 | Unspecified |
| mmu-miR-466q | MIMAT0020631 | Ocln | 18260 | HITS-CLIP | 37986224 | Unspecified |
| mmu-miR-466q | MIMAT0020631 | Orc3 | 50793 | HITS-CLIP | 23597149 | Unspecified |
| mmu-miR-466q | MIMAT0020631 | Ormdl3 | 66612 | HITS-CLIP | 31433990 | Unspecified |
| mmu-miR-466q | MIMAT0020631 | Osbpl11 | 106326 | HITS-CLIP | 23597149 | Unspecified |
| mmu-miR-466q | MIMAT0020631 | Osbpl2 | 228983 | HITS-CLIP | 37986224 | Unspecified |
| mmu-miR-466q | MIMAT0020631 | Ostc | 66357 | HITS-CLIP | 37986224 | Unspecified |
| mmu-miR-466q | MIMAT0020631 | Pag1 | 94212 | HITS-CLIP | 23597149 | Unspecified |
| mmu-miR-466q | MIMAT0020631 | Papss2 | 23972 | HITS-CLIP | 23597149 | Unspecified |
| mmu-miR-466q | MIMAT0020631 | Parp16 | 214424 | HITS-CLIP | 23597149 | Unspecified |
| mmu-miR-466q | MIMAT0020631 | Pbrm1 | 66923 | HITS-CLIP | 23597149 | Unspecified |
| mmu-miR-466q | MIMAT0020631 | Pcgf5 | 76073 | HITS-CLIP | 37986224 | Unspecified |
| mmu-miR-466q | MIMAT0020631 | Pck2 | 74551 | HITS-CLIP | 23597149 | Unspecified |
| mmu-miR-466q | MIMAT0020631 | Pctp | 18559 | HITS-CLIP | 37986224\|23597149 | Unspecified |
| mmu-miR-466q | MIMAT0020631 | Pcx | 18563 | HITS-CLIP | 37986224\|23597149 | Unspecified |
| mmu-miR-466q | MIMAT0020631 | Pcyt1a | 13026 | HITS-CLIP | 37986224 | Unspecified |
| mmu-miR-466q | MIMAT0020631 | Pcyt2 | 68671 | HITS-CLIP | 37986224 | Unspecified |
| mmu-miR-466q | MIMAT0020631 | Pde3b | 18576 | HITS-CLIP | 23597149 | Unspecified |
| mmu-miR-466q | MIMAT0020631 | Pde4dip | 83679 | HITS-CLIP | 37986224 | Unspecified |
| mmu-miR-466q | MIMAT0020631 | Pdk2 | 18604 | HITS-CLIP | 37986224\|23597149 | Unspecified |
| mmu-miR-466q | MIMAT0020631 | Pds5a | 71521 | HITS-CLIP | 37986224 | Unspecified |
| mmu-miR-466q | MIMAT0020631 | Pex13 | 72129 | HITS-CLIP | 37986224 | Unspecified |
| mmu-miR-466q | MIMAT0020631 | Pias1 | 56469 | HITS-CLIP | 31433990 | Unspecified |
| mmu-miR-466q | MIMAT0020631 | Pigx | 72084 | HITS-CLIP | 23597149 | Unspecified |
| mmu-miR-466q | MIMAT0020631 | Pik3ap1 | 83490 | HITS-CLIP | 23597149 | Unspecified |
| mmu-miR-466q | MIMAT0020631 | Pkn1 | 320795 | HITS-CLIP | 37986224 | Unspecified |
| mmu-miR-466q | MIMAT0020631 | Plg | 18815 | HITS-CLIP | 23597149 | Unspecified |
| mmu-miR-466q | MIMAT0020631 | Plin3 | 66905 | HITS-CLIP | 23597149 | Unspecified |
| mmu-miR-466q | MIMAT0020631 | Plxna2 | 18845 | HITS-CLIP | 26203562 | Unspecified |
| mmu-miR-466q | MIMAT0020631 | Pnkd | 56695 | HITS-CLIP | 37986224 | Unspecified |
| mmu-miR-466q | MIMAT0020631 | Pnpla6 | 50767 | HITS-CLIP | 23597149 | Unspecified |

| | | | | | | | |
|---|---|---|---|---|---|---|---|
| mmu-miR-466q | MIMAT0020631 | Poc1b | 382406 | HITS-CLIP | | 37986224 | Unspecified |
| mmu-miR-466q | MIMAT0020631 | Polr1b | 20017 | HITS-CLIP | 37986224\|23597149 | | Unspecified |
| mmu-miR-466q | MIMAT0020631 | Por | 18984 | HITS-CLIP | | 23597149 | Unspecified |
| mmu-miR-466q | MIMAT0020631 | Ppara | 19013 | HITS-CLIP | | 37986224 | Unspecified |
| mmu-miR-466q | MIMAT0020631 | Ppil4 | 67418 | HITS-CLIP | | 37986224 | Unspecified |
| mmu-miR-466q | MIMAT0020631 | Ppp1r15b | 108954 | HITS-CLIP | | 37986224 | Unspecified |
| mmu-miR-466q | MIMAT0020631 | Ppp1r3b | 244416 | HITS-CLIP | | 23597149 | Unspecified |
| mmu-miR-466q | MIMAT0020631 | Ppp1r7 | 66385 | HITS-CLIP | | 23597149 | Unspecified |
| mmu-miR-466q | MIMAT0020631 | Ppp4r4 | 74521 | HITS-CLIP | | 23597149 | Unspecified |
| mmu-miR-466q | MIMAT0020631 | Prim1 | 19075 | HITS-CLIP | | 23597149 | Unspecified |
| mmu-miR-466q | MIMAT0020631 | Prkar1a | 19084 | HITS-CLIP | | 37986224 | Unspecified |
| mmu-miR-466q | MIMAT0020631 | Proc | 19123 | HITS-CLIP | | 37986224 | Unspecified |
| mmu-miR-466q | MIMAT0020631 | Prodh2 | 56189 | HITS-CLIP | | 37986224 | Unspecified |
| mmu-miR-466q | MIMAT0020631 | Proz | 66901 | HITS-CLIP | | 37986224 | Unspecified |
| mmu-miR-466q | MIMAT0020631 | Prpf18 | 67229 | HITS-CLIP | | 23597149 | Unspecified |
| mmu-miR-466q | MIMAT0020631 | Psap | 19156 | HITS-CLIP | | 23597149 | Unspecified |
| mmu-miR-466q | MIMAT0020631 | Ptdss2 | 27388 | HITS-CLIP | 37986224\|23597149 | | Unspecified |
| mmu-miR-466q | MIMAT0020631 | Ptpn18 | 19253 | HITS-CLIP | | 31433990 | Unspecified |
| mmu-miR-466q | MIMAT0020631 | Ptprb | 19263 | HITS-CLIP | | 23597149 | Unspecified |
| mmu-miR-466q | MIMAT0020631 | Ptprd | 19266 | HITS-CLIP | | 23597149 | Unspecified |
| mmu-miR-466q | MIMAT0020631 | Purb | 19291 | HITS-CLIP | | 37986224 | Unspecified |
| mmu-miR-466q | MIMAT0020631 | Pus10 | 74467 | HITS-CLIP | | 23597149 | Unspecified |
| mmu-miR-466q | MIMAT0020631 | Pxn | 19303 | HITS-CLIP | 37986224\|23597149 | | Unspecified |
| mmu-miR-466q | MIMAT0020631 | Pxylp1 | 235534 | HITS-CLIP | | 23597149 | Unspecified |
| mmu-miR-466q | MIMAT0020631 | R3hdm2 | 71750 | HITS-CLIP | | 23597149 | Unspecified |
| mmu-miR-466q | MIMAT0020631 | Rab1a | 5861 | HITS-CLIP | | 23597149 | Unspecified |
| mmu-miR-466q | MIMAT0020631 | Rai14 | 75646 | HITS-CLIP | 37986224\|23597149 | | Unspecified |
| mmu-miR-466q | MIMAT0020631 | Ralgapb | 228850 | HITS-CLIP | 37986224\|23597149 | | Unspecified |
| mmu-miR-466q | MIMAT0020631 | Rap2a | 76108 | HITS-CLIP | | 23597149 | Unspecified |
| mmu-miR-466q | MIMAT0020631 | Rasgef1b | 320292 | HITS-CLIP | | 37986224 | Unspecified |
| mmu-miR-466q | MIMAT0020631 | Rassf3 | 192678 | HITS-CLIP | | 37986224 | Unspecified |
| mmu-miR-466q | MIMAT0020631 | Rbm38 | 56190 | HITS-CLIP | | 23597149 | Unspecified |
| mmu-miR-466q | MIMAT0020631 | Rbm47 | 245945 | HITS-CLIP | | 23597149 | Unspecified |
| mmu-miR-466q | MIMAT0020631 | Rcbtb1 | 71330 | HITS-CLIP | | 37986224 | Unspecified |
| mmu-miR-466q | MIMAT0020631 | Rce1 | 19671 | HITS-CLIP | | 37986224 | Unspecified |
| mmu-miR-466q | MIMAT0020631 | Rcl1 | 59028 | HITS-CLIP | | 37986224 | Unspecified |
| mmu-miR-466q | MIMAT0020631 | Rdh7 | 54150 | HITS-CLIP | | 37986224 | Unspecified |
| mmu-miR-466q | MIMAT0020631 | Reps2 | 194590 | HITS-CLIP | | 23597149 | Unspecified |
| mmu-miR-466q | MIMAT0020631 | Retreg3 | 162427 | HITS-CLIP | | 23597149 | Unspecified |

| | | | | | | | |
|---|---|---|---|---|---|---|---|
| mmu-miR-466q | MIMAT0020631 | Retsat | 67442 | HITS-CLIP | | 37986224 | Unspecified |
| mmu-miR-466q | MIMAT0020631 | Rhot2 | 214952 | HITS-CLIP | | 23597149 | Unspecified |
| mmu-miR-466q | MIMAT0020631 | Rnf111 | 93836 | HITS-CLIP | | 23597149 | Unspecified |
| mmu-miR-466q | MIMAT0020631 | Rnf187 | 108660 | HITS-CLIP | | 37986224 | Unspecified |
| mmu-miR-466q | MIMAT0020631 | Rnft1 | 76892 | HITS-CLIP | | 23597149 | Unspecified |
| mmu-miR-466q | MIMAT0020631 | Sars | 20226 | HITS-CLIP | | 23597149 | Unspecified |
| mmu-miR-466q | MIMAT0020631 | Sc5d | 235293 | HITS-CLIP | | 37986224 | Unspecified |
| mmu-miR-466q | MIMAT0020631 | Scamp1 | 107767 | HITS-CLIP | | 23597149 | Unspecified |
| mmu-miR-466q | MIMAT0020631 | Scamp3 | 24045 | HITS-CLIP | | 37986224 | Unspecified |
| mmu-miR-466q | MIMAT0020631 | Sec14l4 | 103655 | HITS-CLIP | | 37986224 | Unspecified |
| mmu-miR-466q | MIMAT0020631 | Sec16a | 227648 | HITS-CLIP | | 23597149 | Unspecified |
| mmu-miR-466q | MIMAT0020631 | Serpine2 | 20720 | HITS-CLIP | | 37986224 | Unspecified |
| mmu-miR-466q | MIMAT0020631 | Sesn2 | 230784 | HITS-CLIP | | 37986224 | Unspecified |
| mmu-miR-466q | MIMAT0020631 | Set | 56086 | HITS-CLIP | | 37986224 | Unspecified |
| mmu-miR-466q | MIMAT0020631 | Sf3b1 | 81898 | HITS-CLIP | | 37986224 | Unspecified |
| mmu-miR-466q | MIMAT0020631 | Sgk1 | 20393 | HITS-CLIP | | 23597149 | Unspecified |
| mmu-miR-466q | MIMAT0020631 | Sgpp1 | 81535 | HITS-CLIP | | 23597149 | Unspecified |
| mmu-miR-466q | MIMAT0020631 | Sgsm3 | 105835 | HITS-CLIP | | 23597149 | Unspecified |
| mmu-miR-466q | MIMAT0020631 | Sigmar1 | 18391 | HITS-CLIP | | 23597149 | Unspecified |
| mmu-miR-466q | MIMAT0020631 | Slc16a2 | 20502 | HITS-CLIP | | 37986224 | Unspecified |
| mmu-miR-466q | MIMAT0020631 | Slc1a2 | 20511 | HITS-CLIP | | 23597149 | Unspecified |
| mmu-miR-466q | MIMAT0020631 | Slc22a1 | 20517 | HITS-CLIP | | 23597149 | Unspecified |
| mmu-miR-466q | MIMAT0020631 | Slc22a23 | 73102 | HITS-CLIP | | 23597149 | Unspecified |
| mmu-miR-466q | MIMAT0020631 | Slc25a13 | 50799 | HITS-CLIP | | 23597149 | Unspecified |
| mmu-miR-466q | MIMAT0020631 | Slc25a47 | 104910 | HITS-CLIP | | 23597149 | Unspecified |
| mmu-miR-466q | MIMAT0020631 | Slc26a1 | 231583 | HITS-CLIP | 37986224\|23597149 | | Unspecified |
| mmu-miR-466q | MIMAT0020631 | Slc29a1 | 63959 | HITS-CLIP | | 23597149 | Unspecified |
| mmu-miR-466q | MIMAT0020631 | Slc37a3 | 72144 | HITS-CLIP | | 31433990 | Unspecified |
| mmu-miR-466q | MIMAT0020631 | Slc38a3 | 76257 | HITS-CLIP | | 23597149 | Unspecified |
| mmu-miR-466q | MIMAT0020631 | Slc39a13 | 68427 | HITS-CLIP | | 23597149 | Unspecified |
| mmu-miR-466q | MIMAT0020631 | Slc40a1 | 53945 | HITS-CLIP | 37986224\|23597149 | | Unspecified |
| mmu-miR-466q | MIMAT0020631 | Slc6a6 | 21366 | HITS-CLIP | 37986224\|23597149 | | Unspecified |
| mmu-miR-466q | MIMAT0020631 | Slc7a2 | 11988 | HITS-CLIP | 23597149\|28735896 | | Unspecified |
| mmu-miR-466q | MIMAT0020631 | Slco1a1 | 28248 | HITS-CLIP | | 37986224 | Unspecified |
| mmu-miR-466q | MIMAT0020631 | Slco1a4 | 28250 | HITS-CLIP | | 37986224 | Unspecified |
| mmu-miR-466q | MIMAT0020631 | Slco2b1 | 101488 | HITS-CLIP | | 37986224 | Unspecified |
| mmu-miR-466q | MIMAT0020631 | Smad5 | 17129 | HITS-CLIP | | 23597149 | Unspecified |
| mmu-miR-466q | MIMAT0020631 | Smg1 | 233789 | HITS-CLIP | | 37986224 | Unspecified |
| mmu-miR-466q | MIMAT0020631 | Smoc1 | 64075 | HITS-CLIP | | 37986224 | Unspecified |

| | | | | | | | |
|---|---|---|---|---|---|---|---|
| mmu-miR-466q | MIMAT0020631 | Snrk | 20623 | HITS-CLIP | | 37986224 | Unspecified |
| mmu-miR-466q | MIMAT0020631 | Snx5 | 69178 | HITS-CLIP | | 37986224 | Unspecified |
| mmu-miR-466q | MIMAT0020631 | Sod2 | 20656 | HITS-CLIP | | 37986224 | Unspecified |
| mmu-miR-466q | MIMAT0020631 | Sox9 | 20682 | HITS-CLIP | | 37986224 | Unspecified |
| mmu-miR-466q | MIMAT0020631 | Sppl2a | 66552 | HITS-CLIP | | 37986224 | Unspecified |
| mmu-miR-466q | MIMAT0020631 | Spred2 | 114716 | HITS-CLIP | | 37986224 | Unspecified |
| mmu-miR-466q | MIMAT0020631 | Sptbn1 | 20742 | HITS-CLIP | | 26203562 | Unspecified |
| mmu-miR-466q | MIMAT0020631 | Sptlc2 | 20773 | HITS-CLIP | 37986224\|23597149 | | Unspecified |
| mmu-miR-466q | MIMAT0020631 | Ssh2 | 237860 | HITS-CLIP | | 23597149 | Unspecified |
| mmu-miR-466q | MIMAT0020631 | Stbd1 | 52331 | HITS-CLIP | | 37986224 | Unspecified |
| mmu-miR-466q | MIMAT0020631 | Steap4 | 117167 | HITS-CLIP | | 37986224 | Unspecified |
| mmu-miR-466q | MIMAT0020631 | Stxbp3 | 20912 | HITS-CLIP | | 23597149 | Unspecified |
| mmu-miR-466q | MIMAT0020631 | Suox | 211389 | HITS-CLIP | | 37986224 | Unspecified |
| mmu-miR-466q | MIMAT0020631 | Supv3l1 | 338359 | HITS-CLIP | | 37986224 | Unspecified |
| mmu-miR-466q | MIMAT0020631 | Sypl | 19027 | HITS-CLIP | | 37986224 | Unspecified |
| mmu-miR-466q | MIMAT0020631 | Szt2 | 230676 | HITS-CLIP | | 23597149 | Unspecified |
| mmu-miR-466q | MIMAT0020631 | Tax1bp1 | 52440 | HITS-CLIP | | 37986224 | Unspecified |
| mmu-miR-466q | MIMAT0020631 | Tbrg4 | 21379 | HITS-CLIP | | 23597149 | Unspecified |
| mmu-miR-466q | MIMAT0020631 | Tbx3 | 21386 | HITS-CLIP | | 37986224 | Unspecified |
| mmu-miR-466q | MIMAT0020631 | Tcaf1 | 9747 | HITS-CLIP | | 23597149 | Unspecified |
| mmu-miR-466q | MIMAT0020631 | Tctn1 | 654470 | HITS-CLIP | | 23597149 | Unspecified |
| mmu-miR-466q | MIMAT0020631 | Tecr | 106529 | HITS-CLIP | | 37986224 | Unspecified |
| mmu-miR-466q | MIMAT0020631 | Tef | 21685 | HITS-CLIP | | 37986224 | Unspecified |
| mmu-miR-466q | MIMAT0020631 | Tes | 21753 | HITS-CLIP | | 31433990 | Unspecified |
| mmu-miR-466q | MIMAT0020631 | Tgfbr1 | 21812 | HITS-CLIP | | 37986224 | Unspecified |
| mmu-miR-466q | MIMAT0020631 | Thrsp | 21835 | HITS-CLIP | | 23597149 | Unspecified |
| mmu-miR-466q | MIMAT0020631 | Timp3 | 21859 | HITS-CLIP | | 37986224 | Unspecified |
| mmu-miR-466q | MIMAT0020631 | Tm4sf1 | 17112 | HITS-CLIP | | 37986224 | Unspecified |
| mmu-miR-466q | MIMAT0020631 | Tm6sf2 | 107770 | HITS-CLIP | | 37986224 | Unspecified |
| mmu-miR-466q | MIMAT0020631 | Tmbim6 | 110213 | HITS-CLIP | | 37986224 | Unspecified |
| mmu-miR-466q | MIMAT0020631 | Tmed7 | 66676 | HITS-CLIP | | 37986224 | Unspecified |
| mmu-miR-466q | MIMAT0020631 | Tmem170b | 621976 | HITS-CLIP | | 23597149 | Unspecified |
| mmu-miR-466q | MIMAT0020631 | Tmem176a | 66058 | HITS-CLIP | | 37986224 | Unspecified |
| mmu-miR-466q | MIMAT0020631 | Tmem37 | 170706 | HITS-CLIP | 37986224\|23597149 | | Unspecified |
| mmu-miR-466q | MIMAT0020631 | Tmem87b | 72477 | HITS-CLIP | | 37986224 | Unspecified |
| mmu-miR-466q | MIMAT0020631 | Tmx4 | 52837 | HITS-CLIP | | 23597149 | Unspecified |
| mmu-miR-466q | MIMAT0020631 | Tnfaip1 | 21927 | HITS-CLIP | | 23597149 | Unspecified |
| mmu-miR-466q | MIMAT0020631 | Tnks2 | 74493 | HITS-CLIP | 31433990\|26203562 | | Unspecified |
| mmu-miR-466q | MIMAT0020631 | Tnrc6a | 233833 | HITS-CLIP | | 23597149 | Unspecified |

| | | | | | | |
|---|---|---|---|---|---|---|
| mmu-miR-466q | MIMAT0020631 | Tnrc6b | 213988 | HITS-CLIP | 23597149 | Unspecified |
| mmu-miR-466q | MIMAT0020631 | Tnrc6c | 217351 | HITS-CLIP | 37986224 | Unspecified |
| mmu-miR-466q | MIMAT0020631 | Tns2 | 209039 | HITS-CLIP | 23597149 | Unspecified |
| mmu-miR-466q | MIMAT0020631 | Tomm6 | 66119 | HITS-CLIP | 37986224\|23597149 | Unspecified |
| mmu-miR-466q | MIMAT0020631 | Top2a | 21973 | HITS-CLIP | 31433990 | Unspecified |
| mmu-miR-466q | MIMAT0020631 | Trim23 | 81003 | HITS-CLIP | 23597149 | Unspecified |
| mmu-miR-466q | MIMAT0020631 | Trim68 | 101700 | HITS-CLIP | 23597149 | Unspecified |
| mmu-miR-466q | MIMAT0020631 | Trip12 | 14897 | HITS-CLIP | 37986224 | Unspecified |
| mmu-miR-466q | MIMAT0020631 | Trp53bp2 | 209456 | HITS-CLIP | 23597149 | Unspecified |
| mmu-miR-466q | MIMAT0020631 | Trp53inp1 | 60599 | HITS-CLIP | 23597149 | Unspecified |
| mmu-miR-466q | MIMAT0020631 | Trp53inp2 | 68728 | HITS-CLIP | 23597149 | Unspecified |
| mmu-miR-466q | MIMAT0020631 | Tsc22d3 | 14605 | HITS-CLIP | 37986224 | Unspecified |
| mmu-miR-466q | MIMAT0020631 | Ttc36 | 192653 | HITS-CLIP | 37986224 | Unspecified |
| mmu-miR-466q | MIMAT0020631 | Ttc38 | 239570 | HITS-CLIP | 37986224 | Unspecified |
| mmu-miR-466q | MIMAT0020631 | Tusc2 | 80385 | HITS-CLIP | 31433990 | Unspecified |
| mmu-miR-466q | MIMAT0020631 | Tut4 | 23318 | HITS-CLIP | 23597149 | Unspecified |
| mmu-miR-466q | MIMAT0020631 | Txnrd3 | 232223 | HITS-CLIP | 23597149 | Unspecified |
| mmu-miR-466q | MIMAT0020631 | Uap1l1 | 227620 | HITS-CLIP | 37986224 | Unspecified |
| mmu-miR-466q | MIMAT0020631 | Uba1 | 22201 | HITS-CLIP | 23597149 | Unspecified |
| mmu-miR-466q | MIMAT0020631 | Ubr3 | 68795 | HITS-CLIP | 23597149 | Unspecified |
| mmu-miR-466q | MIMAT0020631 | Ubr4 | 69116 | HITS-CLIP | 37986224\|23597149 | Unspecified |
| mmu-miR-466q | MIMAT0020631 | Ubr5 | 70790 | HITS-CLIP | 37986224\|23597149 | Unspecified |
| mmu-miR-466q | MIMAT0020631 | Unc119b | 106840 | HITS-CLIP | 37986224 | Unspecified |
| mmu-miR-466q | MIMAT0020631 | Upp1 | 22271 | HITS-CLIP | 37986224 | Unspecified |
| mmu-miR-466q | MIMAT0020631 | Utrn | 22288 | HITS-CLIP | 23597149 | Unspecified |
| mmu-miR-466q | MIMAT0020631 | Vasn | 246154 | HITS-CLIP | 23597149 | Unspecified |
| mmu-miR-466q | MIMAT0020631 | Vps35 | 65114 | HITS-CLIP | 23597149 | Unspecified |
| mmu-miR-466q | MIMAT0020631 | Vps37b | 330192 | HITS-CLIP | 37986224 | Unspecified |
| mmu-miR-466q | MIMAT0020631 | Vps4b | 20479 | HITS-CLIP | 37986224\|23597149 | Unspecified |
| mmu-miR-466q | MIMAT0020631 | Wasl | 73178 | HITS-CLIP | 23597149 | Unspecified |
| mmu-miR-466q | MIMAT0020631 | Wbp1l | 226178 | HITS-CLIP | 37986224\|23597149\|28735896 | Unspecified |
| mmu-miR-466q | MIMAT0020631 | Wdr45b | 66840 | HITS-CLIP | 23597149 | Unspecified |
| mmu-miR-466q | MIMAT0020631 | Wdr77 | 70465 | HITS-CLIP | 37986224\|23597149 | Unspecified |
| mmu-miR-466q | MIMAT0020631 | Wrnip1 | 78903 | HITS-CLIP | 37986224 | Unspecified |
| mmu-miR-466q | MIMAT0020631 | Wwc1 | 211652 | HITS-CLIP | 37986224 | Unspecified |
| mmu-miR-466q | MIMAT0020631 | Xpo7 | 65246 | HITS-CLIP | 23597149 | Unspecified |
| mmu-miR-466q | MIMAT0020631 | Ywhag | 22628 | HITS-CLIP | 37986224 | Unspecified |
| mmu-miR-466q | MIMAT0020631 | Zbtb20 | 56490 | HITS-CLIP | 37986224\|23597149 | Unspecified |
| mmu-miR-466q | MIMAT0020631 | Zbtb4 | 75580 | HITS-CLIP | 23597149 | Unspecified |

| | | | | | | |
|---|---|---|---|---|---|---|
| mmu-miR-466q | MIMAT0020631 | Zbtb5 | 230119 | HITS-CLIP | 23597149 | Unspecified |
| mmu-miR-466q | MIMAT0020631 | Zbtb7b | 22724 | HITS-CLIP | 25083871\|23597149 | Unspecified |
| mmu-miR-466q | MIMAT0020631 | Zc3h12c | 244871 | HITS-CLIP | 23597149 | Unspecified |
| mmu-miR-466q | MIMAT0020631 | Zcchc2 | 227449 | HITS-CLIP | 37986224 | Unspecified |
| mmu-miR-466q | MIMAT0020631 | Zcchc24 | 71918 | HITS-CLIP | 37986224\|23597149 | Unspecified |
| mmu-miR-466q | MIMAT0020631 | Zdhhc16 | 74168 | HITS-CLIP | 37986224 | Unspecified |
| mmu-miR-466q | MIMAT0020631 | Zer1 | 227693 | HITS-CLIP | 37986224 | Unspecified |
| mmu-miR-466q | MIMAT0020631 | Zfp592 | 233410 | HITS-CLIP | 37986224\|23597149 | Unspecified |
| mmu-miR-466q | MIMAT0020631 | Zfp655 | 72611 | HITS-CLIP | 37986224 | Unspecified |
| mmu-miR-466q | MIMAT0020631 | Zhx3 | 320799 | HITS-CLIP | 37986224 | Unspecified |
| mmu-miR-466q | MIMAT0020631 | Zmiz1 | 328365 | HITS-CLIP | 23597149 | Unspecified |
| mmu-miR-466q | MIMAT0020631 | Znrf2 | 387524 | HITS-CLIP | 37986224 | Unspecified |
| mmu-miR-466q | MIMAT0020631 | Zscan29 | 99334 | HITS-CLIP | 23597149 | Unspecified |
| mmu-miR-466q | MIMAT0020631 | Zyg11b | 414872 | HITS-CLIP | 37986224 | Unspecified |
| mmu-miR-466q | MIMAT0020631 | Zyx | 22793 | HITS-CLIP | 23597149 | Unspecified |
| mmu-miR-344d-3p | MIMAT0014808 | Krt16 | 16666 | HITS-CLIP | 26203562 | Unspecified |
| mmu-miR-344d-3p | MIMAT0014808 | Taf5l | 102162 | HITS-CLIP | 26203562 | Unspecified |
| mmu-miR-3108-5p | MIMAT0014947 | Rspry1 | 67610 | HITS-CLIP | 31433990 | Unspecified |
| mmu-miR-1187 | MIMAT0005837 | 2510039O18Rik | 77034 | HITS-CLIP | 37986224 | Unspecified |
| mmu-miR-1187 | MIMAT0005837 | Aars | 234734 | HITS-CLIP | 37986224 | Unspecified |
| mmu-miR-1187 | MIMAT0005837 | Abca1 | 11303 | HITS-CLIP | 37986224 | Unspecified |
| mmu-miR-1187 | MIMAT0005837 | Abca8a | 217258 | HITS-CLIP | 37986224 | Unspecified |
| mmu-miR-1187 | MIMAT0005837 | Ahnak | 66395 | HITS-CLIP | 37986224 | Unspecified |
| mmu-miR-1187 | MIMAT0005837 | Aldh1a1 | 11668 | HITS-CLIP | 37986224 | Unspecified |
| mmu-miR-1187 | MIMAT0005837 | Apoa4 | 11808 | HITS-CLIP | 37986224 | Unspecified |
| mmu-miR-1187 | MIMAT0005837 | Cat | 12359 | HITS-CLIP | 37986224 | Unspecified |
| mmu-miR-1187 | MIMAT0005837 | Cd84 | 12523 | HITS-CLIP | 37986224 | Unspecified |
| mmu-miR-1187 | MIMAT0005837 | Cebpb | 12608 | HITS-CLIP | 37986224 | Unspecified |
| mmu-miR-1187 | MIMAT0005837 | Chd7 | 320790 | HITS-CLIP | 37986224 | Unspecified |
| mmu-miR-1187 | MIMAT0005837 | Clpx | 270166 | HITS-CLIP | 37986224 | Unspecified |
| mmu-miR-1187 | MIMAT0005837 | Cnot6l | 231464 | HITS-CLIP | 37986224 | Unspecified |
| mmu-miR-1187 | MIMAT0005837 | Coa4 | 68185 | HITS-CLIP | 37986224 | Unspecified |
| mmu-miR-1187 | MIMAT0005837 | Crp | 12944 | HITS-CLIP | 37986224 | Unspecified |
| mmu-miR-1187 | MIMAT0005837 | Ctsd | 13033 | HITS-CLIP | 37986224 | Unspecified |
| mmu-miR-1187 | MIMAT0005837 | Cyb5r3 | 109754 | HITS-CLIP | 37986224 | Unspecified |
| mmu-miR-1187 | MIMAT0005837 | Cyp39a1 | 56050 | HITS-CLIP | 37986224 | Unspecified |
| mmu-miR-1187 | MIMAT0005837 | Ddc | 13195 | HITS-CLIP | 37986224 | Unspecified |
| mmu-miR-1187 | MIMAT0005837 | Dgcr2 | 13356 | HITS-CLIP | 37986224 | Unspecified |
| mmu-miR-1187 | MIMAT0005837 | Epc2 | 227867 | HITS-CLIP | 37986224 | Unspecified |

| | | | | | | |
|---|---|---|---|---|---|---|
| mmu-miR-1187 | MIMAT0005837 | Fbxw8 | 231672 | HITS-CLIP | 37986224 | Unspecified |
| mmu-miR-1187 | MIMAT0005837 | Fndc3b | 72007 | HITS-CLIP | 23597149 | Unspecified |
| mmu-miR-1187 | MIMAT0005837 | Gatad2b | 229542 | HITS-CLIP | 23597149 | Unspecified |
| mmu-miR-1187 | MIMAT0005837 | Gm4799 | 216185 | HITS-CLIP | 37986224 | Unspecified |
| mmu-miR-1187 | MIMAT0005837 | Gnptab | 432486 | HITS-CLIP | 37986224 | Unspecified |
| mmu-miR-1187 | MIMAT0005837 | Gpr146 | 80290 | HITS-CLIP | 37986224 | Unspecified |
| mmu-miR-1187 | MIMAT0005837 | Gsr | 14782 | HITS-CLIP | 37986224 | Unspecified |
| mmu-miR-1187 | MIMAT0005837 | Gxylt1 | 223827 | HITS-CLIP | 37986224 | Unspecified |
| mmu-miR-1187 | MIMAT0005837 | Hectd1 | 207304 | HITS-CLIP | 37986224 | Unspecified |
| mmu-miR-1187 | MIMAT0005837 | Hsd11b1 | 15483 | HITS-CLIP | 37986224 | Unspecified |
| mmu-miR-1187 | MIMAT0005837 | Igf1 | 16000 | HITS-CLIP | 37986224 | Unspecified |
| mmu-miR-1187 | MIMAT0005837 | Il6st | 16195 | HITS-CLIP | 37986224 | Unspecified |
| mmu-miR-1187 | MIMAT0005837 | Iqcc | 230767 | HITS-CLIP | 37986224 | Unspecified |
| mmu-miR-1187 | MIMAT0005837 | Klf6 | 23849 | HITS-CLIP | 37986224 | Unspecified |
| mmu-miR-1187 | MIMAT0005837 | Lamp2 | 16784 | HITS-CLIP | 37986224 | Unspecified |
| mmu-miR-1187 | MIMAT0005837 | Lats1 | 16798 | HITS-CLIP | 37986224 | Unspecified |
| mmu-miR-1187 | MIMAT0005837 | Lcn2 | 16819 | HITS-CLIP | 37986224 | Unspecified |
| mmu-miR-1187 | MIMAT0005837 | Lcp1 | 18826 | HITS-CLIP | 31433990 | Unspecified |
| mmu-miR-1187 | MIMAT0005837 | Lonp1 | 74142 | HITS-CLIP | 37986224 | Unspecified |
| mmu-miR-1187 | MIMAT0005837 | Lpcat3 | 14792 | HITS-CLIP | 37986224 | Unspecified |
| mmu-miR-1187 | MIMAT0005837 | Lpin2 | 64898 | HITS-CLIP | 37986224 | Unspecified |
| mmu-miR-1187 | MIMAT0005837 | Lrp1 | 16971 | HITS-CLIP | 37986224 | Unspecified |
| mmu-miR-1187 | MIMAT0005837 | Maf | 17132 | HITS-CLIP | 23597149 | Unspecified |
| mmu-miR-1187 | MIMAT0005837 | Mcat | 223722 | HITS-CLIP | 37986224 | Unspecified |
| mmu-miR-1187 | MIMAT0005837 | Mettl7b | 71664 | HITS-CLIP | 37986224 | Unspecified |
| mmu-miR-1187 | MIMAT0005837 | Mob1b | 68473 | HITS-CLIP | 37986224 | Unspecified |
| mmu-miR-1187 | MIMAT0005837 | Morc2a | 74522 | HITS-CLIP | 37986224 | Unspecified |
| mmu-miR-1187 | MIMAT0005837 | Mtmr3 | 74302 | HITS-CLIP | 37986224 | Unspecified |
| mmu-miR-1187 | MIMAT0005837 | N4bp2l2 | 381695 | HITS-CLIP | 37986224 | Unspecified |
| mmu-miR-1187 | MIMAT0005837 | Ndufs1 | 227197 | HITS-CLIP | 37986224 | Unspecified |
| mmu-miR-1187 | MIMAT0005837 | Ndufs2 | 226646 | HITS-CLIP | 37986224 | Unspecified |
| mmu-miR-1187 | MIMAT0005837 | Nuak2 | 74137 | HITS-CLIP | 37986224 | Unspecified |
| mmu-miR-1187 | MIMAT0005837 | Nupl1 | 71844 | HITS-CLIP | 37986224 | Unspecified |
| mmu-miR-1187 | MIMAT0005837 | Oxsr1 | 108737 | HITS-CLIP | 37986224 | Unspecified |
| mmu-miR-1187 | MIMAT0005837 | P4hb | 18453 | HITS-CLIP | 37986224 | Unspecified |
| mmu-miR-1187 | MIMAT0005837 | Pafah1b1 | 18472 | HITS-CLIP | 37986224 | Unspecified |
| mmu-miR-1187 | MIMAT0005837 | Parp14 | 547253 | HITS-CLIP | 37986224 | Unspecified |
| mmu-miR-1187 | MIMAT0005837 | Pcna | 18538 | HITS-CLIP | 37986224 | Unspecified |
| mmu-miR-1187 | MIMAT0005837 | Pgm1 | 66681 | HITS-CLIP | 37986224 | Unspecified |

| | | | | | | | |
|---|---|---|---|---|---|---|---|
| mmu-miR-1187 | MIMAT0005837 | Pgrmc1 | 53328 | HITS-CLIP | | 23597149 | Unspecified |
| mmu-miR-1187 | MIMAT0005837 | Pik3ca | 18706 | HITS-CLIP | | 37986224 | Unspecified |
| mmu-miR-1187 | MIMAT0005837 | Pikfyve | 18711 | HITS-CLIP | | 37986224 | Unspecified |
| mmu-miR-1187 | MIMAT0005837 | Ppp6r3 | 52036 | HITS-CLIP | | 37986224 | Unspecified |
| mmu-miR-1187 | MIMAT0005837 | Psme4 | 103554 | HITS-CLIP | | 37986224 | Unspecified |
| mmu-miR-1187 | MIMAT0005837 | Rab7 | 19349 | HITS-CLIP//Western blotting//RNA-seq | 23597149\|34014398 | | Unspecified |
| mmu-miR-1187 | MIMAT0005837 | Rere | 68703 | HITS-CLIP | | 37986224 | Unspecified |
| mmu-miR-1187 | MIMAT0005837 | Rnase4 | 58809 | HITS-CLIP | | 37986224 | Unspecified |
| mmu-miR-1187 | MIMAT0005837 | Rnf44 | 105239 | HITS-CLIP | | 23597149 | Unspecified |
| mmu-miR-1187 | MIMAT0005837 | Sdr42e1 | 74032 | HITS-CLIP | | 37986224 | Unspecified |
| mmu-miR-1187 | MIMAT0005837 | Serpina1c | 20702 | HITS-CLIP | | 37986224 | Unspecified |
| mmu-miR-1187 | MIMAT0005837 | Sigmar1 | 18391 | HITS-CLIP | | 37986224 | Unspecified |
| mmu-miR-1187 | MIMAT0005837 | Slc16a2 | 20502 | HITS-CLIP | | 37986224 | Unspecified |
| mmu-miR-1187 | MIMAT0005837 | Slc25a16 | 73132 | HITS-CLIP | | 37986224 | Unspecified |
| mmu-miR-1187 | MIMAT0005837 | Slc31a1 | 20529 | HITS-CLIP | | 37986224 | Unspecified |
| mmu-miR-1187 | MIMAT0005837 | Slc38a2 | 67760 | HITS-CLIP | | 37986224 | Unspecified |
| mmu-miR-1187 | MIMAT0005837 | Slc43a3 | 58207 | HITS-CLIP | | 37986224 | Unspecified |
| mmu-miR-1187 | MIMAT0005837 | Smarca2 | 67155 | HITS-CLIP | 25083871\|23597149 | | Unspecified |
| mmu-miR-1187 | MIMAT0005837 | Sord | 20322 | HITS-CLIP | | 37986224 | Unspecified |
| mmu-miR-1187 | MIMAT0005837 | Sppl2b | 73218 | HITS-CLIP | | 37986224 | Unspecified |
| mmu-miR-1187 | MIMAT0005837 | Sypl | 19027 | HITS-CLIP | | 37986224 | Unspecified |
| mmu-miR-1187 | MIMAT0005837 | Tmbim6 | 110213 | HITS-CLIP | | 37986224 | Unspecified |
| mmu-miR-1187 | MIMAT0005837 | Trib1 | 211770 | HITS-CLIP | | 37986224 | Unspecified |
| mmu-miR-1187 | MIMAT0005837 | Trpm7 | 58800 | HITS-CLIP | | 37986224 | Unspecified |
| mmu-miR-1187 | MIMAT0005837 | Usp47 | 74996 | HITS-CLIP | | 37986224 | Unspecified |
| mmu-miR-1187 | MIMAT0005837 | Wbp1l | 226178 | HITS-CLIP | | 37986224 | Unspecified |
| mmu-miR-1187 | MIMAT0005837 | Zbtb33 | 56805 | HITS-CLIP | | 37986224 | Unspecified |
| mmu-miR-1187 | MIMAT0005837 | Zfp36l1 | 12192 | HITS-CLIP | | 37986224 | Unspecified |
| mmu-miR-1187 | MIMAT0005837 | Zmiz1 | 328365 | HITS-CLIP | 37986224\|31433990 | | Unspecified |
| mmu-miR-1187 | MIMAT0005837 | Zswim8 | 268721 | HITS-CLIP | | 37986224 | Unspecified |
| mmu-miR-326-3p | MIMAT0000559 | Arrb2 | 216869 | HITS-CLIP | | 29167373 | Unspecified |
| mmu-miR-326-3p | MIMAT0000559 | Atp6v0a1 | 11975 | HITS-CLIP | | 29167373 | Unspecified |
| mmu-miR-326-3p | MIMAT0000559 | Ccndbp1 | 17151 | HITS-CLIP | | 29167373 | Unspecified |
| mmu-miR-326-3p | MIMAT0000559 | Celf2 | 14007 | HITS-CLIP | | 29167373 | Unspecified |
| mmu-miR-326-3p | MIMAT0000559 | Dock10 | 210293 | HITS-CLIP | | 29167373 | Unspecified |
| mmu-miR-326-3p | MIMAT0000559 | Itgb1 | 16412 | HITS-CLIP | | 29167373 | Unspecified |
| mmu-miR-326-3p | MIMAT0000559 | Mcm6 | 17219 | HITS-CLIP | | 29167373 | Unspecified |
| mmu-miR-326-3p | MIMAT0000559 | Mgat2 | 217664 | HITS-CLIP | | 29167373 | Unspecified |
| mmu-miR-326-3p | MIMAT0000559 | Mon2 | 67074 | HITS-CLIP | | 29167373 | Unspecified |

| | | | | | | | |
|---|---|---|---|---|---|---|---|
| mmu-miR-326-3p | MIMAT0000559 | Qrich1 | 69232 | HITS-CLIP | | 29167373 | Unspecified |
| mmu-miR-326-3p | MIMAT0000559 | Rere | 68703 | HITS-CLIP | | 29167373 | Unspecified |
| mmu-miR-326-3p | MIMAT0000559 | Rptor | 74370 | HITS-CLIP | | 29167373 | Unspecified |
| mmu-miR-326-3p | MIMAT0000559 | Ssh2 | 237860 | HITS-CLIP | | 29167373 | Unspecified |
| mmu-miR-326-3p | MIMAT0000559 | Tlr4 | 21898 | HITS-CLIP | | 29167373 | Unspecified |
| mmu-miR-326-3p | MIMAT0000559 | Tmem38b | 52076 | qCLASH | | 31393953 | Unspecified |
| mmu-miR-875-3p | MIMAT0004938 | Klf5 | 12224 | HITS-CLIP | | 26203562 | Unspecified |
| mmu-miR-875-3p | MIMAT0004938 | Lrrcc1 | 71710 | HITS-CLIP | | 31433990 | Unspecified |
| mmu-miR-875-3p | MIMAT0004938 | Yme1l1 | 27377 | HITS-CLIP | | 31433990 | Unspecified |
| mmu-miR-139-5p | MIMAT0000656 | 0610030E20Rik | 68364 | HITS-CLIP | | 31433990 | Unspecified |
| mmu-miR-139-5p | MIMAT0000656 | 1700017B05Rik | 74211 | HITS-CLIP | | 29167373 | Unspecified |
| mmu-miR-139-5p | MIMAT0000656 | 1700123O20Rik | 58248 | HITS-CLIP | | 29167373 | Unspecified |
| mmu-miR-139-5p | MIMAT0000656 | 1810013L24Rik | 69053 | HITS-CLIP | | 29167373 | Unspecified |
| mmu-miR-139-5p | MIMAT0000656 | 2010111I01Rik | 72061 | sRNA-Seq//HITS-CLIP | 33184237\|29167373 | | Unspecified |
| mmu-miR-139-5p | MIMAT0000656 | 2010315B03Rik | 630836 | HITS-CLIP | | 29167373 | Unspecified |
| mmu-miR-139-5p | MIMAT0000656 | 2510009E07Rik | 72190 | sRNA-Seq//HITS-CLIP | 33184237\|29167373 | | Unspecified |
| mmu-miR-139-5p | MIMAT0000656 | 2510039O18Rik | 77034 | HITS-CLIP | | 29167373 | Unspecified |
| mmu-miR-139-5p | MIMAT0000656 | 2610021A01Rik | 668572 | HITS-CLIP | | 29167373 | Unspecified |
| mmu-miR-139-5p | MIMAT0000656 | 2610507B11Rik | 72503 | sRNA-Seq//HITS-CLIP | 33184237\|29167373 | | Unspecified |
| mmu-miR-139-5p | MIMAT0000656 | 4932438A13Rik | 229227 | sRNA-Seq//HITS-CLIP | 33184237\|29167373 | | Unspecified |
| mmu-miR-139-5p | MIMAT0000656 | 5430427O19Rik | 71398 | HITS-CLIP | | 29167373 | Unspecified |
| mmu-miR-139-5p | MIMAT0000656 | AI661453 | 224833 | HITS-CLIP | | 26203562 | Unspecified |
| mmu-miR-139-5p | MIMAT0000656 | AI987944 | 233168 | HITS-CLIP | | 29167373 | Unspecified |
| mmu-miR-139-5p | MIMAT0000656 | AW146154 | 101835 | HITS-CLIP | | 29167373 | Unspecified |
| mmu-miR-139-5p | MIMAT0000656 | AW554918 | 225289 | HITS-CLIP | 26203562\|29167373 | | Unspecified |
| mmu-miR-139-5p | MIMAT0000656 | Aacs | 78894 | sRNA-Seq//HITS-CLIP | 33184237\|29167373 | | Unspecified |
| mmu-miR-139-5p | MIMAT0000656 | Aak1 | 269774 | HITS-CLIP | | 29167373 | Unspecified |
| mmu-miR-139-5p | MIMAT0000656 | Abca1 | 11303 | HITS-CLIP | | 29167373 | Unspecified |
| mmu-miR-139-5p | MIMAT0000656 | Abcb1b | 18669 | HITS-CLIP | | 29167373 | Unspecified |
| mmu-miR-139-5p | MIMAT0000656 | Abcb7 | 11306 | HITS-CLIP | | 31433990 | Unspecified |
| mmu-miR-139-5p | MIMAT0000656 | Abcc1 | 17250 | HITS-CLIP | | 29167373 | Unspecified |
| mmu-miR-139-5p | MIMAT0000656 | Abcc3 | 76408 | HITS-CLIP | | 29167373 | Unspecified |
| mmu-miR-139-5p | MIMAT0000656 | Abce1 | 24015 | sRNA-Seq//HITS-CLIP | 33184237\|29167373 | | Unspecified |
| mmu-miR-139-5p | MIMAT0000656 | Abhd13 | 68904 | HITS-CLIP | | 29167373 | Unspecified |
| mmu-miR-139-5p | MIMAT0000656 | Abhd17b | 226016 | HITS-CLIP | | 29167373 | Unspecified |
| mmu-miR-139-5p | MIMAT0000656 | Abi1 | 11308 | sRNA-Seq//HITS-CLIP | 33184237\|29167373 | | Unspecified |
| mmu-miR-139-5p | MIMAT0000656 | Abl1 | 11350 | HITS-CLIP | | 31433990 | Unspecified |
| mmu-miR-139-5p | MIMAT0000656 | Abl2 | 11352 | HITS-CLIP | | 29167373 | Unspecified |
| mmu-miR-139-5p | MIMAT0000656 | Abr | 109934 | sRNA-Seq//HITS-CLIP | 33184237\|29167373 | | Unspecified |

| | | | | | | | |
|---|---|---|---|---|---|---|---|
| mmu-miR-139-5p | MIMAT0000656 | Acaa2 | 52538 | HITS-CLIP | | 29167373 | Unspecified |
| mmu-miR-139-5p | MIMAT0000656 | Acadm | 11364 | sRNA-Seq//HITS-CLIP | 33184237\|31433990 | | Unspecified |
| mmu-miR-139-5p | MIMAT0000656 | Acadsb | 66885 | HITS-CLIP | | 29167373 | Unspecified |
| mmu-miR-139-5p | MIMAT0000656 | Acap2 | 78618 | HITS-CLIP | | 29167373 | Unspecified |
| mmu-miR-139-5p | MIMAT0000656 | Acbd3 | 170760 | HITS-CLIP | | 29167373 | Unspecified |
| mmu-miR-139-5p | MIMAT0000656 | Acbd6 | 72482 | HITS-CLIP | | 29167373 | Unspecified |
| mmu-miR-139-5p | MIMAT0000656 | Acin1 | 56215 | sRNA-Seq//HITS-CLIP | 33184237\|29167373 | | Unspecified |
| mmu-miR-139-5p | MIMAT0000656 | Acly | 104112 | sRNA-Seq//HITS-CLIP | 33184237\|29167373 | | Unspecified |
| mmu-miR-139-5p | MIMAT0000656 | Acot7 | 70025 | sRNA-Seq//HITS-CLIP | 33184237\|29167373 | | Unspecified |
| mmu-miR-139-5p | MIMAT0000656 | Acox3 | 80911 | HITS-CLIP | | 29167373 | Unspecified |
| mmu-miR-139-5p | MIMAT0000656 | Acp2 | 11432 | HITS-CLIP | | 29167373 | Unspecified |
| mmu-miR-139-5p | MIMAT0000656 | Acsl1 | 14081 | HITS-CLIP | | 29167373 | Unspecified |
| mmu-miR-139-5p | MIMAT0000656 | Acsl4 | 50790 | sRNA-Seq//HITS-CLIP | 33184237\|29167373 | | Unspecified |
| mmu-miR-139-5p | MIMAT0000656 | Actb | 11461 | HITS-CLIP//sRNA-Seq | 31433990\|33184237\|26203562 | | Unspecified |
| mmu-miR-139-5p | MIMAT0000656 | Actn1 | 109711 | sRNA-Seq//HITS-CLIP | 33184237\|29167373 | | Unspecified |
| mmu-miR-139-5p | MIMAT0000656 | Actn4 | 60595 | sRNA-Seq//HITS-CLIP | 33184237\|29167373 | | Unspecified |
| mmu-miR-139-5p | MIMAT0000656 | Actr3 | 74117 | sRNA-Seq//HITS-CLIP | 33184237\|29167373 | | Unspecified |
| mmu-miR-139-5p | MIMAT0000656 | Adam4 | 11498 | HITS-CLIP | | 29167373 | Unspecified |
| mmu-miR-139-5p | MIMAT0000656 | Adam8 | 11501 | HITS-CLIP | | 29167373 | Unspecified |
| mmu-miR-139-5p | MIMAT0000656 | Adam9 | 11502 | HITS-CLIP | | 29167373 | Unspecified |
| mmu-miR-139-5p | MIMAT0000656 | Adamts10 | 224697 | HITS-CLIP | | 29167373 | Unspecified |
| mmu-miR-139-5p | MIMAT0000656 | Adap2 | 216991 | HITS-CLIP | | 29167373 | Unspecified |
| mmu-miR-139-5p | MIMAT0000656 | Adcy6 | 11512 | HITS-CLIP | | 30971389 | Unspecified |
| mmu-miR-139-5p | MIMAT0000656 | Adcy7 | 11513 | HITS-CLIP | | 29167373 | Unspecified |
| mmu-miR-139-5p | MIMAT0000656 | Add1 | 11518 | sRNA-Seq//HITS-CLIP | 33184237\|29167373 | | Unspecified |
| mmu-miR-139-5p | MIMAT0000656 | Adgre1 | 13733 | HITS-CLIP | | 29167373 | Unspecified |
| mmu-miR-139-5p | MIMAT0000656 | Adnp | 11538 | HITS-CLIP | | 29167373 | Unspecified |
| mmu-miR-139-5p | MIMAT0000656 | Adnp2 | 240442 | HITS-CLIP | | 29167373 | Unspecified |
| mmu-miR-139-5p | MIMAT0000656 | Ado | 211488 | HITS-CLIP | | 29167373 | Unspecified |
| mmu-miR-139-5p | MIMAT0000656 | Adrb2 | 11555 | HITS-CLIP | | 29167373 | Unspecified |
| mmu-miR-139-5p | MIMAT0000656 | Aebp2 | 11569 | HITS-CLIP | | 29167373 | Unspecified |
| mmu-miR-139-5p | MIMAT0000656 | Aff1 | 17355 | HITS-CLIP | | 29167373 | Unspecified |
| mmu-miR-139-5p | MIMAT0000656 | Aff4 | 93736 | sRNA-Seq//HITS-CLIP | 33184237\|26203562\|29167373 | | Unspecified |
| mmu-miR-139-5p | MIMAT0000656 | Afg3l2 | 69597 | HITS-CLIP | | 31433990 | Unspecified |
| mmu-miR-139-5p | MIMAT0000656 | Aftph | 216549 | HITS-CLIP | | 29167373 | Unspecified |
| mmu-miR-139-5p | MIMAT0000656 | Agfg1 | 15463 | sRNA-Seq//HITS-CLIP | 33184237\|26203562\|29167373 | | Unspecified |
| mmu-miR-139-5p | MIMAT0000656 | Aggf1 | 66549 | HITS-CLIP | | 29167373 | Unspecified |
| mmu-miR-139-5p | MIMAT0000656 | Ago1 | 236511 | HITS-CLIP | 31433990\|29167373 | | Unspecified |
| mmu-miR-139-5p | MIMAT0000656 | Ago2 | 239528 | sRNA-Seq//HITS-CLIP | 33184237\|31433990\|29167373 | | Unspecified |

| | | | | | | |
|---|---|---|---|---|---|---|
| mmu-miR-139-5p | MIMAT0000656 | Ago3 | 214150 | HITS-CLIP | 31433990\|29167373 | Unspecified |
| mmu-miR-139-5p | MIMAT0000656 | Agpat5 | 52123 | HITS-CLIP | 29167373 | Unspecified |
| mmu-miR-139-5p | MIMAT0000656 | Agps | 228061 | sRNA-Seq//HITS-CLIP | 33184237\|29167373 | Unspecified |
| mmu-miR-139-5p | MIMAT0000656 | Agtpbp1 | 67269 | sRNA-Seq//HITS-CLIP | 33184237\|29167373 | Unspecified |
| mmu-miR-139-5p | MIMAT0000656 | Ahcyl1 | 229709 | sRNA-Seq//HITS-CLIP | 33184237\|29167373 | Unspecified |
| mmu-miR-139-5p | MIMAT0000656 | Ahnak | 66395 | HITS-CLIP | 29167373 | Unspecified |
| mmu-miR-139-5p | MIMAT0000656 | Ahrr | 11624 | HITS-CLIP | 29167373 | Unspecified |
| mmu-miR-139-5p | MIMAT0000656 | Ajuba | 16475 | HITS-CLIP | 21258322\|26203562 | Unspecified |
| mmu-miR-139-5p | MIMAT0000656 | Akap10 | 56697 | HITS-CLIP | 29167373 | Unspecified |
| mmu-miR-139-5p | MIMAT0000656 | Akap11 | 219181 | sRNA-Seq//HITS-CLIP | 33184237\|29167373 | Unspecified |
| mmu-miR-139-5p | MIMAT0000656 | Akap13 | 75547 | HITS-CLIP | 29167373 | Unspecified |
| mmu-miR-139-5p | MIMAT0000656 | Akap9 | 100986 | sRNA-Seq//HITS-CLIP | 33184237\|29167373 | Unspecified |
| mmu-miR-139-5p | MIMAT0000656 | Akna | 100182 | HITS-CLIP | 29167373 | Unspecified |
| mmu-miR-139-5p | MIMAT0000656 | Akr1a1 | 58810 | sRNA-Seq//HITS-CLIP | 33184237\|29167373 | Unspecified |
| mmu-miR-139-5p | MIMAT0000656 | Akt3 | 23797 | sRNA-Seq//HITS-CLIP | 33184237\|29167373 | Unspecified |
| mmu-miR-139-5p | MIMAT0000656 | Alcam | 11658 | sRNA-Seq//HITS-CLIP | 33184237\|31433990 | Unspecified |
| mmu-miR-139-5p | MIMAT0000656 | Aldoa | 11674 | sRNA-Seq//HITS-CLIP | 33184237\|29167373 | Unspecified |
| mmu-miR-139-5p | MIMAT0000656 | Alg10b | 380959 | HITS-CLIP | 29167373 | Unspecified |
| mmu-miR-139-5p | MIMAT0000656 | Alg11 | 207958 | HITS-CLIP | 29167373 | Unspecified |
| mmu-miR-139-5p | MIMAT0000656 | Alms1 | 236266 | HITS-CLIP | 29167373 | Unspecified |
| mmu-miR-139-5p | MIMAT0000656 | Alox5ap | 11690 | HITS-CLIP | 29167373 | Unspecified |
| mmu-miR-139-5p | MIMAT0000656 | Als2 | 74018 | HITS-CLIP | 29167373 | Unspecified |
| mmu-miR-139-5p | MIMAT0000656 | Amdhd2 | 245847 | HITS-CLIP | 29167373 | Unspecified |
| mmu-miR-139-5p | MIMAT0000656 | Amotl2 | 56332 | HITS-CLIP | 26203562 | Unspecified |
| mmu-miR-139-5p | MIMAT0000656 | Anapc11 | 66156 | HITS-CLIP | 29167373 | Unspecified |
| mmu-miR-139-5p | MIMAT0000656 | Anapc5 | 59008 | sRNA-Seq//HITS-CLIP | 33184237\|29167373 | Unspecified |
| mmu-miR-139-5p | MIMAT0000656 | Angel2 | 52477 | HITS-CLIP | 29167373 | Unspecified |
| mmu-miR-139-5p | MIMAT0000656 | Angptl2 | 26360 | HITS-CLIP | 29167373 | Unspecified |
| mmu-miR-139-5p | MIMAT0000656 | Ankfy1 | 11736 | HITS-CLIP | 29167373 | Unspecified |
| mmu-miR-139-5p | MIMAT0000656 | Ankhd1 | 108857 | HITS-CLIP | 29167373 | Unspecified |
| mmu-miR-139-5p | MIMAT0000656 | Ankib1 | 70797 | HITS-CLIP | 29167373 | Unspecified |
| mmu-miR-139-5p | MIMAT0000656 | Ankrd13a | 68420 | sRNA-Seq//HITS-CLIP | 33184237\|29167373 | Unspecified |
| mmu-miR-139-5p | MIMAT0000656 | Ankrd17 | 81702 | sRNA-Seq//HITS-CLIP | 33184237\|29167373 | Unspecified |
| mmu-miR-139-5p | MIMAT0000656 | Ankrd33b | 67434 | HITS-CLIP | 29167373 | Unspecified |
| mmu-miR-139-5p | MIMAT0000656 | Ankrd52 | 237615 | HITS-CLIP | 29167373 | Unspecified |
| mmu-miR-139-5p | MIMAT0000656 | Anln | 68743 | HITS-CLIP | 31433990 | Unspecified |
| mmu-miR-139-5p | MIMAT0000656 | Anxa1 | 16952 | HITS-CLIP | 29167373 | Unspecified |
| mmu-miR-139-5p | MIMAT0000656 | Anxa2 | 12306 | sRNA-Seq//HITS-CLIP | 33184237\|26203562\|29167373 | Unspecified |
| mmu-miR-139-5p | MIMAT0000656 | Ap1b1 | 11764 | sRNA-Seq//HITS-CLIP | 33184237\|29167373 | Unspecified |

| | | | | | | |
|---|---|---|---|---|---|---|
| mmu-miR-139-5p | MIMAT0000656 | Ap1g1 | 11765 | sRNA-Seq//HITS-CLIP | 33184237\|29167373 | Unspecified |
| mmu-miR-139-5p | MIMAT0000656 | Ap2a2 | 11772 | sRNA-Seq//HITS-CLIP | 33184237\|29167373 | Unspecified |
| mmu-miR-139-5p | MIMAT0000656 | Ap2b1 | 71770 | sRNA-Seq//HITS-CLIP | 33184237\|29167373 | Unspecified |
| mmu-miR-139-5p | MIMAT0000656 | Ap3d1 | 11776 | sRNA-Seq//HITS-CLIP | 33184237\|29167373 | Unspecified |
| mmu-miR-139-5p | MIMAT0000656 | Ap4b1 | 67489 | HITS-CLIP | 29167373 | Unspecified |
| mmu-miR-139-5p | MIMAT0000656 | Ap5m1 | 74385 | HITS-CLIP | 29167373 | Unspecified |
| mmu-miR-139-5p | MIMAT0000656 | Apaf1 | 11783 | HITS-CLIP | 29167373 | Unspecified |
| mmu-miR-139-5p | MIMAT0000656 | Apbb1ip | 54519 | HITS-CLIP | 29167373 | Unspecified |
| mmu-miR-139-5p | MIMAT0000656 | Apbb2 | 11787 | sRNA-Seq//HITS-CLIP | 33184237\|29167373 | Unspecified |
| mmu-miR-139-5p | MIMAT0000656 | Apc | 11789 | sRNA-Seq//HITS-CLIP | 33184237\|29167373 | Unspecified |
| mmu-miR-139-5p | MIMAT0000656 | Api5 | 11800 | HITS-CLIP | 23630078 | Unspecified |
| mmu-miR-139-5p | MIMAT0000656 | Aplp2 | 11804 | sRNA-Seq//HITS-CLIP | 33184237\|29167373 | Unspecified |
| mmu-miR-139-5p | MIMAT0000656 | Apobec1 | 11810 | HITS-CLIP | 29167373 | Unspecified |
| mmu-miR-139-5p | MIMAT0000656 | Apobr | 171504 | HITS-CLIP | 29167373 | Unspecified |
| mmu-miR-139-5p | MIMAT0000656 | Apoe | 11816 | HITS-CLIP | 29167373 | Unspecified |
| mmu-miR-139-5p | MIMAT0000656 | App | 11820 | sRNA-Seq//HITS-CLIP | 33184237\|29167373 | Unspecified |
| mmu-miR-139-5p | MIMAT0000656 | Appbp2 | 66884 | sRNA-Seq//HITS-CLIP | 33184237\|29167373 | Unspecified |
| mmu-miR-139-5p | MIMAT0000656 | Arap1 | 69710 | HITS-CLIP | 29167373 | Unspecified |
| mmu-miR-139-5p | MIMAT0000656 | Arap3 | 106952 | HITS-CLIP | 29167373 | Unspecified |
| mmu-miR-139-5p | MIMAT0000656 | Arcn1 | 213827 | sRNA-Seq//HITS-CLIP | 33184237\|29167373 | Unspecified |
| mmu-miR-139-5p | MIMAT0000656 | Arf1 | 11840 | HITS-CLIP//sRNA-Seq | 31433990\|33184237\|29167373 | Unspecified |
| mmu-miR-139-5p | MIMAT0000656 | Arf3 | 11842 | sRNA-Seq//HITS-CLIP | 33184237\|29167373 | Unspecified |
| mmu-miR-139-5p | MIMAT0000656 | Arf6 | 11845 | HITS-CLIP | 29167373 | Unspecified |
| mmu-miR-139-5p | MIMAT0000656 | Arfgef1 | 211673 | sRNA-Seq//HITS-CLIP | 33184237\|29167373 | Unspecified |
| mmu-miR-139-5p | MIMAT0000656 | Arhgap12 | 75415 | sRNA-Seq//HITS-CLIP | 33184237\|23630078 | Unspecified |
| mmu-miR-139-5p | MIMAT0000656 | Arhgap19 | 71085 | HITS-CLIP | 29167373 | Unspecified |
| mmu-miR-139-5p | MIMAT0000656 | Arhgap21 | 71435 | sRNA-Seq//HITS-CLIP | 33184237\|29167373 | Unspecified |
| mmu-miR-139-5p | MIMAT0000656 | Arhgap22 | 239027 | HITS-CLIP | 29167373 | Unspecified |
| mmu-miR-139-5p | MIMAT0000656 | Arhgap30 | 226652 | HITS-CLIP | 29167373 | Unspecified |
| mmu-miR-139-5p | MIMAT0000656 | Arhgap31 | 12549 | HITS-CLIP | 29167373 | Unspecified |
| mmu-miR-139-5p | MIMAT0000656 | Arhgap4 | 393 | HITS-CLIP | 29167373 | Unspecified |
| mmu-miR-139-5p | MIMAT0000656 | Arhgap45 | 23526 | HITS-CLIP | 29167373 | Unspecified |
| mmu-miR-139-5p | MIMAT0000656 | Arhgap5 | 11855 | HITS-CLIP//sRNA-Seq | 29167373\|33184237 | Unspecified |
| mmu-miR-139-5p | MIMAT0000656 | Arhgdia | 192662 | sRNA-Seq//HITS-CLIP | 33184237\|31433990 | Unspecified |
| mmu-miR-139-5p | MIMAT0000656 | Arhgef11 | 213498 | HITS-CLIP | 29167373 | Unspecified |
| mmu-miR-139-5p | MIMAT0000656 | Arhgef12 | 69632 | HITS-CLIP | 31433990 | Unspecified |
| mmu-miR-139-5p | MIMAT0000656 | Arhgef2 | 16800 | HITS-CLIP | 29167373 | Unspecified |
| mmu-miR-139-5p | MIMAT0000656 | Arhgef7 | 54126 | sRNA-Seq//HITS-CLIP | 33184237\|29167373 | Unspecified |
| mmu-miR-139-5p | MIMAT0000656 | Arid1a | 93760 | HITS-CLIP | 29167373 | Unspecified |

| | | | | | | | |
|---|---|---|---|---|---|---|---|
| mmu-miR-139-5p | MIMAT0000656 | Arid1b | 239985 | HITS-CLIP | | 29167373 | Unspecified |
| mmu-miR-139-5p | MIMAT0000656 | Arid4b | 94246 | HITS-CLIP | | 29167373 | Unspecified |
| mmu-miR-139-5p | MIMAT0000656 | Arid5b | 71371 | HITS-CLIP | 31433990\|29167373 | | Unspecified |
| mmu-miR-139-5p | MIMAT0000656 | Arih1 | 23806 | sRNA-Seq//HITS-CLIP | 33184237\|29167373 | | Unspecified |
| mmu-miR-139-5p | MIMAT0000656 | Arl5b | 75869 | HITS-CLIP | | 29167373 | Unspecified |
| mmu-miR-139-5p | MIMAT0000656 | Arl6ip5 | 65106 | sRNA-Seq//HITS-CLIP | 33184237\|29167373 | | Unspecified |
| mmu-miR-139-5p | MIMAT0000656 | Arl8a | 68724 | sRNA-Seq//HITS-CLIP | 33184237\|29167373 | | Unspecified |
| mmu-miR-139-5p | MIMAT0000656 | Arl8b | 67166 | sRNA-Seq//HITS-CLIP | 33184237\|29167373 | | Unspecified |
| mmu-miR-139-5p | MIMAT0000656 | Armc1 | 74252 | HITS-CLIP | | 29167373 | Unspecified |
| mmu-miR-139-5p | MIMAT0000656 | Arnt | 11863 | HITS-CLIP | | 29167373 | Unspecified |
| mmu-miR-139-5p | MIMAT0000656 | Arpc1a | 56443 | HITS-CLIP//sRNA-Seq | 31433990\|33184237 | | Unspecified |
| mmu-miR-139-5p | MIMAT0000656 | Arpc4 | 68089 | sRNA-Seq//HITS-CLIP | 33184237\|29167373 | | Unspecified |
| mmu-miR-139-5p | MIMAT0000656 | Arpc5 | 67771 | sRNA-Seq//HITS-CLIP | 33184237\|29167373 | | Unspecified |
| mmu-miR-139-5p | MIMAT0000656 | Arrb1 | 109689 | HITS-CLIP | | 29167373 | Unspecified |
| mmu-miR-139-5p | MIMAT0000656 | Arrdc1 | 215705 | HITS-CLIP | | 29167373 | Unspecified |
| mmu-miR-139-5p | MIMAT0000656 | Arsb | 11881 | HITS-CLIP | | 29167373 | Unspecified |
| mmu-miR-139-5p | MIMAT0000656 | Asap1 | 13196 | sRNA-Seq//HITS-CLIP | 33184237\|29167373 | | Unspecified |
| mmu-miR-139-5p | MIMAT0000656 | Ascc3 | 77987 | HITS-CLIP | | 29167373 | Unspecified |
| mmu-miR-139-5p | MIMAT0000656 | Asf1b | 66929 | HITS-CLIP | | 31433990 | Unspecified |
| mmu-miR-139-5p | MIMAT0000656 | Ash1l | 192195 | HITS-CLIP | | 29167373 | Unspecified |
| mmu-miR-139-5p | MIMAT0000656 | Asph | 65973 | HITS-CLIP | | 29167373 | Unspecified |
| mmu-miR-139-5p | MIMAT0000656 | Asprv1 | 67855 | HITS-CLIP | | 26203562 | Unspecified |
| mmu-miR-139-5p | MIMAT0000656 | Asxl2 | 75302 | HITS-CLIP | | 29167373 | Unspecified |
| mmu-miR-139-5p | MIMAT0000656 | Atad2 | 70472 | HITS-CLIP | | 31433990 | Unspecified |
| mmu-miR-139-5p | MIMAT0000656 | Atad5 | 237877 | HITS-CLIP | | 29167373 | Unspecified |
| mmu-miR-139-5p | MIMAT0000656 | Atf1 | 11908 | HITS-CLIP | | 29167373 | Unspecified |
| mmu-miR-139-5p | MIMAT0000656 | Atf2 | 11909 | sRNA-Seq//HITS-CLIP | 33184237\|29167373 | | Unspecified |
| mmu-miR-139-5p | MIMAT0000656 | Atf3 | 11910 | HITS-CLIP | | 29167373 | Unspecified |
| mmu-miR-139-5p | MIMAT0000656 | Atf6 | 226641 | HITS-CLIP | | 29167373 | Unspecified |
| mmu-miR-139-5p | MIMAT0000656 | Atf6b | 12915 | HITS-CLIP | | 29167373 | Unspecified |
| mmu-miR-139-5p | MIMAT0000656 | Atf7 | 223922 | HITS-CLIP | | 29167373 | Unspecified |
| mmu-miR-139-5p | MIMAT0000656 | Atf7ip | 54343 | HITS-CLIP | | 29167373 | Unspecified |
| mmu-miR-139-5p | MIMAT0000656 | Atg12 | 67526 | sRNA-Seq//HITS-CLIP | 33184237\|29167373 | | Unspecified |
| mmu-miR-139-5p | MIMAT0000656 | Atg2b | 76559 | HITS-CLIP | | 26203562 | Unspecified |
| mmu-miR-139-5p | MIMAT0000656 | Atm | 11920 | HITS-CLIP | | 29167373 | Unspecified |
| mmu-miR-139-5p | MIMAT0000656 | Atn1 | 13498 | HITS-CLIP | | 29167373 | Unspecified |
| mmu-miR-139-5p | MIMAT0000656 | Atosa | 56204 | HITS-CLIP | | 31433990 | Unspecified |
| mmu-miR-139-5p | MIMAT0000656 | Atox1 | 11927 | sRNA-Seq//HITS-CLIP | 33184237\|29167373 | | Unspecified |
| mmu-miR-139-5p | MIMAT0000656 | Atp11a | 50770 | sRNA-Seq//HITS-CLIP | 33184237\|29167373 | | Unspecified |

| | | | | | | |
|---|---|---|---|---|---|---|
| mmu-miR-139-5p | MIMAT0000656 | Atp13a3 | 224088 | HITS-CLIP | 29167373 | Unspecified |
| mmu-miR-139-5p | MIMAT0000656 | Atp1a1 | 11928 | sRNA-Seq//HITS-CLIP | 33184237\|29167373 | Unspecified |
| mmu-miR-139-5p | MIMAT0000656 | Atp1b3 | 11933 | sRNA-Seq//HITS-CLIP | 33184237\|29167373 | Unspecified |
| mmu-miR-139-5p | MIMAT0000656 | Atp2a2 | 11938 | sRNA-Seq//HITS-CLIP | 33184237\|29167373 | Unspecified |
| mmu-miR-139-5p | MIMAT0000656 | Atp2b1 | 67972 | sRNA-Seq//HITS-CLIP | 33184237\|29167373 | Unspecified |
| mmu-miR-139-5p | MIMAT0000656 | Atp5b | 11947 | sRNA-Seq//HITS-CLIP | 33184237\|29167373 | Unspecified |
| mmu-miR-139-5p | MIMAT0000656 | Atp6v0a1 | 11975 | sRNA-Seq//HITS-CLIP | 33184237\|29167373 | Unspecified |
| mmu-miR-139-5p | MIMAT0000656 | Atp6v0e | 11974 | HITS-CLIP | 29167373 | Unspecified |
| mmu-miR-139-5p | MIMAT0000656 | Atp6v1a | 11964 | sRNA-Seq//HITS-CLIP | 33184237\|29167373 | Unspecified |
| mmu-miR-139-5p | MIMAT0000656 | Atp6v1b2 | 11966 | sRNA-Seq//HITS-CLIP | 33184237\|29167373 | Unspecified |
| mmu-miR-139-5p | MIMAT0000656 | Atp6v1h | 108664 | sRNA-Seq//HITS-CLIP | 33184237\|29167373 | Unspecified |
| mmu-miR-139-5p | MIMAT0000656 | Atp7a | 11977 | HITS-CLIP | 29167373 | Unspecified |
| mmu-miR-139-5p | MIMAT0000656 | Atp8b4 | 241633 | HITS-CLIP | 29167373 | Unspecified |
| mmu-miR-139-5p | MIMAT0000656 | Atpif1 | 11983 | sRNA-Seq//HITS-CLIP | 33184237\|29167373 | Unspecified |
| mmu-miR-139-5p | MIMAT0000656 | Atrn | 11990 | HITS-CLIP | 23142080\|29167373 | Unspecified |
| mmu-miR-139-5p | MIMAT0000656 | Atrx | 22589 | sRNA-Seq//HITS-CLIP | 33184237\|29167373 | Unspecified |
| mmu-miR-139-5p | MIMAT0000656 | Atxn1 | 20238 | HITS-CLIP | 31433990\|29167373 | Unspecified |
| mmu-miR-139-5p | MIMAT0000656 | Atxn10 | 54138 | sRNA-Seq//HITS-CLIP | 33184237\|29167373 | Unspecified |
| mmu-miR-139-5p | MIMAT0000656 | Atxn1l | 52335 | HITS-CLIP | 26203562\|29167373 | Unspecified |
| mmu-miR-139-5p | MIMAT0000656 | Atxn3 | 110616 | HITS-CLIP | 29167373 | Unspecified |
| mmu-miR-139-5p | MIMAT0000656 | Azin1 | 54375 | sRNA-Seq//HITS-CLIP | 33184237\|29167373 | Unspecified |
| mmu-miR-139-5p | MIMAT0000656 | B2m | 12010 | HITS-CLIP | 29167373 | Unspecified |
| mmu-miR-139-5p | MIMAT0000656 | B3gnt2 | 53625 | HITS-CLIP | 29167373 | Unspecified |
| mmu-miR-139-5p | MIMAT0000656 | B430306N03Rik | 320148 | HITS-CLIP | 29167373 | Unspecified |
| mmu-miR-139-5p | MIMAT0000656 | B4galt1 | 14595 | HITS-CLIP | 29167373 | Unspecified |
| mmu-miR-139-5p | MIMAT0000656 | B4galt3 | 57370 | HITS-CLIP | 29167373 | Unspecified |
| mmu-miR-139-5p | MIMAT0000656 | B4galt5 | 56336 | HITS-CLIP | 29167373 | Unspecified |
| mmu-miR-139-5p | MIMAT0000656 | B4galt6 | 56386 | HITS-CLIP | 29167373 | Unspecified |
| mmu-miR-139-5p | MIMAT0000656 | BC005537 | 79555 | sRNA-Seq//HITS-CLIP | 33184237\|29167373 | Unspecified |
| mmu-miR-139-5p | MIMAT0000656 | Bach1 | 12013 | sRNA-Seq//HITS-CLIP | 33184237\|29167373 | Unspecified |
| mmu-miR-139-5p | MIMAT0000656 | Bag6 | 224727 | sRNA-Seq//HITS-CLIP | 33184237\|29167373 | Unspecified |
| mmu-miR-139-5p | MIMAT0000656 | Bahd1 | 228536 | HITS-CLIP | 29167373 | Unspecified |
| mmu-miR-139-5p | MIMAT0000656 | Bak1 | 12018 | HITS-CLIP | 29167373 | Unspecified |
| mmu-miR-139-5p | MIMAT0000656 | Baz1a | 217578 | HITS-CLIP | 29167373 | Unspecified |
| mmu-miR-139-5p | MIMAT0000656 | Baz1b | 22385 | HITS-CLIP | 26203562 | Unspecified |
| mmu-miR-139-5p | MIMAT0000656 | Baz2a | 116848 | HITS-CLIP | 31433990 | Unspecified |
| mmu-miR-139-5p | MIMAT0000656 | Baz2b | 407823 | sRNA-Seq//HITS-CLIP | 33184237\|29167373 | Unspecified |
| mmu-miR-139-5p | MIMAT0000656 | Bcl10 | 12042 | HITS-CLIP | 29167373 | Unspecified |
| mmu-miR-139-5p | MIMAT0000656 | Bcl2 | 12043 | HITS-CLIP | 29167373 | Unspecified |

| | | | | | | |
|---|---|---|---|---|---|---|
| mmu-miR-139-5p | MIMAT0000656 | Bcl2l1 | 12048 | sRNA-Seq//HITS-CLIP | 33184237\|29167373 | Unspecified |
| mmu-miR-139-5p | MIMAT0000656 | Bcl2l11 | 12125 | HITS-CLIP | 29167373 | Unspecified |
| mmu-miR-139-5p | MIMAT0000656 | Bcl2l13 | 94044 | sRNA-Seq//HITS-CLIP | 33184237\|29167373 | Unspecified |
| mmu-miR-139-5p | MIMAT0000656 | Bcl6 | 12053 | HITS-CLIP | 29167373 | Unspecified |
| mmu-miR-139-5p | MIMAT0000656 | Bclaf1 | 72567 | sRNA-Seq//HITS-CLIP | 33184237\|29167373 | Unspecified |
| mmu-miR-139-5p | MIMAT0000656 | Bcorl1 | 320376 | HITS-CLIP | 29167373 | Unspecified |
| mmu-miR-139-5p | MIMAT0000656 | Bdp1 | 544971 | HITS-CLIP | 29167373 | Unspecified |
| mmu-miR-139-5p | MIMAT0000656 | Bend4 | 666938 | HITS-CLIP | 26203562 | Unspecified |
| mmu-miR-139-5p | MIMAT0000656 | Bhlhe41 | 79362 | HITS-CLIP | 29167373 | Unspecified |
| mmu-miR-139-5p | MIMAT0000656 | Birc6 | 12211 | sRNA-Seq//HITS-CLIP | 33184237\|31433990\|29167373 | Unspecified |
| mmu-miR-139-5p | MIMAT0000656 | Blnk | 17060 | HITS-CLIP | 29167373 | Unspecified |
| mmu-miR-139-5p | MIMAT0000656 | Bmf | 171543 | HITS-CLIP | 29167373 | Unspecified |
| mmu-miR-139-5p | MIMAT0000656 | Bmp2k | 140780 | HITS-CLIP | 29167373 | Unspecified |
| mmu-miR-139-5p | MIMAT0000656 | Bmpr2 | 12168 | sRNA-Seq//HITS-CLIP | 33184237\|29167373 | Unspecified |
| mmu-miR-139-5p | MIMAT0000656 | Bmt2 | 101148 | HITS-CLIP | 31433990\|29167373 | Unspecified |
| mmu-miR-139-5p | MIMAT0000656 | Bptf | 207165 | HITS-CLIP | 29167373 | Unspecified |
| mmu-miR-139-5p | MIMAT0000656 | Brd1 | 223770 | HITS-CLIP | 29167373 | Unspecified |
| mmu-miR-139-5p | MIMAT0000656 | Brd2 | 14312 | sRNA-Seq//HITS-CLIP | 33184237\|29167373 | Unspecified |
| mmu-miR-139-5p | MIMAT0000656 | Brd3 | 67382 | HITS-CLIP | 29167373 | Unspecified |
| mmu-miR-139-5p | MIMAT0000656 | Brd4 | 57261 | sRNA-Seq//HITS-CLIP | 33184237\|29167373 | Unspecified |
| mmu-miR-139-5p | MIMAT0000656 | Brd8 | 78656 | HITS-CLIP | 29167373 | Unspecified |
| mmu-miR-139-5p | MIMAT0000656 | Brwd3 | 382236 | HITS-CLIP | 29167373 | Unspecified |
| mmu-miR-139-5p | MIMAT0000656 | Bsn | 12217 | HITS-CLIP | 29167373 | Unspecified |
| mmu-miR-139-5p | MIMAT0000656 | Bst2 | 69550 | HITS-CLIP | 29167373 | Unspecified |
| mmu-miR-139-5p | MIMAT0000656 | Btaf1 | 107182 | HITS-CLIP | 29167373 | Unspecified |
| mmu-miR-139-5p | MIMAT0000656 | Btg1 | 12226 | HITS-CLIP | 29167373 | Unspecified |
| mmu-miR-139-5p | MIMAT0000656 | Btg2 | 12227 | HITS-CLIP | 29167373 | Unspecified |
| mmu-miR-139-5p | MIMAT0000656 | Btk | 12229 | HITS-CLIP | 29167373 | Unspecified |
| mmu-miR-139-5p | MIMAT0000656 | Bub3 | 12237 | HITS-CLIP | 29167373 | Unspecified |
| mmu-miR-139-5p | MIMAT0000656 | Bzw1 | 66882 | sRNA-Seq//HITS-CLIP | 33184237\|29167373 | Unspecified |
| mmu-miR-139-5p | MIMAT0000656 | C1galt1 | 94192 | HITS-CLIP | 29167373 | Unspecified |
| mmu-miR-139-5p | MIMAT0000656 | C1qa | 12259 | HITS-CLIP | 29167373 | Unspecified |
| mmu-miR-139-5p | MIMAT0000656 | C1qb | 12260 | HITS-CLIP | 29167373 | Unspecified |
| mmu-miR-139-5p | MIMAT0000656 | C3ar1 | 12267 | HITS-CLIP | 29167373 | Unspecified |
| mmu-miR-139-5p | MIMAT0000656 | C5ar1 | 12273 | HITS-CLIP | 29167373 | Unspecified |
| mmu-miR-139-5p | MIMAT0000656 | C5ar2 | 319430 | HITS-CLIP | 29167373 | Unspecified |
| mmu-miR-139-5p | MIMAT0000656 | Cab39 | 12283 | sRNA-Seq//HITS-CLIP | 33184237\|29167373 | Unspecified |
| mmu-miR-139-5p | MIMAT0000656 | Cacna1a | 12286 | sRNA-Seq//HITS-CLIP | 33184237\|29167373 | Unspecified |
| mmu-miR-139-5p | MIMAT0000656 | Cacna1d | 12289 | HITS-CLIP | 29167373 | Unspecified |

| | | | | | | |
|---|---|---|---|---|---|---|
| mmu-miR-139-5p | MIMAT0000656 | Cacul1 | 78832 | HITS-CLIP | 29167373 | Unspecified |
| mmu-miR-139-5p | MIMAT0000656 | Calcrl | 54598 | HITS-CLIP | 29167373 | Unspecified |
| mmu-miR-139-5p | MIMAT0000656 | Calm2 | 12314 | sRNA-Seq//HITS-CLIP | 33184237\|29167373 | Unspecified |
| mmu-miR-139-5p | MIMAT0000656 | Calm3 | 12315 | sRNA-Seq//HITS-CLIP | 33184237\|29167373 | Unspecified |
| mmu-miR-139-5p | MIMAT0000656 | Calr | 12317 | sRNA-Seq//HITS-CLIP | 33184237\|29167373 | Unspecified |
| mmu-miR-139-5p | MIMAT0000656 | Camk1 | 52163 | HITS-CLIP | 29167373 | Unspecified |
| mmu-miR-139-5p | MIMAT0000656 | Camta2 | 216874 | HITS-CLIP | 31433990\|29167373 | Unspecified |
| mmu-miR-139-5p | MIMAT0000656 | Canx | 12330 | sRNA-Seq//HITS-CLIP | 33184237\|31433990\|29167373 | Unspecified |
| mmu-miR-139-5p | MIMAT0000656 | Cap1 | 12331 | sRNA-Seq//HITS-CLIP | 33184237\|29167373 | Unspecified |
| mmu-miR-139-5p | MIMAT0000656 | Capg | 12332 | HITS-CLIP | 29167373 | Unspecified |
| mmu-miR-139-5p | MIMAT0000656 | Capza2 | 12343 | sRNA-Seq//HITS-CLIP | 33184237\|31340158\|29167373 | Unspecified |
| mmu-miR-139-5p | MIMAT0000656 | Carhsp1 | 52502 | sRNA-Seq//HITS-CLIP | 33184237\|29167373 | Unspecified |
| mmu-miR-139-5p | MIMAT0000656 | Casd1 | 213819 | HITS-CLIP | 31433990 | Unspecified |
| mmu-miR-139-5p | MIMAT0000656 | Casp2 | 12366 | HITS-CLIP | 29167373 | Unspecified |
| mmu-miR-139-5p | MIMAT0000656 | Castor2 | 729438 | HITS-CLIP | 29167373 | Unspecified |
| mmu-miR-139-5p | MIMAT0000656 | Cav2 | 12390 | HITS-CLIP | 29167373 | Unspecified |
| mmu-miR-139-5p | MIMAT0000656 | Cbl | 12402 | HITS-CLIP | 29167373 | Unspecified |
| mmu-miR-139-5p | MIMAT0000656 | Ccar1 | 67500 | sRNA-Seq//HITS-CLIP | 33184237\|29167373 | Unspecified |
| mmu-miR-139-5p | MIMAT0000656 | Ccdc127 | 67433 | sRNA-Seq//HITS-CLIP | 33184237\|29167373 | Unspecified |
| mmu-miR-139-5p | MIMAT0000656 | Ccdc186 | 55088 | sRNA-Seq//HITS-CLIP | 33184237\|29167373 | Unspecified |
| mmu-miR-139-5p | MIMAT0000656 | Ccdc47 | 67163 | sRNA-Seq//HITS-CLIP | 33184237\|23630078 | Unspecified |
| mmu-miR-139-5p | MIMAT0000656 | Ccdc50 | 67501 | sRNA-Seq//HITS-CLIP | 33184237\|29167373 | Unspecified |
| mmu-miR-139-5p | MIMAT0000656 | Ccdc61 | 232933 | HITS-CLIP | 26203562 | Unspecified |
| mmu-miR-139-5p | MIMAT0000656 | Ccdc82 | 66396 | HITS-CLIP | 29167373 | Unspecified |
| mmu-miR-139-5p | MIMAT0000656 | Ccdc88a | 108686 | sRNA-Seq//HITS-CLIP | 33184237\|29167373 | Unspecified |
| mmu-miR-139-5p | MIMAT0000656 | Ccl2 | 20296 | HITS-CLIP | 29167373 | Unspecified |
| mmu-miR-139-5p | MIMAT0000656 | Ccl9 | 20308 | HITS-CLIP | 29167373 | Unspecified |
| mmu-miR-139-5p | MIMAT0000656 | Ccna2 | 12428 | HITS-CLIP | 31433990 | Unspecified |
| mmu-miR-139-5p | MIMAT0000656 | Ccnd1 | 12443 | HITS-CLIP | 29167373 | Unspecified |
| mmu-miR-139-5p | MIMAT0000656 | Ccnf | 12449 | HITS-CLIP | 31433990 | Unspecified |
| mmu-miR-139-5p | MIMAT0000656 | Ccng1 | 12450 | sRNA-Seq//HITS-CLIP | 33184237\|31433990 | Unspecified |
| mmu-miR-139-5p | MIMAT0000656 | Ccnk | 12454 | HITS-CLIP | 31433990 | Unspecified |
| mmu-miR-139-5p | MIMAT0000656 | Ccnl2 | 56036 | HITS-CLIP | 29167373 | Unspecified |
| mmu-miR-139-5p | MIMAT0000656 | Ccnt2 | 72949 | HITS-CLIP | 29167373 | Unspecified |
| mmu-miR-139-5p | MIMAT0000656 | Ccny | 67974 | HITS-CLIP | 29167373 | Unspecified |
| mmu-miR-139-5p | MIMAT0000656 | Ccr2 | 12772 | HITS-CLIP | 29167373 | Unspecified |
| mmu-miR-139-5p | MIMAT0000656 | Cct5 | 12465 | sRNA-Seq//HITS-CLIP | 33184237\|29167373 | Unspecified |
| mmu-miR-139-5p | MIMAT0000656 | Cd180 | 17079 | HITS-CLIP | 29167373 | Unspecified |
| mmu-miR-139-5p | MIMAT0000656 | Cd28 | 12487 | HITS-CLIP | 29167373 | Unspecified |

| | | | | | | |
|---|---|---|---|---|---|---|
| mmu-miR-139-5p | MIMAT0000656 | Cd2ap | 12488 | HITS-CLIP | 29167373 | Unspecified |
| mmu-miR-139-5p | MIMAT0000656 | Cd300lb | 217304 | HITS-CLIP | 29167373 | Unspecified |
| mmu-miR-139-5p | MIMAT0000656 | Cd300ld | 217305 | HITS-CLIP | 29167373 | Unspecified |
| mmu-miR-139-5p | MIMAT0000656 | Cd36 | 12491 | HITS-CLIP | 29167373 | Unspecified |
| mmu-miR-139-5p | MIMAT0000656 | Cd3e | 12501 | HITS-CLIP | 31433990 | Unspecified |
| mmu-miR-139-5p | MIMAT0000656 | Cd44 | 12505 | HITS-CLIP | 29167373 | Unspecified |
| mmu-miR-139-5p | MIMAT0000656 | Cd47 | 16423 | sRNA-Seq//HITS-CLIP | 33184237\|29167373 | Unspecified |
| mmu-miR-139-5p | MIMAT0000656 | Cd53 | 12508 | HITS-CLIP | 29167373 | Unspecified |
| mmu-miR-139-5p | MIMAT0000656 | Cd74 | 16149 | HITS-CLIP | 29167373 | Unspecified |
| mmu-miR-139-5p | MIMAT0000656 | Cd84 | 12523 | HITS-CLIP | 29167373 | Unspecified |
| mmu-miR-139-5p | MIMAT0000656 | Cd93 | 17064 | HITS-CLIP | 29167373 | Unspecified |
| mmu-miR-139-5p | MIMAT0000656 | Cdc27 | 217232 | sRNA-Seq//HITS-CLIP | 33184237\|29167373 | Unspecified |
| mmu-miR-139-5p | MIMAT0000656 | Cdc42 | 12540 | sRNA-Seq//HITS-CLIP | 33184237\|29167373 | Unspecified |
| mmu-miR-139-5p | MIMAT0000656 | Cdc42bpb | 217866 | sRNA-Seq//HITS-CLIP | 33184237\|29167373 | Unspecified |
| mmu-miR-139-5p | MIMAT0000656 | Cdc42se2 | 72729 | HITS-CLIP | 29167373 | Unspecified |
| mmu-miR-139-5p | MIMAT0000656 | Cdca4 | 71963 | HITS-CLIP | 31433990\|29167373 | Unspecified |
| mmu-miR-139-5p | MIMAT0000656 | Cdk12 | 69131 | HITS-CLIP | 31433990\|29167373 | Unspecified |
| mmu-miR-139-5p | MIMAT0000656 | Cdk14 | 18647 | HITS-CLIP | 29167373 | Unspecified |
| mmu-miR-139-5p | MIMAT0000656 | Cdk17 | 237459 | sRNA-Seq//HITS-CLIP | 33184237\|29167373 | Unspecified |
| mmu-miR-139-5p | MIMAT0000656 | Cdk18 | 18557 | HITS-CLIP | 29167373 | Unspecified |
| mmu-miR-139-5p | MIMAT0000656 | Cdk19 | 78334 | HITS-CLIP | 29167373 | Unspecified |
| mmu-miR-139-5p | MIMAT0000656 | Cdk6 | 12571 | HITS-CLIP | 29167373 | Unspecified |
| mmu-miR-139-5p | MIMAT0000656 | Cdkn1a | 12575 | HITS-CLIP//sRNA-Seq | 29167373\|33184237 | Unspecified |
| mmu-miR-139-5p | MIMAT0000656 | Cdkn1b | 12576 | HITS-CLIP | 29167373 | Unspecified |
| mmu-miR-139-5p | MIMAT0000656 | Cds2 | 110911 | sRNA-Seq//HITS-CLIP | 33184237\|29167373 | Unspecified |
| mmu-miR-139-5p | MIMAT0000656 | Cebpa | 12606 | HITS-CLIP | 26203562 | Unspecified |
| mmu-miR-139-5p | MIMAT0000656 | Cebpb | 12608 | HITS-CLIP | 29167373 | Unspecified |
| mmu-miR-139-5p | MIMAT0000656 | Celf1 | 13046 | HITS-CLIP | 29167373 | Unspecified |
| mmu-miR-139-5p | MIMAT0000656 | Celf2 | 14007 | HITS-CLIP | 29167373 | Unspecified |
| mmu-miR-139-5p | MIMAT0000656 | Cenpe | 229841 | HITS-CLIP | 29167373 | Unspecified |
| mmu-miR-139-5p | MIMAT0000656 | Cep162 | 382090 | HITS-CLIP | 29167373 | Unspecified |
| mmu-miR-139-5p | MIMAT0000656 | Cep170 | 545389 | sRNA-Seq//HITS-CLIP | 33184237\|29167373 | Unspecified |
| mmu-miR-139-5p | MIMAT0000656 | Cep192 | 70799 | HITS-CLIP | 29167373 | Unspecified |
| mmu-miR-139-5p | MIMAT0000656 | Cep250 | 16328 | HITS-CLIP | 29167373 | Unspecified |
| mmu-miR-139-5p | MIMAT0000656 | Cep350 | 74081 | HITS-CLIP | 29167373 | Unspecified |
| mmu-miR-139-5p | MIMAT0000656 | Cep78 | 208518 | HITS-CLIP | 29167373 | Unspecified |
| mmu-miR-139-5p | MIMAT0000656 | Cep83 | 77048 | HITS-CLIP | 29167373 | Unspecified |
| mmu-miR-139-5p | MIMAT0000656 | Cers2 | 76893 | HITS-CLIP | 29167373 | Unspecified |
| mmu-miR-139-5p | MIMAT0000656 | Cers6 | 241447 | sRNA-Seq//HITS-CLIP | 33184237\|29167373 | Unspecified |

| | | | | | | | |
|---|---|---|---|---|---|---|---|
| mmu-miR-139-5p | MIMAT0000656 | Cfap97 | 57587 | HITS-CLIP | | 29167373 | Unspecified |
| mmu-miR-139-5p | MIMAT0000656 | Cfl1 | 12631 | HITS-CLIP//sRNA-Seq | 31433990\|33184237 | | Unspecified |
| mmu-miR-139-5p | MIMAT0000656 | Cfl2 | 12632 | sRNA-Seq//HITS-CLIP | 33184237\|29167373 | | Unspecified |
| mmu-miR-139-5p | MIMAT0000656 | Cflar | 12633 | HITS-CLIP | | 29167373 | Unspecified |
| mmu-miR-139-5p | MIMAT0000656 | Cfp | 18636 | HITS-CLIP | | 29167373 | Unspecified |
| mmu-miR-139-5p | MIMAT0000656 | Cggbp1 | 106143 | HITS-CLIP | | 29167373 | Unspecified |
| mmu-miR-139-5p | MIMAT0000656 | Chd2 | 244059 | HITS-CLIP | | 29167373 | Unspecified |
| mmu-miR-139-5p | MIMAT0000656 | Chd3 | 216848 | sRNA-Seq//HITS-CLIP | 33184237\|29167373 | | Unspecified |
| mmu-miR-139-5p | MIMAT0000656 | Chd4 | 107932 | sRNA-Seq//HITS-CLIP | 33184237\|29167373 | | Unspecified |
| mmu-miR-139-5p | MIMAT0000656 | Chd7 | 320790 | HITS-CLIP | 31433990\|29167373 | | Unspecified |
| mmu-miR-139-5p | MIMAT0000656 | Chd8 | 67772 | sRNA-Seq//HITS-CLIP | 33184237\|29167373 | | Unspecified |
| mmu-miR-139-5p | MIMAT0000656 | Chek2 | 50883 | HITS-CLIP | | 29167373 | Unspecified |
| mmu-miR-139-5p | MIMAT0000656 | Chic2 | 74277 | HITS-CLIP | | 29167373 | Unspecified |
| mmu-miR-139-5p | MIMAT0000656 | Chm | 12662 | HITS-CLIP | | 29167373 | Unspecified |
| mmu-miR-139-5p | MIMAT0000656 | Cic | 71722 | sRNA-Seq//HITS-CLIP | 33184237\|29167373 | | Unspecified |
| mmu-miR-139-5p | MIMAT0000656 | Cipc | 217732 | HITS-CLIP | | 29167373 | Unspecified |
| mmu-miR-139-5p | MIMAT0000656 | Ckap5 | 75786 | sRNA-Seq//HITS-CLIP | 33184237\|29167373 | | Unspecified |
| mmu-miR-139-5p | MIMAT0000656 | Clasp2 | 76499 | sRNA-Seq//HITS-CLIP | 33184237\|29167373 | | Unspecified |
| mmu-miR-139-5p | MIMAT0000656 | Clcn3 | 12725 | sRNA-Seq//HITS-CLIP | 33184237\|29167373 | | Unspecified |
| mmu-miR-139-5p | MIMAT0000656 | Clcn5 | 12728 | sRNA-Seq//HITS-CLIP | 33184237\|29167373 | | Unspecified |
| mmu-miR-139-5p | MIMAT0000656 | Cldnd1 | 224250 | sRNA-Seq//HITS-CLIP | 33184237\|29167373 | | Unspecified |
| mmu-miR-139-5p | MIMAT0000656 | Clec12a | 160364 | HITS-CLIP | | 29167373 | Unspecified |
| mmu-miR-139-5p | MIMAT0000656 | Clec4a1 | 269799 | HITS-CLIP | | 29167373 | Unspecified |
| mmu-miR-139-5p | MIMAT0000656 | Clec4d | 17474 | HITS-CLIP | | 29167373 | Unspecified |
| mmu-miR-139-5p | MIMAT0000656 | Clic1 | 114584 | sRNA-Seq//HITS-CLIP | 33184237\|29167373 | | Unspecified |
| mmu-miR-139-5p | MIMAT0000656 | Clip1 | 56430 | sRNA-Seq//HITS-CLIP | 33184237\|29167373 | | Unspecified |
| mmu-miR-139-5p | MIMAT0000656 | Clip2 | 269713 | sRNA-Seq//HITS-CLIP | 33184237\|29167373 | | Unspecified |
| mmu-miR-139-5p | MIMAT0000656 | Clock | 12753 | HITS-CLIP | | 29167373 | Unspecified |
| mmu-miR-139-5p | MIMAT0000656 | Clta | 12757 | sRNA-Seq//HITS-CLIP | 33184237\|29167373 | | Unspecified |
| mmu-miR-139-5p | MIMAT0000656 | Cltc | 67300 | HITS-CLIP//sRNA-Seq | 31433990\|33184237\|26203562\|29167 | | Unspecified |
| mmu-miR-139-5p | MIMAT0000656 | Cluh | 74148 | HITS-CLIP | | 29167373 | Unspecified |
| mmu-miR-139-5p | MIMAT0000656 | Cmip | 74440 | sRNA-Seq//HITS-CLIP | 33184237\|29167373 | | Unspecified |
| mmu-miR-139-5p | MIMAT0000656 | Cmpk2 | 22169 | HITS-CLIP | | 29167373 | Unspecified |
| mmu-miR-139-5p | MIMAT0000656 | Cmtm6 | 67213 | HITS-CLIP | | 29167373 | Unspecified |
| mmu-miR-139-5p | MIMAT0000656 | Cmtm7 | 102545 | HITS-CLIP | | 31433990 | Unspecified |
| mmu-miR-139-5p | MIMAT0000656 | Cndp2 | 66054 | HITS-CLIP | | 29167373 | Unspecified |
| mmu-miR-139-5p | MIMAT0000656 | Cnih4 | 98417 | HITS-CLIP | | 29167373 | Unspecified |
| mmu-miR-139-5p | MIMAT0000656 | Cnnm3 | 94218 | HITS-CLIP | | 26203562 | Unspecified |
| mmu-miR-139-5p | MIMAT0000656 | Cnot1 | 234594 | sRNA-Seq//HITS-CLIP | 33184237\|29167373 | | Unspecified |

| | | | | | | | |
|---|---|---|---|---|---|---|---|
| mmu-miR-139-5p | MIMAT0000656 | Cnot4 | 53621 | HITS-CLIP | | 29167373 | Unspecified |
| mmu-miR-139-5p | MIMAT0000656 | Cnot6l | 231464 | HITS-CLIP | 26203562\|29167373 | | Unspecified |
| mmu-miR-139-5p | MIMAT0000656 | Cnot7 | 18983 | sRNA-Seq//HITS-CLIP | 33184237\|29167373 | | Unspecified |
| mmu-miR-139-5p | MIMAT0000656 | Cnot9 | 9125 | HITS-CLIP | | 29167373 | Unspecified |
| mmu-miR-139-5p | MIMAT0000656 | Cntrl | 26920 | HITS-CLIP | | 31433990 | Unspecified |
| mmu-miR-139-5p | MIMAT0000656 | Coa5 | 76178 | HITS-CLIP | | 29167373 | Unspecified |
| mmu-miR-139-5p | MIMAT0000656 | Col4a3bp | 68018 | HITS-CLIP | | 29167373 | Unspecified |
| mmu-miR-139-5p | MIMAT0000656 | Colec12 | 140792 | HITS-CLIP | | 29167373 | Unspecified |
| mmu-miR-139-5p | MIMAT0000656 | Commd8 | 27784 | HITS-CLIP//sRNA-Seq | 21258322\|33184237\|29167373\|25083 | | Unspecified |
| mmu-miR-139-5p | MIMAT0000656 | Copa | 12847 | sRNA-Seq//HITS-CLIP | 33184237\|29167373 | | Unspecified |
| mmu-miR-139-5p | MIMAT0000656 | Cops2 | 12848 | sRNA-Seq//HITS-CLIP | 33184237\|29167373 | | Unspecified |
| mmu-miR-139-5p | MIMAT0000656 | Cox14 | 66379 | sRNA-Seq//HITS-CLIP | 33184237\|29167373 | | Unspecified |
| mmu-miR-139-5p | MIMAT0000656 | Cox15 | 226139 | HITS-CLIP | | 29167373 | Unspecified |
| mmu-miR-139-5p | MIMAT0000656 | Cox17 | 12856 | sRNA-Seq//HITS-CLIP | 33184237\|29167373 | | Unspecified |
| mmu-miR-139-5p | MIMAT0000656 | Cox8a | 12868 | HITS-CLIP//sRNA-Seq | 31433990\|33184237 | | Unspecified |
| mmu-miR-139-5p | MIMAT0000656 | Cpeb2 | 231207 | HITS-CLIP | | 29167373 | Unspecified |
| mmu-miR-139-5p | MIMAT0000656 | Cpeb3 | 208922 | HITS-CLIP | | 29167373 | Unspecified |
| mmu-miR-139-5p | MIMAT0000656 | Cpeb4 | 67579 | sRNA-Seq//HITS-CLIP | 33184237\|29167373 | | Unspecified |
| mmu-miR-139-5p | MIMAT0000656 | Cpped1 | 223978 | HITS-CLIP | | 29167373 | Unspecified |
| mmu-miR-139-5p | MIMAT0000656 | Cpt1a | 12894 | HITS-CLIP | | 29167373 | Unspecified |
| mmu-miR-139-5p | MIMAT0000656 | Creb1 | 12912 | HITS-CLIP | | 29167373 | Unspecified |
| mmu-miR-139-5p | MIMAT0000656 | Creb5 | 231991 | HITS-CLIP | | 29167373 | Unspecified |
| mmu-miR-139-5p | MIMAT0000656 | Crebrf | 77128 | HITS-CLIP | | 29167373 | Unspecified |
| mmu-miR-139-5p | MIMAT0000656 | Creg1 | 433375 | HITS-CLIP | | 29167373 | Unspecified |
| mmu-miR-139-5p | MIMAT0000656 | Crim1 | 50766 | HITS-CLIP | | 23630078 | Unspecified |
| mmu-miR-139-5p | MIMAT0000656 | Cript | 56724 | sRNA-Seq//HITS-CLIP | 33184237\|29167373 | | Unspecified |
| mmu-miR-139-5p | MIMAT0000656 | Crk | 12928 | sRNA-Seq//HITS-CLIP | 33184237\|29167373 | | Unspecified |
| mmu-miR-139-5p | MIMAT0000656 | Crtc1 | 382056 | HITS-CLIP | | 29167373 | Unspecified |
| mmu-miR-139-5p | MIMAT0000656 | Cry2 | 12953 | HITS-CLIP | | 26203562 | Unspecified |
| mmu-miR-139-5p | MIMAT0000656 | Crybg3 | 224273 | HITS-CLIP | | 29167373 | Unspecified |
| mmu-miR-139-5p | MIMAT0000656 | Csde1 | 229663 | sRNA-Seq//HITS-CLIP | 33184237\|29167373 | | Unspecified |
| mmu-miR-139-5p | MIMAT0000656 | Csf1r | 12978 | HITS-CLIP | | 29167373 | Unspecified |
| mmu-miR-139-5p | MIMAT0000656 | Csf2ra | 12982 | HITS-CLIP | | 29167373 | Unspecified |
| mmu-miR-139-5p | MIMAT0000656 | Csf2rb | 12983 | HITS-CLIP | | 29167373 | Unspecified |
| mmu-miR-139-5p | MIMAT0000656 | Csk | 12988 | sRNA-Seq//HITS-CLIP | 33184237\|29167373 | | Unspecified |
| mmu-miR-139-5p | MIMAT0000656 | Csnk1a1 | 93687 | sRNA-Seq//HITS-CLIP | 33184237\|31433990\|29167373 | | Unspecified |
| mmu-miR-139-5p | MIMAT0000656 | Csnk1g1 | 214897 | sRNA-Seq//HITS-CLIP | 33184237\|29167373 | | Unspecified |
| mmu-miR-139-5p | MIMAT0000656 | Cspp1 | 211660 | sRNA-Seq//HITS-CLIP | 33184237\|29167373 | | Unspecified |
| mmu-miR-139-5p | MIMAT0000656 | Ctdp1 | 67655 | HITS-CLIP | | 29167373 | Unspecified |

| | | | | | | | |
|---|---|---|---|---|---|---|---|
| mmu-miR-139-5p | MIMAT0000656 | Ctdsp1 | 227292 | HITS-CLIP | | 31433990 | Unspecified |
| mmu-miR-139-5p | MIMAT0000656 | Ctdsp2 | 52468 | HITS-CLIP | 23630078\|31433990 | | Unspecified |
| mmu-miR-139-5p | MIMAT0000656 | Ctdspl2 | 329506 | HITS-CLIP | | 29167373 | Unspecified |
| mmu-miR-139-5p | MIMAT0000656 | Ctnna1 | 12385 | HITS-CLIP | | 31433990 | Unspecified |
| mmu-miR-139-5p | MIMAT0000656 | Ctnnb1 | 12387 | sRNA-Seq//HITS-CLIP | 33184237\|29167373 | | Unspecified |
| mmu-miR-139-5p | MIMAT0000656 | Ctnnd1 | 12388 | sRNA-Seq//HITS-CLIP | 33184237\|29167373 | | Unspecified |
| mmu-miR-139-5p | MIMAT0000656 | Ctnnd2 | 18163 | sRNA-Seq//HITS-CLIP | 33184237\|29167373 | | Unspecified |
| mmu-miR-139-5p | MIMAT0000656 | Ctr9 | 22083 | sRNA-Seq//HITS-CLIP | 33184237\|29167373 | | Unspecified |
| mmu-miR-139-5p | MIMAT0000656 | Ctsa | 19025 | HITS-CLIP | 25083871\|29167373 | | Unspecified |
| mmu-miR-139-5p | MIMAT0000656 | Ctsb | 13030 | sRNA-Seq//HITS-CLIP | 33184237\|29167373 | | Unspecified |
| mmu-miR-139-5p | MIMAT0000656 | Ctsc | 13032 | HITS-CLIP | | 29167373 | Unspecified |
| mmu-miR-139-5p | MIMAT0000656 | Ctsh | 13036 | HITS-CLIP | | 29167373 | Unspecified |
| mmu-miR-139-5p | MIMAT0000656 | Ctso | 229445 | HITS-CLIP | | 29167373 | Unspecified |
| mmu-miR-139-5p | MIMAT0000656 | Ctss | 13040 | HITS-CLIP | | 29167373 | Unspecified |
| mmu-miR-139-5p | MIMAT0000656 | Ctsz | 64138 | HITS-CLIP | | 29167373 | Unspecified |
| mmu-miR-139-5p | MIMAT0000656 | Cttnbp2nl | 80281 | HITS-CLIP | | 29167373 | Unspecified |
| mmu-miR-139-5p | MIMAT0000656 | Cul2 | 71745 | sRNA-Seq//HITS-CLIP | 33184237\|29167373 | | Unspecified |
| mmu-miR-139-5p | MIMAT0000656 | Cul3 | 26554 | sRNA-Seq//HITS-CLIP | 33184237\|29167373 | | Unspecified |
| mmu-miR-139-5p | MIMAT0000656 | Cul4a | 99375 | sRNA-Seq//HITS-CLIP | 33184237\|29167373 | | Unspecified |
| mmu-miR-139-5p | MIMAT0000656 | Cux1 | 13047 | HITS-CLIP | | 29167373 | Unspecified |
| mmu-miR-139-5p | MIMAT0000656 | Cwf19l2 | 244672 | HITS-CLIP | | 29167373 | Unspecified |
| mmu-miR-139-5p | MIMAT0000656 | Cx3cr1 | 13051 | HITS-CLIP | | 29167373 | Unspecified |
| mmu-miR-139-5p | MIMAT0000656 | Cyb561a3 | 225912 | HITS-CLIP | | 29167373 | Unspecified |
| mmu-miR-139-5p | MIMAT0000656 | Cyb5r3 | 109754 | sRNA-Seq//HITS-CLIP | 33184237\|29167373 | | Unspecified |
| mmu-miR-139-5p | MIMAT0000656 | Cybb | 13058 | HITS-CLIP | | 29167373 | Unspecified |
| mmu-miR-139-5p | MIMAT0000656 | Cyld | 74256 | HITS-CLIP | | 29167373 | Unspecified |
| mmu-miR-139-5p | MIMAT0000656 | Cyth1 | 19157 | HITS-CLIP | | 29167373 | Unspecified |
| mmu-miR-139-5p | MIMAT0000656 | D17Wsu92e | 224647 | HITS-CLIP | | 29167373 | Unspecified |
| mmu-miR-139-5p | MIMAT0000656 | D1Ertd622e | 52392 | HITS-CLIP | | 29167373 | Unspecified |
| mmu-miR-139-5p | MIMAT0000656 | Dab2 | 13132 | HITS-CLIP | | 29167373 | Unspecified |
| mmu-miR-139-5p | MIMAT0000656 | Dagla | 269060 | HITS-CLIP | | 29167373 | Unspecified |
| mmu-miR-139-5p | MIMAT0000656 | Dars | 226414 | sRNA-Seq//HITS-CLIP | 33184237\|29167373 | | Unspecified |
| mmu-miR-139-5p | MIMAT0000656 | Dbt | 13171 | HITS-CLIP | | 29167373 | Unspecified |
| mmu-miR-139-5p | MIMAT0000656 | Dcaf12 | 68970 | HITS-CLIP | 31433990\|29167373 | | Unspecified |
| mmu-miR-139-5p | MIMAT0000656 | Dcaf4 | 73828 | HITS-CLIP | | 29167373 | Unspecified |
| mmu-miR-139-5p | MIMAT0000656 | Dcakd | 68087 | HITS-CLIP | | 29167373 | Unspecified |
| mmu-miR-139-5p | MIMAT0000656 | Dclre1c | 227525 | HITS-CLIP | | 29167373 | Unspecified |
| mmu-miR-139-5p | MIMAT0000656 | Dcp1a | 75901 | HITS-CLIP | | 29167373 | Unspecified |
| mmu-miR-139-5p | MIMAT0000656 | Dcp1b | 319618 | HITS-CLIP | | 29167373 | Unspecified |

| | | | | | | | |
|---|---|---|---|---|---|---|---|
| mmu-miR-139-5p | MIMAT0000656 | Dcp2 | 70640 | HITS-CLIP | | 29167373 | Unspecified |
| mmu-miR-139-5p | MIMAT0000656 | Dctn3 | 53598 | sRNA-Seq//HITS-CLIP | 33184237\|29167373 | | Unspecified |
| mmu-miR-139-5p | MIMAT0000656 | Dctn4 | 67665 | sRNA-Seq//HITS-CLIP | 33184237\|29167373 | | Unspecified |
| mmu-miR-139-5p | MIMAT0000656 | Ddb1 | 13194 | sRNA-Seq//HITS-CLIP | 33184237\|29167373 | | Unspecified |
| mmu-miR-139-5p | MIMAT0000656 | Ddi2 | 68817 | HITS-CLIP | | 29167373 | Unspecified |
| mmu-miR-139-5p | MIMAT0000656 | Ddit4 | 74747 | HITS-CLIP | | 26203562 | Unspecified |
| mmu-miR-139-5p | MIMAT0000656 | Ddx17 | 67040 | HITS-CLIP//sRNA-Seq | 29167373\|33184237 | | Unspecified |
| mmu-miR-139-5p | MIMAT0000656 | Ddx24 | 27225 | sRNA-Seq//HITS-CLIP | 33184237\|29167373 | | Unspecified |
| mmu-miR-139-5p | MIMAT0000656 | Ddx3x | 13205 | sRNA-Seq//HITS-CLIP | 33184237\|29167373 | | Unspecified |
| mmu-miR-139-5p | MIMAT0000656 | Ddx3y | 26900 | HITS-CLIP | | 31433990 | Unspecified |
| mmu-miR-139-5p | MIMAT0000656 | Ddx46 | 212880 | HITS-CLIP | | 29167373 | Unspecified |
| mmu-miR-139-5p | MIMAT0000656 | Ddx58 | 230073 | HITS-CLIP | | 29167373 | Unspecified |
| mmu-miR-139-5p | MIMAT0000656 | Ddx6 | 13209 | sRNA-Seq//HITS-CLIP | 33184237\|29167373 | | Unspecified |
| mmu-miR-139-5p | MIMAT0000656 | Degs1 | 13244 | sRNA-Seq//HITS-CLIP | 33184237\|29167373 | | Unspecified |
| mmu-miR-139-5p | MIMAT0000656 | Dennd1a | 227801 | sRNA-Seq//HITS-CLIP | 33184237\|29167373 | | Unspecified |
| mmu-miR-139-5p | MIMAT0000656 | Dennd1b | 329260 | HITS-CLIP | | 31433990 | Unspecified |
| mmu-miR-139-5p | MIMAT0000656 | Dennd4b | 229541 | HITS-CLIP | 31433990\|29167373 | | Unspecified |
| mmu-miR-139-5p | MIMAT0000656 | Dennd5a | 19347 | HITS-CLIP | | 29167373 | Unspecified |
| mmu-miR-139-5p | MIMAT0000656 | Depdc5 | 277854 | HITS-CLIP | | 31433990 | Unspecified |
| mmu-miR-139-5p | MIMAT0000656 | Dgkd | 227333 | HITS-CLIP | | 29167373 | Unspecified |
| mmu-miR-139-5p | MIMAT0000656 | Dgkz | 104418 | sRNA-Seq//HITS-CLIP | 33184237\|29167373 | | Unspecified |
| mmu-miR-139-5p | MIMAT0000656 | Dhrs3 | 20148 | HITS-CLIP | | 29167373 | Unspecified |
| mmu-miR-139-5p | MIMAT0000656 | Dhx15 | 13204 | sRNA-Seq//HITS-CLIP | 33184237\|29167373 | | Unspecified |
| mmu-miR-139-5p | MIMAT0000656 | Dhx33 | 216877 | HITS-CLIP | | 29167373 | Unspecified |
| mmu-miR-139-5p | MIMAT0000656 | Dhx9 | 13211 | sRNA-Seq//HITS-CLIP | 33184237\|29167373 | | Unspecified |
| mmu-miR-139-5p | MIMAT0000656 | Diaph1 | 1729 | HITS-CLIP | | 29167373 | Unspecified |
| mmu-miR-139-5p | MIMAT0000656 | Dido1 | 23856 | HITS-CLIP | | 29167373 | Unspecified |
| mmu-miR-139-5p | MIMAT0000656 | Dis3l | 213550 | HITS-CLIP | | 31433990 | Unspecified |
| mmu-miR-139-5p | MIMAT0000656 | Dlg1 | 13383 | HITS-CLIP | | 29167373 | Unspecified |
| mmu-miR-139-5p | MIMAT0000656 | Dmxl1 | 240283 | HITS-CLIP | | 29167373 | Unspecified |
| mmu-miR-139-5p | MIMAT0000656 | Dnaja2 | 56445 | sRNA-Seq//HITS-CLIP | 33184237\|29167373 | | Unspecified |
| mmu-miR-139-5p | MIMAT0000656 | Dnajb11 | 67838 | HITS-CLIP | | 29167373 | Unspecified |
| mmu-miR-139-5p | MIMAT0000656 | Dnajb14 | 70604 | HITS-CLIP | | 29167373 | Unspecified |
| mmu-miR-139-5p | MIMAT0000656 | Dnajc13 | 235567 | sRNA-Seq//HITS-CLIP | 33184237\|26203562 | | Unspecified |
| mmu-miR-139-5p | MIMAT0000656 | Dnajc16 | 214063 | HITS-CLIP | | 29167373 | Unspecified |
| mmu-miR-139-5p | MIMAT0000656 | Dnajc3 | 100037258 | HITS-CLIP | | 29167373 | Unspecified |
| mmu-miR-139-5p | MIMAT0000656 | Dnajc5 | 13002 | sRNA-Seq//HITS-CLIP | 33184237\|29167373 | | Unspecified |
| mmu-miR-139-5p | MIMAT0000656 | Dnajc7 | 56354 | sRNA-Seq//HITS-CLIP | 33184237\|29167373 | | Unspecified |
| mmu-miR-139-5p | MIMAT0000656 | Dnm2 | 13430 | HITS-CLIP | | 29167373 | Unspecified |

| | | | | | | |
|---|---|---|---|---|---|---|
| mmu-miR-139-5p | MIMAT0000656 | Dnmt1 | 13433 | HITS-CLIP | 29167373 | Unspecified |
| mmu-miR-139-5p | MIMAT0000656 | Dnmt3a | 13435 | HITS-CLIP | 29167373 | Unspecified |
| mmu-miR-139-5p | MIMAT0000656 | Dock10 | 210293 | HITS-CLIP | 31433990\|29167373 | Unspecified |
| mmu-miR-139-5p | MIMAT0000656 | Dock2 | 94176 | HITS-CLIP | 29167373 | Unspecified |
| mmu-miR-139-5p | MIMAT0000656 | Dock4 | 238130 | sRNA-Seq//HITS-CLIP | 33184237\|29167373 | Unspecified |
| mmu-miR-139-5p | MIMAT0000656 | Dock7 | 67299 | sRNA-Seq//HITS-CLIP | 33184237\|29167373 | Unspecified |
| mmu-miR-139-5p | MIMAT0000656 | Dolk | 227697 | HITS-CLIP | 29167373 | Unspecified |
| mmu-miR-139-5p | MIMAT0000656 | Dop1b | 9980 | HITS-CLIP | 29167373 | Unspecified |
| mmu-miR-139-5p | MIMAT0000656 | Dot1l | 208266 | HITS-CLIP | 29167373 | Unspecified |
| mmu-miR-139-5p | MIMAT0000656 | Dram1 | 71712 | HITS-CLIP | 29167373 | Unspecified |
| mmu-miR-139-5p | MIMAT0000656 | Dram2 | 67171 | sRNA-Seq//HITS-CLIP | 33184237\|29167373 | Unspecified |
| mmu-miR-139-5p | MIMAT0000656 | Drg1 | 13494 | sRNA-Seq//HITS-CLIP | 33184237\|29167373 | Unspecified |
| mmu-miR-139-5p | MIMAT0000656 | Dse | 212898 | HITS-CLIP | 29167373 | Unspecified |
| mmu-miR-139-5p | MIMAT0000656 | Dst | 13518 | sRNA-Seq//HITS-CLIP | 33184237\|29167373 | Unspecified |
| mmu-miR-139-5p | MIMAT0000656 | Dstn | 56431 | sRNA-Seq//HITS-CLIP | 33184237\|29167373 | Unspecified |
| mmu-miR-139-5p | MIMAT0000656 | Dtnbp1 | 94245 | HITS-CLIP | 29167373 | Unspecified |
| mmu-miR-139-5p | MIMAT0000656 | Dusp1 | 19252 | HITS-CLIP | 29167373 | Unspecified |
| mmu-miR-139-5p | MIMAT0000656 | Dusp3 | 72349 | HITS-CLIP//sRNA-Seq | 21258322\|33184237\|29167373 | Unspecified |
| mmu-miR-139-5p | MIMAT0000656 | Dusp4 | 319520 | HITS-CLIP | 29167373 | Unspecified |
| mmu-miR-139-5p | MIMAT0000656 | Dusp5 | 240672 | HITS-CLIP | 31433990 | Unspecified |
| mmu-miR-139-5p | MIMAT0000656 | Dync1h1 | 13424 | sRNA-Seq//HITS-CLIP | 33184237\|29167373 | Unspecified |
| mmu-miR-139-5p | MIMAT0000656 | Dync1i2 | 13427 | sRNA-Seq//HITS-CLIP | 33184237\|29167373 | Unspecified |
| mmu-miR-139-5p | MIMAT0000656 | Dync1li2 | 234663 | sRNA-Seq//HITS-CLIP | 33184237\|29167373 | Unspecified |
| mmu-miR-139-5p | MIMAT0000656 | Dynll2 | 68097 | HITS-CLIP//sRNA-Seq | 31433990\|33184237\|29167373 | Unspecified |
| mmu-miR-139-5p | MIMAT0000656 | Dyrk1a | 13548 | HITS-CLIP | 29167373 | Unspecified |
| mmu-miR-139-5p | MIMAT0000656 | Dyrk2 | 69181 | HITS-CLIP | 29167373 | Unspecified |
| mmu-miR-139-5p | MIMAT0000656 | E130311K13Rik | 329659 | HITS-CLIP | 26203562 | Unspecified |
| mmu-miR-139-5p | MIMAT0000656 | Ece1 | 230857 | sRNA-Seq//HITS-CLIP | 33184237\|29167373 | Unspecified |
| mmu-miR-139-5p | MIMAT0000656 | Ecm1 | 13601 | HITS-CLIP | 29167373 | Unspecified |
| mmu-miR-139-5p | MIMAT0000656 | Ecpas | 23392 | sRNA-Seq//HITS-CLIP | 33184237\|29167373 | Unspecified |
| mmu-miR-139-5p | MIMAT0000656 | Edc4 | 234699 | HITS-CLIP | 29167373 | Unspecified |
| mmu-miR-139-5p | MIMAT0000656 | Edem1 | 192193 | HITS-CLIP | 29167373 | Unspecified |
| mmu-miR-139-5p | MIMAT0000656 | Edrf1 | 214764 | HITS-CLIP | 31433990 | Unspecified |
| mmu-miR-139-5p | MIMAT0000656 | Eea1 | 216238 | HITS-CLIP | 31433990\|29167373 | Unspecified |
| mmu-miR-139-5p | MIMAT0000656 | Eef2 | 13629 | sRNA-Seq//HITS-CLIP | 33184237\|29167373 | Unspecified |
| mmu-miR-139-5p | MIMAT0000656 | Efcab14 | 230648 | HITS-CLIP | 29167373 | Unspecified |
| mmu-miR-139-5p | MIMAT0000656 | Efhd2 | 27984 | HITS-CLIP | 31433990\|29167373 | Unspecified |
| mmu-miR-139-5p | MIMAT0000656 | Ehd4 | 98878 | HITS-CLIP | 29167373 | Unspecified |
| mmu-miR-139-5p | MIMAT0000656 | Ehmt1 | 77683 | HITS-CLIP | 29167373 | Unspecified |

| | | | | | | | |
|---|---|---|---|---|---|---|---|
| mmu-miR-139-5p | MIMAT0000656 | Eif2ak1 | 15467 | HITS-CLIP | | 29167373 | Unspecified |
| mmu-miR-139-5p | MIMAT0000656 | Eif2ak2 | 19106 | HITS-CLIP | | 29167373 | Unspecified |
| mmu-miR-139-5p | MIMAT0000656 | Eif2ak4 | 27103 | HITS-CLIP | | 29167373 | Unspecified |
| mmu-miR-139-5p | MIMAT0000656 | Eif3a | 13669 | sRNA-Seq//HITS-CLIP | 33184237\|29167373 | | Unspecified |
| mmu-miR-139-5p | MIMAT0000656 | Eif3b | 27979 | sRNA-Seq//HITS-CLIP | 33184237\|29167373 | | Unspecified |
| mmu-miR-139-5p | MIMAT0000656 | Eif3c | 56347 | sRNA-Seq//HITS-CLIP | 33184237\|29167373 | | Unspecified |
| mmu-miR-139-5p | MIMAT0000656 | Eif3m | 98221 | sRNA-Seq//HITS-CLIP | 33184237\|29167373 | | Unspecified |
| mmu-miR-139-5p | MIMAT0000656 | Eif4a2 | 13682 | sRNA-Seq//HITS-CLIP | 33184237\|29167373 | | Unspecified |
| mmu-miR-139-5p | MIMAT0000656 | Eif4a3 | 192170 | HITS-CLIP//sRNA-Seq | 31433990\|33184237 | | Unspecified |
| mmu-miR-139-5p | MIMAT0000656 | Eif4e | 13684 | sRNA-Seq//HITS-CLIP | 33184237\|29167373 | | Unspecified |
| mmu-miR-139-5p | MIMAT0000656 | Eif4ebp2 | 13688 | HITS-CLIP | | 29167373 | Unspecified |
| mmu-miR-139-5p | MIMAT0000656 | Eif4g1 | 208643 | sRNA-Seq//HITS-CLIP | 33184237\|29167373 | | Unspecified |
| mmu-miR-139-5p | MIMAT0000656 | Eif4g2 | 13690 | sRNA-Seq//HITS-CLIP | 33184237\|29167373 | | Unspecified |
| mmu-miR-139-5p | MIMAT0000656 | Eif4g3 | 230861 | sRNA-Seq//HITS-CLIP | 33184237\|29167373 | | Unspecified |
| mmu-miR-139-5p | MIMAT0000656 | Eif4h | 22384 | sRNA-Seq//HITS-CLIP | 33184237\|29167373 | | Unspecified |
| mmu-miR-139-5p | MIMAT0000656 | Eif5 | 217869 | sRNA-Seq//HITS-CLIP | 33184237\|29167373 | | Unspecified |
| mmu-miR-139-5p | MIMAT0000656 | Eif5a | 276770 | sRNA-Seq//HITS-CLIP | 33184237\|29167373 | | Unspecified |
| mmu-miR-139-5p | MIMAT0000656 | Eif5b | 226982 | sRNA-Seq//HITS-CLIP | 33184237\|29167373 | | Unspecified |
| mmu-miR-139-5p | MIMAT0000656 | Elf2 | 69257 | HITS-CLIP | | 29167373 | Unspecified |
| mmu-miR-139-5p | MIMAT0000656 | Elk3 | 13713 | HITS-CLIP | | 29167373 | Unspecified |
| mmu-miR-139-5p | MIMAT0000656 | Elk4 | 13714 | HITS-CLIP | | 29167373 | Unspecified |
| mmu-miR-139-5p | MIMAT0000656 | Ell | 13716 | HITS-CLIP | | 29167373 | Unspecified |
| mmu-miR-139-5p | MIMAT0000656 | Ell2 | 192657 | HITS-CLIP | | 29167373 | Unspecified |
| mmu-miR-139-5p | MIMAT0000656 | Elmo1 | 140580 | HITS-CLIP | | 29167373 | Unspecified |
| mmu-miR-139-5p | MIMAT0000656 | Elmsan1 | 238317 | HITS-CLIP | | 29167373 | Unspecified |
| mmu-miR-139-5p | MIMAT0000656 | Eloc | 6921 | sRNA-Seq//HITS-CLIP | 33184237\|29167373 | | Unspecified |
| mmu-miR-139-5p | MIMAT0000656 | Elovl1 | 54325 | HITS-CLIP | | 29167373 | Unspecified |
| mmu-miR-139-5p | MIMAT0000656 | Elp3 | 74195 | sRNA-Seq//HITS-CLIP | 33184237\|29167373 | | Unspecified |
| mmu-miR-139-5p | MIMAT0000656 | Emilin2 | 246707 | HITS-CLIP | | 29167373 | Unspecified |
| mmu-miR-139-5p | MIMAT0000656 | Eml4 | 78798 | HITS-CLIP | | 29167373 | Unspecified |
| mmu-miR-139-5p | MIMAT0000656 | Emsy | 56946 | HITS-CLIP | | 29167373 | Unspecified |
| mmu-miR-139-5p | MIMAT0000656 | Enc1 | 13803 | sRNA-Seq//HITS-CLIP | 33184237\|29167373 | | Unspecified |
| mmu-miR-139-5p | MIMAT0000656 | Enoph1 | 67870 | HITS-CLIP | | 29167373 | Unspecified |
| mmu-miR-139-5p | MIMAT0000656 | Ensa | 56205 | sRNA-Seq//HITS-CLIP | 33184237\|29167373 | | Unspecified |
| mmu-miR-139-5p | MIMAT0000656 | Ep300 | 328572 | HITS-CLIP | | 29167373 | Unspecified |
| mmu-miR-139-5p | MIMAT0000656 | Ep400 | 75560 | HITS-CLIP | | 29167373 | Unspecified |
| mmu-miR-139-5p | MIMAT0000656 | Epb41 | 269587 | HITS-CLIP | | 29167373 | Unspecified |
| mmu-miR-139-5p | MIMAT0000656 | Epb41l1 | 13821 | sRNA-Seq//HITS-CLIP | 33184237\|29167373 | | Unspecified |
| mmu-miR-139-5p | MIMAT0000656 | Epb41l2 | 2037 | HITS-CLIP | | 29167373 | Unspecified |

| | | | | | | | |
|---|---|---|---|---|---|---|---|
| mmu-miR-139-5p | MIMAT0000656 | Epc1 | 13831 | HITS-CLIP | | 29167373 | Unspecified |
| mmu-miR-139-5p | MIMAT0000656 | Epg5 | 100502841 | HITS-CLIP | | 29167373 | Unspecified |
| mmu-miR-139-5p | MIMAT0000656 | Eppk1 | 3857 | HITS-CLIP | | 29167373 | Unspecified |
| mmu-miR-139-5p | MIMAT0000656 | Eprs | 107508 | sRNA-Seq//HITS-CLIP | 33184237\|29167373 | | Unspecified |
| mmu-miR-139-5p | MIMAT0000656 | Eps15 | 13858 | sRNA-Seq//HITS-CLIP | 33184237\|29167373 | | Unspecified |
| mmu-miR-139-5p | MIMAT0000656 | Epsti1 | 108670 | HITS-CLIP | | 29167373 | Unspecified |
| mmu-miR-139-5p | MIMAT0000656 | Erap1 | 80898 | HITS-CLIP | | 29167373 | Unspecified |
| mmu-miR-139-5p | MIMAT0000656 | Erbb3 | 13867 | HITS-CLIP | | 26203562 | Unspecified |
| mmu-miR-139-5p | MIMAT0000656 | Ergic2 | 67456 | sRNA-Seq//HITS-CLIP | 33184237\|29167373 | | Unspecified |
| mmu-miR-139-5p | MIMAT0000656 | Erich1 | 234086 | HITS-CLIP | | 31433990 | Unspecified |
| mmu-miR-139-5p | MIMAT0000656 | Erlec1 | 66753 | HITS-CLIP | | 29167373 | Unspecified |
| mmu-miR-139-5p | MIMAT0000656 | Erlin1 | 226144 | HITS-CLIP | | 29167373 | Unspecified |
| mmu-miR-139-5p | MIMAT0000656 | Esco1 | 77805 | HITS-CLIP | | 30971389 | Unspecified |
| mmu-miR-139-5p | MIMAT0000656 | Etnk1 | 75320 | sRNA-Seq//HITS-CLIP | 33184237\|29167373 | | Unspecified |
| mmu-miR-139-5p | MIMAT0000656 | Ets1 | 23871 | HITS-CLIP | | 29167373 | Unspecified |
| mmu-miR-139-5p | MIMAT0000656 | Ets2 | 23872 | HITS-CLIP | | 29167373 | Unspecified |
| mmu-miR-139-5p | MIMAT0000656 | Etv3 | 27049 | HITS-CLIP | | 29167373 | Unspecified |
| mmu-miR-139-5p | MIMAT0000656 | Evi5 | 14020 | HITS-CLIP | | 29167373 | Unspecified |
| mmu-miR-139-5p | MIMAT0000656 | Ewsr1 | 14030 | sRNA-Seq//HITS-CLIP | 33184237\|29167373 | | Unspecified |
| mmu-miR-139-5p | MIMAT0000656 | Exoc5 | 105504 | sRNA-Seq//HITS-CLIP | 33184237\|29167373 | | Unspecified |
| mmu-miR-139-5p | MIMAT0000656 | Exoc8 | 102058 | HITS-CLIP | | 29167373 | Unspecified |
| mmu-miR-139-5p | MIMAT0000656 | Exosc2 | 227715 | HITS-CLIP | | 29167373 | Unspecified |
| mmu-miR-139-5p | MIMAT0000656 | Extl3 | 54616 | HITS-CLIP//sRNA-Seq | 29167373\|33184237 | | Unspecified |
| mmu-miR-139-5p | MIMAT0000656 | F11r | 16456 | HITS-CLIP | | 29167373 | Unspecified |
| mmu-miR-139-5p | MIMAT0000656 | F13a1 | 74145 | HITS-CLIP | | 29167373 | Unspecified |
| mmu-miR-139-5p | MIMAT0000656 | F2r | 14062 | HITS-CLIP//sRNA-Seq | 31433990\|33184237 | | Unspecified |
| mmu-miR-139-5p | MIMAT0000656 | Fam111a | 107373 | HITS-CLIP | | 29167373 | Unspecified |
| mmu-miR-139-5p | MIMAT0000656 | Fam117b | 72750 | HITS-CLIP | | 26203562 | Unspecified |
| mmu-miR-139-5p | MIMAT0000656 | Fam120a | 218236 | HITS-CLIP | | 29167373 | Unspecified |
| mmu-miR-139-5p | MIMAT0000656 | Fam129a | 63913 | HITS-CLIP | | 29167373 | Unspecified |
| mmu-miR-139-5p | MIMAT0000656 | Fam168b | 214469 | sRNA-Seq//HITS-CLIP | 33184237\|29167373 | | Unspecified |
| mmu-miR-139-5p | MIMAT0000656 | Fam193a | 231128 | HITS-CLIP | | 29167373 | Unspecified |
| mmu-miR-139-5p | MIMAT0000656 | Fam208a | 218850 | HITS-CLIP | | 29167373 | Unspecified |
| mmu-miR-139-5p | MIMAT0000656 | Fam208b | 54906 | HITS-CLIP | | 29167373 | Unspecified |
| mmu-miR-139-5p | MIMAT0000656 | Fam214b | 230088 | HITS-CLIP | | 29167373 | Unspecified |
| mmu-miR-139-5p | MIMAT0000656 | Fam49b | 223601 | sRNA-Seq//HITS-CLIP | 33184237\|29167373 | | Unspecified |
| mmu-miR-139-5p | MIMAT0000656 | Fam53c | 66306 | HITS-CLIP | | 29167373 | Unspecified |
| mmu-miR-139-5p | MIMAT0000656 | Fam69a | 388650 | HITS-CLIP | | 29167373 | Unspecified |
| mmu-miR-139-5p | MIMAT0000656 | Fam91a1 | 210998 | HITS-CLIP | | 29167373 | Unspecified |

| | | | | | | |
|---|---|---|---|---|---|---|
| mmu-miR-139-5p | MIMAT0000656 | Fancm | 104806 | HITS-CLIP | 29167373 | Unspecified |
| mmu-miR-139-5p | MIMAT0000656 | Far1 | 67420 | sRNA-Seq//HITS-CLIP | 33184237\|29167373 | Unspecified |
| mmu-miR-139-5p | MIMAT0000656 | Farp2 | 227377 | HITS-CLIP | 29167373 | Unspecified |
| mmu-miR-139-5p | MIMAT0000656 | Farsb | 23874 | sRNA-Seq//HITS-CLIP | 33184237\|29167373 | Unspecified |
| mmu-miR-139-5p | MIMAT0000656 | Fasn | 14104 | sRNA-Seq//HITS-CLIP | 33184237\|31433990 | Unspecified |
| mmu-miR-139-5p | MIMAT0000656 | Fat3 | 270120 | HITS-CLIP | 29167373 | Unspecified |
| mmu-miR-139-5p | MIMAT0000656 | Fbxl14 | 101358 | HITS-CLIP | 29167373 | Unspecified |
| mmu-miR-139-5p | MIMAT0000656 | Fbxo33 | 70611 | HITS-CLIP | 29167373 | Unspecified |
| mmu-miR-139-5p | MIMAT0000656 | Fbxo38 | 107035 | HITS-CLIP | 29167373 | Unspecified |
| mmu-miR-139-5p | MIMAT0000656 | Fbxw11 | 103583 | sRNA-Seq//HITS-CLIP | 33184237\|29167373 | Unspecified |
| mmu-miR-139-5p | MIMAT0000656 | Fcgr1 | 14129 | HITS-CLIP | 29167373 | Unspecified |
| mmu-miR-139-5p | MIMAT0000656 | Fech | 14151 | HITS-CLIP | 29167373 | Unspecified |
| mmu-miR-139-5p | MIMAT0000656 | Fem1b | 14155 | sRNA-Seq//HITS-CLIP | 33184237\|29167373 | Unspecified |
| mmu-miR-139-5p | MIMAT0000656 | Fem1c | 240263 | HITS-CLIP | 29167373 | Unspecified |
| mmu-miR-139-5p | MIMAT0000656 | Fgd4 | 224014 | sRNA-Seq//HITS-CLIP | 33184237\|29167373 | Unspecified |
| mmu-miR-139-5p | MIMAT0000656 | Fgfr1op | 75296 | HITS-CLIP | 29167373 | Unspecified |
| mmu-miR-139-5p | MIMAT0000656 | Fhip2a | 57700 | HITS-CLIP | 31433990 | Unspecified |
| mmu-miR-139-5p | MIMAT0000656 | Filip1l | 78749 | HITS-CLIP | 29167373 | Unspecified |
| mmu-miR-139-5p | MIMAT0000656 | Fiz1 | 23877 | HITS-CLIP | 31433990 | Unspecified |
| mmu-miR-139-5p | MIMAT0000656 | Flii | 14248 | sRNA-Seq//HITS-CLIP | 33184237\|29167373 | Unspecified |
| mmu-miR-139-5p | MIMAT0000656 | Flna | 192176 | HITS-CLIP | 29167373 | Unspecified |
| mmu-miR-139-5p | MIMAT0000656 | Fmn1 | 14260 | HITS-CLIP | 29167373 | Unspecified |
| mmu-miR-139-5p | MIMAT0000656 | Fmnl1 | 57778 | HITS-CLIP | 29167373 | Unspecified |
| mmu-miR-139-5p | MIMAT0000656 | Fmnl2 | 71409 | HITS-CLIP | 26203562\|29167373 | Unspecified |
| mmu-miR-139-5p | MIMAT0000656 | Fnbp1 | 14269 | sRNA-Seq//HITS-CLIP | 33184237\|29167373 | Unspecified |
| mmu-miR-139-5p | MIMAT0000656 | Fndc3a | 319448 | HITS-CLIP | 29167373 | Unspecified |
| mmu-miR-139-5p | MIMAT0000656 | Fndc3b | 72007 | HITS-CLIP | 29167373 | Unspecified |
| mmu-miR-139-5p | MIMAT0000656 | Fnip1 | 216742 | HITS-CLIP | 31433990\|29167373 | Unspecified |
| mmu-miR-139-5p | MIMAT0000656 | Fnip2 | 329679 | HITS-CLIP | 29167373 | Unspecified |
| mmu-miR-139-5p | MIMAT0000656 | Folr2 | 14276 | HITS-CLIP | 29167373 | Unspecified |
| mmu-miR-139-5p | MIMAT0000656 | Fos | 14281 | HITS-CLIP | 29167373 | Unspecified |
| mmu-miR-139-5p | MIMAT0000656 | FoxO1 | 56458 | Western Blot//RT-PCR//Luciferase assay//I | 19883627\|29167373 | Unspecified |
| mmu-miR-139-5p | MIMAT0000656 | Foxk1 | 17425 | HITS-CLIP | 30971389\|31433990 | Unspecified |
| mmu-miR-139-5p | MIMAT0000656 | Foxk2 | 68837 | sRNA-Seq//HITS-CLIP | 33184237\|29167373 | Unspecified |
| mmu-miR-139-5p | MIMAT0000656 | Foxn3 | 71375 | sRNA-Seq//HITS-CLIP | 33184237\|29167373 | Unspecified |
| mmu-miR-139-5p | MIMAT0000656 | Foxo1 | 56458 | Luciferase assay//qRT-PCR//Western blot/ | 19883627\|29167373 | Unspecified |
| mmu-miR-139-5p | MIMAT0000656 | Foxp1 | 108655 | HITS-CLIP | 29167373 | Unspecified |
| mmu-miR-139-5p | MIMAT0000656 | Frmd4a | 209630 | sRNA-Seq//HITS-CLIP | 33184237\|29167373 | Unspecified |
| mmu-miR-139-5p | MIMAT0000656 | Frrs1 | 20321 | HITS-CLIP | 29167373 | Unspecified |

| | | | | | | |
|---|---|---|---|---|---|---|
| mmu-miR-139-5p | MIMAT0000656 | Frs2 | 327826 | HITS-CLIP | 29167373 | Unspecified |
| mmu-miR-139-5p | MIMAT0000656 | Fsbp | 100503583 | HITS-CLIP | 30971389 | Unspecified |
| mmu-miR-139-5p | MIMAT0000656 | Fth1 | 14319 | HITS-CLIP//sRNA-Seq | 29167373\|33184237 | Unspecified |
| mmu-miR-139-5p | MIMAT0000656 | Fubp1 | 51886 | sRNA-Seq//HITS-CLIP | 33184237\|29167373 | Unspecified |
| mmu-miR-139-5p | MIMAT0000656 | Fuca2 | 66848 | HITS-CLIP | 29167373 | Unspecified |
| mmu-miR-139-5p | MIMAT0000656 | Fundc1 | 72018 | HITS-CLIP | 29167373 | Unspecified |
| mmu-miR-139-5p | MIMAT0000656 | Furin | 18550 | HITS-CLIP | 29167373 | Unspecified |
| mmu-miR-139-5p | MIMAT0000656 | Fxyd5 | 18301 | HITS-CLIP | 29167373 | Unspecified |
| mmu-miR-139-5p | MIMAT0000656 | Fyb | 23880 | HITS-CLIP | 29167373 | Unspecified |
| mmu-miR-139-5p | MIMAT0000656 | Fyco1 | 17281 | HITS-CLIP | 31433990 | Unspecified |
| mmu-miR-139-5p | MIMAT0000656 | Fyttd1 | 69823 | HITS-CLIP//sRNA-Seq | 23630078\|33184237 | Unspecified |
| mmu-miR-139-5p | MIMAT0000656 | G2e3 | 217558 | HITS-CLIP | 29167373 | Unspecified |
| mmu-miR-139-5p | MIMAT0000656 | G3bp1 | 27041 | HITS-CLIP | 29167373 | Unspecified |
| mmu-miR-139-5p | MIMAT0000656 | G3bp2 | 23881 | sRNA-Seq//HITS-CLIP | 33184237\|29167373 | Unspecified |
| mmu-miR-139-5p | MIMAT0000656 | Gabpb2 | 213054 | HITS-CLIP | 29167373 | Unspecified |
| mmu-miR-139-5p | MIMAT0000656 | Gak | 231580 | sRNA-Seq//HITS-CLIP | 33184237\|29167373 | Unspecified |
| mmu-miR-139-5p | MIMAT0000656 | Galk1 | 14635 | HITS-CLIP | 31433990 | Unspecified |
| mmu-miR-139-5p | MIMAT0000656 | Galnt7 | 108150 | HITS-CLIP | 29167373 | Unspecified |
| mmu-miR-139-5p | MIMAT0000656 | Gan | 209239 | HITS-CLIP | 29167373 | Unspecified |
| mmu-miR-139-5p | MIMAT0000656 | Gapvd1 | 66691 | sRNA-Seq//HITS-CLIP | 33184237\|29167373 | Unspecified |
| mmu-miR-139-5p | MIMAT0000656 | Gas2l3 | 237436 | HITS-CLIP | 21258322\|29167373\|25083871 | Unspecified |
| mmu-miR-139-5p | MIMAT0000656 | Gas7 | 14457 | HITS-CLIP | 29167373 | Unspecified |
| mmu-miR-139-5p | MIMAT0000656 | Gatad1 | 67210 | sRNA-Seq//HITS-CLIP | 33184237\|29167373 | Unspecified |
| mmu-miR-139-5p | MIMAT0000656 | Gatad2a | 234366 | HITS-CLIP | 31433990 | Unspecified |
| mmu-miR-139-5p | MIMAT0000656 | Gatad2b | 229542 | HITS-CLIP | 29167373 | Unspecified |
| mmu-miR-139-5p | MIMAT0000656 | Gatc | 384281 | HITS-CLIP | 29167373 | Unspecified |
| mmu-miR-139-5p | MIMAT0000656 | Gatm | 67092 | HITS-CLIP | 29167373 | Unspecified |
| mmu-miR-139-5p | MIMAT0000656 | Gcc2 | 70297 | HITS-CLIP | 29167373 | Unspecified |
| mmu-miR-139-5p | MIMAT0000656 | Gclm | 14630 | HITS-CLIP | 29167373 | Unspecified |
| mmu-miR-139-5p | MIMAT0000656 | Gcnt1 | 14537 | HITS-CLIP | 29167373 | Unspecified |
| mmu-miR-139-5p | MIMAT0000656 | Gde1 | 56209 | sRNA-Seq//HITS-CLIP | 33184237\|29167373 | Unspecified |
| mmu-miR-139-5p | MIMAT0000656 | Gdi2 | 14569 | sRNA-Seq//HITS-CLIP | 33184237\|29167373 | Unspecified |
| mmu-miR-139-5p | MIMAT0000656 | Gdpgp1 | 269952 | HITS-CLIP | 21258322\|26203562 | Unspecified |
| mmu-miR-139-5p | MIMAT0000656 | Gfm2 | 320806 | HITS-CLIP | 29167373 | Unspecified |
| mmu-miR-139-5p | MIMAT0000656 | Ggnbp2 | 217039 | sRNA-Seq//HITS-CLIP | 33184237\|29167373 | Unspecified |
| mmu-miR-139-5p | MIMAT0000656 | Ggta1 | 14594 | HITS-CLIP | 26203562 | Unspecified |
| mmu-miR-139-5p | MIMAT0000656 | Ginm1 | 215751 | HITS-CLIP | 29167373 | Unspecified |
| mmu-miR-139-5p | MIMAT0000656 | Git1 | 216963 | HITS-CLIP | 29167373 | Unspecified |
| mmu-miR-139-5p | MIMAT0000656 | Git2 | 26431 | HITS-CLIP | 29167373 | Unspecified |

| | | | | | | |
|---|---|---|---|---|---|---|
| mmu-miR-139-5p | MIMAT0000656 | Gk | 14933 | HITS-CLIP | 29167373 | Unspecified |
| mmu-miR-139-5p | MIMAT0000656 | Glrx3 | 30926 | sRNA-Seq//HITS-CLIP | 33184237\|29167373 | Unspecified |
| mmu-miR-139-5p | MIMAT0000656 | Gls | 14660 | sRNA-Seq//HITS-CLIP | 33184237\|29167373 | Unspecified |
| mmu-miR-139-5p | MIMAT0000656 | Glt8d1 | 76485 | HITS-CLIP | 29167373 | Unspecified |
| mmu-miR-139-5p | MIMAT0000656 | Glud1 | 14661 | sRNA-Seq//HITS-CLIP | 33184237\|29167373 | Unspecified |
| mmu-miR-139-5p | MIMAT0000656 | Gm10521 | 100038548 | HITS-CLIP | 29167373 | Unspecified |
| mmu-miR-139-5p | MIMAT0000656 | Gm14548 | 100038909 | HITS-CLIP | 29167373 | Unspecified |
| mmu-miR-139-5p | MIMAT0000656 | Gm20431 | 6608666 | HITS-CLIP | 29167373 | Unspecified |
| mmu-miR-139-5p | MIMAT0000656 | Gm21969 | 6609957 | HITS-CLIP | 26203562 | Unspecified |
| mmu-miR-139-5p | MIMAT0000656 | Gm340 | 381224 | HITS-CLIP | 29167373 | Unspecified |
| mmu-miR-139-5p | MIMAT0000656 | Gm38394 | 667118 | HITS-CLIP | 29167373 | Unspecified |
| mmu-miR-139-5p | MIMAT0000656 | Gm42674 | | HITS-CLIP | 31433990 | Unspecified |
| mmu-miR-139-5p | MIMAT0000656 | Gm49336 | 127230315 | sRNA-Seq//HITS-CLIP | 33184237\|29167373 | Unspecified |
| mmu-miR-139-5p | MIMAT0000656 | Gm49339 | | HITS-CLIP | 29167373 | Unspecified |
| mmu-miR-139-5p | MIMAT0000656 | Gm49359 | 71508 | HITS-CLIP | 29167373 | Unspecified |
| mmu-miR-139-5p | MIMAT0000656 | Gm49368 | 118261720 | HITS-CLIP | 29167373 | Unspecified |
| mmu-miR-139-5p | MIMAT0000656 | Gmfb | 63985 | sRNA-Seq//HITS-CLIP | 33184237\|29167373 | Unspecified |
| mmu-miR-139-5p | MIMAT0000656 | Gmip | 78816 | HITS-CLIP | 29167373 | Unspecified |
| mmu-miR-139-5p | MIMAT0000656 | Gna12 | 14673 | sRNA-Seq//HITS-CLIP | 33184237\|29167373 | Unspecified |
| mmu-miR-139-5p | MIMAT0000656 | Gna13 | 14674 | HITS-CLIP | 29167373 | Unspecified |
| mmu-miR-139-5p | MIMAT0000656 | Gnai2 | 14678 | sRNA-Seq//HITS-CLIP | 33184237\|31433990\|29167373 | Unspecified |
| mmu-miR-139-5p | MIMAT0000656 | Gnai3 | 14679 | sRNA-Seq//HITS-CLIP | 33184237\|29167373 | Unspecified |
| mmu-miR-139-5p | MIMAT0000656 | Gnaq | 14682 | sRNA-Seq//HITS-CLIP | 33184237\|29167373 | Unspecified |
| mmu-miR-139-5p | MIMAT0000656 | Gnas | 14683 | HITS-CLIP//sRNA-Seq | 31433990\|33184237\|29167373 | Unspecified |
| mmu-miR-139-5p | MIMAT0000656 | Gnb1 | 14688 | sRNA-Seq//HITS-CLIP | 33184237\|29167373 | Unspecified |
| mmu-miR-139-5p | MIMAT0000656 | Gng12 | 14701 | HITS-CLIP | 31433990\|29167373 | Unspecified |
| mmu-miR-139-5p | MIMAT0000656 | Gnptab | 432486 | sRNA-Seq//HITS-CLIP | 33184237\|29167373 | Unspecified |
| mmu-miR-139-5p | MIMAT0000656 | Gns | 75612 | sRNA-Seq//HITS-CLIP | 33184237\|29167373 | Unspecified |
| mmu-miR-139-5p | MIMAT0000656 | Golga3 | 269682 | HITS-CLIP | 29167373 | Unspecified |
| mmu-miR-139-5p | MIMAT0000656 | Golga4 | 54214 | HITS-CLIP | 29167373 | Unspecified |
| mmu-miR-139-5p | MIMAT0000656 | Golgb1 | 224139 | HITS-CLIP | 29167373 | Unspecified |
| mmu-miR-139-5p | MIMAT0000656 | Golm1 | 105348 | HITS-CLIP | 29167373 | Unspecified |
| mmu-miR-139-5p | MIMAT0000656 | Gpatch8 | 237943 | HITS-CLIP | 29167373 | Unspecified |
| mmu-miR-139-5p | MIMAT0000656 | Gpi1 | 14751 | sRNA-Seq//HITS-CLIP | 33184237\|29167373 | Unspecified |
| mmu-miR-139-5p | MIMAT0000656 | Gpnmb | 93695 | HITS-CLIP | 29167373 | Unspecified |
| mmu-miR-139-5p | MIMAT0000656 | Gpr65 | 14744 | HITS-CLIP | 29167373 | Unspecified |
| mmu-miR-139-5p | MIMAT0000656 | Gramd1b | 235283 | HITS-CLIP | 29167373 | Unspecified |
| mmu-miR-139-5p | MIMAT0000656 | Grb2 | 14784 | sRNA-Seq//HITS-CLIP | 33184237\|29167373 | Unspecified |
| mmu-miR-139-5p | MIMAT0000656 | Greb1l | 381157 | sRNA-Seq//HITS-CLIP | 33184237\|23630078 | Unspecified |

| | | | | | | |
|---|---|---|---|---|---|---|
| mmu-miR-139-5p | MIMAT0000656 | Grk5 | 14773 | HITS-CLIP | 31433990 | Unspecified |
| mmu-miR-139-5p | MIMAT0000656 | Grn | 14824 | HITS-CLIP | 29167373 | Unspecified |
| mmu-miR-139-5p | MIMAT0000656 | Gsdmd | 69146 | HITS-CLIP | 29167373 | Unspecified |
| mmu-miR-139-5p | MIMAT0000656 | Gse1 | 382034 | HITS-CLIP | 29167373 | Unspecified |
| mmu-miR-139-5p | MIMAT0000656 | Gsk3a | 606496 | HITS-CLIP | 29167373 | Unspecified |
| mmu-miR-139-5p | MIMAT0000656 | Gsk3b | 56637 | sRNA-Seq//HITS-CLIP | 33184237\|29167373 | Unspecified |
| mmu-miR-139-5p | MIMAT0000656 | Gspt1 | 14852 | sRNA-Seq//HITS-CLIP | 33184237\|29167373 | Unspecified |
| mmu-miR-139-5p | MIMAT0000656 | Gtf3c2 | 71752 | HITS-CLIP | 29167373 | Unspecified |
| mmu-miR-139-5p | MIMAT0000656 | Gxylt1 | 223827 | HITS-CLIP | 29167373 | Unspecified |
| mmu-miR-139-5p | MIMAT0000656 | H13 | 14950 | sRNA-Seq//HITS-CLIP | 33184237\|29167373 | Unspecified |
| mmu-miR-139-5p | MIMAT0000656 | H2-Aa | 14960 | HITS-CLIP | 29167373 | Unspecified |
| mmu-miR-139-5p | MIMAT0000656 | H2ac20 | 8338 | HITS-CLIP | 31433990 | Unspecified |
| mmu-miR-139-5p | MIMAT0000656 | H3c3 | 8350 | HITS-CLIP | 31433990 | Unspecified |
| mmu-miR-139-5p | MIMAT0000656 | H3f3b | 15081 | HITS-CLIP//sRNA-Seq | 31433990\|33184237\|29167373 | Unspecified |
| mmu-miR-139-5p | MIMAT0000656 | H6pd | 100198 | HITS-CLIP | 29167373 | Unspecified |
| mmu-miR-139-5p | MIMAT0000656 | Haus6 | 230376 | HITS-CLIP | 29167373 | Unspecified |
| mmu-miR-139-5p | MIMAT0000656 | Hcfc1 | 15161 | HITS-CLIP | 29167373 | Unspecified |
| mmu-miR-139-5p | MIMAT0000656 | Hcfc2 | 67933 | HITS-CLIP | 29167373 | Unspecified |
| mmu-miR-139-5p | MIMAT0000656 | Hck | 15162 | HITS-CLIP | 29167373 | Unspecified |
| mmu-miR-139-5p | MIMAT0000656 | Hcls1 | 15163 | HITS-CLIP | 29167373 | Unspecified |
| mmu-miR-139-5p | MIMAT0000656 | Hdac4 | 208727 | HITS-CLIP | 29167373 | Unspecified |
| mmu-miR-139-5p | MIMAT0000656 | Hdlbp | 110611 | sRNA-Seq//HITS-CLIP | 33184237\|29167373 | Unspecified |
| mmu-miR-139-5p | MIMAT0000656 | Heatr1 | 217995 | HITS-CLIP | 29167373 | Unspecified |
| mmu-miR-139-5p | MIMAT0000656 | Heatr5a | 320487 | HITS-CLIP | 29167373 | Unspecified |
| mmu-miR-139-5p | MIMAT0000656 | Heca | 380629 | HITS-CLIP | 29167373 | Unspecified |
| mmu-miR-139-5p | MIMAT0000656 | Hectd1 | 207304 | sRNA-Seq//HITS-CLIP | 33184237\|31433990\|29167373 | Unspecified |
| mmu-miR-139-5p | MIMAT0000656 | Hectd2 | 226098 | HITS-CLIP | 29167373 | Unspecified |
| mmu-miR-139-5p | MIMAT0000656 | Hells | 15201 | HITS-CLIP | 29167373 | Unspecified |
| mmu-miR-139-5p | MIMAT0000656 | Helz2 | 229003 | HITS-CLIP | 26203562\|29167373 | Unspecified |
| mmu-miR-139-5p | MIMAT0000656 | Herc1 | 235439 | sRNA-Seq//HITS-CLIP | 33184237\|29167373 | Unspecified |
| mmu-miR-139-5p | MIMAT0000656 | Herc2 | 15204 | sRNA-Seq//HITS-CLIP | 33184237\|29167373 | Unspecified |
| mmu-miR-139-5p | MIMAT0000656 | Herc3 | 73998 | HITS-CLIP | 29167373 | Unspecified |
| mmu-miR-139-5p | MIMAT0000656 | Herpud1 | 64209 | HITS-CLIP | 29167373 | Unspecified |
| mmu-miR-139-5p | MIMAT0000656 | Hexa | 15211 | HITS-CLIP | 29167373 | Unspecified |
| mmu-miR-139-5p | MIMAT0000656 | Hexb | 15212 | sRNA-Seq//HITS-CLIP | 33184237\|29167373 | Unspecified |
| mmu-miR-139-5p | MIMAT0000656 | Hgs | 15239 | HITS-CLIP | 29167373 | Unspecified |
| mmu-miR-139-5p | MIMAT0000656 | Hgsnat | 52120 | HITS-CLIP | 29167373 | Unspecified |
| mmu-miR-139-5p | MIMAT0000656 | Hhex | 15242 | HITS-CLIP | 29167373 | Unspecified |
| mmu-miR-139-5p | MIMAT0000656 | Hic2 | 58180 | HITS-CLIP | 29167373 | Unspecified |

| | | | | | | |
|---|---|---|---|---|---|---|
| mmu-miR-139-5p | MIMAT0000656 | Hif1a | 15251 | HITS-CLIP//sRNA-Seq | 31433990\|33184237\|29167373 | Unspecified |
| mmu-miR-139-5p | MIMAT0000656 | Hip1 | 215114 | sRNA-Seq//HITS-CLIP | 33184237\|29167373 | Unspecified |
| mmu-miR-139-5p | MIMAT0000656 | Hipk1 | 15257 | HITS-CLIP | 31433990\|29167373 | Unspecified |
| mmu-miR-139-5p | MIMAT0000656 | Hipk2 | 15258 | HITS-CLIP | 29167373 | Unspecified |
| mmu-miR-139-5p | MIMAT0000656 | Hipk3 | 15259 | HITS-CLIP | 29167373 | Unspecified |
| mmu-miR-139-5p | MIMAT0000656 | Hist2h3c2 | 97114 | HITS-CLIP | 29167373 | Unspecified |
| mmu-miR-139-5p | MIMAT0000656 | Hk2 | 15277 | HITS-CLIP | 29167373 | Unspecified |
| mmu-miR-139-5p | MIMAT0000656 | Hk3 | 212032 | HITS-CLIP | 29167373 | Unspecified |
| mmu-miR-139-5p | MIMAT0000656 | Hmga2 | 15364 | HITS-CLIP | 29167373 | Unspecified |
| mmu-miR-139-5p | MIMAT0000656 | Hnrnpa1 | 15382 | sRNA-Seq//HITS-CLIP | 33184237\|29167373 | Unspecified |
| mmu-miR-139-5p | MIMAT0000656 | Hnrnpa2b1 | 53379 | sRNA-Seq//HITS-CLIP | 33184237\|29167373 | Unspecified |
| mmu-miR-139-5p | MIMAT0000656 | Hnrnpab | 15384 | HITS-CLIP//sRNA-Seq | 21258322\|33184237\|31433990\|29167 | Unspecified |
| mmu-miR-139-5p | MIMAT0000656 | Hnrnph2 | 56258 | sRNA-Seq//HITS-CLIP | 33184237\|29167373 | Unspecified |
| mmu-miR-139-5p | MIMAT0000656 | Hnrnpk | 15387 | sRNA-Seq//HITS-CLIP | 33184237\|29167373 | Unspecified |
| mmu-miR-139-5p | MIMAT0000656 | Hnrnpl | 15388 | sRNA-Seq//HITS-CLIP | 33184237\|29167373 | Unspecified |
| mmu-miR-139-5p | MIMAT0000656 | Hnrnpll | 72692 | HITS-CLIP | 29167373 | Unspecified |
| mmu-miR-139-5p | MIMAT0000656 | Hnrnpr | 74326 | sRNA-Seq//HITS-CLIP | 33184237\|29167373 | Unspecified |
| mmu-miR-139-5p | MIMAT0000656 | Hnrnpu | 51810 | sRNA-Seq//HITS-CLIP | 33184237\|29167373 | Unspecified |
| mmu-miR-139-5p | MIMAT0000656 | Hnrnpul1 | 232989 | HITS-CLIP//sRNA-Seq | 29167373\|33184237 | Unspecified |
| mmu-miR-139-5p | MIMAT0000656 | Hnrnpul2 | 68693 | sRNA-Seq//HITS-CLIP | 33184237\|29167373 | Unspecified |
| mmu-miR-139-5p | MIMAT0000656 | Hook2 | 170833 | HITS-CLIP | 31433990 | Unspecified |
| mmu-miR-139-5p | MIMAT0000656 | Hps3 | 12807 | HITS-CLIP | 29167373 | Unspecified |
| mmu-miR-139-5p | MIMAT0000656 | Hsd17b12 | 56348 | sRNA-Seq//HITS-CLIP | 33184237\|29167373 | Unspecified |
| mmu-miR-139-5p | MIMAT0000656 | Hsf1 | 15499 | HITS-CLIP | 29167373 | Unspecified |
| mmu-miR-139-5p | MIMAT0000656 | Hsp90ab1 | 15516 | sRNA-Seq//HITS-CLIP | 33184237\|29167373 | Unspecified |
| mmu-miR-139-5p | MIMAT0000656 | Hsp90b1 | 22027 | sRNA-Seq//HITS-CLIP | 33184237\|29167373 | Unspecified |
| mmu-miR-139-5p | MIMAT0000656 | Hspa4 | 15525 | HITS-CLIP//sRNA-Seq | 23630078\|33184237\|29167373 | Unspecified |
| mmu-miR-139-5p | MIMAT0000656 | Hspa5 | 14828 | sRNA-Seq//HITS-CLIP | 33184237\|29167373 | Unspecified |
| mmu-miR-139-5p | MIMAT0000656 | Htatsf1 | 72459 | HITS-CLIP | 29167373 | Unspecified |
| mmu-miR-139-5p | MIMAT0000656 | Htt | 15194 | HITS-CLIP | 29167373 | Unspecified |
| mmu-miR-139-5p | MIMAT0000656 | Huwe1 | 59026 | sRNA-Seq//HITS-CLIP | 33184237\|29167373 | Unspecified |
| mmu-miR-139-5p | MIMAT0000656 | Hyal1 | 15586 | HITS-CLIP | 29167373 | Unspecified |
| mmu-miR-139-5p | MIMAT0000656 | Hycc2 | 285172 | HITS-CLIP | 26203562 | Unspecified |
| mmu-miR-139-5p | MIMAT0000656 | Iars | 105148 | sRNA-Seq//HITS-CLIP | 33184237\|29167373 | Unspecified |
| mmu-miR-139-5p | MIMAT0000656 | Id2 | 15902 | HITS-CLIP | 29167373 | Unspecified |
| mmu-miR-139-5p | MIMAT0000656 | Ide | 15925 | HITS-CLIP | 29167373 | Unspecified |
| mmu-miR-139-5p | MIMAT0000656 | Idh3g | 15929 | sRNA-Seq//HITS-CLIP | 33184237\|29167373 | Unspecified |
| mmu-miR-139-5p | MIMAT0000656 | Ids | 15931 | sRNA-Seq//HITS-CLIP | 33184237\|29167373 | Unspecified |
| mmu-miR-139-5p | MIMAT0000656 | Iffo2 | 212632 | HITS-CLIP | 26203562 | Unspecified |

| | | | | | | |
|---|---|---|---|---|---|---|
| mmu-miR-139-5p | MIMAT0000656 | Ifit2 | 15958 | HITS-CLIP | 29167373 | Unspecified |
| mmu-miR-139-5p | MIMAT0000656 | Ifnar1 | 15975 | sRNA-Seq//HITS-CLIP | 33184237\|29167373 | Unspecified |
| mmu-miR-139-5p | MIMAT0000656 | Ifngr2 | 15980 | HITS-CLIP | 29167373 | Unspecified |
| mmu-miR-139-5p | MIMAT0000656 | Ifrd1 | 15982 | sRNA-Seq//HITS-CLIP | 33184237\|29167373 | Unspecified |
| mmu-miR-139-5p | MIMAT0000656 | Igf1 | 16000 | HITS-CLIP | 29167373 | Unspecified |
| mmu-miR-139-5p | MIMAT0000656 | Igf2r | 16004 | HITS-CLIP | 29167373 | Unspecified |
| mmu-miR-139-5p | MIMAT0000656 | Igfbp2 | 16008 | HITS-CLIP | 26203562 | Unspecified |
| mmu-miR-139-5p | MIMAT0000656 | Iigp1 | 60440 | HITS-CLIP | 31433990 | Unspecified |
| mmu-miR-139-5p | MIMAT0000656 | Ikbke | 56489 | HITS-CLIP | 29167373 | Unspecified |
| mmu-miR-139-5p | MIMAT0000656 | Ikzf1 | 22778 | HITS-CLIP | 31433990 | Unspecified |
| mmu-miR-139-5p | MIMAT0000656 | Il10ra | 16154 | HITS-CLIP | 29167373 | Unspecified |
| mmu-miR-139-5p | MIMAT0000656 | Il21r | 60504 | HITS-CLIP | 29167373 | Unspecified |
| mmu-miR-139-5p | MIMAT0000656 | Il2ra | 16184 | HITS-CLIP | 31433990 | Unspecified |
| mmu-miR-139-5p | MIMAT0000656 | Il2rg | 16186 | HITS-CLIP | 29167373 | Unspecified |
| mmu-miR-139-5p | MIMAT0000656 | Il4ra | 16190 | HITS-CLIP | 31433990 | Unspecified |
| mmu-miR-139-5p | MIMAT0000656 | Il6st | 16195 | sRNA-Seq//HITS-CLIP | 33184237\|29167373 | Unspecified |
| mmu-miR-139-5p | MIMAT0000656 | Il7r | 16197 | HITS-CLIP | 29167373 | Unspecified |
| mmu-miR-139-5p | MIMAT0000656 | Imp4 | 27993 | HITS-CLIP | 31433990 | Unspecified |
| mmu-miR-139-5p | MIMAT0000656 | Impa1 | 55980 | HITS-CLIP | 29167373 | Unspecified |
| mmu-miR-139-5p | MIMAT0000656 | Ing5 | 66262 | HITS-CLIP | 29167373 | Unspecified |
| mmu-miR-139-5p | MIMAT0000656 | Ino80d | 227195 | HITS-CLIP | 29167373 | Unspecified |
| mmu-miR-139-5p | MIMAT0000656 | Inpp5d | 16331 | HITS-CLIP | 29167373 | Unspecified |
| mmu-miR-139-5p | MIMAT0000656 | Inppl1 | 16332 | HITS-CLIP | 29167373 | Unspecified |
| mmu-miR-139-5p | MIMAT0000656 | Ints6 | 18130 | HITS-CLIP | 29167373 | Unspecified |
| mmu-miR-139-5p | MIMAT0000656 | Ints7 | 77065 | HITS-CLIP | 29167373 | Unspecified |
| mmu-miR-139-5p | MIMAT0000656 | Ip6k1 | 27399 | sRNA-Seq//HITS-CLIP | 33184237\|31433990 | Unspecified |
| mmu-miR-139-5p | MIMAT0000656 | Ipo4 | 75751 | sRNA-Seq//HITS-CLIP | 33184237\|29167373 | Unspecified |
| mmu-miR-139-5p | MIMAT0000656 | Ipo5 | 70572 | sRNA-Seq//HITS-CLIP | 33184237\|29167373 | Unspecified |
| mmu-miR-139-5p | MIMAT0000656 | Ipo7 | 233726 | sRNA-Seq//HITS-CLIP | 33184237\|29167373 | Unspecified |
| mmu-miR-139-5p | MIMAT0000656 | Ipo9 | 226432 | sRNA-Seq//HITS-CLIP | 33184237\|29167373 | Unspecified |
| mmu-miR-139-5p | MIMAT0000656 | Ippk | 75678 | HITS-CLIP | 31433990 | Unspecified |
| mmu-miR-139-5p | MIMAT0000656 | Iqgap1 | 29875 | HITS-CLIP | 26203562\|29167373 | Unspecified |
| mmu-miR-139-5p | MIMAT0000656 | Iqgap2 | 544963 | HITS-CLIP | 29167373 | Unspecified |
| mmu-miR-139-5p | MIMAT0000656 | Iqsec1 | 232227 | HITS-CLIP | 29167373 | Unspecified |
| mmu-miR-139-5p | MIMAT0000656 | Irak4 | 266632 | HITS-CLIP | 29167373 | Unspecified |
| mmu-miR-139-5p | MIMAT0000656 | Ireb2 | 64602 | sRNA-Seq//HITS-CLIP | 33184237\|29167373 | Unspecified |
| mmu-miR-139-5p | MIMAT0000656 | Irf1 | 16362 | HITS-CLIP | 29167373 | Unspecified |
| mmu-miR-139-5p | MIMAT0000656 | Irf2 | 16363 | HITS-CLIP | 29167373 | Unspecified |
| mmu-miR-139-5p | MIMAT0000656 | Irf4 | 16364 | HITS-CLIP | 31433990 | Unspecified |

| | | | | | | | |
|---|---|---|---|---|---|---|---|
| mmu-miR-139-5p | MIMAT0000656 | Irf5 | 27056 | HITS-CLIP | | 29167373 | Unspecified |
| mmu-miR-139-5p | MIMAT0000656 | Irgm2 | 54396 | HITS-CLIP | 25083871\|31433990 | | Unspecified |
| mmu-miR-139-5p | MIMAT0000656 | Irs2 | 384783 | HITS-CLIP | | 29167373 | Unspecified |
| mmu-miR-139-5p | MIMAT0000656 | Isoc1 | 66307 | HITS-CLIP | | 29167373 | Unspecified |
| mmu-miR-139-5p | MIMAT0000656 | Ist1 | 71955 | sRNA-Seq//HITS-CLIP | 33184237\|29167373 | | Unspecified |
| mmu-miR-139-5p | MIMAT0000656 | Itga4 | 16401 | HITS-CLIP | | 29167373 | Unspecified |
| mmu-miR-139-5p | MIMAT0000656 | Itga5 | 16402 | HITS-CLIP | | 29167373 | Unspecified |
| mmu-miR-139-5p | MIMAT0000656 | Itga6 | 16403 | sRNA-Seq//HITS-CLIP | 33184237\|29167373 | | Unspecified |
| mmu-miR-139-5p | MIMAT0000656 | Itgam | 16409 | HITS-CLIP | | 29167373 | Unspecified |
| mmu-miR-139-5p | MIMAT0000656 | Itgav | 16410 | HITS-CLIP | | 29167373 | Unspecified |
| mmu-miR-139-5p | MIMAT0000656 | Itgb1 | 16412 | sRNA-Seq//HITS-CLIP | 33184237\|29167373 | | Unspecified |
| mmu-miR-139-5p | MIMAT0000656 | Itm2b | 16432 | sRNA-Seq//HITS-CLIP | 33184237\|29167373 | | Unspecified |
| mmu-miR-139-5p | MIMAT0000656 | Itpkb | 320404 | HITS-CLIP | | 29167373 | Unspecified |
| mmu-miR-139-5p | MIMAT0000656 | Itprid2 | 6744 | HITS-CLIP | | 29167373 | Unspecified |
| mmu-miR-139-5p | MIMAT0000656 | Itpripl2 | 319622 | HITS-CLIP | | 29167373 | Unspecified |
| mmu-miR-139-5p | MIMAT0000656 | Itsn1 | 16443 | sRNA-Seq//HITS-CLIP | 33184237\|29167373 | | Unspecified |
| mmu-miR-139-5p | MIMAT0000656 | Itsn2 | 20403 | HITS-CLIP | | 29167373 | Unspecified |
| mmu-miR-139-5p | MIMAT0000656 | Ivns1abp | 117198 | sRNA-Seq//HITS-CLIP | 33184237\|29167373 | | Unspecified |
| mmu-miR-139-5p | MIMAT0000656 | Iws1 | 73473 | HITS-CLIP | | 29167373 | Unspecified |
| mmu-miR-139-5p | MIMAT0000656 | Jade1 | 269424 | HITS-CLIP | | 29167373 | Unspecified |
| mmu-miR-139-5p | MIMAT0000656 | Jag1 | 16449 | HITS-CLIP | | 29167373 | Unspecified |
| mmu-miR-139-5p | MIMAT0000656 | Jarid2 | 16468 | sRNA-Seq//HITS-CLIP | 33184237\|29167373 | | Unspecified |
| mmu-miR-139-5p | MIMAT0000656 | Jdp2 | 81703 | HITS-CLIP | | 29167373 | Unspecified |
| mmu-miR-139-5p | MIMAT0000656 | Jmjd1c | 108829 | sRNA-Seq//HITS-CLIP | 33184237\|29167373 | | Unspecified |
| mmu-miR-139-5p | MIMAT0000656 | Josd1 | 74158 | HITS-CLIP | | 31433990 | Unspecified |
| mmu-miR-139-5p | MIMAT0000656 | Jun | 16476 | HITS-CLIP | | 29167373 | Unspecified |
| mmu-miR-139-5p | MIMAT0000656 | Kansl1l | 68691 | HITS-CLIP | | 29167373 | Unspecified |
| mmu-miR-139-5p | MIMAT0000656 | Kansl3 | 226976 | HITS-CLIP | | 26203562 | Unspecified |
| mmu-miR-139-5p | MIMAT0000656 | Kat14 | 57325 | HITS-CLIP | | 29167373 | Unspecified |
| mmu-miR-139-5p | MIMAT0000656 | Kat2a | 14534 | HITS-CLIP | | 29167373 | Unspecified |
| mmu-miR-139-5p | MIMAT0000656 | Kat6a | 244349 | HITS-CLIP | | 29167373 | Unspecified |
| mmu-miR-139-5p | MIMAT0000656 | Katnbl1 | 72425 | HITS-CLIP | | 29167373 | Unspecified |
| mmu-miR-139-5p | MIMAT0000656 | Kcnn4 | 16534 | HITS-CLIP | | 29167373 | Unspecified |
| mmu-miR-139-5p | MIMAT0000656 | Kdelr2 | 66913 | HITS-CLIP | | 31433990 | Unspecified |
| mmu-miR-139-5p | MIMAT0000656 | Kdm1b | 218214 | HITS-CLIP | | 29167373 | Unspecified |
| mmu-miR-139-5p | MIMAT0000656 | Kdm2a | 225876 | sRNA-Seq//HITS-CLIP | 33184237\|29167373 | | Unspecified |
| mmu-miR-139-5p | MIMAT0000656 | Kdm3a | 104263 | HITS-CLIP | | 29167373 | Unspecified |
| mmu-miR-139-5p | MIMAT0000656 | Kdm3b | 277250 | HITS-CLIP | | 29167373 | Unspecified |
| mmu-miR-139-5p | MIMAT0000656 | Kdm7a | 338523 | HITS-CLIP | | 29167373 | Unspecified |

| | | | | | | |
|---|---|---|---|---|---|---|
| mmu-miR-139-5p | MIMAT0000656 | Kdr | 16542 | HITS-CLIP | 29167373 | Unspecified |
| mmu-miR-139-5p | MIMAT0000656 | Kif1c | 16562 | HITS-CLIP | 26203562 | Unspecified |
| mmu-miR-139-5p | MIMAT0000656 | Kif20b | 240641 | HITS-CLIP | 29167373 | Unspecified |
| mmu-miR-139-5p | MIMAT0000656 | Kif21b | 16565 | HITS-CLIP | 29167373 | Unspecified |
| mmu-miR-139-5p | MIMAT0000656 | Kif2a | 16563 | sRNA-Seq//HITS-CLIP | 33184237\|29167373 | Unspecified |
| mmu-miR-139-5p | MIMAT0000656 | Kif4 | 16571 | HITS-CLIP | 29167373 | Unspecified |
| mmu-miR-139-5p | MIMAT0000656 | Kif5b | 16573 | HITS-CLIP//sRNA-Seq | 31433990\|33184237\|29167373 | Unspecified |
| mmu-miR-139-5p | MIMAT0000656 | Klc1 | 16593 | sRNA-Seq//HITS-CLIP | 33184237\|29167373 | Unspecified |
| mmu-miR-139-5p | MIMAT0000656 | Klf10 | 21847 | HITS-CLIP | 29167373\|26203562 | Unspecified |
| mmu-miR-139-5p | MIMAT0000656 | Klf13 | 50794 | HITS-CLIP | 29167373 | Unspecified |
| mmu-miR-139-5p | MIMAT0000656 | Klf2 | 16598 | HITS-CLIP | 29167373 | Unspecified |
| mmu-miR-139-5p | MIMAT0000656 | Klf3 | 16599 | HITS-CLIP | 26203562\|29167373 | Unspecified |
| mmu-miR-139-5p | MIMAT0000656 | Klf4 | 16600 | HITS-CLIP | 23630078\|26203562 | Unspecified |
| mmu-miR-139-5p | MIMAT0000656 | Klf6 | 23849 | sRNA-Seq//HITS-CLIP | 33184237\|29167373 | Unspecified |
| mmu-miR-139-5p | MIMAT0000656 | Klhdc2 | 69554 | sRNA-Seq//HITS-CLIP | 33184237\|29167373 | Unspecified |
| mmu-miR-139-5p | MIMAT0000656 | Klhl2 | 77113 | HITS-CLIP | 29167373 | Unspecified |
| mmu-miR-139-5p | MIMAT0000656 | Klhl21 | 242785 | HITS-CLIP | 26203562 | Unspecified |
| mmu-miR-139-5p | MIMAT0000656 | Klhl24 | 75785 | HITS-CLIP | 29167373 | Unspecified |
| mmu-miR-139-5p | MIMAT0000656 | Klhl28 | 66689 | HITS-CLIP | 29167373 | Unspecified |
| mmu-miR-139-5p | MIMAT0000656 | Klhl42 | 232539 | HITS-CLIP | 29167373 | Unspecified |
| mmu-miR-139-5p | MIMAT0000656 | Klhl8 | 246293 | HITS-CLIP | 29167373 | Unspecified |
| mmu-miR-139-5p | MIMAT0000656 | Klhl9 | 242521 | sRNA-Seq//HITS-CLIP | 33184237\|29167373 | Unspecified |
| mmu-miR-139-5p | MIMAT0000656 | Kmt2a | 214162 | HITS-CLIP | 31433990\|29167373 | Unspecified |
| mmu-miR-139-5p | MIMAT0000656 | Kmt2b | 75410 | HITS-CLIP | 29167373 | Unspecified |
| mmu-miR-139-5p | MIMAT0000656 | Kmt2c | 231051 | HITS-CLIP | 29167373 | Unspecified |
| mmu-miR-139-5p | MIMAT0000656 | Kmt2d | 381022 | HITS-CLIP | 26203562\|29167373 | Unspecified |
| mmu-miR-139-5p | MIMAT0000656 | Kmt2e | 69188 | HITS-CLIP | 29167373 | Unspecified |
| mmu-miR-139-5p | MIMAT0000656 | Kmt5b | 225888 | HITS-CLIP | 29167373 | Unspecified |
| mmu-miR-139-5p | MIMAT0000656 | Kntc1 | 208628 | HITS-CLIP | 31433990 | Unspecified |
| mmu-miR-139-5p | MIMAT0000656 | Kpna4 | 16649 | HITS-CLIP | 29167373 | Unspecified |
| mmu-miR-139-5p | MIMAT0000656 | Kpna6 | 16650 | HITS-CLIP | 29167373 | Unspecified |
| mmu-miR-139-5p | MIMAT0000656 | Kpnb1 | 16211 | sRNA-Seq//HITS-CLIP | 33184237\|31433990\|29167373 | Unspecified |
| mmu-miR-139-5p | MIMAT0000656 | Kras | 16653 | sRNA-Seq//HITS-CLIP | 33184237\|29167373 | Unspecified |
| mmu-miR-139-5p | MIMAT0000656 | Kri1 | 215194 | HITS-CLIP | 29167373 | Unspecified |
| mmu-miR-139-5p | MIMAT0000656 | Krit1 | 79264 | HITS-CLIP | 29167373 | Unspecified |
| mmu-miR-139-5p | MIMAT0000656 | L1cam | 16728 | sRNA-Seq//HITS-CLIP | 33184237\|29167373 | Unspecified |
| mmu-miR-139-5p | MIMAT0000656 | L3mbtl2 | 214669 | HITS-CLIP | 26203562 | Unspecified |
| mmu-miR-139-5p | MIMAT0000656 | Lair1 | 52855 | HITS-CLIP | 29167373 | Unspecified |
| mmu-miR-139-5p | MIMAT0000656 | Lamc1 | 226519 | HITS-CLIP | 29167373 | Unspecified |

| | | | | | | |
|---|---|---|---|---|---|---|
| mmu-miR-139-5p | MIMAT0000656 | Laptm4a | 17775 | sRNA-Seq//HITS-CLIP | 33184237\|26203562\|29167373 | Unspecified |
| mmu-miR-139-5p | MIMAT0000656 | Laptm5 | 16792 | HITS-CLIP | 29167373 | Unspecified |
| mmu-miR-139-5p | MIMAT0000656 | Larp1 | 73158 | sRNA-Seq//HITS-CLIP | 33184237\|29167373 | Unspecified |
| mmu-miR-139-5p | MIMAT0000656 | Larp4b | 217980 | sRNA-Seq//HITS-CLIP | 33184237\|29167373 | Unspecified |
| mmu-miR-139-5p | MIMAT0000656 | Lats1 | 16798 | HITS-CLIP | 29167373 | Unspecified |
| mmu-miR-139-5p | MIMAT0000656 | Lcor | 212391 | sRNA-Seq//HITS-CLIP | 33184237\|29167373 | Unspecified |
| mmu-miR-139-5p | MIMAT0000656 | Lcorl | 209707 | HITS-CLIP | 29167373 | Unspecified |
| mmu-miR-139-5p | MIMAT0000656 | Lcp1 | 18826 | HITS-CLIP | 29167373 | Unspecified |
| mmu-miR-139-5p | MIMAT0000656 | Ldah | 38150 | HITS-CLIP | 29167373 | Unspecified |
| mmu-miR-139-5p | MIMAT0000656 | Ldlrap1 | 100017 | HITS-CLIP | 29167373 | Unspecified |
| mmu-miR-139-5p | MIMAT0000656 | Lemd3 | 380664 | HITS-CLIP | 29167373 | Unspecified |
| mmu-miR-139-5p | MIMAT0000656 | Leng8 | 232798 | HITS-CLIP | 29167373 | Unspecified |
| mmu-miR-139-5p | MIMAT0000656 | Lfng | 16848 | HITS-CLIP | 29167373 | Unspecified |
| mmu-miR-139-5p | MIMAT0000656 | Lgals3 | 16854 | HITS-CLIP | 26203562 | Unspecified |
| mmu-miR-139-5p | MIMAT0000656 | Lgals9 | 16859 | HITS-CLIP | 29167373 | Unspecified |
| mmu-miR-139-5p | MIMAT0000656 | Lhfpl2 | 218454 | HITS-CLIP | 29167373 | Unspecified |
| mmu-miR-139-5p | MIMAT0000656 | Lif | 16878 | HITS-CLIP | 31433990 | Unspecified |
| mmu-miR-139-5p | MIMAT0000656 | Lilr4b | 14727 | HITS-CLIP | 29167373 | Unspecified |
| mmu-miR-139-5p | MIMAT0000656 | Lima1 | 65970 | HITS-CLIP | 29167373 | Unspecified |
| mmu-miR-139-5p | MIMAT0000656 | Lims1 | 110829 | sRNA-Seq//HITS-CLIP | 33184237\|29167373 | Unspecified |
| mmu-miR-139-5p | MIMAT0000656 | Lin7c | 22343 | sRNA-Seq//HITS-CLIP | 33184237\|29167373 | Unspecified |
| mmu-miR-139-5p | MIMAT0000656 | Lipa | 16889 | HITS-CLIP | 29167373 | Unspecified |
| mmu-miR-139-5p | MIMAT0000656 | Litaf | 56722 | HITS-CLIP | 29167373 | Unspecified |
| mmu-miR-139-5p | MIMAT0000656 | Lmbrd2 | 320506 | sRNA-Seq//HITS-CLIP | 33184237\|31433990 | Unspecified |
| mmu-miR-139-5p | MIMAT0000656 | Lmf2 | 105847 | HITS-CLIP | 26203562 | Unspecified |
| mmu-miR-139-5p | MIMAT0000656 | Lmnb2 | 16907 | HITS-CLIP | 29167373 | Unspecified |
| mmu-miR-139-5p | MIMAT0000656 | Lmo2 | 16909 | HITS-CLIP | 29167373 | Unspecified |
| mmu-miR-139-5p | MIMAT0000656 | Lmtk2 | 231876 | HITS-CLIP | 29167373 | Unspecified |
| mmu-miR-139-5p | MIMAT0000656 | Lnpep | 240028 | sRNA-Seq//HITS-CLIP | 33184237\|29167373 | Unspecified |
| mmu-miR-139-5p | MIMAT0000656 | Lonrf1 | 244421 | HITS-CLIP | 26203562 | Unspecified |
| mmu-miR-139-5p | MIMAT0000656 | Lpgat1 | 226856 | sRNA-Seq//HITS-CLIP | 33184237\|29167373 | Unspecified |
| mmu-miR-139-5p | MIMAT0000656 | Lpin2 | 64898 | HITS-CLIP | 29167373 | Unspecified |
| mmu-miR-139-5p | MIMAT0000656 | Lpl | 16956 | HITS-CLIP | 29167373 | Unspecified |
| mmu-miR-139-5p | MIMAT0000656 | Lpxn | 107321 | HITS-CLIP | 29167373 | Unspecified |
| mmu-miR-139-5p | MIMAT0000656 | Lrch1 | 380916 | HITS-CLIP | 29167373 | Unspecified |
| mmu-miR-139-5p | MIMAT0000656 | Lrch4 | 231798 | HITS-CLIP | 29167373 | Unspecified |
| mmu-miR-139-5p | MIMAT0000656 | Lrig1 | 16206 | HITS-CLIP | 31433990 | Unspecified |
| mmu-miR-139-5p | MIMAT0000656 | Lrig2 | 269473 | HITS-CLIP | 29167373 | Unspecified |
| mmu-miR-139-5p | MIMAT0000656 | Lrp1 | 16971 | HITS-CLIP | 29167373 | Unspecified |

| | | | | | | |
|---|---|---|---|---|---|---|
| mmu-miR-139-5p | MIMAT0000656 | Lrp10 | 65107 | HITS-CLIP | 29167373 | Unspecified |
| mmu-miR-139-5p | MIMAT0000656 | Lrp12 | 239393 | sRNA-Seq//HITS-CLIP | 33184237\|29167373 | Unspecified |
| mmu-miR-139-5p | MIMAT0000656 | Lrp8 | 16975 | HITS-CLIP//sRNA-Seq | 31433990\|33184237 | Unspecified |
| mmu-miR-139-5p | MIMAT0000656 | Lrpap1 | 16976 | sRNA-Seq//HITS-CLIP | 33184237\|29167373 | Unspecified |
| mmu-miR-139-5p | MIMAT0000656 | Lrrc40 | 67144 | HITS-CLIP | 29167373 | Unspecified |
| mmu-miR-139-5p | MIMAT0000656 | Lrrc58 | 320184 | HITS-CLIP//sRNA-Seq | 25083871\|33184237\|29167373\|2125[illegible] | Unspecified |
| mmu-miR-139-5p | MIMAT0000656 | Lrrc59 | 98238 | HITS-CLIP//sRNA-Seq | 23630078\|31433990\|33184237 | Unspecified |
| mmu-miR-139-5p | MIMAT0000656 | Lrrc8b | 433926 | HITS-CLIP | 29167373 | Unspecified |
| mmu-miR-139-5p | MIMAT0000656 | Lrrfip1 | 16978 | HITS-CLIP | 29167373 | Unspecified |
| mmu-miR-139-5p | MIMAT0000656 | Lrrfip2 | 71268 | HITS-CLIP | 29167373 | Unspecified |
| mmu-miR-139-5p | MIMAT0000656 | Lsg1 | 224092 | HITS-CLIP | 29167373 | Unspecified |
| mmu-miR-139-5p | MIMAT0000656 | Lsm14a | 67070 | HITS-CLIP | 31433990 | Unspecified |
| mmu-miR-139-5p | MIMAT0000656 | Ltn1 | 78913 | HITS-CLIP | 31433990 | Unspecified |
| mmu-miR-139-5p | MIMAT0000656 | Luc7l3 | 67684 | sRNA-Seq//HITS-CLIP | 33184237\|29167373 | Unspecified |
| mmu-miR-139-5p | MIMAT0000656 | Luzp1 | 269593 | HITS-CLIP | 29167373 | Unspecified |
| mmu-miR-139-5p | MIMAT0000656 | Ly6e | 17069 | sRNA-Seq//HITS-CLIP | 33184237\|29167373 | Unspecified |
| mmu-miR-139-5p | MIMAT0000656 | Lyn | 17096 | HITS-CLIP | 31433990\|29167373 | Unspecified |
| mmu-miR-139-5p | MIMAT0000656 | Lyst | 17101 | HITS-CLIP | 31433990\|29167373 | Unspecified |
| mmu-miR-139-5p | MIMAT0000656 | M6pr | 17113 | HITS-CLIP | 29167373 | Unspecified |
| mmu-miR-139-5p | MIMAT0000656 | Macf1 | 11426 | sRNA-Seq//HITS-CLIP | 33184237\|26203562\|29167373 | Unspecified |
| mmu-miR-139-5p | MIMAT0000656 | Madd | 228355 | sRNA-Seq//HITS-CLIP | 33184237\|29167373 | Unspecified |
| mmu-miR-139-5p | MIMAT0000656 | Mafg | 17134 | HITS-CLIP | 29167373 | Unspecified |
| mmu-miR-139-5p | MIMAT0000656 | Man1a2 | 17156 | HITS-CLIP | 29167373 | Unspecified |
| mmu-miR-139-5p | MIMAT0000656 | Man2a1 | 17158 | HITS-CLIP | 29167373 | Unspecified |
| mmu-miR-139-5p | MIMAT0000656 | Map1lc3b | 67443 | sRNA-Seq//HITS-CLIP | 33184237\|29167373 | Unspecified |
| mmu-miR-139-5p | MIMAT0000656 | Map2k4 | 26398 | sRNA-Seq//HITS-CLIP | 33184237\|29167373 | Unspecified |
| mmu-miR-139-5p | MIMAT0000656 | Map3k1 | 26401 | HITS-CLIP | 29167373 | Unspecified |
| mmu-miR-139-5p | MIMAT0000656 | Map3k2 | 26405 | HITS-CLIP | 29167373 | Unspecified |
| mmu-miR-139-5p | MIMAT0000656 | Map3k3 | 26406 | HITS-CLIP | 29167373 | Unspecified |
| mmu-miR-139-5p | MIMAT0000656 | Map3k4 | 26407 | HITS-CLIP | 29167373 | Unspecified |
| mmu-miR-139-5p | MIMAT0000656 | Map3k7 | 26409 | HITS-CLIP | 29167373 | Unspecified |
| mmu-miR-139-5p | MIMAT0000656 | Map3k8 | 26410 | HITS-CLIP | 29167373 | Unspecified |
| mmu-miR-139-5p | MIMAT0000656 | Map4 | 17758 | sRNA-Seq//HITS-CLIP | 33184237\|31433990\|29167373 | Unspecified |
| mmu-miR-139-5p | MIMAT0000656 | Map4k3 | 225028 | HITS-CLIP | 29167373 | Unspecified |
| mmu-miR-139-5p | MIMAT0000656 | Map4k4 | 26921 | sRNA-Seq//HITS-CLIP | 33184237\|29167373 | Unspecified |
| mmu-miR-139-5p | MIMAT0000656 | Mapk1 | 26413 | sRNA-Seq//HITS-CLIP | 33184237\|29167373 | Unspecified |
| mmu-miR-139-5p | MIMAT0000656 | Mapk14 | 26416 | HITS-CLIP | 31433990\|29167373 | Unspecified |
| mmu-miR-139-5p | MIMAT0000656 | Mapk1ip1l | 218975 | HITS-CLIP | 29167373 | Unspecified |
| mmu-miR-139-5p | MIMAT0000656 | Mapk3 | 26417 | sRNA-Seq//HITS-CLIP | 33184237\|26203562 | Unspecified |

| | | | | | | |
|---|---|---|---|---|---|---|
| mmu-miR-139-5p | MIMAT0000656 | Mapk6 | 50772 | sRNA-Seq//HITS-CLIP | 33184237\|29167373 | Unspecified |
| mmu-miR-139-5p | MIMAT0000656 | Mapk9 | 26420 | sRNA-Seq//HITS-CLIP | 33184237\|29167373 | Unspecified |
| mmu-miR-139-5p | MIMAT0000656 | Mapre1 | 13589 | sRNA-Seq//HITS-CLIP | 33184237\|29167373 | Unspecified |
| mmu-miR-139-5p | MIMAT0000656 | March5 | 54708 | sRNA-Seq//HITS-CLIP | 33184237\|29167373 | Unspecified |
| mmu-miR-139-5p | MIMAT0000656 | Marcks | 17118 | sRNA-Seq//HITS-CLIP | 33184237\|26203562\|29167373 | Unspecified |
| mmu-miR-139-5p | MIMAT0000656 | Marf1 | 223989 | HITS-CLIP | 29167373 | Unspecified |
| mmu-miR-139-5p | MIMAT0000656 | Mat2a | 232087 | sRNA-Seq//HITS-CLIP | 33184237\|29167373 | Unspecified |
| mmu-miR-139-5p | MIMAT0000656 | Mat2b | 108645 | HITS-CLIP | 29167373 | Unspecified |
| mmu-miR-139-5p | MIMAT0000656 | Matr3 | 17184 | sRNA-Seq//HITS-CLIP | 33184237\|29167373 | Unspecified |
| mmu-miR-139-5p | MIMAT0000656 | Max | 17187 | sRNA-Seq//HITS-CLIP | 33184237\|29167373 | Unspecified |
| mmu-miR-139-5p | MIMAT0000656 | Mbd5 | 109241 | sRNA-Seq//HITS-CLIP | 33184237\|29167373 | Unspecified |
| mmu-miR-139-5p | MIMAT0000656 | Mbd6 | 110962 | HITS-CLIP | 29167373 | Unspecified |
| mmu-miR-139-5p | MIMAT0000656 | Mbnl1 | 56758 | HITS-CLIP | 29167373 | Unspecified |
| mmu-miR-139-5p | MIMAT0000656 | Mbnl2 | 105559 | HITS-CLIP | 29167373 | Unspecified |
| mmu-miR-139-5p | MIMAT0000656 | Mbtps1 | 56453 | HITS-CLIP | 29167373 | Unspecified |
| mmu-miR-139-5p | MIMAT0000656 | Mbtps2 | 270669 | HITS-CLIP | 29167373 | Unspecified |
| mmu-miR-139-5p | MIMAT0000656 | Mcl1 | 17210 | sRNA-Seq//HITS-CLIP | 33184237\|29167373 | Unspecified |
| mmu-miR-139-5p | MIMAT0000656 | Mcm6 | 17219 | HITS-CLIP | 29167373 | Unspecified |
| mmu-miR-139-5p | MIMAT0000656 | Mcm9 | 71567 | HITS-CLIP | 29167373 | Unspecified |
| mmu-miR-139-5p | MIMAT0000656 | Mctp1 | 78771 | HITS-CLIP | 29167373 | Unspecified |
| mmu-miR-139-5p | MIMAT0000656 | Mdc1 | 240087 | HITS-CLIP | 29167373 | Unspecified |
| mmu-miR-139-5p | MIMAT0000656 | Mdfic | 16543 | HITS-CLIP | 29167373 | Unspecified |
| mmu-miR-139-5p | MIMAT0000656 | Mdh2 | 17448 | sRNA-Seq//HITS-CLIP | 33184237\|31433990 | Unspecified |
| mmu-miR-139-5p | MIMAT0000656 | Mdm4 | 17248 | sRNA-Seq//HITS-CLIP | 33184237\|29167373 | Unspecified |
| mmu-miR-139-5p | MIMAT0000656 | Mdn1 | 100019 | HITS-CLIP | 29167373 | Unspecified |
| mmu-miR-139-5p | MIMAT0000656 | Mecp2 | 17257 | HITS-CLIP | 29167373 | Unspecified |
| mmu-miR-139-5p | MIMAT0000656 | Med1 | 19014 | HITS-CLIP | 29167373 | Unspecified |
| mmu-miR-139-5p | MIMAT0000656 | Med12 | 59024 | HITS-CLIP | 29167373 | Unspecified |
| mmu-miR-139-5p | MIMAT0000656 | Med13 | 327987 | HITS-CLIP | 29167373 | Unspecified |
| mmu-miR-139-5p | MIMAT0000656 | Med13l | 76199 | HITS-CLIP | 29167373 | Unspecified |
| mmu-miR-139-5p | MIMAT0000656 | Med15 | 94112 | HITS-CLIP | 29167373 | Unspecified |
| mmu-miR-139-5p | MIMAT0000656 | Med21 | 108098 | sRNA-Seq//HITS-CLIP | 33184237\|31433990 | Unspecified |
| mmu-miR-139-5p | MIMAT0000656 | Mef2c | 17260 | HITS-CLIP | 29167373 | Unspecified |
| mmu-miR-139-5p | MIMAT0000656 | Metap1 | 75624 | HITS-CLIP | 31433990 | Unspecified |
| mmu-miR-139-5p | MIMAT0000656 | Metrnl | 210029 | HITS-CLIP | 29167373 | Unspecified |
| mmu-miR-139-5p | MIMAT0000656 | Mettl14 | 210529 | HITS-CLIP | 29167373 | Unspecified |
| mmu-miR-139-5p | MIMAT0000656 | Mex3c | 240396 | HITS-CLIP | 29167373 | Unspecified |
| mmu-miR-139-5p | MIMAT0000656 | Mfap3 | 216760 | HITS-CLIP | 29167373 | Unspecified |
| mmu-miR-139-5p | MIMAT0000656 | Mfsd12 | 73822 | HITS-CLIP | 29167373 | Unspecified |

| | | | | | | | |
|---|---|---|---|---|---|---|---|
| mmu-miR-139-5p | MIMAT0000656 | Mfsd14a | 15247 | HITS-CLIP | | 29167373 | Unspecified |
| mmu-miR-139-5p | MIMAT0000656 | Mgea5 | 76055 | HITS-CLIP | | 29167373 | Unspecified |
| mmu-miR-139-5p | MIMAT0000656 | Mia2 | 338320 | HITS-CLIP | | 29167373 | Unspecified |
| mmu-miR-139-5p | MIMAT0000656 | Mia3 | 338366 | sRNA-Seq//HITS-CLIP | 33184237\|29167373 | | Unspecified |
| mmu-miR-139-5p | MIMAT0000656 | Mib1 | 225164 | sRNA-Seq//HITS-CLIP | 33184237\|31433990\|29167373 | | Unspecified |
| mmu-miR-139-5p | MIMAT0000656 | Micall2 | 231830 | HITS-CLIP | | 29167373 | Unspecified |
| mmu-miR-139-5p | MIMAT0000656 | Micu1 | 216001 | HITS-CLIP | | 29167373 | Unspecified |
| mmu-miR-139-5p | MIMAT0000656 | Mideas | 91748 | HITS-CLIP | | 31433990 | Unspecified |
| mmu-miR-139-5p | MIMAT0000656 | Midn | 59090 | HITS-CLIP//sRNA-Seq | 31433990\|33184237 | | Unspecified |
| mmu-miR-139-5p | MIMAT0000656 | Mier2 | 70427 | HITS-CLIP | | 29167373 | Unspecified |
| mmu-miR-139-5p | MIMAT0000656 | Mitf | 17342 | HITS-CLIP | | 29167373 | Unspecified |
| mmu-miR-139-5p | MIMAT0000656 | Mki67 | 17345 | HITS-CLIP | | 26203562 | Unspecified |
| mmu-miR-139-5p | MIMAT0000656 | Mkln1 | 27418 | sRNA-Seq//HITS-CLIP | 33184237\|29167373 | | Unspecified |
| mmu-miR-139-5p | MIMAT0000656 | Mknk2 | 17347 | HITS-CLIP | | 29167373 | Unspecified |
| mmu-miR-139-5p | MIMAT0000656 | Mkrn1 | 54484 | sRNA-Seq//HITS-CLIP | 33184237\|29167373 | | Unspecified |
| mmu-miR-139-5p | MIMAT0000656 | Mlec | 109154 | HITS-CLIP | | 29167373 | Unspecified |
| mmu-miR-139-5p | MIMAT0000656 | Mllt1 | 64144 | HITS-CLIP | | 29167373 | Unspecified |
| mmu-miR-139-5p | MIMAT0000656 | Mllt6 | 246198 | HITS-CLIP | | 29167373 | Unspecified |
| mmu-miR-139-5p | MIMAT0000656 | Mlxip | 208104 | HITS-CLIP | | 29167373 | Unspecified |
| mmu-miR-139-5p | MIMAT0000656 | Mnt | 17428 | HITS-CLIP | | 29167373 | Unspecified |
| mmu-miR-139-5p | MIMAT0000656 | Mob1a | 232157 | HITS-CLIP | | 29167373 | Unspecified |
| mmu-miR-139-5p | MIMAT0000656 | Mob2 | 101513 | HITS-CLIP | | 29167373 | Unspecified |
| mmu-miR-139-5p | MIMAT0000656 | Mob3a | 208228 | HITS-CLIP | | 29167373 | Unspecified |
| mmu-miR-139-5p | MIMAT0000656 | Mocos | 68591 | HITS-CLIP | | 29167373 | Unspecified |
| mmu-miR-139-5p | MIMAT0000656 | Mocs2 | 17434 | sRNA-Seq//HITS-CLIP | 33184237\|29167373 | | Unspecified |
| mmu-miR-139-5p | MIMAT0000656 | Mon2 | 67074 | HITS-CLIP | | 29167373 | Unspecified |
| mmu-miR-139-5p | MIMAT0000656 | Morc2a | 74522 | HITS-CLIP | | 29167373 | Unspecified |
| mmu-miR-139-5p | MIMAT0000656 | Morc3 | 338467 | HITS-CLIP | | 29167373 | Unspecified |
| mmu-miR-139-5p | MIMAT0000656 | Morf4l2 | 56397 | sRNA-Seq//HITS-CLIP | 33184237\|29167373 | | Unspecified |
| mmu-miR-139-5p | MIMAT0000656 | Mpeg1 | 17476 | HITS-CLIP | | 29167373 | Unspecified |
| mmu-miR-139-5p | MIMAT0000656 | Mphosph8 | 75339 | HITS-CLIP | | 29167373 | Unspecified |
| mmu-miR-139-5p | MIMAT0000656 | Mpp1 | 17524 | HITS-CLIP | | 29167373 | Unspecified |
| mmu-miR-139-5p | MIMAT0000656 | Mpv17 | 17527 | HITS-CLIP | | 23630078 | Unspecified |
| mmu-miR-139-5p | MIMAT0000656 | Mpzl2 | 14012 | HITS-CLIP | | 26203562 | Unspecified |
| mmu-miR-139-5p | MIMAT0000656 | Mrm2 | 29960 | HITS-CLIP | | 29167373 | Unspecified |
| mmu-miR-139-5p | MIMAT0000656 | Mroh2a | 100040766 | HITS-CLIP | | 29167373 | Unspecified |
| mmu-miR-139-5p | MIMAT0000656 | Mrpl24 | 67707 | HITS-CLIP | | 29167373 | Unspecified |
| mmu-miR-139-5p | MIMAT0000656 | Mrpl35 | 66223 | HITS-CLIP | | 31433990 | Unspecified |
| mmu-miR-139-5p | MIMAT0000656 | Mrps15 | 66407 | sRNA-Seq//HITS-CLIP | 33184237\|29167373 | | Unspecified |

| | | | | | | |
|---|---|---|---|---|---|---|
| mmu-miR-139-5p | MIMAT0000656 | Mrps2 | 118451 | HITS-CLIP | 29167373 | Unspecified |
| mmu-miR-139-5p | MIMAT0000656 | Mrtfa | 57591 | HITS-CLIP | 29167373 | Unspecified |
| mmu-miR-139-5p | MIMAT0000656 | Mrtfb | 239719 | HITS-CLIP | 29167373 | Unspecified |
| mmu-miR-139-5p | MIMAT0000656 | Ms4a6b | 69774 | HITS-CLIP | 29167373 | Unspecified |
| mmu-miR-139-5p | MIMAT0000656 | Msl2 | 77853 | HITS-CLIP | 29167373 | Unspecified |
| mmu-miR-139-5p | MIMAT0000656 | Msmo1 | 66234 | sRNA-Seq//HITS-CLIP | 33184237\|29167373 | Unspecified |
| mmu-miR-139-5p | MIMAT0000656 | Msn | 17698 | sRNA-Seq//HITS-CLIP | 33184237\|29167373 | Unspecified |
| mmu-miR-139-5p | MIMAT0000656 | Msr1 | 20288 | HITS-CLIP | 29167373 | Unspecified |
| mmu-miR-139-5p | MIMAT0000656 | Mthfr | 17769 | HITS-CLIP | 29167373 | Unspecified |
| mmu-miR-139-5p | MIMAT0000656 | Mthfsl | 100039707 | HITS-CLIP | 29167373 | Unspecified |
| mmu-miR-139-5p | MIMAT0000656 | Mtmr1 | 53332 | HITS-CLIP | 29167373 | Unspecified |
| mmu-miR-139-5p | MIMAT0000656 | Mtmr10 | 233315 | HITS-CLIP | 29167373 | Unspecified |
| mmu-miR-139-5p | MIMAT0000656 | Mtmr2 | 77116 | sRNA-Seq//HITS-CLIP | 33184237\|29167373 | Unspecified |
| mmu-miR-139-5p | MIMAT0000656 | Mtmr3 | 74302 | HITS-CLIP | 31433990\|26203562 | Unspecified |
| mmu-miR-139-5p | MIMAT0000656 | Mtmr6 | 219135 | sRNA-Seq//HITS-CLIP | 33184237\|29167373 | Unspecified |
| mmu-miR-139-5p | MIMAT0000656 | Mtor | 56717 | HITS-CLIP | 29167373 | Unspecified |
| mmu-miR-139-5p | MIMAT0000656 | Mtr | 238505 | HITS-CLIP | 29167373 | Unspecified |
| mmu-miR-139-5p | MIMAT0000656 | Mtss1 | 211401 | HITS-CLIP | 29167373 | Unspecified |
| mmu-miR-139-5p | MIMAT0000656 | Mvp | 78388 | HITS-CLIP | 29167373 | Unspecified |
| mmu-miR-139-5p | MIMAT0000656 | Mx1 | 17857 | HITS-CLIP | 29167373 | Unspecified |
| mmu-miR-139-5p | MIMAT0000656 | Myadm | 50918 | sRNA-Seq//HITS-CLIP | 33184237\|29167373 | Unspecified |
| mmu-miR-139-5p | MIMAT0000656 | Mybbp1a | 18432 | HITS-CLIP | 29167373 | Unspecified |
| mmu-miR-139-5p | MIMAT0000656 | Mycbp2 | 105689 | sRNA-Seq//HITS-CLIP | 33184237\|29167373 | Unspecified |
| mmu-miR-139-5p | MIMAT0000656 | Myg1 | 60315 | HITS-CLIP | 29167373 | Unspecified |
| mmu-miR-139-5p | MIMAT0000656 | Myh9 | 17886 | HITS-CLIP | 26203562\|29167373 | Unspecified |
| mmu-miR-139-5p | MIMAT0000656 | Myo10 | 17909 | HITS-CLIP | 29167373 | Unspecified |
| mmu-miR-139-5p | MIMAT0000656 | Myo18a | 360013 | HITS-CLIP | 29167373 | Unspecified |
| mmu-miR-139-5p | MIMAT0000656 | Myo1c | 17913 | HITS-CLIP | 29167373 | Unspecified |
| mmu-miR-139-5p | MIMAT0000656 | Myo1e | 71602 | HITS-CLIP | 29167373 | Unspecified |
| mmu-miR-139-5p | MIMAT0000656 | Myo1f | 17916 | HITS-CLIP | 29167373 | Unspecified |
| mmu-miR-139-5p | MIMAT0000656 | Myo5a | 17918 | sRNA-Seq//HITS-CLIP | 33184237\|29167373 | Unspecified |
| mmu-miR-139-5p | MIMAT0000656 | Myo9b | 17925 | HITS-CLIP | 29167373 | Unspecified |
| mmu-miR-139-5p | MIMAT0000656 | Myof | 226101 | HITS-CLIP | 29167373 | Unspecified |
| mmu-miR-139-5p | MIMAT0000656 | Mysm1 | 320713 | HITS-CLIP | 29167373 | Unspecified |
| mmu-miR-139-5p | MIMAT0000656 | N4bp1 | 80750 | HITS-CLIP | 29167373 | Unspecified |
| mmu-miR-139-5p | MIMAT0000656 | N4bp2l2 | 381695 | sRNA-Seq//HITS-CLIP | 33184237\|29167373 | Unspecified |
| mmu-miR-139-5p | MIMAT0000656 | Naa15 | 74838 | sRNA-Seq//HITS-CLIP | 33184237\|29167373 | Unspecified |
| mmu-miR-139-5p | MIMAT0000656 | Naa50 | 72117 | HITS-CLIP | 29167373 | Unspecified |
| mmu-miR-139-5p | MIMAT0000656 | Naa60 | 74763 | HITS-CLIP | 29167373 | Unspecified |

| | | | | | | |
|---|---|---|---|---|---|---|
| mmu-miR-139-5p | MIMAT0000656 | Nab1 | 17936 | HITS-CLIP | 29167373 | Unspecified |
| mmu-miR-139-5p | MIMAT0000656 | Naca | 17938 | sRNA-Seq//HITS-CLIP | 33184237\|29167373 | Unspecified |
| mmu-miR-139-5p | MIMAT0000656 | Nadk | 192185 | HITS-CLIP | 29167373 | Unspecified |
| mmu-miR-139-5p | MIMAT0000656 | Nap1l1 | 53605 | sRNA-Seq//HITS-CLIP | 33184237\|29167373 | Unspecified |
| mmu-miR-139-5p | MIMAT0000656 | Napa | 108124 | sRNA-Seq//HITS-CLIP | 33184237\|29167373 | Unspecified |
| mmu-miR-139-5p | MIMAT0000656 | Napg | 108123 | HITS-CLIP | 29167373 | Unspecified |
| mmu-miR-139-5p | MIMAT0000656 | Narf | 67608 | HITS-CLIP | 29167373 | Unspecified |
| mmu-miR-139-5p | MIMAT0000656 | Nav1 | 215690 | sRNA-Seq//HITS-CLIP | 33184237\|29167373 | Unspecified |
| mmu-miR-139-5p | MIMAT0000656 | Nbas | 71169 | HITS-CLIP | 29167373 | Unspecified |
| mmu-miR-139-5p | MIMAT0000656 | Nbeal2 | 235627 | HITS-CLIP | 29167373 | Unspecified |
| mmu-miR-139-5p | MIMAT0000656 | Nbr1 | 17966 | sRNA-Seq//HITS-CLIP | 33184237\|29167373 | Unspecified |
| mmu-miR-139-5p | MIMAT0000656 | Ncald | 52589 | HITS-CLIP | 31433990 | Unspecified |
| mmu-miR-139-5p | MIMAT0000656 | Ncapg2 | 76044 | HITS-CLIP | 29167373 | Unspecified |
| mmu-miR-139-5p | MIMAT0000656 | Ncaph2 | 52683 | HITS-CLIP | 29167373 | Unspecified |
| mmu-miR-139-5p | MIMAT0000656 | Nceh1 | 320024 | HITS-CLIP | 29167373 | Unspecified |
| mmu-miR-139-5p | MIMAT0000656 | Ncf1 | 17969 | HITS-CLIP | 29167373 | Unspecified |
| mmu-miR-139-5p | MIMAT0000656 | Nckap1l | 105855 | HITS-CLIP | 29167373 | Unspecified |
| mmu-miR-139-5p | MIMAT0000656 | Ncl | 17975 | sRNA-Seq//HITS-CLIP | 33184237\|29167373 | Unspecified |
| mmu-miR-139-5p | MIMAT0000656 | Ncoa1 | 17977 | HITS-CLIP | 29167373 | Unspecified |
| mmu-miR-139-5p | MIMAT0000656 | Ncoa2 | 17978 | HITS-CLIP | 29167373 | Unspecified |
| mmu-miR-139-5p | MIMAT0000656 | Ncoa3 | 17979 | HITS-CLIP | 31433990\|29167373 | Unspecified |
| mmu-miR-139-5p | MIMAT0000656 | Ncor1 | 20185 | sRNA-Seq//HITS-CLIP | 33184237\|29167373 | Unspecified |
| mmu-miR-139-5p | MIMAT0000656 | Ncor2 | 20602 | HITS-CLIP | 29167373 | Unspecified |
| mmu-miR-139-5p | MIMAT0000656 | Ndfip2 | 76273 | sRNA-Seq//HITS-CLIP | 33184237\|29167373 | Unspecified |
| mmu-miR-139-5p | MIMAT0000656 | Ndst1 | 15531 | HITS-CLIP | 26203562 | Unspecified |
| mmu-miR-139-5p | MIMAT0000656 | Ndufa4 | 17992 | sRNA-Seq//HITS-CLIP | 33184237\|29167373 | Unspecified |
| mmu-miR-139-5p | MIMAT0000656 | Necap1 | 67602 | sRNA-Seq//HITS-CLIP | 33184237\|29167373 | Unspecified |
| mmu-miR-139-5p | MIMAT0000656 | Nectin1 | 58235 | HITS-CLIP | 29167373 | Unspecified |
| mmu-miR-139-5p | MIMAT0000656 | Nek9 | 217718 | HITS-CLIP | 26203562 | Unspecified |
| mmu-miR-139-5p | MIMAT0000656 | Nemf | 66244 | HITS-CLIP | 29167373 | Unspecified |
| mmu-miR-139-5p | MIMAT0000656 | Nepro | 212547 | HITS-CLIP | 29167373 | Unspecified |
| mmu-miR-139-5p | MIMAT0000656 | Nf1 | 18015 | sRNA-Seq//HITS-CLIP | 33184237\|29167373 | Unspecified |
| mmu-miR-139-5p | MIMAT0000656 | Nfam1 | 74039 | HITS-CLIP | 29167373 | Unspecified |
| mmu-miR-139-5p | MIMAT0000656 | Nfat5 | 54446 | HITS-CLIP | 29167373 | Unspecified |
| mmu-miR-139-5p | MIMAT0000656 | Nfatc1 | 18018 | HITS-CLIP | 29167373 | Unspecified |
| mmu-miR-139-5p | MIMAT0000656 | Nfe2l1 | 18023 | sRNA-Seq//HITS-CLIP | 33184237\|29167373 | Unspecified |
| mmu-miR-139-5p | MIMAT0000656 | Nfkbiz | 80859 | HITS-CLIP | 29167373 | Unspecified |
| mmu-miR-139-5p | MIMAT0000656 | Nfrkb | 235134 | HITS-CLIP | 31433990 | Unspecified |
| mmu-miR-139-5p | MIMAT0000656 | Nfx1 | 74164 | HITS-CLIP | 29167373 | Unspecified |

| | | | | | | |
|---|---|---|---|---|---|---|
| mmu-miR-139-5p | MIMAT0000656 | Nfxl1 | 100978 | HITS-CLIP | 29167373 | Unspecified |
| mmu-miR-139-5p | MIMAT0000656 | Nfya | 18044 | HITS-CLIP | 29167373 | Unspecified |
| mmu-miR-139-5p | MIMAT0000656 | Nhlrc3 | 212114 | HITS-CLIP | 29167373 | Unspecified |
| mmu-miR-139-5p | MIMAT0000656 | Nhp2 | 52530 | HITS-CLIP | 29167373 | Unspecified |
| mmu-miR-139-5p | MIMAT0000656 | Nin | 18080 | HITS-CLIP | 29167373 | Unspecified |
| mmu-miR-139-5p | MIMAT0000656 | Nip7 | 66164 | HITS-CLIP | 21258322\|31433990 | Unspecified |
| mmu-miR-139-5p | MIMAT0000656 | Nipa2 | 93790 | HITS-CLIP | 31433990 | Unspecified |
| mmu-miR-139-5p | MIMAT0000656 | Nipal4 | 214112 | HITS-CLIP | 26203562 | Unspecified |
| mmu-miR-139-5p | MIMAT0000656 | Nipbl | 71175 | HITS-CLIP | 29167373 | Unspecified |
| mmu-miR-139-5p | MIMAT0000656 | Nipsnap3b | 66536 | HITS-CLIP//sRNA-Seq | 31433990\|33184237 | Unspecified |
| mmu-miR-139-5p | MIMAT0000656 | Nisch | 64652 | sRNA-Seq//HITS-CLIP | 33184237\|29167373 | Unspecified |
| mmu-miR-139-5p | MIMAT0000656 | Nktr | 18087 | sRNA-Seq//HITS-CLIP | 33184237\|29167373 | Unspecified |
| mmu-miR-139-5p | MIMAT0000656 | Nlrc5 | 434341 | HITS-CLIP | 29167373 | Unspecified |
| mmu-miR-139-5p | MIMAT0000656 | Nlrp1b | 637515 | HITS-CLIP | 29167373 | Unspecified |
| mmu-miR-139-5p | MIMAT0000656 | Nme7 | 171567 | HITS-CLIP | 31433990 | Unspecified |
| mmu-miR-139-5p | MIMAT0000656 | Nmt2 | 18108 | sRNA-Seq//HITS-CLIP | 33184237\|26203562 | Unspecified |
| mmu-miR-139-5p | MIMAT0000656 | Noc4l | 100608 | HITS-CLIP | 29167373 | Unspecified |
| mmu-miR-139-5p | MIMAT0000656 | Nol4l | 329540 | sRNA-Seq//HITS-CLIP | 33184237\|29167373 | Unspecified |
| mmu-miR-139-5p | MIMAT0000656 | Nom1 | 433864 | HITS-CLIP | 29167373 | Unspecified |
| mmu-miR-139-5p | MIMAT0000656 | Nop53 | 29997 | HITS-CLIP | 31433990 | Unspecified |
| mmu-miR-139-5p | MIMAT0000656 | Nop9 | 67842 | HITS-CLIP | 21258322\|29167373 | Unspecified |
| mmu-miR-139-5p | MIMAT0000656 | Notch2 | 18129 | HITS-CLIP | 31433990\|26203562\|29167373 | Unspecified |
| mmu-miR-139-5p | MIMAT0000656 | Npc1 | 18145 | HITS-CLIP | 29167373 | Unspecified |
| mmu-miR-139-5p | MIMAT0000656 | Npepps | 19155 | sRNA-Seq//HITS-CLIP | 33184237\|31433990 | Unspecified |
| mmu-miR-139-5p | MIMAT0000656 | Nptn | 20320 | sRNA-Seq//HITS-CLIP | 33184237\|29167373 | Unspecified |
| mmu-miR-139-5p | MIMAT0000656 | Nr2f6 | 13864 | HITS-CLIP | 29167373 | Unspecified |
| mmu-miR-139-5p | MIMAT0000656 | Nr3c1 | 14815 | HITS-CLIP | 31433990\|29167373 | Unspecified |
| mmu-miR-139-5p | MIMAT0000656 | Nras | 18176 | sRNA-Seq//HITS-CLIP | 33184237\|29167373 | Unspecified |
| mmu-miR-139-5p | MIMAT0000656 | Nrd1 | 230598 | sRNA-Seq//HITS-CLIP | 33184237\|29167373 | Unspecified |
| mmu-miR-139-5p | MIMAT0000656 | Nrf1 | 18181 | HITS-CLIP | 29167373 | Unspecified |
| mmu-miR-139-5p | MIMAT0000656 | Nrp1 | 18186 | sRNA-Seq//HITS-CLIP | 33184237\|29167373 | Unspecified |
| mmu-miR-139-5p | MIMAT0000656 | Nrp2 | 18187 | HITS-CLIP | 29167373 | Unspecified |
| mmu-miR-139-5p | MIMAT0000656 | Nsd1 | 18193 | sRNA-Seq//HITS-CLIP | 33184237\|29167373 | Unspecified |
| mmu-miR-139-5p | MIMAT0000656 | Nsd2 | 107823 | sRNA-Seq//HITS-CLIP | 33184237\|29167373 | Unspecified |
| mmu-miR-139-5p | MIMAT0000656 | Nsd3 | 234135 | sRNA-Seq//HITS-CLIP | 33184237\|29167373 | Unspecified |
| mmu-miR-139-5p | MIMAT0000656 | Nsrp1 | 237859 | HITS-CLIP | 29167373 | Unspecified |
| mmu-miR-139-5p | MIMAT0000656 | Nsun2 | 28114 | HITS-CLIP | 29167373 | Unspecified |
| mmu-miR-139-5p | MIMAT0000656 | Nt5c2 | 76952 | HITS-CLIP | 29167373 | Unspecified |
| mmu-miR-139-5p | MIMAT0000656 | Nt5dc2 | 70021 | HITS-CLIP | 29167373 | Unspecified |

| | | | | | | |
|---|---|---|---|---|---|---|
| mmu-miR-139-5p | MIMAT0000656 | Nub1 | 53312 | sRNA-Seq//HITS-CLIP | 33184237\|29167373 | Unspecified |
| mmu-miR-139-5p | MIMAT0000656 | Nucks1 | 98415 | sRNA-Seq//HITS-CLIP | 33184237\|29167373 | Unspecified |
| mmu-miR-139-5p | MIMAT0000656 | Nudcd3 | 209586 | HITS-CLIP | 29167373 | Unspecified |
| mmu-miR-139-5p | MIMAT0000656 | Nuf2 | 66977 | HITS-CLIP | 29167373 | Unspecified |
| mmu-miR-139-5p | MIMAT0000656 | Nufip2 | 68564 | sRNA-Seq//HITS-CLIP | 33184237\|29167373 | Unspecified |
| mmu-miR-139-5p | MIMAT0000656 | Numb | 18222 | HITS-CLIP | 31433990 | Unspecified |
| mmu-miR-139-5p | MIMAT0000656 | Nup153 | 218210 | HITS-CLIP | 31433990\|29167373 | Unspecified |
| mmu-miR-139-5p | MIMAT0000656 | Nup205 | 70699 | HITS-CLIP | 29167373 | Unspecified |
| mmu-miR-139-5p | MIMAT0000656 | Nup98 | 269966 | HITS-CLIP | 29167373 | Unspecified |
| mmu-miR-139-5p | MIMAT0000656 | Nus1 | 52014 | HITS-CLIP | 29167373 | Unspecified |
| mmu-miR-139-5p | MIMAT0000656 | Nvl | 67459 | HITS-CLIP | 29167373 | Unspecified |
| mmu-miR-139-5p | MIMAT0000656 | Oas1a | 246730 | HITS-CLIP | 29167373 | Unspecified |
| mmu-miR-139-5p | MIMAT0000656 | Oas1g | 23960 | HITS-CLIP | 29167373 | Unspecified |
| mmu-miR-139-5p | MIMAT0000656 | Oas2 | 246728 | HITS-CLIP | 29167373 | Unspecified |
| mmu-miR-139-5p | MIMAT0000656 | Oasl2 | 23962 | HITS-CLIP | 29167373 | Unspecified |
| mmu-miR-139-5p | MIMAT0000656 | Ogdh | 18293 | sRNA-Seq//HITS-CLIP | 33184237\|29167373 | Unspecified |
| mmu-miR-139-5p | MIMAT0000656 | Olfm1 | 56177 | sRNA-Seq//HITS-CLIP | 33184237\|29167373 | Unspecified |
| mmu-miR-139-5p | MIMAT0000656 | Olfr111 | 545205 | HITS-CLIP | 29167373 | Unspecified |
| mmu-miR-139-5p | MIMAT0000656 | Orc3 | 50793 | HITS-CLIP | 29167373 | Unspecified |
| mmu-miR-139-5p | MIMAT0000656 | Os9 | 216440 | sRNA-Seq//HITS-CLIP | 33184237\|29167373 | Unspecified |
| mmu-miR-139-5p | MIMAT0000656 | Osbpl11 | 106326 | HITS-CLIP | 29167373 | Unspecified |
| mmu-miR-139-5p | MIMAT0000656 | Osbpl3 | 71720 | HITS-CLIP | 29167373 | Unspecified |
| mmu-miR-139-5p | MIMAT0000656 | Osbpl8 | 237542 | HITS-CLIP//sRNA-Seq | 25083871\|33184237\|29167373\|21258 | Unspecified |
| mmu-miR-139-5p | MIMAT0000656 | Osgin2 | 209212 | HITS-CLIP | 29167373 | Unspecified |
| mmu-miR-139-5p | MIMAT0000656 | Ostf1 | 20409 | HITS-CLIP | 21258322\|29167373 | Unspecified |
| mmu-miR-139-5p | MIMAT0000656 | Otulinl | 54491 | HITS-CLIP | 29167373 | Unspecified |
| mmu-miR-139-5p | MIMAT0000656 | Oxsr1 | 108737 | HITS-CLIP | 29167373 | Unspecified |
| mmu-miR-139-5p | MIMAT0000656 | P2rx4 | 18438 | HITS-CLIP | 29167373 | Unspecified |
| mmu-miR-139-5p | MIMAT0000656 | P2ry2 | 18442 | HITS-CLIP | 29167373 | Unspecified |
| mmu-miR-139-5p | MIMAT0000656 | P2ry6 | 233571 | HITS-CLIP | 29167373 | Unspecified |
| mmu-miR-139-5p | MIMAT0000656 | P4hb | 18453 | sRNA-Seq//HITS-CLIP | 33184237\|29167373 | Unspecified |
| mmu-miR-139-5p | MIMAT0000656 | Pacsin2 | 23970 | HITS-CLIP | 29167373 | Unspecified |
| mmu-miR-139-5p | MIMAT0000656 | Pafah1b1 | 18472 | sRNA-Seq//HITS-CLIP | 33184237\|29167373 | Unspecified |
| mmu-miR-139-5p | MIMAT0000656 | Pag1 | 94212 | HITS-CLIP | 31433990 | Unspecified |
| mmu-miR-139-5p | MIMAT0000656 | Paics | 67054 | sRNA-Seq//HITS-CLIP | 33184237\|29167373 | Unspecified |
| mmu-miR-139-5p | MIMAT0000656 | Paip2 | 67869 | HITS-CLIP//sRNA-Seq | 31433990\|33184237 | Unspecified |
| mmu-miR-139-5p | MIMAT0000656 | Pak2 | 224105 | HITS-CLIP | 29167373 | Unspecified |
| mmu-miR-139-5p | MIMAT0000656 | Pak4 | 70584 | HITS-CLIP | 29167373 | Unspecified |
| mmu-miR-139-5p | MIMAT0000656 | Pan3 | 72587 | HITS-CLIP | 29167373 | Unspecified |

| | | | | | | | |
|---|---|---|---|---|---|---|---|
| mmu-miR-139-5p | MIMAT0000656 | Pank2 | 74450 | HITS-CLIP | | 29167373 | Unspecified |
| mmu-miR-139-5p | MIMAT0000656 | Pank3 | 211347 | sRNA-Seq//HITS-CLIP | 33184237\|31433990\|29167373 | | Unspecified |
| mmu-miR-139-5p | MIMAT0000656 | Parg | 26430 | HITS-CLIP | | 29167373 | Unspecified |
| mmu-miR-139-5p | MIMAT0000656 | Parp11 | 101187 | HITS-CLIP | 21258322\|29167373 | | Unspecified |
| mmu-miR-139-5p | MIMAT0000656 | Parp14 | 547253 | HITS-CLIP | 31433990\|29167373 | | Unspecified |
| mmu-miR-139-5p | MIMAT0000656 | Parp4 | 328417 | HITS-CLIP | | 29167373 | Unspecified |
| mmu-miR-139-5p | MIMAT0000656 | Parvb | 170736 | sRNA-Seq//HITS-CLIP | 33184237\|29167373 | | Unspecified |
| mmu-miR-139-5p | MIMAT0000656 | Patl1 | 225929 | HITS-CLIP | | 29167373 | Unspecified |
| mmu-miR-139-5p | MIMAT0000656 | Patz1 | 56218 | HITS-CLIP | | 29167373 | Unspecified |
| mmu-miR-139-5p | MIMAT0000656 | Paxip1 | 55982 | HITS-CLIP | | 29167373 | Unspecified |
| mmu-miR-139-5p | MIMAT0000656 | Pbx2 | 18515 | HITS-CLIP | | 29167373 | Unspecified |
| mmu-miR-139-5p | MIMAT0000656 | Pcbp1 | 23983 | sRNA-Seq//HITS-CLIP | 33184237\|29167373 | | Unspecified |
| mmu-miR-139-5p | MIMAT0000656 | Pcdh7 | 54216 | sRNA-Seq//HITS-CLIP | 33184237\|29167373 | | Unspecified |
| mmu-miR-139-5p | MIMAT0000656 | Pcgf3 | 69587 | HITS-CLIP | | 31433990 | Unspecified |
| mmu-miR-139-5p | MIMAT0000656 | Pcm1 | 18536 | sRNA-Seq//HITS-CLIP | 33184237\|29167373 | | Unspecified |
| mmu-miR-139-5p | MIMAT0000656 | Pcnt | 18541 | HITS-CLIP | | 29167373 | Unspecified |
| mmu-miR-139-5p | MIMAT0000656 | Pcnx3 | 399909 | HITS-CLIP | | 29167373 | Unspecified |
| mmu-miR-139-5p | MIMAT0000656 | Pcolce | 18542 | HITS-CLIP | | 23630078 | Unspecified |
| mmu-miR-139-5p | MIMAT0000656 | Pctp | 18559 | HITS-CLIP | | 29167373 | Unspecified |
| mmu-miR-139-5p | MIMAT0000656 | Pde3b | 18576 | HITS-CLIP | | 29167373 | Unspecified |
| mmu-miR-139-5p | MIMAT0000656 | Pde8a | 18584 | HITS-CLIP | | 29167373 | Unspecified |
| mmu-miR-139-5p | MIMAT0000656 | Pdgfa | 18590 | HITS-CLIP | | 29167373 | Unspecified |
| mmu-miR-139-5p | MIMAT0000656 | Pdhx | 27402 | HITS-CLIP | | 29167373 | Unspecified |
| mmu-miR-139-5p | MIMAT0000656 | Pdia3 | 14827 | sRNA-Seq//HITS-CLIP | 33184237\|29167373 | | Unspecified |
| mmu-miR-139-5p | MIMAT0000656 | Pdia6 | 71853 | sRNA-Seq//HITS-CLIP | 33184237\|29167373 | | Unspecified |
| mmu-miR-139-5p | MIMAT0000656 | Pdlim5 | 56376 | HITS-CLIP | | 29167373 | Unspecified |
| mmu-miR-139-5p | MIMAT0000656 | Pdpk1 | 18607 | sRNA-Seq//HITS-CLIP | 33184237\|29167373 | | Unspecified |
| mmu-miR-139-5p | MIMAT0000656 | Pdxdc1 | 94184 | HITS-CLIP | | 29167373 | Unspecified |
| mmu-miR-139-5p | MIMAT0000656 | Pdxk | 216134 | HITS-CLIP | | 29167373 | Unspecified |
| mmu-miR-139-5p | MIMAT0000656 | Pdzd8 | 107368 | HITS-CLIP | | 29167373 | Unspecified |
| mmu-miR-139-5p | MIMAT0000656 | Pea15a | 18611 | sRNA-Seq//HITS-CLIP | 33184237\|29167373 | | Unspecified |
| mmu-miR-139-5p | MIMAT0000656 | Peak1 | 244895 | HITS-CLIP | | 29167373 | Unspecified |
| mmu-miR-139-5p | MIMAT0000656 | Pfkfb3 | 170768 | HITS-CLIP | | 29167373 | Unspecified |
| mmu-miR-139-5p | MIMAT0000656 | Pfkl | 18641 | HITS-CLIP | 25083871\|29167373\|21258322 | | Unspecified |
| mmu-miR-139-5p | MIMAT0000656 | Pgap1 | 241062 | HITS-CLIP | | 26203562 | Unspecified |
| mmu-miR-139-5p | MIMAT0000656 | Pggt1b | 225467 | HITS-CLIP | | 29167373 | Unspecified |
| mmu-miR-139-5p | MIMAT0000656 | Pgp | 67078 | HITS-CLIP | | 26203562 | Unspecified |
| mmu-miR-139-5p | MIMAT0000656 | Pgrmc2 | 70804 | HITS-CLIP | | 29167373 | Unspecified |
| mmu-miR-139-5p | MIMAT0000656 | Phaf1 | 80262 | HITS-CLIP | | 31433990 | Unspecified |

| | | | | | | |
|---|---|---|---|---|---|---|
| mmu-miR-139-5p | MIMAT0000656 | Phax | 56698 | sRNA-Seq//HITS-CLIP | 33184237\|29167373 | Unspecified |
| mmu-miR-139-5p | MIMAT0000656 | Phc2 | 54383 | sRNA-Seq//HITS-CLIP | 33184237\|31433990\|29167373 | Unspecified |
| mmu-miR-139-5p | MIMAT0000656 | Phc3 | 241915 | HITS-CLIP | 29167373 | Unspecified |
| mmu-miR-139-5p | MIMAT0000656 | Pheta1 | 144717 | HITS-CLIP | 26203562 | Unspecified |
| mmu-miR-139-5p | MIMAT0000656 | Phf14 | 75725 | sRNA-Seq//HITS-CLIP | 33184237\|29167373 | Unspecified |
| mmu-miR-139-5p | MIMAT0000656 | Phf20 | 228829 | sRNA-Seq//HITS-CLIP | 33184237\|29167373 | Unspecified |
| mmu-miR-139-5p | MIMAT0000656 | Phf3 | 213109 | sRNA-Seq//HITS-CLIP | 33184237\|26203562\|29167373 | Unspecified |
| mmu-miR-139-5p | MIMAT0000656 | Phip | 83946 | sRNA-Seq//HITS-CLIP | 33184237\|29167373 | Unspecified |
| mmu-miR-139-5p | MIMAT0000656 | Phka2 | 110094 | HITS-CLIP | 29167373 | Unspecified |
| mmu-miR-139-5p | MIMAT0000656 | Phkb | 102093 | HITS-CLIP | 29167373 | Unspecified |
| mmu-miR-139-5p | MIMAT0000656 | Phrf1 | 101471 | HITS-CLIP | 29167373 | Unspecified |
| mmu-miR-139-5p | MIMAT0000656 | Pi4ka | 224020 | sRNA-Seq//HITS-CLIP | 33184237\|29167373 | Unspecified |
| mmu-miR-139-5p | MIMAT0000656 | Pias1 | 56469 | HITS-CLIP | 29167373 | Unspecified |
| mmu-miR-139-5p | MIMAT0000656 | Pias2 | 17344 | HITS-CLIP | 31433990\|29167373 | Unspecified |
| mmu-miR-139-5p | MIMAT0000656 | Pias3 | 229615 | HITS-CLIP | 29167373 | Unspecified |
| mmu-miR-139-5p | MIMAT0000656 | Pias4 | 59004 | HITS-CLIP | 29167373 | Unspecified |
| mmu-miR-139-5p | MIMAT0000656 | Picalm | 233489 | HITS-CLIP//sRNA-Seq | 29167373\|33184237 | Unspecified |
| mmu-miR-139-5p | MIMAT0000656 | Pid1 | 98496 | sRNA-Seq//HITS-CLIP | 33184237\|29167373 | Unspecified |
| mmu-miR-139-5p | MIMAT0000656 | Piezo1 | 234839 | HITS-CLIP | 29167373 | Unspecified |
| mmu-miR-139-5p | MIMAT0000656 | Pigk | 329777 | sRNA-Seq//HITS-CLIP | 33184237\|29167373 | Unspecified |
| mmu-miR-139-5p | MIMAT0000656 | Pigm | 67556 | HITS-CLIP | 29167373 | Unspecified |
| mmu-miR-139-5p | MIMAT0000656 | Pign | 27392 | HITS-CLIP | 29167373 | Unspecified |
| mmu-miR-139-5p | MIMAT0000656 | Pigx | 72084 | HITS-CLIP | 31433990 | Unspecified |
| mmu-miR-139-5p | MIMAT0000656 | Pik3ap1 | 83490 | HITS-CLIP | 29167373 | Unspecified |
| mmu-miR-139-5p | MIMAT0000656 | Pik3c2a | 18704 | HITS-CLIP | 29167373 | Unspecified |
| mmu-miR-139-5p | MIMAT0000656 | Pik3cb | 74769 | HITS-CLIP | 29167373 | Unspecified |
| mmu-miR-139-5p | MIMAT0000656 | Pik3cg | 30955 | HITS-CLIP | 29167373 | Unspecified |
| mmu-miR-139-5p | MIMAT0000656 | Pik3r1 | 18708 | HITS-CLIP | 29167373 | Unspecified |
| mmu-miR-139-5p | MIMAT0000656 | Pik3r5 | 320207 | HITS-CLIP | 31433990 | Unspecified |
| mmu-miR-139-5p | MIMAT0000656 | Pikfyve | 18711 | HITS-CLIP | 29167373 | Unspecified |
| mmu-miR-139-5p | MIMAT0000656 | Pip4p1 | 90809 | sRNA-Seq//HITS-CLIP | 33184237\|29167373 | Unspecified |
| mmu-miR-139-5p | MIMAT0000656 | Pip5k1c | 18717 | sRNA-Seq//HITS-CLIP | 33184237\|29167373 | Unspecified |
| mmu-miR-139-5p | MIMAT0000656 | Pirb | 18733 | HITS-CLIP | 29167373 | Unspecified |
| mmu-miR-139-5p | MIMAT0000656 | Pisd | 320951 | HITS-CLIP | 29167373 | Unspecified |
| mmu-miR-139-5p | MIMAT0000656 | Pitpna | 18738 | sRNA-Seq//HITS-CLIP | 33184237\|29167373 | Unspecified |
| mmu-miR-139-5p | MIMAT0000656 | Pitpnm1 | 18739 | HITS-CLIP | 29167373 | Unspecified |
| mmu-miR-139-5p | MIMAT0000656 | Pja2 | 224938 | sRNA-Seq//HITS-CLIP | 33184237\|29167373 | Unspecified |
| mmu-miR-139-5p | MIMAT0000656 | Pkd1 | 18763 | HITS-CLIP | 29167373 | Unspecified |
| mmu-miR-139-5p | MIMAT0000656 | Pkm | 18746 | HITS-CLIP//sRNA-Seq | 31433990\|33184237\|29167373 | Unspecified |

| | | | | | | | |
|---|---|---|---|---|---|---|---|
| mmu-miR-139-5p | MIMAT0000656 | Pkn2 | 109333 | HITS-CLIP | | 26203562 | Unspecified |
| mmu-miR-139-5p | MIMAT0000656 | Pkp1 | 18772 | HITS-CLIP | | 26203562 | Unspecified |
| mmu-miR-139-5p | MIMAT0000656 | Plagl2 | 54711 | HITS-CLIP | 31433990\|29167373 | | Unspecified |
| mmu-miR-139-5p | MIMAT0000656 | Plau | 18792 | HITS-CLIP | | 29167373 | Unspecified |
| mmu-miR-139-5p | MIMAT0000656 | Plcb2 | 18796 | HITS-CLIP | | 29167373 | Unspecified |
| mmu-miR-139-5p | MIMAT0000656 | Plcb3 | 18797 | HITS-CLIP | | 29167373 | Unspecified |
| mmu-miR-139-5p | MIMAT0000656 | Plcg2 | 234779 | HITS-CLIP | | 29167373 | Unspecified |
| mmu-miR-139-5p | MIMAT0000656 | Pld1 | 18805 | HITS-CLIP | | 29167373 | Unspecified |
| mmu-miR-139-5p | MIMAT0000656 | Pld4 | 104759 | HITS-CLIP | | 29167373 | Unspecified |
| mmu-miR-139-5p | MIMAT0000656 | Plec | 18810 | sRNA-Seq//HITS-CLIP | 33184237\|29167373 | | Unspecified |
| mmu-miR-139-5p | MIMAT0000656 | Plekhb2 | 226971 | sRNA-Seq//HITS-CLIP | 33184237\|29167373 | | Unspecified |
| mmu-miR-139-5p | MIMAT0000656 | Plekhf2 | 71801 | HITS-CLIP | | 29167373 | Unspecified |
| mmu-miR-139-5p | MIMAT0000656 | Plekhj1 | 78670 | HITS-CLIP | | 29167373 | Unspecified |
| mmu-miR-139-5p | MIMAT0000656 | Plekhm3 | 241075 | HITS-CLIP | | 29167373 | Unspecified |
| mmu-miR-139-5p | MIMAT0000656 | Plekho1 | 67220 | HITS-CLIP | | 29167373 | Unspecified |
| mmu-miR-139-5p | MIMAT0000656 | Plekho2 | 102595 | HITS-CLIP | | 29167373 | Unspecified |
| mmu-miR-139-5p | MIMAT0000656 | Plk2 | 20620 | sRNA-Seq//HITS-CLIP | 33184237\|29167373 | | Unspecified |
| mmu-miR-139-5p | MIMAT0000656 | Plxdc1 | 72324 | HITS-CLIP | | 29167373 | Unspecified |
| mmu-miR-139-5p | MIMAT0000656 | Plxna1 | 18844 | HITS-CLIP | | 29167373 | Unspecified |
| mmu-miR-139-5p | MIMAT0000656 | Plxnb2 | 140570 | HITS-CLIP | | 29167373 | Unspecified |
| mmu-miR-139-5p | MIMAT0000656 | Pmel | 20431 | HITS-CLIP | | 26203562 | Unspecified |
| mmu-miR-139-5p | MIMAT0000656 | Pmm2 | 54128 | HITS-CLIP | | 29167373 | Unspecified |
| mmu-miR-139-5p | MIMAT0000656 | Pmp22 | 18858 | sRNA-Seq//HITS-CLIP | 33184237\|29167373 | | Unspecified |
| mmu-miR-139-5p | MIMAT0000656 | Pnisr | 66625 | HITS-CLIP | | 29167373 | Unspecified |
| mmu-miR-139-5p | MIMAT0000656 | Pnn | 18949 | HITS-CLIP | | 29167373 | Unspecified |
| mmu-miR-139-5p | MIMAT0000656 | Pnpla8 | 67452 | sRNA-Seq//HITS-CLIP | 33184237\|29167373 | | Unspecified |
| mmu-miR-139-5p | MIMAT0000656 | Pnrc1 | 108767 | HITS-CLIP | | 29167373 | Unspecified |
| mmu-miR-139-5p | MIMAT0000656 | Pogz | 229584 | HITS-CLIP | | 29167373 | Unspecified |
| mmu-miR-139-5p | MIMAT0000656 | Pole2 | 18974 | HITS-CLIP | | 31433990 | Unspecified |
| mmu-miR-139-5p | MIMAT0000656 | Polg | 18975 | HITS-CLIP | | 29167373 | Unspecified |
| mmu-miR-139-5p | MIMAT0000656 | Polk | 27015 | HITS-CLIP | | 29167373 | Unspecified |
| mmu-miR-139-5p | MIMAT0000656 | Polq | 77782 | HITS-CLIP | | 31433990 | Unspecified |
| mmu-miR-139-5p | MIMAT0000656 | Polr2a | 20020 | sRNA-Seq//HITS-CLIP | 33184237\|29167373 | | Unspecified |
| mmu-miR-139-5p | MIMAT0000656 | Polr2b | 231329 | HITS-CLIP | | 29167373 | Unspecified |
| mmu-miR-139-5p | MIMAT0000656 | Pon3 | 269823 | HITS-CLIP | | 29167373 | Unspecified |
| mmu-miR-139-5p | MIMAT0000656 | Pot1b | 72836 | HITS-CLIP | | 29167373 | Unspecified |
| mmu-miR-139-5p | MIMAT0000656 | Pou2f2 | 18987 | HITS-CLIP | | 29167373 | Unspecified |
| mmu-miR-139-5p | MIMAT0000656 | Ppfia1 | 233977 | HITS-CLIP | | 29167373 | Unspecified |
| mmu-miR-139-5p | MIMAT0000656 | Ppfibp1 | 67533 | HITS-CLIP | | 29167373 | Unspecified |

| | | | | | | |
|---|---|---|---|---|---|---|
| mmu-miR-139-5p | MIMAT0000656 | Pphln1 | 223828 | HITS-CLIP | 21258322\|29167373 | Unspecified |
| mmu-miR-139-5p | MIMAT0000656 | Ppib | 19035 | sRNA-Seq//HITS-CLIP | 33184237\|29167373 | Unspecified |
| mmu-miR-139-5p | MIMAT0000656 | Ppil4 | 67418 | HITS-CLIP | 31433990 | Unspecified |
| mmu-miR-139-5p | MIMAT0000656 | Ppm1a | 19042 | sRNA-Seq//HITS-CLIP | 33184237\|29167373 | Unspecified |
| mmu-miR-139-5p | MIMAT0000656 | Ppm1g | 14208 | sRNA-Seq//HITS-CLIP | 33184237\|29167373 | Unspecified |
| mmu-miR-139-5p | MIMAT0000656 | Ppp1ca | 19045 | sRNA-Seq//HITS-CLIP | 33184237\|29167373 | Unspecified |
| mmu-miR-139-5p | MIMAT0000656 | Ppp1r10 | 52040 | HITS-CLIP | 29167373 | Unspecified |
| mmu-miR-139-5p | MIMAT0000656 | Ppp1r13b | 21981 | HITS-CLIP | 29167373 | Unspecified |
| mmu-miR-139-5p | MIMAT0000656 | Ppp1r15b | 108954 | HITS-CLIP | 29167373 | Unspecified |
| mmu-miR-139-5p | MIMAT0000656 | Ppp2ca | 19052 | sRNA-Seq//HITS-CLIP | 33184237\|29167373 | Unspecified |
| mmu-miR-139-5p | MIMAT0000656 | Ppp2r2a | 71978 | sRNA-Seq//HITS-CLIP | 33184237\|29167373 | Unspecified |
| mmu-miR-139-5p | MIMAT0000656 | Ppp2r5c | 26931 | sRNA-Seq//HITS-CLIP | 33184237\|29167373 | Unspecified |
| mmu-miR-139-5p | MIMAT0000656 | Ppp2r5e | 26932 | HITS-CLIP | 29167373 | Unspecified |
| mmu-miR-139-5p | MIMAT0000656 | Ppp3cb | 19056 | sRNA-Seq//HITS-CLIP | 33184237\|29167373 | Unspecified |
| mmu-miR-139-5p | MIMAT0000656 | Ppp4c | 56420 | sRNA-Seq//HITS-CLIP | 33184237\|31433990 | Unspecified |
| mmu-miR-139-5p | MIMAT0000656 | Ppp4r1 | 70351 | sRNA-Seq//HITS-CLIP | 33184237\|29167373 | Unspecified |
| mmu-miR-139-5p | MIMAT0000656 | Ppp4r3a | 55671 | HITS-CLIP | 31433990 | Unspecified |
| mmu-miR-139-5p | MIMAT0000656 | Ppp4r3b | 57223 | HITS-CLIP | 29167373 | Unspecified |
| mmu-miR-139-5p | MIMAT0000656 | Ppp6c | 67857 | HITS-CLIP//sRNA-Seq | 31433990\|33184237\|29167373 | Unspecified |
| mmu-miR-139-5p | MIMAT0000656 | Ppp6r3 | 52036 | sRNA-Seq//HITS-CLIP | 33184237\|29167373 | Unspecified |
| mmu-miR-139-5p | MIMAT0000656 | Pprc1 | 226169 | HITS-CLIP | 29167373 | Unspecified |
| mmu-miR-139-5p | MIMAT0000656 | Pptc7 | 320717 | HITS-CLIP | 31433990 | Unspecified |
| mmu-miR-139-5p | MIMAT0000656 | Pqlc3 | 217430 | HITS-CLIP | 29167373 | Unspecified |
| mmu-miR-139-5p | MIMAT0000656 | Prdm15 | 63977 | HITS-CLIP | 26203562\|29167373 | Unspecified |
| mmu-miR-139-5p | MIMAT0000656 | Prdx1 | 18477 | HITS-CLIP//sRNA-Seq | 21258322\|33184237\|29167373 | Unspecified |
| mmu-miR-139-5p | MIMAT0000656 | Prdx6 | 11758 | sRNA-Seq//HITS-CLIP | 33184237\|26203562 | Unspecified |
| mmu-miR-139-5p | MIMAT0000656 | Prex1 | 277360 | HITS-CLIP | 29167373 | Unspecified |
| mmu-miR-139-5p | MIMAT0000656 | Primpol | 408022 | HITS-CLIP | 29167373 | Unspecified |
| mmu-miR-139-5p | MIMAT0000656 | Prkar1a | 19084 | sRNA-Seq//HITS-CLIP | 33184237\|29167373 | Unspecified |
| mmu-miR-139-5p | MIMAT0000656 | Prkcb | 18751 | HITS-CLIP | 29167373 | Unspecified |
| mmu-miR-139-5p | MIMAT0000656 | Prkcd | 18753 | sRNA-Seq//HITS-CLIP | 33184237\|29167373 | Unspecified |
| mmu-miR-139-5p | MIMAT0000656 | Prkch | 18755 | HITS-CLIP | 29167373 | Unspecified |
| mmu-miR-139-5p | MIMAT0000656 | Prkd2 | 101540 | HITS-CLIP | 31433990 | Unspecified |
| mmu-miR-139-5p | MIMAT0000656 | Prkx | 19108 | HITS-CLIP | 31433990 | Unspecified |
| mmu-miR-139-5p | MIMAT0000656 | Prnp | 19122 | sRNA-Seq//HITS-CLIP | 33184237\|29167373 | Unspecified |
| mmu-miR-139-5p | MIMAT0000656 | Prpf40a | 56194 | sRNA-Seq//HITS-CLIP | 33184237\|29167373 | Unspecified |
| mmu-miR-139-5p | MIMAT0000656 | Prpf4b | 19134 | HITS-CLIP | 31433990\|29167373 | Unspecified |
| mmu-miR-139-5p | MIMAT0000656 | Prpf6 | 68879 | HITS-CLIP | 29167373 | Unspecified |
| mmu-miR-139-5p | MIMAT0000656 | Prpf8 | 192159 | sRNA-Seq//HITS-CLIP | 33184237\|29167373 | Unspecified |

| | | | | | | |
|---|---|---|---|---|---|---|
| mmu-miR-139-5p | MIMAT0000656 | Prps1l3 | 328099 | HITS-CLIP | 31433990 | Unspecified |
| mmu-miR-139-5p | MIMAT0000656 | Prr14l | 215476 | HITS-CLIP | 29167373 | Unspecified |
| mmu-miR-139-5p | MIMAT0000656 | Prrc2a | 53761 | sRNA-Seq//HITS-CLIP | 33184237\|29167373 | Unspecified |
| mmu-miR-139-5p | MIMAT0000656 | Prrc2c | 226562 | sRNA-Seq//HITS-CLIP | 33184237\|29167373 | Unspecified |
| mmu-miR-139-5p | MIMAT0000656 | Psap | 19156 | HITS-CLIP//sRNA-Seq | 31433990\|33184237\|26203562\|29167 | Unspecified |
| mmu-miR-139-5p | MIMAT0000656 | Psd4 | 215632 | HITS-CLIP | 29167373 | Unspecified |
| mmu-miR-139-5p | MIMAT0000656 | Pskh1 | 244631 | HITS-CLIP | 26203562 | Unspecified |
| mmu-miR-139-5p | MIMAT0000656 | Psmb2 | 26445 | sRNA-Seq//HITS-CLIP | 33184237\|29167373 | Unspecified |
| mmu-miR-139-5p | MIMAT0000656 | Psmb4 | 19172 | sRNA-Seq//HITS-CLIP | 33184237\|29167373 | Unspecified |
| mmu-miR-139-5p | MIMAT0000656 | Psmd11 | 69077 | sRNA-Seq//HITS-CLIP | 33184237\|29167373 | Unspecified |
| mmu-miR-139-5p | MIMAT0000656 | Psme4 | 103554 | sRNA-Seq//HITS-CLIP | 33184237\|29167373 | Unspecified |
| mmu-miR-139-5p | MIMAT0000656 | Pstpip1 | 19200 | HITS-CLIP | 29167373 | Unspecified |
| mmu-miR-139-5p | MIMAT0000656 | Ptafr | 19204 | HITS-CLIP | 29167373 | Unspecified |
| mmu-miR-139-5p | MIMAT0000656 | Ptbp2 | 56195 | sRNA-Seq//HITS-CLIP | 33184237\|29167373 | Unspecified |
| mmu-miR-139-5p | MIMAT0000656 | Ptbp3 | 230257 | sRNA-Seq//HITS-CLIP | 33184237\|29167373 | Unspecified |
| mmu-miR-139-5p | MIMAT0000656 | Ptcd2 | 68927 | HITS-CLIP | 31433990 | Unspecified |
| mmu-miR-139-5p | MIMAT0000656 | Ptchd1 | 211612 | HITS-CLIP | 29167373 | Unspecified |
| mmu-miR-139-5p | MIMAT0000656 | Ptdss2 | 27388 | HITS-CLIP | 29167373 | Unspecified |
| mmu-miR-139-5p | MIMAT0000656 | Pten | 19211 | sRNA-Seq//HITS-CLIP | 33184237\|29167373 | Unspecified |
| mmu-miR-139-5p | MIMAT0000656 | Ptk2b | 19229 | HITS-CLIP | 29167373 | Unspecified |
| mmu-miR-139-5p | MIMAT0000656 | Ptp4a2 | 19244 | sRNA-Seq//HITS-CLIP | 33184237\|29167373 | Unspecified |
| mmu-miR-139-5p | MIMAT0000656 | Ptpn12 | 19248 | sRNA-Seq//HITS-CLIP | 33184237\|29167373 | Unspecified |
| mmu-miR-139-5p | MIMAT0000656 | Ptpn23 | 104831 | HITS-CLIP | 29167373 | Unspecified |
| mmu-miR-139-5p | MIMAT0000656 | Ptpn6 | 15170 | HITS-CLIP | 29167373 | Unspecified |
| mmu-miR-139-5p | MIMAT0000656 | Ptpn7 | 320139 | HITS-CLIP | 29167373 | Unspecified |
| mmu-miR-139-5p | MIMAT0000656 | Ptpre | 19267 | HITS-CLIP | 29167373 | Unspecified |
| mmu-miR-139-5p | MIMAT0000656 | Ptprj | 19271 | HITS-CLIP | 29167373 | Unspecified |
| mmu-miR-139-5p | MIMAT0000656 | Ptprs | 19280 | sRNA-Seq//HITS-CLIP | 33184237\|29167373 | Unspecified |
| mmu-miR-139-5p | MIMAT0000656 | Pum1 | 80912 | HITS-CLIP | 29167373 | Unspecified |
| mmu-miR-139-5p | MIMAT0000656 | Pum2 | 80913 | sRNA-Seq//HITS-CLIP | 33184237\|29167373 | Unspecified |
| mmu-miR-139-5p | MIMAT0000656 | Pum3 | 52874 | HITS-CLIP | 29167373 | Unspecified |
| mmu-miR-139-5p | MIMAT0000656 | Pura | 19290 | HITS-CLIP | 29167373 | Unspecified |
| mmu-miR-139-5p | MIMAT0000656 | Purb | 19291 | HITS-CLIP | 29167373 | Unspecified |
| mmu-miR-139-5p | MIMAT0000656 | Pwwp2b | 101631 | HITS-CLIP | 29167373 | Unspecified |
| mmu-miR-139-5p | MIMAT0000656 | Qk | 19317 | HITS-CLIP | 29167373 | Unspecified |
| mmu-miR-139-5p | MIMAT0000656 | Qrich1 | 69232 | sRNA-Seq//HITS-CLIP | 33184237\|29167373 | Unspecified |
| mmu-miR-139-5p | MIMAT0000656 | Qser1 | 99003 | HITS-CLIP | 29167373 | Unspecified |
| mmu-miR-139-5p | MIMAT0000656 | R3hcc1l | 52013 | HITS-CLIP | 29167373 | Unspecified |
| mmu-miR-139-5p | MIMAT0000656 | Rab11a | 53869 | sRNA-Seq//HITS-CLIP | 33184237\|29167373 | Unspecified |

| | | | | | | | |
|---|---|---|---|---|---|---|---|
| mmu-miR-139-5p | MIMAT0000656 | Rab11fip5 | 52055 | HITS-CLIP | | 29167373 | Unspecified |
| mmu-miR-139-5p | MIMAT0000656 | Rab12 | 19328 | HITS-CLIP | | 29167373 | Unspecified |
| mmu-miR-139-5p | MIMAT0000656 | Rab18 | 19330 | HITS-CLIP//sRNA-Seq | 29167373\|33184237 | | Unspecified |
| mmu-miR-139-5p | MIMAT0000656 | Rab1a | 5861 | sRNA-Seq//HITS-CLIP | 33184237\|29167373 | | Unspecified |
| mmu-miR-139-5p | MIMAT0000656 | Rab20 | 19332 | HITS-CLIP | | 29167373 | Unspecified |
| mmu-miR-139-5p | MIMAT0000656 | Rab21 | 216344 | HITS-CLIP | | 29167373 | Unspecified |
| mmu-miR-139-5p | MIMAT0000656 | Rab22a | 19334 | HITS-CLIP | | 29167373 | Unspecified |
| mmu-miR-139-5p | MIMAT0000656 | Rab2a | 59021 | HITS-CLIP//sRNA-Seq | 31433990\|33184237 | | Unspecified |
| mmu-miR-139-5p | MIMAT0000656 | Rab3gap2 | 98732 | HITS-CLIP | | 29167373 | Unspecified |
| mmu-miR-139-5p | MIMAT0000656 | Rab3il1 | 74760 | HITS-CLIP | | 29167373 | Unspecified |
| mmu-miR-139-5p | MIMAT0000656 | Rab40c | 224624 | HITS-CLIP | | 29167373 | Unspecified |
| mmu-miR-139-5p | MIMAT0000656 | Rab5c | 19345 | sRNA-Seq//HITS-CLIP | 33184237\|31433990 | | Unspecified |
| mmu-miR-139-5p | MIMAT0000656 | Rab7 | 19349 | sRNA-Seq//HITS-CLIP | 33184237\|29167373 | | Unspecified |
| mmu-miR-139-5p | MIMAT0000656 | Rab7b | 338382 | HITS-CLIP | | 29167373 | Unspecified |
| mmu-miR-139-5p | MIMAT0000656 | Rab8b | 235442 | HITS-CLIP | | 29167373 | Unspecified |
| mmu-miR-139-5p | MIMAT0000656 | Rabep1 | 54189 | sRNA-Seq//HITS-CLIP | 33184237\|29167373 | | Unspecified |
| mmu-miR-139-5p | MIMAT0000656 | Rabgap1 | 227800 | sRNA-Seq//HITS-CLIP | 33184237\|29167373 | | Unspecified |
| mmu-miR-139-5p | MIMAT0000656 | Rabggtb | 19352 | sRNA-Seq//HITS-CLIP | 33184237\|29167373 | | Unspecified |
| mmu-miR-139-5p | MIMAT0000656 | Rad18 | 58186 | HITS-CLIP | | 29167373 | Unspecified |
| mmu-miR-139-5p | MIMAT0000656 | Rad50 | 19360 | HITS-CLIP | | 29167373 | Unspecified |
| mmu-miR-139-5p | MIMAT0000656 | Raf1 | 110157 | sRNA-Seq//HITS-CLIP | 33184237\|29167373 | | Unspecified |
| mmu-miR-139-5p | MIMAT0000656 | Ralb | 64143 | HITS-CLIP | | 29167373 | Unspecified |
| mmu-miR-139-5p | MIMAT0000656 | Ralgapa2 | 241694 | HITS-CLIP | | 29167373 | Unspecified |
| mmu-miR-139-5p | MIMAT0000656 | Ranbp2 | 19386 | sRNA-Seq//HITS-CLIP | 33184237\|29167373 | | Unspecified |
| mmu-miR-139-5p | MIMAT0000656 | Rap1a | 109905 | HITS-CLIP | | 29167373 | Unspecified |
| mmu-miR-139-5p | MIMAT0000656 | Rap1b | 215449 | sRNA-Seq//HITS-CLIP | 33184237\|29167373 | | Unspecified |
| mmu-miR-139-5p | MIMAT0000656 | Rap1gds1 | 229877 | HITS-CLIP//sRNA-Seq | 31433990\|33184237\|29167373 | | Unspecified |
| mmu-miR-139-5p | MIMAT0000656 | Rap2a | 76108 | HITS-CLIP | | 29167373 | Unspecified |
| mmu-miR-139-5p | MIMAT0000656 | Rap2b | 74012 | HITS-CLIP | | 29167373 | Unspecified |
| mmu-miR-139-5p | MIMAT0000656 | Rapgef1 | 107746 | HITS-CLIP | | 29167373 | Unspecified |
| mmu-miR-139-5p | MIMAT0000656 | Rapgef2 | 76089 | HITS-CLIP | | 29167373 | Unspecified |
| mmu-miR-139-5p | MIMAT0000656 | Raph1 | 77300 | sRNA-Seq//HITS-CLIP | 33184237\|29167373 | | Unspecified |
| mmu-miR-139-5p | MIMAT0000656 | Rara | 19401 | HITS-CLIP | | 29167373 | Unspecified |
| mmu-miR-139-5p | MIMAT0000656 | Rasa1 | 218397 | sRNA-Seq//HITS-CLIP | 33184237\|29167373 | | Unspecified |
| mmu-miR-139-5p | MIMAT0000656 | Rasa4 | 54153 | HITS-CLIP | | 29167373 | Unspecified |
| mmu-miR-139-5p | MIMAT0000656 | Rasgef1b | 320292 | HITS-CLIP | | 29167373 | Unspecified |
| mmu-miR-139-5p | MIMAT0000656 | Rasgrp1 | 19419 | HITS-CLIP | | 31433990 | Unspecified |
| mmu-miR-139-5p | MIMAT0000656 | Rassf2 | 215653 | HITS-CLIP | | 29167373 | Unspecified |
| mmu-miR-139-5p | MIMAT0000656 | Rassf4 | 213391 | HITS-CLIP | | 29167373 | Unspecified |

| | | | | | | |
|---|---|---|---|---|---|---|
| mmu-miR-139-5p | MIMAT0000656 | Rb1cc1 | 12421 | sRNA-Seq//HITS-CLIP | 33184237\|29167373 | Unspecified |
| mmu-miR-139-5p | MIMAT0000656 | Rbbp6 | 19647 | HITS-CLIP | 29167373 | Unspecified |
| mmu-miR-139-5p | MIMAT0000656 | Rbl1 | 19650 | HITS-CLIP | 29167373 | Unspecified |
| mmu-miR-139-5p | MIMAT0000656 | Rbm12b2 | 77604 | HITS-CLIP | 29167373 | Unspecified |
| mmu-miR-139-5p | MIMAT0000656 | Rbm15b | 109095 | HITS-CLIP | 31433990 | Unspecified |
| mmu-miR-139-5p | MIMAT0000656 | Rbm25 | 67039 | sRNA-Seq//HITS-CLIP | 33184237\|29167373 | Unspecified |
| mmu-miR-139-5p | MIMAT0000656 | Rbm26 | 74213 | HITS-CLIP | 29167373 | Unspecified |
| mmu-miR-139-5p | MIMAT0000656 | Rbm33 | 381626 | HITS-CLIP | 29167373 | Unspecified |
| mmu-miR-139-5p | MIMAT0000656 | Rbm47 | 245945 | HITS-CLIP | 29167373 | Unspecified |
| mmu-miR-139-5p | MIMAT0000656 | Rbms1 | 56878 | sRNA-Seq//HITS-CLIP | 33184237\|31433990\|29167373 | Unspecified |
| mmu-miR-139-5p | MIMAT0000656 | Rbpj | 19664 | HITS-CLIP | 29167373 | Unspecified |
| mmu-miR-139-5p | MIMAT0000656 | Rc3h2 | 319817 | sRNA-Seq//HITS-CLIP | 33184237\|29167373 | Unspecified |
| mmu-miR-139-5p | MIMAT0000656 | Rcor1 | 217864 | HITS-CLIP | 29167373 | Unspecified |
| mmu-miR-139-5p | MIMAT0000656 | Recql5 | 170472 | HITS-CLIP | 29167373 | Unspecified |
| mmu-miR-139-5p | MIMAT0000656 | Reep3 | 28193 | HITS-CLIP | 26203562\|29167373 | Unspecified |
| mmu-miR-139-5p | MIMAT0000656 | Reep5 | 13476 | sRNA-Seq//HITS-CLIP | 33184237\|29167373 | Unspecified |
| mmu-miR-139-5p | MIMAT0000656 | Relch | 57614 | sRNA-Seq//HITS-CLIP | 33184237\|29167373 | Unspecified |
| mmu-miR-139-5p | MIMAT0000656 | Rere | 68703 | sRNA-Seq//HITS-CLIP | 33184237\|29167373 | Unspecified |
| mmu-miR-139-5p | MIMAT0000656 | Retreg1 | 66270 | HITS-CLIP | 29167373 | Unspecified |
| mmu-miR-139-5p | MIMAT0000656 | Rexo1 | 66932 | HITS-CLIP | 29167373 | Unspecified |
| mmu-miR-139-5p | MIMAT0000656 | Rgs2 | 19735 | HITS-CLIP | 29167373 | Unspecified |
| mmu-miR-139-5p | MIMAT0000656 | Rhoq | 104215 | HITS-CLIP | 31433990 | Unspecified |
| mmu-miR-139-5p | MIMAT0000656 | Rin2 | 74030 | HITS-CLIP | 29167373 | Unspecified |
| mmu-miR-139-5p | MIMAT0000656 | Ripor1 | 75687 | HITS-CLIP | 29167373 | Unspecified |
| mmu-miR-139-5p | MIMAT0000656 | Rlim | 19820 | sRNA-Seq//HITS-CLIP | 33184237\|29167373 | Unspecified |
| mmu-miR-139-5p | MIMAT0000656 | Rmnd5a | 68477 | HITS-CLIP | 29167373 | Unspecified |
| mmu-miR-139-5p | MIMAT0000656 | Rmnd5b | 66089 | HITS-CLIP | 29167373 | Unspecified |
| mmu-miR-139-5p | MIMAT0000656 | Rnf111 | 93836 | HITS-CLIP | 29167373 | Unspecified |
| mmu-miR-139-5p | MIMAT0000656 | Rnf13 | 24017 | HITS-CLIP | 29167373 | Unspecified |
| mmu-miR-139-5p | MIMAT0000656 | Rnf130 | 59044 | sRNA-Seq//HITS-CLIP | 33184237\|29167373 | Unspecified |
| mmu-miR-139-5p | MIMAT0000656 | Rnf14 | 56736 | sRNA-Seq//HITS-CLIP | 33184237\|29167373 | Unspecified |
| mmu-miR-139-5p | MIMAT0000656 | Rnf149 | 67702 | HITS-CLIP | 29167373 | Unspecified |
| mmu-miR-139-5p | MIMAT0000656 | Rnf19a | 30945 | HITS-CLIP | 29167373 | Unspecified |
| mmu-miR-139-5p | MIMAT0000656 | Rnf19b | 75234 | HITS-CLIP | 29167373 | Unspecified |
| mmu-miR-139-5p | MIMAT0000656 | Rnf2 | 19821 | sRNA-Seq//HITS-CLIP | 33184237\|29167373 | Unspecified |
| mmu-miR-139-5p | MIMAT0000656 | Rnf213 | 672511 | HITS-CLIP | 29167373 | Unspecified |
| mmu-miR-139-5p | MIMAT0000656 | Rnf38 | 73469 | sRNA-Seq//HITS-CLIP | 33184237\|29167373 | Unspecified |
| mmu-miR-139-5p | MIMAT0000656 | Rnf4 | 19822 | HITS-CLIP//sRNA-Seq | 31433990\|33184237 | Unspecified |
| mmu-miR-139-5p | MIMAT0000656 | Rnf41 | 67588 | sRNA-Seq//HITS-CLIP | 33184237\|29167373 | Unspecified |

| | | | | | | |
|---|---|---|---|---|---|---|
| mmu-miR-139-5p | MIMAT0000656 | Rnf44 | 105239 | HITS-CLIP | 29167373 | Unspecified |
| mmu-miR-139-5p | MIMAT0000656 | Rngtt | 24018 | HITS-CLIP | 29167373 | Unspecified |
| mmu-miR-139-5p | MIMAT0000656 | Rock1 | 19877 | HITS-CLIP | 29167373 | Unspecified |
| mmu-miR-139-5p | MIMAT0000656 | Rock2 | 19878 | HITS-CLIP | 29167373 | Unspecified |
| mmu-miR-139-5p | MIMAT0000656 | Rpa3 | 68240 | HITS-CLIP | 29167373 | Unspecified |
| mmu-miR-139-5p | MIMAT0000656 | Rpe | 66646 | HITS-CLIP | 29167373 | Unspecified |
| mmu-miR-139-5p | MIMAT0000656 | Rpl10 | 110954 | sRNA-Seq//HITS-CLIP | 33184237\|29167373 | Unspecified |
| mmu-miR-139-5p | MIMAT0000656 | Rpl23 | 65019 | sRNA-Seq//HITS-CLIP | 33184237\|29167373 | Unspecified |
| mmu-miR-139-5p | MIMAT0000656 | Rpl7 | 19989 | sRNA-Seq//HITS-CLIP | 33184237\|29167373 | Unspecified |
| mmu-miR-139-5p | MIMAT0000656 | Rpl7l1 | 66229 | HITS-CLIP | 29167373 | Unspecified |
| mmu-miR-139-5p | MIMAT0000656 | Rprd1b | 70470 | HITS-CLIP | 29167373 | Unspecified |
| mmu-miR-139-5p | MIMAT0000656 | Rps20 | 67427 | sRNA-Seq//HITS-CLIP | 33184237\|31433990 | Unspecified |
| mmu-miR-139-5p | MIMAT0000656 | Rps3 | 27050 | sRNA-Seq//HITS-CLIP | 33184237\|29167373 | Unspecified |
| mmu-miR-139-5p | MIMAT0000656 | Rps6ka3 | 110651 | sRNA-Seq//HITS-CLIP | 33184237\|29167373 | Unspecified |
| mmu-miR-139-5p | MIMAT0000656 | Rps6ka5 | 73086 | HITS-CLIP | 29167373 | Unspecified |
| mmu-miR-139-5p | MIMAT0000656 | Rps9 | 76846 | sRNA-Seq//HITS-CLIP | 33184237\|23630078\|30971389 | Unspecified |
| mmu-miR-139-5p | MIMAT0000656 | Rragc | 54170 | sRNA-Seq//HITS-CLIP | 33184237\|29167373 | Unspecified |
| mmu-miR-139-5p | MIMAT0000656 | Rrbp1 | 81910 | HITS-CLIP | 29167373 | Unspecified |
| mmu-miR-139-5p | MIMAT0000656 | Rreb1 | 68750 | HITS-CLIP | 29167373 | Unspecified |
| mmu-miR-139-5p | MIMAT0000656 | Rrm2 | 20135 | HITS-CLIP | 31433990 | Unspecified |
| mmu-miR-139-5p | MIMAT0000656 | Rrp1 | 18114 | sRNA-Seq//HITS-CLIP | 33184237\|29167373 | Unspecified |
| mmu-miR-139-5p | MIMAT0000656 | Rrp8 | 101867 | HITS-CLIP | 31433990 | Unspecified |
| mmu-miR-139-5p | MIMAT0000656 | Rsad2 | 58185 | HITS-CLIP | 29167373 | Unspecified |
| mmu-miR-139-5p | MIMAT0000656 | Rsbn1l | 242860 | HITS-CLIP | 29167373 | Unspecified |
| mmu-miR-139-5p | MIMAT0000656 | Rsf1 | 233532 | HITS-CLIP | 29167373 | Unspecified |
| mmu-miR-139-5p | MIMAT0000656 | Rsrc2 | 208606 | sRNA-Seq//HITS-CLIP | 33184237\|29167373 | Unspecified |
| mmu-miR-139-5p | MIMAT0000656 | Rsrp1 | 27981 | sRNA-Seq//HITS-CLIP | 33184237\|29167373 | Unspecified |
| mmu-miR-139-5p | MIMAT0000656 | Rtf2 | 51507 | HITS-CLIP | 29167373 | Unspecified |
| mmu-miR-139-5p | MIMAT0000656 | Rtn3 | 20168 | sRNA-Seq//HITS-CLIP | 33184237\|29167373 | Unspecified |
| mmu-miR-139-5p | MIMAT0000656 | Rtn4 | 68585 | sRNA-Seq//HITS-CLIP | 33184237\|29167373 | Unspecified |
| mmu-miR-139-5p | MIMAT0000656 | Rtraf | 51637 | HITS-CLIP//sRNA-Seq | 31433990\|33184237 | Unspecified |
| mmu-miR-139-5p | MIMAT0000656 | Rufy3 | 52822 | sRNA-Seq//HITS-CLIP | 33184237\|29167373 | Unspecified |
| mmu-miR-139-5p | MIMAT0000656 | Runx1 | 12394 | HITS-CLIP | 29167373 | Unspecified |
| mmu-miR-139-5p | MIMAT0000656 | Runx3 | 12399 | HITS-CLIP | 29167373 | Unspecified |
| mmu-miR-139-5p | MIMAT0000656 | Rxra | 20181 | HITS-CLIP | 29167373 | Unspecified |
| mmu-miR-139-5p | MIMAT0000656 | S100a14 | 57402 | HITS-CLIP | 26203562 | Unspecified |
| mmu-miR-139-5p | MIMAT0000656 | S100a4 | 20198 | HITS-CLIP | 31433990 | Unspecified |
| mmu-miR-139-5p | MIMAT0000656 | S1pr1 | 13609 | HITS-CLIP | 29167373 | Unspecified |
| mmu-miR-139-5p | MIMAT0000656 | S1pr2 | 14739 | HITS-CLIP | 29167373 | Unspecified |

| | | | | | | |
|---|---|---|---|---|---|---|
| mmu-miR-139-5p | MIMAT0000656 | Sacs | 50720 | HITS-CLIP | 29167373 | Unspecified |
| mmu-miR-139-5p | MIMAT0000656 | Safb | 224903 | HITS-CLIP | 29167373 | Unspecified |
| mmu-miR-139-5p | MIMAT0000656 | Samd9l | 209086 | HITS-CLIP | 29167373 | Unspecified |
| mmu-miR-139-5p | MIMAT0000656 | Samhd1 | 56045 | HITS-CLIP | 29167373 | Unspecified |
| mmu-miR-139-5p | MIMAT0000656 | Saraf | 51669 | sRNA-Seq//HITS-CLIP | 33184237\|29167373 | Unspecified |
| mmu-miR-139-5p | MIMAT0000656 | Sash1 | 70097 | HITS-CLIP | 29167373 | Unspecified |
| mmu-miR-139-5p | MIMAT0000656 | Sbf2 | 319934 | sRNA-Seq//HITS-CLIP | 33184237\|29167373 | Unspecified |
| mmu-miR-139-5p | MIMAT0000656 | Sbno1 | 243272 | sRNA-Seq//HITS-CLIP | 33184237\|29167373 | Unspecified |
| mmu-miR-139-5p | MIMAT0000656 | Scaf11 | 72193 | HITS-CLIP | 29167373 | Unspecified |
| mmu-miR-139-5p | MIMAT0000656 | Scaf4 | 224432 | HITS-CLIP | 29167373 | Unspecified |
| mmu-miR-139-5p | MIMAT0000656 | Scaf8 | 106583 | HITS-CLIP | 29167373 | Unspecified |
| mmu-miR-139-5p | MIMAT0000656 | Scarf1 | 380713 | HITS-CLIP | 29167373 | Unspecified |
| mmu-miR-139-5p | MIMAT0000656 | Scd2 | 20250 | sRNA-Seq//HITS-CLIP | 33184237\|26203562 | Unspecified |
| mmu-miR-139-5p | MIMAT0000656 | Scfd2 | 212986 | HITS-CLIP | 29167373 | Unspecified |
| mmu-miR-139-5p | MIMAT0000656 | Scyl2 | 213326 | sRNA-Seq//HITS-CLIP | 33184237\|29167373 | Unspecified |
| mmu-miR-139-5p | MIMAT0000656 | Sdad1 | 231452 | HITS-CLIP | 29167373 | Unspecified |
| mmu-miR-139-5p | MIMAT0000656 | Sdc1 | 20969 | HITS-CLIP | 29167373 | Unspecified |
| mmu-miR-139-5p | MIMAT0000656 | Sdc3 | 20970 | sRNA-Seq//HITS-CLIP | 33184237\|26203562\|29167373 | Unspecified |
| mmu-miR-139-5p | MIMAT0000656 | Sdc4 | 20971 | HITS-CLIP | 29167373 | Unspecified |
| mmu-miR-139-5p | MIMAT0000656 | Sdcbp | 53378 | sRNA-Seq//HITS-CLIP | 33184237\|29167373 | Unspecified |
| mmu-miR-139-5p | MIMAT0000656 | Sdhaf2 | 66072 | HITS-CLIP | 29167373 | Unspecified |
| mmu-miR-139-5p | MIMAT0000656 | Sec23a | 20334 | sRNA-Seq//HITS-CLIP | 33184237\|29167373 | Unspecified |
| mmu-miR-139-5p | MIMAT0000656 | Sec24c | 218811 | sRNA-Seq//HITS-CLIP | 33184237\|29167373 | Unspecified |
| mmu-miR-139-5p | MIMAT0000656 | Secisbp2 | 75420 | HITS-CLIP | 29167373 | Unspecified |
| mmu-miR-139-5p | MIMAT0000656 | Secisbp2l | 70354 | HITS-CLIP | 29167373 | Unspecified |
| mmu-miR-139-5p | MIMAT0000656 | Sel1l | 20338 | HITS-CLIP | 29167373 | Unspecified |
| mmu-miR-139-5p | MIMAT0000656 | Selenop | 6414 | HITS-CLIP | 29167373 | Unspecified |
| mmu-miR-139-5p | MIMAT0000656 | Sema4d | 20354 | sRNA-Seq//HITS-CLIP | 33184237\|29167373 | Unspecified |
| mmu-miR-139-5p | MIMAT0000656 | Senp1 | 223870 | HITS-CLIP | 29167373 | Unspecified |
| mmu-miR-139-5p | MIMAT0000656 | Senp5 | 320213 | HITS-CLIP | 29167373 | Unspecified |
| mmu-miR-139-5p | MIMAT0000656 | Senp6 | 215351 | sRNA-Seq//HITS-CLIP | 33184237\|29167373 | Unspecified |
| mmu-miR-139-5p | MIMAT0000656 | Sept2 | 4735 | HITS-CLIP | 29167373 | Unspecified |
| mmu-miR-139-5p | MIMAT0000656 | Sept7 | 235072 | sRNA-Seq//HITS-CLIP | 33184237\|29167373 | Unspecified |
| mmu-miR-139-5p | MIMAT0000656 | Sept8 | 20362 | HITS-CLIP | 29167373 | Unspecified |
| mmu-miR-139-5p | MIMAT0000656 | Serbp1 | 66870 | sRNA-Seq//HITS-CLIP | 33184237\|29167373 | Unspecified |
| mmu-miR-139-5p | MIMAT0000656 | Serf2 | 378702 | sRNA-Seq//HITS-CLIP | 33184237\|29167373 | Unspecified |
| mmu-miR-139-5p | MIMAT0000656 | Serinc1 | 56442 | HITS-CLIP//sRNA-Seq | 31433990\|33184237 | Unspecified |
| mmu-miR-139-5p | MIMAT0000656 | Setd1a | 233904 | HITS-CLIP | 29167373 | Unspecified |
| mmu-miR-139-5p | MIMAT0000656 | Setd1b | 208043 | HITS-CLIP | 29167373 | Unspecified |

| | | | | | | |
|---|---|---|---|---|---|---|
| mmu-miR-139-5p | MIMAT0000656 | Setd3 | 52690 | sRNA-Seq//HITS-CLIP | 33184237\|29167373 | Unspecified |
| mmu-miR-139-5p | MIMAT0000656 | Setx | 269254 | HITS-CLIP | 29167373 | Unspecified |
| mmu-miR-139-5p | MIMAT0000656 | Sf1 | 22668 | sRNA-Seq//HITS-CLIP | 33184237\|26203562\|29167373 | Unspecified |
| mmu-miR-139-5p | MIMAT0000656 | Sf3a3 | 75062 | HITS-CLIP | 29167373 | Unspecified |
| mmu-miR-139-5p | MIMAT0000656 | Sf3b1 | 81898 | sRNA-Seq//HITS-CLIP | 33184237\|31433990\|29167373 | Unspecified |
| mmu-miR-139-5p | MIMAT0000656 | Sfmbt1 | 54650 | HITS-CLIP | 29167373 | Unspecified |
| mmu-miR-139-5p | MIMAT0000656 | Sgms1 | 208449 | HITS-CLIP | 29167373 | Unspecified |
| mmu-miR-139-5p | MIMAT0000656 | Sh2b3 | 16923 | sRNA-Seq//HITS-CLIP | 33184237\|29167373 | Unspecified |
| mmu-miR-139-5p | MIMAT0000656 | Sh3kbp1 | 58194 | sRNA-Seq//HITS-CLIP | 33184237\|29167373 | Unspecified |
| mmu-miR-139-5p | MIMAT0000656 | Sh3pxd2b | 268396 | HITS-CLIP | 29167373 | Unspecified |
| mmu-miR-139-5p | MIMAT0000656 | Sh3tc1 | 231147 | HITS-CLIP | 29167373 | Unspecified |
| mmu-miR-139-5p | MIMAT0000656 | Shoc2 | 56392 | sRNA-Seq//HITS-CLIP | 33184237\|29167373 | Unspecified |
| mmu-miR-139-5p | MIMAT0000656 | Shtn1 | 57698 | sRNA-Seq//HITS-CLIP | 33184237\|29167373 | Unspecified |
| mmu-miR-139-5p | MIMAT0000656 | Sik1 | 17691 | HITS-CLIP | 31433990\|26203562 | Unspecified |
| mmu-miR-139-5p | MIMAT0000656 | Sik2 | 235344 | HITS-CLIP | 31433990 | Unspecified |
| mmu-miR-139-5p | MIMAT0000656 | Sin3b | 20467 | sRNA-Seq//HITS-CLIP | 33184237\|26203562 | Unspecified |
| mmu-miR-139-5p | MIMAT0000656 | Sipa1l2 | 244668 | HITS-CLIP | 29167373 | Unspecified |
| mmu-miR-139-5p | MIMAT0000656 | Sirpa | 19261 | HITS-CLIP | 25083871\|29167373 | Unspecified |
| mmu-miR-139-5p | MIMAT0000656 | Sirt1 | 93759 | HITS-CLIP | 29167373 | Unspecified |
| mmu-miR-139-5p | MIMAT0000656 | Ski | 20481 | HITS-CLIP | 29167373 | Unspecified |
| mmu-miR-139-5p | MIMAT0000656 | Skil | 20482 | HITS-CLIP | 29167373 | Unspecified |
| mmu-miR-139-5p | MIMAT0000656 | Skp1 | 6500 | HITS-CLIP//sRNA-Seq | 31433990\|33184237 | Unspecified |
| mmu-miR-139-5p | MIMAT0000656 | Skp1a | 21402 | sRNA-Seq//HITS-CLIP | 33184237\|29167373 | Unspecified |
| mmu-miR-139-5p | MIMAT0000656 | Skp2 | 27401 | HITS-CLIP | 29167373 | Unspecified |
| mmu-miR-139-5p | MIMAT0000656 | Slamf7 | 75345 | HITS-CLIP | 29167373 | Unspecified |
| mmu-miR-139-5p | MIMAT0000656 | Slbp | 20492 | HITS-CLIP | 29167373 | Unspecified |
| mmu-miR-139-5p | MIMAT0000656 | Slc11a2 | 18174 | HITS-CLIP | 29167373 | Unspecified |
| mmu-miR-139-5p | MIMAT0000656 | Slc12a6 | 107723 | HITS-CLIP | 29167373 | Unspecified |
| mmu-miR-139-5p | MIMAT0000656 | Slc15a3 | 65221 | HITS-CLIP | 29167373 | Unspecified |
| mmu-miR-139-5p | MIMAT0000656 | Slc16a10 | 72472 | HITS-CLIP | 29167373 | Unspecified |
| mmu-miR-139-5p | MIMAT0000656 | Slc16a6 | 104681 | HITS-CLIP | 31433990 | Unspecified |
| mmu-miR-139-5p | MIMAT0000656 | Slc19a2 | 116914 | HITS-CLIP | 26203562 | Unspecified |
| mmu-miR-139-5p | MIMAT0000656 | Slc20a1 | 20515 | sRNA-Seq//HITS-CLIP | 33184237\|29167373 | Unspecified |
| mmu-miR-139-5p | MIMAT0000656 | Slc23a2 | 54338 | HITS-CLIP | 29167373 | Unspecified |
| mmu-miR-139-5p | MIMAT0000656 | Slc25a12 | 78830 | sRNA-Seq//HITS-CLIP | 33184237\|31433990 | Unspecified |
| mmu-miR-139-5p | MIMAT0000656 | Slc25a16 | 73132 | HITS-CLIP | 29167373 | Unspecified |
| mmu-miR-139-5p | MIMAT0000656 | Slc25a24 | 229731 | HITS-CLIP | 29167373 | Unspecified |
| mmu-miR-139-5p | MIMAT0000656 | Slc25a25 | 227731 | HITS-CLIP | 29167373 | Unspecified |
| mmu-miR-139-5p | MIMAT0000656 | Slc25a3 | 18674 | sRNA-Seq//HITS-CLIP | 33184237\|29167373 | Unspecified |

| | | | | | | |
|---|---|---|---|---|---|---|
| mmu-miR-139-5p | MIMAT0000656 | Slc25a30 | 67554 | HITS-CLIP | 29167373 | Unspecified |
| mmu-miR-139-5p | MIMAT0000656 | Slc25a36 | 192287 | sRNA-Seq//HITS-CLIP | 33184237\|29167373 | Unspecified |
| mmu-miR-139-5p | MIMAT0000656 | Slc25a44 | 229517 | HITS-CLIP | 29167373 | Unspecified |
| mmu-miR-139-5p | MIMAT0000656 | Slc30a9 | 109108 | sRNA-Seq//HITS-CLIP | 33184237\|29167373 | Unspecified |
| mmu-miR-139-5p | MIMAT0000656 | Slc31a1 | 20529 | HITS-CLIP | 29167373 | Unspecified |
| mmu-miR-139-5p | MIMAT0000656 | Slc35a4 | 67843 | sRNA-Seq//HITS-CLIP | 33184237\|31433990 | Unspecified |
| mmu-miR-139-5p | MIMAT0000656 | Slc35a5 | 74102 | HITS-CLIP | 29167373 | Unspecified |
| mmu-miR-139-5p | MIMAT0000656 | Slc38a1 | 105727 | sRNA-Seq//HITS-CLIP | 33184237\|29167373 | Unspecified |
| mmu-miR-139-5p | MIMAT0000656 | Slc38a10 | 72055 | HITS-CLIP | 29167373 | Unspecified |
| mmu-miR-139-5p | MIMAT0000656 | Slc38a2 | 67760 | sRNA-Seq//HITS-CLIP | 33184237\|29167373 | Unspecified |
| mmu-miR-139-5p | MIMAT0000656 | Slc39a13 | 68427 | HITS-CLIP | 29167373 | Unspecified |
| mmu-miR-139-5p | MIMAT0000656 | Slc39a6 | 106957 | sRNA-Seq//HITS-CLIP | 33184237\|29167373 | Unspecified |
| mmu-miR-139-5p | MIMAT0000656 | Slc3a2 | 17254 | sRNA-Seq//HITS-CLIP | 33184237\|26203562 | Unspecified |
| mmu-miR-139-5p | MIMAT0000656 | Slc41a2 | 338365 | HITS-CLIP | 29167373 | Unspecified |
| mmu-miR-139-5p | MIMAT0000656 | Slc43a2 | 215113 | HITS-CLIP | 29167373 | Unspecified |
| mmu-miR-139-5p | MIMAT0000656 | Slc4a2 | 20535 | HITS-CLIP | 29167373 | Unspecified |
| mmu-miR-139-5p | MIMAT0000656 | Slc4a7 | 218756 | HITS-CLIP | 29167373 | Unspecified |
| mmu-miR-139-5p | MIMAT0000656 | Slc5a3 | 53881 | HITS-CLIP | 29167373 | Unspecified |
| mmu-miR-139-5p | MIMAT0000656 | Slc6a6 | 21366 | HITS-CLIP | 29167373 | Unspecified |
| mmu-miR-139-5p | MIMAT0000656 | Slc7a8 | 50934 | HITS-CLIP | 29167373 | Unspecified |
| mmu-miR-139-5p | MIMAT0000656 | Slc9a1 | 20544 | HITS-CLIP | 29167373 | Unspecified |
| mmu-miR-139-5p | MIMAT0000656 | Slf2 | 226151 | HITS-CLIP | 29167373 | Unspecified |
| mmu-miR-139-5p | MIMAT0000656 | Slfn2 | 20556 | HITS-CLIP | 31433990\|29167373 | Unspecified |
| mmu-miR-139-5p | MIMAT0000656 | Slfn8 | 276950 | HITS-CLIP | 29167373 | Unspecified |
| mmu-miR-139-5p | MIMAT0000656 | Slfn9 | 237886 | HITS-CLIP | 29167373 | Unspecified |
| mmu-miR-139-5p | MIMAT0000656 | Slk | 20874 | HITS-CLIP | 29167373 | Unspecified |
| mmu-miR-139-5p | MIMAT0000656 | Slmap | 83997 | HITS-CLIP | 29167373 | Unspecified |
| mmu-miR-139-5p | MIMAT0000656 | Sltm | 66660 | HITS-CLIP | 29167373 | Unspecified |
| mmu-miR-139-5p | MIMAT0000656 | Smad2 | 17126 | sRNA-Seq//HITS-CLIP | 33184237\|29167373 | Unspecified |
| mmu-miR-139-5p | MIMAT0000656 | Smad4 | 17128 | HITS-CLIP | 29167373 | Unspecified |
| mmu-miR-139-5p | MIMAT0000656 | Smarca4 | 20586 | sRNA-Seq//HITS-CLIP | 33184237\|31433990\|29167373 | Unspecified |
| mmu-miR-139-5p | MIMAT0000656 | Smarca5 | 93762 | sRNA-Seq//HITS-CLIP | 33184237\|26203562 | Unspecified |
| mmu-miR-139-5p | MIMAT0000656 | Smarcad1 | 13990 | HITS-CLIP | 29167373 | Unspecified |
| mmu-miR-139-5p | MIMAT0000656 | Smarcal1 | 54380 | HITS-CLIP | 29167373 | Unspecified |
| mmu-miR-139-5p | MIMAT0000656 | Smarcd2 | 83796 | HITS-CLIP | 29167373 | Unspecified |
| mmu-miR-139-5p | MIMAT0000656 | Smc2 | 14211 | HITS-CLIP | 29167373 | Unspecified |
| mmu-miR-139-5p | MIMAT0000656 | Smc3 | 13006 | HITS-CLIP | 29167373 | Unspecified |
| mmu-miR-139-5p | MIMAT0000656 | Smc4 | 70099 | HITS-CLIP | 31433990\|29167373 | Unspecified |
| mmu-miR-139-5p | MIMAT0000656 | Smc6 | 67241 | HITS-CLIP | 29167373 | Unspecified |

| | | | | | | |
|---|---|---|---|---|---|---|
| mmu-miR-139-5p | MIMAT0000656 | Smchd1 | 74355 | HITS-CLIP | 26203562\|29167373 | Unspecified |
| mmu-miR-139-5p | MIMAT0000656 | Smcr8 | 237782 | HITS-CLIP | 29167373 | Unspecified |
| mmu-miR-139-5p | MIMAT0000656 | Smg1 | 233789 | HITS-CLIP | 29167373 | Unspecified |
| mmu-miR-139-5p | MIMAT0000656 | Smg7 | 226517 | sRNA-Seq//HITS-CLIP | 33184237\|29167373 | Unspecified |
| mmu-miR-139-5p | MIMAT0000656 | Smox | 228608 | HITS-CLIP | 29167373 | Unspecified |
| mmu-miR-139-5p | MIMAT0000656 | Smurf1 | 75788 | HITS-CLIP | 26203562 | Unspecified |
| mmu-miR-139-5p | MIMAT0000656 | Snap23 | 20619 | HITS-CLIP | 23630078 | Unspecified |
| mmu-miR-139-5p | MIMAT0000656 | Snapc3 | 77634 | HITS-CLIP | 29167373 | Unspecified |
| mmu-miR-139-5p | MIMAT0000656 | Snd1 | 56463 | sRNA-Seq//HITS-CLIP | 33184237\|29167373 | Unspecified |
| mmu-miR-139-5p | MIMAT0000656 | Snrnp200 | 320632 | sRNA-Seq//HITS-CLIP | 33184237\|23630078 | Unspecified |
| mmu-miR-139-5p | MIMAT0000656 | Snrnp70 | 20637 | sRNA-Seq//HITS-CLIP | 33184237\|29167373 | Unspecified |
| mmu-miR-139-5p | MIMAT0000656 | Sntb2 | 20650 | HITS-CLIP | 29167373 | Unspecified |
| mmu-miR-139-5p | MIMAT0000656 | Snx10 | 71982 | sRNA-Seq//HITS-CLIP | 33184237\|29167373 | Unspecified |
| mmu-miR-139-5p | MIMAT0000656 | Snx2 | 67804 | sRNA-Seq//HITS-CLIP | 33184237\|29167373 | Unspecified |
| mmu-miR-139-5p | MIMAT0000656 | Snx20 | 71607 | HITS-CLIP | 29167373 | Unspecified |
| mmu-miR-139-5p | MIMAT0000656 | Snx25 | 102141 | HITS-CLIP | 29167373 | Unspecified |
| mmu-miR-139-5p | MIMAT0000656 | Snx27 | 76742 | HITS-CLIP | 29167373 | Unspecified |
| mmu-miR-139-5p | MIMAT0000656 | Snx29 | 74478 | HITS-CLIP | 29167373 | Unspecified |
| mmu-miR-139-5p | MIMAT0000656 | Snx3 | 54198 | sRNA-Seq//HITS-CLIP | 33184237\|29167373 | Unspecified |
| mmu-miR-139-5p | MIMAT0000656 | Snx30 | 209131 | HITS-CLIP | 31433990\|29167373 | Unspecified |
| mmu-miR-139-5p | MIMAT0000656 | Snx4 | 69150 | sRNA-Seq//HITS-CLIP | 33184237\|29167373 | Unspecified |
| mmu-miR-139-5p | MIMAT0000656 | Snx5 | 69178 | HITS-CLIP | 29167373 | Unspecified |
| mmu-miR-139-5p | MIMAT0000656 | Snx8 | 231834 | HITS-CLIP | 29167373 | Unspecified |
| mmu-miR-139-5p | MIMAT0000656 | Snx9 | 66616 | HITS-CLIP | 29167373 | Unspecified |
| mmu-miR-139-5p | MIMAT0000656 | Soat1 | 20652 | HITS-CLIP | 29167373 | Unspecified |
| mmu-miR-139-5p | MIMAT0000656 | Socs1 | 12703 | HITS-CLIP | 31433990 | Unspecified |
| mmu-miR-139-5p | MIMAT0000656 | Socs3 | 12702 | HITS-CLIP | 31433990 | Unspecified |
| mmu-miR-139-5p | MIMAT0000656 | Socs6 | 54607 | HITS-CLIP | 29167373 | Unspecified |
| mmu-miR-139-5p | MIMAT0000656 | Soga1 | 320706 | HITS-CLIP | 29167373 | Unspecified |
| mmu-miR-139-5p | MIMAT0000656 | Son | 20658 | sRNA-Seq//HITS-CLIP | 33184237\|29167373 | Unspecified |
| mmu-miR-139-5p | MIMAT0000656 | Sp1 | 20683 | HITS-CLIP | 29167373 | Unspecified |
| mmu-miR-139-5p | MIMAT0000656 | Sp100 | 20684 | HITS-CLIP | 29167373 | Unspecified |
| mmu-miR-139-5p | MIMAT0000656 | Spag9 | 70834 | sRNA-Seq//HITS-CLIP | 33184237\|29167373 | Unspecified |
| mmu-miR-139-5p | MIMAT0000656 | Spast | 50850 | sRNA-Seq//HITS-CLIP | 33184237\|29167373 | Unspecified |
| mmu-miR-139-5p | MIMAT0000656 | Spen | 56381 | HITS-CLIP | 29167373 | Unspecified |
| mmu-miR-139-5p | MIMAT0000656 | Spg11 | 214585 | HITS-CLIP | 29167373 | Unspecified |
| mmu-miR-139-5p | MIMAT0000656 | Spg20 | 229285 | HITS-CLIP | 29167373 | Unspecified |
| mmu-miR-139-5p | MIMAT0000656 | Spink5 | 72432 | HITS-CLIP | 26203562 | Unspecified |
| mmu-miR-139-5p | MIMAT0000656 | Spint1 | 20732 | HITS-CLIP | 26203562 | Unspecified |

| | | | | | | |
|---|---|---|---|---|---|---|
| mmu-miR-139-5p | MIMAT0000656 | Spopl | 76857 | HITS-CLIP | 29167373 | Unspecified |
| mmu-miR-139-5p | MIMAT0000656 | Spp1 | 20750 | sRNA-Seq//HITS-CLIP | 33184237\|29167373 | Unspecified |
| mmu-miR-139-5p | MIMAT0000656 | Sppl3 | 74585 | HITS-CLIP | 29167373 | Unspecified |
| mmu-miR-139-5p | MIMAT0000656 | Spred1 | 114715 | HITS-CLIP | 29167373 | Unspecified |
| mmu-miR-139-5p | MIMAT0000656 | Sptan1 | 20740 | sRNA-Seq//HITS-CLIP | 33184237\|29167373 | Unspecified |
| mmu-miR-139-5p | MIMAT0000656 | Sptlc1 | 268656 | HITS-CLIP | 29167373 | Unspecified |
| mmu-miR-139-5p | MIMAT0000656 | Srcap | 100043597 | HITS-CLIP | 29167373 | Unspecified |
| mmu-miR-139-5p | MIMAT0000656 | Srebf2 | 20788 | sRNA-Seq//HITS-CLIP | 33184237\|29167373 | Unspecified |
| mmu-miR-139-5p | MIMAT0000656 | Srek1 | 218543 | HITS-CLIP | 29167373 | Unspecified |
| mmu-miR-139-5p | MIMAT0000656 | Srgap2 | 14270 | HITS-CLIP | 29167373 | Unspecified |
| mmu-miR-139-5p | MIMAT0000656 | Srpr | 67398 | HITS-CLIP | 29167373 | Unspecified |
| mmu-miR-139-5p | MIMAT0000656 | Srrm1 | 51796 | sRNA-Seq//HITS-CLIP | 33184237\|29167373 | Unspecified |
| mmu-miR-139-5p | MIMAT0000656 | Srrm2 | 75956 | HITS-CLIP//sRNA-Seq | 31433990\|33184237\|29167373 | Unspecified |
| mmu-miR-139-5p | MIMAT0000656 | Srsf1 | 110809 | sRNA-Seq//HITS-CLIP | 33184237\|29167373 | Unspecified |
| mmu-miR-139-5p | MIMAT0000656 | Srsf11 | 69207 | sRNA-Seq//HITS-CLIP | 33184237\|29167373 | Unspecified |
| mmu-miR-139-5p | MIMAT0000656 | Srsf6 | 67996 | sRNA-Seq//HITS-CLIP | 33184237\|29167373 | Unspecified |
| mmu-miR-139-5p | MIMAT0000656 | Ssh2 | 237860 | HITS-CLIP | 31433990\|29167373 | Unspecified |
| mmu-miR-139-5p | MIMAT0000656 | Ssh3 | 245857 | HITS-CLIP | 29167373 | Unspecified |
| mmu-miR-139-5p | MIMAT0000656 | Ssr2 | 66256 | sRNA-Seq//HITS-CLIP | 33184237\|29167373 | Unspecified |
| mmu-miR-139-5p | MIMAT0000656 | Ssr3 | 67437 | sRNA-Seq//HITS-CLIP | 33184237\|29167373 | Unspecified |
| mmu-miR-139-5p | MIMAT0000656 | St3gal1 | 20442 | HITS-CLIP | 29167373 | Unspecified |
| mmu-miR-139-5p | MIMAT0000656 | St3gal5 | 20454 | HITS-CLIP | 29167373 | Unspecified |
| mmu-miR-139-5p | MIMAT0000656 | St8sia4 | 20452 | HITS-CLIP | 29167373 | Unspecified |
| mmu-miR-139-5p | MIMAT0000656 | Stag1 | 20842 | HITS-CLIP | 29167373 | Unspecified |
| mmu-miR-139-5p | MIMAT0000656 | Stag2 | 20843 | HITS-CLIP | 29167373 | Unspecified |
| mmu-miR-139-5p | MIMAT0000656 | Stam2 | 56324 | HITS-CLIP | 29167373 | Unspecified |
| mmu-miR-139-5p | MIMAT0000656 | Stard9 | 668880 | HITS-CLIP | 29167373 | Unspecified |
| mmu-miR-139-5p | MIMAT0000656 | Stat3 | 20848 | sRNA-Seq//HITS-CLIP | 33184237\|29167373 | Unspecified |
| mmu-miR-139-5p | MIMAT0000656 | Stil | 20460 | HITS-CLIP | 29167373 | Unspecified |
| mmu-miR-139-5p | MIMAT0000656 | Stim2 | 116873 | HITS-CLIP | 29167373 | Unspecified |
| mmu-miR-139-5p | MIMAT0000656 | Stk10 | 20868 | HITS-CLIP | 29167373 | Unspecified |
| mmu-miR-139-5p | MIMAT0000656 | Stk3 | 56274 | HITS-CLIP | 29167373 | Unspecified |
| mmu-miR-139-5p | MIMAT0000656 | Stk4 | 58231 | HITS-CLIP | 29167373 | Unspecified |
| mmu-miR-139-5p | MIMAT0000656 | Stk40 | 74178 | HITS-CLIP | 26203562 | Unspecified |
| mmu-miR-139-5p | MIMAT0000656 | Stom | 13830 | sRNA-Seq//HITS-CLIP | 33184237\|29167373 | Unspecified |
| mmu-miR-139-5p | MIMAT0000656 | Strn3 | 94186 | HITS-CLIP//sRNA-Seq | 31433990\|33184237 | Unspecified |
| mmu-miR-139-5p | MIMAT0000656 | Stt3b | 68292 | sRNA-Seq//HITS-CLIP | 33184237\|29167373 | Unspecified |
| mmu-miR-139-5p | MIMAT0000656 | Stx12 | 100226 | sRNA-Seq//HITS-CLIP | 33184237\|29167373 | Unspecified |
| mmu-miR-139-5p | MIMAT0000656 | Stx16 | 228960 | HITS-CLIP | 29167373 | Unspecified |

| | | | | | | |
|---|---|---|---|---|---|---|
| mmu-miR-139-5p | MIMAT0000656 | Stx3 | 20908 | HITS-CLIP | 29167373 | Unspecified |
| mmu-miR-139-5p | MIMAT0000656 | Stxbp4 | 20913 | HITS-CLIP | 29167373 | Unspecified |
| mmu-miR-139-5p | MIMAT0000656 | Stxbp5 | 78808 | HITS-CLIP | 29167373 | Unspecified |
| mmu-miR-139-5p | MIMAT0000656 | Suco | 226551 | HITS-CLIP | 29167373 | Unspecified |
| mmu-miR-139-5p | MIMAT0000656 | Sulf2 | 72043 | HITS-CLIP | 26203562\|29167373 | Unspecified |
| mmu-miR-139-5p | MIMAT0000656 | Sun2 | 223697 | HITS-CLIP | 29167373 | Unspecified |
| mmu-miR-139-5p | MIMAT0000656 | Supt16 | 114741 | HITS-CLIP | 29167373 | Unspecified |
| mmu-miR-139-5p | MIMAT0000656 | Susd3 | 66329 | HITS-CLIP | 29167373 | Unspecified |
| mmu-miR-139-5p | MIMAT0000656 | Susd6 | 9766 | HITS-CLIP | 29167373 | Unspecified |
| mmu-miR-139-5p | MIMAT0000656 | Suv39h1 | 20937 | HITS-CLIP | 29167373 | Unspecified |
| mmu-miR-139-5p | MIMAT0000656 | Swap70 | 20947 | HITS-CLIP | 29167373 | Unspecified |
| mmu-miR-139-5p | MIMAT0000656 | Syne1 | 64009 | sRNA-Seq//HITS-CLIP | 33184237\|29167373 | Unspecified |
| mmu-miR-139-5p | MIMAT0000656 | Syngr2 | 20973 | HITS-CLIP | 29167373 | Unspecified |
| mmu-miR-139-5p | MIMAT0000656 | Synj2 | 20975 | HITS-CLIP | 29167373 | Unspecified |
| mmu-miR-139-5p | MIMAT0000656 | Sypl | 19027 | HITS-CLIP | 29167373 | Unspecified |
| mmu-miR-139-5p | MIMAT0000656 | Syt11 | 229521 | sRNA-Seq//HITS-CLIP | 33184237\|31433990 | Unspecified |
| mmu-miR-139-5p | MIMAT0000656 | Syvn1 | 74126 | HITS-CLIP | 26203562 | Unspecified |
| mmu-miR-139-5p | MIMAT0000656 | Szrd1 | 213491 | HITS-CLIP | 29167373 | Unspecified |
| mmu-miR-139-5p | MIMAT0000656 | Tab2 | 68652 | HITS-CLIP | 29167373 | Unspecified |
| mmu-miR-139-5p | MIMAT0000656 | Taf1d | 75316 | sRNA-Seq//HITS-CLIP | 33184237\|29167373 | Unspecified |
| mmu-miR-139-5p | MIMAT0000656 | Taf3 | 209361 | HITS-CLIP | 26203562 | Unspecified |
| mmu-miR-139-5p | MIMAT0000656 | Tank | 21353 | HITS-CLIP | 29167373 | Unspecified |
| mmu-miR-139-5p | MIMAT0000656 | Taok1 | 216965 | HITS-CLIP//sRNA-Seq | 31433990\|33184237\|29167373 | Unspecified |
| mmu-miR-139-5p | MIMAT0000656 | Taok3 | 330177 | HITS-CLIP | 29167373 | Unspecified |
| mmu-miR-139-5p | MIMAT0000656 | Tardbp | 230908 | sRNA-Seq//HITS-CLIP | 33184237\|29167373 | Unspecified |
| mmu-miR-139-5p | MIMAT0000656 | Tatdn2 | 381801 | HITS-CLIP | 29167373 | Unspecified |
| mmu-miR-139-5p | MIMAT0000656 | Tbc1d15 | 66687 | HITS-CLIP | 29167373 | Unspecified |
| mmu-miR-139-5p | MIMAT0000656 | Tbc1d20 | 67231 | HITS-CLIP | 31433990\|29167373 | Unspecified |
| mmu-miR-139-5p | MIMAT0000656 | Tbc1d5 | 72238 | HITS-CLIP | 29167373 | Unspecified |
| mmu-miR-139-5p | MIMAT0000656 | Tbcel | 272589 | HITS-CLIP | 29167373 | Unspecified |
| mmu-miR-139-5p | MIMAT0000656 | Tbk1 | 56480 | HITS-CLIP | 29167373 | Unspecified |
| mmu-miR-139-5p | MIMAT0000656 | Tbl1xr1 | 81004 | HITS-CLIP | 29167373 | Unspecified |
| mmu-miR-139-5p | MIMAT0000656 | Tbrg4 | 21379 | HITS-CLIP | 29167373 | Unspecified |
| mmu-miR-139-5p | MIMAT0000656 | Tbxas1 | 21391 | HITS-CLIP | 29167373 | Unspecified |
| mmu-miR-139-5p | MIMAT0000656 | Tcf20 | 21411 | sRNA-Seq//HITS-CLIP | 33184237\|29167373 | Unspecified |
| mmu-miR-139-5p | MIMAT0000656 | Tcf25 | 66855 | HITS-CLIP//sRNA-Seq | 29167373\|33184237 | Unspecified |
| mmu-miR-139-5p | MIMAT0000656 | Tcirg1 | 27060 | HITS-CLIP | 29167373 | Unspecified |
| mmu-miR-139-5p | MIMAT0000656 | Tdrd7 | 100121 | HITS-CLIP | 29167373 | Unspecified |
| mmu-miR-139-5p | MIMAT0000656 | Tead4 | 21679 | HITS-CLIP | 26203562 | Unspecified |

| | | | | | | |
|---|---|---|---|---|---|---|
| mmu-miR-139-5p | MIMAT0000656 | Tec | 21682 | HITS-CLIP | 29167373 | Unspecified |
| mmu-miR-139-5p | MIMAT0000656 | Tent4b | 64282 | HITS-CLIP | 29167373 | Unspecified |
| mmu-miR-139-5p | MIMAT0000656 | Tent5a | 55603 | HITS-CLIP | 31433990\|29167373 | Unspecified |
| mmu-miR-139-5p | MIMAT0000656 | Tep1 | 21745 | HITS-CLIP | 29167373 | Unspecified |
| mmu-miR-139-5p | MIMAT0000656 | Tet2 | 214133 | HITS-CLIP | 29167373 | Unspecified |
| mmu-miR-139-5p | MIMAT0000656 | Tet3 | 194388 | HITS-CLIP | 29167373 | Unspecified |
| mmu-miR-139-5p | MIMAT0000656 | Tfeb | 21425 | HITS-CLIP | 29167373 | Unspecified |
| mmu-miR-139-5p | MIMAT0000656 | Tfrc | 22042 | HITS-CLIP//sRNA-Seq | 31433990\|33184237 | Unspecified |
| mmu-miR-139-5p | MIMAT0000656 | Tgfbi | 21810 | HITS-CLIP | 29167373 | Unspecified |
| mmu-miR-139-5p | MIMAT0000656 | Tgfbr1 | 21812 | HITS-CLIP | 30971389\|29167373 | Unspecified |
| mmu-miR-139-5p | MIMAT0000656 | Tgs1 | 116940 | HITS-CLIP | 29167373 | Unspecified |
| mmu-miR-139-5p | MIMAT0000656 | Thoc7 | 66231 | sRNA-Seq//HITS-CLIP | 33184237\|29167373 | Unspecified |
| mmu-miR-139-5p | MIMAT0000656 | Tial1 | 21843 | sRNA-Seq//HITS-CLIP | 33184237\|29167373 | Unspecified |
| mmu-miR-139-5p | MIMAT0000656 | Tiam1 | 21844 | HITS-CLIP | 29167373 | Unspecified |
| mmu-miR-139-5p | MIMAT0000656 | Tiparp | 99929 | HITS-CLIP | 29167373 | Unspecified |
| mmu-miR-139-5p | MIMAT0000656 | Tle1 | 21885 | sRNA-Seq//HITS-CLIP | 33184237\|29167373 | Unspecified |
| mmu-miR-139-5p | MIMAT0000656 | Tle4 | 21888 | HITS-CLIP | 29167373 | Unspecified |
| mmu-miR-139-5p | MIMAT0000656 | Tln1 | 21894 | HITS-CLIP | 29167373 | Unspecified |
| mmu-miR-139-5p | MIMAT0000656 | Tlr13 | 279572 | HITS-CLIP | 29167373 | Unspecified |
| mmu-miR-139-5p | MIMAT0000656 | Tlr2 | 24088 | HITS-CLIP | 29167373 | Unspecified |
| mmu-miR-139-5p | MIMAT0000656 | Tlr7 | 170743 | HITS-CLIP | 29167373 | Unspecified |
| mmu-miR-139-5p | MIMAT0000656 | Tm2d2 | 69742 | sRNA-Seq//HITS-CLIP | 33184237\|29167373 | Unspecified |
| mmu-miR-139-5p | MIMAT0000656 | Tm9sf3 | 107358 | sRNA-Seq//HITS-CLIP | 33184237\|29167373 | Unspecified |
| mmu-miR-139-5p | MIMAT0000656 | Tmcc1 | 330401 | HITS-CLIP | 29167373 | Unspecified |
| mmu-miR-139-5p | MIMAT0000656 | Tmed7 | 66676 | sRNA-Seq//HITS-CLIP | 33184237\|29167373 | Unspecified |
| mmu-miR-139-5p | MIMAT0000656 | Tmem104 | 320534 | HITS-CLIP | 29167373 | Unspecified |
| mmu-miR-139-5p | MIMAT0000656 | Tmem106a | 217203 | HITS-CLIP | 29167373 | Unspecified |
| mmu-miR-139-5p | MIMAT0000656 | Tmem106b | 71900 | HITS-CLIP | 26203562 | Unspecified |
| mmu-miR-139-5p | MIMAT0000656 | Tmem127 | 69470 | HITS-CLIP | 29167373 | Unspecified |
| mmu-miR-139-5p | MIMAT0000656 | Tmem128 | 66309 | HITS-CLIP | 29167373 | Unspecified |
| mmu-miR-139-5p | MIMAT0000656 | Tmem14c | 66154 | HITS-CLIP | 29167373 | Unspecified |
| mmu-miR-139-5p | MIMAT0000656 | Tmem170b | 621976 | HITS-CLIP | 31433990\|26203562\|29167373 | Unspecified |
| mmu-miR-139-5p | MIMAT0000656 | Tmem189 | 407243 | HITS-CLIP | 29167373 | Unspecified |
| mmu-miR-139-5p | MIMAT0000656 | Tmem206 | 66950 | HITS-CLIP | 21258322\|29167373\|25083871 | Unspecified |
| mmu-miR-139-5p | MIMAT0000656 | Tmem209 | 72649 | HITS-CLIP | 29167373 | Unspecified |
| mmu-miR-139-5p | MIMAT0000656 | Tmem245 | 242474 | HITS-CLIP | 29167373 | Unspecified |
| mmu-miR-139-5p | MIMAT0000656 | Tmem30a | 69981 | sRNA-Seq//HITS-CLIP | 33184237\|29167373 | Unspecified |
| mmu-miR-139-5p | MIMAT0000656 | Tmem38b | 52076 | HITS-CLIP | 29167373 | Unspecified |
| mmu-miR-139-5p | MIMAT0000656 | Tmem68 | 72098 | HITS-CLIP | 29167373 | Unspecified |

| | | | | | | | |
|---|---|---|---|---|---|---|---|
| mmu-miR-139-5p | MIMAT0000656 | Tmpo | 21917 | HITS-CLIP | | 29167373 | Unspecified |
| mmu-miR-139-5p | MIMAT0000656 | Tmtc3 | 237500 | HITS-CLIP | | 29167373 | Unspecified |
| mmu-miR-139-5p | MIMAT0000656 | Tnfaip8 | 106869 | HITS-CLIP | | 29167373 | Unspecified |
| mmu-miR-139-5p | MIMAT0000656 | Tnfrsf11a | 21934 | HITS-CLIP | | 29167373 | Unspecified |
| mmu-miR-139-5p | MIMAT0000656 | Tnfrsf21 | 94185 | HITS-CLIP | | 29167373 | Unspecified |
| mmu-miR-139-5p | MIMAT0000656 | Tnip3 | 414084 | HITS-CLIP | | 29167373 | Unspecified |
| mmu-miR-139-5p | MIMAT0000656 | Tnks | 21951 | sRNA-Seq//HITS-CLIP | 33184237\|29167373 | | Unspecified |
| mmu-miR-139-5p | MIMAT0000656 | Tnks2 | 74493 | sRNA-Seq//HITS-CLIP | 33184237\|29167373 | | Unspecified |
| mmu-miR-139-5p | MIMAT0000656 | Tnpo1 | 238799 | HITS-CLIP | | 29167373 | Unspecified |
| mmu-miR-139-5p | MIMAT0000656 | Tnpo3 | 320938 | sRNA-Seq//HITS-CLIP | 33184237\|29167373 | | Unspecified |
| mmu-miR-139-5p | MIMAT0000656 | Tnrc6a | 233833 | HITS-CLIP | 31433990\|29167373 | | Unspecified |
| mmu-miR-139-5p | MIMAT0000656 | Tnrc6b | 213988 | HITS-CLIP | | 29167373 | Unspecified |
| mmu-miR-139-5p | MIMAT0000656 | Tnrc6c | 217351 | HITS-CLIP | 26203562\|29167373 | | Unspecified |
| mmu-miR-139-5p | MIMAT0000656 | Tns3 | 319939 | HITS-CLIP | | 29167373 | Unspecified |
| mmu-miR-139-5p | MIMAT0000656 | Tom1l2 | 216810 | sRNA-Seq//HITS-CLIP | 33184237\|29167373 | | Unspecified |
| mmu-miR-139-5p | MIMAT0000656 | Tomm70a | 28185 | sRNA-Seq//HITS-CLIP | 33184237\|29167373 | | Unspecified |
| mmu-miR-139-5p | MIMAT0000656 | Top1 | 21969 | sRNA-Seq//HITS-CLIP | 33184237\|29167373 | | Unspecified |
| mmu-miR-139-5p | MIMAT0000656 | Top2b | 21974 | sRNA-Seq//HITS-CLIP | 33184237\|29167373 | | Unspecified |
| mmu-miR-139-5p | MIMAT0000656 | Topors | 106021 | HITS-CLIP | | 29167373 | Unspecified |
| mmu-miR-139-5p | MIMAT0000656 | Tor1aip1 | 208263 | HITS-CLIP | | 29167373 | Unspecified |
| mmu-miR-139-5p | MIMAT0000656 | Tor1aip2 | 240832 | sRNA-Seq//HITS-CLIP | 33184237\|31433990\|29167373 | | Unspecified |
| mmu-miR-139-5p | MIMAT0000656 | Tor1b | 30934 | HITS-CLIP | 26203562\|29167373 | | Unspecified |
| mmu-miR-139-5p | MIMAT0000656 | Tor4a | 227612 | HITS-CLIP | | 29167373 | Unspecified |
| mmu-miR-139-5p | MIMAT0000656 | Tpcn2 | 233979 | HITS-CLIP | | 29167373 | Unspecified |
| mmu-miR-139-5p | MIMAT0000656 | Tpd52 | 21985 | sRNA-Seq//HITS-CLIP | 33184237\|29167373 | | Unspecified |
| mmu-miR-139-5p | MIMAT0000656 | Tpd52l2 | 66314 | HITS-CLIP//sRNA-Seq | 31433990\|33184237 | | Unspecified |
| mmu-miR-139-5p | MIMAT0000656 | Tpm1 | 22003 | sRNA-Seq//HITS-CLIP | 33184237\|29167373 | | Unspecified |
| mmu-miR-139-5p | MIMAT0000656 | Tpm3 | 59069 | sRNA-Seq//HITS-CLIP | 33184237\|29167373 | | Unspecified |
| mmu-miR-139-5p | MIMAT0000656 | Tpm4 | 326618 | sRNA-Seq//HITS-CLIP | 33184237\|29167373 | | Unspecified |
| mmu-miR-139-5p | MIMAT0000656 | Tpp1 | 12751 | HITS-CLIP | | 29167373 | Unspecified |
| mmu-miR-139-5p | MIMAT0000656 | Tpp2 | 22019 | sRNA-Seq//HITS-CLIP | 33184237\|29167373 | | Unspecified |
| mmu-miR-139-5p | MIMAT0000656 | Tpr | 108989 | sRNA-Seq//HITS-CLIP | 33184237\|29167373 | | Unspecified |
| mmu-miR-139-5p | MIMAT0000656 | Tpra1 | 24100 | HITS-CLIP | | 29167373 | Unspecified |
| mmu-miR-139-5p | MIMAT0000656 | Tpx2 | 72119 | HITS-CLIP | | 29167373 | Unspecified |
| mmu-miR-139-5p | MIMAT0000656 | Tra2a | 101214 | HITS-CLIP | | 29167373 | Unspecified |
| mmu-miR-139-5p | MIMAT0000656 | Trabd | 67976 | HITS-CLIP | | 29167373 | Unspecified |
| mmu-miR-139-5p | MIMAT0000656 | Traf3 | 22031 | HITS-CLIP | | 29167373 | Unspecified |
| mmu-miR-139-5p | MIMAT0000656 | Traf6 | 22034 | HITS-CLIP | | 29167373 | Unspecified |
| mmu-miR-139-5p | MIMAT0000656 | Traf7 | 224619 | HITS-CLIP | | 29167373 | Unspecified |

| | | | | | | | |
|---|---|---|---|---|---|---|---|
| mmu-miR-139-5p | MIMAT0000656 | Trafd1 | 231712 | HITS-CLIP | | 29167373 | Unspecified |
| mmu-miR-139-5p | MIMAT0000656 | Tram1 | 72265 | HITS-CLIP | | 29167373 | Unspecified |
| mmu-miR-139-5p | MIMAT0000656 | Trap1 | 68015 | HITS-CLIP | | 29167373 | Unspecified |
| mmu-miR-139-5p | MIMAT0000656 | Trappc10 | 216131 | HITS-CLIP | | 29167373 | Unspecified |
| mmu-miR-139-5p | MIMAT0000656 | Trappc11 | 320714 | HITS-CLIP | | 29167373 | Unspecified |
| mmu-miR-139-5p | MIMAT0000656 | Trappc12 | 217449 | HITS-CLIP | | 29167373 | Unspecified |
| mmu-miR-139-5p | MIMAT0000656 | Trem2 | 83433 | HITS-CLIP | | 29167373 | Unspecified |
| mmu-miR-139-5p | MIMAT0000656 | Trim25 | 217069 | HITS-CLIP | | 29167373 | Unspecified |
| mmu-miR-139-5p | MIMAT0000656 | Trim30a | 20128 | HITS-CLIP | | 29167373 | Unspecified |
| mmu-miR-139-5p | MIMAT0000656 | Trim30d | 209387 | HITS-CLIP | | 29167373 | Unspecified |
| mmu-miR-139-5p | MIMAT0000656 | Trim44 | 80985 | sRNA-Seq//HITS-CLIP | 33184237\|29167373 | | Unspecified |
| mmu-miR-139-5p | MIMAT0000656 | Trio | 223435 | HITS-CLIP | | 29167373 | Unspecified |
| mmu-miR-139-5p | MIMAT0000656 | Trip11 | 109181 | HITS-CLIP | | 29167373 | Unspecified |
| mmu-miR-139-5p | MIMAT0000656 | Trmt2a | 15547 | HITS-CLIP | | 29167373 | Unspecified |
| mmu-miR-139-5p | MIMAT0000656 | Trp53bp1 | 27223 | sRNA-Seq//HITS-CLIP | 33184237\|29167373 | | Unspecified |
| mmu-miR-139-5p | MIMAT0000656 | Trp53inp1 | 60599 | HITS-CLIP | | 26203562 | Unspecified |
| mmu-miR-139-5p | MIMAT0000656 | Trpc4ap | 56407 | sRNA-Seq//HITS-CLIP | 33184237\|29167373 | | Unspecified |
| mmu-miR-139-5p | MIMAT0000656 | Trpm7 | 58800 | HITS-CLIP | | 29167373 | Unspecified |
| mmu-miR-139-5p | MIMAT0000656 | Trps1 | 83925 | HITS-CLIP | | 29167373 | Unspecified |
| mmu-miR-139-5p | MIMAT0000656 | Trpv3 | 246788 | HITS-CLIP | | 26203562 | Unspecified |
| mmu-miR-139-5p | MIMAT0000656 | Tsc1 | 64930 | HITS-CLIP | | 29167373 | Unspecified |
| mmu-miR-139-5p | MIMAT0000656 | Tsc22d2 | 72033 | HITS-CLIP | | 29167373 | Unspecified |
| mmu-miR-139-5p | MIMAT0000656 | Tsc22d4 | 78829 | HITS-CLIP | | 29167373 | Unspecified |
| mmu-miR-139-5p | MIMAT0000656 | Tsn | 22099 | sRNA-Seq//HITS-CLIP | 33184237\|29167373 | | Unspecified |
| mmu-miR-139-5p | MIMAT0000656 | Tsnax | 53424 | sRNA-Seq//HITS-CLIP | 33184237\|29167373 | | Unspecified |
| mmu-miR-139-5p | MIMAT0000656 | Ttbk2 | 140810 | sRNA-Seq//HITS-CLIP | 33184237\|29167373 | | Unspecified |
| mmu-miR-139-5p | MIMAT0000656 | Ttc13 | 234875 | sRNA-Seq//HITS-CLIP | 33184237\|29167373 | | Unspecified |
| mmu-miR-139-5p | MIMAT0000656 | Ttc14 | 67120 | HITS-CLIP | | 29167373 | Unspecified |
| mmu-miR-139-5p | MIMAT0000656 | Ttc7 | 225049 | HITS-CLIP | | 29167373 | Unspecified |
| mmu-miR-139-5p | MIMAT0000656 | Ttc7b | 104718 | HITS-CLIP | | 29167373 | Unspecified |
| mmu-miR-139-5p | MIMAT0000656 | Tti1 | 75425 | HITS-CLIP | | 29167373 | Unspecified |
| mmu-miR-139-5p | MIMAT0000656 | Tubb2a | 22151 | sRNA-Seq//HITS-CLIP | 33184237\|29167373 | | Unspecified |
| mmu-miR-139-5p | MIMAT0000656 | Tubb5 | 22154 | HITS-CLIP//sRNA-Seq | 31433990\|33184237 | | Unspecified |
| mmu-miR-139-5p | MIMAT0000656 | Tubgcp3 | 259279 | HITS-CLIP | | 29167373 | Unspecified |
| mmu-miR-139-5p | MIMAT0000656 | Tut4 | 23318 | HITS-CLIP | | 29167373 | Unspecified |
| mmu-miR-139-5p | MIMAT0000656 | Txndc15 | 69672 | HITS-CLIP | | 29167373 | Unspecified |
| mmu-miR-139-5p | MIMAT0000656 | Txndc5 | 105245 | HITS-CLIP | | 29167373 | Unspecified |
| mmu-miR-139-5p | MIMAT0000656 | Txnip | 56338 | HITS-CLIP | | 29167373 | Unspecified |
| mmu-miR-139-5p | MIMAT0000656 | Txnrd1 | 50493 | HITS-CLIP//sRNA-Seq | 29167373\|31433990\|33184237 | | Unspecified |

| | | | | | | |
|---|---|---|---|---|---|---|
| mmu-miR-139-5p | MIMAT0000656 | U2af1 | 108121 | sRNA-Seq//HITS-CLIP | 33184237\|29167373 | Unspecified |
| mmu-miR-139-5p | MIMAT0000656 | U2surp | 67958 | sRNA-Seq//HITS-CLIP | 33184237\|29167373 | Unspecified |
| mmu-miR-139-5p | MIMAT0000656 | Uap1l1 | 227620 | HITS-CLIP | 29167373 | Unspecified |
| mmu-miR-139-5p | MIMAT0000656 | Uba1 | 22201 | sRNA-Seq//HITS-CLIP | 33184237\|29167373 | Unspecified |
| mmu-miR-139-5p | MIMAT0000656 | Ubac2 | 68889 | HITS-CLIP | 29167373 | Unspecified |
| mmu-miR-139-5p | MIMAT0000656 | Ubap2 | 68926 | sRNA-Seq//HITS-CLIP | 33184237\|29167373 | Unspecified |
| mmu-miR-139-5p | MIMAT0000656 | Ubap2l | 74383 | sRNA-Seq//HITS-CLIP | 33184237\|29167373 | Unspecified |
| mmu-miR-139-5p | MIMAT0000656 | Ubash3b | 72828 | sRNA-Seq//HITS-CLIP | 33184237\|31433990\|29167373 | Unspecified |
| mmu-miR-139-5p | MIMAT0000656 | Ube2b | 22210 | sRNA-Seq//HITS-CLIP | 33184237\|29167373 | Unspecified |
| mmu-miR-139-5p | MIMAT0000656 | Ube2c | 68612 | HITS-CLIP | 31433990 | Unspecified |
| mmu-miR-139-5p | MIMAT0000656 | Ube2d1 | 216080 | sRNA-Seq//HITS-CLIP | 33184237\|29167373 | Unspecified |
| mmu-miR-139-5p | MIMAT0000656 | Ube2e2 | 218793 | sRNA-Seq//HITS-CLIP | 33184237\|29167373 | Unspecified |
| mmu-miR-139-5p | MIMAT0000656 | Ube2e3 | 22193 | sRNA-Seq//HITS-CLIP | 33184237\|29167373 | Unspecified |
| mmu-miR-139-5p | MIMAT0000656 | Ube2f | 67921 | HITS-CLIP | 29167373 | Unspecified |
| mmu-miR-139-5p | MIMAT0000656 | Ube2g1 | 67128 | HITS-CLIP | 29167373 | Unspecified |
| mmu-miR-139-5p | MIMAT0000656 | Ube2k | 53323 | sRNA-Seq//HITS-CLIP | 33184237\|31433990\|29167373 | Unspecified |
| mmu-miR-139-5p | MIMAT0000656 | Ube2l3 | 22195 | HITS-CLIP//sRNA-Seq | 29167373\|33184237 | Unspecified |
| mmu-miR-139-5p | MIMAT0000656 | Ube2m | 22192 | sRNA-Seq//HITS-CLIP | 33184237\|29167373 | Unspecified |
| mmu-miR-139-5p | MIMAT0000656 | Ube2q1 | 70093 | HITS-CLIP | 29167373 | Unspecified |
| mmu-miR-139-5p | MIMAT0000656 | Ube2s | 77891 | HITS-CLIP//sRNA-Seq | 31433990\|33184237 | Unspecified |
| mmu-miR-139-5p | MIMAT0000656 | Ube2v2 | 70620 | sRNA-Seq//HITS-CLIP | 33184237\|29167373 | Unspecified |
| mmu-miR-139-5p | MIMAT0000656 | Ube2z | 268470 | sRNA-Seq//HITS-CLIP | 33184237\|31433990 | Unspecified |
| mmu-miR-139-5p | MIMAT0000656 | Ube3a | 22215 | sRNA-Seq//HITS-CLIP | 33184237\|29167373 | Unspecified |
| mmu-miR-139-5p | MIMAT0000656 | Ubiad1 | 71707 | HITS-CLIP | 29167373 | Unspecified |
| mmu-miR-139-5p | MIMAT0000656 | Ubl3 | 24109 | sRNA-Seq//HITS-CLIP | 33184237\|29167373 | Unspecified |
| mmu-miR-139-5p | MIMAT0000656 | Ubn1 | 170644 | HITS-CLIP | 29167373 | Unspecified |
| mmu-miR-139-5p | MIMAT0000656 | Ubn2 | 320538 | HITS-CLIP | 29167373 | Unspecified |
| mmu-miR-139-5p | MIMAT0000656 | Ubr2 | 224826 | HITS-CLIP | 29167373 | Unspecified |
| mmu-miR-139-5p | MIMAT0000656 | Ubr3 | 68795 | sRNA-Seq//HITS-CLIP | 33184237\|29167373 | Unspecified |
| mmu-miR-139-5p | MIMAT0000656 | Ubr4 | 69116 | sRNA-Seq//HITS-CLIP | 33184237\|29167373 | Unspecified |
| mmu-miR-139-5p | MIMAT0000656 | Ubr5 | 70790 | sRNA-Seq//HITS-CLIP | 33184237\|29167373 | Unspecified |
| mmu-miR-139-5p | MIMAT0000656 | Ucp2 | 22228 | sRNA-Seq//HITS-CLIP | 33184237\|29167373 | Unspecified |
| mmu-miR-139-5p | MIMAT0000656 | Ugt1a6b | 394435 | HITS-CLIP | 29167373 | Unspecified |
| mmu-miR-139-5p | MIMAT0000656 | Uhmk1 | 16589 | sRNA-Seq//HITS-CLIP | 33184237\|29167373 | Unspecified |
| mmu-miR-139-5p | MIMAT0000656 | Uhrf1bp1 | 224648 | HITS-CLIP | 29167373 | Unspecified |
| mmu-miR-139-5p | MIMAT0000656 | Uhrf2 | 109113 | HITS-CLIP | 29167373 | Unspecified |
| mmu-miR-139-5p | MIMAT0000656 | Umps | 22247 | HITS-CLIP | 29167373 | Unspecified |
| mmu-miR-139-5p | MIMAT0000656 | Unc13d | 70450 | HITS-CLIP | 31433990 | Unspecified |
| mmu-miR-139-5p | MIMAT0000656 | Upf1 | 19704 | HITS-CLIP | 29167373 | Unspecified |

| | | | | | | | |
|---|---|---|---|---|---|---|---|
| mmu-miR-139-5p | MIMAT0000656 | Upf2 | 326622 | HITS-CLIP | | 29167373 | Unspecified |
| mmu-miR-139-5p | MIMAT0000656 | Uqcrb | 67530 | sRNA-Seq//HITS-CLIP | 33184237\|29167373 | | Unspecified |
| mmu-miR-139-5p | MIMAT0000656 | Usf3 | 205717 | HITS-CLIP | | 29167373 | Unspecified |
| mmu-miR-139-5p | MIMAT0000656 | Uso1 | 56041 | HITS-CLIP | | 29167373 | Unspecified |
| mmu-miR-139-5p | MIMAT0000656 | Usp15 | 14479 | HITS-CLIP//sRNA-Seq | 29167373\|33184237 | | Unspecified |
| mmu-miR-139-5p | MIMAT0000656 | Usp19 | 71472 | sRNA-Seq//HITS-CLIP | 33184237\|29167373 | | Unspecified |
| mmu-miR-139-5p | MIMAT0000656 | Usp24 | 329908 | HITS-CLIP | | 29167373 | Unspecified |
| mmu-miR-139-5p | MIMAT0000656 | Usp25 | 30940 | HITS-CLIP | | 31433990 | Unspecified |
| mmu-miR-139-5p | MIMAT0000656 | Usp28 | 235323 | HITS-CLIP | | 29167373 | Unspecified |
| mmu-miR-139-5p | MIMAT0000656 | Usp3 | 235441 | sRNA-Seq//HITS-CLIP | 33184237\|29167373 | | Unspecified |
| mmu-miR-139-5p | MIMAT0000656 | Usp34 | 17847 | sRNA-Seq//HITS-CLIP | 33184237\|29167373 | | Unspecified |
| mmu-miR-139-5p | MIMAT0000656 | Usp36 | 72344 | HITS-CLIP | | 29167373 | Unspecified |
| mmu-miR-139-5p | MIMAT0000656 | Usp45 | 77593 | HITS-CLIP | | 29167373 | Unspecified |
| mmu-miR-139-5p | MIMAT0000656 | Usp47 | 74996 | sRNA-Seq//HITS-CLIP | 33184237\|29167373 | | Unspecified |
| mmu-miR-139-5p | MIMAT0000656 | Usp48 | 170707 | sRNA-Seq//HITS-CLIP | 33184237\|29167373 | | Unspecified |
| mmu-miR-139-5p | MIMAT0000656 | Usp5 | 22225 | sRNA-Seq//HITS-CLIP | 33184237\|31433990 | | Unspecified |
| mmu-miR-139-5p | MIMAT0000656 | Usp7 | 252870 | sRNA-Seq//HITS-CLIP | 33184237\|29167373 | | Unspecified |
| mmu-miR-139-5p | MIMAT0000656 | Usp8 | 84092 | HITS-CLIP | | 29167373 | Unspecified |
| mmu-miR-139-5p | MIMAT0000656 | Usp9x | 22284 | sRNA-Seq//HITS-CLIP | 33184237\|29167373 | | Unspecified |
| mmu-miR-139-5p | MIMAT0000656 | Utp6 | 216987 | HITS-CLIP | | 29167373 | Unspecified |
| mmu-miR-139-5p | MIMAT0000656 | Vac14 | 234729 | HITS-CLIP | | 29167373 | Unspecified |
| mmu-miR-139-5p | MIMAT0000656 | Vamp3 | 22319 | HITS-CLIP | | 29167373 | Unspecified |
| mmu-miR-139-5p | MIMAT0000656 | Vamp4 | 53330 | sRNA-Seq//HITS-CLIP | 33184237\|29167373 | | Unspecified |
| mmu-miR-139-5p | MIMAT0000656 | Vapb | 56491 | sRNA-Seq//HITS-CLIP | 33184237\|29167373 | | Unspecified |
| mmu-miR-139-5p | MIMAT0000656 | Vbp1 | 22327 | sRNA-Seq//HITS-CLIP | 33184237\|29167373 | | Unspecified |
| mmu-miR-139-5p | MIMAT0000656 | Vcpip1 | 70675 | HITS-CLIP | | 29167373 | Unspecified |
| mmu-miR-139-5p | MIMAT0000656 | Vezf1 | 22344 | HITS-CLIP | | 29167373 | Unspecified |
| mmu-miR-139-5p | MIMAT0000656 | Vim | 22352 | sRNA-Seq//HITS-CLIP | 33184237\|29167373 | | Unspecified |
| mmu-miR-139-5p | MIMAT0000656 | Vps13a | 271564 | sRNA-Seq//HITS-CLIP | 33184237\|29167373 | | Unspecified |
| mmu-miR-139-5p | MIMAT0000656 | Vps29 | 56433 | sRNA-Seq//HITS-CLIP | 33184237\|29167373 | | Unspecified |
| mmu-miR-139-5p | MIMAT0000656 | Vps37a | 52348 | HITS-CLIP | | 29167373 | Unspecified |
| mmu-miR-139-5p | MIMAT0000656 | Vps4b | 20479 | sRNA-Seq//HITS-CLIP | 33184237\|29167373 | | Unspecified |
| mmu-miR-139-5p | MIMAT0000656 | Vps50 | 55610 | sRNA-Seq//HITS-CLIP | 33184237\|31433990 | | Unspecified |
| mmu-miR-139-5p | MIMAT0000656 | Vps53 | 68299 | sRNA-Seq//HITS-CLIP | 33184237\|31433990 | | Unspecified |
| mmu-miR-139-5p | MIMAT0000656 | Vps54 | 245944 | HITS-CLIP | | 29167373 | Unspecified |
| mmu-miR-139-5p | MIMAT0000656 | Vps72 | 21427 | HITS-CLIP | | 29167373 | Unspecified |
| mmu-miR-139-5p | MIMAT0000656 | Vrk3 | 101568 | HITS-CLIP | | 29167373 | Unspecified |
| mmu-miR-139-5p | MIMAT0000656 | Vta1 | 66201 | sRNA-Seq//HITS-CLIP | 33184237\|29167373 | | Unspecified |
| mmu-miR-139-5p | MIMAT0000656 | Vwa5a | 67776 | HITS-CLIP | | 29167373 | Unspecified |

| | | | | | | |
|---|---|---|---|---|---|---|
| mmu-miR-139-5p | MIMAT0000656 | Wapl | 23063 | HITS-CLIP | 31433990 | Unspecified |
| mmu-miR-139-5p | MIMAT0000656 | Wasf2 | 242687 | HITS-CLIP | 29167373 | Unspecified |
| mmu-miR-139-5p | MIMAT0000656 | Washc4 | 319277 | HITS-CLIP | 29167373 | Unspecified |
| mmu-miR-139-5p | MIMAT0000656 | Washc5 | 9897 | HITS-CLIP | 29167373 | Unspecified |
| mmu-miR-139-5p | MIMAT0000656 | Wasl | 73178 | HITS-CLIP | 29167373 | Unspecified |
| mmu-miR-139-5p | MIMAT0000656 | Wbp1l | 226178 | HITS-CLIP | 29167373 | Unspecified |
| mmu-miR-139-5p | MIMAT0000656 | Wbp2 | 22378 | sRNA-Seq//HITS-CLIP | 33184237\|26203562\|29167373 | Unspecified |
| mmu-miR-139-5p | MIMAT0000656 | Wdfy3 | 72145 | sRNA-Seq//HITS-CLIP | 33184237\|29167373 | Unspecified |
| mmu-miR-139-5p | MIMAT0000656 | Wdfy4 | 545030 | HITS-CLIP | 29167373 | Unspecified |
| mmu-miR-139-5p | MIMAT0000656 | Wdhd1 | 218973 | HITS-CLIP | 29167373 | Unspecified |
| mmu-miR-139-5p | MIMAT0000656 | Wdr12 | 57750 | HITS-CLIP | 29167373 | Unspecified |
| mmu-miR-139-5p | MIMAT0000656 | Wdr26 | 226757 | sRNA-Seq//HITS-CLIP | 33184237\|29167373 | Unspecified |
| mmu-miR-139-5p | MIMAT0000656 | Wdr36 | 225348 | HITS-CLIP | 29167373 | Unspecified |
| mmu-miR-139-5p | MIMAT0000656 | Wdr43 | 72515 | HITS-CLIP | 29167373 | Unspecified |
| mmu-miR-139-5p | MIMAT0000656 | Wdr5 | 140858 | HITS-CLIP | 29167373 | Unspecified |
| mmu-miR-139-5p | MIMAT0000656 | Wdr62 | 233064 | HITS-CLIP | 29167373 | Unspecified |
| mmu-miR-139-5p | MIMAT0000656 | Wdr7 | 104082 | HITS-CLIP | 29167373 | Unspecified |
| mmu-miR-139-5p | MIMAT0000656 | Wdr82 | 77305 | sRNA-Seq//HITS-CLIP | 33184237\|29167373 | Unspecified |
| mmu-miR-139-5p | MIMAT0000656 | Wdr91 | 101240 | HITS-CLIP | 29167373 | Unspecified |
| mmu-miR-139-5p | MIMAT0000656 | Wee1 | 22390 | HITS-CLIP | 31433990\|26203562 | Unspecified |
| mmu-miR-139-5p | MIMAT0000656 | Wipf1 | 215280 | HITS-CLIP | 29167373 | Unspecified |
| mmu-miR-139-5p | MIMAT0000656 | Wipf2 | 68524 | HITS-CLIP | 29167373 | Unspecified |
| mmu-miR-139-5p | MIMAT0000656 | Wls | 68151 | HITS-CLIP | 29167373 | Unspecified |
| mmu-miR-139-5p | MIMAT0000656 | Wnk1 | 232341 | sRNA-Seq//HITS-CLIP | 33184237\|29167373 | Unspecified |
| mmu-miR-139-5p | MIMAT0000656 | Wsb1 | 78889 | sRNA-Seq//HITS-CLIP | 33184237\|29167373 | Unspecified |
| mmu-miR-139-5p | MIMAT0000656 | Wtap | 60532 | sRNA-Seq//HITS-CLIP | 33184237\|29167373 | Unspecified |
| mmu-miR-139-5p | MIMAT0000656 | Wwp1 | 107568 | HITS-CLIP | 29167373 | Unspecified |
| mmu-miR-139-5p | MIMAT0000656 | Xpo1 | 103573 | HITS-CLIP//sRNA-Seq | 31433990\|33184237\|29167373 | Unspecified |
| mmu-miR-139-5p | MIMAT0000656 | Xpo7 | 65246 | HITS-CLIP | 25083871\|29167373 | Unspecified |
| mmu-miR-139-5p | MIMAT0000656 | Xpr1 | 19775 | sRNA-Seq//HITS-CLIP | 33184237\|29167373 | Unspecified |
| mmu-miR-139-5p | MIMAT0000656 | Xrn1 | 24127 | HITS-CLIP | 29167373 | Unspecified |
| mmu-miR-139-5p | MIMAT0000656 | Xylt2 | 217119 | HITS-CLIP | 29167373 | Unspecified |
| mmu-miR-139-5p | MIMAT0000656 | Yeats2 | 208146 | HITS-CLIP | 29167373 | Unspecified |
| mmu-miR-139-5p | MIMAT0000656 | Yipf6 | 77929 | sRNA-Seq//HITS-CLIP | 33184237\|29167373 | Unspecified |
| mmu-miR-139-5p | MIMAT0000656 | Ylpm1 | 56531 | sRNA-Seq//HITS-CLIP | 33184237\|29167373 | Unspecified |
| mmu-miR-139-5p | MIMAT0000656 | Yod1 | 226418 | HITS-CLIP | 29167373 | Unspecified |
| mmu-miR-139-5p | MIMAT0000656 | Ypel5 | 383295 | HITS-CLIP | 31433990\|29167373 | Unspecified |
| mmu-miR-139-5p | MIMAT0000656 | Ythdf1 | 228994 | sRNA-Seq//HITS-CLIP | 33184237\|29167373 | Unspecified |
| mmu-miR-139-5p | MIMAT0000656 | Ythdf3 | 229096 | sRNA-Seq//HITS-CLIP | 33184237\|29167373 | Unspecified |

| | | | | | | |
|---|---|---|---|---|---|---|
| mmu-miR-139-5p | MIMAT0000656 | Ywhae | 22627 | HITS-CLIP//sRNA-Seq | 31433990\|33184237\|29167373 | Unspecified |
| mmu-miR-139-5p | MIMAT0000656 | Ywhag | 22628 | HITS-CLIP//sRNA-Seq | 31433990\|33184237\|29167373 | Unspecified |
| mmu-miR-139-5p | MIMAT0000656 | Ywhaq | 22630 | sRNA-Seq//HITS-CLIP | 33184237\|29167373 | Unspecified |
| mmu-miR-139-5p | MIMAT0000656 | Yy1 | 22632 | HITS-CLIP | 29167373 | Unspecified |
| mmu-miR-139-5p | MIMAT0000656 | Zap70 | 22637 | HITS-CLIP | 31433990 | Unspecified |
| mmu-miR-139-5p | MIMAT0000656 | Zbtb1 | 268564 | HITS-CLIP | 29167373 | Unspecified |
| mmu-miR-139-5p | MIMAT0000656 | Zbtb20 | 56490 | HITS-CLIP | 29167373 | Unspecified |
| mmu-miR-139-5p | MIMAT0000656 | Zbtb24 | 268294 | HITS-CLIP | 29167373 | Unspecified |
| mmu-miR-139-5p | MIMAT0000656 | Zbtb26 | 320633 | HITS-CLIP | 29167373 | Unspecified |
| mmu-miR-139-5p | MIMAT0000656 | Zbtb34 | 241311 | HITS-CLIP | 29167373 | Unspecified |
| mmu-miR-139-5p | MIMAT0000656 | Zbtb4 | 75580 | HITS-CLIP | 29167373 | Unspecified |
| mmu-miR-139-5p | MIMAT0000656 | Zbtb43 | 71834 | HITS-CLIP | 29167373 | Unspecified |
| mmu-miR-139-5p | MIMAT0000656 | Zbtb44 | 235132 | HITS-CLIP | 29167373 | Unspecified |
| mmu-miR-139-5p | MIMAT0000656 | Zbtb7b | 22724 | HITS-CLIP | 26203562 | Unspecified |
| mmu-miR-139-5p | MIMAT0000656 | Zc3h12c | 244871 | HITS-CLIP | 29167373 | Unspecified |
| mmu-miR-139-5p | MIMAT0000656 | Zc3h7a | 106205 | sRNA-Seq//HITS-CLIP | 33184237\|29167373 | Unspecified |
| mmu-miR-139-5p | MIMAT0000656 | Zcchc14 | 142682 | HITS-CLIP | 29167373 | Unspecified |
| mmu-miR-139-5p | MIMAT0000656 | Zcchc24 | 71918 | HITS-CLIP | 29167373 | Unspecified |
| mmu-miR-139-5p | MIMAT0000656 | Zcchc7 | 319885 | HITS-CLIP | 29167373 | Unspecified |
| mmu-miR-139-5p | MIMAT0000656 | Zdhhc17 | 320150 | sRNA-Seq//HITS-CLIP | 33184237\|29167373 | Unspecified |
| mmu-miR-139-5p | MIMAT0000656 | Zdhhc20 | 75965 | HITS-CLIP | 29167373 | Unspecified |
| mmu-miR-139-5p | MIMAT0000656 | Zdhhc3 | 69035 | HITS-CLIP | 21258322\|29167373\|23597149 | Unspecified |
| mmu-miR-139-5p | MIMAT0000656 | Zeb1 | 21417 | HITS-CLIP | 23630078 | Unspecified |
| mmu-miR-139-5p | MIMAT0000656 | Zeb2 | 24136 | HITS-CLIP | 29167373 | Unspecified |
| mmu-miR-139-5p | MIMAT0000656 | Zfand5 | 22682 | HITS-CLIP//sRNA-Seq | 29167373\|33184237 | Unspecified |
| mmu-miR-139-5p | MIMAT0000656 | Zfat | 57623 | HITS-CLIP | 29167373 | Unspecified |
| mmu-miR-139-5p | MIMAT0000656 | Zfc3h1 | 216345 | HITS-CLIP | 29167373 | Unspecified |
| mmu-miR-139-5p | MIMAT0000656 | Zfhx2 | 239102 | HITS-CLIP | 29167373 | Unspecified |
| mmu-miR-139-5p | MIMAT0000656 | Zfhx3 | 11906 | HITS-CLIP | 29167373 | Unspecified |
| mmu-miR-139-5p | MIMAT0000656 | Zfp1 | 22640 | HITS-CLIP | 29167373 | Unspecified |
| mmu-miR-139-5p | MIMAT0000656 | Zfp101 | 22643 | HITS-CLIP | 29167373 | Unspecified |
| mmu-miR-139-5p | MIMAT0000656 | Zfp106 | 20402 | sRNA-Seq//HITS-CLIP | 33184237\|29167373 | Unspecified |
| mmu-miR-139-5p | MIMAT0000656 | Zfp131 | 72465 | HITS-CLIP | 29167373 | Unspecified |
| mmu-miR-139-5p | MIMAT0000656 | Zfp146 | 26465 | HITS-CLIP | 29167373 | Unspecified |
| mmu-miR-139-5p | MIMAT0000656 | Zfp160 | 224585 | HITS-CLIP | 29167373 | Unspecified |
| mmu-miR-139-5p | MIMAT0000656 | Zfp202 | 80902 | HITS-CLIP | 29167373 | Unspecified |
| mmu-miR-139-5p | MIMAT0000656 | Zfp217 | 228913 | HITS-CLIP | 29167373 | Unspecified |
| mmu-miR-139-5p | MIMAT0000656 | Zfp248 | 72720 | HITS-CLIP | 29167373 | Unspecified |
| mmu-miR-139-5p | MIMAT0000656 | Zfp26 | 22688 | HITS-CLIP | 29167373 | Unspecified |

| | | | | | | |
|---|---|---|---|---|---|---|
| mmu-miR-139-5p | MIMAT0000656 | Zfp273 | 212569 | HITS-CLIP | 29167373 | Unspecified |
| mmu-miR-139-5p | MIMAT0000656 | Zfp280b | 64453 | HITS-CLIP | 31433990 | Unspecified |
| mmu-miR-139-5p | MIMAT0000656 | Zfp280c | 208968 | HITS-CLIP | 26203562 | Unspecified |
| mmu-miR-139-5p | MIMAT0000656 | Zfp292 | 30046 | HITS-CLIP | 31433990\|29167373 | Unspecified |
| mmu-miR-139-5p | MIMAT0000656 | Zfp3 | 193043 | HITS-CLIP | 29167373 | Unspecified |
| mmu-miR-139-5p | MIMAT0000656 | Zfp322a | 218100 | HITS-CLIP | 25083871\|29167373 | Unspecified |
| mmu-miR-139-5p | MIMAT0000656 | Zfp335 | 329559 | HITS-CLIP | 29167373 | Unspecified |
| mmu-miR-139-5p | MIMAT0000656 | Zfp346 | 26919 | HITS-CLIP | 29167373 | Unspecified |
| mmu-miR-139-5p | MIMAT0000656 | Zfp367 | 238673 | HITS-CLIP | 29167373 | Unspecified |
| mmu-miR-139-5p | MIMAT0000656 | Zfp36l1 | 12192 | HITS-CLIP | 29167373 | Unspecified |
| mmu-miR-139-5p | MIMAT0000656 | Zfp384 | 269800 | HITS-CLIP | 29167373 | Unspecified |
| mmu-miR-139-5p | MIMAT0000656 | Zfp386 | 56220 | HITS-CLIP | 29167373 | Unspecified |
| mmu-miR-139-5p | MIMAT0000656 | Zfp397 | 69256 | HITS-CLIP | 29167373 | Unspecified |
| mmu-miR-139-5p | MIMAT0000656 | Zfp426 | 235028 | HITS-CLIP | 29167373 | Unspecified |
| mmu-miR-139-5p | MIMAT0000656 | Zfp445 | 235682 | HITS-CLIP//sRNA-Seq | 29167373\|33184237 | Unspecified |
| mmu-miR-139-5p | MIMAT0000656 | Zfp446 | 269870 | HITS-CLIP | 29167373 | Unspecified |
| mmu-miR-139-5p | MIMAT0000656 | Zfp451 | 98403 | HITS-CLIP | 29167373 | Unspecified |
| mmu-miR-139-5p | MIMAT0000656 | Zfp512 | 269639 | sRNA-Seq//HITS-CLIP | 33184237\|31433990 | Unspecified |
| mmu-miR-139-5p | MIMAT0000656 | Zfp516 | 329003 | HITS-CLIP | 29167373 | Unspecified |
| mmu-miR-139-5p | MIMAT0000656 | Zfp52 | 22710 | HITS-CLIP | 29167373 | Unspecified |
| mmu-miR-139-5p | MIMAT0000656 | Zfp607a | 545938 | HITS-CLIP | 29167373 | Unspecified |
| mmu-miR-139-5p | MIMAT0000656 | Zfp609 | 214812 | HITS-CLIP | 29167373 | Unspecified |
| mmu-miR-139-5p | MIMAT0000656 | Zfp638 | 27332 | sRNA-Seq//HITS-CLIP | 33184237\|29167373 | Unspecified |
| mmu-miR-139-5p | MIMAT0000656 | Zfp644 | 52397 | sRNA-Seq//HITS-CLIP | 33184237\|29167373 | Unspecified |
| mmu-miR-139-5p | MIMAT0000656 | Zfp652 | 268469 | HITS-CLIP | 29167373 | Unspecified |
| mmu-miR-139-5p | MIMAT0000656 | Zfp655 | 72611 | HITS-CLIP | 29167373 | Unspecified |
| mmu-miR-139-5p | MIMAT0000656 | Zfp68 | 24135 | HITS-CLIP | 29167373 | Unspecified |
| mmu-miR-139-5p | MIMAT0000656 | Zfp69 | 381549 | HITS-CLIP | 31433990 | Unspecified |
| mmu-miR-139-5p | MIMAT0000656 | Zfp703 | 353310 | HITS-CLIP | 29167373 | Unspecified |
| mmu-miR-139-5p | MIMAT0000656 | Zfp706 | 68036 | sRNA-Seq//HITS-CLIP | 33184237\|29167373 | Unspecified |
| mmu-miR-139-5p | MIMAT0000656 | Zfp715 | 69930 | HITS-CLIP | 29167373 | Unspecified |
| mmu-miR-139-5p | MIMAT0000656 | Zfp748 | 212276 | HITS-CLIP | 26203562 | Unspecified |
| mmu-miR-139-5p | MIMAT0000656 | Zfp770 | 228491 | HITS-CLIP | 29167373 | Unspecified |
| mmu-miR-139-5p | MIMAT0000656 | Zfp800 | 627049 | HITS-CLIP | 31433990 | Unspecified |
| mmu-miR-139-5p | MIMAT0000656 | Zfp866 | 330788 | HITS-CLIP | 29167373 | Unspecified |
| mmu-miR-139-5p | MIMAT0000656 | Zfp868 | 234362 | HITS-CLIP | 29167373 | Unspecified |
| mmu-miR-139-5p | MIMAT0000656 | Zfp871 | 208292 | sRNA-Seq//HITS-CLIP | 33184237\|29167373 | Unspecified |
| mmu-miR-139-5p | MIMAT0000656 | Zfp91 | 109910 | HITS-CLIP | 29167373 | Unspecified |
| mmu-miR-139-5p | MIMAT0000656 | Zfp933 | 242747 | HITS-CLIP | 26203562 | Unspecified |

| | | | | | | |
|---|---|---|---|---|---|---|
| mmu-miR-139-5p | MIMAT0000656 | Zfp942 | 73233 | HITS-CLIP | 29167373 | Unspecified |
| mmu-miR-139-5p | MIMAT0000656 | Zfp948 | 381066 | HITS-CLIP | 29167373 | Unspecified |
| mmu-miR-139-5p | MIMAT0000656 | Zfp955b | 100043468 | HITS-CLIP | 29167373 | Unspecified |
| mmu-miR-139-5p | MIMAT0000656 | Zfp958 | 233987 | HITS-CLIP | 26203562 | Unspecified |
| mmu-miR-139-5p | MIMAT0000656 | Zfx | 22764 | HITS-CLIP | 26203562 | Unspecified |
| mmu-miR-139-5p | MIMAT0000656 | Zfyve16 | 218441 | HITS-CLIP | 29167373 | Unspecified |
| mmu-miR-139-5p | MIMAT0000656 | Zfyve27 | 319740 | HITS-CLIP | 29167373 | Unspecified |
| mmu-miR-139-5p | MIMAT0000656 | Zhx1 | 22770 | HITS-CLIP | 29167373 | Unspecified |
| mmu-miR-139-5p | MIMAT0000656 | Zmiz1 | 328365 | sRNA-Seq//HITS-CLIP | 33184237\|29167373 | Unspecified |
| mmu-miR-139-5p | MIMAT0000656 | Zmym2 | 76007 | sRNA-Seq//HITS-CLIP | 33184237\|29167373 | Unspecified |
| mmu-miR-139-5p | MIMAT0000656 | Znfx1 | 98999 | HITS-CLIP | 29167373 | Unspecified |
| mmu-miR-139-5p | MIMAT0000656 | Znrf1 | 170737 | HITS-CLIP | 29167373 | Unspecified |
| mmu-miR-139-5p | MIMAT0000656 | Zranb2 | 53861 | sRNA-Seq//HITS-CLIP | 33184237\|29167373 | Unspecified |
| mmu-miR-139-5p | MIMAT0000656 | Zscan26 | 432731 | HITS-CLIP | 29167373 | Unspecified |
| mmu-miR-139-5p | MIMAT0000656 | Zscan29 | 99334 | HITS-CLIP | 29167373 | Unspecified |
| mmu-miR-139-5p | MIMAT0000656 | Zswim3 | 67538 | HITS-CLIP | 31433990 | Unspecified |
| mmu-miR-139-5p | MIMAT0000656 | Zswim4 | 212168 | HITS-CLIP | 29167373 | Unspecified |
| mmu-miR-139-5p | MIMAT0000656 | mt-Nd5 | 4540 | HITS-CLIP//sRNA-Seq | 29167373\|33184237 | Unspecified |
| mmu-miR-3109-3p | MIMAT0014950 | Actr2 | 66713 | HITS-CLIP | 31433990 | Unspecified |
| mmu-miR-3109-3p | MIMAT0014950 | Camsap2 | 67886 | HITS-CLIP | 26203562 | Unspecified |
| mmu-miR-3109-3p | MIMAT0014950 | Coro7 | 78885 | HITS-CLIP | 31433990 | Unspecified |
| mmu-miR-3109-3p | MIMAT0014950 | Diaph1 | 1729 | HITS-CLIP | 31433990 | Unspecified |
| mmu-miR-691 | MIMAT0003470 | Ccn2 | 1490 | HITS-CLIP | 26203562 | Unspecified |
| mmu-miR-691 | MIMAT0003470 | Zfyve26 | 211978 | HITS-CLIP | 31433990 | Unspecified |
| mmu-miR-3095-3p | MIMAT0014912 | Ago2 | 239528 | HITS-CLIP | 37986224 | Unspecified |
| mmu-miR-3095-3p | MIMAT0014912 | Ankrd10 | 102334 | HITS-CLIP | 37986224 | Unspecified |
| mmu-miR-3095-3p | MIMAT0014912 | Bhlhe41 | 79362 | HITS-CLIP | 26203562 | Unspecified |
| mmu-miR-3095-3p | MIMAT0014912 | Cdc42ep4 | 56699 | HITS-CLIP | 37986224 | Unspecified |
| mmu-miR-3095-3p | MIMAT0014912 | Coro2a | 107684 | HITS-CLIP | 31433990 | Unspecified |
| mmu-miR-3095-3p | MIMAT0014912 | Dlgap4 | 228836 | HITS-CLIP | 37986224 | Unspecified |
| mmu-miR-3095-3p | MIMAT0014912 | Gnas | 14683 | HITS-CLIP | 37986224 | Unspecified |
| mmu-miR-3095-3p | MIMAT0014912 | Itpk1 | 217837 | HITS-CLIP | 37986224 | Unspecified |
| mmu-miR-3095-3p | MIMAT0014912 | Nop56 | 67134 | HITS-CLIP | 37986224 | Unspecified |
| mmu-miR-3095-3p | MIMAT0014912 | Rbms2 | 56516 | HITS-CLIP | 37986224 | Unspecified |
| mmu-miR-3095-3p | MIMAT0014912 | Rbpj | 19664 | HITS-CLIP | 37986224 | Unspecified |
| mmu-miR-3095-3p | MIMAT0014912 | Rictor | 78757 | HITS-CLIP | 37986224 | Unspecified |
| mmu-miR-3095-3p | MIMAT0014912 | Rrm1 | 20133 | HITS-CLIP | 37986224 | Unspecified |
| mmu-miR-3095-3p | MIMAT0014912 | Syn3 | 27204 | HITS-CLIP | 37986224 | Unspecified |
| mmu-miR-3095-3p | MIMAT0014912 | Tet1 | 52463 | HITS-CLIP | 37986224 | Unspecified |

| | | | | | | |
|---|---|---|---|---|---|---|
| mmu-miR-3095-3p | MIMAT0014912 | Vps54 | 245944 | HITS-CLIP | 31433990 | Unspecified |
| mmu-miR-669n | MIMAT0009427 | 1810013L24Rik | 69053 | HITS-CLIP | 23142080 | Unspecified |
| mmu-miR-669n | MIMAT0009427 | 4921524J17Rik | 66714 | HITS-CLIP | 23142080 | Unspecified |
| mmu-miR-669n | MIMAT0009427 | 4930503L19Rik | 269033 | HITS-CLIP | 23142080 | Unspecified |
| mmu-miR-669n | MIMAT0009427 | 4931406C07Rik | 70984 | HITS-CLIP | 37986224 | Unspecified |
| mmu-miR-669n | MIMAT0009427 | 4931406P16Rik | 233103 | HITS-CLIP | 37986224 | Unspecified |
| mmu-miR-669n | MIMAT0009427 | 4932438A13Rik | 229227 | HITS-CLIP | 37986224 | Unspecified |
| mmu-miR-669n | MIMAT0009427 | 9930021J03Rik | 240613 | HITS-CLIP | 37986224\|23142080 | Unspecified |
| mmu-miR-669n | MIMAT0009427 | AW549877 | 106064 | HITS-CLIP | 37986224 | Unspecified |
| mmu-miR-669n | MIMAT0009427 | Abcb9 | 56325 | HITS-CLIP | 23142080 | Unspecified |
| mmu-miR-669n | MIMAT0009427 | Abce1 | 24015 | HITS-CLIP | 23142080 | Unspecified |
| mmu-miR-669n | MIMAT0009427 | Abhd10 | 213012 | HITS-CLIP | 37986224\|23142080 | Unspecified |
| mmu-miR-669n | MIMAT0009427 | Abhd17c | 70178 | HITS-CLIP | 23142080 | Unspecified |
| mmu-miR-669n | MIMAT0009427 | Abi1 | 11308 | HITS-CLIP | 23142080 | Unspecified |
| mmu-miR-669n | MIMAT0009427 | Acap2 | 78618 | HITS-CLIP | 23142080 | Unspecified |
| mmu-miR-669n | MIMAT0009427 | Acbd5 | 74159 | HITS-CLIP | 23142080 | Unspecified |
| mmu-miR-669n | MIMAT0009427 | Acp5 | 11433 | HITS-CLIP | 23142080 | Unspecified |
| mmu-miR-669n | MIMAT0009427 | Actb | 11461 | HITS-CLIP | 37986224 | Unspecified |
| mmu-miR-669n | MIMAT0009427 | Actr2 | 66713 | HITS-CLIP | 23142080 | Unspecified |
| mmu-miR-669n | MIMAT0009427 | Adar | 56417 | HITS-CLIP | 37986224\|23142080 | Unspecified |
| mmu-miR-669n | MIMAT0009427 | Add3 | 27360 | HITS-CLIP | 23142080 | Unspecified |
| mmu-miR-669n | MIMAT0009427 | Adgrl2 | 23266 | HITS-CLIP | 23142080 | Unspecified |
| mmu-miR-669n | MIMAT0009427 | Adipor2 | 68465 | HITS-CLIP | 23142080 | Unspecified |
| mmu-miR-669n | MIMAT0009427 | Adnp | 11538 | HITS-CLIP | 37986224\|23142080 | Unspecified |
| mmu-miR-669n | MIMAT0009427 | Adprhl2 | 100206 | HITS-CLIP | 23142080 | Unspecified |
| mmu-miR-669n | MIMAT0009427 | Adrb2 | 11555 | HITS-CLIP | 23142080 | Unspecified |
| mmu-miR-669n | MIMAT0009427 | Adss | 11566 | HITS-CLIP | 23142080 | Unspecified |
| mmu-miR-669n | MIMAT0009427 | Aebp2 | 11569 | HITS-CLIP | 37986224\|23142080 | Unspecified |
| mmu-miR-669n | MIMAT0009427 | Aff3 | 16764 | HITS-CLIP | 23142080 | Unspecified |
| mmu-miR-669n | MIMAT0009427 | Aff4 | 93736 | HITS-CLIP | 37986224 | Unspecified |
| mmu-miR-669n | MIMAT0009427 | Agfg1 | 15463 | HITS-CLIP | 37986224 | Unspecified |
| mmu-miR-669n | MIMAT0009427 | Ago1 | 236511 | HITS-CLIP | 37986224 | Unspecified |
| mmu-miR-669n | MIMAT0009427 | Ago2 | 239528 | HITS-CLIP | 37986224\|23142080 | Unspecified |
| mmu-miR-669n | MIMAT0009427 | Ahctf1 | 226747 | HITS-CLIP | 37986224 | Unspecified |
| mmu-miR-669n | MIMAT0009427 | Akap10 | 56697 | HITS-CLIP | 23142080 | Unspecified |
| mmu-miR-669n | MIMAT0009427 | Akap13 | 75547 | HITS-CLIP | 37986224\|23142080 | Unspecified |
| mmu-miR-669n | MIMAT0009427 | Akirin1 | 68050 | HITS-CLIP | 23142080 | Unspecified |
| mmu-miR-669n | MIMAT0009427 | Alkbh5 | 268420 | HITS-CLIP | 37986224\|23142080 | Unspecified |
| mmu-miR-669n | MIMAT0009427 | Ammecr1l | 225339 | HITS-CLIP | 23142080 | Unspecified |

| | | | | | | | |
|---|---|---|---|---|---|---|---|
| mmu-miR-669n | MIMAT0009427 | Anapc4 | 52206 | HITS-CLIP | | 23142080 | Unspecified |
| mmu-miR-669n | MIMAT0009427 | Angptl2 | 26360 | HITS-CLIP | | 23142080 | Unspecified |
| mmu-miR-669n | MIMAT0009427 | Ankrd11 | 77087 | HITS-CLIP | | 23142080 | Unspecified |
| mmu-miR-669n | MIMAT0009427 | Ankrd12 | 106585 | HITS-CLIP | | 23142080 | Unspecified |
| mmu-miR-669n | MIMAT0009427 | Ankrd17 | 81702 | HITS-CLIP | 37986224\|23142080 | | Unspecified |
| mmu-miR-669n | MIMAT0009427 | Ankrd44 | 329154 | HITS-CLIP | | 37986224 | Unspecified |
| mmu-miR-669n | MIMAT0009427 | Ankrd52 | 237615 | HITS-CLIP | | 37986224 | Unspecified |
| mmu-miR-669n | MIMAT0009427 | Antxr2 | 71914 | HITS-CLIP | 37986224\|23142080 | | Unspecified |
| mmu-miR-669n | MIMAT0009427 | Ap1g1 | 11765 | HITS-CLIP | | 37986224 | Unspecified |
| mmu-miR-669n | MIMAT0009427 | Ap2a2 | 11772 | HITS-CLIP | | 23142080 | Unspecified |
| mmu-miR-669n | MIMAT0009427 | Ap5m1 | 74385 | HITS-CLIP | | 23142080 | Unspecified |
| mmu-miR-669n | MIMAT0009427 | Apaf1 | 11783 | HITS-CLIP | | 23142080 | Unspecified |
| mmu-miR-669n | MIMAT0009427 | Aplp2 | 11804 | HITS-CLIP | | 23142080 | Unspecified |
| mmu-miR-669n | MIMAT0009427 | Appbp2 | 66884 | HITS-CLIP | | 23142080 | Unspecified |
| mmu-miR-669n | MIMAT0009427 | Arap2 | 212285 | HITS-CLIP | | 23142080 | Unspecified |
| mmu-miR-669n | MIMAT0009427 | Arf6 | 11845 | HITS-CLIP | | 23142080 | Unspecified |
| mmu-miR-669n | MIMAT0009427 | Arhgap17 | 70497 | HITS-CLIP | | 23142080 | Unspecified |
| mmu-miR-669n | MIMAT0009427 | Arhgef1 | 16801 | HITS-CLIP | | 23142080 | Unspecified |
| mmu-miR-669n | MIMAT0009427 | Arhgef10 | 234094 | HITS-CLIP | | 37986224 | Unspecified |
| mmu-miR-669n | MIMAT0009427 | Arhgef18 | 102098 | HITS-CLIP | | 23142080 | Unspecified |
| mmu-miR-669n | MIMAT0009427 | Arid4b | 94246 | HITS-CLIP | 37986224\|23142080 | | Unspecified |
| mmu-miR-669n | MIMAT0009427 | Arih1 | 23806 | HITS-CLIP | | 23142080 | Unspecified |
| mmu-miR-669n | MIMAT0009427 | Arl1 | 104303 | HITS-CLIP | | 23142080 | Unspecified |
| mmu-miR-669n | MIMAT0009427 | Arl5b | 75869 | HITS-CLIP | | 37986224 | Unspecified |
| mmu-miR-669n | MIMAT0009427 | Arl8b | 67166 | HITS-CLIP | | 23142080 | Unspecified |
| mmu-miR-669n | MIMAT0009427 | Arpc2 | 76709 | HITS-CLIP | | 23142080 | Unspecified |
| mmu-miR-669n | MIMAT0009427 | Arpp19 | 59046 | HITS-CLIP | | 23142080 | Unspecified |
| mmu-miR-669n | MIMAT0009427 | Asb3 | 65257 | HITS-CLIP | | 37986224 | Unspecified |
| mmu-miR-669n | MIMAT0009427 | Ascc3 | 77987 | HITS-CLIP | | 37986224 | Unspecified |
| mmu-miR-669n | MIMAT0009427 | Ash1l | 192195 | HITS-CLIP | | 37986224 | Unspecified |
| mmu-miR-669n | MIMAT0009427 | Atad2b | 320817 | HITS-CLIP | | 23142080 | Unspecified |
| mmu-miR-669n | MIMAT0009427 | Atad5 | 237877 | HITS-CLIP | | 37986224 | Unspecified |
| mmu-miR-669n | MIMAT0009427 | Atf2 | 11909 | HITS-CLIP | | 37986224 | Unspecified |
| mmu-miR-669n | MIMAT0009427 | Atf7 | 223922 | HITS-CLIP | | 37986224 | Unspecified |
| mmu-miR-669n | MIMAT0009427 | Atg14 | 100504663 | HITS-CLIP | | 23142080 | Unspecified |
| mmu-miR-669n | MIMAT0009427 | Atg3 | 67841 | HITS-CLIP | | 37986224 | Unspecified |
| mmu-miR-669n | MIMAT0009427 | Atp1b3 | 11933 | HITS-CLIP | | 37986224 | Unspecified |
| mmu-miR-669n | MIMAT0009427 | Atp2a2 | 11938 | HITS-CLIP | | 23142080 | Unspecified |
| mmu-miR-669n | MIMAT0009427 | Atp6v1c1 | 66335 | HITS-CLIP | | 23142080 | Unspecified |

| | | | | | | | |
|---|---|---|---|---|---|---|---|
| mmu-miR-669n | MIMAT0009427 | Atp6v1h | 108664 | HITS-CLIP | | 23142080 | Unspecified |
| mmu-miR-669n | MIMAT0009427 | Atr | 245000 | HITS-CLIP | | 23142080 | Unspecified |
| mmu-miR-669n | MIMAT0009427 | Atrn | 11990 | HITS-CLIP | | 23142080 | Unspecified |
| mmu-miR-669n | MIMAT0009427 | Atxn2l | 233871 | HITS-CLIP | | 23142080 | Unspecified |
| mmu-miR-669n | MIMAT0009427 | Atxn7 | 246103 | HITS-CLIP | | 23142080 | Unspecified |
| mmu-miR-669n | MIMAT0009427 | Atxn7l1 | 380753 | HITS-CLIP | | 23142080 | Unspecified |
| mmu-miR-669n | MIMAT0009427 | Aurkb | 20877 | HITS-CLIP | | 23142080 | Unspecified |
| mmu-miR-669n | MIMAT0009427 | B3gnt2 | 53625 | HITS-CLIP | | 23142080 | Unspecified |
| mmu-miR-669n | MIMAT0009427 | B4galt5 | 56336 | HITS-CLIP | 37986224\|23142080 | | Unspecified |
| mmu-miR-669n | MIMAT0009427 | BC005537 | 79555 | HITS-CLIP | | 37986224 | Unspecified |
| mmu-miR-669n | MIMAT0009427 | Bach2 | 12014 | HITS-CLIP | 37986224\|23142080 | | Unspecified |
| mmu-miR-669n | MIMAT0009427 | Baz1a | 217578 | HITS-CLIP | 37986224\|23142080 | | Unspecified |
| mmu-miR-669n | MIMAT0009427 | Bbx | 70508 | HITS-CLIP | 37986224\|23142080 | | Unspecified |
| mmu-miR-669n | MIMAT0009427 | Bcl11b | 58208 | HITS-CLIP | | 23142080 | Unspecified |
| mmu-miR-669n | MIMAT0009427 | Bcl3 | 12051 | HITS-CLIP | | 23142080 | Unspecified |
| mmu-miR-669n | MIMAT0009427 | Bclaf1 | 72567 | HITS-CLIP | | 23142080 | Unspecified |
| mmu-miR-669n | MIMAT0009427 | Bcor | 71458 | HITS-CLIP | | 23142080 | Unspecified |
| mmu-miR-669n | MIMAT0009427 | Bcs1l | 66821 | HITS-CLIP | | 37986224 | Unspecified |
| mmu-miR-669n | MIMAT0009427 | Birc3 | 11796 | HITS-CLIP | | 23142080 | Unspecified |
| mmu-miR-669n | MIMAT0009427 | Birc6 | 12211 | HITS-CLIP | 37986224\|23142080 | | Unspecified |
| mmu-miR-669n | MIMAT0009427 | Blmh | 104184 | HITS-CLIP | | 23142080 | Unspecified |
| mmu-miR-669n | MIMAT0009427 | Blzf1 | 66352 | HITS-CLIP | | 23142080 | Unspecified |
| mmu-miR-669n | MIMAT0009427 | Bmt2 | 101148 | HITS-CLIP | | 23142080 | Unspecified |
| mmu-miR-669n | MIMAT0009427 | Bptf | 207165 | HITS-CLIP | | 23142080 | Unspecified |
| mmu-miR-669n | MIMAT0009427 | Brd1 | 223770 | HITS-CLIP | | 23142080 | Unspecified |
| mmu-miR-669n | MIMAT0009427 | Brd8 | 78656 | HITS-CLIP | | 31433990 | Unspecified |
| mmu-miR-669n | MIMAT0009427 | Brms1l | 52592 | HITS-CLIP | | 23142080 | Unspecified |
| mmu-miR-669n | MIMAT0009427 | Brwd3 | 382236 | HITS-CLIP | | 23142080 | Unspecified |
| mmu-miR-669n | MIMAT0009427 | Btbd8 | 100503185 | HITS-CLIP | | 23142080 | Unspecified |
| mmu-miR-669n | MIMAT0009427 | Btg1 | 12226 | HITS-CLIP | | 23142080 | Unspecified |
| mmu-miR-669n | MIMAT0009427 | C1qbp | 12261 | HITS-CLIP | | 23142080 | Unspecified |
| mmu-miR-669n | MIMAT0009427 | Calhm6 | 441168 | HITS-CLIP | | 23142080 | Unspecified |
| mmu-miR-669n | MIMAT0009427 | Calm1 | 12313 | HITS-CLIP | | 23142080 | Unspecified |
| mmu-miR-669n | MIMAT0009427 | Camsap2 | 67886 | HITS-CLIP | | 23142080 | Unspecified |
| mmu-miR-669n | MIMAT0009427 | Cap1 | 12331 | HITS-CLIP | | 23142080 | Unspecified |
| mmu-miR-669n | MIMAT0009427 | Casc3 | 192160 | HITS-CLIP | | 23142080 | Unspecified |
| mmu-miR-669n | MIMAT0009427 | Casp3 | 12367 | HITS-CLIP | | 23142080 | Unspecified |
| mmu-miR-669n | MIMAT0009427 | Cbl | 12402 | HITS-CLIP | | 23142080 | Unspecified |
| mmu-miR-669n | MIMAT0009427 | Cblb | 208650 | HITS-CLIP | | 37986224 | Unspecified |

| | | | | | | | |
|---|---|---|---|---|---|---|---|
| mmu-miR-669n | MIMAT0009427 | Cbx6 | 494448 | HITS-CLIP | | 23142080 | Unspecified |
| mmu-miR-669n | MIMAT0009427 | Ccdc115 | 69668 | HITS-CLIP | | 23142080 | Unspecified |
| mmu-miR-669n | MIMAT0009427 | Ccdc167 | 68597 | HITS-CLIP | | 23142080 | Unspecified |
| mmu-miR-669n | MIMAT0009427 | Ccdc28a | 215814 | HITS-CLIP | | 37986224 | Unspecified |
| mmu-miR-669n | MIMAT0009427 | Ccna2 | 12428 | HITS-CLIP | 37986224\|23142080 | | Unspecified |
| mmu-miR-669n | MIMAT0009427 | Ccnb2 | 12442 | HITS-CLIP | | 23142080 | Unspecified |
| mmu-miR-669n | MIMAT0009427 | Ccnd2 | 12444 | HITS-CLIP | | 37986224 | Unspecified |
| mmu-miR-669n | MIMAT0009427 | Ccne2 | 12448 | HITS-CLIP | 37986224\|23142080 | | Unspecified |
| mmu-miR-669n | MIMAT0009427 | Ccnk | 12454 | HITS-CLIP | | 37986224 | Unspecified |
| mmu-miR-669n | MIMAT0009427 | Ccnt1 | 12455 | HITS-CLIP | 37986224\|23142080 | | Unspecified |
| mmu-miR-669n | MIMAT0009427 | Ccnt2 | 72949 | HITS-CLIP | | 23142080 | Unspecified |
| mmu-miR-669n | MIMAT0009427 | Ccr4 | 12773 | HITS-CLIP | | 23142080 | Unspecified |
| mmu-miR-669n | MIMAT0009427 | Ccr7 | 12775 | HITS-CLIP | | 37986224 | Unspecified |
| mmu-miR-669n | MIMAT0009427 | Cct3 | 12462 | HITS-CLIP | | 23142080 | Unspecified |
| mmu-miR-669n | MIMAT0009427 | Cct5 | 12465 | HITS-CLIP | 31433990\|23142080 | | Unspecified |
| mmu-miR-669n | MIMAT0009427 | Cd2 | 12481 | HITS-CLIP | | 37986224 | Unspecified |
| mmu-miR-669n | MIMAT0009427 | Cd24a | 12484 | HITS-CLIP | | 23142080 | Unspecified |
| mmu-miR-669n | MIMAT0009427 | Cd274 | 60533 | HITS-CLIP | | 37986224 | Unspecified |
| mmu-miR-669n | MIMAT0009427 | Cd28 | 12487 | HITS-CLIP | 37986224\|23142080 | | Unspecified |
| mmu-miR-669n | MIMAT0009427 | Cd69 | 12515 | HITS-CLIP | | 37986224 | Unspecified |
| mmu-miR-669n | MIMAT0009427 | Cdc25b | 12531 | HITS-CLIP | | 23142080 | Unspecified |
| mmu-miR-669n | MIMAT0009427 | Cdc27 | 217232 | HITS-CLIP | | 23142080 | Unspecified |
| mmu-miR-669n | MIMAT0009427 | Cdc42 | 12540 | HITS-CLIP | | 23142080 | Unspecified |
| mmu-miR-669n | MIMAT0009427 | Cdc42se1 | 57912 | HITS-CLIP | | 37986224 | Unspecified |
| mmu-miR-669n | MIMAT0009427 | Cdc7 | 12545 | HITS-CLIP | | 23142080 | Unspecified |
| mmu-miR-669n | MIMAT0009427 | Cdk1 | 12534 | HITS-CLIP | | 23142080 | Unspecified |
| mmu-miR-669n | MIMAT0009427 | Cdk17 | 237459 | HITS-CLIP | 37986224\|23142080 | | Unspecified |
| mmu-miR-669n | MIMAT0009427 | Cdk19 | 78334 | HITS-CLIP | | 23142080 | Unspecified |
| mmu-miR-669n | MIMAT0009427 | Cdk5r1 | 12569 | HITS-CLIP | | 23142080 | Unspecified |
| mmu-miR-669n | MIMAT0009427 | Cdk8 | 264064 | HITS-CLIP | | 23142080 | Unspecified |
| mmu-miR-669n | MIMAT0009427 | Cdkn1b | 12576 | HITS-CLIP | 37986224\|23142080 | | Unspecified |
| mmu-miR-669n | MIMAT0009427 | Celf1 | 13046 | HITS-CLIP | | 37986224 | Unspecified |
| mmu-miR-669n | MIMAT0009427 | Celf2 | 14007 | HITS-CLIP | | 23142080 | Unspecified |
| mmu-miR-669n | MIMAT0009427 | Cenpe | 229841 | HITS-CLIP | | 23142080 | Unspecified |
| mmu-miR-669n | MIMAT0009427 | Cep120 | 225523 | HITS-CLIP | | 23142080 | Unspecified |
| mmu-miR-669n | MIMAT0009427 | Cep170 | 545389 | HITS-CLIP | | 23142080 | Unspecified |
| mmu-miR-669n | MIMAT0009427 | Cep57 | 74360 | HITS-CLIP | | 37986224 | Unspecified |
| mmu-miR-669n | MIMAT0009427 | Chd2 | 244059 | HITS-CLIP | | 37986224 | Unspecified |
| mmu-miR-669n | MIMAT0009427 | Chd4 | 107932 | HITS-CLIP | | 23142080 | Unspecified |

| | | | | | | | |
|---|---|---|---|---|---|---|---|
| mmu-miR-669n | MIMAT0009427 | Chtop | 66511 | HITS-CLIP | | 37986224 | Unspecified |
| mmu-miR-669n | MIMAT0009427 | Cited2 | 17684 | HITS-CLIP | | 37986224 | Unspecified |
| mmu-miR-669n | MIMAT0009427 | Ckap2l | 70466 | HITS-CLIP | 37986224\|23142080 | | Unspecified |
| mmu-miR-669n | MIMAT0009427 | Ckap5 | 75786 | HITS-CLIP | | 23142080 | Unspecified |
| mmu-miR-669n | MIMAT0009427 | Clcc1 | 229725 | HITS-CLIP | 37986224\|23142080 | | Unspecified |
| mmu-miR-669n | MIMAT0009427 | Clcf1 | 56708 | HITS-CLIP | | 37986224 | Unspecified |
| mmu-miR-669n | MIMAT0009427 | Clcn5 | 12728 | HITS-CLIP | | 23142080 | Unspecified |
| mmu-miR-669n | MIMAT0009427 | Clint1 | 216705 | HITS-CLIP | 37986224\|23142080 | | Unspecified |
| mmu-miR-669n | MIMAT0009427 | Clk4 | 12750 | HITS-CLIP | | 23142080 | Unspecified |
| mmu-miR-669n | MIMAT0009427 | Clock | 12753 | HITS-CLIP | | 23142080 | Unspecified |
| mmu-miR-669n | MIMAT0009427 | Clpb | 20480 | HITS-CLIP | | 23142080 | Unspecified |
| mmu-miR-669n | MIMAT0009427 | Cltc | 67300 | HITS-CLIP | | 23142080 | Unspecified |
| mmu-miR-669n | MIMAT0009427 | Cmpk1 | 66588 | HITS-CLIP | | 37986224 | Unspecified |
| mmu-miR-669n | MIMAT0009427 | Cnot6 | 104625 | HITS-CLIP | | 37986224 | Unspecified |
| mmu-miR-669n | MIMAT0009427 | Cog3 | 338337 | HITS-CLIP | | 23142080 | Unspecified |
| mmu-miR-669n | MIMAT0009427 | Cog8 | 97484 | HITS-CLIP | | 23142080 | Unspecified |
| mmu-miR-669n | MIMAT0009427 | Col4a3bp | 68018 | HITS-CLIP | | 23142080 | Unspecified |
| mmu-miR-669n | MIMAT0009427 | Cops2 | 12848 | HITS-CLIP | | 23142080 | Unspecified |
| mmu-miR-669n | MIMAT0009427 | Cops7a | 26894 | HITS-CLIP | | 37986224 | Unspecified |
| mmu-miR-669n | MIMAT0009427 | Cpeb2 | 231207 | HITS-CLIP | | 23142080 | Unspecified |
| mmu-miR-669n | MIMAT0009427 | Cpsf6 | 432508 | HITS-CLIP | | 23142080 | Unspecified |
| mmu-miR-669n | MIMAT0009427 | Creb1 | 12912 | HITS-CLIP | 37986224\|23142080 | | Unspecified |
| mmu-miR-669n | MIMAT0009427 | Crlf2 | 57914 | HITS-CLIP | | 37986224 | Unspecified |
| mmu-miR-669n | MIMAT0009427 | Cs | 12974 | HITS-CLIP | | 23142080 | Unspecified |
| mmu-miR-669n | MIMAT0009427 | Csde1 | 229663 | HITS-CLIP | | 23142080 | Unspecified |
| mmu-miR-669n | MIMAT0009427 | Cse1l | 110750 | HITS-CLIP | 37986224\|23142080 | | Unspecified |
| mmu-miR-669n | MIMAT0009427 | Csnk1a1 | 93687 | HITS-CLIP | | 23142080 | Unspecified |
| mmu-miR-669n | MIMAT0009427 | Csnk1g3 | 70425 | HITS-CLIP | | 23142080 | Unspecified |
| mmu-miR-669n | MIMAT0009427 | Csnk2b | 13001 | HITS-CLIP | | 23142080 | Unspecified |
| mmu-miR-669n | MIMAT0009427 | Cstf2 | 108062 | HITS-CLIP | | 23142080 | Unspecified |
| mmu-miR-669n | MIMAT0009427 | Ctdp1 | 67655 | HITS-CLIP | | 23142080 | Unspecified |
| mmu-miR-669n | MIMAT0009427 | Cth | 107869 | HITS-CLIP | | 23142080 | Unspecified |
| mmu-miR-669n | MIMAT0009427 | Ctla4 | 12477 | HITS-CLIP | | 37986224 | Unspecified |
| mmu-miR-669n | MIMAT0009427 | Ctnnb1 | 12387 | HITS-CLIP | | 23142080 | Unspecified |
| mmu-miR-669n | MIMAT0009427 | Cul1 | 26965 | HITS-CLIP | | 23142080 | Unspecified |
| mmu-miR-669n | MIMAT0009427 | Cul3 | 26554 | HITS-CLIP | | 23142080 | Unspecified |
| mmu-miR-669n | MIMAT0009427 | Cul4a | 99375 | HITS-CLIP | | 23142080 | Unspecified |
| mmu-miR-669n | MIMAT0009427 | Cul7 | 66515 | HITS-CLIP | | 23142080 | Unspecified |
| mmu-miR-669n | MIMAT0009427 | Cyld | 74256 | HITS-CLIP | 37986224\|23142080 | | Unspecified |

| | | | | | | |
|---|---|---|---|---|---|---|
| mmu-miR-669n | MIMAT0009427 | D230025D16Rik | 234678 | HITS-CLIP | | 23142080 Unspecified |
| mmu-miR-669n | MIMAT0009427 | Dcaf7 | 71833 | HITS-CLIP | | 23142080 Unspecified |
| mmu-miR-669n | MIMAT0009427 | Dcbld1 | 66686 | HITS-CLIP | | 23142080 Unspecified |
| mmu-miR-669n | MIMAT0009427 | Dctn4 | 67665 | HITS-CLIP | | 23142080 Unspecified |
| mmu-miR-669n | MIMAT0009427 | Dctn6 | 22428 | HITS-CLIP | | 23142080 Unspecified |
| mmu-miR-669n | MIMAT0009427 | Ddr1 | 12305 | HITS-CLIP | | 23142080 Unspecified |
| mmu-miR-669n | MIMAT0009427 | Ddx3y | 26900 | HITS-CLIP | | 23142080 Unspecified |
| mmu-miR-669n | MIMAT0009427 | Ddx5 | 13207 | HITS-CLIP | | 37986224 Unspecified |
| mmu-miR-669n | MIMAT0009427 | Dennd4a | 102442 | HITS-CLIP | | 23142080 Unspecified |
| mmu-miR-669n | MIMAT0009427 | Dennd4b | 229541 | HITS-CLIP | | 23142080 Unspecified |
| mmu-miR-669n | MIMAT0009427 | Dennd4c | 329877 | HITS-CLIP | | 23142080 Unspecified |
| mmu-miR-669n | MIMAT0009427 | Dennd6a | 211922 | HITS-CLIP | 37986224\|23142080 | Unspecified |
| mmu-miR-669n | MIMAT0009427 | Dhx15 | 13204 | HITS-CLIP | | 23142080 Unspecified |
| mmu-miR-669n | MIMAT0009427 | Dhx30 | 72831 | HITS-CLIP | | 23142080 Unspecified |
| mmu-miR-669n | MIMAT0009427 | Dhx33 | 216877 | HITS-CLIP | | 23142080 Unspecified |
| mmu-miR-669n | MIMAT0009427 | Dhx36 | 72162 | HITS-CLIP | | 23142080 Unspecified |
| mmu-miR-669n | MIMAT0009427 | Dhx9 | 13211 | HITS-CLIP | | 23142080 Unspecified |
| mmu-miR-669n | MIMAT0009427 | Dmtf1 | 23857 | HITS-CLIP | | 23142080 Unspecified |
| mmu-miR-669n | MIMAT0009427 | Dnaja1 | 15502 | HITS-CLIP | | 37986224 Unspecified |
| mmu-miR-669n | MIMAT0009427 | Dnajc11 | 230935 | HITS-CLIP | | 23142080 Unspecified |
| mmu-miR-669n | MIMAT0009427 | Dnajc16 | 214063 | HITS-CLIP | 37986224\|23142080 | Unspecified |
| mmu-miR-669n | MIMAT0009427 | Dnajc7 | 56354 | HITS-CLIP | 37986224\|23142080 | Unspecified |
| mmu-miR-669n | MIMAT0009427 | Dnm1l | 74006 | HITS-CLIP | | 23142080 Unspecified |
| mmu-miR-669n | MIMAT0009427 | Dnmt3a | 13435 | HITS-CLIP | | 23142080 Unspecified |
| mmu-miR-669n | MIMAT0009427 | Dock11 | 75974 | HITS-CLIP | | 23142080 Unspecified |
| mmu-miR-669n | MIMAT0009427 | Dolk | 227697 | HITS-CLIP | | 23142080 Unspecified |
| mmu-miR-669n | MIMAT0009427 | Dpp4 | 13482 | HITS-CLIP | | 23142080 Unspecified |
| mmu-miR-669n | MIMAT0009427 | Dtl | 76843 | HITS-CLIP | | 37986224 Unspecified |
| mmu-miR-669n | MIMAT0009427 | Dus4l | 71916 | HITS-CLIP | | 37986224 Unspecified |
| mmu-miR-669n | MIMAT0009427 | Dvl1 | 13542 | HITS-CLIP | | 23142080 Unspecified |
| mmu-miR-669n | MIMAT0009427 | Dync1i2 | 13427 | HITS-CLIP | | 23142080 Unspecified |
| mmu-miR-669n | MIMAT0009427 | Dyrk1a | 13548 | HITS-CLIP | | 23142080 Unspecified |
| mmu-miR-669n | MIMAT0009427 | E2f3 | 13557 | HITS-CLIP | 37986224\|23142080 | Unspecified |
| mmu-miR-669n | MIMAT0009427 | E2f8 | 108961 | HITS-CLIP | | 23142080 Unspecified |
| mmu-miR-669n | MIMAT0009427 | Edem1 | 192193 | HITS-CLIP | 37986224\|23142080 | Unspecified |
| mmu-miR-669n | MIMAT0009427 | Eea1 | 216238 | HITS-CLIP | | 37986224 Unspecified |
| mmu-miR-669n | MIMAT0009427 | Eif3a | 13669 | HITS-CLIP | | 23142080 Unspecified |
| mmu-miR-669n | MIMAT0009427 | Eif3c | 56347 | HITS-CLIP | | 23142080 Unspecified |
| mmu-miR-669n | MIMAT0009427 | Eif4g3 | 230861 | HITS-CLIP | | 23142080 Unspecified |

| | | | | | | |
|---|---|---|---|---|---|---|
| mmu-miR-669n | MIMAT0009427 | Elf1 | 13709 | HITS-CLIP | 37986224\|23142080 | Unspecified |
| mmu-miR-669n | MIMAT0009427 | Ell2 | 192657 | HITS-CLIP | 37986224 | Unspecified |
| mmu-miR-669n | MIMAT0009427 | Elmsan1 | 238317 | HITS-CLIP | 37986224\|23142080 | Unspecified |
| mmu-miR-669n | MIMAT0009427 | Enc1 | 13803 | HITS-CLIP | 37986224 | Unspecified |
| mmu-miR-669n | MIMAT0009427 | Ep300 | 328572 | HITS-CLIP | 23142080 | Unspecified |
| mmu-miR-669n | MIMAT0009427 | Ep400 | 75560 | HITS-CLIP | 23142080 | Unspecified |
| mmu-miR-669n | MIMAT0009427 | Epas1 | 13819 | HITS-CLIP | 23142080 | Unspecified |
| mmu-miR-669n | MIMAT0009427 | Erap1 | 80898 | HITS-CLIP | 37986224 | Unspecified |
| mmu-miR-669n | MIMAT0009427 | Ero1l | 50527 | HITS-CLIP | 23142080 | Unspecified |
| mmu-miR-669n | MIMAT0009427 | Esco2 | 71988 | HITS-CLIP | 37986224 | Unspecified |
| mmu-miR-669n | MIMAT0009427 | Etf1 | 225363 | HITS-CLIP | 23142080 | Unspecified |
| mmu-miR-669n | MIMAT0009427 | Etnk1 | 75320 | HITS-CLIP | 37986224 | Unspecified |
| mmu-miR-669n | MIMAT0009427 | Ets1 | 23871 | HITS-CLIP | 37986224 | Unspecified |
| mmu-miR-669n | MIMAT0009427 | Etv6 | 14011 | HITS-CLIP | 23142080 | Unspecified |
| mmu-miR-669n | MIMAT0009427 | Exoc4 | 20336 | HITS-CLIP | 23142080 | Unspecified |
| mmu-miR-669n | MIMAT0009427 | F3 | 14066 | HITS-CLIP | 23142080 | Unspecified |
| mmu-miR-669n | MIMAT0009427 | Faap24 | 91442 | HITS-CLIP | 23142080 | Unspecified |
| mmu-miR-669n | MIMAT0009427 | Fam117a | 215512 | HITS-CLIP | 37986224\|23142080 | Unspecified |
| mmu-miR-669n | MIMAT0009427 | Fam117b | 72750 | HITS-CLIP | 23142080 | Unspecified |
| mmu-miR-669n | MIMAT0009427 | Fam120a | 218236 | HITS-CLIP | 23142080 | Unspecified |
| mmu-miR-669n | MIMAT0009427 | Fam133b | 68152 | HITS-CLIP | 23142080 | Unspecified |
| mmu-miR-669n | MIMAT0009427 | Fam214b | 230088 | HITS-CLIP | 23142080 | Unspecified |
| mmu-miR-669n | MIMAT0009427 | Fam222a | 433940 | HITS-CLIP | 23142080 | Unspecified |
| mmu-miR-669n | MIMAT0009427 | Fam49b | 223601 | HITS-CLIP | 23142080 | Unspecified |
| mmu-miR-669n | MIMAT0009427 | Fam69a | 388650 | HITS-CLIP | 23142080 | Unspecified |
| mmu-miR-669n | MIMAT0009427 | Fam91a1 | 210998 | HITS-CLIP | 23142080 | Unspecified |
| mmu-miR-669n | MIMAT0009427 | Far1 | 67420 | HITS-CLIP | 23142080 | Unspecified |
| mmu-miR-669n | MIMAT0009427 | Fbxl14 | 101358 | HITS-CLIP | 25083871\|23142080 | Unspecified |
| mmu-miR-669n | MIMAT0009427 | Fbxl3 | 50789 | HITS-CLIP | 37986224 | Unspecified |
| mmu-miR-669n | MIMAT0009427 | Fbxo21 | 231670 | HITS-CLIP | 23142080 | Unspecified |
| mmu-miR-669n | MIMAT0009427 | Fbxo3 | 57443 | HITS-CLIP | 37986224\|23142080 | Unspecified |
| mmu-miR-669n | MIMAT0009427 | Fbxo8 | 50753 | HITS-CLIP | 37986224\|23142080 | Unspecified |
| mmu-miR-669n | MIMAT0009427 | Fbxw7 | 50754 | HITS-CLIP | 23142080 | Unspecified |
| mmu-miR-669n | MIMAT0009427 | Fcho2 | 218503 | HITS-CLIP | 23142080 | Unspecified |
| mmu-miR-669n | MIMAT0009427 | Fem1b | 14155 | HITS-CLIP | 37986224 | Unspecified |
| mmu-miR-669n | MIMAT0009427 | Fem1c | 240263 | HITS-CLIP | 23142080 | Unspecified |
| mmu-miR-669n | MIMAT0009427 | Ficd | 231630 | HITS-CLIP | 37986224 | Unspecified |
| mmu-miR-669n | MIMAT0009427 | Filip1l | 78749 | HITS-CLIP | 37986224\|23142080 | Unspecified |
| mmu-miR-669n | MIMAT0009427 | Fip1l1 | 66899 | HITS-CLIP | 23142080 | Unspecified |

| | | | | | | |
|---|---|---|---|---|---|---|
| mmu-miR-669n | MIMAT0009427 | Fkbp5 | 14229 | HITS-CLIP | 23142080 | Unspecified |
| mmu-miR-669n | MIMAT0009427 | Flna | 192176 | HITS-CLIP | 23142080 | Unspecified |
| mmu-miR-669n | MIMAT0009427 | Fmr1 | 14265 | HITS-CLIP | 23142080 | Unspecified |
| mmu-miR-669n | MIMAT0009427 | Foxj3 | 230700 | HITS-CLIP | 23142080 | Unspecified |
| mmu-miR-669n | MIMAT0009427 | Foxo4 | 54601 | HITS-CLIP | 23142080 | Unspecified |
| mmu-miR-669n | MIMAT0009427 | Frrs1 | 20321 | HITS-CLIP | 23142080 | Unspecified |
| mmu-miR-669n | MIMAT0009427 | Fubp1 | 51886 | HITS-CLIP | 23142080 | Unspecified |
| mmu-miR-669n | MIMAT0009427 | Fxr1 | 14359 | HITS-CLIP | 23142080 | Unspecified |
| mmu-miR-669n | MIMAT0009427 | G2e3 | 217558 | HITS-CLIP | 23142080 | Unspecified |
| mmu-miR-669n | MIMAT0009427 | G3bp2 | 23881 | HITS-CLIP | 37986224\|23142080 | Unspecified |
| mmu-miR-669n | MIMAT0009427 | G6pdx | 14381 | HITS-CLIP | 23142080 | Unspecified |
| mmu-miR-669n | MIMAT0009427 | Galnt12 | 230145 | HITS-CLIP | 37986224 | Unspecified |
| mmu-miR-669n | MIMAT0009427 | Galnt3 | 14425 | HITS-CLIP | 23142080 | Unspecified |
| mmu-miR-669n | MIMAT0009427 | Gars | 353172 | HITS-CLIP | 23142080 | Unspecified |
| mmu-miR-669n | MIMAT0009427 | Gart | 14450 | HITS-CLIP | 23142080 | Unspecified |
| mmu-miR-669n | MIMAT0009427 | Gbp4 | 17472 | HITS-CLIP | 23142080 | Unspecified |
| mmu-miR-669n | MIMAT0009427 | Gbp5 | 229898 | HITS-CLIP | 37986224 | Unspecified |
| mmu-miR-669n | MIMAT0009427 | Gcc2 | 70297 | HITS-CLIP | 23142080 | Unspecified |
| mmu-miR-669n | MIMAT0009427 | Ggnbp2 | 217039 | HITS-CLIP | 23142080 | Unspecified |
| mmu-miR-669n | MIMAT0009427 | Gimap4 | 107526 | HITS-CLIP | 37986224 | Unspecified |
| mmu-miR-669n | MIMAT0009427 | Git1 | 216963 | HITS-CLIP | 23142080 | Unspecified |
| mmu-miR-669n | MIMAT0009427 | Glcci1 | 170772 | HITS-CLIP | 23142080 | Unspecified |
| mmu-miR-669n | MIMAT0009427 | Glg1 | 20340 | HITS-CLIP | 23142080 | Unspecified |
| mmu-miR-669n | MIMAT0009427 | Gm10033 | 378466 | HITS-CLIP | 31433990 | Unspecified |
| mmu-miR-669n | MIMAT0009427 | Gmfb | 63985 | HITS-CLIP | 23142080 | Unspecified |
| mmu-miR-669n | MIMAT0009427 | Gnb1 | 14688 | HITS-CLIP | 37986224\|23142080 | Unspecified |
| mmu-miR-669n | MIMAT0009427 | Gng5 | 14707 | HITS-CLIP | 37986224\|23142080 | Unspecified |
| mmu-miR-669n | MIMAT0009427 | Gnpda2 | 67980 | HITS-CLIP | 23142080 | Unspecified |
| mmu-miR-669n | MIMAT0009427 | Gnptab | 432486 | HITS-CLIP | 37986224 | Unspecified |
| mmu-miR-669n | MIMAT0009427 | Golph3 | 66629 | HITS-CLIP | 37986224\|23142080 | Unspecified |
| mmu-miR-669n | MIMAT0009427 | Gpatch8 | 237943 | HITS-CLIP | 23142080 | Unspecified |
| mmu-miR-669n | MIMAT0009427 | Gpbp1 | 73274 | HITS-CLIP | 37986224 | Unspecified |
| mmu-miR-669n | MIMAT0009427 | Gpbp1l1 | 77110 | HITS-CLIP | 23142080 | Unspecified |
| mmu-miR-669n | MIMAT0009427 | Gpd2 | 14571 | HITS-CLIP | 23142080 | Unspecified |
| mmu-miR-669n | MIMAT0009427 | Gpr132 | 56696 | HITS-CLIP | 37986224 | Unspecified |
| mmu-miR-669n | MIMAT0009427 | Gpr171 | 229323 | HITS-CLIP | 23142080 | Unspecified |
| mmu-miR-669n | MIMAT0009427 | Gpr174 | 213439 | HITS-CLIP | 23142080 | Unspecified |
| mmu-miR-669n | MIMAT0009427 | Gprin3 | 243385 | HITS-CLIP | 37986224 | Unspecified |
| mmu-miR-669n | MIMAT0009427 | Gse1 | 382034 | HITS-CLIP | 23142080 | Unspecified |

| | | | | | | |
|---|---|---|---|---|---|---|
| mmu-miR-669n | MIMAT0009427 | H2-K1 | 14972 | HITS-CLIP | 23142080 | Unspecified |
| mmu-miR-669n | MIMAT0009427 | H2-T23 | 15040 | HITS-CLIP | 23142080 | Unspecified |
| mmu-miR-669n | MIMAT0009427 | Hat1 | 107435 | HITS-CLIP | 37986224 | Unspecified |
| mmu-miR-669n | MIMAT0009427 | Hcfc1 | 15161 | HITS-CLIP | 23142080 | Unspecified |
| mmu-miR-669n | MIMAT0009427 | Hcfc2 | 67933 | HITS-CLIP | 37986224\|23142080 | Unspecified |
| mmu-miR-669n | MIMAT0009427 | Hdac4 | 208727 | HITS-CLIP | 23142080 | Unspecified |
| mmu-miR-669n | MIMAT0009427 | Hdlbp | 110611 | HITS-CLIP | 23142080 | Unspecified |
| mmu-miR-669n | MIMAT0009427 | Heatr1 | 217995 | HITS-CLIP | 23142080 | Unspecified |
| mmu-miR-669n | MIMAT0009427 | Herc2 | 15204 | HITS-CLIP | 23142080 | Unspecified |
| mmu-miR-669n | MIMAT0009427 | Herc4 | 67345 | HITS-CLIP | 23142080 | Unspecified |
| mmu-miR-669n | MIMAT0009427 | Herpud2 | 80517 | HITS-CLIP | 23142080 | Unspecified |
| mmu-miR-669n | MIMAT0009427 | Hif1a | 15251 | HITS-CLIP | 37986224 | Unspecified |
| mmu-miR-669n | MIMAT0009427 | Hipk1 | 15257 | HITS-CLIP | 37986224\|23142080 | Unspecified |
| mmu-miR-669n | MIMAT0009427 | Hipk3 | 15259 | HITS-CLIP | 23142080 | Unspecified |
| mmu-miR-669n | MIMAT0009427 | Hist1h2bm | 8342 | HITS-CLIP | 23142080 | Unspecified |
| mmu-miR-669n | MIMAT0009427 | Hivep3 | 16656 | HITS-CLIP | 23142080 | Unspecified |
| mmu-miR-669n | MIMAT0009427 | Hmbox1 | 219150 | HITS-CLIP | 23142080 | Unspecified |
| mmu-miR-669n | MIMAT0009427 | Hmgcr | 15357 | HITS-CLIP | 37986224\|23142080 | Unspecified |
| mmu-miR-669n | MIMAT0009427 | Hmgcs1 | 208715 | HITS-CLIP | 37986224 | Unspecified |
| mmu-miR-669n | MIMAT0009427 | Hnrnpa0 | 77134 | HITS-CLIP | 37986224 | Unspecified |
| mmu-miR-669n | MIMAT0009427 | Hnrnph1 | 59013 | HITS-CLIP | 23142080 | Unspecified |
| mmu-miR-669n | MIMAT0009427 | Hnrnph2 | 56258 | HITS-CLIP | 23142080 | Unspecified |
| mmu-miR-669n | MIMAT0009427 | Hnrnpk | 15387 | HITS-CLIP | 23142080 | Unspecified |
| mmu-miR-669n | MIMAT0009427 | Hp1bp3 | 15441 | HITS-CLIP | 23142080 | Unspecified |
| mmu-miR-669n | MIMAT0009427 | Hs3st3b1 | 54710 | HITS-CLIP | 23142080 | Unspecified |
| mmu-miR-669n | MIMAT0009427 | Hsp90b1 | 22027 | HITS-CLIP | 23142080 | Unspecified |
| mmu-miR-669n | MIMAT0009427 | Hspa4 | 15525 | HITS-CLIP | 23142080 | Unspecified |
| mmu-miR-669n | MIMAT0009427 | Hus1 | 15574 | HITS-CLIP | 23142080 | Unspecified |
| mmu-miR-669n | MIMAT0009427 | Huwe1 | 59026 | HITS-CLIP | 23142080 | Unspecified |
| mmu-miR-669n | MIMAT0009427 | Iars | 105148 | HITS-CLIP | 23142080 | Unspecified |
| mmu-miR-669n | MIMAT0009427 | Ier3 | 15937 | HITS-CLIP | 23142080 | Unspecified |
| mmu-miR-669n | MIMAT0009427 | Ifnar1 | 15975 | HITS-CLIP | 37986224\|23142080 | Unspecified |
| mmu-miR-669n | MIMAT0009427 | Ifngr1 | 15979 | HITS-CLIP | 37986224 | Unspecified |
| mmu-miR-669n | MIMAT0009427 | Ifngr2 | 15980 | HITS-CLIP | 31433990 | Unspecified |
| mmu-miR-669n | MIMAT0009427 | Ikzf1 | 22778 | HITS-CLIP | 37986224 | Unspecified |
| mmu-miR-669n | MIMAT0009427 | Ikzf3 | 22780 | HITS-CLIP | 23142080 | Unspecified |
| mmu-miR-669n | MIMAT0009427 | Il7r | 16197 | HITS-CLIP | 37986224 | Unspecified |
| mmu-miR-669n | MIMAT0009427 | Inhba | 16323 | HITS-CLIP | 23142080 | Unspecified |
| mmu-miR-669n | MIMAT0009427 | Ino80d | 227195 | HITS-CLIP | 23142080 | Unspecified |

| | | | | | | |
|---|---|---|---|---|---|---|
| mmu-miR-669n | MIMAT0009427 | Inpp4b | 234515 | HITS-CLIP | 37986224\|23142080 | Unspecified |
| mmu-miR-669n | MIMAT0009427 | Inpp5a | 212111 | HITS-CLIP | 23142080 | Unspecified |
| mmu-miR-669n | MIMAT0009427 | Inpp5f | 101490 | HITS-CLIP | 23142080 | Unspecified |
| mmu-miR-669n | MIMAT0009427 | Insig2 | 72999 | HITS-CLIP | 23142080 | Unspecified |
| mmu-miR-669n | MIMAT0009427 | Insr | 16337 | HITS-CLIP | 31433990\|26203562 | Unspecified |
| mmu-miR-669n | MIMAT0009427 | Ints1 | 68510 | HITS-CLIP | 23142080 | Unspecified |
| mmu-miR-669n | MIMAT0009427 | Ints6 | 18130 | HITS-CLIP | 21258322\|23142080 | Unspecified |
| mmu-miR-669n | MIMAT0009427 | Ip6k2 | 76500 | HITS-CLIP | 23142080 | Unspecified |
| mmu-miR-669n | MIMAT0009427 | Iqgap1 | 29875 | HITS-CLIP | 23142080 | Unspecified |
| mmu-miR-669n | MIMAT0009427 | Iqgap2 | 544963 | HITS-CLIP | 23142080 | Unspecified |
| mmu-miR-669n | MIMAT0009427 | Irf2bp2 | 270110 | HITS-CLIP | 23142080 | Unspecified |
| mmu-miR-669n | MIMAT0009427 | Irf4 | 16364 | HITS-CLIP | 37986224\|23142080 | Unspecified |
| mmu-miR-669n | MIMAT0009427 | Irgm2 | 54396 | HITS-CLIP | 37986224 | Unspecified |
| mmu-miR-669n | MIMAT0009427 | Itch | 16396 | HITS-CLIP | 23142080 | Unspecified |
| mmu-miR-669n | MIMAT0009427 | Itgb1 | 16412 | HITS-CLIP | 37986224 | Unspecified |
| mmu-miR-669n | MIMAT0009427 | Itpkb | 320404 | HITS-CLIP | 37986224 | Unspecified |
| mmu-miR-669n | MIMAT0009427 | Ivns1abp | 117198 | HITS-CLIP | 23142080 | Unspecified |
| mmu-miR-669n | MIMAT0009427 | Izumo1r | 64931 | HITS-CLIP | 23142080 | Unspecified |
| mmu-miR-669n | MIMAT0009427 | Jade1 | 269424 | HITS-CLIP | 37986224\|23142080 | Unspecified |
| mmu-miR-669n | MIMAT0009427 | Jarid2 | 16468 | HITS-CLIP | 37986224\|23142080 | Unspecified |
| mmu-miR-669n | MIMAT0009427 | Jdp2 | 81703 | HITS-CLIP | 23142080 | Unspecified |
| mmu-miR-669n | MIMAT0009427 | Jmjd8 | 72106 | HITS-CLIP | 23142080 | Unspecified |
| mmu-miR-669n | MIMAT0009427 | Kansl1 | 76719 | HITS-CLIP | 37986224 | Unspecified |
| mmu-miR-669n | MIMAT0009427 | Kansl1l | 68691 | HITS-CLIP | 23142080 | Unspecified |
| mmu-miR-669n | MIMAT0009427 | Kbtbd2 | 210973 | HITS-CLIP | 37986224\|23142080 | Unspecified |
| mmu-miR-669n | MIMAT0009427 | Kbtbd8 | 243574 | HITS-CLIP | 37986224 | Unspecified |
| mmu-miR-669n | MIMAT0009427 | Kctd12 | 239217 | HITS-CLIP | 23142080 | Unspecified |
| mmu-miR-669n | MIMAT0009427 | Kctd9 | 105440 | HITS-CLIP | 23142080 | Unspecified |
| mmu-miR-669n | MIMAT0009427 | Kdelr2 | 66913 | HITS-CLIP | 37986224\|23142080 | Unspecified |
| mmu-miR-669n | MIMAT0009427 | Kdm1a | 99982 | HITS-CLIP | 23142080 | Unspecified |
| mmu-miR-669n | MIMAT0009427 | Kdm2a | 225876 | HITS-CLIP | 23142080 | Unspecified |
| mmu-miR-669n | MIMAT0009427 | Kdm3b | 277250 | HITS-CLIP | 37986224 | Unspecified |
| mmu-miR-669n | MIMAT0009427 | Kdm4b | 193796 | HITS-CLIP | 23142080 | Unspecified |
| mmu-miR-669n | MIMAT0009427 | Kdm4c | 76804 | HITS-CLIP | 37986224 | Unspecified |
| mmu-miR-669n | MIMAT0009427 | Kdm7a | 338523 | HITS-CLIP | 37986224\|23142080 | Unspecified |
| mmu-miR-669n | MIMAT0009427 | Kdsr | 70750 | HITS-CLIP | 23142080 | Unspecified |
| mmu-miR-669n | MIMAT0009427 | Kif11 | 16551 | HITS-CLIP | 37986224 | Unspecified |
| mmu-miR-669n | MIMAT0009427 | Kif1bp | 26128 | HITS-CLIP | 23142080 | Unspecified |
| mmu-miR-669n | MIMAT0009427 | Kif20a | 19348 | HITS-CLIP | 23142080 | Unspecified |

| | | | | | | | |
|---|---|---|---|---|---|---|---|
| mmu-miR-669n | MIMAT0009427 | Kif21b | 16565 | HITS-CLIP | | 37986224 | Unspecified |
| mmu-miR-669n | MIMAT0009427 | Kif5b | 16573 | HITS-CLIP | 37986224\|23142080 | | Unspecified |
| mmu-miR-669n | MIMAT0009427 | Kifap3 | 16579 | HITS-CLIP | | 23142080 | Unspecified |
| mmu-miR-669n | MIMAT0009427 | Klf13 | 50794 | HITS-CLIP | 37986224\|23142080 | | Unspecified |
| mmu-miR-669n | MIMAT0009427 | Klhl9 | 242521 | HITS-CLIP | | 37986224 | Unspecified |
| mmu-miR-669n | MIMAT0009427 | Kmt2a | 214162 | HITS-CLIP | 37986224\|23142080 | | Unspecified |
| mmu-miR-669n | MIMAT0009427 | Kmt2c | 231051 | HITS-CLIP | | 23142080 | Unspecified |
| mmu-miR-669n | MIMAT0009427 | Kmt2d | 381022 | HITS-CLIP | 37986224\|23142080 | | Unspecified |
| mmu-miR-669n | MIMAT0009427 | Kmt2e | 69188 | HITS-CLIP | | 23142080 | Unspecified |
| mmu-miR-669n | MIMAT0009427 | Kmt5b | 225888 | HITS-CLIP | | 23142080 | Unspecified |
| mmu-miR-669n | MIMAT0009427 | Kpna3 | 16648 | HITS-CLIP | | 31433990 | Unspecified |
| mmu-miR-669n | MIMAT0009427 | Kpna4 | 16649 | HITS-CLIP | | 23142080 | Unspecified |
| mmu-miR-669n | MIMAT0009427 | Kpnb1 | 16211 | HITS-CLIP | | 23142080 | Unspecified |
| mmu-miR-669n | MIMAT0009427 | Krit1 | 79264 | HITS-CLIP | | 23142080 | Unspecified |
| mmu-miR-669n | MIMAT0009427 | Ksr1 | 16706 | HITS-CLIP | | 23142080 | Unspecified |
| mmu-miR-669n | MIMAT0009427 | Lamp1 | 16783 | HITS-CLIP | | 23142080 | Unspecified |
| mmu-miR-669n | MIMAT0009427 | Laptm4a | 17775 | HITS-CLIP | | 23142080 | Unspecified |
| mmu-miR-669n | MIMAT0009427 | Lats1 | 16798 | HITS-CLIP | 37986224\|23142080 | | Unspecified |
| mmu-miR-669n | MIMAT0009427 | Lcor | 212391 | HITS-CLIP | | 23142080 | Unspecified |
| mmu-miR-669n | MIMAT0009427 | Lcp1 | 18826 | HITS-CLIP | | 37986224 | Unspecified |
| mmu-miR-669n | MIMAT0009427 | Ldb1 | 16825 | HITS-CLIP | | 23142080 | Unspecified |
| mmu-miR-669n | MIMAT0009427 | Lims1 | 110829 | HITS-CLIP | 37986224\|23142080 | | Unspecified |
| mmu-miR-669n | MIMAT0009427 | Lin54 | 231506 | HITS-CLIP | | 23142080 | Unspecified |
| mmu-miR-669n | MIMAT0009427 | Lipe | 16890 | HITS-CLIP | | 23142080 | Unspecified |
| mmu-miR-669n | MIMAT0009427 | Lnpep | 240028 | HITS-CLIP | | 23142080 | Unspecified |
| mmu-miR-669n | MIMAT0009427 | Lnx2 | 140887 | HITS-CLIP | | 23142080 | Unspecified |
| mmu-miR-669n | MIMAT0009427 | Lonrf1 | 244421 | HITS-CLIP | | 23142080 | Unspecified |
| mmu-miR-669n | MIMAT0009427 | Lonrf3 | 74365 | HITS-CLIP | | 23142080 | Unspecified |
| mmu-miR-669n | MIMAT0009427 | Lpcat3 | 14792 | HITS-CLIP | | 37986224 | Unspecified |
| mmu-miR-669n | MIMAT0009427 | Lpp | 210126 | HITS-CLIP | | 23142080 | Unspecified |
| mmu-miR-669n | MIMAT0009427 | Lrba | 80877 | HITS-CLIP | | 23142080 | Unspecified |
| mmu-miR-669n | MIMAT0009427 | Lrch1 | 380916 | HITS-CLIP | | 23142080 | Unspecified |
| mmu-miR-669n | MIMAT0009427 | Lrig1 | 16206 | HITS-CLIP | | 37986224 | Unspecified |
| mmu-miR-669n | MIMAT0009427 | Lrp6 | 16974 | HITS-CLIP | | 37986224 | Unspecified |
| mmu-miR-669n | MIMAT0009427 | Lrr1 | 69706 | HITS-CLIP | | 23142080 | Unspecified |
| mmu-miR-669n | MIMAT0009427 | Lrrc8c | 100604 | HITS-CLIP | 31433990\|23142080 | | Unspecified |
| mmu-miR-669n | MIMAT0009427 | Lsm12 | 268490 | HITS-CLIP | | 23142080 | Unspecified |
| mmu-miR-669n | MIMAT0009427 | Ltbp3 | 16998 | HITS-CLIP | | 23142080 | Unspecified |
| mmu-miR-669n | MIMAT0009427 | Luc7l3 | 67684 | HITS-CLIP | | 37986224 | Unspecified |

| | | | | | | | |
|---|---|---|---|---|---|---|---|
| mmu-miR-669n | MIMAT0009427 | Ly6a | 110454 | HITS-CLIP | | 23142080 | Unspecified |
| mmu-miR-669n | MIMAT0009427 | Lyst | 17101 | HITS-CLIP | 37986224\|23142080 | | Unspecified |
| mmu-miR-669n | MIMAT0009427 | Macf1 | 11426 | HITS-CLIP | | 23142080 | Unspecified |
| mmu-miR-669n | MIMAT0009427 | Maff | 17133 | HITS-CLIP | 37986224\|23142080 | | Unspecified |
| mmu-miR-669n | MIMAT0009427 | Malt1 | 240354 | HITS-CLIP | | 37986224 | Unspecified |
| mmu-miR-669n | MIMAT0009427 | Map2k3 | 26397 | HITS-CLIP | | 23142080 | Unspecified |
| mmu-miR-669n | MIMAT0009427 | Map3k1 | 26401 | HITS-CLIP | 37986224\|23142080 | | Unspecified |
| mmu-miR-669n | MIMAT0009427 | Map3k2 | 26405 | HITS-CLIP | | 37986224 | Unspecified |
| mmu-miR-669n | MIMAT0009427 | Map4k3 | 225028 | HITS-CLIP | | 23142080 | Unspecified |
| mmu-miR-669n | MIMAT0009427 | Map4k4 | 26921 | HITS-CLIP | | 23142080 | Unspecified |
| mmu-miR-669n | MIMAT0009427 | Mapk1 | 26413 | HITS-CLIP | 37986224\|23142080 | | Unspecified |
| mmu-miR-669n | MIMAT0009427 | Mapk6 | 50772 | HITS-CLIP | | 37986224 | Unspecified |
| mmu-miR-669n | MIMAT0009427 | Mapre2 | 212307 | HITS-CLIP | | 23142080 | Unspecified |
| mmu-miR-669n | MIMAT0009427 | March2 | 51257 | HITS-CLIP | | 23142080 | Unspecified |
| mmu-miR-669n | MIMAT0009427 | Marchf6 | 223455 | HITS-CLIP | 37986224\|23142080 | | Unspecified |
| mmu-miR-669n | MIMAT0009427 | Marcks | 17118 | HITS-CLIP | 37986224\|23142080 | | Unspecified |
| mmu-miR-669n | MIMAT0009427 | Marf1 | 223989 | HITS-CLIP | | 23142080 | Unspecified |
| mmu-miR-669n | MIMAT0009427 | Mast2 | 17776 | HITS-CLIP | | 23142080 | Unspecified |
| mmu-miR-669n | MIMAT0009427 | Mat2b | 108645 | HITS-CLIP | | 23142080 | Unspecified |
| mmu-miR-669n | MIMAT0009427 | Matr3 | 17184 | HITS-CLIP | | 23142080 | Unspecified |
| mmu-miR-669n | MIMAT0009427 | Mbd2 | 17191 | HITS-CLIP | | 37986224 | Unspecified |
| mmu-miR-669n | MIMAT0009427 | Mbd6 | 110962 | HITS-CLIP | | 23142080 | Unspecified |
| mmu-miR-669n | MIMAT0009427 | Mbnl1 | 56758 | HITS-CLIP | 37986224\|23142080 | | Unspecified |
| mmu-miR-669n | MIMAT0009427 | Mcmbp | 210711 | HITS-CLIP | | 37986224 | Unspecified |
| mmu-miR-669n | MIMAT0009427 | Med13 | 327987 | HITS-CLIP | 37986224\|23142080 | | Unspecified |
| mmu-miR-669n | MIMAT0009427 | Med13l | 76199 | HITS-CLIP | | 23142080 | Unspecified |
| mmu-miR-669n | MIMAT0009427 | Med14 | 26896 | HITS-CLIP | | 23142080 | Unspecified |
| mmu-miR-669n | MIMAT0009427 | Mef2a | 17258 | HITS-CLIP | | 23142080 | Unspecified |
| mmu-miR-669n | MIMAT0009427 | Mex3b | 108797 | HITS-CLIP | | 23142080 | Unspecified |
| mmu-miR-669n | MIMAT0009427 | Mex3c | 240396 | HITS-CLIP | 37986224\|23142080 | | Unspecified |
| mmu-miR-669n | MIMAT0009427 | Mga | 29808 | HITS-CLIP | | 23142080 | Unspecified |
| mmu-miR-669n | MIMAT0009427 | Mgat4a | 269181 | HITS-CLIP | | 23142080 | Unspecified |
| mmu-miR-669n | MIMAT0009427 | Mgea5 | 76055 | HITS-CLIP | | 23142080 | Unspecified |
| mmu-miR-669n | MIMAT0009427 | Mib1 | 225164 | HITS-CLIP | | 37986224 | Unspecified |
| mmu-miR-669n | MIMAT0009427 | Mier1 | 71148 | HITS-CLIP | | 23142080 | Unspecified |
| mmu-miR-669n | MIMAT0009427 | Mif4gd | 69674 | HITS-CLIP | | 23142080 | Unspecified |
| mmu-miR-669n | MIMAT0009427 | Mis18bp1 | 217653 | HITS-CLIP | | 23142080 | Unspecified |
| mmu-miR-669n | MIMAT0009427 | Mki67 | 17345 | HITS-CLIP | | 37986224 | Unspecified |
| mmu-miR-669n | MIMAT0009427 | Mkln1 | 27418 | HITS-CLIP | | 23142080 | Unspecified |

| | | | | | | | |
|---|---|---|---|---|---|---|---|
| mmu-miR-669n | MIMAT0009427 | Mmd | 67468 | HITS-CLIP | | 37986224 | Unspecified |
| mmu-miR-669n | MIMAT0009427 | Mob1a | 232157 | HITS-CLIP | | 37986224 | Unspecified |
| mmu-miR-669n | MIMAT0009427 | Mob1b | 68473 | HITS-CLIP | | 37986224 | Unspecified |
| mmu-miR-669n | MIMAT0009427 | Mpp7 | 75739 | HITS-CLIP | | 23142080 | Unspecified |
| mmu-miR-669n | MIMAT0009427 | Mrpl13 | 68537 | HITS-CLIP | | 23142080 | Unspecified |
| mmu-miR-669n | MIMAT0009427 | Mrpl18 | 67681 | HITS-CLIP | | 37986224 | Unspecified |
| mmu-miR-669n | MIMAT0009427 | Mrps10 | 64657 | HITS-CLIP | | 23142080 | Unspecified |
| mmu-miR-669n | MIMAT0009427 | Mrps24 | 64660 | HITS-CLIP | | 23142080 | Unspecified |
| mmu-miR-669n | MIMAT0009427 | Ms4a4b | 60361 | HITS-CLIP | | 37986224 | Unspecified |
| mmu-miR-669n | MIMAT0009427 | Ms4a4c | 64380 | HITS-CLIP | | 23142080 | Unspecified |
| mmu-miR-669n | MIMAT0009427 | Msh6 | 17688 | HITS-CLIP | | 23142080 | Unspecified |
| mmu-miR-669n | MIMAT0009427 | Msi2 | 76626 | HITS-CLIP | | 23142080 | Unspecified |
| mmu-miR-669n | MIMAT0009427 | Mtf2 | 17765 | HITS-CLIP | | 37986224 | Unspecified |
| mmu-miR-669n | MIMAT0009427 | Mtmr10 | 233315 | HITS-CLIP | | 23142080 | Unspecified |
| mmu-miR-669n | MIMAT0009427 | Mtmr14 | 97287 | HITS-CLIP | | 37986224 | Unspecified |
| mmu-miR-669n | MIMAT0009427 | Mtmr3 | 74302 | HITS-CLIP | | 23142080 | Unspecified |
| mmu-miR-669n | MIMAT0009427 | Myb | 17863 | HITS-CLIP | 37986224\|23142080 | | Unspecified |
| mmu-miR-669n | MIMAT0009427 | Mybl1 | 17864 | HITS-CLIP | | 23142080 | Unspecified |
| mmu-miR-669n | MIMAT0009427 | Myc | 17869 | HITS-CLIP | 37986224\|23142080 | | Unspecified |
| mmu-miR-669n | MIMAT0009427 | Myh9 | 17886 | HITS-CLIP | | 23142080 | Unspecified |
| mmu-miR-669n | MIMAT0009427 | Myo10 | 17909 | HITS-CLIP | | 23142080 | Unspecified |
| mmu-miR-669n | MIMAT0009427 | Myo9a | 270163 | HITS-CLIP | | 23142080 | Unspecified |
| mmu-miR-669n | MIMAT0009427 | Myo9b | 17925 | HITS-CLIP | | 23142080 | Unspecified |
| mmu-miR-669n | MIMAT0009427 | Mysm1 | 320713 | HITS-CLIP | | 23142080 | Unspecified |
| mmu-miR-669n | MIMAT0009427 | N4bp2l2 | 381695 | HITS-CLIP | | 23142080 | Unspecified |
| mmu-miR-669n | MIMAT0009427 | Naa25 | 231713 | HITS-CLIP | | 23142080 | Unspecified |
| mmu-miR-669n | MIMAT0009427 | Nap1l4 | 17955 | HITS-CLIP | | 37986224 | Unspecified |
| mmu-miR-669n | MIMAT0009427 | Nbeal2 | 235627 | HITS-CLIP | | 23142080 | Unspecified |
| mmu-miR-669n | MIMAT0009427 | Ncaph | 215387 | HITS-CLIP | | 23142080 | Unspecified |
| mmu-miR-669n | MIMAT0009427 | Nck1 | 17973 | HITS-CLIP | | 23142080 | Unspecified |
| mmu-miR-669n | MIMAT0009427 | Nckap1l | 105855 | HITS-CLIP | | 23142080 | Unspecified |
| mmu-miR-669n | MIMAT0009427 | Ncoa2 | 17978 | HITS-CLIP | 37986224\|23142080 | | Unspecified |
| mmu-miR-669n | MIMAT0009427 | Ncoa6 | 56406 | HITS-CLIP | | 23142080 | Unspecified |
| mmu-miR-669n | MIMAT0009427 | Ncor1 | 20185 | HITS-CLIP | | 23142080 | Unspecified |
| mmu-miR-669n | MIMAT0009427 | Ndufab1 | 70316 | HITS-CLIP | | 23142080 | Unspecified |
| mmu-miR-669n | MIMAT0009427 | Ndufs4 | 17993 | HITS-CLIP | | 23142080 | Unspecified |
| mmu-miR-669n | MIMAT0009427 | Ndufv3 | 78330 | HITS-CLIP | | 23142080 | Unspecified |
| mmu-miR-669n | MIMAT0009427 | Nedd4 | 17999 | HITS-CLIP | | 37986224 | Unspecified |
| mmu-miR-669n | MIMAT0009427 | Nek7 | 59125 | HITS-CLIP | 37986224\|23142080 | | Unspecified |

| | | | | | | | |
|---|---|---|---|---|---|---|---|
| mmu-miR-669n | MIMAT0009427 | Nfatc1 | 18018 | HITS-CLIP | | 23142080 | Unspecified |
| mmu-miR-669n | MIMAT0009427 | Nfatc3 | 18021 | HITS-CLIP | | 23142080 | Unspecified |
| mmu-miR-669n | MIMAT0009427 | Nin | 18080 | HITS-CLIP | | 23142080 | Unspecified |
| mmu-miR-669n | MIMAT0009427 | Nipal4 | 214112 | HITS-CLIP | | 26203562 | Unspecified |
| mmu-miR-669n | MIMAT0009427 | Nop58 | 55989 | HITS-CLIP | | 23142080 | Unspecified |
| mmu-miR-669n | MIMAT0009427 | Npc2 | 67963 | HITS-CLIP | | 23142080 | Unspecified |
| mmu-miR-669n | MIMAT0009427 | Nptn | 20320 | HITS-CLIP | | 23142080 | Unspecified |
| mmu-miR-669n | MIMAT0009427 | Nr3c1 | 14815 | HITS-CLIP | | 37986224 | Unspecified |
| mmu-miR-669n | MIMAT0009427 | Nr4a2 | 18227 | HITS-CLIP | | 23142080 | Unspecified |
| mmu-miR-669n | MIMAT0009427 | Nr4a3 | 18124 | HITS-CLIP | 37986224\|23142080 | | Unspecified |
| mmu-miR-669n | MIMAT0009427 | Nras | 18176 | HITS-CLIP | | 37986224 | Unspecified |
| mmu-miR-669n | MIMAT0009427 | Nrip1 | 268903 | HITS-CLIP | | 23142080 | Unspecified |
| mmu-miR-669n | MIMAT0009427 | Nsd1 | 18193 | HITS-CLIP | 37986224\|23142080 | | Unspecified |
| mmu-miR-669n | MIMAT0009427 | Nsd3 | 234135 | HITS-CLIP | | 23142080 | Unspecified |
| mmu-miR-669n | MIMAT0009427 | Nt5dc1 | 319638 | HITS-CLIP | | 23142080 | Unspecified |
| mmu-miR-669n | MIMAT0009427 | Nt5e | 23959 | HITS-CLIP | | 23142080 | Unspecified |
| mmu-miR-669n | MIMAT0009427 | Nufip2 | 68564 | HITS-CLIP | | 37986224 | Unspecified |
| mmu-miR-669n | MIMAT0009427 | Numa1 | 101706 | HITS-CLIP | | 23142080 | Unspecified |
| mmu-miR-669n | MIMAT0009427 | Nup153 | 218210 | HITS-CLIP | | 37986224 | Unspecified |
| mmu-miR-669n | MIMAT0009427 | Nup93 | 71805 | HITS-CLIP | | 23142080 | Unspecified |
| mmu-miR-669n | MIMAT0009427 | Nup98 | 269966 | HITS-CLIP | | 37986224 | Unspecified |
| mmu-miR-669n | MIMAT0009427 | Nus1 | 52014 | HITS-CLIP | 25083871\|23142080 | | Unspecified |
| mmu-miR-669n | MIMAT0009427 | Orc5 | 26429 | HITS-CLIP | | 23142080 | Unspecified |
| mmu-miR-669n | MIMAT0009427 | Otud7b | 229603 | HITS-CLIP | | 23142080 | Unspecified |
| mmu-miR-669n | MIMAT0009427 | Oxr1 | 170719 | HITS-CLIP | 37986224\|23142080 | | Unspecified |
| mmu-miR-669n | MIMAT0009427 | Pabpc1 | 18458 | HITS-CLIP | | 23142080 | Unspecified |
| mmu-miR-669n | MIMAT0009427 | Pafah1b1 | 18472 | HITS-CLIP | 37986224\|23142080 | | Unspecified |
| mmu-miR-669n | MIMAT0009427 | Paip2 | 67869 | HITS-CLIP | | 37986224 | Unspecified |
| mmu-miR-669n | MIMAT0009427 | Pank3 | 211347 | HITS-CLIP | | 37986224 | Unspecified |
| mmu-miR-669n | MIMAT0009427 | Parp14 | 547253 | HITS-CLIP | | 23142080 | Unspecified |
| mmu-miR-669n | MIMAT0009427 | Patl1 | 225929 | HITS-CLIP | 37986224\|23142080 | | Unspecified |
| mmu-miR-669n | MIMAT0009427 | Paxip1 | 55982 | HITS-CLIP | | 37986224 | Unspecified |
| mmu-miR-669n | MIMAT0009427 | Pbrm1 | 66923 | HITS-CLIP | | 37986224 | Unspecified |
| mmu-miR-669n | MIMAT0009427 | Pcf11 | 74737 | HITS-CLIP | | 23142080 | Unspecified |
| mmu-miR-669n | MIMAT0009427 | Pcmt1 | 18537 | HITS-CLIP | | 23142080 | Unspecified |
| mmu-miR-669n | MIMAT0009427 | Pde4b | 18578 | HITS-CLIP | | 23142080 | Unspecified |
| mmu-miR-669n | MIMAT0009427 | Pde4d | 238871 | HITS-CLIP | | 37986224 | Unspecified |
| mmu-miR-669n | MIMAT0009427 | Pde7a | 18583 | HITS-CLIP | | 37986224 | Unspecified |
| mmu-miR-669n | MIMAT0009427 | Pdzd8 | 107368 | HITS-CLIP | | 23142080 | Unspecified |

| | | | | | | |
|---|---|---|---|---|---|---|
| mmu-miR-669n | MIMAT0009427 | Per1 | 18626 | HITS-CLIP | 23142080 | Unspecified |
| mmu-miR-669n | MIMAT0009427 | Pex3 | 56535 | HITS-CLIP | 23142080 | Unspecified |
| mmu-miR-669n | MIMAT0009427 | Phc3 | 241915 | HITS-CLIP | 37986224 | Unspecified |
| mmu-miR-669n | MIMAT0009427 | Phf20 | 228829 | HITS-CLIP | 23142080 | Unspecified |
| mmu-miR-669n | MIMAT0009427 | Phip | 83946 | HITS-CLIP | 23142080 | Unspecified |
| mmu-miR-669n | MIMAT0009427 | Pik3c2b | 240752 | HITS-CLIP | 23142080 | Unspecified |
| mmu-miR-669n | MIMAT0009427 | Pik3ca | 18706 | HITS-CLIP | 37986224\|23142080 | Unspecified |
| mmu-miR-669n | MIMAT0009427 | Pitpna | 18738 | HITS-CLIP | 37986224 | Unspecified |
| mmu-miR-669n | MIMAT0009427 | Pkn2 | 109333 | HITS-CLIP | 37986224\|23142080 | Unspecified |
| mmu-miR-669n | MIMAT0009427 | Plaa | 18786 | HITS-CLIP | 23142080 | Unspecified |
| mmu-miR-669n | MIMAT0009427 | Plekhm2 | 69582 | HITS-CLIP | 23142080 | Unspecified |
| mmu-miR-669n | MIMAT0009427 | Plgrkt | 67759 | HITS-CLIP | 37986224 | Unspecified |
| mmu-miR-669n | MIMAT0009427 | Plrg1 | 53317 | HITS-CLIP | 23142080 | Unspecified |
| mmu-miR-669n | MIMAT0009427 | Pole | 18973 | HITS-CLIP | 23142080 | Unspecified |
| mmu-miR-669n | MIMAT0009427 | Polr2a | 20020 | HITS-CLIP | 37986224\|23142080 | Unspecified |
| mmu-miR-669n | MIMAT0009427 | Ppfia1 | 233977 | HITS-CLIP | 23142080 | Unspecified |
| mmu-miR-669n | MIMAT0009427 | Ppig | 228005 | HITS-CLIP | 37986224\|23142080 | Unspecified |
| mmu-miR-669n | MIMAT0009427 | Ppil1 | 68816 | HITS-CLIP | 23142080 | Unspecified |
| mmu-miR-669n | MIMAT0009427 | Ppil2 | 66053 | HITS-CLIP | 23142080 | Unspecified |
| mmu-miR-669n | MIMAT0009427 | Ppp1cb | 19046 | HITS-CLIP | 37986224 | Unspecified |
| mmu-miR-669n | MIMAT0009427 | Ppp2ca | 19052 | HITS-CLIP | 23142080 | Unspecified |
| mmu-miR-669n | MIMAT0009427 | Ppp3cb | 19056 | HITS-CLIP | 37986224\|23142080 | Unspecified |
| mmu-miR-669n | MIMAT0009427 | Ppp6c | 67857 | HITS-CLIP | 23142080 | Unspecified |
| mmu-miR-669n | MIMAT0009427 | Ppp6r3 | 52036 | HITS-CLIP | 37986224\|23142080 | Unspecified |
| mmu-miR-669n | MIMAT0009427 | Prc1 | 233406 | HITS-CLIP | 37986224 | Unspecified |
| mmu-miR-669n | MIMAT0009427 | Prdm1 | 12142 | HITS-CLIP | 23142080 | Unspecified |
| mmu-miR-669n | MIMAT0009427 | Prep | 19072 | HITS-CLIP | 23142080 | Unspecified |
| mmu-miR-669n | MIMAT0009427 | Prim2 | 19076 | HITS-CLIP | 23142080 | Unspecified |
| mmu-miR-669n | MIMAT0009427 | Prkag2 | 108099 | HITS-CLIP | 23142080 | Unspecified |
| mmu-miR-669n | MIMAT0009427 | Prkar1a | 19084 | HITS-CLIP | 37986224 | Unspecified |
| mmu-miR-669n | MIMAT0009427 | Prkcd | 18753 | HITS-CLIP | 23142080 | Unspecified |
| mmu-miR-669n | MIMAT0009427 | Prkch | 18755 | HITS-CLIP | 23142080 | Unspecified |
| mmu-miR-669n | MIMAT0009427 | Prkcq | 18761 | HITS-CLIP | 23142080 | Unspecified |
| mmu-miR-669n | MIMAT0009427 | Prkx | 19108 | HITS-CLIP | 23142080 | Unspecified |
| mmu-miR-669n | MIMAT0009427 | Prmt5 | 27374 | HITS-CLIP | 26203562 | Unspecified |
| mmu-miR-669n | MIMAT0009427 | Prpf38b | 66921 | HITS-CLIP | 23142080 | Unspecified |
| mmu-miR-669n | MIMAT0009427 | Prpf40a | 56194 | HITS-CLIP | 23142080\|26203562 | Unspecified |
| mmu-miR-669n | MIMAT0009427 | Prpf4b | 19134 | HITS-CLIP | 37986224 | Unspecified |
| mmu-miR-669n | MIMAT0009427 | Prpf8 | 192159 | HITS-CLIP | 23142080 | Unspecified |

| | | | | | | | |
|---|---|---|---|---|---|---|---|
| mmu-miR-669n | MIMAT0009427 | Psma6 | 26443 | HITS-CLIP | | 23142080 | Unspecified |
| mmu-miR-669n | MIMAT0009427 | Psmc2 | 19181 | HITS-CLIP | | 37986224 | Unspecified |
| mmu-miR-669n | MIMAT0009427 | Psmd1 | 70247 | HITS-CLIP | | 23142080 | Unspecified |
| mmu-miR-669n | MIMAT0009427 | Psmd12 | 66997 | HITS-CLIP | | 23142080 | Unspecified |
| mmu-miR-669n | MIMAT0009427 | Psmd7 | 17463 | HITS-CLIP | | 23142080 | Unspecified |
| mmu-miR-669n | MIMAT0009427 | Ptch1 | 19206 | HITS-CLIP | | 31433990 | Unspecified |
| mmu-miR-669n | MIMAT0009427 | Ptpn4 | 19258 | HITS-CLIP | 37986224\|23142080 | | Unspecified |
| mmu-miR-669n | MIMAT0009427 | Ptprc | 19264 | HITS-CLIP | 37986224\|23142080 | | Unspecified |
| mmu-miR-669n | MIMAT0009427 | Pttg1ip | 108705 | HITS-CLIP | | 23142080 | Unspecified |
| mmu-miR-669n | MIMAT0009427 | Pum1 | 80912 | HITS-CLIP | | 23142080 | Unspecified |
| mmu-miR-669n | MIMAT0009427 | Pum2 | 80913 | HITS-CLIP | | 37986224 | Unspecified |
| mmu-miR-669n | MIMAT0009427 | Pura | 19290 | HITS-CLIP | 37986224\|23142080 | | Unspecified |
| mmu-miR-669n | MIMAT0009427 | Purb | 19291 | HITS-CLIP | | 37986224 | Unspecified |
| mmu-miR-669n | MIMAT0009427 | Pvr | 52118 | HITS-CLIP | | 37986224 | Unspecified |
| mmu-miR-669n | MIMAT0009427 | Qser1 | 99003 | HITS-CLIP | 37986224\|23142080 | | Unspecified |
| mmu-miR-669n | MIMAT0009427 | Rab11a | 53869 | HITS-CLIP | | 37986224 | Unspecified |
| mmu-miR-669n | MIMAT0009427 | Rab11fip2 | 74998 | HITS-CLIP | | 23142080 | Unspecified |
| mmu-miR-669n | MIMAT0009427 | Rab12 | 19328 | HITS-CLIP | | 23142080 | Unspecified |
| mmu-miR-669n | MIMAT0009427 | Rab14 | 68365 | HITS-CLIP | | 37986224 | Unspecified |
| mmu-miR-669n | MIMAT0009427 | Rab33b | 19338 | HITS-CLIP | | 37986224 | Unspecified |
| mmu-miR-669n | MIMAT0009427 | Rab6a | 19346 | HITS-CLIP | | 23142080 | Unspecified |
| mmu-miR-669n | MIMAT0009427 | Rab7 | 19349 | HITS-CLIP | | 23142080 | Unspecified |
| mmu-miR-669n | MIMAT0009427 | Rad21 | 19357 | HITS-CLIP | 37986224\|23142080 | | Unspecified |
| mmu-miR-669n | MIMAT0009427 | Rad50 | 19360 | HITS-CLIP | | 23142080 | Unspecified |
| mmu-miR-669n | MIMAT0009427 | Rapgef2 | 76089 | HITS-CLIP | | 23142080 | Unspecified |
| mmu-miR-669n | MIMAT0009427 | Rapgef6 | 192786 | HITS-CLIP | | 23142080 | Unspecified |
| mmu-miR-669n | MIMAT0009427 | Raph1 | 77300 | HITS-CLIP | | 23142080 | Unspecified |
| mmu-miR-669n | MIMAT0009427 | Rasgrp1 | 19419 | HITS-CLIP | 37986224\|23142080 | | Unspecified |
| mmu-miR-669n | MIMAT0009427 | Rb1cc1 | 12421 | HITS-CLIP | | 23142080 | Unspecified |
| mmu-miR-669n | MIMAT0009427 | Rbbp4 | 19646 | HITS-CLIP | 37986224\|23142080 | | Unspecified |
| mmu-miR-669n | MIMAT0009427 | Rbbp6 | 19647 | HITS-CLIP | | 23142080 | Unspecified |
| mmu-miR-669n | MIMAT0009427 | Rbm26 | 74213 | HITS-CLIP | | 23142080 | Unspecified |
| mmu-miR-669n | MIMAT0009427 | Rbm3 | 19652 | HITS-CLIP | | 23142080 | Unspecified |
| mmu-miR-669n | MIMAT0009427 | Rbm7 | 67010 | HITS-CLIP | | 23142080 | Unspecified |
| mmu-miR-669n | MIMAT0009427 | Rc3h1 | 381305 | HITS-CLIP | | 37986224 | Unspecified |
| mmu-miR-669n | MIMAT0009427 | Rc3h2 | 319817 | HITS-CLIP | | 23142080 | Unspecified |
| mmu-miR-669n | MIMAT0009427 | Rcor1 | 217864 | HITS-CLIP | 37986224\|23142080 | | Unspecified |
| mmu-miR-669n | MIMAT0009427 | Rdx | 19684 | HITS-CLIP | | 37986224 | Unspecified |
| mmu-miR-669n | MIMAT0009427 | Rest | 19712 | HITS-CLIP | 37986224\|23142080 | | Unspecified |

| | | | | | | | |
|---|---|---|---|---|---|---|---|
| mmu-miR-669n | MIMAT0009427 | Rev3l | 19714 | HITS-CLIP | | 23142080 | Unspecified |
| mmu-miR-669n | MIMAT0009427 | Rfx3 | 19726 | HITS-CLIP | 37986224\|23142080 | | Unspecified |
| mmu-miR-669n | MIMAT0009427 | Rfx7 | 319758 | HITS-CLIP | | 23142080 | Unspecified |
| mmu-miR-669n | MIMAT0009427 | Rgs1 | 50778 | HITS-CLIP | 37986224\|23142080 | | Unspecified |
| mmu-miR-669n | MIMAT0009427 | Rhoq | 104215 | HITS-CLIP | 37986224\|23142080 | | Unspecified |
| mmu-miR-669n | MIMAT0009427 | Ric8b | 237422 | HITS-CLIP | | 23142080 | Unspecified |
| mmu-miR-669n | MIMAT0009427 | Rictor | 78757 | HITS-CLIP | 37986224\|23142080 | | Unspecified |
| mmu-miR-669n | MIMAT0009427 | Ripor2 | 193385 | HITS-CLIP | | 23142080 | Unspecified |
| mmu-miR-669n | MIMAT0009427 | Rnf111 | 93836 | HITS-CLIP | | 23142080 | Unspecified |
| mmu-miR-669n | MIMAT0009427 | Rnf181 | 66510 | HITS-CLIP | | 23142080 | Unspecified |
| mmu-miR-669n | MIMAT0009427 | Rnf19a | 30945 | HITS-CLIP | | 37986224 | Unspecified |
| mmu-miR-669n | MIMAT0009427 | Rnf2 | 19821 | HITS-CLIP | | 23142080 | Unspecified |
| mmu-miR-669n | MIMAT0009427 | Rnf214 | 235315 | HITS-CLIP | | 23142080 | Unspecified |
| mmu-miR-669n | MIMAT0009427 | Rnf220 | 66743 | HITS-CLIP | | 23142080 | Unspecified |
| mmu-miR-669n | MIMAT0009427 | Rnf34 | 80751 | HITS-CLIP | | 23142080 | Unspecified |
| mmu-miR-669n | MIMAT0009427 | Rnf38 | 73469 | HITS-CLIP | | 23142080 | Unspecified |
| mmu-miR-669n | MIMAT0009427 | Rnf6 | 74132 | HITS-CLIP | | 23142080 | Unspecified |
| mmu-miR-669n | MIMAT0009427 | Rngtt | 24018 | HITS-CLIP | | 23142080 | Unspecified |
| mmu-miR-669n | MIMAT0009427 | Rock1 | 19877 | HITS-CLIP | | 37986224 | Unspecified |
| mmu-miR-669n | MIMAT0009427 | Rpa2 | 19891 | HITS-CLIP | | 23142080 | Unspecified |
| mmu-miR-669n | MIMAT0009427 | Rpap2 | 231571 | HITS-CLIP | | 23142080 | Unspecified |
| mmu-miR-669n | MIMAT0009427 | Rpl32 | 19951 | HITS-CLIP | | 37986224 | Unspecified |
| mmu-miR-669n | MIMAT0009427 | Rps26 | 27370 | HITS-CLIP | | 31433990 | Unspecified |
| mmu-miR-669n | MIMAT0009427 | Rsf1 | 233532 | HITS-CLIP | | 23142080 | Unspecified |
| mmu-miR-669n | MIMAT0009427 | Rsrp1 | 27981 | HITS-CLIP | | 23142080 | Unspecified |
| mmu-miR-669n | MIMAT0009427 | Rufy2 | 70432 | HITS-CLIP | | 37986224 | Unspecified |
| mmu-miR-669n | MIMAT0009427 | Runx1 | 12394 | HITS-CLIP | | 37986224 | Unspecified |
| mmu-miR-669n | MIMAT0009427 | Sacs | 50720 | HITS-CLIP | 37986224\|23142080 | | Unspecified |
| mmu-miR-669n | MIMAT0009427 | Samhd1 | 56045 | HITS-CLIP | | 37986224 | Unspecified |
| mmu-miR-669n | MIMAT0009427 | Samsn1 | 67742 | HITS-CLIP | | 23142080 | Unspecified |
| mmu-miR-669n | MIMAT0009427 | Sav1 | 64010 | HITS-CLIP | 37986224\|23142080 | | Unspecified |
| mmu-miR-669n | MIMAT0009427 | Sbf2 | 319934 | HITS-CLIP | | 23142080 | Unspecified |
| mmu-miR-669n | MIMAT0009427 | Scaf4 | 224432 | HITS-CLIP | | 23142080 | Unspecified |
| mmu-miR-669n | MIMAT0009427 | Scd2 | 20250 | HITS-CLIP | | 37986224 | Unspecified |
| mmu-miR-669n | MIMAT0009427 | Sec13 | 110379 | HITS-CLIP | | 23142080 | Unspecified |
| mmu-miR-669n | MIMAT0009427 | Sec16a | 227648 | HITS-CLIP | | 37986224 | Unspecified |
| mmu-miR-669n | MIMAT0009427 | Sec22b | 20333 | HITS-CLIP | | 23142080 | Unspecified |
| mmu-miR-669n | MIMAT0009427 | Sec24a | 77371 | HITS-CLIP | | 37986224 | Unspecified |
| mmu-miR-669n | MIMAT0009427 | Selenof | 9403 | HITS-CLIP | | 23142080 | Unspecified |

| | | | | | | | |
|---|---|---|---|---|---|---|---|
| mmu-miR-669n | MIMAT0009427 | Sell | 20343 | HITS-CLIP | | 23142080 | Unspecified |
| mmu-miR-669n | MIMAT0009427 | Sepsecs | 211006 | HITS-CLIP | | 23142080 | Unspecified |
| mmu-miR-669n | MIMAT0009427 | Serbp1 | 66870 | HITS-CLIP | | 23142080 | Unspecified |
| mmu-miR-669n | MIMAT0009427 | Serinc1 | 56442 | HITS-CLIP | | 37986224 | Unspecified |
| mmu-miR-669n | MIMAT0009427 | Serinc3 | 26943 | HITS-CLIP | | 37986224 | Unspecified |
| mmu-miR-669n | MIMAT0009427 | Sertad2 | 58172 | HITS-CLIP | | 23142080 | Unspecified |
| mmu-miR-669n | MIMAT0009427 | Sesn1 | 140742 | HITS-CLIP | | 37986224 | Unspecified |
| mmu-miR-669n | MIMAT0009427 | Sesn3 | 75747 | HITS-CLIP | | 37986224 | Unspecified |
| mmu-miR-669n | MIMAT0009427 | Setbp1 | 240427 | HITS-CLIP | | 23142080 | Unspecified |
| mmu-miR-669n | MIMAT0009427 | Setd2 | 235626 | HITS-CLIP | | 23142080 | Unspecified |
| mmu-miR-669n | MIMAT0009427 | Setd3 | 52690 | HITS-CLIP | | 37986224 | Unspecified |
| mmu-miR-669n | MIMAT0009427 | Setx | 269254 | HITS-CLIP | | 37986224 | Unspecified |
| mmu-miR-669n | MIMAT0009427 | Sf3a3 | 75062 | HITS-CLIP | | 23142080 | Unspecified |
| mmu-miR-669n | MIMAT0009427 | Sfpq | 71514 | HITS-CLIP | | 37986224 | Unspecified |
| mmu-miR-669n | MIMAT0009427 | Sgk1 | 20393 | HITS-CLIP | | 37986224 | Unspecified |
| mmu-miR-669n | MIMAT0009427 | Sh3bp5 | 24056 | HITS-CLIP | | 23142080 | Unspecified |
| mmu-miR-669n | MIMAT0009427 | Shprh | 268281 | HITS-CLIP | | 23142080 | Unspecified |
| mmu-miR-669n | MIMAT0009427 | Sik2 | 235344 | HITS-CLIP | | 37986224 | Unspecified |
| mmu-miR-669n | MIMAT0009427 | Sirt1 | 93759 | HITS-CLIP | | 37986224 | Unspecified |
| mmu-miR-669n | MIMAT0009427 | Skil | 20482 | HITS-CLIP | 37986224\|23142080 | | Unspecified |
| mmu-miR-669n | MIMAT0009427 | Sla | 20491 | HITS-CLIP | | 37986224 | Unspecified |
| mmu-miR-669n | MIMAT0009427 | Slc16a3 | 80879 | HITS-CLIP | | 23142080 | Unspecified |
| mmu-miR-669n | MIMAT0009427 | Slc16a6 | 104681 | HITS-CLIP | | 23142080 | Unspecified |
| mmu-miR-669n | MIMAT0009427 | Slc17a6 | 140919 | HITS-CLIP | 37986224\|23142080 | | Unspecified |
| mmu-miR-669n | MIMAT0009427 | Slc25a46 | 67453 | HITS-CLIP | | 23142080 | Unspecified |
| mmu-miR-669n | MIMAT0009427 | Slc2a1 | 20525 | HITS-CLIP | | 37986224 | Unspecified |
| mmu-miR-669n | MIMAT0009427 | Slc2a3 | 20527 | HITS-CLIP | 37986224\|23142080 | | Unspecified |
| mmu-miR-669n | MIMAT0009427 | Slc35b1 | 110172 | HITS-CLIP | | 37986224 | Unspecified |
| mmu-miR-669n | MIMAT0009427 | Slc38a1 | 105727 | HITS-CLIP | | 23142080 | Unspecified |
| mmu-miR-669n | MIMAT0009427 | Slc41a1 | 98396 | HITS-CLIP | | 37986224 | Unspecified |
| mmu-miR-669n | MIMAT0009427 | Slc4a7 | 218756 | HITS-CLIP | | 37986224 | Unspecified |
| mmu-miR-669n | MIMAT0009427 | Slc5a3 | 53881 | HITS-CLIP | | 37986224 | Unspecified |
| mmu-miR-669n | MIMAT0009427 | Smad3 | 17127 | HITS-CLIP | | 23142080 | Unspecified |
| mmu-miR-669n | MIMAT0009427 | Smarcc1 | 20588 | HITS-CLIP | | 23142080 | Unspecified |
| mmu-miR-669n | MIMAT0009427 | Smarcc2 | 68094 | HITS-CLIP | | 23142080 | Unspecified |
| mmu-miR-669n | MIMAT0009427 | Smc3 | 13006 | HITS-CLIP | | 37986224 | Unspecified |
| mmu-miR-669n | MIMAT0009427 | Smc4 | 70099 | HITS-CLIP | | 37986224 | Unspecified |
| mmu-miR-669n | MIMAT0009427 | Smchd1 | 74355 | HITS-CLIP | 37986224\|23142080 | | Unspecified |
| mmu-miR-669n | MIMAT0009427 | Snrnp200 | 320632 | HITS-CLIP | | 23142080 | Unspecified |

| | | | | | | |
|---|---|---|---|---|---|---|
| mmu-miR-669n | MIMAT0009427 | Snw1 | 66354 | HITS-CLIP | 23142080 | Unspecified |
| mmu-miR-669n | MIMAT0009427 | Snx12 | 55988 | HITS-CLIP | 23142080 | Unspecified |
| mmu-miR-669n | MIMAT0009427 | Snx13 | 217463 | HITS-CLIP | 23142080 | Unspecified |
| mmu-miR-669n | MIMAT0009427 | Snx27 | 76742 | HITS-CLIP | 23142080 | Unspecified |
| mmu-miR-669n | MIMAT0009427 | Socs3 | 12702 | HITS-CLIP | 23142080 | Unspecified |
| mmu-miR-669n | MIMAT0009427 | Son | 20658 | HITS-CLIP | 23142080 | Unspecified |
| mmu-miR-669n | MIMAT0009427 | Sp3 | 20687 | HITS-CLIP | 37986224\|23142080 | Unspecified |
| mmu-miR-669n | MIMAT0009427 | Spag9 | 70834 | HITS-CLIP | 37986224\|23142080 | Unspecified |
| mmu-miR-669n | MIMAT0009427 | Spcs2 | 66624 | HITS-CLIP | 23142080 | Unspecified |
| mmu-miR-669n | MIMAT0009427 | Sphk2 | 56632 | HITS-CLIP | 23142080 | Unspecified |
| mmu-miR-669n | MIMAT0009427 | Spn | 20737 | HITS-CLIP | 37986224 | Unspecified |
| mmu-miR-669n | MIMAT0009427 | Spock2 | 94214 | HITS-CLIP | 23142080 | Unspecified |
| mmu-miR-669n | MIMAT0009427 | Spry1 | 24063 | HITS-CLIP | 23142080 | Unspecified |
| mmu-miR-669n | MIMAT0009427 | Sptbn1 | 20742 | HITS-CLIP | 37986224\|23142080 | Unspecified |
| mmu-miR-669n | MIMAT0009427 | Spty2d1 | 101685 | HITS-CLIP | 23142080 | Unspecified |
| mmu-miR-669n | MIMAT0009427 | Srbd1 | 78586 | HITS-CLIP | 23142080 | Unspecified |
| mmu-miR-669n | MIMAT0009427 | Srebf2 | 20788 | HITS-CLIP | 23142080 | Unspecified |
| mmu-miR-669n | MIMAT0009427 | Srp68 | 217337 | HITS-CLIP | 23142080 | Unspecified |
| mmu-miR-669n | MIMAT0009427 | Srpk1 | 20815 | HITS-CLIP | 23142080 | Unspecified |
| mmu-miR-669n | MIMAT0009427 | Ssbp4 | 76900 | HITS-CLIP | 23142080 | Unspecified |
| mmu-miR-669n | MIMAT0009427 | Stag2 | 20843 | HITS-CLIP | 23142080 | Unspecified |
| mmu-miR-669n | MIMAT0009427 | Stat1 | 20846 | HITS-CLIP | 37986224\|23142080 | Unspecified |
| mmu-miR-669n | MIMAT0009427 | Stat3 | 20848 | HITS-CLIP | 37986224\|23142080 | Unspecified |
| mmu-miR-669n | MIMAT0009427 | Stat4 | 20849 | HITS-CLIP | 23142080 | Unspecified |
| mmu-miR-669n | MIMAT0009427 | Stat5b | 20851 | HITS-CLIP | 37986224 | Unspecified |
| mmu-miR-669n | MIMAT0009427 | Stil | 20460 | HITS-CLIP | 23142080 | Unspecified |
| mmu-miR-669n | MIMAT0009427 | Stk24 | 223255 | HITS-CLIP | 23142080 | Unspecified |
| mmu-miR-669n | MIMAT0009427 | Stk26 | 51765 | HITS-CLIP | 23142080 | Unspecified |
| mmu-miR-669n | MIMAT0009427 | Stk38 | 106504 | HITS-CLIP | 23142080 | Unspecified |
| mmu-miR-669n | MIMAT0009427 | Strn3 | 94186 | HITS-CLIP | 37986224\|23142080 | Unspecified |
| mmu-miR-669n | MIMAT0009427 | Stx16 | 228960 | HITS-CLIP | 23142080 | Unspecified |
| mmu-miR-669n | MIMAT0009427 | Sun2 | 223697 | HITS-CLIP | 37986224\|23142080 | Unspecified |
| mmu-miR-669n | MIMAT0009427 | Surf2 | 20931 | HITS-CLIP | 37986224 | Unspecified |
| mmu-miR-669n | MIMAT0009427 | Suz12 | 52615 | HITS-CLIP | 23142080 | Unspecified |
| mmu-miR-669n | MIMAT0009427 | Syt11 | 229521 | HITS-CLIP | 23142080 | Unspecified |
| mmu-miR-669n | MIMAT0009427 | Tab2 | 68652 | HITS-CLIP | 23142080 | Unspecified |
| mmu-miR-669n | MIMAT0009427 | Taf1 | 270627 | HITS-CLIP | 23142080 | Unspecified |
| mmu-miR-669n | MIMAT0009427 | Taf15 | 70439 | HITS-CLIP | 23142080 | Unspecified |
| mmu-miR-669n | MIMAT0009427 | Taok1 | 216965 | HITS-CLIP | 37986224\|23142080 | Unspecified |

| | | | | | | |
|---|---|---|---|---|---|---|
| mmu-miR-669n | MIMAT0009427 | Taok2 | 381921 | HITS-CLIP | 23142080 | Unspecified |
| mmu-miR-669n | MIMAT0009427 | Tap1 | 21354 | HITS-CLIP | 23142080 | Unspecified |
| mmu-miR-669n | MIMAT0009427 | Tbc1d22b | 381085 | HITS-CLIP | 37986224 | Unspecified |
| mmu-miR-669n | MIMAT0009427 | Tbc1d23 | 67581 | HITS-CLIP | 23142080 | Unspecified |
| mmu-miR-669n | MIMAT0009427 | Tbl1xr1 | 81004 | HITS-CLIP | 37986224\|23142080 | Unspecified |
| mmu-miR-669n | MIMAT0009427 | Tcf12 | 21406 | HITS-CLIP | 23142080 | Unspecified |
| mmu-miR-669n | MIMAT0009427 | Tes | 21753 | HITS-CLIP | 23142080 | Unspecified |
| mmu-miR-669n | MIMAT0009427 | Tet3 | 194388 | HITS-CLIP | 37986224 | Unspecified |
| mmu-miR-669n | MIMAT0009427 | Tex30 | 75623 | HITS-CLIP | 23142080 | Unspecified |
| mmu-miR-669n | MIMAT0009427 | Tfg | 21787 | HITS-CLIP | 37986224 | Unspecified |
| mmu-miR-669n | MIMAT0009427 | Tgfbr2 | 21813 | HITS-CLIP | 23142080 | Unspecified |
| mmu-miR-669n | MIMAT0009427 | Thada | 240174 | HITS-CLIP | 23142080 | Unspecified |
| mmu-miR-669n | MIMAT0009427 | Thoc2 | 331401 | HITS-CLIP | 23142080 | Unspecified |
| mmu-miR-669n | MIMAT0009427 | Tiam1 | 21844 | HITS-CLIP | 37986224\|23142080 | Unspecified |
| mmu-miR-669n | MIMAT0009427 | Ticrr | 77011 | HITS-CLIP | 23142080 | Unspecified |
| mmu-miR-669n | MIMAT0009427 | Timm10 | 30059 | HITS-CLIP | 37986224 | Unspecified |
| mmu-miR-669n | MIMAT0009427 | Tiparp | 99929 | HITS-CLIP | 37986224 | Unspecified |
| mmu-miR-669n | MIMAT0009427 | Tlk1 | 228012 | HITS-CLIP | 23142080 | Unspecified |
| mmu-miR-669n | MIMAT0009427 | Tln1 | 21894 | HITS-CLIP | 37986224\|23142080 | Unspecified |
| mmu-miR-669n | MIMAT0009427 | Tmed5 | 73130 | HITS-CLIP | 37986224 | Unspecified |
| mmu-miR-669n | MIMAT0009427 | Tmed7 | 66676 | HITS-CLIP | 37986224\|23142080 | Unspecified |
| mmu-miR-669n | MIMAT0009427 | Tmem106b | 71900 | HITS-CLIP | 23142080 | Unspecified |
| mmu-miR-669n | MIMAT0009427 | Tmem131 | 56030 | HITS-CLIP | 23142080 | Unspecified |
| mmu-miR-669n | MIMAT0009427 | Tmem170 | 66817 | HITS-CLIP | 37986224 | Unspecified |
| mmu-miR-669n | MIMAT0009427 | Tmem170b | 621976 | HITS-CLIP | 37986224\|23142080 | Unspecified |
| mmu-miR-669n | MIMAT0009427 | Tmem245 | 242474 | HITS-CLIP | 37986224\|23142080 | Unspecified |
| mmu-miR-669n | MIMAT0009427 | Tmem38b | 52076 | HITS-CLIP | 37986224 | Unspecified |
| mmu-miR-669n | MIMAT0009427 | Tmem64 | 100201 | HITS-CLIP | 37986224 | Unspecified |
| mmu-miR-669n | MIMAT0009427 | Tmem70 | 70397 | HITS-CLIP | 23142080 | Unspecified |
| mmu-miR-669n | MIMAT0009427 | Tmem87b | 72477 | HITS-CLIP | 37986224 | Unspecified |
| mmu-miR-669n | MIMAT0009427 | Tmf1 | 232286 | HITS-CLIP | 37986224 | Unspecified |
| mmu-miR-669n | MIMAT0009427 | Tmsb10 | 19240 | HITS-CLIP | 23142080 | Unspecified |
| mmu-miR-669n | MIMAT0009427 | Tmx1 | 72736 | HITS-CLIP | 37986224 | Unspecified |
| mmu-miR-669n | MIMAT0009427 | Tmx3 | 67988 | HITS-CLIP | 23142080 | Unspecified |
| mmu-miR-669n | MIMAT0009427 | Tnfrsf1a | 21937 | HITS-CLIP | 37986224 | Unspecified |
| mmu-miR-669n | MIMAT0009427 | Tnfrsf4 | 22163 | HITS-CLIP | 23142080 | Unspecified |
| mmu-miR-669n | MIMAT0009427 | Tnks2 | 74493 | HITS-CLIP | 23142080 | Unspecified |
| mmu-miR-669n | MIMAT0009427 | Tnpo1 | 238799 | HITS-CLIP | 37986224 | Unspecified |
| mmu-miR-669n | MIMAT0009427 | Tnpo3 | 320938 | HITS-CLIP | 23142080 | Unspecified |

| | | | | | | |
|---|---|---|---|---|---|---|
| mmu-miR-669n | MIMAT0009427 | Tnrc6a | 233833 | HITS-CLIP | 37986224\|23142080 | Unspecified |
| mmu-miR-669n | MIMAT0009427 | Tnrc6b | 213988 | HITS-CLIP | 37986224\|23142080 | Unspecified |
| mmu-miR-669n | MIMAT0009427 | Tnrc6c | 217351 | HITS-CLIP | 23142080 | Unspecified |
| mmu-miR-669n | MIMAT0009427 | Top1 | 21969 | HITS-CLIP | 23142080 | Unspecified |
| mmu-miR-669n | MIMAT0009427 | Topors | 106021 | HITS-CLIP | 37986224 | Unspecified |
| mmu-miR-669n | MIMAT0009427 | Tpk1 | 29807 | HITS-CLIP | 37986224 | Unspecified |
| mmu-miR-669n | MIMAT0009427 | Traf2 | 22030 | HITS-CLIP | 23142080 | Unspecified |
| mmu-miR-669n | MIMAT0009427 | Trak2 | 70827 | HITS-CLIP | 37986224 | Unspecified |
| mmu-miR-669n | MIMAT0009427 | Tram1 | 72265 | HITS-CLIP | 23142080 | Unspecified |
| mmu-miR-669n | MIMAT0009427 | Trib2 | 217410 | HITS-CLIP | 37986224 | Unspecified |
| mmu-miR-669n | MIMAT0009427 | Trim33 | 94093 | HITS-CLIP | 23142080 | Unspecified |
| mmu-miR-669n | MIMAT0009427 | Trip11 | 109181 | HITS-CLIP | 37986224 | Unspecified |
| mmu-miR-669n | MIMAT0009427 | Trip12 | 14897 | HITS-CLIP | 23142080 | Unspecified |
| mmu-miR-669n | MIMAT0009427 | Trp53 | 22059 | HITS-CLIP | 23142080 | Unspecified |
| mmu-miR-669n | MIMAT0009427 | Trp53inp1 | 60599 | HITS-CLIP | 23142080 | Unspecified |
| mmu-miR-669n | MIMAT0009427 | Tshz1 | 110796 | HITS-CLIP | 37986224 | Unspecified |
| mmu-miR-669n | MIMAT0009427 | Tspan5 | 56224 | HITS-CLIP | 23142080 | Unspecified |
| mmu-miR-669n | MIMAT0009427 | Ttc8 | 76260 | HITS-CLIP | 37986224 | Unspecified |
| mmu-miR-669n | MIMAT0009427 | Txlna | 109658 | HITS-CLIP | 23142080 | Unspecified |
| mmu-miR-669n | MIMAT0009427 | Txnip | 56338 | HITS-CLIP | 37986224 | Unspecified |
| mmu-miR-669n | MIMAT0009427 | Ube2d1 | 216080 | HITS-CLIP | 23142080 | Unspecified |
| mmu-miR-669n | MIMAT0009427 | Ube2g2 | 22213 | HITS-CLIP | 23142080 | Unspecified |
| mmu-miR-669n | MIMAT0009427 | Ube2j1 | 56228 | HITS-CLIP | 23142080 | Unspecified |
| mmu-miR-669n | MIMAT0009427 | Ube3a | 22215 | HITS-CLIP | 23142080 | Unspecified |
| mmu-miR-669n | MIMAT0009427 | Ube3c | 100763 | HITS-CLIP | 37986224 | Unspecified |
| mmu-miR-669n | MIMAT0009427 | Ubl3 | 24109 | HITS-CLIP | 23142080 | Unspecified |
| mmu-miR-669n | MIMAT0009427 | Ubqln2 | 54609 | HITS-CLIP | 23142080 | Unspecified |
| mmu-miR-669n | MIMAT0009427 | Ubr4 | 69116 | HITS-CLIP | 23142080 | Unspecified |
| mmu-miR-669n | MIMAT0009427 | Ubxn7 | 224111 | HITS-CLIP | 23142080 | Unspecified |
| mmu-miR-669n | MIMAT0009427 | Uhrf2 | 109113 | HITS-CLIP | 23142080 | Unspecified |
| mmu-miR-669n | MIMAT0009427 | Uqcr10 | 66152 | HITS-CLIP | 23142080 | Unspecified |
| mmu-miR-669n | MIMAT0009427 | Usp15 | 14479 | HITS-CLIP | 37986224\|23142080 | Unspecified |
| mmu-miR-669n | MIMAT0009427 | Usp31 | 76179 | HITS-CLIP | 23142080 | Unspecified |
| mmu-miR-669n | MIMAT0009427 | Usp34 | 17847 | HITS-CLIP | 37986224 | Unspecified |
| mmu-miR-669n | MIMAT0009427 | Usp45 | 77593 | HITS-CLIP | 23142080 | Unspecified |
| mmu-miR-669n | MIMAT0009427 | Usp8 | 84092 | HITS-CLIP | 23142080 | Unspecified |
| mmu-miR-669n | MIMAT0009427 | Uspl1 | 231915 | HITS-CLIP | 37986224 | Unspecified |
| mmu-miR-669n | MIMAT0009427 | Utp11 | 51118 | HITS-CLIP | 23142080 | Unspecified |
| mmu-miR-669n | MIMAT0009427 | Utrn | 22288 | HITS-CLIP | 37986224\|23142080 | Unspecified |

| | | | | | | | |
|---|---|---|---|---|---|---|---|
| mmu-miR-669n | MIMAT0009427 | Vapa | 30960 | HITS-CLIP | | 37986224\|23142080 | Unspecified |
| mmu-miR-669n | MIMAT0009427 | Vapb | 56491 | HITS-CLIP | | 23142080 | Unspecified |
| mmu-miR-669n | MIMAT0009427 | Vezf1 | 22344 | HITS-CLIP | | 23142080 | Unspecified |
| mmu-miR-669n | MIMAT0009427 | Vps29 | 56433 | HITS-CLIP | | 23142080 | Unspecified |
| mmu-miR-669n | MIMAT0009427 | Vps39 | 269338 | HITS-CLIP | | 23142080 | Unspecified |
| mmu-miR-669n | MIMAT0009427 | Vps41 | 218035 | HITS-CLIP | | 31433990 | Unspecified |
| mmu-miR-669n | MIMAT0009427 | Vps54 | 245944 | HITS-CLIP | | 23142080 | Unspecified |
| mmu-miR-669n | MIMAT0009427 | Wac | 225131 | HITS-CLIP | | 37986224 | Unspecified |
| mmu-miR-669n | MIMAT0009427 | Wasl | 73178 | HITS-CLIP | | 37986224\|23142080 | Unspecified |
| mmu-miR-669n | MIMAT0009427 | Wdhd1 | 218973 | HITS-CLIP | | 23142080 | Unspecified |
| mmu-miR-669n | MIMAT0009427 | Wdr1 | 22388 | HITS-CLIP | | 37986224 | Unspecified |
| mmu-miR-669n | MIMAT0009427 | Wdr26 | 226757 | HITS-CLIP | | 37986224 | Unspecified |
| mmu-miR-669n | MIMAT0009427 | Wdr3 | 269470 | HITS-CLIP | | 23142080 | Unspecified |
| mmu-miR-669n | MIMAT0009427 | Wnk1 | 232341 | HITS-CLIP | | 37986224\|23142080 | Unspecified |
| mmu-miR-669n | MIMAT0009427 | Xiap | 11798 | HITS-CLIP | | 23142080 | Unspecified |
| mmu-miR-669n | MIMAT0009427 | Xpo4 | 57258 | HITS-CLIP | | 37986224 | Unspecified |
| mmu-miR-669n | MIMAT0009427 | Xpr1 | 19775 | HITS-CLIP | | 23142080 | Unspecified |
| mmu-miR-669n | MIMAT0009427 | Yme1l1 | 27377 | HITS-CLIP | | 23142080 | Unspecified |
| mmu-miR-669n | MIMAT0009427 | Yod1 | 226418 | HITS-CLIP | | 23142080 | Unspecified |
| mmu-miR-669n | MIMAT0009427 | Ythdf3 | 229096 | HITS-CLIP | | 23142080 | Unspecified |
| mmu-miR-669n | MIMAT0009427 | Zadh2 | 225791 | HITS-CLIP | | 37986224 | Unspecified |
| mmu-miR-669n | MIMAT0009427 | Zbtb14 | 22666 | HITS-CLIP | | 23142080 | Unspecified |
| mmu-miR-669n | MIMAT0009427 | Zbtb25 | 109929 | HITS-CLIP | | 23142080 | Unspecified |
| mmu-miR-669n | MIMAT0009427 | Zbtb33 | 56805 | HITS-CLIP | | 23142080 | Unspecified |
| mmu-miR-669n | MIMAT0009427 | Zbtb34 | 241311 | HITS-CLIP | | 23142080 | Unspecified |
| mmu-miR-669n | MIMAT0009427 | Zbtb38 | 245007 | HITS-CLIP | | 37986224 | Unspecified |
| mmu-miR-669n | MIMAT0009427 | Zbtb7a | 16969 | HITS-CLIP | | 37986224 | Unspecified |
| mmu-miR-669n | MIMAT0009427 | Zc3h12d | 237256 | HITS-CLIP | | 23142080 | Unspecified |
| mmu-miR-669n | MIMAT0009427 | Zc3h14 | 75553 | HITS-CLIP | | 23142080 | Unspecified |
| mmu-miR-669n | MIMAT0009427 | Zc3hav1 | 78781 | HITS-CLIP | | 37986224 | Unspecified |
| mmu-miR-669n | MIMAT0009427 | Zcchc9 | 69085 | HITS-CLIP | | 23142080 | Unspecified |
| mmu-miR-669n | MIMAT0009427 | Zdhhc17 | 320150 | HITS-CLIP | | 23142080 | Unspecified |
| mmu-miR-669n | MIMAT0009427 | Zdhhc21 | 68268 | HITS-CLIP | | 37986224 | Unspecified |
| mmu-miR-669n | MIMAT0009427 | Zer1 | 227693 | HITS-CLIP | | 37986224 | Unspecified |
| mmu-miR-669n | MIMAT0009427 | Zfp148 | 22661 | HITS-CLIP | | 23142080 | Unspecified |
| mmu-miR-669n | MIMAT0009427 | Zfp157 | 72154 | HITS-CLIP | | 23142080 | Unspecified |
| mmu-miR-669n | MIMAT0009427 | Zfp26 | 22688 | HITS-CLIP | | 37986224 | Unspecified |
| mmu-miR-669n | MIMAT0009427 | Zfp260 | 26466 | HITS-CLIP | | 37986224\|23142080 | Unspecified |
| mmu-miR-669n | MIMAT0009427 | Zfp266 | 77519 | HITS-CLIP | | 37986224 | Unspecified |

| | | | | | | |
|---|---|---|---|---|---|---|
| mmu-miR-669n | MIMAT0009427 | Zfp280b | 64453 | HITS-CLIP | 37986224 | Unspecified |
| mmu-miR-669n | MIMAT0009427 | Zfp281 | 226442 | HITS-CLIP | 37986224\|23142080 | Unspecified |
| mmu-miR-669n | MIMAT0009427 | Zfp36l1 | 12192 | HITS-CLIP | 37986224 | Unspecified |
| mmu-miR-669n | MIMAT0009427 | Zfp512b | 269401 | HITS-CLIP | 23142080 | Unspecified |
| mmu-miR-669n | MIMAT0009427 | Zfp518a | 72672 | HITS-CLIP | 37986224 | Unspecified |
| mmu-miR-669n | MIMAT0009427 | Zfp52 | 22710 | HITS-CLIP | 37986224 | Unspecified |
| mmu-miR-669n | MIMAT0009427 | Zfp54 | 22712 | HITS-CLIP | 37986224 | Unspecified |
| mmu-miR-669n | MIMAT0009427 | Zfp592 | 233410 | HITS-CLIP | 23142080 | Unspecified |
| mmu-miR-669n | MIMAT0009427 | Zfp617 | 170938 | HITS-CLIP | 37986224 | Unspecified |
| mmu-miR-669n | MIMAT0009427 | Zfp62 | 22720 | HITS-CLIP | 23142080 | Unspecified |
| mmu-miR-669n | MIMAT0009427 | Zfp644 | 52397 | HITS-CLIP | 23142080 | Unspecified |
| mmu-miR-669n | MIMAT0009427 | Zfp654 | 72020 | HITS-CLIP | 37986224\|23142080 | Unspecified |
| mmu-miR-669n | MIMAT0009427 | Zfp655 | 72611 | HITS-CLIP | 37986224 | Unspecified |
| mmu-miR-669n | MIMAT0009427 | Zfp719 | 210105 | HITS-CLIP | 23142080 | Unspecified |
| mmu-miR-669n | MIMAT0009427 | Zfp729b | 100416706 | HITS-CLIP | 23142080 | Unspecified |
| mmu-miR-669n | MIMAT0009427 | Zfp738 | 408068 | HITS-CLIP | 23142080 | Unspecified |
| mmu-miR-669n | MIMAT0009427 | Zfp763 | 73451 | HITS-CLIP | 23142080 | Unspecified |
| mmu-miR-669n | MIMAT0009427 | Zfp800 | 627049 | HITS-CLIP | 23142080 | Unspecified |
| mmu-miR-669n | MIMAT0009427 | Zfp831 | 100043757 | HITS-CLIP | 23142080 | Unspecified |
| mmu-miR-669n | MIMAT0009427 | Zfp85 | 22746 | HITS-CLIP | 23142080 | Unspecified |
| mmu-miR-669n | MIMAT0009427 | Zfp87 | 170763 | HITS-CLIP | 23142080 | Unspecified |
| mmu-miR-669n | MIMAT0009427 | Zfp871 | 208292 | HITS-CLIP | 23142080 | Unspecified |
| mmu-miR-669n | MIMAT0009427 | Zfp91 | 109910 | HITS-CLIP | 37986224 | Unspecified |
| mmu-miR-669n | MIMAT0009427 | Zfp943 | 74670 | HITS-CLIP | 23142080 | Unspecified |
| mmu-miR-669n | MIMAT0009427 | Zfp949 | 71640 | HITS-CLIP | 23142080 | Unspecified |
| mmu-miR-669n | MIMAT0009427 | Zfpm1 | 22761 | HITS-CLIP | 23142080 | Unspecified |
| mmu-miR-669n | MIMAT0009427 | Zhx2 | 387609 | HITS-CLIP | 23142080 | Unspecified |
| mmu-miR-669n | MIMAT0009427 | Zkscan3 | 72739 | HITS-CLIP | 23142080 | Unspecified |
| mmu-miR-669n | MIMAT0009427 | Zmym4 | 67785 | HITS-CLIP | 23142080 | Unspecified |
| mmu-miR-669n | MIMAT0009427 | Znhit6 | 229937 | HITS-CLIP | 23142080 | Unspecified |
| mmu-miR-669n | MIMAT0009427 | Znrf3 | 407821 | HITS-CLIP | 37986224\|23142080 | Unspecified |
| mmu-miR-669n | MIMAT0009427 | Zswim3 | 67538 | HITS-CLIP | 37986224 | Unspecified |
| mmu-miR-669n | MIMAT0009427 | Zswim6 | 67263 | HITS-CLIP | 37986224 | Unspecified |
| mmu-miR-669n | MIMAT0009427 | mt-Cytb | 19893556 | HITS-CLIP | 23142080 | Unspecified |
| mmu-miR-669n | MIMAT0009427 | mt-Nd2 | 4536 | HITS-CLIP | 23142080 | Unspecified |
| mmu-miR-592-5p | MIMAT0003730 | Bet1l | 54399 | HITS-CLIP | 31433990 | Unspecified |
| mmu-miR-592-5p | MIMAT0003730 | Brwd3 | 382236 | HITS-CLIP | 31433990 | Unspecified |
| mmu-miR-592-5p | MIMAT0003730 | E2f8 | 108961 | HITS-CLIP | 31433990 | Unspecified |
| mmu-miR-592-5p | MIMAT0003730 | Etaa1 | 68145 | HITS-CLIP | 31433990 | Unspecified |

| | | | | | |
|---|---|---|---|---|---|
| mmu-miR-592-5p | MIMAT0003730 | Ezh2 | 14056 | HITS-CLIP | 31433990 Unspecified |
| mmu-miR-592-5p | MIMAT0003730 | Galnt1 | 14423 | HITS-CLIP | 26203562 Unspecified |
| mmu-miR-592-5p | MIMAT0003730 | Gosr2 | 56494 | HITS-CLIP | 31433990 Unspecified |
| mmu-miR-592-5p | MIMAT0003730 | H2bc11 | 8970 | HITS-CLIP | 31433990 Unspecified |
| mmu-miR-592-5p | MIMAT0003730 | Hspa4 | 15525 | HITS-CLIP | 31433990 Unspecified |
| mmu-miR-592-5p | MIMAT0003730 | Ints7 | 77065 | HITS-CLIP | 31433990 Unspecified |
| mmu-miR-592-5p | MIMAT0003730 | Leprotl1 | 68192 | HITS-CLIP | 31433990 Unspecified |
| mmu-miR-592-5p | MIMAT0003730 | Mtfr1 | 67472 | HITS-CLIP | 31433990 Unspecified |
| mmu-miR-592-5p | MIMAT0003730 | Mxd1 | 17119 | HITS-CLIP | 26203562 Unspecified |
| mmu-miR-592-5p | MIMAT0003730 | Nrf1 | 18181 | HITS-CLIP | 31433990 Unspecified |
| mmu-miR-592-5p | MIMAT0003730 | Pcnx4 | 67708 | HITS-CLIP | 31433990 Unspecified |
| mmu-miR-592-5p | MIMAT0003730 | Ptpn23 | 104831 | HITS-CLIP | 31433990 Unspecified |
| mmu-miR-592-5p | MIMAT0003730 | Rdh11 | 17252 | HITS-CLIP | 26203562 Unspecified |
| mmu-miR-592-5p | MIMAT0003730 | Rxrb | 20182 | HITS-CLIP | 31433990 Unspecified |
| mmu-miR-592-5p | MIMAT0003730 | Slc30a7 | 66500 | HITS-CLIP | 31433990 Unspecified |
| mmu-miR-592-5p | MIMAT0003730 | Soat1 | 20652 | HITS-CLIP | 31433990 Unspecified |
| mmu-miR-592-5p | MIMAT0003730 | Sptbn1 | 20742 | HITS-CLIP | 31433990 Unspecified |
| mmu-miR-592-5p | MIMAT0003730 | Tnpo1 | 238799 | HITS-CLIP | 31433990 Unspecified |
| mmu-miR-592-5p | MIMAT0003730 | Top2a | 21973 | HITS-CLIP | 31433990 Unspecified |
| mmu-miR-592-5p | MIMAT0003730 | Trmt10a | 108943 | HITS-CLIP | 31433990 Unspecified |
| mmu-miR-592-5p | MIMAT0003730 | Ttll4 | 67534 | HITS-CLIP | 31433990 Unspecified |
| mmu-miR-592-5p | MIMAT0003730 | Usp34 | 17847 | HITS-CLIP | 31433990 Unspecified |
| mmu-miR-592-5p | MIMAT0003730 | Ywhab | 54401 | HITS-CLIP | 31433990 Unspecified |
| mmu-miR-3086-3p | MIMAT0014881 | Foxo3 | 56484 | HITS-CLIP | 31433990 Unspecified |
| mmu-miR-3086-3p | MIMAT0014881 | Nfil3 | 18030 | HITS-CLIP | 31433990 Unspecified |
| mmu-miR-3086-3p | MIMAT0014881 | Syt11 | 229521 | HITS-CLIP | 31433990 Unspecified |
| mmu-miR-674-5p | MIMAT0003740 | 1810013L24Rik | 69053 | HITS-CLIP | 29167373 Unspecified |
| mmu-miR-674-5p | MIMAT0003740 | 2210016F16Rik | 70153 | qCLASH | 31393953 Unspecified |
| mmu-miR-674-5p | MIMAT0003740 | 4931406P16Rik | 233103 | HITS-CLIP | 29167373 Unspecified |
| mmu-miR-674-5p | MIMAT0003740 | Abcd1 | 11666 | HITS-CLIP | 29167373 Unspecified |
| mmu-miR-674-5p | MIMAT0003740 | Abi1 | 11308 | HITS-CLIP | 29167373 Unspecified |
| mmu-miR-674-5p | MIMAT0003740 | Acsf2 | 264895 | qCLASH | 31393953 Unspecified |
| mmu-miR-674-5p | MIMAT0003740 | Adar | 56417 | HITS-CLIP | 29167373 Unspecified |
| mmu-miR-674-5p | MIMAT0003740 | Adcy7 | 11513 | HITS-CLIP | 29167373 Unspecified |
| mmu-miR-674-5p | MIMAT0003740 | Adnp | 11538 | HITS-CLIP | 25753418 Unspecified |
| mmu-miR-674-5p | MIMAT0003740 | Agps | 228061 | HITS-CLIP | 29167373 Unspecified |
| mmu-miR-674-5p | MIMAT0003740 | Ak1 | 11636 | HITS-CLIP | 25753418 Unspecified |
| mmu-miR-674-5p | MIMAT0003740 | Akt1 | 11651 | HITS-CLIP | 29167373 Unspecified |
| mmu-miR-674-5p | MIMAT0003740 | Anapc1 | 17222 | HITS-CLIP | 29167373 Unspecified |

| | | | | | | |
|---|---|---|---|---|---|---|
| mmu-miR-674-5p | MIMAT0003740 | Ankrd11 | 77087 | HITS-CLIP | 29167373 | Unspecified |
| mmu-miR-674-5p | MIMAT0003740 | Ankrd60 | 70065 | qCLASH | 31393953 | Unspecified |
| mmu-miR-674-5p | MIMAT0003740 | Anp32e | 66471 | qCLASH | 31393953 | Unspecified |
| mmu-miR-674-5p | MIMAT0003740 | Anxa5 | 11747 | HITS-CLIP | 29167373 | Unspecified |
| mmu-miR-674-5p | MIMAT0003740 | Ap2a1 | 11771 | HITS-CLIP | 25753418 | Unspecified |
| mmu-miR-674-5p | MIMAT0003740 | Appbp2 | 66884 | HITS-CLIP | 29167373 | Unspecified |
| mmu-miR-674-5p | MIMAT0003740 | Arhgap30 | 226652 | HITS-CLIP | 29167373 | Unspecified |
| mmu-miR-674-5p | MIMAT0003740 | Arrb2 | 216869 | HITS-CLIP | 29167373 | Unspecified |
| mmu-miR-674-5p | MIMAT0003740 | Atp5a1 | 11946 | HITS-CLIP | 25753418 | Unspecified |
| mmu-miR-674-5p | MIMAT0003740 | Bcl7c | 12055 | HITS-CLIP | 26203562 | Unspecified |
| mmu-miR-674-5p | MIMAT0003740 | Bclaf1 | 72567 | HITS-CLIP | 29167373 | Unspecified |
| mmu-miR-674-5p | MIMAT0003740 | Bmf | 171543 | HITS-CLIP | 29167373 | Unspecified |
| mmu-miR-674-5p | MIMAT0003740 | Bmt2 | 101148 | HITS-CLIP | 25753418 | Unspecified |
| mmu-miR-674-5p | MIMAT0003740 | Cacul1 | 78832 | qCLASH | 31393953 | Unspecified |
| mmu-miR-674-5p | MIMAT0003740 | Camsap2 | 67886 | HITS-CLIP | 25753418 | Unspecified |
| mmu-miR-674-5p | MIMAT0003740 | Cap1 | 12331 | HITS-CLIP | 29167373 | Unspecified |
| mmu-miR-674-5p | MIMAT0003740 | Carmil1 | 55604 | HITS-CLIP | 25753418 | Unspecified |
| mmu-miR-674-5p | MIMAT0003740 | Casd1 | 213819 | HITS-CLIP | 29167373 | Unspecified |
| mmu-miR-674-5p | MIMAT0003740 | Casp6 | 12368 | HITS-CLIP | 25753418 | Unspecified |
| mmu-miR-674-5p | MIMAT0003740 | Cbr4 | 234309 | HITS-CLIP | 25753418 | Unspecified |
| mmu-miR-674-5p | MIMAT0003740 | Ccdc186 | 55088 | HITS-CLIP | 29167373 | Unspecified |
| mmu-miR-674-5p | MIMAT0003740 | Ccdc88a | 108686 | HITS-CLIP | 29167373 | Unspecified |
| mmu-miR-674-5p | MIMAT0003740 | Ccnf | 12449 | qCLASH | 31393953 | Unspecified |
| mmu-miR-674-5p | MIMAT0003740 | Chmp2a | 68953 | qCLASH | 31393953 | Unspecified |
| mmu-miR-674-5p | MIMAT0003740 | Cisd2 | 67006 | HITS-CLIP | 25753418 | Unspecified |
| mmu-miR-674-5p | MIMAT0003740 | Clcf1 | 56708 | HITS-CLIP | 26203562 | Unspecified |
| mmu-miR-674-5p | MIMAT0003740 | Cltc | 67300 | HITS-CLIP | 29167373 | Unspecified |
| mmu-miR-674-5p | MIMAT0003740 | Cog8 | 97484 | HITS-CLIP | 37986224 | Unspecified |
| mmu-miR-674-5p | MIMAT0003740 | Copb1 | 70349 | HITS-CLIP | 29167373 | Unspecified |
| mmu-miR-674-5p | MIMAT0003740 | Copg1 | 54161 | HITS-CLIP | 29167373 | Unspecified |
| mmu-miR-674-5p | MIMAT0003740 | Copg2 | 54160 | HITS-CLIP | 25753418 | Unspecified |
| mmu-miR-674-5p | MIMAT0003740 | Cpsf6 | 432508 | HITS-CLIP | 29167373 | Unspecified |
| mmu-miR-674-5p | MIMAT0003740 | Cpt1a | 12894 | HITS-CLIP | 29167373 | Unspecified |
| mmu-miR-674-5p | MIMAT0003740 | Crebzf | 233490 | HITS-CLIP | 29167373 | Unspecified |
| mmu-miR-674-5p | MIMAT0003740 | Dcaf5 | 320808 | HITS-CLIP | 25753418 | Unspecified |
| mmu-miR-674-5p | MIMAT0003740 | Ddx24 | 27225 | HITS-CLIP | 29167373 | Unspecified |
| mmu-miR-674-5p | MIMAT0003740 | Dennd4b | 229541 | HITS-CLIP | 25753418 | Unspecified |
| mmu-miR-674-5p | MIMAT0003740 | Dhx15 | 13204 | qCLASH | 31393953 | Unspecified |
| mmu-miR-674-5p | MIMAT0003740 | Dnajb6 | 23950 | HITS-CLIP | 37986224 | Unspecified |

| | | | | | | |
|---|---|---|---|---|---|---|
| mmu-miR-674-5p | MIMAT0003740 | Dnajc13 | 235567 | HITS-CLIP | 29167373 | Unspecified |
| mmu-miR-674-5p | MIMAT0003740 | Dst | 13518 | HITS-CLIP | 25753418 | Unspecified |
| mmu-miR-674-5p | MIMAT0003740 | Dynlt1c | 100040563 | qCLASH | 31393953 | Unspecified |
| mmu-miR-674-5p | MIMAT0003740 | Echs1 | 93747 | HITS-CLIP | 25753418 | Unspecified |
| mmu-miR-674-5p | MIMAT0003740 | Ecpas | 23392 | HITS-CLIP | 25753418 | Unspecified |
| mmu-miR-674-5p | MIMAT0003740 | Eef2 | 13629 | qCLASH | 31393953 | Unspecified |
| mmu-miR-674-5p | MIMAT0003740 | Eif3i | 54709 | HITS-CLIP | 25753418 | Unspecified |
| mmu-miR-674-5p | MIMAT0003740 | Eif5 | 217869 | HITS-CLIP | 26203562 | Unspecified |
| mmu-miR-674-5p | MIMAT0003740 | Etf1 | 225363 | HITS-CLIP | 29167373 | Unspecified |
| mmu-miR-674-5p | MIMAT0003740 | Far1 | 67420 | HITS-CLIP | 29167373 | Unspecified |
| mmu-miR-674-5p | MIMAT0003740 | Fgf13 | 14168 | HITS-CLIP | 25753418 | Unspecified |
| mmu-miR-674-5p | MIMAT0003740 | Flna | 192176 | HITS-CLIP | 29167373 | Unspecified |
| mmu-miR-674-5p | MIMAT0003740 | Fndc3a | 319448 | HITS-CLIP | 29167373 | Unspecified |
| mmu-miR-674-5p | MIMAT0003740 | Fndc3b | 72007 | HITS-CLIP | 29167373 | Unspecified |
| mmu-miR-674-5p | MIMAT0003740 | Fstl4 | 320027 | HITS-CLIP | 25753418 | Unspecified |
| mmu-miR-674-5p | MIMAT0003740 | Gabpa | 14390 | HITS-CLIP | 29167373 | Unspecified |
| mmu-miR-674-5p | MIMAT0003740 | Gan | 209239 | HITS-CLIP | 29167373 | Unspecified |
| mmu-miR-674-5p | MIMAT0003740 | Ganab | 14376 | HITS-CLIP | 29167373 | Unspecified |
| mmu-miR-674-5p | MIMAT0003740 | Gatad1 | 67210 | HITS-CLIP | 25753418 | Unspecified |
| mmu-miR-674-5p | MIMAT0003740 | Gigyf1 | 57330 | qCLASH | 31393953 | Unspecified |
| mmu-miR-674-5p | MIMAT0003740 | Gm20716 | 105980075 | HITS-CLIP | 25753418 | Unspecified |
| mmu-miR-674-5p | MIMAT0003740 | Gm21833 | 6609710 | qCLASH | 31393953 | Unspecified |
| mmu-miR-674-5p | MIMAT0003740 | Gm45928 | 105948585 | HITS-CLIP | 26203562 | Unspecified |
| mmu-miR-674-5p | MIMAT0003740 | Gnptab | 432486 | HITS-CLIP | 25753418 | Unspecified |
| mmu-miR-674-5p | MIMAT0003740 | Gpat4 | 137964 | HITS-CLIP | 25753418 | Unspecified |
| mmu-miR-674-5p | MIMAT0003740 | Gramd1b | 235283 | HITS-CLIP | 29167373 | Unspecified |
| mmu-miR-674-5p | MIMAT0003740 | Gtf2f2 | 68705 | qCLASH | 31393953 | Unspecified |
| mmu-miR-674-5p | MIMAT0003740 | Hcfc1 | 15161 | HITS-CLIP | 29167373 | Unspecified |
| mmu-miR-674-5p | MIMAT0003740 | Hivep1 | 110521 | HITS-CLIP | 29167373 | Unspecified |
| mmu-miR-674-5p | MIMAT0003740 | Hnrnpdl | 50926 | HITS-CLIP | 31433990\|25753418 | Unspecified |
| mmu-miR-674-5p | MIMAT0003740 | Hnrnpul1 | 232989 | HITS-CLIP | 29167373 | Unspecified |
| mmu-miR-674-5p | MIMAT0003740 | Hsd17b11 | 114664 | HITS-CLIP | 29167373 | Unspecified |
| mmu-miR-674-5p | MIMAT0003740 | Hsdl2 | 72479 | HITS-CLIP | 25753418 | Unspecified |
| mmu-miR-674-5p | MIMAT0003740 | Huwe1 | 59026 | HITS-CLIP | 25753418 | Unspecified |
| mmu-miR-674-5p | MIMAT0003740 | Id2 | 15902 | HITS-CLIP | 29167373 | Unspecified |
| mmu-miR-674-5p | MIMAT0003740 | Irf2bp2 | 270110 | HITS-CLIP | 37986224 | Unspecified |
| mmu-miR-674-5p | MIMAT0003740 | Irf2bpl | 238330 | HITS-CLIP | 29167373 | Unspecified |
| mmu-miR-674-5p | MIMAT0003740 | Itga8 | 241226 | HITS-CLIP | 25753418 | Unspecified |
| mmu-miR-674-5p | MIMAT0003740 | Ivns1abp | 117198 | HITS-CLIP | 29167373 | Unspecified |

| | | | | | | |
|---|---|---|---|---|---|---|
| mmu-miR-674-5p | MIMAT0003740 | Jmjd1c | 108829 | qCLASH | 31393953 | Unspecified |
| mmu-miR-674-5p | MIMAT0003740 | Kdm6a | 22289 | qCLASH | 31393953 | Unspecified |
| mmu-miR-674-5p | MIMAT0003740 | Kif3b | 16569 | HITS-CLIP | 29167373 | Unspecified |
| mmu-miR-674-5p | MIMAT0003740 | Kif5b | 16573 | HITS-CLIP | 25753418\|29167373 | Unspecified |
| mmu-miR-674-5p | MIMAT0003740 | Klf13 | 50794 | HITS-CLIP | 37986224 | Unspecified |
| mmu-miR-674-5p | MIMAT0003740 | Klhl21 | 242785 | HITS-CLIP | 29167373 | Unspecified |
| mmu-miR-674-5p | MIMAT0003740 | Kmt2c | 231051 | HITS-CLIP | 29167373 | Unspecified |
| mmu-miR-674-5p | MIMAT0003740 | Kmt2d | 381022 | HITS-CLIP | 25753418 | Unspecified |
| mmu-miR-674-5p | MIMAT0003740 | Kpna4 | 16649 | HITS-CLIP | 29167373 | Unspecified |
| mmu-miR-674-5p | MIMAT0003740 | Lair1 | 52855 | HITS-CLIP | 29167373 | Unspecified |
| mmu-miR-674-5p | MIMAT0003740 | Lmbrd2 | 320506 | HITS-CLIP | 29167373 | Unspecified |
| mmu-miR-674-5p | MIMAT0003740 | Lmna | 16905 | HITS-CLIP | 29167373 | Unspecified |
| mmu-miR-674-5p | MIMAT0003740 | Lpgat1 | 226856 | HITS-CLIP | 25753418 | Unspecified |
| mmu-miR-674-5p | MIMAT0003740 | Lrrfip1 | 16978 | HITS-CLIP | 29167373 | Unspecified |
| mmu-miR-674-5p | MIMAT0003740 | Lrtm1 | 319476 | HITS-CLIP | 25753418 | Unspecified |
| mmu-miR-674-5p | MIMAT0003740 | Macf1 | 11426 | HITS-CLIP | 29167373 | Unspecified |
| mmu-miR-674-5p | MIMAT0003740 | Mapk6 | 50772 | HITS-CLIP | 29167373 | Unspecified |
| mmu-miR-674-5p | MIMAT0003740 | Mcl1 | 17210 | HITS-CLIP | 37986224 | Unspecified |
| mmu-miR-674-5p | MIMAT0003740 | Mcm4 | 17217 | HITS-CLIP | 29167373 | Unspecified |
| mmu-miR-674-5p | MIMAT0003740 | Mdh1 | 17449 | HITS-CLIP | 29167373 | Unspecified |
| mmu-miR-674-5p | MIMAT0003740 | Med13 | 327987 | HITS-CLIP | 29167373 | Unspecified |
| mmu-miR-674-5p | MIMAT0003740 | Med13l | 76199 | HITS-CLIP | 29167373 | Unspecified |
| mmu-miR-674-5p | MIMAT0003740 | Mga | 29808 | HITS-CLIP | 25753418 | Unspecified |
| mmu-miR-674-5p | MIMAT0003740 | Mgat5 | 107895 | HITS-CLIP | 29167373 | Unspecified |
| mmu-miR-674-5p | MIMAT0003740 | Mrps18c | 68735 | HITS-CLIP | 25753418 | Unspecified |
| mmu-miR-674-5p | MIMAT0003740 | Mrps5 | 77721 | HITS-CLIP | 25753418 | Unspecified |
| mmu-miR-674-5p | MIMAT0003740 | Msi2 | 76626 | qCLASH | 31393953 | Unspecified |
| mmu-miR-674-5p | MIMAT0003740 | Mtdh | 67154 | qCLASH | 31393953 | Unspecified |
| mmu-miR-674-5p | MIMAT0003740 | Mtfr1l | 76824 | HITS-CLIP | 25753418 | Unspecified |
| mmu-miR-674-5p | MIMAT0003740 | Mycbp2 | 105689 | qCLASH | 31393953 | Unspecified |
| mmu-miR-674-5p | MIMAT0003740 | Myo18a | 360013 | HITS-CLIP | 29167373 | Unspecified |
| mmu-miR-674-5p | MIMAT0003740 | Myo9a | 270163 | HITS-CLIP | 29167373 | Unspecified |
| mmu-miR-674-5p | MIMAT0003740 | Nckap1 | 50884 | qCLASH | 31393953 | Unspecified |
| mmu-miR-674-5p | MIMAT0003740 | Ncoa2 | 17978 | HITS-CLIP | 29167373 | Unspecified |
| mmu-miR-674-5p | MIMAT0003740 | Nelfa | 24116 | qCLASH | 31393953 | Unspecified |
| mmu-miR-674-5p | MIMAT0003740 | Nfatc3 | 18021 | HITS-CLIP | 25753418 | Unspecified |
| mmu-miR-674-5p | MIMAT0003740 | Nfic | 18029 | HITS-CLIP | 29167373 | Unspecified |
| mmu-miR-674-5p | MIMAT0003740 | Nipbl | 71175 | HITS-CLIP | 29167373 | Unspecified |
| mmu-miR-674-5p | MIMAT0003740 | Nol4l | 329540 | HITS-CLIP | 29167373 | Unspecified |

| | | | | | | |
|---|---|---|---|---|---|---|
| mmu-miR-674-5p | MIMAT0003740 | Pbrm1 | 66923 | qCLASH//HITS-CLIP | 31393953\|29167373 | Unspecified |
| mmu-miR-674-5p | MIMAT0003740 | Pcnx3 | 399909 | HITS-CLIP | 29167373 | Unspecified |
| mmu-miR-674-5p | MIMAT0003740 | Pdgfd | 71785 | HITS-CLIP | 25753418 | Unspecified |
| mmu-miR-674-5p | MIMAT0003740 | Pdp1 | 381511 | HITS-CLIP | 25753418 | Unspecified |
| mmu-miR-674-5p | MIMAT0003740 | Phlpp1 | 98432 | HITS-CLIP | 29167373 | Unspecified |
| mmu-miR-674-5p | MIMAT0003740 | Pias2 | 17344 | HITS-CLIP | 37986224 | Unspecified |
| mmu-miR-674-5p | MIMAT0003740 | Pja2 | 224938 | HITS-CLIP | 25753418 | Unspecified |
| mmu-miR-674-5p | MIMAT0003740 | Plag1 | 56711 | HITS-CLIP | 25753418 | Unspecified |
| mmu-miR-674-5p | MIMAT0003740 | Plec | 18810 | HITS-CLIP | 29167373 | Unspecified |
| mmu-miR-674-5p | MIMAT0003740 | Ppil2 | 66053 | qCLASH | 31393953 | Unspecified |
| mmu-miR-674-5p | MIMAT0003740 | Ppm1a | 19042 | HITS-CLIP | 25753418 | Unspecified |
| mmu-miR-674-5p | MIMAT0003740 | Ppp2ca | 19052 | HITS-CLIP | 29167373 | Unspecified |
| mmu-miR-674-5p | MIMAT0003740 | Psmc2 | 19181 | HITS-CLIP | 29167373 | Unspecified |
| mmu-miR-674-5p | MIMAT0003740 | Ptar1 | 72351 | HITS-CLIP | 37986224 | Unspecified |
| mmu-miR-674-5p | MIMAT0003740 | Ptma | 19231 | qCLASH | 31393953 | Unspecified |
| mmu-miR-674-5p | MIMAT0003740 | Pxmp2 | 19301 | HITS-CLIP | 25753418 | Unspecified |
| mmu-miR-674-5p | MIMAT0003740 | Qsox1 | 104009 | HITS-CLIP | 25753418 | Unspecified |
| mmu-miR-674-5p | MIMAT0003740 | Rab10 | 19325 | HITS-CLIP | 25753418 | Unspecified |
| mmu-miR-674-5p | MIMAT0003740 | Ramp2 | 54409 | qCLASH | 31393953 | Unspecified |
| mmu-miR-674-5p | MIMAT0003740 | Rlim | 19820 | HITS-CLIP | 29167373 | Unspecified |
| mmu-miR-674-5p | MIMAT0003740 | Rmnd5b | 66089 | HITS-CLIP | 29167373 | Unspecified |
| mmu-miR-674-5p | MIMAT0003740 | Rnf220 | 66743 | HITS-CLIP | 25753418 | Unspecified |
| mmu-miR-674-5p | MIMAT0003740 | Rpf1 | 70285 | HITS-CLIP | 25753418 | Unspecified |
| mmu-miR-674-5p | MIMAT0003740 | Rpia | 19895 | qCLASH | 31393953 | Unspecified |
| mmu-miR-674-5p | MIMAT0003740 | Rreb1 | 68750 | HITS-CLIP | 29167373 | Unspecified |
| mmu-miR-674-5p | MIMAT0003740 | Scaf4 | 224432 | qCLASH | 31393953 | Unspecified |
| mmu-miR-674-5p | MIMAT0003740 | Sclt1 | 67161 | qCLASH | 31393953 | Unspecified |
| mmu-miR-674-5p | MIMAT0003740 | Sec24c | 218811 | HITS-CLIP | 25753418 | Unspecified |
| mmu-miR-674-5p | MIMAT0003740 | Sh3gl1 | 20405 | HITS-CLIP | 29167373 | Unspecified |
| mmu-miR-674-5p | MIMAT0003740 | Sh3pxd2a | 14218 | HITS-CLIP | 29167373 | Unspecified |
| mmu-miR-674-5p | MIMAT0003740 | Sin3a | 20466 | HITS-CLIP | 29167373 | Unspecified |
| mmu-miR-674-5p | MIMAT0003740 | Slain2 | 75991 | qCLASH | 31393953 | Unspecified |
| mmu-miR-674-5p | MIMAT0003740 | Slc12a6 | 107723 | HITS-CLIP | 29167373 | Unspecified |
| mmu-miR-674-5p | MIMAT0003740 | Slc23a2 | 54338 | HITS-CLIP | 29167373 | Unspecified |
| mmu-miR-674-5p | MIMAT0003740 | Slc25a11 | 67863 | HITS-CLIP | 25753418 | Unspecified |
| mmu-miR-674-5p | MIMAT0003740 | Slc31a2 | 20530 | HITS-CLIP | 29167373 | Unspecified |
| mmu-miR-674-5p | MIMAT0003740 | Slmap | 83997 | HITS-CLIP | 29167373 | Unspecified |
| mmu-miR-674-5p | MIMAT0003740 | Smu1 | 74255 | HITS-CLIP | 25753418 | Unspecified |
| mmu-miR-674-5p | MIMAT0003740 | Snapc4 | 6621 | HITS-CLIP | 29167373 | Unspecified |

| | | | | | | |
|---|---|---|---|---|---|---|
| mmu-miR-674-5p | MIMAT0003740 | Snrnp200 | 320632 | HITS-CLIP | 29167373 | Unspecified |
| mmu-miR-674-5p | MIMAT0003740 | Snrpd2 | 107686 | HITS-CLIP | 29167373 | Unspecified |
| mmu-miR-674-5p | MIMAT0003740 | Sp3 | 20687 | HITS-CLIP | 29167373 | Unspecified |
| mmu-miR-674-5p | MIMAT0003740 | Spen | 56381 | HITS-CLIP | 25753418 | Unspecified |
| mmu-miR-674-5p | MIMAT0003740 | Srrm2 | 75956 | HITS-CLIP | 29167373 | Unspecified |
| mmu-miR-674-5p | MIMAT0003740 | St13 | 70356 | HITS-CLIP | 37986224 | Unspecified |
| mmu-miR-674-5p | MIMAT0003740 | Supt16 | 114741 | HITS-CLIP | 29167373 | Unspecified |
| mmu-miR-674-5p | MIMAT0003740 | Syne1 | 64009 | HITS-CLIP | 25753418 | Unspecified |
| mmu-miR-674-5p | MIMAT0003740 | Tango6 | 272538 | HITS-CLIP | 29167373 | Unspecified |
| mmu-miR-674-5p | MIMAT0003740 | Tbc1d9 | 71310 | HITS-CLIP | 29167373 | Unspecified |
| mmu-miR-674-5p | MIMAT0003740 | Tbcd | 108903 | HITS-CLIP | 25753418 | Unspecified |
| mmu-miR-674-5p | MIMAT0003740 | Tent4b | 64282 | HITS-CLIP | 29167373 | Unspecified |
| mmu-miR-674-5p | MIMAT0003740 | Tln1 | 21894 | HITS-CLIP | 25753418\|29167373 | Unspecified |
| mmu-miR-674-5p | MIMAT0003740 | Tmc7 | 209760 | HITS-CLIP | 25753418 | Unspecified |
| mmu-miR-674-5p | MIMAT0003740 | Tmem192 | 73067 | qCLASH | 31393953 | Unspecified |
| mmu-miR-674-5p | MIMAT0003740 | Topors | 106021 | HITS-CLIP | 25753418 | Unspecified |
| mmu-miR-674-5p | MIMAT0003740 | Trim37 | 68729 | HITS-CLIP | 25753418 | Unspecified |
| mmu-miR-674-5p | MIMAT0003740 | Trim71 | 636931 | HITS-CLIP | 37986224 | Unspecified |
| mmu-miR-674-5p | MIMAT0003740 | Trip11 | 109181 | HITS-CLIP | 25753418 | Unspecified |
| mmu-miR-674-5p | MIMAT0003740 | Tsc1 | 64930 | HITS-CLIP | 29167373 | Unspecified |
| mmu-miR-674-5p | MIMAT0003740 | Ttn | 22138 | HITS-CLIP | 25753418 | Unspecified |
| mmu-miR-674-5p | MIMAT0003740 | Ttyh3 | 78339 | HITS-CLIP | 29167373 | Unspecified |
| mmu-miR-674-5p | MIMAT0003740 | Txndc11 | 106200 | HITS-CLIP | 29167373 | Unspecified |
| mmu-miR-674-5p | MIMAT0003740 | Ubap2l | 74383 | HITS-CLIP | 29167373 | Unspecified |
| mmu-miR-674-5p | MIMAT0003740 | Ube2i | 22196 | HITS-CLIP | 37986224 | Unspecified |
| mmu-miR-674-5p | MIMAT0003740 | Ube2w | 66799 | HITS-CLIP | 29167373 | Unspecified |
| mmu-miR-674-5p | MIMAT0003740 | Ubr4 | 69116 | HITS-CLIP | 29167373 | Unspecified |
| mmu-miR-674-5p | MIMAT0003740 | Ulk1 | 22241 | qCLASH | 31393953 | Unspecified |
| mmu-miR-674-5p | MIMAT0003740 | Vegfb | 22340 | HITS-CLIP | 25753418 | Unspecified |
| mmu-miR-674-5p | MIMAT0003740 | Virma | 25962 | HITS-CLIP | 25753418 | Unspecified |
| mmu-miR-674-5p | MIMAT0003740 | Vps54 | 245944 | HITS-CLIP | 37986224 | Unspecified |
| mmu-miR-674-5p | MIMAT0003740 | Washc4 | 319277 | HITS-CLIP | 29167373\|21258322\|23597149 | Unspecified |
| mmu-miR-674-5p | MIMAT0003740 | Wbp1l | 226178 | HITS-CLIP | 29167373 | Unspecified |
| mmu-miR-674-5p | MIMAT0003740 | Wdr48 | 67561 | HITS-CLIP | 29167373 | Unspecified |
| mmu-miR-674-5p | MIMAT0003740 | Wdr91 | 101240 | HITS-CLIP | 29167373 | Unspecified |
| mmu-miR-674-5p | MIMAT0003740 | Wsb2 | 59043 | HITS-CLIP | 29167373 | Unspecified |
| mmu-miR-674-5p | MIMAT0003740 | Xbp1 | 22433 | HITS-CLIP | 29167373 | Unspecified |
| mmu-miR-674-5p | MIMAT0003740 | Ywhae | 22627 | HITS-CLIP | 37986224\|25753418 | Unspecified |
| mmu-miR-674-5p | MIMAT0003740 | Ywhaz | 22631 | HITS-CLIP | 37986224\|25753418 | Unspecified |

| | | | | | | |
|---|---|---|---|---|---|---|
| mmu-miR-674-5p | MIMAT0003740 | Zeb1 | 21417 | HITS-CLIP | 25753418 | Unspecified |
| mmu-miR-674-5p | MIMAT0003740 | Zfp142 | 77264 | qCLASH | 31393953 | Unspecified |
| mmu-miR-674-5p | MIMAT0003740 | Zfp217 | 228913 | HITS-CLIP | 37986224 | Unspecified |
| mmu-miR-674-5p | MIMAT0003740 | Zfp706 | 68036 | HITS-CLIP | 25753418 | Unspecified |
| mmu-miR-674-5p | MIMAT0003740 | Zfp800 | 627049 | HITS-CLIP | 25753418 | Unspecified |
| mmu-miR-674-5p | MIMAT0003740 | mt-Co1 | 4512 | qCLASH | 31393953 | Unspecified |
| mmu-miR-5114 | MIMAT0020622 | AI467606 | 101602 | HITS-CLIP | 31433990 | Unspecified |
| mmu-miR-5114 | MIMAT0020622 | B2m | 12010 | HITS-CLIP | 31433990 | Unspecified |
| mmu-miR-5114 | MIMAT0020622 | Bcl9l | 80288 | HITS-CLIP | 26203562 | Unspecified |
| mmu-miR-5114 | MIMAT0020622 | Bdp1 | 544971 | HITS-CLIP | 31433990 | Unspecified |
| mmu-miR-5114 | MIMAT0020622 | Cfl2 | 12632 | HITS-CLIP | 26203562 | Unspecified |
| mmu-miR-5114 | MIMAT0020622 | Clic4 | 29876 | HITS-CLIP | 26203562 | Unspecified |
| mmu-miR-5114 | MIMAT0020622 | Creb1 | 12912 | HITS-CLIP | 31433990 | Unspecified |
| mmu-miR-5114 | MIMAT0020622 | Ddx5 | 13207 | HITS-CLIP | 31433990 | Unspecified |
| mmu-miR-5114 | MIMAT0020622 | Dolpp1 | 57170 | HITS-CLIP | 26203562 | Unspecified |
| mmu-miR-5114 | MIMAT0020622 | Eef1g | 67160 | HITS-CLIP | 26203562 | Unspecified |
| mmu-miR-5114 | MIMAT0020622 | Eif5a | 276770 | HITS-CLIP | 26203562 | Unspecified |
| mmu-miR-5114 | MIMAT0020622 | Fam8a1 | 51439 | HITS-CLIP | 31433990 | Unspecified |
| mmu-miR-5114 | MIMAT0020622 | Gna11 | 14672 | HITS-CLIP | 26203562 | Unspecified |
| mmu-miR-5114 | MIMAT0020622 | Gne | 50798 | HITS-CLIP | 31433990 | Unspecified |
| mmu-miR-5114 | MIMAT0020622 | Golga4 | 54214 | HITS-CLIP | 31433990 | Unspecified |
| mmu-miR-5114 | MIMAT0020622 | H13 | 14950 | HITS-CLIP | 31433990 | Unspecified |
| mmu-miR-5114 | MIMAT0020622 | Hdgf | 15191 | HITS-CLIP | 26203562 | Unspecified |
| mmu-miR-5114 | MIMAT0020622 | Hells | 15201 | HITS-CLIP | 31433990 | Unspecified |
| mmu-miR-5114 | MIMAT0020622 | Hspa4 | 15525 | HITS-CLIP | 31433990 | Unspecified |
| mmu-miR-5114 | MIMAT0020622 | Hyou1 | 12282 | HITS-CLIP | 31433990 | Unspecified |
| mmu-miR-5114 | MIMAT0020622 | Ints6 | 18130 | HITS-CLIP | 31433990 | Unspecified |
| mmu-miR-5114 | MIMAT0020622 | Itpkb | 320404 | HITS-CLIP | 31433990 | Unspecified |
| mmu-miR-5114 | MIMAT0020622 | Klf9 | 16601 | HITS-CLIP | 26203562 | Unspecified |
| mmu-miR-5114 | MIMAT0020622 | Lgmn | 19141 | HITS-CLIP | 26203562 | Unspecified |
| mmu-miR-5114 | MIMAT0020622 | Lif | 16878 | HITS-CLIP | 31433990 | Unspecified |
| mmu-miR-5114 | MIMAT0020622 | Lnpep | 240028 | HITS-CLIP | 31433990 | Unspecified |
| mmu-miR-5114 | MIMAT0020622 | Mob4 | 19070 | HITS-CLIP | 31433990 | Unspecified |
| mmu-miR-5114 | MIMAT0020622 | Nfatc2ip | 18020 | HITS-CLIP | 31433990 | Unspecified |
| mmu-miR-5114 | MIMAT0020622 | Nip7 | 66164 | HITS-CLIP | 31433990 | Unspecified |
| mmu-miR-5114 | MIMAT0020622 | Npm1 | 18148 | HITS-CLIP | 31433990 | Unspecified |
| mmu-miR-5114 | MIMAT0020622 | Nup98 | 269966 | HITS-CLIP | 31433990 | Unspecified |
| mmu-miR-5114 | MIMAT0020622 | Pgm2l1 | 70974 | HITS-CLIP | 31433990 | Unspecified |
| mmu-miR-5114 | MIMAT0020622 | Plekha8 | 231999 | HITS-CLIP | 26203562 | Unspecified |

| | | | | | | |
|---|---|---|---|---|---|---|
| mmu-miR-5114 | MIMAT0020622 | Prkar1a | 19084 | HITS-CLIP | 31433990 | Unspecified |
| mmu-miR-5114 | MIMAT0020622 | Prpf8 | 192159 | HITS-CLIP | 31433990 | Unspecified |
| mmu-miR-5114 | MIMAT0020622 | Rpl8 | 26961 | HITS-CLIP | 31433990 | Unspecified |
| mmu-miR-5114 | MIMAT0020622 | Rrm2 | 20135 | HITS-CLIP | 31433990 | Unspecified |
| mmu-miR-5114 | MIMAT0020622 | Setd1b | 208043 | HITS-CLIP | 31433990 | Unspecified |
| mmu-miR-5114 | MIMAT0020622 | Sgk3 | 170755 | HITS-CLIP | 26203562 | Unspecified |
| mmu-miR-5114 | MIMAT0020622 | Srsf2 | 20382 | HITS-CLIP | 31433990 | Unspecified |
| mmu-miR-5114 | MIMAT0020622 | Sumf1 | 58911 | HITS-CLIP | 26203562 | Unspecified |
| mmu-miR-5114 | MIMAT0020622 | Susd6 | 9766 | HITS-CLIP | 26203562 | Unspecified |
| mmu-miR-5114 | MIMAT0020622 | Taok1 | 216965 | HITS-CLIP | 26203562 | Unspecified |
| mmu-miR-5114 | MIMAT0020622 | Tars2 | 71807 | HITS-CLIP | 26203562 | Unspecified |
| mmu-miR-5114 | MIMAT0020622 | Trak2 | 70827 | HITS-CLIP | 31433990 | Unspecified |
| mmu-miR-5114 | MIMAT0020622 | Vps37a | 52348 | HITS-CLIP | 31433990 | Unspecified |
| mmu-miR-5114 | MIMAT0020622 | Ythdc1 | 231386 | HITS-CLIP | 31433990 | Unspecified |
| mmu-miR-5114 | MIMAT0020622 | Zfp217 | 228913 | HITS-CLIP | 26203562 | Unspecified |
| mmu-miR-5114 | MIMAT0020622 | Zfyve1 | 217695 | HITS-CLIP | 31433990 | Unspecified |
| mmu-miR-5114 | MIMAT0020622 | Zfyve26 | 211978 | HITS-CLIP | 31433990 | Unspecified |
| mmu-miR-3106-5p | MIMAT0014817 | Atp5a1 | 11946 | sRNA-Seq//HITS-CLIP | 33184237\|26203562 | Unspecified |
| mmu-miR-3106-5p | MIMAT0014817 | Celf1 | 13046 | HITS-CLIP | 31433990 | Unspecified |
| mmu-miR-3106-5p | MIMAT0014817 | Dnmt1 | 13433 | HITS-CLIP | 31433990 | Unspecified |
| mmu-miR-3106-5p | MIMAT0014817 | Hectd1 | 207304 | sRNA-Seq//HITS-CLIP | 33184237\|26203562 | Unspecified |
| mmu-miR-3106-5p | MIMAT0014817 | Riok1 | 71340 | HITS-CLIP | 31433990 | Unspecified |
| mmu-miR-3106-5p | MIMAT0014817 | Sh3pxd2a | 14218 | HITS-CLIP | 26203562 | Unspecified |
| mmu-miR-3106-5p | MIMAT0014817 | Tnrc6b | 213988 | HITS-CLIP | 31433990 | Unspecified |
| mmu-miR-3106-5p | MIMAT0014817 | Ugcg | 22234 | HITS-CLIP//sRNA-Seq | 31433990\|33184237 | Unspecified |
| mmu-miR-29b-3p | MIMAT0000127 | 0610030E20Rik | 68364 | HITS-CLIP | 37986224\|23597149\|23142080\|25083 | Unspecified |
| mmu-miR-29b-3p | MIMAT0000127 | 1110002E22Rik | 102634333 | HITS-CLIP | 25083871 | Unspecified |
| mmu-miR-29b-3p | MIMAT0000127 | 1110008L16Rik | 66132 | HITS-CLIP | 23142080 | Unspecified |
| mmu-miR-29b-3p | MIMAT0000127 | 1110032A03Rik | 68721 | HITS-CLIP | 23142080 | Unspecified |
| mmu-miR-29b-3p | MIMAT0000127 | 1190002N15Rik | 68861 | HITS-CLIP | 23142080 | Unspecified |
| mmu-miR-29b-3p | MIMAT0000127 | 1500011B03Rik | 66236 | HITS-CLIP//sRNA-Seq | 25083871\|33184237\|23142080 | Unspecified |
| mmu-miR-29b-3p | MIMAT0000127 | 1600012H06Rik | 67912 | HITS-CLIP | 25083871\|23142080\|23597149 | Unspecified |
| mmu-miR-29b-3p | MIMAT0000127 | 1600012P17Rik | 72025 | HITS-CLIP | 37986224 | Unspecified |
| mmu-miR-29b-3p | MIMAT0000127 | 1600014C10Rik | 72244 | HITS-CLIP | 37986224 | Unspecified |
| mmu-miR-29b-3p | MIMAT0000127 | 1700017B05Rik | 74211 | HITS-CLIP | 23597149\|25083871 | Unspecified |
| mmu-miR-29b-3p | MIMAT0000127 | 1700019D03Rik | 67080 | HITS-CLIP | 23142080 | Unspecified |
| mmu-miR-29b-3p | MIMAT0000127 | 1700021F05Rik | 67851 | HITS-CLIP | 23142080 | Unspecified |
| mmu-miR-29b-3p | MIMAT0000127 | 1700030K09Rik | 72254 | HITS-CLIP | 23597149 | Unspecified |
| mmu-miR-29b-3p | MIMAT0000127 | 1700037H04Rik | 67326 | HITS-CLIP//sRNA-Seq | 25083871\|33184237 | Unspecified |

| | | | | | | |
|---|---|---|---|---|---|---|
| mmu-miR-29b-3p | MIMAT0000127 | 1700066M21Rik | 73467 | HITS-CLIP | 23142080 | Unspecified |
| mmu-miR-29b-3p | MIMAT0000127 | 1700123O20Rik | 58248 | HITS-CLIP | 23142080 | Unspecified |
| mmu-miR-29b-3p | MIMAT0000127 | 1810013L24Rik | 69053 | HITS-CLIP | 37986224\|25083871\|23142080 | Unspecified |
| mmu-miR-29b-3p | MIMAT0000127 | 1810055G02Rik | 72056 | HITS-CLIP//sRNA-Seq | 37986224\|23597149\|33184237 | Unspecified |
| mmu-miR-29b-3p | MIMAT0000127 | 2010111I01Rik | 72061 | HITS-CLIP//sRNA-Seq | 25083871\|33184237 | Unspecified |
| mmu-miR-29b-3p | MIMAT0000127 | 2310022B05Rik | 69551 | HITS-CLIP//sRNA-Seq | 37986224\|23597149\|33184237\|23142 | Unspecified |
| mmu-miR-29b-3p | MIMAT0000127 | 2510009E07Rik | 72190 | HITS-CLIP//sRNA-Seq | 37986224\|33184237\|23597149 | Unspecified |
| mmu-miR-29b-3p | MIMAT0000127 | 2510039O18Rik | 77034 | HITS-CLIP | 37986224 | Unspecified |
| mmu-miR-29b-3p | MIMAT0000127 | 2610002M06Rik | 67028 | HITS-CLIP | 25083871\|23597149\|23142080 | Unspecified |
| mmu-miR-29b-3p | MIMAT0000127 | 2610507B11Rik | 72503 | HITS-CLIP//sRNA-Seq | 23597149\|23142080\|33184237\|25083 | Unspecified |
| mmu-miR-29b-3p | MIMAT0000127 | 2700081O15Rik | 108899 | HITS-CLIP | 25083871 | Unspecified |
| mmu-miR-29b-3p | MIMAT0000127 | 4833420G17Rik | 67392 | HITS-CLIP | 23142080\|23597149 | Unspecified |
| mmu-miR-29b-3p | MIMAT0000127 | 4921524J17Rik | 66714 | HITS-CLIP | 37986224\|25083871\|23142080 | Unspecified |
| mmu-miR-29b-3p | MIMAT0000127 | 4930402H24Rik | 228602 | HITS-CLIP//sRNA-Seq | 25083871\|33184237\|23142080 | Unspecified |
| mmu-miR-29b-3p | MIMAT0000127 | 4930503L19Rik | 269033 | HITS-CLIP | 25083871 | Unspecified |
| mmu-miR-29b-3p | MIMAT0000127 | 4930579G24Rik | 75939 | HITS-CLIP | 23142080 | Unspecified |
| mmu-miR-29b-3p | MIMAT0000127 | 4931406C07Rik | 70984 | HITS-CLIP | 23597149 | Unspecified |
| mmu-miR-29b-3p | MIMAT0000127 | 4931406P16Rik | 233103 | HITS-CLIP | 37986224\|25083871\|23597149\|23142 | Unspecified |
| mmu-miR-29b-3p | MIMAT0000127 | 4931428F04Rik | 74356 | HITS-CLIP//sRNA-Seq | 37986224\|25083871\|33184237 | Unspecified |
| mmu-miR-29b-3p | MIMAT0000127 | 4932438A13Rik | 229227 | HITS-CLIP//sRNA-Seq | 25083871\|23597149\|33184237\|23142 | Unspecified |
| mmu-miR-29b-3p | MIMAT0000127 | 8030462N17Rik | 212163 | HITS-CLIP | 25083871\|23142080 | Unspecified |
| mmu-miR-29b-3p | MIMAT0000127 | 9130023H24Rik | 100043133 | HITS-CLIP | 25083871 | Unspecified |
| mmu-miR-29b-3p | MIMAT0000127 | 9530068E07Rik | 213673 | HITS-CLIP//sRNA-Seq | 25083871\|33184237\|23597149\|23142 | Unspecified |
| mmu-miR-29b-3p | MIMAT0000127 | 9930012K11Rik | 268759 | HITS-CLIP | 37986224\|25083871 | Unspecified |
| mmu-miR-29b-3p | MIMAT0000127 | 9930021J03Rik | 240613 | HITS-CLIP | 25083871 | Unspecified |
| mmu-miR-29b-3p | MIMAT0000127 | AI597479 | 98404 | HITS-CLIP | 37986224\|25083871 | Unspecified |
| mmu-miR-29b-3p | MIMAT0000127 | AU018091 | 245128 | HITS-CLIP | 37986224\|25083871 | Unspecified |
| mmu-miR-29b-3p | MIMAT0000127 | AW549877 | 106064 | HITS-CLIP//sRNA-Seq | 37986224\|25083871\|33184237\|23597 | Unspecified |
| mmu-miR-29b-3p | MIMAT0000127 | Aacs | 78894 | HITS-CLIP//sRNA-Seq | 37986224\|25083871\|23597149\|33184 | Unspecified |
| mmu-miR-29b-3p | MIMAT0000127 | Aadac | 67758 | HITS-CLIP | 23597149 | Unspecified |
| mmu-miR-29b-3p | MIMAT0000127 | Aagab | 66939 | HITS-CLIP | 23142080 | Unspecified |
| mmu-miR-29b-3p | MIMAT0000127 | Aak1 | 269774 | HITS-CLIP | 31433990\|23142080 | Unspecified |
| mmu-miR-29b-3p | MIMAT0000127 | Aamp | 227290 | HITS-CLIP//sRNA-Seq | 37986224\|23597149\|33184237 | Unspecified |
| mmu-miR-29b-3p | MIMAT0000127 | Aar2 | 68295 | HITS-CLIP | 23142080 | Unspecified |
| mmu-miR-29b-3p | MIMAT0000127 | Aars | 234734 | HITS-CLIP//sRNA-Seq | 23597149\|25083871\|33184237\|23142 | Unspecified |
| mmu-miR-29b-3p | MIMAT0000127 | Aass | 30956 | HITS-CLIP | 23597149 | Unspecified |
| mmu-miR-29b-3p | MIMAT0000127 | Abat | 268860 | HITS-CLIP | 23597149 | Unspecified |
| mmu-miR-29b-3p | MIMAT0000127 | Abca1 | 11303 | HITS-CLIP | 23597149 | Unspecified |
| mmu-miR-29b-3p | MIMAT0000127 | Abca2 | 11305 | HITS-CLIP | 25083871 | Unspecified |

| | | | | | | |
|---|---|---|---|---|---|---|
| mmu-miR-29b-3p | MIMAT0000127 | Abca3 | 27410 | HITS-CLIP//sRNA-Seq | 23597149\|33184237 | Unspecified |
| mmu-miR-29b-3p | MIMAT0000127 | Abca5 | 217265 | HITS-CLIP | 25083871 | Unspecified |
| mmu-miR-29b-3p | MIMAT0000127 | Abca6 | 76184 | HITS-CLIP | 23597149 | Unspecified |
| mmu-miR-29b-3p | MIMAT0000127 | Abca8b | 27404 | HITS-CLIP | 23597149 | Unspecified |
| mmu-miR-29b-3p | MIMAT0000127 | Abcb11 | 27413 | HITS-CLIP | 37986224\|23597149 | Unspecified |
| mmu-miR-29b-3p | MIMAT0000127 | Abcb1b | 18669 | HITS-CLIP | 23142080\|25083871 | Unspecified |
| mmu-miR-29b-3p | MIMAT0000127 | Abcb6 | 74104 | HITS-CLIP | 37986224\|23597149\|25083871 | Unspecified |
| mmu-miR-29b-3p | MIMAT0000127 | Abcb7 | 11306 | HITS-CLIP | 25083871\|23142080 | Unspecified |
| mmu-miR-29b-3p | MIMAT0000127 | Abcb8 | 74610 | HITS-CLIP | 23142080 | Unspecified |
| mmu-miR-29b-3p | MIMAT0000127 | Abcb9 | 56325 | HITS-CLIP | 23142080 | Unspecified |
| mmu-miR-29b-3p | MIMAT0000127 | Abcc10 | 89845 | HITS-CLIP | 23597149 | Unspecified |
| mmu-miR-29b-3p | MIMAT0000127 | Abcc3 | 76408 | HITS-CLIP | 37986224\|23597149 | Unspecified |
| mmu-miR-29b-3p | MIMAT0000127 | Abcc5 | 27416 | sRNA-Seq//HITS-CLIP | 33184237\|23142080 | Unspecified |
| mmu-miR-29b-3p | MIMAT0000127 | Abcd3 | 19299 | HITS-CLIP//sRNA-Seq | 23597149\|33184237 | Unspecified |
| mmu-miR-29b-3p | MIMAT0000127 | Abce1 | 24015 | HITS-CLIP//sRNA-Seq | 37986224\|25083871\|33184237\|2314[illegible] | Unspecified |
| mmu-miR-29b-3p | MIMAT0000127 | Abcf1 | 224742 | HITS-CLIP//sRNA-Seq | 25083871\|33184237\|23142080 | Unspecified |
| mmu-miR-29b-3p | MIMAT0000127 | Abcf2 | 27407 | HITS-CLIP//sRNA-Seq | 25083871\|33184237\|23142080 | Unspecified |
| mmu-miR-29b-3p | MIMAT0000127 | Abcf3 | 27406 | HITS-CLIP | 37986224\|23597149 | Unspecified |
| mmu-miR-29b-3p | MIMAT0000127 | Abcg2 | 26357 | HITS-CLIP | 25083871\|23597149 | Unspecified |
| mmu-miR-29b-3p | MIMAT0000127 | Abhd13 | 68904 | HITS-CLIP | 37986224\|23142080\|25083871\|2359[illegible] | Unspecified |
| mmu-miR-29b-3p | MIMAT0000127 | Abhd15 | 67477 | HITS-CLIP | 37986224 | Unspecified |
| mmu-miR-29b-3p | MIMAT0000127 | Abhd16a | 193742 | HITS-CLIP//sRNA-Seq | 23597149\|33184237\|23142080 | Unspecified |
| mmu-miR-29b-3p | MIMAT0000127 | Abhd17b | 226016 | HITS-CLIP | 37986224\|25083871\|23597149\|2314[illegible] | Unspecified |
| mmu-miR-29b-3p | MIMAT0000127 | Abhd17c | 70178 | HITS-CLIP | 25083871\|23142080 | Unspecified |
| mmu-miR-29b-3p | MIMAT0000127 | Abhd18 | 269423 | HITS-CLIP | 23142080\|25083871\|23597149 | Unspecified |
| mmu-miR-29b-3p | MIMAT0000127 | Abhd3 | 106861 | HITS-CLIP | 23597149 | Unspecified |
| mmu-miR-29b-3p | MIMAT0000127 | Abhd4 | 105501 | HITS-CLIP//sRNA-Seq | 25083871\|33184237\|23597149\|2314[illegible] | Unspecified |
| mmu-miR-29b-3p | MIMAT0000127 | Abhd5 | 67469 | HITS-CLIP | 25083871 | Unspecified |
| mmu-miR-29b-3p | MIMAT0000127 | Abhd8 | 64296 | HITS-CLIP | 37986224 | Unspecified |
| mmu-miR-29b-3p | MIMAT0000127 | Abl1 | 11350 | HITS-CLIP//sRNA-Seq | 25083871\|33184237\|23597149\|2314[illegible] | Unspecified |
| mmu-miR-29b-3p | MIMAT0000127 | Abl2 | 11352 | HITS-CLIP | 23142080\|25083871 | Unspecified |
| mmu-miR-29b-3p | MIMAT0000127 | Ablim1 | 226251 | HITS-CLIP//sRNA-Seq | 25083871\|33184237\|23142080 | Unspecified |
| mmu-miR-29b-3p | MIMAT0000127 | Ablim3 | 319713 | HITS-CLIP//sRNA-Seq | 25083871\|33184237 | Unspecified |
| mmu-miR-29b-3p | MIMAT0000127 | Abraxas2 | 23172 | HITS-CLIP | 25083871 | Unspecified |
| mmu-miR-29b-3p | MIMAT0000127 | Abtb1 | 80283 | HITS-CLIP | 23142080 | Unspecified |
| mmu-miR-29b-3p | MIMAT0000127 | Abtb2 | 99382 | HITS-CLIP | 23142080 | Unspecified |
| mmu-miR-29b-3p | MIMAT0000127 | Acaca | 107476 | sRNA-Seq//HITS-CLIP | 33184237\|23142080 | Unspecified |
| mmu-miR-29b-3p | MIMAT0000127 | Acacb | 100705 | HITS-CLIP | 23597149 | Unspecified |
| mmu-miR-29b-3p | MIMAT0000127 | Acadsb | 66885 | HITS-CLIP | 23142080\|25083871 | Unspecified |

| | | | | | | |
|---|---|---|---|---|---|---|
| mmu-miR-29b-3p | MIMAT0000127 | Acadvl | 11370 | HITS-CLIP//sRNA-Seq | 37986224\|23597149\|33184237\|23142 | Unspecified |
| mmu-miR-29b-3p | MIMAT0000127 | Acap1 | 216859 | HITS-CLIP | 23142080 | Unspecified |
| mmu-miR-29b-3p | MIMAT0000127 | Acap2 | 78618 | HITS-CLIP | 37986224\|25083871\|23142080 | Unspecified |
| mmu-miR-29b-3p | MIMAT0000127 | Acbd3 | 170760 | HITS-CLIP | 25083871\|23142080 | Unspecified |
| mmu-miR-29b-3p | MIMAT0000127 | Acbd5 | 74159 | HITS-CLIP | 37986224\|23142080\|23597149 | Unspecified |
| mmu-miR-29b-3p | MIMAT0000127 | Acer2 | 230379 | HITS-CLIP | 23597149\|23142080 | Unspecified |
| mmu-miR-29b-3p | MIMAT0000127 | Acer3 | 66190 | HITS-CLIP | 23597149\|23142080 | Unspecified |
| mmu-miR-29b-3p | MIMAT0000127 | Acin1 | 56215 | sRNA-Seq//HITS-CLIP | 33184237\|23142080 | Unspecified |
| mmu-miR-29b-3p | MIMAT0000127 | Acly | 104112 | HITS-CLIP//sRNA-Seq | 25083871\|33184237\|23142080 | Unspecified |
| mmu-miR-29b-3p | MIMAT0000127 | Aco1 | 11428 | HITS-CLIP | 37986224\|25083871\|23597149\|23142 | Unspecified |
| mmu-miR-29b-3p | MIMAT0000127 | Aco2 | 11429 | HITS-CLIP//sRNA-Seq | 23597149\|33184237 | Unspecified |
| mmu-miR-29b-3p | MIMAT0000127 | Acot11 | 329910 | HITS-CLIP | 23142080 | Unspecified |
| mmu-miR-29b-3p | MIMAT0000127 | Acot7 | 70025 | HITS-CLIP//sRNA-Seq | 23142080\|33184237 | Unspecified |
| mmu-miR-29b-3p | MIMAT0000127 | Acot9 | 56360 | sRNA-Seq//HITS-CLIP | 33184237\|23142080 | Unspecified |
| mmu-miR-29b-3p | MIMAT0000127 | Acox2 | 93732 | HITS-CLIP | 37986224\|23597149 | Unspecified |
| mmu-miR-29b-3p | MIMAT0000127 | Acp2 | 11432 | HITS-CLIP | 23142080 | Unspecified |
| mmu-miR-29b-3p | MIMAT0000127 | Acp5 | 11433 | HITS-CLIP | 23142080\|23597149 | Unspecified |
| mmu-miR-29b-3p | MIMAT0000127 | Acp6 | 66659 | HITS-CLIP | 23142080 | Unspecified |
| mmu-miR-29b-3p | MIMAT0000127 | Acsl1 | 14081 | HITS-CLIP | 37986224\|23597149 | Unspecified |
| mmu-miR-29b-3p | MIMAT0000127 | Acsl3 | 74205 | HITS-CLIP//sRNA-Seq | 25083871\|33184237\|23142080 | Unspecified |
| mmu-miR-29b-3p | MIMAT0000127 | Acsl4 | 50790 | HITS-CLIP//sRNA-Seq | 37986224\|25083871\|33184237\|23142 | Unspecified |
| mmu-miR-29b-3p | MIMAT0000127 | Acsm1 | 117147 | HITS-CLIP | 23597149 | Unspecified |
| mmu-miR-29b-3p | MIMAT0000127 | Acsm3 | 20216 | HITS-CLIP | 23597149 | Unspecified |
| mmu-miR-29b-3p | MIMAT0000127 | Acsm5 | 272428 | HITS-CLIP | 23597149 | Unspecified |
| mmu-miR-29b-3p | MIMAT0000127 | Acss1 | 68738 | HITS-CLIP | 23142080 | Unspecified |
| mmu-miR-29b-3p | MIMAT0000127 | Acss2 | 60525 | sRNA-Seq//HITS-CLIP | 33184237\|23142080 | Unspecified |
| mmu-miR-29b-3p | MIMAT0000127 | Actb | 11461 | HITS-CLIP//sRNA-Seq | 31433990\|33184237 | Unspecified |
| mmu-miR-29b-3p | MIMAT0000127 | Actl6a | 56456 | sRNA-Seq//HITS-CLIP | 33184237\|23142080 | Unspecified |
| mmu-miR-29b-3p | MIMAT0000127 | Actn1 | 109711 | sRNA-Seq//HITS-CLIP | 33184237\|23142080\|25083871 | Unspecified |
| mmu-miR-29b-3p | MIMAT0000127 | Actn3 | 11474 | HITS-CLIP | 25083871 | Unspecified |
| mmu-miR-29b-3p | MIMAT0000127 | Actn4 | 60595 | HITS-CLIP//sRNA-Seq | 25083871\|23597149\|33184237\|23142 | Unspecified |
| mmu-miR-29b-3p | MIMAT0000127 | Actr10 | 56444 | HITS-CLIP//sRNA-Seq | 23142080\|25083871\|33184237 | Unspecified |
| mmu-miR-29b-3p | MIMAT0000127 | Actr2 | 66713 | HITS-CLIP//sRNA-Seq | 23597149\|33184237\|23142080\|25083 | Unspecified |
| mmu-miR-29b-3p | MIMAT0000127 | Actr3 | 74117 | HITS-CLIP//sRNA-Seq | 23142080\|33184237 | Unspecified |
| mmu-miR-29b-3p | MIMAT0000127 | Actr6 | 67019 | HITS-CLIP | 23142080 | Unspecified |
| mmu-miR-29b-3p | MIMAT0000127 | Acvr1 | 11477 | HITS-CLIP | 25083871 | Unspecified |
| mmu-miR-29b-3p | MIMAT0000127 | Acvr2a | 11480 | Luciferase assay//Microarray//Northern bl | 19342382\|25083871\|23597149\|23142 | Unspecified |
| mmu-miR-29b-3p | MIMAT0000127 | Ada | 11486 | HITS-CLIP | 23142080 | Unspecified |
| mmu-miR-29b-3p | MIMAT0000127 | Adam10 | 11487 | HITS-CLIP//sRNA-Seq | 25083871\|33184237\|23142080 | Unspecified |

| | | | | | | |
|---|---|---|---|---|---|---|
| mmu-miR-29b-3p | MIMAT0000127 | Adam12 | 11489 | HITS-CLIP//sRNA-Seq | 37986224\|25083871\|33184237 | Unspecified |
| mmu-miR-29b-3p | MIMAT0000127 | Adam17 | 11491 | HITS-CLIP | 25083871\|23597149\|23142080 | Unspecified |
| mmu-miR-29b-3p | MIMAT0000127 | Adam19 | 11492 | HITS-CLIP//sRNA-Seq | 23597149\|23142080\|25083871\|33184 | Unspecified |
| mmu-miR-29b-3p | MIMAT0000127 | Adam1a | 280668 | HITS-CLIP | 37986224\|25083871 | Unspecified |
| mmu-miR-29b-3p | MIMAT0000127 | Adam9 | 11502 | sRNA-Seq//HITS-CLIP | 33184237\|25083871 | Unspecified |
| mmu-miR-29b-3p | MIMAT0000127 | Adamts12 | 239337 | HITS-CLIP | 25083871\|23597149 | Unspecified |
| mmu-miR-29b-3p | MIMAT0000127 | Adamts16 | 271127 | HITS-CLIP | 37986224\|25083871 | Unspecified |
| mmu-miR-29b-3p | MIMAT0000127 | Adamts2 | 216725 | HITS-CLIP | 37986224\|25083871 | Unspecified |
| mmu-miR-29b-3p | MIMAT0000127 | Adamts6 | 108154 | HITS-CLIP | 23142080\|25083871 | Unspecified |
| mmu-miR-29b-3p | MIMAT0000127 | Adamts7 | 108153 | HITS-CLIP | 37986224\|23597149 | Unspecified |
| mmu-miR-29b-3p | MIMAT0000127 | Adamtsl4 | 229595 | HITS-CLIP | 23597149 | Unspecified |
| mmu-miR-29b-3p | MIMAT0000127 | Adar | 56417 | sRNA-Seq//HITS-CLIP | 33184237\|23142080 | Unspecified |
| mmu-miR-29b-3p | MIMAT0000127 | Adarb1 | 110532 | HITS-CLIP | 23142080 | Unspecified |
| mmu-miR-29b-3p | MIMAT0000127 | Adck5 | 268822 | HITS-CLIP | 23142080 | Unspecified |
| mmu-miR-29b-3p | MIMAT0000127 | Adcy3 | 104111 | HITS-CLIP | 23597149 | Unspecified |
| mmu-miR-29b-3p | MIMAT0000127 | Adcy6 | 11512 | HITS-CLIP//sRNA-Seq | 25083871\|33184237\|23142080 | Unspecified |
| mmu-miR-29b-3p | MIMAT0000127 | Adcy7 | 11513 | HITS-CLIP | 25083871\|23142080 | Unspecified |
| mmu-miR-29b-3p | MIMAT0000127 | Adcy9 | 11515 | HITS-CLIP | 37986224 | Unspecified |
| mmu-miR-29b-3p | MIMAT0000127 | Add1 | 11518 | HITS-CLIP//sRNA-Seq | 25083871\|33184237 | Unspecified |
| mmu-miR-29b-3p | MIMAT0000127 | Add3 | 27360 | HITS-CLIP | 25083871\|23597149\|23142080 | Unspecified |
| mmu-miR-29b-3p | MIMAT0000127 | Adgra2 | 25960 | HITS-CLIP//sRNA-Seq | 25083871\|33184237 | Unspecified |
| mmu-miR-29b-3p | MIMAT0000127 | Adgre5 | 976 | HITS-CLIP | 25083871\|23142080 | Unspecified |
| mmu-miR-29b-3p | MIMAT0000127 | Adgrf5 | 224792 | HITS-CLIP | 23597149 | Unspecified |
| mmu-miR-29b-3p | MIMAT0000127 | Adgrl3 | 319387 | HITS-CLIP//sRNA-Seq | 25083871\|33184237 | Unspecified |
| mmu-miR-29b-3p | MIMAT0000127 | Adh1 | 11522 | HITS-CLIP | 23597149 | Unspecified |
| mmu-miR-29b-3p | MIMAT0000127 | Adh5 | 11532 | HITS-CLIP//sRNA-Seq | 23597149\|33184237 | Unspecified |
| mmu-miR-29b-3p | MIMAT0000127 | Adhfe1 | 76187 | HITS-CLIP | 23597149 | Unspecified |
| mmu-miR-29b-3p | MIMAT0000127 | Adipor1 | 72674 | HITS-CLIP//sRNA-Seq | 37986224\|25083871\|33184237\|23597 | Unspecified |
| mmu-miR-29b-3p | MIMAT0000127 | Adipor2 | 68465 | HITS-CLIP//sRNA-Seq | 23597149\|25083871\|33184237\|30971 | Unspecified |
| mmu-miR-29b-3p | MIMAT0000127 | Adnp | 11538 | HITS-CLIP | 37986224\|23597149\|23142080 | Unspecified |
| mmu-miR-29b-3p | MIMAT0000127 | Adnp2 | 240442 | HITS-CLIP | 25083871 | Unspecified |
| mmu-miR-29b-3p | MIMAT0000127 | Ado | 211488 | HITS-CLIP | 25083871\|23597149\|23142080 | Unspecified |
| mmu-miR-29b-3p | MIMAT0000127 | Adora2a | 11540 | HITS-CLIP | 23142080 | Unspecified |
| mmu-miR-29b-3p | MIMAT0000127 | Adora2b | 11541 | HITS-CLIP | 26203562 | Unspecified |
| mmu-miR-29b-3p | MIMAT0000127 | Adprh | 11544 | sRNA-Seq//HITS-CLIP | 33184237\|25083871\|23142080 | Unspecified |
| mmu-miR-29b-3p | MIMAT0000127 | Adrb2 | 11555 | HITS-CLIP | 23142080 | Unspecified |
| mmu-miR-29b-3p | MIMAT0000127 | Adss | 11566 | HITS-CLIP//sRNA-Seq | 23597149\|23142080\|33184237\|25083 | Unspecified |
| mmu-miR-29b-3p | MIMAT0000127 | Adtrp | 109254 | HITS-CLIP | 23597149 | Unspecified |
| mmu-miR-29b-3p | MIMAT0000127 | Aebp1 | 11568 | HITS-CLIP//sRNA-Seq | 37986224\|25083871\|33184237 | Unspecified |

| | | | | | | |
|---|---|---|---|---|---|---|
| mmu-miR-29b-3p | MIMAT0000127 | Aebp2 | 11569 | HITS-CLIP | 25083871\|23142080 | Unspecified |
| mmu-miR-29b-3p | MIMAT0000127 | Afap1 | 70292 | HITS-CLIP | 37986224 | Unspecified |
| mmu-miR-29b-3p | MIMAT0000127 | Afap1l1 | 106877 | HITS-CLIP | 23597149 | Unspecified |
| mmu-miR-29b-3p | MIMAT0000127 | Afdn | 4301 | HITS-CLIP//sRNA-Seq | 25083871\|33184237 | Unspecified |
| mmu-miR-29b-3p | MIMAT0000127 | Aff1 | 17355 | HITS-CLIP | 25083871\|23142080 | Unspecified |
| mmu-miR-29b-3p | MIMAT0000127 | Aff3 | 16764 | sRNA-Seq//HITS-CLIP | 33184237\|23142080 | Unspecified |
| mmu-miR-29b-3p | MIMAT0000127 | Aff4 | 93736 | HITS-CLIP//sRNA-Seq | 37986224\|25083871\|33184237\|23597 | Unspecified |
| mmu-miR-29b-3p | MIMAT0000127 | Afg1l | 246269 | HITS-CLIP | 23142080 | Unspecified |
| mmu-miR-29b-3p | MIMAT0000127 | Afg3l1 | 114896 | HITS-CLIP | 23142080 | Unspecified |
| mmu-miR-29b-3p | MIMAT0000127 | Afg3l2 | 69597 | HITS-CLIP | 25083871 | Unspecified |
| mmu-miR-29b-3p | MIMAT0000127 | Afm | 280662 | HITS-CLIP | 23597149 | Unspecified |
| mmu-miR-29b-3p | MIMAT0000127 | Afmid | 71562 | HITS-CLIP | 23597149 | Unspecified |
| mmu-miR-29b-3p | MIMAT0000127 | Aftph | 216549 | HITS-CLIP | 23142080 | Unspecified |
| mmu-miR-29b-3p | MIMAT0000127 | Agap1 | 347722 | HITS-CLIP//sRNA-Seq | 37986224\|25083871\|33184237 | Unspecified |
| mmu-miR-29b-3p | MIMAT0000127 | Agfg1 | 15463 | HITS-CLIP//sRNA-Seq | 37986224\|25083871\|23142080\|23597 | Unspecified |
| mmu-miR-29b-3p | MIMAT0000127 | Aggf1 | 66549 | HITS-CLIP | 23142080 | Unspecified |
| mmu-miR-29b-3p | MIMAT0000127 | Agl | 77559 | HITS-CLIP | 25083871\|23142080 | Unspecified |
| mmu-miR-29b-3p | MIMAT0000127 | Agmo | 319660 | HITS-CLIP | 23597149 | Unspecified |
| mmu-miR-29b-3p | MIMAT0000127 | Ago1 | 236511 | HITS-CLIP | 37986224\|25083871\|23597149\|23142 | Unspecified |
| mmu-miR-29b-3p | MIMAT0000127 | Ago2 | 239528 | HITS-CLIP//sRNA-Seq | 37986224\|25083871\|33184237\|23597 | Unspecified |
| mmu-miR-29b-3p | MIMAT0000127 | Ago3 | 214150 | HITS-CLIP | 37986224\|25083871\|23142080 | Unspecified |
| mmu-miR-29b-3p | MIMAT0000127 | Agpat1 | 55979 | sRNA-Seq//HITS-CLIP | 33184237\|23142080 | Unspecified |
| mmu-miR-29b-3p | MIMAT0000127 | Agpat3 | 28169 | HITS-CLIP | 23597149\|25083871\|23142080 | Unspecified |
| mmu-miR-29b-3p | MIMAT0000127 | Agpat4 | 68262 | HITS-CLIP//sRNA-Seq | 25083871\|33184237 | Unspecified |
| mmu-miR-29b-3p | MIMAT0000127 | Agpat5 | 52123 | HITS-CLIP | 37986224\|25083871 | Unspecified |
| mmu-miR-29b-3p | MIMAT0000127 | Agps | 228061 | HITS-CLIP//sRNA-Seq | 37986224\|23597149\|33184237\|23142 | Unspecified |
| mmu-miR-29b-3p | MIMAT0000127 | Agrn | 11603 | HITS-CLIP//sRNA-Seq | 25083871\|33184237\|23597149 | Unspecified |
| mmu-miR-29b-3p | MIMAT0000127 | Agxt2 | 268782 | HITS-CLIP | 23597149 | Unspecified |
| mmu-miR-29b-3p | MIMAT0000127 | Ahctf1 | 226747 | HITS-CLIP | 23142080 | Unspecified |
| mmu-miR-29b-3p | MIMAT0000127 | Ahcyl1 | 229709 | HITS-CLIP//sRNA-Seq | 37986224\|23597149\|33184237\|23142 | Unspecified |
| mmu-miR-29b-3p | MIMAT0000127 | Ahcyl2 | 74340 | sRNA-Seq//HITS-CLIP | 33184237\|23142080 | Unspecified |
| mmu-miR-29b-3p | MIMAT0000127 | Ahdc1 | 230793 | HITS-CLIP | 25083871 | Unspecified |
| mmu-miR-29b-3p | MIMAT0000127 | Ahi1 | 52906 | HITS-CLIP | 23597149\|23142080 | Unspecified |
| mmu-miR-29b-3p | MIMAT0000127 | Ahnak | 66395 | HITS-CLIP//sRNA-Seq | 37986224\|25083871\|33184237\|23597 | Unspecified |
| mmu-miR-29b-3p | MIMAT0000127 | Ahr | 11622 | HITS-CLIP | 37986224\|23142080 | Unspecified |
| mmu-miR-29b-3p | MIMAT0000127 | Ahsa1 | 217737 | sRNA-Seq//HITS-CLIP | 33184237\|23142080 | Unspecified |
| mmu-miR-29b-3p | MIMAT0000127 | Ahsa2 | 268390 | HITS-CLIP//sRNA-Seq | 25083871\|33184237\|23597149 | Unspecified |
| mmu-miR-29b-3p | MIMAT0000127 | Aif1l | 108897 | HITS-CLIP | 25083871 | Unspecified |
| mmu-miR-29b-3p | MIMAT0000127 | Aifm1 | 26926 | HITS-CLIP//sRNA-Seq | 37986224\|25083871\|33184237\|23597 | Unspecified |

| | | | | | | |
|---|---|---|---|---|---|---|
| mmu-miR-29b-3p | MIMAT0000127 | Aifm2 | 71361 | HITS-CLIP | 23597149\|23142080 | Unspecified |
| mmu-miR-29b-3p | MIMAT0000127 | Aimp2 | 231872 | HITS-CLIP//sRNA-Seq | 37986224\|23597149\|25083871\|33184 | Unspecified |
| mmu-miR-29b-3p | MIMAT0000127 | Ajuba | 16475 | HITS-CLIP//sRNA-Seq | 25083871\|33184237\|23142080 | Unspecified |
| mmu-miR-29b-3p | MIMAT0000127 | Ak2 | 11637 | HITS-CLIP//sRNA-Seq | 25083871\|23597149\|33184237\|23142 | Unspecified |
| mmu-miR-29b-3p | MIMAT0000127 | Ak3 | 56248 | HITS-CLIP//sRNA-Seq | 37986224\|25083871\|23597149\|23142 | Unspecified |
| mmu-miR-29b-3p | MIMAT0000127 | Ak4 | 11639 | HITS-CLIP | 37986224\|23142080 | Unspecified |
| mmu-miR-29b-3p | MIMAT0000127 | Akap1 | 11640 | HITS-CLIP | 37986224\|23597149 | Unspecified |
| mmu-miR-29b-3p | MIMAT0000127 | Akap10 | 56697 | HITS-CLIP | 25083871\|23142080 | Unspecified |
| mmu-miR-29b-3p | MIMAT0000127 | Akap11 | 219181 | HITS-CLIP//sRNA-Seq | 25083871\|33184237\|23142080 | Unspecified |
| mmu-miR-29b-3p | MIMAT0000127 | Akap13 | 75547 | HITS-CLIP | 37986224\|23142080\|23597149\|25083 | Unspecified |
| mmu-miR-29b-3p | MIMAT0000127 | Akap17b | 338351 | HITS-CLIP | 23142080 | Unspecified |
| mmu-miR-29b-3p | MIMAT0000127 | Akap2 | 11641 | HITS-CLIP | 25083871 | Unspecified |
| mmu-miR-29b-3p | MIMAT0000127 | Akap6 | 238161 | HITS-CLIP//sRNA-Seq | 25083871\|33184237 | Unspecified |
| mmu-miR-29b-3p | MIMAT0000127 | Akap8l | 54194 | HITS-CLIP | 25083871 | Unspecified |
| mmu-miR-29b-3p | MIMAT0000127 | Akap9 | 100986 | HITS-CLIP//sRNA-Seq | 25083871\|23597149\|33184237 | Unspecified |
| mmu-miR-29b-3p | MIMAT0000127 | Akirin1 | 68050 | HITS-CLIP//sRNA-Seq | 25083871\|33184237\|23142080 | Unspecified |
| mmu-miR-29b-3p | MIMAT0000127 | Akna | 100182 | HITS-CLIP | 23142080 | Unspecified |
| mmu-miR-29b-3p | MIMAT0000127 | Akr1a1 | 58810 | HITS-CLIP//sRNA-Seq | 23597149\|33184237\|23142080 | Unspecified |
| mmu-miR-29b-3p | MIMAT0000127 | Akr1c12 | 622402 | HITS-CLIP | 23597149 | Unspecified |
| mmu-miR-29b-3p | MIMAT0000127 | Akr1c18 | 105349 | HITS-CLIP | 23142080 | Unspecified |
| mmu-miR-29b-3p | MIMAT0000127 | Akr1c20 | 116852 | HITS-CLIP | 23597149 | Unspecified |
| mmu-miR-29b-3p | MIMAT0000127 | Akr1c6 | 83702 | HITS-CLIP | 23597149 | Unspecified |
| mmu-miR-29b-3p | MIMAT0000127 | Akt3 | 23797 | Luciferase assay//qRT-PCR//Western blot/ | 23764849\|25083871\|33184237\|23142 | Unspecified |
| mmu-miR-29b-3p | MIMAT0000127 | Alad | 17025 | HITS-CLIP//sRNA-Seq | 23597149\|33184237 | Unspecified |
| mmu-miR-29b-3p | MIMAT0000127 | Alas1 | 11655 | HITS-CLIP//sRNA-Seq | 23142080\|25083871\|23597149\|33184 | Unspecified |
| mmu-miR-29b-3p | MIMAT0000127 | Alas2 | 11656 | HITS-CLIP | 37986224\|23597149 | Unspecified |
| mmu-miR-29b-3p | MIMAT0000127 | Alcam | 11658 | sRNA-Seq//HITS-CLIP | 33184237\|23142080 | Unspecified |
| mmu-miR-29b-3p | MIMAT0000127 | Aldh1l1 | 107747 | HITS-CLIP | 23597149 | Unspecified |
| mmu-miR-29b-3p | MIMAT0000127 | Aldh3a2 | 11671 | HITS-CLIP | 23597149 | Unspecified |
| mmu-miR-29b-3p | MIMAT0000127 | Aldh4a1 | 212647 | HITS-CLIP | 23597149 | Unspecified |
| mmu-miR-29b-3p | MIMAT0000127 | Aldh7a1 | 110695 | HITS-CLIP | 23597149 | Unspecified |
| mmu-miR-29b-3p | MIMAT0000127 | Aldh8a1 | 237320 | HITS-CLIP | 23597149 | Unspecified |
| mmu-miR-29b-3p | MIMAT0000127 | Aldoa | 11674 | HITS-CLIP//sRNA-Seq | 23597149\|33184237\|23142080 | Unspecified |
| mmu-miR-29b-3p | MIMAT0000127 | Aldoart2 | 79459 | HITS-CLIP | 23142080 | Unspecified |
| mmu-miR-29b-3p | MIMAT0000127 | Aldob | 230163 | HITS-CLIP | 23597149 | Unspecified |
| mmu-miR-29b-3p | MIMAT0000127 | Alg10b | 380959 | HITS-CLIP//sRNA-Seq | 25083871\|33184237\|23142080 | Unspecified |
| mmu-miR-29b-3p | MIMAT0000127 | Alg11 | 207958 | HITS-CLIP | 37986224\|25083871\|23142080 | Unspecified |
| mmu-miR-29b-3p | MIMAT0000127 | Alg14 | 66789 | HITS-CLIP | 25083871 | Unspecified |
| mmu-miR-29b-3p | MIMAT0000127 | Alg2 | 56737 | sRNA-Seq//HITS-CLIP | 33184237\|23142080 | Unspecified |

| | | | | | | |
|---|---|---|---|---|---|---|
| mmu-miR-29b-3p | MIMAT0000127 | Alg3 | 208624 | HITS-CLIP | 23142080 | Unspecified |
| mmu-miR-29b-3p | MIMAT0000127 | Alg9 | 102580 | HITS-CLIP | 37986224\|25083871\|23142080 | Unspecified |
| mmu-miR-29b-3p | MIMAT0000127 | Alkbh1 | 211064 | HITS-CLIP | 37986224 | Unspecified |
| mmu-miR-29b-3p | MIMAT0000127 | Alkbh4 | 72041 | HITS-CLIP | 23142080 | Unspecified |
| mmu-miR-29b-3p | MIMAT0000127 | Alkbh5 | 268420 | HITS-CLIP//sRNA-Seq | 37986224\|25083871\|33184237\|23142 | Unspecified |
| mmu-miR-29b-3p | MIMAT0000127 | Alkbh8 | 67667 | HITS-CLIP//sRNA-Seq | 37986224\|23142080\|33184237\|25083 | Unspecified |
| mmu-miR-29b-3p | MIMAT0000127 | Alms1 | 236266 | HITS-CLIP | 23142080\|25083871 | Unspecified |
| mmu-miR-29b-3p | MIMAT0000127 | Alpk3 | 116904 | HITS-CLIP | 25083871 | Unspecified |
| mmu-miR-29b-3p | MIMAT0000127 | Als2 | 74018 | HITS-CLIP | 37986224\|23597149 | Unspecified |
| mmu-miR-29b-3p | MIMAT0000127 | Alyref2 | 56009 | HITS-CLIP | 25083871\|23142080 | Unspecified |
| mmu-miR-29b-3p | MIMAT0000127 | Ambp | 11699 | HITS-CLIP | 23597149 | Unspecified |
| mmu-miR-29b-3p | MIMAT0000127 | Amdhd1 | 71761 | HITS-CLIP | 23597149 | Unspecified |
| mmu-miR-29b-3p | MIMAT0000127 | Amdhd2 | 245847 | HITS-CLIP//sRNA-Seq | 23142080\|33184237 | Unspecified |
| mmu-miR-29b-3p | MIMAT0000127 | Amer1 | 72345 | HITS-CLIP | 37986224\|25083871\|23142080 | Unspecified |
| mmu-miR-29b-3p | MIMAT0000127 | Amfr | 23802 | HITS-CLIP//sRNA-Seq | 37986224\|23597149\|33184237\|23142 | Unspecified |
| mmu-miR-29b-3p | MIMAT0000127 | Ammecr1 | 56068 | HITS-CLIP | 23142080 | Unspecified |
| mmu-miR-29b-3p | MIMAT0000127 | Ammecr1l | 225339 | HITS-CLIP | 37986224\|25083871\|23597149\|23142 | Unspecified |
| mmu-miR-29b-3p | MIMAT0000127 | Amot | 27494 | HITS-CLIP//sRNA-Seq | 25083871\|33184237\|23597149 | Unspecified |
| mmu-miR-29b-3p | MIMAT0000127 | Amotl1 | 75723 | HITS-CLIP//sRNA-Seq | 25083871\|33184237 | Unspecified |
| mmu-miR-29b-3p | MIMAT0000127 | Amotl2 | 56332 | HITS-CLIP//sRNA-Seq | 25083871\|33184237 | Unspecified |
| mmu-miR-29b-3p | MIMAT0000127 | Ampd3 | 11717 | HITS-CLIP | 37986224\|25083871 | Unspecified |
| mmu-miR-29b-3p | MIMAT0000127 | Amy1 | 11722 | HITS-CLIP | 23597149 | Unspecified |
| mmu-miR-29b-3p | MIMAT0000127 | Amz2 | 13929 | HITS-CLIP//sRNA-Seq | 37986224\|33184237\|23142080 | Unspecified |
| mmu-miR-29b-3p | MIMAT0000127 | Anapc1 | 17222 | HITS-CLIP//sRNA-Seq | 37986224\|23597149\|23142080\|25083 | Unspecified |
| mmu-miR-29b-3p | MIMAT0000127 | Anapc11 | 66156 | HITS-CLIP | 23142080 | Unspecified |
| mmu-miR-29b-3p | MIMAT0000127 | Anapc16 | 52717 | HITS-CLIP//sRNA-Seq | 25083871\|33184237 | Unspecified |
| mmu-miR-29b-3p | MIMAT0000127 | Anapc2 | 99152 | HITS-CLIP//sRNA-Seq | 37986224\|23597149\|33184237\|23142 | Unspecified |
| mmu-miR-29b-3p | MIMAT0000127 | Anapc4 | 52206 | HITS-CLIP//sRNA-Seq | 25083871\|33184237\|23142080 | Unspecified |
| mmu-miR-29b-3p | MIMAT0000127 | Anapc5 | 59008 | HITS-CLIP//sRNA-Seq | 37986224\|25083871\|33184237 | Unspecified |
| mmu-miR-29b-3p | MIMAT0000127 | Anapc7 | 56317 | HITS-CLIP | 23142080 | Unspecified |
| mmu-miR-29b-3p | MIMAT0000127 | Angel2 | 52477 | HITS-CLIP | 37986224 | Unspecified |
| mmu-miR-29b-3p | MIMAT0000127 | Angptl4 | 57875 | HITS-CLIP, Luciferase Reporter Assay//HITS | 37986224\|23597149\|23354167 | Unspecified |
| mmu-miR-29b-3p | MIMAT0000127 | Ank | 11732 | HITS-CLIP//sRNA-Seq | 25083871\|33184237 | Unspecified |
| mmu-miR-29b-3p | MIMAT0000127 | Ank1 | 11733 | HITS-CLIP | 25083871 | Unspecified |
| mmu-miR-29b-3p | MIMAT0000127 | Ank2 | 109676 | HITS-CLIP | 37986224 | Unspecified |
| mmu-miR-29b-3p | MIMAT0000127 | Ank3 | 11735 | HITS-CLIP//sRNA-Seq | 37986224\|25083871\|33184237 | Unspecified |
| mmu-miR-29b-3p | MIMAT0000127 | Ankfy1 | 11736 | HITS-CLIP//sRNA-Seq | 25083871\|23142080\|33184237 | Unspecified |
| mmu-miR-29b-3p | MIMAT0000127 | Ankhd1 | 108857 | HITS-CLIP | 23142080\|25083871 | Unspecified |
| mmu-miR-29b-3p | MIMAT0000127 | Ankib1 | 70797 | HITS-CLIP | 37986224\|25083871\|23142080 | Unspecified |

| | | | | | | |
|---|---|---|---|---|---|---|
| mmu-miR-29b-3p | MIMAT0000127 | Ankmy2 | 217473 | HITS-CLIP | 23597149 | Unspecified |
| mmu-miR-29b-3p | MIMAT0000127 | Ankrd10 | 102334 | HITS-CLIP | 37986224\|23142080\|25083871 | Unspecified |
| mmu-miR-29b-3p | MIMAT0000127 | Ankrd11 | 77087 | HITS-CLIP//sRNA-Seq | 25083871\|33184237\|23142080 | Unspecified |
| mmu-miR-29b-3p | MIMAT0000127 | Ankrd13a | 68420 | HITS-CLIP | 37986224 | Unspecified |
| mmu-miR-29b-3p | MIMAT0000127 | Ankrd13b | 268445 | HITS-CLIP | 37986224\|25083871 | Unspecified |
| mmu-miR-29b-3p | MIMAT0000127 | Ankrd13c | 433667 | HITS-CLIP//sRNA-Seq | 23597149\|33184237\|23142080 | Unspecified |
| mmu-miR-29b-3p | MIMAT0000127 | Ankrd17 | 81702 | HITS-CLIP//sRNA-Seq | 37986224\|23597149\|25083871\|23142 | Unspecified |
| mmu-miR-29b-3p | MIMAT0000127 | Ankrd28 | 105522 | HITS-CLIP | 37986224\|23597149\|23142080\|2508 | Unspecified |
| mmu-miR-29b-3p | MIMAT0000127 | Ankrd40 | 71452 | HITS-CLIP//sRNA-Seq | 25083871\|23597149\|33184237\|23142 | Unspecified |
| mmu-miR-29b-3p | MIMAT0000127 | Ankrd44 | 329154 | HITS-CLIP | 37986224\|23142080 | Unspecified |
| mmu-miR-29b-3p | MIMAT0000127 | Ankrd46 | 68839 | sRNA-Seq//HITS-CLIP | 33184237\|23597149\|23142080 | Unspecified |
| mmu-miR-29b-3p | MIMAT0000127 | Ankrd49 | 56503 | HITS-CLIP | 37986224\|23142080\|25083871 | Unspecified |
| mmu-miR-29b-3p | MIMAT0000127 | Ankrd50 | 99696 | HITS-CLIP//sRNA-Seq | 37986224\|25083871\|23142080\|3318 | Unspecified |
| mmu-miR-29b-3p | MIMAT0000127 | Ankrd52 | 237615 | HITS-CLIP | 23597149\|25083871\|23142080 | Unspecified |
| mmu-miR-29b-3p | MIMAT0000127 | Anks4b | 72074 | HITS-CLIP | 37986224\|23597149 | Unspecified |
| mmu-miR-29b-3p | MIMAT0000127 | Ankzf1 | 52231 | HITS-CLIP | 37986224\|23142080 | Unspecified |
| mmu-miR-29b-3p | MIMAT0000127 | Ano10 | 102566 | sRNA-Seq//HITS-CLIP | 33184237\|23597149 | Unspecified |
| mmu-miR-29b-3p | MIMAT0000127 | Ano5 | 233246 | HITS-CLIP | 37986224 | Unspecified |
| mmu-miR-29b-3p | MIMAT0000127 | Anp32a | 11737 | HITS-CLIP//sRNA-Seq | 25083871\|33184237 | Unspecified |
| mmu-miR-29b-3p | MIMAT0000127 | Anpep | 16790 | HITS-CLIP | 23597149 | Unspecified |
| mmu-miR-29b-3p | MIMAT0000127 | Antxr1 | 69538 | HITS-CLIP | 37986224 | Unspecified |
| mmu-miR-29b-3p | MIMAT0000127 | Antxr2 | 71914 | HITS-CLIP | 23597149 | Unspecified |
| mmu-miR-29b-3p | MIMAT0000127 | Anxa2 | 12306 | sRNA-Seq//HITS-CLIP | 33184237\|23142080 | Unspecified |
| mmu-miR-29b-3p | MIMAT0000127 | Anxa5 | 11747 | HITS-CLIP//sRNA-Seq | 25083871\|33184237 | Unspecified |
| mmu-miR-29b-3p | MIMAT0000127 | Anxa7 | 11750 | HITS-CLIP//sRNA-Seq | 23597149\|33184237 | Unspecified |
| mmu-miR-29b-3p | MIMAT0000127 | Aox3 | 71724 | HITS-CLIP | 23597149 | Unspecified |
| mmu-miR-29b-3p | MIMAT0000127 | Ap1b1 | 11764 | HITS-CLIP//sRNA-Seq | 37986224\|23597149\|33184237\|23142 | Unspecified |
| mmu-miR-29b-3p | MIMAT0000127 | Ap1g1 | 11765 | HITS-CLIP//sRNA-Seq | 37986224\|25083871\|23597149\|3318 | Unspecified |
| mmu-miR-29b-3p | MIMAT0000127 | Ap1g2 | 11766 | HITS-CLIP | 23142080 | Unspecified |
| mmu-miR-29b-3p | MIMAT0000127 | Ap1m1 | 11767 | sRNA-Seq//HITS-CLIP | 33184237\|25083871 | Unspecified |
| mmu-miR-29b-3p | MIMAT0000127 | Ap1s2 | 108012 | sRNA-Seq//HITS-CLIP | 33184237\|23142080 | Unspecified |
| mmu-miR-29b-3p | MIMAT0000127 | Ap2a2 | 11772 | HITS-CLIP//sRNA-Seq | 25083871\|33184237 | Unspecified |
| mmu-miR-29b-3p | MIMAT0000127 | Ap2b1 | 71770 | HITS-CLIP//sRNA-Seq | 37986224\|23142080\|25083871\|3318 | Unspecified |
| mmu-miR-29b-3p | MIMAT0000127 | Ap2s1 | 232910 | HITS-CLIP//sRNA-Seq | 37986224\|25083871\|33184237\|2359 | Unspecified |
| mmu-miR-29b-3p | MIMAT0000127 | Ap3b1 | 11774 | sRNA-Seq//HITS-CLIP | 33184237\|23142080\|25083871 | Unspecified |
| mmu-miR-29b-3p | MIMAT0000127 | Ap4b1 | 67489 | HITS-CLIP | 23142080 | Unspecified |
| mmu-miR-29b-3p | MIMAT0000127 | Ap4e1 | 108011 | HITS-CLIP | 37986224\|23142080\|25083871 | Unspecified |
| mmu-miR-29b-3p | MIMAT0000127 | Ap5b1 | 381201 | HITS-CLIP | 25083871\|23142080 | Unspecified |
| mmu-miR-29b-3p | MIMAT0000127 | Apaf1 | 11783 | HITS-CLIP | 37986224\|25083871\|23142080 | Unspecified |

| | | | | | | |
|---|---|---|---|---|---|---|
| mmu-miR-29b-3p | MIMAT0000127 | Apbb1ip | 54519 | HITS-CLIP//sRNA-Seq | 23142080\|33184237\|25083871 | Unspecified |
| mmu-miR-29b-3p | MIMAT0000127 | Apbb2 | 11787 | HITS-CLIP//sRNA-Seq | 25083871\|33184237 | Unspecified |
| mmu-miR-29b-3p | MIMAT0000127 | Apc | 11789 | HITS-CLIP//sRNA-Seq | 37986224\|25083871\|33184237\|23142 | Unspecified |
| mmu-miR-29b-3p | MIMAT0000127 | Apex1 | 11792 | sRNA-Seq//HITS-CLIP | 33184237\|23597149 | Unspecified |
| mmu-miR-29b-3p | MIMAT0000127 | Apex2 | 77622 | HITS-CLIP | 23142080 | Unspecified |
| mmu-miR-29b-3p | MIMAT0000127 | Aplp2 | 11804 | HITS-CLIP//sRNA-Seq | 25083871\|33184237\|23597149\|23142 | Unspecified |
| mmu-miR-29b-3p | MIMAT0000127 | Apmap | 71881 | HITS-CLIP | 25083871 | Unspecified |
| mmu-miR-29b-3p | MIMAT0000127 | Apoa1 | 11806 | HITS-CLIP | 23597149 | Unspecified |
| mmu-miR-29b-3p | MIMAT0000127 | Apoa5 | 66113 | HITS-CLIP | 23597149 | Unspecified |
| mmu-miR-29b-3p | MIMAT0000127 | Apob | 238055 | HITS-CLIP | 37986224\|23597149 | Unspecified |
| mmu-miR-29b-3p | MIMAT0000127 | Apool | 68117 | HITS-CLIP | 23142080 | Unspecified |
| mmu-miR-29b-3p | MIMAT0000127 | App | 11820 | sRNA-Seq//HITS-CLIP | 33184237\|25083871 | Unspecified |
| mmu-miR-29b-3p | MIMAT0000127 | Appbp2 | 66884 | HITS-CLIP//sRNA-Seq | 37986224\|25083871\|33184237\|23142 | Unspecified |
| mmu-miR-29b-3p | MIMAT0000127 | Appl1 | 72993 | HITS-CLIP | 37986224\|25083871\|23142080 | Unspecified |
| mmu-miR-29b-3p | MIMAT0000127 | Appl2 | 216190 | HITS-CLIP | 23142080 | Unspecified |
| mmu-miR-29b-3p | MIMAT0000127 | Aqp11 | 66333 | HITS-CLIP | 23142080\|23597149 | Unspecified |
| mmu-miR-29b-3p | MIMAT0000127 | Araf | 11836 | sRNA-Seq//HITS-CLIP | 33184237\|23142080 | Unspecified |
| mmu-miR-29b-3p | MIMAT0000127 | Arap1 | 69710 | HITS-CLIP | 23142080 | Unspecified |
| mmu-miR-29b-3p | MIMAT0000127 | Arap2 | 212285 | HITS-CLIP | 23142080 | Unspecified |
| mmu-miR-29b-3p | MIMAT0000127 | Arcn1 | 213827 | HITS-CLIP//sRNA-Seq | 25083871\|33184237\|23142080 | Unspecified |
| mmu-miR-29b-3p | MIMAT0000127 | Areg | 11839 | RNA-Seq | 33332475 | Unspecified |
| mmu-miR-29b-3p | MIMAT0000127 | Arel1 | 68497 | HITS-CLIP | 25083871 | Unspecified |
| mmu-miR-29b-3p | MIMAT0000127 | Arf1 | 11840 | HITS-CLIP//sRNA-Seq | 23597149\|33184237 | Unspecified |
| mmu-miR-29b-3p | MIMAT0000127 | Arf2 | 11841 | HITS-CLIP | 37986224\|25083871\|23597149\|23142 | Unspecified |
| mmu-miR-29b-3p | MIMAT0000127 | Arf3 | 11842 | HITS-CLIP//sRNA-Seq | 37986224\|25083871\|23597149\|33184 | Unspecified |
| mmu-miR-29b-3p | MIMAT0000127 | Arf4 | 11843 | HITS-CLIP//sRNA-Seq | 23142080\|33184237 | Unspecified |
| mmu-miR-29b-3p | MIMAT0000127 | Arf5 | 11844 | HITS-CLIP//sRNA-Seq | 37986224\|23597149\|25083871\|33184 | Unspecified |
| mmu-miR-29b-3p | MIMAT0000127 | Arf6 | 11845 | HITS-CLIP//sRNA-Seq | 37986224\|25083871\|33184237\|23142 | Unspecified |
| mmu-miR-29b-3p | MIMAT0000127 | Arfgap1 | 228998 | HITS-CLIP//sRNA-Seq | 23597149\|33184237\|23142080 | Unspecified |
| mmu-miR-29b-3p | MIMAT0000127 | Arfgap3 | 66251 | HITS-CLIP//sRNA-Seq | 25083871\|33184237 | Unspecified |
| mmu-miR-29b-3p | MIMAT0000127 | Arfgef1 | 211673 | HITS-CLIP//sRNA-Seq | 25083871\|33184237\|23142080 | Unspecified |
| mmu-miR-29b-3p | MIMAT0000127 | Arfgef2 | 99371 | HITS-CLIP | 37986224\|25083871\|23597149\|23142 | Unspecified |
| mmu-miR-29b-3p | MIMAT0000127 | Arglu1 | 234023 | HITS-CLIP | 23597149 | Unspecified |
| mmu-miR-29b-3p | MIMAT0000127 | Arhgap11a | 228482 | HITS-CLIP | 37986224\|23142080 | Unspecified |
| mmu-miR-29b-3p | MIMAT0000127 | Arhgap12 | 75415 | HITS-CLIP//sRNA-Seq | 23142080\|33184237 | Unspecified |
| mmu-miR-29b-3p | MIMAT0000127 | Arhgap15 | 76117 | HITS-CLIP | 23142080 | Unspecified |
| mmu-miR-29b-3p | MIMAT0000127 | Arhgap17 | 70497 | HITS-CLIP | 23142080 | Unspecified |
| mmu-miR-29b-3p | MIMAT0000127 | Arhgap19 | 71085 | HITS-CLIP | 23597149\|23142080 | Unspecified |
| mmu-miR-29b-3p | MIMAT0000127 | Arhgap23 | 58996 | HITS-CLIP | 25083871 | Unspecified |

| | | | | | | |
|---|---|---|---|---|---|---|
| mmu-miR-29b-3p | MIMAT0000127 | Arhgap28 | 268970 | HITS-CLIP | 37986224 | Unspecified |
| mmu-miR-29b-3p | MIMAT0000127 | Arhgap29 | 214137 | HITS-CLIP//sRNA-Seq | 23597149\|33184237 | Unspecified |
| mmu-miR-29b-3p | MIMAT0000127 | Arhgap35 | 232906 | sRNA-Seq//HITS-CLIP | 33184237\|23142080 | Unspecified |
| mmu-miR-29b-3p | MIMAT0000127 | Arhgap39 | 223666 | HITS-CLIP | 23597149 | Unspecified |
| mmu-miR-29b-3p | MIMAT0000127 | Arhgap45 | 23526 | HITS-CLIP | 23142080 | Unspecified |
| mmu-miR-29b-3p | MIMAT0000127 | Arhgap5 | 11855 | HITS-CLIP//sRNA-Seq | 37986224\|23142080\|33184237\|23597 | Unspecified |
| mmu-miR-29b-3p | MIMAT0000127 | Arhgap9 | 216445 | HITS-CLIP | 23142080 | Unspecified |
| mmu-miR-29b-3p | MIMAT0000127 | Arhgdia | 192662 | HITS-CLIP//sRNA-Seq | 23597149\|33184237\|23142080 | Unspecified |
| mmu-miR-29b-3p | MIMAT0000127 | Arhgef1 | 16801 | HITS-CLIP | 23142080 | Unspecified |
| mmu-miR-29b-3p | MIMAT0000127 | Arhgef10 | 234094 | HITS-CLIP | 37986224\|23142080 | Unspecified |
| mmu-miR-29b-3p | MIMAT0000127 | Arhgef10l | 72754 | HITS-CLIP | 25083871\|23597149 | Unspecified |
| mmu-miR-29b-3p | MIMAT0000127 | Arhgef12 | 69632 | HITS-CLIP | 25083871\|23142080\|23597149 | Unspecified |
| mmu-miR-29b-3p | MIMAT0000127 | Arhgef18 | 102098 | HITS-CLIP | 37986224 | Unspecified |
| mmu-miR-29b-3p | MIMAT0000127 | Arhgef2 | 16800 | HITS-CLIP//sRNA-Seq | 25083871\|33184237 | Unspecified |
| mmu-miR-29b-3p | MIMAT0000127 | Arhgef6 | 73341 | HITS-CLIP | 23597149 | Unspecified |
| mmu-miR-29b-3p | MIMAT0000127 | Arid1a | 93760 | HITS-CLIP | 23142080\|25083871\|23597149 | Unspecified |
| mmu-miR-29b-3p | MIMAT0000127 | Arid1b | 239985 | HITS-CLIP | 37986224\|25083871\|23142080 | Unspecified |
| mmu-miR-29b-3p | MIMAT0000127 | Arid2 | 77044 | HITS-CLIP | 25083871\|23597149\|23142080 | Unspecified |
| mmu-miR-29b-3p | MIMAT0000127 | Arid4a | 238247 | HITS-CLIP | 25083871\|23142080 | Unspecified |
| mmu-miR-29b-3p | MIMAT0000127 | Arid4b | 94246 | HITS-CLIP | 25083871\|23142080 | Unspecified |
| mmu-miR-29b-3p | MIMAT0000127 | Arid5b | 71371 | HITS-CLIP | 23142080\|23597149\|25083871 | Unspecified |
| mmu-miR-29b-3p | MIMAT0000127 | Arih1 | 23806 | HITS-CLIP//sRNA-Seq | 37986224\|25083871\|23142080\|33184 | Unspecified |
| mmu-miR-29b-3p | MIMAT0000127 | Arih2 | 23807 | HITS-CLIP | 37986224 | Unspecified |
| mmu-miR-29b-3p | MIMAT0000127 | Arl1 | 104303 | HITS-CLIP//sRNA-Seq | 25083871\|33184237 | Unspecified |
| mmu-miR-29b-3p | MIMAT0000127 | Arl10 | 56795 | HITS-CLIP | 37986224 | Unspecified |
| mmu-miR-29b-3p | MIMAT0000127 | Arl14ep | 212772 | HITS-CLIP | 25083871 | Unspecified |
| mmu-miR-29b-3p | MIMAT0000127 | Arl4a | 11861 | HITS-CLIP | 23142080 | Unspecified |
| mmu-miR-29b-3p | MIMAT0000127 | Arl4c | 320982 | HITS-CLIP | 37986224\|25083871 | Unspecified |
| mmu-miR-29b-3p | MIMAT0000127 | Arl5a | 75423 | HITS-CLIP | 37986224\|23142080 | Unspecified |
| mmu-miR-29b-3p | MIMAT0000127 | Arl5b | 75869 | HITS-CLIP | 37986224\|25083871\|23597149\|23142 | Unspecified |
| mmu-miR-29b-3p | MIMAT0000127 | Arl5c | 217151 | HITS-CLIP | 23142080 | Unspecified |
| mmu-miR-29b-3p | MIMAT0000127 | Arl6ip1 | 54208 | HITS-CLIP//sRNA-Seq | 23597149\|25083871\|33184237\|23142 | Unspecified |
| mmu-miR-29b-3p | MIMAT0000127 | Arl6ip5 | 65106 | HITS-CLIP//sRNA-Seq | 25083871\|33184237\|23142080 | Unspecified |
| mmu-miR-29b-3p | MIMAT0000127 | Arl6ip6 | 65103 | HITS-CLIP | 25083871\|23142080 | Unspecified |
| mmu-miR-29b-3p | MIMAT0000127 | Arl8a | 68724 | sRNA-Seq//HITS-CLIP | 33184237\|23142080 | Unspecified |
| mmu-miR-29b-3p | MIMAT0000127 | Arl8b | 67166 | HITS-CLIP//sRNA-Seq | 23597149\|25083871\|33184237 | Unspecified |
| mmu-miR-29b-3p | MIMAT0000127 | Armc1 | 74252 | HITS-CLIP | 37986224 | Unspecified |
| mmu-miR-29b-3p | MIMAT0000127 | Armc8 | 74125 | HITS-CLIP//sRNA-Seq | 37986224\|23142080\|33184237 | Unspecified |
| mmu-miR-29b-3p | MIMAT0000127 | Armcx4 | 100503043 | HITS-CLIP | 23597149\|23142080 | Unspecified |

| | | | | | |
|---|---|---|---|---|---|
| mmu-miR-29b-3p | MIMAT0000127 | Armcx5 | 494468 | HITS-CLIP | 23142080 | Unspecified |
| mmu-miR-29b-3p | MIMAT0000127 | Armh3 | 79591 | HITS-CLIP | 25083871\|23142080 | Unspecified |
| mmu-miR-29b-3p | MIMAT0000127 | Arnt | 11863 | HITS-CLIP//sRNA-Seq | 37986224\|25083871\|23597149\|33184 | Unspecified |
| mmu-miR-29b-3p | MIMAT0000127 | Arntl | 11865 | HITS-CLIP | 23142080\|25083871 | Unspecified |
| mmu-miR-29b-3p | MIMAT0000127 | Arpc1a | 56443 | HITS-CLIP//sRNA-Seq | 25083871\|33184237 | Unspecified |
| mmu-miR-29b-3p | MIMAT0000127 | Arpc2 | 76709 | HITS-CLIP//sRNA-Seq | 25083871\|33184237\|23597149\|23142 | Unspecified |
| mmu-miR-29b-3p | MIMAT0000127 | Arpc5 | 67771 | HITS-CLIP//sRNA-Seq | 37986224\|25083871\|33184237\|23142 | Unspecified |
| mmu-miR-29b-3p | MIMAT0000127 | Arpp19 | 59046 | HITS-CLIP//sRNA-Seq | 37986224\|23597149\|23142080\|33184 | Unspecified |
| mmu-miR-29b-3p | MIMAT0000127 | Arpp21 | 74100 | HITS-CLIP | 25083871 | Unspecified |
| mmu-miR-29b-3p | MIMAT0000127 | Arrb2 | 216869 | HITS-CLIP | 23597149\|23142080 | Unspecified |
| mmu-miR-29b-3p | MIMAT0000127 | Arrdc3 | 105171 | HITS-CLIP | 37986224\|25083871\|23142080 | Unspecified |
| mmu-miR-29b-3p | MIMAT0000127 | Arrdc4 | 66412 | HITS-CLIP | 23597149\|23142080 | Unspecified |
| mmu-miR-29b-3p | MIMAT0000127 | Arvcf | 11877 | HITS-CLIP | 37986224\|25083871\|23597149 | Unspecified |
| mmu-miR-29b-3p | MIMAT0000127 | As3mt | 57344 | HITS-CLIP | 23597149 | Unspecified |
| mmu-miR-29b-3p | MIMAT0000127 | Asah1 | 11886 | HITS-CLIP//sRNA-Seq | 23597149\|33184237 | Unspecified |
| mmu-miR-29b-3p | MIMAT0000127 | Asb2 | 65256 | HITS-CLIP | 23142080 | Unspecified |
| mmu-miR-29b-3p | MIMAT0000127 | Asb3 | 65257 | HITS-CLIP | 37986224\|23142080 | Unspecified |
| mmu-miR-29b-3p | MIMAT0000127 | Asb8 | 78541 | HITS-CLIP | 23142080 | Unspecified |
| mmu-miR-29b-3p | MIMAT0000127 | Asf1a | 66403 | HITS-CLIP | 23142080 | Unspecified |
| mmu-miR-29b-3p | MIMAT0000127 | Asf1b | 66929 | HITS-CLIP | 23142080 | Unspecified |
| mmu-miR-29b-3p | MIMAT0000127 | Ash1l | 192195 | HITS-CLIP | 25083871\|23142080 | Unspecified |
| mmu-miR-29b-3p | MIMAT0000127 | Asph | 65973 | HITS-CLIP | 37986224\|25083871 | Unspecified |
| mmu-miR-29b-3p | MIMAT0000127 | Aspm | 12316 | HITS-CLIP | 23142080 | Unspecified |
| mmu-miR-29b-3p | MIMAT0000127 | Asxl1 | 228790 | HITS-CLIP | 37986224\|23142080\|25083871 | Unspecified |
| mmu-miR-29b-3p | MIMAT0000127 | Atad1 | 67979 | HITS-CLIP//sRNA-Seq | 23142080\|33184237\|25083871 | Unspecified |
| mmu-miR-29b-3p | MIMAT0000127 | Atad2 | 70472 | HITS-CLIP//sRNA-Seq | 37986224\|33184237\|23142080 | Unspecified |
| mmu-miR-29b-3p | MIMAT0000127 | Atad2b | 320817 | HITS-CLIP | 37986224\|25083871\|23597149\|23142 | Unspecified |
| mmu-miR-29b-3p | MIMAT0000127 | Atad5 | 237877 | HITS-CLIP | 23142080 | Unspecified |
| mmu-miR-29b-3p | MIMAT0000127 | Ate1 | 11907 | HITS-CLIP | 23597149 | Unspecified |
| mmu-miR-29b-3p | MIMAT0000127 | Atf2 | 11909 | HITS-CLIP//sRNA-Seq | 25083871\|33184237 | Unspecified |
| mmu-miR-29b-3p | MIMAT0000127 | Atf5 | 107503 | sRNA-Seq//HITS-CLIP | 33184237\|23597149 | Unspecified |
| mmu-miR-29b-3p | MIMAT0000127 | Atf6 | 226641 | HITS-CLIP | 23597149\|25083871\|23142080 | Unspecified |
| mmu-miR-29b-3p | MIMAT0000127 | Atf7 | 223922 | HITS-CLIP | 37986224\|25083871\|23142080 | Unspecified |
| mmu-miR-29b-3p | MIMAT0000127 | Atf7ip | 54343 | HITS-CLIP | 25083871\|23142080 | Unspecified |
| mmu-miR-29b-3p | MIMAT0000127 | Atg12 | 67526 | sRNA-Seq//HITS-CLIP | 33184237\|23142080 | Unspecified |
| mmu-miR-29b-3p | MIMAT0000127 | Atg13 | 51897 | HITS-CLIP | 25083871 | Unspecified |
| mmu-miR-29b-3p | MIMAT0000127 | Atg16l1 | 77040 | HITS-CLIP | 23142080 | Unspecified |
| mmu-miR-29b-3p | MIMAT0000127 | Atg9a | 245860 | HITS-CLIP | 37986224\|23142080\|25083871\|23597 | Unspecified |
| mmu-miR-29b-3p | MIMAT0000127 | Atl2 | 56298 | HITS-CLIP | 25083871\|23142080 | Unspecified |

| | | | | | | |
|---|---|---|---|---|---|---|
| mmu-miR-29b-3p | MIMAT0000127 | Atl3 | 109168 | HITS-CLIP//sRNA-Seq//qCLASH | 37986224\|23142080\|25083871\|33184 | Unspecified |
| mmu-miR-29b-3p | MIMAT0000127 | Atm | 11920 | HITS-CLIP | 23142080 | Unspecified |
| mmu-miR-29b-3p | MIMAT0000127 | Atn1 | 13498 | HITS-CLIP | 25083871 | Unspecified |
| mmu-miR-29b-3p | MIMAT0000127 | Atp10a | 11982 | HITS-CLIP | 23142080 | Unspecified |
| mmu-miR-29b-3p | MIMAT0000127 | Atp11a | 50770 | HITS-CLIP//sRNA-Seq | 25083871\|33184237\|23142080 | Unspecified |
| mmu-miR-29b-3p | MIMAT0000127 | Atp11b | 76295 | HITS-CLIP | 37986224\|25083871\|23142080 | Unspecified |
| mmu-miR-29b-3p | MIMAT0000127 | Atp11c | 320940 | HITS-CLIP | 23597149\|23142080 | Unspecified |
| mmu-miR-29b-3p | MIMAT0000127 | Atp13a1 | 170759 | HITS-CLIP | 23597149 | Unspecified |
| mmu-miR-29b-3p | MIMAT0000127 | Atp13a3 | 224088 | HITS-CLIP//sRNA-Seq | 23597149\|33184237\|23142080 | Unspecified |
| mmu-miR-29b-3p | MIMAT0000127 | Atp1a1 | 11928 | sRNA-Seq//HITS-CLIP | 33184237\|23597149\|23142080 | Unspecified |
| mmu-miR-29b-3p | MIMAT0000127 | Atp1a2 | 98660 | HITS-CLIP | 25083871 | Unspecified |
| mmu-miR-29b-3p | MIMAT0000127 | Atp1b1 | 11931 | HITS-CLIP//sRNA-Seq | 23597149\|25083871\|33184237 | Unspecified |
| mmu-miR-29b-3p | MIMAT0000127 | Atp2a2 | 11938 | HITS-CLIP//sRNA-Seq | 25083871\|23142080\|33184237 | Unspecified |
| mmu-miR-29b-3p | MIMAT0000127 | Atp2a3 | 53313 | HITS-CLIP | 23142080 | Unspecified |
| mmu-miR-29b-3p | MIMAT0000127 | Atp2b1 | 67972 | HITS-CLIP//sRNA-Seq | 37986224\|23142080\|33184237\|25083 | Unspecified |
| mmu-miR-29b-3p | MIMAT0000127 | Atp2b4 | 381290 | HITS-CLIP | 37986224\|25083871\|23142080 | Unspecified |
| mmu-miR-29b-3p | MIMAT0000127 | Atp2c1 | 235574 | sRNA-Seq//HITS-CLIP | 33184237\|23142080 | Unspecified |
| mmu-miR-29b-3p | MIMAT0000127 | Atp5a1 | 11946 | HITS-CLIP//sRNA-Seq | 25083871\|33184237\|23597149\|23142 | Unspecified |
| mmu-miR-29b-3p | MIMAT0000127 | Atp5b | 11947 | HITS-CLIP//qCLASH//sRNA-Seq | 23597149\|25083871\|23142080\|31393 | Unspecified |
| mmu-miR-29b-3p | MIMAT0000127 | Atp5c1 | 11949 | HITS-CLIP//sRNA-Seq | 25083871\|33184237\|23597149\|23142 | Unspecified |
| mmu-miR-29b-3p | MIMAT0000127 | Atp5f1 | 515 | HITS-CLIP//sRNA-Seq | 25083871\|33184237\|23597149\|23142 | Unspecified |
| mmu-miR-29b-3p | MIMAT0000127 | Atp5g3 | 228033 | HITS-CLIP//sRNA-Seq | 37986224\|33184237\|23597149\|23142 | Unspecified |
| mmu-miR-29b-3p | MIMAT0000127 | Atp5j | 11957 | HITS-CLIP//sRNA-Seq | 37986224\|23597149\|23142080\|33184 | Unspecified |
| mmu-miR-29b-3p | MIMAT0000127 | Atp5k | 11958 | HITS-CLIP//sRNA-Seq | 25083871\|33184237\|23597149 | Unspecified |
| mmu-miR-29b-3p | MIMAT0000127 | Atp6ap1 | 54411 | HITS-CLIP//sRNA-Seq | 25083871\|33184237\|23142080 | Unspecified |
| mmu-miR-29b-3p | MIMAT0000127 | Atp6ap2 | 70495 | HITS-CLIP//sRNA-Seq | 23597149\|33184237\|23142080 | Unspecified |
| mmu-miR-29b-3p | MIMAT0000127 | Atp6v0a1 | 11975 | HITS-CLIP//sRNA-Seq | 25083871\|33184237 | Unspecified |
| mmu-miR-29b-3p | MIMAT0000127 | Atp6v0a2 | 21871 | HITS-CLIP | 25083871\|23597149\|23142080 | Unspecified |
| mmu-miR-29b-3p | MIMAT0000127 | Atp6v0b | 114143 | sRNA-Seq//HITS-CLIP | 33184237\|23597149 | Unspecified |
| mmu-miR-29b-3p | MIMAT0000127 | Atp6v0d1 | 11972 | HITS-CLIP//sRNA-Seq | 23597149\|25083871\|33184237\|23142 | Unspecified |
| mmu-miR-29b-3p | MIMAT0000127 | Atp6v0e | 11974 | HITS-CLIP | 25083871 | Unspecified |
| mmu-miR-29b-3p | MIMAT0000127 | Atp6v1a | 11964 | HITS-CLIP//sRNA-Seq | 25083871\|33184237\|23142080 | Unspecified |
| mmu-miR-29b-3p | MIMAT0000127 | Atp6v1b2 | 11966 | HITS-CLIP//sRNA-Seq | 23597149\|25083871\|33184237\|23142 | Unspecified |
| mmu-miR-29b-3p | MIMAT0000127 | Atp6v1c1 | 66335 | HITS-CLIP//sRNA-Seq | 37986224\|23597149\|23142080\|25083 | Unspecified |
| mmu-miR-29b-3p | MIMAT0000127 | Atp6v1h | 108664 | HITS-CLIP//sRNA-Seq | 37986224\|25083871\|33184237\|23142 | Unspecified |
| mmu-miR-29b-3p | MIMAT0000127 | Atp7b | 11979 | HITS-CLIP | 23597149 | Unspecified |
| mmu-miR-29b-3p | MIMAT0000127 | Atp8a1 | 11980 | HITS-CLIP//sRNA-Seq | 23597149\|33184237 | Unspecified |
| mmu-miR-29b-3p | MIMAT0000127 | Atp8b2 | 54667 | HITS-CLIP//sRNA-Seq | 25083871\|33184237\|23142080 | Unspecified |
| mmu-miR-29b-3p | MIMAT0000127 | Atpif1 | 11983 | HITS-CLIP//sRNA-Seq | 25083871\|33184237\|23142080 | Unspecified |

| | | | | | | |
|---|---|---|---|---|---|---|
| mmu-miR-29b-3p | MIMAT0000127 | Atr | 245000 | HITS-CLIP | 23142080 | Unspecified |
| mmu-miR-29b-3p | MIMAT0000127 | Atrn | 11990 | HITS-CLIP | 23597149\|25083871\|23142080 | Unspecified |
| mmu-miR-29b-3p | MIMAT0000127 | Atrx | 22589 | HITS-CLIP//sRNA-Seq | 37986224\|23142080\|25083871\|33184 | Unspecified |
| mmu-miR-29b-3p | MIMAT0000127 | Atxn1 | 20238 | HITS-CLIP | 37986224\|25083871\|23142080 | Unspecified |
| mmu-miR-29b-3p | MIMAT0000127 | Atxn10 | 54138 | HITS-CLIP//sRNA-Seq | 23142080\|25083871\|33184237 | Unspecified |
| mmu-miR-29b-3p | MIMAT0000127 | Atxn1l | 52335 | HITS-CLIP | 37986224\|25083871\|23597149\|23142 | Unspecified |
| mmu-miR-29b-3p | MIMAT0000127 | Atxn2 | 20239 | HITS-CLIP//sRNA-Seq | 25083871\|33184237\|23142080 | Unspecified |
| mmu-miR-29b-3p | MIMAT0000127 | Atxn2l | 233871 | HITS-CLIP//sRNA-Seq | 37986224\|25083871\|23142080\|33184 | Unspecified |
| mmu-miR-29b-3p | MIMAT0000127 | Atxn3 | 110616 | HITS-CLIP//qCLASH | 25083871\|31393953 | Unspecified |
| mmu-miR-29b-3p | MIMAT0000127 | Atxn7 | 246103 | HITS-CLIP | 25083871\|23142080 | Unspecified |
| mmu-miR-29b-3p | MIMAT0000127 | Atxn7l1 | 380753 | HITS-CLIP | 37986224\|23142080 | Unspecified |
| mmu-miR-29b-3p | MIMAT0000127 | Atxn7l3 | 217218 | HITS-CLIP//sRNA-Seq | 25083871\|33184237 | Unspecified |
| mmu-miR-29b-3p | MIMAT0000127 | Atxn7l3b | 382423 | sRNA-Seq//HITS-CLIP | 33184237\|23142080 | Unspecified |
| mmu-miR-29b-3p | MIMAT0000127 | Aurkb | 20877 | HITS-CLIP//sRNA-Seq | 37986224\|23142080\|33184237 | Unspecified |
| mmu-miR-29b-3p | MIMAT0000127 | Auts2 | 319974 | HITS-CLIP | 37986224 | Unspecified |
| mmu-miR-29b-3p | MIMAT0000127 | Avl9 | 78937 | HITS-CLIP | 37986224\|23142080 | Unspecified |
| mmu-miR-29b-3p | MIMAT0000127 | Axl | 26362 | HITS-CLIP//sRNA-Seq | 23597149\|23142080\|33184237\|25083 | Unspecified |
| mmu-miR-29b-3p | MIMAT0000127 | Azgp1 | 12007 | HITS-CLIP | 23597149 | Unspecified |
| mmu-miR-29b-3p | MIMAT0000127 | Azin1 | 54375 | HITS-CLIP//sRNA-Seq | 23597149\|33184237\|23142080 | Unspecified |
| mmu-miR-29b-3p | MIMAT0000127 | B230219D22Rik | 78521 | HITS-CLIP//sRNA-Seq | 25083871\|33184237\|23597149\|23142 | Unspecified |
| mmu-miR-29b-3p | MIMAT0000127 | B2m | 12010 | sRNA-Seq//HITS-CLIP | 33184237\|23597149\|23142080 | Unspecified |
| mmu-miR-29b-3p | MIMAT0000127 | B3galnt1 | 26879 | HITS-CLIP | 37986224 | Unspecified |
| mmu-miR-29b-3p | MIMAT0000127 | B3gat3 | 72727 | sRNA-Seq//HITS-CLIP | 33184237\|23597149\|23142080 | Unspecified |
| mmu-miR-29b-3p | MIMAT0000127 | B3glct | 381694 | HITS-CLIP | 37986224 | Unspecified |
| mmu-miR-29b-3p | MIMAT0000127 | B4galt1 | 14595 | HITS-CLIP//sRNA-Seq | 23597149\|33184237\|26203562 | Unspecified |
| mmu-miR-29b-3p | MIMAT0000127 | B4galt5 | 56336 | HITS-CLIP | 23597149\|23142080 | Unspecified |
| mmu-miR-29b-3p | MIMAT0000127 | B4galt6 | 56386 | HITS-CLIP | 37986224\|25083871 | Unspecified |
| mmu-miR-29b-3p | MIMAT0000127 | B4galt7 | 218271 | HITS-CLIP | 25083871\|23142080 | Unspecified |
| mmu-miR-29b-3p | MIMAT0000127 | BC003965 | 214489 | HITS-CLIP | 23597149 | Unspecified |
| mmu-miR-29b-3p | MIMAT0000127 | BC005537 | 79555 | HITS-CLIP//sRNA-Seq | 23142080\|25083871\|33184237 | Unspecified |
| mmu-miR-29b-3p | MIMAT0000127 | BC005561 | 100042165 | HITS-CLIP | 25083871\|23142080 | Unspecified |
| mmu-miR-29b-3p | MIMAT0000127 | BC016579 | 212998 | HITS-CLIP | 23142080 | Unspecified |
| mmu-miR-29b-3p | MIMAT0000127 | BC030867 | 217216 | HITS-CLIP | 23142080 | Unspecified |
| mmu-miR-29b-3p | MIMAT0000127 | BC055324 | 381306 | HITS-CLIP | 23142080 | Unspecified |
| mmu-miR-29b-3p | MIMAT0000127 | Babam2 | 9577 | HITS-CLIP | 23142080 | Unspecified |
| mmu-miR-29b-3p | MIMAT0000127 | Bace1 | 23821 | Luciferase assay//qRT-PCR//Western blot/ | 22664922\|25083871\|23597149\|33184 | Unspecified |
| mmu-miR-29b-3p | MIMAT0000127 | Bach2 | 12014 | HITS-CLIP | 37986224\|25083871\|23142080\|28735 | Unspecified |
| mmu-miR-29b-3p | MIMAT0000127 | Bag6 | 224727 | HITS-CLIP//sRNA-Seq | 37986224\|23142080\|25083871\|33184 | Unspecified |
| mmu-miR-29b-3p | MIMAT0000127 | Bahcc1 | 268515 | HITS-CLIP | 25083871 | Unspecified |

| | | | | | | |
|---|---|---|---|---|---|---|
| mmu-miR-29b-3p | MIMAT0000127 | Baiap2 | 108100 | HITS-CLIP | 25083871 | Unspecified |
| mmu-miR-29b-3p | MIMAT0000127 | Bak1 | 12018 | Luciferase assay//Microarray//qRT-PCR//V | 21245165\|25083871\|23597149\|33184 | Unspecified |
| mmu-miR-29b-3p | MIMAT0000127 | Banf1 | 23825 | HITS-CLIP//sRNA-Seq | 37986224\|33184237\|23142080 | Unspecified |
| mmu-miR-29b-3p | MIMAT0000127 | Bank1 | 242248 | HITS-CLIP | 37986224\|23597149 | Unspecified |
| mmu-miR-29b-3p | MIMAT0000127 | Banp | 53325 | HITS-CLIP | 23142080 | Unspecified |
| mmu-miR-29b-3p | MIMAT0000127 | Basp1 | 70350 | HITS-CLIP//sRNA-Seq | 37986224\|25083871\|33184237\|23142 | Unspecified |
| mmu-miR-29b-3p | MIMAT0000127 | Bax | 12028 | HITS-CLIP//sRNA-Seq | 23142080\|33184237\|23597149 | Unspecified |
| mmu-miR-29b-3p | MIMAT0000127 | Baz1a | 217578 | HITS-CLIP | 37986224\|25083871\|23142080 | Unspecified |
| mmu-miR-29b-3p | MIMAT0000127 | Baz2a | 116848 | HITS-CLIP | 23142080\|25083871 | Unspecified |
| mmu-miR-29b-3p | MIMAT0000127 | Bbc3 | 170770 | Luciferase assay//Microarray//qRT-PCR//V | 21245165\|23597149\|23142080\|25083 | Unspecified |
| mmu-miR-29b-3p | MIMAT0000127 | Bbip1 | 100503572 | HITS-CLIP | 25083871\|23142080 | Unspecified |
| mmu-miR-29b-3p | MIMAT0000127 | Bbs10 | 71769 | HITS-CLIP | 37986224\|23597149 | Unspecified |
| mmu-miR-29b-3p | MIMAT0000127 | Bbs9 | 319845 | HITS-CLIP | 23142080 | Unspecified |
| mmu-miR-29b-3p | MIMAT0000127 | Bbx | 70508 | HITS-CLIP | 37986224\|23142080\|25083871 | Unspecified |
| mmu-miR-29b-3p | MIMAT0000127 | Bcap29 | 12033 | HITS-CLIP | 23142080 | Unspecified |
| mmu-miR-29b-3p | MIMAT0000127 | Bcap31 | 27061 | HITS-CLIP//sRNA-Seq | 23597149\|33184237 | Unspecified |
| mmu-miR-29b-3p | MIMAT0000127 | Bcas2 | 68183 | sRNA-Seq//HITS-CLIP | 33184237\|26203562 | Unspecified |
| mmu-miR-29b-3p | MIMAT0000127 | Bcas3 | 192197 | HITS-CLIP | 25083871\|23142080 | Unspecified |
| mmu-miR-29b-3p | MIMAT0000127 | Bcat1 | 12035 | HITS-CLIP//sRNA-Seq | 25083871\|33184237 | Unspecified |
| mmu-miR-29b-3p | MIMAT0000127 | Bcat2 | 12036 | HITS-CLIP | 23142080\|23597149 | Unspecified |
| mmu-miR-29b-3p | MIMAT0000127 | Bche | 12038 | HITS-CLIP | 37986224 | Unspecified |
| mmu-miR-29b-3p | MIMAT0000127 | Bcl10 | 12042 | sRNA-Seq//HITS-CLIP | 33184237\|23142080 | Unspecified |
| mmu-miR-29b-3p | MIMAT0000127 | Bcl11a | 14025 | HITS-CLIP | 23142080 | Unspecified |
| mmu-miR-29b-3p | MIMAT0000127 | Bcl11b | 58208 | HITS-CLIP | 37986224\|23142080 | Unspecified |
| mmu-miR-29b-3p | MIMAT0000127 | Bcl2 | 12043 | HITS-CLIP | 23597149\|23142080 | Unspecified |
| mmu-miR-29b-3p | MIMAT0000127 | Bcl2l1 | 12048 | HITS-CLIP//sRNA-Seq | 23597149\|33184237\|23142080 | Unspecified |
| mmu-miR-29b-3p | MIMAT0000127 | Bcl2l11 | 12125 | Luciferase assay//Microarray//qRT-PCR//V | 21245165\|25083871\|23142080 | Unspecified |
| mmu-miR-29b-3p | MIMAT0000127 | Bcl2l13 | 94044 | HITS-CLIP//sRNA-Seq | 23597149\|33184237\|25083871 | Unspecified |
| mmu-miR-29b-3p | MIMAT0000127 | Bcl2l15 | 229672 | HITS-CLIP | 23142080 | Unspecified |
| mmu-miR-29b-3p | MIMAT0000127 | Bcl2l2 | 12050 | HITS-CLIP | 37986224\|25083871\|23142080\|23597 | Unspecified |
| mmu-miR-29b-3p | MIMAT0000127 | Bcl3 | 12051 | HITS-CLIP | 25083871\|23597149\|23142080 | Unspecified |
| mmu-miR-29b-3p | MIMAT0000127 | Bcl7a | 77045 | HITS-CLIP | 37986224\|25083871\|23142080 | Unspecified |
| mmu-miR-29b-3p | MIMAT0000127 | Bcl7b | 12054 | HITS-CLIP | 25083871 | Unspecified |
| mmu-miR-29b-3p | MIMAT0000127 | Bcl9l | 80288 | sRNA-Seq//HITS-CLIP | 33184237\|25083871 | Unspecified |
| mmu-miR-29b-3p | MIMAT0000127 | Bclaf1 | 72567 | HITS-CLIP//sRNA-Seq | 25083871\|33184237\|23142080 | Unspecified |
| mmu-miR-29b-3p | MIMAT0000127 | Bclaf3 | 382252 | HITS-CLIP | 25083871\|23142080 | Unspecified |
| mmu-miR-29b-3p | MIMAT0000127 | Bcor | 71458 | HITS-CLIP | 37986224\|25083871\|23597149\|23142 | Unspecified |
| mmu-miR-29b-3p | MIMAT0000127 | Bcorl1 | 320376 | HITS-CLIP | 37986224\|25083871 | Unspecified |
| mmu-miR-29b-3p | MIMAT0000127 | Bdh1 | 71911 | HITS-CLIP//sRNA-Seq | 37986224\|23597149\|33184237\|23142 | Unspecified |

| | | | | | | |
|---|---|---|---|---|---|---|
| mmu-miR-29b-3p | MIMAT0000127 | Bdp1 | 544971 | HITS-CLIP | 31433990 | Unspecified |
| mmu-miR-29b-3p | MIMAT0000127 | Becn1 | 56208 | HITS-CLIP//sRNA-Seq | 25083871\|33184237 | Unspecified |
| mmu-miR-29b-3p | MIMAT0000127 | Bet1 | 12068 | HITS-CLIP | 25083871 | Unspecified |
| mmu-miR-29b-3p | MIMAT0000127 | Bet1l | 54399 | HITS-CLIP | 37986224\|23597149\|25083871 | Unspecified |
| mmu-miR-29b-3p | MIMAT0000127 | Bfar | 67118 | HITS-CLIP | 23142080 | Unspecified |
| mmu-miR-29b-3p | MIMAT0000127 | Bgn | 12111 | HITS-CLIP//sRNA-Seq | 25083871\|33184237 | Unspecified |
| mmu-miR-29b-3p | MIMAT0000127 | Bhlhe40 | 20893 | HITS-CLIP | 23597149\|25083871\|23142080 | Unspecified |
| mmu-miR-29b-3p | MIMAT0000127 | Bhlhe41 | 79362 | HITS-CLIP | 23142080 | Unspecified |
| mmu-miR-29b-3p | MIMAT0000127 | Bhmt2 | 64918 | HITS-CLIP | 23597149 | Unspecified |
| mmu-miR-29b-3p | MIMAT0000127 | Bicd2 | 76895 | HITS-CLIP | 25083871 | Unspecified |
| mmu-miR-29b-3p | MIMAT0000127 | Bik | 12124 | HITS-CLIP | 23142080 | Unspecified |
| mmu-miR-29b-3p | MIMAT0000127 | Bin1 | 30948 | HITS-CLIP//sRNA-Seq | 25083871\|33184237 | Unspecified |
| mmu-miR-29b-3p | MIMAT0000127 | Birc2 | 11797 | HITS-CLIP | 37986224\|23142080 | Unspecified |
| mmu-miR-29b-3p | MIMAT0000127 | Birc3 | 11796 | HITS-CLIP | 37986224\|23142080 | Unspecified |
| mmu-miR-29b-3p | MIMAT0000127 | Birc6 | 12211 | HITS-CLIP//sRNA-Seq | 37986224\|23142080\|25083871\|2359 | Unspecified |
| mmu-miR-29b-3p | MIMAT0000127 | Bivm | 246229 | HITS-CLIP | 25083871 | Unspecified |
| mmu-miR-29b-3p | MIMAT0000127 | Blcap | 53619 | HITS-CLIP//sRNA-Seq | 37986224\|25083871\|23597149\|3318 | Unspecified |
| mmu-miR-29b-3p | MIMAT0000127 | Blk | 12143 | HITS-CLIP | 23142080 | Unspecified |
| mmu-miR-29b-3p | MIMAT0000127 | Blm | 12144 | HITS-CLIP | 37986224\|23142080\|25083871 | Unspecified |
| mmu-miR-29b-3p | MIMAT0000127 | Blmh | 104184 | HITS-CLIP//sRNA-Seq | 37986224\|23597149\|33184237\|2508 | Unspecified |
| mmu-miR-29b-3p | MIMAT0000127 | Bloc1s6 | 18457 | HITS-CLIP | 37986224 | Unspecified |
| mmu-miR-29b-3p | MIMAT0000127 | Blzf1 | 66352 | HITS-CLIP | 23142080 | Unspecified |
| mmu-miR-29b-3p | MIMAT0000127 | Bmf | 171543 | Luciferase assay//Microarray//qRT-PCR//W | 21245165\|25083871\|23597149\|2314 | Unspecified |
| mmu-miR-29b-3p | MIMAT0000127 | Bmi1 | 12151 | HITS-CLIP | 25083871 | Unspecified |
| mmu-miR-29b-3p | MIMAT0000127 | Bmp2k | 140780 | HITS-CLIP | 37986224\|25083871\|23142080 | Unspecified |
| mmu-miR-29b-3p | MIMAT0000127 | Bmper | 73230 | HITS-CLIP | 25083871 | Unspecified |
| mmu-miR-29b-3p | MIMAT0000127 | Bmpr1a | 12166 | HITS-CLIP//sRNA-Seq | 25083871\|33184237 | Unspecified |
| mmu-miR-29b-3p | MIMAT0000127 | Bmpr2 | 12168 | HITS-CLIP//sRNA-Seq | 25083871\|23597149\|33184237 | Unspecified |
| mmu-miR-29b-3p | MIMAT0000127 | Bms1 | 213895 | HITS-CLIP//sRNA-Seq | 25083871\|33184237\|23142080 | Unspecified |
| mmu-miR-29b-3p | MIMAT0000127 | Bmt2 | 101148 | HITS-CLIP | 23142080\|25083871 | Unspecified |
| mmu-miR-29b-3p | MIMAT0000127 | Bnc2 | 242509 | HITS-CLIP//sRNA-Seq | 25083871\|33184237 | Unspecified |
| mmu-miR-29b-3p | MIMAT0000127 | Bnip2 | 12175 | HITS-CLIP//sRNA-Seq | 23597149\|25083871\|33184237\|2314 | Unspecified |
| mmu-miR-29b-3p | MIMAT0000127 | Bnip3 | 12176 | HITS-CLIP//sRNA-Seq | 23597149\|33184237\|23142080 | Unspecified |
| mmu-miR-29b-3p | MIMAT0000127 | Bod1l | 665775 | HITS-CLIP | 37986224\|23142080 | Unspecified |
| mmu-miR-29b-3p | MIMAT0000127 | Bop1 | 12181 | HITS-CLIP//sRNA-Seq | 25083871\|33184237 | Unspecified |
| mmu-miR-29b-3p | MIMAT0000127 | Bora | 77744 | HITS-CLIP | 37986224\|23597149\|23142080\|2508 | Unspecified |
| mmu-miR-29b-3p | MIMAT0000127 | Borcs6 | 54785 | HITS-CLIP | 23142080 | Unspecified |
| mmu-miR-29b-3p | MIMAT0000127 | Bptf | 207165 | HITS-CLIP | 25083871\|23142080 | Unspecified |
| mmu-miR-29b-3p | MIMAT0000127 | Braf | 109880 | HITS-CLIP | 25083871\|23142080 | Unspecified |

| | | | | | | |
|---|---|---|---|---|---|---|
| mmu-miR-29b-3p | MIMAT0000127 | Brd1 | 223770 | HITS-CLIP//sRNA-Seq | 37986224\|23597149\|33184237 | Unspecified |
| mmu-miR-29b-3p | MIMAT0000127 | Brd3 | 67382 | HITS-CLIP//sRNA-Seq | 37986224\|23597149\|25083871\|33184 | Unspecified |
| mmu-miR-29b-3p | MIMAT0000127 | Brd4 | 57261 | HITS-CLIP//sRNA-Seq | 37986224\|23597149\|25083871\|23142 | Unspecified |
| mmu-miR-29b-3p | MIMAT0000127 | Brd7 | 26992 | HITS-CLIP | 25083871\|23142080 | Unspecified |
| mmu-miR-29b-3p | MIMAT0000127 | Brd8 | 78656 | HITS-CLIP//sRNA-Seq | 25083871\|23142080\|33184237 | Unspecified |
| mmu-miR-29b-3p | MIMAT0000127 | Brdt | 114642 | HITS-CLIP | 23142080 | Unspecified |
| mmu-miR-29b-3p | MIMAT0000127 | Bri3bp | 76809 | HITS-CLIP | 37986224\|25083871\|23597149\|23142 | Unspecified |
| mmu-miR-29b-3p | MIMAT0000127 | Brinp3 | 215378 | HITS-CLIP | 25083871 | Unspecified |
| mmu-miR-29b-3p | MIMAT0000127 | Brip1 | 237911 | HITS-CLIP | 23142080 | Unspecified |
| mmu-miR-29b-3p | MIMAT0000127 | Brms1l | 52592 | HITS-CLIP | 25083871\|23142080 | Unspecified |
| mmu-miR-29b-3p | MIMAT0000127 | Brpf1 | 78783 | HITS-CLIP | 37986224\|25083871 | Unspecified |
| mmu-miR-29b-3p | MIMAT0000127 | Brwd1 | 93871 | HITS-CLIP//sRNA-Seq | 37986224\|25083871\|23142080\|33184 | Unspecified |
| mmu-miR-29b-3p | MIMAT0000127 | Brwd3 | 382236 | HITS-CLIP | 37986224\|25083871\|23142080\|23597 | Unspecified |
| mmu-miR-29b-3p | MIMAT0000127 | Bsdc1 | 100383 | HITS-CLIP | 23142080 | Unspecified |
| mmu-miR-29b-3p | MIMAT0000127 | Bsg | 12215 | HITS-CLIP//sRNA-Seq | 25083871\|33184237\|23597149\|23142 | Unspecified |
| mmu-miR-29b-3p | MIMAT0000127 | Bst2 | 69550 | HITS-CLIP | 37986224 | Unspecified |
| mmu-miR-29b-3p | MIMAT0000127 | Btaf1 | 107182 | HITS-CLIP | 37986224\|25083871\|23597149\|23142 | Unspecified |
| mmu-miR-29b-3p | MIMAT0000127 | Btbd10 | 68815 | HITS-CLIP | 23142080 | Unspecified |
| mmu-miR-29b-3p | MIMAT0000127 | Btbd19 | 78611 | HITS-CLIP | 37986224 | Unspecified |
| mmu-miR-29b-3p | MIMAT0000127 | Btbd6 | 399566 | HITS-CLIP | 23142080\|25083871 | Unspecified |
| mmu-miR-29b-3p | MIMAT0000127 | Btbd7 | 238386 | HITS-CLIP | 23142080 | Unspecified |
| mmu-miR-29b-3p | MIMAT0000127 | Btbd8 | 100503185 | HITS-CLIP | 25083871\|23142080 | Unspecified |
| mmu-miR-29b-3p | MIMAT0000127 | Btbd9 | 224671 | HITS-CLIP | 25083871 | Unspecified |
| mmu-miR-29b-3p | MIMAT0000127 | Btf3 | 218490 | HITS-CLIP//sRNA-Seq | 23597149\|25083871\|33184237\|23142 | Unspecified |
| mmu-miR-29b-3p | MIMAT0000127 | Btg2 | 12227 | HITS-CLIP | 37986224\|23597149\|23142080 | Unspecified |
| mmu-miR-29b-3p | MIMAT0000127 | Btla | 208154 | HITS-CLIP | 23142080 | Unspecified |
| mmu-miR-29b-3p | MIMAT0000127 | Bub3 | 12237 | HITS-CLIP | 23142080 | Unspecified |
| mmu-miR-29b-3p | MIMAT0000127 | Bves | 23828 | HITS-CLIP | 25083871 | Unspecified |
| mmu-miR-29b-3p | MIMAT0000127 | Bzw1 | 66882 | sRNA-Seq//HITS-CLIP | 33184237\|23142080 | Unspecified |
| mmu-miR-29b-3p | MIMAT0000127 | C1d | 57316 | HITS-CLIP//sRNA-Seq | 23142080\|25083871\|33184237 | Unspecified |
| mmu-miR-29b-3p | MIMAT0000127 | C1galt1 | 94192 | HITS-CLIP | 25083871\|23597149\|23142080 | Unspecified |
| mmu-miR-29b-3p | MIMAT0000127 | C1galt1c1 | 59048 | HITS-CLIP | 25083871 | Unspecified |
| mmu-miR-29b-3p | MIMAT0000127 | C1qa | 12259 | HITS-CLIP | 23597149 | Unspecified |
| mmu-miR-29b-3p | MIMAT0000127 | C1qtnf1 | 56745 | HITS-CLIP | 37986224\|23597149 | Unspecified |
| mmu-miR-29b-3p | MIMAT0000127 | C1qtnf3 | 81799 | HITS-CLIP | 37986224\|25083871 | Unspecified |
| mmu-miR-29b-3p | MIMAT0000127 | C1qtnf6 | 72709 | HITS-CLIP//sRNA-Seq | 37986224\|23142080\|33184237 | Unspecified |
| mmu-miR-29b-3p | MIMAT0000127 | C1s1 | 50908 | HITS-CLIP//sRNA-Seq | 23597149\|33184237 | Unspecified |
| mmu-miR-29b-3p | MIMAT0000127 | C2cd2 | 207781 | HITS-CLIP | 23597149 | Unspecified |
| mmu-miR-29b-3p | MIMAT0000127 | C3 | 12266 | HITS-CLIP | 23597149 | Unspecified |

| | | | | | | |
|---|---|---|---|---|---|---|
| mmu-miR-29b-3p | MIMAT0000127 | C4b | 12268 | HITS-CLIP | 23597149 | Unspecified |
| mmu-miR-29b-3p | MIMAT0000127 | C530008M17Rik | 320827 | HITS-CLIP//sRNA-Seq | 25083871\|33184237 | Unspecified |
| mmu-miR-29b-3p | MIMAT0000127 | C77080 | 97130 | HITS-CLIP | 37986224 | Unspecified |
| mmu-miR-29b-3p | MIMAT0000127 | C8a | 230558 | HITS-CLIP | 23597149 | Unspecified |
| mmu-miR-29b-3p | MIMAT0000127 | C9 | 12279 | HITS-CLIP | 37986224\|23597149 | Unspecified |
| mmu-miR-29b-3p | MIMAT0000127 | Caap1 | 67770 | HITS-CLIP | 23142080 | Unspecified |
| mmu-miR-29b-3p | MIMAT0000127 | Cab39 | 12283 | HITS-CLIP | 37986224 | Unspecified |
| mmu-miR-29b-3p | MIMAT0000127 | Cabin1 | 104248 | HITS-CLIP | 23597149 | Unspecified |
| mmu-miR-29b-3p | MIMAT0000127 | Cacfd1 | 381356 | HITS-CLIP | 37986224 | Unspecified |
| mmu-miR-29b-3p | MIMAT0000127 | Cacna1s | 12292 | HITS-CLIP | 25083871 | Unspecified |
| mmu-miR-29b-3p | MIMAT0000127 | Cacna2d1 | 12293 | HITS-CLIP//sRNA-Seq | 25083871\|33184237 | Unspecified |
| mmu-miR-29b-3p | MIMAT0000127 | Cacnb1 | 12295 | HITS-CLIP | 25083871 | Unspecified |
| mmu-miR-29b-3p | MIMAT0000127 | Cacul1 | 78832 | HITS-CLIP | 37986224\|25083871\|23597149\|23142 | Unspecified |
| mmu-miR-29b-3p | MIMAT0000127 | Cad | 69719 | HITS-CLIP | 23142080 | Unspecified |
| mmu-miR-29b-3p | MIMAT0000127 | Calm1 | 12313 | HITS-CLIP//sRNA-Seq | 25083871\|33184237\|23142080 | Unspecified |
| mmu-miR-29b-3p | MIMAT0000127 | Calm3 | 12315 | HITS-CLIP//sRNA-Seq | 37986224\|23142080\|25083871\|33184 | Unspecified |
| mmu-miR-29b-3p | MIMAT0000127 | Calr | 12317 | HITS-CLIP//sRNA-Seq | 23597149\|33184237\|23142080 | Unspecified |
| mmu-miR-29b-3p | MIMAT0000127 | Calu | 12321 | HITS-CLIP//sRNA-Seq | 25083871\|23597149\|33184237\|23142 | Unspecified |
| mmu-miR-29b-3p | MIMAT0000127 | Camk2d | 108058 | HITS-CLIP//sRNA-Seq | 37986224\|23142080\|33184237\|25083 | Unspecified |
| mmu-miR-29b-3p | MIMAT0000127 | Camk2g | 12325 | HITS-CLIP//sRNA-Seq | 37986224\|33184237\|23142080 | Unspecified |
| mmu-miR-29b-3p | MIMAT0000127 | Camk4 | 12326 | HITS-CLIP | 37986224 | Unspecified |
| mmu-miR-29b-3p | MIMAT0000127 | Camkk2 | 207565 | HITS-CLIP | 23142080 | Unspecified |
| mmu-miR-29b-3p | MIMAT0000127 | Camsap1 | 227634 | HITS-CLIP | 37986224 | Unspecified |
| mmu-miR-29b-3p | MIMAT0000127 | Camsap2 | 67886 | HITS-CLIP//sRNA-Seq | 23597149\|33184237\|23142080 | Unspecified |
| mmu-miR-29b-3p | MIMAT0000127 | Camta2 | 216874 | HITS-CLIP | 25083871 | Unspecified |
| mmu-miR-29b-3p | MIMAT0000127 | Cand1 | 71902 | sRNA-Seq//HITS-CLIP | 33184237\|25083871\|23142080 | Unspecified |
| mmu-miR-29b-3p | MIMAT0000127 | Canx | 12330 | HITS-CLIP//sRNA-Seq | 23142080\|25083871\|23597149\|33184 | Unspecified |
| mmu-miR-29b-3p | MIMAT0000127 | Cap1 | 12331 | HITS-CLIP//sRNA-Seq | 25083871\|33184237\|23142080 | Unspecified |
| mmu-miR-29b-3p | MIMAT0000127 | Cap2 | 67252 | HITS-CLIP | 25083871 | Unspecified |
| mmu-miR-29b-3p | MIMAT0000127 | Capn1 | 12333 | HITS-CLIP//sRNA-Seq | 23142080\|33184237 | Unspecified |
| mmu-miR-29b-3p | MIMAT0000127 | Capn11 | 268958 | HITS-CLIP | 23142080 | Unspecified |
| mmu-miR-29b-3p | MIMAT0000127 | Capn6 | 12338 | HITS-CLIP//sRNA-Seq | 37986224\|25083871\|33184237 | Unspecified |
| mmu-miR-29b-3p | MIMAT0000127 | Capn7 | 12339 | HITS-CLIP | 37986224\|23597149\|23142080\|25083 | Unspecified |
| mmu-miR-29b-3p | MIMAT0000127 | Caprin1 | 53872 | HITS-CLIP//sRNA-Seq | 23597149\|25083871\|23142080\|33184 | Unspecified |
| mmu-miR-29b-3p | MIMAT0000127 | Capza2 | 12343 | HITS-CLIP//sRNA-Seq | 37986224\|23142080\|33184237\|25083 | Unspecified |
| mmu-miR-29b-3p | MIMAT0000127 | Capzb | 12345 | HITS-CLIP//sRNA-Seq | 25083871\|23597149\|33184237\|23142 | Unspecified |
| mmu-miR-29b-3p | MIMAT0000127 | Car3 | 12350 | HITS-CLIP | 23597149\|25083871 | Unspecified |
| mmu-miR-29b-3p | MIMAT0000127 | Carmil2 | 146206 | HITS-CLIP | 23142080 | Unspecified |
| mmu-miR-29b-3p | MIMAT0000127 | Cars | 27267 | sRNA-Seq//HITS-CLIP | 33184237\|25083871 | Unspecified |

| | | | | | | |
|---|---|---|---|---|---|---|
| mmu-miR-29b-3p | MIMAT0000127 | Casc4 | 319996 | HITS-CLIP//sRNA-Seq | 25083871\|33184237 | Unspecified |
| mmu-miR-29b-3p | MIMAT0000127 | Casd1 | 213819 | HITS-CLIP | 23142080 | Unspecified |
| mmu-miR-29b-3p | MIMAT0000127 | Casp2 | 12366 | HITS-CLIP//sRNA-Seq | 37986224\|33184237\|23142080 | Unspecified |
| mmu-miR-29b-3p | MIMAT0000127 | Casp3 | 12367 | HITS-CLIP | 37986224 | Unspecified |
| mmu-miR-29b-3p | MIMAT0000127 | Casp6 | 12368 | HITS-CLIP | 23142080 | Unspecified |
| mmu-miR-29b-3p | MIMAT0000127 | Casp8ap2 | 26885 | HITS-CLIP | 25083871 | Unspecified |
| mmu-miR-29b-3p | MIMAT0000127 | Casq1 | 12372 | HITS-CLIP | 25083871 | Unspecified |
| mmu-miR-29b-3p | MIMAT0000127 | Casq2 | 12373 | HITS-CLIP | 25083871 | Unspecified |
| mmu-miR-29b-3p | MIMAT0000127 | Cast | 12380 | HITS-CLIP//sRNA-Seq | 23142080\|33184237\|25083871 | Unspecified |
| mmu-miR-29b-3p | MIMAT0000127 | Castor1 | 652968 | HITS-CLIP | 25083871\|23142080 | Unspecified |
| mmu-miR-29b-3p | MIMAT0000127 | Casz1 | 69743 | HITS-CLIP | 37986224 | Unspecified |
| mmu-miR-29b-3p | MIMAT0000127 | Cat | 12359 | HITS-CLIP//sRNA-Seq | 23597149\|33184237\|23142080 | Unspecified |
| mmu-miR-29b-3p | MIMAT0000127 | Cav1 | 12389 | HITS-CLIP//sRNA-Seq | 25083871\|33184237\|26203562 | Unspecified |
| mmu-miR-29b-3p | MIMAT0000127 | Cav2 | 12390 | reporter_gene//HITS-CLIP//Reporter assay | 37986224\|25083871\|17652184 | Unspecified |
| mmu-miR-29b-3p | MIMAT0000127 | Cav3 | 12391 | HITS-CLIP | 25083871 | Unspecified |
| mmu-miR-29b-3p | MIMAT0000127 | Cavin1 | 284119 | HITS-CLIP//sRNA-Seq | 25083871\|33184237 | Unspecified |
| mmu-miR-29b-3p | MIMAT0000127 | Cbfb | 12400 | HITS-CLIP//sRNA-Seq | 25083871\|33184237\|23597149\|23142 | Unspecified |
| mmu-miR-29b-3p | MIMAT0000127 | Cbl | 12402 | HITS-CLIP | 25083871 | Unspecified |
| mmu-miR-29b-3p | MIMAT0000127 | Cblb | 208650 | HITS-CLIP | 25083871\|23142080 | Unspecified |
| mmu-miR-29b-3p | MIMAT0000127 | Cbll1 | 104836 | HITS-CLIP | 23142080\|25083871 | Unspecified |
| mmu-miR-29b-3p | MIMAT0000127 | Cbs | 12411 | HITS-CLIP | 37986224\|23597149 | Unspecified |
| mmu-miR-29b-3p | MIMAT0000127 | Cbx2 | 12416 | HITS-CLIP | 37986224\|23142080 | Unspecified |
| mmu-miR-29b-3p | MIMAT0000127 | Cbx4 | 12418 | HITS-CLIP | 37986224\|23597149\|23142080 | Unspecified |
| mmu-miR-29b-3p | MIMAT0000127 | Cbx5 | 12419 | sRNA-Seq//HITS-CLIP | 33184237\|23597149\|23142080 | Unspecified |
| mmu-miR-29b-3p | MIMAT0000127 | Cbx6 | 494448 | HITS-CLIP//sRNA-Seq | 37986224\|33184237\|23142080 | Unspecified |
| mmu-miR-29b-3p | MIMAT0000127 | Cbx7 | 52609 | sRNA-Seq//HITS-CLIP | 33184237\|23142080 | Unspecified |
| mmu-miR-29b-3p | MIMAT0000127 | Cc2d1a | 212139 | HITS-CLIP//sRNA-Seq | 23597149\|33184237 | Unspecified |
| mmu-miR-29b-3p | MIMAT0000127 | Cc2d1b | 319965 | HITS-CLIP | 37986224 | Unspecified |
| mmu-miR-29b-3p | MIMAT0000127 | Ccar1 | 67500 | HITS-CLIP//sRNA-Seq | 23597149\|25083871\|23142080\|33184 | Unspecified |
| mmu-miR-29b-3p | MIMAT0000127 | Ccdc115 | 69668 | HITS-CLIP | 25083871\|23142080 | Unspecified |
| mmu-miR-29b-3p | MIMAT0000127 | Ccdc117 | 104479 | HITS-CLIP | 37986224\|25083871\|23142080 | Unspecified |
| mmu-miR-29b-3p | MIMAT0000127 | Ccdc126 | 57895 | HITS-CLIP | 23142080 | Unspecified |
| mmu-miR-29b-3p | MIMAT0000127 | Ccdc127 | 67433 | HITS-CLIP//sRNA-Seq | 23597149\|25083871\|33184237\|23142 | Unspecified |
| mmu-miR-29b-3p | MIMAT0000127 | Ccdc166 | 223648 | HITS-CLIP | 23142080\|25083871 | Unspecified |
| mmu-miR-29b-3p | MIMAT0000127 | Ccdc186 | 55088 | HITS-CLIP//sRNA-Seq | 25083871\|33184237\|23142080 | Unspecified |
| mmu-miR-29b-3p | MIMAT0000127 | Ccdc25 | 67179 | HITS-CLIP | 37986224\|23597149\|25083871 | Unspecified |
| mmu-miR-29b-3p | MIMAT0000127 | Ccdc28b | 66264 | HITS-CLIP//sRNA-Seq | 25083871\|33184237\|23142080 | Unspecified |
| mmu-miR-29b-3p | MIMAT0000127 | Ccdc47 | 67163 | sRNA-Seq//HITS-CLIP | 33184237\|23142080\|25083871 | Unspecified |
| mmu-miR-29b-3p | MIMAT0000127 | Ccdc50 | 67501 | HITS-CLIP//sRNA-Seq | 37986224\|23142080\|33184237 | Unspecified |

| | | | | | | |
|---|---|---|---|---|---|---|
| mmu-miR-29b-3p | MIMAT0000127 | Ccdc6 | 76551 | HITS-CLIP | 23142080 | Unspecified |
| mmu-miR-29b-3p | MIMAT0000127 | Ccdc71l | 72123 | HITS-CLIP | 23142080\|25083871 | Unspecified |
| mmu-miR-29b-3p | MIMAT0000127 | Ccdc8 | 434130 | HITS-CLIP | 37986224\|25083871 | Unspecified |
| mmu-miR-29b-3p | MIMAT0000127 | Ccdc80 | 67896 | HITS-CLIP//sRNA-Seq | 37986224\|25083871\|33184237 | Unspecified |
| mmu-miR-29b-3p | MIMAT0000127 | Ccdc82 | 66396 | HITS-CLIP | 25083871 | Unspecified |
| mmu-miR-29b-3p | MIMAT0000127 | Ccdc88a | 108686 | HITS-CLIP//sRNA-Seq//qCLASH | 37986224\|25083871\|33184237\|3139[illegible] | Unspecified |
| mmu-miR-29b-3p | MIMAT0000127 | Ccdc88b | 78317 | HITS-CLIP | 23142080 | Unspecified |
| mmu-miR-29b-3p | MIMAT0000127 | Ccdc88c | 68339 | HITS-CLIP | 23142080 | Unspecified |
| mmu-miR-29b-3p | MIMAT0000127 | Ccdc90b | 66365 | HITS-CLIP | 23597149\|23142080 | Unspecified |
| mmu-miR-29b-3p | MIMAT0000127 | Cchcr1 | 240084 | HITS-CLIP | 23142080 | Unspecified |
| mmu-miR-29b-3p | MIMAT0000127 | Ccl22 | 20299 | HITS-CLIP | 23142080 | Unspecified |
| mmu-miR-29b-3p | MIMAT0000127 | Ccm2 | 216527 | HITS-CLIP//sRNA-Seq | 23597149\|23142080\|33184237 | Unspecified |
| mmu-miR-29b-3p | MIMAT0000127 | Ccn1 | 3491 | HITS-CLIP | 23597149 | Unspecified |
| mmu-miR-29b-3p | MIMAT0000127 | Ccn2 | 1490 | HITS-CLIP | 25083871\|23597149 | Unspecified |
| mmu-miR-29b-3p | MIMAT0000127 | Ccn4 | 8840 | HITS-CLIP | 25083871\|23142080 | Unspecified |
| mmu-miR-29b-3p | MIMAT0000127 | Ccna2 | 12428 | HITS-CLIP//sRNA-Seq | 37986224\|25083871\|33184237\|2314[illegible] | Unspecified |
| mmu-miR-29b-3p | MIMAT0000127 | Ccnb2 | 12442 | sRNA-Seq//HITS-CLIP | 33184237\|23142080 | Unspecified |
| mmu-miR-29b-3p | MIMAT0000127 | Ccnc | 51813 | HITS-CLIP | 25083871\|23142080 | Unspecified |
| mmu-miR-29b-3p | MIMAT0000127 | Ccnd1 | 12443 | HITS-CLIP | 25083871\|23597149 | Unspecified |
| mmu-miR-29b-3p | MIMAT0000127 | Ccnd2 | 12444 | HITS-CLIP | 37986224\|25083871\|23142080\|2620[illegible] | Unspecified |
| mmu-miR-29b-3p | MIMAT0000127 | Ccne1 | 12447 | HITS-CLIP | 25083871\|23142080 | Unspecified |
| mmu-miR-29b-3p | MIMAT0000127 | Ccne2 | 12448 | HITS-CLIP | 37986224\|25083871\|23142080\|3143[illegible] | Unspecified |
| mmu-miR-29b-3p | MIMAT0000127 | Ccnf | 12449 | HITS-CLIP | 37986224\|25083871\|23597149\|3143[illegible] | Unspecified |
| mmu-miR-29b-3p | MIMAT0000127 | Ccng1 | 12450 | HITS-CLIP//sRNA-Seq | 23142080\|33184237 | Unspecified |
| mmu-miR-29b-3p | MIMAT0000127 | Ccng2 | 12452 | HITS-CLIP | 25083871\|23142080 | Unspecified |
| mmu-miR-29b-3p | MIMAT0000127 | Ccni | 12453 | sRNA-Seq//HITS-CLIP | 33184237\|23142080 | Unspecified |
| mmu-miR-29b-3p | MIMAT0000127 | Ccnj | 240665 | HITS-CLIP | 37986224\|25083871\|23142080 | Unspecified |
| mmu-miR-29b-3p | MIMAT0000127 | Ccnk | 12454 | HITS-CLIP | 23142080\|25083871 | Unspecified |
| mmu-miR-29b-3p | MIMAT0000127 | Ccnl1 | 56706 | HITS-CLIP | 23597149 | Unspecified |
| mmu-miR-29b-3p | MIMAT0000127 | Ccnl2 | 56036 | HITS-CLIP//sRNA-Seq | 37986224\|25083871\|23597149\|3318[illegible] | Unspecified |
| mmu-miR-29b-3p | MIMAT0000127 | Ccnt1 | 12455 | HITS-CLIP | 37986224\|23142080\|23597149 | Unspecified |
| mmu-miR-29b-3p | MIMAT0000127 | Ccnt2 | 72949 | HITS-CLIP | 31433990\|23142080 | Unspecified |
| mmu-miR-29b-3p | MIMAT0000127 | Ccny | 67974 | HITS-CLIP | 23142080 | Unspecified |
| mmu-miR-29b-3p | MIMAT0000127 | Ccnyl1 | 227210 | HITS-CLIP | 37986224\|25083871 | Unspecified |
| mmu-miR-29b-3p | MIMAT0000127 | Ccpg1 | 72278 | HITS-CLIP | 25083871\|23142080 | Unspecified |
| mmu-miR-29b-3p | MIMAT0000127 | Ccr5 | 12774 | HITS-CLIP | 23142080 | Unspecified |
| mmu-miR-29b-3p | MIMAT0000127 | Ccr7 | 12775 | HITS-CLIP | 23142080 | Unspecified |
| mmu-miR-29b-3p | MIMAT0000127 | Ccr8 | 12776 | HITS-CLIP | 23142080 | Unspecified |
| mmu-miR-29b-3p | MIMAT0000127 | Ccs | 12460 | sRNA-Seq//HITS-CLIP | 33184237\|23597149\|23142080 | Unspecified |

| | | | | | | |
|---|---|---|---|---|---|---|
| mmu-miR-29b-3p | MIMAT0000127 | Ccser1 | 232035 | HITS-CLIP | 23142080 | Unspecified |
| mmu-miR-29b-3p | MIMAT0000127 | Ccser2 | 72972 | HITS-CLIP//sRNA-Seq | 37986224\|25083871\|33184237 | Unspecified |
| mmu-miR-29b-3p | MIMAT0000127 | Cct2 | 12461 | sRNA-Seq//HITS-CLIP | 33184237\|23597149\|23142080 | Unspecified |
| mmu-miR-29b-3p | MIMAT0000127 | Cct3 | 12462 | HITS-CLIP//sRNA-Seq | 23597149\|25083871\|33184237\|23142 | Unspecified |
| mmu-miR-29b-3p | MIMAT0000127 | Cct4 | 12464 | HITS-CLIP//sRNA-Seq | 25083871\|23142080\|23597149\|33184 | Unspecified |
| mmu-miR-29b-3p | MIMAT0000127 | Cct5 | 12465 | HITS-CLIP//sRNA-Seq | 23597149\|33184237\|25083871\|23142 | Unspecified |
| mmu-miR-29b-3p | MIMAT0000127 | Cct7 | 12468 | HITS-CLIP//sRNA-Seq | 25083871\|33184237\|23142080 | Unspecified |
| mmu-miR-29b-3p | MIMAT0000127 | Cct8 | 12469 | HITS-CLIP//sRNA-Seq | 23597149\|25083871\|33184237\|23142 | Unspecified |
| mmu-miR-29b-3p | MIMAT0000127 | Ccz1 | 231874 | HITS-CLIP | 37986224 | Unspecified |
| mmu-miR-29b-3p | MIMAT0000127 | Cd101 | 630146 | HITS-CLIP | 23142080 | Unspecified |
| mmu-miR-29b-3p | MIMAT0000127 | Cd151 | 12476 | HITS-CLIP | 37986224 | Unspecified |
| mmu-miR-29b-3p | MIMAT0000127 | Cd163 | 93671 | HITS-CLIP | 23597149 | Unspecified |
| mmu-miR-29b-3p | MIMAT0000127 | Cd164 | 53599 | HITS-CLIP//sRNA-Seq | 23597149\|25083871\|33184237\|23142 | Unspecified |
| mmu-miR-29b-3p | MIMAT0000127 | Cd2 | 12481 | HITS-CLIP | 23142080 | Unspecified |
| mmu-miR-29b-3p | MIMAT0000127 | Cd200 | 17470 | HITS-CLIP//sRNA-Seq | 23142080\|25083871\|33184237 | Unspecified |
| mmu-miR-29b-3p | MIMAT0000127 | Cd226 | 225825 | HITS-CLIP | 23142080 | Unspecified |
| mmu-miR-29b-3p | MIMAT0000127 | Cd274 | 60533 | HITS-CLIP | 23142080 | Unspecified |
| mmu-miR-29b-3p | MIMAT0000127 | Cd276 | 102657 | HITS-CLIP//sRNA-Seq | 37986224\|33184237\|23597149 | Unspecified |
| mmu-miR-29b-3p | MIMAT0000127 | Cd28 | 12487 | HITS-CLIP | 25083871\|23142080 | Unspecified |
| mmu-miR-29b-3p | MIMAT0000127 | Cd37 | 12493 | HITS-CLIP | 23142080 | Unspecified |
| mmu-miR-29b-3p | MIMAT0000127 | Cd38 | 12494 | HITS-CLIP | 23597149\|23142080 | Unspecified |
| mmu-miR-29b-3p | MIMAT0000127 | Cd3d | 12500 | HITS-CLIP | 23142080 | Unspecified |
| mmu-miR-29b-3p | MIMAT0000127 | Cd3e | 12501 | HITS-CLIP | 23597149\|23142080 | Unspecified |
| mmu-miR-29b-3p | MIMAT0000127 | Cd4 | 12504 | HITS-CLIP | 23142080 | Unspecified |
| mmu-miR-29b-3p | MIMAT0000127 | Cd40lg | 959 | HITS-CLIP | 23142080 | Unspecified |
| mmu-miR-29b-3p | MIMAT0000127 | Cd44 | 12505 | HITS-CLIP//sRNA-Seq | 37986224\|23142080\|33184237 | Unspecified |
| mmu-miR-29b-3p | MIMAT0000127 | Cd47 | 16423 | HITS-CLIP//sRNA-Seq | 25083871\|33184237\|23142080 | Unspecified |
| mmu-miR-29b-3p | MIMAT0000127 | Cd48 | 12506 | HITS-CLIP | 23142080 | Unspecified |
| mmu-miR-29b-3p | MIMAT0000127 | Cd5 | 12507 | HITS-CLIP | 23142080 | Unspecified |
| mmu-miR-29b-3p | MIMAT0000127 | Cd52 | 23833 | HITS-CLIP | 23142080 | Unspecified |
| mmu-miR-29b-3p | MIMAT0000127 | Cd53 | 12508 | HITS-CLIP | 37986224 | Unspecified |
| mmu-miR-29b-3p | MIMAT0000127 | Cd6 | 12511 | HITS-CLIP | 23142080 | Unspecified |
| mmu-miR-29b-3p | MIMAT0000127 | Cd69 | 12515 | HITS-CLIP//qCLASH | 37986224\|31393953 | Unspecified |
| mmu-miR-29b-3p | MIMAT0000127 | Cd79a | 12518 | HITS-CLIP | 23142080 | Unspecified |
| mmu-miR-29b-3p | MIMAT0000127 | Cd80 | 12519 | HITS-CLIP | 25083871 | Unspecified |
| mmu-miR-29b-3p | MIMAT0000127 | Cd81 | 12520 | HITS-CLIP//sRNA-Seq | 37986224\|25083871\|33184237\|23597 | Unspecified |
| mmu-miR-29b-3p | MIMAT0000127 | Cd82 | 12521 | HITS-CLIP//sRNA-Seq | 23597149\|25083871\|33184237\|23142 | Unspecified |
| mmu-miR-29b-3p | MIMAT0000127 | Cd99l2 | 171486 | HITS-CLIP | 37986224 | Unspecified |
| mmu-miR-29b-3p | MIMAT0000127 | Cdadc1 | 71891 | HITS-CLIP | 23142080 | Unspecified |

| | | | | | | |
|---|---|---|---|---|---|---|
| mmu-miR-29b-3p | MIMAT0000127 | Cdc14a | 229776 | HITS-CLIP | 23142080 | Unspecified |
| mmu-miR-29b-3p | MIMAT0000127 | Cdc14b | 218294 | HITS-CLIP | 37986224 | Unspecified |
| mmu-miR-29b-3p | MIMAT0000127 | Cdc16 | 69957 | HITS-CLIP//sRNA-Seq | 37986224\|25083871\|33184237\|23597 | Unspecified |
| mmu-miR-29b-3p | MIMAT0000127 | Cdc23 | 52563 | HITS-CLIP | 37986224 | Unspecified |
| mmu-miR-29b-3p | MIMAT0000127 | Cdc25a | 12530 | HITS-CLIP | 25083871 | Unspecified |
| mmu-miR-29b-3p | MIMAT0000127 | Cdc25b | 12531 | HITS-CLIP | 37986224 | Unspecified |
| mmu-miR-29b-3p | MIMAT0000127 | Cdc27 | 217232 | HITS-CLIP//sRNA-Seq | 37986224\|25083871\|33184237\|23142 | Unspecified |
| mmu-miR-29b-3p | MIMAT0000127 | Cdc34 | 216150 | HITS-CLIP//sRNA-Seq | 25083871\|33184237 | Unspecified |
| mmu-miR-29b-3p | MIMAT0000127 | Cdc37 | 12539 | HITS-CLIP//sRNA-Seq | 23142080\|33184237 | Unspecified |
| mmu-miR-29b-3p | MIMAT0000127 | Cdc37l1 | 67072 | HITS-CLIP | 25083871 | Unspecified |
| mmu-miR-29b-3p | MIMAT0000127 | Cdc40 | 71713 | HITS-CLIP | 37986224\|23142080 | Unspecified |
| mmu-miR-29b-3p | MIMAT0000127 | Cdc42 | 12540 | HITS-CLIP//sRNA-Seq | 37986224\|25083871\|33184237\|23597 | Unspecified |
| mmu-miR-29b-3p | MIMAT0000127 | Cdc42bpa | 226751 | HITS-CLIP//sRNA-Seq | 37986224\|25083871\|33184237 | Unspecified |
| mmu-miR-29b-3p | MIMAT0000127 | Cdc42bpb | 217866 | HITS-CLIP//sRNA-Seq | 25083871\|33184237 | Unspecified |
| mmu-miR-29b-3p | MIMAT0000127 | Cdc42se1 | 57912 | HITS-CLIP//sRNA-Seq | 25083871\|33184237 | Unspecified |
| mmu-miR-29b-3p | MIMAT0000127 | Cdc42se2 | 72729 | HITS-CLIP | 25083871\|23142080 | Unspecified |
| mmu-miR-29b-3p | MIMAT0000127 | Cdc7 | 12545 | HITS-CLIP | 37986224\|23142080 | Unspecified |
| mmu-miR-29b-3p | MIMAT0000127 | Cdc73 | 214498 | HITS-CLIP | 25083871\|23142080 | Unspecified |
| mmu-miR-29b-3p | MIMAT0000127 | Cdca2 | 108912 | HITS-CLIP | 23142080 | Unspecified |
| mmu-miR-29b-3p | MIMAT0000127 | Cdca4 | 71963 | HITS-CLIP//sRNA-Seq | 37986224\|33184237\|23142080 | Unspecified |
| mmu-miR-29b-3p | MIMAT0000127 | Cdca7 | 66953 | HITS-CLIP//sRNA-Seq | 37986224\|25083871\|23597149\|33184 | Unspecified |
| mmu-miR-29b-3p | MIMAT0000127 | Cdip1 | 66626 | HITS-CLIP | 37986224 | Unspecified |
| mmu-miR-29b-3p | MIMAT0000127 | Cdipt | 52858 | HITS-CLIP//sRNA-Seq | 23597149\|33184237\|23142080 | Unspecified |
| mmu-miR-29b-3p | MIMAT0000127 | Cdk1 | 12534 | HITS-CLIP//sRNA-Seq | 23142080\|33184237 | Unspecified |
| mmu-miR-29b-3p | MIMAT0000127 | Cdk11b | 12537 | HITS-CLIP//sRNA-Seq | 25083871\|33184237 | Unspecified |
| mmu-miR-29b-3p | MIMAT0000127 | Cdk12 | 69131 | HITS-CLIP | 25083871\|23142080 | Unspecified |
| mmu-miR-29b-3p | MIMAT0000127 | Cdk13 | 69562 | HITS-CLIP | 23142080\|23597149\|25083871 | Unspecified |
| mmu-miR-29b-3p | MIMAT0000127 | Cdk16 | 18555 | HITS-CLIP//sRNA-Seq | 37986224\|25083871\|33184237\|23142 | Unspecified |
| mmu-miR-29b-3p | MIMAT0000127 | Cdk17 | 237459 | HITS-CLIP//sRNA-Seq | 25083871\|33184237\|23142080 | Unspecified |
| mmu-miR-29b-3p | MIMAT0000127 | Cdk19 | 78334 | HITS-CLIP | 25083871 | Unspecified |
| mmu-miR-29b-3p | MIMAT0000127 | Cdk2 | 12566 | HITS-CLIP//sRNA-Seq | 37986224\|25083871\|23142080\|33184 | Unspecified |
| mmu-miR-29b-3p | MIMAT0000127 | Cdk4 | 12567 | HITS-CLIP//sRNA-Seq | 23142080\|33184237 | Unspecified |
| mmu-miR-29b-3p | MIMAT0000127 | Cdk5r1 | 12569 | HITS-CLIP//sRNA-Seq | 25083871\|33184237\|23142080 | Unspecified |
| mmu-miR-29b-3p | MIMAT0000127 | Cdk6 | 12571 | HITS-CLIP//qCLASH | 37986224\|23142080\|25083871\|23597 | Unspecified |
| mmu-miR-29b-3p | MIMAT0000127 | Cdk7 | 12572 | HITS-CLIP | 37986224 | Unspecified |
| mmu-miR-29b-3p | MIMAT0000127 | Cdk8 | 264064 | HITS-CLIP | 37986224\|25083871\|23597149\|23142 | Unspecified |
| mmu-miR-29b-3p | MIMAT0000127 | Cdk9 | 107951 | HITS-CLIP | 37986224\|25083871\|23142080 | Unspecified |
| mmu-miR-29b-3p | MIMAT0000127 | Cdkl2 | 53886 | HITS-CLIP | 37986224 | Unspecified |
| mmu-miR-29b-3p | MIMAT0000127 | Cdkl3 | 213084 | HITS-CLIP | 23142080 | Unspecified |

| | | | | | | |
|---|---|---|---|---|---|---|
| mmu-miR-29b-3p | MIMAT0000127 | Cdkn1a | 12575 | HITS-CLIP//sRNA-Seq | 37986224\|23142080\|33184237\|25083 | Unspecified |
| mmu-miR-29b-3p | MIMAT0000127 | Cdkn1b | 12576 | HITS-CLIP//sRNA-Seq | 25083871\|33184237\|23142080 | Unspecified |
| mmu-miR-29b-3p | MIMAT0000127 | Cdkn2aipnl | 52626 | sRNA-Seq//HITS-CLIP | 33184237\|23142080 | Unspecified |
| mmu-miR-29b-3p | MIMAT0000127 | Cdkn2d | 12581 | HITS-CLIP//sRNA-Seq | 23142080\|33184237 | Unspecified |
| mmu-miR-29b-3p | MIMAT0000127 | Cdo1 | 12583 | HITS-CLIP | 37986224\|23597149 | Unspecified |
| mmu-miR-29b-3p | MIMAT0000127 | Cdon | 57810 | HITS-CLIP//sRNA-Seq | 25083871\|33184237 | Unspecified |
| mmu-miR-29b-3p | MIMAT0000127 | Cdr2 | 12585 | HITS-CLIP | 23142080 | Unspecified |
| mmu-miR-29b-3p | MIMAT0000127 | Cdt1 | 67177 | HITS-CLIP//sRNA-Seq | 23142080\|33184237 | Unspecified |
| mmu-miR-29b-3p | MIMAT0000127 | Cebpa | 12606 | HITS-CLIP | 37986224 | Unspecified |
| mmu-miR-29b-3p | MIMAT0000127 | Cebpg | 12611 | HITS-CLIP | 23597149 | Unspecified |
| mmu-miR-29b-3p | MIMAT0000127 | Celf1 | 13046 | HITS-CLIP | 37986224\|25083871\|23597149\|23142 | Unspecified |
| mmu-miR-29b-3p | MIMAT0000127 | Celf2 | 14007 | HITS-CLIP | 37986224\|23142080\|25083871 | Unspecified |
| mmu-miR-29b-3p | MIMAT0000127 | Cemip | 80982 | HITS-CLIP | 25083871 | Unspecified |
| mmu-miR-29b-3p | MIMAT0000127 | Cemip2 | 23670 | HITS-CLIP | 25083871\|23597149 | Unspecified |
| mmu-miR-29b-3p | MIMAT0000127 | Cenpb | 12616 | HITS-CLIP | 37986224 | Unspecified |
| mmu-miR-29b-3p | MIMAT0000127 | Cenpc1 | 12617 | HITS-CLIP | 23142080 | Unspecified |
| mmu-miR-29b-3p | MIMAT0000127 | Cenpe | 229841 | HITS-CLIP | 25083871\|23597149\|23142080 | Unspecified |
| mmu-miR-29b-3p | MIMAT0000127 | Cenpf | 108000 | HITS-CLIP | 25083871\|23142080 | Unspecified |
| mmu-miR-29b-3p | MIMAT0000127 | Cenpi | 102920 | HITS-CLIP | 23142080 | Unspecified |
| mmu-miR-29b-3p | MIMAT0000127 | Cenpj | 219103 | HITS-CLIP | 23142080 | Unspecified |
| mmu-miR-29b-3p | MIMAT0000127 | Cenpl | 70454 | HITS-CLIP | 23142080 | Unspecified |
| mmu-miR-29b-3p | MIMAT0000127 | Cep120 | 225523 | HITS-CLIP//sRNA-Seq | 25083871\|33184237\|23142080 | Unspecified |
| mmu-miR-29b-3p | MIMAT0000127 | Cep131 | 12009 | HITS-CLIP | 23142080 | Unspecified |
| mmu-miR-29b-3p | MIMAT0000127 | Cep170 | 545389 | HITS-CLIP//sRNA-Seq | 23142080\|33184237\|25083871 | Unspecified |
| mmu-miR-29b-3p | MIMAT0000127 | Cep192 | 70799 | HITS-CLIP | 23597149 | Unspecified |
| mmu-miR-29b-3p | MIMAT0000127 | Cep250 | 16328 | HITS-CLIP | 23142080 | Unspecified |
| mmu-miR-29b-3p | MIMAT0000127 | Cep350 | 74081 | HITS-CLIP | 25083871\|23142080 | Unspecified |
| mmu-miR-29b-3p | MIMAT0000127 | Cep41 | 83922 | HITS-CLIP | 23142080 | Unspecified |
| mmu-miR-29b-3p | MIMAT0000127 | Cep68 | 216543 | HITS-CLIP | 37986224\|25083871\|23142080 | Unspecified |
| mmu-miR-29b-3p | MIMAT0000127 | Cep76 | 225659 | HITS-CLIP | 37986224\|25083871\|23142080 | Unspecified |
| mmu-miR-29b-3p | MIMAT0000127 | Cep83 | 77048 | HITS-CLIP | 23142080 | Unspecified |
| mmu-miR-29b-3p | MIMAT0000127 | Cep85 | 70012 | HITS-CLIP | 25083871 | Unspecified |
| mmu-miR-29b-3p | MIMAT0000127 | Cep97 | 74201 | HITS-CLIP | 37986224\|25083871\|23142080 | Unspecified |
| mmu-miR-29b-3p | MIMAT0000127 | Cept1 | 99712 | sRNA-Seq//HITS-CLIP | 33184237\|23142080 | Unspecified |
| mmu-miR-29b-3p | MIMAT0000127 | Cercam | 99151 | HITS-CLIP | 25083871\|23142080 | Unspecified |
| mmu-miR-29b-3p | MIMAT0000127 | Cerk | 223753 | HITS-CLIP | 25083871\|23142080 | Unspecified |
| mmu-miR-29b-3p | MIMAT0000127 | Cers2 | 76893 | HITS-CLIP//sRNA-Seq | 25083871\|33184237\|23142080 | Unspecified |
| mmu-miR-29b-3p | MIMAT0000127 | Cers5 | 71949 | HITS-CLIP//sRNA-Seq | 37986224\|25083871\|33184237\|23142 | Unspecified |
| mmu-miR-29b-3p | MIMAT0000127 | Ces1b | 382044 | HITS-CLIP | 23597149 | Unspecified |

| | | | | | | |
|---|---|---|---|---|---|---|
| mmu-miR-29b-3p | MIMAT0000127 | Ces1c | 13884 | HITS-CLIP | 23597149 | Unspecified |
| mmu-miR-29b-3p | MIMAT0000127 | Ces1f | 234564 | HITS-CLIP | 23597149 | Unspecified |
| mmu-miR-29b-3p | MIMAT0000127 | Ces2a | 102022 | HITS-CLIP | 23597149 | Unspecified |
| mmu-miR-29b-3p | MIMAT0000127 | Ces2e | 234673 | HITS-CLIP | 23597149 | Unspecified |
| mmu-miR-29b-3p | MIMAT0000127 | Ces3a | 382053 | HITS-CLIP | 23597149 | Unspecified |
| mmu-miR-29b-3p | MIMAT0000127 | Cfap20 | 29105 | HITS-CLIP//sRNA-Seq | 23142080\|33184237\|25083871 | Unspecified |
| mmu-miR-29b-3p | MIMAT0000127 | Cfap36 | 112942 | HITS-CLIP//sRNA-Seq | 23142080\|33184237 | Unspecified |
| mmu-miR-29b-3p | MIMAT0000127 | Cfap97 | 57587 | HITS-CLIP | 23142080 | Unspecified |
| mmu-miR-29b-3p | MIMAT0000127 | Cfh | 12628 | HITS-CLIP | 25083871\|23597149 | Unspecified |
| mmu-miR-29b-3p | MIMAT0000127 | Cfl2 | 12632 | HITS-CLIP | 37986224 | Unspecified |
| mmu-miR-29b-3p | MIMAT0000127 | Cflar | 12633 | HITS-CLIP | 37986224\|25083871\|31433990\|23142 | Unspecified |
| mmu-miR-29b-3p | MIMAT0000127 | Cggbp1 | 106143 | sRNA-Seq//HITS-CLIP | 33184237\|23142080 | Unspecified |
| mmu-miR-29b-3p | MIMAT0000127 | Cgn | 70737 | HITS-CLIP | 23597149 | Unspecified |
| mmu-miR-29b-3p | MIMAT0000127 | Cgnl1 | 68178 | HITS-CLIP | 23597149 | Unspecified |
| mmu-miR-29b-3p | MIMAT0000127 | Chaf1a | 27221 | sRNA-Seq//HITS-CLIP | 33184237\|23142080 | Unspecified |
| mmu-miR-29b-3p | MIMAT0000127 | Chchd1 | 66121 | sRNA-Seq//HITS-CLIP | 33184237\|23142080 | Unspecified |
| mmu-miR-29b-3p | MIMAT0000127 | Chchd3 | 66075 | HITS-CLIP//sRNA-Seq | 37986224\|25083871\|33184237\|23142 | Unspecified |
| mmu-miR-29b-3p | MIMAT0000127 | Chd1 | 12648 | HITS-CLIP | 37986224\|23142080\|25083871 | Unspecified |
| mmu-miR-29b-3p | MIMAT0000127 | Chd2 | 244059 | HITS-CLIP | 25083871\|23142080 | Unspecified |
| mmu-miR-29b-3p | MIMAT0000127 | Chd4 | 107932 | HITS-CLIP//sRNA-Seq | 25083871\|33184237\|23142080 | Unspecified |
| mmu-miR-29b-3p | MIMAT0000127 | Chd6 | 71389 | HITS-CLIP | 25083871 | Unspecified |
| mmu-miR-29b-3p | MIMAT0000127 | Chd7 | 320790 | HITS-CLIP | 37986224\|25083871\|30971389\|23142 | Unspecified |
| mmu-miR-29b-3p | MIMAT0000127 | Chd8 | 67772 | HITS-CLIP//sRNA-Seq | 23142080\|33184237\|23597149\|2508 | Unspecified |
| mmu-miR-29b-3p | MIMAT0000127 | Chd9 | 109151 | HITS-CLIP//sRNA-Seq | 25083871\|23597149\|33184237\|23142 | Unspecified |
| mmu-miR-29b-3p | MIMAT0000127 | Chdh | 218865 | HITS-CLIP | 23597149 | Unspecified |
| mmu-miR-29b-3p | MIMAT0000127 | Chek1 | 12649 | HITS-CLIP | 23142080 | Unspecified |
| mmu-miR-29b-3p | MIMAT0000127 | Cherp | 27967 | HITS-CLIP//sRNA-Seq | 25083871\|33184237\|23142080 | Unspecified |
| mmu-miR-29b-3p | MIMAT0000127 | Chfr | 231600 | HITS-CLIP//sRNA-Seq | 25083871\|33184237\|23142080 | Unspecified |
| mmu-miR-29b-3p | MIMAT0000127 | Chic1 | 12212 | HITS-CLIP | 23142080 | Unspecified |
| mmu-miR-29b-3p | MIMAT0000127 | Chic2 | 74277 | HITS-CLIP | 37986224\|23142080 | Unspecified |
| mmu-miR-29b-3p | MIMAT0000127 | Chm | 12662 | HITS-CLIP | 31433990 | Unspecified |
| mmu-miR-29b-3p | MIMAT0000127 | Chml | 12663 | qCLASH | 31393953 | Unspecified |
| mmu-miR-29b-3p | MIMAT0000127 | Chmp2a | 68953 | HITS-CLIP//sRNA-Seq | 23597149\|23142080\|33184237\|2508 | Unspecified |
| mmu-miR-29b-3p | MIMAT0000127 | Chmp5 | 76959 | HITS-CLIP//sRNA-Seq | 25083871\|33184237 | Unspecified |
| mmu-miR-29b-3p | MIMAT0000127 | Chmp6 | 208092 | HITS-CLIP | 37986224 | Unspecified |
| mmu-miR-29b-3p | MIMAT0000127 | Chmp7 | 105513 | HITS-CLIP | 25083871\|23597149\|23142080 | Unspecified |
| mmu-miR-29b-3p | MIMAT0000127 | Chordc1 | 66917 | HITS-CLIP//sRNA-Seq | 37986224\|33184237\|23142080 | Unspecified |
| mmu-miR-29b-3p | MIMAT0000127 | Chp1 | 56398 | HITS-CLIP//sRNA-Seq | 25083871\|33184237 | Unspecified |
| mmu-miR-29b-3p | MIMAT0000127 | Chpf2 | 100910 | HITS-CLIP | 25083871 | Unspecified |

| | | | | | | |
|---|---|---|---|---|---|---|
| mmu-miR-29b-3p | MIMAT0000127 | Chpt1 | 212862 | HITS-CLIP | 23597149 | Unspecified |
| mmu-miR-29b-3p | MIMAT0000127 | Chrnb1 | 11443 | HITS-CLIP | 25083871 | Unspecified |
| mmu-miR-29b-3p | MIMAT0000127 | Chrng | 11449 | HITS-CLIP | 25083871 | Unspecified |
| mmu-miR-29b-3p | MIMAT0000127 | Chst11 | 58250 | HITS-CLIP | 37986224\|25083871\|23142080 | Unspecified |
| mmu-miR-29b-3p | MIMAT0000127 | Chst13 | 71797 | HITS-CLIP | 23597149 | Unspecified |
| mmu-miR-29b-3p | MIMAT0000127 | Chst14 | 72136 | HITS-CLIP | 23142080 | Unspecified |
| mmu-miR-29b-3p | MIMAT0000127 | Chst2 | 54371 | HITS-CLIP | 37986224\|25083871 | Unspecified |
| mmu-miR-29b-3p | MIMAT0000127 | Chsy1 | 269941 | HITS-CLIP | 37986224\|25083871\|23142080 | Unspecified |
| mmu-miR-29b-3p | MIMAT0000127 | Chtf18 | 214901 | HITS-CLIP | 23142080 | Unspecified |
| mmu-miR-29b-3p | MIMAT0000127 | Chtf8 | 214987 | HITS-CLIP//sRNA-Seq | 23142080\|33184237\|23597149 | Unspecified |
| mmu-miR-29b-3p | MIMAT0000127 | Chtop | 66511 | sRNA-Seq//HITS-CLIP | 33184237\|23142080\|25083871 | Unspecified |
| mmu-miR-29b-3p | MIMAT0000127 | Cib2 | 56506 | HITS-CLIP | 25083871 | Unspecified |
| mmu-miR-29b-3p | MIMAT0000127 | Cic | 71722 | HITS-CLIP//sRNA-Seq | 23142080\|25083871\|33184237\|23597 | Unspecified |
| mmu-miR-29b-3p | MIMAT0000127 | Cideb | 12684 | HITS-CLIP | 37986224\|23597149 | Unspecified |
| mmu-miR-29b-3p | MIMAT0000127 | Cinp | 67236 | HITS-CLIP | 37986224\|23142080 | Unspecified |
| mmu-miR-29b-3p | MIMAT0000127 | Cip2a | 57650 | HITS-CLIP | 23142080 | Unspecified |
| mmu-miR-29b-3p | MIMAT0000127 | Cisd1 | 52637 | HITS-CLIP//sRNA-Seq | 25083871\|33184237\|23142080 | Unspecified |
| mmu-miR-29b-3p | MIMAT0000127 | Cisd2 | 67006 | HITS-CLIP | 21258322\|25083871\|23597149\|23142 | Unspecified |
| mmu-miR-29b-3p | MIMAT0000127 | Cish | 12700 | HITS-CLIP | 23142080 | Unspecified |
| mmu-miR-29b-3p | MIMAT0000127 | Cit | 12704 | HITS-CLIP | 25083871 | Unspecified |
| mmu-miR-29b-3p | MIMAT0000127 | Ckap5 | 75786 | HITS-CLIP//sRNA-Seq | 25083871\|33184237\|23142080 | Unspecified |
| mmu-miR-29b-3p | MIMAT0000127 | Clasp1 | 76707 | HITS-CLIP//sRNA-Seq | 25083871\|33184237 | Unspecified |
| mmu-miR-29b-3p | MIMAT0000127 | Clasp2 | 76499 | sRNA-Seq//HITS-CLIP | 33184237\|23142080 | Unspecified |
| mmu-miR-29b-3p | MIMAT0000127 | Clcf1 | 56708 | HITS-CLIP | 25083871\|23142080 | Unspecified |
| mmu-miR-29b-3p | MIMAT0000127 | Clcn2 | 12724 | HITS-CLIP | 23597149 | Unspecified |
| mmu-miR-29b-3p | MIMAT0000127 | Clcn3 | 12725 | sRNA-Seq//HITS-CLIP | 33184237\|23142080 | Unspecified |
| mmu-miR-29b-3p | MIMAT0000127 | Clcn5 | 12728 | HITS-CLIP//sRNA-Seq | 25083871\|33184237\|23142080 | Unspecified |
| mmu-miR-29b-3p | MIMAT0000127 | Clcn7 | 26373 | HITS-CLIP//sRNA-Seq | 37986224\|25083871\|33184237\|23597 | Unspecified |
| mmu-miR-29b-3p | MIMAT0000127 | Cldn1 | 12737 | HITS-CLIP | 37986224\|23597149 | Unspecified |
| mmu-miR-29b-3p | MIMAT0000127 | Cldn12 | 64945 | HITS-CLIP | 37986224\|25083871\|23597149 | Unspecified |
| mmu-miR-29b-3p | MIMAT0000127 | Cldn2 | 12738 | HITS-CLIP | 23597149 | Unspecified |
| mmu-miR-29b-3p | MIMAT0000127 | Cldn3 | 12739 | HITS-CLIP | 23597149 | Unspecified |
| mmu-miR-29b-3p | MIMAT0000127 | Cldnd1 | 224250 | sRNA-Seq//HITS-CLIP | 33184237\|25083871 | Unspecified |
| mmu-miR-29b-3p | MIMAT0000127 | Clec2d | 93694 | HITS-CLIP | 37986224 | Unspecified |
| mmu-miR-29b-3p | MIMAT0000127 | Clec4f | 51811 | HITS-CLIP | 23597149 | Unspecified |
| mmu-miR-29b-3p | MIMAT0000127 | Clec4g | 75863 | HITS-CLIP | 23597149 | Unspecified |
| mmu-miR-29b-3p | MIMAT0000127 | Clic1 | 114584 | HITS-CLIP//sRNA-Seq | 23597149\|33184237\|25083871\|23142 | Unspecified |
| mmu-miR-29b-3p | MIMAT0000127 | Clic5 | 224796 | HITS-CLIP | 25083871 | Unspecified |
| mmu-miR-29b-3p | MIMAT0000127 | Clint1 | 216705 | HITS-CLIP | 25083871\|23142080 | Unspecified |

| | | | | | | |
|---|---|---|---|---|---|---|
| mmu-miR-29b-3p | MIMAT0000127 | Clip1 | 56430 | HITS-CLIP//sRNA-Seq | 25083871\|33184237\|23142080 | Unspecified |
| mmu-miR-29b-3p | MIMAT0000127 | Clip2 | 269713 | sRNA-Seq//HITS-CLIP | 33184237\|23142080 | Unspecified |
| mmu-miR-29b-3p | MIMAT0000127 | Clk1 | 12747 | sRNA-Seq//HITS-CLIP | 33184237\|23142080 | Unspecified |
| mmu-miR-29b-3p | MIMAT0000127 | Clk2 | 12748 | HITS-CLIP | 25083871\|23142080 | Unspecified |
| mmu-miR-29b-3p | MIMAT0000127 | Clmn | 94040 | HITS-CLIP | 23597149 | Unspecified |
| mmu-miR-29b-3p | MIMAT0000127 | Cln3 | 12752 | HITS-CLIP | 23142080 | Unspecified |
| mmu-miR-29b-3p | MIMAT0000127 | Cln6 | 76524 | HITS-CLIP | 23597149 | Unspecified |
| mmu-miR-29b-3p | MIMAT0000127 | Clns1a | 12729 | sRNA-Seq//HITS-CLIP | 33184237\|23142080 | Unspecified |
| mmu-miR-29b-3p | MIMAT0000127 | Clock | 12753 | HITS-CLIP | 37986224\|23142080\|23597149\|25083 | Unspecified |
| mmu-miR-29b-3p | MIMAT0000127 | Clp1 | 98985 | HITS-CLIP | 23142080\|25083871\|23597149 | Unspecified |
| mmu-miR-29b-3p | MIMAT0000127 | Clptm1 | 56457 | sRNA-Seq//HITS-CLIP | 33184237\|23597149\|23142080 | Unspecified |
| mmu-miR-29b-3p | MIMAT0000127 | Clpx | 270166 | HITS-CLIP | 37986224\|23597149\|23142080\|25083 | Unspecified |
| mmu-miR-29b-3p | MIMAT0000127 | Clspn | 269582 | HITS-CLIP | 23142080 | Unspecified |
| mmu-miR-29b-3p | MIMAT0000127 | Clta | 12757 | HITS-CLIP//sRNA-Seq | 25083871\|33184237\|23142080 | Unspecified |
| mmu-miR-29b-3p | MIMAT0000127 | Cltc | 67300 | HITS-CLIP//sRNA-Seq | 37986224\|25083871\|23597149\|33184 | Unspecified |
| mmu-miR-29b-3p | MIMAT0000127 | Clu | 12759 | HITS-CLIP//sRNA-Seq | 23597149\|33184237 | Unspecified |
| mmu-miR-29b-3p | MIMAT0000127 | Cluh | 74148 | HITS-CLIP | 37986224\|23597149 | Unspecified |
| mmu-miR-29b-3p | MIMAT0000127 | Clybl | 69634 | HITS-CLIP | 23142080 | Unspecified |
| mmu-miR-29b-3p | MIMAT0000127 | Cmah | 12763 | HITS-CLIP | 23597149 | Unspecified |
| mmu-miR-29b-3p | MIMAT0000127 | Cmas | 12764 | HITS-CLIP//sRNA-Seq | 37986224\|33184237\|23142080 | Unspecified |
| mmu-miR-29b-3p | MIMAT0000127 | Cmklr1 | 14747 | HITS-CLIP | 25083871 | Unspecified |
| mmu-miR-29b-3p | MIMAT0000127 | Cmpk1 | 66588 | HITS-CLIP//sRNA-Seq | 37986224\|33184237\|23142080 | Unspecified |
| mmu-miR-29b-3p | MIMAT0000127 | Cmtm3 | 68119 | HITS-CLIP//sRNA-Seq | 37986224\|25083871\|33184237 | Unspecified |
| mmu-miR-29b-3p | MIMAT0000127 | Cmtm4 | 97487 | HITS-CLIP | 23142080 | Unspecified |
| mmu-miR-29b-3p | MIMAT0000127 | Cmtm6 | 67213 | HITS-CLIP | 23142080 | Unspecified |
| mmu-miR-29b-3p | MIMAT0000127 | Cmtm7 | 102545 | HITS-CLIP | 23142080 | Unspecified |
| mmu-miR-29b-3p | MIMAT0000127 | Cmtr1 | 74157 | HITS-CLIP | 23142080 | Unspecified |
| mmu-miR-29b-3p | MIMAT0000127 | Cnbp | 12785 | HITS-CLIP//sRNA-Seq | 23597149\|25083871\|33184237\|23142 | Unspecified |
| mmu-miR-29b-3p | MIMAT0000127 | Cnep1r1 | 382030 | HITS-CLIP | 23142080 | Unspecified |
| mmu-miR-29b-3p | MIMAT0000127 | Cnih1 | 12793 | sRNA-Seq//HITS-CLIP | 33184237\|23142080\|25083871 | Unspecified |
| mmu-miR-29b-3p | MIMAT0000127 | Cnih4 | 98417 | HITS-CLIP | 23142080\|23597149 | Unspecified |
| mmu-miR-29b-3p | MIMAT0000127 | Cnksr3 | 215748 | HITS-CLIP | 25083871\|23142080 | Unspecified |
| mmu-miR-29b-3p | MIMAT0000127 | Cnn2 | 12798 | HITS-CLIP//sRNA-Seq | 23142080\|33184237 | Unspecified |
| mmu-miR-29b-3p | MIMAT0000127 | Cnnm4 | 94220 | HITS-CLIP | 37986224\|23142080 | Unspecified |
| mmu-miR-29b-3p | MIMAT0000127 | Cnot1 | 234594 | HITS-CLIP//sRNA-Seq | 25083871\|33184237\|23597149\|23142 | Unspecified |
| mmu-miR-29b-3p | MIMAT0000127 | Cnot2 | 72068 | sRNA-Seq//HITS-CLIP | 33184237\|23142080 | Unspecified |
| mmu-miR-29b-3p | MIMAT0000127 | Cnot4 | 53621 | HITS-CLIP | 23142080 | Unspecified |
| mmu-miR-29b-3p | MIMAT0000127 | Cnot6 | 104625 | HITS-CLIP//sRNA-Seq | 37986224\|23142080\|33184237 | Unspecified |
| mmu-miR-29b-3p | MIMAT0000127 | Cnot6l | 231464 | HITS-CLIP | 25083871\|23142080\|31433990\|26203 | Unspecified |

| | | | | | | |
|---|---|---|---|---|---|---|
| mmu-miR-29b-3p | MIMAT0000127 | Cnot7 | 18983 | sRNA-Seq//HITS-CLIP | 33184237\|23142080 | Unspecified |
| mmu-miR-29b-3p | MIMAT0000127 | Cnot9 | 9125 | sRNA-Seq//HITS-CLIP | 33184237\|23597149\|23142080 | Unspecified |
| mmu-miR-29b-3p | MIMAT0000127 | Cnp | 12799 | HITS-CLIP | 23597149\|23142080 | Unspecified |
| mmu-miR-29b-3p | MIMAT0000127 | Cnppd1 | 69171 | HITS-CLIP | 23142080\|23597149 | Unspecified |
| mmu-miR-29b-3p | MIMAT0000127 | Cnr2 | 12802 | HITS-CLIP | 23142080 | Unspecified |
| mmu-miR-29b-3p | MIMAT0000127 | Coasy | 71743 | HITS-CLIP | 23597149 | Unspecified |
| mmu-miR-29b-3p | MIMAT0000127 | Cog3 | 338337 | HITS-CLIP | 25083871 | Unspecified |
| mmu-miR-29b-3p | MIMAT0000127 | Cog4 | 102339 | HITS-CLIP//sRNA-Seq | 25083871\|23597149\|33184237 | Unspecified |
| mmu-miR-29b-3p | MIMAT0000127 | Cog5 | 238123 | HITS-CLIP | 30971389 | Unspecified |
| mmu-miR-29b-3p | MIMAT0000127 | Cog6 | 67542 | HITS-CLIP | 25083871\|23597149\|23142080 | Unspecified |
| mmu-miR-29b-3p | MIMAT0000127 | Cog8 | 97484 | HITS-CLIP | 37986224\|23142080 | Unspecified |
| mmu-miR-29b-3p | MIMAT0000127 | Col11a1 | 12814 | HITS-CLIP//sRNA-Seq | 37986224\|25083871\|33184237 | Unspecified |
| mmu-miR-29b-3p | MIMAT0000127 | Col12a1 | 12816 | HITS-CLIP//sRNA-Seq | 25083871\|33184237 | Unspecified |
| mmu-miR-29b-3p | MIMAT0000127 | Col1a1 | 12842 | Luciferase assay//Microarray//Northern bl | 19342382\|23597149\|33184237\|2508 | Unspecified |
| mmu-miR-29b-3p | MIMAT0000127 | Col1a2 | 12843 | Luciferase assay//Microarray//Northern bl | 18723672\|25083871\|23597149\|3318 | Unspecified |
| mmu-miR-29b-3p | MIMAT0000127 | Col25a1 | 77018 | HITS-CLIP | 37986224\|25083871 | Unspecified |
| mmu-miR-29b-3p | MIMAT0000127 | Col3a1 | 12825 | Luciferase assay//Microarray//Northern bl | 18723672\|25083871\|33184237\|2359 | Unspecified |
| mmu-miR-29b-3p | MIMAT0000127 | Col4a1 | 12826 | Luciferase assay//qRT-PCR//Western blot/ | 23139829\|23597149\|25083871\|3318 | Unspecified |
| mmu-miR-29b-3p | MIMAT0000127 | Col4a2 | 12827 | Luciferase assay//qRT-PCR//Western blot/ | 23139829\|25083871\|23597149\|3318 | Unspecified |
| mmu-miR-29b-3p | MIMAT0000127 | Col4a5 | 12830 | Luciferase assay//qRT-PCR//Western blot/ | 23139829\|25083871\|33184237 | Unspecified |
| mmu-miR-29b-3p | MIMAT0000127 | Col5a1 | 12831 | HITS-CLIP//sRNA-Seq | 23597149\|25083871\|33184237\|2314 | Unspecified |
| mmu-miR-29b-3p | MIMAT0000127 | Col5a2 | 12832 | HITS-CLIP//sRNA-Seq | 37986224\|25083871\|33184237 | Unspecified |
| mmu-miR-29b-3p | MIMAT0000127 | Col5a3 | 53867 | Luciferase assay//Microarray//Northern bl | 19342382\|25083871\|33184237 | Unspecified |
| mmu-miR-29b-3p | MIMAT0000127 | Col6a1 | 12833 | HITS-CLIP//sRNA-Seq | 37986224\|23597149\|33184237\|2508 | Unspecified |
| mmu-miR-29b-3p | MIMAT0000127 | Col6a2 | 12834 | HITS-CLIP//sRNA-Seq | 37986224\|25083871\|33184237 | Unspecified |
| mmu-miR-29b-3p | MIMAT0000127 | Col6a3 | 12835 | HITS-CLIP//sRNA-Seq | 37986224\|25083871\|33184237\|2359 | Unspecified |
| mmu-miR-29b-3p | MIMAT0000127 | Col8a1 | 12837 | HITS-CLIP//sRNA-Seq | 37986224\|25083871\|33184237 | Unspecified |
| mmu-miR-29b-3p | MIMAT0000127 | Colgalt1 | 234407 | HITS-CLIP | 37986224 | Unspecified |
| mmu-miR-29b-3p | MIMAT0000127 | Commd1 | 17846 | HITS-CLIP//sRNA-Seq | 25083871\|33184237 | Unspecified |
| mmu-miR-29b-3p | MIMAT0000127 | Commd4 | 66199 | HITS-CLIP | 23142080 | Unspecified |
| mmu-miR-29b-3p | MIMAT0000127 | Comt | 12846 | HITS-CLIP//sRNA-Seq | 25083871\|23142080\|33184237\|2359 | Unspecified |
| mmu-miR-29b-3p | MIMAT0000127 | Copa | 12847 | HITS-CLIP//sRNA-Seq | 37986224\|25083871\|23142080\|3318 | Unspecified |
| mmu-miR-29b-3p | MIMAT0000127 | Copb1 | 70349 | HITS-CLIP//sRNA-Seq | 25083871\|33184237\|23142080 | Unspecified |
| mmu-miR-29b-3p | MIMAT0000127 | Copg1 | 54161 | HITS-CLIP//sRNA-Seq | 25083871\|23597149\|33184237\|2314 | Unspecified |
| mmu-miR-29b-3p | MIMAT0000127 | Copg2 | 54160 | HITS-CLIP | 25083871\|23597149 | Unspecified |
| mmu-miR-29b-3p | MIMAT0000127 | Cops2 | 12848 | HITS-CLIP//sRNA-Seq | 25083871\|23597149\|33184237\|2314 | Unspecified |
| mmu-miR-29b-3p | MIMAT0000127 | Cops4 | 26891 | HITS-CLIP//sRNA-Seq | 25083871\|33184237\|23142080 | Unspecified |
| mmu-miR-29b-3p | MIMAT0000127 | Cops7b | 26895 | HITS-CLIP | 25083871\|23597149 | Unspecified |
| mmu-miR-29b-3p | MIMAT0000127 | Copz1 | 56447 | HITS-CLIP//sRNA-Seq | 23142080\|33184237\|25083871 | Unspecified |

| | | | | | | |
|---|---|---|---|---|---|---|
| mmu-miR-29b-3p | MIMAT0000127 | Coq8a | 56997 | HITS-CLIP | 23597149\|25083871\|23142080 | Unspecified |
| mmu-miR-29b-3p | MIMAT0000127 | Coq8b | 79934 | HITS-CLIP | 23142080 | Unspecified |
| mmu-miR-29b-3p | MIMAT0000127 | Coq9 | 67914 | HITS-CLIP//sRNA-Seq | 23597149\|33184237 | Unspecified |
| mmu-miR-29b-3p | MIMAT0000127 | Coro1a | 12721 | sRNA-Seq//HITS-CLIP | 33184237\|23142080 | Unspecified |
| mmu-miR-29b-3p | MIMAT0000127 | Coro1b | 23789 | HITS-CLIP//sRNA-Seq | 23142080\|25083871\|33184237 | Unspecified |
| mmu-miR-29b-3p | MIMAT0000127 | Coro1c | 23790 | HITS-CLIP//sRNA-Seq | 25083871\|33184237 | Unspecified |
| mmu-miR-29b-3p | MIMAT0000127 | Coro2b | 235431 | HITS-CLIP//sRNA-Seq | 25083871\|33184237 | Unspecified |
| mmu-miR-29b-3p | MIMAT0000127 | Coro6 | 216961 | HITS-CLIP | 25083871 | Unspecified |
| mmu-miR-29b-3p | MIMAT0000127 | Coro7 | 78885 | HITS-CLIP | 23142080 | Unspecified |
| mmu-miR-29b-3p | MIMAT0000127 | Cotl1 | 72042 | sRNA-Seq//HITS-CLIP | 33184237\|23142080 | Unspecified |
| mmu-miR-29b-3p | MIMAT0000127 | Cox11 | 69802 | HITS-CLIP | 37986224\|23597149\|23142080\|25083 | Unspecified |
| mmu-miR-29b-3p | MIMAT0000127 | Cox15 | 226139 | HITS-CLIP | 37986224 | Unspecified |
| mmu-miR-29b-3p | MIMAT0000127 | Cox6a1 | 12861 | HITS-CLIP//sRNA-Seq | 25083871\|33184237\|23597149\|23142 | Unspecified |
| mmu-miR-29b-3p | MIMAT0000127 | Cp | 12870 | HITS-CLIP | 23597149 | Unspecified |
| mmu-miR-29b-3p | MIMAT0000127 | Cpd | 12874 | HITS-CLIP//sRNA-Seq | 23597149\|25083871\|33184237\|23142 | Unspecified |
| mmu-miR-29b-3p | MIMAT0000127 | Cpe | 12876 | HITS-CLIP//sRNA-Seq | 25083871\|33184237 | Unspecified |
| mmu-miR-29b-3p | MIMAT0000127 | Cpeb2 | 231207 | HITS-CLIP | 23142080\|25083871\|31433990 | Unspecified |
| mmu-miR-29b-3p | MIMAT0000127 | Cpeb3 | 208922 | HITS-CLIP | 25083871 | Unspecified |
| mmu-miR-29b-3p | MIMAT0000127 | Cpeb4 | 67579 | HITS-CLIP//sRNA-Seq | 37986224\|25083871\|33184237\|23142 | Unspecified |
| mmu-miR-29b-3p | MIMAT0000127 | Cped1 | 214642 | HITS-CLIP | 23597149 | Unspecified |
| mmu-miR-29b-3p | MIMAT0000127 | Cplane1 | 65250 | HITS-CLIP | 25083871 | Unspecified |
| mmu-miR-29b-3p | MIMAT0000127 | Cplx2 | 12890 | HITS-CLIP//sRNA-Seq | 25083871\|33184237 | Unspecified |
| mmu-miR-29b-3p | MIMAT0000127 | Cpn1 | 93721 | HITS-CLIP | 23597149 | Unspecified |
| mmu-miR-29b-3p | MIMAT0000127 | Cpn2 | 71756 | HITS-CLIP | 23597149 | Unspecified |
| mmu-miR-29b-3p | MIMAT0000127 | Cpne3 | 70568 | HITS-CLIP//sRNA-Seq | 25083871\|23597149\|33184237\|23142 | Unspecified |
| mmu-miR-29b-3p | MIMAT0000127 | Cpq | 54381 | HITS-CLIP | 37986224\|25083871\|23597149 | Unspecified |
| mmu-miR-29b-3p | MIMAT0000127 | Cps1 | 227231 | HITS-CLIP | 23597149 | Unspecified |
| mmu-miR-29b-3p | MIMAT0000127 | Cpsf1 | 94230 | sRNA-Seq//HITS-CLIP | 33184237\|23597149\|23142080 | Unspecified |
| mmu-miR-29b-3p | MIMAT0000127 | Cpsf3 | 54451 | sRNA-Seq//HITS-CLIP | 33184237\|23142080 | Unspecified |
| mmu-miR-29b-3p | MIMAT0000127 | Cpsf6 | 432508 | HITS-CLIP//sRNA-Seq | 25083871\|33184237\|23142080 | Unspecified |
| mmu-miR-29b-3p | MIMAT0000127 | Cpsf7 | 269061 | HITS-CLIP//sRNA-Seq | 37986224\|25083871\|33184237\|23142 | Unspecified |
| mmu-miR-29b-3p | MIMAT0000127 | Cpt1a | 12894 | HITS-CLIP//sRNA-Seq | 23597149\|33184237 | Unspecified |
| mmu-miR-29b-3p | MIMAT0000127 | Cpt2 | 12896 | HITS-CLIP | 23597149\|25083871\|23142080 | Unspecified |
| mmu-miR-29b-3p | MIMAT0000127 | Cramp1l | 57354 | HITS-CLIP | 37986224\|23142080\|25083871 | Unspecified |
| mmu-miR-29b-3p | MIMAT0000127 | Crat | 12908 | HITS-CLIP | 25083871 | Unspecified |
| mmu-miR-29b-3p | MIMAT0000127 | Crbn | 58799 | sRNA-Seq//HITS-CLIP | 33184237\|23597149\|23142080 | Unspecified |
| mmu-miR-29b-3p | MIMAT0000127 | Creb1 | 12912 | HITS-CLIP | 25083871\|23142080 | Unspecified |
| mmu-miR-29b-3p | MIMAT0000127 | Creb3l3 | 208677 | HITS-CLIP | 23597149 | Unspecified |
| mmu-miR-29b-3p | MIMAT0000127 | Creb5 | 231991 | HITS-CLIP | 37986224\|25083871 | Unspecified |

| | | | | | | |
|---|---|---|---|---|---|---|
| mmu-miR-29b-3p | MIMAT0000127 | Crebbp | 12914 | HITS-CLIP | 23597149\|23142080\|25083871 | Unspecified |
| mmu-miR-29b-3p | MIMAT0000127 | Crebl2 | 232430 | HITS-CLIP | 23142080 | Unspecified |
| mmu-miR-29b-3p | MIMAT0000127 | Crebrf | 77128 | HITS-CLIP | 25083871\|23142080 | Unspecified |
| mmu-miR-29b-3p | MIMAT0000127 | Creld1 | 171508 | HITS-CLIP | 23597149\|25083871 | Unspecified |
| mmu-miR-29b-3p | MIMAT0000127 | Crim1 | 50766 | sRNA-Seq//HITS-CLIP | 33184237\|23142080 | Unspecified |
| mmu-miR-29b-3p | MIMAT0000127 | Crip2 | 68337 | HITS-CLIP//sRNA-Seq | 25083871\|33184237\|23597149 | Unspecified |
| mmu-miR-29b-3p | MIMAT0000127 | Cript | 56724 | HITS-CLIP//sRNA-Seq | 23597149\|23142080\|33184237 | Unspecified |
| mmu-miR-29b-3p | MIMAT0000127 | Crk | 12928 | HITS-CLIP//sRNA-Seq | 25083871\|33184237\|23142080 | Unspecified |
| mmu-miR-29b-3p | MIMAT0000127 | Crkl | 12929 | HITS-CLIP//sRNA-Seq | 23142080\|25083871\|33184237 | Unspecified |
| mmu-miR-29b-3p | MIMAT0000127 | Crot | 74114 | HITS-CLIP | 37986224\|23142080 | Unspecified |
| mmu-miR-29b-3p | MIMAT0000127 | Crp | 12944 | HITS-CLIP | 23597149 | Unspecified |
| mmu-miR-29b-3p | MIMAT0000127 | Crybg1 | 202 | HITS-CLIP | 23142080 | Unspecified |
| mmu-miR-29b-3p | MIMAT0000127 | Crybg2 | 55057 | HITS-CLIP | 23142080 | Unspecified |
| mmu-miR-29b-3p | MIMAT0000127 | Cs | 12974 | HITS-CLIP//sRNA-Seq | 23597149\|33184237 | Unspecified |
| mmu-miR-29b-3p | MIMAT0000127 | Csde1 | 229663 | HITS-CLIP//sRNA-Seq | 25083871\|33184237\|23142080 | Unspecified |
| mmu-miR-29b-3p | MIMAT0000127 | Cse1l | 110750 | HITS-CLIP//sRNA-Seq | 37986224\|23597149\|25083871\|33184 | Unspecified |
| mmu-miR-29b-3p | MIMAT0000127 | Csf1r | 12978 | HITS-CLIP | 23597149 | Unspecified |
| mmu-miR-29b-3p | MIMAT0000127 | Csgalnact1 | 234356 | HITS-CLIP | 37986224 | Unspecified |
| mmu-miR-29b-3p | MIMAT0000127 | Csgalnact2 | 78752 | HITS-CLIP | 37986224 | Unspecified |
| mmu-miR-29b-3p | MIMAT0000127 | Csk | 12988 | HITS-CLIP//sRNA-Seq | 37986224\|25083871\|33184237\|23142 | Unspecified |
| mmu-miR-29b-3p | MIMAT0000127 | Csnk1a1 | 93687 | HITS-CLIP//sRNA-Seq | 23142080\|25083871\|33184237 | Unspecified |
| mmu-miR-29b-3p | MIMAT0000127 | Csnk1d | 104318 | HITS-CLIP//sRNA-Seq | 25083871\|33184237 | Unspecified |
| mmu-miR-29b-3p | MIMAT0000127 | Csnk1e | 27373 | HITS-CLIP//sRNA-Seq | 25083871\|33184237 | Unspecified |
| mmu-miR-29b-3p | MIMAT0000127 | Csnk1g1 | 214897 | HITS-CLIP//sRNA-Seq | 37986224\|25083871\|33184237\|23142 | Unspecified |
| mmu-miR-29b-3p | MIMAT0000127 | Csnk1g2 | 103236 | sRNA-Seq//HITS-CLIP | 33184237\|23142080 | Unspecified |
| mmu-miR-29b-3p | MIMAT0000127 | Csnk1g3 | 70425 | HITS-CLIP//sRNA-Seq | 25083871\|33184237\|23142080 | Unspecified |
| mmu-miR-29b-3p | MIMAT0000127 | Csnk2a1 | 12995 | HITS-CLIP//sRNA-Seq | 25083871\|33184237\|23142080 | Unspecified |
| mmu-miR-29b-3p | MIMAT0000127 | Csnk2a2 | 13000 | sRNA-Seq//HITS-CLIP | 33184237\|25083871 | Unspecified |
| mmu-miR-29b-3p | MIMAT0000127 | Csnk2b | 13001 | sRNA-Seq//HITS-CLIP | 33184237\|23142080 | Unspecified |
| mmu-miR-29b-3p | MIMAT0000127 | Cspg4 | 121021 | HITS-CLIP | 37986224\|25083871 | Unspecified |
| mmu-miR-29b-3p | MIMAT0000127 | Cspp1 | 211660 | HITS-CLIP//sRNA-Seq | 25083871\|33184237\|23597149 | Unspecified |
| mmu-miR-29b-3p | MIMAT0000127 | Csrnp1 | 215418 | HITS-CLIP | 23142080 | Unspecified |
| mmu-miR-29b-3p | MIMAT0000127 | Csrnp2 | 207785 | HITS-CLIP | 37986224\|25083871 | Unspecified |
| mmu-miR-29b-3p | MIMAT0000127 | Cst7 | 13011 | HITS-CLIP | 23142080 | Unspecified |
| mmu-miR-29b-3p | MIMAT0000127 | Cstf2 | 108062 | HITS-CLIP | 23142080 | Unspecified |
| mmu-miR-29b-3p | MIMAT0000127 | Cstf2t | 83410 | HITS-CLIP | 25083871 | Unspecified |
| mmu-miR-29b-3p | MIMAT0000127 | Cstf3 | 228410 | HITS-CLIP | 25083871\|23142080 | Unspecified |
| mmu-miR-29b-3p | MIMAT0000127 | Ctbp1 | 13016 | HITS-CLIP//sRNA-Seq | 23142080\|33184237\|23597149 | Unspecified |
| mmu-miR-29b-3p | MIMAT0000127 | Ctcf | 13018 | HITS-CLIP//sRNA-Seq | 23597149\|23142080\|33184237 | Unspecified |

| | | | | | | |
|---|---|---|---|---|---|---|
| mmu-miR-29b-3p | MIMAT0000127 | Ctdnep1 | 67181 | HITS-CLIP//sRNA-Seq | 25083871\|33184237\|23142080 | Unspecified |
| mmu-miR-29b-3p | MIMAT0000127 | Ctdp1 | 67655 | HITS-CLIP | 23597149 | Unspecified |
| mmu-miR-29b-3p | MIMAT0000127 | Ctdsp1 | 227292 | HITS-CLIP//sRNA-Seq | 25083871\|33184237 | Unspecified |
| mmu-miR-29b-3p | MIMAT0000127 | Ctdsp2 | 52468 | HITS-CLIP//sRNA-Seq | 25083871\|33184237 | Unspecified |
| mmu-miR-29b-3p | MIMAT0000127 | Ctdspl2 | 329506 | HITS-CLIP | 37986224\|25083871\|23597149\|23142 | Unspecified |
| mmu-miR-29b-3p | MIMAT0000127 | Cth | 107869 | HITS-CLIP | 23597149 | Unspecified |
| mmu-miR-29b-3p | MIMAT0000127 | Ctla4 | 12477 | HITS-CLIP | 23142080 | Unspecified |
| mmu-miR-29b-3p | MIMAT0000127 | Ctnna1 | 12385 | HITS-CLIP//sRNA-Seq | 25083871\|23597149\|33184237 | Unspecified |
| mmu-miR-29b-3p | MIMAT0000127 | Ctnnb1 | 12387 | HITS-CLIP//sRNA-Seq | 37986224\|23597149\|25083871\|23142 | Unspecified |
| mmu-miR-29b-3p | MIMAT0000127 | Ctnnbip1 | 67087 | Luciferase assay//Microarray//Northern bl | 19342382\|25083871 | Unspecified |
| mmu-miR-29b-3p | MIMAT0000127 | Ctnnbl1 | 66642 | HITS-CLIP | 23597149 | Unspecified |
| mmu-miR-29b-3p | MIMAT0000127 | Ctnnd1 | 12388 | HITS-CLIP//sRNA-Seq | 37986224\|25083871\|23597149\|33184 | Unspecified |
| mmu-miR-29b-3p | MIMAT0000127 | Ctr9 | 22083 | sRNA-Seq//HITS-CLIP | 33184237\|25083871\|23142080 | Unspecified |
| mmu-miR-29b-3p | MIMAT0000127 | Ctsa | 19025 | HITS-CLIP//sRNA-Seq | 37986224\|25083871\|23597149\|33184 | Unspecified |
| mmu-miR-29b-3p | MIMAT0000127 | Ctsc | 13032 | HITS-CLIP | 37986224\|23142080\|23597149 | Unspecified |
| mmu-miR-29b-3p | MIMAT0000127 | Ctsf | 56464 | HITS-CLIP | 37986224\|25083871\|23597149 | Unspecified |
| mmu-miR-29b-3p | MIMAT0000127 | Ctsl | 13039 | HITS-CLIP//sRNA-Seq | 23597149\|33184237\|25083871 | Unspecified |
| mmu-miR-29b-3p | MIMAT0000127 | Ctss | 13040 | HITS-CLIP | 37986224\|23142080 | Unspecified |
| mmu-miR-29b-3p | MIMAT0000127 | Cttnbp2nl | 80281 | HITS-CLIP | 37986224\|23597149 | Unspecified |
| mmu-miR-29b-3p | MIMAT0000127 | Cuedc1 | 103841 | HITS-CLIP | 37986224\|25083871 | Unspecified |
| mmu-miR-29b-3p | MIMAT0000127 | Cul1 | 26965 | sRNA-Seq//HITS-CLIP | 33184237\|23142080 | Unspecified |
| mmu-miR-29b-3p | MIMAT0000127 | Cul2 | 71745 | HITS-CLIP//sRNA-Seq | 25083871\|23142080\|33184237\|23597 | Unspecified |
| mmu-miR-29b-3p | MIMAT0000127 | Cul3 | 26554 | HITS-CLIP//sRNA-Seq | 25083871\|23142080\|33184237\|23597 | Unspecified |
| mmu-miR-29b-3p | MIMAT0000127 | Cul4a | 99375 | sRNA-Seq//HITS-CLIP | 33184237\|25083871\|23142080 | Unspecified |
| mmu-miR-29b-3p | MIMAT0000127 | Cul4b | 72584 | HITS-CLIP//sRNA-Seq | 23142080\|33184237\|25083871 | Unspecified |
| mmu-miR-29b-3p | MIMAT0000127 | Cul5 | 75717 | HITS-CLIP | 25083871\|23142080 | Unspecified |
| mmu-miR-29b-3p | MIMAT0000127 | Cul7 | 66515 | sRNA-Seq//HITS-CLIP | 33184237\|23142080 | Unspecified |
| mmu-miR-29b-3p | MIMAT0000127 | Cux1 | 13047 | HITS-CLIP | 25083871\|23142080 | Unspecified |
| mmu-miR-29b-3p | MIMAT0000127 | Cwc15 | 66070 | sRNA-Seq//HITS-CLIP | 33184237\|23142080\|25083871 | Unspecified |
| mmu-miR-29b-3p | MIMAT0000127 | Cwc22 | 80744 | HITS-CLIP | 23142080 | Unspecified |
| mmu-miR-29b-3p | MIMAT0000127 | Cwc25 | 67480 | HITS-CLIP | 37986224 | Unspecified |
| mmu-miR-29b-3p | MIMAT0000127 | Cwc27 | 67285 | HITS-CLIP | 23142080 | Unspecified |
| mmu-miR-29b-3p | MIMAT0000127 | Cwf19l1 | 72502 | HITS-CLIP | 23142080 | Unspecified |
| mmu-miR-29b-3p | MIMAT0000127 | Cxadr | 13052 | HITS-CLIP | 37986224\|23597149\|23142080 | Unspecified |
| mmu-miR-29b-3p | MIMAT0000127 | Cxcl12 | 20315 | HITS-CLIP//sRNA-Seq | 25083871\|33184237\|23597149 | Unspecified |
| mmu-miR-29b-3p | MIMAT0000127 | Cxcr3 | 12766 | HITS-CLIP | 23142080 | Unspecified |
| mmu-miR-29b-3p | MIMAT0000127 | Cxcr4 | 12767 | HITS-CLIP | 23142080 | Unspecified |
| mmu-miR-29b-3p | MIMAT0000127 | Cxcr5 | 135972058 | HITS-CLIP | 23142080 | Unspecified |
| mmu-miR-29b-3p | MIMAT0000127 | Cxxc5 | 67393 | HITS-CLIP | 23597149\|23142080 | Unspecified |

| | | | | | | |
|---|---|---|---|---|---|---|
| mmu-miR-29b-3p | MIMAT0000127 | Cyb561 | 13056 | HITS-CLIP | 23142080 | Unspecified |
| mmu-miR-29b-3p | MIMAT0000127 | Cyb561a3 | 225912 | HITS-CLIP | 23142080 | Unspecified |
| mmu-miR-29b-3p | MIMAT0000127 | Cyb5a | 109672 | HITS-CLIP//sRNA-Seq | 23142080\|33184237\|23597149 | Unspecified |
| mmu-miR-29b-3p | MIMAT0000127 | Cyb5b | 66427 | HITS-CLIP//sRNA-Seq | 23597149\|33184237 | Unspecified |
| mmu-miR-29b-3p | MIMAT0000127 | Cyb5r3 | 109754 | sRNA-Seq//HITS-CLIP | 33184237\|23142080 | Unspecified |
| mmu-miR-29b-3p | MIMAT0000127 | Cyba | 13057 | sRNA-Seq//HITS-CLIP | 33184237\|23142080 | Unspecified |
| mmu-miR-29b-3p | MIMAT0000127 | Cyc1 | 66445 | HITS-CLIP//sRNA-Seq | 23142080\|33184237 | Unspecified |
| mmu-miR-29b-3p | MIMAT0000127 | Cyfip2 | 76884 | HITS-CLIP//sRNA-Seq | 23142080\|33184237 | Unspecified |
| mmu-miR-29b-3p | MIMAT0000127 | Cyld | 74256 | HITS-CLIP | 25083871\|23142080 | Unspecified |
| mmu-miR-29b-3p | MIMAT0000127 | Cyp11a1 | 13070 | HITS-CLIP | 23142080 | Unspecified |
| mmu-miR-29b-3p | MIMAT0000127 | Cyp1a2 | 13077 | HITS-CLIP | 23597149 | Unspecified |
| mmu-miR-29b-3p | MIMAT0000127 | Cyp2a12 | 13085 | HITS-CLIP | 23597149 | Unspecified |
| mmu-miR-29b-3p | MIMAT0000127 | Cyp2a5 | 13087 | HITS-CLIP | 37986224\|23597149 | Unspecified |
| mmu-miR-29b-3p | MIMAT0000127 | Cyp2c23 | 226143 | HITS-CLIP | 23597149 | Unspecified |
| mmu-miR-29b-3p | MIMAT0000127 | Cyp2c29 | 13095 | HITS-CLIP | 23597149 | Unspecified |
| mmu-miR-29b-3p | MIMAT0000127 | Cyp2c50 | 107141 | HITS-CLIP | 23597149 | Unspecified |
| mmu-miR-29b-3p | MIMAT0000127 | Cyp2c67 | 545288 | HITS-CLIP | 23597149 | Unspecified |
| mmu-miR-29b-3p | MIMAT0000127 | Cyp2c69 | 100043108 | HITS-CLIP | 23597149 | Unspecified |
| mmu-miR-29b-3p | MIMAT0000127 | Cyp2c70 | 226105 | HITS-CLIP | 23597149 | Unspecified |
| mmu-miR-29b-3p | MIMAT0000127 | Cyp2d26 | 76279 | HITS-CLIP | 23597149 | Unspecified |
| mmu-miR-29b-3p | MIMAT0000127 | Cyp2j5 | 13109 | HITS-CLIP | 23597149 | Unspecified |
| mmu-miR-29b-3p | MIMAT0000127 | Cyp3a11 | 13112 | HITS-CLIP | 23597149 | Unspecified |
| mmu-miR-29b-3p | MIMAT0000127 | Cyp3a25 | 56388 | HITS-CLIP | 37986224 | Unspecified |
| mmu-miR-29b-3p | MIMAT0000127 | Cyp4a14 | 13119 | HITS-CLIP | 23597149 | Unspecified |
| mmu-miR-29b-3p | MIMAT0000127 | Cyp4v3 | 102294 | HITS-CLIP | 23597149 | Unspecified |
| mmu-miR-29b-3p | MIMAT0000127 | Cyp51 | 13121 | HITS-CLIP//sRNA-Seq | 37986224\|23597149\|23142080\|3318 | Unspecified |
| mmu-miR-29b-3p | MIMAT0000127 | Cyp7b1 | 13123 | HITS-CLIP | 23597149 | Unspecified |
| mmu-miR-29b-3p | MIMAT0000127 | Cyp8b1 | 13124 | HITS-CLIP | 23597149 | Unspecified |
| mmu-miR-29b-3p | MIMAT0000127 | Cyth1 | 19157 | HITS-CLIP | 37986224\|25083871\|23597149\|2314 | Unspecified |
| mmu-miR-29b-3p | MIMAT0000127 | Cyth3 | 19159 | HITS-CLIP//sRNA-Seq | 37986224\|25083871\|33184237\|2359 | Unspecified |
| mmu-miR-29b-3p | MIMAT0000127 | D17H6S53E | 114585 | HITS-CLIP | 37986224 | Unspecified |
| mmu-miR-29b-3p | MIMAT0000127 | D17Wsu92e | 224647 | HITS-CLIP | 25083871\|23597149\|23142080 | Unspecified |
| mmu-miR-29b-3p | MIMAT0000127 | D1Ertd622e | 52392 | HITS-CLIP | 25083871\|23142080 | Unspecified |
| mmu-miR-29b-3p | MIMAT0000127 | D230025D16Rik | 234678 | HITS-CLIP | 23597149 | Unspecified |
| mmu-miR-29b-3p | MIMAT0000127 | D5Ertd579e | 320661 | HITS-CLIP | 25083871\|23142080 | Unspecified |
| mmu-miR-29b-3p | MIMAT0000127 | D630045J12Rik | 330286 | HITS-CLIP | 25083871 | Unspecified |
| mmu-miR-29b-3p | MIMAT0000127 | Daam1 | 208846 | HITS-CLIP//sRNA-Seq | 37986224\|25083871\|23597149\|3318 | Unspecified |
| mmu-miR-29b-3p | MIMAT0000127 | Daam2 | 76441 | HITS-CLIP | 37986224 | Unspecified |
| mmu-miR-29b-3p | MIMAT0000127 | Dab2 | 13132 | HITS-CLIP//sRNA-Seq | 25083871\|33184237 | Unspecified |

| | | | | | | |
|---|---|---|---|---|---|---|
| mmu-miR-29b-3p | MIMAT0000127 | Dab2ip | 69601 | HITS-CLIP//sRNA-Seq | 37986224\|25083871\|23597149\|33184 | Unspecified |
| mmu-miR-29b-3p | MIMAT0000127 | Dag1 | 13138 | HITS-CLIP//sRNA-Seq | 37986224\|25083871\|23142080\|23597 | Unspecified |
| mmu-miR-29b-3p | MIMAT0000127 | Dagla | 269060 | HITS-CLIP | 25083871 | Unspecified |
| mmu-miR-29b-3p | MIMAT0000127 | Dap3 | 65111 | HITS-CLIP//sRNA-Seq | 23597149\|33184237 | Unspecified |
| mmu-miR-29b-3p | MIMAT0000127 | Dapk1 | 69635 | HITS-CLIP//sRNA-Seq | 23597149\|33184237 | Unspecified |
| mmu-miR-29b-3p | MIMAT0000127 | Dapk2 | 13143 | HITS-CLIP | 25083871 | Unspecified |
| mmu-miR-29b-3p | MIMAT0000127 | Dapk3 | 13144 | HITS-CLIP | 37986224\|23597149 | Unspecified |
| mmu-miR-29b-3p | MIMAT0000127 | Dars | 226414 | sRNA-Seq//HITS-CLIP | 33184237\|23142080 | Unspecified |
| mmu-miR-29b-3p | MIMAT0000127 | Daxx | 13163 | HITS-CLIP//sRNA-Seq | 23142080\|33184237 | Unspecified |
| mmu-miR-29b-3p | MIMAT0000127 | Dazap2 | 23994 | HITS-CLIP//sRNA-Seq | 23597149\|25083871\|23142080\|33184 | Unspecified |
| mmu-miR-29b-3p | MIMAT0000127 | Dbf4 | 27214 | HITS-CLIP//sRNA-Seq | 23142080\|33184237 | Unspecified |
| mmu-miR-29b-3p | MIMAT0000127 | Dbp | 13170 | HITS-CLIP | 25083871 | Unspecified |
| mmu-miR-29b-3p | MIMAT0000127 | Dbr1 | 83703 | HITS-CLIP | 23142080 | Unspecified |
| mmu-miR-29b-3p | MIMAT0000127 | Dbt | 13171 | HITS-CLIP | 37986224\|23597149\|23142080\|25083 | Unspecified |
| mmu-miR-29b-3p | MIMAT0000127 | Dcaf1 | 41836 | HITS-CLIP | 25083871\|23142080 | Unspecified |
| mmu-miR-29b-3p | MIMAT0000127 | Dcaf10 | 242418 | HITS-CLIP | 37986224\|25083871\|23142080\|31433 | Unspecified |
| mmu-miR-29b-3p | MIMAT0000127 | Dcaf11 | 28199 | HITS-CLIP | 23142080\|23597149 | Unspecified |
| mmu-miR-29b-3p | MIMAT0000127 | Dcaf12 | 68970 | HITS-CLIP | 37986224\|25083871\|23142080 | Unspecified |
| mmu-miR-29b-3p | MIMAT0000127 | Dcaf13 | 223499 | HITS-CLIP | 23142080 | Unspecified |
| mmu-miR-29b-3p | MIMAT0000127 | Dcaf15 | 212123 | HITS-CLIP | 25083871\|23142080 | Unspecified |
| mmu-miR-29b-3p | MIMAT0000127 | Dcaf17 | 75763 | HITS-CLIP | 37986224\|23142080\|23597149 | Unspecified |
| mmu-miR-29b-3p | MIMAT0000127 | Dcaf5 | 320808 | HITS-CLIP | 37986224 | Unspecified |
| mmu-miR-29b-3p | MIMAT0000127 | Dcaf6 | 74106 | sRNA-Seq//HITS-CLIP | 33184237\|23142080 | Unspecified |
| mmu-miR-29b-3p | MIMAT0000127 | Dcaf7 | 71833 | HITS-CLIP//sRNA-Seq | 37986224\|25083871\|33184237\|23142 | Unspecified |
| mmu-miR-29b-3p | MIMAT0000127 | Dcakd | 68087 | HITS-CLIP | 37986224 | Unspecified |
| mmu-miR-29b-3p | MIMAT0000127 | Dcbld1 | 66686 | HITS-CLIP | 37986224\|25083871\|23597149\|23142 | Unspecified |
| mmu-miR-29b-3p | MIMAT0000127 | Dcbld2 | 73379 | HITS-CLIP | 25083871\|23597149 | Unspecified |
| mmu-miR-29b-3p | MIMAT0000127 | Dchs2 | 54798 | HITS-CLIP | 25083871 | Unspecified |
| mmu-miR-29b-3p | MIMAT0000127 | Dclk1 | 13175 | HITS-CLIP//sRNA-Seq | 37986224\|25083871\|33184237 | Unspecified |
| mmu-miR-29b-3p | MIMAT0000127 | Dcp1a | 75901 | HITS-CLIP | 37986224\|23142080\|25083871 | Unspecified |
| mmu-miR-29b-3p | MIMAT0000127 | Dcp2 | 70640 | HITS-CLIP | 37986224\|23597149\|23142080\|25083 | Unspecified |
| mmu-miR-29b-3p | MIMAT0000127 | Dctn1 | 13191 | HITS-CLIP//sRNA-Seq | 25083871\|33184237\|23597149\|23142 | Unspecified |
| mmu-miR-29b-3p | MIMAT0000127 | Dctn3 | 53598 | HITS-CLIP//sRNA-Seq | 23597149\|25083871\|33184237\|23142 | Unspecified |
| mmu-miR-29b-3p | MIMAT0000127 | Dctn6 | 22428 | HITS-CLIP//sRNA-Seq | 25083871\|33184237 | Unspecified |
| mmu-miR-29b-3p | MIMAT0000127 | Dcun1d1 | 114893 | sRNA-Seq//HITS-CLIP | 33184237\|31433990\|23142080\|28735 | Unspecified |
| mmu-miR-29b-3p | MIMAT0000127 | Dcun1d3 | 233805 | sRNA-Seq//HITS-CLIP | 33184237\|23142080 | Unspecified |
| mmu-miR-29b-3p | MIMAT0000127 | Dcun1d4 | 100737 | HITS-CLIP//sRNA-Seq | 37986224\|25083871\|33184237\|23597 | Unspecified |
| mmu-miR-29b-3p | MIMAT0000127 | Dcun1d5 | 76863 | HITS-CLIP//sRNA-Seq | 23142080\|33184237\|25083871 | Unspecified |
| mmu-miR-29b-3p | MIMAT0000127 | Dda1 | 66498 | HITS-CLIP//sRNA-Seq | 25083871\|33184237 | Unspecified |

| | | | | | | |
|---|---|---|---|---|---|---|
| mmu-miR-29b-3p | MIMAT0000127 | Ddb1 | 13194 | HITS-CLIP//sRNA-Seq | 23597149\|25083871\|33184237\|23142 | Unspecified |
| mmu-miR-29b-3p | MIMAT0000127 | Ddc | 13195 | HITS-CLIP | 37986224\|23142080\|23597149 | Unspecified |
| mmu-miR-29b-3p | MIMAT0000127 | Ddi2 | 68817 | HITS-CLIP | 37986224\|25083871\|23597149\|23142 | Unspecified |
| mmu-miR-29b-3p | MIMAT0000127 | Ddias | 220042 | HITS-CLIP | 23142080 | Unspecified |
| mmu-miR-29b-3p | MIMAT0000127 | Ddit4 | 74747 | HITS-CLIP//sRNA-Seq | 23597149\|33184237\|23142080\|25083 | Unspecified |
| mmu-miR-29b-3p | MIMAT0000127 | Ddr2 | 18214 | HITS-CLIP//sRNA-Seq | 25083871\|33184237 | Unspecified |
| mmu-miR-29b-3p | MIMAT0000127 | Ddx1 | 104721 | HITS-CLIP//sRNA-Seq | 25083871\|23597149\|33184237\|23142 | Unspecified |
| mmu-miR-29b-3p | MIMAT0000127 | Ddx10 | 77591 | HITS-CLIP | 25083871\|23597149 | Unspecified |
| mmu-miR-29b-3p | MIMAT0000127 | Ddx17 | 67040 | HITS-CLIP//sRNA-Seq | 25083871\|33184237\|23597149\|23142 | Unspecified |
| mmu-miR-29b-3p | MIMAT0000127 | Ddx19a | 13680 | HITS-CLIP//sRNA-Seq | 25083871\|33184237 | Unspecified |
| mmu-miR-29b-3p | MIMAT0000127 | Ddx20 | 53975 | HITS-CLIP | 23142080 | Unspecified |
| mmu-miR-29b-3p | MIMAT0000127 | Ddx21 | 56200 | HITS-CLIP//sRNA-Seq | 25083871\|33184237\|23142080 | Unspecified |
| mmu-miR-29b-3p | MIMAT0000127 | Ddx24 | 27225 | sRNA-Seq//HITS-CLIP | 33184237\|23597149 | Unspecified |
| mmu-miR-29b-3p | MIMAT0000127 | Ddx39b | 53817 | HITS-CLIP | 37986224 | Unspecified |
| mmu-miR-29b-3p | MIMAT0000127 | Ddx3x | 13205 | HITS-CLIP//sRNA-Seq | 37986224\|23597149\|25083871\|23142 | Unspecified |
| mmu-miR-29b-3p | MIMAT0000127 | Ddx3y | 26900 | HITS-CLIP | 37986224\|23597149\|23142080 | Unspecified |
| mmu-miR-29b-3p | MIMAT0000127 | Ddx42 | 72047 | HITS-CLIP//sRNA-Seq | 23142080\|33184237\|25083871 | Unspecified |
| mmu-miR-29b-3p | MIMAT0000127 | Ddx46 | 212880 | HITS-CLIP | 25083871\|23597149 | Unspecified |
| mmu-miR-29b-3p | MIMAT0000127 | Ddx47 | 67755 | HITS-CLIP | 23597149\|23142080 | Unspecified |
| mmu-miR-29b-3p | MIMAT0000127 | Ddx5 | 13207 | HITS-CLIP//sRNA-Seq | 37986224\|23597149\|33184237\|25083 | Unspecified |
| mmu-miR-29b-3p | MIMAT0000127 | Ddx50 | 94213 | sRNA-Seq//HITS-CLIP | 33184237\|23142080 | Unspecified |
| mmu-miR-29b-3p | MIMAT0000127 | Ddx52 | 78394 | sRNA-Seq//HITS-CLIP | 33184237\|23142080 | Unspecified |
| mmu-miR-29b-3p | MIMAT0000127 | Ddx54 | 71990 | HITS-CLIP | 37986224\|23597149\|25083871\|23142 | Unspecified |
| mmu-miR-29b-3p | MIMAT0000127 | Ddx55 | 67848 | HITS-CLIP | 23142080 | Unspecified |
| mmu-miR-29b-3p | MIMAT0000127 | Ddx56 | 52513 | HITS-CLIP//sRNA-Seq | 37986224\|25083871\|33184237 | Unspecified |
| mmu-miR-29b-3p | MIMAT0000127 | Ddx58 | 230073 | sRNA-Seq//HITS-CLIP | 33184237\|23142080 | Unspecified |
| mmu-miR-29b-3p | MIMAT0000127 | Ddx59 | 67997 | HITS-CLIP | 23597149 | Unspecified |
| mmu-miR-29b-3p | MIMAT0000127 | Ddx6 | 13209 | HITS-CLIP//sRNA-Seq | 23142080\|25083871\|33184237 | Unspecified |
| mmu-miR-29b-3p | MIMAT0000127 | Decr2 | 26378 | HITS-CLIP | 23597149 | Unspecified |
| mmu-miR-29b-3p | MIMAT0000127 | Def6 | 23853 | HITS-CLIP | 23142080 | Unspecified |
| mmu-miR-29b-3p | MIMAT0000127 | Degs1 | 13244 | HITS-CLIP//sRNA-Seq | 25083871\|33184237\|23597149\|23142 | Unspecified |
| mmu-miR-29b-3p | MIMAT0000127 | Dek | 110052 | HITS-CLIP | 37986224 | Unspecified |
| mmu-miR-29b-3p | MIMAT0000127 | Dennd1b | 329260 | HITS-CLIP | 37986224\|23142080\|23597149 | Unspecified |
| mmu-miR-29b-3p | MIMAT0000127 | Dennd2c | 329727 | HITS-CLIP | 23142080 | Unspecified |
| mmu-miR-29b-3p | MIMAT0000127 | Dennd2d | 72121 | HITS-CLIP | 23142080 | Unspecified |
| mmu-miR-29b-3p | MIMAT0000127 | Dennd3 | 105841 | HITS-CLIP | 23142080 | Unspecified |
| mmu-miR-29b-3p | MIMAT0000127 | Dennd4a | 102442 | HITS-CLIP | 25083871\|23142080 | Unspecified |
| mmu-miR-29b-3p | MIMAT0000127 | Dennd4b | 229541 | HITS-CLIP | 23142080 | Unspecified |
| mmu-miR-29b-3p | MIMAT0000127 | Dennd4c | 329877 | HITS-CLIP | 25083871 | Unspecified |

| | | | | | | |
|---|---|---|---|---|---|---|
| mmu-miR-29b-3p | MIMAT0000127 | Dennd5a | 19347 | HITS-CLIP//sRNA-Seq | 25083871\|33184237 | Unspecified |
| mmu-miR-29b-3p | MIMAT0000127 | Dennd5b | 320560 | HITS-CLIP | 37986224\|23597149\|23142080 | Unspecified |
| mmu-miR-29b-3p | MIMAT0000127 | Dennd6a | 211922 | HITS-CLIP | 25083871\|23142080 | Unspecified |
| mmu-miR-29b-3p | MIMAT0000127 | Denr | 68184 | sRNA-Seq//HITS-CLIP | 33184237\|23142080 | Unspecified |
| mmu-miR-29b-3p | MIMAT0000127 | Depdc1a | 76131 | HITS-CLIP | 37986224\|25083871\|23142080 | Unspecified |
| mmu-miR-29b-3p | MIMAT0000127 | Depdc5 | 277854 | HITS-CLIP | 23142080 | Unspecified |
| mmu-miR-29b-3p | MIMAT0000127 | Deptor | 97998 | HITS-CLIP | 25083871\|23597149 | Unspecified |
| mmu-miR-29b-3p | MIMAT0000127 | Dera | 232449 | HITS-CLIP | 37986224\|25083871\|23597149\|23142 | Unspecified |
| mmu-miR-29b-3p | MIMAT0000127 | Derl2 | 116891 | HITS-CLIP | 23142080 | Unspecified |
| mmu-miR-29b-3p | MIMAT0000127 | Desi1 | 28075 | HITS-CLIP//sRNA-Seq | 25083871\|23142080\|33184237 | Unspecified |
| mmu-miR-29b-3p | MIMAT0000127 | Desi2 | 78825 | HITS-CLIP | 37986224\|25083871\|23142080 | Unspecified |
| mmu-miR-29b-3p | MIMAT0000127 | Dgat1 | 13350 | HITS-CLIP | 23597149\|23142080 | Unspecified |
| mmu-miR-29b-3p | MIMAT0000127 | Dgat2 | 67800 | HITS-CLIP | 23597149 | Unspecified |
| mmu-miR-29b-3p | MIMAT0000127 | Dgcr2 | 13356 | HITS-CLIP//sRNA-Seq | 23597149\|33184237 | Unspecified |
| mmu-miR-29b-3p | MIMAT0000127 | Dgka | 13139 | HITS-CLIP | 25083871 | Unspecified |
| mmu-miR-29b-3p | MIMAT0000127 | Dgkd | 227333 | HITS-CLIP//qCLASH | 37986224\|25083871\|31393953\|23142 | Unspecified |
| mmu-miR-29b-3p | MIMAT0000127 | Dgkz | 104418 | HITS-CLIP//sRNA-Seq | 25083871\|33184237\|23142080 | Unspecified |
| mmu-miR-29b-3p | MIMAT0000127 | Dguok | 27369 | HITS-CLIP | 37986224\|23597149\|25083871 | Unspecified |
| mmu-miR-29b-3p | MIMAT0000127 | Dhcr24 | 74754 | HITS-CLIP//sRNA-Seq | 37986224\|25083871\|23597149\|33184 | Unspecified |
| mmu-miR-29b-3p | MIMAT0000127 | Dhcr7 | 13360 | HITS-CLIP//sRNA-Seq | 25083871\|33184237 | Unspecified |
| mmu-miR-29b-3p | MIMAT0000127 | Dhps | 330817 | HITS-CLIP | 37986224 | Unspecified |
| mmu-miR-29b-3p | MIMAT0000127 | Dhrs1 | 52585 | HITS-CLIP//sRNA-Seq | 23597149\|23142080\|33184237 | Unspecified |
| mmu-miR-29b-3p | MIMAT0000127 | Dhx15 | 13204 | HITS-CLIP//sRNA-Seq | 23597149\|25083871\|33184237\|23142 | Unspecified |
| mmu-miR-29b-3p | MIMAT0000127 | Dhx16 | 69192 | HITS-CLIP//sRNA-Seq | 37986224\|23597149\|23142080\|2508 | Unspecified |
| mmu-miR-29b-3p | MIMAT0000127 | Dhx30 | 72831 | HITS-CLIP//sRNA-Seq | 23142080\|33184237 | Unspecified |
| mmu-miR-29b-3p | MIMAT0000127 | Dhx32 | 101437 | HITS-CLIP | 23597149\|25083871 | Unspecified |
| mmu-miR-29b-3p | MIMAT0000127 | Dhx33 | 216877 | HITS-CLIP | 25083871\|23142080 | Unspecified |
| mmu-miR-29b-3p | MIMAT0000127 | Dhx36 | 72162 | HITS-CLIP | 25083871\|23142080 | Unspecified |
| mmu-miR-29b-3p | MIMAT0000127 | Dhx37 | 208144 | HITS-CLIP | 23142080 | Unspecified |
| mmu-miR-29b-3p | MIMAT0000127 | Dhx38 | 64340 | HITS-CLIP//sRNA-Seq | 25083871\|33184237\|23142080 | Unspecified |
| mmu-miR-29b-3p | MIMAT0000127 | Dhx40 | 67487 | HITS-CLIP//sRNA-Seq | 23597149\|33184237 | Unspecified |
| mmu-miR-29b-3p | MIMAT0000127 | Dhx58 | 80861 | sRNA-Seq//HITS-CLIP | 33184237\|23142080 | Unspecified |
| mmu-miR-29b-3p | MIMAT0000127 | Dhx8 | 217207 | HITS-CLIP | 37986224\|25083871\|23142080 | Unspecified |
| mmu-miR-29b-3p | MIMAT0000127 | Dhx9 | 13211 | HITS-CLIP//sRNA-Seq | 25083871\|33184237\|23142080 | Unspecified |
| mmu-miR-29b-3p | MIMAT0000127 | Diablo | 66593 | HITS-CLIP//sRNA-Seq | 25083871\|33184237 | Unspecified |
| mmu-miR-29b-3p | MIMAT0000127 | Diaph1 | 1729 | HITS-CLIP | 23142080 | Unspecified |
| mmu-miR-29b-3p | MIMAT0000127 | Diaph2 | 54004 | HITS-CLIP | 25083871\|23142080 | Unspecified |
| mmu-miR-29b-3p | MIMAT0000127 | Dicer1 | 192119 | HITS-CLIP//sRNA-Seq | 37986224\|25083871\|23597149\|33184 | Unspecified |
| mmu-miR-29b-3p | MIMAT0000127 | Dido1 | 23856 | HITS-CLIP | 37986224\|25083871 | Unspecified |

| | | | | | | |
|---|---|---|---|---|---|---|
| mmu-miR-29b-3p | MIMAT0000127 | Diexf | 215193 | HITS-CLIP | 23142080 | Unspecified |
| mmu-miR-29b-3p | MIMAT0000127 | Dimt1 | 66254 | HITS-CLIP | 23142080 | Unspecified |
| mmu-miR-29b-3p | MIMAT0000127 | Dio1 | 13370 | HITS-CLIP | 37986224 | Unspecified |
| mmu-miR-29b-3p | MIMAT0000127 | Dio2 | 13371 | HITS-CLIP | 37986224\|25083871\|23142080 | Unspecified |
| mmu-miR-29b-3p | MIMAT0000127 | Dip2b | 239667 | HITS-CLIP | 37986224\|23142080 | Unspecified |
| mmu-miR-29b-3p | MIMAT0000127 | Dip2c | 208440 | HITS-CLIP//sRNA-Seq | 37986224\|23142080\|25083871\|3318 | Unspecified |
| mmu-miR-29b-3p | MIMAT0000127 | Dis3 | 72662 | HITS-CLIP | 23142080 | Unspecified |
| mmu-miR-29b-3p | MIMAT0000127 | Dkc1 | 245474 | sRNA-Seq//HITS-CLIP | 33184237\|23142080 | Unspecified |
| mmu-miR-29b-3p | MIMAT0000127 | Dlat | 235339 | HITS-CLIP//sRNA-Seq | 25083871\|23597149\|33184237 | Unspecified |
| mmu-miR-29b-3p | MIMAT0000127 | Dlc1 | 50768 | HITS-CLIP | 25083871 | Unspecified |
| mmu-miR-29b-3p | MIMAT0000127 | Dld | 13382 | HITS-CLIP//sRNA-Seq | 25083871\|33184237\|23597149\|2314 | Unspecified |
| mmu-miR-29b-3p | MIMAT0000127 | Dlg1 | 13383 | HITS-CLIP | 25083871 | Unspecified |
| mmu-miR-29b-3p | MIMAT0000127 | Dlg3 | 53310 | HITS-CLIP | 23142080 | Unspecified |
| mmu-miR-29b-3p | MIMAT0000127 | Dlg5 | 71228 | HITS-CLIP | 25083871 | Unspecified |
| mmu-miR-29b-3p | MIMAT0000127 | Dlgap4 | 228836 | HITS-CLIP//sRNA-Seq | 25083871\|33184237 | Unspecified |
| mmu-miR-29b-3p | MIMAT0000127 | Dll1 | 13388 | HITS-CLIP | 23597149 | Unspecified |
| mmu-miR-29b-3p | MIMAT0000127 | Dlst | 78920 | HITS-CLIP//sRNA-Seq | 25083871\|33184237\|23597149 | Unspecified |
| mmu-miR-29b-3p | MIMAT0000127 | Dmap1 | 66233 | HITS-CLIP | 23142080 | Unspecified |
| mmu-miR-29b-3p | MIMAT0000127 | Dmgdh | 74129 | HITS-CLIP | 37986224\|23597149 | Unspecified |
| mmu-miR-29b-3p | MIMAT0000127 | Dmpk | 13400 | HITS-CLIP | 25083871 | Unspecified |
| mmu-miR-29b-3p | MIMAT0000127 | Dmwd | 13401 | HITS-CLIP | 37986224 | Unspecified |
| mmu-miR-29b-3p | MIMAT0000127 | Dmxl1 | 240283 | HITS-CLIP | 37986224\|25083871 | Unspecified |
| mmu-miR-29b-3p | MIMAT0000127 | Dna2 | 327762 | HITS-CLIP | 37986224\|23142080\|23597149 | Unspecified |
| mmu-miR-29b-3p | MIMAT0000127 | Dnaaf5 | 433956 | HITS-CLIP | 23142080 | Unspecified |
| mmu-miR-29b-3p | MIMAT0000127 | Dnaja4 | 58233 | HITS-CLIP | 25083871\|23142080 | Unspecified |
| mmu-miR-29b-3p | MIMAT0000127 | Dnajb1 | 81489 | HITS-CLIP//sRNA-Seq | 25083871\|33184237\|23142080 | Unspecified |
| mmu-miR-29b-3p | MIMAT0000127 | Dnajb11 | 67838 | sRNA-Seq//HITS-CLIP | 33184237\|23142080\|25083871 | Unspecified |
| mmu-miR-29b-3p | MIMAT0000127 | Dnajb12 | 56709 | HITS-CLIP//sRNA-Seq | 23142080\|33184237 | Unspecified |
| mmu-miR-29b-3p | MIMAT0000127 | Dnajb14 | 70604 | HITS-CLIP | 25083871\|23142080 | Unspecified |
| mmu-miR-29b-3p | MIMAT0000127 | Dnajb2 | 56812 | HITS-CLIP//sRNA-Seq | 25083871\|33184237 | Unspecified |
| mmu-miR-29b-3p | MIMAT0000127 | Dnajb4 | 67035 | HITS-CLIP | 25083871\|23597149 | Unspecified |
| mmu-miR-29b-3p | MIMAT0000127 | Dnajb6 | 23950 | HITS-CLIP//sRNA-Seq | 25083871\|33184237\|23142080 | Unspecified |
| mmu-miR-29b-3p | MIMAT0000127 | Dnajb9 | 27362 | HITS-CLIP | 23597149\|25083871 | Unspecified |
| mmu-miR-29b-3p | MIMAT0000127 | Dnajc10 | 66861 | HITS-CLIP//sRNA-Seq | 25083871\|33184237\|23142080 | Unspecified |
| mmu-miR-29b-3p | MIMAT0000127 | Dnajc13 | 235567 | HITS-CLIP//sRNA-Seq | 25083871\|33184237\|23142080 | Unspecified |
| mmu-miR-29b-3p | MIMAT0000127 | Dnajc16 | 214063 | HITS-CLIP | 37986224\|25083871\|23597149 | Unspecified |
| mmu-miR-29b-3p | MIMAT0000127 | Dnajc18 | 76594 | HITS-CLIP | 25083871 | Unspecified |
| mmu-miR-29b-3p | MIMAT0000127 | Dnajc27 | 217378 | HITS-CLIP | 23142080 | Unspecified |
| mmu-miR-29b-3p | MIMAT0000127 | Dnajc30 | 66114 | HITS-CLIP | 25083871\|23142080 | Unspecified |

| | | | | | | |
|---|---|---|---|---|---|---|
| mmu-miR-29b-3p | MIMAT0000127 | Dnajc5 | 13002 | HITS-CLIP//sRNA-Seq | 37986224\|25083871\|23597149\|33184 | Unspecified |
| mmu-miR-29b-3p | MIMAT0000127 | Dnajc8 | 68598 | HITS-CLIP//sRNA-Seq | 25083871\|33184237\|23142080 | Unspecified |
| mmu-miR-29b-3p | MIMAT0000127 | Dnal1 | 105000 | HITS-CLIP | 37986224 | Unspecified |
| mmu-miR-29b-3p | MIMAT0000127 | Dnase1l1 | 69537 | HITS-CLIP | 23142080\|25083871 | Unspecified |
| mmu-miR-29b-3p | MIMAT0000127 | Dnhd1 | 77505 | HITS-CLIP | 23597149 | Unspecified |
| mmu-miR-29b-3p | MIMAT0000127 | Dnm1l | 74006 | HITS-CLIP//sRNA-Seq | 25083871\|33184237 | Unspecified |
| mmu-miR-29b-3p | MIMAT0000127 | Dnmbp | 71972 | HITS-CLIP | 23597149 | Unspecified |
| mmu-miR-29b-3p | MIMAT0000127 | Dnmt1 | 13433 | HITS-CLIP//sRNA-Seq | 23142080\|33184237\|23597149 | Unspecified |
| mmu-miR-29b-3p | MIMAT0000127 | Dnmt3a | 13435 | Luciferase assay//mRAP//Northern blot//H | 19509302\|23597149\|23142080\|25083 | Unspecified |
| mmu-miR-29b-3p | MIMAT0000127 | Dock10 | 210293 | HITS-CLIP | 23597149\|23142080 | Unspecified |
| mmu-miR-29b-3p | MIMAT0000127 | Dock11 | 75974 | sRNA-Seq//HITS-CLIP | 33184237\|23142080 | Unspecified |
| mmu-miR-29b-3p | MIMAT0000127 | Dock2 | 94176 | HITS-CLIP | 23142080 | Unspecified |
| mmu-miR-29b-3p | MIMAT0000127 | Dock5 | 68813 | HITS-CLIP | 37986224\|25083871\|23597149\|23142 | Unspecified |
| mmu-miR-29b-3p | MIMAT0000127 | Dock8 | 76088 | HITS-CLIP | 23142080 | Unspecified |
| mmu-miR-29b-3p | MIMAT0000127 | Dock9 | 105445 | HITS-CLIP | 23142080 | Unspecified |
| mmu-miR-29b-3p | MIMAT0000127 | Dok4 | 114255 | HITS-CLIP | 37986224\|25083871 | Unspecified |
| mmu-miR-29b-3p | MIMAT0000127 | Dolk | 227697 | HITS-CLIP | 37986224\|23597149\|23142080 | Unspecified |
| mmu-miR-29b-3p | MIMAT0000127 | Dop1a | 23033 | HITS-CLIP | 23142080 | Unspecified |
| mmu-miR-29b-3p | MIMAT0000127 | Dop1b | 9980 | HITS-CLIP | 23597149 | Unspecified |
| mmu-miR-29b-3p | MIMAT0000127 | Dot1l | 208266 | HITS-CLIP | 37986224\|25083871\|23597149\|23142 | Unspecified |
| mmu-miR-29b-3p | MIMAT0000127 | Dpm1 | 13480 | HITS-CLIP | 23142080\|25083871\|23597149 | Unspecified |
| mmu-miR-29b-3p | MIMAT0000127 | Dpm3 | 68563 | HITS-CLIP | 23597149\|23142080 | Unspecified |
| mmu-miR-29b-3p | MIMAT0000127 | Dpp3 | 75221 | HITS-CLIP | 37986224 | Unspecified |
| mmu-miR-29b-3p | MIMAT0000127 | Dpp4 | 13482 | HITS-CLIP | 37986224\|23142080 | Unspecified |
| mmu-miR-29b-3p | MIMAT0000127 | Dpp8 | 74388 | HITS-CLIP//sRNA-Seq | 25083871\|33184237\|23142080 | Unspecified |
| mmu-miR-29b-3p | MIMAT0000127 | Dpy19l1 | 244745 | HITS-CLIP | 25083871\|23597149\|23142080 | Unspecified |
| mmu-miR-29b-3p | MIMAT0000127 | Dpy19l3 | 233115 | HITS-CLIP | 37986224\|23142080 | Unspecified |
| mmu-miR-29b-3p | MIMAT0000127 | Dpyd | 99586 | HITS-CLIP | 23597149 | Unspecified |
| mmu-miR-29b-3p | MIMAT0000127 | Dpys | 64705 | HITS-CLIP | 23597149 | Unspecified |
| mmu-miR-29b-3p | MIMAT0000127 | Dpysl2 | 12934 | HITS-CLIP//sRNA-Seq | 37986224\|23142080\|33184237 | Unspecified |
| mmu-miR-29b-3p | MIMAT0000127 | Dpysl3 | 22240 | HITS-CLIP//sRNA-Seq | 37986224\|25083871\|33184237 | Unspecified |
| mmu-miR-29b-3p | MIMAT0000127 | Dr1 | 13486 | HITS-CLIP | 37986224 | Unspecified |
| mmu-miR-29b-3p | MIMAT0000127 | Dram1 | 71712 | HITS-CLIP | 37986224 | Unspecified |
| mmu-miR-29b-3p | MIMAT0000127 | Drg1 | 13494 | sRNA-Seq//HITS-CLIP | 33184237\|23142080\|25083871 | Unspecified |
| mmu-miR-29b-3p | MIMAT0000127 | Drg2 | 13495 | sRNA-Seq//HITS-CLIP | 33184237\|23142080 | Unspecified |
| mmu-miR-29b-3p | MIMAT0000127 | Drosha | 14000 | HITS-CLIP//sRNA-Seq | 37986224\|25083871\|33184237\|23597 | Unspecified |
| mmu-miR-29b-3p | MIMAT0000127 | Drp2 | 13497 | HITS-CLIP | 37986224 | Unspecified |
| mmu-miR-29b-3p | MIMAT0000127 | Dsc2 | 13506 | HITS-CLIP | 37986224 | Unspecified |
| mmu-miR-29b-3p | MIMAT0000127 | Dsel | 319901 | HITS-CLIP | 23142080 | Unspecified |

| | | | | | | |
|---|---|---|---|---|---|---|
| mmu-miR-29b-3p | MIMAT0000127 | Dsp | 109620 | HITS-CLIP | 23597149\|23142080 | Unspecified |
| mmu-miR-29b-3p | MIMAT0000127 | Dst | 13518 | HITS-CLIP//sRNA-Seq | 25083871\|33184237\|23597149 | Unspecified |
| mmu-miR-29b-3p | MIMAT0000127 | Dstyk | 213452 | HITS-CLIP | 37986224\|25083871\|23142080 | Unspecified |
| mmu-miR-29b-3p | MIMAT0000127 | Dtl | 76843 | HITS-CLIP | 23142080 | Unspecified |
| mmu-miR-29b-3p | MIMAT0000127 | Dtna | 13527 | HITS-CLIP | 25083871 | Unspecified |
| mmu-miR-29b-3p | MIMAT0000127 | Dtx4 | 207521 | HITS-CLIP | 37986224\|25083871 | Unspecified |
| mmu-miR-29b-3p | MIMAT0000127 | Dus1l | 68730 | HITS-CLIP | 23597149 | Unspecified |
| mmu-miR-29b-3p | MIMAT0000127 | Dusp10 | 63953 | HITS-CLIP | 37986224\|23142080 | Unspecified |
| mmu-miR-29b-3p | MIMAT0000127 | Dusp11 | 72102 | HITS-CLIP | 37986224\|23142080 | Unspecified |
| mmu-miR-29b-3p | MIMAT0000127 | Dusp18 | 75219 | HITS-CLIP | 37986224\|25083871 | Unspecified |
| mmu-miR-29b-3p | MIMAT0000127 | Dusp2 | 13537 | Luciferase assay//Microarray//Northern bl | 19342382\|25083871\|31393953\|23142 | Unspecified |
| mmu-miR-29b-3p | MIMAT0000127 | Dusp22 | 105352 | HITS-CLIP | 37986224\|25083871\|23142080 | Unspecified |
| mmu-miR-29b-3p | MIMAT0000127 | Dusp3 | 72349 | HITS-CLIP//sRNA-Seq | 25083871\|33184237 | Unspecified |
| mmu-miR-29b-3p | MIMAT0000127 | Dusp4 | 319520 | HITS-CLIP | 23142080 | Unspecified |
| mmu-miR-29b-3p | MIMAT0000127 | Dusp5 | 240672 | HITS-CLIP | 31433990 | Unspecified |
| mmu-miR-29b-3p | MIMAT0000127 | Dusp6 | 67603 | HITS-CLIP//sRNA-Seq | 23597149\|33184237\|23142080\|25083 | Unspecified |
| mmu-miR-29b-3p | MIMAT0000127 | Dvl1 | 13542 | HITS-CLIP | 23142080 | Unspecified |
| mmu-miR-29b-3p | MIMAT0000127 | Dvl2 | 13543 | HITS-CLIP//qCLASH//sRNA-Seq | 25083871\|31393953\|33184237 | Unspecified |
| mmu-miR-29b-3p | MIMAT0000127 | Dvl3 | 13544 | HITS-CLIP//sRNA-Seq | 23142080\|33184237 | Unspecified |
| mmu-miR-29b-3p | MIMAT0000127 | Dym | 69190 | HITS-CLIP | 23142080 | Unspecified |
| mmu-miR-29b-3p | MIMAT0000127 | Dync1h1 | 13424 | HITS-CLIP//sRNA-Seq | 23597149\|25083871\|33184237\|23142 | Unspecified |
| mmu-miR-29b-3p | MIMAT0000127 | Dync1i2 | 13427 | HITS-CLIP//sRNA-Seq | 25083871\|33184237\|23142080 | Unspecified |
| mmu-miR-29b-3p | MIMAT0000127 | Dync1li2 | 234663 | HITS-CLIP//sRNA-Seq | 25083871\|33184237\|23142080 | Unspecified |
| mmu-miR-29b-3p | MIMAT0000127 | Dynll2 | 68097 | HITS-CLIP//sRNA-Seq | 23597149\|23142080\|33184237 | Unspecified |
| mmu-miR-29b-3p | MIMAT0000127 | Dynlt3 | 67117 | HITS-CLIP//sRNA-Seq | 25083871\|33184237\|23142080 | Unspecified |
| mmu-miR-29b-3p | MIMAT0000127 | Dyrk1a | 13548 | HITS-CLIP | 37986224\|25083871\|23597149\|23142 | Unspecified |
| mmu-miR-29b-3p | MIMAT0000127 | Dyrk1b | 13549 | HITS-CLIP | 25083871 | Unspecified |
| mmu-miR-29b-3p | MIMAT0000127 | Dyrk2 | 69181 | HITS-CLIP | 37986224\|23597149\|25083871\|23142 | Unspecified |
| mmu-miR-29b-3p | MIMAT0000127 | Dysf | 26903 | HITS-CLIP | 25083871\|23597149 | Unspecified |
| mmu-miR-29b-3p | MIMAT0000127 | Dzip3 | 224170 | HITS-CLIP//sRNA-Seq | 37986224\|25083871\|33184237\|23142 | Unspecified |
| mmu-miR-29b-3p | MIMAT0000127 | E130308A19Rik | 230259 | HITS-CLIP | 23142080\|26203562 | Unspecified |
| mmu-miR-29b-3p | MIMAT0000127 | E2f1 | 13555 | sRNA-Seq//HITS-CLIP | 33184237\|23142080 | Unspecified |
| mmu-miR-29b-3p | MIMAT0000127 | E2f2 | 242705 | HITS-CLIP | 37986224\|25083871\|23142080 | Unspecified |
| mmu-miR-29b-3p | MIMAT0000127 | E2f3 | 13557 | HITS-CLIP | 25083871\|23142080 | Unspecified |
| mmu-miR-29b-3p | MIMAT0000127 | E2f7 | 52679 | HITS-CLIP | 37986224\|25083871\|23142080 | Unspecified |
| mmu-miR-29b-3p | MIMAT0000127 | E2f8 | 108961 | qCLASH//HITS-CLIP | 31393953\|23597149\|23142080 | Unspecified |
| mmu-miR-29b-3p | MIMAT0000127 | E330009J07Rik | 243780 | HITS-CLIP | 37986224\|25083871\|23597149\|23142 | Unspecified |
| mmu-miR-29b-3p | MIMAT0000127 | Eaf1 | 74427 | HITS-CLIP | 23597149\|25083871 | Unspecified |
| mmu-miR-29b-3p | MIMAT0000127 | Ebf3 | 13593 | HITS-CLIP//sRNA-Seq | 25083871\|33184237 | Unspecified |

| | | | | | | |
|---|---|---|---|---|---|---|
| mmu-miR-29b-3p | MIMAT0000127 | Ebp | 13595 | HITS-CLIP//sRNA-Seq | 23597149\|33184237 | Unspecified |
| mmu-miR-29b-3p | MIMAT0000127 | Ecd | 70601 | HITS-CLIP | 23142080 | Unspecified |
| mmu-miR-29b-3p | MIMAT0000127 | Ece1 | 230857 | HITS-CLIP//sRNA-Seq | 25083871\|33184237\|23142080 | Unspecified |
| mmu-miR-29b-3p | MIMAT0000127 | Ech1 | 51798 | HITS-CLIP//sRNA-Seq | 23597149\|33184237 | Unspecified |
| mmu-miR-29b-3p | MIMAT0000127 | Eci2 | 23986 | HITS-CLIP//sRNA-Seq | 23597149\|33184237\|23142080 | Unspecified |
| mmu-miR-29b-3p | MIMAT0000127 | Ecpas | 23392 | sRNA-Seq//HITS-CLIP | 33184237\|23142080 | Unspecified |
| mmu-miR-29b-3p | MIMAT0000127 | Ecsit | 26940 | HITS-CLIP | 23597149 | Unspecified |
| mmu-miR-29b-3p | MIMAT0000127 | Ect2 | 13605 | sRNA-Seq//HITS-CLIP | 33184237\|23142080 | Unspecified |
| mmu-miR-29b-3p | MIMAT0000127 | Eda2r | 245527 | HITS-CLIP | 37986224 | Unspecified |
| mmu-miR-29b-3p | MIMAT0000127 | Edc3 | 353190 | HITS-CLIP | 37986224\|25083871\|23597149\|23142 | Unspecified |
| mmu-miR-29b-3p | MIMAT0000127 | Edc4 | 234699 | sRNA-Seq//HITS-CLIP | 33184237\|23142080 | Unspecified |
| mmu-miR-29b-3p | MIMAT0000127 | Edem1 | 192193 | HITS-CLIP | 23597149 | Unspecified |
| mmu-miR-29b-3p | MIMAT0000127 | Edem2 | 108687 | HITS-CLIP | 23142080 | Unspecified |
| mmu-miR-29b-3p | MIMAT0000127 | Edem3 | 66967 | HITS-CLIP | 25083871\|23597149\|23142080 | Unspecified |
| mmu-miR-29b-3p | MIMAT0000127 | Ednrb | 13618 | HITS-CLIP | 37986224\|23597149 | Unspecified |
| mmu-miR-29b-3p | MIMAT0000127 | Edrf1 | 214764 | HITS-CLIP | 23142080 | Unspecified |
| mmu-miR-29b-3p | MIMAT0000127 | Eea1 | 216238 | HITS-CLIP | 25083871\|23142080 | Unspecified |
| mmu-miR-29b-3p | MIMAT0000127 | Eed | 13626 | HITS-CLIP | 23142080 | Unspecified |
| mmu-miR-29b-3p | MIMAT0000127 | Eef1a1 | 13627 | sRNA-Seq//HITS-CLIP | 33184237\|23142080 | Unspecified |
| mmu-miR-29b-3p | MIMAT0000127 | Eef1e1 | 66143 | HITS-CLIP | 37986224 | Unspecified |
| mmu-miR-29b-3p | MIMAT0000127 | Eef2 | 13629 | HITS-CLIP//sRNA-Seq | 37986224\|25083871\|33184237\|23597 | Unspecified |
| mmu-miR-29b-3p | MIMAT0000127 | Eefsec | 65967 | HITS-CLIP | 37986224\|23142080 | Unspecified |
| mmu-miR-29b-3p | MIMAT0000127 | Efl1 | 1942 | HITS-CLIP | 23597149\|23142080 | Unspecified |
| mmu-miR-29b-3p | MIMAT0000127 | Efna2 | 13637 | HITS-CLIP | 25083871 | Unspecified |
| mmu-miR-29b-3p | MIMAT0000127 | Efna5 | 13640 | HITS-CLIP//sRNA-Seq | 37986224\|25083871\|33184237\|23142 | Unspecified |
| mmu-miR-29b-3p | MIMAT0000127 | Efr3a | 76740 | sRNA-Seq//HITS-CLIP | 33184237\|25083871 | Unspecified |
| mmu-miR-29b-3p | MIMAT0000127 | Eftud2 | 20624 | HITS-CLIP//sRNA-Seq | 25083871\|33184237\|23597149 | Unspecified |
| mmu-miR-29b-3p | MIMAT0000127 | Egfl7 | 353156 | HITS-CLIP | 23597149 | Unspecified |
| mmu-miR-29b-3p | MIMAT0000127 | Egfr | 13649 | HITS-CLIP | 37986224\|23597149\|25083871\|26203 | Unspecified |
| mmu-miR-29b-3p | MIMAT0000127 | Egln2 | 112406 | HITS-CLIP//sRNA-Seq | 25083871\|33184237 | Unspecified |
| mmu-miR-29b-3p | MIMAT0000127 | Egln3 | 112407 | HITS-CLIP | 23142080 | Unspecified |
| mmu-miR-29b-3p | MIMAT0000127 | Egr1 | 13653 | HITS-CLIP//sRNA-Seq | 25083871\|33184237\|23597149\|23142 | Unspecified |
| mmu-miR-29b-3p | MIMAT0000127 | Egr2 | 13654 | HITS-CLIP | 23142080 | Unspecified |
| mmu-miR-29b-3p | MIMAT0000127 | Egr3 | 13655 | HITS-CLIP | 23142080\|25083871 | Unspecified |
| mmu-miR-29b-3p | MIMAT0000127 | Ehbp1l1 | 114601 | HITS-CLIP | 25083871\|23142080 | Unspecified |
| mmu-miR-29b-3p | MIMAT0000127 | Ehd1 | 13660 | HITS-CLIP//sRNA-Seq | 37986224\|25083871\|23597149\|33184 | Unspecified |
| mmu-miR-29b-3p | MIMAT0000127 | Ehd2 | 259300 | HITS-CLIP//sRNA-Seq | 25083871\|33184237 | Unspecified |
| mmu-miR-29b-3p | MIMAT0000127 | Ehhadh | 74147 | HITS-CLIP | 37986224\|23597149\|23142080 | Unspecified |
| mmu-miR-29b-3p | MIMAT0000127 | Ehmt1 | 77683 | HITS-CLIP | 37986224\|23142080\|25083871 | Unspecified |

| | | | | | | |
|---|---|---|---|---|---|---|
| mmu-miR-29b-3p | MIMAT0000127 | Ehmt2 | 110147 | HITS-CLIP//sRNA-Seq | 23142080\|33184237 | Unspecified |
| mmu-miR-29b-3p | MIMAT0000127 | Ei24 | 13663 | HITS-CLIP//sRNA-Seq | 25083871\|33184237 | Unspecified |
| mmu-miR-29b-3p | MIMAT0000127 | Eif1 | 20918 | HITS-CLIP//sRNA-Seq | 25083871\|33184237\|23142080 | Unspecified |
| mmu-miR-29b-3p | MIMAT0000127 | Eif1a | 13664 | HITS-CLIP//sRNA-Seq | 23597149\|33184237 | Unspecified |
| mmu-miR-29b-3p | MIMAT0000127 | Eif1ad | 69860 | HITS-CLIP//sRNA-Seq | 25083871\|33184237\|23142080 | Unspecified |
| mmu-miR-29b-3p | MIMAT0000127 | Eif1ax | 66235 | HITS-CLIP | 25083871\|23142080 | Unspecified |
| mmu-miR-29b-3p | MIMAT0000127 | Eif2a | 229317 | HITS-CLIP//sRNA-Seq | 25083871\|33184237 | Unspecified |
| mmu-miR-29b-3p | MIMAT0000127 | Eif2ak3 | 13666 | HITS-CLIP | 23597149\|25083871\|23142080 | Unspecified |
| mmu-miR-29b-3p | MIMAT0000127 | Eif2ak4 | 27103 | HITS-CLIP | 23142080 | Unspecified |
| mmu-miR-29b-3p | MIMAT0000127 | Eif2b1 | 209354 | HITS-CLIP | 37986224\|25083871\|23597149\|23142 | Unspecified |
| mmu-miR-29b-3p | MIMAT0000127 | Eif2b2 | 217715 | HITS-CLIP | 23142080 | Unspecified |
| mmu-miR-29b-3p | MIMAT0000127 | Eif2b4 | 13667 | HITS-CLIP | 23597149 | Unspecified |
| mmu-miR-29b-3p | MIMAT0000127 | Eif2b5 | 224045 | sRNA-Seq//HITS-CLIP | 33184237\|23142080 | Unspecified |
| mmu-miR-29b-3p | MIMAT0000127 | Eif2d | 16865 | HITS-CLIP | 25083871\|23142080 | Unspecified |
| mmu-miR-29b-3p | MIMAT0000127 | Eif2s1 | 13665 | HITS-CLIP//sRNA-Seq | 25083871\|33184237\|23142080 | Unspecified |
| mmu-miR-29b-3p | MIMAT0000127 | Eif2s2 | 67204 | HITS-CLIP//sRNA-Seq | 37986224\|33184237\|31433990 | Unspecified |
| mmu-miR-29b-3p | MIMAT0000127 | Eif2s3y | 26908 | HITS-CLIP//sRNA-Seq | 37986224\|33184237\|23597149\|23142 | Unspecified |
| mmu-miR-29b-3p | MIMAT0000127 | Eif3b | 27979 | HITS-CLIP//sRNA-Seq | 25083871\|33184237\|23597149\|23142 | Unspecified |
| mmu-miR-29b-3p | MIMAT0000127 | Eif3c | 56347 | HITS-CLIP//sRNA-Seq | 25083871\|33184237\|23142080 | Unspecified |
| mmu-miR-29b-3p | MIMAT0000127 | Eif3d | 55944 | HITS-CLIP//sRNA-Seq | 25083871\|23597149\|33184237\|23142 | Unspecified |
| mmu-miR-29b-3p | MIMAT0000127 | Eif3i | 54709 | sRNA-Seq//HITS-CLIP | 33184237\|23142080 | Unspecified |
| mmu-miR-29b-3p | MIMAT0000127 | Eif3k | 73830 | HITS-CLIP//sRNA-Seq | 23142080\|33184237 | Unspecified |
| mmu-miR-29b-3p | MIMAT0000127 | Eif3l | 223691 | HITS-CLIP//sRNA-Seq | 25083871\|33184237\|23142080 | Unspecified |
| mmu-miR-29b-3p | MIMAT0000127 | Eif3m | 98221 | sRNA-Seq//HITS-CLIP | 33184237\|23142080 | Unspecified |
| mmu-miR-29b-3p | MIMAT0000127 | Eif4a2 | 13682 | HITS-CLIP//sRNA-Seq | 25083871\|33184237\|23142080 | Unspecified |
| mmu-miR-29b-3p | MIMAT0000127 | Eif4e | 13684 | HITS-CLIP//sRNA-Seq | 25083871\|33184237\|23142080 | Unspecified |
| mmu-miR-29b-3p | MIMAT0000127 | Eif4e2 | 26987 | HITS-CLIP//sRNA-Seq | 37986224\|25083871\|23597149\|33184 | Unspecified |
| mmu-miR-29b-3p | MIMAT0000127 | Eif4ebp2 | 13688 | HITS-CLIP//sRNA-Seq | 25083871\|33184237\|23142080 | Unspecified |
| mmu-miR-29b-3p | MIMAT0000127 | Eif4enif1 | 74203 | HITS-CLIP//sRNA-Seq | 25083871\|33184237 | Unspecified |
| mmu-miR-29b-3p | MIMAT0000127 | Eif4g1 | 208643 | HITS-CLIP//sRNA-Seq | 25083871\|23597149\|33184237\|23142 | Unspecified |
| mmu-miR-29b-3p | MIMAT0000127 | Eif4g2 | 13690 | HITS-CLIP//sRNA-Seq | 23142080\|33184237 | Unspecified |
| mmu-miR-29b-3p | MIMAT0000127 | Eif4h | 22384 | HITS-CLIP//sRNA-Seq | 37986224\|23142080\|33184237 | Unspecified |
| mmu-miR-29b-3p | MIMAT0000127 | Eif5 | 217869 | HITS-CLIP//sRNA-Seq | 25083871\|23142080\|33184237 | Unspecified |
| mmu-miR-29b-3p | MIMAT0000127 | Eif5a | 276770 | HITS-CLIP//sRNA-Seq | 37986224\|25083871\|33184237\|23597 | Unspecified |
| mmu-miR-29b-3p | MIMAT0000127 | Eif5b | 226982 | HITS-CLIP//sRNA-Seq | 25083871\|23597149\|23142080\|30971 | Unspecified |
| mmu-miR-29b-3p | MIMAT0000127 | Eif6 | 16418 | HITS-CLIP//sRNA-Seq | 23597149\|33184237\|23142080 | Unspecified |
| mmu-miR-29b-3p | MIMAT0000127 | Eipr1 | 7260 | HITS-CLIP | 25083871 | Unspecified |
| mmu-miR-29b-3p | MIMAT0000127 | Elac1 | 114615 | HITS-CLIP | 23142080 | Unspecified |
| mmu-miR-29b-3p | MIMAT0000127 | Elavl1 | 15568 | HITS-CLIP | 25083871\|23142080 | Unspecified |

| | | | | | | |
|---|---|---|---|---|---|---|
| mmu-miR-29b-3p | MIMAT0000127 | Elf1 | 13709 | HITS-CLIP | 37986224\|25083871\|23597149\|23142 | Unspecified |
| mmu-miR-29b-3p | MIMAT0000127 | Elf2 | 69257 | HITS-CLIP | 37986224\|23142080 | Unspecified |
| mmu-miR-29b-3p | MIMAT0000127 | Elf4 | 56501 | HITS-CLIP | 37986224\|25083871\|23142080 | Unspecified |
| mmu-miR-29b-3p | MIMAT0000127 | Elk1 | 13712 | HITS-CLIP | 37986224\|25083871 | Unspecified |
| mmu-miR-29b-3p | MIMAT0000127 | Elk4 | 13714 | HITS-CLIP | 25083871\|23597149\|23142080 | Unspecified |
| mmu-miR-29b-3p | MIMAT0000127 | Ell | 13716 | HITS-CLIP | 25083871\|23597149\|23142080 | Unspecified |
| mmu-miR-29b-3p | MIMAT0000127 | Ell2 | 192657 | HITS-CLIP | 23597149 | Unspecified |
| mmu-miR-29b-3p | MIMAT0000127 | Elmod2 | 244548 | HITS-CLIP | 23142080 | Unspecified |
| mmu-miR-29b-3p | MIMAT0000127 | Elmsan1 | 238317 | HITS-CLIP | 37986224\|23597149\|23142080 | Unspecified |
| mmu-miR-29b-3p | MIMAT0000127 | Eloa | 6924 | HITS-CLIP//sRNA-Seq | 25083871\|33184237 | Unspecified |
| mmu-miR-29b-3p | MIMAT0000127 | Eloc | 6921 | HITS-CLIP//sRNA-Seq | 25083871\|23142080\|33184237 | Unspecified |
| mmu-miR-29b-3p | MIMAT0000127 | Elovl1 | 54325 | HITS-CLIP//sRNA-Seq | 25083871\|33184237 | Unspecified |
| mmu-miR-29b-3p | MIMAT0000127 | Elovl2 | 54326 | HITS-CLIP | 23597149 | Unspecified |
| mmu-miR-29b-3p | MIMAT0000127 | Elovl5 | 68801 | HITS-CLIP//sRNA-Seq | 37986224\|25083871\|33184237\|23142 | Unspecified |
| mmu-miR-29b-3p | MIMAT0000127 | Elovl6 | 170439 | HITS-CLIP//sRNA-Seq | 25083871\|33184237\|23597149\|23142 | Unspecified |
| mmu-miR-29b-3p | MIMAT0000127 | Elp1 | 37455 | HITS-CLIP//sRNA-Seq | 25083871\|33184237\|23142080 | Unspecified |
| mmu-miR-29b-3p | MIMAT0000127 | Elp3 | 74195 | HITS-CLIP//sRNA-Seq | 37986224\|33184237\|23142080 | Unspecified |
| mmu-miR-29b-3p | MIMAT0000127 | Elp5 | 54351 | HITS-CLIP//sRNA-Seq | 25083871\|33184237\|23142080 | Unspecified |
| mmu-miR-29b-3p | MIMAT0000127 | Emb | 13723 | sRNA-Seq//HITS-CLIP | 33184237\|23142080 | Unspecified |
| mmu-miR-29b-3p | MIMAT0000127 | Emc1 | 230866 | HITS-CLIP | 23142080 | Unspecified |
| mmu-miR-29b-3p | MIMAT0000127 | Emc10 | 69683 | HITS-CLIP//sRNA-Seq | 25083871\|33184237 | Unspecified |
| mmu-miR-29b-3p | MIMAT0000127 | Emc3 | 66087 | HITS-CLIP | 37986224 | Unspecified |
| mmu-miR-29b-3p | MIMAT0000127 | Emc6 | 66048 | HITS-CLIP | 25083871\|23142080 | Unspecified |
| mmu-miR-29b-3p | MIMAT0000127 | Emc8 | 18117 | HITS-CLIP | 37986224\|23597149\|23142080 | Unspecified |
| mmu-miR-29b-3p | MIMAT0000127 | Emg1 | 14791 | HITS-CLIP | 37986224\|23142080 | Unspecified |
| mmu-miR-29b-3p | MIMAT0000127 | Emilin2 | 246707 | HITS-CLIP | 37986224\|23142080 | Unspecified |
| mmu-miR-29b-3p | MIMAT0000127 | Eml1 | 68519 | HITS-CLIP//sRNA-Seq | 37986224\|25083871\|33184237 | Unspecified |
| mmu-miR-29b-3p | MIMAT0000127 | Eml4 | 78798 | HITS-CLIP//sRNA-Seq | 37986224\|25083871\|33184237\|23597 | Unspecified |
| mmu-miR-29b-3p | MIMAT0000127 | Eml5 | 319670 | HITS-CLIP | 23142080 | Unspecified |
| mmu-miR-29b-3p | MIMAT0000127 | Eml6 | 237711 | HITS-CLIP | 37986224\|25083871 | Unspecified |
| mmu-miR-29b-3p | MIMAT0000127 | Emp1 | 13730 | sRNA-Seq//HITS-CLIP | 33184237\|25083871 | Unspecified |
| mmu-miR-29b-3p | MIMAT0000127 | Emp3 | 2014 | HITS-CLIP//sRNA-Seq | 23142080\|25083871\|33184237 | Unspecified |
| mmu-miR-29b-3p | MIMAT0000127 | Emsy | 56946 | HITS-CLIP | 23142080\|25083871 | Unspecified |
| mmu-miR-29b-3p | MIMAT0000127 | Endog | 13804 | HITS-CLIP | 23597149 | Unspecified |
| mmu-miR-29b-3p | MIMAT0000127 | Eng | 13805 | HITS-CLIP | 23597149 | Unspecified |
| mmu-miR-29b-3p | MIMAT0000127 | Enho | 69638 | HITS-CLIP | 25083871 | Unspecified |
| mmu-miR-29b-3p | MIMAT0000127 | Eno1 | 13806 | HITS-CLIP | 37986224 | Unspecified |
| mmu-miR-29b-3p | MIMAT0000127 | Eno3 | 13808 | HITS-CLIP//sRNA-Seq | 25083871\|33184237 | Unspecified |
| mmu-miR-29b-3p | MIMAT0000127 | Enox2 | 209224 | HITS-CLIP | 23142080 | Unspecified |

| | | | | | | |
|---|---|---|---|---|---|---|
| mmu-miR-29b-3p | MIMAT0000127 | Enpp1 | 18605 | HITS-CLIP | 37986224\|25083871\|23142080 | Unspecified |
| mmu-miR-29b-3p | MIMAT0000127 | Enpp2 | 18606 | HITS-CLIP//sRNA-Seq | 37986224\|33184237\|23597149 | Unspecified |
| mmu-miR-29b-3p | MIMAT0000127 | Enpp3 | 209558 | HITS-CLIP | 23597149 | Unspecified |
| mmu-miR-29b-3p | MIMAT0000127 | Enpp4 | 224794 | HITS-CLIP | 25083871 | Unspecified |
| mmu-miR-29b-3p | MIMAT0000127 | Entpd7 | 93685 | HITS-CLIP | 37986224\|25083871\|23597149\|2314 | Unspecified |
| mmu-miR-29b-3p | MIMAT0000127 | Entr1 | 10807 | sRNA-Seq//HITS-CLIP | 33184237\|23142080 | Unspecified |
| mmu-miR-29b-3p | MIMAT0000127 | Eomes | 13813 | Luciferase assay//Microarray//qRT-PCR//H | 21820330\|23142080 | Unspecified |
| mmu-miR-29b-3p | MIMAT0000127 | Ep300 | 328572 | HITS-CLIP | 25083871\|23142080 | Unspecified |
| mmu-miR-29b-3p | MIMAT0000127 | Ep400 | 75560 | HITS-CLIP | 25083871\|23142080 | Unspecified |
| mmu-miR-29b-3p | MIMAT0000127 | Epas1 | 13819 | HITS-CLIP | 23597149\|23142080 | Unspecified |
| mmu-miR-29b-3p | MIMAT0000127 | Epb41l2 | 2037 | HITS-CLIP//sRNA-Seq | 25083871\|33184237 | Unspecified |
| mmu-miR-29b-3p | MIMAT0000127 | Epb41l5 | 226352 | HITS-CLIP | 23597149 | Unspecified |
| mmu-miR-29b-3p | MIMAT0000127 | Epc1 | 13831 | HITS-CLIP | 25083871\|23597149\|23142080 | Unspecified |
| mmu-miR-29b-3p | MIMAT0000127 | Epc2 | 227867 | HITS-CLIP | 25083871\|23142080 | Unspecified |
| mmu-miR-29b-3p | MIMAT0000127 | Epdr1 | 105298 | HITS-CLIP | 25083871 | Unspecified |
| mmu-miR-29b-3p | MIMAT0000127 | Epg5 | 100502841 | HITS-CLIP | 25083871\|23597149\|23142080 | Unspecified |
| mmu-miR-29b-3p | MIMAT0000127 | Epha2 | 13836 | HITS-CLIP | 23597149\|25083871 | Unspecified |
| mmu-miR-29b-3p | MIMAT0000127 | Ephb2 | 13844 | HITS-CLIP | 25083871 | Unspecified |
| mmu-miR-29b-3p | MIMAT0000127 | Ephb3 | 13845 | HITS-CLIP//sRNA-Seq | 37986224\|25083871\|33184237 | Unspecified |
| mmu-miR-29b-3p | MIMAT0000127 | Ephb6 | 13848 | HITS-CLIP | 25083871 | Unspecified |
| mmu-miR-29b-3p | MIMAT0000127 | Ephx1 | 13849 | sRNA-Seq//HITS-CLIP | 33184237\|23597149\|23142080 | Unspecified |
| mmu-miR-29b-3p | MIMAT0000127 | Ephx2 | 13850 | HITS-CLIP | 23597149 | Unspecified |
| mmu-miR-29b-3p | MIMAT0000127 | Ephx4 | 384214 | HITS-CLIP | 23142080 | Unspecified |
| mmu-miR-29b-3p | MIMAT0000127 | Epm2aip1 | 77781 | sRNA-Seq//HITS-CLIP | 33184237\|23142080 | Unspecified |
| mmu-miR-29b-3p | MIMAT0000127 | Epn1 | 13854 | sRNA-Seq//HITS-CLIP | 33184237\|23142080 | Unspecified |
| mmu-miR-29b-3p | MIMAT0000127 | Eprs | 107508 | HITS-CLIP//sRNA-Seq | 25083871\|23597149\|33184237\|2314 | Unspecified |
| mmu-miR-29b-3p | MIMAT0000127 | Eps15 | 13858 | HITS-CLIP//sRNA-Seq | 21258322\|25083871\|33184237\|2314 | Unspecified |
| mmu-miR-29b-3p | MIMAT0000127 | Eps15l1 | 13859 | sRNA-Seq//HITS-CLIP | 33184237\|25083871 | Unspecified |
| mmu-miR-29b-3p | MIMAT0000127 | Epsti1 | 108670 | HITS-CLIP | 23142080 | Unspecified |
| mmu-miR-29b-3p | MIMAT0000127 | Eral1 | 57837 | HITS-CLIP | 23597149 | Unspecified |
| mmu-miR-29b-3p | MIMAT0000127 | Erap1 | 80898 | HITS-CLIP | 37986224\|25083871\|23142080\|2359 | Unspecified |
| mmu-miR-29b-3p | MIMAT0000127 | Erbb2 | 13866 | HITS-CLIP | 37986224\|25083871 | Unspecified |
| mmu-miR-29b-3p | MIMAT0000127 | Erbb3 | 13867 | HITS-CLIP | 25083871\|23597149 | Unspecified |
| mmu-miR-29b-3p | MIMAT0000127 | Erbin | 55914 | HITS-CLIP//sRNA-Seq | 23597149\|23142080\|25083871\|3318 | Unspecified |
| mmu-miR-29b-3p | MIMAT0000127 | Erc1 | 111173 | HITS-CLIP//sRNA-Seq | 37986224\|25083871\|33184237 | Unspecified |
| mmu-miR-29b-3p | MIMAT0000127 | Ercc2 | 13871 | HITS-CLIP | 23142080 | Unspecified |
| mmu-miR-29b-3p | MIMAT0000127 | Ercc4 | 50505 | HITS-CLIP | 23142080 | Unspecified |
| mmu-miR-29b-3p | MIMAT0000127 | Erf | 13875 | HITS-CLIP | 25083871\|23597149\|23142080 | Unspecified |
| mmu-miR-29b-3p | MIMAT0000127 | Erg28 | 856767 | HITS-CLIP//sRNA-Seq | 23142080\|33184237 | Unspecified |

| | | | | | | |
|---|---|---|---|---|---|---|
| mmu-miR-29b-3p | MIMAT0000127 | Ergic1 | 67458 | HITS-CLIP//sRNA-Seq | 25083871\|33184237 | Unspecified |
| mmu-miR-29b-3p | MIMAT0000127 | Ergic2 | 67456 | HITS-CLIP//sRNA-Seq | 23142080\|33184237 | Unspecified |
| mmu-miR-29b-3p | MIMAT0000127 | Eri3 | 140546 | HITS-CLIP//sRNA-Seq | 25083871\|33184237 | Unspecified |
| mmu-miR-29b-3p | MIMAT0000127 | Erlin1 | 226144 | HITS-CLIP | 23597149\|23142080 | Unspecified |
| mmu-miR-29b-3p | MIMAT0000127 | Erlin2 | 244373 | HITS-CLIP | 37986224\|25083871\|23597149\|23142 | Unspecified |
| mmu-miR-29b-3p | MIMAT0000127 | Ermp1 | 226090 | HITS-CLIP | 37986224\|25083871\|23142080 | Unspecified |
| mmu-miR-29b-3p | MIMAT0000127 | Ern1 | 78943 | HITS-CLIP | 23597149\|25083871 | Unspecified |
| mmu-miR-29b-3p | MIMAT0000127 | Ero1l | 50527 | HITS-CLIP | 23142080 | Unspecified |
| mmu-miR-29b-3p | MIMAT0000127 | Erp29 | 67397 | HITS-CLIP//sRNA-Seq | 37986224\|23142080\|33184237\|23597 | Unspecified |
| mmu-miR-29b-3p | MIMAT0000127 | Erp44 | 76299 | HITS-CLIP//sRNA-Seq | 23597149\|33184237\|23142080 | Unspecified |
| mmu-miR-29b-3p | MIMAT0000127 | Errfi1 | 74155 | HITS-CLIP//sRNA-Seq | 23597149\|25083871\|33184237 | Unspecified |
| mmu-miR-29b-3p | MIMAT0000127 | Esam | 69524 | HITS-CLIP | 37986224\|23597149 | Unspecified |
| mmu-miR-29b-3p | MIMAT0000127 | Esd | 13885 | sRNA-Seq//HITS-CLIP | 33184237\|23142080\|25083871 | Unspecified |
| mmu-miR-29b-3p | MIMAT0000127 | Esf1 | 66580 | HITS-CLIP | 37986224\|25083871\|23142080 | Unspecified |
| mmu-miR-29b-3p | MIMAT0000127 | Espl1 | 105988 | HITS-CLIP | 37986224\|25083871\|23142080 | Unspecified |
| mmu-miR-29b-3p | MIMAT0000127 | Esrp2 | 77411 | HITS-CLIP | 23597149 | Unspecified |
| mmu-miR-29b-3p | MIMAT0000127 | Esyt1 | 23943 | HITS-CLIP//sRNA-Seq | 25083871\|23597149\|33184237\|23142 | Unspecified |
| mmu-miR-29b-3p | MIMAT0000127 | Esyt2 | 52635 | HITS-CLIP//sRNA-Seq | 25083871\|33184237\|23142080 | Unspecified |
| mmu-miR-29b-3p | MIMAT0000127 | Etf1 | 225363 | HITS-CLIP//sRNA-Seq | 23597149\|33184237\|23142080\|25083 | Unspecified |
| mmu-miR-29b-3p | MIMAT0000127 | Etfa | 110842 | HITS-CLIP//sRNA-Seq | 23597149\|33184237 | Unspecified |
| mmu-miR-29b-3p | MIMAT0000127 | Etfb | 110826 | HITS-CLIP | 25083871\|23142080 | Unspecified |
| mmu-miR-29b-3p | MIMAT0000127 | Etfdh | 66841 | HITS-CLIP | 23597149 | Unspecified |
| mmu-miR-29b-3p | MIMAT0000127 | Etnk1 | 75320 | HITS-CLIP//sRNA-Seq | 23597149\|25083871\|33184237\|23142 | Unspecified |
| mmu-miR-29b-3p | MIMAT0000127 | Etnk2 | 214253 | HITS-CLIP | 23597149 | Unspecified |
| mmu-miR-29b-3p | MIMAT0000127 | Ets1 | 23871 | HITS-CLIP | 37986224\|25083871\|23142080 | Unspecified |
| mmu-miR-29b-3p | MIMAT0000127 | Etv5 | 104156 | HITS-CLIP//sRNA-Seq | 25083871\|33184237 | Unspecified |
| mmu-miR-29b-3p | MIMAT0000127 | Etv6 | 14011 | HITS-CLIP | 37986224 | Unspecified |
| mmu-miR-29b-3p | MIMAT0000127 | Eva1a | 232146 | HITS-CLIP | 37986224 | Unspecified |
| mmu-miR-29b-3p | MIMAT0000127 | Eva1b | 230752 | HITS-CLIP//sRNA-Seq | 37986224\|33184237\|23142080 | Unspecified |
| mmu-miR-29b-3p | MIMAT0000127 | Evi2 | 14017 | HITS-CLIP | 23142080 | Unspecified |
| mmu-miR-29b-3p | MIMAT0000127 | Evi5 | 14020 | HITS-CLIP | 25083871 | Unspecified |
| mmu-miR-29b-3p | MIMAT0000127 | Evl | 14026 | sRNA-Seq//HITS-CLIP | 33184237\|23142080 | Unspecified |
| mmu-miR-29b-3p | MIMAT0000127 | Ewsr1 | 14030 | HITS-CLIP//sRNA-Seq | 37986224\|33184237\|23142080 | Unspecified |
| mmu-miR-29b-3p | MIMAT0000127 | Exd2 | 97827 | HITS-CLIP | 25083871\|23142080 | Unspecified |
| mmu-miR-29b-3p | MIMAT0000127 | Exo1 | 26909 | HITS-CLIP | 23142080 | Unspecified |
| mmu-miR-29b-3p | MIMAT0000127 | Exoc2 | 66482 | HITS-CLIP | 23142080 | Unspecified |
| mmu-miR-29b-3p | MIMAT0000127 | Exoc3 | 211446 | HITS-CLIP//sRNA-Seq | 23597149\|33184237\|23142080 | Unspecified |
| mmu-miR-29b-3p | MIMAT0000127 | Exoc4 | 20336 | sRNA-Seq//HITS-CLIP | 33184237\|23142080 | Unspecified |
| mmu-miR-29b-3p | MIMAT0000127 | Exoc6 | 107371 | HITS-CLIP | 23142080 | Unspecified |

| | | | | | | |
|---|---|---|---|---|---|---|
| mmu-miR-29b-3p | MIMAT0000127 | Exosc10 | 50912 | HITS-CLIP//sRNA-Seq | 23597149\|33184237\|25083871\|23142 | Unspecified |
| mmu-miR-29b-3p | MIMAT0000127 | Exosc2 | 227715 | HITS-CLIP//sRNA-Seq | 25083871\|23597149\|33184237\|23142 | Unspecified |
| mmu-miR-29b-3p | MIMAT0000127 | Exosc3 | 66362 | HITS-CLIP | 23142080 | Unspecified |
| mmu-miR-29b-3p | MIMAT0000127 | Exosc4 | 109075 | HITS-CLIP | 23142080 | Unspecified |
| mmu-miR-29b-3p | MIMAT0000127 | Exosc7 | 66446 | HITS-CLIP | 23142080 | Unspecified |
| mmu-miR-29b-3p | MIMAT0000127 | Exosc9 | 50911 | HITS-CLIP | 23142080 | Unspecified |
| mmu-miR-29b-3p | MIMAT0000127 | Ext1 | 14042 | HITS-CLIP | 37986224\|25083871 | Unspecified |
| mmu-miR-29b-3p | MIMAT0000127 | Ext2 | 14043 | HITS-CLIP//sRNA-Seq | 25083871\|33184237\|23142080 | Unspecified |
| mmu-miR-29b-3p | MIMAT0000127 | Extl1 | 56219 | HITS-CLIP | 23597149 | Unspecified |
| mmu-miR-29b-3p | MIMAT0000127 | Extl3 | 54616 | HITS-CLIP//sRNA-Seq | 37986224\|23142080\|23597149\|33184 | Unspecified |
| mmu-miR-29b-3p | MIMAT0000127 | Ezh1 | 14055 | HITS-CLIP | 23142080 | Unspecified |
| mmu-miR-29b-3p | MIMAT0000127 | Ezh2 | 14056 | HITS-CLIP//sRNA-Seq | 37986224\|25083871\|33184237\|23597 | Unspecified |
| mmu-miR-29b-3p | MIMAT0000127 | F12 | 58992 | HITS-CLIP | 37986224\|23597149 | Unspecified |
| mmu-miR-29b-3p | MIMAT0000127 | F13b | 14060 | HITS-CLIP | 23597149 | Unspecified |
| mmu-miR-29b-3p | MIMAT0000127 | F2 | 14061 | HITS-CLIP | 23597149 | Unspecified |
| mmu-miR-29b-3p | MIMAT0000127 | F2r | 14062 | sRNA-Seq//HITS-CLIP | 33184237\|31433990 | Unspecified |
| mmu-miR-29b-3p | MIMAT0000127 | F3 | 14066 | HITS-CLIP//sRNA-Seq | 25083871\|33184237 | Unspecified |
| mmu-miR-29b-3p | MIMAT0000127 | F5 | 14067 | HITS-CLIP | 23597149 | Unspecified |
| mmu-miR-29b-3p | MIMAT0000127 | Faah | 14073 | HITS-CLIP | 37986224\|23597149\|23142080 | Unspecified |
| mmu-miR-29b-3p | MIMAT0000127 | Faap24 | 91442 | HITS-CLIP | 23597149 | Unspecified |
| mmu-miR-29b-3p | MIMAT0000127 | Fads1 | 76267 | HITS-CLIP//sRNA-Seq | 23597149\|25083871\|33184237\|23142 | Unspecified |
| mmu-miR-29b-3p | MIMAT0000127 | Fads6 | 328035 | HITS-CLIP | 23597149 | Unspecified |
| mmu-miR-29b-3p | MIMAT0000127 | Faf2 | 76577 | HITS-CLIP//sRNA-Seq | 37986224\|25083871\|33184237\|23142 | Unspecified |
| mmu-miR-29b-3p | MIMAT0000127 | Fah | 14085 | HITS-CLIP | 23597149 | Unspecified |
| mmu-miR-29b-3p | MIMAT0000127 | Fahd2a | 68126 | HITS-CLIP | 23597149 | Unspecified |
| mmu-miR-29b-3p | MIMAT0000127 | Faim | 23873 | HITS-CLIP | 37986224\|25083871\|23142080 | Unspecified |
| mmu-miR-29b-3p | MIMAT0000127 | Fam102a | 98952 | HITS-CLIP | 23142080 | Unspecified |
| mmu-miR-29b-3p | MIMAT0000127 | Fam117b | 72750 | HITS-CLIP | 23597149\|23142080 | Unspecified |
| mmu-miR-29b-3p | MIMAT0000127 | Fam118a | 73225 | HITS-CLIP | 23142080\|25083871 | Unspecified |
| mmu-miR-29b-3p | MIMAT0000127 | Fam118b | 109229 | HITS-CLIP | 23142080 | Unspecified |
| mmu-miR-29b-3p | MIMAT0000127 | Fam120a | 218236 | HITS-CLIP//sRNA-Seq | 37986224\|25083871\|23597149\|33184 | Unspecified |
| mmu-miR-29b-3p | MIMAT0000127 | Fam120b | 67544 | HITS-CLIP | 23142080 | Unspecified |
| mmu-miR-29b-3p | MIMAT0000127 | Fam126a | 84652 | HITS-CLIP | 37986224\|23597149\|25083871\|23142 | Unspecified |
| mmu-miR-29b-3p | MIMAT0000127 | Fam126b | 213056 | HITS-CLIP | 21258322\|25083871\|23142080 | Unspecified |
| mmu-miR-29b-3p | MIMAT0000127 | Fam129b | 227737 | HITS-CLIP//sRNA-Seq | 25083871\|33184237\|23142080 | Unspecified |
| mmu-miR-29b-3p | MIMAT0000127 | Fam135a | 68187 | HITS-CLIP | 37986224\|25083871 | Unspecified |
| mmu-miR-29b-3p | MIMAT0000127 | Fam136a | 66488 | HITS-CLIP | 37986224\|25083871\|23597149 | Unspecified |
| mmu-miR-29b-3p | MIMAT0000127 | Fam160a1 | 229488 | HITS-CLIP | 23597149 | Unspecified |
| mmu-miR-29b-3p | MIMAT0000127 | Fam160b1 | 226252 | HITS-CLIP | 25083871\|23597149\|23142080 | Unspecified |

| | | | | | | |
|---|---|---|---|---|---|---|
| mmu-miR-29b-3p | MIMAT0000127 | Fam168a | 319604 | HITS-CLIP | 37986224\|25083871 | Unspecified |
| mmu-miR-29b-3p | MIMAT0000127 | Fam168b | 214469 | HITS-CLIP//sRNA-Seq | 37986224\|23142080\|25083871\|33184 | Unspecified |
| mmu-miR-29b-3p | MIMAT0000127 | Fam171a2 | 217219 | HITS-CLIP | 37986224\|25083871\|23597149 | Unspecified |
| mmu-miR-29b-3p | MIMAT0000127 | Fam172a | 68675 | HITS-CLIP | 25083871 | Unspecified |
| mmu-miR-29b-3p | MIMAT0000127 | Fam193a | 231128 | HITS-CLIP | 37986224 | Unspecified |
| mmu-miR-29b-3p | MIMAT0000127 | Fam193b | 212483 | HITS-CLIP | 37986224\|23597149\|25083871 | Unspecified |
| mmu-miR-29b-3p | MIMAT0000127 | Fam199x | 245622 | HITS-CLIP | 25083871\|23142080 | Unspecified |
| mmu-miR-29b-3p | MIMAT0000127 | Fam208a | 218850 | HITS-CLIP | 23597149\|23142080\|25083871 | Unspecified |
| mmu-miR-29b-3p | MIMAT0000127 | Fam214a | 235493 | HITS-CLIP | 37986224\|23142080\|23597149 | Unspecified |
| mmu-miR-29b-3p | MIMAT0000127 | Fam219b | 78323 | HITS-CLIP | 37986224\|25083871\|23597149\|23142 | Unspecified |
| mmu-miR-29b-3p | MIMAT0000127 | Fam234a | 83986 | HITS-CLIP//sRNA-Seq | 25083871\|23597149\|33184237 | Unspecified |
| mmu-miR-29b-3p | MIMAT0000127 | Fam241a | 132720 | HITS-CLIP | 23142080\|25083871 | Unspecified |
| mmu-miR-29b-3p | MIMAT0000127 | Fam32a | 67922 | HITS-CLIP//sRNA-Seq | 23142080\|25083871\|33184237 | Unspecified |
| mmu-miR-29b-3p | MIMAT0000127 | Fam53c | 66306 | HITS-CLIP//sRNA-Seq | 37986224\|33184237\|23142080 | Unspecified |
| mmu-miR-29b-3p | MIMAT0000127 | Fam76b | 72826 | HITS-CLIP | 37986224\|25083871\|23142080 | Unspecified |
| mmu-miR-29b-3p | MIMAT0000127 | Fam84b | 157638 | HITS-CLIP | 23142080 | Unspecified |
| mmu-miR-29b-3p | MIMAT0000127 | Fam91a1 | 210998 | HITS-CLIP//sRNA-Seq | 25083871\|33184237\|23142080 | Unspecified |
| mmu-miR-29b-3p | MIMAT0000127 | Fan1 | 330554 | HITS-CLIP | 37986224\|23142080\|23597149 | Unspecified |
| mmu-miR-29b-3p | MIMAT0000127 | Fancc | 2176 | HITS-CLIP | 23142080 | Unspecified |
| mmu-miR-29b-3p | MIMAT0000127 | Fance | 2178 | HITS-CLIP | 23142080 | Unspecified |
| mmu-miR-29b-3p | MIMAT0000127 | Fancf | 100040608 | HITS-CLIP | 23142080 | Unspecified |
| mmu-miR-29b-3p | MIMAT0000127 | Far1 | 67420 | HITS-CLIP//sRNA-Seq | 25083871\|33184237\|23142080 | Unspecified |
| mmu-miR-29b-3p | MIMAT0000127 | Farp1 | 223254 | HITS-CLIP//sRNA-Seq | 37986224\|25083871\|23597149\|33184 | Unspecified |
| mmu-miR-29b-3p | MIMAT0000127 | Farp2 | 227377 | HITS-CLIP | 37986224\|25083871 | Unspecified |
| mmu-miR-29b-3p | MIMAT0000127 | Fars2 | 69955 | HITS-CLIP | 23597149 | Unspecified |
| mmu-miR-29b-3p | MIMAT0000127 | Farsb | 23874 | sRNA-Seq//HITS-CLIP | 33184237\|23597149 | Unspecified |
| mmu-miR-29b-3p | MIMAT0000127 | Fasl | 14103 | HITS-CLIP | 37986224\|23142080 | Unspecified |
| mmu-miR-29b-3p | MIMAT0000127 | Fasn | 14104 | HITS-CLIP//sRNA-Seq | 37986224\|25083871\|33184237\|23597 | Unspecified |
| mmu-miR-29b-3p | MIMAT0000127 | Fastk | 66587 | HITS-CLIP | 23597149\|23142080 | Unspecified |
| mmu-miR-29b-3p | MIMAT0000127 | Fastkd3 | 69577 | HITS-CLIP | 23142080 | Unspecified |
| mmu-miR-29b-3p | MIMAT0000127 | Fat1 | 14107 | HITS-CLIP | 37986224\|25083871 | Unspecified |
| mmu-miR-29b-3p | MIMAT0000127 | Faxc | 76132 | HITS-CLIP//sRNA-Seq | 25083871\|33184237 | Unspecified |
| mmu-miR-29b-3p | MIMAT0000127 | Fbf1 | 217335 | HITS-CLIP | 23597149 | Unspecified |
| mmu-miR-29b-3p | MIMAT0000127 | Fbh1 | 84893 | HITS-CLIP//sRNA-Seq | 25083871\|33184237\|23142080 | Unspecified |
| mmu-miR-29b-3p | MIMAT0000127 | Fbln2 | 14115 | HITS-CLIP//sRNA-Seq | 25083871\|33184237 | Unspecified |
| mmu-miR-29b-3p | MIMAT0000127 | Fbln5 | 23876 | HITS-CLIP//sRNA-Seq | 25083871\|33184237 | Unspecified |
| mmu-miR-29b-3p | MIMAT0000127 | Fbn1 | 14118 | Luciferase assay//Microarray//Northern bl | 18723672\|25083871\|33184237 | Unspecified |
| mmu-miR-29b-3p | MIMAT0000127 | Fbrs | 14123 | HITS-CLIP | 37986224\|23142080\|23597149\|25083 | Unspecified |
| mmu-miR-29b-3p | MIMAT0000127 | Fbxl14 | 101358 | HITS-CLIP | 23597149\|23142080 | Unspecified |

| | | | | | | |
|---|---|---|---|---|---|---|
| mmu-miR-29b-3p | MIMAT0000127 | Fbxl17 | 50758 | HITS-CLIP | 25083871 | Unspecified |
| mmu-miR-29b-3p | MIMAT0000127 | Fbxl18 | 231863 | HITS-CLIP | 37986224\|23142080 | Unspecified |
| mmu-miR-29b-3p | MIMAT0000127 | Fbxl19 | 233902 | HITS-CLIP//sRNA-Seq | 25083871\|33184237 | Unspecified |
| mmu-miR-29b-3p | MIMAT0000127 | Fbxl20 | 72194 | HITS-CLIP | 37986224 | Unspecified |
| mmu-miR-29b-3p | MIMAT0000127 | Fbxl3 | 50789 | HITS-CLIP | 23142080\|25083871 | Unspecified |
| mmu-miR-29b-3p | MIMAT0000127 | Fbxl5 | 242960 | HITS-CLIP | 25083871\|23142080 | Unspecified |
| mmu-miR-29b-3p | MIMAT0000127 | Fbxl6 | 30840 | HITS-CLIP | 23597149 | Unspecified |
| mmu-miR-29b-3p | MIMAT0000127 | Fbxo10 | 269529 | HITS-CLIP | 37986224 | Unspecified |
| mmu-miR-29b-3p | MIMAT0000127 | Fbxo11 | 225055 | HITS-CLIP//sRNA-Seq | 23597149\|33184237\|25083871\|23142 | Unspecified |
| mmu-miR-29b-3p | MIMAT0000127 | Fbxo21 | 231670 | HITS-CLIP//sRNA-Seq | 37986224\|23142080\|23597149\|33184 | Unspecified |
| mmu-miR-29b-3p | MIMAT0000127 | Fbxo25 | 66822 | HITS-CLIP//sRNA-Seq | 23142080\|33184237 | Unspecified |
| mmu-miR-29b-3p | MIMAT0000127 | Fbxo28 | 67948 | HITS-CLIP | 25083871\|23142080 | Unspecified |
| mmu-miR-29b-3p | MIMAT0000127 | Fbxo3 | 57443 | HITS-CLIP | 37986224\|23597149\|23142080 | Unspecified |
| mmu-miR-29b-3p | MIMAT0000127 | Fbxo30 | 71865 | HITS-CLIP | 25083871\|23142080\|23597149 | Unspecified |
| mmu-miR-29b-3p | MIMAT0000127 | Fbxo32 | 67731 | HITS-CLIP | 37986224\|23142080\|25083871 | Unspecified |
| mmu-miR-29b-3p | MIMAT0000127 | Fbxo33 | 70611 | HITS-CLIP | 23142080 | Unspecified |
| mmu-miR-29b-3p | MIMAT0000127 | Fbxo42 | 213499 | HITS-CLIP | 23142080 | Unspecified |
| mmu-miR-29b-3p | MIMAT0000127 | Fbxo45 | 268882 | HITS-CLIP | 25083871 | Unspecified |
| mmu-miR-29b-3p | MIMAT0000127 | Fbxo7 | 69754 | HITS-CLIP | 23597149 | Unspecified |
| mmu-miR-29b-3p | MIMAT0000127 | Fbxo8 | 50753 | HITS-CLIP | 25083871\|23142080 | Unspecified |
| mmu-miR-29b-3p | MIMAT0000127 | Fbxw11 | 103583 | HITS-CLIP//sRNA-Seq | 25083871\|33184237\|23597149\|23142 | Unspecified |
| mmu-miR-29b-3p | MIMAT0000127 | Fbxw7 | 50754 | HITS-CLIP | 37986224\|25083871\|23142080 | Unspecified |
| mmu-miR-29b-3p | MIMAT0000127 | Fbxw9 | 68628 | HITS-CLIP | 37986224 | Unspecified |
| mmu-miR-29b-3p | MIMAT0000127 | Fcho1 | 23149 | HITS-CLIP | 23142080 | Unspecified |
| mmu-miR-29b-3p | MIMAT0000127 | Fcho2 | 218503 | HITS-CLIP | 23142080 | Unspecified |
| mmu-miR-29b-3p | MIMAT0000127 | Fchsd2 | 207278 | HITS-CLIP//sRNA-Seq | 23597149\|33184237\|28735896\|23142 | Unspecified |
| mmu-miR-29b-3p | MIMAT0000127 | Fdx1l | 112812 | HITS-CLIP | 23142080 | Unspecified |
| mmu-miR-29b-3p | MIMAT0000127 | Fech | 14151 | HITS-CLIP | 23597149 | Unspecified |
| mmu-miR-29b-3p | MIMAT0000127 | Fem1a | 14154 | HITS-CLIP | 25083871 | Unspecified |
| mmu-miR-29b-3p | MIMAT0000127 | Fem1b | 14155 | HITS-CLIP//sRNA-Seq | 37986224\|25083871\|23597149\|33184 | Unspecified |
| mmu-miR-29b-3p | MIMAT0000127 | Fem1c | 240263 | HITS-CLIP | 37986224\|23142080 | Unspecified |
| mmu-miR-29b-3p | MIMAT0000127 | Fer | 14158 | HITS-CLIP | 25083871 | Unspecified |
| mmu-miR-29b-3p | MIMAT0000127 | Fermt2 | 218952 | HITS-CLIP//sRNA-Seq | 37986224\|23597149\|33184237 | Unspecified |
| mmu-miR-29b-3p | MIMAT0000127 | Fermt3 | 108101 | HITS-CLIP | 23142080 | Unspecified |
| mmu-miR-29b-3p | MIMAT0000127 | Fetub | 59083 | HITS-CLIP | 23597149 | Unspecified |
| mmu-miR-29b-3p | MIMAT0000127 | Fga | 14161 | HITS-CLIP | 23597149 | Unspecified |
| mmu-miR-29b-3p | MIMAT0000127 | Fgd1 | 14163 | HITS-CLIP | 25083871 | Unspecified |
| mmu-miR-29b-3p | MIMAT0000127 | Fgd3 | 30938 | HITS-CLIP | 37986224 | Unspecified |
| mmu-miR-29b-3p | MIMAT0000127 | Fgd6 | 13998 | HITS-CLIP | 25083871\|23142080 | Unspecified |

| | | | | | | |
|---|---|---|---|---|---|---|
| mmu-miR-29b-3p | MIMAT0000127 | Fgf2 | 14173 | HITS-CLIP | 23142080\|25083871 | Unspecified |
| mmu-miR-29b-3p | MIMAT0000127 | Fgf7 | 14178 | HITS-CLIP//sRNA-Seq | 25083871\|33184237 | Unspecified |
| mmu-miR-29b-3p | MIMAT0000127 | Fgfr1 | 14182 | HITS-CLIP//sRNA-Seq | 25083871\|33184237 | Unspecified |
| mmu-miR-29b-3p | MIMAT0000127 | Fgfr1op | 75296 | HITS-CLIP | 37986224 | Unspecified |
| mmu-miR-29b-3p | MIMAT0000127 | Fgfr1op2 | 67529 | HITS-CLIP | 23142080 | Unspecified |
| mmu-miR-29b-3p | MIMAT0000127 | Fgfrl1 | 116701 | HITS-CLIP | 37986224\|25083871 | Unspecified |
| mmu-miR-29b-3p | MIMAT0000127 | Fgl1 | 234199 | HITS-CLIP | 23597149 | Unspecified |
| mmu-miR-29b-3p | MIMAT0000127 | Fgl2 | 14190 | HITS-CLIP | 37986224 | Unspecified |
| mmu-miR-29b-3p | MIMAT0000127 | Fh1 | 14194 | HITS-CLIP//sRNA-Seq | 37986224\|25083871\|33184237\|23142 | Unspecified |
| mmu-miR-29b-3p | MIMAT0000127 | Fhod1 | 234686 | HITS-CLIP | 25083871 | Unspecified |
| mmu-miR-29b-3p | MIMAT0000127 | Ficd | 231630 | HITS-CLIP | 25083871\|23142080 | Unspecified |
| mmu-miR-29b-3p | MIMAT0000127 | Fignl1 | 60530 | HITS-CLIP | 23142080 | Unspecified |
| mmu-miR-29b-3p | MIMAT0000127 | Filip1 | 70598 | HITS-CLIP | 25083871 | Unspecified |
| mmu-miR-29b-3p | MIMAT0000127 | Fis1 | 66437 | HITS-CLIP//sRNA-Seq | 25083871\|33184237\|23597149\|23142 | Unspecified |
| mmu-miR-29b-3p | MIMAT0000127 | Fiz1 | 23877 | HITS-CLIP | 25083871 | Unspecified |
| mmu-miR-29b-3p | MIMAT0000127 | Fkbp10 | 14230 | HITS-CLIP//sRNA-Seq | 37986224\|25083871\|33184237 | Unspecified |
| mmu-miR-29b-3p | MIMAT0000127 | Fkbp1a | 14225 | HITS-CLIP//sRNA-Seq | 37986224\|25083871\|23597149\|33184 | Unspecified |
| mmu-miR-29b-3p | MIMAT0000127 | Fkbp1b | 14226 | HITS-CLIP//sRNA-Seq | 37986224\|33184237\|23142080 | Unspecified |
| mmu-miR-29b-3p | MIMAT0000127 | Fkbp4 | 14228 | HITS-CLIP//sRNA-Seq | 37986224\|23597149\|25083871\|33184 | Unspecified |
| mmu-miR-29b-3p | MIMAT0000127 | Fkbp5 | 14229 | sRNA-Seq//HITS-CLIP | 33184237\|23142080 | Unspecified |
| mmu-miR-29b-3p | MIMAT0000127 | Fktn | 246179 | HITS-CLIP | 25083871 | Unspecified |
| mmu-miR-29b-3p | MIMAT0000127 | Flcn | 216805 | HITS-CLIP | 25083871\|23142080 | Unspecified |
| mmu-miR-29b-3p | MIMAT0000127 | Fli1 | 14247 | HITS-CLIP | 23142080 | Unspecified |
| mmu-miR-29b-3p | MIMAT0000127 | Flii | 14248 | HITS-CLIP//sRNA-Seq | 37986224\|23142080\|33184237\|2508З | Unspecified |
| mmu-miR-29b-3p | MIMAT0000127 | Flna | 192176 | HITS-CLIP//sRNA-Seq | 23597149\|33184237\|23142080 | Unspecified |
| mmu-miR-29b-3p | MIMAT0000127 | Flnb | 286940 | HITS-CLIP//sRNA-Seq | 37986224\|25083871\|23597149\|33184 | Unspecified |
| mmu-miR-29b-3p | MIMAT0000127 | Flnc | 68794 | HITS-CLIP//sRNA-Seq | 25083871\|33184237 | Unspecified |
| mmu-miR-29b-3p | MIMAT0000127 | Flt1 | 14254 | HITS-CLIP | 23597149 | Unspecified |
| mmu-miR-29b-3p | MIMAT0000127 | Flt3l | 14256 | HITS-CLIP | 23142080 | Unspecified |
| mmu-miR-29b-3p | MIMAT0000127 | Flywch1 | 224613 | HITS-CLIP//sRNA-Seq | 23142080\|33184237\|25083871 | Unspecified |
| mmu-miR-29b-3p | MIMAT0000127 | Fmnl1 | 57778 | HITS-CLIP | 23142080 | Unspecified |
| mmu-miR-29b-3p | MIMAT0000127 | Fmnl2 | 71409 | HITS-CLIP | 25083871 | Unspecified |
| mmu-miR-29b-3p | MIMAT0000127 | Fmnl3 | 22379 | HITS-CLIP//sRNA-Seq | 25083871\|33184237 | Unspecified |
| mmu-miR-29b-3p | MIMAT0000127 | Fmo1 | 14261 | HITS-CLIP | 25083871 | Unspecified |
| mmu-miR-29b-3p | MIMAT0000127 | Fmo4 | 226564 | HITS-CLIP | 23597149 | Unspecified |
| mmu-miR-29b-3p | MIMAT0000127 | Fmr1 | 14265 | HITS-CLIP//sRNA-Seq | 37986224\|25083871\|33184237\|23142 | Unspecified |
| mmu-miR-29b-3p | MIMAT0000127 | Fn1 | 14268 | HITS-CLIP//sRNA-Seq | 37986224\|25083871\|33184237\|23597 | Unspecified |
| mmu-miR-29b-3p | MIMAT0000127 | Fnbp1 | 14269 | sRNA-Seq//HITS-CLIP | 33184237\|23142080 | Unspecified |
| mmu-miR-29b-3p | MIMAT0000127 | Fnbp1l | 214459 | HITS-CLIP//sRNA-Seq | 25083871\|33184237 | Unspecified |

| | | | | | | |
|---|---|---|---|---|---|---|
| mmu-miR-29b-3p | MIMAT0000127 | Fnbp4 | 55935 | HITS-CLIP | 23142080 | Unspecified |
| mmu-miR-29b-3p | MIMAT0000127 | Fndc3a | 319448 | HITS-CLIP | 23597149\|23142080 | Unspecified |
| mmu-miR-29b-3p | MIMAT0000127 | Fndc3b | 72007 | HITS-CLIP//sRNA-Seq | 23142080\|33184237 | Unspecified |
| mmu-miR-29b-3p | MIMAT0000127 | Fnip1 | 216742 | HITS-CLIP | 23142080 | Unspecified |
| mmu-miR-29b-3p | MIMAT0000127 | Fnip2 | 329679 | HITS-CLIP | 23142080 | Unspecified |
| mmu-miR-29b-3p | MIMAT0000127 | Focad | 230393 | HITS-CLIP | 23597149\|23142080 | Unspecified |
| mmu-miR-29b-3p | MIMAT0000127 | Fos | 14281 | HITS-CLIP | 23597149\|23142080\|25083871 | Unspecified |
| mmu-miR-29b-3p | MIMAT0000127 | Fosl2 | 14284 | sRNA-Seq//HITS-CLIP | 33184237\|23142080 | Unspecified |
| mmu-miR-29b-3p | MIMAT0000127 | Foxj2 | 60611 | HITS-CLIP | 37986224\|23597149\|23142080\|25083 | Unspecified |
| mmu-miR-29b-3p | MIMAT0000127 | Foxj3 | 230700 | HITS-CLIP | 23142080 | Unspecified |
| mmu-miR-29b-3p | MIMAT0000127 | Foxk2 | 68837 | HITS-CLIP//sRNA-Seq | 25083871\|23597149\|33184237 | Unspecified |
| mmu-miR-29b-3p | MIMAT0000127 | Foxn2 | 14236 | HITS-CLIP | 37986224\|25083871\|30971389\|23142 | Unspecified |
| mmu-miR-29b-3p | MIMAT0000127 | Foxo1 | 56458 | HITS-CLIP | 37986224\|23142080 | Unspecified |
| mmu-miR-29b-3p | MIMAT0000127 | Foxo3 | 56484 | HITS-CLIP | 37986224\|25083871\|23597149\|23142 | Unspecified |
| mmu-miR-29b-3p | MIMAT0000127 | Foxo4 | 54601 | HITS-CLIP | 37986224\|25083871\|23142080 | Unspecified |
| mmu-miR-29b-3p | MIMAT0000127 | Foxp1 | 108655 | HITS-CLIP | 23142080 | Unspecified |
| mmu-miR-29b-3p | MIMAT0000127 | Fpgs | 14287 | HITS-CLIP | 23597149 | Unspecified |
| mmu-miR-29b-3p | MIMAT0000127 | Fpgt | 75540 | HITS-CLIP | 23142080 | Unspecified |
| mmu-miR-29b-3p | MIMAT0000127 | Frat2 | 212398 | HITS-CLIP | 37986224\|23142080 | Unspecified |
| mmu-miR-29b-3p | MIMAT0000127 | Frmd4a | 209630 | HITS-CLIP//sRNA-Seq | 37986224\|25083871\|33184237 | Unspecified |
| mmu-miR-29b-3p | MIMAT0000127 | Frmd8 | 67457 | HITS-CLIP | 23142080 | Unspecified |
| mmu-miR-29b-3p | MIMAT0000127 | Frmpd1 | 666060 | HITS-CLIP | 25083871 | Unspecified |
| mmu-miR-29b-3p | MIMAT0000127 | Frrs1 | 20321 | HITS-CLIP | 23142080 | Unspecified |
| mmu-miR-29b-3p | MIMAT0000127 | Frs2 | 327826 | HITS-CLIP | 37986224\|23142080\|25083871 | Unspecified |
| mmu-miR-29b-3p | MIMAT0000127 | Fryl | 72313 | HITS-CLIP | 25083871\|23142080 | Unspecified |
| mmu-miR-29b-3p | MIMAT0000127 | Fscn1 | 14086 | HITS-CLIP//sRNA-Seq | 37986224\|25083871\|33184237 | Unspecified |
| mmu-miR-29b-3p | MIMAT0000127 | Fsd1l | 319636 | HITS-CLIP//sRNA-Seq | 37986224\|25083871\|23142080\|33184 | Unspecified |
| mmu-miR-29b-3p | MIMAT0000127 | Fstl1 | 14314 | HITS-CLIP//sRNA-Seq | 25083871\|33184237 | Unspecified |
| mmu-miR-29b-3p | MIMAT0000127 | Ftcd | 14317 | HITS-CLIP | 37986224\|23597149 | Unspecified |
| mmu-miR-29b-3p | MIMAT0000127 | Fto | 26383 | HITS-CLIP//sRNA-Seq | 37986224\|25083871\|33184237 | Unspecified |
| mmu-miR-29b-3p | MIMAT0000127 | Fubp1 | 51886 | HITS-CLIP//sRNA-Seq | 37986224\|25083871\|23142080\|33184 | Unspecified |
| mmu-miR-29b-3p | MIMAT0000127 | Fubp3 | 320267 | HITS-CLIP//sRNA-Seq | 25083871\|33184237 | Unspecified |
| mmu-miR-29b-3p | MIMAT0000127 | Fuca2 | 66848 | HITS-CLIP | 23597149\|23142080 | Unspecified |
| mmu-miR-29b-3p | MIMAT0000127 | Fuk | 197258 | HITS-CLIP | 23597149 | Unspecified |
| mmu-miR-29b-3p | MIMAT0000127 | Furin | 18550 | HITS-CLIP | 23142080 | Unspecified |
| mmu-miR-29b-3p | MIMAT0000127 | Fut8 | 53618 | HITS-CLIP//sRNA-Seq | 25083871\|33184237\|23597149 | Unspecified |
| mmu-miR-29b-3p | MIMAT0000127 | Fxr2 | 23879 | sRNA-Seq//HITS-CLIP | 33184237\|25083871 | Unspecified |
| mmu-miR-29b-3p | MIMAT0000127 | Fyb2 | 199920 | HITS-CLIP | 23597149 | Unspecified |
| mmu-miR-29b-3p | MIMAT0000127 | Fyco1 | 17281 | HITS-CLIP | 25083871\|23142080 | Unspecified |

| | | | | | | |
|---|---|---|---|---|---|---|
| mmu-miR-29b-3p | MIMAT0000127 | Fyn | 14360 | HITS-CLIP//sRNA-Seq | 37986224\|25083871\|33184237\|23142 | Unspecified |
| mmu-miR-29b-3p | MIMAT0000127 | Fyttd1 | 69823 | sRNA-Seq//HITS-CLIP | 33184237\|25083871 | Unspecified |
| mmu-miR-29b-3p | MIMAT0000127 | Fzd2 | 57265 | HITS-CLIP//sRNA-Seq | 25083871\|33184237 | Unspecified |
| mmu-miR-29b-3p | MIMAT0000127 | Fzd3 | 14365 | HITS-CLIP | 25083871 | Unspecified |
| mmu-miR-29b-3p | MIMAT0000127 | Fzd5 | 14367 | HITS-CLIP | 37986224\|25083871\|23597149 | Unspecified |
| mmu-miR-29b-3p | MIMAT0000127 | Fzd7 | 14369 | HITS-CLIP | 37986224\|25083871 | Unspecified |
| mmu-miR-29b-3p | MIMAT0000127 | Fzr1 | 56371 | HITS-CLIP//sRNA-Seq | 25083871\|23142080\|33184237 | Unspecified |
| mmu-miR-29b-3p | MIMAT0000127 | G2e3 | 217558 | HITS-CLIP | 23142080 | Unspecified |
| mmu-miR-29b-3p | MIMAT0000127 | G3bp1 | 27041 | HITS-CLIP//sRNA-Seq | 23142080\|33184237\|25083871 | Unspecified |
| mmu-miR-29b-3p | MIMAT0000127 | G3bp2 | 23881 | HITS-CLIP//sRNA-Seq | 23142080\|25083871\|33184237\|23597 | Unspecified |
| mmu-miR-29b-3p | MIMAT0000127 | G6pc | 14377 | HITS-CLIP | 37986224\|23597149 | Unspecified |
| mmu-miR-29b-3p | MIMAT0000127 | Gaa | 14387 | HITS-CLIP//sRNA-Seq | 25083871\|23597149\|33184237 | Unspecified |
| mmu-miR-29b-3p | MIMAT0000127 | Gab2 | 14389 | HITS-CLIP | 25083871 | Unspecified |
| mmu-miR-29b-3p | MIMAT0000127 | Gabarap | 56486 | HITS-CLIP//sRNA-Seq | 23597149\|33184237\|25083871\|23142 | Unspecified |
| mmu-miR-29b-3p | MIMAT0000127 | Gabbr1 | 54393 | HITS-CLIP//sRNA-Seq | 25083871\|33184237 | Unspecified |
| mmu-miR-29b-3p | MIMAT0000127 | Gabbr2 | 242425 | HITS-CLIP | 23597149 | Unspecified |
| mmu-miR-29b-3p | MIMAT0000127 | Gabpb1 | 14391 | HITS-CLIP | 37986224\|25083871\|23142080 | Unspecified |
| mmu-miR-29b-3p | MIMAT0000127 | Gabpb2 | 213054 | HITS-CLIP | 37986224\|23142080\|25083871 | Unspecified |
| mmu-miR-29b-3p | MIMAT0000127 | Gadd45g | 23882 | HITS-CLIP | 25083871\|23142080 | Unspecified |
| mmu-miR-29b-3p | MIMAT0000127 | Gak | 231580 | sRNA-Seq//HITS-CLIP | 33184237\|23142080 | Unspecified |
| mmu-miR-29b-3p | MIMAT0000127 | Galc | 14420 | HITS-CLIP | 25083871\|23142080 | Unspecified |
| mmu-miR-29b-3p | MIMAT0000127 | Galk1 | 14635 | HITS-CLIP//sRNA-Seq | 37986224\|23597149\|25083871\|33184 | Unspecified |
| mmu-miR-29b-3p | MIMAT0000127 | Galnt1 | 14423 | HITS-CLIP//sRNA-Seq | 25083871\|33184237 | Unspecified |
| mmu-miR-29b-3p | MIMAT0000127 | Galnt10 | 171212 | HITS-CLIP | 37986224\|25083871 | Unspecified |
| mmu-miR-29b-3p | MIMAT0000127 | Galnt12 | 230145 | HITS-CLIP | 23142080 | Unspecified |
| mmu-miR-29b-3p | MIMAT0000127 | Galnt2 | 108148 | HITS-CLIP | 25083871\|23597149\|23142080 | Unspecified |
| mmu-miR-29b-3p | MIMAT0000127 | Galnt6 | 207839 | HITS-CLIP | 37986224 | Unspecified |
| mmu-miR-29b-3p | MIMAT0000127 | Gamt | 14431 | HITS-CLIP | 25083871 | Unspecified |
| mmu-miR-29b-3p | MIMAT0000127 | Gan | 209239 | HITS-CLIP | 25083871\|23142080 | Unspecified |
| mmu-miR-29b-3p | MIMAT0000127 | Ganab | 14376 | HITS-CLIP//sRNA-Seq | 23597149\|33184237\|23142080 | Unspecified |
| mmu-miR-29b-3p | MIMAT0000127 | Gapvd1 | 66691 | HITS-CLIP//sRNA-Seq | 37986224\|25083871\|33184237 | Unspecified |
| mmu-miR-29b-3p | MIMAT0000127 | Garem1 | 64762 | HITS-CLIP | 23597149 | Unspecified |
| mmu-miR-29b-3p | MIMAT0000127 | Gars | 353172 | HITS-CLIP//sRNA-Seq | 25083871\|33184237 | Unspecified |
| mmu-miR-29b-3p | MIMAT0000127 | Gas1 | 14451 | HITS-CLIP | 37986224 | Unspecified |
| mmu-miR-29b-3p | MIMAT0000127 | Gas2 | 14453 | HITS-CLIP | 23597149 | Unspecified |
| mmu-miR-29b-3p | MIMAT0000127 | Gas2l1 | 78926 | HITS-CLIP | 37986224\|25083871 | Unspecified |
| mmu-miR-29b-3p | MIMAT0000127 | Gas2l3 | 237436 | HITS-CLIP | 23142080 | Unspecified |
| mmu-miR-29b-3p | MIMAT0000127 | Gata2 | 14461 | HITS-CLIP | 25083871 | Unspecified |
| mmu-miR-29b-3p | MIMAT0000127 | Gatad2a | 234366 | HITS-CLIP//sRNA-Seq | 23142080\|33184237 | Unspecified |

| | | | | | | |
|---|---|---|---|---|---|---|
| mmu-miR-29b-3p | MIMAT0000127 | Gatad2b | 229542 | HITS-CLIP | 23142080\|23597149\|25083871 | Unspecified |
| mmu-miR-29b-3p | MIMAT0000127 | Gatd1 | 347862 | HITS-CLIP | 23142080 | Unspecified |
| mmu-miR-29b-3p | MIMAT0000127 | Gatm | 67092 | HITS-CLIP | 25083871\|23142080 | Unspecified |
| mmu-miR-29b-3p | MIMAT0000127 | Gbf1 | 107338 | HITS-CLIP//sRNA-Seq | 25083871\|23597149\|33184237\|23142 | Unspecified |
| mmu-miR-29b-3p | MIMAT0000127 | Gbp2 | 14469 | HITS-CLIP//sRNA-Seq | 23142080\|33184237 | Unspecified |
| mmu-miR-29b-3p | MIMAT0000127 | Gbp5 | 229898 | HITS-CLIP | 23142080 | Unspecified |
| mmu-miR-29b-3p | MIMAT0000127 | Gbp7 | 229900 | HITS-CLIP | 31433990\|23142080 | Unspecified |
| mmu-miR-29b-3p | MIMAT0000127 | Gbp9 | 236573 | HITS-CLIP | 23142080 | Unspecified |
| mmu-miR-29b-3p | MIMAT0000127 | Gc | 14473 | HITS-CLIP | 37986224\|23597149 | Unspecified |
| mmu-miR-29b-3p | MIMAT0000127 | Gcat | 26912 | HITS-CLIP | 23597149 | Unspecified |
| mmu-miR-29b-3p | MIMAT0000127 | Gcc2 | 70297 | HITS-CLIP | 25083871\|23142080 | Unspecified |
| mmu-miR-29b-3p | MIMAT0000127 | Gcgr | 14527 | HITS-CLIP | 23597149 | Unspecified |
| mmu-miR-29b-3p | MIMAT0000127 | Gch1 | 14528 | HITS-CLIP | 23597149 | Unspecified |
| mmu-miR-29b-3p | MIMAT0000127 | Gchfr | 320415 | HITS-CLIP//sRNA-Seq | 23597149\|33184237 | Unspecified |
| mmu-miR-29b-3p | MIMAT0000127 | Gckr | 231103 | HITS-CLIP | 23597149 | Unspecified |
| mmu-miR-29b-3p | MIMAT0000127 | Gclm | 14630 | HITS-CLIP | 23597149 | Unspecified |
| mmu-miR-29b-3p | MIMAT0000127 | Gcn1l1 | 10985 | HITS-CLIP | 23597149\|23142080 | Unspecified |
| mmu-miR-29b-3p | MIMAT0000127 | Gcnt2 | 14538 | HITS-CLIP | 37986224 | Unspecified |
| mmu-miR-29b-3p | MIMAT0000127 | Gcnt4 | 218476 | HITS-CLIP | 37986224 | Unspecified |
| mmu-miR-29b-3p | MIMAT0000127 | Gde1 | 56209 | HITS-CLIP//sRNA-Seq | 37986224\|25083871\|33184237\|23142 | Unspecified |
| mmu-miR-29b-3p | MIMAT0000127 | Gdf15 | 23886 | HITS-CLIP | 23597149 | Unspecified |
| mmu-miR-29b-3p | MIMAT0000127 | Gdi1 | 14567 | HITS-CLIP//sRNA-Seq | 23142080\|33184237\|25083871 | Unspecified |
| mmu-miR-29b-3p | MIMAT0000127 | Gdi2 | 14569 | sRNA-Seq//HITS-CLIP | 33184237\|25083871 | Unspecified |
| mmu-miR-29b-3p | MIMAT0000127 | Gem | 14579 | HITS-CLIP | 23142080 | Unspecified |
| mmu-miR-29b-3p | MIMAT0000127 | Gen1 | 209334 | HITS-CLIP | 23142080 | Unspecified |
| mmu-miR-29b-3p | MIMAT0000127 | Get4 | 67604 | HITS-CLIP//sRNA-Seq | 23142080\|33184237 | Unspecified |
| mmu-miR-29b-3p | MIMAT0000127 | Gfi1 | 14581 | HITS-CLIP | 23142080 | Unspecified |
| mmu-miR-29b-3p | MIMAT0000127 | Gfod1 | 328232 | HITS-CLIP | 37986224\|23142080 | Unspecified |
| mmu-miR-29b-3p | MIMAT0000127 | Gfra1 | 14585 | HITS-CLIP | 23597149 | Unspecified |
| mmu-miR-29b-3p | MIMAT0000127 | Gga3 | 260302 | HITS-CLIP//qCLASH | 23597149\|25083871\|23142080\|31393 | Unspecified |
| mmu-miR-29b-3p | MIMAT0000127 | Ggact | 223267 | HITS-CLIP | 23597149 | Unspecified |
| mmu-miR-29b-3p | MIMAT0000127 | Ggct | 110175 | HITS-CLIP | 23142080 | Unspecified |
| mmu-miR-29b-3p | MIMAT0000127 | Ggcx | 56316 | HITS-CLIP | 25083871\|23597149 | Unspecified |
| mmu-miR-29b-3p | MIMAT0000127 | Ggnbp2 | 217039 | HITS-CLIP//sRNA-Seq | 25083871\|23142080\|33184237\|23597 | Unspecified |
| mmu-miR-29b-3p | MIMAT0000127 | Ggt6 | 71522 | HITS-CLIP | 23597149 | Unspecified |
| mmu-miR-29b-3p | MIMAT0000127 | Ghdc | 80860 | HITS-CLIP | 37986224\|23597149 | Unspecified |
| mmu-miR-29b-3p | MIMAT0000127 | Ghitm | 66092 | HITS-CLIP//sRNA-Seq | 23142080\|33184237 | Unspecified |
| mmu-miR-29b-3p | MIMAT0000127 | Gid8 | 76425 | HITS-CLIP//sRNA-Seq | 37986224\|25083871\|33184237\|23142 | Unspecified |
| mmu-miR-29b-3p | MIMAT0000127 | Gigyf2 | 227331 | HITS-CLIP | 37986224 | Unspecified |

| | | | | | | |
|---|---|---|---|---|---|---|
| mmu-miR-29b-3p | MIMAT0000127 | Gimap1 | 16205 | HITS-CLIP | 23597149\|23142080 | Unspecified |
| mmu-miR-29b-3p | MIMAT0000127 | Gimap3 | 83408 | HITS-CLIP | 23142080 | Unspecified |
| mmu-miR-29b-3p | MIMAT0000127 | Gimap6 | 231931 | HITS-CLIP | 23142080 | Unspecified |
| mmu-miR-29b-3p | MIMAT0000127 | Gimap9 | 317758 | HITS-CLIP | 23142080 | Unspecified |
| mmu-miR-29b-3p | MIMAT0000127 | Gin1 | 252876 | HITS-CLIP | 23142080 | Unspecified |
| mmu-miR-29b-3p | MIMAT0000127 | Gipc1 | 67903 | sRNA-Seq//HITS-CLIP | 33184237\|26203562 | Unspecified |
| mmu-miR-29b-3p | MIMAT0000127 | Git1 | 216963 | HITS-CLIP//sRNA-Seq | 25083871\|33184237 | Unspecified |
| mmu-miR-29b-3p | MIMAT0000127 | Git2 | 26431 | HITS-CLIP//qCLASH | 37986224\|25083871\|23597149\|31393 | Unspecified |
| mmu-miR-29b-3p | MIMAT0000127 | Gja1 | 14609 | HITS-CLIP//sRNA-Seq | 37986224\|33184237\|23597149\|25083 | Unspecified |
| mmu-miR-29b-3p | MIMAT0000127 | Gja4 | 14612 | HITS-CLIP | 37986224 | Unspecified |
| mmu-miR-29b-3p | MIMAT0000127 | Gja5 | 14613 | HITS-CLIP | 25083871 | Unspecified |
| mmu-miR-29b-3p | MIMAT0000127 | Gjb1 | 14618 | HITS-CLIP | 23597149 | Unspecified |
| mmu-miR-29b-3p | MIMAT0000127 | Gjb2 | 14619 | HITS-CLIP | 23597149 | Unspecified |
| mmu-miR-29b-3p | MIMAT0000127 | Gk | 14933 | HITS-CLIP | 25083871\|23597149\|23142080 | Unspecified |
| mmu-miR-29b-3p | MIMAT0000127 | Gk5 | 235533 | HITS-CLIP | 37986224\|23142080 | Unspecified |
| mmu-miR-29b-3p | MIMAT0000127 | Glcci1 | 170772 | HITS-CLIP | 25083871\|23142080 | Unspecified |
| mmu-miR-29b-3p | MIMAT0000127 | Gldc | 104174 | HITS-CLIP | 23597149 | Unspecified |
| mmu-miR-29b-3p | MIMAT0000127 | Glg1 | 20340 | HITS-CLIP//sRNA-Seq | 23142080\|33184237 | Unspecified |
| mmu-miR-29b-3p | MIMAT0000127 | Gli3 | 14634 | HITS-CLIP | 37986224\|25083871 | Unspecified |
| mmu-miR-29b-3p | MIMAT0000127 | Glis2 | 83396 | HITS-CLIP | 37986224\|25083871 | Unspecified |
| mmu-miR-29b-3p | MIMAT0000127 | Glmp | 56700 | sRNA-Seq//HITS-CLIP | 33184237\|26203562 | Unspecified |
| mmu-miR-29b-3p | MIMAT0000127 | Glrx5 | 73046 | HITS-CLIP | 23597149\|23142080 | Unspecified |
| mmu-miR-29b-3p | MIMAT0000127 | Gls | 14660 | sRNA-Seq//HITS-CLIP | 33184237\|23142080 | Unspecified |
| mmu-miR-29b-3p | MIMAT0000127 | Gls2 | 216456 | HITS-CLIP | 23597149 | Unspecified |
| mmu-miR-29b-3p | MIMAT0000127 | Gltp | 56356 | sRNA-Seq//HITS-CLIP | 33184237\|25083871 | Unspecified |
| mmu-miR-29b-3p | MIMAT0000127 | Glud1 | 14661 | sRNA-Seq//HITS-CLIP | 33184237\|25083871\|23142080 | Unspecified |
| mmu-miR-29b-3p | MIMAT0000127 | Glul | 14645 | HITS-CLIP | 37986224\|23597149\|23142080 | Unspecified |
| mmu-miR-29b-3p | MIMAT0000127 | Glyat | 107146 | HITS-CLIP | 23597149 | Unspecified |
| mmu-miR-29b-3p | MIMAT0000127 | Glyr1 | 74022 | HITS-CLIP//sRNA-Seq | 25083871\|33184237\|23597149\|23142 | Unspecified |
| mmu-miR-29b-3p | MIMAT0000127 | Gm12166 | 100039624 | HITS-CLIP | 23142080 | Unspecified |
| mmu-miR-29b-3p | MIMAT0000127 | Gm14085 | 381417 | HITS-CLIP | 23142080 | Unspecified |
| mmu-miR-29b-3p | MIMAT0000127 | Gm16286 | 117903916 | HITS-CLIP//sRNA-Seq | 23597149\|33184237\|23142080 | Unspecified |
| mmu-miR-29b-3p | MIMAT0000127 | Gm20425 | 6608681 | HITS-CLIP | 23597149 | Unspecified |
| mmu-miR-29b-3p | MIMAT0000127 | Gm20547 | 6608472 | HITS-CLIP | 23597149 | Unspecified |
| mmu-miR-29b-3p | MIMAT0000127 | Gm20604 | 100859931 | HITS-CLIP | 37986224\|23142080 | Unspecified |
| mmu-miR-29b-3p | MIMAT0000127 | Gm37240 | | HITS-CLIP | 23142080 | Unspecified |
| mmu-miR-29b-3p | MIMAT0000127 | Gm38394 | 667118 | HITS-CLIP | 23142080\|23597149 | Unspecified |
| mmu-miR-29b-3p | MIMAT0000127 | Gm44502 | 100134861 | RNA-Seq | 33332475 | Unspecified |
| mmu-miR-29b-3p | MIMAT0000127 | Gm49333 | 110599584 | HITS-CLIP | 23142080 | Unspecified |

| | | | | | | |
|---|---|---|---|---|---|---|
| mmu-miR-29b-3p | MIMAT0000127 | Gm49336 | 127230315 | sRNA-Seq//HITS-CLIP | 33184237\|23142080 | Unspecified |
| mmu-miR-29b-3p | MIMAT0000127 | Gm50388 | | RNA-Seq | 33332475 | Unspecified |
| mmu-miR-29b-3p | MIMAT0000127 | Gm8369 | 666926 | HITS-CLIP | 23142080 | Unspecified |
| mmu-miR-29b-3p | MIMAT0000127 | Gmds | 218138 | HITS-CLIP | 23597149 | Unspecified |
| mmu-miR-29b-3p | MIMAT0000127 | Gmeb2 | 229004 | HITS-CLIP | 25083871\|23142080 | Unspecified |
| mmu-miR-29b-3p | MIMAT0000127 | Gmfb | 63985 | HITS-CLIP//sRNA-Seq | 37986224\|25083871\|23597149\|33184 | Unspecified |
| mmu-miR-29b-3p | MIMAT0000127 | Gmip | 78816 | HITS-CLIP | 23142080 | Unspecified |
| mmu-miR-29b-3p | MIMAT0000127 | Gmppb | 331026 | HITS-CLIP | 37986224 | Unspecified |
| mmu-miR-29b-3p | MIMAT0000127 | Gmps | 229363 | HITS-CLIP | 23597149\|23142080 | Unspecified |
| mmu-miR-29b-3p | MIMAT0000127 | Gna12 | 14673 | HITS-CLIP//sRNA-Seq | 23597149\|25083871\|33184237 | Unspecified |
| mmu-miR-29b-3p | MIMAT0000127 | Gna13 | 14674 | HITS-CLIP | 23597149\|23142080 | Unspecified |
| mmu-miR-29b-3p | MIMAT0000127 | Gnai2 | 14678 | HITS-CLIP//sRNA-Seq | 25083871\|23597149\|33184237\|23142 | Unspecified |
| mmu-miR-29b-3p | MIMAT0000127 | Gnai3 | 14679 | HITS-CLIP//sRNA-Seq | 37986224\|25083871\|23142080\|33184 | Unspecified |
| mmu-miR-29b-3p | MIMAT0000127 | Gnaq | 14682 | HITS-CLIP//sRNA-Seq | 25083871\|23142080\|33184237 | Unspecified |
| mmu-miR-29b-3p | MIMAT0000127 | Gnas | 14683 | sRNA-Seq//HITS-CLIP | 33184237\|23142080 | Unspecified |
| mmu-miR-29b-3p | MIMAT0000127 | Gnb1 | 14688 | HITS-CLIP//sRNA-Seq | 23142080\|33184237 | Unspecified |
| mmu-miR-29b-3p | MIMAT0000127 | Gnb4 | 14696 | HITS-CLIP//sRNA-Seq | 23142080\|33184237\|23597149 | Unspecified |
| mmu-miR-29b-3p | MIMAT0000127 | Gne | 50798 | HITS-CLIP | 23597149 | Unspecified |
| mmu-miR-29b-3p | MIMAT0000127 | Gng12 | 14701 | HITS-CLIP//sRNA-Seq | 37986224\|25083871\|33184237\|23597 | Unspecified |
| mmu-miR-29b-3p | MIMAT0000127 | Gng2 | 14702 | HITS-CLIP//sRNA-Seq | 37986224\|25083871\|23142080\|33184 | Unspecified |
| mmu-miR-29b-3p | MIMAT0000127 | Gnl2 | 230737 | sRNA-Seq//HITS-CLIP | 33184237\|23142080 | Unspecified |
| mmu-miR-29b-3p | MIMAT0000127 | Gnl3 | 30877 | HITS-CLIP//sRNA-Seq | 25083871\|33184237 | Unspecified |
| mmu-miR-29b-3p | MIMAT0000127 | Gnmt | 14711 | HITS-CLIP | 23597149 | Unspecified |
| mmu-miR-29b-3p | MIMAT0000127 | Gnpat | 14712 | sRNA-Seq//HITS-CLIP | 33184237\|23142080 | Unspecified |
| mmu-miR-29b-3p | MIMAT0000127 | Gnpda2 | 67980 | HITS-CLIP | 23142080 | Unspecified |
| mmu-miR-29b-3p | MIMAT0000127 | Gnptab | 432486 | HITS-CLIP//qCLASH//sRNA-Seq | 23142080\|31393953\|33184237 | Unspecified |
| mmu-miR-29b-3p | MIMAT0000127 | Golga1 | 76899 | HITS-CLIP | 23142080 | Unspecified |
| mmu-miR-29b-3p | MIMAT0000127 | Golga3 | 269682 | HITS-CLIP | 37986224 | Unspecified |
| mmu-miR-29b-3p | MIMAT0000127 | Golga4 | 54214 | HITS-CLIP | 25083871 | Unspecified |
| mmu-miR-29b-3p | MIMAT0000127 | Golgb1 | 224139 | HITS-CLIP | 25083871 | Unspecified |
| mmu-miR-29b-3p | MIMAT0000127 | Golph3 | 66629 | HITS-CLIP//sRNA-Seq | 23142080\|33184237\|31433990\|25083 | Unspecified |
| mmu-miR-29b-3p | MIMAT0000127 | Golph3l | 229593 | HITS-CLIP | 23142080 | Unspecified |
| mmu-miR-29b-3p | MIMAT0000127 | Gopc | 94221 | HITS-CLIP | 37986224\|25083871\|23597149\|23142 | Unspecified |
| mmu-miR-29b-3p | MIMAT0000127 | Gosr2 | 56494 | HITS-CLIP//sRNA-Seq | 23597149\|33184237\|25083871 | Unspecified |
| mmu-miR-29b-3p | MIMAT0000127 | Gpaa1 | 14731 | HITS-CLIP | 37986224\|23597149 | Unspecified |
| mmu-miR-29b-3p | MIMAT0000127 | Gpam | 14732 | HITS-CLIP | 23142080\|23597149\|25083871 | Unspecified |
| mmu-miR-29b-3p | MIMAT0000127 | Gpat4 | 137964 | HITS-CLIP//sRNA-Seq | 23142080\|25083871\|23597149\|33184 | Unspecified |
| mmu-miR-29b-3p | MIMAT0000127 | Gpatch2 | 67769 | HITS-CLIP | 37986224\|25083871 | Unspecified |
| mmu-miR-29b-3p | MIMAT0000127 | Gpatch2l | 70373 | HITS-CLIP | 23597149 | Unspecified |

| | | | | | | |
|---|---|---|---|---|---|---|
| mmu-miR-29b-3p | MIMAT0000127 | Gpbp1 | 73274 | HITS-CLIP//sRNA-Seq | 37986224\|25083871\|33184237\|23142 | Unspecified |
| mmu-miR-29b-3p | MIMAT0000127 | Gpbp1l1 | 77110 | HITS-CLIP | 25083871 | Unspecified |
| mmu-miR-29b-3p | MIMAT0000127 | Gpc1 | 14733 | HITS-CLIP//sRNA-Seq | 25083871\|33184237 | Unspecified |
| mmu-miR-29b-3p | MIMAT0000127 | Gpc4 | 14735 | HITS-CLIP//sRNA-Seq | 23597149\|33184237 | Unspecified |
| mmu-miR-29b-3p | MIMAT0000127 | Gpcpd1 | 74182 | HITS-CLIP | 37986224\|25083871\|23597149 | Unspecified |
| mmu-miR-29b-3p | MIMAT0000127 | Gpd1l | 333433 | HITS-CLIP | 23142080 | Unspecified |
| mmu-miR-29b-3p | MIMAT0000127 | Gpd2 | 14571 | HITS-CLIP | 23142080 | Unspecified |
| mmu-miR-29b-3p | MIMAT0000127 | Gphn | 268566 | HITS-CLIP | 37986224\|23597149 | Unspecified |
| mmu-miR-29b-3p | MIMAT0000127 | Gpi1 | 14751 | sRNA-Seq//HITS-CLIP | 33184237\|23142080 | Unspecified |
| mmu-miR-29b-3p | MIMAT0000127 | Gpkow | 209416 | HITS-CLIP | 37986224 | Unspecified |
| mmu-miR-29b-3p | MIMAT0000127 | Gpld1 | 14756 | HITS-CLIP | 23597149 | Unspecified |
| mmu-miR-29b-3p | MIMAT0000127 | Gpm6b | 14758 | HITS-CLIP | 23142080 | Unspecified |
| mmu-miR-29b-3p | MIMAT0000127 | Gpn2 | 100210 | HITS-CLIP | 25083871\|23142080 | Unspecified |
| mmu-miR-29b-3p | MIMAT0000127 | Gpr132 | 56696 | HITS-CLIP | 23142080 | Unspecified |
| mmu-miR-29b-3p | MIMAT0000127 | Gpr137 | 107173 | HITS-CLIP | 37986224\|25083871\|23597149 | Unspecified |
| mmu-miR-29b-3p | MIMAT0000127 | Gpr141 | 353346 | HITS-CLIP | 25083871 | Unspecified |
| mmu-miR-29b-3p | MIMAT0000127 | Gpr146 | 80290 | HITS-CLIP | 37986224\|25083871\|23597149\|23142 | Unspecified |
| mmu-miR-29b-3p | MIMAT0000127 | Gpr157 | 269604 | HITS-CLIP | 37986224\|23597149\|23142080 | Unspecified |
| mmu-miR-29b-3p | MIMAT0000127 | Gpr161 | 240888 | HITS-CLIP | 25083871 | Unspecified |
| mmu-miR-29b-3p | MIMAT0000127 | Gpr174 | 213439 | HITS-CLIP | 23142080 | Unspecified |
| mmu-miR-29b-3p | MIMAT0000127 | Gpr3 | 14748 | HITS-CLIP | 23142080 | Unspecified |
| mmu-miR-29b-3p | MIMAT0000127 | Gpr68 | 238377 | HITS-CLIP | 23142080 | Unspecified |
| mmu-miR-29b-3p | MIMAT0000127 | Gpr89 | 67549 | HITS-CLIP | 25083871 | Unspecified |
| mmu-miR-29b-3p | MIMAT0000127 | Gprc5c | 70355 | HITS-CLIP | 37986224\|25083871\|23597149 | Unspecified |
| mmu-miR-29b-3p | MIMAT0000127 | Gprin3 | 243385 | HITS-CLIP | 37986224\|23597149\|23142080 | Unspecified |
| mmu-miR-29b-3p | MIMAT0000127 | Gpt | 76282 | HITS-CLIP | 37986224\|23597149 | Unspecified |
| mmu-miR-29b-3p | MIMAT0000127 | Gpt2 | 108682 | HITS-CLIP//sRNA-Seq | 23597149\|33184237 | Unspecified |
| mmu-miR-29b-3p | MIMAT0000127 | Gpx1 | 14775 | sRNA-Seq//HITS-CLIP | 33184237\|23597149\|23142080 | Unspecified |
| mmu-miR-29b-3p | MIMAT0000127 | Gpx7 | 67305 | HITS-CLIP | 23597149 | Unspecified |
| mmu-miR-29b-3p | MIMAT0000127 | Gramd1b | 235283 | HITS-CLIP | 37986224\|25083871\|23142080 | Unspecified |
| mmu-miR-29b-3p | MIMAT0000127 | Gramd3 | 107022 | HITS-CLIP | 23142080\|23597149\|25083871 | Unspecified |
| mmu-miR-29b-3p | MIMAT0000127 | Gramd4 | 223752 | HITS-CLIP | 23142080 | Unspecified |
| mmu-miR-29b-3p | MIMAT0000127 | Grb14 | 50915 | HITS-CLIP//sRNA-Seq | 23597149\|33184237 | Unspecified |
| mmu-miR-29b-3p | MIMAT0000127 | Grcc10 | 14790 | HITS-CLIP//sRNA-Seq | 23597149\|25083871\|33184237\|23142 | Unspecified |
| mmu-miR-29b-3p | MIMAT0000127 | Grina | 66168 | HITS-CLIP//sRNA-Seq | 37986224\|23597149\|33184237\|25083 | Unspecified |
| mmu-miR-29b-3p | MIMAT0000127 | Grk2 | 156 | sRNA-Seq//HITS-CLIP | 33184237\|23142080 | Unspecified |
| mmu-miR-29b-3p | MIMAT0000127 | Grk6 | 26385 | HITS-CLIP//sRNA-Seq | 37986224\|23142080\|33184237\|25083 | Unspecified |
| mmu-miR-29b-3p | MIMAT0000127 | Grn | 14824 | HITS-CLIP | 37986224 | Unspecified |
| mmu-miR-29b-3p | MIMAT0000127 | Grpel2 | 17714 | HITS-CLIP | 37986224 | Unspecified |

| | | | | | | |
|---|---|---|---|---|---|---|
| mmu-miR-29b-3p | MIMAT0000127 | Grsf1 | 231413 | sRNA-Seq//HITS-CLIP | 33184237\|23142080 | Unspecified |
| mmu-miR-29b-3p | MIMAT0000127 | Gsap | 212167 | HITS-CLIP | 23142080 | Unspecified |
| mmu-miR-29b-3p | MIMAT0000127 | Gsdmd | 69146 | HITS-CLIP | 23142080 | Unspecified |
| mmu-miR-29b-3p | MIMAT0000127 | Gse1 | 382034 | HITS-CLIP | 25083871\|23142080 | Unspecified |
| mmu-miR-29b-3p | MIMAT0000127 | Gsk3b | 56637 | HITS-CLIP, Luciferase Reporter Assay//HITS | 37986224\|23597149\|33184237\|25083 | Unspecified |
| mmu-miR-29b-3p | MIMAT0000127 | Gsn | 227753 | HITS-CLIP//sRNA-Seq | 25083871\|33184237\|23142080 | Unspecified |
| mmu-miR-29b-3p | MIMAT0000127 | Gspt1 | 14852 | HITS-CLIP//sRNA-Seq | 25083871\|23142080\|33184237\|23597 | Unspecified |
| mmu-miR-29b-3p | MIMAT0000127 | Gss | 14854 | HITS-CLIP | 25083871\|23597149\|23142080 | Unspecified |
| mmu-miR-29b-3p | MIMAT0000127 | Gstcd | 67553 | HITS-CLIP | 25083871\|23142080 | Unspecified |
| mmu-miR-29b-3p | MIMAT0000127 | Gstm7 | 68312 | HITS-CLIP | 23597149 | Unspecified |
| mmu-miR-29b-3p | MIMAT0000127 | Gstz1 | 14874 | HITS-CLIP | 37986224\|23597149 | Unspecified |
| mmu-miR-29b-3p | MIMAT0000127 | Gtf2a1 | 83602 | sRNA-Seq//HITS-CLIP | 33184237\|25083871 | Unspecified |
| mmu-miR-29b-3p | MIMAT0000127 | Gtf2b | 229906 | HITS-CLIP//sRNA-Seq | 23597149\|33184237\|23142080 | Unspecified |
| mmu-miR-29b-3p | MIMAT0000127 | Gtf2e2 | 68153 | HITS-CLIP//sRNA-Seq | 37986224\|23142080\|25083871\|33184 | Unspecified |
| mmu-miR-29b-3p | MIMAT0000127 | Gtf2f1 | 98053 | HITS-CLIP//sRNA-Seq | 25083871\|33184237 | Unspecified |
| mmu-miR-29b-3p | MIMAT0000127 | Gtf2f2 | 68705 | HITS-CLIP | 23142080 | Unspecified |
| mmu-miR-29b-3p | MIMAT0000127 | Gtf2h1 | 14884 | HITS-CLIP | 25083871\|23142080 | Unspecified |
| mmu-miR-29b-3p | MIMAT0000127 | Gtf2h3 | 209357 | HITS-CLIP | 23597149 | Unspecified |
| mmu-miR-29b-3p | MIMAT0000127 | Gtf2h5 | 66467 | HITS-CLIP//sRNA-Seq | 23597149\|33184237\|23142080 | Unspecified |
| mmu-miR-29b-3p | MIMAT0000127 | Gtf2i | 14886 | sRNA-Seq//HITS-CLIP | 33184237\|23142080 | Unspecified |
| mmu-miR-29b-3p | MIMAT0000127 | Gtf3c1 | 233863 | HITS-CLIP//sRNA-Seq | 25083871\|23597149\|33184237\|23142 | Unspecified |
| mmu-miR-29b-3p | MIMAT0000127 | Gtf3c2 | 71752 | HITS-CLIP | 25083871\|23597149\|23142080 | Unspecified |
| mmu-miR-29b-3p | MIMAT0000127 | Gtf3c4 | 269252 | HITS-CLIP | 37986224\|23142080\|23597149 | Unspecified |
| mmu-miR-29b-3p | MIMAT0000127 | Gtf3c6 | 67371 | HITS-CLIP//sRNA-Seq | 23142080\|33184237 | Unspecified |
| mmu-miR-29b-3p | MIMAT0000127 | Gtpbp1 | 14904 | sRNA-Seq//HITS-CLIP | 33184237\|23142080 | Unspecified |
| mmu-miR-29b-3p | MIMAT0000127 | Gtpbp2 | 56055 | HITS-CLIP | 37986224\|23597149\|25083871\|23142 | Unspecified |
| mmu-miR-29b-3p | MIMAT0000127 | Gucd1 | 68778 | HITS-CLIP | 37986224\|23597149 | Unspecified |
| mmu-miR-29b-3p | MIMAT0000127 | Guf1 | 231279 | HITS-CLIP | 37986224 | Unspecified |
| mmu-miR-29b-3p | MIMAT0000127 | Gulo | 268756 | HITS-CLIP | 23597149 | Unspecified |
| mmu-miR-29b-3p | MIMAT0000127 | Gxylt2 | 232313 | HITS-CLIP//sRNA-Seq | 37986224\|25083871\|33184237 | Unspecified |
| mmu-miR-29b-3p | MIMAT0000127 | Gyg | 27357 | HITS-CLIP//sRNA-Seq | 25083871\|33184237\|23142080 | Unspecified |
| mmu-miR-29b-3p | MIMAT0000127 | Gys2 | 232493 | HITS-CLIP | 37986224\|23597149 | Unspecified |
| mmu-miR-29b-3p | MIMAT0000127 | Gzmc | 14940 | HITS-CLIP | 23142080 | Unspecified |
| mmu-miR-29b-3p | MIMAT0000127 | H1f0 | 14958 | HITS-CLIP//sRNA-Seq | 23142080\|25083871\|23597149\|33184 | Unspecified |
| mmu-miR-29b-3p | MIMAT0000127 | H2-Ab1 | 14961 | HITS-CLIP | 23142080 | Unspecified |
| mmu-miR-29b-3p | MIMAT0000127 | H2-D1 | 14964 | HITS-CLIP//sRNA-Seq | 23142080\|23597149\|33184237 | Unspecified |
| mmu-miR-29b-3p | MIMAT0000127 | H2-K1 | 14972 | HITS-CLIP//sRNA-Seq | 23597149\|33184237 | Unspecified |
| mmu-miR-29b-3p | MIMAT0000127 | H2-Q10 | 15007 | HITS-CLIP | 23597149 | Unspecified |
| mmu-miR-29b-3p | MIMAT0000127 | H2-Q4 | 15015 | sRNA-Seq//HITS-CLIP | 33184237\|23142080 | Unspecified |

| | | | | | | |
|---|---|---|---|---|---|---|
| mmu-miR-29b-3p | MIMAT0000127 | H2afj | 232440 | HITS-CLIP | 37986224\|23142080 | Unspecified |
| mmu-miR-29b-3p | MIMAT0000127 | H2afx | 15270 | HITS-CLIP | 37986224\|25083871\|23142080 | Unspecified |
| mmu-miR-29b-3p | MIMAT0000127 | H2afy | 9555 | HITS-CLIP//sRNA-Seq | 23142080\|33184237 | Unspecified |
| mmu-miR-29b-3p | MIMAT0000127 | H3f3b | 15081 | HITS-CLIP//sRNA-Seq | 25083871\|33184237\|23142080 | Unspecified |
| mmu-miR-29b-3p | MIMAT0000127 | H6pd | 100198 | HITS-CLIP | 37986224\|25083871\|23597149 | Unspecified |
| mmu-miR-29b-3p | MIMAT0000127 | Habp2 | 226243 | HITS-CLIP | 23597149 | Unspecified |
| mmu-miR-29b-3p | MIMAT0000127 | Habp4 | 56541 | HITS-CLIP | 37986224 | Unspecified |
| mmu-miR-29b-3p | MIMAT0000127 | Hacd3 | 57874 | sRNA-Seq//HITS-CLIP | 33184237\|23142080 | Unspecified |
| mmu-miR-29b-3p | MIMAT0000127 | Hacl1 | 56794 | HITS-CLIP | 37986224\|23597149 | Unspecified |
| mmu-miR-29b-3p | MIMAT0000127 | Hadh | 15107 | HITS-CLIP//sRNA-Seq | 25083871\|33184237\|23142080 | Unspecified |
| mmu-miR-29b-3p | MIMAT0000127 | Hadha | 97212 | HITS-CLIP//sRNA-Seq | 25083871\|23142080\|33184237\|23597 | Unspecified |
| mmu-miR-29b-3p | MIMAT0000127 | Hal | 15109 | HITS-CLIP | 23597149 | Unspecified |
| mmu-miR-29b-3p | MIMAT0000127 | Hao1 | 15112 | HITS-CLIP | 37986224 | Unspecified |
| mmu-miR-29b-3p | MIMAT0000127 | Hapln1 | 12950 | HITS-CLIP | 37986224\|23597149 | Unspecified |
| mmu-miR-29b-3p | MIMAT0000127 | Hars | 15115 | sRNA-Seq//HITS-CLIP | 33184237\|23142080 | Unspecified |
| mmu-miR-29b-3p | MIMAT0000127 | Hars2 | 70791 | HITS-CLIP | 23597149 | Unspecified |
| mmu-miR-29b-3p | MIMAT0000127 | Has2 | 15117 | HITS-CLIP | 37986224\|25083871 | Unspecified |
| mmu-miR-29b-3p | MIMAT0000127 | Hat1 | 107435 | sRNA-Seq//HITS-CLIP | 33184237\|23142080 | Unspecified |
| mmu-miR-29b-3p | MIMAT0000127 | Haus1 | 225745 | HITS-CLIP | 23142080 | Unspecified |
| mmu-miR-29b-3p | MIMAT0000127 | Haus3 | 231123 | HITS-CLIP | 37986224 | Unspecified |
| mmu-miR-29b-3p | MIMAT0000127 | Haus6 | 230376 | HITS-CLIP | 37986224\|25083871\|23142080 | Unspecified |
| mmu-miR-29b-3p | MIMAT0000127 | Hba-a1 | 15122 | HITS-CLIP | 23597149 | Unspecified |
| mmu-miR-29b-3p | MIMAT0000127 | Hbegf | 15200 | HITS-CLIP | 37986224 | Unspecified |
| mmu-miR-29b-3p | MIMAT0000127 | Hbp1 | 73389 | HITS-CLIP//sRNA-Seq | 37986224\|23142080\|25083871\|23597 | Unspecified |
| mmu-miR-29b-3p | MIMAT0000127 | Hbs1l | 56422 | HITS-CLIP//sRNA-Seq | 23142080\|25083871\|33184237 | Unspecified |
| mmu-miR-29b-3p | MIMAT0000127 | Hccs | 15159 | HITS-CLIP | 37986224\|25083871\|23597149\|23142 | Unspecified |
| mmu-miR-29b-3p | MIMAT0000127 | Hcfc1 | 15161 | HITS-CLIP//sRNA-Seq | 25083871\|33184237\|23142080 | Unspecified |
| mmu-miR-29b-3p | MIMAT0000127 | Hcfc2 | 67933 | HITS-CLIP | 37986224\|25083871\|23142080 | Unspecified |
| mmu-miR-29b-3p | MIMAT0000127 | Hcls1 | 15163 | HITS-CLIP | 23142080 | Unspecified |
| mmu-miR-29b-3p | MIMAT0000127 | Hdac10 | 170787 | HITS-CLIP | 23597149 | Unspecified |
| mmu-miR-29b-3p | MIMAT0000127 | Hdac2 | 15182 | HITS-CLIP//sRNA-Seq | 25083871\|33184237\|23142080 | Unspecified |
| mmu-miR-29b-3p | MIMAT0000127 | Hdac4 | 208727 | Luciferase assay//qRT-PCR//Western blot/ | 21324893\|25083871\|23142080\|19342 | Unspecified |
| mmu-miR-29b-3p | MIMAT0000127 | Hdac5 | 15184 | sRNA-Seq//HITS-CLIP | 33184237\|25083871 | Unspecified |
| mmu-miR-29b-3p | MIMAT0000127 | Hdac6 | 15185 | HITS-CLIP | 23142080 | Unspecified |
| mmu-miR-29b-3p | MIMAT0000127 | Hdgf | 15191 | HITS-CLIP//sRNA-Seq | 37986224\|23597149\|23142080\|33184 | Unspecified |
| mmu-miR-29b-3p | MIMAT0000127 | Hdgfl2 | 84717 | sRNA-Seq//HITS-CLIP | 33184237\|25083871 | Unspecified |
| mmu-miR-29b-3p | MIMAT0000127 | Hdhd3 | 72748 | HITS-CLIP | 23597149 | Unspecified |
| mmu-miR-29b-3p | MIMAT0000127 | Hdlbp | 110611 | HITS-CLIP//sRNA-Seq | 23142080\|25083871\|23597149\|33184 | Unspecified |
| mmu-miR-29b-3p | MIMAT0000127 | Heatr1 | 217995 | HITS-CLIP | 23597149 | Unspecified |

| | | | | | | |
|---|---|---|---|---|---|---|
| mmu-miR-29b-3p | MIMAT0000127 | Heatr5a | 320487 | HITS-CLIP | 25083871\|23597149\|23142080 | Unspecified |
| mmu-miR-29b-3p | MIMAT0000127 | Heatr6 | 217026 | HITS-CLIP | 23142080 | Unspecified |
| mmu-miR-29b-3p | MIMAT0000127 | Heca | 380629 | HITS-CLIP | 25083871\|23142080 | Unspecified |
| mmu-miR-29b-3p | MIMAT0000127 | Hectd1 | 207304 | HITS-CLIP//sRNA-Seq | 37986224\|23142080\|25083871\|33184 | Unspecified |
| mmu-miR-29b-3p | MIMAT0000127 | Hectd4 | 283450 | HITS-CLIP//sRNA-Seq | 25083871\|33184237\|23142080 | Unspecified |
| mmu-miR-29b-3p | MIMAT0000127 | Heg1 | 77446 | HITS-CLIP | 25083871\|23597149 | Unspecified |
| mmu-miR-29b-3p | MIMAT0000127 | Helb | 117599 | HITS-CLIP | 37986224\|23142080 | Unspecified |
| mmu-miR-29b-3p | MIMAT0000127 | Helq | 191578 | HITS-CLIP | 23142080 | Unspecified |
| mmu-miR-29b-3p | MIMAT0000127 | Helz | 78455 | HITS-CLIP | 37986224\|25083871\|23142080 | Unspecified |
| mmu-miR-29b-3p | MIMAT0000127 | Helz2 | 229003 | HITS-CLIP | 25083871\|23597149\|26203562 | Unspecified |
| mmu-miR-29b-3p | MIMAT0000127 | Hephl1 | 244698 | HITS-CLIP | 37986224\|25083871 | Unspecified |
| mmu-miR-29b-3p | MIMAT0000127 | Herc1 | 235439 | HITS-CLIP//sRNA-Seq | 37986224\|25083871\|33184237\|23142 | Unspecified |
| mmu-miR-29b-3p | MIMAT0000127 | Herc2 | 15204 | HITS-CLIP//sRNA-Seq | 23142080\|23597149\|31433990\|33184 | Unspecified |
| mmu-miR-29b-3p | MIMAT0000127 | Herc3 | 73998 | HITS-CLIP | 23597149\|23142080 | Unspecified |
| mmu-miR-29b-3p | MIMAT0000127 | Herc6 | 67138 | HITS-CLIP | 23142080 | Unspecified |
| mmu-miR-29b-3p | MIMAT0000127 | Herpud1 | 64209 | HITS-CLIP//sRNA-Seq | 23142080\|33184237\|23597149 | Unspecified |
| mmu-miR-29b-3p | MIMAT0000127 | Hexa | 15211 | sRNA-Seq//HITS-CLIP | 33184237\|23142080 | Unspecified |
| mmu-miR-29b-3p | MIMAT0000127 | Hexim1 | 192231 | HITS-CLIP | 25083871\|23142080 | Unspecified |
| mmu-miR-29b-3p | MIMAT0000127 | Heyl | 56198 | HITS-CLIP | 25083871 | Unspecified |
| mmu-miR-29b-3p | MIMAT0000127 | Hgd | 15233 | HITS-CLIP | 23597149 | Unspecified |
| mmu-miR-29b-3p | MIMAT0000127 | Hgf | 15234 | HITS-CLIP | 23597149 | Unspecified |
| mmu-miR-29b-3p | MIMAT0000127 | Hgfac | 54426 | HITS-CLIP | 23597149 | Unspecified |
| mmu-miR-29b-3p | MIMAT0000127 | Hgs | 15239 | HITS-CLIP | 23597149\|23142080 | Unspecified |
| mmu-miR-29b-3p | MIMAT0000127 | Hic2 | 58180 | HITS-CLIP | 37986224\|25083871\|23142080 | Unspecified |
| mmu-miR-29b-3p | MIMAT0000127 | Hif1a | 15251 | HITS-CLIP//sRNA-Seq | 37986224\|23597149\|23142080\|33184 | Unspecified |
| mmu-miR-29b-3p | MIMAT0000127 | Hif1an | 319594 | HITS-CLIP | 37986224\|25083871\|23597149\|23142 | Unspecified |
| mmu-miR-29b-3p | MIMAT0000127 | Hif3a | 53417 | HITS-CLIP | 21258322\|23142080\|25083871\|23597 | Unspecified |
| mmu-miR-29b-3p | MIMAT0000127 | Higd2a | 67044 | HITS-CLIP//sRNA-Seq | 23597149\|33184237 | Unspecified |
| mmu-miR-29b-3p | MIMAT0000127 | Hinfp | 102423 | HITS-CLIP | 37986224\|25083871 | Unspecified |
| mmu-miR-29b-3p | MIMAT0000127 | Hint1 | 15254 | sRNA-Seq//HITS-CLIP | 33184237\|23142080 | Unspecified |
| mmu-miR-29b-3p | MIMAT0000127 | Hip1 | 215114 | HITS-CLIP//sRNA-Seq | 37986224\|25083871\|33184237\|23142 | Unspecified |
| mmu-miR-29b-3p | MIMAT0000127 | Hipk1 | 15257 | HITS-CLIP | 23597149\|31433990\|23142080 | Unspecified |
| mmu-miR-29b-3p | MIMAT0000127 | Hipk3 | 15259 | HITS-CLIP | 37986224\|23597149\|23142080 | Unspecified |
| mmu-miR-29b-3p | MIMAT0000127 | Hist1h1a | 3024 | HITS-CLIP | 25083871\|23142080 | Unspecified |
| mmu-miR-29b-3p | MIMAT0000127 | Hist1h1b | 3009 | HITS-CLIP | 23142080 | Unspecified |
| mmu-miR-29b-3p | MIMAT0000127 | Hist1h1c | 50708 | HITS-CLIP//sRNA-Seq | 25083871\|23597149\|33184237\|23142 | Unspecified |
| mmu-miR-29b-3p | MIMAT0000127 | Hist1h1d | 14957 | HITS-CLIP | 23142080 | Unspecified |
| mmu-miR-29b-3p | MIMAT0000127 | Hist1h2bc | 3017 | HITS-CLIP | 23597149 | Unspecified |
| mmu-miR-29b-3p | MIMAT0000127 | Hist1h4d | 8360 | sRNA-Seq//HITS-CLIP | 33184237\|23597149\|23142080 | Unspecified |

| | | | | | | |
|---|---|---|---|---|---|---|
| mmu-miR-29b-3p | MIMAT0000127 | Hivep1 | 110521 | HITS-CLIP | 25083871 | Unspecified |
| mmu-miR-29b-3p | MIMAT0000127 | Hivep2 | 15273 | HITS-CLIP | 37986224\|23142080\|25083871 | Unspecified |
| mmu-miR-29b-3p | MIMAT0000127 | Hivep3 | 16656 | HITS-CLIP | 37986224\|23142080 | Unspecified |
| mmu-miR-29b-3p | MIMAT0000127 | Hk1 | 15275 | sRNA-Seq//HITS-CLIP | 33184237\|23142080 | Unspecified |
| mmu-miR-29b-3p | MIMAT0000127 | Hk2 | 15277 | HITS-CLIP | 23142080 | Unspecified |
| mmu-miR-29b-3p | MIMAT0000127 | Hltf | 20585 | HITS-CLIP | 23142080 | Unspecified |
| mmu-miR-29b-3p | MIMAT0000127 | Hlx | 15284 | HITS-CLIP | 23597149 | Unspecified |
| mmu-miR-29b-3p | MIMAT0000127 | Hmbox1 | 219150 | HITS-CLIP//sRNA-Seq | 37986224\|25083871\|33184237 | Unspecified |
| mmu-miR-29b-3p | MIMAT0000127 | Hmgb2 | 97165 | qCLASH | 31393953 | Unspecified |
| mmu-miR-29b-3p | MIMAT0000127 | Hmgcr | 15357 | HITS-CLIP//sRNA-Seq | 37986224\|23597149\|23142080\|33184 | Unspecified |
| mmu-miR-29b-3p | MIMAT0000127 | Hmgcs1 | 208715 | HITS-CLIP//sRNA-Seq | 37986224\|23142080\|23597149\|33184 | Unspecified |
| mmu-miR-29b-3p | MIMAT0000127 | Hmgcs2 | 15360 | HITS-CLIP | 23597149 | Unspecified |
| mmu-miR-29b-3p | MIMAT0000127 | Hmgn3 | 94353 | HITS-CLIP//sRNA-Seq | 37986224\|23142080\|25083871\|33184 | Unspecified |
| mmu-miR-29b-3p | MIMAT0000127 | Hmgxb4 | 70823 | HITS-CLIP | 25083871\|23142080 | Unspecified |
| mmu-miR-29b-3p | MIMAT0000127 | Hmmr | 15366 | HITS-CLIP | 37986224\|23597149\|23142080 | Unspecified |
| mmu-miR-29b-3p | MIMAT0000127 | Hnf4a | 15378 | HITS-CLIP | 37986224\|23597149 | Unspecified |
| mmu-miR-29b-3p | MIMAT0000127 | Hnrnpa2b1 | 53379 | sRNA-Seq//HITS-CLIP | 33184237\|23142080 | Unspecified |
| mmu-miR-29b-3p | MIMAT0000127 | Hnrnpa3 | 229279 | HITS-CLIP | 37986224 | Unspecified |
| mmu-miR-29b-3p | MIMAT0000127 | Hnrnpd | 11991 | HITS-CLIP//sRNA-Seq | 23142080\|33184237 | Unspecified |
| mmu-miR-29b-3p | MIMAT0000127 | Hnrnpdl | 50926 | HITS-CLIP//sRNA-Seq | 23142080\|25083871\|33184237\|23597 | Unspecified |
| mmu-miR-29b-3p | MIMAT0000127 | Hnrnpf | 98758 | HITS-CLIP | 37986224 | Unspecified |
| mmu-miR-29b-3p | MIMAT0000127 | Hnrnph1 | 59013 | HITS-CLIP//sRNA-Seq | 25083871\|33184237\|23142080 | Unspecified |
| mmu-miR-29b-3p | MIMAT0000127 | Hnrnpk | 15387 | sRNA-Seq//HITS-CLIP | 33184237\|23142080 | Unspecified |
| mmu-miR-29b-3p | MIMAT0000127 | Hnrnpl | 15388 | HITS-CLIP//sRNA-Seq | 23142080\|33184237 | Unspecified |
| mmu-miR-29b-3p | MIMAT0000127 | Hnrnpm | 76936 | HITS-CLIP//sRNA-Seq | 23142080\|33184237 | Unspecified |
| mmu-miR-29b-3p | MIMAT0000127 | Hnrnpr | 74326 | HITS-CLIP//sRNA-Seq | 25083871\|23142080\|33184237 | Unspecified |
| mmu-miR-29b-3p | MIMAT0000127 | Hnrnpu | 51810 | HITS-CLIP//sRNA-Seq | 25083871\|33184237\|23597149\|23142 | Unspecified |
| mmu-miR-29b-3p | MIMAT0000127 | Hnrnpul1 | 232989 | HITS-CLIP//sRNA-Seq | 37986224\|23142080\|33184237\|25083 | Unspecified |
| mmu-miR-29b-3p | MIMAT0000127 | Hnrnpul2 | 68693 | HITS-CLIP//sRNA-Seq | 37986224\|25083871\|33184237 | Unspecified |
| mmu-miR-29b-3p | MIMAT0000127 | Hoga1 | 67432 | HITS-CLIP | 37986224 | Unspecified |
| mmu-miR-29b-3p | MIMAT0000127 | Hook1 | 77963 | HITS-CLIP | 37986224\|23597149\|23142080 | Unspecified |
| mmu-miR-29b-3p | MIMAT0000127 | Hoxa11 | 15396 | HITS-CLIP | 25083871 | Unspecified |
| mmu-miR-29b-3p | MIMAT0000127 | Hoxa5 | 15402 | HITS-CLIP | 37986224 | Unspecified |
| mmu-miR-29b-3p | MIMAT0000127 | Hoxc11 | 109663 | HITS-CLIP | 25083871 | Unspecified |
| mmu-miR-29b-3p | MIMAT0000127 | Hoxc13 | 15422 | HITS-CLIP | 25083871 | Unspecified |
| mmu-miR-29b-3p | MIMAT0000127 | Hoxc8 | 15426 | HITS-CLIP | 25083871 | Unspecified |
| mmu-miR-29b-3p | MIMAT0000127 | Hp1bp3 | 15441 | HITS-CLIP//sRNA-Seq | 25083871\|33184237\|23142080 | Unspecified |
| mmu-miR-29b-3p | MIMAT0000127 | Hpf1 | 54969 | HITS-CLIP | 25083871\|23142080 | Unspecified |
| mmu-miR-29b-3p | MIMAT0000127 | Hpgd | 15446 | HITS-CLIP | 23597149 | Unspecified |

| | | | | | | |
|---|---|---|---|---|---|---|
| mmu-miR-29b-3p | MIMAT0000127 | Hpn | 15451 | HITS-CLIP | 23597149 | Unspecified |
| mmu-miR-29b-3p | MIMAT0000127 | Hps4 | 192232 | HITS-CLIP | 37986224\|25083871\|23597149 | Unspecified |
| mmu-miR-29b-3p | MIMAT0000127 | Hrh2 | 15466 | HITS-CLIP | 23142080 | Unspecified |
| mmu-miR-29b-3p | MIMAT0000127 | Hs1bp3 | 58240 | HITS-CLIP | 37986224 | Unspecified |
| mmu-miR-29b-3p | MIMAT0000127 | Hs2st1 | 23908 | HITS-CLIP | 25083871\|23142080 | Unspecified |
| mmu-miR-29b-3p | MIMAT0000127 | Hs3st1 | 15476 | HITS-CLIP | 23142080 | Unspecified |
| mmu-miR-29b-3p | MIMAT0000127 | Hs3st3b1 | 54710 | HITS-CLIP | 37986224\|25083871\|23597149\|23142 | Unspecified |
| mmu-miR-29b-3p | MIMAT0000127 | Hs6st1 | 50785 | HITS-CLIP//sRNA-Seq | 25083871\|23597149\|33184237 | Unspecified |
| mmu-miR-29b-3p | MIMAT0000127 | Hsd11b1 | 15483 | HITS-CLIP | 37986224\|23597149 | Unspecified |
| mmu-miR-29b-3p | MIMAT0000127 | Hsd17b12 | 56348 | HITS-CLIP//sRNA-Seq | 23597149\|25083871\|33184237\|23142 | Unspecified |
| mmu-miR-29b-3p | MIMAT0000127 | Hsd17b13 | 243168 | HITS-CLIP | 23597149 | Unspecified |
| mmu-miR-29b-3p | MIMAT0000127 | Hsd17b6 | 27400 | HITS-CLIP | 23597149 | Unspecified |
| mmu-miR-29b-3p | MIMAT0000127 | Hsd17b7 | 15490 | HITS-CLIP | 37986224 | Unspecified |
| mmu-miR-29b-3p | MIMAT0000127 | Hsd3b3 | 15494 | HITS-CLIP | 23597149 | Unspecified |
| mmu-miR-29b-3p | MIMAT0000127 | Hsf1 | 15499 | HITS-CLIP | 23142080 | Unspecified |
| mmu-miR-29b-3p | MIMAT0000127 | Hspa13 | 110920 | HITS-CLIP | 23597149 | Unspecified |
| mmu-miR-29b-3p | MIMAT0000127 | Hspa14 | 50497 | sRNA-Seq//HITS-CLIP | 33184237\|23142080 | Unspecified |
| mmu-miR-29b-3p | MIMAT0000127 | Hspa2 | 15512 | HITS-CLIP | 23142080\|25083871 | Unspecified |
| mmu-miR-29b-3p | MIMAT0000127 | Hspa4 | 15525 | HITS-CLIP//sRNA-Seq | 23142080\|25083871\|33184237 | Unspecified |
| mmu-miR-29b-3p | MIMAT0000127 | Hspa4l | 18415 | HITS-CLIP//sRNA-Seq | 23142080\|33184237\|25083871 | Unspecified |
| mmu-miR-29b-3p | MIMAT0000127 | Hspa5 | 14828 | HITS-CLIP//sRNA-Seq | 23597149\|33184237\|23142080 | Unspecified |
| mmu-miR-29b-3p | MIMAT0000127 | Hspa9 | 15526 | HITS-CLIP//sRNA-Seq | 23597149\|33184237\|23142080 | Unspecified |
| mmu-miR-29b-3p | MIMAT0000127 | Hspb2 | 69253 | HITS-CLIP | 25083871 | Unspecified |
| mmu-miR-29b-3p | MIMAT0000127 | Hspd1 | 15510 | HITS-CLIP//sRNA-Seq | 25083871\|23597149\|33184237 | Unspecified |
| mmu-miR-29b-3p | MIMAT0000127 | Hspg2 | 15530 | HITS-CLIP//sRNA-Seq | 37986224\|25083871\|33184237 | Unspecified |
| mmu-miR-29b-3p | MIMAT0000127 | Hsph1 | 15505 | sRNA-Seq//HITS-CLIP | 33184237\|23597149 | Unspecified |
| mmu-miR-29b-3p | MIMAT0000127 | Htatsf1 | 72459 | HITS-CLIP | 25083871\|23142080 | Unspecified |
| mmu-miR-29b-3p | MIMAT0000127 | Htr1b | 15551 | HITS-CLIP | 37986224 | Unspecified |
| mmu-miR-29b-3p | MIMAT0000127 | Htt | 15194 | HITS-CLIP | 23597149\|23142080 | Unspecified |
| mmu-miR-29b-3p | MIMAT0000127 | Hus1 | 15574 | HITS-CLIP | 23142080 | Unspecified |
| mmu-miR-29b-3p | MIMAT0000127 | Huwe1 | 59026 | HITS-CLIP//sRNA-Seq | 37986224\|25083871\|23597149\|33184 | Unspecified |
| mmu-miR-29b-3p | MIMAT0000127 | Hycc2 | 285172 | HITS-CLIP | 31433990 | Unspecified |
| mmu-miR-29b-3p | MIMAT0000127 | Hykk | 235386 | HITS-CLIP | 23597149 | Unspecified |
| mmu-miR-29b-3p | MIMAT0000127 | Hyls1 | 76832 | HITS-CLIP | 25083871\|23142080 | Unspecified |
| mmu-miR-29b-3p | MIMAT0000127 | Hyou1 | 12282 | HITS-CLIP//sRNA-Seq | 23142080\|25083871\|33184237\|23597 | Unspecified |
| mmu-miR-29b-3p | MIMAT0000127 | Hypk | 67693 | HITS-CLIP | 25083871\|23597149\|23142080 | Unspecified |
| mmu-miR-29b-3p | MIMAT0000127 | Iars | 105148 | HITS-CLIP//sRNA-Seq | 25083871\|23142080\|33184237 | Unspecified |
| mmu-miR-29b-3p | MIMAT0000127 | Iars2 | 381314 | HITS-CLIP | 23597149 | Unspecified |
| mmu-miR-29b-3p | MIMAT0000127 | Ibtk | 108837 | HITS-CLIP | 25083871\|23142080 | Unspecified |

| | | | | | | |
|---|---|---|---|---|---|---|
| mmu-miR-29b-3p | MIMAT0000127 | Ice1 | 822287 | sRNA-Seq//HITS-CLIP | 33184237\|23142080 | Unspecified |
| mmu-miR-29b-3p | MIMAT0000127 | Ick | 56542 | HITS-CLIP | 37986224 | Unspecified |
| mmu-miR-29b-3p | MIMAT0000127 | Icmt | 57295 | HITS-CLIP | 25083871\|23597149\|23142080 | Unspecified |
| mmu-miR-29b-3p | MIMAT0000127 | Icos | 54167 | HITS-CLIP | 37986224\|23142080 | Unspecified |
| mmu-miR-29b-3p | MIMAT0000127 | Id2 | 15902 | HITS-CLIP | 23142080\|23597149 | Unspecified |
| mmu-miR-29b-3p | MIMAT0000127 | Ide | 15925 | HITS-CLIP | 23142080 | Unspecified |
| mmu-miR-29b-3p | MIMAT0000127 | Idh1 | 15926 | HITS-CLIP//sRNA-Seq | 25083871\|33184237\|23597149 | Unspecified |
| mmu-miR-29b-3p | MIMAT0000127 | Idh2 | 269951 | sRNA-Seq//HITS-CLIP | 33184237\|23142080 | Unspecified |
| mmu-miR-29b-3p | MIMAT0000127 | Idh3b | 170718 | HITS-CLIP//sRNA-Seq | 25083871\|23597149\|33184237 | Unspecified |
| mmu-miR-29b-3p | MIMAT0000127 | Idh3g | 15929 | HITS-CLIP//sRNA-Seq | 23597149\|33184237\|23142080 | Unspecified |
| mmu-miR-29b-3p | MIMAT0000127 | Ido2 | 209176 | HITS-CLIP | 37986224\|23597149 | Unspecified |
| mmu-miR-29b-3p | MIMAT0000127 | Ier2 | 15936 | HITS-CLIP | 37986224 | Unspecified |
| mmu-miR-29b-3p | MIMAT0000127 | Ier3 | 15937 | sRNA-Seq//HITS-CLIP | 33184237\|25083871\|23142080 | Unspecified |
| mmu-miR-29b-3p | MIMAT0000127 | Ier5 | 15939 | HITS-CLIP | 23142080 | Unspecified |
| mmu-miR-29b-3p | MIMAT0000127 | Iffo1 | 320678 | HITS-CLIP | 37986224\|25083871 | Unspecified |
| mmu-miR-29b-3p | MIMAT0000127 | Iffo2 | 212632 | HITS-CLIP | 37986224\|25083871\|23142080 | Unspecified |
| mmu-miR-29b-3p | MIMAT0000127 | Ifi27l2a | 76933 | HITS-CLIP | 23142080 | Unspecified |
| mmu-miR-29b-3p | MIMAT0000127 | Ifi30 | 65972 | HITS-CLIP//qCLASH | 37986224\|31393953 | Unspecified |
| mmu-miR-29b-3p | MIMAT0000127 | Ifng | 15978 | HITS-CLIP | 23142080 | Unspecified |
| mmu-miR-29b-3p | MIMAT0000127 | Ifngr2 | 15980 | HITS-CLIP | 37986224\|23597149\|25083871\|23142 | Unspecified |
| mmu-miR-29b-3p | MIMAT0000127 | Ifrd1 | 15982 | sRNA-Seq//HITS-CLIP | 33184237\|23142080 | Unspecified |
| mmu-miR-29b-3p | MIMAT0000127 | Ift172 | 67661 | HITS-CLIP | 23142080 | Unspecified |
| mmu-miR-29b-3p | MIMAT0000127 | Ift80 | 68259 | HITS-CLIP | 23142080 | Unspecified |
| mmu-miR-29b-3p | MIMAT0000127 | Igdcc4 | 56741 | HITS-CLIP | 25083871 | Unspecified |
| mmu-miR-29b-3p | MIMAT0000127 | Igf1 | 16000 | ELISA//Luciferase assay//qRT-PCR//HITS-C | 23880139\|25083871\|23597149 | Unspecified |
| mmu-miR-29b-3p | MIMAT0000127 | Igf1r | 16001 | HITS-CLIP | 25083871 | Unspecified |
| mmu-miR-29b-3p | MIMAT0000127 | Igf2r | 16004 | HITS-CLIP//sRNA-Seq | 23597149\|23142080\|25083871\|33184 | Unspecified |
| mmu-miR-29b-3p | MIMAT0000127 | Igfals | 16005 | HITS-CLIP | 23597149 | Unspecified |
| mmu-miR-29b-3p | MIMAT0000127 | Igfbp4 | 16010 | HITS-CLIP//sRNA-Seq | 23597149\|33184237 | Unspecified |
| mmu-miR-29b-3p | MIMAT0000127 | Igfbp6 | 16012 | sRNA-Seq//HITS-CLIP | 33184237\|25083871 | Unspecified |
| mmu-miR-29b-3p | MIMAT0000127 | Igfbp7 | 29817 | HITS-CLIP | 25083871\|23597149 | Unspecified |
| mmu-miR-29b-3p | MIMAT0000127 | Igsf3 | 78908 | HITS-CLIP//sRNA-Seq | 37986224\|25083871\|33184237\|23142 | Unspecified |
| mmu-miR-29b-3p | MIMAT0000127 | Igsf8 | 140559 | HITS-CLIP | 37986224 | Unspecified |
| mmu-miR-29b-3p | MIMAT0000127 | Igtp | 16145 | sRNA-Seq//HITS-CLIP | 33184237\|23597149\|23142080 | Unspecified |
| mmu-miR-29b-3p | MIMAT0000127 | Iigp1 | 60440 | HITS-CLIP | 23597149\|23142080 | Unspecified |
| mmu-miR-29b-3p | MIMAT0000127 | Ik | 24010 | HITS-CLIP//sRNA-Seq | 23142080\|33184237 | Unspecified |
| mmu-miR-29b-3p | MIMAT0000127 | Ikbkb | 16150 | HITS-CLIP | 23597149 | Unspecified |
| mmu-miR-29b-3p | MIMAT0000127 | Ikbke | 56489 | HITS-CLIP | 23142080 | Unspecified |
| mmu-miR-29b-3p | MIMAT0000127 | Ikbkg | 16151 | HITS-CLIP | 23597149 | Unspecified |

| | | | | | | |
|---|---|---|---|---|---|---|
| mmu-miR-29b-3p | MIMAT0000127 | Ikzf1 | 22778 | HITS-CLIP | 37986224\|23142080 | Unspecified |
| mmu-miR-29b-3p | MIMAT0000127 | Ikzf2 | 22779 | HITS-CLIP | 25083871\|23142080 | Unspecified |
| mmu-miR-29b-3p | MIMAT0000127 | Il12rb2 | 16162 | HITS-CLIP | 23142080 | Unspecified |
| mmu-miR-29b-3p | MIMAT0000127 | Il13ra1 | 16164 | HITS-CLIP | 25083871 | Unspecified |
| mmu-miR-29b-3p | MIMAT0000127 | Il15ra | 16169 | HITS-CLIP | 23142080 | Unspecified |
| mmu-miR-29b-3p | MIMAT0000127 | Il16 | 16170 | HITS-CLIP | 23142080 | Unspecified |
| mmu-miR-29b-3p | MIMAT0000127 | Il17ra | 16172 | HITS-CLIP | 23597149\|25083871 | Unspecified |
| mmu-miR-29b-3p | MIMAT0000127 | Il1r1 | 16177 | HITS-CLIP | 37986224 | Unspecified |
| mmu-miR-29b-3p | MIMAT0000127 | Il1rap | 16180 | HITS-CLIP | 37986224\|23597149 | Unspecified |
| mmu-miR-29b-3p | MIMAT0000127 | Il1rl2 | 107527 | HITS-CLIP | 23142080 | Unspecified |
| mmu-miR-29b-3p | MIMAT0000127 | Il21 | 60505 | HITS-CLIP | 37986224\|23142080 | Unspecified |
| mmu-miR-29b-3p | MIMAT0000127 | Il21r | 60504 | HITS-CLIP | 23142080 | Unspecified |
| mmu-miR-29b-3p | MIMAT0000127 | Il2ra | 16184 | HITS-CLIP | 37986224\|31433990\|23142080 | Unspecified |
| mmu-miR-29b-3p | MIMAT0000127 | Il2rb | 16185 | HITS-CLIP | 31433990 | Unspecified |
| mmu-miR-29b-3p | MIMAT0000127 | Il2rg | 16186 | HITS-CLIP | 23142080 | Unspecified |
| mmu-miR-29b-3p | MIMAT0000127 | Il3ra | 16188 | HITS-CLIP | 23597149 | Unspecified |
| mmu-miR-29b-3p | MIMAT0000127 | Il4ra | 16190 | HITS-CLIP | 23142080 | Unspecified |
| mmu-miR-29b-3p | MIMAT0000127 | Il5ra | 16192 | HITS-CLIP | 23142080 | Unspecified |
| mmu-miR-29b-3p | MIMAT0000127 | Il6 | 16193 | HITS-CLIP | 25083871 | Unspecified |
| mmu-miR-29b-3p | MIMAT0000127 | Il6ra | 16194 | HITS-CLIP | 23597149\|25083871 | Unspecified |
| mmu-miR-29b-3p | MIMAT0000127 | Il6st | 16195 | HITS-CLIP//sRNA-Seq | 37986224\|23142080\|25083871\|33184 | Unspecified |
| mmu-miR-29b-3p | MIMAT0000127 | Il7r | 16197 | HITS-CLIP | 23142080 | Unspecified |
| mmu-miR-29b-3p | MIMAT0000127 | Ildr2 | 100039795 | HITS-CLIP | 23597149 | Unspecified |
| mmu-miR-29b-3p | MIMAT0000127 | Ilf3 | 16201 | HITS-CLIP//sRNA-Seq | 23142080\|33184237 | Unspecified |
| mmu-miR-29b-3p | MIMAT0000127 | Ilk | 16202 | HITS-CLIP | 37986224 | Unspecified |
| mmu-miR-29b-3p | MIMAT0000127 | Ilkap | 67444 | HITS-CLIP | 25083871 | Unspecified |
| mmu-miR-29b-3p | MIMAT0000127 | Imp3 | 102462 | sRNA-Seq//HITS-CLIP | 33184237\|23142080 | Unspecified |
| mmu-miR-29b-3p | MIMAT0000127 | Impad1 | 242291 | HITS-CLIP | 23597149\|23142080 | Unspecified |
| mmu-miR-29b-3p | MIMAT0000127 | Impdh1 | 23917 | HITS-CLIP//sRNA-Seq | 37986224\|23142080\|33184237 | Unspecified |
| mmu-miR-29b-3p | MIMAT0000127 | Inava | 55765 | HITS-CLIP | 25083871 | Unspecified |
| mmu-miR-29b-3p | MIMAT0000127 | Inf2 | 70435 | sRNA-Seq//HITS-CLIP | 33184237\|23142080 | Unspecified |
| mmu-miR-29b-3p | MIMAT0000127 | Ing3 | 71777 | HITS-CLIP | 25083871\|23142080 | Unspecified |
| mmu-miR-29b-3p | MIMAT0000127 | Ing4 | 28019 | HITS-CLIP | 37986224\|25083871\|23142080 | Unspecified |
| mmu-miR-29b-3p | MIMAT0000127 | Inhba | 16323 | HITS-CLIP | 25083871\|23597149\|23142080 | Unspecified |
| mmu-miR-29b-3p | MIMAT0000127 | Ino80d | 227195 | HITS-CLIP | 37986224\|25083871\|23142080 | Unspecified |
| mmu-miR-29b-3p | MIMAT0000127 | Inpp4a | 269180 | HITS-CLIP//sRNA-Seq | 23597149\|33184237\|23142080 | Unspecified |
| mmu-miR-29b-3p | MIMAT0000127 | Inpp4b | 234515 | HITS-CLIP | 25083871\|23142080 | Unspecified |
| mmu-miR-29b-3p | MIMAT0000127 | Inpp5a | 212111 | HITS-CLIP | 37986224\|25083871\|23142080 | Unspecified |
| mmu-miR-29b-3p | MIMAT0000127 | Inpp5b | 16330 | HITS-CLIP | 23142080 | Unspecified |

| | | | | | | | |
|---|---|---|---|---|---|---|---|
| mmu-miR-29b-3p | MIMAT0000127 | Inpp5d | 16331 | HITS-CLIP | | 23142080 | Unspecified |
| mmu-miR-29b-3p | MIMAT0000127 | Inpp5f | 101490 | HITS-CLIP//sRNA-Seq | 23597149\|33184237\|23142080 | | Unspecified |
| mmu-miR-29b-3p | MIMAT0000127 | Insig1 | 231070 | reporter_gene//sRNA-Seq//HITS-CLIP//Re | 37986224\|33184237\|23597149\|17652 | | Unspecified |
| mmu-miR-29b-3p | MIMAT0000127 | Insig2 | 72999 | HITS-CLIP | | 23597149 | Unspecified |
| mmu-miR-29b-3p | MIMAT0000127 | Insr | 16337 | HITS-CLIP | 25083871\|23597149\|23142080 | | Unspecified |
| mmu-miR-29b-3p | MIMAT0000127 | Ints10 | 70885 | HITS-CLIP | | 23142080 | Unspecified |
| mmu-miR-29b-3p | MIMAT0000127 | Ints11 | 54973 | sRNA-Seq//HITS-CLIP | 33184237\|23142080 | | Unspecified |
| mmu-miR-29b-3p | MIMAT0000127 | Ints12 | 71793 | HITS-CLIP | | 23142080 | Unspecified |
| mmu-miR-29b-3p | MIMAT0000127 | Ints13 | 55726 | HITS-CLIP | | 25083871 | Unspecified |
| mmu-miR-29b-3p | MIMAT0000127 | Ints2 | 70422 | HITS-CLIP | 25083871\|23142080 | | Unspecified |
| mmu-miR-29b-3p | MIMAT0000127 | Ints3 | 229543 | sRNA-Seq//HITS-CLIP | 33184237\|23142080 | | Unspecified |
| mmu-miR-29b-3p | MIMAT0000127 | Ints4 | 101861 | HITS-CLIP | | 23142080 | Unspecified |
| mmu-miR-29b-3p | MIMAT0000127 | Ints5 | 80789 | HITS-CLIP | | 23142080 | Unspecified |
| mmu-miR-29b-3p | MIMAT0000127 | Ints6 | 18130 | HITS-CLIP | | 23142080 | Unspecified |
| mmu-miR-29b-3p | MIMAT0000127 | Ints6l | 236790 | HITS-CLIP | | 23142080 | Unspecified |
| mmu-miR-29b-3p | MIMAT0000127 | Ints7 | 77065 | HITS-CLIP | 37986224\|23142080 | | Unspecified |
| mmu-miR-29b-3p | MIMAT0000127 | Ip6k1 | 27399 | sRNA-Seq//HITS-CLIP | 33184237\|23142080 | | Unspecified |
| mmu-miR-29b-3p | MIMAT0000127 | Ip6k2 | 76500 | HITS-CLIP//sRNA-Seq | 23597149\|33184237\|23142080 | | Unspecified |
| mmu-miR-29b-3p | MIMAT0000127 | Ip6k3 | 271424 | HITS-CLIP | | 25083871 | Unspecified |
| mmu-miR-29b-3p | MIMAT0000127 | Ipcef1 | 320495 | HITS-CLIP | | 23142080 | Unspecified |
| mmu-miR-29b-3p | MIMAT0000127 | Ipmk | 69718 | HITS-CLIP | 37986224\|23142080 | | Unspecified |
| mmu-miR-29b-3p | MIMAT0000127 | Ipo13 | 230673 | HITS-CLIP | 37986224\|25083871\|23597149 | | Unspecified |
| mmu-miR-29b-3p | MIMAT0000127 | Ipo4 | 75751 | HITS-CLIP//sRNA-Seq | 23597149\|33184237 | | Unspecified |
| mmu-miR-29b-3p | MIMAT0000127 | Ipo5 | 70572 | HITS-CLIP//sRNA-Seq | 25083871\|23597149\|33184237\|23142 | | Unspecified |
| mmu-miR-29b-3p | MIMAT0000127 | Ipo7 | 233726 | HITS-CLIP//sRNA-Seq | 25083871\|33184237\|23142080 | | Unspecified |
| mmu-miR-29b-3p | MIMAT0000127 | Ipo9 | 226432 | HITS-CLIP//sRNA-Seq | 37986224\|25083871\|33184237\|23597 | | Unspecified |
| mmu-miR-29b-3p | MIMAT0000127 | Ipp | 16351 | HITS-CLIP | 25083871\|23597149 | | Unspecified |
| mmu-miR-29b-3p | MIMAT0000127 | Ippk | 75678 | HITS-CLIP | 31433990\|25083871 | | Unspecified |
| mmu-miR-29b-3p | MIMAT0000127 | Iqcb1 | 320299 | HITS-CLIP | | 23142080 | Unspecified |
| mmu-miR-29b-3p | MIMAT0000127 | Iqgap2 | 544963 | HITS-CLIP | | 23597149 | Unspecified |
| mmu-miR-29b-3p | MIMAT0000127 | Iqschfp | 100505386 | HITS-CLIP | | 25083871 | Unspecified |
| mmu-miR-29b-3p | MIMAT0000127 | Irak3 | 73914 | HITS-CLIP | | 23142080 | Unspecified |
| mmu-miR-29b-3p | MIMAT0000127 | Ireb2 | 64602 | HITS-CLIP//sRNA-Seq | 37986224\|25083871\|23597149\|33184 | | Unspecified |
| mmu-miR-29b-3p | MIMAT0000127 | Irf1 | 16362 | HITS-CLIP//sRNA-Seq | 25083871\|33184237\|23597149\|23142 | | Unspecified |
| mmu-miR-29b-3p | MIMAT0000127 | Irf2bp1 | 272359 | HITS-CLIP | | 23597149 | Unspecified |
| mmu-miR-29b-3p | MIMAT0000127 | Irf2bp2 | 270110 | HITS-CLIP | 25083871\|23597149\|23142080 | | Unspecified |
| mmu-miR-29b-3p | MIMAT0000127 | Irf4 | 16364 | HITS-CLIP | 23142080\|31433990 | | Unspecified |
| mmu-miR-29b-3p | MIMAT0000127 | Irf8 | 15900 | HITS-CLIP | | 23142080 | Unspecified |
| mmu-miR-29b-3p | MIMAT0000127 | Irgm1 | 15944 | HITS-CLIP//sRNA-Seq | 23597149\|33184237 | | Unspecified |

| | | | | | | |
|---|---|---|---|---|---|---|
| mmu-miR-29b-3p | MIMAT0000127 | Irgm2 | 54396 | HITS-CLIP//sRNA-Seq | 23142080\|33184237 | Unspecified |
| mmu-miR-29b-3p | MIMAT0000127 | Irs1 | 16367 | HITS-CLIP | 37986224\|25083871\|23597149 | Unspecified |
| mmu-miR-29b-3p | MIMAT0000127 | Irs2 | 384783 | HITS-CLIP | 23597149\|23142080 | Unspecified |
| mmu-miR-29b-3p | MIMAT0000127 | Isg20l2 | 229504 | HITS-CLIP | 37986224 | Unspecified |
| mmu-miR-29b-3p | MIMAT0000127 | Islr | 26968 | HITS-CLIP | 37986224\|25083871 | Unspecified |
| mmu-miR-29b-3p | MIMAT0000127 | Isoc2a | 664994 | HITS-CLIP | 23597149 | Unspecified |
| mmu-miR-29b-3p | MIMAT0000127 | Ist1 | 71955 | HITS-CLIP//sRNA-Seq | 25083871\|33184237\|23142080 | Unspecified |
| mmu-miR-29b-3p | MIMAT0000127 | Itch | 16396 | HITS-CLIP | 25083871\|23597149\|23142080 | Unspecified |
| mmu-miR-29b-3p | MIMAT0000127 | Itfg1 | 71927 | HITS-CLIP//sRNA-Seq | 25083871\|33184237\|23142080 | Unspecified |
| mmu-miR-29b-3p | MIMAT0000127 | Itfg2 | 101142 | HITS-CLIP | 23142080 | Unspecified |
| mmu-miR-29b-3p | MIMAT0000127 | Itga3 | 16400 | HITS-CLIP | 25083871\|23142080 | Unspecified |
| mmu-miR-29b-3p | MIMAT0000127 | Itga4 | 16401 | HITS-CLIP | 23142080 | Unspecified |
| mmu-miR-29b-3p | MIMAT0000127 | Itga5 | 16402 | HITS-CLIP//sRNA-Seq | 23597149\|23142080\|25083871\|3318 | Unspecified |
| mmu-miR-29b-3p | MIMAT0000127 | Itga6 | 16403 | sRNA-Seq//HITS-CLIP | 33184237\|23142080 | Unspecified |
| mmu-miR-29b-3p | MIMAT0000127 | Itga7 | 16404 | HITS-CLIP//sRNA-Seq | 23142080\|33184237\|25083871 | Unspecified |
| mmu-miR-29b-3p | MIMAT0000127 | Itgal | 16408 | HITS-CLIP | 23142080 | Unspecified |
| mmu-miR-29b-3p | MIMAT0000127 | Itgav | 16410 | HITS-CLIP//sRNA-Seq | 25083871\|33184237\|23142080 | Unspecified |
| mmu-miR-29b-3p | MIMAT0000127 | Itgb1 | 16412 | HITS-CLIP//sRNA-Seq | 37986224\|25083871\|33184237\|2314 | Unspecified |
| mmu-miR-29b-3p | MIMAT0000127 | Itgb1bp1 | 16413 | HITS-CLIP | 23142080 | Unspecified |
| mmu-miR-29b-3p | MIMAT0000127 | Itgb2 | 16414 | HITS-CLIP | 23142080 | Unspecified |
| mmu-miR-29b-3p | MIMAT0000127 | Itgb3 | 16416 | HITS-CLIP | 37986224\|23142080\|25083871 | Unspecified |
| mmu-miR-29b-3p | MIMAT0000127 | Itgb5 | 16419 | HITS-CLIP//sRNA-Seq | 25083871\|33184237 | Unspecified |
| mmu-miR-29b-3p | MIMAT0000127 | Itgb7 | 16421 | HITS-CLIP | 37986224\|23142080 | Unspecified |
| mmu-miR-29b-3p | MIMAT0000127 | Itgb8 | 320910 | HITS-CLIP | 23142080 | Unspecified |
| mmu-miR-29b-3p | MIMAT0000127 | Itih1 | 16424 | HITS-CLIP | 23597149 | Unspecified |
| mmu-miR-29b-3p | MIMAT0000127 | Itih2 | 16425 | HITS-CLIP | 23597149 | Unspecified |
| mmu-miR-29b-3p | MIMAT0000127 | Itih3 | 16426 | HITS-CLIP | 23597149 | Unspecified |
| mmu-miR-29b-3p | MIMAT0000127 | Itih4 | 16427 | HITS-CLIP | 23597149 | Unspecified |
| mmu-miR-29b-3p | MIMAT0000127 | Itk | 16428 | HITS-CLIP | 23142080 | Unspecified |
| mmu-miR-29b-3p | MIMAT0000127 | Itm2a | 16431 | HITS-CLIP | 23142080 | Unspecified |
| mmu-miR-29b-3p | MIMAT0000127 | Itm2b | 16432 | HITS-CLIP//sRNA-Seq | 23597149\|33184237\|23142080 | Unspecified |
| mmu-miR-29b-3p | MIMAT0000127 | Itm2c | 64294 | sRNA-Seq//HITS-CLIP | 33184237\|23142080 | Unspecified |
| mmu-miR-29b-3p | MIMAT0000127 | Itpk1 | 217837 | HITS-CLIP | 23597149\|23142080 | Unspecified |
| mmu-miR-29b-3p | MIMAT0000127 | Itpkb | 320404 | HITS-CLIP | 23142080 | Unspecified |
| mmu-miR-29b-3p | MIMAT0000127 | Itpr1 | 16438 | HITS-CLIP | 23597149\|23142080 | Unspecified |
| mmu-miR-29b-3p | MIMAT0000127 | Itpr2 | 16439 | HITS-CLIP | 37986224\|23142080 | Unspecified |
| mmu-miR-29b-3p | MIMAT0000127 | Itpr3 | 16440 | HITS-CLIP | 37986224\|25083871\|23142080 | Unspecified |
| mmu-miR-29b-3p | MIMAT0000127 | Itpripl2 | 319622 | HITS-CLIP | 23142080\|25083871 | Unspecified |
| mmu-miR-29b-3p | MIMAT0000127 | Itsn1 | 16443 | HITS-CLIP//sRNA-Seq | 25083871\|33184237\|23597149 | Unspecified |

| | | | | | | |
|---|---|---|---|---|---|---|
| mmu-miR-29b-3p | MIMAT0000127 | Itsn2 | 20403 | HITS-CLIP | 37986224\|23142080\|23597149\|25083 | Unspecified |
| mmu-miR-29b-3p | MIMAT0000127 | Ivd | 56357 | HITS-CLIP | 23597149 | Unspecified |
| mmu-miR-29b-3p | MIMAT0000127 | Ivns1abp | 117198 | HITS-CLIP//sRNA-Seq | 37986224\|25083871\|23597149\|33184 | Unspecified |
| mmu-miR-29b-3p | MIMAT0000127 | Iyd | 70337 | HITS-CLIP | 37986224\|23597149 | Unspecified |
| mmu-miR-29b-3p | MIMAT0000127 | Izumo1r | 64931 | HITS-CLIP | 23142080 | Unspecified |
| mmu-miR-29b-3p | MIMAT0000127 | Jade1 | 269424 | HITS-CLIP | 23142080 | Unspecified |
| mmu-miR-29b-3p | MIMAT0000127 | Jade2 | 76901 | HITS-CLIP | 23597149 | Unspecified |
| mmu-miR-29b-3p | MIMAT0000127 | Jagn1 | 67767 | HITS-CLIP | 37986224\|25083871\|23597149\|23142 | Unspecified |
| mmu-miR-29b-3p | MIMAT0000127 | Jak1 | 16451 | HITS-CLIP//sRNA-Seq | 25083871\|33184237\|23597149\|23142 | Unspecified |
| mmu-miR-29b-3p | MIMAT0000127 | Jak2 | 16452 | HITS-CLIP | 37986224\|25083871\|23597149\|23142 | Unspecified |
| mmu-miR-29b-3p | MIMAT0000127 | Jak3 | 16453 | HITS-CLIP | 23142080 | Unspecified |
| mmu-miR-29b-3p | MIMAT0000127 | Jam2 | 67374 | HITS-CLIP | 25083871 | Unspecified |
| mmu-miR-29b-3p | MIMAT0000127 | Jaml | 120425 | HITS-CLIP | 23142080 | Unspecified |
| mmu-miR-29b-3p | MIMAT0000127 | Jarid2 | 16468 | HITS-CLIP//sRNA-Seq | 37986224\|25083871\|23142080\|33184 | Unspecified |
| mmu-miR-29b-3p | MIMAT0000127 | Jazf1 | 231986 | HITS-CLIP | 37986224\|23142080 | Unspecified |
| mmu-miR-29b-3p | MIMAT0000127 | Jcad | 57608 | HITS-CLIP | 25083871 | Unspecified |
| mmu-miR-29b-3p | MIMAT0000127 | Jchain | 16069 | HITS-CLIP | 23142080 | Unspecified |
| mmu-miR-29b-3p | MIMAT0000127 | Jdp2 | 81703 | HITS-CLIP//sRNA-Seq | 25083871\|23142080\|23597149\|33184 | Unspecified |
| mmu-miR-29b-3p | MIMAT0000127 | Jmjd1c | 108829 | HITS-CLIP//sRNA-Seq | 37986224\|25083871\|23142080\|33184 | Unspecified |
| mmu-miR-29b-3p | MIMAT0000127 | Jmjd6 | 107817 | HITS-CLIP | 37986224\|25083871 | Unspecified |
| mmu-miR-29b-3p | MIMAT0000127 | Jmy | 57748 | HITS-CLIP | 37986224\|25083871\|23142080 | Unspecified |
| mmu-miR-29b-3p | MIMAT0000127 | Josd1 | 74158 | HITS-CLIP//sRNA-Seq | 37986224\|23597149\|33184237\|23142 | Unspecified |
| mmu-miR-29b-3p | MIMAT0000127 | Jph2 | 59091 | HITS-CLIP | 25083871 | Unspecified |
| mmu-miR-29b-3p | MIMAT0000127 | Jpt2 | 90861 | sRNA-Seq//HITS-CLIP | 33184237\|23142080 | Unspecified |
| mmu-miR-29b-3p | MIMAT0000127 | Jrkl | 77532 | HITS-CLIP | 37986224\|23142080 | Unspecified |
| mmu-miR-29b-3p | MIMAT0000127 | Jup | 16480 | HITS-CLIP//sRNA-Seq | 37986224\|23597149\|33184237\|25083 | Unspecified |
| mmu-miR-29b-3p | MIMAT0000127 | Kansl1 | 76719 | HITS-CLIP | 23142080 | Unspecified |
| mmu-miR-29b-3p | MIMAT0000127 | Kansl3 | 226976 | HITS-CLIP | 25083871 | Unspecified |
| mmu-miR-29b-3p | MIMAT0000127 | Kat2b | 18519 | HITS-CLIP | 37986224\|23142080\|25083871 | Unspecified |
| mmu-miR-29b-3p | MIMAT0000127 | Kat6a | 244349 | HITS-CLIP | 25083871\|23142080 | Unspecified |
| mmu-miR-29b-3p | MIMAT0000127 | Kat6b | 54169 | HITS-CLIP | 37986224\|23142080\|25083871 | Unspecified |
| mmu-miR-29b-3p | MIMAT0000127 | Kat7 | 217127 | sRNA-Seq//HITS-CLIP | 33184237\|23142080 | Unspecified |
| mmu-miR-29b-3p | MIMAT0000127 | Katna1 | 23924 | HITS-CLIP//sRNA-Seq | 23142080\|33184237 | Unspecified |
| mmu-miR-29b-3p | MIMAT0000127 | Katnal1 | 231912 | HITS-CLIP | 25083871 | Unspecified |
| mmu-miR-29b-3p | MIMAT0000127 | Katnbl1 | 72425 | HITS-CLIP | 25083871 | Unspecified |
| mmu-miR-29b-3p | MIMAT0000127 | Kbtbd11 | 74901 | HITS-CLIP | 23142080\|28735896 | Unspecified |
| mmu-miR-29b-3p | MIMAT0000127 | Kbtbd2 | 210973 | HITS-CLIP | 23142080 | Unspecified |
| mmu-miR-29b-3p | MIMAT0000127 | Kbtbd3 | 69149 | HITS-CLIP | 23142080 | Unspecified |
| mmu-miR-29b-3p | MIMAT0000127 | Kbtbd8 | 243574 | HITS-CLIP | 25083871 | Unspecified |

| | | | | | | |
|---|---|---|---|---|---|---|
| mmu-miR-29b-3p | MIMAT0000127 | Kcmf1 | 74287 | HITS-CLIP | 37986224\|23142080 | Unspecified |
| mmu-miR-29b-3p | MIMAT0000127 | Kcna3 | 16491 | HITS-CLIP | 23142080 | Unspecified |
| mmu-miR-29b-3p | MIMAT0000127 | Kcnip2 | 80906 | HITS-CLIP | 23142080 | Unspecified |
| mmu-miR-29b-3p | MIMAT0000127 | Kcnj2 | 16518 | HITS-CLIP | 37986224\|25083871 | Unspecified |
| mmu-miR-29b-3p | MIMAT0000127 | Kcnk5 | 16529 | HITS-CLIP | 23597149 | Unspecified |
| mmu-miR-29b-3p | MIMAT0000127 | Kcnk7 | 16530 | HITS-CLIP | 23142080 | Unspecified |
| mmu-miR-29b-3p | MIMAT0000127 | Kcnn3 | 140493 | HITS-CLIP | 37986224\|25083871 | Unspecified |
| mmu-miR-29b-3p | MIMAT0000127 | Kcnn4 | 16534 | HITS-CLIP | 23142080 | Unspecified |
| mmu-miR-29b-3p | MIMAT0000127 | Kcnq4 | 60613 | HITS-CLIP | 25083871 | Unspecified |
| mmu-miR-29b-3p | MIMAT0000127 | Kctd11 | 216858 | HITS-CLIP | 25083871\|23142080 | Unspecified |
| mmu-miR-29b-3p | MIMAT0000127 | Kctd12 | 239217 | HITS-CLIP | 23142080 | Unspecified |
| mmu-miR-29b-3p | MIMAT0000127 | Kctd15 | 233107 | HITS-CLIP | 37986224\|25083871 | Unspecified |
| mmu-miR-29b-3p | MIMAT0000127 | Kctd17 | 72844 | HITS-CLIP//sRNA-Seq | 25083871\|23142080\|33184237\|23597 | Unspecified |
| mmu-miR-29b-3p | MIMAT0000127 | Kctd18 | 51960 | HITS-CLIP | 23142080 | Unspecified |
| mmu-miR-29b-3p | MIMAT0000127 | Kctd2 | 70382 | HITS-CLIP | 25083871 | Unspecified |
| mmu-miR-29b-3p | MIMAT0000127 | Kctd20 | 66989 | HITS-CLIP//sRNA-Seq | 37986224\|33184237\|25083871 | Unspecified |
| mmu-miR-29b-3p | MIMAT0000127 | Kctd21 | 622320 | HITS-CLIP | 37986224\|25083871 | Unspecified |
| mmu-miR-29b-3p | MIMAT0000127 | Kctd3 | 226823 | HITS-CLIP | 37986224\|23597149\|23142080 | Unspecified |
| mmu-miR-29b-3p | MIMAT0000127 | Kctd5 | 69259 | HITS-CLIP//sRNA-Seq | 37986224\|23142080\|33184237\|2508 | Unspecified |
| mmu-miR-29b-3p | MIMAT0000127 | Kctd6 | 71393 | HITS-CLIP | 23597149 | Unspecified |
| mmu-miR-29b-3p | MIMAT0000127 | Kdelc1 | 79070 | HITS-CLIP | 25083871 | Unspecified |
| mmu-miR-29b-3p | MIMAT0000127 | Kdelr1 | 68137 | HITS-CLIP//sRNA-Seq | 23597149\|33184237\|25083871\|2314 | Unspecified |
| mmu-miR-29b-3p | MIMAT0000127 | Kdelr2 | 66913 | HITS-CLIP//sRNA-Seq | 25083871\|33184237\|23597149\|2314 | Unspecified |
| mmu-miR-29b-3p | MIMAT0000127 | Kdelr3 | 105785 | HITS-CLIP//sRNA-Seq | 37986224\|25083871\|33184237 | Unspecified |
| mmu-miR-29b-3p | MIMAT0000127 | Kdm1a | 99982 | HITS-CLIP//sRNA-Seq | 25083871\|33184237\|23142080 | Unspecified |
| mmu-miR-29b-3p | MIMAT0000127 | Kdm2a | 225876 | HITS-CLIP//sRNA-Seq | 25083871\|23597149\|33184237\|2314 | Unspecified |
| mmu-miR-29b-3p | MIMAT0000127 | Kdm3a | 104263 | HITS-CLIP | 25083871\|23142080 | Unspecified |
| mmu-miR-29b-3p | MIMAT0000127 | Kdm3b | 277250 | HITS-CLIP | 37986224 | Unspecified |
| mmu-miR-29b-3p | MIMAT0000127 | Kdm4b | 193796 | HITS-CLIP | 37986224 | Unspecified |
| mmu-miR-29b-3p | MIMAT0000127 | Kdm4c | 76804 | HITS-CLIP | 25083871 | Unspecified |
| mmu-miR-29b-3p | MIMAT0000127 | Kdm5a | 214899 | HITS-CLIP | 37986224\|23597149\|25083871\|2314 | Unspecified |
| mmu-miR-29b-3p | MIMAT0000127 | Kdm5b | 75605 | HITS-CLIP//sRNA-Seq | 37986224\|25083871\|33184237\|2314 | Unspecified |
| mmu-miR-29b-3p | MIMAT0000127 | Kdm5c | 20591 | HITS-CLIP//sRNA-Seq | 37986224\|25083871\|33184237\|23597 | Unspecified |
| mmu-miR-29b-3p | MIMAT0000127 | Kdm6a | 22289 | HITS-CLIP | 37986224\|25083871\|23597149\|2314 | Unspecified |
| mmu-miR-29b-3p | MIMAT0000127 | Kdm6b | 216850 | HITS-CLIP | 37986224\|25083871\|23597149\|2314 | Unspecified |
| mmu-miR-29b-3p | MIMAT0000127 | Kdm7a | 338523 | HITS-CLIP | 23142080\|25083871 | Unspecified |
| mmu-miR-29b-3p | MIMAT0000127 | Kdr | 16542 | HITS-CLIP | 23597149 | Unspecified |
| mmu-miR-29b-3p | MIMAT0000127 | Kdsr | 70750 | HITS-CLIP | 25083871\|23142080 | Unspecified |
| mmu-miR-29b-3p | MIMAT0000127 | Keap1 | 50868 | HITS-CLIP | 25083871 | Unspecified |

| | | | | | | |
|---|---|---|---|---|---|---|
| mmu-miR-29b-3p | MIMAT0000127 | Keg1 | 64697 | HITS-CLIP | 23597149 | Unspecified |
| mmu-miR-29b-3p | MIMAT0000127 | Khdrbs1 | 20218 | sRNA-Seq//HITS-CLIP | 33184237\|23142080 | Unspecified |
| mmu-miR-29b-3p | MIMAT0000127 | Khk | 16548 | HITS-CLIP | 23597149 | Unspecified |
| mmu-miR-29b-3p | MIMAT0000127 | Khnyn | 219094 | HITS-CLIP | 23597149\|25083871\|23142080 | Unspecified |
| mmu-miR-29b-3p | MIMAT0000127 | Khsrp | 16549 | sRNA-Seq//HITS-CLIP | 33184237\|25083871 | Unspecified |
| mmu-miR-29b-3p | MIMAT0000127 | Kidins220 | 77480 | HITS-CLIP//sRNA-Seq | 25083871\|33184237\|23142080 | Unspecified |
| mmu-miR-29b-3p | MIMAT0000127 | Kif11 | 16551 | HITS-CLIP//sRNA-Seq | 37986224\|33184237\|23142080 | Unspecified |
| mmu-miR-29b-3p | MIMAT0000127 | Kif13b | 16554 | HITS-CLIP | 23597149 | Unspecified |
| mmu-miR-29b-3p | MIMAT0000127 | Kif15 | 209737 | HITS-CLIP | 23142080 | Unspecified |
| mmu-miR-29b-3p | MIMAT0000127 | Kif16b | 16558 | HITS-CLIP | 23597149 | Unspecified |
| mmu-miR-29b-3p | MIMAT0000127 | Kif18a | 228421 | HITS-CLIP | 23142080 | Unspecified |
| mmu-miR-29b-3p | MIMAT0000127 | Kif1b | 16561 | HITS-CLIP//sRNA-Seq | 25083871\|23597149\|33184237 | Unspecified |
| mmu-miR-29b-3p | MIMAT0000127 | Kif1bp | 26128 | HITS-CLIP//sRNA-Seq | 25083871\|33184237 | Unspecified |
| mmu-miR-29b-3p | MIMAT0000127 | Kif1c | 16562 | HITS-CLIP//sRNA-Seq | 25083871\|33184237 | Unspecified |
| mmu-miR-29b-3p | MIMAT0000127 | Kif20a | 19348 | HITS-CLIP//sRNA-Seq | 23142080\|33184237\|30971389\|2508: | Unspecified |
| mmu-miR-29b-3p | MIMAT0000127 | Kif20b | 240641 | HITS-CLIP//qCLASH | 37986224\|31393953\|23142080 | Unspecified |
| mmu-miR-29b-3p | MIMAT0000127 | Kif21a | 16564 | HITS-CLIP//sRNA-Seq | 25083871\|33184237 | Unspecified |
| mmu-miR-29b-3p | MIMAT0000127 | Kif21b | 16565 | HITS-CLIP | 25083871 | Unspecified |
| mmu-miR-29b-3p | MIMAT0000127 | Kif22 | 110033 | HITS-CLIP | 37986224\|23142080 | Unspecified |
| mmu-miR-29b-3p | MIMAT0000127 | Kif26b | 269152 | HITS-CLIP | 37986224\|25083871 | Unspecified |
| mmu-miR-29b-3p | MIMAT0000127 | Kif2a | 16563 | sRNA-Seq//HITS-CLIP | 33184237\|23142080 | Unspecified |
| mmu-miR-29b-3p | MIMAT0000127 | Kif2c | 73804 | sRNA-Seq//HITS-CLIP | 33184237\|23142080 | Unspecified |
| mmu-miR-29b-3p | MIMAT0000127 | Kif3b | 16569 | HITS-CLIP | 37986224\|25083871 | Unspecified |
| mmu-miR-29b-3p | MIMAT0000127 | Kif3c | 16570 | HITS-CLIP//sRNA-Seq | 25083871\|23142080\|33184237 | Unspecified |
| mmu-miR-29b-3p | MIMAT0000127 | Kif5b | 16573 | HITS-CLIP//sRNA-Seq | 23597149\|33184237\|23142080 | Unspecified |
| mmu-miR-29b-3p | MIMAT0000127 | Kif7 | 16576 | HITS-CLIP | 37986224\|25083871\|23597149 | Unspecified |
| mmu-miR-29b-3p | MIMAT0000127 | Kifap3 | 16579 | HITS-CLIP//sRNA-Seq | 25083871\|33184237\|23142080 | Unspecified |
| mmu-miR-29b-3p | MIMAT0000127 | Kin | 16588 | HITS-CLIP | 25083871\|23142080 | Unspecified |
| mmu-miR-29b-3p | MIMAT0000127 | Kirrel | 170643 | HITS-CLIP | 37986224 | Unspecified |
| mmu-miR-29b-3p | MIMAT0000127 | Klc2 | 16594 | HITS-CLIP//sRNA-Seq | 25083871\|33184237\|23142080 | Unspecified |
| mmu-miR-29b-3p | MIMAT0000127 | Klf13 | 50794 | HITS-CLIP | 23142080 | Unspecified |
| mmu-miR-29b-3p | MIMAT0000127 | Klf15 | 66277 | HITS-CLIP | 23597149 | Unspecified |
| mmu-miR-29b-3p | MIMAT0000127 | Klf2 | 16598 | HITS-CLIP | 25083871\|23597149\|23142080 | Unspecified |
| mmu-miR-29b-3p | MIMAT0000127 | Klf3 | 16599 | HITS-CLIP//sRNA-Seq | 25083871\|33184237 | Unspecified |
| mmu-miR-29b-3p | MIMAT0000127 | Klf4 | 16600 | HITS-CLIP | 25083871\|23597149\|30971389\|2314: | Unspecified |
| mmu-miR-29b-3p | MIMAT0000127 | Klf5 | 12224 | HITS-CLIP | 37986224 | Unspecified |
| mmu-miR-29b-3p | MIMAT0000127 | Klf6 | 23849 | HITS-CLIP//sRNA-Seq | 23597149\|23142080\|25083871\|3318- | Unspecified |
| mmu-miR-29b-3p | MIMAT0000127 | Klf7 | 93691 | HITS-CLIP//sRNA-Seq | 37986224\|25083871\|33184237\|2314: | Unspecified |
| mmu-miR-29b-3p | MIMAT0000127 | Klf9 | 16601 | HITS-CLIP | 37986224\|23597149\|23142080 | Unspecified |

| | | | | | | |
|---|---|---|---|---|---|---|
| mmu-miR-29b-3p | MIMAT0000127 | Klhdc10 | 76788 | HITS-CLIP//sRNA-Seq | 25083871\|23597149\|33184237\|23142 | Unspecified |
| mmu-miR-29b-3p | MIMAT0000127 | Klhdc2 | 69554 | HITS-CLIP//sRNA-Seq | 25083871\|33184237 | Unspecified |
| mmu-miR-29b-3p | MIMAT0000127 | Klhdc3 | 71765 | HITS-CLIP//sRNA-Seq | 25083871\|23597149\|33184237 | Unspecified |
| mmu-miR-29b-3p | MIMAT0000127 | Klhdc4 | 234825 | HITS-CLIP | 23597149 | Unspecified |
| mmu-miR-29b-3p | MIMAT0000127 | Klhl11 | 217194 | HITS-CLIP | 23142080 | Unspecified |
| mmu-miR-29b-3p | MIMAT0000127 | Klhl12 | 240756 | HITS-CLIP | 23142080 | Unspecified |
| mmu-miR-29b-3p | MIMAT0000127 | Klhl15 | 236904 | HITS-CLIP | 25083871\|23142080 | Unspecified |
| mmu-miR-29b-3p | MIMAT0000127 | Klhl18 | 270201 | HITS-CLIP | 37986224 | Unspecified |
| mmu-miR-29b-3p | MIMAT0000127 | Klhl2 | 77113 | HITS-CLIP | 23142080\|23597149\|25083871 | Unspecified |
| mmu-miR-29b-3p | MIMAT0000127 | Klhl20 | 226541 | HITS-CLIP | 25083871\|23597149\|23142080 | Unspecified |
| mmu-miR-29b-3p | MIMAT0000127 | Klhl21 | 242785 | HITS-CLIP | 23142080 | Unspecified |
| mmu-miR-29b-3p | MIMAT0000127 | Klhl22 | 224023 | sRNA-Seq//HITS-CLIP | 33184237\|23597149 | Unspecified |
| mmu-miR-29b-3p | MIMAT0000127 | Klhl24 | 75785 | HITS-CLIP | 23597149\|23142080\|25083871 | Unspecified |
| mmu-miR-29b-3p | MIMAT0000127 | Klhl25 | 207952 | HITS-CLIP | 37986224\|25083871\|23142080 | Unspecified |
| mmu-miR-29b-3p | MIMAT0000127 | Klhl28 | 66689 | HITS-CLIP | 37986224\|23142080\|23597149\|2508: | Unspecified |
| mmu-miR-29b-3p | MIMAT0000127 | Klhl3 | 100503085 | HITS-CLIP | 23597149 | Unspecified |
| mmu-miR-29b-3p | MIMAT0000127 | Klhl42 | 232539 | HITS-CLIP | 37986224\|23142080 | Unspecified |
| mmu-miR-29b-3p | MIMAT0000127 | Klhl5 | 71778 | HITS-CLIP | 23597149 | Unspecified |
| mmu-miR-29b-3p | MIMAT0000127 | Klhl6 | 239743 | HITS-CLIP | 23142080 | Unspecified |
| mmu-miR-29b-3p | MIMAT0000127 | Klhl8 | 246293 | HITS-CLIP | 37986224\|25083871 | Unspecified |
| mmu-miR-29b-3p | MIMAT0000127 | Klhl9 | 242521 | sRNA-Seq//HITS-CLIP | 33184237\|23142080 | Unspecified |
| mmu-miR-29b-3p | MIMAT0000127 | Kmo | 98256 | HITS-CLIP | 23597149 | Unspecified |
| mmu-miR-29b-3p | MIMAT0000127 | Kmt2a | 214162 | HITS-CLIP | 37986224\|23142080 | Unspecified |
| mmu-miR-29b-3p | MIMAT0000127 | Kmt2b | 75410 | HITS-CLIP | 37986224\|23597149\|23142080\|2508: | Unspecified |
| mmu-miR-29b-3p | MIMAT0000127 | Kmt2c | 231051 | HITS-CLIP | 25083871\|23142080 | Unspecified |
| mmu-miR-29b-3p | MIMAT0000127 | Kmt2d | 381022 | HITS-CLIP | 37986224\|23142080\|25083871 | Unspecified |
| mmu-miR-29b-3p | MIMAT0000127 | Kmt2e | 69188 | HITS-CLIP | 25083871\|23597149\|23142080 | Unspecified |
| mmu-miR-29b-3p | MIMAT0000127 | Kmt5a | 387893 | sRNA-Seq//HITS-CLIP | 33184237\|23142080 | Unspecified |
| mmu-miR-29b-3p | MIMAT0000127 | Kmt5b | 225888 | HITS-CLIP | 25083871\|23142080 | Unspecified |
| mmu-miR-29b-3p | MIMAT0000127 | Kmt5c | 84787 | HITS-CLIP | 25083871\|23142080 | Unspecified |
| mmu-miR-29b-3p | MIMAT0000127 | Knl1 | 57082 | HITS-CLIP | 25083871\|23142080 | Unspecified |
| mmu-miR-29b-3p | MIMAT0000127 | Knop1 | 66356 | HITS-CLIP | 25083871\|23142080 | Unspecified |
| mmu-miR-29b-3p | MIMAT0000127 | Kpna1 | 16646 | HITS-CLIP//sRNA-Seq | 37986224\|25083871\|33184237\|23142 | Unspecified |
| mmu-miR-29b-3p | MIMAT0000127 | Kpna3 | 16648 | HITS-CLIP//sRNA-Seq | 23142080\|25083871\|33184237 | Unspecified |
| mmu-miR-29b-3p | MIMAT0000127 | Kpna4 | 16649 | HITS-CLIP | 37986224\|25083871\|23142080 | Unspecified |
| mmu-miR-29b-3p | MIMAT0000127 | Kpna6 | 16650 | HITS-CLIP//sRNA-Seq | 37986224\|25083871\|33184237\|23142 | Unspecified |
| mmu-miR-29b-3p | MIMAT0000127 | Kpnb1 | 16211 | HITS-CLIP//sRNA-Seq | 37986224\|23597149\|23142080\|33184 | Unspecified |
| mmu-miR-29b-3p | MIMAT0000127 | Kras | 16653 | HITS-CLIP//sRNA-Seq | 25083871\|33184237\|23142080 | Unspecified |
| mmu-miR-29b-3p | MIMAT0000127 | Kremen1 | 84035 | HITS-CLIP | 25083871 | Unspecified |

| | | | | | | |
|---|---|---|---|---|---|---|
| mmu-miR-29b-3p | MIMAT0000127 | Kremen2 | 73016 | HITS-CLIP | 23142080 | Unspecified |
| mmu-miR-29b-3p | MIMAT0000127 | Krit1 | 79264 | HITS-CLIP | 23142080 | Unspecified |
| mmu-miR-29b-3p | MIMAT0000127 | Krr1 | 52705 | HITS-CLIP | 23142080\|25083871 | Unspecified |
| mmu-miR-29b-3p | MIMAT0000127 | Krt6a | 3853 | HITS-CLIP | 26203562 | Unspecified |
| mmu-miR-29b-3p | MIMAT0000127 | Krt80 | 74127 | HITS-CLIP | 25083871 | Unspecified |
| mmu-miR-29b-3p | MIMAT0000127 | Ksr1 | 16706 | HITS-CLIP | 37986224 | Unspecified |
| mmu-miR-29b-3p | MIMAT0000127 | Ksr2 | 333050 | HITS-CLIP | 23597149 | Unspecified |
| mmu-miR-29b-3p | MIMAT0000127 | Ktn1 | 16709 | HITS-CLIP//sRNA-Seq | 25083871\|33184237 | Unspecified |
| mmu-miR-29b-3p | MIMAT0000127 | Ky | 16716 | HITS-CLIP | 23142080 | Unspecified |
| mmu-miR-29b-3p | MIMAT0000127 | Kyat1 | 883 | HITS-CLIP | 23597149 | Unspecified |
| mmu-miR-29b-3p | MIMAT0000127 | Kyat3 | 56267 | HITS-CLIP | 23597149\|23142080 | Unspecified |
| mmu-miR-29b-3p | MIMAT0000127 | Kynu | 70789 | HITS-CLIP | 23597149 | Unspecified |
| mmu-miR-29b-3p | MIMAT0000127 | Lactb2 | 212442 | HITS-CLIP | 21258322\|23597149 | Unspecified |
| mmu-miR-29b-3p | MIMAT0000127 | Lage3 | 66192 | HITS-CLIP | 23142080 | Unspecified |
| mmu-miR-29b-3p | MIMAT0000127 | Lama5 | 16776 | HITS-CLIP | 37986224\|25083871 | Unspecified |
| mmu-miR-29b-3p | MIMAT0000127 | Lamb2 | 16779 | HITS-CLIP//sRNA-Seq | 25083871\|33184237 | Unspecified |
| mmu-miR-29b-3p | MIMAT0000127 | Lamc1 | 226519 | HITS-CLIP//sRNA-Seq | 37986224\|23597149\|33184237\|23142 | Unspecified |
| mmu-miR-29b-3p | MIMAT0000127 | Lamp1 | 16783 | HITS-CLIP//sRNA-Seq | 23597149\|25083871\|33184237\|23142 | Unspecified |
| mmu-miR-29b-3p | MIMAT0000127 | Lamp2 | 16784 | sRNA-Seq//HITS-CLIP | 33184237\|23597149\|23142080 | Unspecified |
| mmu-miR-29b-3p | MIMAT0000127 | Lamtor1 | 66508 | HITS-CLIP//sRNA-Seq | 37986224\|25083871\|33184237 | Unspecified |
| mmu-miR-29b-3p | MIMAT0000127 | Lamtor4 | 66096 | sRNA-Seq//HITS-CLIP | 33184237\|23142080 | Unspecified |
| mmu-miR-29b-3p | MIMAT0000127 | Lancl2 | 71835 | HITS-CLIP | 37986224\|23142080 | Unspecified |
| mmu-miR-29b-3p | MIMAT0000127 | Lancl3 | 236285 | HITS-CLIP | 23142080 | Unspecified |
| mmu-miR-29b-3p | MIMAT0000127 | Lap3 | 66988 | sRNA-Seq//HITS-CLIP | 33184237\|23597149 | Unspecified |
| mmu-miR-29b-3p | MIMAT0000127 | Laptm4a | 17775 | HITS-CLIP//sRNA-Seq | 23142080\|33184237 | Unspecified |
| mmu-miR-29b-3p | MIMAT0000127 | Laptm5 | 16792 | HITS-CLIP | 23142080 | Unspecified |
| mmu-miR-29b-3p | MIMAT0000127 | Larp1 | 73158 | HITS-CLIP//sRNA-Seq | 25083871\|23142080\|23597149\|33184 | Unspecified |
| mmu-miR-29b-3p | MIMAT0000127 | Larp4 | 207214 | HITS-CLIP | 25083871\|23597149\|23142080 | Unspecified |
| mmu-miR-29b-3p | MIMAT0000127 | Larp4b | 217980 | HITS-CLIP//sRNA-Seq | 37986224\|25083871\|23597149\|33184 | Unspecified |
| mmu-miR-29b-3p | MIMAT0000127 | Larp7 | 28036 | sRNA-Seq//HITS-CLIP | 33184237\|23142080 | Unspecified |
| mmu-miR-29b-3p | MIMAT0000127 | Lars | 107045 | HITS-CLIP//sRNA-Seq | 25083871\|33184237\|23142080 | Unspecified |
| mmu-miR-29b-3p | MIMAT0000127 | Lasp1 | 16796 | HITS-CLIP//sRNA-Seq | 37986224\|33184237\|23142080 | Unspecified |
| mmu-miR-29b-3p | MIMAT0000127 | Lats1 | 16798 | HITS-CLIP | 23142080 | Unspecified |
| mmu-miR-29b-3p | MIMAT0000127 | Lbp | 16803 | HITS-CLIP | 23597149 | Unspecified |
| mmu-miR-29b-3p | MIMAT0000127 | Lbr | 98386 | HITS-CLIP | 37986224\|23142080 | Unspecified |
| mmu-miR-29b-3p | MIMAT0000127 | Lcmt1 | 30949 | HITS-CLIP | 23597149 | Unspecified |
| mmu-miR-29b-3p | MIMAT0000127 | Lcor | 212391 | HITS-CLIP//sRNA-Seq | 25083871\|33184237 | Unspecified |
| mmu-miR-29b-3p | MIMAT0000127 | Lcorl | 209707 | HITS-CLIP | 37986224\|23142080 | Unspecified |
| mmu-miR-29b-3p | MIMAT0000127 | Lcp1 | 18826 | HITS-CLIP | 37986224\|23142080\|23597149 | Unspecified |

| | | | | | | |
|---|---|---|---|---|---|---|
| mmu-miR-29b-3p | MIMAT0000127 | Lcp2 | 16822 | HITS-CLIP | 23142080 | Unspecified |
| mmu-miR-29b-3p | MIMAT0000127 | Ldah | 38150 | HITS-CLIP | 23597149 | Unspecified |
| mmu-miR-29b-3p | MIMAT0000127 | Ldha | 16828 | HITS-CLIP//sRNA-Seq | 23142080\|23597149\|33184237\|25083 | Unspecified |
| mmu-miR-29b-3p | MIMAT0000127 | Ldhd | 52815 | HITS-CLIP | 23597149 | Unspecified |
| mmu-miR-29b-3p | MIMAT0000127 | Ldlr | 16835 | HITS-CLIP//sRNA-Seq | 23597149\|33184237\|23142080 | Unspecified |
| mmu-miR-29b-3p | MIMAT0000127 | Ldlrad3 | 241576 | HITS-CLIP | 37986224\|25083871 | Unspecified |
| mmu-miR-29b-3p | MIMAT0000127 | Ldlrad4 | 52662 | HITS-CLIP | 25083871 | Unspecified |
| mmu-miR-29b-3p | MIMAT0000127 | Ldlrap1 | 100017 | HITS-CLIP | 37986224\|25083871\|23142080 | Unspecified |
| mmu-miR-29b-3p | MIMAT0000127 | Lemd2 | 224640 | HITS-CLIP | 37986224 | Unspecified |
| mmu-miR-29b-3p | MIMAT0000127 | Lemd3 | 380664 | HITS-CLIP | 25083871 | Unspecified |
| mmu-miR-29b-3p | MIMAT0000127 | Leng8 | 232798 | HITS-CLIP | 37986224\|23597149 | Unspecified |
| mmu-miR-29b-3p | MIMAT0000127 | Leng9 | 243813 | HITS-CLIP | 23597149 | Unspecified |
| mmu-miR-29b-3p | MIMAT0000127 | Letm1 | 56384 | HITS-CLIP | 23597149\|25083871 | Unspecified |
| mmu-miR-29b-3p | MIMAT0000127 | Letmd1 | 68614 | HITS-CLIP | 25083871\|23597149 | Unspecified |
| mmu-miR-29b-3p | MIMAT0000127 | Lfng | 16848 | HITS-CLIP | 37986224\|23597149\|25083871\|23142 | Unspecified |
| mmu-miR-29b-3p | MIMAT0000127 | Lgals8 | 56048 | HITS-CLIP//sRNA-Seq | 37986224\|25083871\|23597149\|33184 | Unspecified |
| mmu-miR-29b-3p | MIMAT0000127 | Lgalsl | 216551 | HITS-CLIP | 23142080 | Unspecified |
| mmu-miR-29b-3p | MIMAT0000127 | Lgmn | 19141 | HITS-CLIP//sRNA-Seq | 23597149\|33184237 | Unspecified |
| mmu-miR-29b-3p | MIMAT0000127 | Lgr4 | 107515 | HITS-CLIP | 25083871 | Unspecified |
| mmu-miR-29b-3p | MIMAT0000127 | Lhfp | 108927 | HITS-CLIP | 37986224 | Unspecified |
| mmu-miR-29b-3p | MIMAT0000127 | Lhfpl2 | 218454 | HITS-CLIP | 25083871 | Unspecified |
| mmu-miR-29b-3p | MIMAT0000127 | Lif | 16878 | HITS-CLIP | 37986224\|25083871\|31433990\|23142 | Unspecified |
| mmu-miR-29b-3p | MIMAT0000127 | Lifr | 16880 | HITS-CLIP//sRNA-Seq | 25083871\|23597149\|33184237 | Unspecified |
| mmu-miR-29b-3p | MIMAT0000127 | Lig3 | 16882 | HITS-CLIP | 23142080\|25083871 | Unspecified |
| mmu-miR-29b-3p | MIMAT0000127 | Limd1 | 29806 | HITS-CLIP | 37986224\|23597149\|23142080 | Unspecified |
| mmu-miR-29b-3p | MIMAT0000127 | Limk1 | 16885 | sRNA-Seq//HITS-CLIP | 33184237\|25083871 | Unspecified |
| mmu-miR-29b-3p | MIMAT0000127 | Limk2 | 16886 | sRNA-Seq//HITS-CLIP | 33184237\|23142080 | Unspecified |
| mmu-miR-29b-3p | MIMAT0000127 | Lims1 | 110829 | HITS-CLIP//sRNA-Seq | 37986224\|23142080\|25083871\|33184 | Unspecified |
| mmu-miR-29b-3p | MIMAT0000127 | Lims2 | 225341 | HITS-CLIP | 37986224 | Unspecified |
| mmu-miR-29b-3p | MIMAT0000127 | Lin54 | 231506 | HITS-CLIP | 23142080 | Unspecified |
| mmu-miR-29b-3p | MIMAT0000127 | Lin7c | 22343 | HITS-CLIP//sRNA-Seq | 25083871\|23597149\|33184237\|23142 | Unspecified |
| mmu-miR-29b-3p | MIMAT0000127 | Lix1l | 280411 | HITS-CLIP//sRNA-Seq | 25083871\|33184237 | Unspecified |
| mmu-miR-29b-3p | MIMAT0000127 | Llgl1 | 16897 | sRNA-Seq//HITS-CLIP | 33184237\|23142080 | Unspecified |
| mmu-miR-29b-3p | MIMAT0000127 | Lman1 | 70361 | HITS-CLIP//sRNA-Seq | 23142080\|23597149\|33184237\|25083 | Unspecified |
| mmu-miR-29b-3p | MIMAT0000127 | Lmbr1 | 56873 | HITS-CLIP | 23142080 | Unspecified |
| mmu-miR-29b-3p | MIMAT0000127 | Lmbrd1 | 68421 | HITS-CLIP | 37986224\|23142080 | Unspecified |
| mmu-miR-29b-3p | MIMAT0000127 | Lmbrd2 | 320506 | sRNA-Seq//HITS-CLIP | 33184237\|23597149 | Unspecified |
| mmu-miR-29b-3p | MIMAT0000127 | Lmf2 | 105847 | HITS-CLIP//sRNA-Seq | 25083871\|23142080\|33184237 | Unspecified |
| mmu-miR-29b-3p | MIMAT0000127 | Lmln | 239833 | HITS-CLIP | 37986224\|23142080\|25083871 | Unspecified |

| | | | | | | |
|---|---|---|---|---|---|---|
| mmu-miR-29b-3p | MIMAT0000127 | Lmna | 16905 | HITS-CLIP//sRNA-Seq | 25083871\|33184237 | Unspecified |
| mmu-miR-29b-3p | MIMAT0000127 | Lmnb1 | 16906 | HITS-CLIP//sRNA-Seq | 37986224\|23142080\|25083871\|33184 | Unspecified |
| mmu-miR-29b-3p | MIMAT0000127 | Lmo7 | 380928 | HITS-CLIP | 25083871 | Unspecified |
| mmu-miR-29b-3p | MIMAT0000127 | Lnpep | 240028 | HITS-CLIP//sRNA-Seq | 25083871\|33184237\|23142080 | Unspecified |
| mmu-miR-29b-3p | MIMAT0000127 | Lnx2 | 140887 | HITS-CLIP | 23142080 | Unspecified |
| mmu-miR-29b-3p | MIMAT0000127 | Lonrf1 | 244421 | HITS-CLIP | 25083871\|23142080 | Unspecified |
| mmu-miR-29b-3p | MIMAT0000127 | Lox | 16948 | Luciferase Reporter Assay | 37986224 | Unspecified |
| mmu-miR-29b-3p | MIMAT0000127 | Loxl1 | 16949 | HITS-CLIP//sRNA-Seq | 25083871\|33184237 | Unspecified |
| mmu-miR-29b-3p | MIMAT0000127 | Loxl2 | 94352 | HITS-CLIP//sRNA-Seq | 37986224\|33184237\|25083871 | Unspecified |
| mmu-miR-29b-3p | MIMAT0000127 | Loxl3 | 16950 | HITS-CLIP | 37986224 | Unspecified |
| mmu-miR-29b-3p | MIMAT0000127 | Loxl4 | 67573 | Luciferase Reporter Assay | 37986224\|23354167 | Unspecified |
| mmu-miR-29b-3p | MIMAT0000127 | Lpar4 | 78134 | HITS-CLIP | 37986224\|25083871 | Unspecified |
| mmu-miR-29b-3p | MIMAT0000127 | Lpar6 | 67168 | HITS-CLIP | 37986224\|25083871\|23597149\|23142 | Unspecified |
| mmu-miR-29b-3p | MIMAT0000127 | Lpcat1 | 210992 | HITS-CLIP | 25083871 | Unspecified |
| mmu-miR-29b-3p | MIMAT0000127 | Lpcat3 | 14792 | HITS-CLIP//sRNA-Seq | 37986224\|23597149\|25083871\|23142 | Unspecified |
| mmu-miR-29b-3p | MIMAT0000127 | Lpgat1 | 226856 | HITS-CLIP//sRNA-Seq | 25083871\|33184237\|23597149\|23142 | Unspecified |
| mmu-miR-29b-3p | MIMAT0000127 | Lpin1 | 14245 | HITS-CLIP | 23597149\|23142080 | Unspecified |
| mmu-miR-29b-3p | MIMAT0000127 | Lpin2 | 64898 | HITS-CLIP | 25083871\|23142080\|23597149 | Unspecified |
| mmu-miR-29b-3p | MIMAT0000127 | Lpl | 16956 | HITS-CLIP | 37986224\|23597149\|25083871 | Unspecified |
| mmu-miR-29b-3p | MIMAT0000127 | Lpp | 210126 | HITS-CLIP | 25083871 | Unspecified |
| mmu-miR-29b-3p | MIMAT0000127 | Lpxn | 107321 | HITS-CLIP | 23142080 | Unspecified |
| mmu-miR-29b-3p | MIMAT0000127 | Lrba | 80877 | HITS-CLIP | 25083871\|23142080 | Unspecified |
| mmu-miR-29b-3p | MIMAT0000127 | Lrch4 | 231798 | HITS-CLIP | 25083871\|23597149 | Unspecified |
| mmu-miR-29b-3p | MIMAT0000127 | Lrfn4 | 225875 | HITS-CLIP | 37986224 | Unspecified |
| mmu-miR-29b-3p | MIMAT0000127 | Lrif1 | 321000 | HITS-CLIP | 37986224 | Unspecified |
| mmu-miR-29b-3p | MIMAT0000127 | Lrig1 | 16206 | HITS-CLIP//sRNA-Seq | 25083871\|33184237 | Unspecified |
| mmu-miR-29b-3p | MIMAT0000127 | Lrig2 | 269473 | HITS-CLIP | 25083871\|23142080 | Unspecified |
| mmu-miR-29b-3p | MIMAT0000127 | Lrig3 | 320398 | HITS-CLIP | 25083871 | Unspecified |
| mmu-miR-29b-3p | MIMAT0000127 | Lrmp | 16970 | HITS-CLIP | 23142080 | Unspecified |
| mmu-miR-29b-3p | MIMAT0000127 | Lrp1 | 16971 | HITS-CLIP//sRNA-Seq | 37986224\|25083871\|23597149\|33184 | Unspecified |
| mmu-miR-29b-3p | MIMAT0000127 | Lrp10 | 65107 | HITS-CLIP | 37986224\|23597149\|23142080\|25083 | Unspecified |
| mmu-miR-29b-3p | MIMAT0000127 | Lrp12 | 239393 | HITS-CLIP//sRNA-Seq | 25083871\|33184237 | Unspecified |
| mmu-miR-29b-3p | MIMAT0000127 | Lrp4 | 228357 | HITS-CLIP | 37986224\|25083871 | Unspecified |
| mmu-miR-29b-3p | MIMAT0000127 | Lrp6 | 16974 | HITS-CLIP//sRNA-Seq | 37986224\|25083871\|23597149\|33184 | Unspecified |
| mmu-miR-29b-3p | MIMAT0000127 | Lrpprc | 72416 | sRNA-Seq//HITS-CLIP | 33184237\|23597149\|23142080 | Unspecified |
| mmu-miR-29b-3p | MIMAT0000127 | Lrrc14 | 223664 | HITS-CLIP | 26203562 | Unspecified |
| mmu-miR-29b-3p | MIMAT0000127 | Lrrc17 | 74511 | HITS-CLIP//sRNA-Seq | 25083871\|33184237 | Unspecified |
| mmu-miR-29b-3p | MIMAT0000127 | Lrrc3 | 237387 | HITS-CLIP | 23597149 | Unspecified |
| mmu-miR-29b-3p | MIMAT0000127 | Lrrc32 | 434215 | HITS-CLIP | 23142080\|23597149 | Unspecified |

| | | | | | | |
|---|---|---|---|---|---|---|
| mmu-miR-29b-3p | MIMAT0000127 | Lrrc41 | 230654 | HITS-CLIP//sRNA-Seq | 25083871\|33184237\|23142080 | Unspecified |
| mmu-miR-29b-3p | MIMAT0000127 | Lrrc42 | 77809 | HITS-CLIP//sRNA-Seq | 25083871\|33184237\|23142080 | Unspecified |
| mmu-miR-29b-3p | MIMAT0000127 | Lrrc58 | 320184 | HITS-CLIP//sRNA-Seq | 25083871\|23597149\|33184237\|23142 | Unspecified |
| mmu-miR-29b-3p | MIMAT0000127 | Lrrc59 | 98238 | HITS-CLIP//sRNA-Seq | 37986224\|33184237\|25083871\|23142 | Unspecified |
| mmu-miR-29b-3p | MIMAT0000127 | Lrrc8a | 241296 | HITS-CLIP | 37986224\|25083871\|23597149\|23142 | Unspecified |
| mmu-miR-29b-3p | MIMAT0000127 | Lrrc8c | 100604 | HITS-CLIP | 25083871 | Unspecified |
| mmu-miR-29b-3p | MIMAT0000127 | Lrrcc1 | 71710 | HITS-CLIP | 37986224\|25083871\|23142080 | Unspecified |
| mmu-miR-29b-3p | MIMAT0000127 | Lrrfip1 | 16978 | HITS-CLIP//sRNA-Seq | 25083871\|33184237\|23142080 | Unspecified |
| mmu-miR-29b-3p | MIMAT0000127 | Lrrk1 | 233328 | HITS-CLIP | 25083871\|23142080 | Unspecified |
| mmu-miR-29b-3p | MIMAT0000127 | Lsm1 | 67207 | HITS-CLIP | 23142080 | Unspecified |
| mmu-miR-29b-3p | MIMAT0000127 | Lsm11 | 72290 | HITS-CLIP | 37986224\|25083871\|23142080 | Unspecified |
| mmu-miR-29b-3p | MIMAT0000127 | Lsm14a | 67070 | HITS-CLIP//sRNA-Seq | 37986224\|33184237\|23142080 | Unspecified |
| mmu-miR-29b-3p | MIMAT0000127 | Lsm14b | 241846 | HITS-CLIP//sRNA-Seq | 25083871\|33184237 | Unspecified |
| mmu-miR-29b-3p | MIMAT0000127 | Lsm4 | 50783 | sRNA-Seq//HITS-CLIP | 33184237\|23142080 | Unspecified |
| mmu-miR-29b-3p | MIMAT0000127 | Ltbp3 | 16998 | HITS-CLIP//sRNA-Seq | 25083871\|33184237 | Unspecified |
| mmu-miR-29b-3p | MIMAT0000127 | Ltbr | 17000 | HITS-CLIP//sRNA-Seq | 37986224\|25083871\|33184237 | Unspecified |
| mmu-miR-29b-3p | MIMAT0000127 | Ltn1 | 78913 | HITS-CLIP | 37986224\|25083871\|23597149\|23142 | Unspecified |
| mmu-miR-29b-3p | MIMAT0000127 | Luc7l | 66978 | HITS-CLIP//sRNA-Seq | 25083871\|33184237\|23142080 | Unspecified |
| mmu-miR-29b-3p | MIMAT0000127 | Luc7l2 | 192196 | HITS-CLIP//sRNA-Seq | 23142080\|33184237 | Unspecified |
| mmu-miR-29b-3p | MIMAT0000127 | Luc7l3 | 67684 | HITS-CLIP//sRNA-Seq | 37986224\|25083871\|23142080\|33184 | Unspecified |
| mmu-miR-29b-3p | MIMAT0000127 | Lurap1l | 52829 | HITS-CLIP | 25083871 | Unspecified |
| mmu-miR-29b-3p | MIMAT0000127 | Luzp1 | 269593 | HITS-CLIP | 25083871\|23597149 | Unspecified |
| mmu-miR-29b-3p | MIMAT0000127 | Ly6e | 17069 | HITS-CLIP//sRNA-Seq | 25083871\|33184237\|23142080 | Unspecified |
| mmu-miR-29b-3p | MIMAT0000127 | Ly75 | 17076 | HITS-CLIP | 23142080 | Unspecified |
| mmu-miR-29b-3p | MIMAT0000127 | Lyn | 17096 | HITS-CLIP | 31433990\|23597149 | Unspecified |
| mmu-miR-29b-3p | MIMAT0000127 | Lypla1 | 18777 | HITS-CLIP//sRNA-Seq | 37986224\|23597149\|25083871\|33184 | Unspecified |
| mmu-miR-29b-3p | MIMAT0000127 | Lysmd4 | 75099 | HITS-CLIP | 37986224 | Unspecified |
| mmu-miR-29b-3p | MIMAT0000127 | Lyst | 17101 | HITS-CLIP | 25083871\|23142080 | Unspecified |
| mmu-miR-29b-3p | MIMAT0000127 | Macf1 | 11426 | HITS-CLIP//sRNA-Seq | 37986224\|25083871\|23597149\|31433 | Unspecified |
| mmu-miR-29b-3p | MIMAT0000127 | Macrod2 | 72899 | HITS-CLIP | 23142080 | Unspecified |
| mmu-miR-29b-3p | MIMAT0000127 | Mad2l1 | 56150 | HITS-CLIP | 23142080 | Unspecified |
| mmu-miR-29b-3p | MIMAT0000127 | Mad2l1bp | 66591 | HITS-CLIP | 23142080 | Unspecified |
| mmu-miR-29b-3p | MIMAT0000127 | Mad2l2 | 71890 | HITS-CLIP | 23142080 | Unspecified |
| mmu-miR-29b-3p | MIMAT0000127 | Madd | 228355 | HITS-CLIP//sRNA-Seq | 25083871\|33184237\|23142080 | Unspecified |
| mmu-miR-29b-3p | MIMAT0000127 | Maf | 17132 | HITS-CLIP | 37986224\|25083871\|23597149\|23142 | Unspecified |
| mmu-miR-29b-3p | MIMAT0000127 | Mafb | 16658 | HITS-CLIP | 23597149 | Unspecified |
| mmu-miR-29b-3p | MIMAT0000127 | Mafg | 17134 | HITS-CLIP | 37986224\|25083871\|23597149\|23142 | Unspecified |
| mmu-miR-29b-3p | MIMAT0000127 | Maged1 | 94275 | HITS-CLIP//sRNA-Seq | 23597149\|33184237 | Unspecified |
| mmu-miR-29b-3p | MIMAT0000127 | Maged2 | 80884 | HITS-CLIP//sRNA-Seq | 25083871\|33184237 | Unspecified |

| | | | | | | |
|---|---|---|---|---|---|---|
| mmu-miR-29b-3p | MIMAT0000127 | Magi3 | 99470 | HITS-CLIP | 25083871\|23142080 | Unspecified |
| mmu-miR-29b-3p | MIMAT0000127 | Magt1 | 67075 | HITS-CLIP | 37986224\|25083871\|23142080\|23597 | Unspecified |
| mmu-miR-29b-3p | MIMAT0000127 | Malt1 | 240354 | HITS-CLIP | 23142080 | Unspecified |
| mmu-miR-29b-3p | MIMAT0000127 | Maml1 | 103806 | HITS-CLIP | 25083871\|23142080 | Unspecified |
| mmu-miR-29b-3p | MIMAT0000127 | Maml2 | 270118 | HITS-CLIP | 25083871 | Unspecified |
| mmu-miR-29b-3p | MIMAT0000127 | Maml3 | 433586 | HITS-CLIP | 37986224\|25083871\|23142080 | Unspecified |
| mmu-miR-29b-3p | MIMAT0000127 | Man1a | 17155 | HITS-CLIP | 23142080 | Unspecified |
| mmu-miR-29b-3p | MIMAT0000127 | Man1a2 | 17156 | HITS-CLIP | 37986224\|23142080 | Unspecified |
| mmu-miR-29b-3p | MIMAT0000127 | Man2a1 | 17158 | HITS-CLIP//sRNA-Seq | 25083871\|23597149\|33184237\|23142 | Unspecified |
| mmu-miR-29b-3p | MIMAT0000127 | Man2b1 | 17159 | HITS-CLIP//sRNA-Seq | 23597149\|33184237\|23142080 | Unspecified |
| mmu-miR-29b-3p | MIMAT0000127 | Maoa | 17161 | HITS-CLIP//sRNA-Seq | 25083871\|33184237 | Unspecified |
| mmu-miR-29b-3p | MIMAT0000127 | Map1lc3b | 67443 | HITS-CLIP//sRNA-Seq | 37986224\|25083871\|33184237\|23142 | Unspecified |
| mmu-miR-29b-3p | MIMAT0000127 | Map2k3 | 26397 | sRNA-Seq//HITS-CLIP | 33184237\|23142080 | Unspecified |
| mmu-miR-29b-3p | MIMAT0000127 | Map2k4 | 26398 | HITS-CLIP//sRNA-Seq | 37986224\|25083871\|23597149\|33184 | Unspecified |
| mmu-miR-29b-3p | MIMAT0000127 | Map2k6 | 26399 | HITS-CLIP | 37986224\|23597149\|23142080 | Unspecified |
| mmu-miR-29b-3p | MIMAT0000127 | Map2k7 | 26400 | HITS-CLIP | 25083871 | Unspecified |
| mmu-miR-29b-3p | MIMAT0000127 | Map3k1 | 26401 | HITS-CLIP | 23142080 | Unspecified |
| mmu-miR-29b-3p | MIMAT0000127 | Map3k10 | 269881 | HITS-CLIP | 37986224 | Unspecified |
| mmu-miR-29b-3p | MIMAT0000127 | Map3k11 | 26403 | HITS-CLIP | 23142080 | Unspecified |
| mmu-miR-29b-3p | MIMAT0000127 | Map3k12 | 26404 | HITS-CLIP | 23142080 | Unspecified |
| mmu-miR-29b-3p | MIMAT0000127 | Map3k14 | 53859 | HITS-CLIP | 23142080 | Unspecified |
| mmu-miR-29b-3p | MIMAT0000127 | Map3k2 | 26405 | HITS-CLIP | 37986224\|25083871\|23597149\|23142 | Unspecified |
| mmu-miR-29b-3p | MIMAT0000127 | Map3k20 | 65964 | HITS-CLIP | 25083871 | Unspecified |
| mmu-miR-29b-3p | MIMAT0000127 | Map3k7 | 26409 | sRNA-Seq//HITS-CLIP | 33184237\|23142080 | Unspecified |
| mmu-miR-29b-3p | MIMAT0000127 | Map4 | 17758 | HITS-CLIP//sRNA-Seq | 37986224\|25083871\|33184237\|23597 | Unspecified |
| mmu-miR-29b-3p | MIMAT0000127 | Map4k1 | 26411 | HITS-CLIP | 23142080 | Unspecified |
| mmu-miR-29b-3p | MIMAT0000127 | Map4k2 | 26412 | HITS-CLIP | 23142080 | Unspecified |
| mmu-miR-29b-3p | MIMAT0000127 | Map4k3 | 225028 | HITS-CLIP | 25083871\|23597149\|23142080 | Unspecified |
| mmu-miR-29b-3p | MIMAT0000127 | Map4k4 | 26921 | HITS-CLIP//sRNA-Seq | 25083871\|23142080\|33184237\|23597 | Unspecified |
| mmu-miR-29b-3p | MIMAT0000127 | Map6 | 17760 | HITS-CLIP//sRNA-Seq | 25083871\|33184237 | Unspecified |
| mmu-miR-29b-3p | MIMAT0000127 | Map7 | 17761 | HITS-CLIP | 23142080 | Unspecified |
| mmu-miR-29b-3p | MIMAT0000127 | Map7d1 | 245877 | HITS-CLIP//sRNA-Seq | 25083871\|33184237 | Unspecified |
| mmu-miR-29b-3p | MIMAT0000127 | Map9 | 213582 | HITS-CLIP | 37986224\|25083871\|23142080 | Unspecified |
| mmu-miR-29b-3p | MIMAT0000127 | Mapk1 | 26413 | HITS-CLIP//sRNA-Seq | 25083871\|33184237\|23142080 | Unspecified |
| mmu-miR-29b-3p | MIMAT0000127 | Mapk14 | 26416 | HITS-CLIP//sRNA-Seq | 37986224\|25083871\|33184237\|23597 | Unspecified |
| mmu-miR-29b-3p | MIMAT0000127 | Mapk1ip1l | 218975 | HITS-CLIP | 37986224\|23142080\|25083871\|23597 | Unspecified |
| mmu-miR-29b-3p | MIMAT0000127 | Mapk6 | 50772 | HITS-CLIP//sRNA-Seq | 37986224\|23142080\|23597149\|33184 | Unspecified |
| mmu-miR-29b-3p | MIMAT0000127 | Mapk8 | 26419 | HITS-CLIP//sRNA-Seq | 37986224\|25083871\|33184237\|23142 | Unspecified |
| mmu-miR-29b-3p | MIMAT0000127 | Mapkap1 | 227743 | HITS-CLIP//sRNA-Seq | 37986224\|25083871\|33184237 | Unspecified |

| | | | | | | |
|---|---|---|---|---|---|---|
| mmu-miR-29b-3p | MIMAT0000127 | Mapkapk2 | 17164 | HITS-CLIP//sRNA-Seq | 23142080\|33184237 | Unspecified |
| mmu-miR-29b-3p | MIMAT0000127 | Mapre1 | 13589 | HITS-CLIP//sRNA-Seq | 25083871\|33184237\|23142080 | Unspecified |
| mmu-miR-29b-3p | MIMAT0000127 | Mapre2 | 212307 | HITS-CLIP//sRNA-Seq | 37986224\|33184237\|25083871\|23142 | Unspecified |
| mmu-miR-29b-3p | MIMAT0000127 | Marc1 | 66112 | HITS-CLIP | 23597149 | Unspecified |
| mmu-miR-29b-3p | MIMAT0000127 | March7 | 57438 | HITS-CLIP//sRNA-Seq | 37986224\|25083871\|23597149\|33184 | Unspecified |
| mmu-miR-29b-3p | MIMAT0000127 | March9 | 216438 | HITS-CLIP | 23142080 | Unspecified |
| mmu-miR-29b-3p | MIMAT0000127 | Marchf6 | 223455 | HITS-CLIP//sRNA-Seq | 37986224\|25083871\|23597149\|33184 | Unspecified |
| mmu-miR-29b-3p | MIMAT0000127 | Marcks | 17118 | HITS-CLIP//sRNA-Seq | 37986224\|23597149\|33184237\|23142 | Unspecified |
| mmu-miR-29b-3p | MIMAT0000127 | Marf1 | 223989 | HITS-CLIP | 37986224\|25083871\|23597149\|23142 | Unspecified |
| mmu-miR-29b-3p | MIMAT0000127 | Mark2 | 13728 | HITS-CLIP | 37986224\|25083871\|23142080 | Unspecified |
| mmu-miR-29b-3p | MIMAT0000127 | Mark3 | 17169 | HITS-CLIP//sRNA-Seq | 37986224\|25083871\|23597149\|33184 | Unspecified |
| mmu-miR-29b-3p | MIMAT0000127 | Marveld1 | 277010 | HITS-CLIP//sRNA-Seq | 23142080\|25083871\|23597149\|33184 | Unspecified |
| mmu-miR-29b-3p | MIMAT0000127 | Marveld2 | 218518 | HITS-CLIP | 23597149 | Unspecified |
| mmu-miR-29b-3p | MIMAT0000127 | Masp1 | 17174 | HITS-CLIP//sRNA-Seq | 23597149\|33184237 | Unspecified |
| mmu-miR-29b-3p | MIMAT0000127 | Mast2 | 17776 | sRNA-Seq//HITS-CLIP | 33184237\|23142080 | Unspecified |
| mmu-miR-29b-3p | MIMAT0000127 | Mast3 | 546071 | HITS-CLIP | 23597149 | Unspecified |
| mmu-miR-29b-3p | MIMAT0000127 | Mast4 | 328329 | HITS-CLIP | 25083871 | Unspecified |
| mmu-miR-29b-3p | MIMAT0000127 | Mastl | 67121 | HITS-CLIP | 23142080 | Unspecified |
| mmu-miR-29b-3p | MIMAT0000127 | Mat1a | 11720 | HITS-CLIP | 37986224\|23597149 | Unspecified |
| mmu-miR-29b-3p | MIMAT0000127 | Mat2a | 232087 | HITS-CLIP//sRNA-Seq | 23142080\|23597149\|33184237 | Unspecified |
| mmu-miR-29b-3p | MIMAT0000127 | Mat2b | 108645 | HITS-CLIP//sRNA-Seq | 25083871\|33184237 | Unspecified |
| mmu-miR-29b-3p | MIMAT0000127 | Matr3 | 17184 | HITS-CLIP//sRNA-Seq | 23142080\|25083871\|33184237 | Unspecified |
| mmu-miR-29b-3p | MIMAT0000127 | Mau2 | 74549 | HITS-CLIP | 25083871 | Unspecified |
| mmu-miR-29b-3p | MIMAT0000127 | Maz | 17188 | HITS-CLIP//sRNA-Seq | 37986224\|23142080\|23597149\|33184 | Unspecified |
| mmu-miR-29b-3p | MIMAT0000127 | Mb21d2 | 239796 | HITS-CLIP | 23597149 | Unspecified |
| mmu-miR-29b-3p | MIMAT0000127 | Mbd5 | 109241 | sRNA-Seq//HITS-CLIP | 33184237\|23142080 | Unspecified |
| mmu-miR-29b-3p | MIMAT0000127 | Mbd6 | 110962 | HITS-CLIP | 23597149 | Unspecified |
| mmu-miR-29b-3p | MIMAT0000127 | Mbl2 | 17195 | HITS-CLIP | 23597149 | Unspecified |
| mmu-miR-29b-3p | MIMAT0000127 | Mblac2 | 72852 | HITS-CLIP//sRNA-Seq | 37986224\|23597149\|33184237 | Unspecified |
| mmu-miR-29b-3p | MIMAT0000127 | Mbnl3 | 171170 | HITS-CLIP | 23142080 | Unspecified |
| mmu-miR-29b-3p | MIMAT0000127 | Mboat1 | 218121 | HITS-CLIP | 23142080 | Unspecified |
| mmu-miR-29b-3p | MIMAT0000127 | Mboat2 | 67216 | HITS-CLIP | 23142080\|25083871 | Unspecified |
| mmu-miR-29b-3p | MIMAT0000127 | Mbtd1 | 103537 | HITS-CLIP | 37986224\|25083871\|23142080 | Unspecified |
| mmu-miR-29b-3p | MIMAT0000127 | Mbtps1 | 56453 | sRNA-Seq//HITS-CLIP | 33184237\|23142080 | Unspecified |
| mmu-miR-29b-3p | MIMAT0000127 | Mbtps2 | 270669 | HITS-CLIP | 37986224\|25083871\|23142080 | Unspecified |
| mmu-miR-29b-3p | MIMAT0000127 | Mccc2 | 78038 | HITS-CLIP | 23142080 | Unspecified |
| mmu-miR-29b-3p | MIMAT0000127 | Mcl1 | 17210 | HITS-CLIP//sRNA-Seq | 23597149\|25083871\|33184237\|23142 | Unspecified |
| mmu-miR-29b-3p | MIMAT0000127 | Mcm3 | 17215 | sRNA-Seq//HITS-CLIP | 33184237\|23142080 | Unspecified |
| mmu-miR-29b-3p | MIMAT0000127 | Mcm3ap | 54387 | HITS-CLIP | 23142080 | Unspecified |

| | | | | | | |
|---|---|---|---|---|---|---|
| mmu-miR-29b-3p | MIMAT0000127 | Mcm7 | 17220 | HITS-CLIP//sRNA-Seq | 23142080\|33184237 | Unspecified |
| mmu-miR-29b-3p | MIMAT0000127 | Mcm9 | 71567 | HITS-CLIP | 23142080 | Unspecified |
| mmu-miR-29b-3p | MIMAT0000127 | Mcmbp | 210711 | HITS-CLIP | 23142080 | Unspecified |
| mmu-miR-29b-3p | MIMAT0000127 | Mcrip1 | 348262 | HITS-CLIP//sRNA-Seq | 25083871\|33184237\|23142080 | Unspecified |
| mmu-miR-29b-3p | MIMAT0000127 | Mcts1 | 68995 | HITS-CLIP//sRNA-Seq | 25083871\|33184237 | Unspecified |
| mmu-miR-29b-3p | MIMAT0000127 | Mdc1 | 240087 | sRNA-Seq//HITS-CLIP | 33184237\|23597149 | Unspecified |
| mmu-miR-29b-3p | MIMAT0000127 | Mdfic | 16543 | HITS-CLIP | 23142080\|31433990 | Unspecified |
| mmu-miR-29b-3p | MIMAT0000127 | Mdh1 | 17449 | HITS-CLIP//sRNA-Seq | 25083871\|33184237\|23597149\|2314 | Unspecified |
| mmu-miR-29b-3p | MIMAT0000127 | Mdm2 | 17246 | HITS-CLIP//sRNA-Seq | 37986224\|33184237\|23142080 | Unspecified |
| mmu-miR-29b-3p | MIMAT0000127 | Mdm4 | 17248 | HITS-CLIP//sRNA-Seq | 23142080\|25083871\|33184237 | Unspecified |
| mmu-miR-29b-3p | MIMAT0000127 | Mdn1 | 100019 | HITS-CLIP | 25083871\|23142080 | Unspecified |
| mmu-miR-29b-3p | MIMAT0000127 | Mecp2 | 17257 | HITS-CLIP | 25083871 | Unspecified |
| mmu-miR-29b-3p | MIMAT0000127 | Med1 | 19014 | HITS-CLIP | 25083871\|23142080 | Unspecified |
| mmu-miR-29b-3p | MIMAT0000127 | Med12 | 59024 | HITS-CLIP | 23597149\|23142080 | Unspecified |
| mmu-miR-29b-3p | MIMAT0000127 | Med13 | 327987 | HITS-CLIP//qCLASH | 37986224\|25083871\|23597149\|3139 | Unspecified |
| mmu-miR-29b-3p | MIMAT0000127 | Med13l | 76199 | HITS-CLIP | 37986224\|25083871\|23142080 | Unspecified |
| mmu-miR-29b-3p | MIMAT0000127 | Med14 | 26896 | HITS-CLIP | 23142080 | Unspecified |
| mmu-miR-29b-3p | MIMAT0000127 | Med21 | 108098 | HITS-CLIP//sRNA-Seq | 30971389\|33184237 | Unspecified |
| mmu-miR-29b-3p | MIMAT0000127 | Med23 | 70208 | HITS-CLIP | 23142080 | Unspecified |
| mmu-miR-29b-3p | MIMAT0000127 | Med24 | 23989 | HITS-CLIP | 23142080 | Unspecified |
| mmu-miR-29b-3p | MIMAT0000127 | Med25 | 75613 | HITS-CLIP | 23597149 | Unspecified |
| mmu-miR-29b-3p | MIMAT0000127 | Med28 | 66999 | HITS-CLIP//sRNA-Seq | 23142080\|33184237 | Unspecified |
| mmu-miR-29b-3p | MIMAT0000127 | Med7 | 66213 | HITS-CLIP | 25083871\|23142080 | Unspecified |
| mmu-miR-29b-3p | MIMAT0000127 | Med8 | 80509 | HITS-CLIP//sRNA-Seq | 25083871\|33184237\|23597149\|2314 | Unspecified |
| mmu-miR-29b-3p | MIMAT0000127 | Medag | 70717 | HITS-CLIP | 25083871 | Unspecified |
| mmu-miR-29b-3p | MIMAT0000127 | Mef2a | 17258 | HITS-CLIP//sRNA-Seq | 37986224\|23142080\|25083871\|3318 | Unspecified |
| mmu-miR-29b-3p | MIMAT0000127 | Mef2c | 17260 | HITS-CLIP | 23142080\|25083871 | Unspecified |
| mmu-miR-29b-3p | MIMAT0000127 | Mef2d | 17261 | HITS-CLIP | 37986224\|25083871 | Unspecified |
| mmu-miR-29b-3p | MIMAT0000127 | Megf6 | 230971 | HITS-CLIP | 23597149 | Unspecified |
| mmu-miR-29b-3p | MIMAT0000127 | Megf9 | 230316 | HITS-CLIP | 25083871\|23597149 | Unspecified |
| mmu-miR-29b-3p | MIMAT0000127 | Melk | 17279 | HITS-CLIP | 23142080 | Unspecified |
| mmu-miR-29b-3p | MIMAT0000127 | Men1 | 17283 | sRNA-Seq//HITS-CLIP | 33184237\|23142080 | Unspecified |
| mmu-miR-29b-3p | MIMAT0000127 | Mepce | 231803 | HITS-CLIP | 23597149 | Unspecified |
| mmu-miR-29b-3p | MIMAT0000127 | Mest | 17294 | HITS-CLIP//sRNA-Seq | 37986224\|25083871\|33184237 | Unspecified |
| mmu-miR-29b-3p | MIMAT0000127 | Met | 17295 | HITS-CLIP | 25083871\|23597149 | Unspecified |
| mmu-miR-29b-3p | MIMAT0000127 | Metap1 | 75624 | HITS-CLIP | 23142080 | Unspecified |
| mmu-miR-29b-3p | MIMAT0000127 | Metap2 | 56307 | HITS-CLIP//sRNA-Seq | 25083871\|33184237\|23142080 | Unspecified |
| mmu-miR-29b-3p | MIMAT0000127 | Metrnl | 210029 | HITS-CLIP | 37986224\|25083871 | Unspecified |
| mmu-miR-29b-3p | MIMAT0000127 | Mettl13 | 51603 | HITS-CLIP | 23142080 | Unspecified |

| | | | | | | |
|---|---|---|---|---|---|---|
| mmu-miR-29b-3p | MIMAT0000127 | Mettl14 | 210529 | HITS-CLIP | 23597149\|23142080 | Unspecified |
| mmu-miR-29b-3p | MIMAT0000127 | Mettl15 | 76894 | HITS-CLIP | 23597149\|23142080 | Unspecified |
| mmu-miR-29b-3p | MIMAT0000127 | Mettl16 | 67493 | HITS-CLIP | 23142080 | Unspecified |
| mmu-miR-29b-3p | MIMAT0000127 | Mettl21c | 196541 | HITS-CLIP | 23142080 | Unspecified |
| mmu-miR-29b-3p | MIMAT0000127 | Mettl6 | 67011 | HITS-CLIP | 37986224\|23597149 | Unspecified |
| mmu-miR-29b-3p | MIMAT0000127 | Mettl7b | 71664 | HITS-CLIP | 23597149 | Unspecified |
| mmu-miR-29b-3p | MIMAT0000127 | Mettl8 | 228019 | HITS-CLIP | 23142080 | Unspecified |
| mmu-miR-29b-3p | MIMAT0000127 | Mex3b | 108797 | HITS-CLIP | 37986224\|25083871 | Unspecified |
| mmu-miR-29b-3p | MIMAT0000127 | Mex3c | 240396 | HITS-CLIP//sRNA-Seq | 25083871\|33184237\|23142080 | Unspecified |
| mmu-miR-29b-3p | MIMAT0000127 | Mfap3 | 216760 | HITS-CLIP | 23597149\|25083871\|23142080 | Unspecified |
| mmu-miR-29b-3p | MIMAT0000127 | Mfap5 | 50530 | HITS-CLIP | 37986224\|25083871 | Unspecified |
| mmu-miR-29b-3p | MIMAT0000127 | Mff | 75734 | HITS-CLIP//sRNA-Seq | 25083871\|33184237 | Unspecified |
| mmu-miR-29b-3p | MIMAT0000127 | Mfhas1 | 52065 | HITS-CLIP | 23142080 | Unspecified |
| mmu-miR-29b-3p | MIMAT0000127 | Mfn1 | 67414 | HITS-CLIP//sRNA-Seq | 25083871\|33184237 | Unspecified |
| mmu-miR-29b-3p | MIMAT0000127 | Mfn2 | 170731 | HITS-CLIP//sRNA-Seq | 23597149\|33184237 | Unspecified |
| mmu-miR-29b-3p | MIMAT0000127 | Mfsd1 | 66868 | HITS-CLIP//sRNA-Seq | 23597149\|25083871\|33184237\|23142 | Unspecified |
| mmu-miR-29b-3p | MIMAT0000127 | Mfsd10 | 68294 | HITS-CLIP//sRNA-Seq | 25083871\|33184237 | Unspecified |
| mmu-miR-29b-3p | MIMAT0000127 | Mfsd12 | 73822 | HITS-CLIP | 25083871\|23142080 | Unspecified |
| mmu-miR-29b-3p | MIMAT0000127 | Mfsd14a | 15247 | HITS-CLIP | 23142080 | Unspecified |
| mmu-miR-29b-3p | MIMAT0000127 | Mfsd14b | 84641 | HITS-CLIP | 25083871\|23142080\|23597149 | Unspecified |
| mmu-miR-29b-3p | MIMAT0000127 | Mfsd4b1 | 215929 | HITS-CLIP | 23597149 | Unspecified |
| mmu-miR-29b-3p | MIMAT0000127 | Mfsd8 | 72175 | HITS-CLIP | 23142080 | Unspecified |
| mmu-miR-29b-3p | MIMAT0000127 | Mga | 29808 | HITS-CLIP//sRNA-Seq | 37986224\|25083871\|23597149\|33184 | Unspecified |
| mmu-miR-29b-3p | MIMAT0000127 | Mgat1 | 17308 | HITS-CLIP//sRNA-Seq | 37986224\|23142080\|23597149\|33184 | Unspecified |
| mmu-miR-29b-3p | MIMAT0000127 | Mgat2 | 217664 | HITS-CLIP | 25083871 | Unspecified |
| mmu-miR-29b-3p | MIMAT0000127 | Mgat4b | 103534 | HITS-CLIP//sRNA-Seq | 25083871\|33184237\|23142080 | Unspecified |
| mmu-miR-29b-3p | MIMAT0000127 | Mgea5 | 76055 | HITS-CLIP | 23142080 | Unspecified |
| mmu-miR-29b-3p | MIMAT0000127 | Mgp | 17313 | HITS-CLIP//sRNA-Seq | 25083871\|33184237 | Unspecified |
| mmu-miR-29b-3p | MIMAT0000127 | Mgrn1 | 17237 | HITS-CLIP//sRNA-Seq | 25083871\|33184237 | Unspecified |
| mmu-miR-29b-3p | MIMAT0000127 | Mgst3 | 66447 | HITS-CLIP | 37986224 | Unspecified |
| mmu-miR-29b-3p | MIMAT0000127 | Mib1 | 225164 | HITS-CLIP//sRNA-Seq | 37986224\|25083871\|33184237\|23597 | Unspecified |
| mmu-miR-29b-3p | MIMAT0000127 | Mical1 | 171580 | HITS-CLIP | 23142080 | Unspecified |
| mmu-miR-29b-3p | MIMAT0000127 | Mical2 | 320878 | HITS-CLIP | 37986224\|25083871\|23597149 | Unspecified |
| mmu-miR-29b-3p | MIMAT0000127 | Mical3 | 194401 | HITS-CLIP | 25083871 | Unspecified |
| mmu-miR-29b-3p | MIMAT0000127 | Micu1 | 216001 | HITS-CLIP//sRNA-Seq | 25083871\|33184237 | Unspecified |
| mmu-miR-29b-3p | MIMAT0000127 | Micu3 | 78506 | HITS-CLIP//sRNA-Seq | 23142080\|33184237 | Unspecified |
| mmu-miR-29b-3p | MIMAT0000127 | Mid1ip1 | 68041 | sRNA-Seq//HITS-CLIP | 33184237\|23142080 | Unspecified |
| mmu-miR-29b-3p | MIMAT0000127 | Midn | 59090 | HITS-CLIP//sRNA-Seq | 25083871\|23597149\|33184237\|26203 | Unspecified |
| mmu-miR-29b-3p | MIMAT0000127 | Mief1 | 239555 | HITS-CLIP | 37986224\|23597149\|25083871\|23142 | Unspecified |

| | | | | | | |
|---|---|---|---|---|---|---|
| mmu-miR-29b-3p | MIMAT0000127 | Mief2 | 237781 | HITS-CLIP | 37986224\|23597149\|25083871 | Unspecified |
| mmu-miR-29b-3p | MIMAT0000127 | Mien1 | 103742 | sRNA-Seq//HITS-CLIP | 33184237\|23142080 | Unspecified |
| mmu-miR-29b-3p | MIMAT0000127 | Mier2 | 70427 | HITS-CLIP | 23142080\|25083871 | Unspecified |
| mmu-miR-29b-3p | MIMAT0000127 | Mier3 | 218613 | HITS-CLIP | 25083871\|23142080 | Unspecified |
| mmu-miR-29b-3p | MIMAT0000127 | Mif4gd | 69674 | HITS-CLIP | 23597149\|31433990\|23142080 | Unspecified |
| mmu-miR-29b-3p | MIMAT0000127 | Miga1 | 215708 | HITS-CLIP | 23142080 | Unspecified |
| mmu-miR-29b-3p | MIMAT0000127 | Mindy1 | 55793 | HITS-CLIP | 23597149 | Unspecified |
| mmu-miR-29b-3p | MIMAT0000127 | Mindy2 | 235461 | HITS-CLIP | 25083871\|23142080 | Unspecified |
| mmu-miR-29b-3p | MIMAT0000127 | Mink1 | 50932 | HITS-CLIP//sRNA-Seq | 25083871\|33184237\|23142080 | Unspecified |
| mmu-miR-29b-3p | MIMAT0000127 | Minos1 | 440574 | HITS-CLIP//sRNA-Seq | 23142080\|33184237 | Unspecified |
| mmu-miR-29b-3p | MIMAT0000127 | Minpp1 | 17330 | HITS-CLIP | 23597149\|23142080 | Unspecified |
| mmu-miR-29b-3p | MIMAT0000127 | Mios | 252875 | HITS-CLIP | 37986224\|25083871\|23597149\|23142 | Unspecified |
| mmu-miR-29b-3p | MIMAT0000127 | Mis12 | 67139 | HITS-CLIP | 31433990 | Unspecified |
| mmu-miR-29b-3p | MIMAT0000127 | Mitf | 17342 | HITS-CLIP | 25083871 | Unspecified |
| mmu-miR-29b-3p | MIMAT0000127 | Mki67 | 17345 | HITS-CLIP//sRNA-Seq | 37986224\|25083871\|23597149\|33184 | Unspecified |
| mmu-miR-29b-3p | MIMAT0000127 | Mkln1 | 27418 | HITS-CLIP//sRNA-Seq | 37986224\|33184237\|23142080 | Unspecified |
| mmu-miR-29b-3p | MIMAT0000127 | Mknk2 | 17347 | sRNA-Seq//HITS-CLIP | 33184237\|23597149 | Unspecified |
| mmu-miR-29b-3p | MIMAT0000127 | Mkrn1 | 54484 | HITS-CLIP//sRNA-Seq | 37986224\|23597149\|33184237 | Unspecified |
| mmu-miR-29b-3p | MIMAT0000127 | Mkrn2 | 67027 | HITS-CLIP | 23597149\|23142080 | Unspecified |
| mmu-miR-29b-3p | MIMAT0000127 | Mlec | 109154 | HITS-CLIP//sRNA-Seq | 25083871\|23597149\|33184237\|23142 | Unspecified |
| mmu-miR-29b-3p | MIMAT0000127 | Mllt10 | 17354 | HITS-CLIP | 25083871 | Unspecified |
| mmu-miR-29b-3p | MIMAT0000127 | Mllt11 | 56772 | HITS-CLIP//sRNA-Seq | 37986224\|25083871\|33184237\|23142 | Unspecified |
| mmu-miR-29b-3p | MIMAT0000127 | Mllt3 | 70122 | HITS-CLIP | 23142080 | Unspecified |
| mmu-miR-29b-3p | MIMAT0000127 | Mlxip | 208104 | HITS-CLIP | 37986224\|25083871\|23142080 | Unspecified |
| mmu-miR-29b-3p | MIMAT0000127 | Mlxipl | 58805 | HITS-CLIP | 23597149 | Unspecified |
| mmu-miR-29b-3p | MIMAT0000127 | Mmadhc | 109129 | HITS-CLIP//sRNA-Seq | 25083871\|33184237\|23597149\|23142 | Unspecified |
| mmu-miR-29b-3p | MIMAT0000127 | Mmd | 67468 | HITS-CLIP//sRNA-Seq | 23597149\|33184237\|25083871\|23142 | Unspecified |
| mmu-miR-29b-3p | MIMAT0000127 | Mmgt1 | 236792 | HITS-CLIP | 37986224 | Unspecified |
| mmu-miR-29b-3p | MIMAT0000127 | Mmp11 | 17385 | HITS-CLIP | 37986224 | Unspecified |
| mmu-miR-29b-3p | MIMAT0000127 | Mmp14 | 17387 | sRNA-Seq//HITS-CLIP | 33184237\|25083871 | Unspecified |
| mmu-miR-29b-3p | MIMAT0000127 | Mmp15 | 17388 | HITS-CLIP, Luciferase Reporter Assay//HITS | 37986224\|25083871\|23597149\|33184 | Unspecified |
| mmu-miR-29b-3p | MIMAT0000127 | Mmp19 | 58223 | HITS-CLIP | 23597149 | Unspecified |
| mmu-miR-29b-3p | MIMAT0000127 | Mmp2 | 17390 | Luciferase Reporter Assay//HITS-CLIP//sRN | 37986224\|25083871\|33184237 | Unspecified |
| mmu-miR-29b-3p | MIMAT0000127 | Mmp24 | 17391 | Luciferase Reporter Assay | 37986224 | Unspecified |
| mmu-miR-29b-3p | MIMAT0000127 | Mmp9 | 17395 | Luciferase Reporter Assay | 37986224 | Unspecified |
| mmu-miR-29b-3p | MIMAT0000127 | Mmrn2 | 105450 | HITS-CLIP | 37986224 | Unspecified |
| mmu-miR-29b-3p | MIMAT0000127 | Mnd1 | 76915 | HITS-CLIP | 37986224\|25083871\|23142080 | Unspecified |
| mmu-miR-29b-3p | MIMAT0000127 | Mnt | 17428 | HITS-CLIP | 25083871\|23142080 | Unspecified |
| mmu-miR-29b-3p | MIMAT0000127 | Mob1a | 232157 | HITS-CLIP//sRNA-Seq | 37986224\|25083871\|33184237\|23142 | Unspecified |

| | | | | | | |
|---|---|---|---|---|---|---|
| mmu-miR-29b-3p | MIMAT0000127 | Mob1b | 68473 | HITS-CLIP | 25083871\|23597149\|23142080 | Unspecified |
| mmu-miR-29b-3p | MIMAT0000127 | Mob2 | 101513 | HITS-CLIP | 23142080 | Unspecified |
| mmu-miR-29b-3p | MIMAT0000127 | Mob4 | 19070 | HITS-CLIP//sRNA-Seq | 37986224\|23142080\|25083871\|33184 | Unspecified |
| mmu-miR-29b-3p | MIMAT0000127 | Mon2 | 67074 | HITS-CLIP | 23597149\|23142080 | Unspecified |
| mmu-miR-29b-3p | MIMAT0000127 | Morc2a | 74522 | HITS-CLIP | 37986224\|25083871\|23597149\|23142 | Unspecified |
| mmu-miR-29b-3p | MIMAT0000127 | Morc3 | 338467 | HITS-CLIP | 23142080\|25083871 | Unspecified |
| mmu-miR-29b-3p | MIMAT0000127 | Morc4 | 75746 | HITS-CLIP | 37986224 | Unspecified |
| mmu-miR-29b-3p | MIMAT0000127 | Morf4l2 | 56397 | HITS-CLIP//sRNA-Seq | 37986224\|25083871\|33184237\|23142 | Unspecified |
| mmu-miR-29b-3p | MIMAT0000127 | Mosmo | 233812 | HITS-CLIP | 25083871\|23142080 | Unspecified |
| mmu-miR-29b-3p | MIMAT0000127 | Mospd1 | 70380 | HITS-CLIP | 23142080 | Unspecified |
| mmu-miR-29b-3p | MIMAT0000127 | Mospd2 | 76763 | HITS-CLIP | 23142080 | Unspecified |
| mmu-miR-29b-3p | MIMAT0000127 | Mospd3 | 68929 | HITS-CLIP | 25083871\|23597149\|23142080 | Unspecified |
| mmu-miR-29b-3p | MIMAT0000127 | Mov10 | 17454 | sRNA-Seq//HITS-CLIP | 33184237\|23597149 | Unspecified |
| mmu-miR-29b-3p | MIMAT0000127 | Mpc2 | 70456 | HITS-CLIP//sRNA-Seq | 23597149\|33184237 | Unspecified |
| mmu-miR-29b-3p | MIMAT0000127 | Mpdu1 | 24070 | HITS-CLIP | 23597149 | Unspecified |
| mmu-miR-29b-3p | MIMAT0000127 | Mpdz | 17475 | HITS-CLIP | 25083871 | Unspecified |
| mmu-miR-29b-3p | MIMAT0000127 | Mpeg1 | 17476 | HITS-CLIP | 23597149 | Unspecified |
| mmu-miR-29b-3p | MIMAT0000127 | Mphosph10 | 67973 | HITS-CLIP | 26203562 | Unspecified |
| mmu-miR-29b-3p | MIMAT0000127 | Mphosph8 | 75339 | HITS-CLIP | 25083871 | Unspecified |
| mmu-miR-29b-3p | MIMAT0000127 | Mpp5 | 56217 | HITS-CLIP | 23142080 | Unspecified |
| mmu-miR-29b-3p | MIMAT0000127 | Mpp6 | 56524 | HITS-CLIP | 23142080 | Unspecified |
| mmu-miR-29b-3p | MIMAT0000127 | Mprip | 26936 | HITS-CLIP//sRNA-Seq | 37986224\|25083871\|33184237 | Unspecified |
| mmu-miR-29b-3p | MIMAT0000127 | Mpv17 | 17527 | HITS-CLIP//sRNA-Seq | 25083871\|33184237 | Unspecified |
| mmu-miR-29b-3p | MIMAT0000127 | Mpzl1 | 68481 | HITS-CLIP | 37986224 | Unspecified |
| mmu-miR-29b-3p | MIMAT0000127 | Mpzl2 | 14012 | HITS-CLIP | 26203562 | Unspecified |
| mmu-miR-29b-3p | MIMAT0000127 | Mrfap1 | 67568 | HITS-CLIP//sRNA-Seq | 25083871\|33184237\|23142080 | Unspecified |
| mmu-miR-29b-3p | MIMAT0000127 | Mroh1 | 223658 | sRNA-Seq//HITS-CLIP | 33184237\|23597149 | Unspecified |
| mmu-miR-29b-3p | MIMAT0000127 | Mrpl1 | 94061 | HITS-CLIP | 25083871 | Unspecified |
| mmu-miR-29b-3p | MIMAT0000127 | Mrpl10 | 107732 | HITS-CLIP//sRNA-Seq | 25083871\|23142080\|33184237 | Unspecified |
| mmu-miR-29b-3p | MIMAT0000127 | Mrpl12 | 56282 | HITS-CLIP//sRNA-Seq | 25083871\|23597149\|33184237\|23142 | Unspecified |
| mmu-miR-29b-3p | MIMAT0000127 | Mrpl13 | 68537 | HITS-CLIP | 25083871\|23142080\|23597149 | Unspecified |
| mmu-miR-29b-3p | MIMAT0000127 | Mrpl15 | 27395 | HITS-CLIP//sRNA-Seq | 23142080\|33184237 | Unspecified |
| mmu-miR-29b-3p | MIMAT0000127 | Mrpl28 | 68611 | sRNA-Seq//HITS-CLIP | 33184237\|23142080 | Unspecified |
| mmu-miR-29b-3p | MIMAT0000127 | Mrpl32 | 75398 | HITS-CLIP//sRNA-Seq | 37986224\|23142080\|33184237 | Unspecified |
| mmu-miR-29b-3p | MIMAT0000127 | Mrpl37 | 56280 | HITS-CLIP | 23597149\|23142080 | Unspecified |
| mmu-miR-29b-3p | MIMAT0000127 | Mrpl43 | 94067 | sRNA-Seq//HITS-CLIP | 33184237\|23142080 | Unspecified |
| mmu-miR-29b-3p | MIMAT0000127 | Mrpl45 | 67036 | sRNA-Seq//HITS-CLIP | 33184237\|23142080 | Unspecified |
| mmu-miR-29b-3p | MIMAT0000127 | Mrpl50 | 28028 | HITS-CLIP | 37986224 | Unspecified |
| mmu-miR-29b-3p | MIMAT0000127 | Mrpl52 | 68836 | HITS-CLIP//sRNA-Seq | 23142080\|23597149\|33184237\|25083 | Unspecified |

| | | | | | | |
|---|---|---|---|---|---|---|
| mmu-miR-29b-3p | MIMAT0000127 | Mrps14 | 64659 | HITS-CLIP | 23142080 | Unspecified |
| mmu-miR-29b-3p | MIMAT0000127 | Mrps18b | 66973 | sRNA-Seq//HITS-CLIP | 33184237\|23142080 | Unspecified |
| mmu-miR-29b-3p | MIMAT0000127 | Mrps23 | 64656 | HITS-CLIP//sRNA-Seq | 25083871\|33184237 | Unspecified |
| mmu-miR-29b-3p | MIMAT0000127 | Mrps30 | 59054 | HITS-CLIP | 23142080 | Unspecified |
| mmu-miR-29b-3p | MIMAT0000127 | Mrps9 | 69527 | HITS-CLIP | 37986224 | Unspecified |
| mmu-miR-29b-3p | MIMAT0000127 | Mrs2 | 380836 | HITS-CLIP | 25083871 | Unspecified |
| mmu-miR-29b-3p | MIMAT0000127 | Mrtfb | 239719 | HITS-CLIP | 25083871 | Unspecified |
| mmu-miR-29b-3p | MIMAT0000127 | Ms4a4b | 60361 | HITS-CLIP | 23142080 | Unspecified |
| mmu-miR-29b-3p | MIMAT0000127 | Ms4a6b | 69774 | HITS-CLIP | 23142080 | Unspecified |
| mmu-miR-29b-3p | MIMAT0000127 | Msantd2 | 235184 | HITS-CLIP | 23142080 | Unspecified |
| mmu-miR-29b-3p | MIMAT0000127 | Msantd3 | 66665 | HITS-CLIP | 25083871 | Unspecified |
| mmu-miR-29b-3p | MIMAT0000127 | Msantd4 | 78100 | HITS-CLIP | 23142080\|25083871 | Unspecified |
| mmu-miR-29b-3p | MIMAT0000127 | Msc | 17681 | HITS-CLIP | 37986224 | Unspecified |
| mmu-miR-29b-3p | MIMAT0000127 | Msh5 | 4439 | HITS-CLIP | 23597149 | Unspecified |
| mmu-miR-29b-3p | MIMAT0000127 | Msh6 | 17688 | HITS-CLIP//sRNA-Seq | 37986224\|33184237\|25083871\|23142 | Unspecified |
| mmu-miR-29b-3p | MIMAT0000127 | Msi2 | 76626 | HITS-CLIP | 23142080 | Unspecified |
| mmu-miR-29b-3p | MIMAT0000127 | Msl1 | 74026 | HITS-CLIP//sRNA-Seq | 37986224\|25083871\|33184237 | Unspecified |
| mmu-miR-29b-3p | MIMAT0000127 | Msl2 | 77853 | HITS-CLIP | 23142080 | Unspecified |
| mmu-miR-29b-3p | MIMAT0000127 | Msl3 | 17692 | sRNA-Seq//HITS-CLIP | 33184237\|23142080 | Unspecified |
| mmu-miR-29b-3p | MIMAT0000127 | Msn | 17698 | HITS-CLIP//sRNA-Seq | 25083871\|33184237\|23142080 | Unspecified |
| mmu-miR-29b-3p | MIMAT0000127 | Msrb3 | 320183 | HITS-CLIP//sRNA-Seq | 25083871\|33184237 | Unspecified |
| mmu-miR-29b-3p | MIMAT0000127 | Mst1 | 15235 | HITS-CLIP | 23597149 | Unspecified |
| mmu-miR-29b-3p | MIMAT0000127 | Msto1 | 229524 | HITS-CLIP | 23597149 | Unspecified |
| mmu-miR-29b-3p | MIMAT0000127 | Mta1 | 116870 | HITS-CLIP//sRNA-Seq | 25083871\|33184237\|23142080 | Unspecified |
| mmu-miR-29b-3p | MIMAT0000127 | Mta2 | 23942 | sRNA-Seq//HITS-CLIP | 33184237\|23142080 | Unspecified |
| mmu-miR-29b-3p | MIMAT0000127 | Mtbp | 105837 | HITS-CLIP//sRNA-Seq | 23142080\|33184237 | Unspecified |
| mmu-miR-29b-3p | MIMAT0000127 | Mtch1 | 56462 | HITS-CLIP//sRNA-Seq | 25083871\|33184237\|23142080 | Unspecified |
| mmu-miR-29b-3p | MIMAT0000127 | Mtch2 | 56428 | HITS-CLIP//sRNA-Seq | 25083871\|23142080\|33184237 | Unspecified |
| mmu-miR-29b-3p | MIMAT0000127 | Mtdh | 67154 | sRNA-Seq//HITS-CLIP | 33184237\|23142080\|25083871 | Unspecified |
| mmu-miR-29b-3p | MIMAT0000127 | Mtf1 | 17764 | HITS-CLIP | 23142080 | Unspecified |
| mmu-miR-29b-3p | MIMAT0000127 | Mtf2 | 17765 | HITS-CLIP | 25083871\|23142080 | Unspecified |
| mmu-miR-29b-3p | MIMAT0000127 | Mtfmt | 69606 | HITS-CLIP | 23142080 | Unspecified |
| mmu-miR-29b-3p | MIMAT0000127 | Mthfd1 | 108156 | HITS-CLIP//sRNA-Seq | 23597149\|25083871\|23142080\|33184 | Unspecified |
| mmu-miR-29b-3p | MIMAT0000127 | Mthfd1l | 270685 | HITS-CLIP | 23142080 | Unspecified |
| mmu-miR-29b-3p | MIMAT0000127 | Mthfd2 | 17768 | HITS-CLIP//sRNA-Seq | 37986224\|25083871\|33184237 | Unspecified |
| mmu-miR-29b-3p | MIMAT0000127 | Mtif2 | 76784 | HITS-CLIP | 37986224\|25083871 | Unspecified |
| mmu-miR-29b-3p | MIMAT0000127 | Mtmr1 | 53332 | HITS-CLIP | 25083871\|23142080 | Unspecified |
| mmu-miR-29b-3p | MIMAT0000127 | Mtmr12 | 268783 | HITS-CLIP | 23597149 | Unspecified |
| mmu-miR-29b-3p | MIMAT0000127 | Mtmr14 | 97287 | HITS-CLIP | 25083871\|23142080 | Unspecified |

| | | | | | | |
|---|---|---|---|---|---|---|
| mmu-miR-29b-3p | MIMAT0000127 | Mtmr2 | 77116 | HITS-CLIP//sRNA-Seq | 25083871\|33184237\|23142080 | Unspecified |
| mmu-miR-29b-3p | MIMAT0000127 | Mtmr3 | 74302 | sRNA-Seq//HITS-CLIP | 33184237\|23142080 | Unspecified |
| mmu-miR-29b-3p | MIMAT0000127 | Mtmr4 | 170749 | HITS-CLIP//sRNA-Seq | 37986224\|25083871\|33184237\|23142 | Unspecified |
| mmu-miR-29b-3p | MIMAT0000127 | Mtmr6 | 219135 | HITS-CLIP//sRNA-Seq | 25083871\|33184237\|23142080 | Unspecified |
| mmu-miR-29b-3p | MIMAT0000127 | Mtmr9 | 210376 | sRNA-Seq//HITS-CLIP | 33184237\|23142080 | Unspecified |
| mmu-miR-29b-3p | MIMAT0000127 | Mto1 | 68291 | HITS-CLIP | 23142080 | Unspecified |
| mmu-miR-29b-3p | MIMAT0000127 | Mtpap | 67440 | HITS-CLIP | 23142080\|25083871 | Unspecified |
| mmu-miR-29b-3p | MIMAT0000127 | Mtpn | 14489 | HITS-CLIP//sRNA-Seq | 37986224\|25083871\|33184237\|23142 | Unspecified |
| mmu-miR-29b-3p | MIMAT0000127 | Mtr | 238505 | HITS-CLIP | 25083871\|23597149\|23142080 | Unspecified |
| mmu-miR-29b-3p | MIMAT0000127 | Mtrr | 210009 | HITS-CLIP | 23142080 | Unspecified |
| mmu-miR-29b-3p | MIMAT0000127 | Mtss1l | 244654 | HITS-CLIP | 25083871 | Unspecified |
| mmu-miR-29b-3p | MIMAT0000127 | Mtx3 | 382793 | HITS-CLIP | 37986224\|25083871\|23142080 | Unspecified |
| mmu-miR-29b-3p | MIMAT0000127 | Muc2 | 17831 | HITS-CLIP | 23142080 | Unspecified |
| mmu-miR-29b-3p | MIMAT0000127 | Mug1 | 17836 | HITS-CLIP | 23597149 | Unspecified |
| mmu-miR-29b-3p | MIMAT0000127 | Mup20 | 381530 | HITS-CLIP | 23597149 | Unspecified |
| mmu-miR-29b-3p | MIMAT0000127 | Musk | 18198 | HITS-CLIP | 25083871 | Unspecified |
| mmu-miR-29b-3p | MIMAT0000127 | Mut | 4594 | HITS-CLIP | 23597149 | Unspecified |
| mmu-miR-29b-3p | MIMAT0000127 | Mxd1 | 17119 | HITS-CLIP | 37986224\|23597149\|23142080\|25083 | Unspecified |
| mmu-miR-29b-3p | MIMAT0000127 | Mxd3 | 17121 | HITS-CLIP | 23142080 | Unspecified |
| mmu-miR-29b-3p | MIMAT0000127 | Mxd4 | 17122 | HITS-CLIP | 37986224\|25083871\|23597149\|23142 | Unspecified |
| mmu-miR-29b-3p | MIMAT0000127 | Myadm | 50918 | HITS-CLIP//sRNA-Seq | 37986224\|33184237\|23597149\|25083 | Unspecified |
| mmu-miR-29b-3p | MIMAT0000127 | Myb | 17863 | HITS-CLIP | 37986224\|23142080 | Unspecified |
| mmu-miR-29b-3p | MIMAT0000127 | Mybl1 | 17864 | HITS-CLIP | 23142080 | Unspecified |
| mmu-miR-29b-3p | MIMAT0000127 | Mybph | 53311 | HITS-CLIP | 25083871 | Unspecified |
| mmu-miR-29b-3p | MIMAT0000127 | Myc | 17869 | HITS-CLIP//sRNA-Seq | 23142080\|25083871\|33184237 | Unspecified |
| mmu-miR-29b-3p | MIMAT0000127 | Mycbp2 | 105689 | HITS-CLIP//sRNA-Seq | 37986224\|25083871\|23142080\|33184 | Unspecified |
| mmu-miR-29b-3p | MIMAT0000127 | Mycl | 16918 | HITS-CLIP | 25083871 | Unspecified |
| mmu-miR-29b-3p | MIMAT0000127 | Mycn | 18109 | HITS-CLIP | 23142080 | Unspecified |
| mmu-miR-29b-3p | MIMAT0000127 | Myh1 | 17879 | HITS-CLIP | 25083871 | Unspecified |
| mmu-miR-29b-3p | MIMAT0000127 | Myh10 | 77579 | HITS-CLIP//sRNA-Seq | 25083871\|33184237 | Unspecified |
| mmu-miR-29b-3p | MIMAT0000127 | Myh3 | 17883 | HITS-CLIP | 25083871 | Unspecified |
| mmu-miR-29b-3p | MIMAT0000127 | Myh9 | 17886 | HITS-CLIP//sRNA-Seq | 37986224\|23597149\|23142080\|25083 | Unspecified |
| mmu-miR-29b-3p | MIMAT0000127 | Mylk | 107589 | HITS-CLIP | 25083871 | Unspecified |
| mmu-miR-29b-3p | MIMAT0000127 | Mylk4 | 238564 | HITS-CLIP | 25083871 | Unspecified |
| mmu-miR-29b-3p | MIMAT0000127 | Mymk | 66139 | HITS-CLIP | 25083871 | Unspecified |
| mmu-miR-29b-3p | MIMAT0000127 | Mynn | 80732 | HITS-CLIP | 37986224 | Unspecified |
| mmu-miR-29b-3p | MIMAT0000127 | Myo10 | 17909 | HITS-CLIP | 25083871\|23597149 | Unspecified |
| mmu-miR-29b-3p | MIMAT0000127 | Myo18a | 360013 | HITS-CLIP | 25083871\|23597149 | Unspecified |
| mmu-miR-29b-3p | MIMAT0000127 | Myo19 | 66196 | HITS-CLIP | 37986224 | Unspecified |

| | | | | | | |
|---|---|---|---|---|---|---|
| mmu-miR-29b-3p | MIMAT0000127 | Myo1c | 17913 | HITS-CLIP//sRNA-Seq | 37986224\|25083871\|33184237\|23142 | Unspecified |
| mmu-miR-29b-3p | MIMAT0000127 | Myo1e | 71602 | HITS-CLIP | 25083871 | Unspecified |
| mmu-miR-29b-3p | MIMAT0000127 | Myo1g | 246177 | HITS-CLIP | 37986224\|23142080 | Unspecified |
| mmu-miR-29b-3p | MIMAT0000127 | Myo5a | 17918 | HITS-CLIP//sRNA-Seq | 25083871\|33184237\|23142080 | Unspecified |
| mmu-miR-29b-3p | MIMAT0000127 | Myo6 | 17920 | HITS-CLIP | 25083871\|23597149\|23142080 | Unspecified |
| mmu-miR-29b-3p | MIMAT0000127 | Myo9a | 270163 | HITS-CLIP//sRNA-Seq | 25083871\|23142080\|33184237 | Unspecified |
| mmu-miR-29b-3p | MIMAT0000127 | Myo9b | 17925 | HITS-CLIP//sRNA-Seq | 37986224\|25083871\|23142080\|23597 | Unspecified |
| mmu-miR-29b-3p | MIMAT0000127 | Myod1 | 17927 | HITS-CLIP | 25083871 | Unspecified |
| mmu-miR-29b-3p | MIMAT0000127 | Myof | 226101 | HITS-CLIP//sRNA-Seq | 25083871\|33184237 | Unspecified |
| mmu-miR-29b-3p | MIMAT0000127 | Myom2 | 17930 | HITS-CLIP | 25083871 | Unspecified |
| mmu-miR-29b-3p | MIMAT0000127 | Myoz2 | 59006 | HITS-CLIP | 25083871 | Unspecified |
| mmu-miR-29b-3p | MIMAT0000127 | Mysm1 | 320713 | HITS-CLIP | 23142080 | Unspecified |
| mmu-miR-29b-3p | MIMAT0000127 | N4bp1 | 80750 | HITS-CLIP | 25083871\|23142080 | Unspecified |
| mmu-miR-29b-3p | MIMAT0000127 | N4bp2 | 333789 | HITS-CLIP | 37986224\|23142080\|23597149\|25083 | Unspecified |
| mmu-miR-29b-3p | MIMAT0000127 | N4bp2l1 | 100637 | HITS-CLIP | 37986224\|23597149\|23142080 | Unspecified |
| mmu-miR-29b-3p | MIMAT0000127 | N4bp2l2 | 381695 | sRNA-Seq//HITS-CLIP | 33184237\|23142080 | Unspecified |
| mmu-miR-29b-3p | MIMAT0000127 | N6amt1 | 67768 | HITS-CLIP | 23597149 | Unspecified |
| mmu-miR-29b-3p | MIMAT0000127 | Naa15 | 74838 | HITS-CLIP | 37986224 | Unspecified |
| mmu-miR-29b-3p | MIMAT0000127 | Naa25 | 231713 | HITS-CLIP | 25083871\|23597149\|23142080 | Unspecified |
| mmu-miR-29b-3p | MIMAT0000127 | Naa30 | 70646 | HITS-CLIP | 25083871\|23142080 | Unspecified |
| mmu-miR-29b-3p | MIMAT0000127 | Naa40 | 70999 | HITS-CLIP | 37986224\|23142080\|23597149 | Unspecified |
| mmu-miR-29b-3p | MIMAT0000127 | Naa50 | 72117 | HITS-CLIP//sRNA-Seq | 25083871\|33184237 | Unspecified |
| mmu-miR-29b-3p | MIMAT0000127 | Nab1 | 17936 | HITS-CLIP | 23142080 | Unspecified |
| mmu-miR-29b-3p | MIMAT0000127 | Nab2 | 17937 | HITS-CLIP//sRNA-Seq | 37986224\|23597149\|33184237 | Unspecified |
| mmu-miR-29b-3p | MIMAT0000127 | Nacc2 | 67991 | HITS-CLIP | 25083871 | Unspecified |
| mmu-miR-29b-3p | MIMAT0000127 | Nadk | 192185 | HITS-CLIP | 25083871\|23142080 | Unspecified |
| mmu-miR-29b-3p | MIMAT0000127 | Nagpa | 27426 | HITS-CLIP | 23142080 | Unspecified |
| mmu-miR-29b-3p | MIMAT0000127 | Nampt | 59027 | HITS-CLIP//sRNA-Seq | 23597149\|23142080\|33184237 | Unspecified |
| mmu-miR-29b-3p | MIMAT0000127 | Nanos1 | 332397 | HITS-CLIP | 23142080 | Unspecified |
| mmu-miR-29b-3p | MIMAT0000127 | Nanp | 67311 | HITS-CLIP | 25083871\|23142080 | Unspecified |
| mmu-miR-29b-3p | MIMAT0000127 | Nans | 94181 | HITS-CLIP | 37986224 | Unspecified |
| mmu-miR-29b-3p | MIMAT0000127 | Nap1l1 | 53605 | HITS-CLIP//sRNA-Seq | 37986224\|23142080\|33184237\|25083 | Unspecified |
| mmu-miR-29b-3p | MIMAT0000127 | Nap1l3 | 54561 | HITS-CLIP | 23142080 | Unspecified |
| mmu-miR-29b-3p | MIMAT0000127 | Nap1l4 | 17955 | sRNA-Seq//HITS-CLIP | 33184237\|23597149\|23142080 | Unspecified |
| mmu-miR-29b-3p | MIMAT0000127 | Naprt | 93100 | HITS-CLIP | 23597149 | Unspecified |
| mmu-miR-29b-3p | MIMAT0000127 | Narf | 67608 | HITS-CLIP | 37986224\|25083871\|23597149\|23142 | Unspecified |
| mmu-miR-29b-3p | MIMAT0000127 | Nars | 70223 | HITS-CLIP//sRNA-Seq | 23142080\|33184237\|23597149\|25083 | Unspecified |
| mmu-miR-29b-3p | MIMAT0000127 | Nasp | 50927 | HITS-CLIP//sRNA-Seq | 37986224\|25083871\|23597149\|33184 | Unspecified |
| mmu-miR-29b-3p | MIMAT0000127 | Nat8f5 | 69049 | HITS-CLIP | 23597149 | Unspecified |

| | | | | | | |
|---|---|---|---|---|---|---|
| mmu-miR-29b-3p | MIMAT0000127 | Natd1 | 24083 | HITS-CLIP | 25083871\|23142080 | Unspecified |
| mmu-miR-29b-3p | MIMAT0000127 | Nav1 | 215690 | HITS-CLIP//sRNA-Seq | 37986224\|25083871\|33184237\|23597[illegible] | Unspecified |
| mmu-miR-29b-3p | MIMAT0000127 | Nav2 | 78286 | HITS-CLIP | 37986224\|25083871\|23597149 | Unspecified |
| mmu-miR-29b-3p | MIMAT0000127 | Nav3 | 260315 | HITS-CLIP//sRNA-Seq | 37986224\|25083871\|33184237 | Unspecified |
| mmu-miR-29b-3p | MIMAT0000127 | Nbdy | 550643 | HITS-CLIP//sRNA-Seq | 25083871\|33184237 | Unspecified |
| mmu-miR-29b-3p | MIMAT0000127 | Nbeal1 | 269198 | HITS-CLIP | 37986224\|23142080\|25083871 | Unspecified |
| mmu-miR-29b-3p | MIMAT0000127 | Nbeal2 | 235627 | HITS-CLIP | 23597149 | Unspecified |
| mmu-miR-29b-3p | MIMAT0000127 | Nbl1 | 17965 | HITS-CLIP | 25083871 | Unspecified |
| mmu-miR-29b-3p | MIMAT0000127 | Nbr1 | 17966 | HITS-CLIP//sRNA-Seq | 25083871\|33184237\|23142080 | Unspecified |
| mmu-miR-29b-3p | MIMAT0000127 | Ncapd2 | 68298 | sRNA-Seq//HITS-CLIP | 33184237\|23142080 | Unspecified |
| mmu-miR-29b-3p | MIMAT0000127 | Ncapd3 | 78658 | HITS-CLIP | 23142080 | Unspecified |
| mmu-miR-29b-3p | MIMAT0000127 | Ncapg | 54392 | HITS-CLIP | 23142080 | Unspecified |
| mmu-miR-29b-3p | MIMAT0000127 | Ncapg2 | 76044 | HITS-CLIP | 23142080 | Unspecified |
| mmu-miR-29b-3p | MIMAT0000127 | Ncbp1 | 433702 | HITS-CLIP//sRNA-Seq | 25083871\|33184237\|23142080 | Unspecified |
| mmu-miR-29b-3p | MIMAT0000127 | Ncbp3 | 66874 | HITS-CLIP | 25083871 | Unspecified |
| mmu-miR-29b-3p | MIMAT0000127 | Ncdn | 26562 | HITS-CLIP//sRNA-Seq | 23597149\|25083871\|33184237 | Unspecified |
| mmu-miR-29b-3p | MIMAT0000127 | Nceh1 | 320024 | HITS-CLIP | 23597149 | Unspecified |
| mmu-miR-29b-3p | MIMAT0000127 | Ncf1 | 17969 | HITS-CLIP | 23142080 | Unspecified |
| mmu-miR-29b-3p | MIMAT0000127 | Ncf4 | 17972 | HITS-CLIP | 23142080 | Unspecified |
| mmu-miR-29b-3p | MIMAT0000127 | Nck1 | 17973 | sRNA-Seq//HITS-CLIP | 33184237\|23142080 | Unspecified |
| mmu-miR-29b-3p | MIMAT0000127 | Nckap1 | 50884 | HITS-CLIP//sRNA-Seq | 25083871\|33184237 | Unspecified |
| mmu-miR-29b-3p | MIMAT0000127 | Nckap5l | 380969 | HITS-CLIP | 37986224\|23142080\|25083871 | Unspecified |
| mmu-miR-29b-3p | MIMAT0000127 | Ncl | 17975 | HITS-CLIP//sRNA-Seq | 37986224\|25083871\|33184237 | Unspecified |
| mmu-miR-29b-3p | MIMAT0000127 | Ncoa2 | 17978 | HITS-CLIP | 37986224\|23142080\|23597149 | Unspecified |
| mmu-miR-29b-3p | MIMAT0000127 | Ncoa3 | 17979 | HITS-CLIP | 25083871\|23142080 | Unspecified |
| mmu-miR-29b-3p | MIMAT0000127 | Ncoa5 | 228869 | HITS-CLIP | 23142080 | Unspecified |
| mmu-miR-29b-3p | MIMAT0000127 | Ncoa6 | 56406 | HITS-CLIP | 37986224\|23142080 | Unspecified |
| mmu-miR-29b-3p | MIMAT0000127 | Ncoa7 | 211329 | HITS-CLIP//sRNA-Seq | 37986224\|23142080\|33184237 | Unspecified |
| mmu-miR-29b-3p | MIMAT0000127 | Ncor1 | 20185 | sRNA-Seq//HITS-CLIP | 33184237\|25083871\|23142080 | Unspecified |
| mmu-miR-29b-3p | MIMAT0000127 | Ncor2 | 20602 | HITS-CLIP | 37986224\|25083871 | Unspecified |
| mmu-miR-29b-3p | MIMAT0000127 | Ncstn | 59287 | sRNA-Seq//HITS-CLIP | 33184237\|25083871 | Unspecified |
| mmu-miR-29b-3p | MIMAT0000127 | Ndc1 | 72787 | HITS-CLIP | 23142080\|25083871 | Unspecified |
| mmu-miR-29b-3p | MIMAT0000127 | Nde1 | 67203 | sRNA-Seq//HITS-CLIP | 33184237\|23142080 | Unspecified |
| mmu-miR-29b-3p | MIMAT0000127 | Ndfip2 | 76273 | HITS-CLIP//sRNA-Seq | 25083871\|23142080\|33184237 | Unspecified |
| mmu-miR-29b-3p | MIMAT0000127 | Ndrg1 | 17988 | HITS-CLIP//sRNA-Seq | 23142080\|33184237 | Unspecified |
| mmu-miR-29b-3p | MIMAT0000127 | Ndrg2 | 29811 | HITS-CLIP | 23597149\|23142080 | Unspecified |
| mmu-miR-29b-3p | MIMAT0000127 | Ndst1 | 15531 | HITS-CLIP//sRNA-Seq | 37986224\|23142080\|23597149\|3318[illegible] | Unspecified |
| mmu-miR-29b-3p | MIMAT0000127 | Ndst2 | 17423 | HITS-CLIP | 37986224\|25083871 | Unspecified |
| mmu-miR-29b-3p | MIMAT0000127 | Ndufa2 | 17991 | sRNA-Seq//HITS-CLIP | 33184237\|25083871\|23142080 | Unspecified |

| | | | | | | |
|---|---|---|---|---|---|---|
| mmu-miR-29b-3p | MIMAT0000127 | Ndufa8 | 68375 | sRNA-Seq//HITS-CLIP | 33184237\|23597149 | Unspecified |
| mmu-miR-29b-3p | MIMAT0000127 | Ndufa9 | 66108 | sRNA-Seq//HITS-CLIP | 33184237\|23597149 | Unspecified |
| mmu-miR-29b-3p | MIMAT0000127 | Ndufaf7 | 73694 | HITS-CLIP | 25083871 | Unspecified |
| mmu-miR-29b-3p | MIMAT0000127 | Ndufb2 | 68198 | HITS-CLIP//sRNA-Seq | 23142080\|33184237 | Unspecified |
| mmu-miR-29b-3p | MIMAT0000127 | Ndufb6 | 230075 | sRNA-Seq//HITS-CLIP | 33184237\|23142080 | Unspecified |
| mmu-miR-29b-3p | MIMAT0000127 | Ndufb8 | 67264 | sRNA-Seq//HITS-CLIP | 33184237\|23597149 | Unspecified |
| mmu-miR-29b-3p | MIMAT0000127 | Ndufb9 | 66218 | HITS-CLIP//sRNA-Seq | 23597149\|33184237\|25083871\|23142 | Unspecified |
| mmu-miR-29b-3p | MIMAT0000127 | Ndufc2 | 68197 | sRNA-Seq//HITS-CLIP | 33184237\|23142080 | Unspecified |
| mmu-miR-29b-3p | MIMAT0000127 | Ndufs1 | 227197 | HITS-CLIP//sRNA-Seq | 23597149\|33184237\|23142080 | Unspecified |
| mmu-miR-29b-3p | MIMAT0000127 | Ndufs4 | 17993 | HITS-CLIP//sRNA-Seq | 25083871\|33184237\|23142080 | Unspecified |
| mmu-miR-29b-3p | MIMAT0000127 | Ndufv1 | 17995 | HITS-CLIP//sRNA-Seq | 25083871\|23597149\|33184237\|23142 | Unspecified |
| mmu-miR-29b-3p | MIMAT0000127 | Ndufv2 | 72900 | HITS-CLIP//sRNA-Seq | 23142080\|25083871\|33184237\|23597 | Unspecified |
| mmu-miR-29b-3p | MIMAT0000127 | Neb | 17996 | HITS-CLIP | 25083871 | Unspecified |
| mmu-miR-29b-3p | MIMAT0000127 | Nebl | 74103 | sRNA-Seq//HITS-CLIP | 33184237\|23142080 | Unspecified |
| mmu-miR-29b-3p | MIMAT0000127 | Necap1 | 67602 | HITS-CLIP//sRNA-Seq | 25083871\|23597149\|33184237\|23142 | Unspecified |
| mmu-miR-29b-3p | MIMAT0000127 | Nectin1 | 58235 | HITS-CLIP | 23597149 | Unspecified |
| mmu-miR-29b-3p | MIMAT0000127 | Nectin2 | 5819 | HITS-CLIP | 23597149 | Unspecified |
| mmu-miR-29b-3p | MIMAT0000127 | Nedd1 | 17997 | HITS-CLIP | 25083871\|23142080 | Unspecified |
| mmu-miR-29b-3p | MIMAT0000127 | Nedd4 | 17999 | HITS-CLIP//sRNA-Seq | 25083871\|23597149\|33184237 | Unspecified |
| mmu-miR-29b-3p | MIMAT0000127 | Nedd9 | 18003 | HITS-CLIP//sRNA-Seq | 37986224\|33184237\|25083871 | Unspecified |
| mmu-miR-29b-3p | MIMAT0000127 | Neil1 | 72774 | HITS-CLIP | 23142080 | Unspecified |
| mmu-miR-29b-3p | MIMAT0000127 | Nek7 | 59125 | HITS-CLIP | 23142080\|25083871 | Unspecified |
| mmu-miR-29b-3p | MIMAT0000127 | Nek9 | 217718 | sRNA-Seq//HITS-CLIP | 33184237\|23597149 | Unspecified |
| mmu-miR-29b-3p | MIMAT0000127 | Nelfa | 24116 | HITS-CLIP | 25083871\|23142080 | Unspecified |
| mmu-miR-29b-3p | MIMAT0000127 | Nelfb | 58202 | HITS-CLIP//sRNA-Seq | 37986224\|23597149\|33184237\|23142 | Unspecified |
| mmu-miR-29b-3p | MIMAT0000127 | Nelfcd | 57314 | HITS-CLIP//sRNA-Seq | 23597149\|23142080\|33184237 | Unspecified |
| mmu-miR-29b-3p | MIMAT0000127 | Nemf | 66244 | HITS-CLIP | 23142080\|25083871 | Unspecified |
| mmu-miR-29b-3p | MIMAT0000127 | Neo1 | 18007 | HITS-CLIP//sRNA-Seq | 25083871\|23597149\|33184237 | Unspecified |
| mmu-miR-29b-3p | MIMAT0000127 | Nepro | 212547 | HITS-CLIP | 25083871 | Unspecified |
| mmu-miR-29b-3p | MIMAT0000127 | Nes | 18008 | HITS-CLIP//sRNA-Seq | 25083871\|33184237 | Unspecified |
| mmu-miR-29b-3p | MIMAT0000127 | Net1 | 56349 | HITS-CLIP//sRNA-Seq | 25083871\|23597149\|33184237 | Unspecified |
| mmu-miR-29b-3p | MIMAT0000127 | Neu3 | 50877 | HITS-CLIP | 37986224\|23142080 | Unspecified |
| mmu-miR-29b-3p | MIMAT0000127 | Neurl1b | 240055 | HITS-CLIP | 23597149 | Unspecified |
| mmu-miR-29b-3p | MIMAT0000127 | Neurl4 | 216860 | HITS-CLIP | 25083871 | Unspecified |
| mmu-miR-29b-3p | MIMAT0000127 | Nf1 | 18015 | HITS-CLIP//sRNA-Seq | 25083871\|33184237\|23142080 | Unspecified |
| mmu-miR-29b-3p | MIMAT0000127 | Nf2 | 18016 | HITS-CLIP//sRNA-Seq | 37986224\|25083871\|33184237\|23597 | Unspecified |
| mmu-miR-29b-3p | MIMAT0000127 | Nfat5 | 54446 | HITS-CLIP | 37986224\|23142080\|25083871 | Unspecified |
| mmu-miR-29b-3p | MIMAT0000127 | Nfatc1 | 18018 | HITS-CLIP | 23142080 | Unspecified |
| mmu-miR-29b-3p | MIMAT0000127 | Nfatc2 | 18019 | HITS-CLIP | 23142080 | Unspecified |

| | | | | | | |
|---|---|---|---|---|---|---|
| mmu-miR-29b-3p | MIMAT0000127 | Nfatc2ip | 18020 | HITS-CLIP | 37986224\|25083871\|23142080 | Unspecified |
| mmu-miR-29b-3p | MIMAT0000127 | Nfatc3 | 18021 | HITS-CLIP//sRNA-Seq | 25083871\|23597149\|33184237\|23142 | Unspecified |
| mmu-miR-29b-3p | MIMAT0000127 | Nfe2l1 | 18023 | HITS-CLIP//sRNA-Seq | 37986224\|25083871\|23597149\|33184 | Unspecified |
| mmu-miR-29b-3p | MIMAT0000127 | Nfia | 18027 | HITS-CLIP//qCLASH | 37986224\|23597149\|31393953\|23142 | Unspecified |
| mmu-miR-29b-3p | MIMAT0000127 | Nfib | 18028 | HITS-CLIP | 37986224 | Unspecified |
| mmu-miR-29b-3p | MIMAT0000127 | Nfic | 18029 | HITS-CLIP//sRNA-Seq | 37986224\|33184237\|23597149\|23142 | Unspecified |
| mmu-miR-29b-3p | MIMAT0000127 | Nfkb1 | 18033 | HITS-CLIP//sRNA-Seq | 25083871\|33184237 | Unspecified |
| mmu-miR-29b-3p | MIMAT0000127 | Nfrkb | 235134 | HITS-CLIP | 31433990\|23142080 | Unspecified |
| mmu-miR-29b-3p | MIMAT0000127 | Nfs1 | 18041 | HITS-CLIP | 23597149 | Unspecified |
| mmu-miR-29b-3p | MIMAT0000127 | Nfxl1 | 100978 | HITS-CLIP | 37986224\|23597149\|23142080 | Unspecified |
| mmu-miR-29b-3p | MIMAT0000127 | Nfya | 18044 | HITS-CLIP | 23142080\|25083871 | Unspecified |
| mmu-miR-29b-3p | MIMAT0000127 | Ngdn | 68966 | HITS-CLIP//sRNA-Seq | 23597149\|23142080\|33184237 | Unspecified |
| mmu-miR-29b-3p | MIMAT0000127 | Ngly1 | 59007 | HITS-CLIP | 23142080 | Unspecified |
| mmu-miR-29b-3p | MIMAT0000127 | Nhlrc2 | 66866 | HITS-CLIP | 23597149 | Unspecified |
| mmu-miR-29b-3p | MIMAT0000127 | Nhlrc3 | 212114 | HITS-CLIP | 37986224 | Unspecified |
| mmu-miR-29b-3p | MIMAT0000127 | Nhsl1 | 215819 | HITS-CLIP | 37986224 | Unspecified |
| mmu-miR-29b-3p | MIMAT0000127 | Nhsl2 | 100042480 | HITS-CLIP | 23142080 | Unspecified |
| mmu-miR-29b-3p | MIMAT0000127 | Nicn1 | 66257 | HITS-CLIP//sRNA-Seq | 25083871\|33184237\|23142080 | Unspecified |
| mmu-miR-29b-3p | MIMAT0000127 | Nid1 | 18073 | HITS-CLIP//sRNA-Seq | 37986224\|25083871\|33184237 | Unspecified |
| mmu-miR-29b-3p | MIMAT0000127 | Nid2 | 18074 | HITS-CLIP | 37986224 | Unspecified |
| mmu-miR-29b-3p | MIMAT0000127 | Nifk | 67949 | HITS-CLIP | 37986224\|25083871\|23142080 | Unspecified |
| mmu-miR-29b-3p | MIMAT0000127 | Nipal1 | 70701 | HITS-CLIP | 23142080 | Unspecified |
| mmu-miR-29b-3p | MIMAT0000127 | Nipbl | 71175 | HITS-CLIP//sRNA-Seq | 25083871\|33184237\|26203562\|23142 | Unspecified |
| mmu-miR-29b-3p | MIMAT0000127 | Nipsnap2 | 2631 | HITS-CLIP//sRNA-Seq | 25083871\|33184237 | Unspecified |
| mmu-miR-29b-3p | MIMAT0000127 | Nipsnap3b | 66536 | HITS-CLIP//sRNA-Seq | 37986224\|23597149\|33184237\|23142 | Unspecified |
| mmu-miR-29b-3p | MIMAT0000127 | Nisch | 64652 | HITS-CLIP//sRNA-Seq | 37986224\|23597149\|33184237 | Unspecified |
| mmu-miR-29b-3p | MIMAT0000127 | Nit2 | 52633 | HITS-CLIP | 37986224\|23597149\|23142080\|25083 | Unspecified |
| mmu-miR-29b-3p | MIMAT0000127 | Nkap | 67050 | HITS-CLIP | 23142080\|25083871 | Unspecified |
| mmu-miR-29b-3p | MIMAT0000127 | Nkiras2 | 71966 | HITS-CLIP | 37986224\|25083871 | Unspecified |
| mmu-miR-29b-3p | MIMAT0000127 | Nkrf | 77286 | HITS-CLIP | 37986224\|25083871\|23597149\|23142 | Unspecified |
| mmu-miR-29b-3p | MIMAT0000127 | Nktr | 18087 | HITS-CLIP//sRNA-Seq | 37986224\|25083871\|23597149\|33184 | Unspecified |
| mmu-miR-29b-3p | MIMAT0000127 | Nlk | 18099 | HITS-CLIP//sRNA-Seq | 23142080\|25083871\|33184237 | Unspecified |
| mmu-miR-29b-3p | MIMAT0000127 | Nlrc5 | 434341 | HITS-CLIP | 23142080\|23597149 | Unspecified |
| mmu-miR-29b-3p | MIMAT0000127 | Nlrx1 | 270151 | HITS-CLIP | 37986224\|25083871 | Unspecified |
| mmu-miR-29b-3p | MIMAT0000127 | Nmd3 | 97112 | HITS-CLIP//sRNA-Seq | 23597149\|25083871\|33184237\|23142 | Unspecified |
| mmu-miR-29b-3p | MIMAT0000127 | Nme1 | 18102 | HITS-CLIP//sRNA-Seq | 23142080\|33184237 | Unspecified |
| mmu-miR-29b-3p | MIMAT0000127 | Nme3 | 79059 | HITS-CLIP | 23142080 | Unspecified |
| mmu-miR-29b-3p | MIMAT0000127 | Nme7 | 171567 | HITS-CLIP | 25083871 | Unspecified |
| mmu-miR-29b-3p | MIMAT0000127 | Nmrk1 | 225994 | HITS-CLIP | 37986224 | Unspecified |

| | | | | | | |
|---|---|---|---|---|---|---|
| mmu-miR-29b-3p | MIMAT0000127 | Nmt1 | 18107 | HITS-CLIP//sRNA-Seq | 25083871\|23142080\|33184237 | Unspecified |
| mmu-miR-29b-3p | MIMAT0000127 | Nmt2 | 18108 | HITS-CLIP//sRNA-Seq | 25083871\|33184237\|23142080 | Unspecified |
| mmu-miR-29b-3p | MIMAT0000127 | Noc2l | 57741 | HITS-CLIP//sRNA-Seq | 23142080\|25083871\|33184237 | Unspecified |
| mmu-miR-29b-3p | MIMAT0000127 | Noc4l | 100608 | HITS-CLIP | 23597149\|23142080 | Unspecified |
| mmu-miR-29b-3p | MIMAT0000127 | Nod1 | 107607 | HITS-CLIP | 23597149 | Unspecified |
| mmu-miR-29b-3p | MIMAT0000127 | Nod2 | 257632 | HITS-CLIP | 37986224\|23142080 | Unspecified |
| mmu-miR-29b-3p | MIMAT0000127 | Nol10 | 217431 | HITS-CLIP | 23142080 | Unspecified |
| mmu-miR-29b-3p | MIMAT0000127 | Nol4l | 329540 | HITS-CLIP//sRNA-Seq | 37986224\|33184237\|23142080 | Unspecified |
| mmu-miR-29b-3p | MIMAT0000127 | Nol8 | 70930 | HITS-CLIP | 23142080 | Unspecified |
| mmu-miR-29b-3p | MIMAT0000127 | Nom1 | 433864 | HITS-CLIP | 25083871\|23142080 | Unspecified |
| mmu-miR-29b-3p | MIMAT0000127 | Nomo1 | 211548 | HITS-CLIP//sRNA-Seq | 25083871\|33184237 | Unspecified |
| mmu-miR-29b-3p | MIMAT0000127 | Nono | 53610 | HITS-CLIP//sRNA-Seq | 23597149\|25083871\|33184237\|23142 | Unspecified |
| mmu-miR-29b-3p | MIMAT0000127 | Nop10 | 66181 | HITS-CLIP//sRNA-Seq | 23597149\|33184237\|23142080 | Unspecified |
| mmu-miR-29b-3p | MIMAT0000127 | Nop2 | 110109 | sRNA-Seq//HITS-CLIP | 33184237\|23142080 | Unspecified |
| mmu-miR-29b-3p | MIMAT0000127 | Nop53 | 29997 | HITS-CLIP | 23142080 | Unspecified |
| mmu-miR-29b-3p | MIMAT0000127 | Nop56 | 67134 | HITS-CLIP//sRNA-Seq | 37986224\|25083871\|33184237\|23142 | Unspecified |
| mmu-miR-29b-3p | MIMAT0000127 | Nos3 | 18127 | HITS-CLIP | 23597149 | Unspecified |
| mmu-miR-29b-3p | MIMAT0000127 | Notch1 | 18128 | HITS-CLIP | 25083871\|23142080 | Unspecified |
| mmu-miR-29b-3p | MIMAT0000127 | Notch2 | 18129 | HITS-CLIP//sRNA-Seq | 25083871\|23597149\|33184237\|23142 | Unspecified |
| mmu-miR-29b-3p | MIMAT0000127 | Npas3 | 27386 | HITS-CLIP | 37986224\|25083871 | Unspecified |
| mmu-miR-29b-3p | MIMAT0000127 | Npc1 | 18145 | HITS-CLIP | 23597149\|25083871 | Unspecified |
| mmu-miR-29b-3p | MIMAT0000127 | Npc2 | 67963 | sRNA-Seq//HITS-CLIP | 33184237\|25083871\|23142080 | Unspecified |
| mmu-miR-29b-3p | MIMAT0000127 | Npepps | 19155 | HITS-CLIP//sRNA-Seq | 23597149\|33184237\|23142080\|31433 | Unspecified |
| mmu-miR-29b-3p | MIMAT0000127 | Nphp3 | 74025 | HITS-CLIP | 25083871 | Unspecified |
| mmu-miR-29b-3p | MIMAT0000127 | Nploc4 | 217365 | HITS-CLIP//sRNA-Seq | 25083871\|33184237\|23142080 | Unspecified |
| mmu-miR-29b-3p | MIMAT0000127 | Npnt | 114249 | HITS-CLIP | 37986224\|25083871 | Unspecified |
| mmu-miR-29b-3p | MIMAT0000127 | Nptn | 20320 | HITS-CLIP//sRNA-Seq | 37986224\|23142080\|25083871\|33184 | Unspecified |
| mmu-miR-29b-3p | MIMAT0000127 | Nr1d1 | 217166 | HITS-CLIP | 25083871\|23597149 | Unspecified |
| mmu-miR-29b-3p | MIMAT0000127 | Nr1d2 | 353187 | HITS-CLIP | 25083871 | Unspecified |
| mmu-miR-29b-3p | MIMAT0000127 | Nr1h3 | 22259 | HITS-CLIP | 23597149 | Unspecified |
| mmu-miR-29b-3p | MIMAT0000127 | Nr2c2 | 22026 | HITS-CLIP | 37986224\|25083871\|23142080 | Unspecified |
| mmu-miR-29b-3p | MIMAT0000127 | Nr3c1 | 14815 | HITS-CLIP//sRNA-Seq | 25083871\|33184237\|23142080 | Unspecified |
| mmu-miR-29b-3p | MIMAT0000127 | Nr4a1 | 15370 | HITS-CLIP | 23142080 | Unspecified |
| mmu-miR-29b-3p | MIMAT0000127 | Nr4a3 | 18124 | HITS-CLIP | 23142080 | Unspecified |
| mmu-miR-29b-3p | MIMAT0000127 | Nr5a2 | 26424 | HITS-CLIP | 23597149 | Unspecified |
| mmu-miR-29b-3p | MIMAT0000127 | Nras | 18176 | HITS-CLIP//sRNA-Seq | 37986224\|25083871\|23597149\|33184 | Unspecified |
| mmu-miR-29b-3p | MIMAT0000127 | Nrbp1 | 192292 | HITS-CLIP//sRNA-Seq | 23142080\|33184237\|23597149 | Unspecified |
| mmu-miR-29b-3p | MIMAT0000127 | Nrd1 | 230598 | HITS-CLIP//sRNA-Seq | 37986224\|25083871\|33184237\|23597 | Unspecified |
| mmu-miR-29b-3p | MIMAT0000127 | Nrep | 27528 | HITS-CLIP//sRNA-Seq//RNA-Seq | 25083871\|33184237\|33332475\|23597 | Unspecified |

| | | | | | | |
|---|---|---|---|---|---|---|
| mmu-miR-29b-3p | MIMAT0000127 | Nrip1 | 268903 | HITS-CLIP | 37986224\|25083871 | Unspecified |
| mmu-miR-29b-3p | MIMAT0000127 | Nrn1 | 68404 | HITS-CLIP//sRNA-Seq | 23142080\|33184237\|23597149 | Unspecified |
| mmu-miR-29b-3p | MIMAT0000127 | Nrp2 | 18187 | HITS-CLIP//sRNA-Seq | 25083871\|33184237\|23597149 | Unspecified |
| mmu-miR-29b-3p | MIMAT0000127 | Nrros | 224109 | HITS-CLIP | 23142080\|23597149 | Unspecified |
| mmu-miR-29b-3p | MIMAT0000127 | Nsd1 | 18193 | HITS-CLIP//qCLASH//sRNA-Seq | 37986224\|25083871\|23597149\|31393 | Unspecified |
| mmu-miR-29b-3p | MIMAT0000127 | Nsd2 | 107823 | HITS-CLIP//sRNA-Seq | 25083871\|33184237\|23142080 | Unspecified |
| mmu-miR-29b-3p | MIMAT0000127 | Nsd3 | 234135 | HITS-CLIP//sRNA-Seq | 23142080\|25083871\|33184237\|23597 | Unspecified |
| mmu-miR-29b-3p | MIMAT0000127 | Nsf | 18195 | sRNA-Seq//HITS-CLIP | 33184237\|23142080 | Unspecified |
| mmu-miR-29b-3p | MIMAT0000127 | Nsmaf | 18201 | HITS-CLIP | 23142080 | Unspecified |
| mmu-miR-29b-3p | MIMAT0000127 | Nsun2 | 28114 | sRNA-Seq//HITS-CLIP | 33184237\|23142080\|25083871 | Unspecified |
| mmu-miR-29b-3p | MIMAT0000127 | Nsun4 | 72181 | HITS-CLIP | 23142080 | Unspecified |
| mmu-miR-29b-3p | MIMAT0000127 | Nub1 | 53312 | HITS-CLIP//sRNA-Seq | 37986224\|23142080\|23597149\|33184 | Unspecified |
| mmu-miR-29b-3p | MIMAT0000127 | Nucb1 | 18220 | HITS-CLIP//sRNA-Seq | 23142080\|33184237\|23597149\|25083 | Unspecified |
| mmu-miR-29b-3p | MIMAT0000127 | Nucks1 | 98415 | HITS-CLIP//sRNA-Seq | 37986224\|25083871\|33184237\|23597 | Unspecified |
| mmu-miR-29b-3p | MIMAT0000127 | Nudcd2 | 52653 | HITS-CLIP | 37986224\|23142080\|25083871 | Unspecified |
| mmu-miR-29b-3p | MIMAT0000127 | Nudt1 | 17766 | HITS-CLIP | 23142080 | Unspecified |
| mmu-miR-29b-3p | MIMAT0000127 | Nudt18 | 213484 | HITS-CLIP | 25083871\|23597149\|23142080 | Unspecified |
| mmu-miR-29b-3p | MIMAT0000127 | Nudt2 | 66401 | HITS-CLIP | 25083871 | Unspecified |
| mmu-miR-29b-3p | MIMAT0000127 | Nudt3 | 56409 | HITS-CLIP//sRNA-Seq | 23597149\|33184237 | Unspecified |
| mmu-miR-29b-3p | MIMAT0000127 | Nudt9 | 74167 | HITS-CLIP | 37986224 | Unspecified |
| mmu-miR-29b-3p | MIMAT0000127 | Nufip2 | 68564 | HITS-CLIP//sRNA-Seq | 37986224\|25083871\|23597149\|33184 | Unspecified |
| mmu-miR-29b-3p | MIMAT0000127 | Numa1 | 101706 | HITS-CLIP//sRNA-Seq | 37986224\|25083871\|33184237\|23142 | Unspecified |
| mmu-miR-29b-3p | MIMAT0000127 | Numb | 18222 | HITS-CLIP | 25083871\|23142080 | Unspecified |
| mmu-miR-29b-3p | MIMAT0000127 | Nup133 | 234865 | HITS-CLIP | 23597149\|23142080 | Unspecified |
| mmu-miR-29b-3p | MIMAT0000127 | Nup153 | 218210 | HITS-CLIP//sRNA-Seq | 23597149\|25083871\|23142080\|33184 | Unspecified |
| mmu-miR-29b-3p | MIMAT0000127 | Nup155 | 170762 | HITS-CLIP | 23142080 | Unspecified |
| mmu-miR-29b-3p | MIMAT0000127 | Nup160 | 59015 | HITS-CLIP | 37986224\|25083871\|23142080 | Unspecified |
| mmu-miR-29b-3p | MIMAT0000127 | Nup188 | 227699 | sRNA-Seq//HITS-CLIP | 33184237\|23142080 | Unspecified |
| mmu-miR-29b-3p | MIMAT0000127 | Nup205 | 70699 | HITS-CLIP//sRNA-Seq | 25083871\|33184237 | Unspecified |
| mmu-miR-29b-3p | MIMAT0000127 | Nup210 | 54563 | HITS-CLIP | 37986224\|25083871\|23142080 | Unspecified |
| mmu-miR-29b-3p | MIMAT0000127 | Nup214 | 227720 | HITS-CLIP | 37986224\|25083871\|23142080 | Unspecified |
| mmu-miR-29b-3p | MIMAT0000127 | Nup37 | 69736 | HITS-CLIP | 23142080 | Unspecified |
| mmu-miR-29b-3p | MIMAT0000127 | Nup50 | 18141 | HITS-CLIP | 25083871\|23142080 | Unspecified |
| mmu-miR-29b-3p | MIMAT0000127 | Nup62 | 18226 | sRNA-Seq//HITS-CLIP | 33184237\|23142080 | Unspecified |
| mmu-miR-29b-3p | MIMAT0000127 | Nup85 | 445007 | sRNA-Seq//HITS-CLIP | 33184237\|23142080 | Unspecified |
| mmu-miR-29b-3p | MIMAT0000127 | Nup88 | 19069 | sRNA-Seq//HITS-CLIP | 33184237\|23142080 | Unspecified |
| mmu-miR-29b-3p | MIMAT0000127 | Nup93 | 71805 | HITS-CLIP//sRNA-Seq | 23142080\|33184237\|25083871 | Unspecified |
| mmu-miR-29b-3p | MIMAT0000127 | Nup98 | 269966 | HITS-CLIP | 23142080\|23597149\|25083871 | Unspecified |
| mmu-miR-29b-3p | MIMAT0000127 | Nus1 | 52014 | HITS-CLIP | 25083871 | Unspecified |

| | | | | | | |
|---|---|---|---|---|---|---|
| mmu-miR-29b-3p | MIMAT0000127 | Nusap1 | 108907 | sRNA-Seq//HITS-CLIP | 33184237\|23142080 | Unspecified |
| mmu-miR-29b-3p | MIMAT0000127 | Nvl | 67459 | HITS-CLIP | 37986224\|25083871\|23142080 | Unspecified |
| mmu-miR-29b-3p | MIMAT0000127 | Nxf1 | 53319 | HITS-CLIP//sRNA-Seq | 37986224\|25083871\|33184237 | Unspecified |
| mmu-miR-29b-3p | MIMAT0000127 | Nxn | 18230 | HITS-CLIP//sRNA-Seq | 25083871\|33184237 | Unspecified |
| mmu-miR-29b-3p | MIMAT0000127 | Nxpe3 | 385658 | HITS-CLIP | 23142080 | Unspecified |
| mmu-miR-29b-3p | MIMAT0000127 | Nynrin | 277154 | HITS-CLIP | 25083871 | Unspecified |
| mmu-miR-29b-3p | MIMAT0000127 | Oat | 18242 | HITS-CLIP//sRNA-Seq | 37986224\|23597149\|33184237\|23142 | Unspecified |
| mmu-miR-29b-3p | MIMAT0000127 | Obscn | 380698 | HITS-CLIP | 37986224\|25083871\|23597149 | Unspecified |
| mmu-miR-29b-3p | MIMAT0000127 | Obsl1 | 98733 | HITS-CLIP | 37986224\|25083871 | Unspecified |
| mmu-miR-29b-3p | MIMAT0000127 | Ociad2 | 433904 | HITS-CLIP | 23597149 | Unspecified |
| mmu-miR-29b-3p | MIMAT0000127 | Odc1 | 18263 | HITS-CLIP//sRNA-Seq | 25083871\|23142080\|23597149\|33184 | Unspecified |
| mmu-miR-29b-3p | MIMAT0000127 | Ogdh | 18293 | HITS-CLIP//sRNA-Seq | 23597149\|33184237\|25083871\|23142 | Unspecified |
| mmu-miR-29b-3p | MIMAT0000127 | Ogfr | 72075 | HITS-CLIP//sRNA-Seq | 25083871\|33184237\|23142080 | Unspecified |
| mmu-miR-29b-3p | MIMAT0000127 | Ogt | 108155 | HITS-CLIP//sRNA-Seq | 37986224\|25083871\|33184237\|23597 | Unspecified |
| mmu-miR-29b-3p | MIMAT0000127 | Oit3 | 18302 | HITS-CLIP | 23597149 | Unspecified |
| mmu-miR-29b-3p | MIMAT0000127 | Ola1 | 67059 | HITS-CLIP//sRNA-Seq | 25083871\|23597149\|33184237 | Unspecified |
| mmu-miR-29b-3p | MIMAT0000127 | Olfm1 | 56177 | HITS-CLIP//sRNA-Seq | 25083871\|33184237 | Unspecified |
| mmu-miR-29b-3p | MIMAT0000127 | Olfm3 | 229759 | HITS-CLIP | 37986224\|23597149 | Unspecified |
| mmu-miR-29b-3p | MIMAT0000127 | Olfml2b | 320078 | HITS-CLIP//sRNA-Seq | 25083871\|33184237 | Unspecified |
| mmu-miR-29b-3p | MIMAT0000127 | Onecut2 | 225631 | sRNA-Seq//HITS-CLIP | 33184237\|23597149 | Unspecified |
| mmu-miR-29b-3p | MIMAT0000127 | Opa1 | 74143 | HITS-CLIP//sRNA-Seq | 37986224\|23142080\|33184237\|23597 | Unspecified |
| mmu-miR-29b-3p | MIMAT0000127 | Oplah | 75475 | HITS-CLIP | 23597149 | Unspecified |
| mmu-miR-29b-3p | MIMAT0000127 | Orc3 | 50793 | HITS-CLIP | 25083871 | Unspecified |
| mmu-miR-29b-3p | MIMAT0000127 | Osbp | 76303 | HITS-CLIP//sRNA-Seq | 37986224\|25083871\|33184237\|23142 | Unspecified |
| mmu-miR-29b-3p | MIMAT0000127 | Osbpl11 | 106326 | HITS-CLIP | 37986224\|25083871 | Unspecified |
| mmu-miR-29b-3p | MIMAT0000127 | Osbpl1a | 64291 | HITS-CLIP | 37986224 | Unspecified |
| mmu-miR-29b-3p | MIMAT0000127 | Osbpl7 | 71240 | HITS-CLIP | 25083871 | Unspecified |
| mmu-miR-29b-3p | MIMAT0000127 | Osbpl8 | 237542 | HITS-CLIP//sRNA-Seq | 25083871\|33184237\|23142080 | Unspecified |
| mmu-miR-29b-3p | MIMAT0000127 | Osbpl9 | 100273 | HITS-CLIP//sRNA-Seq | 25083871\|33184237\|23597149\|23142 | Unspecified |
| mmu-miR-29b-3p | MIMAT0000127 | Osgin1 | 71839 | HITS-CLIP | 37986224\|23597149 | Unspecified |
| mmu-miR-29b-3p | MIMAT0000127 | Osmr | 18414 | HITS-CLIP | 25083871 | Unspecified |
| mmu-miR-29b-3p | MIMAT0000127 | Ostc | 66357 | HITS-CLIP//sRNA-Seq | 37986224\|25083871\|33184237\|23597 | Unspecified |
| mmu-miR-29b-3p | MIMAT0000127 | Ostm1 | 14628 | HITS-CLIP | 23142080 | Unspecified |
| mmu-miR-29b-3p | MIMAT0000127 | Otc | 18416 | HITS-CLIP | 23597149 | Unspecified |
| mmu-miR-29b-3p | MIMAT0000127 | Otub1 | 107260 | HITS-CLIP//sRNA-Seq | 25083871\|33184237 | Unspecified |
| mmu-miR-29b-3p | MIMAT0000127 | Otub2 | 68149 | HITS-CLIP | 37986224\|25083871 | Unspecified |
| mmu-miR-29b-3p | MIMAT0000127 | Otud1 | 71198 | HITS-CLIP | 25083871\|23597149\|23142080 | Unspecified |
| mmu-miR-29b-3p | MIMAT0000127 | Otud4 | 73945 | HITS-CLIP | 37986224\|25083871\|23142080\|31433 | Unspecified |
| mmu-miR-29b-3p | MIMAT0000127 | Otud7b | 229603 | HITS-CLIP | 37986224\|25083871\|23142080 | Unspecified |

| | | | | | | |
|---|---|---|---|---|---|---|
| mmu-miR-29b-3p | MIMAT0000127 | Otulinl | 54491 | HITS-CLIP | 23142080\|23597149 | Unspecified |
| mmu-miR-29b-3p | MIMAT0000127 | Oxct1 | 67041 | sRNA-Seq//HITS-CLIP | 33184237\|23142080 | Unspecified |
| mmu-miR-29b-3p | MIMAT0000127 | Oxr1 | 170719 | HITS-CLIP//sRNA-Seq | 37986224\|25083871\|23597149\|33184 | Unspecified |
| mmu-miR-29b-3p | MIMAT0000127 | Oxsm | 71147 | HITS-CLIP | 23597149\|23142080 | Unspecified |
| mmu-miR-29b-3p | MIMAT0000127 | Oxsr1 | 108737 | HITS-CLIP | 23142080 | Unspecified |
| mmu-miR-29b-3p | MIMAT0000127 | P2rx1 | 18436 | HITS-CLIP | 30971389 | Unspecified |
| mmu-miR-29b-3p | MIMAT0000127 | P2rx4 | 18438 | HITS-CLIP | 23597149 | Unspecified |
| mmu-miR-29b-3p | MIMAT0000127 | P2rx7 | 18439 | HITS-CLIP | 37986224\|25083871\|23142080 | Unspecified |
| mmu-miR-29b-3p | MIMAT0000127 | P2ry1 | 18441 | HITS-CLIP | 23142080 | Unspecified |
| mmu-miR-29b-3p | MIMAT0000127 | P2ry10 | 78826 | HITS-CLIP | 23142080 | Unspecified |
| mmu-miR-29b-3p | MIMAT0000127 | P2ry12 | 70839 | HITS-CLIP | 23597149 | Unspecified |
| mmu-miR-29b-3p | MIMAT0000127 | P2ry2 | 18442 | HITS-CLIP | 25083871 | Unspecified |
| mmu-miR-29b-3p | MIMAT0000127 | P3h1 | 64175 | HITS-CLIP//sRNA-Seq | 23597149\|25083871\|33184237\|23142 | Unspecified |
| mmu-miR-29b-3p | MIMAT0000127 | P4ha1 | 18451 | sRNA-Seq//HITS-CLIP | 33184237\|23142080\|25083871 | Unspecified |
| mmu-miR-29b-3p | MIMAT0000127 | P4ha2 | 18452 | HITS-CLIP | 25083871\|23142080 | Unspecified |
| mmu-miR-29b-3p | MIMAT0000127 | P4hb | 18453 | HITS-CLIP//sRNA-Seq | 23597149\|25083871\|33184237 | Unspecified |
| mmu-miR-29b-3p | MIMAT0000127 | Pabpc1 | 18458 | qCLASH//sRNA-Seq//HITS-CLIP | 31393953\|33184237\|25083871\|23142 | Unspecified |
| mmu-miR-29b-3p | MIMAT0000127 | Pabpn1 | 54196 | HITS-CLIP//sRNA-Seq | 25083871\|23142080\|33184237 | Unspecified |
| mmu-miR-29b-3p | MIMAT0000127 | Pacs2 | 217893 | HITS-CLIP//sRNA-Seq | 37986224\|25083871\|33184237\|23142 | Unspecified |
| mmu-miR-29b-3p | MIMAT0000127 | Pacsin2 | 23970 | HITS-CLIP//sRNA-Seq | 23142080\|33184237\|23597149 | Unspecified |
| mmu-miR-29b-3p | MIMAT0000127 | Pacsin3 | 80708 | HITS-CLIP | 23597149 | Unspecified |
| mmu-miR-29b-3p | MIMAT0000127 | Paf1 | 54624 | HITS-CLIP | 23142080 | Unspecified |
| mmu-miR-29b-3p | MIMAT0000127 | Pafah1b1 | 18472 | HITS-CLIP//sRNA-Seq | 37986224\|23142080\|33184237\|25083 | Unspecified |
| mmu-miR-29b-3p | MIMAT0000127 | Pafah1b2 | 18475 | HITS-CLIP//sRNA-Seq | 25083871\|23142080\|23597149\|33184 | Unspecified |
| mmu-miR-29b-3p | MIMAT0000127 | Pag1 | 94212 | HITS-CLIP | 23142080\|23597149 | Unspecified |
| mmu-miR-29b-3p | MIMAT0000127 | Pah | 18478 | HITS-CLIP | 23597149 | Unspecified |
| mmu-miR-29b-3p | MIMAT0000127 | Paics | 67054 | HITS-CLIP//sRNA-Seq | 23597149\|25083871\|33184237\|23142 | Unspecified |
| mmu-miR-29b-3p | MIMAT0000127 | Paip1 | 218693 | HITS-CLIP//sRNA-Seq | 25083871\|33184237\|23142080 | Unspecified |
| mmu-miR-29b-3p | MIMAT0000127 | Paip2 | 67869 | HITS-CLIP//sRNA-Seq | 23142080\|23597149\|33184237\|25083 | Unspecified |
| mmu-miR-29b-3p | MIMAT0000127 | Pak1 | 18479 | HITS-CLIP//sRNA-Seq | 25083871\|33184237 | Unspecified |
| mmu-miR-29b-3p | MIMAT0000127 | Pak1ip1 | 68083 | HITS-CLIP | 37986224\|25083871\|23142080 | Unspecified |
| mmu-miR-29b-3p | MIMAT0000127 | Pak2 | 224105 | HITS-CLIP | 23142080\|25083871 | Unspecified |
| mmu-miR-29b-3p | MIMAT0000127 | Pak4 | 70584 | HITS-CLIP | 25083871\|23142080 | Unspecified |
| mmu-miR-29b-3p | MIMAT0000127 | Palm | 18483 | HITS-CLIP | 37986224 | Unspecified |
| mmu-miR-29b-3p | MIMAT0000127 | Palm2 | 242481 | HITS-CLIP | 25083871 | Unspecified |
| mmu-miR-29b-3p | MIMAT0000127 | Pam | 18484 | HITS-CLIP//sRNA-Seq | 25083871\|33184237\|23597149\|23142 | Unspecified |
| mmu-miR-29b-3p | MIMAT0000127 | Pan2 | 103135 | HITS-CLIP | 37986224\|23597149\|23142080 | Unspecified |
| mmu-miR-29b-3p | MIMAT0000127 | Pan3 | 72587 | HITS-CLIP | 25083871\|23597149 | Unspecified |
| mmu-miR-29b-3p | MIMAT0000127 | Pank1 | 75735 | HITS-CLIP | 37986224 | Unspecified |

| | | | | | | |
|---|---|---|---|---|---|---|
| mmu-miR-29b-3p | MIMAT0000127 | Pank3 | 211347 | HITS-CLIP//sRNA-Seq | 23142080\|33184237\|25083871 | Unspecified |
| mmu-miR-29b-3p | MIMAT0000127 | Panx1 | 55991 | HITS-CLIP//sRNA-Seq | 25083871\|23142080\|33184237 | Unspecified |
| mmu-miR-29b-3p | MIMAT0000127 | Papola | 18789 | HITS-CLIP//sRNA-Seq | 37986224\|25083871\|23142080\|33184 | Unspecified |
| mmu-miR-29b-3p | MIMAT0000127 | Papss1 | 23971 | HITS-CLIP//sRNA-Seq | 37986224\|23142080\|33184237 | Unspecified |
| mmu-miR-29b-3p | MIMAT0000127 | Papss2 | 23972 | HITS-CLIP | 23597149 | Unspecified |
| mmu-miR-29b-3p | MIMAT0000127 | Paqr7 | 71904 | HITS-CLIP | 23142080 | Unspecified |
| mmu-miR-29b-3p | MIMAT0000127 | Paqr8 | 74229 | HITS-CLIP | 23142080 | Unspecified |
| mmu-miR-29b-3p | MIMAT0000127 | Paqr9 | 75552 | HITS-CLIP | 37986224\|23597149 | Unspecified |
| mmu-miR-29b-3p | MIMAT0000127 | Pard6b | 58220 | HITS-CLIP | 23142080 | Unspecified |
| mmu-miR-29b-3p | MIMAT0000127 | Parg | 26430 | HITS-CLIP | 25083871\|23142080 | Unspecified |
| mmu-miR-29b-3p | MIMAT0000127 | Parl | 381038 | HITS-CLIP//sRNA-Seq | 23142080\|33184237 | Unspecified |
| mmu-miR-29b-3p | MIMAT0000127 | Parn | 74108 | HITS-CLIP | 23142080 | Unspecified |
| mmu-miR-29b-3p | MIMAT0000127 | Parp11 | 101187 | HITS-CLIP | 23142080 | Unspecified |
| mmu-miR-29b-3p | MIMAT0000127 | Parp14 | 547253 | HITS-CLIP | 37986224\|23142080 | Unspecified |
| mmu-miR-29b-3p | MIMAT0000127 | Parp3 | 235587 | HITS-CLIP//sRNA-Seq | 37986224\|25083871\|33184237\|23142 | Unspecified |
| mmu-miR-29b-3p | MIMAT0000127 | Parp4 | 328417 | HITS-CLIP | 25083871\|23597149 | Unspecified |
| mmu-miR-29b-3p | MIMAT0000127 | Parp6 | 67287 | HITS-CLIP | 37986224 | Unspecified |
| mmu-miR-29b-3p | MIMAT0000127 | Parva | 57342 | HITS-CLIP//sRNA-Seq | 23597149\|33184237 | Unspecified |
| mmu-miR-29b-3p | MIMAT0000127 | Parvb | 170736 | HITS-CLIP//sRNA-Seq | 37986224\|23597149\|33184237\|25083 | Unspecified |
| mmu-miR-29b-3p | MIMAT0000127 | Patl1 | 225929 | HITS-CLIP | 31433990 | Unspecified |
| mmu-miR-29b-3p | MIMAT0000127 | Patz1 | 56218 | HITS-CLIP | 25083871\|23597149\|23142080 | Unspecified |
| mmu-miR-29b-3p | MIMAT0000127 | Pax7 | 18509 | HITS-CLIP | 37986224\|25083871 | Unspecified |
| mmu-miR-29b-3p | MIMAT0000127 | Paxip1 | 55982 | HITS-CLIP | 23142080 | Unspecified |
| mmu-miR-29b-3p | MIMAT0000127 | Pbk | 52033 | HITS-CLIP | 23142080 | Unspecified |
| mmu-miR-29b-3p | MIMAT0000127 | Pbrm1 | 66923 | HITS-CLIP//sRNA-Seq | 25083871\|23597149\|33184237 | Unspecified |
| mmu-miR-29b-3p | MIMAT0000127 | Pbx2 | 18515 | HITS-CLIP | 25083871 | Unspecified |
| mmu-miR-29b-3p | MIMAT0000127 | Pbxip1 | 229534 | HITS-CLIP//sRNA-Seq | 25083871\|33184237 | Unspecified |
| mmu-miR-29b-3p | MIMAT0000127 | Pcbp2 | 18521 | HITS-CLIP//sRNA-Seq | 25083871\|33184237\|23142080 | Unspecified |
| mmu-miR-29b-3p | MIMAT0000127 | Pcca | 110821 | HITS-CLIP | 23142080 | Unspecified |
| mmu-miR-29b-3p | MIMAT0000127 | Pccb | 66904 | HITS-CLIP | 23142080 | Unspecified |
| mmu-miR-29b-3p | MIMAT0000127 | Pcdh1 | 75599 | HITS-CLIP | 23597149 | Unspecified |
| mmu-miR-29b-3p | MIMAT0000127 | Pcdh7 | 54216 | HITS-CLIP//sRNA-Seq | 25083871\|33184237 | Unspecified |
| mmu-miR-29b-3p | MIMAT0000127 | Pcdhga1 | 93709 | RNA-Seq | 33332475 | Unspecified |
| mmu-miR-29b-3p | MIMAT0000127 | Pced1b | 239647 | HITS-CLIP | 23142080 | Unspecified |
| mmu-miR-29b-3p | MIMAT0000127 | Pcf11 | 74737 | HITS-CLIP | 25083871\|23142080 | Unspecified |
| mmu-miR-29b-3p | MIMAT0000127 | Pcgf3 | 69587 | HITS-CLIP | 37986224\|25083871\|23597149\|23142 | Unspecified |
| mmu-miR-29b-3p | MIMAT0000127 | Pcgf5 | 76073 | HITS-CLIP | 23142080\|25083871 | Unspecified |
| mmu-miR-29b-3p | MIMAT0000127 | Pcif1 | 228866 | HITS-CLIP | 23597149 | Unspecified |
| mmu-miR-29b-3p | MIMAT0000127 | Pclaf | 68026 | sRNA-Seq//HITS-CLIP | 33184237\|23142080 | Unspecified |

| | | | | | | |
|---|---|---|---|---|---|---|
| mmu-miR-29b-3p | MIMAT0000127 | Pcm1 | 18536 | HITS-CLIP//sRNA-Seq | 37986224\|25083871\|33184237\|23142 | Unspecified |
| mmu-miR-29b-3p | MIMAT0000127 | Pcmt1 | 18537 | HITS-CLIP | 37986224 | Unspecified |
| mmu-miR-29b-3p | MIMAT0000127 | Pcmtd1 | 319263 | HITS-CLIP | 25083871 | Unspecified |
| mmu-miR-29b-3p | MIMAT0000127 | Pcmtd2 | 245867 | HITS-CLIP | 23597149\|23142080 | Unspecified |
| mmu-miR-29b-3p | MIMAT0000127 | Pcnx | 54604 | HITS-CLIP | 23597149\|25083871 | Unspecified |
| mmu-miR-29b-3p | MIMAT0000127 | Pcnx3 | 399909 | HITS-CLIP | 25083871\|23142080 | Unspecified |
| mmu-miR-29b-3p | MIMAT0000127 | Pcnx4 | 67708 | HITS-CLIP | 25083871\|23597149\|23142080 | Unspecified |
| mmu-miR-29b-3p | MIMAT0000127 | Pcsk9 | 100102 | HITS-CLIP | 23597149 | Unspecified |
| mmu-miR-29b-3p | MIMAT0000127 | Pcx | 18563 | HITS-CLIP//sRNA-Seq | 23597149\|33184237 | Unspecified |
| mmu-miR-29b-3p | MIMAT0000127 | Pcyox1 | 66881 | HITS-CLIP//sRNA-Seq | 25083871\|33184237 | Unspecified |
| mmu-miR-29b-3p | MIMAT0000127 | Pcyt1a | 13026 | HITS-CLIP | 37986224\|25083871\|23597149 | Unspecified |
| mmu-miR-29b-3p | MIMAT0000127 | Pcyt1b | 236899 | HITS-CLIP | 37986224 | Unspecified |
| mmu-miR-29b-3p | MIMAT0000127 | Pcyt2 | 68671 | HITS-CLIP//sRNA-Seq | 23597149\|33184237\|23142080 | Unspecified |
| mmu-miR-29b-3p | MIMAT0000127 | Pdcd10 | 56426 | HITS-CLIP//sRNA-Seq | 25083871\|23142080\|33184237 | Unspecified |
| mmu-miR-29b-3p | MIMAT0000127 | Pdcd11 | 18572 | HITS-CLIP | 23597149 | Unspecified |
| mmu-miR-29b-3p | MIMAT0000127 | Pdcd4 | 18569 | HITS-CLIP | 23142080 | Unspecified |
| mmu-miR-29b-3p | MIMAT0000127 | Pdcd6ip | 18571 | sRNA-Seq//HITS-CLIP | 33184237\|23597149\|23142080 | Unspecified |
| mmu-miR-29b-3p | MIMAT0000127 | Pdcl | 67466 | HITS-CLIP | 37986224\|25083871\|23597149\|23142 | Unspecified |
| mmu-miR-29b-3p | MIMAT0000127 | Pdcl3 | 68833 | HITS-CLIP//sRNA-Seq | 25083871\|33184237 | Unspecified |
| mmu-miR-29b-3p | MIMAT0000127 | Pde12 | 211948 | HITS-CLIP | 23142080 | Unspecified |
| mmu-miR-29b-3p | MIMAT0000127 | Pde3a | 54611 | HITS-CLIP | 25083871 | Unspecified |
| mmu-miR-29b-3p | MIMAT0000127 | Pde3b | 18576 | HITS-CLIP | 37986224\|23597149\|23142080 | Unspecified |
| mmu-miR-29b-3p | MIMAT0000127 | Pde4b | 18578 | HITS-CLIP | 37986224\|25083871\|23597149\|23142 | Unspecified |
| mmu-miR-29b-3p | MIMAT0000127 | Pde4d | 238871 | HITS-CLIP | 23142080 | Unspecified |
| mmu-miR-29b-3p | MIMAT0000127 | Pde4dip | 83679 | HITS-CLIP//sRNA-Seq | 25083871\|33184237\|23597149\|23142 | Unspecified |
| mmu-miR-29b-3p | MIMAT0000127 | Pde5a | 242202 | HITS-CLIP | 25083871 | Unspecified |
| mmu-miR-29b-3p | MIMAT0000127 | Pde7a | 18583 | HITS-CLIP | 37986224\|25083871 | Unspecified |
| mmu-miR-29b-3p | MIMAT0000127 | Pdgfa | 18590 | Luciferase Reporter Assay | 37986224 | Unspecified |
| mmu-miR-29b-3p | MIMAT0000127 | Pdgfb | 18591 | HITS-CLIP, Luciferase Reporter Assay//Luci | 37986224\|23354167 | Unspecified |
| mmu-miR-29b-3p | MIMAT0000127 | Pdgfc | 54635 | HITS-CLIP, Luciferase Reporter Assay//HITS | 37986224\|25083871 | Unspecified |
| mmu-miR-29b-3p | MIMAT0000127 | Pdgfra | 18595 | HITS-CLIP//sRNA-Seq | 37986224\|25083871\|33184237 | Unspecified |
| mmu-miR-29b-3p | MIMAT0000127 | Pdgfrb | 18596 | HITS-CLIP//sRNA-Seq | 37986224\|25083871\|33184237 | Unspecified |
| mmu-miR-29b-3p | MIMAT0000127 | Pdgfrl | 68797 | HITS-CLIP//sRNA-Seq | 25083871\|33184237 | Unspecified |
| mmu-miR-29b-3p | MIMAT0000127 | Pdha1 | 18597 | HITS-CLIP//sRNA-Seq | 23597149\|25083871\|33184237\|23142 | Unspecified |
| mmu-miR-29b-3p | MIMAT0000127 | Pdhb | 68263 | HITS-CLIP//sRNA-Seq | 37986224\|23597149\|25083871\|23142 | Unspecified |
| mmu-miR-29b-3p | MIMAT0000127 | Pdhx | 27402 | HITS-CLIP | 37986224\|23597149\|25083871\|23142 | Unspecified |
| mmu-miR-29b-3p | MIMAT0000127 | Pdia4 | 12304 | HITS-CLIP//sRNA-Seq | 37986224\|23597149\|33184237\|23142 | Unspecified |
| mmu-miR-29b-3p | MIMAT0000127 | Pdia5 | 72599 | HITS-CLIP//sRNA-Seq | 23597149\|33184237 | Unspecified |
| mmu-miR-29b-3p | MIMAT0000127 | Pdia6 | 71853 | sRNA-Seq//HITS-CLIP | 33184237\|23142080 | Unspecified |

| | | | | | | |
|---|---|---|---|---|---|---|
| mmu-miR-29b-3p | MIMAT0000127 | Pdik1l | 230809 | HITS-CLIP | 37986224\|25083871\|23142080 | Unspecified |
| mmu-miR-29b-3p | MIMAT0000127 | Pdk1 | 228026 | HITS-CLIP | 37986224\|23142080 | Unspecified |
| mmu-miR-29b-3p | MIMAT0000127 | Pdk2 | 18604 | HITS-CLIP | 25083871\|23597149 | Unspecified |
| mmu-miR-29b-3p | MIMAT0000127 | Pdk3 | 236900 | sRNA-Seq//HITS-CLIP | 33184237\|23142080 | Unspecified |
| mmu-miR-29b-3p | MIMAT0000127 | Pdk4 | 27273 | HITS-CLIP | 37986224\|25083871 | Unspecified |
| mmu-miR-29b-3p | MIMAT0000127 | Pdlim5 | 56376 | HITS-CLIP//sRNA-Seq | 23597149\|33184237 | Unspecified |
| mmu-miR-29b-3p | MIMAT0000127 | Pdlim7 | 67399 | HITS-CLIP//sRNA-Seq | 37986224\|33184237\|23142080 | Unspecified |
| mmu-miR-29b-3p | MIMAT0000127 | Pdp1 | 381511 | HITS-CLIP | 37986224 | Unspecified |
| mmu-miR-29b-3p | MIMAT0000127 | Pdp2 | 382051 | HITS-CLIP | 25083871\|23597149\|23142080 | Unspecified |
| mmu-miR-29b-3p | MIMAT0000127 | Pdpk1 | 18607 | HITS-CLIP//sRNA-Seq | 37986224\|33184237\|23597149\|23142 | Unspecified |
| mmu-miR-29b-3p | MIMAT0000127 | Pdpr | 319518 | HITS-CLIP | 23597149\|23142080 | Unspecified |
| mmu-miR-29b-3p | MIMAT0000127 | Pds5a | 71521 | HITS-CLIP//sRNA-Seq | 23142080\|33184237 | Unspecified |
| mmu-miR-29b-3p | MIMAT0000127 | Pds5b | 100710 | HITS-CLIP | 37986224\|23142080\|25083871 | Unspecified |
| mmu-miR-29b-3p | MIMAT0000127 | Pdxdc1 | 94184 | HITS-CLIP//sRNA-Seq | 23142080\|33184237\|25083871 | Unspecified |
| mmu-miR-29b-3p | MIMAT0000127 | Pdzd11 | 72621 | HITS-CLIP//sRNA-Seq | 23142080\|33184237 | Unspecified |
| mmu-miR-29b-3p | MIMAT0000127 | Pdzd8 | 107368 | HITS-CLIP | 25083871\|23142080 | Unspecified |
| mmu-miR-29b-3p | MIMAT0000127 | Pdzrn3 | 55983 | HITS-CLIP | 25083871 | Unspecified |
| mmu-miR-29b-3p | MIMAT0000127 | Pea15a | 18611 | HITS-CLIP//sRNA-Seq | 23597149\|33184237 | Unspecified |
| mmu-miR-29b-3p | MIMAT0000127 | Peak1 | 244895 | HITS-CLIP | 37986224\|23142080 | Unspecified |
| mmu-miR-29b-3p | MIMAT0000127 | Peg10 | 170676 | HITS-CLIP//sRNA-Seq | 25083871\|33184237 | Unspecified |
| mmu-miR-29b-3p | MIMAT0000127 | Peg3 | 18616 | HITS-CLIP | 37986224 | Unspecified |
| mmu-miR-29b-3p | MIMAT0000127 | Peli1 | 67245 | HITS-CLIP | 25083871\|23597149\|23142080 | Unspecified |
| mmu-miR-29b-3p | MIMAT0000127 | Pelp1 | 75273 | HITS-CLIP | 23142080 | Unspecified |
| mmu-miR-29b-3p | MIMAT0000127 | Pemt | 18618 | HITS-CLIP | 23597149 | Unspecified |
| mmu-miR-29b-3p | MIMAT0000127 | Pepd | 18624 | HITS-CLIP | 37986224\|23597149 | Unspecified |
| mmu-miR-29b-3p | MIMAT0000127 | Per1 | 18626 | HITS-CLIP | 37986224\|25083871\|23142080 | Unspecified |
| mmu-miR-29b-3p | MIMAT0000127 | Per3 | 18628 | HITS-CLIP | 37986224\|25083871\|23597149 | Unspecified |
| mmu-miR-29b-3p | MIMAT0000127 | Perp | 64058 | HITS-CLIP | 23597149\|23142080 | Unspecified |
| mmu-miR-29b-3p | MIMAT0000127 | Pex1 | 71382 | HITS-CLIP | 23142080 | Unspecified |
| mmu-miR-29b-3p | MIMAT0000127 | Pex14 | 56273 | HITS-CLIP | 23597149 | Unspecified |
| mmu-miR-29b-3p | MIMAT0000127 | Pex16 | 18633 | HITS-CLIP | 25083871\|23597149 | Unspecified |
| mmu-miR-29b-3p | MIMAT0000127 | Pex19 | 19298 | sRNA-Seq//HITS-CLIP | 33184237\|23597149 | Unspecified |
| mmu-miR-29b-3p | MIMAT0000127 | Pex26 | 74043 | HITS-CLIP | 23142080 | Unspecified |
| mmu-miR-29b-3p | MIMAT0000127 | Pex3 | 56535 | HITS-CLIP | 23142080 | Unspecified |
| mmu-miR-29b-3p | MIMAT0000127 | Pex5 | 19305 | HITS-CLIP//sRNA-Seq | 25083871\|33184237\|23597149\|23142 | Unspecified |
| mmu-miR-29b-3p | MIMAT0000127 | Pex6 | 224824 | HITS-CLIP | 23597149 | Unspecified |
| mmu-miR-29b-3p | MIMAT0000127 | Pfas | 237823 | HITS-CLIP | 23597149\|23142080 | Unspecified |
| mmu-miR-29b-3p | MIMAT0000127 | Pfdn2 | 18637 | HITS-CLIP//sRNA-Seq | 23142080\|33184237 | Unspecified |
| mmu-miR-29b-3p | MIMAT0000127 | Pfkfb1 | 18639 | HITS-CLIP | 23597149 | Unspecified |

| | | | | | | |
|---|---|---|---|---|---|---|
| mmu-miR-29b-3p | MIMAT0000127 | Pfkfb2 | 18640 | HITS-CLIP | 23597149 | Unspecified |
| mmu-miR-29b-3p | MIMAT0000127 | Pfkfb3 | 170768 | HITS-CLIP | 23142080 | Unspecified |
| mmu-miR-29b-3p | MIMAT0000127 | Pfkl | 18641 | sRNA-Seq//HITS-CLIP | 33184237\|23142080 | Unspecified |
| mmu-miR-29b-3p | MIMAT0000127 | Pfkm | 18642 | HITS-CLIP//sRNA-Seq | 37986224\|25083871\|33184237\|23142 | Unspecified |
| mmu-miR-29b-3p | MIMAT0000127 | Pgap1 | 241062 | HITS-CLIP | 23142080 | Unspecified |
| mmu-miR-29b-3p | MIMAT0000127 | Pgap2 | 233575 | HITS-CLIP | 37986224\|25083871\|23597149\|23142 | Unspecified |
| mmu-miR-29b-3p | MIMAT0000127 | Pgd | 110208 | sRNA-Seq//HITS-CLIP | 33184237\|23597149 | Unspecified |
| mmu-miR-29b-3p | MIMAT0000127 | Pgls | 66171 | sRNA-Seq//HITS-CLIP | 33184237\|23597149 | Unspecified |
| mmu-miR-29b-3p | MIMAT0000127 | Pglyrp1 | 21946 | HITS-CLIP | 23142080 | Unspecified |
| mmu-miR-29b-3p | MIMAT0000127 | Pglyrp2 | 57757 | HITS-CLIP | 37986224\|23142080 | Unspecified |
| mmu-miR-29b-3p | MIMAT0000127 | Pgm1 | 66681 | HITS-CLIP//sRNA-Seq | 23597149\|33184237\|23142080 | Unspecified |
| mmu-miR-29b-3p | MIMAT0000127 | Pgp | 67078 | HITS-CLIP | 25083871\|23597149\|23142080 | Unspecified |
| mmu-miR-29b-3p | MIMAT0000127 | Pgpep1 | 66522 | HITS-CLIP | 23597149 | Unspecified |
| mmu-miR-29b-3p | MIMAT0000127 | Pgs1 | 74451 | HITS-CLIP | 37986224\|23142080\|23597149 | Unspecified |
| mmu-miR-29b-3p | MIMAT0000127 | Phactr2 | 215789 | HITS-CLIP | 23142080 | Unspecified |
| mmu-miR-29b-3p | MIMAT0000127 | Phactr4 | 100169 | HITS-CLIP | 37986224\|25083871 | Unspecified |
| mmu-miR-29b-3p | MIMAT0000127 | Phb2 | 12034 | HITS-CLIP | 37986224 | Unspecified |
| mmu-miR-29b-3p | MIMAT0000127 | Phc1 | 13619 | HITS-CLIP | 37986224 | Unspecified |
| mmu-miR-29b-3p | MIMAT0000127 | Phc3 | 241915 | HITS-CLIP | 37986224\|23142080\|25083871 | Unspecified |
| mmu-miR-29b-3p | MIMAT0000127 | Phf10 | 72057 | HITS-CLIP//sRNA-Seq | 23597149\|33184237\|25083871\|23142 | Unspecified |
| mmu-miR-29b-3p | MIMAT0000127 | Phf2 | 18676 | HITS-CLIP//sRNA-Seq | 25083871\|23597149\|33184237\|23142 | Unspecified |
| mmu-miR-29b-3p | MIMAT0000127 | Phf20 | 228829 | HITS-CLIP//sRNA-Seq | 25083871\|33184237\|23597149 | Unspecified |
| mmu-miR-29b-3p | MIMAT0000127 | Phf20l1 | 239510 | HITS-CLIP | 37986224\|23142080\|25083871 | Unspecified |
| mmu-miR-29b-3p | MIMAT0000127 | Phf21a | 192285 | HITS-CLIP | 37986224 | Unspecified |
| mmu-miR-29b-3p | MIMAT0000127 | Phf3 | 213109 | HITS-CLIP//sRNA-Seq | 25083871\|23142080\|33184237\|23597 | Unspecified |
| mmu-miR-29b-3p | MIMAT0000127 | Phf5a | 68479 | HITS-CLIP//sRNA-Seq | 37986224\|33184237\|23142080 | Unspecified |
| mmu-miR-29b-3p | MIMAT0000127 | Phf6 | 70998 | HITS-CLIP | 37986224\|25083871\|23142080 | Unspecified |
| mmu-miR-29b-3p | MIMAT0000127 | Phf8 | 320595 | HITS-CLIP | 25083871\|23597149 | Unspecified |
| mmu-miR-29b-3p | MIMAT0000127 | Phip | 83946 | HITS-CLIP//sRNA-Seq | 25083871\|33184237\|23142080 | Unspecified |
| mmu-miR-29b-3p | MIMAT0000127 | Phlda1 | 21664 | HITS-CLIP | 23142080 | Unspecified |
| mmu-miR-29b-3p | MIMAT0000127 | Phldb1 | 102693 | HITS-CLIP//sRNA-Seq | 25083871\|23142080\|23597149\|33184 | Unspecified |
| mmu-miR-29b-3p | MIMAT0000127 | Phldb3 | 232970 | HITS-CLIP | 37986224 | Unspecified |
| mmu-miR-29b-3p | MIMAT0000127 | Phlpp1 | 98432 | HITS-CLIP | 23597149 | Unspecified |
| mmu-miR-29b-3p | MIMAT0000127 | Phrf1 | 101471 | HITS-CLIP | 23142080 | Unspecified |
| mmu-miR-29b-3p | MIMAT0000127 | Phtf1 | 18685 | HITS-CLIP | 23142080 | Unspecified |
| mmu-miR-29b-3p | MIMAT0000127 | Phtf2 | 68770 | HITS-CLIP | 25083871\|23597149\|23142080 | Unspecified |
| mmu-miR-29b-3p | MIMAT0000127 | Phyhd1 | 227696 | HITS-CLIP | 23597149 | Unspecified |
| mmu-miR-29b-3p | MIMAT0000127 | Pi15 | 94227 | HITS-CLIP | 37986224\|25083871 | Unspecified |
| mmu-miR-29b-3p | MIMAT0000127 | Pi4k2b | 67073 | HITS-CLIP | 23142080 | Unspecified |

| | | | | | | |
|---|---|---|---|---|---|---|
| mmu-miR-29b-3p | MIMAT0000127 | Pias1 | 56469 | HITS-CLIP | 23142080 | Unspecified |
| mmu-miR-29b-3p | MIMAT0000127 | Pias2 | 17344 | HITS-CLIP//sRNA-Seq | 25083871\|33184237\|23142080 | Unspecified |
| mmu-miR-29b-3p | MIMAT0000127 | Pias4 | 59004 | HITS-CLIP | 23142080\|25083871 | Unspecified |
| mmu-miR-29b-3p | MIMAT0000127 | Pibf1 | 52023 | HITS-CLIP | 23142080\|25083871 | Unspecified |
| mmu-miR-29b-3p | MIMAT0000127 | Picalm | 233489 | HITS-CLIP//sRNA-Seq | 37986224\|23142080\|33184237\|25083 | Unspecified |
| mmu-miR-29b-3p | MIMAT0000127 | Piezo1 | 234839 | HITS-CLIP//sRNA-Seq | 25083871\|23142080\|33184237 | Unspecified |
| mmu-miR-29b-3p | MIMAT0000127 | Piezo2 | 667742 | HITS-CLIP | 37986224\|25083871 | Unspecified |
| mmu-miR-29b-3p | MIMAT0000127 | Piga | 18700 | HITS-CLIP | 23142080 | Unspecified |
| mmu-miR-29b-3p | MIMAT0000127 | Pigc | 67292 | HITS-CLIP | 37986224\|23142080\|25083871 | Unspecified |
| mmu-miR-29b-3p | MIMAT0000127 | Pigk | 329777 | HITS-CLIP//sRNA-Seq | 25083871\|33184237 | Unspecified |
| mmu-miR-29b-3p | MIMAT0000127 | Pigm | 67556 | HITS-CLIP | 37986224\|25083871\|23142080 | Unspecified |
| mmu-miR-29b-3p | MIMAT0000127 | Pign | 27392 | HITS-CLIP | 37986224\|25083871\|23142080\|23597 | Unspecified |
| mmu-miR-29b-3p | MIMAT0000127 | Pigq | 14755 | HITS-CLIP | 23597149 | Unspecified |
| mmu-miR-29b-3p | MIMAT0000127 | Pigr | 18703 | HITS-CLIP | 37986224\|23597149 | Unspecified |
| mmu-miR-29b-3p | MIMAT0000127 | Pigt | 78928 | HITS-CLIP//sRNA-Seq | 21258322\|33184237\|23142080 | Unspecified |
| mmu-miR-29b-3p | MIMAT0000127 | Pigu | 228812 | HITS-CLIP | 23597149 | Unspecified |
| mmu-miR-29b-3p | MIMAT0000127 | Pik3ap1 | 83490 | HITS-CLIP | 23597149\|23142080 | Unspecified |
| mmu-miR-29b-3p | MIMAT0000127 | Pik3c2a | 18704 | HITS-CLIP | 37986224\|23142080\|25083871\|23597 | Unspecified |
| mmu-miR-29b-3p | MIMAT0000127 | Pik3ca | 18706 | HITS-CLIP | 37986224\|25083871\|23597149 | Unspecified |
| mmu-miR-29b-3p | MIMAT0000127 | Pik3cb | 74769 | HITS-CLIP | 37986224\|25083871\|23597149\|23142 | Unspecified |
| mmu-miR-29b-3p | MIMAT0000127 | Pik3cd | 18707 | sRNA-Seq//HITS-CLIP | 33184237\|23142080 | Unspecified |
| mmu-miR-29b-3p | MIMAT0000127 | Pik3cg | 30955 | HITS-CLIP | 23142080 | Unspecified |
| mmu-miR-29b-3p | MIMAT0000127 | Pik3ip1 | 216505 | HITS-CLIP | 23142080 | Unspecified |
| mmu-miR-29b-3p | MIMAT0000127 | Pik3r1 | 18708 | HITS-CLIP | 37986224\|25083871\|23597149\|23142 | Unspecified |
| mmu-miR-29b-3p | MIMAT0000127 | Pik3r3 | 18710 | HITS-CLIP//sRNA-Seq | 25083871\|33184237 | Unspecified |
| mmu-miR-29b-3p | MIMAT0000127 | Pikfyve | 18711 | HITS-CLIP | 37986224\|25083871\|23597149\|23142 | Unspecified |
| mmu-miR-29b-3p | MIMAT0000127 | Pim1 | 18712 | HITS-CLIP | 23597149\|23142080\|25083871 | Unspecified |
| mmu-miR-29b-3p | MIMAT0000127 | Pim2 | 18715 | HITS-CLIP | 23142080 | Unspecified |
| mmu-miR-29b-3p | MIMAT0000127 | Pim3 | 223775 | HITS-CLIP | 37986224\|23597149\|23142080 | Unspecified |
| mmu-miR-29b-3p | MIMAT0000127 | Pip4k2b | 108083 | sRNA-Seq//HITS-CLIP | 33184237\|25083871 | Unspecified |
| mmu-miR-29b-3p | MIMAT0000127 | Pip4p2 | 55529 | HITS-CLIP | 23142080 | Unspecified |
| mmu-miR-29b-3p | MIMAT0000127 | Pip5k1a | 18720 | sRNA-Seq//HITS-CLIP | 33184237\|23142080 | Unspecified |
| mmu-miR-29b-3p | MIMAT0000127 | Pipox | 19193 | HITS-CLIP | 23597149 | Unspecified |
| mmu-miR-29b-3p | MIMAT0000127 | Pithd1 | 66193 | HITS-CLIP | 25083871 | Unspecified |
| mmu-miR-29b-3p | MIMAT0000127 | Pitpna | 18738 | HITS-CLIP//sRNA-Seq | 37986224\|23142080\|33184237\|23597 | Unspecified |
| mmu-miR-29b-3p | MIMAT0000127 | Pitpnb | 56305 | HITS-CLIP//sRNA-Seq | 25083871\|33184237\|23142080 | Unspecified |
| mmu-miR-29b-3p | MIMAT0000127 | Pitpnm1 | 18739 | HITS-CLIP | 23142080 | Unspecified |
| mmu-miR-29b-3p | MIMAT0000127 | Pitpnm2 | 19679 | HITS-CLIP | 37986224\|23142080 | Unspecified |
| mmu-miR-29b-3p | MIMAT0000127 | Pitrm1 | 69617 | HITS-CLIP | 37986224 | Unspecified |

| | | | | | | |
|---|---|---|---|---|---|---|
| mmu-miR-29b-3p | MIMAT0000127 | Pitx2 | 18741 | HITS-CLIP | 25083871 | Unspecified |
| mmu-miR-29b-3p | MIMAT0000127 | Pja2 | 224938 | HITS-CLIP//sRNA-Seq | 25083871\|23142080\|33184237 | Unspecified |
| mmu-miR-29b-3p | MIMAT0000127 | Pkd1 | 18763 | HITS-CLIP | 37986224\|25083871\|23597149\|23142 | Unspecified |
| mmu-miR-29b-3p | MIMAT0000127 | Pkd2 | 18764 | sRNA-Seq//HITS-CLIP | 33184237\|23597149 | Unspecified |
| mmu-miR-29b-3p | MIMAT0000127 | Pkdcc | 106522 | sRNA-Seq//HITS-CLIP | 33184237\|23597149 | Unspecified |
| mmu-miR-29b-3p | MIMAT0000127 | Pkhd1 | 241035 | HITS-CLIP | 37986224 | Unspecified |
| mmu-miR-29b-3p | MIMAT0000127 | Pkia | 18767 | HITS-CLIP//sRNA-Seq | 25083871\|33184237 | Unspecified |
| mmu-miR-29b-3p | MIMAT0000127 | Pkn1 | 320795 | HITS-CLIP | 23142080\|25083871 | Unspecified |
| mmu-miR-29b-3p | MIMAT0000127 | Pkn2 | 109333 | HITS-CLIP//sRNA-Seq | 37986224\|25083871\|33184237\|23142 | Unspecified |
| mmu-miR-29b-3p | MIMAT0000127 | Pknox2 | 208076 | HITS-CLIP | 37986224 | Unspecified |
| mmu-miR-29b-3p | MIMAT0000127 | Pkp1 | 18772 | HITS-CLIP | 25083871 | Unspecified |
| mmu-miR-29b-3p | MIMAT0000127 | Pkp4 | 227937 | HITS-CLIP | 23142080 | Unspecified |
| mmu-miR-29b-3p | MIMAT0000127 | Pla2g12a | 66350 | HITS-CLIP | 23142080 | Unspecified |
| mmu-miR-29b-3p | MIMAT0000127 | Pla2g12b | 69836 | HITS-CLIP | 23597149 | Unspecified |
| mmu-miR-29b-3p | MIMAT0000127 | Pla2g15 | 192654 | HITS-CLIP | 25083871 | Unspecified |
| mmu-miR-29b-3p | MIMAT0000127 | Pla2g6 | 53357 | HITS-CLIP | 23142080 | Unspecified |
| mmu-miR-29b-3p | MIMAT0000127 | Plaa | 18786 | HITS-CLIP//sRNA-Seq | 37986224\|33184237\|23142080 | Unspecified |
| mmu-miR-29b-3p | MIMAT0000127 | Plag1 | 56711 | HITS-CLIP | 23142080 | Unspecified |
| mmu-miR-29b-3p | MIMAT0000127 | Plagl2 | 54711 | HITS-CLIP | 37986224\|23597149\|23142080 | Unspecified |
| mmu-miR-29b-3p | MIMAT0000127 | Plau | 18792 | HITS-CLIP | 25083871 | Unspecified |
| mmu-miR-29b-3p | MIMAT0000127 | Plcb4 | 18798 | HITS-CLIP | 25083871 | Unspecified |
| mmu-miR-29b-3p | MIMAT0000127 | Plcd1 | 18799 | HITS-CLIP | 23142080 | Unspecified |
| mmu-miR-29b-3p | MIMAT0000127 | Plcg1 | 18803 | HITS-CLIP//sRNA-Seq | 25083871\|33184237\|23142080 | Unspecified |
| mmu-miR-29b-3p | MIMAT0000127 | Plcl2 | 224860 | HITS-CLIP | 23142080 | Unspecified |
| mmu-miR-29b-3p | MIMAT0000127 | Plcxd2 | 433022 | HITS-CLIP | 37986224\|23142080\|23597149 | Unspecified |
| mmu-miR-29b-3p | MIMAT0000127 | Pld3 | 18807 | HITS-CLIP//sRNA-Seq | 25083871\|33184237\|23597149 | Unspecified |
| mmu-miR-29b-3p | MIMAT0000127 | Plec | 18810 | HITS-CLIP//sRNA-Seq | 25083871\|23597149\|33184237 | Unspecified |
| mmu-miR-29b-3p | MIMAT0000127 | Plekha1 | 101476 | HITS-CLIP//sRNA-Seq | 37986224\|25083871\|33184237\|23597 | Unspecified |
| mmu-miR-29b-3p | MIMAT0000127 | Plekha2 | 83436 | HITS-CLIP | 23142080 | Unspecified |
| mmu-miR-29b-3p | MIMAT0000127 | Plekha8 | 231999 | HITS-CLIP | 37986224\|25083871\|23597149 | Unspecified |
| mmu-miR-29b-3p | MIMAT0000127 | Plekhb2 | 226971 | HITS-CLIP//sRNA-Seq | 25083871\|33184237\|23142080 | Unspecified |
| mmu-miR-29b-3p | MIMAT0000127 | Plekhf2 | 71801 | HITS-CLIP | 23142080 | Unspecified |
| mmu-miR-29b-3p | MIMAT0000127 | Plekhg3 | 263406 | HITS-CLIP | 23597149 | Unspecified |
| mmu-miR-29b-3p | MIMAT0000127 | Plekhg5 | 269608 | HITS-CLIP | 23142080 | Unspecified |
| mmu-miR-29b-3p | MIMAT0000127 | Plekhh1 | 211945 | HITS-CLIP | 25083871 | Unspecified |
| mmu-miR-29b-3p | MIMAT0000127 | Plekhh2 | 213556 | HITS-CLIP | 25083871 | Unspecified |
| mmu-miR-29b-3p | MIMAT0000127 | Plekhm1 | 353047 | HITS-CLIP | 25083871\|23142080 | Unspecified |
| mmu-miR-29b-3p | MIMAT0000127 | Plekhm2 | 69582 | HITS-CLIP | 23597149\|23142080 | Unspecified |
| mmu-miR-29b-3p | MIMAT0000127 | Plekho1 | 67220 | HITS-CLIP//sRNA-Seq | 25083871\|33184237 | Unspecified |

| | | | | | | |
|---|---|---|---|---|---|---|
| mmu-miR-29b-3p | MIMAT0000127 | Plg | 18815 | HITS-CLIP | 23597149 | Unspecified |
| mmu-miR-29b-3p | MIMAT0000127 | Plgrkt | 67759 | HITS-CLIP//sRNA-Seq | 37986224\|23142080\|33184237 | Unspecified |
| mmu-miR-29b-3p | MIMAT0000127 | Plin2 | 11520 | HITS-CLIP//sRNA-Seq | 23597149\|33184237\|25083871 | Unspecified |
| mmu-miR-29b-3p | MIMAT0000127 | Plin5 | 66968 | HITS-CLIP | 23597149 | Unspecified |
| mmu-miR-29b-3p | MIMAT0000127 | Plk1 | 18817 | sRNA-Seq//HITS-CLIP | 33184237\|23142080 | Unspecified |
| mmu-miR-29b-3p | MIMAT0000127 | Plk2 | 20620 | HITS-CLIP//sRNA-Seq | 25083871\|23142080\|33184237 | Unspecified |
| mmu-miR-29b-3p | MIMAT0000127 | Plk3 | 12795 | HITS-CLIP | 25083871\|23142080 | Unspecified |
| mmu-miR-29b-3p | MIMAT0000127 | Plk4 | 20873 | HITS-CLIP | 23142080 | Unspecified |
| mmu-miR-29b-3p | MIMAT0000127 | Plod1 | 18822 | HITS-CLIP | 37986224 | Unspecified |
| mmu-miR-29b-3p | MIMAT0000127 | Plod3 | 26433 | HITS-CLIP//sRNA-Seq | 25083871\|23142080\|33184237 | Unspecified |
| mmu-miR-29b-3p | MIMAT0000127 | Plrg1 | 53317 | HITS-CLIP//sRNA-Seq | 23142080\|33184237 | Unspecified |
| mmu-miR-29b-3p | MIMAT0000127 | Pls3 | 102866 | HITS-CLIP//sRNA-Seq | 23597149\|25083871\|33184237 | Unspecified |
| mmu-miR-29b-3p | MIMAT0000127 | Plscr3 | 70310 | HITS-CLIP | 37986224 | Unspecified |
| mmu-miR-29b-3p | MIMAT0000127 | Plxna1 | 18844 | HITS-CLIP//sRNA-Seq | 37986224\|25083871\|23142080\|33184 | Unspecified |
| mmu-miR-29b-3p | MIMAT0000127 | Plxna3 | 18846 | HITS-CLIP | 23597149 | Unspecified |
| mmu-miR-29b-3p | MIMAT0000127 | Plxnb1 | 235611 | HITS-CLIP | 23597149 | Unspecified |
| mmu-miR-29b-3p | MIMAT0000127 | Plxnb2 | 140570 | HITS-CLIP//sRNA-Seq | 23597149\|33184237 | Unspecified |
| mmu-miR-29b-3p | MIMAT0000127 | Plxnd1 | 67784 | HITS-CLIP | 25083871 | Unspecified |
| mmu-miR-29b-3p | MIMAT0000127 | Pmaip1 | 58801 | HITS-CLIP | 23142080\|25083871 | Unspecified |
| mmu-miR-29b-3p | MIMAT0000127 | Pmepa1 | 65112 | HITS-CLIP | 23142080 | Unspecified |
| mmu-miR-29b-3p | MIMAT0000127 | Pml | 18854 | sRNA-Seq//HITS-CLIP | 33184237\|23142080 | Unspecified |
| mmu-miR-29b-3p | MIMAT0000127 | Pmp22 | 18858 | HITS-CLIP//sRNA-Seq | 37986224\|25083871\|33184237 | Unspecified |
| mmu-miR-29b-3p | MIMAT0000127 | Pmpca | 66865 | HITS-CLIP//sRNA-Seq | 37986224\|25083871\|33184237\|23597 | Unspecified |
| mmu-miR-29b-3p | MIMAT0000127 | Pmpcb | 73078 | HITS-CLIP | 23597149\|23142080 | Unspecified |
| mmu-miR-29b-3p | MIMAT0000127 | Pms2 | 18861 | HITS-CLIP | 23142080 | Unspecified |
| mmu-miR-29b-3p | MIMAT0000127 | Pmvk | 68603 | sRNA-Seq//HITS-CLIP | 33184237\|23142080 | Unspecified |
| mmu-miR-29b-3p | MIMAT0000127 | Pnisr | 66625 | HITS-CLIP | 25083871 | Unspecified |
| mmu-miR-29b-3p | MIMAT0000127 | Pnmal2 | 434128 | HITS-CLIP | 25083871 | Unspecified |
| mmu-miR-29b-3p | MIMAT0000127 | Pnn | 18949 | sRNA-Seq//HITS-CLIP | 33184237\|23142080 | Unspecified |
| mmu-miR-29b-3p | MIMAT0000127 | Pno1 | 66249 | HITS-CLIP | 23142080 | Unspecified |
| mmu-miR-29b-3p | MIMAT0000127 | Pnp | 18950 | HITS-CLIP | 23597149 | Unspecified |
| mmu-miR-29b-3p | MIMAT0000127 | Pnpla6 | 50767 | HITS-CLIP | 23597149 | Unspecified |
| mmu-miR-29b-3p | MIMAT0000127 | Pnrc1 | 108767 | HITS-CLIP//sRNA-Seq | 37986224\|33184237\|23597149\|23142 | Unspecified |
| mmu-miR-29b-3p | MIMAT0000127 | Pnrc2 | 52830 | HITS-CLIP//sRNA-Seq | 25083871\|33184237\|23142080 | Unspecified |
| mmu-miR-29b-3p | MIMAT0000127 | Pogk | 71592 | HITS-CLIP | 25083871 | Unspecified |
| mmu-miR-29b-3p | MIMAT0000127 | Pogz | 229584 | HITS-CLIP | 37986224 | Unspecified |
| mmu-miR-29b-3p | MIMAT0000127 | Pola1 | 18968 | sRNA-Seq//HITS-CLIP | 33184237\|23142080 | Unspecified |
| mmu-miR-29b-3p | MIMAT0000127 | Pold1 | 18971 | HITS-CLIP | 23142080 | Unspecified |
| mmu-miR-29b-3p | MIMAT0000127 | Pold3 | 67967 | HITS-CLIP | 25083871\|23142080 | Unspecified |

| | | | | | | |
|---|---|---|---|---|---|---|
| mmu-miR-29b-3p | MIMAT0000127 | Poldip3 | 73826 | HITS-CLIP//sRNA-Seq | 25083871\|33184237\|31433990\|23142 | Unspecified |
| mmu-miR-29b-3p | MIMAT0000127 | Pole | 18973 | HITS-CLIP | 23142080 | Unspecified |
| mmu-miR-29b-3p | MIMAT0000127 | Pole3 | 59001 | HITS-CLIP | 37986224 | Unspecified |
| mmu-miR-29b-3p | MIMAT0000127 | Polh | 80905 | HITS-CLIP | 23142080 | Unspecified |
| mmu-miR-29b-3p | MIMAT0000127 | Polr2a | 20020 | HITS-CLIP//sRNA-Seq | 25083871\|23597149\|33184237\|23142 | Unspecified |
| mmu-miR-29b-3p | MIMAT0000127 | Polr2b | 231329 | HITS-CLIP//sRNA-Seq | 25083871\|33184237\|23142080 | Unspecified |
| mmu-miR-29b-3p | MIMAT0000127 | Polr2d | 69241 | HITS-CLIP | 37986224 | Unspecified |
| mmu-miR-29b-3p | MIMAT0000127 | Polr2g | 67710 | sRNA-Seq//HITS-CLIP | 33184237\|23597149\|23142080 | Unspecified |
| mmu-miR-29b-3p | MIMAT0000127 | Polr3b | 70428 | HITS-CLIP | 25083871\|23142080 | Unspecified |
| mmu-miR-29b-3p | MIMAT0000127 | Polr3f | 70408 | HITS-CLIP | 25083871\|23142080 | Unspecified |
| mmu-miR-29b-3p | MIMAT0000127 | Polrmt | 216151 | HITS-CLIP | 23597149\|23142080 | Unspecified |
| mmu-miR-29b-3p | MIMAT0000127 | Pom121 | 107939 | HITS-CLIP | 25083871\|31433990\|23142080 | Unspecified |
| mmu-miR-29b-3p | MIMAT0000127 | Pomgnt1 | 68273 | HITS-CLIP | 25083871 | Unspecified |
| mmu-miR-29b-3p | MIMAT0000127 | Pomk | 74653 | HITS-CLIP | 37986224\|25083871 | Unspecified |
| mmu-miR-29b-3p | MIMAT0000127 | Pomt2 | 217734 | HITS-CLIP | 23597149 | Unspecified |
| mmu-miR-29b-3p | MIMAT0000127 | Pon2 | 330260 | HITS-CLIP | 23597149 | Unspecified |
| mmu-miR-29b-3p | MIMAT0000127 | Pop1 | 67724 | HITS-CLIP | 23597149 | Unspecified |
| mmu-miR-29b-3p | MIMAT0000127 | Pop4 | 66161 | HITS-CLIP | 23142080 | Unspecified |
| mmu-miR-29b-3p | MIMAT0000127 | Por | 18984 | HITS-CLIP//sRNA-Seq | 23597149\|25083871\|33184237 | Unspecified |
| mmu-miR-29b-3p | MIMAT0000127 | Porcn | 53627 | HITS-CLIP | 25083871 | Unspecified |
| mmu-miR-29b-3p | MIMAT0000127 | Postn | 50706 | HITS-CLIP//sRNA-Seq | 25083871\|33184237 | Unspecified |
| mmu-miR-29b-3p | MIMAT0000127 | Pot1a | 101185 | HITS-CLIP | 37986224\|23142080 | Unspecified |
| mmu-miR-29b-3p | MIMAT0000127 | Pou2f1 | 18986 | HITS-CLIP | 23142080 | Unspecified |
| mmu-miR-29b-3p | MIMAT0000127 | Pou2f2 | 18987 | HITS-CLIP | 23142080 | Unspecified |
| mmu-miR-29b-3p | MIMAT0000127 | Ppard | 19015 | HITS-CLIP | 23142080 | Unspecified |
| mmu-miR-29b-3p | MIMAT0000127 | Ppargc1b | 170826 | HITS-CLIP | 37986224 | Unspecified |
| mmu-miR-29b-3p | MIMAT0000127 | Ppfibp1 | 67533 | HITS-CLIP//sRNA-Seq | 23597149\|33184237\|25083871 | Unspecified |
| mmu-miR-29b-3p | MIMAT0000127 | Ppfibp2 | 19024 | HITS-CLIP | 23597149 | Unspecified |
| mmu-miR-29b-3p | MIMAT0000127 | Ppib | 19035 | sRNA-Seq//HITS-CLIP | 33184237\|23597149 | Unspecified |
| mmu-miR-29b-3p | MIMAT0000127 | Ppic | 19038 | sRNA-Seq//HITS-CLIP//qCLASH | 33184237\|23142080\|31393953 | Unspecified |
| mmu-miR-29b-3p | MIMAT0000127 | Ppid | 67738 | HITS-CLIP | 37986224 | Unspecified |
| mmu-miR-29b-3p | MIMAT0000127 | Ppif | 105675 | HITS-CLIP | 37986224 | Unspecified |
| mmu-miR-29b-3p | MIMAT0000127 | Ppig | 228005 | HITS-CLIP//sRNA-Seq | 25083871\|33184237\|23142080 | Unspecified |
| mmu-miR-29b-3p | MIMAT0000127 | Ppih | 66101 | HITS-CLIP | 23597149 | Unspecified |
| mmu-miR-29b-3p | MIMAT0000127 | Ppil2 | 66053 | HITS-CLIP//sRNA-Seq | 23142080\|33184237\|25083871 | Unspecified |
| mmu-miR-29b-3p | MIMAT0000127 | Ppil3 | 53938 | HITS-CLIP | 23142080 | Unspecified |
| mmu-miR-29b-3p | MIMAT0000127 | Ppil4 | 67418 | HITS-CLIP | 37986224\|23142080\|25083871 | Unspecified |
| mmu-miR-29b-3p | MIMAT0000127 | Ppil6 | 73075 | HITS-CLIP | 23142080 | Unspecified |
| mmu-miR-29b-3p | MIMAT0000127 | Ppip5k1 | 327655 | HITS-CLIP | 23142080 | Unspecified |

| | | | | | | |
|---|---|---|---|---|---|---|
| mmu-miR-29b-3p | MIMAT0000127 | Ppip5k2 | 227399 | HITS-CLIP//sRNA-Seq | 37986224\|25083871\|33184237\|23597 | Unspecified |
| mmu-miR-29b-3p | MIMAT0000127 | Ppl | 19041 | HITS-CLIP | 37986224\|23597149 | Unspecified |
| mmu-miR-29b-3p | MIMAT0000127 | Ppm1a | 19042 | HITS-CLIP//sRNA-Seq | 25083871\|33184237\|23142080 | Unspecified |
| mmu-miR-29b-3p | MIMAT0000127 | Ppm1d | 53892 | HITS-CLIP | 37986224\|23142080 | Unspecified |
| mmu-miR-29b-3p | MIMAT0000127 | Ppm1e | 320472 | HITS-CLIP | 37986224 | Unspecified |
| mmu-miR-29b-3p | MIMAT0000127 | Ppm1g | 14208 | sRNA-Seq//HITS-CLIP | 33184237\|23142080 | Unspecified |
| mmu-miR-29b-3p | MIMAT0000127 | Ppm1k | 243382 | HITS-CLIP | 25083871\|23597149 | Unspecified |
| mmu-miR-29b-3p | MIMAT0000127 | Ppme1 | 72590 | HITS-CLIP//sRNA-Seq | 25083871\|33184237 | Unspecified |
| mmu-miR-29b-3p | MIMAT0000127 | Ppp1ca | 19045 | sRNA-Seq//HITS-CLIP | 33184237\|23597149 | Unspecified |
| mmu-miR-29b-3p | MIMAT0000127 | Ppp1cb | 19046 | HITS-CLIP//sRNA-Seq | 37986224\|23597149\|33184237\|25083 | Unspecified |
| mmu-miR-29b-3p | MIMAT0000127 | Ppp1r10 | 52040 | HITS-CLIP | 37986224\|23142080 | Unspecified |
| mmu-miR-29b-3p | MIMAT0000127 | Ppp1r12a | 17931 | HITS-CLIP//sRNA-Seq | 23142080\|33184237\|25083871 | Unspecified |
| mmu-miR-29b-3p | MIMAT0000127 | Ppp1r12b | 329251 | HITS-CLIP | 37986224 | Unspecified |
| mmu-miR-29b-3p | MIMAT0000127 | Ppp1r12c | 232807 | HITS-CLIP | 23142080 | Unspecified |
| mmu-miR-29b-3p | MIMAT0000127 | Ppp1r13b | 21981 | HITS-CLIP | 37986224\|25083871\|23142080 | Unspecified |
| mmu-miR-29b-3p | MIMAT0000127 | Ppp1r13l | 333654 | HITS-CLIP | 25083871 | Unspecified |
| mmu-miR-29b-3p | MIMAT0000127 | Ppp1r14c | 76142 | HITS-CLIP | 37986224 | Unspecified |
| mmu-miR-29b-3p | MIMAT0000127 | Ppp1r15b | 108954 | HITS-CLIP | 25083871\|23597149\|23142080 | Unspecified |
| mmu-miR-29b-3p | MIMAT0000127 | Ppp1r18 | 76448 | sRNA-Seq//HITS-CLIP | 33184237\|23142080 | Unspecified |
| mmu-miR-29b-3p | MIMAT0000127 | Ppp1r21 | 73825 | HITS-CLIP//sRNA-Seq | 25083871\|33184237\|23142080 | Unspecified |
| mmu-miR-29b-3p | MIMAT0000127 | Ppp1r3f | 54646 | HITS-CLIP | 23142080 | Unspecified |
| mmu-miR-29b-3p | MIMAT0000127 | Ppp1r8 | 100336 | sRNA-Seq//HITS-CLIP | 33184237\|23142080 | Unspecified |
| mmu-miR-29b-3p | MIMAT0000127 | Ppp2ca | 19052 | sRNA-Seq//HITS-CLIP | 33184237\|23142080\|25083871 | Unspecified |
| mmu-miR-29b-3p | MIMAT0000127 | Ppp2cb | 19053 | sRNA-Seq//HITS-CLIP | 33184237\|23142080\|25083871 | Unspecified |
| mmu-miR-29b-3p | MIMAT0000127 | Ppp2r1a | 51792 | HITS-CLIP//sRNA-Seq | 37986224\|23597149\|33184237\|25083 | Unspecified |
| mmu-miR-29b-3p | MIMAT0000127 | Ppp2r1b | 73699 | HITS-CLIP | 25083871\|23142080 | Unspecified |
| mmu-miR-29b-3p | MIMAT0000127 | Ppp2r2a | 71978 | HITS-CLIP//sRNA-Seq | 25083871\|33184237 | Unspecified |
| mmu-miR-29b-3p | MIMAT0000127 | Ppp2r3a | 235542 | HITS-CLIP | 25083871 | Unspecified |
| mmu-miR-29b-3p | MIMAT0000127 | Ppp2r5c | 26931 | HITS-CLIP//sRNA-Seq | 25083871\|33184237\|23142080 | Unspecified |
| mmu-miR-29b-3p | MIMAT0000127 | Ppp2r5e | 26932 | HITS-CLIP | 31433990\|23142080 | Unspecified |
| mmu-miR-29b-3p | MIMAT0000127 | Ppp3ca | 19055 | HITS-CLIP//sRNA-Seq | 25083871\|33184237\|23142080 | Unspecified |
| mmu-miR-29b-3p | MIMAT0000127 | Ppp3cb | 19056 | HITS-CLIP//sRNA-Seq | 37986224\|25083871\|23142080\|33184 | Unspecified |
| mmu-miR-29b-3p | MIMAT0000127 | Ppp3cc | 19057 | HITS-CLIP | 23142080 | Unspecified |
| mmu-miR-29b-3p | MIMAT0000127 | Ppp3r1 | 19058 | HITS-CLIP//sRNA-Seq | 37986224\|33184237\|23142080 | Unspecified |
| mmu-miR-29b-3p | MIMAT0000127 | Ppp4r2 | 232314 | HITS-CLIP//sRNA-Seq | 25083871\|33184237\|23142080 | Unspecified |
| mmu-miR-29b-3p | MIMAT0000127 | Ppp4r3a | 55671 | HITS-CLIP | 23142080 | Unspecified |
| mmu-miR-29b-3p | MIMAT0000127 | Ppp4r3b | 57223 | HITS-CLIP | 23142080\|25083871 | Unspecified |
| mmu-miR-29b-3p | MIMAT0000127 | Ppp6r1 | 243819 | HITS-CLIP//sRNA-Seq | 25083871\|33184237\|23597149\|23142 | Unspecified |
| mmu-miR-29b-3p | MIMAT0000127 | Ppp6r3 | 52036 | HITS-CLIP//sRNA-Seq | 37986224\|25083871\|23597149\|33184 | Unspecified |

| | | | | | | |
|---|---|---|---|---|---|---|
| mmu-miR-29b-3p | MIMAT0000127 | Ppt1 | 19063 | HITS-CLIP//sRNA-Seq | 37986224\|25083871\|33184237\|23142 | Unspecified |
| mmu-miR-29b-3p | MIMAT0000127 | Ppt2 | 54397 | HITS-CLIP | 37986224\|23597149 | Unspecified |
| mmu-miR-29b-3p | MIMAT0000127 | Pptc7 | 320717 | HITS-CLIP | 23142080 | Unspecified |
| mmu-miR-29b-3p | MIMAT0000127 | Pqlc1 | 66943 | HITS-CLIP//sRNA-Seq | 37986224\|33184237\|23597149 | Unspecified |
| mmu-miR-29b-3p | MIMAT0000127 | Pramef8 | 242736 | HITS-CLIP | 23142080 | Unspecified |
| mmu-miR-29b-3p | MIMAT0000127 | Prc1 | 233406 | HITS-CLIP//sRNA-Seq | 23142080\|33184237\|23597149 | Unspecified |
| mmu-miR-29b-3p | MIMAT0000127 | Prcc | 94315 | HITS-CLIP//sRNA-Seq | 37986224\|33184237\|23142080\|25083 | Unspecified |
| mmu-miR-29b-3p | MIMAT0000127 | Prdm1 | 12142 | HITS-CLIP | 23142080 | Unspecified |
| mmu-miR-29b-3p | MIMAT0000127 | Prdm10 | 382066 | HITS-CLIP | 23597149 | Unspecified |
| mmu-miR-29b-3p | MIMAT0000127 | Prdm11 | 100042784 | HITS-CLIP | 23142080 | Unspecified |
| mmu-miR-29b-3p | MIMAT0000127 | Prdm4 | 72843 | HITS-CLIP | 37986224\|23142080\|25083871 | Unspecified |
| mmu-miR-29b-3p | MIMAT0000127 | Prdx1 | 18477 | HITS-CLIP//sRNA-Seq | 23142080\|25083871\|33184237 | Unspecified |
| mmu-miR-29b-3p | MIMAT0000127 | Prdx2 | 21672 | HITS-CLIP//sRNA-Seq | 25083871\|33184237\|23597149\|23142 | Unspecified |
| mmu-miR-29b-3p | MIMAT0000127 | Prdx3 | 11757 | HITS-CLIP//sRNA-Seq | 25083871\|23597149\|33184237\|23142 | Unspecified |
| mmu-miR-29b-3p | MIMAT0000127 | Prdx4 | 53381 | HITS-CLIP//sRNA-Seq | 23597149\|25083871\|23142080\|33184 | Unspecified |
| mmu-miR-29b-3p | MIMAT0000127 | Prelp | 116847 | HITS-CLIP | 37986224\|25083871\|23597149 | Unspecified |
| mmu-miR-29b-3p | MIMAT0000127 | Prep | 19072 | HITS-CLIP | 25083871\|23597149\|23142080 | Unspecified |
| mmu-miR-29b-3p | MIMAT0000127 | Prepl | 213760 | sRNA-Seq//HITS-CLIP | 33184237\|23142080 | Unspecified |
| mmu-miR-29b-3p | MIMAT0000127 | Prg4 | 96875 | HITS-CLIP | 23597149 | Unspecified |
| mmu-miR-29b-3p | MIMAT0000127 | Prkaa1 | 105787 | HITS-CLIP | 25083871\|23597149 | Unspecified |
| mmu-miR-29b-3p | MIMAT0000127 | Prkab1 | 19079 | sRNA-Seq//HITS-CLIP | 33184237\|23142080 | Unspecified |
| mmu-miR-29b-3p | MIMAT0000127 | Prkab2 | 108097 | HITS-CLIP | 37986224\|23142080 | Unspecified |
| mmu-miR-29b-3p | MIMAT0000127 | Prkaca | 18747 | HITS-CLIP//sRNA-Seq | 23142080\|33184237\|25083871 | Unspecified |
| mmu-miR-29b-3p | MIMAT0000127 | Prkacb | 18749 | sRNA-Seq//HITS-CLIP | 33184237\|23597149\|23142080 | Unspecified |
| mmu-miR-29b-3p | MIMAT0000127 | Prkag1 | 19082 | HITS-CLIP//sRNA-Seq | 23142080\|25083871\|33184237 | Unspecified |
| mmu-miR-29b-3p | MIMAT0000127 | Prkag3 | 241113 | HITS-CLIP | 25083871 | Unspecified |
| mmu-miR-29b-3p | MIMAT0000127 | Prkar1a | 19084 | HITS-CLIP//sRNA-Seq | 37986224\|25083871\|23597149\|33184 | Unspecified |
| mmu-miR-29b-3p | MIMAT0000127 | Prkar2a | 19087 | HITS-CLIP//sRNA-Seq | 37986224\|25083871\|33184237\|23142 | Unspecified |
| mmu-miR-29b-3p | MIMAT0000127 | Prkca | 18750 | HITS-CLIP//sRNA-Seq | 37986224\|33184237\|23597149\|23142 | Unspecified |
| mmu-miR-29b-3p | MIMAT0000127 | Prkcd | 18753 | HITS-CLIP//sRNA-Seq | 37986224\|33184237\|23597149\|23142 | Unspecified |
| mmu-miR-29b-3p | MIMAT0000127 | Prkci | 18759 | HITS-CLIP//sRNA-Seq | 37986224\|25083871\|33184237 | Unspecified |
| mmu-miR-29b-3p | MIMAT0000127 | Prkcq | 18761 | HITS-CLIP | 23142080 | Unspecified |
| mmu-miR-29b-3p | MIMAT0000127 | Prkd3 | 75292 | HITS-CLIP//sRNA-Seq | 37986224\|25083871\|33184237 | Unspecified |
| mmu-miR-29b-3p | MIMAT0000127 | Prkra | 23992 | HITS-CLIP//sRNA-Seq | 25083871\|33184237 | Unspecified |
| mmu-miR-29b-3p | MIMAT0000127 | Prkx | 19108 | HITS-CLIP | 25083871\|23142080 | Unspecified |
| mmu-miR-29b-3p | MIMAT0000127 | Prlr | 19116 | HITS-CLIP | 23597149 | Unspecified |
| mmu-miR-29b-3p | MIMAT0000127 | Prmt1 | 15469 | sRNA-Seq//HITS-CLIP | 33184237\|23597149 | Unspecified |
| mmu-miR-29b-3p | MIMAT0000127 | Prmt5 | 27374 | HITS-CLIP//sRNA-Seq | 31433990\|33184237 | Unspecified |
| mmu-miR-29b-3p | MIMAT0000127 | Prnp | 19122 | HITS-CLIP//sRNA-Seq | 25083871\|33184237\|23142080 | Unspecified |

| | | | | | | |
|---|---|---|---|---|---|---|
| mmu-miR-29b-3p | MIMAT0000127 | Pros1 | 19128 | HITS-CLIP | 37986224\|23597149 | Unspecified |
| mmu-miR-29b-3p | MIMAT0000127 | Proser1 | 212127 | HITS-CLIP | 37986224\|25083871\|23597149\|23142 | Unspecified |
| mmu-miR-29b-3p | MIMAT0000127 | Prox1 | 19130 | HITS-CLIP | 23597149 | Unspecified |
| mmu-miR-29b-3p | MIMAT0000127 | Prpf19 | 28000 | sRNA-Seq//HITS-CLIP | 33184237\|23597149 | Unspecified |
| mmu-miR-29b-3p | MIMAT0000127 | Prpf3 | 70767 | HITS-CLIP//sRNA-Seq | 37986224\|23142080\|25083871\|33184 | Unspecified |
| mmu-miR-29b-3p | MIMAT0000127 | Prpf38b | 66921 | sRNA-Seq//HITS-CLIP | 33184237\|23142080 | Unspecified |
| mmu-miR-29b-3p | MIMAT0000127 | Prpf4 | 70052 | HITS-CLIP | 23142080 | Unspecified |
| mmu-miR-29b-3p | MIMAT0000127 | Prpf40a | 56194 | HITS-CLIP//sRNA-Seq | 25083871\|33184237\|23597149\|23142 | Unspecified |
| mmu-miR-29b-3p | MIMAT0000127 | Prpf4b | 19134 | HITS-CLIP | 37986224\|25083871\|23142080 | Unspecified |
| mmu-miR-29b-3p | MIMAT0000127 | Prpf6 | 68879 | HITS-CLIP//sRNA-Seq | 23142080\|33184237 | Unspecified |
| mmu-miR-29b-3p | MIMAT0000127 | Prpf8 | 192159 | HITS-CLIP//sRNA-Seq | 23597149\|23142080\|25083871\|33184 | Unspecified |
| mmu-miR-29b-3p | MIMAT0000127 | Prps1l3 | 328099 | HITS-CLIP | 23142080 | Unspecified |
| mmu-miR-29b-3p | MIMAT0000127 | Prps2 | 110639 | HITS-CLIP | 23142080 | Unspecified |
| mmu-miR-29b-3p | MIMAT0000127 | Prr11 | 270906 | HITS-CLIP | 23142080 | Unspecified |
| mmu-miR-29b-3p | MIMAT0000127 | Prr12 | 233210 | HITS-CLIP | 25083871 | Unspecified |
| mmu-miR-29b-3p | MIMAT0000127 | Prr14 | 233895 | HITS-CLIP | 23142080 | Unspecified |
| mmu-miR-29b-3p | MIMAT0000127 | Prr14l | 215476 | HITS-CLIP | 37986224\|23142080 | Unspecified |
| mmu-miR-29b-3p | MIMAT0000127 | Prr5 | 109270 | HITS-CLIP | 25083871\|23597149 | Unspecified |
| mmu-miR-29b-3p | MIMAT0000127 | Prrc1 | 73137 | HITS-CLIP | 37986224\|25083871\|23142080 | Unspecified |
| mmu-miR-29b-3p | MIMAT0000127 | Prrc2b | 227723 | sRNA-Seq//HITS-CLIP | 33184237\|23597149\|23142080 | Unspecified |
| mmu-miR-29b-3p | MIMAT0000127 | Prrc2c | 226562 | HITS-CLIP//sRNA-Seq | 37986224\|23142080\|25083871\|23597 | Unspecified |
| mmu-miR-29b-3p | MIMAT0000127 | Prrg1 | 546336 | HITS-CLIP | 23142080 | Unspecified |
| mmu-miR-29b-3p | MIMAT0000127 | Prrg3 | 208748 | HITS-CLIP | 23597149 | Unspecified |
| mmu-miR-29b-3p | MIMAT0000127 | Prrx1 | 18933 | HITS-CLIP//sRNA-Seq | 25083871\|33184237 | Unspecified |
| mmu-miR-29b-3p | MIMAT0000127 | Prtg | 235472 | HITS-CLIP | 25083871 | Unspecified |
| mmu-miR-29b-3p | MIMAT0000127 | Prune2 | 353211 | HITS-CLIP//sRNA-Seq | 25083871\|33184237 | Unspecified |
| mmu-miR-29b-3p | MIMAT0000127 | Psap | 19156 | HITS-CLIP//sRNA-Seq | 25083871\|23142080\|33184237\|23597 | Unspecified |
| mmu-miR-29b-3p | MIMAT0000127 | Psat1 | 107272 | HITS-CLIP//sRNA-Seq | 23142080\|33184237 | Unspecified |
| mmu-miR-29b-3p | MIMAT0000127 | Psd3 | 234353 | HITS-CLIP//RNA-Seq//sRNA-Seq | 25083871\|33332475\|33184237 | Unspecified |
| mmu-miR-29b-3p | MIMAT0000127 | Psen1 | 19164 | HITS-CLIP//sRNA-Seq | 25083871\|33184237\|23142080 | Unspecified |
| mmu-miR-29b-3p | MIMAT0000127 | Psip1 | 101739 | HITS-CLIP//sRNA-Seq | 37986224\|25083871\|33184237\|23142 | Unspecified |
| mmu-miR-29b-3p | MIMAT0000127 | Pskh1 | 244631 | HITS-CLIP | 37986224\|25083871\|23597149 | Unspecified |
| mmu-miR-29b-3p | MIMAT0000127 | Psma1 | 26440 | HITS-CLIP//sRNA-Seq | 23597149\|33184237\|23142080 | Unspecified |
| mmu-miR-29b-3p | MIMAT0000127 | Psma3 | 19167 | HITS-CLIP//sRNA-Seq | 37986224\|23142080\|33184237\|23597 | Unspecified |
| mmu-miR-29b-3p | MIMAT0000127 | Psma4 | 26441 | HITS-CLIP//sRNA-Seq | 37986224\|25083871\|33184237\|23142 | Unspecified |
| mmu-miR-29b-3p | MIMAT0000127 | Psmb2 | 26445 | HITS-CLIP//sRNA-Seq | 25083871\|33184237\|23142080 | Unspecified |
| mmu-miR-29b-3p | MIMAT0000127 | Psmb4 | 19172 | HITS-CLIP//sRNA-Seq | 23597149\|33184237\|25083871\|23142 | Unspecified |
| mmu-miR-29b-3p | MIMAT0000127 | Psmb6 | 19175 | sRNA-Seq//HITS-CLIP | 33184237\|23142080\|25083871 | Unspecified |
| mmu-miR-29b-3p | MIMAT0000127 | Psmb7 | 19177 | HITS-CLIP//sRNA-Seq | 25083871\|23597149\|33184237\|23142 | Unspecified |

| | | | | | | |
|---|---|---|---|---|---|---|
| mmu-miR-29b-3p | MIMAT0000127 | Psmb8 | 16913 | HITS-CLIP | 23142080 | Unspecified |
| mmu-miR-29b-3p | MIMAT0000127 | Psmb9 | 16912 | HITS-CLIP | 23142080 | Unspecified |
| mmu-miR-29b-3p | MIMAT0000127 | Psmc1 | 19179 | sRNA-Seq//HITS-CLIP | 33184237\|25083871 | Unspecified |
| mmu-miR-29b-3p | MIMAT0000127 | Psmc6 | 67089 | HITS-CLIP//sRNA-Seq | 23142080\|25083871\|33184237 | Unspecified |
| mmu-miR-29b-3p | MIMAT0000127 | Psmd1 | 70247 | HITS-CLIP//sRNA-Seq | 23142080\|33184237 | Unspecified |
| mmu-miR-29b-3p | MIMAT0000127 | Psmd10 | 53380 | HITS-CLIP | 25083871 | Unspecified |
| mmu-miR-29b-3p | MIMAT0000127 | Psmd11 | 69077 | HITS-CLIP//sRNA-Seq | 23597149\|33184237 | Unspecified |
| mmu-miR-29b-3p | MIMAT0000127 | Psmd12 | 66997 | HITS-CLIP//sRNA-Seq | 23597149\|33184237\|23142080 | Unspecified |
| mmu-miR-29b-3p | MIMAT0000127 | Psmd13 | 23997 | HITS-CLIP//sRNA-Seq | 23597149\|25083871\|23142080\|33184 | Unspecified |
| mmu-miR-29b-3p | MIMAT0000127 | Psmd14 | 59029 | sRNA-Seq//HITS-CLIP | 33184237\|23142080 | Unspecified |
| mmu-miR-29b-3p | MIMAT0000127 | Psmd3 | 22123 | sRNA-Seq//HITS-CLIP | 33184237\|23142080 | Unspecified |
| mmu-miR-29b-3p | MIMAT0000127 | Psmd9 | 67151 | HITS-CLIP | 25083871 | Unspecified |
| mmu-miR-29b-3p | MIMAT0000127 | Psme2b | 621823 | HITS-CLIP | 23142080 | Unspecified |
| mmu-miR-29b-3p | MIMAT0000127 | Psme3 | 19192 | HITS-CLIP//sRNA-Seq | 23597149\|33184237\|23142080 | Unspecified |
| mmu-miR-29b-3p | MIMAT0000127 | Psme4 | 103554 | HITS-CLIP//sRNA-Seq | 25083871\|33184237\|23597149\|23142 | Unspecified |
| mmu-miR-29b-3p | MIMAT0000127 | Psmg2 | 107047 | HITS-CLIP//sRNA-Seq | 23142080\|33184237\|23597149\|25083 | Unspecified |
| mmu-miR-29b-3p | MIMAT0000127 | Pspc1 | 66645 | HITS-CLIP | 25083871 | Unspecified |
| mmu-miR-29b-3p | MIMAT0000127 | Pstk | 214580 | HITS-CLIP | 23142080 | Unspecified |
| mmu-miR-29b-3p | MIMAT0000127 | Pstpip2 | 19201 | HITS-CLIP | 25083871 | Unspecified |
| mmu-miR-29b-3p | MIMAT0000127 | Ptbp2 | 56195 | HITS-CLIP//sRNA-Seq | 23142080\|33184237 | Unspecified |
| mmu-miR-29b-3p | MIMAT0000127 | Ptbp3 | 230257 | HITS-CLIP//sRNA-Seq | 37986224\|23597149\|25083871\|33184 | Unspecified |
| mmu-miR-29b-3p | MIMAT0000127 | Ptch1 | 19206 | HITS-CLIP | 23597149\|23142080 | Unspecified |
| mmu-miR-29b-3p | MIMAT0000127 | Ptdss1 | 19210 | HITS-CLIP | 23142080\|25083871 | Unspecified |
| mmu-miR-29b-3p | MIMAT0000127 | Pten | 19211 | HITS-CLIP//sRNA-Seq | 37986224\|23597149\|25083871\|33184 | Unspecified |
| mmu-miR-29b-3p | MIMAT0000127 | Ptger2 | 19217 | HITS-CLIP | 37986224\|23597149\|23142080 | Unspecified |
| mmu-miR-29b-3p | MIMAT0000127 | Ptgfrn | 19221 | sRNA-Seq//HITS-CLIP | 33184237\|23142080 | Unspecified |
| mmu-miR-29b-3p | MIMAT0000127 | Ptgr1 | 67103 | HITS-CLIP | 25083871 | Unspecified |
| mmu-miR-29b-3p | MIMAT0000127 | Ptk2 | 14083 | sRNA-Seq//HITS-CLIP | 33184237\|25083871 | Unspecified |
| mmu-miR-29b-3p | MIMAT0000127 | Ptp4a1 | 19243 | HITS-CLIP//sRNA-Seq | 37986224\|23142080\|33184237\|23597 | Unspecified |
| mmu-miR-29b-3p | MIMAT0000127 | Ptp4a2 | 19244 | HITS-CLIP//sRNA-Seq | 23597149\|33184237\|23142080 | Unspecified |
| mmu-miR-29b-3p | MIMAT0000127 | Ptpa | 5524 | HITS-CLIP//sRNA-Seq | 25083871\|33184237\|23597149 | Unspecified |
| mmu-miR-29b-3p | MIMAT0000127 | Ptpn11 | 19247 | HITS-CLIP//sRNA-Seq | 37986224\|25083871\|33184237\|23597 | Unspecified |
| mmu-miR-29b-3p | MIMAT0000127 | Ptpn12 | 19248 | HITS-CLIP//sRNA-Seq | 37986224\|25083871\|33184237 | Unspecified |
| mmu-miR-29b-3p | MIMAT0000127 | Ptpn14 | 19250 | HITS-CLIP | 25083871 | Unspecified |
| mmu-miR-29b-3p | MIMAT0000127 | Ptpn18 | 19253 | HITS-CLIP | 31433990 | Unspecified |
| mmu-miR-29b-3p | MIMAT0000127 | Ptpn2 | 19255 | HITS-CLIP | 23142080 | Unspecified |
| mmu-miR-29b-3p | MIMAT0000127 | Ptpn21 | 24000 | HITS-CLIP | 25083871 | Unspecified |
| mmu-miR-29b-3p | MIMAT0000127 | Ptpn22 | 19260 | HITS-CLIP | 23142080 | Unspecified |
| mmu-miR-29b-3p | MIMAT0000127 | Ptpn23 | 104831 | HITS-CLIP//sRNA-Seq | 25083871\|33184237\|23142080 | Unspecified |

| | | | | | | |
|---|---|---|---|---|---|---|
| mmu-miR-29b-3p | MIMAT0000127 | Ptpn3 | 545622 | HITS-CLIP//sRNA-Seq | 25083871\|33184237\|23142080 | Unspecified |
| mmu-miR-29b-3p | MIMAT0000127 | Ptpn4 | 19258 | HITS-CLIP | 25083871\|23142080 | Unspecified |
| mmu-miR-29b-3p | MIMAT0000127 | Ptpn6 | 15170 | HITS-CLIP | 23142080 | Unspecified |
| mmu-miR-29b-3p | MIMAT0000127 | Ptprb | 19263 | HITS-CLIP | 37986224\|23597149 | Unspecified |
| mmu-miR-29b-3p | MIMAT0000127 | Ptprc | 19264 | HITS-CLIP | 23142080 | Unspecified |
| mmu-miR-29b-3p | MIMAT0000127 | Ptprcap | 19265 | HITS-CLIP | 23142080 | Unspecified |
| mmu-miR-29b-3p | MIMAT0000127 | Ptprd | 19266 | HITS-CLIP//sRNA-Seq | 37986224\|25083871\|33184237\|23597 | Unspecified |
| mmu-miR-29b-3p | MIMAT0000127 | Ptprf | 19268 | HITS-CLIP//sRNA-Seq | 37986224\|23597149\|33184237 | Unspecified |
| mmu-miR-29b-3p | MIMAT0000127 | Ptprg | 19270 | HITS-CLIP | 37986224\|23597149\|25083871 | Unspecified |
| mmu-miR-29b-3p | MIMAT0000127 | Ptprk | 19272 | HITS-CLIP//sRNA-Seq | 37986224\|25083871\|33184237 | Unspecified |
| mmu-miR-29b-3p | MIMAT0000127 | Ptprs | 19280 | HITS-CLIP//sRNA-Seq | 25083871\|33184237 | Unspecified |
| mmu-miR-29b-3p | MIMAT0000127 | Ptrh2 | 217057 | HITS-CLIP | 37986224 | Unspecified |
| mmu-miR-29b-3p | MIMAT0000127 | Pttg1ip | 108705 | HITS-CLIP//sRNA-Seq | 23597149\|33184237\|25083871 | Unspecified |
| mmu-miR-29b-3p | MIMAT0000127 | Puf60 | 67959 | HITS-CLIP//sRNA-Seq | 37986224\|33184237\|23597149\|23142 | Unspecified |
| mmu-miR-29b-3p | MIMAT0000127 | Pum1 | 80912 | HITS-CLIP//sRNA-Seq | 25083871\|33184237\|23142080 | Unspecified |
| mmu-miR-29b-3p | MIMAT0000127 | Pum2 | 80913 | HITS-CLIP//sRNA-Seq | 25083871\|23597149\|33184237\|23142 | Unspecified |
| mmu-miR-29b-3p | MIMAT0000127 | Pum3 | 52874 | HITS-CLIP | 23597149\|23142080 | Unspecified |
| mmu-miR-29b-3p | MIMAT0000127 | Pura | 19290 | HITS-CLIP | 37986224\|25083871\|23142080 | Unspecified |
| mmu-miR-29b-3p | MIMAT0000127 | Purb | 19291 | HITS-CLIP | 23142080\|25083871 | Unspecified |
| mmu-miR-29b-3p | MIMAT0000127 | Purg | 75029 | HITS-CLIP//sRNA-Seq | 37986224\|25083871\|33184237\|23597 | Unspecified |
| mmu-miR-29b-3p | MIMAT0000127 | Pus10 | 74467 | HITS-CLIP | 23142080 | Unspecified |
| mmu-miR-29b-3p | MIMAT0000127 | Pxdn | 69675 | HITS-CLIP//sRNA-Seq | 25083871\|33184237\|23597149 | Unspecified |
| mmu-miR-29b-3p | MIMAT0000127 | Pxk | 218699 | HITS-CLIP | 23142080 | Unspecified |
| mmu-miR-29b-3p | MIMAT0000127 | Pxmp2 | 19301 | HITS-CLIP | 37986224 | Unspecified |
| mmu-miR-29b-3p | MIMAT0000127 | Pxmp4 | 59038 | HITS-CLIP | 23597149\|25083871 | Unspecified |
| mmu-miR-29b-3p | MIMAT0000127 | Pxylp1 | 235534 | HITS-CLIP | 25083871 | Unspecified |
| mmu-miR-29b-3p | MIMAT0000127 | Pygb | 110078 | sRNA-Seq//HITS-CLIP | 33184237\|23142080 | Unspecified |
| mmu-miR-29b-3p | MIMAT0000127 | Pyroxd1 | 232491 | HITS-CLIP | 23142080 | Unspecified |
| mmu-miR-29b-3p | MIMAT0000127 | Pzp | 11287 | HITS-CLIP | 23597149 | Unspecified |
| mmu-miR-29b-3p | MIMAT0000127 | Qars | 97541 | HITS-CLIP | 37986224 | Unspecified |
| mmu-miR-29b-3p | MIMAT0000127 | Qk | 19317 | HITS-CLIP | 37986224\|25083871\|23142080\|23597 | Unspecified |
| mmu-miR-29b-3p | MIMAT0000127 | Qser1 | 99003 | HITS-CLIP | 37986224\|23142080\|23597149\|25083 | Unspecified |
| mmu-miR-29b-3p | MIMAT0000127 | Qsox1 | 104009 | HITS-CLIP//sRNA-Seq | 25083871\|33184237 | Unspecified |
| mmu-miR-29b-3p | MIMAT0000127 | R3hcc1l | 52013 | HITS-CLIP | 37986224\|25083871\|23142080 | Unspecified |
| mmu-miR-29b-3p | MIMAT0000127 | R3hdm1 | 226412 | HITS-CLIP | 37986224 | Unspecified |
| mmu-miR-29b-3p | MIMAT0000127 | R3hdm2 | 71750 | HITS-CLIP//sRNA-Seq | 23142080\|25083871\|23597149\|33184 | Unspecified |
| mmu-miR-29b-3p | MIMAT0000127 | R3hdm4 | 109284 | HITS-CLIP//sRNA-Seq | 37986224\|25083871\|33184237\|23142 | Unspecified |
| mmu-miR-29b-3p | MIMAT0000127 | Rab11a | 53869 | HITS-CLIP//sRNA-Seq | 25083871\|23142080\|33184237\|23597 | Unspecified |
| mmu-miR-29b-3p | MIMAT0000127 | Rab11b | 19326 | qCLASH | 31393953 | Unspecified |

| | | | | | | |
|---|---|---|---|---|---|---|
| mmu-miR-29b-3p | MIMAT0000127 | Rab12 | 19328 | HITS-CLIP//sRNA-Seq | 37986224\|25083871\|33184237\|23142 | Unspecified |
| mmu-miR-29b-3p | MIMAT0000127 | Rab14 | 68365 | HITS-CLIP//sRNA-Seq | 23597149\|33184237\|23142080 | Unspecified |
| mmu-miR-29b-3p | MIMAT0000127 | Rab18 | 19330 | HITS-CLIP//sRNA-Seq | 23142080\|33184237 | Unspecified |
| mmu-miR-29b-3p | MIMAT0000127 | Rab1a | 5861 | sRNA-Seq//HITS-CLIP | 33184237\|23142080 | Unspecified |
| mmu-miR-29b-3p | MIMAT0000127 | Rab1b | 76308 | HITS-CLIP//sRNA-Seq | 23597149\|23142080\|33184237 | Unspecified |
| mmu-miR-29b-3p | MIMAT0000127 | Rab21 | 216344 | HITS-CLIP | 23597149\|25083871\|23142080 | Unspecified |
| mmu-miR-29b-3p | MIMAT0000127 | Rab22a | 19334 | HITS-CLIP | 23142080 | Unspecified |
| mmu-miR-29b-3p | MIMAT0000127 | Rab23 | 19335 | HITS-CLIP | 25083871 | Unspecified |
| mmu-miR-29b-3p | MIMAT0000127 | Rab24 | 19336 | HITS-CLIP//sRNA-Seq | 25083871\|33184237 | Unspecified |
| mmu-miR-29b-3p | MIMAT0000127 | Rab2a | 59021 | HITS-CLIP//sRNA-Seq | 25083871\|33184237\|23597149\|31433 | Unspecified |
| mmu-miR-29b-3p | MIMAT0000127 | Rab30 | 75985 | HITS-CLIP | 23142080 | Unspecified |
| mmu-miR-29b-3p | MIMAT0000127 | Rab33b | 19338 | HITS-CLIP | 25083871\|23142080 | Unspecified |
| mmu-miR-29b-3p | MIMAT0000127 | Rab35 | 77407 | sRNA-Seq//HITS-CLIP | 33184237\|23142080 | Unspecified |
| mmu-miR-29b-3p | MIMAT0000127 | Rab38 | 72433 | HITS-CLIP | 25083871 | Unspecified |
| mmu-miR-29b-3p | MIMAT0000127 | Rab3gap1 | 226407 | HITS-CLIP | 25083871\|23597149\|23142080 | Unspecified |
| mmu-miR-29b-3p | MIMAT0000127 | Rab3ip | 216363 | HITS-CLIP | 23142080 | Unspecified |
| mmu-miR-29b-3p | MIMAT0000127 | Rab40c | 224624 | HITS-CLIP | 37986224\|23597149 | Unspecified |
| mmu-miR-29b-3p | MIMAT0000127 | Rab4b | 19342 | HITS-CLIP//sRNA-Seq | 25083871\|33184237 | Unspecified |
| mmu-miR-29b-3p | MIMAT0000127 | Rab5a | 271457 | HITS-CLIP//sRNA-Seq | 23597149\|33184237\|23142080\|25083 | Unspecified |
| mmu-miR-29b-3p | MIMAT0000127 | Rab5c | 19345 | sRNA-Seq//HITS-CLIP | 33184237\|23142080\|25083871 | Unspecified |
| mmu-miR-29b-3p | MIMAT0000127 | Rab6a | 19346 | HITS-CLIP//sRNA-Seq | 25083871\|33184237\|23142080 | Unspecified |
| mmu-miR-29b-3p | MIMAT0000127 | Rab6b | 270192 | sRNA-Seq//HITS-CLIP | 33184237\|23142080 | Unspecified |
| mmu-miR-29b-3p | MIMAT0000127 | Rab8a | 17274 | HITS-CLIP | 25083871\|23142080 | Unspecified |
| mmu-miR-29b-3p | MIMAT0000127 | Rab8b | 235442 | HITS-CLIP | 37986224 | Unspecified |
| mmu-miR-29b-3p | MIMAT0000127 | Rabac1 | 14470 | HITS-CLIP//sRNA-Seq | 23142080\|33184237 | Unspecified |
| mmu-miR-29b-3p | MIMAT0000127 | Rabep1 | 54189 | HITS-CLIP//sRNA-Seq | 37986224\|25083871\|33184237\|23142 | Unspecified |
| mmu-miR-29b-3p | MIMAT0000127 | Rabgap1 | 227800 | HITS-CLIP//sRNA-Seq | 25083871\|33184237 | Unspecified |
| mmu-miR-29b-3p | MIMAT0000127 | Rabgap1l | 29809 | HITS-CLIP//sRNA-Seq | 37986224\|23597149\|33184237\|23142 | Unspecified |
| mmu-miR-29b-3p | MIMAT0000127 | Rabgef1 | 56715 | HITS-CLIP | 25083871 | Unspecified |
| mmu-miR-29b-3p | MIMAT0000127 | Rabggtb | 19352 | HITS-CLIP//sRNA-Seq | 23142080\|33184237\|23597149 | Unspecified |
| mmu-miR-29b-3p | MIMAT0000127 | Rabl6 | 227624 | HITS-CLIP//sRNA-Seq | 25083871\|33184237 | Unspecified |
| mmu-miR-29b-3p | MIMAT0000127 | Rac1 | 19353 | sRNA-Seq//HITS-CLIP | 33184237\|31433990 | Unspecified |
| mmu-miR-29b-3p | MIMAT0000127 | Rack1 | 10399 | HITS-CLIP//sRNA-Seq | 23597149\|25083871\|33184237\|23142 | Unspecified |
| mmu-miR-29b-3p | MIMAT0000127 | Rad18 | 58186 | HITS-CLIP | 23142080 | Unspecified |
| mmu-miR-29b-3p | MIMAT0000127 | Rad23b | 19359 | HITS-CLIP//sRNA-Seq | 23597149\|25083871\|33184237\|23142 | Unspecified |
| mmu-miR-29b-3p | MIMAT0000127 | Rad50 | 19360 | HITS-CLIP | 23142080 | Unspecified |
| mmu-miR-29b-3p | MIMAT0000127 | Rad51 | 19361 | HITS-CLIP | 25083871\|23597149\|23142080 | Unspecified |
| mmu-miR-29b-3p | MIMAT0000127 | Rad54l2 | 81000 | HITS-CLIP | 25083871\|23597149 | Unspecified |
| mmu-miR-29b-3p | MIMAT0000127 | Rad9a | 19367 | HITS-CLIP//sRNA-Seq | 23142080\|33184237 | Unspecified |

| | | | | | | |
|---|---|---|---|---|---|---|
| mmu-miR-29b-3p | MIMAT0000127 | Raf1 | 110157 | sRNA-Seq//HITS-CLIP | 33184237\|23142080 | Unspecified |
| mmu-miR-29b-3p | MIMAT0000127 | Rala | 56044 | HITS-CLIP//sRNA-Seq | 37986224\|23142080\|25083871\|33184 | Unspecified |
| mmu-miR-29b-3p | MIMAT0000127 | Ralbp1 | 19765 | HITS-CLIP | 23142080 | Unspecified |
| mmu-miR-29b-3p | MIMAT0000127 | Ralgapa1 | 56784 | HITS-CLIP//sRNA-Seq | 23142080\|33184237\|25083871 | Unspecified |
| mmu-miR-29b-3p | MIMAT0000127 | Ralgapa2 | 241694 | HITS-CLIP | 23597149\|23142080 | Unspecified |
| mmu-miR-29b-3p | MIMAT0000127 | Ralgapb | 228850 | HITS-CLIP//sRNA-Seq | 25083871\|33184237\|23597149 | Unspecified |
| mmu-miR-29b-3p | MIMAT0000127 | Ralgps2 | 78255 | HITS-CLIP | 37986224\|23142080 | Unspecified |
| mmu-miR-29b-3p | MIMAT0000127 | Raly | 19383 | HITS-CLIP//sRNA-Seq | 25083871\|23597149\|33184237 | Unspecified |
| mmu-miR-29b-3p | MIMAT0000127 | Ranbp2 | 19386 | sRNA-Seq//HITS-CLIP | 33184237\|23142080\|25083871 | Unspecified |
| mmu-miR-29b-3p | MIMAT0000127 | Ranbp3 | 71810 | sRNA-Seq//HITS-CLIP | 33184237\|25083871 | Unspecified |
| mmu-miR-29b-3p | MIMAT0000127 | Ranbp6 | 240614 | HITS-CLIP | 23597149 | Unspecified |
| mmu-miR-29b-3p | MIMAT0000127 | Ranbp9 | 56705 | HITS-CLIP | 23142080\|25083871 | Unspecified |
| mmu-miR-29b-3p | MIMAT0000127 | Rap1b | 215449 | HITS-CLIP//sRNA-Seq | 37986224\|25083871\|33184237\|23142 | Unspecified |
| mmu-miR-29b-3p | MIMAT0000127 | Rap1gds1 | 229877 | HITS-CLIP//sRNA-Seq | 25083871\|33184237\|23597149\|23142 | Unspecified |
| mmu-miR-29b-3p | MIMAT0000127 | Rap2a | 76108 | HITS-CLIP | 25083871\|23142080 | Unspecified |
| mmu-miR-29b-3p | MIMAT0000127 | Rap2b | 74012 | HITS-CLIP | 23142080 | Unspecified |
| mmu-miR-29b-3p | MIMAT0000127 | Rap2c | 72065 | HITS-CLIP | 25083871\|23142080 | Unspecified |
| mmu-miR-29b-3p | MIMAT0000127 | Rapgef1 | 107746 | HITS-CLIP//sRNA-Seq | 25083871\|33184237\|23597149\|23142 | Unspecified |
| mmu-miR-29b-3p | MIMAT0000127 | Rapgef2 | 76089 | HITS-CLIP | 25083871 | Unspecified |
| mmu-miR-29b-3p | MIMAT0000127 | Rapgef6 | 192786 | sRNA-Seq//HITS-CLIP | 33184237\|23142080 | Unspecified |
| mmu-miR-29b-3p | MIMAT0000127 | Rapgefl1 | 268480 | HITS-CLIP | 37986224\|25083871 | Unspecified |
| mmu-miR-29b-3p | MIMAT0000127 | Raph1 | 77300 | HITS-CLIP//sRNA-Seq | 37986224\|23142080\|25083871\|23597 | Unspecified |
| mmu-miR-29b-3p | MIMAT0000127 | Rarg | 19411 | HITS-CLIP | 37986224 | Unspecified |
| mmu-miR-29b-3p | MIMAT0000127 | Rars | 104458 | sRNA-Seq//HITS-CLIP | 33184237\|25083871 | Unspecified |
| mmu-miR-29b-3p | MIMAT0000127 | Rars2 | 109093 | HITS-CLIP | 23142080 | Unspecified |
| mmu-miR-29b-3p | MIMAT0000127 | Rasa1 | 218397 | sRNA-Seq//HITS-CLIP | 33184237\|23142080 | Unspecified |
| mmu-miR-29b-3p | MIMAT0000127 | Rasa2 | 114713 | HITS-CLIP | 25083871\|23142080 | Unspecified |
| mmu-miR-29b-3p | MIMAT0000127 | Rasal2 | 226525 | HITS-CLIP | 37986224 | Unspecified |
| mmu-miR-29b-3p | MIMAT0000127 | Rasal3 | 320484 | HITS-CLIP | 23142080 | Unspecified |
| mmu-miR-29b-3p | MIMAT0000127 | Rasgef1b | 320292 | HITS-CLIP | 23142080 | Unspecified |
| mmu-miR-29b-3p | MIMAT0000127 | Rasgrp1 | 19419 | HITS-CLIP | 37986224\|23142080 | Unspecified |
| mmu-miR-29b-3p | MIMAT0000127 | Rasgrp4 | 233046 | HITS-CLIP | 23142080 | Unspecified |
| mmu-miR-29b-3p | MIMAT0000127 | Rassf5 | 54354 | HITS-CLIP | 37986224\|25083871\|23142080 | Unspecified |
| mmu-miR-29b-3p | MIMAT0000127 | Rb1 | 19645 | HITS-CLIP | 23597149\|23142080\|25083871 | Unspecified |
| mmu-miR-29b-3p | MIMAT0000127 | Rb1cc1 | 12421 | HITS-CLIP//sRNA-Seq | 37986224\|25083871\|33184237\|23142 | Unspecified |
| mmu-miR-29b-3p | MIMAT0000127 | Rbak | 57782 | HITS-CLIP | 23142080 | Unspecified |
| mmu-miR-29b-3p | MIMAT0000127 | Rbbp7 | 245688 | HITS-CLIP//sRNA-Seq | 25083871\|23597149\|33184237\|23142 | Unspecified |
| mmu-miR-29b-3p | MIMAT0000127 | Rbfox2 | 93686 | HITS-CLIP//sRNA-Seq | 25083871\|33184237\|23597149 | Unspecified |
| mmu-miR-29b-3p | MIMAT0000127 | Rbl1 | 19650 | HITS-CLIP | 23142080 | Unspecified |

| | | | | | | |
|---|---|---|---|---|---|---|
| mmu-miR-29b-3p | MIMAT0000127 | Rbl2 | 19651 | HITS-CLIP | 25083871\|23142080 | Unspecified |
| mmu-miR-29b-3p | MIMAT0000127 | Rbm12 | 75710 | HITS-CLIP | 23142080 | Unspecified |
| mmu-miR-29b-3p | MIMAT0000127 | Rbm14 | 56275 | HITS-CLIP | 37986224\|25083871 | Unspecified |
| mmu-miR-29b-3p | MIMAT0000127 | Rbm15 | 229700 | HITS-CLIP | 25083871 | Unspecified |
| mmu-miR-29b-3p | MIMAT0000127 | Rbm15b | 109095 | HITS-CLIP | 37986224 | Unspecified |
| mmu-miR-29b-3p | MIMAT0000127 | Rbm17 | 76938 | HITS-CLIP//sRNA-Seq | 25083871\|33184237\|23142080 | Unspecified |
| mmu-miR-29b-3p | MIMAT0000127 | Rbm18 | 67889 | HITS-CLIP//sRNA-Seq | 23597149\|33184237 | Unspecified |
| mmu-miR-29b-3p | MIMAT0000127 | Rbm22 | 66810 | HITS-CLIP//sRNA-Seq | 25083871\|33184237\|23142080 | Unspecified |
| mmu-miR-29b-3p | MIMAT0000127 | Rbm25 | 67039 | HITS-CLIP//sRNA-Seq | 25083871\|23597149\|33184237\|23142 | Unspecified |
| mmu-miR-29b-3p | MIMAT0000127 | Rbm26 | 74213 | HITS-CLIP | 37986224\|25083871\|23142080 | Unspecified |
| mmu-miR-29b-3p | MIMAT0000127 | Rbm27 | 225432 | HITS-CLIP | 23142080\|25083871 | Unspecified |
| mmu-miR-29b-3p | MIMAT0000127 | Rbm33 | 381626 | HITS-CLIP | 37986224\|23142080 | Unspecified |
| mmu-miR-29b-3p | MIMAT0000127 | Rbm34 | 52202 | HITS-CLIP | 23142080 | Unspecified |
| mmu-miR-29b-3p | MIMAT0000127 | Rbm39 | 170791 | HITS-CLIP//sRNA-Seq | 37986224\|33184237\|23142080 | Unspecified |
| mmu-miR-29b-3p | MIMAT0000127 | Rbm4 | 19653 | HITS-CLIP//sRNA-Seq | 31433990\|33184237 | Unspecified |
| mmu-miR-29b-3p | MIMAT0000127 | Rbm47 | 245945 | HITS-CLIP | 23597149 | Unspecified |
| mmu-miR-29b-3p | MIMAT0000127 | Rbm48 | 269623 | HITS-CLIP | 23597149\|23142080 | Unspecified |
| mmu-miR-29b-3p | MIMAT0000127 | Rbm5 | 83486 | HITS-CLIP//sRNA-Seq | 23142080\|25083871\|33184237 | Unspecified |
| mmu-miR-29b-3p | MIMAT0000127 | Rbm6 | 19654 | sRNA-Seq//HITS-CLIP | 33184237\|23142080 | Unspecified |
| mmu-miR-29b-3p | MIMAT0000127 | Rbm8a | 60365 | HITS-CLIP//sRNA-Seq | 37986224\|23142080\|33184237\|25083 | Unspecified |
| mmu-miR-29b-3p | MIMAT0000127 | Rbms3 | 207181 | HITS-CLIP | 37986224\|25083871 | Unspecified |
| mmu-miR-29b-3p | MIMAT0000127 | Rbsn | 64145 | HITS-CLIP | 25083871 | Unspecified |
| mmu-miR-29b-3p | MIMAT0000127 | Rbx1 | 56438 | HITS-CLIP//sRNA-Seq | 25083871\|33184237\|23142080 | Unspecified |
| mmu-miR-29b-3p | MIMAT0000127 | Rc3h1 | 381305 | HITS-CLIP | 25083871\|23142080 | Unspecified |
| mmu-miR-29b-3p | MIMAT0000127 | Rc3h2 | 319817 | sRNA-Seq//HITS-CLIP | 33184237\|23142080 | Unspecified |
| mmu-miR-29b-3p | MIMAT0000127 | Rcbtb2 | 105670 | HITS-CLIP | 23597149 | Unspecified |
| mmu-miR-29b-3p | MIMAT0000127 | Rcc1 | 100088 | HITS-CLIP | 23142080 | Unspecified |
| mmu-miR-29b-3p | MIMAT0000127 | Rcc2 | 108911 | HITS-CLIP//sRNA-Seq | 37986224\|25083871\|33184237\|23597 | Unspecified |
| mmu-miR-29b-3p | MIMAT0000127 | Rcn2 | 26611 | HITS-CLIP//sRNA-Seq | 25083871\|33184237\|23597149\|23142 | Unspecified |
| mmu-miR-29b-3p | MIMAT0000127 | Rcor1 | 217864 | HITS-CLIP | 37986224\|25083871\|23597149\|23142 | Unspecified |
| mmu-miR-29b-3p | MIMAT0000127 | Rdh11 | 17252 | HITS-CLIP//sRNA-Seq | 25083871\|23142080\|33184237 | Unspecified |
| mmu-miR-29b-3p | MIMAT0000127 | Rdh7 | 54150 | HITS-CLIP | 37986224 | Unspecified |
| mmu-miR-29b-3p | MIMAT0000127 | Rdx | 19684 | HITS-CLIP//sRNA-Seq | 37986224\|23597149\|25083871\|33184 | Unspecified |
| mmu-miR-29b-3p | MIMAT0000127 | Reck | 53614 | HITS-CLIP | 23142080 | Unspecified |
| mmu-miR-29b-3p | MIMAT0000127 | Reep3 | 28193 | HITS-CLIP//sRNA-Seq | 23597149\|25083871\|23142080\|33184 | Unspecified |
| mmu-miR-29b-3p | MIMAT0000127 | Reep4 | 72549 | HITS-CLIP | 26203562 | Unspecified |
| mmu-miR-29b-3p | MIMAT0000127 | Reep5 | 13476 | sRNA-Seq//HITS-CLIP | 33184237\|25083871\|23142080 | Unspecified |
| mmu-miR-29b-3p | MIMAT0000127 | Rel | 19696 | HITS-CLIP | 23142080 | Unspecified |
| mmu-miR-29b-3p | MIMAT0000127 | Relch | 57614 | HITS-CLIP//sRNA-Seq | 25083871\|23597149\|33184237\|23142 | Unspecified |

| | | | | | | |
|---|---|---|---|---|---|---|
| mmu-miR-29b-3p | MIMAT0000127 | Rell1 | 100532 | HITS-CLIP | 23142080 | Unspecified |
| mmu-miR-29b-3p | MIMAT0000127 | Reln | 19699 | HITS-CLIP | 23597149 | Unspecified |
| mmu-miR-29b-3p | MIMAT0000127 | Repin1 | 58887 | HITS-CLIP | 23142080 | Unspecified |
| mmu-miR-29b-3p | MIMAT0000127 | Reps1 | 19707 | HITS-CLIP//sRNA-Seq | 25083871\|33184237\|23142080 | Unspecified |
| mmu-miR-29b-3p | MIMAT0000127 | Rere | 68703 | HITS-CLIP//sRNA-Seq | 37986224\|25083871\|23597149\|33184 | Unspecified |
| mmu-miR-29b-3p | MIMAT0000127 | Resf1 | 55196 | HITS-CLIP | 25083871 | Unspecified |
| mmu-miR-29b-3p | MIMAT0000127 | Rest | 19712 | HITS-CLIP | 37986224\|23142080 | Unspecified |
| mmu-miR-29b-3p | MIMAT0000127 | Retreg2 | 79137 | sRNA-Seq//HITS-CLIP | 33184237\|23142080 | Unspecified |
| mmu-miR-29b-3p | MIMAT0000127 | Retreg3 | 162427 | HITS-CLIP//sRNA-Seq | 25083871\|23597149\|33184237\|23142 | Unspecified |
| mmu-miR-29b-3p | MIMAT0000127 | Rev1 | 56210 | HITS-CLIP | 37986224\|23597149\|23142080 | Unspecified |
| mmu-miR-29b-3p | MIMAT0000127 | Rev3l | 19714 | HITS-CLIP//sRNA-Seq | 37986224\|33184237\|25083871\|23142 | Unspecified |
| mmu-miR-29b-3p | MIMAT0000127 | Rexo1 | 66932 | HITS-CLIP | 37986224\|25083871\|30971389\|23142 | Unspecified |
| mmu-miR-29b-3p | MIMAT0000127 | Rfc1 | 19687 | sRNA-Seq//HITS-CLIP | 33184237\|23597149 | Unspecified |
| mmu-miR-29b-3p | MIMAT0000127 | Rfc2 | 19718 | HITS-CLIP | 23142080 | Unspecified |
| mmu-miR-29b-3p | MIMAT0000127 | Rffl | 67338 | HITS-CLIP | 37986224 | Unspecified |
| mmu-miR-29b-3p | MIMAT0000127 | Rfng | 19719 | HITS-CLIP | 25083871 | Unspecified |
| mmu-miR-29b-3p | MIMAT0000127 | Rftn1 | 76438 | HITS-CLIP | 23142080 | Unspecified |
| mmu-miR-29b-3p | MIMAT0000127 | Rftn2 | 74013 | HITS-CLIP | 25083871 | Unspecified |
| mmu-miR-29b-3p | MIMAT0000127 | Rfwd3 | 234736 | HITS-CLIP | 23597149\|23142080 | Unspecified |
| mmu-miR-29b-3p | MIMAT0000127 | Rfx3 | 19726 | HITS-CLIP | 23142080 | Unspecified |
| mmu-miR-29b-3p | MIMAT0000127 | Rfx5 | 53970 | HITS-CLIP | 23142080 | Unspecified |
| mmu-miR-29b-3p | MIMAT0000127 | Rfx7 | 319758 | HITS-CLIP | 37986224\|25083871\|23142080 | Unspecified |
| mmu-miR-29b-3p | MIMAT0000127 | Rgp1 | 242406 | HITS-CLIP | 25083871\|23142080 | Unspecified |
| mmu-miR-29b-3p | MIMAT0000127 | Rgs10 | 67865 | HITS-CLIP | 37986224\|23142080 | Unspecified |
| mmu-miR-29b-3p | MIMAT0000127 | Rgs12 | 71729 | HITS-CLIP | 23142080 | Unspecified |
| mmu-miR-29b-3p | MIMAT0000127 | Rgs19 | 56470 | HITS-CLIP | 23142080\|25083871 | Unspecified |
| mmu-miR-29b-3p | MIMAT0000127 | Rgs3 | 50780 | HITS-CLIP | 23597149 | Unspecified |
| mmu-miR-29b-3p | MIMAT0000127 | Rhbdd2 | 215160 | HITS-CLIP | 37986224 | Unspecified |
| mmu-miR-29b-3p | MIMAT0000127 | Rhbdf1 | 13650 | HITS-CLIP | 37986224\|25083871 | Unspecified |
| mmu-miR-29b-3p | MIMAT0000127 | Rhbdf2 | 217344 | HITS-CLIP | 37986224 | Unspecified |
| mmu-miR-29b-3p | MIMAT0000127 | Rhob | 11852 | HITS-CLIP | 37986224 | Unspecified |
| mmu-miR-29b-3p | MIMAT0000127 | Rhobtb1 | 69288 | HITS-CLIP | 37986224\|25083871\|23597149 | Unspecified |
| mmu-miR-29b-3p | MIMAT0000127 | Rhobtb3 | 73296 | HITS-CLIP | 25083871 | Unspecified |
| mmu-miR-29b-3p | MIMAT0000127 | Rhoq | 104215 | HITS-CLIP//sRNA-Seq | 23142080\|33184237 | Unspecified |
| mmu-miR-29b-3p | MIMAT0000127 | Rhot1 | 59040 | HITS-CLIP//sRNA-Seq | 25083871\|33184237\|23142080 | Unspecified |
| mmu-miR-29b-3p | MIMAT0000127 | Ric1 | 850728 | HITS-CLIP | 25083871\|23142080 | Unspecified |
| mmu-miR-29b-3p | MIMAT0000127 | Ric8a | 60626 | HITS-CLIP//sRNA-Seq | 25083871\|33184237 | Unspecified |
| mmu-miR-29b-3p | MIMAT0000127 | Rictor | 78757 | HITS-CLIP | 37986224\|25083871\|23142080 | Unspecified |
| mmu-miR-29b-3p | MIMAT0000127 | Rif1 | 51869 | HITS-CLIP | 25083871\|23142080 | Unspecified |

| | | | | | | |
|---|---|---|---|---|---|---|
| mmu-miR-29b-3p | MIMAT0000127 | Rin1 | 225870 | HITS-CLIP | 25083871 | Unspecified |
| mmu-miR-29b-3p | MIMAT0000127 | Rin3 | 217835 | HITS-CLIP | 25083871\|23597149\|23142080 | Unspecified |
| mmu-miR-29b-3p | MIMAT0000127 | Riok3 | 66878 | sRNA-Seq//HITS-CLIP | 33184237\|23142080\|25083871 | Unspecified |
| mmu-miR-29b-3p | MIMAT0000127 | Ripor1 | 75687 | HITS-CLIP//sRNA-Seq | 25083871\|33184237 | Unspecified |
| mmu-miR-29b-3p | MIMAT0000127 | Ripor2 | 193385 | HITS-CLIP | 23142080 | Unspecified |
| mmu-miR-29b-3p | MIMAT0000127 | Rlf | 109263 | HITS-CLIP | 37986224\|23142080\|25083871\|23597 | Unspecified |
| mmu-miR-29b-3p | MIMAT0000127 | Rlim | 19820 | HITS-CLIP//sRNA-Seq | 37986224\|33184237\|23142080 | Unspecified |
| mmu-miR-29b-3p | MIMAT0000127 | Rmc1 | 19775 | HITS-CLIP | 23142080 | Unspecified |
| mmu-miR-29b-3p | MIMAT0000127 | Rmdn2 | 381110 | HITS-CLIP | 23597149 | Unspecified |
| mmu-miR-29b-3p | MIMAT0000127 | Rmdn3 | 67809 | HITS-CLIP | 37986224\|23597149 | Unspecified |
| mmu-miR-29b-3p | MIMAT0000127 | Rmnd5a | 68477 | HITS-CLIP | 37986224\|23597149\|23142080\|25083 | Unspecified |
| mmu-miR-29b-3p | MIMAT0000127 | Rnase4 | 58809 | HITS-CLIP//sRNA-Seq | 23597149\|25083871\|33184237 | Unspecified |
| mmu-miR-29b-3p | MIMAT0000127 | Rnaseh2a | 69724 | HITS-CLIP | 37986224 | Unspecified |
| mmu-miR-29b-3p | MIMAT0000127 | Rnaseh2b | 67153 | HITS-CLIP | 23142080 | Unspecified |
| mmu-miR-29b-3p | MIMAT0000127 | Rnaseh2c | 68209 | HITS-CLIP | 23142080 | Unspecified |
| mmu-miR-29b-3p | MIMAT0000127 | Rnasek | 52898 | sRNA-Seq//HITS-CLIP | 33184237\|23142080 | Unspecified |
| mmu-miR-29b-3p | MIMAT0000127 | Rnasel | 24014 | HITS-CLIP | 23142080 | Unspecified |
| mmu-miR-29b-3p | MIMAT0000127 | Rnd1 | 223881 | HITS-CLIP | 23597149 | Unspecified |
| mmu-miR-29b-3p | MIMAT0000127 | Rnd3 | 74194 | HITS-CLIP//sRNA-Seq | 23142080\|33184237\|23597149 | Unspecified |
| mmu-miR-29b-3p | MIMAT0000127 | Rnf10 | 50849 | sRNA-Seq//HITS-CLIP | 33184237\|23597149 | Unspecified |
| mmu-miR-29b-3p | MIMAT0000127 | Rnf11 | 29864 | HITS-CLIP//sRNA-Seq | 37986224\|25083871\|33184237 | Unspecified |
| mmu-miR-29b-3p | MIMAT0000127 | Rnf111 | 93836 | HITS-CLIP | 23142080\|23597149\|25083871 | Unspecified |
| mmu-miR-29b-3p | MIMAT0000127 | Rnf113a1 | 69942 | HITS-CLIP | 23142080 | Unspecified |
| mmu-miR-29b-3p | MIMAT0000127 | Rnf121 | 75212 | HITS-CLIP | 23142080 | Unspecified |
| mmu-miR-29b-3p | MIMAT0000127 | Rnf123 | 84585 | HITS-CLIP | 23597149 | Unspecified |
| mmu-miR-29b-3p | MIMAT0000127 | Rnf128 | 66889 | HITS-CLIP | 23597149 | Unspecified |
| mmu-miR-29b-3p | MIMAT0000127 | Rnf13 | 24017 | HITS-CLIP | 23142080 | Unspecified |
| mmu-miR-29b-3p | MIMAT0000127 | Rnf130 | 59044 | sRNA-Seq//HITS-CLIP | 33184237\|23142080 | Unspecified |
| mmu-miR-29b-3p | MIMAT0000127 | Rnf138 | 56515 | HITS-CLIP | 37986224\|25083871\|23142080 | Unspecified |
| mmu-miR-29b-3p | MIMAT0000127 | Rnf139 | 75841 | HITS-CLIP | 23142080 | Unspecified |
| mmu-miR-29b-3p | MIMAT0000127 | Rnf144b | 218215 | HITS-CLIP | 23597149 | Unspecified |
| mmu-miR-29b-3p | MIMAT0000127 | Rnf145 | 74315 | HITS-CLIP | 23142080 | Unspecified |
| mmu-miR-29b-3p | MIMAT0000127 | Rnf146 | 68031 | HITS-CLIP//sRNA-Seq | 37986224\|33184237\|23142080 | Unspecified |
| mmu-miR-29b-3p | MIMAT0000127 | Rnf149 | 67702 | HITS-CLIP | 25083871\|23597149\|23142080 | Unspecified |
| mmu-miR-29b-3p | MIMAT0000127 | Rnf150 | 330812 | HITS-CLIP | 37986224\|25083871 | Unspecified |
| mmu-miR-29b-3p | MIMAT0000127 | Rnf165 | 225743 | HITS-CLIP | 37986224\|25083871 | Unspecified |
| mmu-miR-29b-3p | MIMAT0000127 | Rnf167 | 70510 | HITS-CLIP | 25083871\|23142080 | Unspecified |
| mmu-miR-29b-3p | MIMAT0000127 | Rnf168 | 70238 | HITS-CLIP | 23142080 | Unspecified |
| mmu-miR-29b-3p | MIMAT0000127 | Rnf169 | 108937 | HITS-CLIP | 37986224\|25083871\|23597149\|23142 | Unspecified |

| | | | | | | |
|---|---|---|---|---|---|---|
| mmu-miR-29b-3p | MIMAT0000127 | Rnf19a | 30945 | HITS-CLIP | 37986224\|25083871\|23142080\|31433 | Unspecified |
| mmu-miR-29b-3p | MIMAT0000127 | Rnf19b | 75234 | HITS-CLIP//sRNA-Seq | 23142080\|25083871\|33184237\|23597 | Unspecified |
| mmu-miR-29b-3p | MIMAT0000127 | Rnf213 | 672511 | HITS-CLIP//sRNA-Seq | 23142080\|33184237\|23597149 | Unspecified |
| mmu-miR-29b-3p | MIMAT0000127 | Rnf214 | 235315 | HITS-CLIP//sRNA-Seq | 37986224\|25083871\|33184237\|23597 | Unspecified |
| mmu-miR-29b-3p | MIMAT0000127 | Rnf216 | 108086 | HITS-CLIP | 37986224\|25083871 | Unspecified |
| mmu-miR-29b-3p | MIMAT0000127 | Rnf217 | 268291 | HITS-CLIP | 37986224\|25083871\|23597149 | Unspecified |
| mmu-miR-29b-3p | MIMAT0000127 | Rnf34 | 80751 | HITS-CLIP | 25083871\|23142080 | Unspecified |
| mmu-miR-29b-3p | MIMAT0000127 | Rnf38 | 73469 | HITS-CLIP//sRNA-Seq | 37986224\|33184237\|23142080 | Unspecified |
| mmu-miR-29b-3p | MIMAT0000127 | Rnf4 | 19822 | HITS-CLIP//sRNA-Seq | 23142080\|25083871\|33184237 | Unspecified |
| mmu-miR-29b-3p | MIMAT0000127 | Rnf41 | 67588 | HITS-CLIP//sRNA-Seq | 37986224\|23142080\|33184237 | Unspecified |
| mmu-miR-29b-3p | MIMAT0000127 | Rnf44 | 105239 | HITS-CLIP | 23142080 | Unspecified |
| mmu-miR-29b-3p | MIMAT0000127 | Rnf6 | 74132 | HITS-CLIP | 23142080\|23597149 | Unspecified |
| mmu-miR-29b-3p | MIMAT0000127 | Rnft1 | 76892 | HITS-CLIP | 23142080\|25083871 | Unspecified |
| mmu-miR-29b-3p | MIMAT0000127 | Rngtt | 24018 | HITS-CLIP | 23142080 | Unspecified |
| mmu-miR-29b-3p | MIMAT0000127 | Rnh1 | 107702 | HITS-CLIP//sRNA-Seq | 25083871\|33184237\|23597149\|23142 | Unspecified |
| mmu-miR-29b-3p | MIMAT0000127 | Rnls | 67795 | HITS-CLIP | 37986224\|25083871 | Unspecified |
| mmu-miR-29b-3p | MIMAT0000127 | Rnmt | 67897 | HITS-CLIP | 25083871\|23142080 | Unspecified |
| mmu-miR-29b-3p | MIMAT0000127 | Rnpep | 215615 | HITS-CLIP | 25083871\|23597149 | Unspecified |
| mmu-miR-29b-3p | MIMAT0000127 | Rnpepl1 | 108657 | HITS-CLIP | 37986224\|23597149\|25083871\|23142 | Unspecified |
| mmu-miR-29b-3p | MIMAT0000127 | Robo1 | 19876 | HITS-CLIP | 23597149 | Unspecified |
| mmu-miR-29b-3p | MIMAT0000127 | Robo4 | 74144 | HITS-CLIP | 23597149 | Unspecified |
| mmu-miR-29b-3p | MIMAT0000127 | Rock1 | 19877 | HITS-CLIP | 37986224\|25083871\|26203562\|23142 | Unspecified |
| mmu-miR-29b-3p | MIMAT0000127 | Rock2 | 19878 | HITS-CLIP | 23597149\|25083871\|23142080 | Unspecified |
| mmu-miR-29b-3p | MIMAT0000127 | Ror1 | 26563 | HITS-CLIP//RNA-Seq | 23597149\|33332475 | Unspecified |
| mmu-miR-29b-3p | MIMAT0000127 | Rora | 19883 | HITS-CLIP | 37986224\|23597149\|23142080 | Unspecified |
| mmu-miR-29b-3p | MIMAT0000127 | Rp2 | 19889 | HITS-CLIP | 23142080 | Unspecified |
| mmu-miR-29b-3p | MIMAT0000127 | Rpa1 | 68275 | sRNA-Seq//HITS-CLIP | 33184237\|23142080 | Unspecified |
| mmu-miR-29b-3p | MIMAT0000127 | Rpap1 | 68925 | HITS-CLIP | 23142080 | Unspecified |
| mmu-miR-29b-3p | MIMAT0000127 | Rpia | 19895 | HITS-CLIP | 23142080 | Unspecified |
| mmu-miR-29b-3p | MIMAT0000127 | Rpl14 | 67115 | qCLASH | 31393953 | Unspecified |
| mmu-miR-29b-3p | MIMAT0000127 | Rpl15 | 66480 | sRNA-Seq//HITS-CLIP | 33184237\|23142080 | Unspecified |
| mmu-miR-29b-3p | MIMAT0000127 | Rpl17 | 319195 | HITS-CLIP | 37986224 | Unspecified |
| mmu-miR-29b-3p | MIMAT0000127 | Rpl22 | 19934 | HITS-CLIP//sRNA-Seq | 23142080\|33184237 | Unspecified |
| mmu-miR-29b-3p | MIMAT0000127 | Rpl5 | 100503670 | HITS-CLIP//sRNA-Seq | 37986224\|25083871\|33184237 | Unspecified |
| mmu-miR-29b-3p | MIMAT0000127 | Rplp0 | 11837 | HITS-CLIP//sRNA-Seq | 37986224\|25083871\|33184237\|23597 | Unspecified |
| mmu-miR-29b-3p | MIMAT0000127 | Rplp1 | 56040 | HITS-CLIP//sRNA-Seq | 23142080\|33184237 | Unspecified |
| mmu-miR-29b-3p | MIMAT0000127 | Rpn1 | 103963 | HITS-CLIP//sRNA-Seq | 25083871\|33184237 | Unspecified |
| mmu-miR-29b-3p | MIMAT0000127 | Rpn2 | 20014 | HITS-CLIP//sRNA-Seq | 25083871\|33184237\|23142080 | Unspecified |
| mmu-miR-29b-3p | MIMAT0000127 | Rpp38 | 227522 | HITS-CLIP | 23142080 | Unspecified |

| | | | | | | |
|---|---|---|---|---|---|---|
| mmu-miR-29b-3p | MIMAT0000127 | Rprd1a | 225283 | HITS-CLIP | 25083871 | Unspecified |
| mmu-miR-29b-3p | MIMAT0000127 | Rprd1b | 70470 | HITS-CLIP | 37986224\|23142080 | Unspecified |
| mmu-miR-29b-3p | MIMAT0000127 | Rprd2 | 75137 | HITS-CLIP | 25083871\|23142080 | Unspecified |
| mmu-miR-29b-3p | MIMAT0000127 | Rps16 | 20055 | sRNA-Seq//HITS-CLIP | 33184237\|23142080 | Unspecified |
| mmu-miR-29b-3p | MIMAT0000127 | Rps20 | 67427 | HITS-CLIP//sRNA-Seq | 37986224\|23142080\|33184237 | Unspecified |
| mmu-miR-29b-3p | MIMAT0000127 | Rps6ka3 | 110651 | HITS-CLIP//sRNA-Seq | 23597149\|25083871\|33184237\|23142 | Unspecified |
| mmu-miR-29b-3p | MIMAT0000127 | Rps6ka5 | 73086 | HITS-CLIP | 23142080 | Unspecified |
| mmu-miR-29b-3p | MIMAT0000127 | Rps6kb1 | 72508 | HITS-CLIP | 37986224\|25083871\|23142080 | Unspecified |
| mmu-miR-29b-3p | MIMAT0000127 | Rps6kb2 | 58988 | HITS-CLIP | 25083871\|23597149\|23142080 | Unspecified |
| mmu-miR-29b-3p | MIMAT0000127 | Rps6kc1 | 320119 | HITS-CLIP | 25083871 | Unspecified |
| mmu-miR-29b-3p | MIMAT0000127 | Rptor | 74370 | HITS-CLIP | 25083871\|23597149\|23142080 | Unspecified |
| mmu-miR-29b-3p | MIMAT0000127 | Rrad | 56437 | HITS-CLIP | 25083871 | Unspecified |
| mmu-miR-29b-3p | MIMAT0000127 | Rraga | 68441 | HITS-CLIP//sRNA-Seq | 37986224\|25083871\|33184237\|23142 | Unspecified |
| mmu-miR-29b-3p | MIMAT0000127 | Rragb | 245670 | HITS-CLIP | 23142080 | Unspecified |
| mmu-miR-29b-3p | MIMAT0000127 | Rras2 | 66922 | HITS-CLIP | 37986224\|25083871\|23142080 | Unspecified |
| mmu-miR-29b-3p | MIMAT0000127 | Rrbp1 | 81910 | sRNA-Seq//HITS-CLIP | 33184237\|25083871 | Unspecified |
| mmu-miR-29b-3p | MIMAT0000127 | Rreb1 | 68750 | HITS-CLIP | 37986224\|25083871 | Unspecified |
| mmu-miR-29b-3p | MIMAT0000127 | Rrm1 | 20133 | HITS-CLIP//sRNA-Seq//qCLASH | 23597149\|33184237\|31393953\|23142 | Unspecified |
| mmu-miR-29b-3p | MIMAT0000127 | Rrn3 | 106298 | HITS-CLIP//sRNA-Seq | 23142080\|33184237\|25083871 | Unspecified |
| mmu-miR-29b-3p | MIMAT0000127 | Rrnad1 | 229503 | HITS-CLIP | 23142080 | Unspecified |
| mmu-miR-29b-3p | MIMAT0000127 | Rrp15 | 67223 | HITS-CLIP | 23142080 | Unspecified |
| mmu-miR-29b-3p | MIMAT0000127 | Rsbn1 | 229675 | HITS-CLIP | 25083871 | Unspecified |
| mmu-miR-29b-3p | MIMAT0000127 | Rsbn1l | 242860 | HITS-CLIP | 23142080\|25083871 | Unspecified |
| mmu-miR-29b-3p | MIMAT0000127 | Rsf1 | 233532 | HITS-CLIP | 37986224\|23142080 | Unspecified |
| mmu-miR-29b-3p | MIMAT0000127 | Rsrc2 | 208606 | HITS-CLIP//sRNA-Seq | 25083871\|23597149\|33184237\|23142 | Unspecified |
| mmu-miR-29b-3p | MIMAT0000127 | Rsu1 | 20163 | HITS-CLIP//sRNA-Seq | 25083871\|33184237\|23142080 | Unspecified |
| mmu-miR-29b-3p | MIMAT0000127 | Rtel1 | 269400 | HITS-CLIP | 23142080 | Unspecified |
| mmu-miR-29b-3p | MIMAT0000127 | Rtf1 | 76246 | HITS-CLIP | 25083871 | Unspecified |
| mmu-miR-29b-3p | MIMAT0000127 | Rtkn | 20166 | HITS-CLIP | 25083871\|23597149 | Unspecified |
| mmu-miR-29b-3p | MIMAT0000127 | Rtl5 | 331474 | HITS-CLIP | 23142080 | Unspecified |
| mmu-miR-29b-3p | MIMAT0000127 | Rtn3 | 20168 | HITS-CLIP//sRNA-Seq | 25083871\|23597149\|33184237\|23142 | Unspecified |
| mmu-miR-29b-3p | MIMAT0000127 | Rtn4 | 68585 | HITS-CLIP//sRNA-Seq | 25083871\|23597149\|33184237\|23142 | Unspecified |
| mmu-miR-29b-3p | MIMAT0000127 | Rttn | 246102 | HITS-CLIP | 23142080 | Unspecified |
| mmu-miR-29b-3p | MIMAT0000127 | Rubcnl | 80183 | HITS-CLIP | 23142080 | Unspecified |
| mmu-miR-29b-3p | MIMAT0000127 | Rufy2 | 70432 | HITS-CLIP//sRNA-Seq | 37986224\|25083871\|33184237\|23142 | Unspecified |
| mmu-miR-29b-3p | MIMAT0000127 | Rundc1 | 217201 | HITS-CLIP | 37986224\|25083871\|23142080 | Unspecified |
| mmu-miR-29b-3p | MIMAT0000127 | Runx1 | 12394 | HITS-CLIP | 37986224\|23597149\|23142080 | Unspecified |
| mmu-miR-29b-3p | MIMAT0000127 | Runx2 | 12393 | HITS-CLIP | 37986224\|25083871 | Unspecified |
| mmu-miR-29b-3p | MIMAT0000127 | Rusc2 | 100213 | HITS-CLIP | 37986224 | Unspecified |

| | | | | | | |
|---|---|---|---|---|---|---|
| mmu-miR-29b-3p | MIMAT0000127 | Ruvbl2 | 20174 | HITS-CLIP | 23142080 | Unspecified |
| mmu-miR-29b-3p | MIMAT0000127 | Rxylt1 | 10329 | HITS-CLIP | 25083871 | Unspecified |
| mmu-miR-29b-3p | MIMAT0000127 | Rybp | 56353 | HITS-CLIP | 37986224 | Unspecified |
| mmu-miR-29b-3p | MIMAT0000127 | Ryr1 | 20190 | HITS-CLIP | 37986224\|25083871 | Unspecified |
| mmu-miR-29b-3p | MIMAT0000127 | Ryr3 | 20192 | HITS-CLIP | 25083871 | Unspecified |
| mmu-miR-29b-3p | MIMAT0000127 | S100a1 | 20193 | sRNA-Seq//HITS-CLIP | 33184237\|23142080 | Unspecified |
| mmu-miR-29b-3p | MIMAT0000127 | S100a10 | 20194 | HITS-CLIP//sRNA-Seq | 25083871\|33184237\|23142080 | Unspecified |
| mmu-miR-29b-3p | MIMAT0000127 | S100a13 | 20196 | sRNA-Seq//HITS-CLIP | 33184237\|23142080 | Unspecified |
| mmu-miR-29b-3p | MIMAT0000127 | S100pbp | 74648 | HITS-CLIP//sRNA-Seq | 25083871\|33184237\|23142080 | Unspecified |
| mmu-miR-29b-3p | MIMAT0000127 | S1pr1 | 13609 | HITS-CLIP | 23142080 | Unspecified |
| mmu-miR-29b-3p | MIMAT0000127 | S1pr2 | 14739 | HITS-CLIP | 23142080 | Unspecified |
| mmu-miR-29b-3p | MIMAT0000127 | S1pr4 | 13611 | HITS-CLIP | 23142080 | Unspecified |
| mmu-miR-29b-3p | MIMAT0000127 | Saal1 | 78935 | HITS-CLIP | 23142080 | Unspecified |
| mmu-miR-29b-3p | MIMAT0000127 | Sacm1l | 83493 | HITS-CLIP//sRNA-Seq | 23597149\|23142080\|33184237 | Unspecified |
| mmu-miR-29b-3p | MIMAT0000127 | Sacs | 50720 | HITS-CLIP | 37986224\|25083871\|23142080 | Unspecified |
| mmu-miR-29b-3p | MIMAT0000127 | Sall1 | 58198 | HITS-CLIP | 23597149 | Unspecified |
| mmu-miR-29b-3p | MIMAT0000127 | Samd4 | 74480 | HITS-CLIP | 37986224\|25083871 | Unspecified |
| mmu-miR-29b-3p | MIMAT0000127 | Samd4b | 233033 | HITS-CLIP | 23142080 | Unspecified |
| mmu-miR-29b-3p | MIMAT0000127 | Samd8 | 67630 | HITS-CLIP//sRNA-Seq | 25083871\|33184237\|23142080 | Unspecified |
| mmu-miR-29b-3p | MIMAT0000127 | Samhd1 | 56045 | sRNA-Seq//HITS-CLIP | 33184237\|23142080 | Unspecified |
| mmu-miR-29b-3p | MIMAT0000127 | Samsn1 | 67742 | HITS-CLIP | 23142080 | Unspecified |
| mmu-miR-29b-3p | MIMAT0000127 | Sap130 | 269003 | HITS-CLIP | 25083871\|23597149 | Unspecified |
| mmu-miR-29b-3p | MIMAT0000127 | Sap30 | 60406 | HITS-CLIP | 23142080 | Unspecified |
| mmu-miR-29b-3p | MIMAT0000127 | Saraf | 51669 | HITS-CLIP//sRNA-Seq | 23597149\|25083871\|33184237\|23142 | Unspecified |
| mmu-miR-29b-3p | MIMAT0000127 | Sardh | 192166 | HITS-CLIP | 37986224\|23597149 | Unspecified |
| mmu-miR-29b-3p | MIMAT0000127 | Sars | 20226 | HITS-CLIP//sRNA-Seq | 37986224\|25083871\|23597149\|33184 | Unspecified |
| mmu-miR-29b-3p | MIMAT0000127 | Saysd1 | 67509 | HITS-CLIP | 23142080 | Unspecified |
| mmu-miR-29b-3p | MIMAT0000127 | Sbf2 | 319934 | HITS-CLIP//sRNA-Seq | 37986224\|25083871\|33184237 | Unspecified |
| mmu-miR-29b-3p | MIMAT0000127 | Sbk1 | 104175 | HITS-CLIP//sRNA-Seq | 25083871\|33184237 | Unspecified |
| mmu-miR-29b-3p | MIMAT0000127 | Sbno1 | 243272 | HITS-CLIP//sRNA-Seq | 23142080\|33184237\|25083871 | Unspecified |
| mmu-miR-29b-3p | MIMAT0000127 | Sbno2 | 216161 | HITS-CLIP//sRNA-Seq | 37986224\|23142080\|33184237\|25083 | Unspecified |
| mmu-miR-29b-3p | MIMAT0000127 | Sc5d | 235293 | HITS-CLIP//sRNA-Seq | 25083871\|33184237\|23597149\|23142 | Unspecified |
| mmu-miR-29b-3p | MIMAT0000127 | Scaf11 | 72193 | HITS-CLIP//sRNA-Seq | 25083871\|23597149\|33184237\|23142 | Unspecified |
| mmu-miR-29b-3p | MIMAT0000127 | Scai | 320271 | HITS-CLIP | 23142080 | Unspecified |
| mmu-miR-29b-3p | MIMAT0000127 | Scamp2 | 24044 | HITS-CLIP | 23597149 | Unspecified |
| mmu-miR-29b-3p | MIMAT0000127 | Scamp3 | 24045 | sRNA-Seq//HITS-CLIP | 33184237\|23142080 | Unspecified |
| mmu-miR-29b-3p | MIMAT0000127 | Scamp5 | 56807 | HITS-CLIP//sRNA-Seq | 37986224\|25083871\|33184237\|23597 | Unspecified |
| mmu-miR-29b-3p | MIMAT0000127 | Scap | 235623 | sRNA-Seq//HITS-CLIP//Luciferase reporter | 33184237\|23597149\|23142080\|27477 | Unspecified |
| mmu-miR-29b-3p | MIMAT0000127 | Scarb1 | 20778 | HITS-CLIP//sRNA-Seq | 37986224\|33184237\|23142080 | Unspecified |

| | | | | | | |
|---|---|---|---|---|---|---|
| mmu-miR-29b-3p | MIMAT0000127 | Scd1 | 20249 | HITS-CLIP//sRNA-Seq | 37986224\|23597149\|33184237\|23142 | Unspecified |
| mmu-miR-29b-3p | MIMAT0000127 | Scd2 | 20250 | HITS-CLIP//sRNA-Seq | 25083871\|23142080\|33184237 | Unspecified |
| mmu-miR-29b-3p | MIMAT0000127 | Scfd1 | 76983 | HITS-CLIP//sRNA-Seq | 37986224\|23142080\|33184237\|25083 | Unspecified |
| mmu-miR-29b-3p | MIMAT0000127 | Schip1 | 30953 | HITS-CLIP | 37986224 | Unspecified |
| mmu-miR-29b-3p | MIMAT0000127 | Scly | 50880 | HITS-CLIP | 23597149 | Unspecified |
| mmu-miR-29b-3p | MIMAT0000127 | Scml4 | 268297 | HITS-CLIP | 37986224 | Unspecified |
| mmu-miR-29b-3p | MIMAT0000127 | Scn1b | 20266 | HITS-CLIP | 23597149 | Unspecified |
| mmu-miR-29b-3p | MIMAT0000127 | Scp2 | 20280 | HITS-CLIP//sRNA-Seq | 23597149\|33184237 | Unspecified |
| mmu-miR-29b-3p | MIMAT0000127 | Scpep1 | 74617 | HITS-CLIP//sRNA-Seq | 37986224\|23597149\|33184237 | Unspecified |
| mmu-miR-29b-3p | MIMAT0000127 | Scrib | 105782 | HITS-CLIP//sRNA-Seq | 23597149\|33184237\|25083871 | Unspecified |
| mmu-miR-29b-3p | MIMAT0000127 | Scyl1 | 78891 | sRNA-Seq//HITS-CLIP | 33184237\|25083871 | Unspecified |
| mmu-miR-29b-3p | MIMAT0000127 | Scyl2 | 213326 | HITS-CLIP//sRNA-Seq | 23142080\|25083871\|33184237 | Unspecified |
| mmu-miR-29b-3p | MIMAT0000127 | Scyl3 | 240880 | HITS-CLIP | 23142080 | Unspecified |
| mmu-miR-29b-3p | MIMAT0000127 | Sdad1 | 231452 | HITS-CLIP | 37986224\|25083871\|23142080 | Unspecified |
| mmu-miR-29b-3p | MIMAT0000127 | Sdc1 | 20969 | HITS-CLIP//sRNA-Seq | 23597149\|33184237 | Unspecified |
| mmu-miR-29b-3p | MIMAT0000127 | Sdc2 | 15529 | HITS-CLIP//sRNA-Seq | 23597149\|33184237\|25083871 | Unspecified |
| mmu-miR-29b-3p | MIMAT0000127 | Sdc4 | 20971 | HITS-CLIP//sRNA-Seq | 25083871\|33184237 | Unspecified |
| mmu-miR-29b-3p | MIMAT0000127 | Sdcbp | 53378 | HITS-CLIP//sRNA-Seq | 25083871\|33184237\|23142080 | Unspecified |
| mmu-miR-29b-3p | MIMAT0000127 | Sde2 | 208768 | HITS-CLIP | 23142080 | Unspecified |
| mmu-miR-29b-3p | MIMAT0000127 | Sdf4 | 20318 | sRNA-Seq//HITS-CLIP | 33184237\|23142080 | Unspecified |
| mmu-miR-29b-3p | MIMAT0000127 | Sdha | 66945 | HITS-CLIP//sRNA-Seq | 23597149\|33184237 | Unspecified |
| mmu-miR-29b-3p | MIMAT0000127 | Sdhaf2 | 66072 | HITS-CLIP | 37986224 | Unspecified |
| mmu-miR-29b-3p | MIMAT0000127 | Sdhb | 67680 | HITS-CLIP//sRNA-Seq | 23597149\|33184237\|23142080\|25083 | Unspecified |
| mmu-miR-29b-3p | MIMAT0000127 | Sdr9c7 | 70061 | HITS-CLIP | 23597149 | Unspecified |
| mmu-miR-29b-3p | MIMAT0000127 | Sec11a | 56529 | HITS-CLIP | 23142080 | Unspecified |
| mmu-miR-29b-3p | MIMAT0000127 | Sec14l1 | 74136 | HITS-CLIP//sRNA-Seq | 25083871\|33184237 | Unspecified |
| mmu-miR-29b-3p | MIMAT0000127 | Sec14l2 | 67815 | HITS-CLIP | 23597149 | Unspecified |
| mmu-miR-29b-3p | MIMAT0000127 | Sec14l4 | 103655 | HITS-CLIP | 37986224\|23597149 | Unspecified |
| mmu-miR-29b-3p | MIMAT0000127 | Sec16a | 227648 | HITS-CLIP | 23597149 | Unspecified |
| mmu-miR-29b-3p | MIMAT0000127 | Sec16b | 89867 | HITS-CLIP | 23597149 | Unspecified |
| mmu-miR-29b-3p | MIMAT0000127 | Sec22b | 20333 | HITS-CLIP//sRNA-Seq | 37986224\|25083871\|33184237\|23142 | Unspecified |
| mmu-miR-29b-3p | MIMAT0000127 | Sec23a | 20334 | HITS-CLIP//sRNA-Seq | 37986224\|23597149\|33184237 | Unspecified |
| mmu-miR-29b-3p | MIMAT0000127 | Sec23b | 27054 | HITS-CLIP//sRNA-Seq | 37986224\|23597149\|25083871\|33184 | Unspecified |
| mmu-miR-29b-3p | MIMAT0000127 | Sec24a | 77371 | HITS-CLIP | 25083871\|23142080 | Unspecified |
| mmu-miR-29b-3p | MIMAT0000127 | Sec24b | 99683 | HITS-CLIP | 23597149\|23142080 | Unspecified |
| mmu-miR-29b-3p | MIMAT0000127 | Sec24c | 218811 | HITS-CLIP//sRNA-Seq | 25083871\|33184237 | Unspecified |
| mmu-miR-29b-3p | MIMAT0000127 | Sec24d | 69608 | HITS-CLIP//sRNA-Seq | 25083871\|33184237\|23597149\|23142 | Unspecified |
| mmu-miR-29b-3p | MIMAT0000127 | Sec31a | 69162 | HITS-CLIP//sRNA-Seq | 37986224\|25083871\|23597149\|33184 | Unspecified |
| mmu-miR-29b-3p | MIMAT0000127 | Sec61a1 | 53421 | HITS-CLIP//sRNA-Seq | 37986224\|23597149\|25083871\|33184 | Unspecified |

| | | | | | | |
|---|---|---|---|---|---|---|
| mmu-miR-29b-3p | MIMAT0000127 | Sec61a2 | 57743 | HITS-CLIP//sRNA-Seq | 25083871\|33184237 | Unspecified |
| mmu-miR-29b-3p | MIMAT0000127 | Sec63 | 140740 | HITS-CLIP//sRNA-Seq | 37986224\|23597149\|23142080\|25083 | Unspecified |
| mmu-miR-29b-3p | MIMAT0000127 | Secisbp2 | 75420 | HITS-CLIP | 23142080 | Unspecified |
| mmu-miR-29b-3p | MIMAT0000127 | Secisbp2l | 70354 | HITS-CLIP | 23142080 | Unspecified |
| mmu-miR-29b-3p | MIMAT0000127 | Sel1l3 | 231238 | HITS-CLIP//sRNA-Seq | 23597149\|33184237 | Unspecified |
| mmu-miR-29b-3p | MIMAT0000127 | Selenoi | 28042 | HITS-CLIP | 23142080 | Unspecified |
| mmu-miR-29b-3p | MIMAT0000127 | Selenop | 6414 | HITS-CLIP | 23142080\|23597149 | Unspecified |
| mmu-miR-29b-3p | MIMAT0000127 | Selenot | 69227 | HITS-CLIP//sRNA-Seq | 25083871\|33184237\|23142080 | Unspecified |
| mmu-miR-29b-3p | MIMAT0000127 | Selplg | 20345 | HITS-CLIP | 37986224\|23142080 | Unspecified |
| mmu-miR-29b-3p | MIMAT0000127 | Sema3a | 20346 | HITS-CLIP | 37986224 | Unspecified |
| mmu-miR-29b-3p | MIMAT0000127 | Sema3d | 108151 | HITS-CLIP//sRNA-Seq | 25083871\|33184237 | Unspecified |
| mmu-miR-29b-3p | MIMAT0000127 | Sema3e | 20349 | HITS-CLIP | 25083871 | Unspecified |
| mmu-miR-29b-3p | MIMAT0000127 | Sema4b | 20352 | HITS-CLIP | 37986224\|23142080 | Unspecified |
| mmu-miR-29b-3p | MIMAT0000127 | Sema4c | 20353 | HITS-CLIP | 25083871 | Unspecified |
| mmu-miR-29b-3p | MIMAT0000127 | Sema4g | 26456 | HITS-CLIP | 23597149 | Unspecified |
| mmu-miR-29b-3p | MIMAT0000127 | Sema6d | 214968 | HITS-CLIP | 25083871 | Unspecified |
| mmu-miR-29b-3p | MIMAT0000127 | Senp1 | 223870 | HITS-CLIP | 23142080 | Unspecified |
| mmu-miR-29b-3p | MIMAT0000127 | Senp5 | 320213 | HITS-CLIP | 23142080\|25083871 | Unspecified |
| mmu-miR-29b-3p | MIMAT0000127 | Senp6 | 215351 | HITS-CLIP//sRNA-Seq | 25083871\|33184237\|23142080 | Unspecified |
| mmu-miR-29b-3p | MIMAT0000127 | Senp7 | 66315 | HITS-CLIP | 25083871 | Unspecified |
| mmu-miR-29b-3p | MIMAT0000127 | Sephs2 | 20768 | HITS-CLIP | 23142080 | Unspecified |
| mmu-miR-29b-3p | MIMAT0000127 | Sepsecs | 211006 | HITS-CLIP | 25083871\|23142080 | Unspecified |
| mmu-miR-29b-3p | MIMAT0000127 | Sept1 | 54204 | HITS-CLIP | 23142080 | Unspecified |
| mmu-miR-29b-3p | MIMAT0000127 | Sept10 | 103080 | HITS-CLIP | 25083871 | Unspecified |
| mmu-miR-29b-3p | MIMAT0000127 | Sept11 | 52398 | HITS-CLIP//sRNA-Seq | 25083871\|23142080\|33184237 | Unspecified |
| mmu-miR-29b-3p | MIMAT0000127 | Sept6 | 23157 | sRNA-Seq//HITS-CLIP | 33184237\|23142080 | Unspecified |
| mmu-miR-29b-3p | MIMAT0000127 | Sept7 | 235072 | HITS-CLIP//sRNA-Seq | 25083871\|23597149\|33184237\|23142 | Unspecified |
| mmu-miR-29b-3p | MIMAT0000127 | Sept8 | 20362 | HITS-CLIP | 25083871\|23597149 | Unspecified |
| mmu-miR-29b-3p | MIMAT0000127 | Septin2 | 18000 | HITS-CLIP | 37986224\|23597149\|25083871\|23142 | Unspecified |
| mmu-miR-29b-3p | MIMAT0000127 | Serac1 | 321007 | HITS-CLIP | 37986224\|25083871 | Unspecified |
| mmu-miR-29b-3p | MIMAT0000127 | Serbp1 | 66870 | HITS-CLIP//sRNA-Seq | 37986224\|25083871\|23597149\|33184 | Unspecified |
| mmu-miR-29b-3p | MIMAT0000127 | Serf1 | 20365 | HITS-CLIP//sRNA-Seq | 37986224\|33184237\|23142080\|25083 | Unspecified |
| mmu-miR-29b-3p | MIMAT0000127 | Serf2 | 378702 | HITS-CLIP//sRNA-Seq | 25083871\|33184237 | Unspecified |
| mmu-miR-29b-3p | MIMAT0000127 | Serinc1 | 56442 | sRNA-Seq//HITS-CLIP | 33184237\|23142080 | Unspecified |
| mmu-miR-29b-3p | MIMAT0000127 | Serinc3 | 26943 | HITS-CLIP//qCLASH//sRNA-Seq | 37986224\|23597149\|31393953\|25083 | Unspecified |
| mmu-miR-29b-3p | MIMAT0000127 | Serinc5 | 218442 | HITS-CLIP | 37986224\|25083871\|23597149\|23142 | Unspecified |
| mmu-miR-29b-3p | MIMAT0000127 | Serp1 | 28146 | HITS-CLIP//sRNA-Seq | 23597149\|33184237\|23142080 | Unspecified |
| mmu-miR-29b-3p | MIMAT0000127 | Serpina10 | 217847 | HITS-CLIP | 37986224 | Unspecified |
| mmu-miR-29b-3p | MIMAT0000127 | Serpina3g | 20715 | HITS-CLIP | 23142080 | Unspecified |

| | | | | | | |
|---|---|---|---|---|---|---|
| mmu-miR-29b-3p | MIMAT0000127 | Serpina3k | 20714 | HITS-CLIP | 23597149 | Unspecified |
| mmu-miR-29b-3p | MIMAT0000127 | Serpina3m | 20717 | HITS-CLIP | 23597149 | Unspecified |
| mmu-miR-29b-3p | MIMAT0000127 | Serpina6 | 12401 | HITS-CLIP | 23597149 | Unspecified |
| mmu-miR-29b-3p | MIMAT0000127 | Serpinb1a | 66222 | HITS-CLIP | 25083871 | Unspecified |
| mmu-miR-29b-3p | MIMAT0000127 | Serpinb9 | 20723 | HITS-CLIP | 23142080 | Unspecified |
| mmu-miR-29b-3p | MIMAT0000127 | Serpinc1 | 11905 | HITS-CLIP | 37986224\|23597149 | Unspecified |
| mmu-miR-29b-3p | MIMAT0000127 | Serpine1 | 18787 | HITS-CLIP | 23142080 | Unspecified |
| mmu-miR-29b-3p | MIMAT0000127 | Serpinf1 | 20317 | HITS-CLIP//sRNA-Seq | 23597149\|33184237\|25083871 | Unspecified |
| mmu-miR-29b-3p | MIMAT0000127 | Serpinf2 | 18816 | HITS-CLIP | 37986224\|23597149 | Unspecified |
| mmu-miR-29b-3p | MIMAT0000127 | Sertad1 | 55942 | HITS-CLIP | 23142080 | Unspecified |
| mmu-miR-29b-3p | MIMAT0000127 | Sertad2 | 58172 | HITS-CLIP | 25083871\|23597149\|23142080 | Unspecified |
| mmu-miR-29b-3p | MIMAT0000127 | Sesn1 | 140742 | HITS-CLIP | 23142080 | Unspecified |
| mmu-miR-29b-3p | MIMAT0000127 | Sesn2 | 230784 | HITS-CLIP | 25083871 | Unspecified |
| mmu-miR-29b-3p | MIMAT0000127 | Sesn3 | 75747 | HITS-CLIP | 23142080\|25083871 | Unspecified |
| mmu-miR-29b-3p | MIMAT0000127 | Sestd1 | 228071 | HITS-CLIP//sRNA-Seq | 37986224\|25083871\|33184237 | Unspecified |
| mmu-miR-29b-3p | MIMAT0000127 | Setbp1 | 240427 | HITS-CLIP | 25083871\|23142080 | Unspecified |
| mmu-miR-29b-3p | MIMAT0000127 | Setd1b | 208043 | HITS-CLIP | 37986224\|25083871\|23142080 | Unspecified |
| mmu-miR-29b-3p | MIMAT0000127 | Setd2 | 235626 | HITS-CLIP | 25083871\|23142080 | Unspecified |
| mmu-miR-29b-3p | MIMAT0000127 | Setd3 | 52690 | HITS-CLIP | 37986224 | Unspecified |
| mmu-miR-29b-3p | MIMAT0000127 | Setd5 | 72895 | HITS-CLIP//sRNA-Seq | 37986224\|25083871\|33184237\|23142 | Unspecified |
| mmu-miR-29b-3p | MIMAT0000127 | Setd6 | 66083 | HITS-CLIP | 25083871\|23142080 | Unspecified |
| mmu-miR-29b-3p | MIMAT0000127 | Setd7 | 73251 | HITS-CLIP//sRNA-Seq | 37986224\|33184237\|23142080 | Unspecified |
| mmu-miR-29b-3p | MIMAT0000127 | Setdb1 | 84505 | HITS-CLIP | 37986224\|23597149\|23142080\|25083 | Unspecified |
| mmu-miR-29b-3p | MIMAT0000127 | Setx | 269254 | HITS-CLIP | 37986224\|25083871\|23142080 | Unspecified |
| mmu-miR-29b-3p | MIMAT0000127 | Sf1 | 22668 | HITS-CLIP//sRNA-Seq | 23142080\|25083871\|33184237\|23597 | Unspecified |
| mmu-miR-29b-3p | MIMAT0000127 | Sf3a1 | 67465 | HITS-CLIP//sRNA-Seq | 23142080\|33184237 | Unspecified |
| mmu-miR-29b-3p | MIMAT0000127 | Sf3a2 | 20222 | HITS-CLIP//sRNA-Seq | 23142080\|33184237 | Unspecified |
| mmu-miR-29b-3p | MIMAT0000127 | Sf3b1 | 81898 | HITS-CLIP//sRNA-Seq | 37986224\|23597149\|25083871\|23142 | Unspecified |
| mmu-miR-29b-3p | MIMAT0000127 | Sf3b2 | 319322 | HITS-CLIP//sRNA-Seq | 23597149\|33184237 | Unspecified |
| mmu-miR-29b-3p | MIMAT0000127 | Sf3b3 | 101943 | HITS-CLIP//sRNA-Seq | 25083871\|33184237\|23597149\|23142 | Unspecified |
| mmu-miR-29b-3p | MIMAT0000127 | Sf3b6 | 66055 | sRNA-Seq//HITS-CLIP | 33184237\|23142080 | Unspecified |
| mmu-miR-29b-3p | MIMAT0000127 | Sfmbt2 | 353282 | HITS-CLIP | 23142080 | Unspecified |
| mmu-miR-29b-3p | MIMAT0000127 | Sfpq | 71514 | HITS-CLIP//sRNA-Seq | 23597149\|25083871\|33184237\|23142 | Unspecified |
| mmu-miR-29b-3p | MIMAT0000127 | Sfrp2 | 20319 | HITS-CLIP | 25083871 | Unspecified |
| mmu-miR-29b-3p | MIMAT0000127 | Sfswap | 231769 | sRNA-Seq//HITS-CLIP | 33184237\|23142080 | Unspecified |
| mmu-miR-29b-3p | MIMAT0000127 | Sft2d1 | 106489 | HITS-CLIP | 23142080 | Unspecified |
| mmu-miR-29b-3p | MIMAT0000127 | Sft2d2 | 108735 | HITS-CLIP | 37986224\|23142080 | Unspecified |
| mmu-miR-29b-3p | MIMAT0000127 | Sft2d3 | 67158 | HITS-CLIP | 37986224\|25083871\|23597149\|23142 | Unspecified |
| mmu-miR-29b-3p | MIMAT0000127 | Sfxn1 | 14057 | HITS-CLIP//sRNA-Seq | 37986224\|23597149\|33184237\|23142 | Unspecified |

| | | | | | | |
|---|---|---|---|---|---|---|
| mmu-miR-29b-3p | MIMAT0000127 | Sfxn3 | 94280 | HITS-CLIP//sRNA-Seq | 25083871\|33184237\|23142080 | Unspecified |
| mmu-miR-29b-3p | MIMAT0000127 | Sfxn5 | 94282 | HITS-CLIP | 37986224 | Unspecified |
| mmu-miR-29b-3p | MIMAT0000127 | Sgcb | 24051 | HITS-CLIP//sRNA-Seq | 37986224\|33184237\|23142080 | Unspecified |
| mmu-miR-29b-3p | MIMAT0000127 | Sgcg | 24053 | HITS-CLIP | 25083871 | Unspecified |
| mmu-miR-29b-3p | MIMAT0000127 | Sgk1 | 20393 | HITS-CLIP | 25083871\|23142080 | Unspecified |
| mmu-miR-29b-3p | MIMAT0000127 | Sgk3 | 170755 | HITS-CLIP | 23142080 | Unspecified |
| mmu-miR-29b-3p | MIMAT0000127 | Sgms1 | 208449 | HITS-CLIP | 23142080 | Unspecified |
| mmu-miR-29b-3p | MIMAT0000127 | Sgms2 | 74442 | HITS-CLIP | 37986224\|25083871 | Unspecified |
| mmu-miR-29b-3p | MIMAT0000127 | Sgo2a | 68549 | HITS-CLIP | 23142080 | Unspecified |
| mmu-miR-29b-3p | MIMAT0000127 | Sgpl1 | 20397 | sRNA-Seq//HITS-CLIP | 33184237\|23597149 | Unspecified |
| mmu-miR-29b-3p | MIMAT0000127 | Sgpp1 | 81535 | HITS-CLIP | 23597149\|23142080 | Unspecified |
| mmu-miR-29b-3p | MIMAT0000127 | Sgtb | 218544 | HITS-CLIP//sRNA-Seq | 25083871\|33184237 | Unspecified |
| mmu-miR-29b-3p | MIMAT0000127 | Sh2b1 | 20399 | HITS-CLIP | 25083871 | Unspecified |
| mmu-miR-29b-3p | MIMAT0000127 | Sh2b3 | 16923 | HITS-CLIP//sRNA-Seq | 25083871\|33184237 | Unspecified |
| mmu-miR-29b-3p | MIMAT0000127 | Sh2d2a | 27371 | HITS-CLIP | 37986224\|23597149\|23142080 | Unspecified |
| mmu-miR-29b-3p | MIMAT0000127 | Sh3bgrl | 56726 | HITS-CLIP//sRNA-Seq | 37986224\|25083871\|33184237\|2314 | Unspecified |
| mmu-miR-29b-3p | MIMAT0000127 | Sh3bgrl3 | 73723 | HITS-CLIP//sRNA-Seq | 37986224\|33184237\|23142080 | Unspecified |
| mmu-miR-29b-3p | MIMAT0000127 | Sh3bp2 | 24055 | HITS-CLIP | 23597149\|23142080 | Unspecified |
| mmu-miR-29b-3p | MIMAT0000127 | Sh3bp4 | 98402 | HITS-CLIP | 25083871 | Unspecified |
| mmu-miR-29b-3p | MIMAT0000127 | Sh3bp5 | 24056 | HITS-CLIP | 25083871\|23597149\|23142080 | Unspecified |
| mmu-miR-29b-3p | MIMAT0000127 | Sh3bp5l | 79566 | HITS-CLIP | 37986224\|25083871\|23142080 | Unspecified |
| mmu-miR-29b-3p | MIMAT0000127 | Sh3d19 | 27059 | HITS-CLIP | 25083871\|23597149 | Unspecified |
| mmu-miR-29b-3p | MIMAT0000127 | Sh3gl1 | 20405 | HITS-CLIP | 37986224 | Unspecified |
| mmu-miR-29b-3p | MIMAT0000127 | Sh3gl3 | 20408 | HITS-CLIP | 23142080 | Unspecified |
| mmu-miR-29b-3p | MIMAT0000127 | Sh3glb1 | 54673 | HITS-CLIP//sRNA-Seq | 23142080\|33184237 | Unspecified |
| mmu-miR-29b-3p | MIMAT0000127 | Sh3kbp1 | 58194 | HITS-CLIP//sRNA-Seq | 37986224\|25083871\|33184237\|2314 | Unspecified |
| mmu-miR-29b-3p | MIMAT0000127 | Sh3pxd2a | 14218 | HITS-CLIP//sRNA-Seq | 37986224\|25083871\|23597149\|3318 | Unspecified |
| mmu-miR-29b-3p | MIMAT0000127 | Sh3pxd2b | 268396 | HITS-CLIP | 37986224 | Unspecified |
| mmu-miR-29b-3p | MIMAT0000127 | Shank3 | 58234 | HITS-CLIP | 23597149 | Unspecified |
| mmu-miR-29b-3p | MIMAT0000127 | Shb | 230126 | HITS-CLIP | 37986224 | Unspecified |
| mmu-miR-29b-3p | MIMAT0000127 | Shcbp1 | 20419 | HITS-CLIP | 37986224\|23142080 | Unspecified |
| mmu-miR-29b-3p | MIMAT0000127 | Shisa5 | 66940 | HITS-CLIP//sRNA-Seq | 25083871\|33184237\|23142080 | Unspecified |
| mmu-miR-29b-3p | MIMAT0000127 | Shkbp1 | 192192 | HITS-CLIP | 23142080 | Unspecified |
| mmu-miR-29b-3p | MIMAT0000127 | Shld2 | 54537 | HITS-CLIP | 23597149\|23142080 | Unspecified |
| mmu-miR-29b-3p | MIMAT0000127 | Shmt2 | 108037 | HITS-CLIP//sRNA-Seq | 23597149\|33184237 | Unspecified |
| mmu-miR-29b-3p | MIMAT0000127 | Shoc2 | 56392 | HITS-CLIP//sRNA-Seq | 25083871\|33184237\|23142080 | Unspecified |
| mmu-miR-29b-3p | MIMAT0000127 | Shprh | 268281 | HITS-CLIP | 25083871\|23597149\|23142080 | Unspecified |
| mmu-miR-29b-3p | MIMAT0000127 | Shq1 | 72171 | HITS-CLIP | 37986224\|23142080 | Unspecified |
| mmu-miR-29b-3p | MIMAT0000127 | Shroom2 | 110380 | HITS-CLIP | 37986224\|23597149 | Unspecified |

| | | | | | | |
|---|---|---|---|---|---|---|
| mmu-miR-29b-3p | MIMAT0000127 | Shtn1 | 57698 | sRNA-Seq//HITS-CLIP | 33184237\|23142080 | Unspecified |
| mmu-miR-29b-3p | MIMAT0000127 | Siae | 22619 | HITS-CLIP | 23142080\|23597149 | Unspecified |
| mmu-miR-29b-3p | MIMAT0000127 | Siah1b | 20438 | HITS-CLIP | 23142080 | Unspecified |
| mmu-miR-29b-3p | MIMAT0000127 | Siah2 | 20439 | HITS-CLIP | 25083871\|23142080 | Unspecified |
| mmu-miR-29b-3p | MIMAT0000127 | Sidt2 | 214597 | HITS-CLIP | 37986224\|25083871\|23597149\|23142 | Unspecified |
| mmu-miR-29b-3p | MIMAT0000127 | Sigirr | 24058 | HITS-CLIP | 23142080 | Unspecified |
| mmu-miR-29b-3p | MIMAT0000127 | Siglec1 | 20612 | HITS-CLIP | 23597149 | Unspecified |
| mmu-miR-29b-3p | MIMAT0000127 | Sik1 | 17691 | HITS-CLIP | 23597149\|23142080\|26203562 | Unspecified |
| mmu-miR-29b-3p | MIMAT0000127 | Sik3 | 70661 | HITS-CLIP | 25083871\|23142080 | Unspecified |
| mmu-miR-29b-3p | MIMAT0000127 | Sike1 | 66641 | HITS-CLIP | 37986224\|23597149\|25083871\|23142 | Unspecified |
| mmu-miR-29b-3p | MIMAT0000127 | Sim2 | 20465 | HITS-CLIP | 25083871 | Unspecified |
| mmu-miR-29b-3p | MIMAT0000127 | Simc1 | 319719 | HITS-CLIP | 37986224 | Unspecified |
| mmu-miR-29b-3p | MIMAT0000127 | Sin3a | 20466 | HITS-CLIP//sRNA-Seq | 23142080\|25083871\|23597149\|33184 | Unspecified |
| mmu-miR-29b-3p | MIMAT0000127 | Sin3b | 20467 | HITS-CLIP//sRNA-Seq | 23597149\|33184237 | Unspecified |
| mmu-miR-29b-3p | MIMAT0000127 | Sipa1l2 | 244668 | HITS-CLIP//sRNA-Seq | 23142080\|33184237 | Unspecified |
| mmu-miR-29b-3p | MIMAT0000127 | Sipa1l3 | 74206 | HITS-CLIP | 23142080\|23597149 | Unspecified |
| mmu-miR-29b-3p | MIMAT0000127 | Sirt1 | 93759 | Luciferase assay//qRT-PCR//Western blot/ | 24657470\|25083871\|23142080 | Unspecified |
| mmu-miR-29b-3p | MIMAT0000127 | Sirt6 | 50721 | HITS-CLIP | 25083871\|23142080 | Unspecified |
| mmu-miR-29b-3p | MIMAT0000127 | Sirt7 | 209011 | HITS-CLIP | 23142080 | Unspecified |
| mmu-miR-29b-3p | MIMAT0000127 | Sit1 | 54390 | HITS-CLIP | 23142080 | Unspecified |
| mmu-miR-29b-3p | MIMAT0000127 | Six4 | 20474 | HITS-CLIP | 25083871 | Unspecified |
| mmu-miR-29b-3p | MIMAT0000127 | Six5 | 20475 | HITS-CLIP | 37986224\|25083871\|23142080 | Unspecified |
| mmu-miR-29b-3p | MIMAT0000127 | Ska2 | 66140 | HITS-CLIP | 37986224\|23142080 | Unspecified |
| mmu-miR-29b-3p | MIMAT0000127 | Ski | 20481 | HITS-CLIP//sRNA-Seq | 37986224\|25083871\|23597149\|33184 | Unspecified |
| mmu-miR-29b-3p | MIMAT0000127 | Skil | 20482 | HITS-CLIP | 37986224\|25083871\|23142080 | Unspecified |
| mmu-miR-29b-3p | MIMAT0000127 | Skiv2l | 108077 | sRNA-Seq//HITS-CLIP | 33184237\|23597149 | Unspecified |
| mmu-miR-29b-3p | MIMAT0000127 | Skp2 | 27401 | HITS-CLIP | 25083871\|23142080 | Unspecified |
| mmu-miR-29b-3p | MIMAT0000127 | Sla | 20491 | HITS-CLIP | 23142080 | Unspecified |
| mmu-miR-29b-3p | MIMAT0000127 | Slain2 | 75991 | HITS-CLIP | 25083871 | Unspecified |
| mmu-miR-29b-3p | MIMAT0000127 | Slc10a1 | 20493 | HITS-CLIP | 23597149 | Unspecified |
| mmu-miR-29b-3p | MIMAT0000127 | Slc10a2 | 20494 | HITS-CLIP | 37986224\|23597149 | Unspecified |
| mmu-miR-29b-3p | MIMAT0000127 | Slc10a7 | 76775 | HITS-CLIP | 25083871 | Unspecified |
| mmu-miR-29b-3p | MIMAT0000127 | Slc11a2 | 18174 | HITS-CLIP | 25083871 | Unspecified |
| mmu-miR-29b-3p | MIMAT0000127 | Slc12a2 | 20496 | HITS-CLIP | 25083871\|23142080 | Unspecified |
| mmu-miR-29b-3p | MIMAT0000127 | Slc12a6 | 107723 | HITS-CLIP | 23142080 | Unspecified |
| mmu-miR-29b-3p | MIMAT0000127 | Slc12a9 | 83704 | HITS-CLIP | 23142080 | Unspecified |
| mmu-miR-29b-3p | MIMAT0000127 | Slc13a3 | 114644 | HITS-CLIP | 23597149 | Unspecified |
| mmu-miR-29b-3p | MIMAT0000127 | Slc15a3 | 65221 | HITS-CLIP | 23597149 | Unspecified |
| mmu-miR-29b-3p | MIMAT0000127 | Slc15a4 | 100561 | HITS-CLIP | 23597149\|23142080 | Unspecified |

| | | | | | | |
|---|---|---|---|---|---|---|
| mmu-miR-29b-3p | MIMAT0000127 | Slc16a1 | 20501 | HITS-CLIP | 37986224\|25083871\|23597149\|23142 | Unspecified |
| mmu-miR-29b-3p | MIMAT0000127 | Slc16a2 | 20502 | HITS-CLIP | 37986224\|23597149 | Unspecified |
| mmu-miR-29b-3p | MIMAT0000127 | Slc16a3 | 80879 | HITS-CLIP | 37986224\|23142080\|25083871 | Unspecified |
| mmu-miR-29b-3p | MIMAT0000127 | Slc16a6 | 104681 | HITS-CLIP | 25083871 | Unspecified |
| mmu-miR-29b-3p | MIMAT0000127 | Slc16a7 | 20503 | HITS-CLIP//sRNA-Seq | 23597149\|33184237 | Unspecified |
| mmu-miR-29b-3p | MIMAT0000127 | Slc17a2 | 218103 | HITS-CLIP | 23597149 | Unspecified |
| mmu-miR-29b-3p | MIMAT0000127 | Slc17a6 | 140919 | sRNA-Seq//HITS-CLIP | 33184237\|23142080 | Unspecified |
| mmu-miR-29b-3p | MIMAT0000127 | Slc1a2 | 20511 | HITS-CLIP | 23597149 | Unspecified |
| mmu-miR-29b-3p | MIMAT0000127 | Slc1a4 | 55963 | HITS-CLIP | 37986224 | Unspecified |
| mmu-miR-29b-3p | MIMAT0000127 | Slc1a5 | 20514 | HITS-CLIP | 23142080 | Unspecified |
| mmu-miR-29b-3p | MIMAT0000127 | Slc20a1 | 20515 | sRNA-Seq//HITS-CLIP | 33184237\|23142080 | Unspecified |
| mmu-miR-29b-3p | MIMAT0000127 | Slc20a2 | 20516 | HITS-CLIP | 25083871\|23142080 | Unspecified |
| mmu-miR-29b-3p | MIMAT0000127 | Slc22a1 | 20517 | HITS-CLIP | 23597149 | Unspecified |
| mmu-miR-29b-3p | MIMAT0000127 | Slc22a23 | 73102 | HITS-CLIP | 23597149 | Unspecified |
| mmu-miR-29b-3p | MIMAT0000127 | Slc22a5 | 20520 | HITS-CLIP | 23597149 | Unspecified |
| mmu-miR-29b-3p | MIMAT0000127 | Slc22a7 | 108114 | HITS-CLIP | 23597149 | Unspecified |
| mmu-miR-29b-3p | MIMAT0000127 | Slc23a2 | 54338 | HITS-CLIP | 25083871\|23142080 | Unspecified |
| mmu-miR-29b-3p | MIMAT0000127 | Slc24a3 | 94249 | HITS-CLIP | 37986224 | Unspecified |
| mmu-miR-29b-3p | MIMAT0000127 | Slc25a1 | 13358 | HITS-CLIP//sRNA-Seq | 23142080\|33184237 | Unspecified |
| mmu-miR-29b-3p | MIMAT0000127 | Slc25a10 | 27376 | HITS-CLIP | 23597149 | Unspecified |
| mmu-miR-29b-3p | MIMAT0000127 | Slc25a11 | 67863 | sRNA-Seq//HITS-CLIP | 33184237\|23142080 | Unspecified |
| mmu-miR-29b-3p | MIMAT0000127 | Slc25a13 | 50799 | HITS-CLIP | 23142080\|23597149 | Unspecified |
| mmu-miR-29b-3p | MIMAT0000127 | Slc25a16 | 73132 | HITS-CLIP | 25083871\|23597149 | Unspecified |
| mmu-miR-29b-3p | MIMAT0000127 | Slc25a17 | 20524 | sRNA-Seq//HITS-CLIP | 33184237\|25083871 | Unspecified |
| mmu-miR-29b-3p | MIMAT0000127 | Slc25a20 | 57279 | HITS-CLIP | 23597149\|23142080 | Unspecified |
| mmu-miR-29b-3p | MIMAT0000127 | Slc25a22 | 68267 | HITS-CLIP | 37986224\|23597149\|23142080\|25083 | Unspecified |
| mmu-miR-29b-3p | MIMAT0000127 | Slc25a24 | 229731 | HITS-CLIP//sRNA-Seq | 37986224\|25083871\|33184237\|23142 | Unspecified |
| mmu-miR-29b-3p | MIMAT0000127 | Slc25a25 | 227731 | HITS-CLIP | 23597149 | Unspecified |
| mmu-miR-29b-3p | MIMAT0000127 | Slc25a27 | 74011 | HITS-CLIP | 37986224 | Unspecified |
| mmu-miR-29b-3p | MIMAT0000127 | Slc25a3 | 18674 | HITS-CLIP//sRNA-Seq | 37986224\|25083871\|33184237 | Unspecified |
| mmu-miR-29b-3p | MIMAT0000127 | Slc25a32 | 69906 | HITS-CLIP | 25083871\|23142080 | Unspecified |
| mmu-miR-29b-3p | MIMAT0000127 | Slc25a36 | 192287 | HITS-CLIP//sRNA-Seq//qCLASH | 37986224\|25083871\|33184237\|31393 | Unspecified |
| mmu-miR-29b-3p | MIMAT0000127 | Slc25a37 | 67712 | HITS-CLIP | 25083871\|23142080 | Unspecified |
| mmu-miR-29b-3p | MIMAT0000127 | Slc25a4 | 11739 | HITS-CLIP//sRNA-Seq | 37986224\|23597149\|33184237\|23142 | Unspecified |
| mmu-miR-29b-3p | MIMAT0000127 | Slc25a42 | 73095 | HITS-CLIP | 23597149 | Unspecified |
| mmu-miR-29b-3p | MIMAT0000127 | Slc25a45 | 107375 | HITS-CLIP | 23597149 | Unspecified |
| mmu-miR-29b-3p | MIMAT0000127 | Slc25a46 | 67453 | HITS-CLIP | 37986224\|25083871\|23142080 | Unspecified |
| mmu-miR-29b-3p | MIMAT0000127 | Slc25a5 | 11740 | sRNA-Seq//HITS-CLIP | 33184237\|23142080 | Unspecified |
| mmu-miR-29b-3p | MIMAT0000127 | Slc25a51 | 230125 | HITS-CLIP//sRNA-Seq | 37986224\|23597149\|33184237\|23142 | Unspecified |

| | | | | | | |
|---|---|---|---|---|---|---|
| mmu-miR-29b-3p | MIMAT0000127 | Slc26a1 | 231583 | HITS-CLIP | 23597149 | Unspecified |
| mmu-miR-29b-3p | MIMAT0000127 | Slc27a4 | 26569 | HITS-CLIP | 23142080 | Unspecified |
| mmu-miR-29b-3p | MIMAT0000127 | Slc27a5 | 26459 | HITS-CLIP | 23597149 | Unspecified |
| mmu-miR-29b-3p | MIMAT0000127 | Slc29a2 | 13340 | HITS-CLIP | 37986224 | Unspecified |
| mmu-miR-29b-3p | MIMAT0000127 | Slc2a1 | 20525 | HITS-CLIP//sRNA-Seq | 25083871\|33184237\|23142080 | Unspecified |
| mmu-miR-29b-3p | MIMAT0000127 | Slc2a2 | 20526 | HITS-CLIP | 23597149 | Unspecified |
| mmu-miR-29b-3p | MIMAT0000127 | Slc2a3 | 20527 | HITS-CLIP//sRNA-Seq | 37986224\|33184237\|23142080 | Unspecified |
| mmu-miR-29b-3p | MIMAT0000127 | Slc2a6 | 227659 | HITS-CLIP | 23597149 | Unspecified |
| mmu-miR-29b-3p | MIMAT0000127 | Slc30a1 | 22782 | HITS-CLIP | 25083871\|23597149 | Unspecified |
| mmu-miR-29b-3p | MIMAT0000127 | Slc30a4 | 22785 | HITS-CLIP | 25083871 | Unspecified |
| mmu-miR-29b-3p | MIMAT0000127 | Slc30a5 | 69048 | HITS-CLIP | 23142080 | Unspecified |
| mmu-miR-29b-3p | MIMAT0000127 | Slc30a7 | 66500 | HITS-CLIP | 37986224\|23142080 | Unspecified |
| mmu-miR-29b-3p | MIMAT0000127 | Slc31a1 | 20529 | HITS-CLIP | 37986224\|25083871\|23597149\|2314 | Unspecified |
| mmu-miR-29b-3p | MIMAT0000127 | Slc35a1 | 24060 | HITS-CLIP | 23597149\|25083871\|23142080 | Unspecified |
| mmu-miR-29b-3p | MIMAT0000127 | Slc35a4 | 67843 | HITS-CLIP//sRNA-Seq | 25083871\|23142080\|33184237\|2359 | Unspecified |
| mmu-miR-29b-3p | MIMAT0000127 | Slc35b2 | 73836 | HITS-CLIP//sRNA-Seq | 37986224\|25083871\|23597149\|3318 | Unspecified |
| mmu-miR-29b-3p | MIMAT0000127 | Slc35b3 | 108652 | HITS-CLIP | 23142080 | Unspecified |
| mmu-miR-29b-3p | MIMAT0000127 | Slc35b4 | 58246 | HITS-CLIP | 25083871 | Unspecified |
| mmu-miR-29b-3p | MIMAT0000127 | Slc35c1 | 228368 | HITS-CLIP | 37986224\|23142080 | Unspecified |
| mmu-miR-29b-3p | MIMAT0000127 | Slc35d1 | 242585 | HITS-CLIP | 37986224\|23142080\|23597149 | Unspecified |
| mmu-miR-29b-3p | MIMAT0000127 | Slc35e1 | 270066 | HITS-CLIP//sRNA-Seq | 23142080\|33184237 | Unspecified |
| mmu-miR-29b-3p | MIMAT0000127 | Slc35e3 | 215436 | HITS-CLIP | 23597149 | Unspecified |
| mmu-miR-29b-3p | MIMAT0000127 | Slc35f5 | 74150 | HITS-CLIP | 25083871 | Unspecified |
| mmu-miR-29b-3p | MIMAT0000127 | Slc35f6 | 74919 | HITS-CLIP | 25083871\|23597149 | Unspecified |
| mmu-miR-29b-3p | MIMAT0000127 | Slc36a4 | 234967 | HITS-CLIP | 37986224 | Unspecified |
| mmu-miR-29b-3p | MIMAT0000127 | Slc37a3 | 72144 | HITS-CLIP//sRNA-Seq | 37986224\|33184237\|23142080 | Unspecified |
| mmu-miR-29b-3p | MIMAT0000127 | Slc37a4 | 14385 | HITS-CLIP | 37986224\|23597149 | Unspecified |
| mmu-miR-29b-3p | MIMAT0000127 | Slc38a1 | 105727 | HITS-CLIP//sRNA-Seq | 25083871\|33184237\|23142080 | Unspecified |
| mmu-miR-29b-3p | MIMAT0000127 | Slc38a10 | 72055 | HITS-CLIP//sRNA-Seq | 25083871\|23597149\|33184237 | Unspecified |
| mmu-miR-29b-3p | MIMAT0000127 | Slc38a2 | 67760 | HITS-CLIP//sRNA-Seq | 23597149\|33184237\|23142080\|2508 | Unspecified |
| mmu-miR-29b-3p | MIMAT0000127 | Slc38a3 | 76257 | HITS-CLIP | 23597149 | Unspecified |
| mmu-miR-29b-3p | MIMAT0000127 | Slc38a4 | 69354 | HITS-CLIP//sRNA-Seq | 23597149\|33184237 | Unspecified |
| mmu-miR-29b-3p | MIMAT0000127 | Slc38a6 | 625098 | HITS-CLIP | 25083871 | Unspecified |
| mmu-miR-29b-3p | MIMAT0000127 | Slc38a7 | 234595 | HITS-CLIP | 37986224 | Unspecified |
| mmu-miR-29b-3p | MIMAT0000127 | Slc38a9 | 268706 | HITS-CLIP | 23142080 | Unspecified |
| mmu-miR-29b-3p | MIMAT0000127 | Slc39a1 | 30791 | HITS-CLIP//sRNA-Seq | 23597149\|23142080\|33184237\|2508 | Unspecified |
| mmu-miR-29b-3p | MIMAT0000127 | Slc39a10 | 227059 | HITS-CLIP | 23142080 | Unspecified |
| mmu-miR-29b-3p | MIMAT0000127 | Slc39a14 | 213053 | HITS-CLIP | 37986224\|25083871\|23597149 | Unspecified |
| mmu-miR-29b-3p | MIMAT0000127 | Slc39a3 | 106947 | HITS-CLIP | 25083871 | Unspecified |

| | | | | | | |
|---|---|---|---|---|---|---|
| mmu-miR-29b-3p | MIMAT0000127 | Slc39a6 | 106957 | sRNA-Seq//HITS-CLIP | 33184237\|23142080 | Unspecified |
| mmu-miR-29b-3p | MIMAT0000127 | Slc39a7 | 14977 | sRNA-Seq//HITS-CLIP | 33184237\|25083871\|23142080 | Unspecified |
| mmu-miR-29b-3p | MIMAT0000127 | Slc39a9 | 328133 | HITS-CLIP | 37986224\|23142080\|25083871 | Unspecified |
| mmu-miR-29b-3p | MIMAT0000127 | Slc3a2 | 17254 | HITS-CLIP//sRNA-Seq | 25083871\|23142080\|33184237 | Unspecified |
| mmu-miR-29b-3p | MIMAT0000127 | Slc40a1 | 53945 | HITS-CLIP | 37986224\|25083871 | Unspecified |
| mmu-miR-29b-3p | MIMAT0000127 | Slc41a2 | 338365 | HITS-CLIP | 23597149\|23142080 | Unspecified |
| mmu-miR-29b-3p | MIMAT0000127 | Slc43a2 | 215113 | HITS-CLIP | 37986224\|25083871\|23142080\|23597 | Unspecified |
| mmu-miR-29b-3p | MIMAT0000127 | Slc43a3 | 58207 | sRNA-Seq//HITS-CLIP | 33184237\|23142080 | Unspecified |
| mmu-miR-29b-3p | MIMAT0000127 | Slc44a1 | 100434 | HITS-CLIP//sRNA-Seq | 25083871\|33184237 | Unspecified |
| mmu-miR-29b-3p | MIMAT0000127 | Slc44a2 | 68682 | HITS-CLIP//sRNA-Seq | 23142080\|33184237\|25083871 | Unspecified |
| mmu-miR-29b-3p | MIMAT0000127 | Slc46a1 | 52466 | HITS-CLIP | 23597149 | Unspecified |
| mmu-miR-29b-3p | MIMAT0000127 | Slc48a1 | 67739 | HITS-CLIP//sRNA-Seq | 25083871\|33184237 | Unspecified |
| mmu-miR-29b-3p | MIMAT0000127 | Slc4a11 | 269356 | HITS-CLIP | 23142080 | Unspecified |
| mmu-miR-29b-3p | MIMAT0000127 | Slc4a1ap | 20534 | HITS-CLIP | 25083871 | Unspecified |
| mmu-miR-29b-3p | MIMAT0000127 | Slc4a2 | 20535 | HITS-CLIP//sRNA-Seq | 37986224\|33184237\|23597149\|2314 | Unspecified |
| mmu-miR-29b-3p | MIMAT0000127 | Slc4a4 | 54403 | HITS-CLIP | 23597149 | Unspecified |
| mmu-miR-29b-3p | MIMAT0000127 | Slc4a7 | 218756 | HITS-CLIP | 37986224\|25083871\|23142080 | Unspecified |
| mmu-miR-29b-3p | MIMAT0000127 | Slc50a1 | 19729 | sRNA-Seq//HITS-CLIP | 33184237\|23142080 | Unspecified |
| mmu-miR-29b-3p | MIMAT0000127 | Slc5a3 | 53881 | HITS-CLIP | 23142080 | Unspecified |
| mmu-miR-29b-3p | MIMAT0000127 | Slc6a6 | 21366 | HITS-CLIP//sRNA-Seq | 25083871\|33184237\|23142080 | Unspecified |
| mmu-miR-29b-3p | MIMAT0000127 | Slc6a8 | 102857 | HITS-CLIP | 25083871 | Unspecified |
| mmu-miR-29b-3p | MIMAT0000127 | Slc6a9 | 14664 | HITS-CLIP | 25083871\|23597149 | Unspecified |
| mmu-miR-29b-3p | MIMAT0000127 | Slc7a1 | 11987 | HITS-CLIP//sRNA-Seq | 37986224\|25083871\|33184237\|2314 | Unspecified |
| mmu-miR-29b-3p | MIMAT0000127 | Slc7a2 | 11988 | HITS-CLIP | 37986224\|25083871\|23597149 | Unspecified |
| mmu-miR-29b-3p | MIMAT0000127 | Slc7a5 | 20539 | HITS-CLIP//sRNA-Seq//qCLASH | 25083871\|33184237\|31393953\|2314 | Unspecified |
| mmu-miR-29b-3p | MIMAT0000127 | Slc7a6 | 330836 | HITS-CLIP | 37986224\|23142080 | Unspecified |
| mmu-miR-29b-3p | MIMAT0000127 | Slc9a5 | 277973 | HITS-CLIP | 37986224 | Unspecified |
| mmu-miR-29b-3p | MIMAT0000127 | Slc9a6 | 236794 | sRNA-Seq//HITS-CLIP | 33184237\|23142080 | Unspecified |
| mmu-miR-29b-3p | MIMAT0000127 | Slc9a8 | 77031 | HITS-CLIP | 23142080 | Unspecified |
| mmu-miR-29b-3p | MIMAT0000127 | Slc9a9 | 331004 | HITS-CLIP//sRNA-Seq | 25083871\|33184237 | Unspecified |
| mmu-miR-29b-3p | MIMAT0000127 | Slco1b2 | 28253 | HITS-CLIP | 37986224\|23597149 | Unspecified |
| mmu-miR-29b-3p | MIMAT0000127 | Slco3a1 | 108116 | HITS-CLIP | 25083871 | Unspecified |
| mmu-miR-29b-3p | MIMAT0000127 | Slco4a1 | 108115 | HITS-CLIP | 37986224\|23142080 | Unspecified |
| mmu-miR-29b-3p | MIMAT0000127 | Slco5a1 | 240726 | HITS-CLIP | 25083871 | Unspecified |
| mmu-miR-29b-3p | MIMAT0000127 | Slf1 | 852127 | HITS-CLIP | 23142080\|23597149 | Unspecified |
| mmu-miR-29b-3p | MIMAT0000127 | Slf2 | 226151 | HITS-CLIP | 25083871\|23597149\|23142080 | Unspecified |
| mmu-miR-29b-3p | MIMAT0000127 | Slfn1 | 20555 | HITS-CLIP | 23142080 | Unspecified |
| mmu-miR-29b-3p | MIMAT0000127 | Slk | 20874 | HITS-CLIP | 37986224\|25083871\|23142080 | Unspecified |
| mmu-miR-29b-3p | MIMAT0000127 | Slmap | 83997 | HITS-CLIP | 31433990 | Unspecified |

| | | | | | | |
|---|---|---|---|---|---|---|
| mmu-miR-29b-3p | MIMAT0000127 | Slx4ip | 74243 | HITS-CLIP | 37986224 | Unspecified |
| mmu-miR-29b-3p | MIMAT0000127 | Smad1 | 17125 | HITS-CLIP | 25083871\|23597149 | Unspecified |
| mmu-miR-29b-3p | MIMAT0000127 | Smad2 | 17126 | HITS-CLIP//sRNA-Seq | 23142080\|25083871\|33184237 | Unspecified |
| mmu-miR-29b-3p | MIMAT0000127 | Smad3 | 17127 | HITS-CLIP | 25083871\|23597149\|23142080 | Unspecified |
| mmu-miR-29b-3p | MIMAT0000127 | Smad4 | 17128 | HITS-CLIP//sRNA-Seq | 23142080\|25083871\|33184237 | Unspecified |
| mmu-miR-29b-3p | MIMAT0000127 | Smad5 | 17129 | HITS-CLIP | 37986224\|25083871\|23142080 | Unspecified |
| mmu-miR-29b-3p | MIMAT0000127 | Smad7 | 17131 | HITS-CLIP | 25083871\|23142080 | Unspecified |
| mmu-miR-29b-3p | MIMAT0000127 | Smagp | 207818 | HITS-CLIP | 23597149 | Unspecified |
| mmu-miR-29b-3p | MIMAT0000127 | Smap1 | 98366 | HITS-CLIP//sRNA-Seq | 37986224\|25083871\|33184237\|23142 | Unspecified |
| mmu-miR-29b-3p | MIMAT0000127 | Smap2 | 69780 | HITS-CLIP//sRNA-Seq | 23597149\|33184237 | Unspecified |
| mmu-miR-29b-3p | MIMAT0000127 | Smarca4 | 20586 | HITS-CLIP//sRNA-Seq | 25083871\|33184237\|23142080 | Unspecified |
| mmu-miR-29b-3p | MIMAT0000127 | Smarca5 | 93762 | HITS-CLIP//sRNA-Seq | 37986224\|25083871\|33184237\|23142 | Unspecified |
| mmu-miR-29b-3p | MIMAT0000127 | Smarcad1 | 13990 | HITS-CLIP | 37986224\|25083871\|23142080 | Unspecified |
| mmu-miR-29b-3p | MIMAT0000127 | Smarcb1 | 20587 | sRNA-Seq//HITS-CLIP | 33184237\|25083871 | Unspecified |
| mmu-miR-29b-3p | MIMAT0000127 | Smarcc1 | 20588 | HITS-CLIP//sRNA-Seq | 37986224\|23142080\|33184237\|23597 | Unspecified |
| mmu-miR-29b-3p | MIMAT0000127 | Smarcc2 | 68094 | HITS-CLIP//sRNA-Seq | 25083871\|33184237\|23597149\|23142 | Unspecified |
| mmu-miR-29b-3p | MIMAT0000127 | Smarcd2 | 83796 | HITS-CLIP | 23142080\|25083871 | Unspecified |
| mmu-miR-29b-3p | MIMAT0000127 | Smarcd3 | 66993 | HITS-CLIP | 25083871 | Unspecified |
| mmu-miR-29b-3p | MIMAT0000127 | Smarce1 | 57376 | HITS-CLIP//sRNA-Seq | 25083871\|33184237 | Unspecified |
| mmu-miR-29b-3p | MIMAT0000127 | Smc1a | 24061 | sRNA-Seq//HITS-CLIP | 33184237\|23142080 | Unspecified |
| mmu-miR-29b-3p | MIMAT0000127 | Smc2 | 14211 | HITS-CLIP//sRNA-Seq | 37986224\|33184237\|23142080 | Unspecified |
| mmu-miR-29b-3p | MIMAT0000127 | Smc3 | 13006 | HITS-CLIP//sRNA-Seq | 25083871\|33184237\|23142080 | Unspecified |
| mmu-miR-29b-3p | MIMAT0000127 | Smc4 | 70099 | HITS-CLIP//sRNA-Seq | 25083871\|33184237\|23597149\|23142 | Unspecified |
| mmu-miR-29b-3p | MIMAT0000127 | Smchd1 | 74355 | HITS-CLIP//sRNA-Seq | 25083871\|33184237\|23142080 | Unspecified |
| mmu-miR-29b-3p | MIMAT0000127 | Smco4 | 170748 | HITS-CLIP | 37986224\|23597149 | Unspecified |
| mmu-miR-29b-3p | MIMAT0000127 | Smcr8 | 237782 | HITS-CLIP | 37986224\|25083871\|23597149 | Unspecified |
| mmu-miR-29b-3p | MIMAT0000127 | Smg1 | 233789 | HITS-CLIP | 37986224\|25083871\|23142080 | Unspecified |
| mmu-miR-29b-3p | MIMAT0000127 | Smg7 | 226517 | sRNA-Seq//HITS-CLIP | 33184237\|23142080 | Unspecified |
| mmu-miR-29b-3p | MIMAT0000127 | Smg8 | 74133 | HITS-CLIP | 25083871 | Unspecified |
| mmu-miR-29b-3p | MIMAT0000127 | Smim10l2a | 399668 | HITS-CLIP | 23142080 | Unspecified |
| mmu-miR-29b-3p | MIMAT0000127 | Smim12 | 80284 | HITS-CLIP | 23142080 | Unspecified |
| mmu-miR-29b-3p | MIMAT0000127 | Smim13 | 108934 | HITS-CLIP | 25083871 | Unspecified |
| mmu-miR-29b-3p | MIMAT0000127 | Smim14 | 68552 | HITS-CLIP//sRNA-Seq | 37986224\|25083871\|33184237 | Unspecified |
| mmu-miR-29b-3p | MIMAT0000127 | Smim7 | 66818 | HITS-CLIP//sRNA-Seq | 25083871\|33184237 | Unspecified |
| mmu-miR-29b-3p | MIMAT0000127 | Smoc1 | 64075 | HITS-CLIP | 23597149 | Unspecified |
| mmu-miR-29b-3p | MIMAT0000127 | Smpd2 | 20598 | HITS-CLIP | 37986224\|23597149\|23142080 | Unspecified |
| mmu-miR-29b-3p | MIMAT0000127 | Smpdl3a | 57319 | HITS-CLIP | 37986224\|23597149\|23142080 | Unspecified |
| mmu-miR-29b-3p | MIMAT0000127 | Smtnl2 | 276829 | HITS-CLIP | 37986224\|23597149\|23142080 | Unspecified |
| mmu-miR-29b-3p | MIMAT0000127 | Smug1 | 71726 | HITS-CLIP | 37986224\|23142080 | Unspecified |

| | | | | | | |
|---|---|---|---|---|---|---|
| mmu-miR-29b-3p | MIMAT0000127 | Smurf1 | 75788 | HITS-CLIP | 25083871\|23142080 | Unspecified |
| mmu-miR-29b-3p | MIMAT0000127 | Smurf2 | 66313 | HITS-CLIP | 37986224\|25083871\|23597149\|23142 | Unspecified |
| mmu-miR-29b-3p | MIMAT0000127 | Smyd1 | 12180 | HITS-CLIP | 37986224 | Unspecified |
| mmu-miR-29b-3p | MIMAT0000127 | Snai2 | 20583 | HITS-CLIP//sRNA-Seq | 25083871\|33184237 | Unspecified |
| mmu-miR-29b-3p | MIMAT0000127 | Snap29 | 67474 | HITS-CLIP | 23142080\|25083871 | Unspecified |
| mmu-miR-29b-3p | MIMAT0000127 | Snca | 20617 | HITS-CLIP//sRNA-Seq | 23597149\|33184237 | Unspecified |
| mmu-miR-29b-3p | MIMAT0000127 | Snd1 | 56463 | HITS-CLIP//sRNA-Seq | 23597149\|23142080\|25083871\|33184 | Unspecified |
| mmu-miR-29b-3p | MIMAT0000127 | Snn | 20621 | HITS-CLIP | 37986224\|23142080 | Unspecified |
| mmu-miR-29b-3p | MIMAT0000127 | Snrk | 20623 | HITS-CLIP//sRNA-Seq | 37986224\|23597149\|33184237 | Unspecified |
| mmu-miR-29b-3p | MIMAT0000127 | Snrnp200 | 320632 | HITS-CLIP//sRNA-Seq | 25083871\|33184237\|23142080 | Unspecified |
| mmu-miR-29b-3p | MIMAT0000127 | Snrnp27 | 66618 | sRNA-Seq//HITS-CLIP | 33184237\|23142080 | Unspecified |
| mmu-miR-29b-3p | MIMAT0000127 | Snrpa1 | 68981 | HITS-CLIP | 23142080 | Unspecified |
| mmu-miR-29b-3p | MIMAT0000127 | Snrpb | 20638 | sRNA-Seq//HITS-CLIP | 33184237\|23597149\|23142080 | Unspecified |
| mmu-miR-29b-3p | MIMAT0000127 | Snrpd2 | 107686 | sRNA-Seq//HITS-CLIP | 33184237\|23142080 | Unspecified |
| mmu-miR-29b-3p | MIMAT0000127 | Sntb1 | 20649 | HITS-CLIP | 25083871\|23142080 | Unspecified |
| mmu-miR-29b-3p | MIMAT0000127 | Sntb2 | 20650 | HITS-CLIP | 37986224\|25083871 | Unspecified |
| mmu-miR-29b-3p | MIMAT0000127 | Snw1 | 66354 | sRNA-Seq//HITS-CLIP | 33184237\|23142080 | Unspecified |
| mmu-miR-29b-3p | MIMAT0000127 | Snx1 | 56440 | HITS-CLIP//sRNA-Seq | 37986224\|25083871\|33184237\|23142 | Unspecified |
| mmu-miR-29b-3p | MIMAT0000127 | Snx16 | 74718 | HITS-CLIP | 23142080 | Unspecified |
| mmu-miR-29b-3p | MIMAT0000127 | Snx17 | 266781 | HITS-CLIP//sRNA-Seq | 25083871\|33184237 | Unspecified |
| mmu-miR-29b-3p | MIMAT0000127 | Snx18 | 170625 | HITS-CLIP | 25083871\|23142080 | Unspecified |
| mmu-miR-29b-3p | MIMAT0000127 | Snx2 | 67804 | HITS-CLIP//sRNA-Seq | 25083871\|33184237\|23142080 | Unspecified |
| mmu-miR-29b-3p | MIMAT0000127 | Snx20 | 71607 | HITS-CLIP | 23142080 | Unspecified |
| mmu-miR-29b-3p | MIMAT0000127 | Snx24 | 69226 | HITS-CLIP | 37986224 | Unspecified |
| mmu-miR-29b-3p | MIMAT0000127 | Snx25 | 102141 | HITS-CLIP | 23142080 | Unspecified |
| mmu-miR-29b-3p | MIMAT0000127 | Snx33 | 235406 | HITS-CLIP | 37986224 | Unspecified |
| mmu-miR-29b-3p | MIMAT0000127 | Snx4 | 69150 | sRNA-Seq//HITS-CLIP | 33184237\|23142080 | Unspecified |
| mmu-miR-29b-3p | MIMAT0000127 | Snx8 | 231834 | HITS-CLIP//sRNA-Seq | 25083871\|33184237\|23142080 | Unspecified |
| mmu-miR-29b-3p | MIMAT0000127 | Snx9 | 66616 | sRNA-Seq//HITS-CLIP | 33184237\|25083871 | Unspecified |
| mmu-miR-29b-3p | MIMAT0000127 | Soat1 | 20652 | HITS-CLIP | 37986224\|25083871\|23142080 | Unspecified |
| mmu-miR-29b-3p | MIMAT0000127 | Sobp | 109205 | HITS-CLIP | 25083871 | Unspecified |
| mmu-miR-29b-3p | MIMAT0000127 | Socs1 | 12703 | HITS-CLIP | 23142080 | Unspecified |
| mmu-miR-29b-3p | MIMAT0000127 | Socs2 | 216233 | HITS-CLIP | 25083871 | Unspecified |
| mmu-miR-29b-3p | MIMAT0000127 | Socs3 | 12702 | HITS-CLIP | 23597149\|31433990\|23142080 | Unspecified |
| mmu-miR-29b-3p | MIMAT0000127 | Socs4 | 67296 | HITS-CLIP | 23142080 | Unspecified |
| mmu-miR-29b-3p | MIMAT0000127 | Socs5 | 56468 | HITS-CLIP//sRNA-Seq | 37986224\|25083871\|33184237\|23142 | Unspecified |
| mmu-miR-29b-3p | MIMAT0000127 | Socs6 | 54607 | HITS-CLIP | 25083871 | Unspecified |
| mmu-miR-29b-3p | MIMAT0000127 | Socs7 | 192157 | HITS-CLIP | 37986224\|25083871\|23142080 | Unspecified |
| mmu-miR-29b-3p | MIMAT0000127 | Sod1 | 20655 | sRNA-Seq//HITS-CLIP | 33184237\|23142080 | Unspecified |

| | | | | | | |
|---|---|---|---|---|---|---|
| mmu-miR-29b-3p | MIMAT0000127 | Soga1 | 320706 | HITS-CLIP | 25083871 | Unspecified |
| mmu-miR-29b-3p | MIMAT0000127 | Son | 20658 | HITS-CLIP//sRNA-Seq | 37986224\|25083871\|23597149\|33184 | Unspecified |
| mmu-miR-29b-3p | MIMAT0000127 | Sorbs2 | 234214 | HITS-CLIP | 25083871\|23597149 | Unspecified |
| mmu-miR-29b-3p | MIMAT0000127 | Sorcs2 | 81840 | HITS-CLIP | 37986224 | Unspecified |
| mmu-miR-29b-3p | MIMAT0000127 | Sord | 20322 | HITS-CLIP//sRNA-Seq | 23597149\|33184237 | Unspecified |
| mmu-miR-29b-3p | MIMAT0000127 | Sort1 | 20661 | sRNA-Seq//HITS-CLIP | 33184237\|23597149 | Unspecified |
| mmu-miR-29b-3p | MIMAT0000127 | Sos1 | 20662 | HITS-CLIP | 37986224\|25083871\|23597149\|23142 | Unspecified |
| mmu-miR-29b-3p | MIMAT0000127 | Sos2 | 20663 | HITS-CLIP | 37986224\|25083871\|23142080 | Unspecified |
| mmu-miR-29b-3p | MIMAT0000127 | Sowahc | 268301 | HITS-CLIP | 23142080\|25083871 | Unspecified |
| mmu-miR-29b-3p | MIMAT0000127 | Sox9 | 20682 | HITS-CLIP | 25083871 | Unspecified |
| mmu-miR-29b-3p | MIMAT0000127 | Sp1 | 20683 | HITS-CLIP//sRNA-Seq | 37986224\|33184237\|23142080\|25083 | Unspecified |
| mmu-miR-29b-3p | MIMAT0000127 | Sp2 | 78912 | HITS-CLIP | 23142080 | Unspecified |
| mmu-miR-29b-3p | MIMAT0000127 | Sp4 | 20688 | HITS-CLIP | 23142080 | Unspecified |
| mmu-miR-29b-3p | MIMAT0000127 | Spag9 | 70834 | sRNA-Seq//HITS-CLIP | 33184237\|23142080 | Unspecified |
| mmu-miR-29b-3p | MIMAT0000127 | Spast | 50850 | HITS-CLIP//sRNA-Seq | 37986224\|23597149\|25083871\|33184 | Unspecified |
| mmu-miR-29b-3p | MIMAT0000127 | Spata2 | 263876 | HITS-CLIP | 25083871 | Unspecified |
| mmu-miR-29b-3p | MIMAT0000127 | Spata2l | 78779 | HITS-CLIP | 23597149 | Unspecified |
| mmu-miR-29b-3p | MIMAT0000127 | Spata5 | 57815 | HITS-CLIP | 23142080 | Unspecified |
| mmu-miR-29b-3p | MIMAT0000127 | Spata6 | 67946 | HITS-CLIP | 25083871\|23142080 | Unspecified |
| mmu-miR-29b-3p | MIMAT0000127 | Specc1 | 432572 | HITS-CLIP | 37986224 | Unspecified |
| mmu-miR-29b-3p | MIMAT0000127 | Specc1l | 74392 | HITS-CLIP//sRNA-Seq | 25083871\|33184237 | Unspecified |
| mmu-miR-29b-3p | MIMAT0000127 | Speg | 11790 | HITS-CLIP | 25083871 | Unspecified |
| mmu-miR-29b-3p | MIMAT0000127 | Spen | 56381 | HITS-CLIP | 25083871\|23597149\|23142080 | Unspecified |
| mmu-miR-29b-3p | MIMAT0000127 | Spg11 | 214585 | HITS-CLIP | 23142080 | Unspecified |
| mmu-miR-29b-3p | MIMAT0000127 | Spg20 | 229285 | HITS-CLIP//sRNA-Seq | 25083871\|33184237 | Unspecified |
| mmu-miR-29b-3p | MIMAT0000127 | Spg7 | 234847 | HITS-CLIP | 23142080 | Unspecified |
| mmu-miR-29b-3p | MIMAT0000127 | Sphk2 | 56632 | HITS-CLIP | 23597149 | Unspecified |
| mmu-miR-29b-3p | MIMAT0000127 | Spice1 | 212514 | HITS-CLIP | 23142080 | Unspecified |
| mmu-miR-29b-3p | MIMAT0000127 | Spin4 | 270624 | HITS-CLIP | 37986224\|23142080 | Unspecified |
| mmu-miR-29b-3p | MIMAT0000127 | Spint2 | 20733 | HITS-CLIP | 23597149\|23142080 | Unspecified |
| mmu-miR-29b-3p | MIMAT0000127 | Spire1 | 68166 | HITS-CLIP | 25083871 | Unspecified |
| mmu-miR-29b-3p | MIMAT0000127 | Spns1 | 73658 | HITS-CLIP | 37986224\|23597149\|23142080 | Unspecified |
| mmu-miR-29b-3p | MIMAT0000127 | Spock2 | 94214 | HITS-CLIP//sRNA-Seq | 25083871\|33184237 | Unspecified |
| mmu-miR-29b-3p | MIMAT0000127 | Spop | 20747 | HITS-CLIP//sRNA-Seq | 37986224\|25083871\|33184237\|23142 | Unspecified |
| mmu-miR-29b-3p | MIMAT0000127 | Spopl | 76857 | HITS-CLIP | 25083871 | Unspecified |
| mmu-miR-29b-3p | MIMAT0000127 | Spp1 | 20750 | HITS-CLIP//sRNA-Seq | 25083871\|33184237\|23142080 | Unspecified |
| mmu-miR-29b-3p | MIMAT0000127 | Sppl2a | 66552 | HITS-CLIP | 23597149\|25083871\|23142080 | Unspecified |
| mmu-miR-29b-3p | MIMAT0000127 | Sppl3 | 74585 | HITS-CLIP//sRNA-Seq | 25083871\|23597149\|33184237\|23142 | Unspecified |
| mmu-miR-29b-3p | MIMAT0000127 | Spr | 20751 | HITS-CLIP | 23597149 | Unspecified |

| | | | | | | |
|---|---|---|---|---|---|---|
| mmu-miR-29b-3p | MIMAT0000127 | Spred1 | 114715 | HITS-CLIP | 25083871\|23142080 | Unspecified |
| mmu-miR-29b-3p | MIMAT0000127 | Spred2 | 114716 | HITS-CLIP | 23142080 | Unspecified |
| mmu-miR-29b-3p | MIMAT0000127 | Sprtn | 244666 | HITS-CLIP | 23597149\|25083871\|23142080 | Unspecified |
| mmu-miR-29b-3p | MIMAT0000127 | Spry1 | 24063 | HITS-CLIP | 25083871\|23142080 | Unspecified |
| mmu-miR-29b-3p | MIMAT0000127 | Spry2 | 24064 | HITS-CLIP//sRNA-Seq | 23142080\|33184237 | Unspecified |
| mmu-miR-29b-3p | MIMAT0000127 | Spry4 | 24066 | HITS-CLIP//sRNA-Seq | 37986224\|23597149\|33184237 | Unspecified |
| mmu-miR-29b-3p | MIMAT0000127 | Spryd3 | 223918 | HITS-CLIP//sRNA-Seq | 37986224\|23142080\|25083871\|23597 | Unspecified |
| mmu-miR-29b-3p | MIMAT0000127 | Spryd4 | 66701 | HITS-CLIP | 37986224\|23142080 | Unspecified |
| mmu-miR-29b-3p | MIMAT0000127 | Spsb1 | 74646 | HITS-CLIP | 25083871\|23597149\|23142080 | Unspecified |
| mmu-miR-29b-3p | MIMAT0000127 | Spsb4 | 211949 | HITS-CLIP | 37986224 | Unspecified |
| mmu-miR-29b-3p | MIMAT0000127 | Sptan1 | 20740 | HITS-CLIP//sRNA-Seq | 37986224\|25083871\|33184237\|23142 | Unspecified |
| mmu-miR-29b-3p | MIMAT0000127 | Sptbn1 | 20742 | HITS-CLIP//sRNA-Seq | 37986224\|25083871\|23142080\|33184 | Unspecified |
| mmu-miR-29b-3p | MIMAT0000127 | Sptbn2 | 20743 | HITS-CLIP//sRNA-Seq | 23597149\|33184237 | Unspecified |
| mmu-miR-29b-3p | MIMAT0000127 | Sptlc1 | 268656 | HITS-CLIP | 23142080 | Unspecified |
| mmu-miR-29b-3p | MIMAT0000127 | Sptlc2 | 20773 | HITS-CLIP//sRNA-Seq | 37986224\|23597149\|25083871\|33184 | Unspecified |
| mmu-miR-29b-3p | MIMAT0000127 | Spty2d1 | 101685 | HITS-CLIP | 37986224\|25083871\|23597149\|23142 | Unspecified |
| mmu-miR-29b-3p | MIMAT0000127 | Sqle | 20775 | HITS-CLIP//sRNA-Seq | 37986224\|23142080\|33184237\|25083 | Unspecified |
| mmu-miR-29b-3p | MIMAT0000127 | Sqor | 58472 | HITS-CLIP | 23597149 | Unspecified |
| mmu-miR-29b-3p | MIMAT0000127 | Srcap | 100043597 | HITS-CLIP | 37986224\|25083871 | Unspecified |
| mmu-miR-29b-3p | MIMAT0000127 | Srd5a1 | 78925 | HITS-CLIP | 37986224\|23142080 | Unspecified |
| mmu-miR-29b-3p | MIMAT0000127 | Srebf1 | 20787 | sRNA-Seq//HITS-CLIP | 33184237\|23142080 | Unspecified |
| mmu-miR-29b-3p | MIMAT0000127 | Srebf2 | 20788 | HITS-CLIP//sRNA-Seq | 37986224\|23597149\|23142080\|33184 | Unspecified |
| mmu-miR-29b-3p | MIMAT0000127 | Srf | 20807 | HITS-CLIP//sRNA-Seq | 23142080\|33184237 | Unspecified |
| mmu-miR-29b-3p | MIMAT0000127 | Srgap2 | 14270 | HITS-CLIP | 37986224\|23142080 | Unspecified |
| mmu-miR-29b-3p | MIMAT0000127 | Srgap3 | 259302 | HITS-CLIP//sRNA-Seq | 25083871\|33184237 | Unspecified |
| mmu-miR-29b-3p | MIMAT0000127 | Srl | 106393 | HITS-CLIP | 25083871 | Unspecified |
| mmu-miR-29b-3p | MIMAT0000127 | Srp14 | 20813 | sRNA-Seq//HITS-CLIP | 33184237\|25083871 | Unspecified |
| mmu-miR-29b-3p | MIMAT0000127 | Srp68 | 217337 | HITS-CLIP//sRNA-Seq | 23597149\|25083871\|33184237\|23142 | Unspecified |
| mmu-miR-29b-3p | MIMAT0000127 | Srp72 | 66661 | HITS-CLIP//sRNA-Seq | 37986224\|25083871\|23597149\|33184 | Unspecified |
| mmu-miR-29b-3p | MIMAT0000127 | Srpk1 | 20815 | HITS-CLIP//sRNA-Seq | 23142080\|33184237 | Unspecified |
| mmu-miR-29b-3p | MIMAT0000127 | Srpr | 67398 | HITS-CLIP//sRNA-Seq | 23597149\|25083871\|33184237\|23142 | Unspecified |
| mmu-miR-29b-3p | MIMAT0000127 | Srpx2 | 68792 | HITS-CLIP | 25083871 | Unspecified |
| mmu-miR-29b-3p | MIMAT0000127 | Srr | 27364 | HITS-CLIP | 37986224 | Unspecified |
| mmu-miR-29b-3p | MIMAT0000127 | Srrm2 | 75956 | HITS-CLIP//sRNA-Seq | 23142080\|25083871\|33184237\|31433 | Unspecified |
| mmu-miR-29b-3p | MIMAT0000127 | Srsf1 | 110809 | HITS-CLIP//sRNA-Seq | 37986224\|23142080\|25083871\|33184 | Unspecified |
| mmu-miR-29b-3p | MIMAT0000127 | Srsf10 | 14105 | sRNA-Seq//HITS-CLIP | 33184237\|23142080 | Unspecified |
| mmu-miR-29b-3p | MIMAT0000127 | Srsf2 | 20382 | HITS-CLIP//sRNA-Seq | 37986224\|33184237\|23142080 | Unspecified |
| mmu-miR-29b-3p | MIMAT0000127 | Srsf3 | 20383 | HITS-CLIP//sRNA-Seq | 23142080\|33184237 | Unspecified |
| mmu-miR-29b-3p | MIMAT0000127 | Srsf6 | 67996 | HITS-CLIP//sRNA-Seq | 25083871\|33184237 | Unspecified |

| | | | | | | |
|---|---|---|---|---|---|---|
| mmu-miR-29b-3p | MIMAT0000127 | Srsf7 | 225027 | HITS-CLIP//sRNA-Seq | 23142080\|33184237\|25083871 | Unspecified |
| mmu-miR-29b-3p | MIMAT0000127 | Ss18 | 268996 | sRNA-Seq//HITS-CLIP | 33184237\|23597149\|23142080 | Unspecified |
| mmu-miR-29b-3p | MIMAT0000127 | Ss18l1 | 269397 | HITS-CLIP | 37986224\|25083871 | Unspecified |
| mmu-miR-29b-3p | MIMAT0000127 | Ssb | 20823 | HITS-CLIP//sRNA-Seq | 37986224\|23142080\|25083871\|3318[illegible] | Unspecified |
| mmu-miR-29b-3p | MIMAT0000127 | Ssbp2 | 66970 | HITS-CLIP//sRNA-Seq | 25083871\|33184237 | Unspecified |
| mmu-miR-29b-3p | MIMAT0000127 | Ssh1 | 231637 | HITS-CLIP | 23142080 | Unspecified |
| mmu-miR-29b-3p | MIMAT0000127 | Ssh2 | 237860 | HITS-CLIP | 25083871\|23597149\|23142080 | Unspecified |
| mmu-miR-29b-3p | MIMAT0000127 | Ssna1 | 68475 | sRNA-Seq//HITS-CLIP | 33184237\|23142080 | Unspecified |
| mmu-miR-29b-3p | MIMAT0000127 | Sspn | 16651 | HITS-CLIP | 37986224\|25083871 | Unspecified |
| mmu-miR-29b-3p | MIMAT0000127 | Ssr2 | 66256 | HITS-CLIP//sRNA-Seq | 25083871\|33184237\|23597149\|2314[illegible] | Unspecified |
| mmu-miR-29b-3p | MIMAT0000127 | Ssr3 | 67437 | HITS-CLIP//sRNA-Seq | 23597149\|25083871\|33184237\|2314[illegible] | Unspecified |
| mmu-miR-29b-3p | MIMAT0000127 | Ssrp1 | 20833 | sRNA-Seq//HITS-CLIP | 33184237\|23142080 | Unspecified |
| mmu-miR-29b-3p | MIMAT0000127 | Ssx2ip | 99167 | HITS-CLIP | 25083871 | Unspecified |
| mmu-miR-29b-3p | MIMAT0000127 | St3gal1 | 20442 | HITS-CLIP | 23142080 | Unspecified |
| mmu-miR-29b-3p | MIMAT0000127 | St3gal3 | 20441 | HITS-CLIP | 23597149 | Unspecified |
| mmu-miR-29b-3p | MIMAT0000127 | St3gal4 | 20443 | HITS-CLIP | 23597149 | Unspecified |
| mmu-miR-29b-3p | MIMAT0000127 | St3gal6 | 54613 | HITS-CLIP | 37986224 | Unspecified |
| mmu-miR-29b-3p | MIMAT0000127 | St5 | 76954 | HITS-CLIP | 25083871 | Unspecified |
| mmu-miR-29b-3p | MIMAT0000127 | St6galnac6 | 50935 | HITS-CLIP//sRNA-Seq | 23597149\|33184237 | Unspecified |
| mmu-miR-29b-3p | MIMAT0000127 | St8sia1 | 20449 | HITS-CLIP | 23142080 | Unspecified |
| mmu-miR-29b-3p | MIMAT0000127 | St8sia2 | 20450 | HITS-CLIP | 25083871 | Unspecified |
| mmu-miR-29b-3p | MIMAT0000127 | St8sia4 | 20452 | HITS-CLIP | 23142080 | Unspecified |
| mmu-miR-29b-3p | MIMAT0000127 | Stab2 | 192188 | HITS-CLIP | 37986224\|23597149 | Unspecified |
| mmu-miR-29b-3p | MIMAT0000127 | Stag2 | 20843 | HITS-CLIP//sRNA-Seq | 37986224\|25083871\|33184237\|2314[illegible] | Unspecified |
| mmu-miR-29b-3p | MIMAT0000127 | Stam | 20844 | sRNA-Seq//HITS-CLIP | 33184237\|23142080 | Unspecified |
| mmu-miR-29b-3p | MIMAT0000127 | Stambp | 70527 | HITS-CLIP | 25083871 | Unspecified |
| mmu-miR-29b-3p | MIMAT0000127 | Stambpl1 | 76630 | HITS-CLIP | 23142080 | Unspecified |
| mmu-miR-29b-3p | MIMAT0000127 | Stard10 | 56018 | HITS-CLIP | 23597149 | Unspecified |
| mmu-miR-29b-3p | MIMAT0000127 | Stard13 | 243362 | HITS-CLIP | 23597149 | Unspecified |
| mmu-miR-29b-3p | MIMAT0000127 | Stard3 | 59045 | HITS-CLIP | 37986224\|23597149 | Unspecified |
| mmu-miR-29b-3p | MIMAT0000127 | Stard5 | 170460 | HITS-CLIP | 23597149 | Unspecified |
| mmu-miR-29b-3p | MIMAT0000127 | Stard7 | 99138 | HITS-CLIP | 37986224 | Unspecified |
| mmu-miR-29b-3p | MIMAT0000127 | Stard9 | 668880 | HITS-CLIP | 37986224 | Unspecified |
| mmu-miR-29b-3p | MIMAT0000127 | Stat1 | 20846 | HITS-CLIP | 23142080 | Unspecified |
| mmu-miR-29b-3p | MIMAT0000127 | Stat3 | 20848 | HITS-CLIP//sRNA-Seq | 23597149\|33184237\|23142080 | Unspecified |
| mmu-miR-29b-3p | MIMAT0000127 | Stat4 | 20849 | HITS-CLIP | 23142080 | Unspecified |
| mmu-miR-29b-3p | MIMAT0000127 | Stat5a | 20850 | HITS-CLIP | 23142080 | Unspecified |
| mmu-miR-29b-3p | MIMAT0000127 | Stat5b | 20851 | HITS-CLIP | 25083871\|23142080 | Unspecified |
| mmu-miR-29b-3p | MIMAT0000127 | Stat6 | 20852 | sRNA-Seq//HITS-CLIP | 33184237\|23142080 | Unspecified |

| | | | | | | |
|---|---|---|---|---|---|---|
| mmu-miR-29b-3p | MIMAT0000127 | Stau1 | 20853 | HITS-CLIP//sRNA-Seq | 25083871\|23597149\|33184237\|23142 | Unspecified |
| mmu-miR-29b-3p | MIMAT0000127 | Stc2 | 20856 | HITS-CLIP | 23142080 | Unspecified |
| mmu-miR-29b-3p | MIMAT0000127 | Steap4 | 117167 | HITS-CLIP | 23597149 | Unspecified |
| mmu-miR-29b-3p | MIMAT0000127 | Stim1 | 20866 | HITS-CLIP | 37986224\|23142080 | Unspecified |
| mmu-miR-29b-3p | MIMAT0000127 | Stim2 | 116873 | HITS-CLIP | 25083871\|23142080 | Unspecified |
| mmu-miR-29b-3p | MIMAT0000127 | Stk10 | 20868 | HITS-CLIP | 23142080 | Unspecified |
| mmu-miR-29b-3p | MIMAT0000127 | Stk16 | 20872 | HITS-CLIP | 37986224 | Unspecified |
| mmu-miR-29b-3p | MIMAT0000127 | Stk17b | 98267 | HITS-CLIP | 23142080 | Unspecified |
| mmu-miR-29b-3p | MIMAT0000127 | Stk35 | 67333 | HITS-CLIP | 23142080 | Unspecified |
| mmu-miR-29b-3p | MIMAT0000127 | Stk38 | 106504 | HITS-CLIP//sRNA-Seq | 37986224\|33184237\|23142080 | Unspecified |
| mmu-miR-29b-3p | MIMAT0000127 | Stk39 | 53416 | sRNA-Seq//HITS-CLIP | 33184237\|23142080 | Unspecified |
| mmu-miR-29b-3p | MIMAT0000127 | Stk4 | 58231 | HITS-CLIP | 37986224\|23142080 | Unspecified |
| mmu-miR-29b-3p | MIMAT0000127 | Stk40 | 74178 | HITS-CLIP | 25083871\|23597149\|23142080 | Unspecified |
| mmu-miR-29b-3p | MIMAT0000127 | Ston1 | 77057 | HITS-CLIP//sRNA-Seq | 25083871\|33184237 | Unspecified |
| mmu-miR-29b-3p | MIMAT0000127 | Stra6l | 74152 | HITS-CLIP | 23597149 | Unspecified |
| mmu-miR-29b-3p | MIMAT0000127 | Strada | 72149 | HITS-CLIP | 25083871 | Unspecified |
| mmu-miR-29b-3p | MIMAT0000127 | Stradb | 227154 | HITS-CLIP | 25083871\|23597149\|23142080 | Unspecified |
| mmu-miR-29b-3p | MIMAT0000127 | Strap | 20901 | HITS-CLIP//sRNA-Seq | 25083871\|33184237\|23142080 | Unspecified |
| mmu-miR-29b-3p | MIMAT0000127 | Strbp | 20744 | sRNA-Seq//HITS-CLIP | 33184237\|23142080 | Unspecified |
| mmu-miR-29b-3p | MIMAT0000127 | Strn | 268980 | HITS-CLIP | 37986224\|23142080\|25083871 | Unspecified |
| mmu-miR-29b-3p | MIMAT0000127 | Strn3 | 94186 | HITS-CLIP//sRNA-Seq | 37986224\|23597149\|25083871\|23142 | Unspecified |
| mmu-miR-29b-3p | MIMAT0000127 | Strn4 | 97387 | HITS-CLIP//sRNA-Seq | 23142080\|33184237 | Unspecified |
| mmu-miR-29b-3p | MIMAT0000127 | Stt3a | 16430 | HITS-CLIP//sRNA-Seq | 37986224\|25083871\|33184237\|23597 | Unspecified |
| mmu-miR-29b-3p | MIMAT0000127 | Stt3b | 68292 | sRNA-Seq//HITS-CLIP | 33184237\|23142080 | Unspecified |
| mmu-miR-29b-3p | MIMAT0000127 | Stx11 | 74732 | HITS-CLIP | 37986224\|23142080 | Unspecified |
| mmu-miR-29b-3p | MIMAT0000127 | Stx12 | 100226 | HITS-CLIP//sRNA-Seq | 25083871\|33184237\|23142080 | Unspecified |
| mmu-miR-29b-3p | MIMAT0000127 | Stx16 | 228960 | HITS-CLIP | 37986224\|25083871\|23142080 | Unspecified |
| mmu-miR-29b-3p | MIMAT0000127 | Stx17 | 67727 | HITS-CLIP | 37986224\|25083871 | Unspecified |
| mmu-miR-29b-3p | MIMAT0000127 | Stx5a | 56389 | HITS-CLIP//sRNA-Seq | 23597149\|33184237 | Unspecified |
| mmu-miR-29b-3p | MIMAT0000127 | Stx7 | 53331 | sRNA-Seq//HITS-CLIP | 33184237\|25083871 | Unspecified |
| mmu-miR-29b-3p | MIMAT0000127 | Stxbp2 | 20911 | HITS-CLIP | 23142080 | Unspecified |
| mmu-miR-29b-3p | MIMAT0000127 | Stxbp3 | 20912 | HITS-CLIP | 25083871\|23142080 | Unspecified |
| mmu-miR-29b-3p | MIMAT0000127 | Sucla2 | 20916 | HITS-CLIP//sRNA-Seq | 25083871\|23142080\|33184237\|23597 | Unspecified |
| mmu-miR-29b-3p | MIMAT0000127 | Suco | 226551 | HITS-CLIP | 37986224\|25083871\|23142080 | Unspecified |
| mmu-miR-29b-3p | MIMAT0000127 | Suds3 | 71954 | HITS-CLIP | 30971389 | Unspecified |
| mmu-miR-29b-3p | MIMAT0000127 | Sufu | 24069 | HITS-CLIP | 25083871\|23142080 | Unspecified |
| mmu-miR-29b-3p | MIMAT0000127 | Sugp2 | 234373 | HITS-CLIP | 37986224\|25083871\|23142080 | Unspecified |
| mmu-miR-29b-3p | MIMAT0000127 | Sulf1 | 240725 | HITS-CLIP | 25083871 | Unspecified |
| mmu-miR-29b-3p | MIMAT0000127 | Sult2a8 | 76971 | HITS-CLIP | 23597149 | Unspecified |

| | | | | | | |
|---|---|---|---|---|---|---|
| mmu-miR-29b-3p | MIMAT0000127 | Sun1 | 77053 | HITS-CLIP//sRNA-Seq | 25083871\|23597149\|33184237 | Unspecified |
| mmu-miR-29b-3p | MIMAT0000127 | Sun2 | 223697 | sRNA-Seq//HITS-CLIP | 33184237\|23142080 | Unspecified |
| mmu-miR-29b-3p | MIMAT0000127 | Supt16 | 114741 | HITS-CLIP//sRNA-Seq | 25083871\|33184237\|23142080 | Unspecified |
| mmu-miR-29b-3p | MIMAT0000127 | Supt3 | 109115 | HITS-CLIP | 25083871 | Unspecified |
| mmu-miR-29b-3p | MIMAT0000127 | Supt5 | 20924 | HITS-CLIP//sRNA-Seq | 23142080\|33184237\|23597149 | Unspecified |
| mmu-miR-29b-3p | MIMAT0000127 | Supt6 | 20926 | sRNA-Seq//HITS-CLIP | 33184237\|25083871 | Unspecified |
| mmu-miR-29b-3p | MIMAT0000127 | Surf4 | 20932 | HITS-CLIP//sRNA-Seq | 23597149\|33184237\|23142080\|2508: | Unspecified |
| mmu-miR-29b-3p | MIMAT0000127 | Surf6 | 20935 | HITS-CLIP | 37986224 | Unspecified |
| mmu-miR-29b-3p | MIMAT0000127 | Susd6 | 9766 | HITS-CLIP | 25083871 | Unspecified |
| mmu-miR-29b-3p | MIMAT0000127 | Suv39h1 | 20937 | sRNA-Seq//HITS-CLIP | 33184237\|23142080 | Unspecified |
| mmu-miR-29b-3p | MIMAT0000127 | Suz12 | 52615 | HITS-CLIP//sRNA-Seq | 23142080\|33184237\|25083871 | Unspecified |
| mmu-miR-29b-3p | MIMAT0000127 | Svil | 225115 | HITS-CLIP | 37986224\|25083871\|23597149\|2314: | Unspecified |
| mmu-miR-29b-3p | MIMAT0000127 | Swap70 | 20947 | HITS-CLIP | 37986224 | Unspecified |
| mmu-miR-29b-3p | MIMAT0000127 | Swi5 | 72931 | HITS-CLIP//sRNA-Seq | 25083871\|33184237\|23597149\|2314: | Unspecified |
| mmu-miR-29b-3p | MIMAT0000127 | Syncrip | 56403 | HITS-CLIP//sRNA-Seq | 37986224\|25083871\|33184237\|2314: | Unspecified |
| mmu-miR-29b-3p | MIMAT0000127 | Syne1 | 64009 | HITS-CLIP | 37986224 | Unspecified |
| mmu-miR-29b-3p | MIMAT0000127 | Syne2 | 319565 | HITS-CLIP | 23597149 | Unspecified |
| mmu-miR-29b-3p | MIMAT0000127 | Syne3 | 212073 | HITS-CLIP | 23142080 | Unspecified |
| mmu-miR-29b-3p | MIMAT0000127 | Syngr2 | 20973 | HITS-CLIP//sRNA-Seq | 23597149\|33184237\|23142080 | Unspecified |
| mmu-miR-29b-3p | MIMAT0000127 | Synj1 | 104015 | sRNA-Seq//HITS-CLIP | 33184237\|23142080 | Unspecified |
| mmu-miR-29b-3p | MIMAT0000127 | Synj2 | 20975 | HITS-CLIP | 25083871 | Unspecified |
| mmu-miR-29b-3p | MIMAT0000127 | Synrg | 217030 | HITS-CLIP | 23597149 | Unspecified |
| mmu-miR-29b-3p | MIMAT0000127 | Sypl | 19027 | HITS-CLIP | 25083871\|23597149\|23142080 | Unspecified |
| mmu-miR-29b-3p | MIMAT0000127 | Syt1 | 20979 | sRNA-Seq//HITS-CLIP | 33184237\|28735896 | Unspecified |
| mmu-miR-29b-3p | MIMAT0000127 | Sytl2 | 83671 | HITS-CLIP | 25083871 | Unspecified |
| mmu-miR-29b-3p | MIMAT0000127 | Syvn1 | 74126 | HITS-CLIP | 37986224\|23142080\|25083871 | Unspecified |
| mmu-miR-29b-3p | MIMAT0000127 | Szrd1 | 213491 | HITS-CLIP | 37986224 | Unspecified |
| mmu-miR-29b-3p | MIMAT0000127 | Tab1 | 66513 | HITS-CLIP | 37986224\|23142080 | Unspecified |
| mmu-miR-29b-3p | MIMAT0000127 | Tab2 | 68652 | HITS-CLIP//sRNA-Seq | 37986224\|23142080\|25083871\|3318 | Unspecified |
| mmu-miR-29b-3p | MIMAT0000127 | Tacc1 | 320165 | HITS-CLIP | 37986224\|23142080 | Unspecified |
| mmu-miR-29b-3p | MIMAT0000127 | Tacc3 | 21335 | sRNA-Seq//HITS-CLIP | 33184237\|23142080 | Unspecified |
| mmu-miR-29b-3p | MIMAT0000127 | Tada1 | 27878 | HITS-CLIP | 25083871\|23142080 | Unspecified |
| mmu-miR-29b-3p | MIMAT0000127 | Tada2b | 231151 | HITS-CLIP | 23597149 | Unspecified |
| mmu-miR-29b-3p | MIMAT0000127 | Tada3 | 101206 | HITS-CLIP | 25083871 | Unspecified |
| mmu-miR-29b-3p | MIMAT0000127 | Taf1 | 270627 | HITS-CLIP//sRNA-Seq | 25083871\|33184237\|23142080 | Unspecified |
| mmu-miR-29b-3p | MIMAT0000127 | Taf13 | 99730 | HITS-CLIP | 37986224\|25083871\|23142080 | Unspecified |
| mmu-miR-29b-3p | MIMAT0000127 | Taf2 | 319944 | HITS-CLIP | 23142080 | Unspecified |
| mmu-miR-29b-3p | MIMAT0000127 | Taf5 | 226182 | HITS-CLIP | 37986224\|25083871\|23142080 | Unspecified |
| mmu-miR-29b-3p | MIMAT0000127 | Taf6 | 21343 | HITS-CLIP//sRNA-Seq | 25083871\|33184237 | Unspecified |

| | | | | | | |
|---|---|---|---|---|---|---|
| mmu-miR-29b-3p | MIMAT0000127 | Taf7 | 24074 | HITS-CLIP | 37986224\|23142080 | Unspecified |
| mmu-miR-29b-3p | MIMAT0000127 | Taf9 | 108143 | HITS-CLIP | 37986224\|23142080 | Unspecified |
| mmu-miR-29b-3p | MIMAT0000127 | Tanc1 | 66860 | HITS-CLIP//sRNA-Seq | 25083871\|33184237\|23597149\|2314 | Unspecified |
| mmu-miR-29b-3p | MIMAT0000127 | Tanc2 | 77097 | HITS-CLIP//sRNA-Seq | 25083871\|33184237\|23142080 | Unspecified |
| mmu-miR-29b-3p | MIMAT0000127 | Taok1 | 216965 | HITS-CLIP//sRNA-Seq | 37986224\|25083871\|33184237\|2359 | Unspecified |
| mmu-miR-29b-3p | MIMAT0000127 | Taok2 | 381921 | HITS-CLIP//sRNA-Seq | 25083871\|33184237\|23142080 | Unspecified |
| mmu-miR-29b-3p | MIMAT0000127 | Taok3 | 330177 | HITS-CLIP | 23142080 | Unspecified |
| mmu-miR-29b-3p | MIMAT0000127 | Tap1 | 21354 | HITS-CLIP | 23142080 | Unspecified |
| mmu-miR-29b-3p | MIMAT0000127 | Tap2 | 21355 | HITS-CLIP | 37986224\|23597149\|23142080 | Unspecified |
| mmu-miR-29b-3p | MIMAT0000127 | Tapbp | 21356 | HITS-CLIP//sRNA-Seq | 23597149\|25083871\|33184237\|2314 | Unspecified |
| mmu-miR-29b-3p | MIMAT0000127 | Tapt1 | 231225 | HITS-CLIP | 23142080 | Unspecified |
| mmu-miR-29b-3p | MIMAT0000127 | Tarbp1 | 6894 | HITS-CLIP | 23142080\|25083871 | Unspecified |
| mmu-miR-29b-3p | MIMAT0000127 | Tardbp | 230908 | HITS-CLIP//sRNA-Seq | 37986224\|25083871\|33184237\|2314 | Unspecified |
| mmu-miR-29b-3p | MIMAT0000127 | Tars2 | 71807 | HITS-CLIP | 23142080 | Unspecified |
| mmu-miR-29b-3p | MIMAT0000127 | Tasp1 | 75812 | HITS-CLIP | 23142080 | Unspecified |
| mmu-miR-29b-3p | MIMAT0000127 | Tat | 234724 | HITS-CLIP | 23597149 | Unspecified |
| mmu-miR-29b-3p | MIMAT0000127 | Tax1bp1 | 52440 | HITS-CLIP//sRNA-Seq | 25083871\|23597149\|33184237\|2314 | Unspecified |
| mmu-miR-29b-3p | MIMAT0000127 | Tax1bp3 | 76281 | HITS-CLIP | 25083871 | Unspecified |
| mmu-miR-29b-3p | MIMAT0000127 | Tbc1d1 | 57915 | HITS-CLIP | 37986224\|25083871\|23142080 | Unspecified |
| mmu-miR-29b-3p | MIMAT0000127 | Tbc1d10b | 68449 | HITS-CLIP | 23142080 | Unspecified |
| mmu-miR-29b-3p | MIMAT0000127 | Tbc1d10c | 108995 | HITS-CLIP | 23142080 | Unspecified |
| mmu-miR-29b-3p | MIMAT0000127 | Tbc1d13 | 70296 | HITS-CLIP | 37986224 | Unspecified |
| mmu-miR-29b-3p | MIMAT0000127 | Tbc1d14 | 100855 | HITS-CLIP//sRNA-Seq | 23142080\|33184237 | Unspecified |
| mmu-miR-29b-3p | MIMAT0000127 | Tbc1d15 | 66687 | HITS-CLIP | 37986224 | Unspecified |
| mmu-miR-29b-3p | MIMAT0000127 | Tbc1d20 | 67231 | HITS-CLIP//sRNA-Seq | 23597149\|33184237\|23142080 | Unspecified |
| mmu-miR-29b-3p | MIMAT0000127 | Tbc1d22b | 381085 | HITS-CLIP | 37986224\|23142080 | Unspecified |
| mmu-miR-29b-3p | MIMAT0000127 | Tbc1d24 | 224617 | HITS-CLIP | 23597149 | Unspecified |
| mmu-miR-29b-3p | MIMAT0000127 | Tbc1d25 | 209815 | HITS-CLIP | 23142080 | Unspecified |
| mmu-miR-29b-3p | MIMAT0000127 | Tbc1d2b | 67016 | HITS-CLIP | 30971389\|23142080 | Unspecified |
| mmu-miR-29b-3p | MIMAT0000127 | Tbc1d31 | 210544 | HITS-CLIP | 23597149 | Unspecified |
| mmu-miR-29b-3p | MIMAT0000127 | Tbc1d7 | 67046 | HITS-CLIP | 25083871\|23142080\|23597149 | Unspecified |
| mmu-miR-29b-3p | MIMAT0000127 | Tbc1d9b | 76795 | HITS-CLIP//sRNA-Seq | 25083871\|33184237 | Unspecified |
| mmu-miR-29b-3p | MIMAT0000127 | Tbcb | 66411 | HITS-CLIP//sRNA-Seq | 25083871\|33184237 | Unspecified |
| mmu-miR-29b-3p | MIMAT0000127 | Tbccd1 | 70573 | HITS-CLIP | 23142080\|25083871 | Unspecified |
| mmu-miR-29b-3p | MIMAT0000127 | Tbcd | 108903 | HITS-CLIP//sRNA-Seq | 25083871\|23597149\|33184237\|2314 | Unspecified |
| mmu-miR-29b-3p | MIMAT0000127 | Tbce | 70430 | HITS-CLIP | 23597149\|23142080 | Unspecified |
| mmu-miR-29b-3p | MIMAT0000127 | Tbcel | 272589 | HITS-CLIP | 37986224\|25083871\|23597149\|2314 | Unspecified |
| mmu-miR-29b-3p | MIMAT0000127 | Tbk1 | 56480 | HITS-CLIP | 23597149\|31433990\|23142080\|2508 | Unspecified |
| mmu-miR-29b-3p | MIMAT0000127 | Tbl1x | 21372 | HITS-CLIP | 37986224 | Unspecified |

| | | | | | | |
|---|---|---|---|---|---|---|
| mmu-miR-29b-3p | MIMAT0000127 | Tbl1xr1 | 81004 | HITS-CLIP | 37986224\|23142080\|25083871\|23597 | Unspecified |
| mmu-miR-29b-3p | MIMAT0000127 | Tbl2 | 27368 | HITS-CLIP | 37986224\|23142080\|25083871\|23597 | Unspecified |
| mmu-miR-29b-3p | MIMAT0000127 | Tbl3 | 213773 | HITS-CLIP | 25083871\|23142080 | Unspecified |
| mmu-miR-29b-3p | MIMAT0000127 | Tbp | 21374 | HITS-CLIP | 25083871 | Unspecified |
| mmu-miR-29b-3p | MIMAT0000127 | Tbpl1 | 237336 | HITS-CLIP//sRNA-Seq | 25083871\|33184237\|23597149\|23142 | Unspecified |
| mmu-miR-29b-3p | MIMAT0000127 | Tbrg4 | 21379 | HITS-CLIP | 25083871 | Unspecified |
| mmu-miR-29b-3p | MIMAT0000127 | Tbx18 | 76365 | HITS-CLIP | 37986224\|25083871 | Unspecified |
| mmu-miR-29b-3p | MIMAT0000127 | Tbx21 | 57765 | Luciferase assay//Microarray//qRT-PCR//H | 21820330\|23142080 | Unspecified |
| mmu-miR-29b-3p | MIMAT0000127 | Tcaim | 382117 | HITS-CLIP | 23142080 | Unspecified |
| mmu-miR-29b-3p | MIMAT0000127 | Tceanc | 245695 | HITS-CLIP | 23142080 | Unspecified |
| mmu-miR-29b-3p | MIMAT0000127 | Tceanc2 | 66526 | HITS-CLIP | 37986224\|25083871 | Unspecified |
| mmu-miR-29b-3p | MIMAT0000127 | Tcerg1 | 56070 | HITS-CLIP//sRNA-Seq | 23142080\|33184237\|23597149\|25083 | Unspecified |
| mmu-miR-29b-3p | MIMAT0000127 | Tcf12 | 21406 | HITS-CLIP//sRNA-Seq | 23142080\|25083871\|33184237 | Unspecified |
| mmu-miR-29b-3p | MIMAT0000127 | Tcf20 | 21411 | HITS-CLIP//sRNA-Seq | 37986224\|25083871\|33184237 | Unspecified |
| mmu-miR-29b-3p | MIMAT0000127 | Tcf25 | 66855 | HITS-CLIP//sRNA-Seq | 23142080\|23597149\|33184237 | Unspecified |
| mmu-miR-29b-3p | MIMAT0000127 | Tcf4 | 21413 | HITS-CLIP//sRNA-Seq | 37986224\|25083871\|33184237 | Unspecified |
| mmu-miR-29b-3p | MIMAT0000127 | Tcf7l2 | 21416 | HITS-CLIP | 25083871 | Unspecified |
| mmu-miR-29b-3p | MIMAT0000127 | Tcirg1 | 27060 | HITS-CLIP//sRNA-Seq | 37986224\|23597149\|33184237 | Unspecified |
| mmu-miR-29b-3p | MIMAT0000127 | Tcof1 | 21453 | HITS-CLIP//sRNA-Seq | 23142080\|25083871\|33184237 | Unspecified |
| mmu-miR-29b-3p | MIMAT0000127 | Tcp1 | 21454 | HITS-CLIP//sRNA-Seq//qCLASH | 37986224\|25083871\|23142080\|23597 | Unspecified |
| mmu-miR-29b-3p | MIMAT0000127 | Tcp11l1 | 320554 | HITS-CLIP//sRNA-Seq | 37986224\|25083871\|33184237 | Unspecified |
| mmu-miR-29b-3p | MIMAT0000127 | Tcp11l2 | 216198 | HITS-CLIP | 37986224\|23597149\|23142080 | Unspecified |
| mmu-miR-29b-3p | MIMAT0000127 | Tctex1d2 | 66061 | HITS-CLIP | 25083871 | Unspecified |
| mmu-miR-29b-3p | MIMAT0000127 | Tdo2 | 56720 | HITS-CLIP | 37986224\|23597149 | Unspecified |
| mmu-miR-29b-3p | MIMAT0000127 | Tdrd3 | 219249 | HITS-CLIP | 37986224 | Unspecified |
| mmu-miR-29b-3p | MIMAT0000127 | Tdrd7 | 100121 | HITS-CLIP//sRNA-Seq | 37986224\|25083871\|33184237\|23142 | Unspecified |
| mmu-miR-29b-3p | MIMAT0000127 | Tead1 | 21676 | HITS-CLIP | 37986224 | Unspecified |
| mmu-miR-29b-3p | MIMAT0000127 | Tead4 | 21679 | HITS-CLIP | 26203562 | Unspecified |
| mmu-miR-29b-3p | MIMAT0000127 | Tecpr2 | 104859 | HITS-CLIP | 25083871 | Unspecified |
| mmu-miR-29b-3p | MIMAT0000127 | Tecr | 106529 | HITS-CLIP//sRNA-Seq | 23142080\|33184237 | Unspecified |
| mmu-miR-29b-3p | MIMAT0000127 | Tedc1 | 104732 | HITS-CLIP | 23142080 | Unspecified |
| mmu-miR-29b-3p | MIMAT0000127 | Tef | 21685 | HITS-CLIP//sRNA-Seq | 25083871\|33184237\|23142080 | Unspecified |
| mmu-miR-29b-3p | MIMAT0000127 | Tek | 21687 | HITS-CLIP | 23597149 | Unspecified |
| mmu-miR-29b-3p | MIMAT0000127 | Tent4b | 64282 | HITS-CLIP | 25083871\|23597149 | Unspecified |
| mmu-miR-29b-3p | MIMAT0000127 | Tent5a | 55603 | HITS-CLIP | 23142080\|31433990 | Unspecified |
| mmu-miR-29b-3p | MIMAT0000127 | Terf2ip | 57321 | HITS-CLIP | 23142080 | Unspecified |
| mmu-miR-29b-3p | MIMAT0000127 | Tes | 21753 | sRNA-Seq//HITS-CLIP | 33184237\|23142080 | Unspecified |
| mmu-miR-29b-3p | MIMAT0000127 | Tet1 | 52463 | HITS-CLIP | 37986224\|23142080\|25083871 | Unspecified |
| mmu-miR-29b-3p | MIMAT0000127 | Tet2 | 214133 | HITS-CLIP | 37986224\|25083871\|23597149\|23142 | Unspecified |

| | | | | | | |
|---|---|---|---|---|---|---|
| mmu-miR-29b-3p | MIMAT0000127 | Tet3 | 194388 | HITS-CLIP | 37986224\|25083871\|23597149\|23142 | Unspecified |
| mmu-miR-29b-3p | MIMAT0000127 | Tex10 | 269536 | HITS-CLIP | 23142080 | Unspecified |
| mmu-miR-29b-3p | MIMAT0000127 | Tex15 | 104271 | HITS-CLIP | 23142080 | Unspecified |
| mmu-miR-29b-3p | MIMAT0000127 | Tex264 | 21767 | HITS-CLIP//sRNA-Seq | 25083871\|33184237 | Unspecified |
| mmu-miR-29b-3p | MIMAT0000127 | Tex30 | 75623 | HITS-CLIP | 23142080 | Unspecified |
| mmu-miR-29b-3p | MIMAT0000127 | Tfdp2 | 211586 | HITS-CLIP | 23597149 | Unspecified |
| mmu-miR-29b-3p | MIMAT0000127 | Tfeb | 21425 | HITS-CLIP | 37986224\|25083871 | Unspecified |
| mmu-miR-29b-3p | MIMAT0000127 | Tfg | 21787 | HITS-CLIP//sRNA-Seq | 37986224\|25083871\|33184237\|23597 | Unspecified |
| mmu-miR-29b-3p | MIMAT0000127 | Tfrc | 22042 | HITS-CLIP//sRNA-Seq | 23142080\|33184237\|25083871 | Unspecified |
| mmu-miR-29b-3p | MIMAT0000127 | Tgfb3 | 21809 | Luciferase assay//Microarray//Northern bl | 19342382\|25083871\|33184237 | Unspecified |
| mmu-miR-29b-3p | MIMAT0000127 | Tgfbi | 21810 | sRNA-Seq//HITS-CLIP | 33184237\|23597149 | Unspecified |
| mmu-miR-29b-3p | MIMAT0000127 | Tgfbr1 | 21812 | HITS-CLIP | 23597149\|23142080\|25083871 | Unspecified |
| mmu-miR-29b-3p | MIMAT0000127 | Tgfbr3 | 21814 | HITS-CLIP | 25083871 | Unspecified |
| mmu-miR-29b-3p | MIMAT0000127 | Tgfbrap1 | 73122 | HITS-CLIP | 37986224\|25083871 | Unspecified |
| mmu-miR-29b-3p | MIMAT0000127 | Tgif1 | 21815 | HITS-CLIP | 37986224 | Unspecified |
| mmu-miR-29b-3p | MIMAT0000127 | Tgif2 | 228839 | HITS-CLIP | 37986224\|25083871\|23142080 | Unspecified |
| mmu-miR-29b-3p | MIMAT0000127 | Tgm2 | 21817 | HITS-CLIP | 23597149\|23142080 | Unspecified |
| mmu-miR-29b-3p | MIMAT0000127 | Tgoln1 | 22134 | HITS-CLIP | 23597149 | Unspecified |
| mmu-miR-29b-3p | MIMAT0000127 | Thada | 240174 | HITS-CLIP | 23142080 | Unspecified |
| mmu-miR-29b-3p | MIMAT0000127 | Thap11 | 59016 | HITS-CLIP | 25083871 | Unspecified |
| mmu-miR-29b-3p | MIMAT0000127 | Thbs1 | 21825 | HITS-CLIP//sRNA-Seq | 25083871\|33184237 | Unspecified |
| mmu-miR-29b-3p | MIMAT0000127 | Them4 | 75778 | HITS-CLIP | 37986224 | Unspecified |
| mmu-miR-29b-3p | MIMAT0000127 | Themis | 210757 | HITS-CLIP | 37986224\|23142080 | Unspecified |
| mmu-miR-29b-3p | MIMAT0000127 | Thoc2 | 331401 | HITS-CLIP | 25083871\|23142080 | Unspecified |
| mmu-miR-29b-3p | MIMAT0000127 | Thoc5 | 8563 | HITS-CLIP | 23597149\|23142080 | Unspecified |
| mmu-miR-29b-3p | MIMAT0000127 | Thoc7 | 66231 | HITS-CLIP//sRNA-Seq//qCLASH | 37986224\|25083871\|23597149\|3318 | Unspecified |
| mmu-miR-29b-3p | MIMAT0000127 | Thrap3 | 230753 | HITS-CLIP//sRNA-Seq | 25083871\|33184237 | Unspecified |
| mmu-miR-29b-3p | MIMAT0000127 | Thrsp | 21835 | HITS-CLIP | 23597149 | Unspecified |
| mmu-miR-29b-3p | MIMAT0000127 | Thumpd3 | 14911 | HITS-CLIP | 25083871 | Unspecified |
| mmu-miR-29b-3p | MIMAT0000127 | Tia1 | 21841 | HITS-CLIP | 25083871 | Unspecified |
| mmu-miR-29b-3p | MIMAT0000127 | Tiam1 | 21844 | HITS-CLIP | 37986224\|23142080\|25083871 | Unspecified |
| mmu-miR-29b-3p | MIMAT0000127 | Tiam2 | 24001 | HITS-CLIP | 37986224\|25083871\|23597149 | Unspecified |
| mmu-miR-29b-3p | MIMAT0000127 | Tigd2 | 68140 | HITS-CLIP | 37986224\|25083871\|23142080 | Unspecified |
| mmu-miR-29b-3p | MIMAT0000127 | Timd2 | 171284 | HITS-CLIP | 37986224\|23597149 | Unspecified |
| mmu-miR-29b-3p | MIMAT0000127 | Timm17a | 21854 | sRNA-Seq//HITS-CLIP | 33184237\|23142080 | Unspecified |
| mmu-miR-29b-3p | MIMAT0000127 | Timm44 | 21856 | HITS-CLIP | 23597149\|23142080 | Unspecified |
| mmu-miR-29b-3p | MIMAT0000127 | Timm8b | 30057 | HITS-CLIP//sRNA-Seq | 37986224\|33184237\|23142080 | Unspecified |
| mmu-miR-29b-3p | MIMAT0000127 | Timm9 | 30056 | HITS-CLIP//sRNA-Seq | 23597149\|33184237 | Unspecified |
| mmu-miR-29b-3p | MIMAT0000127 | Timp2 | 21858 | HITS-CLIP//sRNA-Seq | 23597149\|33184237 | Unspecified |

| | | | | | | |
|---|---|---|---|---|---|---|
| mmu-miR-29b-3p | MIMAT0000127 | Timp3 | 21859 | HITS-CLIP//sRNA-Seq | 25083871\|33184237 | Unspecified |
| mmu-miR-29b-3p | MIMAT0000127 | Tiparp | 99929 | HITS-CLIP | 23142080 | Unspecified |
| mmu-miR-29b-3p | MIMAT0000127 | Tiprl | 226591 | HITS-CLIP | 23597149\|23142080 | Unspecified |
| mmu-miR-29b-3p | MIMAT0000127 | Tjap1 | 74094 | HITS-CLIP//sRNA-Seq | 23142080\|25083871\|33184237 | Unspecified |
| mmu-miR-29b-3p | MIMAT0000127 | Tjp1 | 21872 | HITS-CLIP | 37986224\|25083871\|23597149 | Unspecified |
| mmu-miR-29b-3p | MIMAT0000127 | Tjp3 | 27375 | HITS-CLIP | 23597149 | Unspecified |
| mmu-miR-29b-3p | MIMAT0000127 | Tkfc | 26007 | HITS-CLIP | 23142080\|23597149 | Unspecified |
| mmu-miR-29b-3p | MIMAT0000127 | Tkt | 21881 | HITS-CLIP//sRNA-Seq | 23142080\|33184237 | Unspecified |
| mmu-miR-29b-3p | MIMAT0000127 | Tlcd1 | 68385 | HITS-CLIP | 23142080 | Unspecified |
| mmu-miR-29b-3p | MIMAT0000127 | Tle3 | 21887 | HITS-CLIP | 25083871\|23142080 | Unspecified |
| mmu-miR-29b-3p | MIMAT0000127 | Tle4 | 21888 | HITS-CLIP | 23142080 | Unspecified |
| mmu-miR-29b-3p | MIMAT0000127 | Tlk1 | 228012 | HITS-CLIP//sRNA-Seq | 25083871\|33184237\|23597149\|23142 | Unspecified |
| mmu-miR-29b-3p | MIMAT0000127 | Tlk2 | 24086 | HITS-CLIP//sRNA-Seq | 25083871\|33184237 | Unspecified |
| mmu-miR-29b-3p | MIMAT0000127 | Tln1 | 21894 | HITS-CLIP//sRNA-Seq | 25083871\|23597149\|33184237\|23142 | Unspecified |
| mmu-miR-29b-3p | MIMAT0000127 | Tln2 | 70549 | HITS-CLIP | 25083871 | Unspecified |
| mmu-miR-29b-3p | MIMAT0000127 | Tlr4 | 21898 | HITS-CLIP | 25083871 | Unspecified |
| mmu-miR-29b-3p | MIMAT0000127 | Tm2d1 | 94043 | HITS-CLIP | 37986224\|25083871\|23142080 | Unspecified |
| mmu-miR-29b-3p | MIMAT0000127 | Tm2d2 | 69742 | HITS-CLIP | 37986224 | Unspecified |
| mmu-miR-29b-3p | MIMAT0000127 | Tm4sf4 | 229302 | HITS-CLIP | 23597149 | Unspecified |
| mmu-miR-29b-3p | MIMAT0000127 | Tm6sf1 | 107769 | HITS-CLIP | 23142080 | Unspecified |
| mmu-miR-29b-3p | MIMAT0000127 | Tm6sf2 | 107770 | HITS-CLIP | 23597149 | Unspecified |
| mmu-miR-29b-3p | MIMAT0000127 | Tm7sf3 | 67623 | sRNA-Seq//HITS-CLIP | 33184237\|23597149 | Unspecified |
| mmu-miR-29b-3p | MIMAT0000127 | Tm9sf1 | 74140 | HITS-CLIP//sRNA-Seq | 37986224\|25083871\|23597149\|33184 | Unspecified |
| mmu-miR-29b-3p | MIMAT0000127 | Tm9sf2 | 68059 | HITS-CLIP//sRNA-Seq | 25083871\|23597149\|23142080\|33184 | Unspecified |
| mmu-miR-29b-3p | MIMAT0000127 | Tm9sf3 | 107358 | HITS-CLIP//sRNA-Seq | 37986224\|23597149\|25083871\|23142 | Unspecified |
| mmu-miR-29b-3p | MIMAT0000127 | Tm9sf4 | 99237 | sRNA-Seq//HITS-CLIP | 33184237\|23142080 | Unspecified |
| mmu-miR-29b-3p | MIMAT0000127 | Tmbim1 | 69660 | HITS-CLIP | 25083871 | Unspecified |
| mmu-miR-29b-3p | MIMAT0000127 | Tmbim4 | 68212 | HITS-CLIP//sRNA-Seq | 23597149\|33184237\|23142080\|25083 | Unspecified |
| mmu-miR-29b-3p | MIMAT0000127 | Tmc6 | 217353 | HITS-CLIP | 23597149\|23142080 | Unspecified |
| mmu-miR-29b-3p | MIMAT0000127 | Tmc8 | 217356 | HITS-CLIP | 23142080 | Unspecified |
| mmu-miR-29b-3p | MIMAT0000127 | Tmcc1 | 330401 | HITS-CLIP | 25083871\|23142080 | Unspecified |
| mmu-miR-29b-3p | MIMAT0000127 | Tmco6 | 71983 | HITS-CLIP | 23142080 | Unspecified |
| mmu-miR-29b-3p | MIMAT0000127 | Tmed1 | 17083 | HITS-CLIP//sRNA-Seq | 23597149\|33184237 | Unspecified |
| mmu-miR-29b-3p | MIMAT0000127 | Tmed4 | 103694 | HITS-CLIP//sRNA-Seq | 37986224\|33184237\|23142080 | Unspecified |
| mmu-miR-29b-3p | MIMAT0000127 | Tmed5 | 73130 | HITS-CLIP | 23142080\|25083871 | Unspecified |
| mmu-miR-29b-3p | MIMAT0000127 | Tmed9 | 67511 | HITS-CLIP//sRNA-Seq | 23142080\|23597149\|33184237\|25083 | Unspecified |
| mmu-miR-29b-3p | MIMAT0000127 | Tmeff1 | 230157 | HITS-CLIP//sRNA-Seq | 37986224\|25083871\|33184237 | Unspecified |
| mmu-miR-29b-3p | MIMAT0000127 | Tmem101 | 76547 | HITS-CLIP | 23597149\|23142080\|25083871 | Unspecified |
| mmu-miR-29b-3p | MIMAT0000127 | Tmem127 | 69470 | HITS-CLIP | 37986224\|25083871\|23142080 | Unspecified |

| | | | | | | |
|---|---|---|---|---|---|---|
| mmu-miR-29b-3p | MIMAT0000127 | Tmem131 | 56030 | HITS-CLIP | 25083871 | Unspecified |
| mmu-miR-29b-3p | MIMAT0000127 | Tmem132a | 98170 | HITS-CLIP//sRNA-Seq | 37986224\|25083871\|33184237 | Unspecified |
| mmu-miR-29b-3p | MIMAT0000127 | Tmem135 | 72759 | HITS-CLIP | 37986224\|25083871\|23597149\|23142 | Unspecified |
| mmu-miR-29b-3p | MIMAT0000127 | Tmem143 | 70209 | HITS-CLIP | 23597149 | Unspecified |
| mmu-miR-29b-3p | MIMAT0000127 | Tmem145 | 330485 | HITS-CLIP | 23142080 | Unspecified |
| mmu-miR-29b-3p | MIMAT0000127 | Tmem147 | 69804 | HITS-CLIP//sRNA-Seq | 23142080\|33184237\|25083871 | Unspecified |
| mmu-miR-29b-3p | MIMAT0000127 | Tmem14c | 66154 | HITS-CLIP//sRNA-Seq | 23597149\|25083871\|33184237 | Unspecified |
| mmu-miR-29b-3p | MIMAT0000127 | Tmem161b | 72745 | HITS-CLIP | 23597149 | Unspecified |
| mmu-miR-29b-3p | MIMAT0000127 | Tmem164 | 209497 | HITS-CLIP | 37986224 | Unspecified |
| mmu-miR-29b-3p | MIMAT0000127 | Tmem167 | 66074 | HITS-CLIP | 37986224\|25083871 | Unspecified |
| mmu-miR-29b-3p | MIMAT0000127 | Tmem167b | 67495 | sRNA-Seq//HITS-CLIP | 33184237\|23142080 | Unspecified |
| mmu-miR-29b-3p | MIMAT0000127 | Tmem168 | 101118 | HITS-CLIP | 37986224\|23142080 | Unspecified |
| mmu-miR-29b-3p | MIMAT0000127 | Tmem170b | 621976 | HITS-CLIP | 37986224\|25083871\|23142080\|2359 | Unspecified |
| mmu-miR-29b-3p | MIMAT0000127 | Tmem173 | 72512 | HITS-CLIP | 23142080\|25083871 | Unspecified |
| mmu-miR-29b-3p | MIMAT0000127 | Tmem179b | 374395 | HITS-CLIP | 23142080\|25083871 | Unspecified |
| mmu-miR-29b-3p | MIMAT0000127 | Tmem182 | 381339 | HITS-CLIP | 25083871 | Unspecified |
| mmu-miR-29b-3p | MIMAT0000127 | Tmem183a | 57439 | HITS-CLIP//sRNA-Seq | 37986224\|25083871\|33184237\|2359 | Unspecified |
| mmu-miR-29b-3p | MIMAT0000127 | Tmem184b | 223693 | HITS-CLIP//sRNA-Seq | 25083871\|33184237\|23597149\|23142 | Unspecified |
| mmu-miR-29b-3p | MIMAT0000127 | Tmem185b | 226351 | HITS-CLIP | 23142080\|25083871 | Unspecified |
| mmu-miR-29b-3p | MIMAT0000127 | Tmem186 | 66690 | HITS-CLIP | 23142080 | Unspecified |
| mmu-miR-29b-3p | MIMAT0000127 | Tmem189 | 407243 | HITS-CLIP | 37986224\|23142080\|25083871 | Unspecified |
| mmu-miR-29b-3p | MIMAT0000127 | Tmem19 | 67226 | HITS-CLIP | 25083871 | Unspecified |
| mmu-miR-29b-3p | MIMAT0000127 | Tmem203 | 227615 | HITS-CLIP | 23142080 | Unspecified |
| mmu-miR-29b-3p | MIMAT0000127 | Tmem204 | 407831 | HITS-CLIP | 23597149 | Unspecified |
| mmu-miR-29b-3p | MIMAT0000127 | Tmem205 | 235043 | HITS-CLIP | 23597149 | Unspecified |
| mmu-miR-29b-3p | MIMAT0000127 | Tmem209 | 72649 | HITS-CLIP | 25083871\|23142080 | Unspecified |
| mmu-miR-29b-3p | MIMAT0000127 | Tmem218 | 66279 | HITS-CLIP | 23142080 | Unspecified |
| mmu-miR-29b-3p | MIMAT0000127 | Tmem223 | 79064 | HITS-CLIP | 23597149 | Unspecified |
| mmu-miR-29b-3p | MIMAT0000127 | Tmem234 | 76799 | HITS-CLIP//sRNA-Seq | 23597149\|33184237 | Unspecified |
| mmu-miR-29b-3p | MIMAT0000127 | Tmem245 | 242474 | HITS-CLIP | 23142080 | Unspecified |
| mmu-miR-29b-3p | MIMAT0000127 | Tmem248 | 71667 | HITS-CLIP | 25083871\|23597149\|23142080 | Unspecified |
| mmu-miR-29b-3p | MIMAT0000127 | Tmem263 | 103266 | HITS-CLIP | 25083871\|23142080 | Unspecified |
| mmu-miR-29b-3p | MIMAT0000127 | Tmem30a | 69981 | sRNA-Seq//HITS-CLIP | 33184237\|23597149\|23142080 | Unspecified |
| mmu-miR-29b-3p | MIMAT0000127 | Tmem33 | 67878 | HITS-CLIP//sRNA-Seq | 25083871\|33184237\|23597149\|23142 | Unspecified |
| mmu-miR-29b-3p | MIMAT0000127 | Tmem38b | 52076 | HITS-CLIP//qCLASH | 23597149\|31393953 | Unspecified |
| mmu-miR-29b-3p | MIMAT0000127 | Tmem39a | 67846 | HITS-CLIP | 23142080 | Unspecified |
| mmu-miR-29b-3p | MIMAT0000127 | Tmem41a | 66664 | HITS-CLIP//sRNA-Seq | 25083871\|33184237 | Unspecified |
| mmu-miR-29b-3p | MIMAT0000127 | Tmem41b | 233724 | HITS-CLIP | 37986224 | Unspecified |
| mmu-miR-29b-3p | MIMAT0000127 | Tmem43 | 74122 | HITS-CLIP//sRNA-Seq | 25083871\|33184237 | Unspecified |

| | | | | | | |
|---|---|---|---|---|---|---|
| mmu-miR-29b-3p | MIMAT0000127 | Tmem50b | 77975 | HITS-CLIP | 23597149\|23142080\|25083871 | Unspecified |
| mmu-miR-29b-3p | MIMAT0000127 | Tmem59 | 56374 | HITS-CLIP//sRNA-Seq | 25083871\|23597149\|33184237\|23142 | Unspecified |
| mmu-miR-29b-3p | MIMAT0000127 | Tmem60 | 212090 | HITS-CLIP | 23142080 | Unspecified |
| mmu-miR-29b-3p | MIMAT0000127 | Tmem62 | 96957 | HITS-CLIP | 25083871 | Unspecified |
| mmu-miR-29b-3p | MIMAT0000127 | Tmem63b | 224807 | HITS-CLIP//sRNA-Seq | 37986224\|25083871\|23597149\|33184 | Unspecified |
| mmu-miR-29b-3p | MIMAT0000127 | Tmem64 | 100201 | HITS-CLIP | 25083871\|23597149\|23142080 | Unspecified |
| mmu-miR-29b-3p | MIMAT0000127 | Tmem65 | 74868 | HITS-CLIP//sRNA-Seq | 37986224\|33184237\|23597149\|23142 | Unspecified |
| mmu-miR-29b-3p | MIMAT0000127 | Tmem86a | 67893 | HITS-CLIP | 37986224\|25083871 | Unspecified |
| mmu-miR-29b-3p | MIMAT0000127 | Tmem87a | 211499 | HITS-CLIP | 25083871\|23142080 | Unspecified |
| mmu-miR-29b-3p | MIMAT0000127 | Tmem87b | 72477 | HITS-CLIP//sRNA-Seq | 37986224\|33184237\|23142080 | Unspecified |
| mmu-miR-29b-3p | MIMAT0000127 | Tmem88b | 320587 | HITS-CLIP | 25083871 | Unspecified |
| mmu-miR-29b-3p | MIMAT0000127 | Tmem9 | 66241 | HITS-CLIP//sRNA-Seq | 25083871\|33184237 | Unspecified |
| mmu-miR-29b-3p | MIMAT0000127 | Tmem94 | 9772 | HITS-CLIP | 25083871 | Unspecified |
| mmu-miR-29b-3p | MIMAT0000127 | Tmem98 | 103743 | HITS-CLIP//sRNA-Seq | 25083871\|33184237\|23597149 | Unspecified |
| mmu-miR-29b-3p | MIMAT0000127 | Tmf1 | 232286 | HITS-CLIP | 25083871\|23597149\|23142080 | Unspecified |
| mmu-miR-29b-3p | MIMAT0000127 | Tmod3 | 50875 | HITS-CLIP//sRNA-Seq | 37986224\|25083871\|33184237\|23142 | Unspecified |
| mmu-miR-29b-3p | MIMAT0000127 | Tmpo | 21917 | HITS-CLIP//sRNA-Seq | 25083871\|23142080\|33184237 | Unspecified |
| mmu-miR-29b-3p | MIMAT0000127 | Tmsb10 | 19240 | sRNA-Seq//HITS-CLIP | 33184237\|23142080 | Unspecified |
| mmu-miR-29b-3p | MIMAT0000127 | Tmtc2 | 278279 | sRNA-Seq//HITS-CLIP | 33184237\|23142080 | Unspecified |
| mmu-miR-29b-3p | MIMAT0000127 | Tmtc3 | 237500 | HITS-CLIP | 37986224\|25083871\|23597149\|23142 | Unspecified |
| mmu-miR-29b-3p | MIMAT0000127 | Tmub2 | 72053 | HITS-CLIP//sRNA-Seq | 37986224\|25083871\|33184237 | Unspecified |
| mmu-miR-29b-3p | MIMAT0000127 | Tmx1 | 72736 | HITS-CLIP//sRNA-Seq | 25083871\|33184237\|23142080\|30971 | Unspecified |
| mmu-miR-29b-3p | MIMAT0000127 | Tmx4 | 52837 | HITS-CLIP//sRNA-Seq | 37986224\|33184237\|23597149 | Unspecified |
| mmu-miR-29b-3p | MIMAT0000127 | Tnc | 21923 | HITS-CLIP//sRNA-Seq | 37986224\|25083871\|33184237 | Unspecified |
| mmu-miR-29b-3p | MIMAT0000127 | Tnf | 21926 | HITS-CLIP | 23142080 | Unspecified |
| mmu-miR-29b-3p | MIMAT0000127 | Tnfaip1 | 21927 | HITS-CLIP//sRNA-Seq | 25083871\|33184237 | Unspecified |
| mmu-miR-29b-3p | MIMAT0000127 | Tnfaip2 | 21928 | HITS-CLIP | 37986224 | Unspecified |
| mmu-miR-29b-3p | MIMAT0000127 | Tnfaip3 | 21929 | Immunoblot//Immunocytochemistry//Luci | 24236890\|23142080\|25083871 | Unspecified |
| mmu-miR-29b-3p | MIMAT0000127 | Tnfrsf18 | 21936 | HITS-CLIP | 23142080 | Unspecified |
| mmu-miR-29b-3p | MIMAT0000127 | Tnfrsf1a | 21937 | sRNA-Seq//HITS-CLIP | 33184237\|23142080 | Unspecified |
| mmu-miR-29b-3p | MIMAT0000127 | Tnfrsf26 | 244237 | HITS-CLIP | 37986224\|23142080 | Unspecified |
| mmu-miR-29b-3p | MIMAT0000127 | Tnfrsf4 | 22163 | HITS-CLIP | 37986224\|23142080 | Unspecified |
| mmu-miR-29b-3p | MIMAT0000127 | Tnfrsf9 | 21942 | HITS-CLIP | 37986224 | Unspecified |
| mmu-miR-29b-3p | MIMAT0000127 | Tnfsf8 | 21949 | HITS-CLIP | 23142080 | Unspecified |
| mmu-miR-29b-3p | MIMAT0000127 | Tnik | 665113 | HITS-CLIP//sRNA-Seq | 23142080\|33184237 | Unspecified |
| mmu-miR-29b-3p | MIMAT0000127 | Tnks | 21951 | HITS-CLIP//sRNA-Seq | 23597149\|25083871\|33184237\|23142 | Unspecified |
| mmu-miR-29b-3p | MIMAT0000127 | Tnks1bp1 | 228140 | HITS-CLIP//sRNA-Seq | 25083871\|33184237 | Unspecified |
| mmu-miR-29b-3p | MIMAT0000127 | Tnks2 | 74493 | HITS-CLIP//sRNA-Seq | 37986224\|25083871\|23142080\|31433 | Unspecified |
| mmu-miR-29b-3p | MIMAT0000127 | Tnni1 | 21952 | HITS-CLIP | 25083871 | Unspecified |

| | | | | | | |
|---|---|---|---|---|---|---|
| mmu-miR-29b-3p | MIMAT0000127 | Tnnt1 | 21955 | HITS-CLIP | 25083871 | Unspecified |
| mmu-miR-29b-3p | MIMAT0000127 | Tnnt2 | 21956 | HITS-CLIP | 25083871 | Unspecified |
| mmu-miR-29b-3p | MIMAT0000127 | Tnpo1 | 238799 | HITS-CLIP | 37986224\|25083871\|23597149\|23142 | Unspecified |
| mmu-miR-29b-3p | MIMAT0000127 | Tnpo2 | 212999 | HITS-CLIP//sRNA-Seq | 23142080\|25083871\|33184237 | Unspecified |
| mmu-miR-29b-3p | MIMAT0000127 | Tnpo3 | 320938 | sRNA-Seq//HITS-CLIP | 33184237\|23142080\|25083871 | Unspecified |
| mmu-miR-29b-3p | MIMAT0000127 | Tnrc18 | 231861 | HITS-CLIP//sRNA-Seq | 25083871\|23142080\|33184237\|23597 | Unspecified |
| mmu-miR-29b-3p | MIMAT0000127 | Tnrc6a | 233833 | HITS-CLIP//qCLASH | 37986224\|25083871\|31393953\|31433 | Unspecified |
| mmu-miR-29b-3p | MIMAT0000127 | Tnrc6b | 213988 | HITS-CLIP | 37986224\|25083871\|23597149\|23142 | Unspecified |
| mmu-miR-29b-3p | MIMAT0000127 | Tnrc6c | 217351 | HITS-CLIP | 37986224\|25083871\|23597149\|23142 | Unspecified |
| mmu-miR-29b-3p | MIMAT0000127 | Tns1 | 21961 | HITS-CLIP | 25083871\|23597149 | Unspecified |
| mmu-miR-29b-3p | MIMAT0000127 | Tns3 | 319939 | HITS-CLIP//sRNA-Seq | 25083871\|33184237 | Unspecified |
| mmu-miR-29b-3p | MIMAT0000127 | Tob1 | 22057 | HITS-CLIP | 25083871 | Unspecified |
| mmu-miR-29b-3p | MIMAT0000127 | Tob2 | 57259 | HITS-CLIP | 37986224\|23142080 | Unspecified |
| mmu-miR-29b-3p | MIMAT0000127 | Toe1 | 68276 | HITS-CLIP | 23142080 | Unspecified |
| mmu-miR-29b-3p | MIMAT0000127 | Togaram1 | 23116 | HITS-CLIP | 23142080\|25083871 | Unspecified |
| mmu-miR-29b-3p | MIMAT0000127 | Tollip | 54473 | HITS-CLIP | 37986224\|23597149\|25083871\|23142 | Unspecified |
| mmu-miR-29b-3p | MIMAT0000127 | Tomm34 | 67145 | HITS-CLIP | 23142080 | Unspecified |
| mmu-miR-29b-3p | MIMAT0000127 | Tomm40l | 641376 | HITS-CLIP | 23597149\|23142080 | Unspecified |
| mmu-miR-29b-3p | MIMAT0000127 | Tomm70a | 28185 | HITS-CLIP//sRNA-Seq | 37986224\|23142080\|25083871\|33184 | Unspecified |
| mmu-miR-29b-3p | MIMAT0000127 | Top2a | 21973 | HITS-CLIP//sRNA-Seq | 37986224\|25083871\|23597149\|33184 | Unspecified |
| mmu-miR-29b-3p | MIMAT0000127 | Top2b | 21974 | HITS-CLIP//sRNA-Seq | 37986224\|23597149\|33184237\|23142 | Unspecified |
| mmu-miR-29b-3p | MIMAT0000127 | Topors | 106021 | HITS-CLIP//sRNA-Seq | 23597149\|33184237\|23142080 | Unspecified |
| mmu-miR-29b-3p | MIMAT0000127 | Tor1aip2 | 240832 | HITS-CLIP//sRNA-Seq | 37986224\|25083871\|23142080\|33184 | Unspecified |
| mmu-miR-29b-3p | MIMAT0000127 | Tor1b | 30934 | HITS-CLIP | 26203562 | Unspecified |
| mmu-miR-29b-3p | MIMAT0000127 | Tor4a | 227612 | HITS-CLIP | 23142080 | Unspecified |
| mmu-miR-29b-3p | MIMAT0000127 | Tox | 252838 | HITS-CLIP | 23142080 | Unspecified |
| mmu-miR-29b-3p | MIMAT0000127 | Tox4 | 268741 | HITS-CLIP | 23142080\|25083871 | Unspecified |
| mmu-miR-29b-3p | MIMAT0000127 | Tpbg | 21983 | HITS-CLIP//sRNA-Seq | 25083871\|33184237 | Unspecified |
| mmu-miR-29b-3p | MIMAT0000127 | Tpcn1 | 252972 | HITS-CLIP | 37986224\|25083871\|23597149\|23142 | Unspecified |
| mmu-miR-29b-3p | MIMAT0000127 | Tpcn2 | 233979 | HITS-CLIP | 23142080 | Unspecified |
| mmu-miR-29b-3p | MIMAT0000127 | Tpk1 | 29807 | HITS-CLIP | 37986224\|23142080 | Unspecified |
| mmu-miR-29b-3p | MIMAT0000127 | Tpm1 | 22003 | sRNA-Seq//HITS-CLIP | 33184237\|23142080 | Unspecified |
| mmu-miR-29b-3p | MIMAT0000127 | Tpm3 | 59069 | sRNA-Seq//HITS-CLIP | 33184237\|23142080 | Unspecified |
| mmu-miR-29b-3p | MIMAT0000127 | Tpm4 | 326618 | sRNA-Seq//HITS-CLIP | 33184237\|25083871\|23142080 | Unspecified |
| mmu-miR-29b-3p | MIMAT0000127 | Tpp1 | 12751 | HITS-CLIP//sRNA-Seq | 37986224\|33184237\|23142080 | Unspecified |
| mmu-miR-29b-3p | MIMAT0000127 | Tpp2 | 22019 | HITS-CLIP//sRNA-Seq | 25083871\|23597149\|33184237\|23142 | Unspecified |
| mmu-miR-29b-3p | MIMAT0000127 | Tppp3 | 67971 | HITS-CLIP//sRNA-Seq | 25083871\|33184237 | Unspecified |
| mmu-miR-29b-3p | MIMAT0000127 | Tpr | 108989 | HITS-CLIP//sRNA-Seq | 25083871\|33184237\|23597149\|23142 | Unspecified |
| mmu-miR-29b-3p | MIMAT0000127 | Tpra1 | 24100 | HITS-CLIP | 23142080 | Unspecified |

| | | | | | | |
|---|---|---|---|---|---|---|
| mmu-miR-29b-3p | MIMAT0000127 | Tprgl | 67808 | HITS-CLIP//sRNA-Seq | 25083871\|33184237\|23142080 | Unspecified |
| mmu-miR-29b-3p | MIMAT0000127 | Tprn | 97031 | HITS-CLIP | 23597149 | Unspecified |
| mmu-miR-29b-3p | MIMAT0000127 | Tpst2 | 22022 | HITS-CLIP | 37986224\|23597149 | Unspecified |
| mmu-miR-29b-3p | MIMAT0000127 | Tpx2 | 72119 | HITS-CLIP//sRNA-Seq | 23142080\|33184237 | Unspecified |
| mmu-miR-29b-3p | MIMAT0000127 | Tra2a | 101214 | sRNA-Seq//HITS-CLIP | 33184237\|23142080 | Unspecified |
| mmu-miR-29b-3p | MIMAT0000127 | Trabd2b | 666048 | HITS-CLIP | 37986224\|25083871 | Unspecified |
| mmu-miR-29b-3p | MIMAT0000127 | Traf3 | 22031 | HITS-CLIP | 37986224\|23597149\|25083871\|23142 | Unspecified |
| mmu-miR-29b-3p | MIMAT0000127 | Traf4 | 22032 | HITS-CLIP//sRNA-Seq | 37986224\|25083871\|33184237 | Unspecified |
| mmu-miR-29b-3p | MIMAT0000127 | Traf6 | 22034 | HITS-CLIP | 23142080\|25083871 | Unspecified |
| mmu-miR-29b-3p | MIMAT0000127 | Trafd1 | 231712 | sRNA-Seq//HITS-CLIP | 33184237\|23597149\|23142080 | Unspecified |
| mmu-miR-29b-3p | MIMAT0000127 | Trak1 | 67095 | HITS-CLIP//sRNA-Seq | 37986224\|23142080\|33184237 | Unspecified |
| mmu-miR-29b-3p | MIMAT0000127 | Trak2 | 70827 | HITS-CLIP//sRNA-Seq | 37986224\|25083871\|33184237 | Unspecified |
| mmu-miR-29b-3p | MIMAT0000127 | Tram1 | 72265 | HITS-CLIP//sRNA-Seq | 25083871\|23597149\|23142080\|33184 | Unspecified |
| mmu-miR-29b-3p | MIMAT0000127 | Tram2 | 170829 | HITS-CLIP | 37986224\|25083871\|23142080 | Unspecified |
| mmu-miR-29b-3p | MIMAT0000127 | Trappc10 | 216131 | HITS-CLIP | 23597149 | Unspecified |
| mmu-miR-29b-3p | MIMAT0000127 | Trappc11 | 320714 | HITS-CLIP | 23142080 | Unspecified |
| mmu-miR-29b-3p | MIMAT0000127 | Trappc13 | 66975 | HITS-CLIP | 23142080\|25083871 | Unspecified |
| mmu-miR-29b-3p | MIMAT0000127 | Trappc2l | 59005 | HITS-CLIP | 23597149\|23142080 | Unspecified |
| mmu-miR-29b-3p | MIMAT0000127 | Trappc8 | 75964 | HITS-CLIP//sRNA-Seq | 37986224\|23142080\|23597149\|33184 | Unspecified |
| mmu-miR-29b-3p | MIMAT0000127 | Treml2 | 328833 | HITS-CLIP | 23142080 | Unspecified |
| mmu-miR-29b-3p | MIMAT0000127 | Trib1 | 211770 | HITS-CLIP | 23597149\|23142080 | Unspecified |
| mmu-miR-29b-3p | MIMAT0000127 | Trib2 | 217410 | HITS-CLIP//sRNA-Seq | 37986224\|23142080\|33184237 | Unspecified |
| mmu-miR-29b-3p | MIMAT0000127 | Trib3 | 228775 | HITS-CLIP//sRNA-Seq | 37986224\|33184237\|23142080 | Unspecified |
| mmu-miR-29b-3p | MIMAT0000127 | Trim2 | 80890 | HITS-CLIP | 37986224 | Unspecified |
| mmu-miR-29b-3p | MIMAT0000127 | Trim21 | 20821 | HITS-CLIP | 37986224\|23597149\|23142080 | Unspecified |
| mmu-miR-29b-3p | MIMAT0000127 | Trim23 | 81003 | HITS-CLIP | 23142080 | Unspecified |
| mmu-miR-29b-3p | MIMAT0000127 | Trim24 | 21848 | HITS-CLIP | 23142080 | Unspecified |
| mmu-miR-29b-3p | MIMAT0000127 | Trim25 | 217069 | HITS-CLIP | 37986224\|23597149\|23142080 | Unspecified |
| mmu-miR-29b-3p | MIMAT0000127 | Trim28 | 21849 | HITS-CLIP//sRNA-Seq | 23597149\|33184237\|23142080 | Unspecified |
| mmu-miR-29b-3p | MIMAT0000127 | Trim3 | 55992 | HITS-CLIP//sRNA-Seq | 23597149\|33184237 | Unspecified |
| mmu-miR-29b-3p | MIMAT0000127 | Trim32 | 69807 | HITS-CLIP//sRNA-Seq | 25083871\|23597149\|33184237\|23142 | Unspecified |
| mmu-miR-29b-3p | MIMAT0000127 | Trim33 | 94093 | HITS-CLIP | 25083871\|23142080 | Unspecified |
| mmu-miR-29b-3p | MIMAT0000127 | Trim35 | 66854 | sRNA-Seq//HITS-CLIP | 33184237\|25083871 | Unspecified |
| mmu-miR-29b-3p | MIMAT0000127 | Trim37 | 68729 | HITS-CLIP//sRNA-Seq | 37986224\|25083871\|33184237\|23142 | Unspecified |
| mmu-miR-29b-3p | MIMAT0000127 | Trim41 | 211007 | HITS-CLIP | 25083871\|23142080 | Unspecified |
| mmu-miR-29b-3p | MIMAT0000127 | Trim44 | 80985 | sRNA-Seq//HITS-CLIP | 33184237\|23142080 | Unspecified |
| mmu-miR-29b-3p | MIMAT0000127 | Trim47 | 217333 | HITS-CLIP | 37986224 | Unspecified |
| mmu-miR-29b-3p | MIMAT0000127 | Trim55 | 381485 | HITS-CLIP | 25083871 | Unspecified |
| mmu-miR-29b-3p | MIMAT0000127 | Trim56 | 384309 | HITS-CLIP | 37986224\|25083871 | Unspecified |

| | | | | | | |
|---|---|---|---|---|---|---|
| mmu-miR-29b-3p | MIMAT0000127 | Trim63 | 433766 | HITS-CLIP | 37986224\|25083871 | Unspecified |
| mmu-miR-29b-3p | MIMAT0000127 | Trim71 | 636931 | HITS-CLIP | 37986224 | Unspecified |
| mmu-miR-29b-3p | MIMAT0000127 | Trio | 223435 | HITS-CLIP | 37986224\|25083871 | Unspecified |
| mmu-miR-29b-3p | MIMAT0000127 | Trip10 | 106628 | HITS-CLIP | 23142080 | Unspecified |
| mmu-miR-29b-3p | MIMAT0000127 | Trip11 | 109181 | HITS-CLIP | 23142080\|25083871 | Unspecified |
| mmu-miR-29b-3p | MIMAT0000127 | Trip12 | 14897 | HITS-CLIP//sRNA-Seq | 37986224\|23597149\|25083871\|23142 | Unspecified |
| mmu-miR-29b-3p | MIMAT0000127 | Trmt1 | 212528 | HITS-CLIP | 25083871 | Unspecified |
| mmu-miR-29b-3p | MIMAT0000127 | Trmt10a | 108943 | HITS-CLIP | 23142080\|25083871 | Unspecified |
| mmu-miR-29b-3p | MIMAT0000127 | Trmt11 | 73681 | HITS-CLIP | 23142080 | Unspecified |
| mmu-miR-29b-3p | MIMAT0000127 | Trmt5 | 76357 | HITS-CLIP | 23142080 | Unspecified |
| mmu-miR-29b-3p | MIMAT0000127 | Trmt6 | 66926 | HITS-CLIP | 23142080 | Unspecified |
| mmu-miR-29b-3p | MIMAT0000127 | Trove2 | 20822 | HITS-CLIP | 23142080\|25083871 | Unspecified |
| mmu-miR-29b-3p | MIMAT0000127 | Trp53inp1 | 60599 | HITS-CLIP | 37986224\|25083871\|23597149\|23142 | Unspecified |
| mmu-miR-29b-3p | MIMAT0000127 | Trp53inp2 | 68728 | HITS-CLIP//sRNA-Seq | 37986224\|25083871\|23597149\|33184 | Unspecified |
| mmu-miR-29b-3p | MIMAT0000127 | Trp53rka | 381406 | HITS-CLIP | 25083871 | Unspecified |
| mmu-miR-29b-3p | MIMAT0000127 | Trpc4ap | 56407 | HITS-CLIP//sRNA-Seq | 37986224\|33184237\|23597149\|23142 | Unspecified |
| mmu-miR-29b-3p | MIMAT0000127 | Trpm7 | 58800 | HITS-CLIP//sRNA-Seq | 25083871\|23597149\|33184237\|26203 | Unspecified |
| mmu-miR-29b-3p | MIMAT0000127 | Trps1 | 83925 | HITS-CLIP | 23142080\|25083871 | Unspecified |
| mmu-miR-29b-3p | MIMAT0000127 | Trrap | 100683 | HITS-CLIP//sRNA-Seq | 23597149\|33184237\|23142080 | Unspecified |
| mmu-miR-29b-3p | MIMAT0000127 | Tsc1 | 64930 | HITS-CLIP | 25083871 | Unspecified |
| mmu-miR-29b-3p | MIMAT0000127 | Tsc22d1 | 21807 | HITS-CLIP//sRNA-Seq | 37986224\|25083871\|33184237 | Unspecified |
| mmu-miR-29b-3p | MIMAT0000127 | Tsc22d2 | 72033 | HITS-CLIP | 37986224\|25083871\|23142080 | Unspecified |
| mmu-miR-29b-3p | MIMAT0000127 | Tsc22d3 | 14605 | HITS-CLIP//sRNA-Seq | 37986224\|25083871\|33184237\|23597 | Unspecified |
| mmu-miR-29b-3p | MIMAT0000127 | Tsg101 | 22088 | sRNA-Seq//HITS-CLIP | 33184237\|25083871 | Unspecified |
| mmu-miR-29b-3p | MIMAT0000127 | Tshz1 | 110796 | HITS-CLIP | 23597149\|23142080 | Unspecified |
| mmu-miR-29b-3p | MIMAT0000127 | Tsku | 244152 | HITS-CLIP | 23597149 | Unspecified |
| mmu-miR-29b-3p | MIMAT0000127 | Tspan13 | 66109 | sRNA-Seq//HITS-CLIP | 33184237\|23142080 | Unspecified |
| mmu-miR-29b-3p | MIMAT0000127 | Tspan14 | 52588 | HITS-CLIP | 37986224\|23142080\|25083871 | Unspecified |
| mmu-miR-29b-3p | MIMAT0000127 | Tspan17 | 74257 | HITS-CLIP//sRNA-Seq | 37986224\|25083871\|33184237 | Unspecified |
| mmu-miR-29b-3p | MIMAT0000127 | Tspan2 | 70747 | HITS-CLIP | 23142080 | Unspecified |
| mmu-miR-29b-3p | MIMAT0000127 | Tspan3 | 56434 | HITS-CLIP//sRNA-Seq | 25083871\|33184237\|23142080 | Unspecified |
| mmu-miR-29b-3p | MIMAT0000127 | Tspan31 | 67125 | sRNA-Seq//HITS-CLIP | 33184237\|23597149 | Unspecified |
| mmu-miR-29b-3p | MIMAT0000127 | Tspan4 | 64540 | HITS-CLIP//sRNA-Seq | 23597149\|25083871\|33184237 | Unspecified |
| mmu-miR-29b-3p | MIMAT0000127 | Tspan5 | 56224 | HITS-CLIP//sRNA-Seq | 25083871\|33184237 | Unspecified |
| mmu-miR-29b-3p | MIMAT0000127 | Tspan8 | 216350 | HITS-CLIP | 37986224\|25083871 | Unspecified |
| mmu-miR-29b-3p | MIMAT0000127 | Tspo | 12257 | HITS-CLIP//sRNA-Seq | 37986224\|25083871\|33184237\|23142 | Unspecified |
| mmu-miR-29b-3p | MIMAT0000127 | Tspyl2 | 52808 | HITS-CLIP | 23142080 | Unspecified |
| mmu-miR-29b-3p | MIMAT0000127 | Tsr3 | 68327 | HITS-CLIP | 25083871 | Unspecified |
| mmu-miR-29b-3p | MIMAT0000127 | Tst | 22117 | HITS-CLIP | 23597149 | Unspecified |

| | | | | | | |
|---|---|---|---|---|---|---|
| mmu-miR-29b-3p | MIMAT0000127 | Tstd2 | 272027 | HITS-CLIP | 37986224 | Unspecified |
| mmu-miR-29b-3p | MIMAT0000127 | Tstd3 | 77032 | HITS-CLIP | 23142080 | Unspecified |
| mmu-miR-29b-3p | MIMAT0000127 | Ttbk2 | 140810 | HITS-CLIP//sRNA-Seq | 37986224\|25083871\|33184237 | Unspecified |
| mmu-miR-29b-3p | MIMAT0000127 | Ttc17 | 74569 | HITS-CLIP | 23142080 | Unspecified |
| mmu-miR-29b-3p | MIMAT0000127 | Ttc19 | 72795 | HITS-CLIP | 25083871 | Unspecified |
| mmu-miR-29b-3p | MIMAT0000127 | Ttc23 | 67009 | HITS-CLIP | 37986224 | Unspecified |
| mmu-miR-29b-3p | MIMAT0000127 | Ttc3 | 22129 | HITS-CLIP//sRNA-Seq | 23142080\|33184237\|25083871 | Unspecified |
| mmu-miR-29b-3p | MIMAT0000127 | Ttc37 | 218343 | HITS-CLIP | 23142080\|25083871 | Unspecified |
| mmu-miR-29b-3p | MIMAT0000127 | Ttc39b | 69863 | HITS-CLIP | 37986224\|25083871\|23142080 | Unspecified |
| mmu-miR-29b-3p | MIMAT0000127 | Ttc39c | 72747 | HITS-CLIP | 23597149 | Unspecified |
| mmu-miR-29b-3p | MIMAT0000127 | Ttc5 | 219022 | HITS-CLIP//sRNA-Seq | 23142080\|33184237 | Unspecified |
| mmu-miR-29b-3p | MIMAT0000127 | Ttc7 | 225049 | HITS-CLIP | 23597149 | Unspecified |
| mmu-miR-29b-3p | MIMAT0000127 | Ttc9c | 70387 | HITS-CLIP | 23142080 | Unspecified |
| mmu-miR-29b-3p | MIMAT0000127 | Tti1 | 75425 | HITS-CLIP | 25083871\|23142080 | Unspecified |
| mmu-miR-29b-3p | MIMAT0000127 | Ttll4 | 67534 | HITS-CLIP | 25083871\|31433990\|23142080 | Unspecified |
| mmu-miR-29b-3p | MIMAT0000127 | Ttn | 22138 | HITS-CLIP | 37986224\|25083871 | Unspecified |
| mmu-miR-29b-3p | MIMAT0000127 | Ttr | 22139 | HITS-CLIP | 23597149 | Unspecified |
| mmu-miR-29b-3p | MIMAT0000127 | Ttyh2 | 117160 | HITS-CLIP | 37986224 | Unspecified |
| mmu-miR-29b-3p | MIMAT0000127 | Ttyh3 | 78339 | HITS-CLIP//sRNA-Seq | 25083871\|33184237\|23142080 | Unspecified |
| mmu-miR-29b-3p | MIMAT0000127 | Tuba1b | 22143 | HITS-CLIP//sRNA-Seq | 25083871\|23597149\|33184237\|23142 | Unspecified |
| mmu-miR-29b-3p | MIMAT0000127 | Tuba4a | 22145 | sRNA-Seq//HITS-CLIP | 33184237\|23597149 | Unspecified |
| mmu-miR-29b-3p | MIMAT0000127 | Tubb2a | 22151 | HITS-CLIP//sRNA-Seq | 37986224\|23597149\|33184237\|23142 | Unspecified |
| mmu-miR-29b-3p | MIMAT0000127 | Tubb5 | 22154 | HITS-CLIP//sRNA-Seq | 37986224\|23597149\|23142080\|3318 | Unspecified |
| mmu-miR-29b-3p | MIMAT0000127 | Tubb6 | 67951 | HITS-CLIP//sRNA-Seq | 37986224\|25083871\|33184237 | Unspecified |
| mmu-miR-29b-3p | MIMAT0000127 | Tubd1 | 56427 | HITS-CLIP | 23142080 | Unspecified |
| mmu-miR-29b-3p | MIMAT0000127 | Tubgcp5 | 233276 | HITS-CLIP | 25083871\|23142080 | Unspecified |
| mmu-miR-29b-3p | MIMAT0000127 | Tusc1 | 69136 | HITS-CLIP | 23142080 | Unspecified |
| mmu-miR-29b-3p | MIMAT0000127 | Tusc2 | 80385 | HITS-CLIP//sRNA-Seq | 25083871\|33184237\|23142080 | Unspecified |
| mmu-miR-29b-3p | MIMAT0000127 | Tut4 | 23318 | HITS-CLIP | 25083871\|23597149\|23142080 | Unspecified |
| mmu-miR-29b-3p | MIMAT0000127 | Tut7 | 79670 | HITS-CLIP | 25083871\|23597149\|23142080 | Unspecified |
| mmu-miR-29b-3p | MIMAT0000127 | Twf1 | 19230 | HITS-CLIP//sRNA-Seq | 23597149\|33184237\|25083871 | Unspecified |
| mmu-miR-29b-3p | MIMAT0000127 | Twsg1 | 65960 | sRNA-Seq//HITS-CLIP | 33184237\|23142080 | Unspecified |
| mmu-miR-29b-3p | MIMAT0000127 | Txlng | 353170 | HITS-CLIP | 25083871\|23142080 | Unspecified |
| mmu-miR-29b-3p | MIMAT0000127 | Txn1 | 22166 | HITS-CLIP//sRNA-Seq | 23597149\|25083871\|33184237\|23142 | Unspecified |
| mmu-miR-29b-3p | MIMAT0000127 | Txndc11 | 106200 | HITS-CLIP | 25083871\|23142080 | Unspecified |
| mmu-miR-29b-3p | MIMAT0000127 | Txndc15 | 69672 | HITS-CLIP | 23597149\|25083871\|23142080 | Unspecified |
| mmu-miR-29b-3p | MIMAT0000127 | Txndc17 | 52700 | HITS-CLIP//sRNA-Seq | 25083871\|33184237\|23142080 | Unspecified |
| mmu-miR-29b-3p | MIMAT0000127 | Txndc5 | 105245 | HITS-CLIP//sRNA-Seq | 25083871\|23597149\|33184237\|23142 | Unspecified |
| mmu-miR-29b-3p | MIMAT0000127 | Txnip | 56338 | HITS-CLIP//sRNA-Seq | 25083871\|23597149\|33184237\|23142 | Unspecified |

| | | | | | | |
|---|---|---|---|---|---|---|
| mmu-miR-29b-3p | MIMAT0000127 | Txnrd1 | 50493 | HITS-CLIP//sRNA-Seq | 23142080\|33184237 | Unspecified |
| mmu-miR-29b-3p | MIMAT0000127 | Txnrd3 | 232223 | HITS-CLIP | 23597149 | Unspecified |
| mmu-miR-29b-3p | MIMAT0000127 | Tysnd1 | 71767 | HITS-CLIP | 23597149 | Unspecified |
| mmu-miR-29b-3p | MIMAT0000127 | U2af2 | 22185 | HITS-CLIP//sRNA-Seq | 23597149\|33184237 | Unspecified |
| mmu-miR-29b-3p | MIMAT0000127 | U2surp | 67958 | HITS-CLIP//sRNA-Seq | 37986224\|33184237\|23142080 | Unspecified |
| mmu-miR-29b-3p | MIMAT0000127 | Uaca | 72565 | HITS-CLIP | 37986224\|25083871\|26203562 | Unspecified |
| mmu-miR-29b-3p | MIMAT0000127 | Uap1 | 107652 | HITS-CLIP//sRNA-Seq | 37986224\|25083871\|33184237\|23142 | Unspecified |
| mmu-miR-29b-3p | MIMAT0000127 | Uap1l1 | 227620 | HITS-CLIP//sRNA-Seq | 25083871\|33184237 | Unspecified |
| mmu-miR-29b-3p | MIMAT0000127 | Uba1 | 22201 | HITS-CLIP//sRNA-Seq | 37986224\|23597149\|33184237\|23142 | Unspecified |
| mmu-miR-29b-3p | MIMAT0000127 | Uba2 | 50995 | HITS-CLIP//sRNA-Seq | 25083871\|33184237\|23142080 | Unspecified |
| mmu-miR-29b-3p | MIMAT0000127 | Uba3 | 22200 | HITS-CLIP//sRNA-Seq | 23142080\|25083871\|33184237 | Unspecified |
| mmu-miR-29b-3p | MIMAT0000127 | Uba6 | 231380 | HITS-CLIP | 23142080 | Unspecified |
| mmu-miR-29b-3p | MIMAT0000127 | Ubac1 | 98766 | HITS-CLIP//sRNA-Seq | 37986224\|25083871\|33184237\|23597 | Unspecified |
| mmu-miR-29b-3p | MIMAT0000127 | Ubap2l | 74383 | HITS-CLIP//sRNA-Seq | 25083871\|33184237\|23142080 | Unspecified |
| mmu-miR-29b-3p | MIMAT0000127 | Ubash3a | 328795 | HITS-CLIP | 23142080 | Unspecified |
| mmu-miR-29b-3p | MIMAT0000127 | Ube2c | 68612 | HITS-CLIP//sRNA-Seq | 23142080\|33184237 | Unspecified |
| mmu-miR-29b-3p | MIMAT0000127 | Ube2d3 | 66105 | HITS-CLIP//sRNA-Seq | 23142080\|33184237\|25083871 | Unspecified |
| mmu-miR-29b-3p | MIMAT0000127 | Ube2f | 67921 | HITS-CLIP | 37986224 | Unspecified |
| mmu-miR-29b-3p | MIMAT0000127 | Ube2h | 22214 | HITS-CLIP//sRNA-Seq | 25083871\|33184237 | Unspecified |
| mmu-miR-29b-3p | MIMAT0000127 | Ube2j1 | 56228 | HITS-CLIP//sRNA-Seq | 37986224\|25083871\|23597149\|33184 | Unspecified |
| mmu-miR-29b-3p | MIMAT0000127 | Ube2k | 53323 | HITS-CLIP | 37986224 | Unspecified |
| mmu-miR-29b-3p | MIMAT0000127 | Ube2m | 22192 | sRNA-Seq//HITS-CLIP | 33184237\|25083871 | Unspecified |
| mmu-miR-29b-3p | MIMAT0000127 | Ube2o | 217342 | HITS-CLIP//sRNA-Seq | 23597149\|33184237\|23142080 | Unspecified |
| mmu-miR-29b-3p | MIMAT0000127 | Ube2q1 | 70093 | HITS-CLIP//sRNA-Seq | 25083871\|23597149\|33184237\|23142 | Unspecified |
| mmu-miR-29b-3p | MIMAT0000127 | Ube2q2 | 109161 | sRNA-Seq//HITS-CLIP | 33184237\|23142080 | Unspecified |
| mmu-miR-29b-3p | MIMAT0000127 | Ube2z | 268470 | sRNA-Seq//HITS-CLIP | 33184237\|23142080 | Unspecified |
| mmu-miR-29b-3p | MIMAT0000127 | Ube3a | 22215 | HITS-CLIP//sRNA-Seq | 23142080\|33184237\|23597149\|25083 | Unspecified |
| mmu-miR-29b-3p | MIMAT0000127 | Ube3b | 117146 | HITS-CLIP//sRNA-Seq | 25083871\|33184237\|23142080 | Unspecified |
| mmu-miR-29b-3p | MIMAT0000127 | Ube3c | 100763 | HITS-CLIP//sRNA-Seq | 25083871\|33184237\|23142080 | Unspecified |
| mmu-miR-29b-3p | MIMAT0000127 | Ube4a | 140630 | HITS-CLIP//sRNA-Seq | 25083871\|33184237\|23597149\|23142 | Unspecified |
| mmu-miR-29b-3p | MIMAT0000127 | Ube4b | 63958 | HITS-CLIP//sRNA-Seq | 25083871\|33184237\|23142080 | Unspecified |
| mmu-miR-29b-3p | MIMAT0000127 | Ubfd1 | 28018 | HITS-CLIP//sRNA-Seq | 37986224\|25083871\|23597149\|33184 | Unspecified |
| mmu-miR-29b-3p | MIMAT0000127 | Ubl5 | 66177 | HITS-CLIP//sRNA-Seq | 23142080\|33184237 | Unspecified |
| mmu-miR-29b-3p | MIMAT0000127 | Ubn1 | 170644 | HITS-CLIP | 37986224\|25083871\|23142080 | Unspecified |
| mmu-miR-29b-3p | MIMAT0000127 | Ubn2 | 320538 | HITS-CLIP | 25083871\|23142080\|23597149 | Unspecified |
| mmu-miR-29b-3p | MIMAT0000127 | Ubp1 | 22221 | HITS-CLIP | 25083871\|23597149\|23142080 | Unspecified |
| mmu-miR-29b-3p | MIMAT0000127 | Ubqln1 | 56085 | HITS-CLIP//sRNA-Seq | 25083871\|33184237 | Unspecified |
| mmu-miR-29b-3p | MIMAT0000127 | Ubqln2 | 54609 | HITS-CLIP//sRNA-Seq | 25083871\|33184237\|23142080 | Unspecified |
| mmu-miR-29b-3p | MIMAT0000127 | Ubr1 | 22222 | HITS-CLIP | 23142080 | Unspecified |

| | | | | | | |
|---|---|---|---|---|---|---|
| mmu-miR-29b-3p | MIMAT0000127 | Ubr2 | 224826 | HITS-CLIP | 25083871\|23597149\|23142080 | Unspecified |
| mmu-miR-29b-3p | MIMAT0000127 | Ubr3 | 68795 | HITS-CLIP//sRNA-Seq | 37986224\|25083871\|33184237\|23597 | Unspecified |
| mmu-miR-29b-3p | MIMAT0000127 | Ubr4 | 69116 | HITS-CLIP//sRNA-Seq | 25083871\|23597149\|33184237\|23142 | Unspecified |
| mmu-miR-29b-3p | MIMAT0000127 | Ubr5 | 70790 | HITS-CLIP//sRNA-Seq | 37986224\|25083871\|33184237\|23142 | Unspecified |
| mmu-miR-29b-3p | MIMAT0000127 | Ubtd1 | 226122 | HITS-CLIP | 25083871 | Unspecified |
| mmu-miR-29b-3p | MIMAT0000127 | Ubtf | 21429 | HITS-CLIP//sRNA-Seq | 37986224\|25083871\|23597149\|33184 | Unspecified |
| mmu-miR-29b-3p | MIMAT0000127 | Ubxn2b | 68053 | HITS-CLIP | 23142080 | Unspecified |
| mmu-miR-29b-3p | MIMAT0000127 | Ubxn4 | 67812 | HITS-CLIP//sRNA-Seq | 25083871\|23597149\|33184237\|23142 | Unspecified |
| mmu-miR-29b-3p | MIMAT0000127 | Ubxn7 | 224111 | HITS-CLIP | 37986224\|23142080 | Unspecified |
| mmu-miR-29b-3p | MIMAT0000127 | Ubxn8 | 108159 | HITS-CLIP | 25083871 | Unspecified |
| mmu-miR-29b-3p | MIMAT0000127 | Uck2 | 80914 | HITS-CLIP//sRNA-Seq | 37986224\|25083871\|33184237\|23142 | Unspecified |
| mmu-miR-29b-3p | MIMAT0000127 | Ucp2 | 22228 | sRNA-Seq//HITS-CLIP | 33184237\|23142080 | Unspecified |
| mmu-miR-29b-3p | MIMAT0000127 | Ufl1 | 67490 | HITS-CLIP | 25083871\|23597149 | Unspecified |
| mmu-miR-29b-3p | MIMAT0000127 | Ufsp2 | 192169 | HITS-CLIP | 23597149 | Unspecified |
| mmu-miR-29b-3p | MIMAT0000127 | Ugcg | 22234 | HITS-CLIP//sRNA-Seq | 25083871\|33184237\|23142080 | Unspecified |
| mmu-miR-29b-3p | MIMAT0000127 | Ugdh | 22235 | HITS-CLIP//sRNA-Seq | 23597149\|25083871\|33184237\|23142 | Unspecified |
| mmu-miR-29b-3p | MIMAT0000127 | Uggt1 | 320011 | HITS-CLIP | 25083871\|23597149\|23142080 | Unspecified |
| mmu-miR-29b-3p | MIMAT0000127 | Ugp2 | 216558 | HITS-CLIP//sRNA-Seq | 23597149\|33184237\|23142080 | Unspecified |
| mmu-miR-29b-3p | MIMAT0000127 | Ugt2b1 | 71773 | HITS-CLIP | 23597149 | Unspecified |
| mmu-miR-29b-3p | MIMAT0000127 | Ugt2b34 | 100727 | HITS-CLIP | 23597149 | Unspecified |
| mmu-miR-29b-3p | MIMAT0000127 | Ugt2b35 | 243085 | HITS-CLIP | 23597149 | Unspecified |
| mmu-miR-29b-3p | MIMAT0000127 | Ugt2b36 | 231396 | HITS-CLIP | 23597149 | Unspecified |
| mmu-miR-29b-3p | MIMAT0000127 | Ugt2b5 | 22238 | HITS-CLIP | 23597149 | Unspecified |
| mmu-miR-29b-3p | MIMAT0000127 | Ugt3a2 | 223337 | HITS-CLIP | 23597149 | Unspecified |
| mmu-miR-29b-3p | MIMAT0000127 | Uhmk1 | 16589 | sRNA-Seq//HITS-CLIP | 33184237\|23142080 | Unspecified |
| mmu-miR-29b-3p | MIMAT0000127 | Uhrf1bp1 | 224648 | HITS-CLIP | 37986224\|25083871 | Unspecified |
| mmu-miR-29b-3p | MIMAT0000127 | Uhrf1bp1l | 75089 | HITS-CLIP | 37986224\|25083871\|23142080 | Unspecified |
| mmu-miR-29b-3p | MIMAT0000127 | Uhrf2 | 109113 | HITS-CLIP//sRNA-Seq | 25083871\|23142080\|33184237 | Unspecified |
| mmu-miR-29b-3p | MIMAT0000127 | Ulk1 | 22241 | HITS-CLIP | 25083871 | Unspecified |
| mmu-miR-29b-3p | MIMAT0000127 | Ulk2 | 29869 | HITS-CLIP//sRNA-Seq | 37986224\|23597149\|23142080\|25083 | Unspecified |
| mmu-miR-29b-3p | MIMAT0000127 | Unc45a | 101869 | HITS-CLIP | 23142080 | Unspecified |
| mmu-miR-29b-3p | MIMAT0000127 | Unc45b | 217012 | HITS-CLIP | 25083871 | Unspecified |
| mmu-miR-29b-3p | MIMAT0000127 | Unc50 | 67387 | HITS-CLIP | 23142080 | Unspecified |
| mmu-miR-29b-3p | MIMAT0000127 | Unc5b | 107449 | HITS-CLIP | 25083871 | Unspecified |
| mmu-miR-29b-3p | MIMAT0000127 | Unc93b1 | 54445 | HITS-CLIP | 23142080 | Unspecified |
| mmu-miR-29b-3p | MIMAT0000127 | Unk | 217331 | HITS-CLIP | 37986224\|25083871 | Unspecified |
| mmu-miR-29b-3p | MIMAT0000127 | Upf1 | 19704 | HITS-CLIP//sRNA-Seq | 25083871\|33184237\|23142080 | Unspecified |
| mmu-miR-29b-3p | MIMAT0000127 | Upf2 | 326622 | HITS-CLIP | 25083871\|23142080 | Unspecified |
| mmu-miR-29b-3p | MIMAT0000127 | Upp1 | 22271 | HITS-CLIP | 23142080 | Unspecified |

| | | | | | | |
|---|---|---|---|---|---|---|
| mmu-miR-29b-3p | MIMAT0000127 | Uqcrc1 | 22273 | HITS-CLIP//sRNA-Seq | 25083871\|23597149\|23142080\|33184 | Unspecified |
| mmu-miR-29b-3p | MIMAT0000127 | Uqcrc2 | 67003 | HITS-CLIP//sRNA-Seq | 37986224\|25083871\|23597149\|33184 | Unspecified |
| mmu-miR-29b-3p | MIMAT0000127 | Uqcrfs1 | 66694 | HITS-CLIP//sRNA-Seq | 25083871\|33184237\|23597149\|23142 | Unspecified |
| mmu-miR-29b-3p | MIMAT0000127 | Urgcp | 72046 | HITS-CLIP | 23597149 | Unspecified |
| mmu-miR-29b-3p | MIMAT0000127 | Uri1 | 19777 | HITS-CLIP | 23142080 | Unspecified |
| mmu-miR-29b-3p | MIMAT0000127 | Urm1 | 68205 | sRNA-Seq//HITS-CLIP | 33184237\|23597149 | Unspecified |
| mmu-miR-29b-3p | MIMAT0000127 | Uroc1 | 243537 | HITS-CLIP | 37986224 | Unspecified |
| mmu-miR-29b-3p | MIMAT0000127 | Urod | 22275 | HITS-CLIP//sRNA-Seq | 23597149\|33184237\|23142080 | Unspecified |
| mmu-miR-29b-3p | MIMAT0000127 | Usb1 | 101985 | HITS-CLIP | 37986224 | Unspecified |
| mmu-miR-29b-3p | MIMAT0000127 | Use1 | 67023 | sRNA-Seq//HITS-CLIP | 33184237\|23142080 | Unspecified |
| mmu-miR-29b-3p | MIMAT0000127 | Usf3 | 205717 | HITS-CLIP | 25083871\|23597149\|23142080 | Unspecified |
| mmu-miR-29b-3p | MIMAT0000127 | Usmg5 | 84833 | HITS-CLIP//sRNA-Seq | 23142080\|33184237\|25083871 | Unspecified |
| mmu-miR-29b-3p | MIMAT0000127 | Uso1 | 56041 | HITS-CLIP//sRNA-Seq | 25083871\|23597149\|33184237\|23142 | Unspecified |
| mmu-miR-29b-3p | MIMAT0000127 | Usp12 | 22217 | HITS-CLIP | 25083871\|23142080 | Unspecified |
| mmu-miR-29b-3p | MIMAT0000127 | Usp14 | 59025 | HITS-CLIP//sRNA-Seq | 37986224\|23597149\|33184237\|23142 | Unspecified |
| mmu-miR-29b-3p | MIMAT0000127 | Usp15 | 14479 | HITS-CLIP//sRNA-Seq | 23142080\|33184237\|25083871 | Unspecified |
| mmu-miR-29b-3p | MIMAT0000127 | Usp16 | 74112 | HITS-CLIP | 23142080 | Unspecified |
| mmu-miR-29b-3p | MIMAT0000127 | Usp19 | 71472 | HITS-CLIP//sRNA-Seq | 23597149\|33184237\|25083871 | Unspecified |
| mmu-miR-29b-3p | MIMAT0000127 | Usp22 | 216825 | HITS-CLIP | 37986224 | Unspecified |
| mmu-miR-29b-3p | MIMAT0000127 | Usp24 | 329908 | HITS-CLIP | 25083871\|23142080\|23597149 | Unspecified |
| mmu-miR-29b-3p | MIMAT0000127 | Usp25 | 30940 | HITS-CLIP | 25083871\|23142080\|26203562 | Unspecified |
| mmu-miR-29b-3p | MIMAT0000127 | Usp28 | 235323 | HITS-CLIP | 37986224\|25083871\|23142080 | Unspecified |
| mmu-miR-29b-3p | MIMAT0000127 | Usp3 | 235441 | HITS-CLIP//sRNA-Seq | 25083871\|23597149\|33184237\|23142 | Unspecified |
| mmu-miR-29b-3p | MIMAT0000127 | Usp31 | 76179 | HITS-CLIP | 37986224 | Unspecified |
| mmu-miR-29b-3p | MIMAT0000127 | Usp32 | 237898 | HITS-CLIP//sRNA-Seq | 25083871\|33184237\|23597149 | Unspecified |
| mmu-miR-29b-3p | MIMAT0000127 | Usp33 | 170822 | HITS-CLIP//sRNA-Seq | 23142080\|33184237\|25083871 | Unspecified |
| mmu-miR-29b-3p | MIMAT0000127 | Usp34 | 17847 | HITS-CLIP//sRNA-Seq | 37986224\|25083871\|23142080\|33184 | Unspecified |
| mmu-miR-29b-3p | MIMAT0000127 | Usp36 | 72344 | HITS-CLIP | 25083871\|23142080 | Unspecified |
| mmu-miR-29b-3p | MIMAT0000127 | Usp37 | 319651 | HITS-CLIP | 37986224\|25083871\|23597149\|23142 | Unspecified |
| mmu-miR-29b-3p | MIMAT0000127 | Usp38 | 74841 | HITS-CLIP | 37986224\|25083871\|23597149\|23142 | Unspecified |
| mmu-miR-29b-3p | MIMAT0000127 | Usp39 | 28035 | HITS-CLIP | 31433990 | Unspecified |
| mmu-miR-29b-3p | MIMAT0000127 | Usp4 | 22258 | HITS-CLIP//sRNA-Seq | 25083871\|33184237\|23597149\|23142 | Unspecified |
| mmu-miR-29b-3p | MIMAT0000127 | Usp40 | 227334 | HITS-CLIP | 23142080 | Unspecified |
| mmu-miR-29b-3p | MIMAT0000127 | Usp42 | 76800 | HITS-CLIP | 37986224\|25083871\|23597149\|23142 | Unspecified |
| mmu-miR-29b-3p | MIMAT0000127 | Usp45 | 77593 | HITS-CLIP | 37986224 | Unspecified |
| mmu-miR-29b-3p | MIMAT0000127 | Usp46 | 69727 | HITS-CLIP | 37986224\|25083871 | Unspecified |
| mmu-miR-29b-3p | MIMAT0000127 | Usp47 | 74996 | HITS-CLIP//sRNA-Seq | 23142080\|33184237\|23597149\|25083 | Unspecified |
| mmu-miR-29b-3p | MIMAT0000127 | Usp48 | 170707 | HITS-CLIP | 37986224 | Unspecified |
| mmu-miR-29b-3p | MIMAT0000127 | Usp49 | 224836 | HITS-CLIP | 25083871\|23142080 | Unspecified |

| | | | | | | |
|---|---|---|---|---|---|---|
| mmu-miR-29b-3p | MIMAT0000127 | Usp7 | 252870 | HITS-CLIP//sRNA-Seq | 25083871\|23142080\|33184237 | Unspecified |
| mmu-miR-29b-3p | MIMAT0000127 | Usp8 | 84092 | sRNA-Seq//HITS-CLIP | 33184237\|23142080 | Unspecified |
| mmu-miR-29b-3p | MIMAT0000127 | Usp9x | 22284 | HITS-CLIP//sRNA-Seq | 37986224\|25083871\|23597149\|3318[illegible] | Unspecified |
| mmu-miR-29b-3p | MIMAT0000127 | Uspl1 | 231915 | HITS-CLIP | 25083871\|23142080 | Unspecified |
| mmu-miR-29b-3p | MIMAT0000127 | Utp14b | 195434 | HITS-CLIP | 37986224\|25083871\|23142080 | Unspecified |
| mmu-miR-29b-3p | MIMAT0000127 | Utp15 | 105372 | HITS-CLIP | 23142080 | Unspecified |
| mmu-miR-29b-3p | MIMAT0000127 | Utp18 | 217109 | HITS-CLIP | 25083871\|23142080 | Unspecified |
| mmu-miR-29b-3p | MIMAT0000127 | Utp23 | 78581 | HITS-CLIP | 23142080 | Unspecified |
| mmu-miR-29b-3p | MIMAT0000127 | Utp3 | 65961 | HITS-CLIP//sRNA-Seq | 37986224\|23597149\|33184237\|2314[illegible] | Unspecified |
| mmu-miR-29b-3p | MIMAT0000127 | Utp6 | 216987 | HITS-CLIP | 37986224 | Unspecified |
| mmu-miR-29b-3p | MIMAT0000127 | Utrn | 22288 | HITS-CLIP//sRNA-Seq | 25083871\|33184237\|23142080 | Unspecified |
| mmu-miR-29b-3p | MIMAT0000127 | Uvrag | 78610 | HITS-CLIP | 37986224\|23597149\|23142080 | Unspecified |
| mmu-miR-29b-3p | MIMAT0000127 | Uvssa | 71101 | HITS-CLIP | 23142080 | Unspecified |
| mmu-miR-29b-3p | MIMAT0000127 | Vamp3 | 22319 | HITS-CLIP//sRNA-Seq | 37986224\|25083871\|23597149\|3318[illegible] | Unspecified |
| mmu-miR-29b-3p | MIMAT0000127 | Vangl2 | 93840 | HITS-CLIP | 37986224\|25083871 | Unspecified |
| mmu-miR-29b-3p | MIMAT0000127 | Vapa | 30960 | HITS-CLIP//sRNA-Seq | 23597149\|23142080\|33184237 | Unspecified |
| mmu-miR-29b-3p | MIMAT0000127 | Vapb | 56491 | HITS-CLIP//sRNA-Seq | 25083871\|33184237 | Unspecified |
| mmu-miR-29b-3p | MIMAT0000127 | Vars | 22321 | HITS-CLIP//sRNA-Seq | 25083871\|33184237 | Unspecified |
| mmu-miR-29b-3p | MIMAT0000127 | Vash1 | 238328 | HITS-CLIP | 25083871 | Unspecified |
| mmu-miR-29b-3p | MIMAT0000127 | Vasn | 246154 | HITS-CLIP//sRNA-Seq | 25083871\|33184237 | Unspecified |
| mmu-miR-29b-3p | MIMAT0000127 | Vav2 | 22325 | HITS-CLIP | 23597149 | Unspecified |
| mmu-miR-29b-3p | MIMAT0000127 | Vcam1 | 22329 | HITS-CLIP//sRNA-Seq | 25083871\|23597149\|33184237 | Unspecified |
| mmu-miR-29b-3p | MIMAT0000127 | Vcan | 13003 | HITS-CLIP//sRNA-Seq | 25083871\|33184237 | Unspecified |
| mmu-miR-29b-3p | MIMAT0000127 | Vcl | 22330 | HITS-CLIP//sRNA-Seq | 25083871\|33184237\|23142080 | Unspecified |
| mmu-miR-29b-3p | MIMAT0000127 | Vcpip1 | 70675 | HITS-CLIP | 37986224\|25083871\|23597149\|2314[illegible] | Unspecified |
| mmu-miR-29b-3p | MIMAT0000127 | Vdac1 | 22333 | HITS-CLIP//sRNA-Seq | 25083871\|33184237\|23142080 | Unspecified |
| mmu-miR-29b-3p | MIMAT0000127 | Vdac2 | 22334 | HITS-CLIP//sRNA-Seq | 23597149\|25083871\|33184237\|2314[illegible] | Unspecified |
| mmu-miR-29b-3p | MIMAT0000127 | Vegfa | 22339 | HITS-CLIP//ELISA//Luciferase reporter ass[illegible] | 37986224\|25083871\|23142080\|2758[illegible] | Unspecified |
| mmu-miR-29b-3p | MIMAT0000127 | Vezt | 215008 | HITS-CLIP//sRNA-Seq | 25083871\|33184237 | Unspecified |
| mmu-miR-29b-3p | MIMAT0000127 | Vgll4 | 232334 | HITS-CLIP//sRNA-Seq | 25083871\|33184237\|23142080 | Unspecified |
| mmu-miR-29b-3p | MIMAT0000127 | Vhl | 22346 | HITS-CLIP | 37986224 | Unspecified |
| mmu-miR-29b-3p | MIMAT0000127 | Vim | 22352 | HITS-CLIP//sRNA-Seq | 25083871\|33184237\|23142080 | Unspecified |
| mmu-miR-29b-3p | MIMAT0000127 | Virma | 25962 | HITS-CLIP | 25083871\|23142080 | Unspecified |
| mmu-miR-29b-3p | MIMAT0000127 | Vkorc1 | 27973 | HITS-CLIP | 23597149 | Unspecified |
| mmu-miR-29b-3p | MIMAT0000127 | Vkorc1l1 | 69568 | HITS-CLIP | 23142080\|25083871 | Unspecified |
| mmu-miR-29b-3p | MIMAT0000127 | Vmn1r55 | 384522 | qCLASH | 31393953 | Unspecified |
| mmu-miR-29b-3p | MIMAT0000127 | Vmn2r107 | 22312 | HITS-CLIP | 25083871 | Unspecified |
| mmu-miR-29b-3p | MIMAT0000127 | Vmp1 | 75909 | HITS-CLIP//sRNA-Seq | 23597149\|33184237\|25083871 | Unspecified |
| mmu-miR-29b-3p | MIMAT0000127 | Vps13a | 271564 | HITS-CLIP//sRNA-Seq | 25083871\|33184237\|23142080 | Unspecified |

| | | | | | | |
|---|---|---|---|---|---|---|
| mmu-miR-29b-3p | MIMAT0000127 | Vps13b | 666173 | HITS-CLIP | 37986224\|25083871\|23597149\|23142 | Unspecified |
| mmu-miR-29b-3p | MIMAT0000127 | Vps13c | 320528 | HITS-CLIP//sRNA-Seq | 25083871\|33184237\|23597149\|23142 | Unspecified |
| mmu-miR-29b-3p | MIMAT0000127 | Vps13d | 230895 | HITS-CLIP | 23597149\|23142080 | Unspecified |
| mmu-miR-29b-3p | MIMAT0000127 | Vps16 | 80743 | sRNA-Seq//HITS-CLIP | 33184237\|23142080 | Unspecified |
| mmu-miR-29b-3p | MIMAT0000127 | Vps18 | 228545 | HITS-CLIP | 23142080 | Unspecified |
| mmu-miR-29b-3p | MIMAT0000127 | Vps25 | 28084 | HITS-CLIP//sRNA-Seq | 37986224\|23597149\|33184237\|23142 | Unspecified |
| mmu-miR-29b-3p | MIMAT0000127 | Vps26a | 30930 | sRNA-Seq//HITS-CLIP | 33184237\|23597149 | Unspecified |
| mmu-miR-29b-3p | MIMAT0000127 | Vps26b | 69091 | HITS-CLIP//sRNA-Seq | 37986224\|23597149\|25083871\|33184 | Unspecified |
| mmu-miR-29b-3p | MIMAT0000127 | Vps33b | 233405 | HITS-CLIP | 23142080 | Unspecified |
| mmu-miR-29b-3p | MIMAT0000127 | Vps35 | 65114 | HITS-CLIP//sRNA-Seq | 23597149\|33184237\|23142080 | Unspecified |
| mmu-miR-29b-3p | MIMAT0000127 | Vps36 | 70160 | HITS-CLIP | 37986224\|25083871\|23142080 | Unspecified |
| mmu-miR-29b-3p | MIMAT0000127 | Vps37c | 107305 | HITS-CLIP | 37986224\|23142080 | Unspecified |
| mmu-miR-29b-3p | MIMAT0000127 | Vps41 | 218035 | sRNA-Seq//HITS-CLIP | 33184237\|23597149 | Unspecified |
| mmu-miR-29b-3p | MIMAT0000127 | Vps4b | 20479 | sRNA-Seq//HITS-CLIP | 33184237\|23597149\|23142080 | Unspecified |
| mmu-miR-29b-3p | MIMAT0000127 | Vps52 | 224705 | HITS-CLIP | 23597149 | Unspecified |
| mmu-miR-29b-3p | MIMAT0000127 | Vps53 | 68299 | HITS-CLIP//sRNA-Seq | 23597149\|33184237 | Unspecified |
| mmu-miR-29b-3p | MIMAT0000127 | Vps54 | 245944 | HITS-CLIP | 25083871\|23142080 | Unspecified |
| mmu-miR-29b-3p | MIMAT0000127 | Vps9d1 | 72325 | HITS-CLIP | 23142080 | Unspecified |
| mmu-miR-29b-3p | MIMAT0000127 | Vrk1 | 22367 | HITS-CLIP | 23142080 | Unspecified |
| mmu-miR-29b-3p | MIMAT0000127 | Vrk2 | 69922 | HITS-CLIP | 23142080 | Unspecified |
| mmu-miR-29b-3p | MIMAT0000127 | Vwa5a | 67776 | HITS-CLIP//sRNA-Seq | 23597149\|33184237 | Unspecified |
| mmu-miR-29b-3p | MIMAT0000127 | Vwa8 | 219189 | HITS-CLIP | 23597149 | Unspecified |
| mmu-miR-29b-3p | MIMAT0000127 | Wac | 225131 | HITS-CLIP//sRNA-Seq | 37986224\|23142080\|33184237 | Unspecified |
| mmu-miR-29b-3p | MIMAT0000127 | Wapl | 23063 | HITS-CLIP | 23597149\|23142080 | Unspecified |
| mmu-miR-29b-3p | MIMAT0000127 | Wars1 | 7453 | HITS-CLIP//sRNA-Seq | 30971389\|33184237 | Unspecified |
| mmu-miR-29b-3p | MIMAT0000127 | Wasf1 | 83767 | HITS-CLIP//sRNA-Seq | 25083871\|33184237 | Unspecified |
| mmu-miR-29b-3p | MIMAT0000127 | Wasf2 | 242687 | HITS-CLIP//sRNA-Seq | 25083871\|33184237 | Unspecified |
| mmu-miR-29b-3p | MIMAT0000127 | Washc3 | 51019 | HITS-CLIP | 23142080 | Unspecified |
| mmu-miR-29b-3p | MIMAT0000127 | Washc4 | 319277 | HITS-CLIP | 23597149\|23142080 | Unspecified |
| mmu-miR-29b-3p | MIMAT0000127 | Washc5 | 9897 | sRNA-Seq//HITS-CLIP | 33184237\|25083871 | Unspecified |
| mmu-miR-29b-3p | MIMAT0000127 | Wasl | 73178 | HITS-CLIP | 37986224\|23597149\|25083871\|23142 | Unspecified |
| mmu-miR-29b-3p | MIMAT0000127 | Wbp11 | 60321 | HITS-CLIP//sRNA-Seq | 25083871\|33184237 | Unspecified |
| mmu-miR-29b-3p | MIMAT0000127 | Wbp1l | 226178 | HITS-CLIP//sRNA-Seq | 37986224\|25083871\|33184237\|23597 | Unspecified |
| mmu-miR-29b-3p | MIMAT0000127 | Wdfy1 | 69368 | HITS-CLIP | 37986224\|25083871\|23142080 | Unspecified |
| mmu-miR-29b-3p | MIMAT0000127 | Wdfy3 | 72145 | HITS-CLIP//sRNA-Seq | 37986224\|25083871\|33184237\|23597 | Unspecified |
| mmu-miR-29b-3p | MIMAT0000127 | Wdr1 | 22388 | sRNA-Seq//HITS-CLIP | 33184237\|23142080 | Unspecified |
| mmu-miR-29b-3p | MIMAT0000127 | Wdr11 | 207425 | HITS-CLIP | 37986224\|23597149\|23142080 | Unspecified |
| mmu-miR-29b-3p | MIMAT0000127 | Wdr13 | 73447 | HITS-CLIP//sRNA-Seq | 37986224\|23142080\|33184237 | Unspecified |
| mmu-miR-29b-3p | MIMAT0000127 | Wdr19 | 213081 | HITS-CLIP | 23142080 | Unspecified |

| | | | | | | |
|---|---|---|---|---|---|---|
| mmu-miR-29b-3p | MIMAT0000127 | Wdr26 | 226757 | HITS-CLIP//sRNA-Seq | 37986224\|23142080\|23597149\|33184 | Unspecified |
| mmu-miR-29b-3p | MIMAT0000127 | Wdr33 | 74320 | sRNA-Seq//HITS-CLIP | 33184237\|23142080 | Unspecified |
| mmu-miR-29b-3p | MIMAT0000127 | Wdr37 | 207615 | HITS-CLIP//sRNA-Seq | 37986224\|25083871\|23142080\|23597 | Unspecified |
| mmu-miR-29b-3p | MIMAT0000127 | Wdr4 | 57773 | HITS-CLIP | 23142080 | Unspecified |
| mmu-miR-29b-3p | MIMAT0000127 | Wdr43 | 72515 | HITS-CLIP//sRNA-Seq | 25083871\|33184237\|23142080 | Unspecified |
| mmu-miR-29b-3p | MIMAT0000127 | Wdr44 | 72404 | HITS-CLIP | 23142080 | Unspecified |
| mmu-miR-29b-3p | MIMAT0000127 | Wdr59 | 319481 | HITS-CLIP | 37986224\|23597149\|23142080 | Unspecified |
| mmu-miR-29b-3p | MIMAT0000127 | Wdr6 | 83669 | sRNA-Seq//HITS-CLIP | 33184237\|23142080 | Unspecified |
| mmu-miR-29b-3p | MIMAT0000127 | Wdr75 | 73674 | HITS-CLIP | 23142080 | Unspecified |
| mmu-miR-29b-3p | MIMAT0000127 | Wdr76 | 241627 | HITS-CLIP | 23142080 | Unspecified |
| mmu-miR-29b-3p | MIMAT0000127 | Wdr77 | 70465 | HITS-CLIP | 25083871\|23142080 | Unspecified |
| mmu-miR-29b-3p | MIMAT0000127 | Wdr82 | 77305 | HITS-CLIP//sRNA-Seq | 23597149\|33184237\|25083871\|23142 | Unspecified |
| mmu-miR-29b-3p | MIMAT0000127 | Wdr89 | 72338 | HITS-CLIP | 23142080 | Unspecified |
| mmu-miR-29b-3p | MIMAT0000127 | Wdr92 | 103784 | HITS-CLIP | 23142080 | Unspecified |
| mmu-miR-29b-3p | MIMAT0000127 | Wdtc1 | 230796 | sRNA-Seq//HITS-CLIP | 33184237\|23597149\|23142080 | Unspecified |
| mmu-miR-29b-3p | MIMAT0000127 | Wdyhv1 | 76773 | HITS-CLIP | 23142080 | Unspecified |
| mmu-miR-29b-3p | MIMAT0000127 | Wee1 | 22390 | HITS-CLIP | 23142080 | Unspecified |
| mmu-miR-29b-3p | MIMAT0000127 | Wipf1 | 215280 | HITS-CLIP | 25083871\|23142080 | Unspecified |
| mmu-miR-29b-3p | MIMAT0000127 | Wipf2 | 68524 | HITS-CLIP | 25083871 | Unspecified |
| mmu-miR-29b-3p | MIMAT0000127 | Wipi1 | 52639 | HITS-CLIP//sRNA-Seq | 25083871\|33184237\|23142080 | Unspecified |
| mmu-miR-29b-3p | MIMAT0000127 | Wiz | 22404 | HITS-CLIP//sRNA-Seq | 23142080\|33184237 | Unspecified |
| mmu-miR-29b-3p | MIMAT0000127 | Wnk1 | 232341 | HITS-CLIP//sRNA-Seq | 37986224\|25083871\|33184237\|23597 | Unspecified |
| mmu-miR-29b-3p | MIMAT0000127 | Wnt9a | 216795 | HITS-CLIP | 25083871 | Unspecified |
| mmu-miR-29b-3p | MIMAT0000127 | Wnt9b | 22412 | HITS-CLIP | 23597149\|28735896 | Unspecified |
| mmu-miR-29b-3p | MIMAT0000127 | Wrb | 71446 | sRNA-Seq//HITS-CLIP | 33184237\|25083871 | Unspecified |
| mmu-miR-29b-3p | MIMAT0000127 | Wrn | 22427 | HITS-CLIP | 23142080 | Unspecified |
| mmu-miR-29b-3p | MIMAT0000127 | Wrnip1 | 78903 | HITS-CLIP | 23597149 | Unspecified |
| mmu-miR-29b-3p | MIMAT0000127 | Wsb1 | 78889 | HITS-CLIP//sRNA-Seq | 23142080\|33184237 | Unspecified |
| mmu-miR-29b-3p | MIMAT0000127 | Wwc1 | 211652 | HITS-CLIP | 23597149 | Unspecified |
| mmu-miR-29b-3p | MIMAT0000127 | Wwc2 | 52357 | HITS-CLIP | 25083871 | Unspecified |
| mmu-miR-29b-3p | MIMAT0000127 | Wwtr1 | 97064 | HITS-CLIP//sRNA-Seq | 37986224\|25083871\|33184237 | Unspecified |
| mmu-miR-29b-3p | MIMAT0000127 | Xbp1 | 22433 | HITS-CLIP//sRNA-Seq | 37986224\|33184237\|23142080 | Unspecified |
| mmu-miR-29b-3p | MIMAT0000127 | Xiap | 11798 | HITS-CLIP | 37986224\|23597149\|25083871\|23142 | Unspecified |
| mmu-miR-29b-3p | MIMAT0000127 | Xirp2 | 241431 | HITS-CLIP | 25083871 | Unspecified |
| mmu-miR-29b-3p | MIMAT0000127 | Xkr8 | 381560 | HITS-CLIP | 23597149 | Unspecified |
| mmu-miR-29b-3p | MIMAT0000127 | Xpc | 22591 | HITS-CLIP | 37986224\|23597149\|23142080 | Unspecified |
| mmu-miR-29b-3p | MIMAT0000127 | Xpnpep1 | 170750 | HITS-CLIP//sRNA-Seq | 37986224\|23597149\|33184237 | Unspecified |
| mmu-miR-29b-3p | MIMAT0000127 | Xpnpep3 | 321003 | HITS-CLIP | 25083871 | Unspecified |
| mmu-miR-29b-3p | MIMAT0000127 | Xpo1 | 103573 | HITS-CLIP//sRNA-Seq | 25083871\|33184237\|23142080 | Unspecified |

| | | | | | | |
|---|---|---|---|---|---|---|
| mmu-miR-29b-3p | MIMAT0000127 | Xpo5 | 72322 | HITS-CLIP | 23597149 | Unspecified |
| mmu-miR-29b-3p | MIMAT0000127 | Xpo7 | 65246 | HITS-CLIP | 23142080\|25083871\|23597149 | Unspecified |
| mmu-miR-29b-3p | MIMAT0000127 | Xpot | 73192 | HITS-CLIP//sRNA-Seq | 23142080\|33184237 | Unspecified |
| mmu-miR-29b-3p | MIMAT0000127 | Xpr1 | 19775 | HITS-CLIP//sRNA-Seq | 23597149\|25083871\|33184237 | Unspecified |
| mmu-miR-29b-3p | MIMAT0000127 | Xrcc1 | 22594 | HITS-CLIP | 23597149 | Unspecified |
| mmu-miR-29b-3p | MIMAT0000127 | Xrcc4 | 108138 | HITS-CLIP | 23142080 | Unspecified |
| mmu-miR-29b-3p | MIMAT0000127 | Xrcc6 | 14375 | HITS-CLIP//sRNA-Seq | 25083871\|33184237\|23142080 | Unspecified |
| mmu-miR-29b-3p | MIMAT0000127 | Xrn1 | 24127 | HITS-CLIP | 25083871\|23142080 | Unspecified |
| mmu-miR-29b-3p | MIMAT0000127 | Xrn2 | 24128 | HITS-CLIP//sRNA-Seq | 23597149\|25083871\|33184237\|2314 | Unspecified |
| mmu-miR-29b-3p | MIMAT0000127 | Xylt1 | 233781 | HITS-CLIP | 23142080 | Unspecified |
| mmu-miR-29b-3p | MIMAT0000127 | Yars | 107271 | HITS-CLIP//sRNA-Seq | 37986224\|25083871\|33184237\|2359 | Unspecified |
| mmu-miR-29b-3p | MIMAT0000127 | Ybey | 216119 | HITS-CLIP | 37986224\|23142080 | Unspecified |
| mmu-miR-29b-3p | MIMAT0000127 | Ybx3 | 56449 | HITS-CLIP//sRNA-Seq | 37986224\|25083871\|33184237\|2314 | Unspecified |
| mmu-miR-29b-3p | MIMAT0000127 | Yes1 | 22612 | HITS-CLIP | 25083871\|23142080 | Unspecified |
| mmu-miR-29b-3p | MIMAT0000127 | Yif1b | 77254 | HITS-CLIP//sRNA-Seq | 37986224\|23142080\|25083871\|2359 | Unspecified |
| mmu-miR-29b-3p | MIMAT0000127 | Yipf2 | 74766 | HITS-CLIP | 23597149\|23142080 | Unspecified |
| mmu-miR-29b-3p | MIMAT0000127 | Yipf3 | 28064 | HITS-CLIP//sRNA-Seq | 23597149\|33184237 | Unspecified |
| mmu-miR-29b-3p | MIMAT0000127 | Yipf5 | 67180 | HITS-CLIP//sRNA-Seq | 37986224\|23597149\|23142080\|3318 | Unspecified |
| mmu-miR-29b-3p | MIMAT0000127 | Yipf6 | 77929 | sRNA-Seq//HITS-CLIP | 33184237\|23142080 | Unspecified |
| mmu-miR-29b-3p | MIMAT0000127 | Ykt6 | 56418 | HITS-CLIP//sRNA-Seq | 23597149\|25083871\|33184237 | Unspecified |
| mmu-miR-29b-3p | MIMAT0000127 | Ylpm1 | 56531 | HITS-CLIP//sRNA-Seq | 23142080\|33184237\|25083871 | Unspecified |
| mmu-miR-29b-3p | MIMAT0000127 | Yod1 | 226418 | HITS-CLIP | 37986224\|25083871\|23597149\|2314 | Unspecified |
| mmu-miR-29b-3p | MIMAT0000127 | Ypel2 | 77864 | HITS-CLIP | 37986224\|23142080\|25083871 | Unspecified |
| mmu-miR-29b-3p | MIMAT0000127 | Ypel3 | 66090 | sRNA-Seq//HITS-CLIP | 33184237\|23142080 | Unspecified |
| mmu-miR-29b-3p | MIMAT0000127 | Ypel5 | 383295 | HITS-CLIP | 23142080 | Unspecified |
| mmu-miR-29b-3p | MIMAT0000127 | Yrdc | 230734 | HITS-CLIP | 23142080 | Unspecified |
| mmu-miR-29b-3p | MIMAT0000127 | Ythdc1 | 231386 | HITS-CLIP | 25083871\|23142080 | Unspecified |
| mmu-miR-29b-3p | MIMAT0000127 | Ythdf1 | 228994 | HITS-CLIP//sRNA-Seq | 37986224\|25083871\|33184237\|2359 | Unspecified |
| mmu-miR-29b-3p | MIMAT0000127 | Ythdf2 | 213541 | HITS-CLIP | 25083871\|23142080 | Unspecified |
| mmu-miR-29b-3p | MIMAT0000127 | Ythdf3 | 229096 | HITS-CLIP//sRNA-Seq | 25083871\|23597149\|33184237\|2314 | Unspecified |
| mmu-miR-29b-3p | MIMAT0000127 | Ywhae | 22627 | HITS-CLIP//sRNA-Seq | 37986224\|25083871\|33184237\|2359 | Unspecified |
| mmu-miR-29b-3p | MIMAT0000127 | Ywhag | 22628 | HITS-CLIP//sRNA-Seq | 25083871\|33184237\|23142080 | Unspecified |
| mmu-miR-29b-3p | MIMAT0000127 | Ywhah | 22629 | HITS-CLIP//sRNA-Seq | 23142080\|25083871\|33184237\|2359 | Unspecified |
| mmu-miR-29b-3p | MIMAT0000127 | Ywhaq | 22630 | HITS-CLIP//sRNA-Seq | 23142080\|25083871\|33184237 | Unspecified |
| mmu-miR-29b-3p | MIMAT0000127 | Yy1 | 22632 | HITS-CLIP//Other | 37986224\|23142080\|25083871\|1897 | Unspecified |
| mmu-miR-29b-3p | MIMAT0000127 | Zadh2 | 225791 | HITS-CLIP//sRNA-Seq | 23597149\|33184237\|23142080 | Unspecified |
| mmu-miR-29b-3p | MIMAT0000127 | Zap70 | 22637 | HITS-CLIP | 23142080 | Unspecified |
| mmu-miR-29b-3p | MIMAT0000127 | Zbed6 | 667118 | HITS-CLIP | 37986224\|26203562 | Unspecified |
| mmu-miR-29b-3p | MIMAT0000127 | Zbtb10 | 229055 | HITS-CLIP | 37986224\|25083871 | Unspecified |

| | | | | | | |
|---|---|---|---|---|---|---|
| mmu-miR-29b-3p | MIMAT0000127 | Zbtb11 | 271377 | HITS-CLIP | 25083871\|23142080 | Unspecified |
| mmu-miR-29b-3p | MIMAT0000127 | Zbtb18 | 30928 | HITS-CLIP | 25083871\|23597149\|23142080 | Unspecified |
| mmu-miR-29b-3p | MIMAT0000127 | Zbtb20 | 56490 | HITS-CLIP | 37986224\|25083871\|23597149\|23142 | Unspecified |
| mmu-miR-29b-3p | MIMAT0000127 | Zbtb26 | 320633 | HITS-CLIP | 25083871 | Unspecified |
| mmu-miR-29b-3p | MIMAT0000127 | Zbtb34 | 241311 | HITS-CLIP | 37986224\|25083871\|23597149\|23142 | Unspecified |
| mmu-miR-29b-3p | MIMAT0000127 | Zbtb37 | 240869 | HITS-CLIP | 25083871 | Unspecified |
| mmu-miR-29b-3p | MIMAT0000127 | Zbtb38 | 245007 | HITS-CLIP | 23142080 | Unspecified |
| mmu-miR-29b-3p | MIMAT0000127 | Zbtb39 | 320080 | HITS-CLIP | 25083871\|23597149\|23142080 | Unspecified |
| mmu-miR-29b-3p | MIMAT0000127 | Zbtb4 | 75580 | HITS-CLIP | 37986224\|25083871 | Unspecified |
| mmu-miR-29b-3p | MIMAT0000127 | Zbtb41 | 226470 | HITS-CLIP | 37986224\|25083871\|23142080 | Unspecified |
| mmu-miR-29b-3p | MIMAT0000127 | Zbtb42 | 382639 | HITS-CLIP | 23142080\|25083871 | Unspecified |
| mmu-miR-29b-3p | MIMAT0000127 | Zbtb43 | 71834 | HITS-CLIP | 37986224\|25083871 | Unspecified |
| mmu-miR-29b-3p | MIMAT0000127 | Zbtb44 | 235132 | HITS-CLIP | 25083871\|23142080 | Unspecified |
| mmu-miR-29b-3p | MIMAT0000127 | Zbtb46 | 72147 | HITS-CLIP | 37986224\|25083871 | Unspecified |
| mmu-miR-29b-3p | MIMAT0000127 | Zbtb5 | 230119 | HITS-CLIP | 23142080\|25083871\|23597149\|28735 | Unspecified |
| mmu-miR-29b-3p | MIMAT0000127 | Zbtb6 | 241322 | HITS-CLIP | 25083871 | Unspecified |
| mmu-miR-29b-3p | MIMAT0000127 | Zbtb7a | 16969 | HITS-CLIP | 23142080 | Unspecified |
| mmu-miR-29b-3p | MIMAT0000127 | Zbtb7b | 22724 | HITS-CLIP | 25083871\|23142080 | Unspecified |
| mmu-miR-29b-3p | MIMAT0000127 | Zbtb9 | 474156 | HITS-CLIP | 25083871\|23597149\|23142080 | Unspecified |
| mmu-miR-29b-3p | MIMAT0000127 | Zc3h11a | 70579 | sRNA-Seq//HITS-CLIP | 33184237\|26203562 | Unspecified |
| mmu-miR-29b-3p | MIMAT0000127 | Zc3h12a | 230738 | HITS-CLIP | 37986224 | Unspecified |
| mmu-miR-29b-3p | MIMAT0000127 | Zc3h12c | 244871 | HITS-CLIP | 23597149 | Unspecified |
| mmu-miR-29b-3p | MIMAT0000127 | Zc3h12d | 237256 | HITS-CLIP | 23142080 | Unspecified |
| mmu-miR-29b-3p | MIMAT0000127 | Zc3h13 | 67302 | HITS-CLIP | 25083871\|23142080 | Unspecified |
| mmu-miR-29b-3p | MIMAT0000127 | Zc3h4 | 330474 | HITS-CLIP | 25083871\|23142080 | Unspecified |
| mmu-miR-29b-3p | MIMAT0000127 | Zc3h6 | 78751 | HITS-CLIP | 23142080 | Unspecified |
| mmu-miR-29b-3p | MIMAT0000127 | Zc3h7b | 20286 | HITS-CLIP//sRNA-Seq | 25083871\|33184237 | Unspecified |
| mmu-miR-29b-3p | MIMAT0000127 | Zc3hav1 | 78781 | HITS-CLIP//sRNA-Seq | 31433990\|33184237 | Unspecified |
| mmu-miR-29b-3p | MIMAT0000127 | Zc3hav1l | 209032 | HITS-CLIP | 37986224\|25083871 | Unspecified |
| mmu-miR-29b-3p | MIMAT0000127 | Zc3hc1 | 51530 | HITS-CLIP | 23142080 | Unspecified |
| mmu-miR-29b-3p | MIMAT0000127 | Zc4h2 | 245522 | HITS-CLIP | 37986224 | Unspecified |
| mmu-miR-29b-3p | MIMAT0000127 | Zcchc14 | 142682 | HITS-CLIP | 25083871\|23597149 | Unspecified |
| mmu-miR-29b-3p | MIMAT0000127 | Zcchc2 | 227449 | HITS-CLIP | 25083871\|23597149\|23142080 | Unspecified |
| mmu-miR-29b-3p | MIMAT0000127 | Zcchc24 | 71918 | HITS-CLIP//sRNA-Seq | 37986224\|23597149\|33184237 | Unspecified |
| mmu-miR-29b-3p | MIMAT0000127 | Zcchc7 | 319885 | HITS-CLIP | 37986224\|25083871\|23142080 | Unspecified |
| mmu-miR-29b-3p | MIMAT0000127 | Zdhhc1 | 70796 | HITS-CLIP | 25083871 | Unspecified |
| mmu-miR-29b-3p | MIMAT0000127 | Zdhhc12 | 66220 | HITS-CLIP | 25083871\|23142080 | Unspecified |
| mmu-miR-29b-3p | MIMAT0000127 | Zdhhc13 | 243983 | HITS-CLIP | 37986224\|25083871\|23142080 | Unspecified |
| mmu-miR-29b-3p | MIMAT0000127 | Zdhhc15 | 108672 | HITS-CLIP | 23142080 | Unspecified |

| | | | | | | |
|---|---|---|---|---|---|---|
| mmu-miR-29b-3p | MIMAT0000127 | Zdhhc17 | 320150 | HITS-CLIP//sRNA-Seq | 25083871\|33184237\|23142080 | Unspecified |
| mmu-miR-29b-3p | MIMAT0000127 | Zdhhc18 | 503610 | HITS-CLIP | 25083871\|23142080 | Unspecified |
| mmu-miR-29b-3p | MIMAT0000127 | Zdhhc2 | 70546 | HITS-CLIP//sRNA-Seq | 25083871\|33184237\|23142080 | Unspecified |
| mmu-miR-29b-3p | MIMAT0000127 | Zdhhc21 | 68268 | HITS-CLIP | 37986224\|23597149\|25083871\|23142 | Unspecified |
| mmu-miR-29b-3p | MIMAT0000127 | Zdhhc24 | 70605 | HITS-CLIP | 23142080 | Unspecified |
| mmu-miR-29b-3p | MIMAT0000127 | Zdhhc4 | 72881 | HITS-CLIP | 23597149 | Unspecified |
| mmu-miR-29b-3p | MIMAT0000127 | Zdhhc5 | 228136 | HITS-CLIP//sRNA-Seq | 37986224\|25083871\|33184237\|23142 | Unspecified |
| mmu-miR-29b-3p | MIMAT0000127 | Zdhhc6 | 66980 | HITS-CLIP | 23142080\|25083871 | Unspecified |
| mmu-miR-29b-3p | MIMAT0000127 | Zdhhc8 | 27801 | HITS-CLIP | 25083871 | Unspecified |
| mmu-miR-29b-3p | MIMAT0000127 | Zdhhc9 | 208884 | HITS-CLIP//sRNA-Seq | 25083871\|33184237\|23142080 | Unspecified |
| mmu-miR-29b-3p | MIMAT0000127 | Zeb1 | 21417 | HITS-CLIP//sRNA-Seq | 25083871\|33184237 | Unspecified |
| mmu-miR-29b-3p | MIMAT0000127 | Zeb2 | 24136 | HITS-CLIP | 23142080 | Unspecified |
| mmu-miR-29b-3p | MIMAT0000127 | Zer1 | 227693 | HITS-CLIP | 23142080 | Unspecified |
| mmu-miR-29b-3p | MIMAT0000127 | Zfand2a | 100494 | HITS-CLIP | 25083871 | Unspecified |
| mmu-miR-29b-3p | MIMAT0000127 | Zfand3 | 21769 | sRNA-Seq//HITS-CLIP | 33184237\|23597149 | Unspecified |
| mmu-miR-29b-3p | MIMAT0000127 | Zfand6 | 65098 | HITS-CLIP | 23597149 | Unspecified |
| mmu-miR-29b-3p | MIMAT0000127 | Zfc3h1 | 216345 | HITS-CLIP | 25083871\|23142080 | Unspecified |
| mmu-miR-29b-3p | MIMAT0000127 | Zfhx3 | 11906 | HITS-CLIP | 37986224\|23597149\|25083871\|26203 | Unspecified |
| mmu-miR-29b-3p | MIMAT0000127 | Zfhx4 | 80892 | HITS-CLIP | 25083871 | Unspecified |
| mmu-miR-29b-3p | MIMAT0000127 | Zfp106 | 20402 | HITS-CLIP//sRNA-Seq | 25083871\|33184237\|23597149\|23142 | Unspecified |
| mmu-miR-29b-3p | MIMAT0000127 | Zfp113 | 56314 | HITS-CLIP | 37986224\|25083871\|23142080 | Unspecified |
| mmu-miR-29b-3p | MIMAT0000127 | Zfp12 | 231866 | HITS-CLIP | 25083871 | Unspecified |
| mmu-miR-29b-3p | MIMAT0000127 | Zfp148 | 22661 | HITS-CLIP | 23142080\|25083871 | Unspecified |
| mmu-miR-29b-3p | MIMAT0000127 | Zfp207 | 22680 | HITS-CLIP//sRNA-Seq | 23142080\|25083871\|33184237 | Unspecified |
| mmu-miR-29b-3p | MIMAT0000127 | Zfp217 | 228913 | HITS-CLIP | 37986224\|25083871\|23597149\|23142 | Unspecified |
| mmu-miR-29b-3p | MIMAT0000127 | Zfp236 | 329002 | HITS-CLIP | 25083871 | Unspecified |
| mmu-miR-29b-3p | MIMAT0000127 | Zfp24 | 59057 | HITS-CLIP | 25083871 | Unspecified |
| mmu-miR-29b-3p | MIMAT0000127 | Zfp248 | 72720 | HITS-CLIP | 25083871 | Unspecified |
| mmu-miR-29b-3p | MIMAT0000127 | Zfp251 | 71591 | HITS-CLIP | 31433990 | Unspecified |
| mmu-miR-29b-3p | MIMAT0000127 | Zfp26 | 22688 | HITS-CLIP | 25083871\|23142080 | Unspecified |
| mmu-miR-29b-3p | MIMAT0000127 | Zfp266 | 77519 | HITS-CLIP | 37986224 | Unspecified |
| mmu-miR-29b-3p | MIMAT0000127 | Zfp275 | 27081 | HITS-CLIP | 23142080 | Unspecified |
| mmu-miR-29b-3p | MIMAT0000127 | Zfp277 | 246196 | HITS-CLIP | 23142080 | Unspecified |
| mmu-miR-29b-3p | MIMAT0000127 | Zfp280b | 64453 | HITS-CLIP | 25083871\|23142080 | Unspecified |
| mmu-miR-29b-3p | MIMAT0000127 | Zfp280c | 208968 | HITS-CLIP | 25083871 | Unspecified |
| mmu-miR-29b-3p | MIMAT0000127 | Zfp280d | 235469 | HITS-CLIP | 25083871\|23142080 | Unspecified |
| mmu-miR-29b-3p | MIMAT0000127 | Zfp281 | 226442 | HITS-CLIP | 25083871\|23142080 | Unspecified |
| mmu-miR-29b-3p | MIMAT0000127 | Zfp282 | 101095 | HITS-CLIP | 37986224\|25083871\|23597149\|23142 | Unspecified |
| mmu-miR-29b-3p | MIMAT0000127 | Zfp292 | 30046 | HITS-CLIP | 37986224\|25083871\|23142080 | Unspecified |

| | | | | | | |
|---|---|---|---|---|---|---|
| mmu-miR-29b-3p | MIMAT0000127 | Zfp318 | 57908 | HITS-CLIP//sRNA-Seq | 37986224\|25083871\|33184237\|23142 | Unspecified |
| mmu-miR-29b-3p | MIMAT0000127 | Zfp319 | 79233 | HITS-CLIP | 25083871 | Unspecified |
| mmu-miR-29b-3p | MIMAT0000127 | Zfp322a | 218100 | HITS-CLIP | 37986224\|25083871 | Unspecified |
| mmu-miR-29b-3p | MIMAT0000127 | Zfp324 | 243834 | HITS-CLIP | 37986224 | Unspecified |
| mmu-miR-29b-3p | MIMAT0000127 | Zfp326 | 54367 | HITS-CLIP | 25083871 | Unspecified |
| mmu-miR-29b-3p | MIMAT0000127 | Zfp334 | 228876 | HITS-CLIP | 25083871 | Unspecified |
| mmu-miR-29b-3p | MIMAT0000127 | Zfp346 | 26919 | HITS-CLIP | 37986224\|25083871 | Unspecified |
| mmu-miR-29b-3p | MIMAT0000127 | Zfp354a | 21408 | HITS-CLIP | 37986224 | Unspecified |
| mmu-miR-29b-3p | MIMAT0000127 | Zfp36 | 22695 | HITS-CLIP | 37986224\|23597149\|31433990\|23142 | Unspecified |
| mmu-miR-29b-3p | MIMAT0000127 | Zfp362 | 230761 | HITS-CLIP | 25083871\|23142080 | Unspecified |
| mmu-miR-29b-3p | MIMAT0000127 | Zfp367 | 238673 | HITS-CLIP | 25083871\|23597149\|23142080 | Unspecified |
| mmu-miR-29b-3p | MIMAT0000127 | Zfp36l1 | 12192 | HITS-CLIP//sRNA-Seq | 37986224\|23597149\|33184237\|25083 | Unspecified |
| mmu-miR-29b-3p | MIMAT0000127 | Zfp36l2 | 12193 | HITS-CLIP//sRNA-Seq | 25083871\|33184237\|23597149 | Unspecified |
| mmu-miR-29b-3p | MIMAT0000127 | Zfp384 | 269800 | HITS-CLIP | 23142080 | Unspecified |
| mmu-miR-29b-3p | MIMAT0000127 | Zfp386 | 56220 | HITS-CLIP | 37986224\|25083871 | Unspecified |
| mmu-miR-29b-3p | MIMAT0000127 | Zfp395 | 380912 | HITS-CLIP | 23142080 | Unspecified |
| mmu-miR-29b-3p | MIMAT0000127 | Zfp40 | 22700 | HITS-CLIP | 25083871 | Unspecified |
| mmu-miR-29b-3p | MIMAT0000127 | Zfp41 | 22701 | HITS-CLIP | 37986224\|25083871\|23142080 | Unspecified |
| mmu-miR-29b-3p | MIMAT0000127 | Zfp445 | 235682 | sRNA-Seq//HITS-CLIP | 33184237\|23597149 | Unspecified |
| mmu-miR-29b-3p | MIMAT0000127 | Zfp451 | 98403 | HITS-CLIP | 23142080 | Unspecified |
| mmu-miR-29b-3p | MIMAT0000127 | Zfp456 | 408065 | HITS-CLIP | 23142080 | Unspecified |
| mmu-miR-29b-3p | MIMAT0000127 | Zfp46 | 22704 | HITS-CLIP | 37986224\|25083871\|23597149\|23142 | Unspecified |
| mmu-miR-29b-3p | MIMAT0000127 | Zfp462 | 242466 | HITS-CLIP | 25083871 | Unspecified |
| mmu-miR-29b-3p | MIMAT0000127 | Zfp507 | 668501 | HITS-CLIP | 37986224 | Unspecified |
| mmu-miR-29b-3p | MIMAT0000127 | Zfp512b | 269401 | HITS-CLIP//sRNA-Seq | 25083871\|33184237\|23597149\|23142 | Unspecified |
| mmu-miR-29b-3p | MIMAT0000127 | Zfp516 | 329003 | HITS-CLIP | 37986224\|25083871 | Unspecified |
| mmu-miR-29b-3p | MIMAT0000127 | Zfp518a | 72672 | HITS-CLIP | 37986224\|23142080 | Unspecified |
| mmu-miR-29b-3p | MIMAT0000127 | Zfp518b | 100515 | HITS-CLIP | 37986224\|25083871 | Unspecified |
| mmu-miR-29b-3p | MIMAT0000127 | Zfp52 | 22710 | HITS-CLIP | 25083871 | Unspecified |
| mmu-miR-29b-3p | MIMAT0000127 | Zfp53 | 24132 | HITS-CLIP | 23142080 | Unspecified |
| mmu-miR-29b-3p | MIMAT0000127 | Zfp532 | 328977 | sRNA-Seq//HITS-CLIP | 33184237\|25083871 | Unspecified |
| mmu-miR-29b-3p | MIMAT0000127 | Zfp563 | 240068 | HITS-CLIP | 23142080 | Unspecified |
| mmu-miR-29b-3p | MIMAT0000127 | Zfp568 | 243905 | HITS-CLIP | 37986224\|25083871\|23142080\|23597 | Unspecified |
| mmu-miR-29b-3p | MIMAT0000127 | Zfp583 | 213011 | HITS-CLIP | 37986224 | Unspecified |
| mmu-miR-29b-3p | MIMAT0000127 | Zfp592 | 233410 | HITS-CLIP | 37986224\|23597149 | Unspecified |
| mmu-miR-29b-3p | MIMAT0000127 | Zfp597 | 71063 | HITS-CLIP | 25083871 | Unspecified |
| mmu-miR-29b-3p | MIMAT0000127 | Zfp609 | 214812 | HITS-CLIP | 25083871\|23142080 | Unspecified |
| mmu-miR-29b-3p | MIMAT0000127 | Zfp622 | 52521 | qCLASH//HITS-CLIP | 31393953\|23142080 | Unspecified |
| mmu-miR-29b-3p | MIMAT0000127 | Zfp623 | 78834 | HITS-CLIP | 37986224\|25083871 | Unspecified |

| | | | | | | |
|---|---|---|---|---|---|---|
| mmu-miR-29b-3p | MIMAT0000127 | Zfp628 | 232816 | HITS-CLIP | 23597149 | Unspecified |
| mmu-miR-29b-3p | MIMAT0000127 | Zfp638 | 27332 | HITS-CLIP//sRNA-Seq | 23142080\|33184237\|25083871 | Unspecified |
| mmu-miR-29b-3p | MIMAT0000127 | Zfp644 | 52397 | HITS-CLIP//sRNA-Seq | 37986224\|33184237\|23142080 | Unspecified |
| mmu-miR-29b-3p | MIMAT0000127 | Zfp646 | 233905 | HITS-CLIP | 23142080 | Unspecified |
| mmu-miR-29b-3p | MIMAT0000127 | Zfp651 | 270210 | HITS-CLIP | 37986224\|25083871 | Unspecified |
| mmu-miR-29b-3p | MIMAT0000127 | Zfp652 | 268469 | HITS-CLIP | 37986224\|25083871\|23597149\|2314 | Unspecified |
| mmu-miR-29b-3p | MIMAT0000127 | Zfp654 | 72020 | HITS-CLIP | 23142080 | Unspecified |
| mmu-miR-29b-3p | MIMAT0000127 | Zfp664 | 269704 | HITS-CLIP//sRNA-Seq | 25083871\|33184237\|23142080 | Unspecified |
| mmu-miR-29b-3p | MIMAT0000127 | Zfp703 | 353310 | HITS-CLIP | 23142080 | Unspecified |
| mmu-miR-29b-3p | MIMAT0000127 | Zfp706 | 68036 | HITS-CLIP//sRNA-Seq | 25083871\|33184237 | Unspecified |
| mmu-miR-29b-3p | MIMAT0000127 | Zfp707 | 69020 | HITS-CLIP | 37986224\|23142080\|23597149 | Unspecified |
| mmu-miR-29b-3p | MIMAT0000127 | Zfp710 | 209225 | HITS-CLIP | 23597149 | Unspecified |
| mmu-miR-29b-3p | MIMAT0000127 | Zfp715 | 69930 | HITS-CLIP | 37986224 | Unspecified |
| mmu-miR-29b-3p | MIMAT0000127 | Zfp729b | 100416706 | HITS-CLIP | 23142080 | Unspecified |
| mmu-miR-29b-3p | MIMAT0000127 | Zfp738 | 408068 | HITS-CLIP | 25083871 | Unspecified |
| mmu-miR-29b-3p | MIMAT0000127 | Zfp74 | 72723 | HITS-CLIP | 37986224 | Unspecified |
| mmu-miR-29b-3p | MIMAT0000127 | Zfp748 | 212276 | HITS-CLIP | 23142080 | Unspecified |
| mmu-miR-29b-3p | MIMAT0000127 | Zfp768 | 233890 | HITS-CLIP | 37986224 | Unspecified |
| mmu-miR-29b-3p | MIMAT0000127 | Zfp770 | 228491 | HITS-CLIP | 37986224\|25083871\|23142080 | Unspecified |
| mmu-miR-29b-3p | MIMAT0000127 | Zfp787 | 67109 | HITS-CLIP | 23142080 | Unspecified |
| mmu-miR-29b-3p | MIMAT0000127 | Zfp800 | 627049 | HITS-CLIP | 23142080 | Unspecified |
| mmu-miR-29b-3p | MIMAT0000127 | Zfp81 | 224694 | HITS-CLIP | 23142080 | Unspecified |
| mmu-miR-29b-3p | MIMAT0000127 | Zfp827 | 622675 | HITS-CLIP | 37986224\|25083871\|23142080 | Unspecified |
| mmu-miR-29b-3p | MIMAT0000127 | Zfp830 | 66983 | HITS-CLIP//qCLASH | 37986224\|31393953 | Unspecified |
| mmu-miR-29b-3p | MIMAT0000127 | Zfp831 | 100043757 | HITS-CLIP | 23142080 | Unspecified |
| mmu-miR-29b-3p | MIMAT0000127 | Zfp871 | 208292 | HITS-CLIP//sRNA-Seq | 23142080\|25083871\|33184237\|2359 | Unspecified |
| mmu-miR-29b-3p | MIMAT0000127 | Zfp91 | 109910 | HITS-CLIP//sRNA-Seq//qCLASH | 37986224\|25083871\|23597149\|3318 | Unspecified |
| mmu-miR-29b-3p | MIMAT0000127 | Zfp933 | 242747 | HITS-CLIP | 37986224 | Unspecified |
| mmu-miR-29b-3p | MIMAT0000127 | Zfp948 | 381066 | HITS-CLIP | 25083871 | Unspecified |
| mmu-miR-29b-3p | MIMAT0000127 | Zfr | 22763 | HITS-CLIP//sRNA-Seq | 25083871\|33184237\|23142080 | Unspecified |
| mmu-miR-29b-3p | MIMAT0000127 | Zfx | 22764 | HITS-CLIP | 37986224\|25083871\|23597149 | Unspecified |
| mmu-miR-29b-3p | MIMAT0000127 | Zfyve1 | 217695 | HITS-CLIP | 23142080\|25083871 | Unspecified |
| mmu-miR-29b-3p | MIMAT0000127 | Zfyve16 | 218441 | HITS-CLIP | 23142080\|25083871 | Unspecified |
| mmu-miR-29b-3p | MIMAT0000127 | Zfyve21 | 68520 | HITS-CLIP | 23597149 | Unspecified |
| mmu-miR-29b-3p | MIMAT0000127 | Zfyve26 | 211978 | HITS-CLIP | 37986224\|23142080 | Unspecified |
| mmu-miR-29b-3p | MIMAT0000127 | Zfyve27 | 319740 | HITS-CLIP | 25083871 | Unspecified |
| mmu-miR-29b-3p | MIMAT0000127 | Zfyve9 | 230597 | HITS-CLIP | 25083871\|23597149 | Unspecified |
| mmu-miR-29b-3p | MIMAT0000127 | Zgpat | 229007 | HITS-CLIP | 23142080 | Unspecified |
| mmu-miR-29b-3p | MIMAT0000127 | Zhx1 | 22770 | HITS-CLIP | 23142080 | Unspecified |

| | | | | | | |
|---|---|---|---|---|---|---|
| mmu-miR-29b-3p | MIMAT0000127 | Zhx2 | 387609 | HITS-CLIP | 37986224\|25083871 | Unspecified |
| mmu-miR-29b-3p | MIMAT0000127 | Zhx3 | 320799 | HITS-CLIP | 37986224\|25083871 | Unspecified |
| mmu-miR-29b-3p | MIMAT0000127 | Zim1 | 22776 | HITS-CLIP | 37986224 | Unspecified |
| mmu-miR-29b-3p | MIMAT0000127 | Zkscan1 | 74570 | HITS-CLIP | 37986224\|23597149\|25083871\|2314 | Unspecified |
| mmu-miR-29b-3p | MIMAT0000127 | Zkscan17 | 268417 | HITS-CLIP | 37986224\|23142080\|25083871 | Unspecified |
| mmu-miR-29b-3p | MIMAT0000127 | Zkscan3 | 72739 | HITS-CLIP | 37986224 | Unspecified |
| mmu-miR-29b-3p | MIMAT0000127 | Zkscan7 | 382118 | HITS-CLIP | 25083871\|23597149\|23142080 | Unspecified |
| mmu-miR-29b-3p | MIMAT0000127 | Zmat3 | 22401 | HITS-CLIP | 25083871\|23142080 | Unspecified |
| mmu-miR-29b-3p | MIMAT0000127 | Zmiz1 | 328365 | HITS-CLIP//sRNA-Seq | 37986224\|23142080\|25083871\|2359 | Unspecified |
| mmu-miR-29b-3p | MIMAT0000127 | Zmiz2 | 52915 | HITS-CLIP | 25083871 | Unspecified |
| mmu-miR-29b-3p | MIMAT0000127 | Zmpste24 | 230709 | HITS-CLIP | 25083871 | Unspecified |
| mmu-miR-29b-3p | MIMAT0000127 | Zmym2 | 76007 | HITS-CLIP//sRNA-Seq | 37986224\|23597149\|23142080\|2508 | Unspecified |
| mmu-miR-29b-3p | MIMAT0000127 | Zmym3 | 56364 | sRNA-Seq//HITS-CLIP | 33184237\|23142080 | Unspecified |
| mmu-miR-29b-3p | MIMAT0000127 | Zmym4 | 67785 | HITS-CLIP//sRNA-Seq | 37986224\|25083871\|33184237\|2314 | Unspecified |
| mmu-miR-29b-3p | MIMAT0000127 | Zmynd11 | 66505 | sRNA-Seq//HITS-CLIP | 33184237\|23142080 | Unspecified |
| mmu-miR-29b-3p | MIMAT0000127 | Zmynd8 | 228880 | sRNA-Seq//HITS-CLIP | 33184237\|23142080 | Unspecified |
| mmu-miR-29b-3p | MIMAT0000127 | Znfx1 | 98999 | HITS-CLIP | 25083871 | Unspecified |
| mmu-miR-29b-3p | MIMAT0000127 | Znhit1 | 70103 | HITS-CLIP//sRNA-Seq | 23597149\|33184237 | Unspecified |
| mmu-miR-29b-3p | MIMAT0000127 | Znrf1 | 170737 | HITS-CLIP | 25083871\|23142080\|23597149 | Unspecified |
| mmu-miR-29b-3p | MIMAT0000127 | Znrf2 | 387524 | HITS-CLIP | 23597149\|23142080 | Unspecified |
| mmu-miR-29b-3p | MIMAT0000127 | Znrf3 | 407821 | HITS-CLIP | 23142080 | Unspecified |
| mmu-miR-29b-3p | MIMAT0000127 | Zranb1 | 360216 | HITS-CLIP | 37986224\|25083871\|23597149\|2314 | Unspecified |
| mmu-miR-29b-3p | MIMAT0000127 | Zscan26 | 432731 | HITS-CLIP | 23142080 | Unspecified |
| mmu-miR-29b-3p | MIMAT0000127 | Zscan29 | 99334 | HITS-CLIP | 37986224\|23142080 | Unspecified |
| mmu-miR-29b-3p | MIMAT0000127 | Zswim1 | 71971 | HITS-CLIP | 25083871 | Unspecified |
| mmu-miR-29b-3p | MIMAT0000127 | Zswim3 | 67538 | HITS-CLIP | 23142080 | Unspecified |
| mmu-miR-29b-3p | MIMAT0000127 | Zswim8 | 268721 | sRNA-Seq//HITS-CLIP | 33184237\|25083871 | Unspecified |
| mmu-miR-29b-3p | MIMAT0000127 | Zwilch | 68014 | HITS-CLIP | 23142080 | Unspecified |
| mmu-miR-29b-3p | MIMAT0000127 | Zxdc | 80292 | HITS-CLIP | 23142080 | Unspecified |
| mmu-miR-29b-3p | MIMAT0000127 | Zyg11b | 414872 | sRNA-Seq//HITS-CLIP | 33184237\|23142080 | Unspecified |
| mmu-miR-29b-3p | MIMAT0000127 | Zyx | 22793 | HITS-CLIP//sRNA-Seq | 23142080\|25083871\|23597149\|3318 | Unspecified |
| mmu-miR-29b-3p | MIMAT0000127 | Zzef1 | 195018 | HITS-CLIP | 23142080\|23597149\|25083871 | Unspecified |
| mmu-miR-29b-3p | MIMAT0000127 | Zzz3 | 108946 | HITS-CLIP | 37986224\|25083871\|23142080 | Unspecified |
| mmu-miR-29b-3p | MIMAT0000127 | mt-Co1 | 4512 | HITS-CLIP//sRNA-Seq | 25083871\|33184237 | Unspecified |
| mmu-miR-29b-3p | MIMAT0000127 | mt-Nd4 | 4538 | sRNA-Seq//HITS-CLIP | 33184237\|23142080 | Unspecified |
| mmu-miR-21a-5p | MIMAT0000530 | 0610010F05Rik | 71675 | HITS-CLIP | 23142080 | Unspecified |
| mmu-miR-21a-5p | MIMAT0000530 | 0610012G03Rik | 106264 | HITS-CLIP | 23142080 | Unspecified |
| mmu-miR-21a-5p | MIMAT0000530 | 0610030E20Rik | 68364 | HITS-CLIP | 23597149\|23142080 | Unspecified |
| mmu-miR-21a-5p | MIMAT0000530 | 1110008P14Rik | 73737 | Microarrays | 37986224 | Unspecified |

| | | | | | | |
|---|---|---|---|---|---|---|
| mmu-miR-21a-5p | MIMAT0000530 | 1110059G10Rik | 66202 | Microarrays | 37986224 | Unspecified |
| mmu-miR-21a-5p | MIMAT0000530 | 1700007K13Rik | 69327 | Microarrays | 37986224 | Unspecified |
| mmu-miR-21a-5p | MIMAT0000530 | 1700010I14Rik | 66931 | Microarrays | 37986224\|25263593 | Unspecified |
| mmu-miR-21a-5p | MIMAT0000530 | 1700017B05Rik | 74211 | Microarrays | 37986224\|25263593 | Unspecified |
| mmu-miR-21a-5p | MIMAT0000530 | 1700025G04Rik | 69399 | HITS-CLIP | 23142080 | Unspecified |
| mmu-miR-21a-5p | MIMAT0000530 | 1700123O20Rik | 58248 | HITS-CLIP | 23142080 | Unspecified |
| mmu-miR-21a-5p | MIMAT0000530 | 1810013L24Rik | 69053 | HITS-CLIP | 23142080 | Unspecified |
| mmu-miR-21a-5p | MIMAT0000530 | 1810030O07Rik | 69155 | HITS-CLIP | 29167373 | Unspecified |
| mmu-miR-21a-5p | MIMAT0000530 | 1810058I24Rik | 67705 | HITS-CLIP | 31433990 | Unspecified |
| mmu-miR-21a-5p | MIMAT0000530 | 2010315B03Rik | 630836 | Microarrays | 37986224 | Unspecified |
| mmu-miR-21a-5p | MIMAT0000530 | 2310022B05Rik | 69551 | HITS-CLIP | 23597149\|23142080 | Unspecified |
| mmu-miR-21a-5p | MIMAT0000530 | 2310057M21Rik | 68277 | HITS-CLIP | 23142080 | Unspecified |
| mmu-miR-21a-5p | MIMAT0000530 | 2310061I04Rik | 69662 | HITS-CLIP | 23142080 | Unspecified |
| mmu-miR-21a-5p | MIMAT0000530 | 2510039O18Rik | 77034 | HITS-CLIP | 23142080 | Unspecified |
| mmu-miR-21a-5p | MIMAT0000530 | 2610002M06Rik | 67028 | HITS-CLIP | 23142080 | Unspecified |
| mmu-miR-21a-5p | MIMAT0000530 | 2610008E11Rik | 72128 | Microarrays | 37986224 | Unspecified |
| mmu-miR-21a-5p | MIMAT0000530 | 2610301B20Rik | 67157 | HITS-CLIP | 23597149 | Unspecified |
| mmu-miR-21a-5p | MIMAT0000530 | 2610318N02Rik | 70458 | Microarrays | 37986224\|25263593 | Unspecified |
| mmu-miR-21a-5p | MIMAT0000530 | 2610507B11Rik | 72503 | HITS-CLIP | 29167373 | Unspecified |
| mmu-miR-21a-5p | MIMAT0000530 | 2700049A03Rik | 76967 | HITS-CLIP | 23142080 | Unspecified |
| mmu-miR-21a-5p | MIMAT0000530 | 2700081O15Rik | 108899 | HITS-CLIP | 23142080 | Unspecified |
| mmu-miR-21a-5p | MIMAT0000530 | 2700097O09Rik | 72658 | HITS-CLIP | 23597149 | Unspecified |
| mmu-miR-21a-5p | MIMAT0000530 | 4833439L19Rik | 97820 | HITS-CLIP | 23142080 | Unspecified |
| mmu-miR-21a-5p | MIMAT0000530 | 4930402H24Rik | 228602 | HITS-CLIP | 23142080 | Unspecified |
| mmu-miR-21a-5p | MIMAT0000530 | 4930503L19Rik | 269033 | Microarrays | 37986224\|25263593 | Unspecified |
| mmu-miR-21a-5p | MIMAT0000530 | 4930523C07Rik | 67647 | HITS-CLIP | 23142080 | Unspecified |
| mmu-miR-21a-5p | MIMAT0000530 | 4930579G24Rik | 75939 | Microarrays | 37986224\|25263593 | Unspecified |
| mmu-miR-21a-5p | MIMAT0000530 | 4931406P16Rik | 233103 | HITS-CLIP | 23142080\|29167373 | Unspecified |
| mmu-miR-21a-5p | MIMAT0000530 | 4932438A13Rik | 229227 | HITS-CLIP | 23597149\|29167373\|23142080 | Unspecified |
| mmu-miR-21a-5p | MIMAT0000530 | 4933430I17Rik | 214106 | Microarrays | 37986224\|25263593 | Unspecified |
| mmu-miR-21a-5p | MIMAT0000530 | 5031439G07Rik | 223739 | HITS-CLIP | 29167373 | Unspecified |
| mmu-miR-21a-5p | MIMAT0000530 | 5830473C10Rik | 622307 | HITS-CLIP | 23597149 | Unspecified |
| mmu-miR-21a-5p | MIMAT0000530 | 6030458C11Rik | 77877 | HITS-CLIP | 19536157 | Unspecified |
| mmu-miR-21a-5p | MIMAT0000530 | 8030462N17Rik | 212163 | HITS-CLIP | 31433990 | Unspecified |
| mmu-miR-21a-5p | MIMAT0000530 | 9930012K11Rik | 268759 | HITS-CLIP | 29167373\|23142080 | Unspecified |
| mmu-miR-21a-5p | MIMAT0000530 | 9930021J03Rik | 240613 | HITS-CLIP | 23142080 | Unspecified |
| mmu-miR-21a-5p | MIMAT0000530 | A430033K04Rik | 243308 | HITS-CLIP | 23142080 | Unspecified |
| mmu-miR-21a-5p | MIMAT0000530 | AA467197 | 433470 | Microarrays | 37986224\|25263593 | Unspecified |
| mmu-miR-21a-5p | MIMAT0000530 | AA986860 | 212439 | HITS-CLIP | 23597149 | Unspecified |

| | | | | | | |
|---|---|---|---|---|---|---|
| mmu-miR-21a-5p | MIMAT0000530 | AB124611 | 382062 | Microarrays | 37986224 | Unspecified |
| mmu-miR-21a-5p | MIMAT0000530 | AI182371 | 98870 | HITS-CLIP | 23597149 | Unspecified |
| mmu-miR-21a-5p | MIMAT0000530 | AW112010 | 107350 | Microarrays | 37986224 | Unspecified |
| mmu-miR-21a-5p | MIMAT0000530 | AW209491 | 105351 | HITS-CLIP | 23142080 | Unspecified |
| mmu-miR-21a-5p | MIMAT0000530 | AW549877 | 106064 | HITS-CLIP | 37986224\|23597149\|23142080\|29167 | Unspecified |
| mmu-miR-21a-5p | MIMAT0000530 | Aacs | 78894 | HITS-CLIP | 23597149 | Unspecified |
| mmu-miR-21a-5p | MIMAT0000530 | Aagab | 66939 | HITS-CLIP | 23142080 | Unspecified |
| mmu-miR-21a-5p | MIMAT0000530 | Aak1 | 269774 | HITS-CLIP | 23597149\|23142080\|29167373 | Unspecified |
| mmu-miR-21a-5p | MIMAT0000530 | Aar2 | 68295 | HITS-CLIP | 23142080 | Unspecified |
| mmu-miR-21a-5p | MIMAT0000530 | Aars | 234734 | Microarrays//HITS-CLIP | 37986224\|23597149\|25263593\|2314 | Unspecified |
| mmu-miR-21a-5p | MIMAT0000530 | Aass | 30956 | HITS-CLIP | 23597149 | Unspecified |
| mmu-miR-21a-5p | MIMAT0000530 | Aatf | 56321 | HITS-CLIP | 23142080 | Unspecified |
| mmu-miR-21a-5p | MIMAT0000530 | Abat | 268860 | HITS-CLIP | 23597149 | Unspecified |
| mmu-miR-21a-5p | MIMAT0000530 | Abca1 | 11303 | Microarrays//HITS-CLIP | 37986224\|23597149\|29167373 | Unspecified |
| mmu-miR-21a-5p | MIMAT0000530 | Abca2 | 11305 | HITS-CLIP | 19536157 | Unspecified |
| mmu-miR-21a-5p | MIMAT0000530 | Abca3 | 27410 | HITS-CLIP | 23597149\|23142080 | Unspecified |
| mmu-miR-21a-5p | MIMAT0000530 | Abca5 | 217265 | HITS-CLIP | 23142080 | Unspecified |
| mmu-miR-21a-5p | MIMAT0000530 | Abca9 | 217262 | Microarrays | 37986224 | Unspecified |
| mmu-miR-21a-5p | MIMAT0000530 | Abcb11 | 27413 | HITS-CLIP | 23597149 | Unspecified |
| mmu-miR-21a-5p | MIMAT0000530 | Abcb1a | 18671 | HITS-CLIP | 37986224 | Unspecified |
| mmu-miR-21a-5p | MIMAT0000530 | Abcb4 | 18670 | HITS-CLIP | 23597149 | Unspecified |
| mmu-miR-21a-5p | MIMAT0000530 | Abcb7 | 11306 | HITS-CLIP | 23597149\|29167373\|23142080 | Unspecified |
| mmu-miR-21a-5p | MIMAT0000530 | Abcc1 | 17250 | HITS-CLIP | 29167373 | Unspecified |
| mmu-miR-21a-5p | MIMAT0000530 | Abcc4 | 239273 | HITS-CLIP | 29167373 | Unspecified |
| mmu-miR-21a-5p | MIMAT0000530 | Abcc5 | 27416 | HITS-CLIP | 23142080 | Unspecified |
| mmu-miR-21a-5p | MIMAT0000530 | Abcd2 | 26874 | HITS-CLIP | 37986224\|23597149 | Unspecified |
| mmu-miR-21a-5p | MIMAT0000530 | Abcd3 | 19299 | HITS-CLIP | 23597149 | Unspecified |
| mmu-miR-21a-5p | MIMAT0000530 | Abce1 | 24015 | HITS-CLIP | 37986224\|23597149\|23142080 | Unspecified |
| mmu-miR-21a-5p | MIMAT0000530 | Abcf2 | 27407 | HITS-CLIP | 23597149\|23142080 | Unspecified |
| mmu-miR-21a-5p | MIMAT0000530 | Abcg1 | 11307 | Microarrays | 37986224 | Unspecified |
| mmu-miR-21a-5p | MIMAT0000530 | Abcg3 | 27405 | Microarrays | 37986224 | Unspecified |
| mmu-miR-21a-5p | MIMAT0000530 | Abhd10 | 213012 | HITS-CLIP | 23142080 | Unspecified |
| mmu-miR-21a-5p | MIMAT0000530 | Abhd13 | 68904 | HITS-CLIP | 23142080 | Unspecified |
| mmu-miR-21a-5p | MIMAT0000530 | Abhd17b | 226016 | HITS-CLIP | 23142080 | Unspecified |
| mmu-miR-21a-5p | MIMAT0000530 | Abhd17c | 70178 | Microarrays | 37986224\|25263593 | Unspecified |
| mmu-miR-21a-5p | MIMAT0000530 | Abhd18 | 269423 | HITS-CLIP | 23142080\|23597149 | Unspecified |
| mmu-miR-21a-5p | MIMAT0000530 | Abi1 | 11308 | HITS-CLIP | 23142080\|31433990\|30971389 | Unspecified |
| mmu-miR-21a-5p | MIMAT0000530 | Abl2 | 11352 | HITS-CLIP//qCLASH | 23142080\|23597149\|31393953\|29167 | Unspecified |
| mmu-miR-21a-5p | MIMAT0000530 | Ablim1 | 226251 | HITS-CLIP | 23142080 | Unspecified |

| | | | | | | |
|---|---|---|---|---|---|---|
| mmu-miR-21a-5p | MIMAT0000530 | Ablim3 | 319713 | HITS-CLIP | 23597149 | Unspecified |
| mmu-miR-21a-5p | MIMAT0000530 | Abracl | 73112 | HITS-CLIP | 23142080 | Unspecified |
| mmu-miR-21a-5p | MIMAT0000530 | Abraxas1 | 70681 | HITS-CLIP | 23142080 | Unspecified |
| mmu-miR-21a-5p | MIMAT0000530 | Abtb1 | 80283 | Microarrays | 37986224 | Unspecified |
| mmu-miR-21a-5p | MIMAT0000530 | Abtb2 | 99382 | Microarrays//HITS-CLIP | 37986224\|25263593\|23597149\|2314 | Unspecified |
| mmu-miR-21a-5p | MIMAT0000530 | Acaa2 | 52538 | HITS-CLIP | 23597149\|23142080\|29167373 | Unspecified |
| mmu-miR-21a-5p | MIMAT0000530 | Acacb | 100705 | HITS-CLIP | 23597149 | Unspecified |
| mmu-miR-21a-5p | MIMAT0000530 | Acadl | 11363 | HITS-CLIP | 23142080 | Unspecified |
| mmu-miR-21a-5p | MIMAT0000530 | Acadm | 11364 | HITS-CLIP | 23597149 | Unspecified |
| mmu-miR-21a-5p | MIMAT0000530 | Acadsb | 66885 | HITS-CLIP | 23142080 | Unspecified |
| mmu-miR-21a-5p | MIMAT0000530 | Acap2 | 78618 | HITS-CLIP | 37986224\|23142080\|29167373 | Unspecified |
| mmu-miR-21a-5p | MIMAT0000530 | Acat1 | 110446 | HITS-CLIP | 37986224\|23142080 | Unspecified |
| mmu-miR-21a-5p | MIMAT0000530 | Acat2 | 110460 | qCLASH | 31393953 | Unspecified |
| mmu-miR-21a-5p | MIMAT0000530 | Acbd3 | 170760 | HITS-CLIP | 31433990\|23142080 | Unspecified |
| mmu-miR-21a-5p | MIMAT0000530 | Acbd4 | 67131 | Microarrays | 37986224\|25263593 | Unspecified |
| mmu-miR-21a-5p | MIMAT0000530 | Acbd5 | 74159 | HITS-CLIP//qCLASH | 23142080\|23597149\|31393953\|2916 | Unspecified |
| mmu-miR-21a-5p | MIMAT0000530 | Acer2 | 230379 | HITS-CLIP | 23142080 | Unspecified |
| mmu-miR-21a-5p | MIMAT0000530 | Ackr3 | 12778 | Microarrays | 37986224 | Unspecified |
| mmu-miR-21a-5p | MIMAT0000530 | Acly | 104112 | HITS-CLIP | 23597149\|23142080 | Unspecified |
| mmu-miR-21a-5p | MIMAT0000530 | Acnat1 | 230161 | HITS-CLIP | 23597149 | Unspecified |
| mmu-miR-21a-5p | MIMAT0000530 | Acnat2 | 209186 | HITS-CLIP | 23597149 | Unspecified |
| mmu-miR-21a-5p | MIMAT0000530 | Aco1 | 11428 | HITS-CLIP | 23597149\|23142080 | Unspecified |
| mmu-miR-21a-5p | MIMAT0000530 | Aco2 | 11429 | HITS-CLIP | 23597149\|19536157\|23142080\|2916 | Unspecified |
| mmu-miR-21a-5p | MIMAT0000530 | Acot7 | 70025 | HITS-CLIP | 23142080 | Unspecified |
| mmu-miR-21a-5p | MIMAT0000530 | Acot8 | 170789 | HITS-CLIP | 23142080 | Unspecified |
| mmu-miR-21a-5p | MIMAT0000530 | Acot9 | 56360 | HITS-CLIP | 23142080\|29167373 | Unspecified |
| mmu-miR-21a-5p | MIMAT0000530 | Acox1 | 11430 | HITS-CLIP | 23597149 | Unspecified |
| mmu-miR-21a-5p | MIMAT0000530 | Acox2 | 93732 | HITS-CLIP | 23597149 | Unspecified |
| mmu-miR-21a-5p | MIMAT0000530 | Acp2 | 11432 | HITS-CLIP | 19536157 | Unspecified |
| mmu-miR-21a-5p | MIMAT0000530 | Acsf2 | 264895 | HITS-CLIP | 31433990 | Unspecified |
| mmu-miR-21a-5p | MIMAT0000530 | Acsl1 | 14081 | HITS-CLIP | 23597149 | Unspecified |
| mmu-miR-21a-5p | MIMAT0000530 | Acsl3 | 74205 | HITS-CLIP, Microarrays//HITS-CLIP | 37986224\|23142080 | Unspecified |
| mmu-miR-21a-5p | MIMAT0000530 | Acsl4 | 50790 | HITS-CLIP | 23142080 | Unspecified |
| mmu-miR-21a-5p | MIMAT0000530 | Acsm3 | 20216 | Microarrays | 37986224 | Unspecified |
| mmu-miR-21a-5p | MIMAT0000530 | Acsm5 | 272428 | HITS-CLIP | 23597149 | Unspecified |
| mmu-miR-21a-5p | MIMAT0000530 | Acss2 | 60525 | Microarrays | 37986224 | Unspecified |
| mmu-miR-21a-5p | MIMAT0000530 | Acta1 | 11459 | HITS-CLIP | 23142080 | Unspecified |
| mmu-miR-21a-5p | MIMAT0000530 | Actb | 11461 | HITS-CLIP | 19536157\|23597149\|23142080 | Unspecified |
| mmu-miR-21a-5p | MIMAT0000530 | Actg2 | 11468 | Microarrays//HITS-CLIP | 37986224\|25263593\|23597149 | Unspecified |

| | | | | | | |
|---|---|---|---|---|---|---|
| mmu-miR-21a-5p | MIMAT0000530 | Actl6a | 56456 | HITS-CLIP | 29167373\|23142080 | Unspecified |
| mmu-miR-21a-5p | MIMAT0000530 | Actn1 | 109711 | HITS-CLIP | 29167373 | Unspecified |
| mmu-miR-21a-5p | MIMAT0000530 | Actn4 | 60595 | HITS-CLIP | 23142080 | Unspecified |
| mmu-miR-21a-5p | MIMAT0000530 | Actr10 | 56444 | HITS-CLIP | 37986224\|23142080 | Unspecified |
| mmu-miR-21a-5p | MIMAT0000530 | Actr1a | 54130 | HITS-CLIP | 23597149\|31433990\|23142080 | Unspecified |
| mmu-miR-21a-5p | MIMAT0000530 | Actr1b | 226977 | HITS-CLIP | 23597149 | Unspecified |
| mmu-miR-21a-5p | MIMAT0000530 | Actr2 | 66713 | HITS-CLIP | 23142080\|29167373 | Unspecified |
| mmu-miR-21a-5p | MIMAT0000530 | Actr3 | 74117 | HITS-CLIP | 23142080 | Unspecified |
| mmu-miR-21a-5p | MIMAT0000530 | Actr6 | 67019 | HITS-CLIP | 23142080 | Unspecified |
| mmu-miR-21a-5p | MIMAT0000530 | Actr8 | 56249 | HITS-CLIP | 23142080 | Unspecified |
| mmu-miR-21a-5p | MIMAT0000530 | Acvr1 | 11477 | HITS-CLIP | 19536157\|29167373 | Unspecified |
| mmu-miR-21a-5p | MIMAT0000530 | Acvr1b | 11479 | HITS-CLIP | 23142080 | Unspecified |
| mmu-miR-21a-5p | MIMAT0000530 | Acvr1c | 269275 | HITS-CLIP | 23597149 | Unspecified |
| mmu-miR-21a-5p | MIMAT0000530 | Acvr2a | 11480 | HITS-CLIP | 19536157\|23142080 | Unspecified |
| mmu-miR-21a-5p | MIMAT0000530 | Acy1 | 109652 | HITS-CLIP | 23597149\|28735896 | Unspecified |
| mmu-miR-21a-5p | MIMAT0000530 | Acy3 | 71670 | Microarrays | 37986224\|25263593 | Unspecified |
| mmu-miR-21a-5p | MIMAT0000530 | Adam10 | 11487 | HITS-CLIP | 37986224\|23597149\|23142080\|29167 | Unspecified |
| mmu-miR-21a-5p | MIMAT0000530 | Adam15 | 11490 | Microarrays | 37986224\|25263593 | Unspecified |
| mmu-miR-21a-5p | MIMAT0000530 | Adam17 | 11491 | HITS-CLIP | 23142080\|29167373 | Unspecified |
| mmu-miR-21a-5p | MIMAT0000530 | Adam19 | 11492 | Microarrays | 37986224 | Unspecified |
| mmu-miR-21a-5p | MIMAT0000530 | Adam1a | 280668 | HITS-CLIP | 23142080 | Unspecified |
| mmu-miR-21a-5p | MIMAT0000530 | Adam22 | 11496 | Microarrays | 37986224 | Unspecified |
| mmu-miR-21a-5p | MIMAT0000530 | Adam8 | 11501 | Microarrays | 37986224\|25263593 | Unspecified |
| mmu-miR-21a-5p | MIMAT0000530 | Adam9 | 11502 | HITS-CLIP | 23142080\|23597149\|19536157\|29167 | Unspecified |
| mmu-miR-21a-5p | MIMAT0000530 | Adamts1 | 11504 | Microarrays | 37986224 | Unspecified |
| mmu-miR-21a-5p | MIMAT0000530 | Adamts6 | 108154 | HITS-CLIP | 30971389 | Unspecified |
| mmu-miR-21a-5p | MIMAT0000530 | Adamtsl5 | 66548 | Microarrays | 37986224\|25263593 | Unspecified |
| mmu-miR-21a-5p | MIMAT0000530 | Adar | 56417 | HITS-CLIP | 23142080 | Unspecified |
| mmu-miR-21a-5p | MIMAT0000530 | Adarb1 | 110532 | HITS-CLIP | 23597149\|23142080 | Unspecified |
| mmu-miR-21a-5p | MIMAT0000530 | Adat2 | 66757 | Microarrays | 37986224 | Unspecified |
| mmu-miR-21a-5p | MIMAT0000530 | Adcy1 | 432530 | HITS-CLIP | 37986224 | Unspecified |
| mmu-miR-21a-5p | MIMAT0000530 | Adcy2 | 210044 | HITS-CLIP | 37986224 | Unspecified |
| mmu-miR-21a-5p | MIMAT0000530 | Adcy7 | 11513 | HITS-CLIP | 29167373 | Unspecified |
| mmu-miR-21a-5p | MIMAT0000530 | Adgb | 215772 | Microarrays | 37986224\|25263593 | Unspecified |
| mmu-miR-21a-5p | MIMAT0000530 | Adgra3 | 166647 | HITS-CLIP | 23597149 | Unspecified |
| mmu-miR-21a-5p | MIMAT0000530 | Adgre1 | 13733 | HITS-CLIP | 29167373 | Unspecified |
| mmu-miR-21a-5p | MIMAT0000530 | Adgre5 | 976 | HITS-CLIP | 23142080 | Unspecified |
| mmu-miR-21a-5p | MIMAT0000530 | Adgrg1 | 9289 | HITS-CLIP | 23597149 | Unspecified |
| mmu-miR-21a-5p | MIMAT0000530 | Adgrg6 | 57211 | HITS-CLIP | 29167373 | Unspecified |

| | | | | | | |
|---|---|---|---|---|---|---|
| mmu-miR-21a-5p | MIMAT0000530 | Adgrl1 | 22859 | HITS-CLIP | 19536157 | Unspecified |
| mmu-miR-21a-5p | MIMAT0000530 | Adh1 | 11522 | HITS-CLIP | 23597149 | Unspecified |
| mmu-miR-21a-5p | MIMAT0000530 | Adipor1 | 72674 | HITS-CLIP | 37986224\|23142080 | Unspecified |
| mmu-miR-21a-5p | MIMAT0000530 | Adipor2 | 68465 | HITS-CLIP | 23142080 | Unspecified |
| mmu-miR-21a-5p | MIMAT0000530 | Adk | 11534 | HITS-CLIP | 23597149 | Unspecified |
| mmu-miR-21a-5p | MIMAT0000530 | Adm | 11535 | HITS-CLIP | 26203562 | Unspecified |
| mmu-miR-21a-5p | MIMAT0000530 | Adnp | 11538 | HITS-CLIP | 37986224\|23142080\|29167373 | Unspecified |
| mmu-miR-21a-5p | MIMAT0000530 | Adnp2 | 240442 | HITS-CLIP | 23142080 | Unspecified |
| mmu-miR-21a-5p | MIMAT0000530 | Ado | 211488 | HITS-CLIP | 37986224\|23142080\|29167373 | Unspecified |
| mmu-miR-21a-5p | MIMAT0000530 | Adora2a | 11540 | HITS-CLIP | 23142080 | Unspecified |
| mmu-miR-21a-5p | MIMAT0000530 | Adprm | 66358 | HITS-CLIP | 23142080 | Unspecified |
| mmu-miR-21a-5p | MIMAT0000530 | Adrb2 | 11555 | Microarrays//HITS-CLIP | 37986224\|29167373 | Unspecified |
| mmu-miR-21a-5p | MIMAT0000530 | Adss | 11566 | HITS-CLIP//qCLASH | 31433990\|23142080\|31393953 | Unspecified |
| mmu-miR-21a-5p | MIMAT0000530 | Adtrp | 109254 | HITS-CLIP | 23597149 | Unspecified |
| mmu-miR-21a-5p | MIMAT0000530 | Aebp2 | 11569 | HITS-CLIP | 37986224\|23597149\|23142080 | Unspecified |
| mmu-miR-21a-5p | MIMAT0000530 | Aen | 68048 | Microarrays | 37986224\|25263593 | Unspecified |
| mmu-miR-21a-5p | MIMAT0000530 | Aes | 14797 | HITS-CLIP | 23597149\|23142080 | Unspecified |
| mmu-miR-21a-5p | MIMAT0000530 | Aff1 | 17355 | HITS-CLIP | 23142080 | Unspecified |
| mmu-miR-21a-5p | MIMAT0000530 | Aff3 | 16764 | HITS-CLIP | 23142080 | Unspecified |
| mmu-miR-21a-5p | MIMAT0000530 | Aff4 | 93736 | HITS-CLIP | 37986224\|31433990\|23597149\|19536 | Unspecified |
| mmu-miR-21a-5p | MIMAT0000530 | Afp | 11576 | Microarrays | 37986224\|25263593 | Unspecified |
| mmu-miR-21a-5p | MIMAT0000530 | Aftph | 216549 | HITS-CLIP | 23142080 | Unspecified |
| mmu-miR-21a-5p | MIMAT0000530 | Agfg1 | 15463 | HITS-CLIP | 23142080\|19536157\|29167373 | Unspecified |
| mmu-miR-21a-5p | MIMAT0000530 | Agfg2 | 231801 | Microarrays | 37986224 | Unspecified |
| mmu-miR-21a-5p | MIMAT0000530 | Agmo | 319660 | Microarrays | 37986224 | Unspecified |
| mmu-miR-21a-5p | MIMAT0000530 | Ago1 | 236511 | HITS-CLIP | 28735896\|23142080\|29167373 | Unspecified |
| mmu-miR-21a-5p | MIMAT0000530 | Ago2 | 239528 | HITS-CLIP//qCLASH | 37986224\|31393953\|23142080\|29167 | Unspecified |
| mmu-miR-21a-5p | MIMAT0000530 | Ago3 | 214150 | HITS-CLIP | 23142080\|29167373 | Unspecified |
| mmu-miR-21a-5p | MIMAT0000530 | Ago4 | 76850 | Microarrays | 37986224 | Unspecified |
| mmu-miR-21a-5p | MIMAT0000530 | Agpat1 | 55979 | HITS-CLIP | 23142080 | Unspecified |
| mmu-miR-21a-5p | MIMAT0000530 | Agpat3 | 28169 | HITS-CLIP | 23142080\|29167373 | Unspecified |
| mmu-miR-21a-5p | MIMAT0000530 | Agpat5 | 52123 | HITS-CLIP | 23142080\|29167373 | Unspecified |
| mmu-miR-21a-5p | MIMAT0000530 | Agps | 228061 | HITS-CLIP//qCLASH | 37986224\|31393953\|29167373 | Unspecified |
| mmu-miR-21a-5p | MIMAT0000530 | Agrn | 11603 | HITS-CLIP | 37986224\|19536157 | Unspecified |
| mmu-miR-21a-5p | MIMAT0000530 | Agtr1a | 11607 | HITS-CLIP | 23597149 | Unspecified |
| mmu-miR-21a-5p | MIMAT0000530 | Agxt | 11611 | HITS-CLIP | 23597149 | Unspecified |
| mmu-miR-21a-5p | MIMAT0000530 | Agxt2 | 268782 | HITS-CLIP | 23597149 | Unspecified |
| mmu-miR-21a-5p | MIMAT0000530 | Ahctf1 | 226747 | HITS-CLIP | 23597149\|19536157\|23142080 | Unspecified |
| mmu-miR-21a-5p | MIMAT0000530 | Ahcyl1 | 229709 | HITS-CLIP | 19536157\|23142080 | Unspecified |

| | | | | | | |
|---|---|---|---|---|---|---|
| mmu-miR-21a-5p | MIMAT0000530 | Ahcyl2 | 74340 | HITS-CLIP | 23142080 | Unspecified |
| mmu-miR-21a-5p | MIMAT0000530 | Ahi1 | 52906 | HITS-CLIP | 23142080 | Unspecified |
| mmu-miR-21a-5p | MIMAT0000530 | Ahnak | 66395 | HITS-CLIP | 37986224\|23142080\|29167373 | Unspecified |
| mmu-miR-21a-5p | MIMAT0000530 | Ahr | 11622 | HITS-CLIP, Microarrays//HITS-CLIP | 37986224\|23597149 | Unspecified |
| mmu-miR-21a-5p | MIMAT0000530 | Ahrr | 11624 | Microarrays | 37986224 | Unspecified |
| mmu-miR-21a-5p | MIMAT0000530 | Ahsa1 | 217737 | Microarrays | 37986224\|25263593 | Unspecified |
| mmu-miR-21a-5p | MIMAT0000530 | Ahsa2 | 268390 | HITS-CLIP | 23142080 | Unspecified |
| mmu-miR-21a-5p | MIMAT0000530 | Aicda | 11628 | qCLASH | 31393953 | Unspecified |
| mmu-miR-21a-5p | MIMAT0000530 | Aif1 | 11629 | Microarrays | 37986224 | Unspecified |
| mmu-miR-21a-5p | MIMAT0000530 | Aifm1 | 26926 | HITS-CLIP | 23142080 | Unspecified |
| mmu-miR-21a-5p | MIMAT0000530 | Aimp1 | 13722 | HITS-CLIP | 23142080 | Unspecified |
| mmu-miR-21a-5p | MIMAT0000530 | Aimp2 | 231872 | HITS-CLIP | 23142080 | Unspecified |
| mmu-miR-21a-5p | MIMAT0000530 | Ak2 | 11637 | HITS-CLIP | 23142080 | Unspecified |
| mmu-miR-21a-5p | MIMAT0000530 | Ak3 | 56248 | HITS-CLIP | 23142080 | Unspecified |
| mmu-miR-21a-5p | MIMAT0000530 | Ak4 | 11639 | HITS-CLIP | 23142080 | Unspecified |
| mmu-miR-21a-5p | MIMAT0000530 | Akap1 | 11640 | HITS-CLIP | 23597149 | Unspecified |
| mmu-miR-21a-5p | MIMAT0000530 | Akap10 | 56697 | HITS-CLIP | 23142080 | Unspecified |
| mmu-miR-21a-5p | MIMAT0000530 | Akap11 | 219181 | HITS-CLIP | 37986224\|23597149\|23142080 | Unspecified |
| mmu-miR-21a-5p | MIMAT0000530 | Akap12 | 83397 | HITS-CLIP | 23597149 | Unspecified |
| mmu-miR-21a-5p | MIMAT0000530 | Akap13 | 75547 | HITS-CLIP | 23142080\|29167373\|26203562 | Unspecified |
| mmu-miR-21a-5p | MIMAT0000530 | Akap17b | 338351 | HITS-CLIP | 23142080 | Unspecified |
| mmu-miR-21a-5p | MIMAT0000530 | Akap2 | 11641 | HITS-CLIP | 23597149 | Unspecified |
| mmu-miR-21a-5p | MIMAT0000530 | Akap6 | 238161 | HITS-CLIP | 37986224 | Unspecified |
| mmu-miR-21a-5p | MIMAT0000530 | Akap8 | 56399 | HITS-CLIP | 23142080 | Unspecified |
| mmu-miR-21a-5p | MIMAT0000530 | Akap9 | 100986 | HITS-CLIP | 23142080\|29167373 | Unspecified |
| mmu-miR-21a-5p | MIMAT0000530 | Akirin1 | 68050 | HITS-CLIP | 29167373 | Unspecified |
| mmu-miR-21a-5p | MIMAT0000530 | Akna | 100182 | HITS-CLIP | 23142080 | Unspecified |
| mmu-miR-21a-5p | MIMAT0000530 | Akr1a1 | 58810 | HITS-CLIP | 26203562 | Unspecified |
| mmu-miR-21a-5p | MIMAT0000530 | Akr1c12 | 622402 | HITS-CLIP | 37986224\|23597149 | Unspecified |
| mmu-miR-21a-5p | MIMAT0000530 | Akr1c13 | 27384 | HITS-CLIP | 37986224 | Unspecified |
| mmu-miR-21a-5p | MIMAT0000530 | Akr1c14 | 105387 | HITS-CLIP | 23597149 | Unspecified |
| mmu-miR-21a-5p | MIMAT0000530 | Akr1c19 | 432720 | HITS-CLIP | 23597149 | Unspecified |
| mmu-miR-21a-5p | MIMAT0000530 | Akr1d1 | 208665 | HITS-CLIP | 23597149 | Unspecified |
| mmu-miR-21a-5p | MIMAT0000530 | Akt1 | 11651 | HITS-CLIP | 23597149 | Unspecified |
| mmu-miR-21a-5p | MIMAT0000530 | Alas1 | 11655 | HITS-CLIP | 23142080\|29167373 | Unspecified |
| mmu-miR-21a-5p | MIMAT0000530 | Alas2 | 11656 | HITS-CLIP | 23597149 | Unspecified |
| mmu-miR-21a-5p | MIMAT0000530 | Alb | 213 | HITS-CLIP | 23597149 | Unspecified |
| mmu-miR-21a-5p | MIMAT0000530 | Alcam | 11658 | HITS-CLIP, Microarrays//Microarrays//HITS | 37986224\|25263593\|29167373 | Unspecified |
| mmu-miR-21a-5p | MIMAT0000530 | Aldh18a1 | 56454 | HITS-CLIP | 23142080 | Unspecified |

| | | | | | | |
|---|---|---|---|---|---|---|
| mmu-miR-21a-5p | MIMAT0000530 | Aldh1a1 | 11668 | HITS-CLIP | 23597149 | Unspecified |
| mmu-miR-21a-5p | MIMAT0000530 | Aldh1l1 | 107747 | HITS-CLIP | 23597149 | Unspecified |
| mmu-miR-21a-5p | MIMAT0000530 | Aldh3a2 | 11671 | HITS-CLIP | 23142080 | Unspecified |
| mmu-miR-21a-5p | MIMAT0000530 | Aldh6a1 | 104776 | HITS-CLIP | 23597149 | Unspecified |
| mmu-miR-21a-5p | MIMAT0000530 | Aldh8a1 | 237320 | HITS-CLIP | 37986224\|23597149 | Unspecified |
| mmu-miR-21a-5p | MIMAT0000530 | Aldoc | 11676 | HITS-CLIP | 23142080 | Unspecified |
| mmu-miR-21a-5p | MIMAT0000530 | Alg10b | 380959 | HITS-CLIP | 19536157\|29167373 | Unspecified |
| mmu-miR-21a-5p | MIMAT0000530 | Alg11 | 207958 | HITS-CLIP | 23142080 | Unspecified |
| mmu-miR-21a-5p | MIMAT0000530 | Alg13 | 67574 | HITS-CLIP | 23142080 | Unspecified |
| mmu-miR-21a-5p | MIMAT0000530 | Alg2 | 56737 | HITS-CLIP | 37986224 | Unspecified |
| mmu-miR-21a-5p | MIMAT0000530 | Alg3 | 208624 | Microarrays | 37986224\|25263593 | Unspecified |
| mmu-miR-21a-5p | MIMAT0000530 | Alg5 | 66248 | HITS-CLIP | 23142080 | Unspecified |
| mmu-miR-21a-5p | MIMAT0000530 | Alkbh2 | 231642 | Microarrays | 37986224\|25263593 | Unspecified |
| mmu-miR-21a-5p | MIMAT0000530 | Alkbh4 | 72041 | HITS-CLIP | 23142080 | Unspecified |
| mmu-miR-21a-5p | MIMAT0000530 | Alkbh5 | 268420 | HITS-CLIP | 23142080 | Unspecified |
| mmu-miR-21a-5p | MIMAT0000530 | Alox5ap | 11690 | HITS-CLIP | 29167373 | Unspecified |
| mmu-miR-21a-5p | MIMAT0000530 | Als2 | 74018 | HITS-CLIP | 23142080 | Unspecified |
| mmu-miR-21a-5p | MIMAT0000530 | Als2cl | 235633 | HITS-CLIP | 23142080 | Unspecified |
| mmu-miR-21a-5p | MIMAT0000530 | Alyref | 21681 | HITS-CLIP | 23142080\|29167373 | Unspecified |
| mmu-miR-21a-5p | MIMAT0000530 | Alyref2 | 56009 | HITS-CLIP | 23597149\|23142080 | Unspecified |
| mmu-miR-21a-5p | MIMAT0000530 | Amacr | 17117 | HITS-CLIP, Microarrays//HITS-CLIP | 37986224\|23597149\|23142080 | Unspecified |
| mmu-miR-21a-5p | MIMAT0000530 | Ambra1 | 228361 | HITS-CLIP | 23142080 | Unspecified |
| mmu-miR-21a-5p | MIMAT0000530 | Amer1 | 72345 | HITS-CLIP | 23142080 | Unspecified |
| mmu-miR-21a-5p | MIMAT0000530 | Amfr | 23802 | HITS-CLIP | 23597149\|23142080 | Unspecified |
| mmu-miR-21a-5p | MIMAT0000530 | Amigo2 | 105827 | HITS-CLIP | 23142080 | Unspecified |
| mmu-miR-21a-5p | MIMAT0000530 | Amn1 | 232566 | HITS-CLIP | 19536157\|23142080\|29167373 | Unspecified |
| mmu-miR-21a-5p | MIMAT0000530 | Amot | 27494 | HITS-CLIP | 23597149 | Unspecified |
| mmu-miR-21a-5p | MIMAT0000530 | Amz1 | 231842 | Microarrays | 37986224\|25263593 | Unspecified |
| mmu-miR-21a-5p | MIMAT0000530 | Amz2 | 13929 | HITS-CLIP | 23142080 | Unspecified |
| mmu-miR-21a-5p | MIMAT0000530 | Anapc1 | 17222 | HITS-CLIP | 23142080\|29167373 | Unspecified |
| mmu-miR-21a-5p | MIMAT0000530 | Anapc10 | 68999 | HITS-CLIP | 23597149 | Unspecified |
| mmu-miR-21a-5p | MIMAT0000530 | Anapc2 | 99152 | HITS-CLIP | 23142080 | Unspecified |
| mmu-miR-21a-5p | MIMAT0000530 | Anapc4 | 52206 | HITS-CLIP | 37986224\|31433990\|23142080 | Unspecified |
| mmu-miR-21a-5p | MIMAT0000530 | Anapc5 | 59008 | HITS-CLIP | 37986224\|23142080 | Unspecified |
| mmu-miR-21a-5p | MIMAT0000530 | Angpt1 | 11600 | Microarrays | 37986224\|25263593 | Unspecified |
| mmu-miR-21a-5p | MIMAT0000530 | Angptl2 | 26360 | HITS-CLIP | 23142080 | Unspecified |
| mmu-miR-21a-5p | MIMAT0000530 | Angptl4 | 57875 | Microarrays | 37986224 | Unspecified |
| mmu-miR-21a-5p | MIMAT0000530 | Ank | 11732 | HITS-CLIP | 19536157\|29167373 | Unspecified |
| mmu-miR-21a-5p | MIMAT0000530 | Ank2 | 109676 | HITS-CLIP | 19536157 | Unspecified |

| | | | | | | |
|---|---|---|---|---|---|---|
| mmu-miR-21a-5p | MIMAT0000530 | Ankfy1 | 11736 | HITS-CLIP | 37986224\|23142080\|29167373 | Unspecified |
| mmu-miR-21a-5p | MIMAT0000530 | Ankhd1 | 108857 | HITS-CLIP | 23142080 | Unspecified |
| mmu-miR-21a-5p | MIMAT0000530 | Ankib1 | 70797 | HITS-CLIP | 23142080 | Unspecified |
| mmu-miR-21a-5p | MIMAT0000530 | Ankle1 | 234396 | Microarrays | 37986224\|25263593 | Unspecified |
| mmu-miR-21a-5p | MIMAT0000530 | Ankle2 | 71782 | HITS-CLIP | 23142080 | Unspecified |
| mmu-miR-21a-5p | MIMAT0000530 | Ankrd11 | 77087 | HITS-CLIP | 23142080 | Unspecified |
| mmu-miR-21a-5p | MIMAT0000530 | Ankrd13a | 68420 | HITS-CLIP | 23597149\|23142080\|29167373 | Unspecified |
| mmu-miR-21a-5p | MIMAT0000530 | Ankrd13b | 268445 | Microarrays | 37986224\|25263593 | Unspecified |
| mmu-miR-21a-5p | MIMAT0000530 | Ankrd13c | 433667 | HITS-CLIP | 23597149\|23142080\|29167373 | Unspecified |
| mmu-miR-21a-5p | MIMAT0000530 | Ankrd17 | 81702 | HITS-CLIP | 23597149\|31433990\|23142080\|29167 | Unspecified |
| mmu-miR-21a-5p | MIMAT0000530 | Ankrd27 | 245886 | HITS-CLIP | 23142080 | Unspecified |
| mmu-miR-21a-5p | MIMAT0000530 | Ankrd28 | 105522 | HITS-CLIP | 23597149\|23142080 | Unspecified |
| mmu-miR-21a-5p | MIMAT0000530 | Ankrd33b | 67434 | Microarrays | 37986224 | Unspecified |
| mmu-miR-21a-5p | MIMAT0000530 | Ankrd37 | 654824 | Microarrays | 37986224 | Unspecified |
| mmu-miR-21a-5p | MIMAT0000530 | Ankrd40 | 71452 | HITS-CLIP | 19536157 | Unspecified |
| mmu-miR-21a-5p | MIMAT0000530 | Ankrd44 | 329154 | HITS-CLIP | 23597149\|23142080\|29167373 | Unspecified |
| mmu-miR-21a-5p | MIMAT0000530 | Ankrd46 | 68839 | HITS-CLIP | 23597149 | Unspecified |
| mmu-miR-21a-5p | MIMAT0000530 | Ankrd49 | 56503 | HITS-CLIP | 37986224\|23142080 | Unspecified |
| mmu-miR-21a-5p | MIMAT0000530 | Ankrd50 | 99696 | HITS-CLIP | 29167373 | Unspecified |
| mmu-miR-21a-5p | MIMAT0000530 | Anks1 | 224650 | HITS-CLIP | 29167373 | Unspecified |
| mmu-miR-21a-5p | MIMAT0000530 | Anln | 68743 | HITS-CLIP, Microarrays//Microarrays//HITS | 37986224\|25263593\|23597149\|23142 | Unspecified |
| mmu-miR-21a-5p | MIMAT0000530 | Ano10 | 102566 | HITS-CLIP | 31433990 | Unspecified |
| mmu-miR-21a-5p | MIMAT0000530 | Anp32a | 11737 | Microarrays//HITS-CLIP | 37986224\|23142080 | Unspecified |
| mmu-miR-21a-5p | MIMAT0000530 | Anp32b | 67628 | HITS-CLIP | 23142080 | Unspecified |
| mmu-miR-21a-5p | MIMAT0000530 | Anp32e | 66471 | HITS-CLIP | 23142080\|23597149\|29167373 | Unspecified |
| mmu-miR-21a-5p | MIMAT0000530 | Antxr2 | 71914 | Microarrays//HITS-CLIP | 37986224\|31433990 | Unspecified |
| mmu-miR-21a-5p | MIMAT0000530 | Anxa1 | 16952 | HITS-CLIP | 29167373 | Unspecified |
| mmu-miR-21a-5p | MIMAT0000530 | Anxa2 | 12306 | HITS-CLIP | 26203562 | Unspecified |
| mmu-miR-21a-5p | MIMAT0000530 | Anxa3 | 11745 | Microarrays | 37986224 | Unspecified |
| mmu-miR-21a-5p | MIMAT0000530 | Anxa5 | 11747 | HITS-CLIP | 23597149 | Unspecified |
| mmu-miR-21a-5p | MIMAT0000530 | Anxa6 | 11749 | HITS-CLIP | 23597149\|23142080 | Unspecified |
| mmu-miR-21a-5p | MIMAT0000530 | Anxa9 | 71790 | Microarrays | 37986224 | Unspecified |
| mmu-miR-21a-5p | MIMAT0000530 | Ap1ar | 211556 | HITS-CLIP | 23142080 | Unspecified |
| mmu-miR-21a-5p | MIMAT0000530 | Ap1b1 | 11764 | HITS-CLIP | 23142080 | Unspecified |
| mmu-miR-21a-5p | MIMAT0000530 | Ap1g1 | 11765 | HITS-CLIP | 37986224\|23142080 | Unspecified |
| mmu-miR-21a-5p | MIMAT0000530 | Ap1s1 | 11769 | Microarrays | 37986224\|25263593 | Unspecified |
| mmu-miR-21a-5p | MIMAT0000530 | Ap1s2 | 108012 | HITS-CLIP | 37986224 | Unspecified |
| mmu-miR-21a-5p | MIMAT0000530 | Ap1s3 | 252903 | Microarrays//HITS-CLIP | 37986224\|23142080\|25263593 | Unspecified |
| mmu-miR-21a-5p | MIMAT0000530 | Ap2a1 | 11771 | HITS-CLIP | 23142080\|23597149\|29167373 | Unspecified |

| | | | | | | |
|---|---|---|---|---|---|---|
| mmu-miR-21a-5p | MIMAT0000530 | Ap2a2 | 11772 | HITS-CLIP, Microarrays//HITS-CLIP | 37986224\|23597149\|19536157 | Unspecified |
| mmu-miR-21a-5p | MIMAT0000530 | Ap2b1 | 71770 | HITS-CLIP | 37986224\|23597149\|19536157\|23142 | Unspecified |
| mmu-miR-21a-5p | MIMAT0000530 | Ap3b1 | 11774 | HITS-CLIP | 23142080 | Unspecified |
| mmu-miR-21a-5p | MIMAT0000530 | Ap3d1 | 11776 | HITS-CLIP | 29167373 | Unspecified |
| mmu-miR-21a-5p | MIMAT0000530 | Ap3m1 | 55946 | HITS-CLIP | 23142080\|23597149 | Unspecified |
| mmu-miR-21a-5p | MIMAT0000530 | Ap4b1 | 67489 | HITS-CLIP | 23142080 | Unspecified |
| mmu-miR-21a-5p | MIMAT0000530 | Ap4e1 | 108011 | HITS-CLIP | 23142080 | Unspecified |
| mmu-miR-21a-5p | MIMAT0000530 | Ap4m1 | 11781 | Microarrays | 37986224 | Unspecified |
| mmu-miR-21a-5p | MIMAT0000530 | Apaf1 | 11783 | HITS-CLIP | 23142080\|29167373 | Unspecified |
| mmu-miR-21a-5p | MIMAT0000530 | Apbb1 | 322 | HITS-CLIP | 23142080 | Unspecified |
| mmu-miR-21a-5p | MIMAT0000530 | Apbb1ip | 54519 | HITS-CLIP | 23142080 | Unspecified |
| mmu-miR-21a-5p | MIMAT0000530 | Apbb2 | 11787 | HITS-CLIP | 23597149\|29167373 | Unspecified |
| mmu-miR-21a-5p | MIMAT0000530 | Apc | 11789 | HITS-CLIP | 19536157\|23142080\|29167373 | Unspecified |
| mmu-miR-21a-5p | MIMAT0000530 | Apcs | 20219 | HITS-CLIP | 23597149 | Unspecified |
| mmu-miR-21a-5p | MIMAT0000530 | Apeh | 235606 | HITS-CLIP | 23142080 | Unspecified |
| mmu-miR-21a-5p | MIMAT0000530 | Apex2 | 77622 | HITS-CLIP | 29167373 | Unspecified |
| mmu-miR-21a-5p | MIMAT0000530 | Aph1b | 208117 | Microarrays | 37986224 | Unspecified |
| mmu-miR-21a-5p | MIMAT0000530 | Aph1c | 68318 | HITS-CLIP | 29167373 | Unspecified |
| mmu-miR-21a-5p | MIMAT0000530 | Api5 | 11800 | HITS-CLIP | 23142080 | Unspecified |
| mmu-miR-21a-5p | MIMAT0000530 | Apln | 30878 | HITS-CLIP | 23142080 | Unspecified |
| mmu-miR-21a-5p | MIMAT0000530 | Aplp1 | 11803 | HITS-CLIP | 19536157 | Unspecified |
| mmu-miR-21a-5p | MIMAT0000530 | Aplp2 | 11804 | HITS-CLIP | 23597149 | Unspecified |
| mmu-miR-21a-5p | MIMAT0000530 | Apoa1 | 11806 | HITS-CLIP | 23597149 | Unspecified |
| mmu-miR-21a-5p | MIMAT0000530 | Apoa5 | 66113 | HITS-CLIP | 23597149 | Unspecified |
| mmu-miR-21a-5p | MIMAT0000530 | Apob | 238055 | HITS-CLIP | 23597149 | Unspecified |
| mmu-miR-21a-5p | MIMAT0000530 | Apobec1 | 11810 | Microarrays | 37986224\|25263593 | Unspecified |
| mmu-miR-21a-5p | MIMAT0000530 | Apobec3 | 80287 | Microarrays//HITS-CLIP | 37986224\|23142080\|29167373 | Unspecified |
| mmu-miR-21a-5p | MIMAT0000530 | Apoc2 | 11813 | Microarrays | 37986224 | Unspecified |
| mmu-miR-21a-5p | MIMAT0000530 | Apoe | 11816 | Microarrays | 37986224 | Unspecified |
| mmu-miR-21a-5p | MIMAT0000530 | App | 11820 | HITS-CLIP | 23597149\|29167373 | Unspecified |
| mmu-miR-21a-5p | MIMAT0000530 | Appbp2 | 66884 | HITS-CLIP | 37986224\|23142080 | Unspecified |
| mmu-miR-21a-5p | MIMAT0000530 | Appl1 | 72993 | HITS-CLIP | 23142080 | Unspecified |
| mmu-miR-21a-5p | MIMAT0000530 | Aptx | 66408 | HITS-CLIP | 23142080 | Unspecified |
| mmu-miR-21a-5p | MIMAT0000530 | Aqp1 | 11826 | Microarrays | 37986224\|25263593 | Unspecified |
| mmu-miR-21a-5p | MIMAT0000530 | Aqp4 | 11829 | HITS-CLIP | 19536157 | Unspecified |
| mmu-miR-21a-5p | MIMAT0000530 | Aqp9 | 64008 | Microarrays | 37986224\|25263593 | Unspecified |
| mmu-miR-21a-5p | MIMAT0000530 | Aqr | 11834 | HITS-CLIP | 23142080 | Unspecified |
| mmu-miR-21a-5p | MIMAT0000530 | Araf | 11836 | HITS-CLIP | 23142080 | Unspecified |
| mmu-miR-21a-5p | MIMAT0000530 | Arap1 | 69710 | HITS-CLIP | 23142080 | Unspecified |

| | | | | | | |
|---|---|---|---|---|---|---|
| mmu-miR-21a-5p | MIMAT0000530 | Arap2 | 212285 | Microarrays//HITS-CLIP | 37986224\|25263593\|23142080 | Unspecified |
| mmu-miR-21a-5p | MIMAT0000530 | Arel1 | 68497 | HITS-CLIP | 23142080 | Unspecified |
| mmu-miR-21a-5p | MIMAT0000530 | Arf1 | 11840 | HITS-CLIP | 23142080\|23597149\|31433990\|29167 | Unspecified |
| mmu-miR-21a-5p | MIMAT0000530 | Arf3 | 11842 | HITS-CLIP | 23597149\|19536157\|23142080 | Unspecified |
| mmu-miR-21a-5p | MIMAT0000530 | Arf4 | 11843 | HITS-CLIP | 23597149\|23142080 | Unspecified |
| mmu-miR-21a-5p | MIMAT0000530 | Arf5 | 11844 | HITS-CLIP | 23597149 | Unspecified |
| mmu-miR-21a-5p | MIMAT0000530 | Arfgef1 | 211673 | HITS-CLIP//qCLASH | 37986224\|31393953\|23142080\|29167 | Unspecified |
| mmu-miR-21a-5p | MIMAT0000530 | Arfrp1 | 76688 | HITS-CLIP | 23142080 | Unspecified |
| mmu-miR-21a-5p | MIMAT0000530 | Arg1 | 11846 | Microarrays//HITS-CLIP | 37986224\|23597149\|25263593 | Unspecified |
| mmu-miR-21a-5p | MIMAT0000530 | Arhgap10 | 78514 | HITS-CLIP | 23142080\|29167373 | Unspecified |
| mmu-miR-21a-5p | MIMAT0000530 | Arhgap11a | 228482 | HITS-CLIP | 23142080 | Unspecified |
| mmu-miR-21a-5p | MIMAT0000530 | Arhgap15 | 76117 | Microarrays//qCLASH//HITS-CLIP | 37986224\|31393953\|23142080 | Unspecified |
| mmu-miR-21a-5p | MIMAT0000530 | Arhgap17 | 70497 | Microarrays | 37986224 | Unspecified |
| mmu-miR-21a-5p | MIMAT0000530 | Arhgap18 | 73910 | Microarrays//HITS-CLIP | 37986224\|23142080\|29167373 | Unspecified |
| mmu-miR-21a-5p | MIMAT0000530 | Arhgap19 | 71085 | HITS-CLIP | 23142080 | Unspecified |
| mmu-miR-21a-5p | MIMAT0000530 | Arhgap24 | 231532 | HITS-CLIP | 23142080 | Unspecified |
| mmu-miR-21a-5p | MIMAT0000530 | Arhgap25 | 232201 | Microarrays//HITS-CLIP | 37986224\|25263593\|31433990\|23142 | Unspecified |
| mmu-miR-21a-5p | MIMAT0000530 | Arhgap26 | 71302 | HITS-CLIP | 23142080 | Unspecified |
| mmu-miR-21a-5p | MIMAT0000530 | Arhgap27 | 544817 | HITS-CLIP | 23142080\|29167373 | Unspecified |
| mmu-miR-21a-5p | MIMAT0000530 | Arhgap29 | 214137 | HITS-CLIP | 23597149 | Unspecified |
| mmu-miR-21a-5p | MIMAT0000530 | Arhgap32 | 330914 | HITS-CLIP | 37986224 | Unspecified |
| mmu-miR-21a-5p | MIMAT0000530 | Arhgap35 | 232906 | HITS-CLIP | 23597149 | Unspecified |
| mmu-miR-21a-5p | MIMAT0000530 | Arhgap45 | 23526 | HITS-CLIP | 23142080 | Unspecified |
| mmu-miR-21a-5p | MIMAT0000530 | Arhgap5 | 11855 | HITS-CLIP | 23142080\|29167373\|19536157 | Unspecified |
| mmu-miR-21a-5p | MIMAT0000530 | Arhgdia | 192662 | HITS-CLIP | 23142080 | Unspecified |
| mmu-miR-21a-5p | MIMAT0000530 | Arhgef1 | 16801 | HITS-CLIP | 23142080 | Unspecified |
| mmu-miR-21a-5p | MIMAT0000530 | Arhgef12 | 69632 | HITS-CLIP | 23142080\|23597149\|29167373 | Unspecified |
| mmu-miR-21a-5p | MIMAT0000530 | Arhgef18 | 102098 | HITS-CLIP | 23142080 | Unspecified |
| mmu-miR-21a-5p | MIMAT0000530 | Arhgef2 | 16800 | HITS-CLIP | 23142080 | Unspecified |
| mmu-miR-21a-5p | MIMAT0000530 | Arhgef26 | 622434 | HITS-CLIP | 23597149 | Unspecified |
| mmu-miR-21a-5p | MIMAT0000530 | Arhgef3 | 71704 | Microarrays | 37986224 | Unspecified |
| mmu-miR-21a-5p | MIMAT0000530 | Arhgef39 | 230098 | Microarrays | 37986224\|25263593 | Unspecified |
| mmu-miR-21a-5p | MIMAT0000530 | Arhgef6 | 73341 | Microarrays//HITS-CLIP | 37986224\|31433990 | Unspecified |
| mmu-miR-21a-5p | MIMAT0000530 | Arhgef7 | 54126 | HITS-CLIP | 23597149\|19536157\|23142080\|29167 | Unspecified |
| mmu-miR-21a-5p | MIMAT0000530 | Arid1a | 93760 | HITS-CLIP | 23142080\|19536157\|29167373 | Unspecified |
| mmu-miR-21a-5p | MIMAT0000530 | Arid1b | 239985 | HITS-CLIP | 23597149 | Unspecified |
| mmu-miR-21a-5p | MIMAT0000530 | Arid2 | 77044 | HITS-CLIP | 37986224\|19536157\|23142080\|29167 | Unspecified |
| mmu-miR-21a-5p | MIMAT0000530 | Arid4a | 238247 | HITS-CLIP | 23142080\|29167373 | Unspecified |
| mmu-miR-21a-5p | MIMAT0000530 | Arid4b | 94246 | HITS-CLIP | 19536157\|23142080\|29167373 | Unspecified |

| | | | | | | |
|---|---|---|---|---|---|---|
| mmu-miR-21a-5p | MIMAT0000530 | Arid5b | 71371 | Microarrays//HITS-CLIP | 37986224\|23597149 | Unspecified |
| mmu-miR-21a-5p | MIMAT0000530 | Arih1 | 23806 | HITS-CLIP | 37986224\|23142080\|23597149 | Unspecified |
| mmu-miR-21a-5p | MIMAT0000530 | Arl10 | 56795 | HITS-CLIP | 37986224 | Unspecified |
| mmu-miR-21a-5p | MIMAT0000530 | Arl15 | 218639 | HITS-CLIP | 19536157 | Unspecified |
| mmu-miR-21a-5p | MIMAT0000530 | Arl2bp | 107566 | HITS-CLIP | 37986224\|19536157\|29167373\|23142 | Unspecified |
| mmu-miR-21a-5p | MIMAT0000530 | Arl3 | 56350 | HITS-CLIP | 23142080\|23597149 | Unspecified |
| mmu-miR-21a-5p | MIMAT0000530 | Arl4a | 11861 | HITS-CLIP | 23142080\|23597149 | Unspecified |
| mmu-miR-21a-5p | MIMAT0000530 | Arl4c | 320982 | Microarrays//HITS-CLIP | 37986224\|23142080\|29167373 | Unspecified |
| mmu-miR-21a-5p | MIMAT0000530 | Arl5a | 75423 | HITS-CLIP | 23142080 | Unspecified |
| mmu-miR-21a-5p | MIMAT0000530 | Arl5b | 75869 | HITS-CLIP//qCLASH | 37986224\|31393953\|23142080 | Unspecified |
| mmu-miR-21a-5p | MIMAT0000530 | Arl5c | 217151 | HITS-CLIP | 23142080 | Unspecified |
| mmu-miR-21a-5p | MIMAT0000530 | Arl6ip1 | 54208 | HITS-CLIP | 37986224\|23597149\|19536157\|23142 | Unspecified |
| mmu-miR-21a-5p | MIMAT0000530 | Arl6ip6 | 65103 | HITS-CLIP | 23142080 | Unspecified |
| mmu-miR-21a-5p | MIMAT0000530 | Arl8a | 68724 | HITS-CLIP | 23597149\|23142080 | Unspecified |
| mmu-miR-21a-5p | MIMAT0000530 | Arl8b | 67166 | HITS-CLIP | 37986224\|23597149\|23142080\|29167 | Unspecified |
| mmu-miR-21a-5p | MIMAT0000530 | Armc6 | 76813 | Microarrays | 37986224\|25263593 | Unspecified |
| mmu-miR-21a-5p | MIMAT0000530 | Armc8 | 74125 | HITS-CLIP | 23142080\|23597149 | Unspecified |
| mmu-miR-21a-5p | MIMAT0000530 | Armcx1 | 78248 | Microarrays//HITS-CLIP | 37986224\|25263593\|23142080 | Unspecified |
| mmu-miR-21a-5p | MIMAT0000530 | Armcx2 | 67416 | HITS-CLIP | 19536157 | Unspecified |
| mmu-miR-21a-5p | MIMAT0000530 | Armcx5 | 494468 | HITS-CLIP | 23142080 | Unspecified |
| mmu-miR-21a-5p | MIMAT0000530 | Armt1 | 79624 | HITS-CLIP | 23142080 | Unspecified |
| mmu-miR-21a-5p | MIMAT0000530 | Arnt | 11863 | HITS-CLIP//qCLASH | 23597149\|31393953\|23142080\|29167 | Unspecified |
| mmu-miR-21a-5p | MIMAT0000530 | Arpc3 | 56378 | HITS-CLIP | 23142080 | Unspecified |
| mmu-miR-21a-5p | MIMAT0000530 | Arpc4 | 68089 | HITS-CLIP | 23142080\|23597149\|19536157\|29167 | Unspecified |
| mmu-miR-21a-5p | MIMAT0000530 | Arpp19 | 59046 | HITS-CLIP | 37986224\|23597149\|23142080 | Unspecified |
| mmu-miR-21a-5p | MIMAT0000530 | Arrb1 | 109689 | HITS-CLIP | 29167373 | Unspecified |
| mmu-miR-21a-5p | MIMAT0000530 | Arrdc1 | 215705 | HITS-CLIP | 23142080 | Unspecified |
| mmu-miR-21a-5p | MIMAT0000530 | Arrdc2 | 70807 | Microarrays//qCLASH | 37986224\|31393953 | Unspecified |
| mmu-miR-21a-5p | MIMAT0000530 | Arrdc3 | 105171 | HITS-CLIP, Microarrays//HITS-CLIP | 37986224\|23597149\|23142080 | Unspecified |
| mmu-miR-21a-5p | MIMAT0000530 | Arrdc4 | 66412 | Microarrays | 37986224 | Unspecified |
| mmu-miR-21a-5p | MIMAT0000530 | Arsk | 77041 | Microarrays | 37986224\|25263593 | Unspecified |
| mmu-miR-21a-5p | MIMAT0000530 | Art2b | 11872 | HITS-CLIP | 23142080 | Unspecified |
| mmu-miR-21a-5p | MIMAT0000530 | Arvcf | 11877 | HITS-CLIP | 23597149 | Unspecified |
| mmu-miR-21a-5p | MIMAT0000530 | Asah1 | 11886 | HITS-CLIP | 23597149\|29167373 | Unspecified |
| mmu-miR-21a-5p | MIMAT0000530 | Asap1 | 13196 | HITS-CLIP | 37986224\|23142080\|23597149 | Unspecified |
| mmu-miR-21a-5p | MIMAT0000530 | Asb1 | 65247 | HITS-CLIP | 23142080 | Unspecified |
| mmu-miR-21a-5p | MIMAT0000530 | Asb3 | 65257 | HITS-CLIP | 23142080 | Unspecified |
| mmu-miR-21a-5p | MIMAT0000530 | Asb4 | 65255 | Microarrays | 37986224 | Unspecified |
| mmu-miR-21a-5p | MIMAT0000530 | Asb7 | 117589 | HITS-CLIP | 23142080 | Unspecified |

| | | | | | | |
|---|---|---|---|---|---|---|
| mmu-miR-21a-5p | MIMAT0000530 | Asb8 | 78541 | HITS-CLIP | 23142080 | Unspecified |
| mmu-miR-21a-5p | MIMAT0000530 | Ascc3 | 77987 | HITS-CLIP | 23142080\|29167373 | Unspecified |
| mmu-miR-21a-5p | MIMAT0000530 | Ascl1 | 17172 | HITS-CLIP | 19536157 | Unspecified |
| mmu-miR-21a-5p | MIMAT0000530 | Asf1a | 66403 | HITS-CLIP | 37986224\|23142080 | Unspecified |
| mmu-miR-21a-5p | MIMAT0000530 | Asf1b | 66929 | Microarrays//HITS-CLIP | 37986224\|25263593\|23142080 | Unspecified |
| mmu-miR-21a-5p | MIMAT0000530 | Asgr1 | 11889 | HITS-CLIP | 23597149 | Unspecified |
| mmu-miR-21a-5p | MIMAT0000530 | Ash1l | 192195 | HITS-CLIP | 23142080\|23597149\|31433990\|29167 | Unspecified |
| mmu-miR-21a-5p | MIMAT0000530 | Ash2l | 23808 | HITS-CLIP | 23142080 | Unspecified |
| mmu-miR-21a-5p | MIMAT0000530 | Asl | 109900 | HITS-CLIP | 23597149 | Unspecified |
| mmu-miR-21a-5p | MIMAT0000530 | Asns | 27053 | Microarrays//HITS-CLIP | 37986224\|25263593\|23142080 | Unspecified |
| mmu-miR-21a-5p | MIMAT0000530 | Aspa | 11484 | HITS-CLIP | 23597149 | Unspecified |
| mmu-miR-21a-5p | MIMAT0000530 | Asph | 65973 | HITS-CLIP | 29167373 | Unspecified |
| mmu-miR-21a-5p | MIMAT0000530 | Aspm | 12316 | HITS-CLIP | 23142080 | Unspecified |
| mmu-miR-21a-5p | MIMAT0000530 | Astn1 | 11899 | Microarrays | 37986224 | Unspecified |
| mmu-miR-21a-5p | MIMAT0000530 | Asxl1 | 228790 | HITS-CLIP | 23142080 | Unspecified |
| mmu-miR-21a-5p | MIMAT0000530 | Asxl2 | 75302 | HITS-CLIP | 37986224\|29167373 | Unspecified |
| mmu-miR-21a-5p | MIMAT0000530 | Atad1 | 67979 | HITS-CLIP | 37986224 | Unspecified |
| mmu-miR-21a-5p | MIMAT0000530 | Atad2 | 70472 | HITS-CLIP | 37986224\|23597149\|23142080 | Unspecified |
| mmu-miR-21a-5p | MIMAT0000530 | Atad2b | 320817 | HITS-CLIP | 23142080\|29167373 | Unspecified |
| mmu-miR-21a-5p | MIMAT0000530 | Atad5 | 237877 | HITS-CLIP | 23142080 | Unspecified |
| mmu-miR-21a-5p | MIMAT0000530 | Atat1 | 73242 | HITS-CLIP | 29167373 | Unspecified |
| mmu-miR-21a-5p | MIMAT0000530 | Atf1 | 11908 | HITS-CLIP | 23597149 | Unspecified |
| mmu-miR-21a-5p | MIMAT0000530 | Atf2 | 11909 | HITS-CLIP | 23142080 | Unspecified |
| mmu-miR-21a-5p | MIMAT0000530 | Atf3 | 11910 | HITS-CLIP | 23597149 | Unspecified |
| mmu-miR-21a-5p | MIMAT0000530 | Atf4 | 11911 | HITS-CLIP | 31433990\|23142080 | Unspecified |
| mmu-miR-21a-5p | MIMAT0000530 | Atf6 | 226641 | HITS-CLIP | 23597149\|23142080 | Unspecified |
| mmu-miR-21a-5p | MIMAT0000530 | Atf7ip2 | 75329 | Microarrays | 37986224\|25263593 | Unspecified |
| mmu-miR-21a-5p | MIMAT0000530 | Atg101 | 68118 | HITS-CLIP | 23142080 | Unspecified |
| mmu-miR-21a-5p | MIMAT0000530 | Atg12 | 67526 | HITS-CLIP | 23142080 | Unspecified |
| mmu-miR-21a-5p | MIMAT0000530 | Atg2b | 76559 | HITS-CLIP | 23597149 | Unspecified |
| mmu-miR-21a-5p | MIMAT0000530 | Atg5 | 11793 | HITS-CLIP | 23142080 | Unspecified |
| mmu-miR-21a-5p | MIMAT0000530 | Atg9a | 245860 | HITS-CLIP | 23142080 | Unspecified |
| mmu-miR-21a-5p | MIMAT0000530 | Atl2 | 56298 | HITS-CLIP | 23597149 | Unspecified |
| mmu-miR-21a-5p | MIMAT0000530 | Atl3 | 109168 | HITS-CLIP | 23142080\|31433990\|30971389 | Unspecified |
| mmu-miR-21a-5p | MIMAT0000530 | Atm | 11920 | Microarrays//HITS-CLIP | 37986224\|25263593\|23597149\|3143 | Unspecified |
| mmu-miR-21a-5p | MIMAT0000530 | Atn1 | 13498 | HITS-CLIP | 23142080 | Unspecified |
| mmu-miR-21a-5p | MIMAT0000530 | Atp10a | 11982 | HITS-CLIP | 23142080 | Unspecified |
| mmu-miR-21a-5p | MIMAT0000530 | Atp11a | 50770 | HITS-CLIP | 37986224\|29167373\|23142080 | Unspecified |
| mmu-miR-21a-5p | MIMAT0000530 | Atp11b | 76295 | HITS-CLIP | 23142080 | Unspecified |

| | | | | | | |
|---|---|---|---|---|---|---|
| mmu-miR-21a-5p | MIMAT0000530 | Atp11c | 320940 | HITS-CLIP | 23142080 | Unspecified |
| mmu-miR-21a-5p | MIMAT0000530 | Atp13a3 | 224088 | HITS-CLIP | 37986224\|23597149\|23142080\|29167 | Unspecified |
| mmu-miR-21a-5p | MIMAT0000530 | Atp1a1 | 11928 | HITS-CLIP | 23597149\|29167373 | Unspecified |
| mmu-miR-21a-5p | MIMAT0000530 | Atp1b1 | 11931 | HITS-CLIP | 23597149\|19536157 | Unspecified |
| mmu-miR-21a-5p | MIMAT0000530 | Atp1b2 | 11932 | HITS-CLIP | 19536157 | Unspecified |
| mmu-miR-21a-5p | MIMAT0000530 | Atp2a2 | 11938 | HITS-CLIP | 37986224\|19536157\|23142080 | Unspecified |
| mmu-miR-21a-5p | MIMAT0000530 | Atp2b1 | 67972 | Microarrays//HITS-CLIP | 37986224\|23597149\|19536157\|23142 | Unspecified |
| mmu-miR-21a-5p | MIMAT0000530 | Atp2b4 | 381290 | HITS-CLIP | 23142080 | Unspecified |
| mmu-miR-21a-5p | MIMAT0000530 | Atp5a1 | 11946 | HITS-CLIP//qCLASH | 37986224\|23597149\|31433990\|31393 | Unspecified |
| mmu-miR-21a-5p | MIMAT0000530 | Atp5b | 11947 | HITS-CLIP | 37986224\|23597149\|23142080\|29167 | Unspecified |
| mmu-miR-21a-5p | MIMAT0000530 | Atp5d | 66043 | HITS-CLIP | 19536157\|23597149\|23142080 | Unspecified |
| mmu-miR-21a-5p | MIMAT0000530 | Atp5j | 11957 | HITS-CLIP | 37986224\|23597149\|23142080\|29167 | Unspecified |
| mmu-miR-21a-5p | MIMAT0000530 | Atp5o | 28080 | HITS-CLIP | 23597149\|23142080 | Unspecified |
| mmu-miR-21a-5p | MIMAT0000530 | Atp6v0a1 | 11975 | HITS-CLIP | 19536157\|23597149 | Unspecified |
| mmu-miR-21a-5p | MIMAT0000530 | Atp6v0d2 | 242341 | Microarrays | 37986224\|25263593 | Unspecified |
| mmu-miR-21a-5p | MIMAT0000530 | Atp6v0e2 | 76252 | HITS-CLIP | 19536157 | Unspecified |
| mmu-miR-21a-5p | MIMAT0000530 | Atp6v1a | 11964 | HITS-CLIP | 31433990\|29167373 | Unspecified |
| mmu-miR-21a-5p | MIMAT0000530 | Atp6v1b2 | 11966 | HITS-CLIP | 37986224\|23597149\|23142080\|26203 | Unspecified |
| mmu-miR-21a-5p | MIMAT0000530 | Atp6v1f | 66144 | HITS-CLIP | 23142080 | Unspecified |
| mmu-miR-21a-5p | MIMAT0000530 | Atp7a | 11977 | HITS-CLIP | 29167373 | Unspecified |
| mmu-miR-21a-5p | MIMAT0000530 | Atp8a1 | 11980 | HITS-CLIP | 23142080 | Unspecified |
| mmu-miR-21a-5p | MIMAT0000530 | Atp9b | 50771 | HITS-CLIP | 23597149 | Unspecified |
| mmu-miR-21a-5p | MIMAT0000530 | Atr | 245000 | HITS-CLIP | 37986224 | Unspecified |
| mmu-miR-21a-5p | MIMAT0000530 | Atrn | 11990 | HITS-CLIP | 26203562\|29167373 | Unspecified |
| mmu-miR-21a-5p | MIMAT0000530 | Atrnl1 | 226255 | Microarrays//HITS-CLIP | 37986224\|19536157 | Unspecified |
| mmu-miR-21a-5p | MIMAT0000530 | Atrx | 22589 | HITS-CLIP | 23142080\|19536157 | Unspecified |
| mmu-miR-21a-5p | MIMAT0000530 | Atxn1 | 20238 | HITS-CLIP | 37986224\|23142080\|29167373 | Unspecified |
| mmu-miR-21a-5p | MIMAT0000530 | Atxn10 | 54138 | HITS-CLIP | 23142080\|29167373 | Unspecified |
| mmu-miR-21a-5p | MIMAT0000530 | Atxn1l | 52335 | HITS-CLIP | 23142080 | Unspecified |
| mmu-miR-21a-5p | MIMAT0000530 | Atxn2 | 20239 | HITS-CLIP | 23142080 | Unspecified |
| mmu-miR-21a-5p | MIMAT0000530 | Atxn2l | 233871 | HITS-CLIP | 23142080\|29167373 | Unspecified |
| mmu-miR-21a-5p | MIMAT0000530 | Atxn3 | 110616 | HITS-CLIP | 23142080 | Unspecified |
| mmu-miR-21a-5p | MIMAT0000530 | Atxn7 | 246103 | HITS-CLIP | 29167373 | Unspecified |
| mmu-miR-21a-5p | MIMAT0000530 | Atxn7l3 | 217218 | HITS-CLIP | 23597149\|23142080 | Unspecified |
| mmu-miR-21a-5p | MIMAT0000530 | Atxn7l3b | 382423 | HITS-CLIP//qCLASH | 37986224\|31393953\|23597149\|23142 | Unspecified |
| mmu-miR-21a-5p | MIMAT0000530 | Aurka | 20878 | Microarrays | 37986224\|25263593 | Unspecified |
| mmu-miR-21a-5p | MIMAT0000530 | Aurkb | 20877 | Microarrays | 37986224\|25263593 | Unspecified |
| mmu-miR-21a-5p | MIMAT0000530 | Avl9 | 78937 | HITS-CLIP | 23142080\|29167373 | Unspecified |
| mmu-miR-21a-5p | MIMAT0000530 | Axl | 26362 | HITS-CLIP | 23142080\|29167373 | Unspecified |

| | | | | | | |
|---|---|---|---|---|---|---|
| mmu-miR-21a-5p | MIMAT0000530 | Azgp1 | 12007 | HITS-CLIP | 23597149 | Unspecified |
| mmu-miR-21a-5p | MIMAT0000530 | Azin1 | 54375 | HITS-CLIP, Microarrays//Microarrays//HITS | 37986224\|25263593\|23597149\|29167 | Unspecified |
| mmu-miR-21a-5p | MIMAT0000530 | B230219D22Rik | 78521 | HITS-CLIP | 19536157\|23142080 | Unspecified |
| mmu-miR-21a-5p | MIMAT0000530 | B2m | 12010 | HITS-CLIP | 23597149\|23142080\|29167373 | Unspecified |
| mmu-miR-21a-5p | MIMAT0000530 | B3gnt2 | 53625 | HITS-CLIP | 23142080 | Unspecified |
| mmu-miR-21a-5p | MIMAT0000530 | B3gnt7 | 227327 | Microarrays | 37986224\|25263593 | Unspecified |
| mmu-miR-21a-5p | MIMAT0000530 | B430306N03Rik | 320148 | Microarrays | 37986224\|25263593 | Unspecified |
| mmu-miR-21a-5p | MIMAT0000530 | B4galt1 | 14595 | HITS-CLIP | 23597149\|23142080 | Unspecified |
| mmu-miR-21a-5p | MIMAT0000530 | B4galt5 | 56336 | HITS-CLIP | 23597149\|23142080 | Unspecified |
| mmu-miR-21a-5p | MIMAT0000530 | B4galt6 | 56386 | HITS-CLIP | 29167373 | Unspecified |
| mmu-miR-21a-5p | MIMAT0000530 | B4galt7 | 218271 | Microarrays | 37986224\|25263593 | Unspecified |
| mmu-miR-21a-5p | MIMAT0000530 | B4gat1 | 135972575 | HITS-CLIP | 23597149 | Unspecified |
| mmu-miR-21a-5p | MIMAT0000530 | BC004004 | 80748 | HITS-CLIP | 23597149 | Unspecified |
| mmu-miR-21a-5p | MIMAT0000530 | BC005537 | 79555 | HITS-CLIP | 29167373 | Unspecified |
| mmu-miR-21a-5p | MIMAT0000530 | BC005561 | 100042165 | HITS-CLIP | 23597149\|23142080 | Unspecified |
| mmu-miR-21a-5p | MIMAT0000530 | BC025446 | | HITS-CLIP | 23597149 | Unspecified |
| mmu-miR-21a-5p | MIMAT0000530 | BC029722 | | HITS-CLIP | 23142080 | Unspecified |
| mmu-miR-21a-5p | MIMAT0000530 | BC035044 | 232406 | Microarrays//HITS-CLIP | 37986224\|23142080 | Unspecified |
| mmu-miR-21a-5p | MIMAT0000530 | BC055324 | 381306 | HITS-CLIP | 23142080 | Unspecified |
| mmu-miR-21a-5p | MIMAT0000530 | Bace1 | 23821 | HITS-CLIP | 23597149 | Unspecified |
| mmu-miR-21a-5p | MIMAT0000530 | Bach1 | 12013 | HITS-CLIP | 23597149\|23142080 | Unspecified |
| mmu-miR-21a-5p | MIMAT0000530 | Bach2 | 12014 | HITS-CLIP | 23597149\|23142080 | Unspecified |
| mmu-miR-21a-5p | MIMAT0000530 | Bag2 | 213539 | HITS-CLIP | 23142080 | Unspecified |
| mmu-miR-21a-5p | MIMAT0000530 | Bag3 | 29810 | HITS-CLIP | 23142080 | Unspecified |
| mmu-miR-21a-5p | MIMAT0000530 | Bahd1 | 228536 | HITS-CLIP | 23597149 | Unspecified |
| mmu-miR-21a-5p | MIMAT0000530 | Baiap2 | 108100 | HITS-CLIP | 23597149 | Unspecified |
| mmu-miR-21a-5p | MIMAT0000530 | Bank1 | 242248 | HITS-CLIP | 23597149 | Unspecified |
| mmu-miR-21a-5p | MIMAT0000530 | Basp1 | 70350 | HITS-CLIP | 23142080 | Unspecified |
| mmu-miR-21a-5p | MIMAT0000530 | Baz1a | 217578 | HITS-CLIP | 37986224\|23142080 | Unspecified |
| mmu-miR-21a-5p | MIMAT0000530 | Baz1b | 22385 | HITS-CLIP | 37986224\|23142080\|29167373 | Unspecified |
| mmu-miR-21a-5p | MIMAT0000530 | Baz2a | 116848 | HITS-CLIP//qCLASH | 19536157\|31393953\|23597149 | Unspecified |
| mmu-miR-21a-5p | MIMAT0000530 | Baz2b | 407823 | HITS-CLIP | 23142080 | Unspecified |
| mmu-miR-21a-5p | MIMAT0000530 | Bbip1 | 100503572 | HITS-CLIP | 23142080 | Unspecified |
| mmu-miR-21a-5p | MIMAT0000530 | Bbln | 79095 | Microarrays | 25263593 | Unspecified |
| mmu-miR-21a-5p | MIMAT0000530 | Bbs2 | 67378 | Microarrays | 37986224 | Unspecified |
| mmu-miR-21a-5p | MIMAT0000530 | Bbx | 70508 | HITS-CLIP | 23142080\|29167373 | Unspecified |
| mmu-miR-21a-5p | MIMAT0000530 | Bcap31 | 27061 | HITS-CLIP | 23597149\|23142080 | Unspecified |
| mmu-miR-21a-5p | MIMAT0000530 | Bcar3 | 29815 | HITS-CLIP | 23597149 | Unspecified |
| mmu-miR-21a-5p | MIMAT0000530 | Bcas2 | 68183 | HITS-CLIP | 23142080 | Unspecified |

| | | | | | | |
|---|---|---|---|---|---|---|
| mmu-miR-21a-5p | MIMAT0000530 | Bccip | 66165 | HITS-CLIP | 23142080 | Unspecified |
| mmu-miR-21a-5p | MIMAT0000530 | Bcl10 | 12042 | HITS-CLIP | 23142080 | Unspecified |
| mmu-miR-21a-5p | MIMAT0000530 | Bcl11a | 14025 | HITS-CLIP | 23142080 | Unspecified |
| mmu-miR-21a-5p | MIMAT0000530 | Bcl11b | 58208 | HITS-CLIP | 37986224\|23142080 | Unspecified |
| mmu-miR-21a-5p | MIMAT0000530 | Bcl2 | 12043 | HITS-CLIP, Microarrays//Microarrays//HITS | 37986224\|25263593\|23142080 | Unspecified |
| mmu-miR-21a-5p | MIMAT0000530 | Bcl2l1 | 12048 | Microarrays | 37986224\|25263593 | Unspecified |
| mmu-miR-21a-5p | MIMAT0000530 | Bcl2l11 | 12125 | Microarrays//qCLASH//HITS-CLIP | 37986224\|31393953\|23142080\|29167 | Unspecified |
| mmu-miR-21a-5p | MIMAT0000530 | Bcl2l13 | 94044 | HITS-CLIP | 23142080 | Unspecified |
| mmu-miR-21a-5p | MIMAT0000530 | Bcl2l2 | 12050 | HITS-CLIP | 23142080 | Unspecified |
| mmu-miR-21a-5p | MIMAT0000530 | Bcl3 | 12051 | Microarrays//HITS-CLIP | 37986224\|23142080 | Unspecified |
| mmu-miR-21a-5p | MIMAT0000530 | Bcl6 | 12053 | HITS-CLIP | 23142080 | Unspecified |
| mmu-miR-21a-5p | MIMAT0000530 | Bcl7a | 77045 | HITS-CLIP//qCLASH | 23597149\|31393953\|23142080 | Unspecified |
| mmu-miR-21a-5p | MIMAT0000530 | Bcl9 | 77578 | HITS-CLIP | 19536157\|23142080 | Unspecified |
| mmu-miR-21a-5p | MIMAT0000530 | Bclaf1 | 72567 | HITS-CLIP | 19536157\|31433990\|23142080 | Unspecified |
| mmu-miR-21a-5p | MIMAT0000530 | Bcor | 71458 | HITS-CLIP | 23142080 | Unspecified |
| mmu-miR-21a-5p | MIMAT0000530 | Bcorl1 | 320376 | HITS-CLIP | 23142080 | Unspecified |
| mmu-miR-21a-5p | MIMAT0000530 | Bdh2 | 69772 | Microarrays | 37986224\|25263593 | Unspecified |
| mmu-miR-21a-5p | MIMAT0000530 | Bdp1 | 544971 | HITS-CLIP | 31433990 | Unspecified |
| mmu-miR-21a-5p | MIMAT0000530 | Bend4 | 666938 | Microarrays | 37986224 | Unspecified |
| mmu-miR-21a-5p | MIMAT0000530 | Bet1 | 12068 | HITS-CLIP | 23597149 | Unspecified |
| mmu-miR-21a-5p | MIMAT0000530 | Bex1 | 19716 | Microarrays | 37986224\|25263593 | Unspecified |
| mmu-miR-21a-5p | MIMAT0000530 | Bex6 | 328660 | Microarrays | 37986224\|25263593 | Unspecified |
| mmu-miR-21a-5p | MIMAT0000530 | Bfar | 67118 | HITS-CLIP | 37986224\|23142080\|29167373 | Unspecified |
| mmu-miR-21a-5p | MIMAT0000530 | Bgn | 12111 | Microarrays | 37986224\|25263593 | Unspecified |
| mmu-miR-21a-5p | MIMAT0000530 | Bhlhe40 | 20893 | HITS-CLIP, Microarrays//Microarrays//HITS | 37986224\|25263593\|23597149\|2314 | Unspecified |
| mmu-miR-21a-5p | MIMAT0000530 | Bhlhe41 | 79362 | HITS-CLIP | 37986224\|23597149\|29167373\|2314 | Unspecified |
| mmu-miR-21a-5p | MIMAT0000530 | Bhmt2 | 64918 | HITS-CLIP | 23597149 | Unspecified |
| mmu-miR-21a-5p | MIMAT0000530 | Bicd1 | 12121 | HITS-CLIP | 37986224 | Unspecified |
| mmu-miR-21a-5p | MIMAT0000530 | Bicd2 | 76895 | HITS-CLIP | 23142080 | Unspecified |
| mmu-miR-21a-5p | MIMAT0000530 | Bicra | 29998 | HITS-CLIP | 23142080 | Unspecified |
| mmu-miR-21a-5p | MIMAT0000530 | Bid | 12122 | Microarrays | 37986224\|25263593 | Unspecified |
| mmu-miR-21a-5p | MIMAT0000530 | Birc2 | 11797 | HITS-CLIP | 23142080 | Unspecified |
| mmu-miR-21a-5p | MIMAT0000530 | Birc3 | 11796 | Microarrays | 37986224 | Unspecified |
| mmu-miR-21a-5p | MIMAT0000530 | Birc5 | 11799 | Microarrays | 37986224\|25263593 | Unspecified |
| mmu-miR-21a-5p | MIMAT0000530 | Birc6 | 12211 | HITS-CLIP | 23142080\|23597149\|29167373 | Unspecified |
| mmu-miR-21a-5p | MIMAT0000530 | Blm | 12144 | HITS-CLIP | 23142080 | Unspecified |
| mmu-miR-21a-5p | MIMAT0000530 | Blmh | 104184 | HITS-CLIP | 23597149\|23142080\|29167373 | Unspecified |
| mmu-miR-21a-5p | MIMAT0000530 | Bloc1s5 | 17828 | HITS-CLIP | 23142080\|29167373 | Unspecified |
| mmu-miR-21a-5p | MIMAT0000530 | Bloc1s6 | 18457 | HITS-CLIP | 23142080 | Unspecified |

| | | | | | | |
|---|---|---|---|---|---|---|
| mmu-miR-21a-5p | MIMAT0000530 | Bmf | 171543 | Microarrays | 37986224 | Unspecified |
| mmu-miR-21a-5p | MIMAT0000530 | Bmp2k | 140780 | HITS-CLIP | 37986224\|23142080\|29167373 | Unspecified |
| mmu-miR-21a-5p | MIMAT0000530 | Bmp8b | 656 | HITS-CLIP | 23597149 | Unspecified |
| mmu-miR-21a-5p | MIMAT0000530 | Bmpr1a | 12166 | HITS-CLIP | 23142080 | Unspecified |
| mmu-miR-21a-5p | MIMAT0000530 | Bmpr2 | 12168 | HITS-CLIP | 37986224\|19536157\|23597149\|23142 | Unspecified |
| mmu-miR-21a-5p | MIMAT0000530 | Bms1 | 213895 | HITS-CLIP | 23142080 | Unspecified |
| mmu-miR-21a-5p | MIMAT0000530 | Bmt2 | 101148 | HITS-CLIP | 23142080\|31433990\|23597149\|28735 | Unspecified |
| mmu-miR-21a-5p | MIMAT0000530 | Bmyc | 107771 | Microarrays | 37986224 | Unspecified |
| mmu-miR-21a-5p | MIMAT0000530 | Bnip1 | 224630 | HITS-CLIP | 23142080 | Unspecified |
| mmu-miR-21a-5p | MIMAT0000530 | Bnip2 | 12175 | HITS-CLIP | 23142080 | Unspecified |
| mmu-miR-21a-5p | MIMAT0000530 | Bod1l | 665775 | qCLASH//HITS-CLIP | 31393953\|23142080 | Unspecified |
| mmu-miR-21a-5p | MIMAT0000530 | Bpifc | 270757 | Microarrays | 37986224\|25263593 | Unspecified |
| mmu-miR-21a-5p | MIMAT0000530 | Bpnt1 | 23827 | Microarrays | 37986224\|25263593 | Unspecified |
| mmu-miR-21a-5p | MIMAT0000530 | Bpnt2 | 54928 | Microarrays//HITS-CLIP | 25263593\|31433990 | Unspecified |
| mmu-miR-21a-5p | MIMAT0000530 | Bptf | 207165 | HITS-CLIP | 23142080\|23597149\|26203562\|29167 | Unspecified |
| mmu-miR-21a-5p | MIMAT0000530 | Braf | 109880 | HITS-CLIP | 37986224\|23142080 | Unspecified |
| mmu-miR-21a-5p | MIMAT0000530 | Brca1 | 12189 | Microarrays//HITS-CLIP | 37986224\|25263593\|23142080 | Unspecified |
| mmu-miR-21a-5p | MIMAT0000530 | Brca2 | 12190 | HITS-CLIP | 23597149\|23142080 | Unspecified |
| mmu-miR-21a-5p | MIMAT0000530 | Brd1 | 223770 | HITS-CLIP | 23142080\|29167373 | Unspecified |
| mmu-miR-21a-5p | MIMAT0000530 | Brd2 | 14312 | HITS-CLIP | 37986224\|23597149\|23142080\|29167 | Unspecified |
| mmu-miR-21a-5p | MIMAT0000530 | Brd3 | 67382 | HITS-CLIP | 23597149\|19536157\|23142080 | Unspecified |
| mmu-miR-21a-5p | MIMAT0000530 | Brd4 | 57261 | HITS-CLIP | 23142080 | Unspecified |
| mmu-miR-21a-5p | MIMAT0000530 | Brd7 | 26992 | HITS-CLIP | 29167373\|23142080 | Unspecified |
| mmu-miR-21a-5p | MIMAT0000530 | Brd8 | 78656 | HITS-CLIP | 23142080 | Unspecified |
| mmu-miR-21a-5p | MIMAT0000530 | Brd9 | 105246 | HITS-CLIP | 23142080 | Unspecified |
| mmu-miR-21a-5p | MIMAT0000530 | Brip1 | 237911 | Microarrays | 37986224\|25263593 | Unspecified |
| mmu-miR-21a-5p | MIMAT0000530 | Brms1l | 52592 | HITS-CLIP | 26203562\|23142080 | Unspecified |
| mmu-miR-21a-5p | MIMAT0000530 | Brpf3 | 268936 | HITS-CLIP | 23597149\|23142080 | Unspecified |
| mmu-miR-21a-5p | MIMAT0000530 | Brwd1 | 93871 | HITS-CLIP | 23142080\|29167373 | Unspecified |
| mmu-miR-21a-5p | MIMAT0000530 | Brwd3 | 382236 | HITS-CLIP | 23142080\|29167373 | Unspecified |
| mmu-miR-21a-5p | MIMAT0000530 | Bscl2 | 14705 | HITS-CLIP | 23142080 | Unspecified |
| mmu-miR-21a-5p | MIMAT0000530 | Bst1 | 12182 | Microarrays | 37986224\|25263593 | Unspecified |
| mmu-miR-21a-5p | MIMAT0000530 | Bst2 | 69550 | Microarrays | 37986224 | Unspecified |
| mmu-miR-21a-5p | MIMAT0000530 | Btaf1 | 107182 | HITS-CLIP | 31433990\|23597149\|23142080\|29167 | Unspecified |
| mmu-miR-21a-5p | MIMAT0000530 | Btbd7 | 238386 | HITS-CLIP | 23142080 | Unspecified |
| mmu-miR-21a-5p | MIMAT0000530 | Btbd8 | 100503185 | HITS-CLIP | 23142080 | Unspecified |
| mmu-miR-21a-5p | MIMAT0000530 | Btf3l4 | 70533 | HITS-CLIP | 23142080 | Unspecified |
| mmu-miR-21a-5p | MIMAT0000530 | Btg1 | 12226 | HITS-CLIP | 29167373 | Unspecified |
| mmu-miR-21a-5p | MIMAT0000530 | Btg2 | 12227 | HITS-CLIP, qPCR//HITS-CLIP//qCLASH//Oth | 37986224\|23597149\|31393953\|29167 | Unspecified |

| | | | | | | |
|---|---|---|---|---|---|---|
| mmu-miR-21a-5p | MIMAT0000530 | Btla | 208154 | HITS-CLIP | 23142080 | Unspecified |
| mmu-miR-21a-5p | MIMAT0000530 | Bub1 | 12235 | Microarrays//HITS-CLIP | 37986224\|25263593\|23142080 | Unspecified |
| mmu-miR-21a-5p | MIMAT0000530 | Bub1b | 12236 | Microarrays//HITS-CLIP | 37986224\|25263593\|23142080 | Unspecified |
| mmu-miR-21a-5p | MIMAT0000530 | Bub3 | 12237 | HITS-CLIP | 23142080 | Unspecified |
| mmu-miR-21a-5p | MIMAT0000530 | Bud13 | 215051 | HITS-CLIP | 23142080 | Unspecified |
| mmu-miR-21a-5p | MIMAT0000530 | Bud31 | 231889 | HITS-CLIP | 23142080 | Unspecified |
| mmu-miR-21a-5p | MIMAT0000530 | Bysl | 53414 | Microarrays//HITS-CLIP | 37986224\|25263593\|23142080 | Unspecified |
| mmu-miR-21a-5p | MIMAT0000530 | Bzw1 | 66882 | HITS-CLIP | 19536157\|29167373 | Unspecified |
| mmu-miR-21a-5p | MIMAT0000530 | C130050O18Rik | 319772 | Microarrays | 37986224 | Unspecified |
| mmu-miR-21a-5p | MIMAT0000530 | C1galt1 | 94192 | HITS-CLIP | 23142080 | Unspecified |
| mmu-miR-21a-5p | MIMAT0000530 | C1qa | 12259 | Microarrays | 37986224 | Unspecified |
| mmu-miR-21a-5p | MIMAT0000530 | C1qbp | 12261 | HITS-CLIP//qCLASH | 31433990\|31393953\|23142080 | Unspecified |
| mmu-miR-21a-5p | MIMAT0000530 | C1ra | 50909 | Microarrays//HITS-CLIP | 37986224\|23597149 | Unspecified |
| mmu-miR-21a-5p | MIMAT0000530 | C2cd2 | 207781 | HITS-CLIP | 23597149 | Unspecified |
| mmu-miR-21a-5p | MIMAT0000530 | C2cd3 | 277939 | HITS-CLIP | 29167373 | Unspecified |
| mmu-miR-21a-5p | MIMAT0000530 | C2cd5 | 74741 | HITS-CLIP | 23142080 | Unspecified |
| mmu-miR-21a-5p | MIMAT0000530 | C3 | 12266 | Microarrays//HITS-CLIP | 37986224\|23597149 | Unspecified |
| mmu-miR-21a-5p | MIMAT0000530 | C330007P06Rik | 77644 | HITS-CLIP | 23142080 | Unspecified |
| mmu-miR-21a-5p | MIMAT0000530 | C330018D20Rik | 77422 | HITS-CLIP | 23597149\|23142080 | Unspecified |
| mmu-miR-21a-5p | MIMAT0000530 | C4b | 12268 | HITS-CLIP | 23597149 | Unspecified |
| mmu-miR-21a-5p | MIMAT0000530 | C6 | 12274 | HITS-CLIP | 23597149 | Unspecified |
| mmu-miR-21a-5p | MIMAT0000530 | C77080 | 97130 | Microarrays//HITS-CLIP | 37986224\|25263593\|23597149 | Unspecified |
| mmu-miR-21a-5p | MIMAT0000530 | C8b | 110382 | HITS-CLIP | 23597149 | Unspecified |
| mmu-miR-21a-5p | MIMAT0000530 | C9 | 12279 | HITS-CLIP | 23597149 | Unspecified |
| mmu-miR-21a-5p | MIMAT0000530 | C9orf72 | 203228 | HITS-CLIP | 23142080\|29167373 | Unspecified |
| mmu-miR-21a-5p | MIMAT0000530 | Caap1 | 67770 | HITS-CLIP | 23142080 | Unspecified |
| mmu-miR-21a-5p | MIMAT0000530 | Cab39 | 12283 | HITS-CLIP | 23142080 | Unspecified |
| mmu-miR-21a-5p | MIMAT0000530 | Cabin1 | 104248 | HITS-CLIP | 23142080 | Unspecified |
| mmu-miR-21a-5p | MIMAT0000530 | Cacna1d | 12289 | HITS-CLIP | 19536157 | Unspecified |
| mmu-miR-21a-5p | MIMAT0000530 | Cacna1e | 12290 | HITS-CLIP | 37986224\|19536157 | Unspecified |
| mmu-miR-21a-5p | MIMAT0000530 | Cacna2d3 | 12294 | HITS-CLIP | 19536157 | Unspecified |
| mmu-miR-21a-5p | MIMAT0000530 | Cacul1 | 78832 | HITS-CLIP | 23142080 | Unspecified |
| mmu-miR-21a-5p | MIMAT0000530 | Cacybp | 12301 | qCLASH | 31393953 | Unspecified |
| mmu-miR-21a-5p | MIMAT0000530 | Cad | 69719 | HITS-CLIP | 23142080 | Unspecified |
| mmu-miR-21a-5p | MIMAT0000530 | Cadm1 | 54725 | HITS-CLIP, Microarrays | 37986224 | Unspecified |
| mmu-miR-21a-5p | MIMAT0000530 | Cadm2 | 239857 | HITS-CLIP | 19536157 | Unspecified |
| mmu-miR-21a-5p | MIMAT0000530 | Cadps | 27062 | Microarrays | 37986224\|25263593 | Unspecified |
| mmu-miR-21a-5p | MIMAT0000530 | Cage1 | 71213 | HITS-CLIP | 23142080 | Unspecified |
| mmu-miR-21a-5p | MIMAT0000530 | Calcb | 116903 | Microarrays | 37986224\|25263593 | Unspecified |

| | | | | | | |
|---|---|---|---|---|---|---|
| mmu-miR-21a-5p | MIMAT0000530 | Calcrl | 54598 | HITS-CLIP | 37986224\|23142080\|29167373 | Unspecified |
| mmu-miR-21a-5p | MIMAT0000530 | Cald1 | 109624 | Microarrays//HITS-CLIP | 37986224\|25263593\|23597149\|28735 | Unspecified |
| mmu-miR-21a-5p | MIMAT0000530 | Calhm2 | 72691 | Microarrays | 37986224 | Unspecified |
| mmu-miR-21a-5p | MIMAT0000530 | Calm1 | 12313 | HITS-CLIP | 37986224\|19536157\|23142080\|29167 | Unspecified |
| mmu-miR-21a-5p | MIMAT0000530 | Calm2 | 12314 | HITS-CLIP | 23142080 | Unspecified |
| mmu-miR-21a-5p | MIMAT0000530 | Calml4 | 75600 | Microarrays | 37986224 | Unspecified |
| mmu-miR-21a-5p | MIMAT0000530 | Calr | 12317 | HITS-CLIP | 23597149\|23142080\|29167373 | Unspecified |
| mmu-miR-21a-5p | MIMAT0000530 | Camk1d | 227541 | qCLASH//HITS-CLIP | 31393953\|23597149 | Unspecified |
| mmu-miR-21a-5p | MIMAT0000530 | Camk2d | 108058 | HITS-CLIP | 37986224\|23142080\|29167373 | Unspecified |
| mmu-miR-21a-5p | MIMAT0000530 | Camk2n1 | 66259 | HITS-CLIP | 23597149 | Unspecified |
| mmu-miR-21a-5p | MIMAT0000530 | Camk2n2 | 73047 | Microarrays | 37986224\|25263593 | Unspecified |
| mmu-miR-21a-5p | MIMAT0000530 | Camk4 | 12326 | HITS-CLIP | 37986224 | Unspecified |
| mmu-miR-21a-5p | MIMAT0000530 | Camkk2 | 207565 | HITS-CLIP | 19536157\|23142080 | Unspecified |
| mmu-miR-21a-5p | MIMAT0000530 | Camkv | 235604 | HITS-CLIP | 37986224\|19536157 | Unspecified |
| mmu-miR-21a-5p | MIMAT0000530 | Camsap2 | 67886 | HITS-CLIP | 19536157\|23142080 | Unspecified |
| mmu-miR-21a-5p | MIMAT0000530 | Cand1 | 71902 | HITS-CLIP | 23142080 | Unspecified |
| mmu-miR-21a-5p | MIMAT0000530 | Cand2 | 67088 | HITS-CLIP | 23142080 | Unspecified |
| mmu-miR-21a-5p | MIMAT0000530 | Cant1 | 76025 | HITS-CLIP | 23597149\|23142080 | Unspecified |
| mmu-miR-21a-5p | MIMAT0000530 | Canx | 12330 | HITS-CLIP | 37986224\|23142080\|19536157\|23597 | Unspecified |
| mmu-miR-21a-5p | MIMAT0000530 | Cap1 | 12331 | Microarrays//HITS-CLIP | 37986224\|23142080 | Unspecified |
| mmu-miR-21a-5p | MIMAT0000530 | Capg | 12332 | HITS-CLIP | 23142080\|29167373 | Unspecified |
| mmu-miR-21a-5p | MIMAT0000530 | Capn1 | 12333 | HITS-CLIP | 23142080 | Unspecified |
| mmu-miR-21a-5p | MIMAT0000530 | Capn10 | 23830 | HITS-CLIP | 23142080 | Unspecified |
| mmu-miR-21a-5p | MIMAT0000530 | Capn2 | 12334 | HITS-CLIP | 23142080 | Unspecified |
| mmu-miR-21a-5p | MIMAT0000530 | Capn3 | 12335 | Microarrays | 37986224\|25263593 | Unspecified |
| mmu-miR-21a-5p | MIMAT0000530 | Capn7 | 12339 | HITS-CLIP | 23142080\|29167373 | Unspecified |
| mmu-miR-21a-5p | MIMAT0000530 | Caprin1 | 53872 | HITS-CLIP | 37986224\|23142080\|23597149\|29167 | Unspecified |
| mmu-miR-21a-5p | MIMAT0000530 | Capsl | 75568 | HITS-CLIP | 23142080 | Unspecified |
| mmu-miR-21a-5p | MIMAT0000530 | Capza2 | 12343 | HITS-CLIP | 23142080 | Unspecified |
| mmu-miR-21a-5p | MIMAT0000530 | Capzb | 12345 | HITS-CLIP | 23142080\|29167373 | Unspecified |
| mmu-miR-21a-5p | MIMAT0000530 | Car14 | 23831 | HITS-CLIP | 23597149 | Unspecified |
| mmu-miR-21a-5p | MIMAT0000530 | Car3 | 12350 | HITS-CLIP | 23597149 | Unspecified |
| mmu-miR-21a-5p | MIMAT0000530 | Card11 | 108723 | HITS-CLIP | 31433990\|23142080 | Unspecified |
| mmu-miR-21a-5p | MIMAT0000530 | Card6 | 239319 | Microarrays | 37986224 | Unspecified |
| mmu-miR-21a-5p | MIMAT0000530 | Carhsp1 | 52502 | HITS-CLIP | 23142080 | Unspecified |
| mmu-miR-21a-5p | MIMAT0000530 | Carmil2 | 146206 | HITS-CLIP | 23142080 | Unspecified |
| mmu-miR-21a-5p | MIMAT0000530 | Casc4 | 319996 | HITS-CLIP | 23142080 | Unspecified |
| mmu-miR-21a-5p | MIMAT0000530 | Casp2 | 12366 | Microarrays//HITS-CLIP | 37986224\|23142080 | Unspecified |
| mmu-miR-21a-5p | MIMAT0000530 | Casp3 | 12367 | Microarrays//HITS-CLIP | 37986224\|23142080\|19536157 | Unspecified |

| | | | | | | |
|---|---|---|---|---|---|---|
| mmu-miR-21a-5p | MIMAT0000530 | Casp6 | 12368 | Microarrays//HITS-CLIP | 37986224\|23142080 | Unspecified |
| mmu-miR-21a-5p | MIMAT0000530 | Casp8 | 12370 | HITS-CLIP | 23142080 | Unspecified |
| mmu-miR-21a-5p | MIMAT0000530 | Casp8ap2 | 26885 | HITS-CLIP | 23142080 | Unspecified |
| mmu-miR-21a-5p | MIMAT0000530 | Cass4 | 320664 | Microarrays//HITS-CLIP | 37986224\|23142080 | Unspecified |
| mmu-miR-21a-5p | MIMAT0000530 | Cast | 12380 | HITS-CLIP | 23597149 | Unspecified |
| mmu-miR-21a-5p | MIMAT0000530 | Casz1 | 69743 | HITS-CLIP | 23142080 | Unspecified |
| mmu-miR-21a-5p | MIMAT0000530 | Cav2 | 12390 | HITS-CLIP | 29167373 | Unspecified |
| mmu-miR-21a-5p | MIMAT0000530 | Cbfa2t2 | 12396 | HITS-CLIP | 29167373 | Unspecified |
| mmu-miR-21a-5p | MIMAT0000530 | Cbfb | 12400 | HITS-CLIP | 23142080 | Unspecified |
| mmu-miR-21a-5p | MIMAT0000530 | Cbl | 12402 | HITS-CLIP | 23142080\|29167373 | Unspecified |
| mmu-miR-21a-5p | MIMAT0000530 | Cblb | 208650 | HITS-CLIP | 23142080\|29167373 | Unspecified |
| mmu-miR-21a-5p | MIMAT0000530 | Cbln1 | 12404 | HITS-CLIP | 19536157 | Unspecified |
| mmu-miR-21a-5p | MIMAT0000530 | Cbx1 | 12412 | HITS-CLIP | 23597149\|23142080 | Unspecified |
| mmu-miR-21a-5p | MIMAT0000530 | Cbx4 | 12418 | HITS-CLIP | 23142080 | Unspecified |
| mmu-miR-21a-5p | MIMAT0000530 | Cbx5 | 12419 | HITS-CLIP | 23142080 | Unspecified |
| mmu-miR-21a-5p | MIMAT0000530 | Cbx7 | 52609 | HITS-CLIP | 37986224\|23142080 | Unspecified |
| mmu-miR-21a-5p | MIMAT0000530 | Ccar1 | 67500 | HITS-CLIP | 23597149 | Unspecified |
| mmu-miR-21a-5p | MIMAT0000530 | Ccdc107 | 622404 | Microarrays | 37986224\|25263593 | Unspecified |
| mmu-miR-21a-5p | MIMAT0000530 | Ccdc117 | 104479 | HITS-CLIP | 23597149\|23142080 | Unspecified |
| mmu-miR-21a-5p | MIMAT0000530 | Ccdc136 | 232664 | HITS-CLIP | 19536157 | Unspecified |
| mmu-miR-21a-5p | MIMAT0000530 | Ccdc137 | 67291 | HITS-CLIP | 37986224 | Unspecified |
| mmu-miR-21a-5p | MIMAT0000530 | Ccdc14 | 239839 | HITS-CLIP | 23142080 | Unspecified |
| mmu-miR-21a-5p | MIMAT0000530 | Ccdc141 | 545428 | HITS-CLIP | 23142080 | Unspecified |
| mmu-miR-21a-5p | MIMAT0000530 | Ccdc163 | 68394 | Microarrays | 37986224 | Unspecified |
| mmu-miR-21a-5p | MIMAT0000530 | Ccdc166 | 223648 | HITS-CLIP | 23142080 | Unspecified |
| mmu-miR-21a-5p | MIMAT0000530 | Ccdc191 | 212153 | HITS-CLIP | 23142080 | Unspecified |
| mmu-miR-21a-5p | MIMAT0000530 | Ccdc25 | 67179 | Microarrays//HITS-CLIP | 37986224\|23142080\|29167373 | Unspecified |
| mmu-miR-21a-5p | MIMAT0000530 | Ccdc43 | 52715 | HITS-CLIP | 23142080 | Unspecified |
| mmu-miR-21a-5p | MIMAT0000530 | Ccdc47 | 67163 | HITS-CLIP | 37986224\|23597149\|23142080 | Unspecified |
| mmu-miR-21a-5p | MIMAT0000530 | Ccdc50 | 67501 | HITS-CLIP | 23142080\|29167373 | Unspecified |
| mmu-miR-21a-5p | MIMAT0000530 | Ccdc59 | 52713 | HITS-CLIP | 23142080 | Unspecified |
| mmu-miR-21a-5p | MIMAT0000530 | Ccdc6 | 76551 | HITS-CLIP | 23142080\|31433990 | Unspecified |
| mmu-miR-21a-5p | MIMAT0000530 | Ccdc71 | 72454 | HITS-CLIP | 29167373\|23142080 | Unspecified |
| mmu-miR-21a-5p | MIMAT0000530 | Ccdc71l | 72123 | HITS-CLIP//qCLASH | 37986224\|23142080\|31393953\|29167 | Unspecified |
| mmu-miR-21a-5p | MIMAT0000530 | Ccdc88a | 108686 | qCLASH//HITS-CLIP | 31393953\|29167373 | Unspecified |
| mmu-miR-21a-5p | MIMAT0000530 | Ccdc88b | 78317 | HITS-CLIP | 23142080\|29167373 | Unspecified |
| mmu-miR-21a-5p | MIMAT0000530 | Ccdc88c | 68339 | HITS-CLIP | 23142080 | Unspecified |
| mmu-miR-21a-5p | MIMAT0000530 | Ccdc90b | 66365 | HITS-CLIP | 23142080 | Unspecified |
| mmu-miR-21a-5p | MIMAT0000530 | Ccl1 | 20290 | HITS-CLIP | 23142080 | Unspecified |

| | | | | | | |
|---|---|---|---|---|---|---|
| mmu-miR-21a-5p | MIMAT0000530 | Ccl12 | 20293 | Microarrays | 37986224\|25263593 | Unspecified |
| mmu-miR-21a-5p | MIMAT0000530 | Ccl17 | 20295 | Microarrays | 37986224\|25263593 | Unspecified |
| mmu-miR-21a-5p | MIMAT0000530 | Ccl2 | 20296 | Microarrays//HITS-CLIP | 37986224\|25263593\|29167373 | Unspecified |
| mmu-miR-21a-5p | MIMAT0000530 | Ccl20 | 6364 | HITS-CLIP | 23142080 | Unspecified |
| mmu-miR-21a-5p | MIMAT0000530 | Ccl22 | 20299 | HITS-CLIP | 23142080 | Unspecified |
| mmu-miR-21a-5p | MIMAT0000530 | Ccl3 | 20302 | Microarrays//HITS-CLIP | 37986224\|25263593\|23142080\|29167 | Unspecified |
| mmu-miR-21a-5p | MIMAT0000530 | Ccl4 | 20303 | Microarrays | 37986224\|25263593 | Unspecified |
| mmu-miR-21a-5p | MIMAT0000530 | Ccl6 | 20305 | Microarrays | 37986224 | Unspecified |
| mmu-miR-21a-5p | MIMAT0000530 | Ccl7 | 20306 | Microarrays | 37986224\|25263593 | Unspecified |
| mmu-miR-21a-5p | MIMAT0000530 | Ccl9 | 20308 | Microarrays | 37986224 | Unspecified |
| mmu-miR-21a-5p | MIMAT0000530 | Ccn1 | 3491 | Microarrays | 25263593 | Unspecified |
| mmu-miR-21a-5p | MIMAT0000530 | Ccn2 | 1490 | Microarrays//HITS-CLIP | 25263593\|23597149 | Unspecified |
| mmu-miR-21a-5p | MIMAT0000530 | Ccna2 | 12428 | Microarrays | 37986224\|25263593 | Unspecified |
| mmu-miR-21a-5p | MIMAT0000530 | Ccnb2 | 12442 | Microarrays//HITS-CLIP | 37986224\|25263593\|23142080 | Unspecified |
| mmu-miR-21a-5p | MIMAT0000530 | Ccnd1 | 12443 | Microarrays | 37986224\|25263593 | Unspecified |
| mmu-miR-21a-5p | MIMAT0000530 | Ccnd2 | 12444 | HITS-CLIP, Microarrays//Microarrays//HITS | 37986224\|25263593\|23142080 | Unspecified |
| mmu-miR-21a-5p | MIMAT0000530 | Ccnd3 | 12445 | HITS-CLIP | 23142080 | Unspecified |
| mmu-miR-21a-5p | MIMAT0000530 | Ccndbp1 | 17151 | HITS-CLIP | 23142080 | Unspecified |
| mmu-miR-21a-5p | MIMAT0000530 | Ccne1 | 12447 | Microarrays//HITS-CLIP | 37986224\|25263593\|23142080 | Unspecified |
| mmu-miR-21a-5p | MIMAT0000530 | Ccne2 | 12448 | HITS-CLIP, Microarrays//HITS-CLIP//Microa | 37986224\|23142080\|25263593\|31433 | Unspecified |
| mmu-miR-21a-5p | MIMAT0000530 | Ccnf | 12449 | Microarrays//HITS-CLIP//qCLASH | 37986224\|25263593\|23597149\|31393 | Unspecified |
| mmu-miR-21a-5p | MIMAT0000530 | Ccng2 | 12452 | HITS-CLIP | 23597149\|19536157\|23142080 | Unspecified |
| mmu-miR-21a-5p | MIMAT0000530 | Ccni | 12453 | HITS-CLIP | 37986224\|19536157\|23142080 | Unspecified |
| mmu-miR-21a-5p | MIMAT0000530 | Ccnj | 240665 | HITS-CLIP | 23142080\|29167373 | Unspecified |
| mmu-miR-21a-5p | MIMAT0000530 | Ccnl1 | 56706 | HITS-CLIP | 29167373\|23142080 | Unspecified |
| mmu-miR-21a-5p | MIMAT0000530 | Ccnl2 | 56036 | HITS-CLIP | 23597149\|29167373\|23142080 | Unspecified |
| mmu-miR-21a-5p | MIMAT0000530 | Ccnt2 | 72949 | HITS-CLIP | 23597149\|29167373\|23142080 | Unspecified |
| mmu-miR-21a-5p | MIMAT0000530 | Ccpg1 | 72278 | Microarrays//HITS-CLIP | 37986224\|23142080 | Unspecified |
| mmu-miR-21a-5p | MIMAT0000530 | Ccr1 | 12768 | Microarrays//HITS-CLIP | 37986224\|25263593\|29167373 | Unspecified |
| mmu-miR-21a-5p | MIMAT0000530 | Ccr2 | 12772 | HITS-CLIP | 29167373 | Unspecified |
| mmu-miR-21a-5p | MIMAT0000530 | Ccr5 | 12774 | HITS-CLIP, Microarrays//Microarrays//HITS | 37986224\|25263593\|23597149\|23142 | Unspecified |
| mmu-miR-21a-5p | MIMAT0000530 | Ccr7 | 12775 | HITS-CLIP | 37986224\|23142080 | Unspecified |
| mmu-miR-21a-5p | MIMAT0000530 | Ccs | 12460 | HITS-CLIP | 23597149\|23142080 | Unspecified |
| mmu-miR-21a-5p | MIMAT0000530 | Ccser2 | 72972 | HITS-CLIP | 23142080 | Unspecified |
| mmu-miR-21a-5p | MIMAT0000530 | Cct2 | 12461 | qCLASH//HITS-CLIP | 31393953\|23142080 | Unspecified |
| mmu-miR-21a-5p | MIMAT0000530 | Cct3 | 12462 | HITS-CLIP | 23597149 | Unspecified |
| mmu-miR-21a-5p | MIMAT0000530 | Cct4 | 12464 | HITS-CLIP//qCLASH | 23142080\|31393953\|23597149\|29167 | Unspecified |
| mmu-miR-21a-5p | MIMAT0000530 | Cct5 | 12465 | HITS-CLIP//qCLASH | 23142080\|31393953 | Unspecified |
| mmu-miR-21a-5p | MIMAT0000530 | Cct6a | 12466 | HITS-CLIP | 31433990 | Unspecified |

| | | | | | | |
|---|---|---|---|---|---|---|
| mmu-miR-21a-5p | MIMAT0000530 | Cct7 | 12468 | HITS-CLIP | 19536157\|23142080 | Unspecified |
| mmu-miR-21a-5p | MIMAT0000530 | Cct8 | 12469 | HITS-CLIP | 26203562 | Unspecified |
| mmu-miR-21a-5p | MIMAT0000530 | Cd109 | 235505 | Microarrays | 37986224\|25263593 | Unspecified |
| mmu-miR-21a-5p | MIMAT0000530 | Cd151 | 12476 | HITS-CLIP | 23142080 | Unspecified |
| mmu-miR-21a-5p | MIMAT0000530 | Cd163 | 93671 | HITS-CLIP | 23597149 | Unspecified |
| mmu-miR-21a-5p | MIMAT0000530 | Cd164 | 53599 | HITS-CLIP | 37986224\|23597149\|23142080 | Unspecified |
| mmu-miR-21a-5p | MIMAT0000530 | Cd180 | 17079 | Microarrays//HITS-CLIP | 37986224\|25263593\|29167373 | Unspecified |
| mmu-miR-21a-5p | MIMAT0000530 | Cd200 | 17470 | Microarrays | 37986224\|25263593 | Unspecified |
| mmu-miR-21a-5p | MIMAT0000530 | Cd209g | 70192 | Microarrays | 37986224\|25263593 | Unspecified |
| mmu-miR-21a-5p | MIMAT0000530 | Cd22 | 12483 | Microarrays | 37986224\|25263593 | Unspecified |
| mmu-miR-21a-5p | MIMAT0000530 | Cd226 | 225825 | HITS-CLIP | 23142080 | Unspecified |
| mmu-miR-21a-5p | MIMAT0000530 | Cd24a | 12484 | Microarrays | 37986224\|25263593 | Unspecified |
| mmu-miR-21a-5p | MIMAT0000530 | Cd28 | 12487 | Microarrays//HITS-CLIP | 37986224\|23142080 | Unspecified |
| mmu-miR-21a-5p | MIMAT0000530 | Cd2ap | 12488 | HITS-CLIP | 23142080 | Unspecified |
| mmu-miR-21a-5p | MIMAT0000530 | Cd300ld | 217305 | Microarrays//HITS-CLIP | 37986224\|29167373 | Unspecified |
| mmu-miR-21a-5p | MIMAT0000530 | Cd300lf | 246746 | Microarrays | 37986224\|25263593 | Unspecified |
| mmu-miR-21a-5p | MIMAT0000530 | Cd302 | 66205 | HITS-CLIP | 23597149 | Unspecified |
| mmu-miR-21a-5p | MIMAT0000530 | Cd34 | 12490 | Microarrays | 37986224\|25263593 | Unspecified |
| mmu-miR-21a-5p | MIMAT0000530 | Cd37 | 12493 | HITS-CLIP | 23142080\|29167373 | Unspecified |
| mmu-miR-21a-5p | MIMAT0000530 | Cd38 | 12494 | HITS-CLIP | 23597149 | Unspecified |
| mmu-miR-21a-5p | MIMAT0000530 | Cd44 | 12505 | HITS-CLIP | 37986224\|23142080 | Unspecified |
| mmu-miR-21a-5p | MIMAT0000530 | Cd47 | 16423 | HITS-CLIP | 37986224\|23142080\|29167373 | Unspecified |
| mmu-miR-21a-5p | MIMAT0000530 | Cd48 | 12506 | HITS-CLIP | 29167373 | Unspecified |
| mmu-miR-21a-5p | MIMAT0000530 | Cd5 | 12507 | HITS-CLIP | 23142080 | Unspecified |
| mmu-miR-21a-5p | MIMAT0000530 | Cd53 | 12508 | HITS-CLIP | 29167373\|23142080 | Unspecified |
| mmu-miR-21a-5p | MIMAT0000530 | Cd59a | 12509 | Microarrays | 37986224\|25263593 | Unspecified |
| mmu-miR-21a-5p | MIMAT0000530 | Cd5l | 11801 | HITS-CLIP | 23597149 | Unspecified |
| mmu-miR-21a-5p | MIMAT0000530 | Cd6 | 12511 | HITS-CLIP | 23142080 | Unspecified |
| mmu-miR-21a-5p | MIMAT0000530 | Cd69 | 12515 | HITS-CLIP | 37986224\|31433990\|23142080 | Unspecified |
| mmu-miR-21a-5p | MIMAT0000530 | Cd72 | 12517 | Microarrays | 37986224\|25263593 | Unspecified |
| mmu-miR-21a-5p | MIMAT0000530 | Cd74 | 16149 | Microarrays//HITS-CLIP | 37986224\|23597149\|23142080\|29167 | Unspecified |
| mmu-miR-21a-5p | MIMAT0000530 | Cd81 | 12520 | HITS-CLIP | 23597149\|19536157\|23142080 | Unspecified |
| mmu-miR-21a-5p | MIMAT0000530 | Cd83 | 12522 | HITS-CLIP, Microarrays//qCLASH//HITS-CLI | 37986224\|31393953\|23142080 | Unspecified |
| mmu-miR-21a-5p | MIMAT0000530 | Cd84 | 12523 | HITS-CLIP | 29167373 | Unspecified |
| mmu-miR-21a-5p | MIMAT0000530 | Cd86 | 12524 | Microarrays | 37986224 | Unspecified |
| mmu-miR-21a-5p | MIMAT0000530 | Cd9 | 12527 | HITS-CLIP | 29167373\|23142080 | Unspecified |
| mmu-miR-21a-5p | MIMAT0000530 | Cd93 | 17064 | HITS-CLIP | 37986224\|29167373 | Unspecified |
| mmu-miR-21a-5p | MIMAT0000530 | Cd99l2 | 171486 | HITS-CLIP | 19536157\|23597149 | Unspecified |
| mmu-miR-21a-5p | MIMAT0000530 | Cda | 72269 | Microarrays | 37986224\|25263593 | Unspecified |

| | | | | | | |
|---|---|---|---|---|---|---|
| mmu-miR-21a-5p | MIMAT0000530 | Cdan1 | 68968 | HITS-CLIP | 23597149\|19536157\|23142080 | Unspecified |
| mmu-miR-21a-5p | MIMAT0000530 | Cdc123 | 98828 | HITS-CLIP | 23142080 | Unspecified |
| mmu-miR-21a-5p | MIMAT0000530 | Cdc14a | 229776 | HITS-CLIP | 23142080 | Unspecified |
| mmu-miR-21a-5p | MIMAT0000530 | Cdc14b | 218294 | HITS-CLIP | 23142080\|29167373 | Unspecified |
| mmu-miR-21a-5p | MIMAT0000530 | Cdc16 | 69957 | qCLASH//HITS-CLIP | 31393953\|23142080 | Unspecified |
| mmu-miR-21a-5p | MIMAT0000530 | Cdc20 | 107995 | Microarrays//HITS-CLIP | 37986224\|25263593\|23597149\|2314 | Unspecified |
| mmu-miR-21a-5p | MIMAT0000530 | Cdc23 | 52563 | HITS-CLIP | 23142080 | Unspecified |
| mmu-miR-21a-5p | MIMAT0000530 | Cdc25a | 12530 | Microarrays//qCLASH//HITS-CLIP | 37986224\|25263593\|31393953\|2314 | Unspecified |
| mmu-miR-21a-5p | MIMAT0000530 | Cdc25c | 12532 | Microarrays//HITS-CLIP | 37986224\|25263593\|23142080 | Unspecified |
| mmu-miR-21a-5p | MIMAT0000530 | Cdc27 | 217232 | HITS-CLIP | 37986224\|23142080 | Unspecified |
| mmu-miR-21a-5p | MIMAT0000530 | Cdc34 | 216150 | HITS-CLIP | 23142080 | Unspecified |
| mmu-miR-21a-5p | MIMAT0000530 | Cdc37l1 | 67072 | HITS-CLIP | 19536157\|29167373 | Unspecified |
| mmu-miR-21a-5p | MIMAT0000530 | Cdc42 | 12540 | HITS-CLIP | 23597149\|23142080 | Unspecified |
| mmu-miR-21a-5p | MIMAT0000530 | Cdc42bpa | 226751 | HITS-CLIP | 19536157 | Unspecified |
| mmu-miR-21a-5p | MIMAT0000530 | Cdc45 | 12544 | Microarrays | 37986224\|25263593 | Unspecified |
| mmu-miR-21a-5p | MIMAT0000530 | Cdc7 | 12545 | Microarrays//HITS-CLIP | 37986224\|25263593\|19536157\|2314 | Unspecified |
| mmu-miR-21a-5p | MIMAT0000530 | Cdca2 | 108912 | Microarrays | 37986224\|25263593 | Unspecified |
| mmu-miR-21a-5p | MIMAT0000530 | Cdca4 | 71963 | HITS-CLIP | 23142080 | Unspecified |
| mmu-miR-21a-5p | MIMAT0000530 | Cdca5 | 67849 | Microarrays | 37986224\|25263593 | Unspecified |
| mmu-miR-21a-5p | MIMAT0000530 | Cdca7 | 66953 | HITS-CLIP | 31433990 | Unspecified |
| mmu-miR-21a-5p | MIMAT0000530 | Cdca7l | 217946 | Microarrays | 37986224\|25263593 | Unspecified |
| mmu-miR-21a-5p | MIMAT0000530 | Cdca8 | 52276 | Microarrays | 37986224\|25263593 | Unspecified |
| mmu-miR-21a-5p | MIMAT0000530 | Cdh1 | 12550 | HITS-CLIP | 23597149 | Unspecified |
| mmu-miR-21a-5p | MIMAT0000530 | Cdh11 | 12552 | Microarrays//HITS-CLIP | 37986224\|25263593\|19536157 | Unspecified |
| mmu-miR-21a-5p | MIMAT0000530 | Cdh2 | 12558 | Microarrays//HITS-CLIP | 37986224\|23597149\|25263593 | Unspecified |
| mmu-miR-21a-5p | MIMAT0000530 | Cdh9 | 12565 | HITS-CLIP | 37986224 | Unspecified |
| mmu-miR-21a-5p | MIMAT0000530 | Cdip1 | 66626 | HITS-CLIP | 37986224\|23597149\|23142080 | Unspecified |
| mmu-miR-21a-5p | MIMAT0000530 | Cdk1 | 12534 | Microarrays | 37986224\|25263593 | Unspecified |
| mmu-miR-21a-5p | MIMAT0000530 | Cdk11b | 12537 | HITS-CLIP | 23142080 | Unspecified |
| mmu-miR-21a-5p | MIMAT0000530 | Cdk13 | 69562 | HITS-CLIP | 23142080\|23597149 | Unspecified |
| mmu-miR-21a-5p | MIMAT0000530 | Cdk14 | 18647 | Microarrays//HITS-CLIP | 37986224\|25263593\|29167373 | Unspecified |
| mmu-miR-21a-5p | MIMAT0000530 | Cdk17 | 237459 | HITS-CLIP | 23142080 | Unspecified |
| mmu-miR-21a-5p | MIMAT0000530 | Cdk18 | 18557 | Microarrays | 37986224\|25263593 | Unspecified |
| mmu-miR-21a-5p | MIMAT0000530 | Cdk19 | 78334 | HITS-CLIP | 23142080\|29167373 | Unspecified |
| mmu-miR-21a-5p | MIMAT0000530 | Cdk2 | 12566 | HITS-CLIP | 23142080 | Unspecified |
| mmu-miR-21a-5p | MIMAT0000530 | Cdk5rap1 | 66971 | Microarrays | 37986224\|25263593 | Unspecified |
| mmu-miR-21a-5p | MIMAT0000530 | Cdk6 | 12571 | Microarrays//HITS-CLIP//qCLASH//Lucifera | 37986224\|23142080\|25263593\|2359 | Unspecified |
| mmu-miR-21a-5p | MIMAT0000530 | Cdk9 | 107951 | HITS-CLIP | 23597149\|23142080 | Unspecified |
| mmu-miR-21a-5p | MIMAT0000530 | Cdkal1 | 68916 | HITS-CLIP | 23142080 | Unspecified |

| | | | | | | |
|---|---|---|---|---|---|---|
| mmu-miR-21a-5p | MIMAT0000530 | Cdkl5 | 382253 | HITS-CLIP | 23142080 | Unspecified |
| mmu-miR-21a-5p | MIMAT0000530 | Cdkn1b | 12576 | HITS-CLIP, Microarrays//HITS-CLIP | 37986224\|23142080\|29167373 | Unspecified |
| mmu-miR-21a-5p | MIMAT0000530 | Cdkn1c | 12577 | Microarrays | 37986224\|25263593 | Unspecified |
| mmu-miR-21a-5p | MIMAT0000530 | Cdkn2a | 12578 | Microarrays | 37986224\|25263593 | Unspecified |
| mmu-miR-21a-5p | MIMAT0000530 | Cdkn2aip | 70925 | HITS-CLIP | 23142080 | Unspecified |
| mmu-miR-21a-5p | MIMAT0000530 | Cdkn2b | 12579 | Microarrays | 37986224\|25263593 | Unspecified |
| mmu-miR-21a-5p | MIMAT0000530 | Cdkn3 | 72391 | Microarrays | 37986224\|25263593 | Unspecified |
| mmu-miR-21a-5p | MIMAT0000530 | Cdon | 57810 | HITS-CLIP | 23142080 | Unspecified |
| mmu-miR-21a-5p | MIMAT0000530 | Cdr2 | 12585 | HITS-CLIP | 23597149\|23142080 | Unspecified |
| mmu-miR-21a-5p | MIMAT0000530 | Cds2 | 110911 | Microarrays//HITS-CLIP | 37986224\|25263593\|23597149\|23142 | Unspecified |
| mmu-miR-21a-5p | MIMAT0000530 | Cdsn | 386463 | HITS-CLIP | 26203562 | Unspecified |
| mmu-miR-21a-5p | MIMAT0000530 | Cdyl | 12593 | HITS-CLIP | 37986224\|23142080 | Unspecified |
| mmu-miR-21a-5p | MIMAT0000530 | Cdyl2 | 75796 | HITS-CLIP | 23142080 | Unspecified |
| mmu-miR-21a-5p | MIMAT0000530 | Ceacam12 | 67315 | Microarrays | 37986224\|25263593 | Unspecified |
| mmu-miR-21a-5p | MIMAT0000530 | Ceacam20 | 71601 | Microarrays | 37986224 | Unspecified |
| mmu-miR-21a-5p | MIMAT0000530 | Cebpb | 12608 | HITS-CLIP | 23597149\|23142080 | Unspecified |
| mmu-miR-21a-5p | MIMAT0000530 | Cebpd | 12609 | HITS-CLIP | 23597149 | Unspecified |
| mmu-miR-21a-5p | MIMAT0000530 | Cebpg | 12611 | HITS-CLIP | 23142080 | Unspecified |
| mmu-miR-21a-5p | MIMAT0000530 | Cebpz | 12607 | HITS-CLIP | 23597149\|23142080 | Unspecified |
| mmu-miR-21a-5p | MIMAT0000530 | Cecr2 | 330409 | HITS-CLIP | 23597149 | Unspecified |
| mmu-miR-21a-5p | MIMAT0000530 | Celf1 | 13046 | HITS-CLIP | 37986224\|23597149\|23142080\|29167 | Unspecified |
| mmu-miR-21a-5p | MIMAT0000530 | Celf2 | 14007 | HITS-CLIP | 23142080\|19536157\|31433990\|29167 | Unspecified |
| mmu-miR-21a-5p | MIMAT0000530 | Cenpa | 12615 | Microarrays//HITS-CLIP | 37986224\|23142080\|23597149 | Unspecified |
| mmu-miR-21a-5p | MIMAT0000530 | Cenpc1 | 12617 | HITS-CLIP | 23142080 | Unspecified |
| mmu-miR-21a-5p | MIMAT0000530 | Cenpe | 229841 | Microarrays//HITS-CLIP | 37986224\|25263593\|23142080 | Unspecified |
| mmu-miR-21a-5p | MIMAT0000530 | Cenpf | 108000 | Microarrays//HITS-CLIP//qCLASH | 37986224\|23142080\|25263593\|31393 | Unspecified |
| mmu-miR-21a-5p | MIMAT0000530 | Cenph | 26886 | Microarrays | 37986224\|25263593 | Unspecified |
| mmu-miR-21a-5p | MIMAT0000530 | Cenpi | 102920 | Microarrays | 37986224\|25263593 | Unspecified |
| mmu-miR-21a-5p | MIMAT0000530 | Cenpk | 60411 | Microarrays | 37986224\|25263593 | Unspecified |
| mmu-miR-21a-5p | MIMAT0000530 | Cenpn | 72155 | Microarrays | 37986224\|25263593 | Unspecified |
| mmu-miR-21a-5p | MIMAT0000530 | Cenpo | 52504 | HITS-CLIP | 23142080 | Unspecified |
| mmu-miR-21a-5p | MIMAT0000530 | Cenpp | 66336 | Microarrays | 37986224\|25263593 | Unspecified |
| mmu-miR-21a-5p | MIMAT0000530 | Cenpq | 83815 | Microarrays | 37986224\|25263593 | Unspecified |
| mmu-miR-21a-5p | MIMAT0000530 | Cenps | 378708 | Microarrays | 25263593 | Unspecified |
| mmu-miR-21a-5p | MIMAT0000530 | Cenpw | 66311 | Microarrays | 37986224\|25263593 | Unspecified |
| mmu-miR-21a-5p | MIMAT0000530 | Cep104 | 230967 | HITS-CLIP | 23597149\|23142080 | Unspecified |
| mmu-miR-21a-5p | MIMAT0000530 | Cep120 | 225523 | HITS-CLIP | 37986224\|23142080 | Unspecified |
| mmu-miR-21a-5p | MIMAT0000530 | Cep162 | 382090 | HITS-CLIP | 23142080 | Unspecified |
| mmu-miR-21a-5p | MIMAT0000530 | Cep170 | 545389 | HITS-CLIP | 37986224\|23142080\|19536157\|29167 | Unspecified |

| | | | | | | |
|---|---|---|---|---|---|---|
| mmu-miR-21a-5p | MIMAT0000530 | Cep170b | 217882 | Microarrays//HITS-CLIP | 37986224\|25263593\|23597149\|23142 | Unspecified |
| mmu-miR-21a-5p | MIMAT0000530 | Cep19 | 66994 | HITS-CLIP | 23142080 | Unspecified |
| mmu-miR-21a-5p | MIMAT0000530 | Cep192 | 70799 | HITS-CLIP | 23142080\|29167373 | Unspecified |
| mmu-miR-21a-5p | MIMAT0000530 | Cep290 | 216274 | HITS-CLIP | 23142080 | Unspecified |
| mmu-miR-21a-5p | MIMAT0000530 | Cep295 | 85459 | HITS-CLIP | 29167373 | Unspecified |
| mmu-miR-21a-5p | MIMAT0000530 | Cep350 | 74081 | HITS-CLIP | 23142080\|19536157\|29167373 | Unspecified |
| mmu-miR-21a-5p | MIMAT0000530 | Cep44 | 382010 | Microarrays//HITS-CLIP | 37986224\|19536157 | Unspecified |
| mmu-miR-21a-5p | MIMAT0000530 | Cep55 | 74107 | Microarrays//HITS-CLIP | 37986224\|23142080\|25263593 | Unspecified |
| mmu-miR-21a-5p | MIMAT0000530 | Cep57 | 74360 | HITS-CLIP | 23142080 | Unspecified |
| mmu-miR-21a-5p | MIMAT0000530 | Cep68 | 216543 | HITS-CLIP | 37986224\|23142080 | Unspecified |
| mmu-miR-21a-5p | MIMAT0000530 | Cep70 | 68121 | HITS-CLIP | 23597149 | Unspecified |
| mmu-miR-21a-5p | MIMAT0000530 | Cep83 | 77048 | HITS-CLIP | 23142080 | Unspecified |
| mmu-miR-21a-5p | MIMAT0000530 | Cep85 | 70012 | HITS-CLIP | 29167373 | Unspecified |
| mmu-miR-21a-5p | MIMAT0000530 | Cep85l | 100038725 | HITS-CLIP | 23597149 | Unspecified |
| mmu-miR-21a-5p | MIMAT0000530 | Cep95 | 320162 | HITS-CLIP | 23142080 | Unspecified |
| mmu-miR-21a-5p | MIMAT0000530 | Cep97 | 74201 | HITS-CLIP | 23142080 | Unspecified |
| mmu-miR-21a-5p | MIMAT0000530 | Cept1 | 99712 | HITS-CLIP | 23142080 | Unspecified |
| mmu-miR-21a-5p | MIMAT0000530 | Cers2 | 76893 | HITS-CLIP | 37986224\|23597149\|29167373 | Unspecified |
| mmu-miR-21a-5p | MIMAT0000530 | Cers4 | 67260 | Microarrays | 37986224\|25263593 | Unspecified |
| mmu-miR-21a-5p | MIMAT0000530 | Ces1f | 234564 | HITS-CLIP | 23597149 | Unspecified |
| mmu-miR-21a-5p | MIMAT0000530 | Ces3b | 13909 | HITS-CLIP | 23597149 | Unspecified |
| mmu-miR-21a-5p | MIMAT0000530 | Cetn3 | 12626 | HITS-CLIP | 37986224\|23597149\|29167373\|23142 | Unspecified |
| mmu-miR-21a-5p | MIMAT0000530 | Cfap36 | 112942 | HITS-CLIP | 23142080 | Unspecified |
| mmu-miR-21a-5p | MIMAT0000530 | Cfdp1 | 23837 | HITS-CLIP | 23142080 | Unspecified |
| mmu-miR-21a-5p | MIMAT0000530 | Cfh | 12628 | Microarrays | 37986224 | Unspecified |
| mmu-miR-21a-5p | MIMAT0000530 | Cfi | 12630 | HITS-CLIP | 23597149 | Unspecified |
| mmu-miR-21a-5p | MIMAT0000530 | Cfl1 | 12631 | qCLASH | 31393953 | Unspecified |
| mmu-miR-21a-5p | MIMAT0000530 | Cfl2 | 12632 | HITS-CLIP | 37986224\|23142080 | Unspecified |
| mmu-miR-21a-5p | MIMAT0000530 | Cfp | 18636 | Microarrays | 37986224\|25263593 | Unspecified |
| mmu-miR-21a-5p | MIMAT0000530 | Cgas | 115004 | HITS-CLIP | 29167373 | Unspecified |
| mmu-miR-21a-5p | MIMAT0000530 | Cggbp1 | 106143 | HITS-CLIP | 23142080\|29167373 | Unspecified |
| mmu-miR-21a-5p | MIMAT0000530 | Ch25h | 12642 | Microarrays//HITS-CLIP | 37986224\|25263593\|29167373 | Unspecified |
| mmu-miR-21a-5p | MIMAT0000530 | Chac1 | 69065 | Microarrays | 37986224\|25263593 | Unspecified |
| mmu-miR-21a-5p | MIMAT0000530 | Chaf1a | 27221 | HITS-CLIP | 31433990 | Unspecified |
| mmu-miR-21a-5p | MIMAT0000530 | Chaf1b | 110749 | Microarrays | 37986224\|25263593 | Unspecified |
| mmu-miR-21a-5p | MIMAT0000530 | Chchd1 | 66121 | HITS-CLIP | 23142080 | Unspecified |
| mmu-miR-21a-5p | MIMAT0000530 | Chchd10 | 103172 | Microarrays | 37986224\|25263593 | Unspecified |
| mmu-miR-21a-5p | MIMAT0000530 | Chchd6 | 66098 | Microarrays | 37986224\|25263593 | Unspecified |
| mmu-miR-21a-5p | MIMAT0000530 | Chd1 | 12648 | HITS-CLIP | 37986224\|23142080 | Unspecified |

| | | | | | | |
|---|---|---|---|---|---|---|
| mmu-miR-21a-5p | MIMAT0000530 | Chd2 | 244059 | HITS-CLIP | 23142080\|23597149\|29167373 | Unspecified |
| mmu-miR-21a-5p | MIMAT0000530 | Chd3 | 216848 | HITS-CLIP | 19536157\|23142080 | Unspecified |
| mmu-miR-21a-5p | MIMAT0000530 | Chd4 | 107932 | qCLASH//HITS-CLIP | 31393953\|23142080 | Unspecified |
| mmu-miR-21a-5p | MIMAT0000530 | Chd6 | 71389 | HITS-CLIP | 23597149 | Unspecified |
| mmu-miR-21a-5p | MIMAT0000530 | Chd7 | 320790 | HITS-CLIP | 37986224\|23597149\|19536157\|31433 | Unspecified |
| mmu-miR-21a-5p | MIMAT0000530 | Chd8 | 67772 | HITS-CLIP | 23142080\|19536157\|23597149 | Unspecified |
| mmu-miR-21a-5p | MIMAT0000530 | Chd9 | 109151 | HITS-CLIP | 23142080 | Unspecified |
| mmu-miR-21a-5p | MIMAT0000530 | Chek1 | 12649 | Microarrays//HITS-CLIP | 37986224\|23142080\|25263593 | Unspecified |
| mmu-miR-21a-5p | MIMAT0000530 | Chfr | 231600 | HITS-CLIP | 19536157\|23142080 | Unspecified |
| mmu-miR-21a-5p | MIMAT0000530 | Chic1 | 12212 | HITS-CLIP//qCLASH | 19536157\|31393953\|23142080 | Unspecified |
| mmu-miR-21a-5p | MIMAT0000530 | Chic2 | 74277 | HITS-CLIP | 23142080 | Unspecified |
| mmu-miR-21a-5p | MIMAT0000530 | Chil3 | 12655 | Microarrays | 37986224\|25263593 | Unspecified |
| mmu-miR-21a-5p | MIMAT0000530 | Chkb | 12651 | HITS-CLIP | 23142080 | Unspecified |
| mmu-miR-21a-5p | MIMAT0000530 | Chm | 12662 | HITS-CLIP | 23597149\|23142080 | Unspecified |
| mmu-miR-21a-5p | MIMAT0000530 | Chmp1b | 67078 | HITS-CLIP | 23142080 | Unspecified |
| mmu-miR-21a-5p | MIMAT0000530 | Chmp2a | 68953 | HITS-CLIP | 23597149 | Unspecified |
| mmu-miR-21a-5p | MIMAT0000530 | Chmp2b | 68942 | HITS-CLIP | 19536157\|23142080 | Unspecified |
| mmu-miR-21a-5p | MIMAT0000530 | Chmp4b | 75608 | HITS-CLIP | 23142080 | Unspecified |
| mmu-miR-21a-5p | MIMAT0000530 | Chmp5 | 76959 | HITS-CLIP | 23142080 | Unspecified |
| mmu-miR-21a-5p | MIMAT0000530 | Chmp6 | 208092 | HITS-CLIP | 23142080 | Unspecified |
| mmu-miR-21a-5p | MIMAT0000530 | Chn2 | 69993 | HITS-CLIP | 23597149 | Unspecified |
| mmu-miR-21a-5p | MIMAT0000530 | Chordc1 | 66917 | HITS-CLIP | 23142080 | Unspecified |
| mmu-miR-21a-5p | MIMAT0000530 | Chp1 | 56398 | HITS-CLIP | 23597149\|23142080 | Unspecified |
| mmu-miR-21a-5p | MIMAT0000530 | Chp2 | 70261 | Microarrays | 37986224 | Unspecified |
| mmu-miR-21a-5p | MIMAT0000530 | Chpt1 | 212862 | HITS-CLIP | 19536157\|29167373 | Unspecified |
| mmu-miR-21a-5p | MIMAT0000530 | Chst11 | 58250 | Microarrays | 37986224\|25263593 | Unspecified |
| mmu-miR-21a-5p | MIMAT0000530 | Chst12 | 59031 | HITS-CLIP | 23142080 | Unspecified |
| mmu-miR-21a-5p | MIMAT0000530 | Chst14 | 72136 | HITS-CLIP | 23142080 | Unspecified |
| mmu-miR-21a-5p | MIMAT0000530 | Chst15 | 77590 | Microarrays | 37986224 | Unspecified |
| mmu-miR-21a-5p | MIMAT0000530 | Chst2 | 54371 | HITS-CLIP | 37986224\|19536157 | Unspecified |
| mmu-miR-21a-5p | MIMAT0000530 | Chst7 | 60322 | Microarrays | 37986224 | Unspecified |
| mmu-miR-21a-5p | MIMAT0000530 | Chsy1 | 269941 | HITS-CLIP | 23142080 | Unspecified |
| mmu-miR-21a-5p | MIMAT0000530 | Chtf18 | 214901 | Microarrays | 37986224\|25263593 | Unspecified |
| mmu-miR-21a-5p | MIMAT0000530 | Chtop | 66511 | HITS-CLIP | 37986224\|23142080 | Unspecified |
| mmu-miR-21a-5p | MIMAT0000530 | Chuk | 12675 | HITS-CLIP | 23142080 | Unspecified |
| mmu-miR-21a-5p | MIMAT0000530 | Ciapin1 | 109006 | Microarrays//HITS-CLIP | 37986224\|25263593\|23597149 | Unspecified |
| mmu-miR-21a-5p | MIMAT0000530 | Cib2 | 56506 | HITS-CLIP | 37986224 | Unspecified |
| mmu-miR-21a-5p | MIMAT0000530 | Cic | 71722 | HITS-CLIP | 23142080 | Unspecified |
| mmu-miR-21a-5p | MIMAT0000530 | Cideb | 12684 | HITS-CLIP | 23597149 | Unspecified |

| | | | | | | |
|---|---|---|---|---|---|---|
| mmu-miR-21a-5p | MIMAT0000530 | Cinp | 67236 | HITS-CLIP | 23142080 | Unspecified |
| mmu-miR-21a-5p | MIMAT0000530 | Cip2a | 57650 | Microarrays//qCLASH//HITS-CLIP | 25263593\|31393953\|23142080 | Unspecified |
| mmu-miR-21a-5p | MIMAT0000530 | Cipc | 217732 | HITS-CLIP | 23597149\|29167373 | Unspecified |
| mmu-miR-21a-5p | MIMAT0000530 | Cisd2 | 67006 | HITS-CLIP | 23142080 | Unspecified |
| mmu-miR-21a-5p | MIMAT0000530 | Cit | 12704 | HITS-CLIP | 23142080 | Unspecified |
| mmu-miR-21a-5p | MIMAT0000530 | Cited2 | 17684 | HITS-CLIP | 23142080 | Unspecified |
| mmu-miR-21a-5p | MIMAT0000530 | Ckap2 | 80986 | Microarrays | 37986224\|25263593 | Unspecified |
| mmu-miR-21a-5p | MIMAT0000530 | Ckap2l | 70466 | Microarrays | 37986224\|25263593 | Unspecified |
| mmu-miR-21a-5p | MIMAT0000530 | Ckap4 | 216197 | Microarrays | 37986224\|25263593 | Unspecified |
| mmu-miR-21a-5p | MIMAT0000530 | Ckap5 | 75786 | HITS-CLIP | 23597149\|23142080 | Unspecified |
| mmu-miR-21a-5p | MIMAT0000530 | Cks1b | 54124 | Microarrays | 37986224\|25263593 | Unspecified |
| mmu-miR-21a-5p | MIMAT0000530 | Cks2 | 66197 | Microarrays | 37986224\|25263593 | Unspecified |
| mmu-miR-21a-5p | MIMAT0000530 | Clasp1 | 76707 | HITS-CLIP | 23142080\|23597149 | Unspecified |
| mmu-miR-21a-5p | MIMAT0000530 | Clasp2 | 76499 | HITS-CLIP | 23142080 | Unspecified |
| mmu-miR-21a-5p | MIMAT0000530 | Clcc1 | 229725 | HITS-CLIP | 23142080\|29167373 | Unspecified |
| mmu-miR-21a-5p | MIMAT0000530 | Clcf1 | 56708 | HITS-CLIP | 23142080 | Unspecified |
| mmu-miR-21a-5p | MIMAT0000530 | Clcn3 | 12725 | HITS-CLIP | 23142080 | Unspecified |
| mmu-miR-21a-5p | MIMAT0000530 | Clcn5 | 12728 | HITS-CLIP | 23142080 | Unspecified |
| mmu-miR-21a-5p | MIMAT0000530 | Clcn7 | 26373 | Microarrays | 37986224\|25263593 | Unspecified |
| mmu-miR-21a-5p | MIMAT0000530 | Cldn3 | 12739 | HITS-CLIP | 23597149 | Unspecified |
| mmu-miR-21a-5p | MIMAT0000530 | Clec10a | 17312 | Microarrays | 37986224 | Unspecified |
| mmu-miR-21a-5p | MIMAT0000530 | Clec1b | 56760 | Microarrays | 37986224 | Unspecified |
| mmu-miR-21a-5p | MIMAT0000530 | Clec2d | 93694 | HITS-CLIP, Microarrays//Microarrays//HITS | 37986224\|25263593\|23142080 | Unspecified |
| mmu-miR-21a-5p | MIMAT0000530 | Clec2i | 93675 | Microarrays//HITS-CLIP | 37986224\|23142080 | Unspecified |
| mmu-miR-21a-5p | MIMAT0000530 | Clec4a2 | 26888 | Microarrays | 37986224 | Unspecified |
| mmu-miR-21a-5p | MIMAT0000530 | Clec4a3 | 73149 | Microarrays | 37986224 | Unspecified |
| mmu-miR-21a-5p | MIMAT0000530 | Clec4b1 | 69810 | Microarrays | 37986224 | Unspecified |
| mmu-miR-21a-5p | MIMAT0000530 | Clec4d | 17474 | Microarrays | 37986224\|25263593 | Unspecified |
| mmu-miR-21a-5p | MIMAT0000530 | Clec4e | 56619 | Microarrays | 37986224 | Unspecified |
| mmu-miR-21a-5p | MIMAT0000530 | Clec4f | 51811 | Microarrays | 37986224 | Unspecified |
| mmu-miR-21a-5p | MIMAT0000530 | Clec4n | 56620 | Microarrays | 37986224 | Unspecified |
| mmu-miR-21a-5p | MIMAT0000530 | Clec5a | 23845 | Microarrays//HITS-CLIP | 37986224\|25263593\|29167373 | Unspecified |
| mmu-miR-21a-5p | MIMAT0000530 | Clec7a | 56644 | Microarrays | 37986224\|25263593 | Unspecified |
| mmu-miR-21a-5p | MIMAT0000530 | Clic1 | 114584 | HITS-CLIP | 23142080\|29167373 | Unspecified |
| mmu-miR-21a-5p | MIMAT0000530 | Clic4 | 29876 | Microarrays | 37986224\|25263593 | Unspecified |
| mmu-miR-21a-5p | MIMAT0000530 | Clint1 | 216705 | HITS-CLIP | 23142080\|29167373 | Unspecified |
| mmu-miR-21a-5p | MIMAT0000530 | Clip1 | 56430 | HITS-CLIP | 23142080\|29167373 | Unspecified |
| mmu-miR-21a-5p | MIMAT0000530 | Clip4 | 78785 | HITS-CLIP | 23597149 | Unspecified |
| mmu-miR-21a-5p | MIMAT0000530 | Clk1 | 12747 | Microarrays//HITS-CLIP | 37986224\|23142080 | Unspecified |

| | | | | | | |
|---|---|---|---|---|---|---|
| mmu-miR-21a-5p | MIMAT0000530 | Clk2 | 12748 | HITS-CLIP | 23142080 | Unspecified |
| mmu-miR-21a-5p | MIMAT0000530 | Clk3 | 102414 | HITS-CLIP | 23142080 | Unspecified |
| mmu-miR-21a-5p | MIMAT0000530 | Clmp | 71566 | Microarrays | 37986224 | Unspecified |
| mmu-miR-21a-5p | MIMAT0000530 | Cln8 | 26889 | HITS-CLIP | 29167373 | Unspecified |
| mmu-miR-21a-5p | MIMAT0000530 | Clns1a | 12729 | HITS-CLIP | 23597149 | Unspecified |
| mmu-miR-21a-5p | MIMAT0000530 | Clock | 12753 | HITS-CLIP | 23597149 | Unspecified |
| mmu-miR-21a-5p | MIMAT0000530 | Clptm1 | 56457 | qCLASH//HITS-CLIP | 31393953\|23597149\|23142080\|29167 | Unspecified |
| mmu-miR-21a-5p | MIMAT0000530 | Clpx | 270166 | Microarrays | 37986224 | Unspecified |
| mmu-miR-21a-5p | MIMAT0000530 | Clspn | 269582 | Microarrays//HITS-CLIP | 37986224\|25263593\|23142080 | Unspecified |
| mmu-miR-21a-5p | MIMAT0000530 | Cltc | 67300 | HITS-CLIP, Microarrays//HITS-CLIP//qCLASI | 37986224\|23597149\|31393953\|23142 | Unspecified |
| mmu-miR-21a-5p | MIMAT0000530 | Cluh | 74148 | HITS-CLIP | 23142080 | Unspecified |
| mmu-miR-21a-5p | MIMAT0000530 | Cmas | 12764 | HITS-CLIP | 23597149\|23142080 | Unspecified |
| mmu-miR-21a-5p | MIMAT0000530 | Cmbl | 69574 | Microarrays | 37986224\|25263593 | Unspecified |
| mmu-miR-21a-5p | MIMAT0000530 | Cmpk1 | 66588 | HITS-CLIP | 23142080 | Unspecified |
| mmu-miR-21a-5p | MIMAT0000530 | Cmtm4 | 97487 | HITS-CLIP | 23142080 | Unspecified |
| mmu-miR-21a-5p | MIMAT0000530 | Cmtm6 | 67213 | HITS-CLIP | 23142080 | Unspecified |
| mmu-miR-21a-5p | MIMAT0000530 | Cmtm7 | 102545 | Microarrays | 37986224 | Unspecified |
| mmu-miR-21a-5p | MIMAT0000530 | Cmtm8 | 70031 | HITS-CLIP | 23597149 | Unspecified |
| mmu-miR-21a-5p | MIMAT0000530 | Cnbp | 12785 | HITS-CLIP | 23597149 | Unspecified |
| mmu-miR-21a-5p | MIMAT0000530 | Cndp2 | 66054 | HITS-CLIP | 29167373 | Unspecified |
| mmu-miR-21a-5p | MIMAT0000530 | Cnep1r1 | 382030 | qCLASH//HITS-CLIP | 31393953\|23142080\|29167373 | Unspecified |
| mmu-miR-21a-5p | MIMAT0000530 | Cnih1 | 12793 | HITS-CLIP | 23597149\|23142080 | Unspecified |
| mmu-miR-21a-5p | MIMAT0000530 | Cnn2 | 12798 | HITS-CLIP | 23142080 | Unspecified |
| mmu-miR-21a-5p | MIMAT0000530 | Cnn3 | 71994 | Microarrays//HITS-CLIP | 37986224\|25263593\|23142080 | Unspecified |
| mmu-miR-21a-5p | MIMAT0000530 | Cnnm3 | 94218 | HITS-CLIP | 23597149\|19536157 | Unspecified |
| mmu-miR-21a-5p | MIMAT0000530 | Cnnm4 | 94220 | HITS-CLIP | 23142080 | Unspecified |
| mmu-miR-21a-5p | MIMAT0000530 | Cnot1 | 234594 | HITS-CLIP | 23597149\|23142080 | Unspecified |
| mmu-miR-21a-5p | MIMAT0000530 | Cnot10 | 78893 | HITS-CLIP | 23142080 | Unspecified |
| mmu-miR-21a-5p | MIMAT0000530 | Cnot2 | 72068 | HITS-CLIP | 23142080 | Unspecified |
| mmu-miR-21a-5p | MIMAT0000530 | Cnot4 | 53621 | HITS-CLIP | 29167373 | Unspecified |
| mmu-miR-21a-5p | MIMAT0000530 | Cnot6 | 104625 | HITS-CLIP | 37986224\|23142080\|23597149\|1953( | Unspecified |
| mmu-miR-21a-5p | MIMAT0000530 | Cnot6l | 231464 | HITS-CLIP | 37986224\|23142080\|29167373 | Unspecified |
| mmu-miR-21a-5p | MIMAT0000530 | Cnot7 | 18983 | HITS-CLIP | 37986224\|23142080 | Unspecified |
| mmu-miR-21a-5p | MIMAT0000530 | Cnot8 | 69125 | HITS-CLIP | 23142080 | Unspecified |
| mmu-miR-21a-5p | MIMAT0000530 | Cnot9 | 9125 | HITS-CLIP | 23142080 | Unspecified |
| mmu-miR-21a-5p | MIMAT0000530 | Cnp | 12799 | HITS-CLIP | 23142080 | Unspecified |
| mmu-miR-21a-5p | MIMAT0000530 | Cnpy2 | 56530 | HITS-CLIP | 37986224\|23597149 | Unspecified |
| mmu-miR-21a-5p | MIMAT0000530 | Cnpy3 | 72029 | Microarrays | 37986224 | Unspecified |
| mmu-miR-21a-5p | MIMAT0000530 | Cnr1 | 12801 | HITS-CLIP | 37986224\|19536157 | Unspecified |

| | | | | | | |
|---|---|---|---|---|---|---|
| mmu-miR-21a-5p | MIMAT0000530 | Cnr2 | 12802 | HITS-CLIP | 23142080 | Unspecified |
| mmu-miR-21a-5p | MIMAT0000530 | Cnst | 226744 | HITS-CLIP | 23142080 | Unspecified |
| mmu-miR-21a-5p | MIMAT0000530 | Cntn1 | 12805 | HITS-CLIP | 37986224 | Unspecified |
| mmu-miR-21a-5p | MIMAT0000530 | Cntrl | 26920 | HITS-CLIP | 23142080 | Unspecified |
| mmu-miR-21a-5p | MIMAT0000530 | Coa4 | 68185 | HITS-CLIP | 23597149 | Unspecified |
| mmu-miR-21a-5p | MIMAT0000530 | Coa5 | 76178 | HITS-CLIP | 29167373 | Unspecified |
| mmu-miR-21a-5p | MIMAT0000530 | Cog1 | 16834 | HITS-CLIP | 23142080 | Unspecified |
| mmu-miR-21a-5p | MIMAT0000530 | Cog4 | 102339 | HITS-CLIP | 23597149\|23142080 | Unspecified |
| mmu-miR-21a-5p | MIMAT0000530 | Cog5 | 238123 | HITS-CLIP | 37986224\|23142080 | Unspecified |
| mmu-miR-21a-5p | MIMAT0000530 | Cog6 | 67542 | HITS-CLIP | 23142080 | Unspecified |
| mmu-miR-21a-5p | MIMAT0000530 | Coil | 12812 | HITS-CLIP | 23597149\|29167373 | Unspecified |
| mmu-miR-21a-5p | MIMAT0000530 | Col1a1 | 12842 | Microarrays | 37986224\|25263593 | Unspecified |
| mmu-miR-21a-5p | MIMAT0000530 | Col1a2 | 12843 | Microarrays | 37986224\|25263593 | Unspecified |
| mmu-miR-21a-5p | MIMAT0000530 | Col4a1 | 12826 | Microarrays | 37986224\|25263593 | Unspecified |
| mmu-miR-21a-5p | MIMAT0000530 | Col4a3bp | 68018 | HITS-CLIP | 37986224\|23142080\|29167373 | Unspecified |
| mmu-miR-21a-5p | MIMAT0000530 | Col5a2 | 12832 | Microarrays | 37986224\|25263593 | Unspecified |
| mmu-miR-21a-5p | MIMAT0000530 | Col6a1 | 12833 | Microarrays | 37986224\|25263593 | Unspecified |
| mmu-miR-21a-5p | MIMAT0000530 | Col6a3 | 12835 | Microarrays | 37986224\|25263593 | Unspecified |
| mmu-miR-21a-5p | MIMAT0000530 | Colec12 | 140792 | Microarrays | 37986224 | Unspecified |
| mmu-miR-21a-5p | MIMAT0000530 | Colgalt1 | 234407 | Microarrays//HITS-CLIP | 37986224\|29167373 | Unspecified |
| mmu-miR-21a-5p | MIMAT0000530 | Commd1 | 17846 | HITS-CLIP | 23142080 | Unspecified |
| mmu-miR-21a-5p | MIMAT0000530 | Commd6 | 66200 | HITS-CLIP | 23142080 | Unspecified |
| mmu-miR-21a-5p | MIMAT0000530 | Commd8 | 27784 | HITS-CLIP | 23142080 | Unspecified |
| mmu-miR-21a-5p | MIMAT0000530 | Comt | 12846 | HITS-CLIP | 23597149 | Unspecified |
| mmu-miR-21a-5p | MIMAT0000530 | Comtd1 | 69156 | Microarrays | 37986224\|25263593 | Unspecified |
| mmu-miR-21a-5p | MIMAT0000530 | Cop1 | 817857 | qCLASH | 31393953 | Unspecified |
| mmu-miR-21a-5p | MIMAT0000530 | Copa | 12847 | HITS-CLIP | 23142080\|23597149 | Unspecified |
| mmu-miR-21a-5p | MIMAT0000530 | Copb1 | 70349 | HITS-CLIP | 23142080 | Unspecified |
| mmu-miR-21a-5p | MIMAT0000530 | Copb2 | 50797 | HITS-CLIP | 23597149\|23142080\|29167373 | Unspecified |
| mmu-miR-21a-5p | MIMAT0000530 | Cope | 59042 | HITS-CLIP | 23597149 | Unspecified |
| mmu-miR-21a-5p | MIMAT0000530 | Copg1 | 54161 | HITS-CLIP | 23142080 | Unspecified |
| mmu-miR-21a-5p | MIMAT0000530 | Copg2 | 54160 | HITS-CLIP | 23142080 | Unspecified |
| mmu-miR-21a-5p | MIMAT0000530 | Cops2 | 12848 | HITS-CLIP | 23142080 | Unspecified |
| mmu-miR-21a-5p | MIMAT0000530 | Cops4 | 26891 | HITS-CLIP | 37986224\|23597149\|23142080 | Unspecified |
| mmu-miR-21a-5p | MIMAT0000530 | Cops6 | 26893 | HITS-CLIP | 23597149 | Unspecified |
| mmu-miR-21a-5p | MIMAT0000530 | Cops7a | 26894 | HITS-CLIP | 23597149\|23142080\|29167373 | Unspecified |
| mmu-miR-21a-5p | MIMAT0000530 | Cops7b | 26895 | HITS-CLIP | 29167373 | Unspecified |
| mmu-miR-21a-5p | MIMAT0000530 | Cops8 | 108679 | HITS-CLIP | 23597149 | Unspecified |
| mmu-miR-21a-5p | MIMAT0000530 | Coq10b | 67876 | HITS-CLIP | 23142080 | Unspecified |

| | | | | | | |
|---|---|---|---|---|---|---|
| mmu-miR-21a-5p | MIMAT0000530 | Coq7 | 12850 | Microarrays | 37986224\|25263593 | Unspecified |
| mmu-miR-21a-5p | MIMAT0000530 | Coq8a | 56997 | HITS-CLIP | 23597149 | Unspecified |
| mmu-miR-21a-5p | MIMAT0000530 | Coq9 | 67914 | HITS-CLIP | 23597149 | Unspecified |
| mmu-miR-21a-5p | MIMAT0000530 | Coro1a | 12721 | HITS-CLIP | 23142080\|29167373 | Unspecified |
| mmu-miR-21a-5p | MIMAT0000530 | Coro1b | 23789 | HITS-CLIP | 23142080\|31433990 | Unspecified |
| mmu-miR-21a-5p | MIMAT0000530 | Coro1c | 23790 | HITS-CLIP | 29167373 | Unspecified |
| mmu-miR-21a-5p | MIMAT0000530 | Cotl1 | 72042 | HITS-CLIP | 29167373 | Unspecified |
| mmu-miR-21a-5p | MIMAT0000530 | Cox10 | 70383 | HITS-CLIP | 23597149 | Unspecified |
| mmu-miR-21a-5p | MIMAT0000530 | Cox11 | 69802 | HITS-CLIP | 23142080 | Unspecified |
| mmu-miR-21a-5p | MIMAT0000530 | Cox16 | 51241 | HITS-CLIP | 29167373 | Unspecified |
| mmu-miR-21a-5p | MIMAT0000530 | Cox4i1 | 12857 | HITS-CLIP | 23142080\|29167373 | Unspecified |
| mmu-miR-21a-5p | MIMAT0000530 | Cox5b | 12859 | HITS-CLIP | 19536157\|23597149 | Unspecified |
| mmu-miR-21a-5p | MIMAT0000530 | Cox5b-ps | 100040340 | HITS-CLIP | 31433990 | Unspecified |
| mmu-miR-21a-5p | MIMAT0000530 | Cox6a1 | 12861 | HITS-CLIP | 37986224 | Unspecified |
| mmu-miR-21a-5p | MIMAT0000530 | Cox6a2 | 12862 | Microarrays | 37986224\|25263593 | Unspecified |
| mmu-miR-21a-5p | MIMAT0000530 | Cox7a1 | 12865 | HITS-CLIP//qCLASH | 23142080\|31393953 | Unspecified |
| mmu-miR-21a-5p | MIMAT0000530 | Cox7a2 | 12866 | HITS-CLIP | 23142080 | Unspecified |
| mmu-miR-21a-5p | MIMAT0000530 | Cox7a2l | 20463 | HITS-CLIP | 23142080 | Unspecified |
| mmu-miR-21a-5p | MIMAT0000530 | Cox8a | 12868 | HITS-CLIP | 23142080 | Unspecified |
| mmu-miR-21a-5p | MIMAT0000530 | Cp | 12870 | Microarrays | 37986224 | Unspecified |
| mmu-miR-21a-5p | MIMAT0000530 | Cpb2 | 56373 | HITS-CLIP | 23597149 | Unspecified |
| mmu-miR-21a-5p | MIMAT0000530 | Cpd | 12874 | Microarrays//HITS-CLIP | 37986224\|23597149\|25263593\|2314 | Unspecified |
| mmu-miR-21a-5p | MIMAT0000530 | Cpe | 12876 | HITS-CLIP | 37986224\|19536157 | Unspecified |
| mmu-miR-21a-5p | MIMAT0000530 | Cpeb1 | 12877 | Microarrays | 37986224\|25263593 | Unspecified |
| mmu-miR-21a-5p | MIMAT0000530 | Cpeb2 | 231207 | qCLASH | 31393953 | Unspecified |
| mmu-miR-21a-5p | MIMAT0000530 | Cpeb4 | 67579 | HITS-CLIP | 23142080 | Unspecified |
| mmu-miR-21a-5p | MIMAT0000530 | Cplx2 | 12890 | HITS-CLIP | 19536157 | Unspecified |
| mmu-miR-21a-5p | MIMAT0000530 | Cpm | 70574 | Microarrays | 37986224\|25263593 | Unspecified |
| mmu-miR-21a-5p | MIMAT0000530 | Cpne1 | 266692 | HITS-CLIP | 23142080 | Unspecified |
| mmu-miR-21a-5p | MIMAT0000530 | Cpne3 | 70568 | HITS-CLIP | 37986224 | Unspecified |
| mmu-miR-21a-5p | MIMAT0000530 | Cpne8 | 66871 | Microarrays | 37986224 | Unspecified |
| mmu-miR-21a-5p | MIMAT0000530 | Cpox | 12892 | HITS-CLIP | 23597149\|23142080 | Unspecified |
| mmu-miR-21a-5p | MIMAT0000530 | Cps1 | 227231 | HITS-CLIP | 23597149 | Unspecified |
| mmu-miR-21a-5p | MIMAT0000530 | Cpsf2 | 51786 | HITS-CLIP | 23142080 | Unspecified |
| mmu-miR-21a-5p | MIMAT0000530 | Cpsf6 | 432508 | HITS-CLIP | 23597149\|23142080 | Unspecified |
| mmu-miR-21a-5p | MIMAT0000530 | Cpt2 | 12896 | HITS-CLIP | 37986224\|23597149\|23142080 | Unspecified |
| mmu-miR-21a-5p | MIMAT0000530 | Cr1l | 12946 | HITS-CLIP | 23597149 | Unspecified |
| mmu-miR-21a-5p | MIMAT0000530 | Crat | 12908 | HITS-CLIP | 23597149 | Unspecified |
| mmu-miR-21a-5p | MIMAT0000530 | Creb1 | 12912 | HITS-CLIP | 23142080 | Unspecified |

| | | | | | | |
|---|---|---|---|---|---|---|
| mmu-miR-21a-5p | MIMAT0000530 | Creb3l1 | 26427 | Microarrays | 37986224\|25263593 | Unspecified |
| mmu-miR-21a-5p | MIMAT0000530 | Creb3l2 | 208647 | Microarrays//HITS-CLIP | 37986224\|23142080 | Unspecified |
| mmu-miR-21a-5p | MIMAT0000530 | Creb3l3 | 208677 | HITS-CLIP | 23597149 | Unspecified |
| mmu-miR-21a-5p | MIMAT0000530 | Creb3l4 | 78284 | Microarrays | 37986224 | Unspecified |
| mmu-miR-21a-5p | MIMAT0000530 | Creb5 | 231991 | HITS-CLIP | 29167373 | Unspecified |
| mmu-miR-21a-5p | MIMAT0000530 | Crebbp | 12914 | HITS-CLIP//qCLASH | 37986224\|23597149\|31393953\|23142 | Unspecified |
| mmu-miR-21a-5p | MIMAT0000530 | Crebl2 | 232430 | HITS-CLIP | 23142080 | Unspecified |
| mmu-miR-21a-5p | MIMAT0000530 | Crebrf | 77128 | Microarrays//HITS-CLIP | 37986224\|23142080\|29167373 | Unspecified |
| mmu-miR-21a-5p | MIMAT0000530 | Crebzf | 233490 | HITS-CLIP | 23142080 | Unspecified |
| mmu-miR-21a-5p | MIMAT0000530 | Creg2 | 263764 | Microarrays | 37986224\|25263593 | Unspecified |
| mmu-miR-21a-5p | MIMAT0000530 | Crem | 12916 | HITS-CLIP | 23142080 | Unspecified |
| mmu-miR-21a-5p | MIMAT0000530 | Crim1 | 50766 | HITS-CLIP | 37986224\|23142080 | Unspecified |
| mmu-miR-21a-5p | MIMAT0000530 | Crip2 | 68337 | Microarrays | 37986224\|25263593 | Unspecified |
| mmu-miR-21a-5p | MIMAT0000530 | Cript | 56724 | HITS-CLIP | 37986224\|23597149\|23142080 | Unspecified |
| mmu-miR-21a-5p | MIMAT0000530 | Crk | 12928 | HITS-CLIP | 37986224\|23142080 | Unspecified |
| mmu-miR-21a-5p | MIMAT0000530 | Crkl | 12929 | HITS-CLIP | 23597149\|23142080 | Unspecified |
| mmu-miR-21a-5p | MIMAT0000530 | Crlf2 | 57914 | Microarrays | 37986224\|25263593 | Unspecified |
| mmu-miR-21a-5p | MIMAT0000530 | Crlf3 | 54394 | Microarrays | 37986224 | Unspecified |
| mmu-miR-21a-5p | MIMAT0000530 | Crls1 | 66586 | HITS-CLIP | 23597149 | Unspecified |
| mmu-miR-21a-5p | MIMAT0000530 | Crnkl1 | 66877 | HITS-CLIP | 23142080 | Unspecified |
| mmu-miR-21a-5p | MIMAT0000530 | Crtc2 | 74343 | HITS-CLIP | 23142080 | Unspecified |
| mmu-miR-21a-5p | MIMAT0000530 | Crybg1 | 202 | HITS-CLIP | 23142080\|29167373 | Unspecified |
| mmu-miR-21a-5p | MIMAT0000530 | Crybg2 | 55057 | HITS-CLIP | 23142080 | Unspecified |
| mmu-miR-21a-5p | MIMAT0000530 | Crybg3 | 224273 | HITS-CLIP | 23597149\|29167373 | Unspecified |
| mmu-miR-21a-5p | MIMAT0000530 | Cs | 12974 | HITS-CLIP | 23142080 | Unspecified |
| mmu-miR-21a-5p | MIMAT0000530 | Csad | 246277 | Microarrays//HITS-CLIP | 37986224\|23142080 | Unspecified |
| mmu-miR-21a-5p | MIMAT0000530 | Csde1 | 229663 | HITS-CLIP | 23597149\|19536157\|29167373\|23142 | Unspecified |
| mmu-miR-21a-5p | MIMAT0000530 | Cse1l | 110750 | HITS-CLIP | 37986224\|23597149\|23142080\|29167 | Unspecified |
| mmu-miR-21a-5p | MIMAT0000530 | Csf1r | 12978 | HITS-CLIP | 29167373 | Unspecified |
| mmu-miR-21a-5p | MIMAT0000530 | Csf2rb | 12983 | Microarrays//HITS-CLIP | 37986224\|23597149 | Unspecified |
| mmu-miR-21a-5p | MIMAT0000530 | Csf2rb2 | 12984 | Microarrays//HITS-CLIP | 37986224\|29167373 | Unspecified |
| mmu-miR-21a-5p | MIMAT0000530 | Csf3r | 12986 | Microarrays | 37986224 | Unspecified |
| mmu-miR-21a-5p | MIMAT0000530 | Csgalnact2 | 78752 | HITS-CLIP | 23142080 | Unspecified |
| mmu-miR-21a-5p | MIMAT0000530 | Csk | 12988 | HITS-CLIP | 23597149\|23142080 | Unspecified |
| mmu-miR-21a-5p | MIMAT0000530 | Csnk1a1 | 93687 | HITS-CLIP//qCLASH | 23142080\|31393953 | Unspecified |
| mmu-miR-21a-5p | MIMAT0000530 | Csnk1d | 104318 | HITS-CLIP | 23142080\|29167373 | Unspecified |
| mmu-miR-21a-5p | MIMAT0000530 | Csnk1g1 | 214897 | HITS-CLIP | 23142080 | Unspecified |
| mmu-miR-21a-5p | MIMAT0000530 | Csnk1g2 | 103236 | HITS-CLIP | 31433990 | Unspecified |
| mmu-miR-21a-5p | MIMAT0000530 | Csnk1g3 | 70425 | HITS-CLIP | 23142080 | Unspecified |

| | | | | | | |
|---|---|---|---|---|---|---|
| mmu-miR-21a-5p | MIMAT0000530 | Csnk2a1 | 12995 | HITS-CLIP | 23142080 | Unspecified |
| mmu-miR-21a-5p | MIMAT0000530 | Csnk2a2 | 13000 | HITS-CLIP | 23142080 | Unspecified |
| mmu-miR-21a-5p | MIMAT0000530 | Cspp1 | 211660 | HITS-CLIP | 19536157\|23142080 | Unspecified |
| mmu-miR-21a-5p | MIMAT0000530 | Csrnp1 | 215418 | HITS-CLIP | 23142080 | Unspecified |
| mmu-miR-21a-5p | MIMAT0000530 | Csrp1 | 13007 | HITS-CLIP | 23142080 | Unspecified |
| mmu-miR-21a-5p | MIMAT0000530 | Cstf2 | 108062 | HITS-CLIP | 23142080 | Unspecified |
| mmu-miR-21a-5p | MIMAT0000530 | Cstf3 | 228410 | HITS-CLIP | 23142080 | Unspecified |
| mmu-miR-21a-5p | MIMAT0000530 | Ctcf | 13018 | HITS-CLIP | 23142080 | Unspecified |
| mmu-miR-21a-5p | MIMAT0000530 | Ctdnep1 | 67181 | HITS-CLIP | 23597149\|23142080 | Unspecified |
| mmu-miR-21a-5p | MIMAT0000530 | Ctdsp1 | 227292 | HITS-CLIP | 23597149 | Unspecified |
| mmu-miR-21a-5p | MIMAT0000530 | Ctdsp2 | 52468 | Microarrays//HITS-CLIP | 37986224\|23597149\|19536157\|23142 | Unspecified |
| mmu-miR-21a-5p | MIMAT0000530 | Ctdspl | 69274 | HITS-CLIP | 37986224 | Unspecified |
| mmu-miR-21a-5p | MIMAT0000530 | Ctdspl2 | 329506 | HITS-CLIP | 23142080 | Unspecified |
| mmu-miR-21a-5p | MIMAT0000530 | Cth | 107869 | HITS-CLIP | 23597149 | Unspecified |
| mmu-miR-21a-5p | MIMAT0000530 | Ctla2a | 13024 | Microarrays | 37986224\|25263593 | Unspecified |
| mmu-miR-21a-5p | MIMAT0000530 | Ctnna1 | 12385 | HITS-CLIP | 23597149\|23142080\|29167373 | Unspecified |
| mmu-miR-21a-5p | MIMAT0000530 | Ctnnal1 | 54366 | HITS-CLIP | 23142080 | Unspecified |
| mmu-miR-21a-5p | MIMAT0000530 | Ctnnb1 | 12387 | HITS-CLIP | 23597149\|23142080\|19536157\|29167 | Unspecified |
| mmu-miR-21a-5p | MIMAT0000530 | Ctnnbl1 | 66642 | HITS-CLIP | 23142080 | Unspecified |
| mmu-miR-21a-5p | MIMAT0000530 | Ctnnd2 | 18163 | Microarrays | 37986224 | Unspecified |
| mmu-miR-21a-5p | MIMAT0000530 | Ctns | 83429 | HITS-CLIP | 23142080 | Unspecified |
| mmu-miR-21a-5p | MIMAT0000530 | Ctps | 51797 | Microarrays | 37986224\|25263593 | Unspecified |
| mmu-miR-21a-5p | MIMAT0000530 | Ctr9 | 22083 | HITS-CLIP | 23142080 | Unspecified |
| mmu-miR-21a-5p | MIMAT0000530 | Ctsa | 19025 | HITS-CLIP | 23597149\|29167373 | Unspecified |
| mmu-miR-21a-5p | MIMAT0000530 | Ctsb | 13030 | HITS-CLIP | 37986224\|23597149\|23142080\|29167 | Unspecified |
| mmu-miR-21a-5p | MIMAT0000530 | Ctsc | 13032 | Microarrays//HITS-CLIP | 37986224\|23597149\|29167373\|23142 | Unspecified |
| mmu-miR-21a-5p | MIMAT0000530 | Ctsd | 13033 | HITS-CLIP | 29167373 | Unspecified |
| mmu-miR-21a-5p | MIMAT0000530 | Ctsk | 13038 | Microarrays | 37986224\|25263593 | Unspecified |
| mmu-miR-21a-5p | MIMAT0000530 | Ctsl | 13039 | Microarrays | 37986224\|25263593 | Unspecified |
| mmu-miR-21a-5p | MIMAT0000530 | Ctso | 229445 | qCLASH | 31393953 | Unspecified |
| mmu-miR-21a-5p | MIMAT0000530 | Cttn | 13043 | Microarrays | 37986224\|25263593 | Unspecified |
| mmu-miR-21a-5p | MIMAT0000530 | Cttnbp2nl | 80281 | HITS-CLIP | 29167373 | Unspecified |
| mmu-miR-21a-5p | MIMAT0000530 | Cuedc2 | 67116 | HITS-CLIP | 23142080 | Unspecified |
| mmu-miR-21a-5p | MIMAT0000530 | Cul1 | 26965 | HITS-CLIP | 23597149\|29167373\|23142080 | Unspecified |
| mmu-miR-21a-5p | MIMAT0000530 | Cul2 | 71745 | HITS-CLIP | 23142080 | Unspecified |
| mmu-miR-21a-5p | MIMAT0000530 | Cul3 | 26554 | HITS-CLIP | 23142080 | Unspecified |
| mmu-miR-21a-5p | MIMAT0000530 | Cul4b | 72584 | HITS-CLIP | 23142080 | Unspecified |
| mmu-miR-21a-5p | MIMAT0000530 | Cul5 | 75717 | HITS-CLIP | 23597149\|23142080 | Unspecified |
| mmu-miR-21a-5p | MIMAT0000530 | Cux1 | 13047 | HITS-CLIP | 23142080\|19536157\|29167373 | Unspecified |

| | | | | | | |
|---|---|---|---|---|---|---|
| mmu-miR-21a-5p | MIMAT0000530 | Cwf19l2 | 244672 | Microarrays | 37986224 | Unspecified |
| mmu-miR-21a-5p | MIMAT0000530 | Cxadr | 13052 | HITS-CLIP | 23597149 | Unspecified |
| mmu-miR-21a-5p | MIMAT0000530 | Cxcl1 | 14825 | Microarrays//HITS-CLIP | 37986224\|23597149 | Unspecified |
| mmu-miR-21a-5p | MIMAT0000530 | Cxcl10 | 15945 | Microarrays//HITS-CLIP | 37986224\|29167373 | Unspecified |
| mmu-miR-21a-5p | MIMAT0000530 | Cxcl12 | 20315 | HITS-CLIP | 23597149 | Unspecified |
| mmu-miR-21a-5p | MIMAT0000530 | Cxcl14 | 57266 | Microarrays | 37986224\|25263593 | Unspecified |
| mmu-miR-21a-5p | MIMAT0000530 | Cxcl16 | 66102 | Microarrays//HITS-CLIP | 37986224\|29167373 | Unspecified |
| mmu-miR-21a-5p | MIMAT0000530 | Cxcl2 | 20310 | Microarrays | 37986224\|25263593 | Unspecified |
| mmu-miR-21a-5p | MIMAT0000530 | Cxcl3 | 330122 | Microarrays | 37986224\|25263593 | Unspecified |
| mmu-miR-21a-5p | MIMAT0000530 | Cxcl5 | 20311 | Microarrays | 37986224\|25263593 | Unspecified |
| mmu-miR-21a-5p | MIMAT0000530 | Cxcr4 | 12767 | Microarrays//qCLASH//HITS-CLIP | 37986224\|31393953\|29167373\|23142 | Unspecified |
| mmu-miR-21a-5p | MIMAT0000530 | Cyb561 | 13056 | Microarrays | 37986224 | Unspecified |
| mmu-miR-21a-5p | MIMAT0000530 | Cyb561d1 | 72023 | HITS-CLIP | 23142080 | Unspecified |
| mmu-miR-21a-5p | MIMAT0000530 | Cyb5a | 109672 | HITS-CLIP | 23142080\|23597149 | Unspecified |
| mmu-miR-21a-5p | MIMAT0000530 | Cyb5d2 | 192986 | Microarrays | 37986224 | Unspecified |
| mmu-miR-21a-5p | MIMAT0000530 | Cyb5r3 | 109754 | HITS-CLIP | 23597149\|29167373 | Unspecified |
| mmu-miR-21a-5p | MIMAT0000530 | Cybb | 13058 | HITS-CLIP | 29167373 | Unspecified |
| mmu-miR-21a-5p | MIMAT0000530 | Cybc1 | 79415 | HITS-CLIP | 23142080 | Unspecified |
| mmu-miR-21a-5p | MIMAT0000530 | Cyc1 | 66445 | HITS-CLIP | 23142080 | Unspecified |
| mmu-miR-21a-5p | MIMAT0000530 | Cycs | 13063 | Microarrays | 37986224\|25263593 | Unspecified |
| mmu-miR-21a-5p | MIMAT0000530 | Cyfip1 | 20430 | HITS-CLIP | 25083871\|23597149\|29167373 | Unspecified |
| mmu-miR-21a-5p | MIMAT0000530 | Cyfip2 | 76884 | Microarrays//HITS-CLIP | 37986224\|23142080\|25263593\|19536 | Unspecified |
| mmu-miR-21a-5p | MIMAT0000530 | Cyld | 74256 | HITS-CLIP | 23142080 | Unspecified |
| mmu-miR-21a-5p | MIMAT0000530 | Cyp1b1 | 13078 | Microarrays | 37986224\|25263593 | Unspecified |
| mmu-miR-21a-5p | MIMAT0000530 | Cyp20a1 | 77951 | HITS-CLIP | 23142080\|29167373 | Unspecified |
| mmu-miR-21a-5p | MIMAT0000530 | Cyp27a1 | 104086 | Microarrays | 37986224 | Unspecified |
| mmu-miR-21a-5p | MIMAT0000530 | Cyp2c23 | 226143 | HITS-CLIP | 23597149 | Unspecified |
| mmu-miR-21a-5p | MIMAT0000530 | Cyp2c37 | 13096 | HITS-CLIP | 23597149 | Unspecified |
| mmu-miR-21a-5p | MIMAT0000530 | Cyp2d22 | 56448 | HITS-CLIP | 37986224 | Unspecified |
| mmu-miR-21a-5p | MIMAT0000530 | Cyp2d26 | 76279 | HITS-CLIP | 23597149 | Unspecified |
| mmu-miR-21a-5p | MIMAT0000530 | Cyp2d40 | 71754 | HITS-CLIP | 23597149 | Unspecified |
| mmu-miR-21a-5p | MIMAT0000530 | Cyp2d9 | 13105 | HITS-CLIP | 23597149 | Unspecified |
| mmu-miR-21a-5p | MIMAT0000530 | Cyp2j5 | 13109 | HITS-CLIP | 23597149 | Unspecified |
| mmu-miR-21a-5p | MIMAT0000530 | Cyp3a13 | 13113 | HITS-CLIP | 23597149 | Unspecified |
| mmu-miR-21a-5p | MIMAT0000530 | Cyp4f14 | 64385 | HITS-CLIP | 28735896 | Unspecified |
| mmu-miR-21a-5p | MIMAT0000530 | Cyp4f18 | 72054 | Microarrays | 37986224 | Unspecified |
| mmu-miR-21a-5p | MIMAT0000530 | Cyp4v3 | 102294 | HITS-CLIP | 23597149 | Unspecified |
| mmu-miR-21a-5p | MIMAT0000530 | Cyp7a1 | 13122 | HITS-CLIP | 23597149 | Unspecified |
| mmu-miR-21a-5p | MIMAT0000530 | Cyp8b1 | 13124 | HITS-CLIP | 23597149 | Unspecified |

| | | | | | | |
|---|---|---|---|---|---|---|
| mmu-miR-21a-5p | MIMAT0000530 | Cysltr1 | 58861 | Microarrays//HITS-CLIP | 37986224\|29167373 | Unspecified |
| mmu-miR-21a-5p | MIMAT0000530 | Cyth1 | 19157 | Microarrays//HITS-CLIP | 37986224\|23597149\|23142080 | Unspecified |
| mmu-miR-21a-5p | MIMAT0000530 | Cyth2 | 19158 | HITS-CLIP | 23597149\|23142080 | Unspecified |
| mmu-miR-21a-5p | MIMAT0000530 | Cytip | 227929 | Microarrays | 37986224 | Unspecified |
| mmu-miR-21a-5p | MIMAT0000530 | Cyyr1 | 224405 | HITS-CLIP | 37986224 | Unspecified |
| mmu-miR-21a-5p | MIMAT0000530 | D030056L22Rik | 225995 | HITS-CLIP | 23142080 | Unspecified |
| mmu-miR-21a-5p | MIMAT0000530 | D10Wsu102e | 28109 | HITS-CLIP | 23142080 | Unspecified |
| mmu-miR-21a-5p | MIMAT0000530 | D16Ertd472e | 67102 | HITS-CLIP | 23142080 | Unspecified |
| mmu-miR-21a-5p | MIMAT0000530 | D1Ertd622e | 52392 | HITS-CLIP | 29167373 | Unspecified |
| mmu-miR-21a-5p | MIMAT0000530 | D5Ertd579e | 320661 | HITS-CLIP | 29167373 | Unspecified |
| mmu-miR-21a-5p | MIMAT0000530 | Daam1 | 208846 | HITS-CLIP | 23142080 | Unspecified |
| mmu-miR-21a-5p | MIMAT0000530 | Dab2 | 13132 | HITS-CLIP | 29167373 | Unspecified |
| mmu-miR-21a-5p | MIMAT0000530 | Dab2ip | 69601 | Microarrays | 37986224\|25263593 | Unspecified |
| mmu-miR-21a-5p | MIMAT0000530 | Dach1 | 13134 | Microarrays | 37986224\|25263593 | Unspecified |
| mmu-miR-21a-5p | MIMAT0000530 | Dag1 | 13138 | HITS-CLIP | 23142080\|23597149\|29167373 | Unspecified |
| mmu-miR-21a-5p | MIMAT0000530 | Dagla | 269060 | Microarrays | 37986224\|25263593 | Unspecified |
| mmu-miR-21a-5p | MIMAT0000530 | Dap | 223453 | HITS-CLIP | 23597149 | Unspecified |
| mmu-miR-21a-5p | MIMAT0000530 | Dap3 | 65111 | HITS-CLIP | 23597149\|23142080 | Unspecified |
| mmu-miR-21a-5p | MIMAT0000530 | Dapk1 | 69635 | HITS-CLIP | 23597149 | Unspecified |
| mmu-miR-21a-5p | MIMAT0000530 | Dapk3 | 13144 | HITS-CLIP | 23142080 | Unspecified |
| mmu-miR-21a-5p | MIMAT0000530 | Dapp1 | 26377 | Microarrays | 37986224 | Unspecified |
| mmu-miR-21a-5p | MIMAT0000530 | Dars | 226414 | HITS-CLIP | 23597149\|23142080 | Unspecified |
| mmu-miR-21a-5p | MIMAT0000530 | Daxx | 13163 | HITS-CLIP | 31433990 | Unspecified |
| mmu-miR-21a-5p | MIMAT0000530 | Dbf4 | 27214 | HITS-CLIP//qCLASH | 23142080\|31393953\|29167373 | Unspecified |
| mmu-miR-21a-5p | MIMAT0000530 | Dbi | 13167 | HITS-CLIP | 23597149\|23142080 | Unspecified |
| mmu-miR-21a-5p | MIMAT0000530 | Dbp | 13170 | Microarrays | 37986224 | Unspecified |
| mmu-miR-21a-5p | MIMAT0000530 | Dbr1 | 83703 | HITS-CLIP | 23142080 | Unspecified |
| mmu-miR-21a-5p | MIMAT0000530 | Dbt | 13171 | HITS-CLIP | 23597149\|23142080 | Unspecified |
| mmu-miR-21a-5p | MIMAT0000530 | Dcaf1 | 41836 | HITS-CLIP | 23142080 | Unspecified |
| mmu-miR-21a-5p | MIMAT0000530 | Dcaf10 | 242418 | HITS-CLIP | 23597149 | Unspecified |
| mmu-miR-21a-5p | MIMAT0000530 | Dcaf11 | 28199 | HITS-CLIP | 37986224\|23142080\|23597149 | Unspecified |
| mmu-miR-21a-5p | MIMAT0000530 | Dcaf12 | 68970 | HITS-CLIP | 23142080 | Unspecified |
| mmu-miR-21a-5p | MIMAT0000530 | Dcaf13 | 223499 | HITS-CLIP | 23142080 | Unspecified |
| mmu-miR-21a-5p | MIMAT0000530 | Dcaf17 | 75763 | HITS-CLIP | 23142080 | Unspecified |
| mmu-miR-21a-5p | MIMAT0000530 | Dcaf5 | 320808 | HITS-CLIP | 23597149\|23142080 | Unspecified |
| mmu-miR-21a-5p | MIMAT0000530 | Dcaf7 | 71833 | HITS-CLIP | 37986224\|23597149\|23142080\|29167 | Unspecified |
| mmu-miR-21a-5p | MIMAT0000530 | Dcaf8 | 98193 | HITS-CLIP | 23142080 | Unspecified |
| mmu-miR-21a-5p | MIMAT0000530 | Dcakd | 68087 | Microarrays | 37986224\|25263593 | Unspecified |
| mmu-miR-21a-5p | MIMAT0000530 | Dcbld1 | 66686 | HITS-CLIP | 23597149 | Unspecified |

| | | | | | | |
|---|---|---|---|---|---|---|
| mmu-miR-21a-5p | MIMAT0000530 | Dck | 13178 | HITS-CLIP | 29167373\|23142080 | Unspecified |
| mmu-miR-21a-5p | MIMAT0000530 | Dclk1 | 13175 | HITS-CLIP | 19536157 | Unspecified |
| mmu-miR-21a-5p | MIMAT0000530 | Dcn | 13179 | Microarrays | 37986224\|25263593 | Unspecified |
| mmu-miR-21a-5p | MIMAT0000530 | Dcp1a | 75901 | HITS-CLIP | 37986224\|23142080\|29167373 | Unspecified |
| mmu-miR-21a-5p | MIMAT0000530 | Dcp1b | 319618 | HITS-CLIP | 23142080 | Unspecified |
| mmu-miR-21a-5p | MIMAT0000530 | Dcp2 | 70640 | HITS-CLIP | 23597149\|23142080\|29167373 | Unspecified |
| mmu-miR-21a-5p | MIMAT0000530 | Dcstamp | 75766 | Microarrays | 37986224\|25263593 | Unspecified |
| mmu-miR-21a-5p | MIMAT0000530 | Dctn2 | 69654 | HITS-CLIP | 23597149\|23142080 | Unspecified |
| mmu-miR-21a-5p | MIMAT0000530 | Dctn3 | 53598 | HITS-CLIP | 23597149 | Unspecified |
| mmu-miR-21a-5p | MIMAT0000530 | Dctn4 | 67665 | HITS-CLIP | 23142080 | Unspecified |
| mmu-miR-21a-5p | MIMAT0000530 | Dctn6 | 22428 | HITS-CLIP | 23142080 | Unspecified |
| mmu-miR-21a-5p | MIMAT0000530 | Dcun1d4 | 100737 | HITS-CLIP | 23142080 | Unspecified |
| mmu-miR-21a-5p | MIMAT0000530 | Ddb1 | 13194 | qCLASH//HITS-CLIP | 31393953\|31433990\|23142080 | Unspecified |
| mmu-miR-21a-5p | MIMAT0000530 | Ddc | 13195 | HITS-CLIP | 23142080\|23597149 | Unspecified |
| mmu-miR-21a-5p | MIMAT0000530 | Ddhd1 | 114874 | HITS-CLIP | 23142080 | Unspecified |
| mmu-miR-21a-5p | MIMAT0000530 | Ddi2 | 68817 | HITS-CLIP | 37986224\|23597149\|23142080\|29167 | Unspecified |
| mmu-miR-21a-5p | MIMAT0000530 | Ddias | 220042 | Microarrays | 25263593 | Unspecified |
| mmu-miR-21a-5p | MIMAT0000530 | Ddit4 | 74747 | HITS-CLIP | 23142080 | Unspecified |
| mmu-miR-21a-5p | MIMAT0000530 | Ddr1 | 12305 | HITS-CLIP | 23142080 | Unspecified |
| mmu-miR-21a-5p | MIMAT0000530 | Ddx1 | 104721 | HITS-CLIP | 23142080 | Unspecified |
| mmu-miR-21a-5p | MIMAT0000530 | Ddx17 | 67040 | HITS-CLIP | 23142080\|19536157\|23597149\|29167 | Unspecified |
| mmu-miR-21a-5p | MIMAT0000530 | Ddx19a | 13680 | HITS-CLIP | 29167373\|23142080 | Unspecified |
| mmu-miR-21a-5p | MIMAT0000530 | Ddx19b | 234733 | HITS-CLIP | 23142080 | Unspecified |
| mmu-miR-21a-5p | MIMAT0000530 | Ddx21 | 56200 | Microarrays//qCLASH//HITS-CLIP | 37986224\|25263593\|31393953\|29167 | Unspecified |
| mmu-miR-21a-5p | MIMAT0000530 | Ddx23 | 74351 | HITS-CLIP | 23142080 | Unspecified |
| mmu-miR-21a-5p | MIMAT0000530 | Ddx24 | 27225 | HITS-CLIP | 23142080 | Unspecified |
| mmu-miR-21a-5p | MIMAT0000530 | Ddx39 | 68278 | Microarrays | 37986224 | Unspecified |
| mmu-miR-21a-5p | MIMAT0000530 | Ddx39a | 10212 | Microarrays | 25263593 | Unspecified |
| mmu-miR-21a-5p | MIMAT0000530 | Ddx39b | 53817 | HITS-CLIP | 23142080 | Unspecified |
| mmu-miR-21a-5p | MIMAT0000530 | Ddx3x | 13205 | qCLASH//HITS-CLIP | 31393953\|23142080\|19536157 | Unspecified |
| mmu-miR-21a-5p | MIMAT0000530 | Ddx3y | 26900 | HITS-CLIP | 23142080 | Unspecified |
| mmu-miR-21a-5p | MIMAT0000530 | Ddx41 | 72935 | HITS-CLIP | 37986224 | Unspecified |
| mmu-miR-21a-5p | MIMAT0000530 | Ddx42 | 72047 | HITS-CLIP | 37986224\|23142080\|29167373 | Unspecified |
| mmu-miR-21a-5p | MIMAT0000530 | Ddx46 | 212880 | HITS-CLIP | 23142080\|29167373 | Unspecified |
| mmu-miR-21a-5p | MIMAT0000530 | Ddx47 | 67755 | qCLASH | 31393953 | Unspecified |
| mmu-miR-21a-5p | MIMAT0000530 | Ddx5 | 13207 | HITS-CLIP//qCLASH | 37986224\|31393953\|23142080 | Unspecified |
| mmu-miR-21a-5p | MIMAT0000530 | Ddx50 | 94213 | HITS-CLIP | 23142080 | Unspecified |
| mmu-miR-21a-5p | MIMAT0000530 | Ddx55 | 67848 | HITS-CLIP | 23142080 | Unspecified |
| mmu-miR-21a-5p | MIMAT0000530 | Ddx58 | 230073 | HITS-CLIP | 23142080 | Unspecified |

| | | | | | | |
|---|---|---|---|---|---|---|
| mmu-miR-21a-5p | MIMAT0000530 | Ddx6 | 13209 | HITS-CLIP | 23142080\|23597149\|29167373 | Unspecified |
| mmu-miR-21a-5p | MIMAT0000530 | Degs1 | 13244 | HITS-CLIP | 37986224 | Unspecified |
| mmu-miR-21a-5p | MIMAT0000530 | Dek | 110052 | HITS-CLIP//qCLASH | 23597149\|23142080\|31393953 | Unspecified |
| mmu-miR-21a-5p | MIMAT0000530 | Dennd1a | 227801 | HITS-CLIP | 37986224 | Unspecified |
| mmu-miR-21a-5p | MIMAT0000530 | Dennd1b | 329260 | HITS-CLIP | 37986224\|23142080\|23597149\|29167 | Unspecified |
| mmu-miR-21a-5p | MIMAT0000530 | Dennd1c | 70785 | HITS-CLIP | 23142080 | Unspecified |
| mmu-miR-21a-5p | MIMAT0000530 | Dennd2c | 329727 | HITS-CLIP | 30971389 | Unspecified |
| mmu-miR-21a-5p | MIMAT0000530 | Dennd4a | 102442 | HITS-CLIP | 23142080 | Unspecified |
| mmu-miR-21a-5p | MIMAT0000530 | Dennd4b | 229541 | HITS-CLIP | 31433990 | Unspecified |
| mmu-miR-21a-5p | MIMAT0000530 | Dennd4c | 329877 | HITS-CLIP | 23597149\|23142080\|29167373 | Unspecified |
| mmu-miR-21a-5p | MIMAT0000530 | Dennd6a | 211922 | HITS-CLIP | 29167373\|23142080 | Unspecified |
| mmu-miR-21a-5p | MIMAT0000530 | Denr | 68184 | HITS-CLIP | 23597149 | Unspecified |
| mmu-miR-21a-5p | MIMAT0000530 | Depdc1a | 76131 | Microarrays//HITS-CLIP | 37986224\|25263593\|23142080 | Unspecified |
| mmu-miR-21a-5p | MIMAT0000530 | Depdc5 | 277854 | HITS-CLIP | 23597149\|23142080 | Unspecified |
| mmu-miR-21a-5p | MIMAT0000530 | Depdc7 | 211896 | HITS-CLIP, Microarrays//HITS-CLIP | 37986224\|23597149 | Unspecified |
| mmu-miR-21a-5p | MIMAT0000530 | Deptor | 97998 | Microarrays | 37986224 | Unspecified |
| mmu-miR-21a-5p | MIMAT0000530 | Derl1 | 67819 | HITS-CLIP | 31433990\|28735896 | Unspecified |
| mmu-miR-21a-5p | MIMAT0000530 | Derl2 | 116891 | HITS-CLIP | 23142080\|23597149 | Unspecified |
| mmu-miR-21a-5p | MIMAT0000530 | Desi2 | 78825 | HITS-CLIP | 23142080 | Unspecified |
| mmu-miR-21a-5p | MIMAT0000530 | Dgat2 | 67800 | HITS-CLIP | 37986224\|23597149\|23142080 | Unspecified |
| mmu-miR-21a-5p | MIMAT0000530 | Dgka | 13139 | HITS-CLIP | 23142080 | Unspecified |
| mmu-miR-21a-5p | MIMAT0000530 | Dgkb | 217480 | HITS-CLIP | 37986224 | Unspecified |
| mmu-miR-21a-5p | MIMAT0000530 | Dgkd | 227333 | HITS-CLIP | 31433990\|23142080\|29167373 | Unspecified |
| mmu-miR-21a-5p | MIMAT0000530 | Dgkg | 110197 | HITS-CLIP | 23142080 | Unspecified |
| mmu-miR-21a-5p | MIMAT0000530 | Dgkq | 110524 | HITS-CLIP | 23142080 | Unspecified |
| mmu-miR-21a-5p | MIMAT0000530 | Dgkz | 104418 | HITS-CLIP | 23142080 | Unspecified |
| mmu-miR-21a-5p | MIMAT0000530 | Dglucy | 217830 | Microarrays//HITS-CLIP | 25263593\|29167373 | Unspecified |
| mmu-miR-21a-5p | MIMAT0000530 | Dhcr24 | 74754 | HITS-CLIP | 37986224\|23597149\|19536157\|23142 | Unspecified |
| mmu-miR-21a-5p | MIMAT0000530 | Dhfr | 13361 | Microarrays//HITS-CLIP | 37986224\|25263593\|23142080 | Unspecified |
| mmu-miR-21a-5p | MIMAT0000530 | Dhrs1 | 52585 | HITS-CLIP | 23597149 | Unspecified |
| mmu-miR-21a-5p | MIMAT0000530 | Dhrs13 | 70451 | Microarrays | 37986224\|25263593 | Unspecified |
| mmu-miR-21a-5p | MIMAT0000530 | Dhrs7 | 66375 | HITS-CLIP | 23142080 | Unspecified |
| mmu-miR-21a-5p | MIMAT0000530 | Dhrs7b | 216820 | HITS-CLIP | 26203562 | Unspecified |
| mmu-miR-21a-5p | MIMAT0000530 | Dhtkd1 | 209692 | HITS-CLIP | 23597149 | Unspecified |
| mmu-miR-21a-5p | MIMAT0000530 | Dhx15 | 13204 | HITS-CLIP | 19536157\|23142080 | Unspecified |
| mmu-miR-21a-5p | MIMAT0000530 | Dhx30 | 72831 | HITS-CLIP | 23142080 | Unspecified |
| mmu-miR-21a-5p | MIMAT0000530 | Dhx33 | 216877 | HITS-CLIP | 23597149\|23142080 | Unspecified |
| mmu-miR-21a-5p | MIMAT0000530 | Dhx36 | 72162 | HITS-CLIP | 37986224\|23142080\|29167373 | Unspecified |
| mmu-miR-21a-5p | MIMAT0000530 | Dhx38 | 64340 | HITS-CLIP | 23142080 | Unspecified |

| | | | | | | |
|---|---|---|---|---|---|---|
| mmu-miR-21a-5p | MIMAT0000530 | Dhx40 | 67487 | Microarrays//HITS-CLIP | 37986224\|23142080 | Unspecified |
| mmu-miR-21a-5p | MIMAT0000530 | Dhx57 | 106794 | Microarrays//HITS-CLIP | 37986224\|29167373 | Unspecified |
| mmu-miR-21a-5p | MIMAT0000530 | Dhx58 | 80861 | HITS-CLIP | 23142080 | Unspecified |
| mmu-miR-21a-5p | MIMAT0000530 | Dhx9 | 13211 | HITS-CLIP | 23597149\|19536157\|23142080 | Unspecified |
| mmu-miR-21a-5p | MIMAT0000530 | Diaph1 | 1729 | HITS-CLIP | 23142080 | Unspecified |
| mmu-miR-21a-5p | MIMAT0000530 | Diaph2 | 54004 | HITS-CLIP | 23142080\|23597149\|29167373 | Unspecified |
| mmu-miR-21a-5p | MIMAT0000530 | Diaph3 | 81624 | HITS-CLIP//Microarrays | 23142080\|25263593 | Unspecified |
| mmu-miR-21a-5p | MIMAT0000530 | Dicer1 | 192119 | HITS-CLIP | 37986224\|23597149\|23142080\|29167 | Unspecified |
| mmu-miR-21a-5p | MIMAT0000530 | Dido1 | 23856 | HITS-CLIP | 23142080\|29167373 | Unspecified |
| mmu-miR-21a-5p | MIMAT0000530 | Diexf | 215193 | HITS-CLIP | 23142080 | Unspecified |
| mmu-miR-21a-5p | MIMAT0000530 | Dio2 | 13371 | Microarrays | 37986224\|25263593 | Unspecified |
| mmu-miR-21a-5p | MIMAT0000530 | Dip2b | 239667 | HITS-CLIP | 19536157\|23142080 | Unspecified |
| mmu-miR-21a-5p | MIMAT0000530 | Dip2c | 208440 | HITS-CLIP | 23597149\|19536157 | Unspecified |
| mmu-miR-21a-5p | MIMAT0000530 | Dirc2 | 224132 | HITS-CLIP | 23142080 | Unspecified |
| mmu-miR-21a-5p | MIMAT0000530 | Dixdc1 | 330938 | HITS-CLIP | 23597149 | Unspecified |
| mmu-miR-21a-5p | MIMAT0000530 | Dkc1 | 245474 | Microarrays//HITS-CLIP | 37986224\|25263593\|23142080\|29167 | Unspecified |
| mmu-miR-21a-5p | MIMAT0000530 | Dkk3 | 50781 | HITS-CLIP | 37986224 | Unspecified |
| mmu-miR-21a-5p | MIMAT0000530 | Dlat | 235339 | HITS-CLIP | 23597149 | Unspecified |
| mmu-miR-21a-5p | MIMAT0000530 | Dld | 13382 | HITS-CLIP | 23142080 | Unspecified |
| mmu-miR-21a-5p | MIMAT0000530 | Dlg1 | 13383 | qCLASH//HITS-CLIP | 31393953\|29167373 | Unspecified |
| mmu-miR-21a-5p | MIMAT0000530 | Dlg3 | 53310 | Microarrays//HITS-CLIP | 37986224\|25263593\|23142080 | Unspecified |
| mmu-miR-21a-5p | MIMAT0000530 | Dlg4 | 13385 | HITS-CLIP | 23142080 | Unspecified |
| mmu-miR-21a-5p | MIMAT0000530 | Dlgap5 | 218977 | Microarrays//HITS-CLIP | 37986224\|25263593\|23142080 | Unspecified |
| mmu-miR-21a-5p | MIMAT0000530 | Dmac1 | 90871 | qCLASH | 31393953 | Unspecified |
| mmu-miR-21a-5p | MIMAT0000530 | Dmap1 | 66233 | HITS-CLIP | 23142080 | Unspecified |
| mmu-miR-21a-5p | MIMAT0000530 | Dmd | 13405 | HITS-CLIP | 19536157\|23597149 | Unspecified |
| mmu-miR-21a-5p | MIMAT0000530 | Dmgdh | 74129 | HITS-CLIP | 23597149 | Unspecified |
| mmu-miR-21a-5p | MIMAT0000530 | Dmpk | 13400 | Microarrays | 37986224\|25263593 | Unspecified |
| mmu-miR-21a-5p | MIMAT0000530 | Dmtf1 | 23857 | HITS-CLIP | 23142080 | Unspecified |
| mmu-miR-21a-5p | MIMAT0000530 | Dmtn | 13829 | HITS-CLIP | 19536157 | Unspecified |
| mmu-miR-21a-5p | MIMAT0000530 | Dmwd | 13401 | Microarrays | 37986224\|25263593 | Unspecified |
| mmu-miR-21a-5p | MIMAT0000530 | Dmxl1 | 240283 | HITS-CLIP | 23597149\|23142080\|29167373 | Unspecified |
| mmu-miR-21a-5p | MIMAT0000530 | Dmxl2 | 235380 | Microarrays//HITS-CLIP | 37986224\|23597149 | Unspecified |
| mmu-miR-21a-5p | MIMAT0000530 | Dnah2 | 327954 | Microarrays | 37986224 | Unspecified |
| mmu-miR-21a-5p | MIMAT0000530 | Dnaja1 | 15502 | qCLASH//HITS-CLIP | 31393953\|29167373 | Unspecified |
| mmu-miR-21a-5p | MIMAT0000530 | Dnaja2 | 56445 | qCLASH//HITS-CLIP | 31393953\|23142080 | Unspecified |
| mmu-miR-21a-5p | MIMAT0000530 | Dnaja4 | 58233 | HITS-CLIP | 23142080 | Unspecified |
| mmu-miR-21a-5p | MIMAT0000530 | Dnajb1 | 81489 | HITS-CLIP | 23142080 | Unspecified |
| mmu-miR-21a-5p | MIMAT0000530 | Dnajb12 | 56709 | HITS-CLIP | 23142080\|19536157\|23597149\|29167 | Unspecified |

| | | | | | | |
|---|---|---|---|---|---|---|
| mmu-miR-21a-5p | MIMAT0000530 | Dnajb14 | 70604 | HITS-CLIP | 23142080\|29167373 | Unspecified |
| mmu-miR-21a-5p | MIMAT0000530 | Dnajb2 | 56812 | HITS-CLIP | 23597149 | Unspecified |
| mmu-miR-21a-5p | MIMAT0000530 | Dnajb4 | 67035 | HITS-CLIP | 23142080 | Unspecified |
| mmu-miR-21a-5p | MIMAT0000530 | Dnajb5 | 56323 | HITS-CLIP | 23142080 | Unspecified |
| mmu-miR-21a-5p | MIMAT0000530 | Dnajb6 | 23950 | HITS-CLIP | 29167373 | Unspecified |
| mmu-miR-21a-5p | MIMAT0000530 | Dnajb9 | 27362 | HITS-CLIP | 37986224\|23142080 | Unspecified |
| mmu-miR-21a-5p | MIMAT0000530 | Dnajc1 | 13418 | HITS-CLIP | 23142080 | Unspecified |
| mmu-miR-21a-5p | MIMAT0000530 | Dnajc10 | 66861 | HITS-CLIP | 23142080\|29167373 | Unspecified |
| mmu-miR-21a-5p | MIMAT0000530 | Dnajc13 | 235567 | Microarrays//HITS-CLIP | 37986224\|23597149\|23142080\|29167 | Unspecified |
| mmu-miR-21a-5p | MIMAT0000530 | Dnajc16 | 214063 | HITS-CLIP | 37986224\|23597149\|23142080\|29167 | Unspecified |
| mmu-miR-21a-5p | MIMAT0000530 | Dnajc2 | 22791 | HITS-CLIP | 23142080 | Unspecified |
| mmu-miR-21a-5p | MIMAT0000530 | Dnajc25 | 72429 | HITS-CLIP | 23142080 | Unspecified |
| mmu-miR-21a-5p | MIMAT0000530 | Dnajc3 | 100037258 | HITS-CLIP | 23597149\|23142080 | Unspecified |
| mmu-miR-21a-5p | MIMAT0000530 | Dnajc5 | 13002 | HITS-CLIP | 37986224\|29167373 | Unspecified |
| mmu-miR-21a-5p | MIMAT0000530 | Dnase1l3 | 13421 | HITS-CLIP | 23597149\|23142080 | Unspecified |
| mmu-miR-21a-5p | MIMAT0000530 | Dnase2a | 13423 | Microarrays | 37986224 | Unspecified |
| mmu-miR-21a-5p | MIMAT0000530 | Dnm1l | 74006 | HITS-CLIP | 23142080 | Unspecified |
| mmu-miR-21a-5p | MIMAT0000530 | Dnm2 | 13430 | HITS-CLIP | 23142080\|23597149 | Unspecified |
| mmu-miR-21a-5p | MIMAT0000530 | Dnmt1 | 13433 | HITS-CLIP, Microarrays//HITS-CLIP//Microa | 37986224\|23142080\|25263593\|31433 | Unspecified |
| mmu-miR-21a-5p | MIMAT0000530 | Dnmt3a | 13435 | HITS-CLIP | 23142080 | Unspecified |
| mmu-miR-21a-5p | MIMAT0000530 | Dnph1 | 381101 | Microarrays | 37986224\|25263593 | Unspecified |
| mmu-miR-21a-5p | MIMAT0000530 | Dntt | 21673 | HITS-CLIP | 23142080 | Unspecified |
| mmu-miR-21a-5p | MIMAT0000530 | Dock1 | 330662 | HITS-CLIP | 23597149 | Unspecified |
| mmu-miR-21a-5p | MIMAT0000530 | Dock10 | 210293 | Microarrays//HITS-CLIP | 37986224\|25263593\|23142080 | Unspecified |
| mmu-miR-21a-5p | MIMAT0000530 | Dock11 | 75974 | HITS-CLIP | 23142080 | Unspecified |
| mmu-miR-21a-5p | MIMAT0000530 | Dock2 | 94176 | HITS-CLIP | 23142080\|29167373 | Unspecified |
| mmu-miR-21a-5p | MIMAT0000530 | Dock4 | 238130 | HITS-CLIP | 29167373 | Unspecified |
| mmu-miR-21a-5p | MIMAT0000530 | Dock7 | 67299 | HITS-CLIP, Microarrays//HITS-CLIP | 37986224\|23597149 | Unspecified |
| mmu-miR-21a-5p | MIMAT0000530 | Dock8 | 76088 | HITS-CLIP | 23142080\|29167373 | Unspecified |
| mmu-miR-21a-5p | MIMAT0000530 | Dok2 | 13449 | Microarrays | 37986224 | Unspecified |
| mmu-miR-21a-5p | MIMAT0000530 | Dop1a | 23033 | HITS-CLIP//Microarrays | 23142080\|25263593 | Unspecified |
| mmu-miR-21a-5p | MIMAT0000530 | Dop1b | 9980 | HITS-CLIP | 23142080\|29167373\|23597149 | Unspecified |
| mmu-miR-21a-5p | MIMAT0000530 | Dot1l | 208266 | HITS-CLIP, Microarrays//Microarrays//HITS | 37986224\|25263593\|23142080\|29167 | Unspecified |
| mmu-miR-21a-5p | MIMAT0000530 | Dph6 | 66632 | HITS-CLIP | 23142080 | Unspecified |
| mmu-miR-21a-5p | MIMAT0000530 | Dpm1 | 13480 | HITS-CLIP | 29167373 | Unspecified |
| mmu-miR-21a-5p | MIMAT0000530 | Dpp10 | 57628 | HITS-CLIP | 19536157 | Unspecified |
| mmu-miR-21a-5p | MIMAT0000530 | Dpp3 | 75221 | HITS-CLIP | 23597149 | Unspecified |
| mmu-miR-21a-5p | MIMAT0000530 | Dpp4 | 13482 | HITS-CLIP | 23597149 | Unspecified |
| mmu-miR-21a-5p | MIMAT0000530 | Dpp6 | 13483 | HITS-CLIP | 37986224 | Unspecified |

| | | | | | | |
|---|---|---|---|---|---|---|
| mmu-miR-21a-5p | MIMAT0000530 | Dpp8 | 74388 | HITS-CLIP | 23142080\|29167373 | Unspecified |
| mmu-miR-21a-5p | MIMAT0000530 | Dpp9 | 224897 | HITS-CLIP | 23597149\|23142080 | Unspecified |
| mmu-miR-21a-5p | MIMAT0000530 | Dpy19l1 | 244745 | HITS-CLIP | 23597149\|23142080\|26203562 | Unspecified |
| mmu-miR-21a-5p | MIMAT0000530 | Dpy19l4 | 381510 | HITS-CLIP | 23142080 | Unspecified |
| mmu-miR-21a-5p | MIMAT0000530 | Dpyd | 99586 | HITS-CLIP | 23597149 | Unspecified |
| mmu-miR-21a-5p | MIMAT0000530 | Dpys | 64705 | Microarrays | 37986224\|25263593 | Unspecified |
| mmu-miR-21a-5p | MIMAT0000530 | Dpysl2 | 12934 | HITS-CLIP | 23142080\|29167373 | Unspecified |
| mmu-miR-21a-5p | MIMAT0000530 | Dr1 | 13486 | HITS-CLIP | 31433990\|29167373\|23142080 | Unspecified |
| mmu-miR-21a-5p | MIMAT0000530 | Dram2 | 67171 | HITS-CLIP | 23597149\|23142080 | Unspecified |
| mmu-miR-21a-5p | MIMAT0000530 | Drd1 | 24316 | HITS-CLIP | 19536157 | Unspecified |
| mmu-miR-21a-5p | MIMAT0000530 | Drg1 | 13494 | HITS-CLIP | 23142080 | Unspecified |
| mmu-miR-21a-5p | MIMAT0000530 | Drg2 | 13495 | HITS-CLIP | 23142080\|29167373 | Unspecified |
| mmu-miR-21a-5p | MIMAT0000530 | Dscc1 | 72107 | Microarrays | 37986224\|25263593 | Unspecified |
| mmu-miR-21a-5p | MIMAT0000530 | Dse | 212898 | Microarrays | 37986224 | Unspecified |
| mmu-miR-21a-5p | MIMAT0000530 | Dst | 13518 | HITS-CLIP | 19536157 | Unspecified |
| mmu-miR-21a-5p | MIMAT0000530 | Dstn | 56431 | HITS-CLIP | 37986224\|23597149\|29167373 | Unspecified |
| mmu-miR-21a-5p | MIMAT0000530 | Dstyk | 213452 | HITS-CLIP | 37986224\|23597149\|29167373\|23142 | Unspecified |
| mmu-miR-21a-5p | MIMAT0000530 | Dtl | 76843 | Microarrays//HITS-CLIP | 37986224\|25263593\|23142080 | Unspecified |
| mmu-miR-21a-5p | MIMAT0000530 | Dtwd1 | 69185 | HITS-CLIP | 23142080 | Unspecified |
| mmu-miR-21a-5p | MIMAT0000530 | Dtwd2 | 68857 | HITS-CLIP | 23142080 | Unspecified |
| mmu-miR-21a-5p | MIMAT0000530 | Dtx3l | 209200 | HITS-CLIP | 23597149\|23142080\|29167373 | Unspecified |
| mmu-miR-21a-5p | MIMAT0000530 | Dus1l | 68730 | HITS-CLIP | 23142080 | Unspecified |
| mmu-miR-21a-5p | MIMAT0000530 | Dusp1 | 19252 | Microarrays//HITS-CLIP | 37986224\|23597149\|25263593 | Unspecified |
| mmu-miR-21a-5p | MIMAT0000530 | Dusp10 | 63953 | HITS-CLIP | 23142080 | Unspecified |
| mmu-miR-21a-5p | MIMAT0000530 | Dusp11 | 72102 | qCLASH//HITS-CLIP | 31393953\|29167373\|23142080 | Unspecified |
| mmu-miR-21a-5p | MIMAT0000530 | Dusp16 | 70686 | Microarrays//HITS-CLIP | 37986224\|25263593\|23142080 | Unspecified |
| mmu-miR-21a-5p | MIMAT0000530 | Dusp2 | 13537 | qCLASH//HITS-CLIP | 31393953\|23142080 | Unspecified |
| mmu-miR-21a-5p | MIMAT0000530 | Dusp22 | 105352 | HITS-CLIP | 23142080 | Unspecified |
| mmu-miR-21a-5p | MIMAT0000530 | Dusp3 | 72349 | Microarrays | 37986224\|25263593 | Unspecified |
| mmu-miR-21a-5p | MIMAT0000530 | Dusp4 | 319520 | Microarrays | 37986224\|25263593 | Unspecified |
| mmu-miR-21a-5p | MIMAT0000530 | Dusp5 | 240672 | HITS-CLIP | 31433990\|23142080 | Unspecified |
| mmu-miR-21a-5p | MIMAT0000530 | Dusp6 | 67603 | Microarrays | 37986224\|25263593 | Unspecified |
| mmu-miR-21a-5p | MIMAT0000530 | Dusp8 | 18218 | HITS-CLIP | 23597149\|19536157 | Unspecified |
| mmu-miR-21a-5p | MIMAT0000530 | Dut | 110074 | HITS-CLIP | 23142080 | Unspecified |
| mmu-miR-21a-5p | MIMAT0000530 | Dvl3 | 13544 | HITS-CLIP | 23142080 | Unspecified |
| mmu-miR-21a-5p | MIMAT0000530 | Dync1h1 | 13424 | HITS-CLIP | 37986224\|23597149\|23142080\|29167 | Unspecified |
| mmu-miR-21a-5p | MIMAT0000530 | Dync1i2 | 13427 | HITS-CLIP | 23142080 | Unspecified |
| mmu-miR-21a-5p | MIMAT0000530 | Dync1li1 | 235661 | HITS-CLIP | 23142080 | Unspecified |
| mmu-miR-21a-5p | MIMAT0000530 | Dync1li2 | 234663 | HITS-CLIP | 29167373 | Unspecified |

| | | | | | | |
|---|---|---|---|---|---|---|
| mmu-miR-21a-5p | MIMAT0000530 | Dync2h1 | 110350 | HITS-CLIP | 23142080 | Unspecified |
| mmu-miR-21a-5p | MIMAT0000530 | Dynll2 | 68097 | Microarrays//HITS-CLIP | 37986224\|25263593\|23597149\|23142 | Unspecified |
| mmu-miR-21a-5p | MIMAT0000530 | Dynlrb1 | 67068 | HITS-CLIP | 23142080 | Unspecified |
| mmu-miR-21a-5p | MIMAT0000530 | Dynlt5 | 200132 | Microarrays | 25263593 | Unspecified |
| mmu-miR-21a-5p | MIMAT0000530 | Dyrk1a | 13548 | HITS-CLIP | 23597149\|23142080 | Unspecified |
| mmu-miR-21a-5p | MIMAT0000530 | Dyrk1b | 13549 | HITS-CLIP | 23597149 | Unspecified |
| mmu-miR-21a-5p | MIMAT0000530 | Dyrk2 | 69181 | HITS-CLIP | 23597149\|23142080 | Unspecified |
| mmu-miR-21a-5p | MIMAT0000530 | Dzank1 | 241688 | HITS-CLIP | 19536157 | Unspecified |
| mmu-miR-21a-5p | MIMAT0000530 | Dzip3 | 224170 | HITS-CLIP | 23142080 | Unspecified |
| mmu-miR-21a-5p | MIMAT0000530 | E030030I06Rik | 319887 | Microarrays | 37986224\|25263593 | Unspecified |
| mmu-miR-21a-5p | MIMAT0000530 | E130308A19Rik | 230259 | HITS-CLIP | 23597149\|23142080 | Unspecified |
| mmu-miR-21a-5p | MIMAT0000530 | E2f1 | 13555 | HITS-CLIP | 31433990 | Unspecified |
| mmu-miR-21a-5p | MIMAT0000530 | E2f2 | 242705 | HITS-CLIP//Microarrays | 23142080\|25263593\|23597149 | Unspecified |
| mmu-miR-21a-5p | MIMAT0000530 | E2f3 | 13557 | HITS-CLIP | 37986224\|23142080\|29167373 | Unspecified |
| mmu-miR-21a-5p | MIMAT0000530 | E2f4 | 104394 | HITS-CLIP | 23142080 | Unspecified |
| mmu-miR-21a-5p | MIMAT0000530 | E2f6 | 50496 | Microarrays//HITS-CLIP | 37986224\|25263593\|23142080 | Unspecified |
| mmu-miR-21a-5p | MIMAT0000530 | E2f7 | 52679 | HITS-CLIP, Microarrays//Microarrays//HITS | 37986224\|25263593\|23142080 | Unspecified |
| mmu-miR-21a-5p | MIMAT0000530 | E2f8 | 108961 | Microarrays//HITS-CLIP | 37986224\|25263593\|23142080 | Unspecified |
| mmu-miR-21a-5p | MIMAT0000530 | Eaf1 | 74427 | HITS-CLIP | 23142080 | Unspecified |
| mmu-miR-21a-5p | MIMAT0000530 | Eapp | 66266 | HITS-CLIP | 23142080 | Unspecified |
| mmu-miR-21a-5p | MIMAT0000530 | Ear1 | 13586 | Microarrays | 37986224 | Unspecified |
| mmu-miR-21a-5p | MIMAT0000530 | Ear2 | 13587 | Microarrays | 37986224 | Unspecified |
| mmu-miR-21a-5p | MIMAT0000530 | Ebi3 | 50498 | Microarrays | 37986224 | Unspecified |
| mmu-miR-21a-5p | MIMAT0000530 | Ebna1bp2 | 69072 | HITS-CLIP | 23597149\|23142080 | Unspecified |
| mmu-miR-21a-5p | MIMAT0000530 | Ece1 | 230857 | HITS-CLIP | 19536157\|23142080 | Unspecified |
| mmu-miR-21a-5p | MIMAT0000530 | Echdc3 | 67856 | Microarrays | 37986224\|25263593 | Unspecified |
| mmu-miR-21a-5p | MIMAT0000530 | Echs1 | 93747 | HITS-CLIP | 23597149 | Unspecified |
| mmu-miR-21a-5p | MIMAT0000530 | Ecpas | 23392 | HITS-CLIP | 31433990\|23142080 | Unspecified |
| mmu-miR-21a-5p | MIMAT0000530 | Ecsit | 26940 | HITS-CLIP | 23597149 | Unspecified |
| mmu-miR-21a-5p | MIMAT0000530 | Ect2 | 13605 | Microarrays//HITS-CLIP | 37986224\|25263593\|23142080 | Unspecified |
| mmu-miR-21a-5p | MIMAT0000530 | Eda2r | 245527 | Microarrays | 37986224\|25263593 | Unspecified |
| mmu-miR-21a-5p | MIMAT0000530 | Edc3 | 353190 | HITS-CLIP | 23142080 | Unspecified |
| mmu-miR-21a-5p | MIMAT0000530 | Edc4 | 234699 | HITS-CLIP | 23597149 | Unspecified |
| mmu-miR-21a-5p | MIMAT0000530 | Edem1 | 192193 | HITS-CLIP | 23142080 | Unspecified |
| mmu-miR-21a-5p | MIMAT0000530 | Edem2 | 108687 | HITS-CLIP | 23142080 | Unspecified |
| mmu-miR-21a-5p | MIMAT0000530 | Edem3 | 66967 | HITS-CLIP | 37986224\|19536157\|23142080\|29167 | Unspecified |
| mmu-miR-21a-5p | MIMAT0000530 | Edil3 | 13612 | Microarrays | 37986224 | Unspecified |
| mmu-miR-21a-5p | MIMAT0000530 | Ednrb | 13618 | Microarrays//HITS-CLIP | 37986224\|25263593\|19536157 | Unspecified |
| mmu-miR-21a-5p | MIMAT0000530 | Edrf1 | 214764 | HITS-CLIP | 23142080 | Unspecified |

| | | | | | | |
|---|---|---|---|---|---|---|
| mmu-miR-21a-5p | MIMAT0000530 | Eea1 | 216238 | HITS-CLIP | 37986224\|29167373\|23142080 | Unspecified |
| mmu-miR-21a-5p | MIMAT0000530 | Eed | 13626 | HITS-CLIP | 23142080 | Unspecified |
| mmu-miR-21a-5p | MIMAT0000530 | Eef1a1 | 13627 | HITS-CLIP | 23142080 | Unspecified |
| mmu-miR-21a-5p | MIMAT0000530 | Eef1akmt2 | 399818 | HITS-CLIP | 23142080 | Unspecified |
| mmu-miR-21a-5p | MIMAT0000530 | Eef1b2 | 55949 | HITS-CLIP//qCLASH | 23142080\|31393953\|23597149 | Unspecified |
| mmu-miR-21a-5p | MIMAT0000530 | Eef1e1 | 66143 | Microarrays//HITS-CLIP | 37986224\|25263593\|23142080 | Unspecified |
| mmu-miR-21a-5p | MIMAT0000530 | Eef2 | 13629 | HITS-CLIP//qCLASH | 23597149\|31393953\|23142080 | Unspecified |
| mmu-miR-21a-5p | MIMAT0000530 | Eef2k | 13631 | HITS-CLIP | 23142080 | Unspecified |
| mmu-miR-21a-5p | MIMAT0000530 | Eepd1 | 67484 | Microarrays | 37986224 | Unspecified |
| mmu-miR-21a-5p | MIMAT0000530 | Efcab11 | 78767 | Microarrays | 37986224\|25263593 | Unspecified |
| mmu-miR-21a-5p | MIMAT0000530 | Efhd2 | 27984 | HITS-CLIP | 29167373 | Unspecified |
| mmu-miR-21a-5p | MIMAT0000530 | Efna1 | 13636 | HITS-CLIP | 26203562 | Unspecified |
| mmu-miR-21a-5p | MIMAT0000530 | Efr3a | 76740 | HITS-CLIP//qCLASH | 23142080\|31393953 | Unspecified |
| mmu-miR-21a-5p | MIMAT0000530 | Eftud2 | 20624 | HITS-CLIP | 23142080 | Unspecified |
| mmu-miR-21a-5p | MIMAT0000530 | Egfr | 13649 | HITS-CLIP | 23597149 | Unspecified |
| mmu-miR-21a-5p | MIMAT0000530 | Egr1 | 13653 | Microarrays | 37986224 | Unspecified |
| mmu-miR-21a-5p | MIMAT0000530 | Egr2 | 13654 | Microarrays | 37986224 | Unspecified |
| mmu-miR-21a-5p | MIMAT0000530 | Egr3 | 13655 | Microarrays//HITS-CLIP | 37986224\|23142080 | Unspecified |
| mmu-miR-21a-5p | MIMAT0000530 | Ehbp1l1 | 114601 | HITS-CLIP | 23142080 | Unspecified |
| mmu-miR-21a-5p | MIMAT0000530 | Ehd1 | 13660 | HITS-CLIP | 23597149\|23142080\|29167373 | Unspecified |
| mmu-miR-21a-5p | MIMAT0000530 | Ehd3 | 57440 | HITS-CLIP | 23597149 | Unspecified |
| mmu-miR-21a-5p | MIMAT0000530 | Ehhadh | 74147 | HITS-CLIP | 23597149 | Unspecified |
| mmu-miR-21a-5p | MIMAT0000530 | Ehmt1 | 77683 | HITS-CLIP | 23142080\|23597149 | Unspecified |
| mmu-miR-21a-5p | MIMAT0000530 | Ehmt2 | 110147 | HITS-CLIP | 23142080 | Unspecified |
| mmu-miR-21a-5p | MIMAT0000530 | Ei24 | 13663 | HITS-CLIP | 23142080 | Unspecified |
| mmu-miR-21a-5p | MIMAT0000530 | Eid1 | 58521 | HITS-CLIP//qCLASH | 23142080\|31393953\|29167373 | Unspecified |
| mmu-miR-21a-5p | MIMAT0000530 | Eif1 | 20918 | HITS-CLIP | 23142080\|29167373 | Unspecified |
| mmu-miR-21a-5p | MIMAT0000530 | Eif1a | 13664 | HITS-CLIP | 37986224\|23142080 | Unspecified |
| mmu-miR-21a-5p | MIMAT0000530 | Eif1ax | 66235 | HITS-CLIP | 23142080\|29167373 | Unspecified |
| mmu-miR-21a-5p | MIMAT0000530 | Eif1b | 68969 | HITS-CLIP | 23142080 | Unspecified |
| mmu-miR-21a-5p | MIMAT0000530 | Eif2a | 229317 | HITS-CLIP | 37986224\|23142080 | Unspecified |
| mmu-miR-21a-5p | MIMAT0000530 | Eif2ak1 | 15467 | HITS-CLIP | 23597149\|23142080 | Unspecified |
| mmu-miR-21a-5p | MIMAT0000530 | Eif2ak3 | 13666 | HITS-CLIP | 23597149\|23142080 | Unspecified |
| mmu-miR-21a-5p | MIMAT0000530 | Eif2ak4 | 27103 | HITS-CLIP | 23142080 | Unspecified |
| mmu-miR-21a-5p | MIMAT0000530 | Eif2b1 | 209354 | HITS-CLIP | 23142080 | Unspecified |
| mmu-miR-21a-5p | MIMAT0000530 | Eif2b2 | 217715 | HITS-CLIP | 23142080 | Unspecified |
| mmu-miR-21a-5p | MIMAT0000530 | Eif2b5 | 224045 | HITS-CLIP | 23597149\|23142080 | Unspecified |
| mmu-miR-21a-5p | MIMAT0000530 | Eif2s3x | 26905 | HITS-CLIP | 23142080 | Unspecified |
| mmu-miR-21a-5p | MIMAT0000530 | Eif2s3y | 26908 | HITS-CLIP | 23142080 | Unspecified |

| | | | | | | |
|---|---|---|---|---|---|---|
| mmu-miR-21a-5p | MIMAT0000530 | Eif3a | 13669 | HITS-CLIP//qCLASH | 23597149\|23142080\|31393953 | Unspecified |
| mmu-miR-21a-5p | MIMAT0000530 | Eif3b | 27979 | Microarrays | 37986224\|25263593 | Unspecified |
| mmu-miR-21a-5p | MIMAT0000530 | Eif3c | 56347 | HITS-CLIP//qCLASH | 23597149\|31393953\|23142080\|29167 | Unspecified |
| mmu-miR-21a-5p | MIMAT0000530 | Eif3d | 55944 | HITS-CLIP | 23597149 | Unspecified |
| mmu-miR-21a-5p | MIMAT0000530 | Eif3f | 66085 | HITS-CLIP | 23142080 | Unspecified |
| mmu-miR-21a-5p | MIMAT0000530 | Eif3i | 54709 | HITS-CLIP | 23142080 | Unspecified |
| mmu-miR-21a-5p | MIMAT0000530 | Eif3l | 223691 | HITS-CLIP | 23597149\|23142080\|29167373 | Unspecified |
| mmu-miR-21a-5p | MIMAT0000530 | Eif4a1 | 13681 | HITS-CLIP | 29167373 | Unspecified |
| mmu-miR-21a-5p | MIMAT0000530 | Eif4a2 | 13682 | HITS-CLIP | 37986224\|23597149\|23142080\|29167 | Unspecified |
| mmu-miR-21a-5p | MIMAT0000530 | Eif4b | 75705 | HITS-CLIP | 23142080\|23597149 | Unspecified |
| mmu-miR-21a-5p | MIMAT0000530 | Eif4e | 13684 | HITS-CLIP | 37986224\|23142080 | Unspecified |
| mmu-miR-21a-5p | MIMAT0000530 | Eif4ebp2 | 13688 | HITS-CLIP | 23597149\|23142080 | Unspecified |
| mmu-miR-21a-5p | MIMAT0000530 | Eif4g1 | 208643 | HITS-CLIP//qCLASH | 23597149\|31393953\|23142080\|29167 | Unspecified |
| mmu-miR-21a-5p | MIMAT0000530 | Eif4g2 | 13690 | HITS-CLIP | 37986224\|23142080\|19536157\|23597 | Unspecified |
| mmu-miR-21a-5p | MIMAT0000530 | Eif4g3 | 230861 | HITS-CLIP | 23142080\|29167373 | Unspecified |
| mmu-miR-21a-5p | MIMAT0000530 | Eif4h | 22384 | HITS-CLIP | 23597149 | Unspecified |
| mmu-miR-21a-5p | MIMAT0000530 | Eif5 | 217869 | HITS-CLIP | 37986224\|23142080\|23597149\|29167 | Unspecified |
| mmu-miR-21a-5p | MIMAT0000530 | Eif5a | 276770 | HITS-CLIP | 23142080 | Unspecified |
| mmu-miR-21a-5p | MIMAT0000530 | Eif5b | 226982 | HITS-CLIP | 37986224\|23597149\|23142080\|29167 | Unspecified |
| mmu-miR-21a-5p | MIMAT0000530 | Eif6 | 16418 | HITS-CLIP | 37986224\|23142080 | Unspecified |
| mmu-miR-21a-5p | MIMAT0000530 | Elac2 | 68626 | HITS-CLIP | 23142080 | Unspecified |
| mmu-miR-21a-5p | MIMAT0000530 | Elavl1 | 15568 | HITS-CLIP | 23142080 | Unspecified |
| mmu-miR-21a-5p | MIMAT0000530 | Elf1 | 13709 | HITS-CLIP | 23142080 | Unspecified |
| mmu-miR-21a-5p | MIMAT0000530 | Elf2 | 69257 | HITS-CLIP | 37986224\|23142080\|29167373 | Unspecified |
| mmu-miR-21a-5p | MIMAT0000530 | Elk3 | 13713 | HITS-CLIP | 23142080\|29167373 | Unspecified |
| mmu-miR-21a-5p | MIMAT0000530 | Elk4 | 13714 | HITS-CLIP | 23142080 | Unspecified |
| mmu-miR-21a-5p | MIMAT0000530 | Ell2 | 192657 | Microarrays//HITS-CLIP | 37986224\|25263593\|23142080\|29167 | Unspecified |
| mmu-miR-21a-5p | MIMAT0000530 | Elmo1 | 140580 | HITS-CLIP | 23142080\|29167373 | Unspecified |
| mmu-miR-21a-5p | MIMAT0000530 | Elmo2 | 140579 | HITS-CLIP | 23142080 | Unspecified |
| mmu-miR-21a-5p | MIMAT0000530 | Elmod2 | 244548 | HITS-CLIP | 23142080 | Unspecified |
| mmu-miR-21a-5p | MIMAT0000530 | Elmsan1 | 238317 | HITS-CLIP | 23142080 | Unspecified |
| mmu-miR-21a-5p | MIMAT0000530 | Eloa | 6924 | HITS-CLIP | 23597149\|23142080 | Unspecified |
| mmu-miR-21a-5p | MIMAT0000530 | Eloc | 6921 | HITS-CLIP | 23597149 | Unspecified |
| mmu-miR-21a-5p | MIMAT0000530 | Elovl1 | 54325 | HITS-CLIP | 23597149\|23142080 | Unspecified |
| mmu-miR-21a-5p | MIMAT0000530 | Elovl2 | 54326 | HITS-CLIP | 23597149 | Unspecified |
| mmu-miR-21a-5p | MIMAT0000530 | Elovl6 | 170439 | HITS-CLIP | 23142080 | Unspecified |
| mmu-miR-21a-5p | MIMAT0000530 | Elp1 | 37455 | HITS-CLIP | 23597149\|23142080 | Unspecified |
| mmu-miR-21a-5p | MIMAT0000530 | Elp3 | 74195 | HITS-CLIP | 23597149\|23142080 | Unspecified |
| mmu-miR-21a-5p | MIMAT0000530 | Elp5 | 54351 | HITS-CLIP | 23597149 | Unspecified |

| | | | | | | |
|---|---|---|---|---|---|---|
| mmu-miR-21a-5p | MIMAT0000530 | Emb | 13723 | Microarrays//HITS-CLIP | 37986224\|23142080 | Unspecified |
| mmu-miR-21a-5p | MIMAT0000530 | Emc2 | 66736 | Microarrays//HITS-CLIP | 37986224\|25263593\|23142080 | Unspecified |
| mmu-miR-21a-5p | MIMAT0000530 | Emc4 | 68032 | Microarrays | 37986224\|25263593 | Unspecified |
| mmu-miR-21a-5p | MIMAT0000530 | Emc7 | 73024 | HITS-CLIP | 23142080 | Unspecified |
| mmu-miR-21a-5p | MIMAT0000530 | Eme2 | 193838 | Microarrays//HITS-CLIP | 37986224\|25263593\|23597149 | Unspecified |
| mmu-miR-21a-5p | MIMAT0000530 | Eml4 | 78798 | HITS-CLIP | 23142080\|29167373 | Unspecified |
| mmu-miR-21a-5p | MIMAT0000530 | Emp1 | 13730 | Microarrays | 37986224\|25263593 | Unspecified |
| mmu-miR-21a-5p | MIMAT0000530 | Emsy | 56946 | HITS-CLIP | 23142080 | Unspecified |
| mmu-miR-21a-5p | MIMAT0000530 | En2 | 13799 | HITS-CLIP | 37986224 | Unspecified |
| mmu-miR-21a-5p | MIMAT0000530 | Enc1 | 13803 | Microarrays//HITS-CLIP | 37986224\|23142080 | Unspecified |
| mmu-miR-21a-5p | MIMAT0000530 | Enpp1 | 18605 | Microarrays//HITS-CLIP | 37986224\|25263593\|23142080 | Unspecified |
| mmu-miR-21a-5p | MIMAT0000530 | Enpp4 | 224794 | Microarrays//HITS-CLIP | 37986224\|19536157 | Unspecified |
| mmu-miR-21a-5p | MIMAT0000530 | Ensa | 56205 | HITS-CLIP | 23597149\|19536157\|29167373 | Unspecified |
| mmu-miR-21a-5p | MIMAT0000530 | Entpd1 | 12495 | Microarrays | 37986224 | Unspecified |
| mmu-miR-21a-5p | MIMAT0000530 | Entpd5 | 12499 | HITS-CLIP | 37986224\|23597149\|23142080 | Unspecified |
| mmu-miR-21a-5p | MIMAT0000530 | Entpd6 | 12497 | HITS-CLIP | 23142080\|29167373 | Unspecified |
| mmu-miR-21a-5p | MIMAT0000530 | Eny2 | 223527 | HITS-CLIP | 23597149 | Unspecified |
| mmu-miR-21a-5p | MIMAT0000530 | Eogt | 101351 | HITS-CLIP | 23142080 | Unspecified |
| mmu-miR-21a-5p | MIMAT0000530 | Eomes | 13813 | HITS-CLIP | 23142080 | Unspecified |
| mmu-miR-21a-5p | MIMAT0000530 | Ep300 | 328572 | HITS-CLIP | 23142080 | Unspecified |
| mmu-miR-21a-5p | MIMAT0000530 | Ep400 | 75560 | HITS-CLIP | 23142080\|29167373 | Unspecified |
| mmu-miR-21a-5p | MIMAT0000530 | Epas1 | 13819 | HITS-CLIP | 31433990 | Unspecified |
| mmu-miR-21a-5p | MIMAT0000530 | Epb41 | 269587 | HITS-CLIP | 29167373 | Unspecified |
| mmu-miR-21a-5p | MIMAT0000530 | Epb41l1 | 13821 | HITS-CLIP | 19536157\|29167373 | Unspecified |
| mmu-miR-21a-5p | MIMAT0000530 | Epb41l3 | 23136 | Microarrays | 25263593 | Unspecified |
| mmu-miR-21a-5p | MIMAT0000530 | Epb41l5 | 226352 | HITS-CLIP | 23597149 | Unspecified |
| mmu-miR-21a-5p | MIMAT0000530 | Epc2 | 227867 | HITS-CLIP | 23142080 | Unspecified |
| mmu-miR-21a-5p | MIMAT0000530 | Epcam | 17075 | HITS-CLIP | 37986224\|23142080 | Unspecified |
| mmu-miR-21a-5p | MIMAT0000530 | Epg5 | 100502841 | HITS-CLIP | 29167373 | Unspecified |
| mmu-miR-21a-5p | MIMAT0000530 | Epha4 | 13838 | HITS-CLIP | 19536157 | Unspecified |
| mmu-miR-21a-5p | MIMAT0000530 | Ephx1 | 13849 | Microarrays | 37986224 | Unspecified |
| mmu-miR-21a-5p | MIMAT0000530 | Ephx2 | 13850 | HITS-CLIP | 23597149 | Unspecified |
| mmu-miR-21a-5p | MIMAT0000530 | Epn1 | 13854 | HITS-CLIP | 23597149 | Unspecified |
| mmu-miR-21a-5p | MIMAT0000530 | Eppk1 | 3857 | HITS-CLIP | 23597149 | Unspecified |
| mmu-miR-21a-5p | MIMAT0000530 | Eprs | 107508 | HITS-CLIP | 23597149\|29167373\|23142080 | Unspecified |
| mmu-miR-21a-5p | MIMAT0000530 | Eps15 | 13858 | HITS-CLIP, Microarrays//HITS-CLIP | 37986224\|19536157\|23142080\|29167 | Unspecified |
| mmu-miR-21a-5p | MIMAT0000530 | Eps8 | 13860 | Microarrays | 37986224 | Unspecified |
| mmu-miR-21a-5p | MIMAT0000530 | Epsti1 | 108670 | Microarrays//HITS-CLIP | 37986224\|23142080 | Unspecified |
| mmu-miR-21a-5p | MIMAT0000530 | Erap1 | 80898 | HITS-CLIP | 23142080 | Unspecified |

| | | | | | | |
|---|---|---|---|---|---|---|
| mmu-miR-21a-5p | MIMAT0000530 | Erbin | 55914 | HITS-CLIP | 19536157\|23142080\|29167373 | Unspecified |
| mmu-miR-21a-5p | MIMAT0000530 | Erc1 | 111173 | HITS-CLIP | 23597149\|29167373 | Unspecified |
| mmu-miR-21a-5p | MIMAT0000530 | Ercc3 | 13872 | HITS-CLIP | 23597149 | Unspecified |
| mmu-miR-21a-5p | MIMAT0000530 | Ercc6 | 319955 | HITS-CLIP | 23142080\|29167373 | Unspecified |
| mmu-miR-21a-5p | MIMAT0000530 | Ercc6l | 236930 | Microarrays//HITS-CLIP | 37986224\|25263593\|23142080 | Unspecified |
| mmu-miR-21a-5p | MIMAT0000530 | Erdr1 | 170942 | Microarrays | 37986224 | Unspecified |
| mmu-miR-21a-5p | MIMAT0000530 | Erg | 13876 | HITS-CLIP | 23597149 | Unspecified |
| mmu-miR-21a-5p | MIMAT0000530 | Ergic1 | 67458 | Microarrays//HITS-CLIP | 37986224\|25263593\|29167373 | Unspecified |
| mmu-miR-21a-5p | MIMAT0000530 | Ergic3 | 66366 | HITS-CLIP | 23142080 | Unspecified |
| mmu-miR-21a-5p | MIMAT0000530 | Eri1 | 67276 | qCLASH//HITS-CLIP | 31393953\|23142080 | Unspecified |
| mmu-miR-21a-5p | MIMAT0000530 | Erlin1 | 226144 | HITS-CLIP | 23597149\|23142080\|29167373 | Unspecified |
| mmu-miR-21a-5p | MIMAT0000530 | Erlin2 | 244373 | HITS-CLIP | 23597149\|23142080 | Unspecified |
| mmu-miR-21a-5p | MIMAT0000530 | Ermard | 381062 | Microarrays//HITS-CLIP | 37986224\|23142080 | Unspecified |
| mmu-miR-21a-5p | MIMAT0000530 | Ern1 | 78943 | HITS-CLIP | 23597149 | Unspecified |
| mmu-miR-21a-5p | MIMAT0000530 | Ero1l | 50527 | HITS-CLIP | 37986224\|23142080 | Unspecified |
| mmu-miR-21a-5p | MIMAT0000530 | Ero1lb | 67475 | HITS-CLIP | 23142080 | Unspecified |
| mmu-miR-21a-5p | MIMAT0000530 | Erp29 | 67397 | HITS-CLIP | 23142080\|23597149\|29167373 | Unspecified |
| mmu-miR-21a-5p | MIMAT0000530 | Erp44 | 76299 | HITS-CLIP | 23142080\|29167373 | Unspecified |
| mmu-miR-21a-5p | MIMAT0000530 | Errfi1 | 74155 | HITS-CLIP | 23597149 | Unspecified |
| mmu-miR-21a-5p | MIMAT0000530 | Esco1 | 77805 | HITS-CLIP | 23142080 | Unspecified |
| mmu-miR-21a-5p | MIMAT0000530 | Esco2 | 71988 | Microarrays | 37986224\|25263593 | Unspecified |
| mmu-miR-21a-5p | MIMAT0000530 | Espl1 | 105988 | Microarrays//HITS-CLIP | 37986224\|25263593\|23142080 | Unspecified |
| mmu-miR-21a-5p | MIMAT0000530 | Espn | 56226 | HITS-CLIP | 23142080 | Unspecified |
| mmu-miR-21a-5p | MIMAT0000530 | Esr1 | 13982 | Microarrays//HITS-CLIP | 37986224\|23597149 | Unspecified |
| mmu-miR-21a-5p | MIMAT0000530 | Esyt1 | 23943 | HITS-CLIP | 23597149 | Unspecified |
| mmu-miR-21a-5p | MIMAT0000530 | Esyt2 | 52635 | HITS-CLIP | 37986224\|23597149\|19536157\|2314 | Unspecified |
| mmu-miR-21a-5p | MIMAT0000530 | Etaa1 | 68145 | HITS-CLIP | 23142080 | Unspecified |
| mmu-miR-21a-5p | MIMAT0000530 | Etf1 | 225363 | HITS-CLIP | 23142080 | Unspecified |
| mmu-miR-21a-5p | MIMAT0000530 | Etfa | 110842 | HITS-CLIP | 29167373 | Unspecified |
| mmu-miR-21a-5p | MIMAT0000530 | Etfb | 110826 | HITS-CLIP | 23597149 | Unspecified |
| mmu-miR-21a-5p | MIMAT0000530 | Etfdh | 66841 | HITS-CLIP | 23142080 | Unspecified |
| mmu-miR-21a-5p | MIMAT0000530 | Etnk1 | 75320 | HITS-CLIP | 37986224\|23597149\|23142080\|2916 | Unspecified |
| mmu-miR-21a-5p | MIMAT0000530 | Ets1 | 23871 | HITS-CLIP | 37986224\|23142080 | Unspecified |
| mmu-miR-21a-5p | MIMAT0000530 | Ets2 | 23872 | Microarrays//HITS-CLIP | 37986224\|25263593\|23597149\|2314 | Unspecified |
| mmu-miR-21a-5p | MIMAT0000530 | Etv3 | 27049 | HITS-CLIP | 23142080 | Unspecified |
| mmu-miR-21a-5p | MIMAT0000530 | Etv5 | 104156 | Microarrays//HITS-CLIP | 37986224\|25263593\|23142080 | Unspecified |
| mmu-miR-21a-5p | MIMAT0000530 | Etv6 | 14011 | HITS-CLIP | 23142080 | Unspecified |
| mmu-miR-21a-5p | MIMAT0000530 | Eva1b | 230752 | Microarrays | 37986224 | Unspecified |
| mmu-miR-21a-5p | MIMAT0000530 | Evi2 | 14017 | HITS-CLIP | 23142080\|29167373 | Unspecified |

| | | | | | | | |
|---|---|---|---|---|---|---|---|
| mmu-miR-21a-5p | MIMAT0000530 | Evi5 | 14020 | HITS-CLIP | | 37986224 | Unspecified |
| mmu-miR-21a-5p | MIMAT0000530 | Evl | 14026 | HITS-CLIP | | 23142080 | Unspecified |
| mmu-miR-21a-5p | MIMAT0000530 | Ewsr1 | 14030 | HITS-CLIP | | 23142080 | Unspecified |
| mmu-miR-21a-5p | MIMAT0000530 | Exd1 | 241624 | Microarrays | | 37986224 | Unspecified |
| mmu-miR-21a-5p | MIMAT0000530 | Exo1 | 26909 | qCLASH | | 31393953 | Unspecified |
| mmu-miR-21a-5p | MIMAT0000530 | Exoc2 | 66482 | HITS-CLIP | | 23142080 | Unspecified |
| mmu-miR-21a-5p | MIMAT0000530 | Exoc3l4 | 74190 | Microarrays | 37986224\|25263593 | | Unspecified |
| mmu-miR-21a-5p | MIMAT0000530 | Exoc4 | 20336 | HITS-CLIP | | 23142080 | Unspecified |
| mmu-miR-21a-5p | MIMAT0000530 | Exoc5 | 105504 | HITS-CLIP | 23142080\|29167373 | | Unspecified |
| mmu-miR-21a-5p | MIMAT0000530 | Exoc6 | 107371 | HITS-CLIP | | 23142080 | Unspecified |
| mmu-miR-21a-5p | MIMAT0000530 | Exoc6b | 75914 | HITS-CLIP | | 23142080 | Unspecified |
| mmu-miR-21a-5p | MIMAT0000530 | Exosc10 | 50912 | HITS-CLIP | 23597149\|23142080 | | Unspecified |
| mmu-miR-21a-5p | MIMAT0000530 | Exosc8 | 69639 | HITS-CLIP | | 23142080 | Unspecified |
| mmu-miR-21a-5p | MIMAT0000530 | Exosc9 | 50911 | HITS-CLIP | | 23142080 | Unspecified |
| mmu-miR-21a-5p | MIMAT0000530 | Eya1 | 14048 | Microarrays | 37986224\|25263593 | | Unspecified |
| mmu-miR-21a-5p | MIMAT0000530 | Ezh1 | 14055 | HITS-CLIP | 23597149\|23142080\|29167373 | | Unspecified |
| mmu-miR-21a-5p | MIMAT0000530 | Ezh2 | 14056 | Microarrays//HITS-CLIP | 37986224\|25263593\|23142080 | | Unspecified |
| mmu-miR-21a-5p | MIMAT0000530 | Ezr | 22350 | HITS-CLIP | 23597149\|23142080 | | Unspecified |
| mmu-miR-21a-5p | MIMAT0000530 | F12 | 58992 | HITS-CLIP | | 23597149 | Unspecified |
| mmu-miR-21a-5p | MIMAT0000530 | F13a1 | 74145 | Microarrays | | 37986224 | Unspecified |
| mmu-miR-21a-5p | MIMAT0000530 | F13b | 14060 | HITS-CLIP | | 23597149 | Unspecified |
| mmu-miR-21a-5p | MIMAT0000530 | F2 | 14061 | HITS-CLIP | | 23597149 | Unspecified |
| mmu-miR-21a-5p | MIMAT0000530 | F2r | 14062 | HITS-CLIP | | 31433990 | Unspecified |
| mmu-miR-21a-5p | MIMAT0000530 | F2rl2 | 14064 | HITS-CLIP | | 23142080 | Unspecified |
| mmu-miR-21a-5p | MIMAT0000530 | F3 | 14066 | Microarrays | 37986224\|25263593 | | Unspecified |
| mmu-miR-21a-5p | MIMAT0000530 | F5 | 14067 | HITS-CLIP | | 23597149 | Unspecified |
| mmu-miR-21a-5p | MIMAT0000530 | F9 | 14071 | Microarrays | 37986224\|25263593 | | Unspecified |
| mmu-miR-21a-5p | MIMAT0000530 | Fabp3 | 14077 | Microarrays | 37986224\|25263593 | | Unspecified |
| mmu-miR-21a-5p | MIMAT0000530 | Fabp7 | 12140 | Microarrays | 37986224\|25263593 | | Unspecified |
| mmu-miR-21a-5p | MIMAT0000530 | Fads1 | 76267 | Microarrays | | 37986224 | Unspecified |
| mmu-miR-21a-5p | MIMAT0000530 | Faf2 | 76577 | HITS-CLIP | 23597149\|23142080 | | Unspecified |
| mmu-miR-21a-5p | MIMAT0000530 | Fah | 14085 | Microarrays//HITS-CLIP | 37986224\|23597149\|25263593\|23142 | | Unspecified |
| mmu-miR-21a-5p | MIMAT0000530 | Fahd1 | 68636 | Microarrays | | 37986224 | Unspecified |
| mmu-miR-21a-5p | MIMAT0000530 | Fam102a | 98952 | Microarrays//HITS-CLIP | 37986224\|25263593\|19536157\|23142 | | Unspecified |
| mmu-miR-21a-5p | MIMAT0000530 | Fam102b | 329739 | Microarrays//HITS-CLIP | 37986224\|23142080 | | Unspecified |
| mmu-miR-21a-5p | MIMAT0000530 | Fam107b | 66540 | HITS-CLIP | 37986224\|23142080\|26203562 | | Unspecified |
| mmu-miR-21a-5p | MIMAT0000530 | Fam111a | 107373 | Microarrays//HITS-CLIP | 37986224\|25263593\|23142080\|29167 | | Unspecified |
| mmu-miR-21a-5p | MIMAT0000530 | Fam114a1 | 68303 | HITS-CLIP | | 23142080 | Unspecified |
| mmu-miR-21a-5p | MIMAT0000530 | Fam114a2 | 67726 | HITS-CLIP | 37986224\|23597149 | | Unspecified |

| | | | | | | |
|---|---|---|---|---|---|---|
| mmu-miR-21a-5p | MIMAT0000530 | Fam117b | 72750 | HITS-CLIP | 23597149 | Unspecified |
| mmu-miR-21a-5p | MIMAT0000530 | Fam120a | 218236 | qCLASH//HITS-CLIP | 31393953\|23142080 | Unspecified |
| mmu-miR-21a-5p | MIMAT0000530 | Fam120b | 67544 | HITS-CLIP | 23142080 | Unspecified |
| mmu-miR-21a-5p | MIMAT0000530 | Fam122b | 78755 | HITS-CLIP | 23142080 | Unspecified |
| mmu-miR-21a-5p | MIMAT0000530 | Fam126a | 84652 | HITS-CLIP | 29167373\|23142080 | Unspecified |
| mmu-miR-21a-5p | MIMAT0000530 | Fam126b | 213056 | HITS-CLIP | 37986224\|23142080\|23597149 | Unspecified |
| mmu-miR-21a-5p | MIMAT0000530 | Fam129a | 63913 | HITS-CLIP | 29167373 | Unspecified |
| mmu-miR-21a-5p | MIMAT0000530 | Fam129b | 227737 | Microarrays//HITS-CLIP | 37986224\|23142080\|29167373 | Unspecified |
| mmu-miR-21a-5p | MIMAT0000530 | Fam129c | 100037278 | Microarrays | 37986224 | Unspecified |
| mmu-miR-21a-5p | MIMAT0000530 | Fam133b | 68152 | HITS-CLIP | 23142080 | Unspecified |
| mmu-miR-21a-5p | MIMAT0000530 | Fam13b | 225358 | Microarrays | 37986224 | Unspecified |
| mmu-miR-21a-5p | MIMAT0000530 | Fam160a1 | 229488 | HITS-CLIP | 23597149 | Unspecified |
| mmu-miR-21a-5p | MIMAT0000530 | Fam160a2 | 74349 | Microarrays | 37986224 | Unspecified |
| mmu-miR-21a-5p | MIMAT0000530 | Fam160b1 | 226252 | HITS-CLIP | 23142080 | Unspecified |
| mmu-miR-21a-5p | MIMAT0000530 | Fam160b2 | 239170 | HITS-CLIP | 23597149 | Unspecified |
| mmu-miR-21a-5p | MIMAT0000530 | Fam168a | 319604 | HITS-CLIP | 37986224\|23142080 | Unspecified |
| mmu-miR-21a-5p | MIMAT0000530 | Fam171a1 | 269233 | Microarrays//HITS-CLIP | 37986224\|25263593\|19536157 | Unspecified |
| mmu-miR-21a-5p | MIMAT0000530 | Fam189b | 68521 | HITS-CLIP | 23142080 | Unspecified |
| mmu-miR-21a-5p | MIMAT0000530 | Fam193a | 231128 | HITS-CLIP | 23142080\|23597149 | Unspecified |
| mmu-miR-21a-5p | MIMAT0000530 | Fam193b | 212483 | HITS-CLIP | 29167373 | Unspecified |
| mmu-miR-21a-5p | MIMAT0000530 | Fam199x | 245622 | HITS-CLIP | 23142080 | Unspecified |
| mmu-miR-21a-5p | MIMAT0000530 | Fam208a | 218850 | HITS-CLIP | 23142080 | Unspecified |
| mmu-miR-21a-5p | MIMAT0000530 | Fam208b | 54906 | HITS-CLIP | 23142080 | Unspecified |
| mmu-miR-21a-5p | MIMAT0000530 | Fam20b | 215015 | HITS-CLIP | 19536157\|23142080\|29167373 | Unspecified |
| mmu-miR-21a-5p | MIMAT0000530 | Fam20c | 80752 | Microarrays//HITS-CLIP | 37986224\|25263593\|29167373 | Unspecified |
| mmu-miR-21a-5p | MIMAT0000530 | Fam210b | 67017 | HITS-CLIP | 23597149 | Unspecified |
| mmu-miR-21a-5p | MIMAT0000530 | Fam214a | 235493 | HITS-CLIP, Microarrays//HITS-CLIP | 37986224\|23142080\|23597149 | Unspecified |
| mmu-miR-21a-5p | MIMAT0000530 | Fam222b | 216971 | HITS-CLIP | 23142080\|23597149 | Unspecified |
| mmu-miR-21a-5p | MIMAT0000530 | Fam234b | 57613 | Microarrays//HITS-CLIP | 25263593\|23142080 | Unspecified |
| mmu-miR-21a-5p | MIMAT0000530 | Fam3c | 27999 | HITS-CLIP | 23142080 | Unspecified |
| mmu-miR-21a-5p | MIMAT0000530 | Fam43a | 224093 | Microarrays | 37986224 | Unspecified |
| mmu-miR-21a-5p | MIMAT0000530 | Fam45a | 67894 | HITS-CLIP | 23142080 | Unspecified |
| mmu-miR-21a-5p | MIMAT0000530 | Fam49b | 223601 | HITS-CLIP | 23597149\|29167373\|23142080 | Unspecified |
| mmu-miR-21a-5p | MIMAT0000530 | Fam57a | 116972 | Microarrays | 37986224 | Unspecified |
| mmu-miR-21a-5p | MIMAT0000530 | Fam72a | 108900 | Microarrays | 37986224\|25263593 | Unspecified |
| mmu-miR-21a-5p | MIMAT0000530 | Fam76b | 72826 | HITS-CLIP | 37986224\|29167373\|23142080 | Unspecified |
| mmu-miR-21a-5p | MIMAT0000530 | Fam78a | 241303 | HITS-CLIP | 23142080 | Unspecified |
| mmu-miR-21a-5p | MIMAT0000530 | Fam83h | 105732 | HITS-CLIP | 23142080 | Unspecified |
| mmu-miR-21a-5p | MIMAT0000530 | Fam84b | 157638 | HITS-CLIP | 23142080 | Unspecified |

| | | | | | | |
|---|---|---|---|---|---|---|
| mmu-miR-21a-5p | MIMAT0000530 | Fam8a1 | 51439 | HITS-CLIP | 23597149\|23142080 | Unspecified |
| mmu-miR-21a-5p | MIMAT0000530 | Fam91a1 | 210998 | HITS-CLIP | 23142080\|29167373 | Unspecified |
| mmu-miR-21a-5p | MIMAT0000530 | Fam98a | 72722 | HITS-CLIP | 23597149\|23142080 | Unspecified |
| mmu-miR-21a-5p | MIMAT0000530 | Fancd2 | 211651 | HITS-CLIP | 23142080 | Unspecified |
| mmu-miR-21a-5p | MIMAT0000530 | Fanci | 208836 | HITS-CLIP | 23142080 | Unspecified |
| mmu-miR-21a-5p | MIMAT0000530 | Fancm | 104806 | HITS-CLIP | 23142080 | Unspecified |
| mmu-miR-21a-5p | MIMAT0000530 | Far1 | 67420 | HITS-CLIP | 23142080\|29167373 | Unspecified |
| mmu-miR-21a-5p | MIMAT0000530 | Far2 | 330450 | HITS-CLIP | 37986224 | Unspecified |
| mmu-miR-21a-5p | MIMAT0000530 | Farp2 | 227377 | HITS-CLIP | 23142080 | Unspecified |
| mmu-miR-21a-5p | MIMAT0000530 | Farsb | 23874 | qCLASH//HITS-CLIP | 31393953\|29167373 | Unspecified |
| mmu-miR-21a-5p | MIMAT0000530 | Fasl | 14103 | Luciferase assay//Western blot//Northern | 20404348\|23142080 | Unspecified |
| mmu-miR-21a-5p | MIMAT0000530 | Fasn | 14104 | HITS-CLIP//qCLASH | 37986224\|23597149\|19536157\|31393 | Unspecified |
| mmu-miR-21a-5p | MIMAT0000530 | Fastkd2 | 75619 | HITS-CLIP | 37986224 | Unspecified |
| mmu-miR-21a-5p | MIMAT0000530 | Fastkd3 | 69577 | HITS-CLIP | 23142080 | Unspecified |
| mmu-miR-21a-5p | MIMAT0000530 | Fastkd5 | 380601 | HITS-CLIP | 23142080 | Unspecified |
| mmu-miR-21a-5p | MIMAT0000530 | Fat1 | 14107 | Microarrays | 37986224\|25263593 | Unspecified |
| mmu-miR-21a-5p | MIMAT0000530 | Fblim1 | 74202 | Microarrays | 37986224\|25263593 | Unspecified |
| mmu-miR-21a-5p | MIMAT0000530 | Fbln2 | 14115 | Microarrays | 37986224\|25263593 | Unspecified |
| mmu-miR-21a-5p | MIMAT0000530 | Fbn1 | 14118 | Microarrays | 37986224\|25263593 | Unspecified |
| mmu-miR-21a-5p | MIMAT0000530 | Fbp1 | 14121 | Microarrays | 37986224 | Unspecified |
| mmu-miR-21a-5p | MIMAT0000530 | Fbxl16 | 214931 | HITS-CLIP | 19536157 | Unspecified |
| mmu-miR-21a-5p | MIMAT0000530 | Fbxl2 | 72179 | HITS-CLIP, Microarrays//Microarrays | 37986224\|25263593 | Unspecified |
| mmu-miR-21a-5p | MIMAT0000530 | Fbxl20 | 72194 | HITS-CLIP | 23597149\|19536157\|23142080 | Unspecified |
| mmu-miR-21a-5p | MIMAT0000530 | Fbxl3 | 50789 | HITS-CLIP | 37986224\|23597149\|23142080 | Unspecified |
| mmu-miR-21a-5p | MIMAT0000530 | Fbxl5 | 242960 | HITS-CLIP | 23597149\|23142080 | Unspecified |
| mmu-miR-21a-5p | MIMAT0000530 | Fbxo11 | 225055 | HITS-CLIP | 37986224\|23142080\|29167373 | Unspecified |
| mmu-miR-21a-5p | MIMAT0000530 | Fbxo21 | 231670 | HITS-CLIP | 23142080 | Unspecified |
| mmu-miR-21a-5p | MIMAT0000530 | Fbxo28 | 67948 | HITS-CLIP | 37986224\|23142080 | Unspecified |
| mmu-miR-21a-5p | MIMAT0000530 | Fbxo3 | 57443 | HITS-CLIP | 23142080\|29167373 | Unspecified |
| mmu-miR-21a-5p | MIMAT0000530 | Fbxo32 | 67731 | HITS-CLIP | 23142080 | Unspecified |
| mmu-miR-21a-5p | MIMAT0000530 | Fbxo33 | 70611 | HITS-CLIP | 23142080\|29167373 | Unspecified |
| mmu-miR-21a-5p | MIMAT0000530 | Fbxo45 | 268882 | HITS-CLIP | 29167373\|23142080 | Unspecified |
| mmu-miR-21a-5p | MIMAT0000530 | Fbxo5 | 67141 | Microarrays | 37986224\|25263593 | Unspecified |
| mmu-miR-21a-5p | MIMAT0000530 | Fbxo8 | 50753 | HITS-CLIP | 23142080 | Unspecified |
| mmu-miR-21a-5p | MIMAT0000530 | Fbxo9 | 71538 | HITS-CLIP | 23597149 | Unspecified |
| mmu-miR-21a-5p | MIMAT0000530 | Fbxw11 | 103583 | HITS-CLIP | 23142080 | Unspecified |
| mmu-miR-21a-5p | MIMAT0000530 | Fbxw7 | 50754 | HITS-CLIP | 23142080\|29167373 | Unspecified |
| mmu-miR-21a-5p | MIMAT0000530 | Fcgr1 | 14129 | Microarrays | 37986224 | Unspecified |
| mmu-miR-21a-5p | MIMAT0000530 | Fcgr2b | 14130 | Microarrays | 37986224 | Unspecified |

| | | | | | | |
|---|---|---|---|---|---|---|
| mmu-miR-21a-5p | MIMAT0000530 | Fcgrt | 14132 | Microarrays | 37986224 | Unspecified |
| mmu-miR-21a-5p | MIMAT0000530 | Fcho2 | 218503 | HITS-CLIP, Microarrays//HITS-CLIP | 37986224\|19536157\|29167373\|23142 | Unspecified |
| mmu-miR-21a-5p | MIMAT0000530 | Fchsd2 | 207278 | HITS-CLIP | 23597149 | Unspecified |
| mmu-miR-21a-5p | MIMAT0000530 | Fcna | 14133 | Microarrays | 37986224\|25263593 | Unspecified |
| mmu-miR-21a-5p | MIMAT0000530 | Fcrls | 80891 | Microarrays | 37986224 | Unspecified |
| mmu-miR-21a-5p | MIMAT0000530 | Fdx1 | 14148 | HITS-CLIP | 23597149\|23142080 | Unspecified |
| mmu-miR-21a-5p | MIMAT0000530 | Fdxr | 14149 | HITS-CLIP | 23142080 | Unspecified |
| mmu-miR-21a-5p | MIMAT0000530 | Fech | 14151 | HITS-CLIP | 23597149 | Unspecified |
| mmu-miR-21a-5p | MIMAT0000530 | Fem1b | 14155 | HITS-CLIP//qCLASH | 37986224\|31393953\|23142080 | Unspecified |
| mmu-miR-21a-5p | MIMAT0000530 | Fem1c | 240263 | Microarrays//HITS-CLIP | 37986224\|23142080\|29167373 | Unspecified |
| mmu-miR-21a-5p | MIMAT0000530 | Fen1 | 14156 | HITS-CLIP | 37986224\|23142080 | Unspecified |
| mmu-miR-21a-5p | MIMAT0000530 | Fer | 14158 | HITS-CLIP | 29167373 | Unspecified |
| mmu-miR-21a-5p | MIMAT0000530 | Fermt2 | 218952 | HITS-CLIP | 37986224\|23597149 | Unspecified |
| mmu-miR-21a-5p | MIMAT0000530 | Fes | 14159 | Microarrays | 37986224 | Unspecified |
| mmu-miR-21a-5p | MIMAT0000530 | Fetub | 59083 | HITS-CLIP | 23597149 | Unspecified |
| mmu-miR-21a-5p | MIMAT0000530 | Fez2 | 225020 | HITS-CLIP | 23142080\|29167373 | Unspecified |
| mmu-miR-21a-5p | MIMAT0000530 | Fga | 14161 | HITS-CLIP | 23597149 | Unspecified |
| mmu-miR-21a-5p | MIMAT0000530 | Fgd2 | 26382 | Microarrays | 37986224 | Unspecified |
| mmu-miR-21a-5p | MIMAT0000530 | Fgd3 | 30938 | HITS-CLIP | 29167373 | Unspecified |
| mmu-miR-21a-5p | MIMAT0000530 | Fgd4 | 224014 | HITS-CLIP | 37986224 | Unspecified |
| mmu-miR-21a-5p | MIMAT0000530 | Fgd6 | 13998 | HITS-CLIP | 29167373 | Unspecified |
| mmu-miR-21a-5p | MIMAT0000530 | Fgf13 | 14168 | HITS-CLIP | 23142080 | Unspecified |
| mmu-miR-21a-5p | MIMAT0000530 | Fgf2 | 14173 | HITS-CLIP | 23142080 | Unspecified |
| mmu-miR-21a-5p | MIMAT0000530 | Fgfbp3 | 72514 | Microarrays | 37986224\|25263593 | Unspecified |
| mmu-miR-21a-5p | MIMAT0000530 | Fgfr1op2 | 67529 | HITS-CLIP | 23142080 | Unspecified |
| mmu-miR-21a-5p | MIMAT0000530 | Fgfr3 | 14184 | HITS-CLIP | 37986224 | Unspecified |
| mmu-miR-21a-5p | MIMAT0000530 | Fgg | 99571 | HITS-CLIP | 23597149 | Unspecified |
| mmu-miR-21a-5p | MIMAT0000530 | Fgl2 | 14190 | Microarrays//HITS-CLIP | 37986224\|23142080 | Unspecified |
| mmu-miR-21a-5p | MIMAT0000530 | Fh1 | 14194 | Microarrays//HITS-CLIP | 37986224\|23142080 | Unspecified |
| mmu-miR-21a-5p | MIMAT0000530 | Fhad1 | 329977 | Microarrays | 37986224\|25263593 | Unspecified |
| mmu-miR-21a-5p | MIMAT0000530 | Fhod1 | 234686 | HITS-CLIP | 29167373 | Unspecified |
| mmu-miR-21a-5p | MIMAT0000530 | Fibp | 58249 | HITS-CLIP | 23597149\|23142080 | Unspecified |
| mmu-miR-21a-5p | MIMAT0000530 | Fignl1 | 60530 | Microarrays//HITS-CLIP | 37986224\|23142080\|25263593 | Unspecified |
| mmu-miR-21a-5p | MIMAT0000530 | Filip1l | 78749 | Microarrays//HITS-CLIP | 37986224\|23142080\|25263593\|29167 | Unspecified |
| mmu-miR-21a-5p | MIMAT0000530 | Fip1l1 | 66899 | HITS-CLIP | 23597149\|23142080 | Unspecified |
| mmu-miR-21a-5p | MIMAT0000530 | Fitm2 | 228859 | HITS-CLIP | 23142080 | Unspecified |
| mmu-miR-21a-5p | MIMAT0000530 | Fiz1 | 23877 | HITS-CLIP | 23142080\|23597149\|29167373 | Unspecified |
| mmu-miR-21a-5p | MIMAT0000530 | Fkbp15 | 338355 | HITS-CLIP | 29167373 | Unspecified |
| mmu-miR-21a-5p | MIMAT0000530 | Fkbp1a | 14225 | HITS-CLIP | 23142080 | Unspecified |

<table>
<tr><td>mmu-miR-21a-5p</td><td>MIMAT0000530</td><td>Fkbp2</td><td>14227</td><td>HITS-CLIP</td><td>23142080</td><td>Unspecified</td></tr>
<tr><td>mmu-miR-21a-5p</td><td>MIMAT0000530</td><td>Fkbp3</td><td>30795</td><td>HITS-CLIP</td><td>23142080</td><td>Unspecified</td></tr>
<tr><td>mmu-miR-21a-5p</td><td>MIMAT0000530</td><td>Fkbp4</td><td>14228</td><td>HITS-CLIP</td><td>23597149</td><td>Unspecified</td></tr>
<tr><td>mmu-miR-21a-5p</td><td>MIMAT0000530</td><td>Fktn</td><td>246179</td><td>HITS-CLIP</td><td>23142080</td><td>Unspecified</td></tr>
<tr><td>mmu-miR-21a-5p</td><td>MIMAT0000530</td><td>Fli1</td><td>14247</td><td>HITS-CLIP</td><td>23142080|29167373</td><td>Unspecified</td></tr>
<tr><td>mmu-miR-21a-5p</td><td>MIMAT0000530</td><td>Flna</td><td>192176</td><td>HITS-CLIP</td><td>23142080|29167373</td><td>Unspecified</td></tr>
<tr><td>mmu-miR-21a-5p</td><td>MIMAT0000530</td><td>Flnb</td><td>286940</td><td>HITS-CLIP</td><td>23597149|23142080</td><td>Unspecified</td></tr>
<tr><td>mmu-miR-21a-5p</td><td>MIMAT0000530</td><td>Flot1</td><td>14251</td><td>HITS-CLIP</td><td>23142080</td><td>Unspecified</td></tr>
<tr><td>mmu-miR-21a-5p</td><td>MIMAT0000530</td><td>Flrt2</td><td>399558</td><td>Microarrays//HITS-CLIP</td><td>37986224|25263593|19536157|29167</td><td>Unspecified</td></tr>
<tr><td>mmu-miR-21a-5p</td><td>MIMAT0000530</td><td>Flt1</td><td>14254</td><td>HITS-CLIP, Microarrays//Microarrays//HITS</td><td>37986224|25263593|19536157</td><td>Unspecified</td></tr>
<tr><td>mmu-miR-21a-5p</td><td>MIMAT0000530</td><td>Flt4</td><td>14257</td><td>HITS-CLIP</td><td>23597149</td><td>Unspecified</td></tr>
<tr><td>mmu-miR-21a-5p</td><td>MIMAT0000530</td><td>Flvcr1</td><td>28982</td><td>HITS-CLIP</td><td>29167373</td><td>Unspecified</td></tr>
<tr><td>mmu-miR-21a-5p</td><td>MIMAT0000530</td><td>Fmn1</td><td>14260</td><td>Microarrays//HITS-CLIP</td><td>37986224|25263593|29167373</td><td>Unspecified</td></tr>
<tr><td>mmu-miR-21a-5p</td><td>MIMAT0000530</td><td>Fmnl1</td><td>57778</td><td>HITS-CLIP</td><td>23142080</td><td>Unspecified</td></tr>
<tr><td>mmu-miR-21a-5p</td><td>MIMAT0000530</td><td>Fmnl2</td><td>71409</td><td>HITS-CLIP</td><td>26203562</td><td>Unspecified</td></tr>
<tr><td>mmu-miR-21a-5p</td><td>MIMAT0000530</td><td>Fmnl3</td><td>22379</td><td>Microarrays</td><td>37986224|25263593</td><td>Unspecified</td></tr>
<tr><td>mmu-miR-21a-5p</td><td>MIMAT0000530</td><td>Fmo5</td><td>14263</td><td>Microarrays</td><td>37986224</td><td>Unspecified</td></tr>
<tr><td>mmu-miR-21a-5p</td><td>MIMAT0000530</td><td>Fmr1</td><td>14265</td><td>qCLASH//HITS-CLIP</td><td>31393953|23142080|29167373</td><td>Unspecified</td></tr>
<tr><td>mmu-miR-21a-5p</td><td>MIMAT0000530</td><td>Fn1</td><td>14268</td><td>Microarrays//HITS-CLIP</td><td>37986224|25263593|23597149</td><td>Unspecified</td></tr>
<tr><td>mmu-miR-21a-5p</td><td>MIMAT0000530</td><td>Fnbp1</td><td>14269</td><td>HITS-CLIP</td><td>29167373</td><td>Unspecified</td></tr>
<tr><td>mmu-miR-21a-5p</td><td>MIMAT0000530</td><td>Fnbp1l</td><td>214459</td><td>HITS-CLIP</td><td>23142080|29167373</td><td>Unspecified</td></tr>
<tr><td>mmu-miR-21a-5p</td><td>MIMAT0000530</td><td>Fnbp4</td><td>55935</td><td>HITS-CLIP</td><td>37986224|23142080</td><td>Unspecified</td></tr>
<tr><td>mmu-miR-21a-5p</td><td>MIMAT0000530</td><td>Fndc3a</td><td>319448</td><td>HITS-CLIP</td><td>23597149|23142080|29167373</td><td>Unspecified</td></tr>
<tr><td>mmu-miR-21a-5p</td><td>MIMAT0000530</td><td>Fndc3b</td><td>72007</td><td>HITS-CLIP</td><td>23142080|29167373</td><td>Unspecified</td></tr>
<tr><td>mmu-miR-21a-5p</td><td>MIMAT0000530</td><td>Fnip1</td><td>216742</td><td>HITS-CLIP</td><td>29167373|23142080</td><td>Unspecified</td></tr>
<tr><td>mmu-miR-21a-5p</td><td>MIMAT0000530</td><td>Fnip2</td><td>329679</td><td>HITS-CLIP</td><td>29167373</td><td>Unspecified</td></tr>
<tr><td>mmu-miR-21a-5p</td><td>MIMAT0000530</td><td>Fnta</td><td>14272</td><td>HITS-CLIP</td><td>23142080</td><td>Unspecified</td></tr>
<tr><td>mmu-miR-21a-5p</td><td>MIMAT0000530</td><td>Focad</td><td>230393</td><td>HITS-CLIP</td><td>23142080</td><td>Unspecified</td></tr>
<tr><td>mmu-miR-21a-5p</td><td>MIMAT0000530</td><td>Folr2</td><td>14276</td><td>Microarrays</td><td>37986224|25263593</td><td>Unspecified</td></tr>
<tr><td>mmu-miR-21a-5p</td><td>MIMAT0000530</td><td>Fopnl</td><td>66086</td><td>HITS-CLIP</td><td>23142080</td><td>Unspecified</td></tr>
<tr><td>mmu-miR-21a-5p</td><td>MIMAT0000530</td><td>Fos</td><td>14281</td><td>HITS-CLIP</td><td>23597149</td><td>Unspecified</td></tr>
<tr><td>mmu-miR-21a-5p</td><td>MIMAT0000530</td><td>Fosb</td><td>14282</td><td>Microarrays</td><td>37986224</td><td>Unspecified</td></tr>
<tr><td>mmu-miR-21a-5p</td><td>MIMAT0000530</td><td>Fosl2</td><td>14284</td><td>HITS-CLIP</td><td>23142080</td><td>Unspecified</td></tr>
<tr><td>mmu-miR-21a-5p</td><td>MIMAT0000530</td><td>Foxj3</td><td>230700</td><td>HITS-CLIP</td><td>23142080</td><td>Unspecified</td></tr>
<tr><td>mmu-miR-21a-5p</td><td>MIMAT0000530</td><td>Foxk1</td><td>17425</td><td>HITS-CLIP</td><td>37986224</td><td>Unspecified</td></tr>
<tr><td>mmu-miR-21a-5p</td><td>MIMAT0000530</td><td>Foxk2</td><td>68837</td><td>HITS-CLIP</td><td>19536157</td><td>Unspecified</td></tr>
<tr><td>mmu-miR-21a-5p</td><td>MIMAT0000530</td><td>Foxn2</td><td>14236</td><td>HITS-CLIP</td><td>23597149|23142080|29167373</td><td>Unspecified</td></tr>
<tr><td>mmu-miR-21a-5p</td><td>MIMAT0000530</td><td>Foxo1</td><td>56458</td><td>HITS-CLIP</td><td>31433990|23142080</td><td>Unspecified</td></tr>
<tr><td>mmu-miR-21a-5p</td><td>MIMAT0000530</td><td>Foxo3</td><td>56484</td><td>HITS-CLIP//qCLASH</td><td>37986224|23597149|31393953|1953</td><td>Unspecified</td></tr>
</table>

| | | | | | | |
|---|---|---|---|---|---|---|
| mmu-miR-21a-5p | MIMAT0000530 | Foxp1 | 108655 | Microarrays//HITS-CLIP | 37986224\|31433990 | Unspecified |
| mmu-miR-21a-5p | MIMAT0000530 | Foxp4 | 74123 | HITS-CLIP | 23142080 | Unspecified |
| mmu-miR-21a-5p | MIMAT0000530 | Foxred1 | 235169 | HITS-CLIP | 23142080 | Unspecified |
| mmu-miR-21a-5p | MIMAT0000530 | Frat1 | 10023 | HITS-CLIP | 23142080 | Unspecified |
| mmu-miR-21a-5p | MIMAT0000530 | Frat2 | 212398 | Microarrays | 37986224 | Unspecified |
| mmu-miR-21a-5p | MIMAT0000530 | Frg1 | 14300 | HITS-CLIP | 23142080 | Unspecified |
| mmu-miR-21a-5p | MIMAT0000530 | Frmd4a | 209630 | Microarrays//HITS-CLIP | 37986224\|23142080 | Unspecified |
| mmu-miR-21a-5p | MIMAT0000530 | Frmd4b | 232288 | Microarrays//HITS-CLIP | 37986224\|23597149 | Unspecified |
| mmu-miR-21a-5p | MIMAT0000530 | Frmd6 | 319710 | Microarrays | 37986224 | Unspecified |
| mmu-miR-21a-5p | MIMAT0000530 | Frmd8 | 67457 | HITS-CLIP | 23142080 | Unspecified |
| mmu-miR-21a-5p | MIMAT0000530 | Frs2 | 327826 | HITS-CLIP | 19536157\|23142080 | Unspecified |
| mmu-miR-21a-5p | MIMAT0000530 | Fry | 320365 | HITS-CLIP | 37986224 | Unspecified |
| mmu-miR-21a-5p | MIMAT0000530 | Fryl | 72313 | HITS-CLIP | 23142080 | Unspecified |
| mmu-miR-21a-5p | MIMAT0000530 | Fscn1 | 14086 | HITS-CLIP, Microarrays//Microarrays | 37986224\|25263593 | Unspecified |
| mmu-miR-21a-5p | MIMAT0000530 | Fsd1l | 319636 | HITS-CLIP | 23142080 | Unspecified |
| mmu-miR-21a-5p | MIMAT0000530 | Fsip1 | 71313 | Microarrays | 37986224\|25263593 | Unspecified |
| mmu-miR-21a-5p | MIMAT0000530 | Fstl1 | 14314 | Microarrays | 37986224\|25263593 | Unspecified |
| mmu-miR-21a-5p | MIMAT0000530 | Fth1 | 14319 | HITS-CLIP | 37986224\|23142080\|23597149\|29167 | Unspecified |
| mmu-miR-21a-5p | MIMAT0000530 | Fto | 26383 | HITS-CLIP | 23597149\|29167373 | Unspecified |
| mmu-miR-21a-5p | MIMAT0000530 | Fubp1 | 51886 | HITS-CLIP | 37986224\|23142080\|29167373 | Unspecified |
| mmu-miR-21a-5p | MIMAT0000530 | Fubp3 | 320267 | HITS-CLIP | 23142080 | Unspecified |
| mmu-miR-21a-5p | MIMAT0000530 | Fuca2 | 66848 | HITS-CLIP | 23597149\|23142080 | Unspecified |
| mmu-miR-21a-5p | MIMAT0000530 | Fundc1 | 72018 | HITS-CLIP | 23142080 | Unspecified |
| mmu-miR-21a-5p | MIMAT0000530 | Furin | 18550 | HITS-CLIP | 23597149\|26203562 | Unspecified |
| mmu-miR-21a-5p | MIMAT0000530 | Fus | 233908 | HITS-CLIP | 37986224\|29167373 | Unspecified |
| mmu-miR-21a-5p | MIMAT0000530 | Fxr1 | 14359 | HITS-CLIP | 37986224\|23142080\|23597149\|29167 | Unspecified |
| mmu-miR-21a-5p | MIMAT0000530 | Fxr2 | 23879 | HITS-CLIP | 23142080 | Unspecified |
| mmu-miR-21a-5p | MIMAT0000530 | Fyb | 23880 | HITS-CLIP | 29167373 | Unspecified |
| mmu-miR-21a-5p | MIMAT0000530 | Fyco1 | 17281 | HITS-CLIP | 23142080\|29167373 | Unspecified |
| mmu-miR-21a-5p | MIMAT0000530 | Fyn | 14360 | HITS-CLIP | 23142080 | Unspecified |
| mmu-miR-21a-5p | MIMAT0000530 | Fzd3 | 14365 | HITS-CLIP | 19536157 | Unspecified |
| mmu-miR-21a-5p | MIMAT0000530 | Fzd7 | 14369 | Microarrays//HITS-CLIP | 37986224\|25263593\|23142080 | Unspecified |
| mmu-miR-21a-5p | MIMAT0000530 | G0s2 | 14373 | HITS-CLIP | 23597149 | Unspecified |
| mmu-miR-21a-5p | MIMAT0000530 | G2e3 | 217558 | Microarrays//HITS-CLIP | 37986224\|25263593\|23142080 | Unspecified |
| mmu-miR-21a-5p | MIMAT0000530 | G3bp2 | 23881 | HITS-CLIP | 23142080\|23597149 | Unspecified |
| mmu-miR-21a-5p | MIMAT0000530 | G6pc | 14377 | HITS-CLIP | 23597149 | Unspecified |
| mmu-miR-21a-5p | MIMAT0000530 | G6pdx | 14381 | HITS-CLIP | 23597149 | Unspecified |
| mmu-miR-21a-5p | MIMAT0000530 | Gab1 | 14388 | HITS-CLIP, Microarrays//HITS-CLIP | 37986224\|23597149 | Unspecified |
| mmu-miR-21a-5p | MIMAT0000530 | Gab2 | 14389 | HITS-CLIP | 29167373 | Unspecified |

| | | | | | | |
|---|---|---|---|---|---|---|
| mmu-miR-21a-5p | MIMAT0000530 | Gabarapl2 | 93739 | HITS-CLIP | 23142080 | Unspecified |
| mmu-miR-21a-5p | MIMAT0000530 | Gabbr2 | 242425 | HITS-CLIP | 19536157 | Unspecified |
| mmu-miR-21a-5p | MIMAT0000530 | Gabpa | 14390 | HITS-CLIP | 37986224\|23597149\|23142080\|29167 | Unspecified |
| mmu-miR-21a-5p | MIMAT0000530 | Gabpb2 | 213054 | HITS-CLIP | 23142080 | Unspecified |
| mmu-miR-21a-5p | MIMAT0000530 | Gabra1 | 14394 | HITS-CLIP | 19536157 | Unspecified |
| mmu-miR-21a-5p | MIMAT0000530 | Gabrb1 | 14400 | HITS-CLIP | 19536157 | Unspecified |
| mmu-miR-21a-5p | MIMAT0000530 | Gad1 | 14415 | HITS-CLIP | 19536157 | Unspecified |
| mmu-miR-21a-5p | MIMAT0000530 | Gadd45g | 23882 | Microarrays | 37986224\|25263593 | Unspecified |
| mmu-miR-21a-5p | MIMAT0000530 | Gak | 231580 | Microarrays//HITS-CLIP | 37986224\|23597149\|29167373 | Unspecified |
| mmu-miR-21a-5p | MIMAT0000530 | Galc | 14420 | Microarrays//HITS-CLIP | 37986224\|25263593\|29167373 | Unspecified |
| mmu-miR-21a-5p | MIMAT0000530 | Gale | 74246 | HITS-CLIP | 23142080\|29167373 | Unspecified |
| mmu-miR-21a-5p | MIMAT0000530 | Galk1 | 14635 | HITS-CLIP | 23142080 | Unspecified |
| mmu-miR-21a-5p | MIMAT0000530 | Galk2 | 69976 | HITS-CLIP | 23142080 | Unspecified |
| mmu-miR-21a-5p | MIMAT0000530 | Galnt12 | 230145 | HITS-CLIP, Microarrays//Microarrays//HITS | 37986224\|25263593\|23142080 | Unspecified |
| mmu-miR-21a-5p | MIMAT0000530 | Galnt16 | 108760 | HITS-CLIP | 37986224 | Unspecified |
| mmu-miR-21a-5p | MIMAT0000530 | Galnt2 | 108148 | HITS-CLIP | 29167373 | Unspecified |
| mmu-miR-21a-5p | MIMAT0000530 | Galnt3 | 14425 | HITS-CLIP | 23142080 | Unspecified |
| mmu-miR-21a-5p | MIMAT0000530 | Galnt4 | 14426 | HITS-CLIP | 29167373 | Unspecified |
| mmu-miR-21a-5p | MIMAT0000530 | Galnt6 | 207839 | HITS-CLIP | 23142080 | Unspecified |
| mmu-miR-21a-5p | MIMAT0000530 | Galnt7 | 108150 | HITS-CLIP | 19536157\|23142080 | Unspecified |
| mmu-miR-21a-5p | MIMAT0000530 | Galnt9 | 231605 | Microarrays | 37986224\|25263593 | Unspecified |
| mmu-miR-21a-5p | MIMAT0000530 | Ganab | 14376 | HITS-CLIP | 23142080 | Unspecified |
| mmu-miR-21a-5p | MIMAT0000530 | Ganc | 76051 | HITS-CLIP | 23142080 | Unspecified |
| mmu-miR-21a-5p | MIMAT0000530 | Gapvd1 | 66691 | HITS-CLIP | 23597149\|23142080\|29167373 | Unspecified |
| mmu-miR-21a-5p | MIMAT0000530 | Gars | 353172 | HITS-CLIP | 23597149\|23142080\|19536157 | Unspecified |
| mmu-miR-21a-5p | MIMAT0000530 | Gart | 14450 | HITS-CLIP | 23597149\|23142080\|29167373 | Unspecified |
| mmu-miR-21a-5p | MIMAT0000530 | Gas2l3 | 237436 | HITS-CLIP | 37986224\|23142080\|29167373 | Unspecified |
| mmu-miR-21a-5p | MIMAT0000530 | Gas7 | 14457 | Microarrays//HITS-CLIP | 37986224\|19536157\|29167373 | Unspecified |
| mmu-miR-21a-5p | MIMAT0000530 | Gask1b | 68659 | Microarrays | 25263593 | Unspecified |
| mmu-miR-21a-5p | MIMAT0000530 | Gata3 | 14462 | qPCR//HITS-CLIP | 37986224\|23416424\|23142080 | Unspecified |
| mmu-miR-21a-5p | MIMAT0000530 | Gatad2a | 234366 | HITS-CLIP | 23142080\|29167373 | Unspecified |
| mmu-miR-21a-5p | MIMAT0000530 | Gatad2b | 229542 | HITS-CLIP | 37986224\|23142080\|29167373 | Unspecified |
| mmu-miR-21a-5p | MIMAT0000530 | Gatd1 | 347862 | HITS-CLIP | 23142080 | Unspecified |
| mmu-miR-21a-5p | MIMAT0000530 | Gatm | 67092 | Microarrays//HITS-CLIP | 37986224\|23142080 | Unspecified |
| mmu-miR-21a-5p | MIMAT0000530 | Gba | 14466 | Microarrays | 37986224\|25263593 | Unspecified |
| mmu-miR-21a-5p | MIMAT0000530 | Gbe1 | 74185 | Microarrays//HITS-CLIP | 37986224\|25263593\|23142080 | Unspecified |
| mmu-miR-21a-5p | MIMAT0000530 | Gbf1 | 107338 | HITS-CLIP | 23142080 | Unspecified |
| mmu-miR-21a-5p | MIMAT0000530 | Gbp2 | 14469 | Microarrays//HITS-CLIP | 37986224\|23142080 | Unspecified |
| mmu-miR-21a-5p | MIMAT0000530 | Gbp2b | 14468 | Microarrays | 25263593 | Unspecified |

| | | | | | | | |
|---|---|---|---|---|---|---|---|
| mmu-miR-21a-5p | MIMAT0000530 | Gbp3 | 55932 | Microarrays | | 37986224 | Unspecified |
| mmu-miR-21a-5p | MIMAT0000530 | Gbp4 | 17472 | HITS-CLIP | 23597149\|23142080 | | Unspecified |
| mmu-miR-21a-5p | MIMAT0000530 | Gbp5 | 229898 | HITS-CLIP, Microarrays//HITS-CLIP | 37986224\|23142080 | | Unspecified |
| mmu-miR-21a-5p | MIMAT0000530 | Gbp6 | 100702 | HITS-CLIP | | 23142080 | Unspecified |
| mmu-miR-21a-5p | MIMAT0000530 | Gbp7 | 229900 | Microarrays//HITS-CLIP | 37986224\|31433990 | | Unspecified |
| mmu-miR-21a-5p | MIMAT0000530 | Gbp8 | 76074 | HITS-CLIP | | 23142080 | Unspecified |
| mmu-miR-21a-5p | MIMAT0000530 | Gbp9 | 236573 | HITS-CLIP | | 23142080 | Unspecified |
| mmu-miR-21a-5p | MIMAT0000530 | Gc | 14473 | HITS-CLIP | | 23597149 | Unspecified |
| mmu-miR-21a-5p | MIMAT0000530 | Gcat | 26912 | HITS-CLIP | | 23142080 | Unspecified |
| mmu-miR-21a-5p | MIMAT0000530 | Gcc2 | 70297 | HITS-CLIP | 19536157\|23142080 | | Unspecified |
| mmu-miR-21a-5p | MIMAT0000530 | Gch1 | 14528 | HITS-CLIP | | 23597149 | Unspecified |
| mmu-miR-21a-5p | MIMAT0000530 | Gchfr | 320415 | Microarrays | | 37986224 | Unspecified |
| mmu-miR-21a-5p | MIMAT0000530 | Gclc | 14629 | Microarrays//HITS-CLIP | 37986224\|25263593\|23597149\|23142 | | Unspecified |
| mmu-miR-21a-5p | MIMAT0000530 | Gclm | 14630 | Microarrays//HITS-CLIP | 37986224\|23597149\|25263593\|1953€ | | Unspecified |
| mmu-miR-21a-5p | MIMAT0000530 | Gcn1l1 | 10985 | HITS-CLIP | 23597149\|29167373 | | Unspecified |
| mmu-miR-21a-5p | MIMAT0000530 | Gcnt1 | 14537 | HITS-CLIP | | 29167373 | Unspecified |
| mmu-miR-21a-5p | MIMAT0000530 | Gdf3 | 14562 | Microarrays//HITS-CLIP | 37986224\|29167373 | | Unspecified |
| mmu-miR-21a-5p | MIMAT0000530 | Gdi2 | 14569 | HITS-CLIP | | 23142080 | Unspecified |
| mmu-miR-21a-5p | MIMAT0000530 | Gdpd1 | 66569 | Microarrays//HITS-CLIP | 37986224\|29167373 | | Unspecified |
| mmu-miR-21a-5p | MIMAT0000530 | Gdpd3 | 68616 | Microarrays | 37986224\|25263593 | | Unspecified |
| mmu-miR-21a-5p | MIMAT0000530 | Gem | 14579 | HITS-CLIP, Microarrays//Microarrays//HITS | 37986224\|25263593\|23142080 | | Unspecified |
| mmu-miR-21a-5p | MIMAT0000530 | Gen1 | 209334 | Microarrays//HITS-CLIP | 37986224\|25263593\|23142080 | | Unspecified |
| mmu-miR-21a-5p | MIMAT0000530 | Gfer | 11692 | HITS-CLIP | | 23142080 | Unspecified |
| mmu-miR-21a-5p | MIMAT0000530 | Gfi1 | 14581 | HITS-CLIP | | 23142080 | Unspecified |
| mmu-miR-21a-5p | MIMAT0000530 | Gfm1 | 28030 | HITS-CLIP | 23597149\|23142080 | | Unspecified |
| mmu-miR-21a-5p | MIMAT0000530 | Gfm2 | 320806 | HITS-CLIP | | 23142080 | Unspecified |
| mmu-miR-21a-5p | MIMAT0000530 | Gfod1 | 328232 | HITS-CLIP | | 23142080 | Unspecified |
| mmu-miR-21a-5p | MIMAT0000530 | Gfpt1 | 14583 | HITS-CLIP | 37986224\|23142080 | | Unspecified |
| mmu-miR-21a-5p | MIMAT0000530 | Gga3 | 260302 | HITS-CLIP | | 31433990 | Unspecified |
| mmu-miR-21a-5p | MIMAT0000530 | Ggact | 223267 | HITS-CLIP | | 23142080 | Unspecified |
| mmu-miR-21a-5p | MIMAT0000530 | Ggnbp2 | 217039 | HITS-CLIP | 23142080\|19536157 | | Unspecified |
| mmu-miR-21a-5p | MIMAT0000530 | Ggps1 | 14593 | HITS-CLIP | | 23142080 | Unspecified |
| mmu-miR-21a-5p | MIMAT0000530 | Ggta1 | 14594 | HITS-CLIP | | 23142080 | Unspecified |
| mmu-miR-21a-5p | MIMAT0000530 | Ghitm | 66092 | HITS-CLIP | 23142080\|23597149 | | Unspecified |
| mmu-miR-21a-5p | MIMAT0000530 | Ghr | 14600 | HITS-CLIP | | 23597149 | Unspecified |
| mmu-miR-21a-5p | MIMAT0000530 | Gid4 | 66771 | HITS-CLIP | 23142080\|23597149 | | Unspecified |
| mmu-miR-21a-5p | MIMAT0000530 | Gid8 | 76425 | HITS-CLIP | | 23142080 | Unspecified |
| mmu-miR-21a-5p | MIMAT0000530 | Gigyf1 | 57330 | qCLASH//HITS-CLIP | 31393953\|23142080 | | Unspecified |
| mmu-miR-21a-5p | MIMAT0000530 | Gigyf2 | 227331 | HITS-CLIP | 23597149\|23142080 | | Unspecified |

| | | | | | | |
|---|---|---|---|---|---|---|
| mmu-miR-21a-5p | MIMAT0000530 | Gimap1 | 16205 | HITS-CLIP | 23142080 | Unspecified |
| mmu-miR-21a-5p | MIMAT0000530 | Gimap4 | 107526 | HITS-CLIP | 31433990\|23142080 | Unspecified |
| mmu-miR-21a-5p | MIMAT0000530 | Gimap6 | 231931 | HITS-CLIP | 23142080 | Unspecified |
| mmu-miR-21a-5p | MIMAT0000530 | Gimap8 | 243374 | HITS-CLIP | 23142080 | Unspecified |
| mmu-miR-21a-5p | MIMAT0000530 | Gimap9 | 317758 | HITS-CLIP | 23142080 | Unspecified |
| mmu-miR-21a-5p | MIMAT0000530 | Gin1 | 252876 | HITS-CLIP | 37986224 | Unspecified |
| mmu-miR-21a-5p | MIMAT0000530 | Ginm1 | 215751 | HITS-CLIP | 23597149\|23142080 | Unspecified |
| mmu-miR-21a-5p | MIMAT0000530 | Gins1 | 69270 | Microarrays//HITS-CLIP | 37986224\|25263593\|23142080 | Unspecified |
| mmu-miR-21a-5p | MIMAT0000530 | Gins2 | 272551 | Microarrays | 37986224\|25263593 | Unspecified |
| mmu-miR-21a-5p | MIMAT0000530 | Gins3 | 78833 | HITS-CLIP | 23142080 | Unspecified |
| mmu-miR-21a-5p | MIMAT0000530 | Git2 | 26431 | Microarrays//HITS-CLIP | 37986224\|29167373 | Unspecified |
| mmu-miR-21a-5p | MIMAT0000530 | Gja1 | 14609 | HITS-CLIP | 37986224\|23142080\|19536157 | Unspecified |
| mmu-miR-21a-5p | MIMAT0000530 | Gk | 14933 | HITS-CLIP//Microarrays | 23597149\|25263593\|29167373 | Unspecified |
| mmu-miR-21a-5p | MIMAT0000530 | Gk5 | 235533 | HITS-CLIP | 23142080 | Unspecified |
| mmu-miR-21a-5p | MIMAT0000530 | Gla | 11605 | qCLASH | 31393953 | Unspecified |
| mmu-miR-21a-5p | MIMAT0000530 | Glcci1 | 170772 | qCLASH//HITS-CLIP | 31393953\|23142080 | Unspecified |
| mmu-miR-21a-5p | MIMAT0000530 | Glce | 93683 | HITS-CLIP | 23597149 | Unspecified |
| mmu-miR-21a-5p | MIMAT0000530 | Gldc | 104174 | HITS-CLIP | 23597149 | Unspecified |
| mmu-miR-21a-5p | MIMAT0000530 | Gle1 | 74412 | HITS-CLIP | 23597149 | Unspecified |
| mmu-miR-21a-5p | MIMAT0000530 | Glg1 | 20340 | HITS-CLIP | 23142080\|29167373 | Unspecified |
| mmu-miR-21a-5p | MIMAT0000530 | Glipr1 | 73690 | HITS-CLIP | 23142080\|29167373 | Unspecified |
| mmu-miR-21a-5p | MIMAT0000530 | Glis2 | 83396 | HITS-CLIP | 23142080 | Unspecified |
| mmu-miR-21a-5p | MIMAT0000530 | Glra4 | 14657 | Microarrays | 37986224\|25263593 | Unspecified |
| mmu-miR-21a-5p | MIMAT0000530 | Glrx | 93692 | Microarrays | 37986224\|25263593 | Unspecified |
| mmu-miR-21a-5p | MIMAT0000530 | Glrx2 | 69367 | HITS-CLIP | 23142080 | Unspecified |
| mmu-miR-21a-5p | MIMAT0000530 | Glrx5 | 73046 | qCLASH | 31393953 | Unspecified |
| mmu-miR-21a-5p | MIMAT0000530 | Gls | 14660 | HITS-CLIP | 37986224\|23142080 | Unspecified |
| mmu-miR-21a-5p | MIMAT0000530 | Gls2 | 216456 | HITS-CLIP | 23597149 | Unspecified |
| mmu-miR-21a-5p | MIMAT0000530 | Glt8d1 | 76485 | HITS-CLIP | 23142080\|19536157\|29167373 | Unspecified |
| mmu-miR-21a-5p | MIMAT0000530 | Gltp | 56356 | HITS-CLIP | 23142080 | Unspecified |
| mmu-miR-21a-5p | MIMAT0000530 | Gltpd2 | 216871 | HITS-CLIP | 23597149 | Unspecified |
| mmu-miR-21a-5p | MIMAT0000530 | Glud1 | 14661 | HITS-CLIP | 23597149\|23142080 | Unspecified |
| mmu-miR-21a-5p | MIMAT0000530 | Glul | 14645 | Microarrays | 37986224 | Unspecified |
| mmu-miR-21a-5p | MIMAT0000530 | Glyr1 | 74022 | HITS-CLIP | 37986224\|23142080 | Unspecified |
| mmu-miR-21a-5p | MIMAT0000530 | Gm10184 | 100043906 | qCLASH | 31393953 | Unspecified |
| mmu-miR-21a-5p | MIMAT0000530 | Gm10521 | 100038548 | HITS-CLIP | 29167373 | Unspecified |
| mmu-miR-21a-5p | MIMAT0000530 | Gm11110 | 100169874 | Microarrays | 37986224\|25263593 | Unspecified |
| mmu-miR-21a-5p | MIMAT0000530 | Gm12166 | 100039624 | HITS-CLIP | 29167373 | Unspecified |
| mmu-miR-21a-5p | MIMAT0000530 | Gm16286 | 117903916 | HITS-CLIP | 23597149\|23142080 | Unspecified |

| | | | | | | |
|---|---|---|---|---|---|---|
| mmu-miR-21a-5p | MIMAT0000530 | Gm17190 | 6614395 | HITS-CLIP | 23142080 | Unspecified |
| mmu-miR-21a-5p | MIMAT0000530 | Gm20425 | 6608681 | HITS-CLIP | 23597149 | Unspecified |
| mmu-miR-21a-5p | MIMAT0000530 | Gm20547 | 6608472 | HITS-CLIP | 23597149 | Unspecified |
| mmu-miR-21a-5p | MIMAT0000530 | Gm21953 | 621580 | Microarrays | 37986224\|25263593 | Unspecified |
| mmu-miR-21a-5p | MIMAT0000530 | Gm2a | 14667 | HITS-CLIP | 23597149\|23142080\|29167373 | Unspecified |
| mmu-miR-21a-5p | MIMAT0000530 | Gm340 | 381224 | HITS-CLIP | 23142080 | Unspecified |
| mmu-miR-21a-5p | MIMAT0000530 | Gm35315 | 102638847 | HITS-CLIP | 23142080 | Unspecified |
| mmu-miR-21a-5p | MIMAT0000530 | Gm37240 | | HITS-CLIP | 23142080\|19536157 | Unspecified |
| mmu-miR-21a-5p | MIMAT0000530 | Gm38394 | 667118 | HITS-CLIP | 23142080\|23597149 | Unspecified |
| mmu-miR-21a-5p | MIMAT0000530 | Gm42372 | 127238614 | HITS-CLIP | 23142080 | Unspecified |
| mmu-miR-21a-5p | MIMAT0000530 | Gm45713 | 103691134 | HITS-CLIP | 31433990 | Unspecified |
| mmu-miR-21a-5p | MIMAT0000530 | Gm47283 | | Microarrays | 25263593 | Unspecified |
| mmu-miR-21a-5p | MIMAT0000530 | Gm4756 | 209183 | HITS-CLIP | 23597149 | Unspecified |
| mmu-miR-21a-5p | MIMAT0000530 | Gm49336 | 127230315 | HITS-CLIP | 23142080 | Unspecified |
| mmu-miR-21a-5p | MIMAT0000530 | Gm4951 | 240327 | Microarrays | 37986224 | Unspecified |
| mmu-miR-21a-5p | MIMAT0000530 | Gm5089 | 328479 | HITS-CLIP | 19536157 | Unspecified |
| mmu-miR-21a-5p | MIMAT0000530 | Gm5617 | 434402 | Microarrays | 37986224 | Unspecified |
| mmu-miR-21a-5p | MIMAT0000530 | Gm7694 | 665574 | HITS-CLIP | 23597149 | Unspecified |
| mmu-miR-21a-5p | MIMAT0000530 | Gm8369 | 666926 | Microarrays//HITS-CLIP | 37986224\|23142080 | Unspecified |
| mmu-miR-21a-5p | MIMAT0000530 | Gmcl1 | 23885 | HITS-CLIP | 23142080 | Unspecified |
| mmu-miR-21a-5p | MIMAT0000530 | Gmfb | 63985 | HITS-CLIP | 23142080 | Unspecified |
| mmu-miR-21a-5p | MIMAT0000530 | Gmfg | 63986 | Microarrays | 37986224 | Unspecified |
| mmu-miR-21a-5p | MIMAT0000530 | Gmip | 78816 | HITS-CLIP | 23142080\|29167373 | Unspecified |
| mmu-miR-21a-5p | MIMAT0000530 | Gmnn | 57441 | Microarrays//HITS-CLIP | 37986224\|25263593\|23142080\|29167 | Unspecified |
| mmu-miR-21a-5p | MIMAT0000530 | Gmppb | 331026 | HITS-CLIP | 23142080 | Unspecified |
| mmu-miR-21a-5p | MIMAT0000530 | Gmpr2 | 105446 | HITS-CLIP | 29167373 | Unspecified |
| mmu-miR-21a-5p | MIMAT0000530 | Gmps | 229363 | HITS-CLIP | 23142080 | Unspecified |
| mmu-miR-21a-5p | MIMAT0000530 | Gna12 | 14673 | Microarrays | 37986224 | Unspecified |
| mmu-miR-21a-5p | MIMAT0000530 | Gna13 | 14674 | HITS-CLIP | 23142080\|29167373 | Unspecified |
| mmu-miR-21a-5p | MIMAT0000530 | Gnai3 | 14679 | HITS-CLIP | 23142080 | Unspecified |
| mmu-miR-21a-5p | MIMAT0000530 | Gnaq | 14682 | HITS-CLIP | 23142080\|29167373 | Unspecified |
| mmu-miR-21a-5p | MIMAT0000530 | Gnb4 | 14696 | Microarrays//HITS-CLIP | 37986224\|25263593\|23142080 | Unspecified |
| mmu-miR-21a-5p | MIMAT0000530 | Gne | 50798 | HITS-CLIP | 23597149 | Unspecified |
| mmu-miR-21a-5p | MIMAT0000530 | Gng12 | 14701 | HITS-CLIP | 37986224\|23597149\|23142080 | Unspecified |
| mmu-miR-21a-5p | MIMAT0000530 | Gng2 | 14702 | HITS-CLIP | 23142080\|29167373 | Unspecified |
| mmu-miR-21a-5p | MIMAT0000530 | Gnl1 | 14670 | HITS-CLIP | 23142080 | Unspecified |
| mmu-miR-21a-5p | MIMAT0000530 | Gnl2 | 230737 | HITS-CLIP | 37986224 | Unspecified |
| mmu-miR-21a-5p | MIMAT0000530 | Gnl3 | 30877 | qCLASH//HITS-CLIP | 31393953\|23142080 | Unspecified |
| mmu-miR-21a-5p | MIMAT0000530 | Gnl3l | 237107 | HITS-CLIP | 23142080 | Unspecified |

| | | | | | | |
|---|---|---|---|---|---|---|
| mmu-miR-21a-5p | MIMAT0000530 | Gnpat | 14712 | HITS-CLIP | 23597149 | Unspecified |
| mmu-miR-21a-5p | MIMAT0000530 | Gnpda2 | 67980 | HITS-CLIP | 23142080 | Unspecified |
| mmu-miR-21a-5p | MIMAT0000530 | Gnptab | 432486 | Microarrays | 37986224\|25263593 | Unspecified |
| mmu-miR-21a-5p | MIMAT0000530 | Gnptg | 214505 | HITS-CLIP | 23142080 | Unspecified |
| mmu-miR-21a-5p | MIMAT0000530 | Gns | 75612 | HITS-CLIP | 23597149 | Unspecified |
| mmu-miR-21a-5p | MIMAT0000530 | Golga1 | 76899 | HITS-CLIP | 23597149\|23142080 | Unspecified |
| mmu-miR-21a-5p | MIMAT0000530 | Golga3 | 269682 | HITS-CLIP | 23142080\|29167373 | Unspecified |
| mmu-miR-21a-5p | MIMAT0000530 | Golga4 | 54214 | HITS-CLIP | 23142080\|29167373 | Unspecified |
| mmu-miR-21a-5p | MIMAT0000530 | Golga7 | 57437 | HITS-CLIP | 23142080 | Unspecified |
| mmu-miR-21a-5p | MIMAT0000530 | Golgb1 | 224139 | HITS-CLIP | 23597149\|29167373 | Unspecified |
| mmu-miR-21a-5p | MIMAT0000530 | Golim4 | 73124 | HITS-CLIP | 23597149 | Unspecified |
| mmu-miR-21a-5p | MIMAT0000530 | Golm1 | 105348 | Microarrays | 37986224\|25263593 | Unspecified |
| mmu-miR-21a-5p | MIMAT0000530 | Golph3 | 66629 | HITS-CLIP | 37986224\|23142080\|31433990\|29167 | Unspecified |
| mmu-miR-21a-5p | MIMAT0000530 | Golph3l | 229593 | Microarrays//HITS-CLIP | 37986224\|23142080 | Unspecified |
| mmu-miR-21a-5p | MIMAT0000530 | Golt1b | 66964 | Microarrays//HITS-CLIP | 37986224\|25263593\|23142080 | Unspecified |
| mmu-miR-21a-5p | MIMAT0000530 | Gopc | 94221 | HITS-CLIP | 23597149\|23142080 | Unspecified |
| mmu-miR-21a-5p | MIMAT0000530 | Gosr2 | 56494 | HITS-CLIP | 23142080\|29167373 | Unspecified |
| mmu-miR-21a-5p | MIMAT0000530 | Got1 | 14718 | HITS-CLIP | 37986224\|23597149\|23142080 | Unspecified |
| mmu-miR-21a-5p | MIMAT0000530 | Gpalpp1 | 67467 | HITS-CLIP | 23142080 | Unspecified |
| mmu-miR-21a-5p | MIMAT0000530 | Gpam | 14732 | HITS-CLIP | 23597149\|19536157 | Unspecified |
| mmu-miR-21a-5p | MIMAT0000530 | Gpatch2 | 67769 | HITS-CLIP | 23142080 | Unspecified |
| mmu-miR-21a-5p | MIMAT0000530 | Gpatch8 | 237943 | HITS-CLIP//qCLASH | 23142080\|31393953\|23597149 | Unspecified |
| mmu-miR-21a-5p | MIMAT0000530 | Gpc3 | 14734 | Microarrays | 37986224 | Unspecified |
| mmu-miR-21a-5p | MIMAT0000530 | Gpc4 | 14735 | HITS-CLIP | 23597149 | Unspecified |
| mmu-miR-21a-5p | MIMAT0000530 | Gpd1 | 14555 | HITS-CLIP | 23597149 | Unspecified |
| mmu-miR-21a-5p | MIMAT0000530 | Gpd1l | 333433 | HITS-CLIP | 19536157 | Unspecified |
| mmu-miR-21a-5p | MIMAT0000530 | Gpkow | 209416 | HITS-CLIP | 23142080 | Unspecified |
| mmu-miR-21a-5p | MIMAT0000530 | Gpld1 | 14756 | HITS-CLIP | 23597149 | Unspecified |
| mmu-miR-21a-5p | MIMAT0000530 | Gpm6a | 234267 | HITS-CLIP | 37986224\|19536157 | Unspecified |
| mmu-miR-21a-5p | MIMAT0000530 | Gpm6b | 14758 | HITS-CLIP | 19536157\|23142080 | Unspecified |
| mmu-miR-21a-5p | MIMAT0000530 | Gpn1 | 74254 | HITS-CLIP | 23142080 | Unspecified |
| mmu-miR-21a-5p | MIMAT0000530 | Gpn2 | 100210 | HITS-CLIP | 23597149\|23142080 | Unspecified |
| mmu-miR-21a-5p | MIMAT0000530 | Gpnmb | 93695 | HITS-CLIP | 29167373 | Unspecified |
| mmu-miR-21a-5p | MIMAT0000530 | Gpr155 | 68526 | HITS-CLIP, Microarrays | 37986224 | Unspecified |
| mmu-miR-21a-5p | MIMAT0000530 | Gpr160 | 71862 | Microarrays | 37986224 | Unspecified |
| mmu-miR-21a-5p | MIMAT0000530 | Gpr162 | 14788 | Microarrays | 37986224 | Unspecified |
| mmu-miR-21a-5p | MIMAT0000530 | Gpr165 | 76206 | HITS-CLIP | 37986224\|19536157 | Unspecified |
| mmu-miR-21a-5p | MIMAT0000530 | Gpr174 | 213439 | HITS-CLIP | 23142080 | Unspecified |
| mmu-miR-21a-5p | MIMAT0000530 | Gpr176 | 381413 | Microarrays | 37986224\|25263593 | Unspecified |

| | | | | | | |
|---|---|---|---|---|---|---|
| mmu-miR-21a-5p | MIMAT0000530 | Gpr18 | 110168 | HITS-CLIP | 23142080 | Unspecified |
| mmu-miR-21a-5p | MIMAT0000530 | Gpr26 | 233919 | HITS-CLIP | 37986224 | Unspecified |
| mmu-miR-21a-5p | MIMAT0000530 | Gpr31b | 436440 | Microarrays | 37986224 | Unspecified |
| mmu-miR-21a-5p | MIMAT0000530 | Gpr34 | 23890 | Microarrays//HITS-CLIP | 37986224\|23142080 | Unspecified |
| mmu-miR-21a-5p | MIMAT0000530 | Gpr65 | 14744 | Microarrays | 37986224 | Unspecified |
| mmu-miR-21a-5p | MIMAT0000530 | Gpr68 | 238377 | HITS-CLIP | 23142080 | Unspecified |
| mmu-miR-21a-5p | MIMAT0000530 | Gpr84 | 80910 | Microarrays | 37986224\|25263593 | Unspecified |
| mmu-miR-21a-5p | MIMAT0000530 | Gpr88 | 64378 | HITS-CLIP | 19536157 | Unspecified |
| mmu-miR-21a-5p | MIMAT0000530 | Gprasp1 | 67298 | HITS-CLIP | 37986224\|19536157 | Unspecified |
| mmu-miR-21a-5p | MIMAT0000530 | Gprasp2 | 245607 | HITS-CLIP | 23142080 | Unspecified |
| mmu-miR-21a-5p | MIMAT0000530 | Gprc5a | 232431 | HITS-CLIP | 23142080 | Unspecified |
| mmu-miR-21a-5p | MIMAT0000530 | Gprc5b | 64297 | Microarrays//HITS-CLIP | 37986224\|25263593\|19536157 | Unspecified |
| mmu-miR-21a-5p | MIMAT0000530 | Gprc5c | 70355 | Microarrays//HITS-CLIP | 37986224\|23597149 | Unspecified |
| mmu-miR-21a-5p | MIMAT0000530 | Gprin2 | 432839 | Microarrays | 37986224\|25263593 | Unspecified |
| mmu-miR-21a-5p | MIMAT0000530 | Gprin3 | 243385 | HITS-CLIP | 23597149 | Unspecified |
| mmu-miR-21a-5p | MIMAT0000530 | Gpt2 | 108682 | HITS-CLIP | 23597149 | Unspecified |
| mmu-miR-21a-5p | MIMAT0000530 | Gpx8 | 69590 | Microarrays | 37986224\|25263593 | Unspecified |
| mmu-miR-21a-5p | MIMAT0000530 | Gramd1b | 235283 | Microarrays | 37986224\|25263593 | Unspecified |
| mmu-miR-21a-5p | MIMAT0000530 | Gramd1c | 207798 | HITS-CLIP | 23142080 | Unspecified |
| mmu-miR-21a-5p | MIMAT0000530 | Gramd3 | 107022 | HITS-CLIP | 37986224\|23597149\|23142080 | Unspecified |
| mmu-miR-21a-5p | MIMAT0000530 | Gramd4 | 223752 | HITS-CLIP | 23142080 | Unspecified |
| mmu-miR-21a-5p | MIMAT0000530 | Grap2 | 17444 | HITS-CLIP | 23142080 | Unspecified |
| mmu-miR-21a-5p | MIMAT0000530 | Grb2 | 14784 | HITS-CLIP | 23142080 | Unspecified |
| mmu-miR-21a-5p | MIMAT0000530 | Grcc10 | 14790 | Microarrays//HITS-CLIP | 37986224\|23597149 | Unspecified |
| mmu-miR-21a-5p | MIMAT0000530 | Grem2 | 23893 | HITS-CLIP | 23597149 | Unspecified |
| mmu-miR-21a-5p | MIMAT0000530 | Grhpr | 76238 | HITS-CLIP | 23597149 | Unspecified |
| mmu-miR-21a-5p | MIMAT0000530 | Gria2 | 14800 | HITS-CLIP | 19536157 | Unspecified |
| mmu-miR-21a-5p | MIMAT0000530 | Gria3 | 53623 | HITS-CLIP | 37986224\|19536157 | Unspecified |
| mmu-miR-21a-5p | MIMAT0000530 | Grk1 | 24013 | qCLASH | 31393953 | Unspecified |
| mmu-miR-21a-5p | MIMAT0000530 | Grk2 | 156 | HITS-CLIP | 23142080 | Unspecified |
| mmu-miR-21a-5p | MIMAT0000530 | Grk6 | 26385 | HITS-CLIP | 23142080 | Unspecified |
| mmu-miR-21a-5p | MIMAT0000530 | Grm5 | 108071 | HITS-CLIP | 19536157 | Unspecified |
| mmu-miR-21a-5p | MIMAT0000530 | Grn | 14824 | HITS-CLIP | 23597149\|29167373 | Unspecified |
| mmu-miR-21a-5p | MIMAT0000530 | Grpel1 | 17713 | HITS-CLIP | 23597149\|23142080 | Unspecified |
| mmu-miR-21a-5p | MIMAT0000530 | Grpel2 | 17714 | HITS-CLIP | 23597149\|23142080 | Unspecified |
| mmu-miR-21a-5p | MIMAT0000530 | Grsf1 | 231413 | HITS-CLIP | 31433990\|23597149\|23142080 | Unspecified |
| mmu-miR-21a-5p | MIMAT0000530 | Grwd1 | 101612 | Microarrays | 37986224\|25263593 | Unspecified |
| mmu-miR-21a-5p | MIMAT0000530 | Gsap | 212167 | HITS-CLIP | 29167373 | Unspecified |
| mmu-miR-21a-5p | MIMAT0000530 | Gsdme | 1687 | Microarrays | 25263593 | Unspecified |

| | | | | | | |
|---|---|---|---|---|---|---|
| mmu-miR-21a-5p | MIMAT0000530 | Gsk3a | 606496 | HITS-CLIP | 23142080 | Unspecified |
| mmu-miR-21a-5p | MIMAT0000530 | Gsk3b | 56637 | HITS-CLIP | 23597149\|23142080\|29167373 | Unspecified |
| mmu-miR-21a-5p | MIMAT0000530 | Gspt1 | 14852 | HITS-CLIP | 23142080\|23597149 | Unspecified |
| mmu-miR-21a-5p | MIMAT0000530 | Gsr | 14782 | Microarrays//HITS-CLIP | 37986224\|25263593\|29167373 | Unspecified |
| mmu-miR-21a-5p | MIMAT0000530 | Gsta3 | 14859 | Microarrays | 37986224\|25263593 | Unspecified |
| mmu-miR-21a-5p | MIMAT0000530 | Gstcd | 67553 | HITS-CLIP | 23142080 | Unspecified |
| mmu-miR-21a-5p | MIMAT0000530 | Gstm2 | 14863 | Microarrays | 37986224\|25263593 | Unspecified |
| mmu-miR-21a-5p | MIMAT0000530 | Gstm7 | 68312 | HITS-CLIP | 23597149 | Unspecified |
| mmu-miR-21a-5p | MIMAT0000530 | Gstp1 | 14870 | Microarrays | 37986224 | Unspecified |
| mmu-miR-21a-5p | MIMAT0000530 | Gstt2 | 14872 | Microarrays | 37986224 | Unspecified |
| mmu-miR-21a-5p | MIMAT0000530 | Gtf2a2 | 235459 | HITS-CLIP | 23597149\|23142080 | Unspecified |
| mmu-miR-21a-5p | MIMAT0000530 | Gtf2b | 229906 | HITS-CLIP | 23597149\|29167373 | Unspecified |
| mmu-miR-21a-5p | MIMAT0000530 | Gtf2e1 | 74197 | HITS-CLIP | 23142080 | Unspecified |
| mmu-miR-21a-5p | MIMAT0000530 | Gtf2e2 | 68153 | HITS-CLIP | 23142080\|29167373 | Unspecified |
| mmu-miR-21a-5p | MIMAT0000530 | Gtf2h1 | 14884 | HITS-CLIP | 23142080 | Unspecified |
| mmu-miR-21a-5p | MIMAT0000530 | Gtf2h2 | 23894 | HITS-CLIP | 23142080 | Unspecified |
| mmu-miR-21a-5p | MIMAT0000530 | Gtf2i | 14886 | HITS-CLIP | 23597149\|23142080 | Unspecified |
| mmu-miR-21a-5p | MIMAT0000530 | Gtf3c1 | 233863 | HITS-CLIP | 23597149\|23142080 | Unspecified |
| mmu-miR-21a-5p | MIMAT0000530 | Gtf3c2 | 71752 | HITS-CLIP | 23142080 | Unspecified |
| mmu-miR-21a-5p | MIMAT0000530 | Gtf3c3 | 98488 | HITS-CLIP | 23142080 | Unspecified |
| mmu-miR-21a-5p | MIMAT0000530 | Gtf3c4 | 269252 | HITS-CLIP | 37986224\|23142080\|23597149 | Unspecified |
| mmu-miR-21a-5p | MIMAT0000530 | Gtpbp1 | 14904 | HITS-CLIP | 23142080 | Unspecified |
| mmu-miR-21a-5p | MIMAT0000530 | Gtpbp2 | 56055 | HITS-CLIP | 23597149\|23142080\|29167373 | Unspecified |
| mmu-miR-21a-5p | MIMAT0000530 | Gtse1 | 29870 | Microarrays | 37986224\|25263593 | Unspecified |
| mmu-miR-21a-5p | MIMAT0000530 | Gucy1a1 | 60596 | HITS-CLIP | 23142080 | Unspecified |
| mmu-miR-21a-5p | MIMAT0000530 | Gulo | 268756 | HITS-CLIP | 23597149 | Unspecified |
| mmu-miR-21a-5p | MIMAT0000530 | Gxylt1 | 223827 | HITS-CLIP | 37986224\|23597149\|23142080\|29167 | Unspecified |
| mmu-miR-21a-5p | MIMAT0000530 | Gykl1 | 14625 | Microarrays | 37986224\|25263593 | Unspecified |
| mmu-miR-21a-5p | MIMAT0000530 | Gys2 | 232493 | HITS-CLIP | 23597149 | Unspecified |
| mmu-miR-21a-5p | MIMAT0000530 | Gzf1 | 74533 | HITS-CLIP | 28735896\|23142080 | Unspecified |
| mmu-miR-21a-5p | MIMAT0000530 | Gzmb | 14939 | HITS-CLIP | 31433990\|23142080 | Unspecified |
| mmu-miR-21a-5p | MIMAT0000530 | Gzmd | 14941 | HITS-CLIP | 23142080 | Unspecified |
| mmu-miR-21a-5p | MIMAT0000530 | H13 | 14950 | HITS-CLIP | 29167373 | Unspecified |
| mmu-miR-21a-5p | MIMAT0000530 | H2-Aa | 14960 | Microarrays | 37986224 | Unspecified |
| mmu-miR-21a-5p | MIMAT0000530 | H2-Ab1 | 14961 | Microarrays | 37986224 | Unspecified |
| mmu-miR-21a-5p | MIMAT0000530 | H2-D1 | 14964 | HITS-CLIP | 23142080\|23597149\|29167373 | Unspecified |
| mmu-miR-21a-5p | MIMAT0000530 | H2-Eb1 | 14969 | Microarrays | 37986224 | Unspecified |
| mmu-miR-21a-5p | MIMAT0000530 | H2-Ob | 15002 | HITS-CLIP | 23142080 | Unspecified |
| mmu-miR-21a-5p | MIMAT0000530 | H2-Q10 | 15007 | HITS-CLIP | 23597149 | Unspecified |

| | | | | | | |
|---|---|---|---|---|---|---|
| mmu-miR-21a-5p | MIMAT0000530 | H2-T24 | 15042 | Microarrays//HITS-CLIP | 37986224\|23142080\|29167373 | Unspecified |
| mmu-miR-21a-5p | MIMAT0000530 | H2afj | 232440 | HITS-CLIP | 37986224\|23142080 | Unspecified |
| mmu-miR-21a-5p | MIMAT0000530 | H2afx | 15270 | Microarrays | 37986224 | Unspecified |
| mmu-miR-21a-5p | MIMAT0000530 | H2afy | 9555 | HITS-CLIP | 23142080\|29167373 | Unspecified |
| mmu-miR-21a-5p | MIMAT0000530 | H2ax | 3014 | Microarrays | 25263593 | Unspecified |
| mmu-miR-21a-5p | MIMAT0000530 | H3c11 | 8350 | HITS-CLIP | 31433990 | Unspecified |
| mmu-miR-21a-5p | MIMAT0000530 | H3c4 | 8350 | Microarrays | 25263593 | Unspecified |
| mmu-miR-21a-5p | MIMAT0000530 | H3f3a | 15078 | Microarrays | 37986224 | Unspecified |
| mmu-miR-21a-5p | MIMAT0000530 | H3f3b | 15081 | Microarrays | 37986224 | Unspecified |
| mmu-miR-21a-5p | MIMAT0000530 | H6pd | 100198 | HITS-CLIP | 23597149\|23142080 | Unspecified |
| mmu-miR-21a-5p | MIMAT0000530 | Habp2 | 226243 | HITS-CLIP | 23597149 | Unspecified |
| mmu-miR-21a-5p | MIMAT0000530 | Habp4 | 56541 | HITS-CLIP | 23597149 | Unspecified |
| mmu-miR-21a-5p | MIMAT0000530 | Hacd2 | 201562 | Microarrays//HITS-CLIP | 25263593\|23142080 | Unspecified |
| mmu-miR-21a-5p | MIMAT0000530 | Hace1 | 209462 | HITS-CLIP | 29167373 | Unspecified |
| mmu-miR-21a-5p | MIMAT0000530 | Hadh | 15107 | HITS-CLIP | 23597149\|23142080 | Unspecified |
| mmu-miR-21a-5p | MIMAT0000530 | Hadha | 97212 | HITS-CLIP | 29167373 | Unspecified |
| mmu-miR-21a-5p | MIMAT0000530 | Hadhb | 231086 | HITS-CLIP | 23597149\|23142080 | Unspecified |
| mmu-miR-21a-5p | MIMAT0000530 | Hagh | 14651 | HITS-CLIP | 23597149 | Unspecified |
| mmu-miR-21a-5p | MIMAT0000530 | Hal | 15109 | Microarrays//HITS-CLIP | 37986224\|25263593\|23597149 | Unspecified |
| mmu-miR-21a-5p | MIMAT0000530 | Hao1 | 15112 | Microarrays//HITS-CLIP | 37986224\|25263593\|23597149 | Unspecified |
| mmu-miR-21a-5p | MIMAT0000530 | Hapln1 | 12950 | HITS-CLIP | 37986224 | Unspecified |
| mmu-miR-21a-5p | MIMAT0000530 | Harbi1 | 241547 | HITS-CLIP | 23142080\|23597149 | Unspecified |
| mmu-miR-21a-5p | MIMAT0000530 | Hat1 | 107435 | HITS-CLIP | 37986224 | Unspecified |
| mmu-miR-21a-5p | MIMAT0000530 | Haus3 | 231123 | HITS-CLIP | 23142080 | Unspecified |
| mmu-miR-21a-5p | MIMAT0000530 | Haus6 | 230376 | Microarrays//HITS-CLIP | 37986224\|25263593\|30971389\|3143 | Unspecified |
| mmu-miR-21a-5p | MIMAT0000530 | Haus7 | 73738 | HITS-CLIP | 23597149\|23142080 | Unspecified |
| mmu-miR-21a-5p | MIMAT0000530 | Havcr2 | 171285 | Microarrays | 37986224\|25263593 | Unspecified |
| mmu-miR-21a-5p | MIMAT0000530 | Hba-a1 | 15122 | HITS-CLIP | 23597149 | Unspecified |
| mmu-miR-21a-5p | MIMAT0000530 | Hba-a2 | 110257 | HITS-CLIP | 23597149 | Unspecified |
| mmu-miR-21a-5p | MIMAT0000530 | Hbegf | 15200 | Microarrays | 37986224\|25263593 | Unspecified |
| mmu-miR-21a-5p | MIMAT0000530 | Hbp1 | 73389 | HITS-CLIP | 37986224\|23142080\|29167373 | Unspecified |
| mmu-miR-21a-5p | MIMAT0000530 | Hbs1l | 56422 | HITS-CLIP | 23597149 | Unspecified |
| mmu-miR-21a-5p | MIMAT0000530 | Hc | 15139 | HITS-CLIP | 23597149 | Unspecified |
| mmu-miR-21a-5p | MIMAT0000530 | Hcar2 | 80885 | Microarrays | 37986224\|25263593 | Unspecified |
| mmu-miR-21a-5p | MIMAT0000530 | Hccs | 15159 | HITS-CLIP | 23142080\|29167373 | Unspecified |
| mmu-miR-21a-5p | MIMAT0000530 | Hcfc2 | 67933 | HITS-CLIP | 37986224\|23142080 | Unspecified |
| mmu-miR-21a-5p | MIMAT0000530 | Hcls1 | 15163 | HITS-CLIP | 23142080\|29167373 | Unspecified |
| mmu-miR-21a-5p | MIMAT0000530 | Hdac4 | 208727 | HITS-CLIP | 23142080 | Unspecified |
| mmu-miR-21a-5p | MIMAT0000530 | Hdac7 | 56233 | HITS-CLIP | 23142080 | Unspecified |

| | | | | | | |
|---|---|---|---|---|---|---|
| mmu-miR-21a-5p | MIMAT0000530 | Hdac9 | 79221 | Microarrays//HITS-CLIP | 37986224\|25263593\|29167373 | Unspecified |
| mmu-miR-21a-5p | MIMAT0000530 | Hddc2 | 69692 | HITS-CLIP | 23142080 | Unspecified |
| mmu-miR-21a-5p | MIMAT0000530 | Hdgfl3 | 29877 | HITS-CLIP | 23142080 | Unspecified |
| mmu-miR-21a-5p | MIMAT0000530 | Hdlbp | 110611 | HITS-CLIP | 23142080\|23597149 | Unspecified |
| mmu-miR-21a-5p | MIMAT0000530 | Heatr3 | 234549 | HITS-CLIP | 23597149\|23142080 | Unspecified |
| mmu-miR-21a-5p | MIMAT0000530 | Heatr5a | 320487 | HITS-CLIP | 23142080\|29167373 | Unspecified |
| mmu-miR-21a-5p | MIMAT0000530 | Heatr5b | 320473 | HITS-CLIP | 23597149\|23142080 | Unspecified |
| mmu-miR-21a-5p | MIMAT0000530 | Hebp1 | 15199 | Microarrays | 37986224\|25263593 | Unspecified |
| mmu-miR-21a-5p | MIMAT0000530 | Heca | 380629 | HITS-CLIP | 23597149\|23142080\|29167373 | Unspecified |
| mmu-miR-21a-5p | MIMAT0000530 | Hectd1 | 207304 | HITS-CLIP | 37986224\|23142080\|23597149 | Unspecified |
| mmu-miR-21a-5p | MIMAT0000530 | Hectd2 | 226098 | HITS-CLIP | 23142080 | Unspecified |
| mmu-miR-21a-5p | MIMAT0000530 | Hectd4 | 283450 | HITS-CLIP | 19536157\|23142080\|29167373 | Unspecified |
| mmu-miR-21a-5p | MIMAT0000530 | Helb | 117599 | HITS-CLIP | 23142080 | Unspecified |
| mmu-miR-21a-5p | MIMAT0000530 | Hells | 15201 | Microarrays//HITS-CLIP | 37986224\|25263593\|23142080 | Unspecified |
| mmu-miR-21a-5p | MIMAT0000530 | Helz | 78455 | HITS-CLIP | 23142080 | Unspecified |
| mmu-miR-21a-5p | MIMAT0000530 | Helz2 | 229003 | Microarrays//HITS-CLIP | 37986224\|25263593\|23597149\|23142 | Unspecified |
| mmu-miR-21a-5p | MIMAT0000530 | Hemgn | 93966 | HITS-CLIP | 23142080 | Unspecified |
| mmu-miR-21a-5p | MIMAT0000530 | Herc1 | 235439 | HITS-CLIP | 37986224\|23597149\|23142080\|29167 | Unspecified |
| mmu-miR-21a-5p | MIMAT0000530 | Herc2 | 15204 | HITS-CLIP | 23142080 | Unspecified |
| mmu-miR-21a-5p | MIMAT0000530 | Herc3 | 73998 | HITS-CLIP | 23142080\|29167373 | Unspecified |
| mmu-miR-21a-5p | MIMAT0000530 | Herc4 | 67345 | Microarrays | 37986224 | Unspecified |
| mmu-miR-21a-5p | MIMAT0000530 | Herc6 | 67138 | HITS-CLIP | 29167373\|23142080 | Unspecified |
| mmu-miR-21a-5p | MIMAT0000530 | Herpud1 | 64209 | Microarrays//HITS-CLIP | 37986224\|23142080\|23597149 | Unspecified |
| mmu-miR-21a-5p | MIMAT0000530 | Herpud2 | 80517 | HITS-CLIP | 37986224\|23142080\|29167373 | Unspecified |
| mmu-miR-21a-5p | MIMAT0000530 | Hexb | 15212 | HITS-CLIP | 23142080 | Unspecified |
| mmu-miR-21a-5p | MIMAT0000530 | Hexim1 | 192231 | Microarrays | 37986224 | Unspecified |
| mmu-miR-21a-5p | MIMAT0000530 | Hfe | 15216 | Microarrays | 37986224 | Unspecified |
| mmu-miR-21a-5p | MIMAT0000530 | Hfe2 | 69585 | HITS-CLIP | 23597149 | Unspecified |
| mmu-miR-21a-5p | MIMAT0000530 | Hgd | 15233 | HITS-CLIP | 23597149 | Unspecified |
| mmu-miR-21a-5p | MIMAT0000530 | Hgs | 15239 | HITS-CLIP | 23597149\|23142080 | Unspecified |
| mmu-miR-21a-5p | MIMAT0000530 | Hibadh | 58875 | HITS-CLIP | 23597149\|23142080 | Unspecified |
| mmu-miR-21a-5p | MIMAT0000530 | Hic2 | 58180 | HITS-CLIP | 23142080 | Unspecified |
| mmu-miR-21a-5p | MIMAT0000530 | Hif1a | 15251 | HITS-CLIP | 37986224\|23142080\|19536157 | Unspecified |
| mmu-miR-21a-5p | MIMAT0000530 | Higd1a | 56295 | Microarrays//HITS-CLIP | 37986224\|25263593\|23142080 | Unspecified |
| mmu-miR-21a-5p | MIMAT0000530 | Hilpda | 69573 | Microarrays | 37986224\|25263593 | Unspecified |
| mmu-miR-21a-5p | MIMAT0000530 | Hip1 | 215114 | Microarrays//HITS-CLIP | 37986224\|25263593\|29167373\|23142 | Unspecified |
| mmu-miR-21a-5p | MIMAT0000530 | Hipk1 | 15257 | HITS-CLIP | 37986224\|23142080\|29167373 | Unspecified |
| mmu-miR-21a-5p | MIMAT0000530 | Hipk2 | 15258 | HITS-CLIP | 23142080 | Unspecified |
| mmu-miR-21a-5p | MIMAT0000530 | Hipk3 | 15259 | HITS-CLIP | 37986224\|19536157\|29167373\|23142 | Unspecified |

| | | | | | | |
|---|---|---|---|---|---|---|
| mmu-miR-21a-5p | MIMAT0000530 | Hira | 15260 | HITS-CLIP | 37986224\|23142080\|29167373 | Unspecified |
| mmu-miR-21a-5p | MIMAT0000530 | Hist1h1c | 50708 | HITS-CLIP | 23142080 | Unspecified |
| mmu-miR-21a-5p | MIMAT0000530 | Hist1h2bb | 3018 | HITS-CLIP | 23597149\|23142080 | Unspecified |
| mmu-miR-21a-5p | MIMAT0000530 | Hist1h2bj | 8970 | HITS-CLIP | 23597149\|23142080 | Unspecified |
| mmu-miR-21a-5p | MIMAT0000530 | Hist1h2bm | 8342 | HITS-CLIP | 23142080 | Unspecified |
| mmu-miR-21a-5p | MIMAT0000530 | Hist1h4f | 8361 | HITS-CLIP | 23142080 | Unspecified |
| mmu-miR-21a-5p | MIMAT0000530 | Hist1h4i | 319158 | HITS-CLIP | 23142080 | Unspecified |
| mmu-miR-21a-5p | MIMAT0000530 | Hist1h4j | 8363 | HITS-CLIP | 23142080 | Unspecified |
| mmu-miR-21a-5p | MIMAT0000530 | Hist2h2bb | 319189 | HITS-CLIP | 23142080\|23597149 | Unspecified |
| mmu-miR-21a-5p | MIMAT0000530 | Hist3h2a | 92815 | HITS-CLIP | 29167373 | Unspecified |
| mmu-miR-21a-5p | MIMAT0000530 | Hivep1 | 110521 | HITS-CLIP | 23142080 | Unspecified |
| mmu-miR-21a-5p | MIMAT0000530 | Hivep2 | 15273 | HITS-CLIP | 37986224\|23142080 | Unspecified |
| mmu-miR-21a-5p | MIMAT0000530 | Hivep3 | 16656 | HITS-CLIP | 23142080 | Unspecified |
| mmu-miR-21a-5p | MIMAT0000530 | Hjurp | 381280 | HITS-CLIP | 23142080 | Unspecified |
| mmu-miR-21a-5p | MIMAT0000530 | Hk1 | 15275 | HITS-CLIP | 23142080 | Unspecified |
| mmu-miR-21a-5p | MIMAT0000530 | Hk2 | 15277 | HITS-CLIP | 23142080 | Unspecified |
| mmu-miR-21a-5p | MIMAT0000530 | Hk3 | 212032 | Microarrays//HITS-CLIP | 37986224\|25263593\|29167373 | Unspecified |
| mmu-miR-21a-5p | MIMAT0000530 | Hlcs | 110948 | Microarrays | 37986224\|25263593 | Unspecified |
| mmu-miR-21a-5p | MIMAT0000530 | Hlx | 15284 | HITS-CLIP | 29167373 | Unspecified |
| mmu-miR-21a-5p | MIMAT0000530 | Hmbox1 | 219150 | HITS-CLIP | 37986224\|23597149\|23142080 | Unspecified |
| mmu-miR-21a-5p | MIMAT0000530 | Hmces | 232210 | HITS-CLIP | 23142080 | Unspecified |
| mmu-miR-21a-5p | MIMAT0000530 | Hmg20a | 66867 | HITS-CLIP | 23142080 | Unspecified |
| mmu-miR-21a-5p | MIMAT0000530 | Hmg20b | 10362 | HITS-CLIP | 23597149 | Unspecified |
| mmu-miR-21a-5p | MIMAT0000530 | Hmga2 | 15364 | Microarrays | 37986224\|25263593 | Unspecified |
| mmu-miR-21a-5p | MIMAT0000530 | Hmgcr | 15357 | HITS-CLIP | 23142080\|29167373 | Unspecified |
| mmu-miR-21a-5p | MIMAT0000530 | Hmgcs1 | 208715 | Microarrays//HITS-CLIP | 37986224\|23142080\|23597149 | Unspecified |
| mmu-miR-21a-5p | MIMAT0000530 | Hmgcs2 | 15360 | HITS-CLIP | 23597149 | Unspecified |
| mmu-miR-21a-5p | MIMAT0000530 | Hmmr | 15366 | Microarrays//HITS-CLIP | 37986224\|25263593\|23142080 | Unspecified |
| mmu-miR-21a-5p | MIMAT0000530 | Hmox1 | 15368 | Microarrays | 37986224\|25263593 | Unspecified |
| mmu-miR-21a-5p | MIMAT0000530 | Hnf4a | 15378 | HITS-CLIP | 37986224\|23597149 | Unspecified |
| mmu-miR-21a-5p | MIMAT0000530 | Hnrnpa0 | 77134 | HITS-CLIP | 23142080\|29167373 | Unspecified |
| mmu-miR-21a-5p | MIMAT0000530 | Hnrnpa1 | 15382 | qCLASH | 31393953 | Unspecified |
| mmu-miR-21a-5p | MIMAT0000530 | Hnrnpa2b1 | 53379 | HITS-CLIP | 37986224\|23597149\|23142080 | Unspecified |
| mmu-miR-21a-5p | MIMAT0000530 | Hnrnpa3 | 229279 | HITS-CLIP | 23142080 | Unspecified |
| mmu-miR-21a-5p | MIMAT0000530 | Hnrnpab | 15384 | HITS-CLIP | 23597149\|23142080 | Unspecified |
| mmu-miR-21a-5p | MIMAT0000530 | Hnrnpdl | 50926 | HITS-CLIP | 23142080\|29167373 | Unspecified |
| mmu-miR-21a-5p | MIMAT0000530 | Hnrnph1 | 59013 | HITS-CLIP | 23142080 | Unspecified |
| mmu-miR-21a-5p | MIMAT0000530 | Hnrnpk | 15387 | HITS-CLIP | 23142080 | Unspecified |
| mmu-miR-21a-5p | MIMAT0000530 | Hnrnpl | 15388 | HITS-CLIP | 23142080 | Unspecified |

| | | | | | | |
|---|---|---|---|---|---|---|
| mmu-miR-21a-5p | MIMAT0000530 | Hnrnpll | 72692 | HITS-CLIP | 37986224 | Unspecified |
| mmu-miR-21a-5p | MIMAT0000530 | Hnrnpm | 76936 | HITS-CLIP | 23142080 | Unspecified |
| mmu-miR-21a-5p | MIMAT0000530 | Hnrnpu | 51810 | HITS-CLIP | 37986224\|23142080\|29167373 | Unspecified |
| mmu-miR-21a-5p | MIMAT0000530 | Hnrnpul1 | 232989 | HITS-CLIP | 23142080\|29167373 | Unspecified |
| mmu-miR-21a-5p | MIMAT0000530 | Hnrnpul2 | 68693 | HITS-CLIP | 37986224\|23142080 | Unspecified |
| mmu-miR-21a-5p | MIMAT0000530 | Homer1 | 26556 | HITS-CLIP | 23597149\|23142080\|29167373 | Unspecified |
| mmu-miR-21a-5p | MIMAT0000530 | Hook1 | 77963 | Microarrays//HITS-CLIP | 37986224\|23142080 | Unspecified |
| mmu-miR-21a-5p | MIMAT0000530 | Hook2 | 170833 | Microarrays//HITS-CLIP | 37986224\|31433990 | Unspecified |
| mmu-miR-21a-5p | MIMAT0000530 | Hook3 | 320191 | HITS-CLIP | 37986224\|23597149\|29167373\|23142 | Unspecified |
| mmu-miR-21a-5p | MIMAT0000530 | Hoxa5 | 15402 | HITS-CLIP | 23597149 | Unspecified |
| mmu-miR-21a-5p | MIMAT0000530 | Hp1bp3 | 15441 | HITS-CLIP | 23597149\|23142080\|29167373 | Unspecified |
| mmu-miR-21a-5p | MIMAT0000530 | Hpgd | 15446 | Microarrays | 37986224 | Unspecified |
| mmu-miR-21a-5p | MIMAT0000530 | Hpgds | 54486 | HITS-CLIP | 37986224 | Unspecified |
| mmu-miR-21a-5p | MIMAT0000530 | Hpn | 15451 | HITS-CLIP | 23597149 | Unspecified |
| mmu-miR-21a-5p | MIMAT0000530 | Hprt | 15452 | HITS-CLIP | 23597149\|23142080\|31433990\|26203 | Unspecified |
| mmu-miR-21a-5p | MIMAT0000530 | Hps1 | 192236 | HITS-CLIP | 23142080 | Unspecified |
| mmu-miR-21a-5p | MIMAT0000530 | Hps3 | 12807 | HITS-CLIP | 23142080 | Unspecified |
| mmu-miR-21a-5p | MIMAT0000530 | Hps6 | 20170 | HITS-CLIP | 23142080 | Unspecified |
| mmu-miR-21a-5p | MIMAT0000530 | Hpse | 15442 | Microarrays | 37986224 | Unspecified |
| mmu-miR-21a-5p | MIMAT0000530 | Hrg | 94175 | HITS-CLIP | 23597149 | Unspecified |
| mmu-miR-21a-5p | MIMAT0000530 | Hrh2 | 15466 | HITS-CLIP | 23142080 | Unspecified |
| mmu-miR-21a-5p | MIMAT0000530 | Hs2st1 | 23908 | HITS-CLIP | 23142080 | Unspecified |
| mmu-miR-21a-5p | MIMAT0000530 | Hs3st1 | 15476 | HITS-CLIP | 23142080 | Unspecified |
| mmu-miR-21a-5p | MIMAT0000530 | Hs3st3b1 | 54710 | HITS-CLIP | 37986224\|23142080 | Unspecified |
| mmu-miR-21a-5p | MIMAT0000530 | Hs3st5 | 319415 | HITS-CLIP | 19536157 | Unspecified |
| mmu-miR-21a-5p | MIMAT0000530 | Hscb | 100900 | Microarrays//HITS-CLIP | 37986224\|23142080 | Unspecified |
| mmu-miR-21a-5p | MIMAT0000530 | Hsd17b11 | 114664 | HITS-CLIP | 23597149\|23142080 | Unspecified |
| mmu-miR-21a-5p | MIMAT0000530 | Hsd17b12 | 56348 | Microarrays//HITS-CLIP | 37986224\|25263593\|23142080 | Unspecified |
| mmu-miR-21a-5p | MIMAT0000530 | Hsd17b6 | 27400 | HITS-CLIP | 23597149 | Unspecified |
| mmu-miR-21a-5p | MIMAT0000530 | Hsd17b7 | 15490 | HITS-CLIP | 23597149\|23142080 | Unspecified |
| mmu-miR-21a-5p | MIMAT0000530 | Hsdl2 | 72479 | HITS-CLIP | 23142080 | Unspecified |
| mmu-miR-21a-5p | MIMAT0000530 | Hsf2bp | 74377 | HITS-CLIP | 23142080 | Unspecified |
| mmu-miR-21a-5p | MIMAT0000530 | Hsp90aa1 | 15519 | Microarrays//HITS-CLIP | 37986224\|25263593\|23142080 | Unspecified |
| mmu-miR-21a-5p | MIMAT0000530 | Hsp90ab1 | 15516 | HITS-CLIP | 37986224\|23597149\|23142080\|19536 | Unspecified |
| mmu-miR-21a-5p | MIMAT0000530 | Hsp90b1 | 22027 | HITS-CLIP | 23597149\|23142080 | Unspecified |
| mmu-miR-21a-5p | MIMAT0000530 | Hspa14 | 50497 | HITS-CLIP | 37986224\|23142080 | Unspecified |
| mmu-miR-21a-5p | MIMAT0000530 | Hspa2 | 15512 | qCLASH | 31393953 | Unspecified |
| mmu-miR-21a-5p | MIMAT0000530 | Hspa4 | 15525 | HITS-CLIP | 37986224\|23597149\|31433990\|23142 | Unspecified |
| mmu-miR-21a-5p | MIMAT0000530 | Hspa4l | 18415 | HITS-CLIP | 37986224\|23142080 | Unspecified |

| | | | | | | |
|---|---|---|---|---|---|---|
| mmu-miR-21a-5p | MIMAT0000530 | Hspa8 | 15481 | HITS-CLIP | 37986224 | Unspecified |
| mmu-miR-21a-5p | MIMAT0000530 | Hspa9 | 15526 | HITS-CLIP | 23597149\|23142080\|29167373 | Unspecified |
| mmu-miR-21a-5p | MIMAT0000530 | Hspbap1 | 66667 | HITS-CLIP | 23142080 | Unspecified |
| mmu-miR-21a-5p | MIMAT0000530 | Hspbp1 | 66245 | HITS-CLIP | 23597149\|23142080 | Unspecified |
| mmu-miR-21a-5p | MIMAT0000530 | Hspd1 | 15510 | HITS-CLIP | 23597149 | Unspecified |
| mmu-miR-21a-5p | MIMAT0000530 | Hsph1 | 15505 | HITS-CLIP | 23142080\|23597149 | Unspecified |
| mmu-miR-21a-5p | MIMAT0000530 | Htr2b | 15559 | Microarrays | 37986224\|25263593 | Unspecified |
| mmu-miR-21a-5p | MIMAT0000530 | Htt | 15194 | HITS-CLIP | 23597149\|23142080 | Unspecified |
| mmu-miR-21a-5p | MIMAT0000530 | Hus1 | 15574 | HITS-CLIP | 23142080 | Unspecified |
| mmu-miR-21a-5p | MIMAT0000530 | Huwe1 | 59026 | HITS-CLIP | 23597149\|19536157\|23142080\|29167 | Unspecified |
| mmu-miR-21a-5p | MIMAT0000530 | Hvcn1 | 74096 | HITS-CLIP | 23142080 | Unspecified |
| mmu-miR-21a-5p | MIMAT0000530 | Hyal1 | 15586 | HITS-CLIP | 23597149 | Unspecified |
| mmu-miR-21a-5p | MIMAT0000530 | Hykk | 235386 | HITS-CLIP | 23597149 | Unspecified |
| mmu-miR-21a-5p | MIMAT0000530 | Hyou1 | 12282 | HITS-CLIP | 23142080\|23597149 | Unspecified |
| mmu-miR-21a-5p | MIMAT0000530 | Iars | 105148 | HITS-CLIP | 23142080 | Unspecified |
| mmu-miR-21a-5p | MIMAT0000530 | Iars2 | 381314 | HITS-CLIP | 23597149\|31433990\|23142080 | Unspecified |
| mmu-miR-21a-5p | MIMAT0000530 | Ibtk | 108837 | HITS-CLIP | 23142080 | Unspecified |
| mmu-miR-21a-5p | MIMAT0000530 | Ica1l | 130026 | HITS-CLIP | 23142080 | Unspecified |
| mmu-miR-21a-5p | MIMAT0000530 | Icam1 | 15894 | HITS-CLIP | 23142080 | Unspecified |
| mmu-miR-21a-5p | MIMAT0000530 | Icam4 | 78369 | Microarrays | 37986224\|25263593 | Unspecified |
| mmu-miR-21a-5p | MIMAT0000530 | Ice1 | 822287 | HITS-CLIP | 23142080 | Unspecified |
| mmu-miR-21a-5p | MIMAT0000530 | Icos | 54167 | HITS-CLIP | 31433990\|23142080 | Unspecified |
| mmu-miR-21a-5p | MIMAT0000530 | Id1 | 15901 | Microarrays | 37986224\|25263593 | Unspecified |
| mmu-miR-21a-5p | MIMAT0000530 | Id2 | 15902 | Microarrays//HITS-CLIP | 37986224\|23142080\|25263593 | Unspecified |
| mmu-miR-21a-5p | MIMAT0000530 | Id3 | 15903 | Microarrays | 37986224\|25263593 | Unspecified |
| mmu-miR-21a-5p | MIMAT0000530 | Id4 | 15904 | HITS-CLIP | 19536157 | Unspecified |
| mmu-miR-21a-5p | MIMAT0000530 | Ide | 15925 | HITS-CLIP | 23597149 | Unspecified |
| mmu-miR-21a-5p | MIMAT0000530 | Idh1 | 15926 | HITS-CLIP | 23597149\|29167373 | Unspecified |
| mmu-miR-21a-5p | MIMAT0000530 | Idh3a | 67834 | HITS-CLIP | 23142080 | Unspecified |
| mmu-miR-21a-5p | MIMAT0000530 | Idh3b | 170718 | HITS-CLIP | 23597149 | Unspecified |
| mmu-miR-21a-5p | MIMAT0000530 | Idh3g | 15929 | HITS-CLIP | 23142080 | Unspecified |
| mmu-miR-21a-5p | MIMAT0000530 | Ids | 15931 | HITS-CLIP | 37986224\|19536157 | Unspecified |
| mmu-miR-21a-5p | MIMAT0000530 | Idua | 15932 | Microarrays//HITS-CLIP | 37986224\|25263593\|29167373 | Unspecified |
| mmu-miR-21a-5p | MIMAT0000530 | Ier3 | 15937 | Microarrays | 37986224 | Unspecified |
| mmu-miR-21a-5p | MIMAT0000530 | Iffo2 | 212632 | HITS-CLIP | 37986224\|29167373 | Unspecified |
| mmu-miR-21a-5p | MIMAT0000530 | Ifi202b | 26388 | Microarrays | 37986224\|25263593 | Unspecified |
| mmu-miR-21a-5p | MIMAT0000530 | Ifi204 | 15951 | Microarrays | 37986224 | Unspecified |
| mmu-miR-21a-5p | MIMAT0000530 | Ifi27l2a | 76933 | Microarrays | 37986224 | Unspecified |
| mmu-miR-21a-5p | MIMAT0000530 | Ifi44 | 99899 | Microarrays | 37986224\|25263593 | Unspecified |

| | | | | | | |
|---|---|---|---|---|---|---|
| mmu-miR-21a-5p | MIMAT0000530 | Ifi44l | 15061 | Microarrays | 37986224\|25263593 | Unspecified |
| mmu-miR-21a-5p | MIMAT0000530 | Ifi47 | 15953 | Microarrays//HITS-CLIP | 37986224\|23142080\|29167373 | Unspecified |
| mmu-miR-21a-5p | MIMAT0000530 | Ifit1 | 15957 | Microarrays//HITS-CLIP | 37986224\|29167373 | Unspecified |
| mmu-miR-21a-5p | MIMAT0000530 | Ifit2 | 15958 | HITS-CLIP | 37986224\|23142080 | Unspecified |
| mmu-miR-21a-5p | MIMAT0000530 | Ifit3 | 15959 | Microarrays | 37986224 | Unspecified |
| mmu-miR-21a-5p | MIMAT0000530 | Ifitm1 | 68713 | Microarrays | 37986224 | Unspecified |
| mmu-miR-21a-5p | MIMAT0000530 | Ifitm2 | 80876 | Microarrays | 37986224 | Unspecified |
| mmu-miR-21a-5p | MIMAT0000530 | Ifitm3 | 66141 | Microarrays | 37986224 | Unspecified |
| mmu-miR-21a-5p | MIMAT0000530 | Ifitm6 | 213002 | Microarrays | 37986224 | Unspecified |
| mmu-miR-21a-5p | MIMAT0000530 | Ifnar1 | 15975 | Microarrays//HITS-CLIP | 37986224\|23142080 | Unspecified |
| mmu-miR-21a-5p | MIMAT0000530 | Ifnar2 | 15976 | HITS-CLIP | 23142080 | Unspecified |
| mmu-miR-21a-5p | MIMAT0000530 | Ifngr1 | 15979 | HITS-CLIP | 37986224\|23142080\|29167373 | Unspecified |
| mmu-miR-21a-5p | MIMAT0000530 | Ifrd1 | 15982 | HITS-CLIP, Microarrays//Microarrays//HITS | 37986224\|25263593\|23142080\|29167 | Unspecified |
| mmu-miR-21a-5p | MIMAT0000530 | Ift172 | 67661 | Microarrays | 37986224 | Unspecified |
| mmu-miR-21a-5p | MIMAT0000530 | Ift80 | 68259 | HITS-CLIP | 23142080 | Unspecified |
| mmu-miR-21a-5p | MIMAT0000530 | Igf1 | 16000 | HITS-CLIP | 23597149\|29167373 | Unspecified |
| mmu-miR-21a-5p | MIMAT0000530 | Igf1r | 16001 | Microarrays//HITS-CLIP | 37986224\|23142080 | Unspecified |
| mmu-miR-21a-5p | MIMAT0000530 | Igf2bp3 | 140488 | Microarrays//HITS-CLIP | 37986224\|25263593\|23142080 | Unspecified |
| mmu-miR-21a-5p | MIMAT0000530 | Igf2r | 16004 | HITS-CLIP | 23597149\|23142080 | Unspecified |
| mmu-miR-21a-5p | MIMAT0000530 | Igfals | 16005 | HITS-CLIP | 23597149 | Unspecified |
| mmu-miR-21a-5p | MIMAT0000530 | Igfbp1 | 16006 | HITS-CLIP | 23597149 | Unspecified |
| mmu-miR-21a-5p | MIMAT0000530 | Igfbp2 | 16008 | HITS-CLIP | 23597149 | Unspecified |
| mmu-miR-21a-5p | MIMAT0000530 | Igfbp4 | 16010 | HITS-CLIP | 23597149 | Unspecified |
| mmu-miR-21a-5p | MIMAT0000530 | Ighmbp2 | 20589 | HITS-CLIP | 23142080 | Unspecified |
| mmu-miR-21a-5p | MIMAT0000530 | Iglon5 | 210094 | HITS-CLIP//qCLASH | 23597149\|31393953 | Unspecified |
| mmu-miR-21a-5p | MIMAT0000530 | Igsf23 | 70080 | HITS-CLIP | 23142080 | Unspecified |
| mmu-miR-21a-5p | MIMAT0000530 | Igsf6 | 80719 | Microarrays | 37986224 | Unspecified |
| mmu-miR-21a-5p | MIMAT0000530 | Igsf8 | 140559 | Microarrays | 37986224\|25263593 | Unspecified |
| mmu-miR-21a-5p | MIMAT0000530 | Iigp1 | 60440 | Microarrays//HITS-CLIP | 37986224\|23597149\|23142080 | Unspecified |
| mmu-miR-21a-5p | MIMAT0000530 | Ik | 24010 | HITS-CLIP | 23597149\|23142080\|29167373 | Unspecified |
| mmu-miR-21a-5p | MIMAT0000530 | Ikbip | 67454 | HITS-CLIP | 23142080 | Unspecified |
| mmu-miR-21a-5p | MIMAT0000530 | Ikbkb | 16150 | HITS-CLIP | 23142080 | Unspecified |
| mmu-miR-21a-5p | MIMAT0000530 | Ikbkg | 16151 | HITS-CLIP | 23597149\|23142080 | Unspecified |
| mmu-miR-21a-5p | MIMAT0000530 | Ikzf1 | 22778 | HITS-CLIP | 37986224\|23142080\|29167373 | Unspecified |
| mmu-miR-21a-5p | MIMAT0000530 | Ikzf2 | 22779 | Microarrays//HITS-CLIP | 37986224\|23142080 | Unspecified |
| mmu-miR-21a-5p | MIMAT0000530 | Ikzf3 | 22780 | HITS-CLIP | 23142080 | Unspecified |
| mmu-miR-21a-5p | MIMAT0000530 | Ikzf4 | 22781 | HITS-CLIP | 23142080 | Unspecified |
| mmu-miR-21a-5p | MIMAT0000530 | Il10 | 16153 | HITS-CLIP | 23142080 | Unspecified |
| mmu-miR-21a-5p | MIMAT0000530 | Il12rb1 | 16161 | Microarrays | 37986224\|25263593 | Unspecified |

| | | | | | | |
|---|---|---|---|---|---|---|
| mmu-miR-21a-5p | MIMAT0000530 | Il12rb2 | 16162 | HITS-CLIP | 23142080 | Unspecified |
| mmu-miR-21a-5p | MIMAT0000530 | Il15 | 16168 | Microarrays | 37986224 | Unspecified |
| mmu-miR-21a-5p | MIMAT0000530 | Il16 | 16170 | HITS-CLIP | 23142080 | Unspecified |
| mmu-miR-21a-5p | MIMAT0000530 | Il18 | 16173 | Microarrays | 37986224 | Unspecified |
| mmu-miR-21a-5p | MIMAT0000530 | Il18bp | 16068 | Microarrays//HITS-CLIP | 37986224\|23597149 | Unspecified |
| mmu-miR-21a-5p | MIMAT0000530 | Il1b | 16176 | Microarrays | 37986224 | Unspecified |
| mmu-miR-21a-5p | MIMAT0000530 | Il1rl2 | 107527 | HITS-CLIP | 23142080 | Unspecified |
| mmu-miR-21a-5p | MIMAT0000530 | Il1rn | 16181 | Microarrays | 37986224\|25263593 | Unspecified |
| mmu-miR-21a-5p | MIMAT0000530 | Il21 | 60505 | HITS-CLIP | 37986224\|23142080 | Unspecified |
| mmu-miR-21a-5p | MIMAT0000530 | Il21r | 60504 | HITS-CLIP | 29167373 | Unspecified |
| mmu-miR-21a-5p | MIMAT0000530 | Il2ra | 16184 | HITS-CLIP | 37986224\|31433990\|23142080 | Unspecified |
| mmu-miR-21a-5p | MIMAT0000530 | Il2rb | 16185 | HITS-CLIP | 23142080 | Unspecified |
| mmu-miR-21a-5p | MIMAT0000530 | Il2rg | 16186 | HITS-CLIP | 23142080 | Unspecified |
| mmu-miR-21a-5p | MIMAT0000530 | Il4ra | 16190 | Microarrays//HITS-CLIP | 37986224\|23597149\|29167373\|2314 | Unspecified |
| mmu-miR-21a-5p | MIMAT0000530 | Il5ra | 16192 | Microarrays | 37986224\|25263593 | Unspecified |
| mmu-miR-21a-5p | MIMAT0000530 | Il6ra | 16194 | Microarrays//HITS-CLIP | 37986224\|23142080\|23597149 | Unspecified |
| mmu-miR-21a-5p | MIMAT0000530 | Il6st | 16195 | HITS-CLIP | 37986224\|23142080\|23597149\|2916 | Unspecified |
| mmu-miR-21a-5p | MIMAT0000530 | Il7r | 16197 | HITS-CLIP | 23142080 | Unspecified |
| mmu-miR-21a-5p | MIMAT0000530 | Ildr2 | 100039795 | HITS-CLIP | 37986224 | Unspecified |
| mmu-miR-21a-5p | MIMAT0000530 | Imp3 | 102462 | HITS-CLIP | 23142080 | Unspecified |
| mmu-miR-21a-5p | MIMAT0000530 | Imp4 | 27993 | Microarrays | 37986224\|25263593 | Unspecified |
| mmu-miR-21a-5p | MIMAT0000530 | Impa1 | 55980 | HITS-CLIP | 23142080 | Unspecified |
| mmu-miR-21a-5p | MIMAT0000530 | Impa2 | 114663 | HITS-CLIP | 23142080 | Unspecified |
| mmu-miR-21a-5p | MIMAT0000530 | Impad1 | 242291 | HITS-CLIP, Microarrays//HITS-CLIP | 37986224\|23597149\|23142080\|2916 | Unspecified |
| mmu-miR-21a-5p | MIMAT0000530 | Impdh1 | 23917 | HITS-CLIP | 23142080 | Unspecified |
| mmu-miR-21a-5p | MIMAT0000530 | Inafm2 | 100505573 | HITS-CLIP | 23142080 | Unspecified |
| mmu-miR-21a-5p | MIMAT0000530 | Incenp | 16319 | HITS-CLIP | 23597149\|23142080 | Unspecified |
| mmu-miR-21a-5p | MIMAT0000530 | Inf2 | 70435 | Microarrays//HITS-CLIP | 37986224\|25263593\|23597149 | Unspecified |
| mmu-miR-21a-5p | MIMAT0000530 | Ing1 | 26356 | HITS-CLIP | 37986224\|23142080 | Unspecified |
| mmu-miR-21a-5p | MIMAT0000530 | Ing2 | 69260 | Microarrays//HITS-CLIP | 37986224\|23142080 | Unspecified |
| mmu-miR-21a-5p | MIMAT0000530 | Ing3 | 71777 | Microarrays//HITS-CLIP | 37986224\|23142080 | Unspecified |
| mmu-miR-21a-5p | MIMAT0000530 | Ing4 | 28019 | HITS-CLIP | 37986224 | Unspecified |
| mmu-miR-21a-5p | MIMAT0000530 | Ing5 | 66262 | HITS-CLIP | 23142080 | Unspecified |
| mmu-miR-21a-5p | MIMAT0000530 | Inhba | 16323 | HITS-CLIP | 19536157\|23142080 | Unspecified |
| mmu-miR-21a-5p | MIMAT0000530 | Inhbb | 16324 | Microarrays | 37986224\|25263593 | Unspecified |
| mmu-miR-21a-5p | MIMAT0000530 | Inip | 66209 | HITS-CLIP | 23142080 | Unspecified |
| mmu-miR-21a-5p | MIMAT0000530 | Ino80d | 227195 | HITS-CLIP | 23142080\|29167373 | Unspecified |
| mmu-miR-21a-5p | MIMAT0000530 | Inpp4b | 234515 | HITS-CLIP, Microarrays//HITS-CLIP//Microa | 37986224\|23142080\|25263593 | Unspecified |
| mmu-miR-21a-5p | MIMAT0000530 | Inpp5a | 212111 | HITS-CLIP | 23142080 | Unspecified |

| | | | | | | |
|---|---|---|---|---|---|---|
| mmu-miR-21a-5p | MIMAT0000530 | Inpp5b | 16330 | HITS-CLIP | 23597149 | Unspecified |
| mmu-miR-21a-5p | MIMAT0000530 | Inpp5d | 16331 | HITS-CLIP | 23142080 | Unspecified |
| mmu-miR-21a-5p | MIMAT0000530 | Inpp5f | 101490 | HITS-CLIP | 37986224\|23142080 | Unspecified |
| mmu-miR-21a-5p | MIMAT0000530 | Inpp5k | 19062 | Microarrays | 37986224 | Unspecified |
| mmu-miR-21a-5p | MIMAT0000530 | Inppl1 | 16332 | HITS-CLIP | 23142080\|29167373 | Unspecified |
| mmu-miR-21a-5p | MIMAT0000530 | Insig1 | 231070 | HITS-CLIP | 23597149 | Unspecified |
| mmu-miR-21a-5p | MIMAT0000530 | Insig2 | 72999 | HITS-CLIP | 37986224\|23597149\|23142080 | Unspecified |
| mmu-miR-21a-5p | MIMAT0000530 | Insr | 16337 | HITS-CLIP | 23597149\|19536157\|23142080 | Unspecified |
| mmu-miR-21a-5p | MIMAT0000530 | Ints1 | 68510 | HITS-CLIP | 29167373 | Unspecified |
| mmu-miR-21a-5p | MIMAT0000530 | Ints10 | 70885 | HITS-CLIP | 23142080 | Unspecified |
| mmu-miR-21a-5p | MIMAT0000530 | Ints11 | 54973 | HITS-CLIP | 23142080 | Unspecified |
| mmu-miR-21a-5p | MIMAT0000530 | Ints12 | 71793 | HITS-CLIP | 23142080 | Unspecified |
| mmu-miR-21a-5p | MIMAT0000530 | Ints2 | 70422 | HITS-CLIP | 23142080 | Unspecified |
| mmu-miR-21a-5p | MIMAT0000530 | Ints6 | 18130 | HITS-CLIP | 23142080 | Unspecified |
| mmu-miR-21a-5p | MIMAT0000530 | Ints7 | 77065 | HITS-CLIP | 23597149 | Unspecified |
| mmu-miR-21a-5p | MIMAT0000530 | Ints8 | 72656 | HITS-CLIP | 31433990\|23142080 | Unspecified |
| mmu-miR-21a-5p | MIMAT0000530 | Ints9 | 210925 | HITS-CLIP | 23142080 | Unspecified |
| mmu-miR-21a-5p | MIMAT0000530 | Ip6k1 | 27399 | HITS-CLIP | 28735896\|23142080 | Unspecified |
| mmu-miR-21a-5p | MIMAT0000530 | Ip6k2 | 76500 | HITS-CLIP | 37986224\|23142080 | Unspecified |
| mmu-miR-21a-5p | MIMAT0000530 | Ipcef1 | 320495 | HITS-CLIP | 37986224 | Unspecified |
| mmu-miR-21a-5p | MIMAT0000530 | Ipo11 | 76582 | Microarrays//HITS-CLIP | 37986224\|23142080 | Unspecified |
| mmu-miR-21a-5p | MIMAT0000530 | Ipo4 | 75751 | HITS-CLIP | 23142080 | Unspecified |
| mmu-miR-21a-5p | MIMAT0000530 | Ipo5 | 70572 | HITS-CLIP | 23597149\|29167373\|23142080 | Unspecified |
| mmu-miR-21a-5p | MIMAT0000530 | Ipo7 | 233726 | HITS-CLIP | 23597149\|31433990\|23142080 | Unspecified |
| mmu-miR-21a-5p | MIMAT0000530 | Ippk | 75678 | HITS-CLIP | 23142080 | Unspecified |
| mmu-miR-21a-5p | MIMAT0000530 | Iqcb1 | 320299 | Microarrays | 37986224 | Unspecified |
| mmu-miR-21a-5p | MIMAT0000530 | Iqce | 74239 | HITS-CLIP | 23142080 | Unspecified |
| mmu-miR-21a-5p | MIMAT0000530 | Iqgap1 | 29875 | HITS-CLIP | 23142080\|23597149\|29167373 | Unspecified |
| mmu-miR-21a-5p | MIMAT0000530 | Iqgap2 | 544963 | HITS-CLIP | 37986224\|23597149\|29167373 | Unspecified |
| mmu-miR-21a-5p | MIMAT0000530 | Iqgap3 | 404710 | HITS-CLIP | 26203562 | Unspecified |
| mmu-miR-21a-5p | MIMAT0000530 | Iqsec1 | 232227 | HITS-CLIP | 23597149 | Unspecified |
| mmu-miR-21a-5p | MIMAT0000530 | Irag2 | 16970 | Microarrays | 25263593 | Unspecified |
| mmu-miR-21a-5p | MIMAT0000530 | Irak2 | 108960 | Microarrays//HITS-CLIP | 37986224\|25263593\|23597149 | Unspecified |
| mmu-miR-21a-5p | MIMAT0000530 | Irak3 | 73914 | HITS-CLIP | 29167373 | Unspecified |
| mmu-miR-21a-5p | MIMAT0000530 | Irak4 | 266632 | Microarrays | 37986224\|25263593 | Unspecified |
| mmu-miR-21a-5p | MIMAT0000530 | Ireb2 | 64602 | HITS-CLIP | 23597149\|29167373\|23142080 | Unspecified |
| mmu-miR-21a-5p | MIMAT0000530 | Irf1 | 16362 | Microarrays//HITS-CLIP | 37986224\|23142080\|29167373 | Unspecified |
| mmu-miR-21a-5p | MIMAT0000530 | Irf2 | 16363 | HITS-CLIP | 23597149\|23142080\|29167373 | Unspecified |
| mmu-miR-21a-5p | MIMAT0000530 | Irf2bp2 | 270110 | Microarrays//HITS-CLIP | 37986224\|23142080\|29167373 | Unspecified |

| | | | | | | |
|---|---|---|---|---|---|---|
| mmu-miR-21a-5p | MIMAT0000530 | Irf2bpl | 238330 | Microarrays//HITS-CLIP | 37986224\|25263593\|23597149 | Unspecified |
| mmu-miR-21a-5p | MIMAT0000530 | Irf4 | 16364 | Microarrays//HITS-CLIP | 37986224\|23142080\|31433990 | Unspecified |
| mmu-miR-21a-5p | MIMAT0000530 | Irf7 | 54123 | Microarrays | 37986224 | Unspecified |
| mmu-miR-21a-5p | MIMAT0000530 | Irf8 | 15900 | Microarrays//HITS-CLIP | 37986224\|23142080 | Unspecified |
| mmu-miR-21a-5p | MIMAT0000530 | Irf9 | 16391 | HITS-CLIP | 23142080 | Unspecified |
| mmu-miR-21a-5p | MIMAT0000530 | Irgm1 | 15944 | HITS-CLIP | 23142080 | Unspecified |
| mmu-miR-21a-5p | MIMAT0000530 | Irgm2 | 54396 | HITS-CLIP | 23142080 | Unspecified |
| mmu-miR-21a-5p | MIMAT0000530 | Irs1 | 16367 | HITS-CLIP | 37986224\|23597149 | Unspecified |
| mmu-miR-21a-5p | MIMAT0000530 | Irs2 | 384783 | HITS-CLIP | 23597149 | Unspecified |
| mmu-miR-21a-5p | MIMAT0000530 | Irs3 | 16369 | Microarrays | 37986224\|25263593 | Unspecified |
| mmu-miR-21a-5p | MIMAT0000530 | Isca2 | 74316 | HITS-CLIP | 23142080 | Unspecified |
| mmu-miR-21a-5p | MIMAT0000530 | Isoc1 | 66307 | HITS-CLIP | 23142080 | Unspecified |
| mmu-miR-21a-5p | MIMAT0000530 | Isoc2a | 664994 | HITS-CLIP | 23597149 | Unspecified |
| mmu-miR-21a-5p | MIMAT0000530 | Ist1 | 71955 | HITS-CLIP | 23142080 | Unspecified |
| mmu-miR-21a-5p | MIMAT0000530 | Isyna1 | 71780 | Microarrays | 37986224\|25263593 | Unspecified |
| mmu-miR-21a-5p | MIMAT0000530 | Itch | 16396 | Microarrays//HITS-CLIP | 37986224\|23597149 | Unspecified |
| mmu-miR-21a-5p | MIMAT0000530 | Itfg1 | 71927 | HITS-CLIP | 19536157 | Unspecified |
| mmu-miR-21a-5p | MIMAT0000530 | Itga4 | 16401 | Microarrays//HITS-CLIP | 37986224\|23142080\|29167373 | Unspecified |
| mmu-miR-21a-5p | MIMAT0000530 | Itga5 | 16402 | HITS-CLIP | 29167373 | Unspecified |
| mmu-miR-21a-5p | MIMAT0000530 | Itga6 | 16403 | HITS-CLIP | 23142080 | Unspecified |
| mmu-miR-21a-5p | MIMAT0000530 | Itga8 | 241226 | Microarrays | 37986224\|25263593 | Unspecified |
| mmu-miR-21a-5p | MIMAT0000530 | Itgal | 16408 | HITS-CLIP | 23142080 | Unspecified |
| mmu-miR-21a-5p | MIMAT0000530 | Itgam | 16409 | HITS-CLIP | 29167373 | Unspecified |
| mmu-miR-21a-5p | MIMAT0000530 | Itgav | 16410 | HITS-CLIP | 37986224\|23597149\|29167373\|23142 | Unspecified |
| mmu-miR-21a-5p | MIMAT0000530 | Itgax | 16411 | Microarrays | 37986224\|25263593 | Unspecified |
| mmu-miR-21a-5p | MIMAT0000530 | Itgb1 | 16412 | HITS-CLIP | 37986224\|23597149\|23142080\|29167 | Unspecified |
| mmu-miR-21a-5p | MIMAT0000530 | Itgb2 | 16414 | HITS-CLIP | 23142080\|29167373 | Unspecified |
| mmu-miR-21a-5p | MIMAT0000530 | Itgb3 | 16416 | Microarrays//HITS-CLIP | 37986224\|25263593\|31433990\|23142 | Unspecified |
| mmu-miR-21a-5p | MIMAT0000530 | Itgb5 | 16419 | HITS-CLIP | 29167373 | Unspecified |
| mmu-miR-21a-5p | MIMAT0000530 | Itgb8 | 320910 | HITS-CLIP | 37986224\|19536157\|23142080 | Unspecified |
| mmu-miR-21a-5p | MIMAT0000530 | Itih2 | 16425 | HITS-CLIP | 23597149 | Unspecified |
| mmu-miR-21a-5p | MIMAT0000530 | Itih4 | 16427 | HITS-CLIP | 23597149 | Unspecified |
| mmu-miR-21a-5p | MIMAT0000530 | Itk | 16428 | HITS-CLIP | 23142080 | Unspecified |
| mmu-miR-21a-5p | MIMAT0000530 | Itm2b | 16432 | HITS-CLIP | 29167373 | Unspecified |
| mmu-miR-21a-5p | MIMAT0000530 | Itpa | 16434 | HITS-CLIP | 23142080 | Unspecified |
| mmu-miR-21a-5p | MIMAT0000530 | Itpkb | 320404 | HITS-CLIP | 23142080 | Unspecified |
| mmu-miR-21a-5p | MIMAT0000530 | Itpr1 | 16438 | HITS-CLIP | 23142080 | Unspecified |
| mmu-miR-21a-5p | MIMAT0000530 | Itpr2 | 16439 | HITS-CLIP | 23142080 | Unspecified |
| mmu-miR-21a-5p | MIMAT0000530 | Itprid2 | 6744 | HITS-CLIP | 19536157\|23142080 | Unspecified |

| | | | | | | |
|---|---|---|---|---|---|---|
| mmu-miR-21a-5p | MIMAT0000530 | Itpripl2 | 319622 | HITS-CLIP | 23142080\|29167373 | Unspecified |
| mmu-miR-21a-5p | MIMAT0000530 | Itsn1 | 16443 | HITS-CLIP | 29167373 | Unspecified |
| mmu-miR-21a-5p | MIMAT0000530 | Itsn2 | 20403 | HITS-CLIP | 37986224\|23142080\|29167373\|23597 | Unspecified |
| mmu-miR-21a-5p | MIMAT0000530 | Ivd | 56357 | Microarrays//HITS-CLIP | 37986224\|25263593\|23597149 | Unspecified |
| mmu-miR-21a-5p | MIMAT0000530 | Ivns1abp | 117198 | HITS-CLIP, Microarrays//HITS-CLIP | 37986224\|23597149\|23142080 | Unspecified |
| mmu-miR-21a-5p | MIMAT0000530 | Iws1 | 73473 | HITS-CLIP | 23142080 | Unspecified |
| mmu-miR-21a-5p | MIMAT0000530 | Iyd | 70337 | HITS-CLIP | 23597149 | Unspecified |
| mmu-miR-21a-5p | MIMAT0000530 | Jade1 | 269424 | HITS-CLIP | 23142080\|23597149 | Unspecified |
| mmu-miR-21a-5p | MIMAT0000530 | Jade2 | 76901 | HITS-CLIP | 23597149 | Unspecified |
| mmu-miR-21a-5p | MIMAT0000530 | Jak1 | 16451 | HITS-CLIP | 23597149\|31433990\|23142080 | Unspecified |
| mmu-miR-21a-5p | MIMAT0000530 | Jak2 | 16452 | HITS-CLIP | 23142080 | Unspecified |
| mmu-miR-21a-5p | MIMAT0000530 | Jak3 | 16453 | HITS-CLIP | 23597149\|23142080 | Unspecified |
| mmu-miR-21a-5p | MIMAT0000530 | Jakmip1 | 76071 | HITS-CLIP | 19536157 | Unspecified |
| mmu-miR-21a-5p | MIMAT0000530 | Jaml | 120425 | HITS-CLIP | 23142080 | Unspecified |
| mmu-miR-21a-5p | MIMAT0000530 | Jarid2 | 16468 | Microarrays//HITS-CLIP//qCLASH | 37986224\|23142080\|31393953\|23597 | Unspecified |
| mmu-miR-21a-5p | MIMAT0000530 | Jazf1 | 231986 | HITS-CLIP | 23142080 | Unspecified |
| mmu-miR-21a-5p | MIMAT0000530 | Jcad | 57608 | Microarrays//HITS-CLIP | 25263593\|31433990 | Unspecified |
| mmu-miR-21a-5p | MIMAT0000530 | Jchain | 16069 | qCLASH | 31393953 | Unspecified |
| mmu-miR-21a-5p | MIMAT0000530 | Jdp2 | 81703 | HITS-CLIP | 23142080\|23597149\|29167373 | Unspecified |
| mmu-miR-21a-5p | MIMAT0000530 | Jmjd1c | 108829 | HITS-CLIP//qCLASH | 37986224\|23142080\|19536157\|31393 | Unspecified |
| mmu-miR-21a-5p | MIMAT0000530 | Jmjd6 | 107817 | HITS-CLIP | 23597149\|23142080 | Unspecified |
| mmu-miR-21a-5p | MIMAT0000530 | Jmy | 57748 | HITS-CLIP | 23142080 | Unspecified |
| mmu-miR-21a-5p | MIMAT0000530 | Josd1 | 74158 | HITS-CLIP | 31433990\|23142080 | Unspecified |
| mmu-miR-21a-5p | MIMAT0000530 | Josd2 | 66124 | HITS-CLIP | 30971389 | Unspecified |
| mmu-miR-21a-5p | MIMAT0000530 | Jph4 | 319984 | HITS-CLIP | 19536157 | Unspecified |
| mmu-miR-21a-5p | MIMAT0000530 | Jpt2 | 90861 | Microarrays | 25263593 | Unspecified |
| mmu-miR-21a-5p | MIMAT0000530 | Jun | 16476 | Microarrays//HITS-CLIP | 37986224\|25263593\|23597149\|23142 | Unspecified |
| mmu-miR-21a-5p | MIMAT0000530 | Jup | 16480 | HITS-CLIP | 26203562 | Unspecified |
| mmu-miR-21a-5p | MIMAT0000530 | Kalrn | 545156 | HITS-CLIP | 19536157 | Unspecified |
| mmu-miR-21a-5p | MIMAT0000530 | Kansl1 | 76719 | HITS-CLIP | 23597149\|23142080\|29167373 | Unspecified |
| mmu-miR-21a-5p | MIMAT0000530 | Kansl1l | 68691 | HITS-CLIP | 29167373 | Unspecified |
| mmu-miR-21a-5p | MIMAT0000530 | Kansl3 | 226976 | HITS-CLIP | 23142080 | Unspecified |
| mmu-miR-21a-5p | MIMAT0000530 | Kantr | 102723508 | HITS-CLIP | 23142080\|19536157\|29167373 | Unspecified |
| mmu-miR-21a-5p | MIMAT0000530 | Kars | 85305 | HITS-CLIP | 23142080 | Unspecified |
| mmu-miR-21a-5p | MIMAT0000530 | Kat14 | 57325 | HITS-CLIP | 23597149 | Unspecified |
| mmu-miR-21a-5p | MIMAT0000530 | Kat2a | 14534 | HITS-CLIP | 19536157\|23597149\|29167373 | Unspecified |
| mmu-miR-21a-5p | MIMAT0000530 | Kat2b | 18519 | HITS-CLIP | 29167373 | Unspecified |
| mmu-miR-21a-5p | MIMAT0000530 | Kat6a | 244349 | HITS-CLIP | 23597149\|23142080 | Unspecified |
| mmu-miR-21a-5p | MIMAT0000530 | Kat6b | 54169 | HITS-CLIP | 23142080 | Unspecified |

| | | | | | | |
|---|---|---|---|---|---|---|
| mmu-miR-21a-5p | MIMAT0000530 | Kat7 | 217127 | HITS-CLIP | 23597149\|23142080\|29167373 | Unspecified |
| mmu-miR-21a-5p | MIMAT0000530 | Kat8 | 67773 | HITS-CLIP | 23142080 | Unspecified |
| mmu-miR-21a-5p | MIMAT0000530 | Katna1 | 23924 | HITS-CLIP | 37986224 | Unspecified |
| mmu-miR-21a-5p | MIMAT0000530 | Kbtbd11 | 74901 | HITS-CLIP | 37986224 | Unspecified |
| mmu-miR-21a-5p | MIMAT0000530 | Kbtbd2 | 210973 | HITS-CLIP | 23142080 | Unspecified |
| mmu-miR-21a-5p | MIMAT0000530 | Kbtbd4 | 67136 | HITS-CLIP | 23142080\|29167373 | Unspecified |
| mmu-miR-21a-5p | MIMAT0000530 | Kbtbd7 | 211255 | HITS-CLIP | 23142080 | Unspecified |
| mmu-miR-21a-5p | MIMAT0000530 | Kcnb1 | 16500 | HITS-CLIP | 19536157 | Unspecified |
| mmu-miR-21a-5p | MIMAT0000530 | Kcnj10 | 16513 | HITS-CLIP | 37986224\|19536157 | Unspecified |
| mmu-miR-21a-5p | MIMAT0000530 | Kcnj2 | 16518 | HITS-CLIP | 29167373 | Unspecified |
| mmu-miR-21a-5p | MIMAT0000530 | Kcnk13 | 56659 | HITS-CLIP | 29167373 | Unspecified |
| mmu-miR-21a-5p | MIMAT0000530 | Kcnn2 | 140492 | HITS-CLIP | 19536157 | Unspecified |
| mmu-miR-21a-5p | MIMAT0000530 | Kcnn4 | 16534 | Microarrays//HITS-CLIP | 37986224\|25263593\|23142080\|29167 | Unspecified |
| mmu-miR-21a-5p | MIMAT0000530 | Kcnq3 | 110862 | HITS-CLIP | 19536157 | Unspecified |
| mmu-miR-21a-5p | MIMAT0000530 | Kcnt1 | 227632 | HITS-CLIP | 19536157 | Unspecified |
| mmu-miR-21a-5p | MIMAT0000530 | Kctd10 | 330171 | HITS-CLIP | 23597149 | Unspecified |
| mmu-miR-21a-5p | MIMAT0000530 | Kctd12 | 239217 | HITS-CLIP | 37986224\|31433990\|23597149 | Unspecified |
| mmu-miR-21a-5p | MIMAT0000530 | Kctd12b | 207474 | Microarrays | 37986224 | Unspecified |
| mmu-miR-21a-5p | MIMAT0000530 | Kctd13 | 233877 | HITS-CLIP | 23142080 | Unspecified |
| mmu-miR-21a-5p | MIMAT0000530 | Kctd17 | 72844 | Microarrays | 37986224\|25263593 | Unspecified |
| mmu-miR-21a-5p | MIMAT0000530 | Kctd5 | 69259 | HITS-CLIP | 23142080 | Unspecified |
| mmu-miR-21a-5p | MIMAT0000530 | Kctd9 | 105440 | HITS-CLIP | 23142080 | Unspecified |
| mmu-miR-21a-5p | MIMAT0000530 | Kdelr2 | 66913 | HITS-CLIP | 23142080 | Unspecified |
| mmu-miR-21a-5p | MIMAT0000530 | Kdm1a | 99982 | qCLASH | 31393953 | Unspecified |
| mmu-miR-21a-5p | MIMAT0000530 | Kdm2a | 225876 | HITS-CLIP | 23597149\|23142080 | Unspecified |
| mmu-miR-21a-5p | MIMAT0000530 | Kdm2b | 30841 | HITS-CLIP | 23142080 | Unspecified |
| mmu-miR-21a-5p | MIMAT0000530 | Kdm3a | 104263 | HITS-CLIP | 23142080\|23597149\|29167373 | Unspecified |
| mmu-miR-21a-5p | MIMAT0000530 | Kdm3b | 277250 | HITS-CLIP | 23142080 | Unspecified |
| mmu-miR-21a-5p | MIMAT0000530 | Kdm5a | 214899 | HITS-CLIP | 23142080 | Unspecified |
| mmu-miR-21a-5p | MIMAT0000530 | Kdm5b | 75605 | HITS-CLIP | 23142080 | Unspecified |
| mmu-miR-21a-5p | MIMAT0000530 | Kdm5c | 20591 | HITS-CLIP | 23597149\|23142080 | Unspecified |
| mmu-miR-21a-5p | MIMAT0000530 | Kdm5d | 20592 | HITS-CLIP | 23142080 | Unspecified |
| mmu-miR-21a-5p | MIMAT0000530 | Kdm6a | 22289 | HITS-CLIP | 23142080\|29167373 | Unspecified |
| mmu-miR-21a-5p | MIMAT0000530 | Kdm6b | 216850 | Microarrays | 37986224 | Unspecified |
| mmu-miR-21a-5p | MIMAT0000530 | Kdm7a | 338523 | HITS-CLIP | 37986224\|23142080\|19536157 | Unspecified |
| mmu-miR-21a-5p | MIMAT0000530 | Kdr | 16542 | HITS-CLIP | 23597149 | Unspecified |
| mmu-miR-21a-5p | MIMAT0000530 | Kdsr | 70750 | HITS-CLIP | 23142080 | Unspecified |
| mmu-miR-21a-5p | MIMAT0000530 | Khdc4 | 22889 | HITS-CLIP | 23142080 | Unspecified |
| mmu-miR-21a-5p | MIMAT0000530 | Khnyn | 219094 | Microarrays//HITS-CLIP | 37986224\|23597149\|23142080\|29167 | Unspecified |

| | | | | | | |
|---|---|---|---|---|---|---|
| mmu-miR-21a-5p | MIMAT0000530 | Kidins220 | 77480 | HITS-CLIP | 37986224\|23597149\|19536157\|23142 | Unspecified |
| mmu-miR-21a-5p | MIMAT0000530 | Kif11 | 16551 | Microarrays//HITS-CLIP | 37986224\|25263593\|23142080 | Unspecified |
| mmu-miR-21a-5p | MIMAT0000530 | Kif13b | 16554 | HITS-CLIP | 29167373 | Unspecified |
| mmu-miR-21a-5p | MIMAT0000530 | Kif14 | 381293 | HITS-CLIP | 23142080 | Unspecified |
| mmu-miR-21a-5p | MIMAT0000530 | Kif15 | 209737 | HITS-CLIP | 23142080 | Unspecified |
| mmu-miR-21a-5p | MIMAT0000530 | Kif16b | 16558 | HITS-CLIP | 23142080 | Unspecified |
| mmu-miR-21a-5p | MIMAT0000530 | Kif18a | 228421 | Microarrays//HITS-CLIP | 37986224\|25263593\|23142080 | Unspecified |
| mmu-miR-21a-5p | MIMAT0000530 | Kif18b | 70218 | Microarrays//HITS-CLIP | 37986224\|25263593\|23142080 | Unspecified |
| mmu-miR-21a-5p | MIMAT0000530 | Kif1a | 16560 | HITS-CLIP | 37986224\|19536157 | Unspecified |
| mmu-miR-21a-5p | MIMAT0000530 | Kif1b | 16561 | HITS-CLIP | 23597149\|29167373\|23142080 | Unspecified |
| mmu-miR-21a-5p | MIMAT0000530 | Kif1c | 16562 | HITS-CLIP | 23597149 | Unspecified |
| mmu-miR-21a-5p | MIMAT0000530 | Kif20a | 19348 | Microarrays//HITS-CLIP | 37986224\|23142080\|25263593 | Unspecified |
| mmu-miR-21a-5p | MIMAT0000530 | Kif20b | 240641 | HITS-CLIP | 23142080 | Unspecified |
| mmu-miR-21a-5p | MIMAT0000530 | Kif21b | 16565 | HITS-CLIP | 19536157\|23142080\|29167373 | Unspecified |
| mmu-miR-21a-5p | MIMAT0000530 | Kif22 | 110033 | Microarrays | 37986224\|25263593 | Unspecified |
| mmu-miR-21a-5p | MIMAT0000530 | Kif23 | 71819 | Microarrays | 37986224\|25263593 | Unspecified |
| mmu-miR-21a-5p | MIMAT0000530 | Kif24 | 109242 | HITS-CLIP | 29167373 | Unspecified |
| mmu-miR-21a-5p | MIMAT0000530 | Kif2a | 16563 | HITS-CLIP | 23142080 | Unspecified |
| mmu-miR-21a-5p | MIMAT0000530 | Kif2c | 73804 | Microarrays//HITS-CLIP | 37986224\|25263593\|23142080 | Unspecified |
| mmu-miR-21a-5p | MIMAT0000530 | Kif3a | 16568 | Microarrays//HITS-CLIP | 37986224\|23142080 | Unspecified |
| mmu-miR-21a-5p | MIMAT0000530 | Kif3b | 16569 | HITS-CLIP | 29167373 | Unspecified |
| mmu-miR-21a-5p | MIMAT0000530 | Kif4 | 16571 | Microarrays//HITS-CLIP | 37986224\|25263593\|23142080 | Unspecified |
| mmu-miR-21a-5p | MIMAT0000530 | Kif5b | 16573 | HITS-CLIP | 23597149\|23142080 | Unspecified |
| mmu-miR-21a-5p | MIMAT0000530 | Kif9 | 16578 | Microarrays | 37986224 | Unspecified |
| mmu-miR-21a-5p | MIMAT0000530 | Kit | 16590 | Microarrays | 37986224\|25263593 | Unspecified |
| mmu-miR-21a-5p | MIMAT0000530 | Kiz | 228730 | Microarrays//HITS-CLIP | 37986224\|23142080 | Unspecified |
| mmu-miR-21a-5p | MIMAT0000530 | Klc1 | 16593 | HITS-CLIP | 29167373 | Unspecified |
| mmu-miR-21a-5p | MIMAT0000530 | Klc2 | 16594 | HITS-CLIP | 23142080 | Unspecified |
| mmu-miR-21a-5p | MIMAT0000530 | Klf10 | 21847 | HITS-CLIP | 23142080\|29167373 | Unspecified |
| mmu-miR-21a-5p | MIMAT0000530 | Klf11 | 194655 | Microarrays//HITS-CLIP | 37986224\|23142080 | Unspecified |
| mmu-miR-21a-5p | MIMAT0000530 | Klf12 | 16597 | HITS-CLIP | 37986224\|23597149 | Unspecified |
| mmu-miR-21a-5p | MIMAT0000530 | Klf13 | 50794 | HITS-CLIP | 29167373\|23142080 | Unspecified |
| mmu-miR-21a-5p | MIMAT0000530 | Klf15 | 66277 | HITS-CLIP | 23597149 | Unspecified |
| mmu-miR-21a-5p | MIMAT0000530 | Klf2 | 16598 | HITS-CLIP | 23142080 | Unspecified |
| mmu-miR-21a-5p | MIMAT0000530 | Klf3 | 16599 | HITS-CLIP | 23142080\|23597149\|29167373 | Unspecified |
| mmu-miR-21a-5p | MIMAT0000530 | Klf6 | 23849 | HITS-CLIP | 23142080\|29167373 | Unspecified |
| mmu-miR-21a-5p | MIMAT0000530 | Klf7 | 93691 | HITS-CLIP | 23142080\|29167373 | Unspecified |
| mmu-miR-21a-5p | MIMAT0000530 | Klf9 | 16601 | HITS-CLIP | 23597149\|23142080 | Unspecified |
| mmu-miR-21a-5p | MIMAT0000530 | Klhdc3 | 71765 | HITS-CLIP | 23142080 | Unspecified |

| | | | | | | |
|---|---|---|---|---|---|---|
| mmu-miR-21a-5p | MIMAT0000530 | Klhl15 | 236904 | HITS-CLIP | 31433990\|23142080 | Unspecified |
| mmu-miR-21a-5p | MIMAT0000530 | Klhl18 | 270201 | HITS-CLIP | 23142080\|23597149 | Unspecified |
| mmu-miR-21a-5p | MIMAT0000530 | Klhl2 | 77113 | Microarrays | 37986224\|25263593 | Unspecified |
| mmu-miR-21a-5p | MIMAT0000530 | Klhl24 | 75785 | Microarrays//HITS-CLIP | 37986224\|23597149\|23142080 | Unspecified |
| mmu-miR-21a-5p | MIMAT0000530 | Klhl28 | 66689 | HITS-CLIP | 37986224\|23142080\|29167373 | Unspecified |
| mmu-miR-21a-5p | MIMAT0000530 | Klhl42 | 232539 | HITS-CLIP | 23142080 | Unspecified |
| mmu-miR-21a-5p | MIMAT0000530 | Klhl6 | 239743 | Microarrays//HITS-CLIP | 37986224\|23142080 | Unspecified |
| mmu-miR-21a-5p | MIMAT0000530 | Klhl7 | 52323 | HITS-CLIP | 23142080 | Unspecified |
| mmu-miR-21a-5p | MIMAT0000530 | Klhl9 | 242521 | HITS-CLIP | 37986224\|19536157\|23597149\|2314 | Unspecified |
| mmu-miR-21a-5p | MIMAT0000530 | Klk1b11 | 16613 | Microarrays | 37986224\|25263593 | Unspecified |
| mmu-miR-21a-5p | MIMAT0000530 | Klra17 | 170733 | Microarrays | 37986224\|25263593 | Unspecified |
| mmu-miR-21a-5p | MIMAT0000530 | Klra2 | 16633 | Microarrays | 37986224\|25263593 | Unspecified |
| mmu-miR-21a-5p | MIMAT0000530 | Klrk1 | 27007 | Microarrays | 37986224 | Unspecified |
| mmu-miR-21a-5p | MIMAT0000530 | Kmo | 98256 | Microarrays | 37986224 | Unspecified |
| mmu-miR-21a-5p | MIMAT0000530 | Kmt2a | 214162 | HITS-CLIP | 19536157\|23142080\|29167373 | Unspecified |
| mmu-miR-21a-5p | MIMAT0000530 | Kmt2c | 231051 | HITS-CLIP | 37986224\|23142080\|19536157\|2359 | Unspecified |
| mmu-miR-21a-5p | MIMAT0000530 | Kmt2d | 381022 | HITS-CLIP | 23597149\|23142080\|26203562\|2916 | Unspecified |
| mmu-miR-21a-5p | MIMAT0000530 | Kmt2e | 69188 | Microarrays//HITS-CLIP | 37986224\|23142080\|29167373 | Unspecified |
| mmu-miR-21a-5p | MIMAT0000530 | Kmt5a | 387893 | HITS-CLIP | 23142080 | Unspecified |
| mmu-miR-21a-5p | MIMAT0000530 | Kng2 | 385643 | HITS-CLIP | 23597149 | Unspecified |
| mmu-miR-21a-5p | MIMAT0000530 | Knl1 | 57082 | HITS-CLIP | 23142080 | Unspecified |
| mmu-miR-21a-5p | MIMAT0000530 | Knop1 | 66356 | HITS-CLIP | 23142080 | Unspecified |
| mmu-miR-21a-5p | MIMAT0000530 | Knstrn | 51944 | Microarrays//HITS-CLIP | 37986224\|25263593\|23142080 | Unspecified |
| mmu-miR-21a-5p | MIMAT0000530 | Kntc1 | 208628 | Microarrays//HITS-CLIP | 37986224\|25263593\|23142080 | Unspecified |
| mmu-miR-21a-5p | MIMAT0000530 | Kpna1 | 16646 | HITS-CLIP | 23142080 | Unspecified |
| mmu-miR-21a-5p | MIMAT0000530 | Kpna2 | 16647 | Microarrays | 37986224\|25263593 | Unspecified |
| mmu-miR-21a-5p | MIMAT0000530 | Kpna3 | 16648 | HITS-CLIP | 23142080\|29167373 | Unspecified |
| mmu-miR-21a-5p | MIMAT0000530 | Kpna4 | 16649 | HITS-CLIP | 29167373\|23142080 | Unspecified |
| mmu-miR-21a-5p | MIMAT0000530 | Kpnb1 | 16211 | HITS-CLIP | 23597149\|23142080\|19536157\|2916 | Unspecified |
| mmu-miR-21a-5p | MIMAT0000530 | Kras | 16653 | HITS-CLIP | 29167373\|23142080 | Unspecified |
| mmu-miR-21a-5p | MIMAT0000530 | Krba1 | 77827 | HITS-CLIP | 23142080 | Unspecified |
| mmu-miR-21a-5p | MIMAT0000530 | Krit1 | 79264 | HITS-CLIP | 37986224\|23142080\|29167373 | Unspecified |
| mmu-miR-21a-5p | MIMAT0000530 | Krt5 | 110308 | HITS-CLIP | 26203562 | Unspecified |
| mmu-miR-21a-5p | MIMAT0000530 | Ktn1 | 16709 | HITS-CLIP | 23142080 | Unspecified |
| mmu-miR-21a-5p | MIMAT0000530 | Kxd1 | 79036 | HITS-CLIP | 23142080 | Unspecified |
| mmu-miR-21a-5p | MIMAT0000530 | Kyat1 | 883 | HITS-CLIP | 23142080 | Unspecified |
| mmu-miR-21a-5p | MIMAT0000530 | Kyat3 | 56267 | HITS-CLIP | 23597149 | Unspecified |
| mmu-miR-21a-5p | MIMAT0000530 | Kynu | 70789 | HITS-CLIP | 23597149 | Unspecified |
| mmu-miR-21a-5p | MIMAT0000530 | L1cam | 16728 | Microarrays | 37986224\|25263593 | Unspecified |

| | | | | | | |
|---|---|---|---|---|---|---|
| mmu-miR-21a-5p | MIMAT0000530 | L2hgdh | 217666 | HITS-CLIP | 23597149\|23142080 | Unspecified |
| mmu-miR-21a-5p | MIMAT0000530 | Lactb | 80907 | Microarrays//HITS-CLIP | 37986224\|25263593\|23597149 | Unspecified |
| mmu-miR-21a-5p | MIMAT0000530 | Lair1 | 52855 | Microarrays | 37986224\|25263593 | Unspecified |
| mmu-miR-21a-5p | MIMAT0000530 | Lamb1 | 16777 | HITS-CLIP | 19536157 | Unspecified |
| mmu-miR-21a-5p | MIMAT0000530 | Lamc1 | 226519 | HITS-CLIP | 23142080 | Unspecified |
| mmu-miR-21a-5p | MIMAT0000530 | Lamp1 | 16783 | HITS-CLIP | 23597149\|29167373\|23142080 | Unspecified |
| mmu-miR-21a-5p | MIMAT0000530 | Lamp2 | 16784 | HITS-CLIP | 26203562 | Unspecified |
| mmu-miR-21a-5p | MIMAT0000530 | Lancl1 | 14768 | HITS-CLIP | 23142080 | Unspecified |
| mmu-miR-21a-5p | MIMAT0000530 | Lancl3 | 236285 | Microarrays | 37986224\|25263593 | Unspecified |
| mmu-miR-21a-5p | MIMAT0000530 | Lap3 | 66988 | HITS-CLIP | 23597149\|23142080 | Unspecified |
| mmu-miR-21a-5p | MIMAT0000530 | Laptm4a | 17775 | HITS-CLIP | 23597149\|23142080 | Unspecified |
| mmu-miR-21a-5p | MIMAT0000530 | Laptm4b | 114128 | Microarrays//HITS-CLIP | 37986224\|25263593\|30971389 | Unspecified |
| mmu-miR-21a-5p | MIMAT0000530 | Larp1 | 73158 | HITS-CLIP | 23142080\|23597149\|29167373 | Unspecified |
| mmu-miR-21a-5p | MIMAT0000530 | Larp4 | 207214 | HITS-CLIP | 23597149\|23142080 | Unspecified |
| mmu-miR-21a-5p | MIMAT0000530 | Lars | 107045 | HITS-CLIP | 23597149\|23142080 | Unspecified |
| mmu-miR-21a-5p | MIMAT0000530 | Lars2 | 102436 | HITS-CLIP | 23597149\|23142080 | Unspecified |
| mmu-miR-21a-5p | MIMAT0000530 | Las1l | 76130 | HITS-CLIP | 23142080 | Unspecified |
| mmu-miR-21a-5p | MIMAT0000530 | Lat | 16797 | Microarrays//HITS-CLIP | 37986224\|25263593\|23142080 | Unspecified |
| mmu-miR-21a-5p | MIMAT0000530 | Lats1 | 16798 | HITS-CLIP | 37986224\|23142080 | Unspecified |
| mmu-miR-21a-5p | MIMAT0000530 | Lax1 | 240754 | HITS-CLIP | 23142080 | Unspecified |
| mmu-miR-21a-5p | MIMAT0000530 | Lbh | 77889 | HITS-CLIP | 31433990 | Unspecified |
| mmu-miR-21a-5p | MIMAT0000530 | Lbhd1 | 79081 | HITS-CLIP | 23142080 | Unspecified |
| mmu-miR-21a-5p | MIMAT0000530 | Lbr | 98386 | HITS-CLIP | 37986224\|23142080 | Unspecified |
| mmu-miR-21a-5p | MIMAT0000530 | Lca5 | 75782 | HITS-CLIP | 23142080 | Unspecified |
| mmu-miR-21a-5p | MIMAT0000530 | Lck | 16818 | HITS-CLIP | 23142080 | Unspecified |
| mmu-miR-21a-5p | MIMAT0000530 | Lclat1 | 225010 | HITS-CLIP | 23142080 | Unspecified |
| mmu-miR-21a-5p | MIMAT0000530 | Lcmt1 | 30949 | HITS-CLIP | 23597149\|23142080 | Unspecified |
| mmu-miR-21a-5p | MIMAT0000530 | Lcn11 | 227630 | Microarrays | 37986224\|25263593 | Unspecified |
| mmu-miR-21a-5p | MIMAT0000530 | Lcn2 | 16819 | HITS-CLIP | 23597149 | Unspecified |
| mmu-miR-21a-5p | MIMAT0000530 | Lcor | 212391 | HITS-CLIP | 23142080 | Unspecified |
| mmu-miR-21a-5p | MIMAT0000530 | Lcorl | 209707 | HITS-CLIP | 19536157\|23142080\|29167373 | Unspecified |
| mmu-miR-21a-5p | MIMAT0000530 | Lcp1 | 18826 | HITS-CLIP | 37986224\|23142080\|29167373 | Unspecified |
| mmu-miR-21a-5p | MIMAT0000530 | Lcp2 | 16822 | HITS-CLIP | 37986224\|23142080 | Unspecified |
| mmu-miR-21a-5p | MIMAT0000530 | Ldb1 | 16825 | HITS-CLIP | 29167373 | Unspecified |
| mmu-miR-21a-5p | MIMAT0000530 | Ldha | 16828 | HITS-CLIP | 23142080\|23597149\|29167373 | Unspecified |
| mmu-miR-21a-5p | MIMAT0000530 | Ldhb | 16832 | HITS-CLIP | 19536157\|23142080 | Unspecified |
| mmu-miR-21a-5p | MIMAT0000530 | Ldhd | 52815 | HITS-CLIP | 23597149 | Unspecified |
| mmu-miR-21a-5p | MIMAT0000530 | Ldlr | 16835 | HITS-CLIP | 23597149\|29167373\|23142080 | Unspecified |
| mmu-miR-21a-5p | MIMAT0000530 | Ldlrad3 | 241576 | Microarrays | 37986224 | Unspecified |

| | | | | | | |
|---|---|---|---|---|---|---|
| mmu-miR-21a-5p | MIMAT0000530 | Ldlrap1 | 100017 | qCLASH//HITS-CLIP | 31393953\|29167373 | Unspecified |
| mmu-miR-21a-5p | MIMAT0000530 | Lemd3 | 380664 | HITS-CLIP | 23142080 | Unspecified |
| mmu-miR-21a-5p | MIMAT0000530 | Leo1 | 235497 | HITS-CLIP | 23142080 | Unspecified |
| mmu-miR-21a-5p | MIMAT0000530 | Leprotl1 | 68192 | HITS-CLIP | 23597149\|23142080 | Unspecified |
| mmu-miR-21a-5p | MIMAT0000530 | Letm1 | 56384 | HITS-CLIP | 23597149 | Unspecified |
| mmu-miR-21a-5p | MIMAT0000530 | Lfng | 16848 | HITS-CLIP | 37986224\|19536157\|23142080\|29167 | Unspecified |
| mmu-miR-21a-5p | MIMAT0000530 | Lgals3 | 16854 | HITS-CLIP | 23142080\|29167373 | Unspecified |
| mmu-miR-21a-5p | MIMAT0000530 | Lgmn | 19141 | Microarrays//HITS-CLIP | 37986224\|23597149 | Unspecified |
| mmu-miR-21a-5p | MIMAT0000530 | Lhfp | 108927 | Microarrays | 37986224\|25263593 | Unspecified |
| mmu-miR-21a-5p | MIMAT0000530 | Lhfpl2 | 218454 | HITS-CLIP | 29167373 | Unspecified |
| mmu-miR-21a-5p | MIMAT0000530 | Lif | 16878 | HITS-CLIP | 23142080 | Unspecified |
| mmu-miR-21a-5p | MIMAT0000530 | Lifr | 16880 | HITS-CLIP, Microarrays//HITS-CLIP | 37986224\|23597149 | Unspecified |
| mmu-miR-21a-5p | MIMAT0000530 | Lig1 | 16881 | Microarrays | 37986224\|25263593 | Unspecified |
| mmu-miR-21a-5p | MIMAT0000530 | Lig3 | 16882 | HITS-CLIP | 23142080 | Unspecified |
| mmu-miR-21a-5p | MIMAT0000530 | Lig4 | 319583 | Microarrays | 37986224\|25263593 | Unspecified |
| mmu-miR-21a-5p | MIMAT0000530 | Limk2 | 16886 | HITS-CLIP | 19536157\|23142080 | Unspecified |
| mmu-miR-21a-5p | MIMAT0000530 | Lims1 | 110829 | HITS-CLIP | 23142080 | Unspecified |
| mmu-miR-21a-5p | MIMAT0000530 | Lims2 | 225341 | HITS-CLIP | 23597149 | Unspecified |
| mmu-miR-21a-5p | MIMAT0000530 | Lin37 | 75660 | HITS-CLIP | 23142080 | Unspecified |
| mmu-miR-21a-5p | MIMAT0000530 | Lin54 | 231506 | HITS-CLIP | 37986224 | Unspecified |
| mmu-miR-21a-5p | MIMAT0000530 | Lin7c | 22343 | HITS-CLIP | 23142080 | Unspecified |
| mmu-miR-21a-5p | MIMAT0000530 | Lin9 | 72568 | HITS-CLIP | 26203562 | Unspecified |
| mmu-miR-21a-5p | MIMAT0000530 | Lipa | 16889 | HITS-CLIP | 23597149\|23142080\|29167373 | Unspecified |
| mmu-miR-21a-5p | MIMAT0000530 | Lipe | 16890 | HITS-CLIP | 23142080 | Unspecified |
| mmu-miR-21a-5p | MIMAT0000530 | Llgl1 | 16897 | HITS-CLIP | 23142080 | Unspecified |
| mmu-miR-21a-5p | MIMAT0000530 | Lman2 | 66890 | HITS-CLIP | 29167373 | Unspecified |
| mmu-miR-21a-5p | MIMAT0000530 | Lmbrd1 | 68421 | HITS-CLIP | 23142080 | Unspecified |
| mmu-miR-21a-5p | MIMAT0000530 | Lmbrd2 | 320506 | HITS-CLIP | 23142080 | Unspecified |
| mmu-miR-21a-5p | MIMAT0000530 | Lmf1 | 76483 | HITS-CLIP | 23597149 | Unspecified |
| mmu-miR-21a-5p | MIMAT0000530 | Lmln | 239833 | HITS-CLIP | 23142080 | Unspecified |
| mmu-miR-21a-5p | MIMAT0000530 | Lmnb1 | 16906 | HITS-CLIP | 37986224\|23142080\|31433990\|30971 | Unspecified |
| mmu-miR-21a-5p | MIMAT0000530 | Lmo7 | 380928 | HITS-CLIP | 23142080 | Unspecified |
| mmu-miR-21a-5p | MIMAT0000530 | Lmtk2 | 231876 | HITS-CLIP | 31433990 | Unspecified |
| mmu-miR-21a-5p | MIMAT0000530 | Lnpep | 240028 | HITS-CLIP | 31433990\|19536157\|23142080\|29167 | Unspecified |
| mmu-miR-21a-5p | MIMAT0000530 | Lnpk | 69605 | HITS-CLIP | 19536157 | Unspecified |
| mmu-miR-21a-5p | MIMAT0000530 | Lnx2 | 140887 | HITS-CLIP | 23597149\|23142080 | Unspecified |
| mmu-miR-21a-5p | MIMAT0000530 | Lonp1 | 74142 | HITS-CLIP | 23597149\|23142080 | Unspecified |
| mmu-miR-21a-5p | MIMAT0000530 | Lonp2 | 66887 | HITS-CLIP | 23597149\|23142080 | Unspecified |
| mmu-miR-21a-5p | MIMAT0000530 | Lonrf3 | 74365 | Microarrays | 37986224\|25263593 | Unspecified |

| | | | | | | |
|---|---|---|---|---|---|---|
| mmu-miR-21a-5p | MIMAT0000530 | Lox | 16948 | Microarrays | 37986224\|25263593 | Unspecified |
| mmu-miR-21a-5p | MIMAT0000530 | Lpar3 | 65086 | HITS-CLIP | 23142080 | Unspecified |
| mmu-miR-21a-5p | MIMAT0000530 | Lpar6 | 67168 | HITS-CLIP, Microarrays//HITS-CLIP | 37986224\|23142080 | Unspecified |
| mmu-miR-21a-5p | MIMAT0000530 | Lpcat1 | 210992 | HITS-CLIP | 29167373 | Unspecified |
| mmu-miR-21a-5p | MIMAT0000530 | Lpcat2 | 270084 | HITS-CLIP | 29167373 | Unspecified |
| mmu-miR-21a-5p | MIMAT0000530 | Lpgat1 | 226856 | HITS-CLIP | 23597149\|19536157\|23142080 | Unspecified |
| mmu-miR-21a-5p | MIMAT0000530 | Lpin1 | 14245 | HITS-CLIP | 37986224\|31433990\|23597149\|2314 | Unspecified |
| mmu-miR-21a-5p | MIMAT0000530 | Lpin2 | 64898 | Microarrays//HITS-CLIP | 37986224\|23142080\|23597149\|2916 | Unspecified |
| mmu-miR-21a-5p | MIMAT0000530 | Lpl | 16956 | HITS-CLIP | 29167373 | Unspecified |
| mmu-miR-21a-5p | MIMAT0000530 | Lpp | 210126 | HITS-CLIP | 23142080 | Unspecified |
| mmu-miR-21a-5p | MIMAT0000530 | Lpxn | 107321 | HITS-CLIP | 23142080\|29167373 | Unspecified |
| mmu-miR-21a-5p | MIMAT0000530 | Lrat | 79235 | HITS-CLIP | 23597149 | Unspecified |
| mmu-miR-21a-5p | MIMAT0000530 | Lrba | 80877 | HITS-CLIP | 23597149\|23142080 | Unspecified |
| mmu-miR-21a-5p | MIMAT0000530 | Lrch3 | 70144 | HITS-CLIP | 37986224\|23597149\|23142080 | Unspecified |
| mmu-miR-21a-5p | MIMAT0000530 | Lrch4 | 231798 | HITS-CLIP | 29167373\|23142080 | Unspecified |
| mmu-miR-21a-5p | MIMAT0000530 | Lrif1 | 321000 | HITS-CLIP | 23597149 | Unspecified |
| mmu-miR-21a-5p | MIMAT0000530 | Lrig1 | 16206 | HITS-CLIP | 23142080 | Unspecified |
| mmu-miR-21a-5p | MIMAT0000530 | Lrig2 | 269473 | HITS-CLIP | 37986224\|23597149 | Unspecified |
| mmu-miR-21a-5p | MIMAT0000530 | Lrmda | 83938 | Microarrays | 25263593 | Unspecified |
| mmu-miR-21a-5p | MIMAT0000530 | Lrmp | 16970 | Microarrays//HITS-CLIP | 37986224\|23142080 | Unspecified |
| mmu-miR-21a-5p | MIMAT0000530 | Lrp1 | 16971 | HITS-CLIP | 23597149\|29167373 | Unspecified |
| mmu-miR-21a-5p | MIMAT0000530 | Lrp10 | 65107 | HITS-CLIP | 23142080 | Unspecified |
| mmu-miR-21a-5p | MIMAT0000530 | Lrp12 | 239393 | Microarrays//HITS-CLIP | 37986224\|23142080 | Unspecified |
| mmu-miR-21a-5p | MIMAT0000530 | Lrp5 | 16973 | HITS-CLIP | 23597149 | Unspecified |
| mmu-miR-21a-5p | MIMAT0000530 | Lrp6 | 16974 | HITS-CLIP, Microarrays//HITS-CLIP | 37986224\|23142080 | Unspecified |
| mmu-miR-21a-5p | MIMAT0000530 | Lrpap1 | 16976 | HITS-CLIP | 29167373\|23142080 | Unspecified |
| mmu-miR-21a-5p | MIMAT0000530 | Lrpprc | 72416 | HITS-CLIP | 37986224\|23597149\|23142080 | Unspecified |
| mmu-miR-21a-5p | MIMAT0000530 | Lrr1 | 69706 | Microarrays//HITS-CLIP | 37986224\|25263593\|23142080 | Unspecified |
| mmu-miR-21a-5p | MIMAT0000530 | Lrrc24 | 378937 | HITS-CLIP | 37986224 | Unspecified |
| mmu-miR-21a-5p | MIMAT0000530 | Lrrc27 | 76612 | Microarrays | 37986224 | Unspecified |
| mmu-miR-21a-5p | MIMAT0000530 | Lrrc39 | 109245 | Microarrays | 37986224\|25263593 | Unspecified |
| mmu-miR-21a-5p | MIMAT0000530 | Lrrc40 | 67144 | HITS-CLIP | 23142080 | Unspecified |
| mmu-miR-21a-5p | MIMAT0000530 | Lrrc42 | 77809 | HITS-CLIP | 23142080 | Unspecified |
| mmu-miR-21a-5p | MIMAT0000530 | Lrrc57 | 66606 | Microarrays | 37986224\|25263593 | Unspecified |
| mmu-miR-21a-5p | MIMAT0000530 | Lrrc58 | 320184 | HITS-CLIP | 23597149\|23142080 | Unspecified |
| mmu-miR-21a-5p | MIMAT0000530 | Lrrc59 | 98238 | HITS-CLIP | 23142080\|19536157 | Unspecified |
| mmu-miR-21a-5p | MIMAT0000530 | Lrrc7 | 242274 | HITS-CLIP | 37986224 | Unspecified |
| mmu-miR-21a-5p | MIMAT0000530 | Lrrc8a | 241296 | HITS-CLIP | 23597149\|23142080 | Unspecified |
| mmu-miR-21a-5p | MIMAT0000530 | Lrrc8b | 433926 | HITS-CLIP | 37986224 | Unspecified |

| | | | | | | |
|---|---|---|---|---|---|---|
| mmu-miR-21a-5p | MIMAT0000530 | Lrrc8c | 100604 | HITS-CLIP | 23142080 | Unspecified |
| mmu-miR-21a-5p | MIMAT0000530 | Lrrc8d | 231549 | HITS-CLIP | 23142080 | Unspecified |
| mmu-miR-21a-5p | MIMAT0000530 | Lrrcc1 | 71710 | HITS-CLIP | 23142080 | Unspecified |
| mmu-miR-21a-5p | MIMAT0000530 | Lrrfip1 | 16978 | HITS-CLIP | 23597149\|29167373 | Unspecified |
| mmu-miR-21a-5p | MIMAT0000530 | Lrrn3 | 16981 | HITS-CLIP | 19536157 | Unspecified |
| mmu-miR-21a-5p | MIMAT0000530 | Lrrtm2 | 107065 | HITS-CLIP | 37986224\|19536157 | Unspecified |
| mmu-miR-21a-5p | MIMAT0000530 | Lrwd1 | 71735 | HITS-CLIP | 23142080 | Unspecified |
| mmu-miR-21a-5p | MIMAT0000530 | Lsamp | 268890 | HITS-CLIP | 37986224 | Unspecified |
| mmu-miR-21a-5p | MIMAT0000530 | Lsg1 | 224092 | HITS-CLIP | 23142080 | Unspecified |
| mmu-miR-21a-5p | MIMAT0000530 | Lsm12 | 268490 | Microarrays | 37986224\|25263593 | Unspecified |
| mmu-miR-21a-5p | MIMAT0000530 | Lsm14a | 67070 | HITS-CLIP | 29167373 | Unspecified |
| mmu-miR-21a-5p | MIMAT0000530 | Lsm14b | 241846 | HITS-CLIP | 30971389 | Unspecified |
| mmu-miR-21a-5p | MIMAT0000530 | Lss | 16987 | HITS-CLIP | 23142080 | Unspecified |
| mmu-miR-21a-5p | MIMAT0000530 | Lst1 | 16988 | Microarrays | 37986224 | Unspecified |
| mmu-miR-21a-5p | MIMAT0000530 | Lta4h | 16993 | HITS-CLIP | 31433990\|23142080 | Unspecified |
| mmu-miR-21a-5p | MIMAT0000530 | Ltbp2 | 16997 | Microarrays | 37986224\|25263593 | Unspecified |
| mmu-miR-21a-5p | MIMAT0000530 | Ltbp4 | 108075 | HITS-CLIP | 37986224 | Unspecified |
| mmu-miR-21a-5p | MIMAT0000530 | Ltn1 | 78913 | HITS-CLIP//qCLASH | 23597149\|31393953\|23142080 | Unspecified |
| mmu-miR-21a-5p | MIMAT0000530 | Ltv1 | 353258 | Microarrays | 37986224\|25263593 | Unspecified |
| mmu-miR-21a-5p | MIMAT0000530 | Luc7l2 | 192196 | HITS-CLIP | 29167373 | Unspecified |
| mmu-miR-21a-5p | MIMAT0000530 | Luc7l3 | 67684 | HITS-CLIP//qCLASH | 23142080\|31393953\|29167373 | Unspecified |
| mmu-miR-21a-5p | MIMAT0000530 | Lurap1 | 68075 | HITS-CLIP | 19536157 | Unspecified |
| mmu-miR-21a-5p | MIMAT0000530 | Luzp1 | 269593 | HITS-CLIP | 23142080\|29167373 | Unspecified |
| mmu-miR-21a-5p | MIMAT0000530 | Luzp2 | 233271 | HITS-CLIP | 37986224 | Unspecified |
| mmu-miR-21a-5p | MIMAT0000530 | Ly6e | 17069 | Microarrays | 37986224 | Unspecified |
| mmu-miR-21a-5p | MIMAT0000530 | Ly6i | 57248 | Microarrays | 37986224 | Unspecified |
| mmu-miR-21a-5p | MIMAT0000530 | Ly86 | 17084 | Microarrays | 37986224 | Unspecified |
| mmu-miR-21a-5p | MIMAT0000530 | Lyl1 | 17095 | Microarrays | 37986224 | Unspecified |
| mmu-miR-21a-5p | MIMAT0000530 | Lypla1 | 18777 | HITS-CLIP | 23597149\|23142080\|19536157 | Unspecified |
| mmu-miR-21a-5p | MIMAT0000530 | Lyplal1 | 226791 | Microarrays | 37986224\|25263593 | Unspecified |
| mmu-miR-21a-5p | MIMAT0000530 | Lysmd3 | 80289 | HITS-CLIP | 23142080 | Unspecified |
| mmu-miR-21a-5p | MIMAT0000530 | Lyst | 17101 | HITS-CLIP | 31433990\|23142080\|29167373 | Unspecified |
| mmu-miR-21a-5p | MIMAT0000530 | Lyz2 | 17105 | HITS-CLIP | 29167373 | Unspecified |
| mmu-miR-21a-5p | MIMAT0000530 | Lztfl1 | 93730 | HITS-CLIP | 23142080 | Unspecified |
| mmu-miR-21a-5p | MIMAT0000530 | Lztr1 | 66863 | HITS-CLIP | 23142080 | Unspecified |
| mmu-miR-21a-5p | MIMAT0000530 | M6pr | 17113 | HITS-CLIP | 23142080\|29167373 | Unspecified |
| mmu-miR-21a-5p | MIMAT0000530 | MAP2K3 | 5606 | HITS-CLIP//Luciferase reporter assay//Flov | 23597149\|23142080\|26996129\|31585 | Unspecified |
| mmu-miR-21a-5p | MIMAT0000530 | Macf1 | 11426 | HITS-CLIP | 37986224\|19536157\|23597149\|23142 | Unspecified |
| mmu-miR-21a-5p | MIMAT0000530 | Mad2l1 | 56150 | Microarrays | 37986224\|25263593 | Unspecified |

| | | | | | | |
|---|---|---|---|---|---|---|
| mmu-miR-21a-5p | MIMAT0000530 | Mad2l1bp | 66591 | HITS-CLIP | 23142080 | Unspecified |
| mmu-miR-21a-5p | MIMAT0000530 | Madd | 228355 | HITS-CLIP | 23597149\|23142080 | Unspecified |
| mmu-miR-21a-5p | MIMAT0000530 | Maea | 59003 | HITS-CLIP | 23142080 | Unspecified |
| mmu-miR-21a-5p | MIMAT0000530 | Maf | 17132 | Microarrays//HITS-CLIP | 37986224\|23142080\|29167373 | Unspecified |
| mmu-miR-21a-5p | MIMAT0000530 | Maf1 | 68877 | HITS-CLIP | 23142080 | Unspecified |
| mmu-miR-21a-5p | MIMAT0000530 | Mafb | 16658 | HITS-CLIP | 29167373 | Unspecified |
| mmu-miR-21a-5p | MIMAT0000530 | Maff | 17133 | Microarrays//HITS-CLIP | 37986224\|25263593\|23142080 | Unspecified |
| mmu-miR-21a-5p | MIMAT0000530 | Mafg | 17134 | HITS-CLIP | 23597149 | Unspecified |
| mmu-miR-21a-5p | MIMAT0000530 | Maged1 | 94275 | HITS-CLIP | 23597149 | Unspecified |
| mmu-miR-21a-5p | MIMAT0000530 | Magee1 | 107528 | HITS-CLIP | 37986224\|19536157\|23142080 | Unspecified |
| mmu-miR-21a-5p | MIMAT0000530 | Magt1 | 67075 | HITS-CLIP | 37986224\|23142080\|23597149\|29167 | Unspecified |
| mmu-miR-21a-5p | MIMAT0000530 | Malsu1 | 75593 | HITS-CLIP | 23142080 | Unspecified |
| mmu-miR-21a-5p | MIMAT0000530 | Malt1 | 240354 | HITS-CLIP | 37986224\|23142080 | Unspecified |
| mmu-miR-21a-5p | MIMAT0000530 | Mamdc2 | 71738 | Microarrays | 37986224 | Unspecified |
| mmu-miR-21a-5p | MIMAT0000530 | Maml1 | 103806 | HITS-CLIP | 37986224\|31433990\|23142080 | Unspecified |
| mmu-miR-21a-5p | MIMAT0000530 | Maml2 | 270118 | HITS-CLIP | 23142080 | Unspecified |
| mmu-miR-21a-5p | MIMAT0000530 | Maml3 | 433586 | HITS-CLIP | 23142080 | Unspecified |
| mmu-miR-21a-5p | MIMAT0000530 | Man1a | 17155 | Microarrays//HITS-CLIP | 37986224\|23142080 | Unspecified |
| mmu-miR-21a-5p | MIMAT0000530 | Man1a2 | 17156 | HITS-CLIP | 23142080\|29167373 | Unspecified |
| mmu-miR-21a-5p | MIMAT0000530 | Man1c1 | 230815 | HITS-CLIP | 23142080 | Unspecified |
| mmu-miR-21a-5p | MIMAT0000530 | Man2a1 | 17158 | HITS-CLIP | 23597149\|23142080\|29167373 | Unspecified |
| mmu-miR-21a-5p | MIMAT0000530 | Man2a2 | 140481 | HITS-CLIP, Microarrays | 37986224 | Unspecified |
| mmu-miR-21a-5p | MIMAT0000530 | Man2b1 | 17159 | Microarrays | 37986224 | Unspecified |
| mmu-miR-21a-5p | MIMAT0000530 | Manea | 242362 | HITS-CLIP | 23142080\|29167373 | Unspecified |
| mmu-miR-21a-5p | MIMAT0000530 | Maneal | 215090 | HITS-CLIP | 37986224 | Unspecified |
| mmu-miR-21a-5p | MIMAT0000530 | Manf | 74840 | HITS-CLIP | 23142080\|23597149 | Unspecified |
| mmu-miR-21a-5p | MIMAT0000530 | Maoa | 17161 | Microarrays | 37986224 | Unspecified |
| mmu-miR-21a-5p | MIMAT0000530 | Maob | 109731 | HITS-CLIP | 23597149 | Unspecified |
| mmu-miR-21a-5p | MIMAT0000530 | Map10 | 74393 | Microarrays | 37986224 | Unspecified |
| mmu-miR-21a-5p | MIMAT0000530 | Map1b | 17755 | HITS-CLIP | 19536157 | Unspecified |
| mmu-miR-21a-5p | MIMAT0000530 | Map1lc3b | 67443 | HITS-CLIP | 29167373 | Unspecified |
| mmu-miR-21a-5p | MIMAT0000530 | Map2 | 17756 | HITS-CLIP | 37986224\|19536157 | Unspecified |
| mmu-miR-21a-5p | MIMAT0000530 | Map2k1 | 26395 | HITS-CLIP | 23142080 | Unspecified |
| mmu-miR-21a-5p | MIMAT0000530 | Map2k4 | 26398 | HITS-CLIP | 19536157\|23597149\|23142080 | Unspecified |
| mmu-miR-21a-5p | MIMAT0000530 | Map2k7 | 26400 | HITS-CLIP | 23597149\|23142080 | Unspecified |
| mmu-miR-21a-5p | MIMAT0000530 | Map3k1 | 26401 | HITS-CLIP | 25083871\|23597149\|19536157\|2314 | Unspecified |
| mmu-miR-21a-5p | MIMAT0000530 | Map3k11 | 26403 | HITS-CLIP | 23597149\|23142080 | Unspecified |
| mmu-miR-21a-5p | MIMAT0000530 | Map3k12 | 26404 | HITS-CLIP | 23142080 | Unspecified |
| mmu-miR-21a-5p | MIMAT0000530 | Map3k13 | 71751 | HITS-CLIP | 23597149 | Unspecified |

| | | | | | | |
|---|---|---|---|---|---|---|
| mmu-miR-21a-5p | MIMAT0000530 | Map3k2 | 26405 | HITS-CLIP | 23597149\|23142080\|29167373 | Unspecified |
| mmu-miR-21a-5p | MIMAT0000530 | Map3k20 | 65964 | HITS-CLIP | 29167373 | Unspecified |
| mmu-miR-21a-5p | MIMAT0000530 | Map3k3 | 26406 | HITS-CLIP | 23597149\|23142080 | Unspecified |
| mmu-miR-21a-5p | MIMAT0000530 | Map3k4 | 26407 | qCLASH//HITS-CLIP | 31393953\|23142080 | Unspecified |
| mmu-miR-21a-5p | MIMAT0000530 | Map3k5 | 26408 | HITS-CLIP | 23597149\|23142080 | Unspecified |
| mmu-miR-21a-5p | MIMAT0000530 | Map3k7 | 26409 | HITS-CLIP | 37986224\|23142080\|29167373 | Unspecified |
| mmu-miR-21a-5p | MIMAT0000530 | Map3k8 | 26410 | Microarrays | 37986224 | Unspecified |
| mmu-miR-21a-5p | MIMAT0000530 | Map4 | 17758 | HITS-CLIP | 19536157\|29167373\|23142080 | Unspecified |
| mmu-miR-21a-5p | MIMAT0000530 | Map4k2 | 26412 | HITS-CLIP | 23142080 | Unspecified |
| mmu-miR-21a-5p | MIMAT0000530 | Map4k3 | 225028 | HITS-CLIP | 23597149\|23142080\|29167373 | Unspecified |
| mmu-miR-21a-5p | MIMAT0000530 | Map4k4 | 26921 | HITS-CLIP | 23142080\|23597149 | Unspecified |
| mmu-miR-21a-5p | MIMAT0000530 | Map9 | 213582 | HITS-CLIP | 23142080 | Unspecified |
| mmu-miR-21a-5p | MIMAT0000530 | Mapk1 | 26413 | HITS-CLIP | 19536157\|23142080\|29167373 | Unspecified |
| mmu-miR-21a-5p | MIMAT0000530 | Mapk14 | 26416 | HITS-CLIP | 23142080\|29167373 | Unspecified |
| mmu-miR-21a-5p | MIMAT0000530 | Mapk15 | 332110 | Microarrays | 37986224 | Unspecified |
| mmu-miR-21a-5p | MIMAT0000530 | Mapk1ip1 | 69546 | Microarrays | 37986224 | Unspecified |
| mmu-miR-21a-5p | MIMAT0000530 | Mapk1ip1l | 218975 | HITS-CLIP | 23142080\|29167373 | Unspecified |
| mmu-miR-21a-5p | MIMAT0000530 | Mapk8 | 26419 | HITS-CLIP | 23142080 | Unspecified |
| mmu-miR-21a-5p | MIMAT0000530 | Mapk8ip3 | 30957 | HITS-CLIP | 23597149\|23142080 | Unspecified |
| mmu-miR-21a-5p | MIMAT0000530 | Mapkap1 | 227743 | HITS-CLIP | 23142080 | Unspecified |
| mmu-miR-21a-5p | MIMAT0000530 | Mapkapk2 | 17164 | HITS-CLIP | 23142080 | Unspecified |
| mmu-miR-21a-5p | MIMAT0000530 | Mapkbp1 | 26390 | HITS-CLIP | 19536157 | Unspecified |
| mmu-miR-21a-5p | MIMAT0000530 | Mapre1 | 13589 | HITS-CLIP | 37986224\|23142080\|29167373 | Unspecified |
| mmu-miR-21a-5p | MIMAT0000530 | Mapre2 | 212307 | HITS-CLIP | 23597149\|23142080\|29167373 | Unspecified |
| mmu-miR-21a-5p | MIMAT0000530 | Marc2 | 67247 | HITS-CLIP | 23597149 | Unspecified |
| mmu-miR-21a-5p | MIMAT0000530 | March5 | 54708 | HITS-CLIP | 23142080 | Unspecified |
| mmu-miR-21a-5p | MIMAT0000530 | March7 | 57438 | HITS-CLIP | 23142080 | Unspecified |
| mmu-miR-21a-5p | MIMAT0000530 | March8 | 71779 | HITS-CLIP | 23597149\|19536157 | Unspecified |
| mmu-miR-21a-5p | MIMAT0000530 | March9 | 216438 | HITS-CLIP | 23142080 | Unspecified |
| mmu-miR-21a-5p | MIMAT0000530 | Marchf6 | 223455 | HITS-CLIP, Microarrays//HITS-CLIP | 37986224\|19536157\|23142080\|29167 | Unspecified |
| mmu-miR-21a-5p | MIMAT0000530 | Marchf8 | 220972 | HITS-CLIP | 26203562 | Unspecified |
| mmu-miR-21a-5p | MIMAT0000530 | Marcks | 17118 | Microarrays//HITS-CLIP | 37986224\|19536157\|23142080 | Unspecified |
| mmu-miR-21a-5p | MIMAT0000530 | Marco | 17167 | Microarrays | 37986224\|25263593 | Unspecified |
| mmu-miR-21a-5p | MIMAT0000530 | Marf1 | 223989 | HITS-CLIP, Microarrays//HITS-CLIP | 37986224\|23597149\|23142080 | Unspecified |
| mmu-miR-21a-5p | MIMAT0000530 | Mark2 | 13728 | HITS-CLIP | 31433990\|23142080 | Unspecified |
| mmu-miR-21a-5p | MIMAT0000530 | Mark3 | 17169 | HITS-CLIP | 19536157\|23597149\|23142080\|29167 | Unspecified |
| mmu-miR-21a-5p | MIMAT0000530 | Masp1 | 17174 | HITS-CLIP | 23597149 | Unspecified |
| mmu-miR-21a-5p | MIMAT0000530 | Mastl | 67121 | Microarrays//HITS-CLIP | 37986224\|25263593\|23142080 | Unspecified |
| mmu-miR-21a-5p | MIMAT0000530 | Mat1a | 11720 | HITS-CLIP | 23597149 | Unspecified |

| | | | | | | |
|---|---|---|---|---|---|---|
| mmu-miR-21a-5p | MIMAT0000530 | Mat2a | 232087 | HITS-CLIP | 23142080\|29167373 | Unspecified |
| mmu-miR-21a-5p | MIMAT0000530 | Matr3 | 17184 | HITS-CLIP | 37986224\|23142080\|19536157 | Unspecified |
| mmu-miR-21a-5p | MIMAT0000530 | Mau2 | 74549 | HITS-CLIP | 23142080\|29167373 | Unspecified |
| mmu-miR-21a-5p | MIMAT0000530 | Mavs | 228607 | HITS-CLIP | 23597149 | Unspecified |
| mmu-miR-21a-5p | MIMAT0000530 | Maz | 17188 | HITS-CLIP | 23142080 | Unspecified |
| mmu-miR-21a-5p | MIMAT0000530 | Mbd1 | 17190 | HITS-CLIP | 23142080 | Unspecified |
| mmu-miR-21a-5p | MIMAT0000530 | Mbd2 | 17191 | HITS-CLIP | 23142080\|29167373 | Unspecified |
| mmu-miR-21a-5p | MIMAT0000530 | Mbd4 | 17193 | Microarrays | 37986224 | Unspecified |
| mmu-miR-21a-5p | MIMAT0000530 | Mbd5 | 109241 | HITS-CLIP | 23142080 | Unspecified |
| mmu-miR-21a-5p | MIMAT0000530 | Mbd6 | 110962 | HITS-CLIP | 23597149\|23142080 | Unspecified |
| mmu-miR-21a-5p | MIMAT0000530 | Mbl1 | 17194 | HITS-CLIP | 23597149 | Unspecified |
| mmu-miR-21a-5p | MIMAT0000530 | Mbnl1 | 56758 | HITS-CLIP | 37986224\|23142080\|29167373 | Unspecified |
| mmu-miR-21a-5p | MIMAT0000530 | Mbnl2 | 105559 | HITS-CLIP | 19536157\|23142080\|29167373 | Unspecified |
| mmu-miR-21a-5p | MIMAT0000530 | Mbnl3 | 171170 | Microarrays//HITS-CLIP | 37986224\|25263593\|23142080 | Unspecified |
| mmu-miR-21a-5p | MIMAT0000530 | Mboat1 | 218121 | HITS-CLIP | 23142080 | Unspecified |
| mmu-miR-21a-5p | MIMAT0000530 | Mboat2 | 67216 | HITS-CLIP | 23142080 | Unspecified |
| mmu-miR-21a-5p | MIMAT0000530 | Mboat7 | 77582 | HITS-CLIP | 23142080\|29167373 | Unspecified |
| mmu-miR-21a-5p | MIMAT0000530 | Mbp | 17196 | HITS-CLIP | 23142080\|29167373 | Unspecified |
| mmu-miR-21a-5p | MIMAT0000530 | Mbtd1 | 103537 | Microarrays//HITS-CLIP | 37986224\|23142080 | Unspecified |
| mmu-miR-21a-5p | MIMAT0000530 | Mbtps1 | 56453 | HITS-CLIP | 23142080\|29167373 | Unspecified |
| mmu-miR-21a-5p | MIMAT0000530 | Mbtps2 | 270669 | HITS-CLIP | 37986224\|23142080 | Unspecified |
| mmu-miR-21a-5p | MIMAT0000530 | Mcat | 223722 | HITS-CLIP | 23597149 | Unspecified |
| mmu-miR-21a-5p | MIMAT0000530 | Mccc2 | 78038 | HITS-CLIP | 23597149 | Unspecified |
| mmu-miR-21a-5p | MIMAT0000530 | Mcf2l | 17207 | HITS-CLIP | 23142080 | Unspecified |
| mmu-miR-21a-5p | MIMAT0000530 | Mcfd2 | 193813 | Microarrays | 37986224\|25263593 | Unspecified |
| mmu-miR-21a-5p | MIMAT0000530 | Mcl1 | 17210 | HITS-CLIP | 37986224\|30971389\|28735896\|2314 | Unspecified |
| mmu-miR-21a-5p | MIMAT0000530 | Mcm10 | 70024 | Microarrays//HITS-CLIP | 37986224\|25263593\|23142080 | Unspecified |
| mmu-miR-21a-5p | MIMAT0000530 | Mcm2 | 17216 | Microarrays | 37986224\|25263593 | Unspecified |
| mmu-miR-21a-5p | MIMAT0000530 | Mcm3 | 17215 | HITS-CLIP | 23142080 | Unspecified |
| mmu-miR-21a-5p | MIMAT0000530 | Mcm4 | 17217 | Microarrays | 37986224\|25263593 | Unspecified |
| mmu-miR-21a-5p | MIMAT0000530 | Mcm5 | 17218 | Microarrays//HITS-CLIP | 37986224\|25263593\|31433990\|2359 | Unspecified |
| mmu-miR-21a-5p | MIMAT0000530 | Mcm6 | 17219 | Microarrays//HITS-CLIP | 37986224\|25263593\|23142080\|2916 | Unspecified |
| mmu-miR-21a-5p | MIMAT0000530 | Mcm7 | 17220 | Microarrays | 37986224\|25263593 | Unspecified |
| mmu-miR-21a-5p | MIMAT0000530 | Mcm8 | 66634 | Microarrays | 37986224\|25263593 | Unspecified |
| mmu-miR-21a-5p | MIMAT0000530 | Mcmbp | 210711 | HITS-CLIP | 29167373\|23142080 | Unspecified |
| mmu-miR-21a-5p | MIMAT0000530 | Mcoln3 | 171166 | Microarrays | 37986224 | Unspecified |
| mmu-miR-21a-5p | MIMAT0000530 | Mctp2 | 244049 | HITS-CLIP | 23142080 | Unspecified |
| mmu-miR-21a-5p | MIMAT0000530 | Mcts1 | 68995 | HITS-CLIP | 23142080 | Unspecified |
| mmu-miR-21a-5p | MIMAT0000530 | Mcub | 11839 | HITS-CLIP | 23142080 | Unspecified |

| | | | | | | |
|---|---|---|---|---|---|---|
| mmu-miR-21a-5p | MIMAT0000530 | Mdfic | 16543 | HITS-CLIP | 23142080\|23597149\|29167373 | Unspecified |
| mmu-miR-21a-5p | MIMAT0000530 | Mdh1 | 17449 | HITS-CLIP | 19536157\|23142080 | Unspecified |
| mmu-miR-21a-5p | MIMAT0000530 | Mdm2 | 17246 | HITS-CLIP | 23142080\|29167373 | Unspecified |
| mmu-miR-21a-5p | MIMAT0000530 | Mdm4 | 17248 | HITS-CLIP | 37986224\|23142080\|23597149\|1953€ | Unspecified |
| mmu-miR-21a-5p | MIMAT0000530 | Mdn1 | 100019 | HITS-CLIP | 23142080\|29167373 | Unspecified |
| mmu-miR-21a-5p | MIMAT0000530 | Me1 | 17436 | Microarrays//HITS-CLIP | 37986224\|25263593\|23597149 | Unspecified |
| mmu-miR-21a-5p | MIMAT0000530 | Me2 | 107029 | HITS-CLIP | 23142080 | Unspecified |
| mmu-miR-21a-5p | MIMAT0000530 | Meaf6 | 70088 | HITS-CLIP | 23142080\|19536157 | Unspecified |
| mmu-miR-21a-5p | MIMAT0000530 | Mecp2 | 17257 | HITS-CLIP | 23142080 | Unspecified |
| mmu-miR-21a-5p | MIMAT0000530 | Med1 | 19014 | HITS-CLIP | 19536157\|23597149\|23142080\|2916 | Unspecified |
| mmu-miR-21a-5p | MIMAT0000530 | Med12l | 329650 | HITS-CLIP | 19536157 | Unspecified |
| mmu-miR-21a-5p | MIMAT0000530 | Med13 | 327987 | HITS-CLIP | 37986224\|23597149\|19536157\|2314 | Unspecified |
| mmu-miR-21a-5p | MIMAT0000530 | Med13l | 76199 | HITS-CLIP | 23142080\|29167373 | Unspecified |
| mmu-miR-21a-5p | MIMAT0000530 | Med14 | 26896 | HITS-CLIP | 23142080 | Unspecified |
| mmu-miR-21a-5p | MIMAT0000530 | Med15 | 94112 | HITS-CLIP | 29167373 | Unspecified |
| mmu-miR-21a-5p | MIMAT0000530 | Med18 | 67219 | HITS-CLIP | 23142080 | Unspecified |
| mmu-miR-21a-5p | MIMAT0000530 | Med19 | 381379 | HITS-CLIP | 23142080 | Unspecified |
| mmu-miR-21a-5p | MIMAT0000530 | Med21 | 108098 | HITS-CLIP | 23142080 | Unspecified |
| mmu-miR-21a-5p | MIMAT0000530 | Med22 | 20933 | HITS-CLIP | 23142080 | Unspecified |
| mmu-miR-21a-5p | MIMAT0000530 | Med23 | 70208 | HITS-CLIP | 23142080 | Unspecified |
| mmu-miR-21a-5p | MIMAT0000530 | Med24 | 23989 | HITS-CLIP | 23142080\|29167373 | Unspecified |
| mmu-miR-21a-5p | MIMAT0000530 | Med25 | 75613 | HITS-CLIP//qCLASH | 23597149\|31393953 | Unspecified |
| mmu-miR-21a-5p | MIMAT0000530 | Med27 | 68975 | HITS-CLIP | 23142080 | Unspecified |
| mmu-miR-21a-5p | MIMAT0000530 | Med6 | 69792 | HITS-CLIP | 23142080\|23597149 | Unspecified |
| mmu-miR-21a-5p | MIMAT0000530 | Mef2a | 17258 | HITS-CLIP | 37986224\|23142080\|29167373 | Unspecified |
| mmu-miR-21a-5p | MIMAT0000530 | Mef2c | 17260 | Microarrays//HITS-CLIP | 37986224\|19536157 | Unspecified |
| mmu-miR-21a-5p | MIMAT0000530 | Mef2d | 17261 | HITS-CLIP | 23597149\|23142080 | Unspecified |
| mmu-miR-21a-5p | MIMAT0000530 | Megf9 | 230316 | Microarrays | 37986224\|25263593 | Unspecified |
| mmu-miR-21a-5p | MIMAT0000530 | Melk | 17279 | Microarrays//HITS-CLIP | 37986224\|25263593\|30971389\|3143 | Unspecified |
| mmu-miR-21a-5p | MIMAT0000530 | Mepce | 231803 | HITS-CLIP | 23142080 | Unspecified |
| mmu-miR-21a-5p | MIMAT0000530 | Mertk | 17289 | Microarrays | 37986224 | Unspecified |
| mmu-miR-21a-5p | MIMAT0000530 | Met | 17295 | HITS-CLIP | 23597149 | Unspecified |
| mmu-miR-21a-5p | MIMAT0000530 | Metap1 | 75624 | HITS-CLIP | 23142080 | Unspecified |
| mmu-miR-21a-5p | MIMAT0000530 | Metap2 | 56307 | HITS-CLIP | 23142080 | Unspecified |
| mmu-miR-21a-5p | MIMAT0000530 | Mettl13 | 51603 | HITS-CLIP | 23142080 | Unspecified |
| mmu-miR-21a-5p | MIMAT0000530 | Mettl14 | 210529 | HITS-CLIP | 29167373 | Unspecified |
| mmu-miR-21a-5p | MIMAT0000530 | Mettl16 | 67493 | HITS-CLIP | 23597149 | Unspecified |
| mmu-miR-21a-5p | MIMAT0000530 | Mettl4 | 76781 | HITS-CLIP | 23142080 | Unspecified |
| mmu-miR-21a-5p | MIMAT0000530 | Mex3a | 72640 | qCLASH | 31393953 | Unspecified |

| | | | | | | |
|---|---|---|---|---|---|---|
| mmu-miR-21a-5p | MIMAT0000530 | Mex3b | 108797 | Microarrays | 37986224 | Unspecified |
| mmu-miR-21a-5p | MIMAT0000530 | Mex3c | 240396 | HITS-CLIP | 23142080 | Unspecified |
| mmu-miR-21a-5p | MIMAT0000530 | Mfap3 | 216760 | HITS-CLIP | 23142080 | Unspecified |
| mmu-miR-21a-5p | MIMAT0000530 | Mfap3l | 71306 | Microarrays | 37986224\|25263593 | Unspecified |
| mmu-miR-21a-5p | MIMAT0000530 | Mfhas1 | 52065 | HITS-CLIP | 23597149\|23142080\|29167373 | Unspecified |
| mmu-miR-21a-5p | MIMAT0000530 | Mfsd1 | 66868 | HITS-CLIP | 23142080 | Unspecified |
| mmu-miR-21a-5p | MIMAT0000530 | Mfsd14a | 15247 | HITS-CLIP | 23597149 | Unspecified |
| mmu-miR-21a-5p | MIMAT0000530 | Mfsd14b | 84641 | HITS-CLIP | 23142080 | Unspecified |
| mmu-miR-21a-5p | MIMAT0000530 | Mfsd5 | 106073 | HITS-CLIP | 29167373 | Unspecified |
| mmu-miR-21a-5p | MIMAT0000530 | Mfsd6 | 98682 | HITS-CLIP | 23142080 | Unspecified |
| mmu-miR-21a-5p | MIMAT0000530 | Mga | 29808 | HITS-CLIP | 23142080\|29167373 | Unspecified |
| mmu-miR-21a-5p | MIMAT0000530 | Mgat1 | 17308 | HITS-CLIP | 23142080 | Unspecified |
| mmu-miR-21a-5p | MIMAT0000530 | Mgat2 | 217664 | HITS-CLIP | 23142080 | Unspecified |
| mmu-miR-21a-5p | MIMAT0000530 | Mgat3 | 17309 | HITS-CLIP | 37986224 | Unspecified |
| mmu-miR-21a-5p | MIMAT0000530 | Mgat4a | 269181 | HITS-CLIP | 23142080 | Unspecified |
| mmu-miR-21a-5p | MIMAT0000530 | Mgat5 | 107895 | Microarrays//HITS-CLIP | 37986224\|25263593\|29167373 | Unspecified |
| mmu-miR-21a-5p | MIMAT0000530 | Mgea5 | 76055 | HITS-CLIP | 23142080 | Unspecified |
| mmu-miR-21a-5p | MIMAT0000530 | Mgl2 | 216864 | Microarrays | 37986224\|25263593 | Unspecified |
| mmu-miR-21a-5p | MIMAT0000530 | Mgll | 23945 | HITS-CLIP | 23142080 | Unspecified |
| mmu-miR-21a-5p | MIMAT0000530 | Mgme1 | 74528 | HITS-CLIP | 37986224\|23142080 | Unspecified |
| mmu-miR-21a-5p | MIMAT0000530 | Mgmt | 17314 | HITS-CLIP | 23142080 | Unspecified |
| mmu-miR-21a-5p | MIMAT0000530 | Mgst2 | 211666 | HITS-CLIP | 23142080 | Unspecified |
| mmu-miR-21a-5p | MIMAT0000530 | Mia2 | 338320 | HITS-CLIP | 23142080 | Unspecified |
| mmu-miR-21a-5p | MIMAT0000530 | Mia3 | 338366 | HITS-CLIP | 23142080\|23597149\|19536157 | Unspecified |
| mmu-miR-21a-5p | MIMAT0000530 | Mib1 | 225164 | HITS-CLIP | 37986224\|23597149\|19536157\|2916 | Unspecified |
| mmu-miR-21a-5p | MIMAT0000530 | Mical2 | 320878 | HITS-CLIP | 29167373 | Unspecified |
| mmu-miR-21a-5p | MIMAT0000530 | Micu1 | 216001 | HITS-CLIP | 23142080\|23597149 | Unspecified |
| mmu-miR-21a-5p | MIMAT0000530 | Micu2 | 68514 | HITS-CLIP | 23142080 | Unspecified |
| mmu-miR-21a-5p | MIMAT0000530 | Micu3 | 78506 | HITS-CLIP | 23142080 | Unspecified |
| mmu-miR-21a-5p | MIMAT0000530 | Mid2 | 23947 | HITS-CLIP | 23142080 | Unspecified |
| mmu-miR-21a-5p | MIMAT0000530 | Midn | 59090 | HITS-CLIP | 23597149\|29167373\|23142080 | Unspecified |
| mmu-miR-21a-5p | MIMAT0000530 | Mier3 | 218613 | HITS-CLIP | 37986224\|23142080 | Unspecified |
| mmu-miR-21a-5p | MIMAT0000530 | Mindy2 | 235461 | HITS-CLIP | 23597149\|31433990\|23142080\|2916 | Unspecified |
| mmu-miR-21a-5p | MIMAT0000530 | Mindy3 | 80013 | HITS-CLIP | 23142080 | Unspecified |
| mmu-miR-21a-5p | MIMAT0000530 | Mis18bp1 | 217653 | HITS-CLIP | 23142080 | Unspecified |
| mmu-miR-21a-5p | MIMAT0000530 | Mitd1 | 69028 | HITS-CLIP | 29167373 | Unspecified |
| mmu-miR-21a-5p | MIMAT0000530 | Mki67 | 17345 | Microarrays//qCLASH//HITS-CLIP | 37986224\|25263593\|31393953\|2314 | Unspecified |
| mmu-miR-21a-5p | MIMAT0000530 | Mkln1 | 27418 | HITS-CLIP | 23142080\|29167373 | Unspecified |
| mmu-miR-21a-5p | MIMAT0000530 | Mknk1 | 17346 | HITS-CLIP | 29167373 | Unspecified |

| | | | | | | |
|---|---|---|---|---|---|---|
| mmu-miR-21a-5p | MIMAT0000530 | Mlec | 109154 | HITS-CLIP | 23142080 | Unspecified |
| mmu-miR-21a-5p | MIMAT0000530 | Mlh1 | 17350 | HITS-CLIP | 23597149 | Unspecified |
| mmu-miR-21a-5p | MIMAT0000530 | Mlkl | 74568 | HITS-CLIP | 23142080 | Unspecified |
| mmu-miR-21a-5p | MIMAT0000530 | Mllt1 | 64144 | HITS-CLIP | 23142080 | Unspecified |
| mmu-miR-21a-5p | MIMAT0000530 | Mllt10 | 17354 | HITS-CLIP | 23142080\|29167373 | Unspecified |
| mmu-miR-21a-5p | MIMAT0000530 | Mllt3 | 70122 | HITS-CLIP | 23142080 | Unspecified |
| mmu-miR-21a-5p | MIMAT0000530 | Mllt6 | 246198 | HITS-CLIP | 23597149\|23142080 | Unspecified |
| mmu-miR-21a-5p | MIMAT0000530 | Mlxip | 208104 | HITS-CLIP | 29167373 | Unspecified |
| mmu-miR-21a-5p | MIMAT0000530 | Mlxipl | 58805 | HITS-CLIP | 37986224\|23597149 | Unspecified |
| mmu-miR-21a-5p | MIMAT0000530 | Mmadhc | 109129 | qCLASH | 31393953 | Unspecified |
| mmu-miR-21a-5p | MIMAT0000530 | Mmel1 | 27390 | Microarrays | 37986224\|25263593 | Unspecified |
| mmu-miR-21a-5p | MIMAT0000530 | Mmgt1 | 236792 | Microarrays//HITS-CLIP | 37986224\|25263593\|23142080\|29167 | Unspecified |
| mmu-miR-21a-5p | MIMAT0000530 | Mmp12 | 17381 | Microarrays | 37986224\|25263593 | Unspecified |
| mmu-miR-21a-5p | MIMAT0000530 | Mmp13 | 17386 | Microarrays | 37986224\|25263593 | Unspecified |
| mmu-miR-21a-5p | MIMAT0000530 | Mmp9 | 17395 | Immunohistochemistry//In situ hybridizati | 23443810\|25263593 | Unspecified |
| mmu-miR-21a-5p | MIMAT0000530 | Mms22l | 212377 | Microarrays//HITS-CLIP | 37986224\|25263593\|23142080 | Unspecified |
| mmu-miR-21a-5p | MIMAT0000530 | Mnat1 | 17420 | HITS-CLIP | 23142080 | Unspecified |
| mmu-miR-21a-5p | MIMAT0000530 | Mnd1 | 76915 | Microarrays | 37986224\|25263593 | Unspecified |
| mmu-miR-21a-5p | MIMAT0000530 | Mndal | 100040462 | HITS-CLIP | 23142080 | Unspecified |
| mmu-miR-21a-5p | MIMAT0000530 | Mob1a | 232157 | HITS-CLIP | 23142080\|29167373 | Unspecified |
| mmu-miR-21a-5p | MIMAT0000530 | Mob1b | 68473 | HITS-CLIP | 19536157\|23142080 | Unspecified |
| mmu-miR-21a-5p | MIMAT0000530 | Mob2 | 101513 | HITS-CLIP | 23142080 | Unspecified |
| mmu-miR-21a-5p | MIMAT0000530 | Mob3a | 208228 | HITS-CLIP | 23142080 | Unspecified |
| mmu-miR-21a-5p | MIMAT0000530 | Mob4 | 19070 | HITS-CLIP | 23142080 | Unspecified |
| mmu-miR-21a-5p | MIMAT0000530 | Mocs3 | 69372 | HITS-CLIP | 23142080 | Unspecified |
| mmu-miR-21a-5p | MIMAT0000530 | Mogs | 57377 | Microarrays//HITS-CLIP | 37986224\|25263593\|23597149\|23142 | Unspecified |
| mmu-miR-21a-5p | MIMAT0000530 | Mon2 | 67074 | HITS-CLIP | 23597149\|23142080 | Unspecified |
| mmu-miR-21a-5p | MIMAT0000530 | Morc3 | 338467 | HITS-CLIP | 23142080\|29167373 | Unspecified |
| mmu-miR-21a-5p | MIMAT0000530 | Morf4l2 | 56397 | HITS-CLIP | 29167373\|23142080 | Unspecified |
| mmu-miR-21a-5p | MIMAT0000530 | Morn1 | 76866 | HITS-CLIP | 23142080 | Unspecified |
| mmu-miR-21a-5p | MIMAT0000530 | Mosmo | 233812 | HITS-CLIP | 19536157\|23142080 | Unspecified |
| mmu-miR-21a-5p | MIMAT0000530 | Mospd2 | 76763 | HITS-CLIP | 23597149 | Unspecified |
| mmu-miR-21a-5p | MIMAT0000530 | Mov10 | 17454 | HITS-CLIP | 23597149\|23142080 | Unspecified |
| mmu-miR-21a-5p | MIMAT0000530 | Mpc1 | 55951 | Microarrays | 37986224\|25263593 | Unspecified |
| mmu-miR-21a-5p | MIMAT0000530 | Mpdu1 | 24070 | HITS-CLIP | 30971389 | Unspecified |
| mmu-miR-21a-5p | MIMAT0000530 | Mpeg1 | 17476 | HITS-CLIP | 23597149\|23142080\|29167373 | Unspecified |
| mmu-miR-21a-5p | MIMAT0000530 | Mphosph10 | 67973 | HITS-CLIP | 23597149\|29167373\|23142080 | Unspecified |
| mmu-miR-21a-5p | MIMAT0000530 | Mphosph6 | 68533 | HITS-CLIP | 23597149 | Unspecified |
| mmu-miR-21a-5p | MIMAT0000530 | Mphosph8 | 75339 | HITS-CLIP | 23142080 | Unspecified |

| | | | | | | |
|---|---|---|---|---|---|---|
| mmu-miR-21a-5p | MIMAT0000530 | Mpp1 | 17524 | Microarrays//HITS-CLIP | 37986224\|29167373 | Unspecified |
| mmu-miR-21a-5p | MIMAT0000530 | Mpp5 | 56217 | HITS-CLIP | 23142080 | Unspecified |
| mmu-miR-21a-5p | MIMAT0000530 | Mprip | 26936 | HITS-CLIP | 23597149\|23142080 | Unspecified |
| mmu-miR-21a-5p | MIMAT0000530 | Mpst | 246221 | HITS-CLIP | 23142080 | Unspecified |
| mmu-miR-21a-5p | MIMAT0000530 | Mpv17l2 | 234384 | Microarrays | 37986224\|25263593 | Unspecified |
| mmu-miR-21a-5p | MIMAT0000530 | Mpzl3 | 319742 | HITS-CLIP | 23142080 | Unspecified |
| mmu-miR-21a-5p | MIMAT0000530 | Mr1 | 15064 | HITS-CLIP | 23142080 | Unspecified |
| mmu-miR-21a-5p | MIMAT0000530 | Mrap | 77037 | HITS-CLIP | 23597149 | Unspecified |
| mmu-miR-21a-5p | MIMAT0000530 | Mrc1 | 17533 | HITS-CLIP | 23597149\|29167373 | Unspecified |
| mmu-miR-21a-5p | MIMAT0000530 | Mreg | 381269 | HITS-CLIP | 37986224\|23142080 | Unspecified |
| mmu-miR-21a-5p | MIMAT0000530 | Mrm2 | 29960 | HITS-CLIP | 29167373 | Unspecified |
| mmu-miR-21a-5p | MIMAT0000530 | Mroh1 | 223658 | HITS-CLIP | 29167373 | Unspecified |
| mmu-miR-21a-5p | MIMAT0000530 | Mrpl10 | 107732 | HITS-CLIP | 23142080 | Unspecified |
| mmu-miR-21a-5p | MIMAT0000530 | Mrpl17 | 27397 | HITS-CLIP | 23142080 | Unspecified |
| mmu-miR-21a-5p | MIMAT0000530 | Mrpl19 | 56284 | HITS-CLIP | 23597149\|23142080 | Unspecified |
| mmu-miR-21a-5p | MIMAT0000530 | Mrpl21 | 353242 | HITS-CLIP | 23142080 | Unspecified |
| mmu-miR-21a-5p | MIMAT0000530 | Mrpl3 | 94062 | HITS-CLIP | 23142080\|23597149 | Unspecified |
| mmu-miR-21a-5p | MIMAT0000530 | Mrpl33 | 66845 | HITS-CLIP | 23142080 | Unspecified |
| mmu-miR-21a-5p | MIMAT0000530 | Mrpl35 | 66223 | qCLASH | 31393953 | Unspecified |
| mmu-miR-21a-5p | MIMAT0000530 | Mrpl37 | 56280 | qCLASH | 31393953 | Unspecified |
| mmu-miR-21a-5p | MIMAT0000530 | Mrpl45 | 67036 | HITS-CLIP | 23142080 | Unspecified |
| mmu-miR-21a-5p | MIMAT0000530 | Mrpl46 | 67308 | HITS-CLIP | 23597149\|23142080 | Unspecified |
| mmu-miR-21a-5p | MIMAT0000530 | Mrpl49 | 18120 | HITS-CLIP | 23142080 | Unspecified |
| mmu-miR-21a-5p | MIMAT0000530 | Mrpl54 | 66047 | HITS-CLIP | 23142080 | Unspecified |
| mmu-miR-21a-5p | MIMAT0000530 | Mrps14 | 64659 | HITS-CLIP | 23142080 | Unspecified |
| mmu-miR-21a-5p | MIMAT0000530 | Mrps18c | 68735 | HITS-CLIP | 23142080 | Unspecified |
| mmu-miR-21a-5p | MIMAT0000530 | Mrps2 | 118451 | HITS-CLIP | 23142080 | Unspecified |
| mmu-miR-21a-5p | MIMAT0000530 | Mrps21 | 66292 | HITS-CLIP | 23142080 | Unspecified |
| mmu-miR-21a-5p | MIMAT0000530 | Mrps22 | 64655 | HITS-CLIP | 23142080 | Unspecified |
| mmu-miR-21a-5p | MIMAT0000530 | Mrps23 | 64656 | HITS-CLIP | 23142080 | Unspecified |
| mmu-miR-21a-5p | MIMAT0000530 | Mrps30 | 59054 | HITS-CLIP | 23142080 | Unspecified |
| mmu-miR-21a-5p | MIMAT0000530 | Mrps35 | 232536 | HITS-CLIP | 23597149\|23142080 | Unspecified |
| mmu-miR-21a-5p | MIMAT0000530 | Mrps7 | 50529 | HITS-CLIP | 23597149\|23142080 | Unspecified |
| mmu-miR-21a-5p | MIMAT0000530 | Mrs2 | 380836 | HITS-CLIP | 19536157 | Unspecified |
| mmu-miR-21a-5p | MIMAT0000530 | Mrtfa | 57591 | HITS-CLIP | 23142080 | Unspecified |
| mmu-miR-21a-5p | MIMAT0000530 | Mrtfb | 239719 | HITS-CLIP | 29167373 | Unspecified |
| mmu-miR-21a-5p | MIMAT0000530 | Ms4a4b | 60361 | Microarrays | 37986224 | Unspecified |
| mmu-miR-21a-5p | MIMAT0000530 | Ms4a4c | 64380 | Microarrays | 37986224 | Unspecified |
| mmu-miR-21a-5p | MIMAT0000530 | Ms4a6b | 69774 | Microarrays | 37986224 | Unspecified |

| | | | | | | |
|---|---|---|---|---|---|---|
| mmu-miR-21a-5p | MIMAT0000530 | Ms4a6c | 73656 | Microarrays | 37986224 | Unspecified |
| mmu-miR-21a-5p | MIMAT0000530 | Ms4a6d | 68774 | HITS-CLIP | 29167373 | Unspecified |
| mmu-miR-21a-5p | MIMAT0000530 | Ms4a7 | 109225 | Microarrays | 37986224 | Unspecified |
| mmu-miR-21a-5p | MIMAT0000530 | Msantd4 | 78100 | HITS-CLIP | 23142080 | Unspecified |
| mmu-miR-21a-5p | MIMAT0000530 | Msh2 | 17685 | HITS-CLIP | 23142080 | Unspecified |
| mmu-miR-21a-5p | MIMAT0000530 | Msh3 | 17686 | HITS-CLIP | 23142080 | Unspecified |
| mmu-miR-21a-5p | MIMAT0000530 | Msh6 | 17688 | HITS-CLIP | 23142080\|19536157 | Unspecified |
| mmu-miR-21a-5p | MIMAT0000530 | Msi2 | 76626 | qCLASH//HITS-CLIP | 31393953\|23142080 | Unspecified |
| mmu-miR-21a-5p | MIMAT0000530 | Msl1 | 74026 | HITS-CLIP | 29167373\|23142080 | Unspecified |
| mmu-miR-21a-5p | MIMAT0000530 | Msl2 | 77853 | HITS-CLIP | 37986224\|23597149\|19536157\|23142 | Unspecified |
| mmu-miR-21a-5p | MIMAT0000530 | Msl3 | 17692 | HITS-CLIP | 29167373 | Unspecified |
| mmu-miR-21a-5p | MIMAT0000530 | Msn | 17698 | HITS-CLIP | 23142080\|29167373 | Unspecified |
| mmu-miR-21a-5p | MIMAT0000530 | Msr1 | 20288 | Microarrays | 37986224\|25263593 | Unspecified |
| mmu-miR-21a-5p | MIMAT0000530 | Msrb1 | 27361 | HITS-CLIP | 23597149\|23142080 | Unspecified |
| mmu-miR-21a-5p | MIMAT0000530 | Mtap | 66902 | Microarrays//HITS-CLIP | 37986224\|25263593\|23142080 | Unspecified |
| mmu-miR-21a-5p | MIMAT0000530 | Mtbp | 105837 | Microarrays//HITS-CLIP | 37986224\|23142080\|25263593 | Unspecified |
| mmu-miR-21a-5p | MIMAT0000530 | Mtch1 | 56462 | HITS-CLIP | 23142080 | Unspecified |
| mmu-miR-21a-5p | MIMAT0000530 | Mtch2 | 56428 | HITS-CLIP | 23597149 | Unspecified |
| mmu-miR-21a-5p | MIMAT0000530 | Mtdh | 67154 | qCLASH//HITS-CLIP | 31393953\|29167373 | Unspecified |
| mmu-miR-21a-5p | MIMAT0000530 | Mtf2 | 17765 | HITS-CLIP | 23142080 | Unspecified |
| mmu-miR-21a-5p | MIMAT0000530 | Mtfr1 | 67472 | HITS-CLIP | 23142080 | Unspecified |
| mmu-miR-21a-5p | MIMAT0000530 | Mtfr2 | 71804 | Microarrays | 37986224\|25263593 | Unspecified |
| mmu-miR-21a-5p | MIMAT0000530 | Mthfd1 | 108156 | Microarrays | 37986224\|25263593 | Unspecified |
| mmu-miR-21a-5p | MIMAT0000530 | Mthfd1l | 270685 | HITS-CLIP | 23142080 | Unspecified |
| mmu-miR-21a-5p | MIMAT0000530 | Mthfd2 | 17768 | Microarrays//HITS-CLIP | 37986224\|25263593\|23597149\|23142 | Unspecified |
| mmu-miR-21a-5p | MIMAT0000530 | Mtln | 67885 | HITS-CLIP | 23142080 | Unspecified |
| mmu-miR-21a-5p | MIMAT0000530 | Mtmr12 | 268783 | HITS-CLIP | 23142080\|29167373 | Unspecified |
| mmu-miR-21a-5p | MIMAT0000530 | Mtmr14 | 97287 | qCLASH | 31393953 | Unspecified |
| mmu-miR-21a-5p | MIMAT0000530 | Mtmr2 | 77116 | HITS-CLIP | 23597149\|23142080\|29167373 | Unspecified |
| mmu-miR-21a-5p | MIMAT0000530 | Mto1 | 68291 | HITS-CLIP | 23142080 | Unspecified |
| mmu-miR-21a-5p | MIMAT0000530 | Mtor | 56717 | qCLASH//HITS-CLIP | 31393953\|31433990\|23142080\|29167 | Unspecified |
| mmu-miR-21a-5p | MIMAT0000530 | Mtpn | 14489 | HITS-CLIP | 37986224\|23142080\|29167373 | Unspecified |
| mmu-miR-21a-5p | MIMAT0000530 | Mtr | 238505 | HITS-CLIP | 23142080 | Unspecified |
| mmu-miR-21a-5p | MIMAT0000530 | Mtrex | 23517 | HITS-CLIP | 23597149\|23142080 | Unspecified |
| mmu-miR-21a-5p | MIMAT0000530 | Mtrf1l | 108853 | HITS-CLIP | 23142080 | Unspecified |
| mmu-miR-21a-5p | MIMAT0000530 | Mtss1 | 211401 | Microarrays//HITS-CLIP//qCLASH | 37986224\|25263593\|23597149\|31393 | Unspecified |
| mmu-miR-21a-5p | MIMAT0000530 | Mttp | 17777 | HITS-CLIP | 23597149 | Unspecified |
| mmu-miR-21a-5p | MIMAT0000530 | Mturn | 68235 | HITS-CLIP | 23597149 | Unspecified |
| mmu-miR-21a-5p | MIMAT0000530 | Mtus1 | 102103 | Microarrays//HITS-CLIP | 37986224\|23597149 | Unspecified |

| | | | | | | | |
|---|---|---|---|---|---|---|---|
| mmu-miR-21a-5p | MIMAT0000530 | Mtx1 | 17827 | HITS-CLIP | | 23597149 | Unspecified |
| mmu-miR-21a-5p | MIMAT0000530 | Mtx2 | 53375 | HITS-CLIP | 23597149\|23142080 | | Unspecified |
| mmu-miR-21a-5p | MIMAT0000530 | Muc2 | 17831 | HITS-CLIP | | 23142080 | Unspecified |
| mmu-miR-21a-5p | MIMAT0000530 | Mug1 | 17836 | HITS-CLIP | | 23597149 | Unspecified |
| mmu-miR-21a-5p | MIMAT0000530 | Mut | 4594 | HITS-CLIP | 23142080\|23597149 | | Unspecified |
| mmu-miR-21a-5p | MIMAT0000530 | Mvd | 192156 | Microarrays | | 37986224 | Unspecified |
| mmu-miR-21a-5p | MIMAT0000530 | Mxd1 | 17119 | HITS-CLIP | | 23142080 | Unspecified |
| mmu-miR-21a-5p | MIMAT0000530 | Mxd4 | 17122 | Microarrays//HITS-CLIP | 37986224\|23142080 | | Unspecified |
| mmu-miR-21a-5p | MIMAT0000530 | Mxi1 | 17859 | Microarrays//HITS-CLIP | 37986224\|30971389\|31433990 | | Unspecified |
| mmu-miR-21a-5p | MIMAT0000530 | Myadm | 50918 | HITS-CLIP | | 29167373 | Unspecified |
| mmu-miR-21a-5p | MIMAT0000530 | Myb | 17863 | Microarrays//HITS-CLIP | 37986224\|23142080\|25263593 | | Unspecified |
| mmu-miR-21a-5p | MIMAT0000530 | Mybl2 | 17865 | Microarrays//HITS-CLIP | 37986224\|25263593\|23142080 | | Unspecified |
| mmu-miR-21a-5p | MIMAT0000530 | Myc | 17869 | Microarrays | 37986224\|25263593 | | Unspecified |
| mmu-miR-21a-5p | MIMAT0000530 | Mycbp | 56309 | HITS-CLIP | 37986224\|23142080 | | Unspecified |
| mmu-miR-21a-5p | MIMAT0000530 | Mycbp2 | 105689 | HITS-CLIP | 23142080\|19536157 | | Unspecified |
| mmu-miR-21a-5p | MIMAT0000530 | Mydgf | 56005 | HITS-CLIP | 23142080\|29167373 | | Unspecified |
| mmu-miR-21a-5p | MIMAT0000530 | Myef2 | 17876 | Microarrays | 37986224\|25263593 | | Unspecified |
| mmu-miR-21a-5p | MIMAT0000530 | Myg1 | 60315 | HITS-CLIP | | 23142080 | Unspecified |
| mmu-miR-21a-5p | MIMAT0000530 | Myh10 | 77579 | HITS-CLIP | | 23142080 | Unspecified |
| mmu-miR-21a-5p | MIMAT0000530 | Myh9 | 17886 | HITS-CLIP | 37986224\|23597149\|23142080\|26203 | | Unspecified |
| mmu-miR-21a-5p | MIMAT0000530 | Myl10 | 59310 | Microarrays | 37986224\|25263593 | | Unspecified |
| mmu-miR-21a-5p | MIMAT0000530 | Myl6 | 17904 | HITS-CLIP | | 29167373 | Unspecified |
| mmu-miR-21a-5p | MIMAT0000530 | Myl9 | 98932 | Microarrays | 37986224\|25263593 | | Unspecified |
| mmu-miR-21a-5p | MIMAT0000530 | Mylip | 218203 | HITS-CLIP | 37986224\|29167373\|23142080 | | Unspecified |
| mmu-miR-21a-5p | MIMAT0000530 | Mynn | 80732 | HITS-CLIP | 37986224\|23142080 | | Unspecified |
| mmu-miR-21a-5p | MIMAT0000530 | Myo10 | 17909 | HITS-CLIP, Microarrays//Microarrays//HITS | 37986224\|25263593\|19536157\|23142 | | Unspecified |
| mmu-miR-21a-5p | MIMAT0000530 | Myo18a | 360013 | HITS-CLIP | 23597149\|23142080 | | Unspecified |
| mmu-miR-21a-5p | MIMAT0000530 | Myo1b | 17912 | HITS-CLIP, Microarrays//Microarrays//HITS | 37986224\|25263593\|23597149 | | Unspecified |
| mmu-miR-21a-5p | MIMAT0000530 | Myo1c | 17913 | HITS-CLIP | | 29167373 | Unspecified |
| mmu-miR-21a-5p | MIMAT0000530 | Myo1d | 338367 | Microarrays | 37986224\|25263593 | | Unspecified |
| mmu-miR-21a-5p | MIMAT0000530 | Myo1e | 71602 | HITS-CLIP | 23597149\|29167373 | | Unspecified |
| mmu-miR-21a-5p | MIMAT0000530 | Myo1f | 17916 | HITS-CLIP | | 29167373 | Unspecified |
| mmu-miR-21a-5p | MIMAT0000530 | Myo5a | 17918 | HITS-CLIP | 23142080\|29167373 | | Unspecified |
| mmu-miR-21a-5p | MIMAT0000530 | Myo6 | 17920 | HITS-CLIP | | 23142080 | Unspecified |
| mmu-miR-21a-5p | MIMAT0000530 | Myo7a | 17921 | Microarrays//HITS-CLIP | 37986224\|25263593\|29167373 | | Unspecified |
| mmu-miR-21a-5p | MIMAT0000530 | Myo9b | 17925 | HITS-CLIP | | 29167373 | Unspecified |
| mmu-miR-21a-5p | MIMAT0000530 | Myorg | 57462 | HITS-CLIP | | 28735896 | Unspecified |
| mmu-miR-21a-5p | MIMAT0000530 | Myrf | 225908 | HITS-CLIP | | 23597149 | Unspecified |
| mmu-miR-21a-5p | MIMAT0000530 | Mysm1 | 320713 | HITS-CLIP | 37986224\|23142080 | | Unspecified |

| | | | | | | |
|---|---|---|---|---|---|---|
| mmu-miR-21a-5p | MIMAT0000530 | Myt1l | 17933 | HITS-CLIP | 37986224 | Unspecified |
| mmu-miR-21a-5p | MIMAT0000530 | Mzt1 | 76789 | HITS-CLIP | 23142080 | Unspecified |
| mmu-miR-21a-5p | MIMAT0000530 | N4bp2 | 333789 | HITS-CLIP | 23142080\|23597149\|29167373 | Unspecified |
| mmu-miR-21a-5p | MIMAT0000530 | N4bp2l1 | 100637 | Microarrays//HITS-CLIP | 37986224\|23597149 | Unspecified |
| mmu-miR-21a-5p | MIMAT0000530 | N4bp2l2 | 381695 | HITS-CLIP | 23142080 | Unspecified |
| mmu-miR-21a-5p | MIMAT0000530 | N4bp3 | 212706 | Microarrays | 37986224 | Unspecified |
| mmu-miR-21a-5p | MIMAT0000530 | Naa15 | 74838 | HITS-CLIP | 37986224\|23142080 | Unspecified |
| mmu-miR-21a-5p | MIMAT0000530 | Naa16 | 66897 | HITS-CLIP | 37986224\|23142080 | Unspecified |
| mmu-miR-21a-5p | MIMAT0000530 | Naa20 | 67877 | Microarrays | 37986224\|25263593 | Unspecified |
| mmu-miR-21a-5p | MIMAT0000530 | Naa25 | 231713 | HITS-CLIP | 29167373\|23142080 | Unspecified |
| mmu-miR-21a-5p | MIMAT0000530 | Naa30 | 70646 | HITS-CLIP | 37986224\|23597149\|23142080 | Unspecified |
| mmu-miR-21a-5p | MIMAT0000530 | Naa50 | 72117 | HITS-CLIP | 37986224\|23142080 | Unspecified |
| mmu-miR-21a-5p | MIMAT0000530 | Naaa | 67111 | Microarrays | 37986224 | Unspecified |
| mmu-miR-21a-5p | MIMAT0000530 | Nab1 | 17936 | HITS-CLIP | 23142080\|29167373 | Unspecified |
| mmu-miR-21a-5p | MIMAT0000530 | Nab2 | 17937 | HITS-CLIP | 23597149 | Unspecified |
| mmu-miR-21a-5p | MIMAT0000530 | Nabp1 | 109019 | HITS-CLIP | 31433990 | Unspecified |
| mmu-miR-21a-5p | MIMAT0000530 | Naca | 17938 | HITS-CLIP | 29167373 | Unspecified |
| mmu-miR-21a-5p | MIMAT0000530 | Nacc1 | 66830 | HITS-CLIP | 31433990 | Unspecified |
| mmu-miR-21a-5p | MIMAT0000530 | Nadk | 192185 | Microarrays | 37986224 | Unspecified |
| mmu-miR-21a-5p | MIMAT0000530 | Naf1 | 234344 | HITS-CLIP | 23142080 | Unspecified |
| mmu-miR-21a-5p | MIMAT0000530 | Naip5 | 17951 | Microarrays | 37986224 | Unspecified |
| mmu-miR-21a-5p | MIMAT0000530 | Naip6 | 191568 | HITS-CLIP | 29167373 | Unspecified |
| mmu-miR-21a-5p | MIMAT0000530 | Nampt | 59027 | HITS-CLIP | 23142080 | Unspecified |
| mmu-miR-21a-5p | MIMAT0000530 | Nanos1 | 332397 | Microarrays | 37986224\|25263593 | Unspecified |
| mmu-miR-21a-5p | MIMAT0000530 | Nap1l4 | 17955 | HITS-CLIP | 23142080 | Unspecified |
| mmu-miR-21a-5p | MIMAT0000530 | Napepld | 242864 | Microarrays | 37986224\|25263593 | Unspecified |
| mmu-miR-21a-5p | MIMAT0000530 | Nars | 70223 | qCLASH | 31393953 | Unspecified |
| mmu-miR-21a-5p | MIMAT0000530 | Nasp | 50927 | HITS-CLIP | 37986224\|23597149\|23142080\|29167 | Unspecified |
| mmu-miR-21a-5p | MIMAT0000530 | Nav1 | 215690 | HITS-CLIP | 29167373 | Unspecified |
| mmu-miR-21a-5p | MIMAT0000530 | Nav2 | 78286 | HITS-CLIP | 23597149\|29167373 | Unspecified |
| mmu-miR-21a-5p | MIMAT0000530 | Nbeal1 | 269198 | HITS-CLIP//qCLASH | 23142080\|31393953\|29167373 | Unspecified |
| mmu-miR-21a-5p | MIMAT0000530 | Nbeal2 | 235627 | HITS-CLIP | 29167373 | Unspecified |
| mmu-miR-21a-5p | MIMAT0000530 | Nbr1 | 17966 | HITS-CLIP | 23597149\|29167373 | Unspecified |
| mmu-miR-21a-5p | MIMAT0000530 | Ncapd2 | 68298 | HITS-CLIP | 23142080 | Unspecified |
| mmu-miR-21a-5p | MIMAT0000530 | Ncapd3 | 78658 | HITS-CLIP | 37986224\|23142080\|29167373 | Unspecified |
| mmu-miR-21a-5p | MIMAT0000530 | Ncapg | 54392 | Microarrays//HITS-CLIP | 37986224\|25263593\|23142080 | Unspecified |
| mmu-miR-21a-5p | MIMAT0000530 | Ncapg2 | 76044 | HITS-CLIP | 23142080 | Unspecified |
| mmu-miR-21a-5p | MIMAT0000530 | Ncaph | 215387 | HITS-CLIP | 23142080 | Unspecified |
| mmu-miR-21a-5p | MIMAT0000530 | Ncaph2 | 52683 | HITS-CLIP | 23142080 | Unspecified |

| | | | | | | |
|---|---|---|---|---|---|---|
| mmu-miR-21a-5p | MIMAT0000530 | Ncbp1 | 433702 | HITS-CLIP | 23597149\|29167373\|23142080 | Unspecified |
| mmu-miR-21a-5p | MIMAT0000530 | Ncbp2 | 68092 | HITS-CLIP | 23142080 | Unspecified |
| mmu-miR-21a-5p | MIMAT0000530 | Ncbp3 | 66874 | HITS-CLIP | 23142080 | Unspecified |
| mmu-miR-21a-5p | MIMAT0000530 | Nceh1 | 320024 | HITS-CLIP | 37986224\|23597149\|29167373 | Unspecified |
| mmu-miR-21a-5p | MIMAT0000530 | Ncf1 | 17969 | HITS-CLIP | 23142080 | Unspecified |
| mmu-miR-21a-5p | MIMAT0000530 | Ncf4 | 17972 | HITS-CLIP | 29167373 | Unspecified |
| mmu-miR-21a-5p | MIMAT0000530 | Nck1 | 17973 | HITS-CLIP | 23597149\|23142080 | Unspecified |
| mmu-miR-21a-5p | MIMAT0000530 | Nckap1 | 50884 | Microarrays//HITS-CLIP | 37986224\|25263593\|23597149\|1953€ | Unspecified |
| mmu-miR-21a-5p | MIMAT0000530 | Nckap1l | 105855 | HITS-CLIP | 23142080\|29167373 | Unspecified |
| mmu-miR-21a-5p | MIMAT0000530 | Ncl | 17975 | HITS-CLIP | 19536157 | Unspecified |
| mmu-miR-21a-5p | MIMAT0000530 | Ncmap | 230822 | Microarrays | 37986224\|25263593 | Unspecified |
| mmu-miR-21a-5p | MIMAT0000530 | Ncoa1 | 17977 | HITS-CLIP | 23142080 | Unspecified |
| mmu-miR-21a-5p | MIMAT0000530 | Ncoa2 | 17978 | HITS-CLIP | 37986224\|26203562\|23142080\|2916 | Unspecified |
| mmu-miR-21a-5p | MIMAT0000530 | Ncoa3 | 17979 | HITS-CLIP | 37986224\|19536157\|23142080\|2916 | Unspecified |
| mmu-miR-21a-5p | MIMAT0000530 | Ncoa4 | 27057 | Microarrays | 37986224 | Unspecified |
| mmu-miR-21a-5p | MIMAT0000530 | Ncoa6 | 56406 | HITS-CLIP | 23142080 | Unspecified |
| mmu-miR-21a-5p | MIMAT0000530 | Ncoa7 | 211329 | Microarrays | 37986224 | Unspecified |
| mmu-miR-21a-5p | MIMAT0000530 | Ncor1 | 20185 | HITS-CLIP | 23142080 | Unspecified |
| mmu-miR-21a-5p | MIMAT0000530 | Ncor2 | 20602 | HITS-CLIP | 28735896\|29167373 | Unspecified |
| mmu-miR-21a-5p | MIMAT0000530 | Ncstn | 59287 | HITS-CLIP | 23142080 | Unspecified |
| mmu-miR-21a-5p | MIMAT0000530 | Ndc1 | 72787 | Microarrays | 37986224\|25263593 | Unspecified |
| mmu-miR-21a-5p | MIMAT0000530 | Ndc80 | 67052 | Microarrays//HITS-CLIP | 37986224\|25263593\|23142080 | Unspecified |
| mmu-miR-21a-5p | MIMAT0000530 | Nde1 | 67203 | HITS-CLIP | 23142080 | Unspecified |
| mmu-miR-21a-5p | MIMAT0000530 | Ndel1 | 83431 | Microarrays | 37986224 | Unspecified |
| mmu-miR-21a-5p | MIMAT0000530 | Ndfip1 | 65113 | HITS-CLIP | 37986224\|23142080\|23597149\|1953€ | Unspecified |
| mmu-miR-21a-5p | MIMAT0000530 | Ndor1 | 78797 | HITS-CLIP | 29167373 | Unspecified |
| mmu-miR-21a-5p | MIMAT0000530 | Ndrg2 | 29811 | HITS-CLIP | 23597149\|19536157\|23142080 | Unspecified |
| mmu-miR-21a-5p | MIMAT0000530 | Ndrg3 | 29812 | HITS-CLIP | 23597149\|23142080 | Unspecified |
| mmu-miR-21a-5p | MIMAT0000530 | Ndrg4 | 234593 | Microarrays | 37986224\|25263593 | Unspecified |
| mmu-miR-21a-5p | MIMAT0000530 | Ndst1 | 15531 | HITS-CLIP | 23142080\|19536157\|23597149 | Unspecified |
| mmu-miR-21a-5p | MIMAT0000530 | Ndst2 | 17423 | HITS-CLIP | 23142080 | Unspecified |
| mmu-miR-21a-5p | MIMAT0000530 | Ndufa10 | 67273 | HITS-CLIP | 23597149 | Unspecified |
| mmu-miR-21a-5p | MIMAT0000530 | Ndufa3 | 66091 | HITS-CLIP | 23142080 | Unspecified |
| mmu-miR-21a-5p | MIMAT0000530 | Ndufa5 | 68202 | HITS-CLIP | 23597149 | Unspecified |
| mmu-miR-21a-5p | MIMAT0000530 | Ndufaf4 | 68493 | HITS-CLIP | 29167373 | Unspecified |
| mmu-miR-21a-5p | MIMAT0000530 | Ndufaf6 | 76947 | HITS-CLIP | 23142080 | Unspecified |
| mmu-miR-21a-5p | MIMAT0000530 | Ndufaf7 | 73694 | HITS-CLIP | 23142080 | Unspecified |
| mmu-miR-21a-5p | MIMAT0000530 | Ndufb11 | 104130 | HITS-CLIP | 23597149\|23142080 | Unspecified |
| mmu-miR-21a-5p | MIMAT0000530 | Ndufb2 | 68198 | HITS-CLIP | 23142080\|23597149 | Unspecified |

| | | | | | | |
|---|---|---|---|---|---|---|
| mmu-miR-21a-5p | MIMAT0000530 | Ndufb5 | 66046 | HITS-CLIP | 23142080 | Unspecified |
| mmu-miR-21a-5p | MIMAT0000530 | Ndufb9 | 66218 | HITS-CLIP | 23597149 | Unspecified |
| mmu-miR-21a-5p | MIMAT0000530 | Ndufs1 | 227197 | HITS-CLIP | 23142080 | Unspecified |
| mmu-miR-21a-5p | MIMAT0000530 | Ndufs2 | 226646 | HITS-CLIP | 23597149\|23142080 | Unspecified |
| mmu-miR-21a-5p | MIMAT0000530 | Ndufs3 | 68349 | HITS-CLIP | 37986224\|23597149\|23142080 | Unspecified |
| mmu-miR-21a-5p | MIMAT0000530 | Ndufs4 | 17993 | HITS-CLIP | 23142080 | Unspecified |
| mmu-miR-21a-5p | MIMAT0000530 | Ndufs8 | 225887 | HITS-CLIP | 23142080 | Unspecified |
| mmu-miR-21a-5p | MIMAT0000530 | Ndufv1 | 17995 | HITS-CLIP | 23597149 | Unspecified |
| mmu-miR-21a-5p | MIMAT0000530 | Ndufv2 | 72900 | HITS-CLIP | 31433990 | Unspecified |
| mmu-miR-21a-5p | MIMAT0000530 | Ndufv3 | 78330 | HITS-CLIP | 19536157\|23142080 | Unspecified |
| mmu-miR-21a-5p | MIMAT0000530 | Nebl | 74103 | HITS-CLIP | 23142080 | Unspecified |
| mmu-miR-21a-5p | MIMAT0000530 | Necap1 | 67602 | HITS-CLIP | 23597149\|29167373\|23142080 | Unspecified |
| mmu-miR-21a-5p | MIMAT0000530 | Necap2 | 66147 | HITS-CLIP | 23142080\|29167373 | Unspecified |
| mmu-miR-21a-5p | MIMAT0000530 | Nectin3 | 58998 | HITS-CLIP | 19536157 | Unspecified |
| mmu-miR-21a-5p | MIMAT0000530 | Nedd4 | 17999 | HITS-CLIP, Microarrays//HITS-CLIP//Microa | 37986224\|23597149\|25263593\|23142 | Unspecified |
| mmu-miR-21a-5p | MIMAT0000530 | Negr1 | 320840 | HITS-CLIP | 19536157 | Unspecified |
| mmu-miR-21a-5p | MIMAT0000530 | Neil1 | 72774 | Microarrays | 37986224 | Unspecified |
| mmu-miR-21a-5p | MIMAT0000530 | Neil3 | 234258 | Microarrays//HITS-CLIP | 37986224\|25263593\|23142080 | Unspecified |
| mmu-miR-21a-5p | MIMAT0000530 | Nek1 | 18004 | HITS-CLIP | 23142080 | Unspecified |
| mmu-miR-21a-5p | MIMAT0000530 | Nek2 | 18005 | Microarrays | 37986224\|25263593 | Unspecified |
| mmu-miR-21a-5p | MIMAT0000530 | Nek3 | 23954 | HITS-CLIP | 23142080 | Unspecified |
| mmu-miR-21a-5p | MIMAT0000530 | Nek6 | 59126 | Microarrays | 37986224\|25263593 | Unspecified |
| mmu-miR-21a-5p | MIMAT0000530 | Nek7 | 59125 | HITS-CLIP | 23142080 | Unspecified |
| mmu-miR-21a-5p | MIMAT0000530 | Nek9 | 217718 | HITS-CLIP | 23597149\|23142080 | Unspecified |
| mmu-miR-21a-5p | MIMAT0000530 | Nelfb | 58202 | HITS-CLIP | 23142080\|29167373 | Unspecified |
| mmu-miR-21a-5p | MIMAT0000530 | Nelfcd | 57314 | HITS-CLIP | 23597149\|23142080\|29167373 | Unspecified |
| mmu-miR-21a-5p | MIMAT0000530 | Nell2 | 54003 | HITS-CLIP | 37986224 | Unspecified |
| mmu-miR-21a-5p | MIMAT0000530 | Nemf | 66244 | HITS-CLIP | 19536157\|23142080 | Unspecified |
| mmu-miR-21a-5p | MIMAT0000530 | Nemp2 | 227094 | HITS-CLIP | 23142080 | Unspecified |
| mmu-miR-21a-5p | MIMAT0000530 | Neo1 | 18007 | Microarrays//HITS-CLIP | 37986224\|23597149 | Unspecified |
| mmu-miR-21a-5p | MIMAT0000530 | Net1 | 56349 | HITS-CLIP | 23142080 | Unspecified |
| mmu-miR-21a-5p | MIMAT0000530 | Neto2 | 74513 | HITS-CLIP | 37986224 | Unspecified |
| mmu-miR-21a-5p | MIMAT0000530 | Neu1 | 18010 | HITS-CLIP | 29167373 | Unspecified |
| mmu-miR-21a-5p | MIMAT0000530 | Neurl1b | 240055 | Microarrays | 37986224\|25263593 | Unspecified |
| mmu-miR-21a-5p | MIMAT0000530 | Neurl3 | 214854 | Microarrays//HITS-CLIP | 37986224\|29167373 | Unspecified |
| mmu-miR-21a-5p | MIMAT0000530 | Neurog2 | 11924 | HITS-CLIP | 23142080 | Unspecified |
| mmu-miR-21a-5p | MIMAT0000530 | Nexmif | 340533 | HITS-CLIP | 19536157 | Unspecified |
| mmu-miR-21a-5p | MIMAT0000530 | Nf1 | 18015 | qCLASH//HITS-CLIP | 31393953\|29167373 | Unspecified |
| mmu-miR-21a-5p | MIMAT0000530 | Nf2 | 18016 | HITS-CLIP | 37986224\|23142080\|29167373 | Unspecified |

| | | | | | | |
|---|---|---|---|---|---|---|
| mmu-miR-21a-5p | MIMAT0000530 | Nfam1 | 74039 | HITS-CLIP | 29167373 | Unspecified |
| mmu-miR-21a-5p | MIMAT0000530 | Nfat5 | 54446 | HITS-CLIP | 37986224\|23142080\|29167373 | Unspecified |
| mmu-miR-21a-5p | MIMAT0000530 | Nfatc1 | 18018 | HITS-CLIP | 23142080 | Unspecified |
| mmu-miR-21a-5p | MIMAT0000530 | Nfatc2 | 18019 | HITS-CLIP | 23142080\|29167373 | Unspecified |
| mmu-miR-21a-5p | MIMAT0000530 | Nfatc2ip | 18020 | HITS-CLIP | 23142080 | Unspecified |
| mmu-miR-21a-5p | MIMAT0000530 | Nfatc3 | 18021 | HITS-CLIP | 37986224\|23142080 | Unspecified |
| mmu-miR-21a-5p | MIMAT0000530 | Nfe2l1 | 18023 | HITS-CLIP | 23597149\|19536157\|23142080 | Unspecified |
| mmu-miR-21a-5p | MIMAT0000530 | Nfe2l2 | 18024 | HITS-CLIP | 23597149\|23142080\|29167373 | Unspecified |
| mmu-miR-21a-5p | MIMAT0000530 | Nfia | 18027 | Microarrays//HITS-CLIP | 37986224\|23142080 | Unspecified |
| mmu-miR-21a-5p | MIMAT0000530 | Nfib | 18028 | HITS-CLIP | 37986224\|23597149\|19536157 | Unspecified |
| mmu-miR-21a-5p | MIMAT0000530 | Nfic | 18029 | HITS-CLIP | 37986224 | Unspecified |
| mmu-miR-21a-5p | MIMAT0000530 | Nfil3 | 18030 | Microarrays//HITS-CLIP | 37986224\|23142080 | Unspecified |
| mmu-miR-21a-5p | MIMAT0000530 | Nfix | 18032 | HITS-CLIP | 37986224\|19536157 | Unspecified |
| mmu-miR-21a-5p | MIMAT0000530 | Nfkb2 | 18034 | HITS-CLIP | 23142080 | Unspecified |
| mmu-miR-21a-5p | MIMAT0000530 | Nfkbia | 18035 | HITS-CLIP | 23597149\|23142080 | Unspecified |
| mmu-miR-21a-5p | MIMAT0000530 | Nfkbib | 18036 | HITS-CLIP | 23142080 | Unspecified |
| mmu-miR-21a-5p | MIMAT0000530 | Nfkbiz | 80859 | HITS-CLIP | 29167373 | Unspecified |
| mmu-miR-21a-5p | MIMAT0000530 | Nfs1 | 18041 | HITS-CLIP | 23597149 | Unspecified |
| mmu-miR-21a-5p | MIMAT0000530 | Nfx1 | 74164 | HITS-CLIP | 30971389\|23142080 | Unspecified |
| mmu-miR-21a-5p | MIMAT0000530 | Nfxl1 | 100978 | Microarrays | 37986224 | Unspecified |
| mmu-miR-21a-5p | MIMAT0000530 | Nfya | 18044 | HITS-CLIP | 23142080\|29167373 | Unspecified |
| mmu-miR-21a-5p | MIMAT0000530 | Nfyb | 18045 | HITS-CLIP | 23142080 | Unspecified |
| mmu-miR-21a-5p | MIMAT0000530 | Ngly1 | 59007 | HITS-CLIP | 23142080 | Unspecified |
| mmu-miR-21a-5p | MIMAT0000530 | Nhp2 | 52530 | Microarrays | 37986224\|25263593 | Unspecified |
| mmu-miR-21a-5p | MIMAT0000530 | Nhsl2 | 100042480 | Microarrays | 37986224 | Unspecified |
| mmu-miR-21a-5p | MIMAT0000530 | Niban2 | 64855 | Microarrays | 25263593 | Unspecified |
| mmu-miR-21a-5p | MIMAT0000530 | Niban3 | 199786 | Microarrays | 25263593 | Unspecified |
| mmu-miR-21a-5p | MIMAT0000530 | Nid1 | 18073 | HITS-CLIP | 19536157 | Unspecified |
| mmu-miR-21a-5p | MIMAT0000530 | Nid2 | 18074 | Microarrays | 37986224\|25263593 | Unspecified |
| mmu-miR-21a-5p | MIMAT0000530 | Nifk | 67949 | Microarrays//qCLASH//HITS-CLIP | 37986224\|25263593\|31393953\|2314[illegible] | Unspecified |
| mmu-miR-21a-5p | MIMAT0000530 | Nim1k | 245269 | Microarrays | 37986224\|25263593 | Unspecified |
| mmu-miR-21a-5p | MIMAT0000530 | Nin | 18080 | HITS-CLIP | 23142080\|29167373 | Unspecified |
| mmu-miR-21a-5p | MIMAT0000530 | Ninj1 | 18081 | Microarrays | 37986224 | Unspecified |
| mmu-miR-21a-5p | MIMAT0000530 | Ninl | 78177 | Microarrays | 37986224\|25263593 | Unspecified |
| mmu-miR-21a-5p | MIMAT0000530 | Nip7 | 66164 | HITS-CLIP | 23142080 | Unspecified |
| mmu-miR-21a-5p | MIMAT0000530 | Nipa1 | 233280 | HITS-CLIP | 37986224 | Unspecified |
| mmu-miR-21a-5p | MIMAT0000530 | Nipa2 | 93790 | HITS-CLIP | 23597149 | Unspecified |
| mmu-miR-21a-5p | MIMAT0000530 | Nipbl | 71175 | HITS-CLIP | 37986224\|23142080\|29167373 | Unspecified |
| mmu-miR-21a-5p | MIMAT0000530 | Nipsnap2 | 2631 | HITS-CLIP | 23142080 | Unspecified |

| | | | | | | |
|---|---|---|---|---|---|---|
| mmu-miR-21a-5p | MIMAT0000530 | Nisch | 64652 | Microarrays//HITS-CLIP | 37986224\|23142080\|19536157 | Unspecified |
| mmu-miR-21a-5p | MIMAT0000530 | Nkain1 | 67149 | Microarrays | 37986224\|25263593 | Unspecified |
| mmu-miR-21a-5p | MIMAT0000530 | Nkap | 67050 | HITS-CLIP | 23142080\|29167373 | Unspecified |
| mmu-miR-21a-5p | MIMAT0000530 | Nkrf | 77286 | HITS-CLIP | 23142080 | Unspecified |
| mmu-miR-21a-5p | MIMAT0000530 | Nktr | 18087 | HITS-CLIP | 23142080 | Unspecified |
| mmu-miR-21a-5p | MIMAT0000530 | Nle1 | 217011 | HITS-CLIP | 23142080 | Unspecified |
| mmu-miR-21a-5p | MIMAT0000530 | Nlgn2 | 216856 | HITS-CLIP | 19536157\|23142080 | Unspecified |
| mmu-miR-21a-5p | MIMAT0000530 | Nln | 75805 | HITS-CLIP | 23597149\|23142080 | Unspecified |
| mmu-miR-21a-5p | MIMAT0000530 | Nlrp1b | 637515 | HITS-CLIP | 29167373 | Unspecified |
| mmu-miR-21a-5p | MIMAT0000530 | Nlrp3 | 216799 | HITS-CLIP | 29167373 | Unspecified |
| mmu-miR-21a-5p | MIMAT0000530 | Nlrx1 | 270151 | HITS-CLIP | 26203562 | Unspecified |
| mmu-miR-21a-5p | MIMAT0000530 | Nmd3 | 97112 | HITS-CLIP | 23142080 | Unspecified |
| mmu-miR-21a-5p | MIMAT0000530 | Nme1 | 18102 | Microarrays | 37986224\|25263593 | Unspecified |
| mmu-miR-21a-5p | MIMAT0000530 | Nme7 | 171567 | HITS-CLIP | 23142080 | Unspecified |
| mmu-miR-21a-5p | MIMAT0000530 | Nmnat1 | 66454 | HITS-CLIP | 23142080 | Unspecified |
| mmu-miR-21a-5p | MIMAT0000530 | Nmrk1 | 225994 | HITS-CLIP | 23597149 | Unspecified |
| mmu-miR-21a-5p | MIMAT0000530 | Nmt1 | 18107 | HITS-CLIP | 23597149\|29167373 | Unspecified |
| mmu-miR-21a-5p | MIMAT0000530 | Nmt2 | 18108 | HITS-CLIP | 37986224\|23142080 | Unspecified |
| mmu-miR-21a-5p | MIMAT0000530 | Nnat | 18111 | HITS-CLIP | 37986224\|19536157 | Unspecified |
| mmu-miR-21a-5p | MIMAT0000530 | Nnmt | 18113 | Microarrays | 37986224 | Unspecified |
| mmu-miR-21a-5p | MIMAT0000530 | Nnt | 18115 | Microarrays | 37986224\|25263593 | Unspecified |
| mmu-miR-21a-5p | MIMAT0000530 | Noct | 12457 | Microarrays//HITS-CLIP | 25263593\|19536157 | Unspecified |
| mmu-miR-21a-5p | MIMAT0000530 | Nod1 | 107607 | Microarrays//HITS-CLIP | 37986224\|23142080 | Unspecified |
| mmu-miR-21a-5p | MIMAT0000530 | Nol10 | 217431 | Microarrays//HITS-CLIP | 37986224\|25263593\|23142080 | Unspecified |
| mmu-miR-21a-5p | MIMAT0000530 | Nol11 | 68979 | HITS-CLIP | 37986224 | Unspecified |
| mmu-miR-21a-5p | MIMAT0000530 | Nol12 | 97961 | HITS-CLIP | 23142080 | Unspecified |
| mmu-miR-21a-5p | MIMAT0000530 | Nol4l | 329540 | HITS-CLIP | 29167373\|23142080 | Unspecified |
| mmu-miR-21a-5p | MIMAT0000530 | Nol7 | 70078 | HITS-CLIP | 23597149\|23142080 | Unspecified |
| mmu-miR-21a-5p | MIMAT0000530 | Nolc1 | 70769 | HITS-CLIP, Microarrays//Microarrays | 37986224\|25263593 | Unspecified |
| mmu-miR-21a-5p | MIMAT0000530 | Nom1 | 433864 | HITS-CLIP | 23142080 | Unspecified |
| mmu-miR-21a-5p | MIMAT0000530 | Nomo1 | 211548 | Microarrays//HITS-CLIP | 37986224\|25263593\|23142080 | Unspecified |
| mmu-miR-21a-5p | MIMAT0000530 | Nono | 53610 | HITS-CLIP | 23142080 | Unspecified |
| mmu-miR-21a-5p | MIMAT0000530 | Nop58 | 55989 | Microarrays | 37986224\|25263593 | Unspecified |
| mmu-miR-21a-5p | MIMAT0000530 | Nopchap1 | 121053 | qCLASH | 31393953 | Unspecified |
| mmu-miR-21a-5p | MIMAT0000530 | Notch1 | 18128 | Microarrays//HITS-CLIP | 37986224\|23597149\|23142080 | Unspecified |
| mmu-miR-21a-5p | MIMAT0000530 | Notch2 | 18129 | HITS-CLIP | 37986224\|23597149 | Unspecified |
| mmu-miR-21a-5p | MIMAT0000530 | Noxo1 | 71893 | Microarrays | 37986224 | Unspecified |
| mmu-miR-21a-5p | MIMAT0000530 | Noxred1 | 71275 | Microarrays | 37986224\|25263593 | Unspecified |
| mmu-miR-21a-5p | MIMAT0000530 | Npat | 244879 | HITS-CLIP | 23142080 | Unspecified |

| | | | | | | |
|---|---|---|---|---|---|---|
| mmu-miR-21a-5p | MIMAT0000530 | Npc1 | 18145 | HITS-CLIP | 23142080 | Unspecified |
| mmu-miR-21a-5p | MIMAT0000530 | Npc2 | 67963 | HITS-CLIP | 23597149\|23142080\|29167373 | Unspecified |
| mmu-miR-21a-5p | MIMAT0000530 | Npdc1 | 18146 | Microarrays | 37986224\|25263593 | Unspecified |
| mmu-miR-21a-5p | MIMAT0000530 | Npepps | 19155 | HITS-CLIP | 37986224\|23597149\|19536157\|2314[illegible] | Unspecified |
| mmu-miR-21a-5p | MIMAT0000530 | Nphp1 | 4867 | HITS-CLIP | 23142080 | Unspecified |
| mmu-miR-21a-5p | MIMAT0000530 | Npr2 | 230103 | HITS-CLIP | 23597149 | Unspecified |
| mmu-miR-21a-5p | MIMAT0000530 | Nptn | 20320 | HITS-CLIP | 23142080\|29167373 | Unspecified |
| mmu-miR-21a-5p | MIMAT0000530 | Nptx1 | 18164 | Microarrays//HITS-CLIP | 37986224\|25263593\|19536157 | Unspecified |
| mmu-miR-21a-5p | MIMAT0000530 | Npy | 109648 | Microarrays | 37986224\|25263593 | Unspecified |
| mmu-miR-21a-5p | MIMAT0000530 | Nqo1 | 18104 | Microarrays | 37986224\|25263593 | Unspecified |
| mmu-miR-21a-5p | MIMAT0000530 | Nr0b2 | 23957 | HITS-CLIP | 23597149 | Unspecified |
| mmu-miR-21a-5p | MIMAT0000530 | Nr1d1 | 217166 | HITS-CLIP | 23597149 | Unspecified |
| mmu-miR-21a-5p | MIMAT0000530 | Nr1h2 | 22260 | HITS-CLIP | 23142080 | Unspecified |
| mmu-miR-21a-5p | MIMAT0000530 | Nr1h4 | 20186 | HITS-CLIP | 23597149 | Unspecified |
| mmu-miR-21a-5p | MIMAT0000530 | Nr2c1 | 22025 | HITS-CLIP | 23597149\|23142080 | Unspecified |
| mmu-miR-21a-5p | MIMAT0000530 | Nr2c2 | 22026 | HITS-CLIP | 23142080 | Unspecified |
| mmu-miR-21a-5p | MIMAT0000530 | Nr3c1 | 14815 | HITS-CLIP | 23142080 | Unspecified |
| mmu-miR-21a-5p | MIMAT0000530 | Nr4a1 | 15370 | HITS-CLIP | 37986224\|23142080 | Unspecified |
| mmu-miR-21a-5p | MIMAT0000530 | Nr4a2 | 18227 | HITS-CLIP | 37986224\|23142080 | Unspecified |
| mmu-miR-21a-5p | MIMAT0000530 | Nr4a3 | 18124 | HITS-CLIP | 23142080 | Unspecified |
| mmu-miR-21a-5p | MIMAT0000530 | Nr5a2 | 26424 | HITS-CLIP | 23597149 | Unspecified |
| mmu-miR-21a-5p | MIMAT0000530 | Nras | 18176 | HITS-CLIP | 23597149\|19536157\|23142080\|29167 | Unspecified |
| mmu-miR-21a-5p | MIMAT0000530 | Nrcam | 319504 | HITS-CLIP | 19536157 | Unspecified |
| mmu-miR-21a-5p | MIMAT0000530 | Nrd1 | 230598 | HITS-CLIP | 23142080 | Unspecified |
| mmu-miR-21a-5p | MIMAT0000530 | Nrf1 | 18181 | HITS-CLIP | 23142080 | Unspecified |
| mmu-miR-21a-5p | MIMAT0000530 | Nrip1 | 268903 | HITS-CLIP | 37986224\|23142080 | Unspecified |
| mmu-miR-21a-5p | MIMAT0000530 | Nrp1 | 18186 | HITS-CLIP | 29167373 | Unspecified |
| mmu-miR-21a-5p | MIMAT0000530 | Nrp2 | 18187 | HITS-CLIP | 29167373 | Unspecified |
| mmu-miR-21a-5p | MIMAT0000530 | Nrxn1 | 18189 | HITS-CLIP | 37986224 | Unspecified |
| mmu-miR-21a-5p | MIMAT0000530 | Nrxn2 | 18190 | HITS-CLIP | 19536157 | Unspecified |
| mmu-miR-21a-5p | MIMAT0000530 | Nsa2 | 59050 | qCLASH | 31393953 | Unspecified |
| mmu-miR-21a-5p | MIMAT0000530 | Nsd1 | 18193 | HITS-CLIP | 23597149\|23142080\|29167373 | Unspecified |
| mmu-miR-21a-5p | MIMAT0000530 | Nsd2 | 107823 | HITS-CLIP | 23142080\|29167373 | Unspecified |
| mmu-miR-21a-5p | MIMAT0000530 | Nsd3 | 234135 | HITS-CLIP | 23142080\|29167373 | Unspecified |
| mmu-miR-21a-5p | MIMAT0000530 | Nsdhl | 18194 | HITS-CLIP | 23597149 | Unspecified |
| mmu-miR-21a-5p | MIMAT0000530 | Nsf | 18195 | HITS-CLIP | 19536157 | Unspecified |
| mmu-miR-21a-5p | MIMAT0000530 | Nsg2 | 18197 | HITS-CLIP | 23142080 | Unspecified |
| mmu-miR-21a-5p | MIMAT0000530 | Nsmaf | 18201 | HITS-CLIP | 23142080 | Unspecified |
| mmu-miR-21a-5p | MIMAT0000530 | Nsmce1 | 67711 | HITS-CLIP | 23142080 | Unspecified |

| | | | | | | |
|---|---|---|---|---|---|---|
| mmu-miR-21a-5p | MIMAT0000530 | Nsrp1 | 237859 | HITS-CLIP | 23142080 | Unspecified |
| mmu-miR-21a-5p | MIMAT0000530 | Nsun3 | 106338 | HITS-CLIP | 23142080 | Unspecified |
| mmu-miR-21a-5p | MIMAT0000530 | Nsun4 | 72181 | HITS-CLIP | 23142080 | Unspecified |
| mmu-miR-21a-5p | MIMAT0000530 | Nt5c2 | 76952 | HITS-CLIP | 23142080 | Unspecified |
| mmu-miR-21a-5p | MIMAT0000530 | Nt5c3 | 107569 | HITS-CLIP | 23142080 | Unspecified |
| mmu-miR-21a-5p | MIMAT0000530 | Nt5dc2 | 70021 | Microarrays | 37986224 | Unspecified |
| mmu-miR-21a-5p | MIMAT0000530 | Nt5dc3 | 103466 | HITS-CLIP | 37986224 | Unspecified |
| mmu-miR-21a-5p | MIMAT0000530 | Nt5e | 23959 | HITS-CLIP | 37986224\|23142080 | Unspecified |
| mmu-miR-21a-5p | MIMAT0000530 | Ntn1 | 18208 | Microarrays//HITS-CLIP | 37986224\|25263593\|23142080 | Unspecified |
| mmu-miR-21a-5p | MIMAT0000530 | Ntrk2 | 18212 | HITS-CLIP | 37986224 | Unspecified |
| mmu-miR-21a-5p | MIMAT0000530 | Nts | 67405 | HITS-CLIP | 19536157 | Unspecified |
| mmu-miR-21a-5p | MIMAT0000530 | Nuak1 | 77976 | Microarrays | 37986224 | Unspecified |
| mmu-miR-21a-5p | MIMAT0000530 | Nuak2 | 74137 | HITS-CLIP | 23597149\|29167373 | Unspecified |
| mmu-miR-21a-5p | MIMAT0000530 | Nub1 | 53312 | HITS-CLIP | 23142080 | Unspecified |
| mmu-miR-21a-5p | MIMAT0000530 | Nubp2 | 26426 | HITS-CLIP | 23597149 | Unspecified |
| mmu-miR-21a-5p | MIMAT0000530 | Nucb1 | 18220 | HITS-CLIP | 23142080\|23597149 | Unspecified |
| mmu-miR-21a-5p | MIMAT0000530 | Nucb2 | 53322 | HITS-CLIP | 23142080 | Unspecified |
| mmu-miR-21a-5p | MIMAT0000530 | Nucks1 | 98415 | HITS-CLIP | 37986224\|23142080 | Unspecified |
| mmu-miR-21a-5p | MIMAT0000530 | Nudcd1 | 67429 | HITS-CLIP | 23142080 | Unspecified |
| mmu-miR-21a-5p | MIMAT0000530 | Nudt12 | 67993 | Microarrays | 37986224\|25263593 | Unspecified |
| mmu-miR-21a-5p | MIMAT0000530 | Nudt16l1 | 66911 | HITS-CLIP | 23142080 | Unspecified |
| mmu-miR-21a-5p | MIMAT0000530 | Nudt5 | 53893 | HITS-CLIP | 23142080 | Unspecified |
| mmu-miR-21a-5p | MIMAT0000530 | Nuf2 | 66977 | Microarrays | 37986224\|25263593 | Unspecified |
| mmu-miR-21a-5p | MIMAT0000530 | Nufip2 | 68564 | HITS-CLIP | 37986224\|23597149\|23142080\|29167 | Unspecified |
| mmu-miR-21a-5p | MIMAT0000530 | Numa1 | 101706 | HITS-CLIP | 23597149\|23142080 | Unspecified |
| mmu-miR-21a-5p | MIMAT0000530 | Numb | 18222 | Microarrays//HITS-CLIP | 37986224\|25263593\|23142080 | Unspecified |
| mmu-miR-21a-5p | MIMAT0000530 | Nup107 | 103468 | HITS-CLIP | 23142080 | Unspecified |
| mmu-miR-21a-5p | MIMAT0000530 | Nup153 | 218210 | HITS-CLIP | 23142080 | Unspecified |
| mmu-miR-21a-5p | MIMAT0000530 | Nup155 | 170762 | HITS-CLIP | 23142080\|29167373 | Unspecified |
| mmu-miR-21a-5p | MIMAT0000530 | Nup160 | 59015 | HITS-CLIP | 23142080 | Unspecified |
| mmu-miR-21a-5p | MIMAT0000530 | Nup188 | 227699 | HITS-CLIP | 23142080 | Unspecified |
| mmu-miR-21a-5p | MIMAT0000530 | Nup205 | 70699 | HITS-CLIP | 23597149\|23142080 | Unspecified |
| mmu-miR-21a-5p | MIMAT0000530 | Nup85 | 445007 | HITS-CLIP | 23142080 | Unspecified |
| mmu-miR-21a-5p | MIMAT0000530 | Nup88 | 19069 | HITS-CLIP | 23142080 | Unspecified |
| mmu-miR-21a-5p | MIMAT0000530 | Nup93 | 71805 | HITS-CLIP | 23142080 | Unspecified |
| mmu-miR-21a-5p | MIMAT0000530 | Nup98 | 269966 | HITS-CLIP | 23142080\|29167373 | Unspecified |
| mmu-miR-21a-5p | MIMAT0000530 | Nus1 | 52014 | HITS-CLIP | 29167373 | Unspecified |
| mmu-miR-21a-5p | MIMAT0000530 | Nusap1 | 108907 | Microarrays//HITS-CLIP | 37986224\|25263593\|23142080 | Unspecified |
| mmu-miR-21a-5p | MIMAT0000530 | Nvl | 67459 | HITS-CLIP | 23142080 | Unspecified |

| | | | | | | |
|---|---|---|---|---|---|---|
| mmu-miR-21a-5p | MIMAT0000530 | Nxpe3 | 385658 | HITS-CLIP | 23142080 | Unspecified |
| mmu-miR-21a-5p | MIMAT0000530 | Nxt2 | 237082 | HITS-CLIP | 23597149\|23142080\|29167373 | Unspecified |
| mmu-miR-21a-5p | MIMAT0000530 | Oas3 | 246727 | Microarrays//HITS-CLIP | 37986224\|25263593\|29167373 | Unspecified |
| mmu-miR-21a-5p | MIMAT0000530 | Oasl1 | 231655 | Microarrays | 37986224 | Unspecified |
| mmu-miR-21a-5p | MIMAT0000530 | Oasl2 | 23962 | HITS-CLIP | 29167373 | Unspecified |
| mmu-miR-21a-5p | MIMAT0000530 | Oat | 18242 | HITS-CLIP | 23597149\|23142080\|29167373 | Unspecified |
| mmu-miR-21a-5p | MIMAT0000530 | Ocel1 | 77090 | Microarrays | 37986224\|25263593 | Unspecified |
| mmu-miR-21a-5p | MIMAT0000530 | Ociad1 | 68095 | HITS-CLIP | 23142080 | Unspecified |
| mmu-miR-21a-5p | MIMAT0000530 | Ocrl | 320634 | Microarrays//HITS-CLIP | 37986224\|23142080 | Unspecified |
| mmu-miR-21a-5p | MIMAT0000530 | Odc1 | 18263 | Microarrays | 37986224\|25263593 | Unspecified |
| mmu-miR-21a-5p | MIMAT0000530 | Ogdh | 18293 | HITS-CLIP | 23142080\|19536157\|31433990\|29167 | Unspecified |
| mmu-miR-21a-5p | MIMAT0000530 | Ogfod1 | 270086 | HITS-CLIP | 37986224\|23142080 | Unspecified |
| mmu-miR-21a-5p | MIMAT0000530 | Ogfod2 | 66627 | HITS-CLIP | 23142080 | Unspecified |
| mmu-miR-21a-5p | MIMAT0000530 | Ogt | 108155 | HITS-CLIP | 23142080 | Unspecified |
| mmu-miR-21a-5p | MIMAT0000530 | Oip5 | 70645 | Microarrays//HITS-CLIP | 37986224\|25263593\|23142080 | Unspecified |
| mmu-miR-21a-5p | MIMAT0000530 | Oit3 | 18302 | Microarrays | 37986224\|25263593 | Unspecified |
| mmu-miR-21a-5p | MIMAT0000530 | Olfm1 | 56177 | Microarrays//HITS-CLIP | 37986224\|19536157 | Unspecified |
| mmu-miR-21a-5p | MIMAT0000530 | Onecut1 | 15379 | HITS-CLIP | 23597149 | Unspecified |
| mmu-miR-21a-5p | MIMAT0000530 | Onecut2 | 225631 | HITS-CLIP | 37986224 | Unspecified |
| mmu-miR-21a-5p | MIMAT0000530 | Opa1 | 74143 | HITS-CLIP | 23142080 | Unspecified |
| mmu-miR-21a-5p | MIMAT0000530 | Oplah | 75475 | HITS-CLIP | 23597149 | Unspecified |
| mmu-miR-21a-5p | MIMAT0000530 | Or5w19 | 258103 | qCLASH | 31393953 | Unspecified |
| mmu-miR-21a-5p | MIMAT0000530 | Orai1 | 109305 | HITS-CLIP | 23142080 | Unspecified |
| mmu-miR-21a-5p | MIMAT0000530 | Orc2 | 18393 | HITS-CLIP | 23142080 | Unspecified |
| mmu-miR-21a-5p | MIMAT0000530 | Orc3 | 50793 | HITS-CLIP | 37986224 | Unspecified |
| mmu-miR-21a-5p | MIMAT0000530 | Orc4 | 26428 | HITS-CLIP | 23142080 | Unspecified |
| mmu-miR-21a-5p | MIMAT0000530 | Orc6 | 56452 | HITS-CLIP | 23142080 | Unspecified |
| mmu-miR-21a-5p | MIMAT0000530 | Ormdl2 | 66844 | HITS-CLIP | 23597149\|23142080 | Unspecified |
| mmu-miR-21a-5p | MIMAT0000530 | Os9 | 216440 | HITS-CLIP//qCLASH | 23597149\|31393953 | Unspecified |
| mmu-miR-21a-5p | MIMAT0000530 | Osbp | 76303 | HITS-CLIP | 23142080 | Unspecified |
| mmu-miR-21a-5p | MIMAT0000530 | Osbpl1a | 64291 | HITS-CLIP | 23597149 | Unspecified |
| mmu-miR-21a-5p | MIMAT0000530 | Osbpl2 | 228983 | HITS-CLIP | 23142080 | Unspecified |
| mmu-miR-21a-5p | MIMAT0000530 | Osbpl8 | 237542 | HITS-CLIP | 23597149\|19536157\|23142080\|29167 | Unspecified |
| mmu-miR-21a-5p | MIMAT0000530 | Osbpl9 | 100273 | Microarrays | 37986224 | Unspecified |
| mmu-miR-21a-5p | MIMAT0000530 | Osgepl1 | 72085 | HITS-CLIP | 23142080 | Unspecified |
| mmu-miR-21a-5p | MIMAT0000530 | Osgin1 | 71839 | HITS-CLIP | 23597149\|29167373 | Unspecified |
| mmu-miR-21a-5p | MIMAT0000530 | Osgin2 | 209212 | HITS-CLIP | 29167373 | Unspecified |
| mmu-miR-21a-5p | MIMAT0000530 | Osm | 18413 | Microarrays | 37986224 | Unspecified |
| mmu-miR-21a-5p | MIMAT0000530 | Otud4 | 73945 | HITS-CLIP | 23142080\|29167373 | Unspecified |

| | | | | | | |
|---|---|---|---|---|---|---|
| mmu-miR-21a-5p | MIMAT0000530 | Otud7b | 229603 | HITS-CLIP | 23142080 | Unspecified |
| mmu-miR-21a-5p | MIMAT0000530 | Otulin | 432940 | HITS-CLIP | 23142080 | Unspecified |
| mmu-miR-21a-5p | MIMAT0000530 | Otulinl | 54491 | HITS-CLIP | 23142080 | Unspecified |
| mmu-miR-21a-5p | MIMAT0000530 | Oxr1 | 170719 | HITS-CLIP | 23142080 | Unspecified |
| mmu-miR-21a-5p | MIMAT0000530 | Oxsr1 | 108737 | HITS-CLIP | 23142080 | Unspecified |
| mmu-miR-21a-5p | MIMAT0000530 | P2rx7 | 18439 | HITS-CLIP | 29167373 | Unspecified |
| mmu-miR-21a-5p | MIMAT0000530 | P2ry1 | 18441 | Microarrays | 37986224\|25263593 | Unspecified |
| mmu-miR-21a-5p | MIMAT0000530 | P2ry10 | 78826 | HITS-CLIP | 23142080 | Unspecified |
| mmu-miR-21a-5p | MIMAT0000530 | P2ry10b | 213438 | Microarrays | 25263593 | Unspecified |
| mmu-miR-21a-5p | MIMAT0000530 | P2ry12 | 70839 | Microarrays | 37986224 | Unspecified |
| mmu-miR-21a-5p | MIMAT0000530 | P2ry14 | 140795 | Microarrays | 37986224 | Unspecified |
| mmu-miR-21a-5p | MIMAT0000530 | P2ry2 | 18442 | Microarrays | 37986224\|25263593 | Unspecified |
| mmu-miR-21a-5p | MIMAT0000530 | P2ry6 | 233571 | HITS-CLIP | 29167373 | Unspecified |
| mmu-miR-21a-5p | MIMAT0000530 | P3h4 | 10609 | Microarrays | 25263593 | Unspecified |
| mmu-miR-21a-5p | MIMAT0000530 | P4ha1 | 18451 | HITS-CLIP | 23142080 | Unspecified |
| mmu-miR-21a-5p | MIMAT0000530 | P4ha2 | 18452 | Microarrays | 37986224\|25263593 | Unspecified |
| mmu-miR-21a-5p | MIMAT0000530 | P4hb | 18453 | HITS-CLIP | 23142080 | Unspecified |
| mmu-miR-21a-5p | MIMAT0000530 | Pa2g4 | 18813 | HITS-CLIP | 23142080 | Unspecified |
| mmu-miR-21a-5p | MIMAT0000530 | Pabpc1 | 18458 | qCLASH//HITS-CLIP | 31393953\|23142080 | Unspecified |
| mmu-miR-21a-5p | MIMAT0000530 | Pabpn1 | 54196 | HITS-CLIP | 37986224\|23142080 | Unspecified |
| mmu-miR-21a-5p | MIMAT0000530 | Pacc1 | 55248 | Microarrays | 25263593 | Unspecified |
| mmu-miR-21a-5p | MIMAT0000530 | Pacrg | 69310 | Microarrays | 37986224\|25263593 | Unspecified |
| mmu-miR-21a-5p | MIMAT0000530 | Pacs2 | 217893 | HITS-CLIP | 37986224\|19536157\|23142080\|29167 | Unspecified |
| mmu-miR-21a-5p | MIMAT0000530 | Pacsin2 | 23970 | HITS-CLIP | 23597149 | Unspecified |
| mmu-miR-21a-5p | MIMAT0000530 | Padi4 | 18602 | Microarrays | 37986224\|25263593 | Unspecified |
| mmu-miR-21a-5p | MIMAT0000530 | Paf1 | 54624 | HITS-CLIP | 23142080 | Unspecified |
| mmu-miR-21a-5p | MIMAT0000530 | Pafah1b2 | 18475 | HITS-CLIP | 37986224\|23142080 | Unspecified |
| mmu-miR-21a-5p | MIMAT0000530 | Pafah2 | 100163 | Microarrays | 37986224\|25263593 | Unspecified |
| mmu-miR-21a-5p | MIMAT0000530 | Pag1 | 94212 | HITS-CLIP | 23142080\|23597149 | Unspecified |
| mmu-miR-21a-5p | MIMAT0000530 | Pah | 18478 | HITS-CLIP | 23597149 | Unspecified |
| mmu-miR-21a-5p | MIMAT0000530 | Paics | 67054 | HITS-CLIP | 23597149 | Unspecified |
| mmu-miR-21a-5p | MIMAT0000530 | Paip1 | 218693 | Microarrays//HITS-CLIP | 37986224\|23142080 | Unspecified |
| mmu-miR-21a-5p | MIMAT0000530 | Paip2 | 67869 | HITS-CLIP | 23142080 | Unspecified |
| mmu-miR-21a-5p | MIMAT0000530 | Paip2b | 232164 | HITS-CLIP | 37986224\|23142080 | Unspecified |
| mmu-miR-21a-5p | MIMAT0000530 | Pak2 | 224105 | HITS-CLIP | 23142080\|29167373 | Unspecified |
| mmu-miR-21a-5p | MIMAT0000530 | Pald1 | 27355 | Microarrays | 37986224\|25263593 | Unspecified |
| mmu-miR-21a-5p | MIMAT0000530 | Pam | 18484 | HITS-CLIP | 23597149 | Unspecified |
| mmu-miR-21a-5p | MIMAT0000530 | Pan3 | 72587 | Microarrays | 37986224 | Unspecified |
| mmu-miR-21a-5p | MIMAT0000530 | Pank1 | 75735 | HITS-CLIP | 23597149\|23142080 | Unspecified |

| | | | | | | |
|---|---|---|---|---|---|---|
| mmu-miR-21a-5p | MIMAT0000530 | Pank3 | 211347 | HITS-CLIP | 37986224\|23142080 | Unspecified |
| mmu-miR-21a-5p | MIMAT0000530 | Pank4 | 55229 | HITS-CLIP | 23142080 | Unspecified |
| mmu-miR-21a-5p | MIMAT0000530 | Panx1 | 55991 | Microarrays//HITS-CLIP | 37986224\|23142080\|25263593 | Unspecified |
| mmu-miR-21a-5p | MIMAT0000530 | Papola | 18789 | HITS-CLIP | 19536157 | Unspecified |
| mmu-miR-21a-5p | MIMAT0000530 | Papolg | 216578 | HITS-CLIP | 23142080 | Unspecified |
| mmu-miR-21a-5p | MIMAT0000530 | Papss2 | 23972 | Microarrays | 37986224\|25263593 | Unspecified |
| mmu-miR-21a-5p | MIMAT0000530 | Paqr8 | 74229 | HITS-CLIP | 37986224 | Unspecified |
| mmu-miR-21a-5p | MIMAT0000530 | Paqr9 | 75552 | HITS-CLIP | 37986224 | Unspecified |
| mmu-miR-21a-5p | MIMAT0000530 | Parg | 26430 | HITS-CLIP | 23142080 | Unspecified |
| mmu-miR-21a-5p | MIMAT0000530 | Parn | 74108 | HITS-CLIP | 23142080 | Unspecified |
| mmu-miR-21a-5p | MIMAT0000530 | Parp1 | 11545 | HITS-CLIP | 29167373 | Unspecified |
| mmu-miR-21a-5p | MIMAT0000530 | Parp11 | 101187 | HITS-CLIP | 29167373 | Unspecified |
| mmu-miR-21a-5p | MIMAT0000530 | Parp14 | 547253 | Microarrays//HITS-CLIP | 37986224\|29167373 | Unspecified |
| mmu-miR-21a-5p | MIMAT0000530 | Parp4 | 328417 | HITS-CLIP | 23142080 | Unspecified |
| mmu-miR-21a-5p | MIMAT0000530 | Parp8 | 52552 | Microarrays//HITS-CLIP | 37986224\|29167373 | Unspecified |
| mmu-miR-21a-5p | MIMAT0000530 | Parpbp | 75317 | Microarrays | 37986224\|25263593 | Unspecified |
| mmu-miR-21a-5p | MIMAT0000530 | Parva | 57342 | HITS-CLIP | 37986224\|19536157 | Unspecified |
| mmu-miR-21a-5p | MIMAT0000530 | Parvg | 64099 | Microarrays//HITS-CLIP | 37986224\|23142080\|25263593 | Unspecified |
| mmu-miR-21a-5p | MIMAT0000530 | Pask | 269224 | HITS-CLIP | 23142080 | Unspecified |
| mmu-miR-21a-5p | MIMAT0000530 | Patl1 | 225929 | HITS-CLIP | 31433990\|29167373 | Unspecified |
| mmu-miR-21a-5p | MIMAT0000530 | Patz1 | 56218 | HITS-CLIP | 23142080\|29167373 | Unspecified |
| mmu-miR-21a-5p | MIMAT0000530 | Paxbp1 | 67367 | HITS-CLIP | 30971389\|23142080 | Unspecified |
| mmu-miR-21a-5p | MIMAT0000530 | Paxip1 | 55982 | HITS-CLIP | 23142080 | Unspecified |
| mmu-miR-21a-5p | MIMAT0000530 | Pbdc1 | 67683 | Microarrays | 37986224\|25263593 | Unspecified |
| mmu-miR-21a-5p | MIMAT0000530 | Pbk | 52033 | HITS-CLIP, Microarrays//HITS-CLIP//Microa | 37986224\|23597149\|25263593\|2314 | Unspecified |
| mmu-miR-21a-5p | MIMAT0000530 | Pbld2 | 67307 | HITS-CLIP | 23597149 | Unspecified |
| mmu-miR-21a-5p | MIMAT0000530 | Pbrm1 | 66923 | HITS-CLIP | 37986224\|19536157\|23142080\|29167 | Unspecified |
| mmu-miR-21a-5p | MIMAT0000530 | Pbx2 | 18515 | HITS-CLIP | 31433990\|26203562 | Unspecified |
| mmu-miR-21a-5p | MIMAT0000530 | Pcbd1 | 13180 | HITS-CLIP | 37986224\|23597149 | Unspecified |
| mmu-miR-21a-5p | MIMAT0000530 | Pcbd2 | 135972048 | HITS-CLIP | 23142080 | Unspecified |
| mmu-miR-21a-5p | MIMAT0000530 | Pcbp1 | 23983 | HITS-CLIP | 23597149\|29167373\|23142080 | Unspecified |
| mmu-miR-21a-5p | MIMAT0000530 | Pcbp2 | 18521 | HITS-CLIP | 37986224\|23142080 | Unspecified |
| mmu-miR-21a-5p | MIMAT0000530 | Pcca | 110821 | HITS-CLIP | 23597149\|23142080 | Unspecified |
| mmu-miR-21a-5p | MIMAT0000530 | Pccb | 66904 | HITS-CLIP | 23597149 | Unspecified |
| mmu-miR-21a-5p | MIMAT0000530 | Pcdh1 | 75599 | HITS-CLIP | 23597149\|19536157 | Unspecified |
| mmu-miR-21a-5p | MIMAT0000530 | Pcdh10 | 18526 | HITS-CLIP | 37986224 | Unspecified |
| mmu-miR-21a-5p | MIMAT0000530 | Pcdh17 | 219228 | HITS-CLIP | 37986224 | Unspecified |
| mmu-miR-21a-5p | MIMAT0000530 | Pcdh7 | 54216 | HITS-CLIP | 19536157\|29167373 | Unspecified |
| mmu-miR-21a-5p | MIMAT0000530 | Pcdh8 | 18530 | HITS-CLIP | 37986224 | Unspecified |

| | | | | | | |
|---|---|---|---|---|---|---|
| mmu-miR-21a-5p | MIMAT0000530 | Pcdhga12 | 93724 | HITS-CLIP | 37986224 | Unspecified |
| mmu-miR-21a-5p | MIMAT0000530 | Pcdhgc3 | 5098 | HITS-CLIP | 19536157 | Unspecified |
| mmu-miR-21a-5p | MIMAT0000530 | Pcf11 | 74737 | HITS-CLIP | 23597149\|23142080\|29167373 | Unspecified |
| mmu-miR-21a-5p | MIMAT0000530 | Pcgf1 | 84759 | HITS-CLIP | 23597149 | Unspecified |
| mmu-miR-21a-5p | MIMAT0000530 | Pcgf3 | 69587 | HITS-CLIP | 23142080\|29167373 | Unspecified |
| mmu-miR-21a-5p | MIMAT0000530 | Pcgf5 | 76073 | HITS-CLIP | 23142080 | Unspecified |
| mmu-miR-21a-5p | MIMAT0000530 | Pcid2 | 234069 | HITS-CLIP | 23142080 | Unspecified |
| mmu-miR-21a-5p | MIMAT0000530 | Pck2 | 74551 | Microarrays | 37986224 | Unspecified |
| mmu-miR-21a-5p | MIMAT0000530 | Pclaf | 68026 | Microarrays | 25263593 | Unspecified |
| mmu-miR-21a-5p | MIMAT0000530 | Pclo | 26875 | HITS-CLIP | 37986224 | Unspecified |
| mmu-miR-21a-5p | MIMAT0000530 | Pcm1 | 18536 | HITS-CLIP | 37986224\|23142080 | Unspecified |
| mmu-miR-21a-5p | MIMAT0000530 | Pcmt1 | 18537 | HITS-CLIP | 37986224\|23142080\|19536157\|29167 | Unspecified |
| mmu-miR-21a-5p | MIMAT0000530 | Pcmtd1 | 319263 | HITS-CLIP | 23142080 | Unspecified |
| mmu-miR-21a-5p | MIMAT0000530 | Pcmtd2 | 245867 | HITS-CLIP, Microarrays//HITS-CLIP | 37986224\|23142080 | Unspecified |
| mmu-miR-21a-5p | MIMAT0000530 | Pcna | 18538 | HITS-CLIP | 23597149\|23142080\|29167373 | Unspecified |
| mmu-miR-21a-5p | MIMAT0000530 | Pcnt | 18541 | HITS-CLIP | 23142080\|26203562 | Unspecified |
| mmu-miR-21a-5p | MIMAT0000530 | Pcnx | 54604 | HITS-CLIP | 23142080 | Unspecified |
| mmu-miR-21a-5p | MIMAT0000530 | Pcsk6 | 18553 | HITS-CLIP | 37986224 | Unspecified |
| mmu-miR-21a-5p | MIMAT0000530 | Pcsk9 | 100102 | HITS-CLIP | 23597149 | Unspecified |
| mmu-miR-21a-5p | MIMAT0000530 | Pcyox1 | 66881 | HITS-CLIP | 23142080 | Unspecified |
| mmu-miR-21a-5p | MIMAT0000530 | Pcyox1l | 240334 | Microarrays | 37986224\|25263593 | Unspecified |
| mmu-miR-21a-5p | MIMAT0000530 | Pcyt1a | 13026 | HITS-CLIP | 23142080 | Unspecified |
| mmu-miR-21a-5p | MIMAT0000530 | Pcyt1b | 236899 | Microarrays | 37986224 | Unspecified |
| mmu-miR-21a-5p | MIMAT0000530 | Pdcd10 | 56426 | Luciferase assay//qRT-PCR//HITS-CLIP//Luc | 24891504\|23142080 | Unspecified |
| mmu-miR-21a-5p | MIMAT0000530 | Pdcd11 | 18572 | HITS-CLIP | 23597149\|23142080 | Unspecified |
| mmu-miR-21a-5p | MIMAT0000530 | Pdcd1lg2 | 58205 | HITS-CLIP | 23142080 | Unspecified |
| mmu-miR-21a-5p | MIMAT0000530 | Pdcd6 | 18570 | HITS-CLIP | 23142080\|23597149\|29167373 | Unspecified |
| mmu-miR-21a-5p | MIMAT0000530 | Pdcd6ip | 18571 | HITS-CLIP | 23597149\|23142080\|29167373 | Unspecified |
| mmu-miR-21a-5p | MIMAT0000530 | Pdcl | 67466 | HITS-CLIP | 23142080\|29167373 | Unspecified |
| mmu-miR-21a-5p | MIMAT0000530 | Pde10a | 23984 | HITS-CLIP | 19536157 | Unspecified |
| mmu-miR-21a-5p | MIMAT0000530 | Pde12 | 211948 | Microarrays//HITS-CLIP | 37986224\|25263593\|23142080 | Unspecified |
| mmu-miR-21a-5p | MIMAT0000530 | Pde1b | 18574 | HITS-CLIP | 19536157 | Unspecified |
| mmu-miR-21a-5p | MIMAT0000530 | Pde2a | 207728 | HITS-CLIP | 23142080 | Unspecified |
| mmu-miR-21a-5p | MIMAT0000530 | Pde3b | 18576 | Microarrays//HITS-CLIP | 37986224\|23142080 | Unspecified |
| mmu-miR-21a-5p | MIMAT0000530 | Pde4a | 18577 | HITS-CLIP | 29167373 | Unspecified |
| mmu-miR-21a-5p | MIMAT0000530 | Pde4b | 18578 | Microarrays//HITS-CLIP | 37986224\|23597149\|23142080\|29167 | Unspecified |
| mmu-miR-21a-5p | MIMAT0000530 | Pde4dip | 83679 | HITS-CLIP | 23597149 | Unspecified |
| mmu-miR-21a-5p | MIMAT0000530 | Pde7a | 18583 | Microarrays//HITS-CLIP | 37986224\|26203562 | Unspecified |
| mmu-miR-21a-5p | MIMAT0000530 | Pde8b | 218461 | Microarrays | 37986224\|25263593 | Unspecified |

| | | | | | | |
|---|---|---|---|---|---|---|
| mmu-miR-21a-5p | MIMAT0000530 | Pdgfb | 18591 | Microarrays | 37986224\|25263593 | Unspecified |
| mmu-miR-21a-5p | MIMAT0000530 | Pdgfra | 18595 | HITS-CLIP | 19536157 | Unspecified |
| mmu-miR-21a-5p | MIMAT0000530 | Pdha1 | 18597 | HITS-CLIP | 23597149\|23142080 | Unspecified |
| mmu-miR-21a-5p | MIMAT0000530 | Pdhb | 68263 | HITS-CLIP | 23597149 | Unspecified |
| mmu-miR-21a-5p | MIMAT0000530 | Pdhx | 27402 | HITS-CLIP | 23597149\|23142080 | Unspecified |
| mmu-miR-21a-5p | MIMAT0000530 | Pdia3 | 14827 | HITS-CLIP//qCLASH | 37986224\|23597149\|19536157\|31393 | Unspecified |
| mmu-miR-21a-5p | MIMAT0000530 | Pdia4 | 12304 | HITS-CLIP | 23597149\|23142080 | Unspecified |
| mmu-miR-21a-5p | MIMAT0000530 | Pdia5 | 72599 | HITS-CLIP | 23597149 | Unspecified |
| mmu-miR-21a-5p | MIMAT0000530 | Pdik1l | 230809 | HITS-CLIP | 23142080 | Unspecified |
| mmu-miR-21a-5p | MIMAT0000530 | Pdk2 | 18604 | Microarrays | 37986224 | Unspecified |
| mmu-miR-21a-5p | MIMAT0000530 | Pdlim5 | 56376 | HITS-CLIP | 23597149\|29167373 | Unspecified |
| mmu-miR-21a-5p | MIMAT0000530 | Pdp2 | 382051 | HITS-CLIP | 23597149 | Unspecified |
| mmu-miR-21a-5p | MIMAT0000530 | Pdpk1 | 18607 | HITS-CLIP | 23142080 | Unspecified |
| mmu-miR-21a-5p | MIMAT0000530 | Pdpn | 14726 | Microarrays | 37986224\|25263593 | Unspecified |
| mmu-miR-21a-5p | MIMAT0000530 | Pds5a | 71521 | HITS-CLIP | 23142080\|19536157 | Unspecified |
| mmu-miR-21a-5p | MIMAT0000530 | Pds5b | 100710 | HITS-CLIP | 23142080\|29167373 | Unspecified |
| mmu-miR-21a-5p | MIMAT0000530 | Pdss1 | 56075 | Microarrays | 37986224\|25263593 | Unspecified |
| mmu-miR-21a-5p | MIMAT0000530 | Pdss2 | 71365 | HITS-CLIP | 23142080 | Unspecified |
| mmu-miR-21a-5p | MIMAT0000530 | Pdxk | 216134 | HITS-CLIP | 29167373 | Unspecified |
| mmu-miR-21a-5p | MIMAT0000530 | Pdzd8 | 107368 | HITS-CLIP//qCLASH | 37986224\|31393953\|23142080 | Unspecified |
| mmu-miR-21a-5p | MIMAT0000530 | Pea15a | 18611 | Microarrays | 37986224\|25263593 | Unspecified |
| mmu-miR-21a-5p | MIMAT0000530 | Peak1 | 244895 | HITS-CLIP | 23142080\|23597149 | Unspecified |
| mmu-miR-21a-5p | MIMAT0000530 | Pear1 | 375033 | HITS-CLIP | 23142080 | Unspecified |
| mmu-miR-21a-5p | MIMAT0000530 | Pebp1 | 23980 | qCLASH | 31393953 | Unspecified |
| mmu-miR-21a-5p | MIMAT0000530 | Peg3 | 18616 | HITS-CLIP | 19536157 | Unspecified |
| mmu-miR-21a-5p | MIMAT0000530 | Peli1 | 67245 | Luciferase assay//Reporter assay//qRT-PCI | 20167875\|23597149\|23142080\|29167 | Unspecified |
| mmu-miR-21a-5p | MIMAT0000530 | Peli2 | 93834 | Microarrays | 37986224 | Unspecified |
| mmu-miR-21a-5p | MIMAT0000530 | Pemt | 18618 | HITS-CLIP | 23597149 | Unspecified |
| mmu-miR-21a-5p | MIMAT0000530 | Pepd | 18624 | HITS-CLIP | 29167373 | Unspecified |
| mmu-miR-21a-5p | MIMAT0000530 | Per1 | 18626 | HITS-CLIP | 23142080 | Unspecified |
| mmu-miR-21a-5p | MIMAT0000530 | Per2 | 18627 | HITS-CLIP | 23142080 | Unspecified |
| mmu-miR-21a-5p | MIMAT0000530 | Per3 | 18628 | Microarrays | 37986224 | Unspecified |
| mmu-miR-21a-5p | MIMAT0000530 | Perp | 64058 | HITS-CLIP | 37986224 | Unspecified |
| mmu-miR-21a-5p | MIMAT0000530 | Pes1 | 64934 | HITS-CLIP | 23142080 | Unspecified |
| mmu-miR-21a-5p | MIMAT0000530 | Pex10 | 668173 | HITS-CLIP | 23597149\|23142080 | Unspecified |
| mmu-miR-21a-5p | MIMAT0000530 | Pex12 | 103737 | HITS-CLIP | 23597149 | Unspecified |
| mmu-miR-21a-5p | MIMAT0000530 | Pex13 | 72129 | HITS-CLIP | 37986224\|23597149\|23142080 | Unspecified |
| mmu-miR-21a-5p | MIMAT0000530 | Pex14 | 56273 | HITS-CLIP | 23142080 | Unspecified |
| mmu-miR-21a-5p | MIMAT0000530 | Pex5 | 19305 | HITS-CLIP | 23142080 | Unspecified |

| | | | | | | |
|---|---|---|---|---|---|---|
| mmu-miR-21a-5p | MIMAT0000530 | Pex7 | 18634 | qCLASH | 31393953 | Unspecified |
| mmu-miR-21a-5p | MIMAT0000530 | Pf4 | 56744 | Microarrays | 37986224 | Unspecified |
| mmu-miR-21a-5p | MIMAT0000530 | Pfdn4 | 109054 | HITS-CLIP | 23142080 | Unspecified |
| mmu-miR-21a-5p | MIMAT0000530 | Pfdn5 | 56612 | HITS-CLIP | 23142080 | Unspecified |
| mmu-miR-21a-5p | MIMAT0000530 | Pfkfb1 | 18639 | HITS-CLIP | 23597149 | Unspecified |
| mmu-miR-21a-5p | MIMAT0000530 | Pfkfb4 | 270198 | Microarrays | 37986224 | Unspecified |
| mmu-miR-21a-5p | MIMAT0000530 | Pfkl | 18641 | HITS-CLIP | 23597149\|23142080 | Unspecified |
| mmu-miR-21a-5p | MIMAT0000530 | Pfkm | 18642 | HITS-CLIP | 37986224\|23142080 | Unspecified |
| mmu-miR-21a-5p | MIMAT0000530 | Pfkp | 56421 | HITS-CLIP | 23142080\|29167373 | Unspecified |
| mmu-miR-21a-5p | MIMAT0000530 | Pfn1 | 18643 | HITS-CLIP | 23142080 | Unspecified |
| mmu-miR-21a-5p | MIMAT0000530 | Pfn2 | 18645 | HITS-CLIP | 23142080 | Unspecified |
| mmu-miR-21a-5p | MIMAT0000530 | Pgap1 | 241062 | Microarrays//HITS-CLIP | 37986224\|26203562\|29167373 | Unspecified |
| mmu-miR-21a-5p | MIMAT0000530 | Pgbd5 | 209966 | Microarrays | 37986224 | Unspecified |
| mmu-miR-21a-5p | MIMAT0000530 | Pglyrp1 | 21946 | HITS-CLIP | 23142080 | Unspecified |
| mmu-miR-21a-5p | MIMAT0000530 | Pgm1 | 66681 | HITS-CLIP | 23597149\|23142080 | Unspecified |
| mmu-miR-21a-5p | MIMAT0000530 | Pgm2 | 72157 | HITS-CLIP | 23597149 | Unspecified |
| mmu-miR-21a-5p | MIMAT0000530 | Pgm2l1 | 70974 | HITS-CLIP | 37986224\|30971389\|23142080 | Unspecified |
| mmu-miR-21a-5p | MIMAT0000530 | Pgpep1 | 66522 | qCLASH | 31393953 | Unspecified |
| mmu-miR-21a-5p | MIMAT0000530 | Pgrmc1 | 53328 | HITS-CLIP | 23597149\|23142080 | Unspecified |
| mmu-miR-21a-5p | MIMAT0000530 | Pgrmc2 | 70804 | HITS-CLIP | 29167373 | Unspecified |
| mmu-miR-21a-5p | MIMAT0000530 | Pgs1 | 74451 | HITS-CLIP | 37986224\|23142080 | Unspecified |
| mmu-miR-21a-5p | MIMAT0000530 | Phactr2 | 215789 | HITS-CLIP | 29167373 | Unspecified |
| mmu-miR-21a-5p | MIMAT0000530 | Phactr4 | 100169 | HITS-CLIP | 29167373 | Unspecified |
| mmu-miR-21a-5p | MIMAT0000530 | Phc2 | 54383 | HITS-CLIP | 29167373 | Unspecified |
| mmu-miR-21a-5p | MIMAT0000530 | Phf10 | 72057 | HITS-CLIP | 23597149 | Unspecified |
| mmu-miR-21a-5p | MIMAT0000530 | Phf12 | 268448 | HITS-CLIP | 23142080 | Unspecified |
| mmu-miR-21a-5p | MIMAT0000530 | Phf14 | 75725 | HITS-CLIP | 23597149 | Unspecified |
| mmu-miR-21a-5p | MIMAT0000530 | Phf20 | 228829 | HITS-CLIP | 23142080 | Unspecified |
| mmu-miR-21a-5p | MIMAT0000530 | Phf20l1 | 239510 | HITS-CLIP | 23142080 | Unspecified |
| mmu-miR-21a-5p | MIMAT0000530 | Phf21a | 192285 | Microarrays//HITS-CLIP | 37986224\|23142080 | Unspecified |
| mmu-miR-21a-5p | MIMAT0000530 | Phf3 | 213109 | HITS-CLIP | 37986224\|23142080\|26203562 | Unspecified |
| mmu-miR-21a-5p | MIMAT0000530 | Phf6 | 70998 | HITS-CLIP | 23142080 | Unspecified |
| mmu-miR-21a-5p | MIMAT0000530 | Phf8 | 320595 | HITS-CLIP | 23142080\|31433990\|29167373 | Unspecified |
| mmu-miR-21a-5p | MIMAT0000530 | Phgdh | 236539 | Microarrays//qCLASH | 37986224\|25263593\|31393953 | Unspecified |
| mmu-miR-21a-5p | MIMAT0000530 | Phip | 83946 | HITS-CLIP | 23597149\|23142080\|29167373 | Unspecified |
| mmu-miR-21a-5p | MIMAT0000530 | Phka2 | 110094 | HITS-CLIP | 23597149 | Unspecified |
| mmu-miR-21a-5p | MIMAT0000530 | Phlda1 | 21664 | Microarrays | 37986224\|25263593 | Unspecified |
| mmu-miR-21a-5p | MIMAT0000530 | Phlda3 | 27280 | Microarrays//HITS-CLIP | 37986224\|25263593\|31433990 | Unspecified |
| mmu-miR-21a-5p | MIMAT0000530 | Phldb2 | 208177 | Microarrays//HITS-CLIP | 37986224\|25263593\|23597149 | Unspecified |

| | | | | | | |
|---|---|---|---|---|---|---|
| mmu-miR-21a-5p | MIMAT0000530 | Phlpp1 | 98432 | HITS-CLIP | 23597149 | Unspecified |
| mmu-miR-21a-5p | MIMAT0000530 | Phlpp2 | 244650 | HITS-CLIP | 23597149\|29167373 | Unspecified |
| mmu-miR-21a-5p | MIMAT0000530 | Phrf1 | 101471 | HITS-CLIP | 23597149\|23142080 | Unspecified |
| mmu-miR-21a-5p | MIMAT0000530 | Phtf2 | 68770 | HITS-CLIP | 23597149\|23142080 | Unspecified |
| mmu-miR-21a-5p | MIMAT0000530 | Pi4k2b | 67073 | HITS-CLIP | 23597149 | Unspecified |
| mmu-miR-21a-5p | MIMAT0000530 | Pi4ka | 224020 | HITS-CLIP | 23142080\|29167373\|19536157 | Unspecified |
| mmu-miR-21a-5p | MIMAT0000530 | Pi4kb | 107650 | HITS-CLIP | 23142080\|29167373 | Unspecified |
| mmu-miR-21a-5p | MIMAT0000530 | Pias1 | 56469 | HITS-CLIP | 23142080 | Unspecified |
| mmu-miR-21a-5p | MIMAT0000530 | Pias2 | 17344 | HITS-CLIP | 37986224\|23142080 | Unspecified |
| mmu-miR-21a-5p | MIMAT0000530 | Picalm | 233489 | Microarrays//HITS-CLIP | 37986224\|23142080 | Unspecified |
| mmu-miR-21a-5p | MIMAT0000530 | Pid1 | 98496 | HITS-CLIP | 29167373 | Unspecified |
| mmu-miR-21a-5p | MIMAT0000530 | Pierce1 | 69327 | Microarrays | 25263593 | Unspecified |
| mmu-miR-21a-5p | MIMAT0000530 | Piezo1 | 234839 | Microarrays//HITS-CLIP | 37986224\|23142080\|25263593 | Unspecified |
| mmu-miR-21a-5p | MIMAT0000530 | Pigg | 433931 | HITS-CLIP | 23142080 | Unspecified |
| mmu-miR-21a-5p | MIMAT0000530 | Pigm | 67556 | HITS-CLIP | 37986224\|23142080 | Unspecified |
| mmu-miR-21a-5p | MIMAT0000530 | Pigq | 14755 | HITS-CLIP | 23597149 | Unspecified |
| mmu-miR-21a-5p | MIMAT0000530 | Pik3ap1 | 83490 | HITS-CLIP | 23597149\|23142080 | Unspecified |
| mmu-miR-21a-5p | MIMAT0000530 | Pik3c2a | 18704 | HITS-CLIP | 23142080\|29167373 | Unspecified |
| mmu-miR-21a-5p | MIMAT0000530 | Pik3c3 | 225326 | HITS-CLIP | 23142080 | Unspecified |
| mmu-miR-21a-5p | MIMAT0000530 | Pik3ca | 18706 | HITS-CLIP | 23142080\|29167373 | Unspecified |
| mmu-miR-21a-5p | MIMAT0000530 | Pik3cb | 74769 | Microarrays//HITS-CLIP | 37986224\|25263593\|23597149 | Unspecified |
| mmu-miR-21a-5p | MIMAT0000530 | Pik3cd | 18707 | HITS-CLIP | 23142080 | Unspecified |
| mmu-miR-21a-5p | MIMAT0000530 | Pik3cg | 30955 | HITS-CLIP | 23142080 | Unspecified |
| mmu-miR-21a-5p | MIMAT0000530 | Pik3ip1 | 216505 | Microarrays | 37986224 | Unspecified |
| mmu-miR-21a-5p | MIMAT0000530 | Pik3r1 | 18708 | HITS-CLIP | 37986224\|23597149\|23142080 | Unspecified |
| mmu-miR-21a-5p | MIMAT0000530 | Pik3r3 | 18710 | Microarrays | 37986224\|25263593 | Unspecified |
| mmu-miR-21a-5p | MIMAT0000530 | Pik3r5 | 320207 | HITS-CLIP | 23142080 | Unspecified |
| mmu-miR-21a-5p | MIMAT0000530 | Pikfyve | 18711 | HITS-CLIP | 23142080\|29167373 | Unspecified |
| mmu-miR-21a-5p | MIMAT0000530 | Pilrb1 | 170741 | Microarrays | 37986224 | Unspecified |
| mmu-miR-21a-5p | MIMAT0000530 | Pim1 | 18712 | qCLASH//HITS-CLIP | 31393953\|23142080\|29167373 | Unspecified |
| mmu-miR-21a-5p | MIMAT0000530 | Pim3 | 223775 | HITS-CLIP | 29167373 | Unspecified |
| mmu-miR-21a-5p | MIMAT0000530 | Pimreg | 54478 | Microarrays | 25263593 | Unspecified |
| mmu-miR-21a-5p | MIMAT0000530 | Pink1 | 68943 | Microarrays | 37986224 | Unspecified |
| mmu-miR-21a-5p | MIMAT0000530 | Pip4k2a | 18718 | HITS-CLIP | 23142080 | Unspecified |
| mmu-miR-21a-5p | MIMAT0000530 | Pip4k2b | 108083 | HITS-CLIP | 23142080\|29167373 | Unspecified |
| mmu-miR-21a-5p | MIMAT0000530 | Pip4k2c | 117150 | HITS-CLIP | 23142080 | Unspecified |
| mmu-miR-21a-5p | MIMAT0000530 | Pip4p1 | 90809 | HITS-CLIP | 29167373 | Unspecified |
| mmu-miR-21a-5p | MIMAT0000530 | Pip5k1c | 18717 | HITS-CLIP | 29167373 | Unspecified |
| mmu-miR-21a-5p | MIMAT0000530 | Pisd | 320951 | HITS-CLIP | 23597149\|23142080 | Unspecified |

| | | | | | | |
|---|---|---|---|---|---|---|
| mmu-miR-21a-5p | MIMAT0000530 | Pitpna | 18738 | HITS-CLIP | 23142080\|29167373 | Unspecified |
| mmu-miR-21a-5p | MIMAT0000530 | Pitpnb | 56305 | HITS-CLIP | 37986224\|23597149\|23142080 | Unspecified |
| mmu-miR-21a-5p | MIMAT0000530 | Pitpnc1 | 71795 | HITS-CLIP | 23142080\|29167373 | Unspecified |
| mmu-miR-21a-5p | MIMAT0000530 | Pitpnm1 | 18739 | HITS-CLIP | 23142080 | Unspecified |
| mmu-miR-21a-5p | MIMAT0000530 | Pitpnm2 | 19679 | HITS-CLIP | 23142080\|19536157 | Unspecified |
| mmu-miR-21a-5p | MIMAT0000530 | Pitrm1 | 69617 | HITS-CLIP | 23597149 | Unspecified |
| mmu-miR-21a-5p | MIMAT0000530 | Pja2 | 224938 | HITS-CLIP | 37986224\|23142080\|19536157 | Unspecified |
| mmu-miR-21a-5p | MIMAT0000530 | Pkd1 | 18763 | HITS-CLIP | 29167373\|23142080 | Unspecified |
| mmu-miR-21a-5p | MIMAT0000530 | Pkib | 18768 | Microarrays | 37986224\|25263593 | Unspecified |
| mmu-miR-21a-5p | MIMAT0000530 | Pkig | 18769 | Microarrays | 37986224 | Unspecified |
| mmu-miR-21a-5p | MIMAT0000530 | Pkm | 18746 | Microarrays | 37986224\|25263593 | Unspecified |
| mmu-miR-21a-5p | MIMAT0000530 | Pkn2 | 109333 | HITS-CLIP | 23597149\|29167373\|23142080 | Unspecified |
| mmu-miR-21a-5p | MIMAT0000530 | Pkp4 | 227937 | HITS-CLIP | 19536157\|23142080\|29167373 | Unspecified |
| mmu-miR-21a-5p | MIMAT0000530 | Pla2g16 | 225845 | HITS-CLIP | 23142080 | Unspecified |
| mmu-miR-21a-5p | MIMAT0000530 | Pla2g2e | 26970 | Microarrays | 37986224\|25263593 | Unspecified |
| mmu-miR-21a-5p | MIMAT0000530 | Pla2g4a | 18783 | Microarrays//HITS-CLIP | 37986224\|25263593\|29167373 | Unspecified |
| mmu-miR-21a-5p | MIMAT0000530 | Pla2g5 | 18784 | Microarrays | 37986224\|25263593 | Unspecified |
| mmu-miR-21a-5p | MIMAT0000530 | Pla2g7 | 27226 | Microarrays | 37986224 | Unspecified |
| mmu-miR-21a-5p | MIMAT0000530 | Plaa | 18786 | HITS-CLIP | 23142080 | Unspecified |
| mmu-miR-21a-5p | MIMAT0000530 | Plag1 | 56711 | HITS-CLIP | 23142080 | Unspecified |
| mmu-miR-21a-5p | MIMAT0000530 | Plagl1 | 22634 | Microarrays//HITS-CLIP | 37986224\|25263593\|23142080 | Unspecified |
| mmu-miR-21a-5p | MIMAT0000530 | Plagl2 | 54711 | HITS-CLIP | 31433990\|29167373 | Unspecified |
| mmu-miR-21a-5p | MIMAT0000530 | Plau | 18792 | Microarrays//HITS-CLIP | 37986224\|25263593\|29167373 | Unspecified |
| mmu-miR-21a-5p | MIMAT0000530 | Plaur | 18793 | Microarrays | 37986224\|25263593 | Unspecified |
| mmu-miR-21a-5p | MIMAT0000530 | Plcb2 | 18796 | HITS-CLIP | 23142080\|29167373 | Unspecified |
| mmu-miR-21a-5p | MIMAT0000530 | Plcb4 | 18798 | HITS-CLIP | 19536157\|23142080 | Unspecified |
| mmu-miR-21a-5p | MIMAT0000530 | Plce1 | 74055 | HITS-CLIP | 23597149 | Unspecified |
| mmu-miR-21a-5p | MIMAT0000530 | Plcg1 | 18803 | HITS-CLIP | 23142080 | Unspecified |
| mmu-miR-21a-5p | MIMAT0000530 | Plcg2 | 234779 | HITS-CLIP | 29167373 | Unspecified |
| mmu-miR-21a-5p | MIMAT0000530 | Plcl1 | 227120 | HITS-CLIP | 23142080 | Unspecified |
| mmu-miR-21a-5p | MIMAT0000530 | Plcl2 | 224860 | Microarrays//HITS-CLIP | 37986224\|23142080 | Unspecified |
| mmu-miR-21a-5p | MIMAT0000530 | Plcxd2 | 433022 | Microarrays//HITS-CLIP | 37986224\|23142080\|25263593\|23597 | Unspecified |
| mmu-miR-21a-5p | MIMAT0000530 | Pld2 | 18806 | Microarrays | 37986224 | Unspecified |
| mmu-miR-21a-5p | MIMAT0000530 | Pld3 | 18807 | HITS-CLIP | 23142080 | Unspecified |
| mmu-miR-21a-5p | MIMAT0000530 | Pld4 | 104759 | Microarrays | 37986224 | Unspecified |
| mmu-miR-21a-5p | MIMAT0000530 | Plec | 18810 | HITS-CLIP | 23597149\|23142080\|29167373 | Unspecified |
| mmu-miR-21a-5p | MIMAT0000530 | Plek | 56193 | Microarrays//HITS-CLIP | 37986224\|25263593\|29167373 | Unspecified |
| mmu-miR-21a-5p | MIMAT0000530 | Plekha1 | 101476 | HITS-CLIP | 23142080 | Unspecified |
| mmu-miR-21a-5p | MIMAT0000530 | Plekha2 | 83436 | HITS-CLIP | 37986224\|23142080\|29167373 | Unspecified |

| | | | | | | |
|---|---|---|---|---|---|---|
| mmu-miR-21a-5p | MIMAT0000530 | Plekha3 | 83435 | HITS-CLIP | 23142080 | Unspecified |
| mmu-miR-21a-5p | MIMAT0000530 | Plekhf2 | 71801 | HITS-CLIP | 23142080\|29167373 | Unspecified |
| mmu-miR-21a-5p | MIMAT0000530 | Plekhg1 | 213783 | Microarrays | 37986224\|25263593 | Unspecified |
| mmu-miR-21a-5p | MIMAT0000530 | Plekhm1 | 353047 | HITS-CLIP | 29167373\|23142080 | Unspecified |
| mmu-miR-21a-5p | MIMAT0000530 | Plekho2 | 102595 | HITS-CLIP | 23142080\|29167373 | Unspecified |
| mmu-miR-21a-5p | MIMAT0000530 | Plgrkt | 67759 | HITS-CLIP | 23142080 | Unspecified |
| mmu-miR-21a-5p | MIMAT0000530 | Plin2 | 11520 | HITS-CLIP | 23597149\|23142080\|29167373 | Unspecified |
| mmu-miR-21a-5p | MIMAT0000530 | Plin3 | 66905 | HITS-CLIP | 23597149 | Unspecified |
| mmu-miR-21a-5p | MIMAT0000530 | Plin5 | 66968 | HITS-CLIP | 23597149 | Unspecified |
| mmu-miR-21a-5p | MIMAT0000530 | Plk1 | 18817 | Microarrays | 37986224\|25263593 | Unspecified |
| mmu-miR-21a-5p | MIMAT0000530 | Plk2 | 20620 | HITS-CLIP | 29167373 | Unspecified |
| mmu-miR-21a-5p | MIMAT0000530 | Plk3 | 12795 | Microarrays | 37986224\|25263593 | Unspecified |
| mmu-miR-21a-5p | MIMAT0000530 | Plk4 | 20873 | HITS-CLIP | 23142080 | Unspecified |
| mmu-miR-21a-5p | MIMAT0000530 | Pllp | 67801 | HITS-CLIP | 23597149 | Unspecified |
| mmu-miR-21a-5p | MIMAT0000530 | Plod2 | 26432 | Microarrays//HITS-CLIP | 37986224\|25263593\|23142080 | Unspecified |
| mmu-miR-21a-5p | MIMAT0000530 | Plod3 | 26433 | HITS-CLIP | 23142080 | Unspecified |
| mmu-miR-21a-5p | MIMAT0000530 | Plp1 | 18823 | HITS-CLIP | 19536157 | Unspecified |
| mmu-miR-21a-5p | MIMAT0000530 | Plp2 | 18824 | HITS-CLIP | 31433990 | Unspecified |
| mmu-miR-21a-5p | MIMAT0000530 | Plpp1 | 8611 | Microarrays | 25263593 | Unspecified |
| mmu-miR-21a-5p | MIMAT0000530 | Plpp2 | 8612 | HITS-CLIP | 23142080 | Unspecified |
| mmu-miR-21a-5p | MIMAT0000530 | Plpp3 | 67916 | Microarrays | 25263593 | Unspecified |
| mmu-miR-21a-5p | MIMAT0000530 | Plppr4 | 229791 | HITS-CLIP | 19536157 | Unspecified |
| mmu-miR-21a-5p | MIMAT0000530 | Pls1 | 102502 | Microarrays | 37986224 | Unspecified |
| mmu-miR-21a-5p | MIMAT0000530 | Pls3 | 102866 | HITS-CLIP, Microarrays//HITS-CLIP//Microa | 37986224\|23597149\|25263593\|29167 | Unspecified |
| mmu-miR-21a-5p | MIMAT0000530 | Pltp | 18830 | Microarrays//HITS-CLIP | 37986224\|23597149 | Unspecified |
| mmu-miR-21a-5p | MIMAT0000530 | Plxdc2 | 67448 | HITS-CLIP, Microarrays | 37986224 | Unspecified |
| mmu-miR-21a-5p | MIMAT0000530 | Plxna1 | 18844 | Microarrays//HITS-CLIP | 37986224\|25263593\|29167373 | Unspecified |
| mmu-miR-21a-5p | MIMAT0000530 | Plxna2 | 18845 | HITS-CLIP | 37986224\|19536157 | Unspecified |
| mmu-miR-21a-5p | MIMAT0000530 | Plxna4 | 243743 | HITS-CLIP | 37986224 | Unspecified |
| mmu-miR-21a-5p | MIMAT0000530 | Plxnb1 | 235611 | HITS-CLIP | 23597149 | Unspecified |
| mmu-miR-21a-5p | MIMAT0000530 | Plxnb2 | 140570 | HITS-CLIP | 23597149\|29167373 | Unspecified |
| mmu-miR-21a-5p | MIMAT0000530 | Plxnc1 | 54712 | Microarrays//HITS-CLIP | 37986224\|31433990\|23142080 | Unspecified |
| mmu-miR-21a-5p | MIMAT0000530 | Plxnd1 | 67784 | Microarrays//HITS-CLIP | 37986224\|23597149\|23142080 | Unspecified |
| mmu-miR-21a-5p | MIMAT0000530 | Pmaip1 | 58801 | Microarrays | 37986224\|25263593 | Unspecified |
| mmu-miR-21a-5p | MIMAT0000530 | Pmepa1 | 65112 | HITS-CLIP | 23142080\|29167373 | Unspecified |
| mmu-miR-21a-5p | MIMAT0000530 | Pmm2 | 54128 | HITS-CLIP | 23597149\|23142080 | Unspecified |
| mmu-miR-21a-5p | MIMAT0000530 | Pmpca | 66865 | HITS-CLIP | 23597149\|23142080 | Unspecified |
| mmu-miR-21a-5p | MIMAT0000530 | Pms2 | 18861 | HITS-CLIP | 23142080 | Unspecified |
| mmu-miR-21a-5p | MIMAT0000530 | Pnisr | 66625 | Microarrays//HITS-CLIP | 37986224\|23142080 | Unspecified |

| | | | | | | |
|---|---|---|---|---|---|---|
| mmu-miR-21a-5p | MIMAT0000530 | Pnmal2 | 434128 | HITS-CLIP | 37986224 | Unspecified |
| mmu-miR-21a-5p | MIMAT0000530 | Pno1 | 66249 | Microarrays | 37986224\|25263593 | Unspecified |
| mmu-miR-21a-5p | MIMAT0000530 | Pnpla2 | 66853 | HITS-CLIP | 23142080\|23597149\|29167373 | Unspecified |
| mmu-miR-21a-5p | MIMAT0000530 | Pnpla7 | 241274 | HITS-CLIP | 23597149 | Unspecified |
| mmu-miR-21a-5p | MIMAT0000530 | Pnpla8 | 67452 | HITS-CLIP | 23597149\|23142080 | Unspecified |
| mmu-miR-21a-5p | MIMAT0000530 | Pnpo | 103711 | HITS-CLIP | 37986224\|23597149 | Unspecified |
| mmu-miR-21a-5p | MIMAT0000530 | Pnrc1 | 108767 | HITS-CLIP | 23142080 | Unspecified |
| mmu-miR-21a-5p | MIMAT0000530 | Pnrc2 | 52830 | HITS-CLIP | 37986224\|23142080 | Unspecified |
| mmu-miR-21a-5p | MIMAT0000530 | Poc1b | 382406 | HITS-CLIP | 23142080 | Unspecified |
| mmu-miR-21a-5p | MIMAT0000530 | Pofut1 | 140484 | HITS-CLIP | 29167373 | Unspecified |
| mmu-miR-21a-5p | MIMAT0000530 | Poglut1 | 224143 | HITS-CLIP | 23142080 | Unspecified |
| mmu-miR-21a-5p | MIMAT0000530 | Pogz | 229584 | HITS-CLIP | 23142080 | Unspecified |
| mmu-miR-21a-5p | MIMAT0000530 | Pola1 | 18968 | HITS-CLIP | 23142080 | Unspecified |
| mmu-miR-21a-5p | MIMAT0000530 | Polb | 18970 | Microarrays | 37986224 | Unspecified |
| mmu-miR-21a-5p | MIMAT0000530 | Pold2 | 18972 | HITS-CLIP | 23142080 | Unspecified |
| mmu-miR-21a-5p | MIMAT0000530 | Pold3 | 67967 | HITS-CLIP | 23142080\|29167373 | Unspecified |
| mmu-miR-21a-5p | MIMAT0000530 | Pole | 18973 | Microarrays//HITS-CLIP | 37986224\|25263593\|23142080 | Unspecified |
| mmu-miR-21a-5p | MIMAT0000530 | Polg | 18975 | HITS-CLIP | 23597149\|23142080\|29167373 | Unspecified |
| mmu-miR-21a-5p | MIMAT0000530 | Polk | 27015 | HITS-CLIP | 23142080 | Unspecified |
| mmu-miR-21a-5p | MIMAT0000530 | Poll | 56626 | HITS-CLIP | 23142080 | Unspecified |
| mmu-miR-21a-5p | MIMAT0000530 | Polr1a | 20019 | HITS-CLIP | 29167373 | Unspecified |
| mmu-miR-21a-5p | MIMAT0000530 | Polr1d | 20018 | HITS-CLIP | 23142080 | Unspecified |
| mmu-miR-21a-5p | MIMAT0000530 | Polr2a | 20020 | HITS-CLIP | 37986224\|23597149\|23142080 | Unspecified |
| mmu-miR-21a-5p | MIMAT0000530 | Polr2b | 231329 | HITS-CLIP | 23142080 | Unspecified |
| mmu-miR-21a-5p | MIMAT0000530 | Polr2m | 28015 | HITS-CLIP | 23142080 | Unspecified |
| mmu-miR-21a-5p | MIMAT0000530 | Polr3a | 218832 | HITS-CLIP | 23142080 | Unspecified |
| mmu-miR-21a-5p | MIMAT0000530 | Polr3b | 70428 | HITS-CLIP | 23142080 | Unspecified |
| mmu-miR-21a-5p | MIMAT0000530 | Polr3e | 26939 | HITS-CLIP | 23597149 | Unspecified |
| mmu-miR-21a-5p | MIMAT0000530 | Polr3f | 70408 | HITS-CLIP | 23142080 | Unspecified |
| mmu-miR-21a-5p | MIMAT0000530 | Polr3gl | 69870 | Microarrays | 37986224 | Unspecified |
| mmu-miR-21a-5p | MIMAT0000530 | Polr3k | 67005 | HITS-CLIP | 23597149\|23142080 | Unspecified |
| mmu-miR-21a-5p | MIMAT0000530 | Polrmt | 216151 | HITS-CLIP | 23597149 | Unspecified |
| mmu-miR-21a-5p | MIMAT0000530 | Pom121 | 107939 | HITS-CLIP | 23597149 | Unspecified |
| mmu-miR-21a-5p | MIMAT0000530 | Pon1 | 18979 | HITS-CLIP | 23597149 | Unspecified |
| mmu-miR-21a-5p | MIMAT0000530 | Pon2 | 330260 | HITS-CLIP | 26203562\|23142080 | Unspecified |
| mmu-miR-21a-5p | MIMAT0000530 | Pon3 | 269823 | HITS-CLIP | 23597149 | Unspecified |
| mmu-miR-21a-5p | MIMAT0000530 | Pop1 | 67724 | HITS-CLIP | 23142080 | Unspecified |
| mmu-miR-21a-5p | MIMAT0000530 | Por | 18984 | HITS-CLIP | 23597149 | Unspecified |
| mmu-miR-21a-5p | MIMAT0000530 | Pot1b | 72836 | Microarrays | 37986224 | Unspecified |

| | | | | | | |
|---|---|---|---|---|---|---|
| mmu-miR-21a-5p | MIMAT0000530 | Pou2af1 | 18985 | qCLASH//HITS-CLIP | 31393953\|23142080 | Unspecified |
| mmu-miR-21a-5p | MIMAT0000530 | Pou2f1 | 18986 | HITS-CLIP | 23142080 | Unspecified |
| mmu-miR-21a-5p | MIMAT0000530 | Pou3f1 | 18991 | Microarrays | 37986224\|25263593 | Unspecified |
| mmu-miR-21a-5p | MIMAT0000530 | Ppa1 | 67895 | Microarrays | 37986224\|25263593 | Unspecified |
| mmu-miR-21a-5p | MIMAT0000530 | Ppa2 | 74776 | HITS-CLIP | 23142080 | Unspecified |
| mmu-miR-21a-5p | MIMAT0000530 | Ppara | 19013 | HITS-CLIP | 23597149 | Unspecified |
| mmu-miR-21a-5p | MIMAT0000530 | Ppard | 19015 | HITS-CLIP | 23597149 | Unspecified |
| mmu-miR-21a-5p | MIMAT0000530 | Ppargc1b | 170826 | HITS-CLIP | 23597149\|23142080 | Unspecified |
| mmu-miR-21a-5p | MIMAT0000530 | Ppat | 231327 | HITS-CLIP | 23142080 | Unspecified |
| mmu-miR-21a-5p | MIMAT0000530 | Ppbp | 57349 | Microarrays | 37986224 | Unspecified |
| mmu-miR-21a-5p | MIMAT0000530 | Ppcdc | 66812 | Microarrays | 37986224\|25263593 | Unspecified |
| mmu-miR-21a-5p | MIMAT0000530 | Ppfia1 | 233977 | HITS-CLIP | 29167373 | Unspecified |
| mmu-miR-21a-5p | MIMAT0000530 | Pphln1 | 223828 | HITS-CLIP | 37986224 | Unspecified |
| mmu-miR-21a-5p | MIMAT0000530 | Ppic | 19038 | Microarrays | 37986224\|25263593 | Unspecified |
| mmu-miR-21a-5p | MIMAT0000530 | Ppid | 67738 | HITS-CLIP | 23597149\|23142080 | Unspecified |
| mmu-miR-21a-5p | MIMAT0000530 | Ppig | 228005 | HITS-CLIP | 37986224\|23142080 | Unspecified |
| mmu-miR-21a-5p | MIMAT0000530 | Ppih | 66101 | HITS-CLIP | 29167373 | Unspecified |
| mmu-miR-21a-5p | MIMAT0000530 | Ppil4 | 67418 | HITS-CLIP | 23142080 | Unspecified |
| mmu-miR-21a-5p | MIMAT0000530 | Ppip5k2 | 227399 | HITS-CLIP | 23142080\|29167373 | Unspecified |
| mmu-miR-21a-5p | MIMAT0000530 | Ppm1d | 53892 | HITS-CLIP | 23142080 | Unspecified |
| mmu-miR-21a-5p | MIMAT0000530 | Ppm1e | 320472 | Microarrays | 37986224\|25263593 | Unspecified |
| mmu-miR-21a-5p | MIMAT0000530 | Ppm1g | 14208 | HITS-CLIP | 29167373 | Unspecified |
| mmu-miR-21a-5p | MIMAT0000530 | Ppm1h | 319468 | HITS-CLIP | 29167373 | Unspecified |
| mmu-miR-21a-5p | MIMAT0000530 | Ppm1j | 71887 | Microarrays | 37986224\|25263593 | Unspecified |
| mmu-miR-21a-5p | MIMAT0000530 | Ppm1k | 243382 | HITS-CLIP | 37986224\|23597149\|23142080 | Unspecified |
| mmu-miR-21a-5p | MIMAT0000530 | Ppm1l | 242083 | HITS-CLIP | 37986224 | Unspecified |
| mmu-miR-21a-5p | MIMAT0000530 | Ppp1ca | 19045 | HITS-CLIP | 31433990\|23142080 | Unspecified |
| mmu-miR-21a-5p | MIMAT0000530 | Ppp1cb | 19046 | HITS-CLIP | 23142080 | Unspecified |
| mmu-miR-21a-5p | MIMAT0000530 | Ppp1r12a | 17931 | HITS-CLIP | 23142080\|19536157\|29167373 | Unspecified |
| mmu-miR-21a-5p | MIMAT0000530 | Ppp1r15b | 108954 | HITS-CLIP | 19536157\|23597149\|23142080 | Unspecified |
| mmu-miR-21a-5p | MIMAT0000530 | Ppp1r16b | 228852 | HITS-CLIP | 23142080 | Unspecified |
| mmu-miR-21a-5p | MIMAT0000530 | Ppp1r21 | 73825 | HITS-CLIP | 37986224\|19536157\|23142080 | Unspecified |
| mmu-miR-21a-5p | MIMAT0000530 | Ppp1r37 | 232947 | HITS-CLIP | 23142080 | Unspecified |
| mmu-miR-21a-5p | MIMAT0000530 | Ppp1r3b | 244416 | HITS-CLIP | 37986224\|23142080\|23597149 | Unspecified |
| mmu-miR-21a-5p | MIMAT0000530 | Ppp1r7 | 66385 | HITS-CLIP | 23142080 | Unspecified |
| mmu-miR-21a-5p | MIMAT0000530 | Ppp1r8 | 100336 | qCLASH//HITS-CLIP | 31393953\|23142080 | Unspecified |
| mmu-miR-21a-5p | MIMAT0000530 | Ppp1r9a | 243725 | HITS-CLIP, Microarrays | 37986224 | Unspecified |
| mmu-miR-21a-5p | MIMAT0000530 | Ppp2ca | 19052 | HITS-CLIP | 31433990\|23142080 | Unspecified |
| mmu-miR-21a-5p | MIMAT0000530 | Ppp2r2a | 71978 | HITS-CLIP | 29167373 | Unspecified |

| | | | | | | |
|---|---|---|---|---|---|---|
| mmu-miR-21a-5p | MIMAT0000530 | Ppp2r2c | 269643 | HITS-CLIP | 37986224\|19536157 | Unspecified |
| mmu-miR-21a-5p | MIMAT0000530 | Ppp2r5a | 226849 | HITS-CLIP | 29167373 | Unspecified |
| mmu-miR-21a-5p | MIMAT0000530 | Ppp2r5b | 225849 | Microarrays | 37986224\|25263593 | Unspecified |
| mmu-miR-21a-5p | MIMAT0000530 | Ppp2r5d | 21770 | HITS-CLIP | 37986224 | Unspecified |
| mmu-miR-21a-5p | MIMAT0000530 | Ppp2r5e | 26932 | HITS-CLIP | 23142080\|29167373 | Unspecified |
| mmu-miR-21a-5p | MIMAT0000530 | Ppp3ca | 19055 | HITS-CLIP | 23142080\|29167373 | Unspecified |
| mmu-miR-21a-5p | MIMAT0000530 | Ppp3cb | 19056 | HITS-CLIP | 23142080\|29167373 | Unspecified |
| mmu-miR-21a-5p | MIMAT0000530 | Ppp3cc | 19057 | HITS-CLIP | 23142080 | Unspecified |
| mmu-miR-21a-5p | MIMAT0000530 | Ppp3r1 | 19058 | HITS-CLIP | 23597149\|23142080 | Unspecified |
| mmu-miR-21a-5p | MIMAT0000530 | Ppp4r1 | 70351 | HITS-CLIP | 23142080\|29167373 | Unspecified |
| mmu-miR-21a-5p | MIMAT0000530 | Ppp4r2 | 232314 | HITS-CLIP | 23142080\|29167373 | Unspecified |
| mmu-miR-21a-5p | MIMAT0000530 | Ppp4r3a | 55671 | HITS-CLIP | 19536157 | Unspecified |
| mmu-miR-21a-5p | MIMAT0000530 | Ppp4r3b | 57223 | HITS-CLIP | 23142080\|19536157\|31433990 | Unspecified |
| mmu-miR-21a-5p | MIMAT0000530 | Ppp5c | 19060 | HITS-CLIP | 23142080 | Unspecified |
| mmu-miR-21a-5p | MIMAT0000530 | Ppp6c | 67857 | HITS-CLIP | 23597149\|23142080 | Unspecified |
| mmu-miR-21a-5p | MIMAT0000530 | Ppp6r1 | 243819 | HITS-CLIP | 23597149\|23142080 | Unspecified |
| mmu-miR-21a-5p | MIMAT0000530 | Ppp6r3 | 52036 | HITS-CLIP | 23142080 | Unspecified |
| mmu-miR-21a-5p | MIMAT0000530 | Pptc7 | 320717 | HITS-CLIP | 37986224\|19536157\|23142080 | Unspecified |
| mmu-miR-21a-5p | MIMAT0000530 | Ppwd1 | 238831 | Microarrays//HITS-CLIP | 37986224\|25263593\|23142080 | Unspecified |
| mmu-miR-21a-5p | MIMAT0000530 | Pqbp1 | 10084 | HITS-CLIP | 23597149\|23142080 | Unspecified |
| mmu-miR-21a-5p | MIMAT0000530 | Pqlc3 | 217430 | Microarrays | 37986224\|25263593 | Unspecified |
| mmu-miR-21a-5p | MIMAT0000530 | Prc1 | 233406 | Microarrays//HITS-CLIP//qCLASH | 37986224\|25263593\|23142080\|3139[illegible] | Unspecified |
| mmu-miR-21a-5p | MIMAT0000530 | Prcp | 72461 | Microarrays | 37986224 | Unspecified |
| mmu-miR-21a-5p | MIMAT0000530 | Prdm1 | 12142 | Microarrays | 37986224 | Unspecified |
| mmu-miR-21a-5p | MIMAT0000530 | Prdm16 | 70673 | HITS-CLIP | 23142080 | Unspecified |
| mmu-miR-21a-5p | MIMAT0000530 | Prdm2 | 110593 | HITS-CLIP | 23597149\|23142080 | Unspecified |
| mmu-miR-21a-5p | MIMAT0000530 | Prdm9 | 213389 | HITS-CLIP | 23597149 | Unspecified |
| mmu-miR-21a-5p | MIMAT0000530 | Prdx2 | 21672 | Microarrays | 37986224 | Unspecified |
| mmu-miR-21a-5p | MIMAT0000530 | Prdx3 | 11757 | qCLASH | 31393953 | Unspecified |
| mmu-miR-21a-5p | MIMAT0000530 | Prdx6 | 11758 | HITS-CLIP | 23597149 | Unspecified |
| mmu-miR-21a-5p | MIMAT0000530 | Prelid3b | 51012 | Microarrays | 25263593 | Unspecified |
| mmu-miR-21a-5p | MIMAT0000530 | Prex1 | 277360 | qCLASH//HITS-CLIP | 31393953\|23142080 | Unspecified |
| mmu-miR-21a-5p | MIMAT0000530 | Prex2 | 109294 | HITS-CLIP | 37986224 | Unspecified |
| mmu-miR-21a-5p | MIMAT0000530 | Prg4 | 96875 | Microarrays | 37986224\|25263593 | Unspecified |
| mmu-miR-21a-5p | MIMAT0000530 | Prim1 | 19075 | Microarrays//HITS-CLIP | 37986224\|25263593\|23597149\|2314[illegible] | Unspecified |
| mmu-miR-21a-5p | MIMAT0000530 | Prim2 | 19076 | HITS-CLIP | 23142080 | Unspecified |
| mmu-miR-21a-5p | MIMAT0000530 | Primpol | 408022 | HITS-CLIP | 23142080 | Unspecified |
| mmu-miR-21a-5p | MIMAT0000530 | Prkaa1 | 105787 | HITS-CLIP | 23142080\|29167373 | Unspecified |
| mmu-miR-21a-5p | MIMAT0000530 | Prkab2 | 108097 | Microarrays | 37986224 | Unspecified |

| | | | | | | | |
|---|---|---|---|---|---|---|---|
| mmu-miR-21a-5p | MIMAT0000530 | Prkaca | 18747 | HITS-CLIP | | 23142080 | Unspecified |
| mmu-miR-21a-5p | MIMAT0000530 | Prkacb | 18749 | HITS-CLIP | 23142080\|29167373 | | Unspecified |
| mmu-miR-21a-5p | MIMAT0000530 | Prkag1 | 19082 | HITS-CLIP | | 23142080 | Unspecified |
| mmu-miR-21a-5p | MIMAT0000530 | Prkag2 | 108099 | HITS-CLIP | | 23142080 | Unspecified |
| mmu-miR-21a-5p | MIMAT0000530 | Prkar1a | 19084 | HITS-CLIP | 37986224\|19536157\|23597149\|23142 | | Unspecified |
| mmu-miR-21a-5p | MIMAT0000530 | Prkar2a | 19087 | qCLASH//HITS-CLIP | 31393953\|23597149 | | Unspecified |
| mmu-miR-21a-5p | MIMAT0000530 | Prkca | 18750 | HITS-CLIP | | 23142080 | Unspecified |
| mmu-miR-21a-5p | MIMAT0000530 | Prkcb | 18751 | HITS-CLIP | 37986224\|19536157 | | Unspecified |
| mmu-miR-21a-5p | MIMAT0000530 | Prkcd | 18753 | HITS-CLIP | 23142080\|29167373 | | Unspecified |
| mmu-miR-21a-5p | MIMAT0000530 | Prkch | 18755 | HITS-CLIP | 23142080\|29167373 | | Unspecified |
| mmu-miR-21a-5p | MIMAT0000530 | Prkci | 18759 | HITS-CLIP | 31433990\|23142080 | | Unspecified |
| mmu-miR-21a-5p | MIMAT0000530 | Prkcsh | 19089 | HITS-CLIP | | 23142080 | Unspecified |
| mmu-miR-21a-5p | MIMAT0000530 | Prkcz | 18762 | HITS-CLIP | | 23142080 | Unspecified |
| mmu-miR-21a-5p | MIMAT0000530 | Prkdc | 19090 | HITS-CLIP | | 23142080 | Unspecified |
| mmu-miR-21a-5p | MIMAT0000530 | Prkg2 | 19092 | HITS-CLIP | | 37986224 | Unspecified |
| mmu-miR-21a-5p | MIMAT0000530 | Prmt9 | 80204 | HITS-CLIP | | 23142080 | Unspecified |
| mmu-miR-21a-5p | MIMAT0000530 | Prnp | 19122 | HITS-CLIP, Microarrays//HITS-CLIP | 37986224\|19536157 | | Unspecified |
| mmu-miR-21a-5p | MIMAT0000530 | Proc | 19123 | HITS-CLIP | | 23597149 | Unspecified |
| mmu-miR-21a-5p | MIMAT0000530 | Procr | 19124 | Microarrays | 37986224\|25263593 | | Unspecified |
| mmu-miR-21a-5p | MIMAT0000530 | Prodh | 19125 | HITS-CLIP | | 23597149 | Unspecified |
| mmu-miR-21a-5p | MIMAT0000530 | Prokr1 | 58182 | Microarrays | 37986224\|25263593 | | Unspecified |
| mmu-miR-21a-5p | MIMAT0000530 | Prorsd1 | 67939 | Microarrays//qCLASH | 37986224\|31393953 | | Unspecified |
| mmu-miR-21a-5p | MIMAT0000530 | Pros1 | 19128 | HITS-CLIP | 23597149\|23142080 | | Unspecified |
| mmu-miR-21a-5p | MIMAT0000530 | Proz | 66901 | HITS-CLIP | | 23597149 | Unspecified |
| mmu-miR-21a-5p | MIMAT0000530 | Prpf31 | 68988 | Microarrays | 37986224\|25263593 | | Unspecified |
| mmu-miR-21a-5p | MIMAT0000530 | Prpf38a | 230596 | HITS-CLIP | | 23142080 | Unspecified |
| mmu-miR-21a-5p | MIMAT0000530 | Prpf40a | 56194 | HITS-CLIP | 37986224\|23142080 | | Unspecified |
| mmu-miR-21a-5p | MIMAT0000530 | Prpf4b | 19134 | HITS-CLIP | 23597149\|29167373 | | Unspecified |
| mmu-miR-21a-5p | MIMAT0000530 | Prpf6 | 68879 | HITS-CLIP | 23597149\|23142080 | | Unspecified |
| mmu-miR-21a-5p | MIMAT0000530 | Prpf8 | 192159 | HITS-CLIP | 37986224\|23597149\|23142080\|29167 | | Unspecified |
| mmu-miR-21a-5p | MIMAT0000530 | Prps1 | 19139 | Microarrays//HITS-CLIP | 37986224\|25263593\|23142080 | | Unspecified |
| mmu-miR-21a-5p | MIMAT0000530 | Prps1l3 | 328099 | HITS-CLIP | 37986224\|23142080\|23597149 | | Unspecified |
| mmu-miR-21a-5p | MIMAT0000530 | Prps2 | 110639 | HITS-CLIP | | 23142080 | Unspecified |
| mmu-miR-21a-5p | MIMAT0000530 | Prr11 | 270906 | Microarrays//HITS-CLIP | 37986224\|25263593\|23142080 | | Unspecified |
| mmu-miR-21a-5p | MIMAT0000530 | Prr13 | 66151 | HITS-CLIP | | 23142080 | Unspecified |
| mmu-miR-21a-5p | MIMAT0000530 | Prr14l | 215476 | HITS-CLIP | | 23597149 | Unspecified |
| mmu-miR-21a-5p | MIMAT0000530 | Prr3 | 75210 | HITS-CLIP | | 23142080 | Unspecified |
| mmu-miR-21a-5p | MIMAT0000530 | Prrc2a | 53761 | HITS-CLIP | 23597149\|29167373 | | Unspecified |
| mmu-miR-21a-5p | MIMAT0000530 | Prrc2b | 227723 | HITS-CLIP | 37986224\|31433990 | | Unspecified |

| | | | | | | |
|---|---|---|---|---|---|---|
| mmu-miR-21a-5p | MIMAT0000530 | Prrc2c | 226562 | HITS-CLIP | 23142080\|29167373 | Unspecified |
| mmu-miR-21a-5p | MIMAT0000530 | Prrg1 | 546336 | Microarrays//HITS-CLIP | 37986224\|23142080 | Unspecified |
| mmu-miR-21a-5p | MIMAT0000530 | Prss16 | 54373 | Microarrays | 37986224\|25263593 | Unspecified |
| mmu-miR-21a-5p | MIMAT0000530 | Prss23 | 76453 | Microarrays | 37986224\|25263593 | Unspecified |
| mmu-miR-21a-5p | MIMAT0000530 | Prss35 | 244954 | HITS-CLIP | 19536157 | Unspecified |
| mmu-miR-21a-5p | MIMAT0000530 | Prune1 | 58497 | HITS-CLIP | 23597149 | Unspecified |
| mmu-miR-21a-5p | MIMAT0000530 | Psap | 19156 | HITS-CLIP | 23142080\|23597149\|19536157\|29167 | Unspecified |
| mmu-miR-21a-5p | MIMAT0000530 | Psat1 | 107272 | Microarrays//HITS-CLIP | 37986224\|25263593\|23142080 | Unspecified |
| mmu-miR-21a-5p | MIMAT0000530 | Psd3 | 234353 | HITS-CLIP | 29167373 | Unspecified |
| mmu-miR-21a-5p | MIMAT0000530 | Psd4 | 215632 | HITS-CLIP | 37986224\|23142080 | Unspecified |
| mmu-miR-21a-5p | MIMAT0000530 | Psen2 | 19165 | HITS-CLIP | 23597149\|29167373 | Unspecified |
| mmu-miR-21a-5p | MIMAT0000530 | Psip1 | 101739 | HITS-CLIP | 37986224\|23142080 | Unspecified |
| mmu-miR-21a-5p | MIMAT0000530 | Pskh1 | 244631 | HITS-CLIP | 23142080 | Unspecified |
| mmu-miR-21a-5p | MIMAT0000530 | Psma1 | 26440 | HITS-CLIP | 23142080 | Unspecified |
| mmu-miR-21a-5p | MIMAT0000530 | Psma2 | 19166 | HITS-CLIP | 23142080 | Unspecified |
| mmu-miR-21a-5p | MIMAT0000530 | Psmb3 | 26446 | HITS-CLIP | 23142080\|29167373 | Unspecified |
| mmu-miR-21a-5p | MIMAT0000530 | Psmb6 | 19175 | HITS-CLIP | 23142080 | Unspecified |
| mmu-miR-21a-5p | MIMAT0000530 | Psmb7 | 19177 | HITS-CLIP | 23597149\|23142080 | Unspecified |
| mmu-miR-21a-5p | MIMAT0000530 | Psmb9 | 16912 | Microarrays//HITS-CLIP | 37986224\|23142080 | Unspecified |
| mmu-miR-21a-5p | MIMAT0000530 | Psmc3 | 19182 | HITS-CLIP | 23142080 | Unspecified |
| mmu-miR-21a-5p | MIMAT0000530 | Psmc3ip | 19183 | Microarrays | 37986224\|25263593 | Unspecified |
| mmu-miR-21a-5p | MIMAT0000530 | Psmc4 | 23996 | HITS-CLIP | 23142080 | Unspecified |
| mmu-miR-21a-5p | MIMAT0000530 | Psmc5 | 19184 | qCLASH | 31393953 | Unspecified |
| mmu-miR-21a-5p | MIMAT0000530 | Psmc6 | 67089 | HITS-CLIP | 23142080\|29167373 | Unspecified |
| mmu-miR-21a-5p | MIMAT0000530 | Psmd1 | 70247 | HITS-CLIP | 23142080 | Unspecified |
| mmu-miR-21a-5p | MIMAT0000530 | Psmd11 | 69077 | HITS-CLIP | 23597149\|19536157\|29167373 | Unspecified |
| mmu-miR-21a-5p | MIMAT0000530 | Psmd14 | 59029 | HITS-CLIP | 23142080 | Unspecified |
| mmu-miR-21a-5p | MIMAT0000530 | Psmd3 | 22123 | HITS-CLIP | 23142080 | Unspecified |
| mmu-miR-21a-5p | MIMAT0000530 | Psmd6 | 66413 | HITS-CLIP | 23597149\|23142080 | Unspecified |
| mmu-miR-21a-5p | MIMAT0000530 | Psmd8 | 57296 | HITS-CLIP | 23142080 | Unspecified |
| mmu-miR-21a-5p | MIMAT0000530 | Psme3 | 19192 | HITS-CLIP | 23142080 | Unspecified |
| mmu-miR-21a-5p | MIMAT0000530 | Psme4 | 103554 | HITS-CLIP//qCLASH | 23597149\|31393953\|23142080\|29167 | Unspecified |
| mmu-miR-21a-5p | MIMAT0000530 | Psmf1 | 9491 | HITS-CLIP | 23597149 | Unspecified |
| mmu-miR-21a-5p | MIMAT0000530 | Psmg1 | 56088 | HITS-CLIP | 23597149\|23142080 | Unspecified |
| mmu-miR-21a-5p | MIMAT0000530 | Psmg2 | 107047 | HITS-CLIP | 23142080 | Unspecified |
| mmu-miR-21a-5p | MIMAT0000530 | Pspc1 | 66645 | HITS-CLIP | 23142080 | Unspecified |
| mmu-miR-21a-5p | MIMAT0000530 | Psph | 100678 | HITS-CLIP | 23142080 | Unspecified |
| mmu-miR-21a-5p | MIMAT0000530 | Psrc1 | 56742 | HITS-CLIP | 37986224\|23142080 | Unspecified |
| mmu-miR-21a-5p | MIMAT0000530 | Pstpip1 | 19200 | Microarrays | 37986224\|25263593 | Unspecified |

| | | | | | | | |
|---|---|---|---|---|---|---|---|
| mmu-miR-21a-5p | MIMAT0000530 | Pstpip2 | 19201 | Microarrays | | 37986224 | Unspecified |
| mmu-miR-21a-5p | MIMAT0000530 | Ptafr | 19204 | HITS-CLIP | | 29167373 | Unspecified |
| mmu-miR-21a-5p | MIMAT0000530 | Ptar1 | 72351 | HITS-CLIP | 23142080\|31433990 | | Unspecified |
| mmu-miR-21a-5p | MIMAT0000530 | Ptbp1 | 19205 | HITS-CLIP | 23142080\|29167373 | | Unspecified |
| mmu-miR-21a-5p | MIMAT0000530 | Ptbp2 | 56195 | HITS-CLIP | | 29167373 | Unspecified |
| mmu-miR-21a-5p | MIMAT0000530 | Ptbp3 | 230257 | HITS-CLIP | 23597149\|23142080 | | Unspecified |
| mmu-miR-21a-5p | MIMAT0000530 | Ptcd2 | 68927 | HITS-CLIP | | 23142080 | Unspecified |
| mmu-miR-21a-5p | MIMAT0000530 | Ptchd4 | 627626 | HITS-CLIP | | 37986224 | Unspecified |
| mmu-miR-21a-5p | MIMAT0000530 | Ptdss1 | 19210 | HITS-CLIP | | 23142080 | Unspecified |
| mmu-miR-21a-5p | MIMAT0000530 | Pten | 19211 | Reporter assay//Western Blot//Luciferase | 20167875\|20404348\|18508928\|26744 | | Unspecified |
| mmu-miR-21a-5p | MIMAT0000530 | Pter | 19212 | Microarrays | 37986224\|25263593 | | Unspecified |
| mmu-miR-21a-5p | MIMAT0000530 | Ptges3l | 73635 | Microarrays | 37986224\|25263593 | | Unspecified |
| mmu-miR-21a-5p | MIMAT0000530 | Ptgfrn | 19221 | HITS-CLIP | 23142080\|26203562 | | Unspecified |
| mmu-miR-21a-5p | MIMAT0000530 | Ptgir | 19222 | Microarrays | 37986224\|25263593 | | Unspecified |
| mmu-miR-21a-5p | MIMAT0000530 | Ptgs1 | 19224 | Microarrays | | 37986224 | Unspecified |
| mmu-miR-21a-5p | MIMAT0000530 | Ptgs2 | 19225 | Microarrays | 37986224\|25263593 | | Unspecified |
| mmu-miR-21a-5p | MIMAT0000530 | Ptk2b | 19229 | HITS-CLIP | 23142080\|29167373 | | Unspecified |
| mmu-miR-21a-5p | MIMAT0000530 | Ptp4a2 | 19244 | HITS-CLIP | | 23142080 | Unspecified |
| mmu-miR-21a-5p | MIMAT0000530 | Ptpn11 | 19247 | HITS-CLIP | | 29167373 | Unspecified |
| mmu-miR-21a-5p | MIMAT0000530 | Ptpn12 | 19248 | HITS-CLIP | | 23142080 | Unspecified |
| mmu-miR-21a-5p | MIMAT0000530 | Ptpn13 | 19249 | HITS-CLIP | | 23142080 | Unspecified |
| mmu-miR-21a-5p | MIMAT0000530 | Ptpn2 | 19255 | HITS-CLIP | 23597149\|23142080 | | Unspecified |
| mmu-miR-21a-5p | MIMAT0000530 | Ptpn22 | 19260 | Microarrays//HITS-CLIP | 37986224\|29167373\|23142080 | | Unspecified |
| mmu-miR-21a-5p | MIMAT0000530 | Ptpn4 | 19258 | HITS-CLIP | 19536157\|23597149\|23142080 | | Unspecified |
| mmu-miR-21a-5p | MIMAT0000530 | Ptpn7 | 320139 | HITS-CLIP | | 23142080 | Unspecified |
| mmu-miR-21a-5p | MIMAT0000530 | Ptpra | 19262 | HITS-CLIP | 29167373\|23142080 | | Unspecified |
| mmu-miR-21a-5p | MIMAT0000530 | Ptprb | 19263 | HITS-CLIP | | 23597149 | Unspecified |
| mmu-miR-21a-5p | MIMAT0000530 | Ptprd | 19266 | HITS-CLIP | | 23597149 | Unspecified |
| mmu-miR-21a-5p | MIMAT0000530 | Ptprf | 19268 | HITS-CLIP | | 23597149 | Unspecified |
| mmu-miR-21a-5p | MIMAT0000530 | Ptprm | 19274 | Microarrays | 37986224\|25263593 | | Unspecified |
| mmu-miR-21a-5p | MIMAT0000530 | Ptpro | 19277 | Microarrays | | 37986224 | Unspecified |
| mmu-miR-21a-5p | MIMAT0000530 | Ptrh1 | 329384 | Microarrays | 37986224\|25263593 | | Unspecified |
| mmu-miR-21a-5p | MIMAT0000530 | Ptrh2 | 217057 | HITS-CLIP | | 23142080 | Unspecified |
| mmu-miR-21a-5p | MIMAT0000530 | Puf60 | 67959 | HITS-CLIP | 19536157\|23142080 | | Unspecified |
| mmu-miR-21a-5p | MIMAT0000530 | Pum1 | 80912 | HITS-CLIP | 37986224\|23597149\|23142080\|29167 | | Unspecified |
| mmu-miR-21a-5p | MIMAT0000530 | Pum2 | 80913 | HITS-CLIP | | 23142080 | Unspecified |
| mmu-miR-21a-5p | MIMAT0000530 | Pum3 | 52874 | HITS-CLIP | | 29167373 | Unspecified |
| mmu-miR-21a-5p | MIMAT0000530 | Pura | 19290 | HITS-CLIP | 37986224\|29167373\|23142080 | | Unspecified |
| mmu-miR-21a-5p | MIMAT0000530 | Purb | 19291 | HITS-CLIP | 23142080\|23597149\|29167373 | | Unspecified |

| | | | | | | |
|---|---|---|---|---|---|---|
| mmu-miR-21a-5p | MIMAT0000530 | Pvr | 52118 | HITS-CLIP | 23597149\|23142080 | Unspecified |
| mmu-miR-21a-5p | MIMAT0000530 | Pwwp2a | 70802 | HITS-CLIP | 37986224\|23142080 | Unspecified |
| mmu-miR-21a-5p | MIMAT0000530 | Pxdc1 | 66895 | HITS-CLIP | 23597149 | Unspecified |
| mmu-miR-21a-5p | MIMAT0000530 | Pxdn | 69675 | HITS-CLIP | 37986224 | Unspecified |
| mmu-miR-21a-5p | MIMAT0000530 | Pxk | 218699 | HITS-CLIP | 23597149\|23142080 | Unspecified |
| mmu-miR-21a-5p | MIMAT0000530 | Pxn | 19303 | Microarrays//HITS-CLIP | 37986224\|23597149 | Unspecified |
| mmu-miR-21a-5p | MIMAT0000530 | Pxylp1 | 235534 | HITS-CLIP | 23142080 | Unspecified |
| mmu-miR-21a-5p | MIMAT0000530 | Pycard | 66824 | HITS-CLIP | 23142080 | Unspecified |
| mmu-miR-21a-5p | MIMAT0000530 | Pycr1 | 209027 | Microarrays | 37986224\|25263593 | Unspecified |
| mmu-miR-21a-5p | MIMAT0000530 | Pygb | 110078 | HITS-CLIP | 23142080 | Unspecified |
| mmu-miR-21a-5p | MIMAT0000530 | Pygl | 110095 | HITS-CLIP | 23597149 | Unspecified |
| mmu-miR-21a-5p | MIMAT0000530 | Pzp | 11287 | HITS-CLIP | 23597149 | Unspecified |
| mmu-miR-21a-5p | MIMAT0000530 | Qars | 97541 | HITS-CLIP | 23142080 | Unspecified |
| mmu-miR-21a-5p | MIMAT0000530 | Qk | 19317 | HITS-CLIP | 37986224\|19536157\|23597149\|2314 | Unspecified |
| mmu-miR-21a-5p | MIMAT0000530 | Qpct | 70536 | Microarrays | 37986224\|25263593 | Unspecified |
| mmu-miR-21a-5p | MIMAT0000530 | Qrich1 | 69232 | HITS-CLIP | 23142080 | Unspecified |
| mmu-miR-21a-5p | MIMAT0000530 | Qrsl1 | 76563 | Microarrays | 37986224\|25263593 | Unspecified |
| mmu-miR-21a-5p | MIMAT0000530 | Qser1 | 99003 | HITS-CLIP | 23142080\|29167373 | Unspecified |
| mmu-miR-21a-5p | MIMAT0000530 | Qsox1 | 104009 | HITS-CLIP | 23597149 | Unspecified |
| mmu-miR-21a-5p | MIMAT0000530 | Qsox2 | 227638 | Microarrays | 37986224\|25263593 | Unspecified |
| mmu-miR-21a-5p | MIMAT0000530 | R3hcc1l | 52013 | HITS-CLIP | 23142080 | Unspecified |
| mmu-miR-21a-5p | MIMAT0000530 | R3hdm1 | 226412 | HITS-CLIP | 23142080\|29167373 | Unspecified |
| mmu-miR-21a-5p | MIMAT0000530 | Rab10 | 19325 | HITS-CLIP | 23597149\|23142080 | Unspecified |
| mmu-miR-21a-5p | MIMAT0000530 | Rab11a | 53869 | HITS-CLIP | 23142080\|29167373 | Unspecified |
| mmu-miR-21a-5p | MIMAT0000530 | Rab11fip2 | 74998 | HITS-CLIP | 23142080 | Unspecified |
| mmu-miR-21a-5p | MIMAT0000530 | Rab11fip5 | 52055 | Microarrays | 37986224 | Unspecified |
| mmu-miR-21a-5p | MIMAT0000530 | Rab12 | 19328 | HITS-CLIP | 23142080\|29167373 | Unspecified |
| mmu-miR-21a-5p | MIMAT0000530 | Rab18 | 19330 | HITS-CLIP | 23142080 | Unspecified |
| mmu-miR-21a-5p | MIMAT0000530 | Rab1a | 5861 | HITS-CLIP | 23142080 | Unspecified |
| mmu-miR-21a-5p | MIMAT0000530 | Rab1b | 76308 | HITS-CLIP | 23597149\|23142080 | Unspecified |
| mmu-miR-21a-5p | MIMAT0000530 | Rab21 | 216344 | HITS-CLIP | 23142080 | Unspecified |
| mmu-miR-21a-5p | MIMAT0000530 | Rab22a | 19334 | HITS-CLIP | 19536157\|23142080\|29167373 | Unspecified |
| mmu-miR-21a-5p | MIMAT0000530 | Rab23 | 19335 | HITS-CLIP | 23142080 | Unspecified |
| mmu-miR-21a-5p | MIMAT0000530 | Rab2a | 59021 | HITS-CLIP | 23142080 | Unspecified |
| mmu-miR-21a-5p | MIMAT0000530 | Rab2b | 76338 | HITS-CLIP | 23142080 | Unspecified |
| mmu-miR-21a-5p | MIMAT0000530 | Rab32 | 67844 | Microarrays | 37986224\|25263593 | Unspecified |
| mmu-miR-21a-5p | MIMAT0000530 | Rab33b | 19338 | HITS-CLIP | 37986224\|29167373\|23142080 | Unspecified |
| mmu-miR-21a-5p | MIMAT0000530 | Rab35 | 77407 | HITS-CLIP | 23142080 | Unspecified |
| mmu-miR-21a-5p | MIMAT0000530 | Rab39 | 270160 | Microarrays | 37986224 | Unspecified |

| | | | | | | |
|---|---|---|---|---|---|---|
| mmu-miR-21a-5p | MIMAT0000530 | Rab3c | 67295 | HITS-CLIP | 19536157 | Unspecified |
| mmu-miR-21a-5p | MIMAT0000530 | Rab3gap1 | 226407 | qCLASH//HITS-CLIP | 31393953\|23142080 | Unspecified |
| mmu-miR-21a-5p | MIMAT0000530 | Rab3ip | 216363 | HITS-CLIP | 23142080 | Unspecified |
| mmu-miR-21a-5p | MIMAT0000530 | Rab40c | 224624 | HITS-CLIP | 29167373 | Unspecified |
| mmu-miR-21a-5p | MIMAT0000530 | Rab4b | 19342 | HITS-CLIP | 23597149 | Unspecified |
| mmu-miR-21a-5p | MIMAT0000530 | Rab5b | 19344 | Microarrays | 37986224 | Unspecified |
| mmu-miR-21a-5p | MIMAT0000530 | Rab5c | 19345 | HITS-CLIP | 23597149\|23142080 | Unspecified |
| mmu-miR-21a-5p | MIMAT0000530 | Rab6a | 19346 | HITS-CLIP | 23142080 | Unspecified |
| mmu-miR-21a-5p | MIMAT0000530 | Rab8b | 235442 | HITS-CLIP | 23142080 | Unspecified |
| mmu-miR-21a-5p | MIMAT0000530 | Rab9b | 319642 | HITS-CLIP | 19536157 | Unspecified |
| mmu-miR-21a-5p | MIMAT0000530 | Rabep1 | 54189 | HITS-CLIP | 37986224\|23142080\|29167373 | Unspecified |
| mmu-miR-21a-5p | MIMAT0000530 | Rabgap1 | 227800 | HITS-CLIP | 37986224\|23142080\|29167373 | Unspecified |
| mmu-miR-21a-5p | MIMAT0000530 | Rabgap1l | 29809 | Microarrays//HITS-CLIP | 37986224\|23142080 | Unspecified |
| mmu-miR-21a-5p | MIMAT0000530 | Rabgef1 | 56715 | HITS-CLIP | 23142080 | Unspecified |
| mmu-miR-21a-5p | MIMAT0000530 | Rabggtb | 19352 | HITS-CLIP | 23142080\|23597149 | Unspecified |
| mmu-miR-21a-5p | MIMAT0000530 | Rabl6 | 227624 | HITS-CLIP | 23597149\|23142080 | Unspecified |
| mmu-miR-21a-5p | MIMAT0000530 | Rac1 | 19353 | HITS-CLIP | 23597149\|23142080 | Unspecified |
| mmu-miR-21a-5p | MIMAT0000530 | Rac2 | 19354 | HITS-CLIP | 29167373\|23142080 | Unspecified |
| mmu-miR-21a-5p | MIMAT0000530 | Racgap1 | 26934 | Microarrays | 37986224\|25263593 | Unspecified |
| mmu-miR-21a-5p | MIMAT0000530 | Rack1 | 10399 | HITS-CLIP | 23142080\|29167373 | Unspecified |
| mmu-miR-21a-5p | MIMAT0000530 | Rad17 | 19356 | HITS-CLIP | 23142080 | Unspecified |
| mmu-miR-21a-5p | MIMAT0000530 | Rad18 | 58186 | HITS-CLIP | 23142080 | Unspecified |
| mmu-miR-21a-5p | MIMAT0000530 | Rad21 | 19357 | HITS-CLIP | 31433990\|23142080 | Unspecified |
| mmu-miR-21a-5p | MIMAT0000530 | Rad23b | 19359 | HITS-CLIP | 37986224\|23142080 | Unspecified |
| mmu-miR-21a-5p | MIMAT0000530 | Rad50 | 19360 | HITS-CLIP | 37986224\|23142080\|29167373 | Unspecified |
| mmu-miR-21a-5p | MIMAT0000530 | Rad51 | 19361 | Microarrays//HITS-CLIP | 37986224\|25263593\|23142080 | Unspecified |
| mmu-miR-21a-5p | MIMAT0000530 | Rad51ap1 | 19362 | Microarrays//HITS-CLIP | 37986224\|25263593\|23142080 | Unspecified |
| mmu-miR-21a-5p | MIMAT0000530 | Rad52 | 19365 | HITS-CLIP | 23142080 | Unspecified |
| mmu-miR-21a-5p | MIMAT0000530 | Rad54b | 623474 | Microarrays | 37986224\|25263593 | Unspecified |
| mmu-miR-21a-5p | MIMAT0000530 | Rad54l2 | 81000 | HITS-CLIP | 23597149\|23142080 | Unspecified |
| mmu-miR-21a-5p | MIMAT0000530 | Rae1 | 66679 | HITS-CLIP | 23142080 | Unspecified |
| mmu-miR-21a-5p | MIMAT0000530 | Raf1 | 110157 | HITS-CLIP | 23142080\|29167373 | Unspecified |
| mmu-miR-21a-5p | MIMAT0000530 | Ralgapa1 | 56784 | HITS-CLIP | 23142080 | Unspecified |
| mmu-miR-21a-5p | MIMAT0000530 | Ralgapa2 | 241694 | Microarrays//HITS-CLIP | 37986224\|25263593\|23142080\|29167 | Unspecified |
| mmu-miR-21a-5p | MIMAT0000530 | Ralgapb | 228850 | HITS-CLIP | 23142080\|29167373 | Unspecified |
| mmu-miR-21a-5p | MIMAT0000530 | Ralgps2 | 78255 | HITS-CLIP | 23142080\|23597149 | Unspecified |
| mmu-miR-21a-5p | MIMAT0000530 | Raly | 19383 | HITS-CLIP | 23142080 | Unspecified |
| mmu-miR-21a-5p | MIMAT0000530 | Ran | 19384 | HITS-CLIP | 19536157 | Unspecified |
| mmu-miR-21a-5p | MIMAT0000530 | Ranbp1 | 19385 | HITS-CLIP | 23597149 | Unspecified |

| | | | | | | |
|---|---|---|---|---|---|---|
| mmu-miR-21a-5p | MIMAT0000530 | Ranbp2 | 19386 | HITS-CLIP | 37986224\|23597149\|23142080\|31433 | Unspecified |
| mmu-miR-21a-5p | MIMAT0000530 | Ranbp9 | 56705 | HITS-CLIP | 23142080 | Unspecified |
| mmu-miR-21a-5p | MIMAT0000530 | Rangrf | 57785 | Microarrays | 37986224\|25263593 | Unspecified |
| mmu-miR-21a-5p | MIMAT0000530 | Rap1a | 109905 | HITS-CLIP | 31433990\|23142080 | Unspecified |
| mmu-miR-21a-5p | MIMAT0000530 | Rap1b | 215449 | HITS-CLIP | 37986224\|19536157\|23597149\|29167 | Unspecified |
| mmu-miR-21a-5p | MIMAT0000530 | Rap1gds1 | 229877 | HITS-CLIP | 23142080 | Unspecified |
| mmu-miR-21a-5p | MIMAT0000530 | Rap2b | 74012 | HITS-CLIP | 26203562\|23142080\|29167373 | Unspecified |
| mmu-miR-21a-5p | MIMAT0000530 | Rapgef2 | 76089 | HITS-CLIP | 29167373 | Unspecified |
| mmu-miR-21a-5p | MIMAT0000530 | Rapgef4 | 56508 | HITS-CLIP | 23142080 | Unspecified |
| mmu-miR-21a-5p | MIMAT0000530 | Rapgef5 | 217944 | Microarrays//HITS-CLIP | 37986224\|25263593\|29167373 | Unspecified |
| mmu-miR-21a-5p | MIMAT0000530 | Rapgef6 | 192786 | HITS-CLIP | 23142080 | Unspecified |
| mmu-miR-21a-5p | MIMAT0000530 | Raph1 | 77300 | HITS-CLIP, Microarrays//HITS-CLIP | 37986224\|23142080\|29167373 | Unspecified |
| mmu-miR-21a-5p | MIMAT0000530 | Rapsn | 19400 | Microarrays | 37986224 | Unspecified |
| mmu-miR-21a-5p | MIMAT0000530 | Rarg | 19411 | Microarrays//HITS-CLIP | 37986224\|23142080 | Unspecified |
| mmu-miR-21a-5p | MIMAT0000530 | Rarres2 | 71660 | HITS-CLIP | 23597149 | Unspecified |
| mmu-miR-21a-5p | MIMAT0000530 | Rars | 104458 | Microarrays//HITS-CLIP | 37986224\|25263593\|23142080 | Unspecified |
| mmu-miR-21a-5p | MIMAT0000530 | Rasa1 | 218397 | HITS-CLIP | 37986224\|19536157\|23142080\|29167 | Unspecified |
| mmu-miR-21a-5p | MIMAT0000530 | Rasa2 | 114713 | HITS-CLIP, Microarrays//Microarrays//HITS | 37986224\|25263593\|23142080 | Unspecified |
| mmu-miR-21a-5p | MIMAT0000530 | Rasal3 | 320484 | HITS-CLIP | 23142080 | Unspecified |
| mmu-miR-21a-5p | MIMAT0000530 | Rasd1 | 19416 | HITS-CLIP | 23597149 | Unspecified |
| mmu-miR-21a-5p | MIMAT0000530 | Rasgef1b | 320292 | Microarrays//HITS-CLIP | 37986224\|23142080\|29167373 | Unspecified |
| mmu-miR-21a-5p | MIMAT0000530 | Rasgrp1 | 19419 | HITS-CLIP | 37986224\|23142080 | Unspecified |
| mmu-miR-21a-5p | MIMAT0000530 | Rasgrp3 | 240168 | Microarrays | 37986224\|25263593 | Unspecified |
| mmu-miR-21a-5p | MIMAT0000530 | Rasl11b | 68939 | HITS-CLIP | 37986224\|23142080 | Unspecified |
| mmu-miR-21a-5p | MIMAT0000530 | Rassf2 | 215653 | Microarrays | 37986224 | Unspecified |
| mmu-miR-21a-5p | MIMAT0000530 | Rassf3 | 192678 | HITS-CLIP//qCLASH | 37986224\|31393953\|23142080\|29167 | Unspecified |
| mmu-miR-21a-5p | MIMAT0000530 | Rassf4 | 213391 | Microarrays//HITS-CLIP | 37986224\|29167373 | Unspecified |
| mmu-miR-21a-5p | MIMAT0000530 | Rb1 | 19645 | Microarrays//HITS-CLIP | 37986224\|23142080\|29167373 | Unspecified |
| mmu-miR-21a-5p | MIMAT0000530 | Rb1cc1 | 12421 | HITS-CLIP | 23597149\|23142080\|29167373 | Unspecified |
| mmu-miR-21a-5p | MIMAT0000530 | Rbak | 57782 | HITS-CLIP | 23142080 | Unspecified |
| mmu-miR-21a-5p | MIMAT0000530 | Rbbp6 | 19647 | HITS-CLIP | 23142080 | Unspecified |
| mmu-miR-21a-5p | MIMAT0000530 | Rbbp7 | 245688 | HITS-CLIP | 23142080 | Unspecified |
| mmu-miR-21a-5p | MIMAT0000530 | Rbbp8 | 225182 | HITS-CLIP | 23142080 | Unspecified |
| mmu-miR-21a-5p | MIMAT0000530 | Rbbp9 | 26450 | HITS-CLIP | 23142080 | Unspecified |
| mmu-miR-21a-5p | MIMAT0000530 | Rbm12 | 75710 | HITS-CLIP | 23142080 | Unspecified |
| mmu-miR-21a-5p | MIMAT0000530 | Rbm12b2 | 77604 | HITS-CLIP | 29167373\|23142080 | Unspecified |
| mmu-miR-21a-5p | MIMAT0000530 | Rbm14 | 56275 | HITS-CLIP | 23142080\|29167373 | Unspecified |
| mmu-miR-21a-5p | MIMAT0000530 | Rbm15b | 109095 | HITS-CLIP | 23597149 | Unspecified |
| mmu-miR-21a-5p | MIMAT0000530 | Rbm17 | 76938 | HITS-CLIP | 23597149 | Unspecified |

| | | | | | | |
|---|---|---|---|---|---|---|
| mmu-miR-21a-5p | MIMAT0000530 | Rbm18 | 67889 | HITS-CLIP | 23142080 | Unspecified |
| mmu-miR-21a-5p | MIMAT0000530 | Rbm24 | 666794 | HITS-CLIP | 23142080 | Unspecified |
| mmu-miR-21a-5p | MIMAT0000530 | Rbm25 | 67039 | HITS-CLIP | 37986224\|29167373\|23142080 | Unspecified |
| mmu-miR-21a-5p | MIMAT0000530 | Rbm26 | 74213 | HITS-CLIP | 23142080\|23597149 | Unspecified |
| mmu-miR-21a-5p | MIMAT0000530 | Rbm27 | 225432 | HITS-CLIP//qCLASH | 23142080\|29167373\|31393953 | Unspecified |
| mmu-miR-21a-5p | MIMAT0000530 | Rbm3 | 19652 | HITS-CLIP | 23142080 | Unspecified |
| mmu-miR-21a-5p | MIMAT0000530 | Rbm33 | 381626 | HITS-CLIP | 23142080 | Unspecified |
| mmu-miR-21a-5p | MIMAT0000530 | Rbm38 | 56190 | HITS-CLIP | 23142080 | Unspecified |
| mmu-miR-21a-5p | MIMAT0000530 | Rbm39 | 170791 | Microarrays | 37986224 | Unspecified |
| mmu-miR-21a-5p | MIMAT0000530 | Rbm43 | 71684 | HITS-CLIP | 23142080 | Unspecified |
| mmu-miR-21a-5p | MIMAT0000530 | Rbm45 | 241490 | Microarrays | 37986224 | Unspecified |
| mmu-miR-21a-5p | MIMAT0000530 | Rbm47 | 245945 | HITS-CLIP | 23597149\|29167373 | Unspecified |
| mmu-miR-21a-5p | MIMAT0000530 | Rbm48 | 269623 | HITS-CLIP | 23142080 | Unspecified |
| mmu-miR-21a-5p | MIMAT0000530 | Rbms1 | 56878 | qCLASH//HITS-CLIP | 31393953\|26203562\|29167373\|23142 | Unspecified |
| mmu-miR-21a-5p | MIMAT0000530 | Rbmx | 19655 | HITS-CLIP | 23142080 | Unspecified |
| mmu-miR-21a-5p | MIMAT0000530 | Rbmxl1 | 19656 | HITS-CLIP | 23142080 | Unspecified |
| mmu-miR-21a-5p | MIMAT0000530 | Rbpms | 19663 | Microarrays | 37986224 | Unspecified |
| mmu-miR-21a-5p | MIMAT0000530 | Rbsn | 64145 | HITS-CLIP | 23142080 | Unspecified |
| mmu-miR-21a-5p | MIMAT0000530 | Rbx1 | 56438 | Microarrays//HITS-CLIP | 37986224\|23142080 | Unspecified |
| mmu-miR-21a-5p | MIMAT0000530 | Rc3h1 | 381305 | HITS-CLIP//In situ hybridization//Luciferase | 29167373\|25697406 | Unspecified |
| mmu-miR-21a-5p | MIMAT0000530 | Rc3h2 | 319817 | HITS-CLIP | 19536157\|23142080\|29167373 | Unspecified |
| mmu-miR-21a-5p | MIMAT0000530 | Rcan1 | 54720 | Microarrays | 37986224\|25263593 | Unspecified |
| mmu-miR-21a-5p | MIMAT0000530 | Rcbtb2 | 105670 | Microarrays | 37986224 | Unspecified |
| mmu-miR-21a-5p | MIMAT0000530 | Rcc2 | 108911 | HITS-CLIP | 31433990\|30971389 | Unspecified |
| mmu-miR-21a-5p | MIMAT0000530 | Rce1 | 19671 | HITS-CLIP | 23142080 | Unspecified |
| mmu-miR-21a-5p | MIMAT0000530 | Rchy1 | 68098 | HITS-CLIP | 23597149 | Unspecified |
| mmu-miR-21a-5p | MIMAT0000530 | Rcn2 | 26611 | HITS-CLIP, Microarrays//Microarrays//HITS | 37986224\|25263593\|19536157\|23142 | Unspecified |
| mmu-miR-21a-5p | MIMAT0000530 | Rcor1 | 217864 | HITS-CLIP//qCLASH | 37986224\|23597149\|31393953\|23142 | Unspecified |
| mmu-miR-21a-5p | MIMAT0000530 | Rcor3 | 214742 | HITS-CLIP | 23142080 | Unspecified |
| mmu-miR-21a-5p | MIMAT0000530 | Rdh7 | 54150 | HITS-CLIP | 23597149 | Unspecified |
| mmu-miR-21a-5p | MIMAT0000530 | Rdm1 | 66599 | Microarrays | 37986224 | Unspecified |
| mmu-miR-21a-5p | MIMAT0000530 | Rdx | 19684 | HITS-CLIP | 37986224\|23597149\|23142080 | Unspecified |
| mmu-miR-21a-5p | MIMAT0000530 | Reck | 53614 | Luciferase assay//HITS-CLIP//Microarrays/ | 18556655\|23597149\|25263593\|31393 | Unspecified |
| mmu-miR-21a-5p | MIMAT0000530 | Reep1 | 52250 | HITS-CLIP | 19536157\|23142080 | Unspecified |
| mmu-miR-21a-5p | MIMAT0000530 | Reep2 | 225362 | HITS-CLIP | 37986224\|23142080 | Unspecified |
| mmu-miR-21a-5p | MIMAT0000530 | Reep3 | 28193 | HITS-CLIP | 23142080\|26203562\|29167373 | Unspecified |
| mmu-miR-21a-5p | MIMAT0000530 | Reep5 | 13476 | HITS-CLIP | 37986224\|23597149\|23142080\|29167 | Unspecified |
| mmu-miR-21a-5p | MIMAT0000530 | Rel | 19696 | HITS-CLIP | 23142080\|29167373 | Unspecified |
| mmu-miR-21a-5p | MIMAT0000530 | Rela | 19697 | HITS-CLIP | 23597149\|23142080 | Unspecified |

| | | | | | | |
|---|---|---|---|---|---|---|
| mmu-miR-21a-5p | MIMAT0000530 | Relch | 57614 | HITS-CLIP | 23597149\|23142080\|29167373 | Unspecified |
| mmu-miR-21a-5p | MIMAT0000530 | Rell1 | 100532 | HITS-CLIP, Microarrays//HITS-CLIP | 37986224\|23142080\|23597149 | Unspecified |
| mmu-miR-21a-5p | MIMAT0000530 | Reln | 19699 | HITS-CLIP | 23597149 | Unspecified |
| mmu-miR-21a-5p | MIMAT0000530 | Relt | 320100 | HITS-CLIP | 31433990 | Unspecified |
| mmu-miR-21a-5p | MIMAT0000530 | Reps1 | 19707 | HITS-CLIP | 23142080 | Unspecified |
| mmu-miR-21a-5p | MIMAT0000530 | Reps2 | 194590 | Microarrays | 37986224 | Unspecified |
| mmu-miR-21a-5p | MIMAT0000530 | Rere | 68703 | HITS-CLIP | 37986224\|23142080 | Unspecified |
| mmu-miR-21a-5p | MIMAT0000530 | Resf1 | 55196 | HITS-CLIP | 23142080\|29167373 | Unspecified |
| mmu-miR-21a-5p | MIMAT0000530 | Rest | 19712 | HITS-CLIP | 23142080 | Unspecified |
| mmu-miR-21a-5p | MIMAT0000530 | Retreg1 | 66270 | HITS-CLIP | 29167373 | Unspecified |
| mmu-miR-21a-5p | MIMAT0000530 | Retreg2 | 79137 | HITS-CLIP | 23597149 | Unspecified |
| mmu-miR-21a-5p | MIMAT0000530 | Retsat | 67442 | HITS-CLIP | 23597149 | Unspecified |
| mmu-miR-21a-5p | MIMAT0000530 | Rev1 | 56210 | HITS-CLIP | 23597149\|23142080 | Unspecified |
| mmu-miR-21a-5p | MIMAT0000530 | Rev3l | 19714 | HITS-CLIP//qCLASH | 23142080\|31393953 | Unspecified |
| mmu-miR-21a-5p | MIMAT0000530 | Rexo2 | 104444 | HITS-CLIP | 23597149\|23142080 | Unspecified |
| mmu-miR-21a-5p | MIMAT0000530 | Rfc4 | 106344 | Microarrays | 37986224\|25263593 | Unspecified |
| mmu-miR-21a-5p | MIMAT0000530 | Rffl | 67338 | HITS-CLIP | 37986224\|23597149\|23142080 | Unspecified |
| mmu-miR-21a-5p | MIMAT0000530 | Rft1 | 328370 | HITS-CLIP | 23142080 | Unspecified |
| mmu-miR-21a-5p | MIMAT0000530 | Rftn1 | 76438 | HITS-CLIP | 23142080 | Unspecified |
| mmu-miR-21a-5p | MIMAT0000530 | Rfx1 | 19724 | HITS-CLIP | 23142080 | Unspecified |
| mmu-miR-21a-5p | MIMAT0000530 | Rfx3 | 19726 | HITS-CLIP | 23142080 | Unspecified |
| mmu-miR-21a-5p | MIMAT0000530 | Rfx7 | 319758 | HITS-CLIP, Microarrays//HITS-CLIP | 37986224\|23142080 | Unspecified |
| mmu-miR-21a-5p | MIMAT0000530 | Rfxank | 19727 | HITS-CLIP | 23597149 | Unspecified |
| mmu-miR-21a-5p | MIMAT0000530 | Rgcc | 66214 | HITS-CLIP, Microarrays//Microarrays//HITS | 37986224\|25263593\|23142080 | Unspecified |
| mmu-miR-21a-5p | MIMAT0000530 | Rgs1 | 50778 | Microarrays | 37986224\|25263593 | Unspecified |
| mmu-miR-21a-5p | MIMAT0000530 | Rgs10 | 67865 | HITS-CLIP | 23142080 | Unspecified |
| mmu-miR-21a-5p | MIMAT0000530 | Rgs12 | 71729 | HITS-CLIP | 29167373 | Unspecified |
| mmu-miR-21a-5p | MIMAT0000530 | Rgs19 | 56470 | HITS-CLIP | 29167373 | Unspecified |
| mmu-miR-21a-5p | MIMAT0000530 | Rgs5 | 19737 | HITS-CLIP | 37986224\|23597149 | Unspecified |
| mmu-miR-21a-5p | MIMAT0000530 | Rgs9bp | 243923 | HITS-CLIP | 23142080 | Unspecified |
| mmu-miR-21a-5p | MIMAT0000530 | Rhbdd1 | 76867 | HITS-CLIP | 23142080 | Unspecified |
| mmu-miR-21a-5p | MIMAT0000530 | Rhbdf2 | 217344 | HITS-CLIP | 23597149\|23142080\|29167373 | Unspecified |
| mmu-miR-21a-5p | MIMAT0000530 | Rhbdl3 | 246104 | HITS-CLIP | 37986224 | Unspecified |
| mmu-miR-21a-5p | MIMAT0000530 | Rhob | 11852 | Microarrays//HITS-CLIP | 37986224\|25263593\|23597149\|1953( | Unspecified |
| mmu-miR-21a-5p | MIMAT0000530 | Rhobtb1 | 69288 | Microarrays//HITS-CLIP | 37986224\|23142080 | Unspecified |
| mmu-miR-21a-5p | MIMAT0000530 | Rhobtb3 | 73296 | HITS-CLIP | 23597149 | Unspecified |
| mmu-miR-21a-5p | MIMAT0000530 | Rhoc | 11853 | Microarrays | 37986224\|25263593 | Unspecified |
| mmu-miR-21a-5p | MIMAT0000530 | Rhoq | 104215 | HITS-CLIP//qCLASH | 37986224\|23142080\|31393953\|29167 | Unspecified |
| mmu-miR-21a-5p | MIMAT0000530 | Rhot1 | 59040 | HITS-CLIP | 23597149\|23142080 | Unspecified |

| | | | | | | |
|---|---|---|---|---|---|---|
| mmu-miR-21a-5p | MIMAT0000530 | Rhou | 69581 | Microarrays | 37986224\|25263593 | Unspecified |
| mmu-miR-21a-5p | MIMAT0000530 | Rhov | 228543 | Microarrays | 37986224\|25263593 | Unspecified |
| mmu-miR-21a-5p | MIMAT0000530 | Rhox4b | 57737 | Microarrays | 37986224\|25263593 | Unspecified |
| mmu-miR-21a-5p | MIMAT0000530 | Ric1 | 850728 | HITS-CLIP | 23597149\|23142080 | Unspecified |
| mmu-miR-21a-5p | MIMAT0000530 | Ric8b | 237422 | HITS-CLIP | 23142080 | Unspecified |
| mmu-miR-21a-5p | MIMAT0000530 | Rictor | 78757 | HITS-CLIP | 23142080 | Unspecified |
| mmu-miR-21a-5p | MIMAT0000530 | Rida | 65151 | Microarrays//HITS-CLIP | 25263593\|23597149\|23142080 | Unspecified |
| mmu-miR-21a-5p | MIMAT0000530 | Rif1 | 51869 | HITS-CLIP | 23142080\|29167373 | Unspecified |
| mmu-miR-21a-5p | MIMAT0000530 | Riiad1 | 66353 | Microarrays | 37986224 | Unspecified |
| mmu-miR-21a-5p | MIMAT0000530 | Rilpl1 | 75695 | HITS-CLIP | 19536157 | Unspecified |
| mmu-miR-21a-5p | MIMAT0000530 | Rims2 | 116838 | Microarrays//HITS-CLIP | 37986224\|25263593\|19536157 | Unspecified |
| mmu-miR-21a-5p | MIMAT0000530 | Rin2 | 74030 | HITS-CLIP | 29167373 | Unspecified |
| mmu-miR-21a-5p | MIMAT0000530 | Ring1 | 19763 | HITS-CLIP | 29167373 | Unspecified |
| mmu-miR-21a-5p | MIMAT0000530 | Rint1 | 72772 | HITS-CLIP | 23142080 | Unspecified |
| mmu-miR-21a-5p | MIMAT0000530 | Riok1 | 71340 | HITS-CLIP | 23142080 | Unspecified |
| mmu-miR-21a-5p | MIMAT0000530 | Riok2 | 67045 | HITS-CLIP | 23597149\|23142080\|29167373 | Unspecified |
| mmu-miR-21a-5p | MIMAT0000530 | Riox1 | 79697 | Microarrays | 25263593 | Unspecified |
| mmu-miR-21a-5p | MIMAT0000530 | Riox2 | 84864 | HITS-CLIP | 23597149 | Unspecified |
| mmu-miR-21a-5p | MIMAT0000530 | Ripk1 | 19766 | HITS-CLIP | 23142080 | Unspecified |
| mmu-miR-21a-5p | MIMAT0000530 | Ripk2 | 192656 | qCLASH | 31393953 | Unspecified |
| mmu-miR-21a-5p | MIMAT0000530 | Ripk3 | 56532 | Microarrays | 37986224\|25263593 | Unspecified |
| mmu-miR-21a-5p | MIMAT0000530 | Ripor2 | 193385 | HITS-CLIP | 23142080 | Unspecified |
| mmu-miR-21a-5p | MIMAT0000530 | Rlf | 109263 | HITS-CLIP | 37986224\|23142080 | Unspecified |
| mmu-miR-21a-5p | MIMAT0000530 | Rlim | 19820 | HITS-CLIP | 23142080\|29167373 | Unspecified |
| mmu-miR-21a-5p | MIMAT0000530 | Rmc1 | 19775 | HITS-CLIP | 23142080 | Unspecified |
| mmu-miR-21a-5p | MIMAT0000530 | Rmdn2 | 381110 | Microarrays | 37986224\|25263593 | Unspecified |
| mmu-miR-21a-5p | MIMAT0000530 | Rmnd1 | 66084 | Microarrays | 37986224 | Unspecified |
| mmu-miR-21a-5p | MIMAT0000530 | Rmnd5a | 68477 | HITS-CLIP | 23597149\|29167373 | Unspecified |
| mmu-miR-21a-5p | MIMAT0000530 | Rnase6 | 78416 | Microarrays | 37986224 | Unspecified |
| mmu-miR-21a-5p | MIMAT0000530 | Rnasel | 24014 | HITS-CLIP | 37986224\|23142080 | Unspecified |
| mmu-miR-21a-5p | MIMAT0000530 | Rnd3 | 74194 | Microarrays | 37986224 | Unspecified |
| mmu-miR-21a-5p | MIMAT0000530 | Rnf103 | 22644 | HITS-CLIP | 37986224\|19536157\|23142080 | Unspecified |
| mmu-miR-21a-5p | MIMAT0000530 | Rnf11 | 29864 | HITS-CLIP | 37986224\|23597149\|19536157\|2314 | Unspecified |
| mmu-miR-21a-5p | MIMAT0000530 | Rnf111 | 93836 | HITS-CLIP | 23142080\|23597149\|29167373 | Unspecified |
| mmu-miR-21a-5p | MIMAT0000530 | Rnf126 | 70294 | HITS-CLIP | 23142080 | Unspecified |
| mmu-miR-21a-5p | MIMAT0000530 | Rnf128 | 66889 | Microarrays | 37986224\|25263593 | Unspecified |
| mmu-miR-21a-5p | MIMAT0000530 | Rnf13 | 24017 | HITS-CLIP | 23142080 | Unspecified |
| mmu-miR-21a-5p | MIMAT0000530 | Rnf135 | 71956 | HITS-CLIP | 23597149 | Unspecified |
| mmu-miR-21a-5p | MIMAT0000530 | Rnf138 | 56515 | HITS-CLIP | 23142080 | Unspecified |

| | | | | | | |
|---|---|---|---|---|---|---|
| mmu-miR-21a-5p | MIMAT0000530 | Rnf139 | 75841 | HITS-CLIP | 23142080 | Unspecified |
| mmu-miR-21a-5p | MIMAT0000530 | Rnf14 | 56736 | HITS-CLIP | 23142080\|23597149 | Unspecified |
| mmu-miR-21a-5p | MIMAT0000530 | Rnf144b | 218215 | Microarrays | 37986224 | Unspecified |
| mmu-miR-21a-5p | MIMAT0000530 | Rnf145 | 74315 | HITS-CLIP, Microarrays//HITS-CLIP | 37986224\|23142080\|19536157\|29167 | Unspecified |
| mmu-miR-21a-5p | MIMAT0000530 | Rnf146 | 68031 | HITS-CLIP | 23142080 | Unspecified |
| mmu-miR-21a-5p | MIMAT0000530 | Rnf149 | 67702 | HITS-CLIP | 23597149\|29167373\|23142080 | Unspecified |
| mmu-miR-21a-5p | MIMAT0000530 | Rnf150 | 330812 | Microarrays | 37986224 | Unspecified |
| mmu-miR-21a-5p | MIMAT0000530 | Rnf152 | 320311 | HITS-CLIP | 23142080 | Unspecified |
| mmu-miR-21a-5p | MIMAT0000530 | Rnf167 | 70510 | HITS-CLIP, Microarrays//Microarrays//qCL | 37986224\|25263593\|31393953\|23597 | Unspecified |
| mmu-miR-21a-5p | MIMAT0000530 | Rnf169 | 108937 | HITS-CLIP | 37986224\|23597149\|23142080 | Unspecified |
| mmu-miR-21a-5p | MIMAT0000530 | Rnf180 | 71816 | HITS-CLIP | 29167373 | Unspecified |
| mmu-miR-21a-5p | MIMAT0000530 | Rnf181 | 66510 | HITS-CLIP | 19536157\|23597149\|23142080 | Unspecified |
| mmu-miR-21a-5p | MIMAT0000530 | Rnf185 | 193670 | HITS-CLIP | 23142080 | Unspecified |
| mmu-miR-21a-5p | MIMAT0000530 | Rnf19a | 30945 | HITS-CLIP, Microarrays//HITS-CLIP | 37986224\|23142080 | Unspecified |
| mmu-miR-21a-5p | MIMAT0000530 | Rnf19b | 75234 | HITS-CLIP | 29167373 | Unspecified |
| mmu-miR-21a-5p | MIMAT0000530 | Rnf2 | 19821 | Microarrays//HITS-CLIP | 37986224\|23142080 | Unspecified |
| mmu-miR-21a-5p | MIMAT0000530 | Rnf20 | 109331 | HITS-CLIP | 23142080 | Unspecified |
| mmu-miR-21a-5p | MIMAT0000530 | Rnf213 | 672511 | HITS-CLIP | 37986224\|23142080\|23597149\|29167 | Unspecified |
| mmu-miR-21a-5p | MIMAT0000530 | Rnf214 | 235315 | HITS-CLIP | 29167373 | Unspecified |
| mmu-miR-21a-5p | MIMAT0000530 | Rnf216 | 108086 | HITS-CLIP | 23597149\|23142080 | Unspecified |
| mmu-miR-21a-5p | MIMAT0000530 | Rnf219 | 72486 | HITS-CLIP | 37986224\|23142080 | Unspecified |
| mmu-miR-21a-5p | MIMAT0000530 | Rnf225 | 381845 | qCLASH | 31393953 | Unspecified |
| mmu-miR-21a-5p | MIMAT0000530 | Rnf24 | 51902 | HITS-CLIP | 23597149 | Unspecified |
| mmu-miR-21a-5p | MIMAT0000530 | Rnf25 | 57751 | HITS-CLIP | 23142080 | Unspecified |
| mmu-miR-21a-5p | MIMAT0000530 | Rnf38 | 73469 | HITS-CLIP | 23597149\|23142080 | Unspecified |
| mmu-miR-21a-5p | MIMAT0000530 | Rnf4 | 19822 | HITS-CLIP | 23142080\|23597149\|29167373 | Unspecified |
| mmu-miR-21a-5p | MIMAT0000530 | Rnf41 | 67588 | HITS-CLIP | 29167373 | Unspecified |
| mmu-miR-21a-5p | MIMAT0000530 | Rnf44 | 105239 | HITS-CLIP | 23597149\|23142080 | Unspecified |
| mmu-miR-21a-5p | MIMAT0000530 | Rnf6 | 74132 | HITS-CLIP | 23142080 | Unspecified |
| mmu-miR-21a-5p | MIMAT0000530 | Rnf8 | 58230 | HITS-CLIP | 23142080 | Unspecified |
| mmu-miR-21a-5p | MIMAT0000530 | Rnft1 | 76892 | HITS-CLIP | 37986224\|23597149\|23142080 | Unspecified |
| mmu-miR-21a-5p | MIMAT0000530 | Rngtt | 24018 | HITS-CLIP | 23142080 | Unspecified |
| mmu-miR-21a-5p | MIMAT0000530 | Rnh1 | 107702 | HITS-CLIP | 23597149\|23142080\|29167373 | Unspecified |
| mmu-miR-21a-5p | MIMAT0000530 | Rnmt | 67897 | HITS-CLIP | 23142080 | Unspecified |
| mmu-miR-21a-5p | MIMAT0000530 | Rnpep | 215615 | HITS-CLIP | 23597149\|23142080 | Unspecified |
| mmu-miR-21a-5p | MIMAT0000530 | Rock1 | 19877 | HITS-CLIP | 23142080\|29167373 | Unspecified |
| mmu-miR-21a-5p | MIMAT0000530 | Rock2 | 19878 | HITS-CLIP | 23142080\|29167373 | Unspecified |
| mmu-miR-21a-5p | MIMAT0000530 | Rora | 19883 | HITS-CLIP | 23597149\|28735896\|23142080 | Unspecified |
| mmu-miR-21a-5p | MIMAT0000530 | Rp2 | 19889 | HITS-CLIP | 23142080 | Unspecified |

| | | | | | | |
|---|---|---|---|---|---|---|
| mmu-miR-21a-5p | MIMAT0000530 | Rpa1 | 68275 | HITS-CLIP | 37986224\|23597149\|30971389\|23142 | Unspecified |
| mmu-miR-21a-5p | MIMAT0000530 | Rpa2 | 19891 | Microarrays//HITS-CLIP | 37986224\|25263593\|23142080 | Unspecified |
| mmu-miR-21a-5p | MIMAT0000530 | Rpa3 | 68240 | HITS-CLIP | 23142080\|31433990 | Unspecified |
| mmu-miR-21a-5p | MIMAT0000530 | Rpain | 69723 | HITS-CLIP | 23142080 | Unspecified |
| mmu-miR-21a-5p | MIMAT0000530 | Rpap2 | 231571 | Microarrays | 37986224\|25263593 | Unspecified |
| mmu-miR-21a-5p | MIMAT0000530 | Rpl12 | 269261 | Microarrays | 37986224\|25263593 | Unspecified |
| mmu-miR-21a-5p | MIMAT0000530 | Rpl13a | 22121 | HITS-CLIP | 23142080\|31433990 | Unspecified |
| mmu-miR-21a-5p | MIMAT0000530 | Rpl14 | 67115 | HITS-CLIP | 23142080\|23597149 | Unspecified |
| mmu-miR-21a-5p | MIMAT0000530 | Rpl15 | 66480 | HITS-CLIP | 23142080\|29167373 | Unspecified |
| mmu-miR-21a-5p | MIMAT0000530 | Rpl18a | 76808 | qCLASH | 31393953 | Unspecified |
| mmu-miR-21a-5p | MIMAT0000530 | Rpl22 | 19934 | HITS-CLIP | 23142080 | Unspecified |
| mmu-miR-21a-5p | MIMAT0000530 | Rpl23a | 268449 | qCLASH//HITS-CLIP | 31393953\|23142080 | Unspecified |
| mmu-miR-21a-5p | MIMAT0000530 | Rpl3 | 27367 | qCLASH | 31393953 | Unspecified |
| mmu-miR-21a-5p | MIMAT0000530 | Rpl39 | 67248 | qCLASH | 31393953 | Unspecified |
| mmu-miR-21a-5p | MIMAT0000530 | Rpl39l | 68172 | Microarrays | 37986224\|25263593 | Unspecified |
| mmu-miR-21a-5p | MIMAT0000530 | Rpl4 | 67891 | HITS-CLIP | 23597149\|23142080 | Unspecified |
| mmu-miR-21a-5p | MIMAT0000530 | Rpl5 | 100503670 | qCLASH//HITS-CLIP | 31393953\|23142080 | Unspecified |
| mmu-miR-21a-5p | MIMAT0000530 | Rpl7l1 | 66229 | HITS-CLIP | 23597149\|23142080 | Unspecified |
| mmu-miR-21a-5p | MIMAT0000530 | Rpl8 | 26961 | HITS-CLIP | 23142080 | Unspecified |
| mmu-miR-21a-5p | MIMAT0000530 | Rpn2 | 20014 | HITS-CLIP | 23597149\|23142080\|29167373 | Unspecified |
| mmu-miR-21a-5p | MIMAT0000530 | Rpp30 | 54364 | HITS-CLIP | 23142080 | Unspecified |
| mmu-miR-21a-5p | MIMAT0000530 | Rprd1b | 70470 | HITS-CLIP | 23142080\|29167373 | Unspecified |
| mmu-miR-21a-5p | MIMAT0000530 | Rps14 | 20044 | HITS-CLIP | 23142080 | Unspecified |
| mmu-miR-21a-5p | MIMAT0000530 | Rps17 | 20068 | qCLASH//HITS-CLIP | 31393953\|23142080 | Unspecified |
| mmu-miR-21a-5p | MIMAT0000530 | Rps19bp1 | 66538 | HITS-CLIP | 23142080 | Unspecified |
| mmu-miR-21a-5p | MIMAT0000530 | Rps25 | 75617 | Microarrays//qCLASH | 37986224\|25263593\|31393953 | Unspecified |
| mmu-miR-21a-5p | MIMAT0000530 | Rps27l | 67941 | HITS-CLIP//qCLASH | 37986224\|23142080\|23597149\|31393 | Unspecified |
| mmu-miR-21a-5p | MIMAT0000530 | Rps5 | 20103 | HITS-CLIP | 31433990\|23142080 | Unspecified |
| mmu-miR-21a-5p | MIMAT0000530 | Rps6 | 20104 | qCLASH | 31393953 | Unspecified |
| mmu-miR-21a-5p | MIMAT0000530 | Rps6ka3 | 110651 | HITS-CLIP | 29167373\|23142080 | Unspecified |
| mmu-miR-21a-5p | MIMAT0000530 | Rps6ka5 | 73086 | HITS-CLIP | 23142080 | Unspecified |
| mmu-miR-21a-5p | MIMAT0000530 | Rps6kb1 | 72508 | HITS-CLIP | 23142080 | Unspecified |
| mmu-miR-21a-5p | MIMAT0000530 | Rps8 | 20116 | qCLASH | 31393953 | Unspecified |
| mmu-miR-21a-5p | MIMAT0000530 | Rps9 | 76846 | Microarrays | 37986224 | Unspecified |
| mmu-miR-21a-5p | MIMAT0000530 | Rrad | 56437 | Microarrays | 37986224 | Unspecified |
| mmu-miR-21a-5p | MIMAT0000530 | Rraga | 68441 | HITS-CLIP | 23142080 | Unspecified |
| mmu-miR-21a-5p | MIMAT0000530 | Rragc | 54170 | HITS-CLIP | 29167373\|23142080 | Unspecified |
| mmu-miR-21a-5p | MIMAT0000530 | Rragd | 52187 | HITS-CLIP | 23142080 | Unspecified |
| mmu-miR-21a-5p | MIMAT0000530 | Rras2 | 66922 | HITS-CLIP | 37986224\|23142080 | Unspecified |

| | | | | | | |
|---|---|---|---|---|---|---|
| mmu-miR-21a-5p | MIMAT0000530 | Rrbp1 | 81910 | HITS-CLIP | 23597149\|29167373 | Unspecified |
| mmu-miR-21a-5p | MIMAT0000530 | Rrm1 | 20133 | HITS-CLIP//qCLASH | 23597149\|31393953\|23142080 | Unspecified |
| mmu-miR-21a-5p | MIMAT0000530 | Rrm2 | 20135 | Microarrays//qCLASH | 37986224\|25263593\|31393953 | Unspecified |
| mmu-miR-21a-5p | MIMAT0000530 | Rrp1 | 18114 | HITS-CLIP | 23142080 | Unspecified |
| mmu-miR-21a-5p | MIMAT0000530 | Rrp12 | 107094 | Microarrays//HITS-CLIP | 37986224\|25263593\|23597149 | Unspecified |
| mmu-miR-21a-5p | MIMAT0000530 | Rrp15 | 67223 | HITS-CLIP | 29167373 | Unspecified |
| mmu-miR-21a-5p | MIMAT0000530 | Rrp1b | 72462 | HITS-CLIP | 23142080 | Unspecified |
| mmu-miR-21a-5p | MIMAT0000530 | Rsad2 | 58185 | Microarrays//HITS-CLIP | 37986224\|25263593\|29167373 | Unspecified |
| mmu-miR-21a-5p | MIMAT0000530 | Rsbn1 | 229675 | HITS-CLIP | 37986224\|23142080 | Unspecified |
| mmu-miR-21a-5p | MIMAT0000530 | Rsbn1l | 242860 | HITS-CLIP | 37986224\|23142080 | Unspecified |
| mmu-miR-21a-5p | MIMAT0000530 | Rsf1 | 233532 | HITS-CLIP | 37986224\|23142080\|29167373 | Unspecified |
| mmu-miR-21a-5p | MIMAT0000530 | Rspo2 | 239405 | HITS-CLIP | 37986224 | Unspecified |
| mmu-miR-21a-5p | MIMAT0000530 | Rsrc2 | 208606 | HITS-CLIP | 23597149\|23142080 | Unspecified |
| mmu-miR-21a-5p | MIMAT0000530 | Rsrp1 | 27981 | Microarrays | 37986224 | Unspecified |
| mmu-miR-21a-5p | MIMAT0000530 | Rtca | 66368 | HITS-CLIP | 37986224 | Unspecified |
| mmu-miR-21a-5p | MIMAT0000530 | Rtcb | 28088 | HITS-CLIP | 23142080 | Unspecified |
| mmu-miR-21a-5p | MIMAT0000530 | Rtf1 | 76246 | HITS-CLIP | 23142080 | Unspecified |
| mmu-miR-21a-5p | MIMAT0000530 | Rtn3 | 20168 | HITS-CLIP | 23597149\|19536157\|29167373\|23142 | Unspecified |
| mmu-miR-21a-5p | MIMAT0000530 | Rtn4 | 68585 | HITS-CLIP | 37986224\|19536157\|23142080\|29167 | Unspecified |
| mmu-miR-21a-5p | MIMAT0000530 | Rtn4rl1 | 237847 | HITS-CLIP | 23142080 | Unspecified |
| mmu-miR-21a-5p | MIMAT0000530 | Rttn | 246102 | HITS-CLIP | 23142080 | Unspecified |
| mmu-miR-21a-5p | MIMAT0000530 | Rubcn | 9711 | HITS-CLIP | 23142080 | Unspecified |
| mmu-miR-21a-5p | MIMAT0000530 | Rufy1 | 216724 | HITS-CLIP | 29167373 | Unspecified |
| mmu-miR-21a-5p | MIMAT0000530 | Rufy2 | 70432 | HITS-CLIP | 23142080 | Unspecified |
| mmu-miR-21a-5p | MIMAT0000530 | Rundc3b | 242819 | HITS-CLIP | 23142080 | Unspecified |
| mmu-miR-21a-5p | MIMAT0000530 | Runx1 | 12394 | HITS-CLIP//qCLASH | 31433990\|31393953 | Unspecified |
| mmu-miR-21a-5p | MIMAT0000530 | Runx3 | 12399 | HITS-CLIP | 23142080 | Unspecified |
| mmu-miR-21a-5p | MIMAT0000530 | Rwdd4a | 192174 | HITS-CLIP//qCLASH | 23597149\|31393953\|29167373 | Unspecified |
| mmu-miR-21a-5p | MIMAT0000530 | Rxylt1 | 10329 | HITS-CLIP | 23142080\|23597149 | Unspecified |
| mmu-miR-21a-5p | MIMAT0000530 | Ryk | 20187 | HITS-CLIP | 37986224\|23142080 | Unspecified |
| mmu-miR-21a-5p | MIMAT0000530 | Ryr2 | 20191 | HITS-CLIP | 19536157 | Unspecified |
| mmu-miR-21a-5p | MIMAT0000530 | S100a10 | 20194 | HITS-CLIP | 26203562 | Unspecified |
| mmu-miR-21a-5p | MIMAT0000530 | S100a6 | 20200 | Microarrays | 37986224\|25263593 | Unspecified |
| mmu-miR-21a-5p | MIMAT0000530 | S100a8 | 20201 | Microarrays | 37986224\|25263593 | Unspecified |
| mmu-miR-21a-5p | MIMAT0000530 | S1pr1 | 13609 | HITS-CLIP | 19536157 | Unspecified |
| mmu-miR-21a-5p | MIMAT0000530 | S1pr4 | 13611 | HITS-CLIP | 23142080 | Unspecified |
| mmu-miR-21a-5p | MIMAT0000530 | SORBS2 | 8470 | HITS-CLIP//Western blot | 19536157\|23597149\|27033308 | Unspecified |
| mmu-miR-21a-5p | MIMAT0000530 | Saa2 | 20209 | HITS-CLIP | 23597149 | Unspecified |
| mmu-miR-21a-5p | MIMAT0000530 | Saa3 | 20210 | Microarrays | 37986224\|25263593 | Unspecified |

| | | | | | | |
|---|---|---|---|---|---|---|
| mmu-miR-21a-5p | MIMAT0000530 | Saal1 | 78935 | HITS-CLIP | 23142080 | Unspecified |
| mmu-miR-21a-5p | MIMAT0000530 | Sacm1l | 83493 | HITS-CLIP | 37986224\|19536157\|23142080 | Unspecified |
| mmu-miR-21a-5p | MIMAT0000530 | Sacs | 50720 | HITS-CLIP | 23142080 | Unspecified |
| mmu-miR-21a-5p | MIMAT0000530 | Sall2 | 50524 | HITS-CLIP | 23142080 | Unspecified |
| mmu-miR-21a-5p | MIMAT0000530 | Samd4 | 74480 | Microarrays//HITS-CLIP | 37986224\|23597149 | Unspecified |
| mmu-miR-21a-5p | MIMAT0000530 | Samd8 | 67630 | HITS-CLIP | 37986224\|23142080 | Unspecified |
| mmu-miR-21a-5p | MIMAT0000530 | Samd9l | 209086 | HITS-CLIP | 23142080\|29167373 | Unspecified |
| mmu-miR-21a-5p | MIMAT0000530 | Samhd1 | 56045 | Microarrays//HITS-CLIP | 37986224\|29167373 | Unspecified |
| mmu-miR-21a-5p | MIMAT0000530 | Sap130 | 269003 | HITS-CLIP | 23142080 | Unspecified |
| mmu-miR-21a-5p | MIMAT0000530 | Sap18 | 20220 | HITS-CLIP | 23142080 | Unspecified |
| mmu-miR-21a-5p | MIMAT0000530 | Sar1a | 20224 | HITS-CLIP | 23597149\|23142080 | Unspecified |
| mmu-miR-21a-5p | MIMAT0000530 | Sars | 20226 | HITS-CLIP | 23142080 | Unspecified |
| mmu-miR-21a-5p | MIMAT0000530 | Sash1 | 70097 | Microarrays//HITS-CLIP | 37986224\|25263593\|23597149 | Unspecified |
| mmu-miR-21a-5p | MIMAT0000530 | Sash3 | 74131 | HITS-CLIP | 23142080 | Unspecified |
| mmu-miR-21a-5p | MIMAT0000530 | Satb1 | 20230 | HITS-CLIP, Microarrays//HITS-CLIP//Micro | 37986224\|23142080\|23416424\|3143 | Unspecified |
| mmu-miR-21a-5p | MIMAT0000530 | Sbf2 | 319934 | qCLASH | 31393953 | Unspecified |
| mmu-miR-21a-5p | MIMAT0000530 | Sbno1 | 243272 | HITS-CLIP | 23142080\|29167373 | Unspecified |
| mmu-miR-21a-5p | MIMAT0000530 | Sc5d | 235293 | HITS-CLIP | 37986224\|23597149\|23142080\|2916 | Unspecified |
| mmu-miR-21a-5p | MIMAT0000530 | Scaf1 | 233208 | HITS-CLIP | 23597149 | Unspecified |
| mmu-miR-21a-5p | MIMAT0000530 | Scaf11 | 72193 | HITS-CLIP | 23142080\|29167373 | Unspecified |
| mmu-miR-21a-5p | MIMAT0000530 | Scaf4 | 224432 | qCLASH | 31393953 | Unspecified |
| mmu-miR-21a-5p | MIMAT0000530 | Scai | 320271 | HITS-CLIP | 23142080 | Unspecified |
| mmu-miR-21a-5p | MIMAT0000530 | Scamp2 | 24044 | Microarrays//HITS-CLIP | 37986224\|23142080 | Unspecified |
| mmu-miR-21a-5p | MIMAT0000530 | Scamp5 | 56807 | HITS-CLIP | 37986224 | Unspecified |
| mmu-miR-21a-5p | MIMAT0000530 | Scarb2 | 12492 | HITS-CLIP | 23597149\|23142080 | Unspecified |
| mmu-miR-21a-5p | MIMAT0000530 | Sccpdh | 109232 | Microarrays | 37986224\|25263593 | Unspecified |
| mmu-miR-21a-5p | MIMAT0000530 | Scd1 | 20249 | Microarrays | 37986224\|25263593 | Unspecified |
| mmu-miR-21a-5p | MIMAT0000530 | Scd2 | 20250 | HITS-CLIP, Microarrays//HITS-CLIP//qCLAS | 37986224\|19536157\|23142080\|3139 | Unspecified |
| mmu-miR-21a-5p | MIMAT0000530 | Scfd1 | 76983 | HITS-CLIP | 37986224\|23142080 | Unspecified |
| mmu-miR-21a-5p | MIMAT0000530 | Scimp | 327957 | HITS-CLIP | 23597149 | Unspecified |
| mmu-miR-21a-5p | MIMAT0000530 | Scly | 50880 | HITS-CLIP | 31433990 | Unspecified |
| mmu-miR-21a-5p | MIMAT0000530 | Scmh1 | 29871 | Microarrays | 37986224 | Unspecified |
| mmu-miR-21a-5p | MIMAT0000530 | Scml4 | 268297 | HITS-CLIP | 23142080 | Unspecified |
| mmu-miR-21a-5p | MIMAT0000530 | Scn2a | 110876 | HITS-CLIP | 19536157 | Unspecified |
| mmu-miR-21a-5p | MIMAT0000530 | Scn3b | 235281 | HITS-CLIP | 37986224 | Unspecified |
| mmu-miR-21a-5p | MIMAT0000530 | Scoc | 56367 | HITS-CLIP | 23142080\|29167373 | Unspecified |
| mmu-miR-21a-5p | MIMAT0000530 | Scrg1 | 20284 | Microarrays | 37986224 | Unspecified |
| mmu-miR-21a-5p | MIMAT0000530 | Scrib | 105782 | HITS-CLIP | 23597149\|23142080 | Unspecified |
| mmu-miR-21a-5p | MIMAT0000530 | Scyl1 | 78891 | HITS-CLIP | 29167373\|23597149 | Unspecified |

| | | | | | | |
|---|---|---|---|---|---|---|
| mmu-miR-21a-5p | MIMAT0000530 | Scyl2 | 213326 | HITS-CLIP | 23142080\|29167373 | Unspecified |
| mmu-miR-21a-5p | MIMAT0000530 | Scyl3 | 240880 | HITS-CLIP | 23142080 | Unspecified |
| mmu-miR-21a-5p | MIMAT0000530 | Sdad1 | 231452 | HITS-CLIP | 37986224\|23142080 | Unspecified |
| mmu-miR-21a-5p | MIMAT0000530 | Sdc3 | 20970 | HITS-CLIP | 23597149\|23142080\|29167373 | Unspecified |
| mmu-miR-21a-5p | MIMAT0000530 | Sdc4 | 20971 | Microarrays//HITS-CLIP | 37986224\|23142080\|29167373 | Unspecified |
| mmu-miR-21a-5p | MIMAT0000530 | Sdcbp | 53378 | HITS-CLIP | 29167373 | Unspecified |
| mmu-miR-21a-5p | MIMAT0000530 | Sde2 | 208768 | HITS-CLIP | 37986224 | Unspecified |
| mmu-miR-21a-5p | MIMAT0000530 | Sdf2 | 20316 | HITS-CLIP | 23597149 | Unspecified |
| mmu-miR-21a-5p | MIMAT0000530 | Sdf4 | 20318 | HITS-CLIP | 23142080 | Unspecified |
| mmu-miR-21a-5p | MIMAT0000530 | Sdhb | 67680 | HITS-CLIP | 23597149 | Unspecified |
| mmu-miR-21a-5p | MIMAT0000530 | Sds | 231691 | HITS-CLIP | 23597149 | Unspecified |
| mmu-miR-21a-5p | MIMAT0000530 | Sec11c | 66286 | HITS-CLIP | 31433990\|23142080 | Unspecified |
| mmu-miR-21a-5p | MIMAT0000530 | Sec13 | 110379 | HITS-CLIP | 31433990\|23142080 | Unspecified |
| mmu-miR-21a-5p | MIMAT0000530 | Sec14l1 | 74136 | HITS-CLIP | 23597149\|29167373 | Unspecified |
| mmu-miR-21a-5p | MIMAT0000530 | Sec16a | 227648 | HITS-CLIP | 23142080\|29167373 | Unspecified |
| mmu-miR-21a-5p | MIMAT0000530 | Sec16b | 89867 | HITS-CLIP | 23597149 | Unspecified |
| mmu-miR-21a-5p | MIMAT0000530 | Sec23a | 20334 | HITS-CLIP | 23597149\|23142080 | Unspecified |
| mmu-miR-21a-5p | MIMAT0000530 | Sec23b | 27054 | HITS-CLIP | 23597149\|23142080 | Unspecified |
| mmu-miR-21a-5p | MIMAT0000530 | Sec24a | 77371 | HITS-CLIP | 23597149\|23142080\|29167373 | Unspecified |
| mmu-miR-21a-5p | MIMAT0000530 | Sec24b | 99683 | HITS-CLIP | 23142080\|31433990 | Unspecified |
| mmu-miR-21a-5p | MIMAT0000530 | Sec24c | 218811 | HITS-CLIP | 37986224\|23142080\|29167373 | Unspecified |
| mmu-miR-21a-5p | MIMAT0000530 | Sec24d | 69608 | HITS-CLIP | 23597149 | Unspecified |
| mmu-miR-21a-5p | MIMAT0000530 | Sec31a | 69162 | HITS-CLIP | 23597149\|23142080 | Unspecified |
| mmu-miR-21a-5p | MIMAT0000530 | Sec62 | 69276 | HITS-CLIP | 19536157 | Unspecified |
| mmu-miR-21a-5p | MIMAT0000530 | Sec63 | 140740 | HITS-CLIP | 23142080\|29167373 | Unspecified |
| mmu-miR-21a-5p | MIMAT0000530 | Secisbp2l | 70354 | HITS-CLIP | 23597149\|31433990\|23142080 | Unspecified |
| mmu-miR-21a-5p | MIMAT0000530 | Seh1l | 72124 | HITS-CLIP | 37986224\|23142080 | Unspecified |
| mmu-miR-21a-5p | MIMAT0000530 | Sel1l | 20338 | HITS-CLIP | 23597149\|23142080 | Unspecified |
| mmu-miR-21a-5p | MIMAT0000530 | Selenbp1 | 20341 | Microarrays | 37986224 | Unspecified |
| mmu-miR-21a-5p | MIMAT0000530 | Selenof | 9403 | HITS-CLIP | 23597149 | Unspecified |
| mmu-miR-21a-5p | MIMAT0000530 | Selenoh | 280636 | Microarrays | 25263593 | Unspecified |
| mmu-miR-21a-5p | MIMAT0000530 | Selenoi | 28042 | HITS-CLIP | 23142080 | Unspecified |
| mmu-miR-21a-5p | MIMAT0000530 | Selenot | 69227 | HITS-CLIP | 23597149\|23142080 | Unspecified |
| mmu-miR-21a-5p | MIMAT0000530 | Selenow | 135964823 | Microarrays | 25263593 | Unspecified |
| mmu-miR-21a-5p | MIMAT0000530 | Sell | 20343 | HITS-CLIP | 37986224\|23142080 | Unspecified |
| mmu-miR-21a-5p | MIMAT0000530 | Selp | 20344 | HITS-CLIP | 23142080 | Unspecified |
| mmu-miR-21a-5p | MIMAT0000530 | Sema3c | 20348 | Microarrays | 37986224\|25263593 | Unspecified |
| mmu-miR-21a-5p | MIMAT0000530 | Sema4a | 20351 | Microarrays//HITS-CLIP | 37986224\|23597149\|23142080 | Unspecified |
| mmu-miR-21a-5p | MIMAT0000530 | Sema4b | 20352 | HITS-CLIP | 31433990\|26203562\|23142080 | Unspecified |

| | | | | | | |
|---|---|---|---|---|---|---|
| mmu-miR-21a-5p | MIMAT0000530 | Sema5a | 20356 | HITS-CLIP | 37986224\|19536157 | Unspecified |
| mmu-miR-21a-5p | MIMAT0000530 | Sema6d | 214968 | HITS-CLIP | 23597149\|23142080 | Unspecified |
| mmu-miR-21a-5p | MIMAT0000530 | Sema7a | 20361 | HITS-CLIP | 37986224\|19536157\|23142080 | Unspecified |
| mmu-miR-21a-5p | MIMAT0000530 | Senp1 | 223870 | HITS-CLIP | 29167373\|23142080 | Unspecified |
| mmu-miR-21a-5p | MIMAT0000530 | Senp3 | 80886 | qCLASH | 31393953 | Unspecified |
| mmu-miR-21a-5p | MIMAT0000530 | Senp6 | 215351 | HITS-CLIP | 23142080\|29167373 | Unspecified |
| mmu-miR-21a-5p | MIMAT0000530 | Senp7 | 66315 | HITS-CLIP | 23142080 | Unspecified |
| mmu-miR-21a-5p | MIMAT0000530 | Sephs1 | 109079 | HITS-CLIP | 31433990\|23142080 | Unspecified |
| mmu-miR-21a-5p | MIMAT0000530 | Sepsecs | 211006 | HITS-CLIP | 23142080 | Unspecified |
| mmu-miR-21a-5p | MIMAT0000530 | Sept1 | 54204 | HITS-CLIP | 23142080 | Unspecified |
| mmu-miR-21a-5p | MIMAT0000530 | Sept10 | 103080 | HITS-CLIP | 37986224\|23597149 | Unspecified |
| mmu-miR-21a-5p | MIMAT0000530 | Sept11 | 52398 | HITS-CLIP | 23142080\|23597149\|29167373 | Unspecified |
| mmu-miR-21a-5p | MIMAT0000530 | Sept2 | 4735 | HITS-CLIP | 23142080 | Unspecified |
| mmu-miR-21a-5p | MIMAT0000530 | Sept6 | 23157 | HITS-CLIP | 23142080 | Unspecified |
| mmu-miR-21a-5p | MIMAT0000530 | Sept7 | 235072 | HITS-CLIP | 37986224\|23142080\|29167373 | Unspecified |
| mmu-miR-21a-5p | MIMAT0000530 | Sept9 | 53860 | HITS-CLIP | 23142080 | Unspecified |
| mmu-miR-21a-5p | MIMAT0000530 | Septin3 | 24050 | HITS-CLIP | 37986224 | Unspecified |
| mmu-miR-21a-5p | MIMAT0000530 | Serbp1 | 66870 | HITS-CLIP | 37986224\|23142080\|29167373 | Unspecified |
| mmu-miR-21a-5p | MIMAT0000530 | Serinc1 | 56442 | HITS-CLIP | 37986224\|19536157\|23142080\|29167 | Unspecified |
| mmu-miR-21a-5p | MIMAT0000530 | Serinc3 | 26943 | Microarrays//HITS-CLIP//qCLASH | 37986224\|23142080\|31393953\|19536 | Unspecified |
| mmu-miR-21a-5p | MIMAT0000530 | Serinc5 | 218442 | HITS-CLIP, Microarrays//HITS-CLIP | 37986224\|23597149\|19536157\|23142 | Unspecified |
| mmu-miR-21a-5p | MIMAT0000530 | Serp1 | 28146 | HITS-CLIP, Microarrays | 37986224 | Unspecified |
| mmu-miR-21a-5p | MIMAT0000530 | Serpina3k | 20714 | HITS-CLIP | 23597149 | Unspecified |
| mmu-miR-21a-5p | MIMAT0000530 | Serpina3n | 20716 | HITS-CLIP | 23597149 | Unspecified |
| mmu-miR-21a-5p | MIMAT0000530 | Serpinb12 | 71869 | Microarrays | 37986224\|25263593 | Unspecified |
| mmu-miR-21a-5p | MIMAT0000530 | Serpinb1a | 66222 | Microarrays | 37986224\|25263593 | Unspecified |
| mmu-miR-21a-5p | MIMAT0000530 | Serpinb1b | 282663 | Microarrays | 37986224\|25263593 | Unspecified |
| mmu-miR-21a-5p | MIMAT0000530 | Serpinb2 | 18788 | Microarrays | 37986224\|25263593 | Unspecified |
| mmu-miR-21a-5p | MIMAT0000530 | Serpinb6a | 20719 | HITS-CLIP | 29167373 | Unspecified |
| mmu-miR-21a-5p | MIMAT0000530 | Serpinb6b | 20708 | Microarrays//HITS-CLIP | 37986224\|25263593\|23142080 | Unspecified |
| mmu-miR-21a-5p | MIMAT0000530 | Serpinb8 | 20725 | HITS-CLIP | 29167373 | Unspecified |
| mmu-miR-21a-5p | MIMAT0000530 | Serpinb9 | 20723 | Microarrays | 37986224\|25263593 | Unspecified |
| mmu-miR-21a-5p | MIMAT0000530 | Serpinb9b | 20706 | Microarrays | 37986224\|25263593 | Unspecified |
| mmu-miR-21a-5p | MIMAT0000530 | Serpinc1 | 11905 | HITS-CLIP | 23597149 | Unspecified |
| mmu-miR-21a-5p | MIMAT0000530 | Serpind1 | 15160 | HITS-CLIP | 23597149 | Unspecified |
| mmu-miR-21a-5p | MIMAT0000530 | Serpine1 | 18787 | Microarrays | 37986224\|25263593 | Unspecified |
| mmu-miR-21a-5p | MIMAT0000530 | Serpine2 | 20720 | Microarrays | 37986224\|25263593 | Unspecified |
| mmu-miR-21a-5p | MIMAT0000530 | Serpinf1 | 20317 | Microarrays//HITS-CLIP | 37986224\|25263593\|23142080 | Unspecified |
| mmu-miR-21a-5p | MIMAT0000530 | Serpinf2 | 18816 | HITS-CLIP | 23597149 | Unspecified |

| | | | | | | |
|---|---|---|---|---|---|---|
| mmu-miR-21a-5p | MIMAT0000530 | Serpinh1 | 12406 | Microarrays | 37986224\|25263593 | Unspecified |
| mmu-miR-21a-5p | MIMAT0000530 | Sertad2 | 58172 | HITS-CLIP | 31433990\|23142080\|29167373 | Unspecified |
| mmu-miR-21a-5p | MIMAT0000530 | Sesn1 | 140742 | Microarrays//HITS-CLIP | 37986224\|23142080 | Unspecified |
| mmu-miR-21a-5p | MIMAT0000530 | Sesn3 | 75747 | HITS-CLIP | 23142080\|19536157 | Unspecified |
| mmu-miR-21a-5p | MIMAT0000530 | Sestd1 | 228071 | HITS-CLIP | 29167373 | Unspecified |
| mmu-miR-21a-5p | MIMAT0000530 | Setd1a | 233904 | HITS-CLIP | 23597149\|23142080 | Unspecified |
| mmu-miR-21a-5p | MIMAT0000530 | Setd1b | 208043 | Microarrays//HITS-CLIP | 37986224\|23142080\|29167373 | Unspecified |
| mmu-miR-21a-5p | MIMAT0000530 | Setd2 | 235626 | HITS-CLIP | 23142080\|23597149\|19536157 | Unspecified |
| mmu-miR-21a-5p | MIMAT0000530 | Setd3 | 52690 | HITS-CLIP | 23142080 | Unspecified |
| mmu-miR-21a-5p | MIMAT0000530 | Setd5 | 72895 | HITS-CLIP | 29167373\|23142080 | Unspecified |
| mmu-miR-21a-5p | MIMAT0000530 | Setd7 | 73251 | HITS-CLIP | 23142080 | Unspecified |
| mmu-miR-21a-5p | MIMAT0000530 | Setdb1 | 84505 | qCLASH | 31393953 | Unspecified |
| mmu-miR-21a-5p | MIMAT0000530 | Setx | 269254 | HITS-CLIP | 37986224\|23597149\|23142080 | Unspecified |
| mmu-miR-21a-5p | MIMAT0000530 | Sez6l2 | 233878 | Microarrays//HITS-CLIP | 37986224\|25263593\|19536157 | Unspecified |
| mmu-miR-21a-5p | MIMAT0000530 | Sf3a3 | 75062 | HITS-CLIP | 23142080 | Unspecified |
| mmu-miR-21a-5p | MIMAT0000530 | Sf3b1 | 81898 | HITS-CLIP | 37986224\|23142080\|29167373 | Unspecified |
| mmu-miR-21a-5p | MIMAT0000530 | Sf3b2 | 319322 | HITS-CLIP | 37986224\|23597149\|23142080 | Unspecified |
| mmu-miR-21a-5p | MIMAT0000530 | Sfi1 | 78887 | Microarrays | 37986224 | Unspecified |
| mmu-miR-21a-5p | MIMAT0000530 | Sfmbt1 | 54650 | HITS-CLIP | 23142080 | Unspecified |
| mmu-miR-21a-5p | MIMAT0000530 | Sfpq | 71514 | HITS-CLIP | 37986224\|23142080\|29167373 | Unspecified |
| mmu-miR-21a-5p | MIMAT0000530 | Sfr1 | 67788 | HITS-CLIP | 23597149\|23142080 | Unspecified |
| mmu-miR-21a-5p | MIMAT0000530 | Sfrp1 | 20377 | HITS-CLIP | 37986224 | Unspecified |
| mmu-miR-21a-5p | MIMAT0000530 | Sft2d1 | 106489 | Microarrays//HITS-CLIP | 37986224\|23142080\|25263593 | Unspecified |
| mmu-miR-21a-5p | MIMAT0000530 | Sft2d2 | 108735 | HITS-CLIP | 29167373\|23142080 | Unspecified |
| mmu-miR-21a-5p | MIMAT0000530 | Sfxn1 | 14057 | Microarrays | 37986224\|25263593 | Unspecified |
| mmu-miR-21a-5p | MIMAT0000530 | Sfxn3 | 94280 | HITS-CLIP | 23142080 | Unspecified |
| mmu-miR-21a-5p | MIMAT0000530 | Sfxn4 | 94281 | Microarrays | 37986224 | Unspecified |
| mmu-miR-21a-5p | MIMAT0000530 | Sgcb | 24051 | HITS-CLIP | 23142080\|29167373 | Unspecified |
| mmu-miR-21a-5p | MIMAT0000530 | Sgk1 | 20393 | HITS-CLIP | 23597149\|19536157\|23142080 | Unspecified |
| mmu-miR-21a-5p | MIMAT0000530 | Sgk2 | 27219 | HITS-CLIP | 23597149 | Unspecified |
| mmu-miR-21a-5p | MIMAT0000530 | Sgk3 | 170755 | HITS-CLIP | 23597149\|29167373 | Unspecified |
| mmu-miR-21a-5p | MIMAT0000530 | Sgms1 | 208449 | HITS-CLIP | 23142080\|29167373 | Unspecified |
| mmu-miR-21a-5p | MIMAT0000530 | Sgms2 | 74442 | Microarrays//HITS-CLIP | 37986224\|25263593\|23597149\|2314 | Unspecified |
| mmu-miR-21a-5p | MIMAT0000530 | Sgo1 | 72415 | Microarrays//HITS-CLIP | 25263593\|23142080 | Unspecified |
| mmu-miR-21a-5p | MIMAT0000530 | Sgo2a | 68549 | Microarrays | 25263593 | Unspecified |
| mmu-miR-21a-5p | MIMAT0000530 | Sgpl1 | 20397 | HITS-CLIP | 23142080 | Unspecified |
| mmu-miR-21a-5p | MIMAT0000530 | Sgpp1 | 81535 | HITS-CLIP | 23597149\|23142080\|19536157 | Unspecified |
| mmu-miR-21a-5p | MIMAT0000530 | Sgsh | 27029 | Microarrays | 37986224 | Unspecified |
| mmu-miR-21a-5p | MIMAT0000530 | Sh2b3 | 16923 | HITS-CLIP | 29167373 | Unspecified |

| | | | | | | |
|---|---|---|---|---|---|---|
| mmu-miR-21a-5p | MIMAT0000530 | Sh2d2a | 27371 | HITS-CLIP | 23142080 | Unspecified |
| mmu-miR-21a-5p | MIMAT0000530 | Sh3bgrl3 | 73723 | HITS-CLIP | 37986224 | Unspecified |
| mmu-miR-21a-5p | MIMAT0000530 | Sh3bp5 | 24056 | HITS-CLIP | 23597149\|23142080 | Unspecified |
| mmu-miR-21a-5p | MIMAT0000530 | Sh3gl3 | 20408 | HITS-CLIP | 23142080 | Unspecified |
| mmu-miR-21a-5p | MIMAT0000530 | Sh3glb1 | 54673 | HITS-CLIP | 23142080\|19536157\|23597149 | Unspecified |
| mmu-miR-21a-5p | MIMAT0000530 | Sh3kbp1 | 58194 | qCLASH//HITS-CLIP | 31393953\|23142080\|29167373 | Unspecified |
| mmu-miR-21a-5p | MIMAT0000530 | Sh3rf1 | 59009 | HITS-CLIP | 23142080 | Unspecified |
| mmu-miR-21a-5p | MIMAT0000530 | Sh3yl1 | 24057 | Microarrays | 37986224\|25263593 | Unspecified |
| mmu-miR-21a-5p | MIMAT0000530 | Shc4 | 271849 | HITS-CLIP | 23142080 | Unspecified |
| mmu-miR-21a-5p | MIMAT0000530 | Shcbp1 | 20419 | Microarrays | 37986224\|25263593 | Unspecified |
| mmu-miR-21a-5p | MIMAT0000530 | Shmt1 | 20425 | HITS-CLIP | 23142080 | Unspecified |
| mmu-miR-21a-5p | MIMAT0000530 | Shmt2 | 108037 | HITS-CLIP | 23142080 | Unspecified |
| mmu-miR-21a-5p | MIMAT0000530 | Shoc2 | 56392 | HITS-CLIP | 23142080 | Unspecified |
| mmu-miR-21a-5p | MIMAT0000530 | Shprh | 268281 | HITS-CLIP | 23142080\|29167373 | Unspecified |
| mmu-miR-21a-5p | MIMAT0000530 | Shtn1 | 57698 | HITS-CLIP | 23142080 | Unspecified |
| mmu-miR-21a-5p | MIMAT0000530 | Siah1a | 20437 | HITS-CLIP | 23142080 | Unspecified |
| mmu-miR-21a-5p | MIMAT0000530 | Siah1b | 20438 | HITS-CLIP | 29167373\|23142080 | Unspecified |
| mmu-miR-21a-5p | MIMAT0000530 | Siah2 | 20439 | HITS-CLIP | 23142080 | Unspecified |
| mmu-miR-21a-5p | MIMAT0000530 | Sidt2 | 214597 | Microarrays//HITS-CLIP | 37986224\|23597149\|23142080 | Unspecified |
| mmu-miR-21a-5p | MIMAT0000530 | Siglecf | 233186 | Microarrays | 25263593 | Unspecified |
| mmu-miR-21a-5p | MIMAT0000530 | Sigmar1 | 18391 | HITS-CLIP | 19536157\|23597149 | Unspecified |
| mmu-miR-21a-5p | MIMAT0000530 | Sik1 | 17691 | HITS-CLIP | 23142080 | Unspecified |
| mmu-miR-21a-5p | MIMAT0000530 | Sik2 | 235344 | HITS-CLIP | 23597149 | Unspecified |
| mmu-miR-21a-5p | MIMAT0000530 | Sil1 | 81500 | HITS-CLIP | 23597149 | Unspecified |
| mmu-miR-21a-5p | MIMAT0000530 | Sin3a | 20466 | HITS-CLIP | 23142080\|29167373 | Unspecified |
| mmu-miR-21a-5p | MIMAT0000530 | Sin3b | 20467 | HITS-CLIP | 23597149\|23142080 | Unspecified |
| mmu-miR-21a-5p | MIMAT0000530 | Sipa1l1 | 217692 | HITS-CLIP | 28735896 | Unspecified |
| mmu-miR-21a-5p | MIMAT0000530 | Sipa1l2 | 244668 | HITS-CLIP | 23142080 | Unspecified |
| mmu-miR-21a-5p | MIMAT0000530 | Sirpa | 19261 | HITS-CLIP | 29167373 | Unspecified |
| mmu-miR-21a-5p | MIMAT0000530 | Sirt1 | 93759 | HITS-CLIP | 37986224\|23142080 | Unspecified |
| mmu-miR-21a-5p | MIMAT0000530 | Sirt3 | 64384 | Microarrays | 37986224 | Unspecified |
| mmu-miR-21a-5p | MIMAT0000530 | Sirt4 | 75387 | Microarrays//HITS-CLIP | 37986224\|23597149 | Unspecified |
| mmu-miR-21a-5p | MIMAT0000530 | Six5 | 20475 | HITS-CLIP | 23142080 | Unspecified |
| mmu-miR-21a-5p | MIMAT0000530 | Ska1 | 66468 | Microarrays | 37986224\|25263593 | Unspecified |
| mmu-miR-21a-5p | MIMAT0000530 | Ska2 | 66140 | Microarrays | 37986224\|25263593 | Unspecified |
| mmu-miR-21a-5p | MIMAT0000530 | Ska3 | 219114 | Microarrays | 37986224\|25263593 | Unspecified |
| mmu-miR-21a-5p | MIMAT0000530 | Skap2 | 54353 | HITS-CLIP | 29167373 | Unspecified |
| mmu-miR-21a-5p | MIMAT0000530 | Ski | 20481 | HITS-CLIP | 37986224\|29167373\|23142080 | Unspecified |
| mmu-miR-21a-5p | MIMAT0000530 | Skida1 | 72668 | HITS-CLIP | 23142080 | Unspecified |

| | | | | | | |
|---|---|---|---|---|---|---|
| mmu-miR-21a-5p | MIMAT0000530 | Skil | 20482 | HITS-CLIP | 23142080\|29167373 | Unspecified |
| mmu-miR-21a-5p | MIMAT0000530 | Skiv2l | 108077 | HITS-CLIP | 23142080 | Unspecified |
| mmu-miR-21a-5p | MIMAT0000530 | Skp1a | 21402 | HITS-CLIP | 23597149\|23142080 | Unspecified |
| mmu-miR-21a-5p | MIMAT0000530 | Skp2 | 27401 | HITS-CLIP, Microarrays//Microarrays//HITS | 37986224\|25263593\|23597149\|23142 | Unspecified |
| mmu-miR-21a-5p | MIMAT0000530 | Sla | 20491 | HITS-CLIP | 23142080 | Unspecified |
| mmu-miR-21a-5p | MIMAT0000530 | Sla2 | 77799 | HITS-CLIP | 23142080 | Unspecified |
| mmu-miR-21a-5p | MIMAT0000530 | Slain2 | 75991 | HITS-CLIP | 37986224\|23142080\|23597149 | Unspecified |
| mmu-miR-21a-5p | MIMAT0000530 | Slamf7 | 75345 | HITS-CLIP | 23142080\|29167373 | Unspecified |
| mmu-miR-21a-5p | MIMAT0000530 | Slamf8 | 74748 | Microarrays | 37986224\|25263593 | Unspecified |
| mmu-miR-21a-5p | MIMAT0000530 | Slamf9 | 98365 | Microarrays | 37986224\|25263593 | Unspecified |
| mmu-miR-21a-5p | MIMAT0000530 | Slc10a2 | 20494 | HITS-CLIP | 23597149 | Unspecified |
| mmu-miR-21a-5p | MIMAT0000530 | Slc11a1 | 18173 | HITS-CLIP | 29167373 | Unspecified |
| mmu-miR-21a-5p | MIMAT0000530 | Slc11a2 | 18174 | HITS-CLIP | 29167373 | Unspecified |
| mmu-miR-21a-5p | MIMAT0000530 | Slc12a2 | 20496 | HITS-CLIP | 37986224\|23142080 | Unspecified |
| mmu-miR-21a-5p | MIMAT0000530 | Slc12a4 | 20498 | HITS-CLIP | 23597149 | Unspecified |
| mmu-miR-21a-5p | MIMAT0000530 | Slc12a6 | 107723 | HITS-CLIP | 37986224\|29167373\|23142080 | Unspecified |
| mmu-miR-21a-5p | MIMAT0000530 | Slc12a9 | 83704 | Microarrays | 37986224 | Unspecified |
| mmu-miR-21a-5p | MIMAT0000530 | Slc13a3 | 114644 | HITS-CLIP | 23142080 | Unspecified |
| mmu-miR-21a-5p | MIMAT0000530 | Slc14a1 | 108052 | Microarrays | 37986224 | Unspecified |
| mmu-miR-21a-5p | MIMAT0000530 | Slc15a2 | 57738 | Microarrays | 37986224 | Unspecified |
| mmu-miR-21a-5p | MIMAT0000530 | Slc15a3 | 65221 | Microarrays | 37986224\|25263593 | Unspecified |
| mmu-miR-21a-5p | MIMAT0000530 | Slc16a1 | 20501 | HITS-CLIP | 19536157 | Unspecified |
| mmu-miR-21a-5p | MIMAT0000530 | Slc16a10 | 72472 | HITS-CLIP | 37986224\|23597149 | Unspecified |
| mmu-miR-21a-5p | MIMAT0000530 | Slc16a12 | 240638 | HITS-CLIP | 23597149 | Unspecified |
| mmu-miR-21a-5p | MIMAT0000530 | Slc16a2 | 20502 | HITS-CLIP | 23597149 | Unspecified |
| mmu-miR-21a-5p | MIMAT0000530 | Slc16a3 | 80879 | HITS-CLIP | 23142080\|29167373 | Unspecified |
| mmu-miR-21a-5p | MIMAT0000530 | Slc16a6 | 104681 | qCLASH | 31393953 | Unspecified |
| mmu-miR-21a-5p | MIMAT0000530 | Slc16a7 | 20503 | Microarrays | 37986224 | Unspecified |
| mmu-miR-21a-5p | MIMAT0000530 | Slc17a6 | 140919 | HITS-CLIP | 37986224 | Unspecified |
| mmu-miR-21a-5p | MIMAT0000530 | Slc19a2 | 116914 | HITS-CLIP | 23597149\|26203562\|23142080 | Unspecified |
| mmu-miR-21a-5p | MIMAT0000530 | Slc1a2 | 20511 | HITS-CLIP | 37986224\|19536157\|23597149 | Unspecified |
| mmu-miR-21a-5p | MIMAT0000530 | Slc1a3 | 20512 | HITS-CLIP | 37986224\|19536157 | Unspecified |
| mmu-miR-21a-5p | MIMAT0000530 | Slc1a4 | 55963 | Microarrays | 37986224\|25263593 | Unspecified |
| mmu-miR-21a-5p | MIMAT0000530 | Slc20a1 | 20515 | HITS-CLIP | 23597149\|19536157\|23142080\|29167 | Unspecified |
| mmu-miR-21a-5p | MIMAT0000530 | Slc22a1 | 20517 | HITS-CLIP | 23597149 | Unspecified |
| mmu-miR-21a-5p | MIMAT0000530 | Slc22a4 | 30805 | Microarrays | 37986224\|25263593 | Unspecified |
| mmu-miR-21a-5p | MIMAT0000530 | Slc23a2 | 54338 | HITS-CLIP | 23142080\|29167373 | Unspecified |
| mmu-miR-21a-5p | MIMAT0000530 | Slc24a3 | 94249 | Microarrays//qCLASH | 37986224\|25263593\|31393953 | Unspecified |
| mmu-miR-21a-5p | MIMAT0000530 | Slc25a12 | 78830 | HITS-CLIP | 23142080\|23597149 | Unspecified |

| | | | | | | |
|---|---|---|---|---|---|---|
| mmu-miR-21a-5p | MIMAT0000530 | Slc25a13 | 50799 | Microarrays//HITS-CLIP | 37986224\|25263593\|23597149 | Unspecified |
| mmu-miR-21a-5p | MIMAT0000530 | Slc25a16 | 73132 | HITS-CLIP | 23142080 | Unspecified |
| mmu-miR-21a-5p | MIMAT0000530 | Slc25a17 | 20524 | HITS-CLIP | 23597149\|23142080 | Unspecified |
| mmu-miR-21a-5p | MIMAT0000530 | Slc25a19 | 67283 | HITS-CLIP | 31433990\|29167373 | Unspecified |
| mmu-miR-21a-5p | MIMAT0000530 | Slc25a20 | 57279 | qCLASH | 31393953 | Unspecified |
| mmu-miR-21a-5p | MIMAT0000530 | Slc25a22 | 68267 | HITS-CLIP | 23142080 | Unspecified |
| mmu-miR-21a-5p | MIMAT0000530 | Slc25a24 | 229731 | HITS-CLIP | 23142080 | Unspecified |
| mmu-miR-21a-5p | MIMAT0000530 | Slc25a25 | 227731 | HITS-CLIP | 37986224\|23597149\|29167373\|2314[illegible] | Unspecified |
| mmu-miR-21a-5p | MIMAT0000530 | Slc25a3 | 18674 | HITS-CLIP | 23142080 | Unspecified |
| mmu-miR-21a-5p | MIMAT0000530 | Slc25a30 | 67554 | HITS-CLIP | 23597149\|23142080 | Unspecified |
| mmu-miR-21a-5p | MIMAT0000530 | Slc25a33 | 70556 | Microarrays | 37986224\|25263593 | Unspecified |
| mmu-miR-21a-5p | MIMAT0000530 | Slc25a36 | 192287 | HITS-CLIP | 37986224\|23142080 | Unspecified |
| mmu-miR-21a-5p | MIMAT0000530 | Slc25a37 | 67712 | Microarrays | 37986224 | Unspecified |
| mmu-miR-21a-5p | MIMAT0000530 | Slc25a39 | 68066 | HITS-CLIP | 23597149 | Unspecified |
| mmu-miR-21a-5p | MIMAT0000530 | Slc25a44 | 229517 | HITS-CLIP | 23597149\|29167373\|23142080 | Unspecified |
| mmu-miR-21a-5p | MIMAT0000530 | Slc25a46 | 67453 | HITS-CLIP | 37986224\|23597149\|23142080 | Unspecified |
| mmu-miR-21a-5p | MIMAT0000530 | Slc25a47 | 104910 | HITS-CLIP | 23597149 | Unspecified |
| mmu-miR-21a-5p | MIMAT0000530 | Slc25a48 | 328258 | HITS-CLIP | 26203562 | Unspecified |
| mmu-miR-21a-5p | MIMAT0000530 | Slc25a5 | 11740 | qCLASH | 31393953 | Unspecified |
| mmu-miR-21a-5p | MIMAT0000530 | Slc25a51 | 230125 | HITS-CLIP | 37986224\|23597149\|23142080\|2916[illegible] | Unspecified |
| mmu-miR-21a-5p | MIMAT0000530 | Slc26a11 | 268512 | Microarrays | 37986224 | Unspecified |
| mmu-miR-21a-5p | MIMAT0000530 | Slc26a2 | 13521 | HITS-CLIP | 23142080 | Unspecified |
| mmu-miR-21a-5p | MIMAT0000530 | Slc27a2 | 26458 | HITS-CLIP | 23597149 | Unspecified |
| mmu-miR-21a-5p | MIMAT0000530 | Slc29a3 | 71279 | HITS-CLIP | 37986224\|23142080\|29167373 | Unspecified |
| mmu-miR-21a-5p | MIMAT0000530 | Slc2a1 | 20525 | HITS-CLIP, Microarrays//Microarrays | 37986224\|25263593 | Unspecified |
| mmu-miR-21a-5p | MIMAT0000530 | Slc2a2 | 20526 | HITS-CLIP | 23597149 | Unspecified |
| mmu-miR-21a-5p | MIMAT0000530 | Slc2a3 | 20527 | HITS-CLIP | 37986224\|23142080 | Unspecified |
| mmu-miR-21a-5p | MIMAT0000530 | Slc30a1 | 22782 | qCLASH//HITS-CLIP | 31393953\|23142080 | Unspecified |
| mmu-miR-21a-5p | MIMAT0000530 | Slc30a10 | 226781 | HITS-CLIP | 37986224 | Unspecified |
| mmu-miR-21a-5p | MIMAT0000530 | Slc30a4 | 22785 | HITS-CLIP | 23142080\|19536157 | Unspecified |
| mmu-miR-21a-5p | MIMAT0000530 | Slc30a6 | 210148 | HITS-CLIP | 23142080 | Unspecified |
| mmu-miR-21a-5p | MIMAT0000530 | Slc30a7 | 66500 | HITS-CLIP | 23142080\|29167373 | Unspecified |
| mmu-miR-21a-5p | MIMAT0000530 | Slc30a9 | 109108 | HITS-CLIP | 23142080 | Unspecified |
| mmu-miR-21a-5p | MIMAT0000530 | Slc31a1 | 20529 | HITS-CLIP | 23597149\|19536157\|29167373 | Unspecified |
| mmu-miR-21a-5p | MIMAT0000530 | Slc33a1 | 11416 | HITS-CLIP | 23142080 | Unspecified |
| mmu-miR-21a-5p | MIMAT0000530 | Slc35a1 | 24060 | HITS-CLIP | 23142080 | Unspecified |
| mmu-miR-21a-5p | MIMAT0000530 | Slc35a3 | 229782 | HITS-CLIP | 23142080\|29167373 | Unspecified |
| mmu-miR-21a-5p | MIMAT0000530 | Slc35a5 | 74102 | HITS-CLIP | 37986224\|23142080\|29167373 | Unspecified |
| mmu-miR-21a-5p | MIMAT0000530 | Slc35b4 | 58246 | HITS-CLIP | 29167373 | Unspecified |

| | | | | | | |
|---|---|---|---|---|---|---|
| mmu-miR-21a-5p | MIMAT0000530 | Slc35d2 | 70484 | HITS-CLIP | 23597149 | Unspecified |
| mmu-miR-21a-5p | MIMAT0000530 | Slc35d3 | 76157 | HITS-CLIP | 37986224 | Unspecified |
| mmu-miR-21a-5p | MIMAT0000530 | Slc36a1 | 215335 | HITS-CLIP | 23597149\|29167373 | Unspecified |
| mmu-miR-21a-5p | MIMAT0000530 | Slc36a2 | 246049 | Microarrays | 37986224\|25263593 | Unspecified |
| mmu-miR-21a-5p | MIMAT0000530 | Slc36a4 | 234967 | HITS-CLIP | 37986224\|23142080 | Unspecified |
| mmu-miR-21a-5p | MIMAT0000530 | Slc37a2 | 56857 | HITS-CLIP | 29167373 | Unspecified |
| mmu-miR-21a-5p | MIMAT0000530 | Slc37a3 | 72144 | HITS-CLIP | 23142080\|29167373 | Unspecified |
| mmu-miR-21a-5p | MIMAT0000530 | Slc38a1 | 105727 | Microarrays//HITS-CLIP | 37986224\|25263593\|23142080 | Unspecified |
| mmu-miR-21a-5p | MIMAT0000530 | Slc38a2 | 67760 | HITS-CLIP | 37986224\|23597149\|23142080\|29167 | Unspecified |
| mmu-miR-21a-5p | MIMAT0000530 | Slc38a4 | 69354 | HITS-CLIP | 37986224\|23597149 | Unspecified |
| mmu-miR-21a-5p | MIMAT0000530 | Slc39a14 | 213053 | HITS-CLIP | 37986224\|23597149 | Unspecified |
| mmu-miR-21a-5p | MIMAT0000530 | Slc39a3 | 106947 | HITS-CLIP | 23597149\|23142080 | Unspecified |
| mmu-miR-21a-5p | MIMAT0000530 | Slc39a7 | 14977 | HITS-CLIP | 23597149\|23142080 | Unspecified |
| mmu-miR-21a-5p | MIMAT0000530 | Slc39a8 | 67547 | Microarrays | 37986224\|25263593 | Unspecified |
| mmu-miR-21a-5p | MIMAT0000530 | Slc39a9 | 328133 | HITS-CLIP | 23142080 | Unspecified |
| mmu-miR-21a-5p | MIMAT0000530 | Slc3a2 | 17254 | HITS-CLIP | 23597149 | Unspecified |
| mmu-miR-21a-5p | MIMAT0000530 | Slc41a1 | 98396 | HITS-CLIP | 23142080 | Unspecified |
| mmu-miR-21a-5p | MIMAT0000530 | Slc41a2 | 338365 | HITS-CLIP | 23597149 | Unspecified |
| mmu-miR-21a-5p | MIMAT0000530 | Slc43a2 | 215113 | HITS-CLIP | 29167373 | Unspecified |
| mmu-miR-21a-5p | MIMAT0000530 | Slc43a3 | 58207 | Microarrays | 37986224\|25263593 | Unspecified |
| mmu-miR-21a-5p | MIMAT0000530 | Slc45a3 | 212980 | Microarrays | 37986224\|25263593 | Unspecified |
| mmu-miR-21a-5p | MIMAT0000530 | Slc45a4 | 106068 | HITS-CLIP | 23142080\|23597149 | Unspecified |
| mmu-miR-21a-5p | MIMAT0000530 | Slc46a1 | 52466 | HITS-CLIP | 23597149 | Unspecified |
| mmu-miR-21a-5p | MIMAT0000530 | Slc46a3 | 71706 | Microarrays//HITS-CLIP | 37986224\|23597149 | Unspecified |
| mmu-miR-21a-5p | MIMAT0000530 | Slc47a1 | 67473 | HITS-CLIP | 23597149 | Unspecified |
| mmu-miR-21a-5p | MIMAT0000530 | Slc4a2 | 20535 | HITS-CLIP | 23142080\|29167373 | Unspecified |
| mmu-miR-21a-5p | MIMAT0000530 | Slc4a4 | 54403 | HITS-CLIP | 37986224 | Unspecified |
| mmu-miR-21a-5p | MIMAT0000530 | Slc4a7 | 218756 | Microarrays//HITS-CLIP | 37986224\|29167373\|23142080 | Unspecified |
| mmu-miR-21a-5p | MIMAT0000530 | Slc5a3 | 53881 | HITS-CLIP | 23142080 | Unspecified |
| mmu-miR-21a-5p | MIMAT0000530 | Slc5a6 | 330064 | HITS-CLIP | 37986224\|23597149\|29167373 | Unspecified |
| mmu-miR-21a-5p | MIMAT0000530 | Slc6a1 | 232333 | HITS-CLIP | 37986224 | Unspecified |
| mmu-miR-21a-5p | MIMAT0000530 | Slc6a11 | 243616 | HITS-CLIP | 19536157 | Unspecified |
| mmu-miR-21a-5p | MIMAT0000530 | Slc6a6 | 21366 | HITS-CLIP | 37986224\|23597149\|19536157\|29167 | Unspecified |
| mmu-miR-21a-5p | MIMAT0000530 | Slc6a8 | 102857 | Microarrays//HITS-CLIP | 37986224\|25263593\|29167373 | Unspecified |
| mmu-miR-21a-5p | MIMAT0000530 | Slc7a1 | 11987 | Microarrays//HITS-CLIP | 37986224\|25263593\|23142080 | Unspecified |
| mmu-miR-21a-5p | MIMAT0000530 | Slc7a11 | 26570 | HITS-CLIP, Microarrays//Microarrays | 37986224\|25263593 | Unspecified |
| mmu-miR-21a-5p | MIMAT0000530 | Slc7a2 | 11988 | Microarrays//HITS-CLIP | 37986224\|25263593\|23597149 | Unspecified |
| mmu-miR-21a-5p | MIMAT0000530 | Slc7a5 | 20539 | Microarrays//HITS-CLIP | 37986224\|25263593\|23142080 | Unspecified |
| mmu-miR-21a-5p | MIMAT0000530 | Slc7a6 | 330836 | Microarrays | 37986224\|25263593 | Unspecified |

| | | | | | | |
|---|---|---|---|---|---|---|
| mmu-miR-21a-5p | MIMAT0000530 | Slc7a8 | 50934 | Microarrays | 37986224\|25263593 | Unspecified |
| mmu-miR-21a-5p | MIMAT0000530 | Slc9a1 | 20544 | qCLASH | 31393953 | Unspecified |
| mmu-miR-21a-5p | MIMAT0000530 | Slc9a3r1 | 26941 | Microarrays//HITS-CLIP | 37986224\|23597149\|23142080 | Unspecified |
| mmu-miR-21a-5p | MIMAT0000530 | Slc9a6 | 236794 | HITS-CLIP | 37986224\|19536157\|23142080\|29167 | Unspecified |
| mmu-miR-21a-5p | MIMAT0000530 | Slc9a9 | 331004 | Microarrays | 37986224 | Unspecified |
| mmu-miR-21a-5p | MIMAT0000530 | Slc9b2 | 97086 | Microarrays | 37986224\|25263593 | Unspecified |
| mmu-miR-21a-5p | MIMAT0000530 | Slco1a1 | 28248 | HITS-CLIP | 23597149 | Unspecified |
| mmu-miR-21a-5p | MIMAT0000530 | Slco1a4 | 28250 | HITS-CLIP | 23597149 | Unspecified |
| mmu-miR-21a-5p | MIMAT0000530 | Slco1b2 | 28253 | HITS-CLIP | 23597149 | Unspecified |
| mmu-miR-21a-5p | MIMAT0000530 | Slco1c1 | 58807 | HITS-CLIP | 37986224 | Unspecified |
| mmu-miR-21a-5p | MIMAT0000530 | Slco2b1 | 101488 | Microarrays | 37986224 | Unspecified |
| mmu-miR-21a-5p | MIMAT0000530 | Slco3a1 | 108116 | Microarrays | 37986224\|25263593 | Unspecified |
| mmu-miR-21a-5p | MIMAT0000530 | Slf2 | 226151 | HITS-CLIP | 23142080 | Unspecified |
| mmu-miR-21a-5p | MIMAT0000530 | Slfn2 | 20556 | HITS-CLIP | 23142080 | Unspecified |
| mmu-miR-21a-5p | MIMAT0000530 | Slfn5 | 327978 | Microarrays//HITS-CLIP | 37986224\|29167373 | Unspecified |
| mmu-miR-21a-5p | MIMAT0000530 | Slfn8 | 276950 | Microarrays | 37986224 | Unspecified |
| mmu-miR-21a-5p | MIMAT0000530 | Slfn9 | 237886 | HITS-CLIP | 23142080 | Unspecified |
| mmu-miR-21a-5p | MIMAT0000530 | Slirp | 380773 | HITS-CLIP | 23142080 | Unspecified |
| mmu-miR-21a-5p | MIMAT0000530 | Slitrk1 | 76965 | HITS-CLIP | 37986224\|19536157 | Unspecified |
| mmu-miR-21a-5p | MIMAT0000530 | Slitrk4 | 245446 | HITS-CLIP | 37986224 | Unspecified |
| mmu-miR-21a-5p | MIMAT0000530 | Slk | 20874 | HITS-CLIP | 23597149\|23142080\|29167373 | Unspecified |
| mmu-miR-21a-5p | MIMAT0000530 | Slmap | 83997 | HITS-CLIP | 23597149\|23142080 | Unspecified |
| mmu-miR-21a-5p | MIMAT0000530 | Slpi | 20568 | Microarrays//HITS-CLIP | 37986224\|25263593\|23142080 | Unspecified |
| mmu-miR-21a-5p | MIMAT0000530 | Sltm | 66660 | HITS-CLIP | 23142080 | Unspecified |
| mmu-miR-21a-5p | MIMAT0000530 | Slx4 | 52864 | HITS-CLIP | 31433990\|23597149 | Unspecified |
| mmu-miR-21a-5p | MIMAT0000530 | Smad1 | 17125 | HITS-CLIP | 23597149 | Unspecified |
| mmu-miR-21a-5p | MIMAT0000530 | Smad2 | 17126 | HITS-CLIP | 23142080 | Unspecified |
| mmu-miR-21a-5p | MIMAT0000530 | Smad3 | 17127 | HITS-CLIP | 31433990\|23142080 | Unspecified |
| mmu-miR-21a-5p | MIMAT0000530 | Smad4 | 17128 | HITS-CLIP | 23142080\|29167373 | Unspecified |
| mmu-miR-21a-5p | MIMAT0000530 | Smad6 | 17130 | Microarrays | 37986224\|25263593 | Unspecified |
| mmu-miR-21a-5p | MIMAT0000530 | Smad7 | 17131 | Luciferase assay//Northern blot//QRTPCR/ | 23784029\|23597149\|25263593\|23142 | Unspecified |
| mmu-miR-21a-5p | MIMAT0000530 | Smarca2 | 67155 | HITS-CLIP | 23142080 | Unspecified |
| mmu-miR-21a-5p | MIMAT0000530 | Smarca5 | 93762 | HITS-CLIP | 23142080 | Unspecified |
| mmu-miR-21a-5p | MIMAT0000530 | Smarcad1 | 13990 | HITS-CLIP | 23142080 | Unspecified |
| mmu-miR-21a-5p | MIMAT0000530 | Smarcal1 | 54380 | HITS-CLIP | 23142080 | Unspecified |
| mmu-miR-21a-5p | MIMAT0000530 | Smarcb1 | 20587 | HITS-CLIP | 23597149 | Unspecified |
| mmu-miR-21a-5p | MIMAT0000530 | Smarcc1 | 20588 | qCLASH//HITS-CLIP | 31393953\|23597149 | Unspecified |
| mmu-miR-21a-5p | MIMAT0000530 | Smarcd2 | 83796 | HITS-CLIP | 23142080 | Unspecified |
| mmu-miR-21a-5p | MIMAT0000530 | Smc1a | 24061 | HITS-CLIP | 23142080 | Unspecified |

| | | | | | | |
|---|---|---|---|---|---|---|
| mmu-miR-21a-5p | MIMAT0000530 | Smc2 | 14211 | Microarrays//HITS-CLIP | 37986224\|25263593\|23142080\|29167 | Unspecified |
| mmu-miR-21a-5p | MIMAT0000530 | Smc3 | 13006 | HITS-CLIP//qCLASH | 37986224\|31393953\|23142080\|29167 | Unspecified |
| mmu-miR-21a-5p | MIMAT0000530 | Smc4 | 70099 | HITS-CLIP | 23597149\|23142080\|29167373 | Unspecified |
| mmu-miR-21a-5p | MIMAT0000530 | Smc5 | 226026 | HITS-CLIP | 37986224\|23597149\|23142080 | Unspecified |
| mmu-miR-21a-5p | MIMAT0000530 | Smc6 | 67241 | HITS-CLIP | 37986224\|23142080\|31433990 | Unspecified |
| mmu-miR-21a-5p | MIMAT0000530 | Smchd1 | 74355 | HITS-CLIP | 23142080\|29167373 | Unspecified |
| mmu-miR-21a-5p | MIMAT0000530 | Smco4 | 170748 | Microarrays | 37986224\|25263593 | Unspecified |
| mmu-miR-21a-5p | MIMAT0000530 | Smcr8 | 237782 | HITS-CLIP | 37986224\|23142080 | Unspecified |
| mmu-miR-21a-5p | MIMAT0000530 | Smg1 | 233789 | HITS-CLIP | 37986224\|23597149\|19536157\|29167 | Unspecified |
| mmu-miR-21a-5p | MIMAT0000530 | Smg7 | 226517 | HITS-CLIP | 29167373\|23142080 | Unspecified |
| mmu-miR-21a-5p | MIMAT0000530 | Smim14 | 68552 | HITS-CLIP | 23142080\|29167373 | Unspecified |
| mmu-miR-21a-5p | MIMAT0000530 | Smim15 | 75616 | HITS-CLIP | 23142080 | Unspecified |
| mmu-miR-21a-5p | MIMAT0000530 | Smim4 | 66487 | HITS-CLIP | 37986224\|23142080 | Unspecified |
| mmu-miR-21a-5p | MIMAT0000530 | Smim5 | 66528 | Microarrays | 37986224\|25263593 | Unspecified |
| mmu-miR-21a-5p | MIMAT0000530 | Smim7 | 66818 | HITS-CLIP | 23142080 | Unspecified |
| mmu-miR-21a-5p | MIMAT0000530 | Smim8 | 66291 | HITS-CLIP | 23142080 | Unspecified |
| mmu-miR-21a-5p | MIMAT0000530 | Smn1 | 20595 | HITS-CLIP | 23142080 | Unspecified |
| mmu-miR-21a-5p | MIMAT0000530 | Smndc1 | 76479 | HITS-CLIP | 23142080 | Unspecified |
| mmu-miR-21a-5p | MIMAT0000530 | Smox | 228608 | HITS-CLIP | 29167373 | Unspecified |
| mmu-miR-21a-5p | MIMAT0000530 | Smpd1 | 20597 | HITS-CLIP | 23597149 | Unspecified |
| mmu-miR-21a-5p | MIMAT0000530 | Smpd4 | 77626 | HITS-CLIP | 23142080 | Unspecified |
| mmu-miR-21a-5p | MIMAT0000530 | Smpdl3a | 57319 | HITS-CLIP | 37986224 | Unspecified |
| mmu-miR-21a-5p | MIMAT0000530 | Smpdl3b | 100340 | Microarrays | 37986224\|25263593 | Unspecified |
| mmu-miR-21a-5p | MIMAT0000530 | Smtn | 29856 | Microarrays | 37986224\|25263593 | Unspecified |
| mmu-miR-21a-5p | MIMAT0000530 | Smtnl2 | 276829 | HITS-CLIP | 23142080 | Unspecified |
| mmu-miR-21a-5p | MIMAT0000530 | Smug1 | 71726 | HITS-CLIP | 23142080 | Unspecified |
| mmu-miR-21a-5p | MIMAT0000530 | Smurf1 | 75788 | HITS-CLIP | 23142080 | Unspecified |
| mmu-miR-21a-5p | MIMAT0000530 | Smurf2 | 66313 | HITS-CLIP | 31433990\|23142080\|29167373 | Unspecified |
| mmu-miR-21a-5p | MIMAT0000530 | Smyd2 | 226830 | HITS-CLIP | 23597149\|23142080 | Unspecified |
| mmu-miR-21a-5p | MIMAT0000530 | Smyd3 | 69726 | HITS-CLIP | 23142080 | Unspecified |
| mmu-miR-21a-5p | MIMAT0000530 | Snap29 | 67474 | HITS-CLIP | 23142080\|28735896 | Unspecified |
| mmu-miR-21a-5p | MIMAT0000530 | Snapc1 | 75627 | Microarrays | 37986224\|25263593 | Unspecified |
| mmu-miR-21a-5p | MIMAT0000530 | Snapin | 20615 | HITS-CLIP | 23142080 | Unspecified |
| mmu-miR-21a-5p | MIMAT0000530 | Sncaip | 67847 | Microarrays | 37986224\|25263593 | Unspecified |
| mmu-miR-21a-5p | MIMAT0000530 | Snd1 | 56463 | HITS-CLIP | 37986224\|23142080\|23597149\|3143 | Unspecified |
| mmu-miR-21a-5p | MIMAT0000530 | Snhg11 | 319317 | HITS-CLIP | 37986224\|19536157 | Unspecified |
| mmu-miR-21a-5p | MIMAT0000530 | Snn | 20621 | HITS-CLIP | 37986224\|19536157\|23142080 | Unspecified |
| mmu-miR-21a-5p | MIMAT0000530 | Snrnp200 | 320632 | qCLASH//HITS-CLIP | 31393953\|23597149\|23142080\|29167 | Unspecified |
| mmu-miR-21a-5p | MIMAT0000530 | Snrnp40 | 66585 | HITS-CLIP | 23142080 | Unspecified |

| | | | | | | |
|---|---|---|---|---|---|---|
| mmu-miR-21a-5p | MIMAT0000530 | Snrnp70 | 20637 | HITS-CLIP | 23142080 | Unspecified |
| mmu-miR-21a-5p | MIMAT0000530 | Snrpa1 | 68981 | Microarrays//HITS-CLIP | 37986224\|25263593\|23142080 | Unspecified |
| mmu-miR-21a-5p | MIMAT0000530 | Snrpb2 | 20639 | Microarrays//HITS-CLIP | 37986224\|23142080 | Unspecified |
| mmu-miR-21a-5p | MIMAT0000530 | Snrpd1 | 20641 | HITS-CLIP | 23142080 | Unspecified |
| mmu-miR-21a-5p | MIMAT0000530 | Snrpd3 | 67332 | HITS-CLIP | 37986224\|23597149 | Unspecified |
| mmu-miR-21a-5p | MIMAT0000530 | Sntb2 | 20650 | HITS-CLIP | 23597149\|29167373 | Unspecified |
| mmu-miR-21a-5p | MIMAT0000530 | Snx11 | 74479 | HITS-CLIP | 23142080 | Unspecified |
| mmu-miR-21a-5p | MIMAT0000530 | Snx13 | 217463 | HITS-CLIP | 23597149\|23142080\|29167373 | Unspecified |
| mmu-miR-21a-5p | MIMAT0000530 | Snx14 | 244962 | HITS-CLIP | 23142080 | Unspecified |
| mmu-miR-21a-5p | MIMAT0000530 | Snx16 | 74718 | HITS-CLIP | 23142080 | Unspecified |
| mmu-miR-21a-5p | MIMAT0000530 | Snx17 | 266781 | HITS-CLIP | 31433990\|23142080 | Unspecified |
| mmu-miR-21a-5p | MIMAT0000530 | Snx18 | 170625 | HITS-CLIP | 23142080 | Unspecified |
| mmu-miR-21a-5p | MIMAT0000530 | Snx27 | 76742 | qCLASH//HITS-CLIP | 31393953\|23142080 | Unspecified |
| mmu-miR-21a-5p | MIMAT0000530 | Snx3 | 54198 | HITS-CLIP | 23142080\|29167373 | Unspecified |
| mmu-miR-21a-5p | MIMAT0000530 | Snx30 | 209131 | HITS-CLIP | 23142080 | Unspecified |
| mmu-miR-21a-5p | MIMAT0000530 | Snx7 | 76561 | Microarrays//HITS-CLIP | 37986224\|25263593\|23597149 | Unspecified |
| mmu-miR-21a-5p | MIMAT0000530 | Snx8 | 231834 | HITS-CLIP | 29167373 | Unspecified |
| mmu-miR-21a-5p | MIMAT0000530 | Snx9 | 66616 | HITS-CLIP | 23597149\|23142080 | Unspecified |
| mmu-miR-21a-5p | MIMAT0000530 | Soat1 | 20652 | HITS-CLIP | 23142080 | Unspecified |
| mmu-miR-21a-5p | MIMAT0000530 | Socs2 | 216233 | HITS-CLIP | 23597149\|23142080 | Unspecified |
| mmu-miR-21a-5p | MIMAT0000530 | Socs3 | 12702 | Microarrays//HITS-CLIP | 37986224\|23597149\|23142080 | Unspecified |
| mmu-miR-21a-5p | MIMAT0000530 | Socs5 | 56468 | HITS-CLIP | 37986224\|31433990\|29167373\|2314 | Unspecified |
| mmu-miR-21a-5p | MIMAT0000530 | Socs6 | 54607 | Microarrays//HITS-CLIP | 37986224\|23142080 | Unspecified |
| mmu-miR-21a-5p | MIMAT0000530 | Socs7 | 192157 | HITS-CLIP | 37986224\|23597149\|23142080 | Unspecified |
| mmu-miR-21a-5p | MIMAT0000530 | Sod2 | 20656 | HITS-CLIP//ChIP-seq//qRT-PCR//Western b | 23597149\|23142080\|26222667 | Unspecified |
| mmu-miR-21a-5p | MIMAT0000530 | Son | 20658 | HITS-CLIP | 23597149\|23142080\|29167373 | Unspecified |
| mmu-miR-21a-5p | MIMAT0000530 | Sorbs3 | 20410 | HITS-CLIP | 23597149 | Unspecified |
| mmu-miR-21a-5p | MIMAT0000530 | Sorl1 | 20660 | Microarrays//HITS-CLIP | 37986224\|19536157\|23142080 | Unspecified |
| mmu-miR-21a-5p | MIMAT0000530 | Sort1 | 20661 | HITS-CLIP | 23597149 | Unspecified |
| mmu-miR-21a-5p | MIMAT0000530 | Sos1 | 20662 | HITS-CLIP | 37986224\|23142080\|29167373 | Unspecified |
| mmu-miR-21a-5p | MIMAT0000530 | Sos2 | 20663 | HITS-CLIP | 37986224\|23142080\|29167373 | Unspecified |
| mmu-miR-21a-5p | MIMAT0000530 | Sowahc | 268301 | HITS-CLIP | 23597149\|29167373 | Unspecified |
| mmu-miR-21a-5p | MIMAT0000530 | Sox5 | 20678 | HITS-CLIP//Luciferase reporter assay//qRT- | 37986224\|23597149\|23142080\|27347 | Unspecified |
| mmu-miR-21a-5p | MIMAT0000530 | Sox7 | 20680 | Microarrays | 37986224\|25263593 | Unspecified |
| mmu-miR-21a-5p | MIMAT0000530 | Sp1 | 20683 | HITS-CLIP | 23142080 | Unspecified |
| mmu-miR-21a-5p | MIMAT0000530 | Sp100 | 20684 | HITS-CLIP | 23142080 | Unspecified |
| mmu-miR-21a-5p | MIMAT0000530 | Sp140 | 434484 | HITS-CLIP | 23142080 | Unspecified |
| mmu-miR-21a-5p | MIMAT0000530 | Sp3 | 20687 | HITS-CLIP | 37986224\|19536157\|23142080\|29167 | Unspecified |
| mmu-miR-21a-5p | MIMAT0000530 | Spaca6 | 75202 | HITS-CLIP | 29167373 | Unspecified |

| | | | | | | |
|---|---|---|---|---|---|---|
| mmu-miR-21a-5p | MIMAT0000530 | Spag5 | 54141 | Microarrays | 37986224\|25263593 | Unspecified |
| mmu-miR-21a-5p | MIMAT0000530 | Spag7 | 216873 | HITS-CLIP | 23142080 | Unspecified |
| mmu-miR-21a-5p | MIMAT0000530 | Spag9 | 70834 | HITS-CLIP//qCLASH | 37986224\|31393953\|19536157\|23142 | Unspecified |
| mmu-miR-21a-5p | MIMAT0000530 | Sparc | 20692 | Microarrays//HITS-CLIP | 37986224\|23597149\|25263593 | Unspecified |
| mmu-miR-21a-5p | MIMAT0000530 | Sparcl1 | 13602 | HITS-CLIP | 37986224 | Unspecified |
| mmu-miR-21a-5p | MIMAT0000530 | Spast | 50850 | HITS-CLIP | 37986224\|23142080 | Unspecified |
| mmu-miR-21a-5p | MIMAT0000530 | Spata13 | 219140 | HITS-CLIP | 23597149\|23142080 | Unspecified |
| mmu-miR-21a-5p | MIMAT0000530 | Spata2 | 263876 | HITS-CLIP | 23597149 | Unspecified |
| mmu-miR-21a-5p | MIMAT0000530 | Spata5 | 57815 | HITS-CLIP | 23142080 | Unspecified |
| mmu-miR-21a-5p | MIMAT0000530 | Spata6 | 67946 | HITS-CLIP | 23142080 | Unspecified |
| mmu-miR-21a-5p | MIMAT0000530 | Spc25 | 66442 | Microarrays//HITS-CLIP | 37986224\|25263593\|23142080 | Unspecified |
| mmu-miR-21a-5p | MIMAT0000530 | Spcs1 | 69019 | HITS-CLIP | 23142080 | Unspecified |
| mmu-miR-21a-5p | MIMAT0000530 | Spcs3 | 76687 | HITS-CLIP | 23142080 | Unspecified |
| mmu-miR-21a-5p | MIMAT0000530 | Spdl1 | 70385 | Microarrays//HITS-CLIP | 37986224\|25263593\|23142080 | Unspecified |
| mmu-miR-21a-5p | MIMAT0000530 | Specc1 | 432572 | HITS-CLIP | 29167373\|23142080 | Unspecified |
| mmu-miR-21a-5p | MIMAT0000530 | Specc1l | 74392 | Microarrays | 37986224 | Unspecified |
| mmu-miR-21a-5p | MIMAT0000530 | Speg | 11790 | Microarrays | 37986224\|25263593 | Unspecified |
| mmu-miR-21a-5p | MIMAT0000530 | Spen | 56381 | HITS-CLIP | 29167373 | Unspecified |
| mmu-miR-21a-5p | MIMAT0000530 | Spg11 | 214585 | HITS-CLIP//qCLASH | 23597149\|31393953 | Unspecified |
| mmu-miR-21a-5p | MIMAT0000530 | Spg20 | 229285 | HITS-CLIP | 37986224\|19536157\|23597149\|23142 | Unspecified |
| mmu-miR-21a-5p | MIMAT0000530 | Spic | 20728 | Microarrays | 37986224 | Unspecified |
| mmu-miR-21a-5p | MIMAT0000530 | Spidr | 224008 | HITS-CLIP | 23142080 | Unspecified |
| mmu-miR-21a-5p | MIMAT0000530 | Spin1 | 20729 | HITS-CLIP | 37986224\|23142080 | Unspecified |
| mmu-miR-21a-5p | MIMAT0000530 | Spin4 | 270624 | HITS-CLIP | 23142080 | Unspecified |
| mmu-miR-21a-5p | MIMAT0000530 | Spn | 20737 | HITS-CLIP | 23142080 | Unspecified |
| mmu-miR-21a-5p | MIMAT0000530 | Spock2 | 94214 | HITS-CLIP | 23142080 | Unspecified |
| mmu-miR-21a-5p | MIMAT0000530 | Spopl | 76857 | HITS-CLIP | 23142080 | Unspecified |
| mmu-miR-21a-5p | MIMAT0000530 | Sppl2a | 66552 | HITS-CLIP | 23142080\|29167373 | Unspecified |
| mmu-miR-21a-5p | MIMAT0000530 | Sppl2b | 73218 | HITS-CLIP | 23597149 | Unspecified |
| mmu-miR-21a-5p | MIMAT0000530 | Sppl3 | 74585 | HITS-CLIP | 23142080 | Unspecified |
| mmu-miR-21a-5p | MIMAT0000530 | Spred1 | 114715 | HITS-CLIP | 19536157\|30971389\|23142080\|29167 | Unspecified |
| mmu-miR-21a-5p | MIMAT0000530 | Spred2 | 114716 | Microarrays//HITS-CLIP | 37986224\|25263593\|23597149\|23142 | Unspecified |
| mmu-miR-21a-5p | MIMAT0000530 | Spry1 | 24063 | Luciferase assay//Western blot//qPCR//Mi | 20216554\|23416424\|23142080\|18508 | Unspecified |
| mmu-miR-21a-5p | MIMAT0000530 | Spry2 | 24064 | Luciferase assay//Western blot//Reporter | 18508928 | Unspecified |
| mmu-miR-21a-5p | MIMAT0000530 | Spry4 | 24066 | qRT-PCR//HITS-CLIP//Other | 19043405\|23597149\|18508928 | Unspecified |
| mmu-miR-21a-5p | MIMAT0000530 | Spryd7 | 66674 | Microarrays | 37986224\|25263593 | Unspecified |
| mmu-miR-21a-5p | MIMAT0000530 | Spsb4 | 211949 | Microarrays | 37986224\|25263593 | Unspecified |
| mmu-miR-21a-5p | MIMAT0000530 | Sptan1 | 20740 | HITS-CLIP | 19536157\|23597149\|23142080\|29167 | Unspecified |
| mmu-miR-21a-5p | MIMAT0000530 | Sptbn1 | 20742 | HITS-CLIP | 19536157\|23597149\|23142080\|31433 | Unspecified |

| | | | | | | |
|---|---|---|---|---|---|---|
| mmu-miR-21a-5p | MIMAT0000530 | Sptbn2 | 20743 | HITS-CLIP | 23597149 | Unspecified |
| mmu-miR-21a-5p | MIMAT0000530 | Sptlc1 | 268656 | HITS-CLIP | 23142080 | Unspecified |
| mmu-miR-21a-5p | MIMAT0000530 | Sptlc2 | 20773 | HITS-CLIP | 23597149\|23142080 | Unspecified |
| mmu-miR-21a-5p | MIMAT0000530 | Sptssa | 104725 | HITS-CLIP | 23142080 | Unspecified |
| mmu-miR-21a-5p | MIMAT0000530 | Sqle | 20775 | Microarrays//HITS-CLIP | 37986224\|23597149\|25263593\|23142 | Unspecified |
| mmu-miR-21a-5p | MIMAT0000530 | Sqstm1 | 18412 | qCLASH//HITS-CLIP | 31393953\|23597149 | Unspecified |
| mmu-miR-21a-5p | MIMAT0000530 | Srcap | 100043597 | HITS-CLIP | 23597149 | Unspecified |
| mmu-miR-21a-5p | MIMAT0000530 | Srd5a1 | 78925 | HITS-CLIP | 23597149\|23142080 | Unspecified |
| mmu-miR-21a-5p | MIMAT0000530 | Srek1 | 218543 | HITS-CLIP | 23142080\|29167373 | Unspecified |
| mmu-miR-21a-5p | MIMAT0000530 | Srfbp1 | 67222 | qCLASH | 31393953 | Unspecified |
| mmu-miR-21a-5p | MIMAT0000530 | Srgap2 | 14270 | HITS-CLIP | 19536157\|23597149\|23142080\|29167 | Unspecified |
| mmu-miR-21a-5p | MIMAT0000530 | Srl | 106393 | Microarrays | 37986224\|25263593 | Unspecified |
| mmu-miR-21a-5p | MIMAT0000530 | Srm | 20810 | Microarrays//HITS-CLIP | 37986224\|23597149\|25263593\|23142 | Unspecified |
| mmu-miR-21a-5p | MIMAT0000530 | Srp14 | 20813 | HITS-CLIP | 23142080 | Unspecified |
| mmu-miR-21a-5p | MIMAT0000530 | Srp19 | 66384 | HITS-CLIP | 23597149\|23142080 | Unspecified |
| mmu-miR-21a-5p | MIMAT0000530 | Srp68 | 217337 | HITS-CLIP | 23142080 | Unspecified |
| mmu-miR-21a-5p | MIMAT0000530 | Srp72 | 66661 | HITS-CLIP | 23142080 | Unspecified |
| mmu-miR-21a-5p | MIMAT0000530 | Srpk1 | 20815 | HITS-CLIP | 23142080 | Unspecified |
| mmu-miR-21a-5p | MIMAT0000530 | Srpk2 | 20817 | HITS-CLIP | 37986224\|23142080 | Unspecified |
| mmu-miR-21a-5p | MIMAT0000530 | Srrm1 | 51796 | HITS-CLIP | 29167373 | Unspecified |
| mmu-miR-21a-5p | MIMAT0000530 | Srrm2 | 75956 | HITS-CLIP, Microarrays//HITS-CLIP | 37986224\|23142080\|23597149\|19536 | Unspecified |
| mmu-miR-21a-5p | MIMAT0000530 | Srrt | 83701 | HITS-CLIP | 29167373 | Unspecified |
| mmu-miR-21a-5p | MIMAT0000530 | Srsf1 | 110809 | HITS-CLIP | 29167373 | Unspecified |
| mmu-miR-21a-5p | MIMAT0000530 | Srsf11 | 69207 | HITS-CLIP | 23142080\|29167373 | Unspecified |
| mmu-miR-21a-5p | MIMAT0000530 | Srsf12 | 272009 | HITS-CLIP | 37986224 | Unspecified |
| mmu-miR-21a-5p | MIMAT0000530 | Srsf2 | 20382 | HITS-CLIP | 23142080\|29167373 | Unspecified |
| mmu-miR-21a-5p | MIMAT0000530 | Srsf3 | 20383 | HITS-CLIP | 29167373 | Unspecified |
| mmu-miR-21a-5p | MIMAT0000530 | Srsf6 | 67996 | HITS-CLIP | 23142080\|29167373 | Unspecified |
| mmu-miR-21a-5p | MIMAT0000530 | Srsf7 | 225027 | HITS-CLIP | 23142080 | Unspecified |
| mmu-miR-21a-5p | MIMAT0000530 | Srsf9 | 108014 | HITS-CLIP | 23142080 | Unspecified |
| mmu-miR-21a-5p | MIMAT0000530 | Ss18l2 | 51188 | Microarrays | 25263593 | Unspecified |
| mmu-miR-21a-5p | MIMAT0000530 | Ssb | 20823 | HITS-CLIP | 37986224\|23142080 | Unspecified |
| mmu-miR-21a-5p | MIMAT0000530 | Ssbp2 | 66970 | HITS-CLIP, Microarrays//HITS-CLIP | 37986224\|23142080 | Unspecified |
| mmu-miR-21a-5p | MIMAT0000530 | Ssh1 | 231637 | HITS-CLIP | 29167373 | Unspecified |
| mmu-miR-21a-5p | MIMAT0000530 | Ssh2 | 237860 | HITS-CLIP | 23142080 | Unspecified |
| mmu-miR-21a-5p | MIMAT0000530 | Ssh3 | 245857 | Microarrays | 37986224 | Unspecified |
| mmu-miR-21a-5p | MIMAT0000530 | Ssna1 | 68475 | HITS-CLIP | 23142080 | Unspecified |
| mmu-miR-21a-5p | MIMAT0000530 | Ssr1 | 107513 | HITS-CLIP | 23142080\|29167373 | Unspecified |
| mmu-miR-21a-5p | MIMAT0000530 | Ssr3 | 67437 | HITS-CLIP | 23142080 | Unspecified |

| | | | | | | |
|---|---|---|---|---|---|---|
| mmu-miR-21a-5p | MIMAT0000530 | Ssrp1 | 20833 | HITS-CLIP | 23142080 | Unspecified |
| mmu-miR-21a-5p | MIMAT0000530 | St13 | 70356 | HITS-CLIP | 23142080 | Unspecified |
| mmu-miR-21a-5p | MIMAT0000530 | St14 | 19143 | Microarrays | 37986224\|25263593 | Unspecified |
| mmu-miR-21a-5p | MIMAT0000530 | St3gal4 | 20443 | Microarrays | 37986224 | Unspecified |
| mmu-miR-21a-5p | MIMAT0000530 | St3gal5 | 20454 | Microarrays//HITS-CLIP | 37986224\|25263593\|23597149 | Unspecified |
| mmu-miR-21a-5p | MIMAT0000530 | St3gal6 | 54613 | Microarrays//HITS-CLIP | 37986224\|23142080\|25263593 | Unspecified |
| mmu-miR-21a-5p | MIMAT0000530 | St6gal1 | 20440 | Microarrays//HITS-CLIP | 37986224\|23142080 | Unspecified |
| mmu-miR-21a-5p | MIMAT0000530 | St8sia1 | 20449 | HITS-CLIP | 37986224\|23142080 | Unspecified |
| mmu-miR-21a-5p | MIMAT0000530 | St8sia4 | 20452 | Microarrays//HITS-CLIP | 37986224\|23142080\|29167373 | Unspecified |
| mmu-miR-21a-5p | MIMAT0000530 | Stab1 | 192187 | HITS-CLIP | 29167373 | Unspecified |
| mmu-miR-21a-5p | MIMAT0000530 | Stab2 | 192188 | Microarrays//HITS-CLIP | 37986224\|23597149 | Unspecified |
| mmu-miR-21a-5p | MIMAT0000530 | Stac2 | 217154 | Microarrays | 37986224\|25263593 | Unspecified |
| mmu-miR-21a-5p | MIMAT0000530 | Stag1 | 20842 | HITS-CLIP | 23142080 | Unspecified |
| mmu-miR-21a-5p | MIMAT0000530 | Stag2 | 20843 | HITS-CLIP | 23597149\|23142080 | Unspecified |
| mmu-miR-21a-5p | MIMAT0000530 | Stag3 | 50878 | HITS-CLIP | 23142080 | Unspecified |
| mmu-miR-21a-5p | MIMAT0000530 | Stam | 20844 | HITS-CLIP | 23142080 | Unspecified |
| mmu-miR-21a-5p | MIMAT0000530 | Stam2 | 56324 | HITS-CLIP | 23142080 | Unspecified |
| mmu-miR-21a-5p | MIMAT0000530 | Stambp | 70527 | HITS-CLIP | 23142080 | Unspecified |
| mmu-miR-21a-5p | MIMAT0000530 | Stambpl1 | 76630 | Microarrays | 37986224 | Unspecified |
| mmu-miR-21a-5p | MIMAT0000530 | Stard13 | 243362 | Microarrays | 37986224 | Unspecified |
| mmu-miR-21a-5p | MIMAT0000530 | Stard4 | 170459 | Microarrays//HITS-CLIP | 37986224\|23142080 | Unspecified |
| mmu-miR-21a-5p | MIMAT0000530 | Stard5 | 170460 | HITS-CLIP | 23597149 | Unspecified |
| mmu-miR-21a-5p | MIMAT0000530 | Stard6 | 170461 | Microarrays//HITS-CLIP | 37986224\|25263593\|23142080 | Unspecified |
| mmu-miR-21a-5p | MIMAT0000530 | Stard7 | 99138 | HITS-CLIP | 19536157 | Unspecified |
| mmu-miR-21a-5p | MIMAT0000530 | Stard8 | 236920 | Microarrays | 37986224 | Unspecified |
| mmu-miR-21a-5p | MIMAT0000530 | Stard9 | 668880 | HITS-CLIP | 29167373 | Unspecified |
| mmu-miR-21a-5p | MIMAT0000530 | Stat1 | 20846 | HITS-CLIP | 29167373 | Unspecified |
| mmu-miR-21a-5p | MIMAT0000530 | Stat3 | 20848 | HITS-CLIP | 37986224\|23597149\|23142080 | Unspecified |
| mmu-miR-21a-5p | MIMAT0000530 | Stat4 | 20849 | HITS-CLIP | 23142080 | Unspecified |
| mmu-miR-21a-5p | MIMAT0000530 | Stat5a | 20850 | HITS-CLIP | 23142080 | Unspecified |
| mmu-miR-21a-5p | MIMAT0000530 | Stat5b | 20851 | HITS-CLIP | 23142080 | Unspecified |
| mmu-miR-21a-5p | MIMAT0000530 | Stat6 | 20852 | HITS-CLIP | 23142080 | Unspecified |
| mmu-miR-21a-5p | MIMAT0000530 | Stau1 | 20853 | HITS-CLIP | 23142080 | Unspecified |
| mmu-miR-21a-5p | MIMAT0000530 | Stau2 | 29819 | Microarrays | 37986224\|25263593 | Unspecified |
| mmu-miR-21a-5p | MIMAT0000530 | Stbd1 | 52331 | Microarrays//HITS-CLIP | 37986224\|25263593\|23597149 | Unspecified |
| mmu-miR-21a-5p | MIMAT0000530 | Stc1 | 20855 | HITS-CLIP | 23142080 | Unspecified |
| mmu-miR-21a-5p | MIMAT0000530 | Steap1 | 70358 | Microarrays | 37986224\|25263593 | Unspecified |
| mmu-miR-21a-5p | MIMAT0000530 | Steap3 | 68428 | Microarrays | 37986224\|25263593 | Unspecified |
| mmu-miR-21a-5p | MIMAT0000530 | Steap4 | 117167 | HITS-CLIP | 37986224\|23597149 | Unspecified |

| | | | | | | |
|---|---|---|---|---|---|---|
| mmu-miR-21a-5p | MIMAT0000530 | Stil | 20460 | Microarrays//HITS-CLIP | 37986224\|25263593\|23142080 | Unspecified |
| mmu-miR-21a-5p | MIMAT0000530 | Stim2 | 116873 | HITS-CLIP | 23142080 | Unspecified |
| mmu-miR-21a-5p | MIMAT0000530 | Stip1 | 20867 | HITS-CLIP | 37986224\|23597149\|23142080 | Unspecified |
| mmu-miR-21a-5p | MIMAT0000530 | Stk10 | 20868 | HITS-CLIP | 23142080 | Unspecified |
| mmu-miR-21a-5p | MIMAT0000530 | Stk11 | 20869 | HITS-CLIP | 23142080 | Unspecified |
| mmu-miR-21a-5p | MIMAT0000530 | Stk17b | 98267 | Microarrays//HITS-CLIP | 37986224\|23142080 | Unspecified |
| mmu-miR-21a-5p | MIMAT0000530 | Stk25 | 59041 | Microarrays | 37986224 | Unspecified |
| mmu-miR-21a-5p | MIMAT0000530 | Stk26 | 51765 | Microarrays//HITS-CLIP | 25263593\|23142080 | Unspecified |
| mmu-miR-21a-5p | MIMAT0000530 | Stk35 | 67333 | HITS-CLIP | 23142080 | Unspecified |
| mmu-miR-21a-5p | MIMAT0000530 | Stk38 | 106504 | HITS-CLIP | 37986224\|23597149\|23142080 | Unspecified |
| mmu-miR-21a-5p | MIMAT0000530 | Stk38l | 232533 | HITS-CLIP | 29167373 | Unspecified |
| mmu-miR-21a-5p | MIMAT0000530 | Stk39 | 53416 | HITS-CLIP | 23142080 | Unspecified |
| mmu-miR-21a-5p | MIMAT0000530 | Stk4 | 58231 | HITS-CLIP | 23142080\|29167373 | Unspecified |
| mmu-miR-21a-5p | MIMAT0000530 | Stk40 | 74178 | HITS-CLIP | 37986224\|23142080\|29167373 | Unspecified |
| mmu-miR-21a-5p | MIMAT0000530 | Stmn1 | 16765 | Microarrays | 37986224\|25263593 | Unspecified |
| mmu-miR-21a-5p | MIMAT0000530 | Ston2 | 108800 | HITS-CLIP | 23142080 | Unspecified |
| mmu-miR-21a-5p | MIMAT0000530 | Stra6l | 74152 | Microarrays//HITS-CLIP | 25263593\|23597149 | Unspecified |
| mmu-miR-21a-5p | MIMAT0000530 | Strada | 72149 | HITS-CLIP | 29167373 | Unspecified |
| mmu-miR-21a-5p | MIMAT0000530 | Stradb | 227154 | HITS-CLIP | 23142080 | Unspecified |
| mmu-miR-21a-5p | MIMAT0000530 | Strn | 268980 | HITS-CLIP | 37986224\|23142080\|23597149 | Unspecified |
| mmu-miR-21a-5p | MIMAT0000530 | Strn3 | 94186 | HITS-CLIP | 23142080 | Unspecified |
| mmu-miR-21a-5p | MIMAT0000530 | Strn4 | 97387 | HITS-CLIP | 23142080\|26203562 | Unspecified |
| mmu-miR-21a-5p | MIMAT0000530 | Stt3a | 16430 | HITS-CLIP | 23142080 | Unspecified |
| mmu-miR-21a-5p | MIMAT0000530 | Stt3b | 68292 | HITS-CLIP | 37986224\|23142080 | Unspecified |
| mmu-miR-21a-5p | MIMAT0000530 | Stub1 | 56424 | HITS-CLIP | 23597149 | Unspecified |
| mmu-miR-21a-5p | MIMAT0000530 | Stx11 | 74732 | HITS-CLIP | 30971389 | Unspecified |
| mmu-miR-21a-5p | MIMAT0000530 | Stx12 | 100226 | HITS-CLIP | 29167373\|23142080 | Unspecified |
| mmu-miR-21a-5p | MIMAT0000530 | Stx16 | 228960 | HITS-CLIP | 23142080 | Unspecified |
| mmu-miR-21a-5p | MIMAT0000530 | Stx18 | 71116 | HITS-CLIP | 23142080\|29167373 | Unspecified |
| mmu-miR-21a-5p | MIMAT0000530 | Stx1b | 56216 | HITS-CLIP | 37986224 | Unspecified |
| mmu-miR-21a-5p | MIMAT0000530 | Stx4a | 20909 | HITS-CLIP | 23142080 | Unspecified |
| mmu-miR-21a-5p | MIMAT0000530 | Stxbp1 | 20910 | HITS-CLIP | 29167373\|23142080 | Unspecified |
| mmu-miR-21a-5p | MIMAT0000530 | Stxbp2 | 20911 | HITS-CLIP | 23597149 | Unspecified |
| mmu-miR-21a-5p | MIMAT0000530 | Stxbp3 | 20912 | HITS-CLIP | 23142080 | Unspecified |
| mmu-miR-21a-5p | MIMAT0000530 | Sub1 | 20024 | Microarrays//HITS-CLIP | 37986224\|23142080\|29167373 | Unspecified |
| mmu-miR-21a-5p | MIMAT0000530 | Sucla2 | 20916 | HITS-CLIP | 23597149 | Unspecified |
| mmu-miR-21a-5p | MIMAT0000530 | Suclg1 | 56451 | HITS-CLIP | 23142080 | Unspecified |
| mmu-miR-21a-5p | MIMAT0000530 | Suclg2 | 20917 | HITS-CLIP | 23142080\|23597149 | Unspecified |
| mmu-miR-21a-5p | MIMAT0000530 | Suco | 226551 | HITS-CLIP | 23597149\|23142080 | Unspecified |

| | | | | | | | |
|---|---|---|---|---|---|---|---|
| mmu-miR-21a-5p | MIMAT0000530 | Sugp1 | 70616 | HITS-CLIP | | 23142080 | Unspecified |
| mmu-miR-21a-5p | MIMAT0000530 | Sugt1 | 67955 | HITS-CLIP | | 23142080 | Unspecified |
| mmu-miR-21a-5p | MIMAT0000530 | Sulf2 | 72043 | Microarrays | | 37986224 | Unspecified |
| mmu-miR-21a-5p | MIMAT0000530 | Sult1a1 | 20887 | Microarrays | | 37986224 | Unspecified |
| mmu-miR-21a-5p | MIMAT0000530 | Sumo3 | 20610 | HITS-CLIP | | 23142080 | Unspecified |
| mmu-miR-21a-5p | MIMAT0000530 | Sun1 | 77053 | HITS-CLIP | | 30971389 | Unspecified |
| mmu-miR-21a-5p | MIMAT0000530 | Sun2 | 223697 | HITS-CLIP | | 23597149 | Unspecified |
| mmu-miR-21a-5p | MIMAT0000530 | Suox | 211389 | HITS-CLIP | 23597149\|23142080 | | Unspecified |
| mmu-miR-21a-5p | MIMAT0000530 | Supt16 | 114741 | HITS-CLIP | 23142080\|29167373 | | Unspecified |
| mmu-miR-21a-5p | MIMAT0000530 | Supt20 | 56790 | HITS-CLIP | | 23142080 | Unspecified |
| mmu-miR-21a-5p | MIMAT0000530 | Supt5 | 20924 | HITS-CLIP | 23142080\|23597149 | | Unspecified |
| mmu-miR-21a-5p | MIMAT0000530 | Supt6 | 20926 | HITS-CLIP | | 23142080 | Unspecified |
| mmu-miR-21a-5p | MIMAT0000530 | Surf1 | 20930 | HITS-CLIP | | 23142080 | Unspecified |
| mmu-miR-21a-5p | MIMAT0000530 | Susd6 | 9766 | HITS-CLIP | 23597149\|23142080\|29167373 | | Unspecified |
| mmu-miR-21a-5p | MIMAT0000530 | Suv39h1 | 20937 | HITS-CLIP | | 23142080 | Unspecified |
| mmu-miR-21a-5p | MIMAT0000530 | Suv39h2 | 64707 | HITS-CLIP | | 23142080 | Unspecified |
| mmu-miR-21a-5p | MIMAT0000530 | Suz12 | 52615 | HITS-CLIP | 37986224\|23597149\|23142080\|29167 | | Unspecified |
| mmu-miR-21a-5p | MIMAT0000530 | Sv2a | 64051 | HITS-CLIP | | 37986224 | Unspecified |
| mmu-miR-21a-5p | MIMAT0000530 | Svip | 75744 | HITS-CLIP, Microarrays//Microarrays//HITS | 37986224\|25263593\|19536157 | | Unspecified |
| mmu-miR-21a-5p | MIMAT0000530 | Swap70 | 20947 | HITS-CLIP | 29167373\|23142080 | | Unspecified |
| mmu-miR-21a-5p | MIMAT0000530 | Swi5 | 72931 | HITS-CLIP | 23597149\|23142080 | | Unspecified |
| mmu-miR-21a-5p | MIMAT0000530 | Swt1 | 66875 | HITS-CLIP | | 23142080 | Unspecified |
| mmu-miR-21a-5p | MIMAT0000530 | Syf2 | 68592 | HITS-CLIP | | 23142080 | Unspecified |
| mmu-miR-21a-5p | MIMAT0000530 | Syk | 20963 | HITS-CLIP | | 29167373 | Unspecified |
| mmu-miR-21a-5p | MIMAT0000530 | Syncrip | 56403 | HITS-CLIP | 37986224\|19536157\|23142080 | | Unspecified |
| mmu-miR-21a-5p | MIMAT0000530 | Syne1 | 64009 | HITS-CLIP | 23142080\|23597149\|19536157 | | Unspecified |
| mmu-miR-21a-5p | MIMAT0000530 | Syne2 | 319565 | HITS-CLIP | 23142080\|23597149 | | Unspecified |
| mmu-miR-21a-5p | MIMAT0000530 | Syne3 | 212073 | HITS-CLIP | 23142080\|29167373 | | Unspecified |
| mmu-miR-21a-5p | MIMAT0000530 | Synj1 | 104015 | HITS-CLIP | 37986224\|19536157\|23142080 | | Unspecified |
| mmu-miR-21a-5p | MIMAT0000530 | Synj2 | 20975 | Microarrays//HITS-CLIP | 37986224\|25263593\|23142080 | | Unspecified |
| mmu-miR-21a-5p | MIMAT0000530 | Sypl | 19027 | HITS-CLIP | | 23142080 | Unspecified |
| mmu-miR-21a-5p | MIMAT0000530 | Syt11 | 229521 | HITS-CLIP | 19536157\|23142080 | | Unspecified |
| mmu-miR-21a-5p | MIMAT0000530 | Syt4 | 20983 | HITS-CLIP | | 19536157 | Unspecified |
| mmu-miR-21a-5p | MIMAT0000530 | Syt5 | 53420 | HITS-CLIP | | 37986224 | Unspecified |
| mmu-miR-21a-5p | MIMAT0000530 | Sytl1 | 269589 | HITS-CLIP | | 23597149 | Unspecified |
| mmu-miR-21a-5p | MIMAT0000530 | Sytl2 | 83671 | HITS-CLIP | | 23142080 | Unspecified |
| mmu-miR-21a-5p | MIMAT0000530 | Syvn1 | 74126 | HITS-CLIP | 23142080\|26203562 | | Unspecified |
| mmu-miR-21a-5p | MIMAT0000530 | Szrd1 | 213491 | HITS-CLIP | | 23142080 | Unspecified |
| mmu-miR-21a-5p | MIMAT0000530 | Tab2 | 68652 | HITS-CLIP | 23142080\|29167373 | | Unspecified |

| | | | | | | |
|---|---|---|---|---|---|---|
| mmu-miR-21a-5p | MIMAT0000530 | Tab3 | 66724 | HITS-CLIP | 23142080 | Unspecified |
| mmu-miR-21a-5p | MIMAT0000530 | Tacc1 | 320165 | HITS-CLIP | 23142080 | Unspecified |
| mmu-miR-21a-5p | MIMAT0000530 | Tacc3 | 21335 | HITS-CLIP | 23142080 | Unspecified |
| mmu-miR-21a-5p | MIMAT0000530 | Taco1 | 70207 | HITS-CLIP | 23597149 | Unspecified |
| mmu-miR-21a-5p | MIMAT0000530 | Tada2b | 231151 | HITS-CLIP | 23142080\|29167373 | Unspecified |
| mmu-miR-21a-5p | MIMAT0000530 | Tada3 | 101206 | HITS-CLIP | 23142080 | Unspecified |
| mmu-miR-21a-5p | MIMAT0000530 | Taf1 | 270627 | HITS-CLIP | 23142080 | Unspecified |
| mmu-miR-21a-5p | MIMAT0000530 | Taf15 | 70439 | Microarrays//qCLASH | 37986224\|31393953 | Unspecified |
| mmu-miR-21a-5p | MIMAT0000530 | Taf1c | 21341 | HITS-CLIP | 23597149 | Unspecified |
| mmu-miR-21a-5p | MIMAT0000530 | Taf1d | 75316 | HITS-CLIP | 29167373 | Unspecified |
| mmu-miR-21a-5p | MIMAT0000530 | Taf2 | 319944 | HITS-CLIP | 23142080 | Unspecified |
| mmu-miR-21a-5p | MIMAT0000530 | Taf3 | 209361 | HITS-CLIP | 23142080 | Unspecified |
| mmu-miR-21a-5p | MIMAT0000530 | Taf4 | 6874 | HITS-CLIP | 23142080 | Unspecified |
| mmu-miR-21a-5p | MIMAT0000530 | Taf4b | 72504 | Microarrays | 37986224\|25263593 | Unspecified |
| mmu-miR-21a-5p | MIMAT0000530 | Taf5 | 226182 | HITS-CLIP | 37986224\|23597149\|23142080 | Unspecified |
| mmu-miR-21a-5p | MIMAT0000530 | Taf5l | 102162 | HITS-CLIP | 19536157\|23597149 | Unspecified |
| mmu-miR-21a-5p | MIMAT0000530 | Taf7 | 24074 | HITS-CLIP, Microarrays//Microarrays//qCL | 37986224\|25263593\|31393953\|23142 | Unspecified |
| mmu-miR-21a-5p | MIMAT0000530 | Taf9 | 108143 | HITS-CLIP | 23142080 | Unspecified |
| mmu-miR-21a-5p | MIMAT0000530 | Taf9b | 407786 | HITS-CLIP | 31433990 | Unspecified |
| mmu-miR-21a-5p | MIMAT0000530 | Tagap | 72536 | HITS-CLIP | 23142080 | Unspecified |
| mmu-miR-21a-5p | MIMAT0000530 | Tagln | 21345 | Microarrays | 37986224\|25263593 | Unspecified |
| mmu-miR-21a-5p | MIMAT0000530 | Tanc1 | 66860 | HITS-CLIP | 23142080 | Unspecified |
| mmu-miR-21a-5p | MIMAT0000530 | Tanc2 | 77097 | Microarrays | 37986224 | Unspecified |
| mmu-miR-21a-5p | MIMAT0000530 | Tango2 | 27883 | HITS-CLIP | 23597149 | Unspecified |
| mmu-miR-21a-5p | MIMAT0000530 | Tango6 | 272538 | HITS-CLIP | 23597149\|23142080 | Unspecified |
| mmu-miR-21a-5p | MIMAT0000530 | Tank | 21353 | HITS-CLIP | 23142080\|29167373 | Unspecified |
| mmu-miR-21a-5p | MIMAT0000530 | Taok1 | 216965 | HITS-CLIP | 19536157\|23142080\|29167373 | Unspecified |
| mmu-miR-21a-5p | MIMAT0000530 | Taok2 | 381921 | HITS-CLIP | 23142080\|29167373 | Unspecified |
| mmu-miR-21a-5p | MIMAT0000530 | Tap1 | 21354 | HITS-CLIP | 23142080 | Unspecified |
| mmu-miR-21a-5p | MIMAT0000530 | Tap2 | 21355 | HITS-CLIP | 23142080 | Unspecified |
| mmu-miR-21a-5p | MIMAT0000530 | Tapt1 | 231225 | HITS-CLIP | 23597149 | Unspecified |
| mmu-miR-21a-5p | MIMAT0000530 | Tarbp1 | 6894 | HITS-CLIP | 23142080 | Unspecified |
| mmu-miR-21a-5p | MIMAT0000530 | Tardbp | 230908 | HITS-CLIP | 23142080\|29167373 | Unspecified |
| mmu-miR-21a-5p | MIMAT0000530 | Tars | 110960 | HITS-CLIP | 23142080 | Unspecified |
| mmu-miR-21a-5p | MIMAT0000530 | Tat | 234724 | HITS-CLIP | 23597149 | Unspecified |
| mmu-miR-21a-5p | MIMAT0000530 | Tatdn2 | 381801 | Microarrays//HITS-CLIP | 37986224\|25263593\|29167373 | Unspecified |
| mmu-miR-21a-5p | MIMAT0000530 | Tax1bp1 | 52440 | HITS-CLIP | 23597149\|23142080 | Unspecified |
| mmu-miR-21a-5p | MIMAT0000530 | Tbc1d1 | 57915 | HITS-CLIP | 23142080 | Unspecified |
| mmu-miR-21a-5p | MIMAT0000530 | Tbc1d15 | 66687 | HITS-CLIP | 37986224\|23142080 | Unspecified |

| | | | | | | |
|---|---|---|---|---|---|---|
| mmu-miR-21a-5p | MIMAT0000530 | Tbc1d16 | 207592 | HITS-CLIP | 37986224\|23597149 | Unspecified |
| mmu-miR-21a-5p | MIMAT0000530 | Tbc1d17 | 233204 | HITS-CLIP | 23142080 | Unspecified |
| mmu-miR-21a-5p | MIMAT0000530 | Tbc1d2 | 381605 | Microarrays | 37986224\|25263593 | Unspecified |
| mmu-miR-21a-5p | MIMAT0000530 | Tbc1d22b | 381085 | HITS-CLIP | 23142080 | Unspecified |
| mmu-miR-21a-5p | MIMAT0000530 | Tbc1d23 | 67581 | HITS-CLIP | 23142080 | Unspecified |
| mmu-miR-21a-5p | MIMAT0000530 | Tbc1d24 | 224617 | HITS-CLIP | 29167373 | Unspecified |
| mmu-miR-21a-5p | MIMAT0000530 | Tbc1d2b | 67016 | HITS-CLIP | 23142080 | Unspecified |
| mmu-miR-21a-5p | MIMAT0000530 | Tbc1d30 | 74694 | HITS-CLIP | 23597149\|23142080 | Unspecified |
| mmu-miR-21a-5p | MIMAT0000530 | Tbc1d31 | 210544 | HITS-CLIP | 23142080 | Unspecified |
| mmu-miR-21a-5p | MIMAT0000530 | Tbc1d4 | 210789 | Microarrays//HITS-CLIP | 37986224\|23142080 | Unspecified |
| mmu-miR-21a-5p | MIMAT0000530 | Tbc1d8 | 54610 | Microarrays | 37986224 | Unspecified |
| mmu-miR-21a-5p | MIMAT0000530 | Tbc1d9b | 76795 | HITS-CLIP | 23142080 | Unspecified |
| mmu-miR-21a-5p | MIMAT0000530 | Tbccd1 | 70573 | HITS-CLIP | 23142080 | Unspecified |
| mmu-miR-21a-5p | MIMAT0000530 | Tbce | 70430 | HITS-CLIP | 23142080 | Unspecified |
| mmu-miR-21a-5p | MIMAT0000530 | Tbcel | 272589 | HITS-CLIP | 23597149\|23142080 | Unspecified |
| mmu-miR-21a-5p | MIMAT0000530 | Tbck | 271981 | HITS-CLIP | 23142080 | Unspecified |
| mmu-miR-21a-5p | MIMAT0000530 | Tbk1 | 56480 | HITS-CLIP | 23142080 | Unspecified |
| mmu-miR-21a-5p | MIMAT0000530 | Tbkbp1 | 73174 | HITS-CLIP | 23142080\|29167373 | Unspecified |
| mmu-miR-21a-5p | MIMAT0000530 | Tbl1xr1 | 81004 | HITS-CLIP | 37986224\|23142080\|29167373 | Unspecified |
| mmu-miR-21a-5p | MIMAT0000530 | Tbl3 | 213773 | HITS-CLIP | 23142080 | Unspecified |
| mmu-miR-21a-5p | MIMAT0000530 | Tbp | 21374 | HITS-CLIP | 23142080 | Unspecified |
| mmu-miR-21a-5p | MIMAT0000530 | Tbpl1 | 237336 | HITS-CLIP | 23597149\|23142080 | Unspecified |
| mmu-miR-21a-5p | MIMAT0000530 | Tbx21 | 57765 | HITS-CLIP | 23142080 | Unspecified |
| mmu-miR-21a-5p | MIMAT0000530 | Tcea1 | 21399 | qCLASH | 31393953 | Unspecified |
| mmu-miR-21a-5p | MIMAT0000530 | Tcea3 | 21401 | HITS-CLIP | 23597149 | Unspecified |
| mmu-miR-21a-5p | MIMAT0000530 | Tceanc | 245695 | Microarrays//HITS-CLIP | 37986224\|23142080 | Unspecified |
| mmu-miR-21a-5p | MIMAT0000530 | Tceanc2 | 66526 | Microarrays | 37986224 | Unspecified |
| mmu-miR-21a-5p | MIMAT0000530 | Tcf20 | 21411 | HITS-CLIP | 23142080\|29167373 | Unspecified |
| mmu-miR-21a-5p | MIMAT0000530 | Tcf25 | 66855 | HITS-CLIP | 23142080\|23597149 | Unspecified |
| mmu-miR-21a-5p | MIMAT0000530 | Tcf4 | 21413 | Microarrays//HITS-CLIP | 37986224\|23142080 | Unspecified |
| mmu-miR-21a-5p | MIMAT0000530 | Tcf7 | 21414 | HITS-CLIP | 23142080 | Unspecified |
| mmu-miR-21a-5p | MIMAT0000530 | Tcf7l1 | 21415 | HITS-CLIP | 23142080 | Unspecified |
| mmu-miR-21a-5p | MIMAT0000530 | Tcp1 | 21454 | HITS-CLIP | 23142080 | Unspecified |
| mmu-miR-21a-5p | MIMAT0000530 | Tcp11l2 | 216198 | Microarrays//HITS-CLIP | 37986224\|23142080 | Unspecified |
| mmu-miR-21a-5p | MIMAT0000530 | Tctex1d1 | 67344 | Microarrays | 37986224 | Unspecified |
| mmu-miR-21a-5p | MIMAT0000530 | Tctex1d2 | 66061 | HITS-CLIP | 37986224\|23597149\|23142080 | Unspecified |
| mmu-miR-21a-5p | MIMAT0000530 | Tdo2 | 56720 | HITS-CLIP | 23597149 | Unspecified |
| mmu-miR-21a-5p | MIMAT0000530 | Tdrd7 | 100121 | HITS-CLIP | 23597149 | Unspecified |
| mmu-miR-21a-5p | MIMAT0000530 | Tdrp | 72148 | HITS-CLIP | 23142080 | Unspecified |

| | | | | | | |
|---|---|---|---|---|---|---|
| mmu-miR-21a-5p | MIMAT0000530 | Tead1 | 21676 | HITS-CLIP | 37986224 | Unspecified |
| mmu-miR-21a-5p | MIMAT0000530 | Tec | 21682 | HITS-CLIP | 23142080 | Unspecified |
| mmu-miR-21a-5p | MIMAT0000530 | Tedc1 | 104732 | HITS-CLIP | 23142080 | Unspecified |
| mmu-miR-21a-5p | MIMAT0000530 | Tef | 21685 | Microarrays | 37986224 | Unspecified |
| mmu-miR-21a-5p | MIMAT0000530 | Tent4a | 11044 | HITS-CLIP | 23142080 | Unspecified |
| mmu-miR-21a-5p | MIMAT0000530 | Tent4b | 64282 | HITS-CLIP | 23142080\|29167373 | Unspecified |
| mmu-miR-21a-5p | MIMAT0000530 | Tent5a | 55603 | HITS-CLIP | 23142080 | Unspecified |
| mmu-miR-21a-5p | MIMAT0000530 | Tent5c | 54855 | Microarrays//HITS-CLIP | 25263593\|23142080 | Unspecified |
| mmu-miR-21a-5p | MIMAT0000530 | Tep1 | 21745 | HITS-CLIP | 23142080\|29167373 | Unspecified |
| mmu-miR-21a-5p | MIMAT0000530 | Terf1 | 21749 | HITS-CLIP | 23142080 | Unspecified |
| mmu-miR-21a-5p | MIMAT0000530 | Tes | 21753 | Microarrays//HITS-CLIP | 37986224\|25263593\|23597149\|23142 | Unspecified |
| mmu-miR-21a-5p | MIMAT0000530 | Tesc | 57816 | Microarrays | 37986224 | Unspecified |
| mmu-miR-21a-5p | MIMAT0000530 | Tesk1 | 21754 | HITS-CLIP | 29167373 | Unspecified |
| mmu-miR-21a-5p | MIMAT0000530 | Tesk2 | 230661 | Microarrays | 37986224 | Unspecified |
| mmu-miR-21a-5p | MIMAT0000530 | Tet1 | 52463 | HITS-CLIP | 37986224\|23142080 | Unspecified |
| mmu-miR-21a-5p | MIMAT0000530 | Tet2 | 214133 | Microarrays//HITS-CLIP | 37986224\|23142080\|29167373 | Unspecified |
| mmu-miR-21a-5p | MIMAT0000530 | Tet3 | 194388 | HITS-CLIP//qCLASH | 37986224\|31393953\|23597149\|23142 | Unspecified |
| mmu-miR-21a-5p | MIMAT0000530 | Tex10 | 269536 | HITS-CLIP | 23142080 | Unspecified |
| mmu-miR-21a-5p | MIMAT0000530 | Tex15 | 104271 | Microarrays//HITS-CLIP | 37986224\|25263593\|23142080 | Unspecified |
| mmu-miR-21a-5p | MIMAT0000530 | Tex2 | 21763 | HITS-CLIP | 37986224\|23142080 | Unspecified |
| mmu-miR-21a-5p | MIMAT0000530 | Tfam | 21780 | HITS-CLIP | 31433990\|23142080 | Unspecified |
| mmu-miR-21a-5p | MIMAT0000530 | Tfap4 | 83383 | HITS-CLIP | 37986224\|23142080 | Unspecified |
| mmu-miR-21a-5p | MIMAT0000530 | Tfdp1 | 21781 | HITS-CLIP | 23142080 | Unspecified |
| mmu-miR-21a-5p | MIMAT0000530 | Tfg | 21787 | HITS-CLIP | 23142080 | Unspecified |
| mmu-miR-21a-5p | MIMAT0000530 | Tfpi2 | 21789 | HITS-CLIP | 23597149 | Unspecified |
| mmu-miR-21a-5p | MIMAT0000530 | Tfr2 | 50765 | HITS-CLIP | 23597149 | Unspecified |
| mmu-miR-21a-5p | MIMAT0000530 | Tfrc | 22042 | HITS-CLIP, Microarrays//Microarrays//HITS | 37986224\|25263593\|23142080 | Unspecified |
| mmu-miR-21a-5p | MIMAT0000530 | Tgds | 76355 | Microarrays | 37986224\|25263593 | Unspecified |
| mmu-miR-21a-5p | MIMAT0000530 | Tgfa | 21802 | HITS-CLIP | 23597149 | Unspecified |
| mmu-miR-21a-5p | MIMAT0000530 | Tgfb3 | 21809 | Microarrays | 37986224 | Unspecified |
| mmu-miR-21a-5p | MIMAT0000530 | Tgfbi | 21810 | ELISA//FACS//Luciferase assay//qRT-PCR// | 21494432\|23597149\|29167373 | Unspecified |
| mmu-miR-21a-5p | MIMAT0000530 | Tgfbr1 | 21812 | HITS-CLIP | 37986224\|23597149\|23142080\|29167 | Unspecified |
| mmu-miR-21a-5p | MIMAT0000530 | Tgfbr2 | 21813 | HITS-CLIP | 23597149\|19536157\|23142080\|29167 | Unspecified |
| mmu-miR-21a-5p | MIMAT0000530 | Tgfbr3 | 21814 | Luciferase assay//qRT-PCR//Western blot/ | 22960625\|23142080 | Unspecified |
| mmu-miR-21a-5p | MIMAT0000530 | Tgif1 | 21815 | HITS-CLIP | 37986224 | Unspecified |
| mmu-miR-21a-5p | MIMAT0000530 | Tgif2 | 228839 | HITS-CLIP | 23142080 | Unspecified |
| mmu-miR-21a-5p | MIMAT0000530 | Tgm2 | 21817 | HITS-CLIP | 37986224\|23142080 | Unspecified |
| mmu-miR-21a-5p | MIMAT0000530 | Tgoln1 | 22134 | HITS-CLIP | 23597149\|23142080 | Unspecified |
| mmu-miR-21a-5p | MIMAT0000530 | Tgs1 | 116940 | HITS-CLIP | 23597149 | Unspecified |

| | | | | | | |
|---|---|---|---|---|---|---|
| mmu-miR-21a-5p | MIMAT0000530 | Thap12 | 72981 | HITS-CLIP | 23142080 | Unspecified |
| mmu-miR-21a-5p | MIMAT0000530 | Thap2 | 66816 | HITS-CLIP | 23142080 | Unspecified |
| mmu-miR-21a-5p | MIMAT0000530 | Thbd | 21824 | HITS-CLIP | 23597149 | Unspecified |
| mmu-miR-21a-5p | MIMAT0000530 | Thbs1 | 21825 | Microarrays//HITS-CLIP | 37986224\|26203562 | Unspecified |
| mmu-miR-21a-5p | MIMAT0000530 | Them6 | 223626 | HITS-CLIP | 37986224 | Unspecified |
| mmu-miR-21a-5p | MIMAT0000530 | Themis | 210757 | HITS-CLIP | 23142080 | Unspecified |
| mmu-miR-21a-5p | MIMAT0000530 | Thnsl2 | 232078 | HITS-CLIP | 23597149 | Unspecified |
| mmu-miR-21a-5p | MIMAT0000530 | Thoc1 | 225160 | HITS-CLIP | 23142080 | Unspecified |
| mmu-miR-21a-5p | MIMAT0000530 | Thoc2 | 331401 | HITS-CLIP | 37986224\|23142080\|29167373 | Unspecified |
| mmu-miR-21a-5p | MIMAT0000530 | Thra | 21833 | HITS-CLIP, Microarrays | 37986224 | Unspecified |
| mmu-miR-21a-5p | MIMAT0000530 | Thrsp | 21835 | HITS-CLIP | 23597149 | Unspecified |
| mmu-miR-21a-5p | MIMAT0000530 | Thumpd3 | 14911 | HITS-CLIP | 23142080 | Unspecified |
| mmu-miR-21a-5p | MIMAT0000530 | Thy1 | 21838 | Microarrays//HITS-CLIP | 37986224\|25263593\|23142080 | Unspecified |
| mmu-miR-21a-5p | MIMAT0000530 | Tiam1 | 21844 | HITS-CLIP | 37986224\|23142080\|29167373 | Unspecified |
| mmu-miR-21a-5p | MIMAT0000530 | Tiam2 | 24001 | HITS-CLIP | 23597149 | Unspecified |
| mmu-miR-21a-5p | MIMAT0000530 | Tifa | 211550 | Microarrays | 37986224 | Unspecified |
| mmu-miR-21a-5p | MIMAT0000530 | Tifab | 212937 | Microarrays | 37986224 | Unspecified |
| mmu-miR-21a-5p | MIMAT0000530 | Tigd2 | 68140 | HITS-CLIP | 23142080 | Unspecified |
| mmu-miR-21a-5p | MIMAT0000530 | Timeless | 21853 | Microarrays | 37986224\|25263593 | Unspecified |
| mmu-miR-21a-5p | MIMAT0000530 | Timm10b | 14356 | HITS-CLIP | 23142080 | Unspecified |
| mmu-miR-21a-5p | MIMAT0000530 | Timm8a1 | 30058 | Microarrays | 37986224\|25263593 | Unspecified |
| mmu-miR-21a-5p | MIMAT0000530 | Timmdc1 | 76916 | Microarrays | 37986224\|25263593 | Unspecified |
| mmu-miR-21a-5p | MIMAT0000530 | Timp1 | 21857 | Microarrays | 37986224\|25263593 | Unspecified |
| mmu-miR-21a-5p | MIMAT0000530 | Timp3 | 21859 | qRT-PCR//HITS-CLIP | 23797704\|19536157 | Unspecified |
| mmu-miR-21a-5p | MIMAT0000530 | Tinf2 | 28113 | HITS-CLIP | 23142080 | Unspecified |
| mmu-miR-21a-5p | MIMAT0000530 | Tiparp | 99929 | HITS-CLIP | 23142080\|29167373 | Unspecified |
| mmu-miR-21a-5p | MIMAT0000530 | Tipin | 66131 | HITS-CLIP, Microarrays//HITS-CLIP//Microa | 37986224\|23142080\|25263593\|23597 | Unspecified |
| mmu-miR-21a-5p | MIMAT0000530 | Tiprl | 226591 | HITS-CLIP | 26203562 | Unspecified |
| mmu-miR-21a-5p | MIMAT0000530 | Tirap | 117149 | HITS-CLIP | 37986224\|23597149 | Unspecified |
| mmu-miR-21a-5p | MIMAT0000530 | Tjp2 | 21873 | HITS-CLIP | 19536157\|23597149 | Unspecified |
| mmu-miR-21a-5p | MIMAT0000530 | Tk1 | 21877 | Microarrays | 37986224\|25263593 | Unspecified |
| mmu-miR-21a-5p | MIMAT0000530 | Tkfc | 26007 | HITS-CLIP | 23597149 | Unspecified |
| mmu-miR-21a-5p | MIMAT0000530 | Tlcd1 | 68385 | HITS-CLIP | 23142080 | Unspecified |
| mmu-miR-21a-5p | MIMAT0000530 | Tlcd2 | 380712 | Microarrays | 37986224\|25263593 | Unspecified |
| mmu-miR-21a-5p | MIMAT0000530 | Tlcd3a | 79850 | Microarrays | 25263593 | Unspecified |
| mmu-miR-21a-5p | MIMAT0000530 | Tle1 | 21885 | HITS-CLIP | 29167373 | Unspecified |
| mmu-miR-21a-5p | MIMAT0000530 | Tle3 | 21887 | HITS-CLIP | 23142080\|23597149 | Unspecified |
| mmu-miR-21a-5p | MIMAT0000530 | Tlk1 | 228012 | HITS-CLIP | 23142080\|29167373 | Unspecified |
| mmu-miR-21a-5p | MIMAT0000530 | Tlk2 | 24086 | HITS-CLIP | 23142080 | Unspecified |

| | | | | | | |
|---|---|---|---|---|---|---|
| mmu-miR-21a-5p | MIMAT0000530 | Tln1 | 21894 | HITS-CLIP | 23597149\|23142080\|29167373 | Unspecified |
| mmu-miR-21a-5p | MIMAT0000530 | Tlnrd1 | 59274 | HITS-CLIP | 23142080 | Unspecified |
| mmu-miR-21a-5p | MIMAT0000530 | Tlr13 | 279572 | HITS-CLIP | 29167373 | Unspecified |
| mmu-miR-21a-5p | MIMAT0000530 | Tlr2 | 24088 | Microarrays | 37986224 | Unspecified |
| mmu-miR-21a-5p | MIMAT0000530 | Tlr4 | 21898 | HITS-CLIP | 29167373 | Unspecified |
| mmu-miR-21a-5p | MIMAT0000530 | Tlr5 | 53791 | HITS-CLIP | 23597149 | Unspecified |
| mmu-miR-21a-5p | MIMAT0000530 | Tlr6 | 21899 | HITS-CLIP | 29167373 | Unspecified |
| mmu-miR-21a-5p | MIMAT0000530 | Tlr7 | 170743 | Microarrays//HITS-CLIP | 37986224\|23142080 | Unspecified |
| mmu-miR-21a-5p | MIMAT0000530 | Tlr8 | 170744 | Microarrays//HITS-CLIP | 37986224\|29167373 | Unspecified |
| mmu-miR-21a-5p | MIMAT0000530 | Tm4sf4 | 229302 | HITS-CLIP | 37986224 | Unspecified |
| mmu-miR-21a-5p | MIMAT0000530 | Tm4sf5 | 75604 | Microarrays | 37986224 | Unspecified |
| mmu-miR-21a-5p | MIMAT0000530 | Tm7sf3 | 67623 | HITS-CLIP | 23142080 | Unspecified |
| mmu-miR-21a-5p | MIMAT0000530 | Tm9sf1 | 74140 | Microarrays | 37986224 | Unspecified |
| mmu-miR-21a-5p | MIMAT0000530 | Tm9sf2 | 68059 | HITS-CLIP | 23597149\|23142080 | Unspecified |
| mmu-miR-21a-5p | MIMAT0000530 | Tm9sf3 | 107358 | HITS-CLIP | 23597149\|23142080 | Unspecified |
| mmu-miR-21a-5p | MIMAT0000530 | Tmbim6 | 110213 | HITS-CLIP | 23597149 | Unspecified |
| mmu-miR-21a-5p | MIMAT0000530 | Tmc7 | 209760 | HITS-CLIP | 37986224 | Unspecified |
| mmu-miR-21a-5p | MIMAT0000530 | Tmcc1 | 330401 | HITS-CLIP | 37986224\|23142080 | Unspecified |
| mmu-miR-21a-5p | MIMAT0000530 | Tmcc3 | 319880 | HITS-CLIP | 23597149 | Unspecified |
| mmu-miR-21a-5p | MIMAT0000530 | Tmco6 | 71983 | HITS-CLIP | 23142080 | Unspecified |
| mmu-miR-21a-5p | MIMAT0000530 | Tmed2 | 10959 | qCLASH | 31393953 | Unspecified |
| mmu-miR-21a-5p | MIMAT0000530 | Tmed4 | 103694 | HITS-CLIP | 23597149\|23142080 | Unspecified |
| mmu-miR-21a-5p | MIMAT0000530 | Tmed5 | 73130 | HITS-CLIP | 23597149\|23142080\|29167373 | Unspecified |
| mmu-miR-21a-5p | MIMAT0000530 | Tmed8 | 382620 | HITS-CLIP | 29167373 | Unspecified |
| mmu-miR-21a-5p | MIMAT0000530 | Tmed9 | 67511 | HITS-CLIP | 23142080\|23597149\|19536157 | Unspecified |
| mmu-miR-21a-5p | MIMAT0000530 | Tmeff1 | 230157 | Microarrays | 37986224\|25263593 | Unspecified |
| mmu-miR-21a-5p | MIMAT0000530 | Tmem104 | 320534 | Microarrays | 37986224 | Unspecified |
| mmu-miR-21a-5p | MIMAT0000530 | Tmem106b | 71900 | HITS-CLIP | 23597149\|29167373\|23142080 | Unspecified |
| mmu-miR-21a-5p | MIMAT0000530 | Tmem120a | 215210 | Microarrays | 37986224\|25263593 | Unspecified |
| mmu-miR-21a-5p | MIMAT0000530 | Tmem123 | 71929 | HITS-CLIP | 23142080 | Unspecified |
| mmu-miR-21a-5p | MIMAT0000530 | Tmem126b | 68472 | HITS-CLIP | 23597149 | Unspecified |
| mmu-miR-21a-5p | MIMAT0000530 | Tmem127 | 69470 | HITS-CLIP | 23142080 | Unspecified |
| mmu-miR-21a-5p | MIMAT0000530 | Tmem128 | 66309 | HITS-CLIP | 23597149 | Unspecified |
| mmu-miR-21a-5p | MIMAT0000530 | Tmem129 | 68366 | HITS-CLIP | 29167373 | Unspecified |
| mmu-miR-21a-5p | MIMAT0000530 | Tmem131 | 56030 | HITS-CLIP | 23142080\|23597149\|19536157 | Unspecified |
| mmu-miR-21a-5p | MIMAT0000530 | Tmem131l | 23240 | HITS-CLIP | 23142080 | Unspecified |
| mmu-miR-21a-5p | MIMAT0000530 | Tmem135 | 72759 | HITS-CLIP | 23597149\|23142080 | Unspecified |
| mmu-miR-21a-5p | MIMAT0000530 | Tmem138 | 72982 | HITS-CLIP | 23142080 | Unspecified |
| mmu-miR-21a-5p | MIMAT0000530 | Tmem150b | 330460 | Microarrays | 37986224 | Unspecified |

| | | | | | | |
|---|---|---|---|---|---|---|
| mmu-miR-21a-5p | MIMAT0000530 | Tmem154 | 320782 | Microarrays//HITS-CLIP | 37986224\|25263593\|23142080 | Unspecified |
| mmu-miR-21a-5p | MIMAT0000530 | Tmem158 | 72309 | Microarrays | 37986224\|25263593 | Unspecified |
| mmu-miR-21a-5p | MIMAT0000530 | Tmem161b | 72745 | HITS-CLIP | 23142080 | Unspecified |
| mmu-miR-21a-5p | MIMAT0000530 | Tmem164 | 209497 | HITS-CLIP | 23142080\|29167373 | Unspecified |
| mmu-miR-21a-5p | MIMAT0000530 | Tmem165 | 21982 | Microarrays | 37986224\|25263593 | Unspecified |
| mmu-miR-21a-5p | MIMAT0000530 | Tmem167b | 67495 | HITS-CLIP | 23142080 | Unspecified |
| mmu-miR-21a-5p | MIMAT0000530 | Tmem168 | 101118 | HITS-CLIP | 29167373\|23142080 | Unspecified |
| mmu-miR-21a-5p | MIMAT0000530 | Tmem170 | 66817 | HITS-CLIP | 37986224\|23142080 | Unspecified |
| mmu-miR-21a-5p | MIMAT0000530 | Tmem170b | 621976 | HITS-CLIP | 23142080\|19536157\|23597149\|29167 | Unspecified |
| mmu-miR-21a-5p | MIMAT0000530 | Tmem171 | 380863 | Microarrays | 37986224\|25263593 | Unspecified |
| mmu-miR-21a-5p | MIMAT0000530 | Tmem173 | 72512 | Microarrays//HITS-CLIP | 37986224\|23142080 | Unspecified |
| mmu-miR-21a-5p | MIMAT0000530 | Tmem176a | 66058 | Microarrays//HITS-CLIP | 37986224\|23597149 | Unspecified |
| mmu-miR-21a-5p | MIMAT0000530 | Tmem176b | 65963 | Microarrays | 37986224 | Unspecified |
| mmu-miR-21a-5p | MIMAT0000530 | Tmem178 | 68027 | HITS-CLIP, Microarrays//Microarrays | 37986224\|25263593 | Unspecified |
| mmu-miR-21a-5p | MIMAT0000530 | Tmem18 | 211986 | HITS-CLIP | 37986224\|23142080 | Unspecified |
| mmu-miR-21a-5p | MIMAT0000530 | Tmem184b | 223693 | HITS-CLIP | 23142080 | Unspecified |
| mmu-miR-21a-5p | MIMAT0000530 | Tmem184c | 234463 | qCLASH//HITS-CLIP | 31393953\|23142080 | Unspecified |
| mmu-miR-21a-5p | MIMAT0000530 | Tmem189 | 407243 | HITS-CLIP | 29167373 | Unspecified |
| mmu-miR-21a-5p | MIMAT0000530 | Tmem206 | 66950 | HITS-CLIP | 29167373\|23142080 | Unspecified |
| mmu-miR-21a-5p | MIMAT0000530 | Tmem209 | 72649 | HITS-CLIP | 37986224\|23597149\|23142080 | Unspecified |
| mmu-miR-21a-5p | MIMAT0000530 | Tmem214 | 68796 | HITS-CLIP | 23142080\|23597149 | Unspecified |
| mmu-miR-21a-5p | MIMAT0000530 | Tmem229b | 268567 | Microarrays//HITS-CLIP | 37986224\|23142080 | Unspecified |
| mmu-miR-21a-5p | MIMAT0000530 | Tmem238 | 664968 | Microarrays | 37986224\|25263593 | Unspecified |
| mmu-miR-21a-5p | MIMAT0000530 | Tmem245 | 242474 | HITS-CLIP | 37986224\|23142080 | Unspecified |
| mmu-miR-21a-5p | MIMAT0000530 | Tmem260 | 218989 | HITS-CLIP | 23142080 | Unspecified |
| mmu-miR-21a-5p | MIMAT0000530 | Tmem263 | 103266 | HITS-CLIP | 23142080 | Unspecified |
| mmu-miR-21a-5p | MIMAT0000530 | Tmem267 | 64417 | Microarrays | 25263593 | Unspecified |
| mmu-miR-21a-5p | MIMAT0000530 | Tmem30a | 69981 | HITS-CLIP | 37986224\|23142080\|29167373 | Unspecified |
| mmu-miR-21a-5p | MIMAT0000530 | Tmem33 | 67878 | HITS-CLIP | 30971389\|23142080 | Unspecified |
| mmu-miR-21a-5p | MIMAT0000530 | Tmem37 | 170706 | Microarrays | 37986224 | Unspecified |
| mmu-miR-21a-5p | MIMAT0000530 | Tmem38a | 74166 | Microarrays | 37986224 | Unspecified |
| mmu-miR-21a-5p | MIMAT0000530 | Tmem38b | 52076 | qCLASH | 31393953 | Unspecified |
| mmu-miR-21a-5p | MIMAT0000530 | Tmem39a | 67846 | HITS-CLIP | 23142080 | Unspecified |
| mmu-miR-21a-5p | MIMAT0000530 | Tmem50a | 71817 | HITS-CLIP | 23142080 | Unspecified |
| mmu-miR-21a-5p | MIMAT0000530 | Tmem50b | 77975 | HITS-CLIP | 23142080 | Unspecified |
| mmu-miR-21a-5p | MIMAT0000530 | Tmem51 | 214359 | Microarrays | 37986224 | Unspecified |
| mmu-miR-21a-5p | MIMAT0000530 | Tmem56 | 99887 | HITS-CLIP | 37986224\|23597149 | Unspecified |
| mmu-miR-21a-5p | MIMAT0000530 | Tmem59 | 56374 | HITS-CLIP | 29167373\|23142080 | Unspecified |
| mmu-miR-21a-5p | MIMAT0000530 | Tmem60 | 212090 | HITS-CLIP | 23142080 | Unspecified |

| | | | | | | | |
|---|---|---|---|---|---|---|---|
| mmu-miR-21a-5p | MIMAT0000530 | Tmem62 | 96957 | HITS-CLIP | | 23597149 | Unspecified |
| mmu-miR-21a-5p | MIMAT0000530 | Tmem64 | 100201 | HITS-CLIP | | 23142080 | Unspecified |
| mmu-miR-21a-5p | MIMAT0000530 | Tmem65 | 74868 | HITS-CLIP | | 23142080 | Unspecified |
| mmu-miR-21a-5p | MIMAT0000530 | Tmem68 | 72098 | HITS-CLIP | | 23142080 | Unspecified |
| mmu-miR-21a-5p | MIMAT0000530 | Tmem71 | 213068 | HITS-CLIP | | 23142080 | Unspecified |
| mmu-miR-21a-5p | MIMAT0000530 | Tmem8 | 60455 | HITS-CLIP | | 23597149 | Unspecified |
| mmu-miR-21a-5p | MIMAT0000530 | Tmem86a | 67893 | Microarrays | | 37986224 | Unspecified |
| mmu-miR-21a-5p | MIMAT0000530 | Tmem87a | 211499 | Microarrays | | 37986224 | Unspecified |
| mmu-miR-21a-5p | MIMAT0000530 | Tmem87b | 72477 | HITS-CLIP | | 23142080 | Unspecified |
| mmu-miR-21a-5p | MIMAT0000530 | Tmem98 | 103743 | Microarrays | 37986224\|25263593 | | Unspecified |
| mmu-miR-21a-5p | MIMAT0000530 | Tmem9b | 56786 | HITS-CLIP | | 23142080 | Unspecified |
| mmu-miR-21a-5p | MIMAT0000530 | Tmf1 | 232286 | HITS-CLIP | | 23142080 | Unspecified |
| mmu-miR-21a-5p | MIMAT0000530 | Tmod2 | 50876 | HITS-CLIP | | 19536157 | Unspecified |
| mmu-miR-21a-5p | MIMAT0000530 | Tmod3 | 50875 | HITS-CLIP | 23142080\|29167373 | | Unspecified |
| mmu-miR-21a-5p | MIMAT0000530 | Tmpo | 21917 | HITS-CLIP | 37986224\|23142080\|29167373 | | Unspecified |
| mmu-miR-21a-5p | MIMAT0000530 | Tmsb4x | 19241 | HITS-CLIP | | 23597149 | Unspecified |
| mmu-miR-21a-5p | MIMAT0000530 | Tmtc3 | 237500 | HITS-CLIP | | 29167373 | Unspecified |
| mmu-miR-21a-5p | MIMAT0000530 | Tmtc4 | 70551 | HITS-CLIP | | 23142080 | Unspecified |
| mmu-miR-21a-5p | MIMAT0000530 | Tmx1 | 72736 | HITS-CLIP | | 23142080 | Unspecified |
| mmu-miR-21a-5p | MIMAT0000530 | Tmx3 | 67988 | HITS-CLIP | 37986224\|23142080\|29167373 | | Unspecified |
| mmu-miR-21a-5p | MIMAT0000530 | Tnf | 21926 | Microarrays//HITS-CLIP | 37986224\|25263593\|23142080\|29167 | | Unspecified |
| mmu-miR-21a-5p | MIMAT0000530 | Tnfaip1 | 21927 | HITS-CLIP | 23597149\|23142080 | | Unspecified |
| mmu-miR-21a-5p | MIMAT0000530 | Tnfaip3 | 21929 | Microarrays//HITS-CLIP | 37986224\|25263593\|23597149\|2314 | | Unspecified |
| mmu-miR-21a-5p | MIMAT0000530 | Tnfaip8l2 | 69769 | ChIP-seq//Flow//Luciferase assay//qRT-PC | 24577093\|23142080 | | Unspecified |
| mmu-miR-21a-5p | MIMAT0000530 | Tnfrsf11a | 21934 | HITS-CLIP | | 29167373 | Unspecified |
| mmu-miR-21a-5p | MIMAT0000530 | Tnfrsf11b | 18383 | HITS-CLIP | 37986224\|23597149 | | Unspecified |
| mmu-miR-21a-5p | MIMAT0000530 | Tnfrsf12a | 27279 | Microarrays | 37986224\|25263593 | | Unspecified |
| mmu-miR-21a-5p | MIMAT0000530 | Tnfrsf17 | 21935 | Microarrays | | 37986224 | Unspecified |
| mmu-miR-21a-5p | MIMAT0000530 | Tnfrsf1a | 21937 | HITS-CLIP | | 37986224 | Unspecified |
| mmu-miR-21a-5p | MIMAT0000530 | Tnfrsf1b | 21938 | Microarrays//HITS-CLIP | 37986224\|23142080 | | Unspecified |
| mmu-miR-21a-5p | MIMAT0000530 | Tnfrsf26 | 244237 | HITS-CLIP | 37986224\|23142080 | | Unspecified |
| mmu-miR-21a-5p | MIMAT0000530 | Tnfrsf4 | 22163 | HITS-CLIP | | 23142080 | Unspecified |
| mmu-miR-21a-5p | MIMAT0000530 | Tnfrsf8 | 21941 | HITS-CLIP | | 23142080 | Unspecified |
| mmu-miR-21a-5p | MIMAT0000530 | Tnfrsf9 | 21942 | HITS-CLIP | | 23142080 | Unspecified |
| mmu-miR-21a-5p | MIMAT0000530 | Tnfsf11 | 21943 | HITS-CLIP | 37986224\|23142080 | | Unspecified |
| mmu-miR-21a-5p | MIMAT0000530 | Tnfsf14 | 50930 | Microarrays//HITS-CLIP | 37986224\|23142080 | | Unspecified |
| mmu-miR-21a-5p | MIMAT0000530 | Tnfsf8 | 21949 | HITS-CLIP | | 23142080 | Unspecified |
| mmu-miR-21a-5p | MIMAT0000530 | Tnfsf9 | 21950 | Microarrays | 37986224\|25263593 | | Unspecified |
| mmu-miR-21a-5p | MIMAT0000530 | Tnik | 665113 | HITS-CLIP | 37986224\|23142080 | | Unspecified |

| | | | | | | |
|---|---|---|---|---|---|---|
| mmu-miR-21a-5p | MIMAT0000530 | Tnip1 | 57783 | HITS-CLIP | 29167373 | Unspecified |
| mmu-miR-21a-5p | MIMAT0000530 | Tnip3 | 414084 | HITS-CLIP | 29167373 | Unspecified |
| mmu-miR-21a-5p | MIMAT0000530 | Tnks | 21951 | HITS-CLIP | 23142080 | Unspecified |
| mmu-miR-21a-5p | MIMAT0000530 | Tnks2 | 74493 | HITS-CLIP | 37986224\|19536157\|23142080 | Unspecified |
| mmu-miR-21a-5p | MIMAT0000530 | Tnpo1 | 238799 | HITS-CLIP//qCLASH | 37986224\|23597149\|31393953\|23142 | Unspecified |
| mmu-miR-21a-5p | MIMAT0000530 | Tnrc6a | 233833 | qCLASH//HITS-CLIP | 31393953\|23142080\|29167373 | Unspecified |
| mmu-miR-21a-5p | MIMAT0000530 | Tnrc6b | 213988 | HITS-CLIP | 37986224\|23597149\|31433990\|19536 | Unspecified |
| mmu-miR-21a-5p | MIMAT0000530 | Tnrc6c | 217351 | HITS-CLIP | 31433990\|23597149\|23142080\|29167 | Unspecified |
| mmu-miR-21a-5p | MIMAT0000530 | Tns1 | 21961 | HITS-CLIP | 23142080\|23597149 | Unspecified |
| mmu-miR-21a-5p | MIMAT0000530 | Tns2 | 209039 | HITS-CLIP | 23597149 | Unspecified |
| mmu-miR-21a-5p | MIMAT0000530 | Tns3 | 319939 | HITS-CLIP | 29167373 | Unspecified |
| mmu-miR-21a-5p | MIMAT0000530 | Tob1 | 22057 | HITS-CLIP | 37986224\|23597149\|23142080\|29167 | Unspecified |
| mmu-miR-21a-5p | MIMAT0000530 | Tom1 | 21968 | Microarrays | 37986224\|25263593 | Unspecified |
| mmu-miR-21a-5p | MIMAT0000530 | Tom1l1 | 71943 | Microarrays | 37986224\|25263593 | Unspecified |
| mmu-miR-21a-5p | MIMAT0000530 | Tomm20 | 67952 | qCLASH | 31393953 | Unspecified |
| mmu-miR-21a-5p | MIMAT0000530 | Tomm70a | 28185 | HITS-CLIP | 37986224\|23142080\|23597149\|29167 | Unspecified |
| mmu-miR-21a-5p | MIMAT0000530 | Top1 | 21969 | HITS-CLIP | 23597149\|23142080\|29167373 | Unspecified |
| mmu-miR-21a-5p | MIMAT0000530 | Top2a | 21973 | Microarrays//qCLASH//HITS-CLIP | 37986224\|25263593\|31393953\|23142 | Unspecified |
| mmu-miR-21a-5p | MIMAT0000530 | Top2b | 21974 | HITS-CLIP | 37986224\|23142080 | Unspecified |
| mmu-miR-21a-5p | MIMAT0000530 | Top3a | 21975 | HITS-CLIP | 23142080 | Unspecified |
| mmu-miR-21a-5p | MIMAT0000530 | Topbp1 | 235559 | HITS-CLIP | 23142080 | Unspecified |
| mmu-miR-21a-5p | MIMAT0000530 | Topors | 106021 | HITS-CLIP | 23597149\|23142080 | Unspecified |
| mmu-miR-21a-5p | MIMAT0000530 | Tor1aip1 | 208263 | HITS-CLIP | 23142080\|29167373 | Unspecified |
| mmu-miR-21a-5p | MIMAT0000530 | Tor1aip2 | 240832 | HITS-CLIP, Microarrays//Microarrays//HITS | 37986224\|25263593\|23142080\|23597 | Unspecified |
| mmu-miR-21a-5p | MIMAT0000530 | Tor1b | 30934 | HITS-CLIP | 23597149 | Unspecified |
| mmu-miR-21a-5p | MIMAT0000530 | Tor4a | 227612 | HITS-CLIP | 31433990\|23142080 | Unspecified |
| mmu-miR-21a-5p | MIMAT0000530 | Tox | 252838 | HITS-CLIP | 23142080 | Unspecified |
| mmu-miR-21a-5p | MIMAT0000530 | Tox4 | 268741 | HITS-CLIP | 23142080 | Unspecified |
| mmu-miR-21a-5p | MIMAT0000530 | Tpcn1 | 252972 | HITS-CLIP | 37986224\|23597149\|23142080 | Unspecified |
| mmu-miR-21a-5p | MIMAT0000530 | Tpcn2 | 233979 | Microarrays | 37986224 | Unspecified |
| mmu-miR-21a-5p | MIMAT0000530 | Tpd52 | 21985 | HITS-CLIP | 29167373 | Unspecified |
| mmu-miR-21a-5p | MIMAT0000530 | Tpgs2 | 66648 | Microarrays | 37986224\|25263593 | Unspecified |
| mmu-miR-21a-5p | MIMAT0000530 | Tpi1 | 21991 | HITS-CLIP | 23142080 | Unspecified |
| mmu-miR-21a-5p | MIMAT0000530 | Tpk1 | 29807 | HITS-CLIP | 23142080 | Unspecified |
| mmu-miR-21a-5p | MIMAT0000530 | Tpm1 | 22003 | HITS-CLIP | 29167373 | Unspecified |
| mmu-miR-21a-5p | MIMAT0000530 | Tpm3 | 59069 | HITS-CLIP | 29167373 | Unspecified |
| mmu-miR-21a-5p | MIMAT0000530 | Tpm4 | 326618 | Microarrays//HITS-CLIP | 37986224\|25263593\|23142080 | Unspecified |
| mmu-miR-21a-5p | MIMAT0000530 | Tppp | 72948 | HITS-CLIP | 23597149 | Unspecified |
| mmu-miR-21a-5p | MIMAT0000530 | Tppp3 | 67971 | HITS-CLIP | 37986224\|19536157 | Unspecified |

| | | | | | | |
|---|---|---|---|---|---|---|
| mmu-miR-21a-5p | MIMAT0000530 | Tpr | 108989 | HITS-CLIP | 37986224\|23597149\|23142080 | Unspecified |
| mmu-miR-21a-5p | MIMAT0000530 | Tprgl | 67808 | HITS-CLIP | 23597149\|23142080 | Unspecified |
| mmu-miR-21a-5p | MIMAT0000530 | Tprkb | 69786 | HITS-CLIP | 23597149 | Unspecified |
| mmu-miR-21a-5p | MIMAT0000530 | Tpx2 | 72119 | Microarrays | 37986224\|25263593 | Unspecified |
| mmu-miR-21a-5p | MIMAT0000530 | Tra2a | 101214 | Microarrays | 37986224 | Unspecified |
| mmu-miR-21a-5p | MIMAT0000530 | Traf3 | 22031 | HITS-CLIP | 23142080 | Unspecified |
| mmu-miR-21a-5p | MIMAT0000530 | Traf6 | 22034 | HITS-CLIP | 23142080\|29167373 | Unspecified |
| mmu-miR-21a-5p | MIMAT0000530 | Trak1 | 67095 | HITS-CLIP | 23597149 | Unspecified |
| mmu-miR-21a-5p | MIMAT0000530 | Trak2 | 70827 | HITS-CLIP | 37986224\|23142080 | Unspecified |
| mmu-miR-21a-5p | MIMAT0000530 | Tram1 | 72265 | HITS-CLIP | 23597149\|23142080\|19536157 | Unspecified |
| mmu-miR-21a-5p | MIMAT0000530 | Tram1l1 | 229801 | HITS-CLIP | 37986224 | Unspecified |
| mmu-miR-21a-5p | MIMAT0000530 | Trap1 | 68015 | HITS-CLIP | 23597149\|23142080\|29167373 | Unspecified |
| mmu-miR-21a-5p | MIMAT0000530 | Trappc10 | 216131 | HITS-CLIP | 23142080 | Unspecified |
| mmu-miR-21a-5p | MIMAT0000530 | Trappc11 | 320714 | HITS-CLIP | 37986224\|19536157\|23142080 | Unspecified |
| mmu-miR-21a-5p | MIMAT0000530 | Trappc2l | 59005 | HITS-CLIP | 23597149\|23142080 | Unspecified |
| mmu-miR-21a-5p | MIMAT0000530 | Trappc4 | 60409 | HITS-CLIP | 23142080 | Unspecified |
| mmu-miR-21a-5p | MIMAT0000530 | Trappc6b | 78232 | HITS-CLIP | 23142080 | Unspecified |
| mmu-miR-21a-5p | MIMAT0000530 | Trappc8 | 75964 | HITS-CLIP | 37986224\|23142080 | Unspecified |
| mmu-miR-21a-5p | MIMAT0000530 | Trappc9 | 76510 | HITS-CLIP | 23142080 | Unspecified |
| mmu-miR-21a-5p | MIMAT0000530 | Trdmt1 | 13434 | HITS-CLIP | 23142080 | Unspecified |
| mmu-miR-21a-5p | MIMAT0000530 | Trem1 | 58217 | Microarrays | 37986224\|25263593 | Unspecified |
| mmu-miR-21a-5p | MIMAT0000530 | Trem3 | 58218 | Microarrays | 37986224\|25263593 | Unspecified |
| mmu-miR-21a-5p | MIMAT0000530 | Treml1 | 71326 | Microarrays | 37986224 | Unspecified |
| mmu-miR-21a-5p | MIMAT0000530 | Treml4 | 224840 | Microarrays | 37986224\|25263593 | Unspecified |
| mmu-miR-21a-5p | MIMAT0000530 | Trib1 | 211770 | HITS-CLIP | 23597149\|23142080\|29167373 | Unspecified |
| mmu-miR-21a-5p | MIMAT0000530 | Trib2 | 217410 | HITS-CLIP | 23142080 | Unspecified |
| mmu-miR-21a-5p | MIMAT0000530 | Trib3 | 228775 | Microarrays | 37986224\|25263593 | Unspecified |
| mmu-miR-21a-5p | MIMAT0000530 | Trim12a | 76681 | Microarrays | 37986224 | Unspecified |
| mmu-miR-21a-5p | MIMAT0000530 | Trim14 | 74735 | HITS-CLIP | 29167373 | Unspecified |
| mmu-miR-21a-5p | MIMAT0000530 | Trim21 | 20821 | Microarrays | 37986224 | Unspecified |
| mmu-miR-21a-5p | MIMAT0000530 | Trim23 | 81003 | HITS-CLIP | 23142080 | Unspecified |
| mmu-miR-21a-5p | MIMAT0000530 | Trim24 | 21848 | HITS-CLIP | 23142080 | Unspecified |
| mmu-miR-21a-5p | MIMAT0000530 | Trim25 | 217069 | HITS-CLIP | 23597149\|23142080\|29167373 | Unspecified |
| mmu-miR-21a-5p | MIMAT0000530 | Trim26 | 22670 | HITS-CLIP | 23142080\|23597149 | Unspecified |
| mmu-miR-21a-5p | MIMAT0000530 | Trim27 | 19720 | HITS-CLIP | 23142080 | Unspecified |
| mmu-miR-21a-5p | MIMAT0000530 | Trim28 | 21849 | HITS-CLIP | 23597149\|23142080 | Unspecified |
| mmu-miR-21a-5p | MIMAT0000530 | Trim30a | 20128 | HITS-CLIP | 29167373 | Unspecified |
| mmu-miR-21a-5p | MIMAT0000530 | Trim30d | 209387 | Microarrays | 37986224 | Unspecified |
| mmu-miR-21a-5p | MIMAT0000530 | Trim32 | 69807 | HITS-CLIP | 23142080 | Unspecified |

| | | | | | | |
|---|---|---|---|---|---|---|
| mmu-miR-21a-5p | MIMAT0000530 | Trim33 | 94093 | HITS-CLIP | 19536157\|23142080 | Unspecified |
| mmu-miR-21a-5p | MIMAT0000530 | Trim34a | 94094 | Microarrays | 37986224 | Unspecified |
| mmu-miR-21a-5p | MIMAT0000530 | Trim35 | 66854 | HITS-CLIP | 29167373 | Unspecified |
| mmu-miR-21a-5p | MIMAT0000530 | Trim37 | 68729 | HITS-CLIP | 23142080 | Unspecified |
| mmu-miR-21a-5p | MIMAT0000530 | Trim44 | 80985 | HITS-CLIP | 19536157\|23142080\|29167373 | Unspecified |
| mmu-miR-21a-5p | MIMAT0000530 | Trim5 | 667823 | Microarrays | 37986224 | Unspecified |
| mmu-miR-21a-5p | MIMAT0000530 | Trim56 | 384309 | HITS-CLIP | 23142080 | Unspecified |
| mmu-miR-21a-5p | MIMAT0000530 | Trim65 | 338364 | Microarrays | 37986224 | Unspecified |
| mmu-miR-21a-5p | MIMAT0000530 | Trim8 | 93679 | HITS-CLIP | 23142080 | Unspecified |
| mmu-miR-21a-5p | MIMAT0000530 | Trip11 | 109181 | HITS-CLIP | 23142080\|29167373 | Unspecified |
| mmu-miR-21a-5p | MIMAT0000530 | Trip12 | 14897 | HITS-CLIP | 23597149\|23142080\|19536157\|29167 | Unspecified |
| mmu-miR-21a-5p | MIMAT0000530 | Trmt10a | 108943 | HITS-CLIP | 23142080 | Unspecified |
| mmu-miR-21a-5p | MIMAT0000530 | Trmt2a | 15547 | HITS-CLIP | 23597149 | Unspecified |
| mmu-miR-21a-5p | MIMAT0000530 | Trmt2b | 215201 | HITS-CLIP | 23142080 | Unspecified |
| mmu-miR-21a-5p | MIMAT0000530 | Trmt5 | 76357 | HITS-CLIP | 23142080 | Unspecified |
| mmu-miR-21a-5p | MIMAT0000530 | Trmt61a | 328162 | Microarrays | 37986224\|25263593 | Unspecified |
| mmu-miR-21a-5p | MIMAT0000530 | Trnt1 | 70047 | HITS-CLIP | 23142080 | Unspecified |
| mmu-miR-21a-5p | MIMAT0000530 | Trp53bp2 | 209456 | HITS-CLIP | 23597149\|23142080 | Unspecified |
| mmu-miR-21a-5p | MIMAT0000530 | Trp53inp1 | 60599 | HITS-CLIP, Microarrays//HITS-CLIP | 37986224\|23597149\|23142080 | Unspecified |
| mmu-miR-21a-5p | MIMAT0000530 | Trp53inp2 | 68728 | HITS-CLIP | 23597149\|23142080 | Unspecified |
| mmu-miR-21a-5p | MIMAT0000530 | Trpc2 | 22064 | Microarrays | 37986224 | Unspecified |
| mmu-miR-21a-5p | MIMAT0000530 | Trpc3 | 22065 | HITS-CLIP | 19536157 | Unspecified |
| mmu-miR-21a-5p | MIMAT0000530 | Trpc4ap | 56407 | HITS-CLIP | 23597149\|23142080 | Unspecified |
| mmu-miR-21a-5p | MIMAT0000530 | Trpm7 | 58800 | qCLASH//HITS-CLIP | 31393953\|23597149\|23142080\|29167 | Unspecified |
| mmu-miR-21a-5p | MIMAT0000530 | Trps1 | 83925 | HITS-CLIP | 29167373 | Unspecified |
| mmu-miR-21a-5p | MIMAT0000530 | Trpv4 | 63873 | Microarrays | 37986224 | Unspecified |
| mmu-miR-21a-5p | MIMAT0000530 | Trrap | 100683 | HITS-CLIP | 23142080 | Unspecified |
| mmu-miR-21a-5p | MIMAT0000530 | Trub1 | 72133 | HITS-CLIP | 23142080 | Unspecified |
| mmu-miR-21a-5p | MIMAT0000530 | Tsc1 | 64930 | HITS-CLIP | 23142080 | Unspecified |
| mmu-miR-21a-5p | MIMAT0000530 | Tsc2 | 22084 | HITS-CLIP | 29167373 | Unspecified |
| mmu-miR-21a-5p | MIMAT0000530 | Tsc22d1 | 21807 | HITS-CLIP | 37986224\|23597149\|23142080 | Unspecified |
| mmu-miR-21a-5p | MIMAT0000530 | Tsc22d2 | 72033 | HITS-CLIP | 31433990\|23142080\|29167373 | Unspecified |
| mmu-miR-21a-5p | MIMAT0000530 | Tsc22d3 | 14605 | Microarrays | 37986224 | Unspecified |
| mmu-miR-21a-5p | MIMAT0000530 | Tsg101 | 22088 | HITS-CLIP | 23142080 | Unspecified |
| mmu-miR-21a-5p | MIMAT0000530 | Tshz1 | 110796 | HITS-CLIP | 37986224\|29167373\|23142080 | Unspecified |
| mmu-miR-21a-5p | MIMAT0000530 | Tslp | 53603 | Microarrays | 37986224 | Unspecified |
| mmu-miR-21a-5p | MIMAT0000530 | Tsn | 22099 | HITS-CLIP | 23142080 | Unspecified |
| mmu-miR-21a-5p | MIMAT0000530 | Tsnax | 53424 | HITS-CLIP | 23142080\|29167373 | Unspecified |
| mmu-miR-21a-5p | MIMAT0000530 | Tspan13 | 66109 | HITS-CLIP | 23142080 | Unspecified |

| | | | | | | |
|---|---|---|---|---|---|---|
| mmu-miR-21a-5p | MIMAT0000530 | Tspan14 | 52588 | HITS-CLIP | 23142080 | Unspecified |
| mmu-miR-21a-5p | MIMAT0000530 | Tspan3 | 56434 | HITS-CLIP | 23142080 | Unspecified |
| mmu-miR-21a-5p | MIMAT0000530 | Tspan31 | 67125 | HITS-CLIP | 23142080 | Unspecified |
| mmu-miR-21a-5p | MIMAT0000530 | Tspyl1 | 22110 | HITS-CLIP | 26203562 | Unspecified |
| mmu-miR-21a-5p | MIMAT0000530 | Tspyl2 | 52808 | HITS-CLIP | 29167373 | Unspecified |
| mmu-miR-21a-5p | MIMAT0000530 | Tst | 22117 | Microarrays | 37986224\|25263593 | Unspecified |
| mmu-miR-21a-5p | MIMAT0000530 | Ttbk2 | 140810 | HITS-CLIP | 19536157 | Unspecified |
| mmu-miR-21a-5p | MIMAT0000530 | Ttc14 | 67120 | HITS-CLIP | 23142080\|29167373 | Unspecified |
| mmu-miR-21a-5p | MIMAT0000530 | Ttc27 | 74196 | Microarrays//HITS-CLIP//qCLASH | 37986224\|25263593\|23597149\|23142 | Unspecified |
| mmu-miR-21a-5p | MIMAT0000530 | Ttc3 | 22129 | HITS-CLIP | 37986224\|23142080 | Unspecified |
| mmu-miR-21a-5p | MIMAT0000530 | Ttc30b | 72421 | HITS-CLIP | 30971389 | Unspecified |
| mmu-miR-21a-5p | MIMAT0000530 | Ttc33 | 67515 | HITS-CLIP | 23597149\|29167373 | Unspecified |
| mmu-miR-21a-5p | MIMAT0000530 | Ttc37 | 218343 | HITS-CLIP | 23142080 | Unspecified |
| mmu-miR-21a-5p | MIMAT0000530 | Ttc39b | 69863 | HITS-CLIP | 37986224\|23142080 | Unspecified |
| mmu-miR-21a-5p | MIMAT0000530 | Ttc39c | 72747 | HITS-CLIP | 23597149 | Unspecified |
| mmu-miR-21a-5p | MIMAT0000530 | Ttc7b | 104718 | Microarrays | 37986224\|25263593 | Unspecified |
| mmu-miR-21a-5p | MIMAT0000530 | Ttf1 | 22130 | HITS-CLIP | 23142080 | Unspecified |
| mmu-miR-21a-5p | MIMAT0000530 | Ttf2 | 74044 | HITS-CLIP | 23597149\|23142080 | Unspecified |
| mmu-miR-21a-5p | MIMAT0000530 | Tti2 | 234138 | HITS-CLIP | 23142080 | Unspecified |
| mmu-miR-21a-5p | MIMAT0000530 | Ttk | 22137 | Microarrays | 37986224\|25263593 | Unspecified |
| mmu-miR-21a-5p | MIMAT0000530 | Ttll12 | 223723 | HITS-CLIP | 23142080 | Unspecified |
| mmu-miR-21a-5p | MIMAT0000530 | Ttll4 | 67534 | HITS-CLIP | 23597149 | Unspecified |
| mmu-miR-21a-5p | MIMAT0000530 | Ttpa | 50500 | HITS-CLIP | 23597149 | Unspecified |
| mmu-miR-21a-5p | MIMAT0000530 | Tuba1a | 22142 | HITS-CLIP | 19536157\|23142080 | Unspecified |
| mmu-miR-21a-5p | MIMAT0000530 | Tuba4a | 22145 | Microarrays | 37986224\|25263593 | Unspecified |
| mmu-miR-21a-5p | MIMAT0000530 | Tubb2b | 73710 | Microarrays//HITS-CLIP | 37986224\|25263593\|19536157 | Unspecified |
| mmu-miR-21a-5p | MIMAT0000530 | Tubb4b | 227613 | Microarrays | 37986224\|25263593 | Unspecified |
| mmu-miR-21a-5p | MIMAT0000530 | Tubd1 | 56427 | HITS-CLIP | 23142080 | Unspecified |
| mmu-miR-21a-5p | MIMAT0000530 | Tubg1 | 103733 | Microarrays//HITS-CLIP | 37986224\|25263593\|23597149\|23142 | Unspecified |
| mmu-miR-21a-5p | MIMAT0000530 | Tubgcp3 | 259279 | HITS-CLIP | 23142080 | Unspecified |
| mmu-miR-21a-5p | MIMAT0000530 | Tubgcp5 | 233276 | HITS-CLIP | 23142080 | Unspecified |
| mmu-miR-21a-5p | MIMAT0000530 | Tubgcp6 | 328580 | HITS-CLIP | 29167373 | Unspecified |
| mmu-miR-21a-5p | MIMAT0000530 | Tulp4 | 68842 | HITS-CLIP | 37986224 | Unspecified |
| mmu-miR-21a-5p | MIMAT0000530 | Tusc2 | 80385 | HITS-CLIP | 37986224 | Unspecified |
| mmu-miR-21a-5p | MIMAT0000530 | Tut4 | 23318 | HITS-CLIP | 23597149\|23142080\|29167373 | Unspecified |
| mmu-miR-21a-5p | MIMAT0000530 | Tut7 | 79670 | HITS-CLIP | 23142080 | Unspecified |
| mmu-miR-21a-5p | MIMAT0000530 | Twf1 | 19230 | HITS-CLIP | 23597149\|23142080 | Unspecified |
| mmu-miR-21a-5p | MIMAT0000530 | Txndc12 | 66073 | HITS-CLIP | 23597149\|23142080 | Unspecified |
| mmu-miR-21a-5p | MIMAT0000530 | Txndc15 | 69672 | HITS-CLIP | 23597149\|23142080\|29167373 | Unspecified |

| | | | | | | |
|---|---|---|---|---|---|---|
| mmu-miR-21a-5p | MIMAT0000530 | Txndc16 | 70561 | Microarrays | 37986224 | Unspecified |
| mmu-miR-21a-5p | MIMAT0000530 | Txndc5 | 105245 | HITS-CLIP | 23597149\|23142080 | Unspecified |
| mmu-miR-21a-5p | MIMAT0000530 | Txnip | 56338 | Microarrays//qCLASH//HITS-CLIP | 37986224\|31393953\|23142080\|29167 | Unspecified |
| mmu-miR-21a-5p | MIMAT0000530 | Txnl1 | 53382 | HITS-CLIP | 23142080\|29167373\|23597149 | Unspecified |
| mmu-miR-21a-5p | MIMAT0000530 | Txnrd1 | 50493 | Microarrays//HITS-CLIP | 37986224\|25263593\|29167373 | Unspecified |
| mmu-miR-21a-5p | MIMAT0000530 | Tyms | 22171 | HITS-CLIP | 30971389 | Unspecified |
| mmu-miR-21a-5p | MIMAT0000530 | U2surp | 67958 | HITS-CLIP | 23142080\|29167373 | Unspecified |
| mmu-miR-21a-5p | MIMAT0000530 | Uap1 | 107652 | HITS-CLIP | 23142080 | Unspecified |
| mmu-miR-21a-5p | MIMAT0000530 | Uba1 | 22201 | HITS-CLIP | 23597149\|23142080\|29167373 | Unspecified |
| mmu-miR-21a-5p | MIMAT0000530 | Uba2 | 50995 | HITS-CLIP | 23142080 | Unspecified |
| mmu-miR-21a-5p | MIMAT0000530 | Uba3 | 22200 | HITS-CLIP//qCLASH | 23142080\|31393953 | Unspecified |
| mmu-miR-21a-5p | MIMAT0000530 | Uba5 | 66663 | HITS-CLIP | 23142080 | Unspecified |
| mmu-miR-21a-5p | MIMAT0000530 | Uba6 | 231380 | HITS-CLIP | 37986224\|23597149\|23142080 | Unspecified |
| mmu-miR-21a-5p | MIMAT0000530 | Uba7 | 74153 | HITS-CLIP | 23142080 | Unspecified |
| mmu-miR-21a-5p | MIMAT0000530 | Ubac2 | 68889 | HITS-CLIP | 23142080 | Unspecified |
| mmu-miR-21a-5p | MIMAT0000530 | Ubap1 | 67123 | HITS-CLIP | 23597149 | Unspecified |
| mmu-miR-21a-5p | MIMAT0000530 | Ubap2 | 68926 | HITS-CLIP | 37986224\|23597149\|23142080 | Unspecified |
| mmu-miR-21a-5p | MIMAT0000530 | Ubap2l | 74383 | HITS-CLIP | 23142080 | Unspecified |
| mmu-miR-21a-5p | MIMAT0000530 | Ubash3a | 328795 | HITS-CLIP | 23142080 | Unspecified |
| mmu-miR-21a-5p | MIMAT0000530 | Ubash3b | 72828 | Microarrays | 37986224\|25263593 | Unspecified |
| mmu-miR-21a-5p | MIMAT0000530 | Ube2a | 22209 | HITS-CLIP | 23142080 | Unspecified |
| mmu-miR-21a-5p | MIMAT0000530 | Ube2b | 22210 | HITS-CLIP | 37986224\|23597149\|23142080 | Unspecified |
| mmu-miR-21a-5p | MIMAT0000530 | Ube2c | 68612 | Microarrays//HITS-CLIP | 37986224\|25263593\|23142080 | Unspecified |
| mmu-miR-21a-5p | MIMAT0000530 | Ube2d2a | 56550 | HITS-CLIP | 29167373 | Unspecified |
| mmu-miR-21a-5p | MIMAT0000530 | Ube2d3 | 66105 | HITS-CLIP | 37986224\|23142080 | Unspecified |
| mmu-miR-21a-5p | MIMAT0000530 | Ube2e1 | 22194 | HITS-CLIP | 23142080 | Unspecified |
| mmu-miR-21a-5p | MIMAT0000530 | Ube2e3 | 22193 | HITS-CLIP | 23597149\|23142080 | Unspecified |
| mmu-miR-21a-5p | MIMAT0000530 | Ube2f | 67921 | HITS-CLIP | 23142080 | Unspecified |
| mmu-miR-21a-5p | MIMAT0000530 | Ube2g1 | 67128 | HITS-CLIP | 23142080 | Unspecified |
| mmu-miR-21a-5p | MIMAT0000530 | Ube2j1 | 56228 | HITS-CLIP | 23142080 | Unspecified |
| mmu-miR-21a-5p | MIMAT0000530 | Ube2k | 53323 | HITS-CLIP | 23142080\|29167373 | Unspecified |
| mmu-miR-21a-5p | MIMAT0000530 | Ube2l3 | 22195 | HITS-CLIP | 23597149 | Unspecified |
| mmu-miR-21a-5p | MIMAT0000530 | Ube2q2 | 109161 | HITS-CLIP | 37986224\|23142080 | Unspecified |
| mmu-miR-21a-5p | MIMAT0000530 | Ube2w | 66799 | HITS-CLIP | 23142080\|29167373 | Unspecified |
| mmu-miR-21a-5p | MIMAT0000530 | Ube2z | 268470 | qCLASH//HITS-CLIP | 31393953\|23142080 | Unspecified |
| mmu-miR-21a-5p | MIMAT0000530 | Ube3a | 22215 | HITS-CLIP | 37986224\|23142080 | Unspecified |
| mmu-miR-21a-5p | MIMAT0000530 | Ube4a | 140630 | HITS-CLIP | 23597149\|29167373 | Unspecified |
| mmu-miR-21a-5p | MIMAT0000530 | Ube4b | 63958 | HITS-CLIP | 19536157\|23142080\|29167373 | Unspecified |
| mmu-miR-21a-5p | MIMAT0000530 | Ubfd1 | 28018 | HITS-CLIP | 23142080 | Unspecified |

| | | | | | | |
|---|---|---|---|---|---|---|
| mmu-miR-21a-5p | MIMAT0000530 | Ubl3 | 24109 | HITS-CLIP | 23142080\|29167373 | Unspecified |
| mmu-miR-21a-5p | MIMAT0000530 | Ubl4a | 8266 | HITS-CLIP | 23142080\|23597149 | Unspecified |
| mmu-miR-21a-5p | MIMAT0000530 | Ubl7 | 69459 | HITS-CLIP | 23142080 | Unspecified |
| mmu-miR-21a-5p | MIMAT0000530 | Ubn1 | 170644 | HITS-CLIP | 29167373\|23142080 | Unspecified |
| mmu-miR-21a-5p | MIMAT0000530 | Ubn2 | 320538 | HITS-CLIP | 23142080\|29167373 | Unspecified |
| mmu-miR-21a-5p | MIMAT0000530 | Ubp1 | 22221 | HITS-CLIP | 23597149\|23142080 | Unspecified |
| mmu-miR-21a-5p | MIMAT0000530 | Ubqln1 | 56085 | HITS-CLIP | 23597149 | Unspecified |
| mmu-miR-21a-5p | MIMAT0000530 | Ubqln2 | 54609 | HITS-CLIP | 23142080 | Unspecified |
| mmu-miR-21a-5p | MIMAT0000530 | Ubr1 | 22222 | HITS-CLIP | 37986224\|23142080 | Unspecified |
| mmu-miR-21a-5p | MIMAT0000530 | Ubr2 | 224826 | HITS-CLIP | 23142080\|29167373 | Unspecified |
| mmu-miR-21a-5p | MIMAT0000530 | Ubr3 | 68795 | HITS-CLIP//qCLASH | 37986224\|31393953\|19536157\|23597 | Unspecified |
| mmu-miR-21a-5p | MIMAT0000530 | Ubr4 | 69116 | HITS-CLIP | 23597149\|23142080 | Unspecified |
| mmu-miR-21a-5p | MIMAT0000530 | Ubr5 | 70790 | HITS-CLIP | 23597149\|23142080\|29167373 | Unspecified |
| mmu-miR-21a-5p | MIMAT0000530 | Ubtd1 | 226122 | HITS-CLIP | 23142080 | Unspecified |
| mmu-miR-21a-5p | MIMAT0000530 | Ubtf | 21429 | HITS-CLIP | 37986224\|23142080\|29167373 | Unspecified |
| mmu-miR-21a-5p | MIMAT0000530 | Ubxn6 | 66530 | HITS-CLIP | 23142080\|29167373 | Unspecified |
| mmu-miR-21a-5p | MIMAT0000530 | Ubxn7 | 224111 | qCLASH//HITS-CLIP | 31393953\|23142080\|29167373 | Unspecified |
| mmu-miR-21a-5p | MIMAT0000530 | Uchl1 | 22223 | Microarrays | 37986224\|25263593 | Unspecified |
| mmu-miR-21a-5p | MIMAT0000530 | Uchl5 | 56207 | Microarrays//HITS-CLIP | 37986224\|25263593\|23597149 | Unspecified |
| mmu-miR-21a-5p | MIMAT0000530 | Ucp2 | 22228 | HITS-CLIP | 23142080 | Unspecified |
| mmu-miR-21a-5p | MIMAT0000530 | Ufd1 | 22230 | HITS-CLIP | 23142080 | Unspecified |
| mmu-miR-21a-5p | MIMAT0000530 | Ufl1 | 67490 | HITS-CLIP | 23597149\|23142080 | Unspecified |
| mmu-miR-21a-5p | MIMAT0000530 | Ufsp2 | 192169 | HITS-CLIP | 23142080 | Unspecified |
| mmu-miR-21a-5p | MIMAT0000530 | Ugcg | 22234 | HITS-CLIP | 37986224\|23597149\|31433990\|23142 | Unspecified |
| mmu-miR-21a-5p | MIMAT0000530 | Uggt1 | 320011 | HITS-CLIP | 23597149\|19536157\|23142080\|29167 | Unspecified |
| mmu-miR-21a-5p | MIMAT0000530 | Ugp2 | 216558 | HITS-CLIP | 23597149\|23142080 | Unspecified |
| mmu-miR-21a-5p | MIMAT0000530 | Ugt2a3 | 72094 | HITS-CLIP | 23597149 | Unspecified |
| mmu-miR-21a-5p | MIMAT0000530 | Ugt2b34 | 100727 | HITS-CLIP | 23597149 | Unspecified |
| mmu-miR-21a-5p | MIMAT0000530 | Ugt3a2 | 223337 | HITS-CLIP | 23597149 | Unspecified |
| mmu-miR-21a-5p | MIMAT0000530 | Ugt8a | 22239 | HITS-CLIP | 37986224\|19536157 | Unspecified |
| mmu-miR-21a-5p | MIMAT0000530 | Uhmk1 | 16589 | HITS-CLIP | 23142080\|29167373 | Unspecified |
| mmu-miR-21a-5p | MIMAT0000530 | Uhrf1 | 18140 | Microarrays//HITS-CLIP | 37986224\|25263593\|23142080\|29167 | Unspecified |
| mmu-miR-21a-5p | MIMAT0000530 | Uhrf1bp1l | 75089 | HITS-CLIP | 37986224\|23142080 | Unspecified |
| mmu-miR-21a-5p | MIMAT0000530 | Ulk2 | 29869 | HITS-CLIP | 23597149\|29167373 | Unspecified |
| mmu-miR-21a-5p | MIMAT0000530 | Ulk3 | 71742 | HITS-CLIP | 29167373 | Unspecified |
| mmu-miR-21a-5p | MIMAT0000530 | Umps | 22247 | HITS-CLIP | 23597149 | Unspecified |
| mmu-miR-21a-5p | MIMAT0000530 | Unc119b | 106840 | HITS-CLIP | 23142080 | Unspecified |
| mmu-miR-21a-5p | MIMAT0000530 | Unc13b | 22249 | HITS-CLIP | 37986224\|23142080\|23597149 | Unspecified |
| mmu-miR-21a-5p | MIMAT0000530 | Unc80 | 329178 | HITS-CLIP | 37986224\|19536157 | Unspecified |

| | | | | | | |
|---|---|---|---|---|---|---|
| mmu-miR-21a-5p | MIMAT0000530 | Unc93b1 | 54445 | Microarrays | 37986224 | Unspecified |
| mmu-miR-21a-5p | MIMAT0000530 | Ung | 22256 | Microarrays | 37986224\|25263593 | Unspecified |
| mmu-miR-21a-5p | MIMAT0000530 | Uox | 22262 | HITS-CLIP | 23597149 | Unspecified |
| mmu-miR-21a-5p | MIMAT0000530 | Upf1 | 19704 | HITS-CLIP | 23597149\|23142080\|29167373 | Unspecified |
| mmu-miR-21a-5p | MIMAT0000530 | Upf2 | 326622 | HITS-CLIP | 23142080 | Unspecified |
| mmu-miR-21a-5p | MIMAT0000530 | Upf3b | 68134 | HITS-CLIP | 23142080 | Unspecified |
| mmu-miR-21a-5p | MIMAT0000530 | Upp2 | 76654 | HITS-CLIP | 23597149 | Unspecified |
| mmu-miR-21a-5p | MIMAT0000530 | Uqcc2 | 67267 | Microarrays | 37986224\|25263593 | Unspecified |
| mmu-miR-21a-5p | MIMAT0000530 | Uqcr10 | 66152 | HITS-CLIP | 23142080 | Unspecified |
| mmu-miR-21a-5p | MIMAT0000530 | Uqcrc2 | 67003 | HITS-CLIP | 23597149\|23142080 | Unspecified |
| mmu-miR-21a-5p | MIMAT0000530 | Uqcrfs1 | 66694 | HITS-CLIP | 23142080 | Unspecified |
| mmu-miR-21a-5p | MIMAT0000530 | Urah | 76974 | HITS-CLIP | 23597149 | Unspecified |
| mmu-miR-21a-5p | MIMAT0000530 | Urb1 | 207932 | HITS-CLIP | 23142080 | Unspecified |
| mmu-miR-21a-5p | MIMAT0000530 | Urb2 | 382038 | HITS-CLIP | 23597149\|23142080 | Unspecified |
| mmu-miR-21a-5p | MIMAT0000530 | Uri1 | 19777 | HITS-CLIP | 37986224\|23142080 | Unspecified |
| mmu-miR-21a-5p | MIMAT0000530 | Uroc1 | 243537 | HITS-CLIP | 37986224\|23597149 | Unspecified |
| mmu-miR-21a-5p | MIMAT0000530 | Use1 | 67023 | Microarrays | 37986224 | Unspecified |
| mmu-miR-21a-5p | MIMAT0000530 | Usf2 | 22282 | HITS-CLIP | 23597149\|23142080 | Unspecified |
| mmu-miR-21a-5p | MIMAT0000530 | Usf3 | 205717 | HITS-CLIP | 23142080\|29167373 | Unspecified |
| mmu-miR-21a-5p | MIMAT0000530 | Uso1 | 56041 | qCLASH//HITS-CLIP | 31393953\|23142080 | Unspecified |
| mmu-miR-21a-5p | MIMAT0000530 | Usp1 | 230484 | HITS-CLIP | 23142080 | Unspecified |
| mmu-miR-21a-5p | MIMAT0000530 | Usp10 | 22224 | Microarrays//HITS-CLIP | 37986224\|25263593\|23597149 | Unspecified |
| mmu-miR-21a-5p | MIMAT0000530 | Usp11 | 236733 | Microarrays | 37986224 | Unspecified |
| mmu-miR-21a-5p | MIMAT0000530 | Usp15 | 14479 | HITS-CLIP | 37986224\|23142080 | Unspecified |
| mmu-miR-21a-5p | MIMAT0000530 | Usp16 | 74112 | HITS-CLIP | 23142080 | Unspecified |
| mmu-miR-21a-5p | MIMAT0000530 | Usp17la | 13531 | Microarrays | 37986224\|25263593 | Unspecified |
| mmu-miR-21a-5p | MIMAT0000530 | Usp18 | 24110 | Microarrays | 37986224\|25263593 | Unspecified |
| mmu-miR-21a-5p | MIMAT0000530 | Usp19 | 71472 | HITS-CLIP | 37986224\|23597149\|23142080\|29167 | Unspecified |
| mmu-miR-21a-5p | MIMAT0000530 | Usp2 | 53376 | HITS-CLIP | 23597149\|23142080 | Unspecified |
| mmu-miR-21a-5p | MIMAT0000530 | Usp21 | 30941 | Microarrays | 37986224 | Unspecified |
| mmu-miR-21a-5p | MIMAT0000530 | Usp24 | 329908 | HITS-CLIP | 23142080\|29167373 | Unspecified |
| mmu-miR-21a-5p | MIMAT0000530 | Usp25 | 30940 | HITS-CLIP | 23597149\|23142080\|29167373 | Unspecified |
| mmu-miR-21a-5p | MIMAT0000530 | Usp28 | 235323 | HITS-CLIP | 23142080 | Unspecified |
| mmu-miR-21a-5p | MIMAT0000530 | Usp31 | 76179 | HITS-CLIP | 26203562 | Unspecified |
| mmu-miR-21a-5p | MIMAT0000530 | Usp32 | 237898 | HITS-CLIP | 23597149\|19536157\|23142080 | Unspecified |
| mmu-miR-21a-5p | MIMAT0000530 | Usp33 | 170822 | HITS-CLIP | 23142080\|29167373 | Unspecified |
| mmu-miR-21a-5p | MIMAT0000530 | Usp34 | 17847 | HITS-CLIP//qCLASH | 37986224\|23142080\|31393953\|1953[illegible] | Unspecified |
| mmu-miR-21a-5p | MIMAT0000530 | Usp37 | 319651 | HITS-CLIP | 37986224\|23142080\|29167373 | Unspecified |
| mmu-miR-21a-5p | MIMAT0000530 | Usp38 | 74841 | HITS-CLIP | 23597149\|23142080 | Unspecified |

| | | | | | | |
|---|---|---|---|---|---|---|
| mmu-miR-21a-5p | MIMAT0000530 | Usp39 | 28035 | HITS-CLIP | 31433990 | Unspecified |
| mmu-miR-21a-5p | MIMAT0000530 | Usp4 | 22258 | HITS-CLIP | 37986224\|23597149\|23142080\|29167 | Unspecified |
| mmu-miR-21a-5p | MIMAT0000530 | Usp42 | 76800 | HITS-CLIP | 37986224\|23597149\|23142080 | Unspecified |
| mmu-miR-21a-5p | MIMAT0000530 | Usp46 | 69727 | Microarrays | 37986224\|25263593 | Unspecified |
| mmu-miR-21a-5p | MIMAT0000530 | Usp47 | 74996 | HITS-CLIP | 23142080\|29167373 | Unspecified |
| mmu-miR-21a-5p | MIMAT0000530 | Usp48 | 170707 | HITS-CLIP | 23142080\|29167373 | Unspecified |
| mmu-miR-21a-5p | MIMAT0000530 | Usp49 | 224836 | HITS-CLIP | 37986224\|23142080 | Unspecified |
| mmu-miR-21a-5p | MIMAT0000530 | Usp5 | 22225 | HITS-CLIP | 31433990\|23142080 | Unspecified |
| mmu-miR-21a-5p | MIMAT0000530 | Usp53 | 99526 | HITS-CLIP | 23142080 | Unspecified |
| mmu-miR-21a-5p | MIMAT0000530 | Usp6nl | 98910 | Microarrays | 37986224\|25263593 | Unspecified |
| mmu-miR-21a-5p | MIMAT0000530 | Usp7 | 252870 | HITS-CLIP | 23142080 | Unspecified |
| mmu-miR-21a-5p | MIMAT0000530 | Usp8 | 84092 | Microarrays//HITS-CLIP | 37986224\|23597149\|23142080 | Unspecified |
| mmu-miR-21a-5p | MIMAT0000530 | Usp9x | 22284 | HITS-CLIP | 23142080\|29167373 | Unspecified |
| mmu-miR-21a-5p | MIMAT0000530 | Uspl1 | 231915 | HITS-CLIP | 23142080 | Unspecified |
| mmu-miR-21a-5p | MIMAT0000530 | Ust | 338362 | HITS-CLIP | 23142080 | Unspecified |
| mmu-miR-21a-5p | MIMAT0000530 | Utp14a | 72554 | HITS-CLIP | 23597149 | Unspecified |
| mmu-miR-21a-5p | MIMAT0000530 | Utp14b | 195434 | HITS-CLIP | 23142080 | Unspecified |
| mmu-miR-21a-5p | MIMAT0000530 | Utp18 | 217109 | HITS-CLIP | 23142080 | Unspecified |
| mmu-miR-21a-5p | MIMAT0000530 | Utp20 | 70683 | HITS-CLIP | 29167373 | Unspecified |
| mmu-miR-21a-5p | MIMAT0000530 | Utp25 | 27042 | HITS-CLIP | 31433990 | Unspecified |
| mmu-miR-21a-5p | MIMAT0000530 | Utp4 | 21771 | qCLASH | 31393953 | Unspecified |
| mmu-miR-21a-5p | MIMAT0000530 | Utp6 | 216987 | HITS-CLIP | 23597149\|23142080 | Unspecified |
| mmu-miR-21a-5p | MIMAT0000530 | Utrn | 22288 | HITS-CLIP | 37986224\|23142080 | Unspecified |
| mmu-miR-21a-5p | MIMAT0000530 | Uty | 22290 | HITS-CLIP | 23142080 | Unspecified |
| mmu-miR-21a-5p | MIMAT0000530 | Uvrag | 78610 | HITS-CLIP | 29167373\|23142080 | Unspecified |
| mmu-miR-21a-5p | MIMAT0000530 | Uvssa | 71101 | HITS-CLIP | 23142080 | Unspecified |
| mmu-miR-21a-5p | MIMAT0000530 | Uxs1 | 67883 | HITS-CLIP | 19536157\|23142080 | Unspecified |
| mmu-miR-21a-5p | MIMAT0000530 | Vac14 | 234729 | HITS-CLIP | 23597149\|23142080 | Unspecified |
| mmu-miR-21a-5p | MIMAT0000530 | Vangl1 | 229658 | HITS-CLIP | 23597149 | Unspecified |
| mmu-miR-21a-5p | MIMAT0000530 | Vapa | 30960 | HITS-CLIP | 37986224\|23597149\|23142080 | Unspecified |
| mmu-miR-21a-5p | MIMAT0000530 | Vasn | 246154 | Microarrays | 37986224\|25263593 | Unspecified |
| mmu-miR-21a-5p | MIMAT0000530 | Vat1 | 26949 | HITS-CLIP | 23597149 | Unspecified |
| mmu-miR-21a-5p | MIMAT0000530 | Vav2 | 22325 | HITS-CLIP | 23597149 | Unspecified |
| mmu-miR-21a-5p | MIMAT0000530 | Vav3 | 57257 | HITS-CLIP | 23142080\|29167373 | Unspecified |
| mmu-miR-21a-5p | MIMAT0000530 | Vbp1 | 22327 | HITS-CLIP | 23142080\|29167373 | Unspecified |
| mmu-miR-21a-5p | MIMAT0000530 | Vcam1 | 22329 | HITS-CLIP | 23597149 | Unspecified |
| mmu-miR-21a-5p | MIMAT0000530 | Vcpip1 | 70675 | HITS-CLIP | 23142080 | Unspecified |
| mmu-miR-21a-5p | MIMAT0000530 | Vdac1 | 22333 | HITS-CLIP | 23597149\|23142080 | Unspecified |
| mmu-miR-21a-5p | MIMAT0000530 | Vegfa | 22339 | Microarrays//HITS-CLIP | 37986224\|23142080 | Unspecified |

| | | | | | | |
|---|---|---|---|---|---|---|
| mmu-miR-21a-5p | MIMAT0000530 | Vezf1 | 22344 | HITS-CLIP | 23597149\|23142080\|29167373 | Unspecified |
| mmu-miR-21a-5p | MIMAT0000530 | Vezt | 215008 | HITS-CLIP | 23597149\|23142080 | Unspecified |
| mmu-miR-21a-5p | MIMAT0000530 | Vgll3 | 73569 | Microarrays | 37986224\|25263593 | Unspecified |
| mmu-miR-21a-5p | MIMAT0000530 | Vhl | 22346 | HITS-CLIP | 23142080 | Unspecified |
| mmu-miR-21a-5p | MIMAT0000530 | Vill | 22351 | Microarrays | 37986224\|25263593 | Unspecified |
| mmu-miR-21a-5p | MIMAT0000530 | Vim | 22352 | HITS-CLIP | 30971389\|31433990\|28735896 | Unspecified |
| mmu-miR-21a-5p | MIMAT0000530 | Virma | 25962 | HITS-CLIP | 23597149\|23142080\|29167373 | Unspecified |
| mmu-miR-21a-5p | MIMAT0000530 | Vldlr | 22359 | HITS-CLIP | 23142080 | Unspecified |
| mmu-miR-21a-5p | MIMAT0000530 | Vma21 | 67048 | HITS-CLIP | 23142080 | Unspecified |
| mmu-miR-21a-5p | MIMAT0000530 | Vmn1r43 | 113847 | qCLASH | 31393953 | Unspecified |
| mmu-miR-21a-5p | MIMAT0000530 | Vmp1 | 75909 | HITS-CLIP | 23597149\|23142080\|31433990\|26203 | Unspecified |
| mmu-miR-21a-5p | MIMAT0000530 | Vps11 | 71732 | HITS-CLIP | 23142080 | Unspecified |
| mmu-miR-21a-5p | MIMAT0000530 | Vps13a | 271564 | HITS-CLIP | 23142080 | Unspecified |
| mmu-miR-21a-5p | MIMAT0000530 | Vps13b | 666173 | HITS-CLIP | 19536157\|23142080\|29167373 | Unspecified |
| mmu-miR-21a-5p | MIMAT0000530 | Vps13c | 320528 | HITS-CLIP//qCLASH | 23597149\|31393953\|23142080\|29167 | Unspecified |
| mmu-miR-21a-5p | MIMAT0000530 | Vps13d | 230895 | HITS-CLIP | 23142080 | Unspecified |
| mmu-miR-21a-5p | MIMAT0000530 | Vps16 | 80743 | HITS-CLIP | 29167373 | Unspecified |
| mmu-miR-21a-5p | MIMAT0000530 | Vps18 | 228545 | HITS-CLIP | 29167373 | Unspecified |
| mmu-miR-21a-5p | MIMAT0000530 | Vps26a | 30930 | HITS-CLIP | 23142080 | Unspecified |
| mmu-miR-21a-5p | MIMAT0000530 | Vps28 | 51160 | HITS-CLIP | 23142080 | Unspecified |
| mmu-miR-21a-5p | MIMAT0000530 | Vps33a | 77573 | HITS-CLIP | 23597149 | Unspecified |
| mmu-miR-21a-5p | MIMAT0000530 | Vps35 | 65114 | HITS-CLIP | 37986224\|23597149\|23142080\|29167 | Unspecified |
| mmu-miR-21a-5p | MIMAT0000530 | Vps36 | 70160 | HITS-CLIP | 37986224\|23142080 | Unspecified |
| mmu-miR-21a-5p | MIMAT0000530 | Vps37a | 52348 | qCLASH//HITS-CLIP | 31393953\|23142080 | Unspecified |
| mmu-miR-21a-5p | MIMAT0000530 | Vps39 | 269338 | HITS-CLIP | 23142080 | Unspecified |
| mmu-miR-21a-5p | MIMAT0000530 | Vps4b | 20479 | HITS-CLIP | 23597149\|23142080\|29167373 | Unspecified |
| mmu-miR-21a-5p | MIMAT0000530 | Vps50 | 55610 | HITS-CLIP | 23142080 | Unspecified |
| mmu-miR-21a-5p | MIMAT0000530 | Vps53 | 68299 | HITS-CLIP | 23142080 | Unspecified |
| mmu-miR-21a-5p | MIMAT0000530 | Vps54 | 245944 | HITS-CLIP | 23142080 | Unspecified |
| mmu-miR-21a-5p | MIMAT0000530 | Vsig10 | 231668 | Microarrays | 37986224 | Unspecified |
| mmu-miR-21a-5p | MIMAT0000530 | Vsir | 64115 | HITS-CLIP | 23142080\|29167373 | Unspecified |
| mmu-miR-21a-5p | MIMAT0000530 | Vta1 | 66201 | HITS-CLIP | 23142080 | Unspecified |
| mmu-miR-21a-5p | MIMAT0000530 | Vtn | 22370 | HITS-CLIP | 23597149 | Unspecified |
| mmu-miR-21a-5p | MIMAT0000530 | Wac | 225131 | HITS-CLIP | 23142080\|23597149\|31433990\|29167 | Unspecified |
| mmu-miR-21a-5p | MIMAT0000530 | Wapl | 23063 | qCLASH//HITS-CLIP | 31393953\|23597149\|23142080\|29167 | Unspecified |
| mmu-miR-21a-5p | MIMAT0000530 | Wasf1 | 83767 | HITS-CLIP | 19536157 | Unspecified |
| mmu-miR-21a-5p | MIMAT0000530 | Wasf2 | 242687 | HITS-CLIP | 23142080 | Unspecified |
| mmu-miR-21a-5p | MIMAT0000530 | Washc2 | 28006 | HITS-CLIP | 29167373 | Unspecified |
| mmu-miR-21a-5p | MIMAT0000530 | Washc4 | 319277 | HITS-CLIP | 23597149\|29167373\|23142080 | Unspecified |

| | | | | | | |
|---|---|---|---|---|---|---|
| mmu-miR-21a-5p | MIMAT0000530 | Washc5 | 9897 | HITS-CLIP | 23142080 | Unspecified |
| mmu-miR-21a-5p | MIMAT0000530 | Wasl | 73178 | HITS-CLIP | 19536157\|23142080 | Unspecified |
| mmu-miR-21a-5p | MIMAT0000530 | Wbp11 | 60321 | HITS-CLIP | 23142080 | Unspecified |
| mmu-miR-21a-5p | MIMAT0000530 | Wbp1l | 226178 | HITS-CLIP | 23597149 | Unspecified |
| mmu-miR-21a-5p | MIMAT0000530 | Wbp2 | 22378 | HITS-CLIP | 23597149\|23142080 | Unspecified |
| mmu-miR-21a-5p | MIMAT0000530 | Wdfy1 | 69368 | HITS-CLIP | 23597149\|23142080 | Unspecified |
| mmu-miR-21a-5p | MIMAT0000530 | Wdfy2 | 268752 | Microarrays | 37986224 | Unspecified |
| mmu-miR-21a-5p | MIMAT0000530 | Wdfy3 | 72145 | HITS-CLIP | 23597149\|19536157\|29167373 | Unspecified |
| mmu-miR-21a-5p | MIMAT0000530 | Wdfy4 | 545030 | HITS-CLIP | 29167373 | Unspecified |
| mmu-miR-21a-5p | MIMAT0000530 | Wdr1 | 22388 | HITS-CLIP | 23142080 | Unspecified |
| mmu-miR-21a-5p | MIMAT0000530 | Wdr11 | 207425 | HITS-CLIP | 23142080 | Unspecified |
| mmu-miR-21a-5p | MIMAT0000530 | Wdr13 | 73447 | HITS-CLIP | 23142080 | Unspecified |
| mmu-miR-21a-5p | MIMAT0000530 | Wdr20 | 69641 | HITS-CLIP | 23142080 | Unspecified |
| mmu-miR-21a-5p | MIMAT0000530 | Wdr26 | 226757 | HITS-CLIP | 37986224\|23142080\|29167373\|23597 | Unspecified |
| mmu-miR-21a-5p | MIMAT0000530 | Wdr3 | 269470 | HITS-CLIP | 23597149 | Unspecified |
| mmu-miR-21a-5p | MIMAT0000530 | Wdr33 | 74320 | HITS-CLIP | 23142080 | Unspecified |
| mmu-miR-21a-5p | MIMAT0000530 | Wdr35 | 74682 | HITS-CLIP | 23142080 | Unspecified |
| mmu-miR-21a-5p | MIMAT0000530 | Wdr41 | 218460 | HITS-CLIP | 23142080 | Unspecified |
| mmu-miR-21a-5p | MIMAT0000530 | Wdr45 | 54636 | Microarrays//HITS-CLIP | 37986224\|23597149\|23142080 | Unspecified |
| mmu-miR-21a-5p | MIMAT0000530 | Wdr45b | 66840 | HITS-CLIP | 23142080 | Unspecified |
| mmu-miR-21a-5p | MIMAT0000530 | Wdr48 | 67561 | HITS-CLIP | 23142080 | Unspecified |
| mmu-miR-21a-5p | MIMAT0000530 | Wdr59 | 319481 | HITS-CLIP | 23142080 | Unspecified |
| mmu-miR-21a-5p | MIMAT0000530 | Wdr6 | 83669 | HITS-CLIP | 23142080 | Unspecified |
| mmu-miR-21a-5p | MIMAT0000530 | Wdr7 | 104082 | HITS-CLIP | 23142080 | Unspecified |
| mmu-miR-21a-5p | MIMAT0000530 | Wdr76 | 241627 | HITS-CLIP | 23142080 | Unspecified |
| mmu-miR-21a-5p | MIMAT0000530 | Wdr77 | 70465 | HITS-CLIP | 23142080 | Unspecified |
| mmu-miR-21a-5p | MIMAT0000530 | Wdr81 | 192652 | HITS-CLIP | 37986224 | Unspecified |
| mmu-miR-21a-5p | MIMAT0000530 | Wdr82 | 77305 | HITS-CLIP | 23142080 | Unspecified |
| mmu-miR-21a-5p | MIMAT0000530 | Wdr89 | 72338 | HITS-CLIP | 29167373\|23142080 | Unspecified |
| mmu-miR-21a-5p | MIMAT0000530 | Wdr92 | 103784 | HITS-CLIP | 23142080 | Unspecified |
| mmu-miR-21a-5p | MIMAT0000530 | Wdsub1 | 72137 | Microarrays | 37986224 | Unspecified |
| mmu-miR-21a-5p | MIMAT0000530 | Wee1 | 22390 | HITS-CLIP | 23597149\|31433990\|23142080\|29167 | Unspecified |
| mmu-miR-21a-5p | MIMAT0000530 | Wfdc17 | 100034251 | Microarrays | 37986224 | Unspecified |
| mmu-miR-21a-5p | MIMAT0000530 | Wfdc18 | 14038 | Microarrays | 37986224 | Unspecified |
| mmu-miR-21a-5p | MIMAT0000530 | Wfikkn2 | 278507 | Microarrays | 37986224 | Unspecified |
| mmu-miR-21a-5p | MIMAT0000530 | Wfs1 | 22393 | HITS-CLIP | 31433990 | Unspecified |
| mmu-miR-21a-5p | MIMAT0000530 | Whamm | 434204 | Microarrays | 37986224 | Unspecified |
| mmu-miR-21a-5p | MIMAT0000530 | Wipf1 | 215280 | HITS-CLIP | 37986224\|23142080 | Unspecified |
| mmu-miR-21a-5p | MIMAT0000530 | Wipi2 | 74781 | HITS-CLIP | 37986224\|23597149\|23142080 | Unspecified |

| | | | | | | |
|---|---|---|---|---|---|---|
| mmu-miR-21a-5p | MIMAT0000530 | Wiz | 22404 | HITS-CLIP | 31433990 | Unspecified |
| mmu-miR-21a-5p | MIMAT0000530 | Wnk1 | 232341 | HITS-CLIP | 37986224\|23597149\|23142080\|29167 | Unspecified |
| mmu-miR-21a-5p | MIMAT0000530 | Wnt7a | 22421 | HITS-CLIP | 37986224 | Unspecified |
| mmu-miR-21a-5p | MIMAT0000530 | Wrn | 22427 | HITS-CLIP, Microarrays//HITS-CLIP | 37986224\|23142080 | Unspecified |
| mmu-miR-21a-5p | MIMAT0000530 | Wsb1 | 78889 | HITS-CLIP | 23142080\|29167373 | Unspecified |
| mmu-miR-21a-5p | MIMAT0000530 | Wwp1 | 107568 | HITS-CLIP, Microarrays//HITS-CLIP | 37986224\|31433990\|23142080\|29167 | Unspecified |
| mmu-miR-21a-5p | MIMAT0000530 | Xaf1 | 327959 | Microarrays | 37986224\|25263593 | Unspecified |
| mmu-miR-21a-5p | MIMAT0000530 | Xbp1 | 22433 | HITS-CLIP | 23142080 | Unspecified |
| mmu-miR-21a-5p | MIMAT0000530 | Xdh | 22436 | HITS-CLIP | 23597149\|23142080 | Unspecified |
| mmu-miR-21a-5p | MIMAT0000530 | Xiap | 11798 | HITS-CLIP | 37986224\|23597149\|23142080 | Unspecified |
| mmu-miR-21a-5p | MIMAT0000530 | Xk | 22439 | HITS-CLIP | 23597149 | Unspecified |
| mmu-miR-21a-5p | MIMAT0000530 | Xkr6 | 219149 | HITS-CLIP | 37986224 | Unspecified |
| mmu-miR-21a-5p | MIMAT0000530 | Xpnpep3 | 321003 | HITS-CLIP | 23142080\|23597149 | Unspecified |
| mmu-miR-21a-5p | MIMAT0000530 | Xpo1 | 103573 | HITS-CLIP | 23142080 | Unspecified |
| mmu-miR-21a-5p | MIMAT0000530 | Xpo4 | 57258 | HITS-CLIP | 23142080\|23597149 | Unspecified |
| mmu-miR-21a-5p | MIMAT0000530 | Xpo5 | 72322 | HITS-CLIP | 23142080 | Unspecified |
| mmu-miR-21a-5p | MIMAT0000530 | Xpo6 | 74204 | qCLASH | 31393953 | Unspecified |
| mmu-miR-21a-5p | MIMAT0000530 | Xpo7 | 65246 | HITS-CLIP | 29167373 | Unspecified |
| mmu-miR-21a-5p | MIMAT0000530 | Xpr1 | 19775 | HITS-CLIP | 23597149\|19536157\|29167373\|23142 | Unspecified |
| mmu-miR-21a-5p | MIMAT0000530 | Xrcc6 | 14375 | HITS-CLIP | 23142080 | Unspecified |
| mmu-miR-21a-5p | MIMAT0000530 | Xrn1 | 24127 | HITS-CLIP | 23142080\|29167373 | Unspecified |
| mmu-miR-21a-5p | MIMAT0000530 | Xrn2 | 24128 | HITS-CLIP | 23142080 | Unspecified |
| mmu-miR-21a-5p | MIMAT0000530 | Yae1d1 | 67008 | Microarrays | 37986224\|25263593 | Unspecified |
| mmu-miR-21a-5p | MIMAT0000530 | Yaf2 | 67057 | HITS-CLIP | 23142080 | Unspecified |
| mmu-miR-21a-5p | MIMAT0000530 | Yap1 | 22601 | HITS-CLIP | 37986224\|23597149\|19536157 | Unspecified |
| mmu-miR-21a-5p | MIMAT0000530 | Yars | 107271 | HITS-CLIP | 23597149\|23142080 | Unspecified |
| mmu-miR-21a-5p | MIMAT0000530 | Yars2 | 70120 | Microarrays | 37986224\|25263593 | Unspecified |
| mmu-miR-21a-5p | MIMAT0000530 | Ybx3 | 56449 | Microarrays | 37986224\|25263593 | Unspecified |
| mmu-miR-21a-5p | MIMAT0000530 | Yeats2 | 208146 | HITS-CLIP | 37986224\|23142080 | Unspecified |
| mmu-miR-21a-5p | MIMAT0000530 | Yipf5 | 67180 | HITS-CLIP | 23597149 | Unspecified |
| mmu-miR-21a-5p | MIMAT0000530 | Yipf6 | 77929 | HITS-CLIP | 37986224\|23597149\|23142080 | Unspecified |
| mmu-miR-21a-5p | MIMAT0000530 | Ylpm1 | 56531 | HITS-CLIP | 23142080\|29167373 | Unspecified |
| mmu-miR-21a-5p | MIMAT0000530 | Yme1l1 | 27377 | HITS-CLIP | 23597149\|23142080 | Unspecified |
| mmu-miR-21a-5p | MIMAT0000530 | Ypel2 | 77864 | Microarrays | 37986224 | Unspecified |
| mmu-miR-21a-5p | MIMAT0000530 | Ypel3 | 66090 | Microarrays | 37986224 | Unspecified |
| mmu-miR-21a-5p | MIMAT0000530 | Yrdc | 230734 | HITS-CLIP | 23142080 | Unspecified |
| mmu-miR-21a-5p | MIMAT0000530 | Ythdc1 | 231386 | HITS-CLIP | 23597149\|23142080 | Unspecified |
| mmu-miR-21a-5p | MIMAT0000530 | Ythdc2 | 240255 | HITS-CLIP | 23142080 | Unspecified |
| mmu-miR-21a-5p | MIMAT0000530 | Ythdf1 | 228994 | HITS-CLIP | 19536157\|23597149\|31433990\|23142 | Unspecified |

| | | | | | | |
|---|---|---|---|---|---|---|
| mmu-miR-21a-5p | MIMAT0000530 | Ythdf2 | 213541 | HITS-CLIP | 23142080 | Unspecified |
| mmu-miR-21a-5p | MIMAT0000530 | Ythdf3 | 229096 | HITS-CLIP | 37986224\|23142080\|29167373 | Unspecified |
| mmu-miR-21a-5p | MIMAT0000530 | Ywhae | 22627 | HITS-CLIP | 37986224\|23142080 | Unspecified |
| mmu-miR-21a-5p | MIMAT0000530 | Ywhah | 22629 | HITS-CLIP | 23597149\|23142080\|29167373 | Unspecified |
| mmu-miR-21a-5p | MIMAT0000530 | Ywhaz | 22631 | HITS-CLIP | 23142080 | Unspecified |
| mmu-miR-21a-5p | MIMAT0000530 | Yy1 | 22632 | Luciferase assay//qRT-PCR//HITS-CLIP//Lu | 24891504\|23597149\|23142080 | Unspecified |
| mmu-miR-21a-5p | MIMAT0000530 | Zbed6 | 667118 | HITS-CLIP | 37986224 | Unspecified |
| mmu-miR-21a-5p | MIMAT0000530 | Zbp1 | 58203 | HITS-CLIP | 29167373 | Unspecified |
| mmu-miR-21a-5p | MIMAT0000530 | Zbtb1 | 268564 | HITS-CLIP | 23142080 | Unspecified |
| mmu-miR-21a-5p | MIMAT0000530 | Zbtb11 | 271377 | HITS-CLIP | 23142080 | Unspecified |
| mmu-miR-21a-5p | MIMAT0000530 | Zbtb18 | 30928 | HITS-CLIP, Microarrays//Microarrays//HITS | 37986224\|25263593\|19536157\|23142 | Unspecified |
| mmu-miR-21a-5p | MIMAT0000530 | Zbtb20 | 56490 | HITS-CLIP | 37986224\|23597149\|19536157\|23142 | Unspecified |
| mmu-miR-21a-5p | MIMAT0000530 | Zbtb21 | 114565 | HITS-CLIP | 23142080\|29167373 | Unspecified |
| mmu-miR-21a-5p | MIMAT0000530 | Zbtb33 | 56805 | HITS-CLIP | 37986224\|23142080 | Unspecified |
| mmu-miR-21a-5p | MIMAT0000530 | Zbtb34 | 241311 | HITS-CLIP | 37986224 | Unspecified |
| mmu-miR-21a-5p | MIMAT0000530 | Zbtb37 | 240869 | HITS-CLIP | 23142080 | Unspecified |
| mmu-miR-21a-5p | MIMAT0000530 | Zbtb38 | 245007 | HITS-CLIP | 23142080\|31433990\|29167373 | Unspecified |
| mmu-miR-21a-5p | MIMAT0000530 | Zbtb40 | 230848 | HITS-CLIP | 23597149 | Unspecified |
| mmu-miR-21a-5p | MIMAT0000530 | Zbtb41 | 226470 | HITS-CLIP//qCLASH | 37986224\|31393953\|23142080 | Unspecified |
| mmu-miR-21a-5p | MIMAT0000530 | Zbtb43 | 71834 | HITS-CLIP | 31433990 | Unspecified |
| mmu-miR-21a-5p | MIMAT0000530 | Zbtb44 | 235132 | HITS-CLIP | 23597149\|23142080 | Unspecified |
| mmu-miR-21a-5p | MIMAT0000530 | Zbtb45 | 232879 | HITS-CLIP | 23142080 | Unspecified |
| mmu-miR-21a-5p | MIMAT0000530 | Zbtb46 | 72147 | HITS-CLIP | 23597149 | Unspecified |
| mmu-miR-21a-5p | MIMAT0000530 | Zbtb49 | 75079 | HITS-CLIP | 23597149 | Unspecified |
| mmu-miR-21a-5p | MIMAT0000530 | Zbtb5 | 230119 | HITS-CLIP | 31433990 | Unspecified |
| mmu-miR-21a-5p | MIMAT0000530 | Zbtb6 | 241322 | HITS-CLIP | 29167373 | Unspecified |
| mmu-miR-21a-5p | MIMAT0000530 | Zbtb7a | 16969 | HITS-CLIP | 37986224\|23597149\|23142080\|29167 | Unspecified |
| mmu-miR-21a-5p | MIMAT0000530 | Zc3h12c | 244871 | HITS-CLIP | 23142080\|29167373 | Unspecified |
| mmu-miR-21a-5p | MIMAT0000530 | Zc3h12d | 237256 | HITS-CLIP | 23142080 | Unspecified |
| mmu-miR-21a-5p | MIMAT0000530 | Zc3h13 | 67302 | HITS-CLIP | 23597149\|23142080 | Unspecified |
| mmu-miR-21a-5p | MIMAT0000530 | Zc3h18 | 76014 | HITS-CLIP | 23597149 | Unspecified |
| mmu-miR-21a-5p | MIMAT0000530 | Zc3h4 | 330474 | Microarrays | 37986224 | Unspecified |
| mmu-miR-21a-5p | MIMAT0000530 | Zc3h6 | 78751 | HITS-CLIP | 19536157\|23142080 | Unspecified |
| mmu-miR-21a-5p | MIMAT0000530 | Zc3h7a | 106205 | HITS-CLIP | 23142080 | Unspecified |
| mmu-miR-21a-5p | MIMAT0000530 | Zc3hav1 | 78781 | HITS-CLIP | 23597149\|23142080\|29167373 | Unspecified |
| mmu-miR-21a-5p | MIMAT0000530 | Zcchc14 | 142682 | HITS-CLIP | 29167373 | Unspecified |
| mmu-miR-21a-5p | MIMAT0000530 | Zcchc2 | 227449 | HITS-CLIP | 23142080\|29167373 | Unspecified |
| mmu-miR-21a-5p | MIMAT0000530 | Zcchc24 | 71918 | HITS-CLIP | 23597149\|29167373 | Unspecified |
| mmu-miR-21a-5p | MIMAT0000530 | Zcchc3 | 67917 | HITS-CLIP | 23142080 | Unspecified |

| | | | | | | |
|---|---|---|---|---|---|---|
| mmu-miR-21a-5p | MIMAT0000530 | Zcchc7 | 319885 | HITS-CLIP | 23142080 | Unspecified |
| mmu-miR-21a-5p | MIMAT0000530 | Zcchc8 | 70650 | HITS-CLIP | 37986224\|23142080\|29167373 | Unspecified |
| mmu-miR-21a-5p | MIMAT0000530 | Zdhhc12 | 66220 | HITS-CLIP | 23142080 | Unspecified |
| mmu-miR-21a-5p | MIMAT0000530 | Zdhhc13 | 243983 | HITS-CLIP | 23142080 | Unspecified |
| mmu-miR-21a-5p | MIMAT0000530 | Zdhhc17 | 320150 | HITS-CLIP | 23142080 | Unspecified |
| mmu-miR-21a-5p | MIMAT0000530 | Zdhhc18 | 503610 | HITS-CLIP | 23142080 | Unspecified |
| mmu-miR-21a-5p | MIMAT0000530 | Zdhhc2 | 70546 | Microarrays//HITS-CLIP | 37986224\|25263593\|23142080 | Unspecified |
| mmu-miR-21a-5p | MIMAT0000530 | Zdhhc21 | 68268 | HITS-CLIP | 37986224\|23142080\|29167373 | Unspecified |
| mmu-miR-21a-5p | MIMAT0000530 | Zdhhc24 | 70605 | HITS-CLIP | 23597149 | Unspecified |
| mmu-miR-21a-5p | MIMAT0000530 | Zdhhc3 | 69035 | HITS-CLIP | 23142080 | Unspecified |
| mmu-miR-21a-5p | MIMAT0000530 | Zdhhc5 | 228136 | HITS-CLIP | 23597149\|29167373 | Unspecified |
| mmu-miR-21a-5p | MIMAT0000530 | Zdhhc7 | 102193 | HITS-CLIP | 23597149 | Unspecified |
| mmu-miR-21a-5p | MIMAT0000530 | Zeb1 | 21417 | HITS-CLIP | 37986224\|23597149\|23142080 | Unspecified |
| mmu-miR-21a-5p | MIMAT0000530 | Zeb2 | 24136 | HITS-CLIP | 23142080\|23597149\|29167373 | Unspecified |
| mmu-miR-21a-5p | MIMAT0000530 | Zer1 | 227693 | HITS-CLIP | 23142080 | Unspecified |
| mmu-miR-21a-5p | MIMAT0000530 | Zfand5 | 22682 | HITS-CLIP | 23142080 | Unspecified |
| mmu-miR-21a-5p | MIMAT0000530 | Zfand6 | 65098 | HITS-CLIP | 23142080 | Unspecified |
| mmu-miR-21a-5p | MIMAT0000530 | Zfat | 57623 | HITS-CLIP | 23142080 | Unspecified |
| mmu-miR-21a-5p | MIMAT0000530 | Zfc3h1 | 216345 | HITS-CLIP | 23142080\|29167373 | Unspecified |
| mmu-miR-21a-5p | MIMAT0000530 | Zfhx3 | 11906 | Microarrays//HITS-CLIP | 37986224\|29167373 | Unspecified |
| mmu-miR-21a-5p | MIMAT0000530 | Zfhx4 | 80892 | HITS-CLIP | 23597149 | Unspecified |
| mmu-miR-21a-5p | MIMAT0000530 | Zfp101 | 22643 | Microarrays//HITS-CLIP | 37986224\|29167373 | Unspecified |
| mmu-miR-21a-5p | MIMAT0000530 | Zfp106 | 20402 | HITS-CLIP | 23142080 | Unspecified |
| mmu-miR-21a-5p | MIMAT0000530 | Zfp110 | 65020 | HITS-CLIP | 23142080 | Unspecified |
| mmu-miR-21a-5p | MIMAT0000530 | Zfp119a | 104349 | Microarrays | 37986224 | Unspecified |
| mmu-miR-21a-5p | MIMAT0000530 | Zfp120 | 104348 | HITS-CLIP | 23142080 | Unspecified |
| mmu-miR-21a-5p | MIMAT0000530 | Zfp131 | 72465 | HITS-CLIP | 23142080 | Unspecified |
| mmu-miR-21a-5p | MIMAT0000530 | Zfp143 | 20841 | HITS-CLIP | 29167373 | Unspecified |
| mmu-miR-21a-5p | MIMAT0000530 | Zfp148 | 22661 | HITS-CLIP | 23142080\|23597149\|29167373 | Unspecified |
| mmu-miR-21a-5p | MIMAT0000530 | Zfp157 | 72154 | Microarrays | 37986224 | Unspecified |
| mmu-miR-21a-5p | MIMAT0000530 | Zfp180 | 210135 | HITS-CLIP | 37986224 | Unspecified |
| mmu-miR-21a-5p | MIMAT0000530 | Zfp207 | 22680 | HITS-CLIP | 23142080 | Unspecified |
| mmu-miR-21a-5p | MIMAT0000530 | Zfp217 | 228913 | Microarrays//HITS-CLIP | 37986224\|23142080\|29167373 | Unspecified |
| mmu-miR-21a-5p | MIMAT0000530 | Zfp236 | 329002 | HITS-CLIP | 23142080 | Unspecified |
| mmu-miR-21a-5p | MIMAT0000530 | Zfp24 | 59057 | HITS-CLIP | 23142080 | Unspecified |
| mmu-miR-21a-5p | MIMAT0000530 | Zfp26 | 22688 | Microarrays | 37986224 | Unspecified |
| mmu-miR-21a-5p | MIMAT0000530 | Zfp260 | 26466 | Microarrays//HITS-CLIP | 37986224\|23142080 | Unspecified |
| mmu-miR-21a-5p | MIMAT0000530 | Zfp263 | 74120 | Microarrays | 37986224 | Unspecified |
| mmu-miR-21a-5p | MIMAT0000530 | Zfp266 | 77519 | HITS-CLIP | 23142080 | Unspecified |

| | | | | | | |
|---|---|---|---|---|---|---|
| mmu-miR-21a-5p | MIMAT0000530 | Zfp27 | 22689 | HITS-CLIP | 23142080 | Unspecified |
| mmu-miR-21a-5p | MIMAT0000530 | Zfp275 | 27081 | HITS-CLIP | 31433990 | Unspecified |
| mmu-miR-21a-5p | MIMAT0000530 | Zfp280b | 64453 | HITS-CLIP | 29167373\|23142080 | Unspecified |
| mmu-miR-21a-5p | MIMAT0000530 | Zfp280c | 208968 | HITS-CLIP | 23597149\|23142080 | Unspecified |
| mmu-miR-21a-5p | MIMAT0000530 | Zfp280d | 235469 | HITS-CLIP | 37986224\|23142080 | Unspecified |
| mmu-miR-21a-5p | MIMAT0000530 | Zfp281 | 226442 | HITS-CLIP | 37986224\|23142080\|29167373 | Unspecified |
| mmu-miR-21a-5p | MIMAT0000530 | Zfp292 | 30046 | HITS-CLIP | 23597149\|31433990\|23142080 | Unspecified |
| mmu-miR-21a-5p | MIMAT0000530 | Zfp318 | 57908 | HITS-CLIP | 19536157 | Unspecified |
| mmu-miR-21a-5p | MIMAT0000530 | Zfp322a | 218100 | HITS-CLIP | 37986224\|23142080 | Unspecified |
| mmu-miR-21a-5p | MIMAT0000530 | Zfp326 | 54367 | HITS-CLIP | 19536157\|29167373\|23142080 | Unspecified |
| mmu-miR-21a-5p | MIMAT0000530 | Zfp335 | 329559 | HITS-CLIP | 23142080 | Unspecified |
| mmu-miR-21a-5p | MIMAT0000530 | Zfp36 | 22695 | Microarrays//HITS-CLIP | 37986224\|23142080 | Unspecified |
| mmu-miR-21a-5p | MIMAT0000530 | Zfp362 | 230761 | HITS-CLIP | 23142080 | Unspecified |
| mmu-miR-21a-5p | MIMAT0000530 | Zfp367 | 238673 | HITS-CLIP, Microarrays//Microarrays//HITS | 37986224\|25263593\|23597149\|23142 | Unspecified |
| mmu-miR-21a-5p | MIMAT0000530 | Zfp36l1 | 12192 | Microarrays//HITS-CLIP | 37986224\|23597149\|23142080\|29167 | Unspecified |
| mmu-miR-21a-5p | MIMAT0000530 | Zfp36l2 | 12193 | HITS-CLIP//qCLASH | 37986224\|31393953\|23142080 | Unspecified |
| mmu-miR-21a-5p | MIMAT0000530 | Zfp39 | 22698 | HITS-CLIP | 23142080 | Unspecified |
| mmu-miR-21a-5p | MIMAT0000530 | Zfp395 | 380912 | Microarrays | 37986224 | Unspecified |
| mmu-miR-21a-5p | MIMAT0000530 | Zfp397 | 69256 | HITS-CLIP | 23597149 | Unspecified |
| mmu-miR-21a-5p | MIMAT0000530 | Zfp40 | 22700 | Microarrays//HITS-CLIP | 37986224\|25263593\|23142080 | Unspecified |
| mmu-miR-21a-5p | MIMAT0000530 | Zfp407 | 240476 | HITS-CLIP | 29167373 | Unspecified |
| mmu-miR-21a-5p | MIMAT0000530 | Zfp408 | 381410 | HITS-CLIP | 23142080 | Unspecified |
| mmu-miR-21a-5p | MIMAT0000530 | Zfp429 | 72807 | Microarrays | 37986224\|25263593 | Unspecified |
| mmu-miR-21a-5p | MIMAT0000530 | Zfp445 | 235682 | HITS-CLIP | 29167373 | Unspecified |
| mmu-miR-21a-5p | MIMAT0000530 | Zfp451 | 98403 | HITS-CLIP | 23142080\|29167373 | Unspecified |
| mmu-miR-21a-5p | MIMAT0000530 | Zfp455 | 218311 | HITS-CLIP | 23142080 | Unspecified |
| mmu-miR-21a-5p | MIMAT0000530 | Zfp46 | 22704 | HITS-CLIP | 37986224\|23597149\|23142080 | Unspecified |
| mmu-miR-21a-5p | MIMAT0000530 | Zfp507 | 668501 | HITS-CLIP | 23142080 | Unspecified |
| mmu-miR-21a-5p | MIMAT0000530 | Zfp512 | 269639 | HITS-CLIP | 37986224\|23142080\|29167373 | Unspecified |
| mmu-miR-21a-5p | MIMAT0000530 | Zfp513 | 101023 | HITS-CLIP | 23142080 | Unspecified |
| mmu-miR-21a-5p | MIMAT0000530 | Zfp516 | 329003 | HITS-CLIP | 23142080 | Unspecified |
| mmu-miR-21a-5p | MIMAT0000530 | Zfp518a | 72672 | HITS-CLIP | 23142080\|29167373 | Unspecified |
| mmu-miR-21a-5p | MIMAT0000530 | Zfp52 | 22710 | Microarrays | 37986224\|25263593 | Unspecified |
| mmu-miR-21a-5p | MIMAT0000530 | Zfp53 | 24132 | Microarrays//HITS-CLIP | 37986224\|23142080 | Unspecified |
| mmu-miR-21a-5p | MIMAT0000530 | Zfp551 | 619331 | Microarrays | 37986224 | Unspecified |
| mmu-miR-21a-5p | MIMAT0000530 | Zfp560 | 434377 | HITS-CLIP | 23142080 | Unspecified |
| mmu-miR-21a-5p | MIMAT0000530 | Zfp563 | 240068 | Microarrays//HITS-CLIP | 37986224\|23142080 | Unspecified |
| mmu-miR-21a-5p | MIMAT0000530 | Zfp568 | 243905 | HITS-CLIP | 23142080 | Unspecified |
| mmu-miR-21a-5p | MIMAT0000530 | Zfp58 | 238693 | Microarrays | 37986224 | Unspecified |

| | | | | | | |
|---|---|---|---|---|---|---|
| mmu-miR-21a-5p | MIMAT0000530 | Zfp592 | 233410 | HITS-CLIP | 23142080 | Unspecified |
| mmu-miR-21a-5p | MIMAT0000530 | Zfp60 | 22718 | HITS-CLIP | 23142080 | Unspecified |
| mmu-miR-21a-5p | MIMAT0000530 | Zfp609 | 214812 | HITS-CLIP | 23142080 | Unspecified |
| mmu-miR-21a-5p | MIMAT0000530 | Zfp617 | 170938 | HITS-CLIP | 23142080 | Unspecified |
| mmu-miR-21a-5p | MIMAT0000530 | Zfp618 | 72701 | HITS-CLIP | 37986224 | Unspecified |
| mmu-miR-21a-5p | MIMAT0000530 | Zfp62 | 22720 | HITS-CLIP | 29167373 | Unspecified |
| mmu-miR-21a-5p | MIMAT0000530 | Zfp623 | 78834 | HITS-CLIP | 23597149\|23142080 | Unspecified |
| mmu-miR-21a-5p | MIMAT0000530 | Zfp626 | 71163 | HITS-CLIP | 23142080\|29167373 | Unspecified |
| mmu-miR-21a-5p | MIMAT0000530 | Zfp638 | 27332 | HITS-CLIP | 23142080\|29167373 | Unspecified |
| mmu-miR-21a-5p | MIMAT0000530 | Zfp644 | 52397 | HITS-CLIP | 23142080\|29167373 | Unspecified |
| mmu-miR-21a-5p | MIMAT0000530 | Zfp646 | 233905 | HITS-CLIP | 23142080 | Unspecified |
| mmu-miR-21a-5p | MIMAT0000530 | Zfp652 | 268469 | HITS-CLIP | 37986224\|23142080\|29167373 | Unspecified |
| mmu-miR-21a-5p | MIMAT0000530 | Zfp654 | 72020 | HITS-CLIP | 37986224\|23142080\|29167373\|19536[illegible] | Unspecified |
| mmu-miR-21a-5p | MIMAT0000530 | Zfp655 | 72611 | HITS-CLIP | 23142080 | Unspecified |
| mmu-miR-21a-5p | MIMAT0000530 | Zfp658 | 210104 | Microarrays | 37986224 | Unspecified |
| mmu-miR-21a-5p | MIMAT0000530 | Zfp667 | 384763 | HITS-CLIP | 23142080 | Unspecified |
| mmu-miR-21a-5p | MIMAT0000530 | Zfp677 | 210503 | HITS-CLIP | 23142080 | Unspecified |
| mmu-miR-21a-5p | MIMAT0000530 | Zfp68 | 24135 | HITS-CLIP | 23142080\|29167373 | Unspecified |
| mmu-miR-21a-5p | MIMAT0000530 | Zfp689 | 71131 | HITS-CLIP | 23142080 | Unspecified |
| mmu-miR-21a-5p | MIMAT0000530 | Zfp704 | 170753 | HITS-CLIP | 37986224\|19536157 | Unspecified |
| mmu-miR-21a-5p | MIMAT0000530 | Zfp707 | 69020 | HITS-CLIP | 37986224\|23142080 | Unspecified |
| mmu-miR-21a-5p | MIMAT0000530 | Zfp708 | 432769 | HITS-CLIP | 23142080 | Unspecified |
| mmu-miR-21a-5p | MIMAT0000530 | Zfp709 | 236193 | HITS-CLIP | 23142080 | Unspecified |
| mmu-miR-21a-5p | MIMAT0000530 | Zfp710 | 209225 | HITS-CLIP | 19536157\|29167373 | Unspecified |
| mmu-miR-21a-5p | MIMAT0000530 | Zfp712 | 78251 | HITS-CLIP | 23142080 | Unspecified |
| mmu-miR-21a-5p | MIMAT0000530 | Zfp715 | 69930 | HITS-CLIP | 23142080 | Unspecified |
| mmu-miR-21a-5p | MIMAT0000530 | Zfp729b | 100416706 | HITS-CLIP | 23142080 | Unspecified |
| mmu-miR-21a-5p | MIMAT0000530 | Zfp740 | 68744 | Microarrays | 37986224 | Unspecified |
| mmu-miR-21a-5p | MIMAT0000530 | Zfp760 | 240034 | Microarrays | 37986224 | Unspecified |
| mmu-miR-21a-5p | MIMAT0000530 | Zfp763 | 73451 | HITS-CLIP | 23142080 | Unspecified |
| mmu-miR-21a-5p | MIMAT0000530 | Zfp770 | 228491 | HITS-CLIP | 23142080 | Unspecified |
| mmu-miR-21a-5p | MIMAT0000530 | Zfp790 | 233056 | Microarrays | 37986224 | Unspecified |
| mmu-miR-21a-5p | MIMAT0000530 | Zfp800 | 627049 | HITS-CLIP | 23142080\|29167373 | Unspecified |
| mmu-miR-21a-5p | MIMAT0000530 | Zfp821 | 75871 | HITS-CLIP | 37986224 | Unspecified |
| mmu-miR-21a-5p | MIMAT0000530 | Zfp830 | 66983 | HITS-CLIP | 23142080 | Unspecified |
| mmu-miR-21a-5p | MIMAT0000530 | Zfp831 | 100043757 | HITS-CLIP | 37986224\|23142080 | Unspecified |
| mmu-miR-21a-5p | MIMAT0000530 | Zfp846 | 244721 | qCLASH | 31393953 | Unspecified |
| mmu-miR-21a-5p | MIMAT0000530 | Zfp850 | 100043772 | HITS-CLIP | 23142080 | Unspecified |
| mmu-miR-21a-5p | MIMAT0000530 | Zfp866 | 330788 | Microarrays//HITS-CLIP | 37986224\|29167373 | Unspecified |

| | | | | | | |
|---|---|---|---|---|---|---|
| mmu-miR-21a-5p | MIMAT0000530 | Zfp869 | 66869 | Microarrays | 37986224 | Unspecified |
| mmu-miR-21a-5p | MIMAT0000530 | Zfp871 | 208292 | HITS-CLIP | 23142080\|23597149 | Unspecified |
| mmu-miR-21a-5p | MIMAT0000530 | Zfp874a | 238692 | Microarrays | 37986224\|25263593 | Unspecified |
| mmu-miR-21a-5p | MIMAT0000530 | Zfp882 | 382019 | HITS-CLIP | 23142080 | Unspecified |
| mmu-miR-21a-5p | MIMAT0000530 | Zfp90 | 22751 | Microarrays | 37986224 | Unspecified |
| mmu-miR-21a-5p | MIMAT0000530 | Zfp91 | 109910 | HITS-CLIP | 23142080\|29167373 | Unspecified |
| mmu-miR-21a-5p | MIMAT0000530 | Zfp943 | 74670 | HITS-CLIP | 29167373 | Unspecified |
| mmu-miR-21a-5p | MIMAT0000530 | Zfp950 | 214368 | HITS-CLIP | 23142080\|29167373 | Unspecified |
| mmu-miR-21a-5p | MIMAT0000530 | Zfp953 | 629016 | Microarrays | 37986224 | Unspecified |
| mmu-miR-21a-5p | MIMAT0000530 | Zfp955a | 77652 | HITS-CLIP | 23142080 | Unspecified |
| mmu-miR-21a-5p | MIMAT0000530 | Zfp955b | 100043468 | HITS-CLIP | 23142080 | Unspecified |
| mmu-miR-21a-5p | MIMAT0000530 | Zfp963 | 620419 | Microarrays | 37986224 | Unspecified |
| mmu-miR-21a-5p | MIMAT0000530 | Zfp994 | 240038 | Microarrays | 25263593 | Unspecified |
| mmu-miR-21a-5p | MIMAT0000530 | Zfpl1 | 81909 | qCLASH | 31393953 | Unspecified |
| mmu-miR-21a-5p | MIMAT0000530 | Zfpm1 | 22761 | HITS-CLIP | 23142080 | Unspecified |
| mmu-miR-21a-5p | MIMAT0000530 | Zfr | 22763 | HITS-CLIP | 37986224\|23142080 | Unspecified |
| mmu-miR-21a-5p | MIMAT0000530 | Zfx | 22764 | HITS-CLIP | 23142080 | Unspecified |
| mmu-miR-21a-5p | MIMAT0000530 | Zfyve1 | 217695 | HITS-CLIP | 23142080\|23597149 | Unspecified |
| mmu-miR-21a-5p | MIMAT0000530 | Zfyve16 | 218441 | HITS-CLIP | 23142080 | Unspecified |
| mmu-miR-21a-5p | MIMAT0000530 | Zfyve26 | 211978 | qCLASH | 31393953 | Unspecified |
| mmu-miR-21a-5p | MIMAT0000530 | Zfyve28 | 231125 | Microarrays | 37986224 | Unspecified |
| mmu-miR-21a-5p | MIMAT0000530 | Zgpat | 229007 | HITS-CLIP | 37986224\|23597149 | Unspecified |
| mmu-miR-21a-5p | MIMAT0000530 | Zhx1 | 22770 | HITS-CLIP | 23142080\|29167373 | Unspecified |
| mmu-miR-21a-5p | MIMAT0000530 | Zhx3 | 320799 | HITS-CLIP | 37986224\|23597149\|19536157\|23142 | Unspecified |
| mmu-miR-21a-5p | MIMAT0000530 | Zik1 | 22775 | HITS-CLIP | 37986224 | Unspecified |
| mmu-miR-21a-5p | MIMAT0000530 | Zkscan1 | 74570 | HITS-CLIP | 37986224\|23597149\|23142080 | Unspecified |
| mmu-miR-21a-5p | MIMAT0000530 | Zkscan17 | 268417 | Microarrays//HITS-CLIP | 37986224\|29167373 | Unspecified |
| mmu-miR-21a-5p | MIMAT0000530 | Zkscan3 | 72739 | Microarrays//HITS-CLIP | 37986224\|31433990 | Unspecified |
| mmu-miR-21a-5p | MIMAT0000530 | Zkscan5 | 22757 | HITS-CLIP | 23142080 | Unspecified |
| mmu-miR-21a-5p | MIMAT0000530 | Zkscan8 | 93681 | Microarrays | 37986224 | Unspecified |
| mmu-miR-21a-5p | MIMAT0000530 | Zmat1 | 215693 | Microarrays | 37986224 | Unspecified |
| mmu-miR-21a-5p | MIMAT0000530 | Zmat3 | 22401 | HITS-CLIP | 37986224\|29167373\|23142080 | Unspecified |
| mmu-miR-21a-5p | MIMAT0000530 | Zmiz1 | 328365 | HITS-CLIP | 23142080\|23597149\|29167373 | Unspecified |
| mmu-miR-21a-5p | MIMAT0000530 | Zmpste24 | 230709 | HITS-CLIP | 23597149 | Unspecified |
| mmu-miR-21a-5p | MIMAT0000530 | Zmym1 | 68310 | HITS-CLIP | 37986224\|23142080 | Unspecified |
| mmu-miR-21a-5p | MIMAT0000530 | Zmym2 | 76007 | HITS-CLIP | 23142080\|29167373 | Unspecified |
| mmu-miR-21a-5p | MIMAT0000530 | Zmym3 | 56364 | HITS-CLIP | 23142080 | Unspecified |
| mmu-miR-21a-5p | MIMAT0000530 | Zmym4 | 67785 | HITS-CLIP | 23142080 | Unspecified |
| mmu-miR-21a-5p | MIMAT0000530 | Zmym5 | 219105 | HITS-CLIP, Microarrays//HITS-CLIP | 37986224\|23142080 | Unspecified |

| | | | | | | |
|---|---|---|---|---|---|---|
| mmu-miR-21a-5p | MIMAT0000530 | Zmym6 | 100177 | HITS-CLIP | 23597149 | Unspecified |
| mmu-miR-21a-5p | MIMAT0000530 | Zmynd11 | 66505 | HITS-CLIP | 23142080\|29167373 | Unspecified |
| mmu-miR-21a-5p | MIMAT0000530 | Zmynd8 | 228880 | HITS-CLIP | 23142080\|29167373 | Unspecified |
| mmu-miR-21a-5p | MIMAT0000530 | Znfx1 | 98999 | HITS-CLIP | 29167373 | Unspecified |
| mmu-miR-21a-5p | MIMAT0000530 | Znrf2 | 387524 | HITS-CLIP | 37986224\|23597149\|23142080\|29167 | Unspecified |
| mmu-miR-21a-5p | MIMAT0000530 | Zpr1 | 22687 | HITS-CLIP | 23142080 | Unspecified |
| mmu-miR-21a-5p | MIMAT0000530 | Zranb2 | 53861 | qCLASH//HITS-CLIP | 31393953\|23597149\|23142080 | Unspecified |
| mmu-miR-21a-5p | MIMAT0000530 | Zranb3 | 226409 | Microarrays | 37986224\|25263593 | Unspecified |
| mmu-miR-21a-5p | MIMAT0000530 | Zrsr1 | 22183 | HITS-CLIP | 23142080 | Unspecified |
| mmu-miR-21a-5p | MIMAT0000530 | Zrsr2 | 22184 | HITS-CLIP | 23142080 | Unspecified |
| mmu-miR-21a-5p | MIMAT0000530 | Zscan26 | 432731 | Microarrays//HITS-CLIP | 37986224\|23142080\|29167373 | Unspecified |
| mmu-miR-21a-5p | MIMAT0000530 | Zscan29 | 99334 | Microarrays//HITS-CLIP | 37986224\|23142080 | Unspecified |
| mmu-miR-21a-5p | MIMAT0000530 | Zswim6 | 67263 | HITS-CLIP | 37986224 | Unspecified |
| mmu-miR-21a-5p | MIMAT0000530 | Zswim8 | 268721 | HITS-CLIP | 23597149\|23142080\|19536157 | Unspecified |
| mmu-miR-21a-5p | MIMAT0000530 | Zwilch | 68014 | Microarrays//HITS-CLIP | 37986224\|25263593\|23142080 | Unspecified |
| mmu-miR-21a-5p | MIMAT0000530 | Zwint | 52696 | HITS-CLIP | 37986224 | Unspecified |
| mmu-miR-21a-5p | MIMAT0000530 | Zyg11b | 414872 | HITS-CLIP | 23142080 | Unspecified |
| mmu-miR-21a-5p | MIMAT0000530 | Zyx | 22793 | HITS-CLIP | 23142080 | Unspecified |
| mmu-miR-21a-5p | MIMAT0000530 | Zzef1 | 195018 | HITS-CLIP | 23142080 | Unspecified |
| mmu-miR-21a-5p | MIMAT0000530 | mt-Atp6 | 4508 | HITS-CLIP | 29167373 | Unspecified |
| mmu-miR-21a-5p | MIMAT0000530 | mt-Co1 | 4512 | qCLASH//HITS-CLIP | 31393953\|19536157\|23597149\|23142 | Unspecified |
| mmu-miR-21a-5p | MIMAT0000530 | mt-Cytb | 19893556 | HITS-CLIP//qCLASH | 19536157\|31393953\|23142080\|29167 | Unspecified |
| mmu-miR-21a-5p | MIMAT0000530 | mt-Nd1 | 4535 | qCLASH | 31393953 | Unspecified |
| mmu-miR-21a-5p | MIMAT0000530 | mt-Nd5 | 4540 | HITS-CLIP | 29167373\|23597149\|19536157\|23142 | Unspecified |
| mmu-miR-3069-3p | MIMAT0014845 | Insr | 16337 | HITS-CLIP | 31433990 | Unspecified |
| mmu-miR-3069-3p | MIMAT0014845 | Irs1 | 16367 | HITS-CLIP | 26203562 | Unspecified |
| mmu-miR-3069-5p | MIMAT0014844 | Socs5 | 56468 | HITS-CLIP | 31433990 | Unspecified |
| mmu-miR-34a-5p | MIMAT0000542 | 0610010F05Rik | 71675 | HITS-CLIP | 23142080 | Unspecified |
| mmu-miR-34a-5p | MIMAT0000542 | 0610012G03Rik | 106264 | HITS-CLIP | 31433990\|23630078 | Unspecified |
| mmu-miR-34a-5p | MIMAT0000542 | 0610030E20Rik | 68364 | HITS-CLIP | 23142080\|29167373 | Unspecified |
| mmu-miR-34a-5p | MIMAT0000542 | 0610037L13Rik | 74098 | HITS-CLIP | 23142080 | Unspecified |
| mmu-miR-34a-5p | MIMAT0000542 | 1110004F10Rik | 56372 | HITS-CLIP | 23142080\|29167373 | Unspecified |
| mmu-miR-34a-5p | MIMAT0000542 | 1110059E24Rik | 66206 | HITS-CLIP | 23142080 | Unspecified |
| mmu-miR-34a-5p | MIMAT0000542 | 1110059G10Rik | 66202 | HITS-CLIP | 23142080 | Unspecified |
| mmu-miR-34a-5p | MIMAT0000542 | 1190002N15Rik | 68861 | HITS-CLIP | 19536157\|23142080\|29167373 | Unspecified |
| mmu-miR-34a-5p | MIMAT0000542 | 1500011B03Rik | 66236 | HITS-CLIP | 23142080 | Unspecified |
| mmu-miR-34a-5p | MIMAT0000542 | 1600012H06Rik | 67912 | HITS-CLIP | 23142080 | Unspecified |
| mmu-miR-34a-5p | MIMAT0000542 | 1600014C10Rik | 72244 | HITS-CLIP | 23142080 | Unspecified |
| mmu-miR-34a-5p | MIMAT0000542 | 1700003E16Rik | 71837 | Microarrays | 37986224 | Unspecified |

| | | | | | | |
|---|---|---|---|---|---|---|
| mmu-miR-34a-5p | MIMAT0000542 | 1700007K13Rik | 69327 | Microarrays | 37986224 | Unspecified |
| mmu-miR-34a-5p | MIMAT0000542 | 1700017B05Rik | 74211 | HITS-CLIP | 31433990\|26203562\|29167373\|23142 | Unspecified |
| mmu-miR-34a-5p | MIMAT0000542 | 1700025G04Rik | 69399 | HITS-CLIP | 29167373 | Unspecified |
| mmu-miR-34a-5p | MIMAT0000542 | 1700037H04Rik | 67326 | Microarrays | 37986224\|23027903 | Unspecified |
| mmu-miR-34a-5p | MIMAT0000542 | 1700066B19Rik | 73449 | HITS-CLIP | 23142080 | Unspecified |
| mmu-miR-34a-5p | MIMAT0000542 | 1700066M21Rik | 73467 | HITS-CLIP | 23142080 | Unspecified |
| mmu-miR-34a-5p | MIMAT0000542 | 1700088E04Rik | 27660 | Microarrays | 37986224 | Unspecified |
| mmu-miR-34a-5p | MIMAT0000542 | 1700123O20Rik | 58248 | HITS-CLIP | 23142080 | Unspecified |
| mmu-miR-34a-5p | MIMAT0000542 | 1810013L24Rik | 69053 | HITS-CLIP | 31433990\|23142080\|19536157\|29167 | Unspecified |
| mmu-miR-34a-5p | MIMAT0000542 | 1810030O07Rik | 69155 | HITS-CLIP | 23142080 | Unspecified |
| mmu-miR-34a-5p | MIMAT0000542 | 1810055G02Rik | 72056 | HITS-CLIP | 23142080\|19536157 | Unspecified |
| mmu-miR-34a-5p | MIMAT0000542 | 2010111I01Rik | 72061 | HITS-CLIP | 29167373 | Unspecified |
| mmu-miR-34a-5p | MIMAT0000542 | 2010315B03Rik | 630836 | HITS-CLIP | 23142080 | Unspecified |
| mmu-miR-34a-5p | MIMAT0000542 | 2300009A05Rik | 69478 | HITS-CLIP | 23142080 | Unspecified |
| mmu-miR-34a-5p | MIMAT0000542 | 2310022A10Rik | 66367 | HITS-CLIP | 30971389 | Unspecified |
| mmu-miR-34a-5p | MIMAT0000542 | 2310022B05Rik | 69551 | HITS-CLIP | 19536157\|23142080 | Unspecified |
| mmu-miR-34a-5p | MIMAT0000542 | 2310057M21Rik | 68277 | HITS-CLIP | 31433990\|23142080 | Unspecified |
| mmu-miR-34a-5p | MIMAT0000542 | 2310061I04Rik | 69662 | HITS-CLIP | 23142080 | Unspecified |
| mmu-miR-34a-5p | MIMAT0000542 | 2410015M20Rik | 224904 | HITS-CLIP | 23142080 | Unspecified |
| mmu-miR-34a-5p | MIMAT0000542 | 2410131K14Rik | 76792 | HITS-CLIP | 23142080 | Unspecified |
| mmu-miR-34a-5p | MIMAT0000542 | 2510002D24Rik | 72307 | HITS-CLIP | 23142080 | Unspecified |
| mmu-miR-34a-5p | MIMAT0000542 | 2510009E07Rik | 72190 | HITS-CLIP | 29167373 | Unspecified |
| mmu-miR-34a-5p | MIMAT0000542 | 2510039O18Rik | 77034 | HITS-CLIP | 31433990\|23142080 | Unspecified |
| mmu-miR-34a-5p | MIMAT0000542 | 2610001J05Rik | 66520 | HITS-CLIP | 29167373 | Unspecified |
| mmu-miR-34a-5p | MIMAT0000542 | 2610002M06Rik | 67028 | HITS-CLIP | 23142080 | Unspecified |
| mmu-miR-34a-5p | MIMAT0000542 | 2610021A01Rik | 668572 | HITS-CLIP | 23142080 | Unspecified |
| mmu-miR-34a-5p | MIMAT0000542 | 2610301B20Rik | 67157 | Microarrays | 37986224 | Unspecified |
| mmu-miR-34a-5p | MIMAT0000542 | 2610507B11Rik | 72503 | HITS-CLIP | 19536157\|23142080\|29167373 | Unspecified |
| mmu-miR-34a-5p | MIMAT0000542 | 2610528A11Rik | 70045 | HITS-CLIP | 23142080 | Unspecified |
| mmu-miR-34a-5p | MIMAT0000542 | 2700049A03Rik | 76967 | HITS-CLIP | 23142080 | Unspecified |
| mmu-miR-34a-5p | MIMAT0000542 | 2810004N23Rik | 66523 | HITS-CLIP | 29167373\|23142080 | Unspecified |
| mmu-miR-34a-5p | MIMAT0000542 | 2810006K23Rik | 72650 | HITS-CLIP | 29167373 | Unspecified |
| mmu-miR-34a-5p | MIMAT0000542 | 2810408A11Rik | 70419 | HITS-CLIP | 23142080 | Unspecified |
| mmu-miR-34a-5p | MIMAT0000542 | 4833420G17Rik | 67392 | HITS-CLIP | 23142080 | Unspecified |
| mmu-miR-34a-5p | MIMAT0000542 | 4833427G06Rik | 235345 | Microarrays | 37986224 | Unspecified |
| mmu-miR-34a-5p | MIMAT0000542 | 4833439L19Rik | 97820 | HITS-CLIP | 29167373 | Unspecified |
| mmu-miR-34a-5p | MIMAT0000542 | 4921524J17Rik | 66714 | HITS-CLIP | 31433990\|23142080 | Unspecified |
| mmu-miR-34a-5p | MIMAT0000542 | 4930402H24Rik | 228602 | HITS-CLIP | 19536157\|23142080\|29167373 | Unspecified |
| mmu-miR-34a-5p | MIMAT0000542 | 4930453N24Rik | 67609 | HITS-CLIP | 23142080 | Unspecified |

| | | | | | | |
|---|---|---|---|---|---|---|
| mmu-miR-34a-5p | MIMAT0000542 | 4930503L19Rik | 269033 | HITS-CLIP | 23142080 | Unspecified |
| mmu-miR-34a-5p | MIMAT0000542 | 4930523C07Rik | 67647 | HITS-CLIP | 23142080 | Unspecified |
| mmu-miR-34a-5p | MIMAT0000542 | 4930539E08Rik | 207819 | HITS-CLIP | 23142080 | Unspecified |
| mmu-miR-34a-5p | MIMAT0000542 | 4930579G24Rik | 75939 | HITS-CLIP | 30971389\|23142080 | Unspecified |
| mmu-miR-34a-5p | MIMAT0000542 | 4931406C07Rik | 70984 | Microarrays | 37986224 | Unspecified |
| mmu-miR-34a-5p | MIMAT0000542 | 4931406P16Rik | 233103 | HITS-CLIP | 23142080 | Unspecified |
| mmu-miR-34a-5p | MIMAT0000542 | 4932438A13Rik | 229227 | HITS-CLIP | 37986224\|19536157\|23142080\|29167 | Unspecified |
| mmu-miR-34a-5p | MIMAT0000542 | 4932438H23Rik | 74387 | HITS-CLIP | 23142080 | Unspecified |
| mmu-miR-34a-5p | MIMAT0000542 | 4933427D14Rik | 74477 | HITS-CLIP | 29167373 | Unspecified |
| mmu-miR-34a-5p | MIMAT0000542 | 5031439G07Rik | 223739 | HITS-CLIP | 19536157\|23630078\|29167373 | Unspecified |
| mmu-miR-34a-5p | MIMAT0000542 | 5430427O19Rik | 71398 | HITS-CLIP | 29167373 | Unspecified |
| mmu-miR-34a-5p | MIMAT0000542 | 5530400C23Rik | 232426 | Microarrays | 37986224 | Unspecified |
| mmu-miR-34a-5p | MIMAT0000542 | 5730455P16Rik | 70591 | HITS-CLIP | 23142080\|29167373 | Unspecified |
| mmu-miR-34a-5p | MIMAT0000542 | 6030458C11Rik | 77877 | Microarrays//HITS-CLIP | 37986224\|23027903\|31433990\|2314 | Unspecified |
| mmu-miR-34a-5p | MIMAT0000542 | 8030462N17Rik | 212163 | HITS-CLIP | 19536157\|23142080\|29167373 | Unspecified |
| mmu-miR-34a-5p | MIMAT0000542 | 9130019O22Rik | 78921 | HITS-CLIP | 23142080 | Unspecified |
| mmu-miR-34a-5p | MIMAT0000542 | 9530068E07Rik | 213673 | HITS-CLIP | 19536157\|23142080 | Unspecified |
| mmu-miR-34a-5p | MIMAT0000542 | 9530077C05Rik | 68283 | HITS-CLIP | 23142080 | Unspecified |
| mmu-miR-34a-5p | MIMAT0000542 | 9930012K11Rik | 268759 | HITS-CLIP | 29167373 | Unspecified |
| mmu-miR-34a-5p | MIMAT0000542 | 9930021J03Rik | 240613 | HITS-CLIP | 23142080\|29167373 | Unspecified |
| mmu-miR-34a-5p | MIMAT0000542 | 9930111J21Rik1 | 667214 | HITS-CLIP | 29167373 | Unspecified |
| mmu-miR-34a-5p | MIMAT0000542 | 9930111J21Rik2 | 245240 | HITS-CLIP | 29167373 | Unspecified |
| mmu-miR-34a-5p | MIMAT0000542 | A130010J15Rik | 319266 | HITS-CLIP | 23142080 | Unspecified |
| mmu-miR-34a-5p | MIMAT0000542 | A2m | 2 | HITS-CLIP | 30971389 | Unspecified |
| mmu-miR-34a-5p | MIMAT0000542 | A630001G21Rik | 319997 | HITS-CLIP | 31433990\|29167373 | Unspecified |
| mmu-miR-34a-5p | MIMAT0000542 | A930004D18Rik | 77940 | HITS-CLIP | 23142080 | Unspecified |
| mmu-miR-34a-5p | MIMAT0000542 | AB124611 | 382062 | HITS-CLIP | 23142080\|29167373 | Unspecified |
| mmu-miR-34a-5p | MIMAT0000542 | AI413582 | 106672 | HITS-CLIP | 23142080 | Unspecified |
| mmu-miR-34a-5p | MIMAT0000542 | AI467606 | 101602 | HITS-CLIP | 23142080 | Unspecified |
| mmu-miR-34a-5p | MIMAT0000542 | AI597479 | 98404 | Microarrays//HITS-CLIP | 37986224\|23027903\|23142080\|3097 | Unspecified |
| mmu-miR-34a-5p | MIMAT0000542 | AI837181 | 107242 | HITS-CLIP | 26203562\|23142080 | Unspecified |
| mmu-miR-34a-5p | MIMAT0000542 | AI987944 | 233168 | HITS-CLIP | 23142080 | Unspecified |
| mmu-miR-34a-5p | MIMAT0000542 | AK157302 | 432732 | HITS-CLIP | 29167373 | Unspecified |
| mmu-miR-34a-5p | MIMAT0000542 | AU022252 | 230696 | HITS-CLIP | 23142080 | Unspecified |
| mmu-miR-34a-5p | MIMAT0000542 | AU040320 | 100317 | HITS-CLIP | 23142080 | Unspecified |
| mmu-miR-34a-5p | MIMAT0000542 | AU041133 | 216177 | HITS-CLIP | 23142080 | Unspecified |
| mmu-miR-34a-5p | MIMAT0000542 | AW549877 | 106064 | Microarrays//HITS-CLIP | 37986224\|19536157\|23142080\|29167 | Unspecified |
| mmu-miR-34a-5p | MIMAT0000542 | Aacs | 78894 | HITS-CLIP | 30971389\|23142080\|29167373 | Unspecified |
| mmu-miR-34a-5p | MIMAT0000542 | Aak1 | 269774 | HITS-CLIP | 19536157\|31433990\|23142080\|29167 | Unspecified |

| | | | | | | |
|---|---|---|---|---|---|---|
| mmu-miR-34a-5p | MIMAT0000542 | Aamp | 227290 | HITS-CLIP | 23142080 | Unspecified |
| mmu-miR-34a-5p | MIMAT0000542 | Aar2 | 68295 | HITS-CLIP | 23142080 | Unspecified |
| mmu-miR-34a-5p | MIMAT0000542 | Aars | 234734 | HITS-CLIP | 31433990\|19536157\|29167373\|23142 | Unspecified |
| mmu-miR-34a-5p | MIMAT0000542 | Aass | 30956 | HITS-CLIP | 30971389 | Unspecified |
| mmu-miR-34a-5p | MIMAT0000542 | Abat | 268860 | HITS-CLIP | 19536157 | Unspecified |
| mmu-miR-34a-5p | MIMAT0000542 | Abca1 | 11303 | HITS-CLIP | 19536157\|30971389\|29167373 | Unspecified |
| mmu-miR-34a-5p | MIMAT0000542 | Abca2 | 11305 | HITS-CLIP | 23142080 | Unspecified |
| mmu-miR-34a-5p | MIMAT0000542 | Abca3 | 27410 | HITS-CLIP | 19536157\|23142080\|29167373 | Unspecified |
| mmu-miR-34a-5p | MIMAT0000542 | Abca8b | 27404 | HITS-CLIP | 29167373 | Unspecified |
| mmu-miR-34a-5p | MIMAT0000542 | Abcb10 | 56199 | HITS-CLIP | 30971389 | Unspecified |
| mmu-miR-34a-5p | MIMAT0000542 | Abcb1b | 18669 | Microarrays//HITS-CLIP | 37986224\|23027903\|23142080\|29167 | Unspecified |
| mmu-miR-34a-5p | MIMAT0000542 | Abcb7 | 11306 | HITS-CLIP | 23142080\|29167373 | Unspecified |
| mmu-miR-34a-5p | MIMAT0000542 | Abcb8 | 74610 | HITS-CLIP | 23142080 | Unspecified |
| mmu-miR-34a-5p | MIMAT0000542 | Abcb9 | 56325 | HITS-CLIP | 23142080 | Unspecified |
| mmu-miR-34a-5p | MIMAT0000542 | Abcc1 | 17250 | HITS-CLIP | 29167373 | Unspecified |
| mmu-miR-34a-5p | MIMAT0000542 | Abcc2 | 12780 | HITS-CLIP | 30971389 | Unspecified |
| mmu-miR-34a-5p | MIMAT0000542 | Abcc3 | 76408 | HITS-CLIP | 29167373 | Unspecified |
| mmu-miR-34a-5p | MIMAT0000542 | Abcc5 | 27416 | HITS-CLIP | 29167373 | Unspecified |
| mmu-miR-34a-5p | MIMAT0000542 | Abcd1 | 11666 | Microarrays//HITS-CLIP | 37986224\|23027903\|23142080\|1953€ | Unspecified |
| mmu-miR-34a-5p | MIMAT0000542 | Abce1 | 24015 | HITS-CLIP | 31433990\|23142080\|29167373 | Unspecified |
| mmu-miR-34a-5p | MIMAT0000542 | Abcf1 | 224742 | HITS-CLIP | 29167373\|23142080 | Unspecified |
| mmu-miR-34a-5p | MIMAT0000542 | Abcf2 | 27407 | Microarrays//HITS-CLIP | 37986224\|23027903\|23142080 | Unspecified |
| mmu-miR-34a-5p | MIMAT0000542 | Abcf3 | 27406 | HITS-CLIP | 19536157 | Unspecified |
| mmu-miR-34a-5p | MIMAT0000542 | Abcg1 | 11307 | HITS-CLIP | 23142080\|29167373 | Unspecified |
| mmu-miR-34a-5p | MIMAT0000542 | Abcg3 | 27405 | HITS-CLIP | 23142080 | Unspecified |
| mmu-miR-34a-5p | MIMAT0000542 | Abhd10 | 213012 | HITS-CLIP | 23142080 | Unspecified |
| mmu-miR-34a-5p | MIMAT0000542 | Abhd12 | 76192 | HITS-CLIP | 19536157\|23142080\|29167373 | Unspecified |
| mmu-miR-34a-5p | MIMAT0000542 | Abhd13 | 68904 | HITS-CLIP | 23142080\|31433990 | Unspecified |
| mmu-miR-34a-5p | MIMAT0000542 | Abhd17b | 226016 | HITS-CLIP | 29167373\|23142080 | Unspecified |
| mmu-miR-34a-5p | MIMAT0000542 | Abhd17c | 70178 | HITS-CLIP | 23142080\|29167373 | Unspecified |
| mmu-miR-34a-5p | MIMAT0000542 | Abhd18 | 269423 | HITS-CLIP | 23142080\|31433990\|29167373 | Unspecified |
| mmu-miR-34a-5p | MIMAT0000542 | Abhd2 | 54608 | HITS-CLIP | 19536157\|30971389 | Unspecified |
| mmu-miR-34a-5p | MIMAT0000542 | Abhd4 | 105501 | HITS-CLIP | 23142080\|31433990 | Unspecified |
| mmu-miR-34a-5p | MIMAT0000542 | Abhd8 | 64296 | HITS-CLIP | 23142080 | Unspecified |
| mmu-miR-34a-5p | MIMAT0000542 | Abi1 | 11308 | HITS-CLIP | 23142080\|31433990 | Unspecified |
| mmu-miR-34a-5p | MIMAT0000542 | Abi2 | 329165 | HITS-CLIP | 19536157\|23142080 | Unspecified |
| mmu-miR-34a-5p | MIMAT0000542 | Abi3 | 66610 | HITS-CLIP | 23142080 | Unspecified |
| mmu-miR-34a-5p | MIMAT0000542 | Abl1 | 11350 | HITS-CLIP | 31433990\|23142080\|29167373 | Unspecified |
| mmu-miR-34a-5p | MIMAT0000542 | Abl2 | 11352 | HITS-CLIP | 23142080\|29167373 | Unspecified |

| | | | | | | |
|---|---|---|---|---|---|---|
| mmu-miR-34a-5p | MIMAT0000542 | Ablim1 | 226251 | HITS-CLIP | 23142080 | Unspecified |
| mmu-miR-34a-5p | MIMAT0000542 | Ablim3 | 319713 | Microarrays | 37986224 | Unspecified |
| mmu-miR-34a-5p | MIMAT0000542 | Abr | 109934 | HITS-CLIP | 19536157\|31433990\|23142080\|29167 | Unspecified |
| mmu-miR-34a-5p | MIMAT0000542 | Abraxas1 | 70681 | HITS-CLIP | 30971389\|23142080 | Unspecified |
| mmu-miR-34a-5p | MIMAT0000542 | Abraxas2 | 23172 | HITS-CLIP | 23142080 | Unspecified |
| mmu-miR-34a-5p | MIMAT0000542 | Abtb2 | 99382 | HITS-CLIP | 30971389\|31433990\|23142080 | Unspecified |
| mmu-miR-34a-5p | MIMAT0000542 | Acaa1a | 113868 | HITS-CLIP | 23142080 | Unspecified |
| mmu-miR-34a-5p | MIMAT0000542 | Acaa2 | 52538 | HITS-CLIP | 23142080 | Unspecified |
| mmu-miR-34a-5p | MIMAT0000542 | Acad11 | 102632 | HITS-CLIP | 23142080 | Unspecified |
| mmu-miR-34a-5p | MIMAT0000542 | Acadl | 11363 | HITS-CLIP | 23142080 | Unspecified |
| mmu-miR-34a-5p | MIMAT0000542 | Acadm | 11364 | HITS-CLIP | 31433990\|23142080 | Unspecified |
| mmu-miR-34a-5p | MIMAT0000542 | Acads | 11409 | HITS-CLIP | 26203562\|23142080 | Unspecified |
| mmu-miR-34a-5p | MIMAT0000542 | Acadsb | 66885 | HITS-CLIP | 23142080\|29167373 | Unspecified |
| mmu-miR-34a-5p | MIMAT0000542 | Acadvl | 11370 | HITS-CLIP | 19536157\|23142080 | Unspecified |
| mmu-miR-34a-5p | MIMAT0000542 | Acap1 | 216859 | HITS-CLIP | 23142080 | Unspecified |
| mmu-miR-34a-5p | MIMAT0000542 | Acap2 | 78618 | HITS-CLIP | 23142080\|29167373 | Unspecified |
| mmu-miR-34a-5p | MIMAT0000542 | Acat1 | 110446 | HITS-CLIP | 23142080 | Unspecified |
| mmu-miR-34a-5p | MIMAT0000542 | Acbd3 | 170760 | HITS-CLIP | 23142080\|29167373 | Unspecified |
| mmu-miR-34a-5p | MIMAT0000542 | Acbd5 | 74159 | HITS-CLIP | 23142080 | Unspecified |
| mmu-miR-34a-5p | MIMAT0000542 | Acer3 | 66190 | HITS-CLIP | 29167373 | Unspecified |
| mmu-miR-34a-5p | MIMAT0000542 | Acin1 | 56215 | HITS-CLIP | 23142080\|29167373 | Unspecified |
| mmu-miR-34a-5p | MIMAT0000542 | Ackr3 | 12778 | HITS-CLIP | 23142080 | Unspecified |
| mmu-miR-34a-5p | MIMAT0000542 | Acly | 104112 | HITS-CLIP | 19536157\|30971389\|26203562\|23142 | Unspecified |
| mmu-miR-34a-5p | MIMAT0000542 | Aco1 | 11428 | HITS-CLIP | 26203562 | Unspecified |
| mmu-miR-34a-5p | MIMAT0000542 | Aco2 | 11429 | HITS-CLIP | 23142080\|30971389\|29167373 | Unspecified |
| mmu-miR-34a-5p | MIMAT0000542 | Acot11 | 329910 | HITS-CLIP | 23142080 | Unspecified |
| mmu-miR-34a-5p | MIMAT0000542 | Acot9 | 56360 | HITS-CLIP | 23142080 | Unspecified |
| mmu-miR-34a-5p | MIMAT0000542 | Acox1 | 11430 | HITS-CLIP | 37986224\|23142080 | Unspecified |
| mmu-miR-34a-5p | MIMAT0000542 | Acox3 | 80911 | HITS-CLIP | 29167373 | Unspecified |
| mmu-miR-34a-5p | MIMAT0000542 | Acp1 | 11431 | Microarrays | 37986224 | Unspecified |
| mmu-miR-34a-5p | MIMAT0000542 | Acp2 | 11432 | HITS-CLIP | 29167373 | Unspecified |
| mmu-miR-34a-5p | MIMAT0000542 | Acp5 | 11433 | HITS-CLIP | 23142080 | Unspecified |
| mmu-miR-34a-5p | MIMAT0000542 | Acp7 | 390928 | HITS-CLIP | 23142080 | Unspecified |
| mmu-miR-34a-5p | MIMAT0000542 | Acsl1 | 14081 | HITS-CLIP | 23142080\|29167373 | Unspecified |
| mmu-miR-34a-5p | MIMAT0000542 | Acsl3 | 74205 | HITS-CLIP | 31433990\|23142080 | Unspecified |
| mmu-miR-34a-5p | MIMAT0000542 | Acsl4 | 50790 | HITS-CLIP | 19536157\|23142080\|29167373 | Unspecified |
| mmu-miR-34a-5p | MIMAT0000542 | Acsl5 | 433256 | HITS-CLIP | 31433990\|23142080 | Unspecified |
| mmu-miR-34a-5p | MIMAT0000542 | Acss1 | 68738 | HITS-CLIP | 23142080 | Unspecified |
| mmu-miR-34a-5p | MIMAT0000542 | Actb | 11461 | Luciferase assay//HITS-CLIP//Luciferase re | 18835850\|19536157\|23142080 | Unspecified |

| | | | | | | |
|---|---|---|---|---|---|---|
| mmu-miR-34a-5p | MIMAT0000542 | Actc1 | 11464 | Microarrays | 37986224 | Unspecified |
| mmu-miR-34a-5p | MIMAT0000542 | Actg1 | 11465 | HITS-CLIP | 31433990\|29167373 | Unspecified |
| mmu-miR-34a-5p | MIMAT0000542 | Actl6a | 56456 | HITS-CLIP | 23142080 | Unspecified |
| mmu-miR-34a-5p | MIMAT0000542 | Actn1 | 109711 | HITS-CLIP | 23142080\|26203562\|29167373 | Unspecified |
| mmu-miR-34a-5p | MIMAT0000542 | Actn4 | 60595 | HITS-CLIP | 19536157\|23142080\|26203562\|29167 | Unspecified |
| mmu-miR-34a-5p | MIMAT0000542 | Actr10 | 56444 | HITS-CLIP | 23142080 | Unspecified |
| mmu-miR-34a-5p | MIMAT0000542 | Actr1a | 54130 | HITS-CLIP | 23142080 | Unspecified |
| mmu-miR-34a-5p | MIMAT0000542 | Actr1b | 226977 | HITS-CLIP | 23142080 | Unspecified |
| mmu-miR-34a-5p | MIMAT0000542 | Actr2 | 66713 | HITS-CLIP | 23142080\|19536157\|29167373 | Unspecified |
| mmu-miR-34a-5p | MIMAT0000542 | Actr3 | 74117 | HITS-CLIP | 29167373 | Unspecified |
| mmu-miR-34a-5p | MIMAT0000542 | Actr8 | 56249 | HITS-CLIP | 23142080 | Unspecified |
| mmu-miR-34a-5p | MIMAT0000542 | Acvr1 | 11477 | HITS-CLIP | 19536157 | Unspecified |
| mmu-miR-34a-5p | MIMAT0000542 | Acvr1b | 11479 | HITS-CLIP | 23142080\|30971389\|29167373 | Unspecified |
| mmu-miR-34a-5p | MIMAT0000542 | Acvr2a | 11480 | Microarrays//HITS-CLIP | 37986224\|19536157\|29167373\|23142 | Unspecified |
| mmu-miR-34a-5p | MIMAT0000542 | Acvr2b | 11481 | HITS-CLIP | 23142080 | Unspecified |
| mmu-miR-34a-5p | MIMAT0000542 | Acvrl1 | 11482 | HITS-CLIP | 23142080 | Unspecified |
| mmu-miR-34a-5p | MIMAT0000542 | Acy1 | 109652 | Microarrays | 37986224\|23027903 | Unspecified |
| mmu-miR-34a-5p | MIMAT0000542 | Adal | 75894 | HITS-CLIP | 30971389 | Unspecified |
| mmu-miR-34a-5p | MIMAT0000542 | Adam10 | 11487 | HITS-CLIP | 19536157\|23142080\|29167373 | Unspecified |
| mmu-miR-34a-5p | MIMAT0000542 | Adam15 | 11490 | HITS-CLIP | 29167373\|23142080 | Unspecified |
| mmu-miR-34a-5p | MIMAT0000542 | Adam19 | 11492 | HITS-CLIP | 23142080 | Unspecified |
| mmu-miR-34a-5p | MIMAT0000542 | Adam1a | 280668 | HITS-CLIP | 23142080 | Unspecified |
| mmu-miR-34a-5p | MIMAT0000542 | Adam22 | 11496 | HITS-CLIP | 19536157 | Unspecified |
| mmu-miR-34a-5p | MIMAT0000542 | Adam33 | 110751 | HITS-CLIP | 29167373 | Unspecified |
| mmu-miR-34a-5p | MIMAT0000542 | Adam8 | 11501 | HITS-CLIP | 29167373 | Unspecified |
| mmu-miR-34a-5p | MIMAT0000542 | Adam9 | 11502 | HITS-CLIP | 23142080\|29167373\|26203562 | Unspecified |
| mmu-miR-34a-5p | MIMAT0000542 | Adamts12 | 239337 | Microarrays | 37986224 | Unspecified |
| mmu-miR-34a-5p | MIMAT0000542 | Adamts6 | 108154 | HITS-CLIP | 23142080\|29167373 | Unspecified |
| mmu-miR-34a-5p | MIMAT0000542 | Adamtsl3 | 269959 | HITS-CLIP | 23142080 | Unspecified |
| mmu-miR-34a-5p | MIMAT0000542 | Adamtsl4 | 229595 | HITS-CLIP | 23142080 | Unspecified |
| mmu-miR-34a-5p | MIMAT0000542 | Adap1 | 231821 | HITS-CLIP | 31433990 | Unspecified |
| mmu-miR-34a-5p | MIMAT0000542 | Adap2 | 216991 | HITS-CLIP | 29167373 | Unspecified |
| mmu-miR-34a-5p | MIMAT0000542 | Adar | 56417 | HITS-CLIP | 23142080\|29167373 | Unspecified |
| mmu-miR-34a-5p | MIMAT0000542 | Adarb1 | 110532 | HITS-CLIP | 31433990\|23142080 | Unspecified |
| mmu-miR-34a-5p | MIMAT0000542 | Adck2 | 57869 | HITS-CLIP | 23142080\|29167373 | Unspecified |
| mmu-miR-34a-5p | MIMAT0000542 | Adck5 | 268822 | HITS-CLIP | 23142080\|29167373 | Unspecified |
| mmu-miR-34a-5p | MIMAT0000542 | Adcy1 | 432530 | HITS-CLIP | 19536157 | Unspecified |
| mmu-miR-34a-5p | MIMAT0000542 | Adcy3 | 104111 | HITS-CLIP | 23142080 | Unspecified |
| mmu-miR-34a-5p | MIMAT0000542 | Adcy6 | 11512 | HITS-CLIP | 23142080 | Unspecified |

| | | | | | | |
|---|---|---|---|---|---|---|
| mmu-miR-34a-5p | MIMAT0000542 | Adcy7 | 11513 | HITS-CLIP | 23142080\|29167373 | Unspecified |
| mmu-miR-34a-5p | MIMAT0000542 | Add1 | 11518 | HITS-CLIP | 30971389\|23142080\|29167373 | Unspecified |
| mmu-miR-34a-5p | MIMAT0000542 | Add2 | 11519 | HITS-CLIP | 19536157 | Unspecified |
| mmu-miR-34a-5p | MIMAT0000542 | Add3 | 27360 | HITS-CLIP | 23142080\|29167373 | Unspecified |
| mmu-miR-34a-5p | MIMAT0000542 | Adgra2 | 25960 | HITS-CLIP | 19536157 | Unspecified |
| mmu-miR-34a-5p | MIMAT0000542 | Adgrb3 | 577 | HITS-CLIP | 19536157 | Unspecified |
| mmu-miR-34a-5p | MIMAT0000542 | Adgre1 | 13733 | HITS-CLIP | 29167373 | Unspecified |
| mmu-miR-34a-5p | MIMAT0000542 | Adgre5 | 976 | HITS-CLIP | 31433990\|23142080\|29167373 | Unspecified |
| mmu-miR-34a-5p | MIMAT0000542 | Adgrf5 | 224792 | HITS-CLIP | 19536157 | Unspecified |
| mmu-miR-34a-5p | MIMAT0000542 | Adgrl1 | 22859 | HITS-CLIP | 23142080\|19536157 | Unspecified |
| mmu-miR-34a-5p | MIMAT0000542 | Adgrl2 | 23266 | HITS-CLIP | 23142080\|29167373 | Unspecified |
| mmu-miR-34a-5p | MIMAT0000542 | Adgrl3 | 319387 | HITS-CLIP | 19536157 | Unspecified |
| mmu-miR-34a-5p | MIMAT0000542 | Adh5 | 11532 | HITS-CLIP | 23142080\|19536157 | Unspecified |
| mmu-miR-34a-5p | MIMAT0000542 | Adi1 | 104923 | Microarrays//HITS-CLIP | 37986224\|23027903\|23142080 | Unspecified |
| mmu-miR-34a-5p | MIMAT0000542 | Adipor1 | 72674 | HITS-CLIP | 37986224\|19536157\|31433990\|2314 | Unspecified |
| mmu-miR-34a-5p | MIMAT0000542 | Adipor2 | 68465 | HITS-CLIP | 23142080\|29167373 | Unspecified |
| mmu-miR-34a-5p | MIMAT0000542 | Adk | 11534 | HITS-CLIP | 23142080 | Unspecified |
| mmu-miR-34a-5p | MIMAT0000542 | Adnp | 11538 | HITS-CLIP | 37986224\|31433990\|19536157\|2314 | Unspecified |
| mmu-miR-34a-5p | MIMAT0000542 | Adnp2 | 240442 | HITS-CLIP | 23142080\|29167373 | Unspecified |
| mmu-miR-34a-5p | MIMAT0000542 | Ado | 211488 | HITS-CLIP | 23142080\|29167373 | Unspecified |
| mmu-miR-34a-5p | MIMAT0000542 | Adora2a | 11540 | HITS-CLIP | 23142080 | Unspecified |
| mmu-miR-34a-5p | MIMAT0000542 | Adora2b | 11541 | HITS-CLIP | 29167373 | Unspecified |
| mmu-miR-34a-5p | MIMAT0000542 | Adpgk | 72141 | HITS-CLIP | 23142080\|29167373 | Unspecified |
| mmu-miR-34a-5p | MIMAT0000542 | Adprh | 11544 | HITS-CLIP | 23142080 | Unspecified |
| mmu-miR-34a-5p | MIMAT0000542 | Adprm | 66358 | HITS-CLIP | 23142080 | Unspecified |
| mmu-miR-34a-5p | MIMAT0000542 | Adra2a | 11551 | HITS-CLIP | 19536157 | Unspecified |
| mmu-miR-34a-5p | MIMAT0000542 | Adrb2 | 11555 | HITS-CLIP | 23142080\|29167373 | Unspecified |
| mmu-miR-34a-5p | MIMAT0000542 | Adsl | 11564 | HITS-CLIP | 23142080 | Unspecified |
| mmu-miR-34a-5p | MIMAT0000542 | Adss | 11566 | HITS-CLIP | 23142080\|29167373 | Unspecified |
| mmu-miR-34a-5p | MIMAT0000542 | Adssl1 | 11565 | Microarrays | 37986224 | Unspecified |
| mmu-miR-34a-5p | MIMAT0000542 | Aebp2 | 11569 | HITS-CLIP | 37986224\|19536157\|23142080\|2916 | Unspecified |
| mmu-miR-34a-5p | MIMAT0000542 | Aes | 14797 | HITS-CLIP | 23142080 | Unspecified |
| mmu-miR-34a-5p | MIMAT0000542 | Afap1 | 70292 | HITS-CLIP | 19536157\|23630078\|26203562 | Unspecified |
| mmu-miR-34a-5p | MIMAT0000542 | Afap1l2 | 226250 | HITS-CLIP | 26203562 | Unspecified |
| mmu-miR-34a-5p | MIMAT0000542 | Afdn | 4301 | HITS-CLIP | 23142080\|19536157\|29167373 | Unspecified |
| mmu-miR-34a-5p | MIMAT0000542 | Aff1 | 17355 | HITS-CLIP | 23142080\|29167373 | Unspecified |
| mmu-miR-34a-5p | MIMAT0000542 | Aff3 | 16764 | HITS-CLIP | 19536157\|23142080 | Unspecified |
| mmu-miR-34a-5p | MIMAT0000542 | Aff4 | 93736 | HITS-CLIP | 37986224\|19536157\|29167373\|2314 | Unspecified |
| mmu-miR-34a-5p | MIMAT0000542 | Afg3l2 | 69597 | HITS-CLIP | 23142080\|31433990 | Unspecified |

| | | | | | | |
|---|---|---|---|---|---|---|
| mmu-miR-34a-5p | MIMAT0000542 | Afmid | 71562 | HITS-CLIP | 23142080 | Unspecified |
| mmu-miR-34a-5p | MIMAT0000542 | Aftph | 216549 | HITS-CLIP | 31433990\|19536157\|23142080\|29167 | Unspecified |
| mmu-miR-34a-5p | MIMAT0000542 | Aga | 11593 | HITS-CLIP | 23142080 | Unspecified |
| mmu-miR-34a-5p | MIMAT0000542 | Agap1 | 347722 | HITS-CLIP | 29167373 | Unspecified |
| mmu-miR-34a-5p | MIMAT0000542 | Agap2 | 216439 | HITS-CLIP | 19536157 | Unspecified |
| mmu-miR-34a-5p | MIMAT0000542 | Agap3 | 213990 | HITS-CLIP | 29167373 | Unspecified |
| mmu-miR-34a-5p | MIMAT0000542 | Agbl3 | 76223 | Microarrays | 37986224 | Unspecified |
| mmu-miR-34a-5p | MIMAT0000542 | Agbl5 | 231093 | HITS-CLIP | 23142080 | Unspecified |
| mmu-miR-34a-5p | MIMAT0000542 | Agfg1 | 15463 | HITS-CLIP | 23142080\|29167373 | Unspecified |
| mmu-miR-34a-5p | MIMAT0000542 | Aggf1 | 66549 | HITS-CLIP | 30971389 | Unspecified |
| mmu-miR-34a-5p | MIMAT0000542 | Agk | 69923 | HITS-CLIP | 23142080 | Unspecified |
| mmu-miR-34a-5p | MIMAT0000542 | Agl | 77559 | HITS-CLIP | 29167373 | Unspecified |
| mmu-miR-34a-5p | MIMAT0000542 | Ago1 | 236511 | HITS-CLIP | 31433990\|23142080\|29167373 | Unspecified |
| mmu-miR-34a-5p | MIMAT0000542 | Ago2 | 239528 | HITS-CLIP | 31433990\|19536157\|23630078\|26203 | Unspecified |
| mmu-miR-34a-5p | MIMAT0000542 | Ago3 | 214150 | HITS-CLIP | 23630078\|19536157\|30971389\|23142 | Unspecified |
| mmu-miR-34a-5p | MIMAT0000542 | Ago4 | 76850 | HITS-CLIP | 23142080 | Unspecified |
| mmu-miR-34a-5p | MIMAT0000542 | Agpat1 | 55979 | HITS-CLIP | 31433990\|23142080 | Unspecified |
| mmu-miR-34a-5p | MIMAT0000542 | Agpat2 | 67512 | HITS-CLIP | 23142080 | Unspecified |
| mmu-miR-34a-5p | MIMAT0000542 | Agpat3 | 28169 | HITS-CLIP | 23142080\|31433990\|29167373 | Unspecified |
| mmu-miR-34a-5p | MIMAT0000542 | Agpat4 | 68262 | HITS-CLIP | 23142080 | Unspecified |
| mmu-miR-34a-5p | MIMAT0000542 | Agpat5 | 52123 | HITS-CLIP | 23142080\|29167373 | Unspecified |
| mmu-miR-34a-5p | MIMAT0000542 | Agps | 228061 | HITS-CLIP | 26203562\|23142080\|29167373 | Unspecified |
| mmu-miR-34a-5p | MIMAT0000542 | Agrn | 11603 | HITS-CLIP | 19536157 | Unspecified |
| mmu-miR-34a-5p | MIMAT0000542 | Agtpbp1 | 67269 | HITS-CLIP | 23142080 | Unspecified |
| mmu-miR-34a-5p | MIMAT0000542 | Agtrap | 11610 | HITS-CLIP | 23142080 | Unspecified |
| mmu-miR-34a-5p | MIMAT0000542 | Ahctf1 | 226747 | HITS-CLIP | 19536157\|23142080 | Unspecified |
| mmu-miR-34a-5p | MIMAT0000542 | Ahcyl1 | 229709 | HITS-CLIP | 19536157\|23142080 | Unspecified |
| mmu-miR-34a-5p | MIMAT0000542 | Ahcyl2 | 74340 | HITS-CLIP | 30971389\|23142080\|29167373 | Unspecified |
| mmu-miR-34a-5p | MIMAT0000542 | Ahdc1 | 230793 | HITS-CLIP | 23142080\|26203562\|29167373 | Unspecified |
| mmu-miR-34a-5p | MIMAT0000542 | Ahi1 | 52906 | HITS-CLIP | 19536157\|23142080 | Unspecified |
| mmu-miR-34a-5p | MIMAT0000542 | Ahnak | 66395 | HITS-CLIP | 37986224\|23142080\|29167373 | Unspecified |
| mmu-miR-34a-5p | MIMAT0000542 | Ahr | 11622 | HITS-CLIP | 23142080 | Unspecified |
| mmu-miR-34a-5p | MIMAT0000542 | Ahrr | 11624 | HITS-CLIP | 29167373 | Unspecified |
| mmu-miR-34a-5p | MIMAT0000542 | Ahsa1 | 217737 | HITS-CLIP | 19536157\|23142080 | Unspecified |
| mmu-miR-34a-5p | MIMAT0000542 | Ahsa2 | 268390 | HITS-CLIP | 23142080 | Unspecified |
| mmu-miR-34a-5p | MIMAT0000542 | Aida | 108909 | HITS-CLIP | 23142080 | Unspecified |
| mmu-miR-34a-5p | MIMAT0000542 | Aifm1 | 26926 | HITS-CLIP | 23142080 | Unspecified |
| mmu-miR-34a-5p | MIMAT0000542 | Aifm2 | 71361 | HITS-CLIP | 29167373 | Unspecified |
| mmu-miR-34a-5p | MIMAT0000542 | Aimp2 | 231872 | HITS-CLIP | 23142080 | Unspecified |

| | | | | | | |
|---|---|---|---|---|---|---|
| mmu-miR-34a-5p | MIMAT0000542 | Aip | 11632 | HITS-CLIP | 31433990\|23142080\|29167373 | Unspecified |
| mmu-miR-34a-5p | MIMAT0000542 | Ajuba | 16475 | HITS-CLIP | 23142080 | Unspecified |
| mmu-miR-34a-5p | MIMAT0000542 | Ak2 | 11637 | HITS-CLIP | 31433990\|19536157\|23142080\|29167 | Unspecified |
| mmu-miR-34a-5p | MIMAT0000542 | Ak3 | 56248 | HITS-CLIP | 19536157\|29167373 | Unspecified |
| mmu-miR-34a-5p | MIMAT0000542 | Ak4 | 11639 | Microarrays//HITS-CLIP | 37986224\|23142080 | Unspecified |
| mmu-miR-34a-5p | MIMAT0000542 | Akap1 | 11640 | HITS-CLIP | 19536157\|30971389 | Unspecified |
| mmu-miR-34a-5p | MIMAT0000542 | Akap10 | 56697 | HITS-CLIP | 23142080\|29167373 | Unspecified |
| mmu-miR-34a-5p | MIMAT0000542 | Akap11 | 219181 | HITS-CLIP | 37986224\|19536157\|30971389\|31433 | Unspecified |
| mmu-miR-34a-5p | MIMAT0000542 | Akap13 | 75547 | HITS-CLIP | 37986224\|23142080\|29167373\|31433 | Unspecified |
| mmu-miR-34a-5p | MIMAT0000542 | Akap2 | 11641 | HITS-CLIP | 19536157 | Unspecified |
| mmu-miR-34a-5p | MIMAT0000542 | Akap6 | 238161 | Microarrays//HITS-CLIP | 37986224\|19536157 | Unspecified |
| mmu-miR-34a-5p | MIMAT0000542 | Akap7 | 432442 | HITS-CLIP | 19536157\|23142080 | Unspecified |
| mmu-miR-34a-5p | MIMAT0000542 | Akap8 | 56399 | HITS-CLIP | 29167373 | Unspecified |
| mmu-miR-34a-5p | MIMAT0000542 | Akap9 | 100986 | HITS-CLIP | 29167373\|23142080 | Unspecified |
| mmu-miR-34a-5p | MIMAT0000542 | Akirin2 | 433693 | HITS-CLIP | 19536157\|23142080 | Unspecified |
| mmu-miR-34a-5p | MIMAT0000542 | Akna | 100182 | HITS-CLIP | 23142080\|29167373 | Unspecified |
| mmu-miR-34a-5p | MIMAT0000542 | Akr1a1 | 58810 | HITS-CLIP | 23142080\|29167373 | Unspecified |
| mmu-miR-34a-5p | MIMAT0000542 | Akr1b8 | 14187 | HITS-CLIP | 29167373 | Unspecified |
| mmu-miR-34a-5p | MIMAT0000542 | Akr1c13 | 27384 | HITS-CLIP | 23142080 | Unspecified |
| mmu-miR-34a-5p | MIMAT0000542 | Akr1e1 | 56043 | HITS-CLIP | 23142080 | Unspecified |
| mmu-miR-34a-5p | MIMAT0000542 | Akt1s1 | 67605 | HITS-CLIP | 23142080 | Unspecified |
| mmu-miR-34a-5p | MIMAT0000542 | Akt2 | 11652 | HITS-CLIP | 23142080 | Unspecified |
| mmu-miR-34a-5p | MIMAT0000542 | Akt3 | 23797 | HITS-CLIP | 19536157\|23142080\|29167373 | Unspecified |
| mmu-miR-34a-5p | MIMAT0000542 | Alas1 | 11655 | HITS-CLIP | 23142080\|29167373 | Unspecified |
| mmu-miR-34a-5p | MIMAT0000542 | Alcam | 11658 | HITS-CLIP | 31433990\|19536157\|23142080\|29167 | Unspecified |
| mmu-miR-34a-5p | MIMAT0000542 | Aldh18a1 | 56454 | HITS-CLIP | 23142080 | Unspecified |
| mmu-miR-34a-5p | MIMAT0000542 | Aldh1b1 | 72535 | HITS-CLIP | 23142080 | Unspecified |
| mmu-miR-34a-5p | MIMAT0000542 | Aldh2 | 11669 | HITS-CLIP | 23142080 | Unspecified |
| mmu-miR-34a-5p | MIMAT0000542 | Aldh3a2 | 11671 | HITS-CLIP | 23142080 | Unspecified |
| mmu-miR-34a-5p | MIMAT0000542 | Aldh4a1 | 212647 | HITS-CLIP | 30971389 | Unspecified |
| mmu-miR-34a-5p | MIMAT0000542 | Aldh5a1 | 214579 | HITS-CLIP | 23142080 | Unspecified |
| mmu-miR-34a-5p | MIMAT0000542 | Aldh6a1 | 104776 | Microarrays//HITS-CLIP | 37986224\|23142080 | Unspecified |
| mmu-miR-34a-5p | MIMAT0000542 | Aldh7a1 | 110695 | HITS-CLIP | 23142080 | Unspecified |
| mmu-miR-34a-5p | MIMAT0000542 | Aldh9a1 | 56752 | HITS-CLIP | 23142080 | Unspecified |
| mmu-miR-34a-5p | MIMAT0000542 | Aldoa | 11674 | HITS-CLIP | 37986224\|19536157\|23142080\|29167 | Unspecified |
| mmu-miR-34a-5p | MIMAT0000542 | Aldoart1 | 353204 | HITS-CLIP | 23142080\|29167373 | Unspecified |
| mmu-miR-34a-5p | MIMAT0000542 | Aldob | 230163 | HITS-CLIP | 30971389 | Unspecified |
| mmu-miR-34a-5p | MIMAT0000542 | Aldoc | 11676 | HITS-CLIP | 23142080\|19536157 | Unspecified |
| mmu-miR-34a-5p | MIMAT0000542 | Alg10b | 380959 | Microarrays//HITS-CLIP | 37986224\|23027903\|23142080\|29167 | Unspecified |

| | | | | | | |
|---|---|---|---|---|---|---|
| mmu-miR-34a-5p | MIMAT0000542 | Alg11 | 207958 | HITS-CLIP | 31433990\|23142080\|29084942 | Unspecified |
| mmu-miR-34a-5p | MIMAT0000542 | Alg14 | 66789 | HITS-CLIP | 23142080 | Unspecified |
| mmu-miR-34a-5p | MIMAT0000542 | Alg2 | 56737 | HITS-CLIP | 19536157\|23142080\|29167373 | Unspecified |
| mmu-miR-34a-5p | MIMAT0000542 | Alg3 | 208624 | HITS-CLIP | 19536157\|23142080\|29167373 | Unspecified |
| mmu-miR-34a-5p | MIMAT0000542 | Alg5 | 66248 | HITS-CLIP | 19536157 | Unspecified |
| mmu-miR-34a-5p | MIMAT0000542 | Alg8 | 381903 | HITS-CLIP | 29167373 | Unspecified |
| mmu-miR-34a-5p | MIMAT0000542 | Alg9 | 102580 | HITS-CLIP | 30971389\|23142080 | Unspecified |
| mmu-miR-34a-5p | MIMAT0000542 | Alkbh1 | 211064 | HITS-CLIP | 31433990 | Unspecified |
| mmu-miR-34a-5p | MIMAT0000542 | Alkbh4 | 72041 | HITS-CLIP | 23142080 | Unspecified |
| mmu-miR-34a-5p | MIMAT0000542 | Alkbh5 | 268420 | HITS-CLIP | 31433990\|23630078\|23142080\|29167 | Unspecified |
| mmu-miR-34a-5p | MIMAT0000542 | Alkbh8 | 67667 | HITS-CLIP | 23142080 | Unspecified |
| mmu-miR-34a-5p | MIMAT0000542 | Alms1 | 236266 | HITS-CLIP | 23142080 | Unspecified |
| mmu-miR-34a-5p | MIMAT0000542 | Alox5ap | 11690 | HITS-CLIP | 29167373 | Unspecified |
| mmu-miR-34a-5p | MIMAT0000542 | Alpk1 | 71481 | HITS-CLIP | 29167373 | Unspecified |
| mmu-miR-34a-5p | MIMAT0000542 | Alpk2 | 225638 | HITS-CLIP | 23142080 | Unspecified |
| mmu-miR-34a-5p | MIMAT0000542 | Alpk3 | 116904 | Microarrays | 37986224 | Unspecified |
| mmu-miR-34a-5p | MIMAT0000542 | Als2 | 74018 | HITS-CLIP | 23142080\|29167373 | Unspecified |
| mmu-miR-34a-5p | MIMAT0000542 | Als2cl | 235633 | HITS-CLIP | 30971389 | Unspecified |
| mmu-miR-34a-5p | MIMAT0000542 | Alyref | 21681 | Microarrays//HITS-CLIP | 37986224\|23027903\|29167373 | Unspecified |
| mmu-miR-34a-5p | MIMAT0000542 | Alyref2 | 56009 | HITS-CLIP | 23142080\|29167373 | Unspecified |
| mmu-miR-34a-5p | MIMAT0000542 | Amacr | 17117 | HITS-CLIP | 30971389 | Unspecified |
| mmu-miR-34a-5p | MIMAT0000542 | Ambra1 | 228361 | HITS-CLIP | 23142080\|29167373 | Unspecified |
| mmu-miR-34a-5p | MIMAT0000542 | Amdhd2 | 245847 | HITS-CLIP | 29167373 | Unspecified |
| mmu-miR-34a-5p | MIMAT0000542 | Amer1 | 72345 | HITS-CLIP | 23142080\|29167373 | Unspecified |
| mmu-miR-34a-5p | MIMAT0000542 | Amfr | 23802 | HITS-CLIP | 29167373\|23142080 | Unspecified |
| mmu-miR-34a-5p | MIMAT0000542 | Amigo2 | 105827 | HITS-CLIP | 23142080 | Unspecified |
| mmu-miR-34a-5p | MIMAT0000542 | Ammecr1 | 56068 | Microarrays//HITS-CLIP | 37986224\|23142080 | Unspecified |
| mmu-miR-34a-5p | MIMAT0000542 | Ammecr1l | 225339 | HITS-CLIP | 23142080 | Unspecified |
| mmu-miR-34a-5p | MIMAT0000542 | Amn | 93835 | HITS-CLIP | 23142080 | Unspecified |
| mmu-miR-34a-5p | MIMAT0000542 | Amn1 | 232566 | HITS-CLIP | 23142080\|29167373 | Unspecified |
| mmu-miR-34a-5p | MIMAT0000542 | Amot | 27494 | HITS-CLIP | 19536157 | Unspecified |
| mmu-miR-34a-5p | MIMAT0000542 | Amotl2 | 56332 | HITS-CLIP | 23630078 | Unspecified |
| mmu-miR-34a-5p | MIMAT0000542 | Ampd2 | 109674 | HITS-CLIP | 23142080\|29167373 | Unspecified |
| mmu-miR-34a-5p | MIMAT0000542 | Ampd3 | 11717 | HITS-CLIP | 29167373 | Unspecified |
| mmu-miR-34a-5p | MIMAT0000542 | Amz2 | 13929 | HITS-CLIP | 23142080 | Unspecified |
| mmu-miR-34a-5p | MIMAT0000542 | Anapc1 | 17222 | HITS-CLIP | 23142080\|29167373 | Unspecified |
| mmu-miR-34a-5p | MIMAT0000542 | Anapc11 | 66156 | HITS-CLIP | 23142080 | Unspecified |
| mmu-miR-34a-5p | MIMAT0000542 | Anapc15 | 75430 | HITS-CLIP | 29167373 | Unspecified |
| mmu-miR-34a-5p | MIMAT0000542 | Anapc16 | 52717 | HITS-CLIP | 23142080 | Unspecified |

| | | | | | | |
|---|---|---|---|---|---|---|
| mmu-miR-34a-5p | MIMAT0000542 | Anapc2 | 99152 | HITS-CLIP | 23142080 | Unspecified |
| mmu-miR-34a-5p | MIMAT0000542 | Anapc4 | 52206 | HITS-CLIP | 23142080 | Unspecified |
| mmu-miR-34a-5p | MIMAT0000542 | Anapc5 | 59008 | HITS-CLIP | 19536157\|23142080\|29167373 | Unspecified |
| mmu-miR-34a-5p | MIMAT0000542 | Anapc7 | 56317 | HITS-CLIP | 23142080 | Unspecified |
| mmu-miR-34a-5p | MIMAT0000542 | Angel2 | 52477 | HITS-CLIP | 30971389\|23142080 | Unspecified |
| mmu-miR-34a-5p | MIMAT0000542 | Angptl2 | 26360 | HITS-CLIP | 23142080\|29167373 | Unspecified |
| mmu-miR-34a-5p | MIMAT0000542 | Angptl4 | 57875 | HITS-CLIP | 23142080 | Unspecified |
| mmu-miR-34a-5p | MIMAT0000542 | Ank | 11732 | HITS-CLIP | 29167373 | Unspecified |
| mmu-miR-34a-5p | MIMAT0000542 | Ank1 | 11733 | HITS-CLIP | 30971389 | Unspecified |
| mmu-miR-34a-5p | MIMAT0000542 | Ank2 | 109676 | HITS-CLIP | 19536157 | Unspecified |
| mmu-miR-34a-5p | MIMAT0000542 | Ank3 | 11735 | HITS-CLIP | 19536157 | Unspecified |
| mmu-miR-34a-5p | MIMAT0000542 | Ankfy1 | 11736 | Microarrays//HITS-CLIP | 37986224\|23142080\|23027903\|2363( | Unspecified |
| mmu-miR-34a-5p | MIMAT0000542 | Ankhd1 | 108857 | HITS-CLIP | 23142080\|31433990\|29167373 | Unspecified |
| mmu-miR-34a-5p | MIMAT0000542 | Ankib1 | 70797 | HITS-CLIP | 23142080 | Unspecified |
| mmu-miR-34a-5p | MIMAT0000542 | Ankle2 | 71782 | HITS-CLIP | 23142080\|29167373 | Unspecified |
| mmu-miR-34a-5p | MIMAT0000542 | Ankmy2 | 217473 | HITS-CLIP | 23142080 | Unspecified |
| mmu-miR-34a-5p | MIMAT0000542 | Ankra2 | 68558 | HITS-CLIP | 23142080 | Unspecified |
| mmu-miR-34a-5p | MIMAT0000542 | Ankrd10 | 102334 | HITS-CLIP | 23142080 | Unspecified |
| mmu-miR-34a-5p | MIMAT0000542 | Ankrd11 | 77087 | HITS-CLIP | 29167373\|23142080 | Unspecified |
| mmu-miR-34a-5p | MIMAT0000542 | Ankrd12 | 106585 | HITS-CLIP | 19536157\|23142080\|29167373 | Unspecified |
| mmu-miR-34a-5p | MIMAT0000542 | Ankrd13a | 68420 | HITS-CLIP | 23142080\|29167373 | Unspecified |
| mmu-miR-34a-5p | MIMAT0000542 | Ankrd13c | 433667 | HITS-CLIP | 19536157 | Unspecified |
| mmu-miR-34a-5p | MIMAT0000542 | Ankrd17 | 81702 | HITS-CLIP | 37986224\|23142080\|19536157\|2916 | Unspecified |
| mmu-miR-34a-5p | MIMAT0000542 | Ankrd2 | 56642 | Microarrays | 37986224 | Unspecified |
| mmu-miR-34a-5p | MIMAT0000542 | Ankrd27 | 245886 | HITS-CLIP | 23142080 | Unspecified |
| mmu-miR-34a-5p | MIMAT0000542 | Ankrd28 | 105522 | HITS-CLIP | 21258322\|23142080\|29167373 | Unspecified |
| mmu-miR-34a-5p | MIMAT0000542 | Ankrd33b | 67434 | HITS-CLIP | 19536157 | Unspecified |
| mmu-miR-34a-5p | MIMAT0000542 | Ankrd40 | 71452 | HITS-CLIP | 19536157 | Unspecified |
| mmu-miR-34a-5p | MIMAT0000542 | Ankrd44 | 329154 | HITS-CLIP | 23142080\|31433990\|29167373 | Unspecified |
| mmu-miR-34a-5p | MIMAT0000542 | Ankrd46 | 68839 | HITS-CLIP | 23142080 | Unspecified |
| mmu-miR-34a-5p | MIMAT0000542 | Ankrd49 | 56503 | HITS-CLIP | 23142080\|31433990 | Unspecified |
| mmu-miR-34a-5p | MIMAT0000542 | Ankrd50 | 99696 | HITS-CLIP | 23142080\|29167373 | Unspecified |
| mmu-miR-34a-5p | MIMAT0000542 | Ankrd52 | 237615 | Microarrays//HITS-CLIP | 37986224\|23027903\|23142080\|1953( | Unspecified |
| mmu-miR-34a-5p | MIMAT0000542 | Ankrd63 | 383787 | HITS-CLIP | 19536157 | Unspecified |
| mmu-miR-34a-5p | MIMAT0000542 | Anks1 | 224650 | HITS-CLIP | 23142080\|29167373 | Unspecified |
| mmu-miR-34a-5p | MIMAT0000542 | Anks1b | 77531 | HITS-CLIP | 19536157 | Unspecified |
| mmu-miR-34a-5p | MIMAT0000542 | Anks3 | 72615 | HITS-CLIP | 23142080 | Unspecified |
| mmu-miR-34a-5p | MIMAT0000542 | Anln | 68743 | HITS-CLIP | 23142080 | Unspecified |
| mmu-miR-34a-5p | MIMAT0000542 | Ano10 | 102566 | HITS-CLIP | 23142080\|29167373 | Unspecified |

| | | | | | | |
|---|---|---|---|---|---|---|
| mmu-miR-34a-5p | MIMAT0000542 | Ano6 | 105722 | HITS-CLIP | 19536157\|23142080\|29167373 | Unspecified |
| mmu-miR-34a-5p | MIMAT0000542 | Anp32a | 11737 | HITS-CLIP | 23142080 | Unspecified |
| mmu-miR-34a-5p | MIMAT0000542 | Anp32b | 67628 | HITS-CLIP | 23630078\|31433990\|23142080\|29167 | Unspecified |
| mmu-miR-34a-5p | MIMAT0000542 | Anp32e | 66471 | Microarrays//HITS-CLIP | 37986224\|23142080\|23027903\|3143 | Unspecified |
| mmu-miR-34a-5p | MIMAT0000542 | Anpep | 16790 | HITS-CLIP | 29167373 | Unspecified |
| mmu-miR-34a-5p | MIMAT0000542 | Antxr1 | 69538 | HITS-CLIP | 26203562 | Unspecified |
| mmu-miR-34a-5p | MIMAT0000542 | Antxr2 | 71914 | HITS-CLIP | 29167373\|23142080 | Unspecified |
| mmu-miR-34a-5p | MIMAT0000542 | Anxa1 | 16952 | HITS-CLIP | 29167373 | Unspecified |
| mmu-miR-34a-5p | MIMAT0000542 | Anxa11 | 11744 | HITS-CLIP | 23142080\|23630078 | Unspecified |
| mmu-miR-34a-5p | MIMAT0000542 | Anxa2 | 12306 | HITS-CLIP | 23630078\|23142080\|29167373 | Unspecified |
| mmu-miR-34a-5p | MIMAT0000542 | Anxa4 | 11746 | HITS-CLIP | 23142080\|29167373 | Unspecified |
| mmu-miR-34a-5p | MIMAT0000542 | Anxa5 | 11747 | HITS-CLIP | 23142080\|29167373 | Unspecified |
| mmu-miR-34a-5p | MIMAT0000542 | Anxa6 | 11749 | HITS-CLIP | 23142080\|29167373 | Unspecified |
| mmu-miR-34a-5p | MIMAT0000542 | Anxa7 | 11750 | HITS-CLIP | 23142080\|29167373 | Unspecified |
| mmu-miR-34a-5p | MIMAT0000542 | Ap1ar | 211556 | HITS-CLIP | 23142080\|29167373 | Unspecified |
| mmu-miR-34a-5p | MIMAT0000542 | Ap1b1 | 11764 | HITS-CLIP | 19536157\|30971389\|23142080\|29167 | Unspecified |
| mmu-miR-34a-5p | MIMAT0000542 | Ap1g1 | 11765 | HITS-CLIP, Microarrays//Microarrays//HITS | 37986224\|23027903\|23142080\|29167 | Unspecified |
| mmu-miR-34a-5p | MIMAT0000542 | Ap1g2 | 11766 | HITS-CLIP | 29167373\|23142080 | Unspecified |
| mmu-miR-34a-5p | MIMAT0000542 | Ap1m1 | 11767 | HITS-CLIP | 23142080\|29167373 | Unspecified |
| mmu-miR-34a-5p | MIMAT0000542 | Ap1s2 | 108012 | HITS-CLIP | 23142080\|29167373 | Unspecified |
| mmu-miR-34a-5p | MIMAT0000542 | Ap1s3 | 252903 | Microarrays//HITS-CLIP | 37986224\|23027903\|23142080 | Unspecified |
| mmu-miR-34a-5p | MIMAT0000542 | Ap2a1 | 11771 | HITS-CLIP | 23142080\|29167373 | Unspecified |
| mmu-miR-34a-5p | MIMAT0000542 | Ap2a2 | 11772 | HITS-CLIP | 31433990\|30971389\|23142080\|29167 | Unspecified |
| mmu-miR-34a-5p | MIMAT0000542 | Ap2b1 | 71770 | HITS-CLIP | 23142080\|31433990\|19536157 | Unspecified |
| mmu-miR-34a-5p | MIMAT0000542 | Ap2m1 | 11773 | HITS-CLIP | 26203562 | Unspecified |
| mmu-miR-34a-5p | MIMAT0000542 | Ap2s1 | 232910 | HITS-CLIP | 23142080\|29167373 | Unspecified |
| mmu-miR-34a-5p | MIMAT0000542 | Ap3b1 | 11774 | HITS-CLIP | 23142080\|29167373 | Unspecified |
| mmu-miR-34a-5p | MIMAT0000542 | Ap3d1 | 11776 | HITS-CLIP | 23142080\|26203562\|29167373 | Unspecified |
| mmu-miR-34a-5p | MIMAT0000542 | Ap3m1 | 55946 | HITS-CLIP | 23142080\|29167373 | Unspecified |
| mmu-miR-34a-5p | MIMAT0000542 | Ap3m2 | 64933 | HITS-CLIP | 19536157 | Unspecified |
| mmu-miR-34a-5p | MIMAT0000542 | Ap3s2 | 11778 | HITS-CLIP | 19536157\|23142080\|29167373 | Unspecified |
| mmu-miR-34a-5p | MIMAT0000542 | Ap4b1 | 67489 | HITS-CLIP | 23142080 | Unspecified |
| mmu-miR-34a-5p | MIMAT0000542 | Ap4e1 | 108011 | HITS-CLIP | 23142080\|19536157\|29167373 | Unspecified |
| mmu-miR-34a-5p | MIMAT0000542 | Ap4m1 | 11781 | HITS-CLIP | 31433990 | Unspecified |
| mmu-miR-34a-5p | MIMAT0000542 | Ap5b1 | 381201 | HITS-CLIP | 23142080\|29167373 | Unspecified |
| mmu-miR-34a-5p | MIMAT0000542 | Ap5m1 | 74385 | HITS-CLIP | 23142080 | Unspecified |
| mmu-miR-34a-5p | MIMAT0000542 | Apaf1 | 11783 | HITS-CLIP | 37986224\|23142080\|31433990\|29167 | Unspecified |
| mmu-miR-34a-5p | MIMAT0000542 | Apba1 | 319924 | HITS-CLIP | 19536157 | Unspecified |
| mmu-miR-34a-5p | MIMAT0000542 | Apba2 | 11784 | HITS-CLIP | 19536157 | Unspecified |

| | | | | | | |
|---|---|---|---|---|---|---|
| mmu-miR-34a-5p | MIMAT0000542 | Apbb1 | 322 | HITS-CLIP | 23142080 | Unspecified |
| mmu-miR-34a-5p | MIMAT0000542 | Apbb1ip | 54519 | HITS-CLIP | 29167373 | Unspecified |
| mmu-miR-34a-5p | MIMAT0000542 | Apbb2 | 11787 | HITS-CLIP | 29167373 | Unspecified |
| mmu-miR-34a-5p | MIMAT0000542 | Apc | 11789 | HITS-CLIP | 19536157\|23142080\|29167373 | Unspecified |
| mmu-miR-34a-5p | MIMAT0000542 | Apc2 | 23805 | HITS-CLIP | 19536157 | Unspecified |
| mmu-miR-34a-5p | MIMAT0000542 | Apcdd1 | 494504 | HITS-CLIP | 19536157 | Unspecified |
| mmu-miR-34a-5p | MIMAT0000542 | Apeh | 235606 | HITS-CLIP | 23142080 | Unspecified |
| mmu-miR-34a-5p | MIMAT0000542 | Apex1 | 11792 | HITS-CLIP | 23142080 | Unspecified |
| mmu-miR-34a-5p | MIMAT0000542 | Apex2 | 77622 | HITS-CLIP | 29167373 | Unspecified |
| mmu-miR-34a-5p | MIMAT0000542 | Aph1c | 68318 | HITS-CLIP | 29167373 | Unspecified |
| mmu-miR-34a-5p | MIMAT0000542 | Api5 | 11800 | HITS-CLIP | 30971389\|23142080\|29167373 | Unspecified |
| mmu-miR-34a-5p | MIMAT0000542 | Apip | 56369 | HITS-CLIP | 23142080 | Unspecified |
| mmu-miR-34a-5p | MIMAT0000542 | Aplf | 72103 | HITS-CLIP | 23142080 | Unspecified |
| mmu-miR-34a-5p | MIMAT0000542 | Aplp1 | 11803 | HITS-CLIP | 23142080\|19536157 | Unspecified |
| mmu-miR-34a-5p | MIMAT0000542 | Aplp2 | 11804 | HITS-CLIP | 19536157\|29167373 | Unspecified |
| mmu-miR-34a-5p | MIMAT0000542 | Apobec1 | 11810 | HITS-CLIP | 29167373 | Unspecified |
| mmu-miR-34a-5p | MIMAT0000542 | Apobec2 | 11811 | Microarrays | 37986224 | Unspecified |
| mmu-miR-34a-5p | MIMAT0000542 | Apobec3 | 80287 | HITS-CLIP | 23142080\|31433990\|29167373 | Unspecified |
| mmu-miR-34a-5p | MIMAT0000542 | Apoe | 11816 | HITS-CLIP | 29167373 | Unspecified |
| mmu-miR-34a-5p | MIMAT0000542 | Apool | 68117 | HITS-CLIP | 23142080 | Unspecified |
| mmu-miR-34a-5p | MIMAT0000542 | App | 11820 | HITS-CLIP | 29167373\|19536157 | Unspecified |
| mmu-miR-34a-5p | MIMAT0000542 | Appbp2 | 66884 | HITS-CLIP | 23142080\|19536157\|29167373 | Unspecified |
| mmu-miR-34a-5p | MIMAT0000542 | Appl1 | 72993 | HITS-CLIP | 29167373\|23142080 | Unspecified |
| mmu-miR-34a-5p | MIMAT0000542 | Appl2 | 216190 | HITS-CLIP | 23142080\|29167373 | Unspecified |
| mmu-miR-34a-5p | MIMAT0000542 | Aprt | 11821 | HITS-CLIP | 23142080 | Unspecified |
| mmu-miR-34a-5p | MIMAT0000542 | Aptx | 66408 | HITS-CLIP | 23142080 | Unspecified |
| mmu-miR-34a-5p | MIMAT0000542 | Aqp11 | 66333 | HITS-CLIP | 23142080 | Unspecified |
| mmu-miR-34a-5p | MIMAT0000542 | Aqp4 | 11829 | HITS-CLIP | 19536157 | Unspecified |
| mmu-miR-34a-5p | MIMAT0000542 | Aqr | 11834 | HITS-CLIP | 30971389\|23142080\|29167373 | Unspecified |
| mmu-miR-34a-5p | MIMAT0000542 | Ar | 11835 | Microarrays//HITS-CLIP | 37986224\|31433990\|23142080 | Unspecified |
| mmu-miR-34a-5p | MIMAT0000542 | Araf | 11836 | HITS-CLIP | 23142080\|29167373 | Unspecified |
| mmu-miR-34a-5p | MIMAT0000542 | Arap1 | 69710 | HITS-CLIP | 23142080\|29167373 | Unspecified |
| mmu-miR-34a-5p | MIMAT0000542 | Arap2 | 212285 | HITS-CLIP | 23142080 | Unspecified |
| mmu-miR-34a-5p | MIMAT0000542 | Arap3 | 106952 | HITS-CLIP | 29167373 | Unspecified |
| mmu-miR-34a-5p | MIMAT0000542 | Arc | 11838 | HITS-CLIP | 19536157\|23142080 | Unspecified |
| mmu-miR-34a-5p | MIMAT0000542 | Arcn1 | 213827 | HITS-CLIP | 23142080 | Unspecified |
| mmu-miR-34a-5p | MIMAT0000542 | Arel1 | 68497 | HITS-CLIP | 19536157\|23142080 | Unspecified |
| mmu-miR-34a-5p | MIMAT0000542 | Arf1 | 11840 | HITS-CLIP | 31433990 | Unspecified |
| mmu-miR-34a-5p | MIMAT0000542 | Arf3 | 11842 | Microarrays//HITS-CLIP | 37986224\|23027903\|19536157\|31433 | Unspecified |

| | | | | | | |
|---|---|---|---|---|---|---|
| mmu-miR-34a-5p | MIMAT0000542 | Arf4 | 11843 | HITS-CLIP | 23142080\|29167373 | Unspecified |
| mmu-miR-34a-5p | MIMAT0000542 | Arf5 | 11844 | HITS-CLIP | 23142080 | Unspecified |
| mmu-miR-34a-5p | MIMAT0000542 | Arf6 | 11845 | HITS-CLIP | 37986224\|29167373\|23142080 | Unspecified |
| mmu-miR-34a-5p | MIMAT0000542 | Arfgap1 | 228998 | HITS-CLIP | 29167373 | Unspecified |
| mmu-miR-34a-5p | MIMAT0000542 | Arfgef1 | 211673 | HITS-CLIP | 23142080\|29167373 | Unspecified |
| mmu-miR-34a-5p | MIMAT0000542 | Arfgef2 | 99371 | HITS-CLIP | 30971389\|29167373 | Unspecified |
| mmu-miR-34a-5p | MIMAT0000542 | Arfgef3 | 57221 | HITS-CLIP | 29167373 | Unspecified |
| mmu-miR-34a-5p | MIMAT0000542 | Arfip2 | 76932 | HITS-CLIP | 23142080 | Unspecified |
| mmu-miR-34a-5p | MIMAT0000542 | Arhgap1 | 228359 | HITS-CLIP | 19536157\|23142080 | Unspecified |
| mmu-miR-34a-5p | MIMAT0000542 | Arhgap10 | 78514 | HITS-CLIP | 29167373 | Unspecified |
| mmu-miR-34a-5p | MIMAT0000542 | Arhgap11a | 228482 | Microarrays//HITS-CLIP | 37986224\|23027903\|23142080\|29167 | Unspecified |
| mmu-miR-34a-5p | MIMAT0000542 | Arhgap12 | 75415 | HITS-CLIP | 37986224\|23142080\|31433990\|23630 | Unspecified |
| mmu-miR-34a-5p | MIMAT0000542 | Arhgap15 | 76117 | HITS-CLIP | 23142080\|29167373 | Unspecified |
| mmu-miR-34a-5p | MIMAT0000542 | Arhgap17 | 70497 | HITS-CLIP | 29167373 | Unspecified |
| mmu-miR-34a-5p | MIMAT0000542 | Arhgap18 | 73910 | HITS-CLIP | 23142080 | Unspecified |
| mmu-miR-34a-5p | MIMAT0000542 | Arhgap19 | 71085 | Microarrays//HITS-CLIP | 37986224\|23027903\|23142080\|26203 | Unspecified |
| mmu-miR-34a-5p | MIMAT0000542 | Arhgap21 | 71435 | HITS-CLIP | 31433990\|30971389\|19536157\|23142 | Unspecified |
| mmu-miR-34a-5p | MIMAT0000542 | Arhgap22 | 239027 | HITS-CLIP | 29167373 | Unspecified |
| mmu-miR-34a-5p | MIMAT0000542 | Arhgap25 | 232201 | HITS-CLIP | 31433990\|29167373 | Unspecified |
| mmu-miR-34a-5p | MIMAT0000542 | Arhgap26 | 71302 | HITS-CLIP | 23142080 | Unspecified |
| mmu-miR-34a-5p | MIMAT0000542 | Arhgap27 | 544817 | HITS-CLIP | 23142080\|29167373 | Unspecified |
| mmu-miR-34a-5p | MIMAT0000542 | Arhgap30 | 226652 | HITS-CLIP | 23142080\|29167373 | Unspecified |
| mmu-miR-34a-5p | MIMAT0000542 | Arhgap31 | 12549 | HITS-CLIP | 29167373 | Unspecified |
| mmu-miR-34a-5p | MIMAT0000542 | Arhgap35 | 232906 | HITS-CLIP | 19536157\|31433990\|23142080\|29167 | Unspecified |
| mmu-miR-34a-5p | MIMAT0000542 | Arhgap4 | 393 | HITS-CLIP | 23142080\|29167373 | Unspecified |
| mmu-miR-34a-5p | MIMAT0000542 | Arhgap45 | 23526 | HITS-CLIP | 23142080\|29167373 | Unspecified |
| mmu-miR-34a-5p | MIMAT0000542 | Arhgap5 | 11855 | HITS-CLIP | 23142080\|29167373 | Unspecified |
| mmu-miR-34a-5p | MIMAT0000542 | Arhgap9 | 216445 | HITS-CLIP | 23142080\|29167373 | Unspecified |
| mmu-miR-34a-5p | MIMAT0000542 | Arhgdia | 192662 | HITS-CLIP | 19536157\|23142080\|29167373 | Unspecified |
| mmu-miR-34a-5p | MIMAT0000542 | Arhgdib | 11857 | Microarrays//HITS-CLIP | 37986224\|31433990\|23142080 | Unspecified |
| mmu-miR-34a-5p | MIMAT0000542 | Arhgef1 | 16801 | HITS-CLIP | 23142080\|29167373 | Unspecified |
| mmu-miR-34a-5p | MIMAT0000542 | Arhgef10 | 234094 | HITS-CLIP | 23142080 | Unspecified |
| mmu-miR-34a-5p | MIMAT0000542 | Arhgef11 | 213498 | HITS-CLIP | 29167373 | Unspecified |
| mmu-miR-34a-5p | MIMAT0000542 | Arhgef12 | 69632 | HITS-CLIP | 23142080\|19536157\|30971389 | Unspecified |
| mmu-miR-34a-5p | MIMAT0000542 | Arhgef17 | 207212 | HITS-CLIP | 30971389\|19536157 | Unspecified |
| mmu-miR-34a-5p | MIMAT0000542 | Arhgef18 | 102098 | HITS-CLIP | 23142080\|31433990 | Unspecified |
| mmu-miR-34a-5p | MIMAT0000542 | Arhgef2 | 16800 | HITS-CLIP | 23142080\|29167373 | Unspecified |
| mmu-miR-34a-5p | MIMAT0000542 | Arhgef3 | 71704 | HITS-CLIP | 23142080 | Unspecified |
| mmu-miR-34a-5p | MIMAT0000542 | Arhgef6 | 73341 | HITS-CLIP | 31433990\|23142080 | Unspecified |

| | | | | | | |
|---|---|---|---|---|---|---|
| mmu-miR-34a-5p | MIMAT0000542 | Arhgef7 | 54126 | Microarrays//HITS-CLIP | 37986224\|23027903\|19536157\|23142 | Unspecified |
| mmu-miR-34a-5p | MIMAT0000542 | Arhgef9 | 236915 | HITS-CLIP | 23142080\|19536157 | Unspecified |
| mmu-miR-34a-5p | MIMAT0000542 | Arid1a | 93760 | HITS-CLIP | 37986224\|23142080\|19536157\|29167 | Unspecified |
| mmu-miR-34a-5p | MIMAT0000542 | Arid1b | 239985 | HITS-CLIP | 23142080 | Unspecified |
| mmu-miR-34a-5p | MIMAT0000542 | Arid2 | 77044 | HITS-CLIP | 31433990\|19536157\|23142080\|29167 | Unspecified |
| mmu-miR-34a-5p | MIMAT0000542 | Arid3a | 13496 | HITS-CLIP | 23142080 | Unspecified |
| mmu-miR-34a-5p | MIMAT0000542 | Arid3b | 56380 | HITS-CLIP | 23142080 | Unspecified |
| mmu-miR-34a-5p | MIMAT0000542 | Arid4a | 238247 | HITS-CLIP | 37986224\|31433990\|26203562\|29167 | Unspecified |
| mmu-miR-34a-5p | MIMAT0000542 | Arid4b | 94246 | HITS-CLIP | 23142080\|29167373 | Unspecified |
| mmu-miR-34a-5p | MIMAT0000542 | Arid5a | 214855 | HITS-CLIP | 23142080 | Unspecified |
| mmu-miR-34a-5p | MIMAT0000542 | Arid5b | 71371 | HITS-CLIP | 23142080\|29167373 | Unspecified |
| mmu-miR-34a-5p | MIMAT0000542 | Arih1 | 23806 | HITS-CLIP | 23142080\|31433990\|19536157\|29167 | Unspecified |
| mmu-miR-34a-5p | MIMAT0000542 | Arih2 | 23807 | HITS-CLIP | 23142080 | Unspecified |
| mmu-miR-34a-5p | MIMAT0000542 | Arl11 | 219144 | HITS-CLIP | 29167373 | Unspecified |
| mmu-miR-34a-5p | MIMAT0000542 | Arl13b | 68146 | HITS-CLIP | 23142080 | Unspecified |
| mmu-miR-34a-5p | MIMAT0000542 | Arl14ep | 212772 | HITS-CLIP | 23630078\|23142080 | Unspecified |
| mmu-miR-34a-5p | MIMAT0000542 | Arl15 | 218639 | HITS-CLIP | 23142080 | Unspecified |
| mmu-miR-34a-5p | MIMAT0000542 | Arl2 | 56327 | HITS-CLIP | 23142080 | Unspecified |
| mmu-miR-34a-5p | MIMAT0000542 | Arl2bp | 107566 | HITS-CLIP | 23630078\|23142080 | Unspecified |
| mmu-miR-34a-5p | MIMAT0000542 | Arl3 | 56350 | HITS-CLIP | 23142080 | Unspecified |
| mmu-miR-34a-5p | MIMAT0000542 | Arl4c | 320982 | HITS-CLIP | 23142080\|29167373 | Unspecified |
| mmu-miR-34a-5p | MIMAT0000542 | Arl5a | 75423 | HITS-CLIP | 23142080 | Unspecified |
| mmu-miR-34a-5p | MIMAT0000542 | Arl5b | 75869 | HITS-CLIP | 37986224\|23142080\|26203562\|29167 | Unspecified |
| mmu-miR-34a-5p | MIMAT0000542 | Arl6ip1 | 54208 | HITS-CLIP | 19536157\|23142080\|29167373 | Unspecified |
| mmu-miR-34a-5p | MIMAT0000542 | Arl6ip4 | 65105 | HITS-CLIP | 30971389 | Unspecified |
| mmu-miR-34a-5p | MIMAT0000542 | Arl6ip5 | 65106 | HITS-CLIP | 23630078\|23142080\|29167373 | Unspecified |
| mmu-miR-34a-5p | MIMAT0000542 | Arl6ip6 | 65103 | HITS-CLIP | 23142080 | Unspecified |
| mmu-miR-34a-5p | MIMAT0000542 | Arl8a | 68724 | HITS-CLIP | 23630078\|23142080 | Unspecified |
| mmu-miR-34a-5p | MIMAT0000542 | Arl8b | 67166 | HITS-CLIP | 23630078\|19536157\|29167373\|23142 | Unspecified |
| mmu-miR-34a-5p | MIMAT0000542 | Armc1 | 74252 | HITS-CLIP | 23142080 | Unspecified |
| mmu-miR-34a-5p | MIMAT0000542 | Armc8 | 74125 | HITS-CLIP | 19536157\|23142080\|29167373 | Unspecified |
| mmu-miR-34a-5p | MIMAT0000542 | Armc9 | 78795 | Microarrays | 37986224 | Unspecified |
| mmu-miR-34a-5p | MIMAT0000542 | Armcx1 | 78248 | HITS-CLIP | 23142080 | Unspecified |
| mmu-miR-34a-5p | MIMAT0000542 | Armcx2 | 67416 | HITS-CLIP | 23142080\|19536157 | Unspecified |
| mmu-miR-34a-5p | MIMAT0000542 | Armcx6 | 278097 | Microarrays | 37986224 | Unspecified |
| mmu-miR-34a-5p | MIMAT0000542 | Armt1 | 79624 | HITS-CLIP | 23142080\|29167373 | Unspecified |
| mmu-miR-34a-5p | MIMAT0000542 | Arnt | 11863 | HITS-CLIP | 23142080 | Unspecified |
| mmu-miR-34a-5p | MIMAT0000542 | Arnt2 | 11864 | HITS-CLIP | 19536157 | Unspecified |
| mmu-miR-34a-5p | MIMAT0000542 | Arntl | 11865 | HITS-CLIP | 23142080 | Unspecified |

| | | | | | | |
|---|---|---|---|---|---|---|
| mmu-miR-34a-5p | MIMAT0000542 | Arpc1a | 56443 | HITS-CLIP | 23142080\|29167373 | Unspecified |
| mmu-miR-34a-5p | MIMAT0000542 | Arpc1b | 11867 | HITS-CLIP | 23142080\|29167373 | Unspecified |
| mmu-miR-34a-5p | MIMAT0000542 | Arpc2 | 76709 | HITS-CLIP | 23630078\|23142080\|29167373 | Unspecified |
| mmu-miR-34a-5p | MIMAT0000542 | Arpc3 | 56378 | Microarrays//HITS-CLIP | 37986224\|23027903\|23142080\|29167 | Unspecified |
| mmu-miR-34a-5p | MIMAT0000542 | Arpc4 | 68089 | HITS-CLIP | 23142080\|31433990\|29167373 | Unspecified |
| mmu-miR-34a-5p | MIMAT0000542 | Arpc5 | 67771 | HITS-CLIP | 23142080\|29167373 | Unspecified |
| mmu-miR-34a-5p | MIMAT0000542 | Arpin | 70420 | Microarrays | 23027903 | Unspecified |
| mmu-miR-34a-5p | MIMAT0000542 | Arpp19 | 59046 | HITS-CLIP | 23142080 | Unspecified |
| mmu-miR-34a-5p | MIMAT0000542 | Arpp21 | 74100 | HITS-CLIP | 19536157 | Unspecified |
| mmu-miR-34a-5p | MIMAT0000542 | Arrb1 | 109689 | HITS-CLIP | 23142080\|19536157\|30971389\|29167 | Unspecified |
| mmu-miR-34a-5p | MIMAT0000542 | Arrb2 | 216869 | HITS-CLIP | 29167373 | Unspecified |
| mmu-miR-34a-5p | MIMAT0000542 | Arrdc1 | 215705 | HITS-CLIP | 29167373 | Unspecified |
| mmu-miR-34a-5p | MIMAT0000542 | Arrdc3 | 105171 | HITS-CLIP | 19536157\|23142080 | Unspecified |
| mmu-miR-34a-5p | MIMAT0000542 | Arrdc4 | 66412 | HITS-CLIP | 23142080 | Unspecified |
| mmu-miR-34a-5p | MIMAT0000542 | Arsb | 11881 | HITS-CLIP | 29167373 | Unspecified |
| mmu-miR-34a-5p | MIMAT0000542 | Art2b | 11872 | HITS-CLIP | 23142080 | Unspecified |
| mmu-miR-34a-5p | MIMAT0000542 | Arv1 | 68865 | HITS-CLIP | 29167373\|23142080 | Unspecified |
| mmu-miR-34a-5p | MIMAT0000542 | Arvcf | 11877 | HITS-CLIP | 29084942 | Unspecified |
| mmu-miR-34a-5p | MIMAT0000542 | Asah1 | 11886 | HITS-CLIP | 29167373\|23142080 | Unspecified |
| mmu-miR-34a-5p | MIMAT0000542 | Asah2 | 54447 | Microarrays | 37986224 | Unspecified |
| mmu-miR-34a-5p | MIMAT0000542 | Asap1 | 13196 | HITS-CLIP | 23142080\|29167373 | Unspecified |
| mmu-miR-34a-5p | MIMAT0000542 | Asb1 | 65247 | HITS-CLIP | 23142080 | Unspecified |
| mmu-miR-34a-5p | MIMAT0000542 | Asb13 | 142688 | HITS-CLIP | 23142080 | Unspecified |
| mmu-miR-34a-5p | MIMAT0000542 | Asb2 | 65256 | Microarrays//HITS-CLIP | 37986224\|31433990 | Unspecified |
| mmu-miR-34a-5p | MIMAT0000542 | Asb3 | 65257 | HITS-CLIP | 23142080 | Unspecified |
| mmu-miR-34a-5p | MIMAT0000542 | Asb4 | 65255 | HITS-CLIP | 30971389 | Unspecified |
| mmu-miR-34a-5p | MIMAT0000542 | Asb7 | 117589 | HITS-CLIP | 23142080\|29167373 | Unspecified |
| mmu-miR-34a-5p | MIMAT0000542 | Asb8 | 78541 | HITS-CLIP | 29167373 | Unspecified |
| mmu-miR-34a-5p | MIMAT0000542 | Ascc2 | 75452 | HITS-CLIP | 29167373 | Unspecified |
| mmu-miR-34a-5p | MIMAT0000542 | Ascc3 | 77987 | HITS-CLIP | 30971389\|19536157\|29167373\|23142 | Unspecified |
| mmu-miR-34a-5p | MIMAT0000542 | Ascl1 | 17172 | HITS-CLIP | 19536157 | Unspecified |
| mmu-miR-34a-5p | MIMAT0000542 | Asf1a | 66403 | HITS-CLIP | 23142080 | Unspecified |
| mmu-miR-34a-5p | MIMAT0000542 | Asf1b | 66929 | HITS-CLIP | 23142080 | Unspecified |
| mmu-miR-34a-5p | MIMAT0000542 | Ash1l | 192195 | HITS-CLIP | 23142080\|19536157\|29167373 | Unspecified |
| mmu-miR-34a-5p | MIMAT0000542 | Ash2l | 23808 | HITS-CLIP | 23630078\|23142080 | Unspecified |
| mmu-miR-34a-5p | MIMAT0000542 | Asic2 | 11418 | HITS-CLIP | 19536157 | Unspecified |
| mmu-miR-34a-5p | MIMAT0000542 | Asl | 109900 | HITS-CLIP | 23142080\|30971389 | Unspecified |
| mmu-miR-34a-5p | MIMAT0000542 | Asna1 | 56495 | HITS-CLIP | 19536157\|23142080 | Unspecified |
| mmu-miR-34a-5p | MIMAT0000542 | Asns | 27053 | HITS-CLIP | 23630078\|31433990\|30971389\|23142 | Unspecified |

| | | | | | | |
|---|---|---|---|---|---|---|
| mmu-miR-34a-5p | MIMAT0000542 | Asph | 65973 | HITS-CLIP | 19536157\|23630078\|23142080\|29167 | Unspecified |
| mmu-miR-34a-5p | MIMAT0000542 | Aspm | 12316 | HITS-CLIP | 23142080 | Unspecified |
| mmu-miR-34a-5p | MIMAT0000542 | Aspn | 66695 | Microarrays | 37986224 | Unspecified |
| mmu-miR-34a-5p | MIMAT0000542 | Aspscr1 | 68938 | HITS-CLIP | 29167373 | Unspecified |
| mmu-miR-34a-5p | MIMAT0000542 | Ass1 | 11898 | HITS-CLIP | 23142080 | Unspecified |
| mmu-miR-34a-5p | MIMAT0000542 | Asxl1 | 228790 | HITS-CLIP | 23142080\|30971389\|29167373 | Unspecified |
| mmu-miR-34a-5p | MIMAT0000542 | Asxl2 | 75302 | HITS-CLIP | 23142080\|29167373 | Unspecified |
| mmu-miR-34a-5p | MIMAT0000542 | Asxl3 | 211961 | HITS-CLIP | 19536157 | Unspecified |
| mmu-miR-34a-5p | MIMAT0000542 | Atad1 | 67979 | HITS-CLIP | 23142080\|29167373\|26203562 | Unspecified |
| mmu-miR-34a-5p | MIMAT0000542 | Atad2 | 70472 | HITS-CLIP | 37986224\|31433990\|23142080 | Unspecified |
| mmu-miR-34a-5p | MIMAT0000542 | Atad2b | 320817 | HITS-CLIP | 30971389\|19536157\|31433990\|26203 | Unspecified |
| mmu-miR-34a-5p | MIMAT0000542 | Atad3a | 108888 | HITS-CLIP | 30971389 | Unspecified |
| mmu-miR-34a-5p | MIMAT0000542 | Atad5 | 237877 | HITS-CLIP | 23142080\|29167373 | Unspecified |
| mmu-miR-34a-5p | MIMAT0000542 | Atat1 | 73242 | HITS-CLIP | 29167373 | Unspecified |
| mmu-miR-34a-5p | MIMAT0000542 | Ate1 | 11907 | HITS-CLIP | 23142080 | Unspecified |
| mmu-miR-34a-5p | MIMAT0000542 | Atf1 | 11908 | HITS-CLIP | 29167373 | Unspecified |
| mmu-miR-34a-5p | MIMAT0000542 | Atf2 | 11909 | HITS-CLIP | 37986224\|23142080\|29167373 | Unspecified |
| mmu-miR-34a-5p | MIMAT0000542 | Atf3 | 11910 | HITS-CLIP | 23142080\|29167373 | Unspecified |
| mmu-miR-34a-5p | MIMAT0000542 | Atf4 | 11911 | HITS-CLIP | 31433990\|23142080\|29167373 | Unspecified |
| mmu-miR-34a-5p | MIMAT0000542 | Atf6 | 226641 | HITS-CLIP | 37986224\|23142080\|29167373 | Unspecified |
| mmu-miR-34a-5p | MIMAT0000542 | Atf6b | 12915 | HITS-CLIP | 29167373 | Unspecified |
| mmu-miR-34a-5p | MIMAT0000542 | Atf7 | 223922 | HITS-CLIP | 23142080\|19536157\|29167373 | Unspecified |
| mmu-miR-34a-5p | MIMAT0000542 | Atf7ip | 54343 | HITS-CLIP | 19536157\|23142080\|29167373 | Unspecified |
| mmu-miR-34a-5p | MIMAT0000542 | Atg12 | 67526 | HITS-CLIP | 23142080 | Unspecified |
| mmu-miR-34a-5p | MIMAT0000542 | Atg13 | 51897 | HITS-CLIP | 31433990\|23142080 | Unspecified |
| mmu-miR-34a-5p | MIMAT0000542 | Atg14 | 100504663 | HITS-CLIP | 23142080 | Unspecified |
| mmu-miR-34a-5p | MIMAT0000542 | Atg16l1 | 77040 | HITS-CLIP | 23142080\|29167373 | Unspecified |
| mmu-miR-34a-5p | MIMAT0000542 | Atg16l2 | 73683 | HITS-CLIP | 23142080 | Unspecified |
| mmu-miR-34a-5p | MIMAT0000542 | Atg2a | 329015 | HITS-CLIP | 23142080 | Unspecified |
| mmu-miR-34a-5p | MIMAT0000542 | Atg2b | 76559 | HITS-CLIP | 19536157\|23142080\|29167373 | Unspecified |
| mmu-miR-34a-5p | MIMAT0000542 | Atg3 | 67841 | HITS-CLIP | 23142080 | Unspecified |
| mmu-miR-34a-5p | MIMAT0000542 | Atg4b | 66615 | HITS-CLIP | 23142080 | Unspecified |
| mmu-miR-34a-5p | MIMAT0000542 | Atg4d | 235040 | HITS-CLIP | 29167373 | Unspecified |
| mmu-miR-34a-5p | MIMAT0000542 | Atg5 | 11793 | HITS-CLIP | 29167373 | Unspecified |
| mmu-miR-34a-5p | MIMAT0000542 | Atg7 | 74244 | HITS-CLIP | 23142080\|30971389 | Unspecified |
| mmu-miR-34a-5p | MIMAT0000542 | Atg9a | 245860 | HITS-CLIP//Immunoblot//Luciferase report | 23142080\|19536157\|26203562\|24973 | Unspecified |
| mmu-miR-34a-5p | MIMAT0000542 | Atic | 108147 | HITS-CLIP | 23142080 | Unspecified |
| mmu-miR-34a-5p | MIMAT0000542 | Atl1 | 73991 | HITS-CLIP | 19536157 | Unspecified |
| mmu-miR-34a-5p | MIMAT0000542 | Atl2 | 56298 | HITS-CLIP | 19536157\|31433990\|23142080\|29167 | Unspecified |

| | | | | | | |
|---|---|---|---|---|---|---|
| mmu-miR-34a-5p | MIMAT0000542 | Atl3 | 109168 | HITS-CLIP | 23142080\|30971389\|29167373 | Unspecified |
| mmu-miR-34a-5p | MIMAT0000542 | Atm | 11920 | HITS-CLIP | 23142080 | Unspecified |
| mmu-miR-34a-5p | MIMAT0000542 | Atmin | 234776 | HITS-CLIP | 26203562 | Unspecified |
| mmu-miR-34a-5p | MIMAT0000542 | Atn1 | 13498 | HITS-CLIP | 23630078\|23142080\|29167373 | Unspecified |
| mmu-miR-34a-5p | MIMAT0000542 | Atoh8 | 71093 | HITS-CLIP | 23142080 | Unspecified |
| mmu-miR-34a-5p | MIMAT0000542 | Atox1 | 11927 | HITS-CLIP | 29167373 | Unspecified |
| mmu-miR-34a-5p | MIMAT0000542 | Atp10a | 11982 | Microarrays//HITS-CLIP | 37986224\|23027903\|23142080 | Unspecified |
| mmu-miR-34a-5p | MIMAT0000542 | Atp11a | 50770 | HITS-CLIP | 23142080\|29167373 | Unspecified |
| mmu-miR-34a-5p | MIMAT0000542 | Atp11b | 76295 | HITS-CLIP | 37986224\|19536157\|23142080 | Unspecified |
| mmu-miR-34a-5p | MIMAT0000542 | Atp11c | 320940 | HITS-CLIP | 23142080\|29167373 | Unspecified |
| mmu-miR-34a-5p | MIMAT0000542 | Atp13a3 | 224088 | HITS-CLIP | 19536157\|23142080 | Unspecified |
| mmu-miR-34a-5p | MIMAT0000542 | Atp1a1 | 11928 | HITS-CLIP | 19536157\|23142080\|29167373 | Unspecified |
| mmu-miR-34a-5p | MIMAT0000542 | Atp1a2 | 98660 | Microarrays//HITS-CLIP | 37986224\|19536157 | Unspecified |
| mmu-miR-34a-5p | MIMAT0000542 | Atp1a3 | 232975 | HITS-CLIP | 19536157 | Unspecified |
| mmu-miR-34a-5p | MIMAT0000542 | Atp1b1 | 11931 | HITS-CLIP | 19536157 | Unspecified |
| mmu-miR-34a-5p | MIMAT0000542 | Atp1b2 | 11932 | HITS-CLIP | 19536157 | Unspecified |
| mmu-miR-34a-5p | MIMAT0000542 | Atp1b3 | 11933 | Microarrays//HITS-CLIP | 37986224\|23027903\|23630078\|23142 | Unspecified |
| mmu-miR-34a-5p | MIMAT0000542 | Atp2a1 | 11937 | HITS-CLIP | 29167373 | Unspecified |
| mmu-miR-34a-5p | MIMAT0000542 | Atp2a2 | 11938 | HITS-CLIP | 37986224\|19536157\|31433990\|29084 | Unspecified |
| mmu-miR-34a-5p | MIMAT0000542 | Atp2a3 | 53313 | HITS-CLIP | 23142080 | Unspecified |
| mmu-miR-34a-5p | MIMAT0000542 | Atp2b1 | 67972 | HITS-CLIP | 23142080\|19536157\|29167373 | Unspecified |
| mmu-miR-34a-5p | MIMAT0000542 | Atp2b2 | 11941 | HITS-CLIP | 19536157 | Unspecified |
| mmu-miR-34a-5p | MIMAT0000542 | Atp2b3 | 320707 | HITS-CLIP | 19536157 | Unspecified |
| mmu-miR-34a-5p | MIMAT0000542 | Atp2b4 | 381290 | HITS-CLIP | 23142080\|19536157 | Unspecified |
| mmu-miR-34a-5p | MIMAT0000542 | Atp2c1 | 235574 | HITS-CLIP | 19536157\|23142080\|29167373 | Unspecified |
| mmu-miR-34a-5p | MIMAT0000542 | Atp5a1 | 11946 | HITS-CLIP | 19536157\|23142080\|29167373 | Unspecified |
| mmu-miR-34a-5p | MIMAT0000542 | Atp5b | 11947 | HITS-CLIP | 23142080\|19536157\|29167373 | Unspecified |
| mmu-miR-34a-5p | MIMAT0000542 | Atp5c1 | 11949 | HITS-CLIP | 23142080 | Unspecified |
| mmu-miR-34a-5p | MIMAT0000542 | Atp5d | 66043 | HITS-CLIP | 23142080 | Unspecified |
| mmu-miR-34a-5p | MIMAT0000542 | Atp5e | 67126 | HITS-CLIP | 23142080 | Unspecified |
| mmu-miR-34a-5p | MIMAT0000542 | Atp5f1 | 515 | HITS-CLIP | 23142080 | Unspecified |
| mmu-miR-34a-5p | MIMAT0000542 | Atp5g3 | 228033 | HITS-CLIP | 37986224\|19536157\|23142080 | Unspecified |
| mmu-miR-34a-5p | MIMAT0000542 | Atp5j2 | 57423 | HITS-CLIP | 23142080 | Unspecified |
| mmu-miR-34a-5p | MIMAT0000542 | Atp5mpl | 9556 | HITS-CLIP | 23142080 | Unspecified |
| mmu-miR-34a-5p | MIMAT0000542 | Atp5o | 28080 | HITS-CLIP | 19536157\|23142080 | Unspecified |
| mmu-miR-34a-5p | MIMAT0000542 | Atp5pb | 515 | HITS-CLIP | 31433990\|23630078 | Unspecified |
| mmu-miR-34a-5p | MIMAT0000542 | Atp5s | 27109 | HITS-CLIP | 23142080 | Unspecified |
| mmu-miR-34a-5p | MIMAT0000542 | Atp6ap1 | 54411 | HITS-CLIP | 23142080\|29167373 | Unspecified |
| mmu-miR-34a-5p | MIMAT0000542 | Atp6ap2 | 70495 | HITS-CLIP | 29167373 | Unspecified |

| | | | | | | |
|---|---|---|---|---|---|---|
| mmu-miR-34a-5p | MIMAT0000542 | Atp6v0a1 | 11975 | HITS-CLIP | 29167373\|23142080 | Unspecified |
| mmu-miR-34a-5p | MIMAT0000542 | Atp6v0a2 | 21871 | HITS-CLIP | 23142080 | Unspecified |
| mmu-miR-34a-5p | MIMAT0000542 | Atp6v0b | 114143 | HITS-CLIP | 30971389\|23142080\|29167373 | Unspecified |
| mmu-miR-34a-5p | MIMAT0000542 | Atp6v0d1 | 11972 | HITS-CLIP | 23142080\|26203562\|29167373 | Unspecified |
| mmu-miR-34a-5p | MIMAT0000542 | Atp6v0d2 | 242341 | HITS-CLIP | 23142080 | Unspecified |
| mmu-miR-34a-5p | MIMAT0000542 | Atp6v0e | 11974 | HITS-CLIP | 23142080\|29167373 | Unspecified |
| mmu-miR-34a-5p | MIMAT0000542 | Atp6v0e2 | 76252 | HITS-CLIP | 19536157 | Unspecified |
| mmu-miR-34a-5p | MIMAT0000542 | Atp6v1a | 11964 | HITS-CLIP | 19536157\|23142080\|29167373 | Unspecified |
| mmu-miR-34a-5p | MIMAT0000542 | Atp6v1b2 | 11966 | HITS-CLIP | 29167373 | Unspecified |
| mmu-miR-34a-5p | MIMAT0000542 | Atp6v1c1 | 66335 | HITS-CLIP | 23142080\|29167373 | Unspecified |
| mmu-miR-34a-5p | MIMAT0000542 | Atp6v1d | 73834 | HITS-CLIP | 23142080\|23630078 | Unspecified |
| mmu-miR-34a-5p | MIMAT0000542 | Atp6v1e1 | 11973 | HITS-CLIP | 23142080 | Unspecified |
| mmu-miR-34a-5p | MIMAT0000542 | Atp6v1f | 66144 | HITS-CLIP | 23142080\|23630078\|31433990 | Unspecified |
| mmu-miR-34a-5p | MIMAT0000542 | Atp6v1g1 | 66290 | Microarrays | 37986224 | Unspecified |
| mmu-miR-34a-5p | MIMAT0000542 | Atp6v1h | 108664 | HITS-CLIP | 23142080\|29167373 | Unspecified |
| mmu-miR-34a-5p | MIMAT0000542 | Atp7a | 11977 | HITS-CLIP | 23142080\|29167373 | Unspecified |
| mmu-miR-34a-5p | MIMAT0000542 | Atp7b | 11979 | HITS-CLIP | 23142080 | Unspecified |
| mmu-miR-34a-5p | MIMAT0000542 | Atp8a1 | 11980 | HITS-CLIP | 19536157\|29167373\|23142080 | Unspecified |
| mmu-miR-34a-5p | MIMAT0000542 | Atp8a2 | 50769 | HITS-CLIP | 23142080 | Unspecified |
| mmu-miR-34a-5p | MIMAT0000542 | Atp8b1 | 54670 | HITS-CLIP | 26203562 | Unspecified |
| mmu-miR-34a-5p | MIMAT0000542 | Atp8b2 | 54667 | HITS-CLIP | 19536157\|23142080\|29167373 | Unspecified |
| mmu-miR-34a-5p | MIMAT0000542 | Atp8b4 | 241633 | HITS-CLIP | 29167373\|23142080 | Unspecified |
| mmu-miR-34a-5p | MIMAT0000542 | Atp9a | 11981 | Microarrays | 37986224 | Unspecified |
| mmu-miR-34a-5p | MIMAT0000542 | Atp9b | 50771 | HITS-CLIP | 19536157\|23142080\|29167373 | Unspecified |
| mmu-miR-34a-5p | MIMAT0000542 | Atpaf2 | 246782 | HITS-CLIP | 23142080 | Unspecified |
| mmu-miR-34a-5p | MIMAT0000542 | Atr | 245000 | HITS-CLIP | 23142080\|29167373 | Unspecified |
| mmu-miR-34a-5p | MIMAT0000542 | Atrn | 11990 | HITS-CLIP | 19536157\|23142080\|29167373 | Unspecified |
| mmu-miR-34a-5p | MIMAT0000542 | Atrnl1 | 226255 | HITS-CLIP | 23142080 | Unspecified |
| mmu-miR-34a-5p | MIMAT0000542 | Atrx | 22589 | HITS-CLIP | 23142080\|19536157\|29167373 | Unspecified |
| mmu-miR-34a-5p | MIMAT0000542 | Atxn1 | 20238 | HITS-CLIP | 37986224\|31433990\|23142080\|29167 | Unspecified |
| mmu-miR-34a-5p | MIMAT0000542 | Atxn10 | 54138 | HITS-CLIP | 29167373 | Unspecified |
| mmu-miR-34a-5p | MIMAT0000542 | Atxn1l | 52335 | Microarrays//HITS-CLIP | 37986224\|23027903\|19536157\|3143 | Unspecified |
| mmu-miR-34a-5p | MIMAT0000542 | Atxn2 | 20239 | HITS-CLIP | 23142080 | Unspecified |
| mmu-miR-34a-5p | MIMAT0000542 | Atxn2l | 233871 | HITS-CLIP | 23142080\|29167373 | Unspecified |
| mmu-miR-34a-5p | MIMAT0000542 | Atxn3 | 110616 | HITS-CLIP | 23142080 | Unspecified |
| mmu-miR-34a-5p | MIMAT0000542 | Atxn7 | 246103 | HITS-CLIP | 23142080 | Unspecified |
| mmu-miR-34a-5p | MIMAT0000542 | Atxn7l1 | 380753 | HITS-CLIP | 23142080\|29167373 | Unspecified |
| mmu-miR-34a-5p | MIMAT0000542 | Atxn7l3 | 217218 | HITS-CLIP | 29167373 | Unspecified |
| mmu-miR-34a-5p | MIMAT0000542 | Atxn7l3b | 382423 | HITS-CLIP | 23142080\|29167373 | Unspecified |

| | | | | | | |
|---|---|---|---|---|---|---|
| mmu-miR-34a-5p | MIMAT0000542 | Auh | 11992 | HITS-CLIP | 23142080\|29167373 | Unspecified |
| mmu-miR-34a-5p | MIMAT0000542 | Aunip | 69885 | HITS-CLIP | 31433990\|23142080\|29167373 | Unspecified |
| mmu-miR-34a-5p | MIMAT0000542 | Aup1 | 11993 | HITS-CLIP | 19536157\|23142080 | Unspecified |
| mmu-miR-34a-5p | MIMAT0000542 | Aurka | 20878 | HITS-CLIP | 23142080 | Unspecified |
| mmu-miR-34a-5p | MIMAT0000542 | Aurkb | 20877 | HITS-CLIP | 23142080\|29167373 | Unspecified |
| mmu-miR-34a-5p | MIMAT0000542 | Avl9 | 78937 | HITS-CLIP | 23142080\|29167373 | Unspecified |
| mmu-miR-34a-5p | MIMAT0000542 | Axin1 | 12005 | HITS-CLIP | 19536157\|23142080 | Unspecified |
| mmu-miR-34a-5p | MIMAT0000542 | Axl | 26362 | Microarrays//HITS-CLIP | 37986224\|23027903\|19536157\|23142 | Unspecified |
| mmu-miR-34a-5p | MIMAT0000542 | Azi2 | 27215 | HITS-CLIP | 29167373\|23142080 | Unspecified |
| mmu-miR-34a-5p | MIMAT0000542 | Azin1 | 54375 | HITS-CLIP | 23630078\|26203562\|29167373\|23142 | Unspecified |
| mmu-miR-34a-5p | MIMAT0000542 | B230219D22Rik | 78521 | HITS-CLIP | 19536157\|29167373\|23142080 | Unspecified |
| mmu-miR-34a-5p | MIMAT0000542 | B2m | 12010 | HITS-CLIP | 31433990\|23142080\|29167373 | Unspecified |
| mmu-miR-34a-5p | MIMAT0000542 | B3galnt1 | 26879 | HITS-CLIP | 19536157 | Unspecified |
| mmu-miR-34a-5p | MIMAT0000542 | B3galnt2 | 97884 | HITS-CLIP | 23142080 | Unspecified |
| mmu-miR-34a-5p | MIMAT0000542 | B3galt2 | 26878 | HITS-CLIP | 19536157 | Unspecified |
| mmu-miR-34a-5p | MIMAT0000542 | B3galt5 | 93961 | Microarrays | 37986224 | Unspecified |
| mmu-miR-34a-5p | MIMAT0000542 | B3gat1 | 76898 | HITS-CLIP | 19536157 | Unspecified |
| mmu-miR-34a-5p | MIMAT0000542 | B3gnt2 | 53625 | HITS-CLIP | 23142080 | Unspecified |
| mmu-miR-34a-5p | MIMAT0000542 | B3gnt3 | 72297 | HITS-CLIP | 23142080 | Unspecified |
| mmu-miR-34a-5p | MIMAT0000542 | B3gnt5 | 108105 | Microarrays | 37986224\|23027903 | Unspecified |
| mmu-miR-34a-5p | MIMAT0000542 | B3gnt7 | 227327 | HITS-CLIP | 23142080 | Unspecified |
| mmu-miR-34a-5p | MIMAT0000542 | B4galnt1 | 14421 | HITS-CLIP | 37986224\|23142080\|29167373 | Unspecified |
| mmu-miR-34a-5p | MIMAT0000542 | B4galt1 | 14595 | HITS-CLIP | 31433990\|23142080\|29167373 | Unspecified |
| mmu-miR-34a-5p | MIMAT0000542 | B4galt3 | 57370 | HITS-CLIP | 29167373 | Unspecified |
| mmu-miR-34a-5p | MIMAT0000542 | B4galt5 | 56336 | HITS-CLIP | 23142080\|29167373 | Unspecified |
| mmu-miR-34a-5p | MIMAT0000542 | B4galt6 | 56386 | HITS-CLIP | 19536157\|29167373 | Unspecified |
| mmu-miR-34a-5p | MIMAT0000542 | B4galt7 | 218271 | HITS-CLIP | 23142080\|29167373 | Unspecified |
| mmu-miR-34a-5p | MIMAT0000542 | B630019K06Rik | 102941 | HITS-CLIP | 19536157 | Unspecified |
| mmu-miR-34a-5p | MIMAT0000542 | B9d1 | 27078 | HITS-CLIP | 23142080 | Unspecified |
| mmu-miR-34a-5p | MIMAT0000542 | BC004004 | 80748 | HITS-CLIP | 29167373 | Unspecified |
| mmu-miR-34a-5p | MIMAT0000542 | BC005537 | 79555 | HITS-CLIP | 23142080\|19536157\|29167373 | Unspecified |
| mmu-miR-34a-5p | MIMAT0000542 | BC005561 | 100042165 | HITS-CLIP | 23142080 | Unspecified |
| mmu-miR-34a-5p | MIMAT0000542 | BC024978 | 414069 | HITS-CLIP | 29167373 | Unspecified |
| mmu-miR-34a-5p | MIMAT0000542 | BC037034 | 231807 | HITS-CLIP | 29167373\|23142080 | Unspecified |
| mmu-miR-34a-5p | MIMAT0000542 | BC052040 | 399568 | HITS-CLIP | 23142080 | Unspecified |
| mmu-miR-34a-5p | MIMAT0000542 | BC055324 | 381306 | HITS-CLIP | 23142080 | Unspecified |
| mmu-miR-34a-5p | MIMAT0000542 | BC067074 | 408066 | Microarrays | 37986224 | Unspecified |
| mmu-miR-34a-5p | MIMAT0000542 | Babam1 | 68251 | HITS-CLIP | 23142080 | Unspecified |
| mmu-miR-34a-5p | MIMAT0000542 | Babam2 | 9577 | HITS-CLIP | 23142080 | Unspecified |

| | | | | | | |
|---|---|---|---|---|---|---|
| mmu-miR-34a-5p | MIMAT0000542 | Bace1 | 23821 | HITS-CLIP | 19536157 | Unspecified |
| mmu-miR-34a-5p | MIMAT0000542 | Bach1 | 12013 | HITS-CLIP | 23142080\|23630078\|29167373 | Unspecified |
| mmu-miR-34a-5p | MIMAT0000542 | Bach2 | 12014 | HITS-CLIP | 37986224\|23630078\|31433990\|23142 | Unspecified |
| mmu-miR-34a-5p | MIMAT0000542 | Bag1 | 12017 | HITS-CLIP | 23142080 | Unspecified |
| mmu-miR-34a-5p | MIMAT0000542 | Bag3 | 29810 | HITS-CLIP | 29167373\|23142080 | Unspecified |
| mmu-miR-34a-5p | MIMAT0000542 | Bag4 | 67384 | HITS-CLIP | 29167373\|23142080 | Unspecified |
| mmu-miR-34a-5p | MIMAT0000542 | Bag5 | 70369 | HITS-CLIP | 23142080 | Unspecified |
| mmu-miR-34a-5p | MIMAT0000542 | Bag6 | 224727 | HITS-CLIP | 23142080\|29167373 | Unspecified |
| mmu-miR-34a-5p | MIMAT0000542 | Bahd1 | 228536 | HITS-CLIP | 23142080 | Unspecified |
| mmu-miR-34a-5p | MIMAT0000542 | Baiap2 | 108100 | HITS-CLIP | 19536157 | Unspecified |
| mmu-miR-34a-5p | MIMAT0000542 | Bank1 | 242248 | HITS-CLIP | 23142080 | Unspecified |
| mmu-miR-34a-5p | MIMAT0000542 | Banp | 53325 | HITS-CLIP | 23142080 | Unspecified |
| mmu-miR-34a-5p | MIMAT0000542 | Bard1 | 12021 | Microarrays//HITS-CLIP | 37986224\|23027903\|23142080\|31433 | Unspecified |
| mmu-miR-34a-5p | MIMAT0000542 | Basp1 | 70350 | HITS-CLIP | 29167373 | Unspecified |
| mmu-miR-34a-5p | MIMAT0000542 | Bax | 12028 | HITS-CLIP | 23142080 | Unspecified |
| mmu-miR-34a-5p | MIMAT0000542 | Baz1a | 217578 | HITS-CLIP | 37986224\|29167373\|23142080 | Unspecified |
| mmu-miR-34a-5p | MIMAT0000542 | Baz1b | 22385 | HITS-CLIP | 31433990\|23142080\|29167373 | Unspecified |
| mmu-miR-34a-5p | MIMAT0000542 | Baz2a | 116848 | HITS-CLIP | 23142080\|19536157\|29167373 | Unspecified |
| mmu-miR-34a-5p | MIMAT0000542 | Baz2b | 407823 | HITS-CLIP | 23142080\|29167373 | Unspecified |
| mmu-miR-34a-5p | MIMAT0000542 | Bbc3 | 170770 | HITS-CLIP | 23142080 | Unspecified |
| mmu-miR-34a-5p | MIMAT0000542 | Bbip1 | 100503572 | HITS-CLIP | 23142080\|29167373 | Unspecified |
| mmu-miR-34a-5p | MIMAT0000542 | Bbs10 | 71769 | HITS-CLIP | 29167373 | Unspecified |
| mmu-miR-34a-5p | MIMAT0000542 | Bbs12 | 241950 | HITS-CLIP | 23142080 | Unspecified |
| mmu-miR-34a-5p | MIMAT0000542 | Bbs2 | 67378 | HITS-CLIP | 30971389 | Unspecified |
| mmu-miR-34a-5p | MIMAT0000542 | Bbs5 | 72569 | Microarrays//HITS-CLIP | 37986224\|23142080 | Unspecified |
| mmu-miR-34a-5p | MIMAT0000542 | Bbs7 | 71492 | Microarrays | 37986224 | Unspecified |
| mmu-miR-34a-5p | MIMAT0000542 | Bbs9 | 319845 | HITS-CLIP | 23142080\|29167373 | Unspecified |
| mmu-miR-34a-5p | MIMAT0000542 | Bbx | 70508 | HITS-CLIP | 23142080\|19536157\|31433990\|29167 | Unspecified |
| mmu-miR-34a-5p | MIMAT0000542 | Bcan | 12032 | HITS-CLIP | 19536157 | Unspecified |
| mmu-miR-34a-5p | MIMAT0000542 | Bcap29 | 12033 | HITS-CLIP | 23142080\|29167373 | Unspecified |
| mmu-miR-34a-5p | MIMAT0000542 | Bcap31 | 27061 | HITS-CLIP | 23142080\|19536157\|29167373 | Unspecified |
| mmu-miR-34a-5p | MIMAT0000542 | Bcar3 | 29815 | HITS-CLIP | 29167373 | Unspecified |
| mmu-miR-34a-5p | MIMAT0000542 | Bcas2 | 68183 | HITS-CLIP | 23142080 | Unspecified |
| mmu-miR-34a-5p | MIMAT0000542 | Bcas3 | 192197 | HITS-CLIP | 23142080\|29167373 | Unspecified |
| mmu-miR-34a-5p | MIMAT0000542 | Bcat1 | 12035 | Microarrays//HITS-CLIP | 37986224\|23027903\|23142080 | Unspecified |
| mmu-miR-34a-5p | MIMAT0000542 | Bcat2 | 12036 | HITS-CLIP | 30971389 | Unspecified |
| mmu-miR-34a-5p | MIMAT0000542 | Bckdha | 12039 | HITS-CLIP | 23142080 | Unspecified |
| mmu-miR-34a-5p | MIMAT0000542 | Bckdk | 12041 | HITS-CLIP | 23142080 | Unspecified |
| mmu-miR-34a-5p | MIMAT0000542 | Bcl10 | 12042 | HITS-CLIP | 29167373 | Unspecified |

| | | | | | | |
|---|---|---|---|---|---|---|
| mmu-miR-34a-5p | MIMAT0000542 | Bcl11a | 14025 | HITS-CLIP | 23142080 | Unspecified |
| mmu-miR-34a-5p | MIMAT0000542 | Bcl11b | 58208 | HITS-CLIP | 19536157\|31433990\|23142080 | Unspecified |
| mmu-miR-34a-5p | MIMAT0000542 | Bcl2 | 12043 | qRT-PCR//Western blot//HITS-CLIP//Other | 19683563\|23142080\|31433990\|29167 | Unspecified |
| mmu-miR-34a-5p | MIMAT0000542 | Bcl2l1 | 12048 | HITS-CLIP | 23142080\|29167373 | Unspecified |
| mmu-miR-34a-5p | MIMAT0000542 | Bcl2l11 | 12125 | HITS-CLIP | 23142080\|29167373 | Unspecified |
| mmu-miR-34a-5p | MIMAT0000542 | Bcl2l13 | 94044 | Microarrays//HITS-CLIP | 37986224\|23027903\|23142080 | Unspecified |
| mmu-miR-34a-5p | MIMAT0000542 | Bcl2l15 | 229672 | HITS-CLIP | 30971389 | Unspecified |
| mmu-miR-34a-5p | MIMAT0000542 | Bcl2l2 | 12050 | HITS-CLIP | 23142080 | Unspecified |
| mmu-miR-34a-5p | MIMAT0000542 | Bcl3 | 12051 | HITS-CLIP | 31433990\|23142080\|26203562 | Unspecified |
| mmu-miR-34a-5p | MIMAT0000542 | Bcl6 | 12053 | Luciferase assay//Northern blot//qRT-PCR | 23047694\|23142080\|29167373 | Unspecified |
| mmu-miR-34a-5p | MIMAT0000542 | Bcl7a | 77045 | HITS-CLIP | 23142080 | Unspecified |
| mmu-miR-34a-5p | MIMAT0000542 | Bcl7b | 12054 | HITS-CLIP | 29167373 | Unspecified |
| mmu-miR-34a-5p | MIMAT0000542 | Bcl9 | 77578 | HITS-CLIP | 23142080 | Unspecified |
| mmu-miR-34a-5p | MIMAT0000542 | Bcl9l | 80288 | HITS-CLIP | 23142080\|19536157 | Unspecified |
| mmu-miR-34a-5p | MIMAT0000542 | Bclaf1 | 72567 | HITS-CLIP | 31433990\|23142080\|29167373 | Unspecified |
| mmu-miR-34a-5p | MIMAT0000542 | Bcor | 71458 | HITS-CLIP | 23142080\|29167373 | Unspecified |
| mmu-miR-34a-5p | MIMAT0000542 | Bcorl1 | 320376 | HITS-CLIP | 23142080 | Unspecified |
| mmu-miR-34a-5p | MIMAT0000542 | Bcr | 110279 | HITS-CLIP | 23142080 | Unspecified |
| mmu-miR-34a-5p | MIMAT0000542 | Bdp1 | 544971 | HITS-CLIP | 19536157\|29167373\|23142080 | Unspecified |
| mmu-miR-34a-5p | MIMAT0000542 | Becn1 | 56208 | HITS-CLIP | 29167373 | Unspecified |
| mmu-miR-34a-5p | MIMAT0000542 | Bend3 | 331623 | HITS-CLIP | 29167373 | Unspecified |
| mmu-miR-34a-5p | MIMAT0000542 | Bend4 | 666938 | HITS-CLIP | 29167373 | Unspecified |
| mmu-miR-34a-5p | MIMAT0000542 | Best1 | 24115 | Microarrays | 37986224 | Unspecified |
| mmu-miR-34a-5p | MIMAT0000542 | Bet1 | 12068 | HITS-CLIP | 26203562\|23142080 | Unspecified |
| mmu-miR-34a-5p | MIMAT0000542 | Bet1l | 54399 | HITS-CLIP | 23142080\|26203562 | Unspecified |
| mmu-miR-34a-5p | MIMAT0000542 | Bfar | 67118 | HITS-CLIP | 23142080\|29167373 | Unspecified |
| mmu-miR-34a-5p | MIMAT0000542 | Bhlhb9 | 70237 | HITS-CLIP | 23142080 | Unspecified |
| mmu-miR-34a-5p | MIMAT0000542 | Bhlhe40 | 20893 | HITS-CLIP | 31433990\|23630078\|23142080 | Unspecified |
| mmu-miR-34a-5p | MIMAT0000542 | Bhlhe41 | 79362 | HITS-CLIP | 29167373\|23142080 | Unspecified |
| mmu-miR-34a-5p | MIMAT0000542 | Bicd1 | 12121 | HITS-CLIP | 19536157 | Unspecified |
| mmu-miR-34a-5p | MIMAT0000542 | Bicd2 | 76895 | HITS-CLIP | 19536157\|29167373\|23142080 | Unspecified |
| mmu-miR-34a-5p | MIMAT0000542 | Bicra | 29998 | HITS-CLIP | 23142080\|29167373 | Unspecified |
| mmu-miR-34a-5p | MIMAT0000542 | Bicral | 23506 | HITS-CLIP | 23142080\|29167373 | Unspecified |
| mmu-miR-34a-5p | MIMAT0000542 | Bid | 12122 | HITS-CLIP | 29167373 | Unspecified |
| mmu-miR-34a-5p | MIMAT0000542 | Bin1 | 30948 | HITS-CLIP | 23142080 | Unspecified |
| mmu-miR-34a-5p | MIMAT0000542 | Bin2 | 668218 | HITS-CLIP | 23142080 | Unspecified |
| mmu-miR-34a-5p | MIMAT0000542 | Bin3 | 57784 | HITS-CLIP | 23142080 | Unspecified |
| mmu-miR-34a-5p | MIMAT0000542 | Birc2 | 11797 | HITS-CLIP | 23142080 | Unspecified |
| mmu-miR-34a-5p | MIMAT0000542 | Birc3 | 11796 | HITS-CLIP | 23142080\|29167373 | Unspecified |

| | | | | | | |
|---|---|---|---|---|---|---|
| mmu-miR-34a-5p | MIMAT0000542 | Birc5 | 11799 | HITS-CLIP | 23142080 | Unspecified |
| mmu-miR-34a-5p | MIMAT0000542 | Birc6 | 12211 | HITS-CLIP | 37986224\|23142080\|19536157\|29167 | Unspecified |
| mmu-miR-34a-5p | MIMAT0000542 | Blcap | 53619 | HITS-CLIP | 23142080 | Unspecified |
| mmu-miR-34a-5p | MIMAT0000542 | Blm | 12144 | HITS-CLIP | 23142080 | Unspecified |
| mmu-miR-34a-5p | MIMAT0000542 | Blmh | 104184 | HITS-CLIP | 23142080\|31433990\|29167373 | Unspecified |
| mmu-miR-34a-5p | MIMAT0000542 | Blnk | 17060 | HITS-CLIP | 29167373 | Unspecified |
| mmu-miR-34a-5p | MIMAT0000542 | Bloc1s3 | 232946 | HITS-CLIP | 21258322\|23142080 | Unspecified |
| mmu-miR-34a-5p | MIMAT0000542 | Bloc1s4 | 117197 | HITS-CLIP | 26203562 | Unspecified |
| mmu-miR-34a-5p | MIMAT0000542 | Bltp2 | 9703 | HITS-CLIP | 30971389\|31433990 | Unspecified |
| mmu-miR-34a-5p | MIMAT0000542 | Blvra | 109778 | HITS-CLIP | 23142080 | Unspecified |
| mmu-miR-34a-5p | MIMAT0000542 | Blzf1 | 66352 | HITS-CLIP | 37986224\|23142080\|29167373 | Unspecified |
| mmu-miR-34a-5p | MIMAT0000542 | Bmi1 | 12151 | HITS-CLIP | 23142080 | Unspecified |
| mmu-miR-34a-5p | MIMAT0000542 | Bmp2k | 140780 | HITS-CLIP | 31433990\|23142080\|29167373 | Unspecified |
| mmu-miR-34a-5p | MIMAT0000542 | Bmpr1a | 12166 | HITS-CLIP | 19536157\|23142080 | Unspecified |
| mmu-miR-34a-5p | MIMAT0000542 | Bmpr2 | 12168 | HITS-CLIP | 19536157\|29167373 | Unspecified |
| mmu-miR-34a-5p | MIMAT0000542 | Bms1 | 213895 | HITS-CLIP | 23142080 | Unspecified |
| mmu-miR-34a-5p | MIMAT0000542 | Bmt2 | 101148 | HITS-CLIP | 23142080\|29167373 | Unspecified |
| mmu-miR-34a-5p | MIMAT0000542 | Bmyc | 107771 | HITS-CLIP | 23142080 | Unspecified |
| mmu-miR-34a-5p | MIMAT0000542 | Bnip3 | 12176 | Microarrays//HITS-CLIP | 37986224\|23142080\|29167373 | Unspecified |
| mmu-miR-34a-5p | MIMAT0000542 | Bnip3l | 12177 | HITS-CLIP | 23142080 | Unspecified |
| mmu-miR-34a-5p | MIMAT0000542 | Bod1 | 69556 | HITS-CLIP | 23142080 | Unspecified |
| mmu-miR-34a-5p | MIMAT0000542 | Bod1l | 665775 | HITS-CLIP | 23142080\|29167373 | Unspecified |
| mmu-miR-34a-5p | MIMAT0000542 | Bora | 77744 | Microarrays//HITS-CLIP | 37986224\|23027903\|23142080 | Unspecified |
| mmu-miR-34a-5p | MIMAT0000542 | Borcs8 | 729991 | HITS-CLIP | 23142080 | Unspecified |
| mmu-miR-34a-5p | MIMAT0000542 | Bpgm | 12183 | HITS-CLIP | 23142080 | Unspecified |
| mmu-miR-34a-5p | MIMAT0000542 | Bphl | 68021 | Microarrays | 37986224 | Unspecified |
| mmu-miR-34a-5p | MIMAT0000542 | Bpnt2 | 54928 | HITS-CLIP | 23630078\|30971389\|26203562 | Unspecified |
| mmu-miR-34a-5p | MIMAT0000542 | Bptf | 207165 | HITS-CLIP | 19536157\|23142080\|29167373 | Unspecified |
| mmu-miR-34a-5p | MIMAT0000542 | Braf | 109880 | HITS-CLIP | 23142080 | Unspecified |
| mmu-miR-34a-5p | MIMAT0000542 | Brap | 72399 | HITS-CLIP | 23142080 | Unspecified |
| mmu-miR-34a-5p | MIMAT0000542 | Brat1 | 231841 | HITS-CLIP | 23142080 | Unspecified |
| mmu-miR-34a-5p | MIMAT0000542 | Brca1 | 12189 | HITS-CLIP//Next Generation Sequencing (N | 23142080\|26403328 | Unspecified |
| mmu-miR-34a-5p | MIMAT0000542 | Brca2 | 12190 | HITS-CLIP | 23142080 | Unspecified |
| mmu-miR-34a-5p | MIMAT0000542 | Brcc3 | 210766 | Microarrays//HITS-CLIP | 37986224\|23027903\|29167373\|23142 | Unspecified |
| mmu-miR-34a-5p | MIMAT0000542 | Brd1 | 223770 | HITS-CLIP | 19536157\|23142080 | Unspecified |
| mmu-miR-34a-5p | MIMAT0000542 | Brd2 | 14312 | HITS-CLIP | 23630078\|23142080\|29167373 | Unspecified |
| mmu-miR-34a-5p | MIMAT0000542 | Brd3 | 67382 | HITS-CLIP | 19536157\|29167373\|23142080 | Unspecified |
| mmu-miR-34a-5p | MIMAT0000542 | Brd4 | 57261 | HITS-CLIP | 23142080\|31433990\|19536157\|29167 | Unspecified |
| mmu-miR-34a-5p | MIMAT0000542 | Brd7 | 26992 | HITS-CLIP | 23142080 | Unspecified |

| | | | | | | |
|---|---|---|---|---|---|---|
| mmu-miR-34a-5p | MIMAT0000542 | Brd8 | 78656 | HITS-CLIP | 23142080\|29167373 | Unspecified |
| mmu-miR-34a-5p | MIMAT0000542 | Bri3bp | 76809 | HITS-CLIP | 19536157\|23142080\|29167373 | Unspecified |
| mmu-miR-34a-5p | MIMAT0000542 | Brinp1 | 56710 | HITS-CLIP | 19536157 | Unspecified |
| mmu-miR-34a-5p | MIMAT0000542 | Brip1 | 237911 | HITS-CLIP | 23142080 | Unspecified |
| mmu-miR-34a-5p | MIMAT0000542 | Brix1 | 67832 | HITS-CLIP | 23142080\|29167373 | Unspecified |
| mmu-miR-34a-5p | MIMAT0000542 | Brk1 | 101314 | HITS-CLIP | 23142080 | Unspecified |
| mmu-miR-34a-5p | MIMAT0000542 | Brms1l | 52592 | HITS-CLIP | 19536157 | Unspecified |
| mmu-miR-34a-5p | MIMAT0000542 | Brox | 71678 | HITS-CLIP | 23142080 | Unspecified |
| mmu-miR-34a-5p | MIMAT0000542 | Brpf1 | 78783 | HITS-CLIP | 23630078\|30971389\|29167373 | Unspecified |
| mmu-miR-34a-5p | MIMAT0000542 | Brwd1 | 93871 | HITS-CLIP | 23142080\|29167373\|23630078\|1953€ | Unspecified |
| mmu-miR-34a-5p | MIMAT0000542 | Brwd3 | 382236 | HITS-CLIP | 23142080\|19536157\|29167373 | Unspecified |
| mmu-miR-34a-5p | MIMAT0000542 | Bscl2 | 14705 | HITS-CLIP | 23142080\|19536157 | Unspecified |
| mmu-miR-34a-5p | MIMAT0000542 | Bsdc1 | 100383 | HITS-CLIP | 23142080 | Unspecified |
| mmu-miR-34a-5p | MIMAT0000542 | Bsg | 12215 | HITS-CLIP | 30971389\|19536157\|23142080 | Unspecified |
| mmu-miR-34a-5p | MIMAT0000542 | Bsn | 12217 | HITS-CLIP | 19536157 | Unspecified |
| mmu-miR-34a-5p | MIMAT0000542 | Bst2 | 69550 | HITS-CLIP | 29167373 | Unspecified |
| mmu-miR-34a-5p | MIMAT0000542 | Btaf1 | 107182 | HITS-CLIP | 31433990\|23142080\|29167373 | Unspecified |
| mmu-miR-34a-5p | MIMAT0000542 | Btbd10 | 68815 | HITS-CLIP | 23142080 | Unspecified |
| mmu-miR-34a-5p | MIMAT0000542 | Btbd11 | 74007 | Microarrays | 37986224\|23027903 | Unspecified |
| mmu-miR-34a-5p | MIMAT0000542 | Btbd19 | 78611 | HITS-CLIP | 29167373 | Unspecified |
| mmu-miR-34a-5p | MIMAT0000542 | Btbd2 | 208198 | HITS-CLIP | 23142080 | Unspecified |
| mmu-miR-34a-5p | MIMAT0000542 | Btbd3 | 228662 | HITS-CLIP | 19536157 | Unspecified |
| mmu-miR-34a-5p | MIMAT0000542 | Btbd6 | 399566 | HITS-CLIP | 23142080 | Unspecified |
| mmu-miR-34a-5p | MIMAT0000542 | Btbd7 | 238386 | HITS-CLIP | 23142080\|29167373 | Unspecified |
| mmu-miR-34a-5p | MIMAT0000542 | Btbd8 | 100503185 | HITS-CLIP | 23142080 | Unspecified |
| mmu-miR-34a-5p | MIMAT0000542 | Btbd9 | 224671 | HITS-CLIP | 23142080 | Unspecified |
| mmu-miR-34a-5p | MIMAT0000542 | Btd | 26363 | HITS-CLIP | 23142080 | Unspecified |
| mmu-miR-34a-5p | MIMAT0000542 | Btf3 | 218490 | HITS-CLIP | 23142080 | Unspecified |
| mmu-miR-34a-5p | MIMAT0000542 | Btf3l4 | 70533 | HITS-CLIP | 23142080 | Unspecified |
| mmu-miR-34a-5p | MIMAT0000542 | Btg1 | 12226 | HITS-CLIP | 29167373\|23142080 | Unspecified |
| mmu-miR-34a-5p | MIMAT0000542 | Btg2 | 12227 | HITS-CLIP, Microarrays//HITS-CLIP | 37986224\|23142080\|19536157\|29167 | Unspecified |
| mmu-miR-34a-5p | MIMAT0000542 | Btk | 12229 | HITS-CLIP | 29167373\|23142080 | Unspecified |
| mmu-miR-34a-5p | MIMAT0000542 | Btla | 208154 | HITS-CLIP | 31433990\|23142080 | Unspecified |
| mmu-miR-34a-5p | MIMAT0000542 | Btrc | 12234 | HITS-CLIP | 23142080 | Unspecified |
| mmu-miR-34a-5p | MIMAT0000542 | Bub1 | 12235 | HITS-CLIP | 23142080 | Unspecified |
| mmu-miR-34a-5p | MIMAT0000542 | Bub1b | 12236 | HITS-CLIP | 31433990\|23630078\|23142080 | Unspecified |
| mmu-miR-34a-5p | MIMAT0000542 | Bub3 | 12237 | HITS-CLIP | 23142080 | Unspecified |
| mmu-miR-34a-5p | MIMAT0000542 | Bud31 | 231889 | HITS-CLIP | 23142080 | Unspecified |
| mmu-miR-34a-5p | MIMAT0000542 | Bysl | 53414 | Microarrays//HITS-CLIP | 37986224\|23027903\|23142080 | Unspecified |

| | | | | | | |
|---|---|---|---|---|---|---|
| mmu-miR-34a-5p | MIMAT0000542 | Bzw1 | 66882 | HITS-CLIP | 31433990\|23142080\|29167373 | Unspecified |
| mmu-miR-34a-5p | MIMAT0000542 | Bzw2 | 66912 | HITS-CLIP | 23142080 | Unspecified |
| mmu-miR-34a-5p | MIMAT0000542 | C030006K11Rik | 223665 | HITS-CLIP | 23142080 | Unspecified |
| mmu-miR-34a-5p | MIMAT0000542 | C1d | 57316 | HITS-CLIP | 23142080 | Unspecified |
| mmu-miR-34a-5p | MIMAT0000542 | C1galt1 | 94192 | HITS-CLIP | 30971389\|26203562\|23142080\|29167 | Unspecified |
| mmu-miR-34a-5p | MIMAT0000542 | C1qa | 12259 | HITS-CLIP | 29167373 | Unspecified |
| mmu-miR-34a-5p | MIMAT0000542 | C1qb | 12260 | HITS-CLIP | 29167373 | Unspecified |
| mmu-miR-34a-5p | MIMAT0000542 | C1qtnf6 | 72709 | HITS-CLIP | 23142080 | Unspecified |
| mmu-miR-34a-5p | MIMAT0000542 | C2cd2l | 71764 | HITS-CLIP | 23142080 | Unspecified |
| mmu-miR-34a-5p | MIMAT0000542 | C2cd3 | 277939 | HITS-CLIP | 29167373 | Unspecified |
| mmu-miR-34a-5p | MIMAT0000542 | C2cd5 | 74741 | HITS-CLIP | 23142080 | Unspecified |
| mmu-miR-34a-5p | MIMAT0000542 | C3 | 12266 | HITS-CLIP | 23142080 | Unspecified |
| mmu-miR-34a-5p | MIMAT0000542 | C330007P06Rik | 77644 | HITS-CLIP | 23142080 | Unspecified |
| mmu-miR-34a-5p | MIMAT0000542 | C3ar1 | 12267 | HITS-CLIP | 29167373 | Unspecified |
| mmu-miR-34a-5p | MIMAT0000542 | C5ar1 | 12273 | HITS-CLIP | 29167373 | Unspecified |
| mmu-miR-34a-5p | MIMAT0000542 | C5ar2 | 319430 | HITS-CLIP | 29167373 | Unspecified |
| mmu-miR-34a-5p | MIMAT0000542 | C9orf72 | 203228 | HITS-CLIP | 23630078\|19536157\|23142080\|29167 | Unspecified |
| mmu-miR-34a-5p | MIMAT0000542 | Caap1 | 67770 | HITS-CLIP | 23142080 | Unspecified |
| mmu-miR-34a-5p | MIMAT0000542 | Cab39 | 12283 | HITS-CLIP | 23630078\|23142080 | Unspecified |
| mmu-miR-34a-5p | MIMAT0000542 | Cabcoco1 | 219621 | HITS-CLIP | 23142080 | Unspecified |
| mmu-miR-34a-5p | MIMAT0000542 | Cabin1 | 104248 | HITS-CLIP | 23142080\|29167373 | Unspecified |
| mmu-miR-34a-5p | MIMAT0000542 | Cables2 | 252966 | HITS-CLIP | 23142080\|29167373 | Unspecified |
| mmu-miR-34a-5p | MIMAT0000542 | Cacfd1 | 381356 | HITS-CLIP | 31433990 | Unspecified |
| mmu-miR-34a-5p | MIMAT0000542 | Cacna1a | 12286 | HITS-CLIP | 19536157\|23142080\|29167373 | Unspecified |
| mmu-miR-34a-5p | MIMAT0000542 | Cacna1b | 12287 | HITS-CLIP | 19536157 | Unspecified |
| mmu-miR-34a-5p | MIMAT0000542 | Cacna1c | 12288 | HITS-CLIP | 19536157 | Unspecified |
| mmu-miR-34a-5p | MIMAT0000542 | Cacna1d | 12289 | HITS-CLIP | 19536157 | Unspecified |
| mmu-miR-34a-5p | MIMAT0000542 | Cacna1e | 12290 | HITS-CLIP | 19536157 | Unspecified |
| mmu-miR-34a-5p | MIMAT0000542 | Cacna2d1 | 12293 | HITS-CLIP | 19536157 | Unspecified |
| mmu-miR-34a-5p | MIMAT0000542 | Cacng4 | 54377 | HITS-CLIP | 19536157 | Unspecified |
| mmu-miR-34a-5p | MIMAT0000542 | Cacul1 | 78832 | HITS-CLIP | 23142080\|29167373 | Unspecified |
| mmu-miR-34a-5p | MIMAT0000542 | Cacybp | 12301 | HITS-CLIP | 23142080 | Unspecified |
| mmu-miR-34a-5p | MIMAT0000542 | Cad | 69719 | HITS-CLIP | 23142080\|29167373 | Unspecified |
| mmu-miR-34a-5p | MIMAT0000542 | Cadm1 | 54725 | HITS-CLIP | 29167373 | Unspecified |
| mmu-miR-34a-5p | MIMAT0000542 | Cadm2 | 239857 | HITS-CLIP | 19536157 | Unspecified |
| mmu-miR-34a-5p | MIMAT0000542 | Cadm3 | 94332 | HITS-CLIP | 19536157 | Unspecified |
| mmu-miR-34a-5p | MIMAT0000542 | Cadps | 27062 | HITS-CLIP | 19536157 | Unspecified |
| mmu-miR-34a-5p | MIMAT0000542 | Cadps2 | 320405 | HITS-CLIP | 19536157 | Unspecified |
| mmu-miR-34a-5p | MIMAT0000542 | Cage1 | 71213 | HITS-CLIP | 23142080 | Unspecified |

| | | | | | | |
|---|---|---|---|---|---|---|
| mmu-miR-34a-5p | MIMAT0000542 | Calb1 | 12307 | HITS-CLIP | 19536157 | Unspecified |
| mmu-miR-34a-5p | MIMAT0000542 | Calcoco1 | 67488 | Microarrays | 37986224 | Unspecified |
| mmu-miR-34a-5p | MIMAT0000542 | Calcrl | 54598 | HITS-CLIP | 37986224\|19536157\|23142080\|29167 | Unspecified |
| mmu-miR-34a-5p | MIMAT0000542 | Cald1 | 109624 | HITS-CLIP | 19536157\|23630078 | Unspecified |
| mmu-miR-34a-5p | MIMAT0000542 | Calm1 | 12313 | HITS-CLIP | 31433990\|19536157\|26203562\|23142 | Unspecified |
| mmu-miR-34a-5p | MIMAT0000542 | Calm2 | 12314 | HITS-CLIP | 23142080\|29167373 | Unspecified |
| mmu-miR-34a-5p | MIMAT0000542 | Calm3 | 12315 | HITS-CLIP | 23142080\|31433990\|19536157\|29167 | Unspecified |
| mmu-miR-34a-5p | MIMAT0000542 | Calr | 12317 | HITS-CLIP | 23142080\|31433990\|19536157\|29167 | Unspecified |
| mmu-miR-34a-5p | MIMAT0000542 | Calu | 12321 | HITS-CLIP | 23142080 | Unspecified |
| mmu-miR-34a-5p | MIMAT0000542 | Camk1d | 227541 | HITS-CLIP | 19536157\|29167373\|23142080 | Unspecified |
| mmu-miR-34a-5p | MIMAT0000542 | Camk2a | 12322 | HITS-CLIP | 19536157 | Unspecified |
| mmu-miR-34a-5p | MIMAT0000542 | Camk2b | 12323 | HITS-CLIP | 23142080 | Unspecified |
| mmu-miR-34a-5p | MIMAT0000542 | Camk2d | 108058 | HITS-CLIP | 23142080\|19536157\|29167373 | Unspecified |
| mmu-miR-34a-5p | MIMAT0000542 | Camk2g | 12325 | HITS-CLIP | 23142080 | Unspecified |
| mmu-miR-34a-5p | MIMAT0000542 | Camk2n1 | 66259 | HITS-CLIP | 19536157 | Unspecified |
| mmu-miR-34a-5p | MIMAT0000542 | Camkk2 | 207565 | HITS-CLIP | 23630078\|29167373 | Unspecified |
| mmu-miR-34a-5p | MIMAT0000542 | Camkv | 235604 | HITS-CLIP | 19536157 | Unspecified |
| mmu-miR-34a-5p | MIMAT0000542 | Caml | 12328 | HITS-CLIP | 23142080 | Unspecified |
| mmu-miR-34a-5p | MIMAT0000542 | Camsap1 | 227634 | HITS-CLIP | 23142080\|29167373 | Unspecified |
| mmu-miR-34a-5p | MIMAT0000542 | Camsap2 | 67886 | HITS-CLIP | 19536157\|31433990\|23142080\|29167 | Unspecified |
| mmu-miR-34a-5p | MIMAT0000542 | Camta1 | 100072 | HITS-CLIP | 23142080 | Unspecified |
| mmu-miR-34a-5p | MIMAT0000542 | Camta2 | 216874 | HITS-CLIP | 23142080\|29167373 | Unspecified |
| mmu-miR-34a-5p | MIMAT0000542 | Cand1 | 71902 | HITS-CLIP | 23142080 | Unspecified |
| mmu-miR-34a-5p | MIMAT0000542 | Cand2 | 67088 | Microarrays//HITS-CLIP | 37986224\|23142080 | Unspecified |
| mmu-miR-34a-5p | MIMAT0000542 | Cant1 | 76025 | HITS-CLIP | 23142080\|29167373 | Unspecified |
| mmu-miR-34a-5p | MIMAT0000542 | Canx | 12330 | HITS-CLIP | 23142080\|19536157\|29167373 | Unspecified |
| mmu-miR-34a-5p | MIMAT0000542 | Cap1 | 12331 | HITS-CLIP | 19536157\|23142080\|29167373 | Unspecified |
| mmu-miR-34a-5p | MIMAT0000542 | Cap2 | 67252 | Microarrays | 37986224 | Unspecified |
| mmu-miR-34a-5p | MIMAT0000542 | Capg | 12332 | HITS-CLIP | 23142080\|29167373 | Unspecified |
| mmu-miR-34a-5p | MIMAT0000542 | Capn1 | 12333 | HITS-CLIP | 23142080 | Unspecified |
| mmu-miR-34a-5p | MIMAT0000542 | Capn10 | 23830 | HITS-CLIP | 23142080 | Unspecified |
| mmu-miR-34a-5p | MIMAT0000542 | Capn15 | 50817 | HITS-CLIP | 23142080\|26203562 | Unspecified |
| mmu-miR-34a-5p | MIMAT0000542 | Capn2 | 12334 | HITS-CLIP | 19536157\|23142080\|29167373 | Unspecified |
| mmu-miR-34a-5p | MIMAT0000542 | Capn5 | 12337 | Microarrays//HITS-CLIP | 37986224\|23027903\|29167373\|23142 | Unspecified |
| mmu-miR-34a-5p | MIMAT0000542 | Capn7 | 12339 | HITS-CLIP | 23142080\|29167373 | Unspecified |
| mmu-miR-34a-5p | MIMAT0000542 | Capns1 | 12336 | HITS-CLIP | 23142080\|29167373 | Unspecified |
| mmu-miR-34a-5p | MIMAT0000542 | Caprin1 | 53872 | HITS-CLIP | 31433990\|23142080\|29167373 | Unspecified |
| mmu-miR-34a-5p | MIMAT0000542 | Capza1 | 12340 | HITS-CLIP | 23142080\|29167373 | Unspecified |
| mmu-miR-34a-5p | MIMAT0000542 | Capza2 | 12343 | HITS-CLIP | 37986224\|23142080\|19536157\|2363C | Unspecified |

| | | | | | | |
|---|---|---|---|---|---|---|
| mmu-miR-34a-5p | MIMAT0000542 | Capzb | 12345 | HITS-CLIP | 31433990\|23142080\|29167373 | Unspecified |
| mmu-miR-34a-5p | MIMAT0000542 | Car13 | 71934 | HITS-CLIP | 29167373 | Unspecified |
| mmu-miR-34a-5p | MIMAT0000542 | Car4 | 12351 | HITS-CLIP | 19536157 | Unspecified |
| mmu-miR-34a-5p | MIMAT0000542 | Car5b | 56078 | HITS-CLIP | 23142080 | Unspecified |
| mmu-miR-34a-5p | MIMAT0000542 | Card11 | 108723 | HITS-CLIP | 23142080 | Unspecified |
| mmu-miR-34a-5p | MIMAT0000542 | Carm1 | 59035 | HITS-CLIP | 23142080\|29167373 | Unspecified |
| mmu-miR-34a-5p | MIMAT0000542 | Carmil1 | 55604 | HITS-CLIP | 23142080 | Unspecified |
| mmu-miR-34a-5p | MIMAT0000542 | Carmil2 | 146206 | HITS-CLIP | 23142080 | Unspecified |
| mmu-miR-34a-5p | MIMAT0000542 | Cars | 27267 | HITS-CLIP | 23142080 | Unspecified |
| mmu-miR-34a-5p | MIMAT0000542 | Casc3 | 192160 | HITS-CLIP | 23142080\|29167373 | Unspecified |
| mmu-miR-34a-5p | MIMAT0000542 | Casc4 | 319996 | HITS-CLIP | 23142080 | Unspecified |
| mmu-miR-34a-5p | MIMAT0000542 | Casd1 | 213819 | HITS-CLIP | 29167373 | Unspecified |
| mmu-miR-34a-5p | MIMAT0000542 | Cask | 12361 | HITS-CLIP | 19536157 | Unspecified |
| mmu-miR-34a-5p | MIMAT0000542 | Casp1 | 12362 | HITS-CLIP | 23142080 | Unspecified |
| mmu-miR-34a-5p | MIMAT0000542 | Casp2 | 12366 | HITS-CLIP, Microarrays//Microarrays//HITS | 37986224\|23027903\|23142080 | Unspecified |
| mmu-miR-34a-5p | MIMAT0000542 | Casp3 | 12367 | Microarrays//HITS-CLIP | 37986224\|23142080\|23027903 | Unspecified |
| mmu-miR-34a-5p | MIMAT0000542 | Casp6 | 12368 | HITS-CLIP | 23630078\|23142080 | Unspecified |
| mmu-miR-34a-5p | MIMAT0000542 | Casp8 | 12370 | HITS-CLIP | 23142080\|29167373 | Unspecified |
| mmu-miR-34a-5p | MIMAT0000542 | Casp8ap2 | 26885 | HITS-CLIP | 31433990\|23142080\|29167373 | Unspecified |
| mmu-miR-34a-5p | MIMAT0000542 | Casq2 | 12373 | Microarrays | 37986224 | Unspecified |
| mmu-miR-34a-5p | MIMAT0000542 | Cast | 12380 | HITS-CLIP | 23142080\|29167373\|26203562 | Unspecified |
| mmu-miR-34a-5p | MIMAT0000542 | Castor1 | 652968 | HITS-CLIP | 23142080 | Unspecified |
| mmu-miR-34a-5p | MIMAT0000542 | Castor2 | 729438 | HITS-CLIP | 19536157 | Unspecified |
| mmu-miR-34a-5p | MIMAT0000542 | Casz1 | 69743 | HITS-CLIP | 23142080 | Unspecified |
| mmu-miR-34a-5p | MIMAT0000542 | Cat | 12359 | Microarrays//HITS-CLIP | 37986224\|30971389\|23142080 | Unspecified |
| mmu-miR-34a-5p | MIMAT0000542 | Catsper2 | 212670 | HITS-CLIP | 29167373 | Unspecified |
| mmu-miR-34a-5p | MIMAT0000542 | Cav1 | 12389 | HITS-CLIP | 19536157\|29167373 | Unspecified |
| mmu-miR-34a-5p | MIMAT0000542 | Cav2 | 12390 | HITS-CLIP | 23630078\|29167373 | Unspecified |
| mmu-miR-34a-5p | MIMAT0000542 | Cavin2 | 8436 | HITS-CLIP | 19536157 | Unspecified |
| mmu-miR-34a-5p | MIMAT0000542 | Cbfa2t2 | 12396 | HITS-CLIP | 30971389\|31433990\|29167373 | Unspecified |
| mmu-miR-34a-5p | MIMAT0000542 | Cbfa2t3 | 12398 | HITS-CLIP | 23142080\|19536157 | Unspecified |
| mmu-miR-34a-5p | MIMAT0000542 | Cbfb | 12400 | HITS-CLIP | 23630078\|31433990\|23142080\|29167 | Unspecified |
| mmu-miR-34a-5p | MIMAT0000542 | Cbl | 12402 | HITS-CLIP | 23142080\|29167373\|19536157 | Unspecified |
| mmu-miR-34a-5p | MIMAT0000542 | Cblb | 208650 | HITS-CLIP | 37986224\|23142080\|19536157\|29167 | Unspecified |
| mmu-miR-34a-5p | MIMAT0000542 | Cbll1 | 104836 | HITS-CLIP | 23142080 | Unspecified |
| mmu-miR-34a-5p | MIMAT0000542 | Cbln4 | 228942 | HITS-CLIP | 19536157 | Unspecified |
| mmu-miR-34a-5p | MIMAT0000542 | Cbr4 | 234309 | HITS-CLIP | 23142080 | Unspecified |
| mmu-miR-34a-5p | MIMAT0000542 | Cbx2 | 12416 | HITS-CLIP | 23142080\|30971389 | Unspecified |
| mmu-miR-34a-5p | MIMAT0000542 | Cbx4 | 12418 | HITS-CLIP | 23142080\|19536157\|31433990 | Unspecified |

| | | | | | | |
|---|---|---|---|---|---|---|
| mmu-miR-34a-5p | MIMAT0000542 | Cbx5 | 12419 | HITS-CLIP | 37986224\|23630078\|19536157\|23142 | Unspecified |
| mmu-miR-34a-5p | MIMAT0000542 | Cbx6 | 494448 | HITS-CLIP | 19536157\|23142080 | Unspecified |
| mmu-miR-34a-5p | MIMAT0000542 | Cbx7 | 52609 | HITS-CLIP | 23142080 | Unspecified |
| mmu-miR-34a-5p | MIMAT0000542 | Cc2d1a | 212139 | HITS-CLIP | 29167373\|23142080 | Unspecified |
| mmu-miR-34a-5p | MIMAT0000542 | Cc2d1b | 319965 | HITS-CLIP | 23142080 | Unspecified |
| mmu-miR-34a-5p | MIMAT0000542 | Ccar1 | 67500 | HITS-CLIP | 23142080\|31433990\|29167373 | Unspecified |
| mmu-miR-34a-5p | MIMAT0000542 | Ccar2 | 219158 | HITS-CLIP | 23142080 | Unspecified |
| mmu-miR-34a-5p | MIMAT0000542 | Ccdc115 | 69668 | HITS-CLIP | 23142080 | Unspecified |
| mmu-miR-34a-5p | MIMAT0000542 | Ccdc117 | 104479 | HITS-CLIP | 31433990 | Unspecified |
| mmu-miR-34a-5p | MIMAT0000542 | Ccdc124 | 234388 | HITS-CLIP | 23142080 | Unspecified |
| mmu-miR-34a-5p | MIMAT0000542 | Ccdc127 | 67433 | HITS-CLIP | 23142080\|30971389\|31433990\|19536 | Unspecified |
| mmu-miR-34a-5p | MIMAT0000542 | Ccdc137 | 67291 | HITS-CLIP | 26203562\|23142080 | Unspecified |
| mmu-miR-34a-5p | MIMAT0000542 | Ccdc14 | 239839 | HITS-CLIP | 23142080 | Unspecified |
| mmu-miR-34a-5p | MIMAT0000542 | Ccdc141 | 545428 | Microarrays//HITS-CLIP | 37986224\|19536157\|23142080 | Unspecified |
| mmu-miR-34a-5p | MIMAT0000542 | Ccdc167 | 68597 | HITS-CLIP | 23142080 | Unspecified |
| mmu-miR-34a-5p | MIMAT0000542 | Ccdc173 | 75051 | Microarrays | 37986224 | Unspecified |
| mmu-miR-34a-5p | MIMAT0000542 | Ccdc174 | 232236 | HITS-CLIP | 23142080 | Unspecified |
| mmu-miR-34a-5p | MIMAT0000542 | Ccdc18 | 73254 | HITS-CLIP | 23142080 | Unspecified |
| mmu-miR-34a-5p | MIMAT0000542 | Ccdc181 | 74895 | HITS-CLIP | 23142080 | Unspecified |
| mmu-miR-34a-5p | MIMAT0000542 | Ccdc186 | 55088 | HITS-CLIP | 23142080\|29167373 | Unspecified |
| mmu-miR-34a-5p | MIMAT0000542 | Ccdc25 | 67179 | HITS-CLIP | 23142080 | Unspecified |
| mmu-miR-34a-5p | MIMAT0000542 | Ccdc28b | 66264 | HITS-CLIP | 23142080 | Unspecified |
| mmu-miR-34a-5p | MIMAT0000542 | Ccdc34 | 68201 | HITS-CLIP | 23142080 | Unspecified |
| mmu-miR-34a-5p | MIMAT0000542 | Ccdc47 | 67163 | HITS-CLIP | 29167373 | Unspecified |
| mmu-miR-34a-5p | MIMAT0000542 | Ccdc50 | 67501 | HITS-CLIP | 23142080\|29167373 | Unspecified |
| mmu-miR-34a-5p | MIMAT0000542 | Ccdc51 | 66658 | HITS-CLIP | 23142080 | Unspecified |
| mmu-miR-34a-5p | MIMAT0000542 | Ccdc59 | 52713 | HITS-CLIP | 23142080\|29167373 | Unspecified |
| mmu-miR-34a-5p | MIMAT0000542 | Ccdc6 | 76551 | HITS-CLIP | 23142080 | Unspecified |
| mmu-miR-34a-5p | MIMAT0000542 | Ccdc65 | 105833 | Microarrays | 37986224 | Unspecified |
| mmu-miR-34a-5p | MIMAT0000542 | Ccdc66 | 320234 | HITS-CLIP | 23142080 | Unspecified |
| mmu-miR-34a-5p | MIMAT0000542 | Ccdc69 | 52570 | HITS-CLIP | 23142080 | Unspecified |
| mmu-miR-34a-5p | MIMAT0000542 | Ccdc71 | 72454 | HITS-CLIP | 23142080 | Unspecified |
| mmu-miR-34a-5p | MIMAT0000542 | Ccdc71l | 72123 | HITS-CLIP | 23142080\|29167373 | Unspecified |
| mmu-miR-34a-5p | MIMAT0000542 | Ccdc82 | 66396 | HITS-CLIP | 23142080 | Unspecified |
| mmu-miR-34a-5p | MIMAT0000542 | Ccdc85c | 668158 | HITS-CLIP | 31433990 | Unspecified |
| mmu-miR-34a-5p | MIMAT0000542 | Ccdc88a | 108686 | HITS-CLIP | 29167373 | Unspecified |
| mmu-miR-34a-5p | MIMAT0000542 | Ccdc88c | 68339 | HITS-CLIP | 31433990\|23142080 | Unspecified |
| mmu-miR-34a-5p | MIMAT0000542 | Ccdc97 | 52132 | HITS-CLIP | 23142080 | Unspecified |
| mmu-miR-34a-5p | MIMAT0000542 | Cchcr1 | 240084 | HITS-CLIP | 23142080 | Unspecified |

| | | | | | | | |
|---|---|---|---|---|---|---|---|
| mmu-miR-34a-5p | MIMAT0000542 | Ccl1 | 20290 | HITS-CLIP | | 23142080 | Unspecified |
| mmu-miR-34a-5p | MIMAT0000542 | Ccl2 | 20296 | HITS-CLIP | | 29167373 | Unspecified |
| mmu-miR-34a-5p | MIMAT0000542 | Ccl22 | 20299 | HITS-CLIP | 37986224\|23142080 | | Unspecified |
| mmu-miR-34a-5p | MIMAT0000542 | Ccl3 | 20302 | HITS-CLIP | 23142080\|29167373 | | Unspecified |
| mmu-miR-34a-5p | MIMAT0000542 | Ccl6 | 20305 | HITS-CLIP | | 23142080 | Unspecified |
| mmu-miR-34a-5p | MIMAT0000542 | Ccl7 | 20306 | HITS-CLIP | | 29167373 | Unspecified |
| mmu-miR-34a-5p | MIMAT0000542 | Ccl9 | 20308 | Microarrays//HITS-CLIP | 37986224\|29167373 | | Unspecified |
| mmu-miR-34a-5p | MIMAT0000542 | Ccm2 | 216527 | HITS-CLIP | | 23142080 | Unspecified |
| mmu-miR-34a-5p | MIMAT0000542 | Ccn2 | 1490 | HITS-CLIP | 23630078\|19536157 | | Unspecified |
| mmu-miR-34a-5p | MIMAT0000542 | Ccn3 | 4856 | HITS-CLIP | | 19536157 | Unspecified |
| mmu-miR-34a-5p | MIMAT0000542 | Ccn4 | 8840 | HITS-CLIP | | 23630078 | Unspecified |
| mmu-miR-34a-5p | MIMAT0000542 | Ccna2 | 12428 | HITS-CLIP | | 23142080 | Unspecified |
| mmu-miR-34a-5p | MIMAT0000542 | Ccnb2 | 12442 | HITS-CLIP | | 23142080 | Unspecified |
| mmu-miR-34a-5p | MIMAT0000542 | Ccnc | 51813 | HITS-CLIP | 23630078\|23142080\|29167373 | | Unspecified |
| mmu-miR-34a-5p | MIMAT0000542 | Ccnd1 | 12443 | Northern blot//qRT-PCR//Western blot//M | 24587330\|23027903\|29167373 | | Unspecified |
| mmu-miR-34a-5p | MIMAT0000542 | Ccnd2 | 12444 | HITS-CLIP | 37986224\|23142080\|19536157\|3143 | | Unspecified |
| mmu-miR-34a-5p | MIMAT0000542 | Ccnd3 | 12445 | HITS-CLIP | 23142080\|31433990 | | Unspecified |
| mmu-miR-34a-5p | MIMAT0000542 | Ccndbp1 | 17151 | HITS-CLIP | | 23142080 | Unspecified |
| mmu-miR-34a-5p | MIMAT0000542 | Ccne1 | 12447 | Microarrays//HITS-CLIP | 37986224\|23027903\|30971389\|3143 | | Unspecified |
| mmu-miR-34a-5p | MIMAT0000542 | Ccne2 | 12448 | Microarrays//HITS-CLIP | 37986224\|23027903\|31433990\|2363 | | Unspecified |
| mmu-miR-34a-5p | MIMAT0000542 | Ccnf | 12449 | HITS-CLIP | 31433990\|23142080 | | Unspecified |
| mmu-miR-34a-5p | MIMAT0000542 | Ccng1 | 12450 | HITS-CLIP | 23142080\|19536157\|29167373 | | Unspecified |
| mmu-miR-34a-5p | MIMAT0000542 | Ccng2 | 12452 | HITS-CLIP | 19536157\|31433990\|23142080 | | Unspecified |
| mmu-miR-34a-5p | MIMAT0000542 | Ccni | 12453 | HITS-CLIP | 23142080\|29167373 | | Unspecified |
| mmu-miR-34a-5p | MIMAT0000542 | Ccnj | 240665 | HITS-CLIP | | 23142080 | Unspecified |
| mmu-miR-34a-5p | MIMAT0000542 | Ccnk | 12454 | HITS-CLIP | | 23142080 | Unspecified |
| mmu-miR-34a-5p | MIMAT0000542 | Ccnl1 | 56706 | HITS-CLIP | 30971389\|29167373\|23142080 | | Unspecified |
| mmu-miR-34a-5p | MIMAT0000542 | Ccnl2 | 56036 | HITS-CLIP | 23142080\|29167373 | | Unspecified |
| mmu-miR-34a-5p | MIMAT0000542 | Ccnt1 | 12455 | HITS-CLIP | 23142080\|29167373 | | Unspecified |
| mmu-miR-34a-5p | MIMAT0000542 | Ccnt2 | 72949 | HITS-CLIP | 23142080\|29167373 | | Unspecified |
| mmu-miR-34a-5p | MIMAT0000542 | Ccny | 67974 | HITS-CLIP | 23630078\|31433990\|23142080 | | Unspecified |
| mmu-miR-34a-5p | MIMAT0000542 | Ccp110 | 101565 | HITS-CLIP | | 23142080 | Unspecified |
| mmu-miR-34a-5p | MIMAT0000542 | Ccpg1 | 72278 | HITS-CLIP | | 23142080 | Unspecified |
| mmu-miR-34a-5p | MIMAT0000542 | Ccpg1os | 546143 | HITS-CLIP | | 23142080 | Unspecified |
| mmu-miR-34a-5p | MIMAT0000542 | Ccr2 | 12772 | HITS-CLIP | | 29167373 | Unspecified |
| mmu-miR-34a-5p | MIMAT0000542 | Ccr4 | 12773 | HITS-CLIP | | 23142080 | Unspecified |
| mmu-miR-34a-5p | MIMAT0000542 | Ccr5 | 12774 | HITS-CLIP | 37986224\|23142080\|29167373 | | Unspecified |
| mmu-miR-34a-5p | MIMAT0000542 | Ccr7 | 12775 | HITS-CLIP | 31433990\|23142080 | | Unspecified |
| mmu-miR-34a-5p | MIMAT0000542 | Ccr8 | 12776 | HITS-CLIP | | 23142080 | Unspecified |

| | | | | | | |
|---|---|---|---|---|---|---|
| mmu-miR-34a-5p | MIMAT0000542 | Ccsap | 73420 | HITS-CLIP | 23142080\|29167373 | Unspecified |
| mmu-miR-34a-5p | MIMAT0000542 | Ccser2 | 72972 | HITS-CLIP | 19536157\|23142080\|29167373 | Unspecified |
| mmu-miR-34a-5p | MIMAT0000542 | Cct2 | 12461 | HITS-CLIP | 23142080 | Unspecified |
| mmu-miR-34a-5p | MIMAT0000542 | Cct3 | 12462 | HITS-CLIP | 23630078\|23142080\|31433990 | Unspecified |
| mmu-miR-34a-5p | MIMAT0000542 | Cct4 | 12464 | HITS-CLIP | 23142080\|29167373 | Unspecified |
| mmu-miR-34a-5p | MIMAT0000542 | Cct5 | 12465 | HITS-CLIP | 23142080 | Unspecified |
| mmu-miR-34a-5p | MIMAT0000542 | Cct7 | 12468 | HITS-CLIP | 23142080 | Unspecified |
| mmu-miR-34a-5p | MIMAT0000542 | Cct8 | 12469 | HITS-CLIP | 23142080\|30971389\|26203562 | Unspecified |
| mmu-miR-34a-5p | MIMAT0000542 | Ccz1 | 231874 | HITS-CLIP | 23142080 | Unspecified |
| mmu-miR-34a-5p | MIMAT0000542 | Cd101 | 630146 | HITS-CLIP | 23142080 | Unspecified |
| mmu-miR-34a-5p | MIMAT0000542 | Cd14 | 12475 | HITS-CLIP | 29167373 | Unspecified |
| mmu-miR-34a-5p | MIMAT0000542 | Cd151 | 12476 | Microarrays//HITS-CLIP | 37986224\|23027903\|19536157\|2314 | Unspecified |
| mmu-miR-34a-5p | MIMAT0000542 | Cd164 | 53599 | HITS-CLIP | 23142080\|19536157\|29167373 | Unspecified |
| mmu-miR-34a-5p | MIMAT0000542 | Cd180 | 17079 | HITS-CLIP | 23142080\|29167373 | Unspecified |
| mmu-miR-34a-5p | MIMAT0000542 | Cd19 | 12478 | HITS-CLIP | 23142080 | Unspecified |
| mmu-miR-34a-5p | MIMAT0000542 | Cd200 | 17470 | HITS-CLIP | 23142080 | Unspecified |
| mmu-miR-34a-5p | MIMAT0000542 | Cd200r1 | 57781 | HITS-CLIP | 29167373 | Unspecified |
| mmu-miR-34a-5p | MIMAT0000542 | Cd200r4 | 239849 | HITS-CLIP | 23142080 | Unspecified |
| mmu-miR-34a-5p | MIMAT0000542 | Cd22 | 12483 | HITS-CLIP | 23142080 | Unspecified |
| mmu-miR-34a-5p | MIMAT0000542 | Cd226 | 225825 | HITS-CLIP | 23142080 | Unspecified |
| mmu-miR-34a-5p | MIMAT0000542 | Cd247 | 12503 | HITS-CLIP | 23142080 | Unspecified |
| mmu-miR-34a-5p | MIMAT0000542 | Cd27 | 21940 | HITS-CLIP | 37986224\|23142080 | Unspecified |
| mmu-miR-34a-5p | MIMAT0000542 | Cd274 | 60533 | HITS-CLIP | 31433990\|23142080\|29167373 | Unspecified |
| mmu-miR-34a-5p | MIMAT0000542 | Cd28 | 12487 | HITS-CLIP, Microarrays//HITS-CLIP | 37986224\|23142080 | Unspecified |
| mmu-miR-34a-5p | MIMAT0000542 | Cd2ap | 12488 | HITS-CLIP | 23142080\|19536157\|29167373 | Unspecified |
| mmu-miR-34a-5p | MIMAT0000542 | Cd2bp2 | 70233 | HITS-CLIP | 23142080\|19536157 | Unspecified |
| mmu-miR-34a-5p | MIMAT0000542 | Cd300lb | 217304 | HITS-CLIP | 29167373 | Unspecified |
| mmu-miR-34a-5p | MIMAT0000542 | Cd300ld | 217305 | HITS-CLIP | 29167373 | Unspecified |
| mmu-miR-34a-5p | MIMAT0000542 | Cd36 | 12491 | HITS-CLIP | 29167373 | Unspecified |
| mmu-miR-34a-5p | MIMAT0000542 | Cd37 | 12493 | HITS-CLIP | 23142080\|29167373 | Unspecified |
| mmu-miR-34a-5p | MIMAT0000542 | Cd3d | 12500 | HITS-CLIP | 23142080 | Unspecified |
| mmu-miR-34a-5p | MIMAT0000542 | Cd3e | 12501 | HITS-CLIP | 23142080 | Unspecified |
| mmu-miR-34a-5p | MIMAT0000542 | Cd4 | 12504 | HITS-CLIP | 23142080\|31433990 | Unspecified |
| mmu-miR-34a-5p | MIMAT0000542 | Cd40lg | 959 | HITS-CLIP | 23142080 | Unspecified |
| mmu-miR-34a-5p | MIMAT0000542 | Cd44 | 12505 | HITS-CLIP | 23142080\|31433990\|29167373 | Unspecified |
| mmu-miR-34a-5p | MIMAT0000542 | Cd47 | 16423 | HITS-CLIP | 19536157\|23142080\|29167373 | Unspecified |
| mmu-miR-34a-5p | MIMAT0000542 | Cd48 | 12506 | HITS-CLIP | 23142080\|29167373 | Unspecified |
| mmu-miR-34a-5p | MIMAT0000542 | Cd5 | 12507 | HITS-CLIP | 23142080 | Unspecified |
| mmu-miR-34a-5p | MIMAT0000542 | Cd52 | 23833 | HITS-CLIP | 23142080\|29167373 | Unspecified |

| | | | | | | |
|---|---|---|---|---|---|---|
| mmu-miR-34a-5p | MIMAT0000542 | Cd53 | 12508 | HITS-CLIP | 37986224\|23142080\|29167373 | Unspecified |
| mmu-miR-34a-5p | MIMAT0000542 | Cd55 | 13136 | HITS-CLIP | 23142080 | Unspecified |
| mmu-miR-34a-5p | MIMAT0000542 | Cd6 | 12511 | HITS-CLIP | 23142080 | Unspecified |
| mmu-miR-34a-5p | MIMAT0000542 | Cd63 | 12512 | HITS-CLIP | 29167373 | Unspecified |
| mmu-miR-34a-5p | MIMAT0000542 | Cd68 | 12514 | HITS-CLIP | 29167373 | Unspecified |
| mmu-miR-34a-5p | MIMAT0000542 | Cd69 | 12515 | HITS-CLIP | 23142080 | Unspecified |
| mmu-miR-34a-5p | MIMAT0000542 | Cd72 | 12517 | HITS-CLIP | 23142080\|29167373 | Unspecified |
| mmu-miR-34a-5p | MIMAT0000542 | Cd74 | 16149 | HITS-CLIP | 23142080\|29167373 | Unspecified |
| mmu-miR-34a-5p | MIMAT0000542 | Cd79a | 12518 | HITS-CLIP | 23142080 | Unspecified |
| mmu-miR-34a-5p | MIMAT0000542 | Cd79b | 15985 | HITS-CLIP | 23142080 | Unspecified |
| mmu-miR-34a-5p | MIMAT0000542 | Cd80 | 12519 | HITS-CLIP | 23142080 | Unspecified |
| mmu-miR-34a-5p | MIMAT0000542 | Cd81 | 12520 | HITS-CLIP | 19536157\|26203562\|29167373\|23142 | Unspecified |
| mmu-miR-34a-5p | MIMAT0000542 | Cd82 | 12521 | HITS-CLIP | 37986224\|23142080 | Unspecified |
| mmu-miR-34a-5p | MIMAT0000542 | Cd83 | 12522 | HITS-CLIP | 31433990\|23142080 | Unspecified |
| mmu-miR-34a-5p | MIMAT0000542 | Cd84 | 12523 | HITS-CLIP | 37986224\|23142080\|29167373 | Unspecified |
| mmu-miR-34a-5p | MIMAT0000542 | Cd8a | 12525 | HITS-CLIP | 23142080 | Unspecified |
| mmu-miR-34a-5p | MIMAT0000542 | Cd9 | 12527 | HITS-CLIP | 29167373 | Unspecified |
| mmu-miR-34a-5p | MIMAT0000542 | Cd93 | 17064 | HITS-CLIP | 21258322\|29167373 | Unspecified |
| mmu-miR-34a-5p | MIMAT0000542 | Cd99l2 | 171486 | HITS-CLIP | 29167373 | Unspecified |
| mmu-miR-34a-5p | MIMAT0000542 | Cdadc1 | 71891 | HITS-CLIP | 29167373 | Unspecified |
| mmu-miR-34a-5p | MIMAT0000542 | Cdan1 | 68968 | HITS-CLIP | 23142080 | Unspecified |
| mmu-miR-34a-5p | MIMAT0000542 | Cdc123 | 98828 | HITS-CLIP | 29167373\|23142080 | Unspecified |
| mmu-miR-34a-5p | MIMAT0000542 | Cdc14a | 229776 | HITS-CLIP | 23142080 | Unspecified |
| mmu-miR-34a-5p | MIMAT0000542 | Cdc14b | 218294 | HITS-CLIP | 29167373\|23142080 | Unspecified |
| mmu-miR-34a-5p | MIMAT0000542 | Cdc16 | 69957 | HITS-CLIP | 23142080\|29167373 | Unspecified |
| mmu-miR-34a-5p | MIMAT0000542 | Cdc20 | 107995 | HITS-CLIP | 23142080\|30971389\|31433990 | Unspecified |
| mmu-miR-34a-5p | MIMAT0000542 | Cdc23 | 52563 | HITS-CLIP | 23142080 | Unspecified |
| mmu-miR-34a-5p | MIMAT0000542 | Cdc25a | 12530 | HITS-CLIP, Microarrays//Microarrays//HITS | 37986224\|23027903\|23142080 | Unspecified |
| mmu-miR-34a-5p | MIMAT0000542 | Cdc25b | 12531 | HITS-CLIP | 23142080 | Unspecified |
| mmu-miR-34a-5p | MIMAT0000542 | Cdc25c | 12532 | HITS-CLIP | 23142080 | Unspecified |
| mmu-miR-34a-5p | MIMAT0000542 | Cdc27 | 217232 | HITS-CLIP | 37986224\|19536157\|23142080\|29167 | Unspecified |
| mmu-miR-34a-5p | MIMAT0000542 | Cdc34 | 216150 | HITS-CLIP | 23142080 | Unspecified |
| mmu-miR-34a-5p | MIMAT0000542 | Cdc37 | 12539 | HITS-CLIP | 23142080\|29167373 | Unspecified |
| mmu-miR-34a-5p | MIMAT0000542 | Cdc37l1 | 67072 | HITS-CLIP | 23142080\|19536157\|29167373 | Unspecified |
| mmu-miR-34a-5p | MIMAT0000542 | Cdc40 | 71713 | HITS-CLIP | 23142080 | Unspecified |
| mmu-miR-34a-5p | MIMAT0000542 | Cdc42 | 12540 | HITS-CLIP | 29167373\|23142080 | Unspecified |
| mmu-miR-34a-5p | MIMAT0000542 | Cdc42bpa | 226751 | HITS-CLIP | 19536157\|23142080 | Unspecified |
| mmu-miR-34a-5p | MIMAT0000542 | Cdc42bpb | 217866 | HITS-CLIP | 29167373 | Unspecified |
| mmu-miR-34a-5p | MIMAT0000542 | Cdc42ep2 | 104252 | HITS-CLIP | 19536157 | Unspecified |

| | | | | | | |
|---|---|---|---|---|---|---|
| mmu-miR-34a-5p | MIMAT0000542 | Cdc42ep3 | 260409 | HITS-CLIP | 23142080\|29167373 | Unspecified |
| mmu-miR-34a-5p | MIMAT0000542 | Cdc42ep4 | 56699 | HITS-CLIP | 19536157\|23142080\|29167373 | Unspecified |
| mmu-miR-34a-5p | MIMAT0000542 | Cdc42se1 | 57912 | HITS-CLIP | 29167373\|23142080 | Unspecified |
| mmu-miR-34a-5p | MIMAT0000542 | Cdc42se2 | 72729 | HITS-CLIP | 29167373\|23142080 | Unspecified |
| mmu-miR-34a-5p | MIMAT0000542 | Cdc45 | 12544 | HITS-CLIP | 23142080 | Unspecified |
| mmu-miR-34a-5p | MIMAT0000542 | Cdc5l | 71702 | HITS-CLIP | 23142080 | Unspecified |
| mmu-miR-34a-5p | MIMAT0000542 | Cdc6 | 23834 | Microarrays//HITS-CLIP | 37986224\|23027903\|23142080 | Unspecified |
| mmu-miR-34a-5p | MIMAT0000542 | Cdc7 | 12545 | Microarrays | 37986224\|23027903 | Unspecified |
| mmu-miR-34a-5p | MIMAT0000542 | Cdc73 | 214498 | HITS-CLIP | 23142080\|29167373 | Unspecified |
| mmu-miR-34a-5p | MIMAT0000542 | Cdca2 | 108912 | HITS-CLIP | 23142080\|30971389 | Unspecified |
| mmu-miR-34a-5p | MIMAT0000542 | Cdca3 | 14793 | HITS-CLIP | 23142080 | Unspecified |
| mmu-miR-34a-5p | MIMAT0000542 | Cdca4 | 71963 | HITS-CLIP | 31433990\|29167373\|23142080 | Unspecified |
| mmu-miR-34a-5p | MIMAT0000542 | Cdca5 | 67849 | Microarrays | 37986224\|23027903 | Unspecified |
| mmu-miR-34a-5p | MIMAT0000542 | Cdca7 | 66953 | Microarrays//HITS-CLIP | 37986224\|23027903\|31433990\|2314 | Unspecified |
| mmu-miR-34a-5p | MIMAT0000542 | Cdca7l | 217946 | HITS-CLIP | 23142080 | Unspecified |
| mmu-miR-34a-5p | MIMAT0000542 | Cdca8 | 52276 | Microarrays//HITS-CLIP | 37986224\|23027903\|31433990 | Unspecified |
| mmu-miR-34a-5p | MIMAT0000542 | Cdh11 | 12552 | HITS-CLIP | 19536157 | Unspecified |
| mmu-miR-34a-5p | MIMAT0000542 | Cdh2 | 12558 | Microarrays//HITS-CLIP | 37986224\|19536157 | Unspecified |
| mmu-miR-34a-5p | MIMAT0000542 | Cdip1 | 66626 | HITS-CLIP | 23142080 | Unspecified |
| mmu-miR-34a-5p | MIMAT0000542 | Cdipt | 52858 | HITS-CLIP | 29167373 | Unspecified |
| mmu-miR-34a-5p | MIMAT0000542 | Cdk1 | 12534 | HITS-CLIP | 23142080 | Unspecified |
| mmu-miR-34a-5p | MIMAT0000542 | Cdk11b | 12537 | HITS-CLIP | 23142080 | Unspecified |
| mmu-miR-34a-5p | MIMAT0000542 | Cdk12 | 69131 | HITS-CLIP | 23142080\|29167373 | Unspecified |
| mmu-miR-34a-5p | MIMAT0000542 | Cdk13 | 69562 | HITS-CLIP | 23142080\|29167373 | Unspecified |
| mmu-miR-34a-5p | MIMAT0000542 | Cdk14 | 18647 | HITS-CLIP | 29167373 | Unspecified |
| mmu-miR-34a-5p | MIMAT0000542 | Cdk16 | 18555 | HITS-CLIP | 31433990\|19536157\|23142080\|2916 | Unspecified |
| mmu-miR-34a-5p | MIMAT0000542 | Cdk17 | 237459 | HITS-CLIP | 31433990\|23142080\|29167373 | Unspecified |
| mmu-miR-34a-5p | MIMAT0000542 | Cdk18 | 18557 | HITS-CLIP | 29167373 | Unspecified |
| mmu-miR-34a-5p | MIMAT0000542 | Cdk19 | 78334 | HITS-CLIP | 19536157\|23142080\|29167373 | Unspecified |
| mmu-miR-34a-5p | MIMAT0000542 | Cdk2 | 12566 | HITS-CLIP | 23142080\|29167373 | Unspecified |
| mmu-miR-34a-5p | MIMAT0000542 | Cdk4 | 12567 | HITS-CLIP | 23142080 | Unspecified |
| mmu-miR-34a-5p | MIMAT0000542 | Cdk5r1 | 12569 | HITS-CLIP | 19536157\|23142080 | Unspecified |
| mmu-miR-34a-5p | MIMAT0000542 | Cdk5r2 | 12570 | HITS-CLIP | 19536157 | Unspecified |
| mmu-miR-34a-5p | MIMAT0000542 | Cdk5rap1 | 66971 | HITS-CLIP | 23142080 | Unspecified |
| mmu-miR-34a-5p | MIMAT0000542 | Cdk5rap3 | 80280 | HITS-CLIP | 23142080 | Unspecified |
| mmu-miR-34a-5p | MIMAT0000542 | Cdk6 | 12571 | Microarrays//HITS-CLIP | 37986224\|23142080\|23027903\|2363 | Unspecified |
| mmu-miR-34a-5p | MIMAT0000542 | Cdk8 | 264064 | HITS-CLIP | 19536157\|23142080 | Unspecified |
| mmu-miR-34a-5p | MIMAT0000542 | Cdk9 | 107951 | HITS-CLIP | 23142080 | Unspecified |
| mmu-miR-34a-5p | MIMAT0000542 | Cdkal1 | 68916 | HITS-CLIP | 23142080\|29167373 | Unspecified |

| | | | | | | |
|---|---|---|---|---|---|---|
| mmu-miR-34a-5p | MIMAT0000542 | Cdkl5 | 382253 | HITS-CLIP | 23142080 | Unspecified |
| mmu-miR-34a-5p | MIMAT0000542 | Cdkn1a | 12575 | HITS-CLIP | 37986224\|23142080\|29167373\|3143 | Unspecified |
| mmu-miR-34a-5p | MIMAT0000542 | Cdkn1b | 12576 | HITS-CLIP | 19536157\|23142080\|29167373 | Unspecified |
| mmu-miR-34a-5p | MIMAT0000542 | Cdkn2aip | 70925 | HITS-CLIP | 23142080 | Unspecified |
| mmu-miR-34a-5p | MIMAT0000542 | Cdkn2aipnl | 52626 | HITS-CLIP | 23142080 | Unspecified |
| mmu-miR-34a-5p | MIMAT0000542 | Cdkn2c | 12580 | HITS-CLIP, Microarrays//Microarrays//HITS | 37986224\|23027903\|23142080 | Unspecified |
| mmu-miR-34a-5p | MIMAT0000542 | Cdkn2d | 12581 | HITS-CLIP | 23142080 | Unspecified |
| mmu-miR-34a-5p | MIMAT0000542 | Cdon | 57810 | HITS-CLIP | 23142080 | Unspecified |
| mmu-miR-34a-5p | MIMAT0000542 | Cdr2l | 237988 | Microarrays//HITS-CLIP | 37986224\|23027903\|19536157 | Unspecified |
| mmu-miR-34a-5p | MIMAT0000542 | Cds2 | 110911 | HITS-CLIP | 37986224\|31433990\|19536157\|2363 | Unspecified |
| mmu-miR-34a-5p | MIMAT0000542 | Cdt1 | 67177 | HITS-CLIP | 23142080 | Unspecified |
| mmu-miR-34a-5p | MIMAT0000542 | Cdv3 | 321022 | HITS-CLIP | 23142080 | Unspecified |
| mmu-miR-34a-5p | MIMAT0000542 | Cdyl | 12593 | HITS-CLIP | 23142080\|31433990\|29167373 | Unspecified |
| mmu-miR-34a-5p | MIMAT0000542 | Cdyl2 | 75796 | HITS-CLIP | 23142080 | Unspecified |
| mmu-miR-34a-5p | MIMAT0000542 | Ceacam1 | 26365 | HITS-CLIP | 23142080\|29167373 | Unspecified |
| mmu-miR-34a-5p | MIMAT0000542 | Cebpa | 12606 | HITS-CLIP | 26203562 | Unspecified |
| mmu-miR-34a-5p | MIMAT0000542 | Cebpb | 12608 | HITS-CLIP | 23142080\|29167373 | Unspecified |
| mmu-miR-34a-5p | MIMAT0000542 | Cebpg | 12611 | HITS-CLIP | 23142080\|29167373 | Unspecified |
| mmu-miR-34a-5p | MIMAT0000542 | Cebpz | 12607 | HITS-CLIP | 23630078 | Unspecified |
| mmu-miR-34a-5p | MIMAT0000542 | Cecr2 | 330409 | HITS-CLIP | 19536157 | Unspecified |
| mmu-miR-34a-5p | MIMAT0000542 | Celf1 | 13046 | HITS-CLIP | 37986224\|23630078\|19536157\|3143 | Unspecified |
| mmu-miR-34a-5p | MIMAT0000542 | Celf2 | 14007 | HITS-CLIP | 23142080\|19536157\|29167373 | Unspecified |
| mmu-miR-34a-5p | MIMAT0000542 | Celf3 | 78784 | HITS-CLIP | 19536157 | Unspecified |
| mmu-miR-34a-5p | MIMAT0000542 | Celf4 | 108013 | HITS-CLIP | 19536157 | Unspecified |
| mmu-miR-34a-5p | MIMAT0000542 | Celf5 | 319586 | HITS-CLIP | 19536157 | Unspecified |
| mmu-miR-34a-5p | MIMAT0000542 | Celsr1 | 12614 | HITS-CLIP | 23142080 | Unspecified |
| mmu-miR-34a-5p | MIMAT0000542 | Celsr2 | 53883 | HITS-CLIP | 19536157 | Unspecified |
| mmu-miR-34a-5p | MIMAT0000542 | Celsr3 | 107934 | HITS-CLIP | 19536157 | Unspecified |
| mmu-miR-34a-5p | MIMAT0000542 | Cemip2 | 23670 | HITS-CLIP | 31433990\|23630078\|23142080 | Unspecified |
| mmu-miR-34a-5p | MIMAT0000542 | Cenpa | 12615 | Microarrays//HITS-CLIP | 37986224\|23142080\|23027903 | Unspecified |
| mmu-miR-34a-5p | MIMAT0000542 | Cenpb | 12616 | HITS-CLIP | 23142080 | Unspecified |
| mmu-miR-34a-5p | MIMAT0000542 | Cenpc1 | 12617 | HITS-CLIP | 30971389\|23142080 | Unspecified |
| mmu-miR-34a-5p | MIMAT0000542 | Cenpe | 229841 | Microarrays//HITS-CLIP | 37986224\|23027903\|23142080 | Unspecified |
| mmu-miR-34a-5p | MIMAT0000542 | Cenpf | 108000 | HITS-CLIP | 23142080 | Unspecified |
| mmu-miR-34a-5p | MIMAT0000542 | Cenph | 26886 | HITS-CLIP | 23142080 | Unspecified |
| mmu-miR-34a-5p | MIMAT0000542 | Cenpi | 102920 | Microarrays//HITS-CLIP | 37986224\|23027903\|23142080 | Unspecified |
| mmu-miR-34a-5p | MIMAT0000542 | Cenpj | 219103 | HITS-CLIP | 23142080 | Unspecified |
| mmu-miR-34a-5p | MIMAT0000542 | Cenpk | 60411 | Microarrays | 37986224\|23027903 | Unspecified |
| mmu-miR-34a-5p | MIMAT0000542 | Cenpl | 70454 | HITS-CLIP//Microarrays | 21258322\|23027903\|23142080 | Unspecified |

| | | | | | | |
|---|---|---|---|---|---|---|
| mmu-miR-34a-5p | MIMAT0000542 | Cenpo | 52504 | HITS-CLIP | 37986224 | Unspecified |
| mmu-miR-34a-5p | MIMAT0000542 | Cenpq | 83815 | Microarrays | 37986224\|23027903 | Unspecified |
| mmu-miR-34a-5p | MIMAT0000542 | Cenpt | 320394 | HITS-CLIP | 23142080 | Unspecified |
| mmu-miR-34a-5p | MIMAT0000542 | Cenpu | 71876 | Microarrays | 37986224\|23027903 | Unspecified |
| mmu-miR-34a-5p | MIMAT0000542 | Cep104 | 230967 | HITS-CLIP | 19536157\|23142080 | Unspecified |
| mmu-miR-34a-5p | MIMAT0000542 | Cep120 | 225523 | HITS-CLIP | 31433990\|23630078\|19536157\|23142 | Unspecified |
| mmu-miR-34a-5p | MIMAT0000542 | Cep131 | 12009 | HITS-CLIP | 23142080 | Unspecified |
| mmu-miR-34a-5p | MIMAT0000542 | Cep135 | 381644 | HITS-CLIP | 23142080 | Unspecified |
| mmu-miR-34a-5p | MIMAT0000542 | Cep162 | 382090 | HITS-CLIP | 23142080\|29167373 | Unspecified |
| mmu-miR-34a-5p | MIMAT0000542 | Cep164 | 214552 | HITS-CLIP | 29167373 | Unspecified |
| mmu-miR-34a-5p | MIMAT0000542 | Cep170 | 545389 | HITS-CLIP | 23142080\|19536157\|29167373 | Unspecified |
| mmu-miR-34a-5p | MIMAT0000542 | Cep170b | 217882 | HITS-CLIP | 23142080 | Unspecified |
| mmu-miR-34a-5p | MIMAT0000542 | Cep19 | 66994 | HITS-CLIP | 23142080 | Unspecified |
| mmu-miR-34a-5p | MIMAT0000542 | Cep192 | 70799 | HITS-CLIP | 37986224\|23142080\|29167373 | Unspecified |
| mmu-miR-34a-5p | MIMAT0000542 | Cep250 | 16328 | HITS-CLIP | 23142080\|29167373 | Unspecified |
| mmu-miR-34a-5p | MIMAT0000542 | Cep295 | 85459 | HITS-CLIP | 23142080\|29167373 | Unspecified |
| mmu-miR-34a-5p | MIMAT0000542 | Cep350 | 74081 | HITS-CLIP | 23142080\|23630078\|29167373 | Unspecified |
| mmu-miR-34a-5p | MIMAT0000542 | Cep41 | 83922 | HITS-CLIP | 23142080 | Unspecified |
| mmu-miR-34a-5p | MIMAT0000542 | Cep44 | 382010 | HITS-CLIP | 19536157 | Unspecified |
| mmu-miR-34a-5p | MIMAT0000542 | Cep55 | 74107 | Microarrays//HITS-CLIP | 37986224\|23142080\|23027903 | Unspecified |
| mmu-miR-34a-5p | MIMAT0000542 | Cep57 | 74360 | HITS-CLIP | 23142080 | Unspecified |
| mmu-miR-34a-5p | MIMAT0000542 | Cep57l1 | 103268 | HITS-CLIP | 23142080 | Unspecified |
| mmu-miR-34a-5p | MIMAT0000542 | Cep68 | 216543 | HITS-CLIP | 23142080 | Unspecified |
| mmu-miR-34a-5p | MIMAT0000542 | Cep72 | 74470 | HITS-CLIP | 23142080 | Unspecified |
| mmu-miR-34a-5p | MIMAT0000542 | Cep76 | 225659 | Microarrays | 37986224\|23027903 | Unspecified |
| mmu-miR-34a-5p | MIMAT0000542 | Cep83 | 77048 | HITS-CLIP | 29167373 | Unspecified |
| mmu-miR-34a-5p | MIMAT0000542 | Cep85 | 70012 | HITS-CLIP | 23142080 | Unspecified |
| mmu-miR-34a-5p | MIMAT0000542 | Cep85l | 100038725 | HITS-CLIP | 23142080 | Unspecified |
| mmu-miR-34a-5p | MIMAT0000542 | Cep95 | 320162 | HITS-CLIP | 23142080 | Unspecified |
| mmu-miR-34a-5p | MIMAT0000542 | Cep97 | 74201 | HITS-CLIP | 23142080 | Unspecified |
| mmu-miR-34a-5p | MIMAT0000542 | Cept1 | 99712 | HITS-CLIP | 37986224\|23142080 | Unspecified |
| mmu-miR-34a-5p | MIMAT0000542 | Cerk | 223753 | HITS-CLIP | 29167373\|23142080 | Unspecified |
| mmu-miR-34a-5p | MIMAT0000542 | Cers2 | 76893 | Microarrays//HITS-CLIP | 37986224\|23027903\|23142080\|29167 | Unspecified |
| mmu-miR-34a-5p | MIMAT0000542 | Cers4 | 67260 | HITS-CLIP | 23142080 | Unspecified |
| mmu-miR-34a-5p | MIMAT0000542 | Cers5 | 71949 | HITS-CLIP | 23630078\|23142080\|29167373 | Unspecified |
| mmu-miR-34a-5p | MIMAT0000542 | Cers6 | 241447 | HITS-CLIP | 19536157\|29167373\|23142080 | Unspecified |
| mmu-miR-34a-5p | MIMAT0000542 | Cert1 | 10087 | HITS-CLIP | 30971389 | Unspecified |
| mmu-miR-34a-5p | MIMAT0000542 | Cetn3 | 12626 | HITS-CLIP | 23142080 | Unspecified |
| mmu-miR-34a-5p | MIMAT0000542 | Cfap20 | 29105 | HITS-CLIP | 23142080 | Unspecified |

| | | | | | | |
|---|---|---|---|---|---|---|
| mmu-miR-34a-5p | MIMAT0000542 | Cfap300 | 85016 | HITS-CLIP | 23630078 | Unspecified |
| mmu-miR-34a-5p | MIMAT0000542 | Cfap36 | 112942 | HITS-CLIP | 23142080 | Unspecified |
| mmu-miR-34a-5p | MIMAT0000542 | Cfap97 | 57587 | HITS-CLIP | 19536157\|23142080\|29167373 | Unspecified |
| mmu-miR-34a-5p | MIMAT0000542 | Cfdp1 | 23837 | HITS-CLIP | 23142080 | Unspecified |
| mmu-miR-34a-5p | MIMAT0000542 | Cfl1 | 12631 | HITS-CLIP | 37986224\|23142080\|29167373\|19536 | Unspecified |
| mmu-miR-34a-5p | MIMAT0000542 | Cfl2 | 12632 | HITS-CLIP | 31433990\|19536157\|23142080\|29167 | Unspecified |
| mmu-miR-34a-5p | MIMAT0000542 | Cflar | 12633 | HITS-CLIP | 31433990\|23142080\|26203562\|29167 | Unspecified |
| mmu-miR-34a-5p | MIMAT0000542 | Cggbp1 | 106143 | HITS-CLIP | 23142080\|29167373 | Unspecified |
| mmu-miR-34a-5p | MIMAT0000542 | Ch25h | 12642 | HITS-CLIP | 29167373 | Unspecified |
| mmu-miR-34a-5p | MIMAT0000542 | Chac1 | 69065 | HITS-CLIP | 23630078\|23142080 | Unspecified |
| mmu-miR-34a-5p | MIMAT0000542 | Chaf1a | 27221 | HITS-CLIP | 23142080\|31433990\|29167373 | Unspecified |
| mmu-miR-34a-5p | MIMAT0000542 | Chaf1b | 110749 | Microarrays | 37986224\|23027903 | Unspecified |
| mmu-miR-34a-5p | MIMAT0000542 | Champ1 | 101994 | HITS-CLIP | 19536157\|29167373\|23142080 | Unspecified |
| mmu-miR-34a-5p | MIMAT0000542 | Chchd3 | 66075 | HITS-CLIP | 29167373 | Unspecified |
| mmu-miR-34a-5p | MIMAT0000542 | Chchd4 | 72170 | HITS-CLIP | 23142080 | Unspecified |
| mmu-miR-34a-5p | MIMAT0000542 | Chchd5 | 66170 | HITS-CLIP | 23142080 | Unspecified |
| mmu-miR-34a-5p | MIMAT0000542 | Chd1 | 12648 | HITS-CLIP | 37986224\|23142080\|19536157\|31433 | Unspecified |
| mmu-miR-34a-5p | MIMAT0000542 | Chd1l | 68058 | HITS-CLIP | 23142080 | Unspecified |
| mmu-miR-34a-5p | MIMAT0000542 | Chd2 | 244059 | HITS-CLIP | 23142080\|31433990\|29167373 | Unspecified |
| mmu-miR-34a-5p | MIMAT0000542 | Chd3 | 216848 | HITS-CLIP | 19536157\|23142080 | Unspecified |
| mmu-miR-34a-5p | MIMAT0000542 | Chd4 | 107932 | HITS-CLIP | 31433990\|19536157\|23142080\|29167 | Unspecified |
| mmu-miR-34a-5p | MIMAT0000542 | Chd7 | 320790 | HITS-CLIP | 37986224\|31433990\|19536157\|23142 | Unspecified |
| mmu-miR-34a-5p | MIMAT0000542 | Chd8 | 67772 | HITS-CLIP | 23142080\|19536157\|29167373 | Unspecified |
| mmu-miR-34a-5p | MIMAT0000542 | Chd9 | 109151 | HITS-CLIP | 19536157\|23142080\|29167373 | Unspecified |
| mmu-miR-34a-5p | MIMAT0000542 | Chek1 | 12649 | Microarrays | 37986224\|23027903 | Unspecified |
| mmu-miR-34a-5p | MIMAT0000542 | Chek2 | 50883 | HITS-CLIP | 23142080 | Unspecified |
| mmu-miR-34a-5p | MIMAT0000542 | Cherp | 27967 | HITS-CLIP | 23142080\|29167373 | Unspecified |
| mmu-miR-34a-5p | MIMAT0000542 | Chfr | 231600 | HITS-CLIP | 19536157\|23142080\|29167373 | Unspecified |
| mmu-miR-34a-5p | MIMAT0000542 | Chgb | 12653 | HITS-CLIP | 19536157 | Unspecified |
| mmu-miR-34a-5p | MIMAT0000542 | Chic1 | 12212 | Microarrays//HITS-CLIP | 37986224\|23142080 | Unspecified |
| mmu-miR-34a-5p | MIMAT0000542 | Chic2 | 74277 | Microarrays//HITS-CLIP | 37986224\|23027903\|23142080 | Unspecified |
| mmu-miR-34a-5p | MIMAT0000542 | Chid1 | 68038 | HITS-CLIP | 23142080\|29167373 | Unspecified |
| mmu-miR-34a-5p | MIMAT0000542 | Chka | 12660 | HITS-CLIP | 23142080 | Unspecified |
| mmu-miR-34a-5p | MIMAT0000542 | Chkb | 12651 | HITS-CLIP | 23142080 | Unspecified |
| mmu-miR-34a-5p | MIMAT0000542 | Chl1 | 12661 | HITS-CLIP | 19536157 | Unspecified |
| mmu-miR-34a-5p | MIMAT0000542 | Chm | 12662 | HITS-CLIP | 23142080 | Unspecified |
| mmu-miR-34a-5p | MIMAT0000542 | Chml | 12663 | HITS-CLIP | 31433990\|23142080 | Unspecified |
| mmu-miR-34a-5p | MIMAT0000542 | Chmp1b | 67078 | HITS-CLIP | 23142080 | Unspecified |
| mmu-miR-34a-5p | MIMAT0000542 | Chmp2b | 68942 | HITS-CLIP | 23142080\|29167373 | Unspecified |

| | | | | | | | |
|---|---|---|---|---|---|---|---|
| mmu-miR-34a-5p | MIMAT0000542 | Chmp3 | 66700 | HITS-CLIP | | 23142080 | Unspecified |
| mmu-miR-34a-5p | MIMAT0000542 | Chmp4b | 75608 | HITS-CLIP | 23142080\|29167373 | | Unspecified |
| mmu-miR-34a-5p | MIMAT0000542 | Chmp5 | 76959 | HITS-CLIP | 23630078\|23142080\|29167373 | | Unspecified |
| mmu-miR-34a-5p | MIMAT0000542 | Chmp6 | 208092 | HITS-CLIP | | 26203562 | Unspecified |
| mmu-miR-34a-5p | MIMAT0000542 | Chmp7 | 105513 | HITS-CLIP | | 23142080 | Unspecified |
| mmu-miR-34a-5p | MIMAT0000542 | Chn1 | 108699 | HITS-CLIP | | 19536157 | Unspecified |
| mmu-miR-34a-5p | MIMAT0000542 | Chn2 | 69993 | Microarrays | 37986224\|23027903 | | Unspecified |
| mmu-miR-34a-5p | MIMAT0000542 | Chordc1 | 66917 | HITS-CLIP | | 23142080 | Unspecified |
| mmu-miR-34a-5p | MIMAT0000542 | Chp1 | 56398 | HITS-CLIP | 19536157\|23142080 | | Unspecified |
| mmu-miR-34a-5p | MIMAT0000542 | Chpf | 74241 | HITS-CLIP | | 23142080 | Unspecified |
| mmu-miR-34a-5p | MIMAT0000542 | Chpt1 | 212862 | HITS-CLIP | 19536157\|23142080 | | Unspecified |
| mmu-miR-34a-5p | MIMAT0000542 | Chrm4 | 12672 | HITS-CLIP | | 23142080 | Unspecified |
| mmu-miR-34a-5p | MIMAT0000542 | Chrna6 | 8973 | HITS-CLIP | | 23142080 | Unspecified |
| mmu-miR-34a-5p | MIMAT0000542 | Chrnd | 11447 | Microarrays | | 37986224 | Unspecified |
| mmu-miR-34a-5p | MIMAT0000542 | Chrng | 11449 | Microarrays | | 37986224 | Unspecified |
| mmu-miR-34a-5p | MIMAT0000542 | Chst12 | 59031 | HITS-CLIP | | 29167373 | Unspecified |
| mmu-miR-34a-5p | MIMAT0000542 | Chst14 | 72136 | HITS-CLIP | 31433990\|26203562\|23142080\|29167 | | Unspecified |
| mmu-miR-34a-5p | MIMAT0000542 | Chst15 | 77590 | Microarrays//HITS-CLIP | 37986224\|23142080 | | Unspecified |
| mmu-miR-34a-5p | MIMAT0000542 | Chst2 | 54371 | HITS-CLIP | | 23142080 | Unspecified |
| mmu-miR-34a-5p | MIMAT0000542 | Chsy1 | 269941 | HITS-CLIP | 23142080\|29167373 | | Unspecified |
| mmu-miR-34a-5p | MIMAT0000542 | Chtf8 | 214987 | HITS-CLIP | | 26203562 | Unspecified |
| mmu-miR-34a-5p | MIMAT0000542 | Chtop | 66511 | HITS-CLIP | | 23142080 | Unspecified |
| mmu-miR-34a-5p | MIMAT0000542 | Chuk | 12675 | HITS-CLIP | 31433990\|23142080\|29167373 | | Unspecified |
| mmu-miR-34a-5p | MIMAT0000542 | Ciao1 | 26371 | HITS-CLIP | | 23142080 | Unspecified |
| mmu-miR-34a-5p | MIMAT0000542 | Ciao2a | 84191 | HITS-CLIP | 30971389\|23142080 | | Unspecified |
| mmu-miR-34a-5p | MIMAT0000542 | Ciao2b | 51647 | HITS-CLIP | | 23142080 | Unspecified |
| mmu-miR-34a-5p | MIMAT0000542 | Ciapin1 | 109006 | HITS-CLIP | | 23142080 | Unspecified |
| mmu-miR-34a-5p | MIMAT0000542 | Ciart | 229599 | HITS-CLIP | | 23142080 | Unspecified |
| mmu-miR-34a-5p | MIMAT0000542 | Cib1 | 23991 | HITS-CLIP | | 23142080 | Unspecified |
| mmu-miR-34a-5p | MIMAT0000542 | Cic | 71722 | HITS-CLIP | 23142080\|19536157\|29167373 | | Unspecified |
| mmu-miR-34a-5p | MIMAT0000542 | Ciita | 12265 | HITS-CLIP | | 23142080 | Unspecified |
| mmu-miR-34a-5p | MIMAT0000542 | Cinp | 67236 | Microarrays//HITS-CLIP | 37986224\|23027903\|31433990\|23142 | | Unspecified |
| mmu-miR-34a-5p | MIMAT0000542 | Cip2a | 57650 | HITS-CLIP | 31433990\|23142080 | | Unspecified |
| mmu-miR-34a-5p | MIMAT0000542 | Cipc | 217732 | HITS-CLIP | 23142080\|29167373 | | Unspecified |
| mmu-miR-34a-5p | MIMAT0000542 | Cisd1 | 52637 | HITS-CLIP | | 23142080 | Unspecified |
| mmu-miR-34a-5p | MIMAT0000542 | Cisd2 | 67006 | HITS-CLIP | 29167373\|23142080 | | Unspecified |
| mmu-miR-34a-5p | MIMAT0000542 | Cish | 12700 | HITS-CLIP | | 23142080 | Unspecified |
| mmu-miR-34a-5p | MIMAT0000542 | Cit | 12704 | HITS-CLIP | 31433990\|23142080 | | Unspecified |
| mmu-miR-34a-5p | MIMAT0000542 | Cited2 | 17684 | HITS-CLIP | 37986224\|23142080\|29167373\|3143 | | Unspecified |

| | | | | | | |
|---|---|---|---|---|---|---|
| mmu-miR-34a-5p | MIMAT0000542 | Ciz1 | 68379 | HITS-CLIP | 19536157 | Unspecified |
| mmu-miR-34a-5p | MIMAT0000542 | Ckap2 | 80986 | HITS-CLIP | 23142080 | Unspecified |
| mmu-miR-34a-5p | MIMAT0000542 | Ckap2l | 70466 | HITS-CLIP | 23142080 | Unspecified |
| mmu-miR-34a-5p | MIMAT0000542 | Ckap5 | 75786 | HITS-CLIP | 30971389\|19536157\|23142080\|2916 | Unspecified |
| mmu-miR-34a-5p | MIMAT0000542 | Ckb | 12709 | HITS-CLIP | 23142080\|29167373 | Unspecified |
| mmu-miR-34a-5p | MIMAT0000542 | Cklf | 75458 | HITS-CLIP | 29167373 | Unspecified |
| mmu-miR-34a-5p | MIMAT0000542 | Ckm | 12715 | Microarrays | 37986224 | Unspecified |
| mmu-miR-34a-5p | MIMAT0000542 | Cks1b | 54124 | HITS-CLIP | 23142080 | Unspecified |
| mmu-miR-34a-5p | MIMAT0000542 | Clasp1 | 76707 | HITS-CLIP | 23142080\|19536157\|29167373 | Unspecified |
| mmu-miR-34a-5p | MIMAT0000542 | Clasp2 | 76499 | HITS-CLIP | 19536157\|23142080\|29167373 | Unspecified |
| mmu-miR-34a-5p | MIMAT0000542 | Clcc1 | 229725 | HITS-CLIP | 23142080\|29167373 | Unspecified |
| mmu-miR-34a-5p | MIMAT0000542 | Clcf1 | 56708 | HITS-CLIP | 23142080 | Unspecified |
| mmu-miR-34a-5p | MIMAT0000542 | Clcn3 | 12725 | HITS-CLIP | 30971389\|19536157\|31433990\|2314 | Unspecified |
| mmu-miR-34a-5p | MIMAT0000542 | Clcn4 | 12727 | HITS-CLIP | 23142080 | Unspecified |
| mmu-miR-34a-5p | MIMAT0000542 | Clcn5 | 12728 | HITS-CLIP | 19536157\|23142080\|29167373 | Unspecified |
| mmu-miR-34a-5p | MIMAT0000542 | Clcn7 | 26373 | HITS-CLIP | 29167373 | Unspecified |
| mmu-miR-34a-5p | MIMAT0000542 | Cldn10 | 58187 | HITS-CLIP | 23142080 | Unspecified |
| mmu-miR-34a-5p | MIMAT0000542 | Cldn4 | 12740 | HITS-CLIP | 26203562 | Unspecified |
| mmu-miR-34a-5p | MIMAT0000542 | Cldnd1 | 224250 | HITS-CLIP | 23142080\|26203562 | Unspecified |
| mmu-miR-34a-5p | MIMAT0000542 | Clec12a | 160364 | HITS-CLIP | 29167373 | Unspecified |
| mmu-miR-34a-5p | MIMAT0000542 | Clec16a | 74374 | HITS-CLIP | 31433990\|23142080 | Unspecified |
| mmu-miR-34a-5p | MIMAT0000542 | Clec2d | 93694 | HITS-CLIP | 23142080\|29167373 | Unspecified |
| mmu-miR-34a-5p | MIMAT0000542 | Clec2i | 93675 | HITS-CLIP | 23142080 | Unspecified |
| mmu-miR-34a-5p | MIMAT0000542 | Clec4d | 17474 | HITS-CLIP | 29167373 | Unspecified |
| mmu-miR-34a-5p | MIMAT0000542 | Clec4e | 56619 | HITS-CLIP | 29167373 | Unspecified |
| mmu-miR-34a-5p | MIMAT0000542 | Clic1 | 114584 | HITS-CLIP | 31433990\|23142080\|29167373 | Unspecified |
| mmu-miR-34a-5p | MIMAT0000542 | Clic4 | 29876 | HITS-CLIP | 19536157\|29167373\|23142080 | Unspecified |
| mmu-miR-34a-5p | MIMAT0000542 | Clint1 | 216705 | HITS-CLIP | 23630078\|23142080\|29167373 | Unspecified |
| mmu-miR-34a-5p | MIMAT0000542 | Clip1 | 56430 | HITS-CLIP | 31433990\|19536157\|23142080\|2916 | Unspecified |
| mmu-miR-34a-5p | MIMAT0000542 | Clip3 | 76686 | HITS-CLIP | 19536157 | Unspecified |
| mmu-miR-34a-5p | MIMAT0000542 | Clk1 | 12747 | HITS-CLIP | 23142080\|29167373 | Unspecified |
| mmu-miR-34a-5p | MIMAT0000542 | Clk2 | 12748 | HITS-CLIP | 23142080 | Unspecified |
| mmu-miR-34a-5p | MIMAT0000542 | Clk3 | 102414 | HITS-CLIP | 23142080 | Unspecified |
| mmu-miR-34a-5p | MIMAT0000542 | Clk4 | 12750 | HITS-CLIP | 23142080\|29167373 | Unspecified |
| mmu-miR-34a-5p | MIMAT0000542 | Clmp | 71566 | HITS-CLIP | 19536157 | Unspecified |
| mmu-miR-34a-5p | MIMAT0000542 | Cln3 | 12752 | HITS-CLIP | 29167373 | Unspecified |
| mmu-miR-34a-5p | MIMAT0000542 | Cln5 | 211286 | HITS-CLIP | 23142080\|29167373 | Unspecified |
| mmu-miR-34a-5p | MIMAT0000542 | Cln6 | 76524 | HITS-CLIP | 29167373\|23142080 | Unspecified |
| mmu-miR-34a-5p | MIMAT0000542 | Cln8 | 26889 | HITS-CLIP | 30971389\|19536157\|29167373 | Unspecified |

| | | | | | | |
|---|---|---|---|---|---|---|
| mmu-miR-34a-5p | MIMAT0000542 | Clock | 12753 | HITS-CLIP | 23142080 | Unspecified |
| mmu-miR-34a-5p | MIMAT0000542 | Clpb | 20480 | HITS-CLIP | 31433990\|29167373 | Unspecified |
| mmu-miR-34a-5p | MIMAT0000542 | Clpp | 53895 | HITS-CLIP | 23142080 | Unspecified |
| mmu-miR-34a-5p | MIMAT0000542 | Clspn | 269582 | Microarrays//HITS-CLIP | 37986224\|23027903\|23142080 | Unspecified |
| mmu-miR-34a-5p | MIMAT0000542 | Clstn1 | 65945 | HITS-CLIP | 19536157 | Unspecified |
| mmu-miR-34a-5p | MIMAT0000542 | Clstn3 | 232370 | HITS-CLIP | 19536157 | Unspecified |
| mmu-miR-34a-5p | MIMAT0000542 | Clta | 12757 | HITS-CLIP | 31433990\|23142080 | Unspecified |
| mmu-miR-34a-5p | MIMAT0000542 | Cltc | 67300 | HITS-CLIP | 37986224\|31433990\|19536157\|2363( | Unspecified |
| mmu-miR-34a-5p | MIMAT0000542 | Clu | 12759 | HITS-CLIP | 19536157 | Unspecified |
| mmu-miR-34a-5p | MIMAT0000542 | Cluap1 | 76779 | HITS-CLIP | 19536157 | Unspecified |
| mmu-miR-34a-5p | MIMAT0000542 | Cluh | 74148 | HITS-CLIP | 23630078 | Unspecified |
| mmu-miR-34a-5p | MIMAT0000542 | Clybl | 69634 | HITS-CLIP | 23142080 | Unspecified |
| mmu-miR-34a-5p | MIMAT0000542 | Cmah | 12763 | Microarrays//HITS-CLIP | 37986224\|23142080 | Unspecified |
| mmu-miR-34a-5p | MIMAT0000542 | Cmas | 12764 | HITS-CLIP | 31433990\|23142080 | Unspecified |
| mmu-miR-34a-5p | MIMAT0000542 | Cmc2 | 66531 | HITS-CLIP | 23142080 | Unspecified |
| mmu-miR-34a-5p | MIMAT0000542 | Cmc4 | 100272147 | HITS-CLIP | 23142080 | Unspecified |
| mmu-miR-34a-5p | MIMAT0000542 | Cmip | 74440 | HITS-CLIP | 23142080\|29167373 | Unspecified |
| mmu-miR-34a-5p | MIMAT0000542 | Cmklr1 | 14747 | HITS-CLIP | 29167373 | Unspecified |
| mmu-miR-34a-5p | MIMAT0000542 | Cmpk1 | 66588 | HITS-CLIP | 23142080\|29167373 | Unspecified |
| mmu-miR-34a-5p | MIMAT0000542 | Cmtm3 | 68119 | Microarrays//HITS-CLIP | 37986224\|29167373\|23142080 | Unspecified |
| mmu-miR-34a-5p | MIMAT0000542 | Cmtm4 | 97487 | HITS-CLIP | 23142080\|31433990\|19536157\|2916 | Unspecified |
| mmu-miR-34a-5p | MIMAT0000542 | Cmtm6 | 67213 | Microarrays//HITS-CLIP | 37986224\|23027903\|31433990\|2314 | Unspecified |
| mmu-miR-34a-5p | MIMAT0000542 | Cmtm7 | 102545 | HITS-CLIP | 23142080 | Unspecified |
| mmu-miR-34a-5p | MIMAT0000542 | Cmtr1 | 74157 | HITS-CLIP | 23142080\|19536157\|29167373 | Unspecified |
| mmu-miR-34a-5p | MIMAT0000542 | Cmtr2 | 234728 | HITS-CLIP | 23142080 | Unspecified |
| mmu-miR-34a-5p | MIMAT0000542 | Cnbp | 12785 | HITS-CLIP | 23142080 | Unspecified |
| mmu-miR-34a-5p | MIMAT0000542 | Cnep1r1 | 382030 | HITS-CLIP | 23142080\|29167373 | Unspecified |
| mmu-miR-34a-5p | MIMAT0000542 | Cnih1 | 12793 | HITS-CLIP | 23142080 | Unspecified |
| mmu-miR-34a-5p | MIMAT0000542 | Cnih2 | 12794 | HITS-CLIP | 23142080 | Unspecified |
| mmu-miR-34a-5p | MIMAT0000542 | Cnih4 | 98417 | HITS-CLIP | 23142080\|29167373 | Unspecified |
| mmu-miR-34a-5p | MIMAT0000542 | Cnksr2 | 245684 | HITS-CLIP | 19536157 | Unspecified |
| mmu-miR-34a-5p | MIMAT0000542 | Cnn2 | 12798 | HITS-CLIP | 23142080 | Unspecified |
| mmu-miR-34a-5p | MIMAT0000542 | Cnn3 | 71994 | HITS-CLIP | 31433990 | Unspecified |
| mmu-miR-34a-5p | MIMAT0000542 | Cnnm3 | 94218 | HITS-CLIP | 21258322\|26203562\|29167373 | Unspecified |
| mmu-miR-34a-5p | MIMAT0000542 | Cnnm4 | 94220 | HITS-CLIP | 23142080 | Unspecified |
| mmu-miR-34a-5p | MIMAT0000542 | Cnot1 | 234594 | HITS-CLIP | 30971389\|23142080\|29167373 | Unspecified |
| mmu-miR-34a-5p | MIMAT0000542 | Cnot10 | 78893 | HITS-CLIP | 23142080 | Unspecified |
| mmu-miR-34a-5p | MIMAT0000542 | Cnot11 | 52846 | HITS-CLIP | 23142080\|19536157 | Unspecified |
| mmu-miR-34a-5p | MIMAT0000542 | Cnot2 | 72068 | HITS-CLIP | 23142080\|29167373 | Unspecified |

| | | | | | | | |
|---|---|---|---|---|---|---|---|
| mmu-miR-34a-5p | MIMAT0000542 | Cnot3 | 232791 | HITS-CLIP | | 23142080 | Unspecified |
| mmu-miR-34a-5p | MIMAT0000542 | Cnot4 | 53621 | HITS-CLIP | 23142080\|29167373 | | Unspecified |
| mmu-miR-34a-5p | MIMAT0000542 | Cnot6 | 104625 | HITS-CLIP | 23142080\|23630078\|29167373 | | Unspecified |
| mmu-miR-34a-5p | MIMAT0000542 | Cnot6l | 231464 | HITS-CLIP | 37986224\|23142080\|23630078\|31433 | | Unspecified |
| mmu-miR-34a-5p | MIMAT0000542 | Cnot7 | 18983 | HITS-CLIP | 23142080\|31433990 | | Unspecified |
| mmu-miR-34a-5p | MIMAT0000542 | Cnot8 | 69125 | HITS-CLIP | | 23142080 | Unspecified |
| mmu-miR-34a-5p | MIMAT0000542 | Cnot9 | 9125 | HITS-CLIP | | 23142080 | Unspecified |
| mmu-miR-34a-5p | MIMAT0000542 | Cnp | 12799 | HITS-CLIP | 23142080\|29167373 | | Unspecified |
| mmu-miR-34a-5p | MIMAT0000542 | Cnppd1 | 69171 | HITS-CLIP | 23142080\|29167373 | | Unspecified |
| mmu-miR-34a-5p | MIMAT0000542 | Cnpy2 | 56530 | HITS-CLIP | | 23142080 | Unspecified |
| mmu-miR-34a-5p | MIMAT0000542 | Cnr1 | 12801 | HITS-CLIP | | 19536157 | Unspecified |
| mmu-miR-34a-5p | MIMAT0000542 | Cnr2 | 12802 | HITS-CLIP | | 23142080 | Unspecified |
| mmu-miR-34a-5p | MIMAT0000542 | Cnst | 226744 | HITS-CLIP | 30971389\|23142080 | | Unspecified |
| mmu-miR-34a-5p | MIMAT0000542 | Cntf | 12803 | HITS-CLIP | | 23142080 | Unspecified |
| mmu-miR-34a-5p | MIMAT0000542 | Cntfr | 12804 | HITS-CLIP | | 19536157 | Unspecified |
| mmu-miR-34a-5p | MIMAT0000542 | Cntn1 | 12805 | HITS-CLIP | | 19536157 | Unspecified |
| mmu-miR-34a-5p | MIMAT0000542 | Cntnap4 | 170571 | HITS-CLIP | | 19536157 | Unspecified |
| mmu-miR-34a-5p | MIMAT0000542 | Cntrl | 26920 | HITS-CLIP | 19536157\|23142080\|29167373 | | Unspecified |
| mmu-miR-34a-5p | MIMAT0000542 | Coa3 | 52469 | HITS-CLIP | | 23142080 | Unspecified |
| mmu-miR-34a-5p | MIMAT0000542 | Coa5 | 76178 | Microarrays//HITS-CLIP | 37986224\|23027903\|23142080\|29167 | | Unspecified |
| mmu-miR-34a-5p | MIMAT0000542 | Coasy | 71743 | HITS-CLIP | | 23142080 | Unspecified |
| mmu-miR-34a-5p | MIMAT0000542 | Cobl | 12808 | HITS-CLIP | | 19536157 | Unspecified |
| mmu-miR-34a-5p | MIMAT0000542 | Cobll1 | 319876 | HITS-CLIP | | 23142080 | Unspecified |
| mmu-miR-34a-5p | MIMAT0000542 | Cog2 | 76332 | HITS-CLIP | | 23142080 | Unspecified |
| mmu-miR-34a-5p | MIMAT0000542 | Cog3 | 338337 | HITS-CLIP | | 23142080 | Unspecified |
| mmu-miR-34a-5p | MIMAT0000542 | Cog4 | 102339 | HITS-CLIP | | 23142080 | Unspecified |
| mmu-miR-34a-5p | MIMAT0000542 | Cog5 | 238123 | HITS-CLIP | 23142080\|29167373 | | Unspecified |
| mmu-miR-34a-5p | MIMAT0000542 | Cog6 | 67542 | HITS-CLIP | | 23142080 | Unspecified |
| mmu-miR-34a-5p | MIMAT0000542 | Cog7 | 233824 | HITS-CLIP | 23142080\|29167373 | | Unspecified |
| mmu-miR-34a-5p | MIMAT0000542 | Cog8 | 97484 | HITS-CLIP | | 23142080 | Unspecified |
| mmu-miR-34a-5p | MIMAT0000542 | Coil | 12812 | HITS-CLIP | | 23142080 | Unspecified |
| mmu-miR-34a-5p | MIMAT0000542 | Col18a1 | 12822 | HITS-CLIP | | 29167373 | Unspecified |
| mmu-miR-34a-5p | MIMAT0000542 | Col1a1 | 12842 | HITS-CLIP | | 23630078 | Unspecified |
| mmu-miR-34a-5p | MIMAT0000542 | Col1a2 | 12843 | HITS-CLIP | | 19536157 | Unspecified |
| mmu-miR-34a-5p | MIMAT0000542 | Col4a1 | 12826 | HITS-CLIP | | 19536157 | Unspecified |
| mmu-miR-34a-5p | MIMAT0000542 | Col4a2 | 12827 | HITS-CLIP | | 19536157 | Unspecified |
| mmu-miR-34a-5p | MIMAT0000542 | Col4a3bp | 68018 | HITS-CLIP | 23142080\|29167373 | | Unspecified |
| mmu-miR-34a-5p | MIMAT0000542 | Col5a1 | 12831 | HITS-CLIP | 19536157\|29167373 | | Unspecified |
| mmu-miR-34a-5p | MIMAT0000542 | Colgalt1 | 234407 | HITS-CLIP | | 29167373 | Unspecified |

| | | | | | | | |
|---|---|---|---|---|---|---|---|
| mmu-miR-34a-5p | MIMAT0000542 | Commd1 | 17846 | HITS-CLIP | | 23142080 | Unspecified |
| mmu-miR-34a-5p | MIMAT0000542 | Commd10 | 69456 | HITS-CLIP | 23142080\|29167373 | | Unspecified |
| mmu-miR-34a-5p | MIMAT0000542 | Commd1b | | HITS-CLIP | | 31433990 | Unspecified |
| mmu-miR-34a-5p | MIMAT0000542 | Commd2 | 52245 | HITS-CLIP | | 23142080 | Unspecified |
| mmu-miR-34a-5p | MIMAT0000542 | Commd3 | 12238 | HITS-CLIP | 23142080\|29167373 | | Unspecified |
| mmu-miR-34a-5p | MIMAT0000542 | Commd4 | 66199 | HITS-CLIP | | 23142080 | Unspecified |
| mmu-miR-34a-5p | MIMAT0000542 | Commd8 | 27784 | HITS-CLIP | | 29167373 | Unspecified |
| mmu-miR-34a-5p | MIMAT0000542 | Commd9 | 76501 | HITS-CLIP | | 29167373 | Unspecified |
| mmu-miR-34a-5p | MIMAT0000542 | Comt | 12846 | HITS-CLIP | 23142080\|19536157\|31433990 | | Unspecified |
| mmu-miR-34a-5p | MIMAT0000542 | Copa | 12847 | HITS-CLIP | 23142080\|29167373 | | Unspecified |
| mmu-miR-34a-5p | MIMAT0000542 | Copb1 | 70349 | HITS-CLIP | 23142080\|29167373 | | Unspecified |
| mmu-miR-34a-5p | MIMAT0000542 | Copb2 | 50797 | HITS-CLIP | 19536157\|23142080\|29167373 | | Unspecified |
| mmu-miR-34a-5p | MIMAT0000542 | Cope | 59042 | HITS-CLIP | 23142080\|29167373 | | Unspecified |
| mmu-miR-34a-5p | MIMAT0000542 | Copg1 | 54161 | HITS-CLIP | 23142080\|29167373 | | Unspecified |
| mmu-miR-34a-5p | MIMAT0000542 | Copg2 | 54160 | HITS-CLIP | | 23142080 | Unspecified |
| mmu-miR-34a-5p | MIMAT0000542 | Cops2 | 12848 | HITS-CLIP | 23142080\|29167373 | | Unspecified |
| mmu-miR-34a-5p | MIMAT0000542 | Cops3 | 26572 | HITS-CLIP | 19536157\|31433990\|23142080 | | Unspecified |
| mmu-miR-34a-5p | MIMAT0000542 | Cops4 | 26891 | HITS-CLIP | 23142080\|29167373 | | Unspecified |
| mmu-miR-34a-5p | MIMAT0000542 | Cops5 | 26754 | HITS-CLIP | | 23142080 | Unspecified |
| mmu-miR-34a-5p | MIMAT0000542 | Cops6 | 26893 | HITS-CLIP | | 23142080 | Unspecified |
| mmu-miR-34a-5p | MIMAT0000542 | Cops7a | 26894 | HITS-CLIP | 30971389\|23142080\|29167373 | | Unspecified |
| mmu-miR-34a-5p | MIMAT0000542 | Cops9 | 150678 | HITS-CLIP | | 29167373 | Unspecified |
| mmu-miR-34a-5p | MIMAT0000542 | Copz1 | 56447 | HITS-CLIP | | 23142080 | Unspecified |
| mmu-miR-34a-5p | MIMAT0000542 | Coq10b | 67876 | HITS-CLIP | | 23142080 | Unspecified |
| mmu-miR-34a-5p | MIMAT0000542 | Coq2 | 71883 | HITS-CLIP | | 29167373 | Unspecified |
| mmu-miR-34a-5p | MIMAT0000542 | Coq9 | 67914 | HITS-CLIP | | 23142080 | Unspecified |
| mmu-miR-34a-5p | MIMAT0000542 | Coro1a | 12721 | HITS-CLIP | 37986224\|19536157\|23142080\|29167 | | Unspecified |
| mmu-miR-34a-5p | MIMAT0000542 | Coro1b | 23789 | HITS-CLIP | | 23142080 | Unspecified |
| mmu-miR-34a-5p | MIMAT0000542 | Coro1c | 23790 | HITS-CLIP | 23142080\|29167373 | | Unspecified |
| mmu-miR-34a-5p | MIMAT0000542 | Coro2a | 107684 | HITS-CLIP | | 23142080 | Unspecified |
| mmu-miR-34a-5p | MIMAT0000542 | Coro2b | 235431 | HITS-CLIP | | 19536157 | Unspecified |
| mmu-miR-34a-5p | MIMAT0000542 | Coro6 | 216961 | Microarrays | | 37986224 | Unspecified |
| mmu-miR-34a-5p | MIMAT0000542 | Coro7 | 78885 | HITS-CLIP | 23142080\|29167373 | | Unspecified |
| mmu-miR-34a-5p | MIMAT0000542 | Cotl1 | 72042 | HITS-CLIP | 29167373\|23142080 | | Unspecified |
| mmu-miR-34a-5p | MIMAT0000542 | Cox10 | 70383 | HITS-CLIP | | 23142080 | Unspecified |
| mmu-miR-34a-5p | MIMAT0000542 | Cox11 | 69802 | HITS-CLIP | | 31433990 | Unspecified |
| mmu-miR-34a-5p | MIMAT0000542 | Cox14 | 66379 | HITS-CLIP | 23630078\|31433990\|23142080 | | Unspecified |
| mmu-miR-34a-5p | MIMAT0000542 | Cox15 | 226139 | HITS-CLIP | | 29167373 | Unspecified |
| mmu-miR-34a-5p | MIMAT0000542 | Cox16 | 51241 | HITS-CLIP | | 23142080 | Unspecified |

| | | | | | | |
|---|---|---|---|---|---|---|
| mmu-miR-34a-5p | MIMAT0000542 | Cox17 | 12856 | HITS-CLIP | 23142080 | Unspecified |
| mmu-miR-34a-5p | MIMAT0000542 | Cox4i1 | 12857 | HITS-CLIP | 23142080\|23630078 | Unspecified |
| mmu-miR-34a-5p | MIMAT0000542 | Cox5a | 12858 | HITS-CLIP | 23142080\|29167373 | Unspecified |
| mmu-miR-34a-5p | MIMAT0000542 | Cox6a1 | 12861 | HITS-CLIP | 23142080 | Unspecified |
| mmu-miR-34a-5p | MIMAT0000542 | Cox6b1 | 110323 | HITS-CLIP | 19536157\|23142080\|29167373 | Unspecified |
| mmu-miR-34a-5p | MIMAT0000542 | Cox7a2 | 12866 | HITS-CLIP | 23142080 | Unspecified |
| mmu-miR-34a-5p | MIMAT0000542 | Cox7a2l | 20463 | HITS-CLIP | 23142080 | Unspecified |
| mmu-miR-34a-5p | MIMAT0000542 | Cox7b | 66142 | HITS-CLIP | 23142080 | Unspecified |
| mmu-miR-34a-5p | MIMAT0000542 | Cox8a | 12868 | HITS-CLIP | 23142080 | Unspecified |
| mmu-miR-34a-5p | MIMAT0000542 | Cox8b | 12869 | Microarrays | 37986224 | Unspecified |
| mmu-miR-34a-5p | MIMAT0000542 | Cp | 12870 | Microarrays | 37986224 | Unspecified |
| mmu-miR-34a-5p | MIMAT0000542 | Cpd | 12874 | HITS-CLIP | 31433990\|23142080\|19536157\|29167 | Unspecified |
| mmu-miR-34a-5p | MIMAT0000542 | Cpe | 12876 | HITS-CLIP | 19536157\|26203562 | Unspecified |
| mmu-miR-34a-5p | MIMAT0000542 | Cpeb1 | 12877 | HITS-CLIP | 19536157 | Unspecified |
| mmu-miR-34a-5p | MIMAT0000542 | Cpeb2 | 231207 | HITS-CLIP | 23142080\|19536157\|29167373 | Unspecified |
| mmu-miR-34a-5p | MIMAT0000542 | Cpeb4 | 67579 | HITS-CLIP | 19536157\|23142080\|29167373 | Unspecified |
| mmu-miR-34a-5p | MIMAT0000542 | Cped1 | 214642 | Microarrays | 37986224\|23027903 | Unspecified |
| mmu-miR-34a-5p | MIMAT0000542 | Cplane1 | 65250 | HITS-CLIP | 30971389\|23142080 | Unspecified |
| mmu-miR-34a-5p | MIMAT0000542 | Cplx2 | 12890 | HITS-CLIP | 19536157 | Unspecified |
| mmu-miR-34a-5p | MIMAT0000542 | Cpm | 70574 | HITS-CLIP | 23142080 | Unspecified |
| mmu-miR-34a-5p | MIMAT0000542 | Cpne1 | 266692 | HITS-CLIP | 23142080\|29167373 | Unspecified |
| mmu-miR-34a-5p | MIMAT0000542 | Cpne2 | 234577 | HITS-CLIP | 29167373 | Unspecified |
| mmu-miR-34a-5p | MIMAT0000542 | Cpne3 | 70568 | HITS-CLIP | 23142080\|29167373 | Unspecified |
| mmu-miR-34a-5p | MIMAT0000542 | Cpox | 12892 | HITS-CLIP | 23142080 | Unspecified |
| mmu-miR-34a-5p | MIMAT0000542 | Cpped1 | 223978 | HITS-CLIP | 31433990\|29167373 | Unspecified |
| mmu-miR-34a-5p | MIMAT0000542 | Cpsf1 | 94230 | HITS-CLIP | 23142080 | Unspecified |
| mmu-miR-34a-5p | MIMAT0000542 | Cpsf2 | 51786 | HITS-CLIP | 29167373\|23142080 | Unspecified |
| mmu-miR-34a-5p | MIMAT0000542 | Cpsf3 | 54451 | HITS-CLIP | 23142080 | Unspecified |
| mmu-miR-34a-5p | MIMAT0000542 | Cpsf4l | 52670 | HITS-CLIP | 29167373 | Unspecified |
| mmu-miR-34a-5p | MIMAT0000542 | Cpsf6 | 432508 | HITS-CLIP | 31433990\|23142080\|29167373 | Unspecified |
| mmu-miR-34a-5p | MIMAT0000542 | Cpsf7 | 269061 | HITS-CLIP | 23630078\|29167373\|23142080 | Unspecified |
| mmu-miR-34a-5p | MIMAT0000542 | Cpt1a | 12894 | HITS-CLIP | 29167373 | Unspecified |
| mmu-miR-34a-5p | MIMAT0000542 | Cpt2 | 12896 | HITS-CLIP | 23142080 | Unspecified |
| mmu-miR-34a-5p | MIMAT0000542 | Cptp | 80772 | HITS-CLIP | 23142080 | Unspecified |
| mmu-miR-34a-5p | MIMAT0000542 | Cr1l | 12946 | HITS-CLIP | 23142080 | Unspecified |
| mmu-miR-34a-5p | MIMAT0000542 | Cramp1 | 57354 | HITS-CLIP | 31433990 | Unspecified |
| mmu-miR-34a-5p | MIMAT0000542 | Cramp1l | 57354 | HITS-CLIP | 23142080\|29167373 | Unspecified |
| mmu-miR-34a-5p | MIMAT0000542 | Crat | 12908 | HITS-CLIP | 19536157\|23142080 | Unspecified |
| mmu-miR-34a-5p | MIMAT0000542 | Crbn | 58799 | HITS-CLIP | 23142080\|29167373 | Unspecified |

| | | | | | | |
|---|---|---|---|---|---|---|
| mmu-miR-34a-5p | MIMAT0000542 | Crcp | 12909 | HITS-CLIP | 31433990 | Unspecified |
| mmu-miR-34a-5p | MIMAT0000542 | Creb1 | 12912 | HITS-CLIP | 31433990\|23142080\|29167373 | Unspecified |
| mmu-miR-34a-5p | MIMAT0000542 | Creb3l2 | 208647 | HITS-CLIP | 23142080 | Unspecified |
| mmu-miR-34a-5p | MIMAT0000542 | Creb3l3 | 208677 | HITS-CLIP | 30971389 | Unspecified |
| mmu-miR-34a-5p | MIMAT0000542 | Creb5 | 231991 | HITS-CLIP | 29167373 | Unspecified |
| mmu-miR-34a-5p | MIMAT0000542 | Crebbp | 12914 | HITS-CLIP | 37986224\|23142080\|31433990\|29167 | Unspecified |
| mmu-miR-34a-5p | MIMAT0000542 | Crebl2 | 232430 | HITS-CLIP | 23142080 | Unspecified |
| mmu-miR-34a-5p | MIMAT0000542 | Crebrf | 77128 | HITS-CLIP | 37986224\|19536157\|23142080\|29167 | Unspecified |
| mmu-miR-34a-5p | MIMAT0000542 | Creg1 | 433375 | HITS-CLIP | 29167373 | Unspecified |
| mmu-miR-34a-5p | MIMAT0000542 | Creld2 | 76737 | HITS-CLIP | 23142080 | Unspecified |
| mmu-miR-34a-5p | MIMAT0000542 | Crem | 12916 | Microarrays//HITS-CLIP | 37986224\|23142080 | Unspecified |
| mmu-miR-34a-5p | MIMAT0000542 | Crim1 | 50766 | HITS-CLIP | 23142080 | Unspecified |
| mmu-miR-34a-5p | MIMAT0000542 | Crip1 | 12925 | HITS-CLIP | 23142080 | Unspecified |
| mmu-miR-34a-5p | MIMAT0000542 | Cript | 56724 | HITS-CLIP | 23142080\|29167373 | Unspecified |
| mmu-miR-34a-5p | MIMAT0000542 | Crk | 12928 | HITS-CLIP | 19536157\|31433990\|23142080\|29167 | Unspecified |
| mmu-miR-34a-5p | MIMAT0000542 | Crkl | 12929 | HITS-CLIP | 23142080\|30971389\|19536157 | Unspecified |
| mmu-miR-34a-5p | MIMAT0000542 | Crlf2 | 57914 | HITS-CLIP | 23142080 | Unspecified |
| mmu-miR-34a-5p | MIMAT0000542 | Crlf3 | 54394 | HITS-CLIP | 29167373\|23142080 | Unspecified |
| mmu-miR-34a-5p | MIMAT0000542 | Crmp1 | 12933 | HITS-CLIP | 19536157 | Unspecified |
| mmu-miR-34a-5p | MIMAT0000542 | Crnkl1 | 66877 | HITS-CLIP | 23142080\|29167373 | Unspecified |
| mmu-miR-34a-5p | MIMAT0000542 | Crot | 74114 | HITS-CLIP | 23630078\|23142080\|30971389 | Unspecified |
| mmu-miR-34a-5p | MIMAT0000542 | Crtc2 | 74343 | HITS-CLIP | 31433990\|23142080 | Unspecified |
| mmu-miR-34a-5p | MIMAT0000542 | Crtc3 | 70461 | HITS-CLIP | 23142080 | Unspecified |
| mmu-miR-34a-5p | MIMAT0000542 | Cry1 | 12952 | HITS-CLIP | 23142080 | Unspecified |
| mmu-miR-34a-5p | MIMAT0000542 | Cry2 | 12953 | HITS-CLIP | 19536157\|31433990\|26203562 | Unspecified |
| mmu-miR-34a-5p | MIMAT0000542 | Crybg1 | 202 | HITS-CLIP | 23142080\|29167373 | Unspecified |
| mmu-miR-34a-5p | MIMAT0000542 | Crybg3 | 224273 | HITS-CLIP | 29167373 | Unspecified |
| mmu-miR-34a-5p | MIMAT0000542 | Cryz | 12972 | HITS-CLIP | 23142080 | Unspecified |
| mmu-miR-34a-5p | MIMAT0000542 | Cryzl1 | 66609 | HITS-CLIP | 29167373 | Unspecified |
| mmu-miR-34a-5p | MIMAT0000542 | Cs | 12974 | HITS-CLIP | 31433990\|23142080\|23630078\|30971 | Unspecified |
| mmu-miR-34a-5p | MIMAT0000542 | Csad | 246277 | HITS-CLIP | 23142080 | Unspecified |
| mmu-miR-34a-5p | MIMAT0000542 | Csde1 | 229663 | HITS-CLIP | 29167373\|23142080 | Unspecified |
| mmu-miR-34a-5p | MIMAT0000542 | Cse1l | 110750 | HITS-CLIP | 23142080\|19536157\|29167373 | Unspecified |
| mmu-miR-34a-5p | MIMAT0000542 | Csf1 | 12977 | HITS-CLIP | 23142080\|31433990\|29167373 | Unspecified |
| mmu-miR-34a-5p | MIMAT0000542 | Csf1r | 12978 | HITS-CLIP//qRT-PCR | 29167373\|32612530 | Unspecified |
| mmu-miR-34a-5p | MIMAT0000542 | Csf2ra | 12982 | HITS-CLIP | 29167373 | Unspecified |
| mmu-miR-34a-5p | MIMAT0000542 | Csf2rb | 12983 | HITS-CLIP | 23142080\|29167373 | Unspecified |
| mmu-miR-34a-5p | MIMAT0000542 | Csf2rb2 | 12984 | HITS-CLIP | 29167373 | Unspecified |
| mmu-miR-34a-5p | MIMAT0000542 | Csf3r | 12986 | HITS-CLIP | 29167373 | Unspecified |

| | | | | | | | |
|---|---|---|---|---|---|---|---|
| mmu-miR-34a-5p | MIMAT0000542 | Csgalnact2 | 78752 | HITS-CLIP | | 23142080 | Unspecified |
| mmu-miR-34a-5p | MIMAT0000542 | Csk | 12988 | HITS-CLIP | 23142080\|29167373 | | Unspecified |
| mmu-miR-34a-5p | MIMAT0000542 | Csmd2 | 329942 | HITS-CLIP | | 19536157 | Unspecified |
| mmu-miR-34a-5p | MIMAT0000542 | Csnk1a1 | 93687 | HITS-CLIP | 23142080\|31433990\|29167373 | | Unspecified |
| mmu-miR-34a-5p | MIMAT0000542 | Csnk1d | 104318 | HITS-CLIP | 19536157\|23142080\|30971389\|31433 | | Unspecified |
| mmu-miR-34a-5p | MIMAT0000542 | Csnk1e | 27373 | HITS-CLIP | | 29167373 | Unspecified |
| mmu-miR-34a-5p | MIMAT0000542 | Csnk1g1 | 214897 | HITS-CLIP | 23142080\|29167373 | | Unspecified |
| mmu-miR-34a-5p | MIMAT0000542 | Csnk1g2 | 103236 | HITS-CLIP | 31433990\|23142080 | | Unspecified |
| mmu-miR-34a-5p | MIMAT0000542 | Csnk1g3 | 70425 | HITS-CLIP | 37986224\|23142080 | | Unspecified |
| mmu-miR-34a-5p | MIMAT0000542 | Csnk2a2 | 13000 | HITS-CLIP | 23142080\|29167373 | | Unspecified |
| mmu-miR-34a-5p | MIMAT0000542 | Csnk2b | 13001 | HITS-CLIP | 23142080\|29167373 | | Unspecified |
| mmu-miR-34a-5p | MIMAT0000542 | Cspp1 | 211660 | HITS-CLIP | | 23142080 | Unspecified |
| mmu-miR-34a-5p | MIMAT0000542 | Csrnp1 | 215418 | HITS-CLIP | | 23142080 | Unspecified |
| mmu-miR-34a-5p | MIMAT0000542 | Csrnp3 | 77771 | HITS-CLIP | | 19536157 | Unspecified |
| mmu-miR-34a-5p | MIMAT0000542 | Cst3 | 13010 | HITS-CLIP | 19536157\|29167373\|23142080 | | Unspecified |
| mmu-miR-34a-5p | MIMAT0000542 | Cst7 | 13011 | HITS-CLIP | | 23142080 | Unspecified |
| mmu-miR-34a-5p | MIMAT0000542 | Cstf1 | 67337 | HITS-CLIP | | 23142080 | Unspecified |
| mmu-miR-34a-5p | MIMAT0000542 | Cstf2 | 108062 | HITS-CLIP | | 23142080 | Unspecified |
| mmu-miR-34a-5p | MIMAT0000542 | Cstf2t | 83410 | HITS-CLIP | | 23142080 | Unspecified |
| mmu-miR-34a-5p | MIMAT0000542 | Cstf3 | 228410 | HITS-CLIP | | 23142080 | Unspecified |
| mmu-miR-34a-5p | MIMAT0000542 | Ctbp1 | 13016 | HITS-CLIP | 23142080\|19536157\|29167373 | | Unspecified |
| mmu-miR-34a-5p | MIMAT0000542 | Ctc1 | 68964 | HITS-CLIP | 31433990\|23142080 | | Unspecified |
| mmu-miR-34a-5p | MIMAT0000542 | Ctcf | 13018 | HITS-CLIP | 23142080\|29167373 | | Unspecified |
| mmu-miR-34a-5p | MIMAT0000542 | Ctdp1 | 67655 | HITS-CLIP | | 29167373 | Unspecified |
| mmu-miR-34a-5p | MIMAT0000542 | Ctdsp1 | 227292 | HITS-CLIP | | 23142080 | Unspecified |
| mmu-miR-34a-5p | MIMAT0000542 | Ctdsp2 | 52468 | Microarrays//HITS-CLIP | 37986224\|23027903\|31433990\|1953€ | | Unspecified |
| mmu-miR-34a-5p | MIMAT0000542 | Ctdspl | 69274 | Microarrays//HITS-CLIP | 37986224\|23027903\|19536157\|23142 | | Unspecified |
| mmu-miR-34a-5p | MIMAT0000542 | Ctdspl2 | 329506 | HITS-CLIP | 23142080\|29167373 | | Unspecified |
| mmu-miR-34a-5p | MIMAT0000542 | Cth | 107869 | HITS-CLIP | 23630078\|23142080 | | Unspecified |
| mmu-miR-34a-5p | MIMAT0000542 | Ctla2a | 13024 | HITS-CLIP | | 23142080 | Unspecified |
| mmu-miR-34a-5p | MIMAT0000542 | Ctla4 | 12477 | HITS-CLIP | | 23142080 | Unspecified |
| mmu-miR-34a-5p | MIMAT0000542 | Ctnna1 | 12385 | HITS-CLIP | 37986224\|23142080\|29167373 | | Unspecified |
| mmu-miR-34a-5p | MIMAT0000542 | Ctnnb1 | 12387 | HITS-CLIP | 37986224\|19536157\|23142080\|29167 | | Unspecified |
| mmu-miR-34a-5p | MIMAT0000542 | Ctnnd1 | 12388 | HITS-CLIP | 19536157\|29167373 | | Unspecified |
| mmu-miR-34a-5p | MIMAT0000542 | Ctnnd2 | 18163 | HITS-CLIP | 19536157\|29167373 | | Unspecified |
| mmu-miR-34a-5p | MIMAT0000542 | Ctns | 83429 | Microarrays//HITS-CLIP | 37986224\|23142080 | | Unspecified |
| mmu-miR-34a-5p | MIMAT0000542 | Ctps | 51797 | HITS-CLIP | | 23142080 | Unspecified |
| mmu-miR-34a-5p | MIMAT0000542 | Ctps2 | 55936 | HITS-CLIP | 23142080\|29167373 | | Unspecified |
| mmu-miR-34a-5p | MIMAT0000542 | Ctr9 | 22083 | HITS-CLIP | | 23142080 | Unspecified |

| | | | | | | |
|---|---|---|---|---|---|---|
| mmu-miR-34a-5p | MIMAT0000542 | Ctrb1 | 66473 | Microarrays | 37986224 | Unspecified |
| mmu-miR-34a-5p | MIMAT0000542 | Ctsa | 19025 | HITS-CLIP | 21258322\|23142080\|29167373 | Unspecified |
| mmu-miR-34a-5p | MIMAT0000542 | Ctsb | 13030 | HITS-CLIP | 23630078\|23142080\|29167373 | Unspecified |
| mmu-miR-34a-5p | MIMAT0000542 | Ctsc | 13032 | HITS-CLIP | 23142080\|19536157\|29167373 | Unspecified |
| mmu-miR-34a-5p | MIMAT0000542 | Ctsd | 13033 | HITS-CLIP | 23142080\|29167373 | Unspecified |
| mmu-miR-34a-5p | MIMAT0000542 | Ctsl | 13039 | HITS-CLIP | 29167373\|19536157 | Unspecified |
| mmu-miR-34a-5p | MIMAT0000542 | Ctso | 229445 | HITS-CLIP | 23142080 | Unspecified |
| mmu-miR-34a-5p | MIMAT0000542 | Ctss | 13040 | HITS-CLIP | 37986224\|23142080\|29167373 | Unspecified |
| mmu-miR-34a-5p | MIMAT0000542 | Ctsz | 64138 | HITS-CLIP | 23142080\|29167373 | Unspecified |
| mmu-miR-34a-5p | MIMAT0000542 | Cttn | 13043 | HITS-CLIP | 30971389 | Unspecified |
| mmu-miR-34a-5p | MIMAT0000542 | Cttnbp2nl | 80281 | HITS-CLIP | 29167373 | Unspecified |
| mmu-miR-34a-5p | MIMAT0000542 | Ctu2 | 66965 | HITS-CLIP | 23142080 | Unspecified |
| mmu-miR-34a-5p | MIMAT0000542 | Cuedc2 | 67116 | HITS-CLIP | 23142080 | Unspecified |
| mmu-miR-34a-5p | MIMAT0000542 | Cul1 | 26965 | HITS-CLIP | 29167373\|23142080 | Unspecified |
| mmu-miR-34a-5p | MIMAT0000542 | Cul2 | 71745 | HITS-CLIP | 23142080\|19536157 | Unspecified |
| mmu-miR-34a-5p | MIMAT0000542 | Cul3 | 26554 | HITS-CLIP | 23142080\|23630078\|31433990 | Unspecified |
| mmu-miR-34a-5p | MIMAT0000542 | Cul4a | 99375 | HITS-CLIP | 23142080 | Unspecified |
| mmu-miR-34a-5p | MIMAT0000542 | Cul4b | 72584 | HITS-CLIP | 23142080\|29167373 | Unspecified |
| mmu-miR-34a-5p | MIMAT0000542 | Cul5 | 75717 | HITS-CLIP | 31433990\|23142080 | Unspecified |
| mmu-miR-34a-5p | MIMAT0000542 | Cul7 | 66515 | HITS-CLIP | 23142080 | Unspecified |
| mmu-miR-34a-5p | MIMAT0000542 | Cuta | 90619386 | HITS-CLIP | 23142080 | Unspecified |
| mmu-miR-34a-5p | MIMAT0000542 | Cux1 | 13047 | HITS-CLIP | 23142080\|26203562\|29167373 | Unspecified |
| mmu-miR-34a-5p | MIMAT0000542 | Cwc15 | 66070 | HITS-CLIP | 37986224\|23142080\|29167373 | Unspecified |
| mmu-miR-34a-5p | MIMAT0000542 | Cwc22 | 80744 | HITS-CLIP | 23142080 | Unspecified |
| mmu-miR-34a-5p | MIMAT0000542 | Cwc25 | 67480 | HITS-CLIP | 23142080 | Unspecified |
| mmu-miR-34a-5p | MIMAT0000542 | Cwf19l1 | 72502 | HITS-CLIP | 29167373 | Unspecified |
| mmu-miR-34a-5p | MIMAT0000542 | Cwf19l2 | 244672 | HITS-CLIP | 23142080 | Unspecified |
| mmu-miR-34a-5p | MIMAT0000542 | Cx3cr1 | 13051 | HITS-CLIP | 29167373 | Unspecified |
| mmu-miR-34a-5p | MIMAT0000542 | Cxcl1 | 14825 | HITS-CLIP | 29167373 | Unspecified |
| mmu-miR-34a-5p | MIMAT0000542 | Cxcl10 | 15945 | HITS-CLIP | 29167373 | Unspecified |
| mmu-miR-34a-5p | MIMAT0000542 | Cxcl12 | 20315 | HITS-CLIP | 23630078 | Unspecified |
| mmu-miR-34a-5p | MIMAT0000542 | Cxcl16 | 66102 | HITS-CLIP | 29167373 | Unspecified |
| mmu-miR-34a-5p | MIMAT0000542 | Cxcr4 | 12767 | Microarrays//HITS-CLIP | 37986224\|29167373\|23142080 | Unspecified |
| mmu-miR-34a-5p | MIMAT0000542 | Cxcr6 | 80901 | HITS-CLIP | 23142080 | Unspecified |
| mmu-miR-34a-5p | MIMAT0000542 | Cxxc1 | 74322 | HITS-CLIP | 23142080 | Unspecified |
| mmu-miR-34a-5p | MIMAT0000542 | Cyb561d1 | 72023 | HITS-CLIP | 23142080 | Unspecified |
| mmu-miR-34a-5p | MIMAT0000542 | Cyb5a | 109672 | HITS-CLIP | 23142080\|19536157\|29167373 | Unspecified |
| mmu-miR-34a-5p | MIMAT0000542 | Cyb5b | 66427 | HITS-CLIP | 23142080 | Unspecified |
| mmu-miR-34a-5p | MIMAT0000542 | Cyb5d2 | 192986 | HITS-CLIP | 23142080 | Unspecified |

| | | | | | | |
|---|---|---|---|---|---|---|
| mmu-miR-34a-5p | MIMAT0000542 | Cyb5r1 | 72017 | HITS-CLIP | 29167373\|23142080 | Unspecified |
| mmu-miR-34a-5p | MIMAT0000542 | Cyb5r3 | 109754 | HITS-CLIP | 37986224\|23142080\|23630078\|29167 | Unspecified |
| mmu-miR-34a-5p | MIMAT0000542 | Cyba | 13057 | HITS-CLIP | 23142080\|29167373 | Unspecified |
| mmu-miR-34a-5p | MIMAT0000542 | Cybb | 13058 | HITS-CLIP | 23142080\|29167373 | Unspecified |
| mmu-miR-34a-5p | MIMAT0000542 | Cybc1 | 79415 | HITS-CLIP | 23142080\|29167373 | Unspecified |
| mmu-miR-34a-5p | MIMAT0000542 | Cyc1 | 66445 | HITS-CLIP | 23142080 | Unspecified |
| mmu-miR-34a-5p | MIMAT0000542 | Cyfip1 | 20430 | HITS-CLIP | 29167373 | Unspecified |
| mmu-miR-34a-5p | MIMAT0000542 | Cyfip2 | 76884 | HITS-CLIP | 23142080\|19536157\|31433990 | Unspecified |
| mmu-miR-34a-5p | MIMAT0000542 | Cyhr1 | 54151 | HITS-CLIP | 23142080\|29167373 | Unspecified |
| mmu-miR-34a-5p | MIMAT0000542 | Cyld | 74256 | HITS-CLIP | 23142080 | Unspecified |
| mmu-miR-34a-5p | MIMAT0000542 | Cyp20a1 | 77951 | HITS-CLIP | 23142080\|29167373 | Unspecified |
| mmu-miR-34a-5p | MIMAT0000542 | Cyp2j6 | 13110 | HITS-CLIP | 26203562 | Unspecified |
| mmu-miR-34a-5p | MIMAT0000542 | Cyp2r1 | 244209 | HITS-CLIP | 23142080 | Unspecified |
| mmu-miR-34a-5p | MIMAT0000542 | Cyp2s1 | 74134 | HITS-CLIP | 23142080 | Unspecified |
| mmu-miR-34a-5p | MIMAT0000542 | Cyp46a1 | 13116 | HITS-CLIP | 19536157 | Unspecified |
| mmu-miR-34a-5p | MIMAT0000542 | Cyp4f13 | 170716 | HITS-CLIP | 23142080 | Unspecified |
| mmu-miR-34a-5p | MIMAT0000542 | Cyp4v3 | 102294 | HITS-CLIP | 23142080 | Unspecified |
| mmu-miR-34a-5p | MIMAT0000542 | Cyp51 | 13121 | HITS-CLIP | 23142080\|19536157 | Unspecified |
| mmu-miR-34a-5p | MIMAT0000542 | Cysltr1 | 58861 | HITS-CLIP | 29167373 | Unspecified |
| mmu-miR-34a-5p | MIMAT0000542 | Cystm1 | 66060 | HITS-CLIP | 23142080\|23630078 | Unspecified |
| mmu-miR-34a-5p | MIMAT0000542 | Cyth1 | 19157 | HITS-CLIP | 23142080\|29167373 | Unspecified |
| mmu-miR-34a-5p | MIMAT0000542 | Cyth2 | 19158 | HITS-CLIP | 23142080 | Unspecified |
| mmu-miR-34a-5p | MIMAT0000542 | Cyth3 | 19159 | HITS-CLIP | 23142080 | Unspecified |
| mmu-miR-34a-5p | MIMAT0000542 | Cyth4 | 72318 | HITS-CLIP | 29167373 | Unspecified |
| mmu-miR-34a-5p | MIMAT0000542 | Cytip | 227929 | HITS-CLIP | 23142080 | Unspecified |
| mmu-miR-34a-5p | MIMAT0000542 | D030056L22Rik | 225995 | HITS-CLIP | 23142080 | Unspecified |
| mmu-miR-34a-5p | MIMAT0000542 | D10Jhu81e | 28295 | HITS-CLIP | 23142080 | Unspecified |
| mmu-miR-34a-5p | MIMAT0000542 | D10Wsu102e | 28109 | Microarrays//HITS-CLIP | 37986224\|23142080\|29167373 | Unspecified |
| mmu-miR-34a-5p | MIMAT0000542 | D11Wsu47e | 276852 | HITS-CLIP | 23142080 | Unspecified |
| mmu-miR-34a-5p | MIMAT0000542 | D130040H23Rik | 211135 | HITS-CLIP | 23142080 | Unspecified |
| mmu-miR-34a-5p | MIMAT0000542 | D16Ertd472e | 67102 | Microarrays//HITS-CLIP | 37986224\|23142080\|23027903 | Unspecified |
| mmu-miR-34a-5p | MIMAT0000542 | D17H6S53E | 114585 | HITS-CLIP | 29167373 | Unspecified |
| mmu-miR-34a-5p | MIMAT0000542 | D1Ertd622e | 52392 | HITS-CLIP | 29167373 | Unspecified |
| mmu-miR-34a-5p | MIMAT0000542 | D230025D16Rik | 234678 | HITS-CLIP | 19536157\|29167373\|23142080 | Unspecified |
| mmu-miR-34a-5p | MIMAT0000542 | D2hgdh | 98314 | HITS-CLIP | 23142080 | Unspecified |
| mmu-miR-34a-5p | MIMAT0000542 | D430041D05Rik | 241589 | HITS-CLIP | 19536157 | Unspecified |
| mmu-miR-34a-5p | MIMAT0000542 | D430042O09Rik | 233865 | HITS-CLIP | 23142080 | Unspecified |
| mmu-miR-34a-5p | MIMAT0000542 | D5Ertd579e | 320661 | HITS-CLIP | 31433990\|19536157\|23142080\|29167 | Unspecified |
| mmu-miR-34a-5p | MIMAT0000542 | D630039A03Rik | 242484 | HITS-CLIP | 23142080 | Unspecified |

| | | | | | | |
|---|---|---|---|---|---|---|
| mmu-miR-34a-5p | MIMAT0000542 | D630045J12Rik | 330286 | HITS-CLIP | 23142080 | Unspecified |
| mmu-miR-34a-5p | MIMAT0000542 | D8Ertd738e | 101966 | HITS-CLIP | 23142080 | Unspecified |
| mmu-miR-34a-5p | MIMAT0000542 | Daam1 | 208846 | HITS-CLIP | 19536157\|23142080 | Unspecified |
| mmu-miR-34a-5p | MIMAT0000542 | Daam2 | 76441 | Microarrays | 37986224 | Unspecified |
| mmu-miR-34a-5p | MIMAT0000542 | Dab1 | 13131 | HITS-CLIP | 19536157 | Unspecified |
| mmu-miR-34a-5p | MIMAT0000542 | Dab2 | 13132 | HITS-CLIP | 19536157\|29167373 | Unspecified |
| mmu-miR-34a-5p | MIMAT0000542 | Dab2ip | 69601 | HITS-CLIP | 19536157 | Unspecified |
| mmu-miR-34a-5p | MIMAT0000542 | Dact1 | 59036 | HITS-CLIP | 19536157 | Unspecified |
| mmu-miR-34a-5p | MIMAT0000542 | Dad1 | 13135 | HITS-CLIP | 23142080 | Unspecified |
| mmu-miR-34a-5p | MIMAT0000542 | Dag1 | 13138 | HITS-CLIP | 23142080\|19536157\|23630078\|29167 | Unspecified |
| mmu-miR-34a-5p | MIMAT0000542 | Daglb | 231871 | HITS-CLIP | 29167373 | Unspecified |
| mmu-miR-34a-5p | MIMAT0000542 | Dand5 | 23863 | HITS-CLIP | 23142080 | Unspecified |
| mmu-miR-34a-5p | MIMAT0000542 | Dap | 223453 | HITS-CLIP | 23142080 | Unspecified |
| mmu-miR-34a-5p | MIMAT0000542 | Dap3 | 65111 | HITS-CLIP | 23142080 | Unspecified |
| mmu-miR-34a-5p | MIMAT0000542 | Dapk1 | 69635 | HITS-CLIP | 29167373 | Unspecified |
| mmu-miR-34a-5p | MIMAT0000542 | Dapk3 | 13144 | HITS-CLIP | 23142080 | Unspecified |
| mmu-miR-34a-5p | MIMAT0000542 | Dapp1 | 26377 | HITS-CLIP | 23142080 | Unspecified |
| mmu-miR-34a-5p | MIMAT0000542 | Dars | 226414 | HITS-CLIP | 23142080 | Unspecified |
| mmu-miR-34a-5p | MIMAT0000542 | Dars2 | 226539 | HITS-CLIP | 23142080 | Unspecified |
| mmu-miR-34a-5p | MIMAT0000542 | Daxx | 13163 | HITS-CLIP | 23142080 | Unspecified |
| mmu-miR-34a-5p | MIMAT0000542 | Dazap2 | 23994 | HITS-CLIP | 23142080\|29167373 | Unspecified |
| mmu-miR-34a-5p | MIMAT0000542 | Dbi | 13167 | HITS-CLIP | 23142080 | Unspecified |
| mmu-miR-34a-5p | MIMAT0000542 | Dbnl | 13169 | HITS-CLIP | 23142080 | Unspecified |
| mmu-miR-34a-5p | MIMAT0000542 | Dbp | 13170 | HITS-CLIP | 23142080 | Unspecified |
| mmu-miR-34a-5p | MIMAT0000542 | Dbt | 13171 | HITS-CLIP | 23142080\|19536157\|29167373 | Unspecified |
| mmu-miR-34a-5p | MIMAT0000542 | Dcaf1 | 41836 | HITS-CLIP | 23142080 | Unspecified |
| mmu-miR-34a-5p | MIMAT0000542 | Dcaf10 | 242418 | HITS-CLIP | 23142080\|31433990 | Unspecified |
| mmu-miR-34a-5p | MIMAT0000542 | Dcaf11 | 28199 | HITS-CLIP | 23142080 | Unspecified |
| mmu-miR-34a-5p | MIMAT0000542 | Dcaf12 | 68970 | HITS-CLIP | 37986224\|31433990\|29167373\|23142 | Unspecified |
| mmu-miR-34a-5p | MIMAT0000542 | Dcaf13 | 223499 | HITS-CLIP | 23142080 | Unspecified |
| mmu-miR-34a-5p | MIMAT0000542 | Dcaf17 | 75763 | HITS-CLIP | 23142080\|29167373 | Unspecified |
| mmu-miR-34a-5p | MIMAT0000542 | Dcaf5 | 320808 | HITS-CLIP | 19536157\|29167373\|23142080 | Unspecified |
| mmu-miR-34a-5p | MIMAT0000542 | Dcaf6 | 74106 | HITS-CLIP | 19536157\|23142080 | Unspecified |
| mmu-miR-34a-5p | MIMAT0000542 | Dcaf7 | 71833 | HITS-CLIP | 23142080 | Unspecified |
| mmu-miR-34a-5p | MIMAT0000542 | Dcaf8 | 98193 | HITS-CLIP | 23142080 | Unspecified |
| mmu-miR-34a-5p | MIMAT0000542 | Dcakd | 68087 | HITS-CLIP | 23142080\|29167373 | Unspecified |
| mmu-miR-34a-5p | MIMAT0000542 | Dcbld2 | 73379 | HITS-CLIP | 19536157\|23142080 | Unspecified |
| mmu-miR-34a-5p | MIMAT0000542 | Dck | 13178 | HITS-CLIP | 37986224\|23630078\|23142080 | Unspecified |
| mmu-miR-34a-5p | MIMAT0000542 | Dclk1 | 13175 | Microarrays//HITS-CLIP | 37986224\|19536157 | Unspecified |

| | | | | | | |
|---|---|---|---|---|---|---|
| mmu-miR-34a-5p | MIMAT0000542 | Dclk2 | 70762 | HITS-CLIP | 19536157 | Unspecified |
| mmu-miR-34a-5p | MIMAT0000542 | Dclre1b | 140917 | Microarrays//HITS-CLIP | 37986224\|23027903\|23142080 | Unspecified |
| mmu-miR-34a-5p | MIMAT0000542 | Dclre1c | 227525 | HITS-CLIP | 29167373 | Unspecified |
| mmu-miR-34a-5p | MIMAT0000542 | Dcn | 13179 | Microarrays | 37986224 | Unspecified |
| mmu-miR-34a-5p | MIMAT0000542 | Dcp1a | 75901 | HITS-CLIP | 37986224\|23142080\|31433990\|29167 | Unspecified |
| mmu-miR-34a-5p | MIMAT0000542 | Dcp1b | 319618 | HITS-CLIP | 23142080 | Unspecified |
| mmu-miR-34a-5p | MIMAT0000542 | Dcp2 | 70640 | Microarrays//HITS-CLIP | 37986224\|23027903\|23142080\|26203 | Unspecified |
| mmu-miR-34a-5p | MIMAT0000542 | Dcps | 69305 | HITS-CLIP | 31433990\|23142080 | Unspecified |
| mmu-miR-34a-5p | MIMAT0000542 | Dctd | 320685 | HITS-CLIP | 23142080 | Unspecified |
| mmu-miR-34a-5p | MIMAT0000542 | Dctn1 | 13191 | HITS-CLIP | 23142080 | Unspecified |
| mmu-miR-34a-5p | MIMAT0000542 | Dctn2 | 69654 | HITS-CLIP | 23142080 | Unspecified |
| mmu-miR-34a-5p | MIMAT0000542 | Dctn3 | 53598 | HITS-CLIP | 31433990\|30971389\|29167373\|23142 | Unspecified |
| mmu-miR-34a-5p | MIMAT0000542 | Dctn4 | 67665 | HITS-CLIP | 23142080\|31433990\|23630078 | Unspecified |
| mmu-miR-34a-5p | MIMAT0000542 | Dctn5 | 59288 | HITS-CLIP | 23142080 | Unspecified |
| mmu-miR-34a-5p | MIMAT0000542 | Dctpp1 | 66422 | Microarrays//HITS-CLIP | 37986224\|23027903\|23142080 | Unspecified |
| mmu-miR-34a-5p | MIMAT0000542 | Dcun1d2 | 102323 | HITS-CLIP | 23142080 | Unspecified |
| mmu-miR-34a-5p | MIMAT0000542 | Dcun1d3 | 233805 | HITS-CLIP | 37986224\|26203562\|23142080 | Unspecified |
| mmu-miR-34a-5p | MIMAT0000542 | Dcun1d4 | 100737 | HITS-CLIP | 23630078\|23142080 | Unspecified |
| mmu-miR-34a-5p | MIMAT0000542 | Dcun1d5 | 76863 | HITS-CLIP | 23142080 | Unspecified |
| mmu-miR-34a-5p | MIMAT0000542 | Dcx | 13193 | HITS-CLIP | 19536157 | Unspecified |
| mmu-miR-34a-5p | MIMAT0000542 | Dda1 | 66498 | HITS-CLIP | 23142080 | Unspecified |
| mmu-miR-34a-5p | MIMAT0000542 | Ddah2 | 51793 | HITS-CLIP | 29084942 | Unspecified |
| mmu-miR-34a-5p | MIMAT0000542 | Ddb1 | 13194 | HITS-CLIP | 23142080\|29167373 | Unspecified |
| mmu-miR-34a-5p | MIMAT0000542 | Ddhd1 | 114874 | HITS-CLIP | 37986224\|23142080\|29167373 | Unspecified |
| mmu-miR-34a-5p | MIMAT0000542 | Ddhd2 | 72108 | HITS-CLIP | 23142080 | Unspecified |
| mmu-miR-34a-5p | MIMAT0000542 | Ddi2 | 68817 | HITS-CLIP | 31433990\|30971389\|19536157\|23142 | Unspecified |
| mmu-miR-34a-5p | MIMAT0000542 | Ddias | 220042 | HITS-CLIP | 26203562\|23142080 | Unspecified |
| mmu-miR-34a-5p | MIMAT0000542 | Ddit3 | 13198 | HITS-CLIP | 29167373 | Unspecified |
| mmu-miR-34a-5p | MIMAT0000542 | Ddit4 | 74747 | HITS-CLIP, Microarrays//HITS-CLIP | 37986224\|23142080 | Unspecified |
| mmu-miR-34a-5p | MIMAT0000542 | Ddit4l | 73284 | Microarrays | 37986224 | Unspecified |
| mmu-miR-34a-5p | MIMAT0000542 | Ddost | 13200 | HITS-CLIP | 26203562 | Unspecified |
| mmu-miR-34a-5p | MIMAT0000542 | Ddr1 | 12305 | HITS-CLIP | 23142080 | Unspecified |
| mmu-miR-34a-5p | MIMAT0000542 | Ddr2 | 18214 | HITS-CLIP | 23630078 | Unspecified |
| mmu-miR-34a-5p | MIMAT0000542 | Ddt | 13202 | HITS-CLIP | 23142080 | Unspecified |
| mmu-miR-34a-5p | MIMAT0000542 | Ddx1 | 104721 | HITS-CLIP | 23142080 | Unspecified |
| mmu-miR-34a-5p | MIMAT0000542 | Ddx10 | 77591 | HITS-CLIP | 23142080\|29167373 | Unspecified |
| mmu-miR-34a-5p | MIMAT0000542 | Ddx11 | 320209 | Microarrays | 37986224\|23027903 | Unspecified |
| mmu-miR-34a-5p | MIMAT0000542 | Ddx17 | 67040 | HITS-CLIP | 37986224\|23142080\|19536157\|29167 | Unspecified |
| mmu-miR-34a-5p | MIMAT0000542 | Ddx19a | 13680 | HITS-CLIP | 23142080 | Unspecified |

| | | | | | | |
|---|---|---|---|---|---|---|
| mmu-miR-34a-5p | MIMAT0000542 | Ddx19b | 234733 | HITS-CLIP | 23142080 | Unspecified |
| mmu-miR-34a-5p | MIMAT0000542 | Ddx20 | 53975 | HITS-CLIP | 23142080 | Unspecified |
| mmu-miR-34a-5p | MIMAT0000542 | Ddx21 | 56200 | HITS-CLIP | 30971389\|31433990\|29167373\|23142 | Unspecified |
| mmu-miR-34a-5p | MIMAT0000542 | Ddx23 | 74351 | HITS-CLIP | 23142080\|29167373 | Unspecified |
| mmu-miR-34a-5p | MIMAT0000542 | Ddx24 | 27225 | HITS-CLIP | 31433990\|23142080 | Unspecified |
| mmu-miR-34a-5p | MIMAT0000542 | Ddx27 | 228889 | HITS-CLIP | 23142080 | Unspecified |
| mmu-miR-34a-5p | MIMAT0000542 | Ddx28 | 71986 | HITS-CLIP | 23142080 | Unspecified |
| mmu-miR-34a-5p | MIMAT0000542 | Ddx31 | 227674 | HITS-CLIP | 23142080\|29167373 | Unspecified |
| mmu-miR-34a-5p | MIMAT0000542 | Ddx39 | 68278 | HITS-CLIP | 23142080\|29167373 | Unspecified |
| mmu-miR-34a-5p | MIMAT0000542 | Ddx39b | 53817 | HITS-CLIP | 23142080\|31433990 | Unspecified |
| mmu-miR-34a-5p | MIMAT0000542 | Ddx3x | 13205 | HITS-CLIP | 19536157\|23142080\|31433990\|29167 | Unspecified |
| mmu-miR-34a-5p | MIMAT0000542 | Ddx3y | 26900 | HITS-CLIP | 31433990\|23142080 | Unspecified |
| mmu-miR-34a-5p | MIMAT0000542 | Ddx41 | 72935 | HITS-CLIP | 23142080 | Unspecified |
| mmu-miR-34a-5p | MIMAT0000542 | Ddx42 | 72047 | HITS-CLIP | 23142080\|29167373 | Unspecified |
| mmu-miR-34a-5p | MIMAT0000542 | Ddx46 | 212880 | HITS-CLIP | 23142080 | Unspecified |
| mmu-miR-34a-5p | MIMAT0000542 | Ddx5 | 13207 | HITS-CLIP | 31433990\|23142080\|19536157\|29167 | Unspecified |
| mmu-miR-34a-5p | MIMAT0000542 | Ddx50 | 94213 | HITS-CLIP | 31433990 | Unspecified |
| mmu-miR-34a-5p | MIMAT0000542 | Ddx54 | 71990 | HITS-CLIP | 23142080 | Unspecified |
| mmu-miR-34a-5p | MIMAT0000542 | Ddx55 | 67848 | HITS-CLIP | 23142080 | Unspecified |
| mmu-miR-34a-5p | MIMAT0000542 | Ddx56 | 52513 | HITS-CLIP | 23142080 | Unspecified |
| mmu-miR-34a-5p | MIMAT0000542 | Ddx58 | 230073 | Microarrays//HITS-CLIP | 37986224\|23142080\|29167373 | Unspecified |
| mmu-miR-34a-5p | MIMAT0000542 | Ddx6 | 13209 | HITS-CLIP | 23142080\|31433990\|29167373 | Unspecified |
| mmu-miR-34a-5p | MIMAT0000542 | Ddx60 | 234311 | HITS-CLIP | 23142080\|29167373 | Unspecified |
| mmu-miR-34a-5p | MIMAT0000542 | Deaf1 | 54006 | HITS-CLIP | 23142080 | Unspecified |
| mmu-miR-34a-5p | MIMAT0000542 | Decr1 | 67460 | HITS-CLIP | 23142080 | Unspecified |
| mmu-miR-34a-5p | MIMAT0000542 | Dedd | 21945 | HITS-CLIP | 23142080 | Unspecified |
| mmu-miR-34a-5p | MIMAT0000542 | Def6 | 23853 | HITS-CLIP | 23142080 | Unspecified |
| mmu-miR-34a-5p | MIMAT0000542 | Def8 | 23854 | Microarrays//HITS-CLIP | 37986224\|23027903\|29167373 | Unspecified |
| mmu-miR-34a-5p | MIMAT0000542 | Degs1 | 13244 | HITS-CLIP | 31433990 | Unspecified |
| mmu-miR-34a-5p | MIMAT0000542 | Dek | 110052 | HITS-CLIP | 23142080\|29167373 | Unspecified |
| mmu-miR-34a-5p | MIMAT0000542 | Dennd11 | 57189 | HITS-CLIP | 31433990 | Unspecified |
| mmu-miR-34a-5p | MIMAT0000542 | Dennd1a | 227801 | HITS-CLIP | 23142080\|29167373 | Unspecified |
| mmu-miR-34a-5p | MIMAT0000542 | Dennd1b | 329260 | HITS-CLIP | 37986224\|23142080\|31433990\|29167 | Unspecified |
| mmu-miR-34a-5p | MIMAT0000542 | Dennd1c | 70785 | HITS-CLIP | 23142080 | Unspecified |
| mmu-miR-34a-5p | MIMAT0000542 | Dennd2d | 72121 | HITS-CLIP | 23142080 | Unspecified |
| mmu-miR-34a-5p | MIMAT0000542 | Dennd3 | 105841 | HITS-CLIP | 31433990\|23142080 | Unspecified |
| mmu-miR-34a-5p | MIMAT0000542 | Dennd4a | 102442 | HITS-CLIP | 30971389\|23142080 | Unspecified |
| mmu-miR-34a-5p | MIMAT0000542 | Dennd4b | 229541 | HITS-CLIP | 23142080\|29167373 | Unspecified |
| mmu-miR-34a-5p | MIMAT0000542 | Dennd4c | 329877 | HITS-CLIP | 23142080\|29167373 | Unspecified |

| | | | | | | |
|---|---|---|---|---|---|---|
| mmu-miR-34a-5p | MIMAT0000542 | Dennd5a | 19347 | HITS-CLIP | 19536157\|23630078\|23142080\|29167 | Unspecified |
| mmu-miR-34a-5p | MIMAT0000542 | Dennd6a | 211922 | Microarrays//HITS-CLIP | 37986224\|31433990\|23142080\|29167 | Unspecified |
| mmu-miR-34a-5p | MIMAT0000542 | Dennd6b | 69440 | HITS-CLIP | 29167373 | Unspecified |
| mmu-miR-34a-5p | MIMAT0000542 | Denr | 68184 | HITS-CLIP | 23142080\|29167373 | Unspecified |
| mmu-miR-34a-5p | MIMAT0000542 | Depdc1a | 76131 | Microarrays//HITS-CLIP | 37986224\|23027903\|23142080 | Unspecified |
| mmu-miR-34a-5p | MIMAT0000542 | Depdc1b | 218581 | HITS-CLIP | 23142080 | Unspecified |
| mmu-miR-34a-5p | MIMAT0000542 | Derl1 | 67819 | HITS-CLIP | 23142080\|31433990\|29167373 | Unspecified |
| mmu-miR-34a-5p | MIMAT0000542 | Derl2 | 116891 | HITS-CLIP | 23142080\|29167373 | Unspecified |
| mmu-miR-34a-5p | MIMAT0000542 | Derpc | 54921 | HITS-CLIP | 26203562 | Unspecified |
| mmu-miR-34a-5p | MIMAT0000542 | Desi1 | 28075 | HITS-CLIP | 23142080 | Unspecified |
| mmu-miR-34a-5p | MIMAT0000542 | Desi2 | 78825 | HITS-CLIP | 29167373\|23142080 | Unspecified |
| mmu-miR-34a-5p | MIMAT0000542 | Det1 | 76375 | HITS-CLIP | 29167373 | Unspecified |
| mmu-miR-34a-5p | MIMAT0000542 | Dgcr2 | 13356 | HITS-CLIP | 19536157\|23142080\|29167373 | Unspecified |
| mmu-miR-34a-5p | MIMAT0000542 | Dgcr8 | 94223 | HITS-CLIP | 23142080\|31433990 | Unspecified |
| mmu-miR-34a-5p | MIMAT0000542 | Dgka | 13139 | Microarrays//HITS-CLIP | 37986224\|23027903\|23142080 | Unspecified |
| mmu-miR-34a-5p | MIMAT0000542 | Dgkd | 227333 | HITS-CLIP | 31433990\|19536157\|23142080\|29167 | Unspecified |
| mmu-miR-34a-5p | MIMAT0000542 | Dgke | 56077 | HITS-CLIP | 19536157\|23142080\|29167373 | Unspecified |
| mmu-miR-34a-5p | MIMAT0000542 | Dgkg | 110197 | HITS-CLIP | 23142080 | Unspecified |
| mmu-miR-34a-5p | MIMAT0000542 | Dgkh | 380921 | HITS-CLIP | 29167373\|19536157 | Unspecified |
| mmu-miR-34a-5p | MIMAT0000542 | Dgkq | 110524 | HITS-CLIP | 23142080 | Unspecified |
| mmu-miR-34a-5p | MIMAT0000542 | Dgkz | 104418 | HITS-CLIP | 23142080\|29167373 | Unspecified |
| mmu-miR-34a-5p | MIMAT0000542 | Dglucy | 217830 | HITS-CLIP | 29167373 | Unspecified |
| mmu-miR-34a-5p | MIMAT0000542 | Dguok | 27369 | Microarrays//HITS-CLIP | 37986224\|23027903\|23142080 | Unspecified |
| mmu-miR-34a-5p | MIMAT0000542 | Dhcr24 | 74754 | HITS-CLIP | 26203562\|23142080 | Unspecified |
| mmu-miR-34a-5p | MIMAT0000542 | Dhcr7 | 13360 | HITS-CLIP | 19536157\|23142080\|29167373 | Unspecified |
| mmu-miR-34a-5p | MIMAT0000542 | Dhdds | 67422 | HITS-CLIP | 23142080 | Unspecified |
| mmu-miR-34a-5p | MIMAT0000542 | Dhdh | 71755 | HITS-CLIP | 23142080 | Unspecified |
| mmu-miR-34a-5p | MIMAT0000542 | Dhfr | 13361 | Microarrays//HITS-CLIP | 37986224\|23027903\|23142080 | Unspecified |
| mmu-miR-34a-5p | MIMAT0000542 | Dhodh | 56749 | HITS-CLIP | 29167373 | Unspecified |
| mmu-miR-34a-5p | MIMAT0000542 | Dhps | 330817 | HITS-CLIP | 23142080 | Unspecified |
| mmu-miR-34a-5p | MIMAT0000542 | Dhrs1 | 52585 | HITS-CLIP | 23142080 | Unspecified |
| mmu-miR-34a-5p | MIMAT0000542 | Dhrs11 | 192970 | Microarrays | 37986224 | Unspecified |
| mmu-miR-34a-5p | MIMAT0000542 | Dhrs3 | 20148 | HITS-CLIP | 29167373 | Unspecified |
| mmu-miR-34a-5p | MIMAT0000542 | Dhrs7b | 216820 | HITS-CLIP | 23630078\|26203562 | Unspecified |
| mmu-miR-34a-5p | MIMAT0000542 | Dhrs9 | 241452 | HITS-CLIP | 29167373 | Unspecified |
| mmu-miR-34a-5p | MIMAT0000542 | Dhx15 | 13204 | HITS-CLIP | 19536157\|23630078\|29167373\|23142 | Unspecified |
| mmu-miR-34a-5p | MIMAT0000542 | Dhx16 | 69192 | HITS-CLIP | 23142080\|23630078 | Unspecified |
| mmu-miR-34a-5p | MIMAT0000542 | Dhx29 | 218629 | HITS-CLIP | 23142080 | Unspecified |
| mmu-miR-34a-5p | MIMAT0000542 | Dhx30 | 72831 | HITS-CLIP | 23142080\|29167373 | Unspecified |

| | | | | | | |
|---|---|---|---|---|---|---|
| mmu-miR-34a-5p | MIMAT0000542 | Dhx32 | 101437 | Microarrays//HITS-CLIP | 37986224\|23142080 | Unspecified |
| mmu-miR-34a-5p | MIMAT0000542 | Dhx33 | 216877 | HITS-CLIP | 31433990\|23142080\|29167373 | Unspecified |
| mmu-miR-34a-5p | MIMAT0000542 | Dhx34 | 71723 | HITS-CLIP | 23142080 | Unspecified |
| mmu-miR-34a-5p | MIMAT0000542 | Dhx35 | 71715 | HITS-CLIP | 23142080\|29167373 | Unspecified |
| mmu-miR-34a-5p | MIMAT0000542 | Dhx36 | 72162 | HITS-CLIP | 19536157\|29167373\|23142080 | Unspecified |
| mmu-miR-34a-5p | MIMAT0000542 | Dhx37 | 208144 | HITS-CLIP | 23142080 | Unspecified |
| mmu-miR-34a-5p | MIMAT0000542 | Dhx40 | 67487 | HITS-CLIP | 23142080 | Unspecified |
| mmu-miR-34a-5p | MIMAT0000542 | Dhx57 | 106794 | HITS-CLIP | 23142080\|29167373 | Unspecified |
| mmu-miR-34a-5p | MIMAT0000542 | Dhx58 | 80861 | HITS-CLIP | 23142080\|29167373 | Unspecified |
| mmu-miR-34a-5p | MIMAT0000542 | Dhx8 | 217207 | HITS-CLIP | 23142080 | Unspecified |
| mmu-miR-34a-5p | MIMAT0000542 | Dhx9 | 13211 | HITS-CLIP | 23142080\|29167373 | Unspecified |
| mmu-miR-34a-5p | MIMAT0000542 | Diablo | 66593 | Microarrays//HITS-CLIP | 37986224\|23027903\|29167373 | Unspecified |
| mmu-miR-34a-5p | MIMAT0000542 | Diaph1 | 1729 | HITS-CLIP | 23142080\|29167373 | Unspecified |
| mmu-miR-34a-5p | MIMAT0000542 | Diaph2 | 54004 | HITS-CLIP | 23142080\|29167373 | Unspecified |
| mmu-miR-34a-5p | MIMAT0000542 | Diaph3 | 81624 | HITS-CLIP | 23142080 | Unspecified |
| mmu-miR-34a-5p | MIMAT0000542 | Dicer1 | 192119 | HITS-CLIP | 19536157\|31433990\|23142080\|29167 | Unspecified |
| mmu-miR-34a-5p | MIMAT0000542 | Dido1 | 23856 | HITS-CLIP | 19536157\|31433990\|23142080\|29167 | Unspecified |
| mmu-miR-34a-5p | MIMAT0000542 | Diexf | 215193 | HITS-CLIP | 23142080 | Unspecified |
| mmu-miR-34a-5p | MIMAT0000542 | Dio2 | 13371 | HITS-CLIP | 19536157 | Unspecified |
| mmu-miR-34a-5p | MIMAT0000542 | Dip2a | 64451 | HITS-CLIP | 31433990 | Unspecified |
| mmu-miR-34a-5p | MIMAT0000542 | Dip2b | 239667 | HITS-CLIP | 19536157\|23142080\|29167373 | Unspecified |
| mmu-miR-34a-5p | MIMAT0000542 | Dipk2a | 205428 | HITS-CLIP | 31433990 | Unspecified |
| mmu-miR-34a-5p | MIMAT0000542 | Dirc2 | 224132 | HITS-CLIP | 23142080 | Unspecified |
| mmu-miR-34a-5p | MIMAT0000542 | Dis3 | 72662 | HITS-CLIP | 23142080 | Unspecified |
| mmu-miR-34a-5p | MIMAT0000542 | Dis3l | 213550 | HITS-CLIP | 29167373 | Unspecified |
| mmu-miR-34a-5p | MIMAT0000542 | Dis3l2 | 208718 | HITS-CLIP | 23142080 | Unspecified |
| mmu-miR-34a-5p | MIMAT0000542 | Disp1 | 68897 | HITS-CLIP | 23142080\|29167373 | Unspecified |
| mmu-miR-34a-5p | MIMAT0000542 | Disp2 | 214240 | HITS-CLIP | 19536157 | Unspecified |
| mmu-miR-34a-5p | MIMAT0000542 | Dixdc1 | 330938 | HITS-CLIP | 23142080 | Unspecified |
| mmu-miR-34a-5p | MIMAT0000542 | Dkc1 | 245474 | HITS-CLIP | 23142080 | Unspecified |
| mmu-miR-34a-5p | MIMAT0000542 | Dkk3 | 50781 | HITS-CLIP | 26203562 | Unspecified |
| mmu-miR-34a-5p | MIMAT0000542 | Dlat | 235339 | HITS-CLIP | 23142080 | Unspecified |
| mmu-miR-34a-5p | MIMAT0000542 | Dld | 13382 | HITS-CLIP | 23142080 | Unspecified |
| mmu-miR-34a-5p | MIMAT0000542 | Dlg1 | 13383 | HITS-CLIP | 29167373 | Unspecified |
| mmu-miR-34a-5p | MIMAT0000542 | Dlg2 | 23859 | HITS-CLIP | 23142080\|19536157 | Unspecified |
| mmu-miR-34a-5p | MIMAT0000542 | Dlg4 | 13385 | HITS-CLIP | 19536157 | Unspecified |
| mmu-miR-34a-5p | MIMAT0000542 | Dlgap1 | 224997 | HITS-CLIP | 19536157 | Unspecified |
| mmu-miR-34a-5p | MIMAT0000542 | Dlgap2 | 244310 | HITS-CLIP | 19536157 | Unspecified |
| mmu-miR-34a-5p | MIMAT0000542 | Dlgap4 | 228836 | HITS-CLIP | 23142080\|29167373 | Unspecified |

| | | | | | | |
|---|---|---|---|---|---|---|
| mmu-miR-34a-5p | MIMAT0000542 | Dlgap5 | 218977 | HITS-CLIP | 23142080 | Unspecified |
| mmu-miR-34a-5p | MIMAT0000542 | Dlk1 | 13386 | HITS-CLIP | 30971389 | Unspecified |
| mmu-miR-34a-5p | MIMAT0000542 | Dlst | 78920 | HITS-CLIP | 29167373\|23142080 | Unspecified |
| mmu-miR-34a-5p | MIMAT0000542 | Dmac1 | 90871 | HITS-CLIP | 23142080 | Unspecified |
| mmu-miR-34a-5p | MIMAT0000542 | Dmac2 | 55101 | HITS-CLIP | 23142080 | Unspecified |
| mmu-miR-34a-5p | MIMAT0000542 | Dmap1 | 66233 | HITS-CLIP | 23142080 | Unspecified |
| mmu-miR-34a-5p | MIMAT0000542 | Dmd | 13405 | HITS-CLIP | 19536157 | Unspecified |
| mmu-miR-34a-5p | MIMAT0000542 | Dmtf1 | 23857 | HITS-CLIP | 23142080\|29167373 | Unspecified |
| mmu-miR-34a-5p | MIMAT0000542 | Dmtn | 13829 | HITS-CLIP | 30971389 | Unspecified |
| mmu-miR-34a-5p | MIMAT0000542 | Dmwd | 13401 | HITS-CLIP | 19536157 | Unspecified |
| mmu-miR-34a-5p | MIMAT0000542 | Dmxl1 | 240283 | HITS-CLIP | 19536157\|23142080\|29167373 | Unspecified |
| mmu-miR-34a-5p | MIMAT0000542 | Dmxl2 | 235380 | HITS-CLIP | 19536157\|29167373 | Unspecified |
| mmu-miR-34a-5p | MIMAT0000542 | Dna2 | 327762 | Microarrays//HITS-CLIP | 37986224\|23027903\|23142080 | Unspecified |
| mmu-miR-34a-5p | MIMAT0000542 | Dnaaf2 | 109065 | HITS-CLIP | 23142080 | Unspecified |
| mmu-miR-34a-5p | MIMAT0000542 | Dnaaf3 | 436022 | Microarrays | 37986224 | Unspecified |
| mmu-miR-34a-5p | MIMAT0000542 | Dnaaf5 | 433956 | HITS-CLIP | 23142080 | Unspecified |
| mmu-miR-34a-5p | MIMAT0000542 | Dnah7a | 627872 | HITS-CLIP | 23142080 | Unspecified |
| mmu-miR-34a-5p | MIMAT0000542 | Dnah8 | 13417 | HITS-CLIP | 23142080\|30971389 | Unspecified |
| mmu-miR-34a-5p | MIMAT0000542 | Dnaic1 | 68922 | HITS-CLIP | 19536157 | Unspecified |
| mmu-miR-34a-5p | MIMAT0000542 | Dnaja2 | 56445 | HITS-CLIP | 23142080\|29167373 | Unspecified |
| mmu-miR-34a-5p | MIMAT0000542 | Dnaja3 | 83945 | HITS-CLIP | 19536157\|29167373 | Unspecified |
| mmu-miR-34a-5p | MIMAT0000542 | Dnaja4 | 58233 | HITS-CLIP | 30971389\|23142080 | Unspecified |
| mmu-miR-34a-5p | MIMAT0000542 | Dnajb1 | 81489 | HITS-CLIP | 23142080 | Unspecified |
| mmu-miR-34a-5p | MIMAT0000542 | Dnajb11 | 67838 | HITS-CLIP | 23142080 | Unspecified |
| mmu-miR-34a-5p | MIMAT0000542 | Dnajb12 | 56709 | HITS-CLIP | 23142080\|30971389 | Unspecified |
| mmu-miR-34a-5p | MIMAT0000542 | Dnajb14 | 70604 | HITS-CLIP | 19536157\|23142080\|29167373 | Unspecified |
| mmu-miR-34a-5p | MIMAT0000542 | Dnajb4 | 67035 | HITS-CLIP | 19536157\|23142080 | Unspecified |
| mmu-miR-34a-5p | MIMAT0000542 | Dnajb5 | 56323 | HITS-CLIP | 31433990\|23142080 | Unspecified |
| mmu-miR-34a-5p | MIMAT0000542 | Dnajb6 | 23950 | HITS-CLIP | 23142080\|29167373 | Unspecified |
| mmu-miR-34a-5p | MIMAT0000542 | Dnajb9 | 27362 | HITS-CLIP | 23142080 | Unspecified |
| mmu-miR-34a-5p | MIMAT0000542 | Dnajc1 | 13418 | HITS-CLIP | 23142080\|29167373 | Unspecified |
| mmu-miR-34a-5p | MIMAT0000542 | Dnajc10 | 66861 | HITS-CLIP | 19536157\|31433990\|29167373\|23142 | Unspecified |
| mmu-miR-34a-5p | MIMAT0000542 | Dnajc13 | 235567 | HITS-CLIP | 19536157\|29167373\|23142080 | Unspecified |
| mmu-miR-34a-5p | MIMAT0000542 | Dnajc16 | 214063 | HITS-CLIP | 19536157\|29167373\|23142080 | Unspecified |
| mmu-miR-34a-5p | MIMAT0000542 | Dnajc2 | 22791 | HITS-CLIP | 23142080 | Unspecified |
| mmu-miR-34a-5p | MIMAT0000542 | Dnajc21 | 78244 | HITS-CLIP | 23142080 | Unspecified |
| mmu-miR-34a-5p | MIMAT0000542 | Dnajc24 | 99349 | Microarrays//HITS-CLIP | 37986224\|23142080 | Unspecified |
| mmu-miR-34a-5p | MIMAT0000542 | Dnajc25 | 72429 | HITS-CLIP | 29167373 | Unspecified |
| mmu-miR-34a-5p | MIMAT0000542 | Dnajc3 | 100037258 | HITS-CLIP | 23142080 | Unspecified |

| | | | | | | |
|---|---|---|---|---|---|---|
| mmu-miR-34a-5p | MIMAT0000542 | Dnajc30 | 66114 | HITS-CLIP | 31433990\|23142080 | Unspecified |
| mmu-miR-34a-5p | MIMAT0000542 | Dnajc4 | 57431 | HITS-CLIP | 23142080 | Unspecified |
| mmu-miR-34a-5p | MIMAT0000542 | Dnajc5 | 13002 | HITS-CLIP | 23142080\|29167373 | Unspecified |
| mmu-miR-34a-5p | MIMAT0000542 | Dnajc6 | 72685 | HITS-CLIP | 19536157 | Unspecified |
| mmu-miR-34a-5p | MIMAT0000542 | Dnajc7 | 56354 | HITS-CLIP | 23142080\|29167373 | Unspecified |
| mmu-miR-34a-5p | MIMAT0000542 | Dnajc8 | 68598 | HITS-CLIP | 23142080 | Unspecified |
| mmu-miR-34a-5p | MIMAT0000542 | Dnal1 | 105000 | HITS-CLIP | 19536157 | Unspecified |
| mmu-miR-34a-5p | MIMAT0000542 | Dnase1l1 | 69537 | HITS-CLIP | 23142080\|29167373 | Unspecified |
| mmu-miR-34a-5p | MIMAT0000542 | Dner | 227325 | HITS-CLIP | 19536157 | Unspecified |
| mmu-miR-34a-5p | MIMAT0000542 | Dnlz | 52838 | HITS-CLIP | 23142080 | Unspecified |
| mmu-miR-34a-5p | MIMAT0000542 | Dnm1 | 13429 | HITS-CLIP | 19536157 | Unspecified |
| mmu-miR-34a-5p | MIMAT0000542 | Dnm1l | 74006 | HITS-CLIP | 37986224\|31433990\|19536157\|23142 | Unspecified |
| mmu-miR-34a-5p | MIMAT0000542 | Dnm2 | 13430 | HITS-CLIP | 23142080 | Unspecified |
| mmu-miR-34a-5p | MIMAT0000542 | Dnmbp | 71972 | HITS-CLIP | 23142080\|19536157 | Unspecified |
| mmu-miR-34a-5p | MIMAT0000542 | Dnmt1 | 13433 | HITS-CLIP | 23142080\|23630078\|31433990\|29167 | Unspecified |
| mmu-miR-34a-5p | MIMAT0000542 | Dnmt3a | 13435 | HITS-CLIP | 23142080\|29167373 | Unspecified |
| mmu-miR-34a-5p | MIMAT0000542 | Dnpep | 13437 | HITS-CLIP | 23142080 | Unspecified |
| mmu-miR-34a-5p | MIMAT0000542 | Dnph1 | 381101 | Microarrays | 37986224\|23027903 | Unspecified |
| mmu-miR-34a-5p | MIMAT0000542 | Dntt | 21673 | HITS-CLIP | 23142080 | Unspecified |
| mmu-miR-34a-5p | MIMAT0000542 | Dnttip1 | 76233 | HITS-CLIP | 23142080 | Unspecified |
| mmu-miR-34a-5p | MIMAT0000542 | Dnttip2 | 99480 | HITS-CLIP | 23142080 | Unspecified |
| mmu-miR-34a-5p | MIMAT0000542 | Dock1 | 330662 | HITS-CLIP | 29167373 | Unspecified |
| mmu-miR-34a-5p | MIMAT0000542 | Dock10 | 210293 | HITS-CLIP | 23142080\|29167373 | Unspecified |
| mmu-miR-34a-5p | MIMAT0000542 | Dock11 | 75974 | HITS-CLIP | 29167373\|23142080 | Unspecified |
| mmu-miR-34a-5p | MIMAT0000542 | Dock2 | 94176 | HITS-CLIP | 23142080\|29167373 | Unspecified |
| mmu-miR-34a-5p | MIMAT0000542 | Dock4 | 238130 | HITS-CLIP | 19536157\|29167373 | Unspecified |
| mmu-miR-34a-5p | MIMAT0000542 | Dock8 | 76088 | HITS-CLIP | 23142080\|31433990\|29167373 | Unspecified |
| mmu-miR-34a-5p | MIMAT0000542 | Dock9 | 105445 | HITS-CLIP | 23142080\|19536157 | Unspecified |
| mmu-miR-34a-5p | MIMAT0000542 | Dok2 | 13449 | HITS-CLIP | 23142080\|29167373 | Unspecified |
| mmu-miR-34a-5p | MIMAT0000542 | Dok4 | 114255 | HITS-CLIP | 23630078 | Unspecified |
| mmu-miR-34a-5p | MIMAT0000542 | Dolpp1 | 57170 | HITS-CLIP | 29084942 | Unspecified |
| mmu-miR-34a-5p | MIMAT0000542 | Dop1a | 23033 | HITS-CLIP | 23142080 | Unspecified |
| mmu-miR-34a-5p | MIMAT0000542 | Dop1b | 9980 | HITS-CLIP | 23142080\|29167373 | Unspecified |
| mmu-miR-34a-5p | MIMAT0000542 | Dot1l | 208266 | HITS-CLIP | 31433990\|23142080\|26203562\|29167 | Unspecified |
| mmu-miR-34a-5p | MIMAT0000542 | Dpagt1 | 13478 | HITS-CLIP | 23142080 | Unspecified |
| mmu-miR-34a-5p | MIMAT0000542 | Dpep1 | 13479 | Microarrays | 37986224 | Unspecified |
| mmu-miR-34a-5p | MIMAT0000542 | Dpf1 | 29861 | HITS-CLIP | 23142080 | Unspecified |
| mmu-miR-34a-5p | MIMAT0000542 | Dpf2 | 19708 | HITS-CLIP | 23142080 | Unspecified |
| mmu-miR-34a-5p | MIMAT0000542 | Dph2 | 67728 | HITS-CLIP | 23142080 | Unspecified |

| | | | | | | |
|---|---|---|---|---|---|---|
| mmu-miR-34a-5p | MIMAT0000542 | Dph6 | 66632 | HITS-CLIP | 23142080 | Unspecified |
| mmu-miR-34a-5p | MIMAT0000542 | Dpm2 | 13481 | HITS-CLIP | 23142080 | Unspecified |
| mmu-miR-34a-5p | MIMAT0000542 | Dpm3 | 68563 | HITS-CLIP | 23142080 | Unspecified |
| mmu-miR-34a-5p | MIMAT0000542 | Dpp3 | 75221 | HITS-CLIP | 23142080\|29167373 | Unspecified |
| mmu-miR-34a-5p | MIMAT0000542 | Dpp4 | 13482 | HITS-CLIP | 23142080 | Unspecified |
| mmu-miR-34a-5p | MIMAT0000542 | Dpp7 | 83768 | HITS-CLIP | 29167373 | Unspecified |
| mmu-miR-34a-5p | MIMAT0000542 | Dpp8 | 74388 | HITS-CLIP | 31433990\|23630078\|26203562\|23142 | Unspecified |
| mmu-miR-34a-5p | MIMAT0000542 | Dpp9 | 224897 | HITS-CLIP | 23142080\|29167373 | Unspecified |
| mmu-miR-34a-5p | MIMAT0000542 | Dpy19l1 | 244745 | HITS-CLIP | 23142080 | Unspecified |
| mmu-miR-34a-5p | MIMAT0000542 | Dpy19l3 | 233115 | HITS-CLIP | 23142080 | Unspecified |
| mmu-miR-34a-5p | MIMAT0000542 | Dpy19l4 | 381510 | HITS-CLIP | 23142080 | Unspecified |
| mmu-miR-34a-5p | MIMAT0000542 | Dpy30 | 66310 | HITS-CLIP | 23142080 | Unspecified |
| mmu-miR-34a-5p | MIMAT0000542 | Dpyd | 99586 | Microarrays | 37986224 | Unspecified |
| mmu-miR-34a-5p | MIMAT0000542 | Dpysl2 | 12934 | HITS-CLIP | 23142080\|19536157\|30971389\|2363( | Unspecified |
| mmu-miR-34a-5p | MIMAT0000542 | Dpysl5 | 65254 | HITS-CLIP | 19536157 | Unspecified |
| mmu-miR-34a-5p | MIMAT0000542 | Dqx1 | 93838 | HITS-CLIP | 30971389 | Unspecified |
| mmu-miR-34a-5p | MIMAT0000542 | Dr1 | 13486 | HITS-CLIP | 31433990\|23142080\|29167373 | Unspecified |
| mmu-miR-34a-5p | MIMAT0000542 | Dram2 | 67171 | HITS-CLIP | 31433990\|23142080\|29167373 | Unspecified |
| mmu-miR-34a-5p | MIMAT0000542 | Drg1 | 13494 | HITS-CLIP | 23142080\|19536157\|29167373 | Unspecified |
| mmu-miR-34a-5p | MIMAT0000542 | Drg2 | 13495 | HITS-CLIP | 23142080\|29167373 | Unspecified |
| mmu-miR-34a-5p | MIMAT0000542 | Drosha | 14000 | HITS-CLIP | 23142080 | Unspecified |
| mmu-miR-34a-5p | MIMAT0000542 | Dscc1 | 72107 | Microarrays | 37986224\|23027903 | Unspecified |
| mmu-miR-34a-5p | MIMAT0000542 | Dsel | 319901 | HITS-CLIP | 23142080 | Unspecified |
| mmu-miR-34a-5p | MIMAT0000542 | Dsn1 | 66934 | HITS-CLIP | 23142080 | Unspecified |
| mmu-miR-34a-5p | MIMAT0000542 | Dst | 13518 | HITS-CLIP | 19536157\|23142080\|29167373 | Unspecified |
| mmu-miR-34a-5p | MIMAT0000542 | Dstn | 56431 | HITS-CLIP | 23142080 | Unspecified |
| mmu-miR-34a-5p | MIMAT0000542 | Dstyk | 213452 | HITS-CLIP | 19536157\|23142080 | Unspecified |
| mmu-miR-34a-5p | MIMAT0000542 | Dtl | 76843 | HITS-CLIP | 23630078\|29167373\|23142080 | Unspecified |
| mmu-miR-34a-5p | MIMAT0000542 | Dtna | 13527 | HITS-CLIP | 19536157 | Unspecified |
| mmu-miR-34a-5p | MIMAT0000542 | Dtnbp1 | 94245 | HITS-CLIP | 29167373 | Unspecified |
| mmu-miR-34a-5p | MIMAT0000542 | Dtwd2 | 68857 | HITS-CLIP | 23142080 | Unspecified |
| mmu-miR-34a-5p | MIMAT0000542 | Dtx3 | 80904 | HITS-CLIP | 23142080 | Unspecified |
| mmu-miR-34a-5p | MIMAT0000542 | Dtx3l | 209200 | HITS-CLIP | 23142080\|29167373 | Unspecified |
| mmu-miR-34a-5p | MIMAT0000542 | Dtx4 | 207521 | HITS-CLIP | 29167373 | Unspecified |
| mmu-miR-34a-5p | MIMAT0000542 | Dtymk | 21915 | HITS-CLIP | 23142080 | Unspecified |
| mmu-miR-34a-5p | MIMAT0000542 | Dus1l | 68730 | HITS-CLIP | 23142080 | Unspecified |
| mmu-miR-34a-5p | MIMAT0000542 | Dus2 | 66369 | HITS-CLIP | 23142080 | Unspecified |
| mmu-miR-34a-5p | MIMAT0000542 | Dusp1 | 19252 | HITS-CLIP | 23142080\|29167373 | Unspecified |
| mmu-miR-34a-5p | MIMAT0000542 | Dusp10 | 63953 | HITS-CLIP | 23142080 | Unspecified |

| | | | | | | |
|---|---|---|---|---|---|---|
| mmu-miR-34a-5p | MIMAT0000542 | Dusp11 | 72102 | HITS-CLIP | 21258322\|29167373\|23142080 | Unspecified |
| mmu-miR-34a-5p | MIMAT0000542 | Dusp13 | 27389 | Microarrays | 37986224 | Unspecified |
| mmu-miR-34a-5p | MIMAT0000542 | Dusp16 | 70686 | HITS-CLIP | 23142080 | Unspecified |
| mmu-miR-34a-5p | MIMAT0000542 | Dusp2 | 13537 | HITS-CLIP | 23142080 | Unspecified |
| mmu-miR-34a-5p | MIMAT0000542 | Dusp22 | 105352 | HITS-CLIP | 19536157\|23142080\|29167373 | Unspecified |
| mmu-miR-34a-5p | MIMAT0000542 | Dusp27 | 240892 | Microarrays | 37986224 | Unspecified |
| mmu-miR-34a-5p | MIMAT0000542 | Dusp3 | 72349 | Microarrays//HITS-CLIP | 37986224\|29167373 | Unspecified |
| mmu-miR-34a-5p | MIMAT0000542 | Dusp4 | 319520 | HITS-CLIP | 23142080\|29167373 | Unspecified |
| mmu-miR-34a-5p | MIMAT0000542 | Dusp5 | 240672 | HITS-CLIP | 19536157\|31433990\|23142080\|29167 | Unspecified |
| mmu-miR-34a-5p | MIMAT0000542 | Dusp6 | 67603 | HITS-CLIP | 23142080 | Unspecified |
| mmu-miR-34a-5p | MIMAT0000542 | Dusp7 | 235584 | HITS-CLIP | 23142080\|26203562\|29167373 | Unspecified |
| mmu-miR-34a-5p | MIMAT0000542 | Dusp8 | 18218 | HITS-CLIP | 19536157 | Unspecified |
| mmu-miR-34a-5p | MIMAT0000542 | Dusp9 | 75590 | HITS-CLIP | 23142080 | Unspecified |
| mmu-miR-34a-5p | MIMAT0000542 | Dut | 110074 | HITS-CLIP | 23142080 | Unspecified |
| mmu-miR-34a-5p | MIMAT0000542 | Dvl1 | 13542 | HITS-CLIP | 23142080 | Unspecified |
| mmu-miR-34a-5p | MIMAT0000542 | Dvl2 | 13543 | HITS-CLIP | 23142080 | Unspecified |
| mmu-miR-34a-5p | MIMAT0000542 | Dvl3 | 13544 | HITS-CLIP | 23142080\|29167373 | Unspecified |
| mmu-miR-34a-5p | MIMAT0000542 | Dym | 69190 | HITS-CLIP | 29167373\|23142080 | Unspecified |
| mmu-miR-34a-5p | MIMAT0000542 | Dync1h1 | 13424 | HITS-CLIP | 23142080\|19536157\|29167373 | Unspecified |
| mmu-miR-34a-5p | MIMAT0000542 | Dync1i2 | 13427 | HITS-CLIP | 30971389\|23142080\|29167373 | Unspecified |
| mmu-miR-34a-5p | MIMAT0000542 | Dync1li1 | 235661 | HITS-CLIP | 23142080 | Unspecified |
| mmu-miR-34a-5p | MIMAT0000542 | Dync1li2 | 234663 | HITS-CLIP | 19536157\|29167373\|23142080 | Unspecified |
| mmu-miR-34a-5p | MIMAT0000542 | Dync2h1 | 110350 | HITS-CLIP | 23142080 | Unspecified |
| mmu-miR-34a-5p | MIMAT0000542 | Dync2li1 | 213575 | Microarrays | 37986224 | Unspecified |
| mmu-miR-34a-5p | MIMAT0000542 | Dynll1 | 56455 | HITS-CLIP | 23142080\|19536157 | Unspecified |
| mmu-miR-34a-5p | MIMAT0000542 | Dynll2 | 68097 | HITS-CLIP | 37986224\|23142080\|19536157\|29167 | Unspecified |
| mmu-miR-34a-5p | MIMAT0000542 | Dynlrb1 | 67068 | HITS-CLIP | 23142080\|29167373 | Unspecified |
| mmu-miR-34a-5p | MIMAT0000542 | Dynlt1a | 100310872 | HITS-CLIP | 23142080\|29167373 | Unspecified |
| mmu-miR-34a-5p | MIMAT0000542 | Dynlt3 | 67117 | HITS-CLIP | 31433990\|29167373 | Unspecified |
| mmu-miR-34a-5p | MIMAT0000542 | Dyrk1a | 13548 | HITS-CLIP | 31433990\|23142080\|29167373 | Unspecified |
| mmu-miR-34a-5p | MIMAT0000542 | Dyrk1b | 13549 | Microarrays//HITS-CLIP | 37986224\|23142080 | Unspecified |
| mmu-miR-34a-5p | MIMAT0000542 | Dyrk2 | 69181 | HITS-CLIP | 23142080 | Unspecified |
| mmu-miR-34a-5p | MIMAT0000542 | Dyrk3 | 226419 | HITS-CLIP | 23142080 | Unspecified |
| mmu-miR-34a-5p | MIMAT0000542 | Dzank1 | 241688 | HITS-CLIP | 19536157 | Unspecified |
| mmu-miR-34a-5p | MIMAT0000542 | Dzip3 | 224170 | HITS-CLIP | 19536157 | Unspecified |
| mmu-miR-34a-5p | MIMAT0000542 | E130309D02Rik | 231868 | HITS-CLIP | 23142080 | Unspecified |
| mmu-miR-34a-5p | MIMAT0000542 | E2f1 | 13555 | HITS-CLIP | 23142080 | Unspecified |
| mmu-miR-34a-5p | MIMAT0000542 | E2f2 | 242705 | HITS-CLIP | 23142080\|29167373 | Unspecified |
| mmu-miR-34a-5p | MIMAT0000542 | E2f3 | 13557 | HITS-CLIP, Microarrays//Microarrays//HIT! | 37986224\|23027903\|19536157\|23142 | Unspecified |

| | | | | | | |
|---|---|---|---|---|---|---|
| mmu-miR-34a-5p | MIMAT0000542 | E2f4 | 104394 | HITS-CLIP | 23142080\|23630078\|29167373 | Unspecified |
| mmu-miR-34a-5p | MIMAT0000542 | E2f5 | 13559 | Microarrays | 37986224\|23027903 | Unspecified |
| mmu-miR-34a-5p | MIMAT0000542 | E2f7 | 52679 | Microarrays//HITS-CLIP | 37986224\|23027903\|31433990\|23142 | Unspecified |
| mmu-miR-34a-5p | MIMAT0000542 | E2f8 | 108961 | HITS-CLIP | 23142080\|29167373 | Unspecified |
| mmu-miR-34a-5p | MIMAT0000542 | E330009J07Rik | 243780 | HITS-CLIP | 23142080 | Unspecified |
| mmu-miR-34a-5p | MIMAT0000542 | Eaf1 | 74427 | HITS-CLIP | 23142080\|29167373 | Unspecified |
| mmu-miR-34a-5p | MIMAT0000542 | Eapp | 66266 | HITS-CLIP | 23142080 | Unspecified |
| mmu-miR-34a-5p | MIMAT0000542 | Ebag9 | 55960 | HITS-CLIP | 23142080 | Unspecified |
| mmu-miR-34a-5p | MIMAT0000542 | Ebi3 | 50498 | HITS-CLIP | 29167373 | Unspecified |
| mmu-miR-34a-5p | MIMAT0000542 | Ebna1bp2 | 69072 | HITS-CLIP | 23142080 | Unspecified |
| mmu-miR-34a-5p | MIMAT0000542 | Ebp | 13595 | HITS-CLIP | 23142080 | Unspecified |
| mmu-miR-34a-5p | MIMAT0000542 | Ecd | 70601 | HITS-CLIP | 19536157\|23142080 | Unspecified |
| mmu-miR-34a-5p | MIMAT0000542 | Ece1 | 230857 | HITS-CLIP | 23142080 | Unspecified |
| mmu-miR-34a-5p | MIMAT0000542 | Ech1 | 51798 | HITS-CLIP | 23142080 | Unspecified |
| mmu-miR-34a-5p | MIMAT0000542 | Echdc1 | 52665 | HITS-CLIP | 23142080\|29167373 | Unspecified |
| mmu-miR-34a-5p | MIMAT0000542 | Echs1 | 93747 | HITS-CLIP | 23142080 | Unspecified |
| mmu-miR-34a-5p | MIMAT0000542 | Eci2 | 23986 | HITS-CLIP | 23142080 | Unspecified |
| mmu-miR-34a-5p | MIMAT0000542 | Ecm1 | 13601 | HITS-CLIP | 23142080 | Unspecified |
| mmu-miR-34a-5p | MIMAT0000542 | Ecpas | 23392 | HITS-CLIP | 30971389\|19536157\|23142080\|29167 | Unspecified |
| mmu-miR-34a-5p | MIMAT0000542 | Ecsit | 26940 | Microarrays | 37986224\|23027903 | Unspecified |
| mmu-miR-34a-5p | MIMAT0000542 | Ect2 | 13605 | HITS-CLIP | 23142080 | Unspecified |
| mmu-miR-34a-5p | MIMAT0000542 | Edc4 | 234699 | HITS-CLIP | 23142080\|29167373 | Unspecified |
| mmu-miR-34a-5p | MIMAT0000542 | Edem1 | 192193 | Microarrays//HITS-CLIP | 37986224\|23027903\|31433990\|23142 | Unspecified |
| mmu-miR-34a-5p | MIMAT0000542 | Edem2 | 108687 | HITS-CLIP | 23142080 | Unspecified |
| mmu-miR-34a-5p | MIMAT0000542 | Edem3 | 66967 | HITS-CLIP | 23142080\|29167373 | Unspecified |
| mmu-miR-34a-5p | MIMAT0000542 | Edf1 | 59022 | HITS-CLIP | 23142080 | Unspecified |
| mmu-miR-34a-5p | MIMAT0000542 | Edn1 | 13614 | HITS-CLIP | 23142080 | Unspecified |
| mmu-miR-34a-5p | MIMAT0000542 | Ednra | 13617 | Microarrays | 37986224 | Unspecified |
| mmu-miR-34a-5p | MIMAT0000542 | Ednrb | 13618 | HITS-CLIP | 19536157 | Unspecified |
| mmu-miR-34a-5p | MIMAT0000542 | Edrf1 | 214764 | HITS-CLIP | 29167373\|23142080 | Unspecified |
| mmu-miR-34a-5p | MIMAT0000542 | Eea1 | 216238 | HITS-CLIP | 23630078\|19536157\|31433990\|23142 | Unspecified |
| mmu-miR-34a-5p | MIMAT0000542 | Eed | 13626 | HITS-CLIP | 31433990\|23142080 | Unspecified |
| mmu-miR-34a-5p | MIMAT0000542 | Eef1a1 | 13627 | HITS-CLIP | 37986224\|19536157\|29167373\|23142 | Unspecified |
| mmu-miR-34a-5p | MIMAT0000542 | Eef1akmt1 | 221143 | HITS-CLIP | 23142080 | Unspecified |
| mmu-miR-34a-5p | MIMAT0000542 | Eef1b2 | 55949 | HITS-CLIP | 23142080 | Unspecified |
| mmu-miR-34a-5p | MIMAT0000542 | Eef1e1 | 66143 | HITS-CLIP | 29167373 | Unspecified |
| mmu-miR-34a-5p | MIMAT0000542 | Eef2 | 13629 | HITS-CLIP | 19536157\|31433990\|23142080\|29167 | Unspecified |
| mmu-miR-34a-5p | MIMAT0000542 | Eef2k | 13631 | HITS-CLIP | 23142080 | Unspecified |
| mmu-miR-34a-5p | MIMAT0000542 | Eefsec | 65967 | HITS-CLIP | 23142080 | Unspecified |

| | | | | | | |
|---|---|---|---|---|---|---|
| mmu-miR-34a-5p | MIMAT0000542 | Eepd1 | 67484 | HITS-CLIP | 29167373 | Unspecified |
| mmu-miR-34a-5p | MIMAT0000542 | Efcab14 | 230648 | HITS-CLIP | 23142080\|29167373 | Unspecified |
| mmu-miR-34a-5p | MIMAT0000542 | Efhd2 | 27984 | HITS-CLIP | 29167373\|23142080 | Unspecified |
| mmu-miR-34a-5p | MIMAT0000542 | Efl1 | 1942 | HITS-CLIP | 31433990\|23142080 | Unspecified |
| mmu-miR-34a-5p | MIMAT0000542 | Efna5 | 13640 | HITS-CLIP | 23142080 | Unspecified |
| mmu-miR-34a-5p | MIMAT0000542 | Efr3a | 76740 | HITS-CLIP | 23142080\|19536157\|31433990 | Unspecified |
| mmu-miR-34a-5p | MIMAT0000542 | Eftud2 | 20624 | HITS-CLIP | 19536157\|23142080\|29167373 | Unspecified |
| mmu-miR-34a-5p | MIMAT0000542 | Egln1 | 112405 | HITS-CLIP | 31433990\|23142080 | Unspecified |
| mmu-miR-34a-5p | MIMAT0000542 | Egln2 | 112406 | HITS-CLIP | 23142080\|29167373 | Unspecified |
| mmu-miR-34a-5p | MIMAT0000542 | Egln3 | 112407 | Microarrays//HITS-CLIP | 37986224\|31433990\|23142080 | Unspecified |
| mmu-miR-34a-5p | MIMAT0000542 | Egr1 | 13653 | HITS-CLIP | 23142080 | Unspecified |
| mmu-miR-34a-5p | MIMAT0000542 | Egr2 | 13654 | HITS-CLIP | 37986224\|23142080 | Unspecified |
| mmu-miR-34a-5p | MIMAT0000542 | Egr3 | 13655 | HITS-CLIP | 23142080 | Unspecified |
| mmu-miR-34a-5p | MIMAT0000542 | Ehbp1l1 | 114601 | HITS-CLIP | 23630078\|30971389\|29167373\|23142 | Unspecified |
| mmu-miR-34a-5p | MIMAT0000542 | Ehd1 | 13660 | HITS-CLIP | 29167373 | Unspecified |
| mmu-miR-34a-5p | MIMAT0000542 | Ehd2 | 259300 | HITS-CLIP | 19536157 | Unspecified |
| mmu-miR-34a-5p | MIMAT0000542 | Ehd3 | 57440 | HITS-CLIP | 23142080 | Unspecified |
| mmu-miR-34a-5p | MIMAT0000542 | Ehd4 | 98878 | HITS-CLIP | 29167373 | Unspecified |
| mmu-miR-34a-5p | MIMAT0000542 | Ehhadh | 74147 | HITS-CLIP | 23142080 | Unspecified |
| mmu-miR-34a-5p | MIMAT0000542 | Ehmt1 | 77683 | HITS-CLIP | 23142080\|29167373 | Unspecified |
| mmu-miR-34a-5p | MIMAT0000542 | Ehmt2 | 110147 | HITS-CLIP | 23142080 | Unspecified |
| mmu-miR-34a-5p | MIMAT0000542 | Ei24 | 13663 | Microarrays//HITS-CLIP | 37986224\|23027903\|23142080 | Unspecified |
| mmu-miR-34a-5p | MIMAT0000542 | Eid1 | 58521 | HITS-CLIP | 23142080\|29167373 | Unspecified |
| mmu-miR-34a-5p | MIMAT0000542 | Eif1 | 20918 | HITS-CLIP | 23142080\|29167373 | Unspecified |
| mmu-miR-34a-5p | MIMAT0000542 | Eif1a | 13664 | HITS-CLIP | 23142080 | Unspecified |
| mmu-miR-34a-5p | MIMAT0000542 | Eif1ad | 69860 | HITS-CLIP | 23142080\|23630078\|26203562 | Unspecified |
| mmu-miR-34a-5p | MIMAT0000542 | Eif1ax | 66235 | HITS-CLIP | 29167373 | Unspecified |
| mmu-miR-34a-5p | MIMAT0000542 | Eif1b | 68969 | HITS-CLIP | 23142080 | Unspecified |
| mmu-miR-34a-5p | MIMAT0000542 | Eif2a | 229317 | HITS-CLIP | 29167373\|23142080 | Unspecified |
| mmu-miR-34a-5p | MIMAT0000542 | Eif2ak1 | 15467 | HITS-CLIP | 23142080 | Unspecified |
| mmu-miR-34a-5p | MIMAT0000542 | Eif2ak2 | 19106 | HITS-CLIP | 29167373 | Unspecified |
| mmu-miR-34a-5p | MIMAT0000542 | Eif2ak4 | 27103 | HITS-CLIP | 23142080\|29167373 | Unspecified |
| mmu-miR-34a-5p | MIMAT0000542 | Eif2b1 | 209354 | HITS-CLIP | 23142080 | Unspecified |
| mmu-miR-34a-5p | MIMAT0000542 | Eif2b4 | 13667 | HITS-CLIP | 23142080 | Unspecified |
| mmu-miR-34a-5p | MIMAT0000542 | Eif2b5 | 224045 | HITS-CLIP | 23142080 | Unspecified |
| mmu-miR-34a-5p | MIMAT0000542 | Eif2s1 | 13665 | HITS-CLIP | 23142080 | Unspecified |
| mmu-miR-34a-5p | MIMAT0000542 | Eif2s2 | 67204 | HITS-CLIP | 23142080 | Unspecified |
| mmu-miR-34a-5p | MIMAT0000542 | Eif3a | 13669 | HITS-CLIP | 23142080\|29167373 | Unspecified |
| mmu-miR-34a-5p | MIMAT0000542 | Eif3b | 27979 | HITS-CLIP | 31433990\|23142080\|29167373 | Unspecified |

| | | | | | | |
|---|---|---|---|---|---|---|
| mmu-miR-34a-5p | MIMAT0000542 | Eif3c | 56347 | HITS-CLIP | 31433990\|19536157\|23142080\|29167 | Unspecified |
| mmu-miR-34a-5p | MIMAT0000542 | Eif3d | 55944 | HITS-CLIP | 23142080 | Unspecified |
| mmu-miR-34a-5p | MIMAT0000542 | Eif3e | 16341 | HITS-CLIP | 23142080\|26203562 | Unspecified |
| mmu-miR-34a-5p | MIMAT0000542 | Eif3f | 66085 | HITS-CLIP | 23142080 | Unspecified |
| mmu-miR-34a-5p | MIMAT0000542 | Eif3g | 8666 | HITS-CLIP | 23142080 | Unspecified |
| mmu-miR-34a-5p | MIMAT0000542 | Eif3h | 68135 | HITS-CLIP | 23142080 | Unspecified |
| mmu-miR-34a-5p | MIMAT0000542 | Eif3i | 54709 | HITS-CLIP | 23142080 | Unspecified |
| mmu-miR-34a-5p | MIMAT0000542 | Eif3k | 73830 | HITS-CLIP | 23142080 | Unspecified |
| mmu-miR-34a-5p | MIMAT0000542 | Eif3l | 223691 | HITS-CLIP | 29167373\|23142080 | Unspecified |
| mmu-miR-34a-5p | MIMAT0000542 | Eif4a1 | 13681 | HITS-CLIP | 31433990 | Unspecified |
| mmu-miR-34a-5p | MIMAT0000542 | Eif4a2 | 13682 | HITS-CLIP | 31433990\|29167373 | Unspecified |
| mmu-miR-34a-5p | MIMAT0000542 | Eif4a3 | 192170 | HITS-CLIP | 23142080\|19536157 | Unspecified |
| mmu-miR-34a-5p | MIMAT0000542 | Eif4b | 75705 | HITS-CLIP | 23142080\|31433990\|19536157\|29167 | Unspecified |
| mmu-miR-34a-5p | MIMAT0000542 | Eif4e | 13684 | HITS-CLIP | 29167373 | Unspecified |
| mmu-miR-34a-5p | MIMAT0000542 | Eif4e2 | 26987 | HITS-CLIP | 30971389\|29167373 | Unspecified |
| mmu-miR-34a-5p | MIMAT0000542 | Eif4e3 | 66892 | Microarrays//HITS-CLIP | 37986224\|23027903\|23142080 | Unspecified |
| mmu-miR-34a-5p | MIMAT0000542 | Eif4ebp1 | 13685 | HITS-CLIP | 23142080 | Unspecified |
| mmu-miR-34a-5p | MIMAT0000542 | Eif4ebp2 | 13688 | Microarrays//HITS-CLIP | 37986224\|23027903\|19536157\|23142 | Unspecified |
| mmu-miR-34a-5p | MIMAT0000542 | Eif4enif1 | 74203 | HITS-CLIP | 23142080\|29167373 | Unspecified |
| mmu-miR-34a-5p | MIMAT0000542 | Eif4g1 | 208643 | HITS-CLIP | 30971389\|31433990\|19536157\|2363C | Unspecified |
| mmu-miR-34a-5p | MIMAT0000542 | Eif4g2 | 13690 | HITS-CLIP | 23142080\|31433990\|19536157\|29167 | Unspecified |
| mmu-miR-34a-5p | MIMAT0000542 | Eif4g3 | 230861 | HITS-CLIP | 19536157\|30971389\|23142080\|29167 | Unspecified |
| mmu-miR-34a-5p | MIMAT0000542 | Eif4h | 22384 | HITS-CLIP | 23142080\|29167373 | Unspecified |
| mmu-miR-34a-5p | MIMAT0000542 | Eif5 | 217869 | HITS-CLIP | 23142080\|19536157\|29167373 | Unspecified |
| mmu-miR-34a-5p | MIMAT0000542 | Eif5a | 276770 | HITS-CLIP | 23142080 | Unspecified |
| mmu-miR-34a-5p | MIMAT0000542 | Eif5a2 | 208691 | Microarrays | 37986224 | Unspecified |
| mmu-miR-34a-5p | MIMAT0000542 | Eif5b | 226982 | HITS-CLIP | 23142080\|29167373 | Unspecified |
| mmu-miR-34a-5p | MIMAT0000542 | Eif6 | 16418 | HITS-CLIP | 23142080 | Unspecified |
| mmu-miR-34a-5p | MIMAT0000542 | Eipr1 | 7260 | HITS-CLIP | 29167373 | Unspecified |
| mmu-miR-34a-5p | MIMAT0000542 | Elac1 | 114615 | HITS-CLIP | 29167373 | Unspecified |
| mmu-miR-34a-5p | MIMAT0000542 | Elavl1 | 15568 | HITS-CLIP | 23142080\|29167373 | Unspecified |
| mmu-miR-34a-5p | MIMAT0000542 | Elavl3 | 15571 | HITS-CLIP | 19536157 | Unspecified |
| mmu-miR-34a-5p | MIMAT0000542 | Elavl4 | 15572 | HITS-CLIP | 19536157 | Unspecified |
| mmu-miR-34a-5p | MIMAT0000542 | Elf1 | 13709 | HITS-CLIP | 31433990\|23142080 | Unspecified |
| mmu-miR-34a-5p | MIMAT0000542 | Elf2 | 69257 | HITS-CLIP | 23142080\|29167373 | Unspecified |
| mmu-miR-34a-5p | MIMAT0000542 | Elf4 | 56501 | HITS-CLIP | 23142080 | Unspecified |
| mmu-miR-34a-5p | MIMAT0000542 | Elk1 | 13712 | HITS-CLIP | 29167373 | Unspecified |
| mmu-miR-34a-5p | MIMAT0000542 | Elk3 | 13713 | HITS-CLIP | 23142080\|29167373 | Unspecified |
| mmu-miR-34a-5p | MIMAT0000542 | Elk4 | 13714 | HITS-CLIP | 19536157\|23142080\|29167373 | Unspecified |

| | | | | | | |
|---|---|---|---|---|---|---|
| mmu-miR-34a-5p | MIMAT0000542 | Ell | 13716 | HITS-CLIP | 23142080\|29167373 | Unspecified |
| mmu-miR-34a-5p | MIMAT0000542 | Ell2 | 192657 | Microarrays//HITS-CLIP | 37986224\|23142080\|31433990\|29167 | Unspecified |
| mmu-miR-34a-5p | MIMAT0000542 | Elmo1 | 140580 | HITS-CLIP | 23142080\|29167373 | Unspecified |
| mmu-miR-34a-5p | MIMAT0000542 | Elmo2 | 140579 | HITS-CLIP | 19536157\|23142080\|29167373 | Unspecified |
| mmu-miR-34a-5p | MIMAT0000542 | Elmod1 | 270162 | HITS-CLIP | 19536157 | Unspecified |
| mmu-miR-34a-5p | MIMAT0000542 | Elmod2 | 244548 | Microarrays//HITS-CLIP | 37986224\|23630078 | Unspecified |
| mmu-miR-34a-5p | MIMAT0000542 | Elmsan1 | 238317 | HITS-CLIP | 23142080\|29167373 | Unspecified |
| mmu-miR-34a-5p | MIMAT0000542 | Eloa | 6924 | HITS-CLIP | 23142080 | Unspecified |
| mmu-miR-34a-5p | MIMAT0000542 | Elof1 | 66126 | HITS-CLIP | 23142080 | Unspecified |
| mmu-miR-34a-5p | MIMAT0000542 | Elovl1 | 54325 | HITS-CLIP | 23142080 | Unspecified |
| mmu-miR-34a-5p | MIMAT0000542 | Elovl4 | 83603 | HITS-CLIP | 26203562 | Unspecified |
| mmu-miR-34a-5p | MIMAT0000542 | Elovl5 | 68801 | HITS-CLIP | 19536157\|23142080 | Unspecified |
| mmu-miR-34a-5p | MIMAT0000542 | Elovl6 | 170439 | HITS-CLIP | 23142080 | Unspecified |
| mmu-miR-34a-5p | MIMAT0000542 | Elp1 | 37455 | HITS-CLIP | 23142080 | Unspecified |
| mmu-miR-34a-5p | MIMAT0000542 | Elp2 | 58523 | HITS-CLIP | 23142080\|26203562 | Unspecified |
| mmu-miR-34a-5p | MIMAT0000542 | Elp3 | 74195 | HITS-CLIP | 23142080 | Unspecified |
| mmu-miR-34a-5p | MIMAT0000542 | Elp4 | 77766 | HITS-CLIP | 37986224\|23142080 | Unspecified |
| mmu-miR-34a-5p | MIMAT0000542 | Elp5 | 54351 | HITS-CLIP | 23142080\|29167373 | Unspecified |
| mmu-miR-34a-5p | MIMAT0000542 | Emb | 13723 | HITS-CLIP | 23142080 | Unspecified |
| mmu-miR-34a-5p | MIMAT0000542 | Emc1 | 230866 | HITS-CLIP | 23142080\|31433990 | Unspecified |
| mmu-miR-34a-5p | MIMAT0000542 | Emc10 | 69683 | HITS-CLIP | 23630078\|19536157\|31433990\|23142 | Unspecified |
| mmu-miR-34a-5p | MIMAT0000542 | Emc2 | 66736 | HITS-CLIP | 19536157\|23142080 | Unspecified |
| mmu-miR-34a-5p | MIMAT0000542 | Emc3 | 66087 | HITS-CLIP | 23142080 | Unspecified |
| mmu-miR-34a-5p | MIMAT0000542 | Emc6 | 66048 | HITS-CLIP | 26203562 | Unspecified |
| mmu-miR-34a-5p | MIMAT0000542 | Emc8 | 18117 | HITS-CLIP | 31433990\|29167373 | Unspecified |
| mmu-miR-34a-5p | MIMAT0000542 | Emd | 13726 | HITS-CLIP | 23142080 | Unspecified |
| mmu-miR-34a-5p | MIMAT0000542 | Eme1 | 268465 | HITS-CLIP | 23142080 | Unspecified |
| mmu-miR-34a-5p | MIMAT0000542 | Emg1 | 14791 | HITS-CLIP | 23142080 | Unspecified |
| mmu-miR-34a-5p | MIMAT0000542 | Emilin2 | 246707 | HITS-CLIP | 23142080\|30971389 | Unspecified |
| mmu-miR-34a-5p | MIMAT0000542 | Eml4 | 78798 | HITS-CLIP | 19536157\|31433990\|23142080\|29167 | Unspecified |
| mmu-miR-34a-5p | MIMAT0000542 | Emp1 | 13730 | HITS-CLIP | 29167373 | Unspecified |
| mmu-miR-34a-5p | MIMAT0000542 | Emp2 | 13731 | HITS-CLIP | 19536157 | Unspecified |
| mmu-miR-34a-5p | MIMAT0000542 | Emp3 | 2014 | HITS-CLIP | 23142080 | Unspecified |
| mmu-miR-34a-5p | MIMAT0000542 | Emsy | 56946 | HITS-CLIP | 23142080\|19536157\|29167373 | Unspecified |
| mmu-miR-34a-5p | MIMAT0000542 | Enah | 13800 | HITS-CLIP | 19536157 | Unspecified |
| mmu-miR-34a-5p | MIMAT0000542 | Enc1 | 13803 | Microarrays//HITS-CLIP | 37986224\|19536157\|31433990\|23142 | Unspecified |
| mmu-miR-34a-5p | MIMAT0000542 | Endod1 | 71946 | HITS-CLIP | 23142080\|30971389\|29167373 | Unspecified |
| mmu-miR-34a-5p | MIMAT0000542 | Eno1 | 13806 | HITS-CLIP | 31433990\|23142080\|29167373 | Unspecified |
| mmu-miR-34a-5p | MIMAT0000542 | Eno2 | 13807 | HITS-CLIP | 19536157 | Unspecified |

| | | | | | | |
|---|---|---|---|---|---|---|
| mmu-miR-34a-5p | MIMAT0000542 | Eno3 | 13808 | Microarrays | 37986224\|23027903 | Unspecified |
| mmu-miR-34a-5p | MIMAT0000542 | Enox2 | 209224 | HITS-CLIP | 23142080 | Unspecified |
| mmu-miR-34a-5p | MIMAT0000542 | Enpp1 | 18605 | HITS-CLIP | 23142080 | Unspecified |
| mmu-miR-34a-5p | MIMAT0000542 | Enpp2 | 18606 | HITS-CLIP | 19536157 | Unspecified |
| mmu-miR-34a-5p | MIMAT0000542 | Enpp4 | 224794 | HITS-CLIP | 23142080\|29167373 | Unspecified |
| mmu-miR-34a-5p | MIMAT0000542 | Enpp5 | 83965 | Microarrays | 37986224 | Unspecified |
| mmu-miR-34a-5p | MIMAT0000542 | Ensa | 56205 | HITS-CLIP | 29167373\|23142080 | Unspecified |
| mmu-miR-34a-5p | MIMAT0000542 | Enthd1 | 383075 | HITS-CLIP | 23142080 | Unspecified |
| mmu-miR-34a-5p | MIMAT0000542 | Entpd3 | 215446 | HITS-CLIP | 19536157 | Unspecified |
| mmu-miR-34a-5p | MIMAT0000542 | Entpd5 | 12499 | Microarrays//HITS-CLIP | 37986224\|23027903\|31433990\|1953( | Unspecified |
| mmu-miR-34a-5p | MIMAT0000542 | Entpd6 | 12497 | HITS-CLIP | 23142080 | Unspecified |
| mmu-miR-34a-5p | MIMAT0000542 | Entpd7 | 93685 | HITS-CLIP | 29167373 | Unspecified |
| mmu-miR-34a-5p | MIMAT0000542 | Entr1 | 10807 | HITS-CLIP | 23142080\|29167373 | Unspecified |
| mmu-miR-34a-5p | MIMAT0000542 | Eny2 | 223527 | HITS-CLIP | 23142080 | Unspecified |
| mmu-miR-34a-5p | MIMAT0000542 | Eogt | 101351 | HITS-CLIP | 26203562 | Unspecified |
| mmu-miR-34a-5p | MIMAT0000542 | Eomes | 13813 | HITS-CLIP | 23142080 | Unspecified |
| mmu-miR-34a-5p | MIMAT0000542 | Ep300 | 328572 | HITS-CLIP | 31433990\|23630078\|23142080\|2916 | Unspecified |
| mmu-miR-34a-5p | MIMAT0000542 | Ep400 | 75560 | HITS-CLIP | 19536157\|23142080\|29167373 | Unspecified |
| mmu-miR-34a-5p | MIMAT0000542 | Epas1 | 13819 | HITS-CLIP | 23142080\|19536157\|31433990 | Unspecified |
| mmu-miR-34a-5p | MIMAT0000542 | Epb41 | 269587 | HITS-CLIP | 23142080\|30971389\|29167373\|1953( | Unspecified |
| mmu-miR-34a-5p | MIMAT0000542 | Epb41l1 | 13821 | HITS-CLIP | 29167373 | Unspecified |
| mmu-miR-34a-5p | MIMAT0000542 | Epb41l2 | 2037 | HITS-CLIP | 31433990\|23142080\|29167373 | Unspecified |
| mmu-miR-34a-5p | MIMAT0000542 | Epb41l3 | 23136 | HITS-CLIP | 19536157 | Unspecified |
| mmu-miR-34a-5p | MIMAT0000542 | Epc1 | 13831 | HITS-CLIP | 23630078\|30971389\|23142080 | Unspecified |
| mmu-miR-34a-5p | MIMAT0000542 | Epc2 | 227867 | HITS-CLIP | 23142080 | Unspecified |
| mmu-miR-34a-5p | MIMAT0000542 | Epcam | 17075 | HITS-CLIP | 23142080 | Unspecified |
| mmu-miR-34a-5p | MIMAT0000542 | Epdr1 | 105298 | HITS-CLIP | 19536157\|23142080 | Unspecified |
| mmu-miR-34a-5p | MIMAT0000542 | Epg5 | 100502841 | HITS-CLIP | 29167373\|23142080 | Unspecified |
| mmu-miR-34a-5p | MIMAT0000542 | Epha4 | 13838 | HITS-CLIP | 19536157 | Unspecified |
| mmu-miR-34a-5p | MIMAT0000542 | Epha5 | 13839 | HITS-CLIP | 19536157 | Unspecified |
| mmu-miR-34a-5p | MIMAT0000542 | Epha6 | 13840 | HITS-CLIP | 19536157 | Unspecified |
| mmu-miR-34a-5p | MIMAT0000542 | Epha7 | 13841 | HITS-CLIP | 19536157 | Unspecified |
| mmu-miR-34a-5p | MIMAT0000542 | Ephb1 | 270190 | HITS-CLIP | 19536157 | Unspecified |
| mmu-miR-34a-5p | MIMAT0000542 | Ephb6 | 13848 | HITS-CLIP | 19536157 | Unspecified |
| mmu-miR-34a-5p | MIMAT0000542 | Ephx1 | 13849 | Microarrays//HITS-CLIP | 37986224\|23142080 | Unspecified |
| mmu-miR-34a-5p | MIMAT0000542 | Epm2aip1 | 77781 | HITS-CLIP | 23142080 | Unspecified |
| mmu-miR-34a-5p | MIMAT0000542 | Eprs | 107508 | HITS-CLIP | 23142080 | Unspecified |
| mmu-miR-34a-5p | MIMAT0000542 | Eps15 | 13858 | HITS-CLIP | 37986224\|23142080\|29167373 | Unspecified |
| mmu-miR-34a-5p | MIMAT0000542 | Eps15l1 | 13859 | HITS-CLIP | 23142080\|29167373 | Unspecified |

| | | | | | | |
|---|---|---|---|---|---|---|
| mmu-miR-34a-5p | MIMAT0000542 | Eps8l2 | 98845 | HITS-CLIP | 26203562 | Unspecified |
| mmu-miR-34a-5p | MIMAT0000542 | Eral1 | 57837 | HITS-CLIP | 23142080 | Unspecified |
| mmu-miR-34a-5p | MIMAT0000542 | Erap1 | 80898 | HITS-CLIP | 23142080\|29167373 | Unspecified |
| mmu-miR-34a-5p | MIMAT0000542 | Erbb3 | 13867 | Microarrays | 37986224 | Unspecified |
| mmu-miR-34a-5p | MIMAT0000542 | Erbin | 55914 | HITS-CLIP | 23142080\|30971389\|19536157\|29167 | Unspecified |
| mmu-miR-34a-5p | MIMAT0000542 | Erc1 | 111173 | Microarrays//HITS-CLIP | 37986224\|23027903\|29167373 | Unspecified |
| mmu-miR-34a-5p | MIMAT0000542 | Erc2 | 238988 | HITS-CLIP | 19536157 | Unspecified |
| mmu-miR-34a-5p | MIMAT0000542 | Ercc1 | 13870 | HITS-CLIP | 23142080 | Unspecified |
| mmu-miR-34a-5p | MIMAT0000542 | Ercc6 | 319955 | HITS-CLIP | 23142080\|29167373 | Unspecified |
| mmu-miR-34a-5p | MIMAT0000542 | Ercc6l | 236930 | HITS-CLIP | 23142080 | Unspecified |
| mmu-miR-34a-5p | MIMAT0000542 | Ercc6l2 | 76251 | HITS-CLIP | 23142080 | Unspecified |
| mmu-miR-34a-5p | MIMAT0000542 | Erg28 | 856767 | HITS-CLIP | 23142080 | Unspecified |
| mmu-miR-34a-5p | MIMAT0000542 | Ergic1 | 67458 | HITS-CLIP | 23142080\|19536157\|26203562\|29167 | Unspecified |
| mmu-miR-34a-5p | MIMAT0000542 | Ergic2 | 67456 | HITS-CLIP | 23142080\|29167373 | Unspecified |
| mmu-miR-34a-5p | MIMAT0000542 | Ergic3 | 66366 | HITS-CLIP | 31433990 | Unspecified |
| mmu-miR-34a-5p | MIMAT0000542 | Eri1 | 67276 | HITS-CLIP | 23142080\|29167373 | Unspecified |
| mmu-miR-34a-5p | MIMAT0000542 | Eri2 | 71151 | HITS-CLIP | 23142080 | Unspecified |
| mmu-miR-34a-5p | MIMAT0000542 | Eri3 | 140546 | HITS-CLIP | 23142080 | Unspecified |
| mmu-miR-34a-5p | MIMAT0000542 | Erlec1 | 66753 | HITS-CLIP | 23142080 | Unspecified |
| mmu-miR-34a-5p | MIMAT0000542 | Erlin1 | 226144 | HITS-CLIP | 23142080\|29167373 | Unspecified |
| mmu-miR-34a-5p | MIMAT0000542 | Erlin2 | 244373 | HITS-CLIP | 23142080\|29167373 | Unspecified |
| mmu-miR-34a-5p | MIMAT0000542 | Ermp1 | 226090 | HITS-CLIP | 23142080\|29167373 | Unspecified |
| mmu-miR-34a-5p | MIMAT0000542 | Ern1 | 78943 | HITS-CLIP | 23142080 | Unspecified |
| mmu-miR-34a-5p | MIMAT0000542 | Ero1a | 30001 | HITS-CLIP | 23630078\|30971389 | Unspecified |
| mmu-miR-34a-5p | MIMAT0000542 | Ero1b | 56605 | Microarrays | 23027903 | Unspecified |
| mmu-miR-34a-5p | MIMAT0000542 | Ero1l | 50527 | HITS-CLIP | 23142080 | Unspecified |
| mmu-miR-34a-5p | MIMAT0000542 | Ero1lb | 67475 | Microarrays | 37986224 | Unspecified |
| mmu-miR-34a-5p | MIMAT0000542 | Erp29 | 67397 | HITS-CLIP | 37986224\|23142080\|29167373 | Unspecified |
| mmu-miR-34a-5p | MIMAT0000542 | Erp44 | 76299 | HITS-CLIP | 23142080 | Unspecified |
| mmu-miR-34a-5p | MIMAT0000542 | Errfi1 | 74155 | HITS-CLIP | 26203562\|23142080 | Unspecified |
| mmu-miR-34a-5p | MIMAT0000542 | Esco1 | 77805 | HITS-CLIP | 23142080 | Unspecified |
| mmu-miR-34a-5p | MIMAT0000542 | Esco2 | 71988 | HITS-CLIP | 23142080 | Unspecified |
| mmu-miR-34a-5p | MIMAT0000542 | Esf1 | 66580 | HITS-CLIP | 31433990\|23142080\|29167373 | Unspecified |
| mmu-miR-34a-5p | MIMAT0000542 | Espl1 | 105988 | Microarrays//HITS-CLIP | 37986224\|23027903\|23142080\|29167 | Unspecified |
| mmu-miR-34a-5p | MIMAT0000542 | Espn | 56226 | HITS-CLIP | 23142080 | Unspecified |
| mmu-miR-34a-5p | MIMAT0000542 | Esrp1 | 54845 | HITS-CLIP | 26203562 | Unspecified |
| mmu-miR-34a-5p | MIMAT0000542 | Esyt1 | 23943 | HITS-CLIP | 23142080\|29167373 | Unspecified |
| mmu-miR-34a-5p | MIMAT0000542 | Esyt2 | 52635 | HITS-CLIP | 19536157\|30971389\|23142080\|29167 | Unspecified |
| mmu-miR-34a-5p | MIMAT0000542 | Etaa1 | 68145 | HITS-CLIP | 23142080 | Unspecified |

| | | | | | | |
|---|---|---|---|---|---|---|
| mmu-miR-34a-5p | MIMAT0000542 | Etf1 | 225363 | HITS-CLIP | 19536157\|23142080\|29167373 | Unspecified |
| mmu-miR-34a-5p | MIMAT0000542 | Etfa | 110842 | HITS-CLIP | 23142080 | Unspecified |
| mmu-miR-34a-5p | MIMAT0000542 | Etfbkmt | 320204 | HITS-CLIP | 23142080 | Unspecified |
| mmu-miR-34a-5p | MIMAT0000542 | Etfdh | 66841 | HITS-CLIP | 23142080 | Unspecified |
| mmu-miR-34a-5p | MIMAT0000542 | Etnk1 | 75320 | HITS-CLIP | 23142080\|29167373 | Unspecified |
| mmu-miR-34a-5p | MIMAT0000542 | Ets1 | 23871 | HITS-CLIP | 37986224\|19536157\|23142080 | Unspecified |
| mmu-miR-34a-5p | MIMAT0000542 | Ets2 | 23872 | HITS-CLIP | 23630078\|23142080\|26203562\|29167 | Unspecified |
| mmu-miR-34a-5p | MIMAT0000542 | Etv1 | 14009 | HITS-CLIP | 23142080 | Unspecified |
| mmu-miR-34a-5p | MIMAT0000542 | Etv3 | 27049 | HITS-CLIP | 23630078\|23142080\|29167373 | Unspecified |
| mmu-miR-34a-5p | MIMAT0000542 | Etv5 | 104156 | HITS-CLIP | 19536157\|23142080\|29167373 | Unspecified |
| mmu-miR-34a-5p | MIMAT0000542 | Etv6 | 14011 | HITS-CLIP | 23142080 | Unspecified |
| mmu-miR-34a-5p | MIMAT0000542 | Eva1c | 70967 | Microarrays | 37986224\|23027903 | Unspecified |
| mmu-miR-34a-5p | MIMAT0000542 | Evi2 | 14017 | HITS-CLIP | 23142080\|29167373 | Unspecified |
| mmu-miR-34a-5p | MIMAT0000542 | Evi2a | 14017 | HITS-CLIP | 29167373 | Unspecified |
| mmu-miR-34a-5p | MIMAT0000542 | Evi5 | 14020 | HITS-CLIP | 31433990\|29167373 | Unspecified |
| mmu-miR-34a-5p | MIMAT0000542 | Evl | 14026 | HITS-CLIP | 23142080\|29167373 | Unspecified |
| mmu-miR-34a-5p | MIMAT0000542 | Ewsr1 | 14030 | HITS-CLIP | 19536157\|23142080\|29167373 | Unspecified |
| mmu-miR-34a-5p | MIMAT0000542 | Exd2 | 97827 | HITS-CLIP | 23142080 | Unspecified |
| mmu-miR-34a-5p | MIMAT0000542 | Exo1 | 26909 | HITS-CLIP | 23142080 | Unspecified |
| mmu-miR-34a-5p | MIMAT0000542 | Exoc1 | 69940 | HITS-CLIP | 23142080\|29167373 | Unspecified |
| mmu-miR-34a-5p | MIMAT0000542 | Exoc2 | 66482 | HITS-CLIP | 23142080\|29167373 | Unspecified |
| mmu-miR-34a-5p | MIMAT0000542 | Exoc3 | 211446 | HITS-CLIP | 23142080\|29167373 | Unspecified |
| mmu-miR-34a-5p | MIMAT0000542 | Exoc4 | 20336 | HITS-CLIP | 37986224\|19536157\|23142080\|29167 | Unspecified |
| mmu-miR-34a-5p | MIMAT0000542 | Exoc5 | 105504 | HITS-CLIP | 23142080\|29167373 | Unspecified |
| mmu-miR-34a-5p | MIMAT0000542 | Exoc6 | 107371 | HITS-CLIP | 23142080 | Unspecified |
| mmu-miR-34a-5p | MIMAT0000542 | Exoc6b | 75914 | HITS-CLIP | 23142080\|19536157\|31433990 | Unspecified |
| mmu-miR-34a-5p | MIMAT0000542 | Exoc8 | 102058 | HITS-CLIP | 29167373 | Unspecified |
| mmu-miR-34a-5p | MIMAT0000542 | Exosc10 | 50912 | HITS-CLIP | 23142080 | Unspecified |
| mmu-miR-34a-5p | MIMAT0000542 | Exosc3 | 66362 | HITS-CLIP | 23142080 | Unspecified |
| mmu-miR-34a-5p | MIMAT0000542 | Exosc4 | 109075 | HITS-CLIP | 23142080 | Unspecified |
| mmu-miR-34a-5p | MIMAT0000542 | Exosc7 | 66446 | HITS-CLIP | 23142080 | Unspecified |
| mmu-miR-34a-5p | MIMAT0000542 | Exosc8 | 69639 | HITS-CLIP | 23142080 | Unspecified |
| mmu-miR-34a-5p | MIMAT0000542 | Exosc9 | 50911 | HITS-CLIP | 23142080 | Unspecified |
| mmu-miR-34a-5p | MIMAT0000542 | Exph5 | 320051 | HITS-CLIP | 23142080 | Unspecified |
| mmu-miR-34a-5p | MIMAT0000542 | Ext1 | 14042 | HITS-CLIP | 23142080 | Unspecified |
| mmu-miR-34a-5p | MIMAT0000542 | Extl2 | 58193 | HITS-CLIP | 23142080 | Unspecified |
| mmu-miR-34a-5p | MIMAT0000542 | Extl3 | 54616 | HITS-CLIP | 23142080\|19536157\|29167373 | Unspecified |
| mmu-miR-34a-5p | MIMAT0000542 | Eya1 | 14048 | HITS-CLIP | 29167373 | Unspecified |
| mmu-miR-34a-5p | MIMAT0000542 | Eya3 | 14050 | HITS-CLIP | 31433990\|19536157\|23142080\|29167 | Unspecified |

| | | | | | | |
|---|---|---|---|---|---|---|
| mmu-miR-34a-5p | MIMAT0000542 | Ezh1 | 14055 | HITS-CLIP | 23142080 | Unspecified |
| mmu-miR-34a-5p | MIMAT0000542 | Ezh2 | 14056 | HITS-CLIP | 23142080\|29167373 | Unspecified |
| mmu-miR-34a-5p | MIMAT0000542 | Ezr | 22350 | HITS-CLIP | 23142080 | Unspecified |
| mmu-miR-34a-5p | MIMAT0000542 | F2r | 14062 | HITS-CLIP | 23142080 | Unspecified |
| mmu-miR-34a-5p | MIMAT0000542 | F2rl1 | 14063 | HITS-CLIP | 23142080 | Unspecified |
| mmu-miR-34a-5p | MIMAT0000542 | F2rl2 | 14064 | HITS-CLIP | 23142080 | Unspecified |
| mmu-miR-34a-5p | MIMAT0000542 | F3 | 14066 | HITS-CLIP | 19536157 | Unspecified |
| mmu-miR-34a-5p | MIMAT0000542 | F8a | 14070 | HITS-CLIP | 23142080 | Unspecified |
| mmu-miR-34a-5p | MIMAT0000542 | Fa2h | 338521 | HITS-CLIP | 19536157 | Unspecified |
| mmu-miR-34a-5p | MIMAT0000542 | Faah | 14073 | HITS-CLIP | 23142080 | Unspecified |
| mmu-miR-34a-5p | MIMAT0000542 | Fabp3 | 14077 | Microarrays | 37986224 | Unspecified |
| mmu-miR-34a-5p | MIMAT0000542 | Fabp7 | 12140 | Microarrays | 37986224 | Unspecified |
| mmu-miR-34a-5p | MIMAT0000542 | Fads1 | 76267 | HITS-CLIP | 23142080\|19536157\|29167373 | Unspecified |
| mmu-miR-34a-5p | MIMAT0000542 | Fads2 | 56473 | HITS-CLIP | 23142080\|19536157\|26203562 | Unspecified |
| mmu-miR-34a-5p | MIMAT0000542 | Faf1 | 14084 | HITS-CLIP | 23630078\|19536157\|23142080\|29167 | Unspecified |
| mmu-miR-34a-5p | MIMAT0000542 | Faf2 | 76577 | HITS-CLIP | 30971389\|23142080\|29167373 | Unspecified |
| mmu-miR-34a-5p | MIMAT0000542 | Faim | 23873 | HITS-CLIP | 23142080 | Unspecified |
| mmu-miR-34a-5p | MIMAT0000542 | Fam102a | 98952 | HITS-CLIP | 37986224\|19536157\|23142080 | Unspecified |
| mmu-miR-34a-5p | MIMAT0000542 | Fam102b | 329739 | HITS-CLIP | 29167373 | Unspecified |
| mmu-miR-34a-5p | MIMAT0000542 | Fam104a | 28081 | HITS-CLIP | 30971389\|23142080 | Unspecified |
| mmu-miR-34a-5p | MIMAT0000542 | Fam107a | 268709 | HITS-CLIP | 19536157 | Unspecified |
| mmu-miR-34a-5p | MIMAT0000542 | Fam107b | 66540 | HITS-CLIP | 37986224\|23142080\|31433990 | Unspecified |
| mmu-miR-34a-5p | MIMAT0000542 | Fam110c | 104943 | HITS-CLIP | 23142080 | Unspecified |
| mmu-miR-34a-5p | MIMAT0000542 | Fam111a | 107373 | Microarrays//HITS-CLIP | 37986224\|23142080\|23027903 | Unspecified |
| mmu-miR-34a-5p | MIMAT0000542 | Fam114a1 | 68303 | HITS-CLIP | 23142080 | Unspecified |
| mmu-miR-34a-5p | MIMAT0000542 | Fam114a2 | 67726 | HITS-CLIP | 23142080\|31433990\|29167373 | Unspecified |
| mmu-miR-34a-5p | MIMAT0000542 | Fam117b | 72750 | HITS-CLIP | 23142080 | Unspecified |
| mmu-miR-34a-5p | MIMAT0000542 | Fam118a | 73225 | HITS-CLIP | 21258322\|23142080 | Unspecified |
| mmu-miR-34a-5p | MIMAT0000542 | Fam118b | 109229 | HITS-CLIP | 30971389\|23142080 | Unspecified |
| mmu-miR-34a-5p | MIMAT0000542 | Fam120a | 218236 | HITS-CLIP | 19536157\|23142080\|29167373 | Unspecified |
| mmu-miR-34a-5p | MIMAT0000542 | Fam120b | 67544 | HITS-CLIP | 23142080 | Unspecified |
| mmu-miR-34a-5p | MIMAT0000542 | Fam122a | 68034 | HITS-CLIP | 23142080 | Unspecified |
| mmu-miR-34a-5p | MIMAT0000542 | Fam122b | 78755 | HITS-CLIP | 23142080\|29167373 | Unspecified |
| mmu-miR-34a-5p | MIMAT0000542 | Fam126a | 84652 | HITS-CLIP | 23142080\|29167373 | Unspecified |
| mmu-miR-34a-5p | MIMAT0000542 | Fam126b | 213056 | HITS-CLIP | 37986224\|23142080\|29167373 | Unspecified |
| mmu-miR-34a-5p | MIMAT0000542 | Fam129a | 63913 | HITS-CLIP | 29167373 | Unspecified |
| mmu-miR-34a-5p | MIMAT0000542 | Fam129b | 227737 | HITS-CLIP | 29167373 | Unspecified |
| mmu-miR-34a-5p | MIMAT0000542 | Fam131b | 76156 | Microarrays//HITS-CLIP | 37986224\|23027903\|19536157 | Unspecified |
| mmu-miR-34a-5p | MIMAT0000542 | Fam133b | 68152 | HITS-CLIP | 29167373 | Unspecified |

| | | | | | | |
|---|---|---|---|---|---|---|
| mmu-miR-34a-5p | MIMAT0000542 | Fam135b | 70363 | HITS-CLIP | 19536157 | Unspecified |
| mmu-miR-34a-5p | MIMAT0000542 | Fam136a | 66488 | Microarrays | 37986224\|23027903 | Unspecified |
| mmu-miR-34a-5p | MIMAT0000542 | Fam13b | 225358 | HITS-CLIP | 29167373\|23142080 | Unspecified |
| mmu-miR-34a-5p | MIMAT0000542 | Fam149b | 105428 | HITS-CLIP | 23142080 | Unspecified |
| mmu-miR-34a-5p | MIMAT0000542 | Fam160a2 | 74349 | HITS-CLIP | 23142080 | Unspecified |
| mmu-miR-34a-5p | MIMAT0000542 | Fam160b1 | 226252 | HITS-CLIP | 19536157\|23142080\|29167373 | Unspecified |
| mmu-miR-34a-5p | MIMAT0000542 | Fam160b2 | 239170 | HITS-CLIP | 23142080 | Unspecified |
| mmu-miR-34a-5p | MIMAT0000542 | Fam162a | 70186 | HITS-CLIP | 23142080 | Unspecified |
| mmu-miR-34a-5p | MIMAT0000542 | Fam168a | 319604 | HITS-CLIP | 23142080 | Unspecified |
| mmu-miR-34a-5p | MIMAT0000542 | Fam168b | 214469 | HITS-CLIP | 23142080\|19536157\|29167373 | Unspecified |
| mmu-miR-34a-5p | MIMAT0000542 | Fam169b | 434197 | HITS-CLIP | 31433990 | Unspecified |
| mmu-miR-34a-5p | MIMAT0000542 | Fam171a1 | 269233 | HITS-CLIP | 19536157 | Unspecified |
| mmu-miR-34a-5p | MIMAT0000542 | Fam171b | 241520 | HITS-CLIP | 19536157 | Unspecified |
| mmu-miR-34a-5p | MIMAT0000542 | Fam172a | 68675 | HITS-CLIP | 23142080 | Unspecified |
| mmu-miR-34a-5p | MIMAT0000542 | Fam173b | 68073 | HITS-CLIP | 23142080 | Unspecified |
| mmu-miR-34a-5p | MIMAT0000542 | Fam174a | 67698 | HITS-CLIP | 23142080 | Unspecified |
| mmu-miR-34a-5p | MIMAT0000542 | Fam189b | 68521 | HITS-CLIP | 19536157 | Unspecified |
| mmu-miR-34a-5p | MIMAT0000542 | Fam193a | 231128 | HITS-CLIP | 29167373 | Unspecified |
| mmu-miR-34a-5p | MIMAT0000542 | Fam199x | 245622 | HITS-CLIP | 23142080 | Unspecified |
| mmu-miR-34a-5p | MIMAT0000542 | Fam19a1 | 320265 | HITS-CLIP | 19536157 | Unspecified |
| mmu-miR-34a-5p | MIMAT0000542 | Fam206a | 230234 | HITS-CLIP | 19536157 | Unspecified |
| mmu-miR-34a-5p | MIMAT0000542 | Fam208a | 218850 | HITS-CLIP | 23142080\|29167373 | Unspecified |
| mmu-miR-34a-5p | MIMAT0000542 | Fam208b | 54906 | HITS-CLIP | 23142080\|29167373 | Unspecified |
| mmu-miR-34a-5p | MIMAT0000542 | Fam20b | 215015 | HITS-CLIP | 19536157\|23142080 | Unspecified |
| mmu-miR-34a-5p | MIMAT0000542 | Fam20c | 80752 | HITS-CLIP | 29167373 | Unspecified |
| mmu-miR-34a-5p | MIMAT0000542 | Fam210a | 108654 | HITS-CLIP | 29167373\|23142080 | Unspecified |
| mmu-miR-34a-5p | MIMAT0000542 | Fam210b | 67017 | HITS-CLIP | 23142080 | Unspecified |
| mmu-miR-34a-5p | MIMAT0000542 | Fam214a | 235493 | Microarrays//HITS-CLIP | 37986224\|23142080 | Unspecified |
| mmu-miR-34a-5p | MIMAT0000542 | Fam214b | 230088 | HITS-CLIP | 23142080\|29167373 | Unspecified |
| mmu-miR-34a-5p | MIMAT0000542 | Fam216a | 68948 | HITS-CLIP | 23142080 | Unspecified |
| mmu-miR-34a-5p | MIMAT0000542 | Fam219a | 71901 | HITS-CLIP | 29167373 | Unspecified |
| mmu-miR-34a-5p | MIMAT0000542 | Fam219b | 78323 | HITS-CLIP | 23142080 | Unspecified |
| mmu-miR-34a-5p | MIMAT0000542 | Fam222b | 216971 | HITS-CLIP | 23142080\|30971389\|29167373 | Unspecified |
| mmu-miR-34a-5p | MIMAT0000542 | Fam234b | 57613 | HITS-CLIP | 23142080 | Unspecified |
| mmu-miR-34a-5p | MIMAT0000542 | Fam241a | 132720 | HITS-CLIP | 29167373 | Unspecified |
| mmu-miR-34a-5p | MIMAT0000542 | Fam32a | 67922 | HITS-CLIP | 23142080 | Unspecified |
| mmu-miR-34a-5p | MIMAT0000542 | Fam3c | 27999 | HITS-CLIP | 23142080 | Unspecified |
| mmu-miR-34a-5p | MIMAT0000542 | Fam45a | 67894 | HITS-CLIP | 23142080\|29167373 | Unspecified |
| mmu-miR-34a-5p | MIMAT0000542 | Fam49a | 76820 | HITS-CLIP | 23142080 | Unspecified |

| | | | | | | |
|---|---|---|---|---|---|---|
| mmu-miR-34a-5p | MIMAT0000542 | Fam49b | 223601 | HITS-CLIP | 19536157\|23142080 | Unspecified |
| mmu-miR-34a-5p | MIMAT0000542 | Fam53b | 77938 | Microarrays//HITS-CLIP | 37986224\|31433990\|23142080\|29167 | Unspecified |
| mmu-miR-34a-5p | MIMAT0000542 | Fam53c | 66306 | HITS-CLIP | 19536157\|29167373 | Unspecified |
| mmu-miR-34a-5p | MIMAT0000542 | Fam57a | 116972 | HITS-CLIP | 37986224\|19536157\|23142080 | Unspecified |
| mmu-miR-34a-5p | MIMAT0000542 | Fam69a | 388650 | HITS-CLIP | 23142080 | Unspecified |
| mmu-miR-34a-5p | MIMAT0000542 | Fam69b | 56279 | HITS-CLIP | 19536157 | Unspecified |
| mmu-miR-34a-5p | MIMAT0000542 | Fam71b | 432552 | HITS-CLIP | 23142080 | Unspecified |
| mmu-miR-34a-5p | MIMAT0000542 | Fam72a | 108900 | Microarrays//HITS-CLIP | 37986224\|23027903\|23142080 | Unspecified |
| mmu-miR-34a-5p | MIMAT0000542 | Fam76b | 72826 | HITS-CLIP | 23630078\|31433990\|23142080\|29167 | Unspecified |
| mmu-miR-34a-5p | MIMAT0000542 | Fam78a | 241303 | Microarrays//HITS-CLIP | 37986224\|23142080 | Unspecified |
| mmu-miR-34a-5p | MIMAT0000542 | Fam83d | 71878 | HITS-CLIP | 30971389\|23142080 | Unspecified |
| mmu-miR-34a-5p | MIMAT0000542 | Fam83h | 105732 | HITS-CLIP | 23142080\|26203562\|29167373 | Unspecified |
| mmu-miR-34a-5p | MIMAT0000542 | Fam84b | 157638 | HITS-CLIP | 23142080\|29167373 | Unspecified |
| mmu-miR-34a-5p | MIMAT0000542 | Fam89b | 17826 | HITS-CLIP | 23142080 | Unspecified |
| mmu-miR-34a-5p | MIMAT0000542 | Fam8a1 | 51439 | HITS-CLIP | 23142080\|29167373 | Unspecified |
| mmu-miR-34a-5p | MIMAT0000542 | Fam91a1 | 210998 | HITS-CLIP | 23142080\|29167373 | Unspecified |
| mmu-miR-34a-5p | MIMAT0000542 | Fam92a | 68099 | HITS-CLIP | 23142080 | Unspecified |
| mmu-miR-34a-5p | MIMAT0000542 | Fam98a | 72722 | HITS-CLIP | 23142080 | Unspecified |
| mmu-miR-34a-5p | MIMAT0000542 | Fam98c | 73833 | HITS-CLIP | 19536157 | Unspecified |
| mmu-miR-34a-5p | MIMAT0000542 | Fanca | 14087 | HITS-CLIP | 29167373 | Unspecified |
| mmu-miR-34a-5p | MIMAT0000542 | Fancb | 237211 | Microarrays//HITS-CLIP | 37986224\|23027903\|23142080 | Unspecified |
| mmu-miR-34a-5p | MIMAT0000542 | Fancf | 100040608 | HITS-CLIP | 31433990 | Unspecified |
| mmu-miR-34a-5p | MIMAT0000542 | Fanci | 208836 | HITS-CLIP | 23142080 | Unspecified |
| mmu-miR-34a-5p | MIMAT0000542 | Fancm | 104806 | HITS-CLIP | 23142080 | Unspecified |
| mmu-miR-34a-5p | MIMAT0000542 | Far1 | 67420 | HITS-CLIP | 23142080 | Unspecified |
| mmu-miR-34a-5p | MIMAT0000542 | Farp2 | 227377 | HITS-CLIP | 29167373 | Unspecified |
| mmu-miR-34a-5p | MIMAT0000542 | Fars2 | 69955 | HITS-CLIP | 23142080 | Unspecified |
| mmu-miR-34a-5p | MIMAT0000542 | Farsb | 23874 | HITS-CLIP | 23142080 | Unspecified |
| mmu-miR-34a-5p | MIMAT0000542 | Fasn | 14104 | HITS-CLIP | 31433990\|30971389\|23142080 | Unspecified |
| mmu-miR-34a-5p | MIMAT0000542 | Fastk | 66587 | HITS-CLIP | 23142080 | Unspecified |
| mmu-miR-34a-5p | MIMAT0000542 | Fastkd2 | 75619 | HITS-CLIP | 23142080 | Unspecified |
| mmu-miR-34a-5p | MIMAT0000542 | Fastkd3 | 69577 | HITS-CLIP | 23142080 | Unspecified |
| mmu-miR-34a-5p | MIMAT0000542 | Fastkd5 | 380601 | HITS-CLIP | 19536157\|23142080\|29167373 | Unspecified |
| mmu-miR-34a-5p | MIMAT0000542 | Fat1 | 14107 | HITS-CLIP | 19536157\|26203562 | Unspecified |
| mmu-miR-34a-5p | MIMAT0000542 | Fat2 | 245827 | HITS-CLIP | 19536157 | Unspecified |
| mmu-miR-34a-5p | MIMAT0000542 | Fat3 | 270120 | HITS-CLIP | 19536157\|29167373 | Unspecified |
| mmu-miR-34a-5p | MIMAT0000542 | Fau | 14109 | HITS-CLIP | 23142080 | Unspecified |
| mmu-miR-34a-5p | MIMAT0000542 | Fbh1 | 84893 | HITS-CLIP | 26203562\|23142080 | Unspecified |
| mmu-miR-34a-5p | MIMAT0000542 | Fblim1 | 74202 | HITS-CLIP | 29167373 | Unspecified |

| | | | | | | |
|---|---|---|---|---|---|---|
| mmu-miR-34a-5p | MIMAT0000542 | Fbrs | 14123 | HITS-CLIP | 23142080\|31433990\|29167373 | Unspecified |
| mmu-miR-34a-5p | MIMAT0000542 | Fbxl14 | 101358 | HITS-CLIP | 29167373\|23142080 | Unspecified |
| mmu-miR-34a-5p | MIMAT0000542 | Fbxl16 | 214931 | HITS-CLIP | 19536157 | Unspecified |
| mmu-miR-34a-5p | MIMAT0000542 | Fbxl17 | 50758 | HITS-CLIP | 23142080\|30971389 | Unspecified |
| mmu-miR-34a-5p | MIMAT0000542 | Fbxl18 | 231863 | HITS-CLIP | 23142080\|29167373 | Unspecified |
| mmu-miR-34a-5p | MIMAT0000542 | Fbxl20 | 72194 | HITS-CLIP | 31433990\|23142080\|29167373 | Unspecified |
| mmu-miR-34a-5p | MIMAT0000542 | Fbxl3 | 50789 | HITS-CLIP | 19536157\|23142080 | Unspecified |
| mmu-miR-34a-5p | MIMAT0000542 | Fbxl4 | 269514 | HITS-CLIP | 23142080 | Unspecified |
| mmu-miR-34a-5p | MIMAT0000542 | Fbxl5 | 242960 | HITS-CLIP | 23142080 | Unspecified |
| mmu-miR-34a-5p | MIMAT0000542 | Fbxl6 | 30840 | HITS-CLIP | 23142080 | Unspecified |
| mmu-miR-34a-5p | MIMAT0000542 | Fbxl8 | 50788 | HITS-CLIP | 23142080 | Unspecified |
| mmu-miR-34a-5p | MIMAT0000542 | Fbxo10 | 269529 | HITS-CLIP | 31433990 | Unspecified |
| mmu-miR-34a-5p | MIMAT0000542 | Fbxo11 | 225055 | HITS-CLIP | 23142080\|31433990 | Unspecified |
| mmu-miR-34a-5p | MIMAT0000542 | Fbxo16 | 50759 | Microarrays | 37986224 | Unspecified |
| mmu-miR-34a-5p | MIMAT0000542 | Fbxo21 | 231670 | HITS-CLIP | 23142080\|31433990 | Unspecified |
| mmu-miR-34a-5p | MIMAT0000542 | Fbxo22 | 71999 | HITS-CLIP | 29167373 | Unspecified |
| mmu-miR-34a-5p | MIMAT0000542 | Fbxo25 | 66822 | HITS-CLIP | 23142080 | Unspecified |
| mmu-miR-34a-5p | MIMAT0000542 | Fbxo28 | 67948 | HITS-CLIP | 23142080 | Unspecified |
| mmu-miR-34a-5p | MIMAT0000542 | Fbxo3 | 57443 | HITS-CLIP | 23142080 | Unspecified |
| mmu-miR-34a-5p | MIMAT0000542 | Fbxo30 | 71865 | HITS-CLIP | 23630078 | Unspecified |
| mmu-miR-34a-5p | MIMAT0000542 | Fbxo31 | 76454 | HITS-CLIP | 29167373 | Unspecified |
| mmu-miR-34a-5p | MIMAT0000542 | Fbxo32 | 67731 | HITS-CLIP | 23142080 | Unspecified |
| mmu-miR-34a-5p | MIMAT0000542 | Fbxo33 | 70611 | HITS-CLIP | 31433990\|23142080\|29167373 | Unspecified |
| mmu-miR-34a-5p | MIMAT0000542 | Fbxo34 | 78938 | HITS-CLIP | 23142080 | Unspecified |
| mmu-miR-34a-5p | MIMAT0000542 | Fbxo36 | 66153 | Microarrays | 37986224 | Unspecified |
| mmu-miR-34a-5p | MIMAT0000542 | Fbxo38 | 107035 | HITS-CLIP | 23142080 | Unspecified |
| mmu-miR-34a-5p | MIMAT0000542 | Fbxo42 | 213499 | HITS-CLIP | 30971389\|29167373 | Unspecified |
| mmu-miR-34a-5p | MIMAT0000542 | Fbxo45 | 268882 | HITS-CLIP | 29167373\|23142080 | Unspecified |
| mmu-miR-34a-5p | MIMAT0000542 | Fbxo46 | 243867 | HITS-CLIP | 23142080 | Unspecified |
| mmu-miR-34a-5p | MIMAT0000542 | Fbxo48 | 319701 | Microarrays | 37986224\|23027903 | Unspecified |
| mmu-miR-34a-5p | MIMAT0000542 | Fbxo5 | 67141 | Microarrays//HITS-CLIP | 37986224\|23027903\|23142080 | Unspecified |
| mmu-miR-34a-5p | MIMAT0000542 | Fbxo7 | 69754 | HITS-CLIP | 29167373 | Unspecified |
| mmu-miR-34a-5p | MIMAT0000542 | Fbxo8 | 50753 | HITS-CLIP | 29167373\|23142080 | Unspecified |
| mmu-miR-34a-5p | MIMAT0000542 | Fbxw11 | 103583 | HITS-CLIP | 23142080\|29167373 | Unspecified |
| mmu-miR-34a-5p | MIMAT0000542 | Fbxw17 | 109082 | HITS-CLIP | 29167373 | Unspecified |
| mmu-miR-34a-5p | MIMAT0000542 | Fbxw2 | 30050 | HITS-CLIP | 23142080 | Unspecified |
| mmu-miR-34a-5p | MIMAT0000542 | Fbxw4 | 30838 | HITS-CLIP | 23142080 | Unspecified |
| mmu-miR-34a-5p | MIMAT0000542 | Fbxw5 | 30839 | HITS-CLIP | 23142080 | Unspecified |
| mmu-miR-34a-5p | MIMAT0000542 | Fbxw7 | 50754 | HITS-CLIP | 23142080\|29167373 | Unspecified |

| | | | | | | |
|---|---|---|---|---|---|---|
| mmu-miR-34a-5p | MIMAT0000542 | Fbxw8 | 231672 | HITS-CLIP | 29167373 | Unspecified |
| mmu-miR-34a-5p | MIMAT0000542 | Fcgr1 | 14129 | HITS-CLIP | 29167373 | Unspecified |
| mmu-miR-34a-5p | MIMAT0000542 | Fcgr4 | 246256 | HITS-CLIP | 29167373 | Unspecified |
| mmu-miR-34a-5p | MIMAT0000542 | Fcho1 | 23149 | HITS-CLIP | 23142080 | Unspecified |
| mmu-miR-34a-5p | MIMAT0000542 | Fcho2 | 218503 | Microarrays//HITS-CLIP | 37986224\|23142080 | Unspecified |
| mmu-miR-34a-5p | MIMAT0000542 | Fdft1 | 14137 | HITS-CLIP | 23142080 | Unspecified |
| mmu-miR-34a-5p | MIMAT0000542 | Fdx1 | 14148 | HITS-CLIP | 23142080 | Unspecified |
| mmu-miR-34a-5p | MIMAT0000542 | Fech | 14151 | HITS-CLIP | 23142080 | Unspecified |
| mmu-miR-34a-5p | MIMAT0000542 | Fem1a | 14154 | HITS-CLIP | 19536157\|30971389\|29167373 | Unspecified |
| mmu-miR-34a-5p | MIMAT0000542 | Fem1b | 14155 | HITS-CLIP | 23142080\|29167373 | Unspecified |
| mmu-miR-34a-5p | MIMAT0000542 | Fem1c | 240263 | HITS-CLIP | 37986224\|19536157\|31433990\|29167 | Unspecified |
| mmu-miR-34a-5p | MIMAT0000542 | Fen1 | 14156 | Microarrays//HITS-CLIP | 37986224\|23027903\|23142080 | Unspecified |
| mmu-miR-34a-5p | MIMAT0000542 | Fermt3 | 108101 | HITS-CLIP | 23142080\|29167373 | Unspecified |
| mmu-miR-34a-5p | MIMAT0000542 | Fes | 14159 | HITS-CLIP | 29167373 | Unspecified |
| mmu-miR-34a-5p | MIMAT0000542 | Fez2 | 225020 | HITS-CLIP | 23142080\|29167373 | Unspecified |
| mmu-miR-34a-5p | MIMAT0000542 | Fezf2 | 54713 | HITS-CLIP | 19536157 | Unspecified |
| mmu-miR-34a-5p | MIMAT0000542 | Fgd3 | 30938 | HITS-CLIP | 23142080\|29167373 | Unspecified |
| mmu-miR-34a-5p | MIMAT0000542 | Fgd4 | 224014 | HITS-CLIP | 29167373 | Unspecified |
| mmu-miR-34a-5p | MIMAT0000542 | Fgd6 | 13998 | HITS-CLIP | 23630078\|29167373 | Unspecified |
| mmu-miR-34a-5p | MIMAT0000542 | Fgf13 | 14168 | HITS-CLIP | 23142080 | Unspecified |
| mmu-miR-34a-5p | MIMAT0000542 | Fgf14 | 14169 | HITS-CLIP | 19536157 | Unspecified |
| mmu-miR-34a-5p | MIMAT0000542 | Fgf2 | 14173 | HITS-CLIP | 23142080 | Unspecified |
| mmu-miR-34a-5p | MIMAT0000542 | Fgf7 | 14178 | Microarrays | 37986224 | Unspecified |
| mmu-miR-34a-5p | MIMAT0000542 | Fgfr1 | 14182 | HITS-CLIP | 19536157 | Unspecified |
| mmu-miR-34a-5p | MIMAT0000542 | Fgfr1op2 | 67529 | HITS-CLIP | 23142080 | Unspecified |
| mmu-miR-34a-5p | MIMAT0000542 | Fgfr2 | 14183 | HITS-CLIP | 19536157 | Unspecified |
| mmu-miR-34a-5p | MIMAT0000542 | Fgl2 | 14190 | HITS-CLIP | 23142080\|29167373 | Unspecified |
| mmu-miR-34a-5p | MIMAT0000542 | Fh1 | 14194 | HITS-CLIP | 19536157\|23142080\|29167373 | Unspecified |
| mmu-miR-34a-5p | MIMAT0000542 | Fhip2a | 57700 | HITS-CLIP | 31433990 | Unspecified |
| mmu-miR-34a-5p | MIMAT0000542 | Fhl3 | 14201 | HITS-CLIP | 23142080 | Unspecified |
| mmu-miR-34a-5p | MIMAT0000542 | Fhod1 | 234686 | HITS-CLIP | 23142080 | Unspecified |
| mmu-miR-34a-5p | MIMAT0000542 | Ficd | 231630 | HITS-CLIP | 30971389\|23142080\|29167373 | Unspecified |
| mmu-miR-34a-5p | MIMAT0000542 | Fignl1 | 60530 | HITS-CLIP | 23142080\|29167373 | Unspecified |
| mmu-miR-34a-5p | MIMAT0000542 | Filip1l | 78749 | HITS-CLIP | 29167373 | Unspecified |
| mmu-miR-34a-5p | MIMAT0000542 | Fis1 | 66437 | HITS-CLIP | 19536157\|23142080 | Unspecified |
| mmu-miR-34a-5p | MIMAT0000542 | Fitm2 | 228859 | HITS-CLIP | 23142080 | Unspecified |
| mmu-miR-34a-5p | MIMAT0000542 | Fjx1 | 14221 | HITS-CLIP | 19536157 | Unspecified |
| mmu-miR-34a-5p | MIMAT0000542 | Fkbp11 | 66120 | HITS-CLIP | 23142080 | Unspecified |
| mmu-miR-34a-5p | MIMAT0000542 | Fkbp15 | 338355 | HITS-CLIP | 29167373 | Unspecified |

| | | | | | | |
|---|---|---|---|---|---|---|
| mmu-miR-34a-5p | MIMAT0000542 | Fkbp1a | 14225 | HITS-CLIP | 19536157\|23142080 | Unspecified |
| mmu-miR-34a-5p | MIMAT0000542 | Fkbp1b | 14226 | HITS-CLIP | 29167373 | Unspecified |
| mmu-miR-34a-5p | MIMAT0000542 | Fkbp2 | 14227 | HITS-CLIP | 23142080\|29167373 | Unspecified |
| mmu-miR-34a-5p | MIMAT0000542 | Fkbp3 | 30795 | HITS-CLIP | 30971389\|23142080 | Unspecified |
| mmu-miR-34a-5p | MIMAT0000542 | Fkbp5 | 14229 | HITS-CLIP | 26203562\|23142080 | Unspecified |
| mmu-miR-34a-5p | MIMAT0000542 | Fkbp8 | 14232 | HITS-CLIP | 23142080 | Unspecified |
| mmu-miR-34a-5p | MIMAT0000542 | Fkbp9 | 27055 | HITS-CLIP | 29167373 | Unspecified |
| mmu-miR-34a-5p | MIMAT0000542 | Fkbpl | 56299 | HITS-CLIP | 23142080 | Unspecified |
| mmu-miR-34a-5p | MIMAT0000542 | Fkrp | 243853 | HITS-CLIP | 23142080 | Unspecified |
| mmu-miR-34a-5p | MIMAT0000542 | Fktn | 246179 | HITS-CLIP | 19536157\|23630078 | Unspecified |
| mmu-miR-34a-5p | MIMAT0000542 | Flcn | 216805 | HITS-CLIP | 23142080\|29167373 | Unspecified |
| mmu-miR-34a-5p | MIMAT0000542 | Fli1 | 14247 | HITS-CLIP | 29167373 | Unspecified |
| mmu-miR-34a-5p | MIMAT0000542 | Flii | 14248 | HITS-CLIP | 23142080\|29167373 | Unspecified |
| mmu-miR-34a-5p | MIMAT0000542 | Flna | 192176 | HITS-CLIP | 19536157\|23142080\|29167373 | Unspecified |
| mmu-miR-34a-5p | MIMAT0000542 | Flnb | 286940 | HITS-CLIP | 26203562\|23142080\|29167373 | Unspecified |
| mmu-miR-34a-5p | MIMAT0000542 | Flnc | 68794 | HITS-CLIP | 23630078 | Unspecified |
| mmu-miR-34a-5p | MIMAT0000542 | Flot1 | 14251 | HITS-CLIP | 23142080 | Unspecified |
| mmu-miR-34a-5p | MIMAT0000542 | Flot2 | 14252 | HITS-CLIP | 19536157\|23630078\|23142080 | Unspecified |
| mmu-miR-34a-5p | MIMAT0000542 | Flrt2 | 399558 | HITS-CLIP | 19536157\|29167373 | Unspecified |
| mmu-miR-34a-5p | MIMAT0000542 | Flvcr1 | 28982 | HITS-CLIP | 29167373 | Unspecified |
| mmu-miR-34a-5p | MIMAT0000542 | Flywch1 | 224613 | HITS-CLIP | 23142080 | Unspecified |
| mmu-miR-34a-5p | MIMAT0000542 | Fmn1 | 14260 | HITS-CLIP | 29167373 | Unspecified |
| mmu-miR-34a-5p | MIMAT0000542 | Fmn2 | 54418 | HITS-CLIP | 19536157 | Unspecified |
| mmu-miR-34a-5p | MIMAT0000542 | Fmnl1 | 57778 | HITS-CLIP | 23142080\|29167373 | Unspecified |
| mmu-miR-34a-5p | MIMAT0000542 | Fmnl2 | 71409 | HITS-CLIP | 19536157\|23142080\|29167373 | Unspecified |
| mmu-miR-34a-5p | MIMAT0000542 | Fmnl3 | 22379 | HITS-CLIP | 29167373 | Unspecified |
| mmu-miR-34a-5p | MIMAT0000542 | Fmo5 | 14263 | HITS-CLIP | 23142080 | Unspecified |
| mmu-miR-34a-5p | MIMAT0000542 | Fmr1 | 14265 | Microarrays//HITS-CLIP | 37986224\|23142080 | Unspecified |
| mmu-miR-34a-5p | MIMAT0000542 | Fnbp1 | 14269 | HITS-CLIP | 37986224\|19536157\|23142080\|29167 | Unspecified |
| mmu-miR-34a-5p | MIMAT0000542 | Fnbp1l | 214459 | HITS-CLIP | 23142080\|23630078\|19536157\|29167 | Unspecified |
| mmu-miR-34a-5p | MIMAT0000542 | Fnbp4 | 55935 | HITS-CLIP | 31433990\|23142080\|30971389\|2363( | Unspecified |
| mmu-miR-34a-5p | MIMAT0000542 | Fndc10 | 230991 | HITS-CLIP | 23142080 | Unspecified |
| mmu-miR-34a-5p | MIMAT0000542 | Fndc3a | 319448 | HITS-CLIP | 37986224\|19536157\|30971389\|3143: | Unspecified |
| mmu-miR-34a-5p | MIMAT0000542 | Fndc3b | 72007 | HITS-CLIP | 23142080\|29167373 | Unspecified |
| mmu-miR-34a-5p | MIMAT0000542 | Fnip1 | 216742 | HITS-CLIP | 31433990\|23142080 | Unspecified |
| mmu-miR-34a-5p | MIMAT0000542 | Fnip2 | 329679 | HITS-CLIP | 23142080\|29167373 | Unspecified |
| mmu-miR-34a-5p | MIMAT0000542 | Fnta | 14272 | HITS-CLIP | 19536157\|23142080 | Unspecified |
| mmu-miR-34a-5p | MIMAT0000542 | Fntb | 110606 | HITS-CLIP | 23142080 | Unspecified |
| mmu-miR-34a-5p | MIMAT0000542 | Focad | 230393 | HITS-CLIP | 31433990\|23142080 | Unspecified |

| | | | | | | |
|---|---|---|---|---|---|---|
| mmu-miR-34a-5p | MIMAT0000542 | Fopnl | 66086 | HITS-CLIP | 23142080\|29167373 | Unspecified |
| mmu-miR-34a-5p | MIMAT0000542 | Fos | 14281 | HITS-CLIP | 23142080\|29167373 | Unspecified |
| mmu-miR-34a-5p | MIMAT0000542 | Fosl1 | 14283 | HITS-CLIP | 23142080 | Unspecified |
| mmu-miR-34a-5p | MIMAT0000542 | Fosl2 | 14284 | HITS-CLIP | 23630078\|31433990\|23142080\|29167 | Unspecified |
| mmu-miR-34a-5p | MIMAT0000542 | Foxj2 | 60611 | Microarrays//HITS-CLIP | 37986224\|23027903\|23142080\|30971 | Unspecified |
| mmu-miR-34a-5p | MIMAT0000542 | Foxj3 | 230700 | HITS-CLIP | 37986224\|23142080\|29167373 | Unspecified |
| mmu-miR-34a-5p | MIMAT0000542 | Foxk1 | 17425 | HITS-CLIP | 31433990\|23142080\|29167373 | Unspecified |
| mmu-miR-34a-5p | MIMAT0000542 | Foxk2 | 68837 | HITS-CLIP | 19536157\|23142080\|29167373 | Unspecified |
| mmu-miR-34a-5p | MIMAT0000542 | Foxn2 | 14236 | HITS-CLIP | 23142080\|29167373 | Unspecified |
| mmu-miR-34a-5p | MIMAT0000542 | Foxn3 | 71375 | HITS-CLIP | 19536157\|23142080\|29167373 | Unspecified |
| mmu-miR-34a-5p | MIMAT0000542 | Foxo1 | 56458 | HITS-CLIP | 37986224\|30971389\|23142080 | Unspecified |
| mmu-miR-34a-5p | MIMAT0000542 | Foxo3 | 56484 | HITS-CLIP | 30971389\|19536157\|23142080\|26203 | Unspecified |
| mmu-miR-34a-5p | MIMAT0000542 | Foxp1 | 108655 | Western blot//Western Blot//HITS-CLIP | 37986224\|20598588\|19536157\|23142 | Unspecified |
| mmu-miR-34a-5p | MIMAT0000542 | Foxq1 | 15220 | HITS-CLIP | 23142080 | Unspecified |
| mmu-miR-34a-5p | MIMAT0000542 | Foxred2 | 239554 | HITS-CLIP | 23142080\|19536157 | Unspecified |
| mmu-miR-34a-5p | MIMAT0000542 | Fpgt | 75540 | HITS-CLIP | 23142080 | Unspecified |
| mmu-miR-34a-5p | MIMAT0000542 | Fras1 | 231470 | Microarrays | 37986224 | Unspecified |
| mmu-miR-34a-5p | MIMAT0000542 | Frg1 | 14300 | HITS-CLIP | 23142080 | Unspecified |
| mmu-miR-34a-5p | MIMAT0000542 | Frmd4a | 209630 | HITS-CLIP | 19536157\|29167373 | Unspecified |
| mmu-miR-34a-5p | MIMAT0000542 | Frmd4b | 232288 | HITS-CLIP | 29167373 | Unspecified |
| mmu-miR-34a-5p | MIMAT0000542 | Frmd6 | 319710 | HITS-CLIP | 23142080 | Unspecified |
| mmu-miR-34a-5p | MIMAT0000542 | Frmd8 | 67457 | HITS-CLIP | 23142080\|19536157\|31433990 | Unspecified |
| mmu-miR-34a-5p | MIMAT0000542 | Frmpd1 | 666060 | Microarrays | 37986224 | Unspecified |
| mmu-miR-34a-5p | MIMAT0000542 | Frrs1 | 20321 | HITS-CLIP | 29167373 | Unspecified |
| mmu-miR-34a-5p | MIMAT0000542 | Frrs1l | 230235 | HITS-CLIP | 19536157\|23142080 | Unspecified |
| mmu-miR-34a-5p | MIMAT0000542 | Frs2 | 327826 | HITS-CLIP | 23142080\|19536157\|31433990 | Unspecified |
| mmu-miR-34a-5p | MIMAT0000542 | Fry | 320365 | HITS-CLIP | 19536157\|23142080 | Unspecified |
| mmu-miR-34a-5p | MIMAT0000542 | Fryl | 72313 | HITS-CLIP | 23142080\|29167373 | Unspecified |
| mmu-miR-34a-5p | MIMAT0000542 | Fscn1 | 14086 | HITS-CLIP | 19536157 | Unspecified |
| mmu-miR-34a-5p | MIMAT0000542 | Fsd1l | 319636 | HITS-CLIP | 23142080 | Unspecified |
| mmu-miR-34a-5p | MIMAT0000542 | Fstl1 | 14314 | HITS-CLIP | 19536157 | Unspecified |
| mmu-miR-34a-5p | MIMAT0000542 | Fth1 | 14319 | HITS-CLIP | 23142080\|19536157\|29167373 | Unspecified |
| mmu-miR-34a-5p | MIMAT0000542 | Fto | 26383 | HITS-CLIP | 29167373 | Unspecified |
| mmu-miR-34a-5p | MIMAT0000542 | Ftsj3 | 56095 | HITS-CLIP | 23630078\|29167373 | Unspecified |
| mmu-miR-34a-5p | MIMAT0000542 | Fubp1 | 51886 | HITS-CLIP | 23142080\|19536157\|29167373 | Unspecified |
| mmu-miR-34a-5p | MIMAT0000542 | Fubp3 | 320267 | Microarrays//HITS-CLIP | 37986224\|23027903\|26203562\|23142 | Unspecified |
| mmu-miR-34a-5p | MIMAT0000542 | Fuca1 | 71665 | HITS-CLIP | 29167373\|23142080 | Unspecified |
| mmu-miR-34a-5p | MIMAT0000542 | Fuca2 | 66848 | HITS-CLIP | 23142080 | Unspecified |
| mmu-miR-34a-5p | MIMAT0000542 | Fuk | 197258 | HITS-CLIP | 23142080 | Unspecified |

| | | | | | | |
|---|---|---|---|---|---|---|
| mmu-miR-34a-5p | MIMAT0000542 | Fundc1 | 72018 | HITS-CLIP | 23142080 | Unspecified |
| mmu-miR-34a-5p | MIMAT0000542 | Furin | 18550 | HITS-CLIP | 23142080\|29167373 | Unspecified |
| mmu-miR-34a-5p | MIMAT0000542 | Fus | 233908 | HITS-CLIP | 23630078\|23142080\|29167373 | Unspecified |
| mmu-miR-34a-5p | MIMAT0000542 | Fut4 | 14345 | HITS-CLIP | 31433990 | Unspecified |
| mmu-miR-34a-5p | MIMAT0000542 | Fut7 | 14347 | HITS-CLIP | 23142080 | Unspecified |
| mmu-miR-34a-5p | MIMAT0000542 | Fut8 | 53618 | HITS-CLIP | 23142080 | Unspecified |
| mmu-miR-34a-5p | MIMAT0000542 | Fuz | 70300 | HITS-CLIP | 23142080 | Unspecified |
| mmu-miR-34a-5p | MIMAT0000542 | Fv1 | 14349 | HITS-CLIP | 31433990\|23142080 | Unspecified |
| mmu-miR-34a-5p | MIMAT0000542 | Fxr1 | 14359 | HITS-CLIP | 23142080\|19536157 | Unspecified |
| mmu-miR-34a-5p | MIMAT0000542 | Fxr2 | 23879 | HITS-CLIP | 23142080 | Unspecified |
| mmu-miR-34a-5p | MIMAT0000542 | Fxyd5 | 18301 | HITS-CLIP | 23142080\|31433990\|29167373 | Unspecified |
| mmu-miR-34a-5p | MIMAT0000542 | Fyb | 23880 | HITS-CLIP | 23142080\|29167373 | Unspecified |
| mmu-miR-34a-5p | MIMAT0000542 | Fyco1 | 17281 | HITS-CLIP | 23142080\|29167373 | Unspecified |
| mmu-miR-34a-5p | MIMAT0000542 | Fyn | 14360 | HITS-CLIP | 19536157\|31433990\|29167373\|23142 | Unspecified |
| mmu-miR-34a-5p | MIMAT0000542 | Fyttd1 | 69823 | HITS-CLIP | 23630078\|19536157\|23142080 | Unspecified |
| mmu-miR-34a-5p | MIMAT0000542 | Fzd3 | 14365 | HITS-CLIP | 19536157 | Unspecified |
| mmu-miR-34a-5p | MIMAT0000542 | Fzd5 | 14367 | HITS-CLIP | 26203562\|29167373 | Unspecified |
| mmu-miR-34a-5p | MIMAT0000542 | Fzd7 | 14369 | HITS-CLIP | 23142080 | Unspecified |
| mmu-miR-34a-5p | MIMAT0000542 | Fzr1 | 56371 | HITS-CLIP | 23142080\|29167373 | Unspecified |
| mmu-miR-34a-5p | MIMAT0000542 | G2e3 | 217558 | HITS-CLIP | 23142080\|29167373 | Unspecified |
| mmu-miR-34a-5p | MIMAT0000542 | G3bp1 | 27041 | HITS-CLIP | 23142080\|31433990\|29167373 | Unspecified |
| mmu-miR-34a-5p | MIMAT0000542 | G3bp2 | 23881 | HITS-CLIP | 23142080\|31433990\|23630078 | Unspecified |
| mmu-miR-34a-5p | MIMAT0000542 | G6pd2 | 14380 | Microarrays//HITS-CLIP | 37986224\|23027903\|29167373 | Unspecified |
| mmu-miR-34a-5p | MIMAT0000542 | G6pdx | 14381 | Microarrays//HITS-CLIP | 37986224\|23027903\|23142080\|29167 | Unspecified |
| mmu-miR-34a-5p | MIMAT0000542 | Gaa | 14387 | HITS-CLIP | 29167373 | Unspecified |
| mmu-miR-34a-5p | MIMAT0000542 | Gab2 | 14389 | HITS-CLIP | 29167373 | Unspecified |
| mmu-miR-34a-5p | MIMAT0000542 | Gab3 | 210710 | HITS-CLIP | 23142080\|29167373 | Unspecified |
| mmu-miR-34a-5p | MIMAT0000542 | Gabarap | 56486 | HITS-CLIP | 37986224\|19536157\|23142080 | Unspecified |
| mmu-miR-34a-5p | MIMAT0000542 | Gabarapl1 | 57436 | HITS-CLIP | 31433990\|29167373 | Unspecified |
| mmu-miR-34a-5p | MIMAT0000542 | Gabbr1 | 54393 | HITS-CLIP | 19536157 | Unspecified |
| mmu-miR-34a-5p | MIMAT0000542 | Gabbr2 | 242425 | HITS-CLIP | 19536157 | Unspecified |
| mmu-miR-34a-5p | MIMAT0000542 | Gabpa | 14390 | HITS-CLIP | 23142080\|29167373 | Unspecified |
| mmu-miR-34a-5p | MIMAT0000542 | Gabpb1 | 14391 | HITS-CLIP | 31433990\|23142080\|29167373 | Unspecified |
| mmu-miR-34a-5p | MIMAT0000542 | Gabpb2 | 213054 | HITS-CLIP | 23142080\|19536157\|26203562\|29167 | Unspecified |
| mmu-miR-34a-5p | MIMAT0000542 | Gabra1 | 14394 | HITS-CLIP | 19536157 | Unspecified |
| mmu-miR-34a-5p | MIMAT0000542 | Gabra2 | 14395 | HITS-CLIP | 19536157 | Unspecified |
| mmu-miR-34a-5p | MIMAT0000542 | Gabra4 | 14397 | HITS-CLIP | 19536157 | Unspecified |
| mmu-miR-34a-5p | MIMAT0000542 | Gabra6 | 14399 | HITS-CLIP | 19536157 | Unspecified |
| mmu-miR-34a-5p | MIMAT0000542 | Gabrb2 | 14401 | HITS-CLIP | 19536157 | Unspecified |

| | | | | | | |
|---|---|---|---|---|---|---|
| mmu-miR-34a-5p | MIMAT0000542 | Gad1 | 14415 | HITS-CLIP | 19536157 | Unspecified |
| mmu-miR-34a-5p | MIMAT0000542 | Gad2 | 14417 | HITS-CLIP | 19536157 | Unspecified |
| mmu-miR-34a-5p | MIMAT0000542 | Gadd45a | 13197 | HITS-CLIP | 23142080 | Unspecified |
| mmu-miR-34a-5p | MIMAT0000542 | Gadd45g | 23882 | HITS-CLIP | 23142080 | Unspecified |
| mmu-miR-34a-5p | MIMAT0000542 | Gak | 231580 | HITS-CLIP | 29167373\|23142080 | Unspecified |
| mmu-miR-34a-5p | MIMAT0000542 | Galc | 14420 | HITS-CLIP | 23142080\|29167373 | Unspecified |
| mmu-miR-34a-5p | MIMAT0000542 | Galk1 | 14635 | HITS-CLIP | 23142080 | Unspecified |
| mmu-miR-34a-5p | MIMAT0000542 | Galk2 | 69976 | HITS-CLIP | 31433990\|23142080\|29167373 | Unspecified |
| mmu-miR-34a-5p | MIMAT0000542 | Galm | 319625 | HITS-CLIP | 23630078 | Unspecified |
| mmu-miR-34a-5p | MIMAT0000542 | Galns | 50917 | HITS-CLIP | 23142080 | Unspecified |
| mmu-miR-34a-5p | MIMAT0000542 | Galnt1 | 14423 | HITS-CLIP | 23142080 | Unspecified |
| mmu-miR-34a-5p | MIMAT0000542 | Galnt11 | 231050 | HITS-CLIP | 19536157 | Unspecified |
| mmu-miR-34a-5p | MIMAT0000542 | Galnt12 | 230145 | HITS-CLIP | 23142080 | Unspecified |
| mmu-miR-34a-5p | MIMAT0000542 | Galnt16 | 108760 | HITS-CLIP | 19536157 | Unspecified |
| mmu-miR-34a-5p | MIMAT0000542 | Galnt2 | 108148 | HITS-CLIP | 23142080\|29167373 | Unspecified |
| mmu-miR-34a-5p | MIMAT0000542 | Galnt3 | 14425 | HITS-CLIP | 23142080 | Unspecified |
| mmu-miR-34a-5p | MIMAT0000542 | Galnt6 | 207839 | HITS-CLIP | 23142080 | Unspecified |
| mmu-miR-34a-5p | MIMAT0000542 | Galnt7 | 108150 | HITS-CLIP | 29167373\|23142080 | Unspecified |
| mmu-miR-34a-5p | MIMAT0000542 | Galnt9 | 231605 | HITS-CLIP | 19536157 | Unspecified |
| mmu-miR-34a-5p | MIMAT0000542 | Galt | 14430 | HITS-CLIP | 31433990\|30971389 | Unspecified |
| mmu-miR-34a-5p | MIMAT0000542 | Gan | 209239 | HITS-CLIP | 23142080\|29167373 | Unspecified |
| mmu-miR-34a-5p | MIMAT0000542 | Ganab | 14376 | HITS-CLIP | 23142080\|29167373 | Unspecified |
| mmu-miR-34a-5p | MIMAT0000542 | Gapvd1 | 66691 | HITS-CLIP | 19536157\|23142080\|29167373 | Unspecified |
| mmu-miR-34a-5p | MIMAT0000542 | Garnl3 | 99326 | HITS-CLIP | 19536157 | Unspecified |
| mmu-miR-34a-5p | MIMAT0000542 | Gars | 353172 | HITS-CLIP | 23142080\|19536157\|29167373 | Unspecified |
| mmu-miR-34a-5p | MIMAT0000542 | Gart | 14450 | Microarrays//HITS-CLIP | 37986224\|23027903\|23142080\|1953 | Unspecified |
| mmu-miR-34a-5p | MIMAT0000542 | Gas1 | 14451 | Immunohistochemistry//Luciferase assay// | 24560136 | Unspecified |
| mmu-miR-34a-5p | MIMAT0000542 | Gas2 | 14453 | HITS-CLIP | 23142080 | Unspecified |
| mmu-miR-34a-5p | MIMAT0000542 | Gas2l3 | 237436 | HITS-CLIP | 23630078\|31433990\|23142080\|2916 | Unspecified |
| mmu-miR-34a-5p | MIMAT0000542 | Gas6 | 14456 | Microarrays | 37986224 | Unspecified |
| mmu-miR-34a-5p | MIMAT0000542 | Gas7 | 14457 | HITS-CLIP | 19536157\|29167373 | Unspecified |
| mmu-miR-34a-5p | MIMAT0000542 | Gata1 | 14460 | HITS-CLIP | 23142080 | Unspecified |
| mmu-miR-34a-5p | MIMAT0000542 | Gata3 | 14462 | HITS-CLIP | 31433990\|23142080\|26203562 | Unspecified |
| mmu-miR-34a-5p | MIMAT0000542 | Gatad2a | 234366 | HITS-CLIP | 23142080 | Unspecified |
| mmu-miR-34a-5p | MIMAT0000542 | Gatad2b | 229542 | HITS-CLIP | 23142080\|31433990\|29167373 | Unspecified |
| mmu-miR-34a-5p | MIMAT0000542 | Gatb | 90603889 | HITS-CLIP | 29167373 | Unspecified |
| mmu-miR-34a-5p | MIMAT0000542 | Gatd1 | 347862 | HITS-CLIP | 23142080 | Unspecified |
| mmu-miR-34a-5p | MIMAT0000542 | Gatd3a | 8209 | HITS-CLIP | 31433990 | Unspecified |
| mmu-miR-34a-5p | MIMAT0000542 | Gatm | 67092 | HITS-CLIP | 29167373 | Unspecified |

| | | | | | | |
|---|---|---|---|---|---|---|
| mmu-miR-34a-5p | MIMAT0000542 | Gba | 14466 | HITS-CLIP | 23142080 | Unspecified |
| mmu-miR-34a-5p | MIMAT0000542 | Gba2 | 230101 | HITS-CLIP | 30971389\|19536157\|23142080 | Unspecified |
| mmu-miR-34a-5p | MIMAT0000542 | Gbe1 | 74185 | HITS-CLIP | 23142080 | Unspecified |
| mmu-miR-34a-5p | MIMAT0000542 | Gbp2 | 14469 | HITS-CLIP | 23142080\|31433990 | Unspecified |
| mmu-miR-34a-5p | MIMAT0000542 | Gbp2b | 14468 | HITS-CLIP | 23142080 | Unspecified |
| mmu-miR-34a-5p | MIMAT0000542 | Gbp3 | 55932 | HITS-CLIP | 23142080\|29167373 | Unspecified |
| mmu-miR-34a-5p | MIMAT0000542 | Gbp4 | 17472 | HITS-CLIP | 23142080 | Unspecified |
| mmu-miR-34a-5p | MIMAT0000542 | Gbp5 | 229898 | HITS-CLIP | 23142080 | Unspecified |
| mmu-miR-34a-5p | MIMAT0000542 | Gbp7 | 229900 | HITS-CLIP | 31433990\|23142080 | Unspecified |
| mmu-miR-34a-5p | MIMAT0000542 | Gbp8 | 76074 | HITS-CLIP | 23142080 | Unspecified |
| mmu-miR-34a-5p | MIMAT0000542 | Gbp9 | 236573 | HITS-CLIP | 23142080\|29167373 | Unspecified |
| mmu-miR-34a-5p | MIMAT0000542 | Gca | 227960 | Microarrays | 37986224 | Unspecified |
| mmu-miR-34a-5p | MIMAT0000542 | Gcat | 26912 | HITS-CLIP | 23142080 | Unspecified |
| mmu-miR-34a-5p | MIMAT0000542 | Gcc1 | 74375 | HITS-CLIP | 29167373 | Unspecified |
| mmu-miR-34a-5p | MIMAT0000542 | Gcc2 | 70297 | HITS-CLIP | 19536157\|23142080 | Unspecified |
| mmu-miR-34a-5p | MIMAT0000542 | Gch1 | 14528 | HITS-CLIP | 23142080\|29167373 | Unspecified |
| mmu-miR-34a-5p | MIMAT0000542 | Gclm | 14630 | HITS-CLIP | 29167373 | Unspecified |
| mmu-miR-34a-5p | MIMAT0000542 | Gcn1l1 | 10985 | HITS-CLIP | 23142080 | Unspecified |
| mmu-miR-34a-5p | MIMAT0000542 | Gdap2 | 14547 | HITS-CLIP | 23142080 | Unspecified |
| mmu-miR-34a-5p | MIMAT0000542 | Gde1 | 56209 | HITS-CLIP | 23142080 | Unspecified |
| mmu-miR-34a-5p | MIMAT0000542 | Gdf11 | 14561 | HITS-CLIP | 23142080 | Unspecified |
| mmu-miR-34a-5p | MIMAT0000542 | Gdi1 | 14567 | HITS-CLIP | 23142080\|19536157\|29167373 | Unspecified |
| mmu-miR-34a-5p | MIMAT0000542 | Gdi2 | 14569 | HITS-CLIP | 23142080\|29167373 | Unspecified |
| mmu-miR-34a-5p | MIMAT0000542 | Gemin5 | 216766 | HITS-CLIP | 23142080 | Unspecified |
| mmu-miR-34a-5p | MIMAT0000542 | Gen1 | 209334 | HITS-CLIP | 23142080 | Unspecified |
| mmu-miR-34a-5p | MIMAT0000542 | Gfer | 11692 | HITS-CLIP | 23142080 | Unspecified |
| mmu-miR-34a-5p | MIMAT0000542 | Gfi1 | 14581 | HITS-CLIP | 23142080 | Unspecified |
| mmu-miR-34a-5p | MIMAT0000542 | Gfm1 | 28030 | HITS-CLIP | 23142080 | Unspecified |
| mmu-miR-34a-5p | MIMAT0000542 | Gfm2 | 320806 | HITS-CLIP | 23142080 | Unspecified |
| mmu-miR-34a-5p | MIMAT0000542 | Gfod1 | 328232 | HITS-CLIP | 23142080 | Unspecified |
| mmu-miR-34a-5p | MIMAT0000542 | Gfpt1 | 14583 | HITS-CLIP | 23142080\|29167373 | Unspecified |
| mmu-miR-34a-5p | MIMAT0000542 | Gga1 | 106039 | HITS-CLIP | 29167373 | Unspecified |
| mmu-miR-34a-5p | MIMAT0000542 | Gga2 | 74105 | HITS-CLIP | 19536157\|29167373 | Unspecified |
| mmu-miR-34a-5p | MIMAT0000542 | Gga3 | 260302 | HITS-CLIP | 37986224\|23142080\|29167373 | Unspecified |
| mmu-miR-34a-5p | MIMAT0000542 | Ggact | 223267 | HITS-CLIP | 23142080 | Unspecified |
| mmu-miR-34a-5p | MIMAT0000542 | Ggn | 243897 | HITS-CLIP | 23142080 | Unspecified |
| mmu-miR-34a-5p | MIMAT0000542 | Ggnbp2 | 217039 | HITS-CLIP | 23142080\|29167373 | Unspecified |
| mmu-miR-34a-5p | MIMAT0000542 | Ggta1 | 14594 | HITS-CLIP | 31433990\|23142080\|29167373 | Unspecified |
| mmu-miR-34a-5p | MIMAT0000542 | Ghitm | 66092 | HITS-CLIP | 37986224\|23142080 | Unspecified |

| | | | | | | |
|---|---|---|---|---|---|---|
| mmu-miR-34a-5p | MIMAT0000542 | Ghr | 14600 | HITS-CLIP | 19536157 | Unspecified |
| mmu-miR-34a-5p | MIMAT0000542 | Gigyf1 | 57330 | HITS-CLIP | 31433990\|23142080 | Unspecified |
| mmu-miR-34a-5p | MIMAT0000542 | Gigyf2 | 227331 | HITS-CLIP | 23142080\|29167373 | Unspecified |
| mmu-miR-34a-5p | MIMAT0000542 | Gimap1 | 16205 | HITS-CLIP | 23142080 | Unspecified |
| mmu-miR-34a-5p | MIMAT0000542 | Gimap3 | 83408 | HITS-CLIP | 23142080\|31433990 | Unspecified |
| mmu-miR-34a-5p | MIMAT0000542 | Gimap4 | 107526 | HITS-CLIP | 23142080 | Unspecified |
| mmu-miR-34a-5p | MIMAT0000542 | Gimap6 | 231931 | HITS-CLIP | 31433990\|23142080 | Unspecified |
| mmu-miR-34a-5p | MIMAT0000542 | Gimap7 | 231932 | HITS-CLIP | 31433990\|23142080 | Unspecified |
| mmu-miR-34a-5p | MIMAT0000542 | Gimap8 | 243374 | HITS-CLIP | 23142080 | Unspecified |
| mmu-miR-34a-5p | MIMAT0000542 | Gimap9 | 317758 | HITS-CLIP | 23142080 | Unspecified |
| mmu-miR-34a-5p | MIMAT0000542 | Ginm1 | 215751 | HITS-CLIP | 29167373 | Unspecified |
| mmu-miR-34a-5p | MIMAT0000542 | Gins1 | 69270 | Microarrays | 37986224\|23027903 | Unspecified |
| mmu-miR-34a-5p | MIMAT0000542 | Git1 | 216963 | HITS-CLIP | 23142080 | Unspecified |
| mmu-miR-34a-5p | MIMAT0000542 | Git2 | 26431 | HITS-CLIP | 23142080\|29167373 | Unspecified |
| mmu-miR-34a-5p | MIMAT0000542 | Gja1 | 14609 | HITS-CLIP | 23142080 | Unspecified |
| mmu-miR-34a-5p | MIMAT0000542 | Gja5 | 14613 | Microarrays | 37986224 | Unspecified |
| mmu-miR-34a-5p | MIMAT0000542 | Gk | 14933 | Microarrays//HITS-CLIP | 23027903\|29167373 | Unspecified |
| mmu-miR-34a-5p | MIMAT0000542 | Gla | 11605 | HITS-CLIP | 29167373\|23142080 | Unspecified |
| mmu-miR-34a-5p | MIMAT0000542 | Glb1 | 12091 | HITS-CLIP | 29167373 | Unspecified |
| mmu-miR-34a-5p | MIMAT0000542 | Glb1l | 74577 | HITS-CLIP | 29167373 | Unspecified |
| mmu-miR-34a-5p | MIMAT0000542 | Glcci1 | 170772 | HITS-CLIP | 31433990\|23142080 | Unspecified |
| mmu-miR-34a-5p | MIMAT0000542 | Glce | 93683 | Microarrays//HITS-CLIP | 37986224\|23142080\|23027903\|1953€ | Unspecified |
| mmu-miR-34a-5p | MIMAT0000542 | Glg1 | 20340 | HITS-CLIP | 23142080\|19536157\|30971389\|3143 | Unspecified |
| mmu-miR-34a-5p | MIMAT0000542 | Glipr1 | 73690 | HITS-CLIP | 23142080 | Unspecified |
| mmu-miR-34a-5p | MIMAT0000542 | Glipr2 | 384009 | HITS-CLIP | 23142080 | Unspecified |
| mmu-miR-34a-5p | MIMAT0000542 | Glis2 | 83396 | HITS-CLIP | 31433990 | Unspecified |
| mmu-miR-34a-5p | MIMAT0000542 | Glis3 | 226075 | Microarrays | 37986224 | Unspecified |
| mmu-miR-34a-5p | MIMAT0000542 | Glo1 | 109801 | HITS-CLIP | 23142080 | Unspecified |
| mmu-miR-34a-5p | MIMAT0000542 | Glod4 | 67201 | HITS-CLIP | 23142080\|29167373 | Unspecified |
| mmu-miR-34a-5p | MIMAT0000542 | Glrb | 14658 | Microarrays | 37986224 | Unspecified |
| mmu-miR-34a-5p | MIMAT0000542 | Glrx | 93692 | HITS-CLIP | 23142080\|19536157\|29167373 | Unspecified |
| mmu-miR-34a-5p | MIMAT0000542 | Glrx2 | 69367 | HITS-CLIP | 23142080\|29167373 | Unspecified |
| mmu-miR-34a-5p | MIMAT0000542 | Glrx3 | 30926 | HITS-CLIP | 29167373\|23142080 | Unspecified |
| mmu-miR-34a-5p | MIMAT0000542 | Gls | 14660 | HITS-CLIP | 31433990\|23142080\|29167373 | Unspecified |
| mmu-miR-34a-5p | MIMAT0000542 | Gls2 | 216456 | HITS-CLIP | 23142080 | Unspecified |
| mmu-miR-34a-5p | MIMAT0000542 | Glt8d1 | 76485 | HITS-CLIP | 23142080 | Unspecified |
| mmu-miR-34a-5p | MIMAT0000542 | Gltp | 56356 | Microarrays//HITS-CLIP | 37986224\|23027903\|23142080\|2620 | Unspecified |
| mmu-miR-34a-5p | MIMAT0000542 | Glud1 | 14661 | HITS-CLIP | 23142080\|19536157\|29167373 | Unspecified |
| mmu-miR-34a-5p | MIMAT0000542 | Glul | 14645 | Microarrays//HITS-CLIP | 37986224\|23027903\|19536157\|2916 | Unspecified |

| | | | | | | |
|---|---|---|---|---|---|---|
| mmu-miR-34a-5p | MIMAT0000542 | Glyr1 | 74022 | HITS-CLIP | 19536157\|31433990\|23142080\|29167 | Unspecified |
| mmu-miR-34a-5p | MIMAT0000542 | Gm10031 | 100039026 | HITS-CLIP | 23142080 | Unspecified |
| mmu-miR-34a-5p | MIMAT0000542 | Gm10337 | 6617069 | HITS-CLIP | 26203562 | Unspecified |
| mmu-miR-34a-5p | MIMAT0000542 | Gm10521 | 100038548 | HITS-CLIP | 29167373 | Unspecified |
| mmu-miR-34a-5p | MIMAT0000542 | Gm11808 | 666586 | HITS-CLIP | 26203562 | Unspecified |
| mmu-miR-34a-5p | MIMAT0000542 | Gm11992 | 626870 | Microarrays | 37986224 | Unspecified |
| mmu-miR-34a-5p | MIMAT0000542 | Gm16286 | 117903916 | HITS-CLIP | 19536157\|23142080 | Unspecified |
| mmu-miR-34a-5p | MIMAT0000542 | Gm2000 | 100038991 | HITS-CLIP | 23142080 | Unspecified |
| mmu-miR-34a-5p | MIMAT0000542 | Gm20431 | 6608666 | HITS-CLIP | 29167373 | Unspecified |
| mmu-miR-34a-5p | MIMAT0000542 | Gm20441 | 6608650 | HITS-CLIP | 23142080 | Unspecified |
| mmu-miR-34a-5p | MIMAT0000542 | Gm20503 | 6608537 | HITS-CLIP | 23142080 | Unspecified |
| mmu-miR-34a-5p | MIMAT0000542 | Gm20604 | 100859931 | HITS-CLIP | 23142080\|19536157\|31433990 | Unspecified |
| mmu-miR-34a-5p | MIMAT0000542 | Gm20716 | 105980075 | HITS-CLIP | 31433990 | Unspecified |
| mmu-miR-34a-5p | MIMAT0000542 | Gm21969 | 6609957 | HITS-CLIP | 30971389 | Unspecified |
| mmu-miR-34a-5p | MIMAT0000542 | Gm21970 | 6609959 | HITS-CLIP | 30971389 | Unspecified |
| mmu-miR-34a-5p | MIMAT0000542 | Gm21983 | 6610009 | HITS-CLIP | 31433990 | Unspecified |
| mmu-miR-34a-5p | MIMAT0000542 | Gm2237 | 100039441 | HITS-CLIP | 23142080 | Unspecified |
| mmu-miR-34a-5p | MIMAT0000542 | Gm28043 | 106014251 | HITS-CLIP | 31433990 | Unspecified |
| mmu-miR-34a-5p | MIMAT0000542 | Gm2a | 14667 | HITS-CLIP | 23142080\|29167373 | Unspecified |
| mmu-miR-34a-5p | MIMAT0000542 | Gm37240 | | HITS-CLIP | 23142080\|19536157 | Unspecified |
| mmu-miR-34a-5p | MIMAT0000542 | Gm38394 | 667118 | HITS-CLIP | 23142080\|19536157\|29167373 | Unspecified |
| mmu-miR-34a-5p | MIMAT0000542 | Gm42674 | | HITS-CLIP | 31433990 | Unspecified |
| mmu-miR-34a-5p | MIMAT0000542 | Gm43064 | | HITS-CLIP | 30971389 | Unspecified |
| mmu-miR-34a-5p | MIMAT0000542 | Gm45062 | | HITS-CLIP | 31433990 | Unspecified |
| mmu-miR-34a-5p | MIMAT0000542 | Gm45234 | | HITS-CLIP | 23630078 | Unspecified |
| mmu-miR-34a-5p | MIMAT0000542 | Gm45713 | 103691134 | HITS-CLIP | 23142080 | Unspecified |
| mmu-miR-34a-5p | MIMAT0000542 | Gm45902 | 119065974 | HITS-CLIP | 31433990\|23142080 | Unspecified |
| mmu-miR-34a-5p | MIMAT0000542 | Gm4787 | 214321 | HITS-CLIP | 29167373 | Unspecified |
| mmu-miR-34a-5p | MIMAT0000542 | Gm4841 | 225594 | HITS-CLIP | 23142080 | Unspecified |
| mmu-miR-34a-5p | MIMAT0000542 | Gm49322 | | HITS-CLIP | 23142080 | Unspecified |
| mmu-miR-34a-5p | MIMAT0000542 | Gm49336 | 127230315 | HITS-CLIP | 30971389\|29167373\|23142080 | Unspecified |
| mmu-miR-34a-5p | MIMAT0000542 | Gm49361 | | HITS-CLIP | 31433990 | Unspecified |
| mmu-miR-34a-5p | MIMAT0000542 | Gm49368 | 118261720 | HITS-CLIP | 30971389\|29167373 | Unspecified |
| mmu-miR-34a-5p | MIMAT0000542 | Gm49380 | | HITS-CLIP | 23142080 | Unspecified |
| mmu-miR-34a-5p | MIMAT0000542 | Gm49486 | | HITS-CLIP | 23142080 | Unspecified |
| mmu-miR-34a-5p | MIMAT0000542 | Gm49510 | | HITS-CLIP | 23142080\|29167373 | Unspecified |
| mmu-miR-34a-5p | MIMAT0000542 | Gm50318 | | HITS-CLIP | 30971389 | Unspecified |
| mmu-miR-34a-5p | MIMAT0000542 | Gm5483 | 433016 | HITS-CLIP | 23142080 | Unspecified |
| mmu-miR-34a-5p | MIMAT0000542 | Gm56303 | | HITS-CLIP | 30971389 | Unspecified |

| | | | | | | |
|---|---|---|---|---|---|---|
| mmu-miR-34a-5p | MIMAT0000542 | Gm7030 | 630294 | HITS-CLIP | 23142080 | Unspecified |
| mmu-miR-34a-5p | MIMAT0000542 | Gm7964 | 666170 | HITS-CLIP | 37986224 | Unspecified |
| mmu-miR-34a-5p | MIMAT0000542 | Gm9844 | 100043712 | HITS-CLIP | 31433990 | Unspecified |
| mmu-miR-34a-5p | MIMAT0000542 | Gmds | 218138 | HITS-CLIP | 23142080 | Unspecified |
| mmu-miR-34a-5p | MIMAT0000542 | Gmeb1 | 56809 | HITS-CLIP | 23142080 | Unspecified |
| mmu-miR-34a-5p | MIMAT0000542 | Gmfb | 63985 | Microarrays//HITS-CLIP | 37986224\|23027903\|23142080\|29167 | Unspecified |
| mmu-miR-34a-5p | MIMAT0000542 | Gmip | 78816 | HITS-CLIP | 23142080\|29167373 | Unspecified |
| mmu-miR-34a-5p | MIMAT0000542 | Gmnn | 57441 | HITS-CLIP | 23142080 | Unspecified |
| mmu-miR-34a-5p | MIMAT0000542 | Gmppa | 69080 | HITS-CLIP | 23142080 | Unspecified |
| mmu-miR-34a-5p | MIMAT0000542 | Gmpr2 | 105446 | HITS-CLIP | 23142080 | Unspecified |
| mmu-miR-34a-5p | MIMAT0000542 | Gmps | 229363 | HITS-CLIP | 23142080 | Unspecified |
| mmu-miR-34a-5p | MIMAT0000542 | Gna11 | 14672 | HITS-CLIP | 26203562\|23142080\|29167373 | Unspecified |
| mmu-miR-34a-5p | MIMAT0000542 | Gna12 | 14673 | HITS-CLIP | 19536157\|29167373 | Unspecified |
| mmu-miR-34a-5p | MIMAT0000542 | Gna13 | 14674 | HITS-CLIP | 31433990\|23142080\|19536157\|29167 | Unspecified |
| mmu-miR-34a-5p | MIMAT0000542 | Gnai2 | 14678 | HITS-CLIP | 31433990\|19536157\|29167373\|23142 | Unspecified |
| mmu-miR-34a-5p | MIMAT0000542 | Gnai3 | 14679 | HITS-CLIP | 23142080\|29167373 | Unspecified |
| mmu-miR-34a-5p | MIMAT0000542 | Gnal | 14680 | HITS-CLIP | 19536157 | Unspecified |
| mmu-miR-34a-5p | MIMAT0000542 | Gnaq | 14682 | HITS-CLIP | 23142080\|19536157\|23630078\|29167 | Unspecified |
| mmu-miR-34a-5p | MIMAT0000542 | Gnas | 14683 | HITS-CLIP | 31433990\|23142080\|29167373 | Unspecified |
| mmu-miR-34a-5p | MIMAT0000542 | Gnb1 | 14688 | HITS-CLIP | 23142080\|19536157\|29167373 | Unspecified |
| mmu-miR-34a-5p | MIMAT0000542 | Gnb2 | 14693 | HITS-CLIP | 31433990\|19536157\|23142080\|29167 | Unspecified |
| mmu-miR-34a-5p | MIMAT0000542 | Gnb5 | 14697 | HITS-CLIP | 31433990 | Unspecified |
| mmu-miR-34a-5p | MIMAT0000542 | Gng12 | 14701 | HITS-CLIP | 23630078\|23142080\|29167373 | Unspecified |
| mmu-miR-34a-5p | MIMAT0000542 | Gng2 | 14702 | HITS-CLIP | 23142080 | Unspecified |
| mmu-miR-34a-5p | MIMAT0000542 | Gnl1 | 14670 | HITS-CLIP | 23142080 | Unspecified |
| mmu-miR-34a-5p | MIMAT0000542 | Gnl2 | 230737 | HITS-CLIP | 23142080 | Unspecified |
| mmu-miR-34a-5p | MIMAT0000542 | Gnl3 | 30877 | HITS-CLIP | 30971389\|29167373\|23142080 | Unspecified |
| mmu-miR-34a-5p | MIMAT0000542 | Gnl3l | 237107 | HITS-CLIP | 23142080 | Unspecified |
| mmu-miR-34a-5p | MIMAT0000542 | Gnpat | 14712 | HITS-CLIP | 29167373\|23142080 | Unspecified |
| mmu-miR-34a-5p | MIMAT0000542 | Gnpda1 | 26384 | HITS-CLIP | 29167373 | Unspecified |
| mmu-miR-34a-5p | MIMAT0000542 | Gnpda2 | 67980 | HITS-CLIP | 23142080 | Unspecified |
| mmu-miR-34a-5p | MIMAT0000542 | Gnptab | 432486 | HITS-CLIP | 23142080\|19536157\|29167373 | Unspecified |
| mmu-miR-34a-5p | MIMAT0000542 | Gns | 75612 | Microarrays//HITS-CLIP | 37986224\|29167373\|23142080 | Unspecified |
| mmu-miR-34a-5p | MIMAT0000542 | Golga1 | 76899 | HITS-CLIP | 23142080\|29167373 | Unspecified |
| mmu-miR-34a-5p | MIMAT0000542 | Golga2 | 99412 | HITS-CLIP | 29167373 | Unspecified |
| mmu-miR-34a-5p | MIMAT0000542 | Golga3 | 269682 | HITS-CLIP | 29167373 | Unspecified |
| mmu-miR-34a-5p | MIMAT0000542 | Golga4 | 54214 | HITS-CLIP | 23142080\|29167373 | Unspecified |
| mmu-miR-34a-5p | MIMAT0000542 | Golgb1 | 224139 | HITS-CLIP | 30971389 | Unspecified |
| mmu-miR-34a-5p | MIMAT0000542 | Golm1 | 105348 | HITS-CLIP | 23142080\|19536157\|31433990 | Unspecified |

| | | | | | | |
|---|---|---|---|---|---|---|
| mmu-miR-34a-5p | MIMAT0000542 | Golph3 | 66629 | HITS-CLIP | 23142080\|29167373 | Unspecified |
| mmu-miR-34a-5p | MIMAT0000542 | Golph3l | 229593 | HITS-CLIP | 23142080 | Unspecified |
| mmu-miR-34a-5p | MIMAT0000542 | Golt1b | 66964 | HITS-CLIP | 31433990\|23142080\|29167373 | Unspecified |
| mmu-miR-34a-5p | MIMAT0000542 | Gon4l | 76022 | HITS-CLIP | 23142080\|29167373 | Unspecified |
| mmu-miR-34a-5p | MIMAT0000542 | Gon7 | 100233175 | HITS-CLIP | 29167373 | Unspecified |
| mmu-miR-34a-5p | MIMAT0000542 | Gopc | 94221 | HITS-CLIP | 37986224\|19536157\|29167373\|23142 | Unspecified |
| mmu-miR-34a-5p | MIMAT0000542 | Gorasp2 | 70231 | HITS-CLIP | 23142080\|29167373 | Unspecified |
| mmu-miR-34a-5p | MIMAT0000542 | Gosr1 | 53334 | HITS-CLIP | 23142080\|29167373 | Unspecified |
| mmu-miR-34a-5p | MIMAT0000542 | Gosr2 | 56494 | HITS-CLIP | 23142080\|29167373 | Unspecified |
| mmu-miR-34a-5p | MIMAT0000542 | Got1 | 14718 | HITS-CLIP | 31433990\|19536157\|23142080 | Unspecified |
| mmu-miR-34a-5p | MIMAT0000542 | Gpaa1 | 14731 | HITS-CLIP | 19536157 | Unspecified |
| mmu-miR-34a-5p | MIMAT0000542 | Gpalpp1 | 67467 | HITS-CLIP | 23142080 | Unspecified |
| mmu-miR-34a-5p | MIMAT0000542 | Gpam | 14732 | HITS-CLIP | 23142080\|19536157 | Unspecified |
| mmu-miR-34a-5p | MIMAT0000542 | Gpank1 | 81845 | HITS-CLIP | 23142080 | Unspecified |
| mmu-miR-34a-5p | MIMAT0000542 | Gpat4 | 137964 | Microarrays | 23027903 | Unspecified |
| mmu-miR-34a-5p | MIMAT0000542 | Gpatch1 | 67471 | HITS-CLIP | 23142080 | Unspecified |
| mmu-miR-34a-5p | MIMAT0000542 | Gpatch11 | 53951 | HITS-CLIP | 23142080 | Unspecified |
| mmu-miR-34a-5p | MIMAT0000542 | Gpatch2 | 67769 | HITS-CLIP | 23142080 | Unspecified |
| mmu-miR-34a-5p | MIMAT0000542 | Gpatch2l | 70373 | HITS-CLIP | 29167373 | Unspecified |
| mmu-miR-34a-5p | MIMAT0000542 | Gpatch8 | 237943 | HITS-CLIP | 23142080\|30971389\|29167373 | Unspecified |
| mmu-miR-34a-5p | MIMAT0000542 | Gpbp1 | 73274 | HITS-CLIP | 23142080\|29167373 | Unspecified |
| mmu-miR-34a-5p | MIMAT0000542 | Gpcpd1 | 74182 | HITS-CLIP | 23142080 | Unspecified |
| mmu-miR-34a-5p | MIMAT0000542 | Gpd2 | 14571 | HITS-CLIP | 31433990\|23142080 | Unspecified |
| mmu-miR-34a-5p | MIMAT0000542 | Gphn | 268566 | HITS-CLIP | 23142080 | Unspecified |
| mmu-miR-34a-5p | MIMAT0000542 | Gpi1 | 14751 | HITS-CLIP | 23142080\|29167373 | Unspecified |
| mmu-miR-34a-5p | MIMAT0000542 | Gpld1 | 14756 | HITS-CLIP | 23142080 | Unspecified |
| mmu-miR-34a-5p | MIMAT0000542 | Gpm6a | 234267 | HITS-CLIP | 19536157 | Unspecified |
| mmu-miR-34a-5p | MIMAT0000542 | Gpm6b | 14758 | Microarrays//HITS-CLIP | 37986224\|19536157\|23142080 | Unspecified |
| mmu-miR-34a-5p | MIMAT0000542 | Gpn2 | 100210 | HITS-CLIP | 23142080 | Unspecified |
| mmu-miR-34a-5p | MIMAT0000542 | Gpnmb | 93695 | HITS-CLIP | 29167373 | Unspecified |
| mmu-miR-34a-5p | MIMAT0000542 | Gpr107 | 277463 | HITS-CLIP | 26203562\|29167373 | Unspecified |
| mmu-miR-34a-5p | MIMAT0000542 | Gpr132 | 56696 | HITS-CLIP | 23142080 | Unspecified |
| mmu-miR-34a-5p | MIMAT0000542 | Gpr137 | 107173 | HITS-CLIP | 23142080 | Unspecified |
| mmu-miR-34a-5p | MIMAT0000542 | Gpr137b | 83924 | HITS-CLIP | 29167373\|31433990 | Unspecified |
| mmu-miR-34a-5p | MIMAT0000542 | Gpr146 | 80290 | HITS-CLIP | 23142080\|29167373 | Unspecified |
| mmu-miR-34a-5p | MIMAT0000542 | Gpr157 | 269604 | HITS-CLIP | 23142080 | Unspecified |
| mmu-miR-34a-5p | MIMAT0000542 | Gpr158 | 241263 | HITS-CLIP | 19536157 | Unspecified |
| mmu-miR-34a-5p | MIMAT0000542 | Gpr171 | 229323 | HITS-CLIP | 23142080 | Unspecified |
| mmu-miR-34a-5p | MIMAT0000542 | Gpr174 | 213439 | HITS-CLIP | 23142080 | Unspecified |

| | | | | | | | |
|---|---|---|---|---|---|---|---|
| mmu-miR-34a-5p | MIMAT0000542 | Gpr18 | 110168 | HITS-CLIP | | 23142080 | Unspecified |
| mmu-miR-34a-5p | MIMAT0000542 | Gpr183 | 321019 | HITS-CLIP | 29167373\|23142080 | | Unspecified |
| mmu-miR-34a-5p | MIMAT0000542 | Gpr19 | 14760 | HITS-CLIP | 19536157\|23142080 | | Unspecified |
| mmu-miR-34a-5p | MIMAT0000542 | Gpr21 | 338346 | HITS-CLIP | | 29167373 | Unspecified |
| mmu-miR-34a-5p | MIMAT0000542 | Gpr26 | 233919 | HITS-CLIP | | 19536157 | Unspecified |
| mmu-miR-34a-5p | MIMAT0000542 | Gpr62 | 118442 | HITS-CLIP | | 23142080 | Unspecified |
| mmu-miR-34a-5p | MIMAT0000542 | Gpr63 | 81006 | HITS-CLIP | | 19536157 | Unspecified |
| mmu-miR-34a-5p | MIMAT0000542 | Gpr65 | 14744 | HITS-CLIP | 23142080\|29167373 | | Unspecified |
| mmu-miR-34a-5p | MIMAT0000542 | Gpr68 | 238377 | HITS-CLIP | | 23142080 | Unspecified |
| mmu-miR-34a-5p | MIMAT0000542 | Gpr85 | 64450 | HITS-CLIP | | 29167373 | Unspecified |
| mmu-miR-34a-5p | MIMAT0000542 | Gpr89 | 67549 | HITS-CLIP | 23142080\|29167373 | | Unspecified |
| mmu-miR-34a-5p | MIMAT0000542 | Gprasp1 | 67298 | HITS-CLIP | | 19536157 | Unspecified |
| mmu-miR-34a-5p | MIMAT0000542 | Gprc5a | 232431 | Microarrays//HITS-CLIP | 37986224\|23027903\|23142080 | | Unspecified |
| mmu-miR-34a-5p | MIMAT0000542 | Gprc5b | 64297 | HITS-CLIP | | 19536157 | Unspecified |
| mmu-miR-34a-5p | MIMAT0000542 | Gprin3 | 243385 | HITS-CLIP | | 23142080 | Unspecified |
| mmu-miR-34a-5p | MIMAT0000542 | Gps1 | 2873 | HITS-CLIP | | 23142080 | Unspecified |
| mmu-miR-34a-5p | MIMAT0000542 | Gpsm2 | 76123 | HITS-CLIP | | 23142080 | Unspecified |
| mmu-miR-34a-5p | MIMAT0000542 | Gpx1 | 14775 | HITS-CLIP | 23142080\|29167373 | | Unspecified |
| mmu-miR-34a-5p | MIMAT0000542 | Gramd1a | 52857 | HITS-CLIP | | 23142080 | Unspecified |
| mmu-miR-34a-5p | MIMAT0000542 | Gramd1b | 235283 | Microarrays//HITS-CLIP | 37986224\|19536157\|23142080\|29167 | | Unspecified |
| mmu-miR-34a-5p | MIMAT0000542 | Gramd3 | 107022 | HITS-CLIP | | 23142080 | Unspecified |
| mmu-miR-34a-5p | MIMAT0000542 | Gramd4 | 223752 | HITS-CLIP | | 23142080 | Unspecified |
| mmu-miR-34a-5p | MIMAT0000542 | Grap | 71520 | HITS-CLIP | | 23142080 | Unspecified |
| mmu-miR-34a-5p | MIMAT0000542 | Grap2 | 17444 | HITS-CLIP | 37986224\|23142080 | | Unspecified |
| mmu-miR-34a-5p | MIMAT0000542 | Grb2 | 14784 | HITS-CLIP | | 29167373 | Unspecified |
| mmu-miR-34a-5p | MIMAT0000542 | Grcc10 | 14790 | HITS-CLIP | | 23142080 | Unspecified |
| mmu-miR-34a-5p | MIMAT0000542 | Grem2 | 23893 | HITS-CLIP | | 29167373 | Unspecified |
| mmu-miR-34a-5p | MIMAT0000542 | Grhpr | 76238 | HITS-CLIP | | 23142080 | Unspecified |
| mmu-miR-34a-5p | MIMAT0000542 | Gria2 | 14800 | HITS-CLIP | | 19536157 | Unspecified |
| mmu-miR-34a-5p | MIMAT0000542 | Gria4 | 14802 | HITS-CLIP | | 19536157 | Unspecified |
| mmu-miR-34a-5p | MIMAT0000542 | Grid1 | 14803 | HITS-CLIP | | 19536157 | Unspecified |
| mmu-miR-34a-5p | MIMAT0000542 | Grik2 | 14806 | HITS-CLIP | | 19536157 | Unspecified |
| mmu-miR-34a-5p | MIMAT0000542 | Grik5 | 14809 | HITS-CLIP | | 19536157 | Unspecified |
| mmu-miR-34a-5p | MIMAT0000542 | Grin1 | 14810 | HITS-CLIP | | 19536157 | Unspecified |
| mmu-miR-34a-5p | MIMAT0000542 | Grin2a | 14811 | HITS-CLIP | | 19536157 | Unspecified |
| mmu-miR-34a-5p | MIMAT0000542 | Grin2b | 14812 | HITS-CLIP | | 19536157 | Unspecified |
| mmu-miR-34a-5p | MIMAT0000542 | Grina | 66168 | HITS-CLIP | | 19536157 | Unspecified |
| mmu-miR-34a-5p | MIMAT0000542 | Gripap1 | 54645 | HITS-CLIP | | 23142080 | Unspecified |
| mmu-miR-34a-5p | MIMAT0000542 | Grk2 | 156 | HITS-CLIP | 23142080\|29167373 | | Unspecified |

| | | | | | | |
|---|---|---|---|---|---|---|
| mmu-miR-34a-5p | MIMAT0000542 | Grk5 | 14773 | HITS-CLIP | 31433990\|29167373 | Unspecified |
| mmu-miR-34a-5p | MIMAT0000542 | Grk6 | 26385 | Microarrays//HITS-CLIP | 37986224\|23027903\|23630078\|3143 | Unspecified |
| mmu-miR-34a-5p | MIMAT0000542 | Grm1 | 14816 | HITS-CLIP | 19536157 | Unspecified |
| mmu-miR-34a-5p | MIMAT0000542 | Grm5 | 108071 | HITS-CLIP | 19536157 | Unspecified |
| mmu-miR-34a-5p | MIMAT0000542 | Grn | 14824 | HITS-CLIP | 19536157\|29167373 | Unspecified |
| mmu-miR-34a-5p | MIMAT0000542 | Grpel1 | 17713 | HITS-CLIP | 31433990\|23142080 | Unspecified |
| mmu-miR-34a-5p | MIMAT0000542 | Grsf1 | 231413 | HITS-CLIP | 23142080 | Unspecified |
| mmu-miR-34a-5p | MIMAT0000542 | Gsap | 212167 | HITS-CLIP | 29167373 | Unspecified |
| mmu-miR-34a-5p | MIMAT0000542 | Gse1 | 382034 | HITS-CLIP | 29167373\|23142080 | Unspecified |
| mmu-miR-34a-5p | MIMAT0000542 | Gsk3a | 606496 | HITS-CLIP | 23142080\|29167373 | Unspecified |
| mmu-miR-34a-5p | MIMAT0000542 | Gsk3b | 56637 | HITS-CLIP | 19536157\|23142080\|31433990\|2916 | Unspecified |
| mmu-miR-34a-5p | MIMAT0000542 | Gskip | 66787 | HITS-CLIP | 23142080 | Unspecified |
| mmu-miR-34a-5p | MIMAT0000542 | Gsn | 227753 | HITS-CLIP | 19536157\|23142080\|29167373 | Unspecified |
| mmu-miR-34a-5p | MIMAT0000542 | Gspt1 | 14852 | HITS-CLIP | 23142080\|26203562\|29167373 | Unspecified |
| mmu-miR-34a-5p | MIMAT0000542 | Gstcd | 67553 | HITS-CLIP | 23142080 | Unspecified |
| mmu-miR-34a-5p | MIMAT0000542 | Gsto1 | 14873 | Microarrays | 37986224\|23027903 | Unspecified |
| mmu-miR-34a-5p | MIMAT0000542 | Gstp1 | 14870 | HITS-CLIP | 23142080 | Unspecified |
| mmu-miR-34a-5p | MIMAT0000542 | Gstp3 | 225884 | HITS-CLIP | 23142080 | Unspecified |
| mmu-miR-34a-5p | MIMAT0000542 | Gstt1 | 14871 | HITS-CLIP | 23142080 | Unspecified |
| mmu-miR-34a-5p | MIMAT0000542 | Gstt3 | 103140 | HITS-CLIP | 23142080 | Unspecified |
| mmu-miR-34a-5p | MIMAT0000542 | Gstz1 | 14874 | HITS-CLIP | 23142080\|29167373 | Unspecified |
| mmu-miR-34a-5p | MIMAT0000542 | Gtdc1 | 227835 | Microarrays | 37986224\|23027903 | Unspecified |
| mmu-miR-34a-5p | MIMAT0000542 | Gtf2a1 | 83602 | HITS-CLIP | 23142080\|29167373 | Unspecified |
| mmu-miR-34a-5p | MIMAT0000542 | Gtf2a2 | 235459 | HITS-CLIP | 31433990 | Unspecified |
| mmu-miR-34a-5p | MIMAT0000542 | Gtf2b | 229906 | Microarrays//HITS-CLIP | 37986224\|23027903\|23142080 | Unspecified |
| mmu-miR-34a-5p | MIMAT0000542 | Gtf2e1 | 74197 | Microarrays//HITS-CLIP | 37986224\|23027903\|23142080 | Unspecified |
| mmu-miR-34a-5p | MIMAT0000542 | Gtf2e2 | 68153 | HITS-CLIP | 23142080\|23630078\|31433990\|30971 | Unspecified |
| mmu-miR-34a-5p | MIMAT0000542 | Gtf2f2 | 68705 | HITS-CLIP | 23142080 | Unspecified |
| mmu-miR-34a-5p | MIMAT0000542 | Gtf2h1 | 14884 | HITS-CLIP | 23142080\|29167373 | Unspecified |
| mmu-miR-34a-5p | MIMAT0000542 | Gtf2h2 | 23894 | HITS-CLIP | 23142080 | Unspecified |
| mmu-miR-34a-5p | MIMAT0000542 | Gtf2h3 | 209357 | HITS-CLIP | 23142080 | Unspecified |
| mmu-miR-34a-5p | MIMAT0000542 | Gtf2i | 14886 | HITS-CLIP | 26203562\|23142080 | Unspecified |
| mmu-miR-34a-5p | MIMAT0000542 | Gtf3a | 66596 | HITS-CLIP | 23142080 | Unspecified |
| mmu-miR-34a-5p | MIMAT0000542 | Gtf3c1 | 233863 | HITS-CLIP | 23142080 | Unspecified |
| mmu-miR-34a-5p | MIMAT0000542 | Gtf3c2 | 71752 | HITS-CLIP | 23142080\|29167373 | Unspecified |
| mmu-miR-34a-5p | MIMAT0000542 | Gtf3c3 | 98488 | HITS-CLIP | 23142080 | Unspecified |
| mmu-miR-34a-5p | MIMAT0000542 | Gtf3c4 | 269252 | HITS-CLIP | 23142080 | Unspecified |
| mmu-miR-34a-5p | MIMAT0000542 | Gtf3c6 | 67371 | HITS-CLIP | 23142080 | Unspecified |
| mmu-miR-34a-5p | MIMAT0000542 | Gtpbp1 | 14904 | HITS-CLIP | 23142080\|29167373 | Unspecified |

| | | | | | | |
|---|---|---|---|---|---|---|
| mmu-miR-34a-5p | MIMAT0000542 | Gtpbp10 | 207704 | HITS-CLIP | 23142080 | Unspecified |
| mmu-miR-34a-5p | MIMAT0000542 | Gtpbp2 | 56055 | HITS-CLIP | 23142080\|29167373 | Unspecified |
| mmu-miR-34a-5p | MIMAT0000542 | Gtpbp4 | 69237 | HITS-CLIP | 23142080 | Unspecified |
| mmu-miR-34a-5p | MIMAT0000542 | Gtpbp8 | 66067 | HITS-CLIP | 19536157 | Unspecified |
| mmu-miR-34a-5p | MIMAT0000542 | Gtse1 | 29870 | HITS-CLIP | 23142080 | Unspecified |
| mmu-miR-34a-5p | MIMAT0000542 | Gucd1 | 68778 | HITS-CLIP | 31433990\|29167373 | Unspecified |
| mmu-miR-34a-5p | MIMAT0000542 | Gucy1a1 | 60596 | HITS-CLIP | 23142080 | Unspecified |
| mmu-miR-34a-5p | MIMAT0000542 | Gucy1b1 | 54195 | HITS-CLIP | 23142080 | Unspecified |
| mmu-miR-34a-5p | MIMAT0000542 | Guf1 | 231279 | HITS-CLIP | 29167373 | Unspecified |
| mmu-miR-34a-5p | MIMAT0000542 | Gusb | 110006 | HITS-CLIP | 29167373 | Unspecified |
| mmu-miR-34a-5p | MIMAT0000542 | Gvin1 | 74558 | HITS-CLIP | 29167373 | Unspecified |
| mmu-miR-34a-5p | MIMAT0000542 | Gxylt1 | 223827 | HITS-CLIP | 31433990\|23142080\|29167373 | Unspecified |
| mmu-miR-34a-5p | MIMAT0000542 | Gyg | 27357 | HITS-CLIP | 23142080\|29167373 | Unspecified |
| mmu-miR-34a-5p | MIMAT0000542 | Gypc | 71683 | HITS-CLIP | 23142080\|30971389 | Unspecified |
| mmu-miR-34a-5p | MIMAT0000542 | Gys1 | 14936 | HITS-CLIP | 23142080 | Unspecified |
| mmu-miR-34a-5p | MIMAT0000542 | Gzf1 | 74533 | HITS-CLIP | 30971389\|23142080 | Unspecified |
| mmu-miR-34a-5p | MIMAT0000542 | Gzma | 14938 | HITS-CLIP | 23142080 | Unspecified |
| mmu-miR-34a-5p | MIMAT0000542 | Gzmb | 14939 | HITS-CLIP | 23142080 | Unspecified |
| mmu-miR-34a-5p | MIMAT0000542 | Gzmc | 14940 | HITS-CLIP | 23142080 | Unspecified |
| mmu-miR-34a-5p | MIMAT0000542 | Gzme | 14942 | Microarrays | 37986224\|23027903 | Unspecified |
| mmu-miR-34a-5p | MIMAT0000542 | H13 | 14950 | HITS-CLIP | 23142080\|29167373 | Unspecified |
| mmu-miR-34a-5p | MIMAT0000542 | H1f0 | 14958 | HITS-CLIP | 23142080\|19536157 | Unspecified |
| mmu-miR-34a-5p | MIMAT0000542 | H1f2 | 3006 | HITS-CLIP | 31433990 | Unspecified |
| mmu-miR-34a-5p | MIMAT0000542 | H2-Aa | 14960 | HITS-CLIP | 23142080 | Unspecified |
| mmu-miR-34a-5p | MIMAT0000542 | H2-Ab1 | 14961 | HITS-CLIP | 23142080 | Unspecified |
| mmu-miR-34a-5p | MIMAT0000542 | H2-D1 | 14964 | HITS-CLIP | 23142080\|29167373 | Unspecified |
| mmu-miR-34a-5p | MIMAT0000542 | H2-DMb1 | 14999 | HITS-CLIP | 23142080 | Unspecified |
| mmu-miR-34a-5p | MIMAT0000542 | H2-Eb1 | 14969 | HITS-CLIP | 23142080 | Unspecified |
| mmu-miR-34a-5p | MIMAT0000542 | H2-K1 | 14972 | HITS-CLIP | 23142080\|29167373 | Unspecified |
| mmu-miR-34a-5p | MIMAT0000542 | H2-M3 | 14991 | HITS-CLIP | 23142080 | Unspecified |
| mmu-miR-34a-5p | MIMAT0000542 | H2-Ob | 15002 | HITS-CLIP | 23142080 | Unspecified |
| mmu-miR-34a-5p | MIMAT0000542 | H2-Q2 | 15013 | HITS-CLIP | 23142080 | Unspecified |
| mmu-miR-34a-5p | MIMAT0000542 | H2-Q4 | 15015 | HITS-CLIP | 23142080 | Unspecified |
| mmu-miR-34a-5p | MIMAT0000542 | H2-Q6 | 110557 | HITS-CLIP | 37986224\|23142080 | Unspecified |
| mmu-miR-34a-5p | MIMAT0000542 | H2-T22 | 15039 | HITS-CLIP | 23142080 | Unspecified |
| mmu-miR-34a-5p | MIMAT0000542 | H2-T23 | 15040 | HITS-CLIP | 23142080 | Unspecified |
| mmu-miR-34a-5p | MIMAT0000542 | H2-T24 | 15042 | HITS-CLIP | 23142080 | Unspecified |
| mmu-miR-34a-5p | MIMAT0000542 | H2ac20 | 8338 | HITS-CLIP | 31433990 | Unspecified |
| mmu-miR-34a-5p | MIMAT0000542 | H2ac4 | 8335 | Microarrays | 23027903 | Unspecified |

| | | | | | | |
|---|---|---|---|---|---|---|
| mmu-miR-34a-5p | MIMAT0000542 | H2ac8 | 8335 | Microarrays | 23027903 | Unspecified |
| mmu-miR-34a-5p | MIMAT0000542 | H2afj | 232440 | HITS-CLIP | 23142080 | Unspecified |
| mmu-miR-34a-5p | MIMAT0000542 | H2afv | 77605 | HITS-CLIP | 23142080 | Unspecified |
| mmu-miR-34a-5p | MIMAT0000542 | H2afx | 15270 | HITS-CLIP | 37986224\|23142080\|29167373 | Unspecified |
| mmu-miR-34a-5p | MIMAT0000542 | H2afy | 9555 | HITS-CLIP | 23142080\|29167373 | Unspecified |
| mmu-miR-34a-5p | MIMAT0000542 | H2ax | 3014 | HITS-CLIP | 31433990 | Unspecified |
| mmu-miR-34a-5p | MIMAT0000542 | H2bc18 | 440689 | HITS-CLIP | 31433990 | Unspecified |
| mmu-miR-34a-5p | MIMAT0000542 | H2bc3 | 3018 | Microarrays | 23027903 | Unspecified |
| mmu-miR-34a-5p | MIMAT0000542 | H3c14 | 126961 | HITS-CLIP | 31433990 | Unspecified |
| mmu-miR-34a-5p | MIMAT0000542 | H3c7 | 8350 | HITS-CLIP | 31433990\|23630078 | Unspecified |
| mmu-miR-34a-5p | MIMAT0000542 | H3c8 | 8350 | HITS-CLIP | 31433990 | Unspecified |
| mmu-miR-34a-5p | MIMAT0000542 | H3f3b | 15081 | HITS-CLIP | 31433990\|23142080\|29167373 | Unspecified |
| mmu-miR-34a-5p | MIMAT0000542 | H4c3 | 8359 | HITS-CLIP | 31433990 | Unspecified |
| mmu-miR-34a-5p | MIMAT0000542 | H6pd | 100198 | HITS-CLIP | 23142080\|29167373 | Unspecified |
| mmu-miR-34a-5p | MIMAT0000542 | Haao | 107766 | HITS-CLIP | 23142080 | Unspecified |
| mmu-miR-34a-5p | MIMAT0000542 | Hacd2 | 201562 | HITS-CLIP | 29167373 | Unspecified |
| mmu-miR-34a-5p | MIMAT0000542 | Hacd3 | 57874 | HITS-CLIP | 26203562\|23142080 | Unspecified |
| mmu-miR-34a-5p | MIMAT0000542 | Hacd4 | 401494 | Microarrays | 23027903 | Unspecified |
| mmu-miR-34a-5p | MIMAT0000542 | Hacl1 | 56794 | HITS-CLIP | 23142080 | Unspecified |
| mmu-miR-34a-5p | MIMAT0000542 | Hadh | 15107 | HITS-CLIP | 19536157\|23142080 | Unspecified |
| mmu-miR-34a-5p | MIMAT0000542 | Hadha | 97212 | HITS-CLIP | 23142080\|19536157 | Unspecified |
| mmu-miR-34a-5p | MIMAT0000542 | Hadhb | 231086 | HITS-CLIP | 23142080 | Unspecified |
| mmu-miR-34a-5p | MIMAT0000542 | Hagh | 14651 | HITS-CLIP | 23142080 | Unspecified |
| mmu-miR-34a-5p | MIMAT0000542 | Harbi1 | 241547 | HITS-CLIP | 23142080 | Unspecified |
| mmu-miR-34a-5p | MIMAT0000542 | Hars | 15115 | HITS-CLIP | 23142080 | Unspecified |
| mmu-miR-34a-5p | MIMAT0000542 | Hars2 | 70791 | HITS-CLIP | 23142080\|30971389 | Unspecified |
| mmu-miR-34a-5p | MIMAT0000542 | Has3 | 15118 | HITS-CLIP | 19536157 | Unspecified |
| mmu-miR-34a-5p | MIMAT0000542 | Hat1 | 107435 | HITS-CLIP | 23142080 | Unspecified |
| mmu-miR-34a-5p | MIMAT0000542 | Haus3 | 231123 | HITS-CLIP | 23142080 | Unspecified |
| mmu-miR-34a-5p | MIMAT0000542 | Haus6 | 230376 | HITS-CLIP | 23142080 | Unspecified |
| mmu-miR-34a-5p | MIMAT0000542 | Haus7 | 73738 | HITS-CLIP | 23142080 | Unspecified |
| mmu-miR-34a-5p | MIMAT0000542 | Hbb-bs | 100503605 | HITS-CLIP | 19536157 | Unspecified |
| mmu-miR-34a-5p | MIMAT0000542 | Hbegf | 15200 | HITS-CLIP | 23142080 | Unspecified |
| mmu-miR-34a-5p | MIMAT0000542 | Hbp1 | 73389 | HITS-CLIP | 23142080\|31433990\|29167373 | Unspecified |
| mmu-miR-34a-5p | MIMAT0000542 | Hbs1l | 56422 | HITS-CLIP | 23142080\|29167373 | Unspecified |
| mmu-miR-34a-5p | MIMAT0000542 | Hc | 15139 | HITS-CLIP | 30971389 | Unspecified |
| mmu-miR-34a-5p | MIMAT0000542 | Hccs | 15159 | Microarrays//HITS-CLIP | 37986224\|23027903\|29167373 | Unspecified |
| mmu-miR-34a-5p | MIMAT0000542 | Hcfc1 | 15161 | HITS-CLIP | 23142080\|29167373 | Unspecified |
| mmu-miR-34a-5p | MIMAT0000542 | Hcfc2 | 67933 | HITS-CLIP | 23142080\|29167373 | Unspecified |

| | | | | | | |
|---|---|---|---|---|---|---|
| mmu-miR-34a-5p | MIMAT0000542 | Hck | 15162 | HITS-CLIP | 29167373 | Unspecified |
| mmu-miR-34a-5p | MIMAT0000542 | Hcls1 | 15163 | HITS-CLIP | 23142080\|29167373 | Unspecified |
| mmu-miR-34a-5p | MIMAT0000542 | Hcn1 | 15165 | HITS-CLIP | 19536157 | Unspecified |
| mmu-miR-34a-5p | MIMAT0000542 | Hdac1 | 433759 | HITS-CLIP | 23142080 | Unspecified |
| mmu-miR-34a-5p | MIMAT0000542 | Hdac10 | 170787 | HITS-CLIP | 23142080 | Unspecified |
| mmu-miR-34a-5p | MIMAT0000542 | Hdac2 | 15182 | HITS-CLIP | 23142080 | Unspecified |
| mmu-miR-34a-5p | MIMAT0000542 | Hdac3 | 15183 | HITS-CLIP | 19536157\|23142080 | Unspecified |
| mmu-miR-34a-5p | MIMAT0000542 | Hdac4 | 208727 | HITS-CLIP | 19536157\|23142080\|29167373 | Unspecified |
| mmu-miR-34a-5p | MIMAT0000542 | Hdac5 | 15184 | HITS-CLIP | 23142080 | Unspecified |
| mmu-miR-34a-5p | MIMAT0000542 | Hdac7 | 56233 | HITS-CLIP | 31433990\|23142080 | Unspecified |
| mmu-miR-34a-5p | MIMAT0000542 | Hdac8 | 70315 | HITS-CLIP | 23142080 | Unspecified |
| mmu-miR-34a-5p | MIMAT0000542 | Hddc2 | 69692 | HITS-CLIP | 23142080 | Unspecified |
| mmu-miR-34a-5p | MIMAT0000542 | Hdgf | 15191 | HITS-CLIP | 23142080 | Unspecified |
| mmu-miR-34a-5p | MIMAT0000542 | Hdhd5 | 27440 | HITS-CLIP | 31433990 | Unspecified |
| mmu-miR-34a-5p | MIMAT0000542 | Hdlbp | 110611 | HITS-CLIP | 23142080\|31433990\|19536157\|29167 | Unspecified |
| mmu-miR-34a-5p | MIMAT0000542 | Heatr1 | 217995 | HITS-CLIP | 23142080\|29167373 | Unspecified |
| mmu-miR-34a-5p | MIMAT0000542 | Heatr3 | 234549 | HITS-CLIP | 23142080 | Unspecified |
| mmu-miR-34a-5p | MIMAT0000542 | Heatr5a | 320487 | HITS-CLIP | 29167373 | Unspecified |
| mmu-miR-34a-5p | MIMAT0000542 | Heatr5b | 320473 | HITS-CLIP | 23142080 | Unspecified |
| mmu-miR-34a-5p | MIMAT0000542 | Heatr6 | 217026 | HITS-CLIP | 29167373 | Unspecified |
| mmu-miR-34a-5p | MIMAT0000542 | Hebp1 | 15199 | Microarrays | 37986224 | Unspecified |
| mmu-miR-34a-5p | MIMAT0000542 | Heca | 380629 | HITS-CLIP | 23142080\|29167373 | Unspecified |
| mmu-miR-34a-5p | MIMAT0000542 | Hectd1 | 207304 | HITS-CLIP | 23142080\|19536157\|31433990\|29167 | Unspecified |
| mmu-miR-34a-5p | MIMAT0000542 | Hectd2 | 226098 | HITS-CLIP | 37986224\|23142080 | Unspecified |
| mmu-miR-34a-5p | MIMAT0000542 | Hectd3 | 76608 | HITS-CLIP | 30971389 | Unspecified |
| mmu-miR-34a-5p | MIMAT0000542 | Hectd4 | 283450 | HITS-CLIP | 31433990\|19536157\|30971389\|23142 | Unspecified |
| mmu-miR-34a-5p | MIMAT0000542 | Heg1 | 77446 | HITS-CLIP | 23142080\|31433990\|19536157 | Unspecified |
| mmu-miR-34a-5p | MIMAT0000542 | Helb | 117599 | HITS-CLIP | 23142080 | Unspecified |
| mmu-miR-34a-5p | MIMAT0000542 | Hells | 15201 | Microarrays//HITS-CLIP | 37986224\|23027903\|23142080 | Unspecified |
| mmu-miR-34a-5p | MIMAT0000542 | Helq | 191578 | HITS-CLIP | 31433990\|23142080 | Unspecified |
| mmu-miR-34a-5p | MIMAT0000542 | Helz | 78455 | HITS-CLIP | 23142080\|29167373 | Unspecified |
| mmu-miR-34a-5p | MIMAT0000542 | Helz2 | 229003 | HITS-CLIP | 30971389\|23142080\|26203562\|29167 | Unspecified |
| mmu-miR-34a-5p | MIMAT0000542 | Hemk1 | 69536 | HITS-CLIP | 23142080 | Unspecified |
| mmu-miR-34a-5p | MIMAT0000542 | Herc1 | 235439 | HITS-CLIP | 19536157\|29167373\|23142080 | Unspecified |
| mmu-miR-34a-5p | MIMAT0000542 | Herc2 | 15204 | HITS-CLIP | 23142080\|19536157\|29167373 | Unspecified |
| mmu-miR-34a-5p | MIMAT0000542 | Herc3 | 73998 | HITS-CLIP | 19536157\|29167373\|23142080 | Unspecified |
| mmu-miR-34a-5p | MIMAT0000542 | Herc4 | 67345 | HITS-CLIP | 23142080\|29167373 | Unspecified |
| mmu-miR-34a-5p | MIMAT0000542 | Herc6 | 67138 | HITS-CLIP | 29167373 | Unspecified |
| mmu-miR-34a-5p | MIMAT0000542 | Herpud1 | 64209 | HITS-CLIP | 23142080\|31433990 | Unspecified |

| | | | | | | | |
|---|---|---|---|---|---|---|---|
| mmu-miR-34a-5p | MIMAT0000542 | Herpud2 | 80517 | HITS-CLIP | | 23142080 | Unspecified |
| mmu-miR-34a-5p | MIMAT0000542 | Hes6 | 55927 | Microarrays//HITS-CLIP | 37986224\|23142080 | | Unspecified |
| mmu-miR-34a-5p | MIMAT0000542 | Hexa | 15211 | HITS-CLIP | | 29167373 | Unspecified |
| mmu-miR-34a-5p | MIMAT0000542 | Hgs | 15239 | HITS-CLIP | | 23142080 | Unspecified |
| mmu-miR-34a-5p | MIMAT0000542 | Hhex | 15242 | HITS-CLIP | | 29167373 | Unspecified |
| mmu-miR-34a-5p | MIMAT0000542 | Hhip | 15245 | Microarrays | 37986224\|23027903 | | Unspecified |
| mmu-miR-34a-5p | MIMAT0000542 | Hibadh | 58875 | HITS-CLIP | | 23142080 | Unspecified |
| mmu-miR-34a-5p | MIMAT0000542 | Hibch | 227095 | HITS-CLIP | 31433990\|23142080 | | Unspecified |
| mmu-miR-34a-5p | MIMAT0000542 | Hic1 | 15248 | HITS-CLIP | | 23142080 | Unspecified |
| mmu-miR-34a-5p | MIMAT0000542 | Hic2 | 58180 | HITS-CLIP | 30971389\|23142080 | | Unspecified |
| mmu-miR-34a-5p | MIMAT0000542 | Hif1a | 15251 | HITS-CLIP | 37986224\|23142080\|29167373 | | Unspecified |
| mmu-miR-34a-5p | MIMAT0000542 | Hif1an | 319594 | Microarrays//HITS-CLIP | 37986224\|23027903\|23630078\|23142 | | Unspecified |
| mmu-miR-34a-5p | MIMAT0000542 | Hif3a | 53417 | HITS-CLIP | | 23142080 | Unspecified |
| mmu-miR-34a-5p | MIMAT0000542 | Higd2a | 67044 | HITS-CLIP | | 23142080 | Unspecified |
| mmu-miR-34a-5p | MIMAT0000542 | Hinfp | 102423 | HITS-CLIP | | 23142080 | Unspecified |
| mmu-miR-34a-5p | MIMAT0000542 | Hint3 | 66847 | Microarrays//HITS-CLIP | 37986224\|23142080\|23027903\|1953€ | | Unspecified |
| mmu-miR-34a-5p | MIMAT0000542 | Hip1 | 215114 | HITS-CLIP | 19536157\|31433990\|23142080\|29167 | | Unspecified |
| mmu-miR-34a-5p | MIMAT0000542 | Hip1r | 29816 | HITS-CLIP | | 26203562 | Unspecified |
| mmu-miR-34a-5p | MIMAT0000542 | Hipk1 | 15257 | HITS-CLIP | 31433990\|23142080\|29167373 | | Unspecified |
| mmu-miR-34a-5p | MIMAT0000542 | Hipk2 | 15258 | HITS-CLIP | 23142080\|30971389\|23630078\|1953€ | | Unspecified |
| mmu-miR-34a-5p | MIMAT0000542 | Hipk3 | 15259 | Microarrays//HITS-CLIP | 37986224\|19536157\|23142080\|29167 | | Unspecified |
| mmu-miR-34a-5p | MIMAT0000542 | Hira | 15260 | HITS-CLIP | 23142080\|29167373 | | Unspecified |
| mmu-miR-34a-5p | MIMAT0000542 | Hist1h1a | 3024 | HITS-CLIP | | 23142080 | Unspecified |
| mmu-miR-34a-5p | MIMAT0000542 | Hist1h1b | 3009 | HITS-CLIP | 23142080\|29167373 | | Unspecified |
| mmu-miR-34a-5p | MIMAT0000542 | Hist1h1c | 50708 | HITS-CLIP | 23142080\|29167373 | | Unspecified |
| mmu-miR-34a-5p | MIMAT0000542 | Hist1h1d | 14957 | HITS-CLIP | | 23142080 | Unspecified |
| mmu-miR-34a-5p | MIMAT0000542 | Hist1h1e | 3008 | HITS-CLIP | 19536157\|23142080 | | Unspecified |
| mmu-miR-34a-5p | MIMAT0000542 | Hist1h2ab | 8335 | HITS-CLIP | | 23142080 | Unspecified |
| mmu-miR-34a-5p | MIMAT0000542 | Hist1h2ag | 8969 | HITS-CLIP | | 23142080 | Unspecified |
| mmu-miR-34a-5p | MIMAT0000542 | Hist1h2ak | 8330 | HITS-CLIP | | 23142080 | Unspecified |
| mmu-miR-34a-5p | MIMAT0000542 | Hist1h2an | 319170 | HITS-CLIP | | 23142080 | Unspecified |
| mmu-miR-34a-5p | MIMAT0000542 | Hist1h2bc | 3017 | HITS-CLIP | | 23142080 | Unspecified |
| mmu-miR-34a-5p | MIMAT0000542 | Hist1h2be | 319179 | HITS-CLIP | | 23142080 | Unspecified |
| mmu-miR-34a-5p | MIMAT0000542 | Hist1h2bf | 3017 | HITS-CLIP | | 23142080 | Unspecified |
| mmu-miR-34a-5p | MIMAT0000542 | Hist1h2bg | 3017 | HITS-CLIP | | 23142080 | Unspecified |
| mmu-miR-34a-5p | MIMAT0000542 | Hist1h2bn | 8341 | HITS-CLIP | | 23142080 | Unspecified |
| mmu-miR-34a-5p | MIMAT0000542 | Hist1h2bp | 319188 | HITS-CLIP | | 23142080 | Unspecified |
| mmu-miR-34a-5p | MIMAT0000542 | Hist1h3a | 8350 | HITS-CLIP | | 29167373 | Unspecified |
| mmu-miR-34a-5p | MIMAT0000542 | Hist1h3d | 8351 | HITS-CLIP | | 23142080 | Unspecified |

| | | | | | | |
|---|---|---|---|---|---|---|
| mmu-miR-34a-5p | MIMAT0000542 | Hist1h3e | 8353 | HITS-CLIP | 23142080 | Unspecified |
| mmu-miR-34a-5p | MIMAT0000542 | Hist1h3f | 260423 | HITS-CLIP | 23142080 | Unspecified |
| mmu-miR-34a-5p | MIMAT0000542 | Hist1h3g | 8355 | HITS-CLIP | 23142080 | Unspecified |
| mmu-miR-34a-5p | MIMAT0000542 | Hist1h4c | 8364 | HITS-CLIP | 23142080 | Unspecified |
| mmu-miR-34a-5p | MIMAT0000542 | Hist1h4i | 319158 | HITS-CLIP | 23142080 | Unspecified |
| mmu-miR-34a-5p | MIMAT0000542 | Hist1h4j | 8363 | HITS-CLIP | 23142080 | Unspecified |
| mmu-miR-34a-5p | MIMAT0000542 | Hist2h2bb | 319189 | HITS-CLIP | 23142080\|29167373 | Unspecified |
| mmu-miR-34a-5p | MIMAT0000542 | Hist2h4 | 8370 | HITS-CLIP | 23142080 | Unspecified |
| mmu-miR-34a-5p | MIMAT0000542 | Hivep1 | 110521 | HITS-CLIP | 19536157\|23142080\|29167373 | Unspecified |
| mmu-miR-34a-5p | MIMAT0000542 | Hivep2 | 15273 | HITS-CLIP | 23142080 | Unspecified |
| mmu-miR-34a-5p | MIMAT0000542 | Hivep3 | 16656 | HITS-CLIP | 37986224\|23142080\|29167373 | Unspecified |
| mmu-miR-34a-5p | MIMAT0000542 | Hjurp | 381280 | HITS-CLIP | 29167373\|23142080 | Unspecified |
| mmu-miR-34a-5p | MIMAT0000542 | Hk1 | 15275 | HITS-CLIP | 23142080\|29167373 | Unspecified |
| mmu-miR-34a-5p | MIMAT0000542 | Hk2 | 15277 | HITS-CLIP | 23142080 | Unspecified |
| mmu-miR-34a-5p | MIMAT0000542 | Hk3 | 212032 | HITS-CLIP | 29167373 | Unspecified |
| mmu-miR-34a-5p | MIMAT0000542 | Hlf | 217082 | HITS-CLIP | 19536157 | Unspecified |
| mmu-miR-34a-5p | MIMAT0000542 | Hltf | 20585 | HITS-CLIP | 23142080 | Unspecified |
| mmu-miR-34a-5p | MIMAT0000542 | Hlx | 15284 | HITS-CLIP | 29167373 | Unspecified |
| mmu-miR-34a-5p | MIMAT0000542 | Hmbox1 | 219150 | HITS-CLIP | 23142080\|29167373 | Unspecified |
| mmu-miR-34a-5p | MIMAT0000542 | Hmbs | 15288 | Microarrays//HITS-CLIP | 37986224\|23027903\|23142080 | Unspecified |
| mmu-miR-34a-5p | MIMAT0000542 | Hmces | 232210 | HITS-CLIP | 23142080 | Unspecified |
| mmu-miR-34a-5p | MIMAT0000542 | Hmga1 | 15361 | HITS-CLIP | 23142080\|30971389 | Unspecified |
| mmu-miR-34a-5p | MIMAT0000542 | Hmga2 | 15364 | HITS-CLIP | 29167373 | Unspecified |
| mmu-miR-34a-5p | MIMAT0000542 | Hmgb2 | 97165 | HITS-CLIP | 23142080 | Unspecified |
| mmu-miR-34a-5p | MIMAT0000542 | Hmgcl | 15356 | HITS-CLIP | 23142080 | Unspecified |
| mmu-miR-34a-5p | MIMAT0000542 | Hmgcll1 | 208982 | Microarrays | 37986224 | Unspecified |
| mmu-miR-34a-5p | MIMAT0000542 | Hmgcr | 15357 | HITS-CLIP | 23142080\|19536157\|30971389\|3143 | Unspecified |
| mmu-miR-34a-5p | MIMAT0000542 | Hmgcs1 | 208715 | HITS-CLIP | 37986224\|23142080\|19536157\|2916 | Unspecified |
| mmu-miR-34a-5p | MIMAT0000542 | Hmgn1 | 15312 | HITS-CLIP | 23630078 | Unspecified |
| mmu-miR-34a-5p | MIMAT0000542 | Hmgn5 | 50887 | Microarrays | 37986224\|23027903 | Unspecified |
| mmu-miR-34a-5p | MIMAT0000542 | Hmgxb3 | 106894 | HITS-CLIP | 23142080\|19536157\|29167373 | Unspecified |
| mmu-miR-34a-5p | MIMAT0000542 | Hmgxb4 | 70823 | HITS-CLIP | 23142080 | Unspecified |
| mmu-miR-34a-5p | MIMAT0000542 | Hmmr | 15366 | HITS-CLIP | 23142080 | Unspecified |
| mmu-miR-34a-5p | MIMAT0000542 | Hmox1 | 15368 | HITS-CLIP | 23142080\|23630078\|29167373 | Unspecified |
| mmu-miR-34a-5p | MIMAT0000542 | Hmox2 | 15369 | HITS-CLIP | 29167373 | Unspecified |
| mmu-miR-34a-5p | MIMAT0000542 | Hnrnpa0 | 77134 | HITS-CLIP | 31433990\|23142080\|23630078\|2916 | Unspecified |
| mmu-miR-34a-5p | MIMAT0000542 | Hnrnpa1 | 15382 | HITS-CLIP | 23142080 | Unspecified |
| mmu-miR-34a-5p | MIMAT0000542 | Hnrnpa2b1 | 53379 | HITS-CLIP | 31433990\|19536157\|23142080\|2916 | Unspecified |
| mmu-miR-34a-5p | MIMAT0000542 | Hnrnpa3 | 229279 | HITS-CLIP | 23142080\|29167373 | Unspecified |

| | | | | | | |
|---|---|---|---|---|---|---|
| mmu-miR-34a-5p | MIMAT0000542 | Hnrnpab | 15384 | HITS-CLIP | 31433990\|23630078\|23142080\|29167 | Unspecified |
| mmu-miR-34a-5p | MIMAT0000542 | Hnrnpc | 15381 | HITS-CLIP | 30971389 | Unspecified |
| mmu-miR-34a-5p | MIMAT0000542 | Hnrnpd | 11991 | HITS-CLIP | 23142080 | Unspecified |
| mmu-miR-34a-5p | MIMAT0000542 | Hnrnpdl | 50926 | HITS-CLIP | 23142080\|31433990\|19536157 | Unspecified |
| mmu-miR-34a-5p | MIMAT0000542 | Hnrnpf | 98758 | HITS-CLIP | 37986224 | Unspecified |
| mmu-miR-34a-5p | MIMAT0000542 | Hnrnph1 | 59013 | HITS-CLIP | 23142080 | Unspecified |
| mmu-miR-34a-5p | MIMAT0000542 | Hnrnpk | 15387 | HITS-CLIP | 19536157\|31433990\|23142080\|29167 | Unspecified |
| mmu-miR-34a-5p | MIMAT0000542 | Hnrnpl | 15388 | HITS-CLIP | 23142080\|31433990\|26203562\|29167 | Unspecified |
| mmu-miR-34a-5p | MIMAT0000542 | Hnrnpll | 72692 | HITS-CLIP | 19536157\|23142080 | Unspecified |
| mmu-miR-34a-5p | MIMAT0000542 | Hnrnpm | 76936 | HITS-CLIP | 23142080\|29167373 | Unspecified |
| mmu-miR-34a-5p | MIMAT0000542 | Hnrnpr | 74326 | HITS-CLIP | 23142080\|29167373 | Unspecified |
| mmu-miR-34a-5p | MIMAT0000542 | Hnrnpu | 51810 | HITS-CLIP | 23142080\|29167373 | Unspecified |
| mmu-miR-34a-5p | MIMAT0000542 | Hnrnpul1 | 232989 | HITS-CLIP | 23142080\|29167373\|31433990 | Unspecified |
| mmu-miR-34a-5p | MIMAT0000542 | Hnrnpul2 | 68693 | HITS-CLIP | 26203562\|23142080\|29167373 | Unspecified |
| mmu-miR-34a-5p | MIMAT0000542 | Homer1 | 26556 | HITS-CLIP | 23142080 | Unspecified |
| mmu-miR-34a-5p | MIMAT0000542 | Homer2 | 26557 | HITS-CLIP | 19536157 | Unspecified |
| mmu-miR-34a-5p | MIMAT0000542 | Hook1 | 77963 | HITS-CLIP | 23142080 | Unspecified |
| mmu-miR-34a-5p | MIMAT0000542 | Hook3 | 320191 | HITS-CLIP | 31433990\|29167373\|23142080 | Unspecified |
| mmu-miR-34a-5p | MIMAT0000542 | Hoxa1 | 15394 | Microarrays | 37986224\|23027903 | Unspecified |
| mmu-miR-34a-5p | MIMAT0000542 | Hp1bp3 | 15441 | HITS-CLIP | 37986224\|23142080 | Unspecified |
| mmu-miR-34a-5p | MIMAT0000542 | Hpcal1 | 53602 | HITS-CLIP | 23142080\|29167373 | Unspecified |
| mmu-miR-34a-5p | MIMAT0000542 | Hpcal4 | 170638 | HITS-CLIP | 19536157 | Unspecified |
| mmu-miR-34a-5p | MIMAT0000542 | Hpdl | 242642 | HITS-CLIP | 23142080 | Unspecified |
| mmu-miR-34a-5p | MIMAT0000542 | Hpgd | 15446 | HITS-CLIP | 29167373 | Unspecified |
| mmu-miR-34a-5p | MIMAT0000542 | Hprt | 15452 | HITS-CLIP | 23142080\|31433990 | Unspecified |
| mmu-miR-34a-5p | MIMAT0000542 | Hps3 | 12807 | HITS-CLIP | 23142080 | Unspecified |
| mmu-miR-34a-5p | MIMAT0000542 | Hps5 | 246694 | HITS-CLIP | 23142080 | Unspecified |
| mmu-miR-34a-5p | MIMAT0000542 | Hps6 | 20170 | HITS-CLIP | 23142080\|29167373 | Unspecified |
| mmu-miR-34a-5p | MIMAT0000542 | Hr | 15460 | HITS-CLIP | 26203562 | Unspecified |
| mmu-miR-34a-5p | MIMAT0000542 | Hrh2 | 15466 | HITS-CLIP | 23142080 | Unspecified |
| mmu-miR-34a-5p | MIMAT0000542 | Hrob | 78995 | HITS-CLIP | 30971389 | Unspecified |
| mmu-miR-34a-5p | MIMAT0000542 | Hs2st1 | 23908 | HITS-CLIP | 26203562\|29167373\|23142080 | Unspecified |
| mmu-miR-34a-5p | MIMAT0000542 | Hs3st1 | 15476 | HITS-CLIP | 23142080 | Unspecified |
| mmu-miR-34a-5p | MIMAT0000542 | Hs3st3b1 | 54710 | HITS-CLIP | 23142080 | Unspecified |
| mmu-miR-34a-5p | MIMAT0000542 | Hs6st1 | 50785 | HITS-CLIP | 23142080\|29167373 | Unspecified |
| mmu-miR-34a-5p | MIMAT0000542 | Hs6st2 | 50786 | HITS-CLIP | 19536157 | Unspecified |
| mmu-miR-34a-5p | MIMAT0000542 | Hsbp1 | 68196 | HITS-CLIP | 19536157\|29167373\|23142080 | Unspecified |
| mmu-miR-34a-5p | MIMAT0000542 | Hsd11b1 | 15483 | HITS-CLIP | 31433990 | Unspecified |
| mmu-miR-34a-5p | MIMAT0000542 | Hsd17b11 | 114664 | HITS-CLIP | 23630078\|23142080\|29167373 | Unspecified |

| | | | | | | |
|---|---|---|---|---|---|---|
| mmu-miR-34a-5p | MIMAT0000542 | Hsd17b12 | 56348 | HITS-CLIP | 23142080 | Unspecified |
| mmu-miR-34a-5p | MIMAT0000542 | Hsd17b4 | 15488 | HITS-CLIP | 23142080\|29167373 | Unspecified |
| mmu-miR-34a-5p | MIMAT0000542 | Hsd17b7 | 15490 | Microarrays//HITS-CLIP | 37986224\|23027903\|29167373\|23142 | Unspecified |
| mmu-miR-34a-5p | MIMAT0000542 | Hsdl1 | 72552 | HITS-CLIP | 23142080 | Unspecified |
| mmu-miR-34a-5p | MIMAT0000542 | Hsdl2 | 72479 | HITS-CLIP | 19536157\|23142080 | Unspecified |
| mmu-miR-34a-5p | MIMAT0000542 | Hsf1 | 15499 | HITS-CLIP | 29167373 | Unspecified |
| mmu-miR-34a-5p | MIMAT0000542 | Hsf2 | 15500 | HITS-CLIP | 23142080 | Unspecified |
| mmu-miR-34a-5p | MIMAT0000542 | Hsh2d | 209488 | HITS-CLIP | 23142080 | Unspecified |
| mmu-miR-34a-5p | MIMAT0000542 | Hsp90aa1 | 15519 | HITS-CLIP | 23142080\|29167373 | Unspecified |
| mmu-miR-34a-5p | MIMAT0000542 | Hsp90ab1 | 15516 | HITS-CLIP | 23142080\|19536157\|31433990\|26203 | Unspecified |
| mmu-miR-34a-5p | MIMAT0000542 | Hsp90b1 | 22027 | HITS-CLIP | 23142080\|29167373 | Unspecified |
| mmu-miR-34a-5p | MIMAT0000542 | Hspa14 | 50497 | HITS-CLIP | 23142080 | Unspecified |
| mmu-miR-34a-5p | MIMAT0000542 | Hspa1b | 15511 | HITS-CLIP | 23142080 | Unspecified |
| mmu-miR-34a-5p | MIMAT0000542 | Hspa2 | 15512 | HITS-CLIP | 23142080 | Unspecified |
| mmu-miR-34a-5p | MIMAT0000542 | Hspa4 | 15525 | HITS-CLIP | 23142080\|23630078\|31433990\|29167 | Unspecified |
| mmu-miR-34a-5p | MIMAT0000542 | Hspa4l | 18415 | HITS-CLIP | 23142080\|30971389 | Unspecified |
| mmu-miR-34a-5p | MIMAT0000542 | Hspa5 | 14828 | HITS-CLIP | 19536157\|23142080\|29167373 | Unspecified |
| mmu-miR-34a-5p | MIMAT0000542 | Hspa8 | 15481 | HITS-CLIP | 23142080 | Unspecified |
| mmu-miR-34a-5p | MIMAT0000542 | Hspa9 | 15526 | HITS-CLIP | 19536157\|23142080\|29167373 | Unspecified |
| mmu-miR-34a-5p | MIMAT0000542 | Hspb6 | 243912 | Microarrays | 37986224\|23027903 | Unspecified |
| mmu-miR-34a-5p | MIMAT0000542 | Hspbp1 | 66245 | HITS-CLIP | 23142080 | Unspecified |
| mmu-miR-34a-5p | MIMAT0000542 | Hspd1 | 15510 | HITS-CLIP | 23142080\|29167373 | Unspecified |
| mmu-miR-34a-5p | MIMAT0000542 | Hspe1 | 15528 | HITS-CLIP | 23142080 | Unspecified |
| mmu-miR-34a-5p | MIMAT0000542 | Hsph1 | 15505 | HITS-CLIP | 23142080\|30971389 | Unspecified |
| mmu-miR-34a-5p | MIMAT0000542 | Htr2b | 15559 | Microarrays | 37986224 | Unspecified |
| mmu-miR-34a-5p | MIMAT0000542 | Htr5a | 15563 | HITS-CLIP | 19536157 | Unspecified |
| mmu-miR-34a-5p | MIMAT0000542 | Htra3 | 78558 | Microarrays | 37986224 | Unspecified |
| mmu-miR-34a-5p | MIMAT0000542 | Htt | 15194 | HITS-CLIP | 31433990\|19536157\|23142080\|29167 | Unspecified |
| mmu-miR-34a-5p | MIMAT0000542 | Hus1 | 15574 | HITS-CLIP | 23142080 | Unspecified |
| mmu-miR-34a-5p | MIMAT0000542 | Huwe1 | 59026 | HITS-CLIP | 23142080\|19536157\|29167373 | Unspecified |
| mmu-miR-34a-5p | MIMAT0000542 | Hvcn1 | 74096 | HITS-CLIP | 23142080 | Unspecified |
| mmu-miR-34a-5p | MIMAT0000542 | Hyal1 | 15586 | HITS-CLIP | 29167373 | Unspecified |
| mmu-miR-34a-5p | MIMAT0000542 | Hyal2 | 15587 | HITS-CLIP | 23142080\|29167373 | Unspecified |
| mmu-miR-34a-5p | MIMAT0000542 | Hycc2 | 285172 | HITS-CLIP | 30971389 | Unspecified |
| mmu-miR-34a-5p | MIMAT0000542 | Hyls1 | 76832 | HITS-CLIP | 23142080 | Unspecified |
| mmu-miR-34a-5p | MIMAT0000542 | Hyou1 | 12282 | HITS-CLIP | 23142080\|31433990 | Unspecified |
| mmu-miR-34a-5p | MIMAT0000542 | Iars | 105148 | HITS-CLIP | 23142080\|29167373 | Unspecified |
| mmu-miR-34a-5p | MIMAT0000542 | Iars2 | 381314 | HITS-CLIP | 29167373\|23142080 | Unspecified |
| mmu-miR-34a-5p | MIMAT0000542 | Ibtk | 108837 | HITS-CLIP | 23142080 | Unspecified |

| | | | | | | |
|---|---|---|---|---|---|---|
| mmu-miR-34a-5p | MIMAT0000542 | Icam1 | 15894 | HITS-CLIP | 29167373 | Unspecified |
| mmu-miR-34a-5p | MIMAT0000542 | Icam2 | 15896 | HITS-CLIP | 30971389 | Unspecified |
| mmu-miR-34a-5p | MIMAT0000542 | Ice1 | 822287 | HITS-CLIP | 23142080 | Unspecified |
| mmu-miR-34a-5p | MIMAT0000542 | Icos | 54167 | HITS-CLIP | 31433990\|23142080 | Unspecified |
| mmu-miR-34a-5p | MIMAT0000542 | Id2 | 15902 | HITS-CLIP | 23142080 | Unspecified |
| mmu-miR-34a-5p | MIMAT0000542 | Id3 | 15903 | HITS-CLIP | 26203562 | Unspecified |
| mmu-miR-34a-5p | MIMAT0000542 | Id4 | 15904 | HITS-CLIP | 19536157 | Unspecified |
| mmu-miR-34a-5p | MIMAT0000542 | Ide | 15925 | HITS-CLIP | 23142080 | Unspecified |
| mmu-miR-34a-5p | MIMAT0000542 | Idh1 | 15926 | HITS-CLIP | 23142080\|29167373 | Unspecified |
| mmu-miR-34a-5p | MIMAT0000542 | Idh2 | 269951 | HITS-CLIP | 23630078 | Unspecified |
| mmu-miR-34a-5p | MIMAT0000542 | Idh3a | 67834 | Microarrays | 37986224\|23027903 | Unspecified |
| mmu-miR-34a-5p | MIMAT0000542 | Idh3b | 170718 | HITS-CLIP | 19536157\|23142080 | Unspecified |
| mmu-miR-34a-5p | MIMAT0000542 | Idh3g | 15929 | HITS-CLIP | 19536157\|23142080 | Unspecified |
| mmu-miR-34a-5p | MIMAT0000542 | Ids | 15931 | HITS-CLIP | 19536157\|31433990\|29167373 | Unspecified |
| mmu-miR-34a-5p | MIMAT0000542 | Idua | 15932 | HITS-CLIP | 29167373 | Unspecified |
| mmu-miR-34a-5p | MIMAT0000542 | Ier2 | 15936 | HITS-CLIP | 23142080\|29167373 | Unspecified |
| mmu-miR-34a-5p | MIMAT0000542 | Ier3 | 15937 | HITS-CLIP | 23142080\|31433990 | Unspecified |
| mmu-miR-34a-5p | MIMAT0000542 | Ier3ip1 | 66191 | HITS-CLIP | 29167373 | Unspecified |
| mmu-miR-34a-5p | MIMAT0000542 | Ier5 | 15939 | HITS-CLIP | 29167373 | Unspecified |
| mmu-miR-34a-5p | MIMAT0000542 | Iffo1 | 320678 | HITS-CLIP | 30971389 | Unspecified |
| mmu-miR-34a-5p | MIMAT0000542 | Iffo2 | 212632 | HITS-CLIP | 23142080\|30971389\|31433990\|26203 | Unspecified |
| mmu-miR-34a-5p | MIMAT0000542 | Ifi204 | 15951 | HITS-CLIP | 29167373 | Unspecified |
| mmu-miR-34a-5p | MIMAT0000542 | Ifi207 | 226691 | HITS-CLIP | 29167373 | Unspecified |
| mmu-miR-34a-5p | MIMAT0000542 | Ifi27 | 52668 | HITS-CLIP | 23142080 | Unspecified |
| mmu-miR-34a-5p | MIMAT0000542 | Ifi27l2a | 76933 | HITS-CLIP | 23142080\|29167373 | Unspecified |
| mmu-miR-34a-5p | MIMAT0000542 | Ifi30 | 65972 | HITS-CLIP | 31433990\|26203562 | Unspecified |
| mmu-miR-34a-5p | MIMAT0000542 | Ifi35 | 70110 | HITS-CLIP | 23142080 | Unspecified |
| mmu-miR-34a-5p | MIMAT0000542 | Ifi44 | 99899 | HITS-CLIP | 29167373 | Unspecified |
| mmu-miR-34a-5p | MIMAT0000542 | Ifi47 | 15953 | HITS-CLIP | 23142080 | Unspecified |
| mmu-miR-34a-5p | MIMAT0000542 | Ifih1 | 71586 | HITS-CLIP | 23142080\|29167373 | Unspecified |
| mmu-miR-34a-5p | MIMAT0000542 | Ifit1 | 15957 | HITS-CLIP | 29167373 | Unspecified |
| mmu-miR-34a-5p | MIMAT0000542 | Ifit1bl1 | 667373 | HITS-CLIP | 23142080 | Unspecified |
| mmu-miR-34a-5p | MIMAT0000542 | Ifit2 | 15958 | HITS-CLIP | 23630078\|23142080\|29167373 | Unspecified |
| mmu-miR-34a-5p | MIMAT0000542 | Ifitm6 | 213002 | HITS-CLIP | 29167373 | Unspecified |
| mmu-miR-34a-5p | MIMAT0000542 | Ifnar1 | 15975 | HITS-CLIP | 23142080\|29167373 | Unspecified |
| mmu-miR-34a-5p | MIMAT0000542 | Ifnar2 | 15976 | HITS-CLIP | 23142080\|30971389\|29167373 | Unspecified |
| mmu-miR-34a-5p | MIMAT0000542 | Ifng | 15978 | HITS-CLIP | 23142080 | Unspecified |
| mmu-miR-34a-5p | MIMAT0000542 | Ifngr1 | 15979 | HITS-CLIP | 19536157\|23142080\|29167373 | Unspecified |
| mmu-miR-34a-5p | MIMAT0000542 | Ifngr2 | 15980 | HITS-CLIP | 23142080\|30971389\|23630078\|19536 | Unspecified |

| | | | | | | | |
|---|---|---|---|---|---|---|---|
| mmu-miR-34a-5p | MIMAT0000542 | Ifrd1 | 15982 | HITS-CLIP | | 23142080 | Unspecified |
| mmu-miR-34a-5p | MIMAT0000542 | Ifrd2 | 15983 | Microarrays | 37986224\|23027903 | | Unspecified |
| mmu-miR-34a-5p | MIMAT0000542 | Ift172 | 67661 | HITS-CLIP | | 23142080 | Unspecified |
| mmu-miR-34a-5p | MIMAT0000542 | Ift22 | 67286 | Microarrays//HITS-CLIP | 37986224\|29167373 | | Unspecified |
| mmu-miR-34a-5p | MIMAT0000542 | Ift46 | 76568 | HITS-CLIP | | 23142080 | Unspecified |
| mmu-miR-34a-5p | MIMAT0000542 | Ift74 | 67694 | Microarrays | | 37986224 | Unspecified |
| mmu-miR-34a-5p | MIMAT0000542 | Ift80 | 68259 | HITS-CLIP | | 23142080 | Unspecified |
| mmu-miR-34a-5p | MIMAT0000542 | Ift88 | 21821 | Microarrays | | 37986224 | Unspecified |
| mmu-miR-34a-5p | MIMAT0000542 | Igbp1 | 18518 | HITS-CLIP | | 23142080 | Unspecified |
| mmu-miR-34a-5p | MIMAT0000542 | Igdcc4 | 56741 | HITS-CLIP | | 19536157 | Unspecified |
| mmu-miR-34a-5p | MIMAT0000542 | Igf1 | 16000 | HITS-CLIP | | 29167373 | Unspecified |
| mmu-miR-34a-5p | MIMAT0000542 | Igf1r | 16001 | HITS-CLIP | 37986224\|23142080\|19536157\|30971 | | Unspecified |
| mmu-miR-34a-5p | MIMAT0000542 | Igf2bp1 | 140486 | HITS-CLIP | | 26203562 | Unspecified |
| mmu-miR-34a-5p | MIMAT0000542 | Igf2bp2 | 319765 | HITS-CLIP | 23142080\|29167373 | | Unspecified |
| mmu-miR-34a-5p | MIMAT0000542 | Igf2r | 16004 | Microarrays//HITS-CLIP | 37986224\|23142080\|31433990\|29167 | | Unspecified |
| mmu-miR-34a-5p | MIMAT0000542 | Igfbp3 | 16009 | HITS-CLIP | | 19536157 | Unspecified |
| mmu-miR-34a-5p | MIMAT0000542 | Igfbp4 | 16010 | HITS-CLIP | | 23142080 | Unspecified |
| mmu-miR-34a-5p | MIMAT0000542 | Ighmbp2 | 20589 | HITS-CLIP | | 23142080 | Unspecified |
| mmu-miR-34a-5p | MIMAT0000542 | Iglon5 | 210094 | HITS-CLIP | | 19536157 | Unspecified |
| mmu-miR-34a-5p | MIMAT0000542 | Igsf23 | 70080 | HITS-CLIP | | 23142080 | Unspecified |
| mmu-miR-34a-5p | MIMAT0000542 | Igsf6 | 80719 | HITS-CLIP | | 29167373 | Unspecified |
| mmu-miR-34a-5p | MIMAT0000542 | Igsf8 | 140559 | HITS-CLIP | 23142080\|29167373 | | Unspecified |
| mmu-miR-34a-5p | MIMAT0000542 | Igtp | 16145 | HITS-CLIP | 37986224\|31433990\|30971389\|23142 | | Unspecified |
| mmu-miR-34a-5p | MIMAT0000542 | Iigp1 | 60440 | HITS-CLIP | 31433990\|23142080 | | Unspecified |
| mmu-miR-34a-5p | MIMAT0000542 | Ik | 24010 | HITS-CLIP | | 23142080 | Unspecified |
| mmu-miR-34a-5p | MIMAT0000542 | Ikbkb | 16150 | HITS-CLIP | 31433990\|23142080\|29167373 | | Unspecified |
| mmu-miR-34a-5p | MIMAT0000542 | Ikbke | 56489 | HITS-CLIP | 29167373\|23142080 | | Unspecified |
| mmu-miR-34a-5p | MIMAT0000542 | Ikbkg | 16151 | HITS-CLIP | | 23142080 | Unspecified |
| mmu-miR-34a-5p | MIMAT0000542 | Ikzf1 | 22778 | HITS-CLIP | 37986224\|31433990\|30971389\|23142 | | Unspecified |
| mmu-miR-34a-5p | MIMAT0000542 | Ikzf2 | 22779 | HITS-CLIP | 23142080\|30971389 | | Unspecified |
| mmu-miR-34a-5p | MIMAT0000542 | Ikzf3 | 22780 | HITS-CLIP | | 23142080 | Unspecified |
| mmu-miR-34a-5p | MIMAT0000542 | Ikzf4 | 22781 | HITS-CLIP | 23142080\|19536157 | | Unspecified |
| mmu-miR-34a-5p | MIMAT0000542 | Ikzf5 | 67143 | HITS-CLIP | 19536157\|31433990 | | Unspecified |
| mmu-miR-34a-5p | MIMAT0000542 | Il10ra | 16154 | HITS-CLIP | 29167373\|23142080 | | Unspecified |
| mmu-miR-34a-5p | MIMAT0000542 | Il10rb | 16155 | HITS-CLIP | 23142080\|29167373 | | Unspecified |
| mmu-miR-34a-5p | MIMAT0000542 | Il12rb2 | 16162 | HITS-CLIP | 37986224\|23142080 | | Unspecified |
| mmu-miR-34a-5p | MIMAT0000542 | Il13 | 16163 | HITS-CLIP | | 23142080 | Unspecified |
| mmu-miR-34a-5p | MIMAT0000542 | Il15 | 16168 | HITS-CLIP | | 29167373 | Unspecified |
| mmu-miR-34a-5p | MIMAT0000542 | Il16 | 16170 | HITS-CLIP | | 23142080 | Unspecified |

| | | | | | | |
|---|---|---|---|---|---|---|
| mmu-miR-34a-5p | MIMAT0000542 | Il17ra | 16172 | HITS-CLIP | 23142080\|23630078\|29167373 | Unspecified |
| mmu-miR-34a-5p | MIMAT0000542 | Il17rb | 50905 | HITS-CLIP | 29167373 | Unspecified |
| mmu-miR-34a-5p | MIMAT0000542 | Il18bp | 16068 | HITS-CLIP | 29167373 | Unspecified |
| mmu-miR-34a-5p | MIMAT0000542 | Il1r1 | 16177 | Microarrays | 37986224\|23027903 | Unspecified |
| mmu-miR-34a-5p | MIMAT0000542 | Il1rap | 16180 | HITS-CLIP | 23142080\|29167373 | Unspecified |
| mmu-miR-34a-5p | MIMAT0000542 | Il1rn | 16181 | HITS-CLIP | 29167373 | Unspecified |
| mmu-miR-34a-5p | MIMAT0000542 | Il21 | 60505 | HITS-CLIP | 23142080 | Unspecified |
| mmu-miR-34a-5p | MIMAT0000542 | Il21r | 60504 | HITS-CLIP | 23142080\|29167373 | Unspecified |
| mmu-miR-34a-5p | MIMAT0000542 | Il23a | 83430 | HITS-CLIP | 23142080 | Unspecified |
| mmu-miR-34a-5p | MIMAT0000542 | Il27ra | 50931 | HITS-CLIP | 31433990 | Unspecified |
| mmu-miR-34a-5p | MIMAT0000542 | Il2ra | 16184 | HITS-CLIP | 31433990\|23142080 | Unspecified |
| mmu-miR-34a-5p | MIMAT0000542 | Il2rb | 16185 | HITS-CLIP | 37986224\|23142080 | Unspecified |
| mmu-miR-34a-5p | MIMAT0000542 | Il2rg | 16186 | HITS-CLIP | 23142080 | Unspecified |
| mmu-miR-34a-5p | MIMAT0000542 | Il3ra | 16188 | HITS-CLIP | 23142080 | Unspecified |
| mmu-miR-34a-5p | MIMAT0000542 | Il4i1 | 14204 | HITS-CLIP | 23142080 | Unspecified |
| mmu-miR-34a-5p | MIMAT0000542 | Il4ra | 16190 | HITS-CLIP | 23142080\|29167373 | Unspecified |
| mmu-miR-34a-5p | MIMAT0000542 | Il6 | 16193 | ELISA, qPCR//ELISA//qPCR | 37986224\|22483937 | Unspecified |
| mmu-miR-34a-5p | MIMAT0000542 | Il6ra | 16194 | HITS-CLIP | 23142080\|23630078\|26203562\|29167 | Unspecified |
| mmu-miR-34a-5p | MIMAT0000542 | Il6st | 16195 | HITS-CLIP, Microarrays//HITS-CLIP//Microa | 37986224\|23142080\|23027903\|19536 | Unspecified |
| mmu-miR-34a-5p | MIMAT0000542 | Il7r | 16197 | HITS-CLIP | 23142080\|29167373 | Unspecified |
| mmu-miR-34a-5p | MIMAT0000542 | Ildr2 | 100039795 | HITS-CLIP | 19536157 | Unspecified |
| mmu-miR-34a-5p | MIMAT0000542 | Ilf3 | 16201 | HITS-CLIP | 23142080\|31433990 | Unspecified |
| mmu-miR-34a-5p | MIMAT0000542 | Ilk | 16202 | HITS-CLIP | 23142080\|29167373 | Unspecified |
| mmu-miR-34a-5p | MIMAT0000542 | Immt | 76614 | HITS-CLIP | 23142080\|29167373 | Unspecified |
| mmu-miR-34a-5p | MIMAT0000542 | Imp3 | 102462 | HITS-CLIP | 23142080 | Unspecified |
| mmu-miR-34a-5p | MIMAT0000542 | Imp4 | 27993 | HITS-CLIP | 23142080\|29167373 | Unspecified |
| mmu-miR-34a-5p | MIMAT0000542 | Impa1 | 55980 | HITS-CLIP | 23142080 | Unspecified |
| mmu-miR-34a-5p | MIMAT0000542 | Impa2 | 114663 | HITS-CLIP | 23142080 | Unspecified |
| mmu-miR-34a-5p | MIMAT0000542 | Impact | 16210 | HITS-CLIP | 23142080\|19536157\|29167373 | Unspecified |
| mmu-miR-34a-5p | MIMAT0000542 | Impad1 | 242291 | HITS-CLIP | 23142080\|29167373 | Unspecified |
| mmu-miR-34a-5p | MIMAT0000542 | Impdh1 | 23917 | HITS-CLIP | 23142080 | Unspecified |
| mmu-miR-34a-5p | MIMAT0000542 | Impdh2 | 23918 | HITS-CLIP | 30971389\|23142080 | Unspecified |
| mmu-miR-34a-5p | MIMAT0000542 | Inafm1 | 66300 | HITS-CLIP | 30971389 | Unspecified |
| mmu-miR-34a-5p | MIMAT0000542 | Incenp | 16319 | HITS-CLIP | 23142080\|29167373 | Unspecified |
| mmu-miR-34a-5p | MIMAT0000542 | Inf2 | 70435 | HITS-CLIP | 29167373 | Unspecified |
| mmu-miR-34a-5p | MIMAT0000542 | Ing2 | 69260 | HITS-CLIP | 30971389 | Unspecified |
| mmu-miR-34a-5p | MIMAT0000542 | Ing3 | 71777 | HITS-CLIP | 23142080 | Unspecified |
| mmu-miR-34a-5p | MIMAT0000542 | Ing4 | 28019 | HITS-CLIP | 29167373\|23142080 | Unspecified |
| mmu-miR-34a-5p | MIMAT0000542 | Ing5 | 66262 | Microarrays//HITS-CLIP | 37986224\|23027903\|23142080 | Unspecified |

| | | | | | | |
|---|---|---|---|---|---|---|
| mmu-miR-34a-5p | MIMAT0000542 | Inhba | 16323 | HITS-CLIP | 31433990\|23142080 | Unspecified |
| mmu-miR-34a-5p | MIMAT0000542 | Inip | 66209 | HITS-CLIP | 23142080 | Unspecified |
| mmu-miR-34a-5p | MIMAT0000542 | Inka2 | 109050 | HITS-CLIP | 19536157 | Unspecified |
| mmu-miR-34a-5p | MIMAT0000542 | Ino80 | 68142 | HITS-CLIP | 26203562\|29167373\|23142080 | Unspecified |
| mmu-miR-34a-5p | MIMAT0000542 | Ino80c | 225280 | HITS-CLIP | 23142080 | Unspecified |
| mmu-miR-34a-5p | MIMAT0000542 | Ino80d | 227195 | HITS-CLIP | 23142080\|29167373 | Unspecified |
| mmu-miR-34a-5p | MIMAT0000542 | Inpp1 | 16329 | HITS-CLIP | 23142080 | Unspecified |
| mmu-miR-34a-5p | MIMAT0000542 | Inpp4a | 269180 | HITS-CLIP | 19536157\|23142080\|31433990\|29167 | Unspecified |
| mmu-miR-34a-5p | MIMAT0000542 | Inpp4b | 234515 | HITS-CLIP, Microarrays//HITS-CLIP | 37986224\|23142080\|31433990\|29167 | Unspecified |
| mmu-miR-34a-5p | MIMAT0000542 | Inpp5a | 212111 | HITS-CLIP | 23142080\|29167373 | Unspecified |
| mmu-miR-34a-5p | MIMAT0000542 | Inpp5b | 16330 | HITS-CLIP | 23142080 | Unspecified |
| mmu-miR-34a-5p | MIMAT0000542 | Inpp5d | 16331 | HITS-CLIP | 23142080\|29167373 | Unspecified |
| mmu-miR-34a-5p | MIMAT0000542 | Inpp5f | 101490 | HITS-CLIP | 29167373\|23142080 | Unspecified |
| mmu-miR-34a-5p | MIMAT0000542 | Inpp5k | 19062 | HITS-CLIP | 23142080\|19536157 | Unspecified |
| mmu-miR-34a-5p | MIMAT0000542 | Inppl1 | 16332 | HITS-CLIP | 29167373 | Unspecified |
| mmu-miR-34a-5p | MIMAT0000542 | Insig1 | 231070 | Microarrays//HITS-CLIP | 37986224\|23142080\|29167373 | Unspecified |
| mmu-miR-34a-5p | MIMAT0000542 | Insig2 | 72999 | HITS-CLIP | 23142080 | Unspecified |
| mmu-miR-34a-5p | MIMAT0000542 | Insr | 16337 | HITS-CLIP | 19536157\|23142080 | Unspecified |
| mmu-miR-34a-5p | MIMAT0000542 | Ints1 | 68510 | HITS-CLIP | 23142080\|29167373 | Unspecified |
| mmu-miR-34a-5p | MIMAT0000542 | Ints10 | 70885 | HITS-CLIP | 23142080 | Unspecified |
| mmu-miR-34a-5p | MIMAT0000542 | Ints11 | 54973 | HITS-CLIP | 23142080 | Unspecified |
| mmu-miR-34a-5p | MIMAT0000542 | Ints12 | 71793 | HITS-CLIP | 23142080 | Unspecified |
| mmu-miR-34a-5p | MIMAT0000542 | Ints13 | 55726 | HITS-CLIP | 23142080 | Unspecified |
| mmu-miR-34a-5p | MIMAT0000542 | Ints14 | 81556 | HITS-CLIP | 23142080 | Unspecified |
| mmu-miR-34a-5p | MIMAT0000542 | Ints2 | 70422 | HITS-CLIP | 23142080\|29167373 | Unspecified |
| mmu-miR-34a-5p | MIMAT0000542 | Ints3 | 229543 | HITS-CLIP | 23142080 | Unspecified |
| mmu-miR-34a-5p | MIMAT0000542 | Ints4 | 101861 | HITS-CLIP | 23142080 | Unspecified |
| mmu-miR-34a-5p | MIMAT0000542 | Ints5 | 80789 | HITS-CLIP | 23142080 | Unspecified |
| mmu-miR-34a-5p | MIMAT0000542 | Ints6 | 18130 | HITS-CLIP | 23142080 | Unspecified |
| mmu-miR-34a-5p | MIMAT0000542 | Ints6l | 236790 | HITS-CLIP | 23142080\|29167373 | Unspecified |
| mmu-miR-34a-5p | MIMAT0000542 | Ints7 | 77065 | Microarrays//HITS-CLIP | 37986224\|23027903\|23142080\|29167 | Unspecified |
| mmu-miR-34a-5p | MIMAT0000542 | Ints8 | 72656 | HITS-CLIP | 23142080 | Unspecified |
| mmu-miR-34a-5p | MIMAT0000542 | Ints9 | 210925 | HITS-CLIP | 23142080 | Unspecified |
| mmu-miR-34a-5p | MIMAT0000542 | Ip6k1 | 27399 | HITS-CLIP | 37986224\|23142080 | Unspecified |
| mmu-miR-34a-5p | MIMAT0000542 | Ip6k2 | 76500 | HITS-CLIP | 23142080 | Unspecified |
| mmu-miR-34a-5p | MIMAT0000542 | Ipcef1 | 320495 | HITS-CLIP | 23142080 | Unspecified |
| mmu-miR-34a-5p | MIMAT0000542 | Ipmk | 69718 | HITS-CLIP | 23142080\|29167373 | Unspecified |
| mmu-miR-34a-5p | MIMAT0000542 | Ipo11 | 76582 | HITS-CLIP | 23142080 | Unspecified |
| mmu-miR-34a-5p | MIMAT0000542 | Ipo13 | 230673 | HITS-CLIP | 29167373\|23142080 | Unspecified |

| | | | | | | |
|---|---|---|---|---|---|---|
| mmu-miR-34a-5p | MIMAT0000542 | Ipo4 | 75751 | HITS-CLIP | 23142080 | Unspecified |
| mmu-miR-34a-5p | MIMAT0000542 | Ipo5 | 70572 | HITS-CLIP | 31433990\|23142080\|29167373 | Unspecified |
| mmu-miR-34a-5p | MIMAT0000542 | Ipo7 | 233726 | HITS-CLIP | 23142080\|29167373 | Unspecified |
| mmu-miR-34a-5p | MIMAT0000542 | Ipo9 | 226432 | HITS-CLIP | 19536157\|29167373\|23142080 | Unspecified |
| mmu-miR-34a-5p | MIMAT0000542 | Ippk | 75678 | HITS-CLIP | 23142080\|31433990 | Unspecified |
| mmu-miR-34a-5p | MIMAT0000542 | Iqcb1 | 320299 | HITS-CLIP | 23142080 | Unspecified |
| mmu-miR-34a-5p | MIMAT0000542 | Iqce | 74239 | HITS-CLIP | 23142080 | Unspecified |
| mmu-miR-34a-5p | MIMAT0000542 | Iqgap1 | 29875 | HITS-CLIP | 23142080\|29167373 | Unspecified |
| mmu-miR-34a-5p | MIMAT0000542 | Iqgap2 | 544963 | HITS-CLIP | 37986224\|31433990\|23142080\|29167 | Unspecified |
| mmu-miR-34a-5p | MIMAT0000542 | Iqgap3 | 404710 | Microarrays//HITS-CLIP | 37986224\|23027903\|23142080 | Unspecified |
| mmu-miR-34a-5p | MIMAT0000542 | Iqsec1 | 232227 | HITS-CLIP | 19536157\|23142080\|29167373 | Unspecified |
| mmu-miR-34a-5p | MIMAT0000542 | Iqsec2 | 245666 | HITS-CLIP | 19536157\|29167373 | Unspecified |
| mmu-miR-34a-5p | MIMAT0000542 | Irak1 | 16179 | HITS-CLIP | 23142080 | Unspecified |
| mmu-miR-34a-5p | MIMAT0000542 | Irak2 | 108960 | HITS-CLIP | 23142080 | Unspecified |
| mmu-miR-34a-5p | MIMAT0000542 | Irak4 | 266632 | HITS-CLIP | 29167373 | Unspecified |
| mmu-miR-34a-5p | MIMAT0000542 | Ireb2 | 64602 | HITS-CLIP | 30971389\|31433990\|29167373\|23142 | Unspecified |
| mmu-miR-34a-5p | MIMAT0000542 | Irf1 | 16362 | HITS-CLIP | 23142080\|29167373 | Unspecified |
| mmu-miR-34a-5p | MIMAT0000542 | Irf2 | 16363 | HITS-CLIP | 23142080\|29167373 | Unspecified |
| mmu-miR-34a-5p | MIMAT0000542 | Irf2bp2 | 270110 | HITS-CLIP | 23142080\|29167373 | Unspecified |
| mmu-miR-34a-5p | MIMAT0000542 | Irf2bpl | 238330 | HITS-CLIP | 23142080\|29167373 | Unspecified |
| mmu-miR-34a-5p | MIMAT0000542 | Irf3 | 54131 | HITS-CLIP | 23142080 | Unspecified |
| mmu-miR-34a-5p | MIMAT0000542 | Irf4 | 16364 | HITS-CLIP | 37986224\|23142080\|31433990 | Unspecified |
| mmu-miR-34a-5p | MIMAT0000542 | Irf5 | 27056 | HITS-CLIP | 30971389\|29167373 | Unspecified |
| mmu-miR-34a-5p | MIMAT0000542 | Irf6 | 54139 | HITS-CLIP | 23142080 | Unspecified |
| mmu-miR-34a-5p | MIMAT0000542 | Irf8 | 15900 | HITS-CLIP | 23142080 | Unspecified |
| mmu-miR-34a-5p | MIMAT0000542 | Irf9 | 16391 | HITS-CLIP | 23142080 | Unspecified |
| mmu-miR-34a-5p | MIMAT0000542 | Irgm1 | 15944 | HITS-CLIP | 29167373\|23142080 | Unspecified |
| mmu-miR-34a-5p | MIMAT0000542 | Irgm2 | 54396 | HITS-CLIP | 23142080\|29167373 | Unspecified |
| mmu-miR-34a-5p | MIMAT0000542 | Irs1 | 16367 | HITS-CLIP | 26203562 | Unspecified |
| mmu-miR-34a-5p | MIMAT0000542 | Irs2 | 384783 | HITS-CLIP | 19536157\|23142080 | Unspecified |
| mmu-miR-34a-5p | MIMAT0000542 | Isca2 | 74316 | HITS-CLIP | 23142080 | Unspecified |
| mmu-miR-34a-5p | MIMAT0000542 | Isg20l2 | 229504 | HITS-CLIP | 23142080 | Unspecified |
| mmu-miR-34a-5p | MIMAT0000542 | Isoc1 | 66307 | HITS-CLIP | 31433990 | Unspecified |
| mmu-miR-34a-5p | MIMAT0000542 | Isoc2a | 664994 | HITS-CLIP | 23142080 | Unspecified |
| mmu-miR-34a-5p | MIMAT0000542 | Ist1 | 71955 | HITS-CLIP | 19536157\|29167373\|23142080 | Unspecified |
| mmu-miR-34a-5p | MIMAT0000542 | Isy1 | 57905 | HITS-CLIP | 23142080 | Unspecified |
| mmu-miR-34a-5p | MIMAT0000542 | Isyna1 | 71780 | HITS-CLIP | 23142080\|29167373 | Unspecified |
| mmu-miR-34a-5p | MIMAT0000542 | Itch | 16396 | HITS-CLIP | 37986224\|23142080\|29167373 | Unspecified |
| mmu-miR-34a-5p | MIMAT0000542 | Itfg1 | 71927 | HITS-CLIP | 23142080\|29167373 | Unspecified |

| | | | | | | |
|---|---|---|---|---|---|---|
| mmu-miR-34a-5p | MIMAT0000542 | Itfg2 | 101142 | Microarrays | 37986224\|23027903 | Unspecified |
| mmu-miR-34a-5p | MIMAT0000542 | Itga11 | 319480 | Microarrays | 37986224 | Unspecified |
| mmu-miR-34a-5p | MIMAT0000542 | Itga3 | 16400 | HITS-CLIP | 26203562 | Unspecified |
| mmu-miR-34a-5p | MIMAT0000542 | Itga4 | 16401 | HITS-CLIP | 23142080\|29167373 | Unspecified |
| mmu-miR-34a-5p | MIMAT0000542 | Itga5 | 16402 | HITS-CLIP | 23142080\|29167373 | Unspecified |
| mmu-miR-34a-5p | MIMAT0000542 | Itga6 | 16403 | HITS-CLIP | 19536157\|23142080\|29167373 | Unspecified |
| mmu-miR-34a-5p | MIMAT0000542 | Itga7 | 16404 | HITS-CLIP | 23142080 | Unspecified |
| mmu-miR-34a-5p | MIMAT0000542 | Itgal | 16408 | HITS-CLIP | 31433990\|23142080 | Unspecified |
| mmu-miR-34a-5p | MIMAT0000542 | Itgam | 16409 | HITS-CLIP | 30971389 | Unspecified |
| mmu-miR-34a-5p | MIMAT0000542 | Itgav | 16410 | HITS-CLIP | 23142080\|29167373 | Unspecified |
| mmu-miR-34a-5p | MIMAT0000542 | Itgb1 | 16412 | HITS-CLIP | 19536157\|23142080\|29167373 | Unspecified |
| mmu-miR-34a-5p | MIMAT0000542 | Itgb1bp1 | 16413 | HITS-CLIP | 23142080 | Unspecified |
| mmu-miR-34a-5p | MIMAT0000542 | Itgb2 | 16414 | HITS-CLIP | 23142080\|30971389\|31433990\|2916 | Unspecified |
| mmu-miR-34a-5p | MIMAT0000542 | Itgb3 | 16416 | HITS-CLIP | 23142080\|31433990 | Unspecified |
| mmu-miR-34a-5p | MIMAT0000542 | Itgb3bp | 67733 | HITS-CLIP | 23142080 | Unspecified |
| mmu-miR-34a-5p | MIMAT0000542 | Itgb7 | 16421 | HITS-CLIP | 37986224\|23142080\|29167373 | Unspecified |
| mmu-miR-34a-5p | MIMAT0000542 | Itgb8 | 320910 | HITS-CLIP | 23142080\|19536157 | Unspecified |
| mmu-miR-34a-5p | MIMAT0000542 | Itih2 | 16425 | HITS-CLIP | 30971389 | Unspecified |
| mmu-miR-34a-5p | MIMAT0000542 | Itih5 | 209378 | HITS-CLIP | 31433990 | Unspecified |
| mmu-miR-34a-5p | MIMAT0000542 | Itk | 16428 | HITS-CLIP | 23142080\|31433990 | Unspecified |
| mmu-miR-34a-5p | MIMAT0000542 | Itm2a | 16431 | HITS-CLIP | 23142080 | Unspecified |
| mmu-miR-34a-5p | MIMAT0000542 | Itm2b | 16432 | HITS-CLIP | 23142080\|19536157\|29167373 | Unspecified |
| mmu-miR-34a-5p | MIMAT0000542 | Itpa | 16434 | Microarrays//HITS-CLIP | 37986224\|23027903\|31433990\|2314 | Unspecified |
| mmu-miR-34a-5p | MIMAT0000542 | Itpk1 | 217837 | HITS-CLIP | 23142080 | Unspecified |
| mmu-miR-34a-5p | MIMAT0000542 | Itpka | 228550 | HITS-CLIP | 23142080 | Unspecified |
| mmu-miR-34a-5p | MIMAT0000542 | Itpkb | 320404 | HITS-CLIP | 31433990\|23142080\|29167373 | Unspecified |
| mmu-miR-34a-5p | MIMAT0000542 | Itpr1 | 16438 | HITS-CLIP | 19536157\|31433990\|23142080 | Unspecified |
| mmu-miR-34a-5p | MIMAT0000542 | Itpr2 | 16439 | HITS-CLIP | 23142080 | Unspecified |
| mmu-miR-34a-5p | MIMAT0000542 | Itpr3 | 16440 | HITS-CLIP | 23142080\|29167373 | Unspecified |
| mmu-miR-34a-5p | MIMAT0000542 | Itprid2 | 6744 | Microarrays//HITS-CLIP | 23027903\|19536157\|23142080\|2916 | Unspecified |
| mmu-miR-34a-5p | MIMAT0000542 | Itprip | 85450 | HITS-CLIP | 23142080 | Unspecified |
| mmu-miR-34a-5p | MIMAT0000542 | Itpripl1 | 73338 | Microarrays//HITS-CLIP | 37986224\|23142080\|23027903\|3097 | Unspecified |
| mmu-miR-34a-5p | MIMAT0000542 | Itpripl2 | 319622 | HITS-CLIP | 23142080\|31433990\|26203562\|2916 | Unspecified |
| mmu-miR-34a-5p | MIMAT0000542 | Itsn1 | 16443 | HITS-CLIP | 19536157\|29167373 | Unspecified |
| mmu-miR-34a-5p | MIMAT0000542 | Itsn2 | 20403 | HITS-CLIP | 37986224\|23142080\|29167373\|3143 | Unspecified |
| mmu-miR-34a-5p | MIMAT0000542 | Ivns1abp | 117198 | HITS-CLIP | 23142080\|19536157 | Unspecified |
| mmu-miR-34a-5p | MIMAT0000542 | Iws1 | 73473 | HITS-CLIP | 23142080\|31433990 | Unspecified |
| mmu-miR-34a-5p | MIMAT0000542 | Izumo1r | 64931 | HITS-CLIP | 23142080 | Unspecified |
| mmu-miR-34a-5p | MIMAT0000542 | Jade1 | 269424 | HITS-CLIP | 23142080\|29167373 | Unspecified |

| | | | | | | |
|---|---|---|---|---|---|---|
| mmu-miR-34a-5p | MIMAT0000542 | Jade2 | 76901 | HITS-CLIP | 23142080\|23630078\|19536157\|29167 | Unspecified |
| mmu-miR-34a-5p | MIMAT0000542 | Jade3 | 382207 | HITS-CLIP | 23142080 | Unspecified |
| mmu-miR-34a-5p | MIMAT0000542 | Jag1 | 16449 | HITS-CLIP | 26203562\|29167373 | Unspecified |
| mmu-miR-34a-5p | MIMAT0000542 | Jag2 | 16450 | HITS-CLIP | 19536157\|26203562\|23142080 | Unspecified |
| mmu-miR-34a-5p | MIMAT0000542 | Jagn1 | 67767 | Microarrays//HITS-CLIP | 37986224\|23027903\|23142080 | Unspecified |
| mmu-miR-34a-5p | MIMAT0000542 | Jak1 | 16451 | HITS-CLIP | 19536157\|23142080\|29167373 | Unspecified |
| mmu-miR-34a-5p | MIMAT0000542 | Jak2 | 16452 | HITS-CLIP | 37986224\|23142080 | Unspecified |
| mmu-miR-34a-5p | MIMAT0000542 | Jak3 | 16453 | HITS-CLIP | 23142080 | Unspecified |
| mmu-miR-34a-5p | MIMAT0000542 | Jakmip1 | 76071 | HITS-CLIP | 19536157\|23142080 | Unspecified |
| mmu-miR-34a-5p | MIMAT0000542 | Jarid2 | 16468 | HITS-CLIP | 37986224\|23142080\|19536157\|31433 | Unspecified |
| mmu-miR-34a-5p | MIMAT0000542 | Jazf1 | 231986 | HITS-CLIP | 23142080 | Unspecified |
| mmu-miR-34a-5p | MIMAT0000542 | Jcad | 57608 | Microarrays//HITS-CLIP | 23027903\|19536157\|31433990\|23142 | Unspecified |
| mmu-miR-34a-5p | MIMAT0000542 | Jdp2 | 81703 | HITS-CLIP | 29167373 | Unspecified |
| mmu-miR-34a-5p | MIMAT0000542 | Jkamp | 104771 | HITS-CLIP | 23142080 | Unspecified |
| mmu-miR-34a-5p | MIMAT0000542 | Jmjd1c | 108829 | HITS-CLIP | 37986224\|23142080\|19536157\|31433 | Unspecified |
| mmu-miR-34a-5p | MIMAT0000542 | Jmjd6 | 107817 | HITS-CLIP | 29167373\|23142080 | Unspecified |
| mmu-miR-34a-5p | MIMAT0000542 | Jmjd8 | 72106 | HITS-CLIP | 23142080 | Unspecified |
| mmu-miR-34a-5p | MIMAT0000542 | Jmy | 57748 | HITS-CLIP | 37986224\|19536157\|23142080 | Unspecified |
| mmu-miR-34a-5p | MIMAT0000542 | Josd1 | 74158 | HITS-CLIP | 23142080 | Unspecified |
| mmu-miR-34a-5p | MIMAT0000542 | Jph4 | 319984 | HITS-CLIP | 19536157 | Unspecified |
| mmu-miR-34a-5p | MIMAT0000542 | Jpt1 | 51155 | HITS-CLIP | 23142080 | Unspecified |
| mmu-miR-34a-5p | MIMAT0000542 | Jpt2 | 90861 | HITS-CLIP | 29167373 | Unspecified |
| mmu-miR-34a-5p | MIMAT0000542 | Jtb | 23922 | HITS-CLIP | 23142080 | Unspecified |
| mmu-miR-34a-5p | MIMAT0000542 | Jun | 16476 | HITS-CLIP | 23142080\|29167373 | Unspecified |
| mmu-miR-34a-5p | MIMAT0000542 | Junb | 16477 | HITS-CLIP | 23142080 | Unspecified |
| mmu-miR-34a-5p | MIMAT0000542 | Jund | 16478 | HITS-CLIP | 31433990\|23142080 | Unspecified |
| mmu-miR-34a-5p | MIMAT0000542 | Jup | 16480 | HITS-CLIP | 23142080\|30971389 | Unspecified |
| mmu-miR-34a-5p | MIMAT0000542 | Kalrn | 545156 | HITS-CLIP | 19536157 | Unspecified |
| mmu-miR-34a-5p | MIMAT0000542 | Kank1 | 107351 | HITS-CLIP | 19536157\|26203562 | Unspecified |
| mmu-miR-34a-5p | MIMAT0000542 | Kansl1 | 76719 | HITS-CLIP | 23142080\|29167373 | Unspecified |
| mmu-miR-34a-5p | MIMAT0000542 | Kansl1l | 68691 | Microarrays//HITS-CLIP | 37986224\|23142080 | Unspecified |
| mmu-miR-34a-5p | MIMAT0000542 | Kansl2 | 69612 | HITS-CLIP | 29167373 | Unspecified |
| mmu-miR-34a-5p | MIMAT0000542 | Kansl3 | 226976 | HITS-CLIP | 29167373 | Unspecified |
| mmu-miR-34a-5p | MIMAT0000542 | Kantr | 102723508 | HITS-CLIP | 23142080\|29167373 | Unspecified |
| mmu-miR-34a-5p | MIMAT0000542 | Kars | 85305 | HITS-CLIP | 23630078 | Unspecified |
| mmu-miR-34a-5p | MIMAT0000542 | Kat14 | 57325 | HITS-CLIP | 29167373\|23142080 | Unspecified |
| mmu-miR-34a-5p | MIMAT0000542 | Kat2b | 18519 | Microarrays//HITS-CLIP | 37986224\|23027903\|23142080\|31433 | Unspecified |
| mmu-miR-34a-5p | MIMAT0000542 | Kat5 | 81601 | HITS-CLIP | 37986224\|23142080\|29167373 | Unspecified |
| mmu-miR-34a-5p | MIMAT0000542 | Kat6a | 244349 | HITS-CLIP | 30971389\|23142080\|29167373 | Unspecified |

| | | | | | | |
|---|---|---|---|---|---|---|
| mmu-miR-34a-5p | MIMAT0000542 | Kat6b | 54169 | HITS-CLIP | 23142080\|19536157\|29167373 | Unspecified |
| mmu-miR-34a-5p | MIMAT0000542 | Kat7 | 217127 | HITS-CLIP | 23142080\|29167373 | Unspecified |
| mmu-miR-34a-5p | MIMAT0000542 | Katna1 | 23924 | HITS-CLIP | 23142080 | Unspecified |
| mmu-miR-34a-5p | MIMAT0000542 | Katnbl1 | 72425 | HITS-CLIP | 31433990\|23630078\|23142080 | Unspecified |
| mmu-miR-34a-5p | MIMAT0000542 | Kbtbd11 | 74901 | HITS-CLIP | 23142080 | Unspecified |
| mmu-miR-34a-5p | MIMAT0000542 | Kbtbd13 | 390594 | HITS-CLIP | 23142080 | Unspecified |
| mmu-miR-34a-5p | MIMAT0000542 | Kbtbd2 | 210973 | HITS-CLIP | 19536157\|23142080\|29167373 | Unspecified |
| mmu-miR-34a-5p | MIMAT0000542 | Kbtbd4 | 67136 | HITS-CLIP | 23142080 | Unspecified |
| mmu-miR-34a-5p | MIMAT0000542 | Kbtbd6 | 89890 | HITS-CLIP | 30971389 | Unspecified |
| mmu-miR-34a-5p | MIMAT0000542 | Kbtbd8 | 243574 | HITS-CLIP | 23142080 | Unspecified |
| mmu-miR-34a-5p | MIMAT0000542 | Kcmf1 | 74287 | HITS-CLIP | 23142080 | Unspecified |
| mmu-miR-34a-5p | MIMAT0000542 | Kcna2 | 16490 | HITS-CLIP | 19536157 | Unspecified |
| mmu-miR-34a-5p | MIMAT0000542 | Kcna3 | 16491 | HITS-CLIP | 29167373 | Unspecified |
| mmu-miR-34a-5p | MIMAT0000542 | Kcnab2 | 16498 | HITS-CLIP | 23142080 | Unspecified |
| mmu-miR-34a-5p | MIMAT0000542 | Kcnb2 | 98741 | HITS-CLIP | 19536157 | Unspecified |
| mmu-miR-34a-5p | MIMAT0000542 | Kcnc1 | 16502 | HITS-CLIP | 19536157 | Unspecified |
| mmu-miR-34a-5p | MIMAT0000542 | Kcnc3 | 16504 | HITS-CLIP | 19536157 | Unspecified |
| mmu-miR-34a-5p | MIMAT0000542 | Kcnd2 | 16508 | HITS-CLIP | 19536157 | Unspecified |
| mmu-miR-34a-5p | MIMAT0000542 | Kcnd3 | 56543 | HITS-CLIP | 19536157 | Unspecified |
| mmu-miR-34a-5p | MIMAT0000542 | Kcnh3 | 16512 | HITS-CLIP | 19536157 | Unspecified |
| mmu-miR-34a-5p | MIMAT0000542 | Kcnh7 | 170738 | HITS-CLIP | 19536157 | Unspecified |
| mmu-miR-34a-5p | MIMAT0000542 | Kcnip4 | 80333 | HITS-CLIP | 19536157 | Unspecified |
| mmu-miR-34a-5p | MIMAT0000542 | Kcnj10 | 16513 | HITS-CLIP | 19536157 | Unspecified |
| mmu-miR-34a-5p | MIMAT0000542 | Kcnj2 | 16518 | HITS-CLIP | 29167373 | Unspecified |
| mmu-miR-34a-5p | MIMAT0000542 | Kcnj6 | 16522 | HITS-CLIP | 19536157 | Unspecified |
| mmu-miR-34a-5p | MIMAT0000542 | Kcnk10 | 72258 | HITS-CLIP | 19536157 | Unspecified |
| mmu-miR-34a-5p | MIMAT0000542 | Kcnk13 | 56659 | HITS-CLIP | 29167373 | Unspecified |
| mmu-miR-34a-5p | MIMAT0000542 | Kcnk3 | 16527 | HITS-CLIP | 19536157 | Unspecified |
| mmu-miR-34a-5p | MIMAT0000542 | Kcnk5 | 16529 | HITS-CLIP, Microarrays//Microarrays//HITS | 37986224\|23027903\|23142080\|3143[illegible] | Unspecified |
| mmu-miR-34a-5p | MIMAT0000542 | Kcnk6 | 52150 | HITS-CLIP | 29167373 | Unspecified |
| mmu-miR-34a-5p | MIMAT0000542 | Kcnk9 | 223604 | HITS-CLIP | 19536157 | Unspecified |
| mmu-miR-34a-5p | MIMAT0000542 | Kcnn4 | 16534 | HITS-CLIP | 23142080\|29167373 | Unspecified |
| mmu-miR-34a-5p | MIMAT0000542 | Kcnq2 | 16536 | HITS-CLIP | 19536157 | Unspecified |
| mmu-miR-34a-5p | MIMAT0000542 | Kcnq4 | 60613 | Microarrays | 37986224 | Unspecified |
| mmu-miR-34a-5p | MIMAT0000542 | Kcnq5 | 226922 | HITS-CLIP | 31433990\|23142080 | Unspecified |
| mmu-miR-34a-5p | MIMAT0000542 | Kctd10 | 330171 | HITS-CLIP | 23142080 | Unspecified |
| mmu-miR-34a-5p | MIMAT0000542 | Kctd12 | 239217 | HITS-CLIP | 23142080\|31433990\|29167373 | Unspecified |
| mmu-miR-34a-5p | MIMAT0000542 | Kctd16 | 383348 | HITS-CLIP | 19536157 | Unspecified |
| mmu-miR-34a-5p | MIMAT0000542 | Kctd17 | 72844 | HITS-CLIP | 23142080\|19536157 | Unspecified |

| | | | | | | |
|---|---|---|---|---|---|---|
| mmu-miR-34a-5p | MIMAT0000542 | Kctd2 | 70382 | Microarrays | 37986224\|23027903 | Unspecified |
| mmu-miR-34a-5p | MIMAT0000542 | Kctd20 | 66989 | HITS-CLIP | 23142080\|29167373 | Unspecified |
| mmu-miR-34a-5p | MIMAT0000542 | Kctd3 | 226823 | HITS-CLIP | 23142080\|29167373 | Unspecified |
| mmu-miR-34a-5p | MIMAT0000542 | Kctd5 | 69259 | HITS-CLIP | 23142080\|29167373 | Unspecified |
| mmu-miR-34a-5p | MIMAT0000542 | Kctd9 | 105440 | HITS-CLIP | 23142080 | Unspecified |
| mmu-miR-34a-5p | MIMAT0000542 | Kdelr1 | 68137 | HITS-CLIP | 37986224\|23142080\|19536157 | Unspecified |
| mmu-miR-34a-5p | MIMAT0000542 | Kdelr2 | 66913 | HITS-CLIP | 31433990\|19536157\|23142080\|2916 | Unspecified |
| mmu-miR-34a-5p | MIMAT0000542 | Kdm1a | 99982 | HITS-CLIP | 23142080 | Unspecified |
| mmu-miR-34a-5p | MIMAT0000542 | Kdm2a | 225876 | HITS-CLIP | 37986224\|19536157\|29167373\|2314 | Unspecified |
| mmu-miR-34a-5p | MIMAT0000542 | Kdm2b | 30841 | HITS-CLIP | 29167373\|23142080 | Unspecified |
| mmu-miR-34a-5p | MIMAT0000542 | Kdm3a | 104263 | HITS-CLIP | 23142080\|29167373 | Unspecified |
| mmu-miR-34a-5p | MIMAT0000542 | Kdm3b | 277250 | HITS-CLIP | 23142080\|29167373 | Unspecified |
| mmu-miR-34a-5p | MIMAT0000542 | Kdm4a | 230674 | HITS-CLIP | 19536157\|23630078\|23142080 | Unspecified |
| mmu-miR-34a-5p | MIMAT0000542 | Kdm4b | 193796 | HITS-CLIP | 23142080 | Unspecified |
| mmu-miR-34a-5p | MIMAT0000542 | Kdm4c | 76804 | HITS-CLIP | 29167373 | Unspecified |
| mmu-miR-34a-5p | MIMAT0000542 | Kdm5a | 214899 | HITS-CLIP | 23142080 | Unspecified |
| mmu-miR-34a-5p | MIMAT0000542 | Kdm5b | 75605 | Microarrays//HITS-CLIP | 37986224\|23142080\|29167373 | Unspecified |
| mmu-miR-34a-5p | MIMAT0000542 | Kdm5c | 20591 | HITS-CLIP | 23142080\|29167373 | Unspecified |
| mmu-miR-34a-5p | MIMAT0000542 | Kdm5d | 20592 | HITS-CLIP | 23142080 | Unspecified |
| mmu-miR-34a-5p | MIMAT0000542 | Kdm6a | 22289 | HITS-CLIP | 19536157\|23142080\|29167373 | Unspecified |
| mmu-miR-34a-5p | MIMAT0000542 | Kdm6b | 216850 | HITS-CLIP | 23630078\|19536157\|23142080\|2620 | Unspecified |
| mmu-miR-34a-5p | MIMAT0000542 | Kdm7a | 338523 | HITS-CLIP | 37986224\|23142080\|29167373 | Unspecified |
| mmu-miR-34a-5p | MIMAT0000542 | Kdm8 | 77035 | HITS-CLIP | 29167373 | Unspecified |
| mmu-miR-34a-5p | MIMAT0000542 | Kdr | 16542 | HITS-CLIP | 29167373 | Unspecified |
| mmu-miR-34a-5p | MIMAT0000542 | Kdsr | 70750 | HITS-CLIP | 23142080\|19536157 | Unspecified |
| mmu-miR-34a-5p | MIMAT0000542 | Keap1 | 50868 | HITS-CLIP | 29167373 | Unspecified |
| mmu-miR-34a-5p | MIMAT0000542 | Khdc4 | 22889 | HITS-CLIP | 23142080 | Unspecified |
| mmu-miR-34a-5p | MIMAT0000542 | Khdrbs1 | 20218 | HITS-CLIP | 23142080\|29167373 | Unspecified |
| mmu-miR-34a-5p | MIMAT0000542 | Khnyn | 219094 | HITS-CLIP | 23142080\|19536157\|29167373 | Unspecified |
| mmu-miR-34a-5p | MIMAT0000542 | Khsrp | 16549 | HITS-CLIP | 31433990\|23142080\|19536157\|2916 | Unspecified |
| mmu-miR-34a-5p | MIMAT0000542 | Kidins220 | 77480 | HITS-CLIP | 23142080\|29167373 | Unspecified |
| mmu-miR-34a-5p | MIMAT0000542 | Kif11 | 16551 | HITS-CLIP | 23142080 | Unspecified |
| mmu-miR-34a-5p | MIMAT0000542 | Kif13a | 16553 | HITS-CLIP | 19536157 | Unspecified |
| mmu-miR-34a-5p | MIMAT0000542 | Kif13b | 16554 | HITS-CLIP | 29167373\|30971389 | Unspecified |
| mmu-miR-34a-5p | MIMAT0000542 | Kif14 | 381293 | HITS-CLIP | 23142080 | Unspecified |
| mmu-miR-34a-5p | MIMAT0000542 | Kif16b | 16558 | HITS-CLIP | 29167373 | Unspecified |
| mmu-miR-34a-5p | MIMAT0000542 | Kif18a | 228421 | HITS-CLIP | 23142080 | Unspecified |
| mmu-miR-34a-5p | MIMAT0000542 | Kif18b | 70218 | Microarrays//HITS-CLIP | 37986224\|23027903\|23142080 | Unspecified |
| mmu-miR-34a-5p | MIMAT0000542 | Kif1a | 16560 | Microarrays//HITS-CLIP | 37986224\|19536157 | Unspecified |

| | | | | | | |
|---|---|---|---|---|---|---|
| mmu-miR-34a-5p | MIMAT0000542 | Kif1b | 16561 | HITS-CLIP | 19536157 | Unspecified |
| mmu-miR-34a-5p | MIMAT0000542 | Kif1bp | 26128 | HITS-CLIP | 23142080 | Unspecified |
| mmu-miR-34a-5p | MIMAT0000542 | Kif1c | 16562 | HITS-CLIP | 19536157\|23142080\|26203562 | Unspecified |
| mmu-miR-34a-5p | MIMAT0000542 | Kif20a | 19348 | HITS-CLIP | 23142080 | Unspecified |
| mmu-miR-34a-5p | MIMAT0000542 | Kif20b | 240641 | HITS-CLIP | 23142080 | Unspecified |
| mmu-miR-34a-5p | MIMAT0000542 | Kif21b | 16565 | HITS-CLIP | 19536157\|23142080\|29167373 | Unspecified |
| mmu-miR-34a-5p | MIMAT0000542 | Kif22 | 110033 | HITS-CLIP | 23142080 | Unspecified |
| mmu-miR-34a-5p | MIMAT0000542 | Kif23 | 71819 | HITS-CLIP | 23142080 | Unspecified |
| mmu-miR-34a-5p | MIMAT0000542 | Kif2a | 16563 | HITS-CLIP | 23142080 | Unspecified |
| mmu-miR-34a-5p | MIMAT0000542 | Kif2c | 73804 | HITS-CLIP | 23142080 | Unspecified |
| mmu-miR-34a-5p | MIMAT0000542 | Kif3a | 16568 | Microarrays//HITS-CLIP | 37986224\|23142080 | Unspecified |
| mmu-miR-34a-5p | MIMAT0000542 | Kif3b | 16569 | HITS-CLIP | 19536157\|23142080 | Unspecified |
| mmu-miR-34a-5p | MIMAT0000542 | Kif4 | 16571 | HITS-CLIP | 23142080 | Unspecified |
| mmu-miR-34a-5p | MIMAT0000542 | Kif5a | 16572 | HITS-CLIP | 19536157 | Unspecified |
| mmu-miR-34a-5p | MIMAT0000542 | Kif5b | 16573 | HITS-CLIP | 31433990\|30971389\|19536157\|23142 | Unspecified |
| mmu-miR-34a-5p | MIMAT0000542 | Kif5c | 16574 | HITS-CLIP | 19536157 | Unspecified |
| mmu-miR-34a-5p | MIMAT0000542 | Kif9 | 16578 | Microarrays | 37986224 | Unspecified |
| mmu-miR-34a-5p | MIMAT0000542 | Kifc2 | 16581 | HITS-CLIP | 19536157 | Unspecified |
| mmu-miR-34a-5p | MIMAT0000542 | Kit | 16590 | HITS-CLIP | 19536157\|30971389 | Unspecified |
| mmu-miR-34a-5p | MIMAT0000542 | Kitl | 17311 | HITS-CLIP | 19536157 | Unspecified |
| mmu-miR-34a-5p | MIMAT0000542 | Klc1 | 16593 | HITS-CLIP | 19536157\|23142080\|29167373 | Unspecified |
| mmu-miR-34a-5p | MIMAT0000542 | Klc2 | 16594 | HITS-CLIP | 23142080 | Unspecified |
| mmu-miR-34a-5p | MIMAT0000542 | Klf10 | 21847 | HITS-CLIP | 23142080\|29167373 | Unspecified |
| mmu-miR-34a-5p | MIMAT0000542 | Klf11 | 194655 | HITS-CLIP | 23142080 | Unspecified |
| mmu-miR-34a-5p | MIMAT0000542 | Klf13 | 50794 | HITS-CLIP | 23630078\|31433990\|29084942\|23142 | Unspecified |
| mmu-miR-34a-5p | MIMAT0000542 | Klf16 | 118445 | HITS-CLIP | 30971389 | Unspecified |
| mmu-miR-34a-5p | MIMAT0000542 | Klf2 | 16598 | HITS-CLIP | 19536157\|23142080\|29167373 | Unspecified |
| mmu-miR-34a-5p | MIMAT0000542 | Klf3 | 16599 | HITS-CLIP | 23142080\|29167373 | Unspecified |
| mmu-miR-34a-5p | MIMAT0000542 | Klf4 | 16600 | HITS-CLIP | 29167373 | Unspecified |
| mmu-miR-34a-5p | MIMAT0000542 | Klf6 | 23849 | HITS-CLIP | 23142080\|29167373 | Unspecified |
| mmu-miR-34a-5p | MIMAT0000542 | Klf7 | 93691 | HITS-CLIP | 19536157\|23142080\|29167373 | Unspecified |
| mmu-miR-34a-5p | MIMAT0000542 | Klf9 | 16601 | HITS-CLIP | 23630078\|19536157\|23142080 | Unspecified |
| mmu-miR-34a-5p | MIMAT0000542 | Klhdc10 | 76788 | HITS-CLIP | 19536157\|30971389 | Unspecified |
| mmu-miR-34a-5p | MIMAT0000542 | Klhdc2 | 69554 | HITS-CLIP | 23630078\|23142080\|31433990 | Unspecified |
| mmu-miR-34a-5p | MIMAT0000542 | Klhdc3 | 71765 | HITS-CLIP | 29167373\|23142080 | Unspecified |
| mmu-miR-34a-5p | MIMAT0000542 | Klhl11 | 217194 | HITS-CLIP | 30971389\|23142080 | Unspecified |
| mmu-miR-34a-5p | MIMAT0000542 | Klhl14 | 57565 | HITS-CLIP | 23142080 | Unspecified |
| mmu-miR-34a-5p | MIMAT0000542 | Klhl15 | 236904 | HITS-CLIP | 23142080 | Unspecified |
| mmu-miR-34a-5p | MIMAT0000542 | Klhl18 | 270201 | HITS-CLIP | 23142080 | Unspecified |

| | | | | | | |
|---|---|---|---|---|---|---|
| mmu-miR-34a-5p | MIMAT0000542 | Klhl2 | 77113 | HITS-CLIP | 23142080\|29167373 | Unspecified |
| mmu-miR-34a-5p | MIMAT0000542 | Klhl20 | 226541 | HITS-CLIP | 23142080 | Unspecified |
| mmu-miR-34a-5p | MIMAT0000542 | Klhl21 | 242785 | HITS-CLIP | 29167373 | Unspecified |
| mmu-miR-34a-5p | MIMAT0000542 | Klhl22 | 224023 | HITS-CLIP | 19536157\|23142080 | Unspecified |
| mmu-miR-34a-5p | MIMAT0000542 | Klhl23 | 277396 | HITS-CLIP | 30971389 | Unspecified |
| mmu-miR-34a-5p | MIMAT0000542 | Klhl24 | 75785 | HITS-CLIP | 23142080\|19536157\|29167373 | Unspecified |
| mmu-miR-34a-5p | MIMAT0000542 | Klhl26 | 234378 | HITS-CLIP | 23142080 | Unspecified |
| mmu-miR-34a-5p | MIMAT0000542 | Klhl28 | 66689 | HITS-CLIP | 23142080\|29167373\|19536157 | Unspecified |
| mmu-miR-34a-5p | MIMAT0000542 | Klhl30 | 70788 | Microarrays | 37986224 | Unspecified |
| mmu-miR-34a-5p | MIMAT0000542 | Klhl31 | 244923 | Microarrays | 37986224 | Unspecified |
| mmu-miR-34a-5p | MIMAT0000542 | Klhl41 | 228003 | Microarrays | 37986224 | Unspecified |
| mmu-miR-34a-5p | MIMAT0000542 | Klhl42 | 232539 | HITS-CLIP | 31433990\|23630078\|23142080 | Unspecified |
| mmu-miR-34a-5p | MIMAT0000542 | Klhl5 | 71778 | Microarrays//HITS-CLIP | 37986224\|19536157\|23142080 | Unspecified |
| mmu-miR-34a-5p | MIMAT0000542 | Klhl6 | 239743 | HITS-CLIP | 23142080 | Unspecified |
| mmu-miR-34a-5p | MIMAT0000542 | Klhl7 | 52323 | HITS-CLIP | 19536157\|23142080 | Unspecified |
| mmu-miR-34a-5p | MIMAT0000542 | Klhl9 | 242521 | HITS-CLIP | 23630078\|30971389\|23142080 | Unspecified |
| mmu-miR-34a-5p | MIMAT0000542 | Kmt2a | 214162 | HITS-CLIP | 37986224\|31433990\|19536157\|30971 | Unspecified |
| mmu-miR-34a-5p | MIMAT0000542 | Kmt2b | 75410 | HITS-CLIP | 23142080\|30971389\|29167373 | Unspecified |
| mmu-miR-34a-5p | MIMAT0000542 | Kmt2c | 231051 | HITS-CLIP | 23142080\|19536157\|26203562\|29167 | Unspecified |
| mmu-miR-34a-5p | MIMAT0000542 | Kmt2d | 381022 | HITS-CLIP | 31433990\|19536157\|23142080\|29167 | Unspecified |
| mmu-miR-34a-5p | MIMAT0000542 | Kmt2e | 69188 | HITS-CLIP | 37986224\|19536157\|31433990\|23142 | Unspecified |
| mmu-miR-34a-5p | MIMAT0000542 | Kmt5b | 225888 | HITS-CLIP | 31433990\|23142080\|29167373 | Unspecified |
| mmu-miR-34a-5p | MIMAT0000542 | Kmt5c | 84787 | HITS-CLIP | 23142080 | Unspecified |
| mmu-miR-34a-5p | MIMAT0000542 | Knl1 | 57082 | HITS-CLIP | 23142080 | Unspecified |
| mmu-miR-34a-5p | MIMAT0000542 | Knop1 | 66356 | HITS-CLIP | 30971389\|31433990\|23142080 | Unspecified |
| mmu-miR-34a-5p | MIMAT0000542 | Knstrn | 51944 | Microarrays//HITS-CLIP | 37986224\|23027903\|23142080\|29167 | Unspecified |
| mmu-miR-34a-5p | MIMAT0000542 | Kntc1 | 208628 | HITS-CLIP | 23142080 | Unspecified |
| mmu-miR-34a-5p | MIMAT0000542 | Kpna1 | 16646 | HITS-CLIP | 23142080\|19536157\|31433990\|29167 | Unspecified |
| mmu-miR-34a-5p | MIMAT0000542 | Kpna2 | 16647 | HITS-CLIP | 23142080 | Unspecified |
| mmu-miR-34a-5p | MIMAT0000542 | Kpna3 | 16648 | HITS-CLIP | 23142080 | Unspecified |
| mmu-miR-34a-5p | MIMAT0000542 | Kpna4 | 16649 | HITS-CLIP | 29167373\|23142080 | Unspecified |
| mmu-miR-34a-5p | MIMAT0000542 | Kpna6 | 16650 | HITS-CLIP | 19536157\|23142080\|29167373 | Unspecified |
| mmu-miR-34a-5p | MIMAT0000542 | Kpnb1 | 16211 | HITS-CLIP | 23142080\|31433990\|19536157\|29167 | Unspecified |
| mmu-miR-34a-5p | MIMAT0000542 | Kras | 16653 | HITS-CLIP | 23142080\|29167373 | Unspecified |
| mmu-miR-34a-5p | MIMAT0000542 | Kri1 | 215194 | HITS-CLIP | 23142080 | Unspecified |
| mmu-miR-34a-5p | MIMAT0000542 | Krit1 | 79264 | HITS-CLIP | 23142080 | Unspecified |
| mmu-miR-34a-5p | MIMAT0000542 | Krt14 | 16664 | HITS-CLIP | 26203562 | Unspecified |
| mmu-miR-34a-5p | MIMAT0000542 | Krt16 | 16666 | HITS-CLIP | 26203562 | Unspecified |
| mmu-miR-34a-5p | MIMAT0000542 | Krt17 | 16667 | HITS-CLIP | 26203562 | Unspecified |

| | | | | | | | |
|---|---|---|---|---|---|---|---|
| mmu-miR-34a-5p | MIMAT0000542 | Krt5 | 110308 | HITS-CLIP | | 26203562 | Unspecified |
| mmu-miR-34a-5p | MIMAT0000542 | Krtcap2 | 66059 | HITS-CLIP | | 23142080 | Unspecified |
| mmu-miR-34a-5p | MIMAT0000542 | Ksr1 | 16706 | HITS-CLIP | 37986224\|23142080 | | Unspecified |
| mmu-miR-34a-5p | MIMAT0000542 | Ksr2 | 333050 | HITS-CLIP | | 19536157 | Unspecified |
| mmu-miR-34a-5p | MIMAT0000542 | Ktn1 | 16709 | HITS-CLIP | 23142080\|31433990\|19536157 | | Unspecified |
| mmu-miR-34a-5p | MIMAT0000542 | L1cam | 16728 | HITS-CLIP | | 19536157 | Unspecified |
| mmu-miR-34a-5p | MIMAT0000542 | L2hgdh | 217666 | HITS-CLIP | 23142080\|30971389 | | Unspecified |
| mmu-miR-34a-5p | MIMAT0000542 | L3mbtl2 | 214669 | HITS-CLIP | | 23142080 | Unspecified |
| mmu-miR-34a-5p | MIMAT0000542 | L3mbtl3 | 237339 | HITS-CLIP | 19536157\|23142080 | | Unspecified |
| mmu-miR-34a-5p | MIMAT0000542 | Lactb2 | 212442 | HITS-CLIP | 23142080\|29167373 | | Unspecified |
| mmu-miR-34a-5p | MIMAT0000542 | Lag3 | 16768 | HITS-CLIP | | 31433990 | Unspecified |
| mmu-miR-34a-5p | MIMAT0000542 | Lair1 | 52855 | HITS-CLIP | | 29167373 | Unspecified |
| mmu-miR-34a-5p | MIMAT0000542 | Lama2 | 16773 | Microarrays | | 37986224 | Unspecified |
| mmu-miR-34a-5p | MIMAT0000542 | Lama5 | 16776 | HITS-CLIP | | 23142080 | Unspecified |
| mmu-miR-34a-5p | MIMAT0000542 | Lamc1 | 226519 | HITS-CLIP | 37986224\|31433990\|23142080\|29167 | | Unspecified |
| mmu-miR-34a-5p | MIMAT0000542 | Lamp1 | 16783 | HITS-CLIP | 29167373\|23142080 | | Unspecified |
| mmu-miR-34a-5p | MIMAT0000542 | Lamp2 | 16784 | HITS-CLIP | 19536157\|23142080\|29167373 | | Unspecified |
| mmu-miR-34a-5p | MIMAT0000542 | Lamp5 | 76161 | HITS-CLIP | | 19536157 | Unspecified |
| mmu-miR-34a-5p | MIMAT0000542 | Lamtor3 | 56692 | HITS-CLIP | 23142080\|29167373 | | Unspecified |
| mmu-miR-34a-5p | MIMAT0000542 | Lancl3 | 236285 | HITS-CLIP | | 23142080 | Unspecified |
| mmu-miR-34a-5p | MIMAT0000542 | Lap3 | 66988 | HITS-CLIP | | 23142080 | Unspecified |
| mmu-miR-34a-5p | MIMAT0000542 | Laptm4a | 17775 | HITS-CLIP | 37986224\|23142080\|29167373 | | Unspecified |
| mmu-miR-34a-5p | MIMAT0000542 | Laptm5 | 16792 | HITS-CLIP | 23142080\|29167373 | | Unspecified |
| mmu-miR-34a-5p | MIMAT0000542 | Larp1 | 73158 | HITS-CLIP | 23142080\|29167373 | | Unspecified |
| mmu-miR-34a-5p | MIMAT0000542 | Larp4 | 207214 | HITS-CLIP | 19536157\|23142080\|29167373 | | Unspecified |
| mmu-miR-34a-5p | MIMAT0000542 | Larp4b | 217980 | HITS-CLIP | 31433990\|23142080\|29167373 | | Unspecified |
| mmu-miR-34a-5p | MIMAT0000542 | Lars | 107045 | HITS-CLIP | | 23142080 | Unspecified |
| mmu-miR-34a-5p | MIMAT0000542 | Lars2 | 102436 | HITS-CLIP | | 23142080 | Unspecified |
| mmu-miR-34a-5p | MIMAT0000542 | Lasp1 | 16796 | HITS-CLIP | 31433990\|29167373 | | Unspecified |
| mmu-miR-34a-5p | MIMAT0000542 | Lat | 16797 | HITS-CLIP | | 23142080 | Unspecified |
| mmu-miR-34a-5p | MIMAT0000542 | Lats1 | 16798 | HITS-CLIP | 23142080\|29167373 | | Unspecified |
| mmu-miR-34a-5p | MIMAT0000542 | Lats2 | 50523 | HITS-CLIP | 23142080\|31433990 | | Unspecified |
| mmu-miR-34a-5p | MIMAT0000542 | Lax1 | 240754 | HITS-CLIP | | 23142080 | Unspecified |
| mmu-miR-34a-5p | MIMAT0000542 | Lbh | 77889 | HITS-CLIP | | 23142080 | Unspecified |
| mmu-miR-34a-5p | MIMAT0000542 | Lbhd1 | 79081 | HITS-CLIP | | 23142080 | Unspecified |
| mmu-miR-34a-5p | MIMAT0000542 | Lbr | 98386 | HITS-CLIP | 23142080\|31433990 | | Unspecified |
| mmu-miR-34a-5p | MIMAT0000542 | Lca5 | 75782 | HITS-CLIP | | 23142080 | Unspecified |
| mmu-miR-34a-5p | MIMAT0000542 | Lck | 16818 | HITS-CLIP | | 23142080 | Unspecified |
| mmu-miR-34a-5p | MIMAT0000542 | Lclat1 | 225010 | HITS-CLIP | 19536157\|23142080\|29167373 | | Unspecified |

| | | | | | | |
|---|---|---|---|---|---|---|
| mmu-miR-34a-5p | MIMAT0000542 | Lcor | 212391 | HITS-CLIP | 23142080 | Unspecified |
| mmu-miR-34a-5p | MIMAT0000542 | Lcorl | 209707 | HITS-CLIP | 19536157\|23142080 | Unspecified |
| mmu-miR-34a-5p | MIMAT0000542 | Lcp1 | 18826 | HITS-CLIP | 23142080\|31433990\|29167373 | Unspecified |
| mmu-miR-34a-5p | MIMAT0000542 | Lcp2 | 16822 | HITS-CLIP | 29167373\|23142080 | Unspecified |
| mmu-miR-34a-5p | MIMAT0000542 | Ldah | 38150 | HITS-CLIP | 23142080\|29167373 | Unspecified |
| mmu-miR-34a-5p | MIMAT0000542 | Ldb1 | 16825 | HITS-CLIP | 23142080 | Unspecified |
| mmu-miR-34a-5p | MIMAT0000542 | Ldha | 16828 | HITS-CLIP | 23142080\|29167373 | Unspecified |
| mmu-miR-34a-5p | MIMAT0000542 | Ldlr | 16835 | HITS-CLIP | 19536157\|31433990\|26203562\|29167 | Unspecified |
| mmu-miR-34a-5p | MIMAT0000542 | Ldlrad3 | 241576 | HITS-CLIP | 19536157 | Unspecified |
| mmu-miR-34a-5p | MIMAT0000542 | Ldlrap1 | 100017 | HITS-CLIP | 31433990\|29167373\|23142080 | Unspecified |
| mmu-miR-34a-5p | MIMAT0000542 | Lef1 | 16842 | HITS-CLIP | 23142080 | Unspecified |
| mmu-miR-34a-5p | MIMAT0000542 | Lemd3 | 380664 | HITS-CLIP | 23142080 | Unspecified |
| mmu-miR-34a-5p | MIMAT0000542 | Leng8 | 232798 | HITS-CLIP | 29167373\|23142080 | Unspecified |
| mmu-miR-34a-5p | MIMAT0000542 | Leo1 | 235497 | HITS-CLIP | 23142080 | Unspecified |
| mmu-miR-34a-5p | MIMAT0000542 | Leprot | 230514 | HITS-CLIP | 23142080 | Unspecified |
| mmu-miR-34a-5p | MIMAT0000542 | Leprotl1 | 68192 | HITS-CLIP | 31433990\|23142080 | Unspecified |
| mmu-miR-34a-5p | MIMAT0000542 | Letm1 | 56384 | HITS-CLIP | 29167373 | Unspecified |
| mmu-miR-34a-5p | MIMAT0000542 | Letm2 | 270035 | HITS-CLIP | 19536157 | Unspecified |
| mmu-miR-34a-5p | MIMAT0000542 | Letmd1 | 68614 | Microarrays//HITS-CLIP | 37986224\|23027903\|23142080 | Unspecified |
| mmu-miR-34a-5p | MIMAT0000542 | Lfng | 16848 | HITS-CLIP | 31433990\|23142080\|29167373 | Unspecified |
| mmu-miR-34a-5p | MIMAT0000542 | Lgals1 | 16852 | HITS-CLIP | 23142080\|29167373 | Unspecified |
| mmu-miR-34a-5p | MIMAT0000542 | Lgals3 | 16854 | HITS-CLIP | 23142080\|29167373 | Unspecified |
| mmu-miR-34a-5p | MIMAT0000542 | Lgals3bp | 19039 | Microarrays//HITS-CLIP | 37986224\|23142080\|29167373 | Unspecified |
| mmu-miR-34a-5p | MIMAT0000542 | Lgals8 | 56048 | HITS-CLIP | 23142080\|29167373 | Unspecified |
| mmu-miR-34a-5p | MIMAT0000542 | Lgals9 | 16859 | HITS-CLIP | 23142080\|29167373 | Unspecified |
| mmu-miR-34a-5p | MIMAT0000542 | Lgalsl | 216551 | HITS-CLIP | 23142080 | Unspecified |
| mmu-miR-34a-5p | MIMAT0000542 | Lgi2 | 246316 | HITS-CLIP | 19536157 | Unspecified |
| mmu-miR-34a-5p | MIMAT0000542 | Lgmn | 19141 | HITS-CLIP | 29167373 | Unspecified |
| mmu-miR-34a-5p | MIMAT0000542 | Lgr4 | 107515 | Microarrays//HITS-CLIP | 37986224\|23027903\|19536157\|23142 | Unspecified |
| mmu-miR-34a-5p | MIMAT0000542 | Lgr5 | 14160 | HITS-CLIP | 23142080 | Unspecified |
| mmu-miR-34a-5p | MIMAT0000542 | Lhfpl2 | 218454 | Microarrays//HITS-CLIP | 37986224\|23630078\|29167373 | Unspecified |
| mmu-miR-34a-5p | MIMAT0000542 | Lhx2 | 16870 | HITS-CLIP | 19536157 | Unspecified |
| mmu-miR-34a-5p | MIMAT0000542 | Lif | 16878 | HITS-CLIP | 31433990\|23142080 | Unspecified |
| mmu-miR-34a-5p | MIMAT0000542 | Lifr | 16880 | Microarrays//HITS-CLIP | 37986224\|29167373 | Unspecified |
| mmu-miR-34a-5p | MIMAT0000542 | Lig1 | 16881 | HITS-CLIP | 23142080\|29167373 | Unspecified |
| mmu-miR-34a-5p | MIMAT0000542 | Lig3 | 16882 | HITS-CLIP | 23142080\|26203562 | Unspecified |
| mmu-miR-34a-5p | MIMAT0000542 | Lilr4b | 14727 | HITS-CLIP | 29167373 | Unspecified |
| mmu-miR-34a-5p | MIMAT0000542 | Lima1 | 65970 | HITS-CLIP | 29167373 | Unspecified |
| mmu-miR-34a-5p | MIMAT0000542 | Limd1 | 29806 | HITS-CLIP | 23142080 | Unspecified |

| | | | | | | |
|---|---|---|---|---|---|---|
| mmu-miR-34a-5p | MIMAT0000542 | Limd2 | 67803 | HITS-CLIP | 31433990\|23142080 | Unspecified |
| mmu-miR-34a-5p | MIMAT0000542 | Lime1 | 72699 | HITS-CLIP | 23142080 | Unspecified |
| mmu-miR-34a-5p | MIMAT0000542 | Limk2 | 16886 | HITS-CLIP | 19536157\|23630078\|23142080 | Unspecified |
| mmu-miR-34a-5p | MIMAT0000542 | Lims1 | 110829 | HITS-CLIP | 37986224\|23142080\|31433990\|2363( | Unspecified |
| mmu-miR-34a-5p | MIMAT0000542 | Lin54 | 231506 | HITS-CLIP | 23142080\|29167373 | Unspecified |
| mmu-miR-34a-5p | MIMAT0000542 | Lin7c | 22343 | Microarrays//HITS-CLIP | 37986224\|31433990\|30971389\|23142 | Unspecified |
| mmu-miR-34a-5p | MIMAT0000542 | Lin9 | 72568 | Microarrays//HITS-CLIP | 37986224\|23142080\|23027903\|29084 | Unspecified |
| mmu-miR-34a-5p | MIMAT0000542 | Lingo1 | 235402 | HITS-CLIP | 19536157 | Unspecified |
| mmu-miR-34a-5p | MIMAT0000542 | Lingo3 | 237403 | HITS-CLIP | 19536157 | Unspecified |
| mmu-miR-34a-5p | MIMAT0000542 | Lins1 | 72635 | HITS-CLIP | 29167373 | Unspecified |
| mmu-miR-34a-5p | MIMAT0000542 | Lipa | 16889 | HITS-CLIP | 29167373 | Unspecified |
| mmu-miR-34a-5p | MIMAT0000542 | Lipe | 16890 | HITS-CLIP | 23142080 | Unspecified |
| mmu-miR-34a-5p | MIMAT0000542 | Lipt2 | 67164 | HITS-CLIP | 31433990 | Unspecified |
| mmu-miR-34a-5p | MIMAT0000542 | Litaf | 56722 | HITS-CLIP | 23142080\|29167373 | Unspecified |
| mmu-miR-34a-5p | MIMAT0000542 | Lix1 | 66643 | HITS-CLIP | 19536157 | Unspecified |
| mmu-miR-34a-5p | MIMAT0000542 | Llgl1 | 16897 | HITS-CLIP | 26203562\|23142080\|29167373 | Unspecified |
| mmu-miR-34a-5p | MIMAT0000542 | Lman1 | 70361 | HITS-CLIP | 23142080\|19536157\|23630078\|29167 | Unspecified |
| mmu-miR-34a-5p | MIMAT0000542 | Lman2 | 66890 | HITS-CLIP | 23142080\|29167373 | Unspecified |
| mmu-miR-34a-5p | MIMAT0000542 | Lmbr1l | 74775 | HITS-CLIP | 23142080 | Unspecified |
| mmu-miR-34a-5p | MIMAT0000542 | Lmbrd1 | 68421 | HITS-CLIP | 23142080\|29167373 | Unspecified |
| mmu-miR-34a-5p | MIMAT0000542 | Lmbrd2 | 320506 | HITS-CLIP | 23142080\|29167373\|19536157 | Unspecified |
| mmu-miR-34a-5p | MIMAT0000542 | Lmf2 | 105847 | HITS-CLIP | 23142080 | Unspecified |
| mmu-miR-34a-5p | MIMAT0000542 | Lmln | 239833 | Microarrays | 37986224 | Unspecified |
| mmu-miR-34a-5p | MIMAT0000542 | Lmna | 16905 | HITS-CLIP | 26203562\|29167373 | Unspecified |
| mmu-miR-34a-5p | MIMAT0000542 | Lmnb1 | 16906 | HITS-CLIP | 23142080\|29167373 | Unspecified |
| mmu-miR-34a-5p | MIMAT0000542 | Lmnb2 | 16907 | HITS-CLIP | 23142080 | Unspecified |
| mmu-miR-34a-5p | MIMAT0000542 | Lmo7 | 380928 | HITS-CLIP | 23142080 | Unspecified |
| mmu-miR-34a-5p | MIMAT0000542 | Lmod2 | 93677 | Microarrays | 37986224 | Unspecified |
| mmu-miR-34a-5p | MIMAT0000542 | Lmod3 | 320502 | Microarrays | 37986224 | Unspecified |
| mmu-miR-34a-5p | MIMAT0000542 | Lmtk2 | 231876 | Microarrays//HITS-CLIP | 37986224\|23027903\|19536157\|23142 | Unspecified |
| mmu-miR-34a-5p | MIMAT0000542 | Lnpep | 240028 | HITS-CLIP | 19536157\|31433990\|29167373\|23142 | Unspecified |
| mmu-miR-34a-5p | MIMAT0000542 | Lnpk | 69605 | HITS-CLIP | 19536157\|29167373 | Unspecified |
| mmu-miR-34a-5p | MIMAT0000542 | Lnx2 | 140887 | HITS-CLIP | 23142080\|19536157 | Unspecified |
| mmu-miR-34a-5p | MIMAT0000542 | Lonp1 | 74142 | HITS-CLIP | 23142080 | Unspecified |
| mmu-miR-34a-5p | MIMAT0000542 | Lonp2 | 66887 | HITS-CLIP | 23142080 | Unspecified |
| mmu-miR-34a-5p | MIMAT0000542 | Lonrf1 | 244421 | HITS-CLIP | 23142080 | Unspecified |
| mmu-miR-34a-5p | MIMAT0000542 | Lonrf2 | 381338 | HITS-CLIP | 19536157 | Unspecified |
| mmu-miR-34a-5p | MIMAT0000542 | Lonrf3 | 74365 | HITS-CLIP | 23142080\|29167373 | Unspecified |
| mmu-miR-34a-5p | MIMAT0000542 | Lpar3 | 65086 | HITS-CLIP | 23142080 | Unspecified |

| | | | | | | |
|---|---|---|---|---|---|---|
| mmu-miR-34a-5p | MIMAT0000542 | Lpar5 | 381810 | HITS-CLIP | 31433990 | Unspecified |
| mmu-miR-34a-5p | MIMAT0000542 | Lpar6 | 67168 | HITS-CLIP | 31433990 | Unspecified |
| mmu-miR-34a-5p | MIMAT0000542 | Lpcat1 | 210992 | Microarrays//HITS-CLIP | 37986224\|23027903\|29167373 | Unspecified |
| mmu-miR-34a-5p | MIMAT0000542 | Lpcat2 | 270084 | HITS-CLIP | 29167373 | Unspecified |
| mmu-miR-34a-5p | MIMAT0000542 | Lpcat3 | 14792 | HITS-CLIP | 37986224\|23142080 | Unspecified |
| mmu-miR-34a-5p | MIMAT0000542 | Lpgat1 | 226856 | HITS-CLIP | 19536157\|23142080\|29167373 | Unspecified |
| mmu-miR-34a-5p | MIMAT0000542 | Lpin1 | 14245 | HITS-CLIP | 37986224\|31433990\|23142080 | Unspecified |
| mmu-miR-34a-5p | MIMAT0000542 | Lpin2 | 64898 | HITS-CLIP | 23142080\|29167373 | Unspecified |
| mmu-miR-34a-5p | MIMAT0000542 | Lpl | 16956 | HITS-CLIP | 29167373 | Unspecified |
| mmu-miR-34a-5p | MIMAT0000542 | Lpp | 210126 | HITS-CLIP | 23630078\|23142080\|29167373 | Unspecified |
| mmu-miR-34a-5p | MIMAT0000542 | Lpxn | 107321 | HITS-CLIP | 23142080\|29167373 | Unspecified |
| mmu-miR-34a-5p | MIMAT0000542 | Lrba | 80877 | HITS-CLIP | 19536157\|23142080 | Unspecified |
| mmu-miR-34a-5p | MIMAT0000542 | Lrch1 | 380916 | HITS-CLIP | 23142080\|29167373 | Unspecified |
| mmu-miR-34a-5p | MIMAT0000542 | Lrch3 | 70144 | HITS-CLIP | 23142080\|29167373 | Unspecified |
| mmu-miR-34a-5p | MIMAT0000542 | Lrch4 | 231798 | HITS-CLIP | 23142080\|29167373 | Unspecified |
| mmu-miR-34a-5p | MIMAT0000542 | Lrig1 | 16206 | HITS-CLIP | 23142080 | Unspecified |
| mmu-miR-34a-5p | MIMAT0000542 | Lrig2 | 269473 | HITS-CLIP | 29167373\|23142080 | Unspecified |
| mmu-miR-34a-5p | MIMAT0000542 | Lrmda | 83938 | HITS-CLIP | 29167373 | Unspecified |
| mmu-miR-34a-5p | MIMAT0000542 | Lrmp | 16970 | HITS-CLIP | 23142080\|29167373 | Unspecified |
| mmu-miR-34a-5p | MIMAT0000542 | Lrp1 | 16971 | HITS-CLIP | 19536157\|29167373 | Unspecified |
| mmu-miR-34a-5p | MIMAT0000542 | Lrp10 | 65107 | HITS-CLIP | 23142080 | Unspecified |
| mmu-miR-34a-5p | MIMAT0000542 | Lrp11 | 237253 | HITS-CLIP | 23142080 | Unspecified |
| mmu-miR-34a-5p | MIMAT0000542 | Lrp12 | 239393 | HITS-CLIP | 23142080\|29167373 | Unspecified |
| mmu-miR-34a-5p | MIMAT0000542 | Lrp5 | 16973 | HITS-CLIP | 30971389 | Unspecified |
| mmu-miR-34a-5p | MIMAT0000542 | Lrp6 | 16974 | HITS-CLIP | 19536157\|23142080\|29167373 | Unspecified |
| mmu-miR-34a-5p | MIMAT0000542 | Lrp8 | 16975 | HITS-CLIP | 19536157\|23142080\|29167373 | Unspecified |
| mmu-miR-34a-5p | MIMAT0000542 | Lrpprc | 72416 | HITS-CLIP | 19536157\|23142080 | Unspecified |
| mmu-miR-34a-5p | MIMAT0000542 | Lrr1 | 69706 | Microarrays//HITS-CLIP | 37986224\|23027903\|23142080 | Unspecified |
| mmu-miR-34a-5p | MIMAT0000542 | Lrrc1 | 214345 | HITS-CLIP | 23142080\|30971389 | Unspecified |
| mmu-miR-34a-5p | MIMAT0000542 | Lrrc32 | 434215 | HITS-CLIP | 23142080 | Unspecified |
| mmu-miR-34a-5p | MIMAT0000542 | Lrrc4 | 192198 | HITS-CLIP | 19536157 | Unspecified |
| mmu-miR-34a-5p | MIMAT0000542 | Lrrc40 | 67144 | HITS-CLIP | 37986224\|23142080\|29167373 | Unspecified |
| mmu-miR-34a-5p | MIMAT0000542 | Lrrc41 | 230654 | HITS-CLIP | 23142080 | Unspecified |
| mmu-miR-34a-5p | MIMAT0000542 | Lrrc42 | 77809 | HITS-CLIP | 23142080 | Unspecified |
| mmu-miR-34a-5p | MIMAT0000542 | Lrrc46 | 69297 | Microarrays | 37986224 | Unspecified |
| mmu-miR-34a-5p | MIMAT0000542 | Lrrc47 | 72946 | HITS-CLIP | 23630078\|30971389\|29167373 | Unspecified |
| mmu-miR-34a-5p | MIMAT0000542 | Lrrc4c | 241568 | HITS-CLIP | 19536157 | Unspecified |
| mmu-miR-34a-5p | MIMAT0000542 | Lrrc55 | 241528 | HITS-CLIP | 19536157 | Unspecified |
| mmu-miR-34a-5p | MIMAT0000542 | Lrrc57 | 66606 | HITS-CLIP | 23142080 | Unspecified |

| | | | | | | |
|---|---|---|---|---|---|---|
| mmu-miR-34a-5p | MIMAT0000542 | Lrrc58 | 320184 | HITS-CLIP | 19536157\|30971389\|23142080\|29167 | Unspecified |
| mmu-miR-34a-5p | MIMAT0000542 | Lrrc59 | 98238 | HITS-CLIP | 23142080 | Unspecified |
| mmu-miR-34a-5p | MIMAT0000542 | Lrrc7 | 242274 | HITS-CLIP | 19536157 | Unspecified |
| mmu-miR-34a-5p | MIMAT0000542 | Lrrc73 | 224813 | Microarrays | 37986224 | Unspecified |
| mmu-miR-34a-5p | MIMAT0000542 | Lrrc8a | 241296 | HITS-CLIP | 19536157\|23142080\|29167373 | Unspecified |
| mmu-miR-34a-5p | MIMAT0000542 | Lrrc8b | 433926 | HITS-CLIP | 19536157\|30971389 | Unspecified |
| mmu-miR-34a-5p | MIMAT0000542 | Lrrc8c | 100604 | HITS-CLIP | 31433990\|23142080\|29167373 | Unspecified |
| mmu-miR-34a-5p | MIMAT0000542 | Lrrc8d | 231549 | HITS-CLIP | 23142080\|29167373 | Unspecified |
| mmu-miR-34a-5p | MIMAT0000542 | Lrrcc1 | 71710 | HITS-CLIP | 23142080\|29167373 | Unspecified |
| mmu-miR-34a-5p | MIMAT0000542 | Lrrfip1 | 16978 | HITS-CLIP | 23142080\|29167373 | Unspecified |
| mmu-miR-34a-5p | MIMAT0000542 | Lrrk1 | 233328 | HITS-CLIP | 23142080\|31433990\|29167373 | Unspecified |
| mmu-miR-34a-5p | MIMAT0000542 | Lrrn1 | 16979 | Microarrays | 37986224 | Unspecified |
| mmu-miR-34a-5p | MIMAT0000542 | Lrwd1 | 71735 | HITS-CLIP | 23142080\|29167373 | Unspecified |
| mmu-miR-34a-5p | MIMAT0000542 | Lsamp | 268890 | HITS-CLIP | 19536157 | Unspecified |
| mmu-miR-34a-5p | MIMAT0000542 | Lsg1 | 224092 | HITS-CLIP | 23142080 | Unspecified |
| mmu-miR-34a-5p | MIMAT0000542 | Lsm1 | 67207 | HITS-CLIP | 31433990\|23142080 | Unspecified |
| mmu-miR-34a-5p | MIMAT0000542 | Lsm14a | 67070 | HITS-CLIP | 31433990\|23142080\|29167373 | Unspecified |
| mmu-miR-34a-5p | MIMAT0000542 | Lsm14b | 241846 | HITS-CLIP | 31433990\|23142080 | Unspecified |
| mmu-miR-34a-5p | MIMAT0000542 | Lsm2 | 27756 | HITS-CLIP | 23142080 | Unspecified |
| mmu-miR-34a-5p | MIMAT0000542 | Lsm4 | 50783 | HITS-CLIP | 30971389 | Unspecified |
| mmu-miR-34a-5p | MIMAT0000542 | Lsm6 | 78651 | HITS-CLIP | 31433990\|29167373 | Unspecified |
| mmu-miR-34a-5p | MIMAT0000542 | Lsm8 | 76522 | HITS-CLIP, Microarrays//Microarrays//HITS | 37986224\|23027903\|23142080 | Unspecified |
| mmu-miR-34a-5p | MIMAT0000542 | Lsr | 54135 | HITS-CLIP | 30971389 | Unspecified |
| mmu-miR-34a-5p | MIMAT0000542 | Lss | 16987 | HITS-CLIP | 23142080\|29167373 | Unspecified |
| mmu-miR-34a-5p | MIMAT0000542 | Lst1 | 16988 | HITS-CLIP | 23142080 | Unspecified |
| mmu-miR-34a-5p | MIMAT0000542 | Lta | 16992 | HITS-CLIP | 23142080 | Unspecified |
| mmu-miR-34a-5p | MIMAT0000542 | Lta4h | 16993 | HITS-CLIP | 23142080 | Unspecified |
| mmu-miR-34a-5p | MIMAT0000542 | Ltb | 16994 | HITS-CLIP | 23142080 | Unspecified |
| mmu-miR-34a-5p | MIMAT0000542 | Ltbp2 | 16997 | HITS-CLIP | 29167373 | Unspecified |
| mmu-miR-34a-5p | MIMAT0000542 | Ltbp3 | 16998 | HITS-CLIP | 23142080 | Unspecified |
| mmu-miR-34a-5p | MIMAT0000542 | Ltbp4 | 108075 | HITS-CLIP | 19536157 | Unspecified |
| mmu-miR-34a-5p | MIMAT0000542 | Ltn1 | 78913 | HITS-CLIP | 23142080 | Unspecified |
| mmu-miR-34a-5p | MIMAT0000542 | Luc7l2 | 192196 | HITS-CLIP | 23142080\|29167373 | Unspecified |
| mmu-miR-34a-5p | MIMAT0000542 | Luc7l3 | 67684 | HITS-CLIP | 23142080\|19536157\|29167373 | Unspecified |
| mmu-miR-34a-5p | MIMAT0000542 | Luzp1 | 269593 | HITS-CLIP | 23142080\|29167373 | Unspecified |
| mmu-miR-34a-5p | MIMAT0000542 | Luzp2 | 233271 | HITS-CLIP | 19536157 | Unspecified |
| mmu-miR-34a-5p | MIMAT0000542 | Lxn | 17035 | HITS-CLIP | 23142080 | Unspecified |
| mmu-miR-34a-5p | MIMAT0000542 | Ly6a | 110454 | HITS-CLIP | 23142080\|29167373 | Unspecified |
| mmu-miR-34a-5p | MIMAT0000542 | Ly6e | 17069 | HITS-CLIP | 37986224\|19536157\|29167373\|23142 | Unspecified |

| | | | | | | |
|---|---|---|---|---|---|---|
| mmu-miR-34a-5p | MIMAT0000542 | Ly6h | 23934 | HITS-CLIP | 19536157 | Unspecified |
| mmu-miR-34a-5p | MIMAT0000542 | Ly75 | 17076 | HITS-CLIP | 23142080 | Unspecified |
| mmu-miR-34a-5p | MIMAT0000542 | Ly9 | 17085 | HITS-CLIP | 23142080\|29167373 | Unspecified |
| mmu-miR-34a-5p | MIMAT0000542 | Ly96 | 17087 | HITS-CLIP | 29167373 | Unspecified |
| mmu-miR-34a-5p | MIMAT0000542 | Lyar | 17089 | HITS-CLIP | 23142080 | Unspecified |
| mmu-miR-34a-5p | MIMAT0000542 | Lyn | 17096 | HITS-CLIP | 29167373\|23142080 | Unspecified |
| mmu-miR-34a-5p | MIMAT0000542 | Lynx1 | 23936 | HITS-CLIP | 19536157 | Unspecified |
| mmu-miR-34a-5p | MIMAT0000542 | Lypla1 | 18777 | HITS-CLIP | 23142080 | Unspecified |
| mmu-miR-34a-5p | MIMAT0000542 | Lypla2 | 26394 | HITS-CLIP | 29167373 | Unspecified |
| mmu-miR-34a-5p | MIMAT0000542 | Lyrm2 | 108755 | HITS-CLIP | 37986224\|23142080 | Unspecified |
| mmu-miR-34a-5p | MIMAT0000542 | Lyrm4 | 380840 | HITS-CLIP | 23142080 | Unspecified |
| mmu-miR-34a-5p | MIMAT0000542 | Lysmd3 | 80289 | HITS-CLIP | 23142080 | Unspecified |
| mmu-miR-34a-5p | MIMAT0000542 | Lyst | 17101 | HITS-CLIP | 23142080\|29167373 | Unspecified |
| mmu-miR-34a-5p | MIMAT0000542 | Lztr1 | 66863 | HITS-CLIP | 23142080\|30971389 | Unspecified |
| mmu-miR-34a-5p | MIMAT0000542 | Lzts1 | 211134 | HITS-CLIP | 23142080 | Unspecified |
| mmu-miR-34a-5p | MIMAT0000542 | Lzts3 | 241638 | HITS-CLIP | 19536157 | Unspecified |
| mmu-miR-34a-5p | MIMAT0000542 | M6pr | 17113 | HITS-CLIP | 23142080\|29167373 | Unspecified |
| mmu-miR-34a-5p | MIMAT0000542 | MSN | 17698 | HITS-CLIP//qRT-PCR//Western blotting | 31433990\|23142080\|29167373\|32319 | Unspecified |
| mmu-miR-34a-5p | MIMAT0000542 | Macf1 | 11426 | HITS-CLIP | 31433990\|19536157\|23142080\|26203 | Unspecified |
| mmu-miR-34a-5p | MIMAT0000542 | Macir | 90355 | HITS-CLIP | 31433990 | Unspecified |
| mmu-miR-34a-5p | MIMAT0000542 | Maco1 | 55219 | HITS-CLIP | 23142080 | Unspecified |
| mmu-miR-34a-5p | MIMAT0000542 | Mad1l1 | 17120 | HITS-CLIP | 23142080 | Unspecified |
| mmu-miR-34a-5p | MIMAT0000542 | Mad2l1 | 56150 | HITS-CLIP | 23142080 | Unspecified |
| mmu-miR-34a-5p | MIMAT0000542 | Mad2l1bp | 66591 | HITS-CLIP | 23142080\|30971389\|29167373 | Unspecified |
| mmu-miR-34a-5p | MIMAT0000542 | Madd | 228355 | HITS-CLIP | 19536157\|23142080 | Unspecified |
| mmu-miR-34a-5p | MIMAT0000542 | Maea | 59003 | HITS-CLIP | 23142080 | Unspecified |
| mmu-miR-34a-5p | MIMAT0000542 | Maf | 17132 | HITS-CLIP | 23142080\|29167373 | Unspecified |
| mmu-miR-34a-5p | MIMAT0000542 | Maf1 | 68877 | HITS-CLIP | 23142080 | Unspecified |
| mmu-miR-34a-5p | MIMAT0000542 | Mafb | 16658 | HITS-CLIP | 29167373 | Unspecified |
| mmu-miR-34a-5p | MIMAT0000542 | Maff | 17133 | HITS-CLIP | 23142080\|29167373 | Unspecified |
| mmu-miR-34a-5p | MIMAT0000542 | Mafg | 17134 | HITS-CLIP | 31433990\|29167373 | Unspecified |
| mmu-miR-34a-5p | MIMAT0000542 | Mafk | 17135 | HITS-CLIP | 23142080\|30971389\|29167373 | Unspecified |
| mmu-miR-34a-5p | MIMAT0000542 | Maged1 | 94275 | HITS-CLIP | 19536157\|23142080\|29167373 | Unspecified |
| mmu-miR-34a-5p | MIMAT0000542 | Maged2 | 80884 | HITS-CLIP | 23142080 | Unspecified |
| mmu-miR-34a-5p | MIMAT0000542 | Magi2 | 50791 | HITS-CLIP | 19536157 | Unspecified |
| mmu-miR-34a-5p | MIMAT0000542 | Magi3 | 99470 | HITS-CLIP | 37986224\|23142080 | Unspecified |
| mmu-miR-34a-5p | MIMAT0000542 | Magoh | 17149 | HITS-CLIP | 23142080 | Unspecified |
| mmu-miR-34a-5p | MIMAT0000542 | Magohb | 66441 | HITS-CLIP | 23142080 | Unspecified |
| mmu-miR-34a-5p | MIMAT0000542 | Magt1 | 67075 | HITS-CLIP | 23142080\|19536157\|29167373 | Unspecified |

| | | | | | | |
|---|---|---|---|---|---|---|
| mmu-miR-34a-5p | MIMAT0000542 | Maip1 | 79568 | HITS-CLIP | 23142080 | Unspecified |
| mmu-miR-34a-5p | MIMAT0000542 | Mak16 | 67920 | HITS-CLIP | 23142080 | Unspecified |
| mmu-miR-34a-5p | MIMAT0000542 | Malsu1 | 75593 | HITS-CLIP | 29167373 | Unspecified |
| mmu-miR-34a-5p | MIMAT0000542 | Malt1 | 240354 | HITS-CLIP | 23142080 | Unspecified |
| mmu-miR-34a-5p | MIMAT0000542 | Mamdc4 | 158056 | HITS-CLIP | 29167373 | Unspecified |
| mmu-miR-34a-5p | MIMAT0000542 | Maml1 | 103806 | HITS-CLIP | 31433990\|23142080 | Unspecified |
| mmu-miR-34a-5p | MIMAT0000542 | Maml2 | 270118 | HITS-CLIP | 23142080 | Unspecified |
| mmu-miR-34a-5p | MIMAT0000542 | Maml3 | 433586 | HITS-CLIP | 23142080 | Unspecified |
| mmu-miR-34a-5p | MIMAT0000542 | Man1a | 17155 | HITS-CLIP | 23142080\|31433990 | Unspecified |
| mmu-miR-34a-5p | MIMAT0000542 | Man1a2 | 17156 | HITS-CLIP | 19536157\|23142080\|26203562\|29167 | Unspecified |
| mmu-miR-34a-5p | MIMAT0000542 | Man1c1 | 230815 | HITS-CLIP | 29167373 | Unspecified |
| mmu-miR-34a-5p | MIMAT0000542 | Man2a1 | 17158 | HITS-CLIP | 37986224\|19536157\|23142080\|29167 | Unspecified |
| mmu-miR-34a-5p | MIMAT0000542 | Man2b1 | 17159 | HITS-CLIP | 29167373\|23142080 | Unspecified |
| mmu-miR-34a-5p | MIMAT0000542 | Man2b2 | 17160 | HITS-CLIP | 29167373\|23142080 | Unspecified |
| mmu-miR-34a-5p | MIMAT0000542 | Man2c1 | 73744 | HITS-CLIP | 23142080\|29167373 | Unspecified |
| mmu-miR-34a-5p | MIMAT0000542 | Manbal | 69161 | HITS-CLIP | 31433990 | Unspecified |
| mmu-miR-34a-5p | MIMAT0000542 | Manea | 242362 | HITS-CLIP | 23142080 | Unspecified |
| mmu-miR-34a-5p | MIMAT0000542 | Manf | 74840 | HITS-CLIP | 23142080\|29167373 | Unspecified |
| mmu-miR-34a-5p | MIMAT0000542 | Map1a | 17754 | Microarrays//HITS-CLIP | 37986224\|23027903\|19536157 | Unspecified |
| mmu-miR-34a-5p | MIMAT0000542 | Map1b | 17755 | HITS-CLIP | 19536157 | Unspecified |
| mmu-miR-34a-5p | MIMAT0000542 | Map1lc3b | 67443 | HITS-CLIP | 23142080\|29167373 | Unspecified |
| mmu-miR-34a-5p | MIMAT0000542 | Map1s | 270058 | HITS-CLIP | 29167373 | Unspecified |
| mmu-miR-34a-5p | MIMAT0000542 | Map2 | 17756 | HITS-CLIP | 19536157 | Unspecified |
| mmu-miR-34a-5p | MIMAT0000542 | Map2k1 | 26395 | Microarrays//HITS-CLIP | 37986224\|23027903\|23142080\|29167 | Unspecified |
| mmu-miR-34a-5p | MIMAT0000542 | Map2k3 | 26397 | HITS-CLIP | 23142080\|26203562\|29167373 | Unspecified |
| mmu-miR-34a-5p | MIMAT0000542 | Map2k4 | 26398 | HITS-CLIP | 23142080\|29167373 | Unspecified |
| mmu-miR-34a-5p | MIMAT0000542 | Map2k5 | 23938 | HITS-CLIP | 23142080 | Unspecified |
| mmu-miR-34a-5p | MIMAT0000542 | Map2k7 | 26400 | HITS-CLIP | 31433990\|23630078\|29167373\|23142 | Unspecified |
| mmu-miR-34a-5p | MIMAT0000542 | Map3k1 | 26401 | HITS-CLIP | 19536157\|26203562\|23142080 | Unspecified |
| mmu-miR-34a-5p | MIMAT0000542 | Map3k11 | 26403 | Microarrays//HITS-CLIP | 37986224\|23027903\|23142080\|29167 | Unspecified |
| mmu-miR-34a-5p | MIMAT0000542 | Map3k12 | 26404 | HITS-CLIP | 23142080\|31433990 | Unspecified |
| mmu-miR-34a-5p | MIMAT0000542 | Map3k14 | 53859 | HITS-CLIP | 23142080\|31433990\|29167373 | Unspecified |
| mmu-miR-34a-5p | MIMAT0000542 | Map3k2 | 26405 | HITS-CLIP | 23142080\|29167373 | Unspecified |
| mmu-miR-34a-5p | MIMAT0000542 | Map3k20 | 65964 | HITS-CLIP | 29167373 | Unspecified |
| mmu-miR-34a-5p | MIMAT0000542 | Map3k3 | 26406 | HITS-CLIP | 19536157\|23142080\|29167373 | Unspecified |
| mmu-miR-34a-5p | MIMAT0000542 | Map3k4 | 26407 | HITS-CLIP | 23142080\|29167373 | Unspecified |
| mmu-miR-34a-5p | MIMAT0000542 | Map3k5 | 26408 | HITS-CLIP | 37986224\|31433990\|23142080 | Unspecified |
| mmu-miR-34a-5p | MIMAT0000542 | Map3k7 | 26409 | HITS-CLIP | 29167373\|23142080 | Unspecified |
| mmu-miR-34a-5p | MIMAT0000542 | Map3k7cl | 224419 | Microarrays | 37986224 | Unspecified |

| | | | | | | | |
|---|---|---|---|---|---|---|---|
| mmu-miR-34a-5p | MIMAT0000542 | Map3k8 | 26410 | HITS-CLIP | | 23142080 | Unspecified |
| mmu-miR-34a-5p | MIMAT0000542 | Map3k9 | 338372 | HITS-CLIP | 19536157\|29167373 | | Unspecified |
| mmu-miR-34a-5p | MIMAT0000542 | Map4 | 17758 | HITS-CLIP | 19536157\|31433990\|23142080\|29167 | | Unspecified |
| mmu-miR-34a-5p | MIMAT0000542 | Map4k1 | 26411 | HITS-CLIP | | 23142080 | Unspecified |
| mmu-miR-34a-5p | MIMAT0000542 | Map4k2 | 26412 | HITS-CLIP | | 23142080 | Unspecified |
| mmu-miR-34a-5p | MIMAT0000542 | Map4k3 | 225028 | HITS-CLIP | | 19536157 | Unspecified |
| mmu-miR-34a-5p | MIMAT0000542 | Map4k4 | 26921 | HITS-CLIP | | 23142080 | Unspecified |
| mmu-miR-34a-5p | MIMAT0000542 | Map4k5 | 399510 | HITS-CLIP | | 19536157 | Unspecified |
| mmu-miR-34a-5p | MIMAT0000542 | Map6 | 17760 | HITS-CLIP | | 19536157 | Unspecified |
| mmu-miR-34a-5p | MIMAT0000542 | Map7 | 17761 | HITS-CLIP | | 23142080 | Unspecified |
| mmu-miR-34a-5p | MIMAT0000542 | Map7d1 | 245877 | HITS-CLIP | 31433990\|29167373 | | Unspecified |
| mmu-miR-34a-5p | MIMAT0000542 | Map9 | 213582 | HITS-CLIP | 19536157\|23142080 | | Unspecified |
| mmu-miR-34a-5p | MIMAT0000542 | Mapk1 | 26413 | HITS-CLIP | 19536157\|23142080\|29167373 | | Unspecified |
| mmu-miR-34a-5p | MIMAT0000542 | Mapk14 | 26416 | Microarrays//HITS-CLIP | 37986224\|23027903\|23142080\|29167 | | Unspecified |
| mmu-miR-34a-5p | MIMAT0000542 | Mapk1ip1l | 218975 | HITS-CLIP, Microarrays//Microarrays//HITS | 37986224\|23027903\|23142080\|29167 | | Unspecified |
| mmu-miR-34a-5p | MIMAT0000542 | Mapk3 | 26417 | HITS-CLIP | | 23142080 | Unspecified |
| mmu-miR-34a-5p | MIMAT0000542 | Mapk6 | 50772 | HITS-CLIP | 23142080\|29167373 | | Unspecified |
| mmu-miR-34a-5p | MIMAT0000542 | Mapk7 | 23939 | HITS-CLIP | | 23142080 | Unspecified |
| mmu-miR-34a-5p | MIMAT0000542 | Mapk8 | 26419 | HITS-CLIP | | 23142080 | Unspecified |
| mmu-miR-34a-5p | MIMAT0000542 | Mapk8ip2 | 60597 | HITS-CLIP | | 19536157 | Unspecified |
| mmu-miR-34a-5p | MIMAT0000542 | Mapk8ip3 | 30957 | HITS-CLIP | 23142080\|29167373 | | Unspecified |
| mmu-miR-34a-5p | MIMAT0000542 | Mapk9 | 26420 | HITS-CLIP | 23142080\|29167373 | | Unspecified |
| mmu-miR-34a-5p | MIMAT0000542 | Mapkapk2 | 17164 | HITS-CLIP | 23142080\|29167373 | | Unspecified |
| mmu-miR-34a-5p | MIMAT0000542 | Mapkapk3 | 102626 | HITS-CLIP | | 23142080 | Unspecified |
| mmu-miR-34a-5p | MIMAT0000542 | Mapkbp1 | 26390 | HITS-CLIP | | 23142080 | Unspecified |
| mmu-miR-34a-5p | MIMAT0000542 | Mapre1 | 13589 | HITS-CLIP | 21258322\|31433990\|23142080\|29167 | | Unspecified |
| mmu-miR-34a-5p | MIMAT0000542 | Mapre2 | 212307 | HITS-CLIP | 23142080\|29167373 | | Unspecified |
| mmu-miR-34a-5p | MIMAT0000542 | Mapre3 | 100732 | HITS-CLIP | | 23142080 | Unspecified |
| mmu-miR-34a-5p | MIMAT0000542 | Mapt | 17762 | HITS-CLIP | | 19536157 | Unspecified |
| mmu-miR-34a-5p | MIMAT0000542 | Marc2 | 67247 | HITS-CLIP | | 23142080 | Unspecified |
| mmu-miR-34a-5p | MIMAT0000542 | March2 | 51257 | HITS-CLIP | 23142080\|29167373 | | Unspecified |
| mmu-miR-34a-5p | MIMAT0000542 | March5 | 54708 | HITS-CLIP | | 29167373 | Unspecified |
| mmu-miR-34a-5p | MIMAT0000542 | March7 | 57438 | HITS-CLIP | 23142080\|29167373 | | Unspecified |
| mmu-miR-34a-5p | MIMAT0000542 | March8 | 71779 | HITS-CLIP | 19536157\|29167373 | | Unspecified |
| mmu-miR-34a-5p | MIMAT0000542 | March9 | 216438 | HITS-CLIP | | 23142080 | Unspecified |
| mmu-miR-34a-5p | MIMAT0000542 | Marchf6 | 223455 | HITS-CLIP | 37986224\|19536157\|23142080\|29167 | | Unspecified |
| mmu-miR-34a-5p | MIMAT0000542 | Marchf8 | 220972 | HITS-CLIP | | 23630078 | Unspecified |
| mmu-miR-34a-5p | MIMAT0000542 | Marcks | 17118 | HITS-CLIP | 19536157\|23142080\|29167373 | | Unspecified |
| mmu-miR-34a-5p | MIMAT0000542 | Marcksl1 | 17357 | HITS-CLIP | 29167373\|23142080 | | Unspecified |

| | | | | | | |
|---|---|---|---|---|---|---|
| mmu-miR-34a-5p | MIMAT0000542 | Marf1 | 223989 | HITS-CLIP | 23142080\|29167373 | Unspecified |
| mmu-miR-34a-5p | MIMAT0000542 | Mark2 | 13728 | HITS-CLIP | 19536157\|23142080\|29167373 | Unspecified |
| mmu-miR-34a-5p | MIMAT0000542 | Mark3 | 17169 | HITS-CLIP | 19536157\|23142080\|26203562 | Unspecified |
| mmu-miR-34a-5p | MIMAT0000542 | Mark4 | 232944 | HITS-CLIP | 23142080 | Unspecified |
| mmu-miR-34a-5p | MIMAT0000542 | Mars | 216443 | HITS-CLIP | 23142080 | Unspecified |
| mmu-miR-34a-5p | MIMAT0000542 | Mars2 | 212679 | HITS-CLIP | 31433990\|23142080 | Unspecified |
| mmu-miR-34a-5p | MIMAT0000542 | Marveld2 | 218518 | HITS-CLIP | 23142080\|31433990 | Unspecified |
| mmu-miR-34a-5p | MIMAT0000542 | Maskbp3 | 404734 | HITS-CLIP | 31433990 | Unspecified |
| mmu-miR-34a-5p | MIMAT0000542 | Mast2 | 17776 | HITS-CLIP | 19536157\|23142080\|29167373 | Unspecified |
| mmu-miR-34a-5p | MIMAT0000542 | Mast3 | 546071 | HITS-CLIP | 29167373 | Unspecified |
| mmu-miR-34a-5p | MIMAT0000542 | Mast4 | 328329 | HITS-CLIP | 23142080 | Unspecified |
| mmu-miR-34a-5p | MIMAT0000542 | Mastl | 67121 | HITS-CLIP | 23142080 | Unspecified |
| mmu-miR-34a-5p | MIMAT0000542 | Mat2a | 232087 | HITS-CLIP | 23142080\|19536157\|29167373 | Unspecified |
| mmu-miR-34a-5p | MIMAT0000542 | Mat2b | 108645 | HITS-CLIP | 23142080 | Unspecified |
| mmu-miR-34a-5p | MIMAT0000542 | Matk | 4145 | HITS-CLIP | 23142080 | Unspecified |
| mmu-miR-34a-5p | MIMAT0000542 | Matr3 | 17184 | HITS-CLIP | 23142080\|19536157\|29167373 | Unspecified |
| mmu-miR-34a-5p | MIMAT0000542 | Mau2 | 74549 | HITS-CLIP | 19536157\|23630078\|23142080\|29167 | Unspecified |
| mmu-miR-34a-5p | MIMAT0000542 | Mavs | 228607 | HITS-CLIP | 30971389\|23142080\|29167373 | Unspecified |
| mmu-miR-34a-5p | MIMAT0000542 | Max | 17187 | HITS-CLIP | 19536157\|23142080 | Unspecified |
| mmu-miR-34a-5p | MIMAT0000542 | Maz | 17188 | HITS-CLIP | 23142080\|29167373 | Unspecified |
| mmu-miR-34a-5p | MIMAT0000542 | Mb | 17189 | Microarrays | 37986224 | Unspecified |
| mmu-miR-34a-5p | MIMAT0000542 | Mb21d2 | 239796 | HITS-CLIP | 19536157 | Unspecified |
| mmu-miR-34a-5p | MIMAT0000542 | Mbd1 | 17190 | HITS-CLIP | 23142080 | Unspecified |
| mmu-miR-34a-5p | MIMAT0000542 | Mbd2 | 17191 | HITS-CLIP | 19536157\|23142080\|29167373 | Unspecified |
| mmu-miR-34a-5p | MIMAT0000542 | Mbd3 | 17192 | HITS-CLIP | 23142080\|29167373 | Unspecified |
| mmu-miR-34a-5p | MIMAT0000542 | Mbd4 | 17193 | HITS-CLIP | 21258322\|23142080 | Unspecified |
| mmu-miR-34a-5p | MIMAT0000542 | Mbd5 | 109241 | HITS-CLIP | 19536157\|23142080\|29167373 | Unspecified |
| mmu-miR-34a-5p | MIMAT0000542 | Mbd6 | 110962 | HITS-CLIP | 23142080\|29167373 | Unspecified |
| mmu-miR-34a-5p | MIMAT0000542 | Mbip | 217588 | HITS-CLIP | 23142080\|29167373 | Unspecified |
| mmu-miR-34a-5p | MIMAT0000542 | Mblac2 | 72852 | HITS-CLIP | 23142080 | Unspecified |
| mmu-miR-34a-5p | MIMAT0000542 | Mbnl1 | 56758 | HITS-CLIP | 37986224\|19536157\|31433990\|23142 | Unspecified |
| mmu-miR-34a-5p | MIMAT0000542 | Mbnl2 | 105559 | HITS-CLIP | 19536157\|23142080\|29167373 | Unspecified |
| mmu-miR-34a-5p | MIMAT0000542 | Mbnl3 | 171170 | HITS-CLIP | 23142080 | Unspecified |
| mmu-miR-34a-5p | MIMAT0000542 | Mboat1 | 218121 | Microarrays//HITS-CLIP | 37986224\|23027903\|23142080 | Unspecified |
| mmu-miR-34a-5p | MIMAT0000542 | Mboat2 | 67216 | HITS-CLIP | 23142080 | Unspecified |
| mmu-miR-34a-5p | MIMAT0000542 | Mbp | 17196 | HITS-CLIP | 19536157\|23142080\|29167373 | Unspecified |
| mmu-miR-34a-5p | MIMAT0000542 | Mbtd1 | 103537 | HITS-CLIP | 23142080 | Unspecified |
| mmu-miR-34a-5p | MIMAT0000542 | Mbtps1 | 56453 | HITS-CLIP | 23142080\|29167373 | Unspecified |
| mmu-miR-34a-5p | MIMAT0000542 | Mbtps2 | 270669 | HITS-CLIP | 29167373\|23142080 | Unspecified |

| | | | | | | | |
|---|---|---|---|---|---|---|---|
| mmu-miR-34a-5p | MIMAT0000542 | Mcam | 84004 | HITS-CLIP | | 26203562 | Unspecified |
| mmu-miR-34a-5p | MIMAT0000542 | Mcat | 223722 | HITS-CLIP | 19536157\|23142080 | | Unspecified |
| mmu-miR-34a-5p | MIMAT0000542 | Mccc2 | 78038 | HITS-CLIP | | 23142080 | Unspecified |
| mmu-miR-34a-5p | MIMAT0000542 | Mcee | 73724 | HITS-CLIP | | 23142080 | Unspecified |
| mmu-miR-34a-5p | MIMAT0000542 | Mcf2l | 17207 | HITS-CLIP | 23142080\|19536157 | | Unspecified |
| mmu-miR-34a-5p | MIMAT0000542 | Mcfd2 | 193813 | HITS-CLIP | | 23142080 | Unspecified |
| mmu-miR-34a-5p | MIMAT0000542 | Mcl1 | 17210 | HITS-CLIP | 23630078\|29167373\|23142080 | | Unspecified |
| mmu-miR-34a-5p | MIMAT0000542 | Mcm10 | 70024 | Microarrays//HITS-CLIP | 37986224\|23027903\|23142080 | | Unspecified |
| mmu-miR-34a-5p | MIMAT0000542 | Mcm2 | 17216 | HITS-CLIP | 23630078\|31433990\|23142080 | | Unspecified |
| mmu-miR-34a-5p | MIMAT0000542 | Mcm3 | 17215 | Microarrays//HITS-CLIP | 37986224\|23027903\|23142080 | | Unspecified |
| mmu-miR-34a-5p | MIMAT0000542 | Mcm3ap | 54387 | HITS-CLIP | | 23142080 | Unspecified |
| mmu-miR-34a-5p | MIMAT0000542 | Mcm4 | 17217 | HITS-CLIP | | 23142080 | Unspecified |
| mmu-miR-34a-5p | MIMAT0000542 | Mcm5 | 17218 | HITS-CLIP, Microarrays//HITS-CLIP//Microa | 37986224\|23142080\|23027903\|29167 | | Unspecified |
| mmu-miR-34a-5p | MIMAT0000542 | Mcm6 | 17219 | HITS-CLIP | 37986224\|23142080\|31433990\|29167 | | Unspecified |
| mmu-miR-34a-5p | MIMAT0000542 | Mcm7 | 17220 | HITS-CLIP | 23630078\|23142080\|30971389 | | Unspecified |
| mmu-miR-34a-5p | MIMAT0000542 | Mcm9 | 71567 | HITS-CLIP | | 29167373 | Unspecified |
| mmu-miR-34a-5p | MIMAT0000542 | Mcmbp | 210711 | HITS-CLIP | 19536157\|23142080\|29167373 | | Unspecified |
| mmu-miR-34a-5p | MIMAT0000542 | Mcph1 | 244329 | HITS-CLIP | | 29167373 | Unspecified |
| mmu-miR-34a-5p | MIMAT0000542 | Mcrip1 | 348262 | HITS-CLIP | 19536157\|23142080 | | Unspecified |
| mmu-miR-34a-5p | MIMAT0000542 | Mctp1 | 78771 | HITS-CLIP | | 23142080 | Unspecified |
| mmu-miR-34a-5p | MIMAT0000542 | Mctp2 | 244049 | HITS-CLIP | | 23142080 | Unspecified |
| mmu-miR-34a-5p | MIMAT0000542 | Mcts1 | 68995 | HITS-CLIP | | 23142080 | Unspecified |
| mmu-miR-34a-5p | MIMAT0000542 | Mcu | 215999 | HITS-CLIP | | 23142080 | Unspecified |
| mmu-miR-34a-5p | MIMAT0000542 | Mcub | 11839 | HITS-CLIP | | 23142080 | Unspecified |
| mmu-miR-34a-5p | MIMAT0000542 | Mdc1 | 240087 | HITS-CLIP | 23142080\|29167373 | | Unspecified |
| mmu-miR-34a-5p | MIMAT0000542 | Mdfic | 16543 | HITS-CLIP | 23142080\|31433990\|26203562\|29167 | | Unspecified |
| mmu-miR-34a-5p | MIMAT0000542 | Mdga2 | 320772 | HITS-CLIP | | 19536157 | Unspecified |
| mmu-miR-34a-5p | MIMAT0000542 | Mdh1 | 17449 | HITS-CLIP | 19536157\|23142080 | | Unspecified |
| mmu-miR-34a-5p | MIMAT0000542 | Mdh2 | 17448 | HITS-CLIP | 23142080\|19536157 | | Unspecified |
| mmu-miR-34a-5p | MIMAT0000542 | Mdm2 | 17246 | HITS-CLIP | 37986224\|23630078\|30971389\|23142 | | Unspecified |
| mmu-miR-34a-5p | MIMAT0000542 | Mdm4 | 17248 | HITS-CLIP, Microarrays//HITS-CLIP//Microa | 37986224\|23142080\|23027903\|29167 | | Unspecified |
| mmu-miR-34a-5p | MIMAT0000542 | Mdn1 | 100019 | HITS-CLIP | 23630078\|31433990\|30971389\|23142 | | Unspecified |
| mmu-miR-34a-5p | MIMAT0000542 | Me2 | 107029 | HITS-CLIP | | 23142080 | Unspecified |
| mmu-miR-34a-5p | MIMAT0000542 | Meaf6 | 70088 | HITS-CLIP | 19536157\|29167373 | | Unspecified |
| mmu-miR-34a-5p | MIMAT0000542 | Mecp2 | 17257 | HITS-CLIP | 31433990\|23142080\|29167373 | | Unspecified |
| mmu-miR-34a-5p | MIMAT0000542 | Med1 | 19014 | HITS-CLIP | 23142080\|29167373 | | Unspecified |
| mmu-miR-34a-5p | MIMAT0000542 | Med10 | 28077 | HITS-CLIP | 23142080\|29167373 | | Unspecified |
| mmu-miR-34a-5p | MIMAT0000542 | Med12 | 59024 | HITS-CLIP | 23142080\|29167373 | | Unspecified |
| mmu-miR-34a-5p | MIMAT0000542 | Med12l | 329650 | HITS-CLIP | | 23142080 | Unspecified |

| | | | | | | |
|---|---|---|---|---|---|---|
| mmu-miR-34a-5p | MIMAT0000542 | Med13 | 327987 | HITS-CLIP | 19536157\|31433990\|29167373\|23142 | Unspecified |
| mmu-miR-34a-5p | MIMAT0000542 | Med13l | 76199 | HITS-CLIP | 31433990\|23142080\|29167373 | Unspecified |
| mmu-miR-34a-5p | MIMAT0000542 | Med14 | 26896 | HITS-CLIP | 19536157\|23142080 | Unspecified |
| mmu-miR-34a-5p | MIMAT0000542 | Med15 | 94112 | HITS-CLIP | 29167373 | Unspecified |
| mmu-miR-34a-5p | MIMAT0000542 | Med16 | 216154 | HITS-CLIP | 23142080 | Unspecified |
| mmu-miR-34a-5p | MIMAT0000542 | Med17 | 234959 | HITS-CLIP | 31433990\|23142080 | Unspecified |
| mmu-miR-34a-5p | MIMAT0000542 | Med18 | 67219 | HITS-CLIP | 23142080 | Unspecified |
| mmu-miR-34a-5p | MIMAT0000542 | Med19 | 381379 | HITS-CLIP | 23142080\|29167373 | Unspecified |
| mmu-miR-34a-5p | MIMAT0000542 | Med20 | 56771 | HITS-CLIP | 23142080\|29167373 | Unspecified |
| mmu-miR-34a-5p | MIMAT0000542 | Med21 | 108098 | HITS-CLIP | 23142080\|29167373 | Unspecified |
| mmu-miR-34a-5p | MIMAT0000542 | Med22 | 20933 | HITS-CLIP//Microarrays | 23142080\|23027903 | Unspecified |
| mmu-miR-34a-5p | MIMAT0000542 | Med24 | 23989 | HITS-CLIP | 23142080 | Unspecified |
| mmu-miR-34a-5p | MIMAT0000542 | Med25 | 75613 | HITS-CLIP | 23142080\|31433990\|29167373 | Unspecified |
| mmu-miR-34a-5p | MIMAT0000542 | Med26 | 70625 | HITS-CLIP | 23142080 | Unspecified |
| mmu-miR-34a-5p | MIMAT0000542 | Med27 | 68975 | HITS-CLIP | 23142080 | Unspecified |
| mmu-miR-34a-5p | MIMAT0000542 | Med28 | 66999 | HITS-CLIP | 29167373 | Unspecified |
| mmu-miR-34a-5p | MIMAT0000542 | Med29 | 67224 | HITS-CLIP | 23142080 | Unspecified |
| mmu-miR-34a-5p | MIMAT0000542 | Med30 | 69790 | HITS-CLIP | 23142080 | Unspecified |
| mmu-miR-34a-5p | MIMAT0000542 | Med31 | 67279 | HITS-CLIP | 23142080 | Unspecified |
| mmu-miR-34a-5p | MIMAT0000542 | Med4 | 67381 | HITS-CLIP | 23142080 | Unspecified |
| mmu-miR-34a-5p | MIMAT0000542 | Med6 | 69792 | HITS-CLIP | 23142080\|29167373 | Unspecified |
| mmu-miR-34a-5p | MIMAT0000542 | Med7 | 66213 | HITS-CLIP | 23142080\|29167373 | Unspecified |
| mmu-miR-34a-5p | MIMAT0000542 | Med8 | 80509 | HITS-CLIP | 23142080 | Unspecified |
| mmu-miR-34a-5p | MIMAT0000542 | Mef2a | 17258 | Microarrays//HITS-CLIP | 37986224\|23142080\|30971389\|1953€ | Unspecified |
| mmu-miR-34a-5p | MIMAT0000542 | Mef2c | 17260 | HITS-CLIP | 23142080\|29167373 | Unspecified |
| mmu-miR-34a-5p | MIMAT0000542 | Mef2d | 17261 | HITS-CLIP | 23142080\|29167373 | Unspecified |
| mmu-miR-34a-5p | MIMAT0000542 | Megf6 | 230971 | HITS-CLIP | 26203562 | Unspecified |
| mmu-miR-34a-5p | MIMAT0000542 | Megf9 | 230316 | HITS-CLIP | 19536157 | Unspecified |
| mmu-miR-34a-5p | MIMAT0000542 | Meig1 | 104362 | Microarrays | 37986224 | Unspecified |
| mmu-miR-34a-5p | MIMAT0000542 | Meis2 | 17536 | HITS-CLIP | 19536157 | Unspecified |
| mmu-miR-34a-5p | MIMAT0000542 | Meis3 | 17537 | HITS-CLIP | 29167373 | Unspecified |
| mmu-miR-34a-5p | MIMAT0000542 | Melk | 17279 | HITS-CLIP | 23142080 | Unspecified |
| mmu-miR-34a-5p | MIMAT0000542 | Men1 | 17283 | HITS-CLIP | 30971389 | Unspecified |
| mmu-miR-34a-5p | MIMAT0000542 | Mepce | 231803 | HITS-CLIP | 23142080 | Unspecified |
| mmu-miR-34a-5p | MIMAT0000542 | Mertk | 17289 | HITS-CLIP | 29167373 | Unspecified |
| mmu-miR-34a-5p | MIMAT0000542 | Mesd | 67943 | HITS-CLIP | 23142080\|29167373 | Unspecified |
| mmu-miR-34a-5p | MIMAT0000542 | Met | 17295 | Microarrays//HITS-CLIP | 37986224\|23027903\|30971389 | Unspecified |
| mmu-miR-34a-5p | MIMAT0000542 | Metap1 | 75624 | HITS-CLIP | 37986224\|23142080\|29167373 | Unspecified |
| mmu-miR-34a-5p | MIMAT0000542 | Metap2 | 56307 | HITS-CLIP | 23142080 | Unspecified |

| | | | | | | |
|---|---|---|---|---|---|---|
| mmu-miR-34a-5p | MIMAT0000542 | Metrnl | 210029 | HITS-CLIP | 29167373 | Unspecified |
| mmu-miR-34a-5p | MIMAT0000542 | Mettl1 | 17299 | HITS-CLIP | 23142080 | Unspecified |
| mmu-miR-34a-5p | MIMAT0000542 | Mettl14 | 210529 | HITS-CLIP | 23142080 | Unspecified |
| mmu-miR-34a-5p | MIMAT0000542 | Mettl21a | 67099 | HITS-CLIP | 23142080 | Unspecified |
| mmu-miR-34a-5p | MIMAT0000542 | Mettl25 | 216292 | HITS-CLIP | 23142080 | Unspecified |
| mmu-miR-34a-5p | MIMAT0000542 | Mettl4 | 76781 | HITS-CLIP | 23142080 | Unspecified |
| mmu-miR-34a-5p | MIMAT0000542 | Mettl5 | 75422 | HITS-CLIP | 23142080 | Unspecified |
| mmu-miR-34a-5p | MIMAT0000542 | Mettl6 | 67011 | HITS-CLIP | 23142080 | Unspecified |
| mmu-miR-34a-5p | MIMAT0000542 | Mettl8 | 228019 | HITS-CLIP | 23142080 | Unspecified |
| mmu-miR-34a-5p | MIMAT0000542 | Mettl9 | 59052 | HITS-CLIP | 30971389\|23142080 | Unspecified |
| mmu-miR-34a-5p | MIMAT0000542 | Mex3a | 72640 | Microarrays | 37986224 | Unspecified |
| mmu-miR-34a-5p | MIMAT0000542 | Mex3b | 108797 | Microarrays//HITS-CLIP | 37986224\|23630078\|29167373 | Unspecified |
| mmu-miR-34a-5p | MIMAT0000542 | Mex3c | 240396 | HITS-CLIP, Microarrays//Microarrays//HIT | 37986224\|23027903\|23142080\|29167 | Unspecified |
| mmu-miR-34a-5p | MIMAT0000542 | Mfap3 | 216760 | HITS-CLIP | 37986224\|19536157\|23142080\|29167 | Unspecified |
| mmu-miR-34a-5p | MIMAT0000542 | Mfap3l | 71306 | HITS-CLIP | 19536157 | Unspecified |
| mmu-miR-34a-5p | MIMAT0000542 | Mff | 75734 | HITS-CLIP | 23142080\|29167373 | Unspecified |
| mmu-miR-34a-5p | MIMAT0000542 | Mfhas1 | 52065 | HITS-CLIP | 23142080\|29167373 | Unspecified |
| mmu-miR-34a-5p | MIMAT0000542 | Mfn1 | 67414 | HITS-CLIP | 23142080 | Unspecified |
| mmu-miR-34a-5p | MIMAT0000542 | Mfn2 | 170731 | HITS-CLIP | 23142080 | Unspecified |
| mmu-miR-34a-5p | MIMAT0000542 | Mfng | 17305 | HITS-CLIP | 23142080 | Unspecified |
| mmu-miR-34a-5p | MIMAT0000542 | Mfsd1 | 66868 | HITS-CLIP | 23142080\|29167373 | Unspecified |
| mmu-miR-34a-5p | MIMAT0000542 | Mfsd10 | 68294 | HITS-CLIP | 23142080 | Unspecified |
| mmu-miR-34a-5p | MIMAT0000542 | Mfsd11 | 69900 | HITS-CLIP | 23142080\|19536157 | Unspecified |
| mmu-miR-34a-5p | MIMAT0000542 | Mfsd12 | 73822 | HITS-CLIP | 29167373 | Unspecified |
| mmu-miR-34a-5p | MIMAT0000542 | Mfsd14a | 15247 | HITS-CLIP | 23142080\|29167373 | Unspecified |
| mmu-miR-34a-5p | MIMAT0000542 | Mfsd14b | 84641 | HITS-CLIP | 23142080\|29167373 | Unspecified |
| mmu-miR-34a-5p | MIMAT0000542 | Mfsd2a | 76574 | HITS-CLIP | 19536157\|31433990\|23142080 | Unspecified |
| mmu-miR-34a-5p | MIMAT0000542 | Mfsd4b4 | 213332 | HITS-CLIP | 23142080\|19536157\|29167373 | Unspecified |
| mmu-miR-34a-5p | MIMAT0000542 | Mfsd5 | 106073 | HITS-CLIP | 29167373 | Unspecified |
| mmu-miR-34a-5p | MIMAT0000542 | Mfsd6 | 98682 | HITS-CLIP | 23142080 | Unspecified |
| mmu-miR-34a-5p | MIMAT0000542 | Mfsd8 | 72175 | HITS-CLIP | 23142080 | Unspecified |
| mmu-miR-34a-5p | MIMAT0000542 | Mga | 29808 | HITS-CLIP | 31433990\|19536157\|23142080\|29167 | Unspecified |
| mmu-miR-34a-5p | MIMAT0000542 | Mgarp | 67749 | HITS-CLIP | 23142080 | Unspecified |
| mmu-miR-34a-5p | MIMAT0000542 | Mgat1 | 17308 | HITS-CLIP | 23142080\|31433990 | Unspecified |
| mmu-miR-34a-5p | MIMAT0000542 | Mgat2 | 217664 | HITS-CLIP | 19536157\|23142080\|29167373 | Unspecified |
| mmu-miR-34a-5p | MIMAT0000542 | Mgat4a | 269181 | HITS-CLIP | 23142080\|29167373 | Unspecified |
| mmu-miR-34a-5p | MIMAT0000542 | Mgat4b | 103534 | HITS-CLIP | 23142080 | Unspecified |
| mmu-miR-34a-5p | MIMAT0000542 | Mgat5 | 107895 | HITS-CLIP | 29167373 | Unspecified |
| mmu-miR-34a-5p | MIMAT0000542 | Mgea5 | 76055 | HITS-CLIP | 23142080\|29167373 | Unspecified |

| | | | | | | |
|---|---|---|---|---|---|---|
| mmu-miR-34a-5p | MIMAT0000542 | Mgll | 23945 | HITS-CLIP | 37986224\|23142080\|31433990\|19536 | Unspecified |
| mmu-miR-34a-5p | MIMAT0000542 | Mgrn1 | 17237 | HITS-CLIP | 23142080\|29167373 | Unspecified |
| mmu-miR-34a-5p | MIMAT0000542 | Mgst3 | 66447 | HITS-CLIP | 19536157 | Unspecified |
| mmu-miR-34a-5p | MIMAT0000542 | Mib1 | 225164 | HITS-CLIP | 31433990\|19536157\|30971389\|29167 | Unspecified |
| mmu-miR-34a-5p | MIMAT0000542 | Mib2 | 76580 | HITS-CLIP | 23142080 | Unspecified |
| mmu-miR-34a-5p | MIMAT0000542 | Mical1 | 171580 | HITS-CLIP | 23142080\|29167373 | Unspecified |
| mmu-miR-34a-5p | MIMAT0000542 | Mical2 | 320878 | HITS-CLIP | 29167373 | Unspecified |
| mmu-miR-34a-5p | MIMAT0000542 | Micall1 | 27008 | HITS-CLIP | 26203562 | Unspecified |
| mmu-miR-34a-5p | MIMAT0000542 | Micall2 | 231830 | HITS-CLIP | 29167373 | Unspecified |
| mmu-miR-34a-5p | MIMAT0000542 | Micu1 | 216001 | HITS-CLIP | 29167373 | Unspecified |
| mmu-miR-34a-5p | MIMAT0000542 | Micu2 | 68514 | HITS-CLIP | 23142080\|29167373 | Unspecified |
| mmu-miR-34a-5p | MIMAT0000542 | Micu3 | 78506 | HITS-CLIP | 23142080 | Unspecified |
| mmu-miR-34a-5p | MIMAT0000542 | Mid1ip1 | 68041 | HITS-CLIP | 23142080 | Unspecified |
| mmu-miR-34a-5p | MIMAT0000542 | Mideas | 91748 | HITS-CLIP | 31433990 | Unspecified |
| mmu-miR-34a-5p | MIMAT0000542 | Midn | 59090 | HITS-CLIP | 23142080\|29167373 | Unspecified |
| mmu-miR-34a-5p | MIMAT0000542 | Mief1 | 239555 | Microarrays//HITS-CLIP | 37986224\|23027903\|23142080\|29167 | Unspecified |
| mmu-miR-34a-5p | MIMAT0000542 | Mier1 | 71148 | HITS-CLIP | 29167373 | Unspecified |
| mmu-miR-34a-5p | MIMAT0000542 | Mier2 | 70427 | HITS-CLIP | 23142080\|31433990\|29167373 | Unspecified |
| mmu-miR-34a-5p | MIMAT0000542 | Mier3 | 218613 | HITS-CLIP | 19536157\|23142080 | Unspecified |
| mmu-miR-34a-5p | MIMAT0000542 | Mif | 17319 | HITS-CLIP | 29167373 | Unspecified |
| mmu-miR-34a-5p | MIMAT0000542 | Mif4gd | 69674 | HITS-CLIP | 30971389\|31433990\|23142080 | Unspecified |
| mmu-miR-34a-5p | MIMAT0000542 | Miga1 | 215708 | HITS-CLIP | 31433990\|23142080 | Unspecified |
| mmu-miR-34a-5p | MIMAT0000542 | Miga2 | 108958 | HITS-CLIP | 29167373 | Unspecified |
| mmu-miR-34a-5p | MIMAT0000542 | Milr1 | 380732 | HITS-CLIP | 23142080\|29167373 | Unspecified |
| mmu-miR-34a-5p | MIMAT0000542 | Mindy2 | 235461 | HITS-CLIP | 19536157\|23142080\|29167373 | Unspecified |
| mmu-miR-34a-5p | MIMAT0000542 | Mindy3 | 80013 | HITS-CLIP | 23142080 | Unspecified |
| mmu-miR-34a-5p | MIMAT0000542 | Mink1 | 50932 | HITS-CLIP | 23142080\|29167373 | Unspecified |
| mmu-miR-34a-5p | MIMAT0000542 | Minos1 | 440574 | HITS-CLIP | 23142080 | Unspecified |
| mmu-miR-34a-5p | MIMAT0000542 | Minpp1 | 17330 | HITS-CLIP | 23142080\|29167373 | Unspecified |
| mmu-miR-34a-5p | MIMAT0000542 | Mios | 252875 | HITS-CLIP | 23142080 | Unspecified |
| mmu-miR-34a-5p | MIMAT0000542 | Mis18bp1 | 217653 | HITS-CLIP | 23142080 | Unspecified |
| mmu-miR-34a-5p | MIMAT0000542 | Mitd1 | 69028 | HITS-CLIP | 23142080 | Unspecified |
| mmu-miR-34a-5p | MIMAT0000542 | Mitf | 17342 | HITS-CLIP | 19536157\|29167373 | Unspecified |
| mmu-miR-34a-5p | MIMAT0000542 | Mki67 | 17345 | HITS-CLIP | 19536157\|31433990\|23142080\|29167 | Unspecified |
| mmu-miR-34a-5p | MIMAT0000542 | Mkks | 59030 | Microarrays//HITS-CLIP | 37986224\|23027903\|23142080 | Unspecified |
| mmu-miR-34a-5p | MIMAT0000542 | Mkln1 | 27418 | HITS-CLIP | 23142080\|29167373 | Unspecified |
| mmu-miR-34a-5p | MIMAT0000542 | Mknk2 | 17347 | HITS-CLIP | 37986224\|23142080\|29167373 | Unspecified |
| mmu-miR-34a-5p | MIMAT0000542 | Mkrn1 | 54484 | HITS-CLIP | 23142080 | Unspecified |
| mmu-miR-34a-5p | MIMAT0000542 | Mkrn2 | 67027 | HITS-CLIP | 31433990 | Unspecified |

| | | | | | | |
|---|---|---|---|---|---|---|
| mmu-miR-34a-5p | MIMAT0000542 | Mlec | 109154 | HITS-CLIP | 19536157\|23630078\|23142080\|29167 | Unspecified |
| mmu-miR-34a-5p | MIMAT0000542 | Mlf2 | 30853 | HITS-CLIP | 23142080 | Unspecified |
| mmu-miR-34a-5p | MIMAT0000542 | Mlh1 | 17350 | HITS-CLIP | 23142080 | Unspecified |
| mmu-miR-34a-5p | MIMAT0000542 | Mlh3 | 217716 | HITS-CLIP | 23142080\|29167373 | Unspecified |
| mmu-miR-34a-5p | MIMAT0000542 | Mlip | 69642 | HITS-CLIP | 19536157 | Unspecified |
| mmu-miR-34a-5p | MIMAT0000542 | Mlkl | 74568 | HITS-CLIP | 23142080 | Unspecified |
| mmu-miR-34a-5p | MIMAT0000542 | Mllt1 | 64144 | HITS-CLIP | 23142080\|26203562 | Unspecified |
| mmu-miR-34a-5p | MIMAT0000542 | Mllt10 | 17354 | HITS-CLIP | 23142080\|29167373 | Unspecified |
| mmu-miR-34a-5p | MIMAT0000542 | Mllt11 | 56772 | HITS-CLIP | 23142080 | Unspecified |
| mmu-miR-34a-5p | MIMAT0000542 | Mllt3 | 70122 | HITS-CLIP | 30971389\|23142080 | Unspecified |
| mmu-miR-34a-5p | MIMAT0000542 | Mllt6 | 246198 | HITS-CLIP | 19536157\|23630078\|31433990\|29167 | Unspecified |
| mmu-miR-34a-5p | MIMAT0000542 | Mlx | 21428 | HITS-CLIP | 23142080\|29167373 | Unspecified |
| mmu-miR-34a-5p | MIMAT0000542 | Mlxip | 208104 | HITS-CLIP | 23142080\|29167373 | Unspecified |
| mmu-miR-34a-5p | MIMAT0000542 | Mmab | 77697 | HITS-CLIP | 30971389 | Unspecified |
| mmu-miR-34a-5p | MIMAT0000542 | Mmachc | 67096 | HITS-CLIP | 23142080 | Unspecified |
| mmu-miR-34a-5p | MIMAT0000542 | Mmd | 67468 | HITS-CLIP | 37986224\|23142080\|19536157\|29167 | Unspecified |
| mmu-miR-34a-5p | MIMAT0000542 | Mmgt1 | 236792 | HITS-CLIP | 19536157\|23142080 | Unspecified |
| mmu-miR-34a-5p | MIMAT0000542 | Mmgt2 | 216829 | HITS-CLIP | 23142080 | Unspecified |
| mmu-miR-34a-5p | MIMAT0000542 | Mmp11 | 17385 | HITS-CLIP | 23142080 | Unspecified |
| mmu-miR-34a-5p | MIMAT0000542 | Mmp12 | 17381 | HITS-CLIP//Microarray//qRT-PCR | 29167373\|32855961 | Unspecified |
| mmu-miR-34a-5p | MIMAT0000542 | Mmp14 | 17387 | HITS-CLIP | 29167373\|26203562 | Unspecified |
| mmu-miR-34a-5p | MIMAT0000542 | Mmp17 | 23948 | HITS-CLIP | 23142080 | Unspecified |
| mmu-miR-34a-5p | MIMAT0000542 | Mms19 | 72199 | HITS-CLIP | 23142080 | Unspecified |
| mmu-miR-34a-5p | MIMAT0000542 | Mms22l | 212377 | HITS-CLIP | 23630078\|30971389\|23142080 | Unspecified |
| mmu-miR-34a-5p | MIMAT0000542 | Mnat1 | 17420 | HITS-CLIP | 23142080 | Unspecified |
| mmu-miR-34a-5p | MIMAT0000542 | Mns1 | 17427 | HITS-CLIP | 30971389 | Unspecified |
| mmu-miR-34a-5p | MIMAT0000542 | Mnt | 17428 | HITS-CLIP | 23142080\|29167373 | Unspecified |
| mmu-miR-34a-5p | MIMAT0000542 | Moap1 | 64113 | HITS-CLIP | 31433990 | Unspecified |
| mmu-miR-34a-5p | MIMAT0000542 | Mob1a | 232157 | HITS-CLIP | 31433990\|23630078\|23142080\|29167 | Unspecified |
| mmu-miR-34a-5p | MIMAT0000542 | Mob1b | 68473 | HITS-CLIP | 23142080\|26203562\|29167373 | Unspecified |
| mmu-miR-34a-5p | MIMAT0000542 | Mob2 | 101513 | HITS-CLIP | 23142080\|26203562 | Unspecified |
| mmu-miR-34a-5p | MIMAT0000542 | Mob3a | 208228 | HITS-CLIP | 23142080 | Unspecified |
| mmu-miR-34a-5p | MIMAT0000542 | Mob4 | 19070 | HITS-CLIP | 37986224\|23142080\|19536157\|29167 | Unspecified |
| mmu-miR-34a-5p | MIMAT0000542 | Mobp | 17433 | HITS-CLIP | 19536157 | Unspecified |
| mmu-miR-34a-5p | MIMAT0000542 | Mocos | 68591 | HITS-CLIP | 29167373 | Unspecified |
| mmu-miR-34a-5p | MIMAT0000542 | Mocs1 | 56738 | HITS-CLIP | 23142080 | Unspecified |
| mmu-miR-34a-5p | MIMAT0000542 | Mocs3 | 69372 | HITS-CLIP | 23142080 | Unspecified |
| mmu-miR-34a-5p | MIMAT0000542 | Mogs | 57377 | HITS-CLIP | 23142080\|29167373 | Unspecified |
| mmu-miR-34a-5p | MIMAT0000542 | Mok | 26448 | Microarrays | 37986224 | Unspecified |

| | | | | | | | |
|---|---|---|---|---|---|---|---|
| mmu-miR-34a-5p | MIMAT0000542 | Mon1b | 270096 | HITS-CLIP | | 23142080 | Unspecified |
| mmu-miR-34a-5p | MIMAT0000542 | Mon2 | 67074 | HITS-CLIP | 31433990\|23142080\|29167373 | | Unspecified |
| mmu-miR-34a-5p | MIMAT0000542 | Morc2a | 74522 | HITS-CLIP | | 23142080 | Unspecified |
| mmu-miR-34a-5p | MIMAT0000542 | Morc3 | 338467 | HITS-CLIP | 23142080\|29167373 | | Unspecified |
| mmu-miR-34a-5p | MIMAT0000542 | Morf4l2 | 56397 | HITS-CLIP | 29167373\|23142080 | | Unspecified |
| mmu-miR-34a-5p | MIMAT0000542 | Morn2 | 378462 | Microarrays | | 37986224 | Unspecified |
| mmu-miR-34a-5p | MIMAT0000542 | Mosmo | 233812 | HITS-CLIP | 23142080\|29167373 | | Unspecified |
| mmu-miR-34a-5p | MIMAT0000542 | Mospd1 | 70380 | HITS-CLIP | | 23142080 | Unspecified |
| mmu-miR-34a-5p | MIMAT0000542 | Mospd2 | 76763 | HITS-CLIP | 23142080\|29167373 | | Unspecified |
| mmu-miR-34a-5p | MIMAT0000542 | Mospd3 | 68929 | Microarrays//HITS-CLIP | 37986224\|23027903\|23142080 | | Unspecified |
| mmu-miR-34a-5p | MIMAT0000542 | Mov10 | 17454 | HITS-CLIP | | 23142080 | Unspecified |
| mmu-miR-34a-5p | MIMAT0000542 | Mpdu1 | 24070 | HITS-CLIP | | 23142080 | Unspecified |
| mmu-miR-34a-5p | MIMAT0000542 | Mpeg1 | 17476 | HITS-CLIP | | 29167373 | Unspecified |
| mmu-miR-34a-5p | MIMAT0000542 | Mphosph9 | 269702 | HITS-CLIP | 19536157\|23142080 | | Unspecified |
| mmu-miR-34a-5p | MIMAT0000542 | Mpi | 110119 | HITS-CLIP | | 23142080 | Unspecified |
| mmu-miR-34a-5p | MIMAT0000542 | Mpp1 | 17524 | HITS-CLIP | | 29167373 | Unspecified |
| mmu-miR-34a-5p | MIMAT0000542 | Mpp2 | 50997 | Microarrays//HITS-CLIP | 37986224\|23027903\|23142080 | | Unspecified |
| mmu-miR-34a-5p | MIMAT0000542 | Mpp3 | 4356 | HITS-CLIP | | 19536157 | Unspecified |
| mmu-miR-34a-5p | MIMAT0000542 | Mpp5 | 56217 | Microarrays//HITS-CLIP | 37986224\|23142080 | | Unspecified |
| mmu-miR-34a-5p | MIMAT0000542 | Mpp6 | 56524 | HITS-CLIP | 23142080\|19536157 | | Unspecified |
| mmu-miR-34a-5p | MIMAT0000542 | Mpp7 | 75739 | HITS-CLIP | | 23142080 | Unspecified |
| mmu-miR-34a-5p | MIMAT0000542 | Mppe1 | 225651 | HITS-CLIP | | 23142080 | Unspecified |
| mmu-miR-34a-5p | MIMAT0000542 | Mprip | 26936 | HITS-CLIP | | 29167373 | Unspecified |
| mmu-miR-34a-5p | MIMAT0000542 | Mpst | 246221 | HITS-CLIP | | 23142080 | Unspecified |
| mmu-miR-34a-5p | MIMAT0000542 | Mpzl2 | 14012 | HITS-CLIP | 26203562\|23142080 | | Unspecified |
| mmu-miR-34a-5p | MIMAT0000542 | Mpzl3 | 319742 | HITS-CLIP | | 23142080 | Unspecified |
| mmu-miR-34a-5p | MIMAT0000542 | Mras | 17532 | HITS-CLIP | | 19536157 | Unspecified |
| mmu-miR-34a-5p | MIMAT0000542 | Mrfap1 | 67568 | HITS-CLIP | | 23142080 | Unspecified |
| mmu-miR-34a-5p | MIMAT0000542 | Mrm2 | 29960 | HITS-CLIP | | 23142080 | Unspecified |
| mmu-miR-34a-5p | MIMAT0000542 | Mroh1 | 223658 | HITS-CLIP | 23142080\|29167373 | | Unspecified |
| mmu-miR-34a-5p | MIMAT0000542 | Mrpl1 | 94061 | HITS-CLIP | 23142080\|29167373 | | Unspecified |
| mmu-miR-34a-5p | MIMAT0000542 | Mrpl10 | 107732 | HITS-CLIP | | 23142080 | Unspecified |
| mmu-miR-34a-5p | MIMAT0000542 | Mrpl13 | 68537 | HITS-CLIP | | 23142080 | Unspecified |
| mmu-miR-34a-5p | MIMAT0000542 | Mrpl15 | 27395 | HITS-CLIP | | 23142080 | Unspecified |
| mmu-miR-34a-5p | MIMAT0000542 | Mrpl16 | 94063 | Microarrays//HITS-CLIP | 37986224\|23142080\|23027903 | | Unspecified |
| mmu-miR-34a-5p | MIMAT0000542 | Mrpl17 | 27397 | HITS-CLIP | | 23142080 | Unspecified |
| mmu-miR-34a-5p | MIMAT0000542 | Mrpl19 | 56284 | HITS-CLIP | | 23142080 | Unspecified |
| mmu-miR-34a-5p | MIMAT0000542 | Mrpl20 | 66448 | HITS-CLIP | 31433990\|23142080 | | Unspecified |
| mmu-miR-34a-5p | MIMAT0000542 | Mrpl27 | 94064 | HITS-CLIP | | 23142080 | Unspecified |

| | | | | | | |
|---|---|---|---|---|---|---|
| mmu-miR-34a-5p | MIMAT0000542 | Mrpl28 | 68611 | HITS-CLIP | 31433990 | Unspecified |
| mmu-miR-34a-5p | MIMAT0000542 | Mrpl30 | 107734 | HITS-CLIP | 23142080 | Unspecified |
| mmu-miR-34a-5p | MIMAT0000542 | Mrpl32 | 75398 | HITS-CLIP | 23142080 | Unspecified |
| mmu-miR-34a-5p | MIMAT0000542 | Mrpl37 | 56280 | HITS-CLIP | 23630078\|30971389 | Unspecified |
| mmu-miR-34a-5p | MIMAT0000542 | Mrpl4 | 66163 | HITS-CLIP | 23142080 | Unspecified |
| mmu-miR-34a-5p | MIMAT0000542 | Mrpl44 | 69163 | HITS-CLIP | 23142080 | Unspecified |
| mmu-miR-34a-5p | MIMAT0000542 | Mrpl47 | 74600 | HITS-CLIP | 23142080 | Unspecified |
| mmu-miR-34a-5p | MIMAT0000542 | Mrpl48 | 52443 | HITS-CLIP | 23142080 | Unspecified |
| mmu-miR-34a-5p | MIMAT0000542 | Mrpl51 | 66493 | HITS-CLIP | 23142080 | Unspecified |
| mmu-miR-34a-5p | MIMAT0000542 | Mrpl52 | 68836 | HITS-CLIP | 23142080 | Unspecified |
| mmu-miR-34a-5p | MIMAT0000542 | Mrpl53 | 68499 | HITS-CLIP | 23142080 | Unspecified |
| mmu-miR-34a-5p | MIMAT0000542 | Mrpl58 | 3396 | HITS-CLIP | 23142080 | Unspecified |
| mmu-miR-34a-5p | MIMAT0000542 | Mrpl9 | 78523 | HITS-CLIP | 23142080 | Unspecified |
| mmu-miR-34a-5p | MIMAT0000542 | Mrps14 | 64659 | HITS-CLIP | 23142080 | Unspecified |
| mmu-miR-34a-5p | MIMAT0000542 | Mrps15 | 66407 | HITS-CLIP | 23142080 | Unspecified |
| mmu-miR-34a-5p | MIMAT0000542 | Mrps16 | 66242 | HITS-CLIP | 23142080 | Unspecified |
| mmu-miR-34a-5p | MIMAT0000542 | Mrps18a | 68565 | HITS-CLIP | 23142080 | Unspecified |
| mmu-miR-34a-5p | MIMAT0000542 | Mrps2 | 118451 | HITS-CLIP | 31433990 | Unspecified |
| mmu-miR-34a-5p | MIMAT0000542 | Mrps35 | 232536 | HITS-CLIP | 23142080 | Unspecified |
| mmu-miR-34a-5p | MIMAT0000542 | Mrps5 | 77721 | HITS-CLIP | 23630078\|23142080 | Unspecified |
| mmu-miR-34a-5p | MIMAT0000542 | Mrps9 | 69527 | HITS-CLIP | 23142080 | Unspecified |
| mmu-miR-34a-5p | MIMAT0000542 | Mrs2 | 380836 | HITS-CLIP | 23142080\|29167373 | Unspecified |
| mmu-miR-34a-5p | MIMAT0000542 | Mrtfa | 57591 | HITS-CLIP | 23142080\|29167373 | Unspecified |
| mmu-miR-34a-5p | MIMAT0000542 | Mrtfb | 239719 | HITS-CLIP | 23630078\|29167373 | Unspecified |
| mmu-miR-34a-5p | MIMAT0000542 | Mrto4 | 69902 | HITS-CLIP | 23142080 | Unspecified |
| mmu-miR-34a-5p | MIMAT0000542 | Ms4a14 | 383435 | HITS-CLIP | 29167373 | Unspecified |
| mmu-miR-34a-5p | MIMAT0000542 | Ms4a4b | 60361 | HITS-CLIP | 23142080 | Unspecified |
| mmu-miR-34a-5p | MIMAT0000542 | Ms4a4c | 64380 | HITS-CLIP | 23142080 | Unspecified |
| mmu-miR-34a-5p | MIMAT0000542 | Ms4a6b | 69774 | HITS-CLIP | 23142080\|29167373 | Unspecified |
| mmu-miR-34a-5p | MIMAT0000542 | Ms4a6c | 73656 | HITS-CLIP | 29167373 | Unspecified |
| mmu-miR-34a-5p | MIMAT0000542 | Ms4a6d | 68774 | HITS-CLIP | 29167373 | Unspecified |
| mmu-miR-34a-5p | MIMAT0000542 | Ms4a7 | 109225 | HITS-CLIP | 29167373 | Unspecified |
| mmu-miR-34a-5p | MIMAT0000542 | Msantd3 | 66665 | HITS-CLIP | 30971389 | Unspecified |
| mmu-miR-34a-5p | MIMAT0000542 | Msantd4 | 78100 | HITS-CLIP | 23142080\|29167373 | Unspecified |
| mmu-miR-34a-5p | MIMAT0000542 | Msh2 | 17685 | Microarrays//HITS-CLIP | 37986224\|23027903\|23142080 | Unspecified |
| mmu-miR-34a-5p | MIMAT0000542 | Msh6 | 17688 | HITS-CLIP | 23142080 | Unspecified |
| mmu-miR-34a-5p | MIMAT0000542 | Msi2 | 76626 | Microarrays//HITS-CLIP | 37986224\|23630078\|30971389\|23142 | Unspecified |
| mmu-miR-34a-5p | MIMAT0000542 | Msl1 | 74026 | HITS-CLIP | 23142080\|29167373 | Unspecified |
| mmu-miR-34a-5p | MIMAT0000542 | Msl2 | 77853 | HITS-CLIP | 31433990\|19536157\|23142080\|29167 | Unspecified |

| | | | | | | |
|---|---|---|---|---|---|---|
| mmu-miR-34a-5p | MIMAT0000542 | Msl3 | 17692 | HITS-CLIP | 23142080 | Unspecified |
| mmu-miR-34a-5p | MIMAT0000542 | Msl3l2 | 73390 | HITS-CLIP | 23142080 | Unspecified |
| mmu-miR-34a-5p | MIMAT0000542 | Msmo1 | 66234 | HITS-CLIP | 19536157\|29167373\|23142080 | Unspecified |
| mmu-miR-34a-5p | MIMAT0000542 | Msr1 | 20288 | HITS-CLIP | 29167373 | Unspecified |
| mmu-miR-34a-5p | MIMAT0000542 | Msra | 110265 | HITS-CLIP | 23142080 | Unspecified |
| mmu-miR-34a-5p | MIMAT0000542 | Msrb1 | 27361 | HITS-CLIP | 29167373 | Unspecified |
| mmu-miR-34a-5p | MIMAT0000542 | Mss51 | 74843 | Microarrays | 37986224 | Unspecified |
| mmu-miR-34a-5p | MIMAT0000542 | Msto1 | 229524 | HITS-CLIP | 23142080 | Unspecified |
| mmu-miR-34a-5p | MIMAT0000542 | Mt1 | 17748 | HITS-CLIP | 23142080\|29167373 | Unspecified |
| mmu-miR-34a-5p | MIMAT0000542 | Mt2 | 17750 | HITS-CLIP | 23142080 | Unspecified |
| mmu-miR-34a-5p | MIMAT0000542 | Mta1 | 116870 | HITS-CLIP | 29167373 | Unspecified |
| mmu-miR-34a-5p | MIMAT0000542 | Mta2 | 23942 | HITS-CLIP | 30971389\|23142080\|29167373 | Unspecified |
| mmu-miR-34a-5p | MIMAT0000542 | Mta3 | 116871 | HITS-CLIP | 23142080 | Unspecified |
| mmu-miR-34a-5p | MIMAT0000542 | Mtap | 66902 | HITS-CLIP | 29167373 | Unspecified |
| mmu-miR-34a-5p | MIMAT0000542 | Mtbp | 105837 | HITS-CLIP | 23142080 | Unspecified |
| mmu-miR-34a-5p | MIMAT0000542 | Mtch1 | 56462 | HITS-CLIP | 23630078\|31433990\|23142080 | Unspecified |
| mmu-miR-34a-5p | MIMAT0000542 | Mtcl1 | 68617 | HITS-CLIP | 19536157 | Unspecified |
| mmu-miR-34a-5p | MIMAT0000542 | Mtcp1 | 17763 | HITS-CLIP | 23142080 | Unspecified |
| mmu-miR-34a-5p | MIMAT0000542 | Mtdh | 67154 | HITS-CLIP | 37986224\|23142080\|19536157\|29167 | Unspecified |
| mmu-miR-34a-5p | MIMAT0000542 | Mterf4 | 69821 | HITS-CLIP | 23142080 | Unspecified |
| mmu-miR-34a-5p | MIMAT0000542 | Mtf1 | 17764 | HITS-CLIP | 23142080\|29167373 | Unspecified |
| mmu-miR-34a-5p | MIMAT0000542 | Mtf2 | 17765 | Microarrays//HITS-CLIP | 37986224\|23142080\|29167373 | Unspecified |
| mmu-miR-34a-5p | MIMAT0000542 | Mtfmt | 69606 | HITS-CLIP | 31433990 | Unspecified |
| mmu-miR-34a-5p | MIMAT0000542 | Mtfr1 | 67472 | HITS-CLIP | 23142080\|30971389\|31433990 | Unspecified |
| mmu-miR-34a-5p | MIMAT0000542 | Mtfr1l | 76824 | HITS-CLIP | 23142080 | Unspecified |
| mmu-miR-34a-5p | MIMAT0000542 | Mtfr2 | 71804 | Microarrays | 37986224\|23027903 | Unspecified |
| mmu-miR-34a-5p | MIMAT0000542 | Mthfd1 | 108156 | Microarrays//HITS-CLIP | 37986224\|23027903\|23142080 | Unspecified |
| mmu-miR-34a-5p | MIMAT0000542 | Mthfd1l | 270685 | HITS-CLIP | 23142080 | Unspecified |
| mmu-miR-34a-5p | MIMAT0000542 | Mthfd2 | 17768 | Microarrays//HITS-CLIP | 37986224\|23027903\|23142080 | Unspecified |
| mmu-miR-34a-5p | MIMAT0000542 | Mthfr | 17769 | HITS-CLIP | 23630078\|29167373 | Unspecified |
| mmu-miR-34a-5p | MIMAT0000542 | Mthfs | 107885 | HITS-CLIP | 29167373 | Unspecified |
| mmu-miR-34a-5p | MIMAT0000542 | Mthfsd | 234814 | HITS-CLIP | 23142080 | Unspecified |
| mmu-miR-34a-5p | MIMAT0000542 | Mtif2 | 76784 | HITS-CLIP | 19536157\|23142080 | Unspecified |
| mmu-miR-34a-5p | MIMAT0000542 | Mtln | 67885 | HITS-CLIP | 23142080 | Unspecified |
| mmu-miR-34a-5p | MIMAT0000542 | Mtm1 | 17772 | HITS-CLIP | 23142080 | Unspecified |
| mmu-miR-34a-5p | MIMAT0000542 | Mtmr1 | 53332 | HITS-CLIP | 31433990\|23142080 | Unspecified |
| mmu-miR-34a-5p | MIMAT0000542 | Mtmr10 | 233315 | HITS-CLIP | 29167373\|23142080 | Unspecified |
| mmu-miR-34a-5p | MIMAT0000542 | Mtmr12 | 268783 | Microarrays//HITS-CLIP | 37986224\|23027903\|23142080\|29167 | Unspecified |
| mmu-miR-34a-5p | MIMAT0000542 | Mtmr14 | 97287 | HITS-CLIP | 23142080\|29167373 | Unspecified |

| | | | | | | |
|---|---|---|---|---|---|---|
| mmu-miR-34a-5p | MIMAT0000542 | Mtmr2 | 77116 | HITS-CLIP | 29167373 | Unspecified |
| mmu-miR-34a-5p | MIMAT0000542 | Mtmr3 | 74302 | HITS-CLIP | 30971389\|23142080\|29167373 | Unspecified |
| mmu-miR-34a-5p | MIMAT0000542 | Mtmr4 | 170749 | Microarrays//HITS-CLIP | 37986224\|23027903\|23142080\|29167 | Unspecified |
| mmu-miR-34a-5p | MIMAT0000542 | Mtmr6 | 219135 | HITS-CLIP | 23142080\|29167373 | Unspecified |
| mmu-miR-34a-5p | MIMAT0000542 | Mtmr7 | 54384 | HITS-CLIP | 23142080\|19536157 | Unspecified |
| mmu-miR-34a-5p | MIMAT0000542 | Mtmr9 | 210376 | HITS-CLIP | 30971389\|23142080 | Unspecified |
| mmu-miR-34a-5p | MIMAT0000542 | Mtor | 56717 | HITS-CLIP | 23142080\|29167373 | Unspecified |
| mmu-miR-34a-5p | MIMAT0000542 | Mtpap | 67440 | HITS-CLIP | 29167373 | Unspecified |
| mmu-miR-34a-5p | MIMAT0000542 | Mtpn | 14489 | HITS-CLIP | 19536157\|23142080\|29167373 | Unspecified |
| mmu-miR-34a-5p | MIMAT0000542 | Mtr | 238505 | HITS-CLIP | 23142080 | Unspecified |
| mmu-miR-34a-5p | MIMAT0000542 | Mtrex | 23517 | HITS-CLIP | 23142080 | Unspecified |
| mmu-miR-34a-5p | MIMAT0000542 | Mtss1 | 211401 | HITS-CLIP | 23142080\|29167373 | Unspecified |
| mmu-miR-34a-5p | MIMAT0000542 | Mtss1l | 244654 | HITS-CLIP | 19536157 | Unspecified |
| mmu-miR-34a-5p | MIMAT0000542 | Mturn | 68235 | HITS-CLIP | 23142080 | Unspecified |
| mmu-miR-34a-5p | MIMAT0000542 | Mtx1 | 17827 | HITS-CLIP | 19536157\|23142080 | Unspecified |
| mmu-miR-34a-5p | MIMAT0000542 | Mtx3 | 382793 | HITS-CLIP | 19536157 | Unspecified |
| mmu-miR-34a-5p | MIMAT0000542 | Muc2 | 17831 | HITS-CLIP | 23142080 | Unspecified |
| mmu-miR-34a-5p | MIMAT0000542 | Mul1 | 68350 | HITS-CLIP | 23142080 | Unspecified |
| mmu-miR-34a-5p | MIMAT0000542 | Mum1 | 3662 | HITS-CLIP | 23142080 | Unspecified |
| mmu-miR-34a-5p | MIMAT0000542 | Mus81 | 71711 | HITS-CLIP | 23142080 | Unspecified |
| mmu-miR-34a-5p | MIMAT0000542 | Musk | 18198 | Microarrays | 37986224 | Unspecified |
| mmu-miR-34a-5p | MIMAT0000542 | Mut | 4594 | HITS-CLIP | 23142080 | Unspecified |
| mmu-miR-34a-5p | MIMAT0000542 | Mutyh | 70603 | HITS-CLIP | 23142080 | Unspecified |
| mmu-miR-34a-5p | MIMAT0000542 | Mvb12a | 73711 | HITS-CLIP | 23142080\|29167373 | Unspecified |
| mmu-miR-34a-5p | MIMAT0000542 | Mvb12b | 72543 | HITS-CLIP | 29167373 | Unspecified |
| mmu-miR-34a-5p | MIMAT0000542 | Mvp | 78388 | HITS-CLIP | 23142080 | Unspecified |
| mmu-miR-34a-5p | MIMAT0000542 | Mxd1 | 17119 | HITS-CLIP | 37986224\|23142080\|31433990\|30971 | Unspecified |
| mmu-miR-34a-5p | MIMAT0000542 | Mxd3 | 17121 | HITS-CLIP | 23142080 | Unspecified |
| mmu-miR-34a-5p | MIMAT0000542 | Mxd4 | 17122 | Microarrays//HITS-CLIP | 37986224\|23630078\|19536157\|23142 | Unspecified |
| mmu-miR-34a-5p | MIMAT0000542 | Mxi1 | 17859 | HITS-CLIP | 23142080 | Unspecified |
| mmu-miR-34a-5p | MIMAT0000542 | Mxra8 | 74761 | Microarrays | 37986224 | Unspecified |
| mmu-miR-34a-5p | MIMAT0000542 | Myadm | 50918 | HITS-CLIP | 29167373 | Unspecified |
| mmu-miR-34a-5p | MIMAT0000542 | Myb | 17863 | Microarrays//HITS-CLIP | 37986224\|23142080\|23027903\|31433 | Unspecified |
| mmu-miR-34a-5p | MIMAT0000542 | Mybbp1a | 18432 | HITS-CLIP | 29167373\|23142080 | Unspecified |
| mmu-miR-34a-5p | MIMAT0000542 | Mybl1 | 17864 | HITS-CLIP | 23142080 | Unspecified |
| mmu-miR-34a-5p | MIMAT0000542 | Mybl2 | 17865 | Microarrays//HITS-CLIP | 37986224\|23027903\|23142080 | Unspecified |
| mmu-miR-34a-5p | MIMAT0000542 | Mybph | 53311 | Microarrays | 37986224 | Unspecified |
| mmu-miR-34a-5p | MIMAT0000542 | Myc | 17869 | HITS-CLIP | 37986224\|23142080\|31433990\|29167 | Unspecified |
| mmu-miR-34a-5p | MIMAT0000542 | Mycbp | 56309 | Microarrays//HITS-CLIP | 37986224\|23027903\|23142080 | Unspecified |

| | | | | | | |
|---|---|---|---|---|---|---|
| mmu-miR-34a-5p | MIMAT0000542 | Mycbp2 | 105689 | HITS-CLIP | 23142080\|19536157\|29167373 | Unspecified |
| mmu-miR-34a-5p | MIMAT0000542 | Mycl | 16918 | Microarrays | 37986224 | Unspecified |
| mmu-miR-34a-5p | MIMAT0000542 | Mycn | 18109 | qPCR, Western Blot//Western Blot//HITS-( | 37986224\|22020437\|23142080 | Unspecified |
| mmu-miR-34a-5p | MIMAT0000542 | Myd88 | 17874 | HITS-CLIP | 29167373\|23142080 | Unspecified |
| mmu-miR-34a-5p | MIMAT0000542 | Mydgf | 56005 | HITS-CLIP | 29167373 | Unspecified |
| mmu-miR-34a-5p | MIMAT0000542 | Myef2 | 17876 | HITS-CLIP | 23142080\|29167373 | Unspecified |
| mmu-miR-34a-5p | MIMAT0000542 | Myh10 | 77579 | HITS-CLIP | 19536157 | Unspecified |
| mmu-miR-34a-5p | MIMAT0000542 | Myh3 | 17883 | Microarrays | 37986224 | Unspecified |
| mmu-miR-34a-5p | MIMAT0000542 | Myh9 | 17886 | HITS-CLIP | 37986224\|23142080\|26203562\|29167 | Unspecified |
| mmu-miR-34a-5p | MIMAT0000542 | Myl12a | 67268 | HITS-CLIP | 23142080\|29167373 | Unspecified |
| mmu-miR-34a-5p | MIMAT0000542 | Myl12b | 67938 | HITS-CLIP | 29167373 | Unspecified |
| mmu-miR-34a-5p | MIMAT0000542 | Myl6 | 17904 | HITS-CLIP | 19536157 | Unspecified |
| mmu-miR-34a-5p | MIMAT0000542 | Mylip | 218203 | HITS-CLIP | 19536157\|23142080\|29167373 | Unspecified |
| mmu-miR-34a-5p | MIMAT0000542 | Mynn | 80732 | HITS-CLIP | 23142080 | Unspecified |
| mmu-miR-34a-5p | MIMAT0000542 | Myo10 | 17909 | HITS-CLIP | 19536157\|23142080\|29167373 | Unspecified |
| mmu-miR-34a-5p | MIMAT0000542 | Myo18a | 360013 | HITS-CLIP | 23142080 | Unspecified |
| mmu-miR-34a-5p | MIMAT0000542 | Myo19 | 66196 | HITS-CLIP | 23142080 | Unspecified |
| mmu-miR-34a-5p | MIMAT0000542 | Myo1c | 17913 | HITS-CLIP | 23142080\|29167373 | Unspecified |
| mmu-miR-34a-5p | MIMAT0000542 | Myo1e | 71602 | HITS-CLIP | 29167373\|23142080 | Unspecified |
| mmu-miR-34a-5p | MIMAT0000542 | Myo1f | 17916 | HITS-CLIP | 31433990\|29167373 | Unspecified |
| mmu-miR-34a-5p | MIMAT0000542 | Myo1g | 246177 | HITS-CLIP | 23142080\|29167373 | Unspecified |
| mmu-miR-34a-5p | MIMAT0000542 | Myo5a | 17918 | HITS-CLIP | 19536157\|23142080\|29167373 | Unspecified |
| mmu-miR-34a-5p | MIMAT0000542 | Myo6 | 17920 | HITS-CLIP | 30971389\|23142080 | Unspecified |
| mmu-miR-34a-5p | MIMAT0000542 | Myo7a | 17921 | HITS-CLIP | 29167373 | Unspecified |
| mmu-miR-34a-5p | MIMAT0000542 | Myo9a | 270163 | HITS-CLIP | 23142080\|19536157 | Unspecified |
| mmu-miR-34a-5p | MIMAT0000542 | Myo9b | 17925 | HITS-CLIP | 23142080\|29167373 | Unspecified |
| mmu-miR-34a-5p | MIMAT0000542 | Myof | 226101 | HITS-CLIP | 29167373 | Unspecified |
| mmu-miR-34a-5p | MIMAT0000542 | Myog | 17928 | Microarrays | 37986224 | Unspecified |
| mmu-miR-34a-5p | MIMAT0000542 | Myom1 | 17929 | Microarrays | 37986224 | Unspecified |
| mmu-miR-34a-5p | MIMAT0000542 | Myom2 | 17930 | Microarrays | 37986224 | Unspecified |
| mmu-miR-34a-5p | MIMAT0000542 | Myoz2 | 59006 | Microarrays | 37986224 | Unspecified |
| mmu-miR-34a-5p | MIMAT0000542 | Myrf | 225908 | HITS-CLIP | 19536157 | Unspecified |
| mmu-miR-34a-5p | MIMAT0000542 | Mysm1 | 320713 | HITS-CLIP | 23142080\|29167373 | Unspecified |
| mmu-miR-34a-5p | MIMAT0000542 | Myt1l | 17933 | HITS-CLIP | 19536157 | Unspecified |
| mmu-miR-34a-5p | MIMAT0000542 | N4bp1 | 80750 | HITS-CLIP | 29167373\|23142080 | Unspecified |
| mmu-miR-34a-5p | MIMAT0000542 | N4bp2 | 333789 | Microarrays//HITS-CLIP | 37986224\|23142080\|19536157 | Unspecified |
| mmu-miR-34a-5p | MIMAT0000542 | N4bp2l2 | 381695 | HITS-CLIP | 23142080\|29167373 | Unspecified |
| mmu-miR-34a-5p | MIMAT0000542 | Naa10 | 56292 | HITS-CLIP | 23142080 | Unspecified |
| mmu-miR-34a-5p | MIMAT0000542 | Naa15 | 74838 | HITS-CLIP | 23142080\|29167373 | Unspecified |

| | | | | | | |
|---|---|---|---|---|---|---|
| mmu-miR-34a-5p | MIMAT0000542 | Naa16 | 66897 | HITS-CLIP | 31433990\|23142080 | Unspecified |
| mmu-miR-34a-5p | MIMAT0000542 | Naa20 | 67877 | HITS-CLIP | 23142080\|29167373 | Unspecified |
| mmu-miR-34a-5p | MIMAT0000542 | Naa25 | 231713 | HITS-CLIP | 23142080\|29167373 | Unspecified |
| mmu-miR-34a-5p | MIMAT0000542 | Naa30 | 70646 | HITS-CLIP | 23142080\|29167373 | Unspecified |
| mmu-miR-34a-5p | MIMAT0000542 | Naa35 | 78689 | HITS-CLIP | 23142080 | Unspecified |
| mmu-miR-34a-5p | MIMAT0000542 | Naa40 | 70999 | HITS-CLIP | 23142080\|19536157 | Unspecified |
| mmu-miR-34a-5p | MIMAT0000542 | Naa50 | 72117 | HITS-CLIP | 23142080\|19536157\|29167373 | Unspecified |
| mmu-miR-34a-5p | MIMAT0000542 | Naa80 | 24142 | HITS-CLIP | 23142080 | Unspecified |
| mmu-miR-34a-5p | MIMAT0000542 | Naaa | 67111 | HITS-CLIP | 23142080 | Unspecified |
| mmu-miR-34a-5p | MIMAT0000542 | Nab1 | 17936 | HITS-CLIP | 23142080\|29167373 | Unspecified |
| mmu-miR-34a-5p | MIMAT0000542 | Nab2 | 17937 | HITS-CLIP | 23142080\|29167373 | Unspecified |
| mmu-miR-34a-5p | MIMAT0000542 | Nabp1 | 109019 | HITS-CLIP | 23142080\|29167373 | Unspecified |
| mmu-miR-34a-5p | MIMAT0000542 | Naca | 17938 | HITS-CLIP | 29167373 | Unspecified |
| mmu-miR-34a-5p | MIMAT0000542 | Nacad | 192950 | HITS-CLIP | 19536157 | Unspecified |
| mmu-miR-34a-5p | MIMAT0000542 | Nacc1 | 66830 | HITS-CLIP | 23142080 | Unspecified |
| mmu-miR-34a-5p | MIMAT0000542 | Nadk2 | 68646 | HITS-CLIP | 30971389\|23630078\|23142080 | Unspecified |
| mmu-miR-34a-5p | MIMAT0000542 | Nadsyn1 | 78914 | HITS-CLIP | 23142080 | Unspecified |
| mmu-miR-34a-5p | MIMAT0000542 | Nae1 | 234664 | HITS-CLIP | 19536157 | Unspecified |
| mmu-miR-34a-5p | MIMAT0000542 | Naga | 17939 | HITS-CLIP | 26203562 | Unspecified |
| mmu-miR-34a-5p | MIMAT0000542 | Nagk | 56174 | Microarrays | 37986224 | Unspecified |
| mmu-miR-34a-5p | MIMAT0000542 | Nagpa | 27426 | HITS-CLIP | 23142080 | Unspecified |
| mmu-miR-34a-5p | MIMAT0000542 | Naif1 | 71254 | HITS-CLIP | 23142080 | Unspecified |
| mmu-miR-34a-5p | MIMAT0000542 | Naip2 | 17948 | HITS-CLIP | 29167373 | Unspecified |
| mmu-miR-34a-5p | MIMAT0000542 | Naip6 | 191568 | HITS-CLIP | 29167373 | Unspecified |
| mmu-miR-34a-5p | MIMAT0000542 | Nampt | 59027 | HITS-CLIP | 23142080\|29167373 | Unspecified |
| mmu-miR-34a-5p | MIMAT0000542 | Nanog | 71950 | Luciferase Reporter Assay, qPCR, Western | 37986224\|22020437 | Unspecified |
| mmu-miR-34a-5p | MIMAT0000542 | Nanos1 | 332397 | HITS-CLIP | 29167373 | Unspecified |
| mmu-miR-34a-5p | MIMAT0000542 | Nans | 94181 | HITS-CLIP | 23630078\|23142080 | Unspecified |
| mmu-miR-34a-5p | MIMAT0000542 | Nap1l1 | 53605 | HITS-CLIP | 23142080 | Unspecified |
| mmu-miR-34a-5p | MIMAT0000542 | Nap1l4 | 17955 | HITS-CLIP | 23142080 | Unspecified |
| mmu-miR-34a-5p | MIMAT0000542 | Nap1l5 | 58243 | HITS-CLIP | 19536157 | Unspecified |
| mmu-miR-34a-5p | MIMAT0000542 | Napa | 108124 | HITS-CLIP | 23142080\|29167373 | Unspecified |
| mmu-miR-34a-5p | MIMAT0000542 | Napb | 17957 | HITS-CLIP | 19536157 | Unspecified |
| mmu-miR-34a-5p | MIMAT0000542 | Narf | 67608 | HITS-CLIP | 23142080\|29167373 | Unspecified |
| mmu-miR-34a-5p | MIMAT0000542 | Narfl | 64428 | HITS-CLIP | 29167373 | Unspecified |
| mmu-miR-34a-5p | MIMAT0000542 | Nars | 70223 | HITS-CLIP | 23142080\|29167373 | Unspecified |
| mmu-miR-34a-5p | MIMAT0000542 | Nars2 | 244141 | HITS-CLIP | 23142080 | Unspecified |
| mmu-miR-34a-5p | MIMAT0000542 | Nasp | 50927 | Microarrays//HITS-CLIP | 37986224\|23027903\|23142080 | Unspecified |
| mmu-miR-34a-5p | MIMAT0000542 | Nat10 | 98956 | HITS-CLIP | 23142080 | Unspecified |

| | | | | | | |
|---|---|---|---|---|---|---|
| mmu-miR-34a-5p | MIMAT0000542 | Nat14 | 269854 | HITS-CLIP | 19536157 | Unspecified |
| mmu-miR-34a-5p | MIMAT0000542 | Natd1 | 24083 | HITS-CLIP | 19536157\|29167373 | Unspecified |
| mmu-miR-34a-5p | MIMAT0000542 | Nav1 | 215690 | Microarrays//HITS-CLIP | 37986224\|23027903\|19536157\|23142 | Unspecified |
| mmu-miR-34a-5p | MIMAT0000542 | Nav2 | 78286 | Microarrays//HITS-CLIP | 37986224\|19536157\|29167373 | Unspecified |
| mmu-miR-34a-5p | MIMAT0000542 | Nav3 | 260315 | HITS-CLIP | 19536157 | Unspecified |
| mmu-miR-34a-5p | MIMAT0000542 | Naxd | 55739 | HITS-CLIP | 23142080 | Unspecified |
| mmu-miR-34a-5p | MIMAT0000542 | Naxe | 128240 | HITS-CLIP | 23142080 | Unspecified |
| mmu-miR-34a-5p | MIMAT0000542 | Nbas | 71169 | HITS-CLIP | 23142080 | Unspecified |
| mmu-miR-34a-5p | MIMAT0000542 | Nbea | 26422 | HITS-CLIP | 19536157 | Unspecified |
| mmu-miR-34a-5p | MIMAT0000542 | Nbeal1 | 269198 | HITS-CLIP | 23142080\|29167373 | Unspecified |
| mmu-miR-34a-5p | MIMAT0000542 | Nbeal2 | 235627 | HITS-CLIP | 23142080\|29167373 | Unspecified |
| mmu-miR-34a-5p | MIMAT0000542 | Nbn | 27354 | HITS-CLIP | 23142080 | Unspecified |
| mmu-miR-34a-5p | MIMAT0000542 | Nbr1 | 17966 | HITS-CLIP | 31433990\|19536157\|30971389\|23142 | Unspecified |
| mmu-miR-34a-5p | MIMAT0000542 | Ncam1 | 17967 | HITS-CLIP | 19536157 | Unspecified |
| mmu-miR-34a-5p | MIMAT0000542 | Ncam2 | 17968 | HITS-CLIP | 19536157 | Unspecified |
| mmu-miR-34a-5p | MIMAT0000542 | Ncan | 13004 | HITS-CLIP | 19536157 | Unspecified |
| mmu-miR-34a-5p | MIMAT0000542 | Ncapd2 | 68298 | HITS-CLIP | 23142080 | Unspecified |
| mmu-miR-34a-5p | MIMAT0000542 | Ncapd3 | 78658 | Microarrays//HITS-CLIP | 37986224\|23027903\|23142080\|29167 | Unspecified |
| mmu-miR-34a-5p | MIMAT0000542 | Ncapg | 54392 | HITS-CLIP | 23142080 | Unspecified |
| mmu-miR-34a-5p | MIMAT0000542 | Ncapg2 | 76044 | Microarrays//HITS-CLIP | 37986224\|23027903\|23142080\|29167 | Unspecified |
| mmu-miR-34a-5p | MIMAT0000542 | Ncaph | 215387 | HITS-CLIP | 23142080 | Unspecified |
| mmu-miR-34a-5p | MIMAT0000542 | Ncaph2 | 52683 | HITS-CLIP | 23142080\|26203562\|29167373 | Unspecified |
| mmu-miR-34a-5p | MIMAT0000542 | Ncbp1 | 433702 | HITS-CLIP | 31433990\|23630078\|23142080 | Unspecified |
| mmu-miR-34a-5p | MIMAT0000542 | Ncbp3 | 66874 | HITS-CLIP | 23142080 | Unspecified |
| mmu-miR-34a-5p | MIMAT0000542 | Ncdn | 26562 | HITS-CLIP | 19536157\|23142080 | Unspecified |
| mmu-miR-34a-5p | MIMAT0000542 | Nceh1 | 320024 | Microarrays//HITS-CLIP | 37986224\|23027903\|19536157\|23142 | Unspecified |
| mmu-miR-34a-5p | MIMAT0000542 | Ncf1 | 17969 | HITS-CLIP | 23142080 | Unspecified |
| mmu-miR-34a-5p | MIMAT0000542 | Ncf4 | 17972 | HITS-CLIP | 23142080\|29167373 | Unspecified |
| mmu-miR-34a-5p | MIMAT0000542 | Nck1 | 17973 | HITS-CLIP | 23142080 | Unspecified |
| mmu-miR-34a-5p | MIMAT0000542 | Nck2 | 17974 | HITS-CLIP | 23142080 | Unspecified |
| mmu-miR-34a-5p | MIMAT0000542 | Nckap1 | 50884 | HITS-CLIP | 19536157\|23142080 | Unspecified |
| mmu-miR-34a-5p | MIMAT0000542 | Nckap1l | 105855 | HITS-CLIP | 29167373\|23142080 | Unspecified |
| mmu-miR-34a-5p | MIMAT0000542 | Nckap5l | 380969 | HITS-CLIP | 23142080 | Unspecified |
| mmu-miR-34a-5p | MIMAT0000542 | Nckipsd | 80987 | HITS-CLIP | 29167373 | Unspecified |
| mmu-miR-34a-5p | MIMAT0000542 | Ncl | 17975 | HITS-CLIP | 23142080\|29167373 | Unspecified |
| mmu-miR-34a-5p | MIMAT0000542 | Ncln | 103425 | HITS-CLIP | 31433990\|29167373 | Unspecified |
| mmu-miR-34a-5p | MIMAT0000542 | Ncoa1 | 17977 | HITS-CLIP | 31433990\|23142080\|29167373 | Unspecified |
| mmu-miR-34a-5p | MIMAT0000542 | Ncoa2 | 17978 | HITS-CLIP | 19536157\|23142080\|29167373 | Unspecified |
| mmu-miR-34a-5p | MIMAT0000542 | Ncoa3 | 17979 | HITS-CLIP | 23142080\|29167373 | Unspecified |

| | | | | | | |
|---|---|---|---|---|---|---|
| mmu-miR-34a-5p | MIMAT0000542 | Ncoa5 | 228869 | HITS-CLIP | 23142080 | Unspecified |
| mmu-miR-34a-5p | MIMAT0000542 | Ncoa6 | 56406 | HITS-CLIP | 31433990\|30971389\|29167373 | Unspecified |
| mmu-miR-34a-5p | MIMAT0000542 | Ncoa7 | 211329 | HITS-CLIP | 23142080\|29167373\|31433990\|30971 | Unspecified |
| mmu-miR-34a-5p | MIMAT0000542 | Ncor1 | 20185 | HITS-CLIP | 19536157\|23142080\|29167373 | Unspecified |
| mmu-miR-34a-5p | MIMAT0000542 | Ncor2 | 20602 | HITS-CLIP | 19536157\|29167373 | Unspecified |
| mmu-miR-34a-5p | MIMAT0000542 | Ncs1 | 14299 | HITS-CLIP | 19536157 | Unspecified |
| mmu-miR-34a-5p | MIMAT0000542 | Ncstn | 59287 | HITS-CLIP | 19536157\|23142080\|29167373 | Unspecified |
| mmu-miR-34a-5p | MIMAT0000542 | Ndc1 | 72787 | HITS-CLIP | 23142080\|19536157\|30971389 | Unspecified |
| mmu-miR-34a-5p | MIMAT0000542 | Ndc80 | 67052 | HITS-CLIP | 23142080 | Unspecified |
| mmu-miR-34a-5p | MIMAT0000542 | Nde1 | 67203 | HITS-CLIP | 23142080\|29167373 | Unspecified |
| mmu-miR-34a-5p | MIMAT0000542 | Ndel1 | 83431 | HITS-CLIP | 23142080 | Unspecified |
| mmu-miR-34a-5p | MIMAT0000542 | Ndfip1 | 65113 | HITS-CLIP | 23142080\|31433990\|23630078 | Unspecified |
| mmu-miR-34a-5p | MIMAT0000542 | Ndfip2 | 76273 | HITS-CLIP | 23142080\|29167373 | Unspecified |
| mmu-miR-34a-5p | MIMAT0000542 | Ndor1 | 78797 | HITS-CLIP | 23142080 | Unspecified |
| mmu-miR-34a-5p | MIMAT0000542 | Ndrg1 | 17988 | HITS-CLIP | 23142080 | Unspecified |
| mmu-miR-34a-5p | MIMAT0000542 | Ndrg2 | 29811 | HITS-CLIP | 31433990\|19536157\|23142080 | Unspecified |
| mmu-miR-34a-5p | MIMAT0000542 | Ndrg3 | 29812 | HITS-CLIP | 23142080 | Unspecified |
| mmu-miR-34a-5p | MIMAT0000542 | Ndrg4 | 234593 | Microarrays//HITS-CLIP | 37986224\|19536157 | Unspecified |
| mmu-miR-34a-5p | MIMAT0000542 | Ndst1 | 15531 | Microarrays//HITS-CLIP | 37986224\|23027903\|23142080\|2363( | Unspecified |
| mmu-miR-34a-5p | MIMAT0000542 | Ndst2 | 17423 | HITS-CLIP | 29167373 | Unspecified |
| mmu-miR-34a-5p | MIMAT0000542 | Ndst3 | 83398 | HITS-CLIP | 19536157 | Unspecified |
| mmu-miR-34a-5p | MIMAT0000542 | Ndst4 | 64580 | Microarrays | 37986224 | Unspecified |
| mmu-miR-34a-5p | MIMAT0000542 | Ndufa1 | 54405 | HITS-CLIP | 23142080 | Unspecified |
| mmu-miR-34a-5p | MIMAT0000542 | Ndufa10 | 67273 | HITS-CLIP | 23142080 | Unspecified |
| mmu-miR-34a-5p | MIMAT0000542 | Ndufa12 | 66414 | HITS-CLIP | 23142080 | Unspecified |
| mmu-miR-34a-5p | MIMAT0000542 | Ndufa13 | 67184 | HITS-CLIP | 23142080 | Unspecified |
| mmu-miR-34a-5p | MIMAT0000542 | Ndufa2 | 17991 | HITS-CLIP | 23142080 | Unspecified |
| mmu-miR-34a-5p | MIMAT0000542 | Ndufa3 | 66091 | HITS-CLIP | 23142080\|29167373 | Unspecified |
| mmu-miR-34a-5p | MIMAT0000542 | Ndufa6 | 67130 | HITS-CLIP | 31433990 | Unspecified |
| mmu-miR-34a-5p | MIMAT0000542 | Ndufa7 | 66416 | HITS-CLIP | 29167373\|23142080 | Unspecified |
| mmu-miR-34a-5p | MIMAT0000542 | Ndufa8 | 68375 | HITS-CLIP | 23142080 | Unspecified |
| mmu-miR-34a-5p | MIMAT0000542 | Ndufa9 | 66108 | HITS-CLIP | 23142080 | Unspecified |
| mmu-miR-34a-5p | MIMAT0000542 | Ndufab1 | 70316 | HITS-CLIP | 23142080 | Unspecified |
| mmu-miR-34a-5p | MIMAT0000542 | Ndufaf3 | 66706 | HITS-CLIP | 23142080 | Unspecified |
| mmu-miR-34a-5p | MIMAT0000542 | Ndufaf4 | 68493 | HITS-CLIP | 30971389\|19536157\|23142080 | Unspecified |
| mmu-miR-34a-5p | MIMAT0000542 | Ndufb10 | 68342 | HITS-CLIP | 23142080 | Unspecified |
| mmu-miR-34a-5p | MIMAT0000542 | Ndufb11 | 104130 | HITS-CLIP | 23142080 | Unspecified |
| mmu-miR-34a-5p | MIMAT0000542 | Ndufb2 | 68198 | HITS-CLIP | 23142080 | Unspecified |
| mmu-miR-34a-5p | MIMAT0000542 | Ndufb7 | 66916 | HITS-CLIP | 23142080 | Unspecified |

| | | | | | | |
|---|---|---|---|---|---|---|
| mmu-miR-34a-5p | MIMAT0000542 | Ndufb8 | 67264 | HITS-CLIP | 23142080 | Unspecified |
| mmu-miR-34a-5p | MIMAT0000542 | Ndufb9 | 66218 | HITS-CLIP | 23142080\|31433990 | Unspecified |
| mmu-miR-34a-5p | MIMAT0000542 | Ndufc1 | 66377 | HITS-CLIP | 23142080 | Unspecified |
| mmu-miR-34a-5p | MIMAT0000542 | Ndufc2 | 68197 | HITS-CLIP | 23142080 | Unspecified |
| mmu-miR-34a-5p | MIMAT0000542 | Ndufs1 | 227197 | HITS-CLIP | 23142080\|19536157\|30971389\|26203 | Unspecified |
| mmu-miR-34a-5p | MIMAT0000542 | Ndufs2 | 226646 | HITS-CLIP | 23142080 | Unspecified |
| mmu-miR-34a-5p | MIMAT0000542 | Ndufs4 | 17993 | HITS-CLIP | 23142080 | Unspecified |
| mmu-miR-34a-5p | MIMAT0000542 | Ndufs6 | 407785 | HITS-CLIP | 23142080 | Unspecified |
| mmu-miR-34a-5p | MIMAT0000542 | Ndufs7 | 75406 | HITS-CLIP | 23142080 | Unspecified |
| mmu-miR-34a-5p | MIMAT0000542 | Ndufs8 | 225887 | HITS-CLIP | 23142080 | Unspecified |
| mmu-miR-34a-5p | MIMAT0000542 | Ndufv1 | 17995 | HITS-CLIP | 23142080 | Unspecified |
| mmu-miR-34a-5p | MIMAT0000542 | Ndufv2 | 72900 | HITS-CLIP | 23142080 | Unspecified |
| mmu-miR-34a-5p | MIMAT0000542 | Ndufv3 | 78330 | HITS-CLIP | 23142080 | Unspecified |
| mmu-miR-34a-5p | MIMAT0000542 | Neb | 17996 | Microarrays//HITS-CLIP | 37986224\|23142080 | Unspecified |
| mmu-miR-34a-5p | MIMAT0000542 | Nebl | 74103 | HITS-CLIP | 23142080 | Unspecified |
| mmu-miR-34a-5p | MIMAT0000542 | Necab3 | 56846 | HITS-CLIP | 23142080 | Unspecified |
| mmu-miR-34a-5p | MIMAT0000542 | Necap1 | 67602 | HITS-CLIP | 29167373\|23142080 | Unspecified |
| mmu-miR-34a-5p | MIMAT0000542 | Necap2 | 66147 | HITS-CLIP | 31433990\|23142080\|29167373 | Unspecified |
| mmu-miR-34a-5p | MIMAT0000542 | Nectin1 | 58235 | Microarrays | 23027903 | Unspecified |
| mmu-miR-34a-5p | MIMAT0000542 | Nectin4 | 71740 | HITS-CLIP | 26203562 | Unspecified |
| mmu-miR-34a-5p | MIMAT0000542 | Nedd1 | 17997 | HITS-CLIP | 23142080 | Unspecified |
| mmu-miR-34a-5p | MIMAT0000542 | Nedd4 | 17999 | HITS-CLIP | 19536157\|23630078\|30971389\|29084 | Unspecified |
| mmu-miR-34a-5p | MIMAT0000542 | Nedd4l | 83814 | HITS-CLIP | 19536157 | Unspecified |
| mmu-miR-34a-5p | MIMAT0000542 | Nedd8 | 18002 | HITS-CLIP | 31433990 | Unspecified |
| mmu-miR-34a-5p | MIMAT0000542 | Nedd9 | 18003 | HITS-CLIP | 23142080\|23630078 | Unspecified |
| mmu-miR-34a-5p | MIMAT0000542 | Neil3 | 234258 | Microarrays//HITS-CLIP | 37986224\|23027903\|23142080 | Unspecified |
| mmu-miR-34a-5p | MIMAT0000542 | Nek1 | 18004 | HITS-CLIP | 23142080 | Unspecified |
| mmu-miR-34a-5p | MIMAT0000542 | Nek2 | 18005 | HITS-CLIP | 23142080 | Unspecified |
| mmu-miR-34a-5p | MIMAT0000542 | Nek4 | 23955 | HITS-CLIP | 23142080 | Unspecified |
| mmu-miR-34a-5p | MIMAT0000542 | Nek6 | 59126 | HITS-CLIP | 29167373 | Unspecified |
| mmu-miR-34a-5p | MIMAT0000542 | Nek7 | 59125 | HITS-CLIP | 23142080\|19536157 | Unspecified |
| mmu-miR-34a-5p | MIMAT0000542 | Nek9 | 217718 | HITS-CLIP | 19536157\|29167373\|23142080 | Unspecified |
| mmu-miR-34a-5p | MIMAT0000542 | Nelfa | 24116 | HITS-CLIP | 23142080 | Unspecified |
| mmu-miR-34a-5p | MIMAT0000542 | Nelfb | 58202 | HITS-CLIP | 23142080\|29167373 | Unspecified |
| mmu-miR-34a-5p | MIMAT0000542 | Nelfcd | 57314 | HITS-CLIP | 23142080 | Unspecified |
| mmu-miR-34a-5p | MIMAT0000542 | Nemf | 66244 | HITS-CLIP | 23142080\|31433990 | Unspecified |
| mmu-miR-34a-5p | MIMAT0000542 | Nemp1 | 23306 | HITS-CLIP | 23142080 | Unspecified |
| mmu-miR-34a-5p | MIMAT0000542 | Nemp2 | 227094 | HITS-CLIP | 23142080 | Unspecified |
| mmu-miR-34a-5p | MIMAT0000542 | Neo1 | 18007 | HITS-CLIP | 19536157 | Unspecified |

| | | | | | | |
|---|---|---|---|---|---|---|
| mmu-miR-34a-5p | MIMAT0000542 | Nepro | 212547 | HITS-CLIP | 23142080 | Unspecified |
| mmu-miR-34a-5p | MIMAT0000542 | Net1 | 56349 | HITS-CLIP | 31433990\|23142080\|29167373 | Unspecified |
| mmu-miR-34a-5p | MIMAT0000542 | Neto1 | 246317 | HITS-CLIP | 19536157 | Unspecified |
| mmu-miR-34a-5p | MIMAT0000542 | Neto2 | 74513 | Microarrays//HITS-CLIP | 37986224\|23027903\|23142080 | Unspecified |
| mmu-miR-34a-5p | MIMAT0000542 | Neu1 | 18010 | HITS-CLIP | 23630078\|29167373 | Unspecified |
| mmu-miR-34a-5p | MIMAT0000542 | Neu3 | 50877 | HITS-CLIP | 31433990\|23142080 | Unspecified |
| mmu-miR-34a-5p | MIMAT0000542 | Neurl3 | 214854 | HITS-CLIP | 29167373\|23142080 | Unspecified |
| mmu-miR-34a-5p | MIMAT0000542 | Neurl4 | 216860 | HITS-CLIP | 23142080 | Unspecified |
| mmu-miR-34a-5p | MIMAT0000542 | Neurod1 | 18012 | HITS-CLIP | 19536157 | Unspecified |
| mmu-miR-34a-5p | MIMAT0000542 | Nexmif | 340533 | HITS-CLIP | 19536157 | Unspecified |
| mmu-miR-34a-5p | MIMAT0000542 | Nf1 | 18015 | HITS-CLIP | 19536157\|23142080\|29167373 | Unspecified |
| mmu-miR-34a-5p | MIMAT0000542 | Nf2 | 18016 | HITS-CLIP | 31433990\|23142080\|29167373 | Unspecified |
| mmu-miR-34a-5p | MIMAT0000542 | Nfam1 | 74039 | HITS-CLIP | 29167373 | Unspecified |
| mmu-miR-34a-5p | MIMAT0000542 | Nfasc | 269116 | HITS-CLIP | 19536157 | Unspecified |
| mmu-miR-34a-5p | MIMAT0000542 | Nfat5 | 54446 | HITS-CLIP | 23142080\|23630078\|29167373 | Unspecified |
| mmu-miR-34a-5p | MIMAT0000542 | Nfatc1 | 18018 | HITS-CLIP | 23142080\|31433990\|29167373 | Unspecified |
| mmu-miR-34a-5p | MIMAT0000542 | Nfatc2 | 18019 | HITS-CLIP | 31433990\|23142080\|29167373 | Unspecified |
| mmu-miR-34a-5p | MIMAT0000542 | Nfatc2ip | 18020 | HITS-CLIP | 31433990\|23142080 | Unspecified |
| mmu-miR-34a-5p | MIMAT0000542 | Nfatc3 | 18021 | HITS-CLIP | 23630078\|23142080\|29167373 | Unspecified |
| mmu-miR-34a-5p | MIMAT0000542 | Nfe2l1 | 18023 | HITS-CLIP | 31433990\|30971389\|19536157\|23142 | Unspecified |
| mmu-miR-34a-5p | MIMAT0000542 | Nfe2l2 | 18024 | HITS-CLIP | 23142080\|29167373 | Unspecified |
| mmu-miR-34a-5p | MIMAT0000542 | Nfia | 18027 | HITS-CLIP | 19536157 | Unspecified |
| mmu-miR-34a-5p | MIMAT0000542 | Nfib | 18028 | HITS-CLIP | 19536157 | Unspecified |
| mmu-miR-34a-5p | MIMAT0000542 | Nfic | 18029 | HITS-CLIP | 23142080\|29167373 | Unspecified |
| mmu-miR-34a-5p | MIMAT0000542 | Nfil3 | 18030 | HITS-CLIP | 23142080 | Unspecified |
| mmu-miR-34a-5p | MIMAT0000542 | Nfkb1 | 18033 | HITS-CLIP | 37986224\|31433990\|19536157\|23142 | Unspecified |
| mmu-miR-34a-5p | MIMAT0000542 | Nfkb2 | 18034 | HITS-CLIP | 23142080\|31433990 | Unspecified |
| mmu-miR-34a-5p | MIMAT0000542 | Nfkbia | 18035 | Western Blot | 37986224 | Unspecified |
| mmu-miR-34a-5p | MIMAT0000542 | Nfkbid | 243910 | HITS-CLIP | 31433990\|23142080\|29167373 | Unspecified |
| mmu-miR-34a-5p | MIMAT0000542 | Nfkbie | 18037 | HITS-CLIP | 31433990\|30971389 | Unspecified |
| mmu-miR-34a-5p | MIMAT0000542 | Nfkbiz | 80859 | HITS-CLIP | 23142080\|29167373 | Unspecified |
| mmu-miR-34a-5p | MIMAT0000542 | Nfrkb | 235134 | HITS-CLIP | 23142080\|29167373 | Unspecified |
| mmu-miR-34a-5p | MIMAT0000542 | Nfs1 | 18041 | HITS-CLIP | 23142080 | Unspecified |
| mmu-miR-34a-5p | MIMAT0000542 | Nfu1 | 56748 | HITS-CLIP | 23142080 | Unspecified |
| mmu-miR-34a-5p | MIMAT0000542 | Nfx1 | 74164 | HITS-CLIP | 30971389\|23142080\|29167373 | Unspecified |
| mmu-miR-34a-5p | MIMAT0000542 | Nfxl1 | 100978 | HITS-CLIP | 23142080\|29167373 | Unspecified |
| mmu-miR-34a-5p | MIMAT0000542 | Nfya | 18044 | HITS-CLIP | 23142080\|31433990\|26203562\|29084 | Unspecified |
| mmu-miR-34a-5p | MIMAT0000542 | Nfyb | 18045 | HITS-CLIP | 23142080 | Unspecified |
| mmu-miR-34a-5p | MIMAT0000542 | Nfyc | 18046 | HITS-CLIP | 23142080 | Unspecified |

| | | | | | | |
|---|---|---|---|---|---|---|
| mmu-miR-34a-5p | MIMAT0000542 | Ngdn | 68966 | HITS-CLIP | 23142080 | Unspecified |
| mmu-miR-34a-5p | MIMAT0000542 | Ngly1 | 59007 | HITS-CLIP | 23142080 | Unspecified |
| mmu-miR-34a-5p | MIMAT0000542 | Nhlrc2 | 66866 | HITS-CLIP | 31433990\|23142080\|29167373 | Unspecified |
| mmu-miR-34a-5p | MIMAT0000542 | Nhlrc3 | 212114 | HITS-CLIP | 29167373 | Unspecified |
| mmu-miR-34a-5p | MIMAT0000542 | Nhp2 | 52530 | HITS-CLIP | 23142080\|29167373 | Unspecified |
| mmu-miR-34a-5p | MIMAT0000542 | Nhsl2 | 100042480 | HITS-CLIP | 19536157\|29167373 | Unspecified |
| mmu-miR-34a-5p | MIMAT0000542 | Nin | 18080 | HITS-CLIP | 23142080\|19536157\|29167373 | Unspecified |
| mmu-miR-34a-5p | MIMAT0000542 | Ninl | 78177 | HITS-CLIP | 30971389 | Unspecified |
| mmu-miR-34a-5p | MIMAT0000542 | Nip7 | 66164 | HITS-CLIP | 23142080 | Unspecified |
| mmu-miR-34a-5p | MIMAT0000542 | Nipa2 | 93790 | HITS-CLIP | 23142080\|29167373 | Unspecified |
| mmu-miR-34a-5p | MIMAT0000542 | Nipbl | 71175 | HITS-CLIP | 19536157\|23142080\|29167373 | Unspecified |
| mmu-miR-34a-5p | MIMAT0000542 | Nipsnap2 | 2631 | HITS-CLIP | 23142080\|29167373 | Unspecified |
| mmu-miR-34a-5p | MIMAT0000542 | Nipsnap3b | 66536 | HITS-CLIP | 29167373 | Unspecified |
| mmu-miR-34a-5p | MIMAT0000542 | Nisch | 64652 | HITS-CLIP | 23142080\|19536157\|29167373 | Unspecified |
| mmu-miR-34a-5p | MIMAT0000542 | Nit1 | 27045 | HITS-CLIP | 23142080 | Unspecified |
| mmu-miR-34a-5p | MIMAT0000542 | Nkap | 67050 | Microarrays//HITS-CLIP | 37986224\|23027903\|23142080 | Unspecified |
| mmu-miR-34a-5p | MIMAT0000542 | Nkiras2 | 71966 | HITS-CLIP | 23142080\|29167373 | Unspecified |
| mmu-miR-34a-5p | MIMAT0000542 | Nkrf | 77286 | Microarrays//HITS-CLIP | 37986224\|23027903\|23142080 | Unspecified |
| mmu-miR-34a-5p | MIMAT0000542 | Nktr | 18087 | HITS-CLIP | 23142080\|29167373 | Unspecified |
| mmu-miR-34a-5p | MIMAT0000542 | Nlgn1 | 192167 | HITS-CLIP | 19536157 | Unspecified |
| mmu-miR-34a-5p | MIMAT0000542 | Nlgn2 | 216856 | HITS-CLIP | 19536157\|23142080 | Unspecified |
| mmu-miR-34a-5p | MIMAT0000542 | Nlk | 18099 | HITS-CLIP | 23142080\|31433990 | Unspecified |
| mmu-miR-34a-5p | MIMAT0000542 | Nln | 75805 | HITS-CLIP | 23142080\|29167373 | Unspecified |
| mmu-miR-34a-5p | MIMAT0000542 | Nlrc5 | 434341 | HITS-CLIP | 23142080 | Unspecified |
| mmu-miR-34a-5p | MIMAT0000542 | Nlrp1b | 637515 | HITS-CLIP | 29167373 | Unspecified |
| mmu-miR-34a-5p | MIMAT0000542 | Nlrp3 | 216799 | HITS-CLIP | 29167373 | Unspecified |
| mmu-miR-34a-5p | MIMAT0000542 | Nlrx1 | 270151 | HITS-CLIP | 29167373 | Unspecified |
| mmu-miR-34a-5p | MIMAT0000542 | Nmd3 | 97112 | HITS-CLIP | 23142080 | Unspecified |
| mmu-miR-34a-5p | MIMAT0000542 | Nme1 | 18102 | HITS-CLIP | 23142080\|19536157 | Unspecified |
| mmu-miR-34a-5p | MIMAT0000542 | Nme4 | 56520 | HITS-CLIP | 23142080 | Unspecified |
| mmu-miR-34a-5p | MIMAT0000542 | Nme5 | 75533 | Microarrays | 37986224 | Unspecified |
| mmu-miR-34a-5p | MIMAT0000542 | Nmnat3 | 74080 | HITS-CLIP | 23142080 | Unspecified |
| mmu-miR-34a-5p | MIMAT0000542 | Nmral1 | 67824 | HITS-CLIP | 23142080 | Unspecified |
| mmu-miR-34a-5p | MIMAT0000542 | Nmrk2 | 69564 | Microarrays | 37986224 | Unspecified |
| mmu-miR-34a-5p | MIMAT0000542 | Nmt1 | 18107 | HITS-CLIP | 23142080\|29167373 | Unspecified |
| mmu-miR-34a-5p | MIMAT0000542 | Nmt2 | 18108 | HITS-CLIP | 23142080\|29167373 | Unspecified |
| mmu-miR-34a-5p | MIMAT0000542 | Noc2l | 57741 | HITS-CLIP | 23142080\|29167373 | Unspecified |
| mmu-miR-34a-5p | MIMAT0000542 | Noc3l | 57753 | HITS-CLIP | 23142080 | Unspecified |
| mmu-miR-34a-5p | MIMAT0000542 | Noct | 12457 | HITS-CLIP | 23142080\|31433990 | Unspecified |

| | | | | | | |
|---|---|---|---|---|---|---|
| mmu-miR-34a-5p | MIMAT0000542 | Nod1 | 107607 | HITS-CLIP | 23142080 | Unspecified |
| mmu-miR-34a-5p | MIMAT0000542 | Nod2 | 257632 | HITS-CLIP | 29167373 | Unspecified |
| mmu-miR-34a-5p | MIMAT0000542 | Nol10 | 217431 | HITS-CLIP | 23142080 | Unspecified |
| mmu-miR-34a-5p | MIMAT0000542 | Nol11 | 68979 | HITS-CLIP | 23142080 | Unspecified |
| mmu-miR-34a-5p | MIMAT0000542 | Nol12 | 97961 | HITS-CLIP | 23142080 | Unspecified |
| mmu-miR-34a-5p | MIMAT0000542 | Nol4l | 329540 | HITS-CLIP | 30971389\|23142080\|29167373 | Unspecified |
| mmu-miR-34a-5p | MIMAT0000542 | Nol8 | 70930 | HITS-CLIP | 29167373 | Unspecified |
| mmu-miR-34a-5p | MIMAT0000542 | Nol9 | 74035 | HITS-CLIP | 19536157\|23142080 | Unspecified |
| mmu-miR-34a-5p | MIMAT0000542 | Nolc1 | 70769 | HITS-CLIP | 23142080\|31433990 | Unspecified |
| mmu-miR-34a-5p | MIMAT0000542 | Nom1 | 433864 | HITS-CLIP | 23142080\|31433990 | Unspecified |
| mmu-miR-34a-5p | MIMAT0000542 | Nomo1 | 211548 | HITS-CLIP | 30971389 | Unspecified |
| mmu-miR-34a-5p | MIMAT0000542 | Nono | 53610 | HITS-CLIP | 23142080 | Unspecified |
| mmu-miR-34a-5p | MIMAT0000542 | Nop10 | 66181 | HITS-CLIP | 23142080 | Unspecified |
| mmu-miR-34a-5p | MIMAT0000542 | Nop14 | 75416 | HITS-CLIP | 23142080 | Unspecified |
| mmu-miR-34a-5p | MIMAT0000542 | Nop56 | 67134 | HITS-CLIP | 23142080 | Unspecified |
| mmu-miR-34a-5p | MIMAT0000542 | Nop58 | 55989 | HITS-CLIP | 23142080\|29167373 | Unspecified |
| mmu-miR-34a-5p | MIMAT0000542 | Nop9 | 67842 | HITS-CLIP | 29167373 | Unspecified |
| mmu-miR-34a-5p | MIMAT0000542 | Nopchap1 | 121053 | Microarrays | 23027903 | Unspecified |
| mmu-miR-34a-5p | MIMAT0000542 | Nosip | 66394 | HITS-CLIP | 23142080 | Unspecified |
| mmu-miR-34a-5p | MIMAT0000542 | Notch1 | 18128 | ELISA//Immunofluorescence//Luciferase a | 22483937\|23142080\|29167373\|24587 | Unspecified |
| mmu-miR-34a-5p | MIMAT0000542 | Notch2 | 18129 | HITS-CLIP | 31433990\|23142080\|26203562\|29167 | Unspecified |
| mmu-miR-34a-5p | MIMAT0000542 | Notch3 | 18131 | HITS-CLIP | 30971389 | Unspecified |
| mmu-miR-34a-5p | MIMAT0000542 | Nox4 | 50490 | Microarrays | 37986224 | Unspecified |
| mmu-miR-34a-5p | MIMAT0000542 | Npas2 | 18143 | HITS-CLIP | 23142080 | Unspecified |
| mmu-miR-34a-5p | MIMAT0000542 | Npat | 244879 | HITS-CLIP | 31433990\|19536157\|23630078\|30971 | Unspecified |
| mmu-miR-34a-5p | MIMAT0000542 | Npc1 | 18145 | HITS-CLIP | 23142080\|29167373 | Unspecified |
| mmu-miR-34a-5p | MIMAT0000542 | Npc2 | 67963 | HITS-CLIP | 23142080\|29167373 | Unspecified |
| mmu-miR-34a-5p | MIMAT0000542 | Npepl1 | 228961 | HITS-CLIP | 23142080 | Unspecified |
| mmu-miR-34a-5p | MIMAT0000542 | Npepps | 19155 | HITS-CLIP | 23142080\|19536157\|23630078\|26203 | Unspecified |
| mmu-miR-34a-5p | MIMAT0000542 | Nphp3 | 74025 | HITS-CLIP | 23142080 | Unspecified |
| mmu-miR-34a-5p | MIMAT0000542 | Nploc4 | 217365 | HITS-CLIP | 23630078\|30971389\|23142080\|29167 | Unspecified |
| mmu-miR-34a-5p | MIMAT0000542 | Npm1 | 18148 | HITS-CLIP | 23142080 | Unspecified |
| mmu-miR-34a-5p | MIMAT0000542 | Nprl2 | 56032 | HITS-CLIP | 23142080 | Unspecified |
| mmu-miR-34a-5p | MIMAT0000542 | Nptn | 20320 | HITS-CLIP | 37986224\|23142080\|19536157\|2363C | Unspecified |
| mmu-miR-34a-5p | MIMAT0000542 | Nptx1 | 18164 | HITS-CLIP | 19536157\|29167373 | Unspecified |
| mmu-miR-34a-5p | MIMAT0000542 | Nptxr | 73340 | HITS-CLIP | 19536157 | Unspecified |
| mmu-miR-34a-5p | MIMAT0000542 | Npy | 109648 | HITS-CLIP | 29167373 | Unspecified |
| mmu-miR-34a-5p | MIMAT0000542 | Npy2r | 18167 | HITS-CLIP | 19536157 | Unspecified |
| mmu-miR-34a-5p | MIMAT0000542 | Nr1d2 | 353187 | HITS-CLIP | 19536157\|23630078 | Unspecified |

| | | | | | | |
|---|---|---|---|---|---|---|
| mmu-miR-34a-5p | MIMAT0000542 | Nr1h2 | 22260 | HITS-CLIP | 23142080 | Unspecified |
| mmu-miR-34a-5p | MIMAT0000542 | Nr2c1 | 22025 | HITS-CLIP | 23142080\|29167373 | Unspecified |
| mmu-miR-34a-5p | MIMAT0000542 | Nr2c2 | 22026 | HITS-CLIP | 37986224\|19536157\|31433990\|23142 | Unspecified |
| mmu-miR-34a-5p | MIMAT0000542 | Nr2f6 | 13864 | HITS-CLIP | 29167373 | Unspecified |
| mmu-miR-34a-5p | MIMAT0000542 | Nr3c1 | 14815 | HITS-CLIP | 19536157\|31433990\|23142080\|29167 | Unspecified |
| mmu-miR-34a-5p | MIMAT0000542 | Nr4a1 | 15370 | HITS-CLIP | 23142080 | Unspecified |
| mmu-miR-34a-5p | MIMAT0000542 | Nr4a2 | 18227 | HITS-CLIP | 37986224\|23142080 | Unspecified |
| mmu-miR-34a-5p | MIMAT0000542 | Nr4a3 | 18124 | HITS-CLIP | 23142080 | Unspecified |
| mmu-miR-34a-5p | MIMAT0000542 | Nr6a1 | 14536 | HITS-CLIP | 23630078\|31433990\|23142080 | Unspecified |
| mmu-miR-34a-5p | MIMAT0000542 | Nras | 18176 | HITS-CLIP | 31433990\|29167373\|23142080 | Unspecified |
| mmu-miR-34a-5p | MIMAT0000542 | Nrbf2 | 641340 | HITS-CLIP | 23142080 | Unspecified |
| mmu-miR-34a-5p | MIMAT0000542 | Nrbp1 | 192292 | HITS-CLIP | 23142080\|29167373 | Unspecified |
| mmu-miR-34a-5p | MIMAT0000542 | Nrbp2 | 223649 | HITS-CLIP | 23142080 | Unspecified |
| mmu-miR-34a-5p | MIMAT0000542 | Nrd1 | 230598 | HITS-CLIP | 23142080\|29167373 | Unspecified |
| mmu-miR-34a-5p | MIMAT0000542 | Nrde2 | 217827 | HITS-CLIP | 23142080\|29167373 | Unspecified |
| mmu-miR-34a-5p | MIMAT0000542 | Nrep | 27528 | HITS-CLIP | 19536157 | Unspecified |
| mmu-miR-34a-5p | MIMAT0000542 | Nrip1 | 268903 | HITS-CLIP | 19536157\|23142080\|29167373 | Unspecified |
| mmu-miR-34a-5p | MIMAT0000542 | Nrn1 | 68404 | HITS-CLIP | 23142080 | Unspecified |
| mmu-miR-34a-5p | MIMAT0000542 | Nrp2 | 18187 | HITS-CLIP | 29167373 | Unspecified |
| mmu-miR-34a-5p | MIMAT0000542 | Nrros | 224109 | HITS-CLIP | 37986224\|23142080\|29167373 | Unspecified |
| mmu-miR-34a-5p | MIMAT0000542 | Nrsn1 | 22360 | HITS-CLIP | 19536157 | Unspecified |
| mmu-miR-34a-5p | MIMAT0000542 | Nrxn1 | 18189 | HITS-CLIP | 19536157 | Unspecified |
| mmu-miR-34a-5p | MIMAT0000542 | Nrxn2 | 18190 | HITS-CLIP | 19536157 | Unspecified |
| mmu-miR-34a-5p | MIMAT0000542 | Nsd1 | 18193 | HITS-CLIP | 37986224\|19536157\|31433990\|23142 | Unspecified |
| mmu-miR-34a-5p | MIMAT0000542 | Nsd2 | 107823 | HITS-CLIP | 23142080\|29167373 | Unspecified |
| mmu-miR-34a-5p | MIMAT0000542 | Nsd3 | 234135 | HITS-CLIP | 23142080\|29167373 | Unspecified |
| mmu-miR-34a-5p | MIMAT0000542 | Nsf | 18195 | HITS-CLIP | 19536157\|23142080 | Unspecified |
| mmu-miR-34a-5p | MIMAT0000542 | Nsg1 | 18196 | HITS-CLIP | 19536157 | Unspecified |
| mmu-miR-34a-5p | MIMAT0000542 | Nsg2 | 18197 | HITS-CLIP | 23142080\|19536157 | Unspecified |
| mmu-miR-34a-5p | MIMAT0000542 | Nsmaf | 18201 | HITS-CLIP | 23142080 | Unspecified |
| mmu-miR-34a-5p | MIMAT0000542 | Nsmce2 | 68501 | HITS-CLIP | 29167373 | Unspecified |
| mmu-miR-34a-5p | MIMAT0000542 | Nsmce4a | 67872 | Microarrays//HITS-CLIP | 37986224\|23027903\|23142080\|29167 | Unspecified |
| mmu-miR-34a-5p | MIMAT0000542 | Nsmf | 56876 | HITS-CLIP | 19536157 | Unspecified |
| mmu-miR-34a-5p | MIMAT0000542 | Nsrp1 | 237859 | HITS-CLIP | 23142080 | Unspecified |
| mmu-miR-34a-5p | MIMAT0000542 | Nsun2 | 28114 | HITS-CLIP | 23142080\|29167373 | Unspecified |
| mmu-miR-34a-5p | MIMAT0000542 | Nsun3 | 106338 | Microarrays | 37986224 | Unspecified |
| mmu-miR-34a-5p | MIMAT0000542 | Nsun4 | 72181 | HITS-CLIP | 23142080 | Unspecified |
| mmu-miR-34a-5p | MIMAT0000542 | Nsun6 | 74455 | HITS-CLIP | 23142080 | Unspecified |
| mmu-miR-34a-5p | MIMAT0000542 | Nt5c2 | 76952 | HITS-CLIP | 29167373 | Unspecified |

| | | | | | | |
|---|---|---|---|---|---|---|
| mmu-miR-34a-5p | MIMAT0000542 | Nt5c3 | 107569 | HITS-CLIP | 23142080 | Unspecified |
| mmu-miR-34a-5p | MIMAT0000542 | Nt5c3b | 68106 | HITS-CLIP | 30971389 | Unspecified |
| mmu-miR-34a-5p | MIMAT0000542 | Nt5dc1 | 319638 | HITS-CLIP | 23142080 | Unspecified |
| mmu-miR-34a-5p | MIMAT0000542 | Nt5dc3 | 103466 | HITS-CLIP | 26203562\|23142080\|29167373 | Unspecified |
| mmu-miR-34a-5p | MIMAT0000542 | Nt5e | 23959 | HITS-CLIP | 23142080 | Unspecified |
| mmu-miR-34a-5p | MIMAT0000542 | Ntan1 | 18203 | HITS-CLIP | 23142080 | Unspecified |
| mmu-miR-34a-5p | MIMAT0000542 | Ntm | 235106 | HITS-CLIP | 19536157 | Unspecified |
| mmu-miR-34a-5p | MIMAT0000542 | Ntn1 | 18208 | Microarrays//HITS-CLIP | 37986224\|23027903\|19536157 | Unspecified |
| mmu-miR-34a-5p | MIMAT0000542 | Ntn4 | 57764 | HITS-CLIP | 19536157 | Unspecified |
| mmu-miR-34a-5p | MIMAT0000542 | Ntng2 | 171171 | HITS-CLIP | 19536157 | Unspecified |
| mmu-miR-34a-5p | MIMAT0000542 | Ntrk2 | 18212 | HITS-CLIP | 19536157 | Unspecified |
| mmu-miR-34a-5p | MIMAT0000542 | Ntrk3 | 18213 | HITS-CLIP | 19536157 | Unspecified |
| mmu-miR-34a-5p | MIMAT0000542 | Nuak1 | 77976 | HITS-CLIP | 26203562 | Unspecified |
| mmu-miR-34a-5p | MIMAT0000542 | Nuak2 | 74137 | HITS-CLIP | 23142080\|29167373 | Unspecified |
| mmu-miR-34a-5p | MIMAT0000542 | Nub1 | 53312 | HITS-CLIP | 23142080\|31433990\|29167373 | Unspecified |
| mmu-miR-34a-5p | MIMAT0000542 | Nubp2 | 26426 | HITS-CLIP | 19536157 | Unspecified |
| mmu-miR-34a-5p | MIMAT0000542 | Nucb1 | 18220 | HITS-CLIP | 23142080 | Unspecified |
| mmu-miR-34a-5p | MIMAT0000542 | Nucks1 | 98415 | HITS-CLIP | 31433990\|30971389\|19536157\|23142 | Unspecified |
| mmu-miR-34a-5p | MIMAT0000542 | Nudcd1 | 67429 | HITS-CLIP | 23142080 | Unspecified |
| mmu-miR-34a-5p | MIMAT0000542 | Nudcd2 | 52653 | HITS-CLIP | 30971389\|23142080 | Unspecified |
| mmu-miR-34a-5p | MIMAT0000542 | Nudcd3 | 209586 | HITS-CLIP | 23142080 | Unspecified |
| mmu-miR-34a-5p | MIMAT0000542 | Nudt1 | 17766 | HITS-CLIP | 23142080 | Unspecified |
| mmu-miR-34a-5p | MIMAT0000542 | Nudt13 | 67725 | HITS-CLIP | 29167373 | Unspecified |
| mmu-miR-34a-5p | MIMAT0000542 | Nudt18 | 213484 | HITS-CLIP | 23142080 | Unspecified |
| mmu-miR-34a-5p | MIMAT0000542 | Nudt19 | 110959 | Microarrays//HITS-CLIP | 37986224\|23027903\|23142080 | Unspecified |
| mmu-miR-34a-5p | MIMAT0000542 | Nudt2 | 66401 | HITS-CLIP | 23142080 | Unspecified |
| mmu-miR-34a-5p | MIMAT0000542 | Nudt21 | 68219 | Microarrays//HITS-CLIP | 37986224\|23027903\|23142080 | Unspecified |
| mmu-miR-34a-5p | MIMAT0000542 | Nudt3 | 56409 | HITS-CLIP | 23142080\|19536157 | Unspecified |
| mmu-miR-34a-5p | MIMAT0000542 | Nudt4 | 71207 | HITS-CLIP | 23142080\|19536157\|29167373 | Unspecified |
| mmu-miR-34a-5p | MIMAT0000542 | Nudt5 | 53893 | HITS-CLIP | 23142080 | Unspecified |
| mmu-miR-34a-5p | MIMAT0000542 | Nudt7 | 67528 | HITS-CLIP | 23142080 | Unspecified |
| mmu-miR-34a-5p | MIMAT0000542 | Nudt8 | 66387 | HITS-CLIP | 23142080 | Unspecified |
| mmu-miR-34a-5p | MIMAT0000542 | Nuf2 | 66977 | HITS-CLIP | 23142080 | Unspecified |
| mmu-miR-34a-5p | MIMAT0000542 | Nufip1 | 27275 | Microarrays | 37986224\|23027903 | Unspecified |
| mmu-miR-34a-5p | MIMAT0000542 | Nufip2 | 68564 | HITS-CLIP | 37986224\|23142080\|29167373 | Unspecified |
| mmu-miR-34a-5p | MIMAT0000542 | Numa1 | 101706 | HITS-CLIP | 23142080\|29167373 | Unspecified |
| mmu-miR-34a-5p | MIMAT0000542 | Numb | 18222 | HITS-CLIP | 23142080\|29167373 | Unspecified |
| mmu-miR-34a-5p | MIMAT0000542 | Nup107 | 103468 | HITS-CLIP | 23142080 | Unspecified |
| mmu-miR-34a-5p | MIMAT0000542 | Nup133 | 234865 | HITS-CLIP | 23142080 | Unspecified |

| | | | | | | |
|---|---|---|---|---|---|---|
| mmu-miR-34a-5p | MIMAT0000542 | Nup153 | 218210 | HITS-CLIP | 23142080\|23630078\|29167373 | Unspecified |
| mmu-miR-34a-5p | MIMAT0000542 | Nup155 | 170762 | HITS-CLIP | 23142080\|31433990\|29167373 | Unspecified |
| mmu-miR-34a-5p | MIMAT0000542 | Nup160 | 59015 | HITS-CLIP | 23142080\|29167373 | Unspecified |
| mmu-miR-34a-5p | MIMAT0000542 | Nup188 | 227699 | HITS-CLIP | 26203562\|29167373\|23142080 | Unspecified |
| mmu-miR-34a-5p | MIMAT0000542 | Nup205 | 70699 | HITS-CLIP | 30971389\|23142080\|29167373 | Unspecified |
| mmu-miR-34a-5p | MIMAT0000542 | Nup210 | 54563 | HITS-CLIP | 31433990\|23142080 | Unspecified |
| mmu-miR-34a-5p | MIMAT0000542 | Nup210l | 77595 | HITS-CLIP | 23142080 | Unspecified |
| mmu-miR-34a-5p | MIMAT0000542 | Nup214 | 227720 | HITS-CLIP | 23142080\|29167373 | Unspecified |
| mmu-miR-34a-5p | MIMAT0000542 | Nup37 | 69736 | HITS-CLIP | 23142080\|31433990 | Unspecified |
| mmu-miR-34a-5p | MIMAT0000542 | Nup43 | 69912 | Microarrays//HITS-CLIP | 37986224\|23027903\|23142080 | Unspecified |
| mmu-miR-34a-5p | MIMAT0000542 | Nup50 | 18141 | HITS-CLIP | 23142080 | Unspecified |
| mmu-miR-34a-5p | MIMAT0000542 | Nup54 | 269113 | HITS-CLIP | 23142080 | Unspecified |
| mmu-miR-34a-5p | MIMAT0000542 | Nup62 | 18226 | HITS-CLIP | 31433990\|23142080\|29167373 | Unspecified |
| mmu-miR-34a-5p | MIMAT0000542 | Nup85 | 445007 | HITS-CLIP | 23142080 | Unspecified |
| mmu-miR-34a-5p | MIMAT0000542 | Nup88 | 19069 | HITS-CLIP | 23142080\|29167373 | Unspecified |
| mmu-miR-34a-5p | MIMAT0000542 | Nup93 | 71805 | HITS-CLIP | 23142080\|29167373 | Unspecified |
| mmu-miR-34a-5p | MIMAT0000542 | Nup98 | 269966 | HITS-CLIP | 23142080 | Unspecified |
| mmu-miR-34a-5p | MIMAT0000542 | Nupl1 | 71844 | HITS-CLIP | 30971389 | Unspecified |
| mmu-miR-34a-5p | MIMAT0000542 | Nupl2 | 231042 | HITS-CLIP | 23142080 | Unspecified |
| mmu-miR-34a-5p | MIMAT0000542 | Nus1 | 52014 | HITS-CLIP | 29167373 | Unspecified |
| mmu-miR-34a-5p | MIMAT0000542 | Nusap1 | 108907 | HITS-CLIP | 31433990\|23142080 | Unspecified |
| mmu-miR-34a-5p | MIMAT0000542 | Nvl | 67459 | HITS-CLIP | 23142080\|29167373 | Unspecified |
| mmu-miR-34a-5p | MIMAT0000542 | Nxf1 | 53319 | HITS-CLIP | 23142080\|23630078\|29167373 | Unspecified |
| mmu-miR-34a-5p | MIMAT0000542 | Nxn | 18230 | HITS-CLIP | 23142080 | Unspecified |
| mmu-miR-34a-5p | MIMAT0000542 | Nxpe2 | 78252 | HITS-CLIP | 23142080 | Unspecified |
| mmu-miR-34a-5p | MIMAT0000542 | Nxpe3 | 385658 | HITS-CLIP | 23142080 | Unspecified |
| mmu-miR-34a-5p | MIMAT0000542 | Nxpe4 | 244853 | HITS-CLIP | 30971389 | Unspecified |
| mmu-miR-34a-5p | MIMAT0000542 | Nxt1 | 56488 | HITS-CLIP | 23142080 | Unspecified |
| mmu-miR-34a-5p | MIMAT0000542 | Nxt2 | 237082 | Microarrays//HITS-CLIP | 37986224\|23027903\|23142080 | Unspecified |
| mmu-miR-34a-5p | MIMAT0000542 | Oaf | 102644 | HITS-CLIP | 19536157 | Unspecified |
| mmu-miR-34a-5p | MIMAT0000542 | Oas1a | 246730 | HITS-CLIP | 29167373 | Unspecified |
| mmu-miR-34a-5p | MIMAT0000542 | Oas1g | 23960 | HITS-CLIP | 29167373 | Unspecified |
| mmu-miR-34a-5p | MIMAT0000542 | Oas2 | 246728 | HITS-CLIP | 29167373 | Unspecified |
| mmu-miR-34a-5p | MIMAT0000542 | Oas3 | 246727 | HITS-CLIP | 23142080\|29167373 | Unspecified |
| mmu-miR-34a-5p | MIMAT0000542 | Oasl2 | 23962 | HITS-CLIP | 29167373 | Unspecified |
| mmu-miR-34a-5p | MIMAT0000542 | Oat | 18242 | HITS-CLIP | 23142080\|29167373 | Unspecified |
| mmu-miR-34a-5p | MIMAT0000542 | Oaz1 | 18245 | HITS-CLIP | 23142080 | Unspecified |
| mmu-miR-34a-5p | MIMAT0000542 | Ociad1 | 68095 | HITS-CLIP | 23142080\|19536157 | Unspecified |
| mmu-miR-34a-5p | MIMAT0000542 | Ocrl | 320634 | HITS-CLIP | 23142080 | Unspecified |

| | | | | | | |
|---|---|---|---|---|---|---|
| mmu-miR-34a-5p | MIMAT0000542 | Odc1 | 18263 | HITS-CLIP | 23142080 | Unspecified |
| mmu-miR-34a-5p | MIMAT0000542 | Odf2l | 52184 | HITS-CLIP | 23142080 | Unspecified |
| mmu-miR-34a-5p | MIMAT0000542 | Odr4 | 54953 | HITS-CLIP | 29167373 | Unspecified |
| mmu-miR-34a-5p | MIMAT0000542 | Ogdh | 18293 | HITS-CLIP | 23142080 | Unspecified |
| mmu-miR-34a-5p | MIMAT0000542 | Ogdhl | 239017 | Microarrays//HITS-CLIP | 37986224\|19536157 | Unspecified |
| mmu-miR-34a-5p | MIMAT0000542 | Ogfod1 | 270086 | Microarrays//HITS-CLIP | 37986224\|23027903\|23142080 | Unspecified |
| mmu-miR-34a-5p | MIMAT0000542 | Ogfod2 | 66627 | HITS-CLIP | 23142080 | Unspecified |
| mmu-miR-34a-5p | MIMAT0000542 | Ogfr | 72075 | HITS-CLIP | 37986224\|29167373\|23142080 | Unspecified |
| mmu-miR-34a-5p | MIMAT0000542 | Ogt | 108155 | HITS-CLIP | 19536157\|23142080\|29167373 | Unspecified |
| mmu-miR-34a-5p | MIMAT0000542 | Ola1 | 67059 | HITS-CLIP | 30971389\|23142080\|29167373 | Unspecified |
| mmu-miR-34a-5p | MIMAT0000542 | Olfm1 | 56177 | HITS-CLIP | 19536157\|29167373 | Unspecified |
| mmu-miR-34a-5p | MIMAT0000542 | Olfr490 | 258491 | HITS-CLIP | 29167373 | Unspecified |
| mmu-miR-34a-5p | MIMAT0000542 | Olfr624 | 258189 | HITS-CLIP | 29167373 | Unspecified |
| mmu-miR-34a-5p | MIMAT0000542 | Oma1 | 67013 | HITS-CLIP | 29167373 | Unspecified |
| mmu-miR-34a-5p | MIMAT0000542 | Opa1 | 74143 | HITS-CLIP | 23142080\|29167373 | Unspecified |
| mmu-miR-34a-5p | MIMAT0000542 | Opcml | 330908 | HITS-CLIP | 19536157 | Unspecified |
| mmu-miR-34a-5p | MIMAT0000542 | Oprl1 | 18389 | HITS-CLIP | 19536157 | Unspecified |
| mmu-miR-34a-5p | MIMAT0000542 | Orai1 | 109305 | HITS-CLIP | 23142080 | Unspecified |
| mmu-miR-34a-5p | MIMAT0000542 | Orai2 | 269717 | HITS-CLIP | 31433990 | Unspecified |
| mmu-miR-34a-5p | MIMAT0000542 | Orai3 | 269999 | HITS-CLIP | 30971389 | Unspecified |
| mmu-miR-34a-5p | MIMAT0000542 | Orc1 | 18392 | Microarrays//HITS-CLIP | 37986224\|23027903\|23142080 | Unspecified |
| mmu-miR-34a-5p | MIMAT0000542 | Orc2 | 18393 | Microarrays//HITS-CLIP | 37986224\|23027903\|19536157 | Unspecified |
| mmu-miR-34a-5p | MIMAT0000542 | Orc3 | 50793 | HITS-CLIP | 23142080\|29167373 | Unspecified |
| mmu-miR-34a-5p | MIMAT0000542 | Orc4 | 26428 | HITS-CLIP | 23142080\|29167373 | Unspecified |
| mmu-miR-34a-5p | MIMAT0000542 | Orc5 | 26429 | HITS-CLIP | 29167373\|23142080 | Unspecified |
| mmu-miR-34a-5p | MIMAT0000542 | Orc6 | 56452 | HITS-CLIP | 23142080 | Unspecified |
| mmu-miR-34a-5p | MIMAT0000542 | Ormdl3 | 66612 | HITS-CLIP | 30971389\|29167373 | Unspecified |
| mmu-miR-34a-5p | MIMAT0000542 | Os9 | 216440 | HITS-CLIP | 29167373 | Unspecified |
| mmu-miR-34a-5p | MIMAT0000542 | Osbp | 76303 | HITS-CLIP | 23142080 | Unspecified |
| mmu-miR-34a-5p | MIMAT0000542 | Osbpl11 | 106326 | HITS-CLIP | 29167373 | Unspecified |
| mmu-miR-34a-5p | MIMAT0000542 | Osbpl1a | 64291 | HITS-CLIP | 19536157 | Unspecified |
| mmu-miR-34a-5p | MIMAT0000542 | Osbpl2 | 228983 | HITS-CLIP | 23142080\|29167373 | Unspecified |
| mmu-miR-34a-5p | MIMAT0000542 | Osbpl3 | 71720 | HITS-CLIP | 23142080 | Unspecified |
| mmu-miR-34a-5p | MIMAT0000542 | Osbpl8 | 237542 | HITS-CLIP | 23142080\|29167373 | Unspecified |
| mmu-miR-34a-5p | MIMAT0000542 | Osbpl9 | 100273 | HITS-CLIP | 23142080 | Unspecified |
| mmu-miR-34a-5p | MIMAT0000542 | Osgepl1 | 72085 | HITS-CLIP | 23142080 | Unspecified |
| mmu-miR-34a-5p | MIMAT0000542 | Osgin2 | 209212 | Microarrays//HITS-CLIP | 37986224\|23027903\|23630078\|3143 | Unspecified |
| mmu-miR-34a-5p | MIMAT0000542 | Osm | 18413 | HITS-CLIP | 23142080 | Unspecified |
| mmu-miR-34a-5p | MIMAT0000542 | Ostf1 | 20409 | HITS-CLIP | 23142080\|29167373 | Unspecified |

| | | | | | | |
|---|---|---|---|---|---|---|
| mmu-miR-34a-5p | MIMAT0000542 | Ostm1 | 14628 | Microarrays//HITS-CLIP | 37986224\|29167373 | Unspecified |
| mmu-miR-34a-5p | MIMAT0000542 | Otub1 | 107260 | HITS-CLIP | 23142080\|29167373 | Unspecified |
| mmu-miR-34a-5p | MIMAT0000542 | Otub2 | 68149 | HITS-CLIP | 23142080 | Unspecified |
| mmu-miR-34a-5p | MIMAT0000542 | Otud1 | 71198 | HITS-CLIP | 23142080 | Unspecified |
| mmu-miR-34a-5p | MIMAT0000542 | Otud4 | 73945 | HITS-CLIP | 23142080\|29167373 | Unspecified |
| mmu-miR-34a-5p | MIMAT0000542 | Otud5 | 54644 | HITS-CLIP | 29167373\|23142080 | Unspecified |
| mmu-miR-34a-5p | MIMAT0000542 | Otud6b | 72201 | HITS-CLIP | 19536157\|23142080\|29167373 | Unspecified |
| mmu-miR-34a-5p | MIMAT0000542 | Otud7b | 229603 | HITS-CLIP | 23142080 | Unspecified |
| mmu-miR-34a-5p | MIMAT0000542 | Otulin | 432940 | HITS-CLIP | 23142080 | Unspecified |
| mmu-miR-34a-5p | MIMAT0000542 | Otulinl | 54491 | HITS-CLIP | 23142080\|29167373 | Unspecified |
| mmu-miR-34a-5p | MIMAT0000542 | Otx2 | 18424 | HITS-CLIP | 29084942 | Unspecified |
| mmu-miR-34a-5p | MIMAT0000542 | Ovca2 | 246257 | HITS-CLIP | 23142080 | Unspecified |
| mmu-miR-34a-5p | MIMAT0000542 | Oxa1l | 69089 | HITS-CLIP | 23630078\|29167373 | Unspecified |
| mmu-miR-34a-5p | MIMAT0000542 | Oxct1 | 67041 | HITS-CLIP | 19536157\|23142080\|29167373 | Unspecified |
| mmu-miR-34a-5p | MIMAT0000542 | Oxr1 | 170719 | HITS-CLIP | 23142080\|29167373 | Unspecified |
| mmu-miR-34a-5p | MIMAT0000542 | Oxsm | 71147 | HITS-CLIP | 23142080 | Unspecified |
| mmu-miR-34a-5p | MIMAT0000542 | Oxsr1 | 108737 | HITS-CLIP | 23630078\|23142080\|29167373 | Unspecified |
| mmu-miR-34a-5p | MIMAT0000542 | P2rx4 | 18438 | HITS-CLIP | 23142080\|29167373 | Unspecified |
| mmu-miR-34a-5p | MIMAT0000542 | P2rx7 | 18439 | HITS-CLIP | 29167373 | Unspecified |
| mmu-miR-34a-5p | MIMAT0000542 | P2ry1 | 18441 | HITS-CLIP | 23142080 | Unspecified |
| mmu-miR-34a-5p | MIMAT0000542 | P2ry10 | 78826 | HITS-CLIP | 23142080 | Unspecified |
| mmu-miR-34a-5p | MIMAT0000542 | P2ry10b | 213438 | HITS-CLIP | 23142080\|29167373 | Unspecified |
| mmu-miR-34a-5p | MIMAT0000542 | P2ry12 | 70839 | HITS-CLIP | 23142080 | Unspecified |
| mmu-miR-34a-5p | MIMAT0000542 | P2ry6 | 233571 | HITS-CLIP | 29167373 | Unspecified |
| mmu-miR-34a-5p | MIMAT0000542 | P3h4 | 10609 | HITS-CLIP | 23630078 | Unspecified |
| mmu-miR-34a-5p | MIMAT0000542 | P4ha1 | 18451 | HITS-CLIP | 23142080 | Unspecified |
| mmu-miR-34a-5p | MIMAT0000542 | P4hb | 18453 | HITS-CLIP | 23142080\|29167373 | Unspecified |
| mmu-miR-34a-5p | MIMAT0000542 | Pabpc1 | 18458 | HITS-CLIP | 23142080\|29167373 | Unspecified |
| mmu-miR-34a-5p | MIMAT0000542 | Pacs1 | 107975 | Microarrays//HITS-CLIP | 37986224\|23027903\|30971389 | Unspecified |
| mmu-miR-34a-5p | MIMAT0000542 | Pacs2 | 217893 | HITS-CLIP | 29167373 | Unspecified |
| mmu-miR-34a-5p | MIMAT0000542 | Pacsin2 | 23970 | HITS-CLIP | 29167373 | Unspecified |
| mmu-miR-34a-5p | MIMAT0000542 | Padi2 | 18600 | HITS-CLIP | 23142080 | Unspecified |
| mmu-miR-34a-5p | MIMAT0000542 | Pafah1b1 | 18472 | HITS-CLIP | 23142080\|19536157 | Unspecified |
| mmu-miR-34a-5p | MIMAT0000542 | Pafah1b2 | 18475 | HITS-CLIP | 23142080\|29167373 | Unspecified |
| mmu-miR-34a-5p | MIMAT0000542 | Pafah1b3 | 18476 | HITS-CLIP | 23142080 | Unspecified |
| mmu-miR-34a-5p | MIMAT0000542 | Pag1 | 94212 | HITS-CLIP | 23142080\|31433990\|19536157\|2363( | Unspecified |
| mmu-miR-34a-5p | MIMAT0000542 | Paics | 67054 | HITS-CLIP | 23142080 | Unspecified |
| mmu-miR-34a-5p | MIMAT0000542 | Paip1 | 218693 | HITS-CLIP | 23142080\|19536157 | Unspecified |
| mmu-miR-34a-5p | MIMAT0000542 | Paip2 | 67869 | HITS-CLIP | 37986224\|23142080\|29167373\|1953( | Unspecified |

| | | | | | | |
|---|---|---|---|---|---|---|
| mmu-miR-34a-5p | MIMAT0000542 | Paip2b | 232164 | HITS-CLIP | 30971389\|23142080 | Unspecified |
| mmu-miR-34a-5p | MIMAT0000542 | Pak2 | 224105 | HITS-CLIP | 23142080\|31433990\|29167373 | Unspecified |
| mmu-miR-34a-5p | MIMAT0000542 | Pak3 | 18481 | Microarrays | 37986224 | Unspecified |
| mmu-miR-34a-5p | MIMAT0000542 | Pak4 | 70584 | HITS-CLIP | 23142080\|29167373 | Unspecified |
| mmu-miR-34a-5p | MIMAT0000542 | Pak6 | 214230 | HITS-CLIP | 26203562 | Unspecified |
| mmu-miR-34a-5p | MIMAT0000542 | Pakap | 677884 | HITS-CLIP | 23630078 | Unspecified |
| mmu-miR-34a-5p | MIMAT0000542 | Palb2 | 233826 | HITS-CLIP | 23142080 | Unspecified |
| mmu-miR-34a-5p | MIMAT0000542 | Palld | 72333 | HITS-CLIP | 23142080 | Unspecified |
| mmu-miR-34a-5p | MIMAT0000542 | Palmd | 114301 | Microarrays | 37986224 | Unspecified |
| mmu-miR-34a-5p | MIMAT0000542 | Pam | 18484 | HITS-CLIP | 29167373 | Unspecified |
| mmu-miR-34a-5p | MIMAT0000542 | Pan2 | 103135 | HITS-CLIP | 23142080\|29167373 | Unspecified |
| mmu-miR-34a-5p | MIMAT0000542 | Pan3 | 72587 | HITS-CLIP | 23142080\|19536157\|29167373 | Unspecified |
| mmu-miR-34a-5p | MIMAT0000542 | Pank1 | 75735 | HITS-CLIP | 23142080 | Unspecified |
| mmu-miR-34a-5p | MIMAT0000542 | Pank2 | 74450 | HITS-CLIP | 29167373 | Unspecified |
| mmu-miR-34a-5p | MIMAT0000542 | Pank3 | 211347 | HITS-CLIP | 37986224\|23142080\|31433990\|1953( | Unspecified |
| mmu-miR-34a-5p | MIMAT0000542 | Pank4 | 55229 | HITS-CLIP | 30971389\|29167373 | Unspecified |
| mmu-miR-34a-5p | MIMAT0000542 | Panx1 | 55991 | HITS-CLIP | 23142080 | Unspecified |
| mmu-miR-34a-5p | MIMAT0000542 | Panx2 | 406218 | HITS-CLIP | 19536157 | Unspecified |
| mmu-miR-34a-5p | MIMAT0000542 | Papola | 18789 | HITS-CLIP | 23142080\|19536157\|29167373 | Unspecified |
| mmu-miR-34a-5p | MIMAT0000542 | Papolg | 216578 | HITS-CLIP | 23142080 | Unspecified |
| mmu-miR-34a-5p | MIMAT0000542 | Pappa | 18491 | Microarrays | 37986224 | Unspecified |
| mmu-miR-34a-5p | MIMAT0000542 | Papss1 | 23971 | HITS-CLIP | 23142080 | Unspecified |
| mmu-miR-34a-5p | MIMAT0000542 | Papss2 | 23972 | Microarrays | 37986224\|23027903 | Unspecified |
| mmu-miR-34a-5p | MIMAT0000542 | Paqr3 | 231474 | HITS-CLIP | 23630078 | Unspecified |
| mmu-miR-34a-5p | MIMAT0000542 | Paqr4 | 76498 | HITS-CLIP | 23142080\|19536157 | Unspecified |
| mmu-miR-34a-5p | MIMAT0000542 | Paqr7 | 71904 | HITS-CLIP | 29167373 | Unspecified |
| mmu-miR-34a-5p | MIMAT0000542 | Paqr8 | 74229 | HITS-CLIP | 23142080 | Unspecified |
| mmu-miR-34a-5p | MIMAT0000542 | Paqr9 | 75552 | HITS-CLIP | 19536157 | Unspecified |
| mmu-miR-34a-5p | MIMAT0000542 | Pard3 | 93742 | HITS-CLIP | 23630078 | Unspecified |
| mmu-miR-34a-5p | MIMAT0000542 | Parg | 26430 | HITS-CLIP | 23142080 | Unspecified |
| mmu-miR-34a-5p | MIMAT0000542 | Park7 | 57320 | HITS-CLIP | 23142080\|29167373 | Unspecified |
| mmu-miR-34a-5p | MIMAT0000542 | Parm1 | 231440 | HITS-CLIP | 19536157 | Unspecified |
| mmu-miR-34a-5p | MIMAT0000542 | Parp1 | 11545 | HITS-CLIP | 23142080\|29167373 | Unspecified |
| mmu-miR-34a-5p | MIMAT0000542 | Parp10 | 671535 | HITS-CLIP | 29167373 | Unspecified |
| mmu-miR-34a-5p | MIMAT0000542 | Parp11 | 101187 | HITS-CLIP | 23142080 | Unspecified |
| mmu-miR-34a-5p | MIMAT0000542 | Parp14 | 547253 | HITS-CLIP | 23142080\|29167373 | Unspecified |
| mmu-miR-34a-5p | MIMAT0000542 | Parp3 | 235587 | Microarrays//HITS-CLIP | 37986224\|23142080 | Unspecified |
| mmu-miR-34a-5p | MIMAT0000542 | Parp4 | 328417 | HITS-CLIP | 23142080\|31433990\|29167373 | Unspecified |
| mmu-miR-34a-5p | MIMAT0000542 | Parp8 | 52552 | HITS-CLIP | 23142080 | Unspecified |

| | | | | | | |
|---|---|---|---|---|---|---|
| mmu-miR-34a-5p | MIMAT0000542 | Parp9 | 80285 | HITS-CLIP | 23142080\|29167373 | Unspecified |
| mmu-miR-34a-5p | MIMAT0000542 | Parpbp | 75317 | HITS-CLIP | 23142080 | Unspecified |
| mmu-miR-34a-5p | MIMAT0000542 | Parva | 57342 | HITS-CLIP | 19536157 | Unspecified |
| mmu-miR-34a-5p | MIMAT0000542 | Parvb | 170736 | HITS-CLIP | 29167373 | Unspecified |
| mmu-miR-34a-5p | MIMAT0000542 | Parvg | 64099 | HITS-CLIP | 23142080\|31433990 | Unspecified |
| mmu-miR-34a-5p | MIMAT0000542 | Pask | 269224 | HITS-CLIP | 23142080 | Unspecified |
| mmu-miR-34a-5p | MIMAT0000542 | Patj | 44100 | HITS-CLIP | 23142080 | Unspecified |
| mmu-miR-34a-5p | MIMAT0000542 | Patl1 | 225929 | HITS-CLIP | 19536157\|23142080\|29167373 | Unspecified |
| mmu-miR-34a-5p | MIMAT0000542 | Pax5 | 5079 | HITS-CLIP | 23142080 | Unspecified |
| mmu-miR-34a-5p | MIMAT0000542 | Paxbp1 | 67367 | HITS-CLIP | 31433990 | Unspecified |
| mmu-miR-34a-5p | MIMAT0000542 | Paxip1 | 55982 | HITS-CLIP | 23142080 | Unspecified |
| mmu-miR-34a-5p | MIMAT0000542 | Pbdc1 | 67683 | HITS-CLIP | 29167373 | Unspecified |
| mmu-miR-34a-5p | MIMAT0000542 | Pbk | 52033 | HITS-CLIP | 23142080 | Unspecified |
| mmu-miR-34a-5p | MIMAT0000542 | Pbrm1 | 66923 | HITS-CLIP | 31433990\|23142080\|29167373 | Unspecified |
| mmu-miR-34a-5p | MIMAT0000542 | Pbx1 | 18514 | HITS-CLIP | 19536157 | Unspecified |
| mmu-miR-34a-5p | MIMAT0000542 | Pbx2 | 18515 | HITS-CLIP | 23142080 | Unspecified |
| mmu-miR-34a-5p | MIMAT0000542 | Pcbp1 | 23983 | HITS-CLIP | 23142080\|29167373 | Unspecified |
| mmu-miR-34a-5p | MIMAT0000542 | Pcbp2 | 18521 | HITS-CLIP | 23630078\|23142080\|29167373 | Unspecified |
| mmu-miR-34a-5p | MIMAT0000542 | Pcca | 110821 | HITS-CLIP | 23142080 | Unspecified |
| mmu-miR-34a-5p | MIMAT0000542 | Pccb | 66904 | HITS-CLIP | 30971389 | Unspecified |
| mmu-miR-34a-5p | MIMAT0000542 | Pcdh10 | 18526 | HITS-CLIP | 19536157 | Unspecified |
| mmu-miR-34a-5p | MIMAT0000542 | Pcdh11x | 245578 | HITS-CLIP | 19536157 | Unspecified |
| mmu-miR-34a-5p | MIMAT0000542 | Pcdh7 | 54216 | HITS-CLIP | 19536157\|29167373 | Unspecified |
| mmu-miR-34a-5p | MIMAT0000542 | Pcdh8 | 18530 | HITS-CLIP | 19536157 | Unspecified |
| mmu-miR-34a-5p | MIMAT0000542 | Pcdh9 | 211712 | HITS-CLIP | 19536157 | Unspecified |
| mmu-miR-34a-5p | MIMAT0000542 | Pcdhac2 | 353237 | HITS-CLIP | 19536157 | Unspecified |
| mmu-miR-34a-5p | MIMAT0000542 | Pcdhb15 | 93886 | Microarrays | 37986224 | Unspecified |
| mmu-miR-34a-5p | MIMAT0000542 | Pcdhb21 | 93892 | Microarrays | 37986224 | Unspecified |
| mmu-miR-34a-5p | MIMAT0000542 | Pcdhb22 | 93893 | Microarrays | 37986224 | Unspecified |
| mmu-miR-34a-5p | MIMAT0000542 | Pcdhga12 | 93724 | HITS-CLIP | 19536157\|23142080 | Unspecified |
| mmu-miR-34a-5p | MIMAT0000542 | Pcdhgc3 | 5098 | HITS-CLIP | 29167373 | Unspecified |
| mmu-miR-34a-5p | MIMAT0000542 | Pced1b | 239647 | HITS-CLIP | 23142080 | Unspecified |
| mmu-miR-34a-5p | MIMAT0000542 | Pcf11 | 74737 | HITS-CLIP | 23142080 | Unspecified |
| mmu-miR-34a-5p | MIMAT0000542 | Pcgf3 | 69587 | HITS-CLIP | 23142080 | Unspecified |
| mmu-miR-34a-5p | MIMAT0000542 | Pcgf5 | 76073 | HITS-CLIP | 23142080\|31433990\|29167373 | Unspecified |
| mmu-miR-34a-5p | MIMAT0000542 | Pcgf6 | 71041 | HITS-CLIP | 31433990\|26203562 | Unspecified |
| mmu-miR-34a-5p | MIMAT0000542 | Pcid2 | 234069 | HITS-CLIP | 23142080 | Unspecified |
| mmu-miR-34a-5p | MIMAT0000542 | Pcif1 | 228866 | HITS-CLIP | 23630078\|29167373\|23142080 | Unspecified |
| mmu-miR-34a-5p | MIMAT0000542 | Pck2 | 74551 | HITS-CLIP | 23142080 | Unspecified |

| | | | | | | |
|---|---|---|---|---|---|---|
| mmu-miR-34a-5p | MIMAT0000542 | Pclaf | 68026 | Microarrays | 23027903 | Unspecified |
| mmu-miR-34a-5p | MIMAT0000542 | Pclo | 26875 | HITS-CLIP | 19536157 | Unspecified |
| mmu-miR-34a-5p | MIMAT0000542 | Pcm1 | 18536 | HITS-CLIP | 19536157\|23142080 | Unspecified |
| mmu-miR-34a-5p | MIMAT0000542 | Pcmt1 | 18537 | HITS-CLIP | 19536157 | Unspecified |
| mmu-miR-34a-5p | MIMAT0000542 | Pcmtd1 | 319263 | HITS-CLIP | 23142080\|29167373 | Unspecified |
| mmu-miR-34a-5p | MIMAT0000542 | Pcmtd2 | 245867 | HITS-CLIP | 31433990\|23142080 | Unspecified |
| mmu-miR-34a-5p | MIMAT0000542 | Pcna | 18538 | HITS-CLIP | 31433990\|23142080\|29167373 | Unspecified |
| mmu-miR-34a-5p | MIMAT0000542 | Pcnt | 18541 | HITS-CLIP | 23142080\|19536157\|29167373 | Unspecified |
| mmu-miR-34a-5p | MIMAT0000542 | Pcnx | 54604 | HITS-CLIP | 19536157\|29167373 | Unspecified |
| mmu-miR-34a-5p | MIMAT0000542 | Pcnx3 | 399909 | HITS-CLIP | 23142080\|29167373 | Unspecified |
| mmu-miR-34a-5p | MIMAT0000542 | Pcp4l1 | 66425 | HITS-CLIP | 19536157 | Unspecified |
| mmu-miR-34a-5p | MIMAT0000542 | Pcsk1n | 30052 | HITS-CLIP | 19536157 | Unspecified |
| mmu-miR-34a-5p | MIMAT0000542 | Pcx | 18563 | HITS-CLIP | 23142080 | Unspecified |
| mmu-miR-34a-5p | MIMAT0000542 | Pcyox1 | 66881 | HITS-CLIP | 23142080\|29167373 | Unspecified |
| mmu-miR-34a-5p | MIMAT0000542 | Pcyt1a | 13026 | HITS-CLIP | 23142080 | Unspecified |
| mmu-miR-34a-5p | MIMAT0000542 | Pcyt2 | 68671 | HITS-CLIP | 23142080 | Unspecified |
| mmu-miR-34a-5p | MIMAT0000542 | Pdcd1 | 18566 | HITS-CLIP | 23142080 | Unspecified |
| mmu-miR-34a-5p | MIMAT0000542 | Pdcd10 | 56426 | HITS-CLIP | 23142080 | Unspecified |
| mmu-miR-34a-5p | MIMAT0000542 | Pdcd1lg2 | 58205 | HITS-CLIP | 23142080 | Unspecified |
| mmu-miR-34a-5p | MIMAT0000542 | Pdcd4 | 18569 | HITS-CLIP | 37986224\|19536157\|30971389\|23142 | Unspecified |
| mmu-miR-34a-5p | MIMAT0000542 | Pdcd6 | 18570 | HITS-CLIP | 23142080\|29167373\|31433990 | Unspecified |
| mmu-miR-34a-5p | MIMAT0000542 | Pdcd6ip | 18571 | HITS-CLIP | 19536157\|23142080\|29167373 | Unspecified |
| mmu-miR-34a-5p | MIMAT0000542 | Pdcd7 | 50996 | HITS-CLIP | 23142080 | Unspecified |
| mmu-miR-34a-5p | MIMAT0000542 | Pdcl | 67466 | HITS-CLIP | 23142080 | Unspecified |
| mmu-miR-34a-5p | MIMAT0000542 | Pde12 | 211948 | HITS-CLIP | 23142080 | Unspecified |
| mmu-miR-34a-5p | MIMAT0000542 | Pde1b | 18574 | HITS-CLIP | 29167373 | Unspecified |
| mmu-miR-34a-5p | MIMAT0000542 | Pde2a | 207728 | Microarrays//HITS-CLIP | 37986224\|29167373 | Unspecified |
| mmu-miR-34a-5p | MIMAT0000542 | Pde3b | 18576 | HITS-CLIP | 23142080 | Unspecified |
| mmu-miR-34a-5p | MIMAT0000542 | Pde4b | 18578 | HITS-CLIP | 37986224\|23142080 | Unspecified |
| mmu-miR-34a-5p | MIMAT0000542 | Pde4d | 238871 | HITS-CLIP | 19536157\|23142080 | Unspecified |
| mmu-miR-34a-5p | MIMAT0000542 | Pde4dip | 83679 | HITS-CLIP | 19536157\|23142080 | Unspecified |
| mmu-miR-34a-5p | MIMAT0000542 | Pde5a | 242202 | HITS-CLIP | 23142080\|19536157 | Unspecified |
| mmu-miR-34a-5p | MIMAT0000542 | Pde7a | 18583 | HITS-CLIP | 23142080 | Unspecified |
| mmu-miR-34a-5p | MIMAT0000542 | Pde7b | 29863 | HITS-CLIP | 19536157 | Unspecified |
| mmu-miR-34a-5p | MIMAT0000542 | Pdgfa | 18590 | HITS-CLIP | 29167373 | Unspecified |
| mmu-miR-34a-5p | MIMAT0000542 | Pdgfb | 18591 | HITS-CLIP | 23142080\|29167373 | Unspecified |
| mmu-miR-34a-5p | MIMAT0000542 | Pdgfd | 71785 | Microarrays | 37986224 | Unspecified |
| mmu-miR-34a-5p | MIMAT0000542 | Pdgfra | 18595 | HITS-CLIP | 19536157 | Unspecified |
| mmu-miR-34a-5p | MIMAT0000542 | Pdha1 | 18597 | HITS-CLIP | 23142080 | Unspecified |

| | | | | | | |
|---|---|---|---|---|---|---|
| mmu-miR-34a-5p | MIMAT0000542 | Pdhb | 68263 | HITS-CLIP | 23142080\|19536157 | Unspecified |
| mmu-miR-34a-5p | MIMAT0000542 | Pdhx | 27402 | HITS-CLIP | 30971389\|29167373 | Unspecified |
| mmu-miR-34a-5p | MIMAT0000542 | Pdia3 | 14827 | HITS-CLIP | 31433990\|19536157\|29167373\|23142 | Unspecified |
| mmu-miR-34a-5p | MIMAT0000542 | Pdia4 | 12304 | HITS-CLIP | 23142080 | Unspecified |
| mmu-miR-34a-5p | MIMAT0000542 | Pdia6 | 71853 | HITS-CLIP | 19536157\|26203562\|23142080\|29167 | Unspecified |
| mmu-miR-34a-5p | MIMAT0000542 | Pdik1l | 230809 | HITS-CLIP | 23142080 | Unspecified |
| mmu-miR-34a-5p | MIMAT0000542 | Pdk1 | 228026 | Microarrays//HITS-CLIP | 37986224\|23142080\|31433990\|29167 | Unspecified |
| mmu-miR-34a-5p | MIMAT0000542 | Pdk2 | 18604 | HITS-CLIP | 23142080 | Unspecified |
| mmu-miR-34a-5p | MIMAT0000542 | Pdk3 | 236900 | HITS-CLIP | 23142080 | Unspecified |
| mmu-miR-34a-5p | MIMAT0000542 | Pdlim1 | 54132 | HITS-CLIP | 23142080 | Unspecified |
| mmu-miR-34a-5p | MIMAT0000542 | Pdlim2 | 213019 | HITS-CLIP | 23142080 | Unspecified |
| mmu-miR-34a-5p | MIMAT0000542 | Pdlim3 | 53318 | Microarrays | 37986224 | Unspecified |
| mmu-miR-34a-5p | MIMAT0000542 | Pdlim5 | 56376 | HITS-CLIP | 29167373\|23142080 | Unspecified |
| mmu-miR-34a-5p | MIMAT0000542 | Pdlim7 | 67399 | HITS-CLIP | 29167373 | Unspecified |
| mmu-miR-34a-5p | MIMAT0000542 | Pdp1 | 381511 | HITS-CLIP | 29167373 | Unspecified |
| mmu-miR-34a-5p | MIMAT0000542 | Pdp2 | 382051 | HITS-CLIP | 23630078\|31433990\|23142080 | Unspecified |
| mmu-miR-34a-5p | MIMAT0000542 | Pdpk1 | 18607 | HITS-CLIP | 23142080\|29167373 | Unspecified |
| mmu-miR-34a-5p | MIMAT0000542 | Pdpn | 14726 | HITS-CLIP | 29167373 | Unspecified |
| mmu-miR-34a-5p | MIMAT0000542 | Pdpr | 319518 | HITS-CLIP | 23142080 | Unspecified |
| mmu-miR-34a-5p | MIMAT0000542 | Pdrg1 | 68559 | HITS-CLIP | 23142080 | Unspecified |
| mmu-miR-34a-5p | MIMAT0000542 | Pds5a | 71521 | HITS-CLIP | 23142080\|19536157\|29167373 | Unspecified |
| mmu-miR-34a-5p | MIMAT0000542 | Pds5b | 100710 | HITS-CLIP | 30971389\|23142080\|19536157 | Unspecified |
| mmu-miR-34a-5p | MIMAT0000542 | Pdss1 | 56075 | Microarrays | 37986224\|23027903 | Unspecified |
| mmu-miR-34a-5p | MIMAT0000542 | Pdxdc1 | 94184 | HITS-CLIP | 23142080\|29167373 | Unspecified |
| mmu-miR-34a-5p | MIMAT0000542 | Pdxk | 216134 | HITS-CLIP | 23142080\|29167373 | Unspecified |
| mmu-miR-34a-5p | MIMAT0000542 | Pdzd11 | 72621 | HITS-CLIP | 23142080 | Unspecified |
| mmu-miR-34a-5p | MIMAT0000542 | Pdzd2 | 68070 | HITS-CLIP | 19536157 | Unspecified |
| mmu-miR-34a-5p | MIMAT0000542 | Pdzd8 | 107368 | HITS-CLIP | 23630078\|23142080\|29167373 | Unspecified |
| mmu-miR-34a-5p | MIMAT0000542 | Pdzrn4 | 239618 | HITS-CLIP | 19536157 | Unspecified |
| mmu-miR-34a-5p | MIMAT0000542 | Pea15a | 18611 | HITS-CLIP | 23142080\|19536157\|29167373 | Unspecified |
| mmu-miR-34a-5p | MIMAT0000542 | Peak1 | 244895 | HITS-CLIP | 23142080\|30971389\|29167373 | Unspecified |
| mmu-miR-34a-5p | MIMAT0000542 | Pear1 | 375033 | HITS-CLIP | 23142080 | Unspecified |
| mmu-miR-34a-5p | MIMAT0000542 | Pebp1 | 23980 | HITS-CLIP | 23142080 | Unspecified |
| mmu-miR-34a-5p | MIMAT0000542 | Pecam1 | 18613 | HITS-CLIP | 23142080 | Unspecified |
| mmu-miR-34a-5p | MIMAT0000542 | Pecr | 111175 | HITS-CLIP | 30971389 | Unspecified |
| mmu-miR-34a-5p | MIMAT0000542 | Pef1 | 67898 | HITS-CLIP | 23142080 | Unspecified |
| mmu-miR-34a-5p | MIMAT0000542 | Peg10 | 170676 | HITS-CLIP | 23630078\|30971389 | Unspecified |
| mmu-miR-34a-5p | MIMAT0000542 | Peg3 | 18616 | Microarrays//HITS-CLIP | 37986224\|19536157\|26203562 | Unspecified |
| mmu-miR-34a-5p | MIMAT0000542 | Peli1 | 67245 | HITS-CLIP | 31433990\|23142080 | Unspecified |

| | | | | | | |
|---|---|---|---|---|---|---|
| mmu-miR-34a-5p | MIMAT0000542 | Pelp1 | 75273 | HITS-CLIP | 31433990 | Unspecified |
| mmu-miR-34a-5p | MIMAT0000542 | Penk | 18619 | HITS-CLIP | 30971389\|23142080 | Unspecified |
| mmu-miR-34a-5p | MIMAT0000542 | Pepd | 18624 | HITS-CLIP | 23142080\|29167373 | Unspecified |
| mmu-miR-34a-5p | MIMAT0000542 | Per1 | 18626 | HITS-CLIP | 23142080\|26203562 | Unspecified |
| mmu-miR-34a-5p | MIMAT0000542 | Per2 | 18627 | HITS-CLIP | 31433990\|23142080 | Unspecified |
| mmu-miR-34a-5p | MIMAT0000542 | Pes1 | 64934 | HITS-CLIP | 23142080 | Unspecified |
| mmu-miR-34a-5p | MIMAT0000542 | Pex1 | 71382 | HITS-CLIP | 19536157 | Unspecified |
| mmu-miR-34a-5p | MIMAT0000542 | Pex10 | 668173 | HITS-CLIP | 23142080 | Unspecified |
| mmu-miR-34a-5p | MIMAT0000542 | Pex11a | 18631 | Microarrays | 37986224 | Unspecified |
| mmu-miR-34a-5p | MIMAT0000542 | Pex11b | 18632 | HITS-CLIP | 23142080 | Unspecified |
| mmu-miR-34a-5p | MIMAT0000542 | Pex13 | 72129 | HITS-CLIP | 23142080\|31433990 | Unspecified |
| mmu-miR-34a-5p | MIMAT0000542 | Pex16 | 18633 | HITS-CLIP | 23142080 | Unspecified |
| mmu-miR-34a-5p | MIMAT0000542 | Pex19 | 19298 | HITS-CLIP | 23142080 | Unspecified |
| mmu-miR-34a-5p | MIMAT0000542 | Pex3 | 56535 | HITS-CLIP | 23142080 | Unspecified |
| mmu-miR-34a-5p | MIMAT0000542 | Pex5 | 19305 | HITS-CLIP | 23142080 | Unspecified |
| mmu-miR-34a-5p | MIMAT0000542 | Pex5l | 58869 | HITS-CLIP | 19536157 | Unspecified |
| mmu-miR-34a-5p | MIMAT0000542 | Pfas | 237823 | Microarrays//HITS-CLIP | 37986224\|23027903\|23142080\|29167 | Unspecified |
| mmu-miR-34a-5p | MIMAT0000542 | Pfdn2 | 18637 | HITS-CLIP | 23142080 | Unspecified |
| mmu-miR-34a-5p | MIMAT0000542 | Pfkfb1 | 18639 | HITS-CLIP | 23142080 | Unspecified |
| mmu-miR-34a-5p | MIMAT0000542 | Pfkfb3 | 170768 | HITS-CLIP | 23142080 | Unspecified |
| mmu-miR-34a-5p | MIMAT0000542 | Pfkfb4 | 270198 | HITS-CLIP | 23142080\|29167373 | Unspecified |
| mmu-miR-34a-5p | MIMAT0000542 | Pfkl | 18641 | HITS-CLIP | 37986224\|23142080\|29167373 | Unspecified |
| mmu-miR-34a-5p | MIMAT0000542 | Pfkm | 18642 | HITS-CLIP | 31433990 | Unspecified |
| mmu-miR-34a-5p | MIMAT0000542 | Pfkp | 56421 | HITS-CLIP | 23142080\|19536157\|29167373 | Unspecified |
| mmu-miR-34a-5p | MIMAT0000542 | Pfn1 | 18643 | HITS-CLIP | 37986224\|23142080\|26203562 | Unspecified |
| mmu-miR-34a-5p | MIMAT0000542 | Pfpl | 56093 | HITS-CLIP | 29167373 | Unspecified |
| mmu-miR-34a-5p | MIMAT0000542 | Pgam1 | 18648 | HITS-CLIP | 29167373 | Unspecified |
| mmu-miR-34a-5p | MIMAT0000542 | Pgap1 | 241062 | Microarrays//HITS-CLIP | 37986224\|23027903\|29167373 | Unspecified |
| mmu-miR-34a-5p | MIMAT0000542 | Pgap2 | 233575 | HITS-CLIP | 23142080 | Unspecified |
| mmu-miR-34a-5p | MIMAT0000542 | Pgap3 | 320655 | HITS-CLIP | 23142080 | Unspecified |
| mmu-miR-34a-5p | MIMAT0000542 | Pgbd5 | 209966 | HITS-CLIP | 19536157 | Unspecified |
| mmu-miR-34a-5p | MIMAT0000542 | Pgd | 110208 | HITS-CLIP | 23142080\|29167373 | Unspecified |
| mmu-miR-34a-5p | MIMAT0000542 | Pggt1b | 225467 | HITS-CLIP | 23142080\|29167373 | Unspecified |
| mmu-miR-34a-5p | MIMAT0000542 | Pglyrp1 | 21946 | HITS-CLIP | 23142080 | Unspecified |
| mmu-miR-34a-5p | MIMAT0000542 | Pglyrp2 | 57757 | HITS-CLIP | 23142080 | Unspecified |
| mmu-miR-34a-5p | MIMAT0000542 | Pgm1 | 66681 | HITS-CLIP | 19536157\|23142080 | Unspecified |
| mmu-miR-34a-5p | MIMAT0000542 | Pgm2 | 72157 | HITS-CLIP | 31433990 | Unspecified |
| mmu-miR-34a-5p | MIMAT0000542 | Pgm2l1 | 70974 | HITS-CLIP | 19536157\|23142080\|29167373 | Unspecified |
| mmu-miR-34a-5p | MIMAT0000542 | Pgp | 67078 | HITS-CLIP | 26203562\|23142080 | Unspecified |

| | | | | | | |
|---|---|---|---|---|---|---|
| mmu-miR-34a-5p | MIMAT0000542 | Pgpep1 | 66522 | HITS-CLIP | 23142080\|29167373 | Unspecified |
| mmu-miR-34a-5p | MIMAT0000542 | Pgrmc1 | 53328 | HITS-CLIP | 19536157\|23142080 | Unspecified |
| mmu-miR-34a-5p | MIMAT0000542 | Pgrmc2 | 70804 | HITS-CLIP | 19536157\|23142080 | Unspecified |
| mmu-miR-34a-5p | MIMAT0000542 | Pgs1 | 74451 | HITS-CLIP | 29167373 | Unspecified |
| mmu-miR-34a-5p | MIMAT0000542 | Phactr1 | 218194 | HITS-CLIP | 19536157 | Unspecified |
| mmu-miR-34a-5p | MIMAT0000542 | Phactr2 | 215789 | HITS-CLIP | 23142080 | Unspecified |
| mmu-miR-34a-5p | MIMAT0000542 | Phactr4 | 100169 | HITS-CLIP | 23142080\|29167373 | Unspecified |
| mmu-miR-34a-5p | MIMAT0000542 | Phb2 | 12034 | HITS-CLIP | 23142080 | Unspecified |
| mmu-miR-34a-5p | MIMAT0000542 | Phc2 | 54383 | HITS-CLIP | 31433990 | Unspecified |
| mmu-miR-34a-5p | MIMAT0000542 | Phc3 | 241915 | HITS-CLIP | 21258322\|23142080\|31433990\|29167 | Unspecified |
| mmu-miR-34a-5p | MIMAT0000542 | Phf10 | 72057 | HITS-CLIP | 23142080 | Unspecified |
| mmu-miR-34a-5p | MIMAT0000542 | Phf11b | 236451 | HITS-CLIP | 23142080 | Unspecified |
| mmu-miR-34a-5p | MIMAT0000542 | Phf11c | 628705 | HITS-CLIP | 29167373 | Unspecified |
| mmu-miR-34a-5p | MIMAT0000542 | Phf12 | 268448 | HITS-CLIP | 23630078\|23142080\|29167373 | Unspecified |
| mmu-miR-34a-5p | MIMAT0000542 | Phf19 | 74016 | HITS-CLIP, Microarrays//Microarrays//HITS | 37986224\|23027903\|23142080 | Unspecified |
| mmu-miR-34a-5p | MIMAT0000542 | Phf2 | 18676 | HITS-CLIP | 29167373\|23142080 | Unspecified |
| mmu-miR-34a-5p | MIMAT0000542 | Phf20 | 228829 | HITS-CLIP | 19536157\|23142080 | Unspecified |
| mmu-miR-34a-5p | MIMAT0000542 | Phf20l1 | 239510 | HITS-CLIP | 23142080\|31433990 | Unspecified |
| mmu-miR-34a-5p | MIMAT0000542 | Phf21a | 192285 | HITS-CLIP | 19536157\|23142080 | Unspecified |
| mmu-miR-34a-5p | MIMAT0000542 | Phf23 | 78246 | HITS-CLIP | 23142080 | Unspecified |
| mmu-miR-34a-5p | MIMAT0000542 | Phf24 | 230085 | HITS-CLIP | 19536157 | Unspecified |
| mmu-miR-34a-5p | MIMAT0000542 | Phf3 | 213109 | HITS-CLIP | 23142080\|19536157\|29167373 | Unspecified |
| mmu-miR-34a-5p | MIMAT0000542 | Phf5a | 68479 | HITS-CLIP | 23142080 | Unspecified |
| mmu-miR-34a-5p | MIMAT0000542 | Phf6 | 70998 | HITS-CLIP | 31433990\|23142080 | Unspecified |
| mmu-miR-34a-5p | MIMAT0000542 | Phf8 | 320595 | HITS-CLIP | 23142080\|29167373 | Unspecified |
| mmu-miR-34a-5p | MIMAT0000542 | Phip | 83946 | HITS-CLIP | 23142080\|29167373 | Unspecified |
| mmu-miR-34a-5p | MIMAT0000542 | Phka2 | 110094 | HITS-CLIP | 23142080\|29167373 | Unspecified |
| mmu-miR-34a-5p | MIMAT0000542 | Phkb | 102093 | HITS-CLIP | 29167373 | Unspecified |
| mmu-miR-34a-5p | MIMAT0000542 | Phkg2 | 68961 | HITS-CLIP | 23142080 | Unspecified |
| mmu-miR-34a-5p | MIMAT0000542 | Phlda1 | 21664 | HITS-CLIP | 23142080 | Unspecified |
| mmu-miR-34a-5p | MIMAT0000542 | Phlda3 | 27280 | HITS-CLIP | 31433990\|26203562 | Unspecified |
| mmu-miR-34a-5p | MIMAT0000542 | Phldb1 | 102693 | HITS-CLIP | 23142080 | Unspecified |
| mmu-miR-34a-5p | MIMAT0000542 | Phlpp2 | 244650 | HITS-CLIP | 31433990 | Unspecified |
| mmu-miR-34a-5p | MIMAT0000542 | Phrf1 | 101471 | HITS-CLIP | 23142080\|29167373 | Unspecified |
| mmu-miR-34a-5p | MIMAT0000542 | Phtf1 | 18685 | HITS-CLIP | 23142080 | Unspecified |
| mmu-miR-34a-5p | MIMAT0000542 | Phtf2 | 68770 | HITS-CLIP | 30971389\|23142080 | Unspecified |
| mmu-miR-34a-5p | MIMAT0000542 | Phyhip | 105653 | HITS-CLIP | 19536157 | Unspecified |
| mmu-miR-34a-5p | MIMAT0000542 | Pi4k2a | 84095 | HITS-CLIP | 23142080 | Unspecified |
| mmu-miR-34a-5p | MIMAT0000542 | Pi4k2b | 67073 | Microarrays | 37986224 | Unspecified |

| | | | | | | |
|---|---|---|---|---|---|---|
| mmu-miR-34a-5p | MIMAT0000542 | Pi4ka | 224020 | HITS-CLIP | 23142080\|29167373 | Unspecified |
| mmu-miR-34a-5p | MIMAT0000542 | Pi4kb | 107650 | HITS-CLIP | 23142080\|29167373 | Unspecified |
| mmu-miR-34a-5p | MIMAT0000542 | Pianp | 319352 | HITS-CLIP | 19536157 | Unspecified |
| mmu-miR-34a-5p | MIMAT0000542 | Pias1 | 56469 | HITS-CLIP | 19536157\|23142080\|29167373 | Unspecified |
| mmu-miR-34a-5p | MIMAT0000542 | Pias2 | 17344 | HITS-CLIP | 31433990\|23142080\|29167373 | Unspecified |
| mmu-miR-34a-5p | MIMAT0000542 | Pias4 | 59004 | HITS-CLIP | 23142080 | Unspecified |
| mmu-miR-34a-5p | MIMAT0000542 | Picalm | 233489 | HITS-CLIP | 23142080\|29167373\|31433990\|2363( | Unspecified |
| mmu-miR-34a-5p | MIMAT0000542 | Pid1 | 98496 | HITS-CLIP | 29167373 | Unspecified |
| mmu-miR-34a-5p | MIMAT0000542 | Piezo1 | 234839 | HITS-CLIP | 23142080\|29167373 | Unspecified |
| mmu-miR-34a-5p | MIMAT0000542 | Piga | 18700 | HITS-CLIP | 23142080 | Unspecified |
| mmu-miR-34a-5p | MIMAT0000542 | Pigc | 67292 | HITS-CLIP | 23142080 | Unspecified |
| mmu-miR-34a-5p | MIMAT0000542 | Pigf | 18701 | HITS-CLIP | 23142080 | Unspecified |
| mmu-miR-34a-5p | MIMAT0000542 | Pigg | 433931 | HITS-CLIP | 23142080 | Unspecified |
| mmu-miR-34a-5p | MIMAT0000542 | Pigh | 110417 | HITS-CLIP | 23142080 | Unspecified |
| mmu-miR-34a-5p | MIMAT0000542 | Pigk | 329777 | HITS-CLIP | 23142080\|29167373 | Unspecified |
| mmu-miR-34a-5p | MIMAT0000542 | Pigm | 67556 | HITS-CLIP | 19536157\|23142080\|29167373 | Unspecified |
| mmu-miR-34a-5p | MIMAT0000542 | Pign | 27392 | HITS-CLIP | 23142080\|31433990\|29167373 | Unspecified |
| mmu-miR-34a-5p | MIMAT0000542 | Pigo | 56703 | HITS-CLIP | 30971389\|23142080\|29167373 | Unspecified |
| mmu-miR-34a-5p | MIMAT0000542 | Pigp | 56176 | HITS-CLIP | 23142080 | Unspecified |
| mmu-miR-34a-5p | MIMAT0000542 | Pigq | 14755 | HITS-CLIP | 19536157\|23142080\|29167373 | Unspecified |
| mmu-miR-34a-5p | MIMAT0000542 | Pigt | 78928 | HITS-CLIP | 23142080\|29167373 | Unspecified |
| mmu-miR-34a-5p | MIMAT0000542 | Pigu | 228812 | HITS-CLIP | 23142080 | Unspecified |
| mmu-miR-34a-5p | MIMAT0000542 | Pigv | 230801 | HITS-CLIP | 23142080 | Unspecified |
| mmu-miR-34a-5p | MIMAT0000542 | Pigx | 72084 | HITS-CLIP | 23142080 | Unspecified |
| mmu-miR-34a-5p | MIMAT0000542 | Pih1d2 | 72614 | Microarrays | 37986224 | Unspecified |
| mmu-miR-34a-5p | MIMAT0000542 | Pik3ap1 | 83490 | HITS-CLIP | 29167373\|23142080 | Unspecified |
| mmu-miR-34a-5p | MIMAT0000542 | Pik3c2a | 18704 | HITS-CLIP | 23142080\|29167373 | Unspecified |
| mmu-miR-34a-5p | MIMAT0000542 | Pik3c2b | 240752 | HITS-CLIP | 23142080 | Unspecified |
| mmu-miR-34a-5p | MIMAT0000542 | Pik3c3 | 225326 | HITS-CLIP | 23142080 | Unspecified |
| mmu-miR-34a-5p | MIMAT0000542 | Pik3ca | 18706 | HITS-CLIP | 19536157\|29167373\|23142080 | Unspecified |
| mmu-miR-34a-5p | MIMAT0000542 | Pik3cb | 74769 | HITS-CLIP | 31433990\|29167373 | Unspecified |
| mmu-miR-34a-5p | MIMAT0000542 | Pik3cd | 18707 | HITS-CLIP | 23142080 | Unspecified |
| mmu-miR-34a-5p | MIMAT0000542 | Pik3cg | 30955 | HITS-CLIP | 30971389\|29167373\|23142080 | Unspecified |
| mmu-miR-34a-5p | MIMAT0000542 | Pik3ip1 | 216505 | HITS-CLIP | 19536157\|23142080 | Unspecified |
| mmu-miR-34a-5p | MIMAT0000542 | Pik3r1 | 18708 | HITS-CLIP | 23142080\|29167373 | Unspecified |
| mmu-miR-34a-5p | MIMAT0000542 | Pik3r5 | 320207 | HITS-CLIP | 23142080\|29167373 | Unspecified |
| mmu-miR-34a-5p | MIMAT0000542 | Pik3r6 | 104709 | HITS-CLIP | 29167373 | Unspecified |
| mmu-miR-34a-5p | MIMAT0000542 | Pikfyve | 18711 | HITS-CLIP | 31433990\|19536157\|23142080\|29167 | Unspecified |
| mmu-miR-34a-5p | MIMAT0000542 | Pilra | 231805 | HITS-CLIP | 29167373 | Unspecified |

| | | | | | | |
|---|---|---|---|---|---|---|
| mmu-miR-34a-5p | MIMAT0000542 | Pim1 | 18712 | HITS-CLIP | 23142080\|31433990 | Unspecified |
| mmu-miR-34a-5p | MIMAT0000542 | Pim3 | 223775 | HITS-CLIP | 23142080 | Unspecified |
| mmu-miR-34a-5p | MIMAT0000542 | Pimreg | 54478 | HITS-CLIP | 23142080 | Unspecified |
| mmu-miR-34a-5p | MIMAT0000542 | Pin1 | 23988 | HITS-CLIP | 21258322\|23142080 | Unspecified |
| mmu-miR-34a-5p | MIMAT0000542 | Pink1 | 68943 | HITS-CLIP | 23142080 | Unspecified |
| mmu-miR-34a-5p | MIMAT0000542 | Pip4k2a | 18718 | HITS-CLIP | 19536157\|23142080\|29167373 | Unspecified |
| mmu-miR-34a-5p | MIMAT0000542 | Pip4k2b | 108083 | HITS-CLIP | 23142080\|19536157\|30971389\|3143 | Unspecified |
| mmu-miR-34a-5p | MIMAT0000542 | Pip4k2c | 117150 | HITS-CLIP | 23142080 | Unspecified |
| mmu-miR-34a-5p | MIMAT0000542 | Pip4p1 | 90809 | HITS-CLIP | 23142080\|29167373 | Unspecified |
| mmu-miR-34a-5p | MIMAT0000542 | Pip4p2 | 55529 | HITS-CLIP | 31433990\|29167373\|23142080 | Unspecified |
| mmu-miR-34a-5p | MIMAT0000542 | Pip5k1a | 18720 | Microarrays//HITS-CLIP | 37986224\|23027903\|23630078\|2620 | Unspecified |
| mmu-miR-34a-5p | MIMAT0000542 | Pip5k1b | 18719 | HITS-CLIP | 23142080 | Unspecified |
| mmu-miR-34a-5p | MIMAT0000542 | Pip5k1c | 18717 | HITS-CLIP | 29167373\|23142080 | Unspecified |
| mmu-miR-34a-5p | MIMAT0000542 | Pirb | 18733 | HITS-CLIP | 29167373 | Unspecified |
| mmu-miR-34a-5p | MIMAT0000542 | Pisd | 320951 | HITS-CLIP | 30971389\|29167373 | Unspecified |
| mmu-miR-34a-5p | MIMAT0000542 | Pitpna | 18738 | HITS-CLIP | 23142080\|19536157\|31433990\|2916 | Unspecified |
| mmu-miR-34a-5p | MIMAT0000542 | Pitpnb | 56305 | HITS-CLIP | 23142080 | Unspecified |
| mmu-miR-34a-5p | MIMAT0000542 | Pitpnc1 | 71795 | HITS-CLIP | 19536157\|23142080 | Unspecified |
| mmu-miR-34a-5p | MIMAT0000542 | Pitpnm1 | 18739 | HITS-CLIP | 31433990\|23142080 | Unspecified |
| mmu-miR-34a-5p | MIMAT0000542 | Pitpnm2 | 19679 | HITS-CLIP | 23142080\|19536157 | Unspecified |
| mmu-miR-34a-5p | MIMAT0000542 | Pitrm1 | 69617 | HITS-CLIP | 23142080 | Unspecified |
| mmu-miR-34a-5p | MIMAT0000542 | Pja2 | 224938 | HITS-CLIP | 23142080\|31433990 | Unspecified |
| mmu-miR-34a-5p | MIMAT0000542 | Pkd1 | 18763 | HITS-CLIP | 31433990\|29167373\|23142080 | Unspecified |
| mmu-miR-34a-5p | MIMAT0000542 | Pkia | 18767 | Microarrays//HITS-CLIP | 37986224\|23027903\|19536157 | Unspecified |
| mmu-miR-34a-5p | MIMAT0000542 | Pkm | 18746 | HITS-CLIP | 29167373\|23142080 | Unspecified |
| mmu-miR-34a-5p | MIMAT0000542 | Pkmyt1 | 268930 | HITS-CLIP | 23142080 | Unspecified |
| mmu-miR-34a-5p | MIMAT0000542 | Pkn1 | 320795 | HITS-CLIP | 23142080\|29167373 | Unspecified |
| mmu-miR-34a-5p | MIMAT0000542 | Pkn2 | 109333 | HITS-CLIP | 31433990\|23142080 | Unspecified |
| mmu-miR-34a-5p | MIMAT0000542 | Pkn3 | 263803 | HITS-CLIP | 23142080 | Unspecified |
| mmu-miR-34a-5p | MIMAT0000542 | Pknox1 | 18771 | HITS-CLIP | 23142080 | Unspecified |
| mmu-miR-34a-5p | MIMAT0000542 | Pkp4 | 227937 | HITS-CLIP | 19536157\|31433990\|23142080 | Unspecified |
| mmu-miR-34a-5p | MIMAT0000542 | Pla2g12a | 66350 | HITS-CLIP | 30971389\|23142080 | Unspecified |
| mmu-miR-34a-5p | MIMAT0000542 | Pla2g15 | 192654 | Microarrays//HITS-CLIP | 37986224\|23027903\|31433990\|2916 | Unspecified |
| mmu-miR-34a-5p | MIMAT0000542 | Pla2g16 | 225845 | HITS-CLIP | 23142080 | Unspecified |
| mmu-miR-34a-5p | MIMAT0000542 | Pla2g2e | 26970 | Microarrays | 37986224 | Unspecified |
| mmu-miR-34a-5p | MIMAT0000542 | Pla2g4a | 18783 | HITS-CLIP | 29167373 | Unspecified |
| mmu-miR-34a-5p | MIMAT0000542 | Plaa | 18786 | HITS-CLIP | 31433990\|30971389\|29167373 | Unspecified |
| mmu-miR-34a-5p | MIMAT0000542 | Plag1 | 56711 | HITS-CLIP | 23142080\|26203562\|29084942 | Unspecified |
| mmu-miR-34a-5p | MIMAT0000542 | Plagl1 | 22634 | HITS-CLIP | 23142080 | Unspecified |

| | | | | | | |
|---|---|---|---|---|---|---|
| mmu-miR-34a-5p | MIMAT0000542 | Plagl2 | 54711 | HITS-CLIP | 37986224\|31433990\|19536157\|23142 | Unspecified |
| mmu-miR-34a-5p | MIMAT0000542 | Plau | 18792 | Microarrays//HITS-CLIP | 37986224\|23027903\|30971389\|29167 | Unspecified |
| mmu-miR-34a-5p | MIMAT0000542 | Plaur | 18793 | HITS-CLIP | 29167373 | Unspecified |
| mmu-miR-34a-5p | MIMAT0000542 | Plbd2 | 71772 | HITS-CLIP | 29167373 | Unspecified |
| mmu-miR-34a-5p | MIMAT0000542 | Plcb2 | 18796 | HITS-CLIP | 23142080\|29167373 | Unspecified |
| mmu-miR-34a-5p | MIMAT0000542 | Plcb3 | 18797 | HITS-CLIP | 23142080 | Unspecified |
| mmu-miR-34a-5p | MIMAT0000542 | Plcb4 | 18798 | HITS-CLIP | 19536157\|23142080 | Unspecified |
| mmu-miR-34a-5p | MIMAT0000542 | Plcg1 | 18803 | HITS-CLIP | 23142080 | Unspecified |
| mmu-miR-34a-5p | MIMAT0000542 | Plcl1 | 227120 | HITS-CLIP | 23142080 | Unspecified |
| mmu-miR-34a-5p | MIMAT0000542 | Plcl2 | 224860 | HITS-CLIP | 23142080\|29167373 | Unspecified |
| mmu-miR-34a-5p | MIMAT0000542 | Plcxd1 | 403178 | HITS-CLIP | 23142080 | Unspecified |
| mmu-miR-34a-5p | MIMAT0000542 | Plcxd2 | 433022 | HITS-CLIP | 23142080 | Unspecified |
| mmu-miR-34a-5p | MIMAT0000542 | Pld2 | 18806 | HITS-CLIP | 29167373 | Unspecified |
| mmu-miR-34a-5p | MIMAT0000542 | Pld3 | 18807 | HITS-CLIP | 23142080\|29167373 | Unspecified |
| mmu-miR-34a-5p | MIMAT0000542 | Pld4 | 104759 | HITS-CLIP | 29167373 | Unspecified |
| mmu-miR-34a-5p | MIMAT0000542 | Plec | 18810 | HITS-CLIP | 31433990\|19536157\|23630078\|23142 | Unspecified |
| mmu-miR-34a-5p | MIMAT0000542 | Plek | 56193 | HITS-CLIP | 23142080\|29167373 | Unspecified |
| mmu-miR-34a-5p | MIMAT0000542 | Plek2 | 27260 | Microarrays//HITS-CLIP | 37986224\|23027903\|30971389 | Unspecified |
| mmu-miR-34a-5p | MIMAT0000542 | Plekha1 | 101476 | HITS-CLIP | 19536157\|29167373 | Unspecified |
| mmu-miR-34a-5p | MIMAT0000542 | Plekha2 | 83436 | HITS-CLIP | 31433990\|23142080 | Unspecified |
| mmu-miR-34a-5p | MIMAT0000542 | Plekha3 | 83435 | Microarrays//HITS-CLIP | 37986224\|23630078\|31433990\|23142 | Unspecified |
| mmu-miR-34a-5p | MIMAT0000542 | Plekha5 | 109135 | HITS-CLIP | 23142080 | Unspecified |
| mmu-miR-34a-5p | MIMAT0000542 | Plekha7 | 233765 | HITS-CLIP | 23142080 | Unspecified |
| mmu-miR-34a-5p | MIMAT0000542 | Plekha8 | 231999 | HITS-CLIP | 23142080 | Unspecified |
| mmu-miR-34a-5p | MIMAT0000542 | Plekhb2 | 226971 | HITS-CLIP | 23142080\|29167373 | Unspecified |
| mmu-miR-34a-5p | MIMAT0000542 | Plekhf2 | 71801 | HITS-CLIP | 29167373 | Unspecified |
| mmu-miR-34a-5p | MIMAT0000542 | Plekhg3 | 263406 | HITS-CLIP | 29167373 | Unspecified |
| mmu-miR-34a-5p | MIMAT0000542 | Plekhg5 | 269608 | HITS-CLIP | 23142080 | Unspecified |
| mmu-miR-34a-5p | MIMAT0000542 | Plekhj1 | 78670 | HITS-CLIP | 23142080 | Unspecified |
| mmu-miR-34a-5p | MIMAT0000542 | Plekhm1 | 353047 | HITS-CLIP | 23142080\|29167373 | Unspecified |
| mmu-miR-34a-5p | MIMAT0000542 | Plekhm2 | 69582 | HITS-CLIP | 29167373 | Unspecified |
| mmu-miR-34a-5p | MIMAT0000542 | Plekhm3 | 241075 | HITS-CLIP | 29167373 | Unspecified |
| mmu-miR-34a-5p | MIMAT0000542 | Plekho2 | 102595 | HITS-CLIP | 23142080\|29167373 | Unspecified |
| mmu-miR-34a-5p | MIMAT0000542 | Plgrkt | 67759 | HITS-CLIP | 23142080 | Unspecified |
| mmu-miR-34a-5p | MIMAT0000542 | Plin2 | 11520 | HITS-CLIP | 23142080\|29167373 | Unspecified |
| mmu-miR-34a-5p | MIMAT0000542 | Plin3 | 66905 | HITS-CLIP | 23142080 | Unspecified |
| mmu-miR-34a-5p | MIMAT0000542 | Plk1 | 18817 | HITS-CLIP | 23142080 | Unspecified |
| mmu-miR-34a-5p | MIMAT0000542 | Plk2 | 20620 | HITS-CLIP | 19536157\|23142080\|29167373 | Unspecified |
| mmu-miR-34a-5p | MIMAT0000542 | Plk3 | 12795 | HITS-CLIP | 23142080 | Unspecified |

| | | | | | | |
|---|---|---|---|---|---|---|
| mmu-miR-34a-5p | MIMAT0000542 | Plk4 | 20873 | HITS-CLIP | 23142080 | Unspecified |
| mmu-miR-34a-5p | MIMAT0000542 | Pllp | 67801 | HITS-CLIP | 19536157 | Unspecified |
| mmu-miR-34a-5p | MIMAT0000542 | Plod1 | 18822 | HITS-CLIP | 29167373 | Unspecified |
| mmu-miR-34a-5p | MIMAT0000542 | Plod2 | 26432 | HITS-CLIP | 23142080\|26203562 | Unspecified |
| mmu-miR-34a-5p | MIMAT0000542 | Plod3 | 26433 | HITS-CLIP | 19536157\|23142080 | Unspecified |
| mmu-miR-34a-5p | MIMAT0000542 | Plp1 | 18823 | HITS-CLIP | 19536157 | Unspecified |
| mmu-miR-34a-5p | MIMAT0000542 | Plpbp | 11212 | HITS-CLIP | 23142080\|29167373 | Unspecified |
| mmu-miR-34a-5p | MIMAT0000542 | Plpp2 | 8612 | HITS-CLIP | 23142080 | Unspecified |
| mmu-miR-34a-5p | MIMAT0000542 | Plpp3 | 67916 | HITS-CLIP | 19536157 | Unspecified |
| mmu-miR-34a-5p | MIMAT0000542 | Plpp5 | 84513 | HITS-CLIP | 23142080\|19536157\|29167373 | Unspecified |
| mmu-miR-34a-5p | MIMAT0000542 | Plpp6 | 403313 | HITS-CLIP | 19536157\|23142080 | Unspecified |
| mmu-miR-34a-5p | MIMAT0000542 | Plrg1 | 53317 | HITS-CLIP | 23142080 | Unspecified |
| mmu-miR-34a-5p | MIMAT0000542 | Plscr3 | 70310 | HITS-CLIP | 23142080\|29167373 | Unspecified |
| mmu-miR-34a-5p | MIMAT0000542 | Plscr4 | 235527 | HITS-CLIP | 23142080 | Unspecified |
| mmu-miR-34a-5p | MIMAT0000542 | Pltp | 18830 | HITS-CLIP | 29167373 | Unspecified |
| mmu-miR-34a-5p | MIMAT0000542 | Plxdc1 | 72324 | HITS-CLIP | 29167373 | Unspecified |
| mmu-miR-34a-5p | MIMAT0000542 | Plxdc2 | 67448 | HITS-CLIP | 23142080\|29167373 | Unspecified |
| mmu-miR-34a-5p | MIMAT0000542 | Plxna1 | 18844 | HITS-CLIP | 23142080\|29167373 | Unspecified |
| mmu-miR-34a-5p | MIMAT0000542 | Plxna2 | 18845 | HITS-CLIP | 19536157 | Unspecified |
| mmu-miR-34a-5p | MIMAT0000542 | Plxna4 | 243743 | HITS-CLIP | 19536157 | Unspecified |
| mmu-miR-34a-5p | MIMAT0000542 | Plxnb2 | 140570 | HITS-CLIP | 29167373 | Unspecified |
| mmu-miR-34a-5p | MIMAT0000542 | Plxnc1 | 54712 | HITS-CLIP | 19536157\|29167373 | Unspecified |
| mmu-miR-34a-5p | MIMAT0000542 | Plxnd1 | 67784 | HITS-CLIP | 19536157\|23142080\|29167373 | Unspecified |
| mmu-miR-34a-5p | MIMAT0000542 | Pmepa1 | 65112 | HITS-CLIP | 23142080 | Unspecified |
| mmu-miR-34a-5p | MIMAT0000542 | Pmf1 | 67037 | Microarrays//HITS-CLIP | 37986224\|23027903\|23142080\|29167 | Unspecified |
| mmu-miR-34a-5p | MIMAT0000542 | Pml | 18854 | HITS-CLIP | 30971389\|23142080\|29167373 | Unspecified |
| mmu-miR-34a-5p | MIMAT0000542 | Pmm1 | 29858 | HITS-CLIP | 23630078 | Unspecified |
| mmu-miR-34a-5p | MIMAT0000542 | Pmm2 | 54128 | HITS-CLIP | 29167373 | Unspecified |
| mmu-miR-34a-5p | MIMAT0000542 | Pmp22 | 18858 | HITS-CLIP | 29167373 | Unspecified |
| mmu-miR-34a-5p | MIMAT0000542 | Pmpca | 66865 | HITS-CLIP | 23142080\|29167373 | Unspecified |
| mmu-miR-34a-5p | MIMAT0000542 | Pmpcb | 73078 | HITS-CLIP | 23142080 | Unspecified |
| mmu-miR-34a-5p | MIMAT0000542 | Pms1 | 227099 | HITS-CLIP | 23142080 | Unspecified |
| mmu-miR-34a-5p | MIMAT0000542 | Pms2 | 18861 | HITS-CLIP | 23142080 | Unspecified |
| mmu-miR-34a-5p | MIMAT0000542 | Pnisr | 66625 | HITS-CLIP | 31433990\|23142080 | Unspecified |
| mmu-miR-34a-5p | MIMAT0000542 | Pnkp | 59047 | HITS-CLIP | 23142080 | Unspecified |
| mmu-miR-34a-5p | MIMAT0000542 | Pnmal2 | 434128 | Microarrays | 37986224 | Unspecified |
| mmu-miR-34a-5p | MIMAT0000542 | Pnn | 18949 | HITS-CLIP | 23142080\|29167373 | Unspecified |
| mmu-miR-34a-5p | MIMAT0000542 | Pnp | 18950 | HITS-CLIP | 23142080\|29167373 | Unspecified |
| mmu-miR-34a-5p | MIMAT0000542 | Pnpla2 | 66853 | HITS-CLIP | 23142080 | Unspecified |

| | | | | | | |
|---|---|---|---|---|---|---|
| mmu-miR-34a-5p | MIMAT0000542 | Pnpla6 | 50767 | HITS-CLIP | 19536157\|23142080 | Unspecified |
| mmu-miR-34a-5p | MIMAT0000542 | Pnpla8 | 67452 | HITS-CLIP | 19536157\|23142080\|29167373 | Unspecified |
| mmu-miR-34a-5p | MIMAT0000542 | Pnpo | 103711 | HITS-CLIP | 23142080 | Unspecified |
| mmu-miR-34a-5p | MIMAT0000542 | Pnpt1 | 71701 | HITS-CLIP | 23142080 | Unspecified |
| mmu-miR-34a-5p | MIMAT0000542 | Pnrc1 | 108767 | Microarrays//HITS-CLIP | 37986224\|19536157\|29167373\|23142 | Unspecified |
| mmu-miR-34a-5p | MIMAT0000542 | Pnrc2 | 52830 | HITS-CLIP | 23142080 | Unspecified |
| mmu-miR-34a-5p | MIMAT0000542 | Poc1a | 70235 | HITS-CLIP | 23142080 | Unspecified |
| mmu-miR-34a-5p | MIMAT0000542 | Poc1b | 382406 | HITS-CLIP | 23142080 | Unspecified |
| mmu-miR-34a-5p | MIMAT0000542 | Podxl2 | 50512 | HITS-CLIP | 19536157 | Unspecified |
| mmu-miR-34a-5p | MIMAT0000542 | Pofut1 | 140484 | Luciferase assay//Northern blot//qRT-PCR | 23047694\|23027903\|23142080\|24587 | Unspecified |
| mmu-miR-34a-5p | MIMAT0000542 | Pogk | 71592 | HITS-CLIP | 23142080 | Unspecified |
| mmu-miR-34a-5p | MIMAT0000542 | Poglut1 | 224143 | HITS-CLIP | 23142080 | Unspecified |
| mmu-miR-34a-5p | MIMAT0000542 | Pogz | 229584 | HITS-CLIP | 23142080\|29167373 | Unspecified |
| mmu-miR-34a-5p | MIMAT0000542 | Pold1 | 18971 | HITS-CLIP | 29167373 | Unspecified |
| mmu-miR-34a-5p | MIMAT0000542 | Pold2 | 18972 | Microarrays | 37986224\|23027903 | Unspecified |
| mmu-miR-34a-5p | MIMAT0000542 | Pold3 | 67967 | HITS-CLIP | 31433990\|23142080 | Unspecified |
| mmu-miR-34a-5p | MIMAT0000542 | Pold4 | 69745 | HITS-CLIP | 23142080 | Unspecified |
| mmu-miR-34a-5p | MIMAT0000542 | Poldip2 | 67811 | HITS-CLIP | 23142080 | Unspecified |
| mmu-miR-34a-5p | MIMAT0000542 | Poldip3 | 73826 | HITS-CLIP | 23142080 | Unspecified |
| mmu-miR-34a-5p | MIMAT0000542 | Pole | 18973 | HITS-CLIP | 23142080 | Unspecified |
| mmu-miR-34a-5p | MIMAT0000542 | Pole2 | 18974 | Microarrays | 37986224\|23027903 | Unspecified |
| mmu-miR-34a-5p | MIMAT0000542 | Pole3 | 59001 | HITS-CLIP | 23142080 | Unspecified |
| mmu-miR-34a-5p | MIMAT0000542 | Polg | 18975 | HITS-CLIP | 30971389\|23142080\|29167373 | Unspecified |
| mmu-miR-34a-5p | MIMAT0000542 | Polh | 80905 | HITS-CLIP | 23142080 | Unspecified |
| mmu-miR-34a-5p | MIMAT0000542 | Poli | 11201 | HITS-CLIP | 23142080 | Unspecified |
| mmu-miR-34a-5p | MIMAT0000542 | Polk | 27015 | HITS-CLIP | 37986224\|23142080 | Unspecified |
| mmu-miR-34a-5p | MIMAT0000542 | Polm | 54125 | HITS-CLIP | 23142080 | Unspecified |
| mmu-miR-34a-5p | MIMAT0000542 | Polq | 77782 | Microarrays//HITS-CLIP | 37986224\|23027903\|30971389 | Unspecified |
| mmu-miR-34a-5p | MIMAT0000542 | Polr1a | 20019 | HITS-CLIP | 23142080 | Unspecified |
| mmu-miR-34a-5p | MIMAT0000542 | Polr1b | 20017 | HITS-CLIP | 30971389 | Unspecified |
| mmu-miR-34a-5p | MIMAT0000542 | Polr1d | 20018 | HITS-CLIP | 23142080 | Unspecified |
| mmu-miR-34a-5p | MIMAT0000542 | Polr1has | 76416 | HITS-CLIP | 31433990 | Unspecified |
| mmu-miR-34a-5p | MIMAT0000542 | Polr2a | 20020 | HITS-CLIP | 31433990\|29167373\|23142080 | Unspecified |
| mmu-miR-34a-5p | MIMAT0000542 | Polr2b | 231329 | HITS-CLIP | 30971389\|23142080\|29167373 | Unspecified |
| mmu-miR-34a-5p | MIMAT0000542 | Polr2d | 69241 | HITS-CLIP | 23142080 | Unspecified |
| mmu-miR-34a-5p | MIMAT0000542 | Polr2h | 245841 | HITS-CLIP | 23142080 | Unspecified |
| mmu-miR-34a-5p | MIMAT0000542 | Polr2i | 69920 | HITS-CLIP | 23142080 | Unspecified |
| mmu-miR-34a-5p | MIMAT0000542 | Polr2j | 20022 | HITS-CLIP | 23142080 | Unspecified |
| mmu-miR-34a-5p | MIMAT0000542 | Polr2m | 28015 | HITS-CLIP | 31433990\|23142080 | Unspecified |

| | | | | | | |
|---|---|---|---|---|---|---|
| mmu-miR-34a-5p | MIMAT0000542 | Polr3a | 218832 | HITS-CLIP | 23142080 | Unspecified |
| mmu-miR-34a-5p | MIMAT0000542 | Polr3b | 70428 | HITS-CLIP | 29167373 | Unspecified |
| mmu-miR-34a-5p | MIMAT0000542 | Polr3c | 74414 | HITS-CLIP | 23142080\|29167373 | Unspecified |
| mmu-miR-34a-5p | MIMAT0000542 | Polr3d | 67065 | HITS-CLIP | 30971389 | Unspecified |
| mmu-miR-34a-5p | MIMAT0000542 | Polr3e | 26939 | HITS-CLIP | 23142080 | Unspecified |
| mmu-miR-34a-5p | MIMAT0000542 | Polr3g | 67486 | HITS-CLIP | 23142080\|29167373 | Unspecified |
| mmu-miR-34a-5p | MIMAT0000542 | Pom121 | 107939 | HITS-CLIP | 31433990\|19536157\|30971389\|23142 | Unspecified |
| mmu-miR-34a-5p | MIMAT0000542 | Pomk | 74653 | HITS-CLIP | 23142080\|29167373 | Unspecified |
| mmu-miR-34a-5p | MIMAT0000542 | Pomp | 66537 | HITS-CLIP | 23142080 | Unspecified |
| mmu-miR-34a-5p | MIMAT0000542 | Pomt1 | 99011 | HITS-CLIP | 23142080 | Unspecified |
| mmu-miR-34a-5p | MIMAT0000542 | Pon2 | 330260 | HITS-CLIP | 23142080 | Unspecified |
| mmu-miR-34a-5p | MIMAT0000542 | Pop1 | 67724 | HITS-CLIP | 23142080 | Unspecified |
| mmu-miR-34a-5p | MIMAT0000542 | Pop5 | 117109 | HITS-CLIP | 23142080 | Unspecified |
| mmu-miR-34a-5p | MIMAT0000542 | Por | 18984 | HITS-CLIP | 19536157 | Unspecified |
| mmu-miR-34a-5p | MIMAT0000542 | Pot1a | 101185 | HITS-CLIP | 23142080 | Unspecified |
| mmu-miR-34a-5p | MIMAT0000542 | Pot1b | 72836 | HITS-CLIP | 29167373 | Unspecified |
| mmu-miR-34a-5p | MIMAT0000542 | Pou2af1 | 18985 | HITS-CLIP | 23142080 | Unspecified |
| mmu-miR-34a-5p | MIMAT0000542 | Pou2f1 | 18986 | HITS-CLIP | 19536157\|23142080\|29167373 | Unspecified |
| mmu-miR-34a-5p | MIMAT0000542 | Pou2f2 | 18987 | HITS-CLIP | 19536157\|23142080\|29167373 | Unspecified |
| mmu-miR-34a-5p | MIMAT0000542 | Pou5f1 | 18999 | qPCR, Western Blot | 37986224 | Unspecified |
| mmu-miR-34a-5p | MIMAT0000542 | Pou6f1 | 19009 | HITS-CLIP | 19536157 | Unspecified |
| mmu-miR-34a-5p | MIMAT0000542 | Ppan | 235036 | HITS-CLIP | 23142080 | Unspecified |
| mmu-miR-34a-5p | MIMAT0000542 | Ppard | 19015 | HITS-CLIP | 23142080 | Unspecified |
| mmu-miR-34a-5p | MIMAT0000542 | Pparg | 19016 | HITS-CLIP | 31433990 | Unspecified |
| mmu-miR-34a-5p | MIMAT0000542 | Ppargc1a | 19017 | Microarrays | 37986224 | Unspecified |
| mmu-miR-34a-5p | MIMAT0000542 | Ppat | 231327 | HITS-CLIP | 23142080 | Unspecified |
| mmu-miR-34a-5p | MIMAT0000542 | Ppfia1 | 233977 | HITS-CLIP | 23142080\|29167373 | Unspecified |
| mmu-miR-34a-5p | MIMAT0000542 | Ppfia4 | 68507 | HITS-CLIP | 19536157 | Unspecified |
| mmu-miR-34a-5p | MIMAT0000542 | Ppfibp2 | 19024 | HITS-CLIP | 19536157 | Unspecified |
| mmu-miR-34a-5p | MIMAT0000542 | Ppia | 268373 | HITS-CLIP | 23142080 | Unspecified |
| mmu-miR-34a-5p | MIMAT0000542 | Ppib | 19035 | HITS-CLIP | 23142080 | Unspecified |
| mmu-miR-34a-5p | MIMAT0000542 | Ppie | 56031 | HITS-CLIP | 23142080 | Unspecified |
| mmu-miR-34a-5p | MIMAT0000542 | Ppif | 105675 | HITS-CLIP | 23142080 | Unspecified |
| mmu-miR-34a-5p | MIMAT0000542 | Ppig | 228005 | HITS-CLIP | 23630078\|23142080 | Unspecified |
| mmu-miR-34a-5p | MIMAT0000542 | Ppil1 | 68816 | HITS-CLIP | 23142080 | Unspecified |
| mmu-miR-34a-5p | MIMAT0000542 | Ppil2 | 66053 | HITS-CLIP | 23142080\|29167373 | Unspecified |
| mmu-miR-34a-5p | MIMAT0000542 | Ppil3 | 53938 | HITS-CLIP | 23142080 | Unspecified |
| mmu-miR-34a-5p | MIMAT0000542 | Ppil4 | 67418 | HITS-CLIP | 23142080\|31433990 | Unspecified |
| mmu-miR-34a-5p | MIMAT0000542 | Ppil6 | 73075 | Microarrays | 37986224 | Unspecified |

| | | | | | | | |
|---|---|---|---|---|---|---|---|
| mmu-miR-34a-5p | MIMAT0000542 | Ppip5k1 | 327655 | HITS-CLIP | | 23142080 | Unspecified |
| mmu-miR-34a-5p | MIMAT0000542 | Ppip5k2 | 227399 | HITS-CLIP | 30971389\|23142080 | | Unspecified |
| mmu-miR-34a-5p | MIMAT0000542 | Ppm1a | 19042 | HITS-CLIP | | 23142080 | Unspecified |
| mmu-miR-34a-5p | MIMAT0000542 | Ppm1b | 19043 | HITS-CLIP | | 23142080 | Unspecified |
| mmu-miR-34a-5p | MIMAT0000542 | Ppm1d | 53892 | HITS-CLIP | | 23142080 | Unspecified |
| mmu-miR-34a-5p | MIMAT0000542 | Ppm1e | 320472 | HITS-CLIP | | 19536157 | Unspecified |
| mmu-miR-34a-5p | MIMAT0000542 | Ppm1f | 68606 | HITS-CLIP | 19536157\|23142080\|31433990 | | Unspecified |
| mmu-miR-34a-5p | MIMAT0000542 | Ppm1h | 319468 | HITS-CLIP | 23142080\|29167373 | | Unspecified |
| mmu-miR-34a-5p | MIMAT0000542 | Ppm1k | 243382 | HITS-CLIP | | 23142080 | Unspecified |
| mmu-miR-34a-5p | MIMAT0000542 | Ppme1 | 72590 | HITS-CLIP | 19536157\|23142080 | | Unspecified |
| mmu-miR-34a-5p | MIMAT0000542 | Ppp1ca | 19045 | HITS-CLIP | 37986224\|19536157\|23142080\|29167 | | Unspecified |
| mmu-miR-34a-5p | MIMAT0000542 | Ppp1cb | 19046 | HITS-CLIP | 19536157\|23142080\|31433990\|29167 | | Unspecified |
| mmu-miR-34a-5p | MIMAT0000542 | Ppp1r10 | 52040 | Northern blot//qRT-PCR//Western blot//H | 24587330\|23142080 | | Unspecified |
| mmu-miR-34a-5p | MIMAT0000542 | Ppp1r11 | 76497 | HITS-CLIP | | 23142080 | Unspecified |
| mmu-miR-34a-5p | MIMAT0000542 | Ppp1r12a | 17931 | HITS-CLIP | 23142080\|29167373\|19536157 | | Unspecified |
| mmu-miR-34a-5p | MIMAT0000542 | Ppp1r12c | 232807 | HITS-CLIP | 23142080\|29167373 | | Unspecified |
| mmu-miR-34a-5p | MIMAT0000542 | Ppp1r13b | 21981 | HITS-CLIP | 31433990\|23630078\|19536157\|23142 | | Unspecified |
| mmu-miR-34a-5p | MIMAT0000542 | Ppp1r15a | 17872 | HITS-CLIP | 23630078\|30971389 | | Unspecified |
| mmu-miR-34a-5p | MIMAT0000542 | Ppp1r15b | 108954 | HITS-CLIP | 31433990\|19536157\|29167373\|23142 | | Unspecified |
| mmu-miR-34a-5p | MIMAT0000542 | Ppp1r16b | 228852 | HITS-CLIP | 31433990\|23142080 | | Unspecified |
| mmu-miR-34a-5p | MIMAT0000542 | Ppp1r2 | 66849 | HITS-CLIP | 30971389\|23142080\|29167373 | | Unspecified |
| mmu-miR-34a-5p | MIMAT0000542 | Ppp1r21 | 73825 | HITS-CLIP | | 23142080 | Unspecified |
| mmu-miR-34a-5p | MIMAT0000542 | Ppp1r3b | 244416 | HITS-CLIP | 23142080\|26203562 | | Unspecified |
| mmu-miR-34a-5p | MIMAT0000542 | Ppp1r7 | 66385 | HITS-CLIP | 19536157\|23142080\|29167373 | | Unspecified |
| mmu-miR-34a-5p | MIMAT0000542 | Ppp1r8 | 100336 | HITS-CLIP | | 23142080 | Unspecified |
| mmu-miR-34a-5p | MIMAT0000542 | Ppp1r9a | 243725 | HITS-CLIP | | 19536157 | Unspecified |
| mmu-miR-34a-5p | MIMAT0000542 | Ppp1r9b | 217124 | HITS-CLIP | | 29167373 | Unspecified |
| mmu-miR-34a-5p | MIMAT0000542 | Ppp2ca | 19052 | HITS-CLIP | 23142080\|19536157\|31433990\|23630 | | Unspecified |
| mmu-miR-34a-5p | MIMAT0000542 | Ppp2cb | 19053 | HITS-CLIP | | 23142080 | Unspecified |
| mmu-miR-34a-5p | MIMAT0000542 | Ppp2r1a | 51792 | HITS-CLIP | 23142080\|19536157\|29167373 | | Unspecified |
| mmu-miR-34a-5p | MIMAT0000542 | Ppp2r1b | 73699 | HITS-CLIP | | 23142080 | Unspecified |
| mmu-miR-34a-5p | MIMAT0000542 | Ppp2r2c | 269643 | HITS-CLIP | | 19536157 | Unspecified |
| mmu-miR-34a-5p | MIMAT0000542 | Ppp2r3a | 235542 | HITS-CLIP | | 23142080 | Unspecified |
| mmu-miR-34a-5p | MIMAT0000542 | Ppp2r5a | 226849 | Microarrays//HITS-CLIP | 37986224\|23027903\|23142080 | | Unspecified |
| mmu-miR-34a-5p | MIMAT0000542 | Ppp2r5c | 26931 | HITS-CLIP | 23142080\|29167373 | | Unspecified |
| mmu-miR-34a-5p | MIMAT0000542 | Ppp2r5d | 21770 | HITS-CLIP | | 19536157 | Unspecified |
| mmu-miR-34a-5p | MIMAT0000542 | Ppp2r5e | 26932 | HITS-CLIP | 23142080\|29167373 | | Unspecified |
| mmu-miR-34a-5p | MIMAT0000542 | Ppp3ca | 19055 | HITS-CLIP | 19536157\|23142080 | | Unspecified |
| mmu-miR-34a-5p | MIMAT0000542 | Ppp3cb | 19056 | HITS-CLIP | | 23142080 | Unspecified |

| | | | | | | |
|---|---|---|---|---|---|---|
| mmu-miR-34a-5p | MIMAT0000542 | Ppp3cc | 19057 | HITS-CLIP | 23142080 | Unspecified |
| mmu-miR-34a-5p | MIMAT0000542 | Ppp3r1 | 19058 | HITS-CLIP | 23630078\|31433990\|23142080 | Unspecified |
| mmu-miR-34a-5p | MIMAT0000542 | Ppp4c | 56420 | HITS-CLIP | 23142080\|29167373 | Unspecified |
| mmu-miR-34a-5p | MIMAT0000542 | Ppp4r1 | 70351 | HITS-CLIP | 29167373 | Unspecified |
| mmu-miR-34a-5p | MIMAT0000542 | Ppp4r2 | 232314 | HITS-CLIP | 23142080 | Unspecified |
| mmu-miR-34a-5p | MIMAT0000542 | Ppp4r3a | 55671 | HITS-CLIP | 23142080\|29167373 | Unspecified |
| mmu-miR-34a-5p | MIMAT0000542 | Ppp4r3b | 57223 | HITS-CLIP | 23142080\|23630078\|29167373 | Unspecified |
| mmu-miR-34a-5p | MIMAT0000542 | Ppp5c | 19060 | HITS-CLIP | 23142080 | Unspecified |
| mmu-miR-34a-5p | MIMAT0000542 | Ppp6c | 67857 | HITS-CLIP | 19536157\|23142080\|29167373 | Unspecified |
| mmu-miR-34a-5p | MIMAT0000542 | Ppp6r1 | 243819 | HITS-CLIP | 29167373\|23142080 | Unspecified |
| mmu-miR-34a-5p | MIMAT0000542 | Ppp6r3 | 52036 | Microarrays//HITS-CLIP | 37986224\|23027903\|23142080\|29167 | Unspecified |
| mmu-miR-34a-5p | MIMAT0000542 | Pprc1 | 226169 | HITS-CLIP | 23142080\|29167373 | Unspecified |
| mmu-miR-34a-5p | MIMAT0000542 | Ppt1 | 19063 | HITS-CLIP | 23142080 | Unspecified |
| mmu-miR-34a-5p | MIMAT0000542 | Ppt2 | 54397 | HITS-CLIP | 23142080 | Unspecified |
| mmu-miR-34a-5p | MIMAT0000542 | Pptc7 | 320717 | HITS-CLIP | 19536157\|23142080 | Unspecified |
| mmu-miR-34a-5p | MIMAT0000542 | Ppwd1 | 238831 | HITS-CLIP | 23142080\|31433990 | Unspecified |
| mmu-miR-34a-5p | MIMAT0000542 | Pqbp1 | 10084 | HITS-CLIP | 23142080 | Unspecified |
| mmu-miR-34a-5p | MIMAT0000542 | Pqlc3 | 217430 | HITS-CLIP | 29167373\|23142080 | Unspecified |
| mmu-miR-34a-5p | MIMAT0000542 | Prag1 | 157285 | HITS-CLIP | 30971389 | Unspecified |
| mmu-miR-34a-5p | MIMAT0000542 | Pramef8 | 242736 | HITS-CLIP | 23142080 | Unspecified |
| mmu-miR-34a-5p | MIMAT0000542 | Prc1 | 233406 | HITS-CLIP | 23142080 | Unspecified |
| mmu-miR-34a-5p | MIMAT0000542 | Prcc | 94315 | HITS-CLIP | 23142080 | Unspecified |
| mmu-miR-34a-5p | MIMAT0000542 | Prcp | 72461 | HITS-CLIP | 30971389 | Unspecified |
| mmu-miR-34a-5p | MIMAT0000542 | Prdm1 | 12142 | HITS-CLIP | 29167373\|23142080 | Unspecified |
| mmu-miR-34a-5p | MIMAT0000542 | Prdm11 | 100042784 | HITS-CLIP | 23142080 | Unspecified |
| mmu-miR-34a-5p | MIMAT0000542 | Prdm15 | 63977 | HITS-CLIP | 23142080 | Unspecified |
| mmu-miR-34a-5p | MIMAT0000542 | Prdm16 | 70673 | HITS-CLIP | 23142080 | Unspecified |
| mmu-miR-34a-5p | MIMAT0000542 | Prdm2 | 110593 | HITS-CLIP | 31433990\|23142080\|29167373 | Unspecified |
| mmu-miR-34a-5p | MIMAT0000542 | Prdm9 | 213389 | HITS-CLIP | 26203562 | Unspecified |
| mmu-miR-34a-5p | MIMAT0000542 | Prdx1 | 18477 | HITS-CLIP | 29167373 | Unspecified |
| mmu-miR-34a-5p | MIMAT0000542 | Prdx2 | 21672 | HITS-CLIP | 23142080 | Unspecified |
| mmu-miR-34a-5p | MIMAT0000542 | Prdx3 | 11757 | HITS-CLIP | 23142080 | Unspecified |
| mmu-miR-34a-5p | MIMAT0000542 | Prdx4 | 53381 | HITS-CLIP | 23142080 | Unspecified |
| mmu-miR-34a-5p | MIMAT0000542 | Prdx6 | 11758 | HITS-CLIP | 19536157\|23142080\|29167373 | Unspecified |
| mmu-miR-34a-5p | MIMAT0000542 | Preb | 50907 | HITS-CLIP | 23142080 | Unspecified |
| mmu-miR-34a-5p | MIMAT0000542 | Prelid1 | 66494 | HITS-CLIP | 29167373 | Unspecified |
| mmu-miR-34a-5p | MIMAT0000542 | Prelid2 | 77619 | HITS-CLIP | 23142080 | Unspecified |
| mmu-miR-34a-5p | MIMAT0000542 | Prelid3b | 51012 | HITS-CLIP | 30971389\|23142080 | Unspecified |
| mmu-miR-34a-5p | MIMAT0000542 | Prelp | 116847 | HITS-CLIP | 19536157 | Unspecified |

| | | | | | | |
|---|---|---|---|---|---|---|
| mmu-miR-34a-5p | MIMAT0000542 | Prep | 19072 | HITS-CLIP | 23142080\|29167373 | Unspecified |
| mmu-miR-34a-5p | MIMAT0000542 | Prepl | 213760 | HITS-CLIP | 23142080 | Unspecified |
| mmu-miR-34a-5p | MIMAT0000542 | Prex1 | 277360 | HITS-CLIP | 23142080\|29167373 | Unspecified |
| mmu-miR-34a-5p | MIMAT0000542 | Prf1 | 18646 | HITS-CLIP | 23142080 | Unspecified |
| mmu-miR-34a-5p | MIMAT0000542 | Prickle2 | 243548 | HITS-CLIP | 19536157 | Unspecified |
| mmu-miR-34a-5p | MIMAT0000542 | Prickle3 | 54630 | HITS-CLIP | 23142080 | Unspecified |
| mmu-miR-34a-5p | MIMAT0000542 | Prim1 | 19075 | Microarrays//HITS-CLIP | 37986224\|23027903\|23142080 | Unspecified |
| mmu-miR-34a-5p | MIMAT0000542 | Primpol | 408022 | HITS-CLIP | 23142080 | Unspecified |
| mmu-miR-34a-5p | MIMAT0000542 | Prkaa1 | 105787 | HITS-CLIP | 23630078\|31433990\|23142080 | Unspecified |
| mmu-miR-34a-5p | MIMAT0000542 | Prkaa2 | 108079 | Microarrays | 37986224 | Unspecified |
| mmu-miR-34a-5p | MIMAT0000542 | Prkab1 | 19079 | HITS-CLIP | 23142080 | Unspecified |
| mmu-miR-34a-5p | MIMAT0000542 | Prkab2 | 108097 | HITS-CLIP | 23142080\|29167373 | Unspecified |
| mmu-miR-34a-5p | MIMAT0000542 | Prkaca | 18747 | HITS-CLIP | 29167373 | Unspecified |
| mmu-miR-34a-5p | MIMAT0000542 | Prkacb | 18749 | Microarrays//HITS-CLIP | 37986224\|23027903\|19536157\|23142 | Unspecified |
| mmu-miR-34a-5p | MIMAT0000542 | Prkag2 | 108099 | HITS-CLIP | 19536157\|23142080 | Unspecified |
| mmu-miR-34a-5p | MIMAT0000542 | Prkar1a | 19084 | HITS-CLIP | 31433990\|23630078\|19536157\|23142 | Unspecified |
| mmu-miR-34a-5p | MIMAT0000542 | Prkar1b | 19085 | HITS-CLIP | 19536157 | Unspecified |
| mmu-miR-34a-5p | MIMAT0000542 | Prkar2a | 19087 | HITS-CLIP | 19536157\|26203562\|29167373\|23142 | Unspecified |
| mmu-miR-34a-5p | MIMAT0000542 | Prkca | 18750 | HITS-CLIP | 37986224\|31433990\|23142080 | Unspecified |
| mmu-miR-34a-5p | MIMAT0000542 | Prkcb | 18751 | HITS-CLIP | 19536157\|29167373\|23142080 | Unspecified |
| mmu-miR-34a-5p | MIMAT0000542 | Prkcd | 18753 | HITS-CLIP | 23142080\|29167373 | Unspecified |
| mmu-miR-34a-5p | MIMAT0000542 | Prkce | 18754 | HITS-CLIP | 23142080 | Unspecified |
| mmu-miR-34a-5p | MIMAT0000542 | Prkch | 18755 | HITS-CLIP | 23142080\|29167373 | Unspecified |
| mmu-miR-34a-5p | MIMAT0000542 | Prkci | 18759 | Microarrays//HITS-CLIP | 37986224\|23027903\|31433990\|23142 | Unspecified |
| mmu-miR-34a-5p | MIMAT0000542 | Prkcq | 18761 | HITS-CLIP | 23142080 | Unspecified |
| mmu-miR-34a-5p | MIMAT0000542 | Prkcsh | 19089 | HITS-CLIP | 23142080 | Unspecified |
| mmu-miR-34a-5p | MIMAT0000542 | Prkcz | 18762 | HITS-CLIP | 23142080 | Unspecified |
| mmu-miR-34a-5p | MIMAT0000542 | Prkd2 | 101540 | HITS-CLIP | 23142080 | Unspecified |
| mmu-miR-34a-5p | MIMAT0000542 | Prkd3 | 75292 | HITS-CLIP | 23630078\|29167373 | Unspecified |
| mmu-miR-34a-5p | MIMAT0000542 | Prkdc | 19090 | HITS-CLIP | 19536157\|23142080\|29167373 | Unspecified |
| mmu-miR-34a-5p | MIMAT0000542 | Prkg1 | 19091 | Microarrays | 37986224 | Unspecified |
| mmu-miR-34a-5p | MIMAT0000542 | Prkrip1 | 66801 | HITS-CLIP | 23142080\|29167373 | Unspecified |
| mmu-miR-34a-5p | MIMAT0000542 | Prkx | 19108 | HITS-CLIP | 23142080\|29167373 | Unspecified |
| mmu-miR-34a-5p | MIMAT0000542 | Prlr | 19116 | HITS-CLIP | 23142080 | Unspecified |
| mmu-miR-34a-5p | MIMAT0000542 | Prmt1 | 15469 | HITS-CLIP | 23142080 | Unspecified |
| mmu-miR-34a-5p | MIMAT0000542 | Prmt2 | 15468 | Microarrays | 37986224 | Unspecified |
| mmu-miR-34a-5p | MIMAT0000542 | Prmt5 | 27374 | HITS-CLIP | 31433990 | Unspecified |
| mmu-miR-34a-5p | MIMAT0000542 | Prmt6 | 99890 | HITS-CLIP | 23142080 | Unspecified |
| mmu-miR-34a-5p | MIMAT0000542 | Prmt7 | 214572 | HITS-CLIP | 23142080 | Unspecified |

| | | | | | | |
|---|---|---|---|---|---|---|
| mmu-miR-34a-5p | MIMAT0000542 | Prmt9 | 80204 | HITS-CLIP | 23142080 | Unspecified |
| mmu-miR-34a-5p | MIMAT0000542 | Prnp | 19122 | HITS-CLIP | 19536157 | Unspecified |
| mmu-miR-34a-5p | MIMAT0000542 | Prom1 | 19126 | HITS-CLIP | 19536157 | Unspecified |
| mmu-miR-34a-5p | MIMAT0000542 | Prorsd1 | 67939 | HITS-CLIP | 23142080 | Unspecified |
| mmu-miR-34a-5p | MIMAT0000542 | Pros1 | 19128 | HITS-CLIP | 23142080 | Unspecified |
| mmu-miR-34a-5p | MIMAT0000542 | Proser1 | 212127 | HITS-CLIP | 23142080\|29167373 | Unspecified |
| mmu-miR-34a-5p | MIMAT0000542 | Proser2 | 227545 | HITS-CLIP | 23142080 | Unspecified |
| mmu-miR-34a-5p | MIMAT0000542 | Prox1 | 19130 | HITS-CLIP | 19536157 | Unspecified |
| mmu-miR-34a-5p | MIMAT0000542 | Prpf18 | 67229 | HITS-CLIP | 23142080 | Unspecified |
| mmu-miR-34a-5p | MIMAT0000542 | Prpf19 | 28000 | HITS-CLIP | 23142080\|19536157 | Unspecified |
| mmu-miR-34a-5p | MIMAT0000542 | Prpf3 | 70767 | HITS-CLIP | 23142080\|19536157\|29167373 | Unspecified |
| mmu-miR-34a-5p | MIMAT0000542 | Prpf38a | 230596 | HITS-CLIP | 23142080 | Unspecified |
| mmu-miR-34a-5p | MIMAT0000542 | Prpf38b | 66921 | HITS-CLIP | 23142080 | Unspecified |
| mmu-miR-34a-5p | MIMAT0000542 | Prpf39 | 328110 | HITS-CLIP | 23142080\|29167373 | Unspecified |
| mmu-miR-34a-5p | MIMAT0000542 | Prpf4 | 70052 | HITS-CLIP | 29167373 | Unspecified |
| mmu-miR-34a-5p | MIMAT0000542 | Prpf40a | 56194 | HITS-CLIP | 31433990\|23142080 | Unspecified |
| mmu-miR-34a-5p | MIMAT0000542 | Prpf4b | 19134 | HITS-CLIP | 31433990\|23142080 | Unspecified |
| mmu-miR-34a-5p | MIMAT0000542 | Prpf6 | 68879 | HITS-CLIP | 23142080 | Unspecified |
| mmu-miR-34a-5p | MIMAT0000542 | Prpf8 | 192159 | HITS-CLIP | 37986224\|23142080\|19536157\|3143[illegible] | Unspecified |
| mmu-miR-34a-5p | MIMAT0000542 | Prps1 | 19139 | HITS-CLIP | 23142080 | Unspecified |
| mmu-miR-34a-5p | MIMAT0000542 | Prps1l3 | 328099 | HITS-CLIP | 23142080\|29167373 | Unspecified |
| mmu-miR-34a-5p | MIMAT0000542 | Prps2 | 110639 | HITS-CLIP | 23142080 | Unspecified |
| mmu-miR-34a-5p | MIMAT0000542 | Prpsap1 | 67763 | HITS-CLIP | 23142080 | Unspecified |
| mmu-miR-34a-5p | MIMAT0000542 | Prr11 | 270906 | HITS-CLIP | 23142080 | Unspecified |
| mmu-miR-34a-5p | MIMAT0000542 | Prr12 | 233210 | HITS-CLIP | 23142080\|29167373 | Unspecified |
| mmu-miR-34a-5p | MIMAT0000542 | Prr13 | 66151 | HITS-CLIP | 23142080\|31433990\|26203562 | Unspecified |
| mmu-miR-34a-5p | MIMAT0000542 | Prr14 | 233895 | HITS-CLIP | 29167373\|23142080 | Unspecified |
| mmu-miR-34a-5p | MIMAT0000542 | Prr14l | 215476 | HITS-CLIP | 30971389\|23142080\|29167373 | Unspecified |
| mmu-miR-34a-5p | MIMAT0000542 | Prr5 | 109270 | Microarrays//HITS-CLIP | 37986224\|23027903\|23142080 | Unspecified |
| mmu-miR-34a-5p | MIMAT0000542 | Prrc1 | 73137 | HITS-CLIP | 23142080 | Unspecified |
| mmu-miR-34a-5p | MIMAT0000542 | Prrc2a | 53761 | HITS-CLIP | 23142080\|26203562\|29167373 | Unspecified |
| mmu-miR-34a-5p | MIMAT0000542 | Prrc2b | 227723 | HITS-CLIP | 19536157\|31433990\|23142080\|29167 | Unspecified |
| mmu-miR-34a-5p | MIMAT0000542 | Prrc2c | 226562 | HITS-CLIP | 23142080\|19536157\|31433990\|30971 | Unspecified |
| mmu-miR-34a-5p | MIMAT0000542 | Prrg1 | 546336 | HITS-CLIP | 23142080 | Unspecified |
| mmu-miR-34a-5p | MIMAT0000542 | Prrg3 | 208748 | HITS-CLIP | 19536157 | Unspecified |
| mmu-miR-34a-5p | MIMAT0000542 | Prrg4 | 228413 | HITS-CLIP | 26203562\|29167373\|23142080 | Unspecified |
| mmu-miR-34a-5p | MIMAT0000542 | Prrt2 | 69017 | HITS-CLIP | 19536157 | Unspecified |
| mmu-miR-34a-5p | MIMAT0000542 | Psap | 19156 | HITS-CLIP | 23142080\|19536157\|29167373 | Unspecified |
| mmu-miR-34a-5p | MIMAT0000542 | Psat1 | 107272 | Microarrays//HITS-CLIP | 37986224\|23027903\|23142080 | Unspecified |

| | | | | | | |
|---|---|---|---|---|---|---|
| mmu-miR-34a-5p | MIMAT0000542 | Psd2 | 74002 | HITS-CLIP | 19536157 | Unspecified |
| mmu-miR-34a-5p | MIMAT0000542 | Psd3 | 234353 | HITS-CLIP | 19536157\|29167373 | Unspecified |
| mmu-miR-34a-5p | MIMAT0000542 | Psd4 | 215632 | HITS-CLIP | 23142080\|29167373 | Unspecified |
| mmu-miR-34a-5p | MIMAT0000542 | Psen1 | 19164 | HITS-CLIP | 23142080\|29167373 | Unspecified |
| mmu-miR-34a-5p | MIMAT0000542 | Psen2 | 19165 | Microarrays//HITS-CLIP | 37986224\|29167373 | Unspecified |
| mmu-miR-34a-5p | MIMAT0000542 | Psip1 | 101739 | HITS-CLIP | 23142080\|29167373 | Unspecified |
| mmu-miR-34a-5p | MIMAT0000542 | Pskh1 | 244631 | HITS-CLIP | 30971389\|19536157\|23142080\|29167 | Unspecified |
| mmu-miR-34a-5p | MIMAT0000542 | Psma1 | 26440 | HITS-CLIP | 23142080 | Unspecified |
| mmu-miR-34a-5p | MIMAT0000542 | Psma2 | 19166 | HITS-CLIP | 23142080 | Unspecified |
| mmu-miR-34a-5p | MIMAT0000542 | Psma4 | 26441 | HITS-CLIP | 23142080 | Unspecified |
| mmu-miR-34a-5p | MIMAT0000542 | Psma5 | 26442 | HITS-CLIP | 26203562 | Unspecified |
| mmu-miR-34a-5p | MIMAT0000542 | Psma6 | 26443 | HITS-CLIP | 23142080\|29167373 | Unspecified |
| mmu-miR-34a-5p | MIMAT0000542 | Psma7 | 26444 | HITS-CLIP | 23142080\|29167373 | Unspecified |
| mmu-miR-34a-5p | MIMAT0000542 | Psmb1 | 19170 | HITS-CLIP | 23142080 | Unspecified |
| mmu-miR-34a-5p | MIMAT0000542 | Psmb10 | 19171 | HITS-CLIP | 23142080 | Unspecified |
| mmu-miR-34a-5p | MIMAT0000542 | Psmb2 | 26445 | HITS-CLIP | 23142080\|29167373 | Unspecified |
| mmu-miR-34a-5p | MIMAT0000542 | Psmb4 | 19172 | HITS-CLIP | 23142080\|23630078 | Unspecified |
| mmu-miR-34a-5p | MIMAT0000542 | Psmb6 | 19175 | HITS-CLIP | 23142080 | Unspecified |
| mmu-miR-34a-5p | MIMAT0000542 | Psmb7 | 19177 | HITS-CLIP | 19536157\|23142080 | Unspecified |
| mmu-miR-34a-5p | MIMAT0000542 | Psmb8 | 16913 | HITS-CLIP | 23142080\|29167373 | Unspecified |
| mmu-miR-34a-5p | MIMAT0000542 | Psmb9 | 16912 | HITS-CLIP | 23142080 | Unspecified |
| mmu-miR-34a-5p | MIMAT0000542 | Psmc1 | 19179 | HITS-CLIP | 23142080\|19536157 | Unspecified |
| mmu-miR-34a-5p | MIMAT0000542 | Psmc2 | 19181 | HITS-CLIP | 19536157\|23142080 | Unspecified |
| mmu-miR-34a-5p | MIMAT0000542 | Psmc3 | 19182 | HITS-CLIP | 23142080 | Unspecified |
| mmu-miR-34a-5p | MIMAT0000542 | Psmc6 | 67089 | HITS-CLIP | 23142080 | Unspecified |
| mmu-miR-34a-5p | MIMAT0000542 | Psmd1 | 70247 | HITS-CLIP | 23142080\|19536157\|29167373 | Unspecified |
| mmu-miR-34a-5p | MIMAT0000542 | Psmd11 | 69077 | HITS-CLIP | 23142080 | Unspecified |
| mmu-miR-34a-5p | MIMAT0000542 | Psmd12 | 66997 | HITS-CLIP | 31433990 | Unspecified |
| mmu-miR-34a-5p | MIMAT0000542 | Psmd13 | 23997 | HITS-CLIP | 23142080 | Unspecified |
| mmu-miR-34a-5p | MIMAT0000542 | Psmd14 | 59029 | HITS-CLIP | 31433990\|23142080 | Unspecified |
| mmu-miR-34a-5p | MIMAT0000542 | Psmd2 | 21762 | HITS-CLIP | 23142080\|29167373 | Unspecified |
| mmu-miR-34a-5p | MIMAT0000542 | Psmd3 | 22123 | HITS-CLIP | 23142080 | Unspecified |
| mmu-miR-34a-5p | MIMAT0000542 | Psmd4 | 19185 | HITS-CLIP | 23142080 | Unspecified |
| mmu-miR-34a-5p | MIMAT0000542 | Psmd5 | 66998 | HITS-CLIP | 23142080\|29167373 | Unspecified |
| mmu-miR-34a-5p | MIMAT0000542 | Psmd6 | 66413 | HITS-CLIP | 23142080 | Unspecified |
| mmu-miR-34a-5p | MIMAT0000542 | Psmd7 | 17463 | HITS-CLIP | 23630078 | Unspecified |
| mmu-miR-34a-5p | MIMAT0000542 | Psmd8 | 57296 | HITS-CLIP | 23142080\|29167373 | Unspecified |
| mmu-miR-34a-5p | MIMAT0000542 | Psme1 | 19186 | HITS-CLIP | 23142080 | Unspecified |
| mmu-miR-34a-5p | MIMAT0000542 | Psme3 | 19192 | HITS-CLIP | 23142080\|29167373 | Unspecified |

| | | | | | | |
|---|---|---|---|---|---|---|
| mmu-miR-34a-5p | MIMAT0000542 | Psme4 | 103554 | HITS-CLIP | 19536157\|23142080\|29167373 | Unspecified |
| mmu-miR-34a-5p | MIMAT0000542 | Psmf1 | 9491 | HITS-CLIP | 23142080 | Unspecified |
| mmu-miR-34a-5p | MIMAT0000542 | Psmg1 | 56088 | HITS-CLIP | 23142080 | Unspecified |
| mmu-miR-34a-5p | MIMAT0000542 | Psmg2 | 107047 | HITS-CLIP | 23142080\|29167373 | Unspecified |
| mmu-miR-34a-5p | MIMAT0000542 | Pspc1 | 66645 | HITS-CLIP | 23142080\|29167373 | Unspecified |
| mmu-miR-34a-5p | MIMAT0000542 | Psph | 100678 | HITS-CLIP | 23142080 | Unspecified |
| mmu-miR-34a-5p | MIMAT0000542 | Psrc1 | 56742 | HITS-CLIP | 23142080 | Unspecified |
| mmu-miR-34a-5p | MIMAT0000542 | Ptafr | 19204 | HITS-CLIP | 29167373 | Unspecified |
| mmu-miR-34a-5p | MIMAT0000542 | Ptar1 | 72351 | HITS-CLIP | 23142080\|31433990 | Unspecified |
| mmu-miR-34a-5p | MIMAT0000542 | Ptbp1 | 19205 | HITS-CLIP | 23142080\|31433990\|29167373 | Unspecified |
| mmu-miR-34a-5p | MIMAT0000542 | Ptbp2 | 56195 | HITS-CLIP | 19536157 | Unspecified |
| mmu-miR-34a-5p | MIMAT0000542 | Ptbp3 | 230257 | Microarrays//HITS-CLIP | 37986224\|23027903\|23630078\|23142 | Unspecified |
| mmu-miR-34a-5p | MIMAT0000542 | Ptcd2 | 68927 | HITS-CLIP | 23142080\|29167373 | Unspecified |
| mmu-miR-34a-5p | MIMAT0000542 | Ptcd3 | 69956 | HITS-CLIP | 23142080 | Unspecified |
| mmu-miR-34a-5p | MIMAT0000542 | Ptch1 | 19206 | HITS-CLIP | 30971389\|23142080\|31433990 | Unspecified |
| mmu-miR-34a-5p | MIMAT0000542 | Ptchd1 | 211612 | HITS-CLIP | 19536157\|29167373 | Unspecified |
| mmu-miR-34a-5p | MIMAT0000542 | Ptdss1 | 19210 | HITS-CLIP | 23142080 | Unspecified |
| mmu-miR-34a-5p | MIMAT0000542 | Pten | 19211 | HITS-CLIP | 30971389\|23142080\|29167373 | Unspecified |
| mmu-miR-34a-5p | MIMAT0000542 | Ptgds | 19215 | HITS-CLIP | 26203562 | Unspecified |
| mmu-miR-34a-5p | MIMAT0000542 | Ptger2 | 19217 | HITS-CLIP | 23142080 | Unspecified |
| mmu-miR-34a-5p | MIMAT0000542 | Ptger4 | 19219 | HITS-CLIP | 23142080\|29167373 | Unspecified |
| mmu-miR-34a-5p | MIMAT0000542 | Ptges3 | 56351 | HITS-CLIP | 29167373 | Unspecified |
| mmu-miR-34a-5p | MIMAT0000542 | Ptgfrn | 19221 | HITS-CLIP | 23630078\|23142080 | Unspecified |
| mmu-miR-34a-5p | MIMAT0000542 | Ptgis | 19223 | Microarrays | 37986224 | Unspecified |
| mmu-miR-34a-5p | MIMAT0000542 | Ptgr2 | 77219 | Microarrays//HITS-CLIP | 37986224\|30971389 | Unspecified |
| mmu-miR-34a-5p | MIMAT0000542 | Ptk2 | 14083 | HITS-CLIP | 23142080 | Unspecified |
| mmu-miR-34a-5p | MIMAT0000542 | Ptk2b | 19229 | HITS-CLIP | 23142080\|29167373 | Unspecified |
| mmu-miR-34a-5p | MIMAT0000542 | Ptn | 19242 | HITS-CLIP | 19536157 | Unspecified |
| mmu-miR-34a-5p | MIMAT0000542 | Ptp4a2 | 19244 | HITS-CLIP | 23142080 | Unspecified |
| mmu-miR-34a-5p | MIMAT0000542 | Ptpa | 5524 | HITS-CLIP | 30971389\|23630078\|29167373\|23142 | Unspecified |
| mmu-miR-34a-5p | MIMAT0000542 | Ptpmt1 | 114971 | HITS-CLIP | 30971389 | Unspecified |
| mmu-miR-34a-5p | MIMAT0000542 | Ptpn1 | 19246 | HITS-CLIP | 19536157\|23142080 | Unspecified |
| mmu-miR-34a-5p | MIMAT0000542 | Ptpn11 | 19247 | HITS-CLIP | 19536157\|31433990\|23142080 | Unspecified |
| mmu-miR-34a-5p | MIMAT0000542 | Ptpn18 | 19253 | HITS-CLIP | 23142080 | Unspecified |
| mmu-miR-34a-5p | MIMAT0000542 | Ptpn2 | 19255 | HITS-CLIP | 37986224\|23142080 | Unspecified |
| mmu-miR-34a-5p | MIMAT0000542 | Ptpn22 | 19260 | HITS-CLIP | 29167373\|23142080 | Unspecified |
| mmu-miR-34a-5p | MIMAT0000542 | Ptpn23 | 104831 | HITS-CLIP | 19536157\|29167373\|23142080 | Unspecified |
| mmu-miR-34a-5p | MIMAT0000542 | Ptpn3 | 545622 | HITS-CLIP | 23142080 | Unspecified |
| mmu-miR-34a-5p | MIMAT0000542 | Ptpn4 | 19258 | HITS-CLIP | 19536157\|29167373\|23142080 | Unspecified |

| | | | | | | | |
|---|---|---|---|---|---|---|---|
| mmu-miR-34a-5p | MIMAT0000542 | Ptpn5 | 19259 | HITS-CLIP | | 19536157 | Unspecified |
| mmu-miR-34a-5p | MIMAT0000542 | Ptpn6 | 15170 | HITS-CLIP | 23142080\|29167373 | | Unspecified |
| mmu-miR-34a-5p | MIMAT0000542 | Ptpn7 | 320139 | HITS-CLIP | 23142080\|29167373 | | Unspecified |
| mmu-miR-34a-5p | MIMAT0000542 | Ptpn9 | 56294 | HITS-CLIP | 19536157\|29167373\|23142080 | | Unspecified |
| mmu-miR-34a-5p | MIMAT0000542 | Ptpra | 19262 | HITS-CLIP | 19536157\|29167373\|23142080 | | Unspecified |
| mmu-miR-34a-5p | MIMAT0000542 | Ptprb | 19263 | HITS-CLIP | | 19536157 | Unspecified |
| mmu-miR-34a-5p | MIMAT0000542 | Ptprc | 19264 | HITS-CLIP | 23142080\|29167373 | | Unspecified |
| mmu-miR-34a-5p | MIMAT0000542 | Ptprcap | 19265 | HITS-CLIP | | 23142080 | Unspecified |
| mmu-miR-34a-5p | MIMAT0000542 | Ptprd | 19266 | Microarrays//HITS-CLIP | 37986224\|19536157 | | Unspecified |
| mmu-miR-34a-5p | MIMAT0000542 | Ptpre | 19267 | HITS-CLIP | | 29167373 | Unspecified |
| mmu-miR-34a-5p | MIMAT0000542 | Ptprg | 19270 | HITS-CLIP | | 19536157 | Unspecified |
| mmu-miR-34a-5p | MIMAT0000542 | Ptprj | 19271 | HITS-CLIP | 19536157\|29167373 | | Unspecified |
| mmu-miR-34a-5p | MIMAT0000542 | Ptprk | 19272 | HITS-CLIP | | 19536157 | Unspecified |
| mmu-miR-34a-5p | MIMAT0000542 | Ptprs | 19280 | HITS-CLIP | 19536157\|23142080\|29167373 | | Unspecified |
| mmu-miR-34a-5p | MIMAT0000542 | Ptpru | 19273 | HITS-CLIP | | 19536157 | Unspecified |
| mmu-miR-34a-5p | MIMAT0000542 | Ptprz1 | 19283 | HITS-CLIP | | 19536157 | Unspecified |
| mmu-miR-34a-5p | MIMAT0000542 | Ptrh1 | 329384 | HITS-CLIP | | 23142080 | Unspecified |
| mmu-miR-34a-5p | MIMAT0000542 | Ptrh2 | 217057 | HITS-CLIP | | 23142080 | Unspecified |
| mmu-miR-34a-5p | MIMAT0000542 | Pttg1ip | 108705 | HITS-CLIP | | 19536157 | Unspecified |
| mmu-miR-34a-5p | MIMAT0000542 | Puf60 | 67959 | HITS-CLIP | 19536157\|23142080 | | Unspecified |
| mmu-miR-34a-5p | MIMAT0000542 | Pum1 | 80912 | HITS-CLIP | 19536157\|23142080\|29167373 | | Unspecified |
| mmu-miR-34a-5p | MIMAT0000542 | Pum2 | 80913 | HITS-CLIP | 31433990\|19536157\|23142080\|29167 | | Unspecified |
| mmu-miR-34a-5p | MIMAT0000542 | Pura | 19290 | HITS-CLIP | | 23142080 | Unspecified |
| mmu-miR-34a-5p | MIMAT0000542 | Purb | 19291 | HITS-CLIP | 23142080\|19536157\|29167373 | | Unspecified |
| mmu-miR-34a-5p | MIMAT0000542 | Purg | 75029 | HITS-CLIP | 19536157\|23142080 | | Unspecified |
| mmu-miR-34a-5p | MIMAT0000542 | Pus1 | 855889 | HITS-CLIP | | 23142080 | Unspecified |
| mmu-miR-34a-5p | MIMAT0000542 | Pus3 | 67049 | HITS-CLIP | 29167373\|23142080 | | Unspecified |
| mmu-miR-34a-5p | MIMAT0000542 | Pus7l | 78895 | Microarrays | 37986224\|23027903 | | Unspecified |
| mmu-miR-34a-5p | MIMAT0000542 | Pvr | 52118 | HITS-CLIP | | 23142080 | Unspecified |
| mmu-miR-34a-5p | MIMAT0000542 | Pwp2 | 110816 | HITS-CLIP | 30971389\|29167373 | | Unspecified |
| mmu-miR-34a-5p | MIMAT0000542 | Pwwp2a | 70802 | HITS-CLIP | | 19536157 | Unspecified |
| mmu-miR-34a-5p | MIMAT0000542 | Pwwp2b | 101631 | HITS-CLIP | | 29167373 | Unspecified |
| mmu-miR-34a-5p | MIMAT0000542 | Pxdn | 69675 | HITS-CLIP | | 19536157 | Unspecified |
| mmu-miR-34a-5p | MIMAT0000542 | Pxk | 218699 | HITS-CLIP | 23142080\|29167373 | | Unspecified |
| mmu-miR-34a-5p | MIMAT0000542 | Pxmp2 | 19301 | HITS-CLIP | | 23142080 | Unspecified |
| mmu-miR-34a-5p | MIMAT0000542 | Pxmp4 | 59038 | HITS-CLIP | | 23142080 | Unspecified |
| mmu-miR-34a-5p | MIMAT0000542 | Pxn | 19303 | HITS-CLIP | 23142080\|29167373 | | Unspecified |
| mmu-miR-34a-5p | MIMAT0000542 | Pxylp1 | 235534 | HITS-CLIP | | 23142080 | Unspecified |
| mmu-miR-34a-5p | MIMAT0000542 | Pycard | 66824 | HITS-CLIP | | 23142080 | Unspecified |

| | | | | | | |
|---|---|---|---|---|---|---|
| mmu-miR-34a-5p | MIMAT0000542 | Pygb | 110078 | HITS-CLIP | 19536157\|23142080 | Unspecified |
| mmu-miR-34a-5p | MIMAT0000542 | Pygl | 110095 | HITS-CLIP | 23142080 | Unspecified |
| mmu-miR-34a-5p | MIMAT0000542 | Pygm | 19309 | Microarrays | 37986224 | Unspecified |
| mmu-miR-34a-5p | MIMAT0000542 | Pygo2 | 68911 | HITS-CLIP | 23142080 | Unspecified |
| mmu-miR-34a-5p | MIMAT0000542 | Pyroxd1 | 232491 | HITS-CLIP | 23142080 | Unspecified |
| mmu-miR-34a-5p | MIMAT0000542 | Qars | 97541 | HITS-CLIP | 23142080 | Unspecified |
| mmu-miR-34a-5p | MIMAT0000542 | Qdpr | 110391 | HITS-CLIP | 23142080 | Unspecified |
| mmu-miR-34a-5p | MIMAT0000542 | Qk | 19317 | HITS-CLIP | 23142080\|29167373\|19536157 | Unspecified |
| mmu-miR-34a-5p | MIMAT0000542 | Qki | 19317 | HITS-CLIP | 31433990 | Unspecified |
| mmu-miR-34a-5p | MIMAT0000542 | Qprt | 67375 | HITS-CLIP | 23142080 | Unspecified |
| mmu-miR-34a-5p | MIMAT0000542 | Qrich1 | 69232 | HITS-CLIP | 19536157\|30971389\|23142080\|29167 | Unspecified |
| mmu-miR-34a-5p | MIMAT0000542 | Qrsl1 | 76563 | HITS-CLIP | 23142080 | Unspecified |
| mmu-miR-34a-5p | MIMAT0000542 | Qser1 | 99003 | HITS-CLIP | 23142080\|31433990 | Unspecified |
| mmu-miR-34a-5p | MIMAT0000542 | Qsox1 | 104009 | HITS-CLIP | 23142080 | Unspecified |
| mmu-miR-34a-5p | MIMAT0000542 | Qtrt1 | 60507 | HITS-CLIP | 23142080 | Unspecified |
| mmu-miR-34a-5p | MIMAT0000542 | Qtrt2 | 79691 | HITS-CLIP | 29167373 | Unspecified |
| mmu-miR-34a-5p | MIMAT0000542 | R3hcc1l | 52013 | HITS-CLIP | 23142080 | Unspecified |
| mmu-miR-34a-5p | MIMAT0000542 | R3hdm1 | 226412 | HITS-CLIP | 37986224\|23142080\|19536157\|31433 | Unspecified |
| mmu-miR-34a-5p | MIMAT0000542 | R3hdm2 | 71750 | HITS-CLIP | 23142080\|19536157 | Unspecified |
| mmu-miR-34a-5p | MIMAT0000542 | Rab10 | 19325 | HITS-CLIP | 23142080 | Unspecified |
| mmu-miR-34a-5p | MIMAT0000542 | Rab11a | 53869 | HITS-CLIP | 23142080\|19536157 | Unspecified |
| mmu-miR-34a-5p | MIMAT0000542 | Rab11b | 19326 | HITS-CLIP | 21258322\|23142080 | Unspecified |
| mmu-miR-34a-5p | MIMAT0000542 | Rab11fip1 | 75767 | HITS-CLIP | 23142080 | Unspecified |
| mmu-miR-34a-5p | MIMAT0000542 | Rab11fip2 | 74998 | HITS-CLIP | 23142080\|29167373 | Unspecified |
| mmu-miR-34a-5p | MIMAT0000542 | Rab11fip3 | 215445 | HITS-CLIP | 23142080 | Unspecified |
| mmu-miR-34a-5p | MIMAT0000542 | Rab11fip5 | 52055 | HITS-CLIP | 29167373 | Unspecified |
| mmu-miR-34a-5p | MIMAT0000542 | Rab12 | 19328 | HITS-CLIP | 23142080\|29167373 | Unspecified |
| mmu-miR-34a-5p | MIMAT0000542 | Rab14 | 68365 | HITS-CLIP | 19536157\|23142080\|29167373 | Unspecified |
| mmu-miR-34a-5p | MIMAT0000542 | Rab18 | 19330 | HITS-CLIP | 23142080 | Unspecified |
| mmu-miR-34a-5p | MIMAT0000542 | Rab1a | 5861 | HITS-CLIP | 23142080\|29167373 | Unspecified |
| mmu-miR-34a-5p | MIMAT0000542 | Rab1b | 76308 | HITS-CLIP | 23142080\|26203562 | Unspecified |
| mmu-miR-34a-5p | MIMAT0000542 | Rab20 | 19332 | HITS-CLIP | 29167373 | Unspecified |
| mmu-miR-34a-5p | MIMAT0000542 | Rab21 | 216344 | HITS-CLIP | 23142080\|19536157\|29167373 | Unspecified |
| mmu-miR-34a-5p | MIMAT0000542 | Rab22a | 19334 | HITS-CLIP | 23142080\|29167373 | Unspecified |
| mmu-miR-34a-5p | MIMAT0000542 | Rab23 | 19335 | HITS-CLIP | 23142080 | Unspecified |
| mmu-miR-34a-5p | MIMAT0000542 | Rab24 | 19336 | HITS-CLIP | 23142080 | Unspecified |
| mmu-miR-34a-5p | MIMAT0000542 | Rab27a | 11891 | HITS-CLIP | 23142080 | Unspecified |
| mmu-miR-34a-5p | MIMAT0000542 | Rab27b | 80718 | HITS-CLIP | 26203562 | Unspecified |
| mmu-miR-34a-5p | MIMAT0000542 | Rab2a | 59021 | HITS-CLIP | 19536157\|23630078\|30971389\|23142 | Unspecified |

| | | | | | | |
|---|---|---|---|---|---|---|
| mmu-miR-34a-5p | MIMAT0000542 | Rab2b | 76338 | HITS-CLIP | 23142080 | Unspecified |
| mmu-miR-34a-5p | MIMAT0000542 | Rab32 | 67844 | HITS-CLIP | 29167373 | Unspecified |
| mmu-miR-34a-5p | MIMAT0000542 | Rab33b | 19338 | HITS-CLIP | 31433990\|23630078\|23142080\|29167 | Unspecified |
| mmu-miR-34a-5p | MIMAT0000542 | Rab35 | 77407 | HITS-CLIP | 23142080 | Unspecified |
| mmu-miR-34a-5p | MIMAT0000542 | Rab39 | 270160 | HITS-CLIP | 30971389 | Unspecified |
| mmu-miR-34a-5p | MIMAT0000542 | Rab3a | 19339 | HITS-CLIP | 19536157 | Unspecified |
| mmu-miR-34a-5p | MIMAT0000542 | Rab3gap1 | 226407 | HITS-CLIP | 23142080\|29167373 | Unspecified |
| mmu-miR-34a-5p | MIMAT0000542 | Rab3gap2 | 98732 | HITS-CLIP | 30971389\|23142080\|29167373 | Unspecified |
| mmu-miR-34a-5p | MIMAT0000542 | Rab3il1 | 74760 | HITS-CLIP | 29167373 | Unspecified |
| mmu-miR-34a-5p | MIMAT0000542 | Rab40c | 224624 | HITS-CLIP | 23142080\|26203562\|29167373 | Unspecified |
| mmu-miR-34a-5p | MIMAT0000542 | Rab4b | 19342 | HITS-CLIP | 31433990\|23142080 | Unspecified |
| mmu-miR-34a-5p | MIMAT0000542 | Rab5b | 19344 | HITS-CLIP | 23142080 | Unspecified |
| mmu-miR-34a-5p | MIMAT0000542 | Rab5c | 19345 | HITS-CLIP | 19536157\|23142080\|29167373 | Unspecified |
| mmu-miR-34a-5p | MIMAT0000542 | Rab5if | 55969 | HITS-CLIP | 23142080 | Unspecified |
| mmu-miR-34a-5p | MIMAT0000542 | Rab6a | 19346 | HITS-CLIP | 29167373\|23142080 | Unspecified |
| mmu-miR-34a-5p | MIMAT0000542 | Rab6b | 270192 | HITS-CLIP | 19536157 | Unspecified |
| mmu-miR-34a-5p | MIMAT0000542 | Rab7 | 19349 | HITS-CLIP | 29167373 | Unspecified |
| mmu-miR-34a-5p | MIMAT0000542 | Rab7b | 338382 | HITS-CLIP | 29167373 | Unspecified |
| mmu-miR-34a-5p | MIMAT0000542 | Rab8a | 17274 | HITS-CLIP | 23142080\|29167373 | Unspecified |
| mmu-miR-34a-5p | MIMAT0000542 | Rab8b | 235442 | HITS-CLIP | 31433990\|30971389\|23142080\|29167 | Unspecified |
| mmu-miR-34a-5p | MIMAT0000542 | Rab9b | 319642 | HITS-CLIP | 19536157 | Unspecified |
| mmu-miR-34a-5p | MIMAT0000542 | Rabac1 | 14470 | HITS-CLIP | 23142080 | Unspecified |
| mmu-miR-34a-5p | MIMAT0000542 | Rabep1 | 54189 | HITS-CLIP | 31433990\|23142080 | Unspecified |
| mmu-miR-34a-5p | MIMAT0000542 | Rabgap1 | 227800 | HITS-CLIP | 19536157\|23142080 | Unspecified |
| mmu-miR-34a-5p | MIMAT0000542 | Rabgap1l | 29809 | HITS-CLIP | 23142080 | Unspecified |
| mmu-miR-34a-5p | MIMAT0000542 | Rabgef1 | 56715 | HITS-CLIP | 23630078 | Unspecified |
| mmu-miR-34a-5p | MIMAT0000542 | Rabggtb | 19352 | HITS-CLIP | 23142080\|29167373 | Unspecified |
| mmu-miR-34a-5p | MIMAT0000542 | Rabif | 98710 | HITS-CLIP | 29167373\|23142080 | Unspecified |
| mmu-miR-34a-5p | MIMAT0000542 | Rabl2 | 68708 | HITS-CLIP | 23142080 | Unspecified |
| mmu-miR-34a-5p | MIMAT0000542 | Rabl3 | 67657 | Microarrays | 37986224\|23027903 | Unspecified |
| mmu-miR-34a-5p | MIMAT0000542 | Rabl6 | 227624 | HITS-CLIP | 29167373 | Unspecified |
| mmu-miR-34a-5p | MIMAT0000542 | Rac1 | 19353 | HITS-CLIP | 31433990\|23142080\|29167373 | Unspecified |
| mmu-miR-34a-5p | MIMAT0000542 | Rac2 | 19354 | HITS-CLIP | 37986224\|31433990\|23142080\|29167 | Unspecified |
| mmu-miR-34a-5p | MIMAT0000542 | Rack1 | 10399 | HITS-CLIP | 23142080\|29167373 | Unspecified |
| mmu-miR-34a-5p | MIMAT0000542 | Rad1 | 19355 | HITS-CLIP | 19536157\|23142080 | Unspecified |
| mmu-miR-34a-5p | MIMAT0000542 | Rad17 | 19356 | HITS-CLIP | 23142080 | Unspecified |
| mmu-miR-34a-5p | MIMAT0000542 | Rad18 | 58186 | HITS-CLIP | 23142080\|29167373 | Unspecified |
| mmu-miR-34a-5p | MIMAT0000542 | Rad21 | 19357 | HITS-CLIP | 23142080\|29167373 | Unspecified |
| mmu-miR-34a-5p | MIMAT0000542 | Rad23b | 19359 | HITS-CLIP | 23630078\|30971389\|29167373\|23142 | Unspecified |

| | | | | | | |
|---|---|---|---|---|---|---|
| mmu-miR-34a-5p | MIMAT0000542 | Rad50 | 19360 | HITS-CLIP | 23142080 | Unspecified |
| mmu-miR-34a-5p | MIMAT0000542 | Rad51 | 19361 | Microarrays | 37986224\|23027903 | Unspecified |
| mmu-miR-34a-5p | MIMAT0000542 | Rad51ap1 | 19362 | Microarrays//HITS-CLIP | 37986224\|23027903\|23142080 | Unspecified |
| mmu-miR-34a-5p | MIMAT0000542 | Rad51c | 114714 | HITS-CLIP | 23142080 | Unspecified |
| mmu-miR-34a-5p | MIMAT0000542 | Rad52 | 19365 | HITS-CLIP | 23142080 | Unspecified |
| mmu-miR-34a-5p | MIMAT0000542 | Rad54l | 19366 | Microarrays//HITS-CLIP | 37986224\|23027903\|31433990 | Unspecified |
| mmu-miR-34a-5p | MIMAT0000542 | Rad54l2 | 81000 | Microarrays//HITS-CLIP | 37986224\|23027903\|26203562\|23142 | Unspecified |
| mmu-miR-34a-5p | MIMAT0000542 | Rad9a | 19367 | HITS-CLIP | 23142080 | Unspecified |
| mmu-miR-34a-5p | MIMAT0000542 | Rad9b | 231724 | HITS-CLIP | 23142080 | Unspecified |
| mmu-miR-34a-5p | MIMAT0000542 | Rae1 | 66679 | HITS-CLIP | 23142080 | Unspecified |
| mmu-miR-34a-5p | MIMAT0000542 | Raf1 | 110157 | HITS-CLIP | 23142080\|26203562 | Unspecified |
| mmu-miR-34a-5p | MIMAT0000542 | Rai1 | 19377 | HITS-CLIP | 23142080 | Unspecified |
| mmu-miR-34a-5p | MIMAT0000542 | Rai14 | 75646 | HITS-CLIP | 31433990 | Unspecified |
| mmu-miR-34a-5p | MIMAT0000542 | Rala | 56044 | HITS-CLIP | 31433990\|29167373 | Unspecified |
| mmu-miR-34a-5p | MIMAT0000542 | Ralb | 64143 | HITS-CLIP | 23142080 | Unspecified |
| mmu-miR-34a-5p | MIMAT0000542 | Ralbp1 | 19765 | HITS-CLIP | 23142080 | Unspecified |
| mmu-miR-34a-5p | MIMAT0000542 | Ralgapa1 | 56784 | HITS-CLIP | 23142080\|19536157\|29167373 | Unspecified |
| mmu-miR-34a-5p | MIMAT0000542 | Ralgapa2 | 241694 | HITS-CLIP | 23142080\|29167373 | Unspecified |
| mmu-miR-34a-5p | MIMAT0000542 | Ralgapb | 228850 | HITS-CLIP | 19536157\|31433990\|23142080\|29167 | Unspecified |
| mmu-miR-34a-5p | MIMAT0000542 | Ralgds | 19730 | HITS-CLIP | 19536157\|23142080\|29167373 | Unspecified |
| mmu-miR-34a-5p | MIMAT0000542 | Ralgps1 | 241308 | HITS-CLIP | 19536157\|23142080\|29167373 | Unspecified |
| mmu-miR-34a-5p | MIMAT0000542 | Ralgps2 | 78255 | HITS-CLIP | 23142080\|19536157 | Unspecified |
| mmu-miR-34a-5p | MIMAT0000542 | Raly | 19383 | HITS-CLIP | 23142080 | Unspecified |
| mmu-miR-34a-5p | MIMAT0000542 | Ramp1 | 51801 | HITS-CLIP | 19536157 | Unspecified |
| mmu-miR-34a-5p | MIMAT0000542 | Ranbp1 | 19385 | HITS-CLIP | 23142080 | Unspecified |
| mmu-miR-34a-5p | MIMAT0000542 | Ranbp10 | 74334 | Microarrays//HITS-CLIP | 37986224\|23027903\|30971389\|29167 | Unspecified |
| mmu-miR-34a-5p | MIMAT0000542 | Ranbp2 | 19386 | HITS-CLIP | 23142080\|30971389\|23630078\|19536 | Unspecified |
| mmu-miR-34a-5p | MIMAT0000542 | Ranbp6 | 240614 | Microarrays//HITS-CLIP | 37986224\|23142080 | Unspecified |
| mmu-miR-34a-5p | MIMAT0000542 | Ranbp9 | 56705 | HITS-CLIP | 23142080\|19536157 | Unspecified |
| mmu-miR-34a-5p | MIMAT0000542 | Rangap1 | 19387 | HITS-CLIP | 23142080 | Unspecified |
| mmu-miR-34a-5p | MIMAT0000542 | Rangrf | 57785 | Microarrays | 37986224\|23027903 | Unspecified |
| mmu-miR-34a-5p | MIMAT0000542 | Rap1a | 109905 | HITS-CLIP | 23142080\|29167373 | Unspecified |
| mmu-miR-34a-5p | MIMAT0000542 | Rap1b | 215449 | HITS-CLIP | 30971389\|31433990\|23142080\|29167 | Unspecified |
| mmu-miR-34a-5p | MIMAT0000542 | Rap1gds1 | 229877 | HITS-CLIP | 19536157\|23142080\|26203562\|29167 | Unspecified |
| mmu-miR-34a-5p | MIMAT0000542 | Rap2a | 76108 | HITS-CLIP | 29167373\|23142080 | Unspecified |
| mmu-miR-34a-5p | MIMAT0000542 | Rap2b | 74012 | Microarrays//HITS-CLIP | 37986224\|26203562\|23142080\|29167 | Unspecified |
| mmu-miR-34a-5p | MIMAT0000542 | Rap2c | 72065 | Microarrays//HITS-CLIP | 37986224\|31433990\|26203562\|23142 | Unspecified |
| mmu-miR-34a-5p | MIMAT0000542 | Rapgef2 | 76089 | HITS-CLIP | 19536157\|26203562 | Unspecified |
| mmu-miR-34a-5p | MIMAT0000542 | Rapgef4 | 56508 | HITS-CLIP | 19536157\|23142080 | Unspecified |

| | | | | | | |
|---|---|---|---|---|---|---|
| mmu-miR-34a-5p | MIMAT0000542 | Rapgef5 | 217944 | HITS-CLIP | 29167373 | Unspecified |
| mmu-miR-34a-5p | MIMAT0000542 | Rapgef6 | 192786 | HITS-CLIP | 23142080\|29167373 | Unspecified |
| mmu-miR-34a-5p | MIMAT0000542 | Rapgefl1 | 268480 | HITS-CLIP | 23142080 | Unspecified |
| mmu-miR-34a-5p | MIMAT0000542 | Raph1 | 77300 | HITS-CLIP | 23142080\|29167373 | Unspecified |
| mmu-miR-34a-5p | MIMAT0000542 | Rapsn | 19400 | Microarrays | 37986224 | Unspecified |
| mmu-miR-34a-5p | MIMAT0000542 | Rara | 19401 | HITS-CLIP | 23142080\|29167373 | Unspecified |
| mmu-miR-34a-5p | MIMAT0000542 | Rarg | 19411 | Microarrays//HITS-CLIP | 37986224\|23027903\|23142080 | Unspecified |
| mmu-miR-34a-5p | MIMAT0000542 | Rars | 104458 | HITS-CLIP | 23142080\|29167373 | Unspecified |
| mmu-miR-34a-5p | MIMAT0000542 | Rasa1 | 218397 | HITS-CLIP | 37986224\|19536157\|23142080\|29167 | Unspecified |
| mmu-miR-34a-5p | MIMAT0000542 | Rasa2 | 114713 | HITS-CLIP | 23142080\|29167373 | Unspecified |
| mmu-miR-34a-5p | MIMAT0000542 | Rasa3 | 19414 | HITS-CLIP | 19536157\|26203562\|23142080\|29167 | Unspecified |
| mmu-miR-34a-5p | MIMAT0000542 | Rasa4 | 54153 | HITS-CLIP | 29167373 | Unspecified |
| mmu-miR-34a-5p | MIMAT0000542 | Rasal1 | 19415 | HITS-CLIP | 23142080 | Unspecified |
| mmu-miR-34a-5p | MIMAT0000542 | Rasal3 | 320484 | HITS-CLIP | 23142080 | Unspecified |
| mmu-miR-34a-5p | MIMAT0000542 | Rasgef1b | 320292 | HITS-CLIP | 29167373 | Unspecified |
| mmu-miR-34a-5p | MIMAT0000542 | Rasgrf2 | 19418 | HITS-CLIP | 19536157 | Unspecified |
| mmu-miR-34a-5p | MIMAT0000542 | Rasgrp1 | 19419 | HITS-CLIP | 31433990\|23142080 | Unspecified |
| mmu-miR-34a-5p | MIMAT0000542 | Rasgrp4 | 233046 | HITS-CLIP | 23142080\|29167373 | Unspecified |
| mmu-miR-34a-5p | MIMAT0000542 | Rasl11b | 68939 | HITS-CLIP | 23142080 | Unspecified |
| mmu-miR-34a-5p | MIMAT0000542 | Rassf2 | 215653 | HITS-CLIP | 23142080\|29167373 | Unspecified |
| mmu-miR-34a-5p | MIMAT0000542 | Rassf3 | 192678 | HITS-CLIP | 23142080\|29167373 | Unspecified |
| mmu-miR-34a-5p | MIMAT0000542 | Rassf4 | 213391 | HITS-CLIP | 29167373 | Unspecified |
| mmu-miR-34a-5p | MIMAT0000542 | Rassf5 | 54354 | HITS-CLIP | 31433990\|23142080\|29167373 | Unspecified |
| mmu-miR-34a-5p | MIMAT0000542 | Rassf7 | 66985 | HITS-CLIP | 23142080\|31433990 | Unspecified |
| mmu-miR-34a-5p | MIMAT0000542 | Rassf8 | 71323 | HITS-CLIP | 29167373 | Unspecified |
| mmu-miR-34a-5p | MIMAT0000542 | Raver1 | 71766 | HITS-CLIP | 23142080\|29167373 | Unspecified |
| mmu-miR-34a-5p | MIMAT0000542 | Raver2 | 242570 | HITS-CLIP | 19536157 | Unspecified |
| mmu-miR-34a-5p | MIMAT0000542 | Rb1 | 19645 | HITS-CLIP | 23142080\|31433990\|29167373 | Unspecified |
| mmu-miR-34a-5p | MIMAT0000542 | Rb1cc1 | 12421 | HITS-CLIP | 23142080 | Unspecified |
| mmu-miR-34a-5p | MIMAT0000542 | Rbak | 57782 | HITS-CLIP | 23142080 | Unspecified |
| mmu-miR-34a-5p | MIMAT0000542 | Rbbp4 | 19646 | HITS-CLIP | 19536157\|23142080 | Unspecified |
| mmu-miR-34a-5p | MIMAT0000542 | Rbbp5 | 213464 | HITS-CLIP | 23142080\|29167373 | Unspecified |
| mmu-miR-34a-5p | MIMAT0000542 | Rbbp6 | 19647 | HITS-CLIP | 19536157\|31433990\|23142080\|29167 | Unspecified |
| mmu-miR-34a-5p | MIMAT0000542 | Rbbp7 | 245688 | HITS-CLIP | 23142080 | Unspecified |
| mmu-miR-34a-5p | MIMAT0000542 | Rbbp8 | 225182 | HITS-CLIP | 23142080\|29167373 | Unspecified |
| mmu-miR-34a-5p | MIMAT0000542 | Rbbp9 | 26450 | HITS-CLIP | 23142080 | Unspecified |
| mmu-miR-34a-5p | MIMAT0000542 | Rbck1 | 24105 | HITS-CLIP | 29167373 | Unspecified |
| mmu-miR-34a-5p | MIMAT0000542 | Rbfox1 | 268859 | Microarrays//HITS-CLIP | 37986224\|19536157 | Unspecified |
| mmu-miR-34a-5p | MIMAT0000542 | Rbl1 | 19650 | HITS-CLIP | 23142080\|29167373 | Unspecified |

| | | | | | | |
|---|---|---|---|---|---|---|
| mmu-miR-34a-5p | MIMAT0000542 | Rbl2 | 19651 | HITS-CLIP | 31433990 | Unspecified |
| mmu-miR-34a-5p | MIMAT0000542 | Rbm10 | 236732 | HITS-CLIP | 23142080 | Unspecified |
| mmu-miR-34a-5p | MIMAT0000542 | Rbm12 | 75710 | HITS-CLIP | 23142080\|29167373 | Unspecified |
| mmu-miR-34a-5p | MIMAT0000542 | Rbm12b2 | 77604 | HITS-CLIP | 29167373 | Unspecified |
| mmu-miR-34a-5p | MIMAT0000542 | Rbm14 | 56275 | HITS-CLIP | 23142080\|29167373 | Unspecified |
| mmu-miR-34a-5p | MIMAT0000542 | Rbm15 | 229700 | HITS-CLIP | 29167373\|23142080 | Unspecified |
| mmu-miR-34a-5p | MIMAT0000542 | Rbm15b | 109095 | HITS-CLIP | 23142080\|29167373 | Unspecified |
| mmu-miR-34a-5p | MIMAT0000542 | Rbm17 | 76938 | HITS-CLIP | 23142080 | Unspecified |
| mmu-miR-34a-5p | MIMAT0000542 | Rbm18 | 67889 | HITS-CLIP | 23142080 | Unspecified |
| mmu-miR-34a-5p | MIMAT0000542 | Rbm25 | 67039 | HITS-CLIP | 23142080\|29167373 | Unspecified |
| mmu-miR-34a-5p | MIMAT0000542 | Rbm26 | 74213 | HITS-CLIP | 23142080\|29167373 | Unspecified |
| mmu-miR-34a-5p | MIMAT0000542 | Rbm27 | 225432 | HITS-CLIP | 23142080\|29167373 | Unspecified |
| mmu-miR-34a-5p | MIMAT0000542 | Rbm28 | 68272 | HITS-CLIP | 23142080 | Unspecified |
| mmu-miR-34a-5p | MIMAT0000542 | Rbm3 | 19652 | HITS-CLIP | 23142080 | Unspecified |
| mmu-miR-34a-5p | MIMAT0000542 | Rbm33 | 381626 | HITS-CLIP | 19536157\|23142080\|29167373 | Unspecified |
| mmu-miR-34a-5p | MIMAT0000542 | Rbm38 | 56190 | HITS-CLIP | 23142080\|29167373 | Unspecified |
| mmu-miR-34a-5p | MIMAT0000542 | Rbm39 | 170791 | HITS-CLIP | 23142080\|29167373 | Unspecified |
| mmu-miR-34a-5p | MIMAT0000542 | Rbm4 | 19653 | HITS-CLIP | 29167373 | Unspecified |
| mmu-miR-34a-5p | MIMAT0000542 | Rbm41 | 237073 | HITS-CLIP | 23142080 | Unspecified |
| mmu-miR-34a-5p | MIMAT0000542 | Rbm43 | 71684 | HITS-CLIP | 23142080 | Unspecified |
| mmu-miR-34a-5p | MIMAT0000542 | Rbm45 | 241490 | HITS-CLIP | 23142080 | Unspecified |
| mmu-miR-34a-5p | MIMAT0000542 | Rbm47 | 245945 | HITS-CLIP | 23142080\|29167373 | Unspecified |
| mmu-miR-34a-5p | MIMAT0000542 | Rbm48 | 269623 | HITS-CLIP | 23142080 | Unspecified |
| mmu-miR-34a-5p | MIMAT0000542 | Rbm5 | 83486 | HITS-CLIP | 23142080\|31433990\|29167373 | Unspecified |
| mmu-miR-34a-5p | MIMAT0000542 | Rbm6 | 19654 | HITS-CLIP | 29167373\|23142080 | Unspecified |
| mmu-miR-34a-5p | MIMAT0000542 | Rbm7 | 67010 | HITS-CLIP | 23142080\|29167373\|19536157 | Unspecified |
| mmu-miR-34a-5p | MIMAT0000542 | Rbm8a | 60365 | HITS-CLIP | 23142080 | Unspecified |
| mmu-miR-34a-5p | MIMAT0000542 | Rbms1 | 56878 | HITS-CLIP | 31433990\|26203562\|23142080\|29167 | Unspecified |
| mmu-miR-34a-5p | MIMAT0000542 | Rbms2 | 56516 | Microarrays//HITS-CLIP | 37986224\|23027903\|23142080 | Unspecified |
| mmu-miR-34a-5p | MIMAT0000542 | Rbmx2 | 209003 | HITS-CLIP | 29167373 | Unspecified |
| mmu-miR-34a-5p | MIMAT0000542 | Rbpj | 19664 | HITS-CLIP | 23142080\|29167373 | Unspecified |
| mmu-miR-34a-5p | MIMAT0000542 | Rbsn | 64145 | HITS-CLIP | 23142080\|29167373 | Unspecified |
| mmu-miR-34a-5p | MIMAT0000542 | Rbx1 | 56438 | HITS-CLIP | 23142080 | Unspecified |
| mmu-miR-34a-5p | MIMAT0000542 | Rc3h1 | 381305 | HITS-CLIP | 23142080\|23630078\|29167373 | Unspecified |
| mmu-miR-34a-5p | MIMAT0000542 | Rc3h2 | 319817 | HITS-CLIP | 31433990\|23142080\|29167373 | Unspecified |
| mmu-miR-34a-5p | MIMAT0000542 | Rcan2 | 53901 | Microarrays | 37986224 | Unspecified |
| mmu-miR-34a-5p | MIMAT0000542 | Rcan3 | 53902 | HITS-CLIP | 29167373\|23142080 | Unspecified |
| mmu-miR-34a-5p | MIMAT0000542 | Rcbtb1 | 71330 | HITS-CLIP | 23142080 | Unspecified |
| mmu-miR-34a-5p | MIMAT0000542 | Rcbtb2 | 105670 | HITS-CLIP | 29167373 | Unspecified |

| | | | | | | |
|---|---|---|---|---|---|---|
| mmu-miR-34a-5p | MIMAT0000542 | Rcc1 | 100088 | HITS-CLIP | 26203562 | Unspecified |
| mmu-miR-34a-5p | MIMAT0000542 | Rcc2 | 108911 | HITS-CLIP | 31433990\|23142080 | Unspecified |
| mmu-miR-34a-5p | MIMAT0000542 | Rce1 | 19671 | HITS-CLIP | 23142080 | Unspecified |
| mmu-miR-34a-5p | MIMAT0000542 | Rchy1 | 68098 | HITS-CLIP | 23142080 | Unspecified |
| mmu-miR-34a-5p | MIMAT0000542 | Rcl1 | 59028 | HITS-CLIP | 23142080 | Unspecified |
| mmu-miR-34a-5p | MIMAT0000542 | Rcn1 | 19672 | HITS-CLIP | 23142080 | Unspecified |
| mmu-miR-34a-5p | MIMAT0000542 | Rcn2 | 26611 | HITS-CLIP | 23142080 | Unspecified |
| mmu-miR-34a-5p | MIMAT0000542 | Rcor1 | 217864 | Microarrays//HITS-CLIP | 37986224\|31433990\|23142080\|29167 | Unspecified |
| mmu-miR-34a-5p | MIMAT0000542 | Rcor3 | 214742 | HITS-CLIP | 19536157 | Unspecified |
| mmu-miR-34a-5p | MIMAT0000542 | Rcsd1 | 226594 | HITS-CLIP | 23142080 | Unspecified |
| mmu-miR-34a-5p | MIMAT0000542 | Rdh11 | 17252 | Microarrays//HITS-CLIP | 37986224\|23142080\|23027903 | Unspecified |
| mmu-miR-34a-5p | MIMAT0000542 | Rdh12 | 77974 | HITS-CLIP | 26203562 | Unspecified |
| mmu-miR-34a-5p | MIMAT0000542 | Rdh14 | 105014 | HITS-CLIP | 23142080 | Unspecified |
| mmu-miR-34a-5p | MIMAT0000542 | Rdh5 | 19682 | HITS-CLIP | 23142080 | Unspecified |
| mmu-miR-34a-5p | MIMAT0000542 | Rdm1 | 66599 | HITS-CLIP | 23142080 | Unspecified |
| mmu-miR-34a-5p | MIMAT0000542 | Rdx | 19684 | HITS-CLIP | 23142080 | Unspecified |
| mmu-miR-34a-5p | MIMAT0000542 | Reck | 53614 | HITS-CLIP | 31433990\|23142080 | Unspecified |
| mmu-miR-34a-5p | MIMAT0000542 | Reep1 | 52250 | HITS-CLIP | 19536157 | Unspecified |
| mmu-miR-34a-5p | MIMAT0000542 | Reep2 | 225362 | HITS-CLIP | 19536157 | Unspecified |
| mmu-miR-34a-5p | MIMAT0000542 | Reep3 | 28193 | HITS-CLIP | 23142080\|29167373 | Unspecified |
| mmu-miR-34a-5p | MIMAT0000542 | Reep4 | 72549 | HITS-CLIP | 23142080 | Unspecified |
| mmu-miR-34a-5p | MIMAT0000542 | Reep5 | 13476 | HITS-CLIP | 23142080\|19536157\|29167373 | Unspecified |
| mmu-miR-34a-5p | MIMAT0000542 | Rel | 19696 | HITS-CLIP | 23142080 | Unspecified |
| mmu-miR-34a-5p | MIMAT0000542 | Rela | 19697 | Immunofluorescence, Western Blot//Immu | 37986224\|22483937\|23142080\|29167 | Unspecified |
| mmu-miR-34a-5p | MIMAT0000542 | Relb | 19698 | HITS-CLIP | 23142080 | Unspecified |
| mmu-miR-34a-5p | MIMAT0000542 | Relch | 57614 | HITS-CLIP | 23142080\|29167373 | Unspecified |
| mmu-miR-34a-5p | MIMAT0000542 | Rell1 | 100532 | HITS-CLIP | 23142080 | Unspecified |
| mmu-miR-34a-5p | MIMAT0000542 | Rem2 | 140743 | Microarrays | 37986224\|23027903 | Unspecified |
| mmu-miR-34a-5p | MIMAT0000542 | Renbp | 19703 | Microarrays//HITS-CLIP | 37986224\|29167373 | Unspecified |
| mmu-miR-34a-5p | MIMAT0000542 | Repin1 | 58887 | HITS-CLIP | 29167373 | Unspecified |
| mmu-miR-34a-5p | MIMAT0000542 | Reps1 | 19707 | HITS-CLIP | 31433990\|23142080 | Unspecified |
| mmu-miR-34a-5p | MIMAT0000542 | Rere | 68703 | HITS-CLIP | 23142080\|29167373 | Unspecified |
| mmu-miR-34a-5p | MIMAT0000542 | Resf1 | 55196 | HITS-CLIP | 23142080\|29167373 | Unspecified |
| mmu-miR-34a-5p | MIMAT0000542 | Rest | 19712 | HITS-CLIP | 23630078\|30971389\|31433990\|23142 | Unspecified |
| mmu-miR-34a-5p | MIMAT0000542 | Retreg1 | 66270 | HITS-CLIP | 23142080\|19536157\|29167373 | Unspecified |
| mmu-miR-34a-5p | MIMAT0000542 | Retreg2 | 79137 | HITS-CLIP | 23142080 | Unspecified |
| mmu-miR-34a-5p | MIMAT0000542 | Retreg3 | 162427 | HITS-CLIP | 23142080 | Unspecified |
| mmu-miR-34a-5p | MIMAT0000542 | Rev1 | 56210 | HITS-CLIP | 23142080\|19536157\|29167373 | Unspecified |
| mmu-miR-34a-5p | MIMAT0000542 | Rev3l | 19714 | HITS-CLIP | 23142080\|19536157\|30971389 | Unspecified |

| | | | | | | |
|---|---|---|---|---|---|---|
| mmu-miR-34a-5p | MIMAT0000542 | Rex1bd | 55049 | HITS-CLIP | 23142080 | Unspecified |
| mmu-miR-34a-5p | MIMAT0000542 | Rexo1 | 66932 | HITS-CLIP | 23142080 | Unspecified |
| mmu-miR-34a-5p | MIMAT0000542 | Rexo2 | 104444 | HITS-CLIP | 23142080 | Unspecified |
| mmu-miR-34a-5p | MIMAT0000542 | Rfc1 | 19687 | Microarrays//HITS-CLIP | 37986224\|23142080\|23027903\|2363( | Unspecified |
| mmu-miR-34a-5p | MIMAT0000542 | Rfc2 | 19718 | HITS-CLIP | 23142080 | Unspecified |
| mmu-miR-34a-5p | MIMAT0000542 | Rfc3 | 69263 | HITS-CLIP | 23142080 | Unspecified |
| mmu-miR-34a-5p | MIMAT0000542 | Rfc4 | 106344 | Microarrays//HITS-CLIP | 37986224\|23027903\|23142080 | Unspecified |
| mmu-miR-34a-5p | MIMAT0000542 | Rfc5 | 72151 | HITS-CLIP | 23142080 | Unspecified |
| mmu-miR-34a-5p | MIMAT0000542 | Rfk | 54391 | HITS-CLIP | 19536157 | Unspecified |
| mmu-miR-34a-5p | MIMAT0000542 | Rflnb | 76566 | HITS-CLIP | 26203562 | Unspecified |
| mmu-miR-34a-5p | MIMAT0000542 | Rftn1 | 76438 | Microarrays//HITS-CLIP | 37986224\|31433990\|23142080 | Unspecified |
| mmu-miR-34a-5p | MIMAT0000542 | Rfwd3 | 234736 | HITS-CLIP | 23142080 | Unspecified |
| mmu-miR-34a-5p | MIMAT0000542 | Rfx1 | 19724 | HITS-CLIP | 31433990\|23142080\|19536157 | Unspecified |
| mmu-miR-34a-5p | MIMAT0000542 | Rfx3 | 19726 | HITS-CLIP | 23142080\|19536157 | Unspecified |
| mmu-miR-34a-5p | MIMAT0000542 | Rfx5 | 53970 | HITS-CLIP | 23630078\|23142080 | Unspecified |
| mmu-miR-34a-5p | MIMAT0000542 | Rfx7 | 319758 | HITS-CLIP | 19536157\|31433990\|30971389\|2314; | Unspecified |
| mmu-miR-34a-5p | MIMAT0000542 | Rfxap | 170767 | HITS-CLIP | 23630078\|23142080 | Unspecified |
| mmu-miR-34a-5p | MIMAT0000542 | Rgcc | 66214 | HITS-CLIP | 23142080 | Unspecified |
| mmu-miR-34a-5p | MIMAT0000542 | Rgl2 | 19732 | HITS-CLIP | 31433990\|19536157\|23142080 | Unspecified |
| mmu-miR-34a-5p | MIMAT0000542 | Rgp1 | 242406 | HITS-CLIP | 23142080\|29167373 | Unspecified |
| mmu-miR-34a-5p | MIMAT0000542 | Rgs1 | 50778 | HITS-CLIP | 23142080\|29167373 | Unspecified |
| mmu-miR-34a-5p | MIMAT0000542 | Rgs10 | 67865 | HITS-CLIP | 19536157\|23142080 | Unspecified |
| mmu-miR-34a-5p | MIMAT0000542 | Rgs12 | 71729 | HITS-CLIP | 29167373 | Unspecified |
| mmu-miR-34a-5p | MIMAT0000542 | Rgs14 | 51791 | HITS-CLIP | 29167373 | Unspecified |
| mmu-miR-34a-5p | MIMAT0000542 | Rgs16 | 19734 | HITS-CLIP | 23142080 | Unspecified |
| mmu-miR-34a-5p | MIMAT0000542 | Rgs18 | 64214 | HITS-CLIP | 29167373 | Unspecified |
| mmu-miR-34a-5p | MIMAT0000542 | Rgs2 | 19735 | HITS-CLIP | 19536157\|23142080\|29167373 | Unspecified |
| mmu-miR-34a-5p | MIMAT0000542 | Rgs3 | 50780 | HITS-CLIP | 23142080 | Unspecified |
| mmu-miR-34a-5p | MIMAT0000542 | Rgs4 | 19736 | HITS-CLIP | 19536157 | Unspecified |
| mmu-miR-34a-5p | MIMAT0000542 | Rgs5 | 19737 | HITS-CLIP | 19536157 | Unspecified |
| mmu-miR-34a-5p | MIMAT0000542 | Rgs7bp | 52882 | HITS-CLIP | 19536157 | Unspecified |
| mmu-miR-34a-5p | MIMAT0000542 | Rgs8 | 67792 | HITS-CLIP | 19536157 | Unspecified |
| mmu-miR-34a-5p | MIMAT0000542 | Rhbdd1 | 76867 | HITS-CLIP | 23142080 | Unspecified |
| mmu-miR-34a-5p | MIMAT0000542 | Rhbdd2 | 215160 | HITS-CLIP | 29167373 | Unspecified |
| mmu-miR-34a-5p | MIMAT0000542 | Rhbdf2 | 217344 | HITS-CLIP | 23142080\|29167373 | Unspecified |
| mmu-miR-34a-5p | MIMAT0000542 | Rhd | 19746 | HITS-CLIP | 30971389 | Unspecified |
| mmu-miR-34a-5p | MIMAT0000542 | Rheb | 19744 | HITS-CLIP | 21258322\|23142080\|29167373 | Unspecified |
| mmu-miR-34a-5p | MIMAT0000542 | Rhebl1 | 69159 | HITS-CLIP | 23142080 | Unspecified |
| mmu-miR-34a-5p | MIMAT0000542 | Rhno1 | 72440 | HITS-CLIP | 23142080 | Unspecified |

| | | | | | | |
|---|---|---|---|---|---|---|
| mmu-miR-34a-5p | MIMAT0000542 | Rhob | 11852 | HITS-CLIP | 29167373 | Unspecified |
| mmu-miR-34a-5p | MIMAT0000542 | Rhobtb1 | 69288 | Microarrays//HITS-CLIP | 37986224\|23142080 | Unspecified |
| mmu-miR-34a-5p | MIMAT0000542 | Rhobtb2 | 246710 | HITS-CLIP | 23142080 | Unspecified |
| mmu-miR-34a-5p | MIMAT0000542 | Rhof | 23912 | HITS-CLIP | 23142080\|31433990 | Unspecified |
| mmu-miR-34a-5p | MIMAT0000542 | Rhog | 56212 | HITS-CLIP | 19536157\|23142080\|29167373 | Unspecified |
| mmu-miR-34a-5p | MIMAT0000542 | Rhoh | 74734 | HITS-CLIP | 23142080 | Unspecified |
| mmu-miR-34a-5p | MIMAT0000542 | Rhoq | 104215 | HITS-CLIP | 23142080\|29167373 | Unspecified |
| mmu-miR-34a-5p | MIMAT0000542 | Rhot1 | 59040 | HITS-CLIP | 23142080 | Unspecified |
| mmu-miR-34a-5p | MIMAT0000542 | Rhot2 | 214952 | HITS-CLIP | 23142080 | Unspecified |
| mmu-miR-34a-5p | MIMAT0000542 | Ric1 | 850728 | HITS-CLIP | 23142080 | Unspecified |
| mmu-miR-34a-5p | MIMAT0000542 | Ric8a | 60626 | HITS-CLIP | 23142080\|29167373 | Unspecified |
| mmu-miR-34a-5p | MIMAT0000542 | Ric8b | 237422 | HITS-CLIP | 29167373\|23142080 | Unspecified |
| mmu-miR-34a-5p | MIMAT0000542 | Rictor | 78757 | HITS-CLIP | 30971389\|31433990\|23142080\|29167 | Unspecified |
| mmu-miR-34a-5p | MIMAT0000542 | Rida | 65151 | HITS-CLIP | 23142080 | Unspecified |
| mmu-miR-34a-5p | MIMAT0000542 | Rif1 | 51869 | HITS-CLIP | 31433990\|23142080\|29167373 | Unspecified |
| mmu-miR-34a-5p | MIMAT0000542 | Rigi | 23586 | Microarrays | 23027903 | Unspecified |
| mmu-miR-34a-5p | MIMAT0000542 | Rilpl2 | 80291 | HITS-CLIP | 23142080\|29167373 | Unspecified |
| mmu-miR-34a-5p | MIMAT0000542 | Rimoc1 | 285636 | Microarrays | 23027903 | Unspecified |
| mmu-miR-34a-5p | MIMAT0000542 | Rims1 | 116837 | HITS-CLIP | 19536157 | Unspecified |
| mmu-miR-34a-5p | MIMAT0000542 | Rin2 | 74030 | HITS-CLIP | 29167373 | Unspecified |
| mmu-miR-34a-5p | MIMAT0000542 | Rin3 | 217835 | HITS-CLIP | 23142080\|29167373 | Unspecified |
| mmu-miR-34a-5p | MIMAT0000542 | Rinl | 320435 | HITS-CLIP | 23142080\|31433990 | Unspecified |
| mmu-miR-34a-5p | MIMAT0000542 | Rint1 | 72772 | HITS-CLIP | 23142080\|29167373 | Unspecified |
| mmu-miR-34a-5p | MIMAT0000542 | Riok1 | 71340 | HITS-CLIP | 23142080 | Unspecified |
| mmu-miR-34a-5p | MIMAT0000542 | Riok2 | 67045 | HITS-CLIP | 23142080 | Unspecified |
| mmu-miR-34a-5p | MIMAT0000542 | Riok3 | 66878 | HITS-CLIP | 23142080\|29167373 | Unspecified |
| mmu-miR-34a-5p | MIMAT0000542 | Riox1 | 79697 | HITS-CLIP | 31433990\|23142080 | Unspecified |
| mmu-miR-34a-5p | MIMAT0000542 | Ripk1 | 19766 | HITS-CLIP | 23142080\|30971389\|29167373 | Unspecified |
| mmu-miR-34a-5p | MIMAT0000542 | Ripk2 | 192656 | HITS-CLIP | 19536157\|29167373\|23142080 | Unspecified |
| mmu-miR-34a-5p | MIMAT0000542 | Ripk3 | 56532 | HITS-CLIP | 23142080 | Unspecified |
| mmu-miR-34a-5p | MIMAT0000542 | Ripor1 | 75687 | HITS-CLIP | 23142080 | Unspecified |
| mmu-miR-34a-5p | MIMAT0000542 | Ripor2 | 193385 | HITS-CLIP | 23142080 | Unspecified |
| mmu-miR-34a-5p | MIMAT0000542 | Rit1 | 19769 | HITS-CLIP | 23142080 | Unspecified |
| mmu-miR-34a-5p | MIMAT0000542 | Rlf | 109263 | HITS-CLIP | 23142080\|31433990\|29167373 | Unspecified |
| mmu-miR-34a-5p | MIMAT0000542 | Rlim | 19820 | HITS-CLIP | 31433990\|23142080\|29167373 | Unspecified |
| mmu-miR-34a-5p | MIMAT0000542 | Rmc1 | 19775 | HITS-CLIP | 23142080\|29167373 | Unspecified |
| mmu-miR-34a-5p | MIMAT0000542 | Rmdn1 | 66302 | HITS-CLIP | 29167373\|23142080 | Unspecified |
| mmu-miR-34a-5p | MIMAT0000542 | Rmdn2 | 381110 | HITS-CLIP | 19536157 | Unspecified |
| mmu-miR-34a-5p | MIMAT0000542 | Rmdn3 | 67809 | HITS-CLIP | 23142080 | Unspecified |

| | | | | | | |
|---|---|---|---|---|---|---|
| mmu-miR-34a-5p | MIMAT0000542 | Rmnd5a | 68477 | HITS-CLIP | 23142080\|31433990\|29167373 | Unspecified |
| mmu-miR-34a-5p | MIMAT0000542 | Rmnd5b | 66089 | HITS-CLIP | 23142080 | Unspecified |
| mmu-miR-34a-5p | MIMAT0000542 | Rnaseh2b | 67153 | HITS-CLIP | 23142080 | Unspecified |
| mmu-miR-34a-5p | MIMAT0000542 | Rnasel | 24014 | HITS-CLIP | 23142080 | Unspecified |
| mmu-miR-34a-5p | MIMAT0000542 | Rnd3 | 74194 | HITS-CLIP | 31433990\|29167373 | Unspecified |
| mmu-miR-34a-5p | MIMAT0000542 | Rnf10 | 50849 | HITS-CLIP | 23142080\|29167373 | Unspecified |
| mmu-miR-34a-5p | MIMAT0000542 | Rnf103 | 22644 | HITS-CLIP | 23142080\|29167373 | Unspecified |
| mmu-miR-34a-5p | MIMAT0000542 | Rnf11 | 29864 | HITS-CLIP | 23142080 | Unspecified |
| mmu-miR-34a-5p | MIMAT0000542 | Rnf111 | 93836 | HITS-CLIP | 23142080\|29167373 | Unspecified |
| mmu-miR-34a-5p | MIMAT0000542 | Rnf113a1 | 69942 | HITS-CLIP | 23142080 | Unspecified |
| mmu-miR-34a-5p | MIMAT0000542 | Rnf113a2 | 66381 | HITS-CLIP | 23142080 | Unspecified |
| mmu-miR-34a-5p | MIMAT0000542 | Rnf114 | 81018 | HITS-CLIP | 23142080 | Unspecified |
| mmu-miR-34a-5p | MIMAT0000542 | Rnf115 | 67845 | HITS-CLIP | 29167373 | Unspecified |
| mmu-miR-34a-5p | MIMAT0000542 | Rnf126 | 70294 | HITS-CLIP | 29167373 | Unspecified |
| mmu-miR-34a-5p | MIMAT0000542 | Rnf128 | 66889 | HITS-CLIP | 29167373 | Unspecified |
| mmu-miR-34a-5p | MIMAT0000542 | Rnf13 | 24017 | HITS-CLIP | 29167373\|23142080 | Unspecified |
| mmu-miR-34a-5p | MIMAT0000542 | Rnf130 | 59044 | HITS-CLIP | 23142080 | Unspecified |
| mmu-miR-34a-5p | MIMAT0000542 | Rnf138 | 56515 | HITS-CLIP | 23142080 | Unspecified |
| mmu-miR-34a-5p | MIMAT0000542 | Rnf139 | 75841 | HITS-CLIP | 19536157\|23142080\|29167373 | Unspecified |
| mmu-miR-34a-5p | MIMAT0000542 | Rnf14 | 56736 | HITS-CLIP | 23142080\|29167373 | Unspecified |
| mmu-miR-34a-5p | MIMAT0000542 | Rnf141 | 67150 | HITS-CLIP | 23142080\|29167373 | Unspecified |
| mmu-miR-34a-5p | MIMAT0000542 | Rnf144a | 108089 | HITS-CLIP | 19536157 | Unspecified |
| mmu-miR-34a-5p | MIMAT0000542 | Rnf145 | 74315 | HITS-CLIP | 31433990\|23142080\|29167373 | Unspecified |
| mmu-miR-34a-5p | MIMAT0000542 | Rnf146 | 68031 | HITS-CLIP | 23142080 | Unspecified |
| mmu-miR-34a-5p | MIMAT0000542 | Rnf148 | 71300 | HITS-CLIP | 19536157 | Unspecified |
| mmu-miR-34a-5p | MIMAT0000542 | Rnf149 | 67702 | HITS-CLIP | 19536157\|29167373\|23142080 | Unspecified |
| mmu-miR-34a-5p | MIMAT0000542 | Rnf152 | 320311 | HITS-CLIP | 31433990\|23142080 | Unspecified |
| mmu-miR-34a-5p | MIMAT0000542 | Rnf157 | 217340 | HITS-CLIP | 31433990 | Unspecified |
| mmu-miR-34a-5p | MIMAT0000542 | Rnf165 | 225743 | HITS-CLIP | 19536157 | Unspecified |
| mmu-miR-34a-5p | MIMAT0000542 | Rnf166 | 68718 | HITS-CLIP | 23142080 | Unspecified |
| mmu-miR-34a-5p | MIMAT0000542 | Rnf167 | 70510 | HITS-CLIP | 31433990\|26203562\|23142080 | Unspecified |
| mmu-miR-34a-5p | MIMAT0000542 | Rnf168 | 70238 | HITS-CLIP | 23142080\|31433990\|29167373 | Unspecified |
| mmu-miR-34a-5p | MIMAT0000542 | Rnf169 | 108937 | HITS-CLIP | 23142080\|29167373 | Unspecified |
| mmu-miR-34a-5p | MIMAT0000542 | Rnf181 | 66510 | HITS-CLIP | 31433990\|23142080 | Unspecified |
| mmu-miR-34a-5p | MIMAT0000542 | Rnf185 | 193670 | HITS-CLIP | 23142080 | Unspecified |
| mmu-miR-34a-5p | MIMAT0000542 | Rnf19a | 30945 | HITS-CLIP | 37986224\|23142080\|23630078\|30971 | Unspecified |
| mmu-miR-34a-5p | MIMAT0000542 | Rnf19b | 75234 | HITS-CLIP | 23142080\|31433990\|29167373 | Unspecified |
| mmu-miR-34a-5p | MIMAT0000542 | Rnf2 | 19821 | HITS-CLIP | 23142080 | Unspecified |
| mmu-miR-34a-5p | MIMAT0000542 | Rnf20 | 109331 | HITS-CLIP | 23142080 | Unspecified |

| | | | | | | |
|---|---|---|---|---|---|---|
| mmu-miR-34a-5p | MIMAT0000542 | Rnf213 | 672511 | HITS-CLIP | 23142080\|29167373 | Unspecified |
| mmu-miR-34a-5p | MIMAT0000542 | Rnf214 | 235315 | HITS-CLIP | 23142080 | Unspecified |
| mmu-miR-34a-5p | MIMAT0000542 | Rnf216 | 108086 | HITS-CLIP | 29167373\|23142080 | Unspecified |
| mmu-miR-34a-5p | MIMAT0000542 | Rnf219 | 72486 | HITS-CLIP | 23142080\|29167373 | Unspecified |
| mmu-miR-34a-5p | MIMAT0000542 | Rnf220 | 66743 | HITS-CLIP | 23142080 | Unspecified |
| mmu-miR-34a-5p | MIMAT0000542 | Rnf24 | 51902 | HITS-CLIP | 23630078 | Unspecified |
| mmu-miR-34a-5p | MIMAT0000542 | Rnf25 | 57751 | HITS-CLIP | 23142080 | Unspecified |
| mmu-miR-34a-5p | MIMAT0000542 | Rnf31 | 268749 | HITS-CLIP | 23142080\|29167373 | Unspecified |
| mmu-miR-34a-5p | MIMAT0000542 | Rnf34 | 80751 | HITS-CLIP | 29167373\|23142080 | Unspecified |
| mmu-miR-34a-5p | MIMAT0000542 | Rnf38 | 73469 | HITS-CLIP | 23142080\|29167373 | Unspecified |
| mmu-miR-34a-5p | MIMAT0000542 | Rnf4 | 19822 | HITS-CLIP | 23142080\|29167373 | Unspecified |
| mmu-miR-34a-5p | MIMAT0000542 | Rnf40 | 233900 | HITS-CLIP | 23142080 | Unspecified |
| mmu-miR-34a-5p | MIMAT0000542 | Rnf41 | 67588 | HITS-CLIP | 29167373 | Unspecified |
| mmu-miR-34a-5p | MIMAT0000542 | Rnf43 | 207742 | HITS-CLIP | 23142080\|30971389 | Unspecified |
| mmu-miR-34a-5p | MIMAT0000542 | Rnf44 | 105239 | HITS-CLIP | 31433990\|23142080\|29167373 | Unspecified |
| mmu-miR-34a-5p | MIMAT0000542 | Rnf5 | 54197 | HITS-CLIP | 23142080 | Unspecified |
| mmu-miR-34a-5p | MIMAT0000542 | Rnf8 | 58230 | HITS-CLIP | 23142080\|31433990 | Unspecified |
| mmu-miR-34a-5p | MIMAT0000542 | Rnft1 | 76892 | HITS-CLIP | 23142080 | Unspecified |
| mmu-miR-34a-5p | MIMAT0000542 | Rnft2 | 269695 | HITS-CLIP | 19536157 | Unspecified |
| mmu-miR-34a-5p | MIMAT0000542 | Rngtt | 24018 | HITS-CLIP | 23142080 | Unspecified |
| mmu-miR-34a-5p | MIMAT0000542 | Rnh1 | 107702 | HITS-CLIP | 23142080\|29167373 | Unspecified |
| mmu-miR-34a-5p | MIMAT0000542 | Rnmt | 67897 | HITS-CLIP | 23142080 | Unspecified |
| mmu-miR-34a-5p | MIMAT0000542 | Rnpep | 215615 | HITS-CLIP | 23142080 | Unspecified |
| mmu-miR-34a-5p | MIMAT0000542 | Rnpepl1 | 108657 | HITS-CLIP | 19536157\|29167373\|23142080 | Unspecified |
| mmu-miR-34a-5p | MIMAT0000542 | Rock1 | 19877 | HITS-CLIP | 23142080 | Unspecified |
| mmu-miR-34a-5p | MIMAT0000542 | Rock2 | 19878 | HITS-CLIP | 23142080\|29167373 | Unspecified |
| mmu-miR-34a-5p | MIMAT0000542 | Ropn1l | 252967 | Microarrays | 37986224 | Unspecified |
| mmu-miR-34a-5p | MIMAT0000542 | Rora | 19883 | Microarrays//HITS-CLIP | 37986224\|23142080 | Unspecified |
| mmu-miR-34a-5p | MIMAT0000542 | Rp2 | 19889 | HITS-CLIP | 23142080 | Unspecified |
| mmu-miR-34a-5p | MIMAT0000542 | Rpa1 | 68275 | HITS-CLIP | 23142080\|29167373 | Unspecified |
| mmu-miR-34a-5p | MIMAT0000542 | Rpa2 | 19891 | Microarrays//HITS-CLIP | 37986224\|23027903\|23142080 | Unspecified |
| mmu-miR-34a-5p | MIMAT0000542 | Rpa3 | 68240 | HITS-CLIP | 23142080 | Unspecified |
| mmu-miR-34a-5p | MIMAT0000542 | Rpain | 69723 | HITS-CLIP | 23142080 | Unspecified |
| mmu-miR-34a-5p | MIMAT0000542 | Rpap1 | 68925 | HITS-CLIP | 23142080 | Unspecified |
| mmu-miR-34a-5p | MIMAT0000542 | Rpap2 | 231571 | HITS-CLIP | 23142080 | Unspecified |
| mmu-miR-34a-5p | MIMAT0000542 | Rpap3 | 71919 | HITS-CLIP | 23142080 | Unspecified |
| mmu-miR-34a-5p | MIMAT0000542 | Rpe | 66646 | HITS-CLIP | 23142080\|29167373 | Unspecified |
| mmu-miR-34a-5p | MIMAT0000542 | Rpf1 | 70285 | HITS-CLIP | 23142080 | Unspecified |
| mmu-miR-34a-5p | MIMAT0000542 | Rpf2 | 67239 | HITS-CLIP | 23142080\|29167373 | Unspecified |

| | | | | | | |
|---|---|---|---|---|---|---|
| mmu-miR-34a-5p | MIMAT0000542 | Rpgr | 19893 | HITS-CLIP | 23142080 | Unspecified |
| mmu-miR-34a-5p | MIMAT0000542 | Rpgrip1l | 244585 | Microarrays//HITS-CLIP | 37986224\|23142080 | Unspecified |
| mmu-miR-34a-5p | MIMAT0000542 | Rpia | 19895 | Microarrays//HITS-CLIP | 37986224\|23027903\|23142080 | Unspecified |
| mmu-miR-34a-5p | MIMAT0000542 | Rpl10a | 19896 | HITS-CLIP | 23142080 | Unspecified |
| mmu-miR-34a-5p | MIMAT0000542 | Rpl12 | 269261 | HITS-CLIP | 23142080 | Unspecified |
| mmu-miR-34a-5p | MIMAT0000542 | Rpl13a | 22121 | HITS-CLIP | 23142080 | Unspecified |
| mmu-miR-34a-5p | MIMAT0000542 | Rpl18 | 6141 | HITS-CLIP | 23142080 | Unspecified |
| mmu-miR-34a-5p | MIMAT0000542 | Rpl18a | 76808 | HITS-CLIP | 23142080 | Unspecified |
| mmu-miR-34a-5p | MIMAT0000542 | Rpl22 | 19934 | HITS-CLIP | 37986224\|23142080 | Unspecified |
| mmu-miR-34a-5p | MIMAT0000542 | Rpl24 | 68193 | HITS-CLIP | 23142080 | Unspecified |
| mmu-miR-34a-5p | MIMAT0000542 | Rpl31 | 114641 | HITS-CLIP | 31433990 | Unspecified |
| mmu-miR-34a-5p | MIMAT0000542 | Rpl32 | 19951 | HITS-CLIP | 23142080 | Unspecified |
| mmu-miR-34a-5p | MIMAT0000542 | Rpl37 | 67281 | HITS-CLIP | 23142080 | Unspecified |
| mmu-miR-34a-5p | MIMAT0000542 | Rpl4 | 67891 | HITS-CLIP | 23142080\|29167373 | Unspecified |
| mmu-miR-34a-5p | MIMAT0000542 | Rpl7 | 19989 | HITS-CLIP | 23142080 | Unspecified |
| mmu-miR-34a-5p | MIMAT0000542 | Rpl7l1 | 66229 | HITS-CLIP | 26203562\|23142080 | Unspecified |
| mmu-miR-34a-5p | MIMAT0000542 | Rpl8 | 26961 | HITS-CLIP | 23142080 | Unspecified |
| mmu-miR-34a-5p | MIMAT0000542 | Rplp0 | 11837 | HITS-CLIP | 31433990\|19536157\|23142080 | Unspecified |
| mmu-miR-34a-5p | MIMAT0000542 | Rplp1 | 56040 | HITS-CLIP | 23142080 | Unspecified |
| mmu-miR-34a-5p | MIMAT0000542 | Rpn1 | 103963 | HITS-CLIP | 23142080\|29167373 | Unspecified |
| mmu-miR-34a-5p | MIMAT0000542 | Rpn2 | 20014 | HITS-CLIP | 23142080 | Unspecified |
| mmu-miR-34a-5p | MIMAT0000542 | Rpp21 | 67676 | HITS-CLIP | 23142080 | Unspecified |
| mmu-miR-34a-5p | MIMAT0000542 | Rpp38 | 227522 | HITS-CLIP | 23142080 | Unspecified |
| mmu-miR-34a-5p | MIMAT0000542 | Rprd1b | 70470 | HITS-CLIP | 23142080 | Unspecified |
| mmu-miR-34a-5p | MIMAT0000542 | Rprd2 | 75137 | HITS-CLIP | 23630078\|19536157\|23142080\|29167 | Unspecified |
| mmu-miR-34a-5p | MIMAT0000542 | Rps14 | 20044 | HITS-CLIP | 29167373 | Unspecified |
| mmu-miR-34a-5p | MIMAT0000542 | Rps15 | 20054 | HITS-CLIP | 23142080 | Unspecified |
| mmu-miR-34a-5p | MIMAT0000542 | Rps15a | 6210 | HITS-CLIP | 31433990\|29167373 | Unspecified |
| mmu-miR-34a-5p | MIMAT0000542 | Rps18 | 20084 | HITS-CLIP | 23142080 | Unspecified |
| mmu-miR-34a-5p | MIMAT0000542 | Rps20 | 67427 | HITS-CLIP | 23142080 | Unspecified |
| mmu-miR-34a-5p | MIMAT0000542 | Rps23 | 66475 | HITS-CLIP | 23142080 | Unspecified |
| mmu-miR-34a-5p | MIMAT0000542 | Rps24 | 20088 | HITS-CLIP | 23142080 | Unspecified |
| mmu-miR-34a-5p | MIMAT0000542 | Rps27l | 67941 | HITS-CLIP | 23142080 | Unspecified |
| mmu-miR-34a-5p | MIMAT0000542 | Rps5 | 20103 | HITS-CLIP | 23142080\|29167373 | Unspecified |
| mmu-miR-34a-5p | MIMAT0000542 | Rps6ka1 | 20111 | HITS-CLIP | 23142080 | Unspecified |
| mmu-miR-34a-5p | MIMAT0000542 | Rps6ka3 | 110651 | HITS-CLIP | 23142080\|29167373 | Unspecified |
| mmu-miR-34a-5p | MIMAT0000542 | Rps6ka4 | 56613 | HITS-CLIP | 23142080 | Unspecified |
| mmu-miR-34a-5p | MIMAT0000542 | Rps6ka5 | 73086 | HITS-CLIP | 23142080\|31433990 | Unspecified |
| mmu-miR-34a-5p | MIMAT0000542 | Rps6kb1 | 72508 | HITS-CLIP | 23142080\|19536157\|29167373 | Unspecified |

| | | | | | | |
|---|---|---|---|---|---|---|
| mmu-miR-34a-5p | MIMAT0000542 | Rps6kc1 | 320119 | HITS-CLIP | 23142080\|29167373 | Unspecified |
| mmu-miR-34a-5p | MIMAT0000542 | Rps8 | 20116 | HITS-CLIP | 31433990\|23630078 | Unspecified |
| mmu-miR-34a-5p | MIMAT0000542 | Rps9 | 76846 | HITS-CLIP | 31433990\|29167373 | Unspecified |
| mmu-miR-34a-5p | MIMAT0000542 | Rpsa | 16785 | HITS-CLIP | 23142080 | Unspecified |
| mmu-miR-34a-5p | MIMAT0000542 | Rptor | 74370 | HITS-CLIP | 23142080\|29167373 | Unspecified |
| mmu-miR-34a-5p | MIMAT0000542 | Rpusd1 | 106707 | HITS-CLIP | 19536157 | Unspecified |
| mmu-miR-34a-5p | MIMAT0000542 | Rpusd2 | 271842 | HITS-CLIP | 23142080 | Unspecified |
| mmu-miR-34a-5p | MIMAT0000542 | Rpusd3 | 101122 | HITS-CLIP | 31433990 | Unspecified |
| mmu-miR-34a-5p | MIMAT0000542 | Rpusd4 | 71989 | HITS-CLIP | 31433990 | Unspecified |
| mmu-miR-34a-5p | MIMAT0000542 | Rraga | 68441 | HITS-CLIP | 23142080 | Unspecified |
| mmu-miR-34a-5p | MIMAT0000542 | Rragc | 54170 | HITS-CLIP | 29167373\|23142080 | Unspecified |
| mmu-miR-34a-5p | MIMAT0000542 | Rragd | 52187 | HITS-CLIP | 31433990\|19536157\|29084942\|23142 | Unspecified |
| mmu-miR-34a-5p | MIMAT0000542 | Rras | 20130 | HITS-CLIP, Microarrays//Microarrays//HITS | 37986224\|23027903\|23142080\|29167 | Unspecified |
| mmu-miR-34a-5p | MIMAT0000542 | Rras2 | 66922 | HITS-CLIP | 23142080 | Unspecified |
| mmu-miR-34a-5p | MIMAT0000542 | Rrbp1 | 81910 | HITS-CLIP | 23142080\|29167373 | Unspecified |
| mmu-miR-34a-5p | MIMAT0000542 | Rreb1 | 68750 | HITS-CLIP | 23142080\|31433990\|23630078\|26203 | Unspecified |
| mmu-miR-34a-5p | MIMAT0000542 | Rrm1 | 20133 | HITS-CLIP | 31433990\|23142080 | Unspecified |
| mmu-miR-34a-5p | MIMAT0000542 | Rrm2 | 20135 | HITS-CLIP | 23142080\|29167373 | Unspecified |
| mmu-miR-34a-5p | MIMAT0000542 | Rrm2b | 382985 | HITS-CLIP | 29167373 | Unspecified |
| mmu-miR-34a-5p | MIMAT0000542 | Rrn3 | 106298 | HITS-CLIP | 23142080\|29167373 | Unspecified |
| mmu-miR-34a-5p | MIMAT0000542 | Rrnad1 | 229503 | HITS-CLIP | 29167373 | Unspecified |
| mmu-miR-34a-5p | MIMAT0000542 | Rrp1 | 18114 | HITS-CLIP | 19536157\|26203562\|23142080\|29167 | Unspecified |
| mmu-miR-34a-5p | MIMAT0000542 | Rrp1b | 72462 | HITS-CLIP | 23142080\|29167373 | Unspecified |
| mmu-miR-34a-5p | MIMAT0000542 | Rrp7a | 74778 | HITS-CLIP | 23142080 | Unspecified |
| mmu-miR-34a-5p | MIMAT0000542 | Rrs1 | 59014 | HITS-CLIP | 31433990\|29167373 | Unspecified |
| mmu-miR-34a-5p | MIMAT0000542 | Rsad2 | 58185 | HITS-CLIP | 29167373 | Unspecified |
| mmu-miR-34a-5p | MIMAT0000542 | Rsbn1 | 229675 | HITS-CLIP | 30971389\|23142080 | Unspecified |
| mmu-miR-34a-5p | MIMAT0000542 | Rsbn1l | 242860 | HITS-CLIP | 23142080\|29167373 | Unspecified |
| mmu-miR-34a-5p | MIMAT0000542 | Rsf1 | 233532 | HITS-CLIP | 31433990\|23142080 | Unspecified |
| mmu-miR-34a-5p | MIMAT0000542 | Rsl1d1 | 66409 | HITS-CLIP | 23630078\|26203562\|23142080 | Unspecified |
| mmu-miR-34a-5p | MIMAT0000542 | Rsph9 | 75564 | Microarrays | 37986224 | Unspecified |
| mmu-miR-34a-5p | MIMAT0000542 | Rspry1 | 67610 | HITS-CLIP | 30971389\|23142080\|29167373 | Unspecified |
| mmu-miR-34a-5p | MIMAT0000542 | Rsrc2 | 208606 | HITS-CLIP | 19536157\|23142080\|29167373 | Unspecified |
| mmu-miR-34a-5p | MIMAT0000542 | Rsrp1 | 27981 | HITS-CLIP | 23142080 | Unspecified |
| mmu-miR-34a-5p | MIMAT0000542 | Rsu1 | 20163 | HITS-CLIP | 29167373 | Unspecified |
| mmu-miR-34a-5p | MIMAT0000542 | Rtca | 66368 | HITS-CLIP | 23142080 | Unspecified |
| mmu-miR-34a-5p | MIMAT0000542 | Rtcb | 28088 | HITS-CLIP | 23142080 | Unspecified |
| mmu-miR-34a-5p | MIMAT0000542 | Rtf1 | 76246 | HITS-CLIP | 29167373\|23142080 | Unspecified |
| mmu-miR-34a-5p | MIMAT0000542 | Rtf2 | 51507 | HITS-CLIP | 30971389\|23142080 | Unspecified |

| | | | | | | |
|---|---|---|---|---|---|---|
| mmu-miR-34a-5p | MIMAT0000542 | Rtl6 | 223732 | HITS-CLIP | 19536157 | Unspecified |
| mmu-miR-34a-5p | MIMAT0000542 | Rtn1 | 104001 | HITS-CLIP | 19536157 | Unspecified |
| mmu-miR-34a-5p | MIMAT0000542 | Rtn3 | 20168 | HITS-CLIP | 19536157\|23142080\|29167373 | Unspecified |
| mmu-miR-34a-5p | MIMAT0000542 | Rtn4 | 68585 | HITS-CLIP | 19536157\|23142080\|29167373 | Unspecified |
| mmu-miR-34a-5p | MIMAT0000542 | Rtn4ip1 | 170728 | HITS-CLIP | 23142080 | Unspecified |
| mmu-miR-34a-5p | MIMAT0000542 | Rtraf | 51637 | HITS-CLIP | 23142080\|29167373 | Unspecified |
| mmu-miR-34a-5p | MIMAT0000542 | Rttn | 246102 | HITS-CLIP | 31433990\|23142080 | Unspecified |
| mmu-miR-34a-5p | MIMAT0000542 | Rubcn | 9711 | HITS-CLIP | 23142080 | Unspecified |
| mmu-miR-34a-5p | MIMAT0000542 | Rufy1 | 216724 | HITS-CLIP | 23142080\|29167373 | Unspecified |
| mmu-miR-34a-5p | MIMAT0000542 | Rufy2 | 70432 | HITS-CLIP | 37986224\|23142080 | Unspecified |
| mmu-miR-34a-5p | MIMAT0000542 | Rufy3 | 52822 | HITS-CLIP | 29167373 | Unspecified |
| mmu-miR-34a-5p | MIMAT0000542 | Rundc1 | 217201 | HITS-CLIP | 31433990\|23142080 | Unspecified |
| mmu-miR-34a-5p | MIMAT0000542 | Rundc3b | 242819 | HITS-CLIP | 23142080 | Unspecified |
| mmu-miR-34a-5p | MIMAT0000542 | Runx1 | 12394 | HITS-CLIP | 23142080\|31433990\|26203562\|29167 | Unspecified |
| mmu-miR-34a-5p | MIMAT0000542 | Runx2 | 12393 | HITS-CLIP | 23142080 | Unspecified |
| mmu-miR-34a-5p | MIMAT0000542 | Runx3 | 12399 | HITS-CLIP | 23142080 | Unspecified |
| mmu-miR-34a-5p | MIMAT0000542 | Rusc2 | 100213 | HITS-CLIP | 19536157 | Unspecified |
| mmu-miR-34a-5p | MIMAT0000542 | Ruvbl2 | 20174 | HITS-CLIP | 23142080\|29167373 | Unspecified |
| mmu-miR-34a-5p | MIMAT0000542 | Rwdd2b | 53858 | HITS-CLIP | 23142080 | Unspecified |
| mmu-miR-34a-5p | MIMAT0000542 | Rxra | 20181 | HITS-CLIP | 29167373 | Unspecified |
| mmu-miR-34a-5p | MIMAT0000542 | Ryr1 | 20190 | HITS-CLIP | 19536157 | Unspecified |
| mmu-miR-34a-5p | MIMAT0000542 | Ryr2 | 20191 | HITS-CLIP | 19536157 | Unspecified |
| mmu-miR-34a-5p | MIMAT0000542 | S100a10 | 20194 | HITS-CLIP | 23142080 | Unspecified |
| mmu-miR-34a-5p | MIMAT0000542 | S100a11 | 20195 | HITS-CLIP | 23142080 | Unspecified |
| mmu-miR-34a-5p | MIMAT0000542 | S100a4 | 20198 | HITS-CLIP | 31433990\|29167373 | Unspecified |
| mmu-miR-34a-5p | MIMAT0000542 | S100pbp | 74648 | HITS-CLIP | 23630078\|31433990\|30971389\|2314 | Unspecified |
| mmu-miR-34a-5p | MIMAT0000542 | S1pr1 | 13609 | HITS-CLIP | 23142080 | Unspecified |
| mmu-miR-34a-5p | MIMAT0000542 | S1pr2 | 14739 | HITS-CLIP | 29167373\|23142080 | Unspecified |
| mmu-miR-34a-5p | MIMAT0000542 | S1pr3 | 13610 | HITS-CLIP | 23142080 | Unspecified |
| mmu-miR-34a-5p | MIMAT0000542 | S1pr4 | 13611 | HITS-CLIP | 23142080 | Unspecified |
| mmu-miR-34a-5p | MIMAT0000542 | Saal1 | 78935 | HITS-CLIP | 23142080 | Unspecified |
| mmu-miR-34a-5p | MIMAT0000542 | Sacm1l | 83493 | HITS-CLIP | 23142080\|29167373 | Unspecified |
| mmu-miR-34a-5p | MIMAT0000542 | Sacs | 50720 | HITS-CLIP | 19536157\|23142080\|29167373 | Unspecified |
| mmu-miR-34a-5p | MIMAT0000542 | Sae1 | 56459 | Microarrays//HITS-CLIP | 37986224\|23027903\|23142080 | Unspecified |
| mmu-miR-34a-5p | MIMAT0000542 | Safb | 224903 | HITS-CLIP | 23142080\|29167373 | Unspecified |
| mmu-miR-34a-5p | MIMAT0000542 | Safb2 | 224902 | HITS-CLIP | 23142080 | Unspecified |
| mmu-miR-34a-5p | MIMAT0000542 | Sall2 | 50524 | HITS-CLIP | 23142080 | Unspecified |
| mmu-miR-34a-5p | MIMAT0000542 | Samd1 | 666704 | HITS-CLIP | 30971389 | Unspecified |
| mmu-miR-34a-5p | MIMAT0000542 | Samd14 | 217125 | HITS-CLIP | 19536157 | Unspecified |

| | | | | | | |
|---|---|---|---|---|---|---|
| mmu-miR-34a-5p | MIMAT0000542 | Samd4b | 233033 | HITS-CLIP | 23142080\|29167373 | Unspecified |
| mmu-miR-34a-5p | MIMAT0000542 | Samd8 | 67630 | HITS-CLIP | 19536157\|23142080 | Unspecified |
| mmu-miR-34a-5p | MIMAT0000542 | Samd9l | 209086 | HITS-CLIP | 23142080\|29167373 | Unspecified |
| mmu-miR-34a-5p | MIMAT0000542 | Samhd1 | 56045 | HITS-CLIP | 23142080\|29167373 | Unspecified |
| mmu-miR-34a-5p | MIMAT0000542 | Samm50 | 68653 | HITS-CLIP | 23142080 | Unspecified |
| mmu-miR-34a-5p | MIMAT0000542 | Samsn1 | 67742 | HITS-CLIP | 23142080\|29167373 | Unspecified |
| mmu-miR-34a-5p | MIMAT0000542 | Sap130 | 269003 | HITS-CLIP | 23142080\|29167373\|19536157 | Unspecified |
| mmu-miR-34a-5p | MIMAT0000542 | Sap18 | 20220 | HITS-CLIP | 31433990 | Unspecified |
| mmu-miR-34a-5p | MIMAT0000542 | Sap25 | 751865 | HITS-CLIP | 29167373 | Unspecified |
| mmu-miR-34a-5p | MIMAT0000542 | Sap30 | 60406 | HITS-CLIP | 23142080 | Unspecified |
| mmu-miR-34a-5p | MIMAT0000542 | Sar1a | 20224 | HITS-CLIP | 23630078\|23142080\|19536157\|31433 | Unspecified |
| mmu-miR-34a-5p | MIMAT0000542 | Sar1b | 66397 | HITS-CLIP | 23142080\|19536157 | Unspecified |
| mmu-miR-34a-5p | MIMAT0000542 | Saraf | 51669 | HITS-CLIP | 19536157\|23142080\|29167373 | Unspecified |
| mmu-miR-34a-5p | MIMAT0000542 | Sars | 20226 | HITS-CLIP | 19536157\|23142080\|29167373 | Unspecified |
| mmu-miR-34a-5p | MIMAT0000542 | Sart1 | 20227 | HITS-CLIP | 29167373\|23142080 | Unspecified |
| mmu-miR-34a-5p | MIMAT0000542 | Sart3 | 53890 | HITS-CLIP | 23142080 | Unspecified |
| mmu-miR-34a-5p | MIMAT0000542 | Sash1 | 70097 | HITS-CLIP | 29167373 | Unspecified |
| mmu-miR-34a-5p | MIMAT0000542 | Sash3 | 74131 | HITS-CLIP | 23142080 | Unspecified |
| mmu-miR-34a-5p | MIMAT0000542 | Sass6 | 72776 | HITS-CLIP | 23142080 | Unspecified |
| mmu-miR-34a-5p | MIMAT0000542 | Sat1 | 20229 | HITS-CLIP | 23142080 | Unspecified |
| mmu-miR-34a-5p | MIMAT0000542 | Satb1 | 20230 | HITS-CLIP | 37986224\|23142080\|31433990 | Unspecified |
| mmu-miR-34a-5p | MIMAT0000542 | Satb2 | 212712 | HITS-CLIP | 23630078 | Unspecified |
| mmu-miR-34a-5p | MIMAT0000542 | Sav1 | 64010 | HITS-CLIP | 23142080\|30971389\|23630078\|29167 | Unspecified |
| mmu-miR-34a-5p | MIMAT0000542 | Sbds | 66711 | HITS-CLIP | 23142080\|29167373 | Unspecified |
| mmu-miR-34a-5p | MIMAT0000542 | Sbf1 | 77980 | HITS-CLIP | 31433990\|23142080\|29167373 | Unspecified |
| mmu-miR-34a-5p | MIMAT0000542 | Sbf2 | 319934 | HITS-CLIP | 19536157\|23142080\|29167373 | Unspecified |
| mmu-miR-34a-5p | MIMAT0000542 | Sbk1 | 104175 | HITS-CLIP | 19536157 | Unspecified |
| mmu-miR-34a-5p | MIMAT0000542 | Sbno1 | 243272 | HITS-CLIP | 23142080\|19536157\|29167373 | Unspecified |
| mmu-miR-34a-5p | MIMAT0000542 | Sbno2 | 216161 | HITS-CLIP | 23142080\|29167373 | Unspecified |
| mmu-miR-34a-5p | MIMAT0000542 | Sc5d | 235293 | HITS-CLIP | 19536157\|23142080\|29167373 | Unspecified |
| mmu-miR-34a-5p | MIMAT0000542 | Scaf11 | 72193 | HITS-CLIP | 29167373\|23142080 | Unspecified |
| mmu-miR-34a-5p | MIMAT0000542 | Scaf4 | 224432 | HITS-CLIP | 23142080\|29167373 | Unspecified |
| mmu-miR-34a-5p | MIMAT0000542 | Scaf8 | 106583 | HITS-CLIP | 23142080 | Unspecified |
| mmu-miR-34a-5p | MIMAT0000542 | Scamp1 | 107767 | HITS-CLIP | 19536157 | Unspecified |
| mmu-miR-34a-5p | MIMAT0000542 | Scamp3 | 24045 | HITS-CLIP | 29167373\|23142080 | Unspecified |
| mmu-miR-34a-5p | MIMAT0000542 | Scamp5 | 56807 | HITS-CLIP | 19536157\|29167373 | Unspecified |
| mmu-miR-34a-5p | MIMAT0000542 | Scaper | 244891 | HITS-CLIP | 23142080 | Unspecified |
| mmu-miR-34a-5p | MIMAT0000542 | Scarb1 | 20778 | HITS-CLIP | 23142080\|29167373 | Unspecified |
| mmu-miR-34a-5p | MIMAT0000542 | Scarb2 | 12492 | HITS-CLIP | 19536157\|23142080\|29167373 | Unspecified |

| | | | | | | |
|---|---|---|---|---|---|---|
| mmu-miR-34a-5p | MIMAT0000542 | Scd1 | 20249 | HITS-CLIP | 19536157\|30971389 | Unspecified |
| mmu-miR-34a-5p | MIMAT0000542 | Scd2 | 20250 | HITS-CLIP | 37986224\|19536157\|23142080\|31433 | Unspecified |
| mmu-miR-34a-5p | MIMAT0000542 | Scfd1 | 76983 | HITS-CLIP | 23142080\|31433990 | Unspecified |
| mmu-miR-34a-5p | MIMAT0000542 | Scfd2 | 212986 | HITS-CLIP | 23142080\|29167373 | Unspecified |
| mmu-miR-34a-5p | MIMAT0000542 | Scg3 | 20255 | HITS-CLIP | 19536157 | Unspecified |
| mmu-miR-34a-5p | MIMAT0000542 | Sclt1 | 67161 | HITS-CLIP | 23630078 | Unspecified |
| mmu-miR-34a-5p | MIMAT0000542 | Scmh1 | 29871 | Microarrays//HITS-CLIP | 37986224\|23027903\|23142080 | Unspecified |
| mmu-miR-34a-5p | MIMAT0000542 | Scml4 | 268297 | HITS-CLIP | 23142080 | Unspecified |
| mmu-miR-34a-5p | MIMAT0000542 | Scn2a | 110876 | HITS-CLIP | 19536157 | Unspecified |
| mmu-miR-34a-5p | MIMAT0000542 | Scn2b | 72821 | HITS-CLIP | 19536157 | Unspecified |
| mmu-miR-34a-5p | MIMAT0000542 | Scn8a | 20273 | HITS-CLIP | 19536157 | Unspecified |
| mmu-miR-34a-5p | MIMAT0000542 | Scnm1 | 69269 | HITS-CLIP | 23142080 | Unspecified |
| mmu-miR-34a-5p | MIMAT0000542 | Sco1 | 52892 | HITS-CLIP | 21258322\|23142080 | Unspecified |
| mmu-miR-34a-5p | MIMAT0000542 | Scoc | 56367 | HITS-CLIP | 29167373 | Unspecified |
| mmu-miR-34a-5p | MIMAT0000542 | Scp2 | 20280 | HITS-CLIP | 31433990 | Unspecified |
| mmu-miR-34a-5p | MIMAT0000542 | Scpep1 | 74617 | HITS-CLIP | 29167373 | Unspecified |
| mmu-miR-34a-5p | MIMAT0000542 | Scrib | 105782 | HITS-CLIP | 23142080\|29167373 | Unspecified |
| mmu-miR-34a-5p | MIMAT0000542 | Scrn1 | 69938 | HITS-CLIP | 19536157 | Unspecified |
| mmu-miR-34a-5p | MIMAT0000542 | Scrn3 | 74616 | HITS-CLIP | 30971389 | Unspecified |
| mmu-miR-34a-5p | MIMAT0000542 | Scrt1 | 170729 | HITS-CLIP | 23142080 | Unspecified |
| mmu-miR-34a-5p | MIMAT0000542 | Scyl2 | 213326 | HITS-CLIP | 23142080\|29167373 | Unspecified |
| mmu-miR-34a-5p | MIMAT0000542 | Sdad1 | 231452 | HITS-CLIP | 31433990\|26203562\|23142080\|29167 | Unspecified |
| mmu-miR-34a-5p | MIMAT0000542 | Sdc2 | 15529 | HITS-CLIP | 19536157\|23630078 | Unspecified |
| mmu-miR-34a-5p | MIMAT0000542 | Sdc3 | 20970 | HITS-CLIP | 29167373 | Unspecified |
| mmu-miR-34a-5p | MIMAT0000542 | Sdc4 | 20971 | HITS-CLIP | 23142080\|29167373 | Unspecified |
| mmu-miR-34a-5p | MIMAT0000542 | Sdcbp | 53378 | HITS-CLIP | 19536157\|29167373\|23142080 | Unspecified |
| mmu-miR-34a-5p | MIMAT0000542 | Sdcbp2 | 228765 | HITS-CLIP | 23142080 | Unspecified |
| mmu-miR-34a-5p | MIMAT0000542 | Sdccag8 | 76816 | HITS-CLIP | 29167373 | Unspecified |
| mmu-miR-34a-5p | MIMAT0000542 | Sde2 | 208768 | HITS-CLIP | 23142080 | Unspecified |
| mmu-miR-34a-5p | MIMAT0000542 | Sdf2 | 20316 | HITS-CLIP | 23142080 | Unspecified |
| mmu-miR-34a-5p | MIMAT0000542 | Sdf2l1 | 64136 | HITS-CLIP | 23142080\|29167373 | Unspecified |
| mmu-miR-34a-5p | MIMAT0000542 | Sdf4 | 20318 | HITS-CLIP | 31433990\|19536157\|23142080\|29167 | Unspecified |
| mmu-miR-34a-5p | MIMAT0000542 | Sdha | 66945 | HITS-CLIP | 23142080\|19536157\|31433990\|29167 | Unspecified |
| mmu-miR-34a-5p | MIMAT0000542 | Sdhaf1 | 68332 | HITS-CLIP | 23142080 | Unspecified |
| mmu-miR-34a-5p | MIMAT0000542 | Sdhaf2 | 66072 | HITS-CLIP | 23142080\|31433990\|29167373 | Unspecified |
| mmu-miR-34a-5p | MIMAT0000542 | Sdhaf4 | 34089 | HITS-CLIP | 23142080 | Unspecified |
| mmu-miR-34a-5p | MIMAT0000542 | Sdhb | 67680 | HITS-CLIP | 23142080 | Unspecified |
| mmu-miR-34a-5p | MIMAT0000542 | Sdhc | 66052 | HITS-CLIP | 23142080\|29167373 | Unspecified |
| mmu-miR-34a-5p | MIMAT0000542 | Sdhd | 66925 | HITS-CLIP | 23142080 | Unspecified |

| | | | | | | |
|---|---|---|---|---|---|---|
| mmu-miR-34a-5p | MIMAT0000542 | Sdr39u1 | 654795 | HITS-CLIP | 23142080 | Unspecified |
| mmu-miR-34a-5p | MIMAT0000542 | Sec11a | 56529 | HITS-CLIP | 23142080 | Unspecified |
| mmu-miR-34a-5p | MIMAT0000542 | Sec11c | 66286 | HITS-CLIP | 23142080 | Unspecified |
| mmu-miR-34a-5p | MIMAT0000542 | Sec13 | 110379 | HITS-CLIP | 31433990\|29167373 | Unspecified |
| mmu-miR-34a-5p | MIMAT0000542 | Sec14l1 | 74136 | HITS-CLIP | 31433990\|23630078\|29167373 | Unspecified |
| mmu-miR-34a-5p | MIMAT0000542 | Sec16a | 227648 | HITS-CLIP | 31433990\|30971389\|23142080\|29167 | Unspecified |
| mmu-miR-34a-5p | MIMAT0000542 | Sec22a | 317717 | HITS-CLIP | 23142080 | Unspecified |
| mmu-miR-34a-5p | MIMAT0000542 | Sec22b | 20333 | HITS-CLIP | 23142080 | Unspecified |
| mmu-miR-34a-5p | MIMAT0000542 | Sec23a | 20334 | HITS-CLIP | 23142080\|19536157\|29167373 | Unspecified |
| mmu-miR-34a-5p | MIMAT0000542 | Sec23b | 27054 | HITS-CLIP | 29167373\|23142080 | Unspecified |
| mmu-miR-34a-5p | MIMAT0000542 | Sec23ip | 207352 | HITS-CLIP | 23142080\|29167373 | Unspecified |
| mmu-miR-34a-5p | MIMAT0000542 | Sec24a | 77371 | HITS-CLIP | 23142080\|29167373 | Unspecified |
| mmu-miR-34a-5p | MIMAT0000542 | Sec24b | 99683 | HITS-CLIP | 23142080 | Unspecified |
| mmu-miR-34a-5p | MIMAT0000542 | Sec24c | 218811 | HITS-CLIP | 37986224\|23630078\|23142080 | Unspecified |
| mmu-miR-34a-5p | MIMAT0000542 | Sec24d | 69608 | HITS-CLIP | 30971389\|29167373\|23142080 | Unspecified |
| mmu-miR-34a-5p | MIMAT0000542 | Sec31a | 69162 | HITS-CLIP | 29167373 | Unspecified |
| mmu-miR-34a-5p | MIMAT0000542 | Sec61a1 | 53421 | HITS-CLIP | 19536157\|23142080 | Unspecified |
| mmu-miR-34a-5p | MIMAT0000542 | Sec61a2 | 57743 | Microarrays | 37986224\|23027903 | Unspecified |
| mmu-miR-34a-5p | MIMAT0000542 | Sec62 | 69276 | HITS-CLIP | 23142080 | Unspecified |
| mmu-miR-34a-5p | MIMAT0000542 | Sec63 | 140740 | HITS-CLIP | 23142080 | Unspecified |
| mmu-miR-34a-5p | MIMAT0000542 | Secisbp2 | 75420 | HITS-CLIP | 23142080 | Unspecified |
| mmu-miR-34a-5p | MIMAT0000542 | Secisbp2l | 70354 | HITS-CLIP | 30971389\|31433990\|19536157\|23142 | Unspecified |
| mmu-miR-34a-5p | MIMAT0000542 | Seh1l | 72124 | HITS-CLIP | 23142080 | Unspecified |
| mmu-miR-34a-5p | MIMAT0000542 | Sel1l | 20338 | HITS-CLIP | 23142080\|19536157\|29167373 | Unspecified |
| mmu-miR-34a-5p | MIMAT0000542 | Selenbp1 | 20341 | HITS-CLIP | 23142080 | Unspecified |
| mmu-miR-34a-5p | MIMAT0000542 | Selenof | 9403 | HITS-CLIP | 31433990 | Unspecified |
| mmu-miR-34a-5p | MIMAT0000542 | Selenoh | 280636 | HITS-CLIP | 23142080 | Unspecified |
| mmu-miR-34a-5p | MIMAT0000542 | Selenoi | 28042 | HITS-CLIP | 26203562 | Unspecified |
| mmu-miR-34a-5p | MIMAT0000542 | Selenok | 58515 | HITS-CLIP | 23142080 | Unspecified |
| mmu-miR-34a-5p | MIMAT0000542 | Selenoo | 83642 | HITS-CLIP | 23142080 | Unspecified |
| mmu-miR-34a-5p | MIMAT0000542 | Selenop | 6414 | HITS-CLIP | 23142080\|19536157\|29167373 | Unspecified |
| mmu-miR-34a-5p | MIMAT0000542 | Selenot | 69227 | HITS-CLIP | 23142080 | Unspecified |
| mmu-miR-34a-5p | MIMAT0000542 | Sell | 20343 | HITS-CLIP | 23142080 | Unspecified |
| mmu-miR-34a-5p | MIMAT0000542 | Selp | 20344 | HITS-CLIP | 23142080 | Unspecified |
| mmu-miR-34a-5p | MIMAT0000542 | Selplg | 20345 | HITS-CLIP | 23142080 | Unspecified |
| mmu-miR-34a-5p | MIMAT0000542 | Sem1 | 7979 | HITS-CLIP | 23142080 | Unspecified |
| mmu-miR-34a-5p | MIMAT0000542 | Sema3c | 20348 | Microarrays | 37986224 | Unspecified |
| mmu-miR-34a-5p | MIMAT0000542 | Sema3d | 108151 | Microarrays | 37986224 | Unspecified |
| mmu-miR-34a-5p | MIMAT0000542 | Sema4a | 20351 | HITS-CLIP | 23142080 | Unspecified |

| | | | | | | |
|---|---|---|---|---|---|---|
| mmu-miR-34a-5p | MIMAT0000542 | Sema4b | 20352 | HITS-CLIP//Microarrays//Luciferase report | 21258322\|23027903\|23142080\|29167 | Unspecified |
| mmu-miR-34a-5p | MIMAT0000542 | Sema4c | 20353 | HITS-CLIP | 23142080 | Unspecified |
| mmu-miR-34a-5p | MIMAT0000542 | Sema4d | 20354 | HITS-CLIP | 23142080\|31433990\|30971389\|29167 | Unspecified |
| mmu-miR-34a-5p | MIMAT0000542 | Sema4f | 20355 | HITS-CLIP | 23142080 | Unspecified |
| mmu-miR-34a-5p | MIMAT0000542 | Sema5b | 20357 | HITS-CLIP | 19536157 | Unspecified |
| mmu-miR-34a-5p | MIMAT0000542 | Sema6d | 214968 | HITS-CLIP | 19536157\|29167373 | Unspecified |
| mmu-miR-34a-5p | MIMAT0000542 | Sema7a | 20361 | HITS-CLIP | 23142080 | Unspecified |
| mmu-miR-34a-5p | MIMAT0000542 | Senp1 | 223870 | HITS-CLIP | 23142080\|29167373 | Unspecified |
| mmu-miR-34a-5p | MIMAT0000542 | Senp2 | 75826 | HITS-CLIP | 23142080\|19536157\|29167373 | Unspecified |
| mmu-miR-34a-5p | MIMAT0000542 | Senp5 | 320213 | HITS-CLIP | 23142080 | Unspecified |
| mmu-miR-34a-5p | MIMAT0000542 | Senp6 | 215351 | HITS-CLIP | 23142080\|29167373 | Unspecified |
| mmu-miR-34a-5p | MIMAT0000542 | Senp7 | 66315 | HITS-CLIP | 30971389\|23142080\|29167373 | Unspecified |
| mmu-miR-34a-5p | MIMAT0000542 | Sephs1 | 109079 | HITS-CLIP | 23142080 | Unspecified |
| mmu-miR-34a-5p | MIMAT0000542 | Sephs2 | 20768 | HITS-CLIP | 31433990\|23142080\|29167373 | Unspecified |
| mmu-miR-34a-5p | MIMAT0000542 | Sepsecs | 211006 | HITS-CLIP | 30971389\|23142080 | Unspecified |
| mmu-miR-34a-5p | MIMAT0000542 | Sept1 | 54204 | HITS-CLIP | 23142080 | Unspecified |
| mmu-miR-34a-5p | MIMAT0000542 | Sept10 | 103080 | HITS-CLIP | 23142080 | Unspecified |
| mmu-miR-34a-5p | MIMAT0000542 | Sept11 | 52398 | HITS-CLIP | 23142080\|19536157 | Unspecified |
| mmu-miR-34a-5p | MIMAT0000542 | Sept2 | 4735 | HITS-CLIP | 23142080\|29167373 | Unspecified |
| mmu-miR-34a-5p | MIMAT0000542 | Sept3 | 55964 | HITS-CLIP | 19536157 | Unspecified |
| mmu-miR-34a-5p | MIMAT0000542 | Sept6 | 23157 | HITS-CLIP | 23142080 | Unspecified |
| mmu-miR-34a-5p | MIMAT0000542 | Sept7 | 235072 | HITS-CLIP | 37986224\|19536157\|23142080\|29167 | Unspecified |
| mmu-miR-34a-5p | MIMAT0000542 | Sept8 | 20362 | HITS-CLIP | 19536157 | Unspecified |
| mmu-miR-34a-5p | MIMAT0000542 | Sept9 | 53860 | HITS-CLIP | 19536157\|23142080 | Unspecified |
| mmu-miR-34a-5p | MIMAT0000542 | Serac1 | 321007 | HITS-CLIP | 29167373 | Unspecified |
| mmu-miR-34a-5p | MIMAT0000542 | Serbp1 | 66870 | HITS-CLIP | 23142080\|29167373 | Unspecified |
| mmu-miR-34a-5p | MIMAT0000542 | Serf1 | 20365 | HITS-CLIP | 23142080 | Unspecified |
| mmu-miR-34a-5p | MIMAT0000542 | Serf2 | 378702 | HITS-CLIP | 23142080\|29167373 | Unspecified |
| mmu-miR-34a-5p | MIMAT0000542 | Serinc1 | 56442 | HITS-CLIP | 37986224\|19536157\|23142080\|29167 | Unspecified |
| mmu-miR-34a-5p | MIMAT0000542 | Serinc3 | 26943 | HITS-CLIP | 23142080\|19536157\|29167373 | Unspecified |
| mmu-miR-34a-5p | MIMAT0000542 | Serinc5 | 218442 | HITS-CLIP | 19536157\|23142080 | Unspecified |
| mmu-miR-34a-5p | MIMAT0000542 | Serpinb6a | 20719 | HITS-CLIP | 23142080 | Unspecified |
| mmu-miR-34a-5p | MIMAT0000542 | Serpinb6b | 20708 | HITS-CLIP | 23142080 | Unspecified |
| mmu-miR-34a-5p | MIMAT0000542 | Serpinb8 | 20725 | HITS-CLIP | 29167373 | Unspecified |
| mmu-miR-34a-5p | MIMAT0000542 | Serpinb9 | 20723 | HITS-CLIP | 23142080 | Unspecified |
| mmu-miR-34a-5p | MIMAT0000542 | Serpine1 | 18787 | HITS-CLIP | 23142080 | Unspecified |
| mmu-miR-34a-5p | MIMAT0000542 | Serpine2 | 20720 | HITS-CLIP | 23142080 | Unspecified |
| mmu-miR-34a-5p | MIMAT0000542 | Serpinf1 | 20317 | HITS-CLIP | 23142080 | Unspecified |
| mmu-miR-34a-5p | MIMAT0000542 | Serpini1 | 20713 | HITS-CLIP | 19536157 | Unspecified |

| | | | | | | |
|---|---|---|---|---|---|---|
| mmu-miR-34a-5p | MIMAT0000542 | Sertad2 | 58172 | HITS-CLIP | 31433990\|23142080 | Unspecified |
| mmu-miR-34a-5p | MIMAT0000542 | Sertad3 | 170742 | HITS-CLIP | 30971389\|23142080 | Unspecified |
| mmu-miR-34a-5p | MIMAT0000542 | Sesn1 | 140742 | HITS-CLIP | 23142080 | Unspecified |
| mmu-miR-34a-5p | MIMAT0000542 | Sesn2 | 230784 | HITS-CLIP | 26203562\|23142080 | Unspecified |
| mmu-miR-34a-5p | MIMAT0000542 | Sesn3 | 75747 | HITS-CLIP | 23142080\|19536157\|31433990\|29084 | Unspecified |
| mmu-miR-34a-5p | MIMAT0000542 | Sestd1 | 228071 | HITS-CLIP | 29167373 | Unspecified |
| mmu-miR-34a-5p | MIMAT0000542 | Setbp1 | 240427 | HITS-CLIP | 23142080 | Unspecified |
| mmu-miR-34a-5p | MIMAT0000542 | Setd1a | 233904 | HITS-CLIP | 29167373 | Unspecified |
| mmu-miR-34a-5p | MIMAT0000542 | Setd1b | 208043 | HITS-CLIP | 23142080\|29167373 | Unspecified |
| mmu-miR-34a-5p | MIMAT0000542 | Setd2 | 235626 | HITS-CLIP | 23142080\|31433990\|30971389\|29167 | Unspecified |
| mmu-miR-34a-5p | MIMAT0000542 | Setd3 | 52690 | HITS-CLIP | 23142080 | Unspecified |
| mmu-miR-34a-5p | MIMAT0000542 | Setd5 | 72895 | HITS-CLIP | 31433990\|19536157\|23142080\|29167 | Unspecified |
| mmu-miR-34a-5p | MIMAT0000542 | Setd7 | 73251 | Microarrays//HITS-CLIP | 37986224\|19536157\|23142080 | Unspecified |
| mmu-miR-34a-5p | MIMAT0000542 | Setdb1 | 84505 | HITS-CLIP | 23142080\|30971389 | Unspecified |
| mmu-miR-34a-5p | MIMAT0000542 | Setdb2 | 83852 | HITS-CLIP | 23142080\|29167373 | Unspecified |
| mmu-miR-34a-5p | MIMAT0000542 | Setx | 269254 | HITS-CLIP | 37986224\|30971389\|23142080\|29167 | Unspecified |
| mmu-miR-34a-5p | MIMAT0000542 | Sez6 | 20370 | HITS-CLIP | 19536157 | Unspecified |
| mmu-miR-34a-5p | MIMAT0000542 | Sez6l | 56747 | HITS-CLIP | 19536157 | Unspecified |
| mmu-miR-34a-5p | MIMAT0000542 | Sf1 | 22668 | HITS-CLIP | 23142080\|29167373 | Unspecified |
| mmu-miR-34a-5p | MIMAT0000542 | Sf3a1 | 67465 | HITS-CLIP | 23142080 | Unspecified |
| mmu-miR-34a-5p | MIMAT0000542 | Sf3a2 | 20222 | HITS-CLIP | 23142080 | Unspecified |
| mmu-miR-34a-5p | MIMAT0000542 | Sf3b1 | 81898 | HITS-CLIP | 37986224\|23142080\|19536157\|29167 | Unspecified |
| mmu-miR-34a-5p | MIMAT0000542 | Sf3b2 | 319322 | HITS-CLIP | 23142080 | Unspecified |
| mmu-miR-34a-5p | MIMAT0000542 | Sf3b3 | 101943 | HITS-CLIP | 31433990\|29167373\|23142080 | Unspecified |
| mmu-miR-34a-5p | MIMAT0000542 | Sf3b5 | 66125 | HITS-CLIP | 23142080 | Unspecified |
| mmu-miR-34a-5p | MIMAT0000542 | Sfi1 | 78887 | HITS-CLIP | 23142080\|29167373 | Unspecified |
| mmu-miR-34a-5p | MIMAT0000542 | Sfmbt1 | 54650 | HITS-CLIP | 29167373 | Unspecified |
| mmu-miR-34a-5p | MIMAT0000542 | Sfmbt2 | 353282 | HITS-CLIP | 23142080 | Unspecified |
| mmu-miR-34a-5p | MIMAT0000542 | Sfpq | 71514 | HITS-CLIP | 23142080 | Unspecified |
| mmu-miR-34a-5p | MIMAT0000542 | Sfr1 | 67788 | HITS-CLIP | 23142080\|29167373 | Unspecified |
| mmu-miR-34a-5p | MIMAT0000542 | Sfrp1 | 20377 | HITS-CLIP | 19536157 | Unspecified |
| mmu-miR-34a-5p | MIMAT0000542 | Sfrp2 | 20319 | Microarrays | 37986224 | Unspecified |
| mmu-miR-34a-5p | MIMAT0000542 | Sfswap | 231769 | HITS-CLIP | 23142080 | Unspecified |
| mmu-miR-34a-5p | MIMAT0000542 | Sft2d2 | 108735 | HITS-CLIP | 31433990\|23142080 | Unspecified |
| mmu-miR-34a-5p | MIMAT0000542 | Sft2d3 | 67158 | HITS-CLIP | 23142080 | Unspecified |
| mmu-miR-34a-5p | MIMAT0000542 | Sfxn1 | 14057 | HITS-CLIP | 23142080 | Unspecified |
| mmu-miR-34a-5p | MIMAT0000542 | Sfxn3 | 94280 | HITS-CLIP | 23142080\|29167373 | Unspecified |
| mmu-miR-34a-5p | MIMAT0000542 | Sfxn5 | 94282 | HITS-CLIP | 29167373 | Unspecified |
| mmu-miR-34a-5p | MIMAT0000542 | Sgcb | 24051 | Microarrays//HITS-CLIP | 37986224\|19536157\|23142080 | Unspecified |

| | | | | | | |
|---|---|---|---|---|---|---|
| mmu-miR-34a-5p | MIMAT0000542 | Sgcg | 24053 | Microarrays | 37986224 | Unspecified |
| mmu-miR-34a-5p | MIMAT0000542 | Sgip1 | 73094 | HITS-CLIP | 19536157 | Unspecified |
| mmu-miR-34a-5p | MIMAT0000542 | Sgk1 | 20393 | HITS-CLIP | 23142080\|29167373 | Unspecified |
| mmu-miR-34a-5p | MIMAT0000542 | Sgk3 | 170755 | HITS-CLIP | 21258322\|29167373 | Unspecified |
| mmu-miR-34a-5p | MIMAT0000542 | Sgms1 | 208449 | HITS-CLIP | 23142080\|31433990\|29167373 | Unspecified |
| mmu-miR-34a-5p | MIMAT0000542 | Sgo1 | 72415 | Microarrays//HITS-CLIP | 23027903\|31433990\|23142080\|29167 | Unspecified |
| mmu-miR-34a-5p | MIMAT0000542 | Sgo2a | 68549 | HITS-CLIP | 30971389 | Unspecified |
| mmu-miR-34a-5p | MIMAT0000542 | Sgpl1 | 20397 | HITS-CLIP | 19536157\|30971389\|23142080\|29167 | Unspecified |
| mmu-miR-34a-5p | MIMAT0000542 | Sgpp1 | 81535 | HITS-CLIP | 23142080 | Unspecified |
| mmu-miR-34a-5p | MIMAT0000542 | Sgsh | 27029 | Microarrays | 37986224 | Unspecified |
| mmu-miR-34a-5p | MIMAT0000542 | Sgsm2 | 97761 | HITS-CLIP | 23142080 | Unspecified |
| mmu-miR-34a-5p | MIMAT0000542 | Sgta | 52551 | HITS-CLIP | 23142080 | Unspecified |
| mmu-miR-34a-5p | MIMAT0000542 | Sh2b1 | 20399 | HITS-CLIP | 29167373\|23142080 | Unspecified |
| mmu-miR-34a-5p | MIMAT0000542 | Sh2b3 | 16923 | HITS-CLIP | 31433990\|23142080\|29167373 | Unspecified |
| mmu-miR-34a-5p | MIMAT0000542 | Sh2d1a | 20400 | HITS-CLIP | 23142080 | Unspecified |
| mmu-miR-34a-5p | MIMAT0000542 | Sh2d2a | 27371 | HITS-CLIP | 23142080 | Unspecified |
| mmu-miR-34a-5p | MIMAT0000542 | Sh2d3c | 27387 | HITS-CLIP | 23142080 | Unspecified |
| mmu-miR-34a-5p | MIMAT0000542 | Sh3bgrl | 56726 | HITS-CLIP | 23142080\|29167373 | Unspecified |
| mmu-miR-34a-5p | MIMAT0000542 | Sh3bgrl2 | 212531 | HITS-CLIP | 29167373 | Unspecified |
| mmu-miR-34a-5p | MIMAT0000542 | Sh3bgrl3 | 73723 | HITS-CLIP | 23142080 | Unspecified |
| mmu-miR-34a-5p | MIMAT0000542 | Sh3bp2 | 24055 | HITS-CLIP | 26203562\|23142080\|29167373 | Unspecified |
| mmu-miR-34a-5p | MIMAT0000542 | Sh3bp5 | 24056 | HITS-CLIP | 23142080\|29167373 | Unspecified |
| mmu-miR-34a-5p | MIMAT0000542 | Sh3bp5l | 79566 | HITS-CLIP | 23142080 | Unspecified |
| mmu-miR-34a-5p | MIMAT0000542 | Sh3d19 | 27059 | HITS-CLIP | 19536157 | Unspecified |
| mmu-miR-34a-5p | MIMAT0000542 | Sh3gl1 | 20405 | HITS-CLIP | 23142080\|29167373 | Unspecified |
| mmu-miR-34a-5p | MIMAT0000542 | Sh3glb1 | 54673 | HITS-CLIP | 23142080\|19536157\|29167373 | Unspecified |
| mmu-miR-34a-5p | MIMAT0000542 | Sh3glb2 | 56904 | HITS-CLIP | 23142080 | Unspecified |
| mmu-miR-34a-5p | MIMAT0000542 | Sh3kbp1 | 58194 | HITS-CLIP | 23142080\|29167373 | Unspecified |
| mmu-miR-34a-5p | MIMAT0000542 | Sh3pxd2a | 14218 | HITS-CLIP | 23142080\|29167373 | Unspecified |
| mmu-miR-34a-5p | MIMAT0000542 | Sh3pxd2b | 268396 | HITS-CLIP | 29167373 | Unspecified |
| mmu-miR-34a-5p | MIMAT0000542 | Sh3rf1 | 59009 | HITS-CLIP | 23142080 | Unspecified |
| mmu-miR-34a-5p | MIMAT0000542 | Sh3tc1 | 231147 | HITS-CLIP | 29167373 | Unspecified |
| mmu-miR-34a-5p | MIMAT0000542 | Shank1 | 243961 | HITS-CLIP | 19536157 | Unspecified |
| mmu-miR-34a-5p | MIMAT0000542 | Shank2 | 210274 | HITS-CLIP | 19536157 | Unspecified |
| mmu-miR-34a-5p | MIMAT0000542 | Shc1 | 20416 | HITS-CLIP | 23142080 | Unspecified |
| mmu-miR-34a-5p | MIMAT0000542 | Shc3 | 53358 | HITS-CLIP | 19536157 | Unspecified |
| mmu-miR-34a-5p | MIMAT0000542 | Shc4 | 271849 | HITS-CLIP | 23142080 | Unspecified |
| mmu-miR-34a-5p | MIMAT0000542 | Shisa5 | 66940 | HITS-CLIP | 23142080\|29167373 | Unspecified |
| mmu-miR-34a-5p | MIMAT0000542 | Shisa6 | 380702 | HITS-CLIP | 19536157 | Unspecified |

| | | | | | | |
|---|---|---|---|---|---|---|
| mmu-miR-34a-5p | MIMAT0000542 | Shkbp1 | 192192 | Microarrays//HITS-CLIP | 37986224\|23027903\|23142080 | Unspecified |
| mmu-miR-34a-5p | MIMAT0000542 | Shld1 | 149840 | HITS-CLIP | 29167373 | Unspecified |
| mmu-miR-34a-5p | MIMAT0000542 | Shmt2 | 108037 | HITS-CLIP | 23142080 | Unspecified |
| mmu-miR-34a-5p | MIMAT0000542 | Shoc2 | 56392 | HITS-CLIP | 23142080 | Unspecified |
| mmu-miR-34a-5p | MIMAT0000542 | Shprh | 268281 | HITS-CLIP | 31433990\|29167373\|23142080 | Unspecified |
| mmu-miR-34a-5p | MIMAT0000542 | Shq1 | 72171 | HITS-CLIP | 23142080 | Unspecified |
| mmu-miR-34a-5p | MIMAT0000542 | Shtn1 | 57698 | HITS-CLIP | 29167373 | Unspecified |
| mmu-miR-34a-5p | MIMAT0000542 | Siae | 22619 | HITS-CLIP | 23142080 | Unspecified |
| mmu-miR-34a-5p | MIMAT0000542 | Siah1a | 20437 | HITS-CLIP | 23142080 | Unspecified |
| mmu-miR-34a-5p | MIMAT0000542 | Siah2 | 20439 | HITS-CLIP | 23142080 | Unspecified |
| mmu-miR-34a-5p | MIMAT0000542 | Sidt1 | 320007 | HITS-CLIP | 19536157 | Unspecified |
| mmu-miR-34a-5p | MIMAT0000542 | Sidt2 | 214597 | Microarrays//HITS-CLIP | 37986224\|23027903\|23142080 | Unspecified |
| mmu-miR-34a-5p | MIMAT0000542 | Sigirr | 24058 | HITS-CLIP | 23142080 | Unspecified |
| mmu-miR-34a-5p | MIMAT0000542 | Siglec1 | 20612 | HITS-CLIP | 29167373 | Unspecified |
| mmu-miR-34a-5p | MIMAT0000542 | Sigmar1 | 18391 | HITS-CLIP | 19536157\|29167373\|23142080 | Unspecified |
| mmu-miR-34a-5p | MIMAT0000542 | Sik1 | 17691 | HITS-CLIP | 23142080\|19536157\|26203562\|29167 | Unspecified |
| mmu-miR-34a-5p | MIMAT0000542 | Sik2 | 235344 | HITS-CLIP | 23142080\|31433990\|30971389 | Unspecified |
| mmu-miR-34a-5p | MIMAT0000542 | Sik3 | 70661 | HITS-CLIP | 23142080\|29167373 | Unspecified |
| mmu-miR-34a-5p | MIMAT0000542 | Sike1 | 66641 | HITS-CLIP | 23142080\|29167373 | Unspecified |
| mmu-miR-34a-5p | MIMAT0000542 | Simc1 | 319719 | HITS-CLIP | 23142080\|29167373 | Unspecified |
| mmu-miR-34a-5p | MIMAT0000542 | Sin3a | 20466 | HITS-CLIP | 23142080\|19536157\|31433990 | Unspecified |
| mmu-miR-34a-5p | MIMAT0000542 | Sin3b | 20467 | HITS-CLIP | 23142080 | Unspecified |
| mmu-miR-34a-5p | MIMAT0000542 | Sipa1 | 20469 | Microarrays//HITS-CLIP | 37986224\|23027903\|23142080\|29167 | Unspecified |
| mmu-miR-34a-5p | MIMAT0000542 | Sipa1l1 | 217692 | HITS-CLIP | 23142080\|23630078 | Unspecified |
| mmu-miR-34a-5p | MIMAT0000542 | Sipa1l2 | 244668 | HITS-CLIP | 23142080\|29167373 | Unspecified |
| mmu-miR-34a-5p | MIMAT0000542 | Sipa1l3 | 74206 | HITS-CLIP | 29167373 | Unspecified |
| mmu-miR-34a-5p | MIMAT0000542 | Sirpa | 19261 | HITS-CLIP | 29167373 | Unspecified |
| mmu-miR-34a-5p | MIMAT0000542 | Sirt7 | 209011 | HITS-CLIP | 23142080 | Unspecified |
| mmu-miR-34a-5p | MIMAT0000542 | Siva1 | 30954 | HITS-CLIP | 31433990 | Unspecified |
| mmu-miR-34a-5p | MIMAT0000542 | Six5 | 20475 | HITS-CLIP | 23142080 | Unspecified |
| mmu-miR-34a-5p | MIMAT0000542 | Ska2 | 66140 | HITS-CLIP | 23142080\|29167373 | Unspecified |
| mmu-miR-34a-5p | MIMAT0000542 | Ska3 | 219114 | HITS-CLIP | 23142080 | Unspecified |
| mmu-miR-34a-5p | MIMAT0000542 | Skap1 | 78473 | HITS-CLIP | 23142080 | Unspecified |
| mmu-miR-34a-5p | MIMAT0000542 | Skap2 | 54353 | HITS-CLIP | 29167373 | Unspecified |
| mmu-miR-34a-5p | MIMAT0000542 | Ski | 20481 | HITS-CLIP | 19536157\|31433990\|26203562\|23142 | Unspecified |
| mmu-miR-34a-5p | MIMAT0000542 | Skil | 20482 | HITS-CLIP | 19536157\|29167373\|23142080 | Unspecified |
| mmu-miR-34a-5p | MIMAT0000542 | Skiv2l | 108077 | HITS-CLIP | 23142080 | Unspecified |
| mmu-miR-34a-5p | MIMAT0000542 | Skp1 | 6500 | HITS-CLIP | 31433990 | Unspecified |
| mmu-miR-34a-5p | MIMAT0000542 | Skp1a | 21402 | HITS-CLIP | 19536157\|23142080\|29167373 | Unspecified |

| | | | | | | |
|---|---|---|---|---|---|---|
| mmu-miR-34a-5p | MIMAT0000542 | Skp2 | 27401 | HITS-CLIP | 23630078\|23142080\|29167373 | Unspecified |
| mmu-miR-34a-5p | MIMAT0000542 | Sla | 20491 | HITS-CLIP | 23142080 | Unspecified |
| mmu-miR-34a-5p | MIMAT0000542 | Slain1 | 105439 | HITS-CLIP | 23142080 | Unspecified |
| mmu-miR-34a-5p | MIMAT0000542 | Slain2 | 75991 | HITS-CLIP | 23142080\|29167373 | Unspecified |
| mmu-miR-34a-5p | MIMAT0000542 | Slamf1 | 27218 | HITS-CLIP | 23142080 | Unspecified |
| mmu-miR-34a-5p | MIMAT0000542 | Slamf6 | 30925 | HITS-CLIP | 23142080\|31433990 | Unspecified |
| mmu-miR-34a-5p | MIMAT0000542 | Slamf7 | 75345 | HITS-CLIP | 23142080\|29167373 | Unspecified |
| mmu-miR-34a-5p | MIMAT0000542 | Slbp | 20492 | HITS-CLIP | 23142080 | Unspecified |
| mmu-miR-34a-5p | MIMAT0000542 | Slc10a3 | 214601 | HITS-CLIP | 29167373 | Unspecified |
| mmu-miR-34a-5p | MIMAT0000542 | Slc10a7 | 76775 | HITS-CLIP | 23142080 | Unspecified |
| mmu-miR-34a-5p | MIMAT0000542 | Slc11a2 | 18174 | HITS-CLIP | 23142080\|29167373 | Unspecified |
| mmu-miR-34a-5p | MIMAT0000542 | Slc12a2 | 20496 | HITS-CLIP | 19536157 | Unspecified |
| mmu-miR-34a-5p | MIMAT0000542 | Slc12a4 | 20498 | HITS-CLIP | 23142080 | Unspecified |
| mmu-miR-34a-5p | MIMAT0000542 | Slc12a5 | 57138 | HITS-CLIP | 19536157 | Unspecified |
| mmu-miR-34a-5p | MIMAT0000542 | Slc12a6 | 107723 | HITS-CLIP | 29167373\|23142080 | Unspecified |
| mmu-miR-34a-5p | MIMAT0000542 | Slc12a7 | 20499 | HITS-CLIP | 23142080 | Unspecified |
| mmu-miR-34a-5p | MIMAT0000542 | Slc12a9 | 83704 | HITS-CLIP | 23142080\|29167373 | Unspecified |
| mmu-miR-34a-5p | MIMAT0000542 | Slc13a3 | 114644 | HITS-CLIP | 23142080 | Unspecified |
| mmu-miR-34a-5p | MIMAT0000542 | Slc14a1 | 108052 | HITS-CLIP | 30971389 | Unspecified |
| mmu-miR-34a-5p | MIMAT0000542 | Slc15a2 | 57738 | HITS-CLIP | 23142080 | Unspecified |
| mmu-miR-34a-5p | MIMAT0000542 | Slc15a3 | 65221 | HITS-CLIP | 29167373 | Unspecified |
| mmu-miR-34a-5p | MIMAT0000542 | Slc15a4 | 100561 | HITS-CLIP | 29167373\|23142080 | Unspecified |
| mmu-miR-34a-5p | MIMAT0000542 | Slc16a1 | 20501 | HITS-CLIP | 19536157\|23142080 | Unspecified |
| mmu-miR-34a-5p | MIMAT0000542 | Slc16a10 | 72472 | HITS-CLIP | 29167373\|31433990\|23142080 | Unspecified |
| mmu-miR-34a-5p | MIMAT0000542 | Slc16a12 | 240638 | Microarrays | 37986224 | Unspecified |
| mmu-miR-34a-5p | MIMAT0000542 | Slc16a2 | 20502 | Microarrays | 37986224 | Unspecified |
| mmu-miR-34a-5p | MIMAT0000542 | Slc16a3 | 80879 | HITS-CLIP | 23142080\|31433990\|29167373 | Unspecified |
| mmu-miR-34a-5p | MIMAT0000542 | Slc16a6 | 104681 | HITS-CLIP | 23142080\|26203562\|29167373 | Unspecified |
| mmu-miR-34a-5p | MIMAT0000542 | Slc16a7 | 20503 | HITS-CLIP | 29167373 | Unspecified |
| mmu-miR-34a-5p | MIMAT0000542 | Slc17a5 | 235504 | HITS-CLIP | 29167373 | Unspecified |
| mmu-miR-34a-5p | MIMAT0000542 | Slc17a6 | 140919 | HITS-CLIP | 23142080 | Unspecified |
| mmu-miR-34a-5p | MIMAT0000542 | Slc17a7 | 72961 | HITS-CLIP | 19536157 | Unspecified |
| mmu-miR-34a-5p | MIMAT0000542 | Slc19a1 | 20509 | HITS-CLIP | 29167373 | Unspecified |
| mmu-miR-34a-5p | MIMAT0000542 | Slc19a2 | 116914 | Microarrays//HITS-CLIP | 37986224\|23142080\|29167373 | Unspecified |
| mmu-miR-34a-5p | MIMAT0000542 | Slc1a1 | 20510 | HITS-CLIP | 19536157 | Unspecified |
| mmu-miR-34a-5p | MIMAT0000542 | Slc1a2 | 20511 | HITS-CLIP | 19536157 | Unspecified |
| mmu-miR-34a-5p | MIMAT0000542 | Slc1a4 | 55963 | HITS-CLIP | 26203562 | Unspecified |
| mmu-miR-34a-5p | MIMAT0000542 | Slc1a5 | 20514 | HITS-CLIP | 23142080 | Unspecified |
| mmu-miR-34a-5p | MIMAT0000542 | Slc20a1 | 20515 | HITS-CLIP | 19536157\|29167373\|23142080 | Unspecified |

| | | | | | | |
|---|---|---|---|---|---|---|
| mmu-miR-34a-5p | MIMAT0000542 | Slc20a2 | 20516 | HITS-CLIP | 23142080\|29167373 | Unspecified |
| mmu-miR-34a-5p | MIMAT0000542 | Slc22a15 | 242126 | HITS-CLIP | 23142080 | Unspecified |
| mmu-miR-34a-5p | MIMAT0000542 | Slc22a21 | 56517 | HITS-CLIP | 23142080 | Unspecified |
| mmu-miR-34a-5p | MIMAT0000542 | Slc22a23 | 73102 | HITS-CLIP | 19536157\|26203562 | Unspecified |
| mmu-miR-34a-5p | MIMAT0000542 | Slc23a2 | 54338 | HITS-CLIP | 29167373\|23142080 | Unspecified |
| mmu-miR-34a-5p | MIMAT0000542 | Slc24a2 | 76376 | HITS-CLIP | 19536157 | Unspecified |
| mmu-miR-34a-5p | MIMAT0000542 | Slc25a1 | 13358 | HITS-CLIP | 23142080 | Unspecified |
| mmu-miR-34a-5p | MIMAT0000542 | Slc25a10 | 27376 | HITS-CLIP | 29167373 | Unspecified |
| mmu-miR-34a-5p | MIMAT0000542 | Slc25a11 | 67863 | HITS-CLIP | 23142080\|29167373 | Unspecified |
| mmu-miR-34a-5p | MIMAT0000542 | Slc25a13 | 50799 | HITS-CLIP | 31433990 | Unspecified |
| mmu-miR-34a-5p | MIMAT0000542 | Slc25a16 | 73132 | HITS-CLIP | 19536157\|31433990\|29167373 | Unspecified |
| mmu-miR-34a-5p | MIMAT0000542 | Slc25a17 | 20524 | HITS-CLIP | 23142080 | Unspecified |
| mmu-miR-34a-5p | MIMAT0000542 | Slc25a19 | 67283 | HITS-CLIP | 23142080 | Unspecified |
| mmu-miR-34a-5p | MIMAT0000542 | Slc25a20 | 57279 | HITS-CLIP | 29167373 | Unspecified |
| mmu-miR-34a-5p | MIMAT0000542 | Slc25a22 | 68267 | HITS-CLIP | 19536157\|23142080 | Unspecified |
| mmu-miR-34a-5p | MIMAT0000542 | Slc25a23 | 66972 | HITS-CLIP | 19536157\|23142080 | Unspecified |
| mmu-miR-34a-5p | MIMAT0000542 | Slc25a24 | 229731 | HITS-CLIP | 29167373\|23142080 | Unspecified |
| mmu-miR-34a-5p | MIMAT0000542 | Slc25a25 | 227731 | HITS-CLIP | 29167373\|23142080 | Unspecified |
| mmu-miR-34a-5p | MIMAT0000542 | Slc25a26 | 67582 | HITS-CLIP | 23142080\|29167373 | Unspecified |
| mmu-miR-34a-5p | MIMAT0000542 | Slc25a28 | 246696 | HITS-CLIP | 29167373 | Unspecified |
| mmu-miR-34a-5p | MIMAT0000542 | Slc25a3 | 18674 | HITS-CLIP | 37986224\|23142080 | Unspecified |
| mmu-miR-34a-5p | MIMAT0000542 | Slc25a30 | 67554 | HITS-CLIP | 23142080 | Unspecified |
| mmu-miR-34a-5p | MIMAT0000542 | Slc25a32 | 69906 | HITS-CLIP | 23142080\|19536157\|29167373 | Unspecified |
| mmu-miR-34a-5p | MIMAT0000542 | Slc25a36 | 192287 | Microarrays//HITS-CLIP | 37986224\|31433990\|23142080\|2916 | Unspecified |
| mmu-miR-34a-5p | MIMAT0000542 | Slc25a37 | 67712 | HITS-CLIP | 23142080 | Unspecified |
| mmu-miR-34a-5p | MIMAT0000542 | Slc25a4 | 11739 | HITS-CLIP | 23142080 | Unspecified |
| mmu-miR-34a-5p | MIMAT0000542 | Slc25a40 | 319653 | HITS-CLIP | 23142080 | Unspecified |
| mmu-miR-34a-5p | MIMAT0000542 | Slc25a43 | 194744 | HITS-CLIP | 29167373 | Unspecified |
| mmu-miR-34a-5p | MIMAT0000542 | Slc25a44 | 229517 | HITS-CLIP | 23142080\|29167373 | Unspecified |
| mmu-miR-34a-5p | MIMAT0000542 | Slc25a46 | 67453 | HITS-CLIP | 37986224\|23142080\|29167373 | Unspecified |
| mmu-miR-34a-5p | MIMAT0000542 | Slc25a5 | 11740 | HITS-CLIP | 23142080 | Unspecified |
| mmu-miR-34a-5p | MIMAT0000542 | Slc25a51 | 230125 | HITS-CLIP | 37986224\|23630078\|30971389\|2314 | Unspecified |
| mmu-miR-34a-5p | MIMAT0000542 | Slc26a2 | 13521 | HITS-CLIP | 23142080\|31433990\|29167373 | Unspecified |
| mmu-miR-34a-5p | MIMAT0000542 | Slc26a4 | 23985 | HITS-CLIP | 29167373 | Unspecified |
| mmu-miR-34a-5p | MIMAT0000542 | Slc27a1 | 26457 | HITS-CLIP | 29167373 | Unspecified |
| mmu-miR-34a-5p | MIMAT0000542 | Slc27a4 | 26569 | Microarrays//HITS-CLIP | 37986224\|23027903\|23142080 | Unspecified |
| mmu-miR-34a-5p | MIMAT0000542 | Slc29a1 | 63959 | HITS-CLIP | 23142080 | Unspecified |
| mmu-miR-34a-5p | MIMAT0000542 | Slc29a3 | 71279 | HITS-CLIP | 23142080\|29167373 | Unspecified |
| mmu-miR-34a-5p | MIMAT0000542 | Slc29a4 | 243328 | HITS-CLIP | 19536157 | Unspecified |

| | | | | | | |
|---|---|---|---|---|---|---|
| mmu-miR-34a-5p | MIMAT0000542 | Slc2a1 | 20525 | HITS-CLIP | 29167373\|23142080 | Unspecified |
| mmu-miR-34a-5p | MIMAT0000542 | Slc2a3 | 20527 | HITS-CLIP | 19536157\|23142080 | Unspecified |
| mmu-miR-34a-5p | MIMAT0000542 | Slc2a6 | 227659 | HITS-CLIP | 23142080 | Unspecified |
| mmu-miR-34a-5p | MIMAT0000542 | Slc30a1 | 22782 | HITS-CLIP | 19536157 | Unspecified |
| mmu-miR-34a-5p | MIMAT0000542 | Slc30a4 | 22785 | HITS-CLIP | 31433990 | Unspecified |
| mmu-miR-34a-5p | MIMAT0000542 | Slc30a5 | 69048 | HITS-CLIP | 31433990\|19536157\|23142080\|29167 | Unspecified |
| mmu-miR-34a-5p | MIMAT0000542 | Slc30a6 | 210148 | HITS-CLIP | 23142080 | Unspecified |
| mmu-miR-34a-5p | MIMAT0000542 | Slc30a7 | 66500 | HITS-CLIP | 23142080\|29167373 | Unspecified |
| mmu-miR-34a-5p | MIMAT0000542 | Slc30a9 | 109108 | HITS-CLIP | 23142080 | Unspecified |
| mmu-miR-34a-5p | MIMAT0000542 | Slc31a1 | 20529 | HITS-CLIP | 19536157\|29167373 | Unspecified |
| mmu-miR-34a-5p | MIMAT0000542 | Slc31a2 | 20530 | HITS-CLIP | 23142080\|29167373 | Unspecified |
| mmu-miR-34a-5p | MIMAT0000542 | Slc33a1 | 11416 | HITS-CLIP | 23142080 | Unspecified |
| mmu-miR-34a-5p | MIMAT0000542 | Slc35a1 | 24060 | HITS-CLIP | 23142080 | Unspecified |
| mmu-miR-34a-5p | MIMAT0000542 | Slc35a2 | 22232 | HITS-CLIP | 23142080 | Unspecified |
| mmu-miR-34a-5p | MIMAT0000542 | Slc35a3 | 229782 | HITS-CLIP | 29167373\|23142080 | Unspecified |
| mmu-miR-34a-5p | MIMAT0000542 | Slc35a4 | 67843 | Microarrays//HITS-CLIP | 37986224\|23027903\|23142080 | Unspecified |
| mmu-miR-34a-5p | MIMAT0000542 | Slc35a5 | 74102 | Microarrays//HITS-CLIP | 37986224\|23142080\|29167373 | Unspecified |
| mmu-miR-34a-5p | MIMAT0000542 | Slc35b1 | 110172 | HITS-CLIP | 23142080\|31433990 | Unspecified |
| mmu-miR-34a-5p | MIMAT0000542 | Slc35b2 | 73836 | HITS-CLIP | 26203562\|29167373 | Unspecified |
| mmu-miR-34a-5p | MIMAT0000542 | Slc35b3 | 108652 | HITS-CLIP | 23142080 | Unspecified |
| mmu-miR-34a-5p | MIMAT0000542 | Slc35b4 | 58246 | HITS-CLIP | 23142080\|29167373 | Unspecified |
| mmu-miR-34a-5p | MIMAT0000542 | Slc35c1 | 228368 | HITS-CLIP | 26203562\|29167373 | Unspecified |
| mmu-miR-34a-5p | MIMAT0000542 | Slc35c2 | 228875 | HITS-CLIP | 19536157 | Unspecified |
| mmu-miR-34a-5p | MIMAT0000542 | Slc35d2 | 70484 | HITS-CLIP | 23142080 | Unspecified |
| mmu-miR-34a-5p | MIMAT0000542 | Slc35d3 | 76157 | HITS-CLIP | 23142080 | Unspecified |
| mmu-miR-34a-5p | MIMAT0000542 | Slc35e1 | 270066 | HITS-CLIP | 23142080\|19536157\|31433990\|29167 | Unspecified |
| mmu-miR-34a-5p | MIMAT0000542 | Slc35e2 | 320541 | HITS-CLIP | 21258322\|23630078\|30971389\|23142 | Unspecified |
| mmu-miR-34a-5p | MIMAT0000542 | Slc35f6 | 74919 | HITS-CLIP | 29167373 | Unspecified |
| mmu-miR-34a-5p | MIMAT0000542 | Slc36a1 | 215335 | HITS-CLIP | 23142080\|26203562\|29167373 | Unspecified |
| mmu-miR-34a-5p | MIMAT0000542 | Slc36a4 | 234967 | HITS-CLIP | 29167373 | Unspecified |
| mmu-miR-34a-5p | MIMAT0000542 | Slc37a1 | 224674 | HITS-CLIP | 23142080 | Unspecified |
| mmu-miR-34a-5p | MIMAT0000542 | Slc37a2 | 56857 | HITS-CLIP | 29167373 | Unspecified |
| mmu-miR-34a-5p | MIMAT0000542 | Slc37a3 | 72144 | HITS-CLIP | 31433990\|23630078\|23142080\|29167 | Unspecified |
| mmu-miR-34a-5p | MIMAT0000542 | Slc37a4 | 14385 | HITS-CLIP | 23142080 | Unspecified |
| mmu-miR-34a-5p | MIMAT0000542 | Slc38a1 | 105727 | Microarrays//HITS-CLIP | 37986224\|23027903\|23142080\|29167 | Unspecified |
| mmu-miR-34a-5p | MIMAT0000542 | Slc38a10 | 72055 | HITS-CLIP | 29167373 | Unspecified |
| mmu-miR-34a-5p | MIMAT0000542 | Slc38a2 | 67760 | HITS-CLIP | 23142080\|30971389\|19536157\|29167 | Unspecified |
| mmu-miR-34a-5p | MIMAT0000542 | Slc38a6 | 625098 | Microarrays | 37986224 | Unspecified |
| mmu-miR-34a-5p | MIMAT0000542 | Slc38a7 | 234595 | HITS-CLIP | 29167373 | Unspecified |

| | | | | | | |
|---|---|---|---|---|---|---|
| mmu-miR-34a-5p | MIMAT0000542 | Slc38a9 | 268706 | HITS-CLIP | 23142080\|29167373 | Unspecified |
| mmu-miR-34a-5p | MIMAT0000542 | Slc39a1 | 30791 | HITS-CLIP | 23142080\|29167373 | Unspecified |
| mmu-miR-34a-5p | MIMAT0000542 | Slc39a10 | 227059 | HITS-CLIP | 23142080\|29167373 | Unspecified |
| mmu-miR-34a-5p | MIMAT0000542 | Slc39a13 | 68427 | HITS-CLIP | 19536157\|29167373 | Unspecified |
| mmu-miR-34a-5p | MIMAT0000542 | Slc39a14 | 213053 | HITS-CLIP | 23630078 | Unspecified |
| mmu-miR-34a-5p | MIMAT0000542 | Slc39a3 | 106947 | HITS-CLIP | 19536157\|23142080 | Unspecified |
| mmu-miR-34a-5p | MIMAT0000542 | Slc39a4 | 72027 | HITS-CLIP | 23142080 | Unspecified |
| mmu-miR-34a-5p | MIMAT0000542 | Slc39a6 | 106957 | HITS-CLIP | 23142080\|29167373 | Unspecified |
| mmu-miR-34a-5p | MIMAT0000542 | Slc39a7 | 14977 | HITS-CLIP | 29167373 | Unspecified |
| mmu-miR-34a-5p | MIMAT0000542 | Slc39a9 | 328133 | HITS-CLIP | 23142080\|29167373\|30971389 | Unspecified |
| mmu-miR-34a-5p | MIMAT0000542 | Slc3a2 | 17254 | HITS-CLIP | 23142080\|29167373 | Unspecified |
| mmu-miR-34a-5p | MIMAT0000542 | Slc41a1 | 98396 | HITS-CLIP | 37986224\|23142080 | Unspecified |
| mmu-miR-34a-5p | MIMAT0000542 | Slc41a2 | 338365 | HITS-CLIP | 23142080\|29167373 | Unspecified |
| mmu-miR-34a-5p | MIMAT0000542 | Slc43a2 | 215113 | HITS-CLIP | 23142080\|30971389\|29167373 | Unspecified |
| mmu-miR-34a-5p | MIMAT0000542 | Slc44a1 | 100434 | HITS-CLIP | 23142080\|29167373 | Unspecified |
| mmu-miR-34a-5p | MIMAT0000542 | Slc44a2 | 68682 | HITS-CLIP | 23142080\|29167373 | Unspecified |
| mmu-miR-34a-5p | MIMAT0000542 | Slc45a4 | 106068 | HITS-CLIP | 29167373 | Unspecified |
| mmu-miR-34a-5p | MIMAT0000542 | Slc46a3 | 71706 | HITS-CLIP | 23142080 | Unspecified |
| mmu-miR-34a-5p | MIMAT0000542 | Slc48a1 | 67739 | HITS-CLIP | 23142080\|29167373 | Unspecified |
| mmu-miR-34a-5p | MIMAT0000542 | Slc4a1ap | 20534 | HITS-CLIP | 30971389 | Unspecified |
| mmu-miR-34a-5p | MIMAT0000542 | Slc4a2 | 20535 | HITS-CLIP | 29167373 | Unspecified |
| mmu-miR-34a-5p | MIMAT0000542 | Slc4a4 | 54403 | HITS-CLIP | 19536157 | Unspecified |
| mmu-miR-34a-5p | MIMAT0000542 | Slc4a7 | 218756 | HITS-CLIP | 23142080\|29167373 | Unspecified |
| mmu-miR-34a-5p | MIMAT0000542 | Slc50a1 | 19729 | HITS-CLIP | 30971389\|23142080 | Unspecified |
| mmu-miR-34a-5p | MIMAT0000542 | Slc52a2 | 52710 | HITS-CLIP | 23142080 | Unspecified |
| mmu-miR-34a-5p | MIMAT0000542 | Slc5a3 | 53881 | HITS-CLIP | 31433990\|19536157\|23142080\|29167 | Unspecified |
| mmu-miR-34a-5p | MIMAT0000542 | Slc6a1 | 232333 | HITS-CLIP | 19536157 | Unspecified |
| mmu-miR-34a-5p | MIMAT0000542 | Slc6a12 | 14411 | HITS-CLIP | 29167373 | Unspecified |
| mmu-miR-34a-5p | MIMAT0000542 | Slc6a17 | 229706 | HITS-CLIP | 19536157 | Unspecified |
| mmu-miR-34a-5p | MIMAT0000542 | Slc6a6 | 21366 | HITS-CLIP | 19536157\|30971389\|23630078\|29167 | Unspecified |
| mmu-miR-34a-5p | MIMAT0000542 | Slc6a8 | 102857 | HITS-CLIP | 29167373 | Unspecified |
| mmu-miR-34a-5p | MIMAT0000542 | Slc7a1 | 11987 | Microarrays//HITS-CLIP | 37986224\|23027903\|30971389\|29167 | Unspecified |
| mmu-miR-34a-5p | MIMAT0000542 | Slc7a4 | 224022 | HITS-CLIP | 19536157 | Unspecified |
| mmu-miR-34a-5p | MIMAT0000542 | Slc7a5 | 20539 | HITS-CLIP | 19536157\|23630078\|23142080 | Unspecified |
| mmu-miR-34a-5p | MIMAT0000542 | Slc7a6 | 330836 | HITS-CLIP | 19536157\|23142080\|29167373 | Unspecified |
| mmu-miR-34a-5p | MIMAT0000542 | Slc7a7 | 20540 | HITS-CLIP | 29167373 | Unspecified |
| mmu-miR-34a-5p | MIMAT0000542 | Slc7a8 | 50934 | HITS-CLIP | 29167373 | Unspecified |
| mmu-miR-34a-5p | MIMAT0000542 | Slc8a1 | 20541 | HITS-CLIP | 29167373\|19536157 | Unspecified |
| mmu-miR-34a-5p | MIMAT0000542 | Slc8a2 | 110891 | HITS-CLIP | 19536157 | Unspecified |

| | | | | | | |
|---|---|---|---|---|---|---|
| mmu-miR-34a-5p | MIMAT0000542 | Slc8b1 | 170756 | HITS-CLIP | 29167373 | Unspecified |
| mmu-miR-34a-5p | MIMAT0000542 | Slc9a1 | 20544 | HITS-CLIP | 23142080\|29167373 | Unspecified |
| mmu-miR-34a-5p | MIMAT0000542 | Slc9a3r1 | 26941 | HITS-CLIP | 23142080 | Unspecified |
| mmu-miR-34a-5p | MIMAT0000542 | Slc9a6 | 236794 | HITS-CLIP | 23142080 | Unspecified |
| mmu-miR-34a-5p | MIMAT0000542 | Slc9a8 | 77031 | HITS-CLIP | 23142080\|29167373 | Unspecified |
| mmu-miR-34a-5p | MIMAT0000542 | Slco1a5 | 108096 | Microarrays | 37986224 | Unspecified |
| mmu-miR-34a-5p | MIMAT0000542 | Slco1c1 | 58807 | HITS-CLIP | 19536157 | Unspecified |
| mmu-miR-34a-5p | MIMAT0000542 | Slco3a1 | 108116 | HITS-CLIP | 31433990 | Unspecified |
| mmu-miR-34a-5p | MIMAT0000542 | Slco4a1 | 108115 | HITS-CLIP | 23142080\|29167373 | Unspecified |
| mmu-miR-34a-5p | MIMAT0000542 | Slf1 | 852127 | HITS-CLIP | 29167373 | Unspecified |
| mmu-miR-34a-5p | MIMAT0000542 | Slf2 | 226151 | HITS-CLIP | 23142080 | Unspecified |
| mmu-miR-34a-5p | MIMAT0000542 | Slfn1 | 20555 | HITS-CLIP | 23142080 | Unspecified |
| mmu-miR-34a-5p | MIMAT0000542 | Slfn2 | 20556 | HITS-CLIP | 29167373\|23142080 | Unspecified |
| mmu-miR-34a-5p | MIMAT0000542 | Slfn3 | 20557 | HITS-CLIP | 23142080 | Unspecified |
| mmu-miR-34a-5p | MIMAT0000542 | Slfn5 | 327978 | HITS-CLIP | 29167373 | Unspecified |
| mmu-miR-34a-5p | MIMAT0000542 | Slit3 | 20564 | HITS-CLIP | 19536157 | Unspecified |
| mmu-miR-34a-5p | MIMAT0000542 | Slitrk2 | 245450 | HITS-CLIP | 19536157 | Unspecified |
| mmu-miR-34a-5p | MIMAT0000542 | Slitrk3 | 386750 | HITS-CLIP | 19536157 | Unspecified |
| mmu-miR-34a-5p | MIMAT0000542 | Slitrk4 | 245446 | HITS-CLIP | 19536157 | Unspecified |
| mmu-miR-34a-5p | MIMAT0000542 | Slk | 20874 | HITS-CLIP | 23142080\|29167373 | Unspecified |
| mmu-miR-34a-5p | MIMAT0000542 | Slmap | 83997 | HITS-CLIP | 19536157\|23142080\|29167373 | Unspecified |
| mmu-miR-34a-5p | MIMAT0000542 | Slpi | 20568 | HITS-CLIP | 23142080 | Unspecified |
| mmu-miR-34a-5p | MIMAT0000542 | Slu7 | 193116 | HITS-CLIP | 23142080 | Unspecified |
| mmu-miR-34a-5p | MIMAT0000542 | Slx1b | 75764 | HITS-CLIP | 23142080 | Unspecified |
| mmu-miR-34a-5p | MIMAT0000542 | Slx4 | 52864 | HITS-CLIP | 31433990\|29167373 | Unspecified |
| mmu-miR-34a-5p | MIMAT0000542 | Slx4ip | 74243 | HITS-CLIP | 23142080 | Unspecified |
| mmu-miR-34a-5p | MIMAT0000542 | Smad2 | 17126 | HITS-CLIP | 23142080\|31433990\|30971389\|29167 | Unspecified |
| mmu-miR-34a-5p | MIMAT0000542 | Smad3 | 17127 | HITS-CLIP | 23142080\|29167373 | Unspecified |
| mmu-miR-34a-5p | MIMAT0000542 | Smad4 | 17128 | HITS-CLIP | 23142080\|29167373 | Unspecified |
| mmu-miR-34a-5p | MIMAT0000542 | Smad5 | 17129 | HITS-CLIP | 19536157\|23142080 | Unspecified |
| mmu-miR-34a-5p | MIMAT0000542 | Smad6 | 17130 | HITS-CLIP | 29167373 | Unspecified |
| mmu-miR-34a-5p | MIMAT0000542 | Smad7 | 17131 | HITS-CLIP | 23142080 | Unspecified |
| mmu-miR-34a-5p | MIMAT0000542 | Smagp | 207818 | HITS-CLIP | 23142080 | Unspecified |
| mmu-miR-34a-5p | MIMAT0000542 | Smap1 | 98366 | HITS-CLIP | 23142080 | Unspecified |
| mmu-miR-34a-5p | MIMAT0000542 | Smap2 | 69780 | HITS-CLIP | 31433990\|23142080 | Unspecified |
| mmu-miR-34a-5p | MIMAT0000542 | Smarca2 | 67155 | HITS-CLIP | 23142080\|29167373 | Unspecified |
| mmu-miR-34a-5p | MIMAT0000542 | Smarca4 | 20586 | HITS-CLIP | 23630078\|23142080\|29167373 | Unspecified |
| mmu-miR-34a-5p | MIMAT0000542 | Smarca5 | 93762 | HITS-CLIP | 23142080\|29167373 | Unspecified |
| mmu-miR-34a-5p | MIMAT0000542 | Smarcad1 | 13990 | HITS-CLIP | 23142080 | Unspecified |

| | | | | | | |
|---|---|---|---|---|---|---|
| mmu-miR-34a-5p | MIMAT0000542 | Smarcal1 | 54380 | HITS-CLIP | 19536157\|23142080 | Unspecified |
| mmu-miR-34a-5p | MIMAT0000542 | Smarcb1 | 20587 | HITS-CLIP | 23142080 | Unspecified |
| mmu-miR-34a-5p | MIMAT0000542 | Smarcc1 | 20588 | HITS-CLIP | 23142080\|31433990\|29167373 | Unspecified |
| mmu-miR-34a-5p | MIMAT0000542 | Smarcc2 | 68094 | HITS-CLIP | 23142080\|29167373 | Unspecified |
| mmu-miR-34a-5p | MIMAT0000542 | Smarcd1 | 83797 | HITS-CLIP | 30971389\|29167373 | Unspecified |
| mmu-miR-34a-5p | MIMAT0000542 | Smarcd2 | 83796 | HITS-CLIP | 23142080 | Unspecified |
| mmu-miR-34a-5p | MIMAT0000542 | Smc1a | 24061 | HITS-CLIP | 23142080\|29167373 | Unspecified |
| mmu-miR-34a-5p | MIMAT0000542 | Smc2 | 14211 | HITS-CLIP | 31433990\|19536157\|23142080\|29167 | Unspecified |
| mmu-miR-34a-5p | MIMAT0000542 | Smc3 | 13006 | HITS-CLIP | 23142080\|30971389 | Unspecified |
| mmu-miR-34a-5p | MIMAT0000542 | Smc4 | 70099 | HITS-CLIP | 23142080\|29167373 | Unspecified |
| mmu-miR-34a-5p | MIMAT0000542 | Smc5 | 226026 | HITS-CLIP | 23142080\|29167373 | Unspecified |
| mmu-miR-34a-5p | MIMAT0000542 | Smc6 | 67241 | HITS-CLIP | 31433990\|23142080\|29167373 | Unspecified |
| mmu-miR-34a-5p | MIMAT0000542 | Smchd1 | 74355 | HITS-CLIP | 23142080\|26203562\|29167373 | Unspecified |
| mmu-miR-34a-5p | MIMAT0000542 | Smco4 | 170748 | HITS-CLIP | 23142080 | Unspecified |
| mmu-miR-34a-5p | MIMAT0000542 | Smcr8 | 237782 | Microarrays//HITS-CLIP | 37986224\|23027903\|29167373\|23142 | Unspecified |
| mmu-miR-34a-5p | MIMAT0000542 | Smg1 | 233789 | HITS-CLIP | 19536157\|31433990\|23142080\|29167 | Unspecified |
| mmu-miR-34a-5p | MIMAT0000542 | Smg6 | 103677 | HITS-CLIP | 23142080 | Unspecified |
| mmu-miR-34a-5p | MIMAT0000542 | Smg7 | 226517 | HITS-CLIP | 23142080\|29167373 | Unspecified |
| mmu-miR-34a-5p | MIMAT0000542 | Smg8 | 74133 | HITS-CLIP | 23142080\|29167373 | Unspecified |
| mmu-miR-34a-5p | MIMAT0000542 | Smg9 | 71997 | HITS-CLIP | 29167373 | Unspecified |
| mmu-miR-34a-5p | MIMAT0000542 | Smim1 | 68859 | HITS-CLIP | 30971389 | Unspecified |
| mmu-miR-34a-5p | MIMAT0000542 | Smim10l1 | 100129361 | HITS-CLIP | 19536157 | Unspecified |
| mmu-miR-34a-5p | MIMAT0000542 | Smim12 | 80284 | HITS-CLIP | 23142080 | Unspecified |
| mmu-miR-34a-5p | MIMAT0000542 | Smim13 | 108934 | HITS-CLIP | 19536157 | Unspecified |
| mmu-miR-34a-5p | MIMAT0000542 | Smim14 | 68552 | HITS-CLIP | 23142080 | Unspecified |
| mmu-miR-34a-5p | MIMAT0000542 | Smim15 | 75616 | HITS-CLIP | 23142080\|29167373 | Unspecified |
| mmu-miR-34a-5p | MIMAT0000542 | Smim20 | 66278 | HITS-CLIP | 23142080 | Unspecified |
| mmu-miR-34a-5p | MIMAT0000542 | Smim26 | 388789 | HITS-CLIP | 23142080 | Unspecified |
| mmu-miR-34a-5p | MIMAT0000542 | Smim5 | 66528 | HITS-CLIP | 30971389 | Unspecified |
| mmu-miR-34a-5p | MIMAT0000542 | Smim7 | 66818 | HITS-CLIP | 23142080\|29167373 | Unspecified |
| mmu-miR-34a-5p | MIMAT0000542 | Smim8 | 66291 | HITS-CLIP | 23142080 | Unspecified |
| mmu-miR-34a-5p | MIMAT0000542 | Smox | 228608 | HITS-CLIP | 29167373 | Unspecified |
| mmu-miR-34a-5p | MIMAT0000542 | Smpd1 | 20597 | HITS-CLIP | 19536157\|23142080 | Unspecified |
| mmu-miR-34a-5p | MIMAT0000542 | Smpd4 | 77626 | HITS-CLIP | 23142080 | Unspecified |
| mmu-miR-34a-5p | MIMAT0000542 | Smpdl3a | 57319 | Microarrays//HITS-CLIP | 37986224\|23142080\|29167373 | Unspecified |
| mmu-miR-34a-5p | MIMAT0000542 | Smpdl3b | 100340 | Microarrays | 37986224\|23027903 | Unspecified |
| mmu-miR-34a-5p | MIMAT0000542 | Smtnl2 | 276829 | HITS-CLIP | 23142080 | Unspecified |
| mmu-miR-34a-5p | MIMAT0000542 | Smu1 | 74255 | HITS-CLIP | 23630078\|23142080 | Unspecified |
| mmu-miR-34a-5p | MIMAT0000542 | Smug1 | 71726 | HITS-CLIP | 23142080 | Unspecified |

| | | | | | | |
|---|---|---|---|---|---|---|
| mmu-miR-34a-5p | MIMAT0000542 | Smurf1 | 75788 | HITS-CLIP | 23630078\|29167373 | Unspecified |
| mmu-miR-34a-5p | MIMAT0000542 | Smurf2 | 66313 | HITS-CLIP | 19536157\|31433990\|23142080\|29167 | Unspecified |
| mmu-miR-34a-5p | MIMAT0000542 | Smyd1 | 12180 | Microarrays | 37986224 | Unspecified |
| mmu-miR-34a-5p | MIMAT0000542 | Smyd3 | 69726 | HITS-CLIP | 23142080 | Unspecified |
| mmu-miR-34a-5p | MIMAT0000542 | Smyd5 | 232187 | HITS-CLIP | 29167373\|23142080 | Unspecified |
| mmu-miR-34a-5p | MIMAT0000542 | Snap23 | 20619 | HITS-CLIP | 23142080\|29167373 | Unspecified |
| mmu-miR-34a-5p | MIMAT0000542 | Snap29 | 67474 | HITS-CLIP | 23142080\|29167373 | Unspecified |
| mmu-miR-34a-5p | MIMAT0000542 | Snap47 | 67826 | HITS-CLIP | 19536157\|23142080\|29167373 | Unspecified |
| mmu-miR-34a-5p | MIMAT0000542 | Snapc1 | 75627 | HITS-CLIP | 23142080 | Unspecified |
| mmu-miR-34a-5p | MIMAT0000542 | Snapc2 | 102209 | HITS-CLIP | 23142080 | Unspecified |
| mmu-miR-34a-5p | MIMAT0000542 | Snapc5 | 330959 | HITS-CLIP | 23142080 | Unspecified |
| mmu-miR-34a-5p | MIMAT0000542 | Sncb | 104069 | HITS-CLIP | 23142080 | Unspecified |
| mmu-miR-34a-5p | MIMAT0000542 | Snd1 | 56463 | HITS-CLIP | 23142080\|30971389\|29167373 | Unspecified |
| mmu-miR-34a-5p | MIMAT0000542 | Snhg11 | 319317 | HITS-CLIP | 19536157 | Unspecified |
| mmu-miR-34a-5p | MIMAT0000542 | Snn | 20621 | HITS-CLIP | 19536157\|23142080 | Unspecified |
| mmu-miR-34a-5p | MIMAT0000542 | Snph | 241727 | HITS-CLIP | 19536157 | Unspecified |
| mmu-miR-34a-5p | MIMAT0000542 | Snrk | 20623 | HITS-CLIP | 23142080 | Unspecified |
| mmu-miR-34a-5p | MIMAT0000542 | Snrnp200 | 320632 | HITS-CLIP | 19536157\|23142080\|29167373 | Unspecified |
| mmu-miR-34a-5p | MIMAT0000542 | Snrnp35 | 76167 | HITS-CLIP | 30971389 | Unspecified |
| mmu-miR-34a-5p | MIMAT0000542 | Snrnp40 | 66585 | HITS-CLIP | 23142080 | Unspecified |
| mmu-miR-34a-5p | MIMAT0000542 | Snrnp70 | 20637 | HITS-CLIP | 23630078\|23142080\|29167373 | Unspecified |
| mmu-miR-34a-5p | MIMAT0000542 | Snrpa | 53607 | HITS-CLIP | 23142080 | Unspecified |
| mmu-miR-34a-5p | MIMAT0000542 | Snrpa1 | 68981 | HITS-CLIP | 23142080 | Unspecified |
| mmu-miR-34a-5p | MIMAT0000542 | Snrpb | 20638 | Microarrays//HITS-CLIP | 37986224\|23027903\|23142080 | Unspecified |
| mmu-miR-34a-5p | MIMAT0000542 | Sntb1 | 20649 | Microarrays//HITS-CLIP | 37986224\|23142080 | Unspecified |
| mmu-miR-34a-5p | MIMAT0000542 | Sntb2 | 20650 | HITS-CLIP | 23142080\|29167373\|30971389 | Unspecified |
| mmu-miR-34a-5p | MIMAT0000542 | Snu13 | 4809 | HITS-CLIP | 23142080 | Unspecified |
| mmu-miR-34a-5p | MIMAT0000542 | Snw1 | 66354 | HITS-CLIP | 23142080 | Unspecified |
| mmu-miR-34a-5p | MIMAT0000542 | Snx1 | 56440 | HITS-CLIP | 23142080\|29167373 | Unspecified |
| mmu-miR-34a-5p | MIMAT0000542 | Snx10 | 71982 | HITS-CLIP | 23630078\|23142080 | Unspecified |
| mmu-miR-34a-5p | MIMAT0000542 | Snx12 | 55988 | Microarrays//HITS-CLIP | 37986224\|23027903\|23142080\|29167 | Unspecified |
| mmu-miR-34a-5p | MIMAT0000542 | Snx13 | 217463 | HITS-CLIP | 23142080 | Unspecified |
| mmu-miR-34a-5p | MIMAT0000542 | Snx15 | 69024 | Microarrays//HITS-CLIP | 37986224\|23027903\|23142080 | Unspecified |
| mmu-miR-34a-5p | MIMAT0000542 | Snx16 | 74718 | HITS-CLIP | 23142080\|29167373 | Unspecified |
| mmu-miR-34a-5p | MIMAT0000542 | Snx17 | 266781 | HITS-CLIP | 23142080\|29167373 | Unspecified |
| mmu-miR-34a-5p | MIMAT0000542 | Snx18 | 170625 | HITS-CLIP | 37986224\|19536157\|23142080\|29167 | Unspecified |
| mmu-miR-34a-5p | MIMAT0000542 | Snx19 | 102607 | HITS-CLIP | 19536157 | Unspecified |
| mmu-miR-34a-5p | MIMAT0000542 | Snx2 | 67804 | HITS-CLIP | 29167373\|23142080 | Unspecified |
| mmu-miR-34a-5p | MIMAT0000542 | Snx20 | 71607 | HITS-CLIP | 23142080 | Unspecified |

| | | | | | | |
|---|---|---|---|---|---|---|
| mmu-miR-34a-5p | MIMAT0000542 | Snx24 | 69226 | HITS-CLIP | 30971389\|29167373 | Unspecified |
| mmu-miR-34a-5p | MIMAT0000542 | Snx25 | 102141 | HITS-CLIP | 23142080 | Unspecified |
| mmu-miR-34a-5p | MIMAT0000542 | Snx27 | 76742 | HITS-CLIP | 23630078\|30971389\|29167373\|23142 | Unspecified |
| mmu-miR-34a-5p | MIMAT0000542 | Snx29 | 74478 | HITS-CLIP | 29167373 | Unspecified |
| mmu-miR-34a-5p | MIMAT0000542 | Snx3 | 54198 | HITS-CLIP | 19536157\|23142080 | Unspecified |
| mmu-miR-34a-5p | MIMAT0000542 | Snx30 | 209131 | HITS-CLIP | 23142080\|29167373 | Unspecified |
| mmu-miR-34a-5p | MIMAT0000542 | Snx32 | 225861 | HITS-CLIP | 29167373 | Unspecified |
| mmu-miR-34a-5p | MIMAT0000542 | Snx4 | 69150 | HITS-CLIP | 31433990\|23142080 | Unspecified |
| mmu-miR-34a-5p | MIMAT0000542 | Snx5 | 69178 | HITS-CLIP | 23142080\|29167373 | Unspecified |
| mmu-miR-34a-5p | MIMAT0000542 | Snx6 | 72183 | HITS-CLIP | 23142080\|29167373 | Unspecified |
| mmu-miR-34a-5p | MIMAT0000542 | Snx7 | 76561 | Microarrays | 37986224 | Unspecified |
| mmu-miR-34a-5p | MIMAT0000542 | Snx8 | 231834 | Microarrays//HITS-CLIP | 37986224\|23027903\|23142080\|29167 | Unspecified |
| mmu-miR-34a-5p | MIMAT0000542 | Snx9 | 66616 | HITS-CLIP | 23142080\|29167373 | Unspecified |
| mmu-miR-34a-5p | MIMAT0000542 | Soat1 | 20652 | HITS-CLIP | 23142080\|26203562\|29167373 | Unspecified |
| mmu-miR-34a-5p | MIMAT0000542 | Socs1 | 12703 | HITS-CLIP | 23142080\|31433990 | Unspecified |
| mmu-miR-34a-5p | MIMAT0000542 | Socs2 | 216233 | HITS-CLIP | 23142080 | Unspecified |
| mmu-miR-34a-5p | MIMAT0000542 | Socs3 | 12702 | HITS-CLIP | 31433990\|26203562\|23142080 | Unspecified |
| mmu-miR-34a-5p | MIMAT0000542 | Socs4 | 67296 | Microarrays//HITS-CLIP | 37986224\|23027903\|31433990\|29167 | Unspecified |
| mmu-miR-34a-5p | MIMAT0000542 | Socs5 | 56468 | HITS-CLIP | 19536157\|29167373\|23142080 | Unspecified |
| mmu-miR-34a-5p | MIMAT0000542 | Socs6 | 54607 | HITS-CLIP | 37986224\|23142080\|29167373 | Unspecified |
| mmu-miR-34a-5p | MIMAT0000542 | Socs7 | 192157 | HITS-CLIP | 23142080 | Unspecified |
| mmu-miR-34a-5p | MIMAT0000542 | Sod2 | 20656 | HITS-CLIP | 19536157\|23142080 | Unspecified |
| mmu-miR-34a-5p | MIMAT0000542 | Soga1 | 320706 | HITS-CLIP | 19536157\|29167373 | Unspecified |
| mmu-miR-34a-5p | MIMAT0000542 | Son | 20658 | HITS-CLIP | 19536157\|23142080\|29167373 | Unspecified |
| mmu-miR-34a-5p | MIMAT0000542 | Sorbs2 | 234214 | HITS-CLIP | 19536157 | Unspecified |
| mmu-miR-34a-5p | MIMAT0000542 | Sorl1 | 20660 | HITS-CLIP | 19536157 | Unspecified |
| mmu-miR-34a-5p | MIMAT0000542 | Sort1 | 20661 | Microarrays | 37986224 | Unspecified |
| mmu-miR-34a-5p | MIMAT0000542 | Sos1 | 20662 | HITS-CLIP | 31433990\|23142080\|29167373 | Unspecified |
| mmu-miR-34a-5p | MIMAT0000542 | Sos2 | 20663 | HITS-CLIP | 23142080\|29167373 | Unspecified |
| mmu-miR-34a-5p | MIMAT0000542 | Sox12 | 20667 | HITS-CLIP | 23142080 | Unspecified |
| mmu-miR-34a-5p | MIMAT0000542 | Sox2 | 20674 | Luciferase Reporter Assay, qPCR, Western | 37986224\|22020437 | Unspecified |
| mmu-miR-34a-5p | MIMAT0000542 | Sox4 | 20677 | HITS-CLIP | 23142080 | Unspecified |
| mmu-miR-34a-5p | MIMAT0000542 | Sox5 | 20678 | HITS-CLIP | 23142080 | Unspecified |
| mmu-miR-34a-5p | MIMAT0000542 | Sox6 | 20679 | HITS-CLIP | 23142080 | Unspecified |
| mmu-miR-34a-5p | MIMAT0000542 | Sp1 | 20683 | HITS-CLIP | 23142080\|19536157\|26203562\|29167 | Unspecified |
| mmu-miR-34a-5p | MIMAT0000542 | Sp100 | 20684 | HITS-CLIP | 23142080\|29167373 | Unspecified |
| mmu-miR-34a-5p | MIMAT0000542 | Sp2 | 78912 | HITS-CLIP | 23142080 | Unspecified |
| mmu-miR-34a-5p | MIMAT0000542 | Sp3 | 20687 | HITS-CLIP | 23142080\|29167373 | Unspecified |
| mmu-miR-34a-5p | MIMAT0000542 | Sp4 | 20688 | HITS-CLIP | 23142080 | Unspecified |

| | | | | | | | |
|---|---|---|---|---|---|---|---|
| mmu-miR-34a-5p | MIMAT0000542 | Spaca6 | 75202 | HITS-CLIP | | 29167373 | Unspecified |
| mmu-miR-34a-5p | MIMAT0000542 | Spag5 | 54141 | HITS-CLIP | | 23142080 | Unspecified |
| mmu-miR-34a-5p | MIMAT0000542 | Spag7 | 216873 | HITS-CLIP | 23142080\|29167373 | | Unspecified |
| mmu-miR-34a-5p | MIMAT0000542 | Spag9 | 70834 | HITS-CLIP | 19536157\|26203562\|29167373\|23142 | | Unspecified |
| mmu-miR-34a-5p | MIMAT0000542 | Sparc | 20692 | HITS-CLIP | | 19536157 | Unspecified |
| mmu-miR-34a-5p | MIMAT0000542 | Sparcl1 | 13602 | HITS-CLIP | | 19536157 | Unspecified |
| mmu-miR-34a-5p | MIMAT0000542 | Spast | 50850 | HITS-CLIP | 23142080\|29167373 | | Unspecified |
| mmu-miR-34a-5p | MIMAT0000542 | Spata13 | 219140 | HITS-CLIP | 23142080\|19536157\|29167373 | | Unspecified |
| mmu-miR-34a-5p | MIMAT0000542 | Spata2 | 263876 | HITS-CLIP | | 23142080 | Unspecified |
| mmu-miR-34a-5p | MIMAT0000542 | Spata24 | 71242 | HITS-CLIP | | 30971389 | Unspecified |
| mmu-miR-34a-5p | MIMAT0000542 | Spata2l | 78779 | HITS-CLIP | | 23142080 | Unspecified |
| mmu-miR-34a-5p | MIMAT0000542 | Spata6 | 67946 | HITS-CLIP | | 31433990 | Unspecified |
| mmu-miR-34a-5p | MIMAT0000542 | Spc25 | 66442 | HITS-CLIP | | 23142080 | Unspecified |
| mmu-miR-34a-5p | MIMAT0000542 | Spcs1 | 69019 | HITS-CLIP | | 23142080 | Unspecified |
| mmu-miR-34a-5p | MIMAT0000542 | Spcs2 | 66624 | HITS-CLIP | | 23142080 | Unspecified |
| mmu-miR-34a-5p | MIMAT0000542 | Spcs3 | 76687 | HITS-CLIP | 19536157\|23142080\|29167373 | | Unspecified |
| mmu-miR-34a-5p | MIMAT0000542 | Specc1 | 432572 | HITS-CLIP | 23142080\|29167373 | | Unspecified |
| mmu-miR-34a-5p | MIMAT0000542 | Specc1l | 74392 | HITS-CLIP | 23142080\|29167373 | | Unspecified |
| mmu-miR-34a-5p | MIMAT0000542 | Speg | 11790 | HITS-CLIP | | 29167373 | Unspecified |
| mmu-miR-34a-5p | MIMAT0000542 | Spen | 56381 | HITS-CLIP | 31433990\|23142080\|29167373 | | Unspecified |
| mmu-miR-34a-5p | MIMAT0000542 | Spg11 | 214585 | HITS-CLIP | 29167373\|23142080 | | Unspecified |
| mmu-miR-34a-5p | MIMAT0000542 | Spg20 | 229285 | HITS-CLIP | 23630078\|23142080\|29167373 | | Unspecified |
| mmu-miR-34a-5p | MIMAT0000542 | Spg7 | 234847 | HITS-CLIP | | 29167373 | Unspecified |
| mmu-miR-34a-5p | MIMAT0000542 | Sphk1 | 20698 | HITS-CLIP | | 30971389 | Unspecified |
| mmu-miR-34a-5p | MIMAT0000542 | Sphk2 | 56632 | HITS-CLIP | 23142080\|29167373 | | Unspecified |
| mmu-miR-34a-5p | MIMAT0000542 | Spi1 | 20375 | HITS-CLIP | | 29167373 | Unspecified |
| mmu-miR-34a-5p | MIMAT0000542 | Spib | 272382 | HITS-CLIP | | 23142080 | Unspecified |
| mmu-miR-34a-5p | MIMAT0000542 | Spice1 | 212514 | HITS-CLIP | | 23142080 | Unspecified |
| mmu-miR-34a-5p | MIMAT0000542 | Spidr | 224008 | HITS-CLIP | | 23142080 | Unspecified |
| mmu-miR-34a-5p | MIMAT0000542 | Spin4 | 270624 | HITS-CLIP | | 23142080 | Unspecified |
| mmu-miR-34a-5p | MIMAT0000542 | Spint2 | 20733 | HITS-CLIP | | 23142080 | Unspecified |
| mmu-miR-34a-5p | MIMAT0000542 | Spire1 | 68166 | HITS-CLIP | | 29167373 | Unspecified |
| mmu-miR-34a-5p | MIMAT0000542 | Spn | 20737 | HITS-CLIP | | 23142080 | Unspecified |
| mmu-miR-34a-5p | MIMAT0000542 | Spns1 | 73658 | HITS-CLIP | 31433990\|23630078\|30971389\|29167 | | Unspecified |
| mmu-miR-34a-5p | MIMAT0000542 | Spock1 | 20745 | HITS-CLIP | | 19536157 | Unspecified |
| mmu-miR-34a-5p | MIMAT0000542 | Spock2 | 94214 | HITS-CLIP | 19536157\|23142080 | | Unspecified |
| mmu-miR-34a-5p | MIMAT0000542 | Spop | 20747 | HITS-CLIP | | 29167373 | Unspecified |
| mmu-miR-34a-5p | MIMAT0000542 | Spopl | 76857 | HITS-CLIP | | 23142080 | Unspecified |
| mmu-miR-34a-5p | MIMAT0000542 | Spp1 | 20750 | HITS-CLIP | 23142080\|29167373 | | Unspecified |

| | | | | | | |
|---|---|---|---|---|---|---|
| mmu-miR-34a-5p | MIMAT0000542 | Sppl2a | 66552 | HITS-CLIP | 23630078\|29167373\|23142080 | Unspecified |
| mmu-miR-34a-5p | MIMAT0000542 | Sppl3 | 74585 | HITS-CLIP | 23142080\|29167373 | Unspecified |
| mmu-miR-34a-5p | MIMAT0000542 | Spred1 | 114715 | HITS-CLIP | 31433990\|23142080\|29167373 | Unspecified |
| mmu-miR-34a-5p | MIMAT0000542 | Spred2 | 114716 | HITS-CLIP | 23142080 | Unspecified |
| mmu-miR-34a-5p | MIMAT0000542 | Sprtn | 244666 | HITS-CLIP | 23142080 | Unspecified |
| mmu-miR-34a-5p | MIMAT0000542 | Spry1 | 24063 | HITS-CLIP | 23142080 | Unspecified |
| mmu-miR-34a-5p | MIMAT0000542 | Spry2 | 24064 | HITS-CLIP | 23142080\|29167373 | Unspecified |
| mmu-miR-34a-5p | MIMAT0000542 | Spry4 | 24066 | HITS-CLIP | 19536157 | Unspecified |
| mmu-miR-34a-5p | MIMAT0000542 | Spryd3 | 223918 | HITS-CLIP | 23142080 | Unspecified |
| mmu-miR-34a-5p | MIMAT0000542 | Spryd4 | 66701 | HITS-CLIP | 23142080 | Unspecified |
| mmu-miR-34a-5p | MIMAT0000542 | Spsb1 | 74646 | Microarrays//HITS-CLIP | 37986224\|23142080 | Unspecified |
| mmu-miR-34a-5p | MIMAT0000542 | Spta1 | 20739 | HITS-CLIP | 30971389 | Unspecified |
| mmu-miR-34a-5p | MIMAT0000542 | Sptan1 | 20740 | HITS-CLIP | 23630078\|19536157\|23142080\|29167 | Unspecified |
| mmu-miR-34a-5p | MIMAT0000542 | Sptbn1 | 20742 | HITS-CLIP | 30971389\|23142080\|19536157\|3143 | Unspecified |
| mmu-miR-34a-5p | MIMAT0000542 | Sptbn2 | 20743 | HITS-CLIP | 19536157\|23142080 | Unspecified |
| mmu-miR-34a-5p | MIMAT0000542 | Sptlc1 | 268656 | HITS-CLIP | 23142080\|31433990\|19536157\|29167 | Unspecified |
| mmu-miR-34a-5p | MIMAT0000542 | Sptlc2 | 20773 | HITS-CLIP | 23142080\|29167373 | Unspecified |
| mmu-miR-34a-5p | MIMAT0000542 | Sptssa | 104725 | HITS-CLIP | 23142080 | Unspecified |
| mmu-miR-34a-5p | MIMAT0000542 | Spty2d1 | 101685 | HITS-CLIP | 23142080 | Unspecified |
| mmu-miR-34a-5p | MIMAT0000542 | Sqle | 20775 | HITS-CLIP | 23142080 | Unspecified |
| mmu-miR-34a-5p | MIMAT0000542 | Sqstm1 | 18412 | HITS-CLIP | 23142080\|26203562 | Unspecified |
| mmu-miR-34a-5p | MIMAT0000542 | Sra1 | 24068 | HITS-CLIP | 23142080 | Unspecified |
| mmu-miR-34a-5p | MIMAT0000542 | Srbd1 | 78586 | HITS-CLIP | 23142080 | Unspecified |
| mmu-miR-34a-5p | MIMAT0000542 | Src | 20779 | HITS-CLIP | 26203562 | Unspecified |
| mmu-miR-34a-5p | MIMAT0000542 | Srcap | 100043597 | HITS-CLIP | 23142080\|29167373 | Unspecified |
| mmu-miR-34a-5p | MIMAT0000542 | Srd5a3 | 57357 | HITS-CLIP | 19536157\|29167373\|23142080 | Unspecified |
| mmu-miR-34a-5p | MIMAT0000542 | Srebf1 | 20787 | HITS-CLIP | 23142080\|31433990\|19536157\|29167 | Unspecified |
| mmu-miR-34a-5p | MIMAT0000542 | Srebf2 | 20788 | HITS-CLIP | 23142080\|29167373 | Unspecified |
| mmu-miR-34a-5p | MIMAT0000542 | Srek1 | 218543 | HITS-CLIP | 23142080 | Unspecified |
| mmu-miR-34a-5p | MIMAT0000542 | Srek1ip1 | 67288 | HITS-CLIP | 23142080 | Unspecified |
| mmu-miR-34a-5p | MIMAT0000542 | Srfbp1 | 67222 | HITS-CLIP | 23142080 | Unspecified |
| mmu-miR-34a-5p | MIMAT0000542 | Srgap2 | 14270 | HITS-CLIP | 23142080\|29167373 | Unspecified |
| mmu-miR-34a-5p | MIMAT0000542 | Srgap3 | 259302 | HITS-CLIP | 23142080\|29167373 | Unspecified |
| mmu-miR-34a-5p | MIMAT0000542 | Srgn | 19073 | HITS-CLIP | 23142080 | Unspecified |
| mmu-miR-34a-5p | MIMAT0000542 | Sri | 109552 | HITS-CLIP | 19536157\|23142080 | Unspecified |
| mmu-miR-34a-5p | MIMAT0000542 | Srl | 106393 | Microarrays | 37986224 | Unspecified |
| mmu-miR-34a-5p | MIMAT0000542 | Srm | 20810 | HITS-CLIP | 23142080 | Unspecified |
| mmu-miR-34a-5p | MIMAT0000542 | Srp14 | 20813 | HITS-CLIP | 31433990 | Unspecified |
| mmu-miR-34a-5p | MIMAT0000542 | Srp19 | 66384 | HITS-CLIP | 23142080 | Unspecified |

| | | | | | | |
|---|---|---|---|---|---|---|
| mmu-miR-34a-5p | MIMAT0000542 | Srp68 | 217337 | HITS-CLIP | 23142080\|29167373 | Unspecified |
| mmu-miR-34a-5p | MIMAT0000542 | Srp72 | 66661 | HITS-CLIP | 19536157\|23142080 | Unspecified |
| mmu-miR-34a-5p | MIMAT0000542 | Srp9 | 27058 | HITS-CLIP | 23142080 | Unspecified |
| mmu-miR-34a-5p | MIMAT0000542 | Srpk1 | 20815 | HITS-CLIP | 23142080\|29167373 | Unspecified |
| mmu-miR-34a-5p | MIMAT0000542 | Srpk2 | 20817 | HITS-CLIP | 23142080 | Unspecified |
| mmu-miR-34a-5p | MIMAT0000542 | Srpr | 67398 | HITS-CLIP | 19536157\|23142080 | Unspecified |
| mmu-miR-34a-5p | MIMAT0000542 | Srr | 27364 | HITS-CLIP | 19536157 | Unspecified |
| mmu-miR-34a-5p | MIMAT0000542 | Srrm1 | 51796 | HITS-CLIP | 23142080\|29167373 | Unspecified |
| mmu-miR-34a-5p | MIMAT0000542 | Srrm2 | 75956 | HITS-CLIP | 23142080\|31433990\|19536157\|29167 | Unspecified |
| mmu-miR-34a-5p | MIMAT0000542 | Srrt | 83701 | HITS-CLIP | 31433990\|23142080 | Unspecified |
| mmu-miR-34a-5p | MIMAT0000542 | Srsf1 | 110809 | HITS-CLIP | 23142080\|29167373 | Unspecified |
| mmu-miR-34a-5p | MIMAT0000542 | Srsf10 | 14105 | HITS-CLIP | 23142080 | Unspecified |
| mmu-miR-34a-5p | MIMAT0000542 | Srsf11 | 69207 | HITS-CLIP | 23142080 | Unspecified |
| mmu-miR-34a-5p | MIMAT0000542 | Srsf2 | 20382 | HITS-CLIP | 23142080\|29167373 | Unspecified |
| mmu-miR-34a-5p | MIMAT0000542 | Srsf4 | 57317 | HITS-CLIP | 31433990\|23142080 | Unspecified |
| mmu-miR-34a-5p | MIMAT0000542 | Srsf5 | 20384 | HITS-CLIP | 29167373 | Unspecified |
| mmu-miR-34a-5p | MIMAT0000542 | Srsf6 | 67996 | HITS-CLIP | 23142080\|29167373 | Unspecified |
| mmu-miR-34a-5p | MIMAT0000542 | Srsf7 | 225027 | HITS-CLIP | 23142080 | Unspecified |
| mmu-miR-34a-5p | MIMAT0000542 | Srxn1 | 76650 | HITS-CLIP | 29167373 | Unspecified |
| mmu-miR-34a-5p | MIMAT0000542 | Ss18 | 268996 | HITS-CLIP | 23142080 | Unspecified |
| mmu-miR-34a-5p | MIMAT0000542 | Ss18l1 | 269397 | HITS-CLIP | 23142080 | Unspecified |
| mmu-miR-34a-5p | MIMAT0000542 | Ssb | 20823 | HITS-CLIP | 23142080\|31433990\|29167373 | Unspecified |
| mmu-miR-34a-5p | MIMAT0000542 | Ssbp1 | 381760 | HITS-CLIP | 29167373 | Unspecified |
| mmu-miR-34a-5p | MIMAT0000542 | Ssbp2 | 66970 | HITS-CLIP | 23142080 | Unspecified |
| mmu-miR-34a-5p | MIMAT0000542 | Ssbp3 | 72475 | HITS-CLIP | 30971389\|23142080 | Unspecified |
| mmu-miR-34a-5p | MIMAT0000542 | Ssc4d | 136853 | HITS-CLIP | 23142080 | Unspecified |
| mmu-miR-34a-5p | MIMAT0000542 | Ssh1 | 231637 | HITS-CLIP | 23142080\|19536157\|29167373 | Unspecified |
| mmu-miR-34a-5p | MIMAT0000542 | Ssh2 | 237860 | HITS-CLIP | 31433990\|23142080\|29167373 | Unspecified |
| mmu-miR-34a-5p | MIMAT0000542 | Sspn | 16651 | Microarrays | 37986224 | Unspecified |
| mmu-miR-34a-5p | MIMAT0000542 | Ssr1 | 107513 | HITS-CLIP | 23142080\|31433990\|19536157\|29167 | Unspecified |
| mmu-miR-34a-5p | MIMAT0000542 | Ssr2 | 66256 | HITS-CLIP | 29167373 | Unspecified |
| mmu-miR-34a-5p | MIMAT0000542 | Ssr3 | 67437 | HITS-CLIP | 19536157\|31433990\|23142080\|29167 | Unspecified |
| mmu-miR-34a-5p | MIMAT0000542 | Ssr4 | 20832 | HITS-CLIP | 23142080\|31433990 | Unspecified |
| mmu-miR-34a-5p | MIMAT0000542 | Ssrp1 | 20833 | HITS-CLIP | 23142080 | Unspecified |
| mmu-miR-34a-5p | MIMAT0000542 | Ssu72 | 68991 | HITS-CLIP | 23142080\|29167373 | Unspecified |
| mmu-miR-34a-5p | MIMAT0000542 | St13 | 70356 | HITS-CLIP | 23142080\|19536157\|29167373 | Unspecified |
| mmu-miR-34a-5p | MIMAT0000542 | St18 | 240690 | HITS-CLIP | 29167373 | Unspecified |
| mmu-miR-34a-5p | MIMAT0000542 | St3gal1 | 20442 | HITS-CLIP | 23142080\|29167373 | Unspecified |
| mmu-miR-34a-5p | MIMAT0000542 | St3gal2 | 20444 | HITS-CLIP | 23142080 | Unspecified |

| | | | | | | |
|---|---|---|---|---|---|---|
| mmu-miR-34a-5p | MIMAT0000542 | St3gal4 | 20443 | HITS-CLIP | 23142080 | Unspecified |
| mmu-miR-34a-5p | MIMAT0000542 | St3gal5 | 20454 | HITS-CLIP | 29167373 | Unspecified |
| mmu-miR-34a-5p | MIMAT0000542 | St3gal6 | 54613 | HITS-CLIP | 23142080 | Unspecified |
| mmu-miR-34a-5p | MIMAT0000542 | St6gal1 | 20440 | HITS-CLIP | 23142080\|29167373 | Unspecified |
| mmu-miR-34a-5p | MIMAT0000542 | St6galnac3 | 20447 | HITS-CLIP | 23142080\|19536157 | Unspecified |
| mmu-miR-34a-5p | MIMAT0000542 | St6galnac5 | 26938 | HITS-CLIP | 19536157 | Unspecified |
| mmu-miR-34a-5p | MIMAT0000542 | St7l | 229681 | HITS-CLIP | 23142080 | Unspecified |
| mmu-miR-34a-5p | MIMAT0000542 | St8sia1 | 20449 | HITS-CLIP | 31433990\|23142080 | Unspecified |
| mmu-miR-34a-5p | MIMAT0000542 | St8sia3 | 20451 | HITS-CLIP | 19536157 | Unspecified |
| mmu-miR-34a-5p | MIMAT0000542 | St8sia4 | 20452 | HITS-CLIP | 23142080\|19536157\|29167373 | Unspecified |
| mmu-miR-34a-5p | MIMAT0000542 | St8sia5 | 225742 | HITS-CLIP | 19536157 | Unspecified |
| mmu-miR-34a-5p | MIMAT0000542 | St8sia6 | 241230 | HITS-CLIP | 23142080 | Unspecified |
| mmu-miR-34a-5p | MIMAT0000542 | Stab1 | 192187 | HITS-CLIP | 29167373 | Unspecified |
| mmu-miR-34a-5p | MIMAT0000542 | Stac2 | 217154 | Microarrays | 37986224\|23027903 | Unspecified |
| mmu-miR-34a-5p | MIMAT0000542 | Stac3 | 237611 | Microarrays | 37986224 | Unspecified |
| mmu-miR-34a-5p | MIMAT0000542 | Stag1 | 20842 | HITS-CLIP | 23142080\|29167373 | Unspecified |
| mmu-miR-34a-5p | MIMAT0000542 | Stag2 | 20843 | HITS-CLIP | 23142080\|29167373 | Unspecified |
| mmu-miR-34a-5p | MIMAT0000542 | Stag3 | 50878 | HITS-CLIP | 23142080 | Unspecified |
| mmu-miR-34a-5p | MIMAT0000542 | Stam | 20844 | HITS-CLIP | 19536157\|23142080\|29167373 | Unspecified |
| mmu-miR-34a-5p | MIMAT0000542 | Stam2 | 56324 | HITS-CLIP | 31433990\|23142080\|29167373 | Unspecified |
| mmu-miR-34a-5p | MIMAT0000542 | Stambp | 70527 | HITS-CLIP | 23142080 | Unspecified |
| mmu-miR-34a-5p | MIMAT0000542 | Stambpl1 | 76630 | HITS-CLIP | 23142080 | Unspecified |
| mmu-miR-34a-5p | MIMAT0000542 | Stap1 | 56792 | HITS-CLIP | 23142080\|29167373 | Unspecified |
| mmu-miR-34a-5p | MIMAT0000542 | Stard4 | 170459 | HITS-CLIP | 23142080 | Unspecified |
| mmu-miR-34a-5p | MIMAT0000542 | Stard5 | 170460 | HITS-CLIP | 23142080 | Unspecified |
| mmu-miR-34a-5p | MIMAT0000542 | Stard6 | 170461 | HITS-CLIP | 23142080 | Unspecified |
| mmu-miR-34a-5p | MIMAT0000542 | Stard7 | 99138 | HITS-CLIP | 31433990\|29167373 | Unspecified |
| mmu-miR-34a-5p | MIMAT0000542 | Stard9 | 668880 | HITS-CLIP | 29167373 | Unspecified |
| mmu-miR-34a-5p | MIMAT0000542 | Stat1 | 20846 | HITS-CLIP | 23142080\|31433990\|29167373 | Unspecified |
| mmu-miR-34a-5p | MIMAT0000542 | Stat2 | 20847 | HITS-CLIP | 23142080\|29167373 | Unspecified |
| mmu-miR-34a-5p | MIMAT0000542 | Stat3 | 20848 | HITS-CLIP | 23142080\|29167373 | Unspecified |
| mmu-miR-34a-5p | MIMAT0000542 | Stat4 | 20849 | HITS-CLIP | 23142080\|31433990 | Unspecified |
| mmu-miR-34a-5p | MIMAT0000542 | Stat5a | 20850 | HITS-CLIP | 23142080 | Unspecified |
| mmu-miR-34a-5p | MIMAT0000542 | Stat5b | 20851 | HITS-CLIP | 23142080 | Unspecified |
| mmu-miR-34a-5p | MIMAT0000542 | Stat6 | 20852 | HITS-CLIP | 23142080\|29167373 | Unspecified |
| mmu-miR-34a-5p | MIMAT0000542 | Stau1 | 20853 | HITS-CLIP | 23142080 | Unspecified |
| mmu-miR-34a-5p | MIMAT0000542 | Stau2 | 29819 | HITS-CLIP | 29167373 | Unspecified |
| mmu-miR-34a-5p | MIMAT0000542 | Stc1 | 20855 | HITS-CLIP | 23142080 | Unspecified |
| mmu-miR-34a-5p | MIMAT0000542 | Stc2 | 20856 | Microarrays//HITS-CLIP | 37986224\|23142080 | Unspecified |

| | | | | | | |
|---|---|---|---|---|---|---|
| mmu-miR-34a-5p | MIMAT0000542 | Stil | 20460 | Microarrays//HITS-CLIP | 37986224\|23027903\|30971389\|23630 | Unspecified |
| mmu-miR-34a-5p | MIMAT0000542 | Stim1 | 20866 | HITS-CLIP | 23142080\|19536157 | Unspecified |
| mmu-miR-34a-5p | MIMAT0000542 | Stim2 | 116873 | HITS-CLIP | 23142080\|29167373 | Unspecified |
| mmu-miR-34a-5p | MIMAT0000542 | Stimate | 375346 | HITS-CLIP | 31433990 | Unspecified |
| mmu-miR-34a-5p | MIMAT0000542 | Stip1 | 20867 | HITS-CLIP | 23142080 | Unspecified |
| mmu-miR-34a-5p | MIMAT0000542 | Stk10 | 20868 | HITS-CLIP | 23142080\|29167373 | Unspecified |
| mmu-miR-34a-5p | MIMAT0000542 | Stk11 | 20869 | HITS-CLIP | 23142080 | Unspecified |
| mmu-miR-34a-5p | MIMAT0000542 | Stk11ip | 71728 | Microarrays | 37986224\|23027903 | Unspecified |
| mmu-miR-34a-5p | MIMAT0000542 | Stk16 | 20872 | HITS-CLIP | 23142080 | Unspecified |
| mmu-miR-34a-5p | MIMAT0000542 | Stk17b | 98267 | HITS-CLIP | 23142080\|29167373 | Unspecified |
| mmu-miR-34a-5p | MIMAT0000542 | Stk19 | 54402 | HITS-CLIP | 23142080 | Unspecified |
| mmu-miR-34a-5p | MIMAT0000542 | Stk24 | 223255 | HITS-CLIP | 23142080\|29167373 | Unspecified |
| mmu-miR-34a-5p | MIMAT0000542 | Stk26 | 51765 | HITS-CLIP | 23142080 | Unspecified |
| mmu-miR-34a-5p | MIMAT0000542 | Stk32c | 282974 | HITS-CLIP | 23142080 | Unspecified |
| mmu-miR-34a-5p | MIMAT0000542 | Stk35 | 67333 | Microarrays//HITS-CLIP | 37986224\|23142080\|23027903\|19536 | Unspecified |
| mmu-miR-34a-5p | MIMAT0000542 | Stk38 | 106504 | HITS-CLIP | 23142080 | Unspecified |
| mmu-miR-34a-5p | MIMAT0000542 | Stk38l | 232533 | HITS-CLIP | 23142080 | Unspecified |
| mmu-miR-34a-5p | MIMAT0000542 | Stk39 | 53416 | HITS-CLIP | 31433990\|23142080 | Unspecified |
| mmu-miR-34a-5p | MIMAT0000542 | Stk4 | 58231 | HITS-CLIP, Microarrays//Microarrays//HITS | 37986224\|23027903\|23142080\|19536 | Unspecified |
| mmu-miR-34a-5p | MIMAT0000542 | Stk40 | 74178 | HITS-CLIP | 31433990\|23142080\|29167373 | Unspecified |
| mmu-miR-34a-5p | MIMAT0000542 | Stmn2 | 20257 | HITS-CLIP | 19536157 | Unspecified |
| mmu-miR-34a-5p | MIMAT0000542 | Stmn3 | 20262 | HITS-CLIP | 19536157 | Unspecified |
| mmu-miR-34a-5p | MIMAT0000542 | Stom | 13830 | HITS-CLIP | 30971389\|29167373 | Unspecified |
| mmu-miR-34a-5p | MIMAT0000542 | Stoml1 | 69106 | HITS-CLIP | 23142080 | Unspecified |
| mmu-miR-34a-5p | MIMAT0000542 | Ston1 | 77057 | HITS-CLIP | 23142080 | Unspecified |
| mmu-miR-34a-5p | MIMAT0000542 | Stra6 | 20897 | HITS-CLIP | 19536157 | Unspecified |
| mmu-miR-34a-5p | MIMAT0000542 | Strada | 72149 | HITS-CLIP | 23142080 | Unspecified |
| mmu-miR-34a-5p | MIMAT0000542 | Stradb | 227154 | HITS-CLIP | 23142080 | Unspecified |
| mmu-miR-34a-5p | MIMAT0000542 | Strap | 20901 | HITS-CLIP | 29167373\|23142080 | Unspecified |
| mmu-miR-34a-5p | MIMAT0000542 | Strbp | 20744 | Microarrays | 37986224 | Unspecified |
| mmu-miR-34a-5p | MIMAT0000542 | Strip1 | 229707 | HITS-CLIP | 23142080 | Unspecified |
| mmu-miR-34a-5p | MIMAT0000542 | Strn | 268980 | HITS-CLIP | 37986224\|23142080\|19536157\|23630 | Unspecified |
| mmu-miR-34a-5p | MIMAT0000542 | Strn3 | 94186 | HITS-CLIP | 37986224\|23142080\|31433990 | Unspecified |
| mmu-miR-34a-5p | MIMAT0000542 | Strn4 | 97387 | HITS-CLIP | 19536157\|23142080 | Unspecified |
| mmu-miR-34a-5p | MIMAT0000542 | Stt3a | 16430 | HITS-CLIP | 23142080\|29167373 | Unspecified |
| mmu-miR-34a-5p | MIMAT0000542 | Stt3b | 68292 | HITS-CLIP | 23142080\|29167373 | Unspecified |
| mmu-miR-34a-5p | MIMAT0000542 | Stub1 | 56424 | HITS-CLIP | 23142080 | Unspecified |
| mmu-miR-34a-5p | MIMAT0000542 | Stx11 | 74732 | HITS-CLIP | 23142080 | Unspecified |
| mmu-miR-34a-5p | MIMAT0000542 | Stx12 | 100226 | HITS-CLIP | 30971389\|23142080\|29167373 | Unspecified |

| | | | | | | |
|---|---|---|---|---|---|---|
| mmu-miR-34a-5p | MIMAT0000542 | Stx16 | 228960 | HITS-CLIP | 19536157\|23142080\|29167373 | Unspecified |
| mmu-miR-34a-5p | MIMAT0000542 | Stx17 | 67727 | HITS-CLIP | 23142080\|29167373 | Unspecified |
| mmu-miR-34a-5p | MIMAT0000542 | Stx18 | 71116 | HITS-CLIP | 23142080 | Unspecified |
| mmu-miR-34a-5p | MIMAT0000542 | Stx2 | 13852 | Microarrays | 37986224\|23027903 | Unspecified |
| mmu-miR-34a-5p | MIMAT0000542 | Stx5a | 56389 | HITS-CLIP | 23142080\|29167373 | Unspecified |
| mmu-miR-34a-5p | MIMAT0000542 | Stx6 | 58244 | HITS-CLIP | 31433990\|23142080 | Unspecified |
| mmu-miR-34a-5p | MIMAT0000542 | Stx7 | 53331 | HITS-CLIP | 23142080\|29167373 | Unspecified |
| mmu-miR-34a-5p | MIMAT0000542 | Stx8 | 55943 | HITS-CLIP | 23142080 | Unspecified |
| mmu-miR-34a-5p | MIMAT0000542 | Stxbp1 | 20910 | HITS-CLIP | 19536157\|23142080 | Unspecified |
| mmu-miR-34a-5p | MIMAT0000542 | Stxbp2 | 20911 | HITS-CLIP | 23142080\|29167373 | Unspecified |
| mmu-miR-34a-5p | MIMAT0000542 | Stxbp3 | 20912 | HITS-CLIP | 23142080\|29167373 | Unspecified |
| mmu-miR-34a-5p | MIMAT0000542 | Stxbp4 | 20913 | HITS-CLIP | 23142080 | Unspecified |
| mmu-miR-34a-5p | MIMAT0000542 | Stxbp5 | 78808 | HITS-CLIP | 23142080 | Unspecified |
| mmu-miR-34a-5p | MIMAT0000542 | Sub1 | 20024 | HITS-CLIP | 23142080\|29167373 | Unspecified |
| mmu-miR-34a-5p | MIMAT0000542 | Sucla2 | 20916 | HITS-CLIP | 23142080\|31433990\|29167373 | Unspecified |
| mmu-miR-34a-5p | MIMAT0000542 | Suclg1 | 56451 | HITS-CLIP | 37986224\|19536157\|23142080 | Unspecified |
| mmu-miR-34a-5p | MIMAT0000542 | Suclg2 | 20917 | HITS-CLIP | 23142080 | Unspecified |
| mmu-miR-34a-5p | MIMAT0000542 | Suco | 226551 | HITS-CLIP | 37986224\|23630078\|30971389\|31433[illegible] | Unspecified |
| mmu-miR-34a-5p | MIMAT0000542 | Sufu | 24069 | HITS-CLIP | 23142080 | Unspecified |
| mmu-miR-34a-5p | MIMAT0000542 | Sugt1 | 67955 | HITS-CLIP | 30971389\|23142080 | Unspecified |
| mmu-miR-34a-5p | MIMAT0000542 | Sulf1 | 240725 | HITS-CLIP | 37986224 | Unspecified |
| mmu-miR-34a-5p | MIMAT0000542 | Sulf2 | 72043 | HITS-CLIP | 29167373 | Unspecified |
| mmu-miR-34a-5p | MIMAT0000542 | Sumf1 | 58911 | HITS-CLIP | 29167373 | Unspecified |
| mmu-miR-34a-5p | MIMAT0000542 | Sumo3 | 20610 | HITS-CLIP | 23630078 | Unspecified |
| mmu-miR-34a-5p | MIMAT0000542 | Sun1 | 77053 | HITS-CLIP | 37986224\|23142080 | Unspecified |
| mmu-miR-34a-5p | MIMAT0000542 | Sun2 | 223697 | HITS-CLIP | 23142080 | Unspecified |
| mmu-miR-34a-5p | MIMAT0000542 | Supt16 | 114741 | HITS-CLIP | 23142080\|29167373 | Unspecified |
| mmu-miR-34a-5p | MIMAT0000542 | Supt20 | 56790 | HITS-CLIP | 23142080\|29167373 | Unspecified |
| mmu-miR-34a-5p | MIMAT0000542 | Supt4a | 20922 | HITS-CLIP | 31433990 | Unspecified |
| mmu-miR-34a-5p | MIMAT0000542 | Supt5 | 20924 | HITS-CLIP | 23142080 | Unspecified |
| mmu-miR-34a-5p | MIMAT0000542 | Supt6 | 20926 | HITS-CLIP | 23142080\|29167373 | Unspecified |
| mmu-miR-34a-5p | MIMAT0000542 | Supt7l | 72195 | HITS-CLIP | 23142080 | Unspecified |
| mmu-miR-34a-5p | MIMAT0000542 | Supv3l1 | 338359 | HITS-CLIP | 23142080 | Unspecified |
| mmu-miR-34a-5p | MIMAT0000542 | Surf4 | 20932 | HITS-CLIP | 19536157 | Unspecified |
| mmu-miR-34a-5p | MIMAT0000542 | Surf6 | 20935 | HITS-CLIP | 23142080 | Unspecified |
| mmu-miR-34a-5p | MIMAT0000542 | Susd3 | 66329 | HITS-CLIP | 29167373 | Unspecified |
| mmu-miR-34a-5p | MIMAT0000542 | Susd6 | 9766 | HITS-CLIP | 19536157\|23142080\|29167373 | Unspecified |
| mmu-miR-34a-5p | MIMAT0000542 | Suv39h1 | 20937 | HITS-CLIP | 23142080 | Unspecified |
| mmu-miR-34a-5p | MIMAT0000542 | Suv39h2 | 64707 | HITS-CLIP | 31433990 | Unspecified |

| | | | | | | |
|---|---|---|---|---|---|---|
| mmu-miR-34a-5p | MIMAT0000542 | Suz12 | 52615 | Microarrays//HITS-CLIP | 37986224\|23027903\|23142080\|31433 | Unspecified |
| mmu-miR-34a-5p | MIMAT0000542 | Sv2a | 64051 | HITS-CLIP | 19536157\|23142080 | Unspecified |
| mmu-miR-34a-5p | MIMAT0000542 | Svil | 225115 | HITS-CLIP | 30971389\|23142080\|29167373 | Unspecified |
| mmu-miR-34a-5p | MIMAT0000542 | Svop | 68666 | HITS-CLIP | 19536157 | Unspecified |
| mmu-miR-34a-5p | MIMAT0000542 | Swap70 | 20947 | HITS-CLIP | 23630078\|31433990\|23142080\|29167 | Unspecified |
| mmu-miR-34a-5p | MIMAT0000542 | Swt1 | 66875 | HITS-CLIP, Microarrays//Microarrays//HITS | 37986224\|23027903\|23142080 | Unspecified |
| mmu-miR-34a-5p | MIMAT0000542 | Syap1 | 67043 | HITS-CLIP | 23142080 | Unspecified |
| mmu-miR-34a-5p | MIMAT0000542 | Sybu | 319613 | HITS-CLIP | 19536157 | Unspecified |
| mmu-miR-34a-5p | MIMAT0000542 | Syce2 | 71846 | HITS-CLIP | 23142080 | Unspecified |
| mmu-miR-34a-5p | MIMAT0000542 | Syf2 | 68592 | HITS-CLIP | 30971389\|23142080 | Unspecified |
| mmu-miR-34a-5p | MIMAT0000542 | Syk | 20963 | HITS-CLIP | 29167373 | Unspecified |
| mmu-miR-34a-5p | MIMAT0000542 | Sympk | 68188 | HITS-CLIP | 23142080 | Unspecified |
| mmu-miR-34a-5p | MIMAT0000542 | Syn1 | 20964 | HITS-CLIP | 19536157 | Unspecified |
| mmu-miR-34a-5p | MIMAT0000542 | Syn2 | 20965 | HITS-CLIP | 19536157 | Unspecified |
| mmu-miR-34a-5p | MIMAT0000542 | Syncrip | 56403 | HITS-CLIP | 31433990\|23142080\|29167373 | Unspecified |
| mmu-miR-34a-5p | MIMAT0000542 | Syne1 | 64009 | HITS-CLIP | 23142080\|19536157\|29167373 | Unspecified |
| mmu-miR-34a-5p | MIMAT0000542 | Syne2 | 319565 | HITS-CLIP | 23142080\|19536157 | Unspecified |
| mmu-miR-34a-5p | MIMAT0000542 | Syne3 | 212073 | HITS-CLIP | 23142080\|29167373 | Unspecified |
| mmu-miR-34a-5p | MIMAT0000542 | Syngap1 | 8831 | HITS-CLIP | 19536157 | Unspecified |
| mmu-miR-34a-5p | MIMAT0000542 | Syngr1 | 20972 | HITS-CLIP | 19536157\|29167373 | Unspecified |
| mmu-miR-34a-5p | MIMAT0000542 | Syngr2 | 20973 | HITS-CLIP | 23142080 | Unspecified |
| mmu-miR-34a-5p | MIMAT0000542 | Syngr3 | 20974 | HITS-CLIP | 19536157 | Unspecified |
| mmu-miR-34a-5p | MIMAT0000542 | Synj1 | 104015 | HITS-CLIP | 31433990\|26203562\|23142080\|29167 | Unspecified |
| mmu-miR-34a-5p | MIMAT0000542 | Synj2 | 20975 | HITS-CLIP | 19536157\|23630078\|23142080 | Unspecified |
| mmu-miR-34a-5p | MIMAT0000542 | Synj2bp | 24071 | HITS-CLIP | 29167373 | Unspecified |
| mmu-miR-34a-5p | MIMAT0000542 | Synpo | 104027 | HITS-CLIP | 23142080\|19536157 | Unspecified |
| mmu-miR-34a-5p | MIMAT0000542 | Synpo2l | 68760 | Microarrays | 37986224 | Unspecified |
| mmu-miR-34a-5p | MIMAT0000542 | Synpr | 72003 | HITS-CLIP | 19536157 | Unspecified |
| mmu-miR-34a-5p | MIMAT0000542 | Synrg | 217030 | HITS-CLIP | 19536157\|23142080\|29167373 | Unspecified |
| mmu-miR-34a-5p | MIMAT0000542 | Syp | 20977 | HITS-CLIP | 19536157 | Unspecified |
| mmu-miR-34a-5p | MIMAT0000542 | Sypl | 19027 | HITS-CLIP | 31433990\|23142080\|29167373 | Unspecified |
| mmu-miR-34a-5p | MIMAT0000542 | Sys1 | 66460 | HITS-CLIP | 23142080 | Unspecified |
| mmu-miR-34a-5p | MIMAT0000542 | Syt1 | 20979 | HITS-CLIP//Immunoblot//Luciferase report | 19536157\|24973144 | Unspecified |
| mmu-miR-34a-5p | MIMAT0000542 | Syt11 | 229521 | HITS-CLIP | 19536157 | Unspecified |
| mmu-miR-34a-5p | MIMAT0000542 | Syt13 | 80976 | Microarrays | 37986224 | Unspecified |
| mmu-miR-34a-5p | MIMAT0000542 | Syt3 | 20981 | HITS-CLIP | 19536157 | Unspecified |
| mmu-miR-34a-5p | MIMAT0000542 | Syt4 | 20983 | HITS-CLIP | 19536157 | Unspecified |
| mmu-miR-34a-5p | MIMAT0000542 | Syt7 | 54525 | HITS-CLIP | 19536157 | Unspecified |
| mmu-miR-34a-5p | MIMAT0000542 | Sytl1 | 269589 | HITS-CLIP | 23142080 | Unspecified |

| | | | | | | |
|---|---|---|---|---|---|---|
| mmu-miR-34a-5p | MIMAT0000542 | Sytl2 | 83671 | HITS-CLIP | 23142080\|19536157 | Unspecified |
| mmu-miR-34a-5p | MIMAT0000542 | Syvn1 | 74126 | HITS-CLIP | 23142080\|23630078 | Unspecified |
| mmu-miR-34a-5p | MIMAT0000542 | Szrd1 | 213491 | HITS-CLIP | 19536157\|23142080\|29167373 | Unspecified |
| mmu-miR-34a-5p | MIMAT0000542 | Szt2 | 230676 | HITS-CLIP | 23142080\|29167373 | Unspecified |
| mmu-miR-34a-5p | MIMAT0000542 | Tab2 | 68652 | HITS-CLIP | 37986224\|23142080\|19536157\|29167 | Unspecified |
| mmu-miR-34a-5p | MIMAT0000542 | Tab3 | 66724 | HITS-CLIP | 31433990\|29167373\|23142080 | Unspecified |
| mmu-miR-34a-5p | MIMAT0000542 | Tacc1 | 320165 | HITS-CLIP | 19536157\|23142080\|29167373 | Unspecified |
| mmu-miR-34a-5p | MIMAT0000542 | Tacc2 | 57752 | HITS-CLIP | 23142080 | Unspecified |
| mmu-miR-34a-5p | MIMAT0000542 | Tacc3 | 21335 | Microarrays//HITS-CLIP | 37986224\|23027903\|23142080 | Unspecified |
| mmu-miR-34a-5p | MIMAT0000542 | Tacstd2 | 56753 | HITS-CLIP | 26203562 | Unspecified |
| mmu-miR-34a-5p | MIMAT0000542 | Tada1 | 27878 | HITS-CLIP | 23142080 | Unspecified |
| mmu-miR-34a-5p | MIMAT0000542 | Tada2b | 231151 | HITS-CLIP | 23142080 | Unspecified |
| mmu-miR-34a-5p | MIMAT0000542 | Taf1 | 270627 | HITS-CLIP | 23142080\|29167373 | Unspecified |
| mmu-miR-34a-5p | MIMAT0000542 | Taf12 | 66464 | HITS-CLIP | 23142080 | Unspecified |
| mmu-miR-34a-5p | MIMAT0000542 | Taf13 | 99730 | HITS-CLIP | 23142080\|19536157 | Unspecified |
| mmu-miR-34a-5p | MIMAT0000542 | Taf15 | 70439 | HITS-CLIP | 23142080 | Unspecified |
| mmu-miR-34a-5p | MIMAT0000542 | Taf1b | 21340 | HITS-CLIP | 29167373 | Unspecified |
| mmu-miR-34a-5p | MIMAT0000542 | Taf2 | 319944 | HITS-CLIP | 23142080\|29167373 | Unspecified |
| mmu-miR-34a-5p | MIMAT0000542 | Taf3 | 209361 | HITS-CLIP | 23142080 | Unspecified |
| mmu-miR-34a-5p | MIMAT0000542 | Taf4 | 6874 | HITS-CLIP | 30971389\|31433990\|23142080 | Unspecified |
| mmu-miR-34a-5p | MIMAT0000542 | Taf4b | 72504 | HITS-CLIP | 23142080 | Unspecified |
| mmu-miR-34a-5p | MIMAT0000542 | Taf5 | 226182 | HITS-CLIP | 23142080 | Unspecified |
| mmu-miR-34a-5p | MIMAT0000542 | Taf5l | 102162 | HITS-CLIP | 37986224\|23142080\|29167373 | Unspecified |
| mmu-miR-34a-5p | MIMAT0000542 | Taf6 | 21343 | HITS-CLIP | 23142080 | Unspecified |
| mmu-miR-34a-5p | MIMAT0000542 | Taf7 | 24074 | HITS-CLIP | 23142080 | Unspecified |
| mmu-miR-34a-5p | MIMAT0000542 | Taf8 | 63856 | HITS-CLIP | 23142080 | Unspecified |
| mmu-miR-34a-5p | MIMAT0000542 | Taf9 | 108143 | HITS-CLIP | 23142080 | Unspecified |
| mmu-miR-34a-5p | MIMAT0000542 | Taf9b | 407786 | Microarrays//HITS-CLIP | 37986224\|23027903\|23142080 | Unspecified |
| mmu-miR-34a-5p | MIMAT0000542 | Tagap | 72536 | HITS-CLIP | 23142080 | Unspecified |
| mmu-miR-34a-5p | MIMAT0000542 | Tagln2 | 21346 | HITS-CLIP | 23142080\|29167373 | Unspecified |
| mmu-miR-34a-5p | MIMAT0000542 | Tal1 | 21349 | HITS-CLIP | 30971389 | Unspecified |
| mmu-miR-34a-5p | MIMAT0000542 | Taldo1 | 21351 | HITS-CLIP | 23142080 | Unspecified |
| mmu-miR-34a-5p | MIMAT0000542 | Tanc2 | 77097 | HITS-CLIP | 19536157\|23142080\|29167373 | Unspecified |
| mmu-miR-34a-5p | MIMAT0000542 | Tango2 | 27883 | HITS-CLIP | 29167373 | Unspecified |
| mmu-miR-34a-5p | MIMAT0000542 | Tank | 21353 | HITS-CLIP | 23630078\|30971389\|23142080\|29167 | Unspecified |
| mmu-miR-34a-5p | MIMAT0000542 | Taok1 | 216965 | HITS-CLIP | 37986224\|23630078\|19536157\|30971 | Unspecified |
| mmu-miR-34a-5p | MIMAT0000542 | Taok2 | 381921 | HITS-CLIP | 23142080 | Unspecified |
| mmu-miR-34a-5p | MIMAT0000542 | Taok3 | 330177 | HITS-CLIP | 23142080\|29167373 | Unspecified |
| mmu-miR-34a-5p | MIMAT0000542 | Tap1 | 21354 | HITS-CLIP | 23142080\|29167373 | Unspecified |

| | | | | | | |
|---|---|---|---|---|---|---|
| mmu-miR-34a-5p | MIMAT0000542 | Tap2 | 21355 | HITS-CLIP | 23142080 | Unspecified |
| mmu-miR-34a-5p | MIMAT0000542 | Tapbp | 21356 | HITS-CLIP | 23142080\|29167373 | Unspecified |
| mmu-miR-34a-5p | MIMAT0000542 | Tapbpl | 213233 | HITS-CLIP | 31433990\|23142080 | Unspecified |
| mmu-miR-34a-5p | MIMAT0000542 | Tapt1 | 231225 | HITS-CLIP | 30971389\|23142080 | Unspecified |
| mmu-miR-34a-5p | MIMAT0000542 | Tarbp1 | 6894 | HITS-CLIP | 23142080 | Unspecified |
| mmu-miR-34a-5p | MIMAT0000542 | Tarbp2 | 21357 | HITS-CLIP | 29167373 | Unspecified |
| mmu-miR-34a-5p | MIMAT0000542 | Tardbp | 230908 | HITS-CLIP | 31433990\|23142080\|29167373 | Unspecified |
| mmu-miR-34a-5p | MIMAT0000542 | Tars | 110960 | HITS-CLIP | 37986224\|23142080\|31433990 | Unspecified |
| mmu-miR-34a-5p | MIMAT0000542 | Tars2 | 71807 | HITS-CLIP | 23142080 | Unspecified |
| mmu-miR-34a-5p | MIMAT0000542 | Tasp1 | 75812 | HITS-CLIP | 23142080 | Unspecified |
| mmu-miR-34a-5p | MIMAT0000542 | Tatdn1 | 69694 | HITS-CLIP | 23142080 | Unspecified |
| mmu-miR-34a-5p | MIMAT0000542 | Tatdn2 | 381801 | HITS-CLIP | 29167373 | Unspecified |
| mmu-miR-34a-5p | MIMAT0000542 | Tatdn3 | 128387 | HITS-CLIP | 23142080 | Unspecified |
| mmu-miR-34a-5p | MIMAT0000542 | Tax1bp1 | 52440 | HITS-CLIP | 19536157\|23142080\|29167373 | Unspecified |
| mmu-miR-34a-5p | MIMAT0000542 | Tbc1d1 | 57915 | HITS-CLIP | 23142080\|29167373 | Unspecified |
| mmu-miR-34a-5p | MIMAT0000542 | Tbc1d10b | 68449 | HITS-CLIP | 23142080\|29167373 | Unspecified |
| mmu-miR-34a-5p | MIMAT0000542 | Tbc1d13 | 70296 | HITS-CLIP | 29167373\|23142080 | Unspecified |
| mmu-miR-34a-5p | MIMAT0000542 | Tbc1d14 | 100855 | HITS-CLIP | 23142080 | Unspecified |
| mmu-miR-34a-5p | MIMAT0000542 | Tbc1d15 | 66687 | HITS-CLIP | 23142080\|31433990\|29167373 | Unspecified |
| mmu-miR-34a-5p | MIMAT0000542 | Tbc1d17 | 233204 | HITS-CLIP | 23142080 | Unspecified |
| mmu-miR-34a-5p | MIMAT0000542 | Tbc1d19 | 67249 | HITS-CLIP | 23142080 | Unspecified |
| mmu-miR-34a-5p | MIMAT0000542 | Tbc1d20 | 67231 | HITS-CLIP | 23630078\|31433990\|23142080 | Unspecified |
| mmu-miR-34a-5p | MIMAT0000542 | Tbc1d22a | 223754 | HITS-CLIP | 29167373\|23142080 | Unspecified |
| mmu-miR-34a-5p | MIMAT0000542 | Tbc1d22b | 381085 | HITS-CLIP | 23142080 | Unspecified |
| mmu-miR-34a-5p | MIMAT0000542 | Tbc1d23 | 67581 | Microarrays//HITS-CLIP | 37986224\|23142080\|29167373 | Unspecified |
| mmu-miR-34a-5p | MIMAT0000542 | Tbc1d24 | 224617 | HITS-CLIP | 19536157\|29167373 | Unspecified |
| mmu-miR-34a-5p | MIMAT0000542 | Tbc1d2b | 67016 | HITS-CLIP | 31433990\|23142080\|29167373 | Unspecified |
| mmu-miR-34a-5p | MIMAT0000542 | Tbc1d30 | 74694 | HITS-CLIP | 23142080\|29167373 | Unspecified |
| mmu-miR-34a-5p | MIMAT0000542 | Tbc1d31 | 210544 | HITS-CLIP | 23142080\|29167373 | Unspecified |
| mmu-miR-34a-5p | MIMAT0000542 | Tbc1d4 | 210789 | Microarrays//HITS-CLIP | 37986224\|23142080\|23027903 | Unspecified |
| mmu-miR-34a-5p | MIMAT0000542 | Tbc1d5 | 72238 | HITS-CLIP | 23142080\|29167373 | Unspecified |
| mmu-miR-34a-5p | MIMAT0000542 | Tbc1d8 | 54610 | Microarrays | 37986224 | Unspecified |
| mmu-miR-34a-5p | MIMAT0000542 | Tbc1d8b | 245638 | HITS-CLIP | 23142080\|29167373 | Unspecified |
| mmu-miR-34a-5p | MIMAT0000542 | Tbc1d9 | 71310 | HITS-CLIP | 29167373 | Unspecified |
| mmu-miR-34a-5p | MIMAT0000542 | Tbc1d9b | 76795 | HITS-CLIP | 23142080 | Unspecified |
| mmu-miR-34a-5p | MIMAT0000542 | Tbca | 21371 | HITS-CLIP | 23142080 | Unspecified |
| mmu-miR-34a-5p | MIMAT0000542 | Tbcb | 66411 | HITS-CLIP | 23142080 | Unspecified |
| mmu-miR-34a-5p | MIMAT0000542 | Tbcc | 72726 | HITS-CLIP | 23142080 | Unspecified |
| mmu-miR-34a-5p | MIMAT0000542 | Tbcd | 108903 | HITS-CLIP | 23142080\|29167373 | Unspecified |

| | | | | | | | |
|---|---|---|---|---|---|---|---|
| mmu-miR-34a-5p | MIMAT0000542 | Tbce | 70430 | HITS-CLIP | | 29167373 | Unspecified |
| mmu-miR-34a-5p | MIMAT0000542 | Tbcel | 272589 | HITS-CLIP | | 30971389 | Unspecified |
| mmu-miR-34a-5p | MIMAT0000542 | Tbck | 271981 | HITS-CLIP | | 23142080 | Unspecified |
| mmu-miR-34a-5p | MIMAT0000542 | Tbk1 | 56480 | HITS-CLIP | 23142080\|29167373 | | Unspecified |
| mmu-miR-34a-5p | MIMAT0000542 | Tbkbp1 | 73174 | HITS-CLIP | | 23142080 | Unspecified |
| mmu-miR-34a-5p | MIMAT0000542 | Tbl1x | 21372 | HITS-CLIP | | 23142080 | Unspecified |
| mmu-miR-34a-5p | MIMAT0000542 | Tbl1xr1 | 81004 | HITS-CLIP | 37986224\|23142080\|31433990\|30971 | | Unspecified |
| mmu-miR-34a-5p | MIMAT0000542 | Tbl2 | 27368 | HITS-CLIP | | 23142080 | Unspecified |
| mmu-miR-34a-5p | MIMAT0000542 | Tbp | 21374 | HITS-CLIP | 23142080\|29167373 | | Unspecified |
| mmu-miR-34a-5p | MIMAT0000542 | Tbpl1 | 237336 | HITS-CLIP | 23142080\|29167373 | | Unspecified |
| mmu-miR-34a-5p | MIMAT0000542 | Tbr1 | 21375 | HITS-CLIP | | 19536157 | Unspecified |
| mmu-miR-34a-5p | MIMAT0000542 | Tbrg1 | 21376 | HITS-CLIP | | 23142080 | Unspecified |
| mmu-miR-34a-5p | MIMAT0000542 | Tbx21 | 57765 | HITS-CLIP | | 23142080 | Unspecified |
| mmu-miR-34a-5p | MIMAT0000542 | Tbxa2r | 21390 | HITS-CLIP | | 23142080 | Unspecified |
| mmu-miR-34a-5p | MIMAT0000542 | Tbxas1 | 21391 | HITS-CLIP | | 29167373 | Unspecified |
| mmu-miR-34a-5p | MIMAT0000542 | Tcaf1 | 9747 | HITS-CLIP | | 19536157 | Unspecified |
| mmu-miR-34a-5p | MIMAT0000542 | Tcaim | 382117 | HITS-CLIP | | 23142080 | Unspecified |
| mmu-miR-34a-5p | MIMAT0000542 | Tcea1 | 21399 | HITS-CLIP | | 31433990 | Unspecified |
| mmu-miR-34a-5p | MIMAT0000542 | Tceal3 | 594844 | HITS-CLIP | | 19536157 | Unspecified |
| mmu-miR-34a-5p | MIMAT0000542 | Tceanc | 245695 | HITS-CLIP | | 23142080 | Unspecified |
| mmu-miR-34a-5p | MIMAT0000542 | Tcerg1 | 56070 | HITS-CLIP | 23142080\|29167373\|30971389\|2363( | | Unspecified |
| mmu-miR-34a-5p | MIMAT0000542 | Tcf12 | 21406 | HITS-CLIP | 23142080\|29167373 | | Unspecified |
| mmu-miR-34a-5p | MIMAT0000542 | Tcf20 | 21411 | HITS-CLIP | 19536157\|23142080\|29167373 | | Unspecified |
| mmu-miR-34a-5p | MIMAT0000542 | Tcf25 | 66855 | HITS-CLIP | 23142080\|29167373\|23630078 | | Unspecified |
| mmu-miR-34a-5p | MIMAT0000542 | Tcf3 | 21423 | HITS-CLIP | 23630078\|29167373 | | Unspecified |
| mmu-miR-34a-5p | MIMAT0000542 | Tcf4 | 21413 | HITS-CLIP | | 29167373 | Unspecified |
| mmu-miR-34a-5p | MIMAT0000542 | Tcf7 | 21414 | HITS-CLIP | | 23142080 | Unspecified |
| mmu-miR-34a-5p | MIMAT0000542 | Tcf7l2 | 21416 | HITS-CLIP | | 29167373 | Unspecified |
| mmu-miR-34a-5p | MIMAT0000542 | Tcirg1 | 27060 | HITS-CLIP | | 29167373 | Unspecified |
| mmu-miR-34a-5p | MIMAT0000542 | Tcn2 | 21452 | Microarrays | | 37986224 | Unspecified |
| mmu-miR-34a-5p | MIMAT0000542 | Tcp1 | 21454 | HITS-CLIP | | 23142080 | Unspecified |
| mmu-miR-34a-5p | MIMAT0000542 | Tcp11l1 | 320554 | HITS-CLIP | | 29167373 | Unspecified |
| mmu-miR-34a-5p | MIMAT0000542 | Tcp11l2 | 216198 | HITS-CLIP | 23142080\|19536157 | | Unspecified |
| mmu-miR-34a-5p | MIMAT0000542 | Tdp1 | 104884 | HITS-CLIP | | 23142080 | Unspecified |
| mmu-miR-34a-5p | MIMAT0000542 | Tdp2 | 56196 | HITS-CLIP | | 23142080 | Unspecified |
| mmu-miR-34a-5p | MIMAT0000542 | Tdrd7 | 100121 | HITS-CLIP | 23142080\|29167373 | | Unspecified |
| mmu-miR-34a-5p | MIMAT0000542 | Tead2 | 21677 | HITS-CLIP | | 23630078 | Unspecified |
| mmu-miR-34a-5p | MIMAT0000542 | Tec | 21682 | HITS-CLIP | | 23142080 | Unspecified |
| mmu-miR-34a-5p | MIMAT0000542 | Tecpr1 | 70381 | HITS-CLIP | 23142080\|29167373 | | Unspecified |

| | | | | | | |
|---|---|---|---|---|---|---|
| mmu-miR-34a-5p | MIMAT0000542 | Tecpr2 | 104859 | HITS-CLIP | 29167373 | Unspecified |
| mmu-miR-34a-5p | MIMAT0000542 | Tecr | 106529 | HITS-CLIP | 23142080 | Unspecified |
| mmu-miR-34a-5p | MIMAT0000542 | Tecrl | 243078 | Microarrays | 37986224 | Unspecified |
| mmu-miR-34a-5p | MIMAT0000542 | Tedc2 | 72016 | HITS-CLIP | 30971389 | Unspecified |
| mmu-miR-34a-5p | MIMAT0000542 | Tef | 21685 | HITS-CLIP | 23630078 | Unspecified |
| mmu-miR-34a-5p | MIMAT0000542 | Ten1 | 69535 | HITS-CLIP | 23142080 | Unspecified |
| mmu-miR-34a-5p | MIMAT0000542 | Tenm1 | 23963 | HITS-CLIP | 19536157 | Unspecified |
| mmu-miR-34a-5p | MIMAT0000542 | Tenm3 | 23965 | HITS-CLIP | 19536157 | Unspecified |
| mmu-miR-34a-5p | MIMAT0000542 | Tenm4 | 23966 | HITS-CLIP | 19536157 | Unspecified |
| mmu-miR-34a-5p | MIMAT0000542 | Tent2 | 167153 | HITS-CLIP | 23142080\|29167373 | Unspecified |
| mmu-miR-34a-5p | MIMAT0000542 | Tent4a | 11044 | HITS-CLIP | 23142080\|29167373 | Unspecified |
| mmu-miR-34a-5p | MIMAT0000542 | Tent4b | 64282 | HITS-CLIP | 19536157\|23142080\|29167373 | Unspecified |
| mmu-miR-34a-5p | MIMAT0000542 | Tent5a | 55603 | HITS-CLIP | 23142080\|31433990\|29167373 | Unspecified |
| mmu-miR-34a-5p | MIMAT0000542 | Tent5c | 54855 | HITS-CLIP | 23142080 | Unspecified |
| mmu-miR-34a-5p | MIMAT0000542 | Tep1 | 21745 | HITS-CLIP | 23142080\|29167373 | Unspecified |
| mmu-miR-34a-5p | MIMAT0000542 | Tepsin | 146705 | HITS-CLIP | 23142080 | Unspecified |
| mmu-miR-34a-5p | MIMAT0000542 | Terf2 | 21750 | HITS-CLIP | 31433990\|23142080 | Unspecified |
| mmu-miR-34a-5p | MIMAT0000542 | Terf2ip | 57321 | HITS-CLIP | 23142080 | Unspecified |
| mmu-miR-34a-5p | MIMAT0000542 | Tes | 21753 | HITS-CLIP | 31433990\|23142080\|29167373 | Unspecified |
| mmu-miR-34a-5p | MIMAT0000542 | Tespa1 | 67596 | HITS-CLIP | 23142080 | Unspecified |
| mmu-miR-34a-5p | MIMAT0000542 | Tet1 | 52463 | HITS-CLIP | 23142080\|19536157 | Unspecified |
| mmu-miR-34a-5p | MIMAT0000542 | Tet2 | 214133 | HITS-CLIP | 23142080\|29167373 | Unspecified |
| mmu-miR-34a-5p | MIMAT0000542 | Tet3 | 194388 | HITS-CLIP | 23142080\|29167373 | Unspecified |
| mmu-miR-34a-5p | MIMAT0000542 | Tex10 | 269536 | HITS-CLIP | 23142080\|31433990 | Unspecified |
| mmu-miR-34a-5p | MIMAT0000542 | Tex15 | 104271 | HITS-CLIP | 23142080 | Unspecified |
| mmu-miR-34a-5p | MIMAT0000542 | Tex2 | 21763 | HITS-CLIP | 23142080 | Unspecified |
| mmu-miR-34a-5p | MIMAT0000542 | Tex261 | 21766 | Microarrays//HITS-CLIP | 37986224\|23027903\|23630078\|23142 | Unspecified |
| mmu-miR-34a-5p | MIMAT0000542 | Tex264 | 21767 | HITS-CLIP | 23142080\|29167373 | Unspecified |
| mmu-miR-34a-5p | MIMAT0000542 | Tex30 | 75623 | HITS-CLIP | 23142080 | Unspecified |
| mmu-miR-34a-5p | MIMAT0000542 | Tfam | 21780 | HITS-CLIP | 23142080 | Unspecified |
| mmu-miR-34a-5p | MIMAT0000542 | Tfap4 | 83383 | HITS-CLIP | 23142080 | Unspecified |
| mmu-miR-34a-5p | MIMAT0000542 | Tfdp1 | 21781 | HITS-CLIP | 23142080\|29167373 | Unspecified |
| mmu-miR-34a-5p | MIMAT0000542 | Tfdp2 | 211586 | HITS-CLIP | 23142080 | Unspecified |
| mmu-miR-34a-5p | MIMAT0000542 | Tfeb | 21425 | HITS-CLIP | 29167373 | Unspecified |
| mmu-miR-34a-5p | MIMAT0000542 | Tfec | 21426 | HITS-CLIP | 30971389 | Unspecified |
| mmu-miR-34a-5p | MIMAT0000542 | Tfg | 21787 | HITS-CLIP | 30971389\|23142080 | Unspecified |
| mmu-miR-34a-5p | MIMAT0000542 | Tfip11 | 54723 | HITS-CLIP | 23142080 | Unspecified |
| mmu-miR-34a-5p | MIMAT0000542 | Tfpi | 21788 | HITS-CLIP | 23142080 | Unspecified |
| mmu-miR-34a-5p | MIMAT0000542 | Tfpt | 69714 | HITS-CLIP | 23142080 | Unspecified |

| | | | | | | |
|---|---|---|---|---|---|---|
| mmu-miR-34a-5p | MIMAT0000542 | Tfrc | 22042 | HITS-CLIP, Microarrays//Microarrays//HITS | 37986224\|23027903\|23142080\|31433 | Unspecified |
| mmu-miR-34a-5p | MIMAT0000542 | Tg | 21819 | Microarrays//HITS-CLIP | 37986224\|31433990\|23142080 | Unspecified |
| mmu-miR-34a-5p | MIMAT0000542 | Tgds | 76355 | HITS-CLIP | 23142080\|19536157 | Unspecified |
| mmu-miR-34a-5p | MIMAT0000542 | Tgfb1 | 21803 | HITS-CLIP | 23142080\|29167373 | Unspecified |
| mmu-miR-34a-5p | MIMAT0000542 | Tgfbi | 21810 | HITS-CLIP | 29167373 | Unspecified |
| mmu-miR-34a-5p | MIMAT0000542 | Tgfbr1 | 21812 | HITS-CLIP | 23142080\|31433990\|23630078\|30971 | Unspecified |
| mmu-miR-34a-5p | MIMAT0000542 | Tgfbr2 | 21813 | Microarrays//HITS-CLIP | 37986224\|19536157\|26203562\|23142 | Unspecified |
| mmu-miR-34a-5p | MIMAT0000542 | Tgfbr3 | 21814 | HITS-CLIP | 31433990\|19536157\|23142080 | Unspecified |
| mmu-miR-34a-5p | MIMAT0000542 | Tgfbrap1 | 73122 | HITS-CLIP | 29167373 | Unspecified |
| mmu-miR-34a-5p | MIMAT0000542 | Tgif1 | 21815 | HITS-CLIP | 23142080\|29167373 | Unspecified |
| mmu-miR-34a-5p | MIMAT0000542 | Tgm2 | 21817 | HITS-CLIP | 23142080 | Unspecified |
| mmu-miR-34a-5p | MIMAT0000542 | Tgoln1 | 22134 | HITS-CLIP | 19536157\|23142080\|29167373 | Unspecified |
| mmu-miR-34a-5p | MIMAT0000542 | Tgs1 | 116940 | HITS-CLIP | 23142080\|29167373 | Unspecified |
| mmu-miR-34a-5p | MIMAT0000542 | Tgtp1 | 21822 | HITS-CLIP | 23142080 | Unspecified |
| mmu-miR-34a-5p | MIMAT0000542 | Tgtp2 | 100039796 | HITS-CLIP | 23142080 | Unspecified |
| mmu-miR-34a-5p | MIMAT0000542 | Thada | 240174 | HITS-CLIP | 23142080\|29167373 | Unspecified |
| mmu-miR-34a-5p | MIMAT0000542 | Thap12 | 72981 | HITS-CLIP | 23142080 | Unspecified |
| mmu-miR-34a-5p | MIMAT0000542 | Thap2 | 66816 | Microarrays | 37986224\|23027903 | Unspecified |
| mmu-miR-34a-5p | MIMAT0000542 | Thap3 | 69876 | HITS-CLIP | 23142080 | Unspecified |
| mmu-miR-34a-5p | MIMAT0000542 | Thap4 | 67026 | HITS-CLIP | 37986224\|23142080 | Unspecified |
| mmu-miR-34a-5p | MIMAT0000542 | Thbd | 21824 | HITS-CLIP | 19536157\|29167373 | Unspecified |
| mmu-miR-34a-5p | MIMAT0000542 | Thbs1 | 21825 | HITS-CLIP | 23142080 | Unspecified |
| mmu-miR-34a-5p | MIMAT0000542 | Them6 | 223626 | HITS-CLIP | 23142080 | Unspecified |
| mmu-miR-34a-5p | MIMAT0000542 | Themis | 210757 | HITS-CLIP | 23142080 | Unspecified |
| mmu-miR-34a-5p | MIMAT0000542 | Thoc2 | 331401 | HITS-CLIP | 23142080 | Unspecified |
| mmu-miR-34a-5p | MIMAT0000542 | Thoc3 | 73666 | HITS-CLIP | 23142080 | Unspecified |
| mmu-miR-34a-5p | MIMAT0000542 | Thoc5 | 8563 | HITS-CLIP | 23142080 | Unspecified |
| mmu-miR-34a-5p | MIMAT0000542 | Thoc7 | 66231 | HITS-CLIP | 23142080 | Unspecified |
| mmu-miR-34a-5p | MIMAT0000542 | Thop1 | 50492 | HITS-CLIP | 23142080\|30971389 | Unspecified |
| mmu-miR-34a-5p | MIMAT0000542 | Thrap3 | 230753 | HITS-CLIP | 29167373 | Unspecified |
| mmu-miR-34a-5p | MIMAT0000542 | Thumpd1 | 233802 | Microarrays//HITS-CLIP | 37986224\|23027903\|23142080 | Unspecified |
| mmu-miR-34a-5p | MIMAT0000542 | Thumpd3 | 14911 | HITS-CLIP | 23142080\|29167373 | Unspecified |
| mmu-miR-34a-5p | MIMAT0000542 | Thy1 | 21838 | HITS-CLIP | 23142080 | Unspecified |
| mmu-miR-34a-5p | MIMAT0000542 | Tia1 | 21841 | HITS-CLIP | 23142080 | Unspecified |
| mmu-miR-34a-5p | MIMAT0000542 | Tial1 | 21843 | HITS-CLIP | 19536157\|26203562\|23142080 | Unspecified |
| mmu-miR-34a-5p | MIMAT0000542 | Tiam1 | 21844 | Microarrays//HITS-CLIP | 37986224\|23027903\|23142080\|29167 | Unspecified |
| mmu-miR-34a-5p | MIMAT0000542 | Ticam1 | 106759 | HITS-CLIP | 23142080\|29167373 | Unspecified |
| mmu-miR-34a-5p | MIMAT0000542 | Ticam2 | 225471 | HITS-CLIP | 29167373 | Unspecified |
| mmu-miR-34a-5p | MIMAT0000542 | Ticrr | 77011 | Microarrays//HITS-CLIP | 37986224\|23027903\|23142080 | Unspecified |

| | | | | | | |
|---|---|---|---|---|---|---|
| mmu-miR-34a-5p | MIMAT0000542 | Tifa | 211550 | Microarrays | 37986224 | Unspecified |
| mmu-miR-34a-5p | MIMAT0000542 | Tifab | 212937 | HITS-CLIP | 29167373 | Unspecified |
| mmu-miR-34a-5p | MIMAT0000542 | Tigar | 319801 | HITS-CLIP | 23142080 | Unspecified |
| mmu-miR-34a-5p | MIMAT0000542 | Tigd2 | 68140 | HITS-CLIP | 23142080 | Unspecified |
| mmu-miR-34a-5p | MIMAT0000542 | Tigit | 100043314 | HITS-CLIP | 23142080 | Unspecified |
| mmu-miR-34a-5p | MIMAT0000542 | Timd2 | 171284 | HITS-CLIP | 23142080 | Unspecified |
| mmu-miR-34a-5p | MIMAT0000542 | Timeless | 21853 | HITS-CLIP | 29167373 | Unspecified |
| mmu-miR-34a-5p | MIMAT0000542 | Timm10 | 30059 | HITS-CLIP | 31433990\|26203562\|23142080 | Unspecified |
| mmu-miR-34a-5p | MIMAT0000542 | Timm10b | 14356 | HITS-CLIP | 29167373 | Unspecified |
| mmu-miR-34a-5p | MIMAT0000542 | Timm44 | 21856 | HITS-CLIP | 23142080 | Unspecified |
| mmu-miR-34a-5p | MIMAT0000542 | Timm50 | 66525 | HITS-CLIP | 23142080 | Unspecified |
| mmu-miR-34a-5p | MIMAT0000542 | Timm8b | 30057 | HITS-CLIP | 23142080 | Unspecified |
| mmu-miR-34a-5p | MIMAT0000542 | Timm9 | 30056 | HITS-CLIP | 23142080 | Unspecified |
| mmu-miR-34a-5p | MIMAT0000542 | Timp2 | 21858 | HITS-CLIP | 23142080 | Unspecified |
| mmu-miR-34a-5p | MIMAT0000542 | Timp3 | 21859 | HITS-CLIP | 26203562 | Unspecified |
| mmu-miR-34a-5p | MIMAT0000542 | Tinf2 | 28113 | Microarrays//HITS-CLIP | 37986224\|23027903\|23142080 | Unspecified |
| mmu-miR-34a-5p | MIMAT0000542 | Tiparp | 99929 | HITS-CLIP | 37986224\|23142080\|29167373 | Unspecified |
| mmu-miR-34a-5p | MIMAT0000542 | Tipin | 66131 | HITS-CLIP | 23142080 | Unspecified |
| mmu-miR-34a-5p | MIMAT0000542 | Tiprl | 226591 | HITS-CLIP | 23142080 | Unspecified |
| mmu-miR-34a-5p | MIMAT0000542 | Tirap | 117149 | HITS-CLIP | 23142080 | Unspecified |
| mmu-miR-34a-5p | MIMAT0000542 | Tjap1 | 74094 | HITS-CLIP | 23142080 | Unspecified |
| mmu-miR-34a-5p | MIMAT0000542 | Tk1 | 21877 | HITS-CLIP | 23142080\|31433990 | Unspecified |
| mmu-miR-34a-5p | MIMAT0000542 | Tk2 | 57813 | HITS-CLIP | 23142080 | Unspecified |
| mmu-miR-34a-5p | MIMAT0000542 | Tkt | 21881 | HITS-CLIP | 23142080\|31433990\|29167373 | Unspecified |
| mmu-miR-34a-5p | MIMAT0000542 | Tlcd1 | 68385 | HITS-CLIP | 23142080 | Unspecified |
| mmu-miR-34a-5p | MIMAT0000542 | Tlcd2 | 380712 | HITS-CLIP | 23142080 | Unspecified |
| mmu-miR-34a-5p | MIMAT0000542 | Tle1 | 21885 | HITS-CLIP | 29167373 | Unspecified |
| mmu-miR-34a-5p | MIMAT0000542 | Tle3 | 21887 | HITS-CLIP | 23142080\|23630078 | Unspecified |
| mmu-miR-34a-5p | MIMAT0000542 | Tle4 | 21888 | HITS-CLIP | 31433990\|23142080 | Unspecified |
| mmu-miR-34a-5p | MIMAT0000542 | Tlk1 | 228012 | HITS-CLIP | 30971389\|31433990\|23142080\|29167 | Unspecified |
| mmu-miR-34a-5p | MIMAT0000542 | Tlk2 | 24086 | HITS-CLIP | 23630078\|23142080\|29167373 | Unspecified |
| mmu-miR-34a-5p | MIMAT0000542 | Tln1 | 21894 | HITS-CLIP | 30971389\|31433990\|23630078\|23142 | Unspecified |
| mmu-miR-34a-5p | MIMAT0000542 | Tlnrd1 | 59274 | HITS-CLIP | 19536157\|23142080\|29167373 | Unspecified |
| mmu-miR-34a-5p | MIMAT0000542 | Tlr13 | 279572 | HITS-CLIP | 29167373 | Unspecified |
| mmu-miR-34a-5p | MIMAT0000542 | Tlr2 | 24088 | HITS-CLIP | 29167373 | Unspecified |
| mmu-miR-34a-5p | MIMAT0000542 | Tlr4 | 21898 | HITS-CLIP | 29167373 | Unspecified |
| mmu-miR-34a-5p | MIMAT0000542 | Tlr6 | 21899 | HITS-CLIP | 23142080 | Unspecified |
| mmu-miR-34a-5p | MIMAT0000542 | Tlr7 | 170743 | HITS-CLIP | 29167373 | Unspecified |
| mmu-miR-34a-5p | MIMAT0000542 | Tlr8 | 170744 | HITS-CLIP | 29167373 | Unspecified |

| | | | | | | |
|---|---|---|---|---|---|---|
| mmu-miR-34a-5p | MIMAT0000542 | Tlr9 | 81897 | HITS-CLIP | 29167373 | Unspecified |
| mmu-miR-34a-5p | MIMAT0000542 | Tm2d2 | 69742 | HITS-CLIP | 23142080 | Unspecified |
| mmu-miR-34a-5p | MIMAT0000542 | Tm6sf1 | 107769 | Microarrays | 37986224 | Unspecified |
| mmu-miR-34a-5p | MIMAT0000542 | Tm7sf2 | 73166 | HITS-CLIP | 23142080 | Unspecified |
| mmu-miR-34a-5p | MIMAT0000542 | Tm9sf1 | 74140 | HITS-CLIP | 23142080\|29167373 | Unspecified |
| mmu-miR-34a-5p | MIMAT0000542 | Tm9sf2 | 68059 | HITS-CLIP | 23142080\|19536157\|29167373 | Unspecified |
| mmu-miR-34a-5p | MIMAT0000542 | Tm9sf3 | 107358 | HITS-CLIP | 31433990\|23142080\|23630078\|29167 | Unspecified |
| mmu-miR-34a-5p | MIMAT0000542 | Tm9sf4 | 99237 | HITS-CLIP | 19536157\|23142080 | Unspecified |
| mmu-miR-34a-5p | MIMAT0000542 | Tma7 | 66167 | HITS-CLIP | 23142080\|19536157 | Unspecified |
| mmu-miR-34a-5p | MIMAT0000542 | Tmbim1 | 69660 | HITS-CLIP | 29167373 | Unspecified |
| mmu-miR-34a-5p | MIMAT0000542 | Tmbim4 | 68212 | HITS-CLIP | 23630078 | Unspecified |
| mmu-miR-34a-5p | MIMAT0000542 | Tmbim6 | 110213 | HITS-CLIP | 23142080\|19536157\|29167373 | Unspecified |
| mmu-miR-34a-5p | MIMAT0000542 | Tmc6 | 217353 | HITS-CLIP | 23142080\|30971389 | Unspecified |
| mmu-miR-34a-5p | MIMAT0000542 | Tmc7 | 209760 | HITS-CLIP | 19536157 | Unspecified |
| mmu-miR-34a-5p | MIMAT0000542 | Tmc8 | 217356 | HITS-CLIP | 23142080 | Unspecified |
| mmu-miR-34a-5p | MIMAT0000542 | Tmcc1 | 330401 | HITS-CLIP | 19536157\|23142080\|29167373 | Unspecified |
| mmu-miR-34a-5p | MIMAT0000542 | Tmcc3 | 319880 | HITS-CLIP | 29167373 | Unspecified |
| mmu-miR-34a-5p | MIMAT0000542 | Tmco1 | 68944 | HITS-CLIP | 23142080 | Unspecified |
| mmu-miR-34a-5p | MIMAT0000542 | Tmco3 | 234076 | HITS-CLIP | 23142080 | Unspecified |
| mmu-miR-34a-5p | MIMAT0000542 | Tmco6 | 71983 | HITS-CLIP | 23142080 | Unspecified |
| mmu-miR-34a-5p | MIMAT0000542 | Tmed3 | 66111 | HITS-CLIP | 23142080 | Unspecified |
| mmu-miR-34a-5p | MIMAT0000542 | Tmed4 | 103694 | HITS-CLIP | 19536157 | Unspecified |
| mmu-miR-34a-5p | MIMAT0000542 | Tmed5 | 73130 | HITS-CLIP | 23142080 | Unspecified |
| mmu-miR-34a-5p | MIMAT0000542 | Tmed7 | 66676 | HITS-CLIP | 23142080\|29167373 | Unspecified |
| mmu-miR-34a-5p | MIMAT0000542 | Tmed8 | 382620 | Microarrays//HITS-CLIP | 37986224\|23027903\|29167373 | Unspecified |
| mmu-miR-34a-5p | MIMAT0000542 | Tmed9 | 67511 | HITS-CLIP | 23142080\|29167373\|19536157 | Unspecified |
| mmu-miR-34a-5p | MIMAT0000542 | Tmeff2 | 56363 | HITS-CLIP | 19536157 | Unspecified |
| mmu-miR-34a-5p | MIMAT0000542 | Tmem101 | 76547 | HITS-CLIP | 23142080 | Unspecified |
| mmu-miR-34a-5p | MIMAT0000542 | Tmem104 | 320534 | HITS-CLIP | 29167373 | Unspecified |
| mmu-miR-34a-5p | MIMAT0000542 | Tmem106a | 217203 | HITS-CLIP | 29167373 | Unspecified |
| mmu-miR-34a-5p | MIMAT0000542 | Tmem106b | 71900 | HITS-CLIP | 31433990\|30971389\|23142080\|29167 | Unspecified |
| mmu-miR-34a-5p | MIMAT0000542 | Tmem108 | 81907 | HITS-CLIP | 19536157\|23142080 | Unspecified |
| mmu-miR-34a-5p | MIMAT0000542 | Tmem109 | 68539 | HITS-CLIP | 31433990\|29167373 | Unspecified |
| mmu-miR-34a-5p | MIMAT0000542 | Tmem11 | 216821 | HITS-CLIP | 23142080\|31433990 | Unspecified |
| mmu-miR-34a-5p | MIMAT0000542 | Tmem110 | 69179 | HITS-CLIP | 23142080\|29167373 | Unspecified |
| mmu-miR-34a-5p | MIMAT0000542 | Tmem123 | 71929 | HITS-CLIP | 37986224\|30971389\|19536157\|29084 | Unspecified |
| mmu-miR-34a-5p | MIMAT0000542 | Tmem126b | 68472 | HITS-CLIP | 23142080 | Unspecified |
| mmu-miR-34a-5p | MIMAT0000542 | Tmem127 | 69470 | HITS-CLIP | 23142080\|29167373 | Unspecified |
| mmu-miR-34a-5p | MIMAT0000542 | Tmem128 | 66309 | HITS-CLIP | 23142080\|29167373 | Unspecified |

| | | | | | | |
|---|---|---|---|---|---|---|
| mmu-miR-34a-5p | MIMAT0000542 | Tmem129 | 68366 | HITS-CLIP | 23142080\|29167373 | Unspecified |
| mmu-miR-34a-5p | MIMAT0000542 | Tmem130 | 243339 | HITS-CLIP | 19536157 | Unspecified |
| mmu-miR-34a-5p | MIMAT0000542 | Tmem131 | 56030 | HITS-CLIP | 23142080\|29167373 | Unspecified |
| mmu-miR-34a-5p | MIMAT0000542 | Tmem131l | 23240 | HITS-CLIP | 23142080 | Unspecified |
| mmu-miR-34a-5p | MIMAT0000542 | Tmem135 | 72759 | HITS-CLIP | 23142080\|29167373 | Unspecified |
| mmu-miR-34a-5p | MIMAT0000542 | Tmem143 | 70209 | HITS-CLIP | 23142080 | Unspecified |
| mmu-miR-34a-5p | MIMAT0000542 | Tmem145 | 330485 | HITS-CLIP | 19536157 | Unspecified |
| mmu-miR-34a-5p | MIMAT0000542 | Tmem147 | 69804 | HITS-CLIP | 19536157 | Unspecified |
| mmu-miR-34a-5p | MIMAT0000542 | Tmem14c | 66154 | HITS-CLIP | 23142080\|29167373 | Unspecified |
| mmu-miR-34a-5p | MIMAT0000542 | Tmem150a | 232086 | HITS-CLIP | 23142080 | Unspecified |
| mmu-miR-34a-5p | MIMAT0000542 | Tmem151b | 210573 | HITS-CLIP | 19536157 | Unspecified |
| mmu-miR-34a-5p | MIMAT0000542 | Tmem154 | 320782 | HITS-CLIP | 23142080 | Unspecified |
| mmu-miR-34a-5p | MIMAT0000542 | Tmem156 | 243025 | HITS-CLIP | 23142080 | Unspecified |
| mmu-miR-34a-5p | MIMAT0000542 | Tmem161a | 234371 | HITS-CLIP | 23142080\|23630078\|31433990 | Unspecified |
| mmu-miR-34a-5p | MIMAT0000542 | Tmem164 | 209497 | HITS-CLIP | 31433990\|23142080\|29167373 | Unspecified |
| mmu-miR-34a-5p | MIMAT0000542 | Tmem165 | 21982 | HITS-CLIP | 31433990\|29167373\|23142080 | Unspecified |
| mmu-miR-34a-5p | MIMAT0000542 | Tmem167 | 66074 | HITS-CLIP | 23142080\|29167373 | Unspecified |
| mmu-miR-34a-5p | MIMAT0000542 | Tmem167b | 67495 | HITS-CLIP | 23142080 | Unspecified |
| mmu-miR-34a-5p | MIMAT0000542 | Tmem168 | 101118 | HITS-CLIP | 23142080 | Unspecified |
| mmu-miR-34a-5p | MIMAT0000542 | Tmem170 | 66817 | HITS-CLIP | 30971389\|26203562\|23142080 | Unspecified |
| mmu-miR-34a-5p | MIMAT0000542 | Tmem170b | 621976 | Microarrays//HITS-CLIP | 37986224\|23142080\|19536157\|29167 | Unspecified |
| mmu-miR-34a-5p | MIMAT0000542 | Tmem173 | 72512 | HITS-CLIP | 23142080\|29167373 | Unspecified |
| mmu-miR-34a-5p | MIMAT0000542 | Tmem175 | 72392 | HITS-CLIP | 23142080 | Unspecified |
| mmu-miR-34a-5p | MIMAT0000542 | Tmem178 | 68027 | HITS-CLIP | 19536157 | Unspecified |
| mmu-miR-34a-5p | MIMAT0000542 | Tmem179b | 374395 | HITS-CLIP | 23142080 | Unspecified |
| mmu-miR-34a-5p | MIMAT0000542 | Tmem18 | 211986 | Microarrays//HITS-CLIP | 37986224\|23027903\|23142080 | Unspecified |
| mmu-miR-34a-5p | MIMAT0000542 | Tmem181a | 77106 | HITS-CLIP | 31433990 | Unspecified |
| mmu-miR-34a-5p | MIMAT0000542 | Tmem182 | 381339 | Microarrays | 37986224 | Unspecified |
| mmu-miR-34a-5p | MIMAT0000542 | Tmem183a | 57439 | HITS-CLIP | 23142080 | Unspecified |
| mmu-miR-34a-5p | MIMAT0000542 | Tmem184a | 231832 | Microarrays | 37986224 | Unspecified |
| mmu-miR-34a-5p | MIMAT0000542 | Tmem184b | 223693 | HITS-CLIP | 23630078\|23142080\|29167373 | Unspecified |
| mmu-miR-34a-5p | MIMAT0000542 | Tmem184c | 234463 | HITS-CLIP | 29167373 | Unspecified |
| mmu-miR-34a-5p | MIMAT0000542 | Tmem185a | 84548 | HITS-CLIP | 19536157 | Unspecified |
| mmu-miR-34a-5p | MIMAT0000542 | Tmem185b | 226351 | HITS-CLIP | 23142080 | Unspecified |
| mmu-miR-34a-5p | MIMAT0000542 | Tmem186 | 66690 | HITS-CLIP | 29167373\|23142080 | Unspecified |
| mmu-miR-34a-5p | MIMAT0000542 | Tmem189 | 407243 | HITS-CLIP | 29167373 | Unspecified |
| mmu-miR-34a-5p | MIMAT0000542 | Tmem19 | 67226 | HITS-CLIP | 23142080 | Unspecified |
| mmu-miR-34a-5p | MIMAT0000542 | Tmem192 | 73067 | HITS-CLIP | 37986224\|23142080\|29167373\|1953[illegible] | Unspecified |
| mmu-miR-34a-5p | MIMAT0000542 | Tmem198b | 73827 | HITS-CLIP | 23142080 | Unspecified |

| | | | | | | |
|---|---|---|---|---|---|---|
| mmu-miR-34a-5p | MIMAT0000542 | Tmem201 | 230917 | HITS-CLIP | 31433990 | Unspecified |
| mmu-miR-34a-5p | MIMAT0000542 | Tmem206 | 66950 | HITS-CLIP | 29167373 | Unspecified |
| mmu-miR-34a-5p | MIMAT0000542 | Tmem209 | 72649 | HITS-CLIP | 23142080 | Unspecified |
| mmu-miR-34a-5p | MIMAT0000542 | Tmem214 | 68796 | HITS-CLIP | 23142080\|30971389 | Unspecified |
| mmu-miR-34a-5p | MIMAT0000542 | Tmem222 | 52174 | HITS-CLIP | 23142080 | Unspecified |
| mmu-miR-34a-5p | MIMAT0000542 | Tmem229a | 319832 | HITS-CLIP | 19536157 | Unspecified |
| mmu-miR-34a-5p | MIMAT0000542 | Tmem229b | 268567 | HITS-CLIP | 31433990\|29167373 | Unspecified |
| mmu-miR-34a-5p | MIMAT0000542 | Tmem230 | 70612 | HITS-CLIP | 31433990\|23142080 | Unspecified |
| mmu-miR-34a-5p | MIMAT0000542 | Tmem231 | 234740 | Microarrays | 37986224 | Unspecified |
| mmu-miR-34a-5p | MIMAT0000542 | Tmem234 | 76799 | HITS-CLIP | 23142080\|29167373 | Unspecified |
| mmu-miR-34a-5p | MIMAT0000542 | Tmem242 | 70544 | HITS-CLIP | 23142080 | Unspecified |
| mmu-miR-34a-5p | MIMAT0000542 | Tmem243 | 652925 | HITS-CLIP | 23142080 | Unspecified |
| mmu-miR-34a-5p | MIMAT0000542 | Tmem245 | 242474 | HITS-CLIP | 19536157\|23142080\|29167373 | Unspecified |
| mmu-miR-34a-5p | MIMAT0000542 | Tmem248 | 71667 | HITS-CLIP | 29167373\|23142080 | Unspecified |
| mmu-miR-34a-5p | MIMAT0000542 | Tmem251 | 320351 | HITS-CLIP | 23142080 | Unspecified |
| mmu-miR-34a-5p | MIMAT0000542 | Tmem256 | 254863 | HITS-CLIP | 23142080 | Unspecified |
| mmu-miR-34a-5p | MIMAT0000542 | Tmem258 | 69038 | HITS-CLIP | 23142080 | Unspecified |
| mmu-miR-34a-5p | MIMAT0000542 | Tmem26 | 327766 | HITS-CLIP | 29167373 | Unspecified |
| mmu-miR-34a-5p | MIMAT0000542 | Tmem260 | 218989 | Microarrays | 37986224 | Unspecified |
| mmu-miR-34a-5p | MIMAT0000542 | Tmem263 | 103266 | HITS-CLIP | 30971389\|19536157 | Unspecified |
| mmu-miR-34a-5p | MIMAT0000542 | Tmem30a | 69981 | HITS-CLIP | 19536157\|23142080\|29167373 | Unspecified |
| mmu-miR-34a-5p | MIMAT0000542 | Tmem33 | 67878 | Microarrays//HITS-CLIP | 37986224\|23027903\|23142080\|29167 | Unspecified |
| mmu-miR-34a-5p | MIMAT0000542 | Tmem35a | 67564 | HITS-CLIP | 19536157 | Unspecified |
| mmu-miR-34a-5p | MIMAT0000542 | Tmem38b | 52076 | HITS-CLIP | 23142080\|29167373 | Unspecified |
| mmu-miR-34a-5p | MIMAT0000542 | Tmem39a | 67846 | HITS-CLIP | 23142080 | Unspecified |
| mmu-miR-34a-5p | MIMAT0000542 | Tmem39b | 230770 | HITS-CLIP | 19536157\|23142080\|29167373 | Unspecified |
| mmu-miR-34a-5p | MIMAT0000542 | Tmem40 | 94346 | HITS-CLIP | 23142080 | Unspecified |
| mmu-miR-34a-5p | MIMAT0000542 | Tmem45a | 56277 | Microarrays//HITS-CLIP | 37986224\|23027903\|23142080 | Unspecified |
| mmu-miR-34a-5p | MIMAT0000542 | Tmem47 | 192216 | HITS-CLIP | 19536157 | Unspecified |
| mmu-miR-34a-5p | MIMAT0000542 | Tmem50b | 77975 | HITS-CLIP | 19536157\|29167373 | Unspecified |
| mmu-miR-34a-5p | MIMAT0000542 | Tmem53 | 68777 | Microarrays | 37986224 | Unspecified |
| mmu-miR-34a-5p | MIMAT0000542 | Tmem56 | 99887 | HITS-CLIP | 19536157 | Unspecified |
| mmu-miR-34a-5p | MIMAT0000542 | Tmem59 | 56374 | HITS-CLIP | 31433990\|23630078\|23142080\|29167 | Unspecified |
| mmu-miR-34a-5p | MIMAT0000542 | Tmem62 | 96957 | Microarrays//HITS-CLIP | 37986224\|23027903\|23142080 | Unspecified |
| mmu-miR-34a-5p | MIMAT0000542 | Tmem63b | 224807 | HITS-CLIP | 23142080 | Unspecified |
| mmu-miR-34a-5p | MIMAT0000542 | Tmem64 | 100201 | Microarrays//HITS-CLIP | 37986224\|23142080\|29167373 | Unspecified |
| mmu-miR-34a-5p | MIMAT0000542 | Tmem65 | 74868 | HITS-CLIP | 37986224\|19536157\|23142080 | Unspecified |
| mmu-miR-34a-5p | MIMAT0000542 | Tmem67 | 329795 | HITS-CLIP | 23142080 | Unspecified |
| mmu-miR-34a-5p | MIMAT0000542 | Tmem68 | 72098 | HITS-CLIP | 23142080 | Unspecified |

| | | | | | | |
|---|---|---|---|---|---|---|
| mmu-miR-34a-5p | MIMAT0000542 | Tmem70 | 70397 | HITS-CLIP | 23142080\|29167373 | Unspecified |
| mmu-miR-34a-5p | MIMAT0000542 | Tmem71 | 213068 | HITS-CLIP | 23142080 | Unspecified |
| mmu-miR-34a-5p | MIMAT0000542 | Tmem86a | 67893 | HITS-CLIP | 29167373 | Unspecified |
| mmu-miR-34a-5p | MIMAT0000542 | Tmem87a | 211499 | HITS-CLIP | 19536157\|23142080\|29167373 | Unspecified |
| mmu-miR-34a-5p | MIMAT0000542 | Tmem87b | 72477 | HITS-CLIP | 23142080\|29167373 | Unspecified |
| mmu-miR-34a-5p | MIMAT0000542 | Tmem94 | 9772 | HITS-CLIP | 19536157\|23142080 | Unspecified |
| mmu-miR-34a-5p | MIMAT0000542 | Tmem97 | 69071 | HITS-CLIP | 23142080 | Unspecified |
| mmu-miR-34a-5p | MIMAT0000542 | Tmem9b | 56786 | HITS-CLIP | 19536157\|23142080\|29167373 | Unspecified |
| mmu-miR-34a-5p | MIMAT0000542 | Tmf1 | 232286 | HITS-CLIP | 23142080\|29167373 | Unspecified |
| mmu-miR-34a-5p | MIMAT0000542 | Tmod1 | 21916 | HITS-CLIP | 29167373 | Unspecified |
| mmu-miR-34a-5p | MIMAT0000542 | Tmod2 | 50876 | HITS-CLIP | 19536157 | Unspecified |
| mmu-miR-34a-5p | MIMAT0000542 | Tmod3 | 50875 | HITS-CLIP | 29167373 | Unspecified |
| mmu-miR-34a-5p | MIMAT0000542 | Tmpo | 21917 | HITS-CLIP | 23142080\|31433990\|26203562\|29167 | Unspecified |
| mmu-miR-34a-5p | MIMAT0000542 | Tmppe | 100504715 | HITS-CLIP | 23142080 | Unspecified |
| mmu-miR-34a-5p | MIMAT0000542 | Tmsb10 | 19240 | HITS-CLIP | 23142080 | Unspecified |
| mmu-miR-34a-5p | MIMAT0000542 | Tmsb15a | 78478 | Microarrays | 37986224 | Unspecified |
| mmu-miR-34a-5p | MIMAT0000542 | Tmsb4x | 19241 | HITS-CLIP | 19536157 | Unspecified |
| mmu-miR-34a-5p | MIMAT0000542 | Tmtc2 | 278279 | HITS-CLIP | 19536157 | Unspecified |
| mmu-miR-34a-5p | MIMAT0000542 | Tmtc3 | 237500 | HITS-CLIP | 23630078\|29167373 | Unspecified |
| mmu-miR-34a-5p | MIMAT0000542 | Tmub1 | 64295 | HITS-CLIP | 23142080 | Unspecified |
| mmu-miR-34a-5p | MIMAT0000542 | Tmub2 | 72053 | HITS-CLIP | 23142080 | Unspecified |
| mmu-miR-34a-5p | MIMAT0000542 | Tmx1 | 72736 | HITS-CLIP | 23142080 | Unspecified |
| mmu-miR-34a-5p | MIMAT0000542 | Tmx3 | 67988 | HITS-CLIP | 23142080\|29167373 | Unspecified |
| mmu-miR-34a-5p | MIMAT0000542 | Tmx4 | 52837 | HITS-CLIP | 37986224\|23142080\|29167373 | Unspecified |
| mmu-miR-34a-5p | MIMAT0000542 | Tnf | 21926 | ELISA, qPCR//qPCR//ELISA//HITS-CLIP | 37986224\|22483937\|23142080 | Unspecified |
| mmu-miR-34a-5p | MIMAT0000542 | Tnfaip1 | 21927 | HITS-CLIP | 19536157\|23142080\|29167373 | Unspecified |
| mmu-miR-34a-5p | MIMAT0000542 | Tnfaip2 | 21928 | HITS-CLIP | 29167373 | Unspecified |
| mmu-miR-34a-5p | MIMAT0000542 | Tnfaip3 | 21929 | HITS-CLIP | 23142080\|26203562\|29167373 | Unspecified |
| mmu-miR-34a-5p | MIMAT0000542 | Tnfaip8 | 106869 | HITS-CLIP | 23142080\|29167373 | Unspecified |
| mmu-miR-34a-5p | MIMAT0000542 | Tnfaip8l1 | 66443 | HITS-CLIP | 23142080 | Unspecified |
| mmu-miR-34a-5p | MIMAT0000542 | Tnfrsf10b | 21933 | HITS-CLIP | 23142080 | Unspecified |
| mmu-miR-34a-5p | MIMAT0000542 | Tnfrsf11a | 21934 | HITS-CLIP | 29167373 | Unspecified |
| mmu-miR-34a-5p | MIMAT0000542 | Tnfrsf18 | 21936 | HITS-CLIP | 37986224\|23142080 | Unspecified |
| mmu-miR-34a-5p | MIMAT0000542 | Tnfrsf19 | 29820 | HITS-CLIP | 19536157 | Unspecified |
| mmu-miR-34a-5p | MIMAT0000542 | Tnfrsf1a | 21937 | HITS-CLIP | 23142080\|29167373 | Unspecified |
| mmu-miR-34a-5p | MIMAT0000542 | Tnfrsf1b | 21938 | HITS-CLIP | 29167373\|23142080 | Unspecified |
| mmu-miR-34a-5p | MIMAT0000542 | Tnfrsf21 | 94185 | HITS-CLIP | 19536157 | Unspecified |
| mmu-miR-34a-5p | MIMAT0000542 | Tnfrsf22 | 79202 | HITS-CLIP | 23142080 | Unspecified |
| mmu-miR-34a-5p | MIMAT0000542 | Tnfrsf26 | 244237 | HITS-CLIP | 23142080 | Unspecified |

| | | | | | | |
|---|---|---|---|---|---|---|
| mmu-miR-34a-5p | MIMAT0000542 | Tnfrsf4 | 22163 | HITS-CLIP | 23142080 | Unspecified |
| mmu-miR-34a-5p | MIMAT0000542 | Tnfrsf8 | 21941 | HITS-CLIP | 23142080 | Unspecified |
| mmu-miR-34a-5p | MIMAT0000542 | Tnfrsf9 | 21942 | HITS-CLIP | 23142080 | Unspecified |
| mmu-miR-34a-5p | MIMAT0000542 | Tnfsf10 | 22035 | HITS-CLIP | 23142080 | Unspecified |
| mmu-miR-34a-5p | MIMAT0000542 | Tnfsf11 | 21943 | HITS-CLIP | 31433990\|23142080 | Unspecified |
| mmu-miR-34a-5p | MIMAT0000542 | Tnfsf12 | 21944 | HITS-CLIP | 29167373 | Unspecified |
| mmu-miR-34a-5p | MIMAT0000542 | Tnfsf4 | 22164 | HITS-CLIP | 31433990 | Unspecified |
| mmu-miR-34a-5p | MIMAT0000542 | Tnfsf8 | 21949 | HITS-CLIP | 23142080 | Unspecified |
| mmu-miR-34a-5p | MIMAT0000542 | Tnfsf9 | 21950 | HITS-CLIP | 23142080\|29167373 | Unspecified |
| mmu-miR-34a-5p | MIMAT0000542 | Tnik | 665113 | HITS-CLIP | 23142080 | Unspecified |
| mmu-miR-34a-5p | MIMAT0000542 | Tnip1 | 57783 | HITS-CLIP | 23142080 | Unspecified |
| mmu-miR-34a-5p | MIMAT0000542 | Tnip2 | 231130 | HITS-CLIP | 29167373 | Unspecified |
| mmu-miR-34a-5p | MIMAT0000542 | Tnip3 | 414084 | HITS-CLIP | 29167373 | Unspecified |
| mmu-miR-34a-5p | MIMAT0000542 | Tnks | 21951 | HITS-CLIP | 19536157\|23142080\|29167373 | Unspecified |
| mmu-miR-34a-5p | MIMAT0000542 | Tnks2 | 74493 | HITS-CLIP | 23142080\|31433990\|19536157 | Unspecified |
| mmu-miR-34a-5p | MIMAT0000542 | Tnnc1 | 21924 | Microarrays | 37986224 | Unspecified |
| mmu-miR-34a-5p | MIMAT0000542 | Tnni1 | 21952 | Microarrays | 37986224 | Unspecified |
| mmu-miR-34a-5p | MIMAT0000542 | Tnnt1 | 21955 | Microarrays | 37986224 | Unspecified |
| mmu-miR-34a-5p | MIMAT0000542 | Tnnt3 | 21957 | Microarrays | 37986224 | Unspecified |
| mmu-miR-34a-5p | MIMAT0000542 | Tnpo1 | 238799 | HITS-CLIP | 19536157\|31433990\|23630078\|30971 | Unspecified |
| mmu-miR-34a-5p | MIMAT0000542 | Tnpo2 | 212999 | HITS-CLIP | 23142080 | Unspecified |
| mmu-miR-34a-5p | MIMAT0000542 | Tnpo3 | 320938 | HITS-CLIP | 23142080\|29167373 | Unspecified |
| mmu-miR-34a-5p | MIMAT0000542 | Tnrc18 | 231861 | HITS-CLIP | 23142080\|29167373 | Unspecified |
| mmu-miR-34a-5p | MIMAT0000542 | Tnrc6a | 233833 | HITS-CLIP, Microarrays//HITS-CLIP | 37986224\|31433990\|19536157\|23142 | Unspecified |
| mmu-miR-34a-5p | MIMAT0000542 | Tnrc6b | 213988 | HITS-CLIP | 37986224\|19536157\|23142080\|29167 | Unspecified |
| mmu-miR-34a-5p | MIMAT0000542 | Tnrc6c | 217351 | HITS-CLIP | 37986224\|31433990\|23142080\|29167 | Unspecified |
| mmu-miR-34a-5p | MIMAT0000542 | Tns1 | 21961 | HITS-CLIP | 23142080\|30971389\|19536157 | Unspecified |
| mmu-miR-34a-5p | MIMAT0000542 | Tns3 | 319939 | HITS-CLIP | 19536157\|29167373 | Unspecified |
| mmu-miR-34a-5p | MIMAT0000542 | Tns4 | 217169 | HITS-CLIP | 23142080 | Unspecified |
| mmu-miR-34a-5p | MIMAT0000542 | Tob1 | 22057 | Microarrays//HITS-CLIP | 37986224\|19536157\|23142080 | Unspecified |
| mmu-miR-34a-5p | MIMAT0000542 | Tob2 | 57259 | HITS-CLIP | 23630078\|23142080 | Unspecified |
| mmu-miR-34a-5p | MIMAT0000542 | Toe1 | 68276 | HITS-CLIP | 31433990 | Unspecified |
| mmu-miR-34a-5p | MIMAT0000542 | Togaram1 | 23116 | HITS-CLIP | 23142080\|29167373 | Unspecified |
| mmu-miR-34a-5p | MIMAT0000542 | Tollip | 54473 | HITS-CLIP | 23142080\|19536157 | Unspecified |
| mmu-miR-34a-5p | MIMAT0000542 | Tom1l2 | 216810 | HITS-CLIP | 19536157\|23142080\|29167373 | Unspecified |
| mmu-miR-34a-5p | MIMAT0000542 | Tomm20 | 67952 | HITS-CLIP | 23142080\|29167373 | Unspecified |
| mmu-miR-34a-5p | MIMAT0000542 | Tomm34 | 67145 | HITS-CLIP | 23142080\|29167373 | Unspecified |
| mmu-miR-34a-5p | MIMAT0000542 | Tomm40 | 53333 | HITS-CLIP | 23142080 | Unspecified |
| mmu-miR-34a-5p | MIMAT0000542 | Tomm70a | 28185 | HITS-CLIP | 23142080\|19536157 | Unspecified |

| | | | | | | |
|---|---|---|---|---|---|---|
| mmu-miR-34a-5p | MIMAT0000542 | Tonsl | 72749 | HITS-CLIP | 23142080 | Unspecified |
| mmu-miR-34a-5p | MIMAT0000542 | Top1 | 21969 | HITS-CLIP | 37986224\|31433990\|23630078\|23142 | Unspecified |
| mmu-miR-34a-5p | MIMAT0000542 | Top2a | 21973 | HITS-CLIP | 37986224\|31433990\|23142080 | Unspecified |
| mmu-miR-34a-5p | MIMAT0000542 | Top2b | 21974 | HITS-CLIP | 23142080\|23630078\|19536157\|29167 | Unspecified |
| mmu-miR-34a-5p | MIMAT0000542 | Top3a | 21975 | HITS-CLIP | 29167373 | Unspecified |
| mmu-miR-34a-5p | MIMAT0000542 | Top3b | 21976 | HITS-CLIP | 23142080\|29167373 | Unspecified |
| mmu-miR-34a-5p | MIMAT0000542 | Topbp1 | 235559 | HITS-CLIP | 31433990\|23142080\|29167373 | Unspecified |
| mmu-miR-34a-5p | MIMAT0000542 | Topors | 106021 | HITS-CLIP | 23630078\|31433990\|23142080\|29167 | Unspecified |
| mmu-miR-34a-5p | MIMAT0000542 | Tor1aip1 | 208263 | HITS-CLIP | 23142080\|29167373 | Unspecified |
| mmu-miR-34a-5p | MIMAT0000542 | Tor1aip2 | 240832 | HITS-CLIP | 37986224\|23142080\|19536157\|29167 | Unspecified |
| mmu-miR-34a-5p | MIMAT0000542 | Tor1b | 30934 | HITS-CLIP | 30971389 | Unspecified |
| mmu-miR-34a-5p | MIMAT0000542 | Tor2a | 30933 | HITS-CLIP | 23142080 | Unspecified |
| mmu-miR-34a-5p | MIMAT0000542 | Tor3a | 30935 | HITS-CLIP | 23630078\|29167373 | Unspecified |
| mmu-miR-34a-5p | MIMAT0000542 | Tor4a | 227612 | HITS-CLIP | 31433990\|23142080\|29167373 | Unspecified |
| mmu-miR-34a-5p | MIMAT0000542 | Tox | 252838 | HITS-CLIP | 23142080 | Unspecified |
| mmu-miR-34a-5p | MIMAT0000542 | Tox4 | 268741 | HITS-CLIP | 23142080 | Unspecified |
| mmu-miR-34a-5p | MIMAT0000542 | Tpbg | 21983 | HITS-CLIP | 23142080 | Unspecified |
| mmu-miR-34a-5p | MIMAT0000542 | Tpcn1 | 252972 | HITS-CLIP | 19536157\|23142080 | Unspecified |
| mmu-miR-34a-5p | MIMAT0000542 | Tpcn2 | 233979 | HITS-CLIP | 29167373 | Unspecified |
| mmu-miR-34a-5p | MIMAT0000542 | Tpd52 | 21985 | Microarrays//HITS-CLIP | 37986224\|23027903\|29167373 | Unspecified |
| mmu-miR-34a-5p | MIMAT0000542 | Tpd52l2 | 66314 | HITS-CLIP | 37986224\|23142080 | Unspecified |
| mmu-miR-34a-5p | MIMAT0000542 | Tpi1 | 21991 | HITS-CLIP | 23142080\|19536157\|29167373 | Unspecified |
| mmu-miR-34a-5p | MIMAT0000542 | Tpk1 | 29807 | HITS-CLIP | 23142080\|29167373 | Unspecified |
| mmu-miR-34a-5p | MIMAT0000542 | Tpm1 | 22003 | HITS-CLIP | 23142080 | Unspecified |
| mmu-miR-34a-5p | MIMAT0000542 | Tpm3 | 59069 | HITS-CLIP | 23142080\|29167373 | Unspecified |
| mmu-miR-34a-5p | MIMAT0000542 | Tpm3-rs7 | 621054 | HITS-CLIP | 26203562 | Unspecified |
| mmu-miR-34a-5p | MIMAT0000542 | Tpm4 | 326618 | HITS-CLIP | 23142080 | Unspecified |
| mmu-miR-34a-5p | MIMAT0000542 | Tpp1 | 12751 | HITS-CLIP | 23142080\|29167373 | Unspecified |
| mmu-miR-34a-5p | MIMAT0000542 | Tpp2 | 22019 | HITS-CLIP | 23142080\|29167373 | Unspecified |
| mmu-miR-34a-5p | MIMAT0000542 | Tpr | 108989 | HITS-CLIP | 23142080\|29167373 | Unspecified |
| mmu-miR-34a-5p | MIMAT0000542 | Tpra1 | 24100 | HITS-CLIP | 31433990 | Unspecified |
| mmu-miR-34a-5p | MIMAT0000542 | Tprkb | 69786 | HITS-CLIP | 19536157\|23142080 | Unspecified |
| mmu-miR-34a-5p | MIMAT0000542 | Tprn | 97031 | HITS-CLIP | 19536157 | Unspecified |
| mmu-miR-34a-5p | MIMAT0000542 | Tpx2 | 72119 | HITS-CLIP | 23142080 | Unspecified |
| mmu-miR-34a-5p | MIMAT0000542 | Tra2a | 101214 | HITS-CLIP | 23142080 | Unspecified |
| mmu-miR-34a-5p | MIMAT0000542 | Tra2b | 20462 | HITS-CLIP | 23142080 | Unspecified |
| mmu-miR-34a-5p | MIMAT0000542 | Trabd | 67976 | HITS-CLIP | 23142080 | Unspecified |
| mmu-miR-34a-5p | MIMAT0000542 | Traf1 | 22029 | HITS-CLIP | 23142080 | Unspecified |
| mmu-miR-34a-5p | MIMAT0000542 | Traf3 | 22031 | HITS-CLIP | 31433990\|23142080\|19536157\|29167 | Unspecified |

| | | | | | | |
|---|---|---|---|---|---|---|
| mmu-miR-34a-5p | MIMAT0000542 | Traf3ip3 | 215243 | HITS-CLIP | 23142080 | Unspecified |
| mmu-miR-34a-5p | MIMAT0000542 | Traf6 | 22034 | HITS-CLIP | 23142080\|19536157\|29167373 | Unspecified |
| mmu-miR-34a-5p | MIMAT0000542 | Traf7 | 224619 | HITS-CLIP | 23142080 | Unspecified |
| mmu-miR-34a-5p | MIMAT0000542 | Trafd1 | 231712 | HITS-CLIP | 29167373\|23142080 | Unspecified |
| mmu-miR-34a-5p | MIMAT0000542 | Trak1 | 67095 | HITS-CLIP | 23142080\|29167373 | Unspecified |
| mmu-miR-34a-5p | MIMAT0000542 | Trak2 | 70827 | Microarrays//HITS-CLIP | 37986224\|23027903\|23142080 | Unspecified |
| mmu-miR-34a-5p | MIMAT0000542 | Tram1 | 72265 | HITS-CLIP | 30971389\|23142080\|29167373 | Unspecified |
| mmu-miR-34a-5p | MIMAT0000542 | Tram2 | 170829 | HITS-CLIP | 29167373\|23142080 | Unspecified |
| mmu-miR-34a-5p | MIMAT0000542 | Trap1 | 68015 | HITS-CLIP | 23142080\|29167373 | Unspecified |
| mmu-miR-34a-5p | MIMAT0000542 | Trappc1 | 245828 | HITS-CLIP | 19536157\|23142080 | Unspecified |
| mmu-miR-34a-5p | MIMAT0000542 | Trappc10 | 216131 | HITS-CLIP | 23142080\|19536157\|29167373 | Unspecified |
| mmu-miR-34a-5p | MIMAT0000542 | Trappc11 | 320714 | HITS-CLIP | 19536157\|23142080 | Unspecified |
| mmu-miR-34a-5p | MIMAT0000542 | Trappc12 | 217449 | HITS-CLIP | 23142080 | Unspecified |
| mmu-miR-34a-5p | MIMAT0000542 | Trappc13 | 66975 | HITS-CLIP | 23142080 | Unspecified |
| mmu-miR-34a-5p | MIMAT0000542 | Trappc4 | 60409 | HITS-CLIP | 23630078\|23142080 | Unspecified |
| mmu-miR-34a-5p | MIMAT0000542 | Trappc5 | 66682 | HITS-CLIP | 23142080 | Unspecified |
| mmu-miR-34a-5p | MIMAT0000542 | Trappc8 | 75964 | HITS-CLIP | 23142080 | Unspecified |
| mmu-miR-34a-5p | MIMAT0000542 | Trappc9 | 76510 | HITS-CLIP | 23142080 | Unspecified |
| mmu-miR-34a-5p | MIMAT0000542 | Treml2 | 328833 | HITS-CLIP | 23142080 | Unspecified |
| mmu-miR-34a-5p | MIMAT0000542 | Trerf1 | 224829 | HITS-CLIP | 31433990\|26203562 | Unspecified |
| mmu-miR-34a-5p | MIMAT0000542 | Trex1 | 22040 | HITS-CLIP | 23142080\|29167373 | Unspecified |
| mmu-miR-34a-5p | MIMAT0000542 | Triap1 | 69076 | HITS-CLIP | 23142080 | Unspecified |
| mmu-miR-34a-5p | MIMAT0000542 | Trib1 | 211770 | HITS-CLIP | 23142080\|29167373 | Unspecified |
| mmu-miR-34a-5p | MIMAT0000542 | Trib2 | 217410 | HITS-CLIP | 23142080 | Unspecified |
| mmu-miR-34a-5p | MIMAT0000542 | Trib3 | 228775 | HITS-CLIP | 23142080 | Unspecified |
| mmu-miR-34a-5p | MIMAT0000542 | Tril | 66873 | HITS-CLIP | 19536157 | Unspecified |
| mmu-miR-34a-5p | MIMAT0000542 | Trim11 | 94091 | HITS-CLIP | 29167373 | Unspecified |
| mmu-miR-34a-5p | MIMAT0000542 | Trim12c | 319236 | HITS-CLIP | 23142080\|29167373 | Unspecified |
| mmu-miR-34a-5p | MIMAT0000542 | Trim14 | 74735 | HITS-CLIP | 29167373 | Unspecified |
| mmu-miR-34a-5p | MIMAT0000542 | Trim2 | 80890 | HITS-CLIP | 19536157 | Unspecified |
| mmu-miR-34a-5p | MIMAT0000542 | Trim21 | 20821 | HITS-CLIP | 31433990\|29167373\|23142080 | Unspecified |
| mmu-miR-34a-5p | MIMAT0000542 | Trim23 | 81003 | HITS-CLIP | 23142080 | Unspecified |
| mmu-miR-34a-5p | MIMAT0000542 | Trim24 | 21848 | HITS-CLIP | 23142080 | Unspecified |
| mmu-miR-34a-5p | MIMAT0000542 | Trim25 | 217069 | HITS-CLIP | 23142080\|29167373 | Unspecified |
| mmu-miR-34a-5p | MIMAT0000542 | Trim26 | 22670 | HITS-CLIP | 23142080\|23630078\|31433990\|2620[illegible] | Unspecified |
| mmu-miR-34a-5p | MIMAT0000542 | Trim27 | 19720 | HITS-CLIP | 23630078\|31433990 | Unspecified |
| mmu-miR-34a-5p | MIMAT0000542 | Trim28 | 21849 | HITS-CLIP | 23142080 | Unspecified |
| mmu-miR-34a-5p | MIMAT0000542 | Trim30a | 20128 | HITS-CLIP | 29167373 | Unspecified |
| mmu-miR-34a-5p | MIMAT0000542 | Trim30d | 209387 | HITS-CLIP | 23142080 | Unspecified |

| | | | | | | |
|---|---|---|---|---|---|---|
| mmu-miR-34a-5p | MIMAT0000542 | Trim32 | 69807 | HITS-CLIP | 23142080 | Unspecified |
| mmu-miR-34a-5p | MIMAT0000542 | Trim33 | 94093 | HITS-CLIP | 29167373\|23142080 | Unspecified |
| mmu-miR-34a-5p | MIMAT0000542 | Trim34a | 94094 | HITS-CLIP | 29167373\|23142080 | Unspecified |
| mmu-miR-34a-5p | MIMAT0000542 | Trim35 | 66854 | HITS-CLIP | 19536157\|23142080\|29167373 | Unspecified |
| mmu-miR-34a-5p | MIMAT0000542 | Trim36 | 28105 | HITS-CLIP | 21258322\|23142080 | Unspecified |
| mmu-miR-34a-5p | MIMAT0000542 | Trim37 | 68729 | HITS-CLIP | 19536157\|30971389\|23142080 | Unspecified |
| mmu-miR-34a-5p | MIMAT0000542 | Trim44 | 80985 | HITS-CLIP | 31433990\|23142080\|26203562 | Unspecified |
| mmu-miR-34a-5p | MIMAT0000542 | Trim46 | 360213 | HITS-CLIP | 23630078 | Unspecified |
| mmu-miR-34a-5p | MIMAT0000542 | Trim55 | 381485 | Microarrays | 37986224 | Unspecified |
| mmu-miR-34a-5p | MIMAT0000542 | Trim56 | 384309 | HITS-CLIP | 29167373 | Unspecified |
| mmu-miR-34a-5p | MIMAT0000542 | Trim59 | 66949 | HITS-CLIP | 23142080 | Unspecified |
| mmu-miR-34a-5p | MIMAT0000542 | Trim62 | 67525 | HITS-CLIP | 19536157 | Unspecified |
| mmu-miR-34a-5p | MIMAT0000542 | Trim65 | 338364 | HITS-CLIP | 23142080 | Unspecified |
| mmu-miR-34a-5p | MIMAT0000542 | Trim72 | 434246 | Microarrays | 37986224 | Unspecified |
| mmu-miR-34a-5p | MIMAT0000542 | Trim8 | 93679 | HITS-CLIP | 23142080\|29167373 | Unspecified |
| mmu-miR-34a-5p | MIMAT0000542 | Trim9 | 94090 | HITS-CLIP | 19536157 | Unspecified |
| mmu-miR-34a-5p | MIMAT0000542 | Trio | 223435 | HITS-CLIP | 30971389\|23630078\|19536157\|29167 | Unspecified |
| mmu-miR-34a-5p | MIMAT0000542 | Triobp | 110253 | HITS-CLIP | 23142080\|29167373 | Unspecified |
| mmu-miR-34a-5p | MIMAT0000542 | Trip10 | 106628 | HITS-CLIP | 23142080 | Unspecified |
| mmu-miR-34a-5p | MIMAT0000542 | Trip11 | 109181 | HITS-CLIP | 23142080 | Unspecified |
| mmu-miR-34a-5p | MIMAT0000542 | Trip12 | 14897 | HITS-CLIP | 23142080\|29167373 | Unspecified |
| mmu-miR-34a-5p | MIMAT0000542 | Trip13 | 69716 | HITS-CLIP | 30971389\|23142080 | Unspecified |
| mmu-miR-34a-5p | MIMAT0000542 | Triqk | 208820 | Microarrays | 37986224 | Unspecified |
| mmu-miR-34a-5p | MIMAT0000542 | Trir | 79002 | HITS-CLIP | 31433990\|23142080 | Unspecified |
| mmu-miR-34a-5p | MIMAT0000542 | Trmt10a | 108943 | HITS-CLIP | 21258322\|23142080 | Unspecified |
| mmu-miR-34a-5p | MIMAT0000542 | Trmt10b | 69934 | HITS-CLIP | 23142080 | Unspecified |
| mmu-miR-34a-5p | MIMAT0000542 | Trmt11 | 73681 | HITS-CLIP | 23142080 | Unspecified |
| mmu-miR-34a-5p | MIMAT0000542 | Trmt2a | 15547 | HITS-CLIP | 29167373 | Unspecified |
| mmu-miR-34a-5p | MIMAT0000542 | Trmt5 | 76357 | HITS-CLIP | 23142080 | Unspecified |
| mmu-miR-34a-5p | MIMAT0000542 | Trmt6 | 66926 | HITS-CLIP | 23142080 | Unspecified |
| mmu-miR-34a-5p | MIMAT0000542 | Trnau1ap | 71787 | HITS-CLIP | 29167373 | Unspecified |
| mmu-miR-34a-5p | MIMAT0000542 | Trnt1 | 70047 | HITS-CLIP | 23142080\|29167373 | Unspecified |
| mmu-miR-34a-5p | MIMAT0000542 | Tro | 7216 | HITS-CLIP | 19536157 | Unspecified |
| mmu-miR-34a-5p | MIMAT0000542 | Trove2 | 20822 | HITS-CLIP | 29167373 | Unspecified |
| mmu-miR-34a-5p | MIMAT0000542 | Trp53 | 22059 | HITS-CLIP | 23142080 | Unspecified |
| mmu-miR-34a-5p | MIMAT0000542 | Trp53bp1 | 27223 | HITS-CLIP | 29167373 | Unspecified |
| mmu-miR-34a-5p | MIMAT0000542 | Trp53bp2 | 209456 | HITS-CLIP | 23142080 | Unspecified |
| mmu-miR-34a-5p | MIMAT0000542 | Trp53i11 | 277414 | Microarrays//HITS-CLIP | 37986224\|23027903\|30971389\|23142 | Unspecified |
| mmu-miR-34a-5p | MIMAT0000542 | Trp53inp1 | 60599 | HITS-CLIP, Microarrays//HITS-CLIP | 37986224\|31433990\|19536157\|23142 | Unspecified |

| | | | | | | |
|---|---|---|---|---|---|---|
| mmu-miR-34a-5p | MIMAT0000542 | Trp53inp2 | 68728 | Microarrays//HITS-CLIP | 37986224\|23027903\|23142080 | Unspecified |
| mmu-miR-34a-5p | MIMAT0000542 | Trpc4ap | 56407 | HITS-CLIP | 23630078\|23142080 | Unspecified |
| mmu-miR-34a-5p | MIMAT0000542 | Trpm3 | 226025 | HITS-CLIP | 19536157 | Unspecified |
| mmu-miR-34a-5p | MIMAT0000542 | Trpm7 | 58800 | HITS-CLIP | 30971389\|19536157\|23142080\|2916 | Unspecified |
| mmu-miR-34a-5p | MIMAT0000542 | Trps1 | 83925 | HITS-CLIP | 23142080\|29167373 | Unspecified |
| mmu-miR-34a-5p | MIMAT0000542 | Trpv2 | 22368 | HITS-CLIP | 23142080 | Unspecified |
| mmu-miR-34a-5p | MIMAT0000542 | Trrap | 100683 | HITS-CLIP | 19536157\|23142080\|29167373 | Unspecified |
| mmu-miR-34a-5p | MIMAT0000542 | Tsc1 | 64930 | HITS-CLIP | 23142080\|29167373 | Unspecified |
| mmu-miR-34a-5p | MIMAT0000542 | Tsc2 | 22084 | HITS-CLIP | 23142080\|29167373 | Unspecified |
| mmu-miR-34a-5p | MIMAT0000542 | Tsc22d1 | 21807 | HITS-CLIP | 19536157 | Unspecified |
| mmu-miR-34a-5p | MIMAT0000542 | Tsc22d2 | 72033 | HITS-CLIP | 23630078\|23142080\|29167373 | Unspecified |
| mmu-miR-34a-5p | MIMAT0000542 | Tsc22d3 | 14605 | HITS-CLIP | 23142080 | Unspecified |
| mmu-miR-34a-5p | MIMAT0000542 | Tsc22d4 | 78829 | HITS-CLIP | 19536157\|23142080 | Unspecified |
| mmu-miR-34a-5p | MIMAT0000542 | Tsen2 | 381802 | HITS-CLIP | 23142080 | Unspecified |
| mmu-miR-34a-5p | MIMAT0000542 | Tshz1 | 110796 | HITS-CLIP | 31433990\|23142080 | Unspecified |
| mmu-miR-34a-5p | MIMAT0000542 | Tshz2 | 228911 | HITS-CLIP | 19536157 | Unspecified |
| mmu-miR-34a-5p | MIMAT0000542 | Tshz3 | 243931 | HITS-CLIP | 19536157 | Unspecified |
| mmu-miR-34a-5p | MIMAT0000542 | Tsku | 244152 | HITS-CLIP | 29167373 | Unspecified |
| mmu-miR-34a-5p | MIMAT0000542 | Tsn | 22099 | HITS-CLIP | 23142080\|29167373 | Unspecified |
| mmu-miR-34a-5p | MIMAT0000542 | Tsnax | 53424 | HITS-CLIP | 30971389\|23142080 | Unspecified |
| mmu-miR-34a-5p | MIMAT0000542 | Tspan13 | 66109 | HITS-CLIP | 23142080\|19536157 | Unspecified |
| mmu-miR-34a-5p | MIMAT0000542 | Tspan14 | 52588 | HITS-CLIP | 23142080\|19536157\|29167373 | Unspecified |
| mmu-miR-34a-5p | MIMAT0000542 | Tspan17 | 74257 | HITS-CLIP | 19536157 | Unspecified |
| mmu-miR-34a-5p | MIMAT0000542 | Tspan3 | 56434 | HITS-CLIP | 23142080 | Unspecified |
| mmu-miR-34a-5p | MIMAT0000542 | Tspan31 | 67125 | HITS-CLIP | 23142080 | Unspecified |
| mmu-miR-34a-5p | MIMAT0000542 | Tspan32 | 27027 | Microarrays//HITS-CLIP | 37986224\|23142080 | Unspecified |
| mmu-miR-34a-5p | MIMAT0000542 | Tspan33 | 232670 | Microarrays//HITS-CLIP | 37986224\|30971389 | Unspecified |
| mmu-miR-34a-5p | MIMAT0000542 | Tspan4 | 64540 | HITS-CLIP | 19536157\|29167373 | Unspecified |
| mmu-miR-34a-5p | MIMAT0000542 | Tspan5 | 56224 | HITS-CLIP | 23142080 | Unspecified |
| mmu-miR-34a-5p | MIMAT0000542 | Tspan7 | 21912 | HITS-CLIP | 19536157 | Unspecified |
| mmu-miR-34a-5p | MIMAT0000542 | Tspo | 12257 | HITS-CLIP | 23142080 | Unspecified |
| mmu-miR-34a-5p | MIMAT0000542 | Tspoap1 | 207777 | HITS-CLIP | 29167373 | Unspecified |
| mmu-miR-34a-5p | MIMAT0000542 | Tspyl1 | 22110 | HITS-CLIP | 37986224\|23142080\|29167373 | Unspecified |
| mmu-miR-34a-5p | MIMAT0000542 | Tspyl2 | 52808 | HITS-CLIP | 23142080 | Unspecified |
| mmu-miR-34a-5p | MIMAT0000542 | Tsr1 | 104662 | HITS-CLIP | 23142080 | Unspecified |
| mmu-miR-34a-5p | MIMAT0000542 | Tsr2 | 69499 | Microarrays | 37986224\|23027903 | Unspecified |
| mmu-miR-34a-5p | MIMAT0000542 | Tsr3 | 68327 | HITS-CLIP | 23142080 | Unspecified |
| mmu-miR-34a-5p | MIMAT0000542 | Tst | 22117 | HITS-CLIP | 19536157 | Unspecified |
| mmu-miR-34a-5p | MIMAT0000542 | Tstd2 | 272027 | HITS-CLIP | 23630078\|23142080 | Unspecified |

| | | | | | | |
|---|---|---|---|---|---|---|
| mmu-miR-34a-5p | MIMAT0000542 | Tstd3 | 77032 | HITS-CLIP | 23142080 | Unspecified |
| mmu-miR-34a-5p | MIMAT0000542 | Ttbk1 | 106763 | HITS-CLIP | 19536157 | Unspecified |
| mmu-miR-34a-5p | MIMAT0000542 | Ttc12 | 235330 | Microarrays | 37986224 | Unspecified |
| mmu-miR-34a-5p | MIMAT0000542 | Ttc13 | 234875 | HITS-CLIP | 23142080 | Unspecified |
| mmu-miR-34a-5p | MIMAT0000542 | Ttc14 | 67120 | HITS-CLIP | 31433990\|23142080\|29167373 | Unspecified |
| mmu-miR-34a-5p | MIMAT0000542 | Ttc17 | 74569 | HITS-CLIP | 23142080\|29167373 | Unspecified |
| mmu-miR-34a-5p | MIMAT0000542 | Ttc19 | 72795 | HITS-CLIP | 23142080 | Unspecified |
| mmu-miR-34a-5p | MIMAT0000542 | Ttc27 | 74196 | HITS-CLIP | 23142080\|29167373 | Unspecified |
| mmu-miR-34a-5p | MIMAT0000542 | Ttc3 | 22129 | HITS-CLIP | 23142080\|19536157 | Unspecified |
| mmu-miR-34a-5p | MIMAT0000542 | Ttc30b | 72421 | Microarrays | 37986224 | Unspecified |
| mmu-miR-34a-5p | MIMAT0000542 | Ttc33 | 67515 | HITS-CLIP | 23142080 | Unspecified |
| mmu-miR-34a-5p | MIMAT0000542 | Ttc37 | 218343 | HITS-CLIP | 23142080 | Unspecified |
| mmu-miR-34a-5p | MIMAT0000542 | Ttc39b | 69863 | HITS-CLIP | 23142080 | Unspecified |
| mmu-miR-34a-5p | MIMAT0000542 | Ttc4 | 72354 | HITS-CLIP | 23142080 | Unspecified |
| mmu-miR-34a-5p | MIMAT0000542 | Ttc7 | 225049 | Microarrays//HITS-CLIP | 37986224\|23027903\|23142080 | Unspecified |
| mmu-miR-34a-5p | MIMAT0000542 | Ttc8 | 76260 | HITS-CLIP | 23142080 | Unspecified |
| mmu-miR-34a-5p | MIMAT0000542 | Ttc9c | 70387 | HITS-CLIP | 23142080\|26203562\|29167373 | Unspecified |
| mmu-miR-34a-5p | MIMAT0000542 | Tti1 | 75425 | HITS-CLIP | 23142080 | Unspecified |
| mmu-miR-34a-5p | MIMAT0000542 | Tti2 | 234138 | HITS-CLIP | 23142080 | Unspecified |
| mmu-miR-34a-5p | MIMAT0000542 | Ttk | 22137 | HITS-CLIP | 31433990\|23142080 | Unspecified |
| mmu-miR-34a-5p | MIMAT0000542 | Ttl | 69737 | HITS-CLIP | 19536157\|23142080 | Unspecified |
| mmu-miR-34a-5p | MIMAT0000542 | Ttll12 | 223723 | HITS-CLIP | 23142080 | Unspecified |
| mmu-miR-34a-5p | MIMAT0000542 | Ttll4 | 67534 | HITS-CLIP | 30971389\|31433990 | Unspecified |
| mmu-miR-34a-5p | MIMAT0000542 | Ttll5 | 320244 | HITS-CLIP | 23142080 | Unspecified |
| mmu-miR-34a-5p | MIMAT0000542 | Ttr | 22139 | HITS-CLIP | 19536157 | Unspecified |
| mmu-miR-34a-5p | MIMAT0000542 | Ttyh1 | 57776 | HITS-CLIP | 19536157 | Unspecified |
| mmu-miR-34a-5p | MIMAT0000542 | Ttyh3 | 78339 | HITS-CLIP | 19536157\|23142080\|29167373 | Unspecified |
| mmu-miR-34a-5p | MIMAT0000542 | Tuba1a | 22142 | HITS-CLIP | 19536157\|23142080 | Unspecified |
| mmu-miR-34a-5p | MIMAT0000542 | Tuba1b | 22143 | HITS-CLIP | 31433990\|23142080 | Unspecified |
| mmu-miR-34a-5p | MIMAT0000542 | Tuba1c | 22146 | HITS-CLIP | 23142080 | Unspecified |
| mmu-miR-34a-5p | MIMAT0000542 | Tuba4a | 22145 | HITS-CLIP | 37986224\|23142080\|19536157 | Unspecified |
| mmu-miR-34a-5p | MIMAT0000542 | Tubb2a | 22151 | HITS-CLIP | 19536157 | Unspecified |
| mmu-miR-34a-5p | MIMAT0000542 | Tubb3 | 22152 | Microarrays//HITS-CLIP | 37986224\|23027903\|19536157 | Unspecified |
| mmu-miR-34a-5p | MIMAT0000542 | Tubb4a | 22153 | HITS-CLIP | 19536157 | Unspecified |
| mmu-miR-34a-5p | MIMAT0000542 | Tubb4b | 227613 | HITS-CLIP | 31433990\|29167373\|23142080 | Unspecified |
| mmu-miR-34a-5p | MIMAT0000542 | Tubb5 | 22154 | HITS-CLIP | 37986224\|23142080\|19536157\|3143[illegible] | Unspecified |
| mmu-miR-34a-5p | MIMAT0000542 | Tubb6 | 67951 | HITS-CLIP | 23142080\|29167373\|26203562 | Unspecified |
| mmu-miR-34a-5p | MIMAT0000542 | Tubd1 | 56427 | HITS-CLIP | 23142080 | Unspecified |
| mmu-miR-34a-5p | MIMAT0000542 | Tube1 | 71924 | HITS-CLIP | 31433990\|23142080 | Unspecified |

| | | | | | | |
|---|---|---|---|---|---|---|
| mmu-miR-34a-5p | MIMAT0000542 | Tubgcp3 | 259279 | HITS-CLIP | 23142080\|31433990\|29167373 | Unspecified |
| mmu-miR-34a-5p | MIMAT0000542 | Tubgcp4 | 51885 | HITS-CLIP | 23142080\|29167373 | Unspecified |
| mmu-miR-34a-5p | MIMAT0000542 | Tubgcp5 | 233276 | HITS-CLIP | 23142080 | Unspecified |
| mmu-miR-34a-5p | MIMAT0000542 | Tubgcp6 | 328580 | HITS-CLIP | 30971389 | Unspecified |
| mmu-miR-34a-5p | MIMAT0000542 | Tuft1 | 22156 | HITS-CLIP | 23142080 | Unspecified |
| mmu-miR-34a-5p | MIMAT0000542 | Tulp3 | 22158 | HITS-CLIP | 19536157 | Unspecified |
| mmu-miR-34a-5p | MIMAT0000542 | Tulp4 | 68842 | HITS-CLIP | 29167373 | Unspecified |
| mmu-miR-34a-5p | MIMAT0000542 | Tusc2 | 80385 | HITS-CLIP | 29167373 | Unspecified |
| mmu-miR-34a-5p | MIMAT0000542 | Tusc3 | 80286 | HITS-CLIP | 23142080 | Unspecified |
| mmu-miR-34a-5p | MIMAT0000542 | Tut1 | 70044 | HITS-CLIP | 29167373 | Unspecified |
| mmu-miR-34a-5p | MIMAT0000542 | Tut4 | 23318 | HITS-CLIP | 23142080\|29167373 | Unspecified |
| mmu-miR-34a-5p | MIMAT0000542 | Tut7 | 79670 | HITS-CLIP | 23142080 | Unspecified |
| mmu-miR-34a-5p | MIMAT0000542 | Twf1 | 19230 | HITS-CLIP | 23142080\|19536157\|29167373 | Unspecified |
| mmu-miR-34a-5p | MIMAT0000542 | Twnk | 56652 | HITS-CLIP | 23142080 | Unspecified |
| mmu-miR-34a-5p | MIMAT0000542 | Txlna | 109658 | HITS-CLIP | 23630078\|30971389 | Unspecified |
| mmu-miR-34a-5p | MIMAT0000542 | Txlnb | 378431 | Microarrays | 37986224 | Unspecified |
| mmu-miR-34a-5p | MIMAT0000542 | Txlng | 353170 | HITS-CLIP | 23142080 | Unspecified |
| mmu-miR-34a-5p | MIMAT0000542 | Txn1 | 22166 | HITS-CLIP | 37986224\|23630078\|23142080\|29167 | Unspecified |
| mmu-miR-34a-5p | MIMAT0000542 | Txn2 | 56551 | HITS-CLIP | 29167373 | Unspecified |
| mmu-miR-34a-5p | MIMAT0000542 | Txndc11 | 106200 | HITS-CLIP | 19536157\|31433990\|29167373 | Unspecified |
| mmu-miR-34a-5p | MIMAT0000542 | Txndc12 | 66073 | HITS-CLIP | 23630078\|19536157\|30971389\|23142 | Unspecified |
| mmu-miR-34a-5p | MIMAT0000542 | Txndc16 | 70561 | HITS-CLIP | 23142080\|19536157 | Unspecified |
| mmu-miR-34a-5p | MIMAT0000542 | Txndc17 | 52700 | HITS-CLIP | 23142080 | Unspecified |
| mmu-miR-34a-5p | MIMAT0000542 | Txndc5 | 105245 | HITS-CLIP | 19536157\|29167373 | Unspecified |
| mmu-miR-34a-5p | MIMAT0000542 | Txndc9 | 98258 | HITS-CLIP | 29167373 | Unspecified |
| mmu-miR-34a-5p | MIMAT0000542 | Txnip | 56338 | Microarrays//HITS-CLIP | 37986224\|23027903\|31433990\|23142 | Unspecified |
| mmu-miR-34a-5p | MIMAT0000542 | Txnl1 | 53382 | HITS-CLIP | 23142080\|19536157 | Unspecified |
| mmu-miR-34a-5p | MIMAT0000542 | Txnl4a | 27366 | HITS-CLIP | 23142080\|29167373 | Unspecified |
| mmu-miR-34a-5p | MIMAT0000542 | Txnrd1 | 50493 | HITS-CLIP | 23142080\|19536157\|29167373 | Unspecified |
| mmu-miR-34a-5p | MIMAT0000542 | Txnrd2 | 26462 | HITS-CLIP | 29167373 | Unspecified |
| mmu-miR-34a-5p | MIMAT0000542 | Tyk2 | 54721 | HITS-CLIP | 23142080 | Unspecified |
| mmu-miR-34a-5p | MIMAT0000542 | Tyms | 22171 | HITS-CLIP | 23142080 | Unspecified |
| mmu-miR-34a-5p | MIMAT0000542 | U2af2 | 22185 | HITS-CLIP | 23142080\|29167373 | Unspecified |
| mmu-miR-34a-5p | MIMAT0000542 | U2surp | 67958 | HITS-CLIP | 23142080\|29167373 | Unspecified |
| mmu-miR-34a-5p | MIMAT0000542 | Uap1 | 107652 | HITS-CLIP | 31433990 | Unspecified |
| mmu-miR-34a-5p | MIMAT0000542 | Uap1l1 | 227620 | HITS-CLIP | 29167373 | Unspecified |
| mmu-miR-34a-5p | MIMAT0000542 | Uba1 | 22201 | HITS-CLIP | 23142080\|19536157\|29167373 | Unspecified |
| mmu-miR-34a-5p | MIMAT0000542 | Uba2 | 50995 | HITS-CLIP | 19536157\|23142080\|29167373 | Unspecified |
| mmu-miR-34a-5p | MIMAT0000542 | Uba3 | 22200 | HITS-CLIP | 37986224\|23142080\|29167373 | Unspecified |

| | | | | | | |
|---|---|---|---|---|---|---|
| mmu-miR-34a-5p | MIMAT0000542 | Uba5 | 66663 | HITS-CLIP | 31433990 | Unspecified |
| mmu-miR-34a-5p | MIMAT0000542 | Uba6 | 231380 | HITS-CLIP | 23142080 | Unspecified |
| mmu-miR-34a-5p | MIMAT0000542 | Uba7 | 74153 | HITS-CLIP | 23142080\|29167373 | Unspecified |
| mmu-miR-34a-5p | MIMAT0000542 | Ubac1 | 98766 | HITS-CLIP | 31433990 | Unspecified |
| mmu-miR-34a-5p | MIMAT0000542 | Ubald1 | 207740 | HITS-CLIP | 29167373 | Unspecified |
| mmu-miR-34a-5p | MIMAT0000542 | Ubap1 | 67123 | HITS-CLIP | 19536157\|23142080 | Unspecified |
| mmu-miR-34a-5p | MIMAT0000542 | Ubap2 | 68926 | HITS-CLIP | 23630078\|31433990\|23142080\|29167 | Unspecified |
| mmu-miR-34a-5p | MIMAT0000542 | Ubap2l | 74383 | HITS-CLIP | 29167373\|23142080 | Unspecified |
| mmu-miR-34a-5p | MIMAT0000542 | Ubash3a | 328795 | HITS-CLIP | 23142080\|31433990 | Unspecified |
| mmu-miR-34a-5p | MIMAT0000542 | Ubash3b | 72828 | HITS-CLIP | 23142080\|29167373 | Unspecified |
| mmu-miR-34a-5p | MIMAT0000542 | Ubb | 22187 | HITS-CLIP | 23142080\|29167373 | Unspecified |
| mmu-miR-34a-5p | MIMAT0000542 | Ube2a | 22209 | HITS-CLIP | 23142080\|29167373 | Unspecified |
| mmu-miR-34a-5p | MIMAT0000542 | Ube2b | 22210 | HITS-CLIP | 29167373 | Unspecified |
| mmu-miR-34a-5p | MIMAT0000542 | Ube2c | 68612 | HITS-CLIP | 23142080 | Unspecified |
| mmu-miR-34a-5p | MIMAT0000542 | Ube2cbp | 70348 | HITS-CLIP | 29167373 | Unspecified |
| mmu-miR-34a-5p | MIMAT0000542 | Ube2d1 | 216080 | HITS-CLIP | 23142080 | Unspecified |
| mmu-miR-34a-5p | MIMAT0000542 | Ube2d2a | 56550 | HITS-CLIP | 23142080 | Unspecified |
| mmu-miR-34a-5p | MIMAT0000542 | Ube2d3 | 66105 | HITS-CLIP | 23142080\|29167373 | Unspecified |
| mmu-miR-34a-5p | MIMAT0000542 | Ube2e1 | 22194 | HITS-CLIP | 23142080 | Unspecified |
| mmu-miR-34a-5p | MIMAT0000542 | Ube2e2 | 218793 | Microarrays//HITS-CLIP | 37986224\|23142080 | Unspecified |
| mmu-miR-34a-5p | MIMAT0000542 | Ube2e3 | 22193 | HITS-CLIP | 29167373\|23142080 | Unspecified |
| mmu-miR-34a-5p | MIMAT0000542 | Ube2f | 67921 | HITS-CLIP | 29167373 | Unspecified |
| mmu-miR-34a-5p | MIMAT0000542 | Ube2g1 | 67128 | HITS-CLIP | 23142080 | Unspecified |
| mmu-miR-34a-5p | MIMAT0000542 | Ube2g2 | 22213 | HITS-CLIP | 23142080\|30971389 | Unspecified |
| mmu-miR-34a-5p | MIMAT0000542 | Ube2h | 22214 | HITS-CLIP | 23142080 | Unspecified |
| mmu-miR-34a-5p | MIMAT0000542 | Ube2i | 22196 | HITS-CLIP | 29167373 | Unspecified |
| mmu-miR-34a-5p | MIMAT0000542 | Ube2j1 | 56228 | HITS-CLIP | 30971389\|23142080 | Unspecified |
| mmu-miR-34a-5p | MIMAT0000542 | Ube2j2 | 140499 | HITS-CLIP | 29167373 | Unspecified |
| mmu-miR-34a-5p | MIMAT0000542 | Ube2k | 53323 | HITS-CLIP | 23142080\|31433990 | Unspecified |
| mmu-miR-34a-5p | MIMAT0000542 | Ube2l3 | 22195 | HITS-CLIP | 23142080 | Unspecified |
| mmu-miR-34a-5p | MIMAT0000542 | Ube2m | 22192 | HITS-CLIP | 23142080\|29167373 | Unspecified |
| mmu-miR-34a-5p | MIMAT0000542 | Ube2n | 93765 | HITS-CLIP | 29167373 | Unspecified |
| mmu-miR-34a-5p | MIMAT0000542 | Ube2o | 217342 | HITS-CLIP | 23142080 | Unspecified |
| mmu-miR-34a-5p | MIMAT0000542 | Ube2q1 | 70093 | HITS-CLIP | 23630078\|29167373\|23142080 | Unspecified |
| mmu-miR-34a-5p | MIMAT0000542 | Ube2r2 | 67615 | HITS-CLIP | 23630078\|30971389\|23142080 | Unspecified |
| mmu-miR-34a-5p | MIMAT0000542 | Ube2t | 67196 | HITS-CLIP | 23142080 | Unspecified |
| mmu-miR-34a-5p | MIMAT0000542 | Ube2v2 | 70620 | HITS-CLIP | 23142080 | Unspecified |
| mmu-miR-34a-5p | MIMAT0000542 | Ube2w | 66799 | HITS-CLIP | 23142080 | Unspecified |
| mmu-miR-34a-5p | MIMAT0000542 | Ube2z | 268470 | Microarrays//HITS-CLIP | 37986224\|23027903\|23142080 | Unspecified |

| | | | | | | |
|---|---|---|---|---|---|---|
| mmu-miR-34a-5p | MIMAT0000542 | Ube3a | 22215 | HITS-CLIP | 23142080\|31433990\|29167373 | Unspecified |
| mmu-miR-34a-5p | MIMAT0000542 | Ube3b | 117146 | HITS-CLIP | 23142080\|26203562 | Unspecified |
| mmu-miR-34a-5p | MIMAT0000542 | Ube3c | 100763 | HITS-CLIP | 19536157\|23142080\|29167373 | Unspecified |
| mmu-miR-34a-5p | MIMAT0000542 | Ube4a | 140630 | HITS-CLIP | 23142080\|29167373 | Unspecified |
| mmu-miR-34a-5p | MIMAT0000542 | Ube4b | 63958 | HITS-CLIP | 19536157\|30971389\|23630078\|23142 | Unspecified |
| mmu-miR-34a-5p | MIMAT0000542 | Ubfd1 | 28018 | Microarrays//HITS-CLIP | 37986224\|23027903\|30971389\|23142 | Unspecified |
| mmu-miR-34a-5p | MIMAT0000542 | Ubl3 | 24109 | HITS-CLIP | 23142080\|29167373 | Unspecified |
| mmu-miR-34a-5p | MIMAT0000542 | Ubl4a | 8266 | HITS-CLIP | 23142080 | Unspecified |
| mmu-miR-34a-5p | MIMAT0000542 | Ubl7 | 69459 | HITS-CLIP | 23142080 | Unspecified |
| mmu-miR-34a-5p | MIMAT0000542 | Ubn1 | 170644 | HITS-CLIP | 31433990\|30971389\|23142080\|29167 | Unspecified |
| mmu-miR-34a-5p | MIMAT0000542 | Ubn2 | 320538 | HITS-CLIP | 23142080\|19536157\|29167373 | Unspecified |
| mmu-miR-34a-5p | MIMAT0000542 | Ubox5 | 140629 | HITS-CLIP | 37986224\|19536157\|29167373\|23142 | Unspecified |
| mmu-miR-34a-5p | MIMAT0000542 | Ubp1 | 22221 | HITS-CLIP | 23142080 | Unspecified |
| mmu-miR-34a-5p | MIMAT0000542 | Ubqln1 | 56085 | HITS-CLIP | 19536157\|23142080\|29167373 | Unspecified |
| mmu-miR-34a-5p | MIMAT0000542 | Ubqln4 | 94232 | HITS-CLIP | 23142080 | Unspecified |
| mmu-miR-34a-5p | MIMAT0000542 | Ubr1 | 22222 | HITS-CLIP | 31433990\|23142080 | Unspecified |
| mmu-miR-34a-5p | MIMAT0000542 | Ubr2 | 224826 | HITS-CLIP | 31433990\|23142080\|26203562\|29167 | Unspecified |
| mmu-miR-34a-5p | MIMAT0000542 | Ubr3 | 68795 | HITS-CLIP | 30971389\|23142080\|29167373 | Unspecified |
| mmu-miR-34a-5p | MIMAT0000542 | Ubr4 | 69116 | HITS-CLIP | 23142080\|29167373 | Unspecified |
| mmu-miR-34a-5p | MIMAT0000542 | Ubr5 | 70790 | HITS-CLIP | 30971389\|19536157\|23142080\|29167 | Unspecified |
| mmu-miR-34a-5p | MIMAT0000542 | Ubr7 | 66622 | HITS-CLIP | 23142080 | Unspecified |
| mmu-miR-34a-5p | MIMAT0000542 | Ubtd1 | 226122 | HITS-CLIP | 23142080 | Unspecified |
| mmu-miR-34a-5p | MIMAT0000542 | Ubtd2 | 327900 | HITS-CLIP | 29167373\|23142080 | Unspecified |
| mmu-miR-34a-5p | MIMAT0000542 | Ubtf | 21429 | HITS-CLIP | 23142080 | Unspecified |
| mmu-miR-34a-5p | MIMAT0000542 | Ubxn1 | 225896 | HITS-CLIP | 23142080\|29167373 | Unspecified |
| mmu-miR-34a-5p | MIMAT0000542 | Ubxn4 | 67812 | HITS-CLIP | 19536157\|23142080\|29167373 | Unspecified |
| mmu-miR-34a-5p | MIMAT0000542 | Ubxn7 | 224111 | HITS-CLIP | 23142080\|29167373 | Unspecified |
| mmu-miR-34a-5p | MIMAT0000542 | Uchl5 | 56207 | HITS-CLIP | 23142080 | Unspecified |
| mmu-miR-34a-5p | MIMAT0000542 | Uck2 | 80914 | HITS-CLIP | 23142080 | Unspecified |
| mmu-miR-34a-5p | MIMAT0000542 | Uckl1 | 68556 | HITS-CLIP | 29167373 | Unspecified |
| mmu-miR-34a-5p | MIMAT0000542 | Ucp2 | 22228 | HITS-CLIP | 23142080\|29167373 | Unspecified |
| mmu-miR-34a-5p | MIMAT0000542 | Uevld | 54122 | HITS-CLIP | 23142080 | Unspecified |
| mmu-miR-34a-5p | MIMAT0000542 | Ufc1 | 66155 | HITS-CLIP | 23142080 | Unspecified |
| mmu-miR-34a-5p | MIMAT0000542 | Ufd1 | 22230 | HITS-CLIP | 23142080 | Unspecified |
| mmu-miR-34a-5p | MIMAT0000542 | Ufl1 | 67490 | HITS-CLIP | 23142080 | Unspecified |
| mmu-miR-34a-5p | MIMAT0000542 | Ufm1 | 67890 | HITS-CLIP | 23142080 | Unspecified |
| mmu-miR-34a-5p | MIMAT0000542 | Ufsp2 | 192169 | HITS-CLIP | 31433990 | Unspecified |
| mmu-miR-34a-5p | MIMAT0000542 | Ugcg | 22234 | HITS-CLIP | 23630078\|29167373\|23142080 | Unspecified |
| mmu-miR-34a-5p | MIMAT0000542 | Ugdh | 22235 | Microarrays//HITS-CLIP | 37986224\|23142080 | Unspecified |

| | | | | | | |
|---|---|---|---|---|---|---|
| mmu-miR-34a-5p | MIMAT0000542 | Uggt1 | 320011 | HITS-CLIP | 29167373\|23142080 | Unspecified |
| mmu-miR-34a-5p | MIMAT0000542 | Ugp2 | 216558 | HITS-CLIP | 19536157 | Unspecified |
| mmu-miR-34a-5p | MIMAT0000542 | Ugt1a6b | 394435 | HITS-CLIP | 29167373 | Unspecified |
| mmu-miR-34a-5p | MIMAT0000542 | Ugt1a7c | 394432 | HITS-CLIP | 29167373 | Unspecified |
| mmu-miR-34a-5p | MIMAT0000542 | Ugt8a | 22239 | HITS-CLIP | 19536157 | Unspecified |
| mmu-miR-34a-5p | MIMAT0000542 | Uhmk1 | 16589 | HITS-CLIP | 31433990\|23630078\|19536157\|23142 | Unspecified |
| mmu-miR-34a-5p | MIMAT0000542 | Uhrf1 | 18140 | Microarrays//HITS-CLIP | 37986224\|23027903\|23142080\|3143 | Unspecified |
| mmu-miR-34a-5p | MIMAT0000542 | Uhrf1bp1 | 224648 | HITS-CLIP | 23142080 | Unspecified |
| mmu-miR-34a-5p | MIMAT0000542 | Uhrf1bp1l | 75089 | HITS-CLIP | 23142080\|29167373 | Unspecified |
| mmu-miR-34a-5p | MIMAT0000542 | Uhrf2 | 109113 | HITS-CLIP | 23142080\|29167373 | Unspecified |
| mmu-miR-34a-5p | MIMAT0000542 | Uimc1 | 20184 | HITS-CLIP | 23142080 | Unspecified |
| mmu-miR-34a-5p | MIMAT0000542 | Ulbp1 | 77777 | HITS-CLIP | 23142080\|29167373 | Unspecified |
| mmu-miR-34a-5p | MIMAT0000542 | Ulk1 | 22241 | HITS-CLIP | 23142080\|29167373 | Unspecified |
| mmu-miR-34a-5p | MIMAT0000542 | Ulk2 | 29869 | HITS-CLIP | 23142080\|19536157\|29167373 | Unspecified |
| mmu-miR-34a-5p | MIMAT0000542 | Ulk3 | 71742 | HITS-CLIP | 23142080\|29167373 | Unspecified |
| mmu-miR-34a-5p | MIMAT0000542 | Umad1 | 729852 | HITS-CLIP | 23142080 | Unspecified |
| mmu-miR-34a-5p | MIMAT0000542 | Umps | 22247 | HITS-CLIP | 23142080 | Unspecified |
| mmu-miR-34a-5p | MIMAT0000542 | Unc119b | 106840 | HITS-CLIP | 23142080 | Unspecified |
| mmu-miR-34a-5p | MIMAT0000542 | Unc13a | 382018 | HITS-CLIP | 23142080 | Unspecified |
| mmu-miR-34a-5p | MIMAT0000542 | Unc13b | 22249 | HITS-CLIP | 23142080 | Unspecified |
| mmu-miR-34a-5p | MIMAT0000542 | Unc13d | 70450 | HITS-CLIP | 23142080 | Unspecified |
| mmu-miR-34a-5p | MIMAT0000542 | Unc45a | 101869 | HITS-CLIP | 23142080\|29167373 | Unspecified |
| mmu-miR-34a-5p | MIMAT0000542 | Unc45b | 217012 | Microarrays | 37986224 | Unspecified |
| mmu-miR-34a-5p | MIMAT0000542 | Unc50 | 67387 | HITS-CLIP | 23142080 | Unspecified |
| mmu-miR-34a-5p | MIMAT0000542 | Unc5b | 107449 | HITS-CLIP | 19536157 | Unspecified |
| mmu-miR-34a-5p | MIMAT0000542 | Unc5cl | 76589 | HITS-CLIP | 23142080 | Unspecified |
| mmu-miR-34a-5p | MIMAT0000542 | Unc79 | 217843 | HITS-CLIP | 19536157 | Unspecified |
| mmu-miR-34a-5p | MIMAT0000542 | Unc80 | 329178 | HITS-CLIP | 19536157 | Unspecified |
| mmu-miR-34a-5p | MIMAT0000542 | Unc93b1 | 54445 | HITS-CLIP | 23142080\|29167373 | Unspecified |
| mmu-miR-34a-5p | MIMAT0000542 | Ung | 22256 | Microarrays//HITS-CLIP | 37986224\|23027903\|23142080 | Unspecified |
| mmu-miR-34a-5p | MIMAT0000542 | Unk | 217331 | HITS-CLIP | 23142080\|29167373 | Unspecified |
| mmu-miR-34a-5p | MIMAT0000542 | Unkl | 74154 | HITS-CLIP | 23142080 | Unspecified |
| mmu-miR-34a-5p | MIMAT0000542 | Upf1 | 19704 | HITS-CLIP | 31433990\|23142080 | Unspecified |
| mmu-miR-34a-5p | MIMAT0000542 | Upf2 | 326622 | HITS-CLIP | 19536157\|31433990\|23142080 | Unspecified |
| mmu-miR-34a-5p | MIMAT0000542 | Upp1 | 22271 | Microarrays | 37986224\|23027903 | Unspecified |
| mmu-miR-34a-5p | MIMAT0000542 | Uqcc1 | 56046 | HITS-CLIP | 31433990\|23142080 | Unspecified |
| mmu-miR-34a-5p | MIMAT0000542 | Uqcc2 | 67267 | HITS-CLIP | 23142080 | Unspecified |
| mmu-miR-34a-5p | MIMAT0000542 | Uqcr10 | 66152 | HITS-CLIP | 23142080 | Unspecified |
| mmu-miR-34a-5p | MIMAT0000542 | Uqcrc1 | 22273 | HITS-CLIP | 23142080 | Unspecified |

| | | | | | | |
|---|---|---|---|---|---|---|
| mmu-miR-34a-5p | MIMAT0000542 | Uqcrc2 | 67003 | HITS-CLIP | 23142080 | Unspecified |
| mmu-miR-34a-5p | MIMAT0000542 | Uqcrfs1 | 66694 | HITS-CLIP | 23142080 | Unspecified |
| mmu-miR-34a-5p | MIMAT0000542 | Uqcrh | 66576 | HITS-CLIP | 23142080 | Unspecified |
| mmu-miR-34a-5p | MIMAT0000542 | Urb1 | 207932 | HITS-CLIP | 23142080 | Unspecified |
| mmu-miR-34a-5p | MIMAT0000542 | Urb2 | 382038 | HITS-CLIP | 23142080\|29167373 | Unspecified |
| mmu-miR-34a-5p | MIMAT0000542 | Urgcp | 72046 | HITS-CLIP | 23142080 | Unspecified |
| mmu-miR-34a-5p | MIMAT0000542 | Uri1 | 19777 | HITS-CLIP | 23142080 | Unspecified |
| mmu-miR-34a-5p | MIMAT0000542 | Urod | 22275 | HITS-CLIP | 23142080 | Unspecified |
| mmu-miR-34a-5p | MIMAT0000542 | Usb1 | 101985 | HITS-CLIP | 29167373 | Unspecified |
| mmu-miR-34a-5p | MIMAT0000542 | Use1 | 67023 | HITS-CLIP | 29167373 | Unspecified |
| mmu-miR-34a-5p | MIMAT0000542 | Usf1 | 22278 | HITS-CLIP | 29167373 | Unspecified |
| mmu-miR-34a-5p | MIMAT0000542 | Usf2 | 22282 | HITS-CLIP | 23142080 | Unspecified |
| mmu-miR-34a-5p | MIMAT0000542 | Usf3 | 205717 | HITS-CLIP | 30971389\|19536157\|23142080\|29167 | Unspecified |
| mmu-miR-34a-5p | MIMAT0000542 | Uso1 | 56041 | HITS-CLIP | 19536157\|29167373\|23142080 | Unspecified |
| mmu-miR-34a-5p | MIMAT0000542 | Usp10 | 22224 | HITS-CLIP | 23142080 | Unspecified |
| mmu-miR-34a-5p | MIMAT0000542 | Usp11 | 236733 | HITS-CLIP | 19536157 | Unspecified |
| mmu-miR-34a-5p | MIMAT0000542 | Usp12 | 22217 | HITS-CLIP | 23142080 | Unspecified |
| mmu-miR-34a-5p | MIMAT0000542 | Usp14 | 59025 | HITS-CLIP | 23142080\|29167373 | Unspecified |
| mmu-miR-34a-5p | MIMAT0000542 | Usp15 | 14479 | HITS-CLIP | 23142080\|29167373 | Unspecified |
| mmu-miR-34a-5p | MIMAT0000542 | Usp16 | 74112 | HITS-CLIP | 23142080 | Unspecified |
| mmu-miR-34a-5p | MIMAT0000542 | Usp18 | 24110 | HITS-CLIP | 23142080\|29167373 | Unspecified |
| mmu-miR-34a-5p | MIMAT0000542 | Usp19 | 71472 | HITS-CLIP | 23142080\|31433990\|29167373 | Unspecified |
| mmu-miR-34a-5p | MIMAT0000542 | Usp22 | 216825 | HITS-CLIP | 19536157\|26203562\|23142080 | Unspecified |
| mmu-miR-34a-5p | MIMAT0000542 | Usp24 | 329908 | HITS-CLIP | 23142080\|31433990\|29167373 | Unspecified |
| mmu-miR-34a-5p | MIMAT0000542 | Usp25 | 30940 | Microarrays//HITS-CLIP | 37986224\|23142080\|29167373 | Unspecified |
| mmu-miR-34a-5p | MIMAT0000542 | Usp28 | 235323 | HITS-CLIP | 19536157 | Unspecified |
| mmu-miR-34a-5p | MIMAT0000542 | Usp3 | 235441 | HITS-CLIP | 23142080\|29167373 | Unspecified |
| mmu-miR-34a-5p | MIMAT0000542 | Usp31 | 76179 | HITS-CLIP | 31433990\|29167373\|23142080 | Unspecified |
| mmu-miR-34a-5p | MIMAT0000542 | Usp32 | 237898 | HITS-CLIP | 19536157\|23142080\|29167373 | Unspecified |
| mmu-miR-34a-5p | MIMAT0000542 | Usp33 | 170822 | HITS-CLIP | 23142080 | Unspecified |
| mmu-miR-34a-5p | MIMAT0000542 | Usp34 | 17847 | HITS-CLIP | 37986224\|23142080\|19536157\|29167 | Unspecified |
| mmu-miR-34a-5p | MIMAT0000542 | Usp36 | 72344 | HITS-CLIP | 23142080 | Unspecified |
| mmu-miR-34a-5p | MIMAT0000542 | Usp37 | 319651 | HITS-CLIP | 23142080\|29167373 | Unspecified |
| mmu-miR-34a-5p | MIMAT0000542 | Usp38 | 74841 | HITS-CLIP | 23630078\|23142080 | Unspecified |
| mmu-miR-34a-5p | MIMAT0000542 | Usp39 | 28035 | HITS-CLIP | 19536157\|23142080 | Unspecified |
| mmu-miR-34a-5p | MIMAT0000542 | Usp4 | 22258 | HITS-CLIP | 19536157\|23142080 | Unspecified |
| mmu-miR-34a-5p | MIMAT0000542 | Usp40 | 227334 | Microarrays//HITS-CLIP | 37986224\|23027903\|23142080 | Unspecified |
| mmu-miR-34a-5p | MIMAT0000542 | Usp42 | 76800 | HITS-CLIP | 37986224\|19536157\|23142080 | Unspecified |
| mmu-miR-34a-5p | MIMAT0000542 | Usp45 | 77593 | HITS-CLIP | 30971389\|29167373\|23142080 | Unspecified |

| | | | | | | |
|---|---|---|---|---|---|---|
| mmu-miR-34a-5p | MIMAT0000542 | Usp47 | 74996 | HITS-CLIP | 23142080\|29167373 | Unspecified |
| mmu-miR-34a-5p | MIMAT0000542 | Usp48 | 170707 | HITS-CLIP | 23142080\|29167373 | Unspecified |
| mmu-miR-34a-5p | MIMAT0000542 | Usp49 | 224836 | HITS-CLIP | 23142080\|29167373 | Unspecified |
| mmu-miR-34a-5p | MIMAT0000542 | Usp5 | 22225 | HITS-CLIP | 31433990\|23142080\|29167373 | Unspecified |
| mmu-miR-34a-5p | MIMAT0000542 | Usp53 | 99526 | HITS-CLIP | 19536157\|23142080 | Unspecified |
| mmu-miR-34a-5p | MIMAT0000542 | Usp6nl | 98910 | HITS-CLIP | 37986224\|23142080\|29167373 | Unspecified |
| mmu-miR-34a-5p | MIMAT0000542 | Usp7 | 252870 | HITS-CLIP | 23142080\|29167373 | Unspecified |
| mmu-miR-34a-5p | MIMAT0000542 | Usp8 | 84092 | HITS-CLIP | 23142080\|29167373 | Unspecified |
| mmu-miR-34a-5p | MIMAT0000542 | Usp9x | 22284 | HITS-CLIP | 23142080\|29167373 | Unspecified |
| mmu-miR-34a-5p | MIMAT0000542 | Uspl1 | 231915 | HITS-CLIP | 23142080\|29084942 | Unspecified |
| mmu-miR-34a-5p | MIMAT0000542 | Ust | 338362 | HITS-CLIP | 23142080 | Unspecified |
| mmu-miR-34a-5p | MIMAT0000542 | Utf1 | 22286 | HITS-CLIP | 23142080 | Unspecified |
| mmu-miR-34a-5p | MIMAT0000542 | Utp14b | 195434 | HITS-CLIP | 23142080 | Unspecified |
| mmu-miR-34a-5p | MIMAT0000542 | Utp15 | 105372 | HITS-CLIP | 30971389\|19536157\|23142080 | Unspecified |
| mmu-miR-34a-5p | MIMAT0000542 | Utp18 | 217109 | HITS-CLIP | 23142080 | Unspecified |
| mmu-miR-34a-5p | MIMAT0000542 | Utp20 | 70683 | HITS-CLIP | 29167373 | Unspecified |
| mmu-miR-34a-5p | MIMAT0000542 | Utp23 | 78581 | HITS-CLIP | 23142080 | Unspecified |
| mmu-miR-34a-5p | MIMAT0000542 | Utp3 | 65961 | HITS-CLIP | 23142080\|29167373 | Unspecified |
| mmu-miR-34a-5p | MIMAT0000542 | Utp4 | 21771 | HITS-CLIP | 23142080 | Unspecified |
| mmu-miR-34a-5p | MIMAT0000542 | Utp6 | 216987 | HITS-CLIP | 23142080 | Unspecified |
| mmu-miR-34a-5p | MIMAT0000542 | Utrn | 22288 | HITS-CLIP | 23142080\|29167373 | Unspecified |
| mmu-miR-34a-5p | MIMAT0000542 | Uty | 22290 | HITS-CLIP | 23142080 | Unspecified |
| mmu-miR-34a-5p | MIMAT0000542 | Uxs1 | 67883 | HITS-CLIP | 23142080 | Unspecified |
| mmu-miR-34a-5p | MIMAT0000542 | Vac14 | 234729 | HITS-CLIP | 29167373 | Unspecified |
| mmu-miR-34a-5p | MIMAT0000542 | Vamp1 | 22317 | HITS-CLIP | 29167373 | Unspecified |
| mmu-miR-34a-5p | MIMAT0000542 | Vamp2 | 22318 | HITS-CLIP | 29167373 | Unspecified |
| mmu-miR-34a-5p | MIMAT0000542 | Vamp3 | 22319 | Microarrays//HITS-CLIP | 37986224\|23027903\|23142080\|29167 | Unspecified |
| mmu-miR-34a-5p | MIMAT0000542 | Vamp4 | 53330 | HITS-CLIP | 29167373 | Unspecified |
| mmu-miR-34a-5p | MIMAT0000542 | Vangl1 | 229658 | HITS-CLIP | 19536157\|23630078\|26203562 | Unspecified |
| mmu-miR-34a-5p | MIMAT0000542 | Vapa | 30960 | HITS-CLIP | 23142080 | Unspecified |
| mmu-miR-34a-5p | MIMAT0000542 | Vapb | 56491 | HITS-CLIP | 23142080 | Unspecified |
| mmu-miR-34a-5p | MIMAT0000542 | Vars | 22321 | HITS-CLIP | 23142080 | Unspecified |
| mmu-miR-34a-5p | MIMAT0000542 | Vash1 | 238328 | HITS-CLIP | 19536157 | Unspecified |
| mmu-miR-34a-5p | MIMAT0000542 | Vash2 | 226841 | Microarrays//HITS-CLIP | 37986224\|19536157 | Unspecified |
| mmu-miR-34a-5p | MIMAT0000542 | Vasp | 22323 | HITS-CLIP | 30971389\|26203562\|29167373\|23142 | Unspecified |
| mmu-miR-34a-5p | MIMAT0000542 | Vat1 | 26949 | HITS-CLIP | 23630078\|30971389\|23142080\|29167 | Unspecified |
| mmu-miR-34a-5p | MIMAT0000542 | Vav1 | 22324 | HITS-CLIP | 23142080\|29167373 | Unspecified |
| mmu-miR-34a-5p | MIMAT0000542 | Vav2 | 22325 | HITS-CLIP | 23142080 | Unspecified |
| mmu-miR-34a-5p | MIMAT0000542 | Vav3 | 57257 | HITS-CLIP | 29167373 | Unspecified |

| | | | | | | |
|---|---|---|---|---|---|---|
| mmu-miR-34a-5p | MIMAT0000542 | Vbp1 | 22327 | HITS-CLIP | 19536157\|29167373\|23142080 | Unspecified |
| mmu-miR-34a-5p | MIMAT0000542 | Vcam1 | 22329 | HITS-CLIP | 23630078 | Unspecified |
| mmu-miR-34a-5p | MIMAT0000542 | Vcan | 13003 | HITS-CLIP | 19536157 | Unspecified |
| mmu-miR-34a-5p | MIMAT0000542 | Vcl | 22330 | Luciferase assay//Northern blot//qRT-PCR | 23047694\|23630078\|23142080\|29167 | Unspecified |
| mmu-miR-34a-5p | MIMAT0000542 | Vcpip1 | 70675 | HITS-CLIP | 19536157\|29167373\|23142080 | Unspecified |
| mmu-miR-34a-5p | MIMAT0000542 | Vcpkmt | 207965 | HITS-CLIP | 23142080 | Unspecified |
| mmu-miR-34a-5p | MIMAT0000542 | Vdac1 | 22333 | HITS-CLIP | 19536157\|31433990\|23142080 | Unspecified |
| mmu-miR-34a-5p | MIMAT0000542 | Vdac2 | 22334 | HITS-CLIP | 23142080\|19536157\|29167373 | Unspecified |
| mmu-miR-34a-5p | MIMAT0000542 | Vdr | 22337 | Microarrays | 37986224\|23027903 | Unspecified |
| mmu-miR-34a-5p | MIMAT0000542 | Vegfa | 22339 | HITS-CLIP | 23142080\|29167373 | Unspecified |
| mmu-miR-34a-5p | MIMAT0000542 | Vegfb | 22340 | Northern blot//qRT-PCR//Western blot//H | 24587330\|23142080 | Unspecified |
| mmu-miR-34a-5p | MIMAT0000542 | Vezf1 | 22344 | HITS-CLIP | 23142080\|29167373 | Unspecified |
| mmu-miR-34a-5p | MIMAT0000542 | Vezt | 215008 | HITS-CLIP | 19536157\|23630078\|23142080 | Unspecified |
| mmu-miR-34a-5p | MIMAT0000542 | Vgll4 | 232334 | HITS-CLIP | 23142080 | Unspecified |
| mmu-miR-34a-5p | MIMAT0000542 | Vhl | 22346 | HITS-CLIP | 23142080 | Unspecified |
| mmu-miR-34a-5p | MIMAT0000542 | Vim | 22352 | HITS-CLIP | 19536157\|31433990\|23630078\|29167 | Unspecified |
| mmu-miR-34a-5p | MIMAT0000542 | Vipas39 | 104799 | HITS-CLIP | 23142080\|29167373 | Unspecified |
| mmu-miR-34a-5p | MIMAT0000542 | Vipr1 | 22354 | HITS-CLIP | 19536157 | Unspecified |
| mmu-miR-34a-5p | MIMAT0000542 | Virma | 25962 | HITS-CLIP | 19536157\|30971389\|31433990\|23142 | Unspecified |
| mmu-miR-34a-5p | MIMAT0000542 | Vkorc1 | 27973 | HITS-CLIP | 19536157 | Unspecified |
| mmu-miR-34a-5p | MIMAT0000542 | Vkorc1l1 | 69568 | HITS-CLIP | 23142080\|29167373 | Unspecified |
| mmu-miR-34a-5p | MIMAT0000542 | Vldlr | 22359 | HITS-CLIP | 23142080\|19536157 | Unspecified |
| mmu-miR-34a-5p | MIMAT0000542 | Vma21 | 67048 | HITS-CLIP | 23142080\|29167373 | Unspecified |
| mmu-miR-34a-5p | MIMAT0000542 | Vmn1r90 | 627280 | HITS-CLIP | 23142080 | Unspecified |
| mmu-miR-34a-5p | MIMAT0000542 | Vmp1 | 75909 | HITS-CLIP | 31433990\|23142080\|29167373 | Unspecified |
| mmu-miR-34a-5p | MIMAT0000542 | Vopp1 | 232023 | HITS-CLIP | 23142080 | Unspecified |
| mmu-miR-34a-5p | MIMAT0000542 | Vps11 | 71732 | HITS-CLIP | 37986224\|23142080 | Unspecified |
| mmu-miR-34a-5p | MIMAT0000542 | Vps13a | 271564 | HITS-CLIP | 23142080\|29167373 | Unspecified |
| mmu-miR-34a-5p | MIMAT0000542 | Vps13b | 666173 | HITS-CLIP | 29167373\|23142080 | Unspecified |
| mmu-miR-34a-5p | MIMAT0000542 | Vps13c | 320528 | HITS-CLIP | 23142080\|29167373 | Unspecified |
| mmu-miR-34a-5p | MIMAT0000542 | Vps13d | 230895 | HITS-CLIP | 31433990\|29167373\|23142080 | Unspecified |
| mmu-miR-34a-5p | MIMAT0000542 | Vps16 | 80743 | HITS-CLIP | 29167373\|23142080 | Unspecified |
| mmu-miR-34a-5p | MIMAT0000542 | Vps18 | 228545 | HITS-CLIP | 23142080 | Unspecified |
| mmu-miR-34a-5p | MIMAT0000542 | Vps25 | 28084 | HITS-CLIP | 23142080 | Unspecified |
| mmu-miR-34a-5p | MIMAT0000542 | Vps26a | 30930 | HITS-CLIP | 23142080\|29167373 | Unspecified |
| mmu-miR-34a-5p | MIMAT0000542 | Vps26c | 135977289 | HITS-CLIP | 23142080\|29167373 | Unspecified |
| mmu-miR-34a-5p | MIMAT0000542 | Vps28 | 51160 | HITS-CLIP | 23142080 | Unspecified |
| mmu-miR-34a-5p | MIMAT0000542 | Vps29 | 56433 | HITS-CLIP | 23142080\|29167373 | Unspecified |
| mmu-miR-34a-5p | MIMAT0000542 | Vps33a | 77573 | HITS-CLIP | 19536157\|23142080\|29167373 | Unspecified |

| | | | | | | |
|---|---|---|---|---|---|---|
| mmu-miR-34a-5p | MIMAT0000542 | Vps33b | 233405 | HITS-CLIP | 23142080 | Unspecified |
| mmu-miR-34a-5p | MIMAT0000542 | Vps35 | 65114 | HITS-CLIP | 19536157\|23142080\|29167373 | Unspecified |
| mmu-miR-34a-5p | MIMAT0000542 | Vps35l | 57020 | HITS-CLIP | 19536157\|23142080 | Unspecified |
| mmu-miR-34a-5p | MIMAT0000542 | Vps36 | 70160 | HITS-CLIP | 29167373 | Unspecified |
| mmu-miR-34a-5p | MIMAT0000542 | Vps37a | 52348 | HITS-CLIP | 23142080\|29167373 | Unspecified |
| mmu-miR-34a-5p | MIMAT0000542 | Vps37b | 330192 | HITS-CLIP | 23142080 | Unspecified |
| mmu-miR-34a-5p | MIMAT0000542 | Vps37c | 107305 | HITS-CLIP | 23142080 | Unspecified |
| mmu-miR-34a-5p | MIMAT0000542 | Vps39 | 269338 | HITS-CLIP | 23142080\|29167373 | Unspecified |
| mmu-miR-34a-5p | MIMAT0000542 | Vps41 | 218035 | HITS-CLIP | 23142080\|29167373 | Unspecified |
| mmu-miR-34a-5p | MIMAT0000542 | Vps4a | 116733 | HITS-CLIP | 23142080 | Unspecified |
| mmu-miR-34a-5p | MIMAT0000542 | Vps4b | 20479 | HITS-CLIP | 30971389\|23142080\|29167373 | Unspecified |
| mmu-miR-34a-5p | MIMAT0000542 | Vps50 | 55610 | HITS-CLIP | 31433990\|29167373 | Unspecified |
| mmu-miR-34a-5p | MIMAT0000542 | Vps51 | 68505 | HITS-CLIP | 23142080\|30971389 | Unspecified |
| mmu-miR-34a-5p | MIMAT0000542 | Vps52 | 224705 | HITS-CLIP | 23142080 | Unspecified |
| mmu-miR-34a-5p | MIMAT0000542 | Vps53 | 68299 | HITS-CLIP | 31433990\|23142080 | Unspecified |
| mmu-miR-34a-5p | MIMAT0000542 | Vps54 | 245944 | HITS-CLIP | 23142080\|31433990\|29167373 | Unspecified |
| mmu-miR-34a-5p | MIMAT0000542 | Vps72 | 21427 | HITS-CLIP | 23142080\|29167373 | Unspecified |
| mmu-miR-34a-5p | MIMAT0000542 | Vps8 | 209018 | HITS-CLIP | 30971389\|23142080 | Unspecified |
| mmu-miR-34a-5p | MIMAT0000542 | Vps9d1 | 72325 | HITS-CLIP | 23142080 | Unspecified |
| mmu-miR-34a-5p | MIMAT0000542 | Vrk1 | 22367 | HITS-CLIP | 23142080 | Unspecified |
| mmu-miR-34a-5p | MIMAT0000542 | Vrk2 | 69922 | Microarrays | 37986224\|23027903 | Unspecified |
| mmu-miR-34a-5p | MIMAT0000542 | Vrk3 | 101568 | HITS-CLIP | 23142080 | Unspecified |
| mmu-miR-34a-5p | MIMAT0000542 | Vsir | 64115 | HITS-CLIP | 23142080\|29167373 | Unspecified |
| mmu-miR-34a-5p | MIMAT0000542 | Vsnl1 | 26950 | HITS-CLIP | 19536157 | Unspecified |
| mmu-miR-34a-5p | MIMAT0000542 | Vstm2a | 211739 | HITS-CLIP | 19536157 | Unspecified |
| mmu-miR-34a-5p | MIMAT0000542 | Vta1 | 66201 | HITS-CLIP | 23142080\|19536157 | Unspecified |
| mmu-miR-34a-5p | MIMAT0000542 | Vti1a | 53611 | HITS-CLIP | 29167373 | Unspecified |
| mmu-miR-34a-5p | MIMAT0000542 | Vwa5a | 67776 | HITS-CLIP | 23142080\|29167373 | Unspecified |
| mmu-miR-34a-5p | MIMAT0000542 | Vwa8 | 219189 | HITS-CLIP | 29167373 | Unspecified |
| mmu-miR-34a-5p | MIMAT0000542 | Vxn | 76982 | HITS-CLIP | 19536157 | Unspecified |
| mmu-miR-34a-5p | MIMAT0000542 | Wac | 225131 | HITS-CLIP | 23142080\|29167373 | Unspecified |
| mmu-miR-34a-5p | MIMAT0000542 | Wapl | 23063 | HITS-CLIP | 31433990\|23142080\|29167373 | Unspecified |
| mmu-miR-34a-5p | MIMAT0000542 | Wars | 22375 | HITS-CLIP | 23142080\|29167373 | Unspecified |
| mmu-miR-34a-5p | MIMAT0000542 | Wars1 | 7453 | HITS-CLIP | 30971389 | Unspecified |
| mmu-miR-34a-5p | MIMAT0000542 | Wars2 | 70560 | HITS-CLIP | 23142080 | Unspecified |
| mmu-miR-34a-5p | MIMAT0000542 | Was | 22376 | HITS-CLIP | 29167373\|23142080 | Unspecified |
| mmu-miR-34a-5p | MIMAT0000542 | Wasf1 | 83767 | HITS-CLIP//Luciferase reporter assay//qRT- | 19536157\|28363996 | Unspecified |
| mmu-miR-34a-5p | MIMAT0000542 | Wasf2 | 242687 | HITS-CLIP | 23142080\|29167373 | Unspecified |
| mmu-miR-34a-5p | MIMAT0000542 | Washc1 | 100287171 | HITS-CLIP | 23142080 | Unspecified |

| | | | | | | |
|---|---|---|---|---|---|---|
| mmu-miR-34a-5p | MIMAT0000542 | Washc2 | 28006 | HITS-CLIP | 29167373 | Unspecified |
| mmu-miR-34a-5p | MIMAT0000542 | Washc3 | 51019 | HITS-CLIP | 23142080 | Unspecified |
| mmu-miR-34a-5p | MIMAT0000542 | Washc4 | 319277 | HITS-CLIP | 31433990\|23142080\|29167373 | Unspecified |
| mmu-miR-34a-5p | MIMAT0000542 | Washc5 | 9897 | HITS-CLIP | 23142080\|29167373 | Unspecified |
| mmu-miR-34a-5p | MIMAT0000542 | Wasl | 73178 | HITS-CLIP | 23142080 | Unspecified |
| mmu-miR-34a-5p | MIMAT0000542 | Wbp11 | 60321 | HITS-CLIP | 23142080\|29167373 | Unspecified |
| mmu-miR-34a-5p | MIMAT0000542 | Wbp1l | 226178 | HITS-CLIP | 19536157\|23142080\|29167373 | Unspecified |
| mmu-miR-34a-5p | MIMAT0000542 | Wbp2 | 22378 | HITS-CLIP | 23142080\|29167373 | Unspecified |
| mmu-miR-34a-5p | MIMAT0000542 | Wbp4 | 22380 | HITS-CLIP | 23142080 | Unspecified |
| mmu-miR-34a-5p | MIMAT0000542 | Wdfy1 | 69368 | HITS-CLIP | 29167373 | Unspecified |
| mmu-miR-34a-5p | MIMAT0000542 | Wdfy2 | 268752 | HITS-CLIP | 23142080 | Unspecified |
| mmu-miR-34a-5p | MIMAT0000542 | Wdfy3 | 72145 | HITS-CLIP | 19536157\|30971389\|29167373 | Unspecified |
| mmu-miR-34a-5p | MIMAT0000542 | Wdfy4 | 545030 | HITS-CLIP | 23142080\|29167373 | Unspecified |
| mmu-miR-34a-5p | MIMAT0000542 | Wdhd1 | 218973 | Microarrays//HITS-CLIP | 37986224\|23142080\|23027903\|2916 | Unspecified |
| mmu-miR-34a-5p | MIMAT0000542 | Wdr1 | 22388 | HITS-CLIP | 23142080\|30971389\|29167373 | Unspecified |
| mmu-miR-34a-5p | MIMAT0000542 | Wdr11 | 207425 | HITS-CLIP | 23142080 | Unspecified |
| mmu-miR-34a-5p | MIMAT0000542 | Wdr12 | 57750 | Microarrays//HITS-CLIP | 37986224\|23027903\|29167373 | Unspecified |
| mmu-miR-34a-5p | MIMAT0000542 | Wdr13 | 73447 | HITS-CLIP | 23142080 | Unspecified |
| mmu-miR-34a-5p | MIMAT0000542 | Wdr19 | 213081 | Microarrays | 37986224 | Unspecified |
| mmu-miR-34a-5p | MIMAT0000542 | Wdr20 | 69641 | HITS-CLIP | 23142080\|29167373 | Unspecified |
| mmu-miR-34a-5p | MIMAT0000542 | Wdr24 | 268933 | HITS-CLIP | 23142080 | Unspecified |
| mmu-miR-34a-5p | MIMAT0000542 | Wdr26 | 226757 | HITS-CLIP | 23142080\|31433990\|29167373 | Unspecified |
| mmu-miR-34a-5p | MIMAT0000542 | Wdr3 | 269470 | HITS-CLIP | 29167373\|23142080 | Unspecified |
| mmu-miR-34a-5p | MIMAT0000542 | Wdr31 | 71354 | Microarrays | 37986224 | Unspecified |
| mmu-miR-34a-5p | MIMAT0000542 | Wdr33 | 74320 | HITS-CLIP | 29167373\|23142080 | Unspecified |
| mmu-miR-34a-5p | MIMAT0000542 | Wdr35 | 74682 | Microarrays | 37986224 | Unspecified |
| mmu-miR-34a-5p | MIMAT0000542 | Wdr36 | 225348 | HITS-CLIP | 31433990\|23142080\|29167373 | Unspecified |
| mmu-miR-34a-5p | MIMAT0000542 | Wdr37 | 207615 | HITS-CLIP | 23142080\|29167373 | Unspecified |
| mmu-miR-34a-5p | MIMAT0000542 | Wdr41 | 218460 | HITS-CLIP | 23142080 | Unspecified |
| mmu-miR-34a-5p | MIMAT0000542 | Wdr43 | 72515 | HITS-CLIP | 29167373 | Unspecified |
| mmu-miR-34a-5p | MIMAT0000542 | Wdr44 | 72404 | HITS-CLIP | 19536157\|23142080 | Unspecified |
| mmu-miR-34a-5p | MIMAT0000542 | Wdr45 | 54636 | HITS-CLIP | 23142080 | Unspecified |
| mmu-miR-34a-5p | MIMAT0000542 | Wdr45b | 66840 | HITS-CLIP | 19536157\|31433990\|23142080 | Unspecified |
| mmu-miR-34a-5p | MIMAT0000542 | Wdr46 | 57315 | HITS-CLIP | 23142080 | Unspecified |
| mmu-miR-34a-5p | MIMAT0000542 | Wdr47 | 99512 | HITS-CLIP | 19536157\|23142080 | Unspecified |
| mmu-miR-34a-5p | MIMAT0000542 | Wdr48 | 67561 | HITS-CLIP | 29167373\|23142080 | Unspecified |
| mmu-miR-34a-5p | MIMAT0000542 | Wdr5 | 140858 | Microarrays//HITS-CLIP | 37986224\|23027903\|26203562\|2314 | Unspecified |
| mmu-miR-34a-5p | MIMAT0000542 | Wdr55 | 67936 | HITS-CLIP | 23142080 | Unspecified |
| mmu-miR-34a-5p | MIMAT0000542 | Wdr59 | 319481 | HITS-CLIP | 31433990\|23142080 | Unspecified |

| | | | | | | |
|---|---|---|---|---|---|---|
| mmu-miR-34a-5p | MIMAT0000542 | Wdr5b | 69544 | HITS-CLIP | 23142080 | Unspecified |
| mmu-miR-34a-5p | MIMAT0000542 | Wdr6 | 83669 | Microarrays//HITS-CLIP | 37986224\|23142080 | Unspecified |
| mmu-miR-34a-5p | MIMAT0000542 | Wdr7 | 104082 | HITS-CLIP | 19536157\|23142080\|29167373 | Unspecified |
| mmu-miR-34a-5p | MIMAT0000542 | Wdr70 | 545085 | HITS-CLIP | 23142080 | Unspecified |
| mmu-miR-34a-5p | MIMAT0000542 | Wdr74 | 107071 | HITS-CLIP | 23142080 | Unspecified |
| mmu-miR-34a-5p | MIMAT0000542 | Wdr75 | 73674 | HITS-CLIP | 23142080 | Unspecified |
| mmu-miR-34a-5p | MIMAT0000542 | Wdr76 | 241627 | Microarrays//HITS-CLIP | 37986224\|23027903\|31433990\|23142 | Unspecified |
| mmu-miR-34a-5p | MIMAT0000542 | Wdr77 | 70465 | HITS-CLIP | 23142080 | Unspecified |
| mmu-miR-34a-5p | MIMAT0000542 | Wdr78 | 242584 | Microarrays | 37986224 | Unspecified |
| mmu-miR-34a-5p | MIMAT0000542 | Wdr81 | 192652 | HITS-CLIP | 23142080 | Unspecified |
| mmu-miR-34a-5p | MIMAT0000542 | Wdr82 | 77305 | HITS-CLIP | 19536157\|23142080\|29167373 | Unspecified |
| mmu-miR-34a-5p | MIMAT0000542 | Wdr83 | 67836 | HITS-CLIP | 23142080 | Unspecified |
| mmu-miR-34a-5p | MIMAT0000542 | Wdr83os | 414077 | HITS-CLIP | 23142080 | Unspecified |
| mmu-miR-34a-5p | MIMAT0000542 | Wdr89 | 72338 | HITS-CLIP | 23142080 | Unspecified |
| mmu-miR-34a-5p | MIMAT0000542 | Wdr90 | 106618 | HITS-CLIP | 23142080 | Unspecified |
| mmu-miR-34a-5p | MIMAT0000542 | Wdr91 | 101240 | HITS-CLIP | 23142080\|29167373 | Unspecified |
| mmu-miR-34a-5p | MIMAT0000542 | Wdr92 | 103784 | HITS-CLIP | 23142080 | Unspecified |
| mmu-miR-34a-5p | MIMAT0000542 | Wdsub1 | 72137 | HITS-CLIP | 31433990\|23142080 | Unspecified |
| mmu-miR-34a-5p | MIMAT0000542 | Wdtc1 | 230796 | HITS-CLIP | 23142080 | Unspecified |
| mmu-miR-34a-5p | MIMAT0000542 | Wdyhv1 | 76773 | HITS-CLIP | 31433990\|23142080 | Unspecified |
| mmu-miR-34a-5p | MIMAT0000542 | Wee1 | 22390 | HITS-CLIP | 31433990\|26203562\|23142080\|29167 | Unspecified |
| mmu-miR-34a-5p | MIMAT0000542 | Wfs1 | 22393 | HITS-CLIP | 23142080 | Unspecified |
| mmu-miR-34a-5p | MIMAT0000542 | Whrn | 73750 | Microarrays | 37986224 | Unspecified |
| mmu-miR-34a-5p | MIMAT0000542 | Wipf1 | 215280 | HITS-CLIP | 37986224\|23142080\|29167373 | Unspecified |
| mmu-miR-34a-5p | MIMAT0000542 | Wipf2 | 68524 | HITS-CLIP | 21258322\|23142080 | Unspecified |
| mmu-miR-34a-5p | MIMAT0000542 | Wipi1 | 52639 | HITS-CLIP | 23630078 | Unspecified |
| mmu-miR-34a-5p | MIMAT0000542 | Wipi2 | 74781 | HITS-CLIP | 31433990\|23142080 | Unspecified |
| mmu-miR-34a-5p | MIMAT0000542 | Wiz | 22404 | HITS-CLIP | 29167373\|30971389\|23142080 | Unspecified |
| mmu-miR-34a-5p | MIMAT0000542 | Wls | 68151 | HITS-CLIP | 29167373 | Unspecified |
| mmu-miR-34a-5p | MIMAT0000542 | Wnk1 | 232341 | HITS-CLIP | 19536157\|31433990\|23142080\|26203 | Unspecified |
| mmu-miR-34a-5p | MIMAT0000542 | Wnk3 | 279561 | HITS-CLIP | 19536157 | Unspecified |
| mmu-miR-34a-5p | MIMAT0000542 | Wnt9a | 216795 | HITS-CLIP | 26203562 | Unspecified |
| mmu-miR-34a-5p | MIMAT0000542 | Wrap73 | 59002 | HITS-CLIP | 31433990 | Unspecified |
| mmu-miR-34a-5p | MIMAT0000542 | Wrb | 71446 | HITS-CLIP | 23142080 | Unspecified |
| mmu-miR-34a-5p | MIMAT0000542 | Wrn | 22427 | HITS-CLIP | 23142080 | Unspecified |
| mmu-miR-34a-5p | MIMAT0000542 | Wrnip1 | 78903 | HITS-CLIP | 23142080 | Unspecified |
| mmu-miR-34a-5p | MIMAT0000542 | Wsb1 | 78889 | HITS-CLIP | 23142080\|29167373 | Unspecified |
| mmu-miR-34a-5p | MIMAT0000542 | Wsb2 | 59043 | Microarrays | 37986224\|23027903 | Unspecified |
| mmu-miR-34a-5p | MIMAT0000542 | Wscd2 | 320916 | HITS-CLIP | 23142080 | Unspecified |

| | | | | | | |
|---|---|---|---|---|---|---|
| mmu-miR-34a-5p | MIMAT0000542 | Wtap | 60532 | HITS-CLIP | 23142080\|31433990\|29167373 | Unspecified |
| mmu-miR-34a-5p | MIMAT0000542 | Wwc1 | 211652 | HITS-CLIP | 23142080\|30971389 | Unspecified |
| mmu-miR-34a-5p | MIMAT0000542 | Wwc2 | 52357 | HITS-CLIP | 23630078 | Unspecified |
| mmu-miR-34a-5p | MIMAT0000542 | Wwp1 | 107568 | HITS-CLIP | 23142080\|29167373 | Unspecified |
| mmu-miR-34a-5p | MIMAT0000542 | Wwp2 | 66894 | HITS-CLIP | 23142080\|29167373 | Unspecified |
| mmu-miR-34a-5p | MIMAT0000542 | Wwtr1 | 97064 | HITS-CLIP | 19536157 | Unspecified |
| mmu-miR-34a-5p | MIMAT0000542 | Xab2 | 67439 | HITS-CLIP | 23142080\|29167373 | Unspecified |
| mmu-miR-34a-5p | MIMAT0000542 | Xaf1 | 327959 | HITS-CLIP | 29167373 | Unspecified |
| mmu-miR-34a-5p | MIMAT0000542 | Xbp1 | 22433 | HITS-CLIP | 23142080\|29167373 | Unspecified |
| mmu-miR-34a-5p | MIMAT0000542 | Xdh | 22436 | HITS-CLIP | 29167373 | Unspecified |
| mmu-miR-34a-5p | MIMAT0000542 | Xiap | 11798 | HITS-CLIP | 23142080\|29167373 | Unspecified |
| mmu-miR-34a-5p | MIMAT0000542 | Xirp1 | 22437 | Microarrays | 37986224 | Unspecified |
| mmu-miR-34a-5p | MIMAT0000542 | Xkr4 | 497097 | HITS-CLIP | 19536157 | Unspecified |
| mmu-miR-34a-5p | MIMAT0000542 | Xkr6 | 219149 | HITS-CLIP | 19536157 | Unspecified |
| mmu-miR-34a-5p | MIMAT0000542 | Xlr4a | 434794 | HITS-CLIP | 23142080 | Unspecified |
| mmu-miR-34a-5p | MIMAT0000542 | Xpnpep1 | 170750 | HITS-CLIP | 23142080 | Unspecified |
| mmu-miR-34a-5p | MIMAT0000542 | Xpo1 | 103573 | HITS-CLIP | 31433990\|19536157\|23142080\|29167 | Unspecified |
| mmu-miR-34a-5p | MIMAT0000542 | Xpo4 | 57258 | Microarrays//HITS-CLIP | 37986224\|23027903\|23142080\|29167 | Unspecified |
| mmu-miR-34a-5p | MIMAT0000542 | Xpo5 | 72322 | HITS-CLIP | 23142080\|29167373 | Unspecified |
| mmu-miR-34a-5p | MIMAT0000542 | Xpo6 | 74204 | HITS-CLIP | 23142080 | Unspecified |
| mmu-miR-34a-5p | MIMAT0000542 | Xpo7 | 65246 | HITS-CLIP | 23142080\|29167373 | Unspecified |
| mmu-miR-34a-5p | MIMAT0000542 | Xpot | 73192 | HITS-CLIP | 23142080\|29167373 | Unspecified |
| mmu-miR-34a-5p | MIMAT0000542 | Xpr1 | 19775 | HITS-CLIP | 30971389\|19536157\|23142080\|29167 | Unspecified |
| mmu-miR-34a-5p | MIMAT0000542 | Xrcc2 | 57434 | Microarrays//HITS-CLIP | 37986224\|23027903\|23142080\|29167 | Unspecified |
| mmu-miR-34a-5p | MIMAT0000542 | Xrcc5 | 22596 | Microarrays//HITS-CLIP | 37986224\|23142080 | Unspecified |
| mmu-miR-34a-5p | MIMAT0000542 | Xrcc6 | 14375 | HITS-CLIP | 23142080 | Unspecified |
| mmu-miR-34a-5p | MIMAT0000542 | Xrn1 | 24127 | HITS-CLIP | 23142080\|29167373 | Unspecified |
| mmu-miR-34a-5p | MIMAT0000542 | Xrn2 | 24128 | HITS-CLIP | 23142080\|29167373 | Unspecified |
| mmu-miR-34a-5p | MIMAT0000542 | Xylt1 | 233781 | HITS-CLIP | 23142080\|29167373 | Unspecified |
| mmu-miR-34a-5p | MIMAT0000542 | Xylt2 | 217119 | HITS-CLIP | 23142080\|29167373 | Unspecified |
| mmu-miR-34a-5p | MIMAT0000542 | Yae1d1 | 67008 | HITS-CLIP | 23142080\|29167373 | Unspecified |
| mmu-miR-34a-5p | MIMAT0000542 | Yaf2 | 67057 | HITS-CLIP | 23142080 | Unspecified |
| mmu-miR-34a-5p | MIMAT0000542 | Yap1 | 22601 | HITS-CLIP | 23630078 | Unspecified |
| mmu-miR-34a-5p | MIMAT0000542 | Yars | 107271 | HITS-CLIP | 29167373 | Unspecified |
| mmu-miR-34a-5p | MIMAT0000542 | Yars2 | 70120 | HITS-CLIP | 19536157 | Unspecified |
| mmu-miR-34a-5p | MIMAT0000542 | Ybey | 216119 | HITS-CLIP | 29167373 | Unspecified |
| mmu-miR-34a-5p | MIMAT0000542 | Ybx1 | 22608 | HITS-CLIP | 23142080 | Unspecified |
| mmu-miR-34a-5p | MIMAT0000542 | Ybx2 | 53422 | HITS-CLIP | 23142080 | Unspecified |
| mmu-miR-34a-5p | MIMAT0000542 | Ybx3 | 56449 | HITS-CLIP | 19536157\|23142080 | Unspecified |

| | | | | | | |
|---|---|---|---|---|---|---|
| mmu-miR-34a-5p | MIMAT0000542 | Yeats2 | 208146 | HITS-CLIP | 23142080\|29167373 | Unspecified |
| mmu-miR-34a-5p | MIMAT0000542 | Yeats4 | 64050 | HITS-CLIP | 23142080\|29167373 | Unspecified |
| mmu-miR-34a-5p | MIMAT0000542 | Yes1 | 22612 | HITS-CLIP | 23142080 | Unspecified |
| mmu-miR-34a-5p | MIMAT0000542 | Yif1a | 68090 | HITS-CLIP | 23142080 | Unspecified |
| mmu-miR-34a-5p | MIMAT0000542 | Yif1b | 77254 | HITS-CLIP | 23142080 | Unspecified |
| mmu-miR-34a-5p | MIMAT0000542 | Yipf1 | 230584 | HITS-CLIP | 23142080 | Unspecified |
| mmu-miR-34a-5p | MIMAT0000542 | Yipf2 | 74766 | HITS-CLIP | 23142080 | Unspecified |
| mmu-miR-34a-5p | MIMAT0000542 | Yipf4 | 67864 | HITS-CLIP | 23142080 | Unspecified |
| mmu-miR-34a-5p | MIMAT0000542 | Yipf5 | 67180 | HITS-CLIP | 29167373 | Unspecified |
| mmu-miR-34a-5p | MIMAT0000542 | Yipf6 | 77929 | HITS-CLIP | 19536157\|23142080 | Unspecified |
| mmu-miR-34a-5p | MIMAT0000542 | Ylpm1 | 56531 | HITS-CLIP | 23142080\|29167373 | Unspecified |
| mmu-miR-34a-5p | MIMAT0000542 | Yme1l1 | 27377 | HITS-CLIP | 29167373\|23142080 | Unspecified |
| mmu-miR-34a-5p | MIMAT0000542 | Yod1 | 226418 | HITS-CLIP | 23142080\|29167373 | Unspecified |
| mmu-miR-34a-5p | MIMAT0000542 | Ypel2 | 77864 | Microarrays//HITS-CLIP | 37986224\|23142080 | Unspecified |
| mmu-miR-34a-5p | MIMAT0000542 | Ypel3 | 66090 | HITS-CLIP | 23142080 | Unspecified |
| mmu-miR-34a-5p | MIMAT0000542 | Ypel4 | 241525 | HITS-CLIP | 23142080 | Unspecified |
| mmu-miR-34a-5p | MIMAT0000542 | Ypel5 | 383295 | HITS-CLIP | 19536157\|23142080\|29167373 | Unspecified |
| mmu-miR-34a-5p | MIMAT0000542 | Ythdc1 | 231386 | HITS-CLIP | 23142080\|29167373 | Unspecified |
| mmu-miR-34a-5p | MIMAT0000542 | Ythdf1 | 228994 | HITS-CLIP | 29167373 | Unspecified |
| mmu-miR-34a-5p | MIMAT0000542 | Ythdf2 | 213541 | HITS-CLIP | 29167373 | Unspecified |
| mmu-miR-34a-5p | MIMAT0000542 | Ythdf3 | 229096 | HITS-CLIP | 23142080\|29167373 | Unspecified |
| mmu-miR-34a-5p | MIMAT0000542 | Ywhab | 54401 | HITS-CLIP | 31433990\|23142080\|19536157 | Unspecified |
| mmu-miR-34a-5p | MIMAT0000542 | Ywhae | 22627 | HITS-CLIP | 19536157\|31433990\|23630078\|30971 | Unspecified |
| mmu-miR-34a-5p | MIMAT0000542 | Ywhag | 22628 | HITS-CLIP | 23142080\|19536157\|29167373 | Unspecified |
| mmu-miR-34a-5p | MIMAT0000542 | Ywhaq | 22630 | HITS-CLIP | 29167373 | Unspecified |
| mmu-miR-34a-5p | MIMAT0000542 | Ywhaz | 22631 | HITS-CLIP | 23142080 | Unspecified |
| mmu-miR-34a-5p | MIMAT0000542 | Yy1 | 22632 | HITS-CLIP | 37986224\|23630078\|23142080\|29167 | Unspecified |
| mmu-miR-34a-5p | MIMAT0000542 | Zadh2 | 225791 | HITS-CLIP | 23142080 | Unspecified |
| mmu-miR-34a-5p | MIMAT0000542 | Zap70 | 22637 | HITS-CLIP | 23142080 | Unspecified |
| mmu-miR-34a-5p | MIMAT0000542 | Zbed6 | 667118 | HITS-CLIP | 37986224 | Unspecified |
| mmu-miR-34a-5p | MIMAT0000542 | Zbp1 | 58203 | HITS-CLIP | 31433990\|23142080\|29167373 | Unspecified |
| mmu-miR-34a-5p | MIMAT0000542 | Zbtb1 | 268564 | HITS-CLIP | 30971389\|23142080\|29167373 | Unspecified |
| mmu-miR-34a-5p | MIMAT0000542 | Zbtb10 | 229055 | HITS-CLIP | 23142080 | Unspecified |
| mmu-miR-34a-5p | MIMAT0000542 | Zbtb11 | 271377 | HITS-CLIP | 23142080 | Unspecified |
| mmu-miR-34a-5p | MIMAT0000542 | Zbtb14 | 22666 | HITS-CLIP | 23142080 | Unspecified |
| mmu-miR-34a-5p | MIMAT0000542 | Zbtb16 | 235320 | HITS-CLIP | 23142080 | Unspecified |
| mmu-miR-34a-5p | MIMAT0000542 | Zbtb18 | 30928 | Microarrays//HITS-CLIP | 37986224\|31433990\|19536157\|2314: | Unspecified |
| mmu-miR-34a-5p | MIMAT0000542 | Zbtb20 | 56490 | HITS-CLIP | 19536157\|23142080\|29167373 | Unspecified |
| mmu-miR-34a-5p | MIMAT0000542 | Zbtb21 | 114565 | HITS-CLIP | 29167373 | Unspecified |

| | | | | | | |
|---|---|---|---|---|---|---|
| mmu-miR-34a-5p | MIMAT0000542 | Zbtb22 | 81630 | HITS-CLIP | 23142080 | Unspecified |
| mmu-miR-34a-5p | MIMAT0000542 | Zbtb24 | 268294 | HITS-CLIP | 23142080 | Unspecified |
| mmu-miR-34a-5p | MIMAT0000542 | Zbtb25 | 109929 | HITS-CLIP | 23142080 | Unspecified |
| mmu-miR-34a-5p | MIMAT0000542 | Zbtb26 | 320633 | HITS-CLIP | 23142080 | Unspecified |
| mmu-miR-34a-5p | MIMAT0000542 | Zbtb3 | 75291 | HITS-CLIP | 23142080 | Unspecified |
| mmu-miR-34a-5p | MIMAT0000542 | Zbtb32 | 58206 | HITS-CLIP | 23142080 | Unspecified |
| mmu-miR-34a-5p | MIMAT0000542 | Zbtb33 | 56805 | HITS-CLIP | 19536157\|23142080 | Unspecified |
| mmu-miR-34a-5p | MIMAT0000542 | Zbtb34 | 241311 | HITS-CLIP | 19536157\|23142080\|29167373 | Unspecified |
| mmu-miR-34a-5p | MIMAT0000542 | Zbtb37 | 240869 | HITS-CLIP | 23142080\|19536157 | Unspecified |
| mmu-miR-34a-5p | MIMAT0000542 | Zbtb38 | 245007 | HITS-CLIP | 23142080\|31433990\|29167373 | Unspecified |
| mmu-miR-34a-5p | MIMAT0000542 | Zbtb39 | 320080 | HITS-CLIP | 23630078\|23142080\|29167373 | Unspecified |
| mmu-miR-34a-5p | MIMAT0000542 | Zbtb4 | 75580 | HITS-CLIP | 23142080 | Unspecified |
| mmu-miR-34a-5p | MIMAT0000542 | Zbtb40 | 230848 | HITS-CLIP | 23142080 | Unspecified |
| mmu-miR-34a-5p | MIMAT0000542 | Zbtb41 | 226470 | HITS-CLIP | 31433990\|23142080 | Unspecified |
| mmu-miR-34a-5p | MIMAT0000542 | Zbtb42 | 382639 | HITS-CLIP | 23142080 | Unspecified |
| mmu-miR-34a-5p | MIMAT0000542 | Zbtb43 | 71834 | HITS-CLIP | 19536157\|29167373 | Unspecified |
| mmu-miR-34a-5p | MIMAT0000542 | Zbtb44 | 235132 | HITS-CLIP | 31433990\|23142080 | Unspecified |
| mmu-miR-34a-5p | MIMAT0000542 | Zbtb45 | 232879 | HITS-CLIP | 23142080 | Unspecified |
| mmu-miR-34a-5p | MIMAT0000542 | Zbtb5 | 230119 | HITS-CLIP | 23142080\|19536157 | Unspecified |
| mmu-miR-34a-5p | MIMAT0000542 | Zbtb6 | 241322 | HITS-CLIP | 31433990\|23142080 | Unspecified |
| mmu-miR-34a-5p | MIMAT0000542 | Zbtb7a | 16969 | HITS-CLIP | 19536157\|23142080\|29167373 | Unspecified |
| mmu-miR-34a-5p | MIMAT0000542 | Zbtb7b | 22724 | HITS-CLIP | 23630078\|26203562\|23142080 | Unspecified |
| mmu-miR-34a-5p | MIMAT0000542 | Zbtb9 | 474156 | HITS-CLIP | 23142080 | Unspecified |
| mmu-miR-34a-5p | MIMAT0000542 | Zc3h10 | 103284 | HITS-CLIP | 30971389 | Unspecified |
| mmu-miR-34a-5p | MIMAT0000542 | Zc3h12a | 230738 | HITS-CLIP | 29167373 | Unspecified |
| mmu-miR-34a-5p | MIMAT0000542 | Zc3h12c | 244871 | HITS-CLIP | 29167373 | Unspecified |
| mmu-miR-34a-5p | MIMAT0000542 | Zc3h12d | 237256 | HITS-CLIP | 23142080 | Unspecified |
| mmu-miR-34a-5p | MIMAT0000542 | Zc3h13 | 67302 | HITS-CLIP | 23142080\|31433990\|29167373 | Unspecified |
| mmu-miR-34a-5p | MIMAT0000542 | Zc3h14 | 75553 | HITS-CLIP | 23142080\|29167373 | Unspecified |
| mmu-miR-34a-5p | MIMAT0000542 | Zc3h15 | 69082 | HITS-CLIP | 31433990\|23630078\|23142080\|29167 | Unspecified |
| mmu-miR-34a-5p | MIMAT0000542 | Zc3h4 | 330474 | HITS-CLIP | 23142080 | Unspecified |
| mmu-miR-34a-5p | MIMAT0000542 | Zc3h6 | 78751 | HITS-CLIP | 23142080\|29167373 | Unspecified |
| mmu-miR-34a-5p | MIMAT0000542 | Zc3h7a | 106205 | HITS-CLIP | 29167373\|23142080 | Unspecified |
| mmu-miR-34a-5p | MIMAT0000542 | Zc3h7b | 20286 | HITS-CLIP | 23142080 | Unspecified |
| mmu-miR-34a-5p | MIMAT0000542 | Zc3hav1 | 78781 | HITS-CLIP | 23142080\|29167373 | Unspecified |
| mmu-miR-34a-5p | MIMAT0000542 | Zc3hav1l | 209032 | HITS-CLIP | 23142080 | Unspecified |
| mmu-miR-34a-5p | MIMAT0000542 | Zc3hc1 | 51530 | HITS-CLIP | 23142080 | Unspecified |
| mmu-miR-34a-5p | MIMAT0000542 | Zcchc14 | 142682 | HITS-CLIP | 19536157 | Unspecified |
| mmu-miR-34a-5p | MIMAT0000542 | Zcchc18 | 66995 | HITS-CLIP | 19536157\|23142080 | Unspecified |

| | | | | | | |
|---|---|---|---|---|---|---|
| mmu-miR-34a-5p | MIMAT0000542 | Zcchc2 | 227449 | HITS-CLIP | 23142080\|29167373 | Unspecified |
| mmu-miR-34a-5p | MIMAT0000542 | Zcchc24 | 71918 | Microarrays//HITS-CLIP | 37986224\|23027903\|23630078\|23142 | Unspecified |
| mmu-miR-34a-5p | MIMAT0000542 | Zcchc4 | 78796 | HITS-CLIP | 23142080 | Unspecified |
| mmu-miR-34a-5p | MIMAT0000542 | Zcchc8 | 70650 | HITS-CLIP | 23142080 | Unspecified |
| mmu-miR-34a-5p | MIMAT0000542 | Zcchc9 | 69085 | HITS-CLIP | 23142080 | Unspecified |
| mmu-miR-34a-5p | MIMAT0000542 | Zdhhc1 | 70796 | Microarrays | 37986224 | Unspecified |
| mmu-miR-34a-5p | MIMAT0000542 | Zdhhc12 | 66220 | HITS-CLIP | 23142080\|29167373 | Unspecified |
| mmu-miR-34a-5p | MIMAT0000542 | Zdhhc13 | 243983 | HITS-CLIP | 23142080\|29167373 | Unspecified |
| mmu-miR-34a-5p | MIMAT0000542 | Zdhhc15 | 108672 | HITS-CLIP | 23142080 | Unspecified |
| mmu-miR-34a-5p | MIMAT0000542 | Zdhhc16 | 74168 | Microarrays//HITS-CLIP | 37986224\|23027903\|23142080 | Unspecified |
| mmu-miR-34a-5p | MIMAT0000542 | Zdhhc17 | 320150 | HITS-CLIP | 19536157\|23142080\|29167373 | Unspecified |
| mmu-miR-34a-5p | MIMAT0000542 | Zdhhc18 | 503610 | Microarrays//HITS-CLIP | 37986224\|23027903\|23142080\|29167 | Unspecified |
| mmu-miR-34a-5p | MIMAT0000542 | Zdhhc2 | 70546 | Microarrays//HITS-CLIP | 37986224\|23027903\|19536157\|23142 | Unspecified |
| mmu-miR-34a-5p | MIMAT0000542 | Zdhhc20 | 75965 | HITS-CLIP | 31433990\|19536157\|23142080\|29167 | Unspecified |
| mmu-miR-34a-5p | MIMAT0000542 | Zdhhc21 | 68268 | HITS-CLIP | 23142080\|29167373 | Unspecified |
| mmu-miR-34a-5p | MIMAT0000542 | Zdhhc24 | 70605 | HITS-CLIP | 23142080\|29167373 | Unspecified |
| mmu-miR-34a-5p | MIMAT0000542 | Zdhhc3 | 69035 | HITS-CLIP | 23142080 | Unspecified |
| mmu-miR-34a-5p | MIMAT0000542 | Zdhhc5 | 228136 | HITS-CLIP | 29167373 | Unspecified |
| mmu-miR-34a-5p | MIMAT0000542 | Zdhhc7 | 102193 | HITS-CLIP | 29167373 | Unspecified |
| mmu-miR-34a-5p | MIMAT0000542 | Zdhhc8 | 27801 | HITS-CLIP | 23142080 | Unspecified |
| mmu-miR-34a-5p | MIMAT0000542 | Zdhhc9 | 208884 | HITS-CLIP | 23142080 | Unspecified |
| mmu-miR-34a-5p | MIMAT0000542 | Zeb1 | 21417 | HITS-CLIP | 23142080\|19536157\|31433990 | Unspecified |
| mmu-miR-34a-5p | MIMAT0000542 | Zeb2 | 24136 | HITS-CLIP | 23142080\|19536157\|29167373 | Unspecified |
| mmu-miR-34a-5p | MIMAT0000542 | Zer1 | 227693 | HITS-CLIP | 23142080 | Unspecified |
| mmu-miR-34a-5p | MIMAT0000542 | Zfand2a | 100494 | HITS-CLIP | 19536157\|23142080\|29167373 | Unspecified |
| mmu-miR-34a-5p | MIMAT0000542 | Zfand2b | 68818 | HITS-CLIP | 23142080 | Unspecified |
| mmu-miR-34a-5p | MIMAT0000542 | Zfand3 | 21769 | HITS-CLIP | 23142080\|29167373\|19536157 | Unspecified |
| mmu-miR-34a-5p | MIMAT0000542 | Zfand5 | 22682 | HITS-CLIP | 23142080\|19536157\|29167373 | Unspecified |
| mmu-miR-34a-5p | MIMAT0000542 | Zfand6 | 65098 | HITS-CLIP | 23142080\|31433990 | Unspecified |
| mmu-miR-34a-5p | MIMAT0000542 | Zfc3h1 | 216345 | HITS-CLIP | 30971389\|19536157\|23142080\|29167 | Unspecified |
| mmu-miR-34a-5p | MIMAT0000542 | Zfhx2 | 239102 | HITS-CLIP | 19536157\|29167373\|23142080 | Unspecified |
| mmu-miR-34a-5p | MIMAT0000542 | Zfhx3 | 11906 | Microarrays//HITS-CLIP | 37986224\|29167373 | Unspecified |
| mmu-miR-34a-5p | MIMAT0000542 | Zfp101 | 22643 | Microarrays//HITS-CLIP | 37986224\|23142080\|23027903\|29167 | Unspecified |
| mmu-miR-34a-5p | MIMAT0000542 | Zfp106 | 20402 | HITS-CLIP | 31433990\|23142080\|29167373 | Unspecified |
| mmu-miR-34a-5p | MIMAT0000542 | Zfp111 | 56707 | HITS-CLIP | 29167373 | Unspecified |
| mmu-miR-34a-5p | MIMAT0000542 | Zfp119a | 104349 | HITS-CLIP | 29167373 | Unspecified |
| mmu-miR-34a-5p | MIMAT0000542 | Zfp12 | 231866 | HITS-CLIP | 23142080 | Unspecified |
| mmu-miR-34a-5p | MIMAT0000542 | Zfp120 | 104348 | HITS-CLIP | 23142080\|29167373 | Unspecified |
| mmu-miR-34a-5p | MIMAT0000542 | Zfp131 | 72465 | HITS-CLIP | 23142080\|29167373 | Unspecified |

| | | | | | | |
|---|---|---|---|---|---|---|
| mmu-miR-34a-5p | MIMAT0000542 | Zfp14 | 243906 | HITS-CLIP | 23142080 | Unspecified |
| mmu-miR-34a-5p | MIMAT0000542 | Zfp141 | 434178 | HITS-CLIP | 23142080 | Unspecified |
| mmu-miR-34a-5p | MIMAT0000542 | Zfp142 | 77264 | HITS-CLIP | 23142080\|29167373 | Unspecified |
| mmu-miR-34a-5p | MIMAT0000542 | Zfp143 | 20841 | HITS-CLIP | 23142080 | Unspecified |
| mmu-miR-34a-5p | MIMAT0000542 | Zfp146 | 26465 | HITS-CLIP | 23142080 | Unspecified |
| mmu-miR-34a-5p | MIMAT0000542 | Zfp148 | 22661 | HITS-CLIP | 23142080\|30971389\|29167373 | Unspecified |
| mmu-miR-34a-5p | MIMAT0000542 | Zfp157 | 72154 | HITS-CLIP | 23142080 | Unspecified |
| mmu-miR-34a-5p | MIMAT0000542 | Zfp189 | 230162 | HITS-CLIP | 23142080 | Unspecified |
| mmu-miR-34a-5p | MIMAT0000542 | Zfp202 | 80902 | HITS-CLIP | 23142080 | Unspecified |
| mmu-miR-34a-5p | MIMAT0000542 | Zfp207 | 22680 | HITS-CLIP | 23142080\|31433990\|29167373 | Unspecified |
| mmu-miR-34a-5p | MIMAT0000542 | Zfp212 | 232784 | HITS-CLIP | 23142080 | Unspecified |
| mmu-miR-34a-5p | MIMAT0000542 | Zfp213 | 449521 | HITS-CLIP | 23142080 | Unspecified |
| mmu-miR-34a-5p | MIMAT0000542 | Zfp217 | 228913 | HITS-CLIP | 31433990\|26203562\|23142080\|29167 | Unspecified |
| mmu-miR-34a-5p | MIMAT0000542 | Zfp219 | 69890 | HITS-CLIP | 29167373 | Unspecified |
| mmu-miR-34a-5p | MIMAT0000542 | Zfp236 | 329002 | HITS-CLIP | 23142080 | Unspecified |
| mmu-miR-34a-5p | MIMAT0000542 | Zfp24 | 59057 | HITS-CLIP | 19536157 | Unspecified |
| mmu-miR-34a-5p | MIMAT0000542 | Zfp248 | 72720 | HITS-CLIP | 23142080 | Unspecified |
| mmu-miR-34a-5p | MIMAT0000542 | Zfp26 | 22688 | HITS-CLIP | 23142080\|29167373 | Unspecified |
| mmu-miR-34a-5p | MIMAT0000542 | Zfp260 | 26466 | HITS-CLIP | 23142080 | Unspecified |
| mmu-miR-34a-5p | MIMAT0000542 | Zfp263 | 74120 | HITS-CLIP | 23142080 | Unspecified |
| mmu-miR-34a-5p | MIMAT0000542 | Zfp266 | 77519 | HITS-CLIP | 23142080\|30971389 | Unspecified |
| mmu-miR-34a-5p | MIMAT0000542 | Zfp27 | 22689 | HITS-CLIP | 23142080 | Unspecified |
| mmu-miR-34a-5p | MIMAT0000542 | Zfp275 | 27081 | HITS-CLIP | 23142080\|29167373 | Unspecified |
| mmu-miR-34a-5p | MIMAT0000542 | Zfp276 | 57247 | HITS-CLIP | 23142080 | Unspecified |
| mmu-miR-34a-5p | MIMAT0000542 | Zfp277 | 246196 | HITS-CLIP | 31433990\|23142080 | Unspecified |
| mmu-miR-34a-5p | MIMAT0000542 | Zfp280b | 64453 | Microarrays//HITS-CLIP | 37986224\|23027903\|23142080 | Unspecified |
| mmu-miR-34a-5p | MIMAT0000542 | Zfp280c | 208968 | HITS-CLIP | 23142080 | Unspecified |
| mmu-miR-34a-5p | MIMAT0000542 | Zfp280d | 235469 | HITS-CLIP | 23142080 | Unspecified |
| mmu-miR-34a-5p | MIMAT0000542 | Zfp281 | 226442 | HITS-CLIP | 37986224\|23142080\|26203562 | Unspecified |
| mmu-miR-34a-5p | MIMAT0000542 | Zfp292 | 30046 | HITS-CLIP | 37986224\|23630078\|23142080\|29167 | Unspecified |
| mmu-miR-34a-5p | MIMAT0000542 | Zfp317 | 244713 | HITS-CLIP | 21258322\|23142080 | Unspecified |
| mmu-miR-34a-5p | MIMAT0000542 | Zfp318 | 57908 | HITS-CLIP | 19536157\|23142080\|29167373 | Unspecified |
| mmu-miR-34a-5p | MIMAT0000542 | Zfp319 | 79233 | HITS-CLIP | 23142080 | Unspecified |
| mmu-miR-34a-5p | MIMAT0000542 | Zfp322a | 218100 | HITS-CLIP | 23142080 | Unspecified |
| mmu-miR-34a-5p | MIMAT0000542 | Zfp326 | 54367 | HITS-CLIP | 19536157\|29167373\|23142080 | Unspecified |
| mmu-miR-34a-5p | MIMAT0000542 | Zfp329 | 67230 | HITS-CLIP | 23142080 | Unspecified |
| mmu-miR-34a-5p | MIMAT0000542 | Zfp330 | 30932 | HITS-CLIP | 23142080 | Unspecified |
| mmu-miR-34a-5p | MIMAT0000542 | Zfp335 | 329559 | HITS-CLIP | 23142080\|29167373 | Unspecified |
| mmu-miR-34a-5p | MIMAT0000542 | Zfp346 | 26919 | HITS-CLIP | 29167373 | Unspecified |

| | | | | | | |
|---|---|---|---|---|---|---|
| mmu-miR-34a-5p | MIMAT0000542 | Zfp35 | 22694 | HITS-CLIP | 23142080 | Unspecified |
| mmu-miR-34a-5p | MIMAT0000542 | Zfp36 | 22695 | HITS-CLIP | 31433990\|23142080\|29167373 | Unspecified |
| mmu-miR-34a-5p | MIMAT0000542 | Zfp362 | 230761 | HITS-CLIP | 31433990\|23142080 | Unspecified |
| mmu-miR-34a-5p | MIMAT0000542 | Zfp367 | 238673 | HITS-CLIP | 31433990\|29167373\|23142080 | Unspecified |
| mmu-miR-34a-5p | MIMAT0000542 | Zfp36l1 | 12192 | Microarrays//HITS-CLIP | 37986224\|23142080\|19536157\|29167 | Unspecified |
| mmu-miR-34a-5p | MIMAT0000542 | Zfp36l2 | 12193 | HITS-CLIP | 31433990\|23142080\|29167373 | Unspecified |
| mmu-miR-34a-5p | MIMAT0000542 | Zfp382 | 233060 | HITS-CLIP | 23142080 | Unspecified |
| mmu-miR-34a-5p | MIMAT0000542 | Zfp384 | 269800 | HITS-CLIP | 23142080\|29167373 | Unspecified |
| mmu-miR-34a-5p | MIMAT0000542 | Zfp385a | 29813 | HITS-CLIP | 29167373 | Unspecified |
| mmu-miR-34a-5p | MIMAT0000542 | Zfp386 | 56220 | HITS-CLIP | 23142080\|29167373 | Unspecified |
| mmu-miR-34a-5p | MIMAT0000542 | Zfp39 | 22698 | HITS-CLIP | 23142080 | Unspecified |
| mmu-miR-34a-5p | MIMAT0000542 | Zfp395 | 380912 | HITS-CLIP | 23142080 | Unspecified |
| mmu-miR-34a-5p | MIMAT0000542 | Zfp397 | 69256 | HITS-CLIP | 23142080\|29167373 | Unspecified |
| mmu-miR-34a-5p | MIMAT0000542 | Zfp407 | 240476 | HITS-CLIP | 23142080 | Unspecified |
| mmu-miR-34a-5p | MIMAT0000542 | Zfp408 | 381410 | HITS-CLIP | 23142080 | Unspecified |
| mmu-miR-34a-5p | MIMAT0000542 | Zfp410 | 52708 | HITS-CLIP | 23142080 | Unspecified |
| mmu-miR-34a-5p | MIMAT0000542 | Zfp426 | 235028 | HITS-CLIP | 23142080 | Unspecified |
| mmu-miR-34a-5p | MIMAT0000542 | Zfp429 | 72807 | HITS-CLIP | 23142080 | Unspecified |
| mmu-miR-34a-5p | MIMAT0000542 | Zfp438 | 240186 | HITS-CLIP | 31433990 | Unspecified |
| mmu-miR-34a-5p | MIMAT0000542 | Zfp445 | 235682 | HITS-CLIP | 23142080\|29167373 | Unspecified |
| mmu-miR-34a-5p | MIMAT0000542 | Zfp449 | 78619 | HITS-CLIP | 23142080 | Unspecified |
| mmu-miR-34a-5p | MIMAT0000542 | Zfp451 | 98403 | HITS-CLIP | 23142080\|29167373 | Unspecified |
| mmu-miR-34a-5p | MIMAT0000542 | Zfp458 | 238690 | HITS-CLIP | 23142080 | Unspecified |
| mmu-miR-34a-5p | MIMAT0000542 | Zfp462 | 242466 | HITS-CLIP | 19536157 | Unspecified |
| mmu-miR-34a-5p | MIMAT0000542 | Zfp467 | 68910 | HITS-CLIP | 23142080\|29167373 | Unspecified |
| mmu-miR-34a-5p | MIMAT0000542 | Zfp472 | 224691 | HITS-CLIP | 23142080 | Unspecified |
| mmu-miR-34a-5p | MIMAT0000542 | Zfp473 | 243963 | HITS-CLIP | 23142080 | Unspecified |
| mmu-miR-34a-5p | MIMAT0000542 | Zfp507 | 668501 | HITS-CLIP | 23142080 | Unspecified |
| mmu-miR-34a-5p | MIMAT0000542 | Zfp512 | 269639 | Microarrays//HITS-CLIP | 37986224\|23027903\|23142080\|29167 | Unspecified |
| mmu-miR-34a-5p | MIMAT0000542 | Zfp512b | 269401 | HITS-CLIP | 23630078\|31433990\|30971389\|23142 | Unspecified |
| mmu-miR-34a-5p | MIMAT0000542 | Zfp516 | 329003 | HITS-CLIP | 23142080\|19536157\|23630078\|29167 | Unspecified |
| mmu-miR-34a-5p | MIMAT0000542 | Zfp518a | 72672 | HITS-CLIP | 23142080 | Unspecified |
| mmu-miR-34a-5p | MIMAT0000542 | Zfp518b | 100515 | HITS-CLIP | 23142080\|29167373 | Unspecified |
| mmu-miR-34a-5p | MIMAT0000542 | Zfp52 | 22710 | HITS-CLIP | 23142080 | Unspecified |
| mmu-miR-34a-5p | MIMAT0000542 | Zfp523 | 7629 | HITS-CLIP | 19536157 | Unspecified |
| mmu-miR-34a-5p | MIMAT0000542 | Zfp526 | 210172 | HITS-CLIP | 23142080 | Unspecified |
| mmu-miR-34a-5p | MIMAT0000542 | Zfp53 | 24132 | HITS-CLIP | 23142080 | Unspecified |
| mmu-miR-34a-5p | MIMAT0000542 | Zfp532 | 328977 | HITS-CLIP | 19536157 | Unspecified |
| mmu-miR-34a-5p | MIMAT0000542 | Zfp54 | 22712 | HITS-CLIP | 23142080\|31433990 | Unspecified |

| | | | | | | |
|---|---|---|---|---|---|---|
| mmu-miR-34a-5p | MIMAT0000542 | Zfp551 | 619331 | HITS-CLIP | 23142080 | Unspecified |
| mmu-miR-34a-5p | MIMAT0000542 | Zfp563 | 240068 | HITS-CLIP | 23142080 | Unspecified |
| mmu-miR-34a-5p | MIMAT0000542 | Zfp568 | 243905 | HITS-CLIP | 23142080\|29167373 | Unspecified |
| mmu-miR-34a-5p | MIMAT0000542 | Zfp58 | 238693 | HITS-CLIP | 23142080 | Unspecified |
| mmu-miR-34a-5p | MIMAT0000542 | Zfp583 | 213011 | HITS-CLIP | 23142080 | Unspecified |
| mmu-miR-34a-5p | MIMAT0000542 | Zfp592 | 233410 | HITS-CLIP | 23142080\|29167373 | Unspecified |
| mmu-miR-34a-5p | MIMAT0000542 | Zfp597 | 71063 | HITS-CLIP | 29167373 | Unspecified |
| mmu-miR-34a-5p | MIMAT0000542 | Zfp598 | 213753 | HITS-CLIP | 23142080 | Unspecified |
| mmu-miR-34a-5p | MIMAT0000542 | Zfp599 | 235048 | HITS-CLIP | 23142080 | Unspecified |
| mmu-miR-34a-5p | MIMAT0000542 | Zfp606 | 67370 | HITS-CLIP | 23142080 | Unspecified |
| mmu-miR-34a-5p | MIMAT0000542 | Zfp607a | 545938 | HITS-CLIP | 23142080 | Unspecified |
| mmu-miR-34a-5p | MIMAT0000542 | Zfp608 | 269023 | HITS-CLIP | 19536157\|23142080 | Unspecified |
| mmu-miR-34a-5p | MIMAT0000542 | Zfp609 | 214812 | HITS-CLIP | 23142080\|29167373 | Unspecified |
| mmu-miR-34a-5p | MIMAT0000542 | Zfp618 | 72701 | HITS-CLIP | 29167373 | Unspecified |
| mmu-miR-34a-5p | MIMAT0000542 | Zfp622 | 52521 | HITS-CLIP | 23142080 | Unspecified |
| mmu-miR-34a-5p | MIMAT0000542 | Zfp629 | 320683 | HITS-CLIP | 23142080 | Unspecified |
| mmu-miR-34a-5p | MIMAT0000542 | Zfp638 | 27332 | HITS-CLIP | 23142080\|19536157\|29167373 | Unspecified |
| mmu-miR-34a-5p | MIMAT0000542 | Zfp644 | 52397 | HITS-CLIP | 19536157\|23142080\|29167373 | Unspecified |
| mmu-miR-34a-5p | MIMAT0000542 | Zfp646 | 233905 | HITS-CLIP | 23142080\|29167373 | Unspecified |
| mmu-miR-34a-5p | MIMAT0000542 | Zfp65 | 235907 | HITS-CLIP | 23142080 | Unspecified |
| mmu-miR-34a-5p | MIMAT0000542 | Zfp652 | 268469 | HITS-CLIP | 23142080\|29167373 | Unspecified |
| mmu-miR-34a-5p | MIMAT0000542 | Zfp653 | 319601 | HITS-CLIP | 23142080 | Unspecified |
| mmu-miR-34a-5p | MIMAT0000542 | Zfp654 | 72020 | HITS-CLIP | 23142080\|19536157\|29167373 | Unspecified |
| mmu-miR-34a-5p | MIMAT0000542 | Zfp655 | 72611 | HITS-CLIP | 23142080\|29167373 | Unspecified |
| mmu-miR-34a-5p | MIMAT0000542 | Zfp664 | 269704 | HITS-CLIP | 23142080 | Unspecified |
| mmu-miR-34a-5p | MIMAT0000542 | Zfp667 | 384763 | Microarrays//HITS-CLIP | 37986224\|23142080 | Unspecified |
| mmu-miR-34a-5p | MIMAT0000542 | Zfp668 | 244219 | HITS-CLIP | 23142080 | Unspecified |
| mmu-miR-34a-5p | MIMAT0000542 | Zfp677 | 210503 | HITS-CLIP | 23142080 | Unspecified |
| mmu-miR-34a-5p | MIMAT0000542 | Zfp687 | 78266 | HITS-CLIP | 23142080\|29167373 | Unspecified |
| mmu-miR-34a-5p | MIMAT0000542 | Zfp689 | 71131 | HITS-CLIP | 23142080 | Unspecified |
| mmu-miR-34a-5p | MIMAT0000542 | Zfp691 | 195522 | HITS-CLIP | 29167373 | Unspecified |
| mmu-miR-34a-5p | MIMAT0000542 | Zfp703 | 353310 | HITS-CLIP | 23142080\|29167373 | Unspecified |
| mmu-miR-34a-5p | MIMAT0000542 | Zfp704 | 170753 | HITS-CLIP | 19536157 | Unspecified |
| mmu-miR-34a-5p | MIMAT0000542 | Zfp706 | 68036 | HITS-CLIP | 29167373 | Unspecified |
| mmu-miR-34a-5p | MIMAT0000542 | Zfp708 | 432769 | HITS-CLIP | 23142080 | Unspecified |
| mmu-miR-34a-5p | MIMAT0000542 | Zfp710 | 209225 | HITS-CLIP | 23142080\|31433990\|29167373 | Unspecified |
| mmu-miR-34a-5p | MIMAT0000542 | Zfp712 | 78251 | HITS-CLIP | 23142080 | Unspecified |
| mmu-miR-34a-5p | MIMAT0000542 | Zfp715 | 69930 | HITS-CLIP | 29167373 | Unspecified |
| mmu-miR-34a-5p | MIMAT0000542 | Zfp719 | 210105 | HITS-CLIP | 23142080 | Unspecified |

| | | | | | | | |
|---|---|---|---|---|---|---|---|
| mmu-miR-34a-5p | MIMAT0000542 | Zfp729a | 212281 | HITS-CLIP | | 23142080 | Unspecified |
| mmu-miR-34a-5p | MIMAT0000542 | Zfp729b | 100416706 | HITS-CLIP | | 23142080 | Unspecified |
| mmu-miR-34a-5p | MIMAT0000542 | Zfp738 | 408068 | HITS-CLIP | | 23142080 | Unspecified |
| mmu-miR-34a-5p | MIMAT0000542 | Zfp740 | 68744 | HITS-CLIP | | 23142080 | Unspecified |
| mmu-miR-34a-5p | MIMAT0000542 | Zfp746 | 69228 | HITS-CLIP | | 23142080 | Unspecified |
| mmu-miR-34a-5p | MIMAT0000542 | Zfp747 | 269997 | HITS-CLIP | | 23142080 | Unspecified |
| mmu-miR-34a-5p | MIMAT0000542 | Zfp748 | 212276 | HITS-CLIP | | 23142080 | Unspecified |
| mmu-miR-34a-5p | MIMAT0000542 | Zfp758 | 224598 | Microarrays | | 37986224 | Unspecified |
| mmu-miR-34a-5p | MIMAT0000542 | Zfp760 | 240034 | HITS-CLIP | 23142080\|29167373 | | Unspecified |
| mmu-miR-34a-5p | MIMAT0000542 | Zfp763 | 73451 | HITS-CLIP | | 23142080 | Unspecified |
| mmu-miR-34a-5p | MIMAT0000542 | Zfp770 | 228491 | HITS-CLIP | | 23142080 | Unspecified |
| mmu-miR-34a-5p | MIMAT0000542 | Zfp772 | 232855 | HITS-CLIP | | 23142080 | Unspecified |
| mmu-miR-34a-5p | MIMAT0000542 | Zfp773 | 76373 | HITS-CLIP | | 23142080 | Unspecified |
| mmu-miR-34a-5p | MIMAT0000542 | Zfp787 | 67109 | HITS-CLIP | 23142080\|29167373 | | Unspecified |
| mmu-miR-34a-5p | MIMAT0000542 | Zfp790 | 233056 | HITS-CLIP | | 23142080 | Unspecified |
| mmu-miR-34a-5p | MIMAT0000542 | Zfp799 | 240064 | HITS-CLIP | | 23142080 | Unspecified |
| mmu-miR-34a-5p | MIMAT0000542 | Zfp800 | 627049 | HITS-CLIP | 31433990\|23142080\|29167373 | | Unspecified |
| mmu-miR-34a-5p | MIMAT0000542 | Zfp809 | 235047 | HITS-CLIP | | 23142080 | Unspecified |
| mmu-miR-34a-5p | MIMAT0000542 | Zfp81 | 224694 | HITS-CLIP | 30971389\|29167373 | | Unspecified |
| mmu-miR-34a-5p | MIMAT0000542 | Zfp827 | 622675 | HITS-CLIP | | 23142080 | Unspecified |
| mmu-miR-34a-5p | MIMAT0000542 | Zfp831 | 100043757 | HITS-CLIP | | 23142080 | Unspecified |
| mmu-miR-34a-5p | MIMAT0000542 | Zfp839 | 72805 | HITS-CLIP | | 23142080 | Unspecified |
| mmu-miR-34a-5p | MIMAT0000542 | Zfp84 | 74352 | HITS-CLIP | | 23142080 | Unspecified |
| mmu-miR-34a-5p | MIMAT0000542 | Zfp850 | 100043772 | HITS-CLIP | | 23142080 | Unspecified |
| mmu-miR-34a-5p | MIMAT0000542 | Zfp865 | 319748 | HITS-CLIP | 23142080\|29167373 | | Unspecified |
| mmu-miR-34a-5p | MIMAT0000542 | Zfp866 | 330788 | HITS-CLIP | | 23142080 | Unspecified |
| mmu-miR-34a-5p | MIMAT0000542 | Zfp869 | 66869 | HITS-CLIP | | 23142080 | Unspecified |
| mmu-miR-34a-5p | MIMAT0000542 | Zfp871 | 208292 | Microarrays//HITS-CLIP | 37986224\|23142080\|23027903\|2363( | | Unspecified |
| mmu-miR-34a-5p | MIMAT0000542 | Zfp874a | 238692 | HITS-CLIP | | 23142080 | Unspecified |
| mmu-miR-34a-5p | MIMAT0000542 | Zfp90 | 22751 | HITS-CLIP | | 29167373 | Unspecified |
| mmu-miR-34a-5p | MIMAT0000542 | Zfp91 | 109910 | HITS-CLIP | 19536157\|23142080\|29167373 | | Unspecified |
| mmu-miR-34a-5p | MIMAT0000542 | Zfp930 | 234358 | Microarrays | 37986224\|23027903 | | Unspecified |
| mmu-miR-34a-5p | MIMAT0000542 | Zfp933 | 242747 | HITS-CLIP | | 30971389 | Unspecified |
| mmu-miR-34a-5p | MIMAT0000542 | Zfp943 | 74670 | HITS-CLIP | | 29167373 | Unspecified |
| mmu-miR-34a-5p | MIMAT0000542 | Zfp949 | 71640 | HITS-CLIP | | 23142080 | Unspecified |
| mmu-miR-34a-5p | MIMAT0000542 | Zfp951 | 626391 | HITS-CLIP | | 23142080 | Unspecified |
| mmu-miR-34a-5p | MIMAT0000542 | Zfp953 | 629016 | HITS-CLIP | | 29167373 | Unspecified |
| mmu-miR-34a-5p | MIMAT0000542 | Zfp955a | 77652 | HITS-CLIP | | 23142080 | Unspecified |
| mmu-miR-34a-5p | MIMAT0000542 | Zfp955b | 100043468 | HITS-CLIP | 31433990\|23142080\|29167373 | | Unspecified |

| | | | | | | |
|---|---|---|---|---|---|---|
| mmu-miR-34a-5p | MIMAT0000542 | Zfp958 | 233987 | HITS-CLIP | 23142080 | Unspecified |
| mmu-miR-34a-5p | MIMAT0000542 | Zfp961 | 234413 | HITS-CLIP | 23142080 | Unspecified |
| mmu-miR-34a-5p | MIMAT0000542 | Zfp971 | 626848 | HITS-CLIP | 23142080 | Unspecified |
| mmu-miR-34a-5p | MIMAT0000542 | Zfp975 | 434179 | HITS-CLIP | 23142080 | Unspecified |
| mmu-miR-34a-5p | MIMAT0000542 | Zfpl1 | 81909 | HITS-CLIP | 23142080 | Unspecified |
| mmu-miR-34a-5p | MIMAT0000542 | Zfpm1 | 22761 | HITS-CLIP | 23142080 | Unspecified |
| mmu-miR-34a-5p | MIMAT0000542 | Zfr | 22763 | HITS-CLIP | 31433990\|19536157\|23142080\|29167 | Unspecified |
| mmu-miR-34a-5p | MIMAT0000542 | Zfr2 | 103406 | HITS-CLIP | 19536157 | Unspecified |
| mmu-miR-34a-5p | MIMAT0000542 | Zfx | 22764 | HITS-CLIP | 23142080\|29167373 | Unspecified |
| mmu-miR-34a-5p | MIMAT0000542 | Zfyve1 | 217695 | HITS-CLIP | 23142080\|31433990\|29167373 | Unspecified |
| mmu-miR-34a-5p | MIMAT0000542 | Zfyve16 | 218441 | HITS-CLIP | 23142080\|29167373 | Unspecified |
| mmu-miR-34a-5p | MIMAT0000542 | Zfyve21 | 68520 | HITS-CLIP | 23142080\|19536157 | Unspecified |
| mmu-miR-34a-5p | MIMAT0000542 | Zfyve26 | 211978 | HITS-CLIP | 23142080\|29167373 | Unspecified |
| mmu-miR-34a-5p | MIMAT0000542 | Zfyve27 | 319740 | HITS-CLIP | 23142080\|29167373 | Unspecified |
| mmu-miR-34a-5p | MIMAT0000542 | Zgpat | 229007 | HITS-CLIP | 23142080 | Unspecified |
| mmu-miR-34a-5p | MIMAT0000542 | Zhx1 | 22770 | HITS-CLIP | 19536157\|23142080\|29167373 | Unspecified |
| mmu-miR-34a-5p | MIMAT0000542 | Zhx2 | 387609 | HITS-CLIP | 19536157\|23142080 | Unspecified |
| mmu-miR-34a-5p | MIMAT0000542 | Zic2 | 22772 | HITS-CLIP | 19536157 | Unspecified |
| mmu-miR-34a-5p | MIMAT0000542 | Zik1 | 22775 | Microarrays//HITS-CLIP | 37986224\|23027903\|23142080 | Unspecified |
| mmu-miR-34a-5p | MIMAT0000542 | Zim1 | 22776 | Microarrays | 37986224 | Unspecified |
| mmu-miR-34a-5p | MIMAT0000542 | Zkscan1 | 74570 | HITS-CLIP | 31433990\|23142080\|26203562\|29167 | Unspecified |
| mmu-miR-34a-5p | MIMAT0000542 | Zkscan3 | 72739 | HITS-CLIP | 23142080\|29167373 | Unspecified |
| mmu-miR-34a-5p | MIMAT0000542 | Zkscan6 | 7566 | HITS-CLIP | 23142080 | Unspecified |
| mmu-miR-34a-5p | MIMAT0000542 | Zkscan8 | 93681 | HITS-CLIP | 23142080\|29167373 | Unspecified |
| mmu-miR-34a-5p | MIMAT0000542 | Zmat1 | 215693 | Microarrays | 37986224 | Unspecified |
| mmu-miR-34a-5p | MIMAT0000542 | Zmat3 | 22401 | HITS-CLIP | 31433990\|29167373\|23142080 | Unspecified |
| mmu-miR-34a-5p | MIMAT0000542 | Zmiz1 | 328365 | HITS-CLIP | 23142080\|31433990\|29167373 | Unspecified |
| mmu-miR-34a-5p | MIMAT0000542 | Zmiz2 | 52915 | HITS-CLIP | 23142080\|29167373 | Unspecified |
| mmu-miR-34a-5p | MIMAT0000542 | Zmpste24 | 230709 | HITS-CLIP | 29167373 | Unspecified |
| mmu-miR-34a-5p | MIMAT0000542 | Zmym1 | 68310 | HITS-CLIP | 23142080 | Unspecified |
| mmu-miR-34a-5p | MIMAT0000542 | Zmym2 | 76007 | HITS-CLIP | 23142080\|19536157\|29167373 | Unspecified |
| mmu-miR-34a-5p | MIMAT0000542 | Zmym3 | 56364 | HITS-CLIP | 23142080 | Unspecified |
| mmu-miR-34a-5p | MIMAT0000542 | Zmym4 | 67785 | HITS-CLIP | 19536157\|23142080\|29167373 | Unspecified |
| mmu-miR-34a-5p | MIMAT0000542 | Zmym5 | 219105 | HITS-CLIP | 23142080 | Unspecified |
| mmu-miR-34a-5p | MIMAT0000542 | Zmym6 | 100177 | HITS-CLIP | 23142080 | Unspecified |
| mmu-miR-34a-5p | MIMAT0000542 | Zmynd11 | 66505 | HITS-CLIP | 31433990\|23142080\|29167373 | Unspecified |
| mmu-miR-34a-5p | MIMAT0000542 | Zmynd19 | 67187 | HITS-CLIP | 29167373\|23142080 | Unspecified |
| mmu-miR-34a-5p | MIMAT0000542 | Zmynd8 | 228880 | HITS-CLIP | 31433990\|23142080 | Unspecified |
| mmu-miR-34a-5p | MIMAT0000542 | Znfx1 | 98999 | HITS-CLIP | 19536157\|29167373\|23142080 | Unspecified |

| | | | | | | |
|---|---|---|---|---|---|---|
| mmu-miR-34a-5p | MIMAT0000542 | Znrd1 | 66136 | HITS-CLIP | 23142080 | Unspecified |
| mmu-miR-34a-5p | MIMAT0000542 | Znrf1 | 170737 | HITS-CLIP | 31433990 | Unspecified |
| mmu-miR-34a-5p | MIMAT0000542 | Znrf2 | 387524 | HITS-CLIP | 31433990\|23142080\|29167373 | Unspecified |
| mmu-miR-34a-5p | MIMAT0000542 | Znrf3 | 407821 | HITS-CLIP | 23142080\|19536157 | Unspecified |
| mmu-miR-34a-5p | MIMAT0000542 | Zpr1 | 22687 | HITS-CLIP | 23142080\|29167373 | Unspecified |
| mmu-miR-34a-5p | MIMAT0000542 | Zranb1 | 360216 | HITS-CLIP | 23142080 | Unspecified |
| mmu-miR-34a-5p | MIMAT0000542 | Zranb2 | 53861 | HITS-CLIP | 23142080 | Unspecified |
| mmu-miR-34a-5p | MIMAT0000542 | Zranb3 | 226409 | HITS-CLIP | 23142080\|29167373 | Unspecified |
| mmu-miR-34a-5p | MIMAT0000542 | Zrsr2 | 22184 | HITS-CLIP | 23142080 | Unspecified |
| mmu-miR-34a-5p | MIMAT0000542 | Zscan12 | 22758 | HITS-CLIP | 23142080 | Unspecified |
| mmu-miR-34a-5p | MIMAT0000542 | Zscan26 | 432731 | HITS-CLIP | 23142080\|29167373 | Unspecified |
| mmu-miR-34a-5p | MIMAT0000542 | Zscan29 | 99334 | HITS-CLIP | 23142080 | Unspecified |
| mmu-miR-34a-5p | MIMAT0000542 | Zswim3 | 67538 | HITS-CLIP | 29167373 | Unspecified |
| mmu-miR-34a-5p | MIMAT0000542 | Zswim4 | 212168 | HITS-CLIP | 29167373 | Unspecified |
| mmu-miR-34a-5p | MIMAT0000542 | Zswim5 | 74464 | HITS-CLIP | 19536157 | Unspecified |
| mmu-miR-34a-5p | MIMAT0000542 | Zswim8 | 268721 | HITS-CLIP | 23142080\|29167373 | Unspecified |
| mmu-miR-34a-5p | MIMAT0000542 | Zup1 | 221302 | HITS-CLIP | 29167373 | Unspecified |
| mmu-miR-34a-5p | MIMAT0000542 | Zwilch | 68014 | HITS-CLIP | 23142080 | Unspecified |
| mmu-miR-34a-5p | MIMAT0000542 | Zwint | 52696 | HITS-CLIP | 23142080 | Unspecified |
| mmu-miR-34a-5p | MIMAT0000542 | Zxdb | 668166 | HITS-CLIP | 23142080 | Unspecified |
| mmu-miR-34a-5p | MIMAT0000542 | Zxdc | 80292 | HITS-CLIP | 23142080\|29167373 | Unspecified |
| mmu-miR-34a-5p | MIMAT0000542 | Zyg11b | 414872 | HITS-CLIP | 37986224\|23630078\|23142080 | Unspecified |
| mmu-miR-34a-5p | MIMAT0000542 | Zyx | 22793 | HITS-CLIP | 23142080 | Unspecified |
| mmu-miR-34a-5p | MIMAT0000542 | Zzef1 | 195018 | HITS-CLIP | 23142080\|29167373 | Unspecified |
| mmu-miR-34a-5p | MIMAT0000542 | Zzz3 | 108946 | HITS-CLIP | 23142080\|29167373 | Unspecified |
| mmu-miR-34a-5p | MIMAT0000542 | mt-Atp6 | 4508 | HITS-CLIP | 29167373 | Unspecified |
| mmu-miR-34a-5p | MIMAT0000542 | mt-Atp8 | 4509 | HITS-CLIP | 29167373 | Unspecified |
| mmu-miR-34a-5p | MIMAT0000542 | mt-Co1 | 4512 | HITS-CLIP | 19536157\|23142080\|29167373 | Unspecified |
| mmu-miR-34a-5p | MIMAT0000542 | mt-Co3 | 4514 | HITS-CLIP | 29167373 | Unspecified |
| mmu-miR-34a-5p | MIMAT0000542 | mt-Cytb | 19893556 | HITS-CLIP | 19536157\|23142080\|29167373 | Unspecified |
| mmu-miR-34a-5p | MIMAT0000542 | mt-Nd1 | 4535 | HITS-CLIP | 19536157\|23142080\|29167373 | Unspecified |
| mmu-miR-34a-5p | MIMAT0000542 | mt-Nd2 | 4536 | HITS-CLIP | 19536157\|23142080\|29167373 | Unspecified |
| mmu-miR-34a-5p | MIMAT0000542 | mt-Nd3 | 4537 | HITS-CLIP | 23142080 | Unspecified |
| mmu-miR-34a-5p | MIMAT0000542 | mt-Nd4 | 4538 | HITS-CLIP | 23142080\|29167373 | Unspecified |
| mmu-miR-34a-5p | MIMAT0000542 | mt-Nd4l | 19893552 | HITS-CLIP | 29167373 | Unspecified |
| mmu-miR-34a-5p | MIMAT0000542 | mt-Nd5 | 4540 | HITS-CLIP | 29167373\|19536157\|23142080 | Unspecified |
| mmu-miR-509-5p | MIMAT0004890 | Macf1 | 11426 | HITS-CLIP | 26203562 | Unspecified |
| mmu-miR-509-5p | MIMAT0004890 | Pom121 | 107939 | HITS-CLIP | 31433990 | Unspecified |
| mmu-miR-509-5p | MIMAT0004890 | Tmem30a | 69981 | HITS-CLIP | 26203562 | Unspecified |

| | | | | | | |
|---|---|---|---|---|---|---|
| mmu-miR-509-5p | MIMAT0004890 | Zbtb7b | 22724 | HITS-CLIP | 26203562 | Unspecified |
| mmu-miR-384-5p | MIMAT0004745 | 4932438A13Rik | 229227 | HITS-CLIP | 19536157 | Unspecified |
| mmu-miR-384-5p | MIMAT0004745 | Abat | 268860 | HITS-CLIP | 19536157 | Unspecified |
| mmu-miR-384-5p | MIMAT0004745 | Abhd17b | 226016 | HITS-CLIP | 19536157 | Unspecified |
| mmu-miR-384-5p | MIMAT0004745 | Abl2 | 11352 | HITS-CLIP | 19536157 | Unspecified |
| mmu-miR-384-5p | MIMAT0004745 | Acp2 | 11432 | HITS-CLIP | 19536157 | Unspecified |
| mmu-miR-384-5p | MIMAT0004745 | Acvr1 | 11477 | HITS-CLIP | 19536157 | Unspecified |
| mmu-miR-384-5p | MIMAT0004745 | Adam9 | 11502 | HITS-CLIP | 19536157 | Unspecified |
| mmu-miR-384-5p | MIMAT0004745 | Adarb1 | 110532 | HITS-CLIP | 19536157 | Unspecified |
| mmu-miR-384-5p | MIMAT0004745 | Adcy1 | 432530 | HITS-CLIP | 19536157 | Unspecified |
| mmu-miR-384-5p | MIMAT0004745 | Aff4 | 93736 | HITS-CLIP | 19536157 | Unspecified |
| mmu-miR-384-5p | MIMAT0004745 | Aggf1 | 66549 | HITS-CLIP | 19536157 | Unspecified |
| mmu-miR-384-5p | MIMAT0004745 | Ago1 | 236511 | HITS-CLIP | 19536157 | Unspecified |
| mmu-miR-384-5p | MIMAT0004745 | Ago3 | 214150 | HITS-CLIP | 19536157 | Unspecified |
| mmu-miR-384-5p | MIMAT0004745 | Ahnak | 66395 | HITS-CLIP | 19536157 | Unspecified |
| mmu-miR-384-5p | MIMAT0004745 | Akap11 | 219181 | HITS-CLIP | 19536157 | Unspecified |
| mmu-miR-384-5p | MIMAT0004745 | Ank2 | 109676 | HITS-CLIP | 19536157 | Unspecified |
| mmu-miR-384-5p | MIMAT0004745 | Ank3 | 11735 | HITS-CLIP | 19536157 | Unspecified |
| mmu-miR-384-5p | MIMAT0004745 | Ankrd17 | 81702 | HITS-CLIP | 19536157 | Unspecified |
| mmu-miR-384-5p | MIMAT0004745 | Ankrd52 | 237615 | HITS-CLIP | 19536157 | Unspecified |
| mmu-miR-384-5p | MIMAT0004745 | Anxa1 | 16952 | HITS-CLIP | 26203562 | Unspecified |
| mmu-miR-384-5p | MIMAT0004745 | Apba1 | 319924 | HITS-CLIP | 19536157 | Unspecified |
| mmu-miR-384-5p | MIMAT0004745 | Apc | 11789 | HITS-CLIP | 19536157 | Unspecified |
| mmu-miR-384-5p | MIMAT0004745 | Arfgef3 | 57221 | HITS-CLIP | 19536157 | Unspecified |
| mmu-miR-384-5p | MIMAT0004745 | Arid4b | 94246 | HITS-CLIP | 19536157 | Unspecified |
| mmu-miR-384-5p | MIMAT0004745 | Arid5b | 71371 | HITS-CLIP | 19536157 | Unspecified |
| mmu-miR-384-5p | MIMAT0004745 | Arl15 | 218639 | HITS-CLIP | 19536157 | Unspecified |
| mmu-miR-384-5p | MIMAT0004745 | Arpc5 | 67771 | HITS-CLIP | 19536157 | Unspecified |
| mmu-miR-384-5p | MIMAT0004745 | Atf4 | 11911 | HITS-CLIP | 31433990 | Unspecified |
| mmu-miR-384-5p | MIMAT0004745 | Atl2 | 56298 | HITS-CLIP | 19536157 | Unspecified |
| mmu-miR-384-5p | MIMAT0004745 | Atl3 | 109168 | HITS-CLIP | 19536157 | Unspecified |
| mmu-miR-384-5p | MIMAT0004745 | Atp2a2 | 11938 | HITS-CLIP | 19536157 | Unspecified |
| mmu-miR-384-5p | MIMAT0004745 | Atp2b2 | 11941 | HITS-CLIP | 19536157 | Unspecified |
| mmu-miR-384-5p | MIMAT0004745 | Atp8a1 | 11980 | HITS-CLIP | 19536157 | Unspecified |
| mmu-miR-384-5p | MIMAT0004745 | Atrx | 22589 | HITS-CLIP | 19536157 | Unspecified |
| mmu-miR-384-5p | MIMAT0004745 | Azin1 | 54375 | HITS-CLIP | 19536157 | Unspecified |
| mmu-miR-384-5p | MIMAT0004745 | B3gat1 | 76898 | HITS-CLIP | 19536157 | Unspecified |
| mmu-miR-384-5p | MIMAT0004745 | Bach2 | 12014 | HITS-CLIP | 31433990 | Unspecified |
| mmu-miR-384-5p | MIMAT0004745 | Brinp2 | 240843 | HITS-CLIP | 19536157 | Unspecified |

| | | | | | | |
|---|---|---|---|---|---|---|
| mmu-miR-384-5p | MIMAT0004745 | Bsn | 12217 | HITS-CLIP | 19536157 | Unspecified |
| mmu-miR-384-5p | MIMAT0004745 | Btbd7 | 238386 | HITS-CLIP | 19536157 | Unspecified |
| mmu-miR-384-5p | MIMAT0004745 | Bzw2 | 66912 | HITS-CLIP | 19536157 | Unspecified |
| mmu-miR-384-5p | MIMAT0004745 | C9orf72 | 203228 | HITS-CLIP | 19536157 | Unspecified |
| mmu-miR-384-5p | MIMAT0004745 | Camk2d | 108058 | HITS-CLIP | 19536157 | Unspecified |
| mmu-miR-384-5p | MIMAT0004745 | Camk2n1 | 66259 | HITS-CLIP | 19536157 | Unspecified |
| mmu-miR-384-5p | MIMAT0004745 | Camkk2 | 207565 | HITS-CLIP | 19536157 | Unspecified |
| mmu-miR-384-5p | MIMAT0004745 | Ccdc71l | 72123 | HITS-CLIP | 19536157 | Unspecified |
| mmu-miR-384-5p | MIMAT0004745 | Ccnd1 | 12443 | HITS-CLIP | 26203562 | Unspecified |
| mmu-miR-384-5p | MIMAT0004745 | Ccne2 | 12448 | HITS-CLIP | 31433990\|19536157\|26203562 | Unspecified |
| mmu-miR-384-5p | MIMAT0004745 | Ccnjl | 380694 | HITS-CLIP | 19536157 | Unspecified |
| mmu-miR-384-5p | MIMAT0004745 | Cd164 | 53599 | HITS-CLIP | 19536157 | Unspecified |
| mmu-miR-384-5p | MIMAT0004745 | Cd81 | 12520 | HITS-CLIP | 19536157 | Unspecified |
| mmu-miR-384-5p | MIMAT0004745 | Cdc123 | 98828 | HITS-CLIP | 19536157 | Unspecified |
| mmu-miR-384-5p | MIMAT0004745 | Cdc42 | 12540 | HITS-CLIP | 19536157 | Unspecified |
| mmu-miR-384-5p | MIMAT0004745 | Cdh2 | 12558 | HITS-CLIP | 19536157 | Unspecified |
| mmu-miR-384-5p | MIMAT0004745 | Cdk19 | 78334 | HITS-CLIP | 19536157 | Unspecified |
| mmu-miR-384-5p | MIMAT0004745 | Celf3 | 78784 | HITS-CLIP | 19536157 | Unspecified |
| mmu-miR-384-5p | MIMAT0004745 | Celf4 | 108013 | HITS-CLIP | 19536157 | Unspecified |
| mmu-miR-384-5p | MIMAT0004745 | Celf5 | 319586 | HITS-CLIP | 19536157 | Unspecified |
| mmu-miR-384-5p | MIMAT0004745 | Cep350 | 74081 | HITS-CLIP | 19536157 | Unspecified |
| mmu-miR-384-5p | MIMAT0004745 | Chd1 | 12648 | HITS-CLIP | 19536157 | Unspecified |
| mmu-miR-384-5p | MIMAT0004745 | Chl1 | 12661 | HITS-CLIP | 19536157 | Unspecified |
| mmu-miR-384-5p | MIMAT0004745 | Chst2 | 54371 | HITS-CLIP | 19536157 | Unspecified |
| mmu-miR-384-5p | MIMAT0004745 | Chsy1 | 269941 | HITS-CLIP | 19536157 | Unspecified |
| mmu-miR-384-5p | MIMAT0004745 | Clasp2 | 76499 | HITS-CLIP | 19536157 | Unspecified |
| mmu-miR-384-5p | MIMAT0004745 | Clcn3 | 12725 | HITS-CLIP | 19536157 | Unspecified |
| mmu-miR-384-5p | MIMAT0004745 | Cldnd1 | 224250 | HITS-CLIP | 19536157 | Unspecified |
| mmu-miR-384-5p | MIMAT0004745 | Clip1 | 56430 | HITS-CLIP | 19536157 | Unspecified |
| mmu-miR-384-5p | MIMAT0004745 | Cnot6l | 231464 | HITS-CLIP | 19536157 | Unspecified |
| mmu-miR-384-5p | MIMAT0004745 | Col9a3 | 12841 | HITS-CLIP | 19536157 | Unspecified |
| mmu-miR-384-5p | MIMAT0004745 | Csnk1g1 | 214897 | HITS-CLIP | 19536157 | Unspecified |
| mmu-miR-384-5p | MIMAT0004745 | Cthrc1 | 68588 | HITS-CLIP | 19536157 | Unspecified |
| mmu-miR-384-5p | MIMAT0004745 | Ctps | 51797 | HITS-CLIP | 26203562 | Unspecified |
| mmu-miR-384-5p | MIMAT0004745 | Cxadr | 13052 | HITS-CLIP | 19536157 | Unspecified |
| mmu-miR-384-5p | MIMAT0004745 | Dab1 | 13131 | HITS-CLIP | 19536157 | Unspecified |
| mmu-miR-384-5p | MIMAT0004745 | Dcaf12 | 68970 | HITS-CLIP | 31433990 | Unspecified |
| mmu-miR-384-5p | MIMAT0004745 | Dcaf7 | 71833 | HITS-CLIP | 19536157 | Unspecified |
| mmu-miR-384-5p | MIMAT0004745 | Dcp1a | 75901 | HITS-CLIP | 19536157 | Unspecified |

| | | | | | | |
|---|---|---|---|---|---|---|
| mmu-miR-384-5p | MIMAT0004745 | Ddit4 | 74747 | HITS-CLIP | 19536157 | Unspecified |
| mmu-miR-384-5p | MIMAT0004745 | Ddx5 | 13207 | HITS-CLIP | 19536157 | Unspecified |
| mmu-miR-384-5p | MIMAT0004745 | Ddx6 | 13209 | HITS-CLIP | 19536157 | Unspecified |
| mmu-miR-384-5p | MIMAT0004745 | Dgkh | 380921 | HITS-CLIP | 19536157 | Unspecified |
| mmu-miR-384-5p | MIMAT0004745 | Dio2 | 13371 | HITS-CLIP | 19536157 | Unspecified |
| mmu-miR-384-5p | MIMAT0004745 | Dmxl2 | 235380 | HITS-CLIP | 19536157 | Unspecified |
| mmu-miR-384-5p | MIMAT0004745 | Dock9 | 105445 | HITS-CLIP | 19536157 | Unspecified |
| mmu-miR-384-5p | MIMAT0004745 | Dpysl2 | 12934 | HITS-CLIP | 19536157 | Unspecified |
| mmu-miR-384-5p | MIMAT0004745 | Dyrk2 | 69181 | HITS-CLIP | 26203562 | Unspecified |
| mmu-miR-384-5p | MIMAT0004745 | Edem3 | 66967 | HITS-CLIP | 19536157 | Unspecified |
| mmu-miR-384-5p | MIMAT0004745 | Ednra | 13617 | HITS-CLIP | 19536157 | Unspecified |
| mmu-miR-384-5p | MIMAT0004745 | Egfr | 13649 | HITS-CLIP | 19536157 | Unspecified |
| mmu-miR-384-5p | MIMAT0004745 | Eif5a2 | 208691 | HITS-CLIP | 19536157 | Unspecified |
| mmu-miR-384-5p | MIMAT0004745 | Elfn2 | 207393 | HITS-CLIP | 19536157 | Unspecified |
| mmu-miR-384-5p | MIMAT0004745 | Elovl5 | 68801 | HITS-CLIP | 19536157 | Unspecified |
| mmu-miR-384-5p | MIMAT0004745 | Elp3 | 74195 | HITS-CLIP | 31433990 | Unspecified |
| mmu-miR-384-5p | MIMAT0004745 | Epha4 | 13838 | HITS-CLIP | 26203562 | Unspecified |
| mmu-miR-384-5p | MIMAT0004745 | Erlin1 | 226144 | HITS-CLIP | 19536157 | Unspecified |
| mmu-miR-384-5p | MIMAT0004745 | Fam117b | 72750 | HITS-CLIP | 19536157 | Unspecified |
| mmu-miR-384-5p | MIMAT0004745 | Fam168b | 214469 | HITS-CLIP | 19536157 | Unspecified |
| mmu-miR-384-5p | MIMAT0004745 | Fam91a1 | 210998 | HITS-CLIP | 19536157 | Unspecified |
| mmu-miR-384-5p | MIMAT0004745 | Fat2 | 245827 | HITS-CLIP | 19536157 | Unspecified |
| mmu-miR-384-5p | MIMAT0004745 | Fbxo42 | 213499 | HITS-CLIP | 19536157 | Unspecified |
| mmu-miR-384-5p | MIMAT0004745 | Fcho2 | 218503 | HITS-CLIP | 19536157 | Unspecified |
| mmu-miR-384-5p | MIMAT0004745 | Fgfr3 | 14184 | HITS-CLIP | 19536157 | Unspecified |
| mmu-miR-384-5p | MIMAT0004745 | Foxn2 | 14236 | HITS-CLIP | 19536157 | Unspecified |
| mmu-miR-384-5p | MIMAT0004745 | Gabra2 | 14395 | HITS-CLIP | 19536157 | Unspecified |
| mmu-miR-384-5p | MIMAT0004745 | Galnt17 | 64409 | HITS-CLIP | 19536157 | Unspecified |
| mmu-miR-384-5p | MIMAT0004745 | Galnt7 | 108150 | HITS-CLIP | 19536157 | Unspecified |
| mmu-miR-384-5p | MIMAT0004745 | Gas7 | 14457 | HITS-CLIP | 19536157 | Unspecified |
| mmu-miR-384-5p | MIMAT0004745 | Gja1 | 14609 | HITS-CLIP | 19536157 | Unspecified |
| mmu-miR-384-5p | MIMAT0004745 | Glce | 93683 | HITS-CLIP | 19536157 | Unspecified |
| mmu-miR-384-5p | MIMAT0004745 | Gnao1 | 14681 | HITS-CLIP | 19536157 | Unspecified |
| mmu-miR-384-5p | MIMAT0004745 | Gpi1 | 14751 | HITS-CLIP | 19536157 | Unspecified |
| mmu-miR-384-5p | MIMAT0004745 | Gpm6a | 234267 | HITS-CLIP | 19536157 | Unspecified |
| mmu-miR-384-5p | MIMAT0004745 | Gpr158 | 241263 | HITS-CLIP | 19536157 | Unspecified |
| mmu-miR-384-5p | MIMAT0004745 | Grm5 | 108071 | HITS-CLIP | 19536157 | Unspecified |
| mmu-miR-384-5p | MIMAT0004745 | Hectd1 | 207304 | HITS-CLIP | 19536157 | Unspecified |
| mmu-miR-384-5p | MIMAT0004745 | Hipk3 | 15259 | HITS-CLIP | 19536157 | Unspecified |

| | | | | | | |
|---|---|---|---|---|---|---|
| mmu-miR-384-5p | MIMAT0004745 | Hnrnpm | 76936 | HITS-CLIP | 19536157 | Unspecified |
| mmu-miR-384-5p | MIMAT0004745 | Hnrnpu | 51810 | HITS-CLIP | 19536157 | Unspecified |
| mmu-miR-384-5p | MIMAT0004745 | Huwe1 | 59026 | HITS-CLIP | 19536157 | Unspecified |
| mmu-miR-384-5p | MIMAT0004745 | Id2 | 15902 | HITS-CLIP | 19536157 | Unspecified |
| mmu-miR-384-5p | MIMAT0004745 | Ier5 | 15939 | HITS-CLIP | 19536157 | Unspecified |
| mmu-miR-384-5p | MIMAT0004745 | Ifnar1 | 15975 | HITS-CLIP | 19536157 | Unspecified |
| mmu-miR-384-5p | MIMAT0004745 | Ifngr2 | 15980 | HITS-CLIP | 19536157 | Unspecified |
| mmu-miR-384-5p | MIMAT0004745 | Igsf9b | 235086 | HITS-CLIP | 19536157 | Unspecified |
| mmu-miR-384-5p | MIMAT0004745 | Il2rb | 16185 | HITS-CLIP | 31433990 | Unspecified |
| mmu-miR-384-5p | MIMAT0004745 | Inhba | 16323 | HITS-CLIP | 19536157 | Unspecified |
| mmu-miR-384-5p | MIMAT0004745 | Inhbb | 16324 | HITS-CLIP | 19536157 | Unspecified |
| mmu-miR-384-5p | MIMAT0004745 | Ints8 | 72656 | HITS-CLIP | 19536157 | Unspecified |
| mmu-miR-384-5p | MIMAT0004745 | Itfg1 | 71927 | HITS-CLIP | 19536157 | Unspecified |
| mmu-miR-384-5p | MIMAT0004745 | Itsn1 | 16443 | HITS-CLIP | 19536157 | Unspecified |
| mmu-miR-384-5p | MIMAT0004745 | Jade3 | 382207 | HITS-CLIP | 19536157 | Unspecified |
| mmu-miR-384-5p | MIMAT0004745 | Jakmip2 | 76217 | HITS-CLIP | 19536157 | Unspecified |
| mmu-miR-384-5p | MIMAT0004745 | Jph4 | 319984 | HITS-CLIP | 19536157 | Unspecified |
| mmu-miR-384-5p | MIMAT0004745 | Kalrn | 545156 | HITS-CLIP | 19536157 | Unspecified |
| mmu-miR-384-5p | MIMAT0004745 | Katnbl1 | 72425 | HITS-CLIP | 19536157 | Unspecified |
| mmu-miR-384-5p | MIMAT0004745 | Kcnk9 | 223604 | HITS-CLIP | 19536157 | Unspecified |
| mmu-miR-384-5p | MIMAT0004745 | Kctd7 | 212919 | HITS-CLIP | 19536157 | Unspecified |
| mmu-miR-384-5p | MIMAT0004745 | Kif1a | 16560 | HITS-CLIP | 19536157 | Unspecified |
| mmu-miR-384-5p | MIMAT0004745 | Kmt2e | 69188 | HITS-CLIP | 19536157 | Unspecified |
| mmu-miR-384-5p | MIMAT0004745 | Lamc1 | 226519 | HITS-CLIP | 19536157 | Unspecified |
| mmu-miR-384-5p | MIMAT0004745 | Large1 | 16795 | HITS-CLIP | 19536157 | Unspecified |
| mmu-miR-384-5p | MIMAT0004745 | Larp1 | 73158 | HITS-CLIP | 19536157 | Unspecified |
| mmu-miR-384-5p | MIMAT0004745 | Ldhb | 16832 | HITS-CLIP | 19536157 | Unspecified |
| mmu-miR-384-5p | MIMAT0004745 | Lepr | 16847 | HITS-CLIP | 19536157 | Unspecified |
| mmu-miR-384-5p | MIMAT0004745 | Lgi1 | 56839 | HITS-CLIP | 19536157 | Unspecified |
| mmu-miR-384-5p | MIMAT0004745 | Lhx1 | 16869 | HITS-CLIP | 19536157 | Unspecified |
| mmu-miR-384-5p | MIMAT0004745 | Lif | 16878 | HITS-CLIP | 31433990 | Unspecified |
| mmu-miR-384-5p | MIMAT0004745 | Lifr | 16880 | HITS-CLIP | 19536157 | Unspecified |
| mmu-miR-384-5p | MIMAT0004745 | Lnpep | 240028 | HITS-CLIP | 19536157 | Unspecified |
| mmu-miR-384-5p | MIMAT0004745 | Lrp8 | 16975 | HITS-CLIP | 19536157 | Unspecified |
| mmu-miR-384-5p | MIMAT0004745 | March6 | 10299 | HITS-CLIP | 19536157 | Unspecified |
| mmu-miR-384-5p | MIMAT0004745 | Marcks | 17118 | HITS-CLIP | 19536157 | Unspecified |
| mmu-miR-384-5p | MIMAT0004745 | Mat2a | 232087 | HITS-CLIP | 19536157 | Unspecified |
| mmu-miR-384-5p | MIMAT0004745 | Mbnl2 | 105559 | HITS-CLIP | 19536157 | Unspecified |
| mmu-miR-384-5p | MIMAT0004745 | Mcf2l | 17207 | HITS-CLIP | 19536157 | Unspecified |

| | | | | | | |
|---|---|---|---|---|---|---|
| mmu-miR-384-5p | MIMAT0004745 | Mecp2 | 17257 | HITS-CLIP | 31433990 | Unspecified |
| mmu-miR-384-5p | MIMAT0004745 | Med13 | 327987 | HITS-CLIP | 19536157 | Unspecified |
| mmu-miR-384-5p | MIMAT0004745 | Mex3b | 108797 | HITS-CLIP | 19536157 | Unspecified |
| mmu-miR-384-5p | MIMAT0004745 | Mia3 | 338366 | HITS-CLIP | 19536157 | Unspecified |
| mmu-miR-384-5p | MIMAT0004745 | Mib1 | 225164 | HITS-CLIP | 19536157 | Unspecified |
| mmu-miR-384-5p | MIMAT0004745 | Mkrn3 | 22652 | HITS-CLIP | 19536157 | Unspecified |
| mmu-miR-384-5p | MIMAT0004745 | Mon2 | 67074 | HITS-CLIP | 19536157 | Unspecified |
| mmu-miR-384-5p | MIMAT0004745 | Mt1 | 17748 | HITS-CLIP | 19536157 | Unspecified |
| mmu-miR-384-5p | MIMAT0004745 | Mtdh | 67154 | HITS-CLIP | 19536157 | Unspecified |
| mmu-miR-384-5p | MIMAT0004745 | Mxd4 | 17122 | HITS-CLIP | 19536157 | Unspecified |
| mmu-miR-384-5p | MIMAT0004745 | Naa25 | 231713 | HITS-CLIP | 19536157 | Unspecified |
| mmu-miR-384-5p | MIMAT0004745 | Nbea | 26422 | HITS-CLIP | 19536157 | Unspecified |
| mmu-miR-384-5p | MIMAT0004745 | Ncam1 | 17967 | HITS-CLIP | 19536157 | Unspecified |
| mmu-miR-384-5p | MIMAT0004745 | Nedd4 | 17999 | HITS-CLIP | 19536157 | Unspecified |
| mmu-miR-384-5p | MIMAT0004745 | Nedd4l | 83814 | HITS-CLIP | 19536157 | Unspecified |
| mmu-miR-384-5p | MIMAT0004745 | Neto2 | 74513 | HITS-CLIP | 19536157 | Unspecified |
| mmu-miR-384-5p | MIMAT0004745 | Nfat5 | 54446 | HITS-CLIP | 19536157 | Unspecified |
| mmu-miR-384-5p | MIMAT0004745 | Nfib | 18028 | HITS-CLIP | 19536157 | Unspecified |
| mmu-miR-384-5p | MIMAT0004745 | Nmt1 | 18107 | HITS-CLIP | 19536157 | Unspecified |
| mmu-miR-384-5p | MIMAT0004745 | Nol4l | 329540 | HITS-CLIP | 19536157 | Unspecified |
| mmu-miR-384-5p | MIMAT0004745 | Npepps | 19155 | HITS-CLIP | 19536157 | Unspecified |
| mmu-miR-384-5p | MIMAT0004745 | Nras | 18176 | HITS-CLIP | 19536157 | Unspecified |
| mmu-miR-384-5p | MIMAT0004745 | Nrep | 27528 | HITS-CLIP | 19536157 | Unspecified |
| mmu-miR-384-5p | MIMAT0004745 | Nrxn3 | 18191 | HITS-CLIP | 19536157 | Unspecified |
| mmu-miR-384-5p | MIMAT0004745 | Nsf | 18195 | HITS-CLIP | 19536157 | Unspecified |
| mmu-miR-384-5p | MIMAT0004745 | Nsg1 | 18196 | HITS-CLIP | 19536157 | Unspecified |
| mmu-miR-384-5p | MIMAT0004745 | P4ha2 | 18452 | HITS-CLIP | 19536157 | Unspecified |
| mmu-miR-384-5p | MIMAT0004745 | Parp6 | 67287 | HITS-CLIP | 19536157 | Unspecified |
| mmu-miR-384-5p | MIMAT0004745 | Pcdh10 | 18526 | HITS-CLIP | 19536157 | Unspecified |
| mmu-miR-384-5p | MIMAT0004745 | Pcgf3 | 69587 | HITS-CLIP | 19536157 | Unspecified |
| mmu-miR-384-5p | MIMAT0004745 | Pclo | 26875 | HITS-CLIP | 19536157 | Unspecified |
| mmu-miR-384-5p | MIMAT0004745 | Pdgfra | 18595 | HITS-CLIP | 19536157 | Unspecified |
| mmu-miR-384-5p | MIMAT0004745 | Pfn2 | 18645 | HITS-CLIP | 19536157 | Unspecified |
| mmu-miR-384-5p | MIMAT0004745 | Pgm2l1 | 70974 | HITS-CLIP | 19536157 | Unspecified |
| mmu-miR-384-5p | MIMAT0004745 | Pheta1 | 144717 | HITS-CLIP | 19536157 | Unspecified |
| mmu-miR-384-5p | MIMAT0004745 | Pik3r1 | 18708 | HITS-CLIP | 19536157 | Unspecified |
| mmu-miR-384-5p | MIMAT0004745 | Pitpnm2 | 19679 | HITS-CLIP | 19536157 | Unspecified |
| mmu-miR-384-5p | MIMAT0004745 | Plekha1 | 101476 | HITS-CLIP | 19536157 | Unspecified |
| mmu-miR-384-5p | MIMAT0004745 | Plp1 | 18823 | HITS-CLIP | 19536157 | Unspecified |

| | | | | | | |
|---|---|---|---|---|---|---|
| mmu-miR-384-5p | MIMAT0004745 | Plxna2 | 18845 | HITS-CLIP | 19536157 | Unspecified |
| mmu-miR-384-5p | MIMAT0004745 | Plxnc1 | 54712 | HITS-CLIP | 19536157 | Unspecified |
| mmu-miR-384-5p | MIMAT0004745 | Pogz | 229584 | HITS-CLIP | 19536157 | Unspecified |
| mmu-miR-384-5p | MIMAT0004745 | Ppp1r12a | 17931 | HITS-CLIP | 19536157 | Unspecified |
| mmu-miR-384-5p | MIMAT0004745 | Ppp3ca | 19055 | HITS-CLIP | 19536157 | Unspecified |
| mmu-miR-384-5p | MIMAT0004745 | Ppp3r1 | 19058 | HITS-CLIP | 19536157 | Unspecified |
| mmu-miR-384-5p | MIMAT0004745 | Prr12 | 233210 | HITS-CLIP | 19536157 | Unspecified |
| mmu-miR-384-5p | MIMAT0004745 | Prrt2 | 69017 | HITS-CLIP | 19536157 | Unspecified |
| mmu-miR-384-5p | MIMAT0004745 | Prune2 | 353211 | HITS-CLIP | 19536157 | Unspecified |
| mmu-miR-384-5p | MIMAT0004745 | Ptprz1 | 19283 | HITS-CLIP | 19536157 | Unspecified |
| mmu-miR-384-5p | MIMAT0004745 | Pura | 19290 | HITS-CLIP | 19536157 | Unspecified |
| mmu-miR-384-5p | MIMAT0004745 | Purb | 19291 | HITS-CLIP | 19536157 | Unspecified |
| mmu-miR-384-5p | MIMAT0004745 | Qk | 19317 | HITS-CLIP | 19536157 | Unspecified |
| mmu-miR-384-5p | MIMAT0004745 | Rab11a | 53869 | HITS-CLIP | 19536157 | Unspecified |
| mmu-miR-384-5p | MIMAT0004745 | Ralgapa1 | 56784 | HITS-CLIP | 19536157 | Unspecified |
| mmu-miR-384-5p | MIMAT0004745 | Rbm12 | 75710 | HITS-CLIP | 19536157 | Unspecified |
| mmu-miR-384-5p | MIMAT0004745 | Rbm7 | 67010 | HITS-CLIP | 19536157 | Unspecified |
| mmu-miR-384-5p | MIMAT0004745 | Rc3h2 | 319817 | HITS-CLIP | 19536157 | Unspecified |
| mmu-miR-384-5p | MIMAT0004745 | Rcor1 | 217864 | HITS-CLIP | 19536157 | Unspecified |
| mmu-miR-384-5p | MIMAT0004745 | Reep1 | 52250 | HITS-CLIP | 19536157 | Unspecified |
| mmu-miR-384-5p | MIMAT0004745 | Reep3 | 28193 | HITS-CLIP | 19536157 | Unspecified |
| mmu-miR-384-5p | MIMAT0004745 | Rev3l | 19714 | HITS-CLIP | 19536157 | Unspecified |
| mmu-miR-384-5p | MIMAT0004745 | Rfx3 | 19726 | HITS-CLIP | 19536157 | Unspecified |
| mmu-miR-384-5p | MIMAT0004745 | Rgs8 | 67792 | HITS-CLIP | 19536157 | Unspecified |
| mmu-miR-384-5p | MIMAT0004745 | Rhob | 11852 | HITS-CLIP | 19536157 | Unspecified |
| mmu-miR-384-5p | MIMAT0004745 | Rictor | 78757 | HITS-CLIP | 19536157 | Unspecified |
| mmu-miR-384-5p | MIMAT0004745 | Rnf128 | 66889 | HITS-CLIP | 19536157 | Unspecified |
| mmu-miR-384-5p | MIMAT0004745 | Rnf44 | 105239 | HITS-CLIP | 19536157 | Unspecified |
| mmu-miR-384-5p | MIMAT0004745 | Rps6ka3 | 110651 | HITS-CLIP | 19536157 | Unspecified |
| mmu-miR-384-5p | MIMAT0004745 | Rps6ka5 | 73086 | HITS-CLIP | 19536157 | Unspecified |
| mmu-miR-384-5p | MIMAT0004745 | Rtn1 | 104001 | HITS-CLIP | 19536157 | Unspecified |
| mmu-miR-384-5p | MIMAT0004745 | Rtn4r | 65079 | HITS-CLIP | 19536157 | Unspecified |
| mmu-miR-384-5p | MIMAT0004745 | Samd4 | 74480 | HITS-CLIP | 19536157 | Unspecified |
| mmu-miR-384-5p | MIMAT0004745 | Samd8 | 67630 | HITS-CLIP | 19536157 | Unspecified |
| mmu-miR-384-5p | MIMAT0004745 | Scaper | 244891 | HITS-CLIP | 19536157 | Unspecified |
| mmu-miR-384-5p | MIMAT0004745 | Scn1a | 20265 | HITS-CLIP | 19536157 | Unspecified |
| mmu-miR-384-5p | MIMAT0004745 | Scn2a | 110876 | HITS-CLIP | 19536157 | Unspecified |
| mmu-miR-384-5p | MIMAT0004745 | Scn3a | 20269 | HITS-CLIP | 19536157 | Unspecified |
| mmu-miR-384-5p | MIMAT0004745 | Scn8a | 20273 | HITS-CLIP | 19536157 | Unspecified |

| | | | | | | |
|---|---|---|---|---|---|---|
| mmu-miR-384-5p | MIMAT0004745 | Scyl2 | 213326 | HITS-CLIP | 19536157 | Unspecified |
| mmu-miR-384-5p | MIMAT0004745 | Sec61a2 | 57743 | HITS-CLIP | 19536157 | Unspecified |
| mmu-miR-384-5p | MIMAT0004745 | Sema3a | 20346 | HITS-CLIP | 19536157 | Unspecified |
| mmu-miR-384-5p | MIMAT0004745 | Sema6d | 214968 | HITS-CLIP | 19536157 | Unspecified |
| mmu-miR-384-5p | MIMAT0004745 | Sept11 | 52398 | HITS-CLIP | 19536157 | Unspecified |
| mmu-miR-384-5p | MIMAT0004745 | Serinc1 | 56442 | HITS-CLIP | 19536157 | Unspecified |
| mmu-miR-384-5p | MIMAT0004745 | Sesn3 | 75747 | HITS-CLIP | 19536157 | Unspecified |
| mmu-miR-384-5p | MIMAT0004745 | Setd2 | 235626 | HITS-CLIP | 19536157 | Unspecified |
| mmu-miR-384-5p | MIMAT0004745 | Sh3pxd2a | 14218 | HITS-CLIP | 19536157 | Unspecified |
| mmu-miR-384-5p | MIMAT0004745 | Sirt1 | 93759 | HITS-CLIP | 19536157 | Unspecified |
| mmu-miR-384-5p | MIMAT0004745 | Skil | 20482 | HITS-CLIP | 19536157 | Unspecified |
| mmu-miR-384-5p | MIMAT0004745 | Skp1a | 21402 | HITS-CLIP | 19536157 | Unspecified |
| mmu-miR-384-5p | MIMAT0004745 | Slc16a1 | 20501 | HITS-CLIP | 23142080\|19536157 | Unspecified |
| mmu-miR-384-5p | MIMAT0004745 | Slc1a2 | 20511 | HITS-CLIP | 19536157 | Unspecified |
| mmu-miR-384-5p | MIMAT0004745 | Slc30a4 | 22785 | HITS-CLIP | 19536157 | Unspecified |
| mmu-miR-384-5p | MIMAT0004745 | Slc30a7 | 66500 | HITS-CLIP | 19536157 | Unspecified |
| mmu-miR-384-5p | MIMAT0004745 | Slc39a10 | 227059 | HITS-CLIP | 19536157 | Unspecified |
| mmu-miR-384-5p | MIMAT0004745 | Slco1c1 | 58807 | HITS-CLIP | 19536157 | Unspecified |
| mmu-miR-384-5p | MIMAT0004745 | Smarca5 | 93762 | HITS-CLIP | 19536157 | Unspecified |
| mmu-miR-384-5p | MIMAT0004745 | Smarcc2 | 68094 | HITS-CLIP | 19536157 | Unspecified |
| mmu-miR-384-5p | MIMAT0004745 | Snapc1 | 75627 | HITS-CLIP | 19536157 | Unspecified |
| mmu-miR-384-5p | MIMAT0004745 | Snx18 | 170625 | HITS-CLIP | 19536157 | Unspecified |
| mmu-miR-384-5p | MIMAT0004745 | Sorbs2 | 234214 | HITS-CLIP | 19536157 | Unspecified |
| mmu-miR-384-5p | MIMAT0004745 | Sparcl1 | 13602 | HITS-CLIP | 19536157 | Unspecified |
| mmu-miR-384-5p | MIMAT0004745 | Srsf7 | 225027 | HITS-CLIP | 19536157 | Unspecified |
| mmu-miR-384-5p | MIMAT0004745 | Ssr3 | 67437 | HITS-CLIP | 19536157 | Unspecified |
| mmu-miR-384-5p | MIMAT0004745 | St8sia4 | 20452 | HITS-CLIP | 19536157 | Unspecified |
| mmu-miR-384-5p | MIMAT0004745 | Steap2 | 74051 | HITS-CLIP | 19536157 | Unspecified |
| mmu-miR-384-5p | MIMAT0004745 | Stox2 | 71069 | HITS-CLIP | 19536157 | Unspecified |
| mmu-miR-384-5p | MIMAT0004745 | Stxbp5 | 78808 | HITS-CLIP | 19536157 | Unspecified |
| mmu-miR-384-5p | MIMAT0004745 | Sugt1 | 67955 | HITS-CLIP | 19536157 | Unspecified |
| mmu-miR-384-5p | MIMAT0004745 | Svop | 68666 | HITS-CLIP | 19536157 | Unspecified |
| mmu-miR-384-5p | MIMAT0004745 | Synpr | 72003 | HITS-CLIP | 19536157 | Unspecified |
| mmu-miR-384-5p | MIMAT0004745 | Tab2 | 68652 | HITS-CLIP | 19536157 | Unspecified |
| mmu-miR-384-5p | MIMAT0004745 | Tcf4 | 21413 | HITS-CLIP | 19536157 | Unspecified |
| mmu-miR-384-5p | MIMAT0004745 | Tenm3 | 23965 | HITS-CLIP | 19536157 | Unspecified |
| mmu-miR-384-5p | MIMAT0004745 | Tfrc | 22042 | HITS-CLIP | 19536157 | Unspecified |
| mmu-miR-384-5p | MIMAT0004745 | Tgfbr1 | 21812 | HITS-CLIP | 19536157 | Unspecified |
| mmu-miR-384-5p | MIMAT0004745 | Timp3 | 21859 | HITS-CLIP | 19536157 | Unspecified |

| | | | | | | |
|---|---|---|---|---|---|---|
| mmu-miR-384-5p | MIMAT0004745 | Tle1 | 21885 | HITS-CLIP | 19536157 | Unspecified |
| mmu-miR-384-5p | MIMAT0004745 | Tmcc1 | 330401 | HITS-CLIP | 19536157 | Unspecified |
| mmu-miR-384-5p | MIMAT0004745 | Tmem108 | 81907 | HITS-CLIP | 19536157 | Unspecified |
| mmu-miR-384-5p | MIMAT0004745 | Tmem151b | 210573 | HITS-CLIP | 19536157 | Unspecified |
| mmu-miR-384-5p | MIMAT0004745 | Tmem62 | 96957 | HITS-CLIP | 19536157 | Unspecified |
| mmu-miR-384-5p | MIMAT0004745 | Tmod2 | 50876 | HITS-CLIP | 19536157 | Unspecified |
| mmu-miR-384-5p | MIMAT0004745 | Tnik | 665113 | HITS-CLIP | 19536157 | Unspecified |
| mmu-miR-384-5p | MIMAT0004745 | Tnrc6a | 233833 | HITS-CLIP | 19536157 | Unspecified |
| mmu-miR-384-5p | MIMAT0004745 | Tprkb | 69786 | HITS-CLIP | 19536157 | Unspecified |
| mmu-miR-384-5p | MIMAT0004745 | Tpst1 | 22021 | HITS-CLIP | 19536157 | Unspecified |
| mmu-miR-384-5p | MIMAT0004745 | Trim37 | 68729 | HITS-CLIP | 19536157 | Unspecified |
| mmu-miR-384-5p | MIMAT0004745 | Trip12 | 14897 | HITS-CLIP | 19536157 | Unspecified |
| mmu-miR-384-5p | MIMAT0004745 | Tusc3 | 80286 | HITS-CLIP | 19536157 | Unspecified |
| mmu-miR-384-5p | MIMAT0004745 | Ubr1 | 22222 | HITS-CLIP | 19536157 | Unspecified |
| mmu-miR-384-5p | MIMAT0004745 | Ugt8a | 22239 | HITS-CLIP | 19536157 | Unspecified |
| mmu-miR-384-5p | MIMAT0004745 | Unc5c | 22253 | HITS-CLIP | 19536157 | Unspecified |
| mmu-miR-384-5p | MIMAT0004745 | Uri1 | 19777 | HITS-CLIP | 31433990 | Unspecified |
| mmu-miR-384-5p | MIMAT0004745 | Usb1 | 101985 | HITS-CLIP | 26203562 | Unspecified |
| mmu-miR-384-5p | MIMAT0004745 | Usp2 | 53376 | HITS-CLIP | 19536157 | Unspecified |
| mmu-miR-384-5p | MIMAT0004745 | Usp48 | 170707 | HITS-CLIP | 19536157 | Unspecified |
| mmu-miR-384-5p | MIMAT0004745 | Vopp1 | 232023 | HITS-CLIP | 19536157 | Unspecified |
| mmu-miR-384-5p | MIMAT0004745 | Vps13c | 320528 | HITS-CLIP | 19536157 | Unspecified |
| mmu-miR-384-5p | MIMAT0004745 | Wipf1 | 215280 | HITS-CLIP | 19536157 | Unspecified |
| mmu-miR-384-5p | MIMAT0004745 | Ywhag | 22628 | HITS-CLIP | 19536157 | Unspecified |
| mmu-miR-384-5p | MIMAT0004745 | Zbtb20 | 56490 | HITS-CLIP | 19536157 | Unspecified |
| mmu-miR-384-5p | MIMAT0004745 | Zcchc14 | 142682 | HITS-CLIP | 19536157 | Unspecified |
| mmu-miR-384-5p | MIMAT0004745 | Zcchc2 | 227449 | HITS-CLIP | 19536157 | Unspecified |
| mmu-miR-384-5p | MIMAT0004745 | Zdhhc20 | 75965 | HITS-CLIP | 19536157 | Unspecified |
| mmu-miR-384-5p | MIMAT0004745 | Zeb1 | 21417 | HITS-CLIP | 19536157 | Unspecified |
| mmu-miR-384-5p | MIMAT0004745 | Zhx1 | 22770 | HITS-CLIP | 19536157 | Unspecified |
| mmu-miR-384-5p | MIMAT0004745 | Zmym4 | 67785 | HITS-CLIP | 19536157 | Unspecified |
| mmu-miR-30c-5p | MIMAT0000514 | 0610010F05Rik | 71675 | HITS-CLIP | 23597149 | Unspecified |
| mmu-miR-30c-5p | MIMAT0000514 | 0610040J01Rik | 76261 | HITS-CLIP | 37986224\|23597149 | Unspecified |
| mmu-miR-30c-5p | MIMAT0000514 | 1110059E24Rik | 66206 | HITS-CLIP | 23142080 | Unspecified |
| mmu-miR-30c-5p | MIMAT0000514 | 1190002N15Rik | 68861 | HITS-CLIP | 23142080 | Unspecified |
| mmu-miR-30c-5p | MIMAT0000514 | 1300017J02Rik | 71775 | HITS-CLIP | 23597149 | Unspecified |
| mmu-miR-30c-5p | MIMAT0000514 | 1600012H06Rik | 67912 | HITS-CLIP | 37986224 | Unspecified |
| mmu-miR-30c-5p | MIMAT0000514 | 1810037I17Rik | 67704 | HITS-CLIP | 23142080 | Unspecified |
| mmu-miR-30c-5p | MIMAT0000514 | 2310022B05Rik | 69551 | HITS-CLIP | 37986224 | Unspecified |

| | | | | | | |
|---|---|---|---|---|---|---|
| mmu-miR-30c-5p | MIMAT0000514 | 2310057M21Rik | 68277 | HITS-CLIP | 37986224 | Unspecified |
| mmu-miR-30c-5p | MIMAT0000514 | 2900026A02Rik | 243219 | HITS-CLIP | 37986224 | Unspecified |
| mmu-miR-30c-5p | MIMAT0000514 | 4921524J17Rik | 66714 | HITS-CLIP | 23597149 | Unspecified |
| mmu-miR-30c-5p | MIMAT0000514 | 4930402H24Rik | 228602 | HITS-CLIP | 37986224 | Unspecified |
| mmu-miR-30c-5p | MIMAT0000514 | 4931406P16Rik | 233103 | HITS-CLIP | 23142080 | Unspecified |
| mmu-miR-30c-5p | MIMAT0000514 | 4932438A13Rik | 229227 | HITS-CLIP | 23142080 | Unspecified |
| mmu-miR-30c-5p | MIMAT0000514 | 4933434E20Rik | 99650 | HITS-CLIP | 37986224 | Unspecified |
| mmu-miR-30c-5p | MIMAT0000514 | 6030458C11Rik | 77877 | HITS-CLIP | 25753418 | Unspecified |
| mmu-miR-30c-5p | MIMAT0000514 | 9130401M01Rik | 75758 | HITS-CLIP | 37986224 | Unspecified |
| mmu-miR-30c-5p | MIMAT0000514 | 9530068E07Rik | 213673 | HITS-CLIP | 25753418 | Unspecified |
| mmu-miR-30c-5p | MIMAT0000514 | 9930021J03Rik | 240613 | HITS-CLIP | 23142080 | Unspecified |
| mmu-miR-30c-5p | MIMAT0000514 | A130010J15Rik | 319266 | HITS-CLIP | 23142080\|23597149 | Unspecified |
| mmu-miR-30c-5p | MIMAT0000514 | A430093F15Rik | 403202 | HITS-CLIP | 37986224 | Unspecified |
| mmu-miR-30c-5p | MIMAT0000514 | AI987944 | 233168 | HITS-CLIP | 37986224 | Unspecified |
| mmu-miR-30c-5p | MIMAT0000514 | AW549877 | 106064 | HITS-CLIP | 23142080 | Unspecified |
| mmu-miR-30c-5p | MIMAT0000514 | Aadac | 67758 | HITS-CLIP | 37986224 | Unspecified |
| mmu-miR-30c-5p | MIMAT0000514 | Aasdhppt | 67618 | HITS-CLIP | 37986224\|25753418 | Unspecified |
| mmu-miR-30c-5p | MIMAT0000514 | Abat | 268860 | HITS-CLIP | 37986224\|19536157\|23597149 | Unspecified |
| mmu-miR-30c-5p | MIMAT0000514 | Abca1 | 11303 | HITS-CLIP | 25753418 | Unspecified |
| mmu-miR-30c-5p | MIMAT0000514 | Abca3 | 27410 | HITS-CLIP | 37986224\|23597149\|23142080 | Unspecified |
| mmu-miR-30c-5p | MIMAT0000514 | Abcb11 | 27413 | HITS-CLIP | 23597149 | Unspecified |
| mmu-miR-30c-5p | MIMAT0000514 | Abcb7 | 11306 | HITS-CLIP | 37986224\|23597149\|23142080 | Unspecified |
| mmu-miR-30c-5p | MIMAT0000514 | Abcc2 | 12780 | HITS-CLIP | 37986224 | Unspecified |
| mmu-miR-30c-5p | MIMAT0000514 | Abcc3 | 76408 | HITS-CLIP | 23597149 | Unspecified |
| mmu-miR-30c-5p | MIMAT0000514 | Abhd16a | 193742 | HITS-CLIP | 23597149 | Unspecified |
| mmu-miR-30c-5p | MIMAT0000514 | Abhd17b | 226016 | HITS-CLIP | 19536157 | Unspecified |
| mmu-miR-30c-5p | MIMAT0000514 | Abi1 | 11308 | HITS-CLIP | 23142080 | Unspecified |
| mmu-miR-30c-5p | MIMAT0000514 | Abl2 | 11352 | HITS-CLIP | 23142080\|23597149 | Unspecified |
| mmu-miR-30c-5p | MIMAT0000514 | Acaca | 107476 | HITS-CLIP | 37986224\|23597149 | Unspecified |
| mmu-miR-30c-5p | MIMAT0000514 | Acad8 | 66948 | HITS-CLIP | 37986224 | Unspecified |
| mmu-miR-30c-5p | MIMAT0000514 | Acad9 | 229211 | HITS-CLIP | 23142080 | Unspecified |
| mmu-miR-30c-5p | MIMAT0000514 | Acap2 | 78618 | HITS-CLIP | 37986224\|23142080 | Unspecified |
| mmu-miR-30c-5p | MIMAT0000514 | Acap3 | 140500 | HITS-CLIP | 37986224 | Unspecified |
| mmu-miR-30c-5p | MIMAT0000514 | Acat1 | 110446 | HITS-CLIP | 23142080 | Unspecified |
| mmu-miR-30c-5p | MIMAT0000514 | Acbd3 | 170760 | HITS-CLIP | 23142080 | Unspecified |
| mmu-miR-30c-5p | MIMAT0000514 | Acbd5 | 74159 | HITS-CLIP | 23142080 | Unspecified |
| mmu-miR-30c-5p | MIMAT0000514 | Acly | 104112 | HITS-CLIP | 37986224 | Unspecified |
| mmu-miR-30c-5p | MIMAT0000514 | Acp2 | 11432 | HITS-CLIP | 37986224 | Unspecified |
| mmu-miR-30c-5p | MIMAT0000514 | Acp6 | 66659 | HITS-CLIP | 23142080 | Unspecified |

| | | | | | | |
|---|---|---|---|---|---|---|
| mmu-miR-30c-5p | MIMAT0000514 | Acsl4 | 50790 | HITS-CLIP | 23142080 | Unspecified |
| mmu-miR-30c-5p | MIMAT0000514 | Acsl6 | 216739 | HITS-CLIP | 37986224 | Unspecified |
| mmu-miR-30c-5p | MIMAT0000514 | Actl6a | 56456 | HITS-CLIP | 23142080 | Unspecified |
| mmu-miR-30c-5p | MIMAT0000514 | Actr1a | 54130 | HITS-CLIP | 23142080 | Unspecified |
| mmu-miR-30c-5p | MIMAT0000514 | Actr2 | 66713 | qCLASH | 31393953 | Unspecified |
| mmu-miR-30c-5p | MIMAT0000514 | Actr3 | 74117 | HITS-CLIP | 25753418 | Unspecified |
| mmu-miR-30c-5p | MIMAT0000514 | Acvr1 | 11477 | HITS-CLIP | 37986224\|19536157\|23597149 | Unspecified |
| mmu-miR-30c-5p | MIMAT0000514 | Adam10 | 11487 | HITS-CLIP | 37986224\|19536157\|23142080 | Unspecified |
| mmu-miR-30c-5p | MIMAT0000514 | Adam19 | 11492 | HITS-CLIP | 23142080 | Unspecified |
| mmu-miR-30c-5p | MIMAT0000514 | Adam9 | 11502 | HITS-CLIP | 37986224\|23142080\|23597149\|19536 | Unspecified |
| mmu-miR-30c-5p | MIMAT0000514 | Adamts16 | 271127 | HITS-CLIP | 37986224 | Unspecified |
| mmu-miR-30c-5p | MIMAT0000514 | Adar | 56417 | HITS-CLIP | 23597149\|19536157\|23142080 | Unspecified |
| mmu-miR-30c-5p | MIMAT0000514 | Adcy1 | 432530 | HITS-CLIP | 19536157 | Unspecified |
| mmu-miR-30c-5p | MIMAT0000514 | Adcy7 | 11513 | HITS-CLIP | 23142080 | Unspecified |
| mmu-miR-30c-5p | MIMAT0000514 | Add3 | 27360 | HITS-CLIP | 23142080 | Unspecified |
| mmu-miR-30c-5p | MIMAT0000514 | Adgra3 | 166647 | HITS-CLIP | 23597149 | Unspecified |
| mmu-miR-30c-5p | MIMAT0000514 | Adgrf5 | 224792 | HITS-CLIP | 23597149\|25753418 | Unspecified |
| mmu-miR-30c-5p | MIMAT0000514 | Adk | 11534 | HITS-CLIP | 23597149 | Unspecified |
| mmu-miR-30c-5p | MIMAT0000514 | Adnp | 11538 | HITS-CLIP | 37986224 | Unspecified |
| mmu-miR-30c-5p | MIMAT0000514 | Ado | 211488 | HITS-CLIP | 23142080 | Unspecified |
| mmu-miR-30c-5p | MIMAT0000514 | Adra2b | 11552 | HITS-CLIP | 37986224 | Unspecified |
| mmu-miR-30c-5p | MIMAT0000514 | Adrb2 | 11555 | HITS-CLIP | 37986224\|23142080 | Unspecified |
| mmu-miR-30c-5p | MIMAT0000514 | Adss | 11566 | HITS-CLIP | 37986224\|23142080 | Unspecified |
| mmu-miR-30c-5p | MIMAT0000514 | Afap1 | 70292 | HITS-CLIP | 37986224 | Unspecified |
| mmu-miR-30c-5p | MIMAT0000514 | Aff2 | 14266 | HITS-CLIP | 37986224 | Unspecified |
| mmu-miR-30c-5p | MIMAT0000514 | Aff4 | 93736 | HITS-CLIP | 37986224\|23597149\|19536157\|23142 | Unspecified |
| mmu-miR-30c-5p | MIMAT0000514 | Afg3l1 | 114896 | HITS-CLIP | 23142080 | Unspecified |
| mmu-miR-30c-5p | MIMAT0000514 | Agfg1 | 15463 | HITS-CLIP | 37986224\|23142080 | Unspecified |
| mmu-miR-30c-5p | MIMAT0000514 | Aggf1 | 66549 | HITS-CLIP | 37986224\|23142080\|19536157 | Unspecified |
| mmu-miR-30c-5p | MIMAT0000514 | Agk | 69923 | HITS-CLIP | 25753418 | Unspecified |
| mmu-miR-30c-5p | MIMAT0000514 | Ago1 | 236511 | HITS-CLIP | 37986224\|19536157\|23597149\|23142 | Unspecified |
| mmu-miR-30c-5p | MIMAT0000514 | Ago2 | 239528 | HITS-CLIP | 37986224\|23142080 | Unspecified |
| mmu-miR-30c-5p | MIMAT0000514 | Ago3 | 214150 | HITS-CLIP | 37986224\|23142080 | Unspecified |
| mmu-miR-30c-5p | MIMAT0000514 | Ago4 | 76850 | HITS-CLIP | 37986224 | Unspecified |
| mmu-miR-30c-5p | MIMAT0000514 | Agpat5 | 52123 | HITS-CLIP | 23142080 | Unspecified |
| mmu-miR-30c-5p | MIMAT0000514 | Agps | 228061 | HITS-CLIP | 23142080 | Unspecified |
| mmu-miR-30c-5p | MIMAT0000514 | Agtr1a | 11607 | HITS-CLIP | 37986224 | Unspecified |
| mmu-miR-30c-5p | MIMAT0000514 | Ahctf1 | 226747 | HITS-CLIP | 37986224\|19536157 | Unspecified |
| mmu-miR-30c-5p | MIMAT0000514 | Ahcyl1 | 229709 | HITS-CLIP | 37986224 | Unspecified |

| | | | | | | |
|---|---|---|---|---|---|---|
| mmu-miR-30c-5p | MIMAT0000514 | Ahi1 | 52906 | HITS-CLIP | 23142080 | Unspecified |
| mmu-miR-30c-5p | MIMAT0000514 | Ahnak | 66395 | HITS-CLIP | 37986224\|23142080 | Unspecified |
| mmu-miR-30c-5p | MIMAT0000514 | Ahr | 11622 | HITS-CLIP | 37986224\|23597149\|19536157\|23142 | Unspecified |
| mmu-miR-30c-5p | MIMAT0000514 | Ahsa1 | 217737 | HITS-CLIP | 37986224\|23597149 | Unspecified |
| mmu-miR-30c-5p | MIMAT0000514 | Ak3 | 56248 | HITS-CLIP | 25753418 | Unspecified |
| mmu-miR-30c-5p | MIMAT0000514 | Akap10 | 56697 | HITS-CLIP | 37986224\|23142080 | Unspecified |
| mmu-miR-30c-5p | MIMAT0000514 | Akap11 | 219181 | HITS-CLIP | 23597149\|23142080 | Unspecified |
| mmu-miR-30c-5p | MIMAT0000514 | Akirin2 | 433693 | HITS-CLIP | 23597149\|23142080\|25753418 | Unspecified |
| mmu-miR-30c-5p | MIMAT0000514 | Akna | 100182 | HITS-CLIP | 37986224\|23142080 | Unspecified |
| mmu-miR-30c-5p | MIMAT0000514 | Akt3 | 23797 | HITS-CLIP | 23142080 | Unspecified |
| mmu-miR-30c-5p | MIMAT0000514 | Alcam | 11658 | HITS-CLIP | 37986224\|23597149\|23142080 | Unspecified |
| mmu-miR-30c-5p | MIMAT0000514 | Aldh1l1 | 107747 | HITS-CLIP | 23597149 | Unspecified |
| mmu-miR-30c-5p | MIMAT0000514 | Alg11 | 207958 | HITS-CLIP | 37986224\|23142080 | Unspecified |
| mmu-miR-30c-5p | MIMAT0000514 | Alg13 | 67574 | HITS-CLIP | 23142080 | Unspecified |
| mmu-miR-30c-5p | MIMAT0000514 | Alg2 | 56737 | HITS-CLIP | 37986224 | Unspecified |
| mmu-miR-30c-5p | MIMAT0000514 | Alkbh1 | 211064 | HITS-CLIP | 23142080 | Unspecified |
| mmu-miR-30c-5p | MIMAT0000514 | Alpk3 | 116904 | HITS-CLIP | 25753418 | Unspecified |
| mmu-miR-30c-5p | MIMAT0000514 | Amer1 | 72345 | HITS-CLIP | 25083871\|23142080 | Unspecified |
| mmu-miR-30c-5p | MIMAT0000514 | Ammecr1 | 56068 | HITS-CLIP | 23142080 | Unspecified |
| mmu-miR-30c-5p | MIMAT0000514 | Amotl2 | 56332 | HITS-CLIP | 37986224\|23597149 | Unspecified |
| mmu-miR-30c-5p | MIMAT0000514 | Amt | 434437 | HITS-CLIP | 23597149 | Unspecified |
| mmu-miR-30c-5p | MIMAT0000514 | Anapc1 | 17222 | HITS-CLIP | 23142080 | Unspecified |
| mmu-miR-30c-5p | MIMAT0000514 | Anapc2 | 99152 | HITS-CLIP | 23142080 | Unspecified |
| mmu-miR-30c-5p | MIMAT0000514 | Anapc7 | 56317 | HITS-CLIP | 37986224\|23597149\|23142080 | Unspecified |
| mmu-miR-30c-5p | MIMAT0000514 | Ankfy1 | 11736 | HITS-CLIP | 23597149 | Unspecified |
| mmu-miR-30c-5p | MIMAT0000514 | Ankhd1 | 108857 | HITS-CLIP | 37986224 | Unspecified |
| mmu-miR-30c-5p | MIMAT0000514 | Ankib1 | 70797 | HITS-CLIP | 37986224 | Unspecified |
| mmu-miR-30c-5p | MIMAT0000514 | Ankle2 | 71782 | HITS-CLIP | 23142080 | Unspecified |
| mmu-miR-30c-5p | MIMAT0000514 | Ankra2 | 68558 | HITS-CLIP | 37986224\|23597149\|23142080 | Unspecified |
| mmu-miR-30c-5p | MIMAT0000514 | Ankrd17 | 81702 | HITS-CLIP | 37986224\|23597149\|23142080\|1953€ | Unspecified |
| mmu-miR-30c-5p | MIMAT0000514 | Ankrd28 | 105522 | HITS-CLIP | 37986224\|23142080 | Unspecified |
| mmu-miR-30c-5p | MIMAT0000514 | Ankrd44 | 329154 | HITS-CLIP | 37986224\|23142080 | Unspecified |
| mmu-miR-30c-5p | MIMAT0000514 | Ankrd52 | 237615 | HITS-CLIP | 37986224 | Unspecified |
| mmu-miR-30c-5p | MIMAT0000514 | Ankrd9 | 74251 | HITS-CLIP | 37986224 | Unspecified |
| mmu-miR-30c-5p | MIMAT0000514 | Anks4b | 72074 | HITS-CLIP | 23597149 | Unspecified |
| mmu-miR-30c-5p | MIMAT0000514 | Anln | 68743 | HITS-CLIP | 23142080 | Unspecified |
| mmu-miR-30c-5p | MIMAT0000514 | Ano10 | 102566 | HITS-CLIP | 37986224 | Unspecified |
| mmu-miR-30c-5p | MIMAT0000514 | Anp32b | 67628 | qCLASH | 31393953 | Unspecified |
| mmu-miR-30c-5p | MIMAT0000514 | Anp32e | 66471 | HITS-CLIP//qCLASH | 37986224\|31393953 | Unspecified |

| | | | | | | |
|---|---|---|---|---|---|---|
| mmu-miR-30c-5p | MIMAT0000514 | Antxr1 | 69538 | HITS-CLIP | 37986224 | Unspecified |
| mmu-miR-30c-5p | MIMAT0000514 | Anxa6 | 11749 | HITS-CLIP | 25753418 | Unspecified |
| mmu-miR-30c-5p | MIMAT0000514 | Aox3 | 71724 | HITS-CLIP | 23597149 | Unspecified |
| mmu-miR-30c-5p | MIMAT0000514 | Ap2a1 | 11771 | HITS-CLIP | 23142080 | Unspecified |
| mmu-miR-30c-5p | MIMAT0000514 | Ap2b1 | 71770 | HITS-CLIP | 37986224\|23597149\|23142080\|1953( | Unspecified |
| mmu-miR-30c-5p | MIMAT0000514 | Ap3s1 | 11777 | HITS-CLIP | 37986224 | Unspecified |
| mmu-miR-30c-5p | MIMAT0000514 | Ap4b1 | 67489 | HITS-CLIP | 23142080 | Unspecified |
| mmu-miR-30c-5p | MIMAT0000514 | Apaf1 | 11783 | HITS-CLIP | 37986224\|23597149\|23142080 | Unspecified |
| mmu-miR-30c-5p | MIMAT0000514 | Apba1 | 319924 | HITS-CLIP | 37986224\|19536157 | Unspecified |
| mmu-miR-30c-5p | MIMAT0000514 | Apbb2 | 11787 | HITS-CLIP | 37986224 | Unspecified |
| mmu-miR-30c-5p | MIMAT0000514 | Apbb3 | 225372 | HITS-CLIP | 23142080 | Unspecified |
| mmu-miR-30c-5p | MIMAT0000514 | Apc | 11789 | HITS-CLIP | 37986224 | Unspecified |
| mmu-miR-30c-5p | MIMAT0000514 | Apeh | 235606 | HITS-CLIP | 23142080 | Unspecified |
| mmu-miR-30c-5p | MIMAT0000514 | Aplp2 | 11804 | HITS-CLIP | 37986224 | Unspecified |
| mmu-miR-30c-5p | MIMAT0000514 | Apob | 238055 | HITS-CLIP | 37986224\|23597149 | Unspecified |
| mmu-miR-30c-5p | MIMAT0000514 | App | 11820 | HITS-CLIP | 37986224 | Unspecified |
| mmu-miR-30c-5p | MIMAT0000514 | Appl2 | 216190 | HITS-CLIP | 37986224\|23597149\|23142080 | Unspecified |
| mmu-miR-30c-5p | MIMAT0000514 | Araf | 11836 | HITS-CLIP | 37986224\|23142080\|25753418 | Unspecified |
| mmu-miR-30c-5p | MIMAT0000514 | Arap2 | 212285 | HITS-CLIP | 37986224\|23142080 | Unspecified |
| mmu-miR-30c-5p | MIMAT0000514 | Arcn1 | 213827 | HITS-CLIP | 25753418 | Unspecified |
| mmu-miR-30c-5p | MIMAT0000514 | Arel1 | 68497 | HITS-CLIP | 25753418\|23142080 | Unspecified |
| mmu-miR-30c-5p | MIMAT0000514 | Arf3 | 11842 | HITS-CLIP | 23142080 | Unspecified |
| mmu-miR-30c-5p | MIMAT0000514 | Arf4 | 11843 | HITS-CLIP | 23597149\|23142080 | Unspecified |
| mmu-miR-30c-5p | MIMAT0000514 | Arfgef1 | 211673 | qCLASH//HITS-CLIP | 31393953\|25753418 | Unspecified |
| mmu-miR-30c-5p | MIMAT0000514 | Arglu1 | 234023 | HITS-CLIP | 23142080 | Unspecified |
| mmu-miR-30c-5p | MIMAT0000514 | Arhgap12 | 75415 | HITS-CLIP | 23142080 | Unspecified |
| mmu-miR-30c-5p | MIMAT0000514 | Arhgap18 | 73910 | HITS-CLIP | 37986224 | Unspecified |
| mmu-miR-30c-5p | MIMAT0000514 | Arhgap19 | 71085 | HITS-CLIP | 23142080 | Unspecified |
| mmu-miR-30c-5p | MIMAT0000514 | Arhgap29 | 214137 | HITS-CLIP | 37986224\|23597149 | Unspecified |
| mmu-miR-30c-5p | MIMAT0000514 | Arhgap5 | 11855 | HITS-CLIP | 37986224 | Unspecified |
| mmu-miR-30c-5p | MIMAT0000514 | Arhgef3 | 71704 | HITS-CLIP | 37986224 | Unspecified |
| mmu-miR-30c-5p | MIMAT0000514 | Arhgef6 | 73341 | HITS-CLIP | 23142080 | Unspecified |
| mmu-miR-30c-5p | MIMAT0000514 | Arid1a | 93760 | HITS-CLIP | 37986224 | Unspecified |
| mmu-miR-30c-5p | MIMAT0000514 | Arid1b | 239985 | HITS-CLIP | 25753418 | Unspecified |
| mmu-miR-30c-5p | MIMAT0000514 | Arid2 | 77044 | HITS-CLIP | 37986224\|23142080 | Unspecified |
| mmu-miR-30c-5p | MIMAT0000514 | Arid4a | 238247 | HITS-CLIP | 37986224\|23142080 | Unspecified |
| mmu-miR-30c-5p | MIMAT0000514 | Arid4b | 94246 | HITS-CLIP | 19536157\|23142080 | Unspecified |
| mmu-miR-30c-5p | MIMAT0000514 | Arid5b | 71371 | HITS-CLIP | 37986224\|23142080\|19536157\|2359 | Unspecified |
| mmu-miR-30c-5p | MIMAT0000514 | Arih1 | 23806 | HITS-CLIP | 37986224\|23142080\|23597149 | Unspecified |

| | | | | | | |
|---|---|---|---|---|---|---|
| mmu-miR-30c-5p | MIMAT0000514 | Arl14ep | 212772 | HITS-CLIP | 23142080 | Unspecified |
| mmu-miR-30c-5p | MIMAT0000514 | Arl4c | 320982 | HITS-CLIP | 23142080 | Unspecified |
| mmu-miR-30c-5p | MIMAT0000514 | Arl6ip1 | 54208 | HITS-CLIP | 37986224 | Unspecified |
| mmu-miR-30c-5p | MIMAT0000514 | Arl6ip5 | 65106 | HITS-CLIP | 37986224\|23142080 | Unspecified |
| mmu-miR-30c-5p | MIMAT0000514 | Arl6ip6 | 65103 | HITS-CLIP | 37986224 | Unspecified |
| mmu-miR-30c-5p | MIMAT0000514 | Arl8b | 67166 | HITS-CLIP | 37986224\|23597149\|23142080 | Unspecified |
| mmu-miR-30c-5p | MIMAT0000514 | Arntl | 11865 | HITS-CLIP | 37986224\|23142080 | Unspecified |
| mmu-miR-30c-5p | MIMAT0000514 | Arpc5 | 67771 | HITS-CLIP | 23142080 | Unspecified |
| mmu-miR-30c-5p | MIMAT0000514 | Arpp19 | 59046 | HITS-CLIP | 37986224\|23142080 | Unspecified |
| mmu-miR-30c-5p | MIMAT0000514 | Arrdc3 | 105171 | HITS-CLIP | 37986224 | Unspecified |
| mmu-miR-30c-5p | MIMAT0000514 | Arrdc4 | 66412 | HITS-CLIP | 23142080 | Unspecified |
| mmu-miR-30c-5p | MIMAT0000514 | Arvcf | 11877 | HITS-CLIP | 23597149 | Unspecified |
| mmu-miR-30c-5p | MIMAT0000514 | As3mt | 57344 | HITS-CLIP | 23597149 | Unspecified |
| mmu-miR-30c-5p | MIMAT0000514 | Asah1 | 11886 | HITS-CLIP | 37986224\|23597149\|23142080 | Unspecified |
| mmu-miR-30c-5p | MIMAT0000514 | Asap1 | 13196 | HITS-CLIP | 23142080 | Unspecified |
| mmu-miR-30c-5p | MIMAT0000514 | Asb3 | 65257 | HITS-CLIP | 37986224\|23597149\|23142080 | Unspecified |
| mmu-miR-30c-5p | MIMAT0000514 | Asb7 | 117589 | HITS-CLIP | 23597149 | Unspecified |
| mmu-miR-30c-5p | MIMAT0000514 | Asf1a | 66403 | HITS-CLIP | 23142080 | Unspecified |
| mmu-miR-30c-5p | MIMAT0000514 | Asf1b | 66929 | qCLASH | 31393953 | Unspecified |
| mmu-miR-30c-5p | MIMAT0000514 | Ash1l | 192195 | HITS-CLIP | 37986224\|23142080\|23597149 | Unspecified |
| mmu-miR-30c-5p | MIMAT0000514 | Aspm | 12316 | HITS-CLIP | 23142080 | Unspecified |
| mmu-miR-30c-5p | MIMAT0000514 | Asxl1 | 228790 | HITS-CLIP | 23142080 | Unspecified |
| mmu-miR-30c-5p | MIMAT0000514 | Atad2 | 70472 | HITS-CLIP | 37986224\|23142080 | Unspecified |
| mmu-miR-30c-5p | MIMAT0000514 | Atad2b | 320817 | HITS-CLIP | 37986224\|19536157\|23597149\|2575[illegible] | Unspecified |
| mmu-miR-30c-5p | MIMAT0000514 | Atad5 | 237877 | HITS-CLIP | 37986224\|23142080 | Unspecified |
| mmu-miR-30c-5p | MIMAT0000514 | Atf1 | 11908 | HITS-CLIP | 23597149\|23142080 | Unspecified |
| mmu-miR-30c-5p | MIMAT0000514 | Atf2 | 11909 | HITS-CLIP | 37986224\|23142080 | Unspecified |
| mmu-miR-30c-5p | MIMAT0000514 | Atf4 | 11911 | HITS-CLIP | 31433990 | Unspecified |
| mmu-miR-30c-5p | MIMAT0000514 | Atf6 | 226641 | HITS-CLIP | 37986224\|23597149\|23142080 | Unspecified |
| mmu-miR-30c-5p | MIMAT0000514 | Atf7ip | 54343 | HITS-CLIP | 37986224\|23142080 | Unspecified |
| mmu-miR-30c-5p | MIMAT0000514 | Atg12 | 67526 | HITS-CLIP | 37986224\|23142080 | Unspecified |
| mmu-miR-30c-5p | MIMAT0000514 | Atg2b | 76559 | HITS-CLIP | 37986224\|23142080 | Unspecified |
| mmu-miR-30c-5p | MIMAT0000514 | Atl2 | 56298 | HITS-CLIP | 37986224\|19536157\|23142080 | Unspecified |
| mmu-miR-30c-5p | MIMAT0000514 | Atp11a | 50770 | HITS-CLIP | 23142080 | Unspecified |
| mmu-miR-30c-5p | MIMAT0000514 | Atp11b | 76295 | HITS-CLIP | 37986224 | Unspecified |
| mmu-miR-30c-5p | MIMAT0000514 | Atp11c | 320940 | HITS-CLIP | 23597149\|23142080 | Unspecified |
| mmu-miR-30c-5p | MIMAT0000514 | Atp13a3 | 224088 | HITS-CLIP | 37986224\|23142080 | Unspecified |
| mmu-miR-30c-5p | MIMAT0000514 | Atp2a2 | 11938 | HITS-CLIP | 37986224\|23597149\|19536157\|2314[illegible] | Unspecified |
| mmu-miR-30c-5p | MIMAT0000514 | Atp2b1 | 67972 | HITS-CLIP | 37986224\|23142080 | Unspecified |

| | | | | | | |
|---|---|---|---|---|---|---|
| mmu-miR-30c-5p | MIMAT0000514 | Atp5a1 | 11946 | HITS-CLIP | 25753418 | Unspecified |
| mmu-miR-30c-5p | MIMAT0000514 | Atp5pb | 515 | HITS-CLIP | 25753418 | Unspecified |
| mmu-miR-30c-5p | MIMAT0000514 | Atp6v0a1 | 11975 | HITS-CLIP | 23597149\|23142080 | Unspecified |
| mmu-miR-30c-5p | MIMAT0000514 | Atp6v0d1 | 11972 | HITS-CLIP | 37986224\|23597149\|23142080 | Unspecified |
| mmu-miR-30c-5p | MIMAT0000514 | Atp6v1c1 | 66335 | HITS-CLIP | 37986224 | Unspecified |
| mmu-miR-30c-5p | MIMAT0000514 | Atp8a1 | 11980 | HITS-CLIP | 23142080 | Unspecified |
| mmu-miR-30c-5p | MIMAT0000514 | Atp8b1 | 54670 | HITS-CLIP | 37986224 | Unspecified |
| mmu-miR-30c-5p | MIMAT0000514 | Atp8b4 | 241633 | HITS-CLIP | 37986224\|23142080 | Unspecified |
| mmu-miR-30c-5p | MIMAT0000514 | Atr | 245000 | HITS-CLIP | 23142080 | Unspecified |
| mmu-miR-30c-5p | MIMAT0000514 | Atrx | 22589 | HITS-CLIP | 23142080 | Unspecified |
| mmu-miR-30c-5p | MIMAT0000514 | Atxn1 | 20238 | HITS-CLIP | 23142080 | Unspecified |
| mmu-miR-30c-5p | MIMAT0000514 | Atxn1l | 52335 | HITS-CLIP | 37986224 | Unspecified |
| mmu-miR-30c-5p | MIMAT0000514 | Atxn2 | 20239 | HITS-CLIP | 25753418 | Unspecified |
| mmu-miR-30c-5p | MIMAT0000514 | Atxn7l1 | 380753 | HITS-CLIP | 37986224 | Unspecified |
| mmu-miR-30c-5p | MIMAT0000514 | Atxn7l3 | 217218 | HITS-CLIP | 37986224 | Unspecified |
| mmu-miR-30c-5p | MIMAT0000514 | Auts2 | 319974 | HITS-CLIP | 37986224 | Unspecified |
| mmu-miR-30c-5p | MIMAT0000514 | Avl9 | 78937 | HITS-CLIP | 37986224\|23597149\|23142080 | Unspecified |
| mmu-miR-30c-5p | MIMAT0000514 | Azin1 | 54375 | HITS-CLIP | 37986224\|19536157\|23597149\|23142 | Unspecified |
| mmu-miR-30c-5p | MIMAT0000514 | B230219D22Rik | 78521 | HITS-CLIP | 23142080 | Unspecified |
| mmu-miR-30c-5p | MIMAT0000514 | B2m | 12010 | HITS-CLIP | 37986224 | Unspecified |
| mmu-miR-30c-5p | MIMAT0000514 | B3galnt1 | 26879 | HITS-CLIP | 37986224 | Unspecified |
| mmu-miR-30c-5p | MIMAT0000514 | B3galt1 | 26877 | HITS-CLIP | 37986224 | Unspecified |
| mmu-miR-30c-5p | MIMAT0000514 | B3gat1 | 76898 | HITS-CLIP | 37986224\|19536157 | Unspecified |
| mmu-miR-30c-5p | MIMAT0000514 | B3gnt2 | 53625 | HITS-CLIP | 23142080 | Unspecified |
| mmu-miR-30c-5p | MIMAT0000514 | B4galt1 | 14595 | HITS-CLIP | 37986224\|23597149\|23142080 | Unspecified |
| mmu-miR-30c-5p | MIMAT0000514 | B4galt5 | 56336 | HITS-CLIP | 37986224\|23142080 | Unspecified |
| mmu-miR-30c-5p | MIMAT0000514 | B4galt6 | 56386 | HITS-CLIP | 37986224 | Unspecified |
| mmu-miR-30c-5p | MIMAT0000514 | BC005561 | 100042165 | HITS-CLIP | 23142080 | Unspecified |
| mmu-miR-30c-5p | MIMAT0000514 | BC031181 | 407819 | HITS-CLIP | 25753418 | Unspecified |
| mmu-miR-30c-5p | MIMAT0000514 | BC055324 | 381306 | HITS-CLIP | 23142080 | Unspecified |
| mmu-miR-30c-5p | MIMAT0000514 | Bach1 | 12013 | HITS-CLIP | 37986224\|23142080 | Unspecified |
| mmu-miR-30c-5p | MIMAT0000514 | Bach2 | 12014 | HITS-CLIP | 37986224\|23142080 | Unspecified |
| mmu-miR-30c-5p | MIMAT0000514 | Banp | 53325 | HITS-CLIP | 23142080 | Unspecified |
| mmu-miR-30c-5p | MIMAT0000514 | Bax | 12028 | qCLASH | 31393953 | Unspecified |
| mmu-miR-30c-5p | MIMAT0000514 | Baz1b | 22385 | HITS-CLIP | 23142080 | Unspecified |
| mmu-miR-30c-5p | MIMAT0000514 | Baz2a | 116848 | HITS-CLIP | 23142080 | Unspecified |
| mmu-miR-30c-5p | MIMAT0000514 | Baz2b | 407823 | HITS-CLIP | 23597149\|23142080 | Unspecified |
| mmu-miR-30c-5p | MIMAT0000514 | Bbs5 | 72569 | HITS-CLIP | 37986224\|23142080 | Unspecified |
| mmu-miR-30c-5p | MIMAT0000514 | Bbx | 70508 | HITS-CLIP | 23142080 | Unspecified |

| | | | | | | |
|---|---|---|---|---|---|---|
| mmu-miR-30c-5p | MIMAT0000514 | Bcl10 | 12042 | HITS-CLIP | 23597149\|23142080 | Unspecified |
| mmu-miR-30c-5p | MIMAT0000514 | Bcl11a | 14025 | HITS-CLIP | 23142080 | Unspecified |
| mmu-miR-30c-5p | MIMAT0000514 | Bcl11b | 58208 | HITS-CLIP | 37986224 | Unspecified |
| mmu-miR-30c-5p | MIMAT0000514 | Bcl2 | 12043 | HITS-CLIP | 37986224\|23142080 | Unspecified |
| mmu-miR-30c-5p | MIMAT0000514 | Bcl2l11 | 12125 | HITS-CLIP | 37986224\|23142080 | Unspecified |
| mmu-miR-30c-5p | MIMAT0000514 | Bcl2l13 | 94044 | HITS-CLIP | 23142080 | Unspecified |
| mmu-miR-30c-5p | MIMAT0000514 | Bcl6b | 12029 | HITS-CLIP | 37986224 | Unspecified |
| mmu-miR-30c-5p | MIMAT0000514 | Bcl9 | 77578 | HITS-CLIP | 37986224\|23142080 | Unspecified |
| mmu-miR-30c-5p | MIMAT0000514 | Bcl9l | 80288 | HITS-CLIP | 23142080 | Unspecified |
| mmu-miR-30c-5p | MIMAT0000514 | Bclaf1 | 72567 | HITS-CLIP | 37986224\|23142080 | Unspecified |
| mmu-miR-30c-5p | MIMAT0000514 | Bcor | 71458 | HITS-CLIP | 23597149\|23142080 | Unspecified |
| mmu-miR-30c-5p | MIMAT0000514 | Bcorl1 | 320376 | HITS-CLIP | 25753418 | Unspecified |
| mmu-miR-30c-5p | MIMAT0000514 | Bdp1 | 544971 | HITS-CLIP | 37986224\|23142080 | Unspecified |
| mmu-miR-30c-5p | MIMAT0000514 | Becn1 | 56208 | HITS-CLIP | 23142080 | Unspecified |
| mmu-miR-30c-5p | MIMAT0000514 | Bend5 | 67621 | HITS-CLIP | 23142080 | Unspecified |
| mmu-miR-30c-5p | MIMAT0000514 | Bhlhe40 | 20893 | HITS-CLIP | 37986224\|25753418\|23142080 | Unspecified |
| mmu-miR-30c-5p | MIMAT0000514 | Bhlhe41 | 79362 | HITS-CLIP | 23142080 | Unspecified |
| mmu-miR-30c-5p | MIMAT0000514 | Bmp2k | 140780 | HITS-CLIP | 37986224\|23597149\|23142080 | Unspecified |
| mmu-miR-30c-5p | MIMAT0000514 | Bmp5 | 12160 | HITS-CLIP | 37986224 | Unspecified |
| mmu-miR-30c-5p | MIMAT0000514 | Bmpr1a | 12166 | HITS-CLIP | 37986224 | Unspecified |
| mmu-miR-30c-5p | MIMAT0000514 | Bmpr2 | 12168 | HITS-CLIP | 37986224\|23597149\|23142080 | Unspecified |
| mmu-miR-30c-5p | MIMAT0000514 | Bmt2 | 101148 | HITS-CLIP | 23142080 | Unspecified |
| mmu-miR-30c-5p | MIMAT0000514 | Bnip3l | 12177 | HITS-CLIP | 23142080 | Unspecified |
| mmu-miR-30c-5p | MIMAT0000514 | Bpnt1 | 23827 | HITS-CLIP | 23597149 | Unspecified |
| mmu-miR-30c-5p | MIMAT0000514 | Bpnt2 | 54928 | HITS-CLIP | 25753418 | Unspecified |
| mmu-miR-30c-5p | MIMAT0000514 | Brap | 72399 | HITS-CLIP | 23597149 | Unspecified |
| mmu-miR-30c-5p | MIMAT0000514 | Brca1 | 12189 | HITS-CLIP | 23142080 | Unspecified |
| mmu-miR-30c-5p | MIMAT0000514 | Brd1 | 223770 | HITS-CLIP | 23597149\|23142080 | Unspecified |
| mmu-miR-30c-5p | MIMAT0000514 | Brd2 | 14312 | HITS-CLIP | 23597149\|23142080 | Unspecified |
| mmu-miR-30c-5p | MIMAT0000514 | Brd3 | 67382 | HITS-CLIP | 37986224\|23597149\|23142080 | Unspecified |
| mmu-miR-30c-5p | MIMAT0000514 | Brd4 | 57261 | HITS-CLIP | 37986224\|23142080\|19536157 | Unspecified |
| mmu-miR-30c-5p | MIMAT0000514 | Brdt | 114642 | HITS-CLIP | 23142080 | Unspecified |
| mmu-miR-30c-5p | MIMAT0000514 | Brinp2 | 240843 | HITS-CLIP | 37986224\|19536157 | Unspecified |
| mmu-miR-30c-5p | MIMAT0000514 | Brix1 | 67832 | HITS-CLIP | 37986224\|23597149\|23142080 | Unspecified |
| mmu-miR-30c-5p | MIMAT0000514 | Brwd1 | 93871 | HITS-CLIP | 37986224\|23142080\|23597149 | Unspecified |
| mmu-miR-30c-5p | MIMAT0000514 | Brwd3 | 382236 | HITS-CLIP | 37986224\|23142080\|19536157 | Unspecified |
| mmu-miR-30c-5p | MIMAT0000514 | Btbd10 | 68815 | HITS-CLIP | 23597149\|23142080 | Unspecified |
| mmu-miR-30c-5p | MIMAT0000514 | Btbd7 | 238386 | HITS-CLIP | 37986224\|23142080\|23597149 | Unspecified |
| mmu-miR-30c-5p | MIMAT0000514 | Btbd8 | 100503185 | HITS-CLIP | 23142080 | Unspecified |

| | | | | | | |
|---|---|---|---|---|---|---|
| mmu-miR-30c-5p | MIMAT0000514 | Btbd9 | 224671 | HITS-CLIP | 23142080 | Unspecified |
| mmu-miR-30c-5p | MIMAT0000514 | Btg2 | 12227 | HITS-CLIP | 23597149\|23142080\|25753418 | Unspecified |
| mmu-miR-30c-5p | MIMAT0000514 | Btrc | 12234 | HITS-CLIP | 37986224\|23597149 | Unspecified |
| mmu-miR-30c-5p | MIMAT0000514 | Bysl | 53414 | HITS-CLIP | 25753418 | Unspecified |
| mmu-miR-30c-5p | MIMAT0000514 | C1d | 57316 | HITS-CLIP | 23142080 | Unspecified |
| mmu-miR-30c-5p | MIMAT0000514 | C1galt1 | 94192 | HITS-CLIP | 23142080 | Unspecified |
| mmu-miR-30c-5p | MIMAT0000514 | C2cd3 | 277939 | HITS-CLIP | 25753418 | Unspecified |
| mmu-miR-30c-5p | MIMAT0000514 | C6 | 12274 | HITS-CLIP | 23597149 | Unspecified |
| mmu-miR-30c-5p | MIMAT0000514 | C77080 | 97130 | HITS-CLIP | 37986224\|23597149 | Unspecified |
| mmu-miR-30c-5p | MIMAT0000514 | C9orf72 | 203228 | HITS-CLIP | 23142080\|25753418 | Unspecified |
| mmu-miR-30c-5p | MIMAT0000514 | Cab39 | 12283 | HITS-CLIP | 37986224 | Unspecified |
| mmu-miR-30c-5p | MIMAT0000514 | Cabin1 | 104248 | HITS-CLIP | 23142080 | Unspecified |
| mmu-miR-30c-5p | MIMAT0000514 | Cacfd1 | 381356 | HITS-CLIP | 23142080 | Unspecified |
| mmu-miR-30c-5p | MIMAT0000514 | Cad | 69719 | HITS-CLIP | 23597149 | Unspecified |
| mmu-miR-30c-5p | MIMAT0000514 | Cadm1 | 54725 | HITS-CLIP | 23597149 | Unspecified |
| mmu-miR-30c-5p | MIMAT0000514 | Calcoco1 | 67488 | HITS-CLIP | 25753418 | Unspecified |
| mmu-miR-30c-5p | MIMAT0000514 | Cald1 | 109624 | HITS-CLIP | 37986224 | Unspecified |
| mmu-miR-30c-5p | MIMAT0000514 | Camk2a | 12322 | HITS-CLIP | 37986224 | Unspecified |
| mmu-miR-30c-5p | MIMAT0000514 | Camk2d | 108058 | HITS-CLIP | 37986224\|25753418 | Unspecified |
| mmu-miR-30c-5p | MIMAT0000514 | Camk2n1 | 66259 | HITS-CLIP | 37986224\|23597149\|19536157\|23142 | Unspecified |
| mmu-miR-30c-5p | MIMAT0000514 | Camk2n2 | 73047 | HITS-CLIP | 37986224 | Unspecified |
| mmu-miR-30c-5p | MIMAT0000514 | Camk4 | 12326 | HITS-CLIP | 23142080 | Unspecified |
| mmu-miR-30c-5p | MIMAT0000514 | Camsap1 | 227634 | HITS-CLIP | 37986224 | Unspecified |
| mmu-miR-30c-5p | MIMAT0000514 | Camsap2 | 67886 | HITS-CLIP | 37986224 | Unspecified |
| mmu-miR-30c-5p | MIMAT0000514 | Cant1 | 76025 | HITS-CLIP | 37986224 | Unspecified |
| mmu-miR-30c-5p | MIMAT0000514 | Canx | 12330 | HITS-CLIP | 37986224\|23142080\|19536157 | Unspecified |
| mmu-miR-30c-5p | MIMAT0000514 | Cap1 | 12331 | HITS-CLIP | 23142080 | Unspecified |
| mmu-miR-30c-5p | MIMAT0000514 | Capn7 | 12339 | HITS-CLIP | 37986224\|23142080\|19536157 | Unspecified |
| mmu-miR-30c-5p | MIMAT0000514 | Capza2 | 12343 | HITS-CLIP | 23597149\|23142080\|25753418 | Unspecified |
| mmu-miR-30c-5p | MIMAT0000514 | Carf | 241066 | HITS-CLIP | 37986224 | Unspecified |
| mmu-miR-30c-5p | MIMAT0000514 | Cars | 27267 | HITS-CLIP | 37986224\|23142080 | Unspecified |
| mmu-miR-30c-5p | MIMAT0000514 | Cask | 12361 | HITS-CLIP | 19536157 | Unspecified |
| mmu-miR-30c-5p | MIMAT0000514 | Casp3 | 12367 | HITS-CLIP | 23142080 | Unspecified |
| mmu-miR-30c-5p | MIMAT0000514 | Casp6 | 12368 | HITS-CLIP | 37986224\|23142080 | Unspecified |
| mmu-miR-30c-5p | MIMAT0000514 | Cast | 12380 | HITS-CLIP | 37986224 | Unspecified |
| mmu-miR-30c-5p | MIMAT0000514 | Cat | 12359 | HITS-CLIP | 37986224 | Unspecified |
| mmu-miR-30c-5p | MIMAT0000514 | Cbfb | 12400 | HITS-CLIP | 37986224\|23597149\|23142080 | Unspecified |
| mmu-miR-30c-5p | MIMAT0000514 | Cblb | 208650 | HITS-CLIP | 37986224\|23597149\|23142080 | Unspecified |
| mmu-miR-30c-5p | MIMAT0000514 | Cbll1 | 104836 | HITS-CLIP | 23142080\|25753418 | Unspecified |

| | | | | | | | |
|---|---|---|---|---|---|---|---|
| mmu-miR-30c-5p | MIMAT0000514 | Cbr4 | 234309 | HITS-CLIP | | 23142080 | Unspecified |
| mmu-miR-30c-5p | MIMAT0000514 | Cbx2 | 12416 | HITS-CLIP | 37986224\|23142080 | | Unspecified |
| mmu-miR-30c-5p | MIMAT0000514 | Cbx5 | 12419 | HITS-CLIP | | 37986224 | Unspecified |
| mmu-miR-30c-5p | MIMAT0000514 | Cbx6 | 494448 | HITS-CLIP//qCLASH | 19536157\|31393953\|23142080 | | Unspecified |
| mmu-miR-30c-5p | MIMAT0000514 | Cc2d1a | 212139 | HITS-CLIP | | 37986224 | Unspecified |
| mmu-miR-30c-5p | MIMAT0000514 | Ccar1 | 67500 | HITS-CLIP | | 37986224 | Unspecified |
| mmu-miR-30c-5p | MIMAT0000514 | Ccdc186 | 55088 | HITS-CLIP | | 23142080 | Unspecified |
| mmu-miR-30c-5p | MIMAT0000514 | Ccdc27 | 381580 | qCLASH | | 31393953 | Unspecified |
| mmu-miR-30c-5p | MIMAT0000514 | Ccdc28a | 215814 | HITS-CLIP | | 25753418 | Unspecified |
| mmu-miR-30c-5p | MIMAT0000514 | Ccdc34 | 68201 | HITS-CLIP | | 23142080 | Unspecified |
| mmu-miR-30c-5p | MIMAT0000514 | Ccdc47 | 67163 | HITS-CLIP | 37986224\|23597149\|23142080 | | Unspecified |
| mmu-miR-30c-5p | MIMAT0000514 | Ccdc6 | 76551 | HITS-CLIP | 37986224\|23142080 | | Unspecified |
| mmu-miR-30c-5p | MIMAT0000514 | Ccdc71l | 72123 | HITS-CLIP | 37986224\|23142080\|23597149\|1953€ | | Unspecified |
| mmu-miR-30c-5p | MIMAT0000514 | Ccdc82 | 66396 | HITS-CLIP | | 23142080 | Unspecified |
| mmu-miR-30c-5p | MIMAT0000514 | Ccdc97 | 52132 | HITS-CLIP | | 23142080 | Unspecified |
| mmu-miR-30c-5p | MIMAT0000514 | Ccl3 | 20302 | HITS-CLIP | | 37986224 | Unspecified |
| mmu-miR-30c-5p | MIMAT0000514 | Ccm2 | 216527 | HITS-CLIP | 37986224\|23142080 | | Unspecified |
| mmu-miR-30c-5p | MIMAT0000514 | Ccn2 | 1490 | HITS-CLIP | | 25753418 | Unspecified |
| mmu-miR-30c-5p | MIMAT0000514 | Ccnb2 | 12442 | HITS-CLIP | | 37986224 | Unspecified |
| mmu-miR-30c-5p | MIMAT0000514 | Ccnd2 | 12444 | HITS-CLIP | 37986224\|23142080\|25753418 | | Unspecified |
| mmu-miR-30c-5p | MIMAT0000514 | Ccne1 | 12447 | HITS-CLIP | | 37986224 | Unspecified |
| mmu-miR-30c-5p | MIMAT0000514 | Ccne2 | 12448 | Luciferase assay//qRT-PCR//HITS-CLIP//Lu | 24555094\|23142080\|23597149\|1953€ | | Unspecified |
| mmu-miR-30c-5p | MIMAT0000514 | Ccnf | 12449 | HITS-CLIP | 37986224\|23597149\|23142080 | | Unspecified |
| mmu-miR-30c-5p | MIMAT0000514 | Ccnjl | 380694 | HITS-CLIP | 37986224\|19536157 | | Unspecified |
| mmu-miR-30c-5p | MIMAT0000514 | Ccnk | 12454 | HITS-CLIP | 37986224\|23142080\|23597149 | | Unspecified |
| mmu-miR-30c-5p | MIMAT0000514 | Ccnl1 | 56706 | HITS-CLIP | | 23142080 | Unspecified |
| mmu-miR-30c-5p | MIMAT0000514 | Ccnl2 | 56036 | qCLASH | | 31393953 | Unspecified |
| mmu-miR-30c-5p | MIMAT0000514 | Ccnt2 | 72949 | HITS-CLIP | 37986224\|23142080 | | Unspecified |
| mmu-miR-30c-5p | MIMAT0000514 | Ccny | 67974 | HITS-CLIP | 25753418\|23142080 | | Unspecified |
| mmu-miR-30c-5p | MIMAT0000514 | Cct5 | 12465 | HITS-CLIP | | 23142080 | Unspecified |
| mmu-miR-30c-5p | MIMAT0000514 | Cct7 | 12468 | HITS-CLIP | 37986224\|23597149 | | Unspecified |
| mmu-miR-30c-5p | MIMAT0000514 | Cct8 | 12469 | HITS-CLIP | | 23142080 | Unspecified |
| mmu-miR-30c-5p | MIMAT0000514 | Cd164 | 53599 | HITS-CLIP | 37986224\|23142080 | | Unspecified |
| mmu-miR-30c-5p | MIMAT0000514 | Cd28 | 12487 | HITS-CLIP | | 23142080 | Unspecified |
| mmu-miR-30c-5p | MIMAT0000514 | Cd34 | 12490 | HITS-CLIP | | 25753418 | Unspecified |
| mmu-miR-30c-5p | MIMAT0000514 | Cd4 | 12504 | HITS-CLIP | | 37986224 | Unspecified |
| mmu-miR-30c-5p | MIMAT0000514 | Cd47 | 16423 | HITS-CLIP | 37986224\|23142080 | | Unspecified |
| mmu-miR-30c-5p | MIMAT0000514 | Cd53 | 12508 | HITS-CLIP | | 23142080 | Unspecified |
| mmu-miR-30c-5p | MIMAT0000514 | Cd69 | 12515 | HITS-CLIP | | 23142080 | Unspecified |

| | | | | | | |
|---|---|---|---|---|---|---|
| mmu-miR-30c-5p | MIMAT0000514 | Cd93 | 17064 | HITS-CLIP | 37986224 | Unspecified |
| mmu-miR-30c-5p | MIMAT0000514 | Cdc123 | 98828 | HITS-CLIP | 23597149\|25753418 | Unspecified |
| mmu-miR-30c-5p | MIMAT0000514 | Cdc16 | 69957 | HITS-CLIP | 23142080 | Unspecified |
| mmu-miR-30c-5p | MIMAT0000514 | Cdc27 | 217232 | HITS-CLIP | 37986224 | Unspecified |
| mmu-miR-30c-5p | MIMAT0000514 | Cdc37l1 | 67072 | HITS-CLIP | 37986224\|23142080 | Unspecified |
| mmu-miR-30c-5p | MIMAT0000514 | Cdc40 | 71713 | HITS-CLIP | 23142080 | Unspecified |
| mmu-miR-30c-5p | MIMAT0000514 | Cdc42 | 12540 | HITS-CLIP | 23142080 | Unspecified |
| mmu-miR-30c-5p | MIMAT0000514 | Cdc42bpb | 217866 | HITS-CLIP | 25753418 | Unspecified |
| mmu-miR-30c-5p | MIMAT0000514 | Cdc5l | 71702 | HITS-CLIP | 37986224 | Unspecified |
| mmu-miR-30c-5p | MIMAT0000514 | Cdc73 | 214498 | HITS-CLIP | 23142080 | Unspecified |
| mmu-miR-30c-5p | MIMAT0000514 | Cdca2 | 108912 | HITS-CLIP | 23142080 | Unspecified |
| mmu-miR-30c-5p | MIMAT0000514 | Cdca3 | 14793 | HITS-CLIP | 23597149 | Unspecified |
| mmu-miR-30c-5p | MIMAT0000514 | Cdh2 | 12558 | HITS-CLIP | 37986224\|23597149\|19536157 | Unspecified |
| mmu-miR-30c-5p | MIMAT0000514 | Cdk1 | 12534 | HITS-CLIP | 37986224 | Unspecified |
| mmu-miR-30c-5p | MIMAT0000514 | Cdk8 | 264064 | qCLASH | 31393953 | Unspecified |
| mmu-miR-30c-5p | MIMAT0000514 | Cds2 | 110911 | HITS-CLIP | 37986224 | Unspecified |
| mmu-miR-30c-5p | MIMAT0000514 | Cecr2 | 330409 | HITS-CLIP | 37986224 | Unspecified |
| mmu-miR-30c-5p | MIMAT0000514 | Celf1 | 13046 | HITS-CLIP | 23597149\|23142080 | Unspecified |
| mmu-miR-30c-5p | MIMAT0000514 | Celf2 | 14007 | HITS-CLIP | 37986224\|23142080\|19536157\|25753 | Unspecified |
| mmu-miR-30c-5p | MIMAT0000514 | Celf3 | 78784 | HITS-CLIP | 37986224\|19536157 | Unspecified |
| mmu-miR-30c-5p | MIMAT0000514 | Celf4 | 108013 | HITS-CLIP | 19536157 | Unspecified |
| mmu-miR-30c-5p | MIMAT0000514 | Cenpb | 12616 | HITS-CLIP | 25753418 | Unspecified |
| mmu-miR-30c-5p | MIMAT0000514 | Cenpq | 83815 | HITS-CLIP | 37986224 | Unspecified |
| mmu-miR-30c-5p | MIMAT0000514 | Cep135 | 381644 | HITS-CLIP | 23142080 | Unspecified |
| mmu-miR-30c-5p | MIMAT0000514 | Cep170 | 545389 | HITS-CLIP | 37986224\|23142080 | Unspecified |
| mmu-miR-30c-5p | MIMAT0000514 | Cep170b | 217882 | HITS-CLIP | 23142080\|23597149\|25083871 | Unspecified |
| mmu-miR-30c-5p | MIMAT0000514 | Cep350 | 74081 | HITS-CLIP | 37986224\|19536157\|23597149\|25753 | Unspecified |
| mmu-miR-30c-5p | MIMAT0000514 | Cep41 | 83922 | HITS-CLIP | 25753418 | Unspecified |
| mmu-miR-30c-5p | MIMAT0000514 | Cep68 | 216543 | HITS-CLIP | 25753418 | Unspecified |
| mmu-miR-30c-5p | MIMAT0000514 | Cep76 | 225659 | HITS-CLIP | 37986224 | Unspecified |
| mmu-miR-30c-5p | MIMAT0000514 | Cept1 | 99712 | HITS-CLIP | 37986224\|23142080 | Unspecified |
| mmu-miR-30c-5p | MIMAT0000514 | Cers5 | 71949 | HITS-CLIP | 23142080 | Unspecified |
| mmu-miR-30c-5p | MIMAT0000514 | Ces1e | 13897 | HITS-CLIP | 37986224 | Unspecified |
| mmu-miR-30c-5p | MIMAT0000514 | Ces1f | 234564 | HITS-CLIP | 23597149 | Unspecified |
| mmu-miR-30c-5p | MIMAT0000514 | Ces2g | 72361 | HITS-CLIP | 37986224 | Unspecified |
| mmu-miR-30c-5p | MIMAT0000514 | Cfh | 12628 | HITS-CLIP | 23597149 | Unspecified |
| mmu-miR-30c-5p | MIMAT0000514 | Cfi | 12630 | HITS-CLIP | 37986224\|23597149 | Unspecified |
| mmu-miR-30c-5p | MIMAT0000514 | Cfl2 | 12632 | HITS-CLIP | 37986224\|23142080 | Unspecified |
| mmu-miR-30c-5p | MIMAT0000514 | Cggbp1 | 106143 | HITS-CLIP | 23142080 | Unspecified |

| | | | | | | |
|---|---|---|---|---|---|---|
| mmu-miR-30c-5p | MIMAT0000514 | Chd1 | 12648 | HITS-CLIP | 37986224\|23597149\|23142080 | Unspecified |
| mmu-miR-30c-5p | MIMAT0000514 | Chd2 | 244059 | HITS-CLIP | 37986224\|23142080\|23597149 | Unspecified |
| mmu-miR-30c-5p | MIMAT0000514 | Chd7 | 320790 | HITS-CLIP | 37986224\|23597149\|31433990\|19536 | Unspecified |
| mmu-miR-30c-5p | MIMAT0000514 | Chd8 | 67772 | HITS-CLIP | 23142080 | Unspecified |
| mmu-miR-30c-5p | MIMAT0000514 | Chd9 | 109151 | HITS-CLIP | 37986224\|23142080 | Unspecified |
| mmu-miR-30c-5p | MIMAT0000514 | Chdh | 218865 | HITS-CLIP | 37986224 | Unspecified |
| mmu-miR-30c-5p | MIMAT0000514 | Chka | 12660 | qCLASH | 31393953 | Unspecified |
| mmu-miR-30c-5p | MIMAT0000514 | Chl1 | 12661 | HITS-CLIP | 37986224\|19536157 | Unspecified |
| mmu-miR-30c-5p | MIMAT0000514 | Chmp4b | 75608 | HITS-CLIP | 23142080 | Unspecified |
| mmu-miR-30c-5p | MIMAT0000514 | Chp1 | 56398 | HITS-CLIP | 37986224 | Unspecified |
| mmu-miR-30c-5p | MIMAT0000514 | Chst2 | 54371 | HITS-CLIP | 37986224\|23597149\|19536157\|23142 | Unspecified |
| mmu-miR-30c-5p | MIMAT0000514 | Chsy1 | 269941 | HITS-CLIP | 37986224\|19536157\|23142080 | Unspecified |
| mmu-miR-30c-5p | MIMAT0000514 | Ciao2b | 51647 | HITS-CLIP | 23142080 | Unspecified |
| mmu-miR-30c-5p | MIMAT0000514 | Cip2a | 57650 | HITS-CLIP | 23142080 | Unspecified |
| mmu-miR-30c-5p | MIMAT0000514 | Cipc | 217732 | HITS-CLIP | 25753418 | Unspecified |
| mmu-miR-30c-5p | MIMAT0000514 | Cited4 | 56222 | HITS-CLIP | 25753418 | Unspecified |
| mmu-miR-30c-5p | MIMAT0000514 | Ckap5 | 75786 | HITS-CLIP | 23142080 | Unspecified |
| mmu-miR-30c-5p | MIMAT0000514 | Clcf1 | 56708 | HITS-CLIP | 23142080 | Unspecified |
| mmu-miR-30c-5p | MIMAT0000514 | Clcn3 | 12725 | HITS-CLIP | 23597149\|19536157\|23142080 | Unspecified |
| mmu-miR-30c-5p | MIMAT0000514 | Cldnd1 | 224250 | HITS-CLIP | 23597149\|19536157\|23142080 | Unspecified |
| mmu-miR-30c-5p | MIMAT0000514 | Clic4 | 29876 | HITS-CLIP | 37986224 | Unspecified |
| mmu-miR-30c-5p | MIMAT0000514 | Clic5 | 224796 | HITS-CLIP | 25753418 | Unspecified |
| mmu-miR-30c-5p | MIMAT0000514 | Clint1 | 216705 | HITS-CLIP | 23142080 | Unspecified |
| mmu-miR-30c-5p | MIMAT0000514 | Clip1 | 56430 | HITS-CLIP | 19536157\|23142080 | Unspecified |
| mmu-miR-30c-5p | MIMAT0000514 | Clip3 | 76686 | HITS-CLIP | 37986224 | Unspecified |
| mmu-miR-30c-5p | MIMAT0000514 | Clptm1l | 218335 | HITS-CLIP | 23597149 | Unspecified |
| mmu-miR-30c-5p | MIMAT0000514 | Clpx | 270166 | HITS-CLIP | 37986224 | Unspecified |
| mmu-miR-30c-5p | MIMAT0000514 | Cltc | 67300 | HITS-CLIP | 25753418 | Unspecified |
| mmu-miR-30c-5p | MIMAT0000514 | Clybl | 69634 | HITS-CLIP | 23597149\|25753418 | Unspecified |
| mmu-miR-30c-5p | MIMAT0000514 | Cmah | 12763 | HITS-CLIP | 37986224 | Unspecified |
| mmu-miR-30c-5p | MIMAT0000514 | Cmas | 12764 | HITS-CLIP | 37986224 | Unspecified |
| mmu-miR-30c-5p | MIMAT0000514 | Cmip | 74440 | HITS-CLIP | 37986224\|23142080\|23597149 | Unspecified |
| mmu-miR-30c-5p | MIMAT0000514 | Cmpk1 | 66588 | qCLASH | 31393953 | Unspecified |
| mmu-miR-30c-5p | MIMAT0000514 | Cmtm4 | 97487 | HITS-CLIP | 37986224\|23142080\|23597149 | Unspecified |
| mmu-miR-30c-5p | MIMAT0000514 | Cmtm6 | 67213 | HITS-CLIP | 37986224 | Unspecified |
| mmu-miR-30c-5p | MIMAT0000514 | Cnksr2 | 245684 | HITS-CLIP | 37986224 | Unspecified |
| mmu-miR-30c-5p | MIMAT0000514 | Cnnm4 | 94220 | HITS-CLIP | 37986224\|23142080 | Unspecified |
| mmu-miR-30c-5p | MIMAT0000514 | Cnot10 | 78893 | HITS-CLIP | 23142080 | Unspecified |
| mmu-miR-30c-5p | MIMAT0000514 | Cnot11 | 52846 | HITS-CLIP | 23142080 | Unspecified |

| | | | | | | |
|---|---|---|---|---|---|---|
| mmu-miR-30c-5p | MIMAT0000514 | Cnot6 | 104625 | HITS-CLIP | 23142080 | Unspecified |
| mmu-miR-30c-5p | MIMAT0000514 | Cnot6l | 231464 | HITS-CLIP | 23142080 | Unspecified |
| mmu-miR-30c-5p | MIMAT0000514 | Cnot9 | 9125 | HITS-CLIP | 23142080 | Unspecified |
| mmu-miR-30c-5p | MIMAT0000514 | Cnr1 | 12801 | HITS-CLIP | 37986224 | Unspecified |
| mmu-miR-30c-5p | MIMAT0000514 | Cntnap1 | 53321 | HITS-CLIP | 37986224 | Unspecified |
| mmu-miR-30c-5p | MIMAT0000514 | Coa4 | 68185 | HITS-CLIP | 37986224\|23597149 | Unspecified |
| mmu-miR-30c-5p | MIMAT0000514 | Coa7 | 69893 | HITS-CLIP | 25753418 | Unspecified |
| mmu-miR-30c-5p | MIMAT0000514 | Cog2 | 76332 | HITS-CLIP | 23142080 | Unspecified |
| mmu-miR-30c-5p | MIMAT0000514 | Cog3 | 338337 | HITS-CLIP | 37986224\|23142080 | Unspecified |
| mmu-miR-30c-5p | MIMAT0000514 | Cog4 | 102339 | HITS-CLIP | 23597149 | Unspecified |
| mmu-miR-30c-5p | MIMAT0000514 | Cog7 | 233824 | HITS-CLIP | 23142080\|25753418 | Unspecified |
| mmu-miR-30c-5p | MIMAT0000514 | Col12a1 | 12816 | HITS-CLIP | 23597149 | Unspecified |
| mmu-miR-30c-5p | MIMAT0000514 | Col14a1 | 12818 | HITS-CLIP | 23597149 | Unspecified |
| mmu-miR-30c-5p | MIMAT0000514 | Col15a1 | 12819 | HITS-CLIP | 25753418 | Unspecified |
| mmu-miR-30c-5p | MIMAT0000514 | Col4a3bp | 68018 | HITS-CLIP | 37986224\|23597149\|23142080 | Unspecified |
| mmu-miR-30c-5p | MIMAT0000514 | Col9a3 | 12841 | HITS-CLIP | 37986224\|19536157 | Unspecified |
| mmu-miR-30c-5p | MIMAT0000514 | Commd3 | 12238 | HITS-CLIP | 25753418 | Unspecified |
| mmu-miR-30c-5p | MIMAT0000514 | Copa | 12847 | HITS-CLIP | 37986224\|23597149\|25753418 | Unspecified |
| mmu-miR-30c-5p | MIMAT0000514 | Copg2 | 54160 | HITS-CLIP | 23142080 | Unspecified |
| mmu-miR-30c-5p | MIMAT0000514 | Cops2 | 12848 | HITS-CLIP | 23142080 | Unspecified |
| mmu-miR-30c-5p | MIMAT0000514 | Cops9 | 150678 | qCLASH | 31393953 | Unspecified |
| mmu-miR-30c-5p | MIMAT0000514 | Coq3 | 230027 | HITS-CLIP | 37986224\|23597149\|25753418\|2314 | Unspecified |
| mmu-miR-30c-5p | MIMAT0000514 | Coq5 | 52064 | HITS-CLIP | 37986224\|19536157\|23142080 | Unspecified |
| mmu-miR-30c-5p | MIMAT0000514 | Corin | 10699 | HITS-CLIP | 25753418 | Unspecified |
| mmu-miR-30c-5p | MIMAT0000514 | Cotl1 | 72042 | HITS-CLIP | 37986224 | Unspecified |
| mmu-miR-30c-5p | MIMAT0000514 | Cox15 | 226139 | HITS-CLIP | 23597149 | Unspecified |
| mmu-miR-30c-5p | MIMAT0000514 | Cpd | 12874 | HITS-CLIP | 37986224 | Unspecified |
| mmu-miR-30c-5p | MIMAT0000514 | Cpeb2 | 231207 | HITS-CLIP | 23142080\|23597149 | Unspecified |
| mmu-miR-30c-5p | MIMAT0000514 | Cpeb4 | 67579 | HITS-CLIP | 25753418 | Unspecified |
| mmu-miR-30c-5p | MIMAT0000514 | Cplx1 | 12889 | HITS-CLIP | 37986224 | Unspecified |
| mmu-miR-30c-5p | MIMAT0000514 | Cpq | 54381 | HITS-CLIP | 23142080 | Unspecified |
| mmu-miR-30c-5p | MIMAT0000514 | Cpsf6 | 432508 | HITS-CLIP | 37986224\|23597149\|23142080 | Unspecified |
| mmu-miR-30c-5p | MIMAT0000514 | Cpt1a | 12894 | HITS-CLIP | 23597149\|23142080 | Unspecified |
| mmu-miR-30c-5p | MIMAT0000514 | Creb1 | 12912 | HITS-CLIP | 37986224 | Unspecified |
| mmu-miR-30c-5p | MIMAT0000514 | Creb3l2 | 208647 | HITS-CLIP | 37986224 | Unspecified |
| mmu-miR-30c-5p | MIMAT0000514 | Crebbp | 12914 | HITS-CLIP | 23597149\|23142080 | Unspecified |
| mmu-miR-30c-5p | MIMAT0000514 | Crebrf | 77128 | HITS-CLIP | 37986224\|23142080 | Unspecified |
| mmu-miR-30c-5p | MIMAT0000514 | Crim1 | 50766 | HITS-CLIP | 23142080 | Unspecified |
| mmu-miR-30c-5p | MIMAT0000514 | Crk | 12928 | HITS-CLIP | 37986224\|23142080 | Unspecified |

| | | | | | | |
|---|---|---|---|---|---|---|
| mmu-miR-30c-5p | MIMAT0000514 | Crkl | 12929 | HITS-CLIP | 23597149\|23142080 | Unspecified |
| mmu-miR-30c-5p | MIMAT0000514 | Crp | 12944 | HITS-CLIP | 23597149 | Unspecified |
| mmu-miR-30c-5p | MIMAT0000514 | Cry2 | 12953 | HITS-CLIP | 37986224 | Unspecified |
| mmu-miR-30c-5p | MIMAT0000514 | Crybg1 | 202 | HITS-CLIP | 23142080 | Unspecified |
| mmu-miR-30c-5p | MIMAT0000514 | Csde1 | 229663 | HITS-CLIP | 37986224\|25753418 | Unspecified |
| mmu-miR-30c-5p | MIMAT0000514 | Csf2rb | 12983 | HITS-CLIP | 37986224\|23597149 | Unspecified |
| mmu-miR-30c-5p | MIMAT0000514 | Csk | 12988 | HITS-CLIP | 37986224\|23142080 | Unspecified |
| mmu-miR-30c-5p | MIMAT0000514 | Csmd1 | 94109 | HITS-CLIP | 23142080 | Unspecified |
| mmu-miR-30c-5p | MIMAT0000514 | Csnk1a1 | 93687 | HITS-CLIP | 37986224\|23142080\|23597149 | Unspecified |
| mmu-miR-30c-5p | MIMAT0000514 | Csnk1g1 | 214897 | HITS-CLIP | 37986224\|19536157\|23142080 | Unspecified |
| mmu-miR-30c-5p | MIMAT0000514 | Csnk1g3 | 70425 | HITS-CLIP | 23142080 | Unspecified |
| mmu-miR-30c-5p | MIMAT0000514 | Csnk2a1 | 12995 | HITS-CLIP | 23142080 | Unspecified |
| mmu-miR-30c-5p | MIMAT0000514 | Cspp1 | 211660 | HITS-CLIP | 37986224\|23142080 | Unspecified |
| mmu-miR-30c-5p | MIMAT0000514 | Csrnp1 | 215418 | HITS-CLIP | 37986224\|23142080 | Unspecified |
| mmu-miR-30c-5p | MIMAT0000514 | Cstf2 | 108062 | HITS-CLIP | 23142080 | Unspecified |
| mmu-miR-30c-5p | MIMAT0000514 | Ctdspl2 | 329506 | HITS-CLIP | 37986224 | Unspecified |
| mmu-miR-30c-5p | MIMAT0000514 | Cth | 107869 | HITS-CLIP | 37986224\|23597149\|25753418 | Unspecified |
| mmu-miR-30c-5p | MIMAT0000514 | Cthrc1 | 68588 | HITS-CLIP | 37986224\|19536157 | Unspecified |
| mmu-miR-30c-5p | MIMAT0000514 | Ctla4 | 12477 | HITS-CLIP | 23142080 | Unspecified |
| mmu-miR-30c-5p | MIMAT0000514 | Ctnna1 | 12385 | HITS-CLIP | 37986224 | Unspecified |
| mmu-miR-30c-5p | MIMAT0000514 | Ctnnb1 | 12387 | HITS-CLIP | 23142080 | Unspecified |
| mmu-miR-30c-5p | MIMAT0000514 | Ctps2 | 55936 | HITS-CLIP | 37986224 | Unspecified |
| mmu-miR-30c-5p | MIMAT0000514 | Cul1 | 26965 | HITS-CLIP | 23142080 | Unspecified |
| mmu-miR-30c-5p | MIMAT0000514 | Cul2 | 71745 | HITS-CLIP | 37986224\|23142080\|23597149\|2575: | Unspecified |
| mmu-miR-30c-5p | MIMAT0000514 | Cul5 | 75717 | HITS-CLIP | 23142080 | Unspecified |
| mmu-miR-30c-5p | MIMAT0000514 | Cux1 | 13047 | HITS-CLIP | 23142080 | Unspecified |
| mmu-miR-30c-5p | MIMAT0000514 | Cwf19l2 | 244672 | HITS-CLIP | 25753418 | Unspecified |
| mmu-miR-30c-5p | MIMAT0000514 | Cxcl12 | 20315 | HITS-CLIP | 23597149 | Unspecified |
| mmu-miR-30c-5p | MIMAT0000514 | Cxcr4 | 12767 | HITS-CLIP | 23142080 | Unspecified |
| mmu-miR-30c-5p | MIMAT0000514 | Cyb5b | 66427 | HITS-CLIP | 23142080 | Unspecified |
| mmu-miR-30c-5p | MIMAT0000514 | Cyb5r4 | 266690 | HITS-CLIP | 37986224 | Unspecified |
| mmu-miR-30c-5p | MIMAT0000514 | Cybc1 | 79415 | HITS-CLIP | 23597149 | Unspecified |
| mmu-miR-30c-5p | MIMAT0000514 | Cyfip1 | 20430 | HITS-CLIP | 23597149\|23142080 | Unspecified |
| mmu-miR-30c-5p | MIMAT0000514 | Cyp2c23 | 226143 | HITS-CLIP | 23597149 | Unspecified |
| mmu-miR-30c-5p | MIMAT0000514 | Cyp2j5 | 13109 | HITS-CLIP | 37986224\|23597149 | Unspecified |
| mmu-miR-30c-5p | MIMAT0000514 | Cyp3a13 | 13113 | HITS-CLIP | 23597149 | Unspecified |
| mmu-miR-30c-5p | MIMAT0000514 | Cyp4v3 | 102294 | HITS-CLIP | 37986224 | Unspecified |
| mmu-miR-30c-5p | MIMAT0000514 | Cyp51 | 13121 | HITS-CLIP | 37986224\|23597149\|23142080 | Unspecified |
| mmu-miR-30c-5p | MIMAT0000514 | Cystm1 | 66060 | HITS-CLIP | 37986224 | Unspecified |

| | | | | | | |
|---|---|---|---|---|---|---|
| mmu-miR-30c-5p | MIMAT0000514 | Cyth3 | 19159 | HITS-CLIP | 23142080 | Unspecified |
| mmu-miR-30c-5p | MIMAT0000514 | D10Wsu102e | 28109 | HITS-CLIP | 23142080 | Unspecified |
| mmu-miR-30c-5p | MIMAT0000514 | Dact1 | 59036 | HITS-CLIP | 23597149 | Unspecified |
| mmu-miR-30c-5p | MIMAT0000514 | Dag1 | 13138 | HITS-CLIP | 37986224\|23142080 | Unspecified |
| mmu-miR-30c-5p | MIMAT0000514 | Dars | 226414 | qCLASH | 31393953 | Unspecified |
| mmu-miR-30c-5p | MIMAT0000514 | Dbt | 13171 | HITS-CLIP | 25753418 | Unspecified |
| mmu-miR-30c-5p | MIMAT0000514 | Dcaf10 | 242418 | HITS-CLIP | 37986224\|25753418 | Unspecified |
| mmu-miR-30c-5p | MIMAT0000514 | Dcaf12 | 68970 | HITS-CLIP | 37986224\|23597149\|25753418\|23142 | Unspecified |
| mmu-miR-30c-5p | MIMAT0000514 | Dcaf17 | 75763 | HITS-CLIP | 23142080\|23597149 | Unspecified |
| mmu-miR-30c-5p | MIMAT0000514 | Dcaf7 | 71833 | HITS-CLIP | 23597149 | Unspecified |
| mmu-miR-30c-5p | MIMAT0000514 | Dcbld1 | 66686 | HITS-CLIP | 23597149 | Unspecified |
| mmu-miR-30c-5p | MIMAT0000514 | Dcp1a | 75901 | HITS-CLIP | 37986224\|19536157\|23142080 | Unspecified |
| mmu-miR-30c-5p | MIMAT0000514 | Dcp2 | 70640 | HITS-CLIP | 37986224 | Unspecified |
| mmu-miR-30c-5p | MIMAT0000514 | Dctn4 | 67665 | HITS-CLIP | 23142080 | Unspecified |
| mmu-miR-30c-5p | MIMAT0000514 | Dctn6 | 22428 | HITS-CLIP | 23142080 | Unspecified |
| mmu-miR-30c-5p | MIMAT0000514 | Dcun1d3 | 233805 | HITS-CLIP | 37986224\|23142080 | Unspecified |
| mmu-miR-30c-5p | MIMAT0000514 | Dda1 | 66498 | HITS-CLIP | 37986224 | Unspecified |
| mmu-miR-30c-5p | MIMAT0000514 | Ddah1 | 69219 | HITS-CLIP | 37986224\|23597149 | Unspecified |
| mmu-miR-30c-5p | MIMAT0000514 | Ddhd1 | 114874 | HITS-CLIP | 23142080 | Unspecified |
| mmu-miR-30c-5p | MIMAT0000514 | Ddit4 | 74747 | HITS-CLIP | 37986224\|23597149\|23142080\|1953€ | Unspecified |
| mmu-miR-30c-5p | MIMAT0000514 | Ddr1 | 12305 | HITS-CLIP | 37986224 | Unspecified |
| mmu-miR-30c-5p | MIMAT0000514 | Ddx17 | 67040 | HITS-CLIP//qCLASH | 23142080\|31393953 | Unspecified |
| mmu-miR-30c-5p | MIMAT0000514 | Ddx19b | 234733 | HITS-CLIP | 23597149\|23142080 | Unspecified |
| mmu-miR-30c-5p | MIMAT0000514 | Ddx24 | 27225 | HITS-CLIP | 23142080\|25753418 | Unspecified |
| mmu-miR-30c-5p | MIMAT0000514 | Ddx3y | 26900 | HITS-CLIP | 23597149\|25753418 | Unspecified |
| mmu-miR-30c-5p | MIMAT0000514 | Ddx46 | 212880 | HITS-CLIP | 23142080 | Unspecified |
| mmu-miR-30c-5p | MIMAT0000514 | Ddx5 | 13207 | HITS-CLIP | 37986224\|23597149\|19536157\|23142 | Unspecified |
| mmu-miR-30c-5p | MIMAT0000514 | Ddx6 | 13209 | HITS-CLIP | 37986224\|19536157 | Unspecified |
| mmu-miR-30c-5p | MIMAT0000514 | Degs1 | 13244 | HITS-CLIP | 37986224 | Unspecified |
| mmu-miR-30c-5p | MIMAT0000514 | Dennd1b | 329260 | HITS-CLIP | 23142080 | Unspecified |
| mmu-miR-30c-5p | MIMAT0000514 | Dennd2c | 329727 | HITS-CLIP | 23142080 | Unspecified |
| mmu-miR-30c-5p | MIMAT0000514 | Dennd4c | 329877 | HITS-CLIP | 23142080 | Unspecified |
| mmu-miR-30c-5p | MIMAT0000514 | Dennd5b | 320560 | HITS-CLIP | 37986224\|23597149\|25753418 | Unspecified |
| mmu-miR-30c-5p | MIMAT0000514 | Dennd6a | 211922 | HITS-CLIP | 23142080 | Unspecified |
| mmu-miR-30c-5p | MIMAT0000514 | Depdc5 | 277854 | qCLASH | 31393953 | Unspecified |
| mmu-miR-30c-5p | MIMAT0000514 | Desi2 | 78825 | HITS-CLIP | 37986224\|23142080 | Unspecified |
| mmu-miR-30c-5p | MIMAT0000514 | Dexi | 58239 | HITS-CLIP | 23597149\|23142080 | Unspecified |
| mmu-miR-30c-5p | MIMAT0000514 | Dgkh | 380921 | HITS-CLIP | 23142080\|19536157\|23597149 | Unspecified |
| mmu-miR-30c-5p | MIMAT0000514 | Dgkq | 110524 | HITS-CLIP | 23142080 | Unspecified |

| | | | | | | |
|---|---|---|---|---|---|---|
| mmu-miR-30c-5p | MIMAT0000514 | Dgkz | 104418 | HITS-CLIP | 37986224\|23142080 | Unspecified |
| mmu-miR-30c-5p | MIMAT0000514 | Dhcr7 | 13360 | HITS-CLIP | 37986224 | Unspecified |
| mmu-miR-30c-5p | MIMAT0000514 | Dhx30 | 72831 | HITS-CLIP | 23142080 | Unspecified |
| mmu-miR-30c-5p | MIMAT0000514 | Dhx36 | 72162 | HITS-CLIP | 23142080 | Unspecified |
| mmu-miR-30c-5p | MIMAT0000514 | Dhx40 | 67487 | HITS-CLIP | 23142080 | Unspecified |
| mmu-miR-30c-5p | MIMAT0000514 | Dhx9 | 13211 | HITS-CLIP | 37986224\|23597149\|23142080 | Unspecified |
| mmu-miR-30c-5p | MIMAT0000514 | Diablo | 66593 | HITS-CLIP | 37986224 | Unspecified |
| mmu-miR-30c-5p | MIMAT0000514 | Diaph3 | 81624 | HITS-CLIP | 23142080 | Unspecified |
| mmu-miR-30c-5p | MIMAT0000514 | Dicer1 | 192119 | HITS-CLIP | 37986224\|23597149 | Unspecified |
| mmu-miR-30c-5p | MIMAT0000514 | Dido1 | 23856 | HITS-CLIP | 37986224 | Unspecified |
| mmu-miR-30c-5p | MIMAT0000514 | Dio2 | 13371 | HITS-CLIP | 37986224\|19536157\|23142080 | Unspecified |
| mmu-miR-30c-5p | MIMAT0000514 | Dip2b | 239667 | HITS-CLIP | 37986224\|23142080 | Unspecified |
| mmu-miR-30c-5p | MIMAT0000514 | Dkc1 | 245474 | HITS-CLIP | 37986224\|23597149\|25753418\|2314 | Unspecified |
| mmu-miR-30c-5p | MIMAT0000514 | Dlg5 | 71228 | HITS-CLIP | 23597149\|25753418 | Unspecified |
| mmu-miR-30c-5p | MIMAT0000514 | Dmtf1 | 23857 | HITS-CLIP | 23142080 | Unspecified |
| mmu-miR-30c-5p | MIMAT0000514 | Dmxl2 | 235380 | HITS-CLIP | 37986224\|19536157 | Unspecified |
| mmu-miR-30c-5p | MIMAT0000514 | Dnaja1 | 15502 | HITS-CLIP | 25753418 | Unspecified |
| mmu-miR-30c-5p | MIMAT0000514 | Dnaja2 | 56445 | HITS-CLIP | 23597149\|23142080 | Unspecified |
| mmu-miR-30c-5p | MIMAT0000514 | Dnajb14 | 70604 | HITS-CLIP | 19536157 | Unspecified |
| mmu-miR-30c-5p | MIMAT0000514 | Dnajc13 | 235567 | HITS-CLIP | 37986224\|23597149\|23142080 | Unspecified |
| mmu-miR-30c-5p | MIMAT0000514 | Dnajc6 | 72685 | HITS-CLIP | 19536157 | Unspecified |
| mmu-miR-30c-5p | MIMAT0000514 | Dnal1 | 105000 | HITS-CLIP | 37986224 | Unspecified |
| mmu-miR-30c-5p | MIMAT0000514 | Dnm1l | 74006 | HITS-CLIP | 23597149\|23142080 | Unspecified |
| mmu-miR-30c-5p | MIMAT0000514 | Dnmbp | 71972 | HITS-CLIP | 23142080\|23597149 | Unspecified |
| mmu-miR-30c-5p | MIMAT0000514 | Dnmt1 | 13433 | HITS-CLIP | 23142080 | Unspecified |
| mmu-miR-30c-5p | MIMAT0000514 | Dnmt3a | 13435 | HITS-CLIP | 37986224\|23142080 | Unspecified |
| mmu-miR-30c-5p | MIMAT0000514 | Dnttip2 | 99480 | HITS-CLIP | 37986224 | Unspecified |
| mmu-miR-30c-5p | MIMAT0000514 | Dock10 | 210293 | HITS-CLIP | 37986224 | Unspecified |
| mmu-miR-30c-5p | MIMAT0000514 | Dock11 | 75974 | HITS-CLIP | 23142080 | Unspecified |
| mmu-miR-30c-5p | MIMAT0000514 | Dock4 | 238130 | HITS-CLIP | 37986224 | Unspecified |
| mmu-miR-30c-5p | MIMAT0000514 | Dock5 | 68813 | HITS-CLIP | 23597149 | Unspecified |
| mmu-miR-30c-5p | MIMAT0000514 | Dock9 | 105445 | HITS-CLIP | 23142080 | Unspecified |
| mmu-miR-30c-5p | MIMAT0000514 | Dolpp1 | 57170 | HITS-CLIP | 23142080 | Unspecified |
| mmu-miR-30c-5p | MIMAT0000514 | Dop1b | 9980 | HITS-CLIP | 23597149 | Unspecified |
| mmu-miR-30c-5p | MIMAT0000514 | Dph5 | 69740 | HITS-CLIP | 23142080 | Unspecified |
| mmu-miR-30c-5p | MIMAT0000514 | Dpm1 | 13480 | HITS-CLIP | 23142080 | Unspecified |
| mmu-miR-30c-5p | MIMAT0000514 | Dpp3 | 75221 | HITS-CLIP | 37986224 | Unspecified |
| mmu-miR-30c-5p | MIMAT0000514 | Dpp8 | 74388 | HITS-CLIP | 37986224 | Unspecified |
| mmu-miR-30c-5p | MIMAT0000514 | Dpy19l1 | 244745 | HITS-CLIP | 25083871\|23142080 | Unspecified |

| | | | | | | |
|---|---|---|---|---|---|---|
| mmu-miR-30c-5p | MIMAT0000514 | Dpy19l4 | 381510 | HITS-CLIP | 37986224 | Unspecified |
| mmu-miR-30c-5p | MIMAT0000514 | Dpyd | 99586 | HITS-CLIP | 23597149 | Unspecified |
| mmu-miR-30c-5p | MIMAT0000514 | Dpysl2 | 12934 | HITS-CLIP | 37986224\|23142080\|19536157 | Unspecified |
| mmu-miR-30c-5p | MIMAT0000514 | Dram2 | 67171 | HITS-CLIP | 37986224 | Unspecified |
| mmu-miR-30c-5p | MIMAT0000514 | Drg2 | 13495 | HITS-CLIP | 23142080 | Unspecified |
| mmu-miR-30c-5p | MIMAT0000514 | Drosha | 14000 | HITS-CLIP | 23142080 | Unspecified |
| mmu-miR-30c-5p | MIMAT0000514 | Dsc2 | 13506 | HITS-CLIP | 37986224\|23597149 | Unspecified |
| mmu-miR-30c-5p | MIMAT0000514 | Dsp | 109620 | HITS-CLIP | 25753418 | Unspecified |
| mmu-miR-30c-5p | MIMAT0000514 | Dst | 13518 | HITS-CLIP | 25753418 | Unspecified |
| mmu-miR-30c-5p | MIMAT0000514 | Dstn | 56431 | HITS-CLIP | 37986224 | Unspecified |
| mmu-miR-30c-5p | MIMAT0000514 | Dstyk | 213452 | HITS-CLIP | 37986224\|23597149 | Unspecified |
| mmu-miR-30c-5p | MIMAT0000514 | Dtx2 | 74198 | HITS-CLIP | 23597149 | Unspecified |
| mmu-miR-30c-5p | MIMAT0000514 | Dusp19 | 68082 | HITS-CLIP | 37986224 | Unspecified |
| mmu-miR-30c-5p | MIMAT0000514 | Dusp5 | 240672 | HITS-CLIP | 23142080 | Unspecified |
| mmu-miR-30c-5p | MIMAT0000514 | Dync1h1 | 13424 | HITS-CLIP | 37986224 | Unspecified |
| mmu-miR-30c-5p | MIMAT0000514 | Dync1li1 | 235661 | HITS-CLIP | 37986224\|23597149\|23142080 | Unspecified |
| mmu-miR-30c-5p | MIMAT0000514 | Dync1li2 | 234663 | HITS-CLIP | 23142080 | Unspecified |
| mmu-miR-30c-5p | MIMAT0000514 | E2f2 | 242705 | HITS-CLIP | 37986224 | Unspecified |
| mmu-miR-30c-5p | MIMAT0000514 | E2f3 | 13557 | HITS-CLIP | 37986224 | Unspecified |
| mmu-miR-30c-5p | MIMAT0000514 | E2f4 | 104394 | HITS-CLIP | 37986224 | Unspecified |
| mmu-miR-30c-5p | MIMAT0000514 | E2f8 | 108961 | HITS-CLIP | 23142080 | Unspecified |
| mmu-miR-30c-5p | MIMAT0000514 | E330009J07Rik | 243780 | HITS-CLIP | 23142080 | Unspecified |
| mmu-miR-30c-5p | MIMAT0000514 | Ebag9 | 55960 | HITS-CLIP | 23597149 | Unspecified |
| mmu-miR-30c-5p | MIMAT0000514 | Echdc1 | 52665 | HITS-CLIP | 37986224 | Unspecified |
| mmu-miR-30c-5p | MIMAT0000514 | Edem1 | 192193 | HITS-CLIP | 23597149\|23142080 | Unspecified |
| mmu-miR-30c-5p | MIMAT0000514 | Edem3 | 66967 | HITS-CLIP | 37986224\|19536157\|23142080 | Unspecified |
| mmu-miR-30c-5p | MIMAT0000514 | Ednra | 13617 | HITS-CLIP | 37986224\|19536157 | Unspecified |
| mmu-miR-30c-5p | MIMAT0000514 | Eea1 | 216238 | HITS-CLIP | 23142080 | Unspecified |
| mmu-miR-30c-5p | MIMAT0000514 | Eed | 13626 | HITS-CLIP | 37986224 | Unspecified |
| mmu-miR-30c-5p | MIMAT0000514 | Eef1b2 | 55949 | qCLASH | 31393953 | Unspecified |
| mmu-miR-30c-5p | MIMAT0000514 | Eef1e1 | 66143 | HITS-CLIP | 23142080 | Unspecified |
| mmu-miR-30c-5p | MIMAT0000514 | Eef2 | 13629 | qCLASH//HITS-CLIP | 31393953\|23597149\|23142080 | Unspecified |
| mmu-miR-30c-5p | MIMAT0000514 | Efna5 | 13640 | HITS-CLIP | 25753418 | Unspecified |
| mmu-miR-30c-5p | MIMAT0000514 | Efr3a | 76740 | HITS-CLIP | 37986224\|23142080\|19536157 | Unspecified |
| mmu-miR-30c-5p | MIMAT0000514 | Eftud2 | 20624 | HITS-CLIP | 23142080 | Unspecified |
| mmu-miR-30c-5p | MIMAT0000514 | Egln3 | 112407 | HITS-CLIP | 37986224 | Unspecified |
| mmu-miR-30c-5p | MIMAT0000514 | Ehd4 | 98878 | HITS-CLIP | 25753418 | Unspecified |
| mmu-miR-30c-5p | MIMAT0000514 | Eif1a | 13664 | HITS-CLIP | 23142080 | Unspecified |
| mmu-miR-30c-5p | MIMAT0000514 | Eif1b | 68969 | HITS-CLIP | 37986224 | Unspecified |

| | | | | | | |
|---|---|---|---|---|---|---|
| mmu-miR-30c-5p | MIMAT0000514 | Eif2b1 | 209354 | HITS-CLIP | 23142080 | Unspecified |
| mmu-miR-30c-5p | MIMAT0000514 | Eif3e | 16341 | HITS-CLIP | 37986224\|23597149\|23142080\|25753 | Unspecified |
| mmu-miR-30c-5p | MIMAT0000514 | Eif3g | 8666 | qCLASH | 31393953 | Unspecified |
| mmu-miR-30c-5p | MIMAT0000514 | Eif4a2 | 13682 | HITS-CLIP | 23142080 | Unspecified |
| mmu-miR-30c-5p | MIMAT0000514 | Eif4b | 75705 | qCLASH | 31393953 | Unspecified |
| mmu-miR-30c-5p | MIMAT0000514 | Eif4g1 | 208643 | HITS-CLIP | 37986224 | Unspecified |
| mmu-miR-30c-5p | MIMAT0000514 | Eif5 | 217869 | HITS-CLIP | 25753418 | Unspecified |
| mmu-miR-30c-5p | MIMAT0000514 | Eif5a2 | 208691 | HITS-CLIP | 19536157\|23142080 | Unspecified |
| mmu-miR-30c-5p | MIMAT0000514 | Eif5b | 226982 | HITS-CLIP//qCLASH | 37986224\|31393953 | Unspecified |
| mmu-miR-30c-5p | MIMAT0000514 | Elavl1 | 15568 | HITS-CLIP | 37986224\|23142080 | Unspecified |
| mmu-miR-30c-5p | MIMAT0000514 | Elf4 | 56501 | HITS-CLIP | 23142080 | Unspecified |
| mmu-miR-30c-5p | MIMAT0000514 | Elfn2 | 207393 | HITS-CLIP | 37986224 | Unspecified |
| mmu-miR-30c-5p | MIMAT0000514 | Ell | 13716 | HITS-CLIP | 37986224\|23142080 | Unspecified |
| mmu-miR-30c-5p | MIMAT0000514 | Ell2 | 192657 | HITS-CLIP | 37986224\|23597149\|23142080 | Unspecified |
| mmu-miR-30c-5p | MIMAT0000514 | Elmo1 | 140580 | HITS-CLIP | 23142080 | Unspecified |
| mmu-miR-30c-5p | MIMAT0000514 | Elmod2 | 244548 | HITS-CLIP | 23142080 | Unspecified |
| mmu-miR-30c-5p | MIMAT0000514 | Elovl3 | 12686 | HITS-CLIP | 37986224 | Unspecified |
| mmu-miR-30c-5p | MIMAT0000514 | Elovl5 | 68801 | HITS-CLIP | 37986224\|23597149\|25753418\|23142 | Unspecified |
| mmu-miR-30c-5p | MIMAT0000514 | Elovl6 | 170439 | HITS-CLIP | 23142080 | Unspecified |
| mmu-miR-30c-5p | MIMAT0000514 | Elovl7 | 74559 | HITS-CLIP | 37986224 | Unspecified |
| mmu-miR-30c-5p | MIMAT0000514 | Emc4 | 68032 | HITS-CLIP | 37986224\|23142080 | Unspecified |
| mmu-miR-30c-5p | MIMAT0000514 | Emd | 13726 | HITS-CLIP | 23142080 | Unspecified |
| mmu-miR-30c-5p | MIMAT0000514 | Emg1 | 14791 | HITS-CLIP | 23142080 | Unspecified |
| mmu-miR-30c-5p | MIMAT0000514 | Eml4 | 78798 | HITS-CLIP | 37986224\|23142080 | Unspecified |
| mmu-miR-30c-5p | MIMAT0000514 | Emsy | 56946 | HITS-CLIP | 23142080 | Unspecified |
| mmu-miR-30c-5p | MIMAT0000514 | En2 | 13799 | HITS-CLIP | 37986224 | Unspecified |
| mmu-miR-30c-5p | MIMAT0000514 | Enc1 | 13803 | HITS-CLIP | 37986224\|23142080 | Unspecified |
| mmu-miR-30c-5p | MIMAT0000514 | Enoph1 | 67870 | HITS-CLIP | 23142080 | Unspecified |
| mmu-miR-30c-5p | MIMAT0000514 | Enox2 | 209224 | HITS-CLIP | 23142080 | Unspecified |
| mmu-miR-30c-5p | MIMAT0000514 | Ensa | 56205 | HITS-CLIP | 37986224\|23142080 | Unspecified |
| mmu-miR-30c-5p | MIMAT0000514 | Entpd5 | 12499 | HITS-CLIP | 37986224\|23597149\|23142080 | Unspecified |
| mmu-miR-30c-5p | MIMAT0000514 | Eny2 | 223527 | HITS-CLIP | 23142080 | Unspecified |
| mmu-miR-30c-5p | MIMAT0000514 | Ep400 | 75560 | HITS-CLIP | 23597149 | Unspecified |
| mmu-miR-30c-5p | MIMAT0000514 | Epas1 | 13819 | HITS-CLIP | 25753418 | Unspecified |
| mmu-miR-30c-5p | MIMAT0000514 | Epb41 | 269587 | HITS-CLIP | 23142080 | Unspecified |
| mmu-miR-30c-5p | MIMAT0000514 | Epb41l4b | 54566 | HITS-CLIP | 23597149 | Unspecified |
| mmu-miR-30c-5p | MIMAT0000514 | Epc1 | 13831 | HITS-CLIP | 25753418 | Unspecified |
| mmu-miR-30c-5p | MIMAT0000514 | Epc2 | 227867 | HITS-CLIP | 37986224\|23142080 | Unspecified |
| mmu-miR-30c-5p | MIMAT0000514 | Epha1 | 13835 | HITS-CLIP | 37986224 | Unspecified |

| | | | | | | |
|---|---|---|---|---|---|---|
| mmu-miR-30c-5p | MIMAT0000514 | Epha7 | 13841 | HITS-CLIP | 37986224 | Unspecified |
| mmu-miR-30c-5p | MIMAT0000514 | Epm2aip1 | 77781 | HITS-CLIP | 23142080 | Unspecified |
| mmu-miR-30c-5p | MIMAT0000514 | Eprs | 107508 | HITS-CLIP | 23597149\|23142080 | Unspecified |
| mmu-miR-30c-5p | MIMAT0000514 | Eps15 | 13858 | HITS-CLIP | 23142080 | Unspecified |
| mmu-miR-30c-5p | MIMAT0000514 | Erbin | 55914 | HITS-CLIP | 23142080 | Unspecified |
| mmu-miR-30c-5p | MIMAT0000514 | Erc1 | 111173 | HITS-CLIP | 25753418 | Unspecified |
| mmu-miR-30c-5p | MIMAT0000514 | Ergic3 | 66366 | HITS-CLIP | 23597149\|23142080 | Unspecified |
| mmu-miR-30c-5p | MIMAT0000514 | Eri1 | 67276 | HITS-CLIP | 23142080 | Unspecified |
| mmu-miR-30c-5p | MIMAT0000514 | Eri2 | 71151 | HITS-CLIP | 23142080 | Unspecified |
| mmu-miR-30c-5p | MIMAT0000514 | Erlin1 | 226144 | HITS-CLIP | 37986224\|23597149\|19536157\|23142 | Unspecified |
| mmu-miR-30c-5p | MIMAT0000514 | Errfi1 | 74155 | HITS-CLIP | 23142080 | Unspecified |
| mmu-miR-30c-5p | MIMAT0000514 | Esf1 | 66580 | HITS-CLIP | 37986224 | Unspecified |
| mmu-miR-30c-5p | MIMAT0000514 | Espn | 56226 | HITS-CLIP | 37986224\|23142080 | Unspecified |
| mmu-miR-30c-5p | MIMAT0000514 | Esyt1 | 23943 | HITS-CLIP | 37986224 | Unspecified |
| mmu-miR-30c-5p | MIMAT0000514 | Etaa1 | 68145 | HITS-CLIP | 37986224\|23142080 | Unspecified |
| mmu-miR-30c-5p | MIMAT0000514 | Etf1 | 225363 | HITS-CLIP | 23597149\|23142080 | Unspecified |
| mmu-miR-30c-5p | MIMAT0000514 | Ethe1 | 66071 | HITS-CLIP | 37986224 | Unspecified |
| mmu-miR-30c-5p | MIMAT0000514 | Etnk1 | 75320 | HITS-CLIP | 23142080 | Unspecified |
| mmu-miR-30c-5p | MIMAT0000514 | Ets1 | 23871 | HITS-CLIP | 37986224\|23142080 | Unspecified |
| mmu-miR-30c-5p | MIMAT0000514 | Etv3 | 27049 | HITS-CLIP | 37986224 | Unspecified |
| mmu-miR-30c-5p | MIMAT0000514 | Etv6 | 14011 | HITS-CLIP | 23142080 | Unspecified |
| mmu-miR-30c-5p | MIMAT0000514 | Evi5 | 14020 | HITS-CLIP | 37986224 | Unspecified |
| mmu-miR-30c-5p | MIMAT0000514 | Exoc2 | 66482 | HITS-CLIP | 23142080 | Unspecified |
| mmu-miR-30c-5p | MIMAT0000514 | Exoc4 | 20336 | HITS-CLIP | 25753418 | Unspecified |
| mmu-miR-30c-5p | MIMAT0000514 | Exoc6 | 107371 | HITS-CLIP | 37986224\|23597149\|23142080 | Unspecified |
| mmu-miR-30c-5p | MIMAT0000514 | Exoc6b | 75914 | HITS-CLIP | 37986224\|23142080 | Unspecified |
| mmu-miR-30c-5p | MIMAT0000514 | Extl2 | 58193 | HITS-CLIP | 37986224\|19536157 | Unspecified |
| mmu-miR-30c-5p | MIMAT0000514 | Ezh2 | 14056 | HITS-CLIP | 23597149\|23142080 | Unspecified |
| mmu-miR-30c-5p | MIMAT0000514 | Fads6 | 328035 | HITS-CLIP | 37986224 | Unspecified |
| mmu-miR-30c-5p | MIMAT0000514 | Faf2 | 76577 | HITS-CLIP | 37986224 | Unspecified |
| mmu-miR-30c-5p | MIMAT0000514 | Fahd1 | 68636 | HITS-CLIP | 37986224 | Unspecified |
| mmu-miR-30c-5p | MIMAT0000514 | Fam117b | 72750 | HITS-CLIP | 37986224\|23142080 | Unspecified |
| mmu-miR-30c-5p | MIMAT0000514 | Fam120a | 218236 | HITS-CLIP | 37986224\|23597149\|23142080 | Unspecified |
| mmu-miR-30c-5p | MIMAT0000514 | Fam124a | 629059 | HITS-CLIP | 37986224\|23597149 | Unspecified |
| mmu-miR-30c-5p | MIMAT0000514 | Fam126b | 213056 | HITS-CLIP | 37986224\|23142080\|23597149 | Unspecified |
| mmu-miR-30c-5p | MIMAT0000514 | Fam133b | 68152 | HITS-CLIP | 23142080 | Unspecified |
| mmu-miR-30c-5p | MIMAT0000514 | Fam136a | 66488 | HITS-CLIP | 37986224 | Unspecified |
| mmu-miR-30c-5p | MIMAT0000514 | Fam160b1 | 226252 | HITS-CLIP | 37986224\|23142080 | Unspecified |
| mmu-miR-30c-5p | MIMAT0000514 | Fam162a | 70186 | HITS-CLIP | 37986224 | Unspecified |

| | | | | | | |
|---|---|---|---|---|---|---|
| mmu-miR-30c-5p | MIMAT0000514 | Fam168b | 214469 | HITS-CLIP | 23142080\|19536157\|23597149 | Unspecified |
| mmu-miR-30c-5p | MIMAT0000514 | Fam199x | 245622 | HITS-CLIP | 37986224\|23597149 | Unspecified |
| mmu-miR-30c-5p | MIMAT0000514 | Fam207a | 108707 | HITS-CLIP | 25753418 | Unspecified |
| mmu-miR-30c-5p | MIMAT0000514 | Fam210a | 108654 | HITS-CLIP | 25753418 | Unspecified |
| mmu-miR-30c-5p | MIMAT0000514 | Fam210b | 67017 | HITS-CLIP | 23597149\|23142080 | Unspecified |
| mmu-miR-30c-5p | MIMAT0000514 | Fam214a | 235493 | HITS-CLIP | 37986224\|23142080 | Unspecified |
| mmu-miR-30c-5p | MIMAT0000514 | Fam234b | 57613 | HITS-CLIP | 23597149\|23142080 | Unspecified |
| mmu-miR-30c-5p | MIMAT0000514 | Fam43a | 224093 | HITS-CLIP | 37986224\|23142080 | Unspecified |
| mmu-miR-30c-5p | MIMAT0000514 | Fam49a | 76820 | HITS-CLIP | 37986224\|23142080 | Unspecified |
| mmu-miR-30c-5p | MIMAT0000514 | Fam49b | 223601 | HITS-CLIP | 37986224\|23142080 | Unspecified |
| mmu-miR-30c-5p | MIMAT0000514 | Fam57a | 116972 | HITS-CLIP | 37986224\|23142080 | Unspecified |
| mmu-miR-30c-5p | MIMAT0000514 | Fam76b | 72826 | HITS-CLIP | 37986224\|23142080 | Unspecified |
| mmu-miR-30c-5p | MIMAT0000514 | Fam83h | 105732 | HITS-CLIP | 23597149 | Unspecified |
| mmu-miR-30c-5p | MIMAT0000514 | Fam91a1 | 210998 | qCLASH//HITS-CLIP | 31393953\|19536157\|23142080 | Unspecified |
| mmu-miR-30c-5p | MIMAT0000514 | Fancf | 100040608 | HITS-CLIP | 37986224 | Unspecified |
| mmu-miR-30c-5p | MIMAT0000514 | Far1 | 67420 | HITS-CLIP | 23142080 | Unspecified |
| mmu-miR-30c-5p | MIMAT0000514 | Fas | 14102 | HITS-CLIP | 23142080 | Unspecified |
| mmu-miR-30c-5p | MIMAT0000514 | Fastkd3 | 69577 | HITS-CLIP | 37986224 | Unspecified |
| mmu-miR-30c-5p | MIMAT0000514 | Fat1 | 14107 | HITS-CLIP | 23597149 | Unspecified |
| mmu-miR-30c-5p | MIMAT0000514 | Fat3 | 270120 | HITS-CLIP | 37986224 | Unspecified |
| mmu-miR-30c-5p | MIMAT0000514 | Fbxl20 | 72194 | HITS-CLIP | 23142080 | Unspecified |
| mmu-miR-30c-5p | MIMAT0000514 | Fbxl3 | 50789 | HITS-CLIP | 23142080 | Unspecified |
| mmu-miR-30c-5p | MIMAT0000514 | Fbxl5 | 242960 | HITS-CLIP | 37986224\|23142080 | Unspecified |
| mmu-miR-30c-5p | MIMAT0000514 | Fbxo11 | 225055 | HITS-CLIP | 37986224\|23142080 | Unspecified |
| mmu-miR-30c-5p | MIMAT0000514 | Fbxo21 | 231670 | HITS-CLIP | 37986224 | Unspecified |
| mmu-miR-30c-5p | MIMAT0000514 | Fbxo28 | 67948 | HITS-CLIP | 37986224\|23142080 | Unspecified |
| mmu-miR-30c-5p | MIMAT0000514 | Fbxo3 | 57443 | HITS-CLIP | 37986224\|23142080\|25753418 | Unspecified |
| mmu-miR-30c-5p | MIMAT0000514 | Fbxo30 | 71865 | HITS-CLIP | 37986224 | Unspecified |
| mmu-miR-30c-5p | MIMAT0000514 | Fbxo32 | 67731 | HITS-CLIP | 23142080 | Unspecified |
| mmu-miR-30c-5p | MIMAT0000514 | Fbxo33 | 70611 | HITS-CLIP | 37986224 | Unspecified |
| mmu-miR-30c-5p | MIMAT0000514 | Fbxo42 | 213499 | HITS-CLIP | 37986224\|23597149\|19536157\|23142 | Unspecified |
| mmu-miR-30c-5p | MIMAT0000514 | Fbxo45 | 268882 | HITS-CLIP | 23597149 | Unspecified |
| mmu-miR-30c-5p | MIMAT0000514 | Fbxo8 | 50753 | HITS-CLIP | 23142080 | Unspecified |
| mmu-miR-30c-5p | MIMAT0000514 | Fbxw11 | 103583 | HITS-CLIP | 23142080 | Unspecified |
| mmu-miR-30c-5p | MIMAT0000514 | Fbxw7 | 50754 | HITS-CLIP | 23597149\|23142080 | Unspecified |
| mmu-miR-30c-5p | MIMAT0000514 | Fchsd2 | 207278 | HITS-CLIP | 37986224 | Unspecified |
| mmu-miR-30c-5p | MIMAT0000514 | Fcmr | 9214 | qCLASH | 31393953 | Unspecified |
| mmu-miR-30c-5p | MIMAT0000514 | Fech | 14151 | HITS-CLIP | 37986224 | Unspecified |
| mmu-miR-30c-5p | MIMAT0000514 | Fem1c | 240263 | HITS-CLIP | 37986224 | Unspecified |

| | | | | | | |
|---|---|---|---|---|---|---|
| mmu-miR-30c-5p | MIMAT0000514 | Fermt2 | 218952 | HITS-CLIP | 25753418 | Unspecified |
| mmu-miR-30c-5p | MIMAT0000514 | Fgd6 | 13998 | HITS-CLIP | 23142080 | Unspecified |
| mmu-miR-30c-5p | MIMAT0000514 | Fgf2 | 14173 | HITS-CLIP | 37986224 | Unspecified |
| mmu-miR-30c-5p | MIMAT0000514 | Fgf4 | 14175 | HITS-CLIP | 37986224 | Unspecified |
| mmu-miR-30c-5p | MIMAT0000514 | Fgfr3 | 14184 | HITS-CLIP | 37986224\|23597149 | Unspecified |
| mmu-miR-30c-5p | MIMAT0000514 | Fgl1 | 234199 | HITS-CLIP | 23597149 | Unspecified |
| mmu-miR-30c-5p | MIMAT0000514 | Fkbp3 | 30795 | HITS-CLIP | 23142080\|25753418 | Unspecified |
| mmu-miR-30c-5p | MIMAT0000514 | Flcn | 216805 | HITS-CLIP | 23142080 | Unspecified |
| mmu-miR-30c-5p | MIMAT0000514 | Flrt3 | 71436 | HITS-CLIP | 37986224 | Unspecified |
| mmu-miR-30c-5p | MIMAT0000514 | Flt1 | 14254 | HITS-CLIP | 37986224 | Unspecified |
| mmu-miR-30c-5p | MIMAT0000514 | Fmnl1 | 57778 | HITS-CLIP | 23142080 | Unspecified |
| mmu-miR-30c-5p | MIMAT0000514 | Fmnl2 | 71409 | HITS-CLIP | 23597149 | Unspecified |
| mmu-miR-30c-5p | MIMAT0000514 | Fmnl3 | 22379 | HITS-CLIP | 23597149 | Unspecified |
| mmu-miR-30c-5p | MIMAT0000514 | Fnbp1l | 214459 | HITS-CLIP | 19536157 | Unspecified |
| mmu-miR-30c-5p | MIMAT0000514 | Fndc3a | 319448 | HITS-CLIP | 37986224\|23142080 | Unspecified |
| mmu-miR-30c-5p | MIMAT0000514 | Fnip1 | 216742 | HITS-CLIP | 37986224 | Unspecified |
| mmu-miR-30c-5p | MIMAT0000514 | Fnip2 | 329679 | HITS-CLIP | 37986224\|23597149 | Unspecified |
| mmu-miR-30c-5p | MIMAT0000514 | Fosl2 | 14284 | HITS-CLIP | 37986224\|25753418\|23142080 | Unspecified |
| mmu-miR-30c-5p | MIMAT0000514 | Foxj3 | 230700 | HITS-CLIP | 23142080 | Unspecified |
| mmu-miR-30c-5p | MIMAT0000514 | Foxn2 | 14236 | HITS-CLIP | 37986224\|19536157 | Unspecified |
| mmu-miR-30c-5p | MIMAT0000514 | Foxo3 | 56484 | HITS-CLIP | 19536157\|23142080 | Unspecified |
| mmu-miR-30c-5p | MIMAT0000514 | Foxp1 | 108655 | HITS-CLIP | 37986224\|23142080 | Unspecified |
| mmu-miR-30c-5p | MIMAT0000514 | Foxp4 | 74123 | HITS-CLIP | 23597149\|23142080 | Unspecified |
| mmu-miR-30c-5p | MIMAT0000514 | Fpgt | 75540 | HITS-CLIP | 23142080 | Unspecified |
| mmu-miR-30c-5p | MIMAT0000514 | Frg1 | 14300 | HITS-CLIP | 23142080 | Unspecified |
| mmu-miR-30c-5p | MIMAT0000514 | Frmd6 | 319710 | HITS-CLIP | 23142080 | Unspecified |
| mmu-miR-30c-5p | MIMAT0000514 | Frrs1l | 230235 | HITS-CLIP | 37986224 | Unspecified |
| mmu-miR-30c-5p | MIMAT0000514 | Frs2 | 327826 | HITS-CLIP | 37986224 | Unspecified |
| mmu-miR-30c-5p | MIMAT0000514 | Fubp1 | 51886 | HITS-CLIP | 23142080 | Unspecified |
| mmu-miR-30c-5p | MIMAT0000514 | Fubp3 | 320267 | HITS-CLIP | 23142080 | Unspecified |
| mmu-miR-30c-5p | MIMAT0000514 | Fundc1 | 72018 | HITS-CLIP | 23142080 | Unspecified |
| mmu-miR-30c-5p | MIMAT0000514 | Fut9 | 14348 | HITS-CLIP | 37986224 | Unspecified |
| mmu-miR-30c-5p | MIMAT0000514 | Fyco1 | 17281 | HITS-CLIP | 37986224 | Unspecified |
| mmu-miR-30c-5p | MIMAT0000514 | Fyn | 14360 | HITS-CLIP | 37986224 | Unspecified |
| mmu-miR-30c-5p | MIMAT0000514 | Fzd3 | 14365 | HITS-CLIP | 37986224\|19536157 | Unspecified |
| mmu-miR-30c-5p | MIMAT0000514 | G2e3 | 217558 | HITS-CLIP | 37986224\|23142080 | Unspecified |
| mmu-miR-30c-5p | MIMAT0000514 | G3bp1 | 27041 | HITS-CLIP | 25753418 | Unspecified |
| mmu-miR-30c-5p | MIMAT0000514 | G3bp2 | 23881 | HITS-CLIP | 23142080 | Unspecified |
| mmu-miR-30c-5p | MIMAT0000514 | Gaa | 14387 | HITS-CLIP | 23597149 | Unspecified |

| | | | | | | |
|---|---|---|---|---|---|---|
| mmu-miR-30c-5p | MIMAT0000514 | Gab2 | 14389 | HITS-CLIP | 23142080 | Unspecified |
| mmu-miR-30c-5p | MIMAT0000514 | Gabra2 | 14395 | HITS-CLIP | 19536157 | Unspecified |
| mmu-miR-30c-5p | MIMAT0000514 | Galnt1 | 14423 | HITS-CLIP | 23142080 | Unspecified |
| mmu-miR-30c-5p | MIMAT0000514 | Galnt16 | 108760 | HITS-CLIP | 37986224 | Unspecified |
| mmu-miR-30c-5p | MIMAT0000514 | Galnt7 | 108150 | HITS-CLIP | 19536157\|23142080 | Unspecified |
| mmu-miR-30c-5p | MIMAT0000514 | Gan | 209239 | HITS-CLIP | 23142080\|23597149 | Unspecified |
| mmu-miR-30c-5p | MIMAT0000514 | Gapvd1 | 66691 | HITS-CLIP | 25753418 | Unspecified |
| mmu-miR-30c-5p | MIMAT0000514 | Gas7 | 14457 | HITS-CLIP | 37986224 | Unspecified |
| mmu-miR-30c-5p | MIMAT0000514 | Gata3 | 14462 | HITS-CLIP | 23142080 | Unspecified |
| mmu-miR-30c-5p | MIMAT0000514 | Gbp7 | 229900 | HITS-CLIP | 37986224 | Unspecified |
| mmu-miR-30c-5p | MIMAT0000514 | Gcdh | 270076 | HITS-CLIP | 23597149 | Unspecified |
| mmu-miR-30c-5p | MIMAT0000514 | Gclc | 14629 | HITS-CLIP | 23597149\|23142080 | Unspecified |
| mmu-miR-30c-5p | MIMAT0000514 | Gcsh | 68133 | HITS-CLIP | 23597149\|23142080 | Unspecified |
| mmu-miR-30c-5p | MIMAT0000514 | Gdap2 | 14547 | HITS-CLIP | 23597149\|23142080 | Unspecified |
| mmu-miR-30c-5p | MIMAT0000514 | Gdi2 | 14569 | HITS-CLIP | 25753418 | Unspecified |
| mmu-miR-30c-5p | MIMAT0000514 | Gem | 14579 | HITS-CLIP | 37986224 | Unspecified |
| mmu-miR-30c-5p | MIMAT0000514 | Get4 | 67604 | HITS-CLIP | 25753418 | Unspecified |
| mmu-miR-30c-5p | MIMAT0000514 | Gfod1 | 328232 | HITS-CLIP | 23142080 | Unspecified |
| mmu-miR-30c-5p | MIMAT0000514 | Gfpt1 | 14583 | HITS-CLIP | 37986224\|23142080 | Unspecified |
| mmu-miR-30c-5p | MIMAT0000514 | Ggcx | 56316 | HITS-CLIP | 23597149 | Unspecified |
| mmu-miR-30c-5p | MIMAT0000514 | Ggnbp2 | 217039 | HITS-CLIP | 37986224\|23142080 | Unspecified |
| mmu-miR-30c-5p | MIMAT0000514 | Gigyf1 | 57330 | HITS-CLIP//qCLASH | 37986224\|23597149\|31393953\|2314; | Unspecified |
| mmu-miR-30c-5p | MIMAT0000514 | Gimap8 | 243374 | HITS-CLIP | 37986224\|23142080 | Unspecified |
| mmu-miR-30c-5p | MIMAT0000514 | Gimap9 | 317758 | HITS-CLIP | 23142080 | Unspecified |
| mmu-miR-30c-5p | MIMAT0000514 | Gin1 | 252876 | HITS-CLIP | 23142080 | Unspecified |
| mmu-miR-30c-5p | MIMAT0000514 | Gins1 | 69270 | HITS-CLIP | 23142080 | Unspecified |
| mmu-miR-30c-5p | MIMAT0000514 | Gja1 | 14609 | HITS-CLIP | 37986224\|23142080\|19536157\|2359; | Unspecified |
| mmu-miR-30c-5p | MIMAT0000514 | Gjb2 | 14619 | HITS-CLIP | 23597149 | Unspecified |
| mmu-miR-30c-5p | MIMAT0000514 | Glce | 93683 | HITS-CLIP | 37986224\|23597149 | Unspecified |
| mmu-miR-30c-5p | MIMAT0000514 | Gldc | 104174 | HITS-CLIP | 37986224\|23597149\|19536157 | Unspecified |
| mmu-miR-30c-5p | MIMAT0000514 | Glg1 | 20340 | HITS-CLIP | 37986224 | Unspecified |
| mmu-miR-30c-5p | MIMAT0000514 | Glipr2 | 384009 | HITS-CLIP | 37986224 | Unspecified |
| mmu-miR-30c-5p | MIMAT0000514 | Gls | 14660 | HITS-CLIP | 37986224 | Unspecified |
| mmu-miR-30c-5p | MIMAT0000514 | Glt1d1 | 319804 | HITS-CLIP | 23597149 | Unspecified |
| mmu-miR-30c-5p | MIMAT0000514 | Glud1 | 14661 | HITS-CLIP | 23142080 | Unspecified |
| mmu-miR-30c-5p | MIMAT0000514 | Glul | 14645 | HITS-CLIP | 37986224 | Unspecified |
| mmu-miR-30c-5p | MIMAT0000514 | Glyr1 | 74022 | HITS-CLIP | 37986224 | Unspecified |
| mmu-miR-30c-5p | MIMAT0000514 | Gm15446 | 100642166 | qCLASH | 31393953 | Unspecified |
| mmu-miR-30c-5p | MIMAT0000514 | Gm20604 | 100859931 | HITS-CLIP | 23597149 | Unspecified |

| | | | | | | |
|---|---|---|---|---|---|---|
| mmu-miR-30c-5p | MIMAT0000514 | Gm38394 | 667118 | HITS-CLIP | 23142080 | Unspecified |
| mmu-miR-30c-5p | MIMAT0000514 | Gm45902 | 119065974 | HITS-CLIP | 23142080 | Unspecified |
| mmu-miR-30c-5p | MIMAT0000514 | Gm49336 | 127230315 | HITS-CLIP | 23142080 | Unspecified |
| mmu-miR-30c-5p | MIMAT0000514 | Gm49510 | | HITS-CLIP | 23142080 | Unspecified |
| mmu-miR-30c-5p | MIMAT0000514 | Gm5434 | 432649 | HITS-CLIP | 37986224 | Unspecified |
| mmu-miR-30c-5p | MIMAT0000514 | Gm7970 | 666186 | qCLASH | 31393953 | Unspecified |
| mmu-miR-30c-5p | MIMAT0000514 | Gm8229 | 666675 | qCLASH | 31393953 | Unspecified |
| mmu-miR-30c-5p | MIMAT0000514 | Gmeb2 | 229004 | HITS-CLIP | 23597149\|23142080 | Unspecified |
| mmu-miR-30c-5p | MIMAT0000514 | Gmfb | 63985 | HITS-CLIP | 23597149\|23142080 | Unspecified |
| mmu-miR-30c-5p | MIMAT0000514 | Gmppa | 69080 | HITS-CLIP | 23597149 | Unspecified |
| mmu-miR-30c-5p | MIMAT0000514 | Gna13 | 14674 | HITS-CLIP | 37986224\|23142080 | Unspecified |
| mmu-miR-30c-5p | MIMAT0000514 | Gnai2 | 14678 | HITS-CLIP | 37986224\|23597149\|19536157\|23142 | Unspecified |
| mmu-miR-30c-5p | MIMAT0000514 | Gnao1 | 14681 | HITS-CLIP | 19536157 | Unspecified |
| mmu-miR-30c-5p | MIMAT0000514 | Gnaq | 14682 | HITS-CLIP | 37986224\|19536157 | Unspecified |
| mmu-miR-30c-5p | MIMAT0000514 | Gnas | 14683 | HITS-CLIP | 23142080 | Unspecified |
| mmu-miR-30c-5p | MIMAT0000514 | Gng12 | 14701 | HITS-CLIP | 23142080 | Unspecified |
| mmu-miR-30c-5p | MIMAT0000514 | Gnl2 | 230737 | HITS-CLIP | 23597149 | Unspecified |
| mmu-miR-30c-5p | MIMAT0000514 | Gnpda1 | 26384 | HITS-CLIP | 23142080 | Unspecified |
| mmu-miR-30c-5p | MIMAT0000514 | Golga1 | 76899 | HITS-CLIP | 23142080 | Unspecified |
| mmu-miR-30c-5p | MIMAT0000514 | Golga4 | 54214 | HITS-CLIP | 37986224\|23142080 | Unspecified |
| mmu-miR-30c-5p | MIMAT0000514 | Golph3 | 66629 | HITS-CLIP | 23142080 | Unspecified |
| mmu-miR-30c-5p | MIMAT0000514 | Gorasp1 | 74498 | HITS-CLIP | 25753418 | Unspecified |
| mmu-miR-30c-5p | MIMAT0000514 | Gpatch2l | 70373 | HITS-CLIP | 23142080 | Unspecified |
| mmu-miR-30c-5p | MIMAT0000514 | Gpatch8 | 237943 | HITS-CLIP | 37986224\|23142080 | Unspecified |
| mmu-miR-30c-5p | MIMAT0000514 | Gpcpd1 | 74182 | HITS-CLIP | 37986224\|23597149\|25753418\|23142 | Unspecified |
| mmu-miR-30c-5p | MIMAT0000514 | Gpd1l | 333433 | HITS-CLIP | 23597149 | Unspecified |
| mmu-miR-30c-5p | MIMAT0000514 | Gphn | 268566 | HITS-CLIP | 23142080 | Unspecified |
| mmu-miR-30c-5p | MIMAT0000514 | Gpkow | 209416 | HITS-CLIP | 23142080 | Unspecified |
| mmu-miR-30c-5p | MIMAT0000514 | Gpm6a | 234267 | HITS-CLIP | 19536157 | Unspecified |
| mmu-miR-30c-5p | MIMAT0000514 | Gpm6b | 14758 | HITS-CLIP | 23142080 | Unspecified |
| mmu-miR-30c-5p | MIMAT0000514 | Gpr158 | 241263 | HITS-CLIP | 37986224 | Unspecified |
| mmu-miR-30c-5p | MIMAT0000514 | Gpr183 | 321019 | HITS-CLIP | 23142080 | Unspecified |
| mmu-miR-30c-5p | MIMAT0000514 | Gpr26 | 233919 | HITS-CLIP | 37986224 | Unspecified |
| mmu-miR-30c-5p | MIMAT0000514 | Gprin3 | 243385 | HITS-CLIP | 23597149\|23142080 | Unspecified |
| mmu-miR-30c-5p | MIMAT0000514 | Gpt2 | 108682 | HITS-CLIP | 37986224\|23597149 | Unspecified |
| mmu-miR-30c-5p | MIMAT0000514 | Gramd3 | 107022 | HITS-CLIP | 37986224\|23142080\|23597149 | Unspecified |
| mmu-miR-30c-5p | MIMAT0000514 | Grb10 | 14783 | HITS-CLIP | 37986224\|19536157 | Unspecified |
| mmu-miR-30c-5p | MIMAT0000514 | Grb14 | 50915 | HITS-CLIP | 23597149 | Unspecified |
| mmu-miR-30c-5p | MIMAT0000514 | Grina | 66168 | HITS-CLIP | 23597149 | Unspecified |

| | | | | | | |
|---|---|---|---|---|---|---|
| mmu-miR-30c-5p | MIMAT0000514 | Grk5 | 14773 | HITS-CLIP | 37986224\|23142080 | Unspecified |
| mmu-miR-30c-5p | MIMAT0000514 | Grk6 | 26385 | HITS-CLIP | 23142080 | Unspecified |
| mmu-miR-30c-5p | MIMAT0000514 | Grm3 | 108069 | HITS-CLIP | 37986224 | Unspecified |
| mmu-miR-30c-5p | MIMAT0000514 | Grm5 | 108071 | HITS-CLIP | 37986224 | Unspecified |
| mmu-miR-30c-5p | MIMAT0000514 | Grpel2 | 17714 | HITS-CLIP | 37986224\|23142080 | Unspecified |
| mmu-miR-30c-5p | MIMAT0000514 | Gskip | 66787 | HITS-CLIP | 25083871\|23142080 | Unspecified |
| mmu-miR-30c-5p | MIMAT0000514 | Gsn | 227753 | HITS-CLIP | 37986224 | Unspecified |
| mmu-miR-30c-5p | MIMAT0000514 | Gstk1 | 76263 | HITS-CLIP | 25753418 | Unspecified |
| mmu-miR-30c-5p | MIMAT0000514 | Gstm3 | 14864 | HITS-CLIP | 37986224 | Unspecified |
| mmu-miR-30c-5p | MIMAT0000514 | Gstm4 | 14865 | HITS-CLIP | 23597149 | Unspecified |
| mmu-miR-30c-5p | MIMAT0000514 | Gtf2a1 | 83602 | HITS-CLIP | 23142080 | Unspecified |
| mmu-miR-30c-5p | MIMAT0000514 | Gtf2e2 | 68153 | HITS-CLIP | 23142080 | Unspecified |
| mmu-miR-30c-5p | MIMAT0000514 | Gtf2f2 | 68705 | HITS-CLIP | 23142080 | Unspecified |
| mmu-miR-30c-5p | MIMAT0000514 | Gtf2h4 | 14885 | HITS-CLIP | 23142080 | Unspecified |
| mmu-miR-30c-5p | MIMAT0000514 | Gtf2i | 14886 | HITS-CLIP | 23142080 | Unspecified |
| mmu-miR-30c-5p | MIMAT0000514 | Gtf3c1 | 233863 | HITS-CLIP | 25753418 | Unspecified |
| mmu-miR-30c-5p | MIMAT0000514 | Gtse1 | 29870 | HITS-CLIP | 37986224 | Unspecified |
| mmu-miR-30c-5p | MIMAT0000514 | Gucd1 | 68778 | HITS-CLIP | 37986224\|23597149\|23142080 | Unspecified |
| mmu-miR-30c-5p | MIMAT0000514 | Gulo | 268756 | HITS-CLIP | 37986224 | Unspecified |
| mmu-miR-30c-5p | MIMAT0000514 | Gzf1 | 74533 | HITS-CLIP | 37986224\|23597149\|23142080 | Unspecified |
| mmu-miR-30c-5p | MIMAT0000514 | Gzmb | 14939 | HITS-CLIP | 37986224\|23142080 | Unspecified |
| mmu-miR-30c-5p | MIMAT0000514 | H2-T23 | 15040 | HITS-CLIP | 31433990 | Unspecified |
| mmu-miR-30c-5p | MIMAT0000514 | H2afy | 9555 | HITS-CLIP | 23142080 | Unspecified |
| mmu-miR-30c-5p | MIMAT0000514 | Hace1 | 209462 | HITS-CLIP | 23142080 | Unspecified |
| mmu-miR-30c-5p | MIMAT0000514 | Hap1 | 15114 | HITS-CLIP | 37986224 | Unspecified |
| mmu-miR-30c-5p | MIMAT0000514 | Haus4 | 219072 | HITS-CLIP | 37986224\|23142080 | Unspecified |
| mmu-miR-30c-5p | MIMAT0000514 | Haus6 | 230376 | HITS-CLIP | 37986224\|23142080 | Unspecified |
| mmu-miR-30c-5p | MIMAT0000514 | Hbp1 | 73389 | HITS-CLIP | 37986224\|23142080\|25753418 | Unspecified |
| mmu-miR-30c-5p | MIMAT0000514 | Hc | 15139 | HITS-CLIP | 23597149 | Unspecified |
| mmu-miR-30c-5p | MIMAT0000514 | Hcfc2 | 67933 | HITS-CLIP | 37986224\|23142080 | Unspecified |
| mmu-miR-30c-5p | MIMAT0000514 | Hdgf | 15191 | HITS-CLIP | 23597149\|23142080 | Unspecified |
| mmu-miR-30c-5p | MIMAT0000514 | Hdgfl3 | 29877 | HITS-CLIP | 23142080 | Unspecified |
| mmu-miR-30c-5p | MIMAT0000514 | Hdlbp | 110611 | HITS-CLIP | 23142080 | Unspecified |
| mmu-miR-30c-5p | MIMAT0000514 | Heatr3 | 234549 | HITS-CLIP | 23597149 | Unspecified |
| mmu-miR-30c-5p | MIMAT0000514 | Heatr5a | 320487 | HITS-CLIP | 23142080 | Unspecified |
| mmu-miR-30c-5p | MIMAT0000514 | Heatr5b | 320473 | HITS-CLIP | 25753418 | Unspecified |
| mmu-miR-30c-5p | MIMAT0000514 | Hectd1 | 207304 | HITS-CLIP | 23142080 | Unspecified |
| mmu-miR-30c-5p | MIMAT0000514 | Helb | 117599 | HITS-CLIP | 23597149 | Unspecified |
| mmu-miR-30c-5p | MIMAT0000514 | Helz | 78455 | HITS-CLIP | 37986224\|23142080 | Unspecified |

| | | | | | | |
|---|---|---|---|---|---|---|
| mmu-miR-30c-5p | MIMAT0000514 | Herc1 | 235439 | HITS-CLIP | 37986224\|25753418\|23142080 | Unspecified |
| mmu-miR-30c-5p | MIMAT0000514 | Herc2 | 15204 | HITS-CLIP | 37986224\|23142080 | Unspecified |
| mmu-miR-30c-5p | MIMAT0000514 | Herpud1 | 64209 | HITS-CLIP | 25753418 | Unspecified |
| mmu-miR-30c-5p | MIMAT0000514 | Hif1a | 15251 | HITS-CLIP | 37986224\|23597149 | Unspecified |
| mmu-miR-30c-5p | MIMAT0000514 | Hint3 | 66847 | HITS-CLIP | 37986224\|23142080 | Unspecified |
| mmu-miR-30c-5p | MIMAT0000514 | Hipk1 | 15257 | HITS-CLIP | 37986224\|23142080 | Unspecified |
| mmu-miR-30c-5p | MIMAT0000514 | Hipk2 | 15258 | HITS-CLIP | 37986224\|25753418 | Unspecified |
| mmu-miR-30c-5p | MIMAT0000514 | Hipk3 | 15259 | HITS-CLIP | 37986224 | Unspecified |
| mmu-miR-30c-5p | MIMAT0000514 | Hirip3 | 233876 | qCLASH | 31393953 | Unspecified |
| mmu-miR-30c-5p | MIMAT0000514 | Hist1h4i | 319158 | HITS-CLIP | 23142080 | Unspecified |
| mmu-miR-30c-5p | MIMAT0000514 | Hist2h2bb | 319189 | HITS-CLIP | 23142080\|23597149 | Unspecified |
| mmu-miR-30c-5p | MIMAT0000514 | Hist2h3c1 | 15077 | HITS-CLIP | 23597149 | Unspecified |
| mmu-miR-30c-5p | MIMAT0000514 | Hivep3 | 16656 | HITS-CLIP | 37986224\|23142080 | Unspecified |
| mmu-miR-30c-5p | MIMAT0000514 | Hmbox1 | 219150 | HITS-CLIP | 23142080 | Unspecified |
| mmu-miR-30c-5p | MIMAT0000514 | Hmbs | 15288 | HITS-CLIP | 23142080 | Unspecified |
| mmu-miR-30c-5p | MIMAT0000514 | Hmgb2 | 97165 | qCLASH | 31393953 | Unspecified |
| mmu-miR-30c-5p | MIMAT0000514 | Hmgcr | 15357 | HITS-CLIP | 37986224 | Unspecified |
| mmu-miR-30c-5p | MIMAT0000514 | Hmgcs1 | 208715 | HITS-CLIP | 37986224 | Unspecified |
| mmu-miR-30c-5p | MIMAT0000514 | Hnrnpa0 | 77134 | HITS-CLIP | 37986224\|23142080 | Unspecified |
| mmu-miR-30c-5p | MIMAT0000514 | Hnrnpa2b1 | 53379 | HITS-CLIP | 23142080 | Unspecified |
| mmu-miR-30c-5p | MIMAT0000514 | Hnrnpab | 15384 | HITS-CLIP | 37986224\|23597149\|23142080 | Unspecified |
| mmu-miR-30c-5p | MIMAT0000514 | Hnrnpd | 11991 | HITS-CLIP | 23142080 | Unspecified |
| mmu-miR-30c-5p | MIMAT0000514 | Hnrnpl | 15388 | HITS-CLIP | 23142080\|23597149 | Unspecified |
| mmu-miR-30c-5p | MIMAT0000514 | Hnrnpm | 76936 | HITS-CLIP | 37986224\|23142080\|19536157 | Unspecified |
| mmu-miR-30c-5p | MIMAT0000514 | Hnrnpu | 51810 | HITS-CLIP | 37986224\|19536157\|23142080 | Unspecified |
| mmu-miR-30c-5p | MIMAT0000514 | Hnrnpul2 | 68693 | HITS-CLIP | 23597149\|23142080 | Unspecified |
| mmu-miR-30c-5p | MIMAT0000514 | Hook1 | 77963 | HITS-CLIP | 23142080 | Unspecified |
| mmu-miR-30c-5p | MIMAT0000514 | Hook3 | 320191 | HITS-CLIP | 37986224 | Unspecified |
| mmu-miR-30c-5p | MIMAT0000514 | Hpcal4 | 170638 | HITS-CLIP | 37986224 | Unspecified |
| mmu-miR-30c-5p | MIMAT0000514 | Hs6st1 | 50785 | HITS-CLIP | 26203562 | Unspecified |
| mmu-miR-30c-5p | MIMAT0000514 | Hsd3b5 | 15496 | HITS-CLIP | 37986224 | Unspecified |
| mmu-miR-30c-5p | MIMAT0000514 | Hsdl2 | 72479 | HITS-CLIP | 25753418\|23142080 | Unspecified |
| mmu-miR-30c-5p | MIMAT0000514 | Hsp90ab1 | 15516 | HITS-CLIP | 23142080 | Unspecified |
| mmu-miR-30c-5p | MIMAT0000514 | Hspa14 | 50497 | HITS-CLIP | 37986224\|23142080 | Unspecified |
| mmu-miR-30c-5p | MIMAT0000514 | Hspa4 | 15525 | HITS-CLIP//qCLASH | 23597149\|31393953\|23142080 | Unspecified |
| mmu-miR-30c-5p | MIMAT0000514 | Hspa4l | 18415 | HITS-CLIP | 25753418 | Unspecified |
| mmu-miR-30c-5p | MIMAT0000514 | Hspa5 | 14828 | HITS-CLIP | 37986224\|23597149\|23142080 | Unspecified |
| mmu-miR-30c-5p | MIMAT0000514 | Hspb1 | 15507 | HITS-CLIP | 25753418 | Unspecified |
| mmu-miR-30c-5p | MIMAT0000514 | Hspd1 | 15510 | qCLASH | 31393953 | Unspecified |

| | | | | | | |
|---|---|---|---|---|---|---|
| mmu-miR-30c-5p | MIMAT0000514 | Hsph1 | 15505 | HITS-CLIP | 23142080 | Unspecified |
| mmu-miR-30c-5p | MIMAT0000514 | Htra2 | 64704 | HITS-CLIP | 23597149\|23142080 | Unspecified |
| mmu-miR-30c-5p | MIMAT0000514 | Hus1 | 15574 | HITS-CLIP | 37986224 | Unspecified |
| mmu-miR-30c-5p | MIMAT0000514 | Huwe1 | 59026 | HITS-CLIP | 19536157\|23142080 | Unspecified |
| mmu-miR-30c-5p | MIMAT0000514 | Hyal2 | 15587 | HITS-CLIP | 23597149\|25753418 | Unspecified |
| mmu-miR-30c-5p | MIMAT0000514 | Iah1 | 67732 | HITS-CLIP | 23597149\|23142080 | Unspecified |
| mmu-miR-30c-5p | MIMAT0000514 | Ibtk | 108837 | HITS-CLIP | 37986224\|23597149\|23142080 | Unspecified |
| mmu-miR-30c-5p | MIMAT0000514 | Icmt | 57295 | qCLASH | 31393953 | Unspecified |
| mmu-miR-30c-5p | MIMAT0000514 | Id2 | 15902 | HITS-CLIP | 23597149 | Unspecified |
| mmu-miR-30c-5p | MIMAT0000514 | Idh1 | 15926 | HITS-CLIP | 23597149 | Unspecified |
| mmu-miR-30c-5p | MIMAT0000514 | Idh3g | 15929 | HITS-CLIP | 23142080 | Unspecified |
| mmu-miR-30c-5p | MIMAT0000514 | Ier2 | 15936 | HITS-CLIP | 23597149\|23142080 | Unspecified |
| mmu-miR-30c-5p | MIMAT0000514 | Ier5 | 15939 | HITS-CLIP | 37986224 | Unspecified |
| mmu-miR-30c-5p | MIMAT0000514 | Iffo2 | 212632 | HITS-CLIP | 23142080\|23597149 | Unspecified |
| mmu-miR-30c-5p | MIMAT0000514 | Ifnar1 | 15975 | HITS-CLIP | 23142080 | Unspecified |
| mmu-miR-30c-5p | MIMAT0000514 | Ifnar2 | 15976 | HITS-CLIP | 23142080 | Unspecified |
| mmu-miR-30c-5p | MIMAT0000514 | Ifngr1 | 15979 | HITS-CLIP | 37986224 | Unspecified |
| mmu-miR-30c-5p | MIMAT0000514 | Ifngr2 | 15980 | HITS-CLIP//qCLASH | 37986224\|23597149\|23142080\|31393 | Unspecified |
| mmu-miR-30c-5p | MIMAT0000514 | Ift172 | 67661 | HITS-CLIP | 25753418 | Unspecified |
| mmu-miR-30c-5p | MIMAT0000514 | Igf1 | 16000 | HITS-CLIP | 37986224\|23597149 | Unspecified |
| mmu-miR-30c-5p | MIMAT0000514 | Igf1r | 16001 | HITS-CLIP | 23142080 | Unspecified |
| mmu-miR-30c-5p | MIMAT0000514 | Igf2r | 16004 | HITS-CLIP | 37986224\|25753418 | Unspecified |
| mmu-miR-30c-5p | MIMAT0000514 | Igfbp4 | 16010 | HITS-CLIP | 23142080 | Unspecified |
| mmu-miR-30c-5p | MIMAT0000514 | Igfbp5 | 16011 | HITS-CLIP | 37986224 | Unspecified |
| mmu-miR-30c-5p | MIMAT0000514 | Igsf3 | 78908 | HITS-CLIP | 37986224 | Unspecified |
| mmu-miR-30c-5p | MIMAT0000514 | Igtp | 16145 | HITS-CLIP | 37986224 | Unspecified |
| mmu-miR-30c-5p | MIMAT0000514 | Ik | 24010 | HITS-CLIP | 23142080 | Unspecified |
| mmu-miR-30c-5p | MIMAT0000514 | Ikzf1 | 22778 | HITS-CLIP | 37986224\|23142080 | Unspecified |
| mmu-miR-30c-5p | MIMAT0000514 | Il10ra | 16154 | HITS-CLIP | 23142080 | Unspecified |
| mmu-miR-30c-5p | MIMAT0000514 | Il12rb2 | 16162 | HITS-CLIP | 37986224\|23142080 | Unspecified |
| mmu-miR-30c-5p | MIMAT0000514 | Il18r1 | 16182 | HITS-CLIP | 25083871\|23142080\|23597149 | Unspecified |
| mmu-miR-30c-5p | MIMAT0000514 | Il2 | 16183 | HITS-CLIP | 23142080 | Unspecified |
| mmu-miR-30c-5p | MIMAT0000514 | Il21r | 60504 | HITS-CLIP | 37986224\|23142080 | Unspecified |
| mmu-miR-30c-5p | MIMAT0000514 | Il2ra | 16184 | HITS-CLIP | 37986224 | Unspecified |
| mmu-miR-30c-5p | MIMAT0000514 | Il4ra | 16190 | HITS-CLIP | 23142080 | Unspecified |
| mmu-miR-30c-5p | MIMAT0000514 | Il6st | 16195 | HITS-CLIP | 37986224\|23142080\|23597149 | Unspecified |
| mmu-miR-30c-5p | MIMAT0000514 | Impact | 16210 | HITS-CLIP | 37986224 | Unspecified |
| mmu-miR-30c-5p | MIMAT0000514 | Ing4 | 28019 | HITS-CLIP | 23597149\|23142080 | Unspecified |
| mmu-miR-30c-5p | MIMAT0000514 | Inhba | 16323 | HITS-CLIP | 19536157\|23142080 | Unspecified |

| | | | | | | |
|---|---|---|---|---|---|---|
| mmu-miR-30c-5p | MIMAT0000514 | Inhbb | 16324 | HITS-CLIP | 19536157 | Unspecified |
| mmu-miR-30c-5p | MIMAT0000514 | Ino80d | 227195 | HITS-CLIP | 37986224\|23142080 | Unspecified |
| mmu-miR-30c-5p | MIMAT0000514 | Inpp4a | 269180 | HITS-CLIP | 37986224\|23142080 | Unspecified |
| mmu-miR-30c-5p | MIMAT0000514 | Inpp5d | 16331 | HITS-CLIP | 23142080 | Unspecified |
| mmu-miR-30c-5p | MIMAT0000514 | Inpp5f | 101490 | HITS-CLIP | 23142080 | Unspecified |
| mmu-miR-30c-5p | MIMAT0000514 | Insig2 | 72999 | HITS-CLIP | 37986224\|23142080 | Unspecified |
| mmu-miR-30c-5p | MIMAT0000514 | Ints2 | 70422 | HITS-CLIP | 23142080 | Unspecified |
| mmu-miR-30c-5p | MIMAT0000514 | Ints3 | 229543 | HITS-CLIP | 23597149 | Unspecified |
| mmu-miR-30c-5p | MIMAT0000514 | Ints6 | 18130 | HITS-CLIP | 23142080 | Unspecified |
| mmu-miR-30c-5p | MIMAT0000514 | Ints8 | 72656 | HITS-CLIP | 19536157\|23597149\|25753418\|23142 | Unspecified |
| mmu-miR-30c-5p | MIMAT0000514 | Ints9 | 210925 | HITS-CLIP | 23597149 | Unspecified |
| mmu-miR-30c-5p | MIMAT0000514 | Ipmk | 69718 | HITS-CLIP | 37986224\|23597149\|23142080 | Unspecified |
| mmu-miR-30c-5p | MIMAT0000514 | Ipo11 | 76582 | HITS-CLIP | 25753418 | Unspecified |
| mmu-miR-30c-5p | MIMAT0000514 | Ipo5 | 70572 | HITS-CLIP | 25753418 | Unspecified |
| mmu-miR-30c-5p | MIMAT0000514 | Ipo7 | 233726 | HITS-CLIP | 37986224 | Unspecified |
| mmu-miR-30c-5p | MIMAT0000514 | Ireb2 | 64602 | HITS-CLIP | 37986224 | Unspecified |
| mmu-miR-30c-5p | MIMAT0000514 | Irf1 | 16362 | HITS-CLIP | 37986224\|23142080 | Unspecified |
| mmu-miR-30c-5p | MIMAT0000514 | Irf2bp2 | 270110 | HITS-CLIP | 37986224\|23142080 | Unspecified |
| mmu-miR-30c-5p | MIMAT0000514 | Irf4 | 16364 | HITS-CLIP | 37986224\|23142080 | Unspecified |
| mmu-miR-30c-5p | MIMAT0000514 | Irs1 | 16367 | HITS-CLIP | 37986224 | Unspecified |
| mmu-miR-30c-5p | MIMAT0000514 | Isca2 | 74316 | HITS-CLIP | 23142080 | Unspecified |
| mmu-miR-30c-5p | MIMAT0000514 | Itch | 16396 | HITS-CLIP | 23597149 | Unspecified |
| mmu-miR-30c-5p | MIMAT0000514 | Itfg1 | 71927 | HITS-CLIP | 37986224\|19536157\|23142080 | Unspecified |
| mmu-miR-30c-5p | MIMAT0000514 | Itgb1 | 16412 | HITS-CLIP | 37986224\|23597149\|23142080 | Unspecified |
| mmu-miR-30c-5p | MIMAT0000514 | Itgb3 | 16416 | HITS-CLIP | 37986224\|23597149\|23142080 | Unspecified |
| mmu-miR-30c-5p | MIMAT0000514 | Itgb8 | 320910 | HITS-CLIP | 37986224\|23142080 | Unspecified |
| mmu-miR-30c-5p | MIMAT0000514 | Itih5 | 209378 | HITS-CLIP | 23142080 | Unspecified |
| mmu-miR-30c-5p | MIMAT0000514 | Itm2c | 64294 | qCLASH | 31393953 | Unspecified |
| mmu-miR-30c-5p | MIMAT0000514 | Itpk1 | 217837 | HITS-CLIP | 23597149\|23142080 | Unspecified |
| mmu-miR-30c-5p | MIMAT0000514 | Itprid2 | 6744 | HITS-CLIP | 23142080 | Unspecified |
| mmu-miR-30c-5p | MIMAT0000514 | Itsn1 | 16443 | HITS-CLIP | 37986224\|19536157 | Unspecified |
| mmu-miR-30c-5p | MIMAT0000514 | Itsn2 | 20403 | HITS-CLIP | 23142080 | Unspecified |
| mmu-miR-30c-5p | MIMAT0000514 | Ivns1abp | 117198 | HITS-CLIP | 23142080 | Unspecified |
| mmu-miR-30c-5p | MIMAT0000514 | Jade1 | 269424 | HITS-CLIP | 37986224 | Unspecified |
| mmu-miR-30c-5p | MIMAT0000514 | Jade3 | 382207 | HITS-CLIP | 37986224\|23597149\|19536157\|23142 | Unspecified |
| mmu-miR-30c-5p | MIMAT0000514 | Jak1 | 16451 | HITS-CLIP | 23142080 | Unspecified |
| mmu-miR-30c-5p | MIMAT0000514 | Jak2 | 16452 | HITS-CLIP | 23142080 | Unspecified |
| mmu-miR-30c-5p | MIMAT0000514 | Jakmip2 | 76217 | HITS-CLIP | 37986224\|19536157 | Unspecified |
| mmu-miR-30c-5p | MIMAT0000514 | Jarid2 | 16468 | HITS-CLIP | 37986224 | Unspecified |

| | | | | | | | |
|---|---|---|---|---|---|---|---|
| mmu-miR-30c-5p | MIMAT0000514 | Jdp2 | 81703 | HITS-CLIP | | 37986224\|23142080\|23597149\|1953[illegible] | Unspecified |
| mmu-miR-30c-5p | MIMAT0000514 | Jmjd1c | 108829 | HITS-CLIP | | 37986224 | Unspecified |
| mmu-miR-30c-5p | MIMAT0000514 | Jmy | 57748 | HITS-CLIP | | 37986224\|23142080 | Unspecified |
| mmu-miR-30c-5p | MIMAT0000514 | Josd1 | 74158 | HITS-CLIP | | 37986224\|23597149\|23142080 | Unspecified |
| mmu-miR-30c-5p | MIMAT0000514 | Jph4 | 319984 | HITS-CLIP | | 37986224\|19536157 | Unspecified |
| mmu-miR-30c-5p | MIMAT0000514 | Jrkl | 77532 | qCLASH | | 31393953 | Unspecified |
| mmu-miR-30c-5p | MIMAT0000514 | Junb | 16477 | HITS-CLIP | | 37986224\|23142080 | Unspecified |
| mmu-miR-30c-5p | MIMAT0000514 | Jup | 16480 | HITS-CLIP | | 23142080 | Unspecified |
| mmu-miR-30c-5p | MIMAT0000514 | Kalrn | 545156 | HITS-CLIP | | 19536157 | Unspecified |
| mmu-miR-30c-5p | MIMAT0000514 | Kansl1 | 76719 | HITS-CLIP | | 23142080 | Unspecified |
| mmu-miR-30c-5p | MIMAT0000514 | Kantr | 102723508 | HITS-CLIP | | 23142080\|19536157 | Unspecified |
| mmu-miR-30c-5p | MIMAT0000514 | Kars | 85305 | HITS-CLIP | | 25753418 | Unspecified |
| mmu-miR-30c-5p | MIMAT0000514 | Kat6a | 244349 | HITS-CLIP | | 23597149 | Unspecified |
| mmu-miR-30c-5p | MIMAT0000514 | Katnbl1 | 72425 | HITS-CLIP | | 37986224 | Unspecified |
| mmu-miR-30c-5p | MIMAT0000514 | Kbtbd2 | 210973 | HITS-CLIP | | 23142080 | Unspecified |
| mmu-miR-30c-5p | MIMAT0000514 | Kbtbd8 | 243574 | HITS-CLIP | | 23142080 | Unspecified |
| mmu-miR-30c-5p | MIMAT0000514 | Kcmf1 | 74287 | HITS-CLIP | | 23597149 | Unspecified |
| mmu-miR-30c-5p | MIMAT0000514 | Kcnh7 | 170738 | HITS-CLIP | | 37986224 | Unspecified |
| mmu-miR-30c-5p | MIMAT0000514 | Kcnj12 | 16515 | HITS-CLIP | | 37986224 | Unspecified |
| mmu-miR-30c-5p | MIMAT0000514 | Kctd20 | 66989 | HITS-CLIP | | 37986224 | Unspecified |
| mmu-miR-30c-5p | MIMAT0000514 | Kctd3 | 226823 | HITS-CLIP | | 37986224 | Unspecified |
| mmu-miR-30c-5p | MIMAT0000514 | Kctd5 | 69259 | HITS-CLIP | | 23597149\|23142080 | Unspecified |
| mmu-miR-30c-5p | MIMAT0000514 | Kctd7 | 212919 | HITS-CLIP | | 23597149\|19536157 | Unspecified |
| mmu-miR-30c-5p | MIMAT0000514 | Kctd8 | 243043 | HITS-CLIP | | 37986224 | Unspecified |
| mmu-miR-30c-5p | MIMAT0000514 | Kdelr1 | 68137 | HITS-CLIP | | 23142080 | Unspecified |
| mmu-miR-30c-5p | MIMAT0000514 | Kdm2a | 225876 | HITS-CLIP | | 37986224 | Unspecified |
| mmu-miR-30c-5p | MIMAT0000514 | Kdm3a | 104263 | HITS-CLIP | | 37986224\|23142080 | Unspecified |
| mmu-miR-30c-5p | MIMAT0000514 | Kdm5a | 214899 | HITS-CLIP | | 37986224\|23142080 | Unspecified |
| mmu-miR-30c-5p | MIMAT0000514 | Kdm5b | 75605 | HITS-CLIP | | 23142080 | Unspecified |
| mmu-miR-30c-5p | MIMAT0000514 | Kdm5c | 20591 | HITS-CLIP | | 37986224 | Unspecified |
| mmu-miR-30c-5p | MIMAT0000514 | Kdm5d | 20592 | HITS-CLIP | | 23142080 | Unspecified |
| mmu-miR-30c-5p | MIMAT0000514 | Kdm6b | 216850 | HITS-CLIP | | 23597149\|23142080 | Unspecified |
| mmu-miR-30c-5p | MIMAT0000514 | Kdm7a | 338523 | HITS-CLIP | | 37986224\|23142080 | Unspecified |
| mmu-miR-30c-5p | MIMAT0000514 | Khnyn | 219094 | HITS-CLIP | | 37986224\|23597149\|23142080 | Unspecified |
| mmu-miR-30c-5p | MIMAT0000514 | Kidins220 | 77480 | HITS-CLIP | | 37986224\|23597149 | Unspecified |
| mmu-miR-30c-5p | MIMAT0000514 | Kif11 | 16551 | HITS-CLIP | | 23142080 | Unspecified |
| mmu-miR-30c-5p | MIMAT0000514 | Kif1a | 16560 | HITS-CLIP | | 37986224\|19536157 | Unspecified |
| mmu-miR-30c-5p | MIMAT0000514 | Kif21b | 16565 | HITS-CLIP | | 37986224\|31433990\|23142080 | Unspecified |
| mmu-miR-30c-5p | MIMAT0000514 | Kif23 | 71819 | HITS-CLIP | | 23142080 | Unspecified |

| | | | | | | |
|---|---|---|---|---|---|---|
| mmu-miR-30c-5p | MIMAT0000514 | Kif3a | 16568 | HITS-CLIP | 37986224 | Unspecified |
| mmu-miR-30c-5p | MIMAT0000514 | Kif5b | 16573 | HITS-CLIP//qCLASH | 37986224\|23597149\|31393953\|23142 | Unspecified |
| mmu-miR-30c-5p | MIMAT0000514 | Kif5c | 16574 | HITS-CLIP | 37986224 | Unspecified |
| mmu-miR-30c-5p | MIMAT0000514 | Klf10 | 21847 | HITS-CLIP | 23597149\|23142080 | Unspecified |
| mmu-miR-30c-5p | MIMAT0000514 | Klf13 | 50794 | HITS-CLIP | 37986224\|23142080 | Unspecified |
| mmu-miR-30c-5p | MIMAT0000514 | Klf6 | 23849 | HITS-CLIP | 37986224 | Unspecified |
| mmu-miR-30c-5p | MIMAT0000514 | Klf7 | 93691 | HITS-CLIP | 37986224 | Unspecified |
| mmu-miR-30c-5p | MIMAT0000514 | Klf9 | 16601 | HITS-CLIP | 37986224 | Unspecified |
| mmu-miR-30c-5p | MIMAT0000514 | Klhdc1 | 271005 | HITS-CLIP | 23142080 | Unspecified |
| mmu-miR-30c-5p | MIMAT0000514 | Klhdc2 | 69554 | HITS-CLIP | 37986224\|23142080 | Unspecified |
| mmu-miR-30c-5p | MIMAT0000514 | Klhl11 | 217194 | HITS-CLIP | 37986224 | Unspecified |
| mmu-miR-30c-5p | MIMAT0000514 | Klhl12 | 240756 | HITS-CLIP | 23597149 | Unspecified |
| mmu-miR-30c-5p | MIMAT0000514 | Klhl20 | 226541 | HITS-CLIP | 23142080 | Unspecified |
| mmu-miR-30c-5p | MIMAT0000514 | Klhl23 | 277396 | HITS-CLIP | 37986224 | Unspecified |
| mmu-miR-30c-5p | MIMAT0000514 | Klhl28 | 66689 | HITS-CLIP | 37986224\|23142080 | Unspecified |
| mmu-miR-30c-5p | MIMAT0000514 | Klhl3 | 100503085 | HITS-CLIP | 23597149 | Unspecified |
| mmu-miR-30c-5p | MIMAT0000514 | Klhl42 | 232539 | HITS-CLIP | 37986224 | Unspecified |
| mmu-miR-30c-5p | MIMAT0000514 | Klhl5 | 71778 | HITS-CLIP | 37986224\|23142080 | Unspecified |
| mmu-miR-30c-5p | MIMAT0000514 | Klhl6 | 239743 | HITS-CLIP | 23142080 | Unspecified |
| mmu-miR-30c-5p | MIMAT0000514 | Klhl9 | 242521 | HITS-CLIP | 37986224 | Unspecified |
| mmu-miR-30c-5p | MIMAT0000514 | Klrd1 | 16643 | HITS-CLIP | 23142080 | Unspecified |
| mmu-miR-30c-5p | MIMAT0000514 | Kmt2a | 214162 | HITS-CLIP | 37986224\|23142080 | Unspecified |
| mmu-miR-30c-5p | MIMAT0000514 | Kmt2b | 75410 | HITS-CLIP | 23142080 | Unspecified |
| mmu-miR-30c-5p | MIMAT0000514 | Kmt2c | 231051 | HITS-CLIP | 23142080 | Unspecified |
| mmu-miR-30c-5p | MIMAT0000514 | Kmt2e | 69188 | HITS-CLIP | 37986224\|23597149\|19536157\|23142 | Unspecified |
| mmu-miR-30c-5p | MIMAT0000514 | Kmt5b | 225888 | HITS-CLIP | 23142080 | Unspecified |
| mmu-miR-30c-5p | MIMAT0000514 | Kpna1 | 16646 | HITS-CLIP | 37986224\|23142080\|25753418 | Unspecified |
| mmu-miR-30c-5p | MIMAT0000514 | Kpna3 | 16648 | HITS-CLIP | 23142080 | Unspecified |
| mmu-miR-30c-5p | MIMAT0000514 | Kpnb1 | 16211 | HITS-CLIP | 23142080 | Unspecified |
| mmu-miR-30c-5p | MIMAT0000514 | Kras | 16653 | HITS-CLIP | 37986224\|23597149\|23142080 | Unspecified |
| mmu-miR-30c-5p | MIMAT0000514 | Kremen1 | 84035 | HITS-CLIP | 23597149 | Unspecified |
| mmu-miR-30c-5p | MIMAT0000514 | Krit1 | 79264 | HITS-CLIP | 23142080 | Unspecified |
| mmu-miR-30c-5p | MIMAT0000514 | Krt1 | 3848 | HITS-CLIP | 26203562 | Unspecified |
| mmu-miR-30c-5p | MIMAT0000514 | Ksr1 | 16706 | HITS-CLIP | 23142080 | Unspecified |
| mmu-miR-30c-5p | MIMAT0000514 | Kyat3 | 56267 | HITS-CLIP | 23597149 | Unspecified |
| mmu-miR-30c-5p | MIMAT0000514 | Lacc1 | 210808 | HITS-CLIP | 23597149 | Unspecified |
| mmu-miR-30c-5p | MIMAT0000514 | Lamp2 | 16784 | HITS-CLIP | 23597149 | Unspecified |
| mmu-miR-30c-5p | MIMAT0000514 | Laptm4a | 17775 | qCLASH//HITS-CLIP | 31393953\|23142080 | Unspecified |
| mmu-miR-30c-5p | MIMAT0000514 | Larp1 | 73158 | HITS-CLIP | 37986224 | Unspecified |

| | | | | | | |
|---|---|---|---|---|---|---|
| mmu-miR-30c-5p | MIMAT0000514 | Larp4b | 217980 | HITS-CLIP | 23142080 | Unspecified |
| mmu-miR-30c-5p | MIMAT0000514 | Larp7 | 28036 | HITS-CLIP | 23142080 | Unspecified |
| mmu-miR-30c-5p | MIMAT0000514 | Lats1 | 16798 | HITS-CLIP | 37986224 | Unspecified |
| mmu-miR-30c-5p | MIMAT0000514 | Lats2 | 50523 | HITS-CLIP | 37986224 | Unspecified |
| mmu-miR-30c-5p | MIMAT0000514 | Lax1 | 240754 | HITS-CLIP | 37986224 | Unspecified |
| mmu-miR-30c-5p | MIMAT0000514 | Lclat1 | 225010 | HITS-CLIP | 23597149 | Unspecified |
| mmu-miR-30c-5p | MIMAT0000514 | Lcmt1 | 30949 | HITS-CLIP | 37986224\|23597149 | Unspecified |
| mmu-miR-30c-5p | MIMAT0000514 | Lcor | 212391 | HITS-CLIP | 23142080 | Unspecified |
| mmu-miR-30c-5p | MIMAT0000514 | Lcorl | 209707 | HITS-CLIP | 23142080 | Unspecified |
| mmu-miR-30c-5p | MIMAT0000514 | Lcp1 | 18826 | HITS-CLIP | 23142080 | Unspecified |
| mmu-miR-30c-5p | MIMAT0000514 | Lcp2 | 16822 | HITS-CLIP | 37986224\|23142080 | Unspecified |
| mmu-miR-30c-5p | MIMAT0000514 | Ldb3 | 24131 | HITS-CLIP | 25753418 | Unspecified |
| mmu-miR-30c-5p | MIMAT0000514 | Ldlrap1 | 100017 | HITS-CLIP | 25753418 | Unspecified |
| mmu-miR-30c-5p | MIMAT0000514 | Lemd3 | 380664 | HITS-CLIP | 23142080 | Unspecified |
| mmu-miR-30c-5p | MIMAT0000514 | Leng8 | 232798 | HITS-CLIP | 25753418 | Unspecified |
| mmu-miR-30c-5p | MIMAT0000514 | Lepr | 16847 | HITS-CLIP | 37986224\|19536157 | Unspecified |
| mmu-miR-30c-5p | MIMAT0000514 | Leprotl1 | 68192 | HITS-CLIP | 23142080 | Unspecified |
| mmu-miR-30c-5p | MIMAT0000514 | Lgals8 | 56048 | HITS-CLIP | 23597149 | Unspecified |
| mmu-miR-30c-5p | MIMAT0000514 | Lgi1 | 56839 | HITS-CLIP | 37986224\|19536157 | Unspecified |
| mmu-miR-30c-5p | MIMAT0000514 | Lhx1 | 16869 | HITS-CLIP | 19536157 | Unspecified |
| mmu-miR-30c-5p | MIMAT0000514 | Lifr | 16880 | HITS-CLIP | 37986224\|23597149\|19536157 | Unspecified |
| mmu-miR-30c-5p | MIMAT0000514 | Lig3 | 16882 | HITS-CLIP | 37986224\|23597149\|25753418 | Unspecified |
| mmu-miR-30c-5p | MIMAT0000514 | Limch1 | 77569 | HITS-CLIP | 37986224\|25753418 | Unspecified |
| mmu-miR-30c-5p | MIMAT0000514 | Limd1 | 29806 | HITS-CLIP | 37986224\|23142080 | Unspecified |
| mmu-miR-30c-5p | MIMAT0000514 | Lims1 | 110829 | HITS-CLIP | 37986224\|23142080\|25753418 | Unspecified |
| mmu-miR-30c-5p | MIMAT0000514 | Lin28a | 83557 | HITS-CLIP | 37986224 | Unspecified |
| mmu-miR-30c-5p | MIMAT0000514 | Lin7c | 22343 | HITS-CLIP | 37986224\|23142080 | Unspecified |
| mmu-miR-30c-5p | MIMAT0000514 | Lipa | 16889 | HITS-CLIP | 37986224 | Unspecified |
| mmu-miR-30c-5p | MIMAT0000514 | Lmbr1l | 74775 | HITS-CLIP//qCLASH | 37986224\|23597149\|31393953\|23142 | Unspecified |
| mmu-miR-30c-5p | MIMAT0000514 | Lmbrd1 | 68421 | HITS-CLIP | 23142080 | Unspecified |
| mmu-miR-30c-5p | MIMAT0000514 | Lmbrd2 | 320506 | HITS-CLIP | 23142080 | Unspecified |
| mmu-miR-30c-5p | MIMAT0000514 | Lmln | 239833 | HITS-CLIP | 23142080 | Unspecified |
| mmu-miR-30c-5p | MIMAT0000514 | Lmnb1 | 16906 | HITS-CLIP | 23142080 | Unspecified |
| mmu-miR-30c-5p | MIMAT0000514 | Lmo7 | 380928 | HITS-CLIP | 25753418 | Unspecified |
| mmu-miR-30c-5p | MIMAT0000514 | Lnpep | 240028 | HITS-CLIP | 37986224\|19536157\|23597149\|23142 | Unspecified |
| mmu-miR-30c-5p | MIMAT0000514 | Lnpk | 69605 | HITS-CLIP | 23597149\|23142080 | Unspecified |
| mmu-miR-30c-5p | MIMAT0000514 | Lnx2 | 140887 | HITS-CLIP | 23597149\|23142080 | Unspecified |
| mmu-miR-30c-5p | MIMAT0000514 | Lonrf1 | 244421 | HITS-CLIP | 23142080 | Unspecified |
| mmu-miR-30c-5p | MIMAT0000514 | Lpar6 | 67168 | HITS-CLIP | 37986224 | Unspecified |

| | | | | | | |
|---|---|---|---|---|---|---|
| mmu-miR-30c-5p | MIMAT0000514 | Lpcat3 | 14792 | HITS-CLIP | 37986224 | Unspecified |
| mmu-miR-30c-5p | MIMAT0000514 | Lpgat1 | 226856 | HITS-CLIP | 37986224\|23597149\|23142080 | Unspecified |
| mmu-miR-30c-5p | MIMAT0000514 | Lpin1 | 14245 | HITS-CLIP | 37986224 | Unspecified |
| mmu-miR-30c-5p | MIMAT0000514 | Lpin2 | 64898 | HITS-CLIP | 23142080\|23597149 | Unspecified |
| mmu-miR-30c-5p | MIMAT0000514 | Lpp | 210126 | HITS-CLIP | 23597149\|23142080 | Unspecified |
| mmu-miR-30c-5p | MIMAT0000514 | Lrba | 80877 | HITS-CLIP | 37986224\|23597149\|23142080 | Unspecified |
| mmu-miR-30c-5p | MIMAT0000514 | Lrch3 | 70144 | HITS-CLIP | 23142080 | Unspecified |
| mmu-miR-30c-5p | MIMAT0000514 | Lrif1 | 321000 | HITS-CLIP | 23142080 | Unspecified |
| mmu-miR-30c-5p | MIMAT0000514 | Lrig1 | 16206 | HITS-CLIP | 23142080 | Unspecified |
| mmu-miR-30c-5p | MIMAT0000514 | Lrig2 | 269473 | HITS-CLIP | 37986224 | Unspecified |
| mmu-miR-30c-5p | MIMAT0000514 | Lrmp | 16970 | HITS-CLIP | 37986224 | Unspecified |
| mmu-miR-30c-5p | MIMAT0000514 | Lrp6 | 16974 | HITS-CLIP | 23142080 | Unspecified |
| mmu-miR-30c-5p | MIMAT0000514 | Lrrc40 | 67144 | HITS-CLIP | 23142080 | Unspecified |
| mmu-miR-30c-5p | MIMAT0000514 | Lrrc4c | 241568 | HITS-CLIP | 37986224 | Unspecified |
| mmu-miR-30c-5p | MIMAT0000514 | Lrrc58 | 320184 | HITS-CLIP | 37986224\|23142080\|25753418 | Unspecified |
| mmu-miR-30c-5p | MIMAT0000514 | Lrrc8d | 231549 | HITS-CLIP | 37986224\|23142080 | Unspecified |
| mmu-miR-30c-5p | MIMAT0000514 | Lrrk2 | 66725 | HITS-CLIP | 23142080 | Unspecified |
| mmu-miR-30c-5p | MIMAT0000514 | Lrrn1 | 16979 | HITS-CLIP | 37986224 | Unspecified |
| mmu-miR-30c-5p | MIMAT0000514 | Lsamp | 268890 | HITS-CLIP | 37986224 | Unspecified |
| mmu-miR-30c-5p | MIMAT0000514 | Lsp1 | 16985 | HITS-CLIP | 23142080 | Unspecified |
| mmu-miR-30c-5p | MIMAT0000514 | Lsr | 54135 | HITS-CLIP | 23597149 | Unspecified |
| mmu-miR-30c-5p | MIMAT0000514 | Lss | 16987 | HITS-CLIP | 23142080 | Unspecified |
| mmu-miR-30c-5p | MIMAT0000514 | Luc7l | 66978 | HITS-CLIP | 37986224\|23597149 | Unspecified |
| mmu-miR-30c-5p | MIMAT0000514 | Luc7l2 | 192196 | HITS-CLIP | 23142080 | Unspecified |
| mmu-miR-30c-5p | MIMAT0000514 | Luc7l3 | 67684 | HITS-CLIP | 23142080 | Unspecified |
| mmu-miR-30c-5p | MIMAT0000514 | Lyst | 17101 | HITS-CLIP | 23597149\|23142080 | Unspecified |
| mmu-miR-30c-5p | MIMAT0000514 | Lztr1 | 66863 | HITS-CLIP | 23142080 | Unspecified |
| mmu-miR-30c-5p | MIMAT0000514 | M6pr | 17113 | HITS-CLIP | 37986224\|23142080 | Unspecified |
| mmu-miR-30c-5p | MIMAT0000514 | Macf1 | 11426 | HITS-CLIP | 37986224\|23142080\|25753418 | Unspecified |
| mmu-miR-30c-5p | MIMAT0000514 | Maco1 | 55219 | HITS-CLIP | 23142080 | Unspecified |
| mmu-miR-30c-5p | MIMAT0000514 | Madd | 228355 | HITS-CLIP | 23142080 | Unspecified |
| mmu-miR-30c-5p | MIMAT0000514 | Maf | 17132 | HITS-CLIP | 37986224\|23142080 | Unspecified |
| mmu-miR-30c-5p | MIMAT0000514 | Mafg | 17134 | HITS-CLIP | 23142080 | Unspecified |
| mmu-miR-30c-5p | MIMAT0000514 | Magi3 | 99470 | HITS-CLIP | 23597149\|23142080 | Unspecified |
| mmu-miR-30c-5p | MIMAT0000514 | Magt1 | 67075 | HITS-CLIP | 37986224\|23142080 | Unspecified |
| mmu-miR-30c-5p | MIMAT0000514 | Maml1 | 103806 | HITS-CLIP | 37986224\|23597149\|23142080 | Unspecified |
| mmu-miR-30c-5p | MIMAT0000514 | Man1a | 17155 | HITS-CLIP | 23597149\|23142080 | Unspecified |
| mmu-miR-30c-5p | MIMAT0000514 | Man1a2 | 17156 | HITS-CLIP | 37986224 | Unspecified |
| mmu-miR-30c-5p | MIMAT0000514 | Man2a1 | 17158 | HITS-CLIP | 37986224\|23597149\|23142080 | Unspecified |

| | | | | | | |
|---|---|---|---|---|---|---|
| mmu-miR-30c-5p | MIMAT0000514 | Manea | 242362 | HITS-CLIP | 37986224 | Unspecified |
| mmu-miR-30c-5p | MIMAT0000514 | Map1b | 17755 | HITS-CLIP | 37986224 | Unspecified |
| mmu-miR-30c-5p | MIMAT0000514 | Map2k1 | 26395 | HITS-CLIP | 23142080 | Unspecified |
| mmu-miR-30c-5p | MIMAT0000514 | Map2k6 | 26399 | HITS-CLIP | 23142080 | Unspecified |
| mmu-miR-30c-5p | MIMAT0000514 | Map3k1 | 26401 | HITS-CLIP | 37986224\|23142080 | Unspecified |
| mmu-miR-30c-5p | MIMAT0000514 | Map3k12 | 26404 | HITS-CLIP | 37986224 | Unspecified |
| mmu-miR-30c-5p | MIMAT0000514 | Map3k2 | 26405 | HITS-CLIP | 37986224 | Unspecified |
| mmu-miR-30c-5p | MIMAT0000514 | Map3k5 | 26408 | HITS-CLIP | 37986224\|23142080 | Unspecified |
| mmu-miR-30c-5p | MIMAT0000514 | Map3k7 | 26409 | HITS-CLIP | 25753418 | Unspecified |
| mmu-miR-30c-5p | MIMAT0000514 | Mapk1 | 26413 | HITS-CLIP | 37986224 | Unspecified |
| mmu-miR-30c-5p | MIMAT0000514 | Mapk14 | 26416 | HITS-CLIP | 37986224 | Unspecified |
| mmu-miR-30c-5p | MIMAT0000514 | Mapk1ip1l | 218975 | HITS-CLIP | 23142080 | Unspecified |
| mmu-miR-30c-5p | MIMAT0000514 | Mapk6 | 50772 | HITS-CLIP | 23142080\|25753418 | Unspecified |
| mmu-miR-30c-5p | MIMAT0000514 | Mapk8 | 26419 | HITS-CLIP | 23142080 | Unspecified |
| mmu-miR-30c-5p | MIMAT0000514 | Mapk9 | 26420 | HITS-CLIP | 37986224 | Unspecified |
| mmu-miR-30c-5p | MIMAT0000514 | Mapkbp1 | 26390 | HITS-CLIP | 23142080 | Unspecified |
| mmu-miR-30c-5p | MIMAT0000514 | Mapre1 | 13589 | HITS-CLIP | 37986224 | Unspecified |
| mmu-miR-30c-5p | MIMAT0000514 | March7 | 57438 | HITS-CLIP | 23142080 | Unspecified |
| mmu-miR-30c-5p | MIMAT0000514 | Marchf6 | 223455 | HITS-CLIP | 37986224\|23597149\|19536157\|23142 | Unspecified |
| mmu-miR-30c-5p | MIMAT0000514 | Marcks | 17118 | HITS-CLIP | 37986224\|19536157\|23142080 | Unspecified |
| mmu-miR-30c-5p | MIMAT0000514 | Marf1 | 223989 | HITS-CLIP | 37986224\|23142080 | Unspecified |
| mmu-miR-30c-5p | MIMAT0000514 | Mark3 | 17169 | HITS-CLIP | 37986224 | Unspecified |
| mmu-miR-30c-5p | MIMAT0000514 | Mars2 | 212679 | HITS-CLIP | 25753418 | Unspecified |
| mmu-miR-30c-5p | MIMAT0000514 | Mast4 | 328329 | HITS-CLIP | 37986224\|23142080\|23597149 | Unspecified |
| mmu-miR-30c-5p | MIMAT0000514 | Mat1a | 11720 | HITS-CLIP | 23597149 | Unspecified |
| mmu-miR-30c-5p | MIMAT0000514 | Mat2a | 232087 | HITS-CLIP | 37986224\|23142080\|19536157\|23597 | Unspecified |
| mmu-miR-30c-5p | MIMAT0000514 | Mat2b | 108645 | HITS-CLIP | 23142080 | Unspecified |
| mmu-miR-30c-5p | MIMAT0000514 | Mbd2 | 17191 | HITS-CLIP | 23142080 | Unspecified |
| mmu-miR-30c-5p | MIMAT0000514 | Mbd5 | 109241 | HITS-CLIP | 37986224\|23142080 | Unspecified |
| mmu-miR-30c-5p | MIMAT0000514 | Mbd6 | 110962 | HITS-CLIP | 23142080 | Unspecified |
| mmu-miR-30c-5p | MIMAT0000514 | Mbnl1 | 56758 | HITS-CLIP | 37986224\|23597149 | Unspecified |
| mmu-miR-30c-5p | MIMAT0000514 | Mboat1 | 218121 | HITS-CLIP | 37986224\|23142080 | Unspecified |
| mmu-miR-30c-5p | MIMAT0000514 | Mbtd1 | 103537 | HITS-CLIP | 23142080 | Unspecified |
| mmu-miR-30c-5p | MIMAT0000514 | Mcf2l | 17207 | HITS-CLIP | 37986224\|23142080\|19536157 | Unspecified |
| mmu-miR-30c-5p | MIMAT0000514 | Mcl1 | 17210 | HITS-CLIP | 37986224 | Unspecified |
| mmu-miR-30c-5p | MIMAT0000514 | Mcm4 | 17217 | HITS-CLIP | 37986224 | Unspecified |
| mmu-miR-30c-5p | MIMAT0000514 | Mcm6 | 17219 | HITS-CLIP | 23142080 | Unspecified |
| mmu-miR-30c-5p | MIMAT0000514 | Mcm7 | 17220 | HITS-CLIP | 23597149\|23142080 | Unspecified |
| mmu-miR-30c-5p | MIMAT0000514 | Mctp2 | 244049 | HITS-CLIP | 23142080 | Unspecified |

| | | | | | | |
|---|---|---|---|---|---|---|
| mmu-miR-30c-5p | MIMAT0000514 | Mcur1 | 76137 | HITS-CLIP | 23597149\|23142080 | Unspecified |
| mmu-miR-30c-5p | MIMAT0000514 | Mdfic | 16543 | HITS-CLIP | 37986224 | Unspecified |
| mmu-miR-30c-5p | MIMAT0000514 | Mdm4 | 17248 | HITS-CLIP | 37986224\|23142080 | Unspecified |
| mmu-miR-30c-5p | MIMAT0000514 | Mdn1 | 100019 | HITS-CLIP | 25753418 | Unspecified |
| mmu-miR-30c-5p | MIMAT0000514 | Me1 | 17436 | HITS-CLIP | 37986224 | Unspecified |
| mmu-miR-30c-5p | MIMAT0000514 | Mecp2 | 17257 | HITS-CLIP | 23142080 | Unspecified |
| mmu-miR-30c-5p | MIMAT0000514 | Med13 | 327987 | HITS-CLIP | 37986224\|19536157\|23142080 | Unspecified |
| mmu-miR-30c-5p | MIMAT0000514 | Med13l | 76199 | HITS-CLIP | 37986224 | Unspecified |
| mmu-miR-30c-5p | MIMAT0000514 | Med17 | 234959 | HITS-CLIP | 23142080 | Unspecified |
| mmu-miR-30c-5p | MIMAT0000514 | Med21 | 108098 | HITS-CLIP | 25753418 | Unspecified |
| mmu-miR-30c-5p | MIMAT0000514 | Med23 | 70208 | HITS-CLIP | 23142080 | Unspecified |
| mmu-miR-30c-5p | MIMAT0000514 | Med26 | 70625 | HITS-CLIP | 37986224 | Unspecified |
| mmu-miR-30c-5p | MIMAT0000514 | Med7 | 66213 | HITS-CLIP | 37986224 | Unspecified |
| mmu-miR-30c-5p | MIMAT0000514 | Mef2c | 17260 | HITS-CLIP | 37986224 | Unspecified |
| mmu-miR-30c-5p | MIMAT0000514 | Mef2d | 17261 | HITS-CLIP | 37986224\|23142080 | Unspecified |
| mmu-miR-30c-5p | MIMAT0000514 | Megf9 | 230316 | HITS-CLIP | 37986224 | Unspecified |
| mmu-miR-30c-5p | MIMAT0000514 | Meis1 | 17268 | HITS-CLIP | 25753418 | Unspecified |
| mmu-miR-30c-5p | MIMAT0000514 | Mertk | 17289 | HITS-CLIP | 37986224 | Unspecified |
| mmu-miR-30c-5p | MIMAT0000514 | Metap1 | 75624 | HITS-CLIP | 37986224\|23597149\|23142080 | Unspecified |
| mmu-miR-30c-5p | MIMAT0000514 | Mettl3 | 56335 | HITS-CLIP | 23142080 | Unspecified |
| mmu-miR-30c-5p | MIMAT0000514 | Mettl4 | 76781 | HITS-CLIP | 23142080 | Unspecified |
| mmu-miR-30c-5p | MIMAT0000514 | Mex3b | 108797 | HITS-CLIP | 19536157 | Unspecified |
| mmu-miR-30c-5p | MIMAT0000514 | Mex3c | 240396 | HITS-CLIP | 37986224\|23142080 | Unspecified |
| mmu-miR-30c-5p | MIMAT0000514 | Mfap3l | 71306 | HITS-CLIP | 37986224 | Unspecified |
| mmu-miR-30c-5p | MIMAT0000514 | Mfsd14a | 15247 | HITS-CLIP | 23142080 | Unspecified |
| mmu-miR-30c-5p | MIMAT0000514 | Mfsd6 | 98682 | HITS-CLIP | 23142080 | Unspecified |
| mmu-miR-30c-5p | MIMAT0000514 | Mfsd8 | 72175 | HITS-CLIP | 37986224\|23142080 | Unspecified |
| mmu-miR-30c-5p | MIMAT0000514 | Mga | 29808 | HITS-CLIP | 23142080 | Unspecified |
| mmu-miR-30c-5p | MIMAT0000514 | Mgea5 | 76055 | HITS-CLIP | 25083871\|23142080 | Unspecified |
| mmu-miR-30c-5p | MIMAT0000514 | Mia3 | 338366 | HITS-CLIP | 37986224\|19536157\|23597149\|23142 | Unspecified |
| mmu-miR-30c-5p | MIMAT0000514 | Mib1 | 225164 | HITS-CLIP | 37986224\|19536157\|23597149\|23142 | Unspecified |
| mmu-miR-30c-5p | MIMAT0000514 | Mical1 | 171580 | HITS-CLIP | 23142080 | Unspecified |
| mmu-miR-30c-5p | MIMAT0000514 | Micu3 | 78506 | HITS-CLIP | 23142080 | Unspecified |
| mmu-miR-30c-5p | MIMAT0000514 | Mid1ip1 | 68041 | HITS-CLIP | 23142080 | Unspecified |
| mmu-miR-30c-5p | MIMAT0000514 | Mier1 | 71148 | HITS-CLIP | 23597149\|23142080 | Unspecified |
| mmu-miR-30c-5p | MIMAT0000514 | Mier2 | 70427 | HITS-CLIP | 23142080 | Unspecified |
| mmu-miR-30c-5p | MIMAT0000514 | Mier3 | 218613 | HITS-CLIP | 37986224\|23142080 | Unspecified |
| mmu-miR-30c-5p | MIMAT0000514 | Mios | 252875 | HITS-CLIP | 23597149\|25753418 | Unspecified |
| mmu-miR-30c-5p | MIMAT0000514 | Mis18bp1 | 217653 | HITS-CLIP | 23142080 | Unspecified |

| | | | | | | |
|---|---|---|---|---|---|---|
| mmu-miR-30c-5p | MIMAT0000514 | Mitd1 | 69028 | HITS-CLIP | 23142080 | Unspecified |
| mmu-miR-30c-5p | MIMAT0000514 | Mkln1 | 27418 | HITS-CLIP | 37986224\|23142080 | Unspecified |
| mmu-miR-30c-5p | MIMAT0000514 | Mkrn3 | 22652 | HITS-CLIP | 37986224\|19536157 | Unspecified |
| mmu-miR-30c-5p | MIMAT0000514 | Mllt10 | 17354 | HITS-CLIP | 37986224 | Unspecified |
| mmu-miR-30c-5p | MIMAT0000514 | Mllt3 | 70122 | HITS-CLIP | 23142080 | Unspecified |
| mmu-miR-30c-5p | MIMAT0000514 | Mllt6 | 246198 | HITS-CLIP | 23142080 | Unspecified |
| mmu-miR-30c-5p | MIMAT0000514 | Mlxip | 208104 | HITS-CLIP | 37986224\|23142080 | Unspecified |
| mmu-miR-30c-5p | MIMAT0000514 | Mmachc | 67096 | HITS-CLIP | 25753418 | Unspecified |
| mmu-miR-30c-5p | MIMAT0000514 | Mmadhc | 109129 | HITS-CLIP | 23142080 | Unspecified |
| mmu-miR-30c-5p | MIMAT0000514 | Mmd | 67468 | HITS-CLIP | 37986224\|23597149\|23142080 | Unspecified |
| mmu-miR-30c-5p | MIMAT0000514 | Mmp14 | 17387 | HITS-CLIP | 37986224 | Unspecified |
| mmu-miR-30c-5p | MIMAT0000514 | Mmp15 | 17388 | HITS-CLIP | 25753418 | Unspecified |
| mmu-miR-30c-5p | MIMAT0000514 | Mmp19 | 58223 | HITS-CLIP | 37986224 | Unspecified |
| mmu-miR-30c-5p | MIMAT0000514 | Mnat1 | 17420 | HITS-CLIP | 23142080 | Unspecified |
| mmu-miR-30c-5p | MIMAT0000514 | Mob1a | 232157 | HITS-CLIP | 23597149\|23142080 | Unspecified |
| mmu-miR-30c-5p | MIMAT0000514 | Mob3a | 208228 | HITS-CLIP | 37986224\|23597149\|23142080 | Unspecified |
| mmu-miR-30c-5p | MIMAT0000514 | Mob3b | 214944 | HITS-CLIP | 37986224 | Unspecified |
| mmu-miR-30c-5p | MIMAT0000514 | Mob3c | 100465 | HITS-CLIP | 23142080 | Unspecified |
| mmu-miR-30c-5p | MIMAT0000514 | Mob4 | 19070 | HITS-CLIP | 37986224\|23142080 | Unspecified |
| mmu-miR-30c-5p | MIMAT0000514 | Mocs2 | 17434 | HITS-CLIP | 37986224 | Unspecified |
| mmu-miR-30c-5p | MIMAT0000514 | Morc3 | 338467 | HITS-CLIP | 37986224\|23142080 | Unspecified |
| mmu-miR-30c-5p | MIMAT0000514 | Morf4l2 | 56397 | HITS-CLIP | 37986224 | Unspecified |
| mmu-miR-30c-5p | MIMAT0000514 | Mrc1 | 17533 | HITS-CLIP | 37986224\|23597149 | Unspecified |
| mmu-miR-30c-5p | MIMAT0000514 | Mrpl1 | 94061 | HITS-CLIP | 23142080 | Unspecified |
| mmu-miR-30c-5p | MIMAT0000514 | Mrpl18 | 67681 | HITS-CLIP | 37986224 | Unspecified |
| mmu-miR-30c-5p | MIMAT0000514 | Mrpl22 | 216767 | HITS-CLIP | 37986224 | Unspecified |
| mmu-miR-30c-5p | MIMAT0000514 | Mrpl38 | 60441 | HITS-CLIP | 25753418 | Unspecified |
| mmu-miR-30c-5p | MIMAT0000514 | Mrps2 | 118451 | HITS-CLIP | 37986224 | Unspecified |
| mmu-miR-30c-5p | MIMAT0000514 | Mrps24 | 64660 | HITS-CLIP | 37986224 | Unspecified |
| mmu-miR-30c-5p | MIMAT0000514 | Mrps31 | 57312 | HITS-CLIP | 25753418 | Unspecified |
| mmu-miR-30c-5p | MIMAT0000514 | Mrs2 | 380836 | HITS-CLIP | 25753418 | Unspecified |
| mmu-miR-30c-5p | MIMAT0000514 | Mrtfb | 239719 | HITS-CLIP | 19536157 | Unspecified |
| mmu-miR-30c-5p | MIMAT0000514 | Msh2 | 17685 | HITS-CLIP | 23142080 | Unspecified |
| mmu-miR-30c-5p | MIMAT0000514 | Msi2 | 76626 | HITS-CLIP | 37986224\|23597149\|23142080 | Unspecified |
| mmu-miR-30c-5p | MIMAT0000514 | Msl2 | 77853 | HITS-CLIP | 23142080 | Unspecified |
| mmu-miR-30c-5p | MIMAT0000514 | Msmo1 | 66234 | HITS-CLIP | 37986224 | Unspecified |
| mmu-miR-30c-5p | MIMAT0000514 | Mt1 | 17748 | HITS-CLIP | 37986224\|23597149 | Unspecified |
| mmu-miR-30c-5p | MIMAT0000514 | Mta3 | 116871 | HITS-CLIP | 23142080 | Unspecified |
| mmu-miR-30c-5p | MIMAT0000514 | Mtbp | 105837 | HITS-CLIP | 37986224 | Unspecified |

| | | | | | | |
|---|---|---|---|---|---|---|
| mmu-miR-30c-5p | MIMAT0000514 | Mtch2 | 56428 | HITS-CLIP | 25753418 | Unspecified |
| mmu-miR-30c-5p | MIMAT0000514 | Mtdh | 67154 | Luciferase assay//qRT-PCR//HITS-CLIP//Luc | 24555094\|23597149\|19536157\|23142 | Unspecified |
| mmu-miR-30c-5p | MIMAT0000514 | Mtf1 | 17764 | HITS-CLIP | 23142080 | Unspecified |
| mmu-miR-30c-5p | MIMAT0000514 | Mtf2 | 17765 | HITS-CLIP | 37986224 | Unspecified |
| mmu-miR-30c-5p | MIMAT0000514 | Mthfd1l | 270685 | HITS-CLIP | 37986224\|23142080 | Unspecified |
| mmu-miR-30c-5p | MIMAT0000514 | Mtmr10 | 233315 | HITS-CLIP | 23597149 | Unspecified |
| mmu-miR-30c-5p | MIMAT0000514 | Mtpn | 14489 | HITS-CLIP | 37986224 | Unspecified |
| mmu-miR-30c-5p | MIMAT0000514 | Mtrex | 23517 | HITS-CLIP | 23142080 | Unspecified |
| mmu-miR-30c-5p | MIMAT0000514 | Mttp | 17777 | HITS-CLIP | 37986224 | Unspecified |
| mmu-miR-30c-5p | MIMAT0000514 | Mtus1 | 102103 | HITS-CLIP | 37986224 | Unspecified |
| mmu-miR-30c-5p | MIMAT0000514 | Mtus2 | 77521 | HITS-CLIP | 37986224 | Unspecified |
| mmu-miR-30c-5p | MIMAT0000514 | Mtx1 | 17827 | HITS-CLIP | 37986224\|23142080 | Unspecified |
| mmu-miR-30c-5p | MIMAT0000514 | Mtx3 | 382793 | HITS-CLIP | 37986224 | Unspecified |
| mmu-miR-30c-5p | MIMAT0000514 | Mug1 | 17836 | HITS-CLIP | 37986224 | Unspecified |
| mmu-miR-30c-5p | MIMAT0000514 | Mul1 | 68350 | HITS-CLIP | 23142080 | Unspecified |
| mmu-miR-30c-5p | MIMAT0000514 | Mxi1 | 17859 | HITS-CLIP | 23142080 | Unspecified |
| mmu-miR-30c-5p | MIMAT0000514 | Myb | 17863 | HITS-CLIP | 23142080 | Unspecified |
| mmu-miR-30c-5p | MIMAT0000514 | Mycbp2 | 105689 | HITS-CLIP | 23142080 | Unspecified |
| mmu-miR-30c-5p | MIMAT0000514 | Myef2 | 17876 | HITS-CLIP | 23597149 | Unspecified |
| mmu-miR-30c-5p | MIMAT0000514 | Myh9 | 17886 | HITS-CLIP | 37986224\|23597149 | Unspecified |
| mmu-miR-30c-5p | MIMAT0000514 | Mylip | 218203 | HITS-CLIP | 23142080 | Unspecified |
| mmu-miR-30c-5p | MIMAT0000514 | Mynn | 80732 | HITS-CLIP | 23142080 | Unspecified |
| mmu-miR-30c-5p | MIMAT0000514 | Myo5a | 17918 | HITS-CLIP | 23142080 | Unspecified |
| mmu-miR-30c-5p | MIMAT0000514 | Myo6 | 17920 | HITS-CLIP | 23142080 | Unspecified |
| mmu-miR-30c-5p | MIMAT0000514 | Mysm1 | 320713 | HITS-CLIP | 37986224 | Unspecified |
| mmu-miR-30c-5p | MIMAT0000514 | N4bp2 | 333789 | HITS-CLIP | 37986224\|23142080 | Unspecified |
| mmu-miR-30c-5p | MIMAT0000514 | Naa15 | 74838 | HITS-CLIP | 23142080 | Unspecified |
| mmu-miR-30c-5p | MIMAT0000514 | Naa25 | 231713 | HITS-CLIP | 37986224\|19536157\|23597149\|23142 | Unspecified |
| mmu-miR-30c-5p | MIMAT0000514 | Naa30 | 70646 | HITS-CLIP | 23142080 | Unspecified |
| mmu-miR-30c-5p | MIMAT0000514 | Nabp1 | 109019 | HITS-CLIP | 37986224\|23142080 | Unspecified |
| mmu-miR-30c-5p | MIMAT0000514 | Nadk | 192185 | HITS-CLIP | 23597149 | Unspecified |
| mmu-miR-30c-5p | MIMAT0000514 | Naf1 | 234344 | HITS-CLIP | 37986224 | Unspecified |
| mmu-miR-30c-5p | MIMAT0000514 | Nagpa | 27426 | HITS-CLIP | 37986224\|23142080\|23597149 | Unspecified |
| mmu-miR-30c-5p | MIMAT0000514 | Nampt | 59027 | HITS-CLIP | 23142080 | Unspecified |
| mmu-miR-30c-5p | MIMAT0000514 | Nanos1 | 332397 | HITS-CLIP | 23142080 | Unspecified |
| mmu-miR-30c-5p | MIMAT0000514 | Nap1l5 | 58243 | HITS-CLIP | 37986224 | Unspecified |
| mmu-miR-30c-5p | MIMAT0000514 | Nasp | 50927 | HITS-CLIP | 23142080 | Unspecified |
| mmu-miR-30c-5p | MIMAT0000514 | Nat1 | 17960 | HITS-CLIP | 23597149 | Unspecified |
| mmu-miR-30c-5p | MIMAT0000514 | Nat2 | 17961 | HITS-CLIP | 37986224 | Unspecified |

| | | | | | | |
|---|---|---|---|---|---|---|
| mmu-miR-30c-5p | MIMAT0000514 | Nbea | 26422 | HITS-CLIP | 37986224\|19536157 | Unspecified |
| mmu-miR-30c-5p | MIMAT0000514 | Nbr1 | 17966 | HITS-CLIP | 37986224 | Unspecified |
| mmu-miR-30c-5p | MIMAT0000514 | Ncam1 | 17967 | HITS-CLIP | 37986224\|19536157 | Unspecified |
| mmu-miR-30c-5p | MIMAT0000514 | Ncan | 13004 | HITS-CLIP | 37986224 | Unspecified |
| mmu-miR-30c-5p | MIMAT0000514 | Ncapd3 | 78658 | HITS-CLIP | 23142080 | Unspecified |
| mmu-miR-30c-5p | MIMAT0000514 | Ncf1 | 17969 | HITS-CLIP | 23142080 | Unspecified |
| mmu-miR-30c-5p | MIMAT0000514 | Nck2 | 17974 | HITS-CLIP | 37986224\|23142080 | Unspecified |
| mmu-miR-30c-5p | MIMAT0000514 | Nckap1 | 50884 | HITS-CLIP | 23597149\|19536157\|23142080\|2575[illegible] | Unspecified |
| mmu-miR-30c-5p | MIMAT0000514 | Ncl | 17975 | qCLASH | 31393953 | Unspecified |
| mmu-miR-30c-5p | MIMAT0000514 | Ncoa2 | 17978 | HITS-CLIP | 23142080 | Unspecified |
| mmu-miR-30c-5p | MIMAT0000514 | Ncoa3 | 17979 | HITS-CLIP | 37986224\|23142080 | Unspecified |
| mmu-miR-30c-5p | MIMAT0000514 | Ncoa7 | 211329 | HITS-CLIP | 37986224\|23142080 | Unspecified |
| mmu-miR-30c-5p | MIMAT0000514 | Ndel1 | 83431 | HITS-CLIP | 37986224\|23142080 | Unspecified |
| mmu-miR-30c-5p | MIMAT0000514 | Ndufs2 | 226646 | HITS-CLIP | 23597149 | Unspecified |
| mmu-miR-30c-5p | MIMAT0000514 | Necap1 | 67602 | HITS-CLIP | 37986224 | Unspecified |
| mmu-miR-30c-5p | MIMAT0000514 | Nedd1 | 17997 | HITS-CLIP | 23142080 | Unspecified |
| mmu-miR-30c-5p | MIMAT0000514 | Nedd4 | 17999 | HITS-CLIP | 37986224\|19536157\|23142080 | Unspecified |
| mmu-miR-30c-5p | MIMAT0000514 | Nedd4l | 83814 | HITS-CLIP | 19536157\|23597149 | Unspecified |
| mmu-miR-30c-5p | MIMAT0000514 | Nefm | 18040 | HITS-CLIP | 37986224 | Unspecified |
| mmu-miR-30c-5p | MIMAT0000514 | Neil3 | 234258 | HITS-CLIP | 23142080 | Unspecified |
| mmu-miR-30c-5p | MIMAT0000514 | Nek7 | 59125 | HITS-CLIP | 37986224\|23142080 | Unspecified |
| mmu-miR-30c-5p | MIMAT0000514 | Nek9 | 217718 | HITS-CLIP | 37986224\|23597149\|23142080 | Unspecified |
| mmu-miR-30c-5p | MIMAT0000514 | Nemp2 | 227094 | HITS-CLIP | 23142080 | Unspecified |
| mmu-miR-30c-5p | MIMAT0000514 | Neto1 | 246317 | HITS-CLIP | 37986224 | Unspecified |
| mmu-miR-30c-5p | MIMAT0000514 | Neurod1 | 18012 | HITS-CLIP | 37986224 | Unspecified |
| mmu-miR-30c-5p | MIMAT0000514 | Nfat5 | 54446 | HITS-CLIP | 37986224\|23142080\|19536157 | Unspecified |
| mmu-miR-30c-5p | MIMAT0000514 | Nfatc2ip | 18020 | HITS-CLIP | 37986224 | Unspecified |
| mmu-miR-30c-5p | MIMAT0000514 | Nfatc3 | 18021 | HITS-CLIP | 25083871\|23142080\|25753418 | Unspecified |
| mmu-miR-30c-5p | MIMAT0000514 | Nfia | 18027 | HITS-CLIP | 37986224\|23142080 | Unspecified |
| mmu-miR-30c-5p | MIMAT0000514 | Nfib | 18028 | HITS-CLIP | 37986224\|23597149\|19536157 | Unspecified |
| mmu-miR-30c-5p | MIMAT0000514 | Nfya | 18044 | HITS-CLIP | 23142080 | Unspecified |
| mmu-miR-30c-5p | MIMAT0000514 | Nid1 | 18073 | HITS-CLIP | 37986224 | Unspecified |
| mmu-miR-30c-5p | MIMAT0000514 | Nif3l1 | 65102 | HITS-CLIP | 23142080 | Unspecified |
| mmu-miR-30c-5p | MIMAT0000514 | Nipa2 | 93790 | HITS-CLIP | 37986224 | Unspecified |
| mmu-miR-30c-5p | MIMAT0000514 | Nipbl | 71175 | HITS-CLIP | 37986224\|23597149\|23142080 | Unspecified |
| mmu-miR-30c-5p | MIMAT0000514 | Nkrf | 77286 | HITS-CLIP | 25753418 | Unspecified |
| mmu-miR-30c-5p | MIMAT0000514 | Nktr | 18087 | HITS-CLIP | 37986224 | Unspecified |
| mmu-miR-30c-5p | MIMAT0000514 | Nmd3 | 97112 | HITS-CLIP | 23142080 | Unspecified |
| mmu-miR-30c-5p | MIMAT0000514 | Nmt1 | 18107 | HITS-CLIP | 37986224\|19536157\|23142080 | Unspecified |

| | | | | | | |
|---|---|---|---|---|---|---|
| mmu-miR-30c-5p | MIMAT0000514 | Nnt | 18115 | HITS-CLIP | 25753418 | Unspecified |
| mmu-miR-30c-5p | MIMAT0000514 | Nob1 | 67619 | HITS-CLIP | 25753418 | Unspecified |
| mmu-miR-30c-5p | MIMAT0000514 | Noct | 12457 | HITS-CLIP | 25753418 | Unspecified |
| mmu-miR-30c-5p | MIMAT0000514 | Nol4l | 329540 | HITS-CLIP | 37986224\|19536157\|23142080 | Unspecified |
| mmu-miR-30c-5p | MIMAT0000514 | Nop58 | 55989 | HITS-CLIP | 23142080\|23597149 | Unspecified |
| mmu-miR-30c-5p | MIMAT0000514 | Notch1 | 18128 | HITS-CLIP | 37986224\|23142080 | Unspecified |
| mmu-miR-30c-5p | MIMAT0000514 | Npat | 244879 | HITS-CLIP | 37986224\|23142080 | Unspecified |
| mmu-miR-30c-5p | MIMAT0000514 | Npc1 | 18145 | HITS-CLIP | 37986224 | Unspecified |
| mmu-miR-30c-5p | MIMAT0000514 | Npepps | 19155 | HITS-CLIP | 37986224\|23597149\|19536157\|23142 | Unspecified |
| mmu-miR-30c-5p | MIMAT0000514 | Nqo1 | 18104 | HITS-CLIP | 23597149 | Unspecified |
| mmu-miR-30c-5p | MIMAT0000514 | Nr2c1 | 22025 | HITS-CLIP | 23597149 | Unspecified |
| mmu-miR-30c-5p | MIMAT0000514 | Nr2c2 | 22026 | HITS-CLIP | 37986224 | Unspecified |
| mmu-miR-30c-5p | MIMAT0000514 | Nr3c1 | 14815 | HITS-CLIP | 37986224\|23142080 | Unspecified |
| mmu-miR-30c-5p | MIMAT0000514 | Nr4a2 | 18227 | HITS-CLIP | 37986224\|23142080 | Unspecified |
| mmu-miR-30c-5p | MIMAT0000514 | Nr4a3 | 18124 | HITS-CLIP | 25753418\|23142080 | Unspecified |
| mmu-miR-30c-5p | MIMAT0000514 | Nr5a2 | 26424 | HITS-CLIP | 23597149 | Unspecified |
| mmu-miR-30c-5p | MIMAT0000514 | Nr6a1 | 14536 | HITS-CLIP | 37986224\|23597149 | Unspecified |
| mmu-miR-30c-5p | MIMAT0000514 | Nrap | 18175 | HITS-CLIP | 25753418 | Unspecified |
| mmu-miR-30c-5p | MIMAT0000514 | Nras | 18176 | HITS-CLIP | 37986224 | Unspecified |
| mmu-miR-30c-5p | MIMAT0000514 | Nrep | 27528 | HITS-CLIP | 37986224\|19536157\|23597149 | Unspecified |
| mmu-miR-30c-5p | MIMAT0000514 | Nrip1 | 268903 | HITS-CLIP | 37986224\|23142080 | Unspecified |
| mmu-miR-30c-5p | MIMAT0000514 | Nrp1 | 18186 | HITS-CLIP | 37986224 | Unspecified |
| mmu-miR-30c-5p | MIMAT0000514 | Nrxn1 | 18189 | HITS-CLIP | 19536157 | Unspecified |
| mmu-miR-30c-5p | MIMAT0000514 | Nrxn3 | 18191 | HITS-CLIP | 19536157 | Unspecified |
| mmu-miR-30c-5p | MIMAT0000514 | Nsa2 | 59050 | HITS-CLIP | 23142080 | Unspecified |
| mmu-miR-30c-5p | MIMAT0000514 | Nsd1 | 18193 | HITS-CLIP | 23597149\|23142080 | Unspecified |
| mmu-miR-30c-5p | MIMAT0000514 | Nsd3 | 234135 | HITS-CLIP | 23142080 | Unspecified |
| mmu-miR-30c-5p | MIMAT0000514 | Nsf | 18195 | HITS-CLIP | 19536157 | Unspecified |
| mmu-miR-30c-5p | MIMAT0000514 | Nsg1 | 18196 | HITS-CLIP | 37986224\|19536157 | Unspecified |
| mmu-miR-30c-5p | MIMAT0000514 | Nsg2 | 18197 | HITS-CLIP | 37986224\|23142080\|19536157 | Unspecified |
| mmu-miR-30c-5p | MIMAT0000514 | Nsmaf | 18201 | HITS-CLIP | 23142080 | Unspecified |
| mmu-miR-30c-5p | MIMAT0000514 | Nsmf | 56876 | HITS-CLIP | 37986224 | Unspecified |
| mmu-miR-30c-5p | MIMAT0000514 | Ntm | 235106 | HITS-CLIP | 37986224 | Unspecified |
| mmu-miR-30c-5p | MIMAT0000514 | Ntrk2 | 18212 | HITS-CLIP | 37986224 | Unspecified |
| mmu-miR-30c-5p | MIMAT0000514 | Nubp2 | 26426 | HITS-CLIP | 37986224 | Unspecified |
| mmu-miR-30c-5p | MIMAT0000514 | Nucks1 | 98415 | HITS-CLIP | 37986224 | Unspecified |
| mmu-miR-30c-5p | MIMAT0000514 | Nudcd1 | 67429 | HITS-CLIP | 23142080 | Unspecified |
| mmu-miR-30c-5p | MIMAT0000514 | Nudcd3 | 209586 | HITS-CLIP | 23142080 | Unspecified |
| mmu-miR-30c-5p | MIMAT0000514 | Nufip2 | 68564 | HITS-CLIP | 37986224\|23597149\|23142080 | Unspecified |

| | | | | | | |
|---|---|---|---|---|---|---|
| mmu-miR-30c-5p | MIMAT0000514 | Numa1 | 101706 | HITS-CLIP | 23142080 | Unspecified |
| mmu-miR-30c-5p | MIMAT0000514 | Nup155 | 170762 | HITS-CLIP | 23142080 | Unspecified |
| mmu-miR-30c-5p | MIMAT0000514 | Nup160 | 59015 | HITS-CLIP | 23142080 | Unspecified |
| mmu-miR-30c-5p | MIMAT0000514 | Nup188 | 227699 | HITS-CLIP | 37986224\|23142080 | Unspecified |
| mmu-miR-30c-5p | MIMAT0000514 | Nup210 | 54563 | HITS-CLIP | 23142080 | Unspecified |
| mmu-miR-30c-5p | MIMAT0000514 | Nup93 | 71805 | HITS-CLIP | 23142080 | Unspecified |
| mmu-miR-30c-5p | MIMAT0000514 | Nup98 | 269966 | HITS-CLIP | 23142080 | Unspecified |
| mmu-miR-30c-5p | MIMAT0000514 | Nus1 | 52014 | HITS-CLIP | 37986224\|23597149\|23142080 | Unspecified |
| mmu-miR-30c-5p | MIMAT0000514 | Nusap1 | 108907 | HITS-CLIP | 37986224 | Unspecified |
| mmu-miR-30c-5p | MIMAT0000514 | Nxpe3 | 385658 | HITS-CLIP | 23142080 | Unspecified |
| mmu-miR-30c-5p | MIMAT0000514 | Nxt2 | 237082 | HITS-CLIP | 23142080 | Unspecified |
| mmu-miR-30c-5p | MIMAT0000514 | Ocrl | 320634 | HITS-CLIP | 23142080 | Unspecified |
| mmu-miR-30c-5p | MIMAT0000514 | Ogdh | 18293 | HITS-CLIP | 37986224\|23597149\|23142080 | Unspecified |
| mmu-miR-30c-5p | MIMAT0000514 | Ogdhl | 239017 | HITS-CLIP | 25753418 | Unspecified |
| mmu-miR-30c-5p | MIMAT0000514 | Ogt | 108155 | HITS-CLIP | 19536157\|23142080 | Unspecified |
| mmu-miR-30c-5p | MIMAT0000514 | Omg | 18377 | HITS-CLIP | 37986224 | Unspecified |
| mmu-miR-30c-5p | MIMAT0000514 | Or5w19 | 258103 | qCLASH | 31393953 | Unspecified |
| mmu-miR-30c-5p | MIMAT0000514 | Orc3 | 50793 | HITS-CLIP | 23142080 | Unspecified |
| mmu-miR-30c-5p | MIMAT0000514 | Orc4 | 26428 | HITS-CLIP | 37986224 | Unspecified |
| mmu-miR-30c-5p | MIMAT0000514 | Orc6 | 56452 | HITS-CLIP | 23142080 | Unspecified |
| mmu-miR-30c-5p | MIMAT0000514 | Ormdl1 | 227102 | HITS-CLIP | 23142080 | Unspecified |
| mmu-miR-30c-5p | MIMAT0000514 | Ormdl2 | 66844 | HITS-CLIP | 37986224 | Unspecified |
| mmu-miR-30c-5p | MIMAT0000514 | Ormdl3 | 66612 | HITS-CLIP | 37986224 | Unspecified |
| mmu-miR-30c-5p | MIMAT0000514 | Osbpl8 | 237542 | HITS-CLIP | 23597149\|23142080 | Unspecified |
| mmu-miR-30c-5p | MIMAT0000514 | Osbpl9 | 100273 | HITS-CLIP | 25753418 | Unspecified |
| mmu-miR-30c-5p | MIMAT0000514 | Osmr | 18414 | HITS-CLIP | 37986224 | Unspecified |
| mmu-miR-30c-5p | MIMAT0000514 | Ostf1 | 20409 | HITS-CLIP | 23597149\|23142080 | Unspecified |
| mmu-miR-30c-5p | MIMAT0000514 | Ostm1 | 14628 | HITS-CLIP | 23142080 | Unspecified |
| mmu-miR-30c-5p | MIMAT0000514 | Otud4 | 73945 | HITS-CLIP | 37986224\|23142080\|19536157 | Unspecified |
| mmu-miR-30c-5p | MIMAT0000514 | Otud5 | 54644 | HITS-CLIP | 25753418 | Unspecified |
| mmu-miR-30c-5p | MIMAT0000514 | Otud6b | 72201 | HITS-CLIP | 25753418 | Unspecified |
| mmu-miR-30c-5p | MIMAT0000514 | Otulinl | 54491 | HITS-CLIP | 23142080 | Unspecified |
| mmu-miR-30c-5p | MIMAT0000514 | Oxct1 | 67041 | HITS-CLIP | 37986224 | Unspecified |
| mmu-miR-30c-5p | MIMAT0000514 | Oxr1 | 170719 | HITS-CLIP | 23597149 | Unspecified |
| mmu-miR-30c-5p | MIMAT0000514 | P4ha1 | 18451 | HITS-CLIP | 23597149\|23142080 | Unspecified |
| mmu-miR-30c-5p | MIMAT0000514 | P4ha2 | 18452 | HITS-CLIP | 37986224\|19536157\|23142080 | Unspecified |
| mmu-miR-30c-5p | MIMAT0000514 | Pabpc1 | 18458 | HITS-CLIP | 37986224\|23142080 | Unspecified |
| mmu-miR-30c-5p | MIMAT0000514 | Paf1 | 54624 | HITS-CLIP | 23142080 | Unspecified |
| mmu-miR-30c-5p | MIMAT0000514 | Pafah1b2 | 18475 | HITS-CLIP | 37986224\|23142080 | Unspecified |

| | | | | | | |
|---|---|---|---|---|---|---|
| mmu-miR-30c-5p | MIMAT0000514 | Pag1 | 94212 | HITS-CLIP | 37986224 | Unspecified |
| mmu-miR-30c-5p | MIMAT0000514 | Pak1ip1 | 68083 | HITS-CLIP | 37986224 | Unspecified |
| mmu-miR-30c-5p | MIMAT0000514 | Pak2 | 224105 | HITS-CLIP | 37986224 | Unspecified |
| mmu-miR-30c-5p | MIMAT0000514 | Pakap | 677884 | HITS-CLIP | 25753418 | Unspecified |
| mmu-miR-30c-5p | MIMAT0000514 | Palb2 | 233826 | HITS-CLIP | 23142080 | Unspecified |
| mmu-miR-30c-5p | MIMAT0000514 | Pan3 | 72587 | HITS-CLIP | 37986224\|23142080 | Unspecified |
| mmu-miR-30c-5p | MIMAT0000514 | Pank3 | 211347 | HITS-CLIP | 37986224\|23142080 | Unspecified |
| mmu-miR-30c-5p | MIMAT0000514 | Papola | 18789 | HITS-CLIP | 37986224\|23142080 | Unspecified |
| mmu-miR-30c-5p | MIMAT0000514 | Papolg | 216578 | HITS-CLIP | 37986224 | Unspecified |
| mmu-miR-30c-5p | MIMAT0000514 | Paqr7 | 71904 | HITS-CLIP | 23597149 | Unspecified |
| mmu-miR-30c-5p | MIMAT0000514 | Paqr9 | 75552 | HITS-CLIP | 37986224\|23597149 | Unspecified |
| mmu-miR-30c-5p | MIMAT0000514 | Parp16 | 214424 | HITS-CLIP | 37986224\|23597149 | Unspecified |
| mmu-miR-30c-5p | MIMAT0000514 | Parp2 | 11546 | HITS-CLIP | 23142080 | Unspecified |
| mmu-miR-30c-5p | MIMAT0000514 | Parp6 | 67287 | HITS-CLIP | 23142080\|19536157 | Unspecified |
| mmu-miR-30c-5p | MIMAT0000514 | Parpbp | 75317 | HITS-CLIP | 23142080 | Unspecified |
| mmu-miR-30c-5p | MIMAT0000514 | Pax5 | 5079 | qCLASH | 31393953 | Unspecified |
| mmu-miR-30c-5p | MIMAT0000514 | Paxip1 | 55982 | HITS-CLIP | 23142080 | Unspecified |
| mmu-miR-30c-5p | MIMAT0000514 | Pbld1 | 68371 | HITS-CLIP | 23597149 | Unspecified |
| mmu-miR-30c-5p | MIMAT0000514 | Pbrm1 | 66923 | HITS-CLIP | 37986224\|25753418\|23142080 | Unspecified |
| mmu-miR-30c-5p | MIMAT0000514 | Pcbp2 | 18521 | HITS-CLIP | 37986224 | Unspecified |
| mmu-miR-30c-5p | MIMAT0000514 | Pcdh10 | 18526 | HITS-CLIP | 37986224\|19536157 | Unspecified |
| mmu-miR-30c-5p | MIMAT0000514 | Pcdh17 | 219228 | HITS-CLIP | 37986224 | Unspecified |
| mmu-miR-30c-5p | MIMAT0000514 | Pcdhga12 | 93724 | HITS-CLIP | 37986224 | Unspecified |
| mmu-miR-30c-5p | MIMAT0000514 | Pcdhgc3 | 5098 | HITS-CLIP | 19536157\|23597149 | Unspecified |
| mmu-miR-30c-5p | MIMAT0000514 | Pcgf3 | 69587 | HITS-CLIP | 37986224\|23597149\|19536157\|2314 | Unspecified |
| mmu-miR-30c-5p | MIMAT0000514 | Pcgf5 | 76073 | HITS-CLIP | 37986224\|23142080 | Unspecified |
| mmu-miR-30c-5p | MIMAT0000514 | Pck1 | 18534 | HITS-CLIP | 23597149 | Unspecified |
| mmu-miR-30c-5p | MIMAT0000514 | Pcm1 | 18536 | HITS-CLIP | 37986224 | Unspecified |
| mmu-miR-30c-5p | MIMAT0000514 | Pcmtd2 | 245867 | HITS-CLIP | 23142080 | Unspecified |
| mmu-miR-30c-5p | MIMAT0000514 | Pcnx | 54604 | HITS-CLIP | 37986224 | Unspecified |
| mmu-miR-30c-5p | MIMAT0000514 | Pcsk2 | 18549 | HITS-CLIP | 37986224 | Unspecified |
| mmu-miR-30c-5p | MIMAT0000514 | Pctp | 18559 | HITS-CLIP | 37986224\|23597149\|23142080 | Unspecified |
| mmu-miR-30c-5p | MIMAT0000514 | Pcyt1a | 13026 | HITS-CLIP | 23142080 | Unspecified |
| mmu-miR-30c-5p | MIMAT0000514 | Pcyt2 | 68671 | HITS-CLIP | 23597149 | Unspecified |
| mmu-miR-30c-5p | MIMAT0000514 | Pdap1 | 231887 | qCLASH | 31393953 | Unspecified |
| mmu-miR-30c-5p | MIMAT0000514 | Pdcd4 | 18569 | HITS-CLIP | 23597149\|23142080 | Unspecified |
| mmu-miR-30c-5p | MIMAT0000514 | Pdcl | 67466 | HITS-CLIP | 23142080 | Unspecified |
| mmu-miR-30c-5p | MIMAT0000514 | Pde3b | 18576 | qCLASH | 31393953 | Unspecified |
| mmu-miR-30c-5p | MIMAT0000514 | Pde4b | 18578 | HITS-CLIP | 19536157\|23142080 | Unspecified |

| | | | | | | |
|---|---|---|---|---|---|---|
| mmu-miR-30c-5p | MIMAT0000514 | Pde4d | 238871 | HITS-CLIP | 37986224 | Unspecified |
| mmu-miR-30c-5p | MIMAT0000514 | Pde7a | 18583 | HITS-CLIP | 37986224\|23142080 | Unspecified |
| mmu-miR-30c-5p | MIMAT0000514 | Pdgfrb | 18596 | HITS-CLIP | 37986224 | Unspecified |
| mmu-miR-30c-5p | MIMAT0000514 | Pdha1 | 18597 | HITS-CLIP | 23142080 | Unspecified |
| mmu-miR-30c-5p | MIMAT0000514 | Pdk1 | 228026 | HITS-CLIP | 37986224\|23142080 | Unspecified |
| mmu-miR-30c-5p | MIMAT0000514 | Pds5b | 100710 | HITS-CLIP | 23142080 | Unspecified |
| mmu-miR-30c-5p | MIMAT0000514 | Pdss1 | 56075 | HITS-CLIP | 23142080 | Unspecified |
| mmu-miR-30c-5p | MIMAT0000514 | Pdzd8 | 107368 | HITS-CLIP | 37986224 | Unspecified |
| mmu-miR-30c-5p | MIMAT0000514 | Pdzrn3 | 55983 | HITS-CLIP | 23597149\|25753418 | Unspecified |
| mmu-miR-30c-5p | MIMAT0000514 | Peak1 | 244895 | HITS-CLIP | 37986224\|23142080 | Unspecified |
| mmu-miR-30c-5p | MIMAT0000514 | Pecr | 111175 | HITS-CLIP | 23597149 | Unspecified |
| mmu-miR-30c-5p | MIMAT0000514 | Peg3 | 18616 | HITS-CLIP | 37986224 | Unspecified |
| mmu-miR-30c-5p | MIMAT0000514 | Peli1 | 67245 | HITS-CLIP | 37986224 | Unspecified |
| mmu-miR-30c-5p | MIMAT0000514 | Per2 | 18627 | HITS-CLIP | 37986224\|23597149\|23142080 | Unspecified |
| mmu-miR-30c-5p | MIMAT0000514 | Pex11a | 18631 | HITS-CLIP | 37986224 | Unspecified |
| mmu-miR-30c-5p | MIMAT0000514 | Pex5 | 19305 | HITS-CLIP | 23597149 | Unspecified |
| mmu-miR-30c-5p | MIMAT0000514 | Pfkm | 18642 | HITS-CLIP | 25753418 | Unspecified |
| mmu-miR-30c-5p | MIMAT0000514 | Pfkp | 56421 | HITS-CLIP | 23142080 | Unspecified |
| mmu-miR-30c-5p | MIMAT0000514 | Pfn2 | 18645 | HITS-CLIP | 23597149\|19536157 | Unspecified |
| mmu-miR-30c-5p | MIMAT0000514 | Pgam5 | 72542 | HITS-CLIP | 37986224 | Unspecified |
| mmu-miR-30c-5p | MIMAT0000514 | Pgap1 | 241062 | HITS-CLIP | 37986224 | Unspecified |
| mmu-miR-30c-5p | MIMAT0000514 | Pgap4 | 84302 | HITS-CLIP | 25753418 | Unspecified |
| mmu-miR-30c-5p | MIMAT0000514 | Pgm1 | 66681 | HITS-CLIP | 23597149\|23142080 | Unspecified |
| mmu-miR-30c-5p | MIMAT0000514 | Pgm2l1 | 70974 | HITS-CLIP | 37986224\|19536157\|23142080 | Unspecified |
| mmu-miR-30c-5p | MIMAT0000514 | Pgp | 67078 | HITS-CLIP | 37986224\|23142080 | Unspecified |
| mmu-miR-30c-5p | MIMAT0000514 | Pgrmc1 | 53328 | HITS-CLIP | 37986224 | Unspecified |
| mmu-miR-30c-5p | MIMAT0000514 | Phactr1 | 218194 | HITS-CLIP | 37986224 | Unspecified |
| mmu-miR-30c-5p | MIMAT0000514 | Phactr2 | 215789 | HITS-CLIP | 23142080 | Unspecified |
| mmu-miR-30c-5p | MIMAT0000514 | Phax | 56698 | HITS-CLIP | 23142080 | Unspecified |
| mmu-miR-30c-5p | MIMAT0000514 | Phc3 | 241915 | HITS-CLIP | 37986224\|23142080 | Unspecified |
| mmu-miR-30c-5p | MIMAT0000514 | Pheta1 | 144717 | HITS-CLIP | 19536157\|23597149 | Unspecified |
| mmu-miR-30c-5p | MIMAT0000514 | Phf10 | 72057 | HITS-CLIP | 37986224 | Unspecified |
| mmu-miR-30c-5p | MIMAT0000514 | Phf13 | 230936 | HITS-CLIP | 23142080 | Unspecified |
| mmu-miR-30c-5p | MIMAT0000514 | Phf14 | 75725 | HITS-CLIP | 37986224 | Unspecified |
| mmu-miR-30c-5p | MIMAT0000514 | Phf19 | 74016 | HITS-CLIP | 37986224\|23142080 | Unspecified |
| mmu-miR-30c-5p | MIMAT0000514 | Phf20 | 228829 | HITS-CLIP | 37986224\|23142080 | Unspecified |
| mmu-miR-30c-5p | MIMAT0000514 | Phf6 | 70998 | HITS-CLIP | 23142080 | Unspecified |
| mmu-miR-30c-5p | MIMAT0000514 | Phip | 83946 | HITS-CLIP | 37986224\|23142080 | Unspecified |
| mmu-miR-30c-5p | MIMAT0000514 | Phtf2 | 68770 | HITS-CLIP | 37986224\|23142080 | Unspecified |

| | | | | | | |
|---|---|---|---|---|---|---|
| mmu-miR-30c-5p | MIMAT0000514 | Picalm | 233489 | HITS-CLIP | 37986224\|23142080 | Unspecified |
| mmu-miR-30c-5p | MIMAT0000514 | Pid1 | 98496 | HITS-CLIP | 23597149 | Unspecified |
| mmu-miR-30c-5p | MIMAT0000514 | Piga | 18700 | HITS-CLIP | 23142080 | Unspecified |
| mmu-miR-30c-5p | MIMAT0000514 | Pigb | 9488 | HITS-CLIP | 25753418 | Unspecified |
| mmu-miR-30c-5p | MIMAT0000514 | Pign | 27392 | HITS-CLIP | 37986224\|23142080\|23597149 | Unspecified |
| mmu-miR-30c-5p | MIMAT0000514 | Pigs | 276846 | HITS-CLIP | 37986224 | Unspecified |
| mmu-miR-30c-5p | MIMAT0000514 | Pigw | 70325 | HITS-CLIP | 23142080 | Unspecified |
| mmu-miR-30c-5p | MIMAT0000514 | Pik3c2a | 18704 | HITS-CLIP | 23142080 | Unspecified |
| mmu-miR-30c-5p | MIMAT0000514 | Pik3c2b | 240752 | HITS-CLIP | 23597149 | Unspecified |
| mmu-miR-30c-5p | MIMAT0000514 | Pik3c3 | 225326 | HITS-CLIP | 23597149\|23142080 | Unspecified |
| mmu-miR-30c-5p | MIMAT0000514 | Pik3ca | 18706 | HITS-CLIP | 37986224\|23597149\|23142080 | Unspecified |
| mmu-miR-30c-5p | MIMAT0000514 | Pik3cb | 74769 | HITS-CLIP | 23597149 | Unspecified |
| mmu-miR-30c-5p | MIMAT0000514 | Pik3cd | 18707 | HITS-CLIP | 37986224\|23142080 | Unspecified |
| mmu-miR-30c-5p | MIMAT0000514 | Pik3r1 | 18708 | HITS-CLIP | 37986224\|19536157\|23597149\|23142 | Unspecified |
| mmu-miR-30c-5p | MIMAT0000514 | Pin1 | 23988 | HITS-CLIP | 25753418 | Unspecified |
| mmu-miR-30c-5p | MIMAT0000514 | Pip4k2a | 18718 | HITS-CLIP | 37986224\|23142080 | Unspecified |
| mmu-miR-30c-5p | MIMAT0000514 | Pip4k2b | 108083 | HITS-CLIP | 23597149\|23142080 | Unspecified |
| mmu-miR-30c-5p | MIMAT0000514 | Pip4p2 | 55529 | HITS-CLIP | 23597149 | Unspecified |
| mmu-miR-30c-5p | MIMAT0000514 | Pitpna | 18738 | HITS-CLIP | 23142080\|23597149 | Unspecified |
| mmu-miR-30c-5p | MIMAT0000514 | Pitpnc1 | 71795 | HITS-CLIP | 23142080 | Unspecified |
| mmu-miR-30c-5p | MIMAT0000514 | Pitpnm2 | 19679 | HITS-CLIP | 23142080\|19536157 | Unspecified |
| mmu-miR-30c-5p | MIMAT0000514 | Pkn2 | 109333 | HITS-CLIP | 37986224\|23597149\|23142080 | Unspecified |
| mmu-miR-30c-5p | MIMAT0000514 | Pkp2 | 67451 | HITS-CLIP | 25753418 | Unspecified |
| mmu-miR-30c-5p | MIMAT0000514 | Pla2g6 | 53357 | HITS-CLIP | 23142080 | Unspecified |
| mmu-miR-30c-5p | MIMAT0000514 | Plaa | 18786 | HITS-CLIP | 23142080 | Unspecified |
| mmu-miR-30c-5p | MIMAT0000514 | Plagl2 | 54711 | HITS-CLIP | 37986224\|23597149\|23142080 | Unspecified |
| mmu-miR-30c-5p | MIMAT0000514 | Plcxd2 | 433022 | HITS-CLIP | 37986224 | Unspecified |
| mmu-miR-30c-5p | MIMAT0000514 | Plekha1 | 101476 | HITS-CLIP | 37986224 | Unspecified |
| mmu-miR-30c-5p | MIMAT0000514 | Plekha2 | 83436 | HITS-CLIP | 23142080 | Unspecified |
| mmu-miR-30c-5p | MIMAT0000514 | Plekhb2 | 226971 | HITS-CLIP | 23142080 | Unspecified |
| mmu-miR-30c-5p | MIMAT0000514 | Plekhm1 | 353047 | HITS-CLIP | 25753418 | Unspecified |
| mmu-miR-30c-5p | MIMAT0000514 | Plekhm3 | 241075 | HITS-CLIP | 37986224\|23142080 | Unspecified |
| mmu-miR-30c-5p | MIMAT0000514 | Plgrkt | 67759 | HITS-CLIP | 37986224\|23142080 | Unspecified |
| mmu-miR-30c-5p | MIMAT0000514 | Plk1 | 18817 | HITS-CLIP | 37986224 | Unspecified |
| mmu-miR-30c-5p | MIMAT0000514 | Plk3 | 12795 | HITS-CLIP | 23142080 | Unspecified |
| mmu-miR-30c-5p | MIMAT0000514 | Plp1 | 18823 | HITS-CLIP | 37986224 | Unspecified |
| mmu-miR-30c-5p | MIMAT0000514 | Plrg1 | 53317 | HITS-CLIP | 23142080 | Unspecified |
| mmu-miR-30c-5p | MIMAT0000514 | Pls3 | 102866 | HITS-CLIP | 23597149 | Unspecified |
| mmu-miR-30c-5p | MIMAT0000514 | Plxna2 | 18845 | HITS-CLIP | 37986224\|23597149\|19536157 | Unspecified |

| | | | | | | |
|---|---|---|---|---|---|---|
| mmu-miR-30c-5p | MIMAT0000514 | Plxnc1 | 54712 | HITS-CLIP | 37986224\|23597149 | Unspecified |
| mmu-miR-30c-5p | MIMAT0000514 | Pmaip1 | 58801 | HITS-CLIP | 37986224 | Unspecified |
| mmu-miR-30c-5p | MIMAT0000514 | Pmepa1 | 65112 | HITS-CLIP | 23142080 | Unspecified |
| mmu-miR-30c-5p | MIMAT0000514 | Pnisr | 66625 | HITS-CLIP | 25753418 | Unspecified |
| mmu-miR-30c-5p | MIMAT0000514 | Pnkd | 56695 | HITS-CLIP | 37986224\|19536157 | Unspecified |
| mmu-miR-30c-5p | MIMAT0000514 | Pnpla8 | 67452 | HITS-CLIP | 37986224\|23142080 | Unspecified |
| mmu-miR-30c-5p | MIMAT0000514 | Pnpo | 103711 | HITS-CLIP | 37986224 | Unspecified |
| mmu-miR-30c-5p | MIMAT0000514 | Pogz | 229584 | HITS-CLIP | 23142080 | Unspecified |
| mmu-miR-30c-5p | MIMAT0000514 | Pola1 | 18968 | HITS-CLIP | 23142080 | Unspecified |
| mmu-miR-30c-5p | MIMAT0000514 | Pold3 | 67967 | HITS-CLIP | 37986224 | Unspecified |
| mmu-miR-30c-5p | MIMAT0000514 | Poldip3 | 73826 | HITS-CLIP | 23597149\|23142080 | Unspecified |
| mmu-miR-30c-5p | MIMAT0000514 | Poli | 11201 | HITS-CLIP | 23142080 | Unspecified |
| mmu-miR-30c-5p | MIMAT0000514 | Polr2g | 67710 | HITS-CLIP | 23142080 | Unspecified |
| mmu-miR-30c-5p | MIMAT0000514 | Polr3e | 26939 | HITS-CLIP | 37986224\|23597149\|23142080 | Unspecified |
| mmu-miR-30c-5p | MIMAT0000514 | Polr3g | 67486 | HITS-CLIP | 37986224\|23142080 | Unspecified |
| mmu-miR-30c-5p | MIMAT0000514 | Pon2 | 330260 | HITS-CLIP | 37986224\|25753418 | Unspecified |
| mmu-miR-30c-5p | MIMAT0000514 | Pou2af1 | 18985 | qCLASH | 31393953 | Unspecified |
| mmu-miR-30c-5p | MIMAT0000514 | Pou2f1 | 18986 | HITS-CLIP | 23597149\|23142080 | Unspecified |
| mmu-miR-30c-5p | MIMAT0000514 | Pou6f1 | 19009 | HITS-CLIP | 37986224 | Unspecified |
| mmu-miR-30c-5p | MIMAT0000514 | Ppa1 | 67895 | HITS-CLIP | 37986224\|23597149\|23142080 | Unspecified |
| mmu-miR-30c-5p | MIMAT0000514 | Ppargc1b | 170826 | HITS-CLIP | 37986224\|23597149\|23142080 | Unspecified |
| mmu-miR-30c-5p | MIMAT0000514 | Ppfia1 | 233977 | HITS-CLIP | 25753418 | Unspecified |
| mmu-miR-30c-5p | MIMAT0000514 | Ppid | 67738 | HITS-CLIP | 37986224\|23142080 | Unspecified |
| mmu-miR-30c-5p | MIMAT0000514 | Ppig | 228005 | HITS-CLIP | 37986224 | Unspecified |
| mmu-miR-30c-5p | MIMAT0000514 | Ppih | 66101 | HITS-CLIP | 23142080 | Unspecified |
| mmu-miR-30c-5p | MIMAT0000514 | Ppil4 | 67418 | HITS-CLIP//qCLASH | 37986224\|31393953\|23142080 | Unspecified |
| mmu-miR-30c-5p | MIMAT0000514 | Ppip5k2 | 227399 | HITS-CLIP | 37986224\|23597149 | Unspecified |
| mmu-miR-30c-5p | MIMAT0000514 | Ppm1a | 19042 | HITS-CLIP | 23597149\|19536157 | Unspecified |
| mmu-miR-30c-5p | MIMAT0000514 | Ppm1b | 19043 | HITS-CLIP | 23142080 | Unspecified |
| mmu-miR-30c-5p | MIMAT0000514 | Ppm1g | 14208 | qCLASH | 31393953 | Unspecified |
| mmu-miR-30c-5p | MIMAT0000514 | Ppm1l | 242083 | HITS-CLIP | 37986224 | Unspecified |
| mmu-miR-30c-5p | MIMAT0000514 | Ppp1cb | 19046 | HITS-CLIP | 23142080 | Unspecified |
| mmu-miR-30c-5p | MIMAT0000514 | Ppp1r12a | 17931 | HITS-CLIP | 37986224\|23142080\|23597149\|19536 | Unspecified |
| mmu-miR-30c-5p | MIMAT0000514 | Ppp1r13l | 333654 | HITS-CLIP | 23142080 | Unspecified |
| mmu-miR-30c-5p | MIMAT0000514 | Ppp1r15b | 108954 | HITS-CLIP | 23142080 | Unspecified |
| mmu-miR-30c-5p | MIMAT0000514 | Ppp1r18 | 76448 | HITS-CLIP | 37986224\|23142080 | Unspecified |
| mmu-miR-30c-5p | MIMAT0000514 | Ppp1r3a | 140491 | HITS-CLIP | 25753418 | Unspecified |
| mmu-miR-30c-5p | MIMAT0000514 | Ppp1r3c | 53412 | HITS-CLIP | 37986224\|23597149 | Unspecified |
| mmu-miR-30c-5p | MIMAT0000514 | Ppp1r8 | 100336 | qCLASH | 31393953 | Unspecified |

| | | | | | | |
|---|---|---|---|---|---|---|
| mmu-miR-30c-5p | MIMAT0000514 | Ppp2r1a | 51792 | HITS-CLIP | 25753418\|26203562 | Unspecified |
| mmu-miR-30c-5p | MIMAT0000514 | Ppp2r2a | 71978 | HITS-CLIP | 23142080 | Unspecified |
| mmu-miR-30c-5p | MIMAT0000514 | Ppp2r2c | 269643 | HITS-CLIP | 19536157 | Unspecified |
| mmu-miR-30c-5p | MIMAT0000514 | Ppp2r2d | 52432 | HITS-CLIP | 23597149 | Unspecified |
| mmu-miR-30c-5p | MIMAT0000514 | Ppp3ca | 19055 | HITS-CLIP | 37986224 | Unspecified |
| mmu-miR-30c-5p | MIMAT0000514 | Ppp3cb | 19056 | HITS-CLIP | 37986224 | Unspecified |
| mmu-miR-30c-5p | MIMAT0000514 | Ppp3r1 | 19058 | HITS-CLIP | 37986224\|19536157\|23142080 | Unspecified |
| mmu-miR-30c-5p | MIMAT0000514 | Ppp4r3a | 55671 | HITS-CLIP | 23597149\|23142080 | Unspecified |
| mmu-miR-30c-5p | MIMAT0000514 | Ppp4r3b | 57223 | HITS-CLIP | 23142080 | Unspecified |
| mmu-miR-30c-5p | MIMAT0000514 | Ppp6r1 | 243819 | HITS-CLIP | 23142080 | Unspecified |
| mmu-miR-30c-5p | MIMAT0000514 | Ppp6r3 | 52036 | HITS-CLIP | 23142080 | Unspecified |
| mmu-miR-30c-5p | MIMAT0000514 | Pptc7 | 320717 | HITS-CLIP//qCLASH | 37986224\|31393953\|23142080 | Unspecified |
| mmu-miR-30c-5p | MIMAT0000514 | Prc1 | 233406 | HITS-CLIP | 23142080 | Unspecified |
| mmu-miR-30c-5p | MIMAT0000514 | Prdm1 | 12142 | HITS-CLIP | 37986224\|23142080 | Unspecified |
| mmu-miR-30c-5p | MIMAT0000514 | Prelid1 | 66494 | HITS-CLIP | 23597149\|23142080 | Unspecified |
| mmu-miR-30c-5p | MIMAT0000514 | Prkab2 | 108097 | HITS-CLIP | 37986224\|23142080 | Unspecified |
| mmu-miR-30c-5p | MIMAT0000514 | Prkaca | 18747 | HITS-CLIP | 23597149 | Unspecified |
| mmu-miR-30c-5p | MIMAT0000514 | Prkar1a | 19084 | HITS-CLIP | 37986224\|23597149\|23142080 | Unspecified |
| mmu-miR-30c-5p | MIMAT0000514 | Prkar2a | 19087 | HITS-CLIP | 23142080 | Unspecified |
| mmu-miR-30c-5p | MIMAT0000514 | Prkca | 18750 | HITS-CLIP | 37986224\|23142080 | Unspecified |
| mmu-miR-30c-5p | MIMAT0000514 | Prkch | 18755 | HITS-CLIP | 23142080 | Unspecified |
| mmu-miR-30c-5p | MIMAT0000514 | Prkcq | 18761 | HITS-CLIP | 23142080 | Unspecified |
| mmu-miR-30c-5p | MIMAT0000514 | Prkd3 | 75292 | HITS-CLIP | 23597149 | Unspecified |
| mmu-miR-30c-5p | MIMAT0000514 | Prkdc | 19090 | HITS-CLIP | 23142080 | Unspecified |
| mmu-miR-30c-5p | MIMAT0000514 | Prlr | 19116 | HITS-CLIP | 37986224\|23597149\|23142080 | Unspecified |
| mmu-miR-30c-5p | MIMAT0000514 | Prmt1 | 15469 | HITS-CLIP | 37986224 | Unspecified |
| mmu-miR-30c-5p | MIMAT0000514 | Prodh | 19125 | HITS-CLIP | 23597149 | Unspecified |
| mmu-miR-30c-5p | MIMAT0000514 | Pros1 | 19128 | HITS-CLIP | 23142080 | Unspecified |
| mmu-miR-30c-5p | MIMAT0000514 | Proser1 | 212127 | HITS-CLIP | 37986224\|23597149 | Unspecified |
| mmu-miR-30c-5p | MIMAT0000514 | Prpf4 | 70052 | HITS-CLIP | 37986224 | Unspecified |
| mmu-miR-30c-5p | MIMAT0000514 | Prpf4b | 19134 | qCLASH | 31393953 | Unspecified |
| mmu-miR-30c-5p | MIMAT0000514 | Prr11 | 270906 | HITS-CLIP | 37986224\|23597149\|23142080 | Unspecified |
| mmu-miR-30c-5p | MIMAT0000514 | Prr36 | 80164 | HITS-CLIP | 23142080 | Unspecified |
| mmu-miR-30c-5p | MIMAT0000514 | Prrc2a | 53761 | HITS-CLIP | 37986224 | Unspecified |
| mmu-miR-30c-5p | MIMAT0000514 | Prrc2b | 227723 | HITS-CLIP | 37986224\|23142080 | Unspecified |
| mmu-miR-30c-5p | MIMAT0000514 | Prrt2 | 69017 | HITS-CLIP | 19536157 | Unspecified |
| mmu-miR-30c-5p | MIMAT0000514 | Prune1 | 58497 | HITS-CLIP | 23142080 | Unspecified |
| mmu-miR-30c-5p | MIMAT0000514 | Prune2 | 353211 | HITS-CLIP | 25753418 | Unspecified |
| mmu-miR-30c-5p | MIMAT0000514 | Psd3 | 234353 | HITS-CLIP | 37986224\|23597149 | Unspecified |

| | | | | | | |
|---|---|---|---|---|---|---|
| mmu-miR-30c-5p | MIMAT0000514 | Psen1 | 19164 | HITS-CLIP | 25753418 | Unspecified |
| mmu-miR-30c-5p | MIMAT0000514 | Psen2 | 19165 | HITS-CLIP | 23142080 | Unspecified |
| mmu-miR-30c-5p | MIMAT0000514 | Psma2 | 19166 | HITS-CLIP | 37986224 | Unspecified |
| mmu-miR-30c-5p | MIMAT0000514 | Psmb1 | 19170 | HITS-CLIP | 23142080 | Unspecified |
| mmu-miR-30c-5p | MIMAT0000514 | Psmb2 | 26445 | HITS-CLIP | 23142080 | Unspecified |
| mmu-miR-30c-5p | MIMAT0000514 | Psmb8 | 16913 | HITS-CLIP | 37986224 | Unspecified |
| mmu-miR-30c-5p | MIMAT0000514 | Psmc3 | 19182 | HITS-CLIP | 23597149 | Unspecified |
| mmu-miR-30c-5p | MIMAT0000514 | Psmd12 | 66997 | HITS-CLIP | 23142080 | Unspecified |
| mmu-miR-30c-5p | MIMAT0000514 | Psmd5 | 66998 | HITS-CLIP | 23142080 | Unspecified |
| mmu-miR-30c-5p | MIMAT0000514 | Psmd6 | 66413 | HITS-CLIP | 23142080 | Unspecified |
| mmu-miR-30c-5p | MIMAT0000514 | Psme3 | 19192 | HITS-CLIP | 23142080 | Unspecified |
| mmu-miR-30c-5p | MIMAT0000514 | Psme4 | 103554 | HITS-CLIP | 37986224\|23597149\|25753418 | Unspecified |
| mmu-miR-30c-5p | MIMAT0000514 | Ptar1 | 72351 | HITS-CLIP | 23142080 | Unspecified |
| mmu-miR-30c-5p | MIMAT0000514 | Ptbp3 | 230257 | HITS-CLIP | 37986224\|23142080 | Unspecified |
| mmu-miR-30c-5p | MIMAT0000514 | Ptgfrn | 19221 | HITS-CLIP | 37986224\|23142080 | Unspecified |
| mmu-miR-30c-5p | MIMAT0000514 | Ptk7 | 71461 | HITS-CLIP | 26203562 | Unspecified |
| mmu-miR-30c-5p | MIMAT0000514 | Ptp4a2 | 19244 | HITS-CLIP | 23142080 | Unspecified |
| mmu-miR-30c-5p | MIMAT0000514 | Ptpa | 5524 | HITS-CLIP | 23597149\|25753418\|23142080 | Unspecified |
| mmu-miR-30c-5p | MIMAT0000514 | Ptpn14 | 19250 | HITS-CLIP | 37986224 | Unspecified |
| mmu-miR-30c-5p | MIMAT0000514 | Ptpn2 | 19255 | HITS-CLIP | 37986224\|23597149\|23142080 | Unspecified |
| mmu-miR-30c-5p | MIMAT0000514 | Ptpn21 | 24000 | HITS-CLIP | 37986224\|23597149 | Unspecified |
| mmu-miR-30c-5p | MIMAT0000514 | Ptpn22 | 19260 | HITS-CLIP | 23142080 | Unspecified |
| mmu-miR-30c-5p | MIMAT0000514 | Ptpn3 | 545622 | HITS-CLIP | 23597149\|25753418 | Unspecified |
| mmu-miR-30c-5p | MIMAT0000514 | Ptpn4 | 19258 | HITS-CLIP | 37986224\|23597149\|23142080 | Unspecified |
| mmu-miR-30c-5p | MIMAT0000514 | Ptprb | 19263 | HITS-CLIP | 37986224 | Unspecified |
| mmu-miR-30c-5p | MIMAT0000514 | Ptprc | 19264 | HITS-CLIP | 37986224\|23142080 | Unspecified |
| mmu-miR-30c-5p | MIMAT0000514 | Ptpre | 19267 | HITS-CLIP | 23142080 | Unspecified |
| mmu-miR-30c-5p | MIMAT0000514 | Ptprf | 19268 | HITS-CLIP | 23597149 | Unspecified |
| mmu-miR-30c-5p | MIMAT0000514 | Ptprg | 19270 | HITS-CLIP | 37986224 | Unspecified |
| mmu-miR-30c-5p | MIMAT0000514 | Ptprz1 | 19283 | HITS-CLIP | 37986224\|19536157 | Unspecified |
| mmu-miR-30c-5p | MIMAT0000514 | Pttg1ip | 108705 | HITS-CLIP | 23597149\|23142080 | Unspecified |
| mmu-miR-30c-5p | MIMAT0000514 | Pum1 | 80912 | HITS-CLIP | 37986224\|23142080 | Unspecified |
| mmu-miR-30c-5p | MIMAT0000514 | Pum2 | 80913 | HITS-CLIP | 37986224 | Unspecified |
| mmu-miR-30c-5p | MIMAT0000514 | Pum3 | 52874 | HITS-CLIP | 23142080 | Unspecified |
| mmu-miR-30c-5p | MIMAT0000514 | Pura | 19290 | HITS-CLIP | 37986224\|19536157\|23142080 | Unspecified |
| mmu-miR-30c-5p | MIMAT0000514 | Purb | 19291 | HITS-CLIP | 37986224\|19536157 | Unspecified |
| mmu-miR-30c-5p | MIMAT0000514 | Purg | 75029 | HITS-CLIP | 23142080 | Unspecified |
| mmu-miR-30c-5p | MIMAT0000514 | Pus10 | 74467 | HITS-CLIP | 23597149\|23142080 | Unspecified |
| mmu-miR-30c-5p | MIMAT0000514 | Pwwp2a | 70802 | HITS-CLIP | 37986224 | Unspecified |

| | | | | | | |
|---|---|---|---|---|---|---|
| mmu-miR-30c-5p | MIMAT0000514 | Pxmp4 | 59038 | HITS-CLIP | 23597149 | Unspecified |
| mmu-miR-30c-5p | MIMAT0000514 | Pxn | 19303 | HITS-CLIP | 23597149\|23142080 | Unspecified |
| mmu-miR-30c-5p | MIMAT0000514 | Pycard | 66824 | HITS-CLIP | 23142080 | Unspecified |
| mmu-miR-30c-5p | MIMAT0000514 | Qk | 19317 | HITS-CLIP | 37986224\|23142080\|19536157 | Unspecified |
| mmu-miR-30c-5p | MIMAT0000514 | Qki | 19317 | qCLASH | 31393953 | Unspecified |
| mmu-miR-30c-5p | MIMAT0000514 | Qrich1 | 69232 | HITS-CLIP | 23142080 | Unspecified |
| mmu-miR-30c-5p | MIMAT0000514 | Qser1 | 99003 | HITS-CLIP | 37986224\|23142080\|19536157 | Unspecified |
| mmu-miR-30c-5p | MIMAT0000514 | R3hdm1 | 226412 | HITS-CLIP | 37986224\|23597149\|23142080 | Unspecified |
| mmu-miR-30c-5p | MIMAT0000514 | Rab10 | 19325 | HITS-CLIP | 23142080 | Unspecified |
| mmu-miR-30c-5p | MIMAT0000514 | Rab11a | 53869 | HITS-CLIP | 19536157 | Unspecified |
| mmu-miR-30c-5p | MIMAT0000514 | Rab11b | 19326 | HITS-CLIP | 37986224 | Unspecified |
| mmu-miR-30c-5p | MIMAT0000514 | Rab11fip2 | 74998 | HITS-CLIP | 23142080 | Unspecified |
| mmu-miR-30c-5p | MIMAT0000514 | Rab18 | 19330 | HITS-CLIP | 23142080\|23597149 | Unspecified |
| mmu-miR-30c-5p | MIMAT0000514 | Rab1a | 5861 | HITS-CLIP | 23142080 | Unspecified |
| mmu-miR-30c-5p | MIMAT0000514 | Rab22a | 19334 | HITS-CLIP | 37986224\|23142080 | Unspecified |
| mmu-miR-30c-5p | MIMAT0000514 | Rab23 | 19335 | HITS-CLIP | 23142080 | Unspecified |
| mmu-miR-30c-5p | MIMAT0000514 | Rab2a | 59021 | HITS-CLIP | 23597149\|25753418\|23142080 | Unspecified |
| mmu-miR-30c-5p | MIMAT0000514 | Rab2b | 76338 | HITS-CLIP | 23142080 | Unspecified |
| mmu-miR-30c-5p | MIMAT0000514 | Rab32 | 67844 | HITS-CLIP | 37986224 | Unspecified |
| mmu-miR-30c-5p | MIMAT0000514 | Rab33b | 19338 | HITS-CLIP | 23142080 | Unspecified |
| mmu-miR-30c-5p | MIMAT0000514 | Rab35 | 77407 | HITS-CLIP | 23597149\|23142080 | Unspecified |
| mmu-miR-30c-5p | MIMAT0000514 | Rab4b | 19342 | HITS-CLIP | 37986224\|23597149\|23142080 | Unspecified |
| mmu-miR-30c-5p | MIMAT0000514 | Rab5if | 55969 | HITS-CLIP | 23597149 | Unspecified |
| mmu-miR-30c-5p | MIMAT0000514 | Rab6a | 19346 | HITS-CLIP | 23142080 | Unspecified |
| mmu-miR-30c-5p | MIMAT0000514 | Rab8a | 17274 | HITS-CLIP | 23142080 | Unspecified |
| mmu-miR-30c-5p | MIMAT0000514 | Rab8b | 235442 | HITS-CLIP | 37986224\|23142080 | Unspecified |
| mmu-miR-30c-5p | MIMAT0000514 | Rabep1 | 54189 | HITS-CLIP | 23142080 | Unspecified |
| mmu-miR-30c-5p | MIMAT0000514 | Rabgap1l | 29809 | HITS-CLIP | 37986224\|23142080 | Unspecified |
| mmu-miR-30c-5p | MIMAT0000514 | Rabif | 98710 | qCLASH | 31393953 | Unspecified |
| mmu-miR-30c-5p | MIMAT0000514 | Rac1 | 19353 | HITS-CLIP | 37986224 | Unspecified |
| mmu-miR-30c-5p | MIMAT0000514 | Rad18 | 58186 | HITS-CLIP | 23597149\|23142080 | Unspecified |
| mmu-miR-30c-5p | MIMAT0000514 | Rad23b | 19359 | HITS-CLIP | 37986224\|23142080 | Unspecified |
| mmu-miR-30c-5p | MIMAT0000514 | Rad51 | 19361 | HITS-CLIP | 23142080 | Unspecified |
| mmu-miR-30c-5p | MIMAT0000514 | Rad51ap1 | 19362 | HITS-CLIP | 37986224 | Unspecified |
| mmu-miR-30c-5p | MIMAT0000514 | Rad54l2 | 81000 | HITS-CLIP | 23597149 | Unspecified |
| mmu-miR-30c-5p | MIMAT0000514 | Ralbp1 | 19765 | HITS-CLIP | 23597149\|23142080 | Unspecified |
| mmu-miR-30c-5p | MIMAT0000514 | Ralgapa1 | 56784 | HITS-CLIP | 19536157 | Unspecified |
| mmu-miR-30c-5p | MIMAT0000514 | Ralgds | 19730 | HITS-CLIP | 37986224 | Unspecified |
| mmu-miR-30c-5p | MIMAT0000514 | Ranbp2 | 19386 | HITS-CLIP | 37986224\|25753418 | Unspecified |

| | | | | | | |
|---|---|---|---|---|---|---|
| mmu-miR-30c-5p | MIMAT0000514 | Ranbp6 | 240614 | HITS-CLIP | 37986224\|23142080\|25753418 | Unspecified |
| mmu-miR-30c-5p | MIMAT0000514 | Ranbp9 | 56705 | HITS-CLIP | 37986224\|23142080 | Unspecified |
| mmu-miR-30c-5p | MIMAT0000514 | Rap1b | 215449 | HITS-CLIP | 37986224\|23597149\|23142080 | Unspecified |
| mmu-miR-30c-5p | MIMAT0000514 | Rap1gds1 | 229877 | HITS-CLIP | 23142080 | Unspecified |
| mmu-miR-30c-5p | MIMAT0000514 | Rap2c | 72065 | HITS-CLIP | 37986224\|19536157\|23142080 | Unspecified |
| mmu-miR-30c-5p | MIMAT0000514 | Rapgef2 | 76089 | HITS-CLIP | 37986224\|23142080\|23597149 | Unspecified |
| mmu-miR-30c-5p | MIMAT0000514 | Rapgef6 | 192786 | HITS-CLIP | 37986224\|23142080 | Unspecified |
| mmu-miR-30c-5p | MIMAT0000514 | Rapgefl1 | 268480 | qCLASH | 31393953 | Unspecified |
| mmu-miR-30c-5p | MIMAT0000514 | Raph1 | 77300 | HITS-CLIP | 23142080\|23597149 | Unspecified |
| mmu-miR-30c-5p | MIMAT0000514 | Rarb | 218772 | HITS-CLIP | 37986224 | Unspecified |
| mmu-miR-30c-5p | MIMAT0000514 | Rarg | 19411 | HITS-CLIP | 23597149\|23142080 | Unspecified |
| mmu-miR-30c-5p | MIMAT0000514 | Rars2 | 109093 | HITS-CLIP | 23597149\|25753418\|23142080 | Unspecified |
| mmu-miR-30c-5p | MIMAT0000514 | Rasa1 | 218397 | HITS-CLIP | 37986224\|23597149\|23142080 | Unspecified |
| mmu-miR-30c-5p | MIMAT0000514 | Rasa2 | 114713 | HITS-CLIP | 37986224\|23142080 | Unspecified |
| mmu-miR-30c-5p | MIMAT0000514 | Rasa3 | 19414 | HITS-CLIP | 23142080 | Unspecified |
| mmu-miR-30c-5p | MIMAT0000514 | Rasd1 | 19416 | HITS-CLIP | 37986224\|23142080 | Unspecified |
| mmu-miR-30c-5p | MIMAT0000514 | Rasgef1a | 70727 | HITS-CLIP | 23142080\|19536157 | Unspecified |
| mmu-miR-30c-5p | MIMAT0000514 | Rasgef1b | 320292 | HITS-CLIP | 23142080 | Unspecified |
| mmu-miR-30c-5p | MIMAT0000514 | Rasgrp1 | 19419 | HITS-CLIP | 37986224\|23142080 | Unspecified |
| mmu-miR-30c-5p | MIMAT0000514 | Rassf8 | 71323 | HITS-CLIP | 37986224\|23597149 | Unspecified |
| mmu-miR-30c-5p | MIMAT0000514 | Raver1 | 71766 | HITS-CLIP | 25753418 | Unspecified |
| mmu-miR-30c-5p | MIMAT0000514 | Rbbp6 | 19647 | HITS-CLIP | 37986224\|23142080 | Unspecified |
| mmu-miR-30c-5p | MIMAT0000514 | Rbm12 | 75710 | HITS-CLIP | 19536157 | Unspecified |
| mmu-miR-30c-5p | MIMAT0000514 | Rbm15 | 229700 | HITS-CLIP | 23597149\|23142080 | Unspecified |
| mmu-miR-30c-5p | MIMAT0000514 | Rbm25 | 67039 | HITS-CLIP | 25753418\|23142080 | Unspecified |
| mmu-miR-30c-5p | MIMAT0000514 | Rbm26 | 74213 | HITS-CLIP | 37986224\|23142080\|23597149 | Unspecified |
| mmu-miR-30c-5p | MIMAT0000514 | Rbm39 | 170791 | HITS-CLIP | 37986224 | Unspecified |
| mmu-miR-30c-5p | MIMAT0000514 | Rbm47 | 245945 | HITS-CLIP | 23142080 | Unspecified |
| mmu-miR-30c-5p | MIMAT0000514 | Rbm6 | 19654 | HITS-CLIP | 23142080 | Unspecified |
| mmu-miR-30c-5p | MIMAT0000514 | Rbm7 | 67010 | HITS-CLIP | 37986224\|23142080\|23597149 | Unspecified |
| mmu-miR-30c-5p | MIMAT0000514 | Rbms1 | 56878 | HITS-CLIP | 37986224\|23597149\|23142080 | Unspecified |
| mmu-miR-30c-5p | MIMAT0000514 | Rbpj | 19664 | HITS-CLIP | 37986224\|23142080 | Unspecified |
| mmu-miR-30c-5p | MIMAT0000514 | Rc3h1 | 381305 | HITS-CLIP | 25753418 | Unspecified |
| mmu-miR-30c-5p | MIMAT0000514 | Rc3h2 | 319817 | HITS-CLIP | 37986224\|23142080 | Unspecified |
| mmu-miR-30c-5p | MIMAT0000514 | Rcbtb1 | 71330 | HITS-CLIP | 23597149 | Unspecified |
| mmu-miR-30c-5p | MIMAT0000514 | Rcc2 | 108911 | HITS-CLIP | 23142080 | Unspecified |
| mmu-miR-30c-5p | MIMAT0000514 | Rcor1 | 217864 | HITS-CLIP | 37986224\|23597149\|19536157\|2314[illegible] | Unspecified |
| mmu-miR-30c-5p | MIMAT0000514 | Rcor3 | 214742 | HITS-CLIP | 37986224 | Unspecified |
| mmu-miR-30c-5p | MIMAT0000514 | Rdx | 19684 | HITS-CLIP | 23142080\|19536157 | Unspecified |

| | | | | | | |
|---|---|---|---|---|---|---|
| mmu-miR-30c-5p | MIMAT0000514 | Reep1 | 52250 | HITS-CLIP | 19536157\|23142080 | Unspecified |
| mmu-miR-30c-5p | MIMAT0000514 | Reep3 | 28193 | HITS-CLIP | 37986224\|19536157 | Unspecified |
| mmu-miR-30c-5p | MIMAT0000514 | Reep4 | 72549 | HITS-CLIP | 23597149\|23142080\|25083871 | Unspecified |
| mmu-miR-30c-5p | MIMAT0000514 | Relch | 57614 | HITS-CLIP | 23142080 | Unspecified |
| mmu-miR-30c-5p | MIMAT0000514 | Rest | 19712 | HITS-CLIP | 37986224\|23142080 | Unspecified |
| mmu-miR-30c-5p | MIMAT0000514 | Retreg1 | 66270 | HITS-CLIP | 23142080\|25753418 | Unspecified |
| mmu-miR-30c-5p | MIMAT0000514 | Rev3l | 19714 | HITS-CLIP | 37986224\|23597149\|23142080 | Unspecified |
| mmu-miR-30c-5p | MIMAT0000514 | Rfc3 | 69263 | HITS-CLIP | 23142080 | Unspecified |
| mmu-miR-30c-5p | MIMAT0000514 | Rffl | 67338 | HITS-CLIP | 23142080 | Unspecified |
| mmu-miR-30c-5p | MIMAT0000514 | Rfx1 | 19724 | HITS-CLIP | 23142080 | Unspecified |
| mmu-miR-30c-5p | MIMAT0000514 | Rfx3 | 19726 | HITS-CLIP | 37986224\|23142080\|19536157 | Unspecified |
| mmu-miR-30c-5p | MIMAT0000514 | Rfx7 | 319758 | HITS-CLIP | 23597149 | Unspecified |
| mmu-miR-30c-5p | MIMAT0000514 | Rfxap | 170767 | HITS-CLIP | 37986224 | Unspecified |
| mmu-miR-30c-5p | MIMAT0000514 | Rgcc | 66214 | HITS-CLIP | 37986224 | Unspecified |
| mmu-miR-30c-5p | MIMAT0000514 | Rgl1 | 19731 | HITS-CLIP | 37986224 | Unspecified |
| mmu-miR-30c-5p | MIMAT0000514 | Rgl2 | 19732 | HITS-CLIP | 23597149\|19536157\|23142080 | Unspecified |
| mmu-miR-30c-5p | MIMAT0000514 | Rgs12 | 71729 | HITS-CLIP | 37986224\|23597149 | Unspecified |
| mmu-miR-30c-5p | MIMAT0000514 | Rgs17 | 56533 | HITS-CLIP | 19536157 | Unspecified |
| mmu-miR-30c-5p | MIMAT0000514 | Rgs2 | 19735 | HITS-CLIP | 37986224 | Unspecified |
| mmu-miR-30c-5p | MIMAT0000514 | Rgs8 | 67792 | HITS-CLIP | 37986224\|19536157 | Unspecified |
| mmu-miR-30c-5p | MIMAT0000514 | Rhebl1 | 69159 | HITS-CLIP | 37986224\|23142080 | Unspecified |
| mmu-miR-30c-5p | MIMAT0000514 | Rhob | 11852 | HITS-CLIP | 37986224\|19536157\|23597149\|2314 | Unspecified |
| mmu-miR-30c-5p | MIMAT0000514 | Rhobtb1 | 69288 | HITS-CLIP | 37986224 | Unspecified |
| mmu-miR-30c-5p | MIMAT0000514 | Ric8b | 237422 | HITS-CLIP | 23142080 | Unspecified |
| mmu-miR-30c-5p | MIMAT0000514 | Rictor | 78757 | HITS-CLIP | 37986224\|19536157\|23142080 | Unspecified |
| mmu-miR-30c-5p | MIMAT0000514 | Rif1 | 51869 | HITS-CLIP | 23142080 | Unspecified |
| mmu-miR-30c-5p | MIMAT0000514 | Rimbp2 | 231760 | HITS-CLIP | 37986224 | Unspecified |
| mmu-miR-30c-5p | MIMAT0000514 | Rint1 | 72772 | HITS-CLIP | 23597149 | Unspecified |
| mmu-miR-30c-5p | MIMAT0000514 | Riok1 | 71340 | HITS-CLIP | 23142080 | Unspecified |
| mmu-miR-30c-5p | MIMAT0000514 | Riok3 | 66878 | HITS-CLIP | 37986224 | Unspecified |
| mmu-miR-30c-5p | MIMAT0000514 | Rlf | 109263 | HITS-CLIP | 23142080 | Unspecified |
| mmu-miR-30c-5p | MIMAT0000514 | Rlim | 19820 | HITS-CLIP | 37986224\|23597149\|23142080 | Unspecified |
| mmu-miR-30c-5p | MIMAT0000514 | Rmdn1 | 66302 | HITS-CLIP | 25753418 | Unspecified |
| mmu-miR-30c-5p | MIMAT0000514 | Rmnd5a | 68477 | HITS-CLIP | 37986224\|23142080 | Unspecified |
| mmu-miR-30c-5p | MIMAT0000514 | Rnf10 | 50849 | HITS-CLIP | 37986224 | Unspecified |
| mmu-miR-30c-5p | MIMAT0000514 | Rnf103 | 22644 | HITS-CLIP | 37986224\|23597149 | Unspecified |
| mmu-miR-30c-5p | MIMAT0000514 | Rnf11 | 29864 | HITS-CLIP | 37986224\|23597149\|19536157\|2314 | Unspecified |
| mmu-miR-30c-5p | MIMAT0000514 | Rnf113a2 | 66381 | HITS-CLIP | 23142080 | Unspecified |
| mmu-miR-30c-5p | MIMAT0000514 | Rnf115 | 67845 | HITS-CLIP | 23597149 | Unspecified |

| | | | | | | |
|---|---|---|---|---|---|---|
| mmu-miR-30c-5p | MIMAT0000514 | Rnf138 | 56515 | HITS-CLIP | 37986224\|23142080 | Unspecified |
| mmu-miR-30c-5p | MIMAT0000514 | Rnf139 | 75841 | HITS-CLIP | 37986224\|23142080 | Unspecified |
| mmu-miR-30c-5p | MIMAT0000514 | Rnf145 | 74315 | HITS-CLIP | 37986224\|23142080 | Unspecified |
| mmu-miR-30c-5p | MIMAT0000514 | Rnf146 | 68031 | HITS-CLIP | 23142080 | Unspecified |
| mmu-miR-30c-5p | MIMAT0000514 | Rnf157 | 217340 | HITS-CLIP | 37986224\|23142080 | Unspecified |
| mmu-miR-30c-5p | MIMAT0000514 | Rnf167 | 70510 | qCLASH | 31393953 | Unspecified |
| mmu-miR-30c-5p | MIMAT0000514 | Rnf169 | 108937 | HITS-CLIP | 37986224\|23142080 | Unspecified |
| mmu-miR-30c-5p | MIMAT0000514 | Rnf19a | 30945 | HITS-CLIP | 37986224\|23142080\|25753418 | Unspecified |
| mmu-miR-30c-5p | MIMAT0000514 | Rnf19b | 75234 | HITS-CLIP | 23142080\|23597149 | Unspecified |
| mmu-miR-30c-5p | MIMAT0000514 | Rnf2 | 19821 | HITS-CLIP | 37986224\|23142080 | Unspecified |
| mmu-miR-30c-5p | MIMAT0000514 | Rnf213 | 672511 | HITS-CLIP | 23142080 | Unspecified |
| mmu-miR-30c-5p | MIMAT0000514 | Rnf219 | 72486 | HITS-CLIP | 23142080 | Unspecified |
| mmu-miR-30c-5p | MIMAT0000514 | Rnf220 | 66743 | HITS-CLIP | 37986224\|23597149\|19536157\|2314 | Unspecified |
| mmu-miR-30c-5p | MIMAT0000514 | Rnf38 | 73469 | HITS-CLIP | 23142080 | Unspecified |
| mmu-miR-30c-5p | MIMAT0000514 | Rnf44 | 105239 | HITS-CLIP | 23597149\|19536157 | Unspecified |
| mmu-miR-30c-5p | MIMAT0000514 | Rnf6 | 74132 | HITS-CLIP | 37986224\|23142080\|23597149\|1953 | Unspecified |
| mmu-miR-30c-5p | MIMAT0000514 | Rnf7 | 19823 | qCLASH | 31393953 | Unspecified |
| mmu-miR-30c-5p | MIMAT0000514 | Rnf8 | 58230 | HITS-CLIP | 23142080 | Unspecified |
| mmu-miR-30c-5p | MIMAT0000514 | Rnft1 | 76892 | HITS-CLIP | 23142080 | Unspecified |
| mmu-miR-30c-5p | MIMAT0000514 | Rnmt | 67897 | HITS-CLIP | 23597149\|23142080 | Unspecified |
| mmu-miR-30c-5p | MIMAT0000514 | Robo2 | 268902 | HITS-CLIP | 37986224 | Unspecified |
| mmu-miR-30c-5p | MIMAT0000514 | Rock2 | 19878 | HITS-CLIP | 23142080 | Unspecified |
| mmu-miR-30c-5p | MIMAT0000514 | Rogdi | 66049 | HITS-CLIP | 23597149 | Unspecified |
| mmu-miR-30c-5p | MIMAT0000514 | Ror1 | 26563 | HITS-CLIP | 37986224 | Unspecified |
| mmu-miR-30c-5p | MIMAT0000514 | Rora | 19883 | HITS-CLIP | 37986224\|23142080 | Unspecified |
| mmu-miR-30c-5p | MIMAT0000514 | Rpa2 | 19891 | HITS-CLIP | 37986224\|23597149\|23142080 | Unspecified |
| mmu-miR-30c-5p | MIMAT0000514 | Rpf2 | 67239 | HITS-CLIP | 37986224 | Unspecified |
| mmu-miR-30c-5p | MIMAT0000514 | Rpl22 | 19934 | qCLASH | 31393953 | Unspecified |
| mmu-miR-30c-5p | MIMAT0000514 | Rpl27a | 26451 | qCLASH | 31393953 | Unspecified |
| mmu-miR-30c-5p | MIMAT0000514 | Rpl32 | 19951 | HITS-CLIP | 37986224 | Unspecified |
| mmu-miR-30c-5p | MIMAT0000514 | Rpl4 | 67891 | HITS-CLIP | 23142080 | Unspecified |
| mmu-miR-30c-5p | MIMAT0000514 | Rplp2 | 67186 | qCLASH | 31393953 | Unspecified |
| mmu-miR-30c-5p | MIMAT0000514 | Rpn2 | 20014 | HITS-CLIP | 37986224\|23142080 | Unspecified |
| mmu-miR-30c-5p | MIMAT0000514 | Rpp14 | 67053 | HITS-CLIP | 23597149\|23142080 | Unspecified |
| mmu-miR-30c-5p | MIMAT0000514 | Rprm | 67874 | HITS-CLIP | 37986224 | Unspecified |
| mmu-miR-30c-5p | MIMAT0000514 | Rps17 | 20068 | qCLASH | 31393953 | Unspecified |
| mmu-miR-30c-5p | MIMAT0000514 | Rps3 | 27050 | qCLASH | 31393953 | Unspecified |
| mmu-miR-30c-5p | MIMAT0000514 | Rps6ka1 | 20111 | HITS-CLIP | 23597149\|23142080 | Unspecified |
| mmu-miR-30c-5p | MIMAT0000514 | Rps6ka3 | 110651 | HITS-CLIP | 19536157\|23142080 | Unspecified |

| | | | | | | |
|---|---|---|---|---|---|---|
| mmu-miR-30c-5p | MIMAT0000514 | Rps6ka5 | 73086 | HITS-CLIP | 37986224\|23142080\|23597149\|19536 | Unspecified |
| mmu-miR-30c-5p | MIMAT0000514 | Rps6kb1 | 72508 | HITS-CLIP | 37986224 | Unspecified |
| mmu-miR-30c-5p | MIMAT0000514 | Rps6kc1 | 320119 | HITS-CLIP | 37986224 | Unspecified |
| mmu-miR-30c-5p | MIMAT0000514 | Rrad | 56437 | HITS-CLIP | 25083871\|23142080 | Unspecified |
| mmu-miR-30c-5p | MIMAT0000514 | Rragd | 52187 | HITS-CLIP | 25753418 | Unspecified |
| mmu-miR-30c-5p | MIMAT0000514 | Rras2 | 66922 | HITS-CLIP | 37986224 | Unspecified |
| mmu-miR-30c-5p | MIMAT0000514 | Rrbp1 | 81910 | HITS-CLIP | 25753418 | Unspecified |
| mmu-miR-30c-5p | MIMAT0000514 | Rreb1 | 68750 | HITS-CLIP | 23142080 | Unspecified |
| mmu-miR-30c-5p | MIMAT0000514 | Rrm1 | 20133 | qCLASH | 31393953 | Unspecified |
| mmu-miR-30c-5p | MIMAT0000514 | Rrm2 | 20135 | HITS-CLIP | 23142080 | Unspecified |
| mmu-miR-30c-5p | MIMAT0000514 | Rsbn1 | 229675 | HITS-CLIP | 23142080 | Unspecified |
| mmu-miR-30c-5p | MIMAT0000514 | Rsbn1l | 242860 | HITS-CLIP | 37986224\|23142080 | Unspecified |
| mmu-miR-30c-5p | MIMAT0000514 | Rspry1 | 67610 | HITS-CLIP | 23597149\|23142080 | Unspecified |
| mmu-miR-30c-5p | MIMAT0000514 | Rtf1 | 76246 | HITS-CLIP | 25753418 | Unspecified |
| mmu-miR-30c-5p | MIMAT0000514 | Rtkn | 20166 | HITS-CLIP | 23597149 | Unspecified |
| mmu-miR-30c-5p | MIMAT0000514 | Rtn1 | 104001 | HITS-CLIP | 19536157 | Unspecified |
| mmu-miR-30c-5p | MIMAT0000514 | Rtn4 | 68585 | HITS-CLIP | 37986224 | Unspecified |
| mmu-miR-30c-5p | MIMAT0000514 | Rtn4r | 65079 | HITS-CLIP | 37986224\|19536157 | Unspecified |
| mmu-miR-30c-5p | MIMAT0000514 | Rtp3 | 235636 | HITS-CLIP | 23597149 | Unspecified |
| mmu-miR-30c-5p | MIMAT0000514 | Rttn | 246102 | HITS-CLIP | 23142080 | Unspecified |
| mmu-miR-30c-5p | MIMAT0000514 | Rundc3b | 242819 | HITS-CLIP | 23142080 | Unspecified |
| mmu-miR-30c-5p | MIMAT0000514 | Runx2 | 12393 | Western Blot//HITS-CLIP | 37986224\|22253433\|23142080 | Unspecified |
| mmu-miR-30c-5p | MIMAT0000514 | S100a10 | 20194 | HITS-CLIP | 37986224\|23142080 | Unspecified |
| mmu-miR-30c-5p | MIMAT0000514 | Sacm1l | 83493 | HITS-CLIP | 37986224 | Unspecified |
| mmu-miR-30c-5p | MIMAT0000514 | Sacs | 50720 | HITS-CLIP | 37986224\|23142080 | Unspecified |
| mmu-miR-30c-5p | MIMAT0000514 | Samd1 | 666704 | qCLASH | 31393953 | Unspecified |
| mmu-miR-30c-5p | MIMAT0000514 | Samd4 | 74480 | HITS-CLIP | 37986224\|19536157\|25753418 | Unspecified |
| mmu-miR-30c-5p | MIMAT0000514 | Samd4b | 233033 | HITS-CLIP | 23142080 | Unspecified |
| mmu-miR-30c-5p | MIMAT0000514 | Samd8 | 67630 | HITS-CLIP | 37986224\|23597149\|19536157\|25753 | Unspecified |
| mmu-miR-30c-5p | MIMAT0000514 | Sap30 | 60406 | HITS-CLIP | 37986224\|23142080 | Unspecified |
| mmu-miR-30c-5p | MIMAT0000514 | Sar1a | 20224 | HITS-CLIP | 23142080 | Unspecified |
| mmu-miR-30c-5p | MIMAT0000514 | Sar1b | 66397 | HITS-CLIP | 23597149\|23142080 | Unspecified |
| mmu-miR-30c-5p | MIMAT0000514 | Saraf | 51669 | HITS-CLIP | 25753418 | Unspecified |
| mmu-miR-30c-5p | MIMAT0000514 | Satb1 | 20230 | HITS-CLIP | 37986224\|23142080 | Unspecified |
| mmu-miR-30c-5p | MIMAT0000514 | Sav1 | 64010 | HITS-CLIP | 23142080 | Unspecified |
| mmu-miR-30c-5p | MIMAT0000514 | Sbf1 | 77980 | HITS-CLIP | 37986224\|23597149\|23142080 | Unspecified |
| mmu-miR-30c-5p | MIMAT0000514 | Sbk1 | 104175 | HITS-CLIP | 37986224\|23597149\|19536157 | Unspecified |
| mmu-miR-30c-5p | MIMAT0000514 | Sbno1 | 243272 | HITS-CLIP | 23142080 | Unspecified |
| mmu-miR-30c-5p | MIMAT0000514 | Sc5d | 235293 | HITS-CLIP | 23597149 | Unspecified |

| | | | | | | |
|---|---|---|---|---|---|---|
| mmu-miR-30c-5p | MIMAT0000514 | Scamp1 | 107767 | HITS-CLIP | 37986224\|23142080 | Unspecified |
| mmu-miR-30c-5p | MIMAT0000514 | Scamp5 | 56807 | HITS-CLIP | 25753418 | Unspecified |
| mmu-miR-30c-5p | MIMAT0000514 | Scaper | 244891 | HITS-CLIP | 37986224\|19536157 | Unspecified |
| mmu-miR-30c-5p | MIMAT0000514 | Scara5 | 71145 | HITS-CLIP | 37986224\|23597149 | Unspecified |
| mmu-miR-30c-5p | MIMAT0000514 | Scarb2 | 12492 | HITS-CLIP | 23142080 | Unspecified |
| mmu-miR-30c-5p | MIMAT0000514 | Scg3 | 20255 | HITS-CLIP | 37986224 | Unspecified |
| mmu-miR-30c-5p | MIMAT0000514 | Scn1a | 20265 | HITS-CLIP | 19536157\|23142080 | Unspecified |
| mmu-miR-30c-5p | MIMAT0000514 | Scn2a | 110876 | HITS-CLIP | 19536157 | Unspecified |
| mmu-miR-30c-5p | MIMAT0000514 | Scn3a | 20269 | HITS-CLIP | 19536157 | Unspecified |
| mmu-miR-30c-5p | MIMAT0000514 | Scn8a | 20273 | HITS-CLIP | 19536157 | Unspecified |
| mmu-miR-30c-5p | MIMAT0000514 | Scyl2 | 213326 | HITS-CLIP | 23142080\|19536157\|23597149 | Unspecified |
| mmu-miR-30c-5p | MIMAT0000514 | Sdad1 | 231452 | HITS-CLIP | 23142080 | Unspecified |
| mmu-miR-30c-5p | MIMAT0000514 | Sdcbp | 53378 | HITS-CLIP | 37986224 | Unspecified |
| mmu-miR-30c-5p | MIMAT0000514 | Sdf4 | 20318 | HITS-CLIP | 37986224 | Unspecified |
| mmu-miR-30c-5p | MIMAT0000514 | Sec14l2 | 67815 | HITS-CLIP | 37986224 | Unspecified |
| mmu-miR-30c-5p | MIMAT0000514 | Sec16a | 227648 | HITS-CLIP | 23597149\|23142080\|25753418 | Unspecified |
| mmu-miR-30c-5p | MIMAT0000514 | Sec23a | 20334 | HITS-CLIP | 37986224\|23597149\|23142080 | Unspecified |
| mmu-miR-30c-5p | MIMAT0000514 | Sec23b | 27054 | HITS-CLIP | 23142080 | Unspecified |
| mmu-miR-30c-5p | MIMAT0000514 | Sec24a | 77371 | HITS-CLIP | 37986224\|23597149\|23142080 | Unspecified |
| mmu-miR-30c-5p | MIMAT0000514 | Sec24b | 99683 | HITS-CLIP | 37986224 | Unspecified |
| mmu-miR-30c-5p | MIMAT0000514 | Sec24c | 218811 | HITS-CLIP | 23597149\|23142080 | Unspecified |
| mmu-miR-30c-5p | MIMAT0000514 | Sec61a2 | 57743 | HITS-CLIP | 37986224 | Unspecified |
| mmu-miR-30c-5p | MIMAT0000514 | Sec63 | 140740 | HITS-CLIP | 37986224 | Unspecified |
| mmu-miR-30c-5p | MIMAT0000514 | Secisbp2l | 70354 | HITS-CLIP | 25753418\|23142080 | Unspecified |
| mmu-miR-30c-5p | MIMAT0000514 | Seh1l | 72124 | HITS-CLIP | 37986224 | Unspecified |
| mmu-miR-30c-5p | MIMAT0000514 | Sel1l | 20338 | HITS-CLIP | 23142080 | Unspecified |
| mmu-miR-30c-5p | MIMAT0000514 | Sel1l3 | 231238 | HITS-CLIP | 37986224\|23597149 | Unspecified |
| mmu-miR-30c-5p | MIMAT0000514 | Sele | 20339 | HITS-CLIP | 25753418 | Unspecified |
| mmu-miR-30c-5p | MIMAT0000514 | Selenop | 6414 | HITS-CLIP | 23597149 | Unspecified |
| mmu-miR-30c-5p | MIMAT0000514 | Sema3a | 20346 | HITS-CLIP | 37986224\|23597149\|19536157 | Unspecified |
| mmu-miR-30c-5p | MIMAT0000514 | Sema4d | 20354 | HITS-CLIP | 37986224\|23142080 | Unspecified |
| mmu-miR-30c-5p | MIMAT0000514 | Sema6a | 20358 | HITS-CLIP | 37986224 | Unspecified |
| mmu-miR-30c-5p | MIMAT0000514 | Sema6d | 214968 | HITS-CLIP | 37986224\|23597149\|19536157 | Unspecified |
| mmu-miR-30c-5p | MIMAT0000514 | Senp1 | 223870 | HITS-CLIP | 23142080 | Unspecified |
| mmu-miR-30c-5p | MIMAT0000514 | Senp2 | 75826 | HITS-CLIP | 37986224\|23142080 | Unspecified |
| mmu-miR-30c-5p | MIMAT0000514 | Senp3 | 80886 | HITS-CLIP | 25753418 | Unspecified |
| mmu-miR-30c-5p | MIMAT0000514 | Senp6 | 215351 | HITS-CLIP | 37986224 | Unspecified |
| mmu-miR-30c-5p | MIMAT0000514 | Sept11 | 52398 | HITS-CLIP | 19536157 | Unspecified |
| mmu-miR-30c-5p | MIMAT0000514 | Sept6 | 23157 | HITS-CLIP | 23142080 | Unspecified |

| | | | | | | |
|---|---|---|---|---|---|---|
| mmu-miR-30c-5p | MIMAT0000514 | Sept7 | 235072 | HITS-CLIP | 37986224\|23142080 | Unspecified |
| mmu-miR-30c-5p | MIMAT0000514 | Sept8 | 20362 | HITS-CLIP | 37986224 | Unspecified |
| mmu-miR-30c-5p | MIMAT0000514 | Septin2 | 18000 | HITS-CLIP | 37986224\|23142080 | Unspecified |
| mmu-miR-30c-5p | MIMAT0000514 | Serbp1 | 66870 | HITS-CLIP | 37986224 | Unspecified |
| mmu-miR-30c-5p | MIMAT0000514 | Serinc1 | 56442 | HITS-CLIP | 37986224 | Unspecified |
| mmu-miR-30c-5p | MIMAT0000514 | Serp1 | 28146 | HITS-CLIP | 37986224 | Unspecified |
| mmu-miR-30c-5p | MIMAT0000514 | Serpina3m | 20717 | HITS-CLIP | 37986224\|23597149 | Unspecified |
| mmu-miR-30c-5p | MIMAT0000514 | Serpinb6b | 20708 | HITS-CLIP | 37986224 | Unspecified |
| mmu-miR-30c-5p | MIMAT0000514 | Serpinf1 | 20317 | HITS-CLIP | 23142080 | Unspecified |
| mmu-miR-30c-5p | MIMAT0000514 | Sertad2 | 58172 | HITS-CLIP | 37986224 | Unspecified |
| mmu-miR-30c-5p | MIMAT0000514 | Sesn1 | 140742 | HITS-CLIP | 37986224 | Unspecified |
| mmu-miR-30c-5p | MIMAT0000514 | Sesn3 | 75747 | HITS-CLIP | 37986224\|23142080 | Unspecified |
| mmu-miR-30c-5p | MIMAT0000514 | Setd1b | 208043 | HITS-CLIP | 23142080 | Unspecified |
| mmu-miR-30c-5p | MIMAT0000514 | Setd2 | 235626 | HITS-CLIP | 37986224\|23142080\|19536157\|23597 | Unspecified |
| mmu-miR-30c-5p | MIMAT0000514 | Setd5 | 72895 | HITS-CLIP | 37986224\|23142080 | Unspecified |
| mmu-miR-30c-5p | MIMAT0000514 | Setd7 | 73251 | HITS-CLIP | 37986224\|23597149\|23142080 | Unspecified |
| mmu-miR-30c-5p | MIMAT0000514 | Setdb1 | 84505 | HITS-CLIP | 25753418 | Unspecified |
| mmu-miR-30c-5p | MIMAT0000514 | Setmar | 6419 | HITS-CLIP | 25753418 | Unspecified |
| mmu-miR-30c-5p | MIMAT0000514 | Setx | 269254 | qCLASH//HITS-CLIP | 31393953\|23142080 | Unspecified |
| mmu-miR-30c-5p | MIMAT0000514 | Sez6l | 56747 | qCLASH | 31393953 | Unspecified |
| mmu-miR-30c-5p | MIMAT0000514 | Sf1 | 22668 | HITS-CLIP | 25753418 | Unspecified |
| mmu-miR-30c-5p | MIMAT0000514 | Sf3b1 | 81898 | HITS-CLIP | 23597149\|23142080 | Unspecified |
| mmu-miR-30c-5p | MIMAT0000514 | Sfr1 | 67788 | HITS-CLIP | 23142080 | Unspecified |
| mmu-miR-30c-5p | MIMAT0000514 | Sgcb | 24051 | HITS-CLIP | 23142080 | Unspecified |
| mmu-miR-30c-5p | MIMAT0000514 | Sgk3 | 170755 | HITS-CLIP | 25753418 | Unspecified |
| mmu-miR-30c-5p | MIMAT0000514 | Sgms2 | 74442 | HITS-CLIP | 37986224 | Unspecified |
| mmu-miR-30c-5p | MIMAT0000514 | Sgpp1 | 81535 | HITS-CLIP | 37986224\|23597149 | Unspecified |
| mmu-miR-30c-5p | MIMAT0000514 | Sh2b3 | 16923 | HITS-CLIP | 23142080 | Unspecified |
| mmu-miR-30c-5p | MIMAT0000514 | Sh3pxd2a | 14218 | HITS-CLIP//qCLASH | 37986224\|19536157\|31393953 | Unspecified |
| mmu-miR-30c-5p | MIMAT0000514 | Sh3rf1 | 59009 | HITS-CLIP | 37986224\|23142080 | Unspecified |
| mmu-miR-30c-5p | MIMAT0000514 | Shank2 | 210274 | HITS-CLIP | 37986224 | Unspecified |
| mmu-miR-30c-5p | MIMAT0000514 | Shank3 | 58234 | HITS-CLIP | 25753418 | Unspecified |
| mmu-miR-30c-5p | MIMAT0000514 | Shmt1 | 20425 | HITS-CLIP | 37986224 | Unspecified |
| mmu-miR-30c-5p | MIMAT0000514 | Shoc2 | 56392 | HITS-CLIP | 23142080 | Unspecified |
| mmu-miR-30c-5p | MIMAT0000514 | Shroom4 | 208431 | HITS-CLIP | 25753418 | Unspecified |
| mmu-miR-30c-5p | MIMAT0000514 | Siah2 | 20439 | HITS-CLIP | 37986224\|23597149\|23142080 | Unspecified |
| mmu-miR-30c-5p | MIMAT0000514 | Sidt2 | 214597 | HITS-CLIP | 37986224 | Unspecified |
| mmu-miR-30c-5p | MIMAT0000514 | Sik1 | 17691 | HITS-CLIP | 25753418 | Unspecified |
| mmu-miR-30c-5p | MIMAT0000514 | Sik3 | 70661 | HITS-CLIP | 23597149\|23142080 | Unspecified |

| | | | | | | |
|---|---|---|---|---|---|---|
| mmu-miR-30c-5p | MIMAT0000514 | Simc1 | 319719 | HITS-CLIP | 23142080 | Unspecified |
| mmu-miR-30c-5p | MIMAT0000514 | Sin3a | 20466 | HITS-CLIP | 25753418 | Unspecified |
| mmu-miR-30c-5p | MIMAT0000514 | Sirt1 | 93759 | HITS-CLIP | 19536157 | Unspecified |
| mmu-miR-30c-5p | MIMAT0000514 | Sirt3 | 64384 | HITS-CLIP | 37986224 | Unspecified |
| mmu-miR-30c-5p | MIMAT0000514 | Ski | 20481 | HITS-CLIP | 37986224\|23142080 | Unspecified |
| mmu-miR-30c-5p | MIMAT0000514 | Skil | 20482 | HITS-CLIP | 37986224 | Unspecified |
| mmu-miR-30c-5p | MIMAT0000514 | Skp1a | 21402 | HITS-CLIP | 23142080 | Unspecified |
| mmu-miR-30c-5p | MIMAT0000514 | Skp2 | 27401 | HITS-CLIP | 37986224\|23597149\|23142080 | Unspecified |
| mmu-miR-30c-5p | MIMAT0000514 | Slamf7 | 75345 | HITS-CLIP | 23142080 | Unspecified |
| mmu-miR-30c-5p | MIMAT0000514 | Slc10a7 | 76775 | HITS-CLIP | 23142080 | Unspecified |
| mmu-miR-30c-5p | MIMAT0000514 | Slc11a2 | 18174 | HITS-CLIP | 37986224\|23142080\|23597149 | Unspecified |
| mmu-miR-30c-5p | MIMAT0000514 | Slc16a3 | 80879 | HITS-CLIP | 23142080 | Unspecified |
| mmu-miR-30c-5p | MIMAT0000514 | Slc16a6 | 104681 | HITS-CLIP | 37986224 | Unspecified |
| mmu-miR-30c-5p | MIMAT0000514 | Slc1a2 | 20511 | HITS-CLIP | 37986224\|19536157 | Unspecified |
| mmu-miR-30c-5p | MIMAT0000514 | Slc20a1 | 20515 | HITS-CLIP | 37986224 | Unspecified |
| mmu-miR-30c-5p | MIMAT0000514 | Slc20a2 | 20516 | HITS-CLIP | 37986224 | Unspecified |
| mmu-miR-30c-5p | MIMAT0000514 | Slc22a23 | 73102 | HITS-CLIP | 37986224\|23597149 | Unspecified |
| mmu-miR-30c-5p | MIMAT0000514 | Slc25a40 | 319653 | HITS-CLIP | 23142080 | Unspecified |
| mmu-miR-30c-5p | MIMAT0000514 | Slc25a46 | 67453 | HITS-CLIP | 37986224 | Unspecified |
| mmu-miR-30c-5p | MIMAT0000514 | Slc26a2 | 13521 | HITS-CLIP | 37986224 | Unspecified |
| mmu-miR-30c-5p | MIMAT0000514 | Slc29a3 | 71279 | HITS-CLIP | 23142080 | Unspecified |
| mmu-miR-30c-5p | MIMAT0000514 | Slc2a3 | 20527 | HITS-CLIP | 23142080 | Unspecified |
| mmu-miR-30c-5p | MIMAT0000514 | Slc30a4 | 22785 | HITS-CLIP | 23597149\|23142080\|19536157 | Unspecified |
| mmu-miR-30c-5p | MIMAT0000514 | Slc30a5 | 69048 | HITS-CLIP | 23142080 | Unspecified |
| mmu-miR-30c-5p | MIMAT0000514 | Slc30a7 | 66500 | HITS-CLIP | 37986224\|23597149\|23142080 | Unspecified |
| mmu-miR-30c-5p | MIMAT0000514 | Slc35a5 | 74102 | HITS-CLIP//qCLASH | 37986224\|23142080\|31393953 | Unspecified |
| mmu-miR-30c-5p | MIMAT0000514 | Slc35b2 | 73836 | HITS-CLIP | 23597149 | Unspecified |
| mmu-miR-30c-5p | MIMAT0000514 | Slc35b3 | 108652 | HITS-CLIP | 23142080 | Unspecified |
| mmu-miR-30c-5p | MIMAT0000514 | Slc35b4 | 58246 | HITS-CLIP | 23142080 | Unspecified |
| mmu-miR-30c-5p | MIMAT0000514 | Slc35c1 | 228368 | HITS-CLIP | 37986224\|23597149 | Unspecified |
| mmu-miR-30c-5p | MIMAT0000514 | Slc35d1 | 242585 | HITS-CLIP | 37986224\|23142080\|23597149 | Unspecified |
| mmu-miR-30c-5p | MIMAT0000514 | Slc35d3 | 76157 | HITS-CLIP | 37986224\|23142080 | Unspecified |
| mmu-miR-30c-5p | MIMAT0000514 | Slc35f2 | 72022 | qCLASH | 31393953 | Unspecified |
| mmu-miR-30c-5p | MIMAT0000514 | Slc36a1 | 215335 | HITS-CLIP | 37986224\|23597149\|23142080 | Unspecified |
| mmu-miR-30c-5p | MIMAT0000514 | Slc38a2 | 67760 | HITS-CLIP | 37986224\|23597149\|23142080 | Unspecified |
| mmu-miR-30c-5p | MIMAT0000514 | Slc38a4 | 69354 | HITS-CLIP | 37986224\|23597149 | Unspecified |
| mmu-miR-30c-5p | MIMAT0000514 | Slc38a7 | 234595 | HITS-CLIP | 23597149\|23142080 | Unspecified |
| mmu-miR-30c-5p | MIMAT0000514 | Slc38a9 | 268706 | HITS-CLIP | 23597149\|23142080 | Unspecified |
| mmu-miR-30c-5p | MIMAT0000514 | Slc39a10 | 227059 | HITS-CLIP | 37986224\|23142080\|19536157 | Unspecified |

| | | | | | | |
|---|---|---|---|---|---|---|
| mmu-miR-30c-5p | MIMAT0000514 | Slc40a1 | 53945 | HITS-CLIP | 23597149 | Unspecified |
| mmu-miR-30c-5p | MIMAT0000514 | Slc41a2 | 338365 | HITS-CLIP | 37986224\|23597149\|23142080 | Unspecified |
| mmu-miR-30c-5p | MIMAT0000514 | Slc44a1 | 100434 | HITS-CLIP | 37986224 | Unspecified |
| mmu-miR-30c-5p | MIMAT0000514 | Slc4a1ap | 20534 | HITS-CLIP | 23142080 | Unspecified |
| mmu-miR-30c-5p | MIMAT0000514 | Slc4a4 | 54403 | HITS-CLIP | 37986224 | Unspecified |
| mmu-miR-30c-5p | MIMAT0000514 | Slc4a7 | 218756 | HITS-CLIP | 37986224\|23142080 | Unspecified |
| mmu-miR-30c-5p | MIMAT0000514 | Slc5a3 | 53881 | HITS-CLIP | 37986224\|23142080 | Unspecified |
| mmu-miR-30c-5p | MIMAT0000514 | Slc6a6 | 21366 | HITS-CLIP | 37986224\|23142080 | Unspecified |
| mmu-miR-30c-5p | MIMAT0000514 | Slc6a9 | 14664 | HITS-CLIP | 23597149 | Unspecified |
| mmu-miR-30c-5p | MIMAT0000514 | Slc7a1 | 11987 | HITS-CLIP | 37986224 | Unspecified |
| mmu-miR-30c-5p | MIMAT0000514 | Slc7a11 | 26570 | HITS-CLIP | 37986224 | Unspecified |
| mmu-miR-30c-5p | MIMAT0000514 | Slc7a14 | 241919 | HITS-CLIP | 37986224 | Unspecified |
| mmu-miR-30c-5p | MIMAT0000514 | Slc7a5 | 20539 | HITS-CLIP | 37986224 | Unspecified |
| mmu-miR-30c-5p | MIMAT0000514 | Slc9a8 | 77031 | HITS-CLIP | 37986224\|23597149\|23142080 | Unspecified |
| mmu-miR-30c-5p | MIMAT0000514 | Slco1a1 | 28248 | HITS-CLIP | 23597149 | Unspecified |
| mmu-miR-30c-5p | MIMAT0000514 | Slco1c1 | 58807 | HITS-CLIP | 37986224\|19536157 | Unspecified |
| mmu-miR-30c-5p | MIMAT0000514 | Slco4a1 | 108115 | HITS-CLIP | 23142080 | Unspecified |
| mmu-miR-30c-5p | MIMAT0000514 | Slco5a1 | 240726 | HITS-CLIP | 25753418 | Unspecified |
| mmu-miR-30c-5p | MIMAT0000514 | Slf1 | 852127 | HITS-CLIP | 25753418 | Unspecified |
| mmu-miR-30c-5p | MIMAT0000514 | Slf2 | 226151 | HITS-CLIP | 23142080 | Unspecified |
| mmu-miR-30c-5p | MIMAT0000514 | Slitrk2 | 245450 | HITS-CLIP | 37986224 | Unspecified |
| mmu-miR-30c-5p | MIMAT0000514 | Slitrk5 | 75409 | HITS-CLIP | 37986224 | Unspecified |
| mmu-miR-30c-5p | MIMAT0000514 | Slk | 20874 | HITS-CLIP | 37986224\|23142080 | Unspecified |
| mmu-miR-30c-5p | MIMAT0000514 | Smad1 | 17125 | Luciferase assay//qRT-PCR//Western Blot/ | 24555094\|22253433 | Unspecified |
| mmu-miR-30c-5p | MIMAT0000514 | Smad3 | 17127 | HITS-CLIP | 23142080 | Unspecified |
| mmu-miR-30c-5p | MIMAT0000514 | Smad4 | 17128 | HITS-CLIP | 23142080 | Unspecified |
| mmu-miR-30c-5p | MIMAT0000514 | Smap1 | 98366 | HITS-CLIP | 37986224\|23142080 | Unspecified |
| mmu-miR-30c-5p | MIMAT0000514 | Smarcb1 | 20587 | HITS-CLIP | 37986224\|23597149 | Unspecified |
| mmu-miR-30c-5p | MIMAT0000514 | Smarcc2 | 68094 | HITS-CLIP | 19536157\|23142080 | Unspecified |
| mmu-miR-30c-5p | MIMAT0000514 | Smarcd2 | 83796 | HITS-CLIP | 37986224\|23597149\|23142080 | Unspecified |
| mmu-miR-30c-5p | MIMAT0000514 | Smc1a | 24061 | HITS-CLIP | 23142080 | Unspecified |
| mmu-miR-30c-5p | MIMAT0000514 | Smc2 | 14211 | HITS-CLIP | 37986224\|23142080 | Unspecified |
| mmu-miR-30c-5p | MIMAT0000514 | Smg7 | 226517 | HITS-CLIP | 23142080 | Unspecified |
| mmu-miR-30c-5p | MIMAT0000514 | Smim14 | 68552 | HITS-CLIP | 23142080 | Unspecified |
| mmu-miR-30c-5p | MIMAT0000514 | Smoc1 | 64075 | HITS-CLIP | 23597149 | Unspecified |
| mmu-miR-30c-5p | MIMAT0000514 | Smu1 | 74255 | HITS-CLIP | 37986224\|23142080 | Unspecified |
| mmu-miR-30c-5p | MIMAT0000514 | Smurf1 | 75788 | HITS-CLIP | 37986224 | Unspecified |
| mmu-miR-30c-5p | MIMAT0000514 | Snai2 | 20583 | HITS-CLIP | 23597149 | Unspecified |
| mmu-miR-30c-5p | MIMAT0000514 | Snapc1 | 75627 | HITS-CLIP | 23597149\|19536157\|23142080 | Unspecified |

| | | | | | | |
|---|---|---|---|---|---|---|
| mmu-miR-30c-5p | MIMAT0000514 | Snd1 | 56463 | HITS-CLIP | 23597149 | Unspecified |
| mmu-miR-30c-5p | MIMAT0000514 | Sned1 | 208777 | HITS-CLIP | 37986224 | Unspecified |
| mmu-miR-30c-5p | MIMAT0000514 | Snw1 | 66354 | HITS-CLIP | 23142080 | Unspecified |
| mmu-miR-30c-5p | MIMAT0000514 | Snx10 | 71982 | HITS-CLIP | 37986224 | Unspecified |
| mmu-miR-30c-5p | MIMAT0000514 | Snx12 | 55988 | HITS-CLIP | 23142080 | Unspecified |
| mmu-miR-30c-5p | MIMAT0000514 | Snx13 | 217463 | HITS-CLIP | 37986224 | Unspecified |
| mmu-miR-30c-5p | MIMAT0000514 | Snx17 | 266781 | HITS-CLIP | 25753418 | Unspecified |
| mmu-miR-30c-5p | MIMAT0000514 | Snx18 | 170625 | HITS-CLIP | 37986224\|19536157\|23597149\|23142 | Unspecified |
| mmu-miR-30c-5p | MIMAT0000514 | Snx2 | 67804 | HITS-CLIP | 37986224\|23142080 | Unspecified |
| mmu-miR-30c-5p | MIMAT0000514 | Snx27 | 76742 | HITS-CLIP | 37986224\|19536157\|23142080 | Unspecified |
| mmu-miR-30c-5p | MIMAT0000514 | Snx3 | 54198 | HITS-CLIP | 23142080 | Unspecified |
| mmu-miR-30c-5p | MIMAT0000514 | Snx30 | 209131 | HITS-CLIP | 25753418\|23142080 | Unspecified |
| mmu-miR-30c-5p | MIMAT0000514 | Snx33 | 235406 | HITS-CLIP | 23597149 | Unspecified |
| mmu-miR-30c-5p | MIMAT0000514 | Snx4 | 69150 | HITS-CLIP | 23142080 | Unspecified |
| mmu-miR-30c-5p | MIMAT0000514 | Snx5 | 69178 | HITS-CLIP | 37986224\|23597149\|23142080 | Unspecified |
| mmu-miR-30c-5p | MIMAT0000514 | Snx6 | 72183 | HITS-CLIP | 19536157\|23142080 | Unspecified |
| mmu-miR-30c-5p | MIMAT0000514 | Snx8 | 231834 | HITS-CLIP | 23142080 | Unspecified |
| mmu-miR-30c-5p | MIMAT0000514 | Soat1 | 20652 | HITS-CLIP | 23142080 | Unspecified |
| mmu-miR-30c-5p | MIMAT0000514 | Sobp | 109205 | HITS-CLIP | 37986224\|19536157 | Unspecified |
| mmu-miR-30c-5p | MIMAT0000514 | Socs1 | 12703 | HITS-CLIP | 37986224\|23142080 | Unspecified |
| mmu-miR-30c-5p | MIMAT0000514 | Socs3 | 12702 | HITS-CLIP | 37986224\|23597149\|23142080 | Unspecified |
| mmu-miR-30c-5p | MIMAT0000514 | Socs4 | 67296 | HITS-CLIP | 23597149\|23142080 | Unspecified |
| mmu-miR-30c-5p | MIMAT0000514 | Socs5 | 56468 | HITS-CLIP | 37986224\|23142080 | Unspecified |
| mmu-miR-30c-5p | MIMAT0000514 | Son | 20658 | HITS-CLIP | 37986224 | Unspecified |
| mmu-miR-30c-5p | MIMAT0000514 | Sorbs2 | 234214 | HITS-CLIP | 37986224\|25753418 | Unspecified |
| mmu-miR-30c-5p | MIMAT0000514 | Sorl1 | 20660 | HITS-CLIP | 23142080 | Unspecified |
| mmu-miR-30c-5p | MIMAT0000514 | Sos1 | 20662 | HITS-CLIP | 23597149\|23142080 | Unspecified |
| mmu-miR-30c-5p | MIMAT0000514 | Sox12 | 20667 | HITS-CLIP | 23597149 | Unspecified |
| mmu-miR-30c-5p | MIMAT0000514 | Sox4 | 20677 | HITS-CLIP | 37986224\|23142080 | Unspecified |
| mmu-miR-30c-5p | MIMAT0000514 | Sp3 | 20687 | HITS-CLIP | 37986224 | Unspecified |
| mmu-miR-30c-5p | MIMAT0000514 | Sp4 | 20688 | HITS-CLIP | 37986224 | Unspecified |
| mmu-miR-30c-5p | MIMAT0000514 | Spag9 | 70834 | HITS-CLIP | 37986224\|23597149\|23142080 | Unspecified |
| mmu-miR-30c-5p | MIMAT0000514 | Spast | 50850 | HITS-CLIP | 37986224\|23597149\|23142080\|1953€ | Unspecified |
| mmu-miR-30c-5p | MIMAT0000514 | Spata2 | 263876 | HITS-CLIP | 23597149\|23142080 | Unspecified |
| mmu-miR-30c-5p | MIMAT0000514 | Spcs3 | 76687 | HITS-CLIP | 37986224\|23597149\|23142080 | Unspecified |
| mmu-miR-30c-5p | MIMAT0000514 | Spdl1 | 70385 | HITS-CLIP | 23142080 | Unspecified |
| mmu-miR-30c-5p | MIMAT0000514 | Spen | 56381 | HITS-CLIP | 37986224\|19536157\|25753418\|23142 | Unspecified |
| mmu-miR-30c-5p | MIMAT0000514 | Spin4 | 270624 | HITS-CLIP | 23142080 | Unspecified |
| mmu-miR-30c-5p | MIMAT0000514 | Spop | 20747 | HITS-CLIP | 23597149\|23142080 | Unspecified |

| | | | | | | | |
|---|---|---|---|---|---|---|---|
| mmu-miR-30c-5p | MIMAT0000514 | Spopl | 76857 | HITS-CLIP | | 23142080 | Unspecified |
| mmu-miR-30c-5p | MIMAT0000514 | Sppl2a | 66552 | HITS-CLIP | | 37986224 | Unspecified |
| mmu-miR-30c-5p | MIMAT0000514 | Sppl3 | 74585 | HITS-CLIP | | 37986224 | Unspecified |
| mmu-miR-30c-5p | MIMAT0000514 | Spryd3 | 223918 | HITS-CLIP | | 23142080 | Unspecified |
| mmu-miR-30c-5p | MIMAT0000514 | Sptan1 | 20740 | HITS-CLIP | | 25753418 | Unspecified |
| mmu-miR-30c-5p | MIMAT0000514 | Sptb | 20741 | HITS-CLIP | | 25753418 | Unspecified |
| mmu-miR-30c-5p | MIMAT0000514 | Sptlc2 | 20773 | HITS-CLIP | | 23142080 | Unspecified |
| mmu-miR-30c-5p | MIMAT0000514 | Spty2d1 | 101685 | HITS-CLIP | | 37986224 | Unspecified |
| mmu-miR-30c-5p | MIMAT0000514 | Sqle | 20775 | HITS-CLIP | | 37986224 | Unspecified |
| mmu-miR-30c-5p | MIMAT0000514 | Srcap | 100043597 | HITS-CLIP | | 23597149 | Unspecified |
| mmu-miR-30c-5p | MIMAT0000514 | Srf | 20807 | HITS-CLIP | | 37986224 | Unspecified |
| mmu-miR-30c-5p | MIMAT0000514 | Srfbp1 | 67222 | HITS-CLIP | | 37986224 | Unspecified |
| mmu-miR-30c-5p | MIMAT0000514 | Srgap3 | 259302 | HITS-CLIP | | 23142080 | Unspecified |
| mmu-miR-30c-5p | MIMAT0000514 | Srsf1 | 110809 | HITS-CLIP | 37986224\|23142080 | | Unspecified |
| mmu-miR-30c-5p | MIMAT0000514 | Srsf10 | 14105 | HITS-CLIP | 23597149\|23142080 | | Unspecified |
| mmu-miR-30c-5p | MIMAT0000514 | Srsf4 | 57317 | HITS-CLIP | | 25753418 | Unspecified |
| mmu-miR-30c-5p | MIMAT0000514 | Srsf7 | 225027 | HITS-CLIP | 37986224\|23142080\|19536157 | | Unspecified |
| mmu-miR-30c-5p | MIMAT0000514 | Ss18 | 268996 | HITS-CLIP | | 23142080 | Unspecified |
| mmu-miR-30c-5p | MIMAT0000514 | Ssh1 | 231637 | HITS-CLIP | | 37986224 | Unspecified |
| mmu-miR-30c-5p | MIMAT0000514 | Ssh2 | 237860 | HITS-CLIP | | 37986224 | Unspecified |
| mmu-miR-30c-5p | MIMAT0000514 | Ssr1 | 107513 | HITS-CLIP | | 37986224 | Unspecified |
| mmu-miR-30c-5p | MIMAT0000514 | Ssr3 | 67437 | HITS-CLIP | 23597149\|19536157 | | Unspecified |
| mmu-miR-30c-5p | MIMAT0000514 | St6gal2 | 240119 | HITS-CLIP | | 37986224 | Unspecified |
| mmu-miR-30c-5p | MIMAT0000514 | St8sia4 | 20452 | HITS-CLIP | 37986224\|19536157\|23142080 | | Unspecified |
| mmu-miR-30c-5p | MIMAT0000514 | Stag1 | 20842 | HITS-CLIP | | 23142080 | Unspecified |
| mmu-miR-30c-5p | MIMAT0000514 | Stag2 | 20843 | HITS-CLIP | | 23142080 | Unspecified |
| mmu-miR-30c-5p | MIMAT0000514 | Stam2 | 56324 | HITS-CLIP | | 23142080 | Unspecified |
| mmu-miR-30c-5p | MIMAT0000514 | Stambpl1 | 76630 | HITS-CLIP | 37986224\|23142080 | | Unspecified |
| mmu-miR-30c-5p | MIMAT0000514 | Stat5b | 20851 | HITS-CLIP | | 37986224 | Unspecified |
| mmu-miR-30c-5p | MIMAT0000514 | Stat6 | 20852 | HITS-CLIP | 37986224\|23142080 | | Unspecified |
| mmu-miR-30c-5p | MIMAT0000514 | Stbd1 | 52331 | HITS-CLIP | | 37986224 | Unspecified |
| mmu-miR-30c-5p | MIMAT0000514 | Stim2 | 116873 | HITS-CLIP | 37986224\|23597149\|23142080 | | Unspecified |
| mmu-miR-30c-5p | MIMAT0000514 | Sting1 | 72512 | HITS-CLIP | | 25753418 | Unspecified |
| mmu-miR-30c-5p | MIMAT0000514 | Stk17b | 98267 | HITS-CLIP | 37986224\|23142080 | | Unspecified |
| mmu-miR-30c-5p | MIMAT0000514 | Stk35 | 67333 | HITS-CLIP | 37986224\|23142080\|23597149 | | Unspecified |
| mmu-miR-30c-5p | MIMAT0000514 | Stk39 | 53416 | HITS-CLIP | | 23142080 | Unspecified |
| mmu-miR-30c-5p | MIMAT0000514 | Stradb | 227154 | HITS-CLIP | | 23142080 | Unspecified |
| mmu-miR-30c-5p | MIMAT0000514 | Strip1 | 229707 | HITS-CLIP | 25083871\|23597149\|23142080 | | Unspecified |
| mmu-miR-30c-5p | MIMAT0000514 | Stt3a | 16430 | HITS-CLIP | | 23142080 | Unspecified |

| | | | | | | |
|---|---|---|---|---|---|---|
| mmu-miR-30c-5p | MIMAT0000514 | Stx16 | 228960 | HITS-CLIP | 37986224 | Unspecified |
| mmu-miR-30c-5p | MIMAT0000514 | Stxbp1 | 20910 | HITS-CLIP | 23142080 | Unspecified |
| mmu-miR-30c-5p | MIMAT0000514 | Stxbp5 | 78808 | HITS-CLIP | 37986224\|19536157\|23142080 | Unspecified |
| mmu-miR-30c-5p | MIMAT0000514 | Stxbp6 | 217517 | HITS-CLIP | 23597149 | Unspecified |
| mmu-miR-30c-5p | MIMAT0000514 | Sub1 | 20024 | HITS-CLIP | 37986224 | Unspecified |
| mmu-miR-30c-5p | MIMAT0000514 | Sucla2 | 20916 | HITS-CLIP | 23142080 | Unspecified |
| mmu-miR-30c-5p | MIMAT0000514 | Suco | 226551 | HITS-CLIP | 37986224 | Unspecified |
| mmu-miR-30c-5p | MIMAT0000514 | Sugt1 | 67955 | HITS-CLIP | 37986224\|19536157 | Unspecified |
| mmu-miR-30c-5p | MIMAT0000514 | Supt16 | 114741 | HITS-CLIP | 23142080 | Unspecified |
| mmu-miR-30c-5p | MIMAT0000514 | Surf2 | 20931 | HITS-CLIP | 23142080 | Unspecified |
| mmu-miR-30c-5p | MIMAT0000514 | Susd6 | 9766 | HITS-CLIP | 23597149 | Unspecified |
| mmu-miR-30c-5p | MIMAT0000514 | Suz12 | 52615 | qCLASH | 31393953 | Unspecified |
| mmu-miR-30c-5p | MIMAT0000514 | Sv2a | 64051 | HITS-CLIP | 37986224 | Unspecified |
| mmu-miR-30c-5p | MIMAT0000514 | Svil | 225115 | HITS-CLIP | 25753418 | Unspecified |
| mmu-miR-30c-5p | MIMAT0000514 | Svop | 68666 | HITS-CLIP | 37986224\|19536157 | Unspecified |
| mmu-miR-30c-5p | MIMAT0000514 | Swt1 | 66875 | HITS-CLIP | 37986224\|23142080 | Unspecified |
| mmu-miR-30c-5p | MIMAT0000514 | Syap1 | 67043 | HITS-CLIP | 23142080 | Unspecified |
| mmu-miR-30c-5p | MIMAT0000514 | Syngr3 | 20974 | HITS-CLIP | 37986224\|19536157 | Unspecified |
| mmu-miR-30c-5p | MIMAT0000514 | Synj1 | 104015 | HITS-CLIP | 37986224\|23142080 | Unspecified |
| mmu-miR-30c-5p | MIMAT0000514 | Synpr | 72003 | HITS-CLIP | 19536157 | Unspecified |
| mmu-miR-30c-5p | MIMAT0000514 | Sypl | 19027 | HITS-CLIP | 37986224\|23142080 | Unspecified |
| mmu-miR-30c-5p | MIMAT0000514 | Tab2 | 68652 | HITS-CLIP | 37986224\|23142080\|23597149\|19536 | Unspecified |
| mmu-miR-30c-5p | MIMAT0000514 | Tab3 | 66724 | HITS-CLIP | 23142080 | Unspecified |
| mmu-miR-30c-5p | MIMAT0000514 | Tada2a | 217031 | HITS-CLIP | 25753418 | Unspecified |
| mmu-miR-30c-5p | MIMAT0000514 | Taf5 | 226182 | HITS-CLIP | 23142080 | Unspecified |
| mmu-miR-30c-5p | MIMAT0000514 | Taok1 | 216965 | HITS-CLIP | 37986224\|23597149\|23142080 | Unspecified |
| mmu-miR-30c-5p | MIMAT0000514 | Tap2 | 21355 | HITS-CLIP | 25753418 | Unspecified |
| mmu-miR-30c-5p | MIMAT0000514 | Tardbp | 230908 | HITS-CLIP | 23142080 | Unspecified |
| mmu-miR-30c-5p | MIMAT0000514 | Tars | 110960 | HITS-CLIP | 23142080 | Unspecified |
| mmu-miR-30c-5p | MIMAT0000514 | Tasp1 | 75812 | HITS-CLIP | 23142080 | Unspecified |
| mmu-miR-30c-5p | MIMAT0000514 | Tax1bp1 | 52440 | HITS-CLIP | 23142080 | Unspecified |
| mmu-miR-30c-5p | MIMAT0000514 | Tbc1d10b | 68449 | HITS-CLIP | 37986224\|23597149\|23142080 | Unspecified |
| mmu-miR-30c-5p | MIMAT0000514 | Tbc1d15 | 66687 | HITS-CLIP | 37986224\|23142080 | Unspecified |
| mmu-miR-30c-5p | MIMAT0000514 | Tbc1d22b | 381085 | HITS-CLIP | 23597149\|23142080 | Unspecified |
| mmu-miR-30c-5p | MIMAT0000514 | Tbc1d23 | 67581 | HITS-CLIP | 23142080 | Unspecified |
| mmu-miR-30c-5p | MIMAT0000514 | Tbc1d4 | 210789 | HITS-CLIP | 25753418 | Unspecified |
| mmu-miR-30c-5p | MIMAT0000514 | Tbc1d8 | 54610 | HITS-CLIP | 37986224 | Unspecified |
| mmu-miR-30c-5p | MIMAT0000514 | Tbc1d8b | 245638 | HITS-CLIP | 23142080 | Unspecified |
| mmu-miR-30c-5p | MIMAT0000514 | Tbcel | 272589 | HITS-CLIP | 37986224\|23597149 | Unspecified |

| | | | | | | |
|---|---|---|---|---|---|---|
| mmu-miR-30c-5p | MIMAT0000514 | Tbl1xr1 | 81004 | HITS-CLIP | 37986224\|23142080\|23597149 | Unspecified |
| mmu-miR-30c-5p | MIMAT0000514 | Tbpl1 | 237336 | HITS-CLIP | 37986224 | Unspecified |
| mmu-miR-30c-5p | MIMAT0000514 | Tbx15 | 21384 | HITS-CLIP | 37986224 | Unspecified |
| mmu-miR-30c-5p | MIMAT0000514 | Tcaim | 382117 | HITS-CLIP | 37986224 | Unspecified |
| mmu-miR-30c-5p | MIMAT0000514 | Tceanc2 | 66526 | HITS-CLIP | 23142080 | Unspecified |
| mmu-miR-30c-5p | MIMAT0000514 | Tcerg1 | 56070 | HITS-CLIP | 37986224\|23142080 | Unspecified |
| mmu-miR-30c-5p | MIMAT0000514 | Tcf4 | 21413 | HITS-CLIP | 19536157\|23142080 | Unspecified |
| mmu-miR-30c-5p | MIMAT0000514 | Tcf7 | 21414 | HITS-CLIP | 37986224\|23142080 | Unspecified |
| mmu-miR-30c-5p | MIMAT0000514 | Tcf7l2 | 21416 | HITS-CLIP | 23597149 | Unspecified |
| mmu-miR-30c-5p | MIMAT0000514 | Tcfl5 | 277353 | HITS-CLIP | 37986224 | Unspecified |
| mmu-miR-30c-5p | MIMAT0000514 | Tcp11l2 | 216198 | HITS-CLIP | 23142080\|25753418 | Unspecified |
| mmu-miR-30c-5p | MIMAT0000514 | Tdgf1 | 21667 | HITS-CLIP | 37986224 | Unspecified |
| mmu-miR-30c-5p | MIMAT0000514 | Tdo2 | 56720 | HITS-CLIP | 37986224\|23597149 | Unspecified |
| mmu-miR-30c-5p | MIMAT0000514 | Tek | 21687 | HITS-CLIP | 37986224 | Unspecified |
| mmu-miR-30c-5p | MIMAT0000514 | Tenm3 | 23965 | HITS-CLIP | 19536157 | Unspecified |
| mmu-miR-30c-5p | MIMAT0000514 | Tent4a | 11044 | HITS-CLIP | 23142080 | Unspecified |
| mmu-miR-30c-5p | MIMAT0000514 | Tent4b | 64282 | HITS-CLIP | 23142080 | Unspecified |
| mmu-miR-30c-5p | MIMAT0000514 | Tent5a | 55603 | HITS-CLIP | 23142080 | Unspecified |
| mmu-miR-30c-5p | MIMAT0000514 | Tepsin | 146705 | HITS-CLIP | 23142080 | Unspecified |
| mmu-miR-30c-5p | MIMAT0000514 | Terf1 | 21749 | HITS-CLIP | 37986224 | Unspecified |
| mmu-miR-30c-5p | MIMAT0000514 | Tet1 | 52463 | HITS-CLIP | 37986224\|23142080 | Unspecified |
| mmu-miR-30c-5p | MIMAT0000514 | Tet2 | 214133 | HITS-CLIP | 37986224\|23142080 | Unspecified |
| mmu-miR-30c-5p | MIMAT0000514 | Tet3 | 194388 | HITS-CLIP | 23142080 | Unspecified |
| mmu-miR-30c-5p | MIMAT0000514 | Tex10 | 269536 | HITS-CLIP | 37986224\|23142080\|25753418 | Unspecified |
| mmu-miR-30c-5p | MIMAT0000514 | Tex2 | 21763 | HITS-CLIP | 37986224\|25753418 | Unspecified |
| mmu-miR-30c-5p | MIMAT0000514 | Tfdp1 | 21781 | HITS-CLIP | 23142080 | Unspecified |
| mmu-miR-30c-5p | MIMAT0000514 | Tfrc | 22042 | HITS-CLIP | 37986224\|23142080\|19536157 | Unspecified |
| mmu-miR-30c-5p | MIMAT0000514 | Tgds | 76355 | HITS-CLIP | 23142080 | Unspecified |
| mmu-miR-30c-5p | MIMAT0000514 | Tgfbr1 | 21812 | HITS-CLIP | 23142080\|25753418 | Unspecified |
| mmu-miR-30c-5p | MIMAT0000514 | Tgm2 | 21817 | HITS-CLIP | 23597149 | Unspecified |
| mmu-miR-30c-5p | MIMAT0000514 | Themis | 210757 | HITS-CLIP | 23142080 | Unspecified |
| mmu-miR-30c-5p | MIMAT0000514 | Thoc7 | 66231 | HITS-CLIP | 23597149 | Unspecified |
| mmu-miR-30c-5p | MIMAT0000514 | Timm22 | 56322 | HITS-CLIP | 25753418 | Unspecified |
| mmu-miR-30c-5p | MIMAT0000514 | Tiparp | 99929 | HITS-CLIP | 37986224\|23142080 | Unspecified |
| mmu-miR-30c-5p | MIMAT0000514 | Tlcd2 | 380712 | HITS-CLIP | 37986224 | Unspecified |
| mmu-miR-30c-5p | MIMAT0000514 | Tle1 | 21885 | HITS-CLIP | 37986224 | Unspecified |
| mmu-miR-30c-5p | MIMAT0000514 | Tlk1 | 228012 | HITS-CLIP | 23142080 | Unspecified |
| mmu-miR-30c-5p | MIMAT0000514 | Tlk2 | 24086 | HITS-CLIP | 23597149\|23142080 | Unspecified |
| mmu-miR-30c-5p | MIMAT0000514 | Tm9sf2 | 68059 | HITS-CLIP | 23142080 | Unspecified |

| | | | | | | |
|---|---|---|---|---|---|---|
| mmu-miR-30c-5p | MIMAT0000514 | Tmcc1 | 330401 | HITS-CLIP | 37986224\|23597149\|19536157\|23142 | Unspecified |
| mmu-miR-30c-5p | MIMAT0000514 | Tmed3 | 66111 | HITS-CLIP | 23142080 | Unspecified |
| mmu-miR-30c-5p | MIMAT0000514 | Tmed7 | 66676 | HITS-CLIP | 37986224\|23142080 | Unspecified |
| mmu-miR-30c-5p | MIMAT0000514 | Tmeff1 | 230157 | HITS-CLIP | 25753418 | Unspecified |
| mmu-miR-30c-5p | MIMAT0000514 | Tmem106b | 71900 | HITS-CLIP | 25753418 | Unspecified |
| mmu-miR-30c-5p | MIMAT0000514 | Tmem108 | 81907 | HITS-CLIP | 19536157 | Unspecified |
| mmu-miR-30c-5p | MIMAT0000514 | Tmem109 | 68539 | HITS-CLIP | 25753418 | Unspecified |
| mmu-miR-30c-5p | MIMAT0000514 | Tmem117 | 320709 | HITS-CLIP | 25753418 | Unspecified |
| mmu-miR-30c-5p | MIMAT0000514 | Tmem131 | 56030 | HITS-CLIP | 23597149 | Unspecified |
| mmu-miR-30c-5p | MIMAT0000514 | Tmem131l | 23240 | HITS-CLIP | 23142080 | Unspecified |
| mmu-miR-30c-5p | MIMAT0000514 | Tmem135 | 72759 | HITS-CLIP | 37986224\|23597149\|23142080 | Unspecified |
| mmu-miR-30c-5p | MIMAT0000514 | Tmem165 | 21982 | HITS-CLIP | 23142080 | Unspecified |
| mmu-miR-30c-5p | MIMAT0000514 | Tmem167b | 67495 | HITS-CLIP | 23142080 | Unspecified |
| mmu-miR-30c-5p | MIMAT0000514 | Tmem170b | 621976 | HITS-CLIP | 37986224\|23142080\|23597149\|1953€ | Unspecified |
| mmu-miR-30c-5p | MIMAT0000514 | Tmem178 | 68027 | HITS-CLIP | 37986224 | Unspecified |
| mmu-miR-30c-5p | MIMAT0000514 | Tmem181a | 77106 | HITS-CLIP | 23142080 | Unspecified |
| mmu-miR-30c-5p | MIMAT0000514 | Tmem183a | 57439 | HITS-CLIP | 37986224 | Unspecified |
| mmu-miR-30c-5p | MIMAT0000514 | Tmem220 | 338369 | HITS-CLIP | 37986224\|23597149 | Unspecified |
| mmu-miR-30c-5p | MIMAT0000514 | Tmem245 | 242474 | HITS-CLIP//qCLASH | 37986224\|31393953\|25753418\|23142 | Unspecified |
| mmu-miR-30c-5p | MIMAT0000514 | Tmem258 | 69038 | HITS-CLIP | 37986224\|23142080 | Unspecified |
| mmu-miR-30c-5p | MIMAT0000514 | Tmem26 | 327766 | HITS-CLIP | 23597149 | Unspecified |
| mmu-miR-30c-5p | MIMAT0000514 | Tmem260 | 218989 | HITS-CLIP | 23142080 | Unspecified |
| mmu-miR-30c-5p | MIMAT0000514 | Tmem30a | 69981 | HITS-CLIP | 23142080 | Unspecified |
| mmu-miR-30c-5p | MIMAT0000514 | Tmem38a | 74166 | HITS-CLIP | 25753418 | Unspecified |
| mmu-miR-30c-5p | MIMAT0000514 | Tmem38b | 52076 | qCLASH | 31393953 | Unspecified |
| mmu-miR-30c-5p | MIMAT0000514 | Tmem41a | 66664 | HITS-CLIP | 23597149 | Unspecified |
| mmu-miR-30c-5p | MIMAT0000514 | Tmem50b | 77975 | HITS-CLIP | 23142080 | Unspecified |
| mmu-miR-30c-5p | MIMAT0000514 | Tmem56 | 99887 | HITS-CLIP | 37986224 | Unspecified |
| mmu-miR-30c-5p | MIMAT0000514 | Tmem59 | 56374 | HITS-CLIP | 23597149\|23142080 | Unspecified |
| mmu-miR-30c-5p | MIMAT0000514 | Tmem60 | 212090 | HITS-CLIP | 23142080 | Unspecified |
| mmu-miR-30c-5p | MIMAT0000514 | Tmem62 | 96957 | HITS-CLIP | 37986224\|23597149\|19536157 | Unspecified |
| mmu-miR-30c-5p | MIMAT0000514 | Tmem64 | 100201 | HITS-CLIP | 37986224\|23597149\|25753418\|23142 | Unspecified |
| mmu-miR-30c-5p | MIMAT0000514 | Tmem68 | 72098 | HITS-CLIP | 25753418 | Unspecified |
| mmu-miR-30c-5p | MIMAT0000514 | Tmem87a | 211499 | HITS-CLIP | 37986224\|23597149\|23142080 | Unspecified |
| mmu-miR-30c-5p | MIMAT0000514 | Tmem87b | 72477 | HITS-CLIP | 23142080 | Unspecified |
| mmu-miR-30c-5p | MIMAT0000514 | Tmf1 | 232286 | qCLASH | 31393953 | Unspecified |
| mmu-miR-30c-5p | MIMAT0000514 | Tmod3 | 50875 | HITS-CLIP | 37986224\|25753418 | Unspecified |
| mmu-miR-30c-5p | MIMAT0000514 | Tmpo | 21917 | HITS-CLIP | 37986224\|23142080 | Unspecified |
| mmu-miR-30c-5p | MIMAT0000514 | Tmprss13 | 214531 | HITS-CLIP | 25753418 | Unspecified |

| | | | | | | |
|---|---|---|---|---|---|---|
| mmu-miR-30c-5p | MIMAT0000514 | Tmx1 | 72736 | HITS-CLIP | 23142080 | Unspecified |
| mmu-miR-30c-5p | MIMAT0000514 | Tmx3 | 67988 | HITS-CLIP | 37986224 | Unspecified |
| mmu-miR-30c-5p | MIMAT0000514 | Tmx4 | 52837 | qCLASH | 31393953 | Unspecified |
| mmu-miR-30c-5p | MIMAT0000514 | Tnf | 21926 | HITS-CLIP | 37986224 | Unspecified |
| mmu-miR-30c-5p | MIMAT0000514 | Tnfsf8 | 21949 | HITS-CLIP | 37986224 | Unspecified |
| mmu-miR-30c-5p | MIMAT0000514 | Tnik | 665113 | HITS-CLIP | 37986224\|19536157 | Unspecified |
| mmu-miR-30c-5p | MIMAT0000514 | Tnip1 | 57783 | HITS-CLIP | 23142080 | Unspecified |
| mmu-miR-30c-5p | MIMAT0000514 | Tnks2 | 74493 | HITS-CLIP | 37986224 | Unspecified |
| mmu-miR-30c-5p | MIMAT0000514 | Tnpo1 | 238799 | HITS-CLIP | 37986224\|23597149\|23142080 | Unspecified |
| mmu-miR-30c-5p | MIMAT0000514 | Tnr | 21960 | HITS-CLIP | 37986224 | Unspecified |
| mmu-miR-30c-5p | MIMAT0000514 | Tnrc6a | 233833 | HITS-CLIP | 37986224\|23597149\|19536157\|23142 | Unspecified |
| mmu-miR-30c-5p | MIMAT0000514 | Tnrc6b | 213988 | HITS-CLIP | 25083871\|23597149\|23142080\|19536 | Unspecified |
| mmu-miR-30c-5p | MIMAT0000514 | Tnrc6c | 217351 | HITS-CLIP | 37986224\|23597149\|23142080 | Unspecified |
| mmu-miR-30c-5p | MIMAT0000514 | Tns3 | 319939 | HITS-CLIP | 37986224 | Unspecified |
| mmu-miR-30c-5p | MIMAT0000514 | Tob2 | 57259 | HITS-CLIP | 37986224 | Unspecified |
| mmu-miR-30c-5p | MIMAT0000514 | Tollip | 54473 | HITS-CLIP | 23142080 | Unspecified |
| mmu-miR-30c-5p | MIMAT0000514 | Tom1l2 | 216810 | HITS-CLIP | 23142080 | Unspecified |
| mmu-miR-30c-5p | MIMAT0000514 | Tomm70a | 28185 | HITS-CLIP | 25083871\|23142080 | Unspecified |
| mmu-miR-30c-5p | MIMAT0000514 | Top1 | 21969 | HITS-CLIP//qCLASH | 37986224\|31393953\|23597149 | Unspecified |
| mmu-miR-30c-5p | MIMAT0000514 | Top2a | 21973 | HITS-CLIP | 23142080 | Unspecified |
| mmu-miR-30c-5p | MIMAT0000514 | Top2b | 21974 | HITS-CLIP | 37986224\|23142080 | Unspecified |
| mmu-miR-30c-5p | MIMAT0000514 | Topors | 106021 | HITS-CLIP | 23142080 | Unspecified |
| mmu-miR-30c-5p | MIMAT0000514 | Tor1aip2 | 240832 | HITS-CLIP | 37986224 | Unspecified |
| mmu-miR-30c-5p | MIMAT0000514 | Tor1b | 30934 | HITS-CLIP | 37986224\|23597149 | Unspecified |
| mmu-miR-30c-5p | MIMAT0000514 | Tor3a | 30935 | HITS-CLIP | 25753418 | Unspecified |
| mmu-miR-30c-5p | MIMAT0000514 | Tox | 252838 | HITS-CLIP | 23142080 | Unspecified |
| mmu-miR-30c-5p | MIMAT0000514 | Tox3 | 244579 | HITS-CLIP | 25753418 | Unspecified |
| mmu-miR-30c-5p | MIMAT0000514 | Tpp2 | 22019 | HITS-CLIP | 37986224 | Unspecified |
| mmu-miR-30c-5p | MIMAT0000514 | Tprkb | 69786 | HITS-CLIP | 37986224\|19536157\|23597149\|23142 | Unspecified |
| mmu-miR-30c-5p | MIMAT0000514 | Traf3ip2 | 103213 | HITS-CLIP | 23142080 | Unspecified |
| mmu-miR-30c-5p | MIMAT0000514 | Tram1 | 72265 | HITS-CLIP | 37986224\|23142080 | Unspecified |
| mmu-miR-30c-5p | MIMAT0000514 | Trappc1 | 245828 | HITS-CLIP | 23142080 | Unspecified |
| mmu-miR-30c-5p | MIMAT0000514 | Trappc8 | 75964 | HITS-CLIP | 37986224\|19536157 | Unspecified |
| mmu-miR-30c-5p | MIMAT0000514 | Trim23 | 81003 | HITS-CLIP | 37986224 | Unspecified |
| mmu-miR-30c-5p | MIMAT0000514 | Trim24 | 21848 | HITS-CLIP | 23142080 | Unspecified |
| mmu-miR-30c-5p | MIMAT0000514 | Trim35 | 66854 | HITS-CLIP | 23142080 | Unspecified |
| mmu-miR-30c-5p | MIMAT0000514 | Trim36 | 28105 | HITS-CLIP | 23597149 | Unspecified |
| mmu-miR-30c-5p | MIMAT0000514 | Trim37 | 68729 | HITS-CLIP | 37986224\|19536157\|23597149 | Unspecified |
| mmu-miR-30c-5p | MIMAT0000514 | Trim44 | 80985 | HITS-CLIP | 37986224 | Unspecified |

| | | | | | | |
|---|---|---|---|---|---|---|
| mmu-miR-30c-5p | MIMAT0000514 | Trio | 223435 | HITS-CLIP | 23142080 | Unspecified |
| mmu-miR-30c-5p | MIMAT0000514 | Trip11 | 109181 | HITS-CLIP | 37986224 | Unspecified |
| mmu-miR-30c-5p | MIMAT0000514 | Trip12 | 14897 | HITS-CLIP | 23597149\|19536157\|23142080 | Unspecified |
| mmu-miR-30c-5p | MIMAT0000514 | Trip13 | 69716 | HITS-CLIP | 23142080 | Unspecified |
| mmu-miR-30c-5p | MIMAT0000514 | Trip4 | 56404 | HITS-CLIP | 23597149 | Unspecified |
| mmu-miR-30c-5p | MIMAT0000514 | Trmt10a | 108943 | HITS-CLIP | 37986224 | Unspecified |
| mmu-miR-30c-5p | MIMAT0000514 | Trnt1 | 70047 | HITS-CLIP | 37986224\|23597149\|23142080 | Unspecified |
| mmu-miR-30c-5p | MIMAT0000514 | Trp53bp1 | 27223 | HITS-CLIP | 23142080 | Unspecified |
| mmu-miR-30c-5p | MIMAT0000514 | Trp53inp1 | 60599 | HITS-CLIP | 37986224\|23597149\|25753418\|23142 | Unspecified |
| mmu-miR-30c-5p | MIMAT0000514 | Trpm7 | 58800 | HITS-CLIP | 23142080 | Unspecified |
| mmu-miR-30c-5p | MIMAT0000514 | Trps1 | 83925 | HITS-CLIP | 23142080 | Unspecified |
| mmu-miR-30c-5p | MIMAT0000514 | Trrap | 100683 | HITS-CLIP | 23142080 | Unspecified |
| mmu-miR-30c-5p | MIMAT0000514 | Trub1 | 72133 | HITS-CLIP | 23142080 | Unspecified |
| mmu-miR-30c-5p | MIMAT0000514 | Tsc22d4 | 78829 | HITS-CLIP | 37986224\|23597149\|23142080 | Unspecified |
| mmu-miR-30c-5p | MIMAT0000514 | Tspan12 | 269831 | HITS-CLIP | 23597149\|25753418 | Unspecified |
| mmu-miR-30c-5p | MIMAT0000514 | Tspyl1 | 22110 | HITS-CLIP | 37986224\|23142080 | Unspecified |
| mmu-miR-30c-5p | MIMAT0000514 | Tspyl3 | 241732 | HITS-CLIP | 37986224 | Unspecified |
| mmu-miR-30c-5p | MIMAT0000514 | Tspyl4 | 72480 | HITS-CLIP | 37986224\|25753418 | Unspecified |
| mmu-miR-30c-5p | MIMAT0000514 | Ttc14 | 67120 | HITS-CLIP | 37986224\|23142080 | Unspecified |
| mmu-miR-30c-5p | MIMAT0000514 | Ttc3 | 22129 | HITS-CLIP | 37986224 | Unspecified |
| mmu-miR-30c-5p | MIMAT0000514 | Ttc39b | 69863 | HITS-CLIP | 37986224 | Unspecified |
| mmu-miR-30c-5p | MIMAT0000514 | Ttll4 | 67534 | HITS-CLIP | 23142080 | Unspecified |
| mmu-miR-30c-5p | MIMAT0000514 | Ttn | 22138 | HITS-CLIP | 25753418 | Unspecified |
| mmu-miR-30c-5p | MIMAT0000514 | Ttyh3 | 78339 | HITS-CLIP | 23142080 | Unspecified |
| mmu-miR-30c-5p | MIMAT0000514 | Tuba1a | 22142 | HITS-CLIP | 23142080 | Unspecified |
| mmu-miR-30c-5p | MIMAT0000514 | Tuba4a | 22145 | HITS-CLIP | 23142080 | Unspecified |
| mmu-miR-30c-5p | MIMAT0000514 | Tubb5 | 22154 | HITS-CLIP | 37986224\|23597149\|23142080 | Unspecified |
| mmu-miR-30c-5p | MIMAT0000514 | Tubgcp2 | 74237 | HITS-CLIP | 23142080\|23597149 | Unspecified |
| mmu-miR-30c-5p | MIMAT0000514 | Tubgcp4 | 51885 | HITS-CLIP | 37986224\|23597149 | Unspecified |
| mmu-miR-30c-5p | MIMAT0000514 | Tulp3 | 22158 | HITS-CLIP | 37986224 | Unspecified |
| mmu-miR-30c-5p | MIMAT0000514 | Tusc2 | 80385 | HITS-CLIP | 37986224 | Unspecified |
| mmu-miR-30c-5p | MIMAT0000514 | Tusc3 | 80286 | HITS-CLIP | 37986224\|23142080\|19536157 | Unspecified |
| mmu-miR-30c-5p | MIMAT0000514 | Tut4 | 23318 | HITS-CLIP | 23142080 | Unspecified |
| mmu-miR-30c-5p | MIMAT0000514 | Tut7 | 79670 | HITS-CLIP | 23142080 | Unspecified |
| mmu-miR-30c-5p | MIMAT0000514 | Tvp23b | 67510 | HITS-CLIP | 37986224 | Unspecified |
| mmu-miR-30c-5p | MIMAT0000514 | Twf1 | 19230 | Luciferase assay//qRT-PCR//HITS-CLIP//Lu | 24555094\|23597149\|23142080 | Unspecified |
| mmu-miR-30c-5p | MIMAT0000514 | Txk | 22165 | HITS-CLIP | 23142080 | Unspecified |
| mmu-miR-30c-5p | MIMAT0000514 | Txndc11 | 106200 | HITS-CLIP | 23597149\|23142080 | Unspecified |
| mmu-miR-30c-5p | MIMAT0000514 | Txndc16 | 70561 | HITS-CLIP | 23142080\|23597149 | Unspecified |

| | | | | | | |
|---|---|---|---|---|---|---|
| mmu-miR-30c-5p | MIMAT0000514 | Txnip | 56338 | HITS-CLIP | 37986224 | Unspecified |
| mmu-miR-30c-5p | MIMAT0000514 | Txnl1 | 53382 | HITS-CLIP | 37986224\|23142080\|23597149 | Unspecified |
| mmu-miR-30c-5p | MIMAT0000514 | U2surp | 67958 | HITS-CLIP | 23142080 | Unspecified |
| mmu-miR-30c-5p | MIMAT0000514 | Uba6 | 231380 | HITS-CLIP | 37986224\|23142080 | Unspecified |
| mmu-miR-30c-5p | MIMAT0000514 | Ube2d3 | 66105 | HITS-CLIP | 37986224 | Unspecified |
| mmu-miR-30c-5p | MIMAT0000514 | Ube2g1 | 67128 | HITS-CLIP | 23142080 | Unspecified |
| mmu-miR-30c-5p | MIMAT0000514 | Ube2j1 | 56228 | HITS-CLIP | 37986224\|23142080\|25753418 | Unspecified |
| mmu-miR-30c-5p | MIMAT0000514 | Ube2k | 53323 | HITS-CLIP | 23142080 | Unspecified |
| mmu-miR-30c-5p | MIMAT0000514 | Ube2w | 66799 | HITS-CLIP | 37986224\|23142080 | Unspecified |
| mmu-miR-30c-5p | MIMAT0000514 | Ube3a | 22215 | HITS-CLIP | 23142080\|19536157 | Unspecified |
| mmu-miR-30c-5p | MIMAT0000514 | Ube3c | 100763 | HITS-CLIP | 37986224\|23597149\|23142080 | Unspecified |
| mmu-miR-30c-5p | MIMAT0000514 | Ube4b | 63958 | HITS-CLIP | 37986224\|23597149\|23142080 | Unspecified |
| mmu-miR-30c-5p | MIMAT0000514 | Ubfd1 | 28018 | HITS-CLIP | 23142080 | Unspecified |
| mmu-miR-30c-5p | MIMAT0000514 | Ubn1 | 170644 | HITS-CLIP | 37986224\|23597149\|23142080 | Unspecified |
| mmu-miR-30c-5p | MIMAT0000514 | Ubn2 | 320538 | HITS-CLIP | 37986224\|23142080\|23597149 | Unspecified |
| mmu-miR-30c-5p | MIMAT0000514 | Ubp1 | 22221 | HITS-CLIP | 37986224\|25753418 | Unspecified |
| mmu-miR-30c-5p | MIMAT0000514 | Ubr1 | 22222 | HITS-CLIP | 37986224\|19536157 | Unspecified |
| mmu-miR-30c-5p | MIMAT0000514 | Ubr2 | 224826 | HITS-CLIP | 23597149 | Unspecified |
| mmu-miR-30c-5p | MIMAT0000514 | Ubr3 | 68795 | HITS-CLIP | 37986224\|23597149 | Unspecified |
| mmu-miR-30c-5p | MIMAT0000514 | Ubr4 | 69116 | HITS-CLIP | 23597149\|23142080 | Unspecified |
| mmu-miR-30c-5p | MIMAT0000514 | Ubr5 | 70790 | HITS-CLIP | 23597149\|23142080 | Unspecified |
| mmu-miR-30c-5p | MIMAT0000514 | Ubxn7 | 224111 | HITS-CLIP | 37986224\|23142080 | Unspecified |
| mmu-miR-30c-5p | MIMAT0000514 | Uchl1 | 22223 | HITS-CLIP | 37986224 | Unspecified |
| mmu-miR-30c-5p | MIMAT0000514 | Ufsp2 | 192169 | HITS-CLIP | 23597149 | Unspecified |
| mmu-miR-30c-5p | MIMAT0000514 | Ugdh | 22235 | HITS-CLIP | 23597149 | Unspecified |
| mmu-miR-30c-5p | MIMAT0000514 | Ugt2b35 | 243085 | HITS-CLIP | 23597149 | Unspecified |
| mmu-miR-30c-5p | MIMAT0000514 | Uhrf1bp1l | 75089 | HITS-CLIP | 37986224\|23142080 | Unspecified |
| mmu-miR-30c-5p | MIMAT0000514 | Ulk2 | 29869 | HITS-CLIP | 23142080 | Unspecified |
| mmu-miR-30c-5p | MIMAT0000514 | Umps | 22247 | HITS-CLIP | 23597149 | Unspecified |
| mmu-miR-30c-5p | MIMAT0000514 | Unc13c | 208898 | HITS-CLIP | 37986224 | Unspecified |
| mmu-miR-30c-5p | MIMAT0000514 | Unc50 | 67387 | HITS-CLIP | 23142080 | Unspecified |
| mmu-miR-30c-5p | MIMAT0000514 | Unc5c | 22253 | HITS-CLIP | 37986224\|19536157 | Unspecified |
| mmu-miR-30c-5p | MIMAT0000514 | Unc93b1 | 54445 | HITS-CLIP | 37986224 | Unspecified |
| mmu-miR-30c-5p | MIMAT0000514 | Ung | 22256 | HITS-CLIP | 37986224\|23597149 | Unspecified |
| mmu-miR-30c-5p | MIMAT0000514 | Unk | 217331 | HITS-CLIP | 23142080 | Unspecified |
| mmu-miR-30c-5p | MIMAT0000514 | Upf1 | 19704 | HITS-CLIP | 23142080 | Unspecified |
| mmu-miR-30c-5p | MIMAT0000514 | Usb1 | 101985 | HITS-CLIP | 26203562 | Unspecified |
| mmu-miR-30c-5p | MIMAT0000514 | Uso1 | 56041 | HITS-CLIP | 37986224\|23597149\|23142080 | Unspecified |
| mmu-miR-30c-5p | MIMAT0000514 | Usp14 | 59025 | HITS-CLIP | 23142080\|25753418 | Unspecified |

| | | | | | | |
|---|---|---|---|---|---|---|
| mmu-miR-30c-5p | MIMAT0000514 | Usp15 | 14479 | HITS-CLIP | 37986224\|23142080 | Unspecified |
| mmu-miR-30c-5p | MIMAT0000514 | Usp2 | 53376 | HITS-CLIP | 23597149\|19536157 | Unspecified |
| mmu-miR-30c-5p | MIMAT0000514 | Usp21 | 30941 | HITS-CLIP | 23597149 | Unspecified |
| mmu-miR-30c-5p | MIMAT0000514 | Usp24 | 329908 | HITS-CLIP | 37986224 | Unspecified |
| mmu-miR-30c-5p | MIMAT0000514 | Usp25 | 30940 | HITS-CLIP | 37986224\|23142080 | Unspecified |
| mmu-miR-30c-5p | MIMAT0000514 | Usp28 | 235323 | HITS-CLIP | 25753418\|23142080 | Unspecified |
| mmu-miR-30c-5p | MIMAT0000514 | Usp32 | 237898 | HITS-CLIP | 23142080 | Unspecified |
| mmu-miR-30c-5p | MIMAT0000514 | Usp33 | 170822 | HITS-CLIP | 23142080 | Unspecified |
| mmu-miR-30c-5p | MIMAT0000514 | Usp34 | 17847 | HITS-CLIP | 23142080\|25753418 | Unspecified |
| mmu-miR-30c-5p | MIMAT0000514 | Usp37 | 319651 | HITS-CLIP | 37986224\|23142080 | Unspecified |
| mmu-miR-30c-5p | MIMAT0000514 | Usp38 | 74841 | HITS-CLIP | 37986224\|23597149\|23142080 | Unspecified |
| mmu-miR-30c-5p | MIMAT0000514 | Usp45 | 77593 | HITS-CLIP | 37986224\|19536157\|23597149\|25753 | Unspecified |
| mmu-miR-30c-5p | MIMAT0000514 | Usp47 | 74996 | HITS-CLIP | 37986224\|23142080 | Unspecified |
| mmu-miR-30c-5p | MIMAT0000514 | Usp48 | 170707 | HITS-CLIP | 37986224\|23597149\|19536157 | Unspecified |
| mmu-miR-30c-5p | MIMAT0000514 | Usp9x | 22284 | HITS-CLIP | 37986224\|23597149\|23142080 | Unspecified |
| mmu-miR-30c-5p | MIMAT0000514 | Uspl1 | 231915 | HITS-CLIP | 37986224 | Unspecified |
| mmu-miR-30c-5p | MIMAT0000514 | Utp15 | 105372 | HITS-CLIP | 23142080 | Unspecified |
| mmu-miR-30c-5p | MIMAT0000514 | Utp20 | 70683 | HITS-CLIP | 37986224\|23142080 | Unspecified |
| mmu-miR-30c-5p | MIMAT0000514 | Uty | 22290 | HITS-CLIP | 25753418 | Unspecified |
| mmu-miR-30c-5p | MIMAT0000514 | Vamp3 | 22319 | HITS-CLIP | 37986224 | Unspecified |
| mmu-miR-30c-5p | MIMAT0000514 | Vapa | 30960 | HITS-CLIP | 23597149\|23142080 | Unspecified |
| mmu-miR-30c-5p | MIMAT0000514 | Vash2 | 226841 | HITS-CLIP | 25753418 | Unspecified |
| mmu-miR-30c-5p | MIMAT0000514 | Vat1 | 26949 | HITS-CLIP | 23142080 | Unspecified |
| mmu-miR-30c-5p | MIMAT0000514 | Vat1l | 270097 | HITS-CLIP | 37986224 | Unspecified |
| mmu-miR-30c-5p | MIMAT0000514 | Vcl | 22330 | HITS-CLIP | 19536157\|23142080 | Unspecified |
| mmu-miR-30c-5p | MIMAT0000514 | Vcp | 269523 | HITS-CLIP | 23142080 | Unspecified |
| mmu-miR-30c-5p | MIMAT0000514 | Vcpip1 | 70675 | HITS-CLIP | 37986224\|23142080 | Unspecified |
| mmu-miR-30c-5p | MIMAT0000514 | Vdac1 | 22333 | HITS-CLIP | 37986224\|23597149\|19536157\|23142 | Unspecified |
| mmu-miR-30c-5p | MIMAT0000514 | Vegfa | 22339 | HITS-CLIP | 19536157 | Unspecified |
| mmu-miR-30c-5p | MIMAT0000514 | Vezf1 | 22344 | HITS-CLIP | 37986224 | Unspecified |
| mmu-miR-30c-5p | MIMAT0000514 | Vim | 22352 | HITS-CLIP | 23142080 | Unspecified |
| mmu-miR-30c-5p | MIMAT0000514 | Vopp1 | 232023 | HITS-CLIP | 37986224\|19536157 | Unspecified |
| mmu-miR-30c-5p | MIMAT0000514 | Vps13a | 271564 | HITS-CLIP | 23597149\|23142080 | Unspecified |
| mmu-miR-30c-5p | MIMAT0000514 | Vps13c | 320528 | HITS-CLIP | 37986224\|19536157\|25753418 | Unspecified |
| mmu-miR-30c-5p | MIMAT0000514 | Vps18 | 228545 | HITS-CLIP | 25753418 | Unspecified |
| mmu-miR-30c-5p | MIMAT0000514 | Vps26b | 69091 | HITS-CLIP | 23142080 | Unspecified |
| mmu-miR-30c-5p | MIMAT0000514 | Vps37a | 52348 | HITS-CLIP | 23142080 | Unspecified |
| mmu-miR-30c-5p | MIMAT0000514 | Vps4b | 20479 | HITS-CLIP | 37986224\|23142080 | Unspecified |
| mmu-miR-30c-5p | MIMAT0000514 | Vps8 | 209018 | HITS-CLIP | 23142080 | Unspecified |

| | | | | | | |
|---|---|---|---|---|---|---|
| mmu-miR-30c-5p | MIMAT0000514 | Vrk2 | 69922 | HITS-CLIP | 23142080 | Unspecified |
| mmu-miR-30c-5p | MIMAT0000514 | Wac | 225131 | HITS-CLIP | 23142080 | Unspecified |
| mmu-miR-30c-5p | MIMAT0000514 | Wapl | 23063 | HITS-CLIP | 23142080 | Unspecified |
| mmu-miR-30c-5p | MIMAT0000514 | Washc4 | 319277 | HITS-CLIP | 19536157 | Unspecified |
| mmu-miR-30c-5p | MIMAT0000514 | Wasl | 73178 | HITS-CLIP | 37986224\|23597149\|23142080 | Unspecified |
| mmu-miR-30c-5p | MIMAT0000514 | Wbp11 | 60321 | HITS-CLIP | 37986224 | Unspecified |
| mmu-miR-30c-5p | MIMAT0000514 | Wbp1l | 226178 | HITS-CLIP | 23142080 | Unspecified |
| mmu-miR-30c-5p | MIMAT0000514 | Wbp4 | 22380 | HITS-CLIP | 23142080 | Unspecified |
| mmu-miR-30c-5p | MIMAT0000514 | Wdfy3 | 72145 | HITS-CLIP | 37986224\|23597149 | Unspecified |
| mmu-miR-30c-5p | MIMAT0000514 | Wdr11 | 207425 | qCLASH | 31393953 | Unspecified |
| mmu-miR-30c-5p | MIMAT0000514 | Wdr18 | 216156 | HITS-CLIP | 23597149 | Unspecified |
| mmu-miR-30c-5p | MIMAT0000514 | Wdr19 | 213081 | HITS-CLIP | 37986224 | Unspecified |
| mmu-miR-30c-5p | MIMAT0000514 | Wdr26 | 226757 | HITS-CLIP | 23142080 | Unspecified |
| mmu-miR-30c-5p | MIMAT0000514 | Wdr7 | 104082 | HITS-CLIP | 37986224\|23597149\|23142080 | Unspecified |
| mmu-miR-30c-5p | MIMAT0000514 | Wdr75 | 73674 | HITS-CLIP | 23142080 | Unspecified |
| mmu-miR-30c-5p | MIMAT0000514 | Wdr82 | 77305 | HITS-CLIP | 37986224\|23597149\|23142080 | Unspecified |
| mmu-miR-30c-5p | MIMAT0000514 | Wee1 | 22390 | HITS-CLIP | 23142080 | Unspecified |
| mmu-miR-30c-5p | MIMAT0000514 | Wipf1 | 215280 | HITS-CLIP | 37986224\|19536157\|23142080 | Unspecified |
| mmu-miR-30c-5p | MIMAT0000514 | Wipf3 | 644150 | HITS-CLIP | 23597149 | Unspecified |
| mmu-miR-30c-5p | MIMAT0000514 | Wnk1 | 232341 | HITS-CLIP | 23142080 | Unspecified |
| mmu-miR-30c-5p | MIMAT0000514 | Wnt5a | 22418 | HITS-CLIP | 37986224 | Unspecified |
| mmu-miR-30c-5p | MIMAT0000514 | Wrn | 22427 | HITS-CLIP | 37986224\|23142080 | Unspecified |
| mmu-miR-30c-5p | MIMAT0000514 | Wwp1 | 107568 | HITS-CLIP | 25753418 | Unspecified |
| mmu-miR-30c-5p | MIMAT0000514 | Xirp1 | 22437 | HITS-CLIP | 25753418 | Unspecified |
| mmu-miR-30c-5p | MIMAT0000514 | Xkr5 | 319581 | HITS-CLIP | 23142080 | Unspecified |
| mmu-miR-30c-5p | MIMAT0000514 | Xkr9 | 381246 | HITS-CLIP | 37986224 | Unspecified |
| mmu-miR-30c-5p | MIMAT0000514 | Xpnpep3 | 321003 | HITS-CLIP | 23142080\|23597149 | Unspecified |
| mmu-miR-30c-5p | MIMAT0000514 | Xpo1 | 103573 | HITS-CLIP | 37986224\|23597149\|23142080 | Unspecified |
| mmu-miR-30c-5p | MIMAT0000514 | Xpo4 | 57258 | HITS-CLIP | 23142080 | Unspecified |
| mmu-miR-30c-5p | MIMAT0000514 | Xpo5 | 72322 | HITS-CLIP | 23142080 | Unspecified |
| mmu-miR-30c-5p | MIMAT0000514 | Xpo6 | 74204 | HITS-CLIP | 37986224\|23142080\|23597149 | Unspecified |
| mmu-miR-30c-5p | MIMAT0000514 | Xpo7 | 65246 | HITS-CLIP | 23142080 | Unspecified |
| mmu-miR-30c-5p | MIMAT0000514 | Xpr1 | 19775 | HITS-CLIP | 23142080 | Unspecified |
| mmu-miR-30c-5p | MIMAT0000514 | Ybx3 | 56449 | HITS-CLIP | 19536157\|23142080 | Unspecified |
| mmu-miR-30c-5p | MIMAT0000514 | Yeats4 | 64050 | HITS-CLIP | 37986224 | Unspecified |
| mmu-miR-30c-5p | MIMAT0000514 | Yod1 | 226418 | HITS-CLIP | 37986224\|23597149\|19536157\|23142 | Unspecified |
| mmu-miR-30c-5p | MIMAT0000514 | Ypel2 | 77864 | HITS-CLIP | 37986224\|23597149\|23142080\|2575: | Unspecified |
| mmu-miR-30c-5p | MIMAT0000514 | Ypel5 | 383295 | HITS-CLIP | 37986224\|23142080 | Unspecified |
| mmu-miR-30c-5p | MIMAT0000514 | Ythdc1 | 231386 | HITS-CLIP | 37986224\|23142080 | Unspecified |

| | | | | | | |
|---|---|---|---|---|---|---|
| mmu-miR-30c-5p | MIMAT0000514 | Ythdc2 | 240255 | HITS-CLIP | 23142080 | Unspecified |
| mmu-miR-30c-5p | MIMAT0000514 | Ythdf3 | 229096 | HITS-CLIP | 37986224\|23597149\|23142080 | Unspecified |
| mmu-miR-30c-5p | MIMAT0000514 | Ywhab | 54401 | HITS-CLIP | 37986224 | Unspecified |
| mmu-miR-30c-5p | MIMAT0000514 | Ywhag | 22628 | HITS-CLIP | 19536157\|23142080 | Unspecified |
| mmu-miR-30c-5p | MIMAT0000514 | Yy1 | 22632 | HITS-CLIP | 23597149 | Unspecified |
| mmu-miR-30c-5p | MIMAT0000514 | Zadh2 | 225791 | HITS-CLIP | 37986224\|23142080 | Unspecified |
| mmu-miR-30c-5p | MIMAT0000514 | Zbed6 | 667118 | HITS-CLIP | 37986224 | Unspecified |
| mmu-miR-30c-5p | MIMAT0000514 | Zbtb1 | 268564 | HITS-CLIP | 23142080 | Unspecified |
| mmu-miR-30c-5p | MIMAT0000514 | Zbtb11 | 271377 | HITS-CLIP | 37986224\|23142080 | Unspecified |
| mmu-miR-30c-5p | MIMAT0000514 | Zbtb16 | 235320 | HITS-CLIP | 25753418 | Unspecified |
| mmu-miR-30c-5p | MIMAT0000514 | Zbtb18 | 30928 | HITS-CLIP | 37986224\|23597149\|23142080 | Unspecified |
| mmu-miR-30c-5p | MIMAT0000514 | Zbtb20 | 56490 | HITS-CLIP | 23597149\|19536157 | Unspecified |
| mmu-miR-30c-5p | MIMAT0000514 | Zbtb21 | 114565 | HITS-CLIP | 23142080 | Unspecified |
| mmu-miR-30c-5p | MIMAT0000514 | Zbtb25 | 109929 | HITS-CLIP | 23142080 | Unspecified |
| mmu-miR-30c-5p | MIMAT0000514 | Zbtb34 | 241311 | HITS-CLIP | 19536157\|23142080 | Unspecified |
| mmu-miR-30c-5p | MIMAT0000514 | Zbtb37 | 240869 | HITS-CLIP | 23142080 | Unspecified |
| mmu-miR-30c-5p | MIMAT0000514 | Zbtb38 | 245007 | HITS-CLIP | 37986224\|23142080 | Unspecified |
| mmu-miR-30c-5p | MIMAT0000514 | Zbtb39 | 320080 | HITS-CLIP | 37986224\|23142080 | Unspecified |
| mmu-miR-30c-5p | MIMAT0000514 | Zbtb41 | 226470 | HITS-CLIP | 23142080 | Unspecified |
| mmu-miR-30c-5p | MIMAT0000514 | Zbtb44 | 235132 | HITS-CLIP | 23597149\|23142080 | Unspecified |
| mmu-miR-30c-5p | MIMAT0000514 | Zbtb6 | 241322 | HITS-CLIP | 23142080 | Unspecified |
| mmu-miR-30c-5p | MIMAT0000514 | Zbtb7a | 16969 | HITS-CLIP | 37986224\|23597149\|23142080 | Unspecified |
| mmu-miR-30c-5p | MIMAT0000514 | Zc3h15 | 69082 | HITS-CLIP | 37986224\|23597149\|23142080 | Unspecified |
| mmu-miR-30c-5p | MIMAT0000514 | Zcchc14 | 142682 | HITS-CLIP | 37986224\|23597149\|19536157 | Unspecified |
| mmu-miR-30c-5p | MIMAT0000514 | Zcchc2 | 227449 | HITS-CLIP | 37986224\|19536157\|23597149\|2314[illegible] | Unspecified |
| mmu-miR-30c-5p | MIMAT0000514 | Zcchc24 | 71918 | HITS-CLIP | 37986224\|23597149\|23142080 | Unspecified |
| mmu-miR-30c-5p | MIMAT0000514 | Zdhhc17 | 320150 | HITS-CLIP | 23597149 | Unspecified |
| mmu-miR-30c-5p | MIMAT0000514 | Zdhhc20 | 75965 | HITS-CLIP | 37986224\|23597149\|19536157\|2314[illegible] | Unspecified |
| mmu-miR-30c-5p | MIMAT0000514 | Zdhhc21 | 68268 | HITS-CLIP | 37986224\|23142080 | Unspecified |
| mmu-miR-30c-5p | MIMAT0000514 | Zdhhc6 | 66980 | HITS-CLIP | 23142080\|23597149 | Unspecified |
| mmu-miR-30c-5p | MIMAT0000514 | Zeb1 | 21417 | HITS-CLIP | 37986224 | Unspecified |
| mmu-miR-30c-5p | MIMAT0000514 | Zeb2 | 24136 | HITS-CLIP | 23597149 | Unspecified |
| mmu-miR-30c-5p | MIMAT0000514 | Zfand4 | 67492 | HITS-CLIP | 37986224 | Unspecified |
| mmu-miR-30c-5p | MIMAT0000514 | Zfand5 | 22682 | HITS-CLIP | 37986224\|19536157\|25753418 | Unspecified |
| mmu-miR-30c-5p | MIMAT0000514 | Zfand6 | 65098 | HITS-CLIP | 23142080\|23597149 | Unspecified |
| mmu-miR-30c-5p | MIMAT0000514 | Zfat | 57623 | HITS-CLIP | 23142080 | Unspecified |
| mmu-miR-30c-5p | MIMAT0000514 | Zfc3h1 | 216345 | HITS-CLIP | 37986224\|23142080 | Unspecified |
| mmu-miR-30c-5p | MIMAT0000514 | Zfp106 | 20402 | HITS-CLIP | 23142080 | Unspecified |
| mmu-miR-30c-5p | MIMAT0000514 | Zfp142 | 77264 | HITS-CLIP | 25753418 | Unspecified |

| | | | | | | |
|---|---|---|---|---|---|---|
| mmu-miR-30c-5p | MIMAT0000514 | Zfp143 | 20841 | HITS-CLIP | 23142080 | Unspecified |
| mmu-miR-30c-5p | MIMAT0000514 | Zfp148 | 22661 | HITS-CLIP | 37986224\|23142080 | Unspecified |
| mmu-miR-30c-5p | MIMAT0000514 | Zfp207 | 22680 | HITS-CLIP | 23142080\|19536157 | Unspecified |
| mmu-miR-30c-5p | MIMAT0000514 | Zfp26 | 22688 | HITS-CLIP | 37986224 | Unspecified |
| mmu-miR-30c-5p | MIMAT0000514 | Zfp280b | 64453 | HITS-CLIP | 37986224 | Unspecified |
| mmu-miR-30c-5p | MIMAT0000514 | Zfp281 | 226442 | HITS-CLIP | 37986224\|23142080 | Unspecified |
| mmu-miR-30c-5p | MIMAT0000514 | Zfp292 | 30046 | HITS-CLIP | 37986224\|25753418\|23142080 | Unspecified |
| mmu-miR-30c-5p | MIMAT0000514 | Zfp36l1 | 12192 | HITS-CLIP | 37986224 | Unspecified |
| mmu-miR-30c-5p | MIMAT0000514 | Zfp397 | 69256 | HITS-CLIP | 37986224 | Unspecified |
| mmu-miR-30c-5p | MIMAT0000514 | Zfp407 | 240476 | HITS-CLIP | 23142080 | Unspecified |
| mmu-miR-30c-5p | MIMAT0000514 | Zfp422 | 67255 | HITS-CLIP | 23142080 | Unspecified |
| mmu-miR-30c-5p | MIMAT0000514 | Zfp445 | 235682 | HITS-CLIP | 23142080 | Unspecified |
| mmu-miR-30c-5p | MIMAT0000514 | Zfp451 | 98403 | HITS-CLIP | 23142080 | Unspecified |
| mmu-miR-30c-5p | MIMAT0000514 | Zfp46 | 22704 | HITS-CLIP | 23142080 | Unspecified |
| mmu-miR-30c-5p | MIMAT0000514 | Zfp462 | 242466 | HITS-CLIP | 37986224 | Unspecified |
| mmu-miR-30c-5p | MIMAT0000514 | Zfp507 | 668501 | HITS-CLIP | 37986224\|23597149\|23142080 | Unspecified |
| mmu-miR-30c-5p | MIMAT0000514 | Zfp516 | 329003 | HITS-CLIP | 23142080\|23597149 | Unspecified |
| mmu-miR-30c-5p | MIMAT0000514 | Zfp518a | 72672 | HITS-CLIP | 37986224\|23142080 | Unspecified |
| mmu-miR-30c-5p | MIMAT0000514 | Zfp521 | 225207 | HITS-CLIP | 37986224\|23597149 | Unspecified |
| mmu-miR-30c-5p | MIMAT0000514 | Zfp593 | 68040 | qCLASH | 31393953 | Unspecified |
| mmu-miR-30c-5p | MIMAT0000514 | Zfp644 | 52397 | HITS-CLIP | 37986224\|23142080 | Unspecified |
| mmu-miR-30c-5p | MIMAT0000514 | Zfp652 | 268469 | HITS-CLIP | 37986224 | Unspecified |
| mmu-miR-30c-5p | MIMAT0000514 | Zfp654 | 72020 | HITS-CLIP | 23142080\|25753418 | Unspecified |
| mmu-miR-30c-5p | MIMAT0000514 | Zfp655 | 72611 | HITS-CLIP | 37986224 | Unspecified |
| mmu-miR-30c-5p | MIMAT0000514 | Zfp687 | 78266 | HITS-CLIP | 37986224 | Unspecified |
| mmu-miR-30c-5p | MIMAT0000514 | Zfp710 | 209225 | HITS-CLIP | 23142080 | Unspecified |
| mmu-miR-30c-5p | MIMAT0000514 | Zfp729a | 212281 | HITS-CLIP | 23142080 | Unspecified |
| mmu-miR-30c-5p | MIMAT0000514 | Zfp746 | 69228 | HITS-CLIP | 37986224\|23597149\|23142080 | Unspecified |
| mmu-miR-30c-5p | MIMAT0000514 | Zfp790 | 233056 | HITS-CLIP | 25753418 | Unspecified |
| mmu-miR-30c-5p | MIMAT0000514 | Zfp800 | 627049 | HITS-CLIP | 23142080 | Unspecified |
| mmu-miR-30c-5p | MIMAT0000514 | Zfp827 | 622675 | HITS-CLIP | 23142080 | Unspecified |
| mmu-miR-30c-5p | MIMAT0000514 | Zfp831 | 100043757 | HITS-CLIP | 23142080 | Unspecified |
| mmu-miR-30c-5p | MIMAT0000514 | Zfp871 | 208292 | HITS-CLIP | 37986224\|23142080\|23597149 | Unspecified |
| mmu-miR-30c-5p | MIMAT0000514 | Zfp930 | 234358 | HITS-CLIP | 37986224 | Unspecified |
| mmu-miR-30c-5p | MIMAT0000514 | Zfp948 | 381066 | HITS-CLIP | 37986224 | Unspecified |
| mmu-miR-30c-5p | MIMAT0000514 | Zfr | 22763 | HITS-CLIP | 23142080 | Unspecified |
| mmu-miR-30c-5p | MIMAT0000514 | Zfx | 22764 | HITS-CLIP | 23142080 | Unspecified |
| mmu-miR-30c-5p | MIMAT0000514 | Zfyve26 | 211978 | HITS-CLIP | 23142080 | Unspecified |
| mmu-miR-30c-5p | MIMAT0000514 | Zhx1 | 22770 | HITS-CLIP | 37986224\|19536157\|23142080 | Unspecified |

| | | | | | | | |
|---|---|---|---|---|---|---|---|
| mmu-miR-30c-5p | MIMAT0000514 | Zik1 | 22775 | HITS-CLIP | | 37986224 | Unspecified |
| mmu-miR-30c-5p | MIMAT0000514 | Zkscan3 | 72739 | HITS-CLIP | | 37986224 | Unspecified |
| mmu-miR-30c-5p | MIMAT0000514 | Zmiz1 | 328365 | HITS-CLIP | | 23142080 | Unspecified |
| mmu-miR-30c-5p | MIMAT0000514 | Zmym1 | 68310 | HITS-CLIP | 23597149\|23142080 | | Unspecified |
| mmu-miR-30c-5p | MIMAT0000514 | Zmym2 | 76007 | HITS-CLIP | | 37986224 | Unspecified |
| mmu-miR-30c-5p | MIMAT0000514 | Zmym3 | 56364 | HITS-CLIP | | 23142080 | Unspecified |
| mmu-miR-30c-5p | MIMAT0000514 | Zmym4 | 67785 | HITS-CLIP | 23597149\|19536157\|23142080 | | Unspecified |
| mmu-miR-30c-5p | MIMAT0000514 | Zmynd8 | 228880 | HITS-CLIP | 25753418\|23142080 | | Unspecified |
| mmu-miR-30c-5p | MIMAT0000514 | Znrd1 | 66136 | HITS-CLIP | | 23142080 | Unspecified |
| mmu-miR-30c-5p | MIMAT0000514 | Znrf1 | 170737 | HITS-CLIP | 37986224\|23142080\|23597149 | | Unspecified |
| mmu-miR-30c-5p | MIMAT0000514 | Zranb1 | 360216 | HITS-CLIP | 37986224\|23597149\|19536157\|23142 | | Unspecified |
| mmu-miR-30c-5p | MIMAT0000514 | Zranb3 | 226409 | HITS-CLIP | | 23142080 | Unspecified |
| mmu-miR-30c-5p | MIMAT0000514 | Zrsr1 | 22183 | HITS-CLIP | | 23597149 | Unspecified |
| mmu-miR-30c-5p | MIMAT0000514 | Zrsr2 | 22184 | HITS-CLIP | | 23142080 | Unspecified |
| mmu-miR-30c-5p | MIMAT0000514 | Zswim3 | 67538 | HITS-CLIP | | 23142080 | Unspecified |
| mmu-miR-30c-5p | MIMAT0000514 | Zswim6 | 67263 | HITS-CLIP | | 37986224 | Unspecified |
| mmu-miR-30c-5p | MIMAT0000514 | Zzef1 | 195018 | HITS-CLIP | 23142080\|25753418 | | Unspecified |
| mmu-miR-30c-5p | MIMAT0000514 | Zzz3 | 108946 | HITS-CLIP | | 23142080 | Unspecified |
| mmu-miR-30c-5p | MIMAT0000514 | mt-Co1 | 4512 | qCLASH | | 31393953 | Unspecified |
| mmu-miR-30c-5p | MIMAT0000514 | mt-Nd5 | 4540 | HITS-CLIP | | 23142080 | Unspecified |
| mmu-miR-139-3p | MIMAT0004662 | Aff4 | 93736 | HITS-CLIP | | 31433990 | Unspecified |
| mmu-miR-139-3p | MIMAT0004662 | B4galt1 | 14595 | HITS-CLIP | | 31433990 | Unspecified |
| mmu-miR-139-3p | MIMAT0004662 | Canx | 12330 | HITS-CLIP | | 31433990 | Unspecified |
| mmu-miR-139-3p | MIMAT0004662 | Dnajb5 | 56323 | HITS-CLIP | | 31433990 | Unspecified |
| mmu-miR-139-3p | MIMAT0004662 | H4c14 | 8359 | HITS-CLIP | | 31433990 | Unspecified |
| mmu-miR-139-3p | MIMAT0004662 | Pgm2l1 | 70974 | HITS-CLIP | | 31433990 | Unspecified |
| mmu-miR-139-3p | MIMAT0004662 | Pofut2 | 80294 | HITS-CLIP | | 31433990 | Unspecified |
| mmu-miR-139-3p | MIMAT0004662 | Stx6 | 58244 | HITS-CLIP | | 31433990 | Unspecified |
| mmu-miR-139-3p | MIMAT0004662 | Zfp598 | 213753 | HITS-CLIP | | 26203562 | Unspecified |
| mmu-miR-152-3p | MIMAT0000162 | 1190002N15Rik | 68861 | HITS-CLIP | | 23597149 | Unspecified |
| mmu-miR-152-3p | MIMAT0000162 | 1810013L24Rik | 69053 | HITS-CLIP | | 37986224 | Unspecified |
| mmu-miR-152-3p | MIMAT0000162 | 2010111I01Rik | 72061 | HITS-CLIP | 25083871\|29167373 | | Unspecified |
| mmu-miR-152-3p | MIMAT0000162 | 2510009E07Rik | 72190 | HITS-CLIP | 23597149\|29167373 | | Unspecified |
| mmu-miR-152-3p | MIMAT0000162 | 2610008E11Rik | 72128 | HITS-CLIP | | 37986224 | Unspecified |
| mmu-miR-152-3p | MIMAT0000162 | 2610507B11Rik | 72503 | HITS-CLIP | 37986224\|29167373 | | Unspecified |
| mmu-miR-152-3p | MIMAT0000162 | 2700081O15Rik | 108899 | HITS-CLIP | 37986224\|25083871 | | Unspecified |
| mmu-miR-152-3p | MIMAT0000162 | 2900026A02Rik | 243219 | HITS-CLIP | 37986224\|23597149 | | Unspecified |
| mmu-miR-152-3p | MIMAT0000162 | 4833420G17Rik | 67392 | HITS-CLIP | | 23597149 | Unspecified |
| mmu-miR-152-3p | MIMAT0000162 | 4921524J17Rik | 66714 | HITS-CLIP | | 25083871 | Unspecified |

| | | | | | | |
|---|---|---|---|---|---|---|
| mmu-miR-152-3p | MIMAT0000162 | 4930402H24Rik | 228602 | HITS-CLIP | 23597149\|29167373 | Unspecified |
| mmu-miR-152-3p | MIMAT0000162 | 4931406C07Rik | 70984 | HITS-CLIP | 23597149 | Unspecified |
| mmu-miR-152-3p | MIMAT0000162 | 4932438A13Rik | 229227 | HITS-CLIP | 25083871 | Unspecified |
| mmu-miR-152-3p | MIMAT0000162 | 5730480H06Rik | 70592 | HITS-CLIP | 25083871 | Unspecified |
| mmu-miR-152-3p | MIMAT0000162 | 6030458C11Rik | 77877 | HITS-CLIP | 37986224\|25083871 | Unspecified |
| mmu-miR-152-3p | MIMAT0000162 | 8030462N17Rik | 212163 | HITS-CLIP | 25083871 | Unspecified |
| mmu-miR-152-3p | MIMAT0000162 | 9930021J03Rik | 240613 | HITS-CLIP | 25083871 | Unspecified |
| mmu-miR-152-3p | MIMAT0000162 | 9930111J21Rik2 | 245240 | HITS-CLIP | 29167373 | Unspecified |
| mmu-miR-152-3p | MIMAT0000162 | A1cf | 69865 | HITS-CLIP | 37986224\|23597149 | Unspecified |
| mmu-miR-152-3p | MIMAT0000162 | A930033H14Rik | 320700 | HITS-CLIP | 37986224 | Unspecified |
| mmu-miR-152-3p | MIMAT0000162 | AI467606 | 101602 | HITS-CLIP | 25083871 | Unspecified |
| mmu-miR-152-3p | MIMAT0000162 | AI606181 | 226118 | HITS-CLIP | 37986224 | Unspecified |
| mmu-miR-152-3p | MIMAT0000162 | AI837181 | 107242 | HITS-CLIP | 23597149 | Unspecified |
| mmu-miR-152-3p | MIMAT0000162 | AI987944 | 233168 | HITS-CLIP | 37986224\|25083871 | Unspecified |
| mmu-miR-152-3p | MIMAT0000162 | AU040320 | 100317 | HITS-CLIP | 25083871\|29167373 | Unspecified |
| mmu-miR-152-3p | MIMAT0000162 | AW549877 | 106064 | HITS-CLIP | 29167373 | Unspecified |
| mmu-miR-152-3p | MIMAT0000162 | Aacs | 78894 | HITS-CLIP | 23597149 | Unspecified |
| mmu-miR-152-3p | MIMAT0000162 | Aak1 | 269774 | HITS-CLIP | 25083871\|23597149 | Unspecified |
| mmu-miR-152-3p | MIMAT0000162 | Abat | 268860 | HITS-CLIP | 23597149 | Unspecified |
| mmu-miR-152-3p | MIMAT0000162 | Abca1 | 11303 | HITS-CLIP | 23597149 | Unspecified |
| mmu-miR-152-3p | MIMAT0000162 | Abcb11 | 27413 | HITS-CLIP | 37986224\|23597149 | Unspecified |
| mmu-miR-152-3p | MIMAT0000162 | Abcb6 | 74104 | HITS-CLIP | 23597149 | Unspecified |
| mmu-miR-152-3p | MIMAT0000162 | Abcb7 | 11306 | HITS-CLIP | 37986224\|25083871\|23597149\|29167 | Unspecified |
| mmu-miR-152-3p | MIMAT0000162 | Abcc1 | 17250 | HITS-CLIP | 29167373 | Unspecified |
| mmu-miR-152-3p | MIMAT0000162 | Abcc2 | 12780 | HITS-CLIP | 23597149 | Unspecified |
| mmu-miR-152-3p | MIMAT0000162 | Abcc5 | 27416 | HITS-CLIP | 37986224\|25083871\|29167373 | Unspecified |
| mmu-miR-152-3p | MIMAT0000162 | Abce1 | 24015 | HITS-CLIP | 25083871 | Unspecified |
| mmu-miR-152-3p | MIMAT0000162 | Abhd13 | 68904 | HITS-CLIP | 25083871 | Unspecified |
| mmu-miR-152-3p | MIMAT0000162 | Abhd17b | 226016 | HITS-CLIP | 25083871\|23597149 | Unspecified |
| mmu-miR-152-3p | MIMAT0000162 | Abhd3 | 106861 | HITS-CLIP | 25083871 | Unspecified |
| mmu-miR-152-3p | MIMAT0000162 | Abi1 | 11308 | HITS-CLIP | 25083871 | Unspecified |
| mmu-miR-152-3p | MIMAT0000162 | Abi2 | 329165 | HITS-CLIP | 37986224\|25083871 | Unspecified |
| mmu-miR-152-3p | MIMAT0000162 | Abl2 | 11352 | HITS-CLIP | 25083871\|23597149\|29167373 | Unspecified |
| mmu-miR-152-3p | MIMAT0000162 | Abraxas2 | 23172 | HITS-CLIP | 25083871\|23597149 | Unspecified |
| mmu-miR-152-3p | MIMAT0000162 | Acaa2 | 52538 | HITS-CLIP | 37986224\|23597149 | Unspecified |
| mmu-miR-152-3p | MIMAT0000162 | Acad10 | 71985 | HITS-CLIP | 37986224 | Unspecified |
| mmu-miR-152-3p | MIMAT0000162 | Acap2 | 78618 | HITS-CLIP | 37986224 | Unspecified |
| mmu-miR-152-3p | MIMAT0000162 | Ackr3 | 12778 | HITS-CLIP | 37986224 | Unspecified |
| mmu-miR-152-3p | MIMAT0000162 | Aco1 | 11428 | HITS-CLIP | 25083871 | Unspecified |

| | | | | | | |
|---|---|---|---|---|---|---|
| mmu-miR-152-3p | MIMAT0000162 | Aco2 | 11429 | HITS-CLIP | 37986224\|23597149\|25083871 | Unspecified |
| mmu-miR-152-3p | MIMAT0000162 | Acod1 | 16365 | HITS-CLIP | 29167373 | Unspecified |
| mmu-miR-152-3p | MIMAT0000162 | Acot8 | 170789 | HITS-CLIP | 37986224\|23597149 | Unspecified |
| mmu-miR-152-3p | MIMAT0000162 | Acox1 | 11430 | HITS-CLIP | 37986224 | Unspecified |
| mmu-miR-152-3p | MIMAT0000162 | Acox2 | 93732 | HITS-CLIP | 23597149 | Unspecified |
| mmu-miR-152-3p | MIMAT0000162 | Acsl1 | 14081 | HITS-CLIP | 25083871 | Unspecified |
| mmu-miR-152-3p | MIMAT0000162 | Acsl4 | 50790 | HITS-CLIP | 37986224\|25083871\|23597149\|29167 | Unspecified |
| mmu-miR-152-3p | MIMAT0000162 | Acsm5 | 272428 | HITS-CLIP | 23597149 | Unspecified |
| mmu-miR-152-3p | MIMAT0000162 | Actb | 11461 | HITS-CLIP | 37986224 | Unspecified |
| mmu-miR-152-3p | MIMAT0000162 | Actn4 | 60595 | HITS-CLIP | 25083871 | Unspecified |
| mmu-miR-152-3p | MIMAT0000162 | Acvr1 | 11477 | HITS-CLIP | 25083871 | Unspecified |
| mmu-miR-152-3p | MIMAT0000162 | Acvr2a | 11480 | HITS-CLIP | 25083871\|29167373 | Unspecified |
| mmu-miR-152-3p | MIMAT0000162 | Adam10 | 11487 | HITS-CLIP | 37986224\|25083871\|23597149\|29167 | Unspecified |
| mmu-miR-152-3p | MIMAT0000162 | Adam12 | 11489 | HITS-CLIP | 25083871 | Unspecified |
| mmu-miR-152-3p | MIMAT0000162 | Adam17 | 11491 | HITS-CLIP | 37986224 | Unspecified |
| mmu-miR-152-3p | MIMAT0000162 | Adam19 | 11492 | HITS-CLIP | 25083871 | Unspecified |
| mmu-miR-152-3p | MIMAT0000162 | Adamts1 | 11504 | HITS-CLIP | 37986224 | Unspecified |
| mmu-miR-152-3p | MIMAT0000162 | Adamts5 | 23794 | HITS-CLIP | 25083871 | Unspecified |
| mmu-miR-152-3p | MIMAT0000162 | Adamts6 | 108154 | HITS-CLIP | 25083871 | Unspecified |
| mmu-miR-152-3p | MIMAT0000162 | Adarb1 | 110532 | HITS-CLIP | 25083871 | Unspecified |
| mmu-miR-152-3p | MIMAT0000162 | Adck5 | 268822 | HITS-CLIP | 23597149 | Unspecified |
| mmu-miR-152-3p | MIMAT0000162 | Adcy7 | 11513 | HITS-CLIP | 37986224\|25083871\|29167373 | Unspecified |
| mmu-miR-152-3p | MIMAT0000162 | Adcy9 | 11515 | HITS-CLIP | 25083871 | Unspecified |
| mmu-miR-152-3p | MIMAT0000162 | Add1 | 11518 | HITS-CLIP | 37986224\|25083871 | Unspecified |
| mmu-miR-152-3p | MIMAT0000162 | Adgre1 | 13733 | HITS-CLIP | 29167373 | Unspecified |
| mmu-miR-152-3p | MIMAT0000162 | Adgrg1 | 9289 | HITS-CLIP | 25083871 | Unspecified |
| mmu-miR-152-3p | MIMAT0000162 | Adgrl2 | 23266 | HITS-CLIP | 25083871 | Unspecified |
| mmu-miR-152-3p | MIMAT0000162 | Adh4 | 26876 | HITS-CLIP | 23597149 | Unspecified |
| mmu-miR-152-3p | MIMAT0000162 | Adh5 | 11532 | HITS-CLIP | 23597149\|25083871 | Unspecified |
| mmu-miR-152-3p | MIMAT0000162 | Adipor2 | 68465 | HITS-CLIP | 23597149 | Unspecified |
| mmu-miR-152-3p | MIMAT0000162 | Adk | 11534 | HITS-CLIP | 23597149 | Unspecified |
| mmu-miR-152-3p | MIMAT0000162 | Adnp | 11538 | HITS-CLIP | 37986224 | Unspecified |
| mmu-miR-152-3p | MIMAT0000162 | Adra1a | 148 | HITS-CLIP | 29167373 | Unspecified |
| mmu-miR-152-3p | MIMAT0000162 | Adrb2 | 11555 | HITS-CLIP | 37986224\|25083871\|23597149 | Unspecified |
| mmu-miR-152-3p | MIMAT0000162 | Adsl | 11564 | HITS-CLIP | 23597149 | Unspecified |
| mmu-miR-152-3p | MIMAT0000162 | Aebp2 | 11569 | HITS-CLIP | 37986224 | Unspecified |
| mmu-miR-152-3p | MIMAT0000162 | Afap1l1 | 106877 | HITS-CLIP | 37986224 | Unspecified |
| mmu-miR-152-3p | MIMAT0000162 | Afdn | 4301 | HITS-CLIP | 29167373 | Unspecified |
| mmu-miR-152-3p | MIMAT0000162 | Aff1 | 17355 | HITS-CLIP | 37986224\|23597149 | Unspecified |

| | | | | | | |
|---|---|---|---|---|---|---|
| mmu-miR-152-3p | MIMAT0000162 | Aff4 | 93736 | HITS-CLIP | 25083871\|23597149\|29167373 | Unspecified |
| mmu-miR-152-3p | MIMAT0000162 | Afg3l2 | 69597 | HITS-CLIP | 37986224 | Unspecified |
| mmu-miR-152-3p | MIMAT0000162 | Aftph | 216549 | HITS-CLIP | 25083871 | Unspecified |
| mmu-miR-152-3p | MIMAT0000162 | Agfg2 | 231801 | HITS-CLIP | 23597149 | Unspecified |
| mmu-miR-152-3p | MIMAT0000162 | Ago1 | 236511 | HITS-CLIP | 37986224\|25083871\|23597149\|29167 | Unspecified |
| mmu-miR-152-3p | MIMAT0000162 | Ago2 | 239528 | HITS-CLIP | 37986224\|25083871\|29167373 | Unspecified |
| mmu-miR-152-3p | MIMAT0000162 | Ago3 | 214150 | HITS-CLIP | 37986224\|25083871 | Unspecified |
| mmu-miR-152-3p | MIMAT0000162 | Ago4 | 76850 | HITS-CLIP | 37986224 | Unspecified |
| mmu-miR-152-3p | MIMAT0000162 | Agt | 11606 | HITS-CLIP | 23597149 | Unspecified |
| mmu-miR-152-3p | MIMAT0000162 | Ahcyl1 | 229709 | HITS-CLIP | 23597149\|25083871 | Unspecified |
| mmu-miR-152-3p | MIMAT0000162 | Ahdc1 | 230793 | HITS-CLIP | 37986224\|25083871 | Unspecified |
| mmu-miR-152-3p | MIMAT0000162 | Ahnak | 66395 | HITS-CLIP | 29167373 | Unspecified |
| mmu-miR-152-3p | MIMAT0000162 | Ajuba | 16475 | HITS-CLIP | 37986224 | Unspecified |
| mmu-miR-152-3p | MIMAT0000162 | Ak2 | 11637 | HITS-CLIP | 23597149 | Unspecified |
| mmu-miR-152-3p | MIMAT0000162 | Akap1 | 11640 | HITS-CLIP | 25083871 | Unspecified |
| mmu-miR-152-3p | MIMAT0000162 | Akap10 | 56697 | HITS-CLIP | 25083871 | Unspecified |
| mmu-miR-152-3p | MIMAT0000162 | Akap11 | 219181 | HITS-CLIP | 25083871 | Unspecified |
| mmu-miR-152-3p | MIMAT0000162 | Akap13 | 75547 | HITS-CLIP | 37986224\|29167373\|25083871 | Unspecified |
| mmu-miR-152-3p | MIMAT0000162 | Akap6 | 238161 | HITS-CLIP | 37986224\|25083871 | Unspecified |
| mmu-miR-152-3p | MIMAT0000162 | Akirin2 | 433693 | HITS-CLIP | 25083871 | Unspecified |
| mmu-miR-152-3p | MIMAT0000162 | Akr1a1 | 58810 | HITS-CLIP | 25083871 | Unspecified |
| mmu-miR-152-3p | MIMAT0000162 | Akr1c6 | 83702 | HITS-CLIP | 23597149 | Unspecified |
| mmu-miR-152-3p | MIMAT0000162 | Akr7a5 | 110198 | HITS-CLIP | 23597149 | Unspecified |
| mmu-miR-152-3p | MIMAT0000162 | Akt2 | 11652 | HITS-CLIP | 29167373 | Unspecified |
| mmu-miR-152-3p | MIMAT0000162 | Aldh1a1 | 11668 | HITS-CLIP | 23597149 | Unspecified |
| mmu-miR-152-3p | MIMAT0000162 | Aldh3a2 | 11671 | HITS-CLIP | 37986224\|23597149 | Unspecified |
| mmu-miR-152-3p | MIMAT0000162 | Aldh4a1 | 212647 | HITS-CLIP | 37986224 | Unspecified |
| mmu-miR-152-3p | MIMAT0000162 | Alg11 | 207958 | HITS-CLIP | 37986224 | Unspecified |
| mmu-miR-152-3p | MIMAT0000162 | Alg2 | 56737 | HITS-CLIP | 37986224 | Unspecified |
| mmu-miR-152-3p | MIMAT0000162 | Alg6 | 320438 | HITS-CLIP | 25083871 | Unspecified |
| mmu-miR-152-3p | MIMAT0000162 | Alkbh5 | 268420 | HITS-CLIP | 25083871 | Unspecified |
| mmu-miR-152-3p | MIMAT0000162 | Alox5 | 11689 | HITS-CLIP | 25083871 | Unspecified |
| mmu-miR-152-3p | MIMAT0000162 | Alpk3 | 116904 | HITS-CLIP | 25083871 | Unspecified |
| mmu-miR-152-3p | MIMAT0000162 | Amer1 | 72345 | HITS-CLIP | 37986224 | Unspecified |
| mmu-miR-152-3p | MIMAT0000162 | Ammecr1 | 56068 | HITS-CLIP | 37986224 | Unspecified |
| mmu-miR-152-3p | MIMAT0000162 | Amn1 | 232566 | HITS-CLIP | 25083871 | Unspecified |
| mmu-miR-152-3p | MIMAT0000162 | Amot | 27494 | HITS-CLIP | 37986224 | Unspecified |
| mmu-miR-152-3p | MIMAT0000162 | Amotl1 | 75723 | HITS-CLIP | 25083871 | Unspecified |
| mmu-miR-152-3p | MIMAT0000162 | Anapc1 | 17222 | HITS-CLIP | 37986224\|25083871 | Unspecified |

| | | | | | | |
|---|---|---|---|---|---|---|
| mmu-miR-152-3p | MIMAT0000162 | Anapc11 | 66156 | HITS-CLIP | 25083871 | Unspecified |
| mmu-miR-152-3p | MIMAT0000162 | Anapc2 | 99152 | HITS-CLIP | 25083871 | Unspecified |
| mmu-miR-152-3p | MIMAT0000162 | Anapc5 | 59008 | HITS-CLIP | 37986224\|23597149 | Unspecified |
| mmu-miR-152-3p | MIMAT0000162 | Angptl2 | 26360 | HITS-CLIP | 25083871\|29167373 | Unspecified |
| mmu-miR-152-3p | MIMAT0000162 | Angptl4 | 57875 | HITS-CLIP | 37986224 | Unspecified |
| mmu-miR-152-3p | MIMAT0000162 | Ankfy1 | 11736 | HITS-CLIP | 25083871\|23597149\|29167373 | Unspecified |
| mmu-miR-152-3p | MIMAT0000162 | Ankhd1 | 108857 | HITS-CLIP | 25083871 | Unspecified |
| mmu-miR-152-3p | MIMAT0000162 | Ankib1 | 70797 | HITS-CLIP | 37986224\|25083871 | Unspecified |
| mmu-miR-152-3p | MIMAT0000162 | Ankrd10 | 102334 | HITS-CLIP | 37986224\|25083871 | Unspecified |
| mmu-miR-152-3p | MIMAT0000162 | Ankrd11 | 77087 | HITS-CLIP | 37986224\|29167373 | Unspecified |
| mmu-miR-152-3p | MIMAT0000162 | Ankrd12 | 106585 | HITS-CLIP | 25083871 | Unspecified |
| mmu-miR-152-3p | MIMAT0000162 | Ankrd17 | 81702 | HITS-CLIP | 25083871\|29167373 | Unspecified |
| mmu-miR-152-3p | MIMAT0000162 | Ankrd33b | 67434 | HITS-CLIP | 37986224 | Unspecified |
| mmu-miR-152-3p | MIMAT0000162 | Ankrd40 | 71452 | HITS-CLIP | 37986224\|29167373 | Unspecified |
| mmu-miR-152-3p | MIMAT0000162 | Ankrd44 | 329154 | HITS-CLIP | 25083871\|23597149 | Unspecified |
| mmu-miR-152-3p | MIMAT0000162 | Ankrd50 | 99696 | HITS-CLIP | 37986224 | Unspecified |
| mmu-miR-152-3p | MIMAT0000162 | Ankrd52 | 237615 | HITS-CLIP | 37986224\|23597149\|25083871\|29167 | Unspecified |
| mmu-miR-152-3p | MIMAT0000162 | Anpep | 16790 | HITS-CLIP | 29167373 | Unspecified |
| mmu-miR-152-3p | MIMAT0000162 | Antxr2 | 71914 | HITS-CLIP | 25083871\|29167373 | Unspecified |
| mmu-miR-152-3p | MIMAT0000162 | Anxa2 | 12306 | HITS-CLIP | 37986224 | Unspecified |
| mmu-miR-152-3p | MIMAT0000162 | Aox3 | 71724 | HITS-CLIP | 23597149 | Unspecified |
| mmu-miR-152-3p | MIMAT0000162 | Ap1b1 | 11764 | HITS-CLIP | 37986224\|23597149 | Unspecified |
| mmu-miR-152-3p | MIMAT0000162 | Ap1g1 | 11765 | HITS-CLIP | 37986224\|25083871\|23597149 | Unspecified |
| mmu-miR-152-3p | MIMAT0000162 | Ap2a2 | 11772 | HITS-CLIP | 37986224 | Unspecified |
| mmu-miR-152-3p | MIMAT0000162 | Ap3b1 | 11774 | HITS-CLIP | 23597149\|25083871\|29167373 | Unspecified |
| mmu-miR-152-3p | MIMAT0000162 | Ap3d1 | 11776 | HITS-CLIP | 25083871\|29167373 | Unspecified |
| mmu-miR-152-3p | MIMAT0000162 | Ap3s2 | 11778 | HITS-CLIP | 25083871\|23597149 | Unspecified |
| mmu-miR-152-3p | MIMAT0000162 | Apbb2 | 11787 | HITS-CLIP | 29167373 | Unspecified |
| mmu-miR-152-3p | MIMAT0000162 | Apc | 11789 | HITS-CLIP | 25083871 | Unspecified |
| mmu-miR-152-3p | MIMAT0000162 | Aph1a | 226548 | HITS-CLIP | 25083871 | Unspecified |
| mmu-miR-152-3p | MIMAT0000162 | Api5 | 11800 | HITS-CLIP | 25083871 | Unspecified |
| mmu-miR-152-3p | MIMAT0000162 | Apip | 56369 | HITS-CLIP | 23597149 | Unspecified |
| mmu-miR-152-3p | MIMAT0000162 | Aplp2 | 11804 | HITS-CLIP | 37986224\|23597149 | Unspecified |
| mmu-miR-152-3p | MIMAT0000162 | Apob | 238055 | HITS-CLIP | 37986224\|23597149 | Unspecified |
| mmu-miR-152-3p | MIMAT0000162 | Apobec2 | 11811 | HITS-CLIP | 25083871 | Unspecified |
| mmu-miR-152-3p | MIMAT0000162 | App | 11820 | HITS-CLIP | 25083871 | Unspecified |
| mmu-miR-152-3p | MIMAT0000162 | Appl1 | 72993 | HITS-CLIP | 37986224\|25083871\|31433990 | Unspecified |
| mmu-miR-152-3p | MIMAT0000162 | Appl2 | 216190 | HITS-CLIP | 29167373 | Unspecified |
| mmu-miR-152-3p | MIMAT0000162 | Aqp9 | 64008 | HITS-CLIP | 23597149 | Unspecified |

| | | | | | | |
|---|---|---|---|---|---|---|
| mmu-miR-152-3p | MIMAT0000162 | Ar | 11835 | HITS-CLIP | 25083871 | Unspecified |
| mmu-miR-152-3p | MIMAT0000162 | Arcn1 | 213827 | HITS-CLIP | 25083871 | Unspecified |
| mmu-miR-152-3p | MIMAT0000162 | Arel1 | 68497 | HITS-CLIP | 37986224 | Unspecified |
| mmu-miR-152-3p | MIMAT0000162 | Arf1 | 11840 | HITS-CLIP | 25083871\|23597149 | Unspecified |
| mmu-miR-152-3p | MIMAT0000162 | Arf3 | 11842 | HITS-CLIP | 37986224\|25083871 | Unspecified |
| mmu-miR-152-3p | MIMAT0000162 | Arf4 | 11843 | HITS-CLIP | 23597149\|25083871 | Unspecified |
| mmu-miR-152-3p | MIMAT0000162 | Arf6 | 11845 | HITS-CLIP | 37986224 | Unspecified |
| mmu-miR-152-3p | MIMAT0000162 | Arfgef1 | 211673 | HITS-CLIP | 25083871\|23597149\|29167373 | Unspecified |
| mmu-miR-152-3p | MIMAT0000162 | Arfgef2 | 99371 | HITS-CLIP | 37986224 | Unspecified |
| mmu-miR-152-3p | MIMAT0000162 | Arfip1 | 99889 | HITS-CLIP | 37986224 | Unspecified |
| mmu-miR-152-3p | MIMAT0000162 | Arg1 | 11846 | HITS-CLIP | 23597149 | Unspecified |
| mmu-miR-152-3p | MIMAT0000162 | Arhgap11a | 228482 | HITS-CLIP | 25083871 | Unspecified |
| mmu-miR-152-3p | MIMAT0000162 | Arhgap23 | 58996 | HITS-CLIP | 37986224 | Unspecified |
| mmu-miR-152-3p | MIMAT0000162 | Arhgap24 | 231532 | HITS-CLIP | 25083871 | Unspecified |
| mmu-miR-152-3p | MIMAT0000162 | Arhgap28 | 268970 | HITS-CLIP | 37986224 | Unspecified |
| mmu-miR-152-3p | MIMAT0000162 | Arhgap29 | 214137 | HITS-CLIP | 37986224\|25083871\|23597149 | Unspecified |
| mmu-miR-152-3p | MIMAT0000162 | Arhgap32 | 330914 | HITS-CLIP | 23597149 | Unspecified |
| mmu-miR-152-3p | MIMAT0000162 | Arhgap42 | 71544 | HITS-CLIP | 37986224 | Unspecified |
| mmu-miR-152-3p | MIMAT0000162 | Arhgap5 | 11855 | HITS-CLIP | 23597149\|25083871 | Unspecified |
| mmu-miR-152-3p | MIMAT0000162 | Arhgef1 | 16801 | HITS-CLIP | 23597149 | Unspecified |
| mmu-miR-152-3p | MIMAT0000162 | Arhgef11 | 213498 | HITS-CLIP | 37986224 | Unspecified |
| mmu-miR-152-3p | MIMAT0000162 | Arhgef12 | 69632 | HITS-CLIP | 37986224 | Unspecified |
| mmu-miR-152-3p | MIMAT0000162 | Arhgef17 | 207212 | HITS-CLIP | 37986224 | Unspecified |
| mmu-miR-152-3p | MIMAT0000162 | Arhgef7 | 54126 | HITS-CLIP | 37986224\|25083871\|29167373 | Unspecified |
| mmu-miR-152-3p | MIMAT0000162 | Arhgef9 | 236915 | HITS-CLIP | 37986224 | Unspecified |
| mmu-miR-152-3p | MIMAT0000162 | Arid1a | 93760 | HITS-CLIP | 37986224\|25083871\|23597149 | Unspecified |
| mmu-miR-152-3p | MIMAT0000162 | Arid1b | 239985 | HITS-CLIP | 25083871 | Unspecified |
| mmu-miR-152-3p | MIMAT0000162 | Arid2 | 77044 | HITS-CLIP | 37986224\|25083871 | Unspecified |
| mmu-miR-152-3p | MIMAT0000162 | Arid4a | 238247 | HITS-CLIP | 25083871 | Unspecified |
| mmu-miR-152-3p | MIMAT0000162 | Arid5b | 71371 | HITS-CLIP | 23597149 | Unspecified |
| mmu-miR-152-3p | MIMAT0000162 | Arl15 | 218639 | HITS-CLIP | 25083871 | Unspecified |
| mmu-miR-152-3p | MIMAT0000162 | Arl6ip1 | 54208 | HITS-CLIP | 37986224 | Unspecified |
| mmu-miR-152-3p | MIMAT0000162 | Arl8b | 67166 | HITS-CLIP | 37986224\|25083871 | Unspecified |
| mmu-miR-152-3p | MIMAT0000162 | Armc8 | 74125 | HITS-CLIP | 29167373 | Unspecified |
| mmu-miR-152-3p | MIMAT0000162 | Armcx4 | 100503043 | HITS-CLIP | 37986224 | Unspecified |
| mmu-miR-152-3p | MIMAT0000162 | Arpc1a | 56443 | HITS-CLIP | 23597149 | Unspecified |
| mmu-miR-152-3p | MIMAT0000162 | Arpc2 | 76709 | HITS-CLIP | 25083871 | Unspecified |
| mmu-miR-152-3p | MIMAT0000162 | Arpc4 | 68089 | HITS-CLIP | 37986224 | Unspecified |
| mmu-miR-152-3p | MIMAT0000162 | Arpp19 | 59046 | HITS-CLIP | 37986224 | Unspecified |

| | | | | | | |
|---|---|---|---|---|---|---|
| mmu-miR-152-3p | MIMAT0000162 | Arrb1 | 109689 | HITS-CLIP | 29167373 | Unspecified |
| mmu-miR-152-3p | MIMAT0000162 | Arrb2 | 216869 | HITS-CLIP | 29167373 | Unspecified |
| mmu-miR-152-3p | MIMAT0000162 | Arsa | 11883 | HITS-CLIP | 23597149 | Unspecified |
| mmu-miR-152-3p | MIMAT0000162 | Asap1 | 13196 | HITS-CLIP | 25083871\|23597149 | Unspecified |
| mmu-miR-152-3p | MIMAT0000162 | Ascc3 | 77987 | HITS-CLIP | 25083871 | Unspecified |
| mmu-miR-152-3p | MIMAT0000162 | Ash1l | 192195 | HITS-CLIP | 25083871\|23597149 | Unspecified |
| mmu-miR-152-3p | MIMAT0000162 | Asph | 65973 | HITS-CLIP | 37986224\|25083871\|23597149\|29167 | Unspecified |
| mmu-miR-152-3p | MIMAT0000162 | Asxl1 | 228790 | HITS-CLIP | 29167373 | Unspecified |
| mmu-miR-152-3p | MIMAT0000162 | Asxl2 | 75302 | HITS-CLIP | 25083871 | Unspecified |
| mmu-miR-152-3p | MIMAT0000162 | Atad2 | 70472 | HITS-CLIP | 25083871 | Unspecified |
| mmu-miR-152-3p | MIMAT0000162 | Atad2b | 320817 | HITS-CLIP | 37986224\|25083871 | Unspecified |
| mmu-miR-152-3p | MIMAT0000162 | Atf2 | 11909 | HITS-CLIP | 25083871\|23597149 | Unspecified |
| mmu-miR-152-3p | MIMAT0000162 | Atf7 | 223922 | HITS-CLIP | 25083871 | Unspecified |
| mmu-miR-152-3p | MIMAT0000162 | Atg101 | 68118 | HITS-CLIP | 23597149 | Unspecified |
| mmu-miR-152-3p | MIMAT0000162 | Atg12 | 67526 | HITS-CLIP | 37986224 | Unspecified |
| mmu-miR-152-3p | MIMAT0000162 | Atg13 | 51897 | HITS-CLIP | 23597149 | Unspecified |
| mmu-miR-152-3p | MIMAT0000162 | Atg14 | 100504663 | HITS-CLIP | 37986224 | Unspecified |
| mmu-miR-152-3p | MIMAT0000162 | Atg5 | 11793 | HITS-CLIP | 23597149 | Unspecified |
| mmu-miR-152-3p | MIMAT0000162 | Atl2 | 56298 | HITS-CLIP | 37986224 | Unspecified |
| mmu-miR-152-3p | MIMAT0000162 | Atl3 | 109168 | HITS-CLIP | 29167373 | Unspecified |
| mmu-miR-152-3p | MIMAT0000162 | Atp11a | 50770 | HITS-CLIP | 25083871 | Unspecified |
| mmu-miR-152-3p | MIMAT0000162 | Atp11b | 76295 | HITS-CLIP | 25083871 | Unspecified |
| mmu-miR-152-3p | MIMAT0000162 | Atp11c | 320940 | HITS-CLIP | 23597149 | Unspecified |
| mmu-miR-152-3p | MIMAT0000162 | Atp1b1 | 11931 | HITS-CLIP | 25083871 | Unspecified |
| mmu-miR-152-3p | MIMAT0000162 | Atp1b3 | 11933 | HITS-CLIP | 25083871 | Unspecified |
| mmu-miR-152-3p | MIMAT0000162 | Atp2a1 | 11937 | HITS-CLIP | 25083871 | Unspecified |
| mmu-miR-152-3p | MIMAT0000162 | Atp2a2 | 11938 | HITS-CLIP | 37986224\|25083871\|23597149 | Unspecified |
| mmu-miR-152-3p | MIMAT0000162 | Atp2b1 | 67972 | HITS-CLIP | 25083871 | Unspecified |
| mmu-miR-152-3p | MIMAT0000162 | Atp2b4 | 381290 | HITS-CLIP | 37986224 | Unspecified |
| mmu-miR-152-3p | MIMAT0000162 | Atp2c1 | 235574 | HITS-CLIP | 29167373 | Unspecified |
| mmu-miR-152-3p | MIMAT0000162 | Atp5b | 11947 | HITS-CLIP | 37986224 | Unspecified |
| mmu-miR-152-3p | MIMAT0000162 | Atp6ap1 | 54411 | HITS-CLIP | 37986224 | Unspecified |
| mmu-miR-152-3p | MIMAT0000162 | Atp6ap2 | 70495 | HITS-CLIP | 25083871\|23597149 | Unspecified |
| mmu-miR-152-3p | MIMAT0000162 | Atp6v0a2 | 21871 | HITS-CLIP | 25083871\|23597149 | Unspecified |
| mmu-miR-152-3p | MIMAT0000162 | Atp6v1b2 | 11966 | HITS-CLIP | 25083871 | Unspecified |
| mmu-miR-152-3p | MIMAT0000162 | Atp6v1c1 | 66335 | HITS-CLIP | 25083871\|29167373 | Unspecified |
| mmu-miR-152-3p | MIMAT0000162 | Atp6v1h | 108664 | HITS-CLIP | 25083871 | Unspecified |
| mmu-miR-152-3p | MIMAT0000162 | Atp7a | 11977 | HITS-CLIP | 37986224\|25083871\|29167373 | Unspecified |
| mmu-miR-152-3p | MIMAT0000162 | Atp8b1 | 54670 | HITS-CLIP | 23597149 | Unspecified |

| | | | | | | |
|---|---|---|---|---|---|---|
| mmu-miR-152-3p | MIMAT0000162 | Atp8b2 | 54667 | HITS-CLIP | 37986224 | Unspecified |
| mmu-miR-152-3p | MIMAT0000162 | Atp9b | 50771 | HITS-CLIP | 23597149 | Unspecified |
| mmu-miR-152-3p | MIMAT0000162 | Atrx | 22589 | HITS-CLIP | 37986224\|25083871 | Unspecified |
| mmu-miR-152-3p | MIMAT0000162 | Atxn1 | 20238 | HITS-CLIP | 37986224\|25083871 | Unspecified |
| mmu-miR-152-3p | MIMAT0000162 | Atxn10 | 54138 | HITS-CLIP | 37986224 | Unspecified |
| mmu-miR-152-3p | MIMAT0000162 | Atxn1l | 52335 | HITS-CLIP | 37986224 | Unspecified |
| mmu-miR-152-3p | MIMAT0000162 | Atxn2 | 20239 | HITS-CLIP | 25083871 | Unspecified |
| mmu-miR-152-3p | MIMAT0000162 | Atxn2l | 233871 | HITS-CLIP | 25083871 | Unspecified |
| mmu-miR-152-3p | MIMAT0000162 | Atxn3 | 110616 | HITS-CLIP | 37986224\|25083871 | Unspecified |
| mmu-miR-152-3p | MIMAT0000162 | Atxn7l1 | 380753 | HITS-CLIP | 29167373 | Unspecified |
| mmu-miR-152-3p | MIMAT0000162 | Avl9 | 78937 | HITS-CLIP | 25083871 | Unspecified |
| mmu-miR-152-3p | MIMAT0000162 | Axin1 | 12005 | HITS-CLIP | 25083871\|29167373 | Unspecified |
| mmu-miR-152-3p | MIMAT0000162 | Axl | 26362 | HITS-CLIP | 37986224\|25083871 | Unspecified |
| mmu-miR-152-3p | MIMAT0000162 | B230219D22Rik | 78521 | HITS-CLIP | 25083871\|21258322\|29167373 | Unspecified |
| mmu-miR-152-3p | MIMAT0000162 | B3galnt1 | 26879 | HITS-CLIP | 37986224 | Unspecified |
| mmu-miR-152-3p | MIMAT0000162 | B3galt1 | 26877 | HITS-CLIP | 23597149 | Unspecified |
| mmu-miR-152-3p | MIMAT0000162 | B4galt1 | 14595 | HITS-CLIP | 37986224 | Unspecified |
| mmu-miR-152-3p | MIMAT0000162 | B4galt5 | 56336 | HITS-CLIP | 37986224\|25083871\|23597149\|29167 | Unspecified |
| mmu-miR-152-3p | MIMAT0000162 | BC004004 | 80748 | HITS-CLIP | 29167373\|23597149\|25083871 | Unspecified |
| mmu-miR-152-3p | MIMAT0000162 | BC005537 | 79555 | HITS-CLIP | 23597149\|25083871\|29167373 | Unspecified |
| mmu-miR-152-3p | MIMAT0000162 | BC005561 | 100042165 | HITS-CLIP | 25083871 | Unspecified |
| mmu-miR-152-3p | MIMAT0000162 | Baat | 12012 | HITS-CLIP | 37986224\|23597149 | Unspecified |
| mmu-miR-152-3p | MIMAT0000162 | Bace1 | 23821 | HITS-CLIP | 37986224\|25083871 | Unspecified |
| mmu-miR-152-3p | MIMAT0000162 | Bag4 | 67384 | HITS-CLIP | 37986224 | Unspecified |
| mmu-miR-152-3p | MIMAT0000162 | Bahcc1 | 268515 | HITS-CLIP | 25083871 | Unspecified |
| mmu-miR-152-3p | MIMAT0000162 | Bahd1 | 228536 | HITS-CLIP | 25083871 | Unspecified |
| mmu-miR-152-3p | MIMAT0000162 | Baiap2 | 108100 | HITS-CLIP | 37986224 | Unspecified |
| mmu-miR-152-3p | MIMAT0000162 | Baiap2l1 | 66898 | HITS-CLIP | 37986224 | Unspecified |
| mmu-miR-152-3p | MIMAT0000162 | Bap1 | 104416 | HITS-CLIP | 23597149 | Unspecified |
| mmu-miR-152-3p | MIMAT0000162 | Basp1 | 70350 | HITS-CLIP | 25083871 | Unspecified |
| mmu-miR-152-3p | MIMAT0000162 | Baz1a | 217578 | HITS-CLIP | 29167373 | Unspecified |
| mmu-miR-152-3p | MIMAT0000162 | Baz1b | 22385 | HITS-CLIP | 25083871\|29167373 | Unspecified |
| mmu-miR-152-3p | MIMAT0000162 | Baz2b | 407823 | HITS-CLIP | 25083871 | Unspecified |
| mmu-miR-152-3p | MIMAT0000162 | Bbc3 | 170770 | HITS-CLIP | 37986224\|25083871 | Unspecified |
| mmu-miR-152-3p | MIMAT0000162 | Bbs10 | 71769 | HITS-CLIP | 29167373 | Unspecified |
| mmu-miR-152-3p | MIMAT0000162 | Bbx | 70508 | HITS-CLIP | 29167373 | Unspecified |
| mmu-miR-152-3p | MIMAT0000162 | Bcap29 | 12033 | HITS-CLIP | 37986224 | Unspecified |
| mmu-miR-152-3p | MIMAT0000162 | Bckdhb | 12040 | HITS-CLIP | 37986224 | Unspecified |
| mmu-miR-152-3p | MIMAT0000162 | Bcl2l11 | 12125 | HITS-CLIP | 37986224 | Unspecified |

| | | | | | | |
|---|---|---|---|---|---|---|
| mmu-miR-152-3p | MIMAT0000162 | Bcl6 | 12053 | HITS-CLIP | 25083871\|29167373 | Unspecified |
| mmu-miR-152-3p | MIMAT0000162 | Bcl7a | 77045 | HITS-CLIP | 37986224\|25083871 | Unspecified |
| mmu-miR-152-3p | MIMAT0000162 | Bcl7b | 12054 | HITS-CLIP | 37986224\|25083871 | Unspecified |
| mmu-miR-152-3p | MIMAT0000162 | Bcl9l | 80288 | HITS-CLIP | 25083871 | Unspecified |
| mmu-miR-152-3p | MIMAT0000162 | Bclaf1 | 72567 | HITS-CLIP | 37986224\|25083871\|23597149\|29167 | Unspecified |
| mmu-miR-152-3p | MIMAT0000162 | Bcor | 71458 | HITS-CLIP | 37986224 | Unspecified |
| mmu-miR-152-3p | MIMAT0000162 | Bdnf | 12064 | HITS-CLIP | 25083871 | Unspecified |
| mmu-miR-152-3p | MIMAT0000162 | Bdp1 | 544971 | HITS-CLIP | 37986224\|23597149 | Unspecified |
| mmu-miR-152-3p | MIMAT0000162 | Becn1 | 56208 | HITS-CLIP | 25083871 | Unspecified |
| mmu-miR-152-3p | MIMAT0000162 | Bet1l | 54399 | HITS-CLIP | 37986224 | Unspecified |
| mmu-miR-152-3p | MIMAT0000162 | Bfar | 67118 | HITS-CLIP | 25083871 | Unspecified |
| mmu-miR-152-3p | MIMAT0000162 | Bgn | 12111 | HITS-CLIP | 25083871 | Unspecified |
| mmu-miR-152-3p | MIMAT0000162 | Bhlhe40 | 20893 | HITS-CLIP | 23597149\|25083871 | Unspecified |
| mmu-miR-152-3p | MIMAT0000162 | Bhlhe41 | 79362 | HITS-CLIP | 25083871 | Unspecified |
| mmu-miR-152-3p | MIMAT0000162 | Bicc1 | 83675 | HITS-CLIP | 37986224\|25083871 | Unspecified |
| mmu-miR-152-3p | MIMAT0000162 | Bicd2 | 76895 | HITS-CLIP | 25083871 | Unspecified |
| mmu-miR-152-3p | MIMAT0000162 | Birc6 | 12211 | HITS-CLIP | 25083871\|23597149 | Unspecified |
| mmu-miR-152-3p | MIMAT0000162 | Blvra | 109778 | HITS-CLIP | 37986224 | Unspecified |
| mmu-miR-152-3p | MIMAT0000162 | Bmp1 | 12153 | HITS-CLIP | 25083871 | Unspecified |
| mmu-miR-152-3p | MIMAT0000162 | Bmp2k | 140780 | HITS-CLIP | 37986224\|25083871 | Unspecified |
| mmu-miR-152-3p | MIMAT0000162 | Bmper | 73230 | HITS-CLIP | 37986224 | Unspecified |
| mmu-miR-152-3p | MIMAT0000162 | Bmpr2 | 12168 | HITS-CLIP | 37986224\|25083871\|23597149\|29167 | Unspecified |
| mmu-miR-152-3p | MIMAT0000162 | Bmt2 | 101148 | HITS-CLIP | 25083871\|29167373 | Unspecified |
| mmu-miR-152-3p | MIMAT0000162 | Bnip2 | 12175 | HITS-CLIP | 25083871 | Unspecified |
| mmu-miR-152-3p | MIMAT0000162 | Bod1l | 665775 | HITS-CLIP | 37986224\|25083871 | Unspecified |
| mmu-miR-152-3p | MIMAT0000162 | Bpnt1 | 23827 | HITS-CLIP | 37986224 | Unspecified |
| mmu-miR-152-3p | MIMAT0000162 | Braf | 109880 | HITS-CLIP | 37986224 | Unspecified |
| mmu-miR-152-3p | MIMAT0000162 | Brd4 | 57261 | HITS-CLIP | 37986224 | Unspecified |
| mmu-miR-152-3p | MIMAT0000162 | Brd7 | 26992 | HITS-CLIP | 25083871 | Unspecified |
| mmu-miR-152-3p | MIMAT0000162 | Brinp3 | 215378 | HITS-CLIP | 25083871 | Unspecified |
| mmu-miR-152-3p | MIMAT0000162 | Brms1l | 52592 | HITS-CLIP | 25083871 | Unspecified |
| mmu-miR-152-3p | MIMAT0000162 | Brpf1 | 78783 | HITS-CLIP | 37986224\|25083871\|23597149 | Unspecified |
| mmu-miR-152-3p | MIMAT0000162 | Brwd1 | 93871 | HITS-CLIP | 25083871 | Unspecified |
| mmu-miR-152-3p | MIMAT0000162 | Brwd3 | 382236 | HITS-CLIP | 37986224\|25083871 | Unspecified |
| mmu-miR-152-3p | MIMAT0000162 | Btaf1 | 107182 | HITS-CLIP | 25083871 | Unspecified |
| mmu-miR-152-3p | MIMAT0000162 | Btbd10 | 68815 | HITS-CLIP | 37986224 | Unspecified |
| mmu-miR-152-3p | MIMAT0000162 | Btbd2 | 208198 | HITS-CLIP | 37986224 | Unspecified |
| mmu-miR-152-3p | MIMAT0000162 | Btbd7 | 238386 | HITS-CLIP | 37986224\|23597149\|25083871 | Unspecified |
| mmu-miR-152-3p | MIMAT0000162 | Bzw1 | 66882 | HITS-CLIP | 25083871 | Unspecified |

| | | | | | | |
|---|---|---|---|---|---|---|
| mmu-miR-152-3p | MIMAT0000162 | C1galt1 | 94192 | HITS-CLIP | 25083871\|23597149 | Unspecified |
| mmu-miR-152-3p | MIMAT0000162 | C1galt1c1 | 59048 | HITS-CLIP | 37986224 | Unspecified |
| mmu-miR-152-3p | MIMAT0000162 | C330018D20Rik | 77422 | HITS-CLIP | 37986224 | Unspecified |
| mmu-miR-152-3p | MIMAT0000162 | C5ar2 | 319430 | HITS-CLIP | 29167373 | Unspecified |
| mmu-miR-152-3p | MIMAT0000162 | C6 | 12274 | HITS-CLIP | 37986224 | Unspecified |
| mmu-miR-152-3p | MIMAT0000162 | C8a | 230558 | HITS-CLIP | 37986224 | Unspecified |
| mmu-miR-152-3p | MIMAT0000162 | C8b | 110382 | HITS-CLIP | 37986224 | Unspecified |
| mmu-miR-152-3p | MIMAT0000162 | CT030661.1 | | HITS-CLIP | 25083871 | Unspecified |
| mmu-miR-152-3p | MIMAT0000162 | Cables2 | 252966 | HITS-CLIP | 23597149 | Unspecified |
| mmu-miR-152-3p | MIMAT0000162 | Cacfd1 | 381356 | HITS-CLIP | 25083871 | Unspecified |
| mmu-miR-152-3p | MIMAT0000162 | Cacna2d1 | 12293 | HITS-CLIP | 37986224\|25083871 | Unspecified |
| mmu-miR-152-3p | MIMAT0000162 | Cadm1 | 54725 | HITS-CLIP | 37986224 | Unspecified |
| mmu-miR-152-3p | MIMAT0000162 | Calm1 | 12313 | HITS-CLIP | 25083871 | Unspecified |
| mmu-miR-152-3p | MIMAT0000162 | Calm2 | 12314 | HITS-CLIP | 25083871 | Unspecified |
| mmu-miR-152-3p | MIMAT0000162 | Calr | 12317 | HITS-CLIP | 37986224\|23597149 | Unspecified |
| mmu-miR-152-3p | MIMAT0000162 | Camk1d | 227541 | HITS-CLIP | 29167373 | Unspecified |
| mmu-miR-152-3p | MIMAT0000162 | Camk2d | 108058 | HITS-CLIP | 37986224\|25083871\|29167373 | Unspecified |
| mmu-miR-152-3p | MIMAT0000162 | Camsap2 | 67886 | HITS-CLIP | 37986224\|25083871 | Unspecified |
| mmu-miR-152-3p | MIMAT0000162 | Camsap3 | 69697 | HITS-CLIP | 23597149 | Unspecified |
| mmu-miR-152-3p | MIMAT0000162 | Cand1 | 71902 | HITS-CLIP | 37986224\|23597149\|29167373 | Unspecified |
| mmu-miR-152-3p | MIMAT0000162 | Cant1 | 76025 | HITS-CLIP | 23597149\|25083871 | Unspecified |
| mmu-miR-152-3p | MIMAT0000162 | Canx | 12330 | HITS-CLIP | 37986224 | Unspecified |
| mmu-miR-152-3p | MIMAT0000162 | Cap1 | 12331 | HITS-CLIP | 25083871\|29167373 | Unspecified |
| mmu-miR-152-3p | MIMAT0000162 | Capn15 | 50817 | HITS-CLIP | 25083871 | Unspecified |
| mmu-miR-152-3p | MIMAT0000162 | Capza1 | 12340 | HITS-CLIP | 29167373 | Unspecified |
| mmu-miR-152-3p | MIMAT0000162 | Capza2 | 12343 | HITS-CLIP | 23597149\|25083871 | Unspecified |
| mmu-miR-152-3p | MIMAT0000162 | Car12 | 76459 | HITS-CLIP | 25083871 | Unspecified |
| mmu-miR-152-3p | MIMAT0000162 | Car9 | 230099 | HITS-CLIP | 25083871 | Unspecified |
| mmu-miR-152-3p | MIMAT0000162 | Carm1 | 59035 | HITS-CLIP | 37986224 | Unspecified |
| mmu-miR-152-3p | MIMAT0000162 | Casd1 | 213819 | HITS-CLIP | 25083871\|29167373 | Unspecified |
| mmu-miR-152-3p | MIMAT0000162 | Casp8 | 12370 | HITS-CLIP | 37986224 | Unspecified |
| mmu-miR-152-3p | MIMAT0000162 | Cast | 12380 | HITS-CLIP | 25083871 | Unspecified |
| mmu-miR-152-3p | MIMAT0000162 | Casz1 | 69743 | HITS-CLIP | 37986224 | Unspecified |
| mmu-miR-152-3p | MIMAT0000162 | Cbl | 12402 | HITS-CLIP | 29167373\|25083871 | Unspecified |
| mmu-miR-152-3p | MIMAT0000162 | Cblb | 208650 | HITS-CLIP | 37986224 | Unspecified |
| mmu-miR-152-3p | MIMAT0000162 | Cbll1 | 104836 | HITS-CLIP | 37986224\|25083871 | Unspecified |
| mmu-miR-152-3p | MIMAT0000162 | Cbx5 | 12419 | HITS-CLIP | 37986224 | Unspecified |
| mmu-miR-152-3p | MIMAT0000162 | Cc2d1b | 319965 | HITS-CLIP | 29167373 | Unspecified |
| mmu-miR-152-3p | MIMAT0000162 | Ccar1 | 67500 | HITS-CLIP | 25083871\|29167373 | Unspecified |

| | | | | | | | |
|---|---|---|---|---|---|---|---|
| mmu-miR-152-3p | MIMAT0000162 | Ccdc186 | 55088 | HITS-CLIP | | 29167373 | Unspecified |
| mmu-miR-152-3p | MIMAT0000162 | Ccdc47 | 67163 | HITS-CLIP | 37986224\|25083871 | | Unspecified |
| mmu-miR-152-3p | MIMAT0000162 | Ccdc50 | 67501 | HITS-CLIP | | 25083871 | Unspecified |
| mmu-miR-152-3p | MIMAT0000162 | Ccdc71 | 72454 | HITS-CLIP | | 25083871 | Unspecified |
| mmu-miR-152-3p | MIMAT0000162 | Ccdc80 | 67896 | HITS-CLIP | | 25083871 | Unspecified |
| mmu-miR-152-3p | MIMAT0000162 | Ccdc88a | 108686 | HITS-CLIP | | 37986224 | Unspecified |
| mmu-miR-152-3p | MIMAT0000162 | Ccn2 | 1490 | HITS-CLIP | | 23597149 | Unspecified |
| mmu-miR-152-3p | MIMAT0000162 | Ccn4 | 8840 | HITS-CLIP | | 25083871 | Unspecified |
| mmu-miR-152-3p | MIMAT0000162 | Ccna2 | 12428 | HITS-CLIP | | 25083871 | Unspecified |
| mmu-miR-152-3p | MIMAT0000162 | Ccnd2 | 12444 | HITS-CLIP | | 25083871 | Unspecified |
| mmu-miR-152-3p | MIMAT0000162 | Ccne2 | 12448 | HITS-CLIP | | 25083871 | Unspecified |
| mmu-miR-152-3p | MIMAT0000162 | Ccnf | 12449 | HITS-CLIP | | 37986224 | Unspecified |
| mmu-miR-152-3p | MIMAT0000162 | Ccng1 | 12450 | HITS-CLIP | 23597149\|25083871 | | Unspecified |
| mmu-miR-152-3p | MIMAT0000162 | Ccnj | 240665 | HITS-CLIP | | 25083871 | Unspecified |
| mmu-miR-152-3p | MIMAT0000162 | Ccnt2 | 72949 | HITS-CLIP | | 25083871 | Unspecified |
| mmu-miR-152-3p | MIMAT0000162 | Cct3 | 12462 | HITS-CLIP | 37986224\|25083871 | | Unspecified |
| mmu-miR-152-3p | MIMAT0000162 | Cd2ap | 12488 | HITS-CLIP | | 25083871 | Unspecified |
| mmu-miR-152-3p | MIMAT0000162 | Cd302 | 66205 | HITS-CLIP | | 23597149 | Unspecified |
| mmu-miR-152-3p | MIMAT0000162 | Cd36 | 12491 | HITS-CLIP | | 29167373 | Unspecified |
| mmu-miR-152-3p | MIMAT0000162 | Cd44 | 12505 | HITS-CLIP | 25083871\|29167373 | | Unspecified |
| mmu-miR-152-3p | MIMAT0000162 | Cdc123 | 98828 | HITS-CLIP | | 23597149 | Unspecified |
| mmu-miR-152-3p | MIMAT0000162 | Cdc14a | 229776 | HITS-CLIP | 37986224\|25083871 | | Unspecified |
| mmu-miR-152-3p | MIMAT0000162 | Cdc25b | 12531 | HITS-CLIP | | 37986224 | Unspecified |
| mmu-miR-152-3p | MIMAT0000162 | Cdc27 | 217232 | HITS-CLIP | | 29167373 | Unspecified |
| mmu-miR-152-3p | MIMAT0000162 | Cdc37 | 12539 | HITS-CLIP | | 23597149 | Unspecified |
| mmu-miR-152-3p | MIMAT0000162 | Cdc42 | 12540 | HITS-CLIP | 37986224\|25083871 | | Unspecified |
| mmu-miR-152-3p | MIMAT0000162 | Cdc42bpa | 226751 | HITS-CLIP | | 25083871 | Unspecified |
| mmu-miR-152-3p | MIMAT0000162 | Cdc42se2 | 72729 | HITS-CLIP | | 25083871 | Unspecified |
| mmu-miR-152-3p | MIMAT0000162 | Cdh11 | 12552 | HITS-CLIP | | 37986224 | Unspecified |
| mmu-miR-152-3p | MIMAT0000162 | Cdh15 | 12555 | HITS-CLIP | | 25083871 | Unspecified |
| mmu-miR-152-3p | MIMAT0000162 | Cdip1 | 66626 | HITS-CLIP | | 37986224 | Unspecified |
| mmu-miR-152-3p | MIMAT0000162 | Cdk12 | 69131 | HITS-CLIP | | 25083871 | Unspecified |
| mmu-miR-152-3p | MIMAT0000162 | Cdk13 | 69562 | HITS-CLIP | | 37986224 | Unspecified |
| mmu-miR-152-3p | MIMAT0000162 | Cdk14 | 18647 | HITS-CLIP | 37986224\|25083871 | | Unspecified |
| mmu-miR-152-3p | MIMAT0000162 | Cdk17 | 237459 | HITS-CLIP | | 25083871 | Unspecified |
| mmu-miR-152-3p | MIMAT0000162 | Cdk19 | 78334 | HITS-CLIP | 37986224\|25083871 | | Unspecified |
| mmu-miR-152-3p | MIMAT0000162 | Cdk2 | 12566 | HITS-CLIP | | 37986224 | Unspecified |
| mmu-miR-152-3p | MIMAT0000162 | Cdk5 | 12568 | HITS-CLIP | | 37986224 | Unspecified |
| mmu-miR-152-3p | MIMAT0000162 | Cdk5r1 | 12569 | HITS-CLIP | | 37986224 | Unspecified |

| | | | | | | |
|---|---|---|---|---|---|---|
| mmu-miR-152-3p | MIMAT0000162 | Cdk6 | 12571 | HITS-CLIP | 25083871\|29167373 | Unspecified |
| mmu-miR-152-3p | MIMAT0000162 | Cdk7 | 12572 | HITS-CLIP | 37986224 | Unspecified |
| mmu-miR-152-3p | MIMAT0000162 | Cdk8 | 264064 | HITS-CLIP | 25083871 | Unspecified |
| mmu-miR-152-3p | MIMAT0000162 | Cdkn1b | 12576 | HITS-CLIP | 37986224 | Unspecified |
| mmu-miR-152-3p | MIMAT0000162 | Cdkn2aip | 70925 | HITS-CLIP | 37986224\|25083871\|23597149 | Unspecified |
| mmu-miR-152-3p | MIMAT0000162 | Cebpa | 12606 | HITS-CLIP | 23597149 | Unspecified |
| mmu-miR-152-3p | MIMAT0000162 | Cebpd | 12609 | HITS-CLIP | 25083871 | Unspecified |
| mmu-miR-152-3p | MIMAT0000162 | Cebpzos | 68554 | HITS-CLIP | 23597149 | Unspecified |
| mmu-miR-152-3p | MIMAT0000162 | Celf1 | 13046 | HITS-CLIP | 25083871\|23597149\|29167373 | Unspecified |
| mmu-miR-152-3p | MIMAT0000162 | Celf2 | 14007 | HITS-CLIP | 37986224\|25083871 | Unspecified |
| mmu-miR-152-3p | MIMAT0000162 | Cemip2 | 23670 | HITS-CLIP | 25083871\|23597149 | Unspecified |
| mmu-miR-152-3p | MIMAT0000162 | Cenpb | 12616 | HITS-CLIP | 25083871 | Unspecified |
| mmu-miR-152-3p | MIMAT0000162 | Cenpf | 108000 | HITS-CLIP | 37986224 | Unspecified |
| mmu-miR-152-3p | MIMAT0000162 | Cep135 | 381644 | HITS-CLIP | 25083871 | Unspecified |
| mmu-miR-152-3p | MIMAT0000162 | Cep162 | 382090 | HITS-CLIP | 37986224 | Unspecified |
| mmu-miR-152-3p | MIMAT0000162 | Cep164 | 214552 | HITS-CLIP | 37986224 | Unspecified |
| mmu-miR-152-3p | MIMAT0000162 | Cep170 | 545389 | HITS-CLIP | 25083871\|29167373 | Unspecified |
| mmu-miR-152-3p | MIMAT0000162 | Cep192 | 70799 | HITS-CLIP | 25083871 | Unspecified |
| mmu-miR-152-3p | MIMAT0000162 | Cep295 | 85459 | HITS-CLIP | 29167373 | Unspecified |
| mmu-miR-152-3p | MIMAT0000162 | Cep350 | 74081 | HITS-CLIP | 37986224 | Unspecified |
| mmu-miR-152-3p | MIMAT0000162 | Cep55 | 74107 | HITS-CLIP | 25083871 | Unspecified |
| mmu-miR-152-3p | MIMAT0000162 | Cep83 | 77048 | HITS-CLIP | 37986224 | Unspecified |
| mmu-miR-152-3p | MIMAT0000162 | Cep85 | 70012 | HITS-CLIP | 23597149 | Unspecified |
| mmu-miR-152-3p | MIMAT0000162 | Cep85l | 100038725 | HITS-CLIP | 23597149 | Unspecified |
| mmu-miR-152-3p | MIMAT0000162 | Cerk | 223753 | HITS-CLIP | 37986224\|29167373 | Unspecified |
| mmu-miR-152-3p | MIMAT0000162 | Cers5 | 71949 | HITS-CLIP | 37986224 | Unspecified |
| mmu-miR-152-3p | MIMAT0000162 | Cers6 | 241447 | HITS-CLIP | 25083871 | Unspecified |
| mmu-miR-152-3p | MIMAT0000162 | Ces1c | 13884 | HITS-CLIP | 23597149 | Unspecified |
| mmu-miR-152-3p | MIMAT0000162 | Ces1f | 234564 | HITS-CLIP | 23597149 | Unspecified |
| mmu-miR-152-3p | MIMAT0000162 | Ces2e | 234673 | HITS-CLIP | 23597149 | Unspecified |
| mmu-miR-152-3p | MIMAT0000162 | Ces4a | 234677 | HITS-CLIP | 37986224 | Unspecified |
| mmu-miR-152-3p | MIMAT0000162 | Cfh | 12628 | HITS-CLIP | 37986224 | Unspecified |
| mmu-miR-152-3p | MIMAT0000162 | Cfl1 | 12631 | HITS-CLIP | 29167373\|23597149\|25083871 | Unspecified |
| mmu-miR-152-3p | MIMAT0000162 | Cflar | 12633 | HITS-CLIP | 37986224\|25083871\|23597149\|29167 | Unspecified |
| mmu-miR-152-3p | MIMAT0000162 | Cfp | 18636 | HITS-CLIP | 29167373 | Unspecified |
| mmu-miR-152-3p | MIMAT0000162 | Cggbp1 | 106143 | HITS-CLIP | 37986224 | Unspecified |
| mmu-miR-152-3p | MIMAT0000162 | Chac1 | 69065 | HITS-CLIP | 25083871 | Unspecified |
| mmu-miR-152-3p | MIMAT0000162 | Chchd4 | 72170 | HITS-CLIP | 37986224\|25083871\|23597149 | Unspecified |
| mmu-miR-152-3p | MIMAT0000162 | Chd1 | 12648 | HITS-CLIP | 25083871 | Unspecified |

| | | | | | | | |
|---|---|---|---|---|---|---|---|
| mmu-miR-152-3p | MIMAT0000162 | Chd2 | 244059 | HITS-CLIP | | 37986224 | Unspecified |
| mmu-miR-152-3p | MIMAT0000162 | Chd3 | 216848 | HITS-CLIP | | 37986224 | Unspecified |
| mmu-miR-152-3p | MIMAT0000162 | Chd4 | 107932 | HITS-CLIP | 25083871\|29167373 | | Unspecified |
| mmu-miR-152-3p | MIMAT0000162 | Chd6 | 71389 | HITS-CLIP | | 37986224 | Unspecified |
| mmu-miR-152-3p | MIMAT0000162 | Chd7 | 320790 | HITS-CLIP | 37986224\|25083871 | | Unspecified |
| mmu-miR-152-3p | MIMAT0000162 | Chd8 | 67772 | HITS-CLIP | 37986224\|25083871 | | Unspecified |
| mmu-miR-152-3p | MIMAT0000162 | Chd9 | 109151 | HITS-CLIP | | 25083871 | Unspecified |
| mmu-miR-152-3p | MIMAT0000162 | Chfr | 231600 | HITS-CLIP | | 25083871 | Unspecified |
| mmu-miR-152-3p | MIMAT0000162 | Chic2 | 74277 | HITS-CLIP | 25083871\|23597149 | | Unspecified |
| mmu-miR-152-3p | MIMAT0000162 | Chmp2b | 68942 | HITS-CLIP | | 25083871 | Unspecified |
| mmu-miR-152-3p | MIMAT0000162 | Chmp4b | 75608 | HITS-CLIP | | 25083871 | Unspecified |
| mmu-miR-152-3p | MIMAT0000162 | Chpf2 | 100910 | HITS-CLIP | | 37986224 | Unspecified |
| mmu-miR-152-3p | MIMAT0000162 | Chrnd | 11447 | HITS-CLIP | | 25083871 | Unspecified |
| mmu-miR-152-3p | MIMAT0000162 | Chst3 | 53374 | HITS-CLIP | | 37986224 | Unspecified |
| mmu-miR-152-3p | MIMAT0000162 | Chsy1 | 269941 | HITS-CLIP | 37986224\|25083871 | | Unspecified |
| mmu-miR-152-3p | MIMAT0000162 | Chuk | 12675 | HITS-CLIP | 37986224\|25083871 | | Unspecified |
| mmu-miR-152-3p | MIMAT0000162 | Cic | 71722 | HITS-CLIP | | 23597149 | Unspecified |
| mmu-miR-152-3p | MIMAT0000162 | Cip2a | 57650 | HITS-CLIP | | 31433990 | Unspecified |
| mmu-miR-152-3p | MIMAT0000162 | Cit | 12704 | HITS-CLIP | | 25083871 | Unspecified |
| mmu-miR-152-3p | MIMAT0000162 | Cited2 | 17684 | HITS-CLIP | | 25083871 | Unspecified |
| mmu-miR-152-3p | MIMAT0000162 | Ciz1 | 68379 | HITS-CLIP | | 25083871 | Unspecified |
| mmu-miR-152-3p | MIMAT0000162 | Ckap4 | 216197 | HITS-CLIP | | 25083871 | Unspecified |
| mmu-miR-152-3p | MIMAT0000162 | Ckap5 | 75786 | HITS-CLIP | 25083871\|23597149 | | Unspecified |
| mmu-miR-152-3p | MIMAT0000162 | Clasp2 | 76499 | HITS-CLIP | 37986224\|25083871\|29167373 | | Unspecified |
| mmu-miR-152-3p | MIMAT0000162 | Cldn12 | 64945 | HITS-CLIP | | 25083871 | Unspecified |
| mmu-miR-152-3p | MIMAT0000162 | Cldn4 | 12740 | HITS-CLIP | | 26203562 | Unspecified |
| mmu-miR-152-3p | MIMAT0000162 | Clec12a | 160364 | HITS-CLIP | | 29167373 | Unspecified |
| mmu-miR-152-3p | MIMAT0000162 | Clec1b | 56760 | HITS-CLIP | | 37986224 | Unspecified |
| mmu-miR-152-3p | MIMAT0000162 | Clec4f | 51811 | HITS-CLIP | | 37986224 | Unspecified |
| mmu-miR-152-3p | MIMAT0000162 | Clock | 12753 | HITS-CLIP | | 23597149 | Unspecified |
| mmu-miR-152-3p | MIMAT0000162 | Clptm1 | 56457 | HITS-CLIP | | 25083871 | Unspecified |
| mmu-miR-152-3p | MIMAT0000162 | Clta | 12757 | HITS-CLIP | 37986224\|25083871\|29167373 | | Unspecified |
| mmu-miR-152-3p | MIMAT0000162 | Cltc | 67300 | HITS-CLIP | 37986224\|25083871\|23597149\|21258 | | Unspecified |
| mmu-miR-152-3p | MIMAT0000162 | Cluh | 74148 | HITS-CLIP | | 25083871 | Unspecified |
| mmu-miR-152-3p | MIMAT0000162 | Cmah | 12763 | HITS-CLIP | | 23597149 | Unspecified |
| mmu-miR-152-3p | MIMAT0000162 | Cmc1 | 67899 | HITS-CLIP | 37986224\|23597149 | | Unspecified |
| mmu-miR-152-3p | MIMAT0000162 | Cmtm3 | 68119 | HITS-CLIP | 25083871\|29167373 | | Unspecified |
| mmu-miR-152-3p | MIMAT0000162 | Cmtm4 | 97487 | HITS-CLIP | | 37986224 | Unspecified |
| mmu-miR-152-3p | MIMAT0000162 | Cnih1 | 12793 | HITS-CLIP | | 25083871 | Unspecified |

| | | | | | | |
|---|---|---|---|---|---|---|
| mmu-miR-152-3p | MIMAT0000162 | Cnih4 | 98417 | HITS-CLIP | 37986224\|25083871 | Unspecified |
| mmu-miR-152-3p | MIMAT0000162 | Cnn3 | 71994 | HITS-CLIP | 25083871 | Unspecified |
| mmu-miR-152-3p | MIMAT0000162 | Cnot1 | 234594 | HITS-CLIP | 37986224\|25083871\|23597149\|29167 | Unspecified |
| mmu-miR-152-3p | MIMAT0000162 | Cnot11 | 52846 | HITS-CLIP | 25083871 | Unspecified |
| mmu-miR-152-3p | MIMAT0000162 | Cnot2 | 72068 | HITS-CLIP | 25083871 | Unspecified |
| mmu-miR-152-3p | MIMAT0000162 | Cnot4 | 53621 | HITS-CLIP | 37986224\|29167373 | Unspecified |
| mmu-miR-152-3p | MIMAT0000162 | Cnot6 | 104625 | HITS-CLIP | 37986224\|25083871\|29167373 | Unspecified |
| mmu-miR-152-3p | MIMAT0000162 | Cnot6l | 231464 | HITS-CLIP | 37986224\|25083871 | Unspecified |
| mmu-miR-152-3p | MIMAT0000162 | Coa5 | 76178 | HITS-CLIP | 37986224\|29167373 | Unspecified |
| mmu-miR-152-3p | MIMAT0000162 | Cog6 | 67542 | HITS-CLIP | 23597149 | Unspecified |
| mmu-miR-152-3p | MIMAT0000162 | Cog7 | 233824 | HITS-CLIP | 29167373 | Unspecified |
| mmu-miR-152-3p | MIMAT0000162 | Col18a1 | 12822 | HITS-CLIP | 23597149 | Unspecified |
| mmu-miR-152-3p | MIMAT0000162 | Col4a1 | 12826 | HITS-CLIP | 25083871\|23597149 | Unspecified |
| mmu-miR-152-3p | MIMAT0000162 | Col4a3bp | 68018 | HITS-CLIP | 25083871 | Unspecified |
| mmu-miR-152-3p | MIMAT0000162 | Col8a1 | 12837 | HITS-CLIP | 37986224 | Unspecified |
| mmu-miR-152-3p | MIMAT0000162 | Colgalt1 | 234407 | HITS-CLIP | 37986224 | Unspecified |
| mmu-miR-152-3p | MIMAT0000162 | Commd10 | 69456 | HITS-CLIP | 37986224\|25083871 | Unspecified |
| mmu-miR-152-3p | MIMAT0000162 | Commd3 | 12238 | HITS-CLIP | 25083871 | Unspecified |
| mmu-miR-152-3p | MIMAT0000162 | Comt | 12846 | HITS-CLIP | 37986224\|25083871 | Unspecified |
| mmu-miR-152-3p | MIMAT0000162 | Copb1 | 70349 | HITS-CLIP | 37986224\|23597149 | Unspecified |
| mmu-miR-152-3p | MIMAT0000162 | Copg1 | 54161 | HITS-CLIP | 25083871 | Unspecified |
| mmu-miR-152-3p | MIMAT0000162 | Cops2 | 12848 | HITS-CLIP | 29167373 | Unspecified |
| mmu-miR-152-3p | MIMAT0000162 | Cops4 | 26891 | HITS-CLIP | 25083871\|23597149 | Unspecified |
| mmu-miR-152-3p | MIMAT0000162 | Cops5 | 26754 | HITS-CLIP | 25083871\|23597149 | Unspecified |
| mmu-miR-152-3p | MIMAT0000162 | Cops7b | 26895 | HITS-CLIP | 23597149 | Unspecified |
| mmu-miR-152-3p | MIMAT0000162 | Copz1 | 56447 | HITS-CLIP | 37986224\|23597149 | Unspecified |
| mmu-miR-152-3p | MIMAT0000162 | Coq10a | 210582 | HITS-CLIP | 37986224 | Unspecified |
| mmu-miR-152-3p | MIMAT0000162 | Coq2 | 71883 | HITS-CLIP | 37986224 | Unspecified |
| mmu-miR-152-3p | MIMAT0000162 | Coq6 | 217707 | HITS-CLIP | 23597149 | Unspecified |
| mmu-miR-152-3p | MIMAT0000162 | Coq8a | 56997 | HITS-CLIP | 23597149 | Unspecified |
| mmu-miR-152-3p | MIMAT0000162 | Coq9 | 67914 | HITS-CLIP | 37986224\|25083871 | Unspecified |
| mmu-miR-152-3p | MIMAT0000162 | Coro1a | 12721 | HITS-CLIP | 23597149 | Unspecified |
| mmu-miR-152-3p | MIMAT0000162 | Coro1b | 23789 | HITS-CLIP | 25083871 | Unspecified |
| mmu-miR-152-3p | MIMAT0000162 | Coro6 | 216961 | HITS-CLIP | 25083871 | Unspecified |
| mmu-miR-152-3p | MIMAT0000162 | Cp | 12870 | HITS-CLIP | 37986224\|23597149 | Unspecified |
| mmu-miR-152-3p | MIMAT0000162 | Cpb2 | 56373 | HITS-CLIP | 23597149 | Unspecified |
| mmu-miR-152-3p | MIMAT0000162 | Cpd | 12874 | HITS-CLIP | 37986224\|25083871 | Unspecified |
| mmu-miR-152-3p | MIMAT0000162 | Cpe | 12876 | HITS-CLIP | 25083871 | Unspecified |
| mmu-miR-152-3p | MIMAT0000162 | Cpeb2 | 231207 | HITS-CLIP | 25083871\|29167373 | Unspecified |

| | | | | | | |
|---|---|---|---|---|---|---|
| mmu-miR-152-3p | MIMAT0000162 | Cpeb3 | 208922 | HITS-CLIP | 37986224\|25083871\|29167373 | Unspecified |
| mmu-miR-152-3p | MIMAT0000162 | Cpeb4 | 67579 | HITS-CLIP | 37986224\|25083871\|23597149\|29167 | Unspecified |
| mmu-miR-152-3p | MIMAT0000162 | Cpne3 | 70568 | HITS-CLIP | 23597149\|29167373 | Unspecified |
| mmu-miR-152-3p | MIMAT0000162 | Cpox | 12892 | HITS-CLIP | 37986224\|23597149 | Unspecified |
| mmu-miR-152-3p | MIMAT0000162 | Cps1 | 227231 | HITS-CLIP | 23597149 | Unspecified |
| mmu-miR-152-3p | MIMAT0000162 | Cpsf1 | 94230 | HITS-CLIP | 25083871 | Unspecified |
| mmu-miR-152-3p | MIMAT0000162 | Cpsf3 | 54451 | HITS-CLIP | 29167373 | Unspecified |
| mmu-miR-152-3p | MIMAT0000162 | Cpt1a | 12894 | HITS-CLIP | 23597149 | Unspecified |
| mmu-miR-152-3p | MIMAT0000162 | Cpt2 | 12896 | HITS-CLIP | 23597149\|25083871 | Unspecified |
| mmu-miR-152-3p | MIMAT0000162 | Cptp | 80772 | HITS-CLIP | 25083871 | Unspecified |
| mmu-miR-152-3p | MIMAT0000162 | Cramp1l | 57354 | HITS-CLIP | 37986224 | Unspecified |
| mmu-miR-152-3p | MIMAT0000162 | Crat | 12908 | HITS-CLIP | 37986224\|25083871 | Unspecified |
| mmu-miR-152-3p | MIMAT0000162 | Creb1 | 12912 | HITS-CLIP | 25083871 | Unspecified |
| mmu-miR-152-3p | MIMAT0000162 | Creb3l2 | 208647 | HITS-CLIP | 37986224 | Unspecified |
| mmu-miR-152-3p | MIMAT0000162 | Creb5 | 231991 | HITS-CLIP | 29167373 | Unspecified |
| mmu-miR-152-3p | MIMAT0000162 | Crebbp | 12914 | HITS-CLIP | 37986224\|25083871 | Unspecified |
| mmu-miR-152-3p | MIMAT0000162 | Crebl2 | 232430 | HITS-CLIP | 23597149 | Unspecified |
| mmu-miR-152-3p | MIMAT0000162 | Crebrf | 77128 | HITS-CLIP | 25083871 | Unspecified |
| mmu-miR-152-3p | MIMAT0000162 | Creg1 | 433375 | HITS-CLIP | 29167373 | Unspecified |
| mmu-miR-152-3p | MIMAT0000162 | Cript | 56724 | HITS-CLIP | 37986224\|25083871 | Unspecified |
| mmu-miR-152-3p | MIMAT0000162 | Crkl | 12929 | HITS-CLIP | 23597149\|29167373 | Unspecified |
| mmu-miR-152-3p | MIMAT0000162 | Crlf3 | 54394 | HITS-CLIP | 29167373 | Unspecified |
| mmu-miR-152-3p | MIMAT0000162 | Crls1 | 66586 | HITS-CLIP | 25083871 | Unspecified |
| mmu-miR-152-3p | MIMAT0000162 | Crp | 12944 | HITS-CLIP | 37986224 | Unspecified |
| mmu-miR-152-3p | MIMAT0000162 | Csde1 | 229663 | HITS-CLIP | 25083871 | Unspecified |
| mmu-miR-152-3p | MIMAT0000162 | Csnk1a1 | 93687 | HITS-CLIP | 37986224 | Unspecified |
| mmu-miR-152-3p | MIMAT0000162 | Csnk1d | 104318 | HITS-CLIP | 23597149 | Unspecified |
| mmu-miR-152-3p | MIMAT0000162 | Csnk1g1 | 214897 | HITS-CLIP | 37986224\|23597149 | Unspecified |
| mmu-miR-152-3p | MIMAT0000162 | Csnk1g3 | 70425 | HITS-CLIP | 37986224 | Unspecified |
| mmu-miR-152-3p | MIMAT0000162 | Csnk2a1 | 12995 | HITS-CLIP | 37986224\|25083871\|29167373 | Unspecified |
| mmu-miR-152-3p | MIMAT0000162 | Cspp1 | 211660 | HITS-CLIP | 29167373 | Unspecified |
| mmu-miR-152-3p | MIMAT0000162 | Csrnp1 | 215418 | HITS-CLIP | 37986224\|23597149 | Unspecified |
| mmu-miR-152-3p | MIMAT0000162 | Cstf2 | 108062 | HITS-CLIP | 25083871 | Unspecified |
| mmu-miR-152-3p | MIMAT0000162 | Ctbp1 | 13016 | HITS-CLIP | 25083871 | Unspecified |
| mmu-miR-152-3p | MIMAT0000162 | Ctbp2 | 13017 | HITS-CLIP | 37986224 | Unspecified |
| mmu-miR-152-3p | MIMAT0000162 | Ctcf | 13018 | HITS-CLIP | 37986224 | Unspecified |
| mmu-miR-152-3p | MIMAT0000162 | Ctdsp1 | 227292 | HITS-CLIP | 37986224 | Unspecified |
| mmu-miR-152-3p | MIMAT0000162 | Ctif | 269037 | HITS-CLIP | 37986224 | Unspecified |
| mmu-miR-152-3p | MIMAT0000162 | Ctnna1 | 12385 | HITS-CLIP | 25083871\|23597149 | Unspecified |

| | | | | | | |
|---|---|---|---|---|---|---|
| mmu-miR-152-3p | MIMAT0000162 | Ctnnbl1 | 66642 | HITS-CLIP | 23597149 | Unspecified |
| mmu-miR-152-3p | MIMAT0000162 | Ctnnd1 | 12388 | HITS-CLIP | 29167373 | Unspecified |
| mmu-miR-152-3p | MIMAT0000162 | Ctr9 | 22083 | HITS-CLIP | 23597149\|25083871 | Unspecified |
| mmu-miR-152-3p | MIMAT0000162 | Ctsz | 64138 | HITS-CLIP | 29167373 | Unspecified |
| mmu-miR-152-3p | MIMAT0000162 | Cttnbp2nl | 80281 | HITS-CLIP | 25083871 | Unspecified |
| mmu-miR-152-3p | MIMAT0000162 | Cul3 | 26554 | HITS-CLIP | 37986224 | Unspecified |
| mmu-miR-152-3p | MIMAT0000162 | Cul7 | 66515 | HITS-CLIP | 37986224 | Unspecified |
| mmu-miR-152-3p | MIMAT0000162 | Cux1 | 13047 | HITS-CLIP | 37986224\|25083871 | Unspecified |
| mmu-miR-152-3p | MIMAT0000162 | Cwc25 | 67480 | HITS-CLIP | 37986224 | Unspecified |
| mmu-miR-152-3p | MIMAT0000162 | Cxcl12 | 20315 | HITS-CLIP | 25083871\|23597149 | Unspecified |
| mmu-miR-152-3p | MIMAT0000162 | Cxcl14 | 57266 | HITS-CLIP | 29167373 | Unspecified |
| mmu-miR-152-3p | MIMAT0000162 | Cxcr4 | 12767 | HITS-CLIP | 29167373 | Unspecified |
| mmu-miR-152-3p | MIMAT0000162 | Cxxc1 | 74322 | HITS-CLIP | 23597149 | Unspecified |
| mmu-miR-152-3p | MIMAT0000162 | Cyb5b | 66427 | HITS-CLIP | 23597149 | Unspecified |
| mmu-miR-152-3p | MIMAT0000162 | Cyb5r4 | 266690 | HITS-CLIP | 25083871 | Unspecified |
| mmu-miR-152-3p | MIMAT0000162 | Cyhr1 | 54151 | HITS-CLIP | 37986224 | Unspecified |
| mmu-miR-152-3p | MIMAT0000162 | Cyld | 74256 | HITS-CLIP | 25083871 | Unspecified |
| mmu-miR-152-3p | MIMAT0000162 | Cyp27a1 | 104086 | HITS-CLIP | 23597149 | Unspecified |
| mmu-miR-152-3p | MIMAT0000162 | Cyp2c29 | 13095 | HITS-CLIP | 23597149 | Unspecified |
| mmu-miR-152-3p | MIMAT0000162 | Cyp2c68 | 433247 | HITS-CLIP | 23597149 | Unspecified |
| mmu-miR-152-3p | MIMAT0000162 | Cyp2c70 | 226105 | HITS-CLIP | 37986224\|23597149 | Unspecified |
| mmu-miR-152-3p | MIMAT0000162 | Cyp2d22 | 56448 | HITS-CLIP | 23597149 | Unspecified |
| mmu-miR-152-3p | MIMAT0000162 | Cyp2e1 | 13106 | HITS-CLIP | 23597149 | Unspecified |
| mmu-miR-152-3p | MIMAT0000162 | Cyp2j5 | 13109 | HITS-CLIP | 37986224\|23597149 | Unspecified |
| mmu-miR-152-3p | MIMAT0000162 | Cyp39a1 | 56050 | HITS-CLIP | 23597149 | Unspecified |
| mmu-miR-152-3p | MIMAT0000162 | Cyp3a13 | 13113 | HITS-CLIP | 23597149 | Unspecified |
| mmu-miR-152-3p | MIMAT0000162 | Cyp4a14 | 13119 | HITS-CLIP | 37986224 | Unspecified |
| mmu-miR-152-3p | MIMAT0000162 | Cyp51 | 13121 | HITS-CLIP | 25083871 | Unspecified |
| mmu-miR-152-3p | MIMAT0000162 | Cyp7a1 | 13122 | HITS-CLIP | 23597149 | Unspecified |
| mmu-miR-152-3p | MIMAT0000162 | Cyth3 | 19159 | HITS-CLIP | 37986224 | Unspecified |
| mmu-miR-152-3p | MIMAT0000162 | D5Ertd579e | 320661 | HITS-CLIP | 37986224 | Unspecified |
| mmu-miR-152-3p | MIMAT0000162 | Daam1 | 208846 | HITS-CLIP | 25083871\|23597149 | Unspecified |
| mmu-miR-152-3p | MIMAT0000162 | Dab2ip | 69601 | HITS-CLIP | 25083871 | Unspecified |
| mmu-miR-152-3p | MIMAT0000162 | Dag1 | 13138 | HITS-CLIP | 23597149 | Unspecified |
| mmu-miR-152-3p | MIMAT0000162 | Dagla | 269060 | HITS-CLIP | 29167373 | Unspecified |
| mmu-miR-152-3p | MIMAT0000162 | Dapk3 | 13144 | HITS-CLIP | 23597149 | Unspecified |
| mmu-miR-152-3p | MIMAT0000162 | Dars2 | 226539 | HITS-CLIP | 37986224\|23597149 | Unspecified |
| mmu-miR-152-3p | MIMAT0000162 | Daxx | 13163 | HITS-CLIP | 25083871 | Unspecified |
| mmu-miR-152-3p | MIMAT0000162 | Dazap2 | 23994 | HITS-CLIP | 25083871 | Unspecified |

| | | | | | | |
|---|---|---|---|---|---|---|
| mmu-miR-152-3p | MIMAT0000162 | Dcaf10 | 242418 | HITS-CLIP | 25083871 | Unspecified |
| mmu-miR-152-3p | MIMAT0000162 | Dcaf11 | 28199 | HITS-CLIP | 23597149 | Unspecified |
| mmu-miR-152-3p | MIMAT0000162 | Dcaf15 | 212123 | HITS-CLIP | 25083871\|23597149 | Unspecified |
| mmu-miR-152-3p | MIMAT0000162 | Dcaf8 | 98193 | HITS-CLIP | 25083871\|23597149 | Unspecified |
| mmu-miR-152-3p | MIMAT0000162 | Dclk1 | 13175 | HITS-CLIP | 25083871 | Unspecified |
| mmu-miR-152-3p | MIMAT0000162 | Dclre1c | 227525 | HITS-CLIP | 23597149 | Unspecified |
| mmu-miR-152-3p | MIMAT0000162 | Dcp1a | 75901 | HITS-CLIP | 37986224\|25083871 | Unspecified |
| mmu-miR-152-3p | MIMAT0000162 | Dcp2 | 70640 | HITS-CLIP | 37986224\|23597149 | Unspecified |
| mmu-miR-152-3p | MIMAT0000162 | Dctn6 | 22428 | HITS-CLIP | 25083871 | Unspecified |
| mmu-miR-152-3p | MIMAT0000162 | Dcun1d3 | 233805 | HITS-CLIP | 37986224 | Unspecified |
| mmu-miR-152-3p | MIMAT0000162 | Dcun1d5 | 76863 | HITS-CLIP | 25083871 | Unspecified |
| mmu-miR-152-3p | MIMAT0000162 | Ddr2 | 18214 | HITS-CLIP | 37986224\|25083871 | Unspecified |
| mmu-miR-152-3p | MIMAT0000162 | Ddx31 | 227674 | HITS-CLIP | 23597149 | Unspecified |
| mmu-miR-152-3p | MIMAT0000162 | Ddx3x | 13205 | HITS-CLIP | 29167373 | Unspecified |
| mmu-miR-152-3p | MIMAT0000162 | Ddx46 | 212880 | HITS-CLIP | 37986224 | Unspecified |
| mmu-miR-152-3p | MIMAT0000162 | Ddx49 | 234374 | HITS-CLIP | 23597149 | Unspecified |
| mmu-miR-152-3p | MIMAT0000162 | Ddx5 | 13207 | HITS-CLIP | 25083871\|29167373 | Unspecified |
| mmu-miR-152-3p | MIMAT0000162 | Ddx58 | 230073 | HITS-CLIP | 29167373 | Unspecified |
| mmu-miR-152-3p | MIMAT0000162 | Ddx6 | 13209 | HITS-CLIP | 25083871\|21258322 | Unspecified |
| mmu-miR-152-3p | MIMAT0000162 | Ddx60 | 234311 | HITS-CLIP | 29167373 | Unspecified |
| mmu-miR-152-3p | MIMAT0000162 | Dedd | 21945 | HITS-CLIP | 37986224 | Unspecified |
| mmu-miR-152-3p | MIMAT0000162 | Dennd1b | 329260 | HITS-CLIP | 25083871 | Unspecified |
| mmu-miR-152-3p | MIMAT0000162 | Dennd4a | 102442 | HITS-CLIP | 37986224 | Unspecified |
| mmu-miR-152-3p | MIMAT0000162 | Dennd4c | 329877 | HITS-CLIP | 29167373 | Unspecified |
| mmu-miR-152-3p | MIMAT0000162 | Dennd5a | 19347 | HITS-CLIP | 25083871 | Unspecified |
| mmu-miR-152-3p | MIMAT0000162 | Dennd6a | 211922 | HITS-CLIP | 37986224 | Unspecified |
| mmu-miR-152-3p | MIMAT0000162 | Deptor | 97998 | HITS-CLIP | 37986224\|25083871\|23597149 | Unspecified |
| mmu-miR-152-3p | MIMAT0000162 | Derl2 | 116891 | HITS-CLIP | 25083871 | Unspecified |
| mmu-miR-152-3p | MIMAT0000162 | Desi2 | 78825 | HITS-CLIP | 29167373 | Unspecified |
| mmu-miR-152-3p | MIMAT0000162 | Dhcr24 | 74754 | HITS-CLIP | 37986224 | Unspecified |
| mmu-miR-152-3p | MIMAT0000162 | Dhrs11 | 192970 | HITS-CLIP | 23597149 | Unspecified |
| mmu-miR-152-3p | MIMAT0000162 | Dhrs3 | 20148 | HITS-CLIP | 23597149 | Unspecified |
| mmu-miR-152-3p | MIMAT0000162 | Dhx36 | 72162 | HITS-CLIP | 23597149 | Unspecified |
| mmu-miR-152-3p | MIMAT0000162 | Dhx40 | 67487 | HITS-CLIP | 37986224 | Unspecified |
| mmu-miR-152-3p | MIMAT0000162 | Dhx9 | 13211 | HITS-CLIP | 25083871 | Unspecified |
| mmu-miR-152-3p | MIMAT0000162 | Diaph1 | 1729 | HITS-CLIP | 23597149 | Unspecified |
| mmu-miR-152-3p | MIMAT0000162 | Dicer1 | 192119 | HITS-CLIP | 37986224\|25083871 | Unspecified |
| mmu-miR-152-3p | MIMAT0000162 | Dido1 | 23856 | HITS-CLIP | 25083871\|29167373 | Unspecified |
| mmu-miR-152-3p | MIMAT0000162 | Dio1 | 13370 | HITS-CLIP | 23597149 | Unspecified |

| | | | | | | |
|---|---|---|---|---|---|---|
| mmu-miR-152-3p | MIMAT0000162 | Dip2a | 64451 | HITS-CLIP | 37986224 | Unspecified |
| mmu-miR-152-3p | MIMAT0000162 | Dip2c | 208440 | HITS-CLIP | 25083871 | Unspecified |
| mmu-miR-152-3p | MIMAT0000162 | Dkc1 | 245474 | HITS-CLIP | 23597149 | Unspecified |
| mmu-miR-152-3p | MIMAT0000162 | Dlc1 | 50768 | HITS-CLIP | 25083871 | Unspecified |
| mmu-miR-152-3p | MIMAT0000162 | Dlg5 | 71228 | HITS-CLIP | 25083871 | Unspecified |
| mmu-miR-152-3p | MIMAT0000162 | Dlgap5 | 218977 | HITS-CLIP | 23597149 | Unspecified |
| mmu-miR-152-3p | MIMAT0000162 | Dll1 | 13388 | HITS-CLIP | 25083871 | Unspecified |
| mmu-miR-152-3p | MIMAT0000162 | Dll4 | 54485 | HITS-CLIP | 23597149 | Unspecified |
| mmu-miR-152-3p | MIMAT0000162 | Dlst | 78920 | HITS-CLIP | 25083871 | Unspecified |
| mmu-miR-152-3p | MIMAT0000162 | Dmpk | 13400 | HITS-CLIP | 25083871 | Unspecified |
| mmu-miR-152-3p | MIMAT0000162 | Dmrta2 | 242620 | HITS-CLIP | 37986224\|25083871 | Unspecified |
| mmu-miR-152-3p | MIMAT0000162 | Dmxl1 | 240283 | HITS-CLIP | 37986224\|25083871\|23597149 | Unspecified |
| mmu-miR-152-3p | MIMAT0000162 | Dnaaf5 | 433956 | HITS-CLIP | 23597149 | Unspecified |
| mmu-miR-152-3p | MIMAT0000162 | Dnajb1 | 81489 | HITS-CLIP | 25083871 | Unspecified |
| mmu-miR-152-3p | MIMAT0000162 | Dnajb14 | 70604 | HITS-CLIP | 37986224 | Unspecified |
| mmu-miR-152-3p | MIMAT0000162 | Dnajb4 | 67035 | HITS-CLIP | 29167373 | Unspecified |
| mmu-miR-152-3p | MIMAT0000162 | Dnajb6 | 23950 | HITS-CLIP | 37986224 | Unspecified |
| mmu-miR-152-3p | MIMAT0000162 | Dnajb9 | 27362 | HITS-CLIP | 37986224\|23597149\|25083871 | Unspecified |
| mmu-miR-152-3p | MIMAT0000162 | Dnajc10 | 66861 | HITS-CLIP | 25083871 | Unspecified |
| mmu-miR-152-3p | MIMAT0000162 | Dnajc11 | 230935 | HITS-CLIP | 25083871\|23597149 | Unspecified |
| mmu-miR-152-3p | MIMAT0000162 | Dnajc13 | 235567 | HITS-CLIP | 25083871\|29167373 | Unspecified |
| mmu-miR-152-3p | MIMAT0000162 | Dnajc3 | 100037258 | HITS-CLIP | 25083871 | Unspecified |
| mmu-miR-152-3p | MIMAT0000162 | Dnase2a | 13423 | HITS-CLIP | 25083871 | Unspecified |
| mmu-miR-152-3p | MIMAT0000162 | Dnm2 | 13430 | HITS-CLIP | 25083871 | Unspecified |
| mmu-miR-152-3p | MIMAT0000162 | Dnmt3a | 13435 | HITS-CLIP | 37986224\|23597149\|25083871 | Unspecified |
| mmu-miR-152-3p | MIMAT0000162 | Dock1 | 330662 | HITS-CLIP | 23597149 | Unspecified |
| mmu-miR-152-3p | MIMAT0000162 | Dock11 | 75974 | HITS-CLIP | 25083871 | Unspecified |
| mmu-miR-152-3p | MIMAT0000162 | Dock4 | 238130 | HITS-CLIP | 23597149 | Unspecified |
| mmu-miR-152-3p | MIMAT0000162 | Dock5 | 68813 | HITS-CLIP | 37986224 | Unspecified |
| mmu-miR-152-3p | MIMAT0000162 | Dock6 | 319899 | HITS-CLIP | 37986224 | Unspecified |
| mmu-miR-152-3p | MIMAT0000162 | Dock7 | 67299 | HITS-CLIP | 25083871 | Unspecified |
| mmu-miR-152-3p | MIMAT0000162 | Dok7 | 231134 | HITS-CLIP | 37986224 | Unspecified |
| mmu-miR-152-3p | MIMAT0000162 | Dop1a | 23033 | HITS-CLIP | 25083871\|29167373 | Unspecified |
| mmu-miR-152-3p | MIMAT0000162 | Dop1b | 9980 | HITS-CLIP | 23597149 | Unspecified |
| mmu-miR-152-3p | MIMAT0000162 | Dot1l | 208266 | HITS-CLIP | 25083871\|23597149\|29167373 | Unspecified |
| mmu-miR-152-3p | MIMAT0000162 | Dpf2 | 19708 | HITS-CLIP | 37986224 | Unspecified |
| mmu-miR-152-3p | MIMAT0000162 | Dpyd | 99586 | HITS-CLIP | 23597149 | Unspecified |
| mmu-miR-152-3p | MIMAT0000162 | Dpysl3 | 22240 | HITS-CLIP | 37986224 | Unspecified |
| mmu-miR-152-3p | MIMAT0000162 | Dram2 | 67171 | HITS-CLIP | 25083871\|23597149\|29167373 | Unspecified |

| | | | | | | |
|---|---|---|---|---|---|---|
| mmu-miR-152-3p | MIMAT0000162 | Dsel | 319901 | HITS-CLIP | 37986224\|25083871 | Unspecified |
| mmu-miR-152-3p | MIMAT0000162 | Dst | 13518 | HITS-CLIP | 37986224\|25083871 | Unspecified |
| mmu-miR-152-3p | MIMAT0000162 | Dstyk | 213452 | HITS-CLIP | 25083871 | Unspecified |
| mmu-miR-152-3p | MIMAT0000162 | Dtna | 13527 | HITS-CLIP | 37986224\|25083871 | Unspecified |
| mmu-miR-152-3p | MIMAT0000162 | Dus3l | 224907 | HITS-CLIP | 23597149 | Unspecified |
| mmu-miR-152-3p | MIMAT0000162 | Dusp10 | 63953 | HITS-CLIP | 37986224 | Unspecified |
| mmu-miR-152-3p | MIMAT0000162 | Dusp11 | 72102 | HITS-CLIP | 23597149 | Unspecified |
| mmu-miR-152-3p | MIMAT0000162 | Dusp16 | 70686 | HITS-CLIP | 37986224 | Unspecified |
| mmu-miR-152-3p | MIMAT0000162 | Dusp3 | 72349 | HITS-CLIP | 37986224 | Unspecified |
| mmu-miR-152-3p | MIMAT0000162 | Dusp7 | 235584 | HITS-CLIP | 25083871 | Unspecified |
| mmu-miR-152-3p | MIMAT0000162 | Dvl1 | 13542 | HITS-CLIP | 37986224 | Unspecified |
| mmu-miR-152-3p | MIMAT0000162 | Dynap | 75577 | HITS-CLIP | 37986224 | Unspecified |
| mmu-miR-152-3p | MIMAT0000162 | Dync1h1 | 13424 | HITS-CLIP | 37986224\|23597149\|25083871 | Unspecified |
| mmu-miR-152-3p | MIMAT0000162 | Dync1i2 | 13427 | HITS-CLIP | 25083871 | Unspecified |
| mmu-miR-152-3p | MIMAT0000162 | Dync1li2 | 234663 | HITS-CLIP | 37986224 | Unspecified |
| mmu-miR-152-3p | MIMAT0000162 | Dyrk1a | 13548 | HITS-CLIP | 37986224\|25083871\|29167373 | Unspecified |
| mmu-miR-152-3p | MIMAT0000162 | Dyrk2 | 69181 | HITS-CLIP | 37986224 | Unspecified |
| mmu-miR-152-3p | MIMAT0000162 | Dysf | 26903 | HITS-CLIP | 25083871 | Unspecified |
| mmu-miR-152-3p | MIMAT0000162 | E2f2 | 242705 | HITS-CLIP | 37986224 | Unspecified |
| mmu-miR-152-3p | MIMAT0000162 | E2f3 | 13557 | HITS-CLIP | 25083871 | Unspecified |
| mmu-miR-152-3p | MIMAT0000162 | E2f6 | 50496 | HITS-CLIP | 25083871 | Unspecified |
| mmu-miR-152-3p | MIMAT0000162 | Ebag9 | 55960 | HITS-CLIP | 37986224\|25083871 | Unspecified |
| mmu-miR-152-3p | MIMAT0000162 | Ebf1 | 13591 | HITS-CLIP | 25083871 | Unspecified |
| mmu-miR-152-3p | MIMAT0000162 | Echs1 | 93747 | HITS-CLIP | 25083871 | Unspecified |
| mmu-miR-152-3p | MIMAT0000162 | Eci2 | 23986 | HITS-CLIP | 23597149\|25083871 | Unspecified |
| mmu-miR-152-3p | MIMAT0000162 | Edc3 | 353190 | HITS-CLIP | 23597149 | Unspecified |
| mmu-miR-152-3p | MIMAT0000162 | Edem3 | 66967 | HITS-CLIP | 25083871 | Unspecified |
| mmu-miR-152-3p | MIMAT0000162 | Edn1 | 13614 | HITS-CLIP | 37986224 | Unspecified |
| mmu-miR-152-3p | MIMAT0000162 | Eepd1 | 67484 | HITS-CLIP | 23597149 | Unspecified |
| mmu-miR-152-3p | MIMAT0000162 | Efcab14 | 230648 | HITS-CLIP | 37986224\|25083871 | Unspecified |
| mmu-miR-152-3p | MIMAT0000162 | Efr3a | 76740 | HITS-CLIP | 37986224\|25083871 | Unspecified |
| mmu-miR-152-3p | MIMAT0000162 | Efr3b | 668212 | HITS-CLIP | 23597149 | Unspecified |
| mmu-miR-152-3p | MIMAT0000162 | Egfr | 13649 | HITS-CLIP | 23597149 | Unspecified |
| mmu-miR-152-3p | MIMAT0000162 | Egr3 | 13655 | HITS-CLIP | 25083871 | Unspecified |
| mmu-miR-152-3p | MIMAT0000162 | Ehbp1l1 | 114601 | HITS-CLIP | 25083871 | Unspecified |
| mmu-miR-152-3p | MIMAT0000162 | Ehd2 | 259300 | HITS-CLIP | 25083871 | Unspecified |
| mmu-miR-152-3p | MIMAT0000162 | Ehmt1 | 77683 | HITS-CLIP | 25083871\|23597149 | Unspecified |
| mmu-miR-152-3p | MIMAT0000162 | Ehmt2 | 110147 | HITS-CLIP | 23597149 | Unspecified |
| mmu-miR-152-3p | MIMAT0000162 | Eif2a | 229317 | HITS-CLIP | 23597149\|29167373 | Unspecified |

| | | | | | | |
|---|---|---|---|---|---|---|
| mmu-miR-152-3p | MIMAT0000162 | Eif2ak1 | 15467 | HITS-CLIP | 37986224 | Unspecified |
| mmu-miR-152-3p | MIMAT0000162 | Eif2d | 16865 | HITS-CLIP | 23597149 | Unspecified |
| mmu-miR-152-3p | MIMAT0000162 | Eif3a | 13669 | HITS-CLIP | 29167373 | Unspecified |
| mmu-miR-152-3p | MIMAT0000162 | Eif3b | 27979 | HITS-CLIP | 25083871 | Unspecified |
| mmu-miR-152-3p | MIMAT0000162 | Eif3f | 66085 | HITS-CLIP | 25083871 | Unspecified |
| mmu-miR-152-3p | MIMAT0000162 | Eif3l | 223691 | HITS-CLIP | 23597149 | Unspecified |
| mmu-miR-152-3p | MIMAT0000162 | Eif4a2 | 13682 | HITS-CLIP | 25083871\|23597149 | Unspecified |
| mmu-miR-152-3p | MIMAT0000162 | Eif4b | 75705 | HITS-CLIP | 37986224\|25083871\|29167373 | Unspecified |
| mmu-miR-152-3p | MIMAT0000162 | Eif4g1 | 208643 | HITS-CLIP | 25083871 | Unspecified |
| mmu-miR-152-3p | MIMAT0000162 | Eif4g2 | 13690 | HITS-CLIP | 25083871\|29167373 | Unspecified |
| mmu-miR-152-3p | MIMAT0000162 | Eif4g3 | 230861 | HITS-CLIP | 25083871\|29167373 | Unspecified |
| mmu-miR-152-3p | MIMAT0000162 | Eif4h | 22384 | HITS-CLIP | 37986224\|25083871 | Unspecified |
| mmu-miR-152-3p | MIMAT0000162 | Elac1 | 114615 | HITS-CLIP | 37986224 | Unspecified |
| mmu-miR-152-3p | MIMAT0000162 | Elavl1 | 15568 | HITS-CLIP | 25083871 | Unspecified |
| mmu-miR-152-3p | MIMAT0000162 | Elf4 | 56501 | HITS-CLIP | 37986224\|25083871 | Unspecified |
| mmu-miR-152-3p | MIMAT0000162 | Elk3 | 13713 | HITS-CLIP | 25083871 | Unspecified |
| mmu-miR-152-3p | MIMAT0000162 | Elk4 | 13714 | HITS-CLIP | 25083871\|23597149 | Unspecified |
| mmu-miR-152-3p | MIMAT0000162 | Ell | 13716 | HITS-CLIP | 23597149 | Unspecified |
| mmu-miR-152-3p | MIMAT0000162 | Eloc | 6921 | HITS-CLIP | 25083871 | Unspecified |
| mmu-miR-152-3p | MIMAT0000162 | Elp2 | 58523 | HITS-CLIP | 29167373 | Unspecified |
| mmu-miR-152-3p | MIMAT0000162 | Emc1 | 230866 | HITS-CLIP | 23597149 | Unspecified |
| mmu-miR-152-3p | MIMAT0000162 | Emc2 | 66736 | HITS-CLIP | 23597149 | Unspecified |
| mmu-miR-152-3p | MIMAT0000162 | Emilin2 | 246707 | HITS-CLIP | 37986224 | Unspecified |
| mmu-miR-152-3p | MIMAT0000162 | Eml4 | 78798 | HITS-CLIP | 25083871\|29167373 | Unspecified |
| mmu-miR-152-3p | MIMAT0000162 | Enah | 13800 | HITS-CLIP | 37986224 | Unspecified |
| mmu-miR-152-3p | MIMAT0000162 | Eno3 | 13808 | HITS-CLIP | 25083871 | Unspecified |
| mmu-miR-152-3p | MIMAT0000162 | Entpd6 | 12497 | HITS-CLIP | 23597149 | Unspecified |
| mmu-miR-152-3p | MIMAT0000162 | Eogt | 101351 | HITS-CLIP | 37986224\|25083871 | Unspecified |
| mmu-miR-152-3p | MIMAT0000162 | Ep300 | 328572 | HITS-CLIP | 37986224\|25083871\|23597149\|29167 | Unspecified |
| mmu-miR-152-3p | MIMAT0000162 | Ep400 | 75560 | HITS-CLIP | 25083871 | Unspecified |
| mmu-miR-152-3p | MIMAT0000162 | Epb41l5 | 226352 | HITS-CLIP | 25083871 | Unspecified |
| mmu-miR-152-3p | MIMAT0000162 | Epc1 | 13831 | HITS-CLIP | 37986224 | Unspecified |
| mmu-miR-152-3p | MIMAT0000162 | Epc2 | 227867 | HITS-CLIP | 25083871\|23597149 | Unspecified |
| mmu-miR-152-3p | MIMAT0000162 | Epdr1 | 105298 | HITS-CLIP | 25083871 | Unspecified |
| mmu-miR-152-3p | MIMAT0000162 | Epg5 | 100502841 | HITS-CLIP | 25083871 | Unspecified |
| mmu-miR-152-3p | MIMAT0000162 | Epha1 | 13835 | HITS-CLIP | 37986224\|23597149 | Unspecified |
| mmu-miR-152-3p | MIMAT0000162 | Epha4 | 13838 | HITS-CLIP | 37986224 | Unspecified |
| mmu-miR-152-3p | MIMAT0000162 | Ephb6 | 13848 | HITS-CLIP | 37986224 | Unspecified |
| mmu-miR-152-3p | MIMAT0000162 | Epn2 | 13855 | HITS-CLIP | 37986224 | Unspecified |

| | | | | | | |
|---|---|---|---|---|---|---|
| mmu-miR-152-3p | MIMAT0000162 | Eppk1 | 3857 | HITS-CLIP | 23597149 | Unspecified |
| mmu-miR-152-3p | MIMAT0000162 | Eprs | 107508 | HITS-CLIP | 25083871 | Unspecified |
| mmu-miR-152-3p | MIMAT0000162 | Eps15 | 13858 | HITS-CLIP | 37986224\|25083871\|23597149 | Unspecified |
| mmu-miR-152-3p | MIMAT0000162 | Erbb2 | 13866 | HITS-CLIP | 25083871 | Unspecified |
| mmu-miR-152-3p | MIMAT0000162 | Erbb3 | 13867 | HITS-CLIP | 37986224 | Unspecified |
| mmu-miR-152-3p | MIMAT0000162 | Erbin | 55914 | HITS-CLIP | 25083871 | Unspecified |
| mmu-miR-152-3p | MIMAT0000162 | Erc1 | 111173 | HITS-CLIP | 25083871 | Unspecified |
| mmu-miR-152-3p | MIMAT0000162 | Ercc2 | 13871 | HITS-CLIP | 23597149 | Unspecified |
| mmu-miR-152-3p | MIMAT0000162 | Ercc6 | 319955 | HITS-CLIP | 29167373 | Unspecified |
| mmu-miR-152-3p | MIMAT0000162 | Ergic1 | 67458 | HITS-CLIP | 25083871 | Unspecified |
| mmu-miR-152-3p | MIMAT0000162 | Erlin1 | 226144 | HITS-CLIP | 37986224\|29167373 | Unspecified |
| mmu-miR-152-3p | MIMAT0000162 | Ermp1 | 226090 | HITS-CLIP | 37986224 | Unspecified |
| mmu-miR-152-3p | MIMAT0000162 | Ern1 | 78943 | HITS-CLIP | 25083871 | Unspecified |
| mmu-miR-152-3p | MIMAT0000162 | Ero1l | 50527 | HITS-CLIP | 25083871 | Unspecified |
| mmu-miR-152-3p | MIMAT0000162 | Erp44 | 76299 | HITS-CLIP | 25083871 | Unspecified |
| mmu-miR-152-3p | MIMAT0000162 | Errfi1 | 74155 | HITS-CLIP | 37986224\|25083871\|23597149 | Unspecified |
| mmu-miR-152-3p | MIMAT0000162 | Esrp2 | 77411 | HITS-CLIP | 37986224 | Unspecified |
| mmu-miR-152-3p | MIMAT0000162 | Esyt2 | 52635 | HITS-CLIP | 25083871 | Unspecified |
| mmu-miR-152-3p | MIMAT0000162 | Etf1 | 225363 | HITS-CLIP | 23597149 | Unspecified |
| mmu-miR-152-3p | MIMAT0000162 | Etfb | 110826 | HITS-CLIP | 23597149\|25083871 | Unspecified |
| mmu-miR-152-3p | MIMAT0000162 | Etfdh | 66841 | HITS-CLIP | 37986224\|25083871\|23597149 | Unspecified |
| mmu-miR-152-3p | MIMAT0000162 | Etnk1 | 75320 | HITS-CLIP | 37986224\|29167373 | Unspecified |
| mmu-miR-152-3p | MIMAT0000162 | Etnk2 | 214253 | HITS-CLIP | 23597149 | Unspecified |
| mmu-miR-152-3p | MIMAT0000162 | Ets1 | 23871 | HITS-CLIP | 23597149\|25083871 | Unspecified |
| mmu-miR-152-3p | MIMAT0000162 | Evi5 | 14020 | HITS-CLIP | 37986224\|25083871\|29167373 | Unspecified |
| mmu-miR-152-3p | MIMAT0000162 | Exd2 | 97827 | HITS-CLIP | 25083871 | Unspecified |
| mmu-miR-152-3p | MIMAT0000162 | Exo1 | 26909 | HITS-CLIP | 29167373 | Unspecified |
| mmu-miR-152-3p | MIMAT0000162 | Exoc1 | 69940 | HITS-CLIP | 37986224\|25083871 | Unspecified |
| mmu-miR-152-3p | MIMAT0000162 | Exoc2 | 66482 | HITS-CLIP | 37986224\|23597149\|29167373 | Unspecified |
| mmu-miR-152-3p | MIMAT0000162 | Exoc4 | 20336 | HITS-CLIP | 29167373 | Unspecified |
| mmu-miR-152-3p | MIMAT0000162 | Exoc5 | 105504 | HITS-CLIP | 25083871\|23597149 | Unspecified |
| mmu-miR-152-3p | MIMAT0000162 | Exoc6b | 75914 | HITS-CLIP | 25083871\|23597149 | Unspecified |
| mmu-miR-152-3p | MIMAT0000162 | Exosc3 | 66362 | HITS-CLIP | 23597149 | Unspecified |
| mmu-miR-152-3p | MIMAT0000162 | Exph5 | 320051 | HITS-CLIP | 37986224 | Unspecified |
| mmu-miR-152-3p | MIMAT0000162 | Eya1 | 14048 | HITS-CLIP | 25083871 | Unspecified |
| mmu-miR-152-3p | MIMAT0000162 | Eya3 | 14050 | HITS-CLIP | 25083871\|29167373 | Unspecified |
| mmu-miR-152-3p | MIMAT0000162 | Ezh1 | 14055 | HITS-CLIP | 37986224\|25083871 | Unspecified |
| mmu-miR-152-3p | MIMAT0000162 | F11 | 109821 | HITS-CLIP | 23597149 | Unspecified |
| mmu-miR-152-3p | MIMAT0000162 | F2r | 14062 | HITS-CLIP | 25083871 | Unspecified |

| | | | | | | |
|---|---|---|---|---|---|---|
| mmu-miR-152-3p | MIMAT0000162 | F9 | 14071 | HITS-CLIP | 23597149 | Unspecified |
| mmu-miR-152-3p | MIMAT0000162 | Faah | 14073 | HITS-CLIP | 23597149 | Unspecified |
| mmu-miR-152-3p | MIMAT0000162 | Fads2 | 56473 | HITS-CLIP | 23597149 | Unspecified |
| mmu-miR-152-3p | MIMAT0000162 | Faf1 | 14084 | HITS-CLIP | 37986224 | Unspecified |
| mmu-miR-152-3p | MIMAT0000162 | Fam102a | 98952 | HITS-CLIP | 25083871 | Unspecified |
| mmu-miR-152-3p | MIMAT0000162 | Fam104a | 28081 | HITS-CLIP | 23597149 | Unspecified |
| mmu-miR-152-3p | MIMAT0000162 | Fam114a1 | 68303 | HITS-CLIP | 25083871 | Unspecified |
| mmu-miR-152-3p | MIMAT0000162 | Fam117b | 72750 | HITS-CLIP | 25083871\|23597149 | Unspecified |
| mmu-miR-152-3p | MIMAT0000162 | Fam120a | 218236 | HITS-CLIP | 37986224\|25083871\|23597149 | Unspecified |
| mmu-miR-152-3p | MIMAT0000162 | Fam126a | 84652 | HITS-CLIP | 25083871\|23597149 | Unspecified |
| mmu-miR-152-3p | MIMAT0000162 | Fam126b | 213056 | HITS-CLIP | 37986224 | Unspecified |
| mmu-miR-152-3p | MIMAT0000162 | Fam135a | 68187 | HITS-CLIP | 37986224 | Unspecified |
| mmu-miR-152-3p | MIMAT0000162 | Fam13c | 71721 | HITS-CLIP | 25083871 | Unspecified |
| mmu-miR-152-3p | MIMAT0000162 | Fam160a1 | 229488 | HITS-CLIP | 23597149 | Unspecified |
| mmu-miR-152-3p | MIMAT0000162 | Fam168a | 319604 | HITS-CLIP | 37986224 | Unspecified |
| mmu-miR-152-3p | MIMAT0000162 | Fam168b | 214469 | HITS-CLIP | 25083871\|23597149 | Unspecified |
| mmu-miR-152-3p | MIMAT0000162 | Fam171a1 | 269233 | HITS-CLIP | 25083871 | Unspecified |
| mmu-miR-152-3p | MIMAT0000162 | Fam193a | 231128 | HITS-CLIP | 37986224\|29167373 | Unspecified |
| mmu-miR-152-3p | MIMAT0000162 | Fam193b | 212483 | HITS-CLIP | 25083871 | Unspecified |
| mmu-miR-152-3p | MIMAT0000162 | Fam199x | 245622 | HITS-CLIP | 37986224 | Unspecified |
| mmu-miR-152-3p | MIMAT0000162 | Fam208b | 54906 | HITS-CLIP | 29167373 | Unspecified |
| mmu-miR-152-3p | MIMAT0000162 | Fam20b | 215015 | HITS-CLIP | 25083871 | Unspecified |
| mmu-miR-152-3p | MIMAT0000162 | Fam210a | 108654 | HITS-CLIP | 23597149 | Unspecified |
| mmu-miR-152-3p | MIMAT0000162 | Fam214a | 235493 | HITS-CLIP | 23597149 | Unspecified |
| mmu-miR-152-3p | MIMAT0000162 | Fam43a | 224093 | HITS-CLIP | 25083871 | Unspecified |
| mmu-miR-152-3p | MIMAT0000162 | Fam45a | 67894 | HITS-CLIP | 25083871 | Unspecified |
| mmu-miR-152-3p | MIMAT0000162 | Fam49a | 76820 | HITS-CLIP | 29167373 | Unspecified |
| mmu-miR-152-3p | MIMAT0000162 | Fam49b | 223601 | HITS-CLIP | 25083871 | Unspecified |
| mmu-miR-152-3p | MIMAT0000162 | Fam53b | 77938 | HITS-CLIP | 25083871 | Unspecified |
| mmu-miR-152-3p | MIMAT0000162 | Fam57a | 116972 | HITS-CLIP | 25083871 | Unspecified |
| mmu-miR-152-3p | MIMAT0000162 | Fam89b | 17826 | HITS-CLIP | 25083871 | Unspecified |
| mmu-miR-152-3p | MIMAT0000162 | Fam8a1 | 51439 | HITS-CLIP | 25083871 | Unspecified |
| mmu-miR-152-3p | MIMAT0000162 | Fam91a1 | 210998 | HITS-CLIP | 25083871\|29167373 | Unspecified |
| mmu-miR-152-3p | MIMAT0000162 | Fancb | 237211 | HITS-CLIP | 25083871 | Unspecified |
| mmu-miR-152-3p | MIMAT0000162 | Far1 | 67420 | HITS-CLIP | 25083871 | Unspecified |
| mmu-miR-152-3p | MIMAT0000162 | Farsa | 66590 | HITS-CLIP | 25083871 | Unspecified |
| mmu-miR-152-3p | MIMAT0000162 | Fastkd1 | 320720 | HITS-CLIP | 37986224 | Unspecified |
| mmu-miR-152-3p | MIMAT0000162 | Fastkd3 | 69577 | HITS-CLIP | 37986224 | Unspecified |
| mmu-miR-152-3p | MIMAT0000162 | Fat1 | 14107 | HITS-CLIP | 25083871 | Unspecified |

| | | | | | | |
|---|---|---|---|---|---|---|
| mmu-miR-152-3p | MIMAT0000162 | Fbf1 | 217335 | HITS-CLIP | 37986224 | Unspecified |
| mmu-miR-152-3p | MIMAT0000162 | Fbn1 | 14118 | HITS-CLIP | 37986224\|25083871 | Unspecified |
| mmu-miR-152-3p | MIMAT0000162 | Fbxl16 | 214931 | HITS-CLIP | 25083871 | Unspecified |
| mmu-miR-152-3p | MIMAT0000162 | Fbxl19 | 233902 | HITS-CLIP | 37986224\|25083871\|23597149 | Unspecified |
| mmu-miR-152-3p | MIMAT0000162 | Fbxl20 | 72194 | HITS-CLIP | 37986224 | Unspecified |
| mmu-miR-152-3p | MIMAT0000162 | Fbxo11 | 225055 | HITS-CLIP | 37986224 | Unspecified |
| mmu-miR-152-3p | MIMAT0000162 | Fbxo28 | 67948 | HITS-CLIP | 37986224 | Unspecified |
| mmu-miR-152-3p | MIMAT0000162 | Fbxo3 | 57443 | HITS-CLIP | 37986224 | Unspecified |
| mmu-miR-152-3p | MIMAT0000162 | Fbxo33 | 70611 | HITS-CLIP | 37986224\|25083871 | Unspecified |
| mmu-miR-152-3p | MIMAT0000162 | Fbxo38 | 107035 | HITS-CLIP | 25083871 | Unspecified |
| mmu-miR-152-3p | MIMAT0000162 | Fbxo45 | 268882 | HITS-CLIP | 37986224 | Unspecified |
| mmu-miR-152-3p | MIMAT0000162 | Fbxo8 | 50753 | HITS-CLIP | 25083871 | Unspecified |
| mmu-miR-152-3p | MIMAT0000162 | Fbxw11 | 103583 | HITS-CLIP | 37986224 | Unspecified |
| mmu-miR-152-3p | MIMAT0000162 | Fbxw2 | 30050 | HITS-CLIP | 21258322 | Unspecified |
| mmu-miR-152-3p | MIMAT0000162 | Fbxw8 | 231672 | HITS-CLIP | 25083871 | Unspecified |
| mmu-miR-152-3p | MIMAT0000162 | Fech | 14151 | HITS-CLIP | 23597149 | Unspecified |
| mmu-miR-152-3p | MIMAT0000162 | Fem1a | 14154 | HITS-CLIP | 37986224 | Unspecified |
| mmu-miR-152-3p | MIMAT0000162 | Fem1b | 14155 | HITS-CLIP | 25083871 | Unspecified |
| mmu-miR-152-3p | MIMAT0000162 | Fem1c | 240263 | HITS-CLIP | 37986224\|25083871 | Unspecified |
| mmu-miR-152-3p | MIMAT0000162 | Fer | 14158 | HITS-CLIP | 37986224\|29167373 | Unspecified |
| mmu-miR-152-3p | MIMAT0000162 | Fermt2 | 218952 | HITS-CLIP | 23597149 | Unspecified |
| mmu-miR-152-3p | MIMAT0000162 | Fermt3 | 108101 | HITS-CLIP | 29167373 | Unspecified |
| mmu-miR-152-3p | MIMAT0000162 | Fgb | 110135 | HITS-CLIP | 23597149 | Unspecified |
| mmu-miR-152-3p | MIMAT0000162 | Fgd6 | 13998 | HITS-CLIP | 29167373 | Unspecified |
| mmu-miR-152-3p | MIMAT0000162 | Fgf11 | 14166 | HITS-CLIP | 37986224 | Unspecified |
| mmu-miR-152-3p | MIMAT0000162 | Fgfr1 | 14182 | HITS-CLIP | 25083871 | Unspecified |
| mmu-miR-152-3p | MIMAT0000162 | Fgfr3 | 14184 | HITS-CLIP | 37986224 | Unspecified |
| mmu-miR-152-3p | MIMAT0000162 | Fgfrl1 | 116701 | HITS-CLIP | 25083871 | Unspecified |
| mmu-miR-152-3p | MIMAT0000162 | Fgl2 | 14190 | HITS-CLIP | 29167373 | Unspecified |
| mmu-miR-152-3p | MIMAT0000162 | Filip1l | 78749 | HITS-CLIP | 25083871 | Unspecified |
| mmu-miR-152-3p | MIMAT0000162 | Fip1l1 | 66899 | HITS-CLIP | 25083871\|29167373 | Unspecified |
| mmu-miR-152-3p | MIMAT0000162 | Fkbp10 | 14230 | HITS-CLIP | 25083871 | Unspecified |
| mmu-miR-152-3p | MIMAT0000162 | Fkbp14 | 231997 | HITS-CLIP | 37986224 | Unspecified |
| mmu-miR-152-3p | MIMAT0000162 | Fkbp1a | 14225 | HITS-CLIP | 25083871\|23597149 | Unspecified |
| mmu-miR-152-3p | MIMAT0000162 | Fkbp1b | 14226 | HITS-CLIP | 25083871 | Unspecified |
| mmu-miR-152-3p | MIMAT0000162 | Fkbp2 | 14227 | HITS-CLIP | 29167373\|23597149 | Unspecified |
| mmu-miR-152-3p | MIMAT0000162 | Fkbp4 | 14228 | HITS-CLIP | 25083871 | Unspecified |
| mmu-miR-152-3p | MIMAT0000162 | Fkbp9 | 27055 | HITS-CLIP | 37986224 | Unspecified |
| mmu-miR-152-3p | MIMAT0000162 | Fktn | 246179 | HITS-CLIP | 25083871 | Unspecified |

| | | | | | | |
|---|---|---|---|---|---|---|
| mmu-miR-152-3p | MIMAT0000162 | Flcn | 216805 | HITS-CLIP | 23597149 | Unspecified |
| mmu-miR-152-3p | MIMAT0000162 | Flna | 192176 | HITS-CLIP | 25083871\|23597149\|29167373 | Unspecified |
| mmu-miR-152-3p | MIMAT0000162 | Flnc | 68794 | HITS-CLIP | 37986224\|25083871 | Unspecified |
| mmu-miR-152-3p | MIMAT0000162 | Flot2 | 14252 | HITS-CLIP | 25083871 | Unspecified |
| mmu-miR-152-3p | MIMAT0000162 | Flrt2 | 399558 | HITS-CLIP | 37986224 | Unspecified |
| mmu-miR-152-3p | MIMAT0000162 | Fmnl3 | 22379 | HITS-CLIP | 25083871\|29167373 | Unspecified |
| mmu-miR-152-3p | MIMAT0000162 | Fmod | 14264 | HITS-CLIP | 37986224\|25083871 | Unspecified |
| mmu-miR-152-3p | MIMAT0000162 | Fmr1 | 14265 | HITS-CLIP | 37986224\|25083871\|23597149 | Unspecified |
| mmu-miR-152-3p | MIMAT0000162 | Fn1 | 14268 | HITS-CLIP | 25083871\|23597149 | Unspecified |
| mmu-miR-152-3p | MIMAT0000162 | Fnbp4 | 55935 | HITS-CLIP | 25083871 | Unspecified |
| mmu-miR-152-3p | MIMAT0000162 | Fndc1 | 68655 | HITS-CLIP | 25083871 | Unspecified |
| mmu-miR-152-3p | MIMAT0000162 | Fndc3a | 319448 | HITS-CLIP | 29167373 | Unspecified |
| mmu-miR-152-3p | MIMAT0000162 | Fndc3b | 72007 | HITS-CLIP | 23597149\|25083871 | Unspecified |
| mmu-miR-152-3p | MIMAT0000162 | Fntb | 110606 | HITS-CLIP | 25083871 | Unspecified |
| mmu-miR-152-3p | MIMAT0000162 | Fos | 14281 | HITS-CLIP | 23597149\|29167373 | Unspecified |
| mmu-miR-152-3p | MIMAT0000162 | Fosl2 | 14284 | HITS-CLIP | 23597149\|25083871 | Unspecified |
| mmu-miR-152-3p | MIMAT0000162 | Foxf2 | 14238 | HITS-CLIP | 25083871 | Unspecified |
| mmu-miR-152-3p | MIMAT0000162 | Foxj3 | 230700 | HITS-CLIP | 23597149 | Unspecified |
| mmu-miR-152-3p | MIMAT0000162 | Foxk2 | 68837 | HITS-CLIP | 37986224 | Unspecified |
| mmu-miR-152-3p | MIMAT0000162 | Foxo4 | 54601 | HITS-CLIP | 25083871 | Unspecified |
| mmu-miR-152-3p | MIMAT0000162 | Frg1 | 14300 | HITS-CLIP | 37986224 | Unspecified |
| mmu-miR-152-3p | MIMAT0000162 | Frk | 14302 | HITS-CLIP | 25083871 | Unspecified |
| mmu-miR-152-3p | MIMAT0000162 | Frmd4a | 209630 | HITS-CLIP | 37986224\|25083871\|23597149\|29167 | Unspecified |
| mmu-miR-152-3p | MIMAT0000162 | Frmd6 | 319710 | HITS-CLIP | 37986224 | Unspecified |
| mmu-miR-152-3p | MIMAT0000162 | Fryl | 72313 | HITS-CLIP | 23597149 | Unspecified |
| mmu-miR-152-3p | MIMAT0000162 | Fst | 14313 | HITS-CLIP | 25083871 | Unspecified |
| mmu-miR-152-3p | MIMAT0000162 | Fstl1 | 14314 | HITS-CLIP | 25083871 | Unspecified |
| mmu-miR-152-3p | MIMAT0000162 | Fubp1 | 51886 | HITS-CLIP | 29167373 | Unspecified |
| mmu-miR-152-3p | MIMAT0000162 | Furin | 18550 | HITS-CLIP | 23597149 | Unspecified |
| mmu-miR-152-3p | MIMAT0000162 | Fyco1 | 17281 | HITS-CLIP | 37986224\|25083871 | Unspecified |
| mmu-miR-152-3p | MIMAT0000162 | Fzd5 | 14367 | HITS-CLIP | 37986224 | Unspecified |
| mmu-miR-152-3p | MIMAT0000162 | G3bp2 | 23881 | HITS-CLIP | 37986224\|21258322 | Unspecified |
| mmu-miR-152-3p | MIMAT0000162 | G6pdx | 14381 | HITS-CLIP | 21258322 | Unspecified |
| mmu-miR-152-3p | MIMAT0000162 | Gab1 | 14388 | HITS-CLIP | 37986224 | Unspecified |
| mmu-miR-152-3p | MIMAT0000162 | Gabarap | 56486 | HITS-CLIP | 25083871 | Unspecified |
| mmu-miR-152-3p | MIMAT0000162 | Gabarapl1 | 57436 | HITS-CLIP | 25083871 | Unspecified |
| mmu-miR-152-3p | MIMAT0000162 | Gadd45g | 23882 | HITS-CLIP | 37986224 | Unspecified |
| mmu-miR-152-3p | MIMAT0000162 | Gadd45gip1 | 102060 | HITS-CLIP | 26203562 | Unspecified |
| mmu-miR-152-3p | MIMAT0000162 | Galc | 14420 | HITS-CLIP | 25083871 | Unspecified |

| | | | | | | |
|---|---|---|---|---|---|---|
| mmu-miR-152-3p | MIMAT0000162 | Gale | 74246 | HITS-CLIP | 25083871 | Unspecified |
| mmu-miR-152-3p | MIMAT0000162 | Galk1 | 14635 | HITS-CLIP | 25083871 | Unspecified |
| mmu-miR-152-3p | MIMAT0000162 | Galnt2 | 108148 | HITS-CLIP | 37986224 | Unspecified |
| mmu-miR-152-3p | MIMAT0000162 | Galnt7 | 108150 | HITS-CLIP | 29167373 | Unspecified |
| mmu-miR-152-3p | MIMAT0000162 | Gamt | 14431 | HITS-CLIP | 37986224 | Unspecified |
| mmu-miR-152-3p | MIMAT0000162 | Gapvd1 | 66691 | HITS-CLIP | 37986224 | Unspecified |
| mmu-miR-152-3p | MIMAT0000162 | Gatad2b | 229542 | HITS-CLIP | 25083871 | Unspecified |
| mmu-miR-152-3p | MIMAT0000162 | Gatb | 90603889 | HITS-CLIP | 23597149 | Unspecified |
| mmu-miR-152-3p | MIMAT0000162 | Gbf1 | 107338 | HITS-CLIP | 25083871 | Unspecified |
| mmu-miR-152-3p | MIMAT0000162 | Gca | 227960 | HITS-CLIP | 23597149 | Unspecified |
| mmu-miR-152-3p | MIMAT0000162 | Gcc2 | 70297 | HITS-CLIP | 37986224 | Unspecified |
| mmu-miR-152-3p | MIMAT0000162 | Gcdh | 270076 | HITS-CLIP | 23597149 | Unspecified |
| mmu-miR-152-3p | MIMAT0000162 | Gcn1l1 | 10985 | HITS-CLIP | 23597149 | Unspecified |
| mmu-miR-152-3p | MIMAT0000162 | Gcnt2 | 14538 | HITS-CLIP | 37986224\|23597149 | Unspecified |
| mmu-miR-152-3p | MIMAT0000162 | Gcnt4 | 218476 | HITS-CLIP | 37986224\|25083871\|23597149 | Unspecified |
| mmu-miR-152-3p | MIMAT0000162 | Gdap2 | 14547 | HITS-CLIP | 37986224 | Unspecified |
| mmu-miR-152-3p | MIMAT0000162 | Gde1 | 56209 | HITS-CLIP | 25083871 | Unspecified |
| mmu-miR-152-3p | MIMAT0000162 | Gdi1 | 14567 | HITS-CLIP | 37986224 | Unspecified |
| mmu-miR-152-3p | MIMAT0000162 | Gdi2 | 14569 | HITS-CLIP | 25083871 | Unspecified |
| mmu-miR-152-3p | MIMAT0000162 | Gdnf | 14573 | HITS-CLIP | 37986224 | Unspecified |
| mmu-miR-152-3p | MIMAT0000162 | Gfpt1 | 14583 | HITS-CLIP | 37986224\|25083871 | Unspecified |
| mmu-miR-152-3p | MIMAT0000162 | Gfra2 | 14586 | HITS-CLIP | 37986224 | Unspecified |
| mmu-miR-152-3p | MIMAT0000162 | Gga2 | 74105 | HITS-CLIP | 23597149 | Unspecified |
| mmu-miR-152-3p | MIMAT0000162 | Ggcx | 56316 | HITS-CLIP | 37986224\|23597149 | Unspecified |
| mmu-miR-152-3p | MIMAT0000162 | Ggnbp2 | 217039 | HITS-CLIP | 25083871 | Unspecified |
| mmu-miR-152-3p | MIMAT0000162 | Ghr | 14600 | HITS-CLIP | 37986224 | Unspecified |
| mmu-miR-152-3p | MIMAT0000162 | Gigyf1 | 57330 | HITS-CLIP | 37986224 | Unspecified |
| mmu-miR-152-3p | MIMAT0000162 | Git2 | 26431 | HITS-CLIP | 25083871 | Unspecified |
| mmu-miR-152-3p | MIMAT0000162 | Gk5 | 235533 | HITS-CLIP | 37986224 | Unspecified |
| mmu-miR-152-3p | MIMAT0000162 | Glcci1 | 170772 | HITS-CLIP | 25083871 | Unspecified |
| mmu-miR-152-3p | MIMAT0000162 | Glg1 | 20340 | HITS-CLIP | 37986224 | Unspecified |
| mmu-miR-152-3p | MIMAT0000162 | Glis2 | 83396 | HITS-CLIP | 37986224\|25083871 | Unspecified |
| mmu-miR-152-3p | MIMAT0000162 | Glrx5 | 73046 | HITS-CLIP | 37986224 | Unspecified |
| mmu-miR-152-3p | MIMAT0000162 | Gls2 | 216456 | HITS-CLIP | 37986224 | Unspecified |
| mmu-miR-152-3p | MIMAT0000162 | Glud1 | 14661 | HITS-CLIP | 23597149 | Unspecified |
| mmu-miR-152-3p | MIMAT0000162 | Glyctk | 235582 | HITS-CLIP | 37986224\|23597149 | Unspecified |
| mmu-miR-152-3p | MIMAT0000162 | Gm16286 | 117903916 | HITS-CLIP | 25083871\|23597149 | Unspecified |
| mmu-miR-152-3p | MIMAT0000162 | Gm38394 | 667118 | HITS-CLIP | 25083871 | Unspecified |
| mmu-miR-152-3p | MIMAT0000162 | Gm527 | 217648 | HITS-CLIP | 25083871 | Unspecified |

| | | | | | | |
|---|---|---|---|---|---|---|
| mmu-miR-152-3p | MIMAT0000162 | Gm5878 | 545861 | HITS-CLIP | 37986224 | Unspecified |
| mmu-miR-152-3p | MIMAT0000162 | Gmfb | 63985 | HITS-CLIP | 37986224 | Unspecified |
| mmu-miR-152-3p | MIMAT0000162 | Gmppa | 69080 | HITS-CLIP | 29167373 | Unspecified |
| mmu-miR-152-3p | MIMAT0000162 | Gna11 | 14672 | HITS-CLIP | 25083871\|23597149\|29167373 | Unspecified |
| mmu-miR-152-3p | MIMAT0000162 | Gnai2 | 14678 | HITS-CLIP | 31433990 | Unspecified |
| mmu-miR-152-3p | MIMAT0000162 | Gnao1 | 14681 | HITS-CLIP | 25083871 | Unspecified |
| mmu-miR-152-3p | MIMAT0000162 | Gnaq | 14682 | HITS-CLIP | 25083871\|29167373 | Unspecified |
| mmu-miR-152-3p | MIMAT0000162 | Gnas | 14683 | HITS-CLIP | 29167373 | Unspecified |
| mmu-miR-152-3p | MIMAT0000162 | Gnb1 | 14688 | HITS-CLIP | 29167373 | Unspecified |
| mmu-miR-152-3p | MIMAT0000162 | Gnb2 | 14693 | HITS-CLIP | 25083871\|23597149 | Unspecified |
| mmu-miR-152-3p | MIMAT0000162 | Gng5 | 14707 | HITS-CLIP | 37986224 | Unspecified |
| mmu-miR-152-3p | MIMAT0000162 | Gnl2 | 230737 | HITS-CLIP | 25083871 | Unspecified |
| mmu-miR-152-3p | MIMAT0000162 | Gnl3 | 30877 | HITS-CLIP | 25083871 | Unspecified |
| mmu-miR-152-3p | MIMAT0000162 | Gnpat | 14712 | HITS-CLIP | 37986224 | Unspecified |
| mmu-miR-152-3p | MIMAT0000162 | Gns | 75612 | HITS-CLIP | 25083871 | Unspecified |
| mmu-miR-152-3p | MIMAT0000162 | Golga1 | 76899 | HITS-CLIP | 37986224 | Unspecified |
| mmu-miR-152-3p | MIMAT0000162 | Golga4 | 54214 | HITS-CLIP | 25083871\|23597149\|29167373 | Unspecified |
| mmu-miR-152-3p | MIMAT0000162 | Golgb1 | 224139 | HITS-CLIP | 37986224 | Unspecified |
| mmu-miR-152-3p | MIMAT0000162 | Golph3 | 66629 | HITS-CLIP | 25083871 | Unspecified |
| mmu-miR-152-3p | MIMAT0000162 | Golt1b | 66964 | HITS-CLIP | 25083871 | Unspecified |
| mmu-miR-152-3p | MIMAT0000162 | Gon4l | 76022 | HITS-CLIP | 25083871 | Unspecified |
| mmu-miR-152-3p | MIMAT0000162 | Gosr1 | 53334 | HITS-CLIP | 25083871 | Unspecified |
| mmu-miR-152-3p | MIMAT0000162 | Gpam | 14732 | HITS-CLIP | 23597149 | Unspecified |
| mmu-miR-152-3p | MIMAT0000162 | Gpatch2 | 67769 | HITS-CLIP | 25083871 | Unspecified |
| mmu-miR-152-3p | MIMAT0000162 | Gpatch8 | 237943 | HITS-CLIP | 37986224 | Unspecified |
| mmu-miR-152-3p | MIMAT0000162 | Gpcpd1 | 74182 | HITS-CLIP | 37986224\|25083871\|23597149 | Unspecified |
| mmu-miR-152-3p | MIMAT0000162 | Gpd2 | 14571 | HITS-CLIP | 25083871 | Unspecified |
| mmu-miR-152-3p | MIMAT0000162 | Gpi1 | 14751 | HITS-CLIP | 37986224 | Unspecified |
| mmu-miR-152-3p | MIMAT0000162 | Gpihbp1 | 68453 | HITS-CLIP | 23597149 | Unspecified |
| mmu-miR-152-3p | MIMAT0000162 | Gpkow | 209416 | HITS-CLIP | 37986224 | Unspecified |
| mmu-miR-152-3p | MIMAT0000162 | Gpr107 | 277463 | HITS-CLIP | 37986224 | Unspecified |
| mmu-miR-152-3p | MIMAT0000162 | Gpr108 | 78308 | HITS-CLIP | 29167373 | Unspecified |
| mmu-miR-152-3p | MIMAT0000162 | Gpr146 | 80290 | HITS-CLIP | 37986224\|25083871\|29167373 | Unspecified |
| mmu-miR-152-3p | MIMAT0000162 | Gpr63 | 81006 | HITS-CLIP | 37986224 | Unspecified |
| mmu-miR-152-3p | MIMAT0000162 | Gprc5b | 64297 | HITS-CLIP | 37986224 | Unspecified |
| mmu-miR-152-3p | MIMAT0000162 | Gprin3 | 243385 | HITS-CLIP | 25083871 | Unspecified |
| mmu-miR-152-3p | MIMAT0000162 | Gpt | 76282 | HITS-CLIP | 25083871 | Unspecified |
| mmu-miR-152-3p | MIMAT0000162 | Gramd1b | 235283 | HITS-CLIP | 25083871 | Unspecified |
| mmu-miR-152-3p | MIMAT0000162 | Grem2 | 23893 | HITS-CLIP | 37986224 | Unspecified |

| | | | | | | |
|---|---|---|---|---|---|---|
| mmu-miR-152-3p | MIMAT0000162 | Gripap1 | 54645 | HITS-CLIP | 23597149 | Unspecified |
| mmu-miR-152-3p | MIMAT0000162 | Grn | 14824 | HITS-CLIP | 37986224 | Unspecified |
| mmu-miR-152-3p | MIMAT0000162 | Gse1 | 382034 | HITS-CLIP | 25083871 | Unspecified |
| mmu-miR-152-3p | MIMAT0000162 | Gsk3b | 56637 | HITS-CLIP | 29167373 | Unspecified |
| mmu-miR-152-3p | MIMAT0000162 | Gskip | 66787 | HITS-CLIP | 37986224 | Unspecified |
| mmu-miR-152-3p | MIMAT0000162 | Gspt1 | 14852 | HITS-CLIP | 37986224\|25083871\|23597149\|29167 | Unspecified |
| mmu-miR-152-3p | MIMAT0000162 | Gstm3 | 14864 | HITS-CLIP | 37986224 | Unspecified |
| mmu-miR-152-3p | MIMAT0000162 | Gstm6 | 14867 | HITS-CLIP | 23597149 | Unspecified |
| mmu-miR-152-3p | MIMAT0000162 | Gstz1 | 14874 | HITS-CLIP | 23597149 | Unspecified |
| mmu-miR-152-3p | MIMAT0000162 | Gtf3c1 | 233863 | HITS-CLIP | 25083871 | Unspecified |
| mmu-miR-152-3p | MIMAT0000162 | Gtf3c4 | 269252 | HITS-CLIP | 37986224\|25083871 | Unspecified |
| mmu-miR-152-3p | MIMAT0000162 | Gxylt2 | 232313 | HITS-CLIP | 25083871 | Unspecified |
| mmu-miR-152-3p | MIMAT0000162 | Gyg | 27357 | HITS-CLIP | 25083871 | Unspecified |
| mmu-miR-152-3p | MIMAT0000162 | Gzf1 | 74533 | HITS-CLIP | 37986224 | Unspecified |
| mmu-miR-152-3p | MIMAT0000162 | H2afy | 9555 | HITS-CLIP | 25083871 | Unspecified |
| mmu-miR-152-3p | MIMAT0000162 | H2bc18 | 440689 | HITS-CLIP | 31433990 | Unspecified |
| mmu-miR-152-3p | MIMAT0000162 | Hal | 15109 | HITS-CLIP | 23597149 | Unspecified |
| mmu-miR-152-3p | MIMAT0000162 | Hars2 | 70791 | HITS-CLIP | 25083871 | Unspecified |
| mmu-miR-152-3p | MIMAT0000162 | Haus6 | 230376 | HITS-CLIP | 37986224\|25083871 | Unspecified |
| mmu-miR-152-3p | MIMAT0000162 | Hbp1 | 73389 | HITS-CLIP | 37986224\|25083871\|23597149 | Unspecified |
| mmu-miR-152-3p | MIMAT0000162 | Hc | 15139 | HITS-CLIP | 23597149 | Unspecified |
| mmu-miR-152-3p | MIMAT0000162 | Hcfc1 | 15161 | HITS-CLIP | 25083871 | Unspecified |
| mmu-miR-152-3p | MIMAT0000162 | Hcfc2 | 67933 | HITS-CLIP | 25083871 | Unspecified |
| mmu-miR-152-3p | MIMAT0000162 | Hdlbp | 110611 | HITS-CLIP | 25083871 | Unspecified |
| mmu-miR-152-3p | MIMAT0000162 | Heatr1 | 217995 | HITS-CLIP | 25083871 | Unspecified |
| mmu-miR-152-3p | MIMAT0000162 | Heatr5a | 320487 | HITS-CLIP | 37986224 | Unspecified |
| mmu-miR-152-3p | MIMAT0000162 | Heca | 380629 | HITS-CLIP | 37986224\|25083871 | Unspecified |
| mmu-miR-152-3p | MIMAT0000162 | Hectd1 | 207304 | HITS-CLIP | 37986224\|25083871\|23597149\|29167 | Unspecified |
| mmu-miR-152-3p | MIMAT0000162 | Hectd2 | 226098 | HITS-CLIP | 37986224\|25083871 | Unspecified |
| mmu-miR-152-3p | MIMAT0000162 | Hectd4 | 283450 | HITS-CLIP | 25083871 | Unspecified |
| mmu-miR-152-3p | MIMAT0000162 | Heg1 | 77446 | HITS-CLIP | 37986224\|25083871 | Unspecified |
| mmu-miR-152-3p | MIMAT0000162 | Herc1 | 235439 | HITS-CLIP | 37986224\|25083871 | Unspecified |
| mmu-miR-152-3p | MIMAT0000162 | Herc2 | 15204 | HITS-CLIP | 25083871\|23597149 | Unspecified |
| mmu-miR-152-3p | MIMAT0000162 | Herc3 | 73998 | HITS-CLIP | 25083871 | Unspecified |
| mmu-miR-152-3p | MIMAT0000162 | Hes1 | 15205 | HITS-CLIP | 37986224 | Unspecified |
| mmu-miR-152-3p | MIMAT0000162 | Hgfac | 54426 | HITS-CLIP | 23597149 | Unspecified |
| mmu-miR-152-3p | MIMAT0000162 | Hgsnat | 52120 | HITS-CLIP | 29167373 | Unspecified |
| mmu-miR-152-3p | MIMAT0000162 | Hhex | 15242 | HITS-CLIP | 23597149 | Unspecified |
| mmu-miR-152-3p | MIMAT0000162 | Hibch | 227095 | HITS-CLIP | 23597149 | Unspecified |

| | | | | | | |
|---|---|---|---|---|---|---|
| mmu-miR-152-3p | MIMAT0000162 | Hif1a | 15251 | HITS-CLIP | 25083871 | Unspecified |
| mmu-miR-152-3p | MIMAT0000162 | Hif1an | 319594 | HITS-CLIP | 37986224\|25083871 | Unspecified |
| mmu-miR-152-3p | MIMAT0000162 | Hint2 | 68917 | HITS-CLIP | 25083871 | Unspecified |
| mmu-miR-152-3p | MIMAT0000162 | Hipk1 | 15257 | HITS-CLIP | 25083871\|29167373 | Unspecified |
| mmu-miR-152-3p | MIMAT0000162 | Hipk2 | 15258 | HITS-CLIP | 37986224\|25083871\|29167373 | Unspecified |
| mmu-miR-152-3p | MIMAT0000162 | Hira | 15260 | HITS-CLIP | 37986224 | Unspecified |
| mmu-miR-152-3p | MIMAT0000162 | Hivep2 | 15273 | HITS-CLIP | 25083871 | Unspecified |
| mmu-miR-152-3p | MIMAT0000162 | Hmg20a | 66867 | HITS-CLIP | 37986224\|25083871\|23597149 | Unspecified |
| mmu-miR-152-3p | MIMAT0000162 | Hmgcr | 15357 | HITS-CLIP | 37986224 | Unspecified |
| mmu-miR-152-3p | MIMAT0000162 | Hmgcs1 | 208715 | HITS-CLIP | 25083871 | Unspecified |
| mmu-miR-152-3p | MIMAT0000162 | Hmgxb4 | 70823 | HITS-CLIP | 37986224 | Unspecified |
| mmu-miR-152-3p | MIMAT0000162 | Hnf4a | 15378 | HITS-CLIP | 37986224 | Unspecified |
| mmu-miR-152-3p | MIMAT0000162 | Hnrnpa2b1 | 53379 | HITS-CLIP | 37986224 | Unspecified |
| mmu-miR-152-3p | MIMAT0000162 | Hnrnpc | 15381 | HITS-CLIP | 37986224 | Unspecified |
| mmu-miR-152-3p | MIMAT0000162 | Hnrnpd | 11991 | HITS-CLIP | 25083871 | Unspecified |
| mmu-miR-152-3p | MIMAT0000162 | Hnrnph3 | 432467 | HITS-CLIP | 23597149 | Unspecified |
| mmu-miR-152-3p | MIMAT0000162 | Hnrnpk | 15387 | HITS-CLIP | 29167373 | Unspecified |
| mmu-miR-152-3p | MIMAT0000162 | Hnrnpl | 15388 | HITS-CLIP | 25083871\|23597149\|29167373 | Unspecified |
| mmu-miR-152-3p | MIMAT0000162 | Hnrnpll | 72692 | HITS-CLIP | 37986224 | Unspecified |
| mmu-miR-152-3p | MIMAT0000162 | Hnrnpm | 76936 | HITS-CLIP | 25083871\|29167373 | Unspecified |
| mmu-miR-152-3p | MIMAT0000162 | Hnrnpr | 74326 | HITS-CLIP | 29167373 | Unspecified |
| mmu-miR-152-3p | MIMAT0000162 | Hnrnpu | 51810 | HITS-CLIP | 37986224 | Unspecified |
| mmu-miR-152-3p | MIMAT0000162 | Hnrnpul1 | 232989 | HITS-CLIP | 25083871 | Unspecified |
| mmu-miR-152-3p | MIMAT0000162 | Hnrnpul2 | 68693 | HITS-CLIP | 29167373 | Unspecified |
| mmu-miR-152-3p | MIMAT0000162 | Homer1 | 26556 | HITS-CLIP | 37986224 | Unspecified |
| mmu-miR-152-3p | MIMAT0000162 | Hook3 | 320191 | HITS-CLIP | 37986224\|25083871 | Unspecified |
| mmu-miR-152-3p | MIMAT0000162 | Hoxa3 | 15400 | HITS-CLIP | 37986224 | Unspecified |
| mmu-miR-152-3p | MIMAT0000162 | Hoxc5 | 15424 | HITS-CLIP | 25083871 | Unspecified |
| mmu-miR-152-3p | MIMAT0000162 | Hoxc8 | 15426 | HITS-CLIP | 37986224 | Unspecified |
| mmu-miR-152-3p | MIMAT0000162 | Hprt | 15452 | HITS-CLIP | 25083871\|23597149 | Unspecified |
| mmu-miR-152-3p | MIMAT0000162 | Hps5 | 246694 | HITS-CLIP | 37986224 | Unspecified |
| mmu-miR-152-3p | MIMAT0000162 | Hrc | 15464 | HITS-CLIP | 25083871 | Unspecified |
| mmu-miR-152-3p | MIMAT0000162 | Hs3st1 | 15476 | HITS-CLIP | 37986224 | Unspecified |
| mmu-miR-152-3p | MIMAT0000162 | Hs6st1 | 50785 | HITS-CLIP | 29167373 | Unspecified |
| mmu-miR-152-3p | MIMAT0000162 | Hsd17b12 | 56348 | HITS-CLIP | 23597149 | Unspecified |
| mmu-miR-152-3p | MIMAT0000162 | Hsf1 | 15499 | HITS-CLIP | 37986224 | Unspecified |
| mmu-miR-152-3p | MIMAT0000162 | Hsp90b1 | 22027 | HITS-CLIP | 37986224\|23597149\|26203562 | Unspecified |
| mmu-miR-152-3p | MIMAT0000162 | Hspa1b | 15511 | HITS-CLIP | 25083871 | Unspecified |
| mmu-miR-152-3p | MIMAT0000162 | Hspa4 | 15525 | HITS-CLIP | 23597149 | Unspecified |

| | | | | | | |
|---|---|---|---|---|---|---|
| mmu-miR-152-3p | MIMAT0000162 | Hspa4l | 18415 | HITS-CLIP | 37986224\|25083871 | Unspecified |
| mmu-miR-152-3p | MIMAT0000162 | Hspa5 | 14828 | HITS-CLIP | 25083871 | Unspecified |
| mmu-miR-152-3p | MIMAT0000162 | Hspa9 | 15526 | HITS-CLIP | 21258322\|25083871 | Unspecified |
| mmu-miR-152-3p | MIMAT0000162 | Hspg2 | 15530 | HITS-CLIP | 25083871 | Unspecified |
| mmu-miR-152-3p | MIMAT0000162 | Htr1b | 15551 | HITS-CLIP | 25083871 | Unspecified |
| mmu-miR-152-3p | MIMAT0000162 | Htra1 | 56213 | HITS-CLIP | 25083871 | Unspecified |
| mmu-miR-152-3p | MIMAT0000162 | Htt | 15194 | HITS-CLIP | 25083871 | Unspecified |
| mmu-miR-152-3p | MIMAT0000162 | Huwe1 | 59026 | HITS-CLIP | 25083871\|23597149 | Unspecified |
| mmu-miR-152-3p | MIMAT0000162 | Hyou1 | 12282 | HITS-CLIP | 25083871 | Unspecified |
| mmu-miR-152-3p | MIMAT0000162 | Iars | 105148 | HITS-CLIP | 25083871 | Unspecified |
| mmu-miR-152-3p | MIMAT0000162 | Icmt | 57295 | HITS-CLIP | 37986224 | Unspecified |
| mmu-miR-152-3p | MIMAT0000162 | Ier3ip1 | 66191 | HITS-CLIP | 25083871 | Unspecified |
| mmu-miR-152-3p | MIMAT0000162 | Iffo1 | 320678 | HITS-CLIP | 25083871 | Unspecified |
| mmu-miR-152-3p | MIMAT0000162 | Iffo2 | 212632 | HITS-CLIP | 25083871\|23597149\|29167373 | Unspecified |
| mmu-miR-152-3p | MIMAT0000162 | Ifit2 | 15958 | HITS-CLIP | 23597149\|29167373 | Unspecified |
| mmu-miR-152-3p | MIMAT0000162 | Ifnar1 | 15975 | HITS-CLIP | 37986224 | Unspecified |
| mmu-miR-152-3p | MIMAT0000162 | Ifnar2 | 15976 | HITS-CLIP | 25083871 | Unspecified |
| mmu-miR-152-3p | MIMAT0000162 | Ifngr1 | 15979 | HITS-CLIP | 37986224 | Unspecified |
| mmu-miR-152-3p | MIMAT0000162 | Ifrd2 | 15983 | HITS-CLIP | 25083871 | Unspecified |
| mmu-miR-152-3p | MIMAT0000162 | Ift46 | 76568 | HITS-CLIP | 23597149 | Unspecified |
| mmu-miR-152-3p | MIMAT0000162 | Igbp1 | 18518 | HITS-CLIP | 23597149 | Unspecified |
| mmu-miR-152-3p | MIMAT0000162 | Igdcc4 | 56741 | HITS-CLIP | 37986224 | Unspecified |
| mmu-miR-152-3p | MIMAT0000162 | Igf1 | 16000 | HITS-CLIP | 37986224 | Unspecified |
| mmu-miR-152-3p | MIMAT0000162 | Igf1r | 16001 | HITS-CLIP | 25083871 | Unspecified |
| mmu-miR-152-3p | MIMAT0000162 | Igsf3 | 78908 | HITS-CLIP | 37986224 | Unspecified |
| mmu-miR-152-3p | MIMAT0000162 | Igsf5 | 72058 | HITS-CLIP | 23597149 | Unspecified |
| mmu-miR-152-3p | MIMAT0000162 | Ikzf2 | 22779 | HITS-CLIP | 25083871 | Unspecified |
| mmu-miR-152-3p | MIMAT0000162 | Il15 | 16168 | HITS-CLIP | 29167373 | Unspecified |
| mmu-miR-152-3p | MIMAT0000162 | Il17rd | 171463 | HITS-CLIP | 37986224 | Unspecified |
| mmu-miR-152-3p | MIMAT0000162 | Il1r1 | 16177 | HITS-CLIP | 25083871 | Unspecified |
| mmu-miR-152-3p | MIMAT0000162 | Il1rap | 16180 | HITS-CLIP | 23597149 | Unspecified |
| mmu-miR-152-3p | MIMAT0000162 | Il6st | 16195 | HITS-CLIP | 37986224\|25083871 | Unspecified |
| mmu-miR-152-3p | MIMAT0000162 | Ilf3 | 16201 | HITS-CLIP | 37986224 | Unspecified |
| mmu-miR-152-3p | MIMAT0000162 | Ilk | 16202 | HITS-CLIP | 37986224\|23597149\|25083871 | Unspecified |
| mmu-miR-152-3p | MIMAT0000162 | Impa1 | 55980 | HITS-CLIP | 37986224 | Unspecified |
| mmu-miR-152-3p | MIMAT0000162 | Inafm2 | 100505573 | HITS-CLIP | 25083871\|23597149 | Unspecified |
| mmu-miR-152-3p | MIMAT0000162 | Incenp | 16319 | HITS-CLIP | 37986224\|29167373 | Unspecified |
| mmu-miR-152-3p | MIMAT0000162 | Inf2 | 70435 | HITS-CLIP | 25083871 | Unspecified |
| mmu-miR-152-3p | MIMAT0000162 | Ing3 | 71777 | HITS-CLIP | 25083871\|23597149 | Unspecified |

| | | | | | | |
|---|---|---|---|---|---|---|
| mmu-miR-152-3p | MIMAT0000162 | Ing5 | 66262 | HITS-CLIP | 37986224 | Unspecified |
| mmu-miR-152-3p | MIMAT0000162 | Inhba | 16323 | HITS-CLIP | 25083871 | Unspecified |
| mmu-miR-152-3p | MIMAT0000162 | Inhbb | 16324 | HITS-CLIP | 37986224 | Unspecified |
| mmu-miR-152-3p | MIMAT0000162 | Ino80 | 68142 | HITS-CLIP | 37986224\|25083871\|23597149 | Unspecified |
| mmu-miR-152-3p | MIMAT0000162 | Ino80d | 227195 | HITS-CLIP | 37986224\|25083871\|29167373 | Unspecified |
| mmu-miR-152-3p | MIMAT0000162 | Inpp4a | 269180 | HITS-CLIP | 25083871 | Unspecified |
| mmu-miR-152-3p | MIMAT0000162 | Inpp4b | 234515 | HITS-CLIP | 25083871 | Unspecified |
| mmu-miR-152-3p | MIMAT0000162 | Inpp5d | 16331 | HITS-CLIP | 29167373 | Unspecified |
| mmu-miR-152-3p | MIMAT0000162 | Insig1 | 231070 | HITS-CLIP | 37986224 | Unspecified |
| mmu-miR-152-3p | MIMAT0000162 | Insig2 | 72999 | HITS-CLIP | 37986224\|23597149\|25083871 | Unspecified |
| mmu-miR-152-3p | MIMAT0000162 | Ints13 | 55726 | HITS-CLIP | 25083871 | Unspecified |
| mmu-miR-152-3p | MIMAT0000162 | Ints6 | 18130 | HITS-CLIP | 37986224\|25083871 | Unspecified |
| mmu-miR-152-3p | MIMAT0000162 | Invs | 16348 | HITS-CLIP | 23597149\|25083871 | Unspecified |
| mmu-miR-152-3p | MIMAT0000162 | Ipo5 | 70572 | HITS-CLIP | 25083871 | Unspecified |
| mmu-miR-152-3p | MIMAT0000162 | Ipo7 | 233726 | HITS-CLIP | 25083871\|29167373 | Unspecified |
| mmu-miR-152-3p | MIMAT0000162 | Ipo8 | 320727 | HITS-CLIP | 37986224\|25083871 | Unspecified |
| mmu-miR-152-3p | MIMAT0000162 | Ippk | 75678 | HITS-CLIP | 37986224 | Unspecified |
| mmu-miR-152-3p | MIMAT0000162 | Iqgap1 | 29875 | HITS-CLIP | 37986224\|25083871 | Unspecified |
| mmu-miR-152-3p | MIMAT0000162 | Iqgap2 | 544963 | HITS-CLIP | 23597149 | Unspecified |
| mmu-miR-152-3p | MIMAT0000162 | Iqsec1 | 232227 | HITS-CLIP | 23597149 | Unspecified |
| mmu-miR-152-3p | MIMAT0000162 | Iqsec2 | 245666 | HITS-CLIP | 23597149 | Unspecified |
| mmu-miR-152-3p | MIMAT0000162 | Ireb2 | 64602 | HITS-CLIP | 37986224\|23597149\|25083871 | Unspecified |
| mmu-miR-152-3p | MIMAT0000162 | Irf2bp2 | 270110 | HITS-CLIP | 37986224 | Unspecified |
| mmu-miR-152-3p | MIMAT0000162 | Irf2bpl | 238330 | HITS-CLIP | 37986224 | Unspecified |
| mmu-miR-152-3p | MIMAT0000162 | Itch | 16396 | HITS-CLIP | 25083871\|23597149 | Unspecified |
| mmu-miR-152-3p | MIMAT0000162 | Itga5 | 16402 | HITS-CLIP | 37986224\|23597149\|25083871 | Unspecified |
| mmu-miR-152-3p | MIMAT0000162 | Itgav | 16410 | HITS-CLIP | 25083871 | Unspecified |
| mmu-miR-152-3p | MIMAT0000162 | Itgb2 | 16414 | HITS-CLIP | 29167373 | Unspecified |
| mmu-miR-152-3p | MIMAT0000162 | Itgb5 | 16419 | HITS-CLIP | 25083871 | Unspecified |
| mmu-miR-152-3p | MIMAT0000162 | Itgb6 | 16420 | HITS-CLIP | 25083871 | Unspecified |
| mmu-miR-152-3p | MIMAT0000162 | Itgb8 | 320910 | HITS-CLIP | 37986224 | Unspecified |
| mmu-miR-152-3p | MIMAT0000162 | Itpr3 | 16440 | HITS-CLIP | 25083871 | Unspecified |
| mmu-miR-152-3p | MIMAT0000162 | Itpripl1 | 73338 | HITS-CLIP | 37986224 | Unspecified |
| mmu-miR-152-3p | MIMAT0000162 | Itpripl2 | 319622 | HITS-CLIP | 26203562 | Unspecified |
| mmu-miR-152-3p | MIMAT0000162 | Itsn2 | 20403 | HITS-CLIP | 37986224 | Unspecified |
| mmu-miR-152-3p | MIMAT0000162 | Jade2 | 76901 | HITS-CLIP | 23597149 | Unspecified |
| mmu-miR-152-3p | MIMAT0000162 | Jagn1 | 67767 | HITS-CLIP | 23597149 | Unspecified |
| mmu-miR-152-3p | MIMAT0000162 | Jak1 | 16451 | HITS-CLIP | 37986224\|25083871\|23597149\|29167 | Unspecified |
| mmu-miR-152-3p | MIMAT0000162 | Jam2 | 67374 | HITS-CLIP | 25083871 | Unspecified |

| | | | | | | |
|---|---|---|---|---|---|---|
| mmu-miR-152-3p | MIMAT0000162 | Jarid2 | 16468 | HITS-CLIP | 37986224\|25083871\|23597149\|29167 | Unspecified |
| mmu-miR-152-3p | MIMAT0000162 | Jcad | 57608 | HITS-CLIP | 25083871 | Unspecified |
| mmu-miR-152-3p | MIMAT0000162 | Jdp2 | 81703 | HITS-CLIP | 25083871\|23597149 | Unspecified |
| mmu-miR-152-3p | MIMAT0000162 | Jkamp | 104771 | HITS-CLIP | 25083871\|23597149 | Unspecified |
| mmu-miR-152-3p | MIMAT0000162 | Jmjd1c | 108829 | HITS-CLIP | 23597149\|29167373 | Unspecified |
| mmu-miR-152-3p | MIMAT0000162 | Jmy | 57748 | HITS-CLIP | 37986224\|25083871 | Unspecified |
| mmu-miR-152-3p | MIMAT0000162 | Jph1 | 57339 | HITS-CLIP | 37986224\|25083871 | Unspecified |
| mmu-miR-152-3p | MIMAT0000162 | Jph2 | 59091 | HITS-CLIP | 37986224 | Unspecified |
| mmu-miR-152-3p | MIMAT0000162 | Kalrn | 545156 | HITS-CLIP | 37986224\|23597149 | Unspecified |
| mmu-miR-152-3p | MIMAT0000162 | Kansl1 | 76719 | HITS-CLIP | 37986224\|29167373 | Unspecified |
| mmu-miR-152-3p | MIMAT0000162 | Kat5 | 81601 | HITS-CLIP | 23597149 | Unspecified |
| mmu-miR-152-3p | MIMAT0000162 | Kat6a | 244349 | HITS-CLIP | 37986224 | Unspecified |
| mmu-miR-152-3p | MIMAT0000162 | Kat6b | 54169 | HITS-CLIP | 25083871 | Unspecified |
| mmu-miR-152-3p | MIMAT0000162 | Kat7 | 217127 | HITS-CLIP | 37986224 | Unspecified |
| mmu-miR-152-3p | MIMAT0000162 | Katnal1 | 231912 | HITS-CLIP | 37986224 | Unspecified |
| mmu-miR-152-3p | MIMAT0000162 | Kbtbd2 | 210973 | HITS-CLIP | 37986224 | Unspecified |
| mmu-miR-152-3p | MIMAT0000162 | Kbtbd7 | 211255 | HITS-CLIP | 37986224 | Unspecified |
| mmu-miR-152-3p | MIMAT0000162 | Kcnj2 | 16518 | HITS-CLIP | 25083871 | Unspecified |
| mmu-miR-152-3p | MIMAT0000162 | Kctd1 | 106931 | HITS-CLIP | 25083871 | Unspecified |
| mmu-miR-152-3p | MIMAT0000162 | Kctd11 | 216858 | HITS-CLIP | 37986224\|25083871 | Unspecified |
| mmu-miR-152-3p | MIMAT0000162 | Kctd5 | 69259 | HITS-CLIP | 37986224\|23597149\|25083871 | Unspecified |
| mmu-miR-152-3p | MIMAT0000162 | Kctd9 | 105440 | HITS-CLIP | 25083871 | Unspecified |
| mmu-miR-152-3p | MIMAT0000162 | Kdelr1 | 68137 | HITS-CLIP | 25083871\|29167373 | Unspecified |
| mmu-miR-152-3p | MIMAT0000162 | Kdm2a | 225876 | HITS-CLIP | 25083871\|29167373 | Unspecified |
| mmu-miR-152-3p | MIMAT0000162 | Kdm3a | 104263 | HITS-CLIP | 37986224 | Unspecified |
| mmu-miR-152-3p | MIMAT0000162 | Kdm4a | 230674 | HITS-CLIP | 25083871 | Unspecified |
| mmu-miR-152-3p | MIMAT0000162 | Kdm4b | 193796 | HITS-CLIP | 25083871 | Unspecified |
| mmu-miR-152-3p | MIMAT0000162 | Kdm4c | 76804 | HITS-CLIP | 23597149\|25083871 | Unspecified |
| mmu-miR-152-3p | MIMAT0000162 | Kdm5b | 75605 | HITS-CLIP | 29167373 | Unspecified |
| mmu-miR-152-3p | MIMAT0000162 | Kdm5c | 20591 | HITS-CLIP | 37986224 | Unspecified |
| mmu-miR-152-3p | MIMAT0000162 | Kdm6a | 22289 | HITS-CLIP | 25083871 | Unspecified |
| mmu-miR-152-3p | MIMAT0000162 | Kdm6b | 216850 | HITS-CLIP | 37986224 | Unspecified |
| mmu-miR-152-3p | MIMAT0000162 | Kdm7a | 338523 | HITS-CLIP | 25083871 | Unspecified |
| mmu-miR-152-3p | MIMAT0000162 | Keg1 | 64697 | HITS-CLIP | 23597149 | Unspecified |
| mmu-miR-152-3p | MIMAT0000162 | Khnyn | 219094 | HITS-CLIP | 37986224 | Unspecified |
| mmu-miR-152-3p | MIMAT0000162 | Kidins220 | 77480 | HITS-CLIP | 31433990 | Unspecified |
| mmu-miR-152-3p | MIMAT0000162 | Kif13a | 16553 | HITS-CLIP | 37986224 | Unspecified |
| mmu-miR-152-3p | MIMAT0000162 | Kif13b | 16554 | HITS-CLIP | 23597149 | Unspecified |
| mmu-miR-152-3p | MIMAT0000162 | Kif5b | 16573 | HITS-CLIP | 25083871 | Unspecified |

| | | | | | | |
|---|---|---|---|---|---|---|
| mmu-miR-152-3p | MIMAT0000162 | Kin | 16588 | HITS-CLIP | 37986224 | Unspecified |
| mmu-miR-152-3p | MIMAT0000162 | Klb | 83379 | HITS-CLIP | 37986224 | Unspecified |
| mmu-miR-152-3p | MIMAT0000162 | Klc1 | 16593 | HITS-CLIP | 29167373 | Unspecified |
| mmu-miR-152-3p | MIMAT0000162 | Klc2 | 16594 | HITS-CLIP | 23597149 | Unspecified |
| mmu-miR-152-3p | MIMAT0000162 | Klf13 | 50794 | HITS-CLIP | 23597149 | Unspecified |
| mmu-miR-152-3p | MIMAT0000162 | Klf15 | 66277 | HITS-CLIP | 37986224 | Unspecified |
| mmu-miR-152-3p | MIMAT0000162 | Klf3 | 16599 | HITS-CLIP | 25083871 | Unspecified |
| mmu-miR-152-3p | MIMAT0000162 | Klf4 | 16600 | HITS-CLIP | 37986224 | Unspecified |
| mmu-miR-152-3p | MIMAT0000162 | Klf6 | 23849 | HITS-CLIP | 37986224\|25083871 | Unspecified |
| mmu-miR-152-3p | MIMAT0000162 | Klf7 | 93691 | HITS-CLIP | 37986224\|25083871 | Unspecified |
| mmu-miR-152-3p | MIMAT0000162 | Klf9 | 16601 | HITS-CLIP | 25083871 | Unspecified |
| mmu-miR-152-3p | MIMAT0000162 | Klhdc10 | 76788 | HITS-CLIP | 25083871\|23597149 | Unspecified |
| mmu-miR-152-3p | MIMAT0000162 | Klhdc4 | 234825 | HITS-CLIP | 23597149 | Unspecified |
| mmu-miR-152-3p | MIMAT0000162 | Klhl12 | 240756 | HITS-CLIP | 37986224 | Unspecified |
| mmu-miR-152-3p | MIMAT0000162 | Klhl15 | 236904 | HITS-CLIP | 37986224\|25083871 | Unspecified |
| mmu-miR-152-3p | MIMAT0000162 | Klhl2 | 77113 | HITS-CLIP | 37986224 | Unspecified |
| mmu-miR-152-3p | MIMAT0000162 | Klhl24 | 75785 | HITS-CLIP | 25083871 | Unspecified |
| mmu-miR-152-3p | MIMAT0000162 | Klhl28 | 66689 | HITS-CLIP | 37986224 | Unspecified |
| mmu-miR-152-3p | MIMAT0000162 | Klhl31 | 244923 | HITS-CLIP | 37986224\|25083871 | Unspecified |
| mmu-miR-152-3p | MIMAT0000162 | Klhl42 | 232539 | HITS-CLIP | 37986224\|25083871 | Unspecified |
| mmu-miR-152-3p | MIMAT0000162 | Klhl9 | 242521 | HITS-CLIP | 37986224\|25083871 | Unspecified |
| mmu-miR-152-3p | MIMAT0000162 | Kmt2a | 214162 | HITS-CLIP | 37986224\|25083871\|23597149 | Unspecified |
| mmu-miR-152-3p | MIMAT0000162 | Kmt2b | 75410 | HITS-CLIP | 25083871 | Unspecified |
| mmu-miR-152-3p | MIMAT0000162 | Kmt2c | 231051 | HITS-CLIP | 23597149\|29167373 | Unspecified |
| mmu-miR-152-3p | MIMAT0000162 | Kmt2d | 381022 | HITS-CLIP | 37986224\|25083871\|29167373 | Unspecified |
| mmu-miR-152-3p | MIMAT0000162 | Kmt2e | 69188 | HITS-CLIP | 25083871 | Unspecified |
| mmu-miR-152-3p | MIMAT0000162 | Kmt5b | 225888 | HITS-CLIP | 25083871 | Unspecified |
| mmu-miR-152-3p | MIMAT0000162 | Kpna1 | 16646 | HITS-CLIP | 37986224\|25083871 | Unspecified |
| mmu-miR-152-3p | MIMAT0000162 | Kpna4 | 16649 | HITS-CLIP | 37986224 | Unspecified |
| mmu-miR-152-3p | MIMAT0000162 | Kpna6 | 16650 | HITS-CLIP | 25083871 | Unspecified |
| mmu-miR-152-3p | MIMAT0000162 | Kpnb1 | 16211 | HITS-CLIP | 37986224\|29167373 | Unspecified |
| mmu-miR-152-3p | MIMAT0000162 | Kras | 16653 | HITS-CLIP | 37986224 | Unspecified |
| mmu-miR-152-3p | MIMAT0000162 | Kremen1 | 84035 | HITS-CLIP | 25083871 | Unspecified |
| mmu-miR-152-3p | MIMAT0000162 | L3mbtl3 | 237339 | HITS-CLIP | 25083871 | Unspecified |
| mmu-miR-152-3p | MIMAT0000162 | Lama5 | 16776 | HITS-CLIP | 25083871 | Unspecified |
| mmu-miR-152-3p | MIMAT0000162 | Lamc1 | 226519 | HITS-CLIP | 25083871 | Unspecified |
| mmu-miR-152-3p | MIMAT0000162 | Lamp1 | 16783 | HITS-CLIP | 37986224\|23597149 | Unspecified |
| mmu-miR-152-3p | MIMAT0000162 | Lamp2 | 16784 | HITS-CLIP | 37986224\|25083871 | Unspecified |
| mmu-miR-152-3p | MIMAT0000162 | Laptm4b | 114128 | HITS-CLIP | 37986224\|23597149\|25083871 | Unspecified |

| | | | | | | |
|---|---|---|---|---|---|---|
| mmu-miR-152-3p | MIMAT0000162 | Large1 | 16795 | HITS-CLIP | 25083871 | Unspecified |
| mmu-miR-152-3p | MIMAT0000162 | Larp4 | 207214 | HITS-CLIP | 25083871\|23597149 | Unspecified |
| mmu-miR-152-3p | MIMAT0000162 | Larp4b | 217980 | HITS-CLIP | 25083871 | Unspecified |
| mmu-miR-152-3p | MIMAT0000162 | Lars | 107045 | HITS-CLIP | 25083871 | Unspecified |
| mmu-miR-152-3p | MIMAT0000162 | Lats1 | 16798 | HITS-CLIP | 37986224\|29167373 | Unspecified |
| mmu-miR-152-3p | MIMAT0000162 | Lbr | 98386 | HITS-CLIP | 37986224\|23597149\|25083871 | Unspecified |
| mmu-miR-152-3p | MIMAT0000162 | Lcor | 212391 | HITS-CLIP | 25083871 | Unspecified |
| mmu-miR-152-3p | MIMAT0000162 | Lcorl | 209707 | HITS-CLIP | 25083871\|29167373 | Unspecified |
| mmu-miR-152-3p | MIMAT0000162 | Ldha | 16828 | HITS-CLIP | 37986224\|23597149\|25083871 | Unspecified |
| mmu-miR-152-3p | MIMAT0000162 | Ldlr | 16835 | HITS-CLIP | 23597149\|31433990\|29167373 | Unspecified |
| mmu-miR-152-3p | MIMAT0000162 | Ldlrad4 | 52662 | HITS-CLIP | 25083871 | Unspecified |
| mmu-miR-152-3p | MIMAT0000162 | Lefty1 | 13590 | HITS-CLIP | 37986224 | Unspecified |
| mmu-miR-152-3p | MIMAT0000162 | Leng8 | 232798 | HITS-CLIP | 37986224\|25083871 | Unspecified |
| mmu-miR-152-3p | MIMAT0000162 | Lgals3bp | 19039 | HITS-CLIP | 23597149 | Unspecified |
| mmu-miR-152-3p | MIMAT0000162 | Lgals8 | 56048 | HITS-CLIP | 23597149 | Unspecified |
| mmu-miR-152-3p | MIMAT0000162 | Lgalsl | 216551 | HITS-CLIP | 37986224 | Unspecified |
| mmu-miR-152-3p | MIMAT0000162 | Lhfpl2 | 218454 | HITS-CLIP | 25083871 | Unspecified |
| mmu-miR-152-3p | MIMAT0000162 | Lhpp | 76429 | HITS-CLIP | 37986224 | Unspecified |
| mmu-miR-152-3p | MIMAT0000162 | Lifr | 16880 | HITS-CLIP | 25083871 | Unspecified |
| mmu-miR-152-3p | MIMAT0000162 | Limd2 | 67803 | HITS-CLIP | 31433990 | Unspecified |
| mmu-miR-152-3p | MIMAT0000162 | Lin28a | 83557 | HITS-CLIP | 37986224 | Unspecified |
| mmu-miR-152-3p | MIMAT0000162 | Lin54 | 231506 | HITS-CLIP | 29167373 | Unspecified |
| mmu-miR-152-3p | MIMAT0000162 | Lipa | 16889 | HITS-CLIP | 37986224 | Unspecified |
| mmu-miR-152-3p | MIMAT0000162 | Lipg | 16891 | HITS-CLIP | 23597149 | Unspecified |
| mmu-miR-152-3p | MIMAT0000162 | Llgl2 | 217325 | HITS-CLIP | 23597149 | Unspecified |
| mmu-miR-152-3p | MIMAT0000162 | Lman2 | 66890 | HITS-CLIP | 37986224 | Unspecified |
| mmu-miR-152-3p | MIMAT0000162 | Lmbr1 | 56873 | HITS-CLIP | 37986224 | Unspecified |
| mmu-miR-152-3p | MIMAT0000162 | Lmbr1l | 74775 | HITS-CLIP | 37986224 | Unspecified |
| mmu-miR-152-3p | MIMAT0000162 | Lmbrd1 | 68421 | HITS-CLIP | 25083871\|29167373 | Unspecified |
| mmu-miR-152-3p | MIMAT0000162 | Lmln | 239833 | HITS-CLIP | 23597149\|25083871 | Unspecified |
| mmu-miR-152-3p | MIMAT0000162 | Lmnb1 | 16906 | HITS-CLIP | 23597149 | Unspecified |
| mmu-miR-152-3p | MIMAT0000162 | Lmtk2 | 231876 | HITS-CLIP | 37986224 | Unspecified |
| mmu-miR-152-3p | MIMAT0000162 | Lnpep | 240028 | HITS-CLIP | 29167373 | Unspecified |
| mmu-miR-152-3p | MIMAT0000162 | Lonp2 | 66887 | HITS-CLIP | 23597149 | Unspecified |
| mmu-miR-152-3p | MIMAT0000162 | Lonrf1 | 244421 | HITS-CLIP | 25083871 | Unspecified |
| mmu-miR-152-3p | MIMAT0000162 | Lonrf3 | 74365 | HITS-CLIP | 29167373 | Unspecified |
| mmu-miR-152-3p | MIMAT0000162 | Lox | 16948 | HITS-CLIP | 25083871 | Unspecified |
| mmu-miR-152-3p | MIMAT0000162 | Lpcat3 | 14792 | HITS-CLIP | 37986224\|25083871 | Unspecified |
| mmu-miR-152-3p | MIMAT0000162 | Lpgat1 | 226856 | HITS-CLIP | 25083871\|23597149 | Unspecified |

| | | | | | | |
|---|---|---|---|---|---|---|
| mmu-miR-152-3p | MIMAT0000162 | Lpin2 | 64898 | HITS-CLIP | 37986224\|23597149 | Unspecified |
| mmu-miR-152-3p | MIMAT0000162 | Lpl | 16956 | HITS-CLIP | 25083871\|29167373 | Unspecified |
| mmu-miR-152-3p | MIMAT0000162 | Lpp | 210126 | HITS-CLIP | 25083871\|23597149 | Unspecified |
| mmu-miR-152-3p | MIMAT0000162 | Lrba | 80877 | HITS-CLIP | 25083871\|23597149 | Unspecified |
| mmu-miR-152-3p | MIMAT0000162 | Lrch3 | 70144 | HITS-CLIP | 37986224\|25083871 | Unspecified |
| mmu-miR-152-3p | MIMAT0000162 | Lrig2 | 269473 | HITS-CLIP | 25083871 | Unspecified |
| mmu-miR-152-3p | MIMAT0000162 | Lrp1 | 16971 | HITS-CLIP | 37986224\|25083871\|23597149\|29167 | Unspecified |
| mmu-miR-152-3p | MIMAT0000162 | Lrp10 | 65107 | HITS-CLIP | 23597149 | Unspecified |
| mmu-miR-152-3p | MIMAT0000162 | Lrp4 | 228357 | HITS-CLIP | 37986224\|25083871 | Unspecified |
| mmu-miR-152-3p | MIMAT0000162 | Lrp6 | 16974 | HITS-CLIP | 37986224 | Unspecified |
| mmu-miR-152-3p | MIMAT0000162 | Lrrc17 | 74511 | HITS-CLIP | 25083871 | Unspecified |
| mmu-miR-152-3p | MIMAT0000162 | Lrrc41 | 230654 | HITS-CLIP | 37986224\|25083871\|23597149 | Unspecified |
| mmu-miR-152-3p | MIMAT0000162 | Lrrc47 | 72946 | HITS-CLIP | 23597149 | Unspecified |
| mmu-miR-152-3p | MIMAT0000162 | Lrrc58 | 320184 | HITS-CLIP | 25083871 | Unspecified |
| mmu-miR-152-3p | MIMAT0000162 | Lrrc8a | 241296 | HITS-CLIP | 25083871\|23597149 | Unspecified |
| mmu-miR-152-3p | MIMAT0000162 | Lrrcc1 | 71710 | HITS-CLIP | 25083871 | Unspecified |
| mmu-miR-152-3p | MIMAT0000162 | Lrrfip1 | 16978 | HITS-CLIP | 25083871 | Unspecified |
| mmu-miR-152-3p | MIMAT0000162 | Lrrk1 | 233328 | HITS-CLIP | 23597149\|25083871 | Unspecified |
| mmu-miR-152-3p | MIMAT0000162 | Lss | 16987 | HITS-CLIP | 37986224\|23597149 | Unspecified |
| mmu-miR-152-3p | MIMAT0000162 | Ltbp1 | 268977 | HITS-CLIP | 37986224 | Unspecified |
| mmu-miR-152-3p | MIMAT0000162 | Ltbr | 17000 | HITS-CLIP | 25083871\|29167373 | Unspecified |
| mmu-miR-152-3p | MIMAT0000162 | Luzp1 | 269593 | HITS-CLIP | 37986224\|25083871 | Unspecified |
| mmu-miR-152-3p | MIMAT0000162 | Lyl1 | 17095 | HITS-CLIP | 29167373 | Unspecified |
| mmu-miR-152-3p | MIMAT0000162 | Lynx1 | 23936 | HITS-CLIP | 37986224 | Unspecified |
| mmu-miR-152-3p | MIMAT0000162 | Lypla1 | 18777 | HITS-CLIP | 23597149\|25083871 | Unspecified |
| mmu-miR-152-3p | MIMAT0000162 | Lyst | 17101 | HITS-CLIP | 37986224\|29167373 | Unspecified |
| mmu-miR-152-3p | MIMAT0000162 | Lztfl1 | 93730 | HITS-CLIP | 37986224 | Unspecified |
| mmu-miR-152-3p | MIMAT0000162 | Lztr1 | 66863 | HITS-CLIP | 37986224 | Unspecified |
| mmu-miR-152-3p | MIMAT0000162 | Macf1 | 11426 | HITS-CLIP | 25083871 | Unspecified |
| mmu-miR-152-3p | MIMAT0000162 | Maf | 17132 | HITS-CLIP | 37986224 | Unspecified |
| mmu-miR-152-3p | MIMAT0000162 | Maf1 | 68877 | HITS-CLIP | 37986224\|23597149 | Unspecified |
| mmu-miR-152-3p | MIMAT0000162 | Mafb | 16658 | HITS-CLIP | 29167373 | Unspecified |
| mmu-miR-152-3p | MIMAT0000162 | Magi3 | 99470 | HITS-CLIP | 37986224\|23597149 | Unspecified |
| mmu-miR-152-3p | MIMAT0000162 | Maip1 | 79568 | HITS-CLIP | 23597149\|25083871 | Unspecified |
| mmu-miR-152-3p | MIMAT0000162 | Maml2 | 270118 | HITS-CLIP | 25083871 | Unspecified |
| mmu-miR-152-3p | MIMAT0000162 | Man1a2 | 17156 | HITS-CLIP | 25083871 | Unspecified |
| mmu-miR-152-3p | MIMAT0000162 | Man1c1 | 230815 | HITS-CLIP | 29167373 | Unspecified |
| mmu-miR-152-3p | MIMAT0000162 | Man2a1 | 17158 | HITS-CLIP | 25083871\|23597149\|29167373 | Unspecified |
| mmu-miR-152-3p | MIMAT0000162 | Manea | 242362 | HITS-CLIP | 37986224 | Unspecified |

| | | | | | | |
|---|---|---|---|---|---|---|
| mmu-miR-152-3p | MIMAT0000162 | Manf | 74840 | HITS-CLIP | 37986224 | Unspecified |
| mmu-miR-152-3p | MIMAT0000162 | Map1lc3b | 67443 | HITS-CLIP | 37986224\|23597149\|25083871 | Unspecified |
| mmu-miR-152-3p | MIMAT0000162 | Map1s | 270058 | HITS-CLIP | 29167373 | Unspecified |
| mmu-miR-152-3p | MIMAT0000162 | Map2k1 | 26395 | HITS-CLIP | 37986224 | Unspecified |
| mmu-miR-152-3p | MIMAT0000162 | Map2k4 | 26398 | HITS-CLIP | 25083871 | Unspecified |
| mmu-miR-152-3p | MIMAT0000162 | Map3k2 | 26405 | HITS-CLIP | 37986224\|25083871 | Unspecified |
| mmu-miR-152-3p | MIMAT0000162 | Map3k20 | 65964 | HITS-CLIP | 25083871 | Unspecified |
| mmu-miR-152-3p | MIMAT0000162 | Map3k3 | 26406 | HITS-CLIP | 37986224 | Unspecified |
| mmu-miR-152-3p | MIMAT0000162 | Map3k4 | 26407 | HITS-CLIP | 23597149 | Unspecified |
| mmu-miR-152-3p | MIMAT0000162 | Map3k5 | 26408 | HITS-CLIP | 29167373 | Unspecified |
| mmu-miR-152-3p | MIMAT0000162 | Map3k9 | 338372 | HITS-CLIP | 29167373 | Unspecified |
| mmu-miR-152-3p | MIMAT0000162 | Map4 | 17758 | HITS-CLIP | 37986224\|25083871 | Unspecified |
| mmu-miR-152-3p | MIMAT0000162 | Map4k4 | 26921 | HITS-CLIP | 37986224\|25083871 | Unspecified |
| mmu-miR-152-3p | MIMAT0000162 | Mapk1 | 26413 | HITS-CLIP | 37986224\|25083871 | Unspecified |
| mmu-miR-152-3p | MIMAT0000162 | Mapk8 | 26419 | HITS-CLIP | 25083871 | Unspecified |
| mmu-miR-152-3p | MIMAT0000162 | Mapkap1 | 227743 | HITS-CLIP | 25083871 | Unspecified |
| mmu-miR-152-3p | MIMAT0000162 | Mapre1 | 13589 | HITS-CLIP | 23597149 | Unspecified |
| mmu-miR-152-3p | MIMAT0000162 | March3 | 320253 | HITS-CLIP | 37986224 | Unspecified |
| mmu-miR-152-3p | MIMAT0000162 | March5 | 54708 | HITS-CLIP | 25083871 | Unspecified |
| mmu-miR-152-3p | MIMAT0000162 | March7 | 57438 | HITS-CLIP | 25083871 | Unspecified |
| mmu-miR-152-3p | MIMAT0000162 | March8 | 71779 | HITS-CLIP | 25083871 | Unspecified |
| mmu-miR-152-3p | MIMAT0000162 | Marchf6 | 223455 | HITS-CLIP | 37986224\|23597149 | Unspecified |
| mmu-miR-152-3p | MIMAT0000162 | Mark2 | 13728 | HITS-CLIP | 25083871\|23597149\|29167373 | Unspecified |
| mmu-miR-152-3p | MIMAT0000162 | Mark3 | 17169 | HITS-CLIP | 25083871 | Unspecified |
| mmu-miR-152-3p | MIMAT0000162 | Masp1 | 17174 | HITS-CLIP | 23597149 | Unspecified |
| mmu-miR-152-3p | MIMAT0000162 | Masp2 | 17175 | HITS-CLIP | 23597149 | Unspecified |
| mmu-miR-152-3p | MIMAT0000162 | Mast2 | 17776 | HITS-CLIP | 37986224\|25083871 | Unspecified |
| mmu-miR-152-3p | MIMAT0000162 | Mast3 | 546071 | HITS-CLIP | 37986224 | Unspecified |
| mmu-miR-152-3p | MIMAT0000162 | Mast4 | 328329 | HITS-CLIP | 25083871 | Unspecified |
| mmu-miR-152-3p | MIMAT0000162 | Mastl | 67121 | HITS-CLIP | 25083871 | Unspecified |
| mmu-miR-152-3p | MIMAT0000162 | Mat2a | 232087 | HITS-CLIP | 25083871\|29167373 | Unspecified |
| mmu-miR-152-3p | MIMAT0000162 | Mat2b | 108645 | HITS-CLIP | 21258322 | Unspecified |
| mmu-miR-152-3p | MIMAT0000162 | Matr3 | 17184 | HITS-CLIP | 25083871 | Unspecified |
| mmu-miR-152-3p | MIMAT0000162 | Max | 17187 | HITS-CLIP | 25083871 | Unspecified |
| mmu-miR-152-3p | MIMAT0000162 | Maz | 17188 | HITS-CLIP | 25083871 | Unspecified |
| mmu-miR-152-3p | MIMAT0000162 | Mbd5 | 109241 | HITS-CLIP | 25083871 | Unspecified |
| mmu-miR-152-3p | MIMAT0000162 | Mbnl1 | 56758 | HITS-CLIP | 25083871\|29167373 | Unspecified |
| mmu-miR-152-3p | MIMAT0000162 | Mbnl2 | 105559 | HITS-CLIP | 25083871\|23597149 | Unspecified |
| mmu-miR-152-3p | MIMAT0000162 | Mbtd1 | 103537 | HITS-CLIP | 25083871 | Unspecified |

| | | | | | | |
|---|---|---|---|---|---|---|
| mmu-miR-152-3p | MIMAT0000162 | Mcm3ap | 54387 | HITS-CLIP | 25083871 | Unspecified |
| mmu-miR-152-3p | MIMAT0000162 | Mcmbp | 210711 | HITS-CLIP | 37986224 | Unspecified |
| mmu-miR-152-3p | MIMAT0000162 | Mcur1 | 76137 | HITS-CLIP | 25083871 | Unspecified |
| mmu-miR-152-3p | MIMAT0000162 | Mdc1 | 240087 | HITS-CLIP | 23597149 | Unspecified |
| mmu-miR-152-3p | MIMAT0000162 | Mdfic | 16543 | HITS-CLIP | 25083871 | Unspecified |
| mmu-miR-152-3p | MIMAT0000162 | Mdh2 | 17448 | HITS-CLIP | 23597149 | Unspecified |
| mmu-miR-152-3p | MIMAT0000162 | Mdn1 | 100019 | HITS-CLIP | 25083871 | Unspecified |
| mmu-miR-152-3p | MIMAT0000162 | Mecp2 | 17257 | HITS-CLIP | 37986224\|25083871\|23597149 | Unspecified |
| mmu-miR-152-3p | MIMAT0000162 | Med1 | 19014 | HITS-CLIP | 37986224\|25083871\|29167373 | Unspecified |
| mmu-miR-152-3p | MIMAT0000162 | Med12 | 59024 | HITS-CLIP | 25083871 | Unspecified |
| mmu-miR-152-3p | MIMAT0000162 | Med13 | 327987 | HITS-CLIP | 37986224\|25083871\|23597149 | Unspecified |
| mmu-miR-152-3p | MIMAT0000162 | Med13l | 76199 | HITS-CLIP | 25083871 | Unspecified |
| mmu-miR-152-3p | MIMAT0000162 | Med14 | 26896 | HITS-CLIP | 37986224\|25083871\|29167373 | Unspecified |
| mmu-miR-152-3p | MIMAT0000162 | Med15 | 94112 | HITS-CLIP | 25083871\|29167373 | Unspecified |
| mmu-miR-152-3p | MIMAT0000162 | Med24 | 23989 | HITS-CLIP | 23597149 | Unspecified |
| mmu-miR-152-3p | MIMAT0000162 | Mef2a | 17258 | HITS-CLIP | 25083871\|29167373 | Unspecified |
| mmu-miR-152-3p | MIMAT0000162 | Mef2c | 17260 | HITS-CLIP | 25083871 | Unspecified |
| mmu-miR-152-3p | MIMAT0000162 | Megf10 | 70417 | HITS-CLIP | 25083871 | Unspecified |
| mmu-miR-152-3p | MIMAT0000162 | Meis1 | 17268 | HITS-CLIP | 37986224 | Unspecified |
| mmu-miR-152-3p | MIMAT0000162 | Memo1 | 76890 | HITS-CLIP | 25083871 | Unspecified |
| mmu-miR-152-3p | MIMAT0000162 | Mepce | 231803 | HITS-CLIP | 37986224 | Unspecified |
| mmu-miR-152-3p | MIMAT0000162 | Mest | 17294 | HITS-CLIP | 25083871 | Unspecified |
| mmu-miR-152-3p | MIMAT0000162 | Met | 17295 | HITS-CLIP | 37986224\|25083871 | Unspecified |
| mmu-miR-152-3p | MIMAT0000162 | Mettl6 | 67011 | HITS-CLIP | 37986224 | Unspecified |
| mmu-miR-152-3p | MIMAT0000162 | Mex3b | 108797 | HITS-CLIP | 25083871 | Unspecified |
| mmu-miR-152-3p | MIMAT0000162 | Mex3c | 240396 | HITS-CLIP | 37986224\|25083871 | Unspecified |
| mmu-miR-152-3p | MIMAT0000162 | Mfap3l | 71306 | HITS-CLIP | 25083871 | Unspecified |
| mmu-miR-152-3p | MIMAT0000162 | Mfhas1 | 52065 | HITS-CLIP | 23597149 | Unspecified |
| mmu-miR-152-3p | MIMAT0000162 | Mfn1 | 67414 | HITS-CLIP | 23597149 | Unspecified |
| mmu-miR-152-3p | MIMAT0000162 | Mfn2 | 170731 | HITS-CLIP | 37986224\|25083871 | Unspecified |
| mmu-miR-152-3p | MIMAT0000162 | Mfsd1 | 66868 | HITS-CLIP | 25083871 | Unspecified |
| mmu-miR-152-3p | MIMAT0000162 | Mfsd12 | 73822 | HITS-CLIP | 25083871 | Unspecified |
| mmu-miR-152-3p | MIMAT0000162 | Mfsd14b | 84641 | HITS-CLIP | 29167373 | Unspecified |
| mmu-miR-152-3p | MIMAT0000162 | Mfsd2a | 76574 | HITS-CLIP | 37986224\|23597149\|25083871 | Unspecified |
| mmu-miR-152-3p | MIMAT0000162 | Mfsd3 | 69572 | HITS-CLIP | 37986224\|23597149 | Unspecified |
| mmu-miR-152-3p | MIMAT0000162 | Mgam | 232714 | HITS-CLIP | 37986224 | Unspecified |
| mmu-miR-152-3p | MIMAT0000162 | Mgat5 | 107895 | HITS-CLIP | 37986224 | Unspecified |
| mmu-miR-152-3p | MIMAT0000162 | Mgea5 | 76055 | HITS-CLIP | 23597149\|25083871 | Unspecified |
| mmu-miR-152-3p | MIMAT0000162 | Mgst1 | 56615 | HITS-CLIP | 23597149 | Unspecified |

| | | | | | | |
|---|---|---|---|---|---|---|
| mmu-miR-152-3p | MIMAT0000162 | Mia2 | 338320 | HITS-CLIP | 23597149 | Unspecified |
| mmu-miR-152-3p | MIMAT0000162 | Mib1 | 225164 | HITS-CLIP | 25083871\|23597149\|29167373 | Unspecified |
| mmu-miR-152-3p | MIMAT0000162 | Mid2 | 23947 | HITS-CLIP | 37986224 | Unspecified |
| mmu-miR-152-3p | MIMAT0000162 | Mief1 | 239555 | HITS-CLIP | 37986224\|25083871 | Unspecified |
| mmu-miR-152-3p | MIMAT0000162 | Mier2 | 70427 | HITS-CLIP | 25083871 | Unspecified |
| mmu-miR-152-3p | MIMAT0000162 | Mier3 | 218613 | HITS-CLIP | 37986224\|25083871 | Unspecified |
| mmu-miR-152-3p | MIMAT0000162 | Mindy2 | 235461 | HITS-CLIP | 25083871\|23597149\|29167373 | Unspecified |
| mmu-miR-152-3p | MIMAT0000162 | Mitf | 17342 | HITS-CLIP | 37986224 | Unspecified |
| mmu-miR-152-3p | MIMAT0000162 | Mkln1 | 27418 | HITS-CLIP | 37986224\|25083871 | Unspecified |
| mmu-miR-152-3p | MIMAT0000162 | Mknk1 | 17346 | HITS-CLIP | 29167373 | Unspecified |
| mmu-miR-152-3p | MIMAT0000162 | Mkrn1 | 54484 | HITS-CLIP | 37986224\|21258322\|29167373 | Unspecified |
| mmu-miR-152-3p | MIMAT0000162 | Mlec | 109154 | HITS-CLIP | 37986224 | Unspecified |
| mmu-miR-152-3p | MIMAT0000162 | Mllt6 | 246198 | HITS-CLIP | 25083871 | Unspecified |
| mmu-miR-152-3p | MIMAT0000162 | Mmd | 67468 | HITS-CLIP | 37986224 | Unspecified |
| mmu-miR-152-3p | MIMAT0000162 | Mmgt1 | 236792 | HITS-CLIP | 25083871 | Unspecified |
| mmu-miR-152-3p | MIMAT0000162 | Mmgt2 | 216829 | HITS-CLIP | 37986224 | Unspecified |
| mmu-miR-152-3p | MIMAT0000162 | Mmp12 | 17381 | HITS-CLIP | 29167373 | Unspecified |
| mmu-miR-152-3p | MIMAT0000162 | Mmp15 | 17388 | HITS-CLIP | 25083871 | Unspecified |
| mmu-miR-152-3p | MIMAT0000162 | Mmp19 | 58223 | HITS-CLIP | 37986224\|25083871 | Unspecified |
| mmu-miR-152-3p | MIMAT0000162 | Mnt | 17428 | HITS-CLIP | 37986224 | Unspecified |
| mmu-miR-152-3p | MIMAT0000162 | Mob1a | 232157 | HITS-CLIP | 25083871 | Unspecified |
| mmu-miR-152-3p | MIMAT0000162 | Mob3b | 214944 | HITS-CLIP | 37986224\|23597149 | Unspecified |
| mmu-miR-152-3p | MIMAT0000162 | Mogat2 | 233549 | HITS-CLIP | 37986224 | Unspecified |
| mmu-miR-152-3p | MIMAT0000162 | Mogs | 57377 | HITS-CLIP | 37986224\|25083871 | Unspecified |
| mmu-miR-152-3p | MIMAT0000162 | Mon2 | 67074 | HITS-CLIP | 25083871 | Unspecified |
| mmu-miR-152-3p | MIMAT0000162 | Mosmo | 233812 | HITS-CLIP | 25083871\|29167373 | Unspecified |
| mmu-miR-152-3p | MIMAT0000162 | Mpdu1 | 24070 | HITS-CLIP | 23597149\|25083871 | Unspecified |
| mmu-miR-152-3p | MIMAT0000162 | Mpdz | 17475 | HITS-CLIP | 25083871 | Unspecified |
| mmu-miR-152-3p | MIMAT0000162 | Mpeg1 | 17476 | HITS-CLIP | 23597149\|29167373 | Unspecified |
| mmu-miR-152-3p | MIMAT0000162 | Mphosph8 | 75339 | HITS-CLIP | 25083871 | Unspecified |
| mmu-miR-152-3p | MIMAT0000162 | Mpp5 | 56217 | HITS-CLIP | 25083871 | Unspecified |
| mmu-miR-152-3p | MIMAT0000162 | Mpp6 | 56524 | HITS-CLIP | 23597149\|25083871\|29167373 | Unspecified |
| mmu-miR-152-3p | MIMAT0000162 | Mprip | 26936 | HITS-CLIP | 25083871\|23597149 | Unspecified |
| mmu-miR-152-3p | MIMAT0000162 | Mrps18a | 68565 | HITS-CLIP | 23597149 | Unspecified |
| mmu-miR-152-3p | MIMAT0000162 | Mrps31 | 57312 | HITS-CLIP | 25083871 | Unspecified |
| mmu-miR-152-3p | MIMAT0000162 | Mrrf | 67871 | HITS-CLIP | 37986224 | Unspecified |
| mmu-miR-152-3p | MIMAT0000162 | Msantd4 | 78100 | HITS-CLIP | 37986224 | Unspecified |
| mmu-miR-152-3p | MIMAT0000162 | Msi2 | 76626 | HITS-CLIP | 25083871 | Unspecified |
| mmu-miR-152-3p | MIMAT0000162 | Msl2 | 77853 | HITS-CLIP | 25083871\|29167373 | Unspecified |

| | | | | | | |
|---|---|---|---|---|---|---|
| mmu-miR-152-3p | MIMAT0000162 | Msl3 | 17692 | HITS-CLIP | 37986224 | Unspecified |
| mmu-miR-152-3p | MIMAT0000162 | Msn | 17698 | HITS-CLIP | 37986224\|25083871 | Unspecified |
| mmu-miR-152-3p | MIMAT0000162 | Msrb3 | 320183 | HITS-CLIP | 25083871\|23597149 | Unspecified |
| mmu-miR-152-3p | MIMAT0000162 | Mtcl1 | 68617 | HITS-CLIP | 25083871 | Unspecified |
| mmu-miR-152-3p | MIMAT0000162 | Mtdh | 67154 | HITS-CLIP | 23597149\|25083871\|29167373 | Unspecified |
| mmu-miR-152-3p | MIMAT0000162 | Mtf1 | 17764 | HITS-CLIP | 37986224 | Unspecified |
| mmu-miR-152-3p | MIMAT0000162 | Mtf2 | 17765 | HITS-CLIP | 25083871 | Unspecified |
| mmu-miR-152-3p | MIMAT0000162 | Mtfr1l | 76824 | HITS-CLIP | 37986224\|25083871 | Unspecified |
| mmu-miR-152-3p | MIMAT0000162 | Mthfd2 | 17768 | HITS-CLIP | 25083871 | Unspecified |
| mmu-miR-152-3p | MIMAT0000162 | Mthfr | 17769 | HITS-CLIP | 25083871 | Unspecified |
| mmu-miR-152-3p | MIMAT0000162 | Mtmr10 | 233315 | HITS-CLIP | 37986224 | Unspecified |
| mmu-miR-152-3p | MIMAT0000162 | Mtmr12 | 268783 | HITS-CLIP | 37986224\|25083871\|29167373 | Unspecified |
| mmu-miR-152-3p | MIMAT0000162 | Mtmr3 | 74302 | HITS-CLIP | 25083871 | Unspecified |
| mmu-miR-152-3p | MIMAT0000162 | Mtmr4 | 170749 | HITS-CLIP | 23597149 | Unspecified |
| mmu-miR-152-3p | MIMAT0000162 | Mtpn | 14489 | HITS-CLIP | 37986224 | Unspecified |
| mmu-miR-152-3p | MIMAT0000162 | Mtss1 | 211401 | HITS-CLIP | 37986224\|25083871 | Unspecified |
| mmu-miR-152-3p | MIMAT0000162 | Mtss1l | 244654 | HITS-CLIP | 25083871 | Unspecified |
| mmu-miR-152-3p | MIMAT0000162 | Mttp | 17777 | HITS-CLIP | 23597149 | Unspecified |
| mmu-miR-152-3p | MIMAT0000162 | Mug1 | 17836 | HITS-CLIP | 23597149 | Unspecified |
| mmu-miR-152-3p | MIMAT0000162 | Mup3 | 17842 | HITS-CLIP | 23597149 | Unspecified |
| mmu-miR-152-3p | MIMAT0000162 | Musk | 18198 | HITS-CLIP | 25083871 | Unspecified |
| mmu-miR-152-3p | MIMAT0000162 | Mut | 4594 | HITS-CLIP | 23597149 | Unspecified |
| mmu-miR-152-3p | MIMAT0000162 | Mxd4 | 17122 | HITS-CLIP | 25083871 | Unspecified |
| mmu-miR-152-3p | MIMAT0000162 | Mxi1 | 17859 | HITS-CLIP | 25083871 | Unspecified |
| mmu-miR-152-3p | MIMAT0000162 | Myadm | 50918 | HITS-CLIP | 25083871\|29167373 | Unspecified |
| mmu-miR-152-3p | MIMAT0000162 | Mybl1 | 17864 | HITS-CLIP | 25083871\|23597149 | Unspecified |
| mmu-miR-152-3p | MIMAT0000162 | Mycbp2 | 105689 | HITS-CLIP | 25083871\|23597149\|29167373 | Unspecified |
| mmu-miR-152-3p | MIMAT0000162 | Mycl | 16918 | HITS-CLIP | 25083871 | Unspecified |
| mmu-miR-152-3p | MIMAT0000162 | Myh10 | 77579 | HITS-CLIP | 25083871 | Unspecified |
| mmu-miR-152-3p | MIMAT0000162 | Myh3 | 17883 | HITS-CLIP | 25083871 | Unspecified |
| mmu-miR-152-3p | MIMAT0000162 | Myh9 | 17886 | HITS-CLIP | 37986224 | Unspecified |
| mmu-miR-152-3p | MIMAT0000162 | Myl12b | 67938 | HITS-CLIP | 37986224 | Unspecified |
| mmu-miR-152-3p | MIMAT0000162 | Myo10 | 17909 | HITS-CLIP | 25083871\|23597149 | Unspecified |
| mmu-miR-152-3p | MIMAT0000162 | Myo9a | 270163 | HITS-CLIP | 37986224\|25083871 | Unspecified |
| mmu-miR-152-3p | MIMAT0000162 | Myo9b | 17925 | HITS-CLIP | 23597149 | Unspecified |
| mmu-miR-152-3p | MIMAT0000162 | Myod1 | 17927 | HITS-CLIP | 25083871 | Unspecified |
| mmu-miR-152-3p | MIMAT0000162 | Myog | 17928 | HITS-CLIP | 25083871 | Unspecified |
| mmu-miR-152-3p | MIMAT0000162 | N4bp1 | 80750 | HITS-CLIP | 37986224\|25083871 | Unspecified |
| mmu-miR-152-3p | MIMAT0000162 | Naa15 | 74838 | HITS-CLIP | 37986224\|25083871 | Unspecified |

| | | | | | | |
|---|---|---|---|---|---|---|
| mmu-miR-152-3p | MIMAT0000162 | Naa25 | 231713 | HITS-CLIP | 25083871\|23597149\|29167373 | Unspecified |
| mmu-miR-152-3p | MIMAT0000162 | Naa38 | 78304 | HITS-CLIP | 25083871 | Unspecified |
| mmu-miR-152-3p | MIMAT0000162 | Naa40 | 70999 | HITS-CLIP | 25083871 | Unspecified |
| mmu-miR-152-3p | MIMAT0000162 | Nab2 | 17937 | HITS-CLIP | 37986224\|25083871 | Unspecified |
| mmu-miR-152-3p | MIMAT0000162 | Nacc2 | 67991 | HITS-CLIP | 25083871 | Unspecified |
| mmu-miR-152-3p | MIMAT0000162 | Nadk | 192185 | HITS-CLIP | 37986224 | Unspecified |
| mmu-miR-152-3p | MIMAT0000162 | Nadk2 | 68646 | HITS-CLIP | 25083871 | Unspecified |
| mmu-miR-152-3p | MIMAT0000162 | Naga | 17939 | HITS-CLIP | 37986224 | Unspecified |
| mmu-miR-152-3p | MIMAT0000162 | Nagk | 56174 | HITS-CLIP | 25083871 | Unspecified |
| mmu-miR-152-3p | MIMAT0000162 | Nampt | 59027 | HITS-CLIP | 29167373 | Unspecified |
| mmu-miR-152-3p | MIMAT0000162 | Nap1l1 | 53605 | HITS-CLIP | 37986224 | Unspecified |
| mmu-miR-152-3p | MIMAT0000162 | Nap1l4 | 17955 | HITS-CLIP | 37986224 | Unspecified |
| mmu-miR-152-3p | MIMAT0000162 | Nasp | 50927 | HITS-CLIP | 37986224 | Unspecified |
| mmu-miR-152-3p | MIMAT0000162 | Nav1 | 215690 | HITS-CLIP | 25083871 | Unspecified |
| mmu-miR-152-3p | MIMAT0000162 | Nav2 | 78286 | HITS-CLIP | 37986224\|25083871 | Unspecified |
| mmu-miR-152-3p | MIMAT0000162 | Nbeal1 | 269198 | HITS-CLIP | 25083871 | Unspecified |
| mmu-miR-152-3p | MIMAT0000162 | Nbeal2 | 235627 | HITS-CLIP | 25083871\|23597149 | Unspecified |
| mmu-miR-152-3p | MIMAT0000162 | Ncam1 | 17967 | HITS-CLIP | 25083871 | Unspecified |
| mmu-miR-152-3p | MIMAT0000162 | Ncapg2 | 76044 | HITS-CLIP | 23597149 | Unspecified |
| mmu-miR-152-3p | MIMAT0000162 | Ncaph2 | 52683 | HITS-CLIP | 25083871 | Unspecified |
| mmu-miR-152-3p | MIMAT0000162 | Ncbp1 | 433702 | HITS-CLIP | 25083871 | Unspecified |
| mmu-miR-152-3p | MIMAT0000162 | Ncdn | 26562 | HITS-CLIP | 23597149\|25083871 | Unspecified |
| mmu-miR-152-3p | MIMAT0000162 | Nceh1 | 320024 | HITS-CLIP | 37986224\|25083871 | Unspecified |
| mmu-miR-152-3p | MIMAT0000162 | Nck1 | 17973 | HITS-CLIP | 37986224 | Unspecified |
| mmu-miR-152-3p | MIMAT0000162 | Nckap1 | 50884 | HITS-CLIP | 37986224\|25083871\|23597149 | Unspecified |
| mmu-miR-152-3p | MIMAT0000162 | Nckap5l | 380969 | HITS-CLIP | 37986224 | Unspecified |
| mmu-miR-152-3p | MIMAT0000162 | Ncoa2 | 17978 | HITS-CLIP | 37986224\|25083871\|23597149\|29167 | Unspecified |
| mmu-miR-152-3p | MIMAT0000162 | Ncoa3 | 17979 | HITS-CLIP | 25083871\|29167373 | Unspecified |
| mmu-miR-152-3p | MIMAT0000162 | Ncor1 | 20185 | HITS-CLIP | 37986224 | Unspecified |
| mmu-miR-152-3p | MIMAT0000162 | Ncs1 | 14299 | HITS-CLIP | 37986224 | Unspecified |
| mmu-miR-152-3p | MIMAT0000162 | Ndfip1 | 65113 | HITS-CLIP | 25083871 | Unspecified |
| mmu-miR-152-3p | MIMAT0000162 | Ndrg2 | 29811 | HITS-CLIP | 23597149 | Unspecified |
| mmu-miR-152-3p | MIMAT0000162 | Ndst1 | 15531 | HITS-CLIP | 25083871 | Unspecified |
| mmu-miR-152-3p | MIMAT0000162 | Ndst2 | 17423 | HITS-CLIP | 37986224 | Unspecified |
| mmu-miR-152-3p | MIMAT0000162 | Ndufs1 | 227197 | HITS-CLIP | 37986224\|23597149\|25083871 | Unspecified |
| mmu-miR-152-3p | MIMAT0000162 | Ndufs8 | 225887 | HITS-CLIP | 37986224 | Unspecified |
| mmu-miR-152-3p | MIMAT0000162 | Ndufv1 | 17995 | HITS-CLIP | 37986224\|25083871\|23597149 | Unspecified |
| mmu-miR-152-3p | MIMAT0000162 | Neb | 17996 | HITS-CLIP | 25083871 | Unspecified |
| mmu-miR-152-3p | MIMAT0000162 | Necap2 | 66147 | HITS-CLIP | 23597149\|25083871\|29167373 | Unspecified |

| | | | | | | |
|---|---|---|---|---|---|---|
| mmu-miR-152-3p | MIMAT0000162 | Nectin1 | 58235 | HITS-CLIP | 23597149 | Unspecified |
| mmu-miR-152-3p | MIMAT0000162 | Nectin2 | 5819 | HITS-CLIP | 25083871 | Unspecified |
| mmu-miR-152-3p | MIMAT0000162 | Nedd1 | 17997 | HITS-CLIP | 37986224\|23597149 | Unspecified |
| mmu-miR-152-3p | MIMAT0000162 | Nedd4 | 17999 | HITS-CLIP | 37986224 | Unspecified |
| mmu-miR-152-3p | MIMAT0000162 | Nek7 | 59125 | HITS-CLIP | 25083871 | Unspecified |
| mmu-miR-152-3p | MIMAT0000162 | Nek9 | 217718 | HITS-CLIP | 25083871\|23597149 | Unspecified |
| mmu-miR-152-3p | MIMAT0000162 | Nemf | 66244 | HITS-CLIP | 23597149\|25083871 | Unspecified |
| mmu-miR-152-3p | MIMAT0000162 | Net1 | 56349 | HITS-CLIP | 25083871 | Unspecified |
| mmu-miR-152-3p | MIMAT0000162 | Neurl4 | 216860 | HITS-CLIP | 37986224 | Unspecified |
| mmu-miR-152-3p | MIMAT0000162 | Nf1 | 18015 | HITS-CLIP | 25083871 | Unspecified |
| mmu-miR-152-3p | MIMAT0000162 | Nfam1 | 74039 | HITS-CLIP | 29167373 | Unspecified |
| mmu-miR-152-3p | MIMAT0000162 | Nfat5 | 54446 | HITS-CLIP | 37986224\|25083871 | Unspecified |
| mmu-miR-152-3p | MIMAT0000162 | Nfe2l1 | 18023 | HITS-CLIP | 37986224 | Unspecified |
| mmu-miR-152-3p | MIMAT0000162 | Nfib | 18028 | HITS-CLIP | 37986224 | Unspecified |
| mmu-miR-152-3p | MIMAT0000162 | Nfic | 18029 | HITS-CLIP | 23597149\|29167373 | Unspecified |
| mmu-miR-152-3p | MIMAT0000162 | Nfkbia | 18035 | HITS-CLIP | 29167373 | Unspecified |
| mmu-miR-152-3p | MIMAT0000162 | Nfx1 | 74164 | HITS-CLIP | 23597149 | Unspecified |
| mmu-miR-152-3p | MIMAT0000162 | Nfxl1 | 100978 | HITS-CLIP | 37986224 | Unspecified |
| mmu-miR-152-3p | MIMAT0000162 | Nfya | 18044 | HITS-CLIP | 25083871\|23597149\|29167373 | Unspecified |
| mmu-miR-152-3p | MIMAT0000162 | Nfyb | 18045 | HITS-CLIP | 37986224 | Unspecified |
| mmu-miR-152-3p | MIMAT0000162 | Nhlrc3 | 212114 | HITS-CLIP | 37986224 | Unspecified |
| mmu-miR-152-3p | MIMAT0000162 | Nhs | 195727 | HITS-CLIP | 37986224\|25083871 | Unspecified |
| mmu-miR-152-3p | MIMAT0000162 | Nid2 | 18074 | HITS-CLIP | 37986224 | Unspecified |
| mmu-miR-152-3p | MIMAT0000162 | Nin | 18080 | HITS-CLIP | 37986224\|29167373\|25083871 | Unspecified |
| mmu-miR-152-3p | MIMAT0000162 | Nip7 | 66164 | HITS-CLIP | 37986224 | Unspecified |
| mmu-miR-152-3p | MIMAT0000162 | Nipa2 | 93790 | HITS-CLIP | 37986224\|25083871\|29167373 | Unspecified |
| mmu-miR-152-3p | MIMAT0000162 | Nisch | 64652 | HITS-CLIP | 23597149\|25083871 | Unspecified |
| mmu-miR-152-3p | MIMAT0000162 | Nkiras1 | 69721 | HITS-CLIP | 25083871 | Unspecified |
| mmu-miR-152-3p | MIMAT0000162 | Nlk | 18099 | HITS-CLIP | 37986224\|25083871 | Unspecified |
| mmu-miR-152-3p | MIMAT0000162 | Nme1 | 18102 | HITS-CLIP | 25083871 | Unspecified |
| mmu-miR-152-3p | MIMAT0000162 | Nmt1 | 18107 | HITS-CLIP | 37986224\|25083871 | Unspecified |
| mmu-miR-152-3p | MIMAT0000162 | Nod2 | 257632 | HITS-CLIP | 37986224 | Unspecified |
| mmu-miR-152-3p | MIMAT0000162 | Nol6 | 230082 | HITS-CLIP | 23597149 | Unspecified |
| mmu-miR-152-3p | MIMAT0000162 | Nom1 | 433864 | HITS-CLIP | 37986224 | Unspecified |
| mmu-miR-152-3p | MIMAT0000162 | Notch1 | 18128 | HITS-CLIP | 23597149 | Unspecified |
| mmu-miR-152-3p | MIMAT0000162 | Notch2 | 18129 | HITS-CLIP | 37986224 | Unspecified |
| mmu-miR-152-3p | MIMAT0000162 | Notch3 | 18131 | HITS-CLIP | 37986224 | Unspecified |
| mmu-miR-152-3p | MIMAT0000162 | Npat | 244879 | HITS-CLIP | 37986224\|23597149 | Unspecified |
| mmu-miR-152-3p | MIMAT0000162 | Npepl1 | 228961 | HITS-CLIP | 37986224 | Unspecified |

| | | | | | | |
|---|---|---|---|---|---|---|
| mmu-miR-152-3p | MIMAT0000162 | Nploc4 | 217365 | HITS-CLIP | 37986224 | Unspecified |
| mmu-miR-152-3p | MIMAT0000162 | Npnt | 114249 | HITS-CLIP | 25083871 | Unspecified |
| mmu-miR-152-3p | MIMAT0000162 | Nptn | 20320 | HITS-CLIP | 37986224\|25083871\|29167373 | Unspecified |
| mmu-miR-152-3p | MIMAT0000162 | Nptx1 | 18164 | HITS-CLIP | 37986224\|25083871 | Unspecified |
| mmu-miR-152-3p | MIMAT0000162 | Nr1d2 | 353187 | HITS-CLIP | 37986224\|25083871 | Unspecified |
| mmu-miR-152-3p | MIMAT0000162 | Nr2c1 | 22025 | HITS-CLIP | 37986224\|23597149 | Unspecified |
| mmu-miR-152-3p | MIMAT0000162 | Nrarp | 67122 | HITS-CLIP | 37986224\|25083871 | Unspecified |
| mmu-miR-152-3p | MIMAT0000162 | Nras | 18176 | HITS-CLIP | 25083871 | Unspecified |
| mmu-miR-152-3p | MIMAT0000162 | Nrbp1 | 192292 | HITS-CLIP | 25083871 | Unspecified |
| mmu-miR-152-3p | MIMAT0000162 | Nrd1 | 230598 | HITS-CLIP | 25083871 | Unspecified |
| mmu-miR-152-3p | MIMAT0000162 | Nrep | 27528 | HITS-CLIP | 37986224\|23597149 | Unspecified |
| mmu-miR-152-3p | MIMAT0000162 | Nrf1 | 18181 | HITS-CLIP | 25083871 | Unspecified |
| mmu-miR-152-3p | MIMAT0000162 | Nrip1 | 268903 | HITS-CLIP | 37986224 | Unspecified |
| mmu-miR-152-3p | MIMAT0000162 | Nrp1 | 18186 | HITS-CLIP | 37986224 | Unspecified |
| mmu-miR-152-3p | MIMAT0000162 | Nrp2 | 18187 | HITS-CLIP | 37986224\|25083871\|23597149\|29167 | Unspecified |
| mmu-miR-152-3p | MIMAT0000162 | Nsd1 | 18193 | HITS-CLIP | 37986224\|25083871\|29167373 | Unspecified |
| mmu-miR-152-3p | MIMAT0000162 | Nsd2 | 107823 | HITS-CLIP | 29167373 | Unspecified |
| mmu-miR-152-3p | MIMAT0000162 | Nsd3 | 234135 | HITS-CLIP | 25083871 | Unspecified |
| mmu-miR-152-3p | MIMAT0000162 | Nsg1 | 18196 | HITS-CLIP | 37986224 | Unspecified |
| mmu-miR-152-3p | MIMAT0000162 | Nsmce2 | 68501 | HITS-CLIP | 29167373 | Unspecified |
| mmu-miR-152-3p | MIMAT0000162 | Nsun2 | 28114 | HITS-CLIP | 25083871 | Unspecified |
| mmu-miR-152-3p | MIMAT0000162 | Nt5dc2 | 70021 | HITS-CLIP | 37986224\|25083871 | Unspecified |
| mmu-miR-152-3p | MIMAT0000162 | Ntn4 | 57764 | HITS-CLIP | 25083871 | Unspecified |
| mmu-miR-152-3p | MIMAT0000162 | Nuak1 | 77976 | HITS-CLIP | 25083871 | Unspecified |
| mmu-miR-152-3p | MIMAT0000162 | Nucks1 | 98415 | HITS-CLIP | 25083871\|29167373 | Unspecified |
| mmu-miR-152-3p | MIMAT0000162 | Nudt19 | 110959 | HITS-CLIP | 37986224 | Unspecified |
| mmu-miR-152-3p | MIMAT0000162 | Nudt21 | 68219 | HITS-CLIP | 37986224 | Unspecified |
| mmu-miR-152-3p | MIMAT0000162 | Nudt7 | 67528 | HITS-CLIP | 37986224\|23597149 | Unspecified |
| mmu-miR-152-3p | MIMAT0000162 | Nuf2 | 66977 | HITS-CLIP | 37986224 | Unspecified |
| mmu-miR-152-3p | MIMAT0000162 | Numa1 | 101706 | HITS-CLIP | 37986224\|25083871 | Unspecified |
| mmu-miR-152-3p | MIMAT0000162 | Numb | 18222 | HITS-CLIP | 37986224\|25083871 | Unspecified |
| mmu-miR-152-3p | MIMAT0000162 | Nup153 | 218210 | HITS-CLIP | 37986224\|25083871 | Unspecified |
| mmu-miR-152-3p | MIMAT0000162 | Nup214 | 227720 | HITS-CLIP | 23597149 | Unspecified |
| mmu-miR-152-3p | MIMAT0000162 | Nup62 | 18226 | HITS-CLIP | 25083871 | Unspecified |
| mmu-miR-152-3p | MIMAT0000162 | Nup93 | 71805 | HITS-CLIP | 37986224 | Unspecified |
| mmu-miR-152-3p | MIMAT0000162 | Nus1 | 52014 | HITS-CLIP | 37986224 | Unspecified |
| mmu-miR-152-3p | MIMAT0000162 | Nusap1 | 108907 | HITS-CLIP | 23597149 | Unspecified |
| mmu-miR-152-3p | MIMAT0000162 | Nxt2 | 237082 | HITS-CLIP | 37986224 | Unspecified |
| mmu-miR-152-3p | MIMAT0000162 | Oas2 | 246728 | HITS-CLIP | 29167373 | Unspecified |

| | | | | | | | |
|---|---|---|---|---|---|---|---|
| mmu-miR-152-3p | MIMAT0000162 | Odr4 | 54953 | HITS-CLIP | | 25083871 | Unspecified |
| mmu-miR-152-3p | MIMAT0000162 | Ogdh | 18293 | HITS-CLIP | | 37986224\|23597149\|25083871\|29167 | Unspecified |
| mmu-miR-152-3p | MIMAT0000162 | Ogt | 108155 | HITS-CLIP | | 25083871 | Unspecified |
| mmu-miR-152-3p | MIMAT0000162 | Olfm1 | 56177 | HITS-CLIP | | 25083871 | Unspecified |
| mmu-miR-152-3p | MIMAT0000162 | Olfr111 | 545205 | HITS-CLIP | | 29167373 | Unspecified |
| mmu-miR-152-3p | MIMAT0000162 | Omd | 27047 | HITS-CLIP | | 25083871 | Unspecified |
| mmu-miR-152-3p | MIMAT0000162 | Orai3 | 269999 | HITS-CLIP | | 37986224 | Unspecified |
| mmu-miR-152-3p | MIMAT0000162 | Orc3 | 50793 | HITS-CLIP | | 25083871 | Unspecified |
| mmu-miR-152-3p | MIMAT0000162 | Ormdl2 | 66844 | HITS-CLIP | | 25083871 | Unspecified |
| mmu-miR-152-3p | MIMAT0000162 | Osbpl11 | 106326 | HITS-CLIP | | 37986224\|25083871 | Unspecified |
| mmu-miR-152-3p | MIMAT0000162 | Osbpl1a | 64291 | HITS-CLIP | | 23597149 | Unspecified |
| mmu-miR-152-3p | MIMAT0000162 | Osbpl3 | 71720 | HITS-CLIP | | 37986224 | Unspecified |
| mmu-miR-152-3p | MIMAT0000162 | Osgin2 | 209212 | HITS-CLIP | | 37986224\|25083871 | Unspecified |
| mmu-miR-152-3p | MIMAT0000162 | Osmr | 18414 | HITS-CLIP | | 23597149 | Unspecified |
| mmu-miR-152-3p | MIMAT0000162 | Ostc | 66357 | HITS-CLIP | | 25083871 | Unspecified |
| mmu-miR-152-3p | MIMAT0000162 | Ostf1 | 20409 | HITS-CLIP | | 37986224 | Unspecified |
| mmu-miR-152-3p | MIMAT0000162 | Otc | 18416 | HITS-CLIP | | 23597149 | Unspecified |
| mmu-miR-152-3p | MIMAT0000162 | Otud4 | 73945 | HITS-CLIP | | 37986224\|25083871\|29167373 | Unspecified |
| mmu-miR-152-3p | MIMAT0000162 | Otud7b | 229603 | HITS-CLIP | | 25083871 | Unspecified |
| mmu-miR-152-3p | MIMAT0000162 | Otulin | 432940 | HITS-CLIP | | 37986224\|25083871 | Unspecified |
| mmu-miR-152-3p | MIMAT0000162 | Oxct1 | 67041 | HITS-CLIP | | 25083871 | Unspecified |
| mmu-miR-152-3p | MIMAT0000162 | Oxsr1 | 108737 | HITS-CLIP | | 37986224 | Unspecified |
| mmu-miR-152-3p | MIMAT0000162 | Pacs2 | 217893 | HITS-CLIP | | 25083871 | Unspecified |
| mmu-miR-152-3p | MIMAT0000162 | Pafah1b1 | 18472 | HITS-CLIP | | 25083871 | Unspecified |
| mmu-miR-152-3p | MIMAT0000162 | Pafah1b2 | 18475 | HITS-CLIP | | 37986224 | Unspecified |
| mmu-miR-152-3p | MIMAT0000162 | Paip2 | 67869 | HITS-CLIP | | 37986224 | Unspecified |
| mmu-miR-152-3p | MIMAT0000162 | Pak1 | 18479 | HITS-CLIP | | 37986224 | Unspecified |
| mmu-miR-152-3p | MIMAT0000162 | Pak2 | 224105 | HITS-CLIP | | 37986224 | Unspecified |
| mmu-miR-152-3p | MIMAT0000162 | Pald1 | 27355 | HITS-CLIP | | 23597149 | Unspecified |
| mmu-miR-152-3p | MIMAT0000162 | Palld | 72333 | HITS-CLIP | | 25083871 | Unspecified |
| mmu-miR-152-3p | MIMAT0000162 | Pan3 | 72587 | HITS-CLIP | | 37986224 | Unspecified |
| mmu-miR-152-3p | MIMAT0000162 | Pank1 | 75735 | HITS-CLIP | | 23597149 | Unspecified |
| mmu-miR-152-3p | MIMAT0000162 | Pank3 | 211347 | HITS-CLIP | | 23597149\|25083871 | Unspecified |
| mmu-miR-152-3p | MIMAT0000162 | Pank4 | 55229 | HITS-CLIP | | 23597149 | Unspecified |
| mmu-miR-152-3p | MIMAT0000162 | Papola | 18789 | HITS-CLIP | | 25083871 | Unspecified |
| mmu-miR-152-3p | MIMAT0000162 | Paqr9 | 75552 | HITS-CLIP | | 37986224 | Unspecified |
| mmu-miR-152-3p | MIMAT0000162 | Pard3 | 93742 | HITS-CLIP | | 37986224 | Unspecified |
| mmu-miR-152-3p | MIMAT0000162 | Parp11 | 101187 | HITS-CLIP | | 29167373 | Unspecified |
| mmu-miR-152-3p | MIMAT0000162 | Parva | 57342 | HITS-CLIP | | 37986224 | Unspecified |

| | | | | | | |
|---|---|---|---|---|---|---|
| mmu-miR-152-3p | MIMAT0000162 | Parvb | 170736 | HITS-CLIP | 25083871 | Unspecified |
| mmu-miR-152-3p | MIMAT0000162 | Patl1 | 225929 | HITS-CLIP | 25083871 | Unspecified |
| mmu-miR-152-3p | MIMAT0000162 | Pbxip1 | 229534 | HITS-CLIP | 37986224 | Unspecified |
| mmu-miR-152-3p | MIMAT0000162 | Pcbp1 | 23983 | HITS-CLIP | 37986224\|23597149 | Unspecified |
| mmu-miR-152-3p | MIMAT0000162 | Pcca | 110821 | HITS-CLIP | 23597149 | Unspecified |
| mmu-miR-152-3p | MIMAT0000162 | Pcdh18 | 73173 | HITS-CLIP | 37986224 | Unspecified |
| mmu-miR-152-3p | MIMAT0000162 | Pcdh19 | 279653 | HITS-CLIP | 37986224 | Unspecified |
| mmu-miR-152-3p | MIMAT0000162 | Pcdhga12 | 93724 | HITS-CLIP | 25083871 | Unspecified |
| mmu-miR-152-3p | MIMAT0000162 | Pcdhgc3 | 5098 | HITS-CLIP | 25083871 | Unspecified |
| mmu-miR-152-3p | MIMAT0000162 | Pcgf3 | 69587 | HITS-CLIP | 25083871 | Unspecified |
| mmu-miR-152-3p | MIMAT0000162 | Pck1 | 18534 | HITS-CLIP | 23597149 | Unspecified |
| mmu-miR-152-3p | MIMAT0000162 | Pcm1 | 18536 | HITS-CLIP | 25083871 | Unspecified |
| mmu-miR-152-3p | MIMAT0000162 | Pcnx | 54604 | HITS-CLIP | 37986224\|25083871 | Unspecified |
| mmu-miR-152-3p | MIMAT0000162 | Pcolce2 | 76477 | HITS-CLIP | 37986224 | Unspecified |
| mmu-miR-152-3p | MIMAT0000162 | Pcsk6 | 18553 | HITS-CLIP | 23597149 | Unspecified |
| mmu-miR-152-3p | MIMAT0000162 | Pcsk9 | 100102 | HITS-CLIP | 37986224 | Unspecified |
| mmu-miR-152-3p | MIMAT0000162 | Pcx | 18563 | HITS-CLIP | 23597149 | Unspecified |
| mmu-miR-152-3p | MIMAT0000162 | Pcyox1 | 66881 | HITS-CLIP | 25083871 | Unspecified |
| mmu-miR-152-3p | MIMAT0000162 | Pdcd2l | 68079 | HITS-CLIP | 23597149\|25083871 | Unspecified |
| mmu-miR-152-3p | MIMAT0000162 | Pdcd6ip | 18571 | HITS-CLIP | 25083871 | Unspecified |
| mmu-miR-152-3p | MIMAT0000162 | Pde3a | 54611 | HITS-CLIP | 37986224 | Unspecified |
| mmu-miR-152-3p | MIMAT0000162 | Pde3b | 18576 | HITS-CLIP | 25083871 | Unspecified |
| mmu-miR-152-3p | MIMAT0000162 | Pde4b | 18578 | HITS-CLIP | 25083871\|29167373 | Unspecified |
| mmu-miR-152-3p | MIMAT0000162 | Pde4d | 238871 | HITS-CLIP | 25083871 | Unspecified |
| mmu-miR-152-3p | MIMAT0000162 | Pde4dip | 83679 | HITS-CLIP | 37986224\|25083871\|23597149 | Unspecified |
| mmu-miR-152-3p | MIMAT0000162 | Pdgfa | 18590 | HITS-CLIP | 25083871 | Unspecified |
| mmu-miR-152-3p | MIMAT0000162 | Pdgfb | 18591 | HITS-CLIP | 37986224 | Unspecified |
| mmu-miR-152-3p | MIMAT0000162 | Pdgfrb | 18596 | HITS-CLIP | 25083871 | Unspecified |
| mmu-miR-152-3p | MIMAT0000162 | Pdha1 | 18597 | HITS-CLIP | 25083871 | Unspecified |
| mmu-miR-152-3p | MIMAT0000162 | Pdhb | 68263 | HITS-CLIP | 23597149 | Unspecified |
| mmu-miR-152-3p | MIMAT0000162 | Pdia3 | 14827 | HITS-CLIP | 37986224\|25083871\|29167373 | Unspecified |
| mmu-miR-152-3p | MIMAT0000162 | Pdia4 | 12304 | HITS-CLIP | 37986224 | Unspecified |
| mmu-miR-152-3p | MIMAT0000162 | Pdk1 | 228026 | HITS-CLIP | 25083871 | Unspecified |
| mmu-miR-152-3p | MIMAT0000162 | Pdk4 | 27273 | HITS-CLIP | 37986224 | Unspecified |
| mmu-miR-152-3p | MIMAT0000162 | Pdp1 | 381511 | HITS-CLIP | 25083871 | Unspecified |
| mmu-miR-152-3p | MIMAT0000162 | Pds5a | 71521 | HITS-CLIP | 29167373 | Unspecified |
| mmu-miR-152-3p | MIMAT0000162 | Pdxk | 216134 | HITS-CLIP | 29167373 | Unspecified |
| mmu-miR-152-3p | MIMAT0000162 | Pdzd8 | 107368 | HITS-CLIP | 37986224\|25083871\|23597149\|29167 | Unspecified |
| mmu-miR-152-3p | MIMAT0000162 | Peak1 | 244895 | HITS-CLIP | 37986224\|29167373 | Unspecified |

| | | | | | | |
|---|---|---|---|---|---|---|
| mmu-miR-152-3p | MIMAT0000162 | Peg3 | 18616 | HITS-CLIP | 37986224\|25083871 | Unspecified |
| mmu-miR-152-3p | MIMAT0000162 | Pemt | 18618 | HITS-CLIP | 37986224\|23597149 | Unspecified |
| mmu-miR-152-3p | MIMAT0000162 | Per1 | 18626 | HITS-CLIP | 23597149 | Unspecified |
| mmu-miR-152-3p | MIMAT0000162 | Pex11a | 18631 | HITS-CLIP | 25083871 | Unspecified |
| mmu-miR-152-3p | MIMAT0000162 | Pex19 | 19298 | HITS-CLIP | 25083871\|21258322 | Unspecified |
| mmu-miR-152-3p | MIMAT0000162 | Pfas | 237823 | HITS-CLIP | 37986224 | Unspecified |
| mmu-miR-152-3p | MIMAT0000162 | Pfkfb1 | 18639 | HITS-CLIP | 23597149 | Unspecified |
| mmu-miR-152-3p | MIMAT0000162 | Pfkm | 18642 | HITS-CLIP | 25083871 | Unspecified |
| mmu-miR-152-3p | MIMAT0000162 | Pgap1 | 241062 | HITS-CLIP | 29167373 | Unspecified |
| mmu-miR-152-3p | MIMAT0000162 | Pglyrp2 | 57757 | HITS-CLIP | 37986224 | Unspecified |
| mmu-miR-152-3p | MIMAT0000162 | Pgm1 | 66681 | HITS-CLIP | 23597149 | Unspecified |
| mmu-miR-152-3p | MIMAT0000162 | Pgp | 67078 | HITS-CLIP | 25083871 | Unspecified |
| mmu-miR-152-3p | MIMAT0000162 | Pgrmc1 | 53328 | HITS-CLIP | 37986224 | Unspecified |
| mmu-miR-152-3p | MIMAT0000162 | Pgs1 | 74451 | HITS-CLIP | 29167373 | Unspecified |
| mmu-miR-152-3p | MIMAT0000162 | Phc2 | 54383 | HITS-CLIP | 25083871 | Unspecified |
| mmu-miR-152-3p | MIMAT0000162 | Phc3 | 241915 | HITS-CLIP | 37986224\|29167373 | Unspecified |
| mmu-miR-152-3p | MIMAT0000162 | Phf20 | 228829 | HITS-CLIP | 25083871\|29167373 | Unspecified |
| mmu-miR-152-3p | MIMAT0000162 | Phf21a | 192285 | HITS-CLIP | 25083871 | Unspecified |
| mmu-miR-152-3p | MIMAT0000162 | Phkb | 102093 | HITS-CLIP | 25083871 | Unspecified |
| mmu-miR-152-3p | MIMAT0000162 | Phldb2 | 208177 | HITS-CLIP | 37986224 | Unspecified |
| mmu-miR-152-3p | MIMAT0000162 | Phyh | 16922 | HITS-CLIP | 23597149 | Unspecified |
| mmu-miR-152-3p | MIMAT0000162 | Phyhd1 | 227696 | HITS-CLIP | 37986224 | Unspecified |
| mmu-miR-152-3p | MIMAT0000162 | Pi4k2a | 84095 | HITS-CLIP | 37986224 | Unspecified |
| mmu-miR-152-3p | MIMAT0000162 | Pias1 | 56469 | HITS-CLIP | 29167373 | Unspecified |
| mmu-miR-152-3p | MIMAT0000162 | Picalm | 233489 | HITS-CLIP | 37986224\|25083871 | Unspecified |
| mmu-miR-152-3p | MIMAT0000162 | Piezo1 | 234839 | HITS-CLIP | 25083871 | Unspecified |
| mmu-miR-152-3p | MIMAT0000162 | Piezo2 | 667742 | HITS-CLIP | 25083871 | Unspecified |
| mmu-miR-152-3p | MIMAT0000162 | Pigc | 67292 | HITS-CLIP | 23597149 | Unspecified |
| mmu-miR-152-3p | MIMAT0000162 | Pigg | 433931 | HITS-CLIP | 37986224 | Unspecified |
| mmu-miR-152-3p | MIMAT0000162 | Pigk | 329777 | HITS-CLIP | 25083871 | Unspecified |
| mmu-miR-152-3p | MIMAT0000162 | Pign | 27392 | HITS-CLIP | 37986224 | Unspecified |
| mmu-miR-152-3p | MIMAT0000162 | Pigq | 14755 | HITS-CLIP | 23597149 | Unspecified |
| mmu-miR-152-3p | MIMAT0000162 | Pigx | 72084 | HITS-CLIP | 25083871\|23597149 | Unspecified |
| mmu-miR-152-3p | MIMAT0000162 | Pik3ca | 18706 | HITS-CLIP | 37986224 | Unspecified |
| mmu-miR-152-3p | MIMAT0000162 | Pik3r1 | 18708 | HITS-CLIP | 37986224\|25083871 | Unspecified |
| mmu-miR-152-3p | MIMAT0000162 | Pik3r3 | 18710 | HITS-CLIP | 37986224 | Unspecified |
| mmu-miR-152-3p | MIMAT0000162 | Pink1 | 68943 | HITS-CLIP | 23597149 | Unspecified |
| mmu-miR-152-3p | MIMAT0000162 | Pip4k2b | 108083 | HITS-CLIP | 37986224 | Unspecified |
| mmu-miR-152-3p | MIMAT0000162 | Pirb | 18733 | HITS-CLIP | 29167373 | Unspecified |

| | | | | | | |
|---|---|---|---|---|---|---|
| mmu-miR-152-3p | MIMAT0000162 | Pitpnb | 56305 | HITS-CLIP | 25083871 | Unspecified |
| mmu-miR-152-3p | MIMAT0000162 | Pja1 | 18744 | HITS-CLIP | 37986224 | Unspecified |
| mmu-miR-152-3p | MIMAT0000162 | Pja2 | 224938 | HITS-CLIP | 25083871\|23597149 | Unspecified |
| mmu-miR-152-3p | MIMAT0000162 | Pkd1 | 18763 | HITS-CLIP | 37986224\|25083871\|21258322\|23597 | Unspecified |
| mmu-miR-152-3p | MIMAT0000162 | Pkia | 18767 | HITS-CLIP | 25083871 | Unspecified |
| mmu-miR-152-3p | MIMAT0000162 | Pkm | 18746 | HITS-CLIP | 25083871\|29167373 | Unspecified |
| mmu-miR-152-3p | MIMAT0000162 | Pknox1 | 18771 | HITS-CLIP | 37986224\|25083871 | Unspecified |
| mmu-miR-152-3p | MIMAT0000162 | Pla2g15 | 192654 | HITS-CLIP | 37986224 | Unspecified |
| mmu-miR-152-3p | MIMAT0000162 | Plaa | 18786 | HITS-CLIP | 37986224\|25083871\|23597149 | Unspecified |
| mmu-miR-152-3p | MIMAT0000162 | Plcb1 | 18795 | HITS-CLIP | 25083871 | Unspecified |
| mmu-miR-152-3p | MIMAT0000162 | Plec | 18810 | HITS-CLIP | 37986224\|25083871\|29167373 | Unspecified |
| mmu-miR-152-3p | MIMAT0000162 | Plekha3 | 83435 | HITS-CLIP | 25083871 | Unspecified |
| mmu-miR-152-3p | MIMAT0000162 | Plekha5 | 109135 | HITS-CLIP | 25083871\|23597149 | Unspecified |
| mmu-miR-152-3p | MIMAT0000162 | Plekhf2 | 71801 | HITS-CLIP | 29167373 | Unspecified |
| mmu-miR-152-3p | MIMAT0000162 | Plekhg5 | 269608 | HITS-CLIP | 25083871 | Unspecified |
| mmu-miR-152-3p | MIMAT0000162 | Plekhh1 | 211945 | HITS-CLIP | 25083871 | Unspecified |
| mmu-miR-152-3p | MIMAT0000162 | Plekhm1 | 353047 | HITS-CLIP | 23597149 | Unspecified |
| mmu-miR-152-3p | MIMAT0000162 | Plekho1 | 67220 | HITS-CLIP | 37986224 | Unspecified |
| mmu-miR-152-3p | MIMAT0000162 | Plk2 | 20620 | HITS-CLIP | 37986224 | Unspecified |
| mmu-miR-152-3p | MIMAT0000162 | Plk3 | 12795 | HITS-CLIP | 23597149 | Unspecified |
| mmu-miR-152-3p | MIMAT0000162 | Plod2 | 26432 | HITS-CLIP | 25083871 | Unspecified |
| mmu-miR-152-3p | MIMAT0000162 | Plpp1 | 8611 | HITS-CLIP | 25083871 | Unspecified |
| mmu-miR-152-3p | MIMAT0000162 | Pmpcb | 73078 | HITS-CLIP | 23597149\|25083871 | Unspecified |
| mmu-miR-152-3p | MIMAT0000162 | Pnisr | 66625 | HITS-CLIP | 25083871\|29167373 | Unspecified |
| mmu-miR-152-3p | MIMAT0000162 | Pnpla6 | 50767 | HITS-CLIP | 37986224\|29167373 | Unspecified |
| mmu-miR-152-3p | MIMAT0000162 | Pnpla8 | 67452 | HITS-CLIP | 25083871 | Unspecified |
| mmu-miR-152-3p | MIMAT0000162 | Pofut2 | 80294 | HITS-CLIP | 37986224 | Unspecified |
| mmu-miR-152-3p | MIMAT0000162 | Pogz | 229584 | HITS-CLIP | 25083871 | Unspecified |
| mmu-miR-152-3p | MIMAT0000162 | Polg | 18975 | HITS-CLIP | 37986224\|23597149 | Unspecified |
| mmu-miR-152-3p | MIMAT0000162 | Polr2a | 20020 | HITS-CLIP | 25083871\|23597149\|29167373 | Unspecified |
| mmu-miR-152-3p | MIMAT0000162 | Polr2m | 28015 | HITS-CLIP | 37986224 | Unspecified |
| mmu-miR-152-3p | MIMAT0000162 | Polr3b | 70428 | HITS-CLIP | 29167373 | Unspecified |
| mmu-miR-152-3p | MIMAT0000162 | Polr3k | 67005 | HITS-CLIP | 23597149 | Unspecified |
| mmu-miR-152-3p | MIMAT0000162 | Pou2f1 | 18986 | HITS-CLIP | 25083871 | Unspecified |
| mmu-miR-152-3p | MIMAT0000162 | Ppan | 235036 | HITS-CLIP | 31433990 | Unspecified |
| mmu-miR-152-3p | MIMAT0000162 | Ppara | 19013 | HITS-CLIP | 23597149 | Unspecified |
| mmu-miR-152-3p | MIMAT0000162 | Ppard | 19015 | HITS-CLIP | 37986224 | Unspecified |
| mmu-miR-152-3p | MIMAT0000162 | Ppfibp1 | 67533 | HITS-CLIP | 25083871 | Unspecified |
| mmu-miR-152-3p | MIMAT0000162 | Ppl | 19041 | HITS-CLIP | 23597149 | Unspecified |

| | | | | | | | |
|---|---|---|---|---|---|---|---|
| mmu-miR-152-3p | MIMAT0000162 | Ppme1 | 72590 | HITS-CLIP | | 25083871 | Unspecified |
| mmu-miR-152-3p | MIMAT0000162 | Ppp1cb | 19046 | HITS-CLIP | | 23597149 | Unspecified |
| mmu-miR-152-3p | MIMAT0000162 | Ppp1cc | 19047 | HITS-CLIP | | 29167373 | Unspecified |
| mmu-miR-152-3p | MIMAT0000162 | Ppp1r12a | 17931 | HITS-CLIP | 25083871\|23597149 | | Unspecified |
| mmu-miR-152-3p | MIMAT0000162 | Ppp1r15a | 17872 | HITS-CLIP | | 29167373 | Unspecified |
| mmu-miR-152-3p | MIMAT0000162 | Ppp1r15b | 108954 | HITS-CLIP | | 25083871 | Unspecified |
| mmu-miR-152-3p | MIMAT0000162 | Ppp1r16a | 73062 | HITS-CLIP | | 23597149 | Unspecified |
| mmu-miR-152-3p | MIMAT0000162 | Ppp1r2 | 66849 | HITS-CLIP | | 25083871 | Unspecified |
| mmu-miR-152-3p | MIMAT0000162 | Ppp1r9b | 217124 | HITS-CLIP | | 25083871 | Unspecified |
| mmu-miR-152-3p | MIMAT0000162 | Ppp2cb | 19053 | HITS-CLIP | | 37986224 | Unspecified |
| mmu-miR-152-3p | MIMAT0000162 | Ppp2r1a | 51792 | HITS-CLIP | 25083871\|29167373 | | Unspecified |
| mmu-miR-152-3p | MIMAT0000162 | Ppp2r2a | 71978 | HITS-CLIP | | 37986224 | Unspecified |
| mmu-miR-152-3p | MIMAT0000162 | Ppp2r3a | 235542 | HITS-CLIP | 37986224\|25083871 | | Unspecified |
| mmu-miR-152-3p | MIMAT0000162 | Ppp2r5a | 226849 | HITS-CLIP | | 23597149 | Unspecified |
| mmu-miR-152-3p | MIMAT0000162 | Ppp2r5d | 21770 | HITS-CLIP | | 37986224 | Unspecified |
| mmu-miR-152-3p | MIMAT0000162 | Ppp3ca | 19055 | HITS-CLIP | | 25083871 | Unspecified |
| mmu-miR-152-3p | MIMAT0000162 | Ppp3cb | 19056 | HITS-CLIP | 37986224\|25083871 | | Unspecified |
| mmu-miR-152-3p | MIMAT0000162 | Ppp5c | 19060 | HITS-CLIP | 37986224\|29167373 | | Unspecified |
| mmu-miR-152-3p | MIMAT0000162 | Ppp6c | 67857 | HITS-CLIP | 37986224\|23597149\|25083871\|29167 | | Unspecified |
| mmu-miR-152-3p | MIMAT0000162 | Ppp6r1 | 243819 | HITS-CLIP | | 37986224 | Unspecified |
| mmu-miR-152-3p | MIMAT0000162 | Ppp6r3 | 52036 | HITS-CLIP | | 25083871 | Unspecified |
| mmu-miR-152-3p | MIMAT0000162 | Pprc1 | 226169 | HITS-CLIP | | 37986224 | Unspecified |
| mmu-miR-152-3p | MIMAT0000162 | Ppt2 | 54397 | HITS-CLIP | | 37986224 | Unspecified |
| mmu-miR-152-3p | MIMAT0000162 | Pptc7 | 320717 | HITS-CLIP | 25083871\|29167373 | | Unspecified |
| mmu-miR-152-3p | MIMAT0000162 | Prdm2 | 110593 | HITS-CLIP | | 25083871 | Unspecified |
| mmu-miR-152-3p | MIMAT0000162 | Prdm4 | 72843 | HITS-CLIP | | 37986224 | Unspecified |
| mmu-miR-152-3p | MIMAT0000162 | Prkaa1 | 105787 | HITS-CLIP | 37986224\|25083871 | | Unspecified |
| mmu-miR-152-3p | MIMAT0000162 | Prkab1 | 19079 | HITS-CLIP | | 23597149 | Unspecified |
| mmu-miR-152-3p | MIMAT0000162 | Prkab2 | 108097 | HITS-CLIP | | 37986224 | Unspecified |
| mmu-miR-152-3p | MIMAT0000162 | Prkaca | 18747 | HITS-CLIP | 23597149\|25083871 | | Unspecified |
| mmu-miR-152-3p | MIMAT0000162 | Prkar2a | 19087 | HITS-CLIP | 25083871\|23597149 | | Unspecified |
| mmu-miR-152-3p | MIMAT0000162 | Prkci | 18759 | HITS-CLIP | 37986224\|25083871 | | Unspecified |
| mmu-miR-152-3p | MIMAT0000162 | Prkd3 | 75292 | HITS-CLIP | 25083871\|23597149 | | Unspecified |
| mmu-miR-152-3p | MIMAT0000162 | Prkra | 23992 | HITS-CLIP | | 23597149 | Unspecified |
| mmu-miR-152-3p | MIMAT0000162 | Prlr | 19116 | HITS-CLIP | | 23597149 | Unspecified |
| mmu-miR-152-3p | MIMAT0000162 | Prmt9 | 80204 | HITS-CLIP | | 23597149 | Unspecified |
| mmu-miR-152-3p | MIMAT0000162 | Prnp | 19122 | HITS-CLIP | 37986224\|29167373 | | Unspecified |
| mmu-miR-152-3p | MIMAT0000162 | Prodh | 19125 | HITS-CLIP | | 23597149 | Unspecified |
| mmu-miR-152-3p | MIMAT0000162 | Prox1 | 19130 | HITS-CLIP | | 23597149 | Unspecified |

| | | | | | | |
|---|---|---|---|---|---|---|
| mmu-miR-152-3p | MIMAT0000162 | Prpf31 | 68988 | HITS-CLIP | 21258322 | Unspecified |
| mmu-miR-152-3p | MIMAT0000162 | Prpf38b | 66921 | HITS-CLIP | 37986224\|25083871 | Unspecified |
| mmu-miR-152-3p | MIMAT0000162 | Prpf6 | 68879 | HITS-CLIP | 37986224 | Unspecified |
| mmu-miR-152-3p | MIMAT0000162 | Prpf8 | 192159 | HITS-CLIP | 37986224\|23597149\|25083871 | Unspecified |
| mmu-miR-152-3p | MIMAT0000162 | Prr14l | 215476 | HITS-CLIP | 37986224\|25083871 | Unspecified |
| mmu-miR-152-3p | MIMAT0000162 | Prrc1 | 73137 | HITS-CLIP | 37986224 | Unspecified |
| mmu-miR-152-3p | MIMAT0000162 | Prune2 | 353211 | HITS-CLIP | 37986224\|25083871 | Unspecified |
| mmu-miR-152-3p | MIMAT0000162 | Psd3 | 234353 | HITS-CLIP | 37986224\|29167373 | Unspecified |
| mmu-miR-152-3p | MIMAT0000162 | Psmb6 | 19175 | HITS-CLIP | 25083871 | Unspecified |
| mmu-miR-152-3p | MIMAT0000162 | Psmc3 | 19182 | HITS-CLIP | 25083871 | Unspecified |
| mmu-miR-152-3p | MIMAT0000162 | Psmd1 | 70247 | HITS-CLIP | 37986224\|23597149 | Unspecified |
| mmu-miR-152-3p | MIMAT0000162 | Psmd11 | 69077 | HITS-CLIP | 37986224\|25083871\|29167373 | Unspecified |
| mmu-miR-152-3p | MIMAT0000162 | Psmd13 | 23997 | HITS-CLIP | 37986224\|23597149\|25083871 | Unspecified |
| mmu-miR-152-3p | MIMAT0000162 | Psmd2 | 21762 | HITS-CLIP | 37986224 | Unspecified |
| mmu-miR-152-3p | MIMAT0000162 | Psmd3 | 22123 | HITS-CLIP | 23597149 | Unspecified |
| mmu-miR-152-3p | MIMAT0000162 | Psmd4 | 19185 | HITS-CLIP | 25083871 | Unspecified |
| mmu-miR-152-3p | MIMAT0000162 | Psme4 | 103554 | HITS-CLIP | 25083871 | Unspecified |
| mmu-miR-152-3p | MIMAT0000162 | Pstk | 214580 | HITS-CLIP | 25083871 | Unspecified |
| mmu-miR-152-3p | MIMAT0000162 | Pstpip2 | 19201 | HITS-CLIP | 25083871\|29167373 | Unspecified |
| mmu-miR-152-3p | MIMAT0000162 | Ptar1 | 72351 | HITS-CLIP | 25083871 | Unspecified |
| mmu-miR-152-3p | MIMAT0000162 | Ptbp3 | 230257 | HITS-CLIP | 37986224 | Unspecified |
| mmu-miR-152-3p | MIMAT0000162 | Ptcd2 | 68927 | HITS-CLIP | 23597149 | Unspecified |
| mmu-miR-152-3p | MIMAT0000162 | Pten | 19211 | HITS-CLIP//Luciferase reporter assay//Wes | 25083871\|26996529 | Unspecified |
| mmu-miR-152-3p | MIMAT0000162 | Ptk7 | 71461 | HITS-CLIP | 37986224 | Unspecified |
| mmu-miR-152-3p | MIMAT0000162 | Ptp4a2 | 19244 | HITS-CLIP | 25083871\|23597149 | Unspecified |
| mmu-miR-152-3p | MIMAT0000162 | Ptpn1 | 19246 | HITS-CLIP | 37986224\|25083871 | Unspecified |
| mmu-miR-152-3p | MIMAT0000162 | Ptpn11 | 19247 | HITS-CLIP | 25083871 | Unspecified |
| mmu-miR-152-3p | MIMAT0000162 | Ptpn14 | 19250 | HITS-CLIP | 37986224 | Unspecified |
| mmu-miR-152-3p | MIMAT0000162 | Ptpn4 | 19258 | HITS-CLIP | 25083871 | Unspecified |
| mmu-miR-152-3p | MIMAT0000162 | Ptpra | 19262 | HITS-CLIP | 37986224\|25083871\|29167373 | Unspecified |
| mmu-miR-152-3p | MIMAT0000162 | Ptprd | 19266 | HITS-CLIP | 37986224 | Unspecified |
| mmu-miR-152-3p | MIMAT0000162 | Ptprg | 19270 | HITS-CLIP | 25083871 | Unspecified |
| mmu-miR-152-3p | MIMAT0000162 | Ptprk | 19272 | HITS-CLIP | 37986224\|23597149 | Unspecified |
| mmu-miR-152-3p | MIMAT0000162 | Pttg1ip | 108705 | HITS-CLIP | 23597149 | Unspecified |
| mmu-miR-152-3p | MIMAT0000162 | Pum1 | 80912 | HITS-CLIP | 25083871 | Unspecified |
| mmu-miR-152-3p | MIMAT0000162 | Pum2 | 80913 | HITS-CLIP | 37986224\|25083871\|23597149 | Unspecified |
| mmu-miR-152-3p | MIMAT0000162 | Pura | 19290 | HITS-CLIP | 25083871 | Unspecified |
| mmu-miR-152-3p | MIMAT0000162 | Purb | 19291 | HITS-CLIP | 25083871\|29167373 | Unspecified |
| mmu-miR-152-3p | MIMAT0000162 | Pus1 | 855889 | HITS-CLIP | 25083871 | Unspecified |

| | | | | | | |
|---|---|---|---|---|---|---|
| mmu-miR-152-3p | MIMAT0000162 | Pvr | 52118 | HITS-CLIP | 23597149 | Unspecified |
| mmu-miR-152-3p | MIMAT0000162 | Pxdn | 69675 | HITS-CLIP | 37986224 | Unspecified |
| mmu-miR-152-3p | MIMAT0000162 | Pxmp4 | 59038 | HITS-CLIP | 37986224\|23597149 | Unspecified |
| mmu-miR-152-3p | MIMAT0000162 | Pxylp1 | 235534 | HITS-CLIP | 25083871 | Unspecified |
| mmu-miR-152-3p | MIMAT0000162 | Pygm | 19309 | HITS-CLIP | 25083871 | Unspecified |
| mmu-miR-152-3p | MIMAT0000162 | Pygo2 | 68911 | HITS-CLIP | 25083871 | Unspecified |
| mmu-miR-152-3p | MIMAT0000162 | Pzp | 11287 | HITS-CLIP | 23597149 | Unspecified |
| mmu-miR-152-3p | MIMAT0000162 | Qk | 19317 | HITS-CLIP | 37986224\|25083871\|23597149\|29167 | Unspecified |
| mmu-miR-152-3p | MIMAT0000162 | Qrich1 | 69232 | HITS-CLIP | 25083871 | Unspecified |
| mmu-miR-152-3p | MIMAT0000162 | Qser1 | 99003 | HITS-CLIP | 37986224\|25083871 | Unspecified |
| mmu-miR-152-3p | MIMAT0000162 | Qsox1 | 104009 | HITS-CLIP | 37986224 | Unspecified |
| mmu-miR-152-3p | MIMAT0000162 | R3hdm1 | 226412 | HITS-CLIP | 37986224 | Unspecified |
| mmu-miR-152-3p | MIMAT0000162 | Rab10 | 19325 | HITS-CLIP | 37986224\|23597149\|25083871\|29167 | Unspecified |
| mmu-miR-152-3p | MIMAT0000162 | Rab11b | 19326 | HITS-CLIP | 37986224 | Unspecified |
| mmu-miR-152-3p | MIMAT0000162 | Rab12 | 19328 | HITS-CLIP | 37986224 | Unspecified |
| mmu-miR-152-3p | MIMAT0000162 | Rab2b | 76338 | HITS-CLIP | 25083871 | Unspecified |
| mmu-miR-152-3p | MIMAT0000162 | Rab30 | 75985 | HITS-CLIP | 23597149 | Unspecified |
| mmu-miR-152-3p | MIMAT0000162 | Rab34 | 19376 | HITS-CLIP | 25083871 | Unspecified |
| mmu-miR-152-3p | MIMAT0000162 | Rab35 | 77407 | HITS-CLIP | 23597149 | Unspecified |
| mmu-miR-152-3p | MIMAT0000162 | Rab3gap2 | 98732 | HITS-CLIP | 23597149\|29167373 | Unspecified |
| mmu-miR-152-3p | MIMAT0000162 | Rab43 | 69834 | HITS-CLIP | 25083871\|23597149 | Unspecified |
| mmu-miR-152-3p | MIMAT0000162 | Rab44 | 442827 | HITS-CLIP | 25083871 | Unspecified |
| mmu-miR-152-3p | MIMAT0000162 | Rab6a | 19346 | HITS-CLIP | 37986224\|23597149 | Unspecified |
| mmu-miR-152-3p | MIMAT0000162 | Rab8b | 235442 | HITS-CLIP | 37986224 | Unspecified |
| mmu-miR-152-3p | MIMAT0000162 | Rabep1 | 54189 | HITS-CLIP | 25083871 | Unspecified |
| mmu-miR-152-3p | MIMAT0000162 | Rabggta | 56187 | HITS-CLIP | 23597149 | Unspecified |
| mmu-miR-152-3p | MIMAT0000162 | Rack1 | 10399 | HITS-CLIP | 23597149\|29167373 | Unspecified |
| mmu-miR-152-3p | MIMAT0000162 | Rad54l2 | 81000 | HITS-CLIP | 25083871 | Unspecified |
| mmu-miR-152-3p | MIMAT0000162 | Rai1 | 19377 | HITS-CLIP | 37986224 | Unspecified |
| mmu-miR-152-3p | MIMAT0000162 | Rai14 | 75646 | HITS-CLIP | 37986224 | Unspecified |
| mmu-miR-152-3p | MIMAT0000162 | Ralgapa1 | 56784 | HITS-CLIP | 37986224\|23597149\|25083871 | Unspecified |
| mmu-miR-152-3p | MIMAT0000162 | Ralgapb | 228850 | HITS-CLIP | 37986224\|25083871 | Unspecified |
| mmu-miR-152-3p | MIMAT0000162 | Ranbp2 | 19386 | HITS-CLIP | 23597149\|25083871\|29167373 | Unspecified |
| mmu-miR-152-3p | MIMAT0000162 | Ranbp3 | 71810 | HITS-CLIP | 25083871 | Unspecified |
| mmu-miR-152-3p | MIMAT0000162 | Ranbp9 | 56705 | HITS-CLIP | 25083871 | Unspecified |
| mmu-miR-152-3p | MIMAT0000162 | Rangap1 | 19387 | HITS-CLIP | 37986224 | Unspecified |
| mmu-miR-152-3p | MIMAT0000162 | Rap1b | 215449 | HITS-CLIP | 23597149 | Unspecified |
| mmu-miR-152-3p | MIMAT0000162 | Rap1gds1 | 229877 | HITS-CLIP | 25083871\|23597149 | Unspecified |
| mmu-miR-152-3p | MIMAT0000162 | Rap2a | 76108 | HITS-CLIP | 25083871 | Unspecified |

| | | | | | | |
|---|---|---|---|---|---|---|
| mmu-miR-152-3p | MIMAT0000162 | Rap2c | 72065 | HITS-CLIP | 37986224\|25083871 | Unspecified |
| mmu-miR-152-3p | MIMAT0000162 | Rapgef1 | 107746 | HITS-CLIP | 25083871 | Unspecified |
| mmu-miR-152-3p | MIMAT0000162 | Rapgef2 | 76089 | HITS-CLIP | 23597149\|25083871 | Unspecified |
| mmu-miR-152-3p | MIMAT0000162 | Rapgef4 | 56508 | HITS-CLIP | 37986224 | Unspecified |
| mmu-miR-152-3p | MIMAT0000162 | Rapgef6 | 192786 | HITS-CLIP | 29167373 | Unspecified |
| mmu-miR-152-3p | MIMAT0000162 | Rara | 19401 | HITS-CLIP | 37986224 | Unspecified |
| mmu-miR-152-3p | MIMAT0000162 | Rars | 104458 | HITS-CLIP | 25083871 | Unspecified |
| mmu-miR-152-3p | MIMAT0000162 | Rars2 | 109093 | HITS-CLIP | 23597149 | Unspecified |
| mmu-miR-152-3p | MIMAT0000162 | Rasal2 | 226525 | HITS-CLIP | 25083871 | Unspecified |
| mmu-miR-152-3p | MIMAT0000162 | Rassf2 | 215653 | HITS-CLIP | 29167373 | Unspecified |
| mmu-miR-152-3p | MIMAT0000162 | Rassf5 | 54354 | HITS-CLIP | 23597149 | Unspecified |
| mmu-miR-152-3p | MIMAT0000162 | Rassf7 | 66985 | HITS-CLIP | 37986224 | Unspecified |
| mmu-miR-152-3p | MIMAT0000162 | Rassf8 | 71323 | HITS-CLIP | 37986224\|25083871 | Unspecified |
| mmu-miR-152-3p | MIMAT0000162 | Raver1 | 71766 | HITS-CLIP | 37986224 | Unspecified |
| mmu-miR-152-3p | MIMAT0000162 | Rb1 | 19645 | HITS-CLIP | 25083871 | Unspecified |
| mmu-miR-152-3p | MIMAT0000162 | Rb1cc1 | 12421 | HITS-CLIP | 37986224\|25083871 | Unspecified |
| mmu-miR-152-3p | MIMAT0000162 | Rbbp4 | 19646 | HITS-CLIP | 25083871\|23597149 | Unspecified |
| mmu-miR-152-3p | MIMAT0000162 | Rbbp7 | 245688 | HITS-CLIP | 23597149\|25083871 | Unspecified |
| mmu-miR-152-3p | MIMAT0000162 | Rbl2 | 19651 | HITS-CLIP | 25083871 | Unspecified |
| mmu-miR-152-3p | MIMAT0000162 | Rbm15 | 229700 | HITS-CLIP | 25083871\|29167373 | Unspecified |
| mmu-miR-152-3p | MIMAT0000162 | Rbm18 | 67889 | HITS-CLIP | 25083871\|29167373 | Unspecified |
| mmu-miR-152-3p | MIMAT0000162 | Rbm26 | 74213 | HITS-CLIP | 37986224\|25083871 | Unspecified |
| mmu-miR-152-3p | MIMAT0000162 | Rbm33 | 381626 | HITS-CLIP | 37986224\|25083871 | Unspecified |
| mmu-miR-152-3p | MIMAT0000162 | Rbm39 | 170791 | HITS-CLIP | 25083871 | Unspecified |
| mmu-miR-152-3p | MIMAT0000162 | Rbm47 | 245945 | HITS-CLIP | 37986224\|23597149 | Unspecified |
| mmu-miR-152-3p | MIMAT0000162 | Rbm7 | 67010 | HITS-CLIP | 37986224 | Unspecified |
| mmu-miR-152-3p | MIMAT0000162 | Rbms2 | 56516 | HITS-CLIP | 37986224 | Unspecified |
| mmu-miR-152-3p | MIMAT0000162 | Rbpj | 19664 | HITS-CLIP | 37986224\|29167373 | Unspecified |
| mmu-miR-152-3p | MIMAT0000162 | Rbpms | 19663 | HITS-CLIP | 23597149 | Unspecified |
| mmu-miR-152-3p | MIMAT0000162 | Rbpms2 | 71973 | HITS-CLIP | 25083871 | Unspecified |
| mmu-miR-152-3p | MIMAT0000162 | Rbsn | 64145 | HITS-CLIP | 23597149\|25083871 | Unspecified |
| mmu-miR-152-3p | MIMAT0000162 | Rc3h1 | 381305 | HITS-CLIP | 37986224 | Unspecified |
| mmu-miR-152-3p | MIMAT0000162 | Rc3h2 | 319817 | HITS-CLIP | 37986224\|25083871\|29167373 | Unspecified |
| mmu-miR-152-3p | MIMAT0000162 | Rcc2 | 108911 | HITS-CLIP | 25083871\|29167373 | Unspecified |
| mmu-miR-152-3p | MIMAT0000162 | Rcor1 | 217864 | HITS-CLIP | 25083871 | Unspecified |
| mmu-miR-152-3p | MIMAT0000162 | Rdh16 | 19683 | HITS-CLIP | 37986224 | Unspecified |
| mmu-miR-152-3p | MIMAT0000162 | Rdx | 19684 | HITS-CLIP | 25083871\|29167373 | Unspecified |
| mmu-miR-152-3p | MIMAT0000162 | Reep3 | 28193 | HITS-CLIP | 37986224 | Unspecified |
| mmu-miR-152-3p | MIMAT0000162 | Rere | 68703 | HITS-CLIP | 37986224 | Unspecified |

| | | | | | | |
|---|---|---|---|---|---|---|
| mmu-miR-152-3p | MIMAT0000162 | Resf1 | 55196 | HITS-CLIP | 25083871\|29167373 | Unspecified |
| mmu-miR-152-3p | MIMAT0000162 | Retreg1 | 66270 | HITS-CLIP | 25083871 | Unspecified |
| mmu-miR-152-3p | MIMAT0000162 | Retreg3 | 162427 | HITS-CLIP | 25083871 | Unspecified |
| mmu-miR-152-3p | MIMAT0000162 | Rfx5 | 53970 | HITS-CLIP | 29167373 | Unspecified |
| mmu-miR-152-3p | MIMAT0000162 | Rfx7 | 319758 | HITS-CLIP | 25083871\|23597149 | Unspecified |
| mmu-miR-152-3p | MIMAT0000162 | Rgl1 | 19731 | HITS-CLIP | 37986224 | Unspecified |
| mmu-miR-152-3p | MIMAT0000162 | Rgma | 244058 | HITS-CLIP | 37986224\|25083871 | Unspecified |
| mmu-miR-152-3p | MIMAT0000162 | Rgmb | 68799 | HITS-CLIP | 25083871 | Unspecified |
| mmu-miR-152-3p | MIMAT0000162 | Rgs1 | 50778 | HITS-CLIP | 29167373 | Unspecified |
| mmu-miR-152-3p | MIMAT0000162 | Rhebl1 | 69159 | HITS-CLIP | 37986224 | Unspecified |
| mmu-miR-152-3p | MIMAT0000162 | Rhot1 | 59040 | HITS-CLIP | 25083871 | Unspecified |
| mmu-miR-152-3p | MIMAT0000162 | Ric8b | 237422 | HITS-CLIP | 37986224 | Unspecified |
| mmu-miR-152-3p | MIMAT0000162 | Rictor | 78757 | HITS-CLIP | 37986224\|25083871\|29167373 | Unspecified |
| mmu-miR-152-3p | MIMAT0000162 | Rif1 | 51869 | HITS-CLIP | 25083871 | Unspecified |
| mmu-miR-152-3p | MIMAT0000162 | Rin2 | 74030 | HITS-CLIP | 23597149 | Unspecified |
| mmu-miR-152-3p | MIMAT0000162 | Ripk2 | 192656 | HITS-CLIP | 25083871 | Unspecified |
| mmu-miR-152-3p | MIMAT0000162 | Ripor1 | 75687 | HITS-CLIP | 25083871 | Unspecified |
| mmu-miR-152-3p | MIMAT0000162 | Rlim | 19820 | HITS-CLIP | 37986224 | Unspecified |
| mmu-miR-152-3p | MIMAT0000162 | Rnf103 | 22644 | HITS-CLIP | 37986224\|25083871\|23597149 | Unspecified |
| mmu-miR-152-3p | MIMAT0000162 | Rnf11 | 29864 | HITS-CLIP | 23597149\|29167373 | Unspecified |
| mmu-miR-152-3p | MIMAT0000162 | Rnf111 | 93836 | HITS-CLIP | 25083871 | Unspecified |
| mmu-miR-152-3p | MIMAT0000162 | Rnf128 | 66889 | HITS-CLIP | 29167373 | Unspecified |
| mmu-miR-152-3p | MIMAT0000162 | Rnf13 | 24017 | HITS-CLIP | 25083871 | Unspecified |
| mmu-miR-152-3p | MIMAT0000162 | Rnf139 | 75841 | HITS-CLIP | 37986224\|25083871 | Unspecified |
| mmu-miR-152-3p | MIMAT0000162 | Rnf150 | 330812 | HITS-CLIP | 29167373 | Unspecified |
| mmu-miR-152-3p | MIMAT0000162 | Rnf157 | 217340 | HITS-CLIP | 29167373 | Unspecified |
| mmu-miR-152-3p | MIMAT0000162 | Rnf165 | 225743 | HITS-CLIP | 37986224\|25083871 | Unspecified |
| mmu-miR-152-3p | MIMAT0000162 | Rnf19a | 30945 | HITS-CLIP | 37986224\|25083871\|29167373 | Unspecified |
| mmu-miR-152-3p | MIMAT0000162 | Rnf2 | 19821 | HITS-CLIP | 25083871 | Unspecified |
| mmu-miR-152-3p | MIMAT0000162 | Rnf213 | 672511 | HITS-CLIP | 37986224\|23597149\|29167373 | Unspecified |
| mmu-miR-152-3p | MIMAT0000162 | Rnf214 | 235315 | HITS-CLIP | 37986224 | Unspecified |
| mmu-miR-152-3p | MIMAT0000162 | Rnf216 | 108086 | HITS-CLIP | 25083871 | Unspecified |
| mmu-miR-152-3p | MIMAT0000162 | Rnf217 | 268291 | HITS-CLIP | 25083871 | Unspecified |
| mmu-miR-152-3p | MIMAT0000162 | Rnf219 | 72486 | HITS-CLIP | 37986224 | Unspecified |
| mmu-miR-152-3p | MIMAT0000162 | Rnf38 | 73469 | HITS-CLIP | 25083871\|23597149 | Unspecified |
| mmu-miR-152-3p | MIMAT0000162 | Rnf44 | 105239 | HITS-CLIP | 37986224 | Unspecified |
| mmu-miR-152-3p | MIMAT0000162 | Rnf7 | 19823 | HITS-CLIP | 25083871 | Unspecified |
| mmu-miR-152-3p | MIMAT0000162 | Rngtt | 24018 | HITS-CLIP | 23597149 | Unspecified |
| mmu-miR-152-3p | MIMAT0000162 | Rnpepl1 | 108657 | HITS-CLIP | 37986224 | Unspecified |

| | | | | | | |
|---|---|---|---|---|---|---|
| mmu-miR-152-3p | MIMAT0000162 | Rock1 | 19877 | HITS-CLIP | 37986224\|29167373 | Unspecified |
| mmu-miR-152-3p | MIMAT0000162 | Rock2 | 19878 | HITS-CLIP | 25083871 | Unspecified |
| mmu-miR-152-3p | MIMAT0000162 | Rpf2 | 67239 | HITS-CLIP | 37986224\|23597149 | Unspecified |
| mmu-miR-152-3p | MIMAT0000162 | Rpl22 | 19934 | HITS-CLIP | 25083871 | Unspecified |
| mmu-miR-152-3p | MIMAT0000162 | Rpl4 | 67891 | HITS-CLIP | 25083871\|23597149\|29167373 | Unspecified |
| mmu-miR-152-3p | MIMAT0000162 | Rpp38 | 227522 | HITS-CLIP | 37986224 | Unspecified |
| mmu-miR-152-3p | MIMAT0000162 | Rprd2 | 75137 | HITS-CLIP | 23597149 | Unspecified |
| mmu-miR-152-3p | MIMAT0000162 | Rps5 | 20103 | HITS-CLIP | 25083871\|29167373 | Unspecified |
| mmu-miR-152-3p | MIMAT0000162 | Rptor | 74370 | HITS-CLIP | 23597149\|25083871 | Unspecified |
| mmu-miR-152-3p | MIMAT0000162 | Rragc | 54170 | HITS-CLIP | 25083871 | Unspecified |
| mmu-miR-152-3p | MIMAT0000162 | Rrm1 | 20133 | HITS-CLIP | 37986224 | Unspecified |
| mmu-miR-152-3p | MIMAT0000162 | Rrm2 | 20135 | HITS-CLIP | 21258322 | Unspecified |
| mmu-miR-152-3p | MIMAT0000162 | Rrn3 | 106298 | HITS-CLIP | 23597149\|25083871 | Unspecified |
| mmu-miR-152-3p | MIMAT0000162 | Rsbn1l | 242860 | HITS-CLIP | 37986224 | Unspecified |
| mmu-miR-152-3p | MIMAT0000162 | Rspry1 | 67610 | HITS-CLIP | 37986224\|23597149 | Unspecified |
| mmu-miR-152-3p | MIMAT0000162 | Rsrc2 | 208606 | HITS-CLIP | 25083871 | Unspecified |
| mmu-miR-152-3p | MIMAT0000162 | Rsu1 | 20163 | HITS-CLIP | 25083871 | Unspecified |
| mmu-miR-152-3p | MIMAT0000162 | Rtca | 66368 | HITS-CLIP | 37986224\|25083871 | Unspecified |
| mmu-miR-152-3p | MIMAT0000162 | Rtkn | 20166 | HITS-CLIP | 25083871 | Unspecified |
| mmu-miR-152-3p | MIMAT0000162 | Rtn3 | 20168 | HITS-CLIP | 37986224\|23597149 | Unspecified |
| mmu-miR-152-3p | MIMAT0000162 | Rtn4 | 68585 | HITS-CLIP | 37986224\|25083871 | Unspecified |
| mmu-miR-152-3p | MIMAT0000162 | Rttn | 246102 | HITS-CLIP | 23597149 | Unspecified |
| mmu-miR-152-3p | MIMAT0000162 | Rubcn | 9711 | HITS-CLIP | 25083871 | Unspecified |
| mmu-miR-152-3p | MIMAT0000162 | Runx1 | 12394 | HITS-CLIP | 37986224\|29167373 | Unspecified |
| mmu-miR-152-3p | MIMAT0000162 | Runx2 | 12393 | HITS-CLIP | 25083871 | Unspecified |
| mmu-miR-152-3p | MIMAT0000162 | Rxra | 20181 | HITS-CLIP | 23597149 | Unspecified |
| mmu-miR-152-3p | MIMAT0000162 | Ryk | 20187 | HITS-CLIP | 25083871 | Unspecified |
| mmu-miR-152-3p | MIMAT0000162 | Ryr1 | 20190 | HITS-CLIP | 25083871 | Unspecified |
| mmu-miR-152-3p | MIMAT0000162 | S100a10 | 20194 | HITS-CLIP | 25083871 | Unspecified |
| mmu-miR-152-3p | MIMAT0000162 | S1pr2 | 14739 | HITS-CLIP | 37986224 | Unspecified |
| mmu-miR-152-3p | MIMAT0000162 | Sacs | 50720 | HITS-CLIP | 37986224\|25083871 | Unspecified |
| mmu-miR-152-3p | MIMAT0000162 | Samd4b | 233033 | HITS-CLIP | 23597149\|25083871 | Unspecified |
| mmu-miR-152-3p | MIMAT0000162 | Samd9l | 209086 | HITS-CLIP | 29167373 | Unspecified |
| mmu-miR-152-3p | MIMAT0000162 | Sap30 | 60406 | HITS-CLIP | 25083871 | Unspecified |
| mmu-miR-152-3p | MIMAT0000162 | Sar1a | 20224 | HITS-CLIP | 25083871 | Unspecified |
| mmu-miR-152-3p | MIMAT0000162 | Sar1b | 66397 | HITS-CLIP | 25083871 | Unspecified |
| mmu-miR-152-3p | MIMAT0000162 | Sars | 20226 | HITS-CLIP | 37986224\|23597149 | Unspecified |
| mmu-miR-152-3p | MIMAT0000162 | Sash1 | 70097 | HITS-CLIP | 29167373 | Unspecified |
| mmu-miR-152-3p | MIMAT0000162 | Sat1 | 20229 | HITS-CLIP | 29167373 | Unspecified |

| | | | | | | |
|---|---|---|---|---|---|---|
| mmu-miR-152-3p | MIMAT0000162 | Sbno1 | 243272 | HITS-CLIP | 37986224\|23597149\|25083871\|29167 | Unspecified |
| mmu-miR-152-3p | MIMAT0000162 | Sbno2 | 216161 | HITS-CLIP | 25083871 | Unspecified |
| mmu-miR-152-3p | MIMAT0000162 | Scaf11 | 72193 | HITS-CLIP | 25083871\|29167373 | Unspecified |
| mmu-miR-152-3p | MIMAT0000162 | Scai | 320271 | HITS-CLIP | 37986224 | Unspecified |
| mmu-miR-152-3p | MIMAT0000162 | Scamp2 | 24044 | HITS-CLIP | 23597149 | Unspecified |
| mmu-miR-152-3p | MIMAT0000162 | Scamp5 | 56807 | HITS-CLIP | 37986224 | Unspecified |
| mmu-miR-152-3p | MIMAT0000162 | Scd1 | 20249 | HITS-CLIP | 37986224\|25083871\|23597149 | Unspecified |
| mmu-miR-152-3p | MIMAT0000162 | Scd2 | 20250 | HITS-CLIP | 21258322 | Unspecified |
| mmu-miR-152-3p | MIMAT0000162 | Scmh1 | 29871 | HITS-CLIP | 31433990 | Unspecified |
| mmu-miR-152-3p | MIMAT0000162 | Scml4 | 268297 | HITS-CLIP | 21258322\|25083871 | Unspecified |
| mmu-miR-152-3p | MIMAT0000162 | Scn5a | 20271 | HITS-CLIP | 25083871 | Unspecified |
| mmu-miR-152-3p | MIMAT0000162 | Sdc4 | 20971 | HITS-CLIP | 23597149\|25083871 | Unspecified |
| mmu-miR-152-3p | MIMAT0000162 | Sec16a | 227648 | HITS-CLIP | 23597149 | Unspecified |
| mmu-miR-152-3p | MIMAT0000162 | Sec16b | 89867 | HITS-CLIP | 23597149 | Unspecified |
| mmu-miR-152-3p | MIMAT0000162 | Sec22b | 20333 | HITS-CLIP | 23597149\|25083871 | Unspecified |
| mmu-miR-152-3p | MIMAT0000162 | Sec23b | 27054 | HITS-CLIP | 25083871 | Unspecified |
| mmu-miR-152-3p | MIMAT0000162 | Sec24a | 77371 | HITS-CLIP | 25083871\|29167373 | Unspecified |
| mmu-miR-152-3p | MIMAT0000162 | Sec24c | 218811 | HITS-CLIP | 25083871\|23597149 | Unspecified |
| mmu-miR-152-3p | MIMAT0000162 | Sec31a | 69162 | HITS-CLIP | 37986224\|25083871\|23597149 | Unspecified |
| mmu-miR-152-3p | MIMAT0000162 | Sec63 | 140740 | HITS-CLIP | 25083871 | Unspecified |
| mmu-miR-152-3p | MIMAT0000162 | Secisbp2l | 70354 | HITS-CLIP | 37986224 | Unspecified |
| mmu-miR-152-3p | MIMAT0000162 | Sel1l | 20338 | HITS-CLIP | 25083871 | Unspecified |
| mmu-miR-152-3p | MIMAT0000162 | Sema3e | 20349 | HITS-CLIP | 37986224 | Unspecified |
| mmu-miR-152-3p | MIMAT0000162 | Sema4b | 20352 | HITS-CLIP | 25083871 | Unspecified |
| mmu-miR-152-3p | MIMAT0000162 | Sema4g | 26456 | HITS-CLIP | 23597149 | Unspecified |
| mmu-miR-152-3p | MIMAT0000162 | Sema5a | 20356 | HITS-CLIP | 37986224 | Unspecified |
| mmu-miR-152-3p | MIMAT0000162 | Sema6a | 20358 | HITS-CLIP | 25083871 | Unspecified |
| mmu-miR-152-3p | MIMAT0000162 | Sema7a | 20361 | HITS-CLIP | 25083871 | Unspecified |
| mmu-miR-152-3p | MIMAT0000162 | Senp2 | 75826 | HITS-CLIP | 37986224 | Unspecified |
| mmu-miR-152-3p | MIMAT0000162 | Senp3 | 80886 | HITS-CLIP | 23597149\|25083871 | Unspecified |
| mmu-miR-152-3p | MIMAT0000162 | Senp6 | 215351 | HITS-CLIP | 25083871 | Unspecified |
| mmu-miR-152-3p | MIMAT0000162 | Senp7 | 66315 | HITS-CLIP | 37986224\|25083871 | Unspecified |
| mmu-miR-152-3p | MIMAT0000162 | Sept6 | 23157 | HITS-CLIP | 23597149 | Unspecified |
| mmu-miR-152-3p | MIMAT0000162 | Serinc5 | 218442 | HITS-CLIP | 37986224 | Unspecified |
| mmu-miR-152-3p | MIMAT0000162 | Serpinc1 | 11905 | HITS-CLIP | 23597149 | Unspecified |
| mmu-miR-152-3p | MIMAT0000162 | Serpind1 | 15160 | HITS-CLIP | 23597149 | Unspecified |
| mmu-miR-152-3p | MIMAT0000162 | Serpine1 | 18787 | HITS-CLIP | 37986224 | Unspecified |
| mmu-miR-152-3p | MIMAT0000162 | Sesn1 | 140742 | HITS-CLIP | 25083871\|23597149 | Unspecified |
| mmu-miR-152-3p | MIMAT0000162 | Sesn2 | 230784 | HITS-CLIP | 26203562 | Unspecified |

| | | | | | | |
|---|---|---|---|---|---|---|
| mmu-miR-152-3p | MIMAT0000162 | Sesn3 | 75747 | HITS-CLIP | 25083871 | Unspecified |
| mmu-miR-152-3p | MIMAT0000162 | Sestd1 | 228071 | HITS-CLIP | 25083871 | Unspecified |
| mmu-miR-152-3p | MIMAT0000162 | Setd1a | 233904 | HITS-CLIP | 37986224\|23597149 | Unspecified |
| mmu-miR-152-3p | MIMAT0000162 | Setd2 | 235626 | HITS-CLIP | 25083871\|23597149 | Unspecified |
| mmu-miR-152-3p | MIMAT0000162 | Setd5 | 72895 | HITS-CLIP | 25083871\|21258322 | Unspecified |
| mmu-miR-152-3p | MIMAT0000162 | Setd7 | 73251 | HITS-CLIP | 25083871 | Unspecified |
| mmu-miR-152-3p | MIMAT0000162 | Sf3b1 | 81898 | HITS-CLIP | 25083871 | Unspecified |
| mmu-miR-152-3p | MIMAT0000162 | Sfpq | 71514 | HITS-CLIP | 37986224\|25083871 | Unspecified |
| mmu-miR-152-3p | MIMAT0000162 | Sft2d2 | 108735 | HITS-CLIP | 37986224 | Unspecified |
| mmu-miR-152-3p | MIMAT0000162 | Sfxn3 | 94280 | HITS-CLIP | 25083871 | Unspecified |
| mmu-miR-152-3p | MIMAT0000162 | Sgcb | 24051 | HITS-CLIP | 37986224\|29167373 | Unspecified |
| mmu-miR-152-3p | MIMAT0000162 | Sgcg | 24053 | HITS-CLIP | 25083871 | Unspecified |
| mmu-miR-152-3p | MIMAT0000162 | Sgms1 | 208449 | HITS-CLIP | 37986224\|25083871 | Unspecified |
| mmu-miR-152-3p | MIMAT0000162 | Sgms2 | 74442 | HITS-CLIP | 37986224\|25083871 | Unspecified |
| mmu-miR-152-3p | MIMAT0000162 | Sgpl1 | 20397 | HITS-CLIP | 29167373 | Unspecified |
| mmu-miR-152-3p | MIMAT0000162 | Sgpp1 | 81535 | HITS-CLIP | 23597149\|25083871 | Unspecified |
| mmu-miR-152-3p | MIMAT0000162 | Sh2b3 | 16923 | HITS-CLIP | 37986224 | Unspecified |
| mmu-miR-152-3p | MIMAT0000162 | Sh3glb1 | 54673 | HITS-CLIP | 23597149 | Unspecified |
| mmu-miR-152-3p | MIMAT0000162 | Sh3pxd2a | 14218 | HITS-CLIP | 25083871 | Unspecified |
| mmu-miR-152-3p | MIMAT0000162 | Sh3rf1 | 59009 | HITS-CLIP | 25083871 | Unspecified |
| mmu-miR-152-3p | MIMAT0000162 | Sharpin | 106025 | HITS-CLIP | 23597149 | Unspecified |
| mmu-miR-152-3p | MIMAT0000162 | Shc1 | 20416 | HITS-CLIP | 25083871 | Unspecified |
| mmu-miR-152-3p | MIMAT0000162 | Shmt1 | 20425 | HITS-CLIP | 23597149 | Unspecified |
| mmu-miR-152-3p | MIMAT0000162 | Shpk | 74637 | HITS-CLIP | 23597149 | Unspecified |
| mmu-miR-152-3p | MIMAT0000162 | Shprh | 268281 | HITS-CLIP | 37986224 | Unspecified |
| mmu-miR-152-3p | MIMAT0000162 | Sidt2 | 214597 | HITS-CLIP | 23597149 | Unspecified |
| mmu-miR-152-3p | MIMAT0000162 | Sigmar1 | 18391 | HITS-CLIP | 37986224\|25083871 | Unspecified |
| mmu-miR-152-3p | MIMAT0000162 | Sik1 | 17691 | HITS-CLIP | 37986224 | Unspecified |
| mmu-miR-152-3p | MIMAT0000162 | Sik2 | 235344 | HITS-CLIP | 31433990 | Unspecified |
| mmu-miR-152-3p | MIMAT0000162 | Sil1 | 81500 | HITS-CLIP | 25083871 | Unspecified |
| mmu-miR-152-3p | MIMAT0000162 | Sipa1 | 20469 | HITS-CLIP | 37986224\|25083871\|29167373 | Unspecified |
| mmu-miR-152-3p | MIMAT0000162 | Sirt4 | 75387 | HITS-CLIP | 23597149 | Unspecified |
| mmu-miR-152-3p | MIMAT0000162 | Sirt7 | 209011 | HITS-CLIP | 37986224\|25083871 | Unspecified |
| mmu-miR-152-3p | MIMAT0000162 | Six4 | 20474 | HITS-CLIP | 37986224\|25083871 | Unspecified |
| mmu-miR-152-3p | MIMAT0000162 | Skap2 | 54353 | HITS-CLIP | 25083871 | Unspecified |
| mmu-miR-152-3p | MIMAT0000162 | Ski | 20481 | HITS-CLIP | 25083871 | Unspecified |
| mmu-miR-152-3p | MIMAT0000162 | Skiv2l | 108077 | HITS-CLIP | 23597149 | Unspecified |
| mmu-miR-152-3p | MIMAT0000162 | Skp1a | 21402 | HITS-CLIP | 25083871 | Unspecified |
| mmu-miR-152-3p | MIMAT0000162 | Slain2 | 75991 | HITS-CLIP | 37986224 | Unspecified |

| | | | | | | |
|---|---|---|---|---|---|---|
| mmu-miR-152-3p | MIMAT0000162 | Slbp | 20492 | HITS-CLIP | 29167373 | Unspecified |
| mmu-miR-152-3p | MIMAT0000162 | Slc10a3 | 214601 | HITS-CLIP | 37986224\|25083871\|29167373 | Unspecified |
| mmu-miR-152-3p | MIMAT0000162 | Slc10a5 | 241877 | HITS-CLIP | 23597149 | Unspecified |
| mmu-miR-152-3p | MIMAT0000162 | Slc12a2 | 20496 | HITS-CLIP | 25083871 | Unspecified |
| mmu-miR-152-3p | MIMAT0000162 | Slc12a9 | 83704 | HITS-CLIP | 23597149 | Unspecified |
| mmu-miR-152-3p | MIMAT0000162 | Slc19a2 | 116914 | HITS-CLIP | 23597149 | Unspecified |
| mmu-miR-152-3p | MIMAT0000162 | Slc20a1 | 20515 | HITS-CLIP | 25083871 | Unspecified |
| mmu-miR-152-3p | MIMAT0000162 | Slc23a2 | 54338 | HITS-CLIP | 37986224\|25083871 | Unspecified |
| mmu-miR-152-3p | MIMAT0000162 | Slc25a1 | 13358 | HITS-CLIP | 23597149\|25083871 | Unspecified |
| mmu-miR-152-3p | MIMAT0000162 | Slc25a12 | 78830 | HITS-CLIP | 23597149 | Unspecified |
| mmu-miR-152-3p | MIMAT0000162 | Slc25a13 | 50799 | HITS-CLIP | 37986224 | Unspecified |
| mmu-miR-152-3p | MIMAT0000162 | Slc25a15 | 18408 | HITS-CLIP | 37986224 | Unspecified |
| mmu-miR-152-3p | MIMAT0000162 | Slc25a3 | 18674 | HITS-CLIP | 37986224\|23597149\|21258322 | Unspecified |
| mmu-miR-152-3p | MIMAT0000162 | Slc25a32 | 69906 | HITS-CLIP | 23597149\|25083871 | Unspecified |
| mmu-miR-152-3p | MIMAT0000162 | Slc25a36 | 192287 | HITS-CLIP | 25083871 | Unspecified |
| mmu-miR-152-3p | MIMAT0000162 | Slc25a4 | 11739 | HITS-CLIP | 23597149 | Unspecified |
| mmu-miR-152-3p | MIMAT0000162 | Slc26a1 | 231583 | HITS-CLIP | 37986224 | Unspecified |
| mmu-miR-152-3p | MIMAT0000162 | Slc27a2 | 26458 | HITS-CLIP | 23597149 | Unspecified |
| mmu-miR-152-3p | MIMAT0000162 | Slc2a2 | 20526 | HITS-CLIP | 23597149 | Unspecified |
| mmu-miR-152-3p | MIMAT0000162 | Slc30a1 | 22782 | HITS-CLIP | 25083871 | Unspecified |
| mmu-miR-152-3p | MIMAT0000162 | Slc30a4 | 22785 | HITS-CLIP | 37986224\|25083871\|29167373 | Unspecified |
| mmu-miR-152-3p | MIMAT0000162 | Slc30a6 | 210148 | HITS-CLIP | 25083871 | Unspecified |
| mmu-miR-152-3p | MIMAT0000162 | Slc30a7 | 66500 | HITS-CLIP | 25083871 | Unspecified |
| mmu-miR-152-3p | MIMAT0000162 | Slc35a2 | 22232 | HITS-CLIP | 25083871\|23597149 | Unspecified |
| mmu-miR-152-3p | MIMAT0000162 | Slc35a4 | 67843 | HITS-CLIP | 37986224\|25083871\|23597149 | Unspecified |
| mmu-miR-152-3p | MIMAT0000162 | Slc35b4 | 58246 | HITS-CLIP | 25083871 | Unspecified |
| mmu-miR-152-3p | MIMAT0000162 | Slc35f5 | 74150 | HITS-CLIP | 25083871 | Unspecified |
| mmu-miR-152-3p | MIMAT0000162 | Slc37a2 | 56857 | HITS-CLIP | 29167373 | Unspecified |
| mmu-miR-152-3p | MIMAT0000162 | Slc38a3 | 76257 | HITS-CLIP | 23597149 | Unspecified |
| mmu-miR-152-3p | MIMAT0000162 | Slc38a9 | 268706 | HITS-CLIP | 29167373 | Unspecified |
| mmu-miR-152-3p | MIMAT0000162 | Slc39a10 | 227059 | HITS-CLIP | 29167373 | Unspecified |
| mmu-miR-152-3p | MIMAT0000162 | Slc39a13 | 68427 | HITS-CLIP | 37986224 | Unspecified |
| mmu-miR-152-3p | MIMAT0000162 | Slc39a14 | 213053 | HITS-CLIP | 37986224 | Unspecified |
| mmu-miR-152-3p | MIMAT0000162 | Slc3a2 | 17254 | HITS-CLIP | 37986224 | Unspecified |
| mmu-miR-152-3p | MIMAT0000162 | Slc41a2 | 338365 | HITS-CLIP | 37986224 | Unspecified |
| mmu-miR-152-3p | MIMAT0000162 | Slc44a1 | 100434 | HITS-CLIP | 25083871 | Unspecified |
| mmu-miR-152-3p | MIMAT0000162 | Slc4a7 | 218756 | HITS-CLIP | 25083871 | Unspecified |
| mmu-miR-152-3p | MIMAT0000162 | Slc6a9 | 14664 | HITS-CLIP | 25083871 | Unspecified |
| mmu-miR-152-3p | MIMAT0000162 | Slc9a9 | 331004 | HITS-CLIP | 37986224\|25083871 | Unspecified |

| | | | | | | |
|---|---|---|---|---|---|---|
| mmu-miR-152-3p | MIMAT0000162 | Slf1 | 852127 | HITS-CLIP | 25083871 | Unspecified |
| mmu-miR-152-3p | MIMAT0000162 | Slf2 | 226151 | HITS-CLIP | 25083871\|23597149 | Unspecified |
| mmu-miR-152-3p | MIMAT0000162 | Slirp | 380773 | HITS-CLIP | 25083871\|23597149\|29167373 | Unspecified |
| mmu-miR-152-3p | MIMAT0000162 | Slmap | 83997 | HITS-CLIP | 37986224\|25083871 | Unspecified |
| mmu-miR-152-3p | MIMAT0000162 | Sltm | 66660 | HITS-CLIP | 29167373 | Unspecified |
| mmu-miR-152-3p | MIMAT0000162 | Smad1 | 17125 | HITS-CLIP | 25083871 | Unspecified |
| mmu-miR-152-3p | MIMAT0000162 | Smad2 | 17126 | HITS-CLIP | 25083871 | Unspecified |
| mmu-miR-152-3p | MIMAT0000162 | Smad7 | 17131 | HITS-CLIP | 25083871\|21258322 | Unspecified |
| mmu-miR-152-3p | MIMAT0000162 | Smarca4 | 20586 | HITS-CLIP | 29167373 | Unspecified |
| mmu-miR-152-3p | MIMAT0000162 | Smarcc1 | 20588 | HITS-CLIP | 23597149\|25083871 | Unspecified |
| mmu-miR-152-3p | MIMAT0000162 | Smarcc2 | 68094 | HITS-CLIP | 25083871 | Unspecified |
| mmu-miR-152-3p | MIMAT0000162 | Smarcd1 | 83797 | HITS-CLIP | 37986224\|25083871 | Unspecified |
| mmu-miR-152-3p | MIMAT0000162 | Smarcd2 | 83796 | HITS-CLIP | 29167373 | Unspecified |
| mmu-miR-152-3p | MIMAT0000162 | Smc4 | 70099 | HITS-CLIP | 25083871 | Unspecified |
| mmu-miR-152-3p | MIMAT0000162 | Smcr8 | 237782 | HITS-CLIP | 25083871 | Unspecified |
| mmu-miR-152-3p | MIMAT0000162 | Smg1 | 233789 | HITS-CLIP | 37986224\|25083871\|29167373 | Unspecified |
| mmu-miR-152-3p | MIMAT0000162 | Smim12 | 80284 | HITS-CLIP | 37986224 | Unspecified |
| mmu-miR-152-3p | MIMAT0000162 | Smim15 | 75616 | HITS-CLIP | 37986224\|25083871 | Unspecified |
| mmu-miR-152-3p | MIMAT0000162 | Smim4 | 66487 | HITS-CLIP | 37986224 | Unspecified |
| mmu-miR-152-3p | MIMAT0000162 | Smim7 | 66818 | HITS-CLIP | 25083871 | Unspecified |
| mmu-miR-152-3p | MIMAT0000162 | Smurf2 | 66313 | HITS-CLIP | 25083871 | Unspecified |
| mmu-miR-152-3p | MIMAT0000162 | Smyd1 | 12180 | HITS-CLIP | 37986224 | Unspecified |
| mmu-miR-152-3p | MIMAT0000162 | Snap47 | 67826 | HITS-CLIP | 37986224\|23597149 | Unspecified |
| mmu-miR-152-3p | MIMAT0000162 | Snd1 | 56463 | HITS-CLIP | 23597149 | Unspecified |
| mmu-miR-152-3p | MIMAT0000162 | Snip1 | 76793 | HITS-CLIP | 37986224 | Unspecified |
| mmu-miR-152-3p | MIMAT0000162 | Snn | 20621 | HITS-CLIP | 37986224 | Unspecified |
| mmu-miR-152-3p | MIMAT0000162 | Snrpa1 | 68981 | HITS-CLIP | 25083871 | Unspecified |
| mmu-miR-152-3p | MIMAT0000162 | Sntb2 | 20650 | HITS-CLIP | 37986224 | Unspecified |
| mmu-miR-152-3p | MIMAT0000162 | Snx10 | 71982 | HITS-CLIP | 29167373 | Unspecified |
| mmu-miR-152-3p | MIMAT0000162 | Snx13 | 217463 | HITS-CLIP | 23597149 | Unspecified |
| mmu-miR-152-3p | MIMAT0000162 | Snx19 | 102607 | HITS-CLIP | 37986224 | Unspecified |
| mmu-miR-152-3p | MIMAT0000162 | Snx2 | 67804 | HITS-CLIP | 25083871\|29167373 | Unspecified |
| mmu-miR-152-3p | MIMAT0000162 | Snx27 | 76742 | HITS-CLIP | 25083871\|23597149 | Unspecified |
| mmu-miR-152-3p | MIMAT0000162 | Snx30 | 209131 | HITS-CLIP | 29167373 | Unspecified |
| mmu-miR-152-3p | MIMAT0000162 | Snx33 | 235406 | HITS-CLIP | 37986224 | Unspecified |
| mmu-miR-152-3p | MIMAT0000162 | Snx5 | 69178 | HITS-CLIP | 23597149 | Unspecified |
| mmu-miR-152-3p | MIMAT0000162 | Soat1 | 20652 | HITS-CLIP | 37986224\|25083871 | Unspecified |
| mmu-miR-152-3p | MIMAT0000162 | Socs2 | 216233 | HITS-CLIP | 25083871 | Unspecified |
| mmu-miR-152-3p | MIMAT0000162 | Socs5 | 56468 | HITS-CLIP | 25083871 | Unspecified |

| | | | | | | |
|---|---|---|---|---|---|---|
| mmu-miR-152-3p | MIMAT0000162 | Socs6 | 54607 | HITS-CLIP | 37986224\|29167373 | Unspecified |
| mmu-miR-152-3p | MIMAT0000162 | Socs7 | 192157 | HITS-CLIP | 25083871 | Unspecified |
| mmu-miR-152-3p | MIMAT0000162 | Sod2 | 20656 | HITS-CLIP | 23597149 | Unspecified |
| mmu-miR-152-3p | MIMAT0000162 | Soga1 | 320706 | HITS-CLIP | 25083871 | Unspecified |
| mmu-miR-152-3p | MIMAT0000162 | Son | 20658 | HITS-CLIP | 37986224\|25083871\|29167373 | Unspecified |
| mmu-miR-152-3p | MIMAT0000162 | Sox11 | 20666 | HITS-CLIP | 37986224 | Unspecified |
| mmu-miR-152-3p | MIMAT0000162 | Sox9 | 20682 | HITS-CLIP | 37986224 | Unspecified |
| mmu-miR-152-3p | MIMAT0000162 | Sp1 | 20683 | HITS-CLIP | 25083871 | Unspecified |
| mmu-miR-152-3p | MIMAT0000162 | Sp2 | 78912 | HITS-CLIP | 23597149 | Unspecified |
| mmu-miR-152-3p | MIMAT0000162 | Sp3 | 20687 | HITS-CLIP | 25083871 | Unspecified |
| mmu-miR-152-3p | MIMAT0000162 | Spag9 | 70834 | HITS-CLIP | 25083871 | Unspecified |
| mmu-miR-152-3p | MIMAT0000162 | Spata5 | 57815 | HITS-CLIP | 25083871 | Unspecified |
| mmu-miR-152-3p | MIMAT0000162 | Spata6 | 67946 | HITS-CLIP | 23597149 | Unspecified |
| mmu-miR-152-3p | MIMAT0000162 | Spen | 56381 | HITS-CLIP | 37986224\|25083871\|29167373 | Unspecified |
| mmu-miR-152-3p | MIMAT0000162 | Spg11 | 214585 | HITS-CLIP | 37986224 | Unspecified |
| mmu-miR-152-3p | MIMAT0000162 | Spg20 | 229285 | HITS-CLIP | 25083871\|23597149 | Unspecified |
| mmu-miR-152-3p | MIMAT0000162 | Spock2 | 94214 | HITS-CLIP | 25083871 | Unspecified |
| mmu-miR-152-3p | MIMAT0000162 | Spred1 | 114715 | HITS-CLIP | 25083871\|29167373 | Unspecified |
| mmu-miR-152-3p | MIMAT0000162 | Spry4 | 24066 | HITS-CLIP | 25083871 | Unspecified |
| mmu-miR-152-3p | MIMAT0000162 | Spryd7 | 66674 | HITS-CLIP | 23597149 | Unspecified |
| mmu-miR-152-3p | MIMAT0000162 | Spsb1 | 74646 | HITS-CLIP | 25083871 | Unspecified |
| mmu-miR-152-3p | MIMAT0000162 | Sptbn1 | 20742 | HITS-CLIP | 37986224\|25083871\|23597149\|29167 | Unspecified |
| mmu-miR-152-3p | MIMAT0000162 | Sptbn2 | 20743 | HITS-CLIP | 23597149 | Unspecified |
| mmu-miR-152-3p | MIMAT0000162 | Srebf1 | 20787 | HITS-CLIP | 23597149 | Unspecified |
| mmu-miR-152-3p | MIMAT0000162 | Srebf2 | 20788 | HITS-CLIP | 23597149 | Unspecified |
| mmu-miR-152-3p | MIMAT0000162 | Srgap2 | 14270 | HITS-CLIP | 37986224\|29167373 | Unspecified |
| mmu-miR-152-3p | MIMAT0000162 | Srp68 | 217337 | HITS-CLIP | 25083871 | Unspecified |
| mmu-miR-152-3p | MIMAT0000162 | Srp72 | 66661 | HITS-CLIP | 25083871 | Unspecified |
| mmu-miR-152-3p | MIMAT0000162 | Srpk1 | 20815 | HITS-CLIP | 37986224 | Unspecified |
| mmu-miR-152-3p | MIMAT0000162 | Srrm2 | 75956 | HITS-CLIP | 37986224\|29167373 | Unspecified |
| mmu-miR-152-3p | MIMAT0000162 | Srsf1 | 110809 | HITS-CLIP | 25083871\|29167373 | Unspecified |
| mmu-miR-152-3p | MIMAT0000162 | Srsf11 | 69207 | HITS-CLIP | 37986224\|25083871 | Unspecified |
| mmu-miR-152-3p | MIMAT0000162 | Srsf6 | 67996 | HITS-CLIP | 25083871 | Unspecified |
| mmu-miR-152-3p | MIMAT0000162 | Ss18 | 268996 | HITS-CLIP | 25083871 | Unspecified |
| mmu-miR-152-3p | MIMAT0000162 | Ssb | 20823 | HITS-CLIP | 25083871\|29167373 | Unspecified |
| mmu-miR-152-3p | MIMAT0000162 | Ssh1 | 231637 | HITS-CLIP | 37986224 | Unspecified |
| mmu-miR-152-3p | MIMAT0000162 | Ssh2 | 237860 | HITS-CLIP | 37986224\|25083871\|23597149 | Unspecified |
| mmu-miR-152-3p | MIMAT0000162 | Stag1 | 20842 | HITS-CLIP | 37986224\|25083871 | Unspecified |
| mmu-miR-152-3p | MIMAT0000162 | Stag2 | 20843 | HITS-CLIP | 25083871\|23597149 | Unspecified |

| | | | | | | |
|---|---|---|---|---|---|---|
| mmu-miR-152-3p | MIMAT0000162 | Stam | 20844 | HITS-CLIP | 37986224\|29167373 | Unspecified |
| mmu-miR-152-3p | MIMAT0000162 | Stam2 | 56324 | HITS-CLIP | 25083871 | Unspecified |
| mmu-miR-152-3p | MIMAT0000162 | Stap1 | 56792 | HITS-CLIP | 29167373 | Unspecified |
| mmu-miR-152-3p | MIMAT0000162 | Stard13 | 243362 | HITS-CLIP | 37986224\|25083871\|23597149 | Unspecified |
| mmu-miR-152-3p | MIMAT0000162 | Stau1 | 20853 | HITS-CLIP | 37986224\|25083871 | Unspecified |
| mmu-miR-152-3p | MIMAT0000162 | Stbd1 | 52331 | HITS-CLIP | 37986224\|25083871\|23597149 | Unspecified |
| mmu-miR-152-3p | MIMAT0000162 | Steap2 | 74051 | HITS-CLIP | 37986224\|25083871 | Unspecified |
| mmu-miR-152-3p | MIMAT0000162 | Stil | 20460 | HITS-CLIP | 25083871 | Unspecified |
| mmu-miR-152-3p | MIMAT0000162 | Stim2 | 116873 | HITS-CLIP | 25083871\|23597149 | Unspecified |
| mmu-miR-152-3p | MIMAT0000162 | Stk10 | 20868 | HITS-CLIP | 25083871 | Unspecified |
| mmu-miR-152-3p | MIMAT0000162 | Stk35 | 67333 | HITS-CLIP | 25083871 | Unspecified |
| mmu-miR-152-3p | MIMAT0000162 | Stk38l | 232533 | HITS-CLIP | 37986224\|25083871 | Unspecified |
| mmu-miR-152-3p | MIMAT0000162 | Stk4 | 58231 | HITS-CLIP | 37986224\|25083871 | Unspecified |
| mmu-miR-152-3p | MIMAT0000162 | Stk40 | 74178 | HITS-CLIP | 37986224 | Unspecified |
| mmu-miR-152-3p | MIMAT0000162 | Stom | 13830 | HITS-CLIP | 25083871 | Unspecified |
| mmu-miR-152-3p | MIMAT0000162 | Stradb | 227154 | HITS-CLIP | 23597149 | Unspecified |
| mmu-miR-152-3p | MIMAT0000162 | Strn3 | 94186 | HITS-CLIP | 37986224\|23597149\|25083871\|29167 | Unspecified |
| mmu-miR-152-3p | MIMAT0000162 | Stt3a | 16430 | HITS-CLIP | 37986224 | Unspecified |
| mmu-miR-152-3p | MIMAT0000162 | Stt3b | 68292 | HITS-CLIP | 37986224\|25083871\|23597149\|29167 | Unspecified |
| mmu-miR-152-3p | MIMAT0000162 | Stx18 | 71116 | HITS-CLIP | 29167373 | Unspecified |
| mmu-miR-152-3p | MIMAT0000162 | Stx6 | 58244 | HITS-CLIP | 25083871 | Unspecified |
| mmu-miR-152-3p | MIMAT0000162 | Stx7 | 53331 | HITS-CLIP | 37986224\|25083871 | Unspecified |
| mmu-miR-152-3p | MIMAT0000162 | Stxbp5 | 78808 | HITS-CLIP | 37986224\|25083871\|23597149 | Unspecified |
| mmu-miR-152-3p | MIMAT0000162 | Sugt1 | 67955 | HITS-CLIP | 25083871 | Unspecified |
| mmu-miR-152-3p | MIMAT0000162 | Sumf1 | 58911 | HITS-CLIP | 25083871 | Unspecified |
| mmu-miR-152-3p | MIMAT0000162 | Sun2 | 223697 | HITS-CLIP | 25083871 | Unspecified |
| mmu-miR-152-3p | MIMAT0000162 | Susd6 | 9766 | HITS-CLIP | 25083871 | Unspecified |
| mmu-miR-152-3p | MIMAT0000162 | Suz12 | 52615 | HITS-CLIP | 25083871\|31433990 | Unspecified |
| mmu-miR-152-3p | MIMAT0000162 | Svil | 225115 | HITS-CLIP | 25083871 | Unspecified |
| mmu-miR-152-3p | MIMAT0000162 | Syap1 | 67043 | HITS-CLIP | 23597149 | Unspecified |
| mmu-miR-152-3p | MIMAT0000162 | Syde1 | 71709 | HITS-CLIP | 25083871 | Unspecified |
| mmu-miR-152-3p | MIMAT0000162 | Sympk | 68188 | HITS-CLIP | 23597149 | Unspecified |
| mmu-miR-152-3p | MIMAT0000162 | Syne2 | 319565 | HITS-CLIP | 23597149 | Unspecified |
| mmu-miR-152-3p | MIMAT0000162 | Syne3 | 212073 | HITS-CLIP | 37986224 | Unspecified |
| mmu-miR-152-3p | MIMAT0000162 | Syngr2 | 20973 | HITS-CLIP | 37986224 | Unspecified |
| mmu-miR-152-3p | MIMAT0000162 | Synj1 | 104015 | HITS-CLIP | 37986224\|25083871 | Unspecified |
| mmu-miR-152-3p | MIMAT0000162 | Sypl | 19027 | HITS-CLIP | 25083871 | Unspecified |
| mmu-miR-152-3p | MIMAT0000162 | Sytl5 | 236643 | HITS-CLIP | 25083871 | Unspecified |
| mmu-miR-152-3p | MIMAT0000162 | Syvn1 | 74126 | HITS-CLIP | 23597149 | Unspecified |

| | | | | | | | |
|---|---|---|---|---|---|---|---|
| mmu-miR-152-3p | MIMAT0000162 | Szrd1 | 213491 | HITS-CLIP | | 37986224 | Unspecified |
| mmu-miR-152-3p | MIMAT0000162 | Szt2 | 230676 | HITS-CLIP | | 25083871 | Unspecified |
| mmu-miR-152-3p | MIMAT0000162 | Tacc2 | 57752 | HITS-CLIP | | 25083871 | Unspecified |
| mmu-miR-152-3p | MIMAT0000162 | Taf10 | 24075 | HITS-CLIP | | 37986224 | Unspecified |
| mmu-miR-152-3p | MIMAT0000162 | Taf15 | 70439 | HITS-CLIP | | 23597149 | Unspecified |
| mmu-miR-152-3p | MIMAT0000162 | Taf9 | 108143 | HITS-CLIP | 37986224\|25083871 | | Unspecified |
| mmu-miR-152-3p | MIMAT0000162 | Tanc2 | 77097 | HITS-CLIP | | 25083871 | Unspecified |
| mmu-miR-152-3p | MIMAT0000162 | Taok1 | 216965 | HITS-CLIP | 37986224\|25083871 | | Unspecified |
| mmu-miR-152-3p | MIMAT0000162 | Taok2 | 381921 | HITS-CLIP | | 25083871 | Unspecified |
| mmu-miR-152-3p | MIMAT0000162 | Tapt1 | 231225 | HITS-CLIP | | 25083871 | Unspecified |
| mmu-miR-152-3p | MIMAT0000162 | Tardbp | 230908 | HITS-CLIP | 37986224\|25083871\|29167373 | | Unspecified |
| mmu-miR-152-3p | MIMAT0000162 | Tatdn1 | 69694 | HITS-CLIP | 37986224\|25083871 | | Unspecified |
| mmu-miR-152-3p | MIMAT0000162 | Tbc1d1 | 57915 | HITS-CLIP | 25083871\|23597149 | | Unspecified |
| mmu-miR-152-3p | MIMAT0000162 | Tbc1d15 | 66687 | HITS-CLIP | | 25083871 | Unspecified |
| mmu-miR-152-3p | MIMAT0000162 | Tbc1d23 | 67581 | HITS-CLIP | 37986224\|23597149\|29167373 | | Unspecified |
| mmu-miR-152-3p | MIMAT0000162 | Tbc1d31 | 210544 | HITS-CLIP | | 29167373 | Unspecified |
| mmu-miR-152-3p | MIMAT0000162 | Tbc1d5 | 72238 | HITS-CLIP | | 25083871 | Unspecified |
| mmu-miR-152-3p | MIMAT0000162 | Tbc1d9 | 71310 | HITS-CLIP | | 29167373 | Unspecified |
| mmu-miR-152-3p | MIMAT0000162 | Tbcel | 272589 | HITS-CLIP | | 25083871 | Unspecified |
| mmu-miR-152-3p | MIMAT0000162 | Tbck | 271981 | HITS-CLIP | | 37986224 | Unspecified |
| mmu-miR-152-3p | MIMAT0000162 | Tbl1x | 21372 | HITS-CLIP | | 25083871 | Unspecified |
| mmu-miR-152-3p | MIMAT0000162 | Tbl1xr1 | 81004 | HITS-CLIP | 37986224\|25083871 | | Unspecified |
| mmu-miR-152-3p | MIMAT0000162 | Tbp | 21374 | HITS-CLIP | | 37986224 | Unspecified |
| mmu-miR-152-3p | MIMAT0000162 | Tbpl1 | 237336 | HITS-CLIP | | 25083871 | Unspecified |
| mmu-miR-152-3p | MIMAT0000162 | Tbx15 | 21384 | HITS-CLIP | 37986224\|25083871 | | Unspecified |
| mmu-miR-152-3p | MIMAT0000162 | Tbx18 | 76365 | HITS-CLIP | 37986224\|25083871 | | Unspecified |
| mmu-miR-152-3p | MIMAT0000162 | Tbx3 | 21386 | HITS-CLIP | | 25083871 | Unspecified |
| mmu-miR-152-3p | MIMAT0000162 | Tcaf1 | 9747 | HITS-CLIP | | 23597149 | Unspecified |
| mmu-miR-152-3p | MIMAT0000162 | Tceal8 | 66684 | HITS-CLIP | 37986224\|25083871 | | Unspecified |
| mmu-miR-152-3p | MIMAT0000162 | Tcf25 | 66855 | HITS-CLIP | | 37986224 | Unspecified |
| mmu-miR-152-3p | MIMAT0000162 | Tcf4 | 21413 | HITS-CLIP | 37986224\|25083871\|29167373 | | Unspecified |
| mmu-miR-152-3p | MIMAT0000162 | Tcf7l1 | 21415 | HITS-CLIP | | 37986224 | Unspecified |
| mmu-miR-152-3p | MIMAT0000162 | Tcirg1 | 27060 | HITS-CLIP | 31433990\|29167373 | | Unspecified |
| mmu-miR-152-3p | MIMAT0000162 | Tcp11l1 | 320554 | HITS-CLIP | | 37986224 | Unspecified |
| mmu-miR-152-3p | MIMAT0000162 | Tead1 | 21676 | HITS-CLIP | 37986224\|25083871 | | Unspecified |
| mmu-miR-152-3p | MIMAT0000162 | Tead4 | 21679 | HITS-CLIP | | 37986224 | Unspecified |
| mmu-miR-152-3p | MIMAT0000162 | Tef | 21685 | HITS-CLIP | | 25083871 | Unspecified |
| mmu-miR-152-3p | MIMAT0000162 | Tenm3 | 23965 | HITS-CLIP | 37986224\|25083871 | | Unspecified |
| mmu-miR-152-3p | MIMAT0000162 | Tent4b | 64282 | HITS-CLIP | | 25083871 | Unspecified |

| | | | | | | |
|---|---|---|---|---|---|---|
| mmu-miR-152-3p | MIMAT0000162 | Tep1 | 21745 | HITS-CLIP | 37986224\|23597149 | Unspecified |
| mmu-miR-152-3p | MIMAT0000162 | Tes | 21753 | HITS-CLIP | 25083871 | Unspecified |
| mmu-miR-152-3p | MIMAT0000162 | Tet2 | 214133 | HITS-CLIP | 25083871 | Unspecified |
| mmu-miR-152-3p | MIMAT0000162 | Tex10 | 269536 | HITS-CLIP | 37986224 | Unspecified |
| mmu-miR-152-3p | MIMAT0000162 | Tex2 | 21763 | HITS-CLIP | 23597149 | Unspecified |
| mmu-miR-152-3p | MIMAT0000162 | Tex261 | 21766 | HITS-CLIP | 23597149\|25083871 | Unspecified |
| mmu-miR-152-3p | MIMAT0000162 | Tfdp2 | 211586 | HITS-CLIP | 37986224 | Unspecified |
| mmu-miR-152-3p | MIMAT0000162 | Tfr2 | 50765 | HITS-CLIP | 23597149 | Unspecified |
| mmu-miR-152-3p | MIMAT0000162 | Tfrc | 22042 | HITS-CLIP | 37986224\|25083871 | Unspecified |
| mmu-miR-152-3p | MIMAT0000162 | Tgfb2 | 21808 | HITS-CLIP | 25083871 | Unspecified |
| mmu-miR-152-3p | MIMAT0000162 | Tgfb3 | 21809 | HITS-CLIP | 25083871 | Unspecified |
| mmu-miR-152-3p | MIMAT0000162 | Tgfbr1 | 21812 | HITS-CLIP | 37986224\|23597149\|25083871\|29167 | Unspecified |
| mmu-miR-152-3p | MIMAT0000162 | Tgfbr2 | 21813 | HITS-CLIP | 25083871 | Unspecified |
| mmu-miR-152-3p | MIMAT0000162 | Tgfbr3 | 21814 | HITS-CLIP | 23597149 | Unspecified |
| mmu-miR-152-3p | MIMAT0000162 | Tgoln1 | 22134 | HITS-CLIP | 37986224\|23597149 | Unspecified |
| mmu-miR-152-3p | MIMAT0000162 | Tgs1 | 116940 | HITS-CLIP | 37986224\|25083871 | Unspecified |
| mmu-miR-152-3p | MIMAT0000162 | Thbd | 21824 | HITS-CLIP | 37986224 | Unspecified |
| mmu-miR-152-3p | MIMAT0000162 | Thbs2 | 21826 | HITS-CLIP | 25083871 | Unspecified |
| mmu-miR-152-3p | MIMAT0000162 | Thoc2 | 331401 | HITS-CLIP | 25083871 | Unspecified |
| mmu-miR-152-3p | MIMAT0000162 | Thra | 21833 | HITS-CLIP | 25083871 | Unspecified |
| mmu-miR-152-3p | MIMAT0000162 | Thrap3 | 230753 | HITS-CLIP | 29167373 | Unspecified |
| mmu-miR-152-3p | MIMAT0000162 | Tia1 | 21841 | HITS-CLIP | 25083871 | Unspecified |
| mmu-miR-152-3p | MIMAT0000162 | Tiam2 | 24001 | HITS-CLIP | 37986224 | Unspecified |
| mmu-miR-152-3p | MIMAT0000162 | Timm22 | 56322 | HITS-CLIP | 23597149 | Unspecified |
| mmu-miR-152-3p | MIMAT0000162 | Tiparp | 99929 | HITS-CLIP | 37986224\|25083871\|29167373 | Unspecified |
| mmu-miR-152-3p | MIMAT0000162 | Tipin | 66131 | HITS-CLIP | 37986224\|25083871 | Unspecified |
| mmu-miR-152-3p | MIMAT0000162 | Tiprl | 226591 | HITS-CLIP | 37986224 | Unspecified |
| mmu-miR-152-3p | MIMAT0000162 | Tirap | 117149 | HITS-CLIP | 25083871 | Unspecified |
| mmu-miR-152-3p | MIMAT0000162 | Tkt | 21881 | HITS-CLIP | 37986224 | Unspecified |
| mmu-miR-152-3p | MIMAT0000162 | Tle4 | 21888 | HITS-CLIP | 29167373 | Unspecified |
| mmu-miR-152-3p | MIMAT0000162 | Tlk2 | 24086 | HITS-CLIP | 25083871 | Unspecified |
| mmu-miR-152-3p | MIMAT0000162 | Tln1 | 21894 | HITS-CLIP | 25083871\|23597149\|29167373 | Unspecified |
| mmu-miR-152-3p | MIMAT0000162 | Tln2 | 70549 | HITS-CLIP | 25083871 | Unspecified |
| mmu-miR-152-3p | MIMAT0000162 | Tlnrd1 | 59274 | HITS-CLIP | 25083871\|29167373 | Unspecified |
| mmu-miR-152-3p | MIMAT0000162 | Tlr13 | 279572 | HITS-CLIP | 29167373 | Unspecified |
| mmu-miR-152-3p | MIMAT0000162 | Tlr4 | 21898 | HITS-CLIP | 37986224 | Unspecified |
| mmu-miR-152-3p | MIMAT0000162 | Tm2d3 | 68634 | HITS-CLIP | 25083871 | Unspecified |
| mmu-miR-152-3p | MIMAT0000162 | Tm4sf1 | 17112 | HITS-CLIP | 25083871 | Unspecified |
| mmu-miR-152-3p | MIMAT0000162 | Tm6sf2 | 107770 | HITS-CLIP | 23597149 | Unspecified |

| | | | | | | |
|---|---|---|---|---|---|---|
| mmu-miR-152-3p | MIMAT0000162 | Tm9sf1 | 74140 | HITS-CLIP | 37986224 | Unspecified |
| mmu-miR-152-3p | MIMAT0000162 | Tmbim4 | 68212 | HITS-CLIP | 37986224\|23597149\|25083871 | Unspecified |
| mmu-miR-152-3p | MIMAT0000162 | Tmbim6 | 110213 | HITS-CLIP | 23597149\|25083871\|29167373 | Unspecified |
| mmu-miR-152-3p | MIMAT0000162 | Tmcc1 | 330401 | HITS-CLIP | 37986224 | Unspecified |
| mmu-miR-152-3p | MIMAT0000162 | Tmed4 | 103694 | HITS-CLIP | 21258322 | Unspecified |
| mmu-miR-152-3p | MIMAT0000162 | Tmed8 | 382620 | HITS-CLIP | 37986224 | Unspecified |
| mmu-miR-152-3p | MIMAT0000162 | Tmeff1 | 230157 | HITS-CLIP | 25083871 | Unspecified |
| mmu-miR-152-3p | MIMAT0000162 | Tmem106b | 71900 | HITS-CLIP | 23597149 | Unspecified |
| mmu-miR-152-3p | MIMAT0000162 | Tmem115 | 56395 | HITS-CLIP | 37986224 | Unspecified |
| mmu-miR-152-3p | MIMAT0000162 | Tmem120a | 215210 | HITS-CLIP | 23597149 | Unspecified |
| mmu-miR-152-3p | MIMAT0000162 | Tmem127 | 69470 | HITS-CLIP | 37986224\|25083871 | Unspecified |
| mmu-miR-152-3p | MIMAT0000162 | Tmem164 | 209497 | HITS-CLIP | 37986224 | Unspecified |
| mmu-miR-152-3p | MIMAT0000162 | Tmem165 | 21982 | HITS-CLIP | 25083871 | Unspecified |
| mmu-miR-152-3p | MIMAT0000162 | Tmem168 | 101118 | HITS-CLIP | 25083871 | Unspecified |
| mmu-miR-152-3p | MIMAT0000162 | Tmem170b | 621976 | HITS-CLIP | 37986224 | Unspecified |
| mmu-miR-152-3p | MIMAT0000162 | Tmem19 | 67226 | HITS-CLIP | 37986224 | Unspecified |
| mmu-miR-152-3p | MIMAT0000162 | Tmem201 | 230917 | HITS-CLIP | 37986224 | Unspecified |
| mmu-miR-152-3p | MIMAT0000162 | Tmem245 | 242474 | HITS-CLIP | 37986224\|25083871\|23597149 | Unspecified |
| mmu-miR-152-3p | MIMAT0000162 | Tmem47 | 192216 | HITS-CLIP | 37986224 | Unspecified |
| mmu-miR-152-3p | MIMAT0000162 | Tmem87b | 72477 | HITS-CLIP | 25083871\|23597149 | Unspecified |
| mmu-miR-152-3p | MIMAT0000162 | Tmem9b | 56786 | HITS-CLIP | 37986224\|25083871\|23597149\|29167 | Unspecified |
| mmu-miR-152-3p | MIMAT0000162 | Tmod1 | 21916 | HITS-CLIP | 25083871 | Unspecified |
| mmu-miR-152-3p | MIMAT0000162 | Tmod3 | 50875 | HITS-CLIP | 25083871 | Unspecified |
| mmu-miR-152-3p | MIMAT0000162 | Tmpo | 21917 | HITS-CLIP | 37986224 | Unspecified |
| mmu-miR-152-3p | MIMAT0000162 | Tmsb10 | 19240 | HITS-CLIP | 37986224 | Unspecified |
| mmu-miR-152-3p | MIMAT0000162 | Tmsb4x | 19241 | HITS-CLIP | 37986224 | Unspecified |
| mmu-miR-152-3p | MIMAT0000162 | Tmx3 | 67988 | HITS-CLIP | 37986224 | Unspecified |
| mmu-miR-152-3p | MIMAT0000162 | Tnc | 21923 | HITS-CLIP | 25083871 | Unspecified |
| mmu-miR-152-3p | MIMAT0000162 | Tnfaip1 | 21927 | HITS-CLIP | 37986224 | Unspecified |
| mmu-miR-152-3p | MIMAT0000162 | Tnfaip3 | 21929 | HITS-CLIP | 25083871 | Unspecified |
| mmu-miR-152-3p | MIMAT0000162 | Tnfrsf1b | 21938 | HITS-CLIP | 37986224 | Unspecified |
| mmu-miR-152-3p | MIMAT0000162 | Tnks | 21951 | HITS-CLIP | 25083871\|29167373 | Unspecified |
| mmu-miR-152-3p | MIMAT0000162 | Tnks1bp1 | 228140 | HITS-CLIP | 25083871 | Unspecified |
| mmu-miR-152-3p | MIMAT0000162 | Tnks2 | 74493 | HITS-CLIP | 25083871 | Unspecified |
| mmu-miR-152-3p | MIMAT0000162 | Tnnt1 | 21955 | HITS-CLIP | 25083871 | Unspecified |
| mmu-miR-152-3p | MIMAT0000162 | Tnnt2 | 21956 | HITS-CLIP | 25083871 | Unspecified |
| mmu-miR-152-3p | MIMAT0000162 | Tnpo2 | 212999 | HITS-CLIP | 25083871 | Unspecified |
| mmu-miR-152-3p | MIMAT0000162 | Tnpo3 | 320938 | HITS-CLIP | 25083871 | Unspecified |
| mmu-miR-152-3p | MIMAT0000162 | Tnrc18 | 231861 | HITS-CLIP | 25083871 | Unspecified |

| | | | | | | |
|---|---|---|---|---|---|---|
| mmu-miR-152-3p | MIMAT0000162 | Tnrc6a | 233833 | HITS-CLIP | 37986224\|25083871 | Unspecified |
| mmu-miR-152-3p | MIMAT0000162 | Tnrc6b | 213988 | HITS-CLIP | 37986224\|25083871\|29167373 | Unspecified |
| mmu-miR-152-3p | MIMAT0000162 | Tnrc6c | 217351 | HITS-CLIP | 37986224\|25083871\|23597149\|29167 | Unspecified |
| mmu-miR-152-3p | MIMAT0000162 | Tns1 | 21961 | HITS-CLIP | 37986224 | Unspecified |
| mmu-miR-152-3p | MIMAT0000162 | Tns2 | 209039 | HITS-CLIP | 25083871 | Unspecified |
| mmu-miR-152-3p | MIMAT0000162 | Tns3 | 319939 | HITS-CLIP | 25083871 | Unspecified |
| mmu-miR-152-3p | MIMAT0000162 | Tob1 | 22057 | HITS-CLIP | 25083871\|23597149 | Unspecified |
| mmu-miR-152-3p | MIMAT0000162 | Tob2 | 57259 | HITS-CLIP | 37986224\|25083871\|23597149 | Unspecified |
| mmu-miR-152-3p | MIMAT0000162 | Togaram1 | 23116 | HITS-CLIP | 25083871 | Unspecified |
| mmu-miR-152-3p | MIMAT0000162 | Tollip | 54473 | HITS-CLIP | 37986224 | Unspecified |
| mmu-miR-152-3p | MIMAT0000162 | Tomm70a | 28185 | HITS-CLIP | 37986224\|25083871\|23597149 | Unspecified |
| mmu-miR-152-3p | MIMAT0000162 | Top1 | 21969 | HITS-CLIP | 25083871 | Unspecified |
| mmu-miR-152-3p | MIMAT0000162 | Top2a | 21973 | HITS-CLIP | 25083871 | Unspecified |
| mmu-miR-152-3p | MIMAT0000162 | Top2b | 21974 | HITS-CLIP | 25083871 | Unspecified |
| mmu-miR-152-3p | MIMAT0000162 | Tox4 | 268741 | HITS-CLIP | 37986224 | Unspecified |
| mmu-miR-152-3p | MIMAT0000162 | Tpm1 | 22003 | HITS-CLIP | 25083871 | Unspecified |
| mmu-miR-152-3p | MIMAT0000162 | Tpm3 | 59069 | HITS-CLIP | 25083871 | Unspecified |
| mmu-miR-152-3p | MIMAT0000162 | Tpp1 | 12751 | HITS-CLIP | 37986224\|25083871 | Unspecified |
| mmu-miR-152-3p | MIMAT0000162 | Tpp2 | 22019 | HITS-CLIP | 25083871 | Unspecified |
| mmu-miR-152-3p | MIMAT0000162 | Tpr | 108989 | HITS-CLIP | 25083871 | Unspecified |
| mmu-miR-152-3p | MIMAT0000162 | Trak2 | 70827 | HITS-CLIP | 37986224 | Unspecified |
| mmu-miR-152-3p | MIMAT0000162 | Tram1 | 72265 | HITS-CLIP | 23597149\|25083871 | Unspecified |
| mmu-miR-152-3p | MIMAT0000162 | Tram2 | 170829 | HITS-CLIP | 23597149 | Unspecified |
| mmu-miR-152-3p | MIMAT0000162 | Trap1 | 68015 | HITS-CLIP | 23597149 | Unspecified |
| mmu-miR-152-3p | MIMAT0000162 | Trappc1 | 245828 | HITS-CLIP | 29167373 | Unspecified |
| mmu-miR-152-3p | MIMAT0000162 | Trappc10 | 216131 | HITS-CLIP | 23597149\|25083871 | Unspecified |
| mmu-miR-152-3p | MIMAT0000162 | Trappc11 | 320714 | HITS-CLIP | 25083871 | Unspecified |
| mmu-miR-152-3p | MIMAT0000162 | Trib1 | 211770 | HITS-CLIP | 37986224\|23597149 | Unspecified |
| mmu-miR-152-3p | MIMAT0000162 | Trim11 | 94091 | HITS-CLIP | 37986224 | Unspecified |
| mmu-miR-152-3p | MIMAT0000162 | Trim16 | 94092 | HITS-CLIP | 37986224 | Unspecified |
| mmu-miR-152-3p | MIMAT0000162 | Trim2 | 80890 | HITS-CLIP | 37986224 | Unspecified |
| mmu-miR-152-3p | MIMAT0000162 | Trim23 | 81003 | HITS-CLIP | 23597149 | Unspecified |
| mmu-miR-152-3p | MIMAT0000162 | Trim25 | 217069 | HITS-CLIP | 37986224\|29167373 | Unspecified |
| mmu-miR-152-3p | MIMAT0000162 | Trim27 | 19720 | HITS-CLIP | 25083871 | Unspecified |
| mmu-miR-152-3p | MIMAT0000162 | Trim33 | 94093 | HITS-CLIP | 37986224\|23597149\|25083871 | Unspecified |
| mmu-miR-152-3p | MIMAT0000162 | Trim44 | 80985 | HITS-CLIP | 25083871 | Unspecified |
| mmu-miR-152-3p | MIMAT0000162 | Trim68 | 101700 | HITS-CLIP | 23597149 | Unspecified |
| mmu-miR-152-3p | MIMAT0000162 | Trim8 | 93679 | HITS-CLIP | 25083871 | Unspecified |
| mmu-miR-152-3p | MIMAT0000162 | Trip11 | 109181 | HITS-CLIP | 37986224\|25083871\|29167373 | Unspecified |

| | | | | | | |
|---|---|---|---|---|---|---|
| mmu-miR-152-3p | MIMAT0000162 | Trip12 | 14897 | HITS-CLIP | 37986224\|25083871\|23597149 | Unspecified |
| mmu-miR-152-3p | MIMAT0000162 | Trmt1 | 212528 | HITS-CLIP | 37986224 | Unspecified |
| mmu-miR-152-3p | MIMAT0000162 | Trove2 | 20822 | HITS-CLIP | 25083871 | Unspecified |
| mmu-miR-152-3p | MIMAT0000162 | Trp53inp1 | 60599 | HITS-CLIP | 37986224\|23597149\|25083871 | Unspecified |
| mmu-miR-152-3p | MIMAT0000162 | Trp53rkb | 76367 | HITS-CLIP | 25083871 | Unspecified |
| mmu-miR-152-3p | MIMAT0000162 | Trpc4ap | 56407 | HITS-CLIP | 37986224 | Unspecified |
| mmu-miR-152-3p | MIMAT0000162 | Trps1 | 83925 | HITS-CLIP | 37986224\|25083871\|29167373 | Unspecified |
| mmu-miR-152-3p | MIMAT0000162 | Tsc22d3 | 14605 | HITS-CLIP | 23597149 | Unspecified |
| mmu-miR-152-3p | MIMAT0000162 | Tsg101 | 22088 | HITS-CLIP | 25083871 | Unspecified |
| mmu-miR-152-3p | MIMAT0000162 | Tshz1 | 110796 | HITS-CLIP | 37986224\|25083871 | Unspecified |
| mmu-miR-152-3p | MIMAT0000162 | Tsn | 22099 | HITS-CLIP | 25083871 | Unspecified |
| mmu-miR-152-3p | MIMAT0000162 | Tspan17 | 74257 | HITS-CLIP | 25083871 | Unspecified |
| mmu-miR-152-3p | MIMAT0000162 | Tspan4 | 64540 | HITS-CLIP | 25083871 | Unspecified |
| mmu-miR-152-3p | MIMAT0000162 | Tstd1 | 100131187 | HITS-CLIP | 23597149 | Unspecified |
| mmu-miR-152-3p | MIMAT0000162 | Ttc28 | 209683 | HITS-CLIP | 37986224\|25083871 | Unspecified |
| mmu-miR-152-3p | MIMAT0000162 | Ttc3 | 22129 | HITS-CLIP | 25083871 | Unspecified |
| mmu-miR-152-3p | MIMAT0000162 | Ttc33 | 67515 | HITS-CLIP | 37986224 | Unspecified |
| mmu-miR-152-3p | MIMAT0000162 | Ttc37 | 218343 | HITS-CLIP | 23597149 | Unspecified |
| mmu-miR-152-3p | MIMAT0000162 | Ttc39b | 69863 | HITS-CLIP | 37986224 | Unspecified |
| mmu-miR-152-3p | MIMAT0000162 | Ttc39c | 72747 | HITS-CLIP | 23597149 | Unspecified |
| mmu-miR-152-3p | MIMAT0000162 | Ttn | 22138 | HITS-CLIP | 37986224\|25083871 | Unspecified |
| mmu-miR-152-3p | MIMAT0000162 | Ttyh3 | 78339 | HITS-CLIP | 37986224 | Unspecified |
| mmu-miR-152-3p | MIMAT0000162 | Tubb2a | 22151 | HITS-CLIP | 37986224 | Unspecified |
| mmu-miR-152-3p | MIMAT0000162 | Tubb4b | 227613 | HITS-CLIP | 23597149 | Unspecified |
| mmu-miR-152-3p | MIMAT0000162 | Tubb6 | 67951 | HITS-CLIP | 37986224 | Unspecified |
| mmu-miR-152-3p | MIMAT0000162 | Tubgcp3 | 259279 | HITS-CLIP | 29167373 | Unspecified |
| mmu-miR-152-3p | MIMAT0000162 | Tubgcp5 | 233276 | HITS-CLIP | 23597149 | Unspecified |
| mmu-miR-152-3p | MIMAT0000162 | Tulp4 | 68842 | HITS-CLIP | 37986224 | Unspecified |
| mmu-miR-152-3p | MIMAT0000162 | Tut4 | 23318 | HITS-CLIP | 29167373 | Unspecified |
| mmu-miR-152-3p | MIMAT0000162 | Twnk | 56652 | HITS-CLIP | 25083871 | Unspecified |
| mmu-miR-152-3p | MIMAT0000162 | Txlna | 109658 | HITS-CLIP | 25083871 | Unspecified |
| mmu-miR-152-3p | MIMAT0000162 | Txndc11 | 106200 | HITS-CLIP | 25083871\|29167373 | Unspecified |
| mmu-miR-152-3p | MIMAT0000162 | Txndc15 | 69672 | HITS-CLIP | 25083871 | Unspecified |
| mmu-miR-152-3p | MIMAT0000162 | Txnip | 56338 | HITS-CLIP | 37986224 | Unspecified |
| mmu-miR-152-3p | MIMAT0000162 | Uba1 | 22201 | HITS-CLIP | 23597149 | Unspecified |
| mmu-miR-152-3p | MIMAT0000162 | Uba3 | 22200 | HITS-CLIP | 23597149 | Unspecified |
| mmu-miR-152-3p | MIMAT0000162 | Uba6 | 231380 | HITS-CLIP | 25083871\|29167373 | Unspecified |
| mmu-miR-152-3p | MIMAT0000162 | Ubac1 | 98766 | HITS-CLIP | 37986224\|25083871 | Unspecified |
| mmu-miR-152-3p | MIMAT0000162 | Ubac2 | 68889 | HITS-CLIP | 37986224\|25083871\|23597149 | Unspecified |

| | | | | | | |
|---|---|---|---|---|---|---|
| mmu-miR-152-3p | MIMAT0000162 | Ubald1 | 207740 | HITS-CLIP | 23597149 | Unspecified |
| mmu-miR-152-3p | MIMAT0000162 | Ubap2 | 68926 | HITS-CLIP | 25083871\|31433990\|29167373 | Unspecified |
| mmu-miR-152-3p | MIMAT0000162 | Ubap2l | 74383 | HITS-CLIP | 25083871 | Unspecified |
| mmu-miR-152-3p | MIMAT0000162 | Ubash3b | 72828 | HITS-CLIP | 37986224 | Unspecified |
| mmu-miR-152-3p | MIMAT0000162 | Ubc | 22190 | HITS-CLIP | 23597149 | Unspecified |
| mmu-miR-152-3p | MIMAT0000162 | Ube2d1 | 216080 | HITS-CLIP | 29167373 | Unspecified |
| mmu-miR-152-3p | MIMAT0000162 | Ube2d3 | 66105 | HITS-CLIP | 25083871 | Unspecified |
| mmu-miR-152-3p | MIMAT0000162 | Ube2h | 22214 | HITS-CLIP | 25083871 | Unspecified |
| mmu-miR-152-3p | MIMAT0000162 | Ube2j1 | 56228 | HITS-CLIP | 37986224\|25083871 | Unspecified |
| mmu-miR-152-3p | MIMAT0000162 | Ube2w | 66799 | HITS-CLIP | 37986224 | Unspecified |
| mmu-miR-152-3p | MIMAT0000162 | Ube3a | 22215 | HITS-CLIP | 25083871\|29167373 | Unspecified |
| mmu-miR-152-3p | MIMAT0000162 | Ube3b | 117146 | HITS-CLIP | 37986224 | Unspecified |
| mmu-miR-152-3p | MIMAT0000162 | Ube3c | 100763 | HITS-CLIP | 25083871 | Unspecified |
| mmu-miR-152-3p | MIMAT0000162 | Ube4b | 63958 | HITS-CLIP | 25083871 | Unspecified |
| mmu-miR-152-3p | MIMAT0000162 | Ubiad1 | 71707 | HITS-CLIP | 25083871 | Unspecified |
| mmu-miR-152-3p | MIMAT0000162 | Ubl3 | 24109 | HITS-CLIP | 23597149\|29167373 | Unspecified |
| mmu-miR-152-3p | MIMAT0000162 | Ubn1 | 170644 | HITS-CLIP | 29167373 | Unspecified |
| mmu-miR-152-3p | MIMAT0000162 | Ubn2 | 320538 | HITS-CLIP | 25083871\|23597149 | Unspecified |
| mmu-miR-152-3p | MIMAT0000162 | Ubp1 | 22221 | HITS-CLIP | 25083871 | Unspecified |
| mmu-miR-152-3p | MIMAT0000162 | Ubr1 | 22222 | HITS-CLIP | 23597149 | Unspecified |
| mmu-miR-152-3p | MIMAT0000162 | Ubr2 | 224826 | HITS-CLIP | 25083871\|23597149 | Unspecified |
| mmu-miR-152-3p | MIMAT0000162 | Ubr4 | 69116 | HITS-CLIP | 25083871\|23597149\|29167373 | Unspecified |
| mmu-miR-152-3p | MIMAT0000162 | Ubr5 | 70790 | HITS-CLIP | 25083871 | Unspecified |
| mmu-miR-152-3p | MIMAT0000162 | Ubxn4 | 67812 | HITS-CLIP | 25083871\|29167373 | Unspecified |
| mmu-miR-152-3p | MIMAT0000162 | Ubxn7 | 224111 | HITS-CLIP | 25083871 | Unspecified |
| mmu-miR-152-3p | MIMAT0000162 | Ufl1 | 67490 | HITS-CLIP | 37986224 | Unspecified |
| mmu-miR-152-3p | MIMAT0000162 | Ugdh | 22235 | HITS-CLIP | 37986224\|23597149\|25083871 | Unspecified |
| mmu-miR-152-3p | MIMAT0000162 | Ugt2b34 | 100727 | HITS-CLIP | 37986224\|23597149 | Unspecified |
| mmu-miR-152-3p | MIMAT0000162 | Ugt2b35 | 243085 | HITS-CLIP | 37986224\|23597149 | Unspecified |
| mmu-miR-152-3p | MIMAT0000162 | Ugt2b36 | 231396 | HITS-CLIP | 37986224 | Unspecified |
| mmu-miR-152-3p | MIMAT0000162 | Ugt2b5 | 22238 | HITS-CLIP | 23597149 | Unspecified |
| mmu-miR-152-3p | MIMAT0000162 | Uhmk1 | 16589 | HITS-CLIP | 29167373 | Unspecified |
| mmu-miR-152-3p | MIMAT0000162 | Uhrf1bp1 | 224648 | HITS-CLIP | 37986224 | Unspecified |
| mmu-miR-152-3p | MIMAT0000162 | Uhrf1bp1l | 75089 | HITS-CLIP | 25083871 | Unspecified |
| mmu-miR-152-3p | MIMAT0000162 | Ulk3 | 71742 | HITS-CLIP | 25083871 | Unspecified |
| mmu-miR-152-3p | MIMAT0000162 | Unc119b | 106840 | HITS-CLIP | 37986224 | Unspecified |
| mmu-miR-152-3p | MIMAT0000162 | Unc45b | 217012 | HITS-CLIP | 25083871 | Unspecified |
| mmu-miR-152-3p | MIMAT0000162 | Unc5b | 107449 | HITS-CLIP | 25083871 | Unspecified |
| mmu-miR-152-3p | MIMAT0000162 | Unk | 217331 | HITS-CLIP | 37986224 | Unspecified |

| | | | | | | |
|---|---|---|---|---|---|---|
| mmu-miR-152-3p | MIMAT0000162 | Unkl | 74154 | HITS-CLIP | | 25083871 | Unspecified |
| mmu-miR-152-3p | MIMAT0000162 | Uox | 22262 | HITS-CLIP | | 23597149 | Unspecified |
| mmu-miR-152-3p | MIMAT0000162 | Upf2 | 326622 | HITS-CLIP | 25083871\|29167373 | | Unspecified |
| mmu-miR-152-3p | MIMAT0000162 | Upf3a | 67031 | HITS-CLIP | | 25083871 | Unspecified |
| mmu-miR-152-3p | MIMAT0000162 | Uqcrc1 | 22273 | HITS-CLIP | | 25083871 | Unspecified |
| mmu-miR-152-3p | MIMAT0000162 | Urgcp | 72046 | HITS-CLIP | 25083871\|23597149 | | Unspecified |
| mmu-miR-152-3p | MIMAT0000162 | Uroc1 | 243537 | HITS-CLIP | 37986224\|23597149 | | Unspecified |
| mmu-miR-152-3p | MIMAT0000162 | Usf3 | 205717 | HITS-CLIP | | 25083871 | Unspecified |
| mmu-miR-152-3p | MIMAT0000162 | Usp1 | 230484 | HITS-CLIP | | 25083871 | Unspecified |
| mmu-miR-152-3p | MIMAT0000162 | Usp15 | 14479 | HITS-CLIP | 29167373\|23597149\|25083871 | | Unspecified |
| mmu-miR-152-3p | MIMAT0000162 | Usp20 | 74270 | HITS-CLIP | 37986224\|25083871 | | Unspecified |
| mmu-miR-152-3p | MIMAT0000162 | Usp24 | 329908 | HITS-CLIP | | 25083871 | Unspecified |
| mmu-miR-152-3p | MIMAT0000162 | Usp25 | 30940 | HITS-CLIP | 23597149\|29167373 | | Unspecified |
| mmu-miR-152-3p | MIMAT0000162 | Usp31 | 76179 | HITS-CLIP | 37986224\|25083871 | | Unspecified |
| mmu-miR-152-3p | MIMAT0000162 | Usp32 | 237898 | HITS-CLIP | | 37986224 | Unspecified |
| mmu-miR-152-3p | MIMAT0000162 | Usp33 | 170822 | HITS-CLIP | 23597149\|25083871 | | Unspecified |
| mmu-miR-152-3p | MIMAT0000162 | Usp34 | 17847 | HITS-CLIP | | 25083871 | Unspecified |
| mmu-miR-152-3p | MIMAT0000162 | Usp37 | 319651 | HITS-CLIP | | 25083871 | Unspecified |
| mmu-miR-152-3p | MIMAT0000162 | Usp38 | 74841 | HITS-CLIP | | 29167373 | Unspecified |
| mmu-miR-152-3p | MIMAT0000162 | Usp4 | 22258 | HITS-CLIP | | 37986224 | Unspecified |
| mmu-miR-152-3p | MIMAT0000162 | Usp47 | 74996 | HITS-CLIP | 37986224\|25083871 | | Unspecified |
| mmu-miR-152-3p | MIMAT0000162 | Usp48 | 170707 | HITS-CLIP | 37986224\|25083871 | | Unspecified |
| mmu-miR-152-3p | MIMAT0000162 | Usp49 | 224836 | HITS-CLIP | | 25083871 | Unspecified |
| mmu-miR-152-3p | MIMAT0000162 | Usp6nl | 98910 | HITS-CLIP | | 37986224 | Unspecified |
| mmu-miR-152-3p | MIMAT0000162 | Usp7 | 252870 | HITS-CLIP | 37986224\|25083871 | | Unspecified |
| mmu-miR-152-3p | MIMAT0000162 | Usp8 | 84092 | HITS-CLIP | | 25083871 | Unspecified |
| mmu-miR-152-3p | MIMAT0000162 | Usp9x | 22284 | HITS-CLIP | 37986224\|25083871\|23597149 | | Unspecified |
| mmu-miR-152-3p | MIMAT0000162 | Utp15 | 105372 | HITS-CLIP | | 29167373 | Unspecified |
| mmu-miR-152-3p | MIMAT0000162 | Vamp3 | 22319 | HITS-CLIP | | 25083871 | Unspecified |
| mmu-miR-152-3p | MIMAT0000162 | Vamp4 | 53330 | HITS-CLIP | | 29167373 | Unspecified |
| mmu-miR-152-3p | MIMAT0000162 | Vasn | 246154 | HITS-CLIP | | 23597149 | Unspecified |
| mmu-miR-152-3p | MIMAT0000162 | Vav2 | 22325 | HITS-CLIP | | 23597149 | Unspecified |
| mmu-miR-152-3p | MIMAT0000162 | Vcam1 | 22329 | HITS-CLIP | 25083871\|23597149 | | Unspecified |
| mmu-miR-152-3p | MIMAT0000162 | Vcpip1 | 70675 | HITS-CLIP | 37986224\|29167373 | | Unspecified |
| mmu-miR-152-3p | MIMAT0000162 | Vdac2 | 22334 | HITS-CLIP | | 37986224 | Unspecified |
| mmu-miR-152-3p | MIMAT0000162 | Vezf1 | 22344 | HITS-CLIP | | 25083871 | Unspecified |
| mmu-miR-152-3p | MIMAT0000162 | Virma | 25962 | HITS-CLIP | | 25083871 | Unspecified |
| mmu-miR-152-3p | MIMAT0000162 | Vkorc1l1 | 69568 | HITS-CLIP | | 25083871 | Unspecified |
| mmu-miR-152-3p | MIMAT0000162 | Vps13b | 666173 | HITS-CLIP | | 25083871 | Unspecified |

| | | | | | | |
|---|---|---|---|---|---|---|
| mmu-miR-152-3p | MIMAT0000162 | Vps13c | 320528 | HITS-CLIP | 25083871 | Unspecified |
| mmu-miR-152-3p | MIMAT0000162 | Vps13d | 230895 | HITS-CLIP | 37986224\|23597149\|29167373 | Unspecified |
| mmu-miR-152-3p | MIMAT0000162 | Vps25 | 28084 | HITS-CLIP | 37986224 | Unspecified |
| mmu-miR-152-3p | MIMAT0000162 | Vps35 | 65114 | HITS-CLIP | 29167373 | Unspecified |
| mmu-miR-152-3p | MIMAT0000162 | Vps36 | 70160 | HITS-CLIP | 37986224 | Unspecified |
| mmu-miR-152-3p | MIMAT0000162 | Vps37a | 52348 | HITS-CLIP | 25083871 | Unspecified |
| mmu-miR-152-3p | MIMAT0000162 | Vps41 | 218035 | HITS-CLIP | 25083871\|23597149\|29167373 | Unspecified |
| mmu-miR-152-3p | MIMAT0000162 | Vps4b | 20479 | HITS-CLIP | 37986224\|25083871 | Unspecified |
| mmu-miR-152-3p | MIMAT0000162 | Vps54 | 245944 | HITS-CLIP | 23597149\|25083871\|29167373 | Unspecified |
| mmu-miR-152-3p | MIMAT0000162 | Vrk2 | 69922 | HITS-CLIP | 23597149 | Unspecified |
| mmu-miR-152-3p | MIMAT0000162 | Vtn | 22370 | HITS-CLIP | 23597149 | Unspecified |
| mmu-miR-152-3p | MIMAT0000162 | Vwa8 | 219189 | HITS-CLIP | 23597149 | Unspecified |
| mmu-miR-152-3p | MIMAT0000162 | Wac | 225131 | HITS-CLIP | 25083871 | Unspecified |
| mmu-miR-152-3p | MIMAT0000162 | Wapl | 23063 | HITS-CLIP | 25083871 | Unspecified |
| mmu-miR-152-3p | MIMAT0000162 | Wars2 | 70560 | HITS-CLIP | 25083871 | Unspecified |
| mmu-miR-152-3p | MIMAT0000162 | Washc4 | 319277 | HITS-CLIP | 25083871 | Unspecified |
| mmu-miR-152-3p | MIMAT0000162 | Wasl | 73178 | HITS-CLIP | 25083871 | Unspecified |
| mmu-miR-152-3p | MIMAT0000162 | Wbp1l | 226178 | HITS-CLIP | 25083871\|23597149 | Unspecified |
| mmu-miR-152-3p | MIMAT0000162 | Wdfy4 | 545030 | HITS-CLIP | 29167373 | Unspecified |
| mmu-miR-152-3p | MIMAT0000162 | Wdr1 | 22388 | HITS-CLIP | 23597149\|25083871\|29167373 | Unspecified |
| mmu-miR-152-3p | MIMAT0000162 | Wdr13 | 73447 | HITS-CLIP | 37986224 | Unspecified |
| mmu-miR-152-3p | MIMAT0000162 | Wdr37 | 207615 | HITS-CLIP | 37986224\|29167373 | Unspecified |
| mmu-miR-152-3p | MIMAT0000162 | Wdr44 | 72404 | HITS-CLIP | 37986224 | Unspecified |
| mmu-miR-152-3p | MIMAT0000162 | Wdr47 | 99512 | HITS-CLIP | 25083871\|29167373 | Unspecified |
| mmu-miR-152-3p | MIMAT0000162 | Wdr55 | 67936 | HITS-CLIP | 23597149 | Unspecified |
| mmu-miR-152-3p | MIMAT0000162 | Wee1 | 22390 | HITS-CLIP | 25083871 | Unspecified |
| mmu-miR-152-3p | MIMAT0000162 | Wfs1 | 22393 | HITS-CLIP | 25083871 | Unspecified |
| mmu-miR-152-3p | MIMAT0000162 | Wls | 68151 | HITS-CLIP | 25083871 | Unspecified |
| mmu-miR-152-3p | MIMAT0000162 | Wrn | 22427 | HITS-CLIP | 25083871 | Unspecified |
| mmu-miR-152-3p | MIMAT0000162 | Wrnip1 | 78903 | HITS-CLIP | 25083871 | Unspecified |
| mmu-miR-152-3p | MIMAT0000162 | Wsb2 | 59043 | HITS-CLIP | 25083871 | Unspecified |
| mmu-miR-152-3p | MIMAT0000162 | Wwc2 | 52357 | HITS-CLIP | 25083871 | Unspecified |
| mmu-miR-152-3p | MIMAT0000162 | Wwp1 | 107568 | HITS-CLIP | 37986224 | Unspecified |
| mmu-miR-152-3p | MIMAT0000162 | Wwtr1 | 97064 | HITS-CLIP | 37986224 | Unspecified |
| mmu-miR-152-3p | MIMAT0000162 | Xdh | 22436 | HITS-CLIP | 23597149 | Unspecified |
| mmu-miR-152-3p | MIMAT0000162 | Xiap | 11798 | HITS-CLIP | 25083871\|29167373 | Unspecified |
| mmu-miR-152-3p | MIMAT0000162 | Xirp1 | 22437 | HITS-CLIP | 37986224 | Unspecified |
| mmu-miR-152-3p | MIMAT0000162 | Xpo1 | 103573 | HITS-CLIP | 37986224 | Unspecified |
| mmu-miR-152-3p | MIMAT0000162 | Xpo4 | 57258 | HITS-CLIP | 37986224\|25083871 | Unspecified |

| | | | | | | |
|---|---|---|---|---|---|---|
| mmu-miR-152-3p | MIMAT0000162 | Xpo5 | 72322 | HITS-CLIP | 29167373 | Unspecified |
| mmu-miR-152-3p | MIMAT0000162 | Xrn2 | 24128 | HITS-CLIP | 25083871 | Unspecified |
| mmu-miR-152-3p | MIMAT0000162 | Yaf2 | 67057 | HITS-CLIP | 37986224 | Unspecified |
| mmu-miR-152-3p | MIMAT0000162 | Yap1 | 22601 | HITS-CLIP | 37986224 | Unspecified |
| mmu-miR-152-3p | MIMAT0000162 | Yes1 | 22612 | HITS-CLIP | 25083871\|21258322 | Unspecified |
| mmu-miR-152-3p | MIMAT0000162 | Yif1b | 77254 | HITS-CLIP | 37986224\|23597149 | Unspecified |
| mmu-miR-152-3p | MIMAT0000162 | Yipf3 | 28064 | HITS-CLIP | 25083871 | Unspecified |
| mmu-miR-152-3p | MIMAT0000162 | Yipf6 | 77929 | HITS-CLIP | 25083871\|23597149 | Unspecified |
| mmu-miR-152-3p | MIMAT0000162 | Yme1l1 | 27377 | HITS-CLIP | 23597149\|25083871 | Unspecified |
| mmu-miR-152-3p | MIMAT0000162 | Yod1 | 226418 | HITS-CLIP | 37986224\|25083871\|23597149\|29167 | Unspecified |
| mmu-miR-152-3p | MIMAT0000162 | Ypel3 | 66090 | HITS-CLIP | 37986224\|25083871 | Unspecified |
| mmu-miR-152-3p | MIMAT0000162 | Ypel5 | 383295 | HITS-CLIP | 25083871\|29167373 | Unspecified |
| mmu-miR-152-3p | MIMAT0000162 | Ythdc2 | 240255 | HITS-CLIP | 37986224 | Unspecified |
| mmu-miR-152-3p | MIMAT0000162 | Ythdf2 | 213541 | HITS-CLIP | 25083871\|23597149 | Unspecified |
| mmu-miR-152-3p | MIMAT0000162 | Ythdf3 | 229096 | HITS-CLIP | 25083871 | Unspecified |
| mmu-miR-152-3p | MIMAT0000162 | Ywhae | 22627 | HITS-CLIP | 25083871\|23597149 | Unspecified |
| mmu-miR-152-3p | MIMAT0000162 | Ywhaq | 22630 | HITS-CLIP | 25083871\|29167373 | Unspecified |
| mmu-miR-152-3p | MIMAT0000162 | Yy1 | 22632 | HITS-CLIP | 25083871 | Unspecified |
| mmu-miR-152-3p | MIMAT0000162 | Zadh2 | 225791 | HITS-CLIP | 37986224 | Unspecified |
| mmu-miR-152-3p | MIMAT0000162 | Zbed3 | 72114 | HITS-CLIP | 37986224\|23597149\|25083871 | Unspecified |
| mmu-miR-152-3p | MIMAT0000162 | Zbed4 | 223773 | HITS-CLIP | 25083871 | Unspecified |
| mmu-miR-152-3p | MIMAT0000162 | Zbtb1 | 268564 | HITS-CLIP | 37986224 | Unspecified |
| mmu-miR-152-3p | MIMAT0000162 | Zbtb10 | 229055 | HITS-CLIP | 25083871 | Unspecified |
| mmu-miR-152-3p | MIMAT0000162 | Zbtb16 | 235320 | HITS-CLIP | 23597149 | Unspecified |
| mmu-miR-152-3p | MIMAT0000162 | Zbtb18 | 30928 | HITS-CLIP | 37986224\|25083871 | Unspecified |
| mmu-miR-152-3p | MIMAT0000162 | Zbtb20 | 56490 | HITS-CLIP | 37986224\|25083871\|23597149 | Unspecified |
| mmu-miR-152-3p | MIMAT0000162 | Zbtb22 | 81630 | HITS-CLIP | 37986224 | Unspecified |
| mmu-miR-152-3p | MIMAT0000162 | Zbtb34 | 241311 | HITS-CLIP | 25083871 | Unspecified |
| mmu-miR-152-3p | MIMAT0000162 | Zbtb37 | 240869 | HITS-CLIP | 25083871 | Unspecified |
| mmu-miR-152-3p | MIMAT0000162 | Zbtb38 | 245007 | HITS-CLIP | 37986224 | Unspecified |
| mmu-miR-152-3p | MIMAT0000162 | Zbtb4 | 75580 | HITS-CLIP | 25083871 | Unspecified |
| mmu-miR-152-3p | MIMAT0000162 | Zbtb41 | 226470 | HITS-CLIP | 37986224 | Unspecified |
| mmu-miR-152-3p | MIMAT0000162 | Zbtb43 | 71834 | HITS-CLIP | 37986224 | Unspecified |
| mmu-miR-152-3p | MIMAT0000162 | Zbtb44 | 235132 | HITS-CLIP | 37986224\|25083871\|23597149 | Unspecified |
| mmu-miR-152-3p | MIMAT0000162 | Zbtb6 | 241322 | HITS-CLIP | 37986224 | Unspecified |
| mmu-miR-152-3p | MIMAT0000162 | Zbtb7a | 16969 | HITS-CLIP | 37986224\|25083871 | Unspecified |
| mmu-miR-152-3p | MIMAT0000162 | Zc3h12c | 244871 | HITS-CLIP | 37986224 | Unspecified |
| mmu-miR-152-3p | MIMAT0000162 | Zc3h13 | 67302 | HITS-CLIP | 37986224 | Unspecified |
| mmu-miR-152-3p | MIMAT0000162 | Zc3h18 | 76014 | HITS-CLIP | 37986224 | Unspecified |

| | | | | | | |
|---|---|---|---|---|---|---|
| mmu-miR-152-3p | MIMAT0000162 | Zc3h4 | 330474 | HITS-CLIP | 23597149 | Unspecified |
| mmu-miR-152-3p | MIMAT0000162 | Zc3h7a | 106205 | HITS-CLIP | 25083871 | Unspecified |
| mmu-miR-152-3p | MIMAT0000162 | Zcchc2 | 227449 | HITS-CLIP | 37986224\|25083871\|23597149 | Unspecified |
| mmu-miR-152-3p | MIMAT0000162 | Zcchc24 | 71918 | HITS-CLIP | 23597149 | Unspecified |
| mmu-miR-152-3p | MIMAT0000162 | Zdhhc1 | 70796 | HITS-CLIP | 25083871 | Unspecified |
| mmu-miR-152-3p | MIMAT0000162 | Zdhhc21 | 68268 | HITS-CLIP | 37986224 | Unspecified |
| mmu-miR-152-3p | MIMAT0000162 | Zdhhc3 | 69035 | HITS-CLIP | 23597149 | Unspecified |
| mmu-miR-152-3p | MIMAT0000162 | Zdhhc6 | 66980 | HITS-CLIP | 25083871 | Unspecified |
| mmu-miR-152-3p | MIMAT0000162 | Zdhhc8 | 27801 | HITS-CLIP | 37986224\|25083871 | Unspecified |
| mmu-miR-152-3p | MIMAT0000162 | Zdhhc9 | 208884 | HITS-CLIP | 37986224 | Unspecified |
| mmu-miR-152-3p | MIMAT0000162 | Zeb1 | 21417 | HITS-CLIP | 25083871 | Unspecified |
| mmu-miR-152-3p | MIMAT0000162 | Zeb2 | 24136 | HITS-CLIP | 37986224\|25083871\|23597149\|29167 | Unspecified |
| mmu-miR-152-3p | MIMAT0000162 | Zer1 | 227693 | HITS-CLIP | 25083871\|23597149 | Unspecified |
| mmu-miR-152-3p | MIMAT0000162 | Zfand4 | 67492 | HITS-CLIP | 25083871 | Unspecified |
| mmu-miR-152-3p | MIMAT0000162 | Zfc3h1 | 216345 | HITS-CLIP | 37986224\|25083871 | Unspecified |
| mmu-miR-152-3p | MIMAT0000162 | Zfhx3 | 11906 | HITS-CLIP | 37986224\|25083871 | Unspecified |
| mmu-miR-152-3p | MIMAT0000162 | Zfhx4 | 80892 | HITS-CLIP | 25083871 | Unspecified |
| mmu-miR-152-3p | MIMAT0000162 | Zfp106 | 20402 | HITS-CLIP | 25083871 | Unspecified |
| mmu-miR-152-3p | MIMAT0000162 | Zfp110 | 65020 | HITS-CLIP | 37986224\|25083871 | Unspecified |
| mmu-miR-152-3p | MIMAT0000162 | Zfp113 | 56314 | HITS-CLIP | 37986224 | Unspecified |
| mmu-miR-152-3p | MIMAT0000162 | Zfp12 | 231866 | HITS-CLIP | 37986224 | Unspecified |
| mmu-miR-152-3p | MIMAT0000162 | Zfp142 | 77264 | HITS-CLIP | 37986224 | Unspecified |
| mmu-miR-152-3p | MIMAT0000162 | Zfp148 | 22661 | HITS-CLIP | 37986224\|29167373 | Unspecified |
| mmu-miR-152-3p | MIMAT0000162 | Zfp236 | 329002 | HITS-CLIP | 25083871 | Unspecified |
| mmu-miR-152-3p | MIMAT0000162 | Zfp24 | 59057 | HITS-CLIP | 25083871 | Unspecified |
| mmu-miR-152-3p | MIMAT0000162 | Zfp26 | 22688 | HITS-CLIP | 23597149 | Unspecified |
| mmu-miR-152-3p | MIMAT0000162 | Zfp266 | 77519 | HITS-CLIP | 37986224 | Unspecified |
| mmu-miR-152-3p | MIMAT0000162 | Zfp280b | 64453 | HITS-CLIP | 31433990 | Unspecified |
| mmu-miR-152-3p | MIMAT0000162 | Zfp280c | 208968 | HITS-CLIP | 37986224\|25083871 | Unspecified |
| mmu-miR-152-3p | MIMAT0000162 | Zfp281 | 226442 | HITS-CLIP | 37986224\|25083871\|29167373 | Unspecified |
| mmu-miR-152-3p | MIMAT0000162 | Zfp292 | 30046 | HITS-CLIP | 37986224\|25083871 | Unspecified |
| mmu-miR-152-3p | MIMAT0000162 | Zfp319 | 79233 | HITS-CLIP | 37986224 | Unspecified |
| mmu-miR-152-3p | MIMAT0000162 | Zfp365 | 216049 | HITS-CLIP | 25083871 | Unspecified |
| mmu-miR-152-3p | MIMAT0000162 | Zfp369 | 170936 | HITS-CLIP | 37986224 | Unspecified |
| mmu-miR-152-3p | MIMAT0000162 | Zfp385a | 29813 | HITS-CLIP | 25083871 | Unspecified |
| mmu-miR-152-3p | MIMAT0000162 | Zfp386 | 56220 | HITS-CLIP | 37986224 | Unspecified |
| mmu-miR-152-3p | MIMAT0000162 | Zfp41 | 22701 | HITS-CLIP | 37986224 | Unspecified |
| mmu-miR-152-3p | MIMAT0000162 | Zfp426 | 235028 | HITS-CLIP | 23597149 | Unspecified |
| mmu-miR-152-3p | MIMAT0000162 | Zfp451 | 98403 | HITS-CLIP | 25083871 | Unspecified |

| | | | | | | |
|---|---|---|---|---|---|---|
| mmu-miR-152-3p | MIMAT0000162 | Zfp462 | 242466 | HITS-CLIP | 37986224 | Unspecified |
| mmu-miR-152-3p | MIMAT0000162 | Zfp467 | 68910 | HITS-CLIP | 23597149 | Unspecified |
| mmu-miR-152-3p | MIMAT0000162 | Zfp472 | 224691 | HITS-CLIP | 29167373 | Unspecified |
| mmu-miR-152-3p | MIMAT0000162 | Zfp516 | 329003 | HITS-CLIP | 25083871 | Unspecified |
| mmu-miR-152-3p | MIMAT0000162 | Zfp532 | 328977 | HITS-CLIP | 37986224\|25083871 | Unspecified |
| mmu-miR-152-3p | MIMAT0000162 | Zfp568 | 243905 | HITS-CLIP | 37986224\|25083871\|23597149 | Unspecified |
| mmu-miR-152-3p | MIMAT0000162 | Zfp606 | 67370 | HITS-CLIP | 37986224\|25083871\|29167373 | Unspecified |
| mmu-miR-152-3p | MIMAT0000162 | Zfp644 | 52397 | HITS-CLIP | 25083871 | Unspecified |
| mmu-miR-152-3p | MIMAT0000162 | Zfp654 | 72020 | HITS-CLIP | 37986224 | Unspecified |
| mmu-miR-152-3p | MIMAT0000162 | Zfp672 | 319475 | HITS-CLIP | 37986224 | Unspecified |
| mmu-miR-152-3p | MIMAT0000162 | Zfp710 | 209225 | HITS-CLIP | 29167373 | Unspecified |
| mmu-miR-152-3p | MIMAT0000162 | Zfp746 | 69228 | HITS-CLIP | 25083871 | Unspecified |
| mmu-miR-152-3p | MIMAT0000162 | Zfp800 | 627049 | HITS-CLIP | 23597149 | Unspecified |
| mmu-miR-152-3p | MIMAT0000162 | Zfp81 | 224694 | HITS-CLIP | 37986224 | Unspecified |
| mmu-miR-152-3p | MIMAT0000162 | Zfp827 | 622675 | HITS-CLIP | 37986224\|25083871 | Unspecified |
| mmu-miR-152-3p | MIMAT0000162 | Zfp865 | 319748 | HITS-CLIP | 23597149 | Unspecified |
| mmu-miR-152-3p | MIMAT0000162 | Zfp866 | 330788 | HITS-CLIP | 37986224 | Unspecified |
| mmu-miR-152-3p | MIMAT0000162 | Zfx | 22764 | HITS-CLIP | 29167373 | Unspecified |
| mmu-miR-152-3p | MIMAT0000162 | Zfyve21 | 68520 | HITS-CLIP | 23597149 | Unspecified |
| mmu-miR-152-3p | MIMAT0000162 | Zfyve26 | 211978 | HITS-CLIP | 37986224 | Unspecified |
| mmu-miR-152-3p | MIMAT0000162 | Zfyve9 | 230597 | HITS-CLIP | 37986224 | Unspecified |
| mmu-miR-152-3p | MIMAT0000162 | Zhx1 | 22770 | HITS-CLIP | 37986224\|25083871 | Unspecified |
| mmu-miR-152-3p | MIMAT0000162 | Zhx3 | 320799 | HITS-CLIP | 37986224 | Unspecified |
| mmu-miR-152-3p | MIMAT0000162 | Zkscan1 | 74570 | HITS-CLIP | 37986224\|25083871 | Unspecified |
| mmu-miR-152-3p | MIMAT0000162 | Zmat3 | 22401 | HITS-CLIP | 37986224\|25083871\|23597149 | Unspecified |
| mmu-miR-152-3p | MIMAT0000162 | Zmiz1 | 328365 | HITS-CLIP | 25083871\|29167373 | Unspecified |
| mmu-miR-152-3p | MIMAT0000162 | Zmpste24 | 230709 | HITS-CLIP | 25083871 | Unspecified |
| mmu-miR-152-3p | MIMAT0000162 | Zmynd8 | 228880 | HITS-CLIP | 25083871 | Unspecified |
| mmu-miR-152-3p | MIMAT0000162 | Znfx1 | 98999 | HITS-CLIP | 25083871\|29167373 | Unspecified |
| mmu-miR-152-3p | MIMAT0000162 | Znrf1 | 170737 | HITS-CLIP | 29167373 | Unspecified |
| mmu-miR-152-3p | MIMAT0000162 | Znrf2 | 387524 | HITS-CLIP | 25083871 | Unspecified |
| mmu-miR-152-3p | MIMAT0000162 | Zranb1 | 360216 | HITS-CLIP | 23597149 | Unspecified |
| mmu-miR-152-3p | MIMAT0000162 | Zrsr1 | 22183 | HITS-CLIP | 37986224 | Unspecified |
| mmu-miR-152-3p | MIMAT0000162 | Zswim4 | 212168 | HITS-CLIP | 37986224 | Unspecified |
| mmu-miR-152-3p | MIMAT0000162 | Zswim6 | 67263 | HITS-CLIP | 37986224 | Unspecified |
| mmu-miR-152-3p | MIMAT0000162 | Zswim8 | 268721 | HITS-CLIP | 37986224\|29167373 | Unspecified |
| mmu-miR-152-3p | MIMAT0000162 | Zup1 | 221302 | HITS-CLIP | 29167373 | Unspecified |
| mmu-miR-152-3p | MIMAT0000162 | Zwilch | 68014 | HITS-CLIP | 25083871 | Unspecified |
| mmu-miR-152-3p | MIMAT0000162 | Zyg11a | 230590 | HITS-CLIP | 23597149 | Unspecified |

| | | | | | | |
|---|---|---|---|---|---|---|
| mmu-miR-152-3p | MIMAT0000162 | Zzz3 | 108946 | HITS-CLIP | 37986224\|25083871 | Unspecified |
| mmu-miR-1946a | MIMAT0009412 | Sbno1 | 243272 | HITS-CLIP | 31433990 | Unspecified |
| mmu-miR-1249-3p | MIMAT0010560 | Acadm | 11364 | qCLASH | 31393953 | Unspecified |
| mmu-miR-1249-3p | MIMAT0010560 | Actr2 | 66713 | HITS-CLIP | 29167373 | Unspecified |
| mmu-miR-1249-3p | MIMAT0010560 | Anp32b | 67628 | qCLASH | 31393953 | Unspecified |
| mmu-miR-1249-3p | MIMAT0010560 | Anxa1 | 16952 | HITS-CLIP | 29167373 | Unspecified |
| mmu-miR-1249-3p | MIMAT0010560 | C1qa | 12259 | HITS-CLIP | 29167373 | Unspecified |
| mmu-miR-1249-3p | MIMAT0010560 | Ccdc127 | 67433 | HITS-CLIP | 29167373 | Unspecified |
| mmu-miR-1249-3p | MIMAT0010560 | Cltc | 67300 | HITS-CLIP | 29167373 | Unspecified |
| mmu-miR-1249-3p | MIMAT0010560 | Cpsf3 | 54451 | qCLASH | 31393953 | Unspecified |
| mmu-miR-1249-3p | MIMAT0010560 | Ddx18 | 66942 | qCLASH | 31393953 | Unspecified |
| mmu-miR-1249-3p | MIMAT0010560 | Fau | 14109 | qCLASH | 31393953 | Unspecified |
| mmu-miR-1249-3p | MIMAT0010560 | Fbxw11 | 103583 | HITS-CLIP | 29167373 | Unspecified |
| mmu-miR-1249-3p | MIMAT0010560 | Grk1 | 24013 | qCLASH | 31393953 | Unspecified |
| mmu-miR-1249-3p | MIMAT0010560 | Hmgn1 | 15312 | qCLASH | 31393953 | Unspecified |
| mmu-miR-1249-3p | MIMAT0010560 | Hnrnpa0 | 77134 | qCLASH | 31393953 | Unspecified |
| mmu-miR-1249-3p | MIMAT0010560 | Hnrnpul2 | 68693 | qCLASH | 31393953 | Unspecified |
| mmu-miR-1249-3p | MIMAT0010560 | Iqgap1 | 29875 | HITS-CLIP | 29167373 | Unspecified |
| mmu-miR-1249-3p | MIMAT0010560 | Kmt2d | 381022 | qCLASH | 31393953 | Unspecified |
| mmu-miR-1249-3p | MIMAT0010560 | Laptm4a | 17775 | qCLASH | 31393953 | Unspecified |
| mmu-miR-1249-3p | MIMAT0010560 | Lcp1 | 18826 | HITS-CLIP | 29167373 | Unspecified |
| mmu-miR-1249-3p | MIMAT0010560 | Mbd2 | 17191 | qCLASH | 31393953 | Unspecified |
| mmu-miR-1249-3p | MIMAT0010560 | Mtrex | 23517 | qCLASH | 31393953 | Unspecified |
| mmu-miR-1249-3p | MIMAT0010560 | Pafah1b1 | 18472 | HITS-CLIP | 29167373 | Unspecified |
| mmu-miR-1249-3p | MIMAT0010560 | Prkaa1 | 105787 | qCLASH | 31393953 | Unspecified |
| mmu-miR-1249-3p | MIMAT0010560 | Psip1 | 101739 | qCLASH | 31393953 | Unspecified |
| mmu-miR-1249-3p | MIMAT0010560 | Rpl22 | 19934 | qCLASH | 31393953 | Unspecified |
| mmu-miR-1249-3p | MIMAT0010560 | Rps16 | 20055 | qCLASH | 31393953 | Unspecified |
| mmu-miR-1249-3p | MIMAT0010560 | Rps25 | 75617 | qCLASH | 31393953 | Unspecified |
| mmu-miR-1249-3p | MIMAT0010560 | Rps6 | 20104 | qCLASH | 31393953 | Unspecified |
| mmu-miR-1249-3p | MIMAT0010560 | Sez6l | 56747 | qCLASH | 31393953 | Unspecified |
| mmu-miR-1249-3p | MIMAT0010560 | Srsf6 | 67996 | HITS-CLIP | 29167373 | Unspecified |
| mmu-miR-1249-3p | MIMAT0010560 | Syncrip | 56403 | HITS-CLIP | 29167373 | Unspecified |
| mmu-miR-1249-3p | MIMAT0010560 | Tmem245 | 242474 | qCLASH | 31393953 | Unspecified |
| mmu-miR-1249-3p | MIMAT0010560 | Tmem38b | 52076 | qCLASH | 31393953 | Unspecified |
| mmu-miR-1249-3p | MIMAT0010560 | Tnpo1 | 238799 | qCLASH | 31393953 | Unspecified |
| mmu-miR-1249-3p | MIMAT0010560 | Tomm20 | 67952 | qCLASH | 31393953 | Unspecified |
| mmu-miR-1249-3p | MIMAT0010560 | Uri1 | 19777 | HITS-CLIP | 29167373 | Unspecified |
| mmu-miR-1249-3p | MIMAT0010560 | Wwp1 | 107568 | HITS-CLIP | 29167373 | Unspecified |

| | | | | | | |
|---|---|---|---|---|---|---|
| mmu-let-7i-5p | MIMAT0000122 | 0610010F05Rik | 71675 | HITS-CLIP | 19536157 | Unspecified |
| mmu-let-7i-5p | MIMAT0000122 | 0610010K14Rik | 104457 | HITS-CLIP | 23142080 | Unspecified |
| mmu-let-7i-5p | MIMAT0000122 | 0610030E20Rik | 68364 | HITS-CLIP | 23597149\|23142080 | Unspecified |
| mmu-let-7i-5p | MIMAT0000122 | 0610040J01Rik | 76261 | HITS-CLIP | 23597149 | Unspecified |
| mmu-let-7i-5p | MIMAT0000122 | 1110059E24Rik | 66206 | HITS-CLIP | 23142080 | Unspecified |
| mmu-let-7i-5p | MIMAT0000122 | 1110059G10Rik | 66202 | HITS-CLIP | 23142080 | Unspecified |
| mmu-let-7i-5p | MIMAT0000122 | 1190002N15Rik | 68861 | HITS-CLIP | 23142080 | Unspecified |
| mmu-let-7i-5p | MIMAT0000122 | 1300017J02Rik | 71775 | HITS-CLIP | 23597149 | Unspecified |
| mmu-let-7i-5p | MIMAT0000122 | 1600012H06Rik | 67912 | HITS-CLIP | 37986224\|23142080\|19536157 | Unspecified |
| mmu-let-7i-5p | MIMAT0000122 | 1600014C10Rik | 72244 | HITS-CLIP | 37986224\|23142080\|23597149 | Unspecified |
| mmu-let-7i-5p | MIMAT0000122 | 1700017B05Rik | 74211 | HITS-CLIP | 37986224\|23597149\|23142080 | Unspecified |
| mmu-let-7i-5p | MIMAT0000122 | 1700066B19Rik | 73449 | HITS-CLIP | 23142080 | Unspecified |
| mmu-let-7i-5p | MIMAT0000122 | 1700066M21Rik | 73467 | HITS-CLIP | 19536157 | Unspecified |
| mmu-let-7i-5p | MIMAT0000122 | 1810013L24Rik | 69053 | HITS-CLIP | 37986224\|23597149\|23142080\|29167 | Unspecified |
| mmu-let-7i-5p | MIMAT0000122 | 1810037I17Rik | 67704 | HITS-CLIP | 37986224 | Unspecified |
| mmu-let-7i-5p | MIMAT0000122 | 1810055G02Rik | 72056 | HITS-CLIP | 23142080\|23597149 | Unspecified |
| mmu-let-7i-5p | MIMAT0000122 | 2010003K11Rik | 69861 | HITS-CLIP | 23597149 | Unspecified |
| mmu-let-7i-5p | MIMAT0000122 | 2310022A10Rik | 66367 | HITS-CLIP | 23142080 | Unspecified |
| mmu-let-7i-5p | MIMAT0000122 | 2310022B05Rik | 69551 | HITS-CLIP | 23597149\|19536157\|23142080 | Unspecified |
| mmu-let-7i-5p | MIMAT0000122 | 2410015M20Rik | 224904 | HITS-CLIP | 23597149 | Unspecified |
| mmu-let-7i-5p | MIMAT0000122 | 2510009E07Rik | 72190 | HITS-CLIP | 23597149\|29167373 | Unspecified |
| mmu-let-7i-5p | MIMAT0000122 | 2510039O18Rik | 77034 | HITS-CLIP | 37986224\|23597149\|19536157\|23142 | Unspecified |
| mmu-let-7i-5p | MIMAT0000122 | 2610002M06Rik | 67028 | HITS-CLIP | 23142080 | Unspecified |
| mmu-let-7i-5p | MIMAT0000122 | 2610008E11Rik | 72128 | HITS-CLIP | 37986224 | Unspecified |
| mmu-let-7i-5p | MIMAT0000122 | 2610021A01Rik | 668572 | HITS-CLIP | 37986224\|19536157 | Unspecified |
| mmu-let-7i-5p | MIMAT0000122 | 2610507B11Rik | 72503 | HITS-CLIP | 37986224\|23597149\|19536157\|23142 | Unspecified |
| mmu-let-7i-5p | MIMAT0000122 | 2700049A03Rik | 76967 | HITS-CLIP | 23142080\|19536157 | Unspecified |
| mmu-let-7i-5p | MIMAT0000122 | 2810004N23Rik | 66523 | HITS-CLIP | 19536157 | Unspecified |
| mmu-let-7i-5p | MIMAT0000122 | 2810459M11Rik | 72792 | HITS-CLIP | 37986224\|23597149 | Unspecified |
| mmu-let-7i-5p | MIMAT0000122 | 2900026A02Rik | 243219 | HITS-CLIP | 23597149\|29167373 | Unspecified |
| mmu-let-7i-5p | MIMAT0000122 | 4833420G17Rik | 67392 | HITS-CLIP | 23142080 | Unspecified |
| mmu-let-7i-5p | MIMAT0000122 | 4921524J17Rik | 66714 | HITS-CLIP | 23142080 | Unspecified |
| mmu-let-7i-5p | MIMAT0000122 | 4930402H24Rik | 228602 | HITS-CLIP | 37986224\|23597149\|19536157\|23142 | Unspecified |
| mmu-let-7i-5p | MIMAT0000122 | 4930430F08Rik | 68281 | HITS-CLIP | 23142080 | Unspecified |
| mmu-let-7i-5p | MIMAT0000122 | 4930453N24Rik | 67609 | HITS-CLIP | 23142080 | Unspecified |
| mmu-let-7i-5p | MIMAT0000122 | 4930503L19Rik | 269033 | HITS-CLIP | 23142080 | Unspecified |
| mmu-let-7i-5p | MIMAT0000122 | 4930523C07Rik | 67647 | HITS-CLIP | 37986224\|23142080 | Unspecified |
| mmu-let-7i-5p | MIMAT0000122 | 4931406C07Rik | 70984 | HITS-CLIP | 23142080\|23597149 | Unspecified |
| mmu-let-7i-5p | MIMAT0000122 | 4931406P16Rik | 233103 | HITS-CLIP | 37986224\|23142080 | Unspecified |

| | | | | | | |
|---|---|---|---|---|---|---|
| mmu-let-7i-5p | MIMAT0000122 | 4931428F04Rik | 74356 | HITS-CLIP | 19536157 | Unspecified |
| mmu-let-7i-5p | MIMAT0000122 | 4932438A13Rik | 229227 | HITS-CLIP | 23597149\|19536157\|23142080\|29167 | Unspecified |
| mmu-let-7i-5p | MIMAT0000122 | 5031439G07Rik | 223739 | HITS-CLIP | 29167373 | Unspecified |
| mmu-let-7i-5p | MIMAT0000122 | 6030458C11Rik | 77877 | HITS-CLIP | 23142080 | Unspecified |
| mmu-let-7i-5p | MIMAT0000122 | 6430548M08Rik | 234797 | HITS-CLIP | 23597149 | Unspecified |
| mmu-let-7i-5p | MIMAT0000122 | 6430571L13Rik | 235599 | HITS-CLIP | 23142080 | Unspecified |
| mmu-let-7i-5p | MIMAT0000122 | 8030462N17Rik | 212163 | HITS-CLIP | 23142080 | Unspecified |
| mmu-let-7i-5p | MIMAT0000122 | 9430015G10Rik | 230996 | HITS-CLIP | 23142080\|23597149 | Unspecified |
| mmu-let-7i-5p | MIMAT0000122 | 9930012K11Rik | 268759 | HITS-CLIP | 29167373\|23142080 | Unspecified |
| mmu-let-7i-5p | MIMAT0000122 | 9930021J03Rik | 240613 | HITS-CLIP | 23142080\|23597149 | Unspecified |
| mmu-let-7i-5p | MIMAT0000122 | 9930111J21Rik1 | 667214 | HITS-CLIP | 29167373 | Unspecified |
| mmu-let-7i-5p | MIMAT0000122 | 9930111J21Rik2 | 245240 | HITS-CLIP | 29167373 | Unspecified |
| mmu-let-7i-5p | MIMAT0000122 | A130010J15Rik | 319266 | HITS-CLIP | 23142080 | Unspecified |
| mmu-let-7i-5p | MIMAT0000122 | A1cf | 69865 | HITS-CLIP | 23597149 | Unspecified |
| mmu-let-7i-5p | MIMAT0000122 | A430033K04Rik | 243308 | HITS-CLIP | 23142080 | Unspecified |
| mmu-let-7i-5p | MIMAT0000122 | A630001G21Rik | 319997 | HITS-CLIP | 23142080 | Unspecified |
| mmu-let-7i-5p | MIMAT0000122 | AB124611 | 382062 | HITS-CLIP | 29167373 | Unspecified |
| mmu-let-7i-5p | MIMAT0000122 | AC115631.1 | | HITS-CLIP | 23142080 | Unspecified |
| mmu-let-7i-5p | MIMAT0000122 | AC170998.1 | | HITS-CLIP | 23142080 | Unspecified |
| mmu-let-7i-5p | MIMAT0000122 | AI464131 | 329828 | HITS-CLIP | 23597149 | Unspecified |
| mmu-let-7i-5p | MIMAT0000122 | AI593442 | 330941 | HITS-CLIP | 19536157 | Unspecified |
| mmu-let-7i-5p | MIMAT0000122 | AI597479 | 98404 | HITS-CLIP | 23142080 | Unspecified |
| mmu-let-7i-5p | MIMAT0000122 | AI837181 | 107242 | HITS-CLIP | 23142080 | Unspecified |
| mmu-let-7i-5p | MIMAT0000122 | AI987944 | 233168 | HITS-CLIP | 23142080 | Unspecified |
| mmu-let-7i-5p | MIMAT0000122 | AU021092 | 239691 | HITS-CLIP | 23597149 | Unspecified |
| mmu-let-7i-5p | MIMAT0000122 | AU022252 | 230696 | HITS-CLIP | 37986224\|23142080\|23597149 | Unspecified |
| mmu-let-7i-5p | MIMAT0000122 | AU041133 | 216177 | HITS-CLIP | 37986224 | Unspecified |
| mmu-let-7i-5p | MIMAT0000122 | AW549877 | 106064 | HITS-CLIP | 37986224\|19536157\|23142080 | Unspecified |
| mmu-let-7i-5p | MIMAT0000122 | Aaas | 223921 | HITS-CLIP | 23142080 | Unspecified |
| mmu-let-7i-5p | MIMAT0000122 | Aacs | 78894 | HITS-CLIP | 23597149 | Unspecified |
| mmu-let-7i-5p | MIMAT0000122 | Aadac | 67758 | HITS-CLIP | 23597149 | Unspecified |
| mmu-let-7i-5p | MIMAT0000122 | Aadat | 23923 | HITS-CLIP | 23597149 | Unspecified |
| mmu-let-7i-5p | MIMAT0000122 | Aagab | 66939 | HITS-CLIP | 29167373 | Unspecified |
| mmu-let-7i-5p | MIMAT0000122 | Aak1 | 269774 | HITS-CLIP | 23597149\|19536157\|23142080\|29167 | Unspecified |
| mmu-let-7i-5p | MIMAT0000122 | Aar2 | 68295 | HITS-CLIP | 23597149\|23142080 | Unspecified |
| mmu-let-7i-5p | MIMAT0000122 | Aars | 234734 | HITS-CLIP | 37986224\|23597149\|23142080 | Unspecified |
| mmu-let-7i-5p | MIMAT0000122 | Aass | 30956 | HITS-CLIP | 23597149 | Unspecified |
| mmu-let-7i-5p | MIMAT0000122 | Aatf | 56321 | HITS-CLIP | 30971389 | Unspecified |
| mmu-let-7i-5p | MIMAT0000122 | Aatk | 11302 | HITS-CLIP | 23597149 | Unspecified |

| | | | | | | |
|---|---|---|---|---|---|---|
| mmu-let-7i-5p | MIMAT0000122 | Abat | 268860 | HITS-CLIP | 23597149 | Unspecified |
| mmu-let-7i-5p | MIMAT0000122 | Abca1 | 11303 | HITS-CLIP | 37986224\|23597149\|29167373\|23142 | Unspecified |
| mmu-let-7i-5p | MIMAT0000122 | Abca2 | 11305 | HITS-CLIP | 37986224\|23597149\|23142080 | Unspecified |
| mmu-let-7i-5p | MIMAT0000122 | Abca3 | 27410 | HITS-CLIP | 37986224\|23597149\|23142080 | Unspecified |
| mmu-let-7i-5p | MIMAT0000122 | Abca6 | 76184 | HITS-CLIP | 23597149 | Unspecified |
| mmu-let-7i-5p | MIMAT0000122 | Abca7 | 27403 | HITS-CLIP | 23142080 | Unspecified |
| mmu-let-7i-5p | MIMAT0000122 | Abcb10 | 56199 | HITS-CLIP | 37986224\|23597149\|23142080 | Unspecified |
| mmu-let-7i-5p | MIMAT0000122 | Abcb11 | 27413 | HITS-CLIP | 37986224\|23597149 | Unspecified |
| mmu-let-7i-5p | MIMAT0000122 | Abcb4 | 18670 | HITS-CLIP | 37986224\|23597149 | Unspecified |
| mmu-let-7i-5p | MIMAT0000122 | Abcb7 | 11306 | HITS-CLIP | 37986224\|23597149\|23142080 | Unspecified |
| mmu-let-7i-5p | MIMAT0000122 | Abcb9 | 56325 | HITS-CLIP | 37986224\|23142080 | Unspecified |
| mmu-let-7i-5p | MIMAT0000122 | Abcc1 | 17250 | HITS-CLIP | 29167373 | Unspecified |
| mmu-let-7i-5p | MIMAT0000122 | Abcc10 | 89845 | HITS-CLIP | 23142080 | Unspecified |
| mmu-let-7i-5p | MIMAT0000122 | Abcc2 | 12780 | HITS-CLIP | 23597149 | Unspecified |
| mmu-let-7i-5p | MIMAT0000122 | Abcc5 | 27416 | HITS-CLIP | 37986224\|19536157\|23597149\|23142 | Unspecified |
| mmu-let-7i-5p | MIMAT0000122 | Abcc6 | 27421 | HITS-CLIP | 37986224\|23597149 | Unspecified |
| mmu-let-7i-5p | MIMAT0000122 | Abcc8 | 20927 | HITS-CLIP | 37986224\|19536157 | Unspecified |
| mmu-let-7i-5p | MIMAT0000122 | Abcd1 | 11666 | HITS-CLIP | 37986224\|23597149\|23142080 | Unspecified |
| mmu-let-7i-5p | MIMAT0000122 | Abcd3 | 19299 | HITS-CLIP | 23597149 | Unspecified |
| mmu-let-7i-5p | MIMAT0000122 | Abcd4 | 19300 | HITS-CLIP | 23142080\|23597149 | Unspecified |
| mmu-let-7i-5p | MIMAT0000122 | Abce1 | 24015 | HITS-CLIP | 37986224\|23142080 | Unspecified |
| mmu-let-7i-5p | MIMAT0000122 | Abcf1 | 224742 | HITS-CLIP | 23142080 | Unspecified |
| mmu-let-7i-5p | MIMAT0000122 | Abcf3 | 27406 | HITS-CLIP | 23597149 | Unspecified |
| mmu-let-7i-5p | MIMAT0000122 | Abcg1 | 11307 | HITS-CLIP | 37986224\|19536157\|23142080 | Unspecified |
| mmu-let-7i-5p | MIMAT0000122 | Abcg2 | 26357 | HITS-CLIP | 23597149\|19536157 | Unspecified |
| mmu-let-7i-5p | MIMAT0000122 | Abcg3 | 27405 | HITS-CLIP | 23142080 | Unspecified |
| mmu-let-7i-5p | MIMAT0000122 | Abcg8 | 67470 | HITS-CLIP | 37986224\|23597149 | Unspecified |
| mmu-let-7i-5p | MIMAT0000122 | Abhd12 | 76192 | HITS-CLIP | 37986224\|23597149\|19536157 | Unspecified |
| mmu-let-7i-5p | MIMAT0000122 | Abhd13 | 68904 | HITS-CLIP | 23142080 | Unspecified |
| mmu-let-7i-5p | MIMAT0000122 | Abhd15 | 67477 | HITS-CLIP | 23597149 | Unspecified |
| mmu-let-7i-5p | MIMAT0000122 | Abhd16a | 193742 | HITS-CLIP | 23142080 | Unspecified |
| mmu-let-7i-5p | MIMAT0000122 | Abhd17b | 226016 | HITS-CLIP | 23142080 | Unspecified |
| mmu-let-7i-5p | MIMAT0000122 | Abhd17c | 70178 | HITS-CLIP | 23597149\|19536157\|23142080 | Unspecified |
| mmu-let-7i-5p | MIMAT0000122 | Abhd18 | 269423 | HITS-CLIP | 23142080 | Unspecified |
| mmu-let-7i-5p | MIMAT0000122 | Abhd2 | 54608 | HITS-CLIP | 37986224\|23597149\|19536157 | Unspecified |
| mmu-let-7i-5p | MIMAT0000122 | Abhd3 | 106861 | HITS-CLIP | 23597149 | Unspecified |
| mmu-let-7i-5p | MIMAT0000122 | Abhd5 | 67469 | HITS-CLIP | 23597149 | Unspecified |
| mmu-let-7i-5p | MIMAT0000122 | Abi1 | 11308 | HITS-CLIP | 23142080\|29167373 | Unspecified |
| mmu-let-7i-5p | MIMAT0000122 | Abi3 | 66610 | HITS-CLIP | 23142080 | Unspecified |

| | | | | | | |
|---|---|---|---|---|---|---|
| mmu-let-7i-5p | MIMAT0000122 | Abl1 | 11350 | HITS-CLIP | 23142080\|29167373 | Unspecified |
| mmu-let-7i-5p | MIMAT0000122 | Abl2 | 11352 | HITS-CLIP | 37986224\|23142080\|23597149\|29167 | Unspecified |
| mmu-let-7i-5p | MIMAT0000122 | Ablim1 | 226251 | HITS-CLIP | 37986224\|23597149\|19536157\|23142 | Unspecified |
| mmu-let-7i-5p | MIMAT0000122 | Abr | 109934 | HITS-CLIP | 19536157\|23142080\|29167373 | Unspecified |
| mmu-let-7i-5p | MIMAT0000122 | Abtb1 | 80283 | HITS-CLIP | 23142080 | Unspecified |
| mmu-let-7i-5p | MIMAT0000122 | Abtb2 | 99382 | HITS-CLIP | 23597149\|23142080 | Unspecified |
| mmu-let-7i-5p | MIMAT0000122 | Acaa1b | 235674 | HITS-CLIP | 23597149 | Unspecified |
| mmu-let-7i-5p | MIMAT0000122 | Acaa2 | 52538 | HITS-CLIP | 23597149 | Unspecified |
| mmu-let-7i-5p | MIMAT0000122 | Acacb | 100705 | HITS-CLIP | 37986224\|23597149 | Unspecified |
| mmu-let-7i-5p | MIMAT0000122 | Acad8 | 66948 | HITS-CLIP | 23597149\|23142080 | Unspecified |
| mmu-let-7i-5p | MIMAT0000122 | Acad9 | 229211 | HITS-CLIP | 23597149 | Unspecified |
| mmu-let-7i-5p | MIMAT0000122 | Acadl | 11363 | HITS-CLIP | 23597149\|23142080 | Unspecified |
| mmu-let-7i-5p | MIMAT0000122 | Acadm | 11364 | HITS-CLIP | 23597149 | Unspecified |
| mmu-let-7i-5p | MIMAT0000122 | Acap1 | 216859 | HITS-CLIP | 23142080 | Unspecified |
| mmu-let-7i-5p | MIMAT0000122 | Acap2 | 78618 | HITS-CLIP | 23597149\|23142080\|29167373 | Unspecified |
| mmu-let-7i-5p | MIMAT0000122 | Acat1 | 110446 | HITS-CLIP | 37986224\|23597149\|23142080 | Unspecified |
| mmu-let-7i-5p | MIMAT0000122 | Acbd3 | 170760 | HITS-CLIP | 23142080\|29167373 | Unspecified |
| mmu-let-7i-5p | MIMAT0000122 | Acbd5 | 74159 | HITS-CLIP | 37986224\|23142080\|23597149 | Unspecified |
| mmu-let-7i-5p | MIMAT0000122 | Acer2 | 230379 | HITS-CLIP | 37986224\|23597149\|23142080 | Unspecified |
| mmu-let-7i-5p | MIMAT0000122 | Acer3 | 66190 | HITS-CLIP | 37986224\|23597149\|19536157\|23142 | Unspecified |
| mmu-let-7i-5p | MIMAT0000122 | Acin1 | 56215 | HITS-CLIP | 23597149\|23142080 | Unspecified |
| mmu-let-7i-5p | MIMAT0000122 | Ackr4 | 252837 | HITS-CLIP | 23597149 | Unspecified |
| mmu-let-7i-5p | MIMAT0000122 | Acly | 104112 | HITS-CLIP | 23597149\|23142080\|26203562 | Unspecified |
| mmu-let-7i-5p | MIMAT0000122 | Acnat1 | 230161 | HITS-CLIP | 23597149 | Unspecified |
| mmu-let-7i-5p | MIMAT0000122 | Acnat2 | 209186 | HITS-CLIP | 23597149 | Unspecified |
| mmu-let-7i-5p | MIMAT0000122 | Aco1 | 11428 | HITS-CLIP | 23597149\|23142080 | Unspecified |
| mmu-let-7i-5p | MIMAT0000122 | Aco2 | 11429 | HITS-CLIP | 37986224\|23597149 | Unspecified |
| mmu-let-7i-5p | MIMAT0000122 | Acot11 | 329910 | HITS-CLIP | 37986224\|19536157\|23142080 | Unspecified |
| mmu-let-7i-5p | MIMAT0000122 | Acot13 | 66834 | HITS-CLIP | 23142080\|29167373 | Unspecified |
| mmu-let-7i-5p | MIMAT0000122 | Acot7 | 70025 | HITS-CLIP | 19536157\|29167373 | Unspecified |
| mmu-let-7i-5p | MIMAT0000122 | Acox1 | 11430 | HITS-CLIP | 23597149\|29167373 | Unspecified |
| mmu-let-7i-5p | MIMAT0000122 | Acox2 | 93732 | HITS-CLIP | 23597149 | Unspecified |
| mmu-let-7i-5p | MIMAT0000122 | Acp2 | 11432 | HITS-CLIP | 23597149\|19536157\|29167373 | Unspecified |
| mmu-let-7i-5p | MIMAT0000122 | Acp5 | 11433 | HITS-CLIP | 23142080\|23597149 | Unspecified |
| mmu-let-7i-5p | MIMAT0000122 | Acp6 | 66659 | HITS-CLIP | 23597149\|23142080 | Unspecified |
| mmu-let-7i-5p | MIMAT0000122 | Acsf2 | 264895 | HITS-CLIP | 23597149 | Unspecified |
| mmu-let-7i-5p | MIMAT0000122 | Acsl1 | 14081 | HITS-CLIP | 37986224\|23597149\|29167373 | Unspecified |
| mmu-let-7i-5p | MIMAT0000122 | Acsl3 | 74205 | HITS-CLIP | 31433990\|23142080 | Unspecified |
| mmu-let-7i-5p | MIMAT0000122 | Acsl4 | 50790 | HITS-CLIP | 23597149\|23142080\|29167373 | Unspecified |

| | | | | | | |
|---|---|---|---|---|---|---|
| mmu-let-7i-5p | MIMAT0000122 | Acsl5 | 433256 | HITS-CLIP | 23597149 | Unspecified |
| mmu-let-7i-5p | MIMAT0000122 | Acsm1 | 117147 | HITS-CLIP | 23597149 | Unspecified |
| mmu-let-7i-5p | MIMAT0000122 | Acsm5 | 272428 | HITS-CLIP | 37986224 | Unspecified |
| mmu-let-7i-5p | MIMAT0000122 | Acss1 | 68738 | HITS-CLIP | 37986224\|23142080 | Unspecified |
| mmu-let-7i-5p | MIMAT0000122 | Acta2 | 11475 | HITS-CLIP | 37986224 | Unspecified |
| mmu-let-7i-5p | MIMAT0000122 | Actb | 11461 | HITS-CLIP | 23597149\|19536157\|31433990\|23142 | Unspecified |
| mmu-let-7i-5p | MIMAT0000122 | Actg2 | 11468 | HITS-CLIP | 23142080 | Unspecified |
| mmu-let-7i-5p | MIMAT0000122 | Actl6a | 56456 | HITS-CLIP | 23142080 | Unspecified |
| mmu-let-7i-5p | MIMAT0000122 | Actn1 | 109711 | HITS-CLIP | 23597149\|29167373 | Unspecified |
| mmu-let-7i-5p | MIMAT0000122 | Actn4 | 60595 | HITS-CLIP | 23597149\|23142080 | Unspecified |
| mmu-let-7i-5p | MIMAT0000122 | Actr10 | 56444 | HITS-CLIP | 23142080 | Unspecified |
| mmu-let-7i-5p | MIMAT0000122 | Actr1a | 54130 | qCLASH//HITS-CLIP | 31393953\|19536157\|23142080 | Unspecified |
| mmu-let-7i-5p | MIMAT0000122 | Actr1b | 226977 | HITS-CLIP | 23142080 | Unspecified |
| mmu-let-7i-5p | MIMAT0000122 | Actr2 | 66713 | HITS-CLIP | 23142080\|29167373 | Unspecified |
| mmu-let-7i-5p | MIMAT0000122 | Actr3 | 74117 | HITS-CLIP | 23142080\|29167373 | Unspecified |
| mmu-let-7i-5p | MIMAT0000122 | Actr6 | 67019 | HITS-CLIP | 23597149\|23142080 | Unspecified |
| mmu-let-7i-5p | MIMAT0000122 | Actr8 | 56249 | HITS-CLIP | 23142080 | Unspecified |
| mmu-let-7i-5p | MIMAT0000122 | Acvr1 | 11477 | HITS-CLIP | 29167373 | Unspecified |
| mmu-let-7i-5p | MIMAT0000122 | Acvr1b | 11479 | HITS-CLIP//qCLASH | 37986224\|23597149\|31393953\|1953( | Unspecified |
| mmu-let-7i-5p | MIMAT0000122 | Acvr1c | 269275 | HITS-CLIP | 23142080 | Unspecified |
| mmu-let-7i-5p | MIMAT0000122 | Acvr2a | 11480 | HITS-CLIP | 19536157\|23597149\|23142080 | Unspecified |
| mmu-let-7i-5p | MIMAT0000122 | Acvr2b | 11481 | HITS-CLIP | 23142080 | Unspecified |
| mmu-let-7i-5p | MIMAT0000122 | Acy3 | 71670 | HITS-CLIP | 23597149 | Unspecified |
| mmu-let-7i-5p | MIMAT0000122 | Adam10 | 11487 | HITS-CLIP | 37986224\|23597149\|19536157\|23142 | Unspecified |
| mmu-let-7i-5p | MIMAT0000122 | Adam15 | 11490 | HITS-CLIP | 23142080\|29167373 | Unspecified |
| mmu-let-7i-5p | MIMAT0000122 | Adam17 | 11491 | HITS-CLIP | 23142080\|29167373 | Unspecified |
| mmu-let-7i-5p | MIMAT0000122 | Adam19 | 11492 | HITS-CLIP | 23142080 | Unspecified |
| mmu-let-7i-5p | MIMAT0000122 | Adam1a | 280668 | HITS-CLIP | 23142080\|23597149 | Unspecified |
| mmu-let-7i-5p | MIMAT0000122 | Adam8 | 11501 | HITS-CLIP | 29167373 | Unspecified |
| mmu-let-7i-5p | MIMAT0000122 | Adam9 | 11502 | HITS-CLIP | 29167373\|26203562 | Unspecified |
| mmu-let-7i-5p | MIMAT0000122 | Adamts1 | 11504 | HITS-CLIP | 26203562 | Unspecified |
| mmu-let-7i-5p | MIMAT0000122 | Adamts15 | 235130 | HITS-CLIP | 37986224\|19536157 | Unspecified |
| mmu-let-7i-5p | MIMAT0000122 | Adamts6 | 108154 | HITS-CLIP | 23142080 | Unspecified |
| mmu-let-7i-5p | MIMAT0000122 | Adamtsl3 | 269959 | HITS-CLIP | 23142080 | Unspecified |
| mmu-let-7i-5p | MIMAT0000122 | Adap2 | 216991 | HITS-CLIP | 23597149\|19536157\|29167373 | Unspecified |
| mmu-let-7i-5p | MIMAT0000122 | Adar | 56417 | HITS-CLIP | 23597149\|23142080\|29167373 | Unspecified |
| mmu-let-7i-5p | MIMAT0000122 | Adarb1 | 110532 | HITS-CLIP | 23597149\|23142080 | Unspecified |
| mmu-let-7i-5p | MIMAT0000122 | Adarb2 | 94191 | HITS-CLIP | 37986224 | Unspecified |
| mmu-let-7i-5p | MIMAT0000122 | Adck1 | 72113 | HITS-CLIP | 23142080\|23597149 | Unspecified |

| | | | | | | | |
|---|---|---|---|---|---|---|---|
| mmu-let-7i-5p | MIMAT0000122 | Adcy1 | 432530 | HITS-CLIP | | 19536157 | Unspecified |
| mmu-let-7i-5p | MIMAT0000122 | Adcy6 | 11512 | HITS-CLIP | | 23597149 | Unspecified |
| mmu-let-7i-5p | MIMAT0000122 | Adcy7 | 11513 | HITS-CLIP | | 23142080 | Unspecified |
| mmu-let-7i-5p | MIMAT0000122 | Adcyap1r1 | 11517 | HITS-CLIP | | 37986224\|19536157 | Unspecified |
| mmu-let-7i-5p | MIMAT0000122 | Add1 | 11518 | HITS-CLIP | | 23597149\|23142080 | Unspecified |
| mmu-let-7i-5p | MIMAT0000122 | Add2 | 11519 | HITS-CLIP | | 19536157 | Unspecified |
| mmu-let-7i-5p | MIMAT0000122 | Add3 | 27360 | HITS-CLIP | | 23142080 | Unspecified |
| mmu-let-7i-5p | MIMAT0000122 | Adgra2 | 25960 | HITS-CLIP | | 23597149 | Unspecified |
| mmu-let-7i-5p | MIMAT0000122 | Adgre1 | 13733 | HITS-CLIP | | 29167373 | Unspecified |
| mmu-let-7i-5p | MIMAT0000122 | Adgre4 | 52614 | HITS-CLIP | | 30971389 | Unspecified |
| mmu-let-7i-5p | MIMAT0000122 | Adgrf5 | 224792 | HITS-CLIP | | 19536157\|23597149 | Unspecified |
| mmu-let-7i-5p | MIMAT0000122 | Adgrg1 | 9289 | HITS-CLIP | | 19536157 | Unspecified |
| mmu-let-7i-5p | MIMAT0000122 | Adgrl1 | 22859 | HITS-CLIP | | 23142080\|19536157 | Unspecified |
| mmu-let-7i-5p | MIMAT0000122 | Adgrl2 | 23266 | HITS-CLIP | | 23597149 | Unspecified |
| mmu-let-7i-5p | MIMAT0000122 | Adgrl4 | 64123 | HITS-CLIP | | 23597149 | Unspecified |
| mmu-let-7i-5p | MIMAT0000122 | Adh1 | 11522 | HITS-CLIP | | 23597149 | Unspecified |
| mmu-let-7i-5p | MIMAT0000122 | Adh4 | 26876 | HITS-CLIP | | 37986224\|23597149 | Unspecified |
| mmu-let-7i-5p | MIMAT0000122 | Adhfe1 | 76187 | HITS-CLIP | | 23597149 | Unspecified |
| mmu-let-7i-5p | MIMAT0000122 | Adi1 | 104923 | HITS-CLIP | | 23597149\|23142080 | Unspecified |
| mmu-let-7i-5p | MIMAT0000122 | Adipor1 | 72674 | HITS-CLIP | | 23597149\|19536157\|23142080 | Unspecified |
| mmu-let-7i-5p | MIMAT0000122 | Adipor2 | 68465 | HITS-CLIP | | 37986224\|23597149\|19536157\|23142 | Unspecified |
| mmu-let-7i-5p | MIMAT0000122 | Adk | 11534 | HITS-CLIP | | 23142080 | Unspecified |
| mmu-let-7i-5p | MIMAT0000122 | Adnp | 11538 | HITS-CLIP | | 37986224\|19536157\|23597149\|23142 | Unspecified |
| mmu-let-7i-5p | MIMAT0000122 | Adnp2 | 240442 | HITS-CLIP | | 23142080 | Unspecified |
| mmu-let-7i-5p | MIMAT0000122 | Ado | 211488 | HITS-CLIP | | 23142080 | Unspecified |
| mmu-let-7i-5p | MIMAT0000122 | Adora1 | 11539 | HITS-CLIP | | 23597149 | Unspecified |
| mmu-let-7i-5p | MIMAT0000122 | Adora2a | 11540 | HITS-CLIP | | 23142080 | Unspecified |
| mmu-let-7i-5p | MIMAT0000122 | Adora2b | 11541 | HITS-CLIP | | 19536157\|23142080\|26203562 | Unspecified |
| mmu-let-7i-5p | MIMAT0000122 | Adprhl2 | 100206 | HITS-CLIP | | 23142080 | Unspecified |
| mmu-let-7i-5p | MIMAT0000122 | Adra1b | 11548 | HITS-CLIP | | 23597149 | Unspecified |
| mmu-let-7i-5p | MIMAT0000122 | Adra2a | 11551 | HITS-CLIP | | 37986224\|19536157 | Unspecified |
| mmu-let-7i-5p | MIMAT0000122 | Adrb1 | 11554 | HITS-CLIP | | 37986224\|23597149 | Unspecified |
| mmu-let-7i-5p | MIMAT0000122 | Adrb2 | 11555 | HITS-CLIP | | 37986224\|19536157\|29167373\|23142 | Unspecified |
| mmu-let-7i-5p | MIMAT0000122 | Adrb3 | 11556 | HITS-CLIP | | 37986224\|23597149 | Unspecified |
| mmu-let-7i-5p | MIMAT0000122 | Adss | 11566 | HITS-CLIP | | 23597149\|23142080\|29167373 | Unspecified |
| mmu-let-7i-5p | MIMAT0000122 | Aebp2 | 11569 | HITS-CLIP | | 23142080 | Unspecified |
| mmu-let-7i-5p | MIMAT0000122 | Aes | 14797 | HITS-CLIP | | 23597149\|23142080 | Unspecified |
| mmu-let-7i-5p | MIMAT0000122 | Afdn | 4301 | HITS-CLIP | | 23630078 | Unspecified |
| mmu-let-7i-5p | MIMAT0000122 | Aff1 | 17355 | HITS-CLIP | | 23597149\|29167373\|23142080 | Unspecified |

| | | | | | | |
|---|---|---|---|---|---|---|
| mmu-let-7i-5p | MIMAT0000122 | Aff2 | 14266 | HITS-CLIP | 37986224\|19536157 | Unspecified |
| mmu-let-7i-5p | MIMAT0000122 | Aff3 | 16764 | HITS-CLIP | 23142080 | Unspecified |
| mmu-let-7i-5p | MIMAT0000122 | Aff4 | 93736 | HITS-CLIP | 37986224\|19536157\|23597149\|29167 | Unspecified |
| mmu-let-7i-5p | MIMAT0000122 | Afg1l | 246269 | HITS-CLIP | 23597149 | Unspecified |
| mmu-let-7i-5p | MIMAT0000122 | Afg3l1 | 114896 | HITS-CLIP | 37986224\|29167373 | Unspecified |
| mmu-let-7i-5p | MIMAT0000122 | Afm | 280662 | HITS-CLIP | 23597149 | Unspecified |
| mmu-let-7i-5p | MIMAT0000122 | Afmid | 71562 | HITS-CLIP | 23597149 | Unspecified |
| mmu-let-7i-5p | MIMAT0000122 | Aftph | 216549 | HITS-CLIP | 29167373 | Unspecified |
| mmu-let-7i-5p | MIMAT0000122 | Aga | 11593 | HITS-CLIP | 23597149 | Unspecified |
| mmu-let-7i-5p | MIMAT0000122 | Agap1 | 347722 | HITS-CLIP | 23597149\|19536157\|29167373 | Unspecified |
| mmu-let-7i-5p | MIMAT0000122 | Agap2 | 216439 | HITS-CLIP | 37986224\|19536157\|23142080 | Unspecified |
| mmu-let-7i-5p | MIMAT0000122 | Agbl5 | 231093 | HITS-CLIP | 23142080 | Unspecified |
| mmu-let-7i-5p | MIMAT0000122 | Agfg1 | 15463 | HITS-CLIP | 23142080\|23597149\|29167373 | Unspecified |
| mmu-let-7i-5p | MIMAT0000122 | Agfg2 | 231801 | HITS-CLIP | 23597149 | Unspecified |
| mmu-let-7i-5p | MIMAT0000122 | Aggf1 | 66549 | HITS-CLIP | 23597149\|23142080 | Unspecified |
| mmu-let-7i-5p | MIMAT0000122 | Agk | 69923 | HITS-CLIP | 23142080 | Unspecified |
| mmu-let-7i-5p | MIMAT0000122 | Agl | 77559 | HITS-CLIP | 37986224\|23597149 | Unspecified |
| mmu-let-7i-5p | MIMAT0000122 | Agmo | 319660 | HITS-CLIP | 23597149 | Unspecified |
| mmu-let-7i-5p | MIMAT0000122 | Ago1 | 236511 | HITS-CLIP | 37986224\|19536157\|23597149\|23142 | Unspecified |
| mmu-let-7i-5p | MIMAT0000122 | Ago2 | 239528 | HITS-CLIP | 37986224\|19536157\|23597149\|26203 | Unspecified |
| mmu-let-7i-5p | MIMAT0000122 | Ago3 | 214150 | HITS-CLIP | 37986224\|23597149\|19536157\|23142 | Unspecified |
| mmu-let-7i-5p | MIMAT0000122 | Ago4 | 76850 | HITS-CLIP | 29167373 | Unspecified |
| mmu-let-7i-5p | MIMAT0000122 | Agpat1 | 55979 | HITS-CLIP | 19536157\|23142080 | Unspecified |
| mmu-let-7i-5p | MIMAT0000122 | Agpat3 | 28169 | HITS-CLIP | 37986224\|23597149\|19536157\|29167 | Unspecified |
| mmu-let-7i-5p | MIMAT0000122 | Agpat5 | 52123 | HITS-CLIP | 23142080 | Unspecified |
| mmu-let-7i-5p | MIMAT0000122 | Agps | 228061 | HITS-CLIP | 23597149\|23142080\|29167373 | Unspecified |
| mmu-let-7i-5p | MIMAT0000122 | Agrn | 11603 | HITS-CLIP | 23597149 | Unspecified |
| mmu-let-7i-5p | MIMAT0000122 | Agt | 11606 | HITS-CLIP | 23597149 | Unspecified |
| mmu-let-7i-5p | MIMAT0000122 | Agtrap | 11610 | HITS-CLIP | 23597149 | Unspecified |
| mmu-let-7i-5p | MIMAT0000122 | Agxt2 | 268782 | HITS-CLIP | 37986224\|23597149 | Unspecified |
| mmu-let-7i-5p | MIMAT0000122 | Ahctf1 | 226747 | HITS-CLIP | 37986224\|23597149\|19536157\|29167 | Unspecified |
| mmu-let-7i-5p | MIMAT0000122 | Ahcyl1 | 229709 | HITS-CLIP | 19536157\|23142080\|29167373 | Unspecified |
| mmu-let-7i-5p | MIMAT0000122 | Ahcyl2 | 74340 | HITS-CLIP | 23142080 | Unspecified |
| mmu-let-7i-5p | MIMAT0000122 | Ahdc1 | 230793 | HITS-CLIP | 23142080 | Unspecified |
| mmu-let-7i-5p | MIMAT0000122 | Ahi1 | 52906 | HITS-CLIP | 37986224\|23142080 | Unspecified |
| mmu-let-7i-5p | MIMAT0000122 | Ahnak | 66395 | HITS-CLIP | 37986224\|23597149\|23142080\|29167 | Unspecified |
| mmu-let-7i-5p | MIMAT0000122 | Ahr | 11622 | HITS-CLIP | 23597149\|23142080 | Unspecified |
| mmu-let-7i-5p | MIMAT0000122 | Ahsa1 | 217737 | HITS-CLIP | 23597149\|19536157\|23142080\|29167 | Unspecified |
| mmu-let-7i-5p | MIMAT0000122 | Ahsg | 11625 | HITS-CLIP | 23597149 | Unspecified |

| | | | | | | |
|---|---|---|---|---|---|---|
| mmu-let-7i-5p | MIMAT0000122 | Aifm1 | 26926 | HITS-CLIP | 37986224\|23597149\|23142080 | Unspecified |
| mmu-let-7i-5p | MIMAT0000122 | Aim2 | 383619 | HITS-CLIP | 23142080 | Unspecified |
| mmu-let-7i-5p | MIMAT0000122 | Aip | 11632 | HITS-CLIP | 23142080 | Unspecified |
| mmu-let-7i-5p | MIMAT0000122 | Ajuba | 16475 | HITS-CLIP | 23142080 | Unspecified |
| mmu-let-7i-5p | MIMAT0000122 | Ak2 | 11637 | HITS-CLIP | 37986224 | Unspecified |
| mmu-let-7i-5p | MIMAT0000122 | Ak3 | 56248 | HITS-CLIP | 19536157 | Unspecified |
| mmu-let-7i-5p | MIMAT0000122 | Ak4 | 11639 | HITS-CLIP | 23142080 | Unspecified |
| mmu-let-7i-5p | MIMAT0000122 | Akap1 | 11640 | HITS-CLIP | 37986224 | Unspecified |
| mmu-let-7i-5p | MIMAT0000122 | Akap10 | 56697 | HITS-CLIP | 23597149 | Unspecified |
| mmu-let-7i-5p | MIMAT0000122 | Akap11 | 219181 | HITS-CLIP | 23597149\|19536157\|23142080\|29167 | Unspecified |
| mmu-let-7i-5p | MIMAT0000122 | Akap12 | 83397 | HITS-CLIP | 23597149 | Unspecified |
| mmu-let-7i-5p | MIMAT0000122 | Akap13 | 75547 | HITS-CLIP | 23142080\|23597149\|29167373 | Unspecified |
| mmu-let-7i-5p | MIMAT0000122 | Akap2 | 11641 | HITS-CLIP | 23597149 | Unspecified |
| mmu-let-7i-5p | MIMAT0000122 | Akap6 | 238161 | HITS-CLIP | 37986224 | Unspecified |
| mmu-let-7i-5p | MIMAT0000122 | Akap7 | 432442 | HITS-CLIP | 23142080 | Unspecified |
| mmu-let-7i-5p | MIMAT0000122 | Akap8 | 56399 | HITS-CLIP | 37986224 | Unspecified |
| mmu-let-7i-5p | MIMAT0000122 | Akap9 | 100986 | HITS-CLIP | 37986224\|23142080 | Unspecified |
| mmu-let-7i-5p | MIMAT0000122 | Akirin1 | 68050 | HITS-CLIP | 23142080 | Unspecified |
| mmu-let-7i-5p | MIMAT0000122 | Akirin2 | 433693 | HITS-CLIP | 23142080 | Unspecified |
| mmu-let-7i-5p | MIMAT0000122 | Akna | 100182 | HITS-CLIP | 37986224\|23142080 | Unspecified |
| mmu-let-7i-5p | MIMAT0000122 | Akr1b10 | 67861 | HITS-CLIP | 23142080 | Unspecified |
| mmu-let-7i-5p | MIMAT0000122 | Akr1b8 | 14187 | HITS-CLIP | 29167373 | Unspecified |
| mmu-let-7i-5p | MIMAT0000122 | Akr1c18 | 105349 | HITS-CLIP | 23142080 | Unspecified |
| mmu-let-7i-5p | MIMAT0000122 | Akr1c20 | 116852 | HITS-CLIP | 23597149 | Unspecified |
| mmu-let-7i-5p | MIMAT0000122 | Akr1c6 | 83702 | HITS-CLIP | 23597149 | Unspecified |
| mmu-let-7i-5p | MIMAT0000122 | Akr1d1 | 208665 | HITS-CLIP | 23597149 | Unspecified |
| mmu-let-7i-5p | MIMAT0000122 | Akr1e1 | 56043 | HITS-CLIP | 37986224 | Unspecified |
| mmu-let-7i-5p | MIMAT0000122 | Akr7a5 | 110198 | HITS-CLIP | 23597149 | Unspecified |
| mmu-let-7i-5p | MIMAT0000122 | Akt1s1 | 67605 | HITS-CLIP | 23142080 | Unspecified |
| mmu-let-7i-5p | MIMAT0000122 | Akt3 | 23797 | HITS-CLIP | 23142080\|29167373 | Unspecified |
| mmu-let-7i-5p | MIMAT0000122 | Alad | 17025 | HITS-CLIP | 23597149\|23142080 | Unspecified |
| mmu-let-7i-5p | MIMAT0000122 | Alas1 | 11655 | HITS-CLIP | 23597149 | Unspecified |
| mmu-let-7i-5p | MIMAT0000122 | Alas2 | 11656 | HITS-CLIP | 23597149 | Unspecified |
| mmu-let-7i-5p | MIMAT0000122 | Alb | 213 | HITS-CLIP | 23597149 | Unspecified |
| mmu-let-7i-5p | MIMAT0000122 | Alcam | 11658 | HITS-CLIP | 37986224\|23142080\|31433990\|29167 | Unspecified |
| mmu-let-7i-5p | MIMAT0000122 | Aldh16a1 | 69748 | HITS-CLIP | 23142080\|23597149 | Unspecified |
| mmu-let-7i-5p | MIMAT0000122 | Aldh1a1 | 11668 | HITS-CLIP | 23597149 | Unspecified |
| mmu-let-7i-5p | MIMAT0000122 | Aldh1a7 | 26358 | HITS-CLIP | 23597149 | Unspecified |
| mmu-let-7i-5p | MIMAT0000122 | Aldh1b1 | 72535 | HITS-CLIP | 23597149 | Unspecified |

| | | | | | | |
|---|---|---|---|---|---|---|
| mmu-let-7i-5p | MIMAT0000122 | Aldh1l1 | 107747 | HITS-CLIP | 23597149 | Unspecified |
| mmu-let-7i-5p | MIMAT0000122 | Aldh2 | 11669 | HITS-CLIP | 23597149 | Unspecified |
| mmu-let-7i-5p | MIMAT0000122 | Aldh4a1 | 212647 | HITS-CLIP | 23597149 | Unspecified |
| mmu-let-7i-5p | MIMAT0000122 | Aldh6a1 | 104776 | HITS-CLIP | 37986224\|23597149 | Unspecified |
| mmu-let-7i-5p | MIMAT0000122 | Aldh8a1 | 237320 | HITS-CLIP | 23597149 | Unspecified |
| mmu-let-7i-5p | MIMAT0000122 | Aldoa | 11674 | HITS-CLIP | 23142080\|29167373 | Unspecified |
| mmu-let-7i-5p | MIMAT0000122 | Aldoart1 | 353204 | HITS-CLIP | 29167373 | Unspecified |
| mmu-let-7i-5p | MIMAT0000122 | Aldoart2 | 79459 | HITS-CLIP | 23142080 | Unspecified |
| mmu-let-7i-5p | MIMAT0000122 | Aldoc | 11676 | HITS-CLIP | 23142080 | Unspecified |
| mmu-let-7i-5p | MIMAT0000122 | Alg10b | 380959 | HITS-CLIP | 37986224\|29167373 | Unspecified |
| mmu-let-7i-5p | MIMAT0000122 | Alg11 | 207958 | HITS-CLIP | 23142080 | Unspecified |
| mmu-let-7i-5p | MIMAT0000122 | Alg14 | 66789 | HITS-CLIP | 23597149 | Unspecified |
| mmu-let-7i-5p | MIMAT0000122 | Alg3 | 208624 | HITS-CLIP | 37986224\|23597149\|19536157\|29167 | Unspecified |
| mmu-let-7i-5p | MIMAT0000122 | Alg8 | 381903 | HITS-CLIP | 23597149\|23142080 | Unspecified |
| mmu-let-7i-5p | MIMAT0000122 | Alg9 | 102580 | qCLASH | 31393953 | Unspecified |
| mmu-let-7i-5p | MIMAT0000122 | Alkbh4 | 72041 | HITS-CLIP | 37986224 | Unspecified |
| mmu-let-7i-5p | MIMAT0000122 | Alkbh5 | 268420 | HITS-CLIP | 23142080 | Unspecified |
| mmu-let-7i-5p | MIMAT0000122 | Alkbh6 | 233065 | HITS-CLIP | 23142080 | Unspecified |
| mmu-let-7i-5p | MIMAT0000122 | Alkbh7 | 66400 | HITS-CLIP | 23142080 | Unspecified |
| mmu-let-7i-5p | MIMAT0000122 | Alkbh8 | 67667 | HITS-CLIP | 23142080\|31433990 | Unspecified |
| mmu-let-7i-5p | MIMAT0000122 | Alox5ap | 11690 | HITS-CLIP | 23597149\|29167373 | Unspecified |
| mmu-let-7i-5p | MIMAT0000122 | Alpk1 | 71481 | HITS-CLIP | 23597149\|29167373 | Unspecified |
| mmu-let-7i-5p | MIMAT0000122 | Als2 | 74018 | HITS-CLIP | 23597149\|23142080 | Unspecified |
| mmu-let-7i-5p | MIMAT0000122 | Alyref | 21681 | HITS-CLIP | 31433990 | Unspecified |
| mmu-let-7i-5p | MIMAT0000122 | Ambra1 | 228361 | HITS-CLIP | 19536157\|23142080 | Unspecified |
| mmu-let-7i-5p | MIMAT0000122 | Amer1 | 72345 | HITS-CLIP | 23142080 | Unspecified |
| mmu-let-7i-5p | MIMAT0000122 | Amfr | 23802 | HITS-CLIP | 37986224\|23597149\|29167373 | Unspecified |
| mmu-let-7i-5p | MIMAT0000122 | Ammecr1 | 56068 | HITS-CLIP | 23142080 | Unspecified |
| mmu-let-7i-5p | MIMAT0000122 | Amn | 93835 | HITS-CLIP | 23142080 | Unspecified |
| mmu-let-7i-5p | MIMAT0000122 | Amot | 27494 | HITS-CLIP | 37986224\|23597149\|19536157 | Unspecified |
| mmu-let-7i-5p | MIMAT0000122 | Amotl1 | 75723 | HITS-CLIP | 23597149 | Unspecified |
| mmu-let-7i-5p | MIMAT0000122 | Ampd3 | 11717 | HITS-CLIP | 29167373 | Unspecified |
| mmu-let-7i-5p | MIMAT0000122 | Amz2 | 13929 | HITS-CLIP | 23142080 | Unspecified |
| mmu-let-7i-5p | MIMAT0000122 | Anapc1 | 17222 | HITS-CLIP | 23597149\|23142080 | Unspecified |
| mmu-let-7i-5p | MIMAT0000122 | Anapc4 | 52206 | HITS-CLIP | 23142080 | Unspecified |
| mmu-let-7i-5p | MIMAT0000122 | Anapc5 | 59008 | HITS-CLIP | 23597149\|19536157 | Unspecified |
| mmu-let-7i-5p | MIMAT0000122 | Anapc7 | 56317 | HITS-CLIP | 23597149 | Unspecified |
| mmu-let-7i-5p | MIMAT0000122 | Ang | 11727 | HITS-CLIP | 23597149 | Unspecified |
| mmu-let-7i-5p | MIMAT0000122 | Angptl2 | 26360 | HITS-CLIP | 37986224\|23597149\|23142080\|29167 | Unspecified |

| | | | | | | |
|---|---|---|---|---|---|---|
| mmu-let-7i-5p | MIMAT0000122 | Angptl3 | 30924 | HITS-CLIP | 23597149 | Unspecified |
| mmu-let-7i-5p | MIMAT0000122 | Angptl4 | 57875 | HITS-CLIP | 23142080 | Unspecified |
| mmu-let-7i-5p | MIMAT0000122 | Angptl6 | 70726 | HITS-CLIP | 23142080 | Unspecified |
| mmu-let-7i-5p | MIMAT0000122 | Ank | 11732 | HITS-CLIP | 37986224\|19536157 | Unspecified |
| mmu-let-7i-5p | MIMAT0000122 | Ank2 | 109676 | HITS-CLIP | 19536157 | Unspecified |
| mmu-let-7i-5p | MIMAT0000122 | Ank3 | 11735 | HITS-CLIP | 37986224\|19536157 | Unspecified |
| mmu-let-7i-5p | MIMAT0000122 | Ankfy1 | 11736 | HITS-CLIP | 37986224\|23142080\|29167373 | Unspecified |
| mmu-let-7i-5p | MIMAT0000122 | Ankhd1 | 108857 | HITS-CLIP | 37986224\|23142080\|23597149\|29167 | Unspecified |
| mmu-let-7i-5p | MIMAT0000122 | Ankib1 | 70797 | HITS-CLIP | 23597149\|23142080 | Unspecified |
| mmu-let-7i-5p | MIMAT0000122 | Ankmy2 | 217473 | HITS-CLIP | 23142080 | Unspecified |
| mmu-let-7i-5p | MIMAT0000122 | Ankra2 | 68558 | HITS-CLIP | 23142080 | Unspecified |
| mmu-let-7i-5p | MIMAT0000122 | Ankrd11 | 77087 | HITS-CLIP | 19536157\|23597149\|23142080\|29167 | Unspecified |
| mmu-let-7i-5p | MIMAT0000122 | Ankrd12 | 106585 | HITS-CLIP | 37986224\|19536157\|23142080 | Unspecified |
| mmu-let-7i-5p | MIMAT0000122 | Ankrd13c | 433667 | HITS-CLIP | 29167373 | Unspecified |
| mmu-let-7i-5p | MIMAT0000122 | Ankrd17 | 81702 | HITS-CLIP | 23597149\|23142080\|26203562\|29167 | Unspecified |
| mmu-let-7i-5p | MIMAT0000122 | Ankrd27 | 245886 | HITS-CLIP | 23142080 | Unspecified |
| mmu-let-7i-5p | MIMAT0000122 | Ankrd28 | 105522 | HITS-CLIP | 37986224\|23597149\|23142080\|29167 | Unspecified |
| mmu-let-7i-5p | MIMAT0000122 | Ankrd34b | 218440 | HITS-CLIP | 37986224 | Unspecified |
| mmu-let-7i-5p | MIMAT0000122 | Ankrd37 | 654824 | HITS-CLIP | 23142080 | Unspecified |
| mmu-let-7i-5p | MIMAT0000122 | Ankrd40 | 71452 | HITS-CLIP | 37986224 | Unspecified |
| mmu-let-7i-5p | MIMAT0000122 | Ankrd44 | 329154 | HITS-CLIP | 37986224\|23142080 | Unspecified |
| mmu-let-7i-5p | MIMAT0000122 | Ankrd49 | 56503 | HITS-CLIP | 37986224\|23142080\|31433990\|29167 | Unspecified |
| mmu-let-7i-5p | MIMAT0000122 | Ankrd50 | 99696 | HITS-CLIP | 37986224\|29167373 | Unspecified |
| mmu-let-7i-5p | MIMAT0000122 | Ankrd52 | 237615 | HITS-CLIP | 23597149\|23142080\|19536157 | Unspecified |
| mmu-let-7i-5p | MIMAT0000122 | Ankrd9 | 74251 | HITS-CLIP | 23142080 | Unspecified |
| mmu-let-7i-5p | MIMAT0000122 | Anks1 | 224650 | HITS-CLIP | 23597149\|19536157\|23142080\|29167 | Unspecified |
| mmu-let-7i-5p | MIMAT0000122 | Anks4b | 72074 | HITS-CLIP | 23597149 | Unspecified |
| mmu-let-7i-5p | MIMAT0000122 | Ankzf1 | 52231 | HITS-CLIP | 23597149 | Unspecified |
| mmu-let-7i-5p | MIMAT0000122 | Anln | 68743 | HITS-CLIP | 31433990\|23142080 | Unspecified |
| mmu-let-7i-5p | MIMAT0000122 | Ano10 | 102566 | HITS-CLIP | 37986224 | Unspecified |
| mmu-let-7i-5p | MIMAT0000122 | Ano6 | 105722 | HITS-CLIP | 23142080\|29167373 | Unspecified |
| mmu-let-7i-5p | MIMAT0000122 | Ano8 | 382014 | HITS-CLIP | 37986224\|23142080 | Unspecified |
| mmu-let-7i-5p | MIMAT0000122 | Anp32a | 11737 | HITS-CLIP | 37986224\|23142080 | Unspecified |
| mmu-let-7i-5p | MIMAT0000122 | Anp32b | 67628 | HITS-CLIP | 23142080 | Unspecified |
| mmu-let-7i-5p | MIMAT0000122 | Anp32e | 66471 | HITS-CLIP//qCLASH | 37986224\|23142080\|31393953\|23597 | Unspecified |
| mmu-let-7i-5p | MIMAT0000122 | Anpep | 16790 | HITS-CLIP | 37986224\|23597149\|29167373 | Unspecified |
| mmu-let-7i-5p | MIMAT0000122 | Antxr2 | 71914 | HITS-CLIP | 29167373 | Unspecified |
| mmu-let-7i-5p | MIMAT0000122 | Anxa1 | 16952 | HITS-CLIP | 29167373 | Unspecified |
| mmu-let-7i-5p | MIMAT0000122 | Anxa11 | 11744 | HITS-CLIP | 37986224\|23597149 | Unspecified |

| | | | | | | |
|---|---|---|---|---|---|---|
| mmu-let-7i-5p | MIMAT0000122 | Anxa2 | 12306 | HITS-CLIP | 23142080 | Unspecified |
| mmu-let-7i-5p | MIMAT0000122 | Anxa4 | 11746 | HITS-CLIP | 23142080 | Unspecified |
| mmu-let-7i-5p | MIMAT0000122 | Anxa5 | 11747 | HITS-CLIP | 23597149\|29167373\|23142080 | Unspecified |
| mmu-let-7i-5p | MIMAT0000122 | Aox1 | 11761 | HITS-CLIP | 23597149 | Unspecified |
| mmu-let-7i-5p | MIMAT0000122 | Aox3 | 71724 | HITS-CLIP | 23597149 | Unspecified |
| mmu-let-7i-5p | MIMAT0000122 | Ap1b1 | 11764 | HITS-CLIP | 23597149\|23142080\|29167373 | Unspecified |
| mmu-let-7i-5p | MIMAT0000122 | Ap1g1 | 11765 | HITS-CLIP | 23597149\|23142080 | Unspecified |
| mmu-let-7i-5p | MIMAT0000122 | Ap1m1 | 11767 | HITS-CLIP | 23597149\|23142080 | Unspecified |
| mmu-let-7i-5p | MIMAT0000122 | Ap1s2 | 108012 | HITS-CLIP | 23142080 | Unspecified |
| mmu-let-7i-5p | MIMAT0000122 | Ap2a1 | 11771 | HITS-CLIP | 23142080 | Unspecified |
| mmu-let-7i-5p | MIMAT0000122 | Ap2a2 | 11772 | HITS-CLIP | 23597149 | Unspecified |
| mmu-let-7i-5p | MIMAT0000122 | Ap2b1 | 71770 | HITS-CLIP | 37986224\|23597149\|23142080\|3143 | Unspecified |
| mmu-let-7i-5p | MIMAT0000122 | Ap3b1 | 11774 | HITS-CLIP | 23142080 | Unspecified |
| mmu-let-7i-5p | MIMAT0000122 | Ap3d1 | 11776 | HITS-CLIP | 23597149\|23142080\|29167373 | Unspecified |
| mmu-let-7i-5p | MIMAT0000122 | Ap3m1 | 55946 | HITS-CLIP | 37986224\|23142080 | Unspecified |
| mmu-let-7i-5p | MIMAT0000122 | Ap3s2 | 11778 | HITS-CLIP | 37986224 | Unspecified |
| mmu-let-7i-5p | MIMAT0000122 | Ap4b1 | 67489 | HITS-CLIP | 37986224\|30971389\|23142080 | Unspecified |
| mmu-let-7i-5p | MIMAT0000122 | Ap4e1 | 108011 | HITS-CLIP | 23142080\|19536157 | Unspecified |
| mmu-let-7i-5p | MIMAT0000122 | Ap5b1 | 381201 | HITS-CLIP | 23142080 | Unspecified |
| mmu-let-7i-5p | MIMAT0000122 | Ap5m1 | 74385 | HITS-CLIP | 23142080 | Unspecified |
| mmu-let-7i-5p | MIMAT0000122 | Ap5z1 | 231855 | HITS-CLIP | 23597149 | Unspecified |
| mmu-let-7i-5p | MIMAT0000122 | Apaf1 | 11783 | HITS-CLIP | 37986224\|23142080\|31433990\|29167 | Unspecified |
| mmu-let-7i-5p | MIMAT0000122 | Apba1 | 319924 | HITS-CLIP | 19536157 | Unspecified |
| mmu-let-7i-5p | MIMAT0000122 | Apbb2 | 11787 | HITS-CLIP | 29167373 | Unspecified |
| mmu-let-7i-5p | MIMAT0000122 | Apbb3 | 225372 | HITS-CLIP | 19536157\|23142080 | Unspecified |
| mmu-let-7i-5p | MIMAT0000122 | Apc | 11789 | HITS-CLIP | 37986224\|19536157\|23597149\|23142 | Unspecified |
| mmu-let-7i-5p | MIMAT0000122 | Apc2 | 23805 | HITS-CLIP | 25083871\|19536157\|23142080\|2125 | Unspecified |
| mmu-let-7i-5p | MIMAT0000122 | Apcdd1 | 494504 | HITS-CLIP | 37986224\|19536157 | Unspecified |
| mmu-let-7i-5p | MIMAT0000122 | Apeh | 235606 | HITS-CLIP | 23597149 | Unspecified |
| mmu-let-7i-5p | MIMAT0000122 | Apex1 | 11792 | HITS-CLIP | 23142080 | Unspecified |
| mmu-let-7i-5p | MIMAT0000122 | Apex2 | 77622 | HITS-CLIP | 29167373 | Unspecified |
| mmu-let-7i-5p | MIMAT0000122 | Api5 | 11800 | HITS-CLIP | 23142080\|29167373 | Unspecified |
| mmu-let-7i-5p | MIMAT0000122 | Apip | 56369 | HITS-CLIP | 37986224\|23597149 | Unspecified |
| mmu-let-7i-5p | MIMAT0000122 | Aplp2 | 11804 | HITS-CLIP | 37986224\|23597149\|29167373 | Unspecified |
| mmu-let-7i-5p | MIMAT0000122 | Apmap | 71881 | HITS-CLIP | 23597149 | Unspecified |
| mmu-let-7i-5p | MIMAT0000122 | Apoa2 | 11807 | HITS-CLIP | 30971389\|23597149 | Unspecified |
| mmu-let-7i-5p | MIMAT0000122 | Apoa5 | 66113 | HITS-CLIP | 23597149 | Unspecified |
| mmu-let-7i-5p | MIMAT0000122 | Apob | 238055 | HITS-CLIP | 23597149 | Unspecified |
| mmu-let-7i-5p | MIMAT0000122 | Apobec3 | 80287 | HITS-CLIP | 23142080\|23597149\|29167373 | Unspecified |

| | | | | | | | |
|---|---|---|---|---|---|---|---|
| mmu-let-7i-5p | MIMAT0000122 | Apobr | 171504 | HITS-CLIP | | 29167373 | Unspecified |
| mmu-let-7i-5p | MIMAT0000122 | Apoc1 | 11812 | HITS-CLIP | | 23597149 | Unspecified |
| mmu-let-7i-5p | MIMAT0000122 | Apoc3 | 345 | HITS-CLIP | | 23597149 | Unspecified |
| mmu-let-7i-5p | MIMAT0000122 | Apoc4 | 11425 | HITS-CLIP | | 23597149 | Unspecified |
| mmu-let-7i-5p | MIMAT0000122 | Apof | 103161 | HITS-CLIP | | 23597149 | Unspecified |
| mmu-let-7i-5p | MIMAT0000122 | Apol7a | 75761 | HITS-CLIP | | 23597149 | Unspecified |
| mmu-let-7i-5p | MIMAT0000122 | Apom | 55938 | HITS-CLIP | | 23597149 | Unspecified |
| mmu-let-7i-5p | MIMAT0000122 | Apool | 68117 | HITS-CLIP | 23597149\|23142080 | | Unspecified |
| mmu-let-7i-5p | MIMAT0000122 | App | 11820 | HITS-CLIP | | 19536157 | Unspecified |
| mmu-let-7i-5p | MIMAT0000122 | Appbp2 | 66884 | HITS-CLIP | 23597149\|23142080\|19536157\|29167 | | Unspecified |
| mmu-let-7i-5p | MIMAT0000122 | Appl2 | 216190 | HITS-CLIP | | 23142080 | Unspecified |
| mmu-let-7i-5p | MIMAT0000122 | Aptx | 66408 | HITS-CLIP | | 23142080 | Unspecified |
| mmu-let-7i-5p | MIMAT0000122 | Aqp1 | 11826 | HITS-CLIP | | 23597149 | Unspecified |
| mmu-let-7i-5p | MIMAT0000122 | Aqp4 | 11829 | HITS-CLIP | 37986224\|19536157 | | Unspecified |
| mmu-let-7i-5p | MIMAT0000122 | Aqp9 | 64008 | HITS-CLIP | | 23597149 | Unspecified |
| mmu-let-7i-5p | MIMAT0000122 | Aqr | 11834 | HITS-CLIP | | 23597149 | Unspecified |
| mmu-let-7i-5p | MIMAT0000122 | Ar | 11835 | HITS-CLIP | | 23142080 | Unspecified |
| mmu-let-7i-5p | MIMAT0000122 | Araf | 11836 | HITS-CLIP | 37986224\|23142080\|29167373 | | Unspecified |
| mmu-let-7i-5p | MIMAT0000122 | Arap1 | 69710 | HITS-CLIP | 37986224\|23597149\|29167373 | | Unspecified |
| mmu-let-7i-5p | MIMAT0000122 | Arap2 | 212285 | HITS-CLIP | | 23142080 | Unspecified |
| mmu-let-7i-5p | MIMAT0000122 | Arap3 | 106952 | HITS-CLIP | | 23597149 | Unspecified |
| mmu-let-7i-5p | MIMAT0000122 | Arc | 11838 | HITS-CLIP | 23142080\|19536157 | | Unspecified |
| mmu-let-7i-5p | MIMAT0000122 | Arcn1 | 213827 | HITS-CLIP | | 23142080 | Unspecified |
| mmu-let-7i-5p | MIMAT0000122 | Arel1 | 68497 | HITS-CLIP | 37986224\|23597149\|23142080 | | Unspecified |
| mmu-let-7i-5p | MIMAT0000122 | Arf2 | 11841 | HITS-CLIP | | 23597149 | Unspecified |
| mmu-let-7i-5p | MIMAT0000122 | Arf3 | 11842 | HITS-CLIP | 37986224\|23597149\|19536157\|23142 | | Unspecified |
| mmu-let-7i-5p | MIMAT0000122 | Arfgap1 | 228998 | HITS-CLIP | 29167373\|23142080 | | Unspecified |
| mmu-let-7i-5p | MIMAT0000122 | Arfgef1 | 211673 | HITS-CLIP | 19536157\|23597149\|23142080\|29167 | | Unspecified |
| mmu-let-7i-5p | MIMAT0000122 | Arfgef2 | 99371 | HITS-CLIP | 37986224\|23597149\|23142080\|29167 | | Unspecified |
| mmu-let-7i-5p | MIMAT0000122 | Arfgef3 | 57221 | HITS-CLIP | | 19536157 | Unspecified |
| mmu-let-7i-5p | MIMAT0000122 | Arfip2 | 76932 | HITS-CLIP | | 23597149 | Unspecified |
| mmu-let-7i-5p | MIMAT0000122 | Arg1 | 11846 | HITS-CLIP | | 23597149 | Unspecified |
| mmu-let-7i-5p | MIMAT0000122 | Arglu1 | 234023 | HITS-CLIP | 23142080\|29167373 | | Unspecified |
| mmu-let-7i-5p | MIMAT0000122 | Arhgap1 | 228359 | HITS-CLIP | 37986224\|23630078\|31433990\|23597 | | Unspecified |
| mmu-let-7i-5p | MIMAT0000122 | Arhgap10 | 78514 | HITS-CLIP | 23142080\|29167373 | | Unspecified |
| mmu-let-7i-5p | MIMAT0000122 | Arhgap12 | 75415 | HITS-CLIP | 37986224\|23142080\|29167373 | | Unspecified |
| mmu-let-7i-5p | MIMAT0000122 | Arhgap15 | 76117 | HITS-CLIP | | 23142080 | Unspecified |
| mmu-let-7i-5p | MIMAT0000122 | Arhgap17 | 70497 | HITS-CLIP | | 29167373 | Unspecified |
| mmu-let-7i-5p | MIMAT0000122 | Arhgap21 | 71435 | HITS-CLIP | | 19536157 | Unspecified |

| | | | | | | | |
|---|---|---|---|---|---|---|---|
| mmu-let-7i-5p | MIMAT0000122 | Arhgap22 | 239027 | HITS-CLIP | | 29167373 | Unspecified |
| mmu-let-7i-5p | MIMAT0000122 | Arhgap23 | 58996 | HITS-CLIP | | 19536157 | Unspecified |
| mmu-let-7i-5p | MIMAT0000122 | Arhgap26 | 71302 | HITS-CLIP | | 23597149\|23142080 | Unspecified |
| mmu-let-7i-5p | MIMAT0000122 | Arhgap28 | 268970 | HITS-CLIP | | 37986224 | Unspecified |
| mmu-let-7i-5p | MIMAT0000122 | Arhgap29 | 214137 | HITS-CLIP | | 37986224 | Unspecified |
| mmu-let-7i-5p | MIMAT0000122 | Arhgap31 | 12549 | HITS-CLIP | | 23597149 | Unspecified |
| mmu-let-7i-5p | MIMAT0000122 | Arhgap32 | 330914 | HITS-CLIP | | 37986224\|19536157\|23597149 | Unspecified |
| mmu-let-7i-5p | MIMAT0000122 | Arhgap33 | 233071 | HITS-CLIP | | 37986224 | Unspecified |
| mmu-let-7i-5p | MIMAT0000122 | Arhgap35 | 232906 | HITS-CLIP | | 23597149\|19536157 | Unspecified |
| mmu-let-7i-5p | MIMAT0000122 | Arhgap39 | 223666 | HITS-CLIP | | 19536157 | Unspecified |
| mmu-let-7i-5p | MIMAT0000122 | Arhgap4 | 393 | HITS-CLIP | | 23142080\|29167373 | Unspecified |
| mmu-let-7i-5p | MIMAT0000122 | Arhgap42 | 71544 | HITS-CLIP | | 23597149 | Unspecified |
| mmu-let-7i-5p | MIMAT0000122 | Arhgap45 | 23526 | HITS-CLIP | | 31433990\|23142080\|29167373 | Unspecified |
| mmu-let-7i-5p | MIMAT0000122 | Arhgap5 | 11855 | HITS-CLIP | | 37986224\|23142080\|29167373\|1953( | Unspecified |
| mmu-let-7i-5p | MIMAT0000122 | Arhgap9 | 216445 | HITS-CLIP | | 23142080 | Unspecified |
| mmu-let-7i-5p | MIMAT0000122 | Arhgdia | 192662 | HITS-CLIP | | 23597149\|23142080 | Unspecified |
| mmu-let-7i-5p | MIMAT0000122 | Arhgdib | 11857 | HITS-CLIP | | 29167373\|23142080 | Unspecified |
| mmu-let-7i-5p | MIMAT0000122 | Arhgef1 | 16801 | HITS-CLIP | | 19536157\|23142080 | Unspecified |
| mmu-let-7i-5p | MIMAT0000122 | Arhgef11 | 213498 | HITS-CLIP | | 29167373 | Unspecified |
| mmu-let-7i-5p | MIMAT0000122 | Arhgef12 | 69632 | HITS-CLIP | | 37986224\|23142080\|23597149 | Unspecified |
| mmu-let-7i-5p | MIMAT0000122 | Arhgef15 | 442801 | HITS-CLIP | | 37986224\|23597149 | Unspecified |
| mmu-let-7i-5p | MIMAT0000122 | Arhgef2 | 16800 | HITS-CLIP | | 23142080 | Unspecified |
| mmu-let-7i-5p | MIMAT0000122 | Arhgef3 | 71704 | HITS-CLIP | | 37986224\|23142080\|23597149\|3143: | Unspecified |
| mmu-let-7i-5p | MIMAT0000122 | Arhgef39 | 230098 | HITS-CLIP | | 23142080 | Unspecified |
| mmu-let-7i-5p | MIMAT0000122 | Arhgef5 | 54324 | HITS-CLIP | | 23597149 | Unspecified |
| mmu-let-7i-5p | MIMAT0000122 | Arhgef6 | 73341 | HITS-CLIP | | 23142080 | Unspecified |
| mmu-let-7i-5p | MIMAT0000122 | Arhgef7 | 54126 | HITS-CLIP | | 29167373 | Unspecified |
| mmu-let-7i-5p | MIMAT0000122 | Arid1a | 93760 | HITS-CLIP | | 23142080\|23597149\|19536157\|2916i | Unspecified |
| mmu-let-7i-5p | MIMAT0000122 | Arid1b | 239985 | HITS-CLIP | | 37986224\|23597149\|23142080 | Unspecified |
| mmu-let-7i-5p | MIMAT0000122 | Arid2 | 77044 | HITS-CLIP | | 37986224\|19536157\|23597149\|3143: | Unspecified |
| mmu-let-7i-5p | MIMAT0000122 | Arid3a | 13496 | HITS-CLIP | | 37986224\|23142080 | Unspecified |
| mmu-let-7i-5p | MIMAT0000122 | Arid3b | 56380 | HITS-CLIP | | 37986224\|23142080\|19536157\|2359i | Unspecified |
| mmu-let-7i-5p | MIMAT0000122 | Arid4a | 238247 | HITS-CLIP | | 23142080\|29167373 | Unspecified |
| mmu-let-7i-5p | MIMAT0000122 | Arid4b | 94246 | HITS-CLIP | | 23142080\|29167373 | Unspecified |
| mmu-let-7i-5p | MIMAT0000122 | Arid5b | 71371 | HITS-CLIP | | 23142080\|23597149\|29167373 | Unspecified |
| mmu-let-7i-5p | MIMAT0000122 | Arih1 | 23806 | HITS-CLIP | | 37986224\|23142080\|23597149\|2916i | Unspecified |
| mmu-let-7i-5p | MIMAT0000122 | Arih2 | 23807 | HITS-CLIP | | 23597149\|23142080 | Unspecified |
| mmu-let-7i-5p | MIMAT0000122 | Arl14ep | 212772 | HITS-CLIP | | 23142080\|29167373 | Unspecified |
| mmu-let-7i-5p | MIMAT0000122 | Arl2 | 56327 | HITS-CLIP | | 31433990 | Unspecified |

| | | | | | | |
|---|---|---|---|---|---|---|
| mmu-let-7i-5p | MIMAT0000122 | Arl4c | 320982 | HITS-CLIP | 23142080\|29167373 | Unspecified |
| mmu-let-7i-5p | MIMAT0000122 | Arl4d | 80981 | HITS-CLIP | 23597149 | Unspecified |
| mmu-let-7i-5p | MIMAT0000122 | Arl5a | 75423 | HITS-CLIP | 37986224\|19536157\|23597149\|23142 | Unspecified |
| mmu-let-7i-5p | MIMAT0000122 | Arl5b | 75869 | HITS-CLIP | 37986224\|23142080\|29167373 | Unspecified |
| mmu-let-7i-5p | MIMAT0000122 | Arl6ip1 | 54208 | HITS-CLIP | 23597149\|23142080 | Unspecified |
| mmu-let-7i-5p | MIMAT0000122 | Arl6ip5 | 65106 | HITS-CLIP | 23142080 | Unspecified |
| mmu-let-7i-5p | MIMAT0000122 | Arl6ip6 | 65103 | HITS-CLIP | 23142080 | Unspecified |
| mmu-let-7i-5p | MIMAT0000122 | Arl8b | 67166 | HITS-CLIP | 23597149\|23142080 | Unspecified |
| mmu-let-7i-5p | MIMAT0000122 | Armc8 | 74125 | HITS-CLIP | 19536157\|23142080\|29167373 | Unspecified |
| mmu-let-7i-5p | MIMAT0000122 | Armcx2 | 67416 | HITS-CLIP | 19536157 | Unspecified |
| mmu-let-7i-5p | MIMAT0000122 | Armt1 | 79624 | HITS-CLIP | 19536157\|23597149\|23142080\|29167 | Unspecified |
| mmu-let-7i-5p | MIMAT0000122 | Arnt | 11863 | HITS-CLIP | 23142080 | Unspecified |
| mmu-let-7i-5p | MIMAT0000122 | Arnt2 | 11864 | HITS-CLIP | 37986224 | Unspecified |
| mmu-let-7i-5p | MIMAT0000122 | Arntl | 11865 | HITS-CLIP | 23142080\|29167373 | Unspecified |
| mmu-let-7i-5p | MIMAT0000122 | Arpc1a | 56443 | HITS-CLIP | 23142080 | Unspecified |
| mmu-let-7i-5p | MIMAT0000122 | Arpc2 | 76709 | HITS-CLIP | 23142080 | Unspecified |
| mmu-let-7i-5p | MIMAT0000122 | Arpc3 | 56378 | HITS-CLIP | 19536157\|23597149\|23142080\|29167 | Unspecified |
| mmu-let-7i-5p | MIMAT0000122 | Arpc4 | 68089 | HITS-CLIP | 23142080\|23597149\|29167373 | Unspecified |
| mmu-let-7i-5p | MIMAT0000122 | Arpc5l | 74192 | HITS-CLIP | 23142080 | Unspecified |
| mmu-let-7i-5p | MIMAT0000122 | Arpp19 | 59046 | HITS-CLIP | 37986224\|23597149\|23142080\|1953€ | Unspecified |
| mmu-let-7i-5p | MIMAT0000122 | Arrb1 | 109689 | HITS-CLIP | 29167373 | Unspecified |
| mmu-let-7i-5p | MIMAT0000122 | Arrdc1 | 215705 | HITS-CLIP | 23142080 | Unspecified |
| mmu-let-7i-5p | MIMAT0000122 | Arrdc3 | 105171 | qCLASH//HITS-CLIP | 31393953\|23597149\|23142080 | Unspecified |
| mmu-let-7i-5p | MIMAT0000122 | Arrdc4 | 66412 | HITS-CLIP | 37986224\|23597149 | Unspecified |
| mmu-let-7i-5p | MIMAT0000122 | Arsb | 11881 | HITS-CLIP | 37986224\|23597149\|23142080 | Unspecified |
| mmu-let-7i-5p | MIMAT0000122 | Arsg | 74008 | HITS-CLIP | 23597149 | Unspecified |
| mmu-let-7i-5p | MIMAT0000122 | Art4 | 109978 | HITS-CLIP | 23597149 | Unspecified |
| mmu-let-7i-5p | MIMAT0000122 | Arvcf | 11877 | HITS-CLIP | 23597149 | Unspecified |
| mmu-let-7i-5p | MIMAT0000122 | Asah1 | 11886 | HITS-CLIP | 23597149\|29167373\|23142080 | Unspecified |
| mmu-let-7i-5p | MIMAT0000122 | Asap1 | 13196 | HITS-CLIP | 37986224\|23142080\|23597149\|29167 | Unspecified |
| mmu-let-7i-5p | MIMAT0000122 | Asb13 | 142688 | HITS-CLIP | 37986224 | Unspecified |
| mmu-let-7i-5p | MIMAT0000122 | Asb8 | 78541 | HITS-CLIP | 37986224\|23142080\|23597149\|29167 | Unspecified |
| mmu-let-7i-5p | MIMAT0000122 | Ascc2 | 75452 | HITS-CLIP | 23597149 | Unspecified |
| mmu-let-7i-5p | MIMAT0000122 | Ascc3 | 77987 | HITS-CLIP | 37986224\|23597149\|19536157\|23142 | Unspecified |
| mmu-let-7i-5p | MIMAT0000122 | Asf1a | 66403 | HITS-CLIP | 23142080 | Unspecified |
| mmu-let-7i-5p | MIMAT0000122 | Asf1b | 66929 | HITS-CLIP | 23142080 | Unspecified |
| mmu-let-7i-5p | MIMAT0000122 | Asgr2 | 11890 | HITS-CLIP | 23597149 | Unspecified |
| mmu-let-7i-5p | MIMAT0000122 | Ash1l | 192195 | HITS-CLIP | 37986224\|23142080\|23597149\|29167 | Unspecified |
| mmu-let-7i-5p | MIMAT0000122 | Asic1 | 11419 | HITS-CLIP | 37986224\|19536157 | Unspecified |

| | | | | | | |
|---|---|---|---|---|---|---|
| mmu-let-7i-5p | MIMAT0000122 | Asl | 109900 | HITS-CLIP | 23597149 | Unspecified |
| mmu-let-7i-5p | MIMAT0000122 | Asna1 | 56495 | HITS-CLIP | 37986224\|23597149\|23142080 | Unspecified |
| mmu-let-7i-5p | MIMAT0000122 | Asns | 27053 | HITS-CLIP | 23597149\|23142080 | Unspecified |
| mmu-let-7i-5p | MIMAT0000122 | Asnsd1 | 70396 | HITS-CLIP | 19536157 | Unspecified |
| mmu-let-7i-5p | MIMAT0000122 | Asph | 65973 | HITS-CLIP | 29167373 | Unspecified |
| mmu-let-7i-5p | MIMAT0000122 | Aspm | 12316 | HITS-CLIP | 23142080 | Unspecified |
| mmu-let-7i-5p | MIMAT0000122 | Aste1 | 66595 | HITS-CLIP | 23142080 | Unspecified |
| mmu-let-7i-5p | MIMAT0000122 | Astn2 | 56079 | HITS-CLIP | 19536157 | Unspecified |
| mmu-let-7i-5p | MIMAT0000122 | Asxl1 | 228790 | HITS-CLIP | 37986224\|23142080\|29167373\|23597 | Unspecified |
| mmu-let-7i-5p | MIMAT0000122 | Atad2 | 70472 | HITS-CLIP | 31433990\|23142080 | Unspecified |
| mmu-let-7i-5p | MIMAT0000122 | Atad2b | 320817 | HITS-CLIP | 37986224\|23597149\|19536157\|23142 | Unspecified |
| mmu-let-7i-5p | MIMAT0000122 | Atad5 | 237877 | HITS-CLIP | 23142080 | Unspecified |
| mmu-let-7i-5p | MIMAT0000122 | Atcay | 16467 | HITS-CLIP | 23142080 | Unspecified |
| mmu-let-7i-5p | MIMAT0000122 | Ate1 | 11907 | HITS-CLIP | 29167373 | Unspecified |
| mmu-let-7i-5p | MIMAT0000122 | Atf1 | 11908 | HITS-CLIP | 23142080 | Unspecified |
| mmu-let-7i-5p | MIMAT0000122 | Atf2 | 11909 | HITS-CLIP | 37986224\|23142080\|29167373 | Unspecified |
| mmu-let-7i-5p | MIMAT0000122 | Atf3 | 11910 | HITS-CLIP | 29167373 | Unspecified |
| mmu-let-7i-5p | MIMAT0000122 | Atf4 | 11911 | HITS-CLIP | 23597149 | Unspecified |
| mmu-let-7i-5p | MIMAT0000122 | Atf6 | 226641 | HITS-CLIP | 23597149\|23142080\|29167373 | Unspecified |
| mmu-let-7i-5p | MIMAT0000122 | Atf6b | 12915 | HITS-CLIP//qCLASH | 37986224\|23142080\|31393953 | Unspecified |
| mmu-let-7i-5p | MIMAT0000122 | Atf7 | 223922 | HITS-CLIP | 37986224\|23142080 | Unspecified |
| mmu-let-7i-5p | MIMAT0000122 | Atf7ip | 54343 | HITS-CLIP | 37986224\|23597149\|19536157\|23142 | Unspecified |
| mmu-let-7i-5p | MIMAT0000122 | Atg10 | 66795 | HITS-CLIP | 23142080 | Unspecified |
| mmu-let-7i-5p | MIMAT0000122 | Atg12 | 67526 | HITS-CLIP | 31433990 | Unspecified |
| mmu-let-7i-5p | MIMAT0000122 | Atg14 | 100504663 | HITS-CLIP | 23597149 | Unspecified |
| mmu-let-7i-5p | MIMAT0000122 | Atg16l1 | 77040 | HITS-CLIP | 37986224\|23597149\|19536157\|29167 | Unspecified |
| mmu-let-7i-5p | MIMAT0000122 | Atg2a | 329015 | HITS-CLIP | 23142080 | Unspecified |
| mmu-let-7i-5p | MIMAT0000122 | Atg2b | 76559 | HITS-CLIP | 37986224\|23597149\|23142080 | Unspecified |
| mmu-let-7i-5p | MIMAT0000122 | Atg4c | 242557 | HITS-CLIP | 23142080 | Unspecified |
| mmu-let-7i-5p | MIMAT0000122 | Atg4d | 235040 | HITS-CLIP | 23142080 | Unspecified |
| mmu-let-7i-5p | MIMAT0000122 | Atg7 | 74244 | HITS-CLIP | 23597149 | Unspecified |
| mmu-let-7i-5p | MIMAT0000122 | Atg9a | 245860 | HITS-CLIP | 23142080 | Unspecified |
| mmu-let-7i-5p | MIMAT0000122 | Atic | 108147 | HITS-CLIP | 23142080 | Unspecified |
| mmu-let-7i-5p | MIMAT0000122 | Atl1 | 73991 | HITS-CLIP | 37986224\|19536157 | Unspecified |
| mmu-let-7i-5p | MIMAT0000122 | Atl2 | 56298 | HITS-CLIP | 23597149\|23142080 | Unspecified |
| mmu-let-7i-5p | MIMAT0000122 | Atl3 | 109168 | HITS-CLIP | 23142080\|23597149 | Unspecified |
| mmu-let-7i-5p | MIMAT0000122 | Atm | 11920 | HITS-CLIP//qCLASH | 37986224\|31393953\|23142080 | Unspecified |
| mmu-let-7i-5p | MIMAT0000122 | Atmin | 234776 | HITS-CLIP | 23142080 | Unspecified |
| mmu-let-7i-5p | MIMAT0000122 | Atn1 | 13498 | HITS-CLIP | 23597149\|19536157\|29167373 | Unspecified |

| | | | | | | |
|---|---|---|---|---|---|---|
| mmu-let-7i-5p | MIMAT0000122 | Atosa | 56204 | qCLASH | 31393953 | Unspecified |
| mmu-let-7i-5p | MIMAT0000122 | Atosb | 80256 | HITS-CLIP | 30971389\|23630078 | Unspecified |
| mmu-let-7i-5p | MIMAT0000122 | Atox1 | 11927 | HITS-CLIP | 23597149 | Unspecified |
| mmu-let-7i-5p | MIMAT0000122 | Atp10a | 11982 | HITS-CLIP | 23142080 | Unspecified |
| mmu-let-7i-5p | MIMAT0000122 | Atp11a | 50770 | HITS-CLIP | 23142080\|29167373 | Unspecified |
| mmu-let-7i-5p | MIMAT0000122 | Atp11b | 76295 | HITS-CLIP | 23142080 | Unspecified |
| mmu-let-7i-5p | MIMAT0000122 | Atp13a2 | 23400 | HITS-CLIP | 29167373 | Unspecified |
| mmu-let-7i-5p | MIMAT0000122 | Atp13a3 | 224088 | HITS-CLIP | 37986224\|23597149\|19536157\|31433 | Unspecified |
| mmu-let-7i-5p | MIMAT0000122 | Atp1a1 | 11928 | HITS-CLIP | 23597149\|23142080 | Unspecified |
| mmu-let-7i-5p | MIMAT0000122 | Atp1a2 | 98660 | HITS-CLIP | 19536157 | Unspecified |
| mmu-let-7i-5p | MIMAT0000122 | Atp1a3 | 232975 | HITS-CLIP | 19536157 | Unspecified |
| mmu-let-7i-5p | MIMAT0000122 | Atp1b1 | 11931 | HITS-CLIP | 37986224\|23597149\|19536157 | Unspecified |
| mmu-let-7i-5p | MIMAT0000122 | Atp1b2 | 11932 | HITS-CLIP | 19536157 | Unspecified |
| mmu-let-7i-5p | MIMAT0000122 | Atp1b3 | 11933 | HITS-CLIP | 23142080\|29167373 | Unspecified |
| mmu-let-7i-5p | MIMAT0000122 | Atp2a2 | 11938 | HITS-CLIP//qCLASH | 37986224\|23597149\|19536157\|31393 | Unspecified |
| mmu-let-7i-5p | MIMAT0000122 | Atp2a3 | 53313 | HITS-CLIP | 31433990 | Unspecified |
| mmu-let-7i-5p | MIMAT0000122 | Atp2b1 | 67972 | HITS-CLIP | 37986224\|23142080\|19536157\|29167 | Unspecified |
| mmu-let-7i-5p | MIMAT0000122 | Atp2b2 | 11941 | HITS-CLIP | 21258322\|19536157 | Unspecified |
| mmu-let-7i-5p | MIMAT0000122 | Atp2b3 | 320707 | HITS-CLIP | 19536157 | Unspecified |
| mmu-let-7i-5p | MIMAT0000122 | Atp2b4 | 381290 | HITS-CLIP | 23142080\|19536157 | Unspecified |
| mmu-let-7i-5p | MIMAT0000122 | Atp2c1 | 235574 | HITS-CLIP | 19536157\|23142080 | Unspecified |
| mmu-let-7i-5p | MIMAT0000122 | Atp5a1 | 11946 | HITS-CLIP | 23597149\|23142080\|29167373 | Unspecified |
| mmu-let-7i-5p | MIMAT0000122 | Atp5d | 66043 | HITS-CLIP | 23597149 | Unspecified |
| mmu-let-7i-5p | MIMAT0000122 | Atp5f1 | 515 | HITS-CLIP | 23597149\|23142080 | Unspecified |
| mmu-let-7i-5p | MIMAT0000122 | Atp5g3 | 228033 | HITS-CLIP | 23597149\|23142080 | Unspecified |
| mmu-let-7i-5p | MIMAT0000122 | Atp5s | 27109 | HITS-CLIP | 23597149 | Unspecified |
| mmu-let-7i-5p | MIMAT0000122 | Atp6v0a2 | 21871 | HITS-CLIP | 23597149 | Unspecified |
| mmu-let-7i-5p | MIMAT0000122 | Atp6v0b | 114143 | HITS-CLIP | 37986224\|19536157\|23597149\|23142 | Unspecified |
| mmu-let-7i-5p | MIMAT0000122 | Atp6v0d1 | 11972 | HITS-CLIP | 37986224\|23597149\|19536157 | Unspecified |
| mmu-let-7i-5p | MIMAT0000122 | Atp6v0e | 11974 | HITS-CLIP | 23597149\|23142080 | Unspecified |
| mmu-let-7i-5p | MIMAT0000122 | Atp6v0e2 | 76252 | HITS-CLIP | 37986224 | Unspecified |
| mmu-let-7i-5p | MIMAT0000122 | Atp6v1a | 11964 | HITS-CLIP | 23142080\|29167373 | Unspecified |
| mmu-let-7i-5p | MIMAT0000122 | Atp6v1b2 | 11966 | HITS-CLIP | 37986224\|23597149\|19536157\|23142 | Unspecified |
| mmu-let-7i-5p | MIMAT0000122 | Atp6v1c1 | 66335 | HITS-CLIP | 23142080 | Unspecified |
| mmu-let-7i-5p | MIMAT0000122 | Atp6v1e1 | 11973 | HITS-CLIP | 37986224\|23142080 | Unspecified |
| mmu-let-7i-5p | MIMAT0000122 | Atp6v1h | 108664 | HITS-CLIP | 23142080 | Unspecified |
| mmu-let-7i-5p | MIMAT0000122 | Atp7a | 11977 | HITS-CLIP | 23597149\|23142080 | Unspecified |
| mmu-let-7i-5p | MIMAT0000122 | Atp8a1 | 11980 | HITS-CLIP | 19536157 | Unspecified |
| mmu-let-7i-5p | MIMAT0000122 | Atp8a2 | 50769 | HITS-CLIP | 19536157 | Unspecified |

| | | | | | | |
|---|---|---|---|---|---|---|
| mmu-let-7i-5p | MIMAT0000122 | Atp8b4 | 241633 | HITS-CLIP | 29167373\|23142080 | Unspecified |
| mmu-let-7i-5p | MIMAT0000122 | Atp9b | 50771 | HITS-CLIP | 37986224\|19536157\|23597149\|23142 | Unspecified |
| mmu-let-7i-5p | MIMAT0000122 | Atr | 245000 | HITS-CLIP//qCLASH | 37986224\|23142080\|19536157\|31393 | Unspecified |
| mmu-let-7i-5p | MIMAT0000122 | Atrn | 11990 | HITS-CLIP | 23597149\|19536157 | Unspecified |
| mmu-let-7i-5p | MIMAT0000122 | Atrx | 22589 | HITS-CLIP | 37986224\|19536157\|23142080\|29167 | Unspecified |
| mmu-let-7i-5p | MIMAT0000122 | Atxn1 | 20238 | HITS-CLIP | 23597149\|23142080 | Unspecified |
| mmu-let-7i-5p | MIMAT0000122 | Atxn10 | 54138 | HITS-CLIP | 37986224\|23142080\|23597149 | Unspecified |
| mmu-let-7i-5p | MIMAT0000122 | Atxn1l | 52335 | HITS-CLIP | 19536157\|23142080\|29167373 | Unspecified |
| mmu-let-7i-5p | MIMAT0000122 | Atxn2 | 20239 | HITS-CLIP | 23597149 | Unspecified |
| mmu-let-7i-5p | MIMAT0000122 | Atxn2l | 233871 | HITS-CLIP | 23142080\|23597149\|19536157\|29167 | Unspecified |
| mmu-let-7i-5p | MIMAT0000122 | Atxn7 | 246103 | HITS-CLIP | 29167373 | Unspecified |
| mmu-let-7i-5p | MIMAT0000122 | Atxn7l1 | 380753 | HITS-CLIP | 23142080\|23597149\|29167373 | Unspecified |
| mmu-let-7i-5p | MIMAT0000122 | Atxn7l2 | 72522 | HITS-CLIP | 19536157\|23142080 | Unspecified |
| mmu-let-7i-5p | MIMAT0000122 | Atxn7l3 | 217218 | HITS-CLIP | 19536157\|23597149\|23142080 | Unspecified |
| mmu-let-7i-5p | MIMAT0000122 | Atxn7l3b | 382423 | HITS-CLIP | 37986224\|23597149\|19536157\|23142 | Unspecified |
| mmu-let-7i-5p | MIMAT0000122 | Auh | 11992 | HITS-CLIP | 23142080\|29167373 | Unspecified |
| mmu-let-7i-5p | MIMAT0000122 | Aurkb | 20877 | HITS-CLIP | 23142080 | Unspecified |
| mmu-let-7i-5p | MIMAT0000122 | Aven | 74268 | HITS-CLIP | 23142080 | Unspecified |
| mmu-let-7i-5p | MIMAT0000122 | Avl9 | 78937 | HITS-CLIP | 37986224\|23597149\|19536157\|31433 | Unspecified |
| mmu-let-7i-5p | MIMAT0000122 | Axl | 26362 | HITS-CLIP | 23142080\|29167373 | Unspecified |
| mmu-let-7i-5p | MIMAT0000122 | Azgp1 | 12007 | HITS-CLIP | 23597149 | Unspecified |
| mmu-let-7i-5p | MIMAT0000122 | Azi2 | 27215 | HITS-CLIP | 29167373\|23142080 | Unspecified |
| mmu-let-7i-5p | MIMAT0000122 | Azin1 | 54375 | HITS-CLIP | 23142080 | Unspecified |
| mmu-let-7i-5p | MIMAT0000122 | B230118H07Rik | 68170 | HITS-CLIP | 23597149 | Unspecified |
| mmu-let-7i-5p | MIMAT0000122 | B230219D22Rik | 78521 | HITS-CLIP | 23142080 | Unspecified |
| mmu-let-7i-5p | MIMAT0000122 | B230307C23Rik | 245305 | HITS-CLIP | 23142080 | Unspecified |
| mmu-let-7i-5p | MIMAT0000122 | B2m | 12010 | HITS-CLIP | 23597149\|23142080\|29167373 | Unspecified |
| mmu-let-7i-5p | MIMAT0000122 | B3galnt1 | 26879 | HITS-CLIP | 37986224\|29167373 | Unspecified |
| mmu-let-7i-5p | MIMAT0000122 | B3galt1 | 26877 | HITS-CLIP | 37986224\|23597149 | Unspecified |
| mmu-let-7i-5p | MIMAT0000122 | B3galt2 | 26878 | HITS-CLIP | 19536157 | Unspecified |
| mmu-let-7i-5p | MIMAT0000122 | B3galt4 | 54218 | HITS-CLIP | 23142080 | Unspecified |
| mmu-let-7i-5p | MIMAT0000122 | B3gat1 | 76898 | HITS-CLIP | 37986224 | Unspecified |
| mmu-let-7i-5p | MIMAT0000122 | B3gat3 | 72727 | HITS-CLIP | 23597149\|23142080 | Unspecified |
| mmu-let-7i-5p | MIMAT0000122 | B3gnt2 | 53625 | HITS-CLIP | 37986224 | Unspecified |
| mmu-let-7i-5p | MIMAT0000122 | B3gntl1 | 210004 | qCLASH | 31393953 | Unspecified |
| mmu-let-7i-5p | MIMAT0000122 | B4galnt1 | 14421 | HITS-CLIP | 23142080 | Unspecified |
| mmu-let-7i-5p | MIMAT0000122 | B4galt1 | 14595 | HITS-CLIP | 23597149\|31433990\|23142080\|29167 | Unspecified |
| mmu-let-7i-5p | MIMAT0000122 | B4galt3 | 57370 | HITS-CLIP | 23142080 | Unspecified |
| mmu-let-7i-5p | MIMAT0000122 | B4galt5 | 56336 | HITS-CLIP | 23597149\|23142080\|29167373 | Unspecified |

| | | | | | | |
|---|---|---|---|---|---|---|
| mmu-let-7i-5p | MIMAT0000122 | B4galt6 | 56386 | HITS-CLIP | 37986224\|19536157\|29167373 | Unspecified |
| mmu-let-7i-5p | MIMAT0000122 | B4gat1 | 135972575 | HITS-CLIP | 19536157 | Unspecified |
| mmu-let-7i-5p | MIMAT0000122 | BC003965 | 214489 | HITS-CLIP | 23597149 | Unspecified |
| mmu-let-7i-5p | MIMAT0000122 | BC005537 | 79555 | HITS-CLIP | 37986224\|23597149\|23142080\|1953( | Unspecified |
| mmu-let-7i-5p | MIMAT0000122 | BC037034 | 231807 | HITS-CLIP | 23142080 | Unspecified |
| mmu-let-7i-5p | MIMAT0000122 | BC052040 | 399568 | HITS-CLIP | 37986224\|23142080\|19536157 | Unspecified |
| mmu-let-7i-5p | MIMAT0000122 | BC055324 | 381306 | HITS-CLIP | 23142080 | Unspecified |
| mmu-let-7i-5p | MIMAT0000122 | Babam2 | 9577 | HITS-CLIP | 23142080\|23597149\|29167373 | Unspecified |
| mmu-let-7i-5p | MIMAT0000122 | Bace1 | 23821 | HITS-CLIP | 23597149\|19536157 | Unspecified |
| mmu-let-7i-5p | MIMAT0000122 | Bace2 | 56175 | HITS-CLIP | 23597149 | Unspecified |
| mmu-let-7i-5p | MIMAT0000122 | Bach1 | 12013 | HITS-CLIP | 37986224\|23597149\|23142080\|1953( | Unspecified |
| mmu-let-7i-5p | MIMAT0000122 | Bach2 | 12014 | HITS-CLIP | 37986224\|23597149\|23630078\|1953( | Unspecified |
| mmu-let-7i-5p | MIMAT0000122 | Bag4 | 67384 | HITS-CLIP | 19536157\|29167373 | Unspecified |
| mmu-let-7i-5p | MIMAT0000122 | Bag6 | 224727 | HITS-CLIP | 37986224\|23597149\|23142080 | Unspecified |
| mmu-let-7i-5p | MIMAT0000122 | Bahcc1 | 268515 | HITS-CLIP | 23597149 | Unspecified |
| mmu-let-7i-5p | MIMAT0000122 | Bahd1 | 228536 | HITS-CLIP | 23597149\|23142080 | Unspecified |
| mmu-let-7i-5p | MIMAT0000122 | Baiap2 | 108100 | HITS-CLIP | 23597149 | Unspecified |
| mmu-let-7i-5p | MIMAT0000122 | Bap1 | 104416 | HITS-CLIP | 23142080 | Unspecified |
| mmu-let-7i-5p | MIMAT0000122 | Bard1 | 12021 | HITS-CLIP | 23142080 | Unspecified |
| mmu-let-7i-5p | MIMAT0000122 | Basp1 | 70350 | HITS-CLIP | 19536157\|29167373 | Unspecified |
| mmu-let-7i-5p | MIMAT0000122 | Baz1a | 217578 | HITS-CLIP | 37986224\|23597149\|23142080 | Unspecified |
| mmu-let-7i-5p | MIMAT0000122 | Baz1b | 22385 | HITS-CLIP | 23597149\|23142080\|29167373 | Unspecified |
| mmu-let-7i-5p | MIMAT0000122 | Baz2a | 116848 | HITS-CLIP//qCLASH | 37986224\|23142080\|31393953\|29167 | Unspecified |
| mmu-let-7i-5p | MIMAT0000122 | Baz2b | 407823 | HITS-CLIP | 19536157\|23597149\|23142080\|29167 | Unspecified |
| mmu-let-7i-5p | MIMAT0000122 | Bbc3 | 170770 | HITS-CLIP | 23597149 | Unspecified |
| mmu-let-7i-5p | MIMAT0000122 | Bbox1 | 170442 | HITS-CLIP | 23597149 | Unspecified |
| mmu-let-7i-5p | MIMAT0000122 | Bbs10 | 71769 | HITS-CLIP | 29167373\|23142080 | Unspecified |
| mmu-let-7i-5p | MIMAT0000122 | Bbs5 | 72569 | HITS-CLIP | 23142080 | Unspecified |
| mmu-let-7i-5p | MIMAT0000122 | Bbx | 70508 | HITS-CLIP | 37986224\|23142080 | Unspecified |
| mmu-let-7i-5p | MIMAT0000122 | Bcan | 12032 | HITS-CLIP | 19536157 | Unspecified |
| mmu-let-7i-5p | MIMAT0000122 | Bcap29 | 12033 | HITS-CLIP | 37986224\|23142080\|23597149 | Unspecified |
| mmu-let-7i-5p | MIMAT0000122 | Bcar1 | 12927 | HITS-CLIP | 37986224\|23597149 | Unspecified |
| mmu-let-7i-5p | MIMAT0000122 | Bcar3 | 29815 | HITS-CLIP | 37986224\|29167373 | Unspecified |
| mmu-let-7i-5p | MIMAT0000122 | Bcat1 | 12035 | HITS-CLIP | 19536157\|23142080 | Unspecified |
| mmu-let-7i-5p | MIMAT0000122 | Bcat2 | 12036 | HITS-CLIP | 23142080\|23597149 | Unspecified |
| mmu-let-7i-5p | MIMAT0000122 | Bccip | 66165 | HITS-CLIP | 23142080 | Unspecified |
| mmu-let-7i-5p | MIMAT0000122 | Bche | 12038 | HITS-CLIP | 23597149 | Unspecified |
| mmu-let-7i-5p | MIMAT0000122 | Bckdha | 12039 | HITS-CLIP | 23597149 | Unspecified |
| mmu-let-7i-5p | MIMAT0000122 | Bckdhb | 12040 | HITS-CLIP | 23597149 | Unspecified |

| | | | | | | |
|---|---|---|---|---|---|---|
| mmu-let-7i-5p | MIMAT0000122 | Bcl11a | 14025 | HITS-CLIP | 23142080 | Unspecified |
| mmu-let-7i-5p | MIMAT0000122 | Bcl11b | 58208 | HITS-CLIP | 19536157\|23142080 | Unspecified |
| mmu-let-7i-5p | MIMAT0000122 | Bcl2 | 12043 | HITS-CLIP | 37986224\|31433990\|23142080 | Unspecified |
| mmu-let-7i-5p | MIMAT0000122 | Bcl2a1d | 12047 | HITS-CLIP | 23142080 | Unspecified |
| mmu-let-7i-5p | MIMAT0000122 | Bcl2l11 | 12125 | HITS-CLIP | 37986224\|23597149\|23142080\|29167 | Unspecified |
| mmu-let-7i-5p | MIMAT0000122 | Bcl2l2 | 12050 | HITS-CLIP | 37986224\|19536157 | Unspecified |
| mmu-let-7i-5p | MIMAT0000122 | Bcl6 | 12053 | HITS-CLIP | 23142080\|29167373 | Unspecified |
| mmu-let-7i-5p | MIMAT0000122 | Bcl7a | 77045 | HITS-CLIP | 23597149\|23142080 | Unspecified |
| mmu-let-7i-5p | MIMAT0000122 | Bcl7b | 12054 | HITS-CLIP | 23597149 | Unspecified |
| mmu-let-7i-5p | MIMAT0000122 | Bcl9 | 77578 | HITS-CLIP | 37986224\|23142080 | Unspecified |
| mmu-let-7i-5p | MIMAT0000122 | Bcl9l | 80288 | HITS-CLIP | 23142080\|19536157 | Unspecified |
| mmu-let-7i-5p | MIMAT0000122 | Bclaf1 | 72567 | HITS-CLIP | 19536157\|29167373\|23142080 | Unspecified |
| mmu-let-7i-5p | MIMAT0000122 | Bcor | 71458 | HITS-CLIP | 23597149\|23142080 | Unspecified |
| mmu-let-7i-5p | MIMAT0000122 | Bcorl1 | 320376 | HITS-CLIP | 23142080 | Unspecified |
| mmu-let-7i-5p | MIMAT0000122 | Bcr | 110279 | HITS-CLIP | 37986224 | Unspecified |
| mmu-let-7i-5p | MIMAT0000122 | Bdh1 | 71911 | HITS-CLIP | 23142080 | Unspecified |
| mmu-let-7i-5p | MIMAT0000122 | Bdp1 | 544971 | HITS-CLIP | 31433990\|23597149\|23142080\|29167 | Unspecified |
| mmu-let-7i-5p | MIMAT0000122 | Bend3 | 331623 | HITS-CLIP | 29167373 | Unspecified |
| mmu-let-7i-5p | MIMAT0000122 | Bet1 | 12068 | HITS-CLIP | 23142080 | Unspecified |
| mmu-let-7i-5p | MIMAT0000122 | Bfar | 67118 | HITS-CLIP | 23142080 | Unspecified |
| mmu-let-7i-5p | MIMAT0000122 | Bgn | 12111 | HITS-CLIP | 23597149 | Unspecified |
| mmu-let-7i-5p | MIMAT0000122 | Bhlhe40 | 20893 | HITS-CLIP | 23597149\|29167373\|23142080 | Unspecified |
| mmu-let-7i-5p | MIMAT0000122 | Bhlhe41 | 79362 | HITS-CLIP | 37986224\|23597149\|19536157\|29167 | Unspecified |
| mmu-let-7i-5p | MIMAT0000122 | Bicc1 | 83675 | HITS-CLIP | 23597149 | Unspecified |
| mmu-let-7i-5p | MIMAT0000122 | Bicra | 29998 | HITS-CLIP | 23142080\|23597149 | Unspecified |
| mmu-let-7i-5p | MIMAT0000122 | Bicral | 23506 | HITS-CLIP | 23597149 | Unspecified |
| mmu-let-7i-5p | MIMAT0000122 | Birc2 | 11797 | HITS-CLIP | 23142080 | Unspecified |
| mmu-let-7i-5p | MIMAT0000122 | Birc3 | 11796 | HITS-CLIP | 37986224\|23142080\|29167373 | Unspecified |
| mmu-let-7i-5p | MIMAT0000122 | Birc6 | 12211 | HITS-CLIP | 37986224\|23142080\|23597149\|19536 | Unspecified |
| mmu-let-7i-5p | MIMAT0000122 | Blcap | 53619 | HITS-CLIP | 23597149\|23142080\|29167373 | Unspecified |
| mmu-let-7i-5p | MIMAT0000122 | Blm | 12144 | HITS-CLIP | 23142080 | Unspecified |
| mmu-let-7i-5p | MIMAT0000122 | Blmh | 104184 | HITS-CLIP | 23142080 | Unspecified |
| mmu-let-7i-5p | MIMAT0000122 | Bloc1s6 | 18457 | HITS-CLIP | 23142080 | Unspecified |
| mmu-let-7i-5p | MIMAT0000122 | Bltp2 | 9703 | HITS-CLIP | 31433990 | Unspecified |
| mmu-let-7i-5p | MIMAT0000122 | Blzf1 | 66352 | HITS-CLIP | 23142080 | Unspecified |
| mmu-let-7i-5p | MIMAT0000122 | Bmp2k | 140780 | HITS-CLIP | 23597149\|23142080\|29167373 | Unspecified |
| mmu-let-7i-5p | MIMAT0000122 | Bmp4 | 12159 | HITS-CLIP | 23597149 | Unspecified |
| mmu-let-7i-5p | MIMAT0000122 | Bmpr2 | 12168 | HITS-CLIP | 37986224\|23597149\|19536157\|23142 | Unspecified |
| mmu-let-7i-5p | MIMAT0000122 | Bmt2 | 101148 | HITS-CLIP | 23142080\|23597149\|29167373 | Unspecified |

| | | | | | | | |
|---|---|---|---|---|---|---|---|
| mmu-let-7i-5p | MIMAT0000122 | Bmyc | 107771 | HITS-CLIP | | 23142080 | Unspecified |
| mmu-let-7i-5p | MIMAT0000122 | Bod1l | 665775 | HITS-CLIP | | 23142080 | Unspecified |
| mmu-let-7i-5p | MIMAT0000122 | Bola1 | 51027 | HITS-CLIP | | 23142080 | Unspecified |
| mmu-let-7i-5p | MIMAT0000122 | Bora | 77744 | HITS-CLIP | | 23142080 | Unspecified |
| mmu-let-7i-5p | MIMAT0000122 | Bphl | 68021 | HITS-CLIP | | 23597149 | Unspecified |
| mmu-let-7i-5p | MIMAT0000122 | Bpnt2 | 54928 | HITS-CLIP | | 31433990 | Unspecified |
| mmu-let-7i-5p | MIMAT0000122 | Bptf | 207165 | HITS-CLIP | | 19536157\|23142080 | Unspecified |
| mmu-let-7i-5p | MIMAT0000122 | Braf | 109880 | HITS-CLIP | | 23142080 | Unspecified |
| mmu-let-7i-5p | MIMAT0000122 | Brca2 | 12190 | HITS-CLIP | | 23142080 | Unspecified |
| mmu-let-7i-5p | MIMAT0000122 | Brcc3 | 210766 | HITS-CLIP | | 37986224\|23597149 | Unspecified |
| mmu-let-7i-5p | MIMAT0000122 | Brd1 | 223770 | HITS-CLIP | | 23597149\|23142080 | Unspecified |
| mmu-let-7i-5p | MIMAT0000122 | Brd2 | 14312 | HITS-CLIP | | 37986224\|23597149\|30971389\|23142 | Unspecified |
| mmu-let-7i-5p | MIMAT0000122 | Brd3 | 67382 | HITS-CLIP | | 37986224\|23597149\|23142080 | Unspecified |
| mmu-let-7i-5p | MIMAT0000122 | Brd4 | 57261 | HITS-CLIP | | 23142080\|29167373 | Unspecified |
| mmu-let-7i-5p | MIMAT0000122 | Brd8 | 78656 | HITS-CLIP | | 29167373 | Unspecified |
| mmu-let-7i-5p | MIMAT0000122 | Brinp2 | 240843 | HITS-CLIP | | 19536157 | Unspecified |
| mmu-let-7i-5p | MIMAT0000122 | Brip1 | 237911 | HITS-CLIP | | 23142080 | Unspecified |
| mmu-let-7i-5p | MIMAT0000122 | Brix1 | 67832 | HITS-CLIP | | 23142080 | Unspecified |
| mmu-let-7i-5p | MIMAT0000122 | Brpf1 | 78783 | HITS-CLIP | | 23597149 | Unspecified |
| mmu-let-7i-5p | MIMAT0000122 | Brwd1 | 93871 | HITS-CLIP | | 37986224\|23142080\|19536157\|23597 | Unspecified |
| mmu-let-7i-5p | MIMAT0000122 | Brwd3 | 382236 | HITS-CLIP | | 37986224\|23142080\|19536157\|23597 | Unspecified |
| mmu-let-7i-5p | MIMAT0000122 | Bscl2 | 14705 | HITS-CLIP | | 23142080\|19536157 | Unspecified |
| mmu-let-7i-5p | MIMAT0000122 | Bsg | 12215 | HITS-CLIP | | 37986224\|23142080\|23597149\|19536 | Unspecified |
| mmu-let-7i-5p | MIMAT0000122 | Bsn | 12217 | HITS-CLIP | | 37986224\|19536157\|23142080 | Unspecified |
| mmu-let-7i-5p | MIMAT0000122 | Btaf1 | 107182 | HITS-CLIP | | 23142080 | Unspecified |
| mmu-let-7i-5p | MIMAT0000122 | Btbd10 | 68815 | HITS-CLIP | | 23142080 | Unspecified |
| mmu-let-7i-5p | MIMAT0000122 | Btbd19 | 78611 | HITS-CLIP | | 23142080 | Unspecified |
| mmu-let-7i-5p | MIMAT0000122 | Btbd3 | 228662 | HITS-CLIP | | 19536157 | Unspecified |
| mmu-let-7i-5p | MIMAT0000122 | Btbd7 | 238386 | HITS-CLIP | | 23142080\|29167373 | Unspecified |
| mmu-let-7i-5p | MIMAT0000122 | Btbd9 | 224671 | HITS-CLIP | | 23142080\|23597149\|29167373 | Unspecified |
| mmu-let-7i-5p | MIMAT0000122 | Btd | 26363 | HITS-CLIP | | 23142080 | Unspecified |
| mmu-let-7i-5p | MIMAT0000122 | Btg1 | 12226 | HITS-CLIP | | 29167373 | Unspecified |
| mmu-let-7i-5p | MIMAT0000122 | Btg2 | 12227 | HITS-CLIP//qCLASH | | 37986224\|23597149\|23142080\|31393 | Unspecified |
| mmu-let-7i-5p | MIMAT0000122 | Btla | 208154 | HITS-CLIP | | 23142080 | Unspecified |
| mmu-let-7i-5p | MIMAT0000122 | Btrc | 12234 | HITS-CLIP | | 23597149\|23142080 | Unspecified |
| mmu-let-7i-5p | MIMAT0000122 | Bub1 | 12235 | HITS-CLIP | | 23142080 | Unspecified |
| mmu-let-7i-5p | MIMAT0000122 | Bub1b | 12236 | HITS-CLIP | | 23142080 | Unspecified |
| mmu-let-7i-5p | MIMAT0000122 | Bub3 | 12237 | HITS-CLIP | | 23142080 | Unspecified |
| mmu-let-7i-5p | MIMAT0000122 | Bud13 | 215051 | HITS-CLIP | | 23142080 | Unspecified |

| | | | | | | |
|---|---|---|---|---|---|---|
| mmu-let-7i-5p | MIMAT0000122 | Bud23 | 114049 | HITS-CLIP | 23142080 | Unspecified |
| mmu-let-7i-5p | MIMAT0000122 | Bysl | 53414 | HITS-CLIP | 23142080\|29167373 | Unspecified |
| mmu-let-7i-5p | MIMAT0000122 | Bzw1 | 66882 | qCLASH//HITS-CLIP | 31393953\|29167373\|23142080 | Unspecified |
| mmu-let-7i-5p | MIMAT0000122 | Bzw2 | 66912 | HITS-CLIP | 23142080 | Unspecified |
| mmu-let-7i-5p | MIMAT0000122 | C030006K11Rik | 223665 | HITS-CLIP | 23142080 | Unspecified |
| mmu-let-7i-5p | MIMAT0000122 | C1galt1 | 94192 | HITS-CLIP | 31433990\|23597149\|19536157\|2363( | Unspecified |
| mmu-let-7i-5p | MIMAT0000122 | C1galt1c1 | 59048 | HITS-CLIP | 23142080 | Unspecified |
| mmu-let-7i-5p | MIMAT0000122 | C1qbp | 12261 | HITS-CLIP | 23142080 | Unspecified |
| mmu-let-7i-5p | MIMAT0000122 | C1qtnf1 | 56745 | HITS-CLIP | 23597149 | Unspecified |
| mmu-let-7i-5p | MIMAT0000122 | C1s1 | 50908 | HITS-CLIP | 23597149 | Unspecified |
| mmu-let-7i-5p | MIMAT0000122 | C2cd2 | 207781 | HITS-CLIP | 37986224\|23597149 | Unspecified |
| mmu-let-7i-5p | MIMAT0000122 | C2cd2l | 71764 | HITS-CLIP | 23597149\|23142080 | Unspecified |
| mmu-let-7i-5p | MIMAT0000122 | C2cd3 | 277939 | HITS-CLIP | 23597149 | Unspecified |
| mmu-let-7i-5p | MIMAT0000122 | C2cd5 | 74741 | HITS-CLIP | 23142080\|31433990 | Unspecified |
| mmu-let-7i-5p | MIMAT0000122 | C3 | 12266 | HITS-CLIP | 37986224\|23597149\|23142080 | Unspecified |
| mmu-let-7i-5p | MIMAT0000122 | C330007P06Rik | 77644 | HITS-CLIP | 23597149\|23142080 | Unspecified |
| mmu-let-7i-5p | MIMAT0000122 | C330018D20Rik | 77422 | HITS-CLIP | 23142080 | Unspecified |
| mmu-let-7i-5p | MIMAT0000122 | C3ar1 | 12267 | HITS-CLIP | 29167373 | Unspecified |
| mmu-let-7i-5p | MIMAT0000122 | C4b | 12268 | HITS-CLIP | 23597149 | Unspecified |
| mmu-let-7i-5p | MIMAT0000122 | C4bp | 12269 | HITS-CLIP | 23597149 | Unspecified |
| mmu-let-7i-5p | MIMAT0000122 | C5ar1 | 12273 | HITS-CLIP | 29167373 | Unspecified |
| mmu-let-7i-5p | MIMAT0000122 | C6 | 12274 | HITS-CLIP | 23597149 | Unspecified |
| mmu-let-7i-5p | MIMAT0000122 | C8a | 230558 | HITS-CLIP | 23597149 | Unspecified |
| mmu-let-7i-5p | MIMAT0000122 | C9 | 12279 | HITS-CLIP | 23597149 | Unspecified |
| mmu-let-7i-5p | MIMAT0000122 | C9orf72 | 203228 | HITS-CLIP | 23597149\|23142080\|29167373 | Unspecified |
| mmu-let-7i-5p | MIMAT0000122 | Cab39 | 12283 | HITS-CLIP | 23142080 | Unspecified |
| mmu-let-7i-5p | MIMAT0000122 | Cabin1 | 104248 | HITS-CLIP | 23142080\|29167373 | Unspecified |
| mmu-let-7i-5p | MIMAT0000122 | Cables2 | 252966 | HITS-CLIP | 23142080 | Unspecified |
| mmu-let-7i-5p | MIMAT0000122 | Cacfd1 | 381356 | HITS-CLIP | 37986224\|23597149\|23142080 | Unspecified |
| mmu-let-7i-5p | MIMAT0000122 | Cacna1a | 12286 | HITS-CLIP | 29167373 | Unspecified |
| mmu-let-7i-5p | MIMAT0000122 | Cacna1c | 12288 | HITS-CLIP | 19536157 | Unspecified |
| mmu-let-7i-5p | MIMAT0000122 | Cacna1d | 12289 | HITS-CLIP | 19536157 | Unspecified |
| mmu-let-7i-5p | MIMAT0000122 | Cacna2d1 | 12293 | HITS-CLIP | 23597149 | Unspecified |
| mmu-let-7i-5p | MIMAT0000122 | Cacna2d3 | 12294 | HITS-CLIP | 37986224\|19536157 | Unspecified |
| mmu-let-7i-5p | MIMAT0000122 | Cacng4 | 54377 | HITS-CLIP | 37986224\|19536157 | Unspecified |
| mmu-let-7i-5p | MIMAT0000122 | Cactin | 70312 | HITS-CLIP | 23142080 | Unspecified |
| mmu-let-7i-5p | MIMAT0000122 | Cacul1 | 78832 | qCLASH//HITS-CLIP | 31393953\|23597149\|23142080\|29167 | Unspecified |
| mmu-let-7i-5p | MIMAT0000122 | Cacybp | 12301 | HITS-CLIP | 23142080 | Unspecified |
| mmu-let-7i-5p | MIMAT0000122 | Cad | 69719 | HITS-CLIP | 23142080 | Unspecified |

| | | | | | | |
|---|---|---|---|---|---|---|
| mmu-let-7i-5p | MIMAT0000122 | Cadm1 | 54725 | HITS-CLIP | 23597149 | Unspecified |
| mmu-let-7i-5p | MIMAT0000122 | Cadm2 | 239857 | HITS-CLIP | 37986224\|19536157 | Unspecified |
| mmu-let-7i-5p | MIMAT0000122 | Cadm3 | 94332 | HITS-CLIP | 37986224\|19536157 | Unspecified |
| mmu-let-7i-5p | MIMAT0000122 | Cadps | 27062 | HITS-CLIP | 19536157 | Unspecified |
| mmu-let-7i-5p | MIMAT0000122 | Calcrl | 54598 | HITS-CLIP | 23142080\|29167373 | Unspecified |
| mmu-let-7i-5p | MIMAT0000122 | Cald1 | 109624 | HITS-CLIP | 37986224\|23597149\|19536157 | Unspecified |
| mmu-let-7i-5p | MIMAT0000122 | Calm1 | 12313 | HITS-CLIP | 37986224\|19536157\|23597149\|2314 | Unspecified |
| mmu-let-7i-5p | MIMAT0000122 | Calm2 | 12314 | HITS-CLIP | 23597149\|29167373\|23142080 | Unspecified |
| mmu-let-7i-5p | MIMAT0000122 | Calm3 | 12315 | HITS-CLIP | 23142080 | Unspecified |
| mmu-let-7i-5p | MIMAT0000122 | Calu | 12321 | HITS-CLIP | 31433990 | Unspecified |
| mmu-let-7i-5p | MIMAT0000122 | Camk1d | 227541 | HITS-CLIP | 23597149\|23142080 | Unspecified |
| mmu-let-7i-5p | MIMAT0000122 | Camk2a | 12322 | HITS-CLIP | 37986224\|19536157 | Unspecified |
| mmu-let-7i-5p | MIMAT0000122 | Camk2d | 108058 | HITS-CLIP | 23142080\|19536157 | Unspecified |
| mmu-let-7i-5p | MIMAT0000122 | Camk4 | 12326 | HITS-CLIP | 37986224\|23142080 | Unspecified |
| mmu-let-7i-5p | MIMAT0000122 | Camkv | 235604 | HITS-CLIP | 37986224 | Unspecified |
| mmu-let-7i-5p | MIMAT0000122 | Camsap2 | 67886 | HITS-CLIP | 19536157\|23142080 | Unspecified |
| mmu-let-7i-5p | MIMAT0000122 | Camta1 | 100072 | HITS-CLIP | 19536157 | Unspecified |
| mmu-let-7i-5p | MIMAT0000122 | Cand1 | 71902 | HITS-CLIP | 23597149\|23142080 | Unspecified |
| mmu-let-7i-5p | MIMAT0000122 | Cand2 | 67088 | HITS-CLIP | 37986224 | Unspecified |
| mmu-let-7i-5p | MIMAT0000122 | Canx | 12330 | HITS-CLIP | 23142080\|23597149\|29167373 | Unspecified |
| mmu-let-7i-5p | MIMAT0000122 | Cap1 | 12331 | HITS-CLIP | 19536157\|23142080 | Unspecified |
| mmu-let-7i-5p | MIMAT0000122 | Cap2 | 67252 | HITS-CLIP | 37986224\|19536157\|23142080 | Unspecified |
| mmu-let-7i-5p | MIMAT0000122 | Capg | 12332 | HITS-CLIP | 23142080\|29167373 | Unspecified |
| mmu-let-7i-5p | MIMAT0000122 | Capn2 | 12334 | HITS-CLIP | 19536157\|23142080 | Unspecified |
| mmu-let-7i-5p | MIMAT0000122 | Capn5 | 12337 | HITS-CLIP | 23142080 | Unspecified |
| mmu-let-7i-5p | MIMAT0000122 | Capn7 | 12339 | HITS-CLIP | 23142080 | Unspecified |
| mmu-let-7i-5p | MIMAT0000122 | Caprin1 | 53872 | HITS-CLIP | 23597149\|19536157\|23142080 | Unspecified |
| mmu-let-7i-5p | MIMAT0000122 | Caprin2 | 232560 | HITS-CLIP | 23142080 | Unspecified |
| mmu-let-7i-5p | MIMAT0000122 | Capza2 | 12343 | HITS-CLIP | 23142080\|19536157\|31433990\|29167 | Unspecified |
| mmu-let-7i-5p | MIMAT0000122 | Capzb | 12345 | HITS-CLIP | 37986224\|19536157\|23597149\|2314 | Unspecified |
| mmu-let-7i-5p | MIMAT0000122 | Car10 | 72605 | HITS-CLIP | 37986224\|19536157 | Unspecified |
| mmu-let-7i-5p | MIMAT0000122 | Car12 | 76459 | HITS-CLIP | 37986224 | Unspecified |
| mmu-let-7i-5p | MIMAT0000122 | Car2 | 12349 | HITS-CLIP | 37986224\|19536157 | Unspecified |
| mmu-let-7i-5p | MIMAT0000122 | Card11 | 108723 | HITS-CLIP | 23142080 | Unspecified |
| mmu-let-7i-5p | MIMAT0000122 | Card6 | 239319 | HITS-CLIP | 23142080 | Unspecified |
| mmu-let-7i-5p | MIMAT0000122 | Carm1 | 59035 | HITS-CLIP | 23597149\|23142080 | Unspecified |
| mmu-let-7i-5p | MIMAT0000122 | Carmil2 | 146206 | HITS-CLIP | 23142080 | Unspecified |
| mmu-let-7i-5p | MIMAT0000122 | Cars | 27267 | HITS-CLIP | 23142080 | Unspecified |
| mmu-let-7i-5p | MIMAT0000122 | Casc3 | 192160 | HITS-CLIP | 23597149\|23142080 | Unspecified |

| | | | | | | |
|---|---|---|---|---|---|---|
| mmu-let-7i-5p | MIMAT0000122 | Cask | 12361 | HITS-CLIP | 23142080 | Unspecified |
| mmu-let-7i-5p | MIMAT0000122 | Casp1 | 12362 | HITS-CLIP | 29167373 | Unspecified |
| mmu-let-7i-5p | MIMAT0000122 | Casp2 | 12366 | HITS-CLIP | 23597149\|23142080 | Unspecified |
| mmu-let-7i-5p | MIMAT0000122 | Casp3 | 12367 | HITS-CLIP | 37986224\|23142080 | Unspecified |
| mmu-let-7i-5p | MIMAT0000122 | Casp4 | 12363 | HITS-CLIP | 23630078\|23142080\|29167373 | Unspecified |
| mmu-let-7i-5p | MIMAT0000122 | Casp6 | 12368 | HITS-CLIP | 23142080 | Unspecified |
| mmu-let-7i-5p | MIMAT0000122 | Casp7 | 12369 | HITS-CLIP | 23597149 | Unspecified |
| mmu-let-7i-5p | MIMAT0000122 | Casp8 | 12370 | HITS-CLIP | 23597149\|23142080\|29167373 | Unspecified |
| mmu-let-7i-5p | MIMAT0000122 | Casp8ap2 | 26885 | HITS-CLIP | 29167373\|23142080 | Unspecified |
| mmu-let-7i-5p | MIMAT0000122 | Castor1 | 652968 | HITS-CLIP | 23142080 | Unspecified |
| mmu-let-7i-5p | MIMAT0000122 | Casz1 | 69743 | HITS-CLIP | 26203562 | Unspecified |
| mmu-let-7i-5p | MIMAT0000122 | Cat | 12359 | HITS-CLIP | 23597149\|23142080 | Unspecified |
| mmu-let-7i-5p | MIMAT0000122 | Cbfa2t3 | 12398 | HITS-CLIP | 37986224\|19536157 | Unspecified |
| mmu-let-7i-5p | MIMAT0000122 | Cbfb | 12400 | HITS-CLIP | 23597149\|23142080 | Unspecified |
| mmu-let-7i-5p | MIMAT0000122 | Cbl | 12402 | HITS-CLIP | 23142080\|29167373\|23597149 | Unspecified |
| mmu-let-7i-5p | MIMAT0000122 | Cblb | 208650 | HITS-CLIP | 37986224\|23142080\|29167373 | Unspecified |
| mmu-let-7i-5p | MIMAT0000122 | Cbll1 | 104836 | HITS-CLIP | 23142080\|23597149 | Unspecified |
| mmu-let-7i-5p | MIMAT0000122 | Cbln1 | 12404 | HITS-CLIP | 19536157 | Unspecified |
| mmu-let-7i-5p | MIMAT0000122 | Cbln2 | 12405 | HITS-CLIP | 37986224 | Unspecified |
| mmu-let-7i-5p | MIMAT0000122 | Cbr1 | 12408 | HITS-CLIP | 23142080 | Unspecified |
| mmu-let-7i-5p | MIMAT0000122 | Cbx2 | 12416 | HITS-CLIP | 23142080 | Unspecified |
| mmu-let-7i-5p | MIMAT0000122 | Cbx5 | 12419 | HITS-CLIP | 37986224\|23597149\|19536157\|23142 | Unspecified |
| mmu-let-7i-5p | MIMAT0000122 | Cbx6 | 494448 | HITS-CLIP | 23630078 | Unspecified |
| mmu-let-7i-5p | MIMAT0000122 | Cc2d1a | 212139 | HITS-CLIP | 23597149\|23142080 | Unspecified |
| mmu-let-7i-5p | MIMAT0000122 | Cc2d1b | 319965 | HITS-CLIP | 23142080 | Unspecified |
| mmu-let-7i-5p | MIMAT0000122 | Ccar1 | 67500 | HITS-CLIP | 37986224\|23142080\|29167373 | Unspecified |
| mmu-let-7i-5p | MIMAT0000122 | Ccar2 | 219158 | HITS-CLIP | 23142080 | Unspecified |
| mmu-let-7i-5p | MIMAT0000122 | Ccdc115 | 69668 | HITS-CLIP | 23142080 | Unspecified |
| mmu-let-7i-5p | MIMAT0000122 | Ccdc117 | 104479 | HITS-CLIP | 23142080 | Unspecified |
| mmu-let-7i-5p | MIMAT0000122 | Ccdc126 | 57895 | HITS-CLIP | 23142080 | Unspecified |
| mmu-let-7i-5p | MIMAT0000122 | Ccdc127 | 67433 | HITS-CLIP | 23142080 | Unspecified |
| mmu-let-7i-5p | MIMAT0000122 | Ccdc14 | 239839 | HITS-CLIP | 23142080 | Unspecified |
| mmu-let-7i-5p | MIMAT0000122 | Ccdc141 | 545428 | HITS-CLIP | 37986224\|23142080 | Unspecified |
| mmu-let-7i-5p | MIMAT0000122 | Ccdc174 | 232236 | HITS-CLIP | 23597149 | Unspecified |
| mmu-let-7i-5p | MIMAT0000122 | Ccdc186 | 55088 | HITS-CLIP | 23142080\|29167373 | Unspecified |
| mmu-let-7i-5p | MIMAT0000122 | Ccdc25 | 67179 | HITS-CLIP | 23142080 | Unspecified |
| mmu-let-7i-5p | MIMAT0000122 | Ccdc47 | 67163 | HITS-CLIP | 23142080\|19536157 | Unspecified |
| mmu-let-7i-5p | MIMAT0000122 | Ccdc50 | 67501 | HITS-CLIP | 37986224\|23142080 | Unspecified |
| mmu-let-7i-5p | MIMAT0000122 | Ccdc6 | 76551 | HITS-CLIP | 23142080 | Unspecified |

| | | | | | | |
|---|---|---|---|---|---|---|
| mmu-let-7i-5p | MIMAT0000122 | Ccdc71l | 72123 | HITS-CLIP | 37986224\|23142080\|23597149 | Unspecified |
| mmu-let-7i-5p | MIMAT0000122 | Ccdc80 | 67896 | HITS-CLIP | 23597149 | Unspecified |
| mmu-let-7i-5p | MIMAT0000122 | Ccdc82 | 66396 | HITS-CLIP | 23142080 | Unspecified |
| mmu-let-7i-5p | MIMAT0000122 | Ccdc85b | 240514 | HITS-CLIP | 23597149 | Unspecified |
| mmu-let-7i-5p | MIMAT0000122 | Ccdc88a | 108686 | HITS-CLIP | 37986224\|19536157\|29167373 | Unspecified |
| mmu-let-7i-5p | MIMAT0000122 | Ccdc88b | 78317 | HITS-CLIP | 29167373 | Unspecified |
| mmu-let-7i-5p | MIMAT0000122 | Ccdc92 | 215707 | HITS-CLIP | 37986224 | Unspecified |
| mmu-let-7i-5p | MIMAT0000122 | Ccdc93 | 70829 | HITS-CLIP | 23142080\|29167373 | Unspecified |
| mmu-let-7i-5p | MIMAT0000122 | Ccdc97 | 52132 | HITS-CLIP | 23142080 | Unspecified |
| mmu-let-7i-5p | MIMAT0000122 | Ccl1 | 20290 | HITS-CLIP | 23142080 | Unspecified |
| mmu-let-7i-5p | MIMAT0000122 | Ccl2 | 20296 | HITS-CLIP | 37986224\|29167373 | Unspecified |
| mmu-let-7i-5p | MIMAT0000122 | Ccm2 | 216527 | HITS-CLIP | 23142080\|29167373 | Unspecified |
| mmu-let-7i-5p | MIMAT0000122 | Ccn2 | 1490 | HITS-CLIP | 23597149\|26203562 | Unspecified |
| mmu-let-7i-5p | MIMAT0000122 | Ccna2 | 12428 | HITS-CLIP | 37986224\|23597149\|30971389\|23142 | Unspecified |
| mmu-let-7i-5p | MIMAT0000122 | Ccnd1 | 12443 | HITS-CLIP | 37986224\|23597149\|19536157\|2620 | Unspecified |
| mmu-let-7i-5p | MIMAT0000122 | Ccnd2 | 12444 | HITS-CLIP | 37986224\|23142080\|31433990\|1953 | Unspecified |
| mmu-let-7i-5p | MIMAT0000122 | Ccnd3 | 12445 | HITS-CLIP | 23142080 | Unspecified |
| mmu-let-7i-5p | MIMAT0000122 | Ccndbp1 | 17151 | HITS-CLIP | 23597149 | Unspecified |
| mmu-let-7i-5p | MIMAT0000122 | Ccne2 | 12448 | HITS-CLIP | 37986224\|23142080\|23630078 | Unspecified |
| mmu-let-7i-5p | MIMAT0000122 | Ccnf | 12449 | HITS-CLIP | 37986224\|23142080 | Unspecified |
| mmu-let-7i-5p | MIMAT0000122 | Ccng1 | 12450 | HITS-CLIP | 23597149\|23142080 | Unspecified |
| mmu-let-7i-5p | MIMAT0000122 | Ccng2 | 12452 | HITS-CLIP | 19536157\|23142080 | Unspecified |
| mmu-let-7i-5p | MIMAT0000122 | Ccnh | 66671 | HITS-CLIP | 37986224\|23142080\|19536157 | Unspecified |
| mmu-let-7i-5p | MIMAT0000122 | Ccnj | 240665 | HITS-CLIP | 37986224\|23142080 | Unspecified |
| mmu-let-7i-5p | MIMAT0000122 | Ccnjl | 380694 | HITS-CLIP | 19536157 | Unspecified |
| mmu-let-7i-5p | MIMAT0000122 | Ccnl1 | 56706 | HITS-CLIP | 23142080\|29167373 | Unspecified |
| mmu-let-7i-5p | MIMAT0000122 | Ccnl2 | 56036 | HITS-CLIP | 23142080\|29167373 | Unspecified |
| mmu-let-7i-5p | MIMAT0000122 | Ccnt1 | 12455 | HITS-CLIP | 23142080\|29167373 | Unspecified |
| mmu-let-7i-5p | MIMAT0000122 | Ccnt2 | 72949 | HITS-CLIP | 37986224\|23597149\|23142080 | Unspecified |
| mmu-let-7i-5p | MIMAT0000122 | Ccny | 67974 | HITS-CLIP | 37986224\|23630078\|31433990\|1953 | Unspecified |
| mmu-let-7i-5p | MIMAT0000122 | Ccpg1 | 72278 | HITS-CLIP | 23142080 | Unspecified |
| mmu-let-7i-5p | MIMAT0000122 | Ccpg1os | 546143 | HITS-CLIP | 23142080 | Unspecified |
| mmu-let-7i-5p | MIMAT0000122 | Ccr2 | 12772 | HITS-CLIP | 29167373 | Unspecified |
| mmu-let-7i-5p | MIMAT0000122 | Ccr5 | 12774 | HITS-CLIP | 23142080\|29167373 | Unspecified |
| mmu-let-7i-5p | MIMAT0000122 | Ccr7 | 12775 | HITS-CLIP | 37986224\|23142080 | Unspecified |
| mmu-let-7i-5p | MIMAT0000122 | Ccr8 | 12776 | HITS-CLIP | 37986224\|23142080 | Unspecified |
| mmu-let-7i-5p | MIMAT0000122 | Ccser2 | 72972 | HITS-CLIP | 23142080 | Unspecified |
| mmu-let-7i-5p | MIMAT0000122 | Cct2 | 12461 | HITS-CLIP | 37986224\|23597149\|19536157\|23142 | Unspecified |
| mmu-let-7i-5p | MIMAT0000122 | Cct3 | 12462 | HITS-CLIP | 23597149\|23142080 | Unspecified |

| | | | | | | |
|---|---|---|---|---|---|---|
| mmu-let-7i-5p | MIMAT0000122 | Cct4 | 12464 | HITS-CLIP | 23142080\|23597149\|29167373 | Unspecified |
| mmu-let-7i-5p | MIMAT0000122 | Cct5 | 12465 | HITS-CLIP | 23597149\|23142080\|29167373 | Unspecified |
| mmu-let-7i-5p | MIMAT0000122 | Cct7 | 12468 | HITS-CLIP | 23597149\|29167373\|23142080 | Unspecified |
| mmu-let-7i-5p | MIMAT0000122 | Cct8 | 12469 | HITS-CLIP | 23597149\|19536157\|23142080\|29167 | Unspecified |
| mmu-let-7i-5p | MIMAT0000122 | Ccz1 | 231874 | HITS-CLIP | 23142080 | Unspecified |
| mmu-let-7i-5p | MIMAT0000122 | Cd151 | 12476 | HITS-CLIP | 19536157\|23142080\|29167373 | Unspecified |
| mmu-let-7i-5p | MIMAT0000122 | Cd163 | 93671 | HITS-CLIP | 23597149 | Unspecified |
| mmu-let-7i-5p | MIMAT0000122 | Cd164 | 53599 | HITS-CLIP | 37986224\|23597149\|23142080\|1953€ | Unspecified |
| mmu-let-7i-5p | MIMAT0000122 | Cd180 | 17079 | HITS-CLIP | 29167373 | Unspecified |
| mmu-let-7i-5p | MIMAT0000122 | Cd200 | 17470 | HITS-CLIP | 23142080 | Unspecified |
| mmu-let-7i-5p | MIMAT0000122 | Cd200r1 | 57781 | HITS-CLIP | 29167373\|23142080 | Unspecified |
| mmu-let-7i-5p | MIMAT0000122 | Cd226 | 225825 | HITS-CLIP | 37986224\|23142080 | Unspecified |
| mmu-let-7i-5p | MIMAT0000122 | Cd247 | 12503 | HITS-CLIP | 23142080 | Unspecified |
| mmu-let-7i-5p | MIMAT0000122 | Cd24a | 12484 | HITS-CLIP | 23142080 | Unspecified |
| mmu-let-7i-5p | MIMAT0000122 | Cd27 | 21940 | HITS-CLIP | 23142080 | Unspecified |
| mmu-let-7i-5p | MIMAT0000122 | Cd274 | 60533 | HITS-CLIP | 37986224\|23142080\|29167373 | Unspecified |
| mmu-let-7i-5p | MIMAT0000122 | Cd28 | 12487 | HITS-CLIP | 23142080 | Unspecified |
| mmu-let-7i-5p | MIMAT0000122 | Cd2ap | 12488 | HITS-CLIP | 23142080 | Unspecified |
| mmu-let-7i-5p | MIMAT0000122 | Cd300lb | 217304 | HITS-CLIP | 29167373 | Unspecified |
| mmu-let-7i-5p | MIMAT0000122 | Cd34 | 12490 | HITS-CLIP | 30971389 | Unspecified |
| mmu-let-7i-5p | MIMAT0000122 | Cd37 | 12493 | HITS-CLIP | 23142080 | Unspecified |
| mmu-let-7i-5p | MIMAT0000122 | Cd3d | 12500 | HITS-CLIP | 23142080 | Unspecified |
| mmu-let-7i-5p | MIMAT0000122 | Cd3e | 12501 | HITS-CLIP | 37986224\|23142080 | Unspecified |
| mmu-let-7i-5p | MIMAT0000122 | Cd4 | 12504 | HITS-CLIP | 23142080 | Unspecified |
| mmu-let-7i-5p | MIMAT0000122 | Cd40lg | 959 | HITS-CLIP | 23142080 | Unspecified |
| mmu-let-7i-5p | MIMAT0000122 | Cd44 | 12505 | HITS-CLIP | 23142080\|29167373 | Unspecified |
| mmu-let-7i-5p | MIMAT0000122 | Cd47 | 16423 | HITS-CLIP | 23142080\|29167373 | Unspecified |
| mmu-let-7i-5p | MIMAT0000122 | Cd5 | 12507 | HITS-CLIP | 23142080 | Unspecified |
| mmu-let-7i-5p | MIMAT0000122 | Cd52 | 23833 | HITS-CLIP | 37986224\|23142080 | Unspecified |
| mmu-let-7i-5p | MIMAT0000122 | Cd53 | 12508 | HITS-CLIP | 23142080 | Unspecified |
| mmu-let-7i-5p | MIMAT0000122 | Cd5l | 11801 | HITS-CLIP | 23597149 | Unspecified |
| mmu-let-7i-5p | MIMAT0000122 | Cd68 | 12514 | HITS-CLIP | 29167373 | Unspecified |
| mmu-let-7i-5p | MIMAT0000122 | Cd69 | 12515 | qCLASH//HITS-CLIP | 31393953\|23142080 | Unspecified |
| mmu-let-7i-5p | MIMAT0000122 | Cd74 | 16149 | HITS-CLIP | 23142080 | Unspecified |
| mmu-let-7i-5p | MIMAT0000122 | Cd79a | 12518 | HITS-CLIP | 23142080 | Unspecified |
| mmu-let-7i-5p | MIMAT0000122 | Cd80 | 12519 | HITS-CLIP | 23142080 | Unspecified |
| mmu-let-7i-5p | MIMAT0000122 | Cd81 | 12520 | HITS-CLIP | 37986224\|23597149\|19536157\|2314 | Unspecified |
| mmu-let-7i-5p | MIMAT0000122 | Cd83 | 12522 | HITS-CLIP | 23142080 | Unspecified |
| mmu-let-7i-5p | MIMAT0000122 | Cd84 | 12523 | HITS-CLIP | 23142080\|29167373 | Unspecified |

| | | | | | | |
|---|---|---|---|---|---|---|
| mmu-let-7i-5p | MIMAT0000122 | Cd93 | 17064 | HITS-CLIP | 37986224\|29167373 | Unspecified |
| mmu-let-7i-5p | MIMAT0000122 | Cdan1 | 68968 | HITS-CLIP | 23142080 | Unspecified |
| mmu-let-7i-5p | MIMAT0000122 | Cdc14a | 229776 | HITS-CLIP | 23142080 | Unspecified |
| mmu-let-7i-5p | MIMAT0000122 | Cdc14b | 218294 | HITS-CLIP | 23597149\|23142080 | Unspecified |
| mmu-let-7i-5p | MIMAT0000122 | Cdc16 | 69957 | HITS-CLIP | 37986224\|23597149\|19536157\|2916 | Unspecified |
| mmu-let-7i-5p | MIMAT0000122 | Cdc20 | 107995 | HITS-CLIP | 23597149\|23142080 | Unspecified |
| mmu-let-7i-5p | MIMAT0000122 | Cdc25a | 12530 | qCLASH//HITS-CLIP | 31393953\|31433990\|23597149\|2314 | Unspecified |
| mmu-let-7i-5p | MIMAT0000122 | Cdc25b | 12531 | HITS-CLIP | 23142080\|29167373 | Unspecified |
| mmu-let-7i-5p | MIMAT0000122 | Cdc27 | 217232 | HITS-CLIP | 19536157\|23142080\|29167373 | Unspecified |
| mmu-let-7i-5p | MIMAT0000122 | Cdc37 | 12539 | HITS-CLIP | 23142080\|29167373 | Unspecified |
| mmu-let-7i-5p | MIMAT0000122 | Cdc40 | 71713 | HITS-CLIP | 37986224\|23597149\|23142080 | Unspecified |
| mmu-let-7i-5p | MIMAT0000122 | Cdc42ep5 | 58804 | HITS-CLIP | 23597149 | Unspecified |
| mmu-let-7i-5p | MIMAT0000122 | Cdc42se1 | 57912 | HITS-CLIP | 19536157\|26203562\|23142080 | Unspecified |
| mmu-let-7i-5p | MIMAT0000122 | Cdc42se2 | 72729 | HITS-CLIP | 23597149 | Unspecified |
| mmu-let-7i-5p | MIMAT0000122 | Cdc45 | 12544 | HITS-CLIP | 23142080 | Unspecified |
| mmu-let-7i-5p | MIMAT0000122 | Cdc5l | 71702 | HITS-CLIP | 23142080 | Unspecified |
| mmu-let-7i-5p | MIMAT0000122 | Cdc6 | 23834 | HITS-CLIP | 23142080 | Unspecified |
| mmu-let-7i-5p | MIMAT0000122 | Cdc73 | 214498 | HITS-CLIP | 23142080 | Unspecified |
| mmu-let-7i-5p | MIMAT0000122 | Cdca2 | 108912 | HITS-CLIP | 23142080 | Unspecified |
| mmu-let-7i-5p | MIMAT0000122 | Cdh2 | 12558 | HITS-CLIP | 23597149\|26203562 | Unspecified |
| mmu-let-7i-5p | MIMAT0000122 | Cdh5 | 12562 | HITS-CLIP | 37986224 | Unspecified |
| mmu-let-7i-5p | MIMAT0000122 | Cdh8 | 12564 | HITS-CLIP | 37986224\|19536157 | Unspecified |
| mmu-let-7i-5p | MIMAT0000122 | Cdhr5 | 72040 | HITS-CLIP | 23597149 | Unspecified |
| mmu-let-7i-5p | MIMAT0000122 | Cdip1 | 66626 | HITS-CLIP | 23597149 | Unspecified |
| mmu-let-7i-5p | MIMAT0000122 | Cdk11b | 12537 | HITS-CLIP | 23597149 | Unspecified |
| mmu-let-7i-5p | MIMAT0000122 | Cdk12 | 69131 | HITS-CLIP | 37986224\|19536157\|23142080\|2916 | Unspecified |
| mmu-let-7i-5p | MIMAT0000122 | Cdk13 | 69562 | HITS-CLIP | 23142080\|19536157\|23597149 | Unspecified |
| mmu-let-7i-5p | MIMAT0000122 | Cdk14 | 18647 | HITS-CLIP | 29167373 | Unspecified |
| mmu-let-7i-5p | MIMAT0000122 | Cdk16 | 18555 | HITS-CLIP | 23142080\|29167373 | Unspecified |
| mmu-let-7i-5p | MIMAT0000122 | Cdk17 | 237459 | HITS-CLIP | 23142080 | Unspecified |
| mmu-let-7i-5p | MIMAT0000122 | Cdk18 | 18557 | HITS-CLIP | 29167373 | Unspecified |
| mmu-let-7i-5p | MIMAT0000122 | Cdk19 | 78334 | HITS-CLIP | 23597149\|19536157\|29167373\|2314 | Unspecified |
| mmu-let-7i-5p | MIMAT0000122 | Cdk2 | 12566 | HITS-CLIP | 23597149 | Unspecified |
| mmu-let-7i-5p | MIMAT0000122 | Cdk2ap2 | 52004 | HITS-CLIP | 23142080 | Unspecified |
| mmu-let-7i-5p | MIMAT0000122 | Cdk4 | 12567 | HITS-CLIP | 23597149 | Unspecified |
| mmu-let-7i-5p | MIMAT0000122 | Cdk5r1 | 12569 | HITS-CLIP | 37986224\|19536157 | Unspecified |
| mmu-let-7i-5p | MIMAT0000122 | Cdk5r2 | 12570 | HITS-CLIP | 37986224\|19536157 | Unspecified |
| mmu-let-7i-5p | MIMAT0000122 | Cdk5rap1 | 66971 | HITS-CLIP | 29167373 | Unspecified |
| mmu-let-7i-5p | MIMAT0000122 | Cdk6 | 12571 | HITS-CLIP | 23142080\|29167373 | Unspecified |

| | | | | | | |
|---|---|---|---|---|---|---|
| mmu-let-7i-5p | MIMAT0000122 | Cdk8 | 264064 | HITS-CLIP | 37986224\|23597149\|23142080 | Unspecified |
| mmu-let-7i-5p | MIMAT0000122 | Cdkn1a | 12575 | HITS-CLIP | 23142080 | Unspecified |
| mmu-let-7i-5p | MIMAT0000122 | Cdkn1b | 12576 | HITS-CLIP | 19536157\|23142080 | Unspecified |
| mmu-let-7i-5p | MIMAT0000122 | Cdkn2aip | 70925 | HITS-CLIP | 37986224\|19536157\|23142080 | Unspecified |
| mmu-let-7i-5p | MIMAT0000122 | Cdkn2aipnl | 52626 | HITS-CLIP | 37986224 | Unspecified |
| mmu-let-7i-5p | MIMAT0000122 | Cdkn2c | 12580 | HITS-CLIP | 23142080 | Unspecified |
| mmu-let-7i-5p | MIMAT0000122 | Cdr2 | 12585 | HITS-CLIP | 23142080 | Unspecified |
| mmu-let-7i-5p | MIMAT0000122 | Cdr2l | 237988 | HITS-CLIP | 37986224 | Unspecified |
| mmu-let-7i-5p | MIMAT0000122 | Cds2 | 110911 | qCLASH | 31393953 | Unspecified |
| mmu-let-7i-5p | MIMAT0000122 | Cdt1 | 67177 | HITS-CLIP | 23142080 | Unspecified |
| mmu-let-7i-5p | MIMAT0000122 | Cdv3 | 321022 | HITS-CLIP | 23597149\|29167373 | Unspecified |
| mmu-let-7i-5p | MIMAT0000122 | Cebpd | 12609 | HITS-CLIP | 23597149 | Unspecified |
| mmu-let-7i-5p | MIMAT0000122 | Cebpg | 12611 | HITS-CLIP | 23142080\|29167373 | Unspecified |
| mmu-let-7i-5p | MIMAT0000122 | Cebpz | 12607 | HITS-CLIP | 23142080 | Unspecified |
| mmu-let-7i-5p | MIMAT0000122 | Cecr2 | 330409 | HITS-CLIP | 23597149\|26203562 | Unspecified |
| mmu-let-7i-5p | MIMAT0000122 | Celf1 | 13046 | HITS-CLIP | 37986224\|23597149\|23142080\|29167 | Unspecified |
| mmu-let-7i-5p | MIMAT0000122 | Celf2 | 14007 | HITS-CLIP | 37986224\|23142080\|19536157\|31433 | Unspecified |
| mmu-let-7i-5p | MIMAT0000122 | Celf3 | 78784 | HITS-CLIP | 37986224\|19536157 | Unspecified |
| mmu-let-7i-5p | MIMAT0000122 | Celsr1 | 12614 | HITS-CLIP | 23142080 | Unspecified |
| mmu-let-7i-5p | MIMAT0000122 | Celsr3 | 107934 | HITS-CLIP | 37986224\|19536157 | Unspecified |
| mmu-let-7i-5p | MIMAT0000122 | Cemip2 | 23670 | HITS-CLIP | 19536157\|23597149\|23142080\|29167 | Unspecified |
| mmu-let-7i-5p | MIMAT0000122 | Cend1 | 57754 | HITS-CLIP | 19536157 | Unspecified |
| mmu-let-7i-5p | MIMAT0000122 | Cenpa | 12615 | HITS-CLIP | 23142080 | Unspecified |
| mmu-let-7i-5p | MIMAT0000122 | Cenpb | 12616 | HITS-CLIP | 37986224\|23597149\|19536157\|23142 | Unspecified |
| mmu-let-7i-5p | MIMAT0000122 | Cenpq | 83815 | HITS-CLIP | 37986224\|23142080 | Unspecified |
| mmu-let-7i-5p | MIMAT0000122 | Cenpv | 73139 | HITS-CLIP | 23597149 | Unspecified |
| mmu-let-7i-5p | MIMAT0000122 | Cenpw | 66311 | HITS-CLIP | 23142080 | Unspecified |
| mmu-let-7i-5p | MIMAT0000122 | Cenpx | 201254 | HITS-CLIP | 23597149 | Unspecified |
| mmu-let-7i-5p | MIMAT0000122 | Cep120 | 225523 | HITS-CLIP | 37986224\|19536157\|23597149\|23142 | Unspecified |
| mmu-let-7i-5p | MIMAT0000122 | Cep135 | 381644 | HITS-CLIP | 37986224\|23142080 | Unspecified |
| mmu-let-7i-5p | MIMAT0000122 | Cep162 | 382090 | HITS-CLIP | 23142080 | Unspecified |
| mmu-let-7i-5p | MIMAT0000122 | Cep170 | 545389 | HITS-CLIP | 37986224\|23142080\|19536157\|23597 | Unspecified |
| mmu-let-7i-5p | MIMAT0000122 | Cep170b | 217882 | HITS-CLIP | 23142080 | Unspecified |
| mmu-let-7i-5p | MIMAT0000122 | Cep192 | 70799 | HITS-CLIP | 29167373 | Unspecified |
| mmu-let-7i-5p | MIMAT0000122 | Cep295 | 85459 | HITS-CLIP | 23142080\|23597149 | Unspecified |
| mmu-let-7i-5p | MIMAT0000122 | Cep350 | 74081 | HITS-CLIP | 37986224\|23142080\|23597149\|31433 | Unspecified |
| mmu-let-7i-5p | MIMAT0000122 | Cep55 | 74107 | HITS-CLIP | 37986224\|23142080\|29167373 | Unspecified |
| mmu-let-7i-5p | MIMAT0000122 | Cep57 | 74360 | HITS-CLIP | 23142080 | Unspecified |
| mmu-let-7i-5p | MIMAT0000122 | Cep68 | 216543 | HITS-CLIP | 37986224\|23597149\|19536157\|23142 | Unspecified |

| | | | | | | |
|---|---|---|---|---|---|---|
| mmu-let-7i-5p | MIMAT0000122 | Cep85 | 70012 | HITS-CLIP | 23597149\|29167373 | Unspecified |
| mmu-let-7i-5p | MIMAT0000122 | Cep85l | 100038725 | HITS-CLIP | 23142080\|19536157\|23597149 | Unspecified |
| mmu-let-7i-5p | MIMAT0000122 | Cep97 | 74201 | HITS-CLIP | 37986224\|23142080\|19536157 | Unspecified |
| mmu-let-7i-5p | MIMAT0000122 | Cept1 | 99712 | HITS-CLIP | 23142080 | Unspecified |
| mmu-let-7i-5p | MIMAT0000122 | Cerk | 223753 | HITS-CLIP | 19536157 | Unspecified |
| mmu-let-7i-5p | MIMAT0000122 | Cers2 | 76893 | HITS-CLIP | 23597149\|23142080\|29167373 | Unspecified |
| mmu-let-7i-5p | MIMAT0000122 | Cers4 | 67260 | HITS-CLIP | 23142080 | Unspecified |
| mmu-let-7i-5p | MIMAT0000122 | Cers5 | 71949 | HITS-CLIP | 23142080 | Unspecified |
| mmu-let-7i-5p | MIMAT0000122 | Cers6 | 241447 | HITS-CLIP | 23142080\|29167373 | Unspecified |
| mmu-let-7i-5p | MIMAT0000122 | Ces1d | 104158 | HITS-CLIP | 23597149 | Unspecified |
| mmu-let-7i-5p | MIMAT0000122 | Ces1e | 13897 | HITS-CLIP | 23597149 | Unspecified |
| mmu-let-7i-5p | MIMAT0000122 | Ces1f | 234564 | HITS-CLIP | 23597149 | Unspecified |
| mmu-let-7i-5p | MIMAT0000122 | Ces1g | 12623 | HITS-CLIP | 23597149 | Unspecified |
| mmu-let-7i-5p | MIMAT0000122 | Ces2a | 102022 | HITS-CLIP | 23597149 | Unspecified |
| mmu-let-7i-5p | MIMAT0000122 | Ces3a | 382053 | HITS-CLIP | 23597149 | Unspecified |
| mmu-let-7i-5p | MIMAT0000122 | Ces3b | 13909 | HITS-CLIP | 23597149 | Unspecified |
| mmu-let-7i-5p | MIMAT0000122 | Cfap36 | 112942 | HITS-CLIP | 23142080 | Unspecified |
| mmu-let-7i-5p | MIMAT0000122 | Cfap97 | 57587 | HITS-CLIP | 23142080 | Unspecified |
| mmu-let-7i-5p | MIMAT0000122 | Cfh | 12628 | HITS-CLIP | 23597149 | Unspecified |
| mmu-let-7i-5p | MIMAT0000122 | Cfhr1 | 50702 | HITS-CLIP | 23597149 | Unspecified |
| mmu-let-7i-5p | MIMAT0000122 | Cfhr2 | 545366 | HITS-CLIP | 23597149 | Unspecified |
| mmu-let-7i-5p | MIMAT0000122 | Cfi | 12630 | HITS-CLIP | 23597149 | Unspecified |
| mmu-let-7i-5p | MIMAT0000122 | Cfl1 | 12631 | HITS-CLIP | 23142080 | Unspecified |
| mmu-let-7i-5p | MIMAT0000122 | Cfl2 | 12632 | HITS-CLIP | 23142080 | Unspecified |
| mmu-let-7i-5p | MIMAT0000122 | Cflar | 12633 | HITS-CLIP | 37986224\|23597149\|23142080\|29167 | Unspecified |
| mmu-let-7i-5p | MIMAT0000122 | Cggbp1 | 106143 | HITS-CLIP | 37986224\|23142080 | Unspecified |
| mmu-let-7i-5p | MIMAT0000122 | Cgnl1 | 68178 | HITS-CLIP | 19536157\|23597149\|29167373 | Unspecified |
| mmu-let-7i-5p | MIMAT0000122 | Chac1 | 69065 | HITS-CLIP | 23142080 | Unspecified |
| mmu-let-7i-5p | MIMAT0000122 | Chac2 | 68044 | HITS-CLIP | 23597149 | Unspecified |
| mmu-let-7i-5p | MIMAT0000122 | Chd1 | 12648 | HITS-CLIP//qCLASH | 37986224\|23597149\|19536157\|23142 | Unspecified |
| mmu-let-7i-5p | MIMAT0000122 | Chd2 | 244059 | HITS-CLIP | 23142080\|29167373 | Unspecified |
| mmu-let-7i-5p | MIMAT0000122 | Chd3 | 216848 | HITS-CLIP | 37986224\|23597149\|19536157\|23142 | Unspecified |
| mmu-let-7i-5p | MIMAT0000122 | Chd4 | 107932 | HITS-CLIP | 19536157\|23142080\|29167373 | Unspecified |
| mmu-let-7i-5p | MIMAT0000122 | Chd7 | 320790 | HITS-CLIP | 37986224\|19536157\|23630078\|3143 | Unspecified |
| mmu-let-7i-5p | MIMAT0000122 | Chd8 | 67772 | HITS-CLIP | 23142080\|23597149\|31433990\|29167 | Unspecified |
| mmu-let-7i-5p | MIMAT0000122 | Chd9 | 109151 | HITS-CLIP | 37986224\|19536157\|23142080 | Unspecified |
| mmu-let-7i-5p | MIMAT0000122 | Chdh | 218865 | HITS-CLIP | 23597149 | Unspecified |
| mmu-let-7i-5p | MIMAT0000122 | Chek1 | 12649 | HITS-CLIP | 23142080 | Unspecified |
| mmu-let-7i-5p | MIMAT0000122 | Chfr | 231600 | HITS-CLIP | 23142080\|29167373 | Unspecified |

| | | | | | | |
|---|---|---|---|---|---|---|
| mmu-let-7i-5p | MIMAT0000122 | Chgb | 12653 | HITS-CLIP | 19536157 | Unspecified |
| mmu-let-7i-5p | MIMAT0000122 | Chic1 | 12212 | qCLASH//HITS-CLIP | 31393953\|19536157 | Unspecified |
| mmu-let-7i-5p | MIMAT0000122 | Chic2 | 74277 | HITS-CLIP | 23142080\|29167373 | Unspecified |
| mmu-let-7i-5p | MIMAT0000122 | Chid1 | 68038 | HITS-CLIP | 23142080\|29167373 | Unspecified |
| mmu-let-7i-5p | MIMAT0000122 | Chkb | 12651 | HITS-CLIP | 23597149\|23142080 | Unspecified |
| mmu-let-7i-5p | MIMAT0000122 | Chl1 | 12661 | HITS-CLIP | 37986224\|19536157 | Unspecified |
| mmu-let-7i-5p | MIMAT0000122 | Chml | 12663 | HITS-CLIP | 31433990\|23142080 | Unspecified |
| mmu-let-7i-5p | MIMAT0000122 | Chmp1b | 67078 | HITS-CLIP | 23142080 | Unspecified |
| mmu-let-7i-5p | MIMAT0000122 | Chmp2b | 68942 | HITS-CLIP | 19536157 | Unspecified |
| mmu-let-7i-5p | MIMAT0000122 | Chmp3 | 66700 | HITS-CLIP | 23597149 | Unspecified |
| mmu-let-7i-5p | MIMAT0000122 | Chmp4b | 75608 | HITS-CLIP | 23142080 | Unspecified |
| mmu-let-7i-5p | MIMAT0000122 | Chmp7 | 105513 | HITS-CLIP | 37986224\|23597149\|23142080 | Unspecified |
| mmu-let-7i-5p | MIMAT0000122 | Chp1 | 56398 | HITS-CLIP | 37986224\|23597149\|23142080 | Unspecified |
| mmu-let-7i-5p | MIMAT0000122 | Chrac1 | 93696 | HITS-CLIP | 23142080 | Unspecified |
| mmu-let-7i-5p | MIMAT0000122 | Chrm1 | 12669 | HITS-CLIP | 37986224 | Unspecified |
| mmu-let-7i-5p | MIMAT0000122 | Chrm3 | 12671 | HITS-CLIP | 19536157 | Unspecified |
| mmu-let-7i-5p | MIMAT0000122 | Chrnb2 | 11444 | HITS-CLIP | 37986224\|19536157 | Unspecified |
| mmu-let-7i-5p | MIMAT0000122 | Chst1 | 76969 | HITS-CLIP | 19536157 | Unspecified |
| mmu-let-7i-5p | MIMAT0000122 | Chst14 | 72136 | HITS-CLIP | 37986224\|23142080 | Unspecified |
| mmu-let-7i-5p | MIMAT0000122 | Chst15 | 77590 | HITS-CLIP | 23597149 | Unspecified |
| mmu-let-7i-5p | MIMAT0000122 | Chst2 | 54371 | HITS-CLIP | 37986224 | Unspecified |
| mmu-let-7i-5p | MIMAT0000122 | Chsy1 | 269941 | HITS-CLIP | 23142080 | Unspecified |
| mmu-let-7i-5p | MIMAT0000122 | Chuk | 12675 | HITS-CLIP | 37986224\|23597149\|23142080 | Unspecified |
| mmu-let-7i-5p | MIMAT0000122 | Ciao2a | 84191 | HITS-CLIP | 23597149\|23142080 | Unspecified |
| mmu-let-7i-5p | MIMAT0000122 | Ciapin1 | 109006 | HITS-CLIP | 23597149 | Unspecified |
| mmu-let-7i-5p | MIMAT0000122 | Cic | 71722 | HITS-CLIP | 29167373 | Unspecified |
| mmu-let-7i-5p | MIMAT0000122 | Cideb | 12684 | HITS-CLIP | 23597149 | Unspecified |
| mmu-let-7i-5p | MIMAT0000122 | Cinp | 67236 | HITS-CLIP | 23142080 | Unspecified |
| mmu-let-7i-5p | MIMAT0000122 | Cip2a | 57650 | HITS-CLIP | 23142080 | Unspecified |
| mmu-let-7i-5p | MIMAT0000122 | Cisd2 | 67006 | HITS-CLIP | 23142080 | Unspecified |
| mmu-let-7i-5p | MIMAT0000122 | Cish | 12700 | HITS-CLIP | 37986224\|23142080 | Unspecified |
| mmu-let-7i-5p | MIMAT0000122 | Cit | 12704 | HITS-CLIP | 23142080 | Unspecified |
| mmu-let-7i-5p | MIMAT0000122 | Cited2 | 17684 | HITS-CLIP | 23142080 | Unspecified |
| mmu-let-7i-5p | MIMAT0000122 | Ckap4 | 216197 | HITS-CLIP | 19536157\|23597149 | Unspecified |
| mmu-let-7i-5p | MIMAT0000122 | Ckap5 | 75786 | HITS-CLIP | 23597149\|19536157\|23142080\|29167 | Unspecified |
| mmu-let-7i-5p | MIMAT0000122 | Cklf | 75458 | HITS-CLIP | 29167373 | Unspecified |
| mmu-let-7i-5p | MIMAT0000122 | Cks2 | 66197 | HITS-CLIP | 23142080 | Unspecified |
| mmu-let-7i-5p | MIMAT0000122 | Clasp1 | 76707 | HITS-CLIP | 37986224\|23142080\|19536157\|23597 | Unspecified |
| mmu-let-7i-5p | MIMAT0000122 | Clasp2 | 76499 | HITS-CLIP | 23142080 | Unspecified |

| | | | | | | |
|---|---|---|---|---|---|---|
| mmu-let-7i-5p | MIMAT0000122 | Clcc1 | 229725 | HITS-CLIP | 37986224 | Unspecified |
| mmu-let-7i-5p | MIMAT0000122 | Clcf1 | 56708 | HITS-CLIP | 23142080 | Unspecified |
| mmu-let-7i-5p | MIMAT0000122 | Clcn3 | 12725 | HITS-CLIP | 37986224\|19536157\|29167373\|23142 | Unspecified |
| mmu-let-7i-5p | MIMAT0000122 | Clcn5 | 12728 | HITS-CLIP | 37986224\|23142080\|19536157\|29167 | Unspecified |
| mmu-let-7i-5p | MIMAT0000122 | Clcn7 | 26373 | HITS-CLIP | 37986224\|23597149 | Unspecified |
| mmu-let-7i-5p | MIMAT0000122 | Cldn1 | 12737 | HITS-CLIP | 23597149 | Unspecified |
| mmu-let-7i-5p | MIMAT0000122 | Cldn10 | 58187 | HITS-CLIP | 23142080 | Unspecified |
| mmu-let-7i-5p | MIMAT0000122 | Cldn12 | 64945 | HITS-CLIP | 37986224\|23142080\|19536157\|23597 | Unspecified |
| mmu-let-7i-5p | MIMAT0000122 | Clec16a | 74374 | HITS-CLIP | 23142080 | Unspecified |
| mmu-let-7i-5p | MIMAT0000122 | Clec1b | 56760 | HITS-CLIP | 23597149 | Unspecified |
| mmu-let-7i-5p | MIMAT0000122 | Clec2d | 93694 | HITS-CLIP | 29167373\|23142080 | Unspecified |
| mmu-let-7i-5p | MIMAT0000122 | Clec4a1 | 269799 | HITS-CLIP | 29167373 | Unspecified |
| mmu-let-7i-5p | MIMAT0000122 | Clic1 | 114584 | HITS-CLIP | 23142080 | Unspecified |
| mmu-let-7i-5p | MIMAT0000122 | Clic4 | 29876 | HITS-CLIP | 37986224\|23597149\|23142080\|29167 | Unspecified |
| mmu-let-7i-5p | MIMAT0000122 | Clint1 | 216705 | HITS-CLIP | 37986224\|23597149\|23142080 | Unspecified |
| mmu-let-7i-5p | MIMAT0000122 | Clip1 | 56430 | HITS-CLIP | 23597149\|31433990\|23142080\|29167 | Unspecified |
| mmu-let-7i-5p | MIMAT0000122 | Clip3 | 76686 | HITS-CLIP | 19536157 | Unspecified |
| mmu-let-7i-5p | MIMAT0000122 | Clk1 | 12747 | HITS-CLIP | 23142080 | Unspecified |
| mmu-let-7i-5p | MIMAT0000122 | Clk2 | 12748 | HITS-CLIP | 23142080 | Unspecified |
| mmu-let-7i-5p | MIMAT0000122 | Clk4 | 12750 | HITS-CLIP | 19536157\|23142080 | Unspecified |
| mmu-let-7i-5p | MIMAT0000122 | Clmn | 94040 | HITS-CLIP | 23597149 | Unspecified |
| mmu-let-7i-5p | MIMAT0000122 | Cln3 | 12752 | HITS-CLIP | 23597149\|23142080 | Unspecified |
| mmu-let-7i-5p | MIMAT0000122 | Cln5 | 211286 | HITS-CLIP | 23142080 | Unspecified |
| mmu-let-7i-5p | MIMAT0000122 | Clock | 12753 | HITS-CLIP | 23142080 | Unspecified |
| mmu-let-7i-5p | MIMAT0000122 | Clp1 | 98985 | HITS-CLIP | 23142080\|23597149 | Unspecified |
| mmu-let-7i-5p | MIMAT0000122 | Clptm1 | 56457 | HITS-CLIP | 19536157\|23142080 | Unspecified |
| mmu-let-7i-5p | MIMAT0000122 | Clptm1l | 218335 | HITS-CLIP | 37986224\|23142080 | Unspecified |
| mmu-let-7i-5p | MIMAT0000122 | Clpx | 270166 | HITS-CLIP | 23597149\|23142080\|29167373 | Unspecified |
| mmu-let-7i-5p | MIMAT0000122 | Clspn | 269582 | HITS-CLIP | 23142080 | Unspecified |
| mmu-let-7i-5p | MIMAT0000122 | Clstn1 | 65945 | HITS-CLIP | 19536157 | Unspecified |
| mmu-let-7i-5p | MIMAT0000122 | Cltb | 74325 | HITS-CLIP | 31433990 | Unspecified |
| mmu-let-7i-5p | MIMAT0000122 | Cltc | 67300 | HITS-CLIP | 23597149\|19536157\|23142080 | Unspecified |
| mmu-let-7i-5p | MIMAT0000122 | Clu | 12759 | HITS-CLIP | 23597149 | Unspecified |
| mmu-let-7i-5p | MIMAT0000122 | Cluh | 74148 | HITS-CLIP | 23597149\|19536157\|23142080 | Unspecified |
| mmu-let-7i-5p | MIMAT0000122 | Cmah | 12763 | HITS-CLIP | 37986224\|23597149\|23142080 | Unspecified |
| mmu-let-7i-5p | MIMAT0000122 | Cmas | 12764 | HITS-CLIP | 37986224\|23597149\|19536157\|23142 | Unspecified |
| mmu-let-7i-5p | MIMAT0000122 | Cmc2 | 66531 | HITS-CLIP | 23142080 | Unspecified |
| mmu-let-7i-5p | MIMAT0000122 | Cmip | 74440 | HITS-CLIP | 23142080\|23597149\|31433990\|29167 | Unspecified |
| mmu-let-7i-5p | MIMAT0000122 | Cmpk1 | 66588 | HITS-CLIP | 23142080 | Unspecified |

| | | | | | | |
|---|---|---|---|---|---|---|
| mmu-let-7i-5p | MIMAT0000122 | Cmtm4 | 97487 | HITS-CLIP | 37986224\|23142080 | Unspecified |
| mmu-let-7i-5p | MIMAT0000122 | Cmtm6 | 67213 | HITS-CLIP | 37986224\|23597149\|23142080\|29167 | Unspecified |
| mmu-let-7i-5p | MIMAT0000122 | Cmtm7 | 102545 | HITS-CLIP | 23142080 | Unspecified |
| mmu-let-7i-5p | MIMAT0000122 | Cmtr1 | 74157 | HITS-CLIP | 23597149\|23142080 | Unspecified |
| mmu-let-7i-5p | MIMAT0000122 | Cmtr2 | 234728 | HITS-CLIP | 23142080 | Unspecified |
| mmu-let-7i-5p | MIMAT0000122 | Cnbp | 12785 | HITS-CLIP | 37986224\|23142080\|29167373 | Unspecified |
| mmu-let-7i-5p | MIMAT0000122 | Cnih1 | 12793 | HITS-CLIP | 37986224\|23597149\|23142080\|19536 | Unspecified |
| mmu-let-7i-5p | MIMAT0000122 | Cnksr2 | 245684 | HITS-CLIP | 19536157 | Unspecified |
| mmu-let-7i-5p | MIMAT0000122 | Cnn2 | 12798 | HITS-CLIP | 23142080 | Unspecified |
| mmu-let-7i-5p | MIMAT0000122 | Cnnm1 | 83674 | HITS-CLIP | 37986224 | Unspecified |
| mmu-let-7i-5p | MIMAT0000122 | Cnnm2 | 94219 | HITS-CLIP | 19536157 | Unspecified |
| mmu-let-7i-5p | MIMAT0000122 | Cnnm3 | 94218 | HITS-CLIP | 23597149\|23142080 | Unspecified |
| mmu-let-7i-5p | MIMAT0000122 | Cnnm4 | 94220 | HITS-CLIP | 23142080\|23597149 | Unspecified |
| mmu-let-7i-5p | MIMAT0000122 | Cnot1 | 234594 | HITS-CLIP | 23597149\|23142080\|26203562 | Unspecified |
| mmu-let-7i-5p | MIMAT0000122 | Cnot10 | 78893 | HITS-CLIP | 29167373 | Unspecified |
| mmu-let-7i-5p | MIMAT0000122 | Cnot11 | 52846 | HITS-CLIP | 23142080\|19536157 | Unspecified |
| mmu-let-7i-5p | MIMAT0000122 | Cnot2 | 72068 | HITS-CLIP | 23142080 | Unspecified |
| mmu-let-7i-5p | MIMAT0000122 | Cnot4 | 53621 | HITS-CLIP | 23142080 | Unspecified |
| mmu-let-7i-5p | MIMAT0000122 | Cnot6 | 104625 | HITS-CLIP | 23597149 | Unspecified |
| mmu-let-7i-5p | MIMAT0000122 | Cnot6l | 231464 | HITS-CLIP | 37986224\|23142080\|23597149 | Unspecified |
| mmu-let-7i-5p | MIMAT0000122 | Cnot7 | 18983 | HITS-CLIP | 23142080 | Unspecified |
| mmu-let-7i-5p | MIMAT0000122 | Cnot9 | 9125 | HITS-CLIP | 23142080 | Unspecified |
| mmu-let-7i-5p | MIMAT0000122 | Cnp | 12799 | HITS-CLIP | 23597149\|19536157\|23142080 | Unspecified |
| mmu-let-7i-5p | MIMAT0000122 | Cnppd1 | 69171 | HITS-CLIP | 23597149\|29167373 | Unspecified |
| mmu-let-7i-5p | MIMAT0000122 | Cnpy2 | 56530 | HITS-CLIP | 23142080 | Unspecified |
| mmu-let-7i-5p | MIMAT0000122 | Cnst | 226744 | HITS-CLIP | 23142080 | Unspecified |
| mmu-let-7i-5p | MIMAT0000122 | Cntf | 12803 | HITS-CLIP | 23142080 | Unspecified |
| mmu-let-7i-5p | MIMAT0000122 | Cntn2 | 21367 | HITS-CLIP | 19536157 | Unspecified |
| mmu-let-7i-5p | MIMAT0000122 | Cntnap1 | 53321 | HITS-CLIP | 37986224\|23142080 | Unspecified |
| mmu-let-7i-5p | MIMAT0000122 | Cntnap4 | 170571 | HITS-CLIP | 19536157 | Unspecified |
| mmu-let-7i-5p | MIMAT0000122 | Cntrl | 26920 | HITS-CLIP | 37986224\|23597149\|19536157\|29167 | Unspecified |
| mmu-let-7i-5p | MIMAT0000122 | Coa4 | 68185 | HITS-CLIP | 37986224 | Unspecified |
| mmu-let-7i-5p | MIMAT0000122 | Coa5 | 76178 | HITS-CLIP | 37986224\|23142080 | Unspecified |
| mmu-let-7i-5p | MIMAT0000122 | Coa7 | 69893 | HITS-CLIP | 37986224\|19536157 | Unspecified |
| mmu-let-7i-5p | MIMAT0000122 | Cobll1 | 319876 | HITS-CLIP | 23142080 | Unspecified |
| mmu-let-7i-5p | MIMAT0000122 | Cog1 | 16834 | HITS-CLIP | 23597149\|19536157 | Unspecified |
| mmu-let-7i-5p | MIMAT0000122 | Cog3 | 338337 | HITS-CLIP | 37986224\|31433990\|23597149\|23142 | Unspecified |
| mmu-let-7i-5p | MIMAT0000122 | Cog4 | 102339 | HITS-CLIP | 23597149\|19536157\|23142080 | Unspecified |
| mmu-let-7i-5p | MIMAT0000122 | Cog5 | 238123 | HITS-CLIP | 23142080 | Unspecified |

| | | | | | | |
|---|---|---|---|---|---|---|
| mmu-let-7i-5p | MIMAT0000122 | Cog6 | 67542 | HITS-CLIP | 37986224\|23597149 | Unspecified |
| mmu-let-7i-5p | MIMAT0000122 | Cog7 | 233824 | HITS-CLIP | 37986224\|23142080\|23597149\|29167 | Unspecified |
| mmu-let-7i-5p | MIMAT0000122 | Coil | 12812 | HITS-CLIP | 23142080 | Unspecified |
| mmu-let-7i-5p | MIMAT0000122 | Col15a1 | 12819 | HITS-CLIP | 23597149 | Unspecified |
| mmu-let-7i-5p | MIMAT0000122 | Col18a1 | 12822 | HITS-CLIP | 23597149\|29167373 | Unspecified |
| mmu-let-7i-5p | MIMAT0000122 | Col1a1 | 12842 | HITS-CLIP | 37986224 | Unspecified |
| mmu-let-7i-5p | MIMAT0000122 | Col1a2 | 12843 | HITS-CLIP//Immunohistochemistry//Lucife | 37986224\|19536157\|23630078\|2625 | Unspecified |
| mmu-let-7i-5p | MIMAT0000122 | Col27a1 | 373864 | HITS-CLIP | 23597149 | Unspecified |
| mmu-let-7i-5p | MIMAT0000122 | Col4a1 | 12826 | HITS-CLIP | 37986224\|19536157 | Unspecified |
| mmu-let-7i-5p | MIMAT0000122 | Col4a2 | 12827 | HITS-CLIP | 37986224\|23597149\|19536157 | Unspecified |
| mmu-let-7i-5p | MIMAT0000122 | Col4a3bp | 68018 | HITS-CLIP | 23142080 | Unspecified |
| mmu-let-7i-5p | MIMAT0000122 | Col5a1 | 12831 | HITS-CLIP | 23597149 | Unspecified |
| mmu-let-7i-5p | MIMAT0000122 | Col6a3 | 12835 | HITS-CLIP | 23597149 | Unspecified |
| mmu-let-7i-5p | MIMAT0000122 | Colec12 | 140792 | HITS-CLIP | 29167373 | Unspecified |
| mmu-let-7i-5p | MIMAT0000122 | Colgalt1 | 234407 | HITS-CLIP | 29167373 | Unspecified |
| mmu-let-7i-5p | MIMAT0000122 | Colq | 382864 | HITS-CLIP | 23142080 | Unspecified |
| mmu-let-7i-5p | MIMAT0000122 | Commd2 | 52245 | HITS-CLIP | 23142080 | Unspecified |
| mmu-let-7i-5p | MIMAT0000122 | Comt | 12846 | HITS-CLIP | 23597149\|29167373 | Unspecified |
| mmu-let-7i-5p | MIMAT0000122 | Copa | 12847 | HITS-CLIP | 23142080\|23597149 | Unspecified |
| mmu-let-7i-5p | MIMAT0000122 | Copb1 | 70349 | HITS-CLIP | 23142080 | Unspecified |
| mmu-let-7i-5p | MIMAT0000122 | Copb2 | 50797 | HITS-CLIP | 23142080 | Unspecified |
| mmu-let-7i-5p | MIMAT0000122 | Cope | 59042 | HITS-CLIP | 23597149\|23142080\|29167373 | Unspecified |
| mmu-let-7i-5p | MIMAT0000122 | Copg1 | 54161 | HITS-CLIP | 37986224\|19536157\|23597149\|2314 | Unspecified |
| mmu-let-7i-5p | MIMAT0000122 | Copg2 | 54160 | HITS-CLIP | 23597149\|23142080 | Unspecified |
| mmu-let-7i-5p | MIMAT0000122 | Cops2 | 12848 | HITS-CLIP | 19536157\|23142080\|29167373 | Unspecified |
| mmu-let-7i-5p | MIMAT0000122 | Cops3 | 26572 | HITS-CLIP | 37986224 | Unspecified |
| mmu-let-7i-5p | MIMAT0000122 | Cops4 | 26891 | HITS-CLIP | 23142080 | Unspecified |
| mmu-let-7i-5p | MIMAT0000122 | Cops5 | 26754 | HITS-CLIP | 23142080 | Unspecified |
| mmu-let-7i-5p | MIMAT0000122 | Cops6 | 26893 | HITS-CLIP | 23142080 | Unspecified |
| mmu-let-7i-5p | MIMAT0000122 | Cops7a | 26894 | HITS-CLIP | 23142080 | Unspecified |
| mmu-let-7i-5p | MIMAT0000122 | Copz1 | 56447 | HITS-CLIP | 23142080 | Unspecified |
| mmu-let-7i-5p | MIMAT0000122 | Coq10b | 67876 | HITS-CLIP | 23597149 | Unspecified |
| mmu-let-7i-5p | MIMAT0000122 | Coq3 | 230027 | HITS-CLIP | 23142080 | Unspecified |
| mmu-let-7i-5p | MIMAT0000122 | Coq5 | 52064 | HITS-CLIP | 23142080 | Unspecified |
| mmu-let-7i-5p | MIMAT0000122 | Coq8a | 56997 | HITS-CLIP | 23597149 | Unspecified |
| mmu-let-7i-5p | MIMAT0000122 | Coq8b | 79934 | HITS-CLIP | 23597149 | Unspecified |
| mmu-let-7i-5p | MIMAT0000122 | Coq9 | 67914 | HITS-CLIP | 23142080 | Unspecified |
| mmu-let-7i-5p | MIMAT0000122 | Coro1a | 12721 | HITS-CLIP | 19536157\|23142080\|29167373 | Unspecified |
| mmu-let-7i-5p | MIMAT0000122 | Coro1b | 23789 | HITS-CLIP | 23142080 | Unspecified |

| | | | | | | |
|---|---|---|---|---|---|---|
| mmu-let-7i-5p | MIMAT0000122 | Coro1c | 23790 | HITS-CLIP | 37986224\|23142080\|29167373 | Unspecified |
| mmu-let-7i-5p | MIMAT0000122 | Coro2b | 235431 | HITS-CLIP | 37986224\|23597149\|19536157 | Unspecified |
| mmu-let-7i-5p | MIMAT0000122 | Cotl1 | 72042 | HITS-CLIP | 30971389 | Unspecified |
| mmu-let-7i-5p | MIMAT0000122 | Cox10 | 70383 | HITS-CLIP | 23597149\|19536157\|23142080 | Unspecified |
| mmu-let-7i-5p | MIMAT0000122 | Cox14 | 66379 | HITS-CLIP | 37986224\|23142080 | Unspecified |
| mmu-let-7i-5p | MIMAT0000122 | Cox16 | 51241 | HITS-CLIP | 23142080 | Unspecified |
| mmu-let-7i-5p | MIMAT0000122 | Cox17 | 12856 | HITS-CLIP | 23142080 | Unspecified |
| mmu-let-7i-5p | MIMAT0000122 | Cox4i1 | 12857 | HITS-CLIP | 23142080\|19536157\|23597149 | Unspecified |
| mmu-let-7i-5p | MIMAT0000122 | Cox5b | 12859 | HITS-CLIP | 23142080\|23597149\|19536157\|29167 | Unspecified |
| mmu-let-7i-5p | MIMAT0000122 | Cox6b1 | 110323 | HITS-CLIP | 37986224 | Unspecified |
| mmu-let-7i-5p | MIMAT0000122 | Cp | 12870 | HITS-CLIP | 23597149 | Unspecified |
| mmu-let-7i-5p | MIMAT0000122 | Cpa4 | 71791 | HITS-CLIP | 26203562 | Unspecified |
| mmu-let-7i-5p | MIMAT0000122 | Cpd | 12874 | HITS-CLIP | 37986224\|23597149\|31433990\|29167 | Unspecified |
| mmu-let-7i-5p | MIMAT0000122 | Cpe | 12876 | HITS-CLIP | 19536157 | Unspecified |
| mmu-let-7i-5p | MIMAT0000122 | Cpeb2 | 231207 | HITS-CLIP | 23142080\|23597149\|29167373 | Unspecified |
| mmu-let-7i-5p | MIMAT0000122 | Cpeb4 | 67579 | HITS-CLIP | 37986224\|23597149\|19536157\|23142 | Unspecified |
| mmu-let-7i-5p | MIMAT0000122 | Cplane1 | 65250 | HITS-CLIP | 23142080 | Unspecified |
| mmu-let-7i-5p | MIMAT0000122 | Cpm | 70574 | HITS-CLIP | 37986224\|23142080 | Unspecified |
| mmu-let-7i-5p | MIMAT0000122 | Cpn1 | 93721 | HITS-CLIP | 23597149 | Unspecified |
| mmu-let-7i-5p | MIMAT0000122 | Cpn2 | 71756 | HITS-CLIP | 23597149 | Unspecified |
| mmu-let-7i-5p | MIMAT0000122 | Cpne2 | 234577 | HITS-CLIP | 23597149 | Unspecified |
| mmu-let-7i-5p | MIMAT0000122 | Cpne3 | 70568 | HITS-CLIP | 23142080 | Unspecified |
| mmu-let-7i-5p | MIMAT0000122 | Cpox | 12892 | HITS-CLIP | 37986224\|23597149\|23142080 | Unspecified |
| mmu-let-7i-5p | MIMAT0000122 | Cpped1 | 223978 | HITS-CLIP | 37986224\|23597149\|23142080\|29167 | Unspecified |
| mmu-let-7i-5p | MIMAT0000122 | Cps1 | 227231 | HITS-CLIP | 23597149 | Unspecified |
| mmu-let-7i-5p | MIMAT0000122 | Cpsf1 | 94230 | HITS-CLIP | 23597149 | Unspecified |
| mmu-let-7i-5p | MIMAT0000122 | Cpsf3 | 54451 | HITS-CLIP | 23142080 | Unspecified |
| mmu-let-7i-5p | MIMAT0000122 | Cpsf6 | 432508 | HITS-CLIP | 23630078 | Unspecified |
| mmu-let-7i-5p | MIMAT0000122 | Cpsf7 | 269061 | HITS-CLIP | 29167373 | Unspecified |
| mmu-let-7i-5p | MIMAT0000122 | Cpt1a | 12894 | HITS-CLIP | 23597149 | Unspecified |
| mmu-let-7i-5p | MIMAT0000122 | Cpt2 | 12896 | HITS-CLIP | 23597149\|23142080 | Unspecified |
| mmu-let-7i-5p | MIMAT0000122 | Cptp | 80772 | HITS-CLIP | 23597149\|23142080 | Unspecified |
| mmu-let-7i-5p | MIMAT0000122 | Cramp1l | 57354 | HITS-CLIP | 37986224\|23142080\|23597149 | Unspecified |
| mmu-let-7i-5p | MIMAT0000122 | Crat | 12908 | HITS-CLIP | 23597149\|23142080 | Unspecified |
| mmu-let-7i-5p | MIMAT0000122 | Crb3 | 224912 | HITS-CLIP | 23597149 | Unspecified |
| mmu-let-7i-5p | MIMAT0000122 | Crbn | 58799 | HITS-CLIP | 37986224\|23597149\|23142080 | Unspecified |
| mmu-let-7i-5p | MIMAT0000122 | Creb1 | 12912 | HITS-CLIP | 23597149\|23142080 | Unspecified |
| mmu-let-7i-5p | MIMAT0000122 | Creb3 | 12913 | HITS-CLIP | 23597149 | Unspecified |
| mmu-let-7i-5p | MIMAT0000122 | Creb3l3 | 208677 | HITS-CLIP | 23597149 | Unspecified |

| | | | | | | |
|---|---|---|---|---|---|---|
| mmu-let-7i-5p | MIMAT0000122 | Creb5 | 231991 | HITS-CLIP | 29167373 | Unspecified |
| mmu-let-7i-5p | MIMAT0000122 | Crebbp | 12914 | HITS-CLIP | 37986224\|23597149\|23142080\|19536 | Unspecified |
| mmu-let-7i-5p | MIMAT0000122 | Crebrf | 77128 | HITS-CLIP | 23142080 | Unspecified |
| mmu-let-7i-5p | MIMAT0000122 | Creg1 | 433375 | HITS-CLIP | 23597149 | Unspecified |
| mmu-let-7i-5p | MIMAT0000122 | Crem | 12916 | HITS-CLIP | 37986224\|23142080 | Unspecified |
| mmu-let-7i-5p | MIMAT0000122 | Crim1 | 50766 | HITS-CLIP | 23142080 | Unspecified |
| mmu-let-7i-5p | MIMAT0000122 | Crip1 | 12925 | HITS-CLIP | 23142080\|23597149 | Unspecified |
| mmu-let-7i-5p | MIMAT0000122 | Crip2 | 68337 | HITS-CLIP | 23597149 | Unspecified |
| mmu-let-7i-5p | MIMAT0000122 | Crk | 12928 | HITS-CLIP | 37986224\|19536157\|31433990\|23142 | Unspecified |
| mmu-let-7i-5p | MIMAT0000122 | Crkl | 12929 | HITS-CLIP | 37986224\|23597149\|23142080 | Unspecified |
| mmu-let-7i-5p | MIMAT0000122 | Crlf3 | 54394 | HITS-CLIP | 29167373 | Unspecified |
| mmu-let-7i-5p | MIMAT0000122 | Crmp1 | 12933 | HITS-CLIP | 37986224\|19536157 | Unspecified |
| mmu-let-7i-5p | MIMAT0000122 | Crnkl1 | 66877 | HITS-CLIP | 23142080 | Unspecified |
| mmu-let-7i-5p | MIMAT0000122 | Crot | 74114 | HITS-CLIP | 23597149\|23142080 | Unspecified |
| mmu-let-7i-5p | MIMAT0000122 | Crp | 12944 | HITS-CLIP | 37986224\|23597149 | Unspecified |
| mmu-let-7i-5p | MIMAT0000122 | Crtam | 54698 | HITS-CLIP | 37986224\|23142080\|19536157 | Unspecified |
| mmu-let-7i-5p | MIMAT0000122 | Crtap | 56693 | HITS-CLIP | 29167373 | Unspecified |
| mmu-let-7i-5p | MIMAT0000122 | Cry1 | 12952 | HITS-CLIP | 23142080 | Unspecified |
| mmu-let-7i-5p | MIMAT0000122 | Cry2 | 12953 | HITS-CLIP | 37986224\|23597149\|23142080 | Unspecified |
| mmu-let-7i-5p | MIMAT0000122 | Cryab | 12955 | HITS-CLIP | 37986224 | Unspecified |
| mmu-let-7i-5p | MIMAT0000122 | Crybg3 | 224273 | HITS-CLIP | 23142080 | Unspecified |
| mmu-let-7i-5p | MIMAT0000122 | Cryz | 12972 | HITS-CLIP | 23597149 | Unspecified |
| mmu-let-7i-5p | MIMAT0000122 | Cryzl1 | 66609 | HITS-CLIP | 29167373 | Unspecified |
| mmu-let-7i-5p | MIMAT0000122 | Cs | 12974 | HITS-CLIP | 37986224\|23597149\|23142080\|19536 | Unspecified |
| mmu-let-7i-5p | MIMAT0000122 | Csad | 246277 | HITS-CLIP | 23597149 | Unspecified |
| mmu-let-7i-5p | MIMAT0000122 | Csde1 | 229663 | HITS-CLIP | 23597149\|19536157\|29167373\|23142 | Unspecified |
| mmu-let-7i-5p | MIMAT0000122 | Cse1l | 110750 | HITS-CLIP | 23597149\|19536157\|29167373 | Unspecified |
| mmu-let-7i-5p | MIMAT0000122 | Csf1 | 12977 | HITS-CLIP | 23142080 | Unspecified |
| mmu-let-7i-5p | MIMAT0000122 | Csf1r | 12978 | HITS-CLIP | 29167373 | Unspecified |
| mmu-let-7i-5p | MIMAT0000122 | Csf2rb | 12983 | HITS-CLIP | 23142080\|29167373 | Unspecified |
| mmu-let-7i-5p | MIMAT0000122 | Csk | 12988 | HITS-CLIP | 23597149\|23142080\|29167373 | Unspecified |
| mmu-let-7i-5p | MIMAT0000122 | Csmd1 | 94109 | HITS-CLIP | 37986224 | Unspecified |
| mmu-let-7i-5p | MIMAT0000122 | Csmd2 | 329942 | HITS-CLIP | 37986224 | Unspecified |
| mmu-let-7i-5p | MIMAT0000122 | Csnk1a1 | 93687 | HITS-CLIP | 37986224\|23142080 | Unspecified |
| mmu-let-7i-5p | MIMAT0000122 | Csnk1d | 104318 | HITS-CLIP | 23597149\|23142080\|19536157 | Unspecified |
| mmu-let-7i-5p | MIMAT0000122 | Csnk1g1 | 214897 | HITS-CLIP | 37986224\|23597149\|23142080 | Unspecified |
| mmu-let-7i-5p | MIMAT0000122 | Csnk1g2 | 103236 | HITS-CLIP | 23142080 | Unspecified |
| mmu-let-7i-5p | MIMAT0000122 | Csnk2a1 | 12995 | HITS-CLIP | 37986224\|23597149\|23142080 | Unspecified |
| mmu-let-7i-5p | MIMAT0000122 | Csnk2a2 | 13000 | HITS-CLIP | 23142080\|23597149 | Unspecified |

| | | | | | | |
|---|---|---|---|---|---|---|
| mmu-let-7i-5p | MIMAT0000122 | Csnk2b | 13001 | HITS-CLIP | 23597149\|23142080\|29167373 | Unspecified |
| mmu-let-7i-5p | MIMAT0000122 | Cspg5 | 29873 | HITS-CLIP | 37986224 | Unspecified |
| mmu-let-7i-5p | MIMAT0000122 | Cspp1 | 211660 | HITS-CLIP | 23142080 | Unspecified |
| mmu-let-7i-5p | MIMAT0000122 | Csrnp1 | 215418 | HITS-CLIP | 23142080\|23597149 | Unspecified |
| mmu-let-7i-5p | MIMAT0000122 | Csrnp2 | 207785 | HITS-CLIP | 37986224\|19536157\|23142080 | Unspecified |
| mmu-let-7i-5p | MIMAT0000122 | Csrp1 | 13007 | HITS-CLIP | 23142080 | Unspecified |
| mmu-let-7i-5p | MIMAT0000122 | Cstf1 | 67337 | HITS-CLIP | 23142080 | Unspecified |
| mmu-let-7i-5p | MIMAT0000122 | Cstf2 | 108062 | HITS-CLIP | 29167373 | Unspecified |
| mmu-let-7i-5p | MIMAT0000122 | Ctbs | 74245 | HITS-CLIP | 29167373\|23142080 | Unspecified |
| mmu-let-7i-5p | MIMAT0000122 | Ctdnep1 | 67181 | HITS-CLIP | 23597149 | Unspecified |
| mmu-let-7i-5p | MIMAT0000122 | Ctdp1 | 67655 | HITS-CLIP | 37986224\|23597149\|29167373\|23142 | Unspecified |
| mmu-let-7i-5p | MIMAT0000122 | Ctdsp2 | 52468 | HITS-CLIP | 23597149\|19536157 | Unspecified |
| mmu-let-7i-5p | MIMAT0000122 | Ctdspl | 69274 | HITS-CLIP | 23597149 | Unspecified |
| mmu-let-7i-5p | MIMAT0000122 | Ctdspl2 | 329506 | HITS-CLIP | 23142080 | Unspecified |
| mmu-let-7i-5p | MIMAT0000122 | Cth | 107869 | HITS-CLIP | 23597149 | Unspecified |
| mmu-let-7i-5p | MIMAT0000122 | Ctla4 | 12477 | HITS-CLIP | 37986224 | Unspecified |
| mmu-let-7i-5p | MIMAT0000122 | Ctnna1 | 12385 | HITS-CLIP | 23597149\|19536157\|23142080 | Unspecified |
| mmu-let-7i-5p | MIMAT0000122 | Ctnnb1 | 12387 | HITS-CLIP | 23597149\|23142080\|31433990\|1953€ | Unspecified |
| mmu-let-7i-5p | MIMAT0000122 | Ctnnbl1 | 66642 | HITS-CLIP | 23142080 | Unspecified |
| mmu-let-7i-5p | MIMAT0000122 | Ctnnd1 | 12388 | HITS-CLIP | 37986224 | Unspecified |
| mmu-let-7i-5p | MIMAT0000122 | Ctnnd2 | 18163 | HITS-CLIP | 37986224\|19536157\|29167373 | Unspecified |
| mmu-let-7i-5p | MIMAT0000122 | Ctns | 83429 | HITS-CLIP | 23597149 | Unspecified |
| mmu-let-7i-5p | MIMAT0000122 | Ctr9 | 22083 | HITS-CLIP | 37986224\|23597149\|23142080 | Unspecified |
| mmu-let-7i-5p | MIMAT0000122 | Ctsa | 19025 | HITS-CLIP | 23597149\|23142080\|29167373 | Unspecified |
| mmu-let-7i-5p | MIMAT0000122 | Ctsb | 13030 | HITS-CLIP | 37986224\|23597149\|19536157\|29167 | Unspecified |
| mmu-let-7i-5p | MIMAT0000122 | Ctsc | 13032 | HITS-CLIP | 37986224\|23597149\|23142080\|29167 | Unspecified |
| mmu-let-7i-5p | MIMAT0000122 | Ctsd | 13033 | HITS-CLIP | 23597149 | Unspecified |
| mmu-let-7i-5p | MIMAT0000122 | Ctsh | 13036 | HITS-CLIP | 23597149 | Unspecified |
| mmu-let-7i-5p | MIMAT0000122 | Ctsl | 13039 | HITS-CLIP | 37986224\|23142080\|23597149\|29167 | Unspecified |
| mmu-let-7i-5p | MIMAT0000122 | Ctso | 229445 | qCLASH | 31393953 | Unspecified |
| mmu-let-7i-5p | MIMAT0000122 | Cul1 | 26965 | HITS-CLIP | 23597149\|19536157\|29167373\|23142 | Unspecified |
| mmu-let-7i-5p | MIMAT0000122 | Cul2 | 71745 | HITS-CLIP | 37986224\|23142080\|30971389\|23597 | Unspecified |
| mmu-let-7i-5p | MIMAT0000122 | Cul3 | 26554 | HITS-CLIP | 23142080\|23597149\|31433990 | Unspecified |
| mmu-let-7i-5p | MIMAT0000122 | Cul4a | 99375 | HITS-CLIP | 29167373 | Unspecified |
| mmu-let-7i-5p | MIMAT0000122 | Cul4b | 72584 | HITS-CLIP | 23142080 | Unspecified |
| mmu-let-7i-5p | MIMAT0000122 | Cul5 | 75717 | HITS-CLIP | 23142080 | Unspecified |
| mmu-let-7i-5p | MIMAT0000122 | Cul7 | 66515 | HITS-CLIP | 23142080 | Unspecified |
| mmu-let-7i-5p | MIMAT0000122 | Cux1 | 13047 | HITS-CLIP | 23142080\|29167373 | Unspecified |
| mmu-let-7i-5p | MIMAT0000122 | Cwf19l2 | 244672 | HITS-CLIP | 23597149\|23142080 | Unspecified |

| | | | | | | |
|---|---|---|---|---|---|---|
| mmu-let-7i-5p | MIMAT0000122 | Cx3cl1 | 20312 | HITS-CLIP | 37986224 | Unspecified |
| mmu-let-7i-5p | MIMAT0000122 | Cx3cr1 | 13051 | HITS-CLIP | 29167373 | Unspecified |
| mmu-let-7i-5p | MIMAT0000122 | Cxadr | 13052 | HITS-CLIP | 37986224\|19536157 | Unspecified |
| mmu-let-7i-5p | MIMAT0000122 | Cxcl1 | 14825 | HITS-CLIP | 23597149 | Unspecified |
| mmu-let-7i-5p | MIMAT0000122 | Cxcl12 | 20315 | HITS-CLIP | 23597149 | Unspecified |
| mmu-let-7i-5p | MIMAT0000122 | Cxcr2 | 12765 | HITS-CLIP | 37986224 | Unspecified |
| mmu-let-7i-5p | MIMAT0000122 | Cxcr4 | 12767 | HITS-CLIP | 23142080 | Unspecified |
| mmu-let-7i-5p | MIMAT0000122 | Cxxc5 | 67393 | HITS-CLIP | 37986224\|23597149\|19536157 | Unspecified |
| mmu-let-7i-5p | MIMAT0000122 | Cyb561d1 | 72023 | HITS-CLIP | 23142080 | Unspecified |
| mmu-let-7i-5p | MIMAT0000122 | Cyb5b | 66427 | HITS-CLIP | 23142080 | Unspecified |
| mmu-let-7i-5p | MIMAT0000122 | Cyb5r3 | 109754 | HITS-CLIP | 23142080\|23597149\|19536157 | Unspecified |
| mmu-let-7i-5p | MIMAT0000122 | Cyb5r4 | 266690 | HITS-CLIP | 37986224\|23142080\|23597149\|29167 | Unspecified |
| mmu-let-7i-5p | MIMAT0000122 | Cyb5rl | 230582 | HITS-CLIP | 23597149 | Unspecified |
| mmu-let-7i-5p | MIMAT0000122 | Cyba | 13057 | HITS-CLIP | 23142080 | Unspecified |
| mmu-let-7i-5p | MIMAT0000122 | Cybb | 13058 | HITS-CLIP | 29167373 | Unspecified |
| mmu-let-7i-5p | MIMAT0000122 | Cyc1 | 66445 | HITS-CLIP | 23597149\|23142080 | Unspecified |
| mmu-let-7i-5p | MIMAT0000122 | Cyfip1 | 20430 | HITS-CLIP | 23597149\|29167373 | Unspecified |
| mmu-let-7i-5p | MIMAT0000122 | Cyfip2 | 76884 | HITS-CLIP | 23142080 | Unspecified |
| mmu-let-7i-5p | MIMAT0000122 | Cyhr1 | 54151 | HITS-CLIP | 23597149 | Unspecified |
| mmu-let-7i-5p | MIMAT0000122 | Cyld | 74256 | HITS-CLIP | 23142080\|29167373 | Unspecified |
| mmu-let-7i-5p | MIMAT0000122 | Cyp11a1 | 13070 | HITS-CLIP | 23142080 | Unspecified |
| mmu-let-7i-5p | MIMAT0000122 | Cyp1a2 | 13077 | HITS-CLIP | 23597149 | Unspecified |
| mmu-let-7i-5p | MIMAT0000122 | Cyp1b1 | 13078 | HITS-CLIP | 19536157 | Unspecified |
| mmu-let-7i-5p | MIMAT0000122 | Cyp2c29 | 13095 | HITS-CLIP | 23597149 | Unspecified |
| mmu-let-7i-5p | MIMAT0000122 | Cyp2c37 | 13096 | HITS-CLIP | 23597149 | Unspecified |
| mmu-let-7i-5p | MIMAT0000122 | Cyp2c70 | 226105 | HITS-CLIP | 37986224\|23597149 | Unspecified |
| mmu-let-7i-5p | MIMAT0000122 | Cyp2d10 | 13101 | HITS-CLIP | 23597149 | Unspecified |
| mmu-let-7i-5p | MIMAT0000122 | Cyp2d26 | 76279 | HITS-CLIP | 23597149 | Unspecified |
| mmu-let-7i-5p | MIMAT0000122 | Cyp2e1 | 13106 | HITS-CLIP | 23597149 | Unspecified |
| mmu-let-7i-5p | MIMAT0000122 | Cyp2f2 | 13107 | HITS-CLIP | 23597149 | Unspecified |
| mmu-let-7i-5p | MIMAT0000122 | Cyp2r1 | 244209 | HITS-CLIP | 37986224\|23597149 | Unspecified |
| mmu-let-7i-5p | MIMAT0000122 | Cyp2s1 | 74134 | HITS-CLIP | 23142080 | Unspecified |
| mmu-let-7i-5p | MIMAT0000122 | Cyp2u1 | 71519 | HITS-CLIP | 23597149 | Unspecified |
| mmu-let-7i-5p | MIMAT0000122 | Cyp3a11 | 13112 | HITS-CLIP | 23597149 | Unspecified |
| mmu-let-7i-5p | MIMAT0000122 | Cyp3a13 | 13113 | HITS-CLIP | 23597149 | Unspecified |
| mmu-let-7i-5p | MIMAT0000122 | Cyp46a1 | 13116 | HITS-CLIP | 19536157 | Unspecified |
| mmu-let-7i-5p | MIMAT0000122 | Cyp4a10 | 13117 | HITS-CLIP | 37986224\|23597149 | Unspecified |
| mmu-let-7i-5p | MIMAT0000122 | Cyp4b1 | 13120 | HITS-CLIP | 23597149 | Unspecified |
| mmu-let-7i-5p | MIMAT0000122 | Cyp4f13 | 170716 | HITS-CLIP | 23597149\|23142080 | Unspecified |

| | | | | | | |
|---|---|---|---|---|---|---|
| mmu-let-7i-5p | MIMAT0000122 | Cyp4v3 | 102294 | HITS-CLIP | 23597149 | Unspecified |
| mmu-let-7i-5p | MIMAT0000122 | Cyp51 | 13121 | HITS-CLIP | 37986224\|23597149\|23142080 | Unspecified |
| mmu-let-7i-5p | MIMAT0000122 | Cyp7a1 | 13122 | HITS-CLIP | 23597149 | Unspecified |
| mmu-let-7i-5p | MIMAT0000122 | Cystm1 | 66060 | HITS-CLIP | 23142080 | Unspecified |
| mmu-let-7i-5p | MIMAT0000122 | Cyth2 | 19158 | HITS-CLIP | 23142080 | Unspecified |
| mmu-let-7i-5p | MIMAT0000122 | Cyth3 | 19159 | HITS-CLIP | 37986224\|23597149\|23142080 | Unspecified |
| mmu-let-7i-5p | MIMAT0000122 | Cyth4 | 72318 | HITS-CLIP | 29167373\|23142080 | Unspecified |
| mmu-let-7i-5p | MIMAT0000122 | Cytip | 227929 | HITS-CLIP | 23142080\|29167373 | Unspecified |
| mmu-let-7i-5p | MIMAT0000122 | D10Wsu102e | 28109 | HITS-CLIP | 23142080\|29167373 | Unspecified |
| mmu-let-7i-5p | MIMAT0000122 | D16Ertd472e | 67102 | HITS-CLIP | 23142080 | Unspecified |
| mmu-let-7i-5p | MIMAT0000122 | D17H6S53E | 114585 | HITS-CLIP | 23142080 | Unspecified |
| mmu-let-7i-5p | MIMAT0000122 | D1Ertd622e | 52392 | HITS-CLIP | 23597149\|29167373 | Unspecified |
| mmu-let-7i-5p | MIMAT0000122 | D230025D16Rik | 234678 | HITS-CLIP | 19536157\|23142080\|29167373 | Unspecified |
| mmu-let-7i-5p | MIMAT0000122 | D430019H16Rik | 268595 | HITS-CLIP | 37986224 | Unspecified |
| mmu-let-7i-5p | MIMAT0000122 | D430041D05Rik | 241589 | HITS-CLIP | 19536157 | Unspecified |
| mmu-let-7i-5p | MIMAT0000122 | D5Ertd579e | 320661 | HITS-CLIP | 23597149\|26203562\|23142080 | Unspecified |
| mmu-let-7i-5p | MIMAT0000122 | D630045J12Rik | 330286 | HITS-CLIP | 37986224\|19536157 | Unspecified |
| mmu-let-7i-5p | MIMAT0000122 | D6Wsu163e | 28040 | HITS-CLIP | 23142080 | Unspecified |
| mmu-let-7i-5p | MIMAT0000122 | Daam1 | 208846 | HITS-CLIP | 19536157\|29167373 | Unspecified |
| mmu-let-7i-5p | MIMAT0000122 | Dab1 | 13131 | HITS-CLIP | 37986224\|19536157 | Unspecified |
| mmu-let-7i-5p | MIMAT0000122 | Dab2 | 13132 | HITS-CLIP | 23597149\|29167373 | Unspecified |
| mmu-let-7i-5p | MIMAT0000122 | Dach1 | 13134 | HITS-CLIP | 19536157 | Unspecified |
| mmu-let-7i-5p | MIMAT0000122 | Dact3 | 629378 | HITS-CLIP | 19536157 | Unspecified |
| mmu-let-7i-5p | MIMAT0000122 | Dad1 | 13135 | HITS-CLIP | 23142080 | Unspecified |
| mmu-let-7i-5p | MIMAT0000122 | Dag1 | 13138 | HITS-CLIP | 23142080\|23597149\|29167373 | Unspecified |
| mmu-let-7i-5p | MIMAT0000122 | Dagla | 269060 | HITS-CLIP | 23597149 | Unspecified |
| mmu-let-7i-5p | MIMAT0000122 | Dalrd3 | 67789 | HITS-CLIP | 23597149 | Unspecified |
| mmu-let-7i-5p | MIMAT0000122 | Dap | 223453 | HITS-CLIP | 37986224\|23142080 | Unspecified |
| mmu-let-7i-5p | MIMAT0000122 | Dapk1 | 69635 | HITS-CLIP | 23597149\|19536157\|29167373 | Unspecified |
| mmu-let-7i-5p | MIMAT0000122 | Dapk3 | 13144 | HITS-CLIP | 23142080\|23597149 | Unspecified |
| mmu-let-7i-5p | MIMAT0000122 | Dars | 226414 | HITS-CLIP | 23597149\|23142080 | Unspecified |
| mmu-let-7i-5p | MIMAT0000122 | Daxx | 13163 | HITS-CLIP | 23142080\|23597149\|29167373 | Unspecified |
| mmu-let-7i-5p | MIMAT0000122 | Dazap2 | 23994 | HITS-CLIP | 23630078 | Unspecified |
| mmu-let-7i-5p | MIMAT0000122 | Dbf4 | 27214 | HITS-CLIP | 23142080 | Unspecified |
| mmu-let-7i-5p | MIMAT0000122 | Dbt | 13171 | HITS-CLIP | 23597149\|23142080 | Unspecified |
| mmu-let-7i-5p | MIMAT0000122 | Dcaf1 | 41836 | HITS-CLIP | 23142080\|23597149\|19536157 | Unspecified |
| mmu-let-7i-5p | MIMAT0000122 | Dcaf11 | 28199 | HITS-CLIP | 23142080\|23597149 | Unspecified |
| mmu-let-7i-5p | MIMAT0000122 | Dcaf12 | 68970 | HITS-CLIP//qCLASH | 23597149\|31393953\|23142080 | Unspecified |
| mmu-let-7i-5p | MIMAT0000122 | Dcaf15 | 212123 | HITS-CLIP | 29167373 | Unspecified |

| | | | | | | |
|---|---|---|---|---|---|---|
| mmu-let-7i-5p | MIMAT0000122 | Dcaf17 | 75763 | HITS-CLIP | 23142080\|23597149 | Unspecified |
| mmu-let-7i-5p | MIMAT0000122 | Dcaf5 | 320808 | HITS-CLIP | 37986224\|23597149\|19536157 | Unspecified |
| mmu-let-7i-5p | MIMAT0000122 | Dcaf6 | 74106 | HITS-CLIP | 23142080 | Unspecified |
| mmu-let-7i-5p | MIMAT0000122 | Dcaf7 | 71833 | HITS-CLIP | 23597149\|23142080 | Unspecified |
| mmu-let-7i-5p | MIMAT0000122 | Dcaf8 | 98193 | HITS-CLIP | 23597149\|23142080 | Unspecified |
| mmu-let-7i-5p | MIMAT0000122 | Dcbld1 | 66686 | HITS-CLIP | 23597149 | Unspecified |
| mmu-let-7i-5p | MIMAT0000122 | Dcc | 13176 | HITS-CLIP | 19536157 | Unspecified |
| mmu-let-7i-5p | MIMAT0000122 | Dck | 13178 | HITS-CLIP | 23142080 | Unspecified |
| mmu-let-7i-5p | MIMAT0000122 | Dclk1 | 13175 | HITS-CLIP | 23142080\|19536157 | Unspecified |
| mmu-let-7i-5p | MIMAT0000122 | Dclre1b | 140917 | HITS-CLIP | 23142080 | Unspecified |
| mmu-let-7i-5p | MIMAT0000122 | Dclre1c | 227525 | HITS-CLIP | 29167373 | Unspecified |
| mmu-let-7i-5p | MIMAT0000122 | Dcp1a | 75901 | HITS-CLIP | 37986224\|29167373 | Unspecified |
| mmu-let-7i-5p | MIMAT0000122 | Dcp1b | 319618 | HITS-CLIP | 23142080 | Unspecified |
| mmu-let-7i-5p | MIMAT0000122 | Dcp2 | 70640 | HITS-CLIP | 23142080\|19536157 | Unspecified |
| mmu-let-7i-5p | MIMAT0000122 | Dctn1 | 13191 | HITS-CLIP | 23142080 | Unspecified |
| mmu-let-7i-5p | MIMAT0000122 | Dctn4 | 67665 | HITS-CLIP | 23142080 | Unspecified |
| mmu-let-7i-5p | MIMAT0000122 | Dctpp1 | 66422 | HITS-CLIP | 23597149\|23142080 | Unspecified |
| mmu-let-7i-5p | MIMAT0000122 | Dcun1d1 | 114893 | HITS-CLIP | 23142080 | Unspecified |
| mmu-let-7i-5p | MIMAT0000122 | Dcun1d2 | 102323 | HITS-CLIP | 23142080 | Unspecified |
| mmu-let-7i-5p | MIMAT0000122 | Dcun1d3 | 233805 | HITS-CLIP | 19536157\|23142080 | Unspecified |
| mmu-let-7i-5p | MIMAT0000122 | Dcxr | 67880 | HITS-CLIP | 23597149 | Unspecified |
| mmu-let-7i-5p | MIMAT0000122 | Ddah1 | 69219 | HITS-CLIP | 23597149 | Unspecified |
| mmu-let-7i-5p | MIMAT0000122 | Ddb1 | 13194 | HITS-CLIP | 37986224\|23597149\|23142080\|1953( | Unspecified |
| mmu-let-7i-5p | MIMAT0000122 | Ddc | 13195 | HITS-CLIP | 23142080\|23597149 | Unspecified |
| mmu-let-7i-5p | MIMAT0000122 | Ddhd1 | 114874 | HITS-CLIP | 37986224\|23597149\|19536157\|2314 | Unspecified |
| mmu-let-7i-5p | MIMAT0000122 | Ddhd2 | 72108 | HITS-CLIP | 23142080 | Unspecified |
| mmu-let-7i-5p | MIMAT0000122 | Ddi2 | 68817 | HITS-CLIP | 37986224\|23597149\|19536157\|2314 | Unspecified |
| mmu-let-7i-5p | MIMAT0000122 | Ddias | 220042 | HITS-CLIP | 23142080 | Unspecified |
| mmu-let-7i-5p | MIMAT0000122 | Ddit4 | 74747 | HITS-CLIP | 23597149\|26203562 | Unspecified |
| mmu-let-7i-5p | MIMAT0000122 | Ddn | 13199 | HITS-CLIP | 37986224\|19536157 | Unspecified |
| mmu-let-7i-5p | MIMAT0000122 | Ddost | 13200 | HITS-CLIP | 37986224\|23597149\|23142080 | Unspecified |
| mmu-let-7i-5p | MIMAT0000122 | Ddr1 | 12305 | HITS-CLIP | 26203562\|23142080 | Unspecified |
| mmu-let-7i-5p | MIMAT0000122 | Ddx1 | 104721 | HITS-CLIP | 23142080 | Unspecified |
| mmu-let-7i-5p | MIMAT0000122 | Ddx10 | 77591 | HITS-CLIP | 37986224\|19536157\|23142080 | Unspecified |
| mmu-let-7i-5p | MIMAT0000122 | Ddx17 | 67040 | HITS-CLIP | 23597149\|23142080\|29167373 | Unspecified |
| mmu-let-7i-5p | MIMAT0000122 | Ddx19a | 13680 | HITS-CLIP | 23142080 | Unspecified |
| mmu-let-7i-5p | MIMAT0000122 | Ddx20 | 53975 | HITS-CLIP | 30971389\|31433990\|23142080 | Unspecified |
| mmu-let-7i-5p | MIMAT0000122 | Ddx21 | 56200 | HITS-CLIP | 31433990 | Unspecified |
| mmu-let-7i-5p | MIMAT0000122 | Ddx24 | 27225 | HITS-CLIP | 23597149\|23142080 | Unspecified |

| | | | | | | |
|---|---|---|---|---|---|---|
| mmu-let-7i-5p | MIMAT0000122 | Ddx39 | 68278 | HITS-CLIP | 29167373 | Unspecified |
| mmu-let-7i-5p | MIMAT0000122 | Ddx39b | 53817 | HITS-CLIP | 23142080 | Unspecified |
| mmu-let-7i-5p | MIMAT0000122 | Ddx3x | 13205 | HITS-CLIP | 23597149\|23142080 | Unspecified |
| mmu-let-7i-5p | MIMAT0000122 | Ddx3y | 26900 | HITS-CLIP | 23597149\|23142080 | Unspecified |
| mmu-let-7i-5p | MIMAT0000122 | Ddx41 | 72935 | HITS-CLIP | 23142080 | Unspecified |
| mmu-let-7i-5p | MIMAT0000122 | Ddx42 | 72047 | HITS-CLIP | 23597149\|23142080 | Unspecified |
| mmu-let-7i-5p | MIMAT0000122 | Ddx47 | 67755 | HITS-CLIP | 23142080 | Unspecified |
| mmu-let-7i-5p | MIMAT0000122 | Ddx5 | 13207 | HITS-CLIP | 23597149\|23142080\|19536157\|29167 | Unspecified |
| mmu-let-7i-5p | MIMAT0000122 | Ddx54 | 71990 | HITS-CLIP | 37986224\|23597149\|23142080 | Unspecified |
| mmu-let-7i-5p | MIMAT0000122 | Ddx56 | 52513 | HITS-CLIP | 23597149 | Unspecified |
| mmu-let-7i-5p | MIMAT0000122 | Ddx58 | 230073 | HITS-CLIP | 37986224\|23597149\|23142080\|29167 | Unspecified |
| mmu-let-7i-5p | MIMAT0000122 | Ddx6 | 13209 | HITS-CLIP | 23142080\|19536157\|23597149\|29167 | Unspecified |
| mmu-let-7i-5p | MIMAT0000122 | Dedd | 21945 | HITS-CLIP | 37986224\|23597149\|23142080\|29167 | Unspecified |
| mmu-let-7i-5p | MIMAT0000122 | Def6 | 23853 | HITS-CLIP | 37986224\|23142080 | Unspecified |
| mmu-let-7i-5p | MIMAT0000122 | Degs1 | 13244 | HITS-CLIP | 23597149\|29167373\|23142080 | Unspecified |
| mmu-let-7i-5p | MIMAT0000122 | Dele1 | 9812 | HITS-CLIP | 23597149 | Unspecified |
| mmu-let-7i-5p | MIMAT0000122 | Dennd1a | 227801 | HITS-CLIP | 23597149\|29167373 | Unspecified |
| mmu-let-7i-5p | MIMAT0000122 | Dennd1b | 329260 | HITS-CLIP | 37986224\|23142080\|23597149\|31433 | Unspecified |
| mmu-let-7i-5p | MIMAT0000122 | Dennd2c | 329727 | HITS-CLIP | 23142080 | Unspecified |
| mmu-let-7i-5p | MIMAT0000122 | Dennd2d | 72121 | HITS-CLIP | 23142080 | Unspecified |
| mmu-let-7i-5p | MIMAT0000122 | Dennd4a | 102442 | HITS-CLIP | 23142080 | Unspecified |
| mmu-let-7i-5p | MIMAT0000122 | Dennd4b | 229541 | HITS-CLIP | 23142080 | Unspecified |
| mmu-let-7i-5p | MIMAT0000122 | Dennd4c | 329877 | HITS-CLIP | 23597149\|23142080\|29167373 | Unspecified |
| mmu-let-7i-5p | MIMAT0000122 | Dennd5a | 19347 | HITS-CLIP | 23597149 | Unspecified |
| mmu-let-7i-5p | MIMAT0000122 | Dennd5b | 320560 | HITS-CLIP | 37986224\|19536157\|23597149 | Unspecified |
| mmu-let-7i-5p | MIMAT0000122 | Dennd6a | 211922 | HITS-CLIP | 29167373\|23142080 | Unspecified |
| mmu-let-7i-5p | MIMAT0000122 | Depp1 | 11067 | HITS-CLIP | 23597149 | Unspecified |
| mmu-let-7i-5p | MIMAT0000122 | Dera | 232449 | HITS-CLIP | 23142080\|29167373 | Unspecified |
| mmu-let-7i-5p | MIMAT0000122 | Derl2 | 116891 | HITS-CLIP | 37986224 | Unspecified |
| mmu-let-7i-5p | MIMAT0000122 | Desi2 | 78825 | HITS-CLIP | 23597149\|23142080 | Unspecified |
| mmu-let-7i-5p | MIMAT0000122 | Dffb | 13368 | HITS-CLIP | 23142080 | Unspecified |
| mmu-let-7i-5p | MIMAT0000122 | Dgcr2 | 13356 | HITS-CLIP | 37986224\|23597149\|23142080\|19536 | Unspecified |
| mmu-let-7i-5p | MIMAT0000122 | Dgcr8 | 94223 | HITS-CLIP | 30971389\|29167373 | Unspecified |
| mmu-let-7i-5p | MIMAT0000122 | Dgka | 13139 | HITS-CLIP | 37986224\|23597149\|23142080 | Unspecified |
| mmu-let-7i-5p | MIMAT0000122 | Dgkd | 227333 | HITS-CLIP | 31433990\|29167373 | Unspecified |
| mmu-let-7i-5p | MIMAT0000122 | Dgke | 56077 | HITS-CLIP | 23142080\|26203562 | Unspecified |
| mmu-let-7i-5p | MIMAT0000122 | Dgkh | 380921 | HITS-CLIP | 29167373 | Unspecified |
| mmu-let-7i-5p | MIMAT0000122 | Dgkq | 110524 | HITS-CLIP | 37986224\|23597149 | Unspecified |
| mmu-let-7i-5p | MIMAT0000122 | Dgkz | 104418 | HITS-CLIP | 23142080\|29167373 | Unspecified |

| | | | | | | |
|---|---|---|---|---|---|---|
| mmu-let-7i-5p | MIMAT0000122 | Dglucy | 217830 | HITS-CLIP | 23597149\|29167373 | Unspecified |
| mmu-let-7i-5p | MIMAT0000122 | Dhcr24 | 74754 | HITS-CLIP | 37986224\|23597149\|26203562\|23142 | Unspecified |
| mmu-let-7i-5p | MIMAT0000122 | Dhcr7 | 13360 | HITS-CLIP | 23597149 | Unspecified |
| mmu-let-7i-5p | MIMAT0000122 | Dhdh | 71755 | HITS-CLIP | 23597149\|23142080 | Unspecified |
| mmu-let-7i-5p | MIMAT0000122 | Dhfr | 13361 | HITS-CLIP | 37986224\|23142080 | Unspecified |
| mmu-let-7i-5p | MIMAT0000122 | Dhrs13 | 70451 | HITS-CLIP | 23142080 | Unspecified |
| mmu-let-7i-5p | MIMAT0000122 | Dhrs3 | 20148 | HITS-CLIP | 23597149\|29167373 | Unspecified |
| mmu-let-7i-5p | MIMAT0000122 | Dhrs7 | 66375 | HITS-CLIP | 23597149 | Unspecified |
| mmu-let-7i-5p | MIMAT0000122 | Dhrs9 | 241452 | HITS-CLIP | 23597149 | Unspecified |
| mmu-let-7i-5p | MIMAT0000122 | Dhtkd1 | 209692 | HITS-CLIP | 23597149 | Unspecified |
| mmu-let-7i-5p | MIMAT0000122 | Dhx15 | 13204 | HITS-CLIP | 23597149\|29167373\|23142080 | Unspecified |
| mmu-let-7i-5p | MIMAT0000122 | Dhx16 | 69192 | HITS-CLIP | 23597149\|23142080 | Unspecified |
| mmu-let-7i-5p | MIMAT0000122 | Dhx30 | 72831 | HITS-CLIP | 23597149 | Unspecified |
| mmu-let-7i-5p | MIMAT0000122 | Dhx32 | 101437 | HITS-CLIP | 23142080 | Unspecified |
| mmu-let-7i-5p | MIMAT0000122 | Dhx34 | 71723 | HITS-CLIP | 23597149\|23142080 | Unspecified |
| mmu-let-7i-5p | MIMAT0000122 | Dhx35 | 71715 | HITS-CLIP | 29167373 | Unspecified |
| mmu-let-7i-5p | MIMAT0000122 | Dhx36 | 72162 | HITS-CLIP | 23597149\|23142080 | Unspecified |
| mmu-let-7i-5p | MIMAT0000122 | Dhx38 | 64340 | HITS-CLIP | 23597149\|23142080 | Unspecified |
| mmu-let-7i-5p | MIMAT0000122 | Dhx40 | 67487 | HITS-CLIP | 23597149\|23142080 | Unspecified |
| mmu-let-7i-5p | MIMAT0000122 | Dhx57 | 106794 | HITS-CLIP | 23142080 | Unspecified |
| mmu-let-7i-5p | MIMAT0000122 | Dhx8 | 217207 | HITS-CLIP | 37986224\|23597149\|23142080 | Unspecified |
| mmu-let-7i-5p | MIMAT0000122 | Dhx9 | 13211 | HITS-CLIP | 23597149\|23142080 | Unspecified |
| mmu-let-7i-5p | MIMAT0000122 | Diablo | 66593 | HITS-CLIP | 23142080 | Unspecified |
| mmu-let-7i-5p | MIMAT0000122 | Diaph1 | 1729 | HITS-CLIP | 23142080\|29167373 | Unspecified |
| mmu-let-7i-5p | MIMAT0000122 | Diaph2 | 54004 | HITS-CLIP | 29167373 | Unspecified |
| mmu-let-7i-5p | MIMAT0000122 | Diaph3 | 81624 | HITS-CLIP | 23142080 | Unspecified |
| mmu-let-7i-5p | MIMAT0000122 | Dicer1 | 192119 | HITS-CLIP | 37986224\|19536157\|23597149\|26203 | Unspecified |
| mmu-let-7i-5p | MIMAT0000122 | Diexf | 215193 | HITS-CLIP | 23142080\|29167373 | Unspecified |
| mmu-let-7i-5p | MIMAT0000122 | Dio2 | 13371 | HITS-CLIP | 19536157 | Unspecified |
| mmu-let-7i-5p | MIMAT0000122 | Dip2a | 64451 | HITS-CLIP | 23142080 | Unspecified |
| mmu-let-7i-5p | MIMAT0000122 | Dip2b | 239667 | HITS-CLIP | 37986224\|19536157\|23142080\|29167 | Unspecified |
| mmu-let-7i-5p | MIMAT0000122 | Dirc2 | 224132 | HITS-CLIP | 23597149\|19536157 | Unspecified |
| mmu-let-7i-5p | MIMAT0000122 | Dis3l | 213550 | HITS-CLIP | 23597149 | Unspecified |
| mmu-let-7i-5p | MIMAT0000122 | Dkc1 | 245474 | HITS-CLIP | 23142080 | Unspecified |
| mmu-let-7i-5p | MIMAT0000122 | Dkk3 | 50781 | HITS-CLIP | 37986224 | Unspecified |
| mmu-let-7i-5p | MIMAT0000122 | Dlat | 235339 | HITS-CLIP | 23142080 | Unspecified |
| mmu-let-7i-5p | MIMAT0000122 | Dlc1 | 50768 | HITS-CLIP | 23597149 | Unspecified |
| mmu-let-7i-5p | MIMAT0000122 | Dlg4 | 13385 | HITS-CLIP | 29167373 | Unspecified |
| mmu-let-7i-5p | MIMAT0000122 | Dlgap1 | 224997 | HITS-CLIP | 19536157 | Unspecified |

| | | | | | | |
|---|---|---|---|---|---|---|
| mmu-let-7i-5p | MIMAT0000122 | Dlgap4 | 228836 | HITS-CLIP | 23142080 | Unspecified |
| mmu-let-7i-5p | MIMAT0000122 | Dlgap5 | 218977 | HITS-CLIP | 23142080 | Unspecified |
| mmu-let-7i-5p | MIMAT0000122 | Dlst | 78920 | HITS-CLIP | 23597149\|23142080 | Unspecified |
| mmu-let-7i-5p | MIMAT0000122 | Dmd | 13405 | HITS-CLIP | 37986224\|23597149\|19536157 | Unspecified |
| mmu-let-7i-5p | MIMAT0000122 | Dmgdh | 74129 | HITS-CLIP | 23597149 | Unspecified |
| mmu-let-7i-5p | MIMAT0000122 | Dmtf1 | 23857 | HITS-CLIP | 23142080\|29167373 | Unspecified |
| mmu-let-7i-5p | MIMAT0000122 | Dmwd | 13401 | HITS-CLIP | 19536157 | Unspecified |
| mmu-let-7i-5p | MIMAT0000122 | Dmxl1 | 240283 | HITS-CLIP | 23597149\|29167373\|23142080 | Unspecified |
| mmu-let-7i-5p | MIMAT0000122 | Dmxl2 | 235380 | HITS-CLIP | 29167373 | Unspecified |
| mmu-let-7i-5p | MIMAT0000122 | Dna2 | 327762 | HITS-CLIP | 37986224\|23142080 | Unspecified |
| mmu-let-7i-5p | MIMAT0000122 | Dnaic1 | 68922 | HITS-CLIP | 19536157 | Unspecified |
| mmu-let-7i-5p | MIMAT0000122 | Dnaja1 | 15502 | HITS-CLIP | 19536157\|23597149\|29167373 | Unspecified |
| mmu-let-7i-5p | MIMAT0000122 | Dnaja2 | 56445 | HITS-CLIP | 37986224\|23142080\|29167373 | Unspecified |
| mmu-let-7i-5p | MIMAT0000122 | Dnaja3 | 83945 | HITS-CLIP | 23597149 | Unspecified |
| mmu-let-7i-5p | MIMAT0000122 | Dnajb1 | 81489 | HITS-CLIP | 23142080 | Unspecified |
| mmu-let-7i-5p | MIMAT0000122 | Dnajb11 | 67838 | HITS-CLIP | 23597149\|23142080 | Unspecified |
| mmu-let-7i-5p | MIMAT0000122 | Dnajb14 | 70604 | HITS-CLIP | 37986224\|19536157\|29167373 | Unspecified |
| mmu-let-7i-5p | MIMAT0000122 | Dnajb4 | 67035 | HITS-CLIP | 29167373 | Unspecified |
| mmu-let-7i-5p | MIMAT0000122 | Dnajb9 | 27362 | HITS-CLIP | 37986224\|23597149\|19536157 | Unspecified |
| mmu-let-7i-5p | MIMAT0000122 | Dnajc1 | 13418 | HITS-CLIP | 19536157 | Unspecified |
| mmu-let-7i-5p | MIMAT0000122 | Dnajc10 | 66861 | HITS-CLIP | 37986224\|19536157\|23142080 | Unspecified |
| mmu-let-7i-5p | MIMAT0000122 | Dnajc13 | 235567 | HITS-CLIP | 37986224\|23597149\|26203562\|23142 | Unspecified |
| mmu-let-7i-5p | MIMAT0000122 | Dnajc14 | 74330 | HITS-CLIP | 37986224\|23597149 | Unspecified |
| mmu-let-7i-5p | MIMAT0000122 | Dnajc16 | 214063 | HITS-CLIP | 23597149 | Unspecified |
| mmu-let-7i-5p | MIMAT0000122 | Dnajc2 | 22791 | HITS-CLIP | 23142080 | Unspecified |
| mmu-let-7i-5p | MIMAT0000122 | Dnajc22 | 72778 | HITS-CLIP | 23597149 | Unspecified |
| mmu-let-7i-5p | MIMAT0000122 | Dnajc25 | 72429 | HITS-CLIP | 23597149 | Unspecified |
| mmu-let-7i-5p | MIMAT0000122 | Dnajc27 | 217378 | HITS-CLIP | 37986224\|23142080 | Unspecified |
| mmu-let-7i-5p | MIMAT0000122 | Dnajc30 | 66114 | HITS-CLIP | 23142080 | Unspecified |
| mmu-let-7i-5p | MIMAT0000122 | Dnajc5 | 13002 | HITS-CLIP | 23597149\|19536157\|23142080\|29167 | Unspecified |
| mmu-let-7i-5p | MIMAT0000122 | Dnajc7 | 56354 | HITS-CLIP | 23142080 | Unspecified |
| mmu-let-7i-5p | MIMAT0000122 | Dnm1l | 74006 | HITS-CLIP | 23142080 | Unspecified |
| mmu-let-7i-5p | MIMAT0000122 | Dnm2 | 13430 | HITS-CLIP | 37986224\|23142080\|30971389\|29167 | Unspecified |
| mmu-let-7i-5p | MIMAT0000122 | Dnmbp | 71972 | HITS-CLIP | 37986224\|23142080\|23597149\|1953 | Unspecified |
| mmu-let-7i-5p | MIMAT0000122 | Dnmt1 | 13433 | HITS-CLIP | 37986224\|23142080\|23597149\|29167 | Unspecified |
| mmu-let-7i-5p | MIMAT0000122 | Dnmt3a | 13435 | HITS-CLIP | 37986224\|23142080\|19536157\|29167 | Unspecified |
| mmu-let-7i-5p | MIMAT0000122 | Dnmt3b | 13436 | HITS-CLIP | 23142080 | Unspecified |
| mmu-let-7i-5p | MIMAT0000122 | Dnpep | 13437 | HITS-CLIP//qCLASH | 23142080\|31393953\|23597149 | Unspecified |
| mmu-let-7i-5p | MIMAT0000122 | Dntt | 21673 | HITS-CLIP | 23142080 | Unspecified |

| | | | | | | |
|---|---|---|---|---|---|---|
| mmu-let-7i-5p | MIMAT0000122 | Dnttip2 | 99480 | HITS-CLIP | 23142080 | Unspecified |
| mmu-let-7i-5p | MIMAT0000122 | Dock10 | 210293 | HITS-CLIP | 19536157\|23142080\|29167373 | Unspecified |
| mmu-let-7i-5p | MIMAT0000122 | Dock11 | 75974 | HITS-CLIP | 23142080\|29167373 | Unspecified |
| mmu-let-7i-5p | MIMAT0000122 | Dock2 | 94176 | HITS-CLIP | 23142080\|29167373 | Unspecified |
| mmu-let-7i-5p | MIMAT0000122 | Dock3 | 208869 | HITS-CLIP | 19536157 | Unspecified |
| mmu-let-7i-5p | MIMAT0000122 | Dock4 | 238130 | HITS-CLIP | 37986224\|19536157\|23597149\|29167 | Unspecified |
| mmu-let-7i-5p | MIMAT0000122 | Dock6 | 319899 | HITS-CLIP | 37986224\|23142080 | Unspecified |
| mmu-let-7i-5p | MIMAT0000122 | Dock8 | 76088 | HITS-CLIP | 37986224\|23597149\|23142080\|30971 | Unspecified |
| mmu-let-7i-5p | MIMAT0000122 | Dock9 | 105445 | HITS-CLIP | 19536157 | Unspecified |
| mmu-let-7i-5p | MIMAT0000122 | Dok2 | 13449 | HITS-CLIP | 23142080 | Unspecified |
| mmu-let-7i-5p | MIMAT0000122 | Dolk | 227697 | HITS-CLIP | 23142080 | Unspecified |
| mmu-let-7i-5p | MIMAT0000122 | Donson | 60364 | HITS-CLIP | 23142080 | Unspecified |
| mmu-let-7i-5p | MIMAT0000122 | Dop1a | 23033 | HITS-CLIP | 23142080\|23597149 | Unspecified |
| mmu-let-7i-5p | MIMAT0000122 | Dop1b | 9980 | HITS-CLIP | 29167373\|23597149 | Unspecified |
| mmu-let-7i-5p | MIMAT0000122 | Dot1l | 208266 | HITS-CLIP//qCLASH | 23597149\|31393953\|19536157\|23142 | Unspecified |
| mmu-let-7i-5p | MIMAT0000122 | Dpagt1 | 13478 | HITS-CLIP | 37986224\|23597149\|23142080 | Unspecified |
| mmu-let-7i-5p | MIMAT0000122 | Dpf2 | 19708 | HITS-CLIP | 23142080 | Unspecified |
| mmu-let-7i-5p | MIMAT0000122 | Dpm1 | 13480 | HITS-CLIP | 23142080 | Unspecified |
| mmu-let-7i-5p | MIMAT0000122 | Dpm2 | 13481 | HITS-CLIP | 23597149 | Unspecified |
| mmu-let-7i-5p | MIMAT0000122 | Dpm3 | 68563 | HITS-CLIP | 23142080 | Unspecified |
| mmu-let-7i-5p | MIMAT0000122 | Dpp3 | 75221 | HITS-CLIP | 23597149\|23142080\|29167373 | Unspecified |
| mmu-let-7i-5p | MIMAT0000122 | Dpp4 | 13482 | HITS-CLIP | 23597149\|23142080 | Unspecified |
| mmu-let-7i-5p | MIMAT0000122 | Dpp6 | 13483 | HITS-CLIP | 19536157 | Unspecified |
| mmu-let-7i-5p | MIMAT0000122 | Dpp7 | 83768 | HITS-CLIP | 29167373 | Unspecified |
| mmu-let-7i-5p | MIMAT0000122 | Dpp8 | 74388 | HITS-CLIP | 29167373\|23142080 | Unspecified |
| mmu-let-7i-5p | MIMAT0000122 | Dpp9 | 224897 | HITS-CLIP | 23597149\|29167373 | Unspecified |
| mmu-let-7i-5p | MIMAT0000122 | Dpy19l1 | 244745 | HITS-CLIP | 23597149\|19536157\|23142080 | Unspecified |
| mmu-let-7i-5p | MIMAT0000122 | Dpy19l4 | 381510 | HITS-CLIP | 19536157\|23597149\|23142080 | Unspecified |
| mmu-let-7i-5p | MIMAT0000122 | Dpyd | 99586 | HITS-CLIP | 23597149 | Unspecified |
| mmu-let-7i-5p | MIMAT0000122 | Dpys | 64705 | HITS-CLIP | 23597149 | Unspecified |
| mmu-let-7i-5p | MIMAT0000122 | Dpysl2 | 12934 | HITS-CLIP | 37986224\|23142080 | Unspecified |
| mmu-let-7i-5p | MIMAT0000122 | Dpysl3 | 22240 | HITS-CLIP | 19536157 | Unspecified |
| mmu-let-7i-5p | MIMAT0000122 | Dpysl5 | 65254 | HITS-CLIP | 19536157 | Unspecified |
| mmu-let-7i-5p | MIMAT0000122 | Dr1 | 13486 | HITS-CLIP | 23142080 | Unspecified |
| mmu-let-7i-5p | MIMAT0000122 | Dram2 | 67171 | HITS-CLIP | 23597149\|23142080 | Unspecified |
| mmu-let-7i-5p | MIMAT0000122 | Drg1 | 13494 | HITS-CLIP | 23142080 | Unspecified |
| mmu-let-7i-5p | MIMAT0000122 | Dse | 212898 | HITS-CLIP | 29167373 | Unspecified |
| mmu-let-7i-5p | MIMAT0000122 | Dsel | 319901 | HITS-CLIP | 37986224\|23597149\|23142080 | Unspecified |
| mmu-let-7i-5p | MIMAT0000122 | Dst | 13518 | HITS-CLIP | 37986224\|23597149\|19536157\|23142 | Unspecified |

| | | | | | | |
|---|---|---|---|---|---|---|
| mmu-let-7i-5p | MIMAT0000122 | Dstn | 56431 | HITS-CLIP | 23142080 | Unspecified |
| mmu-let-7i-5p | MIMAT0000122 | Dtl | 76843 | HITS-CLIP | 23142080 | Unspecified |
| mmu-let-7i-5p | MIMAT0000122 | Dtna | 13527 | HITS-CLIP | 37986224 | Unspecified |
| mmu-let-7i-5p | MIMAT0000122 | Dtnb | 13528 | HITS-CLIP | 37986224 | Unspecified |
| mmu-let-7i-5p | MIMAT0000122 | Dtx2 | 74198 | HITS-CLIP | 23142080 | Unspecified |
| mmu-let-7i-5p | MIMAT0000122 | Dtx3l | 209200 | HITS-CLIP | 23597149\|23142080\|29167373 | Unspecified |
| mmu-let-7i-5p | MIMAT0000122 | Dtx4 | 207521 | HITS-CLIP | 29167373 | Unspecified |
| mmu-let-7i-5p | MIMAT0000122 | Dus1l | 68730 | HITS-CLIP | 23597149\|23142080 | Unspecified |
| mmu-let-7i-5p | MIMAT0000122 | Dusp1 | 19252 | HITS-CLIP | 37986224\|23597149\|23142080\|19536 | Unspecified |
| mmu-let-7i-5p | MIMAT0000122 | Dusp10 | 63953 | HITS-CLIP | 23142080 | Unspecified |
| mmu-let-7i-5p | MIMAT0000122 | Dusp11 | 72102 | HITS-CLIP | 23597149\|23142080 | Unspecified |
| mmu-let-7i-5p | MIMAT0000122 | Dusp16 | 70686 | HITS-CLIP | 23142080 | Unspecified |
| mmu-let-7i-5p | MIMAT0000122 | Dusp2 | 13537 | qCLASH//HITS-CLIP | 31393953\|23142080 | Unspecified |
| mmu-let-7i-5p | MIMAT0000122 | Dusp22 | 105352 | HITS-CLIP | 23597149 | Unspecified |
| mmu-let-7i-5p | MIMAT0000122 | Dusp3 | 72349 | HITS-CLIP | 37986224\|23597149 | Unspecified |
| mmu-let-7i-5p | MIMAT0000122 | Dusp4 | 319520 | HITS-CLIP | 37986224\|23142080 | Unspecified |
| mmu-let-7i-5p | MIMAT0000122 | Dusp5 | 240672 | HITS-CLIP//qCLASH | 37986224\|31433990\|31393953\|29167 | Unspecified |
| mmu-let-7i-5p | MIMAT0000122 | Dusp6 | 67603 | HITS-CLIP | 37986224\|23597149\|23142080\|19536 | Unspecified |
| mmu-let-7i-5p | MIMAT0000122 | Dusp7 | 235584 | HITS-CLIP | 37986224\|23142080\|23597149\|19536 | Unspecified |
| mmu-let-7i-5p | MIMAT0000122 | Dusp8 | 18218 | HITS-CLIP | 37986224\|19536157 | Unspecified |
| mmu-let-7i-5p | MIMAT0000122 | Dvl3 | 13544 | HITS-CLIP | 23142080\|23597149 | Unspecified |
| mmu-let-7i-5p | MIMAT0000122 | Dync1h1 | 13424 | HITS-CLIP | 37986224\|23597149\|23142080\|19536 | Unspecified |
| mmu-let-7i-5p | MIMAT0000122 | Dync1i2 | 13427 | HITS-CLIP | 37986224\|23597149\|19536157\|23142 | Unspecified |
| mmu-let-7i-5p | MIMAT0000122 | Dync1li2 | 234663 | HITS-CLIP | 37986224\|23142080\|29167373 | Unspecified |
| mmu-let-7i-5p | MIMAT0000122 | Dynlrb1 | 67068 | HITS-CLIP | 23142080\|23597149\|19536157 | Unspecified |
| mmu-let-7i-5p | MIMAT0000122 | Dynlt3 | 67117 | HITS-CLIP | 23142080 | Unspecified |
| mmu-let-7i-5p | MIMAT0000122 | Dyrk1a | 13548 | HITS-CLIP | 37986224\|23597149\|23142080\|29167 | Unspecified |
| mmu-let-7i-5p | MIMAT0000122 | Dyrk1b | 13549 | HITS-CLIP | 23142080 | Unspecified |
| mmu-let-7i-5p | MIMAT0000122 | Dyrk2 | 69181 | HITS-CLIP | 37986224\|23597149\|23142080 | Unspecified |
| mmu-let-7i-5p | MIMAT0000122 | Dzank1 | 241688 | HITS-CLIP | 19536157 | Unspecified |
| mmu-let-7i-5p | MIMAT0000122 | Dzip3 | 224170 | HITS-CLIP | 23142080\|29167373 | Unspecified |
| mmu-let-7i-5p | MIMAT0000122 | E130308A19Rik | 230259 | HITS-CLIP | 37986224\|23597149 | Unspecified |
| mmu-let-7i-5p | MIMAT0000122 | E2f2 | 242705 | HITS-CLIP | 23142080 | Unspecified |
| mmu-let-7i-5p | MIMAT0000122 | E2f3 | 13557 | HITS-CLIP | 37986224\|23142080 | Unspecified |
| mmu-let-7i-5p | MIMAT0000122 | E2f4 | 104394 | HITS-CLIP | 23142080\|19536157 | Unspecified |
| mmu-let-7i-5p | MIMAT0000122 | E2f5 | 13559 | HITS-CLIP | 23142080 | Unspecified |
| mmu-let-7i-5p | MIMAT0000122 | E2f6 | 50496 | HITS-CLIP | 37986224\|23597149\|23142080 | Unspecified |
| mmu-let-7i-5p | MIMAT0000122 | E2f7 | 52679 | HITS-CLIP | 23142080 | Unspecified |
| mmu-let-7i-5p | MIMAT0000122 | E2f8 | 108961 | HITS-CLIP | 29167373\|23142080 | Unspecified |

| | | | | | | |
|---|---|---|---|---|---|---|
| mmu-let-7i-5p | MIMAT0000122 | E330009J07Rik | 243780 | HITS-CLIP | 37986224\|19536157\|23597149\|23142 | Unspecified |
| mmu-let-7i-5p | MIMAT0000122 | Eaf1 | 74427 | HITS-CLIP | 23142080 | Unspecified |
| mmu-let-7i-5p | MIMAT0000122 | Ebag9 | 55960 | HITS-CLIP | 23142080\|23597149 | Unspecified |
| mmu-let-7i-5p | MIMAT0000122 | Ecd | 70601 | HITS-CLIP | 23142080 | Unspecified |
| mmu-let-7i-5p | MIMAT0000122 | Ece1 | 230857 | HITS-CLIP | 23142080 | Unspecified |
| mmu-let-7i-5p | MIMAT0000122 | Ech1 | 51798 | HITS-CLIP | 23597149\|23142080 | Unspecified |
| mmu-let-7i-5p | MIMAT0000122 | Echdc1 | 52665 | HITS-CLIP | 23597149 | Unspecified |
| mmu-let-7i-5p | MIMAT0000122 | Echs1 | 93747 | HITS-CLIP | 23597149 | Unspecified |
| mmu-let-7i-5p | MIMAT0000122 | Eci2 | 23986 | HITS-CLIP | 23597149 | Unspecified |
| mmu-let-7i-5p | MIMAT0000122 | Ecm1 | 13601 | HITS-CLIP | 23597149\|23142080 | Unspecified |
| mmu-let-7i-5p | MIMAT0000122 | Ecpas | 23392 | HITS-CLIP | 23597149\|23142080\|29167373 | Unspecified |
| mmu-let-7i-5p | MIMAT0000122 | Ecsit | 26940 | HITS-CLIP | 23597149 | Unspecified |
| mmu-let-7i-5p | MIMAT0000122 | Ect2 | 13605 | HITS-CLIP | 23142080 | Unspecified |
| mmu-let-7i-5p | MIMAT0000122 | Eda2r | 245527 | HITS-CLIP | 37986224 | Unspecified |
| mmu-let-7i-5p | MIMAT0000122 | Edc3 | 353190 | HITS-CLIP | 23597149\|23142080 | Unspecified |
| mmu-let-7i-5p | MIMAT0000122 | Edc4 | 234699 | qCLASH//HITS-CLIP | 31393953\|23142080 | Unspecified |
| mmu-let-7i-5p | MIMAT0000122 | Edem1 | 192193 | HITS-CLIP | 23597149\|19536157\|23142080\|29167 | Unspecified |
| mmu-let-7i-5p | MIMAT0000122 | Edem2 | 108687 | HITS-CLIP | 23597149\|23142080 | Unspecified |
| mmu-let-7i-5p | MIMAT0000122 | Edem3 | 66967 | HITS-CLIP | 37986224\|23597149\|19536157\|23142 | Unspecified |
| mmu-let-7i-5p | MIMAT0000122 | Edf1 | 59022 | HITS-CLIP | 23142080 | Unspecified |
| mmu-let-7i-5p | MIMAT0000122 | Edn1 | 13614 | HITS-CLIP | 37986224\|23597149\|23142080 | Unspecified |
| mmu-let-7i-5p | MIMAT0000122 | Edrf1 | 214764 | HITS-CLIP | 23597149\|23142080 | Unspecified |
| mmu-let-7i-5p | MIMAT0000122 | Eea1 | 216238 | HITS-CLIP | 37986224\|31433990\|23597149\|23142 | Unspecified |
| mmu-let-7i-5p | MIMAT0000122 | Eed | 13626 | HITS-CLIP | 23142080 | Unspecified |
| mmu-let-7i-5p | MIMAT0000122 | Eef1a1 | 13627 | HITS-CLIP | 23597149\|19536157\|23142080\|29167 | Unspecified |
| mmu-let-7i-5p | MIMAT0000122 | Eef1akmt2 | 399818 | HITS-CLIP | 23142080 | Unspecified |
| mmu-let-7i-5p | MIMAT0000122 | Eef1e1 | 66143 | HITS-CLIP | 23142080 | Unspecified |
| mmu-let-7i-5p | MIMAT0000122 | Eef2 | 13629 | HITS-CLIP//qCLASH | 37986224\|23597149\|31393953\|23142 | Unspecified |
| mmu-let-7i-5p | MIMAT0000122 | Eef2k | 13631 | HITS-CLIP | 37986224\|19536157\|23597149\|23142 | Unspecified |
| mmu-let-7i-5p | MIMAT0000122 | Eefsec | 65967 | HITS-CLIP | 23597149 | Unspecified |
| mmu-let-7i-5p | MIMAT0000122 | Efcab14 | 230648 | HITS-CLIP | 23142080\|29167373 | Unspecified |
| mmu-let-7i-5p | MIMAT0000122 | Efhd2 | 27984 | HITS-CLIP | 37986224\|23597149\|23142080\|29167 | Unspecified |
| mmu-let-7i-5p | MIMAT0000122 | Efna1 | 13636 | HITS-CLIP | 23597149 | Unspecified |
| mmu-let-7i-5p | MIMAT0000122 | Efna5 | 13640 | HITS-CLIP | 23142080 | Unspecified |
| mmu-let-7i-5p | MIMAT0000122 | Efnb1 | 13641 | HITS-CLIP | 26203562 | Unspecified |
| mmu-let-7i-5p | MIMAT0000122 | Efr3a | 76740 | HITS-CLIP | 23142080 | Unspecified |
| mmu-let-7i-5p | MIMAT0000122 | Eftud2 | 20624 | HITS-CLIP | 23597149\|23142080 | Unspecified |
| mmu-let-7i-5p | MIMAT0000122 | Egfl7 | 353156 | HITS-CLIP | 23597149 | Unspecified |
| mmu-let-7i-5p | MIMAT0000122 | Egfr | 13649 | HITS-CLIP | 37986224\|23597149 | Unspecified |

| | | | | | | |
|---|---|---|---|---|---|---|
| mmu-let-7i-5p | MIMAT0000122 | Egln1 | 112405 | HITS-CLIP | 23142080 | Unspecified |
| mmu-let-7i-5p | MIMAT0000122 | Egln2 | 112406 | HITS-CLIP | 37986224\|23597149\|23142080\|29167 | Unspecified |
| mmu-let-7i-5p | MIMAT0000122 | Egln3 | 112407 | HITS-CLIP | 23142080 | Unspecified |
| mmu-let-7i-5p | MIMAT0000122 | Egr1 | 13653 | HITS-CLIP | 37986224\|23142080 | Unspecified |
| mmu-let-7i-5p | MIMAT0000122 | Egr2 | 13654 | HITS-CLIP | 23142080 | Unspecified |
| mmu-let-7i-5p | MIMAT0000122 | Egr3 | 13655 | HITS-CLIP | 23142080\|23597149 | Unspecified |
| mmu-let-7i-5p | MIMAT0000122 | Ehd3 | 57440 | HITS-CLIP | 23597149\|23142080 | Unspecified |
| mmu-let-7i-5p | MIMAT0000122 | Ehd4 | 98878 | HITS-CLIP | 37986224\|23142080 | Unspecified |
| mmu-let-7i-5p | MIMAT0000122 | Ehhadh | 74147 | HITS-CLIP | 37986224\|23597149\|23142080 | Unspecified |
| mmu-let-7i-5p | MIMAT0000122 | Ehmt1 | 77683 | HITS-CLIP | 37986224\|23597149\|29167373 | Unspecified |
| mmu-let-7i-5p | MIMAT0000122 | Ehmt2 | 110147 | HITS-CLIP | 23597149\|23142080 | Unspecified |
| mmu-let-7i-5p | MIMAT0000122 | Ei24 | 13663 | HITS-CLIP | 23142080\|19536157\|23597149\|26203 | Unspecified |
| mmu-let-7i-5p | MIMAT0000122 | Eif1a | 13664 | HITS-CLIP | 23597149\|23142080\|29167373 | Unspecified |
| mmu-let-7i-5p | MIMAT0000122 | Eif1ad | 69860 | HITS-CLIP | 23142080 | Unspecified |
| mmu-let-7i-5p | MIMAT0000122 | Eif1ax | 66235 | HITS-CLIP | 37986224\|23142080 | Unspecified |
| mmu-let-7i-5p | MIMAT0000122 | Eif1b | 68969 | HITS-CLIP | 23597149\|23142080 | Unspecified |
| mmu-let-7i-5p | MIMAT0000122 | Eif2a | 229317 | HITS-CLIP | 31433990\|23142080\|29167373 | Unspecified |
| mmu-let-7i-5p | MIMAT0000122 | Eif2ak1 | 15467 | HITS-CLIP | 23597149\|23142080 | Unspecified |
| mmu-let-7i-5p | MIMAT0000122 | Eif2ak3 | 13666 | HITS-CLIP | 23597149\|23142080 | Unspecified |
| mmu-let-7i-5p | MIMAT0000122 | Eif2ak4 | 27103 | HITS-CLIP | 23142080 | Unspecified |
| mmu-let-7i-5p | MIMAT0000122 | Eif2b3 | 108067 | HITS-CLIP | 23142080 | Unspecified |
| mmu-let-7i-5p | MIMAT0000122 | Eif2b4 | 13667 | HITS-CLIP | 23597149\|23142080 | Unspecified |
| mmu-let-7i-5p | MIMAT0000122 | Eif2b5 | 224045 | HITS-CLIP | 23597149\|23142080 | Unspecified |
| mmu-let-7i-5p | MIMAT0000122 | Eif2s2 | 67204 | HITS-CLIP | 23142080 | Unspecified |
| mmu-let-7i-5p | MIMAT0000122 | Eif2s3x | 26905 | HITS-CLIP | 23142080 | Unspecified |
| mmu-let-7i-5p | MIMAT0000122 | Eif2s3y | 26908 | HITS-CLIP | 23597149\|23142080 | Unspecified |
| mmu-let-7i-5p | MIMAT0000122 | Eif3a | 13669 | HITS-CLIP | 29167373 | Unspecified |
| mmu-let-7i-5p | MIMAT0000122 | Eif3c | 56347 | HITS-CLIP | 23597149\|23142080\|29167373 | Unspecified |
| mmu-let-7i-5p | MIMAT0000122 | Eif3d | 55944 | HITS-CLIP | 37986224\|23597149\|23142080 | Unspecified |
| mmu-let-7i-5p | MIMAT0000122 | Eif3e | 16341 | HITS-CLIP | 23142080 | Unspecified |
| mmu-let-7i-5p | MIMAT0000122 | Eif3f | 66085 | HITS-CLIP | 23142080 | Unspecified |
| mmu-let-7i-5p | MIMAT0000122 | Eif3i | 54709 | HITS-CLIP | 37986224\|23142080 | Unspecified |
| mmu-let-7i-5p | MIMAT0000122 | Eif3k | 73830 | HITS-CLIP | 23142080\|23597149 | Unspecified |
| mmu-let-7i-5p | MIMAT0000122 | Eif3l | 223691 | HITS-CLIP | 23597149\|23142080 | Unspecified |
| mmu-let-7i-5p | MIMAT0000122 | Eif3m | 98221 | HITS-CLIP | 23597149\|23142080 | Unspecified |
| mmu-let-7i-5p | MIMAT0000122 | Eif4a1 | 13681 | HITS-CLIP | 23142080\|31433990 | Unspecified |
| mmu-let-7i-5p | MIMAT0000122 | Eif4a2 | 13682 | HITS-CLIP | 23597149\|29167373 | Unspecified |
| mmu-let-7i-5p | MIMAT0000122 | Eif4b | 75705 | HITS-CLIP | 23142080\|23597149 | Unspecified |
| mmu-let-7i-5p | MIMAT0000122 | Eif4e | 13684 | HITS-CLIP | 23142080 | Unspecified |

| | | | | | | |
|---|---|---|---|---|---|---|
| mmu-let-7i-5p | MIMAT0000122 | Eif4e2 | 26987 | HITS-CLIP | 23597149\|23142080\|29167373 | Unspecified |
| mmu-let-7i-5p | MIMAT0000122 | Eif4ebp1 | 13685 | HITS-CLIP | 37986224\|23597149\|23142080 | Unspecified |
| mmu-let-7i-5p | MIMAT0000122 | Eif4ebp2 | 13688 | HITS-CLIP | 37986224\|23597149\|23142080 | Unspecified |
| mmu-let-7i-5p | MIMAT0000122 | Eif4enif1 | 74203 | HITS-CLIP | 23142080\|29167373 | Unspecified |
| mmu-let-7i-5p | MIMAT0000122 | Eif4g1 | 208643 | HITS-CLIP | 37986224\|23597149\|23142080\|29167 | Unspecified |
| mmu-let-7i-5p | MIMAT0000122 | Eif4g2 | 13690 | HITS-CLIP | 37986224\|23142080\|19536157\|23597 | Unspecified |
| mmu-let-7i-5p | MIMAT0000122 | Eif4g3 | 230861 | HITS-CLIP | 23597149\|23142080\|29167373 | Unspecified |
| mmu-let-7i-5p | MIMAT0000122 | Eif5 | 217869 | HITS-CLIP | 23142080\|29167373 | Unspecified |
| mmu-let-7i-5p | MIMAT0000122 | Eif5b | 226982 | HITS-CLIP | 37986224\|23597149\|23142080\|29167 | Unspecified |
| mmu-let-7i-5p | MIMAT0000122 | Eif6 | 16418 | HITS-CLIP | 23142080 | Unspecified |
| mmu-let-7i-5p | MIMAT0000122 | Eipr1 | 7260 | HITS-CLIP | 23142080 | Unspecified |
| mmu-let-7i-5p | MIMAT0000122 | Elac1 | 114615 | HITS-CLIP | 23142080 | Unspecified |
| mmu-let-7i-5p | MIMAT0000122 | Elavl1 | 15568 | HITS-CLIP | 37986224\|23597149\|23142080\|29167 | Unspecified |
| mmu-let-7i-5p | MIMAT0000122 | Elavl3 | 15571 | HITS-CLIP | 37986224 | Unspecified |
| mmu-let-7i-5p | MIMAT0000122 | Elf1 | 13709 | HITS-CLIP | 23142080 | Unspecified |
| mmu-let-7i-5p | MIMAT0000122 | Elf2 | 69257 | HITS-CLIP | 37986224\|23142080\|29167373 | Unspecified |
| mmu-let-7i-5p | MIMAT0000122 | Elf4 | 56501 | HITS-CLIP | 23597149\|23142080 | Unspecified |
| mmu-let-7i-5p | MIMAT0000122 | Elk3 | 13713 | HITS-CLIP | 23142080\|29167373 | Unspecified |
| mmu-let-7i-5p | MIMAT0000122 | Elk4 | 13714 | HITS-CLIP | 37986224\|23597149\|23142080\|29167 | Unspecified |
| mmu-let-7i-5p | MIMAT0000122 | Ell2 | 192657 | HITS-CLIP | 37986224\|23597149\|23142080\|29167 | Unspecified |
| mmu-let-7i-5p | MIMAT0000122 | Elmo1 | 140580 | HITS-CLIP | 23142080 | Unspecified |
| mmu-let-7i-5p | MIMAT0000122 | Elmo2 | 140579 | HITS-CLIP | 31433990\|23142080\|29167373 | Unspecified |
| mmu-let-7i-5p | MIMAT0000122 | Elmod2 | 244548 | HITS-CLIP | 23142080 | Unspecified |
| mmu-let-7i-5p | MIMAT0000122 | Elmsan1 | 238317 | HITS-CLIP | 37986224\|23597149\|23142080\|29167 | Unspecified |
| mmu-let-7i-5p | MIMAT0000122 | Eln | 13717 | HITS-CLIP | 19536157 | Unspecified |
| mmu-let-7i-5p | MIMAT0000122 | Eloa | 6924 | HITS-CLIP | 23597149\|29167373\|23142080 | Unspecified |
| mmu-let-7i-5p | MIMAT0000122 | Eloc | 6921 | HITS-CLIP | 23142080\|23597149 | Unspecified |
| mmu-let-7i-5p | MIMAT0000122 | Elovl1 | 54325 | HITS-CLIP | 23597149\|23142080 | Unspecified |
| mmu-let-7i-5p | MIMAT0000122 | Elovl2 | 54326 | HITS-CLIP | 37986224\|23597149 | Unspecified |
| mmu-let-7i-5p | MIMAT0000122 | Elovl3 | 12686 | HITS-CLIP | 23597149 | Unspecified |
| mmu-let-7i-5p | MIMAT0000122 | Elovl4 | 83603 | HITS-CLIP | 37986224\|19536157 | Unspecified |
| mmu-let-7i-5p | MIMAT0000122 | Elovl5 | 68801 | HITS-CLIP | 19536157\|23142080\|29167373 | Unspecified |
| mmu-let-7i-5p | MIMAT0000122 | Elp1 | 37455 | HITS-CLIP | 23597149\|23142080 | Unspecified |
| mmu-let-7i-5p | MIMAT0000122 | Elp2 | 58523 | HITS-CLIP | 37986224\|23142080 | Unspecified |
| mmu-let-7i-5p | MIMAT0000122 | Emc1 | 230866 | HITS-CLIP | 23597149 | Unspecified |
| mmu-let-7i-5p | MIMAT0000122 | Emc4 | 68032 | HITS-CLIP | 37986224\|23142080 | Unspecified |
| mmu-let-7i-5p | MIMAT0000122 | Eme1 | 268465 | HITS-CLIP | 23142080 | Unspecified |
| mmu-let-7i-5p | MIMAT0000122 | Emg1 | 14791 | HITS-CLIP | 23142080 | Unspecified |
| mmu-let-7i-5p | MIMAT0000122 | Eml1 | 68519 | HITS-CLIP | 37986224 | Unspecified |

| | | | | | | |
|---|---|---|---|---|---|---|
| mmu-let-7i-5p | MIMAT0000122 | Eml3 | 225898 | HITS-CLIP | 23142080 | Unspecified |
| mmu-let-7i-5p | MIMAT0000122 | Eml4 | 78798 | HITS-CLIP | 23142080\|29167373 | Unspecified |
| mmu-let-7i-5p | MIMAT0000122 | Eml5 | 319670 | HITS-CLIP | 37986224\|23142080 | Unspecified |
| mmu-let-7i-5p | MIMAT0000122 | Eml6 | 237711 | HITS-CLIP | 23597149 | Unspecified |
| mmu-let-7i-5p | MIMAT0000122 | Emp1 | 13730 | HITS-CLIP | 29167373 | Unspecified |
| mmu-let-7i-5p | MIMAT0000122 | Emp2 | 13731 | HITS-CLIP | 29167373 | Unspecified |
| mmu-let-7i-5p | MIMAT0000122 | Emsy | 56946 | HITS-CLIP | 23142080\|23597149\|29167373 | Unspecified |
| mmu-let-7i-5p | MIMAT0000122 | Enc1 | 13803 | HITS-CLIP | 37986224\|19536157\|23142080 | Unspecified |
| mmu-let-7i-5p | MIMAT0000122 | Endod1 | 71946 | HITS-CLIP | 23142080 | Unspecified |
| mmu-let-7i-5p | MIMAT0000122 | Eng | 13805 | HITS-CLIP | 37986224\|23597149 | Unspecified |
| mmu-let-7i-5p | MIMAT0000122 | Enkur | 71233 | HITS-CLIP | 19536157 | Unspecified |
| mmu-let-7i-5p | MIMAT0000122 | Eno2 | 13807 | HITS-CLIP | 19536157 | Unspecified |
| mmu-let-7i-5p | MIMAT0000122 | Eno3 | 13808 | HITS-CLIP | 23142080 | Unspecified |
| mmu-let-7i-5p | MIMAT0000122 | Enox2 | 209224 | HITS-CLIP | 23142080 | Unspecified |
| mmu-let-7i-5p | MIMAT0000122 | Enpp1 | 18605 | HITS-CLIP | 23597149\|23142080 | Unspecified |
| mmu-let-7i-5p | MIMAT0000122 | Entpd5 | 12499 | HITS-CLIP | 37986224\|23597149\|23142080 | Unspecified |
| mmu-let-7i-5p | MIMAT0000122 | Entpd7 | 93685 | HITS-CLIP | 37986224\|31433990\|23142080 | Unspecified |
| mmu-let-7i-5p | MIMAT0000122 | Entpd8 | 72090 | HITS-CLIP | 23597149 | Unspecified |
| mmu-let-7i-5p | MIMAT0000122 | Eogt | 101351 | HITS-CLIP | 37986224\|29167373\|23142080 | Unspecified |
| mmu-let-7i-5p | MIMAT0000122 | Eomes | 13813 | HITS-CLIP | 23142080 | Unspecified |
| mmu-let-7i-5p | MIMAT0000122 | Ep300 | 328572 | HITS-CLIP | 23597149\|31433990\|23142080\|29167 | Unspecified |
| mmu-let-7i-5p | MIMAT0000122 | Ep400 | 75560 | HITS-CLIP | 23142080\|29167373 | Unspecified |
| mmu-let-7i-5p | MIMAT0000122 | Epb41 | 269587 | HITS-CLIP | 23597149\|23142080\|19536157\|29167 | Unspecified |
| mmu-let-7i-5p | MIMAT0000122 | Epb41l1 | 13821 | HITS-CLIP | 19536157 | Unspecified |
| mmu-let-7i-5p | MIMAT0000122 | Epb41l2 | 2037 | HITS-CLIP | 23142080\|29167373 | Unspecified |
| mmu-let-7i-5p | MIMAT0000122 | Epb41l3 | 23136 | HITS-CLIP | 19536157 | Unspecified |
| mmu-let-7i-5p | MIMAT0000122 | Epb41l4b | 54566 | HITS-CLIP | 23597149 | Unspecified |
| mmu-let-7i-5p | MIMAT0000122 | Epb41l5 | 226352 | HITS-CLIP | 19536157\|23597149 | Unspecified |
| mmu-let-7i-5p | MIMAT0000122 | Epc1 | 13831 | HITS-CLIP | 37986224\|23597149\|31433990\|26203 | Unspecified |
| mmu-let-7i-5p | MIMAT0000122 | Epc2 | 227867 | HITS-CLIP//qCLASH | 37986224\|31393953\|23142080 | Unspecified |
| mmu-let-7i-5p | MIMAT0000122 | Epdr1 | 105298 | HITS-CLIP | 19536157 | Unspecified |
| mmu-let-7i-5p | MIMAT0000122 | Epg5 | 100502841 | HITS-CLIP | 23142080 | Unspecified |
| mmu-let-7i-5p | MIMAT0000122 | Epha4 | 13838 | HITS-CLIP | 19536157 | Unspecified |
| mmu-let-7i-5p | MIMAT0000122 | Epha6 | 13840 | HITS-CLIP | 19536157 | Unspecified |
| mmu-let-7i-5p | MIMAT0000122 | Epha7 | 13841 | HITS-CLIP | 19536157 | Unspecified |
| mmu-let-7i-5p | MIMAT0000122 | Ephb2 | 13844 | HITS-CLIP | 19536157 | Unspecified |
| mmu-let-7i-5p | MIMAT0000122 | Ephb3 | 13845 | HITS-CLIP | 37986224 | Unspecified |
| mmu-let-7i-5p | MIMAT0000122 | Ephx1 | 13849 | HITS-CLIP | 23597149 | Unspecified |
| mmu-let-7i-5p | MIMAT0000122 | Ephx2 | 13850 | HITS-CLIP | 23597149 | Unspecified |

| | | | | | | |
|---|---|---|---|---|---|---|
| mmu-let-7i-5p | MIMAT0000122 | Epm2aip1 | 77781 | HITS-CLIP | 23142080\|29167373 | Unspecified |
| mmu-let-7i-5p | MIMAT0000122 | Epn1 | 13854 | HITS-CLIP | 23597149\|19536157\|23142080 | Unspecified |
| mmu-let-7i-5p | MIMAT0000122 | Epn2 | 13855 | HITS-CLIP | 23597149 | Unspecified |
| mmu-let-7i-5p | MIMAT0000122 | Eppk1 | 3857 | HITS-CLIP | 23597149 | Unspecified |
| mmu-let-7i-5p | MIMAT0000122 | Eprs | 107508 | HITS-CLIP | 23142080 | Unspecified |
| mmu-let-7i-5p | MIMAT0000122 | Eps15 | 13858 | HITS-CLIP | 19536157\|23142080\|29167373 | Unspecified |
| mmu-let-7i-5p | MIMAT0000122 | Eps15l1 | 13859 | HITS-CLIP | 37986224\|23142080 | Unspecified |
| mmu-let-7i-5p | MIMAT0000122 | Eps8 | 13860 | HITS-CLIP | 29167373 | Unspecified |
| mmu-let-7i-5p | MIMAT0000122 | Erap1 | 80898 | HITS-CLIP | 23597149\|29167373 | Unspecified |
| mmu-let-7i-5p | MIMAT0000122 | Erbb3 | 13867 | HITS-CLIP | 23597149\|23142080 | Unspecified |
| mmu-let-7i-5p | MIMAT0000122 | Erbb4 | 13869 | HITS-CLIP | 19536157 | Unspecified |
| mmu-let-7i-5p | MIMAT0000122 | Erbin | 55914 | HITS-CLIP | 29167373 | Unspecified |
| mmu-let-7i-5p | MIMAT0000122 | Erc1 | 111173 | HITS-CLIP | 37986224 | Unspecified |
| mmu-let-7i-5p | MIMAT0000122 | Erc2 | 238988 | HITS-CLIP | 37986224 | Unspecified |
| mmu-let-7i-5p | MIMAT0000122 | Ercc2 | 13871 | HITS-CLIP | 37986224\|19536157\|23597149\|2916 | Unspecified |
| mmu-let-7i-5p | MIMAT0000122 | Ercc3 | 13872 | HITS-CLIP | 23142080 | Unspecified |
| mmu-let-7i-5p | MIMAT0000122 | Ercc6 | 319955 | HITS-CLIP | 37986224\|19536157\|23597149\|2314 | Unspecified |
| mmu-let-7i-5p | MIMAT0000122 | Ercc6l | 236930 | HITS-CLIP | 23142080 | Unspecified |
| mmu-let-7i-5p | MIMAT0000122 | Ercc6l2 | 76251 | HITS-CLIP | 37986224 | Unspecified |
| mmu-let-7i-5p | MIMAT0000122 | Erf | 13875 | HITS-CLIP | 37986224\|23597149 | Unspecified |
| mmu-let-7i-5p | MIMAT0000122 | Erg | 13876 | HITS-CLIP | 23597149 | Unspecified |
| mmu-let-7i-5p | MIMAT0000122 | Erg28 | 856767 | HITS-CLIP | 23597149\|23142080\|29167373 | Unspecified |
| mmu-let-7i-5p | MIMAT0000122 | Ergic1 | 67458 | HITS-CLIP | 37986224\|23142080\|23597149 | Unspecified |
| mmu-let-7i-5p | MIMAT0000122 | Ergic2 | 67456 | HITS-CLIP | 37986224\|23142080\|19536157\|2359 | Unspecified |
| mmu-let-7i-5p | MIMAT0000122 | Ergic3 | 66366 | HITS-CLIP | 23142080 | Unspecified |
| mmu-let-7i-5p | MIMAT0000122 | Eri1 | 67276 | HITS-CLIP | 23142080 | Unspecified |
| mmu-let-7i-5p | MIMAT0000122 | Eri3 | 140546 | HITS-CLIP | 23142080 | Unspecified |
| mmu-let-7i-5p | MIMAT0000122 | Erlec1 | 66753 | HITS-CLIP | 23142080 | Unspecified |
| mmu-let-7i-5p | MIMAT0000122 | Erlin1 | 226144 | HITS-CLIP | 23597149 | Unspecified |
| mmu-let-7i-5p | MIMAT0000122 | Erlin2 | 244373 | HITS-CLIP | 23142080 | Unspecified |
| mmu-let-7i-5p | MIMAT0000122 | Ermn | 77767 | HITS-CLIP | 37986224 | Unspecified |
| mmu-let-7i-5p | MIMAT0000122 | Ermp1 | 226090 | HITS-CLIP | 37986224\|23142080 | Unspecified |
| mmu-let-7i-5p | MIMAT0000122 | Ern1 | 78943 | HITS-CLIP | 23597149\|23142080\|29167373 | Unspecified |
| mmu-let-7i-5p | MIMAT0000122 | Ero1a | 30001 | qCLASH | 31393953 | Unspecified |
| mmu-let-7i-5p | MIMAT0000122 | Ero1l | 50527 | HITS-CLIP | 23142080 | Unspecified |
| mmu-let-7i-5p | MIMAT0000122 | Erp29 | 67397 | HITS-CLIP | 23597149 | Unspecified |
| mmu-let-7i-5p | MIMAT0000122 | Erp44 | 76299 | HITS-CLIP | 37986224 | Unspecified |
| mmu-let-7i-5p | MIMAT0000122 | Errfi1 | 74155 | HITS-CLIP | 37986224\|23597149\|29167373\|2314 | Unspecified |
| mmu-let-7i-5p | MIMAT0000122 | Esco2 | 71988 | HITS-CLIP | 37986224\|23597149\|23142080 | Unspecified |

| | | | | | | |
|---|---|---|---|---|---|---|
| mmu-let-7i-5p | MIMAT0000122 | Esd | 13885 | HITS-CLIP | 23597149\|23142080\|29167373 | Unspecified |
| mmu-let-7i-5p | MIMAT0000122 | Esf1 | 66580 | HITS-CLIP | 37986224 | Unspecified |
| mmu-let-7i-5p | MIMAT0000122 | Espl1 | 105988 | HITS-CLIP | 23142080 | Unspecified |
| mmu-let-7i-5p | MIMAT0000122 | Esrp2 | 77411 | HITS-CLIP | 23597149 | Unspecified |
| mmu-let-7i-5p | MIMAT0000122 | Esrrg | 26381 | HITS-CLIP | 19536157 | Unspecified |
| mmu-let-7i-5p | MIMAT0000122 | Esyt1 | 23943 | HITS-CLIP | 23142080 | Unspecified |
| mmu-let-7i-5p | MIMAT0000122 | Esyt2 | 52635 | HITS-CLIP | 37986224\|19536157\|23597149\|23142 | Unspecified |
| mmu-let-7i-5p | MIMAT0000122 | Etaa1 | 68145 | HITS-CLIP | 37986224\|23597149\|23142080 | Unspecified |
| mmu-let-7i-5p | MIMAT0000122 | Etf1 | 225363 | HITS-CLIP | 29167373 | Unspecified |
| mmu-let-7i-5p | MIMAT0000122 | Etfdh | 66841 | HITS-CLIP | 23142080 | Unspecified |
| mmu-let-7i-5p | MIMAT0000122 | Etl4 | 208618 | HITS-CLIP | 37986224\|19536157 | Unspecified |
| mmu-let-7i-5p | MIMAT0000122 | Etnk1 | 75320 | HITS-CLIP | 37986224\|23142080\|29167373 | Unspecified |
| mmu-let-7i-5p | MIMAT0000122 | Etnk2 | 214253 | HITS-CLIP | 37986224\|23597149 | Unspecified |
| mmu-let-7i-5p | MIMAT0000122 | Etnppl | 71760 | HITS-CLIP | 23597149 | Unspecified |
| mmu-let-7i-5p | MIMAT0000122 | Ets1 | 23871 | HITS-CLIP | 37986224\|23597149\|19536157\|23142 | Unspecified |
| mmu-let-7i-5p | MIMAT0000122 | Ets2 | 23872 | HITS-CLIP | 23142080\|29167373 | Unspecified |
| mmu-let-7i-5p | MIMAT0000122 | Etv3 | 27049 | HITS-CLIP | 23142080\|29167373 | Unspecified |
| mmu-let-7i-5p | MIMAT0000122 | Etv5 | 104156 | HITS-CLIP | 19536157 | Unspecified |
| mmu-let-7i-5p | MIMAT0000122 | Etv6 | 14011 | HITS-CLIP | 23142080 | Unspecified |
| mmu-let-7i-5p | MIMAT0000122 | Eva1a | 232146 | HITS-CLIP | 23597149 | Unspecified |
| mmu-let-7i-5p | MIMAT0000122 | Eva1b | 230752 | HITS-CLIP | 23142080 | Unspecified |
| mmu-let-7i-5p | MIMAT0000122 | Evi2 | 14017 | HITS-CLIP | 23142080 | Unspecified |
| mmu-let-7i-5p | MIMAT0000122 | Evi2a | 14017 | HITS-CLIP | 29167373 | Unspecified |
| mmu-let-7i-5p | MIMAT0000122 | Evi5 | 14020 | HITS-CLIP | 23597149 | Unspecified |
| mmu-let-7i-5p | MIMAT0000122 | Ewsr1 | 14030 | HITS-CLIP | 23597149\|23142080\|29167373 | Unspecified |
| mmu-let-7i-5p | MIMAT0000122 | Exd2 | 97827 | HITS-CLIP | 23142080 | Unspecified |
| mmu-let-7i-5p | MIMAT0000122 | Exo1 | 26909 | HITS-CLIP//qCLASH | 37986224\|31393953\|23142080 | Unspecified |
| mmu-let-7i-5p | MIMAT0000122 | Exoc1 | 69940 | HITS-CLIP | 23597149\|23142080\|29167373 | Unspecified |
| mmu-let-7i-5p | MIMAT0000122 | Exoc2 | 66482 | HITS-CLIP | 23142080 | Unspecified |
| mmu-let-7i-5p | MIMAT0000122 | Exoc4 | 20336 | HITS-CLIP | 37986224\|23597149\|29167373\|23142 | Unspecified |
| mmu-let-7i-5p | MIMAT0000122 | Exoc6b | 75914 | HITS-CLIP | 37986224\|23142080\|29167373 | Unspecified |
| mmu-let-7i-5p | MIMAT0000122 | Exoc8 | 102058 | HITS-CLIP | 23142080 | Unspecified |
| mmu-let-7i-5p | MIMAT0000122 | Exog | 208194 | HITS-CLIP | 23142080 | Unspecified |
| mmu-let-7i-5p | MIMAT0000122 | Exosc10 | 50912 | HITS-CLIP | 23142080 | Unspecified |
| mmu-let-7i-5p | MIMAT0000122 | Exosc2 | 227715 | HITS-CLIP | 23142080 | Unspecified |
| mmu-let-7i-5p | MIMAT0000122 | Exosc8 | 69639 | HITS-CLIP | 23142080 | Unspecified |
| mmu-let-7i-5p | MIMAT0000122 | Exph5 | 320051 | HITS-CLIP | 23597149 | Unspecified |
| mmu-let-7i-5p | MIMAT0000122 | Ext2 | 14043 | HITS-CLIP | 23142080 | Unspecified |
| mmu-let-7i-5p | MIMAT0000122 | Extl3 | 54616 | HITS-CLIP | 37986224\|23142080\|29167373\|23597 | Unspecified |

| | | | | | | |
|---|---|---|---|---|---|---|
| mmu-let-7i-5p | MIMAT0000122 | Eya1 | 14048 | HITS-CLIP | 29167373 | Unspecified |
| mmu-let-7i-5p | MIMAT0000122 | Eya3 | 14050 | HITS-CLIP | 37986224\|23142080\|23597149 | Unspecified |
| mmu-let-7i-5p | MIMAT0000122 | Ezh2 | 14056 | HITS-CLIP | 23142080\|29167373 | Unspecified |
| mmu-let-7i-5p | MIMAT0000122 | Ezr | 22350 | HITS-CLIP | 37986224\|23142080 | Unspecified |
| mmu-let-7i-5p | MIMAT0000122 | F11 | 109821 | HITS-CLIP | 23597149 | Unspecified |
| mmu-let-7i-5p | MIMAT0000122 | F11r | 16456 | HITS-CLIP | 23597149 | Unspecified |
| mmu-let-7i-5p | MIMAT0000122 | F12 | 58992 | HITS-CLIP | 23597149 | Unspecified |
| mmu-let-7i-5p | MIMAT0000122 | F13b | 14060 | HITS-CLIP | 23597149 | Unspecified |
| mmu-let-7i-5p | MIMAT0000122 | F2r | 14062 | HITS-CLIP | 23597149\|23142080 | Unspecified |
| mmu-let-7i-5p | MIMAT0000122 | F5 | 14067 | HITS-CLIP | 23597149 | Unspecified |
| mmu-let-7i-5p | MIMAT0000122 | F8 | 14069 | HITS-CLIP | 23597149 | Unspecified |
| mmu-let-7i-5p | MIMAT0000122 | F9 | 14071 | HITS-CLIP | 23597149 | Unspecified |
| mmu-let-7i-5p | MIMAT0000122 | Fa2h | 338521 | HITS-CLIP | 37986224 | Unspecified |
| mmu-let-7i-5p | MIMAT0000122 | Faah | 14073 | HITS-CLIP | 23597149\|23142080 | Unspecified |
| mmu-let-7i-5p | MIMAT0000122 | Fabp1 | 14080 | HITS-CLIP | 23597149 | Unspecified |
| mmu-let-7i-5p | MIMAT0000122 | Fadd | 14082 | HITS-CLIP | 23142080 | Unspecified |
| mmu-let-7i-5p | MIMAT0000122 | Fads1 | 76267 | HITS-CLIP | 23597149\|29167373 | Unspecified |
| mmu-let-7i-5p | MIMAT0000122 | Fads2 | 56473 | HITS-CLIP | 37986224\|23597149 | Unspecified |
| mmu-let-7i-5p | MIMAT0000122 | Fads3 | 60527 | HITS-CLIP | 37986224\|23597149\|23142080 | Unspecified |
| mmu-let-7i-5p | MIMAT0000122 | Faf1 | 14084 | HITS-CLIP | 19536157\|23597149\|23142080 | Unspecified |
| mmu-let-7i-5p | MIMAT0000122 | Fah | 14085 | HITS-CLIP | 23597149 | Unspecified |
| mmu-let-7i-5p | MIMAT0000122 | Faim2 | 72393 | HITS-CLIP | 19536157 | Unspecified |
| mmu-let-7i-5p | MIMAT0000122 | Fam102a | 98952 | HITS-CLIP | 23597149\|23142080 | Unspecified |
| mmu-let-7i-5p | MIMAT0000122 | Fam117b | 72750 | HITS-CLIP | 23597149\|23142080\|29167373 | Unspecified |
| mmu-let-7i-5p | MIMAT0000122 | Fam118a | 73225 | HITS-CLIP | 23142080\|29167373 | Unspecified |
| mmu-let-7i-5p | MIMAT0000122 | Fam120a | 218236 | qCLASH//HITS-CLIP | 31393953\|23597149\|23142080\|29167 | Unspecified |
| mmu-let-7i-5p | MIMAT0000122 | Fam120b | 67544 | HITS-CLIP | 37986224\|23597149\|23142080 | Unspecified |
| mmu-let-7i-5p | MIMAT0000122 | Fam120c | 207375 | HITS-CLIP | 23597149 | Unspecified |
| mmu-let-7i-5p | MIMAT0000122 | Fam126a | 84652 | HITS-CLIP | 23597149\|23142080 | Unspecified |
| mmu-let-7i-5p | MIMAT0000122 | Fam126b | 213056 | HITS-CLIP | 23142080\|19536157\|23597149 | Unspecified |
| mmu-let-7i-5p | MIMAT0000122 | Fam131b | 76156 | HITS-CLIP | 19536157 | Unspecified |
| mmu-let-7i-5p | MIMAT0000122 | Fam135a | 68187 | HITS-CLIP | 37986224\|23597149\|23142080\|1953€ | Unspecified |
| mmu-let-7i-5p | MIMAT0000122 | Fam135b | 70363 | HITS-CLIP | 19536157 | Unspecified |
| mmu-let-7i-5p | MIMAT0000122 | Fam13b | 225358 | HITS-CLIP | 23142080 | Unspecified |
| mmu-let-7i-5p | MIMAT0000122 | Fam160a1 | 229488 | HITS-CLIP | 23597149 | Unspecified |
| mmu-let-7i-5p | MIMAT0000122 | Fam160a2 | 74349 | HITS-CLIP | 37986224\|23597149 | Unspecified |
| mmu-let-7i-5p | MIMAT0000122 | Fam160b1 | 226252 | HITS-CLIP | 37986224\|23597149\|23142080 | Unspecified |
| mmu-let-7i-5p | MIMAT0000122 | Fam160b2 | 239170 | HITS-CLIP | 23142080\|19536157 | Unspecified |
| mmu-let-7i-5p | MIMAT0000122 | Fam168a | 319604 | HITS-CLIP | 37986224 | Unspecified |

| | | | | | | |
|---|---|---|---|---|---|---|
| mmu-let-7i-5p | MIMAT0000122 | Fam168b | 214469 | HITS-CLIP | 23142080\|19536157\|29167373 | Unspecified |
| mmu-let-7i-5p | MIMAT0000122 | Fam169b | 434197 | HITS-CLIP | 23142080 | Unspecified |
| mmu-let-7i-5p | MIMAT0000122 | Fam171a1 | 269233 | HITS-CLIP | 37986224\|23597149 | Unspecified |
| mmu-let-7i-5p | MIMAT0000122 | Fam171b | 241520 | HITS-CLIP | 19536157 | Unspecified |
| mmu-let-7i-5p | MIMAT0000122 | Fam172a | 68675 | HITS-CLIP | 23142080 | Unspecified |
| mmu-let-7i-5p | MIMAT0000122 | Fam174a | 67698 | HITS-CLIP | 37986224\|19536157\|23142080 | Unspecified |
| mmu-let-7i-5p | MIMAT0000122 | Fam189a2 | 381217 | HITS-CLIP | 19536157\|23597149 | Unspecified |
| mmu-let-7i-5p | MIMAT0000122 | Fam189b | 68521 | HITS-CLIP | 37986224\|23142080 | Unspecified |
| mmu-let-7i-5p | MIMAT0000122 | Fam193a | 231128 | HITS-CLIP | 23142080\|23597149 | Unspecified |
| mmu-let-7i-5p | MIMAT0000122 | Fam198a | 245050 | HITS-CLIP | 23597149 | Unspecified |
| mmu-let-7i-5p | MIMAT0000122 | Fam199x | 245622 | HITS-CLIP | 23142080 | Unspecified |
| mmu-let-7i-5p | MIMAT0000122 | Fam204a | 76539 | HITS-CLIP | 23597149 | Unspecified |
| mmu-let-7i-5p | MIMAT0000122 | Fam207a | 108707 | HITS-CLIP | 23142080 | Unspecified |
| mmu-let-7i-5p | MIMAT0000122 | Fam208a | 218850 | HITS-CLIP | 23142080 | Unspecified |
| mmu-let-7i-5p | MIMAT0000122 | Fam208b | 54906 | HITS-CLIP | 23142080 | Unspecified |
| mmu-let-7i-5p | MIMAT0000122 | Fam20a | 208659 | HITS-CLIP | 23597149 | Unspecified |
| mmu-let-7i-5p | MIMAT0000122 | Fam20b | 215015 | HITS-CLIP | 19536157 | Unspecified |
| mmu-let-7i-5p | MIMAT0000122 | Fam210a | 108654 | HITS-CLIP | 23597149\|29167373 | Unspecified |
| mmu-let-7i-5p | MIMAT0000122 | Fam214a | 235493 | HITS-CLIP | 23142080\|23597149 | Unspecified |
| mmu-let-7i-5p | MIMAT0000122 | Fam216a | 68948 | HITS-CLIP | 23142080 | Unspecified |
| mmu-let-7i-5p | MIMAT0000122 | Fam222b | 216971 | HITS-CLIP | 37986224\|23142080\|23597149\|29167 | Unspecified |
| mmu-let-7i-5p | MIMAT0000122 | Fam32a | 67922 | HITS-CLIP | 23597149 | Unspecified |
| mmu-let-7i-5p | MIMAT0000122 | Fam3c | 27999 | HITS-CLIP | 23142080 | Unspecified |
| mmu-let-7i-5p | MIMAT0000122 | Fam43a | 224093 | HITS-CLIP | 23597149\|23142080 | Unspecified |
| mmu-let-7i-5p | MIMAT0000122 | Fam45a | 67894 | HITS-CLIP | 23142080 | Unspecified |
| mmu-let-7i-5p | MIMAT0000122 | Fam49a | 76820 | HITS-CLIP | 37986224\|23597149\|19536157\|23142 | Unspecified |
| mmu-let-7i-5p | MIMAT0000122 | Fam49b | 223601 | HITS-CLIP | 37986224\|19536157\|23597149\|29167 | Unspecified |
| mmu-let-7i-5p | MIMAT0000122 | Fam53b | 77938 | HITS-CLIP | 23597149\|29167373 | Unspecified |
| mmu-let-7i-5p | MIMAT0000122 | Fam53c | 66306 | HITS-CLIP | 29167373\|23142080 | Unspecified |
| mmu-let-7i-5p | MIMAT0000122 | Fam57a | 116972 | HITS-CLIP | 19536157\|23142080 | Unspecified |
| mmu-let-7i-5p | MIMAT0000122 | Fam69a | 388650 | HITS-CLIP | 23597149\|23142080 | Unspecified |
| mmu-let-7i-5p | MIMAT0000122 | Fam76a | 230789 | HITS-CLIP | 19536157 | Unspecified |
| mmu-let-7i-5p | MIMAT0000122 | Fam76b | 72826 | HITS-CLIP | 23142080 | Unspecified |
| mmu-let-7i-5p | MIMAT0000122 | Fam78a | 241303 | HITS-CLIP | 23142080 | Unspecified |
| mmu-let-7i-5p | MIMAT0000122 | Fam81a | 76886 | HITS-CLIP | 19536157 | Unspecified |
| mmu-let-7i-5p | MIMAT0000122 | Fam83d | 71878 | HITS-CLIP | 23142080 | Unspecified |
| mmu-let-7i-5p | MIMAT0000122 | Fam84a | 105005 | HITS-CLIP | 23597149 | Unspecified |
| mmu-let-7i-5p | MIMAT0000122 | Fam84b | 157638 | HITS-CLIP | 23142080 | Unspecified |
| mmu-let-7i-5p | MIMAT0000122 | Fam8a1 | 51439 | qCLASH//HITS-CLIP | 31393953\|23142080 | Unspecified |

| | | | | | | |
|---|---|---|---|---|---|---|
| mmu-let-7i-5p | MIMAT0000122 | Fam91a1 | 210998 | HITS-CLIP | 29167373\|23142080 | Unspecified |
| mmu-let-7i-5p | MIMAT0000122 | Fam92a | 68099 | HITS-CLIP | 23142080 | Unspecified |
| mmu-let-7i-5p | MIMAT0000122 | Fanca | 14087 | HITS-CLIP | 23597149\|23142080 | Unspecified |
| mmu-let-7i-5p | MIMAT0000122 | Fancd2 | 211651 | HITS-CLIP | 23142080 | Unspecified |
| mmu-let-7i-5p | MIMAT0000122 | Fance | 2178 | HITS-CLIP | 23597149 | Unspecified |
| mmu-let-7i-5p | MIMAT0000122 | Fancf | 100040608 | HITS-CLIP | 23142080\|23597149 | Unspecified |
| mmu-let-7i-5p | MIMAT0000122 | Far1 | 67420 | HITS-CLIP | 37986224\|23142080 | Unspecified |
| mmu-let-7i-5p | MIMAT0000122 | Farp1 | 223254 | HITS-CLIP | 37986224\|23142080\|19536157 | Unspecified |
| mmu-let-7i-5p | MIMAT0000122 | Farp2 | 227377 | HITS-CLIP | 19536157 | Unspecified |
| mmu-let-7i-5p | MIMAT0000122 | Fars2 | 69955 | HITS-CLIP | 23142080 | Unspecified |
| mmu-let-7i-5p | MIMAT0000122 | Farsb | 23874 | HITS-CLIP | 29167373 | Unspecified |
| mmu-let-7i-5p | MIMAT0000122 | Fas | 14102 | HITS-CLIP | 23142080 | Unspecified |
| mmu-let-7i-5p | MIMAT0000122 | Fasl | 14103 | HITS-CLIP | 23142080 | Unspecified |
| mmu-let-7i-5p | MIMAT0000122 | Fasn | 14104 | HITS-CLIP | 23597149\|19536157\|23142080 | Unspecified |
| mmu-let-7i-5p | MIMAT0000122 | Fastk | 66587 | HITS-CLIP | 19536157\|23142080 | Unspecified |
| mmu-let-7i-5p | MIMAT0000122 | Fastkd1 | 320720 | HITS-CLIP | 23597149\|23142080 | Unspecified |
| mmu-let-7i-5p | MIMAT0000122 | Fastkd2 | 75619 | HITS-CLIP | 23597149\|23142080\|19536157 | Unspecified |
| mmu-let-7i-5p | MIMAT0000122 | Fastkd3 | 69577 | HITS-CLIP | 29167373 | Unspecified |
| mmu-let-7i-5p | MIMAT0000122 | Fastkd5 | 380601 | HITS-CLIP | 37986224 | Unspecified |
| mmu-let-7i-5p | MIMAT0000122 | Fat2 | 245827 | HITS-CLIP | 19536157 | Unspecified |
| mmu-let-7i-5p | MIMAT0000122 | Fat3 | 270120 | HITS-CLIP | 37986224\|19536157\|29167373 | Unspecified |
| mmu-let-7i-5p | MIMAT0000122 | Fat4 | 329628 | HITS-CLIP | 37986224 | Unspecified |
| mmu-let-7i-5p | MIMAT0000122 | Fbh1 | 84893 | HITS-CLIP | 23142080 | Unspecified |
| mmu-let-7i-5p | MIMAT0000122 | Fbn1 | 14118 | HITS-CLIP | 23597149 | Unspecified |
| mmu-let-7i-5p | MIMAT0000122 | Fbp1 | 14121 | HITS-CLIP | 23597149 | Unspecified |
| mmu-let-7i-5p | MIMAT0000122 | Fbrsl1 | 381668 | HITS-CLIP | 29167373 | Unspecified |
| mmu-let-7i-5p | MIMAT0000122 | Fbxl12 | 30843 | HITS-CLIP | 23142080\|29167373 | Unspecified |
| mmu-let-7i-5p | MIMAT0000122 | Fbxl14 | 101358 | HITS-CLIP | 25083871\|29167373\|23142080 | Unspecified |
| mmu-let-7i-5p | MIMAT0000122 | Fbxl16 | 214931 | HITS-CLIP | 37986224\|19536157 | Unspecified |
| mmu-let-7i-5p | MIMAT0000122 | Fbxl17 | 50758 | HITS-CLIP | 23142080 | Unspecified |
| mmu-let-7i-5p | MIMAT0000122 | Fbxl19 | 233902 | HITS-CLIP | 23142080 | Unspecified |
| mmu-let-7i-5p | MIMAT0000122 | Fbxl2 | 72179 | HITS-CLIP | 29167373 | Unspecified |
| mmu-let-7i-5p | MIMAT0000122 | Fbxl20 | 72194 | HITS-CLIP | 23142080 | Unspecified |
| mmu-let-7i-5p | MIMAT0000122 | Fbxl3 | 50789 | HITS-CLIP | 23597149\|23142080 | Unspecified |
| mmu-let-7i-5p | MIMAT0000122 | Fbxl5 | 242960 | HITS-CLIP | 23597149\|23142080 | Unspecified |
| mmu-let-7i-5p | MIMAT0000122 | Fbxo10 | 269529 | HITS-CLIP | 37986224 | Unspecified |
| mmu-let-7i-5p | MIMAT0000122 | Fbxo11 | 225055 | HITS-CLIP | 23142080\|19536157\|31433990\|29167 | Unspecified |
| mmu-let-7i-5p | MIMAT0000122 | Fbxo21 | 231670 | HITS-CLIP | 23142080\|23597149\|29167373 | Unspecified |
| mmu-let-7i-5p | MIMAT0000122 | Fbxo22 | 71999 | HITS-CLIP | 23597149\|23142080 | Unspecified |

| | | | | | | | |
|---|---|---|---|---|---|---|---|
| mmu-let-7i-5p | MIMAT0000122 | Fbxo28 | 67948 | HITS-CLIP | | 23142080 | Unspecified |
| mmu-let-7i-5p | MIMAT0000122 | Fbxo3 | 57443 | HITS-CLIP | 23142080\|29167373 | | Unspecified |
| mmu-let-7i-5p | MIMAT0000122 | Fbxo31 | 76454 | HITS-CLIP | | 37986224 | Unspecified |
| mmu-let-7i-5p | MIMAT0000122 | Fbxo32 | 67731 | HITS-CLIP | 37986224\|23597149\|23142080\|1953( | | Unspecified |
| mmu-let-7i-5p | MIMAT0000122 | Fbxo33 | 70611 | HITS-CLIP | 37986224\|19536157\|23597149\|23142 | | Unspecified |
| mmu-let-7i-5p | MIMAT0000122 | Fbxo38 | 107035 | HITS-CLIP | 37986224\|23142080 | | Unspecified |
| mmu-let-7i-5p | MIMAT0000122 | Fbxo42 | 213499 | HITS-CLIP | | 19536157 | Unspecified |
| mmu-let-7i-5p | MIMAT0000122 | Fbxo45 | 268882 | HITS-CLIP | | 23597149 | Unspecified |
| mmu-let-7i-5p | MIMAT0000122 | Fbxo7 | 69754 | HITS-CLIP | | 23597149 | Unspecified |
| mmu-let-7i-5p | MIMAT0000122 | Fbxo8 | 50753 | HITS-CLIP | 37986224\|23597149\|19536157\|23142 | | Unspecified |
| mmu-let-7i-5p | MIMAT0000122 | Fbxw11 | 103583 | HITS-CLIP | | 23142080 | Unspecified |
| mmu-let-7i-5p | MIMAT0000122 | Fbxw17 | 109082 | HITS-CLIP | | 23142080 | Unspecified |
| mmu-let-7i-5p | MIMAT0000122 | Fbxw2 | 30050 | HITS-CLIP | | 23142080 | Unspecified |
| mmu-let-7i-5p | MIMAT0000122 | Fbxw7 | 50754 | HITS-CLIP | | 23142080 | Unspecified |
| mmu-let-7i-5p | MIMAT0000122 | Fbxw8 | 231672 | HITS-CLIP | 23597149\|29167373 | | Unspecified |
| mmu-let-7i-5p | MIMAT0000122 | Fcgr3 | 14131 | HITS-CLIP | | 29167373 | Unspecified |
| mmu-let-7i-5p | MIMAT0000122 | Fcgrt | 14132 | HITS-CLIP | | 23597149 | Unspecified |
| mmu-let-7i-5p | MIMAT0000122 | Fcho2 | 218503 | HITS-CLIP | 37986224\|23597149\|19536157\|23142 | | Unspecified |
| mmu-let-7i-5p | MIMAT0000122 | Fchsd2 | 207278 | HITS-CLIP | | 23142080 | Unspecified |
| mmu-let-7i-5p | MIMAT0000122 | Fem1b | 14155 | HITS-CLIP | 23597149\|23142080\|29167373 | | Unspecified |
| mmu-let-7i-5p | MIMAT0000122 | Fem1c | 240263 | qCLASH//HITS-CLIP | 31393953\|23142080 | | Unspecified |
| mmu-let-7i-5p | MIMAT0000122 | Fen1 | 14156 | HITS-CLIP | | 23142080 | Unspecified |
| mmu-let-7i-5p | MIMAT0000122 | Fermt3 | 108101 | HITS-CLIP | | 23142080 | Unspecified |
| mmu-let-7i-5p | MIMAT0000122 | Fetub | 59083 | HITS-CLIP | | 23597149 | Unspecified |
| mmu-let-7i-5p | MIMAT0000122 | Fez1 | 235180 | HITS-CLIP | | 19536157 | Unspecified |
| mmu-let-7i-5p | MIMAT0000122 | Fezf2 | 54713 | HITS-CLIP | | 19536157 | Unspecified |
| mmu-let-7i-5p | MIMAT0000122 | Fga | 14161 | HITS-CLIP | | 23597149 | Unspecified |
| mmu-let-7i-5p | MIMAT0000122 | Fgd2 | 26382 | HITS-CLIP | | 23597149 | Unspecified |
| mmu-let-7i-5p | MIMAT0000122 | Fgd3 | 30938 | HITS-CLIP | | 29167373 | Unspecified |
| mmu-let-7i-5p | MIMAT0000122 | Fgd4 | 224014 | HITS-CLIP | 37986224\|29167373 | | Unspecified |
| mmu-let-7i-5p | MIMAT0000122 | Fgd6 | 13998 | HITS-CLIP | 37986224\|23597149\|23142080 | | Unspecified |
| mmu-let-7i-5p | MIMAT0000122 | Fgf1 | 14164 | HITS-CLIP | | 23597149 | Unspecified |
| mmu-let-7i-5p | MIMAT0000122 | Fgf13 | 14168 | HITS-CLIP | | 23142080 | Unspecified |
| mmu-let-7i-5p | MIMAT0000122 | Fgf2 | 14173 | HITS-CLIP | 37986224\|23142080 | | Unspecified |
| mmu-let-7i-5p | MIMAT0000122 | Fgfbp1 | 14181 | HITS-CLIP | | 37986224 | Unspecified |
| mmu-let-7i-5p | MIMAT0000122 | Fgfrl1 | 116701 | HITS-CLIP | 37986224\|23597149 | | Unspecified |
| mmu-let-7i-5p | MIMAT0000122 | Fgg | 99571 | HITS-CLIP | | 23597149 | Unspecified |
| mmu-let-7i-5p | MIMAT0000122 | Fh1 | 14194 | HITS-CLIP | | 23597149 | Unspecified |
| mmu-let-7i-5p | MIMAT0000122 | Fhip2b | 64760 | qCLASH//HITS-CLIP | 31393953\|31433990 | | Unspecified |

| | | | | | | |
|---|---|---|---|---|---|---|
| mmu-let-7i-5p | MIMAT0000122 | Fhl3 | 14201 | HITS-CLIP | 23142080 | Unspecified |
| mmu-let-7i-5p | MIMAT0000122 | Ficd | 231630 | HITS-CLIP | 23142080 | Unspecified |
| mmu-let-7i-5p | MIMAT0000122 | Fig4 | 103199 | HITS-CLIP | 29167373 | Unspecified |
| mmu-let-7i-5p | MIMAT0000122 | Fign | 60344 | HITS-CLIP | 37986224\|23597149 | Unspecified |
| mmu-let-7i-5p | MIMAT0000122 | Fignl1 | 60530 | HITS-CLIP | 23142080\|29167373 | Unspecified |
| mmu-let-7i-5p | MIMAT0000122 | Fignl2 | 668225 | HITS-CLIP | 37986224\|23142080\|19536157\|2359 | Unspecified |
| mmu-let-7i-5p | MIMAT0000122 | Filip1l | 78749 | HITS-CLIP | 37986224\|23142080\|29167373 | Unspecified |
| mmu-let-7i-5p | MIMAT0000122 | Fis1 | 66437 | HITS-CLIP | 23142080 | Unspecified |
| mmu-let-7i-5p | MIMAT0000122 | Fitm1 | 68680 | HITS-CLIP | 23597149 | Unspecified |
| mmu-let-7i-5p | MIMAT0000122 | Fkbp15 | 338355 | HITS-CLIP | 23597149\|23142080\|29167373 | Unspecified |
| mmu-let-7i-5p | MIMAT0000122 | Fkbp1a | 14225 | HITS-CLIP | 23597149\|23142080 | Unspecified |
| mmu-let-7i-5p | MIMAT0000122 | Fkbp1b | 14226 | HITS-CLIP | 19536157 | Unspecified |
| mmu-let-7i-5p | MIMAT0000122 | Fkbp9 | 27055 | HITS-CLIP | 23597149 | Unspecified |
| mmu-let-7i-5p | MIMAT0000122 | Fktn | 246179 | HITS-CLIP | 23142080 | Unspecified |
| mmu-let-7i-5p | MIMAT0000122 | Flad1 | 319945 | HITS-CLIP | 23597149 | Unspecified |
| mmu-let-7i-5p | MIMAT0000122 | Flcn | 216805 | HITS-CLIP | 23597149\|23142080\|29167373 | Unspecified |
| mmu-let-7i-5p | MIMAT0000122 | Fli1 | 14247 | HITS-CLIP | 23597149\|19536157\|23142080\|2916 | Unspecified |
| mmu-let-7i-5p | MIMAT0000122 | Flii | 14248 | HITS-CLIP | 37986224\|23597149\|23142080 | Unspecified |
| mmu-let-7i-5p | MIMAT0000122 | Flna | 192176 | HITS-CLIP | 23142080\|29167373 | Unspecified |
| mmu-let-7i-5p | MIMAT0000122 | Flnb | 286940 | HITS-CLIP | 23597149 | Unspecified |
| mmu-let-7i-5p | MIMAT0000122 | Flot1 | 14251 | HITS-CLIP | 23142080 | Unspecified |
| mmu-let-7i-5p | MIMAT0000122 | Flot2 | 14252 | HITS-CLIP | 23630078 | Unspecified |
| mmu-let-7i-5p | MIMAT0000122 | Flrt2 | 399558 | HITS-CLIP | 37986224\|19536157\|29167373 | Unspecified |
| mmu-let-7i-5p | MIMAT0000122 | Flrt3 | 71436 | HITS-CLIP | 19536157 | Unspecified |
| mmu-let-7i-5p | MIMAT0000122 | Flt1 | 14254 | HITS-CLIP | 37986224\|23597149\|19536157 | Unspecified |
| mmu-let-7i-5p | MIMAT0000122 | Flt4 | 14257 | HITS-CLIP | 23597149 | Unspecified |
| mmu-let-7i-5p | MIMAT0000122 | Fmn1 | 14260 | HITS-CLIP | 29167373 | Unspecified |
| mmu-let-7i-5p | MIMAT0000122 | Fmnl1 | 57778 | HITS-CLIP | 23142080\|29167373 | Unspecified |
| mmu-let-7i-5p | MIMAT0000122 | Fmnl2 | 71409 | HITS-CLIP | 23597149 | Unspecified |
| mmu-let-7i-5p | MIMAT0000122 | Fmo4 | 226564 | HITS-CLIP | 23597149 | Unspecified |
| mmu-let-7i-5p | MIMAT0000122 | Fmo5 | 14263 | HITS-CLIP | 23597149 | Unspecified |
| mmu-let-7i-5p | MIMAT0000122 | Fmod | 14264 | HITS-CLIP | 37986224\|19536157 | Unspecified |
| mmu-let-7i-5p | MIMAT0000122 | Fmr1 | 14265 | HITS-CLIP | 23142080 | Unspecified |
| mmu-let-7i-5p | MIMAT0000122 | Fn1 | 14268 | HITS-CLIP | 23597149 | Unspecified |
| mmu-let-7i-5p | MIMAT0000122 | Fnbp1 | 14269 | HITS-CLIP | 19536157\|23597149\|23142080\|2916 | Unspecified |
| mmu-let-7i-5p | MIMAT0000122 | Fnbp1l | 214459 | HITS-CLIP | 37986224 | Unspecified |
| mmu-let-7i-5p | MIMAT0000122 | Fnbp4 | 55935 | HITS-CLIP | 23142080\|29167373 | Unspecified |
| mmu-let-7i-5p | MIMAT0000122 | Fndc1 | 68655 | HITS-CLIP | 23597149 | Unspecified |
| mmu-let-7i-5p | MIMAT0000122 | Fndc3a | 319448 | HITS-CLIP | 37986224\|19536157\|23597149\|2314 | Unspecified |

| | | | | | | |
|---|---|---|---|---|---|---|
| mmu-let-7i-5p | MIMAT0000122 | Fndc3b | 72007 | HITS-CLIP | 23597149\|29167373 | Unspecified |
| mmu-let-7i-5p | MIMAT0000122 | Fnip1 | 216742 | HITS-CLIP//qCLASH | 37986224\|31393953\|19536157\|23597 | Unspecified |
| mmu-let-7i-5p | MIMAT0000122 | Fnip2 | 329679 | HITS-CLIP | 37986224\|23142080\|19536157\|23597 | Unspecified |
| mmu-let-7i-5p | MIMAT0000122 | Fntb | 110606 | HITS-CLIP | 37986224\|23597149\|23142080 | Unspecified |
| mmu-let-7i-5p | MIMAT0000122 | Fopnl | 66086 | HITS-CLIP | 23142080 | Unspecified |
| mmu-let-7i-5p | MIMAT0000122 | Fosl1 | 14283 | HITS-CLIP | 37986224 | Unspecified |
| mmu-let-7i-5p | MIMAT0000122 | Fosl2 | 14284 | HITS-CLIP | 37986224\|23597149\|31433990\|23142 | Unspecified |
| mmu-let-7i-5p | MIMAT0000122 | Foxj2 | 60611 | HITS-CLIP | 23597149 | Unspecified |
| mmu-let-7i-5p | MIMAT0000122 | Foxj3 | 230700 | HITS-CLIP | 23597149\|23142080 | Unspecified |
| mmu-let-7i-5p | MIMAT0000122 | Foxk1 | 17425 | HITS-CLIP | 37986224\|23142080 | Unspecified |
| mmu-let-7i-5p | MIMAT0000122 | Foxk2 | 68837 | HITS-CLIP | 23597149\|23142080 | Unspecified |
| mmu-let-7i-5p | MIMAT0000122 | Foxm1 | 14235 | HITS-CLIP | 23142080 | Unspecified |
| mmu-let-7i-5p | MIMAT0000122 | Foxn2 | 14236 | HITS-CLIP | 23142080 | Unspecified |
| mmu-let-7i-5p | MIMAT0000122 | Foxn3 | 71375 | HITS-CLIP | 19536157\|23142080 | Unspecified |
| mmu-let-7i-5p | MIMAT0000122 | Foxo1 | 56458 | HITS-CLIP//qCLASH | 37986224\|31393953\|23597149\|23142 | Unspecified |
| mmu-let-7i-5p | MIMAT0000122 | Foxo3 | 56484 | HITS-CLIP | 23597149\|19536157\|23142080\|29167 | Unspecified |
| mmu-let-7i-5p | MIMAT0000122 | Foxp1 | 108655 | HITS-CLIP | 37986224\|23597149\|23142080 | Unspecified |
| mmu-let-7i-5p | MIMAT0000122 | Foxp2 | 114142 | HITS-CLIP | 37986224\|23597149\|19536157 | Unspecified |
| mmu-let-7i-5p | MIMAT0000122 | Foxq1 | 15220 | HITS-CLIP | 23597149 | Unspecified |
| mmu-let-7i-5p | MIMAT0000122 | Fpgs | 14287 | HITS-CLIP | 23597149 | Unspecified |
| mmu-let-7i-5p | MIMAT0000122 | Fpgt | 75540 | HITS-CLIP | 23142080 | Unspecified |
| mmu-let-7i-5p | MIMAT0000122 | Fras1 | 231470 | HITS-CLIP | 37986224\|19536157 | Unspecified |
| mmu-let-7i-5p | MIMAT0000122 | Frem2 | 242022 | HITS-CLIP | 37986224 | Unspecified |
| mmu-let-7i-5p | MIMAT0000122 | Frmd4a | 209630 | HITS-CLIP | 19536157\|29167373 | Unspecified |
| mmu-let-7i-5p | MIMAT0000122 | Frmd4b | 232288 | HITS-CLIP | 29167373 | Unspecified |
| mmu-let-7i-5p | MIMAT0000122 | Frmd6 | 319710 | HITS-CLIP | 23142080 | Unspecified |
| mmu-let-7i-5p | MIMAT0000122 | Frmd8 | 67457 | HITS-CLIP | 23597149\|23142080 | Unspecified |
| mmu-let-7i-5p | MIMAT0000122 | Frrs1 | 20321 | HITS-CLIP | 29167373 | Unspecified |
| mmu-let-7i-5p | MIMAT0000122 | Frs2 | 327826 | HITS-CLIP | 23142080 | Unspecified |
| mmu-let-7i-5p | MIMAT0000122 | Fry | 320365 | HITS-CLIP | 37986224\|19536157\|23142080 | Unspecified |
| mmu-let-7i-5p | MIMAT0000122 | Fryl | 72313 | HITS-CLIP | 23142080 | Unspecified |
| mmu-let-7i-5p | MIMAT0000122 | Fsbp | 100503583 | HITS-CLIP | 23142080 | Unspecified |
| mmu-let-7i-5p | MIMAT0000122 | Fscn1 | 14086 | HITS-CLIP | 37986224 | Unspecified |
| mmu-let-7i-5p | MIMAT0000122 | Fsd1 | 240121 | HITS-CLIP | 37986224 | Unspecified |
| mmu-let-7i-5p | MIMAT0000122 | Fstl1 | 14314 | HITS-CLIP | 37986224\|19536157\|23597149 | Unspecified |
| mmu-let-7i-5p | MIMAT0000122 | Ftcd | 14317 | HITS-CLIP | 23597149 | Unspecified |
| mmu-let-7i-5p | MIMAT0000122 | Fth1 | 14319 | HITS-CLIP | 29167373 | Unspecified |
| mmu-let-7i-5p | MIMAT0000122 | Ftl1 | 14325 | qCLASH | 31393953 | Unspecified |
| mmu-let-7i-5p | MIMAT0000122 | Fto | 26383 | HITS-CLIP | 37986224\|23142080\|23597149 | Unspecified |

| | | | | | | |
|---|---|---|---|---|---|---|
| mmu-let-7i-5p | MIMAT0000122 | Fubp1 | 51886 | HITS-CLIP | 23142080\|19536157\|29167373 | Unspecified |
| mmu-let-7i-5p | MIMAT0000122 | Fubp3 | 320267 | HITS-CLIP | 23142080 | Unspecified |
| mmu-let-7i-5p | MIMAT0000122 | Fuca2 | 66848 | HITS-CLIP | 23597149\|23142080\|29167373 | Unspecified |
| mmu-let-7i-5p | MIMAT0000122 | Fuk | 197258 | HITS-CLIP | 23142080 | Unspecified |
| mmu-let-7i-5p | MIMAT0000122 | Fundc1 | 72018 | HITS-CLIP | 23597149\|23142080 | Unspecified |
| mmu-let-7i-5p | MIMAT0000122 | Fundc2 | 67391 | HITS-CLIP | 23597149 | Unspecified |
| mmu-let-7i-5p | MIMAT0000122 | Fuom | 69064 | HITS-CLIP | 23597149 | Unspecified |
| mmu-let-7i-5p | MIMAT0000122 | Furin | 18550 | HITS-CLIP | 23597149\|29167373 | Unspecified |
| mmu-let-7i-5p | MIMAT0000122 | Fus | 233908 | HITS-CLIP | 31433990 | Unspecified |
| mmu-let-7i-5p | MIMAT0000122 | Fut11 | 73068 | HITS-CLIP | 23142080 | Unspecified |
| mmu-let-7i-5p | MIMAT0000122 | Fut9 | 14348 | HITS-CLIP | 37986224 | Unspecified |
| mmu-let-7i-5p | MIMAT0000122 | Fuz | 70300 | HITS-CLIP | 23142080 | Unspecified |
| mmu-let-7i-5p | MIMAT0000122 | Fxr1 | 14359 | HITS-CLIP | 23142080\|23597149 | Unspecified |
| mmu-let-7i-5p | MIMAT0000122 | Fxyd5 | 18301 | HITS-CLIP | 29167373 | Unspecified |
| mmu-let-7i-5p | MIMAT0000122 | Fxyd6 | 59095 | HITS-CLIP | 37986224 | Unspecified |
| mmu-let-7i-5p | MIMAT0000122 | Fyb | 23880 | HITS-CLIP | 23142080 | Unspecified |
| mmu-let-7i-5p | MIMAT0000122 | Fyco1 | 17281 | HITS-CLIP | 37986224\|23142080 | Unspecified |
| mmu-let-7i-5p | MIMAT0000122 | Fyn | 14360 | HITS-CLIP | 23142080 | Unspecified |
| mmu-let-7i-5p | MIMAT0000122 | Fzd3 | 14365 | HITS-CLIP | 19536157 | Unspecified |
| mmu-let-7i-5p | MIMAT0000122 | Fzd5 | 14367 | HITS-CLIP | 23597149 | Unspecified |
| mmu-let-7i-5p | MIMAT0000122 | Fzd7 | 14369 | HITS-CLIP | 23142080 | Unspecified |
| mmu-let-7i-5p | MIMAT0000122 | Fzd8 | 14370 | HITS-CLIP | 23630078 | Unspecified |
| mmu-let-7i-5p | MIMAT0000122 | G2e3 | 217558 | HITS-CLIP | 37986224\|23597149\|23142080 | Unspecified |
| mmu-let-7i-5p | MIMAT0000122 | G3bp1 | 27041 | HITS-CLIP | 37986224\|23597149\|23142080\|1953 | Unspecified |
| mmu-let-7i-5p | MIMAT0000122 | G3bp2 | 23881 | HITS-CLIP | 23142080\|23597149\|29167373 | Unspecified |
| mmu-let-7i-5p | MIMAT0000122 | G6pc | 14377 | HITS-CLIP | 23597149 | Unspecified |
| mmu-let-7i-5p | MIMAT0000122 | G6pdx | 14381 | HITS-CLIP | 23142080 | Unspecified |
| mmu-let-7i-5p | MIMAT0000122 | Gab1 | 14388 | HITS-CLIP | 37986224 | Unspecified |
| mmu-let-7i-5p | MIMAT0000122 | Gab2 | 14389 | HITS-CLIP | 29167373 | Unspecified |
| mmu-let-7i-5p | MIMAT0000122 | Gabarapl1 | 57436 | HITS-CLIP | 37986224\|23597149 | Unspecified |
| mmu-let-7i-5p | MIMAT0000122 | Gabbr1 | 54393 | HITS-CLIP | 19536157 | Unspecified |
| mmu-let-7i-5p | MIMAT0000122 | Gabbr2 | 242425 | HITS-CLIP | 37986224\|19536157 | Unspecified |
| mmu-let-7i-5p | MIMAT0000122 | Gabpa | 14390 | HITS-CLIP | 23142080 | Unspecified |
| mmu-let-7i-5p | MIMAT0000122 | Gabpb1 | 14391 | HITS-CLIP | 23142080 | Unspecified |
| mmu-let-7i-5p | MIMAT0000122 | Gabpb2 | 213054 | HITS-CLIP | 23142080\|29167373 | Unspecified |
| mmu-let-7i-5p | MIMAT0000122 | Gabra6 | 14399 | HITS-CLIP | 19536157 | Unspecified |
| mmu-let-7i-5p | MIMAT0000122 | Gabrb2 | 14401 | HITS-CLIP | 37986224 | Unspecified |
| mmu-let-7i-5p | MIMAT0000122 | Gabrb3 | 14402 | HITS-CLIP | 19536157 | Unspecified |
| mmu-let-7i-5p | MIMAT0000122 | Gabrg2 | 14406 | HITS-CLIP | 19536157 | Unspecified |

| | | | | | | |
|---|---|---|---|---|---|---|
| mmu-let-7i-5p | MIMAT0000122 | Gad1 | 14415 | HITS-CLIP | 37986224 | Unspecified |
| mmu-let-7i-5p | MIMAT0000122 | Gadd45gip1 | 102060 | HITS-CLIP | 23597149\|26203562\|23142080 | Unspecified |
| mmu-let-7i-5p | MIMAT0000122 | Gak | 231580 | HITS-CLIP | 23597149 | Unspecified |
| mmu-let-7i-5p | MIMAT0000122 | Galc | 14420 | HITS-CLIP | 29167373 | Unspecified |
| mmu-let-7i-5p | MIMAT0000122 | Gale | 74246 | HITS-CLIP | 23597149 | Unspecified |
| mmu-let-7i-5p | MIMAT0000122 | Galk1 | 14635 | HITS-CLIP | 23597149\|23142080 | Unspecified |
| mmu-let-7i-5p | MIMAT0000122 | Galk2 | 69976 | HITS-CLIP | 23142080 | Unspecified |
| mmu-let-7i-5p | MIMAT0000122 | Galm | 319625 | HITS-CLIP | 23597149 | Unspecified |
| mmu-let-7i-5p | MIMAT0000122 | Galnt1 | 14423 | HITS-CLIP | 23142080\|29167373 | Unspecified |
| mmu-let-7i-5p | MIMAT0000122 | Galnt12 | 230145 | HITS-CLIP | 23142080 | Unspecified |
| mmu-let-7i-5p | MIMAT0000122 | Galnt17 | 64409 | HITS-CLIP | 19536157 | Unspecified |
| mmu-let-7i-5p | MIMAT0000122 | Galnt2 | 108148 | HITS-CLIP | 37986224\|23597149\|19536157\|23142 | Unspecified |
| mmu-let-7i-5p | MIMAT0000122 | Galnt3 | 14425 | HITS-CLIP | 23142080 | Unspecified |
| mmu-let-7i-5p | MIMAT0000122 | Galnt4 | 14426 | HITS-CLIP | 23142080\|23597149 | Unspecified |
| mmu-let-7i-5p | MIMAT0000122 | Galnt7 | 108150 | HITS-CLIP | 23142080 | Unspecified |
| mmu-let-7i-5p | MIMAT0000122 | Gamt | 14431 | HITS-CLIP | 23597149 | Unspecified |
| mmu-let-7i-5p | MIMAT0000122 | Gan | 209239 | HITS-CLIP | 23142080\|29167373 | Unspecified |
| mmu-let-7i-5p | MIMAT0000122 | Ganab | 14376 | HITS-CLIP | 23142080 | Unspecified |
| mmu-let-7i-5p | MIMAT0000122 | Ganc | 76051 | HITS-CLIP | 23597149 | Unspecified |
| mmu-let-7i-5p | MIMAT0000122 | Gapvd1 | 66691 | HITS-CLIP | 37986224\|23597149\|23142080 | Unspecified |
| mmu-let-7i-5p | MIMAT0000122 | Gars | 353172 | HITS-CLIP//qCLASH | 37986224\|23597149\|31393953\|23142 | Unspecified |
| mmu-let-7i-5p | MIMAT0000122 | Gas1 | 14451 | HITS-CLIP | 23597149 | Unspecified |
| mmu-let-7i-5p | MIMAT0000122 | Gas7 | 14457 | HITS-CLIP | 37986224\|29167373 | Unspecified |
| mmu-let-7i-5p | MIMAT0000122 | Gas8 | 104346 | HITS-CLIP | 37986224 | Unspecified |
| mmu-let-7i-5p | MIMAT0000122 | Gata3 | 14462 | HITS-CLIP | 23142080 | Unspecified |
| mmu-let-7i-5p | MIMAT0000122 | Gata6 | 14465 | HITS-CLIP | 23597149 | Unspecified |
| mmu-let-7i-5p | MIMAT0000122 | Gatad1 | 67210 | HITS-CLIP | 23597149 | Unspecified |
| mmu-let-7i-5p | MIMAT0000122 | Gatad2b | 229542 | HITS-CLIP | 23142080\|29167373 | Unspecified |
| mmu-let-7i-5p | MIMAT0000122 | Gatc | 384281 | HITS-CLIP | 29167373 | Unspecified |
| mmu-let-7i-5p | MIMAT0000122 | Gatm | 67092 | HITS-CLIP | 37986224\|23597149\|19536157\|23142 | Unspecified |
| mmu-let-7i-5p | MIMAT0000122 | Gba2 | 230101 | HITS-CLIP | 19536157 | Unspecified |
| mmu-let-7i-5p | MIMAT0000122 | Gbe1 | 74185 | HITS-CLIP | 23142080\|23597149 | Unspecified |
| mmu-let-7i-5p | MIMAT0000122 | Gbf1 | 107338 | HITS-CLIP | 37986224\|23597149\|23142080\|29167 | Unspecified |
| mmu-let-7i-5p | MIMAT0000122 | Gbp2 | 14469 | HITS-CLIP | 37986224\|23142080 | Unspecified |
| mmu-let-7i-5p | MIMAT0000122 | Gbp3 | 55932 | HITS-CLIP | 23142080 | Unspecified |
| mmu-let-7i-5p | MIMAT0000122 | Gbp4 | 17472 | HITS-CLIP | 23597149\|23142080 | Unspecified |
| mmu-let-7i-5p | MIMAT0000122 | Gbp5 | 229898 | HITS-CLIP | 23142080 | Unspecified |
| mmu-let-7i-5p | MIMAT0000122 | Gbp7 | 229900 | HITS-CLIP | 37986224\|23597149\|23142080 | Unspecified |
| mmu-let-7i-5p | MIMAT0000122 | Gc | 14473 | HITS-CLIP | 23597149 | Unspecified |

| | | | | | | |
|---|---|---|---|---|---|---|
| mmu-let-7i-5p | MIMAT0000122 | Gcat | 26912 | HITS-CLIP | 23597149 | Unspecified |
| mmu-let-7i-5p | MIMAT0000122 | Gcdh | 270076 | HITS-CLIP | 23597149 | Unspecified |
| mmu-let-7i-5p | MIMAT0000122 | Gchfr | 320415 | HITS-CLIP | 23597149 | Unspecified |
| mmu-let-7i-5p | MIMAT0000122 | Gckr | 231103 | HITS-CLIP | 37986224\|23597149 | Unspecified |
| mmu-let-7i-5p | MIMAT0000122 | Gclc | 14629 | HITS-CLIP | 23597149\|19536157\|23142080 | Unspecified |
| mmu-let-7i-5p | MIMAT0000122 | Gclm | 14630 | HITS-CLIP | 29167373\|23142080 | Unspecified |
| mmu-let-7i-5p | MIMAT0000122 | Gcnt1 | 14537 | HITS-CLIP | 23142080 | Unspecified |
| mmu-let-7i-5p | MIMAT0000122 | Gcnt2 | 14538 | HITS-CLIP | 23597149 | Unspecified |
| mmu-let-7i-5p | MIMAT0000122 | Gcnt4 | 218476 | HITS-CLIP | 23597149 | Unspecified |
| mmu-let-7i-5p | MIMAT0000122 | Gcsh | 68133 | HITS-CLIP | 23597149 | Unspecified |
| mmu-let-7i-5p | MIMAT0000122 | Gdap2 | 14547 | HITS-CLIP | 37986224\|23597149 | Unspecified |
| mmu-let-7i-5p | MIMAT0000122 | Gde1 | 56209 | HITS-CLIP | 23597149\|23142080 | Unspecified |
| mmu-let-7i-5p | MIMAT0000122 | Gdf11 | 14561 | HITS-CLIP | 23142080 | Unspecified |
| mmu-let-7i-5p | MIMAT0000122 | Gdf15 | 23886 | HITS-CLIP | 23597149 | Unspecified |
| mmu-let-7i-5p | MIMAT0000122 | Gdpd5 | 233552 | HITS-CLIP | 23142080 | Unspecified |
| mmu-let-7i-5p | MIMAT0000122 | Gdpgp1 | 269952 | HITS-CLIP | 29167373\|23142080 | Unspecified |
| mmu-let-7i-5p | MIMAT0000122 | Gemin5 | 216766 | HITS-CLIP | 23142080\|31433990 | Unspecified |
| mmu-let-7i-5p | MIMAT0000122 | Gemin7 | 79760 | HITS-CLIP | 23597149 | Unspecified |
| mmu-let-7i-5p | MIMAT0000122 | Gen1 | 209334 | HITS-CLIP | 23142080 | Unspecified |
| mmu-let-7i-5p | MIMAT0000122 | Gfap | 14580 | HITS-CLIP | 19536157 | Unspecified |
| mmu-let-7i-5p | MIMAT0000122 | Gfi1 | 14581 | HITS-CLIP | 23142080 | Unspecified |
| mmu-let-7i-5p | MIMAT0000122 | Gfm2 | 320806 | HITS-CLIP | 37986224\|23597149 | Unspecified |
| mmu-let-7i-5p | MIMAT0000122 | Gfod1 | 328232 | HITS-CLIP | 23142080 | Unspecified |
| mmu-let-7i-5p | MIMAT0000122 | Gfpt1 | 14583 | HITS-CLIP | 26203562 | Unspecified |
| mmu-let-7i-5p | MIMAT0000122 | Gfra1 | 14585 | HITS-CLIP | 23597149 | Unspecified |
| mmu-let-7i-5p | MIMAT0000122 | Gga3 | 260302 | HITS-CLIP | 37986224\|23142080\|31433990 | Unspecified |
| mmu-let-7i-5p | MIMAT0000122 | Ggact | 223267 | HITS-CLIP | 23597149\|23142080 | Unspecified |
| mmu-let-7i-5p | MIMAT0000122 | Ggnbp2 | 217039 | HITS-CLIP | 23142080 | Unspecified |
| mmu-let-7i-5p | MIMAT0000122 | Ggt7 | 207182 | HITS-CLIP | 37986224\|23142080 | Unspecified |
| mmu-let-7i-5p | MIMAT0000122 | Ggta1 | 14594 | HITS-CLIP | 23142080\|29167373 | Unspecified |
| mmu-let-7i-5p | MIMAT0000122 | Ghitm | 66092 | HITS-CLIP | 23142080\|23597149 | Unspecified |
| mmu-let-7i-5p | MIMAT0000122 | Ghr | 14600 | HITS-CLIP | 37986224\|23597149 | Unspecified |
| mmu-let-7i-5p | MIMAT0000122 | Gid8 | 76425 | HITS-CLIP | 37986224\|23597149\|19536157\|2314 | Unspecified |
| mmu-let-7i-5p | MIMAT0000122 | Gigyf1 | 57330 | HITS-CLIP | 37986224\|23597149\|19536157\|2916 | Unspecified |
| mmu-let-7i-5p | MIMAT0000122 | Gigyf2 | 227331 | HITS-CLIP | 26203562 | Unspecified |
| mmu-let-7i-5p | MIMAT0000122 | Gimap3 | 83408 | HITS-CLIP | 23142080 | Unspecified |
| mmu-let-7i-5p | MIMAT0000122 | Gimap4 | 107526 | HITS-CLIP | 23597149\|23142080 | Unspecified |
| mmu-let-7i-5p | MIMAT0000122 | Gimap7 | 231932 | HITS-CLIP | 23142080 | Unspecified |
| mmu-let-7i-5p | MIMAT0000122 | Gimap8 | 243374 | HITS-CLIP | 23142080 | Unspecified |

| | | | | | | | |
|---|---|---|---|---|---|---|---|
| mmu-let-7i-5p | MIMAT0000122 | Ginm1 | 215751 | HITS-CLIP | | 23142080 | Unspecified |
| mmu-let-7i-5p | MIMAT0000122 | Git1 | 216963 | HITS-CLIP | | 23142080 | Unspecified |
| mmu-let-7i-5p | MIMAT0000122 | Git2 | 26431 | HITS-CLIP | 23597149\|23142080 | | Unspecified |
| mmu-let-7i-5p | MIMAT0000122 | Gja1 | 14609 | HITS-CLIP | 23142080\|19536157 | | Unspecified |
| mmu-let-7i-5p | MIMAT0000122 | Gja4 | 14612 | HITS-CLIP | | 23597149 | Unspecified |
| mmu-let-7i-5p | MIMAT0000122 | Gjb1 | 14618 | HITS-CLIP | | 37986224 | Unspecified |
| mmu-let-7i-5p | MIMAT0000122 | Gla | 11605 | HITS-CLIP | 37986224\|19536157\|29167373\|23142 | | Unspecified |
| mmu-let-7i-5p | MIMAT0000122 | Glb1 | 12091 | HITS-CLIP | 37986224\|23597149 | | Unspecified |
| mmu-let-7i-5p | MIMAT0000122 | Glce | 93683 | HITS-CLIP | 37986224\|19536157\|23597149 | | Unspecified |
| mmu-let-7i-5p | MIMAT0000122 | Gldc | 104174 | HITS-CLIP | 23597149\|23142080 | | Unspecified |
| mmu-let-7i-5p | MIMAT0000122 | Gle1 | 74412 | HITS-CLIP | 37986224\|23142080 | | Unspecified |
| mmu-let-7i-5p | MIMAT0000122 | Glg1 | 20340 | HITS-CLIP | | 23142080 | Unspecified |
| mmu-let-7i-5p | MIMAT0000122 | Glipr1 | 73690 | HITS-CLIP | | 23142080 | Unspecified |
| mmu-let-7i-5p | MIMAT0000122 | Glipr2 | 384009 | HITS-CLIP | | 23142080 | Unspecified |
| mmu-let-7i-5p | MIMAT0000122 | Glmn | 170823 | HITS-CLIP | | 23142080 | Unspecified |
| mmu-let-7i-5p | MIMAT0000122 | Glo1 | 109801 | HITS-CLIP | | 23142080 | Unspecified |
| mmu-let-7i-5p | MIMAT0000122 | Glod4 | 67201 | HITS-CLIP | 23142080\|29167373 | | Unspecified |
| mmu-let-7i-5p | MIMAT0000122 | Glrb | 14658 | HITS-CLIP | | 19536157 | Unspecified |
| mmu-let-7i-5p | MIMAT0000122 | Glrx | 93692 | HITS-CLIP | 37986224\|19536157\|23142080\|29167 | | Unspecified |
| mmu-let-7i-5p | MIMAT0000122 | Glrx2 | 69367 | HITS-CLIP | 23142080\|29167373 | | Unspecified |
| mmu-let-7i-5p | MIMAT0000122 | Glrx3 | 30926 | HITS-CLIP | | 23142080 | Unspecified |
| mmu-let-7i-5p | MIMAT0000122 | Glrx5 | 73046 | HITS-CLIP | | 23597149 | Unspecified |
| mmu-let-7i-5p | MIMAT0000122 | Gls | 14660 | HITS-CLIP | 37986224\|19536157\|23142080 | | Unspecified |
| mmu-let-7i-5p | MIMAT0000122 | Gls2 | 216456 | HITS-CLIP | 37986224\|23597149 | | Unspecified |
| mmu-let-7i-5p | MIMAT0000122 | Glt8d1 | 76485 | HITS-CLIP | | 19536157 | Unspecified |
| mmu-let-7i-5p | MIMAT0000122 | Gltp | 56356 | HITS-CLIP | 19536157\|23142080\|31433990\|29167 | | Unspecified |
| mmu-let-7i-5p | MIMAT0000122 | Glud1 | 14661 | HITS-CLIP | 23597149\|23142080 | | Unspecified |
| mmu-let-7i-5p | MIMAT0000122 | Glul | 14645 | HITS-CLIP | | 23597149 | Unspecified |
| mmu-let-7i-5p | MIMAT0000122 | Glyctk | 235582 | HITS-CLIP | | 23597149 | Unspecified |
| mmu-let-7i-5p | MIMAT0000122 | Glyr1 | 74022 | HITS-CLIP | | 23597149 | Unspecified |
| mmu-let-7i-5p | MIMAT0000122 | Gm10030 | 791282 | HITS-CLIP | | 37986224 | Unspecified |
| mmu-let-7i-5p | MIMAT0000122 | Gm10184 | 100043906 | qCLASH | | 31393953 | Unspecified |
| mmu-let-7i-5p | MIMAT0000122 | Gm11273 | 100040340 | HITS-CLIP | 23597149\|29167373 | | Unspecified |
| mmu-let-7i-5p | MIMAT0000122 | Gm16286 | 117903916 | HITS-CLIP | | 23597149 | Unspecified |
| mmu-let-7i-5p | MIMAT0000122 | Gm2000 | 100038991 | HITS-CLIP | | 26203562 | Unspecified |
| mmu-let-7i-5p | MIMAT0000122 | Gm20390 | 127232040 | HITS-CLIP | | 31433990 | Unspecified |
| mmu-let-7i-5p | MIMAT0000122 | Gm20441 | 6608650 | HITS-CLIP | | 23597149 | Unspecified |
| mmu-let-7i-5p | MIMAT0000122 | Gm20498 | 105940408 | HITS-CLIP | | 29167373 | Unspecified |
| mmu-let-7i-5p | MIMAT0000122 | Gm20503 | 6608537 | HITS-CLIP | | 23142080 | Unspecified |

| | | | | | | |
|---|---|---|---|---|---|---|
| mmu-let-7i-5p | MIMAT0000122 | Gm20517 | 112694759 | HITS-CLIP | 31433990 | Unspecified |
| mmu-let-7i-5p | MIMAT0000122 | Gm20547 | 6608472 | HITS-CLIP | 23597149 | Unspecified |
| mmu-let-7i-5p | MIMAT0000122 | Gm20605 | 100316903 | qCLASH | 31393953 | Unspecified |
| mmu-let-7i-5p | MIMAT0000122 | Gm2a | 14667 | HITS-CLIP | 23142080\|29167373 | Unspecified |
| mmu-let-7i-5p | MIMAT0000122 | Gm35315 | 102638847 | HITS-CLIP | 23142080 | Unspecified |
| mmu-let-7i-5p | MIMAT0000122 | Gm37240 | | HITS-CLIP | 23597149 | Unspecified |
| mmu-let-7i-5p | MIMAT0000122 | Gm38394 | 667118 | HITS-CLIP | 23142080\|23597149\|19536157 | Unspecified |
| mmu-let-7i-5p | MIMAT0000122 | Gm4070 | 100042856 | HITS-CLIP | 29167373 | Unspecified |
| mmu-let-7i-5p | MIMAT0000122 | Gm4353 | 100043313 | HITS-CLIP | 23142080 | Unspecified |
| mmu-let-7i-5p | MIMAT0000122 | Gm44805 | 105886299 | HITS-CLIP | 23597149\|29167373 | Unspecified |
| mmu-let-7i-5p | MIMAT0000122 | Gm45713 | 103691134 | HITS-CLIP | 31433990 | Unspecified |
| mmu-let-7i-5p | MIMAT0000122 | Gm4756 | 209183 | HITS-CLIP | 23597149 | Unspecified |
| mmu-let-7i-5p | MIMAT0000122 | Gm4788 | 214403 | HITS-CLIP | 23597149 | Unspecified |
| mmu-let-7i-5p | MIMAT0000122 | Gm49336 | 127230315 | HITS-CLIP | 23597149\|23142080\|29167373 | Unspecified |
| mmu-let-7i-5p | MIMAT0000122 | Gm49339 | | HITS-CLIP | 29167373 | Unspecified |
| mmu-let-7i-5p | MIMAT0000122 | Gm49368 | 118261720 | HITS-CLIP | 29167373 | Unspecified |
| mmu-let-7i-5p | MIMAT0000122 | Gm49380 | | HITS-CLIP | 23142080 | Unspecified |
| mmu-let-7i-5p | MIMAT0000122 | Gm4951 | 240327 | HITS-CLIP | 23597149 | Unspecified |
| mmu-let-7i-5p | MIMAT0000122 | Gm49510 | | HITS-CLIP | 23142080 | Unspecified |
| mmu-let-7i-5p | MIMAT0000122 | Gm4952 | 240549 | HITS-CLIP | 23597149 | Unspecified |
| mmu-let-7i-5p | MIMAT0000122 | Gm5096 | 329008 | HITS-CLIP | 23597149 | Unspecified |
| mmu-let-7i-5p | MIMAT0000122 | Gm6377 | 622976 | HITS-CLIP | 23142080 | Unspecified |
| mmu-let-7i-5p | MIMAT0000122 | Gm7694 | 665574 | HITS-CLIP | 23597149 | Unspecified |
| mmu-let-7i-5p | MIMAT0000122 | Gmip | 78816 | HITS-CLIP | 23142080 | Unspecified |
| mmu-let-7i-5p | MIMAT0000122 | Gmppa | 69080 | HITS-CLIP | 23597149 | Unspecified |
| mmu-let-7i-5p | MIMAT0000122 | Gmppb | 331026 | HITS-CLIP | 23142080 | Unspecified |
| mmu-let-7i-5p | MIMAT0000122 | Gmps | 229363 | HITS-CLIP | 29167373\|23142080 | Unspecified |
| mmu-let-7i-5p | MIMAT0000122 | Gna12 | 14673 | HITS-CLIP | 23597149 | Unspecified |
| mmu-let-7i-5p | MIMAT0000122 | Gna13 | 14674 | HITS-CLIP | 23142080\|31433990 | Unspecified |
| mmu-let-7i-5p | MIMAT0000122 | Gnai2 | 14678 | HITS-CLIP | 23142080 | Unspecified |
| mmu-let-7i-5p | MIMAT0000122 | Gnai3 | 14679 | HITS-CLIP | 23597149\|23142080\|29167373 | Unspecified |
| mmu-let-7i-5p | MIMAT0000122 | Gnal | 14680 | HITS-CLIP | 19536157 | Unspecified |
| mmu-let-7i-5p | MIMAT0000122 | Gnao1 | 14681 | HITS-CLIP | 19536157 | Unspecified |
| mmu-let-7i-5p | MIMAT0000122 | Gnaq | 14682 | HITS-CLIP | 37986224\|23142080\|19536157\|29167 | Unspecified |
| mmu-let-7i-5p | MIMAT0000122 | Gnas | 14683 | HITS-CLIP | 23597149\|23142080 | Unspecified |
| mmu-let-7i-5p | MIMAT0000122 | Gnb1 | 14688 | HITS-CLIP | 26203562 | Unspecified |
| mmu-let-7i-5p | MIMAT0000122 | Gnb2 | 14693 | HITS-CLIP | 23597149\|19536157\|23142080\|29167 | Unspecified |
| mmu-let-7i-5p | MIMAT0000122 | Gne | 50798 | HITS-CLIP | 23597149 | Unspecified |
| mmu-let-7i-5p | MIMAT0000122 | Gng2 | 14702 | HITS-CLIP | 29167373 | Unspecified |

| | | | | | | |
|---|---|---|---|---|---|---|
| mmu-let-7i-5p | MIMAT0000122 | Gng5 | 14707 | qCLASH | 31393953 | Unspecified |
| mmu-let-7i-5p | MIMAT0000122 | Gnl3 | 30877 | HITS-CLIP | 23597149 | Unspecified |
| mmu-let-7i-5p | MIMAT0000122 | Gnpat | 14712 | HITS-CLIP | 29167373 | Unspecified |
| mmu-let-7i-5p | MIMAT0000122 | Gnpda1 | 26384 | HITS-CLIP | 29167373 | Unspecified |
| mmu-let-7i-5p | MIMAT0000122 | Gnptab | 432486 | HITS-CLIP | 37986224\|19536157\|23142080\|29167 | Unspecified |
| mmu-let-7i-5p | MIMAT0000122 | Gns | 75612 | HITS-CLIP | 37986224\|23597149\|23142080\|29167 | Unspecified |
| mmu-let-7i-5p | MIMAT0000122 | Golga2 | 99412 | HITS-CLIP | 23142080 | Unspecified |
| mmu-let-7i-5p | MIMAT0000122 | Golga4 | 54214 | HITS-CLIP | 37986224\|23597149\|23142080\|29167 | Unspecified |
| mmu-let-7i-5p | MIMAT0000122 | Golga7 | 57437 | HITS-CLIP | 37986224\|23597149\|23142080 | Unspecified |
| mmu-let-7i-5p | MIMAT0000122 | Golgb1 | 224139 | HITS-CLIP | 37986224\|23597149\|23142080 | Unspecified |
| mmu-let-7i-5p | MIMAT0000122 | Golim4 | 73124 | HITS-CLIP | 23142080 | Unspecified |
| mmu-let-7i-5p | MIMAT0000122 | Golm1 | 105348 | HITS-CLIP | 23142080 | Unspecified |
| mmu-let-7i-5p | MIMAT0000122 | Golm2 | 113201 | HITS-CLIP | 31433990 | Unspecified |
| mmu-let-7i-5p | MIMAT0000122 | Golph3 | 66629 | HITS-CLIP | 37986224\|23142080\|23597149 | Unspecified |
| mmu-let-7i-5p | MIMAT0000122 | Golph3l | 229593 | HITS-CLIP | 23597149\|29167373 | Unspecified |
| mmu-let-7i-5p | MIMAT0000122 | Golt1b | 66964 | HITS-CLIP | 37986224\|23597149\|23142080\|29167 | Unspecified |
| mmu-let-7i-5p | MIMAT0000122 | Gon4l | 76022 | HITS-CLIP | 23597149\|29167373 | Unspecified |
| mmu-let-7i-5p | MIMAT0000122 | Gopc | 94221 | HITS-CLIP | 23597149 | Unspecified |
| mmu-let-7i-5p | MIMAT0000122 | Gorab | 98376 | HITS-CLIP | 23142080 | Unspecified |
| mmu-let-7i-5p | MIMAT0000122 | Gorasp2 | 70231 | HITS-CLIP | 23142080 | Unspecified |
| mmu-let-7i-5p | MIMAT0000122 | Gosr1 | 53334 | HITS-CLIP | 23597149\|23142080 | Unspecified |
| mmu-let-7i-5p | MIMAT0000122 | Gosr2 | 56494 | HITS-CLIP | 23597149\|23142080 | Unspecified |
| mmu-let-7i-5p | MIMAT0000122 | Got1 | 14718 | HITS-CLIP | 37986224\|23597149\|23142080\|19536 | Unspecified |
| mmu-let-7i-5p | MIMAT0000122 | Got2 | 14719 | HITS-CLIP | 23597149 | Unspecified |
| mmu-let-7i-5p | MIMAT0000122 | Gpaa1 | 14731 | HITS-CLIP | 23597149\|23142080 | Unspecified |
| mmu-let-7i-5p | MIMAT0000122 | Gpalpp1 | 67467 | HITS-CLIP | 23142080 | Unspecified |
| mmu-let-7i-5p | MIMAT0000122 | Gpam | 14732 | HITS-CLIP | 37986224\|23142080\|23597149\|19536 | Unspecified |
| mmu-let-7i-5p | MIMAT0000122 | Gpat2 | 215456 | HITS-CLIP | 23597149 | Unspecified |
| mmu-let-7i-5p | MIMAT0000122 | Gpat4 | 137964 | HITS-CLIP | 23142080\|23597149\|29167373 | Unspecified |
| mmu-let-7i-5p | MIMAT0000122 | Gpatch1 | 67471 | HITS-CLIP | 23142080 | Unspecified |
| mmu-let-7i-5p | MIMAT0000122 | Gpatch2 | 67769 | HITS-CLIP//qCLASH | 37986224\|19536157\|23597149\|31393 | Unspecified |
| mmu-let-7i-5p | MIMAT0000122 | Gpatch3 | 242691 | HITS-CLIP | 37986224\|23142080 | Unspecified |
| mmu-let-7i-5p | MIMAT0000122 | Gpatch8 | 237943 | HITS-CLIP | 37986224\|23142080\|23597149\|29167 | Unspecified |
| mmu-let-7i-5p | MIMAT0000122 | Gpbp1 | 73274 | HITS-CLIP | 23142080 | Unspecified |
| mmu-let-7i-5p | MIMAT0000122 | Gpbp1l1 | 77110 | HITS-CLIP | 23142080 | Unspecified |
| mmu-let-7i-5p | MIMAT0000122 | Gpc1 | 14733 | HITS-CLIP | 26203562 | Unspecified |
| mmu-let-7i-5p | MIMAT0000122 | Gpc4 | 14735 | HITS-CLIP | 23597149 | Unspecified |
| mmu-let-7i-5p | MIMAT0000122 | Gpcpd1 | 74182 | HITS-CLIP | 37986224\|19536157\|23142080 | Unspecified |
| mmu-let-7i-5p | MIMAT0000122 | Gpd1 | 14555 | HITS-CLIP | 23597149 | Unspecified |

| | | | | | | |
|---|---|---|---|---|---|---|
| mmu-let-7i-5p | MIMAT0000122 | Gpd2 | 14571 | HITS-CLIP | 23597149\|23142080\|29167373 | Unspecified |
| mmu-let-7i-5p | MIMAT0000122 | Gphn | 268566 | HITS-CLIP | 37986224\|19536157\|23597149\|23142 | Unspecified |
| mmu-let-7i-5p | MIMAT0000122 | Gpld1 | 14756 | HITS-CLIP | 23597149 | Unspecified |
| mmu-let-7i-5p | MIMAT0000122 | Gpm6a | 234267 | HITS-CLIP | 37986224\|19536157 | Unspecified |
| mmu-let-7i-5p | MIMAT0000122 | Gpm6b | 14758 | HITS-CLIP | 19536157\|23142080 | Unspecified |
| mmu-let-7i-5p | MIMAT0000122 | Gpn2 | 100210 | HITS-CLIP | 23142080 | Unspecified |
| mmu-let-7i-5p | MIMAT0000122 | Gpnmb | 93695 | HITS-CLIP | 29167373 | Unspecified |
| mmu-let-7i-5p | MIMAT0000122 | Gpr108 | 78308 | HITS-CLIP | 23142080 | Unspecified |
| mmu-let-7i-5p | MIMAT0000122 | Gpr132 | 56696 | HITS-CLIP | 37986224\|23142080 | Unspecified |
| mmu-let-7i-5p | MIMAT0000122 | Gpr137b | 83924 | HITS-CLIP | 37986224\|29167373 | Unspecified |
| mmu-let-7i-5p | MIMAT0000122 | Gpr137c | 70713 | HITS-CLIP | 37986224 | Unspecified |
| mmu-let-7i-5p | MIMAT0000122 | Gpr146 | 80290 | HITS-CLIP | 23597149\|31433990\|23142080\|29167 | Unspecified |
| mmu-let-7i-5p | MIMAT0000122 | Gpr155 | 68526 | HITS-CLIP | 23142080 | Unspecified |
| mmu-let-7i-5p | MIMAT0000122 | Gpr157 | 269604 | HITS-CLIP | 23142080 | Unspecified |
| mmu-let-7i-5p | MIMAT0000122 | Gpr158 | 241263 | HITS-CLIP | 19536157 | Unspecified |
| mmu-let-7i-5p | MIMAT0000122 | Gpr17 | 574402 | HITS-CLIP | 19536157 | Unspecified |
| mmu-let-7i-5p | MIMAT0000122 | Gpr171 | 229323 | HITS-CLIP | 23142080 | Unspecified |
| mmu-let-7i-5p | MIMAT0000122 | Gpr174 | 213439 | HITS-CLIP | 23142080 | Unspecified |
| mmu-let-7i-5p | MIMAT0000122 | Gpr183 | 321019 | HITS-CLIP | 23142080 | Unspecified |
| mmu-let-7i-5p | MIMAT0000122 | Gpr22 | 73010 | HITS-CLIP | 37986224\|19536157 | Unspecified |
| mmu-let-7i-5p | MIMAT0000122 | Gpr26 | 233919 | HITS-CLIP | 37986224\|19536157 | Unspecified |
| mmu-let-7i-5p | MIMAT0000122 | Gpr34 | 23890 | HITS-CLIP | 19536157\|23142080 | Unspecified |
| mmu-let-7i-5p | MIMAT0000122 | Gpr35 | 64095 | HITS-CLIP | 23597149 | Unspecified |
| mmu-let-7i-5p | MIMAT0000122 | Gpr37l1 | 171469 | HITS-CLIP | 37986224 | Unspecified |
| mmu-let-7i-5p | MIMAT0000122 | Gpr63 | 81006 | HITS-CLIP | 19536157 | Unspecified |
| mmu-let-7i-5p | MIMAT0000122 | Gpr85 | 64450 | HITS-CLIP | 37986224\|19536157 | Unspecified |
| mmu-let-7i-5p | MIMAT0000122 | Gprc5b | 64297 | HITS-CLIP | 19536157 | Unspecified |
| mmu-let-7i-5p | MIMAT0000122 | Gprc5c | 70355 | HITS-CLIP | 37986224\|23597149 | Unspecified |
| mmu-let-7i-5p | MIMAT0000122 | Gprin3 | 243385 | HITS-CLIP | 23142080 | Unspecified |
| mmu-let-7i-5p | MIMAT0000122 | Gps2 | 56310 | HITS-CLIP | 37986224\|23597149 | Unspecified |
| mmu-let-7i-5p | MIMAT0000122 | Gpsm2 | 76123 | HITS-CLIP | 23142080 | Unspecified |
| mmu-let-7i-5p | MIMAT0000122 | Gpt | 76282 | HITS-CLIP | 23597149 | Unspecified |
| mmu-let-7i-5p | MIMAT0000122 | Gpt2 | 108682 | HITS-CLIP | 23597149 | Unspecified |
| mmu-let-7i-5p | MIMAT0000122 | Gpx1 | 14775 | HITS-CLIP | 23597149\|23142080 | Unspecified |
| mmu-let-7i-5p | MIMAT0000122 | Gramd1b | 235283 | HITS-CLIP | 37986224\|19536157\|31433990\|23597 | Unspecified |
| mmu-let-7i-5p | MIMAT0000122 | Gramd1c | 207798 | HITS-CLIP | 37986224\|23597149\|23142080 | Unspecified |
| mmu-let-7i-5p | MIMAT0000122 | Gramd3 | 107022 | HITS-CLIP | 23142080\|23597149 | Unspecified |
| mmu-let-7i-5p | MIMAT0000122 | Gramd4 | 223752 | HITS-CLIP | 23142080 | Unspecified |
| mmu-let-7i-5p | MIMAT0000122 | Grb14 | 50915 | HITS-CLIP | 23597149 | Unspecified |

| | | | | | | |
|---|---|---|---|---|---|---|
| mmu-let-7i-5p | MIMAT0000122 | Grb2 | 14784 | HITS-CLIP | 23142080\|19536157\|29167373 | Unspecified |
| mmu-let-7i-5p | MIMAT0000122 | Grb7 | 14786 | HITS-CLIP | 23597149 | Unspecified |
| mmu-let-7i-5p | MIMAT0000122 | Greb1l | 381157 | HITS-CLIP | 23630078 | Unspecified |
| mmu-let-7i-5p | MIMAT0000122 | Grem2 | 23893 | HITS-CLIP | 37986224 | Unspecified |
| mmu-let-7i-5p | MIMAT0000122 | Grhpr | 76238 | HITS-CLIP | 23597149 | Unspecified |
| mmu-let-7i-5p | MIMAT0000122 | Gria1 | 14799 | HITS-CLIP | 19536157 | Unspecified |
| mmu-let-7i-5p | MIMAT0000122 | Gria2 | 14800 | HITS-CLIP | 19536157 | Unspecified |
| mmu-let-7i-5p | MIMAT0000122 | Grik2 | 14806 | HITS-CLIP | 37986224\|19536157 | Unspecified |
| mmu-let-7i-5p | MIMAT0000122 | Grin1 | 14810 | HITS-CLIP | 37986224 | Unspecified |
| mmu-let-7i-5p | MIMAT0000122 | Grin2a | 14811 | HITS-CLIP | 19536157 | Unspecified |
| mmu-let-7i-5p | MIMAT0000122 | Grin2b | 14812 | HITS-CLIP | 37986224\|19536157 | Unspecified |
| mmu-let-7i-5p | MIMAT0000122 | Grina | 66168 | HITS-CLIP | 23597149\|23142080 | Unspecified |
| mmu-let-7i-5p | MIMAT0000122 | Gripap1 | 54645 | HITS-CLIP | 29167373 | Unspecified |
| mmu-let-7i-5p | MIMAT0000122 | Grk2 | 156 | HITS-CLIP | 23142080\|29167373 | Unspecified |
| mmu-let-7i-5p | MIMAT0000122 | Grk3 | 320129 | HITS-CLIP | 29167373 | Unspecified |
| mmu-let-7i-5p | MIMAT0000122 | Grk6 | 26385 | HITS-CLIP | 23142080 | Unspecified |
| mmu-let-7i-5p | MIMAT0000122 | Grm1 | 14816 | HITS-CLIP | 19536157 | Unspecified |
| mmu-let-7i-5p | MIMAT0000122 | Grm3 | 108069 | HITS-CLIP | 37986224\|19536157 | Unspecified |
| mmu-let-7i-5p | MIMAT0000122 | Grn | 14824 | HITS-CLIP | 23597149\|29167373\|23142080 | Unspecified |
| mmu-let-7i-5p | MIMAT0000122 | Grpel1 | 17713 | HITS-CLIP | 23597149\|29167373\|23142080 | Unspecified |
| mmu-let-7i-5p | MIMAT0000122 | Grsf1 | 231413 | HITS-CLIP | 23142080 | Unspecified |
| mmu-let-7i-5p | MIMAT0000122 | Gsap | 212167 | HITS-CLIP | 23597149\|29167373 | Unspecified |
| mmu-let-7i-5p | MIMAT0000122 | Gse1 | 382034 | HITS-CLIP | 23142080 | Unspecified |
| mmu-let-7i-5p | MIMAT0000122 | Gsk3a | 606496 | HITS-CLIP | 23597149\|23142080 | Unspecified |
| mmu-let-7i-5p | MIMAT0000122 | Gsk3b | 56637 | HITS-CLIP | 23597149\|23142080 | Unspecified |
| mmu-let-7i-5p | MIMAT0000122 | Gsn | 227753 | HITS-CLIP | 23142080 | Unspecified |
| mmu-let-7i-5p | MIMAT0000122 | Gsr | 14782 | HITS-CLIP | 23597149\|23142080 | Unspecified |
| mmu-let-7i-5p | MIMAT0000122 | Gss | 14854 | HITS-CLIP | 23597149 | Unspecified |
| mmu-let-7i-5p | MIMAT0000122 | Gsta3 | 14859 | HITS-CLIP | 23597149 | Unspecified |
| mmu-let-7i-5p | MIMAT0000122 | Gsta4 | 14860 | HITS-CLIP | 23597149 | Unspecified |
| mmu-let-7i-5p | MIMAT0000122 | Gstm1 | 14862 | HITS-CLIP | 23597149 | Unspecified |
| mmu-let-7i-5p | MIMAT0000122 | Gstm5 | 14866 | HITS-CLIP | 19536157 | Unspecified |
| mmu-let-7i-5p | MIMAT0000122 | Gstm7 | 68312 | HITS-CLIP | 23597149 | Unspecified |
| mmu-let-7i-5p | MIMAT0000122 | Gstp3 | 225884 | HITS-CLIP | 23142080 | Unspecified |
| mmu-let-7i-5p | MIMAT0000122 | Gstt1 | 14871 | HITS-CLIP | 23597149\|23142080 | Unspecified |
| mmu-let-7i-5p | MIMAT0000122 | Gstt3 | 103140 | HITS-CLIP | 23597149 | Unspecified |
| mmu-let-7i-5p | MIMAT0000122 | Gstz1 | 14874 | HITS-CLIP | 23597149\|23142080 | Unspecified |
| mmu-let-7i-5p | MIMAT0000122 | Gtf2a1 | 83602 | HITS-CLIP | 23597149\|23142080\|29167373 | Unspecified |
| mmu-let-7i-5p | MIMAT0000122 | Gtf2a2 | 235459 | HITS-CLIP | 37986224\|23142080 | Unspecified |

| | | | | | | |
|---|---|---|---|---|---|---|
| mmu-let-7i-5p | MIMAT0000122 | Gtf2f1 | 98053 | HITS-CLIP | 23142080 | Unspecified |
| mmu-let-7i-5p | MIMAT0000122 | Gtf2f2 | 68705 | HITS-CLIP | 23142080 | Unspecified |
| mmu-let-7i-5p | MIMAT0000122 | Gtf2h1 | 14884 | HITS-CLIP | 23142080 | Unspecified |
| mmu-let-7i-5p | MIMAT0000122 | Gtf2i | 14886 | HITS-CLIP | 23597149\|23142080 | Unspecified |
| mmu-let-7i-5p | MIMAT0000122 | Gtf3a | 66596 | HITS-CLIP | 23142080 | Unspecified |
| mmu-let-7i-5p | MIMAT0000122 | Gtf3c2 | 71752 | HITS-CLIP | 37986224\|23597149\|23142080 | Unspecified |
| mmu-let-7i-5p | MIMAT0000122 | Gtf3c3 | 98488 | HITS-CLIP | 23142080 | Unspecified |
| mmu-let-7i-5p | MIMAT0000122 | Gtf3c4 | 269252 | HITS-CLIP | 30971389 | Unspecified |
| mmu-let-7i-5p | MIMAT0000122 | Gtpbp8 | 66067 | HITS-CLIP | 23597149 | Unspecified |
| mmu-let-7i-5p | MIMAT0000122 | Gtse1 | 29870 | HITS-CLIP | 23142080\|23597149 | Unspecified |
| mmu-let-7i-5p | MIMAT0000122 | Gucd1 | 68778 | HITS-CLIP | 23597149\|19536157\|29167373 | Unspecified |
| mmu-let-7i-5p | MIMAT0000122 | Gucy1a1 | 60596 | HITS-CLIP | 23142080 | Unspecified |
| mmu-let-7i-5p | MIMAT0000122 | Gucy2e | 14919 | HITS-CLIP | 23142080 | Unspecified |
| mmu-let-7i-5p | MIMAT0000122 | Gulo | 268756 | HITS-CLIP | 23597149 | Unspecified |
| mmu-let-7i-5p | MIMAT0000122 | Gusb | 110006 | HITS-CLIP | 37986224\|23597149\|23142080\|29167 | Unspecified |
| mmu-let-7i-5p | MIMAT0000122 | Gxylt1 | 223827 | HITS-CLIP | 37986224\|23597149\|23142080\|29167 | Unspecified |
| mmu-let-7i-5p | MIMAT0000122 | Gyg | 27357 | HITS-CLIP | 23142080 | Unspecified |
| mmu-let-7i-5p | MIMAT0000122 | Gypc | 71683 | HITS-CLIP | 23597149 | Unspecified |
| mmu-let-7i-5p | MIMAT0000122 | Gys1 | 14936 | HITS-CLIP | 23142080 | Unspecified |
| mmu-let-7i-5p | MIMAT0000122 | Gzmb | 14939 | HITS-CLIP | 23142080 | Unspecified |
| mmu-let-7i-5p | MIMAT0000122 | H13 | 14950 | HITS-CLIP | 37986224\|23597149\|23142080 | Unspecified |
| mmu-let-7i-5p | MIMAT0000122 | H1f0 | 14958 | HITS-CLIP | 37986224\|19536157 | Unspecified |
| mmu-let-7i-5p | MIMAT0000122 | H1f4 | 3008 | qCLASH | 31393953 | Unspecified |
| mmu-let-7i-5p | MIMAT0000122 | H2-Ab1 | 14961 | HITS-CLIP | 23597149\|23142080\|29167373 | Unspecified |
| mmu-let-7i-5p | MIMAT0000122 | H2-DMa | 14998 | HITS-CLIP | 23142080 | Unspecified |
| mmu-let-7i-5p | MIMAT0000122 | H2-K1 | 14972 | HITS-CLIP | 23597149 | Unspecified |
| mmu-let-7i-5p | MIMAT0000122 | H2-Ob | 15002 | HITS-CLIP | 23142080 | Unspecified |
| mmu-let-7i-5p | MIMAT0000122 | H2-Q4 | 15015 | HITS-CLIP | 23142080 | Unspecified |
| mmu-let-7i-5p | MIMAT0000122 | H2-Q6 | 110557 | HITS-CLIP | 23142080 | Unspecified |
| mmu-let-7i-5p | MIMAT0000122 | H2ac20 | 8338 | HITS-CLIP | 31433990 | Unspecified |
| mmu-let-7i-5p | MIMAT0000122 | H2afv | 77605 | HITS-CLIP | 23142080 | Unspecified |
| mmu-let-7i-5p | MIMAT0000122 | H2afx | 15270 | HITS-CLIP | 37986224\|23142080 | Unspecified |
| mmu-let-7i-5p | MIMAT0000122 | H2afy | 9555 | HITS-CLIP | 23142080\|29167373 | Unspecified |
| mmu-let-7i-5p | MIMAT0000122 | H2bc18 | 440689 | HITS-CLIP | 31433990 | Unspecified |
| mmu-let-7i-5p | MIMAT0000122 | H2bc3 | 3018 | HITS-CLIP | 31433990\|23630078 | Unspecified |
| mmu-let-7i-5p | MIMAT0000122 | H3f3b | 15081 | HITS-CLIP | 37986224\|19536157\|23597149\|23142 | Unspecified |
| mmu-let-7i-5p | MIMAT0000122 | H6pd | 100198 | HITS-CLIP | 37986224\|23597149\|23142080\|29167 | Unspecified |
| mmu-let-7i-5p | MIMAT0000122 | Habp2 | 226243 | HITS-CLIP | 23597149 | Unspecified |
| mmu-let-7i-5p | MIMAT0000122 | Habp4 | 56541 | HITS-CLIP | 23597149\|23142080 | Unspecified |

| | | | | | | |
|---|---|---|---|---|---|---|
| mmu-let-7i-5p | MIMAT0000122 | Hacd3 | 57874 | HITS-CLIP | 19536157\|23597149\|29167373\|23142 | Unspecified |
| mmu-let-7i-5p | MIMAT0000122 | Hadha | 97212 | HITS-CLIP | 23597149 | Unspecified |
| mmu-let-7i-5p | MIMAT0000122 | Hagh | 14651 | HITS-CLIP | 23597149 | Unspecified |
| mmu-let-7i-5p | MIMAT0000122 | Hal | 15109 | HITS-CLIP | 23597149 | Unspecified |
| mmu-let-7i-5p | MIMAT0000122 | Hamp | 84506 | HITS-CLIP | 23597149 | Unspecified |
| mmu-let-7i-5p | MIMAT0000122 | Hao1 | 15112 | HITS-CLIP | 37986224\|23597149 | Unspecified |
| mmu-let-7i-5p | MIMAT0000122 | Hapln1 | 12950 | HITS-CLIP | 37986224\|19536157 | Unspecified |
| mmu-let-7i-5p | MIMAT0000122 | Harbi1 | 241547 | HITS-CLIP | 23597149 | Unspecified |
| mmu-let-7i-5p | MIMAT0000122 | Haspin | 83903 | HITS-CLIP | 30971389\|23142080 | Unspecified |
| mmu-let-7i-5p | MIMAT0000122 | Hat1 | 107435 | HITS-CLIP | 23142080 | Unspecified |
| mmu-let-7i-5p | MIMAT0000122 | Haus3 | 231123 | HITS-CLIP | 23142080 | Unspecified |
| mmu-let-7i-5p | MIMAT0000122 | Haus4 | 219072 | HITS-CLIP | 23142080 | Unspecified |
| mmu-let-7i-5p | MIMAT0000122 | Haus6 | 230376 | HITS-CLIP | 37986224\|23142080\|29167373 | Unspecified |
| mmu-let-7i-5p | MIMAT0000122 | Hba-a1 | 15122 | HITS-CLIP | 19536157\|23597149 | Unspecified |
| mmu-let-7i-5p | MIMAT0000122 | Hba-a2 | 110257 | HITS-CLIP | 23597149 | Unspecified |
| mmu-let-7i-5p | MIMAT0000122 | Hbp1 | 73389 | HITS-CLIP//qCLASH | 37986224\|23142080\|31393953\|23597 | Unspecified |
| mmu-let-7i-5p | MIMAT0000122 | Hc | 15139 | HITS-CLIP | 23597149 | Unspecified |
| mmu-let-7i-5p | MIMAT0000122 | Hccs | 15159 | HITS-CLIP | 23597149\|23142080 | Unspecified |
| mmu-let-7i-5p | MIMAT0000122 | Hcfc1 | 15161 | HITS-CLIP | 23597149\|23142080\|29167373 | Unspecified |
| mmu-let-7i-5p | MIMAT0000122 | Hcfc2 | 67933 | HITS-CLIP | 37986224\|23142080\|19536157\|23597 | Unspecified |
| mmu-let-7i-5p | MIMAT0000122 | Hcls1 | 15163 | HITS-CLIP | 23142080 | Unspecified |
| mmu-let-7i-5p | MIMAT0000122 | Hcn1 | 15165 | HITS-CLIP | 19536157 | Unspecified |
| mmu-let-7i-5p | MIMAT0000122 | Hcn3 | 15168 | qCLASH | 31393953 | Unspecified |
| mmu-let-7i-5p | MIMAT0000122 | Hcst | 23900 | HITS-CLIP | 23142080 | Unspecified |
| mmu-let-7i-5p | MIMAT0000122 | Hdac3 | 15183 | HITS-CLIP | 37986224\|19536157\|23597149\|23142 | Unspecified |
| mmu-let-7i-5p | MIMAT0000122 | Hdac4 | 208727 | HITS-CLIP | 23142080 | Unspecified |
| mmu-let-7i-5p | MIMAT0000122 | Hdac5 | 15184 | HITS-CLIP | 23142080 | Unspecified |
| mmu-let-7i-5p | MIMAT0000122 | Hdac6 | 15185 | HITS-CLIP | 23142080 | Unspecified |
| mmu-let-7i-5p | MIMAT0000122 | Hdac7 | 56233 | HITS-CLIP | 23142080 | Unspecified |
| mmu-let-7i-5p | MIMAT0000122 | Hdc | 15186 | HITS-CLIP | 23142080 | Unspecified |
| mmu-let-7i-5p | MIMAT0000122 | Hddc2 | 69692 | HITS-CLIP | 37986224 | Unspecified |
| mmu-let-7i-5p | MIMAT0000122 | Hdhd2 | 76987 | HITS-CLIP | 23597149 | Unspecified |
| mmu-let-7i-5p | MIMAT0000122 | Hdlbp | 110611 | HITS-CLIP | 37986224\|23142080\|23597149 | Unspecified |
| mmu-let-7i-5p | MIMAT0000122 | Heatr5a | 320487 | HITS-CLIP | 29167373 | Unspecified |
| mmu-let-7i-5p | MIMAT0000122 | Heatr5b | 320473 | HITS-CLIP//qCLASH | 37986224\|31393953\|23142080 | Unspecified |
| mmu-let-7i-5p | MIMAT0000122 | Heca | 380629 | HITS-CLIP | 23142080 | Unspecified |
| mmu-let-7i-5p | MIMAT0000122 | Hectd1 | 207304 | HITS-CLIP | 37986224\|23142080\|19536157\|23597 | Unspecified |
| mmu-let-7i-5p | MIMAT0000122 | Hectd2 | 226098 | HITS-CLIP | 23142080 | Unspecified |
| mmu-let-7i-5p | MIMAT0000122 | Hectd4 | 283450 | HITS-CLIP | 19536157\|23142080 | Unspecified |

| | | | | | | |
|---|---|---|---|---|---|---|
| mmu-let-7i-5p | MIMAT0000122 | Heg1 | 77446 | HITS-CLIP | 37986224\|23142080\|23597149 | Unspecified |
| mmu-let-7i-5p | MIMAT0000122 | Helb | 117599 | HITS-CLIP | 37986224\|23142080 | Unspecified |
| mmu-let-7i-5p | MIMAT0000122 | Hells | 15201 | HITS-CLIP | 23142080\|29167373 | Unspecified |
| mmu-let-7i-5p | MIMAT0000122 | Helz | 78455 | HITS-CLIP | 19536157\|23142080\|29167373 | Unspecified |
| mmu-let-7i-5p | MIMAT0000122 | Helz2 | 229003 | HITS-CLIP | 23597149\|23142080 | Unspecified |
| mmu-let-7i-5p | MIMAT0000122 | Herc1 | 235439 | HITS-CLIP | 37986224\|23597149\|19536157\|29167 | Unspecified |
| mmu-let-7i-5p | MIMAT0000122 | Herc2 | 15204 | HITS-CLIP | 23142080\|23597149\|29167373 | Unspecified |
| mmu-let-7i-5p | MIMAT0000122 | Herc3 | 73998 | HITS-CLIP | 23142080 | Unspecified |
| mmu-let-7i-5p | MIMAT0000122 | Herc4 | 67345 | HITS-CLIP | 23142080\|23597149\|19536157 | Unspecified |
| mmu-let-7i-5p | MIMAT0000122 | Herpud1 | 64209 | HITS-CLIP//qCLASH | 23142080\|23597149\|31393953 | Unspecified |
| mmu-let-7i-5p | MIMAT0000122 | Herpud2 | 80517 | HITS-CLIP | 37986224\|23142080 | Unspecified |
| mmu-let-7i-5p | MIMAT0000122 | Hexim1 | 192231 | HITS-CLIP | 23142080 | Unspecified |
| mmu-let-7i-5p | MIMAT0000122 | Hgf | 15234 | HITS-CLIP | 23597149 | Unspecified |
| mmu-let-7i-5p | MIMAT0000122 | Hgfac | 54426 | HITS-CLIP | 23597149 | Unspecified |
| mmu-let-7i-5p | MIMAT0000122 | Hgs | 15239 | HITS-CLIP | 23142080 | Unspecified |
| mmu-let-7i-5p | MIMAT0000122 | Hhex | 15242 | HITS-CLIP | 23597149 | Unspecified |
| mmu-let-7i-5p | MIMAT0000122 | Hic1 | 15248 | HITS-CLIP | 23142080 | Unspecified |
| mmu-let-7i-5p | MIMAT0000122 | Hic2 | 58180 | HITS-CLIP | 37986224\|23597149\|23142080 | Unspecified |
| mmu-let-7i-5p | MIMAT0000122 | Hid1 | 217310 | HITS-CLIP | 19536157 | Unspecified |
| mmu-let-7i-5p | MIMAT0000122 | Hif1a | 15251 | HITS-CLIP | 37986224\|23597149\|23142080\|19536 | Unspecified |
| mmu-let-7i-5p | MIMAT0000122 | Hif1an | 319594 | HITS-CLIP//qCLASH | 37986224\|23597149\|19536157\|31393 | Unspecified |
| mmu-let-7i-5p | MIMAT0000122 | Hif3a | 53417 | HITS-CLIP | 23142080 | Unspecified |
| mmu-let-7i-5p | MIMAT0000122 | Hinfp | 102423 | HITS-CLIP | 31433990 | Unspecified |
| mmu-let-7i-5p | MIMAT0000122 | Hip1 | 215114 | HITS-CLIP | 37986224\|23597149\|19536157\|29167 | Unspecified |
| mmu-let-7i-5p | MIMAT0000122 | Hip1r | 29816 | HITS-CLIP | 37986224\|23597149 | Unspecified |
| mmu-let-7i-5p | MIMAT0000122 | Hipk1 | 15257 | HITS-CLIP | 23597149\|23142080\|29167373 | Unspecified |
| mmu-let-7i-5p | MIMAT0000122 | Hipk2 | 15258 | HITS-CLIP | 23142080\|23597149\|29167373 | Unspecified |
| mmu-let-7i-5p | MIMAT0000122 | Hipk3 | 15259 | HITS-CLIP | 37986224\|23597149\|19536157\|29167 | Unspecified |
| mmu-let-7i-5p | MIMAT0000122 | Hira | 15260 | HITS-CLIP | 23597149 | Unspecified |
| mmu-let-7i-5p | MIMAT0000122 | Hist1h1b | 3009 | HITS-CLIP | 23597149\|23142080 | Unspecified |
| mmu-let-7i-5p | MIMAT0000122 | Hist1h1c | 50708 | HITS-CLIP | 23142080 | Unspecified |
| mmu-let-7i-5p | MIMAT0000122 | Hist1h3a | 8350 | HITS-CLIP | 23142080 | Unspecified |
| mmu-let-7i-5p | MIMAT0000122 | Hist1h3i | 8354 | HITS-CLIP | 23142080 | Unspecified |
| mmu-let-7i-5p | MIMAT0000122 | Hivep1 | 110521 | HITS-CLIP | 19536157\|23597149 | Unspecified |
| mmu-let-7i-5p | MIMAT0000122 | Hivep2 | 15273 | HITS-CLIP | 37986224\|23142080\|23597149\|19536 | Unspecified |
| mmu-let-7i-5p | MIMAT0000122 | Hivep3 | 16656 | HITS-CLIP | 23142080 | Unspecified |
| mmu-let-7i-5p | MIMAT0000122 | Hjurp | 381280 | HITS-CLIP | 23597149\|23142080 | Unspecified |
| mmu-let-7i-5p | MIMAT0000122 | Hk1 | 15275 | HITS-CLIP | 23142080 | Unspecified |
| mmu-let-7i-5p | MIMAT0000122 | Hk2 | 15277 | HITS-CLIP | 23142080 | Unspecified |

| | | | | | | |
|---|---|---|---|---|---|---|
| mmu-let-7i-5p | MIMAT0000122 | Hlf | 217082 | HITS-CLIP | 37986224 | Unspecified |
| mmu-let-7i-5p | MIMAT0000122 | Hmbox1 | 219150 | HITS-CLIP | 23142080\|29167373 | Unspecified |
| mmu-let-7i-5p | MIMAT0000122 | Hmbs | 15288 | HITS-CLIP | 23142080 | Unspecified |
| mmu-let-7i-5p | MIMAT0000122 | Hmg20b | 10362 | HITS-CLIP | 23142080 | Unspecified |
| mmu-let-7i-5p | MIMAT0000122 | Hmga1 | 15361 | HITS-CLIP | 23142080\|29167373 | Unspecified |
| mmu-let-7i-5p | MIMAT0000122 | Hmga1b | 15361 | HITS-CLIP | 23142080 | Unspecified |
| mmu-let-7i-5p | MIMAT0000122 | Hmga2 | 15364 | HITS-CLIP | 29167373 | Unspecified |
| mmu-let-7i-5p | MIMAT0000122 | Hmgcl | 15356 | HITS-CLIP | 23597149\|23142080 | Unspecified |
| mmu-let-7i-5p | MIMAT0000122 | Hmgcr | 15357 | HITS-CLIP | 23597149\|23142080\|19536157 | Unspecified |
| mmu-let-7i-5p | MIMAT0000122 | Hmgcs1 | 208715 | HITS-CLIP | 37986224\|23142080\|19536157\|23597 | Unspecified |
| mmu-let-7i-5p | MIMAT0000122 | Hmgcs2 | 15360 | HITS-CLIP | 23597149\|19536157 | Unspecified |
| mmu-let-7i-5p | MIMAT0000122 | Hmgxb3 | 106894 | HITS-CLIP | 37986224 | Unspecified |
| mmu-let-7i-5p | MIMAT0000122 | Hmgxb4 | 70823 | HITS-CLIP | 23142080 | Unspecified |
| mmu-let-7i-5p | MIMAT0000122 | Hmmr | 15366 | HITS-CLIP | 23142080 | Unspecified |
| mmu-let-7i-5p | MIMAT0000122 | Hmox1 | 15368 | HITS-CLIP | 23597149\|29167373 | Unspecified |
| mmu-let-7i-5p | MIMAT0000122 | Hmox2 | 15369 | HITS-CLIP | 23597149\|23142080 | Unspecified |
| mmu-let-7i-5p | MIMAT0000122 | Hnf1a | 21405 | HITS-CLIP | 23597149 | Unspecified |
| mmu-let-7i-5p | MIMAT0000122 | Hnf4a | 15378 | HITS-CLIP | 37986224\|23597149 | Unspecified |
| mmu-let-7i-5p | MIMAT0000122 | Hnrnpa0 | 77134 | HITS-CLIP | 23142080\|29167373 | Unspecified |
| mmu-let-7i-5p | MIMAT0000122 | Hnrnpa2b1 | 53379 | HITS-CLIP | 31433990\|19536157\|23142080\|29167 | Unspecified |
| mmu-let-7i-5p | MIMAT0000122 | Hnrnpd | 11991 | HITS-CLIP | 23142080 | Unspecified |
| mmu-let-7i-5p | MIMAT0000122 | Hnrnph2 | 56258 | HITS-CLIP | 37986224 | Unspecified |
| mmu-let-7i-5p | MIMAT0000122 | Hnrnph3 | 432467 | HITS-CLIP | 23597149 | Unspecified |
| mmu-let-7i-5p | MIMAT0000122 | Hnrnpk | 15387 | HITS-CLIP | 23142080 | Unspecified |
| mmu-let-7i-5p | MIMAT0000122 | Hnrnpl | 15388 | HITS-CLIP | 23142080\|19536157\|29167373 | Unspecified |
| mmu-let-7i-5p | MIMAT0000122 | Hnrnpll | 72692 | HITS-CLIP | 23142080 | Unspecified |
| mmu-let-7i-5p | MIMAT0000122 | Hnrnpr | 74326 | HITS-CLIP | 29167373 | Unspecified |
| mmu-let-7i-5p | MIMAT0000122 | Hnrnpu | 51810 | HITS-CLIP | 37986224\|23597149\|23142080\|29167 | Unspecified |
| mmu-let-7i-5p | MIMAT0000122 | Hnrnpul1 | 232989 | HITS-CLIP | 23142080\|29167373 | Unspecified |
| mmu-let-7i-5p | MIMAT0000122 | Hnrnpul2 | 68693 | HITS-CLIP | 29167373 | Unspecified |
| mmu-let-7i-5p | MIMAT0000122 | Hoga1 | 67432 | HITS-CLIP | 23597149 | Unspecified |
| mmu-let-7i-5p | MIMAT0000122 | Homer1 | 26556 | HITS-CLIP | 23597149\|19536157\|23142080 | Unspecified |
| mmu-let-7i-5p | MIMAT0000122 | Hook1 | 77963 | HITS-CLIP | 19536157\|23142080 | Unspecified |
| mmu-let-7i-5p | MIMAT0000122 | Hook2 | 170833 | HITS-CLIP | 23597149 | Unspecified |
| mmu-let-7i-5p | MIMAT0000122 | Hp1bp3 | 15441 | HITS-CLIP | 23142080 | Unspecified |
| mmu-let-7i-5p | MIMAT0000122 | Hpca | 15444 | HITS-CLIP | 19536157 | Unspecified |
| mmu-let-7i-5p | MIMAT0000122 | Hpd | 15445 | HITS-CLIP | 23597149 | Unspecified |
| mmu-let-7i-5p | MIMAT0000122 | Hpgd | 15446 | HITS-CLIP | 23597149 | Unspecified |
| mmu-let-7i-5p | MIMAT0000122 | Hpn | 15451 | HITS-CLIP | 23597149 | Unspecified |

| | | | | | | |
|---|---|---|---|---|---|---|
| mmu-let-7i-5p | MIMAT0000122 | Hps1 | 192236 | HITS-CLIP | 23142080 | Unspecified |
| mmu-let-7i-5p | MIMAT0000122 | Hps3 | 12807 | HITS-CLIP | 23142080\|29167373 | Unspecified |
| mmu-let-7i-5p | MIMAT0000122 | Hps4 | 192232 | HITS-CLIP | 23597149 | Unspecified |
| mmu-let-7i-5p | MIMAT0000122 | Hps5 | 246694 | HITS-CLIP | 23142080 | Unspecified |
| mmu-let-7i-5p | MIMAT0000122 | Hr | 15460 | HITS-CLIP | 19536157 | Unspecified |
| mmu-let-7i-5p | MIMAT0000122 | Hrh3 | 99296 | HITS-CLIP | 37986224\|19536157 | Unspecified |
| mmu-let-7i-5p | MIMAT0000122 | Hs1bp3 | 58240 | HITS-CLIP | 37986224\|19536157\|23142080 | Unspecified |
| mmu-let-7i-5p | MIMAT0000122 | Hs6st1 | 50785 | HITS-CLIP | 23597149 | Unspecified |
| mmu-let-7i-5p | MIMAT0000122 | Hsbp1 | 68196 | HITS-CLIP | 37986224\|23597149\|19536157\|23142 | Unspecified |
| mmu-let-7i-5p | MIMAT0000122 | Hsd11b1 | 15483 | HITS-CLIP | 37986224\|23597149 | Unspecified |
| mmu-let-7i-5p | MIMAT0000122 | Hsd17b13 | 243168 | HITS-CLIP | 23597149 | Unspecified |
| mmu-let-7i-5p | MIMAT0000122 | Hsd17b2 | 15486 | HITS-CLIP | 37986224\|30971389 | Unspecified |
| mmu-let-7i-5p | MIMAT0000122 | Hsd17b4 | 15488 | HITS-CLIP | 23597149 | Unspecified |
| mmu-let-7i-5p | MIMAT0000122 | Hsd17b7 | 15490 | HITS-CLIP | 23142080 | Unspecified |
| mmu-let-7i-5p | MIMAT0000122 | Hsd3b2 | 15493 | HITS-CLIP | 23597149 | Unspecified |
| mmu-let-7i-5p | MIMAT0000122 | Hsd3b3 | 15494 | HITS-CLIP | 37986224\|23597149 | Unspecified |
| mmu-let-7i-5p | MIMAT0000122 | Hsd3b7 | 101502 | HITS-CLIP | 37986224\|23597149 | Unspecified |
| mmu-let-7i-5p | MIMAT0000122 | Hsdl1 | 72552 | HITS-CLIP | 23142080 | Unspecified |
| mmu-let-7i-5p | MIMAT0000122 | Hsf1 | 15499 | HITS-CLIP | 37986224 | Unspecified |
| mmu-let-7i-5p | MIMAT0000122 | Hsf2 | 15500 | HITS-CLIP | 23597149 | Unspecified |
| mmu-let-7i-5p | MIMAT0000122 | Hsh2d | 209488 | HITS-CLIP | 23142080 | Unspecified |
| mmu-let-7i-5p | MIMAT0000122 | Hsp90ab1 | 15516 | HITS-CLIP | 37986224\|23597149\|23142080\|3143 | Unspecified |
| mmu-let-7i-5p | MIMAT0000122 | Hsp90b1 | 22027 | HITS-CLIP | 23597149\|19536157\|23142080 | Unspecified |
| mmu-let-7i-5p | MIMAT0000122 | Hspa12a | 73442 | HITS-CLIP | 37986224 | Unspecified |
| mmu-let-7i-5p | MIMAT0000122 | Hspa2 | 15512 | HITS-CLIP | 23142080 | Unspecified |
| mmu-let-7i-5p | MIMAT0000122 | Hspa4 | 15525 | HITS-CLIP | 23142080\|29167373 | Unspecified |
| mmu-let-7i-5p | MIMAT0000122 | Hspa4l | 18415 | HITS-CLIP | 23142080 | Unspecified |
| mmu-let-7i-5p | MIMAT0000122 | Hspa5 | 14828 | HITS-CLIP | 23142080 | Unspecified |
| mmu-let-7i-5p | MIMAT0000122 | Hspa9 | 15526 | HITS-CLIP | 23142080\|29167373 | Unspecified |
| mmu-let-7i-5p | MIMAT0000122 | Hspb6 | 243912 | HITS-CLIP | 23597149 | Unspecified |
| mmu-let-7i-5p | MIMAT0000122 | Hspb8 | 80888 | HITS-CLIP | 23597149 | Unspecified |
| mmu-let-7i-5p | MIMAT0000122 | Hspd1 | 15510 | HITS-CLIP | 23142080\|29167373 | Unspecified |
| mmu-let-7i-5p | MIMAT0000122 | Hspg2 | 15530 | HITS-CLIP | 23597149 | Unspecified |
| mmu-let-7i-5p | MIMAT0000122 | Hsph1 | 15505 | HITS-CLIP | 37986224\|23142080\|23597149\|1953 | Unspecified |
| mmu-let-7i-5p | MIMAT0000122 | Htr2c | 15560 | HITS-CLIP | 19536157 | Unspecified |
| mmu-let-7i-5p | MIMAT0000122 | Htt | 15194 | HITS-CLIP | 37986224\|19536157\|23142080\|2916 | Unspecified |
| mmu-let-7i-5p | MIMAT0000122 | Hus1 | 15574 | HITS-CLIP | 19536157\|23142080 | Unspecified |
| mmu-let-7i-5p | MIMAT0000122 | Huwe1 | 59026 | HITS-CLIP | 37986224\|23597149\|23142080\|1953 | Unspecified |
| mmu-let-7i-5p | MIMAT0000122 | Hyal1 | 15586 | HITS-CLIP | 23597149\|26203562\|29167373 | Unspecified |

| | | | | | | |
|---|---|---|---|---|---|---|
| mmu-let-7i-5p | MIMAT0000122 | Hyal2 | 15587 | HITS-CLIP | 23142080\|23597149\|29167373 | Unspecified |
| mmu-let-7i-5p | MIMAT0000122 | Hyal6 | 74409 | HITS-CLIP | 23142080 | Unspecified |
| mmu-let-7i-5p | MIMAT0000122 | Hykk | 235386 | HITS-CLIP | 37986224\|23597149 | Unspecified |
| mmu-let-7i-5p | MIMAT0000122 | Hyls1 | 76832 | HITS-CLIP | 23142080 | Unspecified |
| mmu-let-7i-5p | MIMAT0000122 | Hyou1 | 12282 | HITS-CLIP | 23597149 | Unspecified |
| mmu-let-7i-5p | MIMAT0000122 | Iars | 105148 | HITS-CLIP | 23142080\|23597149\|29167373 | Unspecified |
| mmu-let-7i-5p | MIMAT0000122 | Iars2 | 381314 | HITS-CLIP | 19536157 | Unspecified |
| mmu-let-7i-5p | MIMAT0000122 | Ibtk | 108837 | HITS-CLIP | 23142080 | Unspecified |
| mmu-let-7i-5p | MIMAT0000122 | Icam1 | 15894 | HITS-CLIP | 23597149\|29167373 | Unspecified |
| mmu-let-7i-5p | MIMAT0000122 | Ice1 | 822287 | HITS-CLIP | 19536157\|23142080\|29167373 | Unspecified |
| mmu-let-7i-5p | MIMAT0000122 | Ice2 | 93697 | HITS-CLIP | 23597149\|23142080 | Unspecified |
| mmu-let-7i-5p | MIMAT0000122 | Id3 | 15903 | HITS-CLIP | 23142080 | Unspecified |
| mmu-let-7i-5p | MIMAT0000122 | Ide | 15925 | HITS-CLIP | 37986224\|23142080\|23597149 | Unspecified |
| mmu-let-7i-5p | MIMAT0000122 | Idh1 | 15926 | HITS-CLIP | 23142080 | Unspecified |
| mmu-let-7i-5p | MIMAT0000122 | Idh2 | 269951 | HITS-CLIP | 37986224\|23597149\|23142080\|29167 | Unspecified |
| mmu-let-7i-5p | MIMAT0000122 | Idh3b | 170718 | HITS-CLIP | 23597149\|19536157 | Unspecified |
| mmu-let-7i-5p | MIMAT0000122 | Idh3g | 15929 | HITS-CLIP | 23142080 | Unspecified |
| mmu-let-7i-5p | MIMAT0000122 | Ids | 15931 | HITS-CLIP | 37986224 | Unspecified |
| mmu-let-7i-5p | MIMAT0000122 | Idua | 15932 | HITS-CLIP | 23142080 | Unspecified |
| mmu-let-7i-5p | MIMAT0000122 | Iffo2 | 212632 | HITS-CLIP | 23597149\|26203562 | Unspecified |
| mmu-let-7i-5p | MIMAT0000122 | Ifi203 | 15950 | HITS-CLIP | 23142080 | Unspecified |
| mmu-let-7i-5p | MIMAT0000122 | Ifi204 | 15951 | HITS-CLIP | 29167373 | Unspecified |
| mmu-let-7i-5p | MIMAT0000122 | Ifi27l2a | 76933 | HITS-CLIP | 23142080 | Unspecified |
| mmu-let-7i-5p | MIMAT0000122 | Ifi44 | 99899 | HITS-CLIP | 29167373 | Unspecified |
| mmu-let-7i-5p | MIMAT0000122 | Ifih1 | 71586 | HITS-CLIP | 23597149\|23142080\|29167373 | Unspecified |
| mmu-let-7i-5p | MIMAT0000122 | Ifit2 | 15958 | HITS-CLIP | 29167373 | Unspecified |
| mmu-let-7i-5p | MIMAT0000122 | Ifitm2 | 80876 | HITS-CLIP | 23142080 | Unspecified |
| mmu-let-7i-5p | MIMAT0000122 | Ifnar1 | 15975 | HITS-CLIP | 21258322\|23142080\|29167373 | Unspecified |
| mmu-let-7i-5p | MIMAT0000122 | Ifnar2 | 15976 | HITS-CLIP | 37986224\|23142080 | Unspecified |
| mmu-let-7i-5p | MIMAT0000122 | Ifng | 15978 | HITS-CLIP | 23142080 | Unspecified |
| mmu-let-7i-5p | MIMAT0000122 | Ifngr2 | 15980 | HITS-CLIP | 23142080 | Unspecified |
| mmu-let-7i-5p | MIMAT0000122 | Ift74 | 67694 | HITS-CLIP | 37986224 | Unspecified |
| mmu-let-7i-5p | MIMAT0000122 | Igdcc3 | 19289 | HITS-CLIP | 19536157 | Unspecified |
| mmu-let-7i-5p | MIMAT0000122 | Igdcc4 | 56741 | HITS-CLIP | 19536157 | Unspecified |
| mmu-let-7i-5p | MIMAT0000122 | Igf1 | 16000 | HITS-CLIP | 37986224\|23597149\|29167373 | Unspecified |
| mmu-let-7i-5p | MIMAT0000122 | Igf1r | 16001 | HITS-CLIP//qCLASH | 37986224\|19536157\|31393953\|23142 | Unspecified |
| mmu-let-7i-5p | MIMAT0000122 | Igf2 | 16002 | HITS-CLIP | 37986224 | Unspecified |
| mmu-let-7i-5p | MIMAT0000122 | Igf2bp3 | 140488 | HITS-CLIP | 23142080 | Unspecified |
| mmu-let-7i-5p | MIMAT0000122 | Igf2r | 16004 | HITS-CLIP | 37986224\|23597149\|23142080\|29167 | Unspecified |

| | | | | | | |
|---|---|---|---|---|---|---|
| mmu-let-7i-5p | MIMAT0000122 | Igfals | 16005 | HITS-CLIP | 37986224\|23597149 | Unspecified |
| mmu-let-7i-5p | MIMAT0000122 | Igfbp4 | 16010 | HITS-CLIP | 23597149 | Unspecified |
| mmu-let-7i-5p | MIMAT0000122 | Igfbp5 | 16011 | HITS-CLIP | 37986224\|19536157 | Unspecified |
| mmu-let-7i-5p | MIMAT0000122 | Iglon5 | 210094 | HITS-CLIP | 37986224 | Unspecified |
| mmu-let-7i-5p | MIMAT0000122 | Igsf5 | 72058 | HITS-CLIP | 23597149 | Unspecified |
| mmu-let-7i-5p | MIMAT0000122 | Igsf6 | 80719 | HITS-CLIP | 29167373 | Unspecified |
| mmu-let-7i-5p | MIMAT0000122 | Igsf9 | 93842 | HITS-CLIP | 29167373 | Unspecified |
| mmu-let-7i-5p | MIMAT0000122 | Iigp1 | 60440 | HITS-CLIP | 23597149\|23142080 | Unspecified |
| mmu-let-7i-5p | MIMAT0000122 | Ik | 24010 | HITS-CLIP | 23142080\|29167373 | Unspecified |
| mmu-let-7i-5p | MIMAT0000122 | Ikbke | 56489 | HITS-CLIP | 23597149\|23142080\|29167373 | Unspecified |
| mmu-let-7i-5p | MIMAT0000122 | Ikbkg | 16151 | HITS-CLIP | 37986224\|23597149 | Unspecified |
| mmu-let-7i-5p | MIMAT0000122 | Ikzf1 | 22778 | HITS-CLIP | 23142080\|29167373 | Unspecified |
| mmu-let-7i-5p | MIMAT0000122 | Ikzf3 | 22780 | HITS-CLIP | 23142080 | Unspecified |
| mmu-let-7i-5p | MIMAT0000122 | Ikzf4 | 22781 | HITS-CLIP | 23142080\|19536157\|23597149 | Unspecified |
| mmu-let-7i-5p | MIMAT0000122 | Il10ra | 16154 | HITS-CLIP | 23142080 | Unspecified |
| mmu-let-7i-5p | MIMAT0000122 | Il10rb | 16155 | HITS-CLIP | 29167373 | Unspecified |
| mmu-let-7i-5p | MIMAT0000122 | Il12rb2 | 16162 | HITS-CLIP | 37986224\|23142080 | Unspecified |
| mmu-let-7i-5p | MIMAT0000122 | Il13 | 16163 | HITS-CLIP | 37986224\|23142080 | Unspecified |
| mmu-let-7i-5p | MIMAT0000122 | Il17ra | 16172 | HITS-CLIP | 23142080 | Unspecified |
| mmu-let-7i-5p | MIMAT0000122 | Il18r1 | 16182 | HITS-CLIP | 23142080 | Unspecified |
| mmu-let-7i-5p | MIMAT0000122 | Il18rap | 16174 | HITS-CLIP | 23142080 | Unspecified |
| mmu-let-7i-5p | MIMAT0000122 | Il1a | 16175 | HITS-CLIP | 37986224\|23597149 | Unspecified |
| mmu-let-7i-5p | MIMAT0000122 | Il1rap | 16180 | HITS-CLIP | 23597149 | Unspecified |
| mmu-let-7i-5p | MIMAT0000122 | Il1rn | 16181 | HITS-CLIP | 23597149 | Unspecified |
| mmu-let-7i-5p | MIMAT0000122 | Il2 | 16183 | HITS-CLIP | 23142080 | Unspecified |
| mmu-let-7i-5p | MIMAT0000122 | Il21 | 60505 | HITS-CLIP | 23142080 | Unspecified |
| mmu-let-7i-5p | MIMAT0000122 | Il21r | 60504 | HITS-CLIP | 23142080 | Unspecified |
| mmu-let-7i-5p | MIMAT0000122 | Il22ra1 | 230828 | HITS-CLIP | 23597149 | Unspecified |
| mmu-let-7i-5p | MIMAT0000122 | Il24 | 93672 | HITS-CLIP | 23142080 | Unspecified |
| mmu-let-7i-5p | MIMAT0000122 | Il27ra | 50931 | HITS-CLIP | 23142080 | Unspecified |
| mmu-let-7i-5p | MIMAT0000122 | Il2ra | 16184 | HITS-CLIP | 23142080 | Unspecified |
| mmu-let-7i-5p | MIMAT0000122 | Il2rb | 16185 | HITS-CLIP | 37986224\|23142080 | Unspecified |
| mmu-let-7i-5p | MIMAT0000122 | Il2rg | 16186 | HITS-CLIP | 23142080 | Unspecified |
| mmu-let-7i-5p | MIMAT0000122 | Il33 | 77125 | HITS-CLIP | 23597149 | Unspecified |
| mmu-let-7i-5p | MIMAT0000122 | Il4ra | 16190 | HITS-CLIP | 23142080 | Unspecified |
| mmu-let-7i-5p | MIMAT0000122 | Il6ra | 16194 | HITS-CLIP | 23142080\|23597149\|29167373 | Unspecified |
| mmu-let-7i-5p | MIMAT0000122 | Il6st | 16195 | HITS-CLIP | 23142080\|23597149\|19536157\|29167 | Unspecified |
| mmu-let-7i-5p | MIMAT0000122 | Il7r | 16197 | HITS-CLIP | 23142080\|29167373 | Unspecified |
| mmu-let-7i-5p | MIMAT0000122 | Il9r | 3581 | qCLASH | 31393953 | Unspecified |

| | | | | | | |
|---|---|---|---|---|---|---|
| mmu-let-7i-5p | MIMAT0000122 | Ilf3 | 16201 | HITS-CLIP | 31433990 | Unspecified |
| mmu-let-7i-5p | MIMAT0000122 | Immt | 76614 | HITS-CLIP | 23597149 | Unspecified |
| mmu-let-7i-5p | MIMAT0000122 | Imp3 | 102462 | HITS-CLIP | 23142080 | Unspecified |
| mmu-let-7i-5p | MIMAT0000122 | Imp4 | 27993 | HITS-CLIP | 37986224\|23597149\|23142080 | Unspecified |
| mmu-let-7i-5p | MIMAT0000122 | Impa1 | 55980 | HITS-CLIP | 29167373 | Unspecified |
| mmu-let-7i-5p | MIMAT0000122 | Impa2 | 114663 | HITS-CLIP | 23142080 | Unspecified |
| mmu-let-7i-5p | MIMAT0000122 | Impad1 | 242291 | HITS-CLIP | 23597149\|23142080 | Unspecified |
| mmu-let-7i-5p | MIMAT0000122 | Impdh1 | 23917 | HITS-CLIP | 23142080 | Unspecified |
| mmu-let-7i-5p | MIMAT0000122 | Incenp | 16319 | HITS-CLIP | 23142080 | Unspecified |
| mmu-let-7i-5p | MIMAT0000122 | Inf2 | 70435 | HITS-CLIP | 23142080 | Unspecified |
| mmu-let-7i-5p | MIMAT0000122 | Ing4 | 28019 | HITS-CLIP | 23142080 | Unspecified |
| mmu-let-7i-5p | MIMAT0000122 | Inhbe | 16326 | HITS-CLIP | 23597149 | Unspecified |
| mmu-let-7i-5p | MIMAT0000122 | Inip | 66209 | HITS-CLIP | 23142080 | Unspecified |
| mmu-let-7i-5p | MIMAT0000122 | Inka1 | 389119 | HITS-CLIP | 23142080 | Unspecified |
| mmu-let-7i-5p | MIMAT0000122 | Ino80 | 68142 | HITS-CLIP | 37986224\|29167373\|23142080 | Unspecified |
| mmu-let-7i-5p | MIMAT0000122 | Ino80c | 225280 | HITS-CLIP | 23142080 | Unspecified |
| mmu-let-7i-5p | MIMAT0000122 | Ino80d | 227195 | HITS-CLIP | 37986224\|23597149\|23142080\|29167 | Unspecified |
| mmu-let-7i-5p | MIMAT0000122 | Ino80e | 233875 | HITS-CLIP | 37986224\|23142080 | Unspecified |
| mmu-let-7i-5p | MIMAT0000122 | Inpp4a | 269180 | HITS-CLIP | 23142080 | Unspecified |
| mmu-let-7i-5p | MIMAT0000122 | Inpp4b | 234515 | HITS-CLIP | 23142080\|29167373 | Unspecified |
| mmu-let-7i-5p | MIMAT0000122 | Inpp5a | 212111 | HITS-CLIP | 37986224\|23142080 | Unspecified |
| mmu-let-7i-5p | MIMAT0000122 | Inpp5b | 16330 | HITS-CLIP | 23597149 | Unspecified |
| mmu-let-7i-5p | MIMAT0000122 | Inpp5d | 16331 | HITS-CLIP | 23142080 | Unspecified |
| mmu-let-7i-5p | MIMAT0000122 | Inpp5f | 101490 | HITS-CLIP | 37986224\|23597149\|23142080 | Unspecified |
| mmu-let-7i-5p | MIMAT0000122 | Inpp5k | 19062 | HITS-CLIP | 23142080 | Unspecified |
| mmu-let-7i-5p | MIMAT0000122 | Insig2 | 72999 | HITS-CLIP | 23597149 | Unspecified |
| mmu-let-7i-5p | MIMAT0000122 | Insr | 16337 | HITS-CLIP | 37986224\|23597149\|23142080 | Unspecified |
| mmu-let-7i-5p | MIMAT0000122 | Ints1 | 68510 | HITS-CLIP | 37986224\|23597149 | Unspecified |
| mmu-let-7i-5p | MIMAT0000122 | Ints11 | 54973 | HITS-CLIP | 23142080 | Unspecified |
| mmu-let-7i-5p | MIMAT0000122 | Ints12 | 71793 | HITS-CLIP | 23142080 | Unspecified |
| mmu-let-7i-5p | MIMAT0000122 | Ints2 | 70422 | HITS-CLIP | 23597149\|23142080 | Unspecified |
| mmu-let-7i-5p | MIMAT0000122 | Ints3 | 229543 | HITS-CLIP | 23597149\|23142080 | Unspecified |
| mmu-let-7i-5p | MIMAT0000122 | Ints4 | 101861 | HITS-CLIP | 37986224 | Unspecified |
| mmu-let-7i-5p | MIMAT0000122 | Ints5 | 80789 | HITS-CLIP | 23142080 | Unspecified |
| mmu-let-7i-5p | MIMAT0000122 | Ints6 | 18130 | HITS-CLIP | 23142080 | Unspecified |
| mmu-let-7i-5p | MIMAT0000122 | Ints6l | 236790 | HITS-CLIP | 23142080 | Unspecified |
| mmu-let-7i-5p | MIMAT0000122 | Ints7 | 77065 | HITS-CLIP | 23142080 | Unspecified |
| mmu-let-7i-5p | MIMAT0000122 | Ints8 | 72656 | HITS-CLIP//qCLASH | 23597149\|31393953\|31433990\|23630 | Unspecified |
| mmu-let-7i-5p | MIMAT0000122 | Ints9 | 210925 | HITS-CLIP | 23142080 | Unspecified |

| | | | | | | |
|---|---|---|---|---|---|---|
| mmu-let-7i-5p | MIMAT0000122 | Invs | 16348 | HITS-CLIP | 23597149 | Unspecified |
| mmu-let-7i-5p | MIMAT0000122 | Ip6k1 | 27399 | HITS-CLIP | 23597149\|23142080 | Unspecified |
| mmu-let-7i-5p | MIMAT0000122 | Ipcef1 | 320495 | HITS-CLIP | 23142080 | Unspecified |
| mmu-let-7i-5p | MIMAT0000122 | Ipmk | 69718 | HITS-CLIP | 23597149 | Unspecified |
| mmu-let-7i-5p | MIMAT0000122 | Ipo11 | 76582 | HITS-CLIP | 19536157\|23142080 | Unspecified |
| mmu-let-7i-5p | MIMAT0000122 | Ipo5 | 70572 | HITS-CLIP | 23142080\|29167373 | Unspecified |
| mmu-let-7i-5p | MIMAT0000122 | Ipo7 | 233726 | HITS-CLIP | 23597149\|23142080\|29167373 | Unspecified |
| mmu-let-7i-5p | MIMAT0000122 | Ipo8 | 320727 | HITS-CLIP | 23142080 | Unspecified |
| mmu-let-7i-5p | MIMAT0000122 | Ippk | 75678 | HITS-CLIP | 23142080 | Unspecified |
| mmu-let-7i-5p | MIMAT0000122 | Iqcb1 | 320299 | HITS-CLIP | 23142080 | Unspecified |
| mmu-let-7i-5p | MIMAT0000122 | Iqgap1 | 29875 | HITS-CLIP | 23142080\|29167373 | Unspecified |
| mmu-let-7i-5p | MIMAT0000122 | Iqgap2 | 544963 | HITS-CLIP | 23597149\|23142080 | Unspecified |
| mmu-let-7i-5p | MIMAT0000122 | Iqgap3 | 404710 | HITS-CLIP | 23597149 | Unspecified |
| mmu-let-7i-5p | MIMAT0000122 | Iqsec2 | 245666 | HITS-CLIP | 23597149\|19536157\|29167373 | Unspecified |
| mmu-let-7i-5p | MIMAT0000122 | Irak3 | 73914 | HITS-CLIP | 23597149 | Unspecified |
| mmu-let-7i-5p | MIMAT0000122 | Ireb2 | 64602 | HITS-CLIP | 37986224\|19536157\|23142080 | Unspecified |
| mmu-let-7i-5p | MIMAT0000122 | Irf1 | 16362 | HITS-CLIP | 23142080 | Unspecified |
| mmu-let-7i-5p | MIMAT0000122 | Irf2 | 16363 | HITS-CLIP | 23142080 | Unspecified |
| mmu-let-7i-5p | MIMAT0000122 | Irf2bp2 | 270110 | HITS-CLIP//qCLASH | 37986224\|31393953\|23597149\|29167 | Unspecified |
| mmu-let-7i-5p | MIMAT0000122 | Irf3 | 54131 | HITS-CLIP | 23597149 | Unspecified |
| mmu-let-7i-5p | MIMAT0000122 | Irf4 | 16364 | HITS-CLIP | 37986224\|23142080 | Unspecified |
| mmu-let-7i-5p | MIMAT0000122 | Irf8 | 15900 | HITS-CLIP | 29167373\|23142080 | Unspecified |
| mmu-let-7i-5p | MIMAT0000122 | Irf9 | 16391 | HITS-CLIP | 23142080 | Unspecified |
| mmu-let-7i-5p | MIMAT0000122 | Irs1 | 16367 | HITS-CLIP | 37986224\|23597149 | Unspecified |
| mmu-let-7i-5p | MIMAT0000122 | Irs2 | 384783 | HITS-CLIP | 37986224\|23597149\|19536157\|23142 | Unspecified |
| mmu-let-7i-5p | MIMAT0000122 | Isca2 | 74316 | HITS-CLIP | 37986224\|23142080\|29167373 | Unspecified |
| mmu-let-7i-5p | MIMAT0000122 | Isg20l2 | 229504 | HITS-CLIP | 23142080 | Unspecified |
| mmu-let-7i-5p | MIMAT0000122 | Islr | 26968 | HITS-CLIP | 19536157\|23597149 | Unspecified |
| mmu-let-7i-5p | MIMAT0000122 | Isoc2a | 664994 | HITS-CLIP | 23142080 | Unspecified |
| mmu-let-7i-5p | MIMAT0000122 | Isoc2b | 67441 | HITS-CLIP | 23597149 | Unspecified |
| mmu-let-7i-5p | MIMAT0000122 | Ist1 | 71955 | HITS-CLIP | 23597149\|29167373\|23142080 | Unspecified |
| mmu-let-7i-5p | MIMAT0000122 | Isyna1 | 71780 | HITS-CLIP | 23597149 | Unspecified |
| mmu-let-7i-5p | MIMAT0000122 | Itch | 16396 | HITS-CLIP | 23597149\|23142080 | Unspecified |
| mmu-let-7i-5p | MIMAT0000122 | Itga4 | 16401 | HITS-CLIP | 23142080\|29167373 | Unspecified |
| mmu-let-7i-5p | MIMAT0000122 | Itga5 | 16402 | HITS-CLIP | 29167373 | Unspecified |
| mmu-let-7i-5p | MIMAT0000122 | Itga6 | 16403 | HITS-CLIP | 19536157 | Unspecified |
| mmu-let-7i-5p | MIMAT0000122 | Itga7 | 16404 | HITS-CLIP | 23142080 | Unspecified |
| mmu-let-7i-5p | MIMAT0000122 | Itgal | 16408 | HITS-CLIP | 37986224\|23597149\|23142080\|29167 | Unspecified |
| mmu-let-7i-5p | MIMAT0000122 | Itgav | 16410 | HITS-CLIP | 23142080\|29167373 | Unspecified |

| | | | | | | |
|---|---|---|---|---|---|---|
| mmu-let-7i-5p | MIMAT0000122 | Itgb1 | 16412 | HITS-CLIP | 23142080\|29167373 | Unspecified |
| mmu-let-7i-5p | MIMAT0000122 | Itgb2 | 16414 | HITS-CLIP | 37986224\|23597149\|23142080\|29167 | Unspecified |
| mmu-let-7i-5p | MIMAT0000122 | Itgb3 | 16416 | HITS-CLIP | 37986224\|23597149\|23142080 | Unspecified |
| mmu-let-7i-5p | MIMAT0000122 | Itgb5 | 16419 | HITS-CLIP | 29167373 | Unspecified |
| mmu-let-7i-5p | MIMAT0000122 | Itgb7 | 16421 | HITS-CLIP | 23142080 | Unspecified |
| mmu-let-7i-5p | MIMAT0000122 | Itgb8 | 320910 | HITS-CLIP | 23142080 | Unspecified |
| mmu-let-7i-5p | MIMAT0000122 | Itih1 | 16424 | HITS-CLIP | 23597149 | Unspecified |
| mmu-let-7i-5p | MIMAT0000122 | Itih2 | 16425 | HITS-CLIP | 23597149 | Unspecified |
| mmu-let-7i-5p | MIMAT0000122 | Itih3 | 16426 | HITS-CLIP | 23597149 | Unspecified |
| mmu-let-7i-5p | MIMAT0000122 | Itih4 | 16427 | HITS-CLIP | 23597149 | Unspecified |
| mmu-let-7i-5p | MIMAT0000122 | Itih5 | 209378 | HITS-CLIP | 23142080 | Unspecified |
| mmu-let-7i-5p | MIMAT0000122 | Itk | 16428 | HITS-CLIP | 23142080 | Unspecified |
| mmu-let-7i-5p | MIMAT0000122 | Itm2a | 16431 | HITS-CLIP | 23142080 | Unspecified |
| mmu-let-7i-5p | MIMAT0000122 | Itm2b | 16432 | HITS-CLIP | 23597149\|23142080\|29167373 | Unspecified |
| mmu-let-7i-5p | MIMAT0000122 | Itm2c | 64294 | HITS-CLIP | 19536157\|23142080 | Unspecified |
| mmu-let-7i-5p | MIMAT0000122 | Itpkb | 320404 | HITS-CLIP | 37986224\|23597149\|23142080 | Unspecified |
| mmu-let-7i-5p | MIMAT0000122 | Itpkc | 233011 | HITS-CLIP | 23597149 | Unspecified |
| mmu-let-7i-5p | MIMAT0000122 | Itpr2 | 16439 | HITS-CLIP | 23142080 | Unspecified |
| mmu-let-7i-5p | MIMAT0000122 | Itpr3 | 16440 | HITS-CLIP | 23142080\|29167373 | Unspecified |
| mmu-let-7i-5p | MIMAT0000122 | Itprid2 | 6744 | HITS-CLIP | 23142080\|29167373 | Unspecified |
| mmu-let-7i-5p | MIMAT0000122 | Itpripl2 | 319622 | HITS-CLIP | 23597149 | Unspecified |
| mmu-let-7i-5p | MIMAT0000122 | Itsn1 | 16443 | HITS-CLIP | 23597149\|29167373 | Unspecified |
| mmu-let-7i-5p | MIMAT0000122 | Itsn2 | 20403 | HITS-CLIP | 37986224\|23142080\|29167373\|19536 | Unspecified |
| mmu-let-7i-5p | MIMAT0000122 | Ivd | 56357 | HITS-CLIP | 23597149 | Unspecified |
| mmu-let-7i-5p | MIMAT0000122 | Ivns1abp | 117198 | HITS-CLIP | 23142080 | Unspecified |
| mmu-let-7i-5p | MIMAT0000122 | Iws1 | 73473 | HITS-CLIP | 23597149\|23142080 | Unspecified |
| mmu-let-7i-5p | MIMAT0000122 | Iyd | 70337 | HITS-CLIP | 23597149 | Unspecified |
| mmu-let-7i-5p | MIMAT0000122 | Izumo1r | 64931 | HITS-CLIP | 23142080 | Unspecified |
| mmu-let-7i-5p | MIMAT0000122 | Jade1 | 269424 | HITS-CLIP | 23142080 | Unspecified |
| mmu-let-7i-5p | MIMAT0000122 | Jade3 | 382207 | HITS-CLIP | 37986224\|23597149\|23142080 | Unspecified |
| mmu-let-7i-5p | MIMAT0000122 | Jag1 | 16449 | HITS-CLIP | 29167373 | Unspecified |
| mmu-let-7i-5p | MIMAT0000122 | Jagn1 | 67767 | HITS-CLIP | 23597149 | Unspecified |
| mmu-let-7i-5p | MIMAT0000122 | Jak1 | 16451 | HITS-CLIP | 37986224\|23597149\|23142080\|29167 | Unspecified |
| mmu-let-7i-5p | MIMAT0000122 | Jak2 | 16452 | HITS-CLIP | 29167373\|23142080 | Unspecified |
| mmu-let-7i-5p | MIMAT0000122 | Jakmip1 | 76071 | HITS-CLIP | 19536157 | Unspecified |
| mmu-let-7i-5p | MIMAT0000122 | Jakmip2 | 76217 | HITS-CLIP | 37986224 | Unspecified |
| mmu-let-7i-5p | MIMAT0000122 | Jam2 | 67374 | HITS-CLIP | 37986224\|19536157\|23597149 | Unspecified |
| mmu-let-7i-5p | MIMAT0000122 | Jarid2 | 16468 | HITS-CLIP | 37986224\|23142080\|23597149\|19536 | Unspecified |
| mmu-let-7i-5p | MIMAT0000122 | Jcad | 57608 | HITS-CLIP | 19536157 | Unspecified |

| | | | | | | |
|---|---|---|---|---|---|---|
| mmu-let-7i-5p | MIMAT0000122 | Jmjd1c | 108829 | HITS-CLIP//qCLASH | 37986224\|23142080\|19536157\|23597 | Unspecified |
| mmu-let-7i-5p | MIMAT0000122 | Jmjd4 | 194952 | HITS-CLIP | 37986224 | Unspecified |
| mmu-let-7i-5p | MIMAT0000122 | Jmjd6 | 107817 | HITS-CLIP | 23142080 | Unspecified |
| mmu-let-7i-5p | MIMAT0000122 | Jmjd7 | 433466 | HITS-CLIP | 23142080 | Unspecified |
| mmu-let-7i-5p | MIMAT0000122 | Jmjd8 | 72106 | HITS-CLIP | 23142080 | Unspecified |
| mmu-let-7i-5p | MIMAT0000122 | Jmy | 57748 | HITS-CLIP | 37986224\|19536157\|23597149\|23142 | Unspecified |
| mmu-let-7i-5p | MIMAT0000122 | Jph1 | 57339 | HITS-CLIP | 23597149 | Unspecified |
| mmu-let-7i-5p | MIMAT0000122 | Jph4 | 319984 | HITS-CLIP | 37986224 | Unspecified |
| mmu-let-7i-5p | MIMAT0000122 | Jpt1 | 51155 | HITS-CLIP | 23142080 | Unspecified |
| mmu-let-7i-5p | MIMAT0000122 | Jrkl | 77532 | HITS-CLIP | 23142080 | Unspecified |
| mmu-let-7i-5p | MIMAT0000122 | Jun | 16476 | HITS-CLIP | 23597149\|23142080\|29167373 | Unspecified |
| mmu-let-7i-5p | MIMAT0000122 | Junb | 16477 | HITS-CLIP | 23597149\|23142080\|29167373 | Unspecified |
| mmu-let-7i-5p | MIMAT0000122 | Jund | 16478 | HITS-CLIP | 23597149\|23142080 | Unspecified |
| mmu-let-7i-5p | MIMAT0000122 | Jup | 16480 | HITS-CLIP | 23597149 | Unspecified |
| mmu-let-7i-5p | MIMAT0000122 | Kalrn | 545156 | HITS-CLIP | 37986224\|19536157\|23597149 | Unspecified |
| mmu-let-7i-5p | MIMAT0000122 | Kank1 | 107351 | HITS-CLIP | 23597149 | Unspecified |
| mmu-let-7i-5p | MIMAT0000122 | Kank2 | 235041 | HITS-CLIP | 30971389 | Unspecified |
| mmu-let-7i-5p | MIMAT0000122 | Kansl1 | 76719 | HITS-CLIP | 23597149\|23142080 | Unspecified |
| mmu-let-7i-5p | MIMAT0000122 | Kansl1l | 68691 | HITS-CLIP | 23142080 | Unspecified |
| mmu-let-7i-5p | MIMAT0000122 | Kansl3 | 226976 | HITS-CLIP | 23142080\|29167373 | Unspecified |
| mmu-let-7i-5p | MIMAT0000122 | Kantr | 102723508 | HITS-CLIP | 23142080 | Unspecified |
| mmu-let-7i-5p | MIMAT0000122 | Kars | 85305 | HITS-CLIP//qCLASH | 37986224\|23142080\|23597149\|31393 | Unspecified |
| mmu-let-7i-5p | MIMAT0000122 | Kat2a | 14534 | HITS-CLIP | 19536157\|23142080\|29167373 | Unspecified |
| mmu-let-7i-5p | MIMAT0000122 | Kat5 | 81601 | HITS-CLIP | 23597149\|23142080 | Unspecified |
| mmu-let-7i-5p | MIMAT0000122 | Kat6a | 244349 | HITS-CLIP | 37986224\|23597149\|29167373\|23142 | Unspecified |
| mmu-let-7i-5p | MIMAT0000122 | Kat6b | 54169 | HITS-CLIP | 23142080 | Unspecified |
| mmu-let-7i-5p | MIMAT0000122 | Kat7 | 217127 | HITS-CLIP | 23597149\|23142080 | Unspecified |
| mmu-let-7i-5p | MIMAT0000122 | Katnbl1 | 72425 | HITS-CLIP | 37986224\|23142080 | Unspecified |
| mmu-let-7i-5p | MIMAT0000122 | Kbtbd11 | 74901 | HITS-CLIP | 31433990 | Unspecified |
| mmu-let-7i-5p | MIMAT0000122 | Kbtbd2 | 210973 | HITS-CLIP | 37986224\|23597149\|23142080\|29167 | Unspecified |
| mmu-let-7i-5p | MIMAT0000122 | Kbtbd4 | 67136 | HITS-CLIP | 23142080 | Unspecified |
| mmu-let-7i-5p | MIMAT0000122 | Kbtbd8 | 243574 | HITS-CLIP | 23142080\|29167373 | Unspecified |
| mmu-let-7i-5p | MIMAT0000122 | Kcmf1 | 74287 | qCLASH | 31393953 | Unspecified |
| mmu-let-7i-5p | MIMAT0000122 | Kcna1 | 16485 | HITS-CLIP | 37986224 | Unspecified |
| mmu-let-7i-5p | MIMAT0000122 | Kcnb1 | 16500 | HITS-CLIP | 23142080 | Unspecified |
| mmu-let-7i-5p | MIMAT0000122 | Kcnb2 | 98741 | HITS-CLIP | 19536157 | Unspecified |
| mmu-let-7i-5p | MIMAT0000122 | Kcnc1 | 16502 | HITS-CLIP | 37986224 | Unspecified |
| mmu-let-7i-5p | MIMAT0000122 | Kcnc4 | 99738 | HITS-CLIP | 19536157 | Unspecified |
| mmu-let-7i-5p | MIMAT0000122 | Kcnf1 | 382571 | HITS-CLIP | 37986224 | Unspecified |

| | | | | | | | |
|---|---|---|---|---|---|---|---|
| mmu-let-7i-5p | MIMAT0000122 | Kcnh1 | 16510 | HITS-CLIP | | 19536157 | Unspecified |
| mmu-let-7i-5p | MIMAT0000122 | Kcnip2 | 80906 | HITS-CLIP | | 23142080 | Unspecified |
| mmu-let-7i-5p | MIMAT0000122 | Kcnj10 | 16513 | HITS-CLIP | | 37986224\|19536157 | Unspecified |
| mmu-let-7i-5p | MIMAT0000122 | Kcnj2 | 16518 | HITS-CLIP | | 37986224\|19536157 | Unspecified |
| mmu-let-7i-5p | MIMAT0000122 | Kcnk1 | 16525 | HITS-CLIP | | 37986224 | Unspecified |
| mmu-let-7i-5p | MIMAT0000122 | Kcnk10 | 72258 | HITS-CLIP | | 37986224 | Unspecified |
| mmu-let-7i-5p | MIMAT0000122 | Kcnk3 | 16527 | HITS-CLIP | | 19536157 | Unspecified |
| mmu-let-7i-5p | MIMAT0000122 | Kcnk5 | 16529 | HITS-CLIP | | 23142080 | Unspecified |
| mmu-let-7i-5p | MIMAT0000122 | Kcnk9 | 223604 | HITS-CLIP | | 19536157 | Unspecified |
| mmu-let-7i-5p | MIMAT0000122 | Kcnn2 | 140492 | HITS-CLIP | | 19536157 | Unspecified |
| mmu-let-7i-5p | MIMAT0000122 | Kcnn4 | 16534 | HITS-CLIP | | 23142080\|29167373 | Unspecified |
| mmu-let-7i-5p | MIMAT0000122 | Kcnq3 | 110862 | HITS-CLIP | | 37986224 | Unspecified |
| mmu-let-7i-5p | MIMAT0000122 | Kcnt2 | 240776 | HITS-CLIP | | 37986224 | Unspecified |
| mmu-let-7i-5p | MIMAT0000122 | Kctd1 | 106931 | HITS-CLIP | | 23142080 | Unspecified |
| mmu-let-7i-5p | MIMAT0000122 | Kctd10 | 330171 | HITS-CLIP | | 23597149\|23142080 | Unspecified |
| mmu-let-7i-5p | MIMAT0000122 | Kctd12 | 239217 | HITS-CLIP | | 30971389\|29167373 | Unspecified |
| mmu-let-7i-5p | MIMAT0000122 | Kctd12b | 207474 | HITS-CLIP | | 19536157 | Unspecified |
| mmu-let-7i-5p | MIMAT0000122 | Kctd16 | 383348 | HITS-CLIP | | 19536157 | Unspecified |
| mmu-let-7i-5p | MIMAT0000122 | Kctd18 | 51960 | HITS-CLIP | | 23142080 | Unspecified |
| mmu-let-7i-5p | MIMAT0000122 | Kctd2 | 70382 | HITS-CLIP | | 19536157 | Unspecified |
| mmu-let-7i-5p | MIMAT0000122 | Kctd20 | 66989 | HITS-CLIP | | 37986224\|23142080\|29167373\|23597 | Unspecified |
| mmu-let-7i-5p | MIMAT0000122 | Kctd21 | 622320 | HITS-CLIP | | 23142080\|29167373 | Unspecified |
| mmu-let-7i-5p | MIMAT0000122 | Kctd3 | 226823 | HITS-CLIP | | 37986224\|23597149\|29167373 | Unspecified |
| mmu-let-7i-5p | MIMAT0000122 | Kctd5 | 69259 | HITS-CLIP | | 37986224\|23142080\|19536157\|2916 | Unspecified |
| mmu-let-7i-5p | MIMAT0000122 | Kctd8 | 243043 | HITS-CLIP | | 37986224 | Unspecified |
| mmu-let-7i-5p | MIMAT0000122 | Kctd9 | 105440 | HITS-CLIP | | 23142080 | Unspecified |
| mmu-let-7i-5p | MIMAT0000122 | Kdelr1 | 68137 | HITS-CLIP | | 23597149\|19536157\|23142080 | Unspecified |
| mmu-let-7i-5p | MIMAT0000122 | Kdelr2 | 66913 | HITS-CLIP//qCLASH | | 37986224\|23597149\|31393953\|2314 | Unspecified |
| mmu-let-7i-5p | MIMAT0000122 | Kdm1a | 99982 | HITS-CLIP | | 23142080 | Unspecified |
| mmu-let-7i-5p | MIMAT0000122 | Kdm2a | 225876 | HITS-CLIP | | 19536157\|23142080\|29167373 | Unspecified |
| mmu-let-7i-5p | MIMAT0000122 | Kdm2b | 30841 | HITS-CLIP | | 37986224\|23597149\|23142080 | Unspecified |
| mmu-let-7i-5p | MIMAT0000122 | Kdm3a | 104263 | HITS-CLIP | | 37986224\|23142080 | Unspecified |
| mmu-let-7i-5p | MIMAT0000122 | Kdm3b | 277250 | HITS-CLIP | | 19536157\|29167373\|23142080 | Unspecified |
| mmu-let-7i-5p | MIMAT0000122 | Kdm4a | 230674 | HITS-CLIP | | 23142080 | Unspecified |
| mmu-let-7i-5p | MIMAT0000122 | Kdm4b | 193796 | HITS-CLIP | | 23142080 | Unspecified |
| mmu-let-7i-5p | MIMAT0000122 | Kdm4c | 76804 | HITS-CLIP | | 23142080 | Unspecified |
| mmu-let-7i-5p | MIMAT0000122 | Kdm5a | 214899 | HITS-CLIP | | 23597149\|23142080\|29167373 | Unspecified |
| mmu-let-7i-5p | MIMAT0000122 | Kdm5b | 75605 | HITS-CLIP | | 23597149\|23142080 | Unspecified |
| mmu-let-7i-5p | MIMAT0000122 | Kdm5c | 20591 | HITS-CLIP | | 23597149\|19536157\|23142080 | Unspecified |

| | | | | | | |
|---|---|---|---|---|---|---|
| mmu-let-7i-5p | MIMAT0000122 | Kdm5d | 20592 | HITS-CLIP | 23142080 | Unspecified |
| mmu-let-7i-5p | MIMAT0000122 | Kdm6a | 22289 | HITS-CLIP//qCLASH | 37986224\|23597149\|31393953\|23142 | Unspecified |
| mmu-let-7i-5p | MIMAT0000122 | Kdm6b | 216850 | HITS-CLIP | 23597149\|19536157\|23142080 | Unspecified |
| mmu-let-7i-5p | MIMAT0000122 | Kdm7a | 338523 | HITS-CLIP | 37986224\|23142080 | Unspecified |
| mmu-let-7i-5p | MIMAT0000122 | Kdr | 16542 | HITS-CLIP | 37986224\|23597149\|19536157 | Unspecified |
| mmu-let-7i-5p | MIMAT0000122 | Keg1 | 64697 | HITS-CLIP | 23597149 | Unspecified |
| mmu-let-7i-5p | MIMAT0000122 | Khdc4 | 22889 | HITS-CLIP | 29167373 | Unspecified |
| mmu-let-7i-5p | MIMAT0000122 | Khdrbs1 | 20218 | HITS-CLIP | 23597149 | Unspecified |
| mmu-let-7i-5p | MIMAT0000122 | Khdrbs3 | 13992 | HITS-CLIP | 23597149 | Unspecified |
| mmu-let-7i-5p | MIMAT0000122 | Khk | 16548 | HITS-CLIP | 23597149 | Unspecified |
| mmu-let-7i-5p | MIMAT0000122 | Khnyn | 219094 | HITS-CLIP | 23597149\|23142080 | Unspecified |
| mmu-let-7i-5p | MIMAT0000122 | Khsrp | 16549 | HITS-CLIP//qCLASH | 37986224\|23597149\|19536157\|23142 | Unspecified |
| mmu-let-7i-5p | MIMAT0000122 | Kidins220 | 77480 | HITS-CLIP | 37986224\|23142080 | Unspecified |
| mmu-let-7i-5p | MIMAT0000122 | Kif11 | 16551 | HITS-CLIP | 23142080 | Unspecified |
| mmu-let-7i-5p | MIMAT0000122 | Kif13b | 16554 | HITS-CLIP | 23142080\|29167373 | Unspecified |
| mmu-let-7i-5p | MIMAT0000122 | Kif14 | 381293 | HITS-CLIP | 23142080 | Unspecified |
| mmu-let-7i-5p | MIMAT0000122 | Kif1a | 16560 | HITS-CLIP | 37986224 | Unspecified |
| mmu-let-7i-5p | MIMAT0000122 | Kif1b | 16561 | HITS-CLIP | 37986224\|23597149\|19536157\|23142 | Unspecified |
| mmu-let-7i-5p | MIMAT0000122 | Kif1bp | 26128 | HITS-CLIP | 23142080 | Unspecified |
| mmu-let-7i-5p | MIMAT0000122 | Kif1c | 16562 | HITS-CLIP | 23597149\|29167373 | Unspecified |
| mmu-let-7i-5p | MIMAT0000122 | Kif20b | 240641 | HITS-CLIP | 23142080 | Unspecified |
| mmu-let-7i-5p | MIMAT0000122 | Kif21a | 16564 | HITS-CLIP | 19536157 | Unspecified |
| mmu-let-7i-5p | MIMAT0000122 | Kif21b | 16565 | HITS-CLIP | 37986224\|19536157\|23142080\|29167 | Unspecified |
| mmu-let-7i-5p | MIMAT0000122 | Kif22 | 110033 | HITS-CLIP | 23142080 | Unspecified |
| mmu-let-7i-5p | MIMAT0000122 | Kif23 | 71819 | HITS-CLIP | 23142080 | Unspecified |
| mmu-let-7i-5p | MIMAT0000122 | Kif24 | 109242 | HITS-CLIP | 23630078\|23142080 | Unspecified |
| mmu-let-7i-5p | MIMAT0000122 | Kif2a | 16563 | HITS-CLIP | 37986224\|23597149\|23142080 | Unspecified |
| mmu-let-7i-5p | MIMAT0000122 | Kif3a | 16568 | HITS-CLIP | 23142080 | Unspecified |
| mmu-let-7i-5p | MIMAT0000122 | Kif3b | 16569 | HITS-CLIP | 19536157\|29167373 | Unspecified |
| mmu-let-7i-5p | MIMAT0000122 | Kif5a | 16572 | HITS-CLIP | 37986224 | Unspecified |
| mmu-let-7i-5p | MIMAT0000122 | Kif5b | 16573 | HITS-CLIP//qCLASH | 31433990\|23142080\|31393953\|29167 | Unspecified |
| mmu-let-7i-5p | MIMAT0000122 | Kif5c | 16574 | HITS-CLIP | 19536157 | Unspecified |
| mmu-let-7i-5p | MIMAT0000122 | Kifap3 | 16579 | HITS-CLIP | 19536157 | Unspecified |
| mmu-let-7i-5p | MIMAT0000122 | Kit | 16590 | HITS-CLIP | 23597149 | Unspecified |
| mmu-let-7i-5p | MIMAT0000122 | Klc1 | 16593 | HITS-CLIP | 19536157\|23597149\|23142080\|29167 | Unspecified |
| mmu-let-7i-5p | MIMAT0000122 | Klc4 | 74764 | HITS-CLIP | 23142080 | Unspecified |
| mmu-let-7i-5p | MIMAT0000122 | Klf10 | 21847 | HITS-CLIP | 37986224\|23597149 | Unspecified |
| mmu-let-7i-5p | MIMAT0000122 | Klf12 | 16597 | HITS-CLIP | 23597149 | Unspecified |
| mmu-let-7i-5p | MIMAT0000122 | Klf13 | 50794 | HITS-CLIP | 23597149\|19536157\|29167373\|23142 | Unspecified |

| | | | | | | |
|---|---|---|---|---|---|---|
| mmu-let-7i-5p | MIMAT0000122 | Klf15 | 66277 | HITS-CLIP | 23597149 | Unspecified |
| mmu-let-7i-5p | MIMAT0000122 | Klf3 | 16599 | HITS-CLIP | 23142080\|29167373 | Unspecified |
| mmu-let-7i-5p | MIMAT0000122 | Klf4 | 16600 | HITS-CLIP | 23597149 | Unspecified |
| mmu-let-7i-5p | MIMAT0000122 | Klf6 | 23849 | HITS-CLIP | 37986224\|23142080\|19536157\|29167 | Unspecified |
| mmu-let-7i-5p | MIMAT0000122 | Klf7 | 93691 | HITS-CLIP | 37986224\|23597149\|19536157 | Unspecified |
| mmu-let-7i-5p | MIMAT0000122 | Klf8 | 245671 | HITS-CLIP | 23142080 | Unspecified |
| mmu-let-7i-5p | MIMAT0000122 | Klf9 | 16601 | HITS-CLIP | 37986224\|19536157\|23597149\|23142 | Unspecified |
| mmu-let-7i-5p | MIMAT0000122 | Klhdc1 | 271005 | HITS-CLIP | 23142080 | Unspecified |
| mmu-let-7i-5p | MIMAT0000122 | Klhdc10 | 76788 | HITS-CLIP | 23597149\|19536157\|30971389\|23142 | Unspecified |
| mmu-let-7i-5p | MIMAT0000122 | Klhdc2 | 69554 | HITS-CLIP | 23142080 | Unspecified |
| mmu-let-7i-5p | MIMAT0000122 | Klhdc3 | 71765 | HITS-CLIP | 23142080 | Unspecified |
| mmu-let-7i-5p | MIMAT0000122 | Klhdc7a | 242721 | HITS-CLIP | 37986224 | Unspecified |
| mmu-let-7i-5p | MIMAT0000122 | Klhdc8a | 213417 | HITS-CLIP | 23597149 | Unspecified |
| mmu-let-7i-5p | MIMAT0000122 | Klhl11 | 217194 | HITS-CLIP | 37986224\|19536157 | Unspecified |
| mmu-let-7i-5p | MIMAT0000122 | Klhl15 | 236904 | HITS-CLIP | 23597149\|23142080 | Unspecified |
| mmu-let-7i-5p | MIMAT0000122 | Klhl2 | 77113 | HITS-CLIP | 37986224\|23142080\|23597149 | Unspecified |
| mmu-let-7i-5p | MIMAT0000122 | Klhl20 | 226541 | HITS-CLIP | 23597149\|23142080 | Unspecified |
| mmu-let-7i-5p | MIMAT0000122 | Klhl21 | 242785 | HITS-CLIP | 23597149 | Unspecified |
| mmu-let-7i-5p | MIMAT0000122 | Klhl24 | 75785 | HITS-CLIP | 37986224\|23597149\|19536157\|23142 | Unspecified |
| mmu-let-7i-5p | MIMAT0000122 | Klhl28 | 66689 | HITS-CLIP | 23142080 | Unspecified |
| mmu-let-7i-5p | MIMAT0000122 | Klhl42 | 232539 | HITS-CLIP | 19536157\|23142080\|29167373 | Unspecified |
| mmu-let-7i-5p | MIMAT0000122 | Klhl6 | 239743 | HITS-CLIP | 23597149\|23142080 | Unspecified |
| mmu-let-7i-5p | MIMAT0000122 | Klhl9 | 242521 | HITS-CLIP | 23597149\|23142080 | Unspecified |
| mmu-let-7i-5p | MIMAT0000122 | Klkb1 | 16621 | HITS-CLIP | 37986224\|23597149 | Unspecified |
| mmu-let-7i-5p | MIMAT0000122 | Kmo | 98256 | HITS-CLIP | 23597149 | Unspecified |
| mmu-let-7i-5p | MIMAT0000122 | Kmt2a | 214162 | HITS-CLIP | 37986224\|23597149\|19536157\|23142 | Unspecified |
| mmu-let-7i-5p | MIMAT0000122 | Kmt2b | 75410 | HITS-CLIP//qCLASH | 37986224\|31393953\|23142080 | Unspecified |
| mmu-let-7i-5p | MIMAT0000122 | Kmt2c | 231051 | HITS-CLIP | 23597149\|23142080 | Unspecified |
| mmu-let-7i-5p | MIMAT0000122 | Kmt2d | 381022 | HITS-CLIP//qCLASH | 37986224\|23597149\|23142080\|31393 | Unspecified |
| mmu-let-7i-5p | MIMAT0000122 | Kmt2e | 69188 | HITS-CLIP | 37986224\|23597149\|19536157\|23142 | Unspecified |
| mmu-let-7i-5p | MIMAT0000122 | Kmt5a | 387893 | HITS-CLIP | 23142080 | Unspecified |
| mmu-let-7i-5p | MIMAT0000122 | Kmt5b | 225888 | HITS-CLIP | 23597149\|19536157\|23142080\|29167 | Unspecified |
| mmu-let-7i-5p | MIMAT0000122 | Kmt5c | 84787 | HITS-CLIP | 23597149\|23142080 | Unspecified |
| mmu-let-7i-5p | MIMAT0000122 | Kng1 | 16644 | HITS-CLIP | 23597149 | Unspecified |
| mmu-let-7i-5p | MIMAT0000122 | Knl1 | 57082 | HITS-CLIP | 23142080 | Unspecified |
| mmu-let-7i-5p | MIMAT0000122 | Knop1 | 66356 | HITS-CLIP | 23142080 | Unspecified |
| mmu-let-7i-5p | MIMAT0000122 | Knstrn | 51944 | HITS-CLIP | 23142080 | Unspecified |
| mmu-let-7i-5p | MIMAT0000122 | Kntc1 | 208628 | HITS-CLIP | 23142080 | Unspecified |
| mmu-let-7i-5p | MIMAT0000122 | Kpna1 | 16646 | HITS-CLIP | 37986224\|19536157\|23142080\|29167 | Unspecified |

| | | | | | | |
|---|---|---|---|---|---|---|
| mmu-let-7i-5p | MIMAT0000122 | Kpna3 | 16648 | HITS-CLIP | 23142080 | Unspecified |
| mmu-let-7i-5p | MIMAT0000122 | Kpna4 | 16649 | HITS-CLIP | 37986224\|23597149\|23142080\|29167 | Unspecified |
| mmu-let-7i-5p | MIMAT0000122 | Kpna6 | 16650 | HITS-CLIP | 23597149\|23142080 | Unspecified |
| mmu-let-7i-5p | MIMAT0000122 | Kpnb1 | 16211 | HITS-CLIP | 19536157\|23142080\|29167373 | Unspecified |
| mmu-let-7i-5p | MIMAT0000122 | Kras | 16653 | HITS-CLIP | 23142080\|29167373 | Unspecified |
| mmu-let-7i-5p | MIMAT0000122 | Krba1 | 77827 | HITS-CLIP | 23142080 | Unspecified |
| mmu-let-7i-5p | MIMAT0000122 | Krcc1 | 57896 | HITS-CLIP | 23142080 | Unspecified |
| mmu-let-7i-5p | MIMAT0000122 | Kremen1 | 84035 | HITS-CLIP | 23597149 | Unspecified |
| mmu-let-7i-5p | MIMAT0000122 | Kremen2 | 73016 | HITS-CLIP | 23142080 | Unspecified |
| mmu-let-7i-5p | MIMAT0000122 | Kri1 | 215194 | HITS-CLIP | 23142080 | Unspecified |
| mmu-let-7i-5p | MIMAT0000122 | Krit1 | 79264 | HITS-CLIP | 23142080 | Unspecified |
| mmu-let-7i-5p | MIMAT0000122 | Krt7 | 110310 | HITS-CLIP | 26203562 | Unspecified |
| mmu-let-7i-5p | MIMAT0000122 | Ksr1 | 16706 | HITS-CLIP | 23142080 | Unspecified |
| mmu-let-7i-5p | MIMAT0000122 | Kti12 | 100087 | HITS-CLIP | 23142080 | Unspecified |
| mmu-let-7i-5p | MIMAT0000122 | Ktn1 | 16709 | HITS-CLIP | 23142080 | Unspecified |
| mmu-let-7i-5p | MIMAT0000122 | Kxd1 | 79036 | HITS-CLIP | 23142080 | Unspecified |
| mmu-let-7i-5p | MIMAT0000122 | Kyat1 | 883 | HITS-CLIP | 23597149 | Unspecified |
| mmu-let-7i-5p | MIMAT0000122 | Kyat3 | 56267 | HITS-CLIP | 23597149 | Unspecified |
| mmu-let-7i-5p | MIMAT0000122 | L1cam | 16728 | HITS-CLIP | 19536157\|29167373 | Unspecified |
| mmu-let-7i-5p | MIMAT0000122 | L2hgdh | 217666 | HITS-CLIP | 23142080 | Unspecified |
| mmu-let-7i-5p | MIMAT0000122 | L3mbtl3 | 237339 | HITS-CLIP | 23142080 | Unspecified |
| mmu-let-7i-5p | MIMAT0000122 | Lactb | 80907 | HITS-CLIP | 19536157 | Unspecified |
| mmu-let-7i-5p | MIMAT0000122 | Lair1 | 52855 | HITS-CLIP | 29167373 | Unspecified |
| mmu-let-7i-5p | MIMAT0000122 | Lama1 | 16772 | HITS-CLIP | 37986224\|19536157 | Unspecified |
| mmu-let-7i-5p | MIMAT0000122 | Lamb1 | 16777 | HITS-CLIP | 37986224\|19536157 | Unspecified |
| mmu-let-7i-5p | MIMAT0000122 | Lamc1 | 226519 | HITS-CLIP | 23597149\|23142080 | Unspecified |
| mmu-let-7i-5p | MIMAT0000122 | Lamp1 | 16783 | HITS-CLIP | 23597149\|19536157\|29167373\|23142 | Unspecified |
| mmu-let-7i-5p | MIMAT0000122 | Lamp2 | 16784 | HITS-CLIP | 37986224\|19536157\|23597149\|23142 | Unspecified |
| mmu-let-7i-5p | MIMAT0000122 | Lamp5 | 76161 | HITS-CLIP | 19536157 | Unspecified |
| mmu-let-7i-5p | MIMAT0000122 | Lancl1 | 14768 | HITS-CLIP | 23142080 | Unspecified |
| mmu-let-7i-5p | MIMAT0000122 | Lap3 | 66988 | HITS-CLIP | 37986224\|23597149\|23142080 | Unspecified |
| mmu-let-7i-5p | MIMAT0000122 | Laptm4a | 17775 | HITS-CLIP | 23597149\|19536157\|23142080\|26203 | Unspecified |
| mmu-let-7i-5p | MIMAT0000122 | Laptm4b | 114128 | HITS-CLIP | 23597149 | Unspecified |
| mmu-let-7i-5p | MIMAT0000122 | Laptm5 | 16792 | HITS-CLIP//qCLASH | 23142080\|31393953\|29167373 | Unspecified |
| mmu-let-7i-5p | MIMAT0000122 | Larp1 | 73158 | HITS-CLIP | 37986224\|23142080\|23597149\|29167 | Unspecified |
| mmu-let-7i-5p | MIMAT0000122 | Larp4 | 207214 | HITS-CLIP | 29167373 | Unspecified |
| mmu-let-7i-5p | MIMAT0000122 | Larp4b | 217980 | HITS-CLIP | 37986224\|19536157\|23142080 | Unspecified |
| mmu-let-7i-5p | MIMAT0000122 | Lars2 | 102436 | HITS-CLIP | 23597149\|29167373 | Unspecified |
| mmu-let-7i-5p | MIMAT0000122 | Lasp1 | 16796 | HITS-CLIP | 23597149\|29167373 | Unspecified |

| | | | | | | |
|---|---|---|---|---|---|---|
| mmu-let-7i-5p | MIMAT0000122 | Lats1 | 16798 | HITS-CLIP | 23142080\|19536157\|23597149\|29167 | Unspecified |
| mmu-let-7i-5p | MIMAT0000122 | Lax1 | 240754 | HITS-CLIP | 23142080 | Unspecified |
| mmu-let-7i-5p | MIMAT0000122 | Lbh | 77889 | HITS-CLIP | 37986224\|23142080 | Unspecified |
| mmu-let-7i-5p | MIMAT0000122 | Lbhd1 | 79081 | HITS-CLIP | 23142080 | Unspecified |
| mmu-let-7i-5p | MIMAT0000122 | Lbr | 98386 | HITS-CLIP | 37986224\|23597149\|23142080 | Unspecified |
| mmu-let-7i-5p | MIMAT0000122 | Lcat | 16816 | HITS-CLIP | 23597149 | Unspecified |
| mmu-let-7i-5p | MIMAT0000122 | Lck | 16818 | HITS-CLIP | 23142080 | Unspecified |
| mmu-let-7i-5p | MIMAT0000122 | Lcmt1 | 30949 | HITS-CLIP | 23597149 | Unspecified |
| mmu-let-7i-5p | MIMAT0000122 | Lcn2 | 16819 | HITS-CLIP | 23597149 | Unspecified |
| mmu-let-7i-5p | MIMAT0000122 | Lcor | 212391 | HITS-CLIP | 37986224\|23597149\|23142080\|29167 | Unspecified |
| mmu-let-7i-5p | MIMAT0000122 | Lcorl | 209707 | HITS-CLIP | 23142080 | Unspecified |
| mmu-let-7i-5p | MIMAT0000122 | Lcp1 | 18826 | HITS-CLIP | 23142080\|23597149\|29167373 | Unspecified |
| mmu-let-7i-5p | MIMAT0000122 | Lcp2 | 16822 | HITS-CLIP | 23142080 | Unspecified |
| mmu-let-7i-5p | MIMAT0000122 | Ldah | 38150 | HITS-CLIP | 23597149\|23142080 | Unspecified |
| mmu-let-7i-5p | MIMAT0000122 | Ldb1 | 16825 | HITS-CLIP | 23142080\|23597149 | Unspecified |
| mmu-let-7i-5p | MIMAT0000122 | Ldb2 | 16826 | HITS-CLIP | 37986224\|23597149 | Unspecified |
| mmu-let-7i-5p | MIMAT0000122 | Ldlr | 16835 | HITS-CLIP | 37986224\|23597149\|31433990\|2363( | Unspecified |
| mmu-let-7i-5p | MIMAT0000122 | Ldlrap1 | 100017 | HITS-CLIP | 29167373 | Unspecified |
| mmu-let-7i-5p | MIMAT0000122 | Lef1 | 16842 | HITS-CLIP | 23142080 | Unspecified |
| mmu-let-7i-5p | MIMAT0000122 | Lemd3 | 380664 | HITS-CLIP | 23597149\|23142080\|29167373 | Unspecified |
| mmu-let-7i-5p | MIMAT0000122 | Leng8 | 232798 | HITS-CLIP | 29167373\|23142080 | Unspecified |
| mmu-let-7i-5p | MIMAT0000122 | Leprotl1 | 68192 | HITS-CLIP | 37986224\|23597149\|19536157 | Unspecified |
| mmu-let-7i-5p | MIMAT0000122 | Letm1 | 56384 | HITS-CLIP | 37986224\|23597149 | Unspecified |
| mmu-let-7i-5p | MIMAT0000122 | Letmd1 | 68614 | HITS-CLIP | 37986224\|23597149\|19536157\|23142 | Unspecified |
| mmu-let-7i-5p | MIMAT0000122 | Lfng | 16848 | HITS-CLIP | 29167373 | Unspecified |
| mmu-let-7i-5p | MIMAT0000122 | Lgals1 | 16852 | HITS-CLIP | 23142080\|29167373 | Unspecified |
| mmu-let-7i-5p | MIMAT0000122 | Lgals3bp | 19039 | HITS-CLIP | 37986224\|23597149\|29167373\|23142 | Unspecified |
| mmu-let-7i-5p | MIMAT0000122 | Lgals8 | 56048 | HITS-CLIP | 37986224\|23142080\|29167373 | Unspecified |
| mmu-let-7i-5p | MIMAT0000122 | Lgals9 | 16859 | HITS-CLIP | 23597149 | Unspecified |
| mmu-let-7i-5p | MIMAT0000122 | Lgmn | 19141 | HITS-CLIP | 23597149 | Unspecified |
| mmu-let-7i-5p | MIMAT0000122 | Lgr4 | 107515 | HITS-CLIP | 23597149\|19536157 | Unspecified |
| mmu-let-7i-5p | MIMAT0000122 | Lhfpl2 | 218454 | HITS-CLIP | 29167373 | Unspecified |
| mmu-let-7i-5p | MIMAT0000122 | Lhx1 | 16869 | HITS-CLIP | 37986224 | Unspecified |
| mmu-let-7i-5p | MIMAT0000122 | Lif | 16878 | HITS-CLIP | 31433990 | Unspecified |
| mmu-let-7i-5p | MIMAT0000122 | Lifr | 16880 | HITS-CLIP | 37986224\|23597149 | Unspecified |
| mmu-let-7i-5p | MIMAT0000122 | Lig1 | 16881 | HITS-CLIP | 37986224\|23142080\|23597149 | Unspecified |
| mmu-let-7i-5p | MIMAT0000122 | Limd1 | 29806 | HITS-CLIP | 23597149\|23142080 | Unspecified |
| mmu-let-7i-5p | MIMAT0000122 | Limd2 | 67803 | HITS-CLIP | 37986224\|23142080 | Unspecified |
| mmu-let-7i-5p | MIMAT0000122 | Limk2 | 16886 | HITS-CLIP | 23142080 | Unspecified |

| | | | | | | |
|---|---|---|---|---|---|---|
| mmu-let-7i-5p | MIMAT0000122 | Lims1 | 110829 | HITS-CLIP | 23142080\|29167373 | Unspecified |
| mmu-let-7i-5p | MIMAT0000122 | Lims2 | 225341 | HITS-CLIP | 23597149 | Unspecified |
| mmu-let-7i-5p | MIMAT0000122 | Lin54 | 231506 | HITS-CLIP | 23142080 | Unspecified |
| mmu-let-7i-5p | MIMAT0000122 | Lin7c | 22343 | HITS-CLIP | 37986224\|23142080 | Unspecified |
| mmu-let-7i-5p | MIMAT0000122 | Lingo1 | 235402 | HITS-CLIP | 37986224\|19536157 | Unspecified |
| mmu-let-7i-5p | MIMAT0000122 | Lins1 | 72635 | HITS-CLIP | 23142080\|25083871 | Unspecified |
| mmu-let-7i-5p | MIMAT0000122 | Lipa | 16889 | HITS-CLIP | 23597149\|29167373 | Unspecified |
| mmu-let-7i-5p | MIMAT0000122 | Lipe | 16890 | HITS-CLIP | 23142080 | Unspecified |
| mmu-let-7i-5p | MIMAT0000122 | Lipt2 | 67164 | HITS-CLIP | 37986224\|23597149\|23142080 | Unspecified |
| mmu-let-7i-5p | MIMAT0000122 | Litaf | 56722 | HITS-CLIP | 23142080 | Unspecified |
| mmu-let-7i-5p | MIMAT0000122 | Llgl1 | 16897 | HITS-CLIP | 23142080 | Unspecified |
| mmu-let-7i-5p | MIMAT0000122 | Llgl2 | 217325 | HITS-CLIP | 37986224 | Unspecified |
| mmu-let-7i-5p | MIMAT0000122 | Lman1 | 70361 | HITS-CLIP | 23142080\|23597149 | Unspecified |
| mmu-let-7i-5p | MIMAT0000122 | Lman2 | 66890 | HITS-CLIP | 23597149\|23142080\|29167373 | Unspecified |
| mmu-let-7i-5p | MIMAT0000122 | Lmbr1l | 74775 | HITS-CLIP//qCLASH | 37986224\|23597149\|31393953\|23142 | Unspecified |
| mmu-let-7i-5p | MIMAT0000122 | Lmbrd1 | 68421 | HITS-CLIP | 23142080 | Unspecified |
| mmu-let-7i-5p | MIMAT0000122 | Lmbrd2 | 320506 | HITS-CLIP | 37986224\|23142080\|19536157\|29167 | Unspecified |
| mmu-let-7i-5p | MIMAT0000122 | Lmna | 16905 | HITS-CLIP | 23597149\|29167373 | Unspecified |
| mmu-let-7i-5p | MIMAT0000122 | Lmnb1 | 16906 | HITS-CLIP | 37986224\|23597149\|23142080\|19536 | Unspecified |
| mmu-let-7i-5p | MIMAT0000122 | Lmo4 | 16911 | HITS-CLIP | 23597149\|23142080 | Unspecified |
| mmu-let-7i-5p | MIMAT0000122 | Lmtk2 | 231876 | HITS-CLIP | 23142080 | Unspecified |
| mmu-let-7i-5p | MIMAT0000122 | Lnpep | 240028 | HITS-CLIP | 23597149\|19536157\|23142080\|29167 | Unspecified |
| mmu-let-7i-5p | MIMAT0000122 | Lnpk | 69605 | HITS-CLIP | 19536157 | Unspecified |
| mmu-let-7i-5p | MIMAT0000122 | Lonp1 | 74142 | HITS-CLIP | 23142080 | Unspecified |
| mmu-let-7i-5p | MIMAT0000122 | Lonp2 | 66887 | HITS-CLIP | 23597149\|19536157\|23142080 | Unspecified |
| mmu-let-7i-5p | MIMAT0000122 | Lonrf1 | 244421 | HITS-CLIP | 23142080 | Unspecified |
| mmu-let-7i-5p | MIMAT0000122 | Lonrf3 | 74365 | HITS-CLIP | 29167373 | Unspecified |
| mmu-let-7i-5p | MIMAT0000122 | Loxl4 | 67573 | HITS-CLIP | 37986224 | Unspecified |
| mmu-let-7i-5p | MIMAT0000122 | Lpar6 | 67168 | HITS-CLIP | 37986224\|23597149\|29167373\|23142 | Unspecified |
| mmu-let-7i-5p | MIMAT0000122 | Lpcat1 | 210992 | HITS-CLIP//qCLASH | 23142080\|31393953\|29167373 | Unspecified |
| mmu-let-7i-5p | MIMAT0000122 | Lpcat3 | 14792 | HITS-CLIP | 37986224\|23142080 | Unspecified |
| mmu-let-7i-5p | MIMAT0000122 | Lpgat1 | 226856 | HITS-CLIP | 37986224\|23597149\|19536157\|23142 | Unspecified |
| mmu-let-7i-5p | MIMAT0000122 | Lpin1 | 14245 | HITS-CLIP | 23597149\|23142080 | Unspecified |
| mmu-let-7i-5p | MIMAT0000122 | Lpin2 | 64898 | HITS-CLIP | 37986224\|23597149\|31433990\|29167 | Unspecified |
| mmu-let-7i-5p | MIMAT0000122 | Lpl | 16956 | HITS-CLIP | 23597149\|29167373 | Unspecified |
| mmu-let-7i-5p | MIMAT0000122 | Lpp | 210126 | HITS-CLIP | 19536157\|23597149\|29167373\|23142 | Unspecified |
| mmu-let-7i-5p | MIMAT0000122 | Lpxn | 107321 | HITS-CLIP | 23142080\|29167373 | Unspecified |
| mmu-let-7i-5p | MIMAT0000122 | Lrba | 80877 | HITS-CLIP | 37986224\|23597149\|23142080 | Unspecified |
| mmu-let-7i-5p | MIMAT0000122 | Lrch1 | 380916 | HITS-CLIP | 29167373 | Unspecified |

| | | | | | | |
|---|---|---|---|---|---|---|
| mmu-let-7i-5p | MIMAT0000122 | Lrch3 | 70144 | HITS-CLIP | 23142080 | Unspecified |
| mmu-let-7i-5p | MIMAT0000122 | Lrch4 | 231798 | HITS-CLIP | 37986224\|23597149\|19536157\|23142 | Unspecified |
| mmu-let-7i-5p | MIMAT0000122 | Lrfn5 | 238205 | HITS-CLIP | 19536157 | Unspecified |
| mmu-let-7i-5p | MIMAT0000122 | Lrig1 | 16206 | HITS-CLIP | 23597149\|23142080 | Unspecified |
| mmu-let-7i-5p | MIMAT0000122 | Lrig2 | 269473 | HITS-CLIP | 37986224\|23597149\|19536157\|29167 | Unspecified |
| mmu-let-7i-5p | MIMAT0000122 | Lrig3 | 320398 | HITS-CLIP | 23597149 | Unspecified |
| mmu-let-7i-5p | MIMAT0000122 | Lrmda | 83938 | HITS-CLIP | 29167373 | Unspecified |
| mmu-let-7i-5p | MIMAT0000122 | Lrp1 | 16971 | HITS-CLIP | 37986224\|23597149\|19536157\|29167 | Unspecified |
| mmu-let-7i-5p | MIMAT0000122 | Lrp10 | 65107 | HITS-CLIP | 23597149 | Unspecified |
| mmu-let-7i-5p | MIMAT0000122 | Lrp11 | 237253 | HITS-CLIP | 37986224 | Unspecified |
| mmu-let-7i-5p | MIMAT0000122 | Lrp12 | 239393 | HITS-CLIP | 23142080\|29167373 | Unspecified |
| mmu-let-7i-5p | MIMAT0000122 | Lrp6 | 16974 | HITS-CLIP | 23597149\|19536157\|23142080 | Unspecified |
| mmu-let-7i-5p | MIMAT0000122 | Lrp8 | 16975 | HITS-CLIP | 23142080 | Unspecified |
| mmu-let-7i-5p | MIMAT0000122 | Lrpprc | 72416 | HITS-CLIP | 23597149 | Unspecified |
| mmu-let-7i-5p | MIMAT0000122 | Lrr1 | 69706 | HITS-CLIP | 23142080 | Unspecified |
| mmu-let-7i-5p | MIMAT0000122 | Lrrc1 | 214345 | HITS-CLIP | 23142080 | Unspecified |
| mmu-let-7i-5p | MIMAT0000122 | Lrrc14 | 223664 | HITS-CLIP | 23142080 | Unspecified |
| mmu-let-7i-5p | MIMAT0000122 | Lrrc28 | 67867 | HITS-CLIP | 23597149 | Unspecified |
| mmu-let-7i-5p | MIMAT0000122 | Lrrc32 | 434215 | HITS-CLIP | 37986224\|23142080 | Unspecified |
| mmu-let-7i-5p | MIMAT0000122 | Lrrc41 | 230654 | HITS-CLIP | 23597149\|23142080 | Unspecified |
| mmu-let-7i-5p | MIMAT0000122 | Lrrc42 | 77809 | HITS-CLIP | 23142080 | Unspecified |
| mmu-let-7i-5p | MIMAT0000122 | Lrrc4b | 272381 | HITS-CLIP | 37986224 | Unspecified |
| mmu-let-7i-5p | MIMAT0000122 | Lrrc4c | 241568 | HITS-CLIP | 19536157 | Unspecified |
| mmu-let-7i-5p | MIMAT0000122 | Lrrc58 | 320184 | HITS-CLIP | 37986224\|23597149\|19536157\|23142 | Unspecified |
| mmu-let-7i-5p | MIMAT0000122 | Lrrc59 | 98238 | HITS-CLIP | 23597149\|23142080 | Unspecified |
| mmu-let-7i-5p | MIMAT0000122 | Lrrc7 | 242274 | HITS-CLIP | 37986224\|19536157 | Unspecified |
| mmu-let-7i-5p | MIMAT0000122 | Lrrc75a | 192976 | HITS-CLIP | 29167373 | Unspecified |
| mmu-let-7i-5p | MIMAT0000122 | Lrrc8a | 241296 | HITS-CLIP | 23597149\|19536157\|23142080 | Unspecified |
| mmu-let-7i-5p | MIMAT0000122 | Lrrc8d | 231549 | HITS-CLIP | 23597149 | Unspecified |
| mmu-let-7i-5p | MIMAT0000122 | Lrrcc1 | 71710 | HITS-CLIP | 23142080 | Unspecified |
| mmu-let-7i-5p | MIMAT0000122 | Lrrfip1 | 16978 | HITS-CLIP | 37986224\|23597149\|23142080\|29167 | Unspecified |
| mmu-let-7i-5p | MIMAT0000122 | Lrrn1 | 16979 | HITS-CLIP | 37986224\|19536157 | Unspecified |
| mmu-let-7i-5p | MIMAT0000122 | Lrrn3 | 16981 | HITS-CLIP | 37986224\|19536157 | Unspecified |
| mmu-let-7i-5p | MIMAT0000122 | Lrwd1 | 71735 | HITS-CLIP//qCLASH | 37986224\|23597149\|31393953\|23142 | Unspecified |
| mmu-let-7i-5p | MIMAT0000122 | Lsamp | 268890 | HITS-CLIP | 37986224 | Unspecified |
| mmu-let-7i-5p | MIMAT0000122 | Lsg1 | 224092 | HITS-CLIP | 23597149\|23142080 | Unspecified |
| mmu-let-7i-5p | MIMAT0000122 | Lsm12 | 268490 | HITS-CLIP | 23142080\|29167373 | Unspecified |
| mmu-let-7i-5p | MIMAT0000122 | Lsm14a | 67070 | HITS-CLIP | 23142080\|26203562\|29167373 | Unspecified |
| mmu-let-7i-5p | MIMAT0000122 | Lsm2 | 27756 | HITS-CLIP | 23142080 | Unspecified |

| | | | | | | |
|---|---|---|---|---|---|---|
| mmu-let-7i-5p | MIMAT0000122 | Lsm3 | 67678 | HITS-CLIP | 30971389 | Unspecified |
| mmu-let-7i-5p | MIMAT0000122 | Lsr | 54135 | HITS-CLIP | 23597149 | Unspecified |
| mmu-let-7i-5p | MIMAT0000122 | Lss | 16987 | HITS-CLIP | 23597149\|23142080 | Unspecified |
| mmu-let-7i-5p | MIMAT0000122 | Lta | 16992 | HITS-CLIP | 23142080 | Unspecified |
| mmu-let-7i-5p | MIMAT0000122 | Ltb | 16994 | HITS-CLIP | 37986224\|23597149\|23142080 | Unspecified |
| mmu-let-7i-5p | MIMAT0000122 | Ltbp4 | 108075 | HITS-CLIP | 23597149 | Unspecified |
| mmu-let-7i-5p | MIMAT0000122 | Ltn1 | 78913 | HITS-CLIP | 37986224\|19536157\|23142080\|29167 | Unspecified |
| mmu-let-7i-5p | MIMAT0000122 | Ltv1 | 353258 | HITS-CLIP | 19536157 | Unspecified |
| mmu-let-7i-5p | MIMAT0000122 | Luc7l2 | 192196 | HITS-CLIP | 23142080 | Unspecified |
| mmu-let-7i-5p | MIMAT0000122 | Luc7l3 | 67684 | HITS-CLIP | 29167373 | Unspecified |
| mmu-let-7i-5p | MIMAT0000122 | Luzp1 | 269593 | HITS-CLIP | 23597149\|29167373 | Unspecified |
| mmu-let-7i-5p | MIMAT0000122 | Ly6g6g | 78725 | qCLASH | 31393953 | Unspecified |
| mmu-let-7i-5p | MIMAT0000122 | Ly75 | 17076 | HITS-CLIP | 23597149 | Unspecified |
| mmu-let-7i-5p | MIMAT0000122 | Lyn | 17096 | HITS-CLIP | 23597149\|29167373 | Unspecified |
| mmu-let-7i-5p | MIMAT0000122 | Lynx1 | 23936 | HITS-CLIP | 37986224\|19536157 | Unspecified |
| mmu-let-7i-5p | MIMAT0000122 | Lypla1 | 18777 | HITS-CLIP | 23142080 | Unspecified |
| mmu-let-7i-5p | MIMAT0000122 | Lypla2 | 26394 | HITS-CLIP | 23597149 | Unspecified |
| mmu-let-7i-5p | MIMAT0000122 | Lyrm1 | 57149 | HITS-CLIP | 23597149 | Unspecified |
| mmu-let-7i-5p | MIMAT0000122 | Lysmd3 | 80289 | HITS-CLIP | 23142080\|19536157\|23597149 | Unspecified |
| mmu-let-7i-5p | MIMAT0000122 | Lysmd4 | 75099 | HITS-CLIP | 23142080 | Unspecified |
| mmu-let-7i-5p | MIMAT0000122 | Lyst | 17101 | HITS-CLIP | 37986224\|23597149\|23142080\|29167 | Unspecified |
| mmu-let-7i-5p | MIMAT0000122 | Lzts3 | 241638 | HITS-CLIP | 37986224\|23597149 | Unspecified |
| mmu-let-7i-5p | MIMAT0000122 | M6pr | 17113 | HITS-CLIP | 23142080\|29167373 | Unspecified |
| mmu-let-7i-5p | MIMAT0000122 | Macf1 | 11426 | HITS-CLIP//qCLASH | 37986224\|19536157\|23597149\|3139 | Unspecified |
| mmu-let-7i-5p | MIMAT0000122 | Mad1l1 | 17120 | HITS-CLIP | 23142080 | Unspecified |
| mmu-let-7i-5p | MIMAT0000122 | Madd | 228355 | HITS-CLIP | 23142080 | Unspecified |
| mmu-let-7i-5p | MIMAT0000122 | Maea | 59003 | HITS-CLIP | 23597149\|23142080 | Unspecified |
| mmu-let-7i-5p | MIMAT0000122 | Maf | 17132 | HITS-CLIP | 23597149\|23142080\|29167373 | Unspecified |
| mmu-let-7i-5p | MIMAT0000122 | Maf1 | 68877 | HITS-CLIP | 23142080 | Unspecified |
| mmu-let-7i-5p | MIMAT0000122 | Mafg | 17134 | HITS-CLIP | 23630078\|23597149\|26203562\|29167 | Unspecified |
| mmu-let-7i-5p | MIMAT0000122 | Maged1 | 94275 | HITS-CLIP | 23597149 | Unspecified |
| mmu-let-7i-5p | MIMAT0000122 | Magi1 | 14924 | HITS-CLIP | 23597149 | Unspecified |
| mmu-let-7i-5p | MIMAT0000122 | Magi2 | 50791 | HITS-CLIP | 37986224 | Unspecified |
| mmu-let-7i-5p | MIMAT0000122 | Magi3 | 99470 | HITS-CLIP | 23597149 | Unspecified |
| mmu-let-7i-5p | MIMAT0000122 | Magohb | 66441 | HITS-CLIP | 23142080 | Unspecified |
| mmu-let-7i-5p | MIMAT0000122 | Magt1 | 67075 | HITS-CLIP | 37986224\|23142080\|23597149\|1953 | Unspecified |
| mmu-let-7i-5p | MIMAT0000122 | Maip1 | 79568 | HITS-CLIP | 23142080 | Unspecified |
| mmu-let-7i-5p | MIMAT0000122 | Mak16 | 67920 | HITS-CLIP | 23142080 | Unspecified |
| mmu-let-7i-5p | MIMAT0000122 | Mal2 | 105853 | HITS-CLIP | 37986224\|23597149 | Unspecified |

| | | | | | | |
|---|---|---|---|---|---|---|
| mmu-let-7i-5p | MIMAT0000122 | Malrd1 | 340895 | HITS-CLIP | 23142080 | Unspecified |
| mmu-let-7i-5p | MIMAT0000122 | Malsu1 | 75593 | HITS-CLIP | 23142080 | Unspecified |
| mmu-let-7i-5p | MIMAT0000122 | Malt1 | 240354 | HITS-CLIP | 23142080 | Unspecified |
| mmu-let-7i-5p | MIMAT0000122 | Maml1 | 103806 | HITS-CLIP | 23597149\|23142080\|29167373 | Unspecified |
| mmu-let-7i-5p | MIMAT0000122 | Maml2 | 270118 | HITS-CLIP | 31433990 | Unspecified |
| mmu-let-7i-5p | MIMAT0000122 | Maml3 | 433586 | HITS-CLIP | 19536157\|23142080 | Unspecified |
| mmu-let-7i-5p | MIMAT0000122 | Man1a | 17155 | HITS-CLIP | 37986224\|23597149\|23142080 | Unspecified |
| mmu-let-7i-5p | MIMAT0000122 | Man1a2 | 17156 | HITS-CLIP | 23597149\|19536157\|23142080\|29167 | Unspecified |
| mmu-let-7i-5p | MIMAT0000122 | Man1b1 | 227619 | HITS-CLIP | 23597149 | Unspecified |
| mmu-let-7i-5p | MIMAT0000122 | Man2a1 | 17158 | HITS-CLIP | 37986224\|23597149\|23142080 | Unspecified |
| mmu-let-7i-5p | MIMAT0000122 | Man2a2 | 140481 | HITS-CLIP | 37986224 | Unspecified |
| mmu-let-7i-5p | MIMAT0000122 | Man2b1 | 17159 | HITS-CLIP | 23597149\|29167373\|23142080 | Unspecified |
| mmu-let-7i-5p | MIMAT0000122 | Man2c1 | 73744 | HITS-CLIP | 37986224\|23597149 | Unspecified |
| mmu-let-7i-5p | MIMAT0000122 | Maneal | 215090 | HITS-CLIP | 19536157 | Unspecified |
| mmu-let-7i-5p | MIMAT0000122 | Maob | 109731 | HITS-CLIP | 23597149 | Unspecified |
| mmu-let-7i-5p | MIMAT0000122 | Map1a | 17754 | HITS-CLIP | 19536157 | Unspecified |
| mmu-let-7i-5p | MIMAT0000122 | Map1b | 17755 | HITS-CLIP | 37986224\|19536157 | Unspecified |
| mmu-let-7i-5p | MIMAT0000122 | Map1lc3b | 67443 | HITS-CLIP | 23142080 | Unspecified |
| mmu-let-7i-5p | MIMAT0000122 | Map2 | 17756 | HITS-CLIP | 37986224\|19536157 | Unspecified |
| mmu-let-7i-5p | MIMAT0000122 | Map2k1 | 26395 | HITS-CLIP | 23142080 | Unspecified |
| mmu-let-7i-5p | MIMAT0000122 | Map2k2 | 26396 | HITS-CLIP | 23597149 | Unspecified |
| mmu-let-7i-5p | MIMAT0000122 | Map2k3 | 26397 | HITS-CLIP | 23142080 | Unspecified |
| mmu-let-7i-5p | MIMAT0000122 | Map2k4 | 26398 | HITS-CLIP | 37986224\|23597149\|19536157\|23142 | Unspecified |
| mmu-let-7i-5p | MIMAT0000122 | Map2k6 | 26399 | HITS-CLIP | 23142080 | Unspecified |
| mmu-let-7i-5p | MIMAT0000122 | Map3k1 | 26401 | HITS-CLIP | 37986224\|23597149\|19536157\|23142 | Unspecified |
| mmu-let-7i-5p | MIMAT0000122 | Map3k11 | 26403 | HITS-CLIP | 23142080 | Unspecified |
| mmu-let-7i-5p | MIMAT0000122 | Map3k13 | 71751 | HITS-CLIP | 23597149\|19536157 | Unspecified |
| mmu-let-7i-5p | MIMAT0000122 | Map3k14 | 53859 | HITS-CLIP | 37986224 | Unspecified |
| mmu-let-7i-5p | MIMAT0000122 | Map3k2 | 26405 | HITS-CLIP | 19536157\|23597149\|23142080\|29167 | Unspecified |
| mmu-let-7i-5p | MIMAT0000122 | Map3k3 | 26406 | HITS-CLIP | 37986224\|23142080\|29167373 | Unspecified |
| mmu-let-7i-5p | MIMAT0000122 | Map3k4 | 26407 | HITS-CLIP | 37986224\|23597149\|23142080 | Unspecified |
| mmu-let-7i-5p | MIMAT0000122 | Map3k5 | 26408 | HITS-CLIP | 23597149\|23142080 | Unspecified |
| mmu-let-7i-5p | MIMAT0000122 | Map3k7 | 26409 | HITS-CLIP | 23630078\|31433990\|30971389\|23142 | Unspecified |
| mmu-let-7i-5p | MIMAT0000122 | Map3k8 | 26410 | HITS-CLIP | 23597149\|29167373 | Unspecified |
| mmu-let-7i-5p | MIMAT0000122 | Map3k9 | 338372 | HITS-CLIP | 37986224\|19536157 | Unspecified |
| mmu-let-7i-5p | MIMAT0000122 | Map4 | 17758 | HITS-CLIP | 29167373\|23142080 | Unspecified |
| mmu-let-7i-5p | MIMAT0000122 | Map4k1 | 26411 | HITS-CLIP | 23142080 | Unspecified |
| mmu-let-7i-5p | MIMAT0000122 | Map4k2 | 26412 | HITS-CLIP | 23142080 | Unspecified |
| mmu-let-7i-5p | MIMAT0000122 | Map4k3 | 225028 | HITS-CLIP | 37986224\|19536157\|23597149\|23142 | Unspecified |

| | | | | | | |
|---|---|---|---|---|---|---|
| mmu-let-7i-5p | MIMAT0000122 | Map4k4 | 26921 | HITS-CLIP | 23597149 | Unspecified |
| mmu-let-7i-5p | MIMAT0000122 | Map4k5 | 399510 | HITS-CLIP | 23597149 | Unspecified |
| mmu-let-7i-5p | MIMAT0000122 | Map6 | 17760 | HITS-CLIP | 37986224 | Unspecified |
| mmu-let-7i-5p | MIMAT0000122 | Map7 | 17761 | HITS-CLIP | 23142080 | Unspecified |
| mmu-let-7i-5p | MIMAT0000122 | Map7d1 | 245877 | HITS-CLIP | 19536157\|29167373\|23142080 | Unspecified |
| mmu-let-7i-5p | MIMAT0000122 | Map9 | 213582 | HITS-CLIP | 37986224\|19536157 | Unspecified |
| mmu-let-7i-5p | MIMAT0000122 | Mapk1 | 26413 | HITS-CLIP | 37986224\|23142080 | Unspecified |
| mmu-let-7i-5p | MIMAT0000122 | Mapk11 | 19094 | HITS-CLIP | 37986224 | Unspecified |
| mmu-let-7i-5p | MIMAT0000122 | Mapk14 | 26416 | HITS-CLIP | 31433990\|23597149\|19536157\|29167 | Unspecified |
| mmu-let-7i-5p | MIMAT0000122 | Mapk1ip1l | 218975 | HITS-CLIP | 23142080\|23597149\|29167373 | Unspecified |
| mmu-let-7i-5p | MIMAT0000122 | Mapk3 | 26417 | HITS-CLIP | 23597149\|23142080 | Unspecified |
| mmu-let-7i-5p | MIMAT0000122 | Mapk4 | 225724 | HITS-CLIP | 37986224 | Unspecified |
| mmu-let-7i-5p | MIMAT0000122 | Mapk6 | 50772 | HITS-CLIP | 37986224\|23142080\|19536157\|23597 | Unspecified |
| mmu-let-7i-5p | MIMAT0000122 | Mapk7 | 23939 | HITS-CLIP | 23142080 | Unspecified |
| mmu-let-7i-5p | MIMAT0000122 | Mapk8 | 26419 | HITS-CLIP | 23142080 | Unspecified |
| mmu-let-7i-5p | MIMAT0000122 | Mapk8ip3 | 30957 | HITS-CLIP | 23142080 | Unspecified |
| mmu-let-7i-5p | MIMAT0000122 | Mapk9 | 26420 | HITS-CLIP | 37986224\|23597149\|23142080 | Unspecified |
| mmu-let-7i-5p | MIMAT0000122 | Mapkap1 | 227743 | HITS-CLIP | 37986224\|23142080\|19536157\|23597 | Unspecified |
| mmu-let-7i-5p | MIMAT0000122 | Mapkapk2 | 17164 | HITS-CLIP | 23142080 | Unspecified |
| mmu-let-7i-5p | MIMAT0000122 | Mapkbp1 | 26390 | HITS-CLIP | 37986224 | Unspecified |
| mmu-let-7i-5p | MIMAT0000122 | Mapre1 | 13589 | HITS-CLIP | 19536157\|23142080\|29167373 | Unspecified |
| mmu-let-7i-5p | MIMAT0000122 | Mapre2 | 212307 | HITS-CLIP | 37986224\|23142080\|19536157 | Unspecified |
| mmu-let-7i-5p | MIMAT0000122 | Mapre3 | 100732 | HITS-CLIP | 23597149 | Unspecified |
| mmu-let-7i-5p | MIMAT0000122 | Mapt | 17762 | HITS-CLIP | 37986224\|19536157 | Unspecified |
| mmu-let-7i-5p | MIMAT0000122 | Marc1 | 66112 | HITS-CLIP | 23597149 | Unspecified |
| mmu-let-7i-5p | MIMAT0000122 | Marc2 | 67247 | HITS-CLIP | 23597149 | Unspecified |
| mmu-let-7i-5p | MIMAT0000122 | March4 | 381270 | HITS-CLIP | 37986224 | Unspecified |
| mmu-let-7i-5p | MIMAT0000122 | March6 | 10299 | HITS-CLIP | 23597149\|19536157\|23142080 | Unspecified |
| mmu-let-7i-5p | MIMAT0000122 | March7 | 57438 | HITS-CLIP | 37986224\|23597149\|19536157\|23142 | Unspecified |
| mmu-let-7i-5p | MIMAT0000122 | March8 | 71779 | HITS-CLIP | 29167373 | Unspecified |
| mmu-let-7i-5p | MIMAT0000122 | March9 | 216438 | HITS-CLIP | 37986224\|19536157\|23142080 | Unspecified |
| mmu-let-7i-5p | MIMAT0000122 | Marcks | 17118 | HITS-CLIP | 37986224\|23142080\|29167373 | Unspecified |
| mmu-let-7i-5p | MIMAT0000122 | Marcksl1 | 17357 | HITS-CLIP | 23597149\|29167373 | Unspecified |
| mmu-let-7i-5p | MIMAT0000122 | Marf1 | 223989 | HITS-CLIP | 23597149\|23142080\|29167373 | Unspecified |
| mmu-let-7i-5p | MIMAT0000122 | Mark2 | 13728 | HITS-CLIP | 23597149\|19536157\|23142080 | Unspecified |
| mmu-let-7i-5p | MIMAT0000122 | Mark3 | 17169 | HITS-CLIP | 37986224 | Unspecified |
| mmu-let-7i-5p | MIMAT0000122 | Mark4 | 232944 | HITS-CLIP | 23142080 | Unspecified |
| mmu-let-7i-5p | MIMAT0000122 | Mars | 216443 | HITS-CLIP | 23142080\|23597149 | Unspecified |
| mmu-let-7i-5p | MIMAT0000122 | Mars2 | 212679 | HITS-CLIP | 23597149\|23142080 | Unspecified |

| | | | | | | |
|---|---|---|---|---|---|---|
| mmu-let-7i-5p | MIMAT0000122 | Marveld2 | 218518 | HITS-CLIP | 23142080\|23597149 | Unspecified |
| mmu-let-7i-5p | MIMAT0000122 | Masp1 | 17174 | HITS-CLIP | 37986224\|23597149 | Unspecified |
| mmu-let-7i-5p | MIMAT0000122 | Masp2 | 17175 | HITS-CLIP | 23597149 | Unspecified |
| mmu-let-7i-5p | MIMAT0000122 | Mast2 | 17776 | qCLASH | 31393953 | Unspecified |
| mmu-let-7i-5p | MIMAT0000122 | Mast4 | 328329 | HITS-CLIP | 23142080\|19536157 | Unspecified |
| mmu-let-7i-5p | MIMAT0000122 | Mastl | 67121 | HITS-CLIP | 23142080 | Unspecified |
| mmu-let-7i-5p | MIMAT0000122 | Mat1a | 11720 | HITS-CLIP | 23597149 | Unspecified |
| mmu-let-7i-5p | MIMAT0000122 | Mat2a | 232087 | HITS-CLIP | 23142080 | Unspecified |
| mmu-let-7i-5p | MIMAT0000122 | Mat2b | 108645 | HITS-CLIP | 23597149\|23142080 | Unspecified |
| mmu-let-7i-5p | MIMAT0000122 | Matr3 | 17184 | HITS-CLIP | 37986224\|23142080\|23597149\|1953€ | Unspecified |
| mmu-let-7i-5p | MIMAT0000122 | Mau2 | 74549 | HITS-CLIP | 23597149\|23142080\|29167373 | Unspecified |
| mmu-let-7i-5p | MIMAT0000122 | Mavs | 228607 | HITS-CLIP | 23597149\|29167373 | Unspecified |
| mmu-let-7i-5p | MIMAT0000122 | Max | 17187 | HITS-CLIP | 37986224\|31433990 | Unspecified |
| mmu-let-7i-5p | MIMAT0000122 | Maz | 17188 | HITS-CLIP | 23142080 | Unspecified |
| mmu-let-7i-5p | MIMAT0000122 | Mbd1 | 17190 | HITS-CLIP | 23142080 | Unspecified |
| mmu-let-7i-5p | MIMAT0000122 | Mbd2 | 17191 | HITS-CLIP | 23142080 | Unspecified |
| mmu-let-7i-5p | MIMAT0000122 | Mbd3 | 17192 | HITS-CLIP | 37986224 | Unspecified |
| mmu-let-7i-5p | MIMAT0000122 | Mbd5 | 109241 | HITS-CLIP | 19536157\|23142080 | Unspecified |
| mmu-let-7i-5p | MIMAT0000122 | Mbd6 | 110962 | HITS-CLIP | 23597149\|19536157\|23142080\|2916 | Unspecified |
| mmu-let-7i-5p | MIMAT0000122 | Mbnl1 | 56758 | HITS-CLIP | 23630078\|23142080\|29167373 | Unspecified |
| mmu-let-7i-5p | MIMAT0000122 | Mbnl2 | 105559 | HITS-CLIP | 19536157\|23142080\|29167373 | Unspecified |
| mmu-let-7i-5p | MIMAT0000122 | Mbnl3 | 171170 | HITS-CLIP | 31433990 | Unspecified |
| mmu-let-7i-5p | MIMAT0000122 | Mbp | 17196 | HITS-CLIP | 23142080 | Unspecified |
| mmu-let-7i-5p | MIMAT0000122 | Mbtd1 | 103537 | HITS-CLIP | 23597149\|23142080\|29167373 | Unspecified |
| mmu-let-7i-5p | MIMAT0000122 | Mcat | 223722 | HITS-CLIP | 19536157\|23597149\|23142080 | Unspecified |
| mmu-let-7i-5p | MIMAT0000122 | Mcee | 73724 | HITS-CLIP | 37986224 | Unspecified |
| mmu-let-7i-5p | MIMAT0000122 | Mcf2l | 17207 | HITS-CLIP | 37986224\|19536157 | Unspecified |
| mmu-let-7i-5p | MIMAT0000122 | Mcfd2 | 193813 | HITS-CLIP | 23142080 | Unspecified |
| mmu-let-7i-5p | MIMAT0000122 | Mcl1 | 17210 | HITS-CLIP | 31433990 | Unspecified |
| mmu-let-7i-5p | MIMAT0000122 | Mcm10 | 70024 | HITS-CLIP | 23142080 | Unspecified |
| mmu-let-7i-5p | MIMAT0000122 | Mcm2 | 17216 | HITS-CLIP | 29167373\|23142080 | Unspecified |
| mmu-let-7i-5p | MIMAT0000122 | Mcm3 | 17215 | HITS-CLIP | 23142080 | Unspecified |
| mmu-let-7i-5p | MIMAT0000122 | Mcm3ap | 54387 | HITS-CLIP | 23142080\|23597149 | Unspecified |
| mmu-let-7i-5p | MIMAT0000122 | Mcm4 | 17217 | HITS-CLIP | 23142080\|29167373 | Unspecified |
| mmu-let-7i-5p | MIMAT0000122 | Mcm5 | 17218 | HITS-CLIP | 23142080\|23597149 | Unspecified |
| mmu-let-7i-5p | MIMAT0000122 | Mcm6 | 17219 | HITS-CLIP | 23142080\|23597149 | Unspecified |
| mmu-let-7i-5p | MIMAT0000122 | Mcm7 | 17220 | HITS-CLIP | 23142080 | Unspecified |
| mmu-let-7i-5p | MIMAT0000122 | Mcm8 | 66634 | HITS-CLIP | 23142080 | Unspecified |
| mmu-let-7i-5p | MIMAT0000122 | Mcmbp | 210711 | HITS-CLIP | 23142080 | Unspecified |

| | | | | | | |
|---|---|---|---|---|---|---|
| mmu-let-7i-5p | MIMAT0000122 | Mcoln1 | 94178 | HITS-CLIP | 23597149 | Unspecified |
| mmu-let-7i-5p | MIMAT0000122 | Mcph1 | 244329 | HITS-CLIP | 23142080 | Unspecified |
| mmu-let-7i-5p | MIMAT0000122 | Mcrip2 | 84331 | HITS-CLIP | 23597149 | Unspecified |
| mmu-let-7i-5p | MIMAT0000122 | Mctp1 | 78771 | HITS-CLIP | 19536157 | Unspecified |
| mmu-let-7i-5p | MIMAT0000122 | Mdc1 | 240087 | HITS-CLIP | 23142080 | Unspecified |
| mmu-let-7i-5p | MIMAT0000122 | Mdfic | 16543 | HITS-CLIP | 23142080\|29167373 | Unspecified |
| mmu-let-7i-5p | MIMAT0000122 | Mdga2 | 320772 | HITS-CLIP | 19536157 | Unspecified |
| mmu-let-7i-5p | MIMAT0000122 | Mdh1 | 17449 | HITS-CLIP | 19536157\|23597149\|23142080 | Unspecified |
| mmu-let-7i-5p | MIMAT0000122 | Mdh2 | 17448 | HITS-CLIP | 23597149\|23142080\|19536157 | Unspecified |
| mmu-let-7i-5p | MIMAT0000122 | Mdm2 | 17246 | HITS-CLIP | 37986224\|23597149\|23142080 | Unspecified |
| mmu-let-7i-5p | MIMAT0000122 | Mdm4 | 17248 | HITS-CLIP | 37986224\|23142080\|19536157\|23597 | Unspecified |
| mmu-let-7i-5p | MIMAT0000122 | Mdn1 | 100019 | HITS-CLIP | 23597149\|23142080\|29167373 | Unspecified |
| mmu-let-7i-5p | MIMAT0000122 | Me2 | 107029 | HITS-CLIP | 23142080 | Unspecified |
| mmu-let-7i-5p | MIMAT0000122 | Meaf6 | 70088 | HITS-CLIP | 37986224\|23142080\|19536157 | Unspecified |
| mmu-let-7i-5p | MIMAT0000122 | Mecp2 | 17257 | HITS-CLIP | 23142080\|29167373 | Unspecified |
| mmu-let-7i-5p | MIMAT0000122 | Med1 | 19014 | HITS-CLIP | 23597149\|23142080 | Unspecified |
| mmu-let-7i-5p | MIMAT0000122 | Med10 | 28077 | HITS-CLIP | 23142080 | Unspecified |
| mmu-let-7i-5p | MIMAT0000122 | Med12 | 59024 | HITS-CLIP | 23597149\|23142080 | Unspecified |
| mmu-let-7i-5p | MIMAT0000122 | Med13 | 327987 | HITS-CLIP | 23597149\|29167373\|23142080 | Unspecified |
| mmu-let-7i-5p | MIMAT0000122 | Med13l | 76199 | HITS-CLIP | 23142080\|29167373 | Unspecified |
| mmu-let-7i-5p | MIMAT0000122 | Med14 | 26896 | HITS-CLIP | 37986224\|19536157\|23597149\|2314 | Unspecified |
| mmu-let-7i-5p | MIMAT0000122 | Med15 | 94112 | HITS-CLIP | 23597149\|29167373 | Unspecified |
| mmu-let-7i-5p | MIMAT0000122 | Med16 | 216154 | HITS-CLIP | 23597149\|23142080 | Unspecified |
| mmu-let-7i-5p | MIMAT0000122 | Med19 | 381379 | HITS-CLIP | 23142080 | Unspecified |
| mmu-let-7i-5p | MIMAT0000122 | Med20 | 56771 | HITS-CLIP | 37986224\|23142080\|29167373 | Unspecified |
| mmu-let-7i-5p | MIMAT0000122 | Med23 | 70208 | HITS-CLIP | 23142080\|29167373 | Unspecified |
| mmu-let-7i-5p | MIMAT0000122 | Med25 | 75613 | qCLASH//HITS-CLIP | 31393953\|23597149 | Unspecified |
| mmu-let-7i-5p | MIMAT0000122 | Med28 | 66999 | HITS-CLIP | 23142080\|23597149 | Unspecified |
| mmu-let-7i-5p | MIMAT0000122 | Med8 | 80509 | HITS-CLIP | 23597149\|23142080 | Unspecified |
| mmu-let-7i-5p | MIMAT0000122 | Mef2a | 17258 | HITS-CLIP | 23142080\|19536157\|29167373 | Unspecified |
| mmu-let-7i-5p | MIMAT0000122 | Mef2c | 17260 | HITS-CLIP | 23142080\|29167373 | Unspecified |
| mmu-let-7i-5p | MIMAT0000122 | Mef2d | 17261 | HITS-CLIP | 23597149\|23142080\|19536157\|29167 | Unspecified |
| mmu-let-7i-5p | MIMAT0000122 | Megf9 | 230316 | HITS-CLIP | 37986224\|23597149 | Unspecified |
| mmu-let-7i-5p | MIMAT0000122 | Meis3 | 17537 | HITS-CLIP | 29167373 | Unspecified |
| mmu-let-7i-5p | MIMAT0000122 | Men1 | 17283 | HITS-CLIP | 23597149\|23142080 | Unspecified |
| mmu-let-7i-5p | MIMAT0000122 | Mepce | 231803 | HITS-CLIP | 29167373 | Unspecified |
| mmu-let-7i-5p | MIMAT0000122 | Mertk | 17289 | HITS-CLIP | 37986224 | Unspecified |
| mmu-let-7i-5p | MIMAT0000122 | Mest | 17294 | HITS-CLIP | 37986224\|23597149 | Unspecified |
| mmu-let-7i-5p | MIMAT0000122 | Met | 17295 | HITS-CLIP | 37986224\|23597149 | Unspecified |

| | | | | | | |
|---|---|---|---|---|---|---|
| mmu-let-7i-5p | MIMAT0000122 | Metap1 | 75624 | HITS-CLIP | 37986224\|23597149\|19536157\|23142 | Unspecified |
| mmu-let-7i-5p | MIMAT0000122 | Mettl1 | 17299 | HITS-CLIP | 23142080 | Unspecified |
| mmu-let-7i-5p | MIMAT0000122 | Mettl14 | 210529 | HITS-CLIP | 29167373 | Unspecified |
| mmu-let-7i-5p | MIMAT0000122 | Mettl21a | 67099 | HITS-CLIP | 23142080 | Unspecified |
| mmu-let-7i-5p | MIMAT0000122 | Mettl25 | 216292 | HITS-CLIP | 23142080 | Unspecified |
| mmu-let-7i-5p | MIMAT0000122 | Mettl27 | 155368 | HITS-CLIP | 23597149 | Unspecified |
| mmu-let-7i-5p | MIMAT0000122 | Mettl3 | 56335 | HITS-CLIP | 23142080 | Unspecified |
| mmu-let-7i-5p | MIMAT0000122 | Mettl4 | 76781 | HITS-CLIP | 23142080 | Unspecified |
| mmu-let-7i-5p | MIMAT0000122 | Mettl6 | 67011 | HITS-CLIP | 23597149\|29167373\|23142080 | Unspecified |
| mmu-let-7i-5p | MIMAT0000122 | Mettl7b | 71664 | HITS-CLIP | 23597149 | Unspecified |
| mmu-let-7i-5p | MIMAT0000122 | Mex3a | 72640 | HITS-CLIP | 37986224\|23142080\|19536157 | Unspecified |
| mmu-let-7i-5p | MIMAT0000122 | Mex3c | 240396 | HITS-CLIP | 23597149\|23142080\|29167373 | Unspecified |
| mmu-let-7i-5p | MIMAT0000122 | Mfap3 | 216760 | HITS-CLIP | 23597149\|23142080 | Unspecified |
| mmu-let-7i-5p | MIMAT0000122 | Mfhas1 | 52065 | HITS-CLIP | 37986224\|23142080 | Unspecified |
| mmu-let-7i-5p | MIMAT0000122 | Mfn1 | 67414 | HITS-CLIP | 23142080 | Unspecified |
| mmu-let-7i-5p | MIMAT0000122 | Mfsd10 | 68294 | HITS-CLIP | 23142080\|29167373 | Unspecified |
| mmu-let-7i-5p | MIMAT0000122 | Mfsd11 | 69900 | HITS-CLIP | 37986224\|19536157 | Unspecified |
| mmu-let-7i-5p | MIMAT0000122 | Mfsd12 | 73822 | HITS-CLIP | 29167373 | Unspecified |
| mmu-let-7i-5p | MIMAT0000122 | Mfsd14a | 15247 | HITS-CLIP | 23142080 | Unspecified |
| mmu-let-7i-5p | MIMAT0000122 | Mfsd14b | 84641 | HITS-CLIP | 23142080\|29167373 | Unspecified |
| mmu-let-7i-5p | MIMAT0000122 | Mfsd2a | 76574 | HITS-CLIP | 23597149\|23142080 | Unspecified |
| mmu-let-7i-5p | MIMAT0000122 | Mfsd5 | 106073 | HITS-CLIP | 23597149\|29167373\|23142080 | Unspecified |
| mmu-let-7i-5p | MIMAT0000122 | Mfsd6 | 98682 | HITS-CLIP | 19536157 | Unspecified |
| mmu-let-7i-5p | MIMAT0000122 | Mfsd8 | 72175 | HITS-CLIP | 23597149\|23142080 | Unspecified |
| mmu-let-7i-5p | MIMAT0000122 | Mfsd9 | 211798 | HITS-CLIP | 23597149 | Unspecified |
| mmu-let-7i-5p | MIMAT0000122 | Mga | 29808 | HITS-CLIP | 37986224\|23597149\|19536157\|23142 | Unspecified |
| mmu-let-7i-5p | MIMAT0000122 | Mgam | 232714 | HITS-CLIP | 37986224\|23597149 | Unspecified |
| mmu-let-7i-5p | MIMAT0000122 | Mgat1 | 17308 | HITS-CLIP | 23142080 | Unspecified |
| mmu-let-7i-5p | MIMAT0000122 | Mgat2 | 217664 | HITS-CLIP | 37986224\|23142080 | Unspecified |
| mmu-let-7i-5p | MIMAT0000122 | Mgat4a | 269181 | HITS-CLIP | 23142080 | Unspecified |
| mmu-let-7i-5p | MIMAT0000122 | Mgat5 | 107895 | HITS-CLIP | 37986224\|29167373 | Unspecified |
| mmu-let-7i-5p | MIMAT0000122 | Mgea5 | 76055 | HITS-CLIP | 23142080 | Unspecified |
| mmu-let-7i-5p | MIMAT0000122 | Mgll | 23945 | HITS-CLIP | 23142080\|23597149 | Unspecified |
| mmu-let-7i-5p | MIMAT0000122 | Mgrn1 | 17237 | HITS-CLIP | 23597149\|23142080 | Unspecified |
| mmu-let-7i-5p | MIMAT0000122 | Mia2 | 338320 | HITS-CLIP | 23142080\|23597149\|29167373 | Unspecified |
| mmu-let-7i-5p | MIMAT0000122 | Mia3 | 338366 | HITS-CLIP | 31433990\|23597149\|23142080 | Unspecified |
| mmu-let-7i-5p | MIMAT0000122 | Mib1 | 225164 | HITS-CLIP | 37986224\|19536157\|23597149\|23142 | Unspecified |
| mmu-let-7i-5p | MIMAT0000122 | Mical1 | 171580 | HITS-CLIP | 23142080 | Unspecified |
| mmu-let-7i-5p | MIMAT0000122 | Mical2 | 320878 | HITS-CLIP | 37986224\|23597149\|29167373 | Unspecified |

| | | | | | | |
|---|---|---|---|---|---|---|
| mmu-let-7i-5p | MIMAT0000122 | Mical3 | 194401 | HITS-CLIP | 19536157 | Unspecified |
| mmu-let-7i-5p | MIMAT0000122 | Micall2 | 231830 | HITS-CLIP | 19536157\|29167373 | Unspecified |
| mmu-let-7i-5p | MIMAT0000122 | Micu3 | 78506 | HITS-CLIP | 23142080 | Unspecified |
| mmu-let-7i-5p | MIMAT0000122 | Midn | 59090 | HITS-CLIP | 37986224\|23597149\|23142080 | Unspecified |
| mmu-let-7i-5p | MIMAT0000122 | Mief1 | 239555 | HITS-CLIP | 37986224\|23597149\|23142080 | Unspecified |
| mmu-let-7i-5p | MIMAT0000122 | Mier1 | 71148 | HITS-CLIP | 23142080\|29167373 | Unspecified |
| mmu-let-7i-5p | MIMAT0000122 | Mier2 | 70427 | HITS-CLIP | 37986224\|23142080 | Unspecified |
| mmu-let-7i-5p | MIMAT0000122 | Mier3 | 218613 | HITS-CLIP | 23142080 | Unspecified |
| mmu-let-7i-5p | MIMAT0000122 | Mif4gd | 69674 | HITS-CLIP | 23597149\|23142080 | Unspecified |
| mmu-let-7i-5p | MIMAT0000122 | Milr1 | 380732 | HITS-CLIP | 29167373 | Unspecified |
| mmu-let-7i-5p | MIMAT0000122 | Mindy1 | 55793 | HITS-CLIP | 23597149 | Unspecified |
| mmu-let-7i-5p | MIMAT0000122 | Mindy2 | 235461 | HITS-CLIP | 23142080\|29167373 | Unspecified |
| mmu-let-7i-5p | MIMAT0000122 | Mindy3 | 80013 | HITS-CLIP | 23597149\|23142080 | Unspecified |
| mmu-let-7i-5p | MIMAT0000122 | Minos1 | 440574 | HITS-CLIP | 23142080 | Unspecified |
| mmu-let-7i-5p | MIMAT0000122 | Mis18a | 66578 | HITS-CLIP | 23597149 | Unspecified |
| mmu-let-7i-5p | MIMAT0000122 | Mis18bp1 | 217653 | HITS-CLIP | 23142080 | Unspecified |
| mmu-let-7i-5p | MIMAT0000122 | Mitf | 17342 | HITS-CLIP | 29167373 | Unspecified |
| mmu-let-7i-5p | MIMAT0000122 | Mki67 | 17345 | HITS-CLIP | 37986224\|23597149\|23142080 | Unspecified |
| mmu-let-7i-5p | MIMAT0000122 | Mkln1 | 27418 | HITS-CLIP | 37986224\|23142080\|19536157\|29167 | Unspecified |
| mmu-let-7i-5p | MIMAT0000122 | Mknk2 | 17347 | HITS-CLIP | 37986224\|23142080\|23597149 | Unspecified |
| mmu-let-7i-5p | MIMAT0000122 | Mkrn2 | 67027 | HITS-CLIP | 23142080 | Unspecified |
| mmu-let-7i-5p | MIMAT0000122 | Mkx | 210719 | HITS-CLIP | 19536157 | Unspecified |
| mmu-let-7i-5p | MIMAT0000122 | Mlec | 109154 | HITS-CLIP | 23597149\|23142080 | Unspecified |
| mmu-let-7i-5p | MIMAT0000122 | Mlf2 | 30853 | HITS-CLIP | 23142080 | Unspecified |
| mmu-let-7i-5p | MIMAT0000122 | Mlh1 | 17350 | HITS-CLIP | 23597149\|29167373\|23142080 | Unspecified |
| mmu-let-7i-5p | MIMAT0000122 | Mllt10 | 17354 | HITS-CLIP | 37986224\|23142080\|29167373 | Unspecified |
| mmu-let-7i-5p | MIMAT0000122 | Mlst8 | 56716 | HITS-CLIP | 23142080 | Unspecified |
| mmu-let-7i-5p | MIMAT0000122 | Mlxip | 208104 | HITS-CLIP | 37986224\|23142080 | Unspecified |
| mmu-let-7i-5p | MIMAT0000122 | Mlxipl | 58805 | HITS-CLIP | 23597149 | Unspecified |
| mmu-let-7i-5p | MIMAT0000122 | Mmd | 67468 | HITS-CLIP | 23597149\|23142080\|29167373 | Unspecified |
| mmu-let-7i-5p | MIMAT0000122 | Mmgt1 | 236792 | HITS-CLIP | 37986224 | Unspecified |
| mmu-let-7i-5p | MIMAT0000122 | Mmp11 | 17385 | HITS-CLIP | 23142080 | Unspecified |
| mmu-let-7i-5p | MIMAT0000122 | Mmp14 | 17387 | HITS-CLIP | 37986224\|23597149 | Unspecified |
| mmu-let-7i-5p | MIMAT0000122 | Mmp15 | 17388 | HITS-CLIP | 23597149 | Unspecified |
| mmu-let-7i-5p | MIMAT0000122 | Mmp16 | 17389 | HITS-CLIP | 37986224 | Unspecified |
| mmu-let-7i-5p | MIMAT0000122 | Mms19 | 72199 | HITS-CLIP | 23142080 | Unspecified |
| mmu-let-7i-5p | MIMAT0000122 | Mms22l | 212377 | HITS-CLIP | 23142080\|31433990\|23597149 | Unspecified |
| mmu-let-7i-5p | MIMAT0000122 | Mn1 | 433938 | HITS-CLIP | 23597149 | Unspecified |
| mmu-let-7i-5p | MIMAT0000122 | Mnt | 17428 | HITS-CLIP | 37986224\|23597149\|23142080 | Unspecified |

| | | | | | | |
|---|---|---|---|---|---|---|
| mmu-let-7i-5p | MIMAT0000122 | Mob1a | 232157 | HITS-CLIP | 37986224\|23142080\|29167373 | Unspecified |
| mmu-let-7i-5p | MIMAT0000122 | Mob1b | 68473 | HITS-CLIP | 23597149\|23142080 | Unspecified |
| mmu-let-7i-5p | MIMAT0000122 | Mob3a | 208228 | HITS-CLIP | 23597149\|23142080\|29167373 | Unspecified |
| mmu-let-7i-5p | MIMAT0000122 | Mob3b | 214944 | HITS-CLIP | 37986224\|23597149 | Unspecified |
| mmu-let-7i-5p | MIMAT0000122 | Mob3c | 100465 | HITS-CLIP | 23597149 | Unspecified |
| mmu-let-7i-5p | MIMAT0000122 | Mob4 | 19070 | HITS-CLIP | 23142080\|29167373 | Unspecified |
| mmu-let-7i-5p | MIMAT0000122 | Mobp | 17433 | HITS-CLIP | 19536157 | Unspecified |
| mmu-let-7i-5p | MIMAT0000122 | Mocos | 68591 | HITS-CLIP | 37986224\|23597149\|29167373 | Unspecified |
| mmu-let-7i-5p | MIMAT0000122 | Mogs | 57377 | HITS-CLIP | 23142080 | Unspecified |
| mmu-let-7i-5p | MIMAT0000122 | Mon1a | 72825 | HITS-CLIP | 23142080 | Unspecified |
| mmu-let-7i-5p | MIMAT0000122 | Mon1b | 270096 | HITS-CLIP | 23597149 | Unspecified |
| mmu-let-7i-5p | MIMAT0000122 | Mon2 | 67074 | HITS-CLIP | 23597149\|19536157\|29167373 | Unspecified |
| mmu-let-7i-5p | MIMAT0000122 | Morc3 | 338467 | HITS-CLIP | 23142080\|29167373 | Unspecified |
| mmu-let-7i-5p | MIMAT0000122 | Morf4l2 | 56397 | HITS-CLIP | 23142080 | Unspecified |
| mmu-let-7i-5p | MIMAT0000122 | Mosmo | 233812 | HITS-CLIP | 23597149\|23142080 | Unspecified |
| mmu-let-7i-5p | MIMAT0000122 | Mospd2 | 76763 | HITS-CLIP | 29167373 | Unspecified |
| mmu-let-7i-5p | MIMAT0000122 | Mospd3 | 68929 | HITS-CLIP | 37986224\|23597149 | Unspecified |
| mmu-let-7i-5p | MIMAT0000122 | Mov10 | 17454 | HITS-CLIP | 23142080\|23597149 | Unspecified |
| mmu-let-7i-5p | MIMAT0000122 | Mpdu1 | 24070 | HITS-CLIP | 23597149\|23142080 | Unspecified |
| mmu-let-7i-5p | MIMAT0000122 | Mpeg1 | 17476 | HITS-CLIP | 23597149\|23142080\|29167373 | Unspecified |
| mmu-let-7i-5p | MIMAT0000122 | Mpi | 110119 | HITS-CLIP | 23597149 | Unspecified |
| mmu-let-7i-5p | MIMAT0000122 | Mpp1 | 17524 | HITS-CLIP | 23142080 | Unspecified |
| mmu-let-7i-5p | MIMAT0000122 | Mpp2 | 50997 | HITS-CLIP | 23142080 | Unspecified |
| mmu-let-7i-5p | MIMAT0000122 | Mpp5 | 56217 | HITS-CLIP | 37986224\|19536157\|23142080 | Unspecified |
| mmu-let-7i-5p | MIMAT0000122 | Mpp6 | 56524 | HITS-CLIP | 23142080\|19536157 | Unspecified |
| mmu-let-7i-5p | MIMAT0000122 | Mpp7 | 75739 | HITS-CLIP | 37986224\|19536157\|23142080 | Unspecified |
| mmu-let-7i-5p | MIMAT0000122 | Mppe1 | 225651 | HITS-CLIP | 37986224 | Unspecified |
| mmu-let-7i-5p | MIMAT0000122 | Mprip | 26936 | HITS-CLIP | 19536157\|23142080 | Unspecified |
| mmu-let-7i-5p | MIMAT0000122 | Mpv17 | 17527 | HITS-CLIP | 23142080 | Unspecified |
| mmu-let-7i-5p | MIMAT0000122 | Mrc1 | 17533 | HITS-CLIP | 23597149\|29167373 | Unspecified |
| mmu-let-7i-5p | MIMAT0000122 | Mrfap1 | 67568 | HITS-CLIP | 23142080 | Unspecified |
| mmu-let-7i-5p | MIMAT0000122 | Mrgbp | 73247 | HITS-CLIP | 23142080 | Unspecified |
| mmu-let-7i-5p | MIMAT0000122 | Mrm1 | 217038 | HITS-CLIP | 23597149 | Unspecified |
| mmu-let-7i-5p | MIMAT0000122 | Mroh1 | 223658 | HITS-CLIP | 23597149 | Unspecified |
| mmu-let-7i-5p | MIMAT0000122 | Mroh8 | 629499 | HITS-CLIP | 23142080 | Unspecified |
| mmu-let-7i-5p | MIMAT0000122 | Mrpl10 | 107732 | HITS-CLIP | 23142080 | Unspecified |
| mmu-let-7i-5p | MIMAT0000122 | Mrpl12 | 56282 | HITS-CLIP | 37986224 | Unspecified |
| mmu-let-7i-5p | MIMAT0000122 | Mrpl13 | 68537 | HITS-CLIP | 23597149 | Unspecified |
| mmu-let-7i-5p | MIMAT0000122 | Mrpl17 | 27397 | HITS-CLIP | 23597149 | Unspecified |

| | | | | | | |
|---|---|---|---|---|---|---|
| mmu-let-7i-5p | MIMAT0000122 | Mrpl19 | 56284 | HITS-CLIP | 31433990 | Unspecified |
| mmu-let-7i-5p | MIMAT0000122 | Mrpl20 | 66448 | HITS-CLIP | 23142080\|29167373 | Unspecified |
| mmu-let-7i-5p | MIMAT0000122 | Mrpl21 | 353242 | HITS-CLIP | 23597149\|23142080 | Unspecified |
| mmu-let-7i-5p | MIMAT0000122 | Mrpl22 | 216767 | HITS-CLIP | 23597149 | Unspecified |
| mmu-let-7i-5p | MIMAT0000122 | Mrpl28 | 68611 | qCLASH//HITS-CLIP | 31393953\|23142080 | Unspecified |
| mmu-let-7i-5p | MIMAT0000122 | Mrpl33 | 66845 | HITS-CLIP | 23142080 | Unspecified |
| mmu-let-7i-5p | MIMAT0000122 | Mrpl39 | 27393 | HITS-CLIP | 23142080 | Unspecified |
| mmu-let-7i-5p | MIMAT0000122 | Mrpl4 | 66163 | HITS-CLIP | 23597149 | Unspecified |
| mmu-let-7i-5p | MIMAT0000122 | Mrpl40 | 18100 | HITS-CLIP | 23597149\|23142080 | Unspecified |
| mmu-let-7i-5p | MIMAT0000122 | Mrpl44 | 69163 | HITS-CLIP | 23142080 | Unspecified |
| mmu-let-7i-5p | MIMAT0000122 | Mrpl46 | 67308 | HITS-CLIP | 23597149 | Unspecified |
| mmu-let-7i-5p | MIMAT0000122 | Mrpl50 | 28028 | HITS-CLIP | 23597149\|23142080 | Unspecified |
| mmu-let-7i-5p | MIMAT0000122 | Mrpl58 | 3396 | HITS-CLIP | 23142080\|19536157 | Unspecified |
| mmu-let-7i-5p | MIMAT0000122 | Mrps16 | 66242 | HITS-CLIP | 23597149\|23142080\|29167373 | Unspecified |
| mmu-let-7i-5p | MIMAT0000122 | Mrps2 | 118451 | HITS-CLIP | 19536157\|23142080 | Unspecified |
| mmu-let-7i-5p | MIMAT0000122 | Mrps24 | 64660 | HITS-CLIP | 23142080 | Unspecified |
| mmu-let-7i-5p | MIMAT0000122 | Mrps26 | 99045 | HITS-CLIP | 23597149 | Unspecified |
| mmu-let-7i-5p | MIMAT0000122 | Mrps35 | 232536 | HITS-CLIP | 23597149\|23142080 | Unspecified |
| mmu-let-7i-5p | MIMAT0000122 | Mrps5 | 77721 | HITS-CLIP | 23597149 | Unspecified |
| mmu-let-7i-5p | MIMAT0000122 | Mrps7 | 50529 | HITS-CLIP | 23597149 | Unspecified |
| mmu-let-7i-5p | MIMAT0000122 | Mrs2 | 380836 | HITS-CLIP | 19536157 | Unspecified |
| mmu-let-7i-5p | MIMAT0000122 | Mrtfa | 57591 | HITS-CLIP | 23142080 | Unspecified |
| mmu-let-7i-5p | MIMAT0000122 | Mrtfb | 239719 | HITS-CLIP | 23142080\|19536157\|23597149\|29167 | Unspecified |
| mmu-let-7i-5p | MIMAT0000122 | Ms4a14 | 383435 | HITS-CLIP | 29167373 | Unspecified |
| mmu-let-7i-5p | MIMAT0000122 | Ms4a4b | 60361 | HITS-CLIP | 23142080 | Unspecified |
| mmu-let-7i-5p | MIMAT0000122 | Ms4a4c | 64380 | HITS-CLIP | 29167373 | Unspecified |
| mmu-let-7i-5p | MIMAT0000122 | Ms4a6b | 69774 | HITS-CLIP | 37986224\|23142080\|29167373 | Unspecified |
| mmu-let-7i-5p | MIMAT0000122 | Ms4a6d | 68774 | HITS-CLIP | 23142080 | Unspecified |
| mmu-let-7i-5p | MIMAT0000122 | Msh6 | 17688 | HITS-CLIP | 23597149\|23142080 | Unspecified |
| mmu-let-7i-5p | MIMAT0000122 | Msi2 | 76626 | HITS-CLIP//qCLASH | 25083871\|31393953\|23142080 | Unspecified |
| mmu-let-7i-5p | MIMAT0000122 | Msl1 | 74026 | HITS-CLIP | 23597149\|19536157\|23142080 | Unspecified |
| mmu-let-7i-5p | MIMAT0000122 | Msl2 | 77853 | HITS-CLIP | 23597149\|23142080 | Unspecified |
| mmu-let-7i-5p | MIMAT0000122 | Msmo1 | 66234 | HITS-CLIP | 23597149\|29167373\|23142080 | Unspecified |
| mmu-let-7i-5p | MIMAT0000122 | Msn | 17698 | HITS-CLIP | 23142080\|29167373 | Unspecified |
| mmu-let-7i-5p | MIMAT0000122 | Msr1 | 20288 | HITS-CLIP | 23597149\|29167373 | Unspecified |
| mmu-let-7i-5p | MIMAT0000122 | Msrb1 | 27361 | HITS-CLIP | 23597149\|23142080 | Unspecified |
| mmu-let-7i-5p | MIMAT0000122 | Mst1 | 15235 | HITS-CLIP | 23597149 | Unspecified |
| mmu-let-7i-5p | MIMAT0000122 | Mta1 | 116870 | HITS-CLIP | 23142080 | Unspecified |
| mmu-let-7i-5p | MIMAT0000122 | Mta3 | 116871 | HITS-CLIP | 23142080 | Unspecified |

| | | | | | | |
|---|---|---|---|---|---|---|
| mmu-let-7i-5p | MIMAT0000122 | Mtap | 66902 | HITS-CLIP | 23597149\|23142080 | Unspecified |
| mmu-let-7i-5p | MIMAT0000122 | Mtbp | 105837 | HITS-CLIP | 23597149 | Unspecified |
| mmu-let-7i-5p | MIMAT0000122 | Mtch1 | 56462 | HITS-CLIP | 23597149\|23142080 | Unspecified |
| mmu-let-7i-5p | MIMAT0000122 | Mtch2 | 56428 | HITS-CLIP | 23142080 | Unspecified |
| mmu-let-7i-5p | MIMAT0000122 | Mtcl1 | 68617 | HITS-CLIP | 19536157 | Unspecified |
| mmu-let-7i-5p | MIMAT0000122 | Mtdh | 67154 | HITS-CLIP | 37986224\|23142080\|19536157\|29167 | Unspecified |
| mmu-let-7i-5p | MIMAT0000122 | Mterf1a | 545725 | HITS-CLIP | 23142080 | Unspecified |
| mmu-let-7i-5p | MIMAT0000122 | Mterf2 | 80298 | HITS-CLIP | 23142080 | Unspecified |
| mmu-let-7i-5p | MIMAT0000122 | Mterf3 | 51001 | HITS-CLIP | 23142080 | Unspecified |
| mmu-let-7i-5p | MIMAT0000122 | Mterf4 | 69821 | HITS-CLIP | 23597149 | Unspecified |
| mmu-let-7i-5p | MIMAT0000122 | Mtf1 | 17764 | HITS-CLIP | 31433990\|23630078\|29167373\|23142 | Unspecified |
| mmu-let-7i-5p | MIMAT0000122 | Mtf2 | 17765 | HITS-CLIP//qCLASH | 37986224\|31393953\|23142080 | Unspecified |
| mmu-let-7i-5p | MIMAT0000122 | Mtfr1 | 67472 | HITS-CLIP | 31433990 | Unspecified |
| mmu-let-7i-5p | MIMAT0000122 | Mtfr1l | 76824 | HITS-CLIP | 23597149 | Unspecified |
| mmu-let-7i-5p | MIMAT0000122 | Mtg1 | 212508 | HITS-CLIP | 23597149 | Unspecified |
| mmu-let-7i-5p | MIMAT0000122 | Mthfd1 | 108156 | HITS-CLIP | 23597149 | Unspecified |
| mmu-let-7i-5p | MIMAT0000122 | Mthfr | 17769 | HITS-CLIP | 23597149\|29167373 | Unspecified |
| mmu-let-7i-5p | MIMAT0000122 | Mthfsd | 234814 | HITS-CLIP | 23597149\|23142080 | Unspecified |
| mmu-let-7i-5p | MIMAT0000122 | Mthfsl | 100039707 | HITS-CLIP | 23142080 | Unspecified |
| mmu-let-7i-5p | MIMAT0000122 | Mtif3 | 76366 | HITS-CLIP | 23597149 | Unspecified |
| mmu-let-7i-5p | MIMAT0000122 | Mtmr1 | 53332 | HITS-CLIP | 23142080\|29167373 | Unspecified |
| mmu-let-7i-5p | MIMAT0000122 | Mtmr10 | 233315 | HITS-CLIP | 29167373 | Unspecified |
| mmu-let-7i-5p | MIMAT0000122 | Mtmr12 | 268783 | HITS-CLIP | 37986224\|23142080 | Unspecified |
| mmu-let-7i-5p | MIMAT0000122 | Mtmr14 | 97287 | HITS-CLIP | 23142080 | Unspecified |
| mmu-let-7i-5p | MIMAT0000122 | Mtmr3 | 74302 | HITS-CLIP | 23142080 | Unspecified |
| mmu-let-7i-5p | MIMAT0000122 | Mtmr4 | 170749 | HITS-CLIP | 19536157\|23597149\|23142080\|29167 | Unspecified |
| mmu-let-7i-5p | MIMAT0000122 | Mtmr6 | 219135 | HITS-CLIP | 23142080 | Unspecified |
| mmu-let-7i-5p | MIMAT0000122 | Mtmr7 | 54384 | HITS-CLIP | 23142080 | Unspecified |
| mmu-let-7i-5p | MIMAT0000122 | Mtmr9 | 210376 | HITS-CLIP | 23142080 | Unspecified |
| mmu-let-7i-5p | MIMAT0000122 | Mtor | 56717 | HITS-CLIP//qCLASH | 37986224\|31393953 | Unspecified |
| mmu-let-7i-5p | MIMAT0000122 | Mtpn | 14489 | qCLASH//HITS-CLIP | 31393953\|23142080\|29167373 | Unspecified |
| mmu-let-7i-5p | MIMAT0000122 | Mtr | 238505 | HITS-CLIP | 37986224\|23597149\|23142080\|29167 | Unspecified |
| mmu-let-7i-5p | MIMAT0000122 | Mtrex | 23517 | HITS-CLIP | 23597149\|23142080 | Unspecified |
| mmu-let-7i-5p | MIMAT0000122 | Mtss1 | 211401 | HITS-CLIP | 23597149\|29167373 | Unspecified |
| mmu-let-7i-5p | MIMAT0000122 | Mttp | 17777 | HITS-CLIP | 23597149 | Unspecified |
| mmu-let-7i-5p | MIMAT0000122 | Mtus2 | 77521 | HITS-CLIP | 37986224\|19536157 | Unspecified |
| mmu-let-7i-5p | MIMAT0000122 | Mtx2 | 53375 | HITS-CLIP | 23142080\|29167373 | Unspecified |
| mmu-let-7i-5p | MIMAT0000122 | Muc2 | 17831 | HITS-CLIP | 23142080 | Unspecified |
| mmu-let-7i-5p | MIMAT0000122 | Mug1 | 17836 | HITS-CLIP | 23597149 | Unspecified |

| | | | | | | | |
|---|---|---|---|---|---|---|---|
| mmu-let-7i-5p | MIMAT0000122 | Mup17 | 100039206 | HITS-CLIP | | 23597149 | Unspecified |
| mmu-let-7i-5p | MIMAT0000122 | Mup3 | 17842 | HITS-CLIP | | 23597149 | Unspecified |
| mmu-let-7i-5p | MIMAT0000122 | Mup4 | 17843 | HITS-CLIP | | 23597149 | Unspecified |
| mmu-let-7i-5p | MIMAT0000122 | Mvd | 192156 | HITS-CLIP | | 37986224\|23142080 | Unspecified |
| mmu-let-7i-5p | MIMAT0000122 | Mvk | 17855 | HITS-CLIP | | 23142080 | Unspecified |
| mmu-let-7i-5p | MIMAT0000122 | Mvp | 78388 | HITS-CLIP | | 23142080 | Unspecified |
| mmu-let-7i-5p | MIMAT0000122 | Mxd1 | 17119 | HITS-CLIP | | 37986224\|23142080 | Unspecified |
| mmu-let-7i-5p | MIMAT0000122 | Mxd4 | 17122 | HITS-CLIP | | 37986224\|23597149\|23142080 | Unspecified |
| mmu-let-7i-5p | MIMAT0000122 | Myadm | 50918 | HITS-CLIP | | 26203562\|29167373 | Unspecified |
| mmu-let-7i-5p | MIMAT0000122 | Myb | 17863 | HITS-CLIP | | 37986224\|23142080\|31433990 | Unspecified |
| mmu-let-7i-5p | MIMAT0000122 | Myc | 17869 | HITS-CLIP | | 23597149\|23142080\|29167373 | Unspecified |
| mmu-let-7i-5p | MIMAT0000122 | Mycbp | 56309 | HITS-CLIP | | 23597149\|23142080 | Unspecified |
| mmu-let-7i-5p | MIMAT0000122 | Mycbp2 | 105689 | HITS-CLIP | | 37986224\|23142080\|19536157\|23597 | Unspecified |
| mmu-let-7i-5p | MIMAT0000122 | Mydgf | 56005 | HITS-CLIP | | 29167373 | Unspecified |
| mmu-let-7i-5p | MIMAT0000122 | Myg1 | 60315 | qCLASH//HITS-CLIP | | 31393953\|23142080 | Unspecified |
| mmu-let-7i-5p | MIMAT0000122 | Myh9 | 17886 | HITS-CLIP | | 37986224\|23597149\|23142080\|29167 | Unspecified |
| mmu-let-7i-5p | MIMAT0000122 | Mylip | 218203 | HITS-CLIP | | 37986224\|23597149\|23142080 | Unspecified |
| mmu-let-7i-5p | MIMAT0000122 | Mynn | 80732 | HITS-CLIP | | 23142080 | Unspecified |
| mmu-let-7i-5p | MIMAT0000122 | Myo10 | 17909 | HITS-CLIP | | 23142080 | Unspecified |
| mmu-let-7i-5p | MIMAT0000122 | Myo18a | 360013 | HITS-CLIP | | 23597149\|23142080 | Unspecified |
| mmu-let-7i-5p | MIMAT0000122 | Myo1b | 17912 | HITS-CLIP | | 23597149 | Unspecified |
| mmu-let-7i-5p | MIMAT0000122 | Myo1c | 17913 | HITS-CLIP | | 23142080 | Unspecified |
| mmu-let-7i-5p | MIMAT0000122 | Myo1e | 71602 | HITS-CLIP | | 37986224\|23597149\|23142080\|29167 | Unspecified |
| mmu-let-7i-5p | MIMAT0000122 | Myo1f | 17916 | HITS-CLIP | | 23142080\|29167373 | Unspecified |
| mmu-let-7i-5p | MIMAT0000122 | Myo1g | 246177 | HITS-CLIP | | 23142080 | Unspecified |
| mmu-let-7i-5p | MIMAT0000122 | Myo5a | 17918 | HITS-CLIP | | 19536157\|29167373 | Unspecified |
| mmu-let-7i-5p | MIMAT0000122 | Myo7a | 17921 | HITS-CLIP | | 37986224 | Unspecified |
| mmu-let-7i-5p | MIMAT0000122 | Myo9a | 270163 | HITS-CLIP | | 19536157 | Unspecified |
| mmu-let-7i-5p | MIMAT0000122 | Myo9b | 17925 | HITS-CLIP | | 37986224\|23142080\|23597149 | Unspecified |
| mmu-let-7i-5p | MIMAT0000122 | Myof | 226101 | HITS-CLIP | | 29167373 | Unspecified |
| mmu-let-7i-5p | MIMAT0000122 | Mypop | 232934 | HITS-CLIP | | 19536157 | Unspecified |
| mmu-let-7i-5p | MIMAT0000122 | Mysm1 | 320713 | HITS-CLIP | | 23142080 | Unspecified |
| mmu-let-7i-5p | MIMAT0000122 | N4bp1 | 80750 | HITS-CLIP | | 37986224\|23142080\|29167373 | Unspecified |
| mmu-let-7i-5p | MIMAT0000122 | N4bp2 | 333789 | HITS-CLIP | | 23142080\|23597149 | Unspecified |
| mmu-let-7i-5p | MIMAT0000122 | N4bp2l1 | 100637 | HITS-CLIP | | 23597149\|23142080\|29167373 | Unspecified |
| mmu-let-7i-5p | MIMAT0000122 | N4bp2l2 | 381695 | HITS-CLIP | | 23597149\|23142080 | Unspecified |
| mmu-let-7i-5p | MIMAT0000122 | Naa10 | 56292 | HITS-CLIP | | 23142080 | Unspecified |
| mmu-let-7i-5p | MIMAT0000122 | Naa15 | 74838 | HITS-CLIP | | 23142080 | Unspecified |
| mmu-let-7i-5p | MIMAT0000122 | Naa16 | 66897 | HITS-CLIP | | 23142080 | Unspecified |

| | | | | | | |
|---|---|---|---|---|---|---|
| mmu-let-7i-5p | MIMAT0000122 | Naa20 | 67877 | HITS-CLIP | 23142080 | Unspecified |
| mmu-let-7i-5p | MIMAT0000122 | Naa25 | 231713 | HITS-CLIP | 23597149\|23142080 | Unspecified |
| mmu-let-7i-5p | MIMAT0000122 | Naa30 | 70646 | HITS-CLIP | 23142080 | Unspecified |
| mmu-let-7i-5p | MIMAT0000122 | Naa35 | 78689 | HITS-CLIP | 23597149\|23142080 | Unspecified |
| mmu-let-7i-5p | MIMAT0000122 | Naa38 | 78304 | HITS-CLIP | 23142080 | Unspecified |
| mmu-let-7i-5p | MIMAT0000122 | Naa40 | 70999 | HITS-CLIP | 37986224 | Unspecified |
| mmu-let-7i-5p | MIMAT0000122 | Naa50 | 72117 | HITS-CLIP | 23142080\|26203562 | Unspecified |
| mmu-let-7i-5p | MIMAT0000122 | Naa60 | 74763 | HITS-CLIP | 23142080 | Unspecified |
| mmu-let-7i-5p | MIMAT0000122 | Nab1 | 17936 | HITS-CLIP | 37986224\|23597149\|23142080\|29167 | Unspecified |
| mmu-let-7i-5p | MIMAT0000122 | Nab2 | 17937 | HITS-CLIP | 23597149 | Unspecified |
| mmu-let-7i-5p | MIMAT0000122 | Nabp1 | 109019 | HITS-CLIP | 23142080 | Unspecified |
| mmu-let-7i-5p | MIMAT0000122 | Naca | 17938 | HITS-CLIP | 37986224\|29167373 | Unspecified |
| mmu-let-7i-5p | MIMAT0000122 | Nacad | 192950 | HITS-CLIP | 19536157 | Unspecified |
| mmu-let-7i-5p | MIMAT0000122 | Nacc1 | 66830 | HITS-CLIP | 37986224\|23142080\|19536157\|29167 | Unspecified |
| mmu-let-7i-5p | MIMAT0000122 | Nacc2 | 67991 | HITS-CLIP | 37986224 | Unspecified |
| mmu-let-7i-5p | MIMAT0000122 | Nadk | 192185 | HITS-CLIP | 23597149\|23142080\|29167373 | Unspecified |
| mmu-let-7i-5p | MIMAT0000122 | Naf1 | 234344 | HITS-CLIP | 23142080 | Unspecified |
| mmu-let-7i-5p | MIMAT0000122 | Nagpa | 27426 | HITS-CLIP | 23597149\|29167373 | Unspecified |
| mmu-let-7i-5p | MIMAT0000122 | Naif1 | 71254 | HITS-CLIP | 23597149 | Unspecified |
| mmu-let-7i-5p | MIMAT0000122 | Nampt | 59027 | HITS-CLIP | 23142080 | Unspecified |
| mmu-let-7i-5p | MIMAT0000122 | Nap1l4 | 17955 | HITS-CLIP | 23142080 | Unspecified |
| mmu-let-7i-5p | MIMAT0000122 | Naprt | 93100 | HITS-CLIP | 23597149 | Unspecified |
| mmu-let-7i-5p | MIMAT0000122 | Narf | 67608 | HITS-CLIP | 37986224\|19536157\|23597149\|23142 | Unspecified |
| mmu-let-7i-5p | MIMAT0000122 | Nars2 | 244141 | HITS-CLIP | 23597149\|23142080 | Unspecified |
| mmu-let-7i-5p | MIMAT0000122 | Nasp | 50927 | HITS-CLIP | 23597149\|23142080 | Unspecified |
| mmu-let-7i-5p | MIMAT0000122 | Nat2 | 17961 | HITS-CLIP | 23597149 | Unspecified |
| mmu-let-7i-5p | MIMAT0000122 | Nat8f1 | 66116 | HITS-CLIP | 23597149 | Unspecified |
| mmu-let-7i-5p | MIMAT0000122 | Nat8f4 | 75541 | HITS-CLIP | 23597149 | Unspecified |
| mmu-let-7i-5p | MIMAT0000122 | Nav1 | 215690 | HITS-CLIP | 19536157\|23597149\|29167373 | Unspecified |
| mmu-let-7i-5p | MIMAT0000122 | Nav2 | 78286 | HITS-CLIP | 37986224\|23142080\|29167373 | Unspecified |
| mmu-let-7i-5p | MIMAT0000122 | Nav3 | 260315 | HITS-CLIP | 19536157 | Unspecified |
| mmu-let-7i-5p | MIMAT0000122 | Nbas | 71169 | HITS-CLIP | 37986224\|23597149\|19536157 | Unspecified |
| mmu-let-7i-5p | MIMAT0000122 | Nbea | 26422 | HITS-CLIP | 37986224\|23597149\|19536157 | Unspecified |
| mmu-let-7i-5p | MIMAT0000122 | Nbeal1 | 269198 | HITS-CLIP | 37986224\|23142080\|19536157\|23597 | Unspecified |
| mmu-let-7i-5p | MIMAT0000122 | Nbeal2 | 235627 | HITS-CLIP | 37986224\|23142080\|23597149\|29167 | Unspecified |
| mmu-let-7i-5p | MIMAT0000122 | Nbn | 27354 | HITS-CLIP | 31433990\|23142080 | Unspecified |
| mmu-let-7i-5p | MIMAT0000122 | Nbr1 | 17966 | HITS-CLIP | 23597149\|26203562\|23142080\|29167 | Unspecified |
| mmu-let-7i-5p | MIMAT0000122 | Ncam1 | 17967 | HITS-CLIP | 37986224\|19536157 | Unspecified |
| mmu-let-7i-5p | MIMAT0000122 | Ncam2 | 17968 | HITS-CLIP | 37986224\|19536157 | Unspecified |

| | | | | | | |
|---|---|---|---|---|---|---|
| mmu-let-7i-5p | MIMAT0000122 | Ncan | 13004 | HITS-CLIP | 37986224\|19536157 | Unspecified |
| mmu-let-7i-5p | MIMAT0000122 | Ncapd2 | 68298 | HITS-CLIP | 23142080 | Unspecified |
| mmu-let-7i-5p | MIMAT0000122 | Ncapg | 54392 | HITS-CLIP | 23142080 | Unspecified |
| mmu-let-7i-5p | MIMAT0000122 | Ncapg2 | 76044 | HITS-CLIP | 23142080\|29167373 | Unspecified |
| mmu-let-7i-5p | MIMAT0000122 | Ncaph | 215387 | HITS-CLIP | 23142080\|29167373 | Unspecified |
| mmu-let-7i-5p | MIMAT0000122 | Ncaph2 | 52683 | HITS-CLIP | 23597149 | Unspecified |
| mmu-let-7i-5p | MIMAT0000122 | Ncbp1 | 433702 | HITS-CLIP | 23630078 | Unspecified |
| mmu-let-7i-5p | MIMAT0000122 | Nceh1 | 320024 | HITS-CLIP | 23597149\|23142080\|29167373 | Unspecified |
| mmu-let-7i-5p | MIMAT0000122 | Ncf4 | 17972 | HITS-CLIP | 23142080\|29167373 | Unspecified |
| mmu-let-7i-5p | MIMAT0000122 | Nck1 | 17973 | HITS-CLIP | 23597149\|23142080 | Unspecified |
| mmu-let-7i-5p | MIMAT0000122 | Nckap1 | 50884 | HITS-CLIP | 37986224\|23597149\|23142080 | Unspecified |
| mmu-let-7i-5p | MIMAT0000122 | Nckap1l | 105855 | HITS-CLIP | 23142080\|29167373 | Unspecified |
| mmu-let-7i-5p | MIMAT0000122 | Nckipsd | 80987 | HITS-CLIP | 23142080 | Unspecified |
| mmu-let-7i-5p | MIMAT0000122 | Ncl | 17975 | HITS-CLIP | 23597149\|26203562\|23142080 | Unspecified |
| mmu-let-7i-5p | MIMAT0000122 | Ncln | 103425 | HITS-CLIP | 23597149 | Unspecified |
| mmu-let-7i-5p | MIMAT0000122 | Ncoa1 | 17977 | HITS-CLIP | 23597149\|19536157\|29167373 | Unspecified |
| mmu-let-7i-5p | MIMAT0000122 | Ncoa2 | 17978 | HITS-CLIP | 37986224\|23597149\|23142080\|26203 | Unspecified |
| mmu-let-7i-5p | MIMAT0000122 | Ncoa3 | 17979 | HITS-CLIP | 37986224\|23597149\|23142080\|29167 | Unspecified |
| mmu-let-7i-5p | MIMAT0000122 | Ncoa6 | 56406 | HITS-CLIP | 37986224\|23597149\|23142080\|29167 | Unspecified |
| mmu-let-7i-5p | MIMAT0000122 | Ncoa7 | 211329 | HITS-CLIP | 23142080\|29167373 | Unspecified |
| mmu-let-7i-5p | MIMAT0000122 | Ncor1 | 20185 | HITS-CLIP | 37986224\|23597149\|23142080\|19536 | Unspecified |
| mmu-let-7i-5p | MIMAT0000122 | Ncor2 | 20602 | HITS-CLIP | 23597149\|31433990\|23142080\|29167 | Unspecified |
| mmu-let-7i-5p | MIMAT0000122 | Ncs1 | 14299 | HITS-CLIP | 23142080 | Unspecified |
| mmu-let-7i-5p | MIMAT0000122 | Ncstn | 59287 | HITS-CLIP | 23142080\|29167373 | Unspecified |
| mmu-let-7i-5p | MIMAT0000122 | Ndc1 | 72787 | HITS-CLIP | 23142080\|23597149\|30971389\|19536 | Unspecified |
| mmu-let-7i-5p | MIMAT0000122 | Ndel1 | 83431 | HITS-CLIP | 37986224\|23597149\|23142080 | Unspecified |
| mmu-let-7i-5p | MIMAT0000122 | Ndfip1 | 65113 | HITS-CLIP | 23142080\|19536157 | Unspecified |
| mmu-let-7i-5p | MIMAT0000122 | Ndfip2 | 76273 | HITS-CLIP | 23142080 | Unspecified |
| mmu-let-7i-5p | MIMAT0000122 | Ndn | 17984 | HITS-CLIP | 19536157 | Unspecified |
| mmu-let-7i-5p | MIMAT0000122 | Ndor1 | 78797 | HITS-CLIP | 23597149 | Unspecified |
| mmu-let-7i-5p | MIMAT0000122 | Ndrg1 | 17988 | HITS-CLIP | 23142080 | Unspecified |
| mmu-let-7i-5p | MIMAT0000122 | Ndrg2 | 29811 | HITS-CLIP | 37986224\|19536157\|23597149\|23142 | Unspecified |
| mmu-let-7i-5p | MIMAT0000122 | Ndrg3 | 29812 | HITS-CLIP | 23597149\|23142080\|29167373 | Unspecified |
| mmu-let-7i-5p | MIMAT0000122 | Ndst1 | 15531 | HITS-CLIP | 23597149 | Unspecified |
| mmu-let-7i-5p | MIMAT0000122 | Ndst2 | 17423 | HITS-CLIP | 29167373 | Unspecified |
| mmu-let-7i-5p | MIMAT0000122 | Ndst3 | 83398 | HITS-CLIP | 23142080\|19536157 | Unspecified |
| mmu-let-7i-5p | MIMAT0000122 | Ndufa10 | 67273 | HITS-CLIP | 37986224\|23597149\|19536157\|23142 | Unspecified |
| mmu-let-7i-5p | MIMAT0000122 | Ndufa11 | 69875 | HITS-CLIP | 23142080\|23597149 | Unspecified |
| mmu-let-7i-5p | MIMAT0000122 | Ndufa3 | 66091 | HITS-CLIP | 23142080 | Unspecified |

| | | | | | | |
|---|---|---|---|---|---|---|
| mmu-let-7i-5p | MIMAT0000122 | Ndufa5 | 68202 | HITS-CLIP | 23142080\|31433990 | Unspecified |
| mmu-let-7i-5p | MIMAT0000122 | Ndufa6 | 67130 | HITS-CLIP | 23597149 | Unspecified |
| mmu-let-7i-5p | MIMAT0000122 | Ndufa9 | 66108 | HITS-CLIP | 23142080\|23597149 | Unspecified |
| mmu-let-7i-5p | MIMAT0000122 | Ndufaf1 | 69702 | HITS-CLIP | 37986224\|23597149 | Unspecified |
| mmu-let-7i-5p | MIMAT0000122 | Ndufaf4 | 68493 | HITS-CLIP | 23597149\|23142080 | Unspecified |
| mmu-let-7i-5p | MIMAT0000122 | Ndufaf7 | 73694 | HITS-CLIP | 23142080 | Unspecified |
| mmu-let-7i-5p | MIMAT0000122 | Ndufb10 | 68342 | HITS-CLIP | 23597149\|23142080 | Unspecified |
| mmu-let-7i-5p | MIMAT0000122 | Ndufb5 | 66046 | HITS-CLIP | 23597149\|23142080 | Unspecified |
| mmu-let-7i-5p | MIMAT0000122 | Ndufb7 | 66916 | HITS-CLIP | 37986224\|19536157\|23597149\|29167 | Unspecified |
| mmu-let-7i-5p | MIMAT0000122 | Ndufb9 | 66218 | HITS-CLIP | 23142080 | Unspecified |
| mmu-let-7i-5p | MIMAT0000122 | Ndufs1 | 227197 | HITS-CLIP | 23597149\|23142080 | Unspecified |
| mmu-let-7i-5p | MIMAT0000122 | Ndufv2 | 72900 | HITS-CLIP | 23142080\|23597149 | Unspecified |
| mmu-let-7i-5p | MIMAT0000122 | Ndufv3 | 78330 | HITS-CLIP | 23142080 | Unspecified |
| mmu-let-7i-5p | MIMAT0000122 | Neb | 17996 | HITS-CLIP | 23142080 | Unspecified |
| mmu-let-7i-5p | MIMAT0000122 | Necap1 | 67602 | HITS-CLIP | 23142080 | Unspecified |
| mmu-let-7i-5p | MIMAT0000122 | Necap2 | 66147 | HITS-CLIP | 23142080 | Unspecified |
| mmu-let-7i-5p | MIMAT0000122 | Nectin3 | 58998 | HITS-CLIP | 19536157 | Unspecified |
| mmu-let-7i-5p | MIMAT0000122 | Nedd4 | 17999 | HITS-CLIP | 37986224\|19536157 | Unspecified |
| mmu-let-7i-5p | MIMAT0000122 | Nedd4l | 83814 | HITS-CLIP | 19536157 | Unspecified |
| mmu-let-7i-5p | MIMAT0000122 | Nefl | 18039 | HITS-CLIP | 37986224 | Unspecified |
| mmu-let-7i-5p | MIMAT0000122 | Nefm | 18040 | HITS-CLIP | 37986224\|19536157 | Unspecified |
| mmu-let-7i-5p | MIMAT0000122 | Neil1 | 72774 | HITS-CLIP | 29167373 | Unspecified |
| mmu-let-7i-5p | MIMAT0000122 | Nek1 | 18004 | HITS-CLIP | 23142080 | Unspecified |
| mmu-let-7i-5p | MIMAT0000122 | Nek6 | 59126 | HITS-CLIP | 29167373 | Unspecified |
| mmu-let-7i-5p | MIMAT0000122 | Nek7 | 59125 | HITS-CLIP | 23142080\|23597149\|19536157 | Unspecified |
| mmu-let-7i-5p | MIMAT0000122 | Nek9 | 217718 | HITS-CLIP | 37986224\|23597149 | Unspecified |
| mmu-let-7i-5p | MIMAT0000122 | Nelfa | 24116 | HITS-CLIP//qCLASH | 23597149\|31393953\|23142080 | Unspecified |
| mmu-let-7i-5p | MIMAT0000122 | Nell2 | 54003 | HITS-CLIP | 19536157 | Unspecified |
| mmu-let-7i-5p | MIMAT0000122 | Nemp1 | 23306 | HITS-CLIP | 23597149\|19536157\|23142080 | Unspecified |
| mmu-let-7i-5p | MIMAT0000122 | Nemp2 | 227094 | HITS-CLIP | 23142080 | Unspecified |
| mmu-let-7i-5p | MIMAT0000122 | Neo1 | 18007 | HITS-CLIP | 23597149 | Unspecified |
| mmu-let-7i-5p | MIMAT0000122 | Net1 | 56349 | HITS-CLIP | 23597149\|23142080 | Unspecified |
| mmu-let-7i-5p | MIMAT0000122 | Neu3 | 50877 | HITS-CLIP | 37986224\|23142080 | Unspecified |
| mmu-let-7i-5p | MIMAT0000122 | Neurl1b | 240055 | HITS-CLIP | 23142080 | Unspecified |
| mmu-let-7i-5p | MIMAT0000122 | Neurl4 | 216860 | HITS-CLIP | 37986224\|23597149\|23142080 | Unspecified |
| mmu-let-7i-5p | MIMAT0000122 | Neurod1 | 18012 | HITS-CLIP | 19536157 | Unspecified |
| mmu-let-7i-5p | MIMAT0000122 | Nexmif | 340533 | HITS-CLIP | 19536157 | Unspecified |
| mmu-let-7i-5p | MIMAT0000122 | Nf1 | 18015 | HITS-CLIP | 23597149\|19536157\|23142080\|29167 | Unspecified |
| mmu-let-7i-5p | MIMAT0000122 | Nf2 | 18016 | HITS-CLIP | 23597149\|19536157\|29167373\|25083 | Unspecified |

| | | | | | | |
|---|---|---|---|---|---|---|
| mmu-let-7i-5p | MIMAT0000122 | Nfam1 | 74039 | HITS-CLIP | 29167373\|23597149 | Unspecified |
| mmu-let-7i-5p | MIMAT0000122 | Nfasc | 269116 | HITS-CLIP | 37986224 | Unspecified |
| mmu-let-7i-5p | MIMAT0000122 | Nfat5 | 54446 | HITS-CLIP | 37986224\|23142080\|23597149\|1953 | Unspecified |
| mmu-let-7i-5p | MIMAT0000122 | Nfatc1 | 18018 | HITS-CLIP | 23142080 | Unspecified |
| mmu-let-7i-5p | MIMAT0000122 | Nfatc2 | 18019 | HITS-CLIP | 31433990\|23142080\|29167373 | Unspecified |
| mmu-let-7i-5p | MIMAT0000122 | Nfatc2ip | 18020 | HITS-CLIP | 23142080\|29167373 | Unspecified |
| mmu-let-7i-5p | MIMAT0000122 | Nfatc3 | 18021 | HITS-CLIP//qCLASH | 37986224\|31393953\|19536157\|2314 | Unspecified |
| mmu-let-7i-5p | MIMAT0000122 | Nfe2l1 | 18023 | HITS-CLIP | 23597149\|19536157\|29167373 | Unspecified |
| mmu-let-7i-5p | MIMAT0000122 | Nfe2l2 | 18024 | HITS-CLIP | 23142080 | Unspecified |
| mmu-let-7i-5p | MIMAT0000122 | Nfib | 18028 | HITS-CLIP | 19536157 | Unspecified |
| mmu-let-7i-5p | MIMAT0000122 | Nfic | 18029 | HITS-CLIP | 19536157\|23142080\|29167373 | Unspecified |
| mmu-let-7i-5p | MIMAT0000122 | Nfil3 | 18030 | HITS-CLIP | 23142080 | Unspecified |
| mmu-let-7i-5p | MIMAT0000122 | Nfix | 18032 | HITS-CLIP | 37986224\|19536157 | Unspecified |
| mmu-let-7i-5p | MIMAT0000122 | Nfkb1 | 18033 | HITS-CLIP | 23142080 | Unspecified |
| mmu-let-7i-5p | MIMAT0000122 | Nfkbia | 18035 | HITS-CLIP | 29167373 | Unspecified |
| mmu-let-7i-5p | MIMAT0000122 | Nfkbib | 18036 | HITS-CLIP | 23597149\|23142080 | Unspecified |
| mmu-let-7i-5p | MIMAT0000122 | Nfrkb | 235134 | HITS-CLIP | 29167373 | Unspecified |
| mmu-let-7i-5p | MIMAT0000122 | Nfx1 | 74164 | HITS-CLIP | 23597149\|23142080 | Unspecified |
| mmu-let-7i-5p | MIMAT0000122 | Nfxl1 | 100978 | HITS-CLIP | 37986224\|19536157\|23597149\|2314 | Unspecified |
| mmu-let-7i-5p | MIMAT0000122 | Nfya | 18044 | HITS-CLIP | 23142080\|29167373 | Unspecified |
| mmu-let-7i-5p | MIMAT0000122 | Nfyb | 18045 | HITS-CLIP | 23142080\|23597149 | Unspecified |
| mmu-let-7i-5p | MIMAT0000122 | Ngly1 | 59007 | HITS-CLIP | 23142080 | Unspecified |
| mmu-let-7i-5p | MIMAT0000122 | Ngrn | 83485 | HITS-CLIP | 37986224 | Unspecified |
| mmu-let-7i-5p | MIMAT0000122 | Nhlh2 | 18072 | HITS-CLIP | 19536157 | Unspecified |
| mmu-let-7i-5p | MIMAT0000122 | Nhlrc3 | 212114 | HITS-CLIP | 37986224\|23597149\|23142080\|2916 | Unspecified |
| mmu-let-7i-5p | MIMAT0000122 | Nhp2 | 52530 | HITS-CLIP | 29167373 | Unspecified |
| mmu-let-7i-5p | MIMAT0000122 | Nhsl2 | 100042480 | HITS-CLIP | 19536157\|29167373 | Unspecified |
| mmu-let-7i-5p | MIMAT0000122 | Niban2 | 64855 | HITS-CLIP | 26203562 | Unspecified |
| mmu-let-7i-5p | MIMAT0000122 | Nid1 | 18073 | HITS-CLIP | 37986224\|19536157 | Unspecified |
| mmu-let-7i-5p | MIMAT0000122 | Nifk | 67949 | HITS-CLIP | 23142080 | Unspecified |
| mmu-let-7i-5p | MIMAT0000122 | Nin | 18080 | HITS-CLIP | 23142080\|23597149 | Unspecified |
| mmu-let-7i-5p | MIMAT0000122 | Nipa2 | 93790 | HITS-CLIP | 37986224\|23142080 | Unspecified |
| mmu-let-7i-5p | MIMAT0000122 | Nipal1 | 70701 | HITS-CLIP | 23597149 | Unspecified |
| mmu-let-7i-5p | MIMAT0000122 | Nipbl | 71175 | HITS-CLIP | 37986224\|23597149\|19536157\|2314 | Unspecified |
| mmu-let-7i-5p | MIMAT0000122 | Nipsnap1 | 18082 | HITS-CLIP | 37986224\|23597149 | Unspecified |
| mmu-let-7i-5p | MIMAT0000122 | Nipsnap2 | 2631 | HITS-CLIP | 19536157\|23142080 | Unspecified |
| mmu-let-7i-5p | MIMAT0000122 | Nipsnap3b | 66536 | HITS-CLIP | 23597149 | Unspecified |
| mmu-let-7i-5p | MIMAT0000122 | Nisch | 64652 | HITS-CLIP | 23597149\|23142080\|29167373 | Unspecified |
| mmu-let-7i-5p | MIMAT0000122 | Nit1 | 27045 | HITS-CLIP | 23142080 | Unspecified |

| | | | | | | |
|---|---|---|---|---|---|---|
| mmu-let-7i-5p | MIMAT0000122 | Nkap | 67050 | HITS-CLIP | 23142080 | Unspecified |
| mmu-let-7i-5p | MIMAT0000122 | Nkiras2 | 71966 | HITS-CLIP | 23142080 | Unspecified |
| mmu-let-7i-5p | MIMAT0000122 | Nktr | 18087 | HITS-CLIP | 23142080 | Unspecified |
| mmu-let-7i-5p | MIMAT0000122 | Nlgn2 | 216856 | HITS-CLIP | 19536157 | Unspecified |
| mmu-let-7i-5p | MIMAT0000122 | Nlk | 18099 | HITS-CLIP | 37986224\|23142080\|31433990\|23597 | Unspecified |
| mmu-let-7i-5p | MIMAT0000122 | Nlrc3 | 268857 | HITS-CLIP | 23142080 | Unspecified |
| mmu-let-7i-5p | MIMAT0000122 | Nlrc5 | 434341 | HITS-CLIP | 37986224\|23142080\|29167373\|23597 | Unspecified |
| mmu-let-7i-5p | MIMAT0000122 | Nlrp3 | 216799 | HITS-CLIP | 29167373 | Unspecified |
| mmu-let-7i-5p | MIMAT0000122 | Nme2 | 18103 | HITS-CLIP | 23597149\|31433990 | Unspecified |
| mmu-let-7i-5p | MIMAT0000122 | Nme6 | 54369 | HITS-CLIP | 23597149\|23142080 | Unspecified |
| mmu-let-7i-5p | MIMAT0000122 | Nmnat1 | 66454 | HITS-CLIP | 23142080 | Unspecified |
| mmu-let-7i-5p | MIMAT0000122 | Nmrk1 | 225994 | HITS-CLIP | 23597149 | Unspecified |
| mmu-let-7i-5p | MIMAT0000122 | Nmt1 | 18107 | HITS-CLIP | 23142080 | Unspecified |
| mmu-let-7i-5p | MIMAT0000122 | Nmt2 | 18108 | HITS-CLIP | 19536157\|23142080 | Unspecified |
| mmu-let-7i-5p | MIMAT0000122 | Nnt | 18115 | HITS-CLIP | 23597149 | Unspecified |
| mmu-let-7i-5p | MIMAT0000122 | Noc2l | 57741 | HITS-CLIP | 29167373 | Unspecified |
| mmu-let-7i-5p | MIMAT0000122 | Noct | 12457 | HITS-CLIP | 23142080 | Unspecified |
| mmu-let-7i-5p | MIMAT0000122 | Nod1 | 107607 | HITS-CLIP | 23597149 | Unspecified |
| mmu-let-7i-5p | MIMAT0000122 | Nol4l | 329540 | HITS-CLIP | 23142080 | Unspecified |
| mmu-let-7i-5p | MIMAT0000122 | Nol9 | 74035 | HITS-CLIP | 19536157\|23142080\|29167373 | Unspecified |
| mmu-let-7i-5p | MIMAT0000122 | Nono | 53610 | HITS-CLIP | 23142080 | Unspecified |
| mmu-let-7i-5p | MIMAT0000122 | Nop10 | 66181 | HITS-CLIP | 37986224\|23597149\|19536157\|23142 | Unspecified |
| mmu-let-7i-5p | MIMAT0000122 | Nop2 | 110109 | HITS-CLIP | 23142080 | Unspecified |
| mmu-let-7i-5p | MIMAT0000122 | Nop9 | 67842 | HITS-CLIP | 23597149\|19536157\|23142080\|29167 | Unspecified |
| mmu-let-7i-5p | MIMAT0000122 | Notch1 | 18128 | HITS-CLIP | 23597149\|23142080 | Unspecified |
| mmu-let-7i-5p | MIMAT0000122 | Notch2 | 18129 | HITS-CLIP | 37986224\|23597149\|23142080\|29167 | Unspecified |
| mmu-let-7i-5p | MIMAT0000122 | Notch4 | 18132 | HITS-CLIP | 23597149 | Unspecified |
| mmu-let-7i-5p | MIMAT0000122 | Nova1 | 664883 | HITS-CLIP//qCLASH | 19536157\|31393953 | Unspecified |
| mmu-let-7i-5p | MIMAT0000122 | Npas4 | 225872 | HITS-CLIP | 23142080 | Unspecified |
| mmu-let-7i-5p | MIMAT0000122 | Npat | 244879 | HITS-CLIP | 19536157\|23597149\|23142080\|29167 | Unspecified |
| mmu-let-7i-5p | MIMAT0000122 | Npc2 | 67963 | HITS-CLIP | 37986224\|23142080 | Unspecified |
| mmu-let-7i-5p | MIMAT0000122 | Npepl1 | 228961 | HITS-CLIP | 23597149\|23142080 | Unspecified |
| mmu-let-7i-5p | MIMAT0000122 | Npepps | 19155 | HITS-CLIP | 23597149\|23142080\|29167373 | Unspecified |
| mmu-let-7i-5p | MIMAT0000122 | Nphp3 | 74025 | HITS-CLIP | 37986224\|23142080 | Unspecified |
| mmu-let-7i-5p | MIMAT0000122 | Nploc4 | 217365 | HITS-CLIP | 37986224\|23597149 | Unspecified |
| mmu-let-7i-5p | MIMAT0000122 | Npm1 | 18148 | HITS-CLIP | 23142080 | Unspecified |
| mmu-let-7i-5p | MIMAT0000122 | Nr1d2 | 353187 | HITS-CLIP | 23597149 | Unspecified |
| mmu-let-7i-5p | MIMAT0000122 | Nr2c2 | 22026 | HITS-CLIP | 23597149\|23142080 | Unspecified |
| mmu-let-7i-5p | MIMAT0000122 | Nr2f2 | 11819 | HITS-CLIP | 23597149 | Unspecified |

| | | | | | | |
|---|---|---|---|---|---|---|
| mmu-let-7i-5p | MIMAT0000122 | Nr3c1 | 14815 | HITS-CLIP | 31433990\|23142080\|29167373 | Unspecified |
| mmu-let-7i-5p | MIMAT0000122 | Nr4a1 | 15370 | HITS-CLIP | 23142080\|23597149 | Unspecified |
| mmu-let-7i-5p | MIMAT0000122 | Nr4a2 | 18227 | HITS-CLIP | 23142080 | Unspecified |
| mmu-let-7i-5p | MIMAT0000122 | Nr4a3 | 18124 | HITS-CLIP | 23142080 | Unspecified |
| mmu-let-7i-5p | MIMAT0000122 | Nr5a2 | 26424 | HITS-CLIP | 23597149 | Unspecified |
| mmu-let-7i-5p | MIMAT0000122 | Nr6a1 | 14536 | HITS-CLIP | 37986224\|23597149\|19536157\|23142 | Unspecified |
| mmu-let-7i-5p | MIMAT0000122 | Nras | 18176 | HITS-CLIP | 37986224\|19536157\|23142080\|29167 | Unspecified |
| mmu-let-7i-5p | MIMAT0000122 | Nrbp1 | 192292 | HITS-CLIP | 23142080\|23597149 | Unspecified |
| mmu-let-7i-5p | MIMAT0000122 | Nrd1 | 230598 | HITS-CLIP | 37986224\|23597149\|19536157\|29167 | Unspecified |
| mmu-let-7i-5p | MIMAT0000122 | Nrde2 | 217827 | HITS-CLIP | 23142080 | Unspecified |
| mmu-let-7i-5p | MIMAT0000122 | Nrf1 | 18181 | HITS-CLIP | 31433990\|23142080 | Unspecified |
| mmu-let-7i-5p | MIMAT0000122 | Nrip1 | 268903 | HITS-CLIP | 37986224\|23597149\|23142080 | Unspecified |
| mmu-let-7i-5p | MIMAT0000122 | Nrm | 106582 | HITS-CLIP | 23142080 | Unspecified |
| mmu-let-7i-5p | MIMAT0000122 | Nrn1 | 68404 | HITS-CLIP | 37986224\|23597149 | Unspecified |
| mmu-let-7i-5p | MIMAT0000122 | Nrp1 | 18186 | HITS-CLIP | 29167373 | Unspecified |
| mmu-let-7i-5p | MIMAT0000122 | Nrp2 | 18187 | HITS-CLIP | 37986224\|23597149\|19536157\|29167 | Unspecified |
| mmu-let-7i-5p | MIMAT0000122 | Nrros | 224109 | HITS-CLIP | 37986224\|23142080\|23597149\|29167 | Unspecified |
| mmu-let-7i-5p | MIMAT0000122 | Nrxn1 | 18189 | HITS-CLIP | 19536157 | Unspecified |
| mmu-let-7i-5p | MIMAT0000122 | Nsd1 | 18193 | HITS-CLIP | 37986224\|23597149\|19536157\|23142 | Unspecified |
| mmu-let-7i-5p | MIMAT0000122 | Nsd2 | 107823 | HITS-CLIP | 23597149\|23142080\|29167373 | Unspecified |
| mmu-let-7i-5p | MIMAT0000122 | Nsd3 | 234135 | HITS-CLIP | 23142080\|23597149\|29167373 | Unspecified |
| mmu-let-7i-5p | MIMAT0000122 | Nsf | 18195 | HITS-CLIP | 23142080 | Unspecified |
| mmu-let-7i-5p | MIMAT0000122 | Nsfl1c | 386649 | HITS-CLIP | 23142080 | Unspecified |
| mmu-let-7i-5p | MIMAT0000122 | Nsg2 | 18197 | HITS-CLIP | 23142080 | Unspecified |
| mmu-let-7i-5p | MIMAT0000122 | Nsmaf | 18201 | HITS-CLIP | 23142080\|29167373 | Unspecified |
| mmu-let-7i-5p | MIMAT0000122 | Nsmce1 | 67711 | HITS-CLIP | 23142080 | Unspecified |
| mmu-let-7i-5p | MIMAT0000122 | Nsmf | 56876 | HITS-CLIP | 37986224 | Unspecified |
| mmu-let-7i-5p | MIMAT0000122 | Nsun3 | 106338 | HITS-CLIP | 23597149\|29167373\|23142080 | Unspecified |
| mmu-let-7i-5p | MIMAT0000122 | Nsun4 | 72181 | HITS-CLIP | 37986224\|23142080\|23597149 | Unspecified |
| mmu-let-7i-5p | MIMAT0000122 | Nsun5 | 100609 | HITS-CLIP | 23142080\|29167373 | Unspecified |
| mmu-let-7i-5p | MIMAT0000122 | Nsun6 | 74455 | HITS-CLIP | 23142080 | Unspecified |
| mmu-let-7i-5p | MIMAT0000122 | Nt5c2 | 76952 | HITS-CLIP | 23142080 | Unspecified |
| mmu-let-7i-5p | MIMAT0000122 | Nt5dc1 | 319638 | HITS-CLIP | 23142080 | Unspecified |
| mmu-let-7i-5p | MIMAT0000122 | Nt5e | 23959 | HITS-CLIP | 37986224\|23142080 | Unspecified |
| mmu-let-7i-5p | MIMAT0000122 | Ntm | 235106 | HITS-CLIP | 19536157 | Unspecified |
| mmu-let-7i-5p | MIMAT0000122 | Ntn1 | 18208 | HITS-CLIP | 23597149 | Unspecified |
| mmu-let-7i-5p | MIMAT0000122 | Ntng2 | 171171 | HITS-CLIP | 19536157 | Unspecified |
| mmu-let-7i-5p | MIMAT0000122 | Ntrk2 | 18212 | HITS-CLIP | 37986224\|19536157 | Unspecified |
| mmu-let-7i-5p | MIMAT0000122 | Ntrk3 | 18213 | HITS-CLIP | 19536157 | Unspecified |

| | | | | | | |
|---|---|---|---|---|---|---|
| mmu-let-7i-5p | MIMAT0000122 | Nuak2 | 74137 | HITS-CLIP | 23597149\|29167373 | Unspecified |
| mmu-let-7i-5p | MIMAT0000122 | Nub1 | 53312 | HITS-CLIP | 29167373 | Unspecified |
| mmu-let-7i-5p | MIMAT0000122 | Nubp1 | 26425 | HITS-CLIP | 29167373 | Unspecified |
| mmu-let-7i-5p | MIMAT0000122 | Nucb1 | 18220 | HITS-CLIP | 23142080\|23597149\|19536157 | Unspecified |
| mmu-let-7i-5p | MIMAT0000122 | Nucks1 | 98415 | HITS-CLIP | 31433990\|19536157\|23142080 | Unspecified |
| mmu-let-7i-5p | MIMAT0000122 | Nudcd3 | 209586 | HITS-CLIP | 29167373 | Unspecified |
| mmu-let-7i-5p | MIMAT0000122 | Nudt18 | 213484 | HITS-CLIP | 23142080 | Unspecified |
| mmu-let-7i-5p | MIMAT0000122 | Nudt3 | 56409 | HITS-CLIP | 23597149 | Unspecified |
| mmu-let-7i-5p | MIMAT0000122 | Nudt4 | 71207 | HITS-CLIP | 29167373 | Unspecified |
| mmu-let-7i-5p | MIMAT0000122 | Nudt5 | 53893 | HITS-CLIP | 23597149\|23142080\|29167373 | Unspecified |
| mmu-let-7i-5p | MIMAT0000122 | Nudt7 | 67528 | HITS-CLIP | 37986224 | Unspecified |
| mmu-let-7i-5p | MIMAT0000122 | Nufip2 | 68564 | HITS-CLIP | 23597149\|23142080 | Unspecified |
| mmu-let-7i-5p | MIMAT0000122 | Numa1 | 101706 | HITS-CLIP | 23597149\|23142080 | Unspecified |
| mmu-let-7i-5p | MIMAT0000122 | Nup107 | 103468 | HITS-CLIP | 23142080 | Unspecified |
| mmu-let-7i-5p | MIMAT0000122 | Nup133 | 234865 | HITS-CLIP | 23142080 | Unspecified |
| mmu-let-7i-5p | MIMAT0000122 | Nup153 | 218210 | HITS-CLIP | 37986224\|23142080\|19536157\|29167 | Unspecified |
| mmu-let-7i-5p | MIMAT0000122 | Nup155 | 170762 | HITS-CLIP | 23142080\|23597149 | Unspecified |
| mmu-let-7i-5p | MIMAT0000122 | Nup160 | 59015 | HITS-CLIP | 23597149\|23142080 | Unspecified |
| mmu-let-7i-5p | MIMAT0000122 | Nup188 | 227699 | HITS-CLIP | 23597149\|23142080 | Unspecified |
| mmu-let-7i-5p | MIMAT0000122 | Nup205 | 70699 | HITS-CLIP | 37986224\|23142080 | Unspecified |
| mmu-let-7i-5p | MIMAT0000122 | Nup210 | 54563 | HITS-CLIP | 23597149\|23142080 | Unspecified |
| mmu-let-7i-5p | MIMAT0000122 | Nup214 | 227720 | HITS-CLIP | 21258322\|23142080 | Unspecified |
| mmu-let-7i-5p | MIMAT0000122 | Nup37 | 69736 | HITS-CLIP | 23142080 | Unspecified |
| mmu-let-7i-5p | MIMAT0000122 | Nup54 | 269113 | HITS-CLIP | 23597149 | Unspecified |
| mmu-let-7i-5p | MIMAT0000122 | Nup62 | 18226 | HITS-CLIP | 23597149\|19536157\|23142080\|29167 | Unspecified |
| mmu-let-7i-5p | MIMAT0000122 | Nup85 | 445007 | HITS-CLIP | 23142080 | Unspecified |
| mmu-let-7i-5p | MIMAT0000122 | Nup88 | 19069 | HITS-CLIP | 37986224\|23597149\|23142080 | Unspecified |
| mmu-let-7i-5p | MIMAT0000122 | Nup93 | 71805 | HITS-CLIP | 23142080\|23597149 | Unspecified |
| mmu-let-7i-5p | MIMAT0000122 | Nup98 | 269966 | HITS-CLIP | 23142080\|23597149\|29167373\|1953€ | Unspecified |
| mmu-let-7i-5p | MIMAT0000122 | Nupl1 | 71844 | HITS-CLIP | 37986224 | Unspecified |
| mmu-let-7i-5p | MIMAT0000122 | Nusap1 | 108907 | HITS-CLIP | 23142080 | Unspecified |
| mmu-let-7i-5p | MIMAT0000122 | Nwd2 | 319807 | HITS-CLIP | 19536157 | Unspecified |
| mmu-let-7i-5p | MIMAT0000122 | Nxf1 | 53319 | HITS-CLIP | 23142080\|23597149\|29167373 | Unspecified |
| mmu-let-7i-5p | MIMAT0000122 | Nxpe2 | 78252 | HITS-CLIP | 23597149 | Unspecified |
| mmu-let-7i-5p | MIMAT0000122 | Nxpe3 | 385658 | HITS-CLIP | 23142080 | Unspecified |
| mmu-let-7i-5p | MIMAT0000122 | Oaf | 102644 | HITS-CLIP | 37986224\|23597149 | Unspecified |
| mmu-let-7i-5p | MIMAT0000122 | Oas3 | 246727 | HITS-CLIP | 29167373 | Unspecified |
| mmu-let-7i-5p | MIMAT0000122 | Oat | 18242 | HITS-CLIP | 37986224 | Unspecified |
| mmu-let-7i-5p | MIMAT0000122 | Ociad1 | 68095 | HITS-CLIP | 23142080 | Unspecified |

| | | | | | | |
|---|---|---|---|---|---|---|
| mmu-let-7i-5p | MIMAT0000122 | Ocln | 18260 | HITS-CLIP | 23597149 | Unspecified |
| mmu-let-7i-5p | MIMAT0000122 | Ocrl | 320634 | HITS-CLIP | 19536157 | Unspecified |
| mmu-let-7i-5p | MIMAT0000122 | Odc1 | 18263 | HITS-CLIP | 23142080\|29167373 | Unspecified |
| mmu-let-7i-5p | MIMAT0000122 | Odf2l | 52184 | HITS-CLIP | 23142080 | Unspecified |
| mmu-let-7i-5p | MIMAT0000122 | Odr4 | 54953 | HITS-CLIP | 23142080 | Unspecified |
| mmu-let-7i-5p | MIMAT0000122 | Ogdh | 18293 | HITS-CLIP | 23597149\|29167373 | Unspecified |
| mmu-let-7i-5p | MIMAT0000122 | Ogfr | 72075 | HITS-CLIP | 23142080 | Unspecified |
| mmu-let-7i-5p | MIMAT0000122 | Ogt | 108155 | HITS-CLIP | 23142080 | Unspecified |
| mmu-let-7i-5p | MIMAT0000122 | Oit3 | 18302 | HITS-CLIP | 37986224\|23597149\|23142080 | Unspecified |
| mmu-let-7i-5p | MIMAT0000122 | Olfm1 | 56177 | HITS-CLIP | 37986224 | Unspecified |
| mmu-let-7i-5p | MIMAT0000122 | Olfr111 | 545205 | HITS-CLIP | 29167373 | Unspecified |
| mmu-let-7i-5p | MIMAT0000122 | Oma1 | 67013 | HITS-CLIP | 19536157 | Unspecified |
| mmu-let-7i-5p | MIMAT0000122 | Omg | 18377 | HITS-CLIP | 37986224 | Unspecified |
| mmu-let-7i-5p | MIMAT0000122 | Onecut1 | 15379 | HITS-CLIP | 23597149 | Unspecified |
| mmu-let-7i-5p | MIMAT0000122 | Onecut2 | 225631 | HITS-CLIP | 37986224\|23597149 | Unspecified |
| mmu-let-7i-5p | MIMAT0000122 | Opa1 | 74143 | HITS-CLIP | 23142080\|23597149\|29167373 | Unspecified |
| mmu-let-7i-5p | MIMAT0000122 | Oplah | 75475 | HITS-CLIP | 23597149 | Unspecified |
| mmu-let-7i-5p | MIMAT0000122 | Orai1 | 109305 | HITS-CLIP | 23142080 | Unspecified |
| mmu-let-7i-5p | MIMAT0000122 | Orai3 | 269999 | qCLASH//HITS-CLIP | 31393953\|23597149\|23142080 | Unspecified |
| mmu-let-7i-5p | MIMAT0000122 | Orc2 | 18393 | HITS-CLIP | 23142080 | Unspecified |
| mmu-let-7i-5p | MIMAT0000122 | Orc3 | 50793 | HITS-CLIP | 31433990\|23630078\|23597149\|23142 | Unspecified |
| mmu-let-7i-5p | MIMAT0000122 | Orc4 | 26428 | HITS-CLIP | 37986224\|29167373\|23142080 | Unspecified |
| mmu-let-7i-5p | MIMAT0000122 | Orc5 | 26429 | HITS-CLIP | 23142080 | Unspecified |
| mmu-let-7i-5p | MIMAT0000122 | Orc6 | 56452 | HITS-CLIP | 23142080 | Unspecified |
| mmu-let-7i-5p | MIMAT0000122 | Orm1 | 18405 | HITS-CLIP | 23597149 | Unspecified |
| mmu-let-7i-5p | MIMAT0000122 | Orm2 | 18406 | HITS-CLIP | 23597149 | Unspecified |
| mmu-let-7i-5p | MIMAT0000122 | Ormdl1 | 227102 | HITS-CLIP | 23142080 | Unspecified |
| mmu-let-7i-5p | MIMAT0000122 | Ormdl2 | 66844 | HITS-CLIP | 23597149\|23142080 | Unspecified |
| mmu-let-7i-5p | MIMAT0000122 | Os9 | 216440 | HITS-CLIP | 23142080\|23597149 | Unspecified |
| mmu-let-7i-5p | MIMAT0000122 | Osbp | 76303 | HITS-CLIP | 23597149\|23142080 | Unspecified |
| mmu-let-7i-5p | MIMAT0000122 | Osbp2 | 74309 | HITS-CLIP | 19536157 | Unspecified |
| mmu-let-7i-5p | MIMAT0000122 | Osbpl11 | 106326 | HITS-CLIP | 23597149\|23142080 | Unspecified |
| mmu-let-7i-5p | MIMAT0000122 | Osbpl1a | 64291 | HITS-CLIP | 23597149\|19536157 | Unspecified |
| mmu-let-7i-5p | MIMAT0000122 | Osbpl2 | 228983 | HITS-CLIP | 23597149\|23142080 | Unspecified |
| mmu-let-7i-5p | MIMAT0000122 | Osbpl3 | 71720 | HITS-CLIP | 23142080 | Unspecified |
| mmu-let-7i-5p | MIMAT0000122 | Osbpl8 | 237542 | HITS-CLIP | 23597149\|23142080\|29167373 | Unspecified |
| mmu-let-7i-5p | MIMAT0000122 | Osbpl9 | 100273 | HITS-CLIP | 23597149 | Unspecified |
| mmu-let-7i-5p | MIMAT0000122 | Oser1 | 51526 | HITS-CLIP | 23142080 | Unspecified |
| mmu-let-7i-5p | MIMAT0000122 | Osm | 18413 | HITS-CLIP | 23142080 | Unspecified |

| | | | | | | |
|---|---|---|---|---|---|---|
| mmu-let-7i-5p | MIMAT0000122 | Ostc | 66357 | HITS-CLIP | 23142080 | Unspecified |
| mmu-let-7i-5p | MIMAT0000122 | Ostf1 | 20409 | HITS-CLIP | 23142080 | Unspecified |
| mmu-let-7i-5p | MIMAT0000122 | Ostm1 | 14628 | HITS-CLIP | 37986224\|23597149\|23142080 | Unspecified |
| mmu-let-7i-5p | MIMAT0000122 | Otc | 18416 | HITS-CLIP | 23597149 | Unspecified |
| mmu-let-7i-5p | MIMAT0000122 | Otub1 | 107260 | HITS-CLIP | 23142080\|29167373 | Unspecified |
| mmu-let-7i-5p | MIMAT0000122 | Otud1 | 71198 | HITS-CLIP | 23142080 | Unspecified |
| mmu-let-7i-5p | MIMAT0000122 | Otud4 | 73945 | HITS-CLIP | 23142080\|23597149\|29167373 | Unspecified |
| mmu-let-7i-5p | MIMAT0000122 | Otud5 | 54644 | HITS-CLIP | 23597149\|23142080 | Unspecified |
| mmu-let-7i-5p | MIMAT0000122 | Otx1 | 18423 | HITS-CLIP | 23142080 | Unspecified |
| mmu-let-7i-5p | MIMAT0000122 | Oxa1l | 69089 | HITS-CLIP | 23597149\|23142080 | Unspecified |
| mmu-let-7i-5p | MIMAT0000122 | Oxct1 | 67041 | HITS-CLIP | 23142080 | Unspecified |
| mmu-let-7i-5p | MIMAT0000122 | Oxr1 | 170719 | HITS-CLIP | 23142080 | Unspecified |
| mmu-let-7i-5p | MIMAT0000122 | Oxsm | 71147 | HITS-CLIP | 23597149 | Unspecified |
| mmu-let-7i-5p | MIMAT0000122 | Oxsr1 | 108737 | HITS-CLIP | 37986224\|23597149\|23142080\|2916 | Unspecified |
| mmu-let-7i-5p | MIMAT0000122 | P2rx4 | 18438 | HITS-CLIP | 29167373\|23142080 | Unspecified |
| mmu-let-7i-5p | MIMAT0000122 | P2ry10 | 78826 | HITS-CLIP | 23142080 | Unspecified |
| mmu-let-7i-5p | MIMAT0000122 | P2ry10b | 213438 | HITS-CLIP | 23142080 | Unspecified |
| mmu-let-7i-5p | MIMAT0000122 | P2ry13 | 74191 | HITS-CLIP | 37986224\|23597149 | Unspecified |
| mmu-let-7i-5p | MIMAT0000122 | P2ry2 | 18442 | HITS-CLIP | 23597149\|29167373 | Unspecified |
| mmu-let-7i-5p | MIMAT0000122 | P3h1 | 64175 | HITS-CLIP | 23142080 | Unspecified |
| mmu-let-7i-5p | MIMAT0000122 | P3h4 | 10609 | HITS-CLIP | 23597149 | Unspecified |
| mmu-let-7i-5p | MIMAT0000122 | P4hb | 18453 | HITS-CLIP | 23597149\|23142080 | Unspecified |
| mmu-let-7i-5p | MIMAT0000122 | Pa2g4 | 18813 | HITS-CLIP | 23630078 | Unspecified |
| mmu-let-7i-5p | MIMAT0000122 | Pabpc1 | 18458 | HITS-CLIP//qCLASH | 23142080\|31393953\|29167373 | Unspecified |
| mmu-let-7i-5p | MIMAT0000122 | Pabpc4l | 241989 | HITS-CLIP | 23597149 | Unspecified |
| mmu-let-7i-5p | MIMAT0000122 | Pacs1 | 107975 | HITS-CLIP | 23142080 | Unspecified |
| mmu-let-7i-5p | MIMAT0000122 | Pacs2 | 217893 | HITS-CLIP | 37986224\|23597149\|23142080 | Unspecified |
| mmu-let-7i-5p | MIMAT0000122 | Padi2 | 18600 | HITS-CLIP | 23142080 | Unspecified |
| mmu-let-7i-5p | MIMAT0000122 | Padi3 | 18601 | HITS-CLIP | 23142080 | Unspecified |
| mmu-let-7i-5p | MIMAT0000122 | Paf1 | 54624 | qCLASH | 31393953 | Unspecified |
| mmu-let-7i-5p | MIMAT0000122 | Pafah1b1 | 18472 | HITS-CLIP | 23597149\|23142080 | Unspecified |
| mmu-let-7i-5p | MIMAT0000122 | Pafah1b2 | 18475 | HITS-CLIP | 23597149\|29167373 | Unspecified |
| mmu-let-7i-5p | MIMAT0000122 | Pafah1b3 | 18476 | HITS-CLIP | 23142080 | Unspecified |
| mmu-let-7i-5p | MIMAT0000122 | Pag1 | 94212 | HITS-CLIP | 37986224\|23142080\|23597149\|1953 | Unspecified |
| mmu-let-7i-5p | MIMAT0000122 | Pah | 18478 | HITS-CLIP | 23597149 | Unspecified |
| mmu-let-7i-5p | MIMAT0000122 | Paip1 | 218693 | HITS-CLIP//qCLASH | 23597149\|23142080\|31393953 | Unspecified |
| mmu-let-7i-5p | MIMAT0000122 | Pak1 | 18479 | HITS-CLIP | 37986224 | Unspecified |
| mmu-let-7i-5p | MIMAT0000122 | Pak2 | 224105 | HITS-CLIP | 23142080 | Unspecified |
| mmu-let-7i-5p | MIMAT0000122 | Pak4 | 70584 | HITS-CLIP | 37986224 | Unspecified |

| | | | | | | |
|---|---|---|---|---|---|---|
| mmu-let-7i-5p | MIMAT0000122 | Pak6 | 214230 | HITS-CLIP | 37986224\|19536157 | Unspecified |
| mmu-let-7i-5p | MIMAT0000122 | Palb2 | 233826 | HITS-CLIP | 23142080 | Unspecified |
| mmu-let-7i-5p | MIMAT0000122 | Palld | 72333 | HITS-CLIP | 23597149\|29167373 | Unspecified |
| mmu-let-7i-5p | MIMAT0000122 | Palmd | 114301 | HITS-CLIP | 23597149 | Unspecified |
| mmu-let-7i-5p | MIMAT0000122 | Pan3 | 72587 | HITS-CLIP | 37986224\|23142080 | Unspecified |
| mmu-let-7i-5p | MIMAT0000122 | Pank1 | 75735 | HITS-CLIP | 19536157\|23597149 | Unspecified |
| mmu-let-7i-5p | MIMAT0000122 | Pank3 | 211347 | HITS-CLIP | 23597149\|23142080\|19536157\|29167 | Unspecified |
| mmu-let-7i-5p | MIMAT0000122 | Pank4 | 55229 | qCLASH | 31393953 | Unspecified |
| mmu-let-7i-5p | MIMAT0000122 | Panx1 | 55991 | HITS-CLIP | 23142080\|23597149 | Unspecified |
| mmu-let-7i-5p | MIMAT0000122 | Panx2 | 406218 | HITS-CLIP | 37986224\|19536157 | Unspecified |
| mmu-let-7i-5p | MIMAT0000122 | Papola | 18789 | HITS-CLIP | 19536157\|23597149\|23142080\|29167 | Unspecified |
| mmu-let-7i-5p | MIMAT0000122 | Papolg | 216578 | HITS-CLIP | 23142080 | Unspecified |
| mmu-let-7i-5p | MIMAT0000122 | Papss2 | 23972 | HITS-CLIP | 23597149 | Unspecified |
| mmu-let-7i-5p | MIMAT0000122 | Paqr4 | 76498 | HITS-CLIP | 23142080 | Unspecified |
| mmu-let-7i-5p | MIMAT0000122 | Paqr7 | 71904 | HITS-CLIP | 23597149\|23142080 | Unspecified |
| mmu-let-7i-5p | MIMAT0000122 | Paqr9 | 75552 | HITS-CLIP | 37986224\|23597149 | Unspecified |
| mmu-let-7i-5p | MIMAT0000122 | Pard3 | 93742 | HITS-CLIP | 37986224\|23597149 | Unspecified |
| mmu-let-7i-5p | MIMAT0000122 | Pard6b | 58220 | HITS-CLIP | 23597149 | Unspecified |
| mmu-let-7i-5p | MIMAT0000122 | Parg | 26430 | HITS-CLIP | 23142080 | Unspecified |
| mmu-let-7i-5p | MIMAT0000122 | Park7 | 57320 | HITS-CLIP | 23597149\|23142080 | Unspecified |
| mmu-let-7i-5p | MIMAT0000122 | Parm1 | 231440 | HITS-CLIP | 37986224 | Unspecified |
| mmu-let-7i-5p | MIMAT0000122 | Parp1 | 11545 | HITS-CLIP | 23597149\|23142080 | Unspecified |
| mmu-let-7i-5p | MIMAT0000122 | Parp11 | 101187 | HITS-CLIP | 23142080 | Unspecified |
| mmu-let-7i-5p | MIMAT0000122 | Parp12 | 243771 | HITS-CLIP | 37986224\|23597149 | Unspecified |
| mmu-let-7i-5p | MIMAT0000122 | Parp14 | 547253 | HITS-CLIP | 23597149\|23142080\|29167373 | Unspecified |
| mmu-let-7i-5p | MIMAT0000122 | Parp16 | 214424 | HITS-CLIP | 37986224\|23142080\|23597149 | Unspecified |
| mmu-let-7i-5p | MIMAT0000122 | Parp2 | 11546 | HITS-CLIP | 23142080 | Unspecified |
| mmu-let-7i-5p | MIMAT0000122 | Parp4 | 328417 | HITS-CLIP | 23142080 | Unspecified |
| mmu-let-7i-5p | MIMAT0000122 | Parp6 | 67287 | HITS-CLIP | 23142080 | Unspecified |
| mmu-let-7i-5p | MIMAT0000122 | Parp9 | 80285 | HITS-CLIP | 23142080\|29167373 | Unspecified |
| mmu-let-7i-5p | MIMAT0000122 | Pars2 | 230577 | HITS-CLIP | 23597149 | Unspecified |
| mmu-let-7i-5p | MIMAT0000122 | Parva | 57342 | HITS-CLIP | 23597149 | Unspecified |
| mmu-let-7i-5p | MIMAT0000122 | Parvb | 170736 | HITS-CLIP | 29167373 | Unspecified |
| mmu-let-7i-5p | MIMAT0000122 | Parvg | 64099 | HITS-CLIP | 23142080 | Unspecified |
| mmu-let-7i-5p | MIMAT0000122 | Pask | 269224 | HITS-CLIP | 23142080 | Unspecified |
| mmu-let-7i-5p | MIMAT0000122 | Patj | 44100 | HITS-CLIP | 23142080 | Unspecified |
| mmu-let-7i-5p | MIMAT0000122 | Patl1 | 225929 | HITS-CLIP//qCLASH | 37986224\|19536157\|23597149\|3139 | Unspecified |
| mmu-let-7i-5p | MIMAT0000122 | Paxbp1 | 67367 | HITS-CLIP | 29167373 | Unspecified |
| mmu-let-7i-5p | MIMAT0000122 | Pbld2 | 67307 | HITS-CLIP | 23597149 | Unspecified |

| | | | | | | |
|---|---|---|---|---|---|---|
| mmu-let-7i-5p | MIMAT0000122 | Pbrm1 | 66923 | HITS-CLIP | 37986224\|23597149\|19536157\|23142 | Unspecified |
| mmu-let-7i-5p | MIMAT0000122 | Pbx1 | 18514 | HITS-CLIP | 37986224\|19536157 | Unspecified |
| mmu-let-7i-5p | MIMAT0000122 | Pbx2 | 18515 | HITS-CLIP | 37986224\|31433990\|19536157\|23597 | Unspecified |
| mmu-let-7i-5p | MIMAT0000122 | Pbx3 | 18516 | HITS-CLIP | 19536157\|23142080 | Unspecified |
| mmu-let-7i-5p | MIMAT0000122 | Pcbp2 | 18521 | HITS-CLIP | 23142080\|29167373 | Unspecified |
| mmu-let-7i-5p | MIMAT0000122 | Pcca | 110821 | HITS-CLIP | 37986224\|23597149 | Unspecified |
| mmu-let-7i-5p | MIMAT0000122 | Pcdh1 | 75599 | HITS-CLIP | 37986224\|19536157\|23597149 | Unspecified |
| mmu-let-7i-5p | MIMAT0000122 | Pcdh10 | 18526 | HITS-CLIP | 19536157 | Unspecified |
| mmu-let-7i-5p | MIMAT0000122 | Pcdh19 | 279653 | HITS-CLIP | 19536157 | Unspecified |
| mmu-let-7i-5p | MIMAT0000122 | Pcdh7 | 54216 | HITS-CLIP | 37986224\|19536157\|29167373 | Unspecified |
| mmu-let-7i-5p | MIMAT0000122 | Pcdh9 | 211712 | HITS-CLIP | 37986224\|19536157 | Unspecified |
| mmu-let-7i-5p | MIMAT0000122 | Pcdha10 | 12943 | HITS-CLIP | 37986224 | Unspecified |
| mmu-let-7i-5p | MIMAT0000122 | Pcdhac2 | 353237 | HITS-CLIP | 19536157 | Unspecified |
| mmu-let-7i-5p | MIMAT0000122 | Pcdhga12 | 93724 | HITS-CLIP | 37986224 | Unspecified |
| mmu-let-7i-5p | MIMAT0000122 | Pcdhgc3 | 5098 | HITS-CLIP | 19536157 | Unspecified |
| mmu-let-7i-5p | MIMAT0000122 | Pced1b | 239647 | HITS-CLIP | 23142080 | Unspecified |
| mmu-let-7i-5p | MIMAT0000122 | Pcf11 | 74737 | HITS-CLIP | 23142080\|29167373 | Unspecified |
| mmu-let-7i-5p | MIMAT0000122 | Pcgf3 | 69587 | HITS-CLIP | 37986224\|23597149\|19536157\|29167 | Unspecified |
| mmu-let-7i-5p | MIMAT0000122 | Pcgf6 | 71041 | HITS-CLIP | 23142080 | Unspecified |
| mmu-let-7i-5p | MIMAT0000122 | Pcif1 | 228866 | HITS-CLIP | 23597149 | Unspecified |
| mmu-let-7i-5p | MIMAT0000122 | Pck1 | 18534 | HITS-CLIP | 23597149 | Unspecified |
| mmu-let-7i-5p | MIMAT0000122 | Pck2 | 74551 | HITS-CLIP | 23597149 | Unspecified |
| mmu-let-7i-5p | MIMAT0000122 | Pclaf | 68026 | HITS-CLIP | 23142080 | Unspecified |
| mmu-let-7i-5p | MIMAT0000122 | Pclo | 26875 | HITS-CLIP | 19536157 | Unspecified |
| mmu-let-7i-5p | MIMAT0000122 | Pcm1 | 18536 | HITS-CLIP | 23597149\|19536157\|29167373\|23142 | Unspecified |
| mmu-let-7i-5p | MIMAT0000122 | Pcmt1 | 18537 | HITS-CLIP | 23142080 | Unspecified |
| mmu-let-7i-5p | MIMAT0000122 | Pcmtd1 | 319263 | HITS-CLIP | 23597149\|23142080 | Unspecified |
| mmu-let-7i-5p | MIMAT0000122 | Pcnx | 54604 | HITS-CLIP | 37986224\|23597149\|23142080\|19536 | Unspecified |
| mmu-let-7i-5p | MIMAT0000122 | Pcnx3 | 399909 | HITS-CLIP | 23597149\|23142080\|29167373 | Unspecified |
| mmu-let-7i-5p | MIMAT0000122 | Pcp4l1 | 66425 | HITS-CLIP | 37986224\|19536157 | Unspecified |
| mmu-let-7i-5p | MIMAT0000122 | Pcsk2 | 18549 | HITS-CLIP | 37986224 | Unspecified |
| mmu-let-7i-5p | MIMAT0000122 | Pcsk4 | 18551 | HITS-CLIP | 23142080 | Unspecified |
| mmu-let-7i-5p | MIMAT0000122 | Pcsk7 | 18554 | HITS-CLIP | 23597149 | Unspecified |
| mmu-let-7i-5p | MIMAT0000122 | Pcx | 18563 | HITS-CLIP | 23597149 | Unspecified |
| mmu-let-7i-5p | MIMAT0000122 | Pcyox1 | 66881 | HITS-CLIP | 37986224\|23597149\|23142080\|29167 | Unspecified |
| mmu-let-7i-5p | MIMAT0000122 | Pcyt1a | 13026 | HITS-CLIP | 37986224 | Unspecified |
| mmu-let-7i-5p | MIMAT0000122 | Pcyt2 | 68671 | HITS-CLIP | 23597149 | Unspecified |
| mmu-let-7i-5p | MIMAT0000122 | Pdcd1 | 18566 | HITS-CLIP | 23142080 | Unspecified |
| mmu-let-7i-5p | MIMAT0000122 | Pdcd11 | 18572 | HITS-CLIP | 23142080 | Unspecified |

| | | | | | | |
|---|---|---|---|---|---|---|
| mmu-let-7i-5p | MIMAT0000122 | Pdcd4 | 18569 | HITS-CLIP | 23597149\|23142080 | Unspecified |
| mmu-let-7i-5p | MIMAT0000122 | Pdcd6ip | 18571 | HITS-CLIP | 19536157\|23142080\|29167373 | Unspecified |
| mmu-let-7i-5p | MIMAT0000122 | Pdcd7 | 50996 | HITS-CLIP | 23597149\|23142080 | Unspecified |
| mmu-let-7i-5p | MIMAT0000122 | Pdcl3 | 68833 | HITS-CLIP | 23142080 | Unspecified |
| mmu-let-7i-5p | MIMAT0000122 | Pde12 | 211948 | HITS-CLIP | 23142080 | Unspecified |
| mmu-let-7i-5p | MIMAT0000122 | Pde2a | 207728 | HITS-CLIP | 23597149\|19536157 | Unspecified |
| mmu-let-7i-5p | MIMAT0000122 | Pde3b | 18576 | HITS-CLIP | 37986224\|23597149\|19536157\|23142 | Unspecified |
| mmu-let-7i-5p | MIMAT0000122 | Pde4a | 18577 | HITS-CLIP | 37986224 | Unspecified |
| mmu-let-7i-5p | MIMAT0000122 | Pde4b | 18578 | HITS-CLIP | 23142080 | Unspecified |
| mmu-let-7i-5p | MIMAT0000122 | Pde4d | 238871 | HITS-CLIP | 19536157\|23142080 | Unspecified |
| mmu-let-7i-5p | MIMAT0000122 | Pde4dip | 83679 | HITS-CLIP | 23597149\|19536157\|23142080 | Unspecified |
| mmu-let-7i-5p | MIMAT0000122 | Pde7a | 18583 | HITS-CLIP | 19536157\|23597149\|23142080 | Unspecified |
| mmu-let-7i-5p | MIMAT0000122 | Pdgfb | 18591 | HITS-CLIP | 23142080\|29167373 | Unspecified |
| mmu-let-7i-5p | MIMAT0000122 | Pdgfrb | 18596 | HITS-CLIP | 37986224 | Unspecified |
| mmu-let-7i-5p | MIMAT0000122 | Pdhx | 27402 | HITS-CLIP | 23597149 | Unspecified |
| mmu-let-7i-5p | MIMAT0000122 | Pdia3 | 14827 | HITS-CLIP | 23597149\|23142080\|29167373 | Unspecified |
| mmu-let-7i-5p | MIMAT0000122 | Pdia4 | 12304 | HITS-CLIP | 23597149\|23142080\|29167373 | Unspecified |
| mmu-let-7i-5p | MIMAT0000122 | Pdia6 | 71853 | HITS-CLIP | 23597149\|29167373 | Unspecified |
| mmu-let-7i-5p | MIMAT0000122 | Pdik1l | 230809 | HITS-CLIP | 30971389 | Unspecified |
| mmu-let-7i-5p | MIMAT0000122 | Pdk1 | 228026 | HITS-CLIP | 23142080 | Unspecified |
| mmu-let-7i-5p | MIMAT0000122 | Pdk2 | 18604 | HITS-CLIP | 37986224\|19536157\|23597149\|23142 | Unspecified |
| mmu-let-7i-5p | MIMAT0000122 | Pdlim1 | 54132 | HITS-CLIP | 23597149\|23142080 | Unspecified |
| mmu-let-7i-5p | MIMAT0000122 | Pdp1 | 381511 | HITS-CLIP | 23630078\|19536157\|23142080 | Unspecified |
| mmu-let-7i-5p | MIMAT0000122 | Pdp2 | 382051 | HITS-CLIP | 23597149\|23142080 | Unspecified |
| mmu-let-7i-5p | MIMAT0000122 | Pdpk1 | 18607 | HITS-CLIP | 37986224\|23597149\|23142080 | Unspecified |
| mmu-let-7i-5p | MIMAT0000122 | Pdpr | 319518 | HITS-CLIP | 37986224\|23597149\|23142080 | Unspecified |
| mmu-let-7i-5p | MIMAT0000122 | Pds5a | 71521 | HITS-CLIP//qCLASH | 23142080\|31393953\|19536157\|29167 | Unspecified |
| mmu-let-7i-5p | MIMAT0000122 | Pds5b | 100710 | HITS-CLIP | 23597149\|30971389\|23142080 | Unspecified |
| mmu-let-7i-5p | MIMAT0000122 | Pdss1 | 56075 | HITS-CLIP | 29167373 | Unspecified |
| mmu-let-7i-5p | MIMAT0000122 | Pdxdc1 | 94184 | HITS-CLIP | 23597149\|23142080\|19536157 | Unspecified |
| mmu-let-7i-5p | MIMAT0000122 | Pdxk | 216134 | HITS-CLIP | 19536157\|29167373 | Unspecified |
| mmu-let-7i-5p | MIMAT0000122 | Pdzd11 | 72621 | HITS-CLIP | 31433990 | Unspecified |
| mmu-let-7i-5p | MIMAT0000122 | Pdzd2 | 68070 | HITS-CLIP | 37986224\|19536157 | Unspecified |
| mmu-let-7i-5p | MIMAT0000122 | Pdzd8 | 107368 | HITS-CLIP | 31433990\|23142080 | Unspecified |
| mmu-let-7i-5p | MIMAT0000122 | Pdzk1 | 59020 | HITS-CLIP | 37986224\|23597149 | Unspecified |
| mmu-let-7i-5p | MIMAT0000122 | Pdzrn4 | 239618 | HITS-CLIP | 19536157 | Unspecified |
| mmu-let-7i-5p | MIMAT0000122 | Peak1 | 244895 | HITS-CLIP | 37986224\|23142080\|23597149\|1953€ | Unspecified |
| mmu-let-7i-5p | MIMAT0000122 | Pear1 | 375033 | HITS-CLIP | 23142080 | Unspecified |
| mmu-let-7i-5p | MIMAT0000122 | Pecam1 | 18613 | HITS-CLIP | 23142080 | Unspecified |

| | | | | | | |
|---|---|---|---|---|---|---|
| mmu-let-7i-5p | MIMAT0000122 | Pecr | 111175 | HITS-CLIP | 23597149 | Unspecified |
| mmu-let-7i-5p | MIMAT0000122 | Pef1 | 67898 | HITS-CLIP | 23597149 | Unspecified |
| mmu-let-7i-5p | MIMAT0000122 | Peg10 | 170676 | HITS-CLIP | 23630078 | Unspecified |
| mmu-let-7i-5p | MIMAT0000122 | Peg3 | 18616 | HITS-CLIP | 37986224\|19536157 | Unspecified |
| mmu-let-7i-5p | MIMAT0000122 | Peli1 | 67245 | HITS-CLIP | 37986224\|23142080 | Unspecified |
| mmu-let-7i-5p | MIMAT0000122 | Pemt | 18618 | HITS-CLIP | 23597149 | Unspecified |
| mmu-let-7i-5p | MIMAT0000122 | Penk | 18619 | HITS-CLIP | 37986224\|31433990\|23142080 | Unspecified |
| mmu-let-7i-5p | MIMAT0000122 | Pepd | 18624 | HITS-CLIP | 23597149\|29167373 | Unspecified |
| mmu-let-7i-5p | MIMAT0000122 | Per1 | 18626 | HITS-CLIP | 23597149\|19536157\|23142080 | Unspecified |
| mmu-let-7i-5p | MIMAT0000122 | Per2 | 18627 | HITS-CLIP | 23142080 | Unspecified |
| mmu-let-7i-5p | MIMAT0000122 | Per3 | 18628 | HITS-CLIP | 23597149 | Unspecified |
| mmu-let-7i-5p | MIMAT0000122 | Pes1 | 64934 | HITS-CLIP | 23142080 | Unspecified |
| mmu-let-7i-5p | MIMAT0000122 | Pet100 | 100503890 | HITS-CLIP | 23142080 | Unspecified |
| mmu-let-7i-5p | MIMAT0000122 | Pex11a | 18631 | HITS-CLIP | 23597149 | Unspecified |
| mmu-let-7i-5p | MIMAT0000122 | Pex11b | 18632 | HITS-CLIP | 23597149 | Unspecified |
| mmu-let-7i-5p | MIMAT0000122 | Pex12 | 103737 | HITS-CLIP | 23597149\|23142080 | Unspecified |
| mmu-let-7i-5p | MIMAT0000122 | Pex13 | 72129 | HITS-CLIP | 23597149\|23142080 | Unspecified |
| mmu-let-7i-5p | MIMAT0000122 | Pex14 | 56273 | HITS-CLIP | 23597149\|23142080 | Unspecified |
| mmu-let-7i-5p | MIMAT0000122 | Pex16 | 18633 | HITS-CLIP | 23597149 | Unspecified |
| mmu-let-7i-5p | MIMAT0000122 | Pex26 | 74043 | HITS-CLIP | 23597149 | Unspecified |
| mmu-let-7i-5p | MIMAT0000122 | Pex3 | 56535 | HITS-CLIP | 23142080 | Unspecified |
| mmu-let-7i-5p | MIMAT0000122 | Pex5 | 19305 | HITS-CLIP | 37986224\|23597149\|23142080 | Unspecified |
| mmu-let-7i-5p | MIMAT0000122 | Pex5l | 58869 | HITS-CLIP | 19536157 | Unspecified |
| mmu-let-7i-5p | MIMAT0000122 | Pex6 | 224824 | HITS-CLIP | 23597149 | Unspecified |
| mmu-let-7i-5p | MIMAT0000122 | Pfas | 237823 | HITS-CLIP | 29167373 | Unspecified |
| mmu-let-7i-5p | MIMAT0000122 | Pfdn1 | 67199 | HITS-CLIP | 23142080 | Unspecified |
| mmu-let-7i-5p | MIMAT0000122 | Pfdn2 | 18637 | HITS-CLIP | 23142080 | Unspecified |
| mmu-let-7i-5p | MIMAT0000122 | Pfkfb1 | 18639 | HITS-CLIP | 23597149\|23142080 | Unspecified |
| mmu-let-7i-5p | MIMAT0000122 | Pfkfb2 | 18640 | HITS-CLIP | 37986224\|23597149\|19536157 | Unspecified |
| mmu-let-7i-5p | MIMAT0000122 | Pfkfb3 | 170768 | HITS-CLIP | 37986224\|23142080\|23597149\|29167 | Unspecified |
| mmu-let-7i-5p | MIMAT0000122 | Pfkfb4 | 270198 | HITS-CLIP | 23142080 | Unspecified |
| mmu-let-7i-5p | MIMAT0000122 | Pfkl | 18641 | HITS-CLIP | 37986224\|23142080 | Unspecified |
| mmu-let-7i-5p | MIMAT0000122 | Pfkm | 18642 | HITS-CLIP | 23142080 | Unspecified |
| mmu-let-7i-5p | MIMAT0000122 | Pfkp | 56421 | HITS-CLIP | 37986224\|23142080 | Unspecified |
| mmu-let-7i-5p | MIMAT0000122 | Pfn1 | 18643 | HITS-CLIP | 31433990\|23142080\|26203562 | Unspecified |
| mmu-let-7i-5p | MIMAT0000122 | Pgap2 | 233575 | HITS-CLIP | 37986224\|23597149\|23142080 | Unspecified |
| mmu-let-7i-5p | MIMAT0000122 | Pgap3 | 320655 | HITS-CLIP | 23142080 | Unspecified |
| mmu-let-7i-5p | MIMAT0000122 | Pggt1b | 225467 | HITS-CLIP | 23142080\|23597149 | Unspecified |
| mmu-let-7i-5p | MIMAT0000122 | Pgk1 | 18655 | HITS-CLIP | 23142080 | Unspecified |

| | | | | | | |
|---|---|---|---|---|---|---|
| mmu-let-7i-5p | MIMAT0000122 | Pglyrp2 | 57757 | HITS-CLIP | 23597149 | Unspecified |
| mmu-let-7i-5p | MIMAT0000122 | Pgm1 | 66681 | HITS-CLIP | 37986224 | Unspecified |
| mmu-let-7i-5p | MIMAT0000122 | Pgm2 | 72157 | HITS-CLIP | 19536157\|23142080 | Unspecified |
| mmu-let-7i-5p | MIMAT0000122 | Pgm2l1 | 70974 | HITS-CLIP | 23142080\|29167373 | Unspecified |
| mmu-let-7i-5p | MIMAT0000122 | Pgm3 | 109785 | HITS-CLIP | 23597149\|23142080 | Unspecified |
| mmu-let-7i-5p | MIMAT0000122 | Pgpep1 | 66522 | HITS-CLIP | 37986224\|23142080 | Unspecified |
| mmu-let-7i-5p | MIMAT0000122 | Pgrmc1 | 53328 | HITS-CLIP | 37986224\|19536157\|23597149\|23142 | Unspecified |
| mmu-let-7i-5p | MIMAT0000122 | Pgs1 | 74451 | HITS-CLIP | 23597149 | Unspecified |
| mmu-let-7i-5p | MIMAT0000122 | Phactr1 | 218194 | HITS-CLIP | 19536157 | Unspecified |
| mmu-let-7i-5p | MIMAT0000122 | Phactr2 | 215789 | HITS-CLIP | 37986224\|23142080 | Unspecified |
| mmu-let-7i-5p | MIMAT0000122 | Phactr4 | 100169 | HITS-CLIP | 37986224\|23597149\|23142080 | Unspecified |
| mmu-let-7i-5p | MIMAT0000122 | Phb2 | 12034 | HITS-CLIP | 37986224\|23142080 | Unspecified |
| mmu-let-7i-5p | MIMAT0000122 | Phc2 | 54383 | HITS-CLIP | 37986224\|23597149\|23142080 | Unspecified |
| mmu-let-7i-5p | MIMAT0000122 | Phc3 | 241915 | HITS-CLIP | 23142080\|29167373 | Unspecified |
| mmu-let-7i-5p | MIMAT0000122 | Phf1 | 21652 | HITS-CLIP | 23142080 | Unspecified |
| mmu-let-7i-5p | MIMAT0000122 | Phf10 | 72057 | HITS-CLIP | 23142080 | Unspecified |
| mmu-let-7i-5p | MIMAT0000122 | Phf12 | 268448 | HITS-CLIP | 29167373 | Unspecified |
| mmu-let-7i-5p | MIMAT0000122 | Phf13 | 230936 | HITS-CLIP | 23142080 | Unspecified |
| mmu-let-7i-5p | MIMAT0000122 | Phf14 | 75725 | HITS-CLIP | 23597149 | Unspecified |
| mmu-let-7i-5p | MIMAT0000122 | Phf19 | 74016 | HITS-CLIP | 23142080 | Unspecified |
| mmu-let-7i-5p | MIMAT0000122 | Phf2 | 18676 | HITS-CLIP | 23597149 | Unspecified |
| mmu-let-7i-5p | MIMAT0000122 | Phf21a | 192285 | HITS-CLIP | 37986224\|23142080\|29167373 | Unspecified |
| mmu-let-7i-5p | MIMAT0000122 | Phf3 | 213109 | HITS-CLIP | 37986224\|23142080\|23597149\|19536 | Unspecified |
| mmu-let-7i-5p | MIMAT0000122 | Phf5a | 68479 | HITS-CLIP | 23142080 | Unspecified |
| mmu-let-7i-5p | MIMAT0000122 | Phf6 | 70998 | HITS-CLIP | 23142080 | Unspecified |
| mmu-let-7i-5p | MIMAT0000122 | Phf8 | 320595 | HITS-CLIP | 23142080\|23597149\|29167373 | Unspecified |
| mmu-let-7i-5p | MIMAT0000122 | Phip | 83946 | HITS-CLIP | 23597149\|19536157\|23142080\|29167 | Unspecified |
| mmu-let-7i-5p | MIMAT0000122 | Phka1 | 18679 | HITS-CLIP | 23597149 | Unspecified |
| mmu-let-7i-5p | MIMAT0000122 | Phkg2 | 68961 | HITS-CLIP | 23597149 | Unspecified |
| mmu-let-7i-5p | MIMAT0000122 | Phldb2 | 208177 | HITS-CLIP | 37986224\|23597149\|19536157\|26203 | Unspecified |
| mmu-let-7i-5p | MIMAT0000122 | Phlpp1 | 98432 | HITS-CLIP | 37986224\|19536157 | Unspecified |
| mmu-let-7i-5p | MIMAT0000122 | Phlpp2 | 244650 | HITS-CLIP | 37986224\|23142080\|19536157 | Unspecified |
| mmu-let-7i-5p | MIMAT0000122 | Phospho2 | 73373 | HITS-CLIP | 37986224 | Unspecified |
| mmu-let-7i-5p | MIMAT0000122 | Phrf1 | 101471 | HITS-CLIP | 23597149\|19536157\|23142080 | Unspecified |
| mmu-let-7i-5p | MIMAT0000122 | Phtf1 | 18685 | HITS-CLIP | 23142080 | Unspecified |
| mmu-let-7i-5p | MIMAT0000122 | Phtf2 | 68770 | HITS-CLIP | 37986224\|19536157\|23597149\|23142 | Unspecified |
| mmu-let-7i-5p | MIMAT0000122 | Phyh | 16922 | HITS-CLIP | 23597149 | Unspecified |
| mmu-let-7i-5p | MIMAT0000122 | Phyhd1 | 227696 | HITS-CLIP | 37986224\|23597149 | Unspecified |
| mmu-let-7i-5p | MIMAT0000122 | Phyhip | 105653 | HITS-CLIP | 37986224 | Unspecified |

| | | | | | | |
|---|---|---|---|---|---|---|
| mmu-let-7i-5p | MIMAT0000122 | Phyhipl | 70911 | HITS-CLIP | 37986224 | Unspecified |
| mmu-let-7i-5p | MIMAT0000122 | Pi4k2a | 84095 | HITS-CLIP | 23597149 | Unspecified |
| mmu-let-7i-5p | MIMAT0000122 | Pi4ka | 224020 | HITS-CLIP | 23142080\|29167373 | Unspecified |
| mmu-let-7i-5p | MIMAT0000122 | Pi4kb | 107650 | HITS-CLIP | 19536157\|29167373 | Unspecified |
| mmu-let-7i-5p | MIMAT0000122 | Pianp | 319352 | HITS-CLIP | 19536157 | Unspecified |
| mmu-let-7i-5p | MIMAT0000122 | Pias1 | 56469 | HITS-CLIP | 37986224 | Unspecified |
| mmu-let-7i-5p | MIMAT0000122 | Pias2 | 17344 | HITS-CLIP | 23142080 | Unspecified |
| mmu-let-7i-5p | MIMAT0000122 | Pias3 | 229615 | HITS-CLIP | 23597149 | Unspecified |
| mmu-let-7i-5p | MIMAT0000122 | Pias4 | 59004 | HITS-CLIP | 37986224\|23597149\|23142080 | Unspecified |
| mmu-let-7i-5p | MIMAT0000122 | Pibf1 | 52023 | HITS-CLIP | 23142080 | Unspecified |
| mmu-let-7i-5p | MIMAT0000122 | Picalm | 233489 | HITS-CLIP//qCLASH | 37986224\|23142080\|29167373\|31393 | Unspecified |
| mmu-let-7i-5p | MIMAT0000122 | Pidd1 | 57913 | HITS-CLIP | 23597149 | Unspecified |
| mmu-let-7i-5p | MIMAT0000122 | Piezo1 | 234839 | HITS-CLIP | 23142080\|29167373 | Unspecified |
| mmu-let-7i-5p | MIMAT0000122 | Pif1 | 208084 | HITS-CLIP | 23142080 | Unspecified |
| mmu-let-7i-5p | MIMAT0000122 | Piga | 18700 | HITS-CLIP | 37986224\|23597149\|23142080 | Unspecified |
| mmu-let-7i-5p | MIMAT0000122 | Pigc | 67292 | HITS-CLIP | 37986224\|23142080 | Unspecified |
| mmu-let-7i-5p | MIMAT0000122 | Pigf | 18701 | HITS-CLIP | 23142080 | Unspecified |
| mmu-let-7i-5p | MIMAT0000122 | Pigm | 67556 | HITS-CLIP | 37986224\|23597149\|23142080 | Unspecified |
| mmu-let-7i-5p | MIMAT0000122 | Pign | 27392 | HITS-CLIP | 23142080 | Unspecified |
| mmu-let-7i-5p | MIMAT0000122 | Pigq | 14755 | HITS-CLIP | 23597149 | Unspecified |
| mmu-let-7i-5p | MIMAT0000122 | Pigr | 18703 | HITS-CLIP | 23597149 | Unspecified |
| mmu-let-7i-5p | MIMAT0000122 | Pigs | 276846 | HITS-CLIP | 37986224\|19536157\|23597149\|23142 | Unspecified |
| mmu-let-7i-5p | MIMAT0000122 | Pigt | 78928 | HITS-CLIP | 23597149\|23142080 | Unspecified |
| mmu-let-7i-5p | MIMAT0000122 | Pigu | 228812 | HITS-CLIP | 23597149\|23142080 | Unspecified |
| mmu-let-7i-5p | MIMAT0000122 | Pigv | 230801 | HITS-CLIP | 23142080 | Unspecified |
| mmu-let-7i-5p | MIMAT0000122 | Pigw | 70325 | HITS-CLIP | 23597149 | Unspecified |
| mmu-let-7i-5p | MIMAT0000122 | Pik3ap1 | 83490 | HITS-CLIP | 23597149\|29167373\|23142080 | Unspecified |
| mmu-let-7i-5p | MIMAT0000122 | Pik3c2a | 18704 | HITS-CLIP | 23142080\|29167373 | Unspecified |
| mmu-let-7i-5p | MIMAT0000122 | Pik3c2b | 240752 | HITS-CLIP | 23597149\|23142080 | Unspecified |
| mmu-let-7i-5p | MIMAT0000122 | Pik3c3 | 225326 | HITS-CLIP | 23142080 | Unspecified |
| mmu-let-7i-5p | MIMAT0000122 | Pik3ca | 18706 | HITS-CLIP | 23142080\|29167373 | Unspecified |
| mmu-let-7i-5p | MIMAT0000122 | Pik3cb | 74769 | HITS-CLIP | 29167373\|23142080 | Unspecified |
| mmu-let-7i-5p | MIMAT0000122 | Pik3cd | 18707 | HITS-CLIP | 23142080 | Unspecified |
| mmu-let-7i-5p | MIMAT0000122 | Pik3cg | 30955 | HITS-CLIP | 29167373 | Unspecified |
| mmu-let-7i-5p | MIMAT0000122 | Pik3ip1 | 216505 | HITS-CLIP | 37986224\|19536157\|23142080 | Unspecified |
| mmu-let-7i-5p | MIMAT0000122 | Pik3r1 | 18708 | HITS-CLIP | 37986224\|23597149\|19536157\|23142 | Unspecified |
| mmu-let-7i-5p | MIMAT0000122 | Pik3r5 | 320207 | HITS-CLIP | 23597149\|23142080\|29167373 | Unspecified |
| mmu-let-7i-5p | MIMAT0000122 | Pikfyve | 18711 | HITS-CLIP | 23142080\|29167373 | Unspecified |
| mmu-let-7i-5p | MIMAT0000122 | Pim1 | 18712 | HITS-CLIP//qCLASH | 23142080\|31393953 | Unspecified |

| | | | | | | |
|---|---|---|---|---|---|---|
| mmu-let-7i-5p | MIMAT0000122 | Pim3 | 223775 | HITS-CLIP | 23142080 | Unspecified |
| mmu-let-7i-5p | MIMAT0000122 | Pink1 | 68943 | HITS-CLIP | 23597149\|23142080 | Unspecified |
| mmu-let-7i-5p | MIMAT0000122 | Pip4k2b | 108083 | HITS-CLIP | 23597149\|23142080\|29167373 | Unspecified |
| mmu-let-7i-5p | MIMAT0000122 | Pip4k2c | 117150 | HITS-CLIP | 37986224 | Unspecified |
| mmu-let-7i-5p | MIMAT0000122 | Pip4p1 | 90809 | HITS-CLIP | 23142080\|23597149 | Unspecified |
| mmu-let-7i-5p | MIMAT0000122 | Pip4p2 | 55529 | HITS-CLIP | 23142080\|29167373 | Unspecified |
| mmu-let-7i-5p | MIMAT0000122 | Pip5k1a | 18720 | HITS-CLIP | 23142080 | Unspecified |
| mmu-let-7i-5p | MIMAT0000122 | Pip5k1c | 18717 | HITS-CLIP | 37986224\|23597149\|19536157\|23142 | Unspecified |
| mmu-let-7i-5p | MIMAT0000122 | Pirb | 18733 | HITS-CLIP | 29167373 | Unspecified |
| mmu-let-7i-5p | MIMAT0000122 | Pitpna | 18738 | HITS-CLIP | 29167373 | Unspecified |
| mmu-let-7i-5p | MIMAT0000122 | Pitpnb | 56305 | HITS-CLIP | 23142080 | Unspecified |
| mmu-let-7i-5p | MIMAT0000122 | Pitpnm2 | 19679 | HITS-CLIP | 23597149 | Unspecified |
| mmu-let-7i-5p | MIMAT0000122 | Pitrm1 | 69617 | HITS-CLIP | 23142080\|23597149 | Unspecified |
| mmu-let-7i-5p | MIMAT0000122 | Piwil2 | 57746 | HITS-CLIP | 23142080 | Unspecified |
| mmu-let-7i-5p | MIMAT0000122 | Pja2 | 224938 | HITS-CLIP | 23142080 | Unspecified |
| mmu-let-7i-5p | MIMAT0000122 | Pkd1 | 18763 | HITS-CLIP | 23142080\|29167373 | Unspecified |
| mmu-let-7i-5p | MIMAT0000122 | Pkib | 18768 | HITS-CLIP | 23142080 | Unspecified |
| mmu-let-7i-5p | MIMAT0000122 | Pkm | 18746 | HITS-CLIP | 23142080\|29167373 | Unspecified |
| mmu-let-7i-5p | MIMAT0000122 | Pkn1 | 320795 | HITS-CLIP | 29167373 | Unspecified |
| mmu-let-7i-5p | MIMAT0000122 | Pkn2 | 109333 | HITS-CLIP//qCLASH | 37986224\|31393953\|23597149\|1953€ | Unspecified |
| mmu-let-7i-5p | MIMAT0000122 | Pkp4 | 227937 | HITS-CLIP | 23597149\|23142080 | Unspecified |
| mmu-let-7i-5p | MIMAT0000122 | Pla2g12a | 66350 | HITS-CLIP | 23142080 | Unspecified |
| mmu-let-7i-5p | MIMAT0000122 | Pla2g12b | 69836 | HITS-CLIP | 23597149 | Unspecified |
| mmu-let-7i-5p | MIMAT0000122 | Pla2g15 | 192654 | HITS-CLIP | 37986224\|23142080 | Unspecified |
| mmu-let-7i-5p | MIMAT0000122 | Pla2g16 | 225845 | HITS-CLIP | 23142080 | Unspecified |
| mmu-let-7i-5p | MIMAT0000122 | Pla2g4a | 18783 | HITS-CLIP | 29167373 | Unspecified |
| mmu-let-7i-5p | MIMAT0000122 | Pla2g6 | 53357 | HITS-CLIP | 23142080 | Unspecified |
| mmu-let-7i-5p | MIMAT0000122 | Pla2r1 | 18779 | HITS-CLIP | 23597149 | Unspecified |
| mmu-let-7i-5p | MIMAT0000122 | Plaa | 18786 | HITS-CLIP | 29167373 | Unspecified |
| mmu-let-7i-5p | MIMAT0000122 | Plag1 | 56711 | HITS-CLIP | 23142080\|30971389 | Unspecified |
| mmu-let-7i-5p | MIMAT0000122 | Plagl1 | 22634 | HITS-CLIP | 23142080 | Unspecified |
| mmu-let-7i-5p | MIMAT0000122 | Plagl2 | 54711 | HITS-CLIP | 37986224\|19536157\|23597149\|3143: | Unspecified |
| mmu-let-7i-5p | MIMAT0000122 | Plat | 18791 | HITS-CLIP | 37986224 | Unspecified |
| mmu-let-7i-5p | MIMAT0000122 | Plaur | 18793 | HITS-CLIP | 29167373 | Unspecified |
| mmu-let-7i-5p | MIMAT0000122 | Plbd1 | 66857 | HITS-CLIP | 23597149 | Unspecified |
| mmu-let-7i-5p | MIMAT0000122 | Plbd2 | 71772 | HITS-CLIP | 37986224\|23597149\|19536157\|29167 | Unspecified |
| mmu-let-7i-5p | MIMAT0000122 | Plcb1 | 18795 | HITS-CLIP | 23597149\|19536157 | Unspecified |
| mmu-let-7i-5p | MIMAT0000122 | Plcb4 | 18798 | HITS-CLIP | 19536157\|23142080\|29167373 | Unspecified |
| mmu-let-7i-5p | MIMAT0000122 | Plcd1 | 18799 | HITS-CLIP | 23142080 | Unspecified |

| | | | | | | |
|---|---|---|---|---|---|---|
| mmu-let-7i-5p | MIMAT0000122 | Plce1 | 74055 | HITS-CLIP | 19536157\|23597149 | Unspecified |
| mmu-let-7i-5p | MIMAT0000122 | Plcg1 | 18803 | HITS-CLIP | 37986224\|23597149 | Unspecified |
| mmu-let-7i-5p | MIMAT0000122 | Plcg2 | 234779 | HITS-CLIP | 29167373 | Unspecified |
| mmu-let-7i-5p | MIMAT0000122 | Plch2 | 269615 | HITS-CLIP | 19536157 | Unspecified |
| mmu-let-7i-5p | MIMAT0000122 | Plcl2 | 224860 | HITS-CLIP | 37986224\|29167373 | Unspecified |
| mmu-let-7i-5p | MIMAT0000122 | Plcxd1 | 403178 | HITS-CLIP | 23142080 | Unspecified |
| mmu-let-7i-5p | MIMAT0000122 | Plcxd2 | 433022 | HITS-CLIP | 37986224\|23142080\|23597149 | Unspecified |
| mmu-let-7i-5p | MIMAT0000122 | Pld2 | 18806 | HITS-CLIP | 23597149 | Unspecified |
| mmu-let-7i-5p | MIMAT0000122 | Pld3 | 18807 | HITS-CLIP | 23597149\|23142080\|29167373 | Unspecified |
| mmu-let-7i-5p | MIMAT0000122 | Plec | 18810 | HITS-CLIP | 37986224\|23597149\|23142080\|29167 | Unspecified |
| mmu-let-7i-5p | MIMAT0000122 | Plek | 56193 | HITS-CLIP | 23142080\|29167373 | Unspecified |
| mmu-let-7i-5p | MIMAT0000122 | Plekha1 | 101476 | HITS-CLIP | 23142080 | Unspecified |
| mmu-let-7i-5p | MIMAT0000122 | Plekha2 | 83436 | HITS-CLIP | 37986224\|23597149\|23142080 | Unspecified |
| mmu-let-7i-5p | MIMAT0000122 | Plekha5 | 109135 | HITS-CLIP | 23142080 | Unspecified |
| mmu-let-7i-5p | MIMAT0000122 | Plekha6 | 240753 | HITS-CLIP | 23597149 | Unspecified |
| mmu-let-7i-5p | MIMAT0000122 | Plekha8 | 231999 | HITS-CLIP | 37986224\|23597149 | Unspecified |
| mmu-let-7i-5p | MIMAT0000122 | Plekhb2 | 226971 | HITS-CLIP | 37986224 | Unspecified |
| mmu-let-7i-5p | MIMAT0000122 | Plekhg3 | 263406 | HITS-CLIP | 23597149 | Unspecified |
| mmu-let-7i-5p | MIMAT0000122 | Plekhm1 | 353047 | HITS-CLIP | 23597149\|29167373\|23142080 | Unspecified |
| mmu-let-7i-5p | MIMAT0000122 | Plekhm3 | 241075 | HITS-CLIP | 29167373 | Unspecified |
| mmu-let-7i-5p | MIMAT0000122 | Plekho1 | 67220 | HITS-CLIP | 37986224\|23597149\|23142080\|29167 | Unspecified |
| mmu-let-7i-5p | MIMAT0000122 | Plekho2 | 102595 | HITS-CLIP | 23142080 | Unspecified |
| mmu-let-7i-5p | MIMAT0000122 | Plin2 | 11520 | HITS-CLIP | 23142080 | Unspecified |
| mmu-let-7i-5p | MIMAT0000122 | Plin3 | 66905 | HITS-CLIP | 23597149\|23142080 | Unspecified |
| mmu-let-7i-5p | MIMAT0000122 | Plk1 | 18817 | HITS-CLIP | 23142080 | Unspecified |
| mmu-let-7i-5p | MIMAT0000122 | Plk2 | 20620 | HITS-CLIP | 37986224\|23597149\|23142080\|29167 | Unspecified |
| mmu-let-7i-5p | MIMAT0000122 | Plk3 | 12795 | HITS-CLIP | 23597149\|23142080 | Unspecified |
| mmu-let-7i-5p | MIMAT0000122 | Plk4 | 20873 | HITS-CLIP | 23142080 | Unspecified |
| mmu-let-7i-5p | MIMAT0000122 | Pllp | 67801 | HITS-CLIP | 19536157 | Unspecified |
| mmu-let-7i-5p | MIMAT0000122 | Plod1 | 18822 | HITS-CLIP | 23597149 | Unspecified |
| mmu-let-7i-5p | MIMAT0000122 | Plod2 | 26432 | HITS-CLIP | 23142080\|19536157 | Unspecified |
| mmu-let-7i-5p | MIMAT0000122 | Plp1 | 18823 | HITS-CLIP | 37986224\|19536157 | Unspecified |
| mmu-let-7i-5p | MIMAT0000122 | Plpp2 | 8612 | HITS-CLIP | 23142080 | Unspecified |
| mmu-let-7i-5p | MIMAT0000122 | Plpp3 | 67916 | HITS-CLIP | 23597149\|19536157 | Unspecified |
| mmu-let-7i-5p | MIMAT0000122 | Plpp5 | 84513 | HITS-CLIP | 23142080\|23597149\|29167373 | Unspecified |
| mmu-let-7i-5p | MIMAT0000122 | Plppr4 | 229791 | HITS-CLIP | 19536157 | Unspecified |
| mmu-let-7i-5p | MIMAT0000122 | Plrg1 | 53317 | HITS-CLIP | 23142080 | Unspecified |
| mmu-let-7i-5p | MIMAT0000122 | Pls3 | 102866 | HITS-CLIP | 23597149\|19536157\|29167373 | Unspecified |
| mmu-let-7i-5p | MIMAT0000122 | Plvap | 84094 | HITS-CLIP | 23597149 | Unspecified |

| | | | | | | |
|---|---|---|---|---|---|---|
| mmu-let-7i-5p | MIMAT0000122 | Plxna1 | 18844 | HITS-CLIP | 23142080\|23597149\|29167373 | Unspecified |
| mmu-let-7i-5p | MIMAT0000122 | Plxna2 | 18845 | HITS-CLIP | 37986224\|23597149\|19536157\|29167 | Unspecified |
| mmu-let-7i-5p | MIMAT0000122 | Plxna4 | 243743 | HITS-CLIP | 19536157 | Unspecified |
| mmu-let-7i-5p | MIMAT0000122 | Plxnb2 | 140570 | HITS-CLIP | 37986224\|23597149\|29167373 | Unspecified |
| mmu-let-7i-5p | MIMAT0000122 | Plxnc1 | 54712 | HITS-CLIP | 37986224\|19536157\|23142080 | Unspecified |
| mmu-let-7i-5p | MIMAT0000122 | Plxnd1 | 67784 | HITS-CLIP | 37986224\|23597149\|23142080 | Unspecified |
| mmu-let-7i-5p | MIMAT0000122 | Pmaip1 | 58801 | HITS-CLIP | 37986224\|23142080 | Unspecified |
| mmu-let-7i-5p | MIMAT0000122 | Pmepa1 | 65112 | HITS-CLIP | 23630078\|23142080 | Unspecified |
| mmu-let-7i-5p | MIMAT0000122 | Pml | 18854 | HITS-CLIP | 23142080 | Unspecified |
| mmu-let-7i-5p | MIMAT0000122 | Pmm2 | 54128 | HITS-CLIP | 23597149 | Unspecified |
| mmu-let-7i-5p | MIMAT0000122 | Pmp22 | 18858 | HITS-CLIP | 29167373 | Unspecified |
| mmu-let-7i-5p | MIMAT0000122 | Pmpca | 66865 | HITS-CLIP | 37986224\|23597149\|23142080\|29167 | Unspecified |
| mmu-let-7i-5p | MIMAT0000122 | Pms1 | 227099 | HITS-CLIP | 23142080 | Unspecified |
| mmu-let-7i-5p | MIMAT0000122 | Pms2 | 18861 | HITS-CLIP | 37986224\|23142080\|19536157 | Unspecified |
| mmu-let-7i-5p | MIMAT0000122 | Pnisr | 66625 | HITS-CLIP | 23142080 | Unspecified |
| mmu-let-7i-5p | MIMAT0000122 | Pnkd | 56695 | HITS-CLIP | 37986224\|23597149 | Unspecified |
| mmu-let-7i-5p | MIMAT0000122 | Pnn | 18949 | HITS-CLIP | 37986224\|23597149\|23142080\|19536 | Unspecified |
| mmu-let-7i-5p | MIMAT0000122 | Pnpla6 | 50767 | HITS-CLIP | 23597149\|23142080 | Unspecified |
| mmu-let-7i-5p | MIMAT0000122 | Pnpla7 | 241274 | HITS-CLIP | 29167373 | Unspecified |
| mmu-let-7i-5p | MIMAT0000122 | Pnpo | 103711 | HITS-CLIP | 37986224\|23597149\|23142080 | Unspecified |
| mmu-let-7i-5p | MIMAT0000122 | Pnpt1 | 71701 | HITS-CLIP | 23142080 | Unspecified |
| mmu-let-7i-5p | MIMAT0000122 | Pnrc2 | 52830 | HITS-CLIP | 31433990 | Unspecified |
| mmu-let-7i-5p | MIMAT0000122 | Poc1a | 70235 | HITS-CLIP | 23597149 | Unspecified |
| mmu-let-7i-5p | MIMAT0000122 | Pofut1 | 140484 | HITS-CLIP | 37986224\|23142080 | Unspecified |
| mmu-let-7i-5p | MIMAT0000122 | Pogk | 71592 | HITS-CLIP | 23142080 | Unspecified |
| mmu-let-7i-5p | MIMAT0000122 | Poglut1 | 224143 | HITS-CLIP | 23597149\|23142080 | Unspecified |
| mmu-let-7i-5p | MIMAT0000122 | Pogz | 229584 | HITS-CLIP | 37986224\|23597149\|23142080 | Unspecified |
| mmu-let-7i-5p | MIMAT0000122 | Polb | 18970 | HITS-CLIP | 37986224\|19536157\|29167373 | Unspecified |
| mmu-let-7i-5p | MIMAT0000122 | Pold2 | 18972 | HITS-CLIP | 23597149 | Unspecified |
| mmu-let-7i-5p | MIMAT0000122 | Pold3 | 67967 | HITS-CLIP | 37986224\|23597149\|23142080 | Unspecified |
| mmu-let-7i-5p | MIMAT0000122 | Poldip2 | 67811 | HITS-CLIP | 23597149\|23142080 | Unspecified |
| mmu-let-7i-5p | MIMAT0000122 | Poldip3 | 73826 | HITS-CLIP | 23142080 | Unspecified |
| mmu-let-7i-5p | MIMAT0000122 | Pole | 18973 | HITS-CLIP//qCLASH | 23597149\|31393953\|23142080 | Unspecified |
| mmu-let-7i-5p | MIMAT0000122 | Pole2 | 18974 | HITS-CLIP | 23142080 | Unspecified |
| mmu-let-7i-5p | MIMAT0000122 | Poli | 11201 | HITS-CLIP | 23142080 | Unspecified |
| mmu-let-7i-5p | MIMAT0000122 | Polq | 77782 | HITS-CLIP | 23142080 | Unspecified |
| mmu-let-7i-5p | MIMAT0000122 | Polr1a | 20019 | HITS-CLIP | 37986224\|23597149\|29167373 | Unspecified |
| mmu-let-7i-5p | MIMAT0000122 | Polr1b | 20017 | HITS-CLIP | 29167373 | Unspecified |
| mmu-let-7i-5p | MIMAT0000122 | Polr1d | 20018 | HITS-CLIP | 37986224 | Unspecified |

| | | | | | | |
|---|---|---|---|---|---|---|
| mmu-let-7i-5p | MIMAT0000122 | Polr2a | 20020 | HITS-CLIP | 23597149\|23142080\|29167373 | Unspecified |
| mmu-let-7i-5p | MIMAT0000122 | Polr2b | 231329 | HITS-CLIP | 23142080 | Unspecified |
| mmu-let-7i-5p | MIMAT0000122 | Polr2f | 69833 | HITS-CLIP | 23597149 | Unspecified |
| mmu-let-7i-5p | MIMAT0000122 | Polr2i | 69920 | HITS-CLIP | 23142080 | Unspecified |
| mmu-let-7i-5p | MIMAT0000122 | Polr3a | 218832 | HITS-CLIP | 23142080 | Unspecified |
| mmu-let-7i-5p | MIMAT0000122 | Polr3b | 70428 | HITS-CLIP | 23597149 | Unspecified |
| mmu-let-7i-5p | MIMAT0000122 | Polr3d | 67065 | HITS-CLIP | 23597149 | Unspecified |
| mmu-let-7i-5p | MIMAT0000122 | Polr3e | 26939 | HITS-CLIP | 29167373 | Unspecified |
| mmu-let-7i-5p | MIMAT0000122 | Polr3g | 67486 | HITS-CLIP | 37986224\|23142080 | Unspecified |
| mmu-let-7i-5p | MIMAT0000122 | Polrmt | 216151 | HITS-CLIP | 37986224\|23597149 | Unspecified |
| mmu-let-7i-5p | MIMAT0000122 | Pom121 | 107939 | HITS-CLIP | 37986224\|23597149\|19536157\|2363( | Unspecified |
| mmu-let-7i-5p | MIMAT0000122 | Pomk | 74653 | HITS-CLIP | 23142080 | Unspecified |
| mmu-let-7i-5p | MIMAT0000122 | Pomt2 | 217734 | HITS-CLIP | 37986224 | Unspecified |
| mmu-let-7i-5p | MIMAT0000122 | Pon1 | 18979 | HITS-CLIP | 23597149 | Unspecified |
| mmu-let-7i-5p | MIMAT0000122 | Pon3 | 269823 | HITS-CLIP | 23597149 | Unspecified |
| mmu-let-7i-5p | MIMAT0000122 | Pop1 | 67724 | HITS-CLIP | 23597149 | Unspecified |
| mmu-let-7i-5p | MIMAT0000122 | Pop4 | 66161 | HITS-CLIP | 23142080 | Unspecified |
| mmu-let-7i-5p | MIMAT0000122 | Pop5 | 117109 | HITS-CLIP | 37986224 | Unspecified |
| mmu-let-7i-5p | MIMAT0000122 | Por | 18984 | HITS-CLIP | 23142080 | Unspecified |
| mmu-let-7i-5p | MIMAT0000122 | Postn | 50706 | HITS-CLIP | 23597149 | Unspecified |
| mmu-let-7i-5p | MIMAT0000122 | Pot1b | 72836 | HITS-CLIP | 23142080 | Unspecified |
| mmu-let-7i-5p | MIMAT0000122 | Pou2f1 | 18986 | HITS-CLIP//qCLASH | 19536157\|31433990\|31393953\|2314: | Unspecified |
| mmu-let-7i-5p | MIMAT0000122 | Pou2f2 | 18987 | HITS-CLIP | 37986224\|23142080\|29167373 | Unspecified |
| mmu-let-7i-5p | MIMAT0000122 | Pou3f3 | 18993 | HITS-CLIP | 19536157 | Unspecified |
| mmu-let-7i-5p | MIMAT0000122 | Ppan | 235036 | HITS-CLIP | 23597149\|23142080 | Unspecified |
| mmu-let-7i-5p | MIMAT0000122 | Ppara | 19013 | HITS-CLIP | 23597149 | Unspecified |
| mmu-let-7i-5p | MIMAT0000122 | Ppard | 19015 | HITS-CLIP | 23142080 | Unspecified |
| mmu-let-7i-5p | MIMAT0000122 | Ppargc1a | 19017 | HITS-CLIP | 23597149 | Unspecified |
| mmu-let-7i-5p | MIMAT0000122 | Ppargc1b | 170826 | HITS-CLIP | 23597149\|23142080 | Unspecified |
| mmu-let-7i-5p | MIMAT0000122 | Ppat | 231327 | HITS-CLIP | 23142080 | Unspecified |
| mmu-let-7i-5p | MIMAT0000122 | Ppfia1 | 233977 | HITS-CLIP | 37986224\|23142080\|19536157\|2359; | Unspecified |
| mmu-let-7i-5p | MIMAT0000122 | Ppfibp2 | 19024 | HITS-CLIP | 23597149 | Unspecified |
| mmu-let-7i-5p | MIMAT0000122 | Ppig | 228005 | HITS-CLIP | 23142080 | Unspecified |
| mmu-let-7i-5p | MIMAT0000122 | Ppil1 | 68816 | qCLASH | 31393953 | Unspecified |
| mmu-let-7i-5p | MIMAT0000122 | Ppil4 | 67418 | HITS-CLIP | 23142080 | Unspecified |
| mmu-let-7i-5p | MIMAT0000122 | Ppip5k2 | 227399 | HITS-CLIP | 37986224\|23597149\|23142080 | Unspecified |
| mmu-let-7i-5p | MIMAT0000122 | Ppm1d | 53892 | HITS-CLIP | 23142080 | Unspecified |
| mmu-let-7i-5p | MIMAT0000122 | Ppm1e | 320472 | HITS-CLIP | 37986224 | Unspecified |
| mmu-let-7i-5p | MIMAT0000122 | Ppm1h | 319468 | HITS-CLIP | 29167373 | Unspecified |

| | | | | | | |
|---|---|---|---|---|---|---|
| mmu-let-7i-5p | MIMAT0000122 | Ppm1k | 243382 | HITS-CLIP | 23597149\|23142080 | Unspecified |
| mmu-let-7i-5p | MIMAT0000122 | Ppme1 | 72590 | HITS-CLIP | 19536157\|23142080 | Unspecified |
| mmu-let-7i-5p | MIMAT0000122 | Ppp1ca | 19045 | HITS-CLIP | 23597149\|31433990\|23142080 | Unspecified |
| mmu-let-7i-5p | MIMAT0000122 | Ppp1cb | 19046 | HITS-CLIP | 31433990\|23142080\|29167373 | Unspecified |
| mmu-let-7i-5p | MIMAT0000122 | Ppp1r10 | 52040 | HITS-CLIP | 29167373 | Unspecified |
| mmu-let-7i-5p | MIMAT0000122 | Ppp1r12a | 17931 | HITS-CLIP//qCLASH | 37986224\|23142080\|23597149\|29167 | Unspecified |
| mmu-let-7i-5p | MIMAT0000122 | Ppp1r13b | 21981 | HITS-CLIP | 37986224\|23142080 | Unspecified |
| mmu-let-7i-5p | MIMAT0000122 | Ppp1r15a | 17872 | HITS-CLIP | 23142080 | Unspecified |
| mmu-let-7i-5p | MIMAT0000122 | Ppp1r15b | 108954 | HITS-CLIP//qCLASH | 37986224\|31393953\|23597149\|26203 | Unspecified |
| mmu-let-7i-5p | MIMAT0000122 | Ppp1r16b | 228852 | HITS-CLIP | 23142080 | Unspecified |
| mmu-let-7i-5p | MIMAT0000122 | Ppp1r18 | 76448 | HITS-CLIP | 23142080 | Unspecified |
| mmu-let-7i-5p | MIMAT0000122 | Ppp1r2 | 66849 | HITS-CLIP | 19536157\|23142080 | Unspecified |
| mmu-let-7i-5p | MIMAT0000122 | Ppp1r37 | 232947 | HITS-CLIP | 23142080 | Unspecified |
| mmu-let-7i-5p | MIMAT0000122 | Ppp1r3b | 244416 | HITS-CLIP | 23142080\|23597149 | Unspecified |
| mmu-let-7i-5p | MIMAT0000122 | Ppp1r3d | 228966 | HITS-CLIP | 19536157 | Unspecified |
| mmu-let-7i-5p | MIMAT0000122 | Ppp1r7 | 66385 | HITS-CLIP | 29167373 | Unspecified |
| mmu-let-7i-5p | MIMAT0000122 | Ppp1r8 | 100336 | qCLASH | 31393953 | Unspecified |
| mmu-let-7i-5p | MIMAT0000122 | Ppp1r9a | 243725 | HITS-CLIP | 37986224\|23597149 | Unspecified |
| mmu-let-7i-5p | MIMAT0000122 | Ppp2ca | 19052 | HITS-CLIP | 23597149\|23142080\|19536157 | Unspecified |
| mmu-let-7i-5p | MIMAT0000122 | Ppp2r1a | 51792 | HITS-CLIP | 23597149\|23142080 | Unspecified |
| mmu-let-7i-5p | MIMAT0000122 | Ppp2r1b | 73699 | HITS-CLIP | 37986224\|23142080 | Unspecified |
| mmu-let-7i-5p | MIMAT0000122 | Ppp2r2a | 71978 | HITS-CLIP | 23142080\|29167373 | Unspecified |
| mmu-let-7i-5p | MIMAT0000122 | Ppp2r2c | 269643 | HITS-CLIP | 19536157 | Unspecified |
| mmu-let-7i-5p | MIMAT0000122 | Ppp2r2d | 52432 | HITS-CLIP | 19536157\|23597149\|23142080 | Unspecified |
| mmu-let-7i-5p | MIMAT0000122 | Ppp2r3c | 59032 | HITS-CLIP | 23142080\|29167373 | Unspecified |
| mmu-let-7i-5p | MIMAT0000122 | Ppp2r5d | 21770 | HITS-CLIP | 23597149 | Unspecified |
| mmu-let-7i-5p | MIMAT0000122 | Ppp2r5e | 26932 | HITS-CLIP | 23597149\|23142080 | Unspecified |
| mmu-let-7i-5p | MIMAT0000122 | Ppp3ca | 19055 | HITS-CLIP | 37986224\|19536157\|23597149\|23142 | Unspecified |
| mmu-let-7i-5p | MIMAT0000122 | Ppp3cb | 19056 | HITS-CLIP | 23142080 | Unspecified |
| mmu-let-7i-5p | MIMAT0000122 | Ppp3cc | 19057 | HITS-CLIP | 23142080 | Unspecified |
| mmu-let-7i-5p | MIMAT0000122 | Ppp3r1 | 19058 | HITS-CLIP | 19536157\|23142080 | Unspecified |
| mmu-let-7i-5p | MIMAT0000122 | Ppp4c | 56420 | HITS-CLIP | 37986224\|23142080\|23597149\|19536 | Unspecified |
| mmu-let-7i-5p | MIMAT0000122 | Ppp4r3a | 55671 | HITS-CLIP | 23597149\|23142080\|29167373 | Unspecified |
| mmu-let-7i-5p | MIMAT0000122 | Ppp4r3b | 57223 | HITS-CLIP | 31433990\|30971389\|23142080 | Unspecified |
| mmu-let-7i-5p | MIMAT0000122 | Ppp6r1 | 243819 | HITS-CLIP | 37986224\|23597149\|19536157\|23142 | Unspecified |
| mmu-let-7i-5p | MIMAT0000122 | Ppp6r2 | 71474 | HITS-CLIP | 23597149 | Unspecified |
| mmu-let-7i-5p | MIMAT0000122 | Ppp6r3 | 52036 | HITS-CLIP | 23597149\|19536157\|23142080 | Unspecified |
| mmu-let-7i-5p | MIMAT0000122 | Ppt2 | 54397 | HITS-CLIP | 23597149 | Unspecified |
| mmu-let-7i-5p | MIMAT0000122 | Pptc7 | 320717 | HITS-CLIP | 37986224\|23597149\|23142080 | Unspecified |

| | | | | | | |
|---|---|---|---|---|---|---|
| mmu-let-7i-5p | MIMAT0000122 | Ppwd1 | 238831 | HITS-CLIP | 31433990 | Unspecified |
| mmu-let-7i-5p | MIMAT0000122 | Pqlc1 | 66943 | HITS-CLIP | 23597149 | Unspecified |
| mmu-let-7i-5p | MIMAT0000122 | Pqlc2 | 212555 | HITS-CLIP | 37986224\|23597149\|23142080 | Unspecified |
| mmu-let-7i-5p | MIMAT0000122 | Pradc1 | 73327 | HITS-CLIP | 23142080 | Unspecified |
| mmu-let-7i-5p | MIMAT0000122 | Prag1 | 157285 | HITS-CLIP | 29167373 | Unspecified |
| mmu-let-7i-5p | MIMAT0000122 | Prc1 | 233406 | HITS-CLIP//qCLASH | 23142080\|31393953 | Unspecified |
| mmu-let-7i-5p | MIMAT0000122 | Prcc | 94315 | HITS-CLIP | 23142080 | Unspecified |
| mmu-let-7i-5p | MIMAT0000122 | Prdm1 | 12142 | HITS-CLIP | 37986224\|23597149\|19536157\|2314 | Unspecified |
| mmu-let-7i-5p | MIMAT0000122 | Prdm10 | 382066 | HITS-CLIP | 23142080\|23597149 | Unspecified |
| mmu-let-7i-5p | MIMAT0000122 | Prdm15 | 63977 | HITS-CLIP | 23597149 | Unspecified |
| mmu-let-7i-5p | MIMAT0000122 | Prdm2 | 110593 | HITS-CLIP | 29167373 | Unspecified |
| mmu-let-7i-5p | MIMAT0000122 | Prdm4 | 72843 | HITS-CLIP | 23142080\|23597149 | Unspecified |
| mmu-let-7i-5p | MIMAT0000122 | Prdx2 | 21672 | HITS-CLIP | 23142080 | Unspecified |
| mmu-let-7i-5p | MIMAT0000122 | Prdx3 | 11757 | HITS-CLIP//qCLASH | 23597149\|31393953\|19536157 | Unspecified |
| mmu-let-7i-5p | MIMAT0000122 | Prdx5 | 54683 | HITS-CLIP | 23597149\|29167373 | Unspecified |
| mmu-let-7i-5p | MIMAT0000122 | Prdx6 | 11758 | HITS-CLIP | 23597149\|23142080 | Unspecified |
| mmu-let-7i-5p | MIMAT0000122 | Preb | 50907 | HITS-CLIP | 23597149 | Unspecified |
| mmu-let-7i-5p | MIMAT0000122 | Prelid2 | 77619 | HITS-CLIP | 23142080 | Unspecified |
| mmu-let-7i-5p | MIMAT0000122 | Prep | 19072 | HITS-CLIP | 37986224\|23597149\|23142080 | Unspecified |
| mmu-let-7i-5p | MIMAT0000122 | Prex1 | 277360 | HITS-CLIP | 29167373\|23142080 | Unspecified |
| mmu-let-7i-5p | MIMAT0000122 | Prex2 | 109294 | HITS-CLIP | 37986224 | Unspecified |
| mmu-let-7i-5p | MIMAT0000122 | Prim1 | 19075 | HITS-CLIP | 23597149\|23142080 | Unspecified |
| mmu-let-7i-5p | MIMAT0000122 | Prim2 | 19076 | HITS-CLIP | 23142080 | Unspecified |
| mmu-let-7i-5p | MIMAT0000122 | Primpol | 408022 | HITS-CLIP | 23142080 | Unspecified |
| mmu-let-7i-5p | MIMAT0000122 | Prkaa2 | 108079 | HITS-CLIP | 37986224\|23597149\|19536157 | Unspecified |
| mmu-let-7i-5p | MIMAT0000122 | Prkab1 | 19079 | HITS-CLIP | 23142080 | Unspecified |
| mmu-let-7i-5p | MIMAT0000122 | Prkacb | 18749 | HITS-CLIP | 31433990\|23597149\|23142080\|29167 | Unspecified |
| mmu-let-7i-5p | MIMAT0000122 | Prkag1 | 19082 | HITS-CLIP | 23142080 | Unspecified |
| mmu-let-7i-5p | MIMAT0000122 | Prkag2 | 108099 | HITS-CLIP | 37986224\|23597149 | Unspecified |
| mmu-let-7i-5p | MIMAT0000122 | Prkar1a | 19084 | HITS-CLIP | 23142080 | Unspecified |
| mmu-let-7i-5p | MIMAT0000122 | Prkar2a | 19087 | HITS-CLIP | 23597149\|19536157\|23142080\|29167 | Unspecified |
| mmu-let-7i-5p | MIMAT0000122 | Prkca | 18750 | HITS-CLIP | 23597149\|19536157\|23142080 | Unspecified |
| mmu-let-7i-5p | MIMAT0000122 | Prkcb | 18751 | HITS-CLIP | 19536157\|23142080 | Unspecified |
| mmu-let-7i-5p | MIMAT0000122 | Prkcd | 18753 | HITS-CLIP | 23142080 | Unspecified |
| mmu-let-7i-5p | MIMAT0000122 | Prkcq | 18761 | HITS-CLIP | 37986224\|23142080 | Unspecified |
| mmu-let-7i-5p | MIMAT0000122 | Prkcsh | 19089 | HITS-CLIP | 37986224\|23597149 | Unspecified |
| mmu-let-7i-5p | MIMAT0000122 | Prkd2 | 101540 | HITS-CLIP | 23142080 | Unspecified |
| mmu-let-7i-5p | MIMAT0000122 | Prkd3 | 75292 | HITS-CLIP | 23142080\|23597149 | Unspecified |
| mmu-let-7i-5p | MIMAT0000122 | Prkdc | 19090 | HITS-CLIP | 23142080 | Unspecified |

| | | | | | | |
|---|---|---|---|---|---|---|
| mmu-let-7i-5p | MIMAT0000122 | Prlr | 19116 | HITS-CLIP | 23597149\|23142080 | Unspecified |
| mmu-let-7i-5p | MIMAT0000122 | Prmt1 | 15469 | HITS-CLIP | 23142080\|23597149 | Unspecified |
| mmu-let-7i-5p | MIMAT0000122 | Prmt5 | 27374 | HITS-CLIP | 23597149\|23142080 | Unspecified |
| mmu-let-7i-5p | MIMAT0000122 | Prmt6 | 99890 | HITS-CLIP | 23142080\|29167373 | Unspecified |
| mmu-let-7i-5p | MIMAT0000122 | Prmt7 | 214572 | HITS-CLIP | 23142080 | Unspecified |
| mmu-let-7i-5p | MIMAT0000122 | Prnp | 19122 | HITS-CLIP | 37986224\|19536157 | Unspecified |
| mmu-let-7i-5p | MIMAT0000122 | Prob1 | 381148 | HITS-CLIP | 23597149 | Unspecified |
| mmu-let-7i-5p | MIMAT0000122 | Proc | 19123 | HITS-CLIP | 37986224\|23597149 | Unspecified |
| mmu-let-7i-5p | MIMAT0000122 | Prodh | 19125 | HITS-CLIP | 23597149 | Unspecified |
| mmu-let-7i-5p | MIMAT0000122 | Pros1 | 19128 | HITS-CLIP | 23597149\|23142080\|19536157 | Unspecified |
| mmu-let-7i-5p | MIMAT0000122 | Prox1 | 19130 | HITS-CLIP | 23597149\|19536157 | Unspecified |
| mmu-let-7i-5p | MIMAT0000122 | Proz | 66901 | HITS-CLIP | 23597149 | Unspecified |
| mmu-let-7i-5p | MIMAT0000122 | Prpf18 | 67229 | HITS-CLIP | 23597149 | Unspecified |
| mmu-let-7i-5p | MIMAT0000122 | Prpf19 | 28000 | HITS-CLIP | 23597149 | Unspecified |
| mmu-let-7i-5p | MIMAT0000122 | Prpf3 | 70767 | HITS-CLIP | 29167373 | Unspecified |
| mmu-let-7i-5p | MIMAT0000122 | Prpf31 | 68988 | HITS-CLIP | 37986224\|23597149\|23142080 | Unspecified |
| mmu-let-7i-5p | MIMAT0000122 | Prpf38b | 66921 | HITS-CLIP | 23142080 | Unspecified |
| mmu-let-7i-5p | MIMAT0000122 | Prpf39 | 328110 | HITS-CLIP | 37986224\|19536157\|23597149\|23142 | Unspecified |
| mmu-let-7i-5p | MIMAT0000122 | Prpf40a | 56194 | HITS-CLIP | 23142080\|29167373 | Unspecified |
| mmu-let-7i-5p | MIMAT0000122 | Prpf4b | 19134 | HITS-CLIP | 31433990\|23597149\|23142080\|29167 | Unspecified |
| mmu-let-7i-5p | MIMAT0000122 | Prpf8 | 192159 | HITS-CLIP | 37986224\|23597149\|23142080\|29167 | Unspecified |
| mmu-let-7i-5p | MIMAT0000122 | Prps1 | 19139 | HITS-CLIP | 23142080\|29167373 | Unspecified |
| mmu-let-7i-5p | MIMAT0000122 | Prps1l3 | 328099 | HITS-CLIP | 37986224\|23142080\|23597149\|29167 | Unspecified |
| mmu-let-7i-5p | MIMAT0000122 | Prr11 | 270906 | HITS-CLIP | 23142080 | Unspecified |
| mmu-let-7i-5p | MIMAT0000122 | Prr12 | 233210 | HITS-CLIP | 19536157\|31433990\|29167373 | Unspecified |
| mmu-let-7i-5p | MIMAT0000122 | Prr14l | 215476 | HITS-CLIP | 23142080 | Unspecified |
| mmu-let-7i-5p | MIMAT0000122 | Prr36 | 80164 | HITS-CLIP | 23142080 | Unspecified |
| mmu-let-7i-5p | MIMAT0000122 | Prr5 | 109270 | HITS-CLIP | 23142080 | Unspecified |
| mmu-let-7i-5p | MIMAT0000122 | Prrc2a | 53761 | HITS-CLIP | 23597149\|23142080\|29167373 | Unspecified |
| mmu-let-7i-5p | MIMAT0000122 | Prrc2b | 227723 | HITS-CLIP | 37986224\|19536157\|26203562\|23142 | Unspecified |
| mmu-let-7i-5p | MIMAT0000122 | Prrc2c | 226562 | HITS-CLIP | 37986224\|23142080\|19536157\|23597 | Unspecified |
| mmu-let-7i-5p | MIMAT0000122 | Prrg1 | 546336 | HITS-CLIP | 23142080 | Unspecified |
| mmu-let-7i-5p | MIMAT0000122 | Prrt1 | 260297 | HITS-CLIP | 23142080\|19536157 | Unspecified |
| mmu-let-7i-5p | MIMAT0000122 | Prss36 | 77613 | HITS-CLIP | 23142080 | Unspecified |
| mmu-let-7i-5p | MIMAT0000122 | Prtg | 235472 | HITS-CLIP | 37986224\|19536157 | Unspecified |
| mmu-let-7i-5p | MIMAT0000122 | Psap | 19156 | HITS-CLIP | 29167373 | Unspecified |
| mmu-let-7i-5p | MIMAT0000122 | Psd3 | 234353 | HITS-CLIP | 37986224\|19536157\|23597149\|29167 | Unspecified |
| mmu-let-7i-5p | MIMAT0000122 | Psd4 | 215632 | HITS-CLIP | 23142080\|29167373 | Unspecified |
| mmu-let-7i-5p | MIMAT0000122 | Psen2 | 19165 | HITS-CLIP | 23597149\|23142080 | Unspecified |

| | | | | | | |
|---|---|---|---|---|---|---|
| mmu-let-7i-5p | MIMAT0000122 | Psip1 | 101739 | HITS-CLIP | 23142080 | Unspecified |
| mmu-let-7i-5p | MIMAT0000122 | Pskh1 | 244631 | HITS-CLIP | 23597149\|23142080 | Unspecified |
| mmu-let-7i-5p | MIMAT0000122 | Psma1 | 26440 | HITS-CLIP | 23142080\|29167373 | Unspecified |
| mmu-let-7i-5p | MIMAT0000122 | Psma2 | 19166 | HITS-CLIP | 23142080\|23597149 | Unspecified |
| mmu-let-7i-5p | MIMAT0000122 | Psma6 | 26443 | HITS-CLIP | 23597149\|23142080 | Unspecified |
| mmu-let-7i-5p | MIMAT0000122 | Psma7 | 26444 | HITS-CLIP | 23142080 | Unspecified |
| mmu-let-7i-5p | MIMAT0000122 | Psmb2 | 26445 | HITS-CLIP | 37986224\|23597149\|23142080 | Unspecified |
| mmu-let-7i-5p | MIMAT0000122 | Psmb5 | 19173 | HITS-CLIP | 23142080\|29167373 | Unspecified |
| mmu-let-7i-5p | MIMAT0000122 | Psmb7 | 19177 | HITS-CLIP | 23597149\|23142080 | Unspecified |
| mmu-let-7i-5p | MIMAT0000122 | Psmb8 | 16913 | HITS-CLIP | 23142080 | Unspecified |
| mmu-let-7i-5p | MIMAT0000122 | Psmb9 | 16912 | HITS-CLIP | 23142080 | Unspecified |
| mmu-let-7i-5p | MIMAT0000122 | Psmc1 | 19179 | HITS-CLIP | 23142080\|29167373 | Unspecified |
| mmu-let-7i-5p | MIMAT0000122 | Psmc2 | 19181 | HITS-CLIP | 37986224\|23597149\|19536157\|29167 | Unspecified |
| mmu-let-7i-5p | MIMAT0000122 | Psmc3 | 19182 | HITS-CLIP | 23597149 | Unspecified |
| mmu-let-7i-5p | MIMAT0000122 | Psmc4 | 23996 | HITS-CLIP | 23597149\|19536157\|23142080 | Unspecified |
| mmu-let-7i-5p | MIMAT0000122 | Psmc5 | 19184 | HITS-CLIP | 23597149\|23142080 | Unspecified |
| mmu-let-7i-5p | MIMAT0000122 | Psmc6 | 67089 | HITS-CLIP | 23597149\|23142080 | Unspecified |
| mmu-let-7i-5p | MIMAT0000122 | Psmd1 | 70247 | HITS-CLIP | 37986224\|23597149\|23142080 | Unspecified |
| mmu-let-7i-5p | MIMAT0000122 | Psmd12 | 66997 | HITS-CLIP | 23597149\|23142080 | Unspecified |
| mmu-let-7i-5p | MIMAT0000122 | Psmd14 | 59029 | HITS-CLIP | 23142080 | Unspecified |
| mmu-let-7i-5p | MIMAT0000122 | Psmd2 | 21762 | HITS-CLIP | 23597149\|23142080 | Unspecified |
| mmu-let-7i-5p | MIMAT0000122 | Psmd3 | 22123 | HITS-CLIP | 23597149\|23142080 | Unspecified |
| mmu-let-7i-5p | MIMAT0000122 | Psmd5 | 66998 | HITS-CLIP | 23142080 | Unspecified |
| mmu-let-7i-5p | MIMAT0000122 | Psmd6 | 66413 | HITS-CLIP | 23597149\|19536157\|23142080 | Unspecified |
| mmu-let-7i-5p | MIMAT0000122 | Psmd7 | 17463 | HITS-CLIP | 23142080 | Unspecified |
| mmu-let-7i-5p | MIMAT0000122 | Psmd8 | 57296 | HITS-CLIP | 23142080\|23597149\|19536157 | Unspecified |
| mmu-let-7i-5p | MIMAT0000122 | Psme2b | 621823 | HITS-CLIP | 23597149 | Unspecified |
| mmu-let-7i-5p | MIMAT0000122 | Psme3 | 19192 | HITS-CLIP | 23142080 | Unspecified |
| mmu-let-7i-5p | MIMAT0000122 | Psme4 | 103554 | HITS-CLIP | 37986224\|23597149\|19536157\|23142 | Unspecified |
| mmu-let-7i-5p | MIMAT0000122 | Pspc1 | 66645 | HITS-CLIP | 37986224\|29167373 | Unspecified |
| mmu-let-7i-5p | MIMAT0000122 | Pstk | 214580 | HITS-CLIP | 23597149 | Unspecified |
| mmu-let-7i-5p | MIMAT0000122 | Pstpip2 | 19201 | HITS-CLIP | 23597149\|29167373 | Unspecified |
| mmu-let-7i-5p | MIMAT0000122 | Ptafr | 19204 | HITS-CLIP | 23597149\|29167373 | Unspecified |
| mmu-let-7i-5p | MIMAT0000122 | Ptar1 | 72351 | HITS-CLIP | 37986224\|23142080\|19536157\|23597 | Unspecified |
| mmu-let-7i-5p | MIMAT0000122 | Ptbp1 | 19205 | HITS-CLIP | 23142080\|19536157\|29167373 | Unspecified |
| mmu-let-7i-5p | MIMAT0000122 | Ptbp2 | 56195 | HITS-CLIP | 37986224\|19536157 | Unspecified |
| mmu-let-7i-5p | MIMAT0000122 | Ptbp3 | 230257 | HITS-CLIP | 23597149\|23142080\|29167373 | Unspecified |
| mmu-let-7i-5p | MIMAT0000122 | Ptcd3 | 69956 | HITS-CLIP | 23142080 | Unspecified |
| mmu-let-7i-5p | MIMAT0000122 | Ptch1 | 19206 | HITS-CLIP | 37986224\|23597149\|23142080 | Unspecified |

| | | | | | | |
|---|---|---|---|---|---|---|
| mmu-let-7i-5p | MIMAT0000122 | Ptchd1 | 211612 | HITS-CLIP | 19536157\|29167373 | Unspecified |
| mmu-let-7i-5p | MIMAT0000122 | Ptdss1 | 19210 | HITS-CLIP | 37986224\|23597149\|19536157 | Unspecified |
| mmu-let-7i-5p | MIMAT0000122 | Pten | 19211 | HITS-CLIP | 23142080 | Unspecified |
| mmu-let-7i-5p | MIMAT0000122 | Ptgds | 19215 | HITS-CLIP | 19536157 | Unspecified |
| mmu-let-7i-5p | MIMAT0000122 | Ptger2 | 19217 | HITS-CLIP | 23142080 | Unspecified |
| mmu-let-7i-5p | MIMAT0000122 | Ptger4 | 19219 | HITS-CLIP | 23142080\|29167373 | Unspecified |
| mmu-let-7i-5p | MIMAT0000122 | Ptk2 | 14083 | HITS-CLIP | 23142080\|19536157 | Unspecified |
| mmu-let-7i-5p | MIMAT0000122 | Ptk2b | 19229 | HITS-CLIP | 23142080\|29167373 | Unspecified |
| mmu-let-7i-5p | MIMAT0000122 | Ptn | 19242 | HITS-CLIP | 37986224 | Unspecified |
| mmu-let-7i-5p | MIMAT0000122 | Ptov1 | 84113 | HITS-CLIP | 37986224\|23142080 | Unspecified |
| mmu-let-7i-5p | MIMAT0000122 | Ptp4a2 | 19244 | HITS-CLIP | 23142080 | Unspecified |
| mmu-let-7i-5p | MIMAT0000122 | Ptpa | 5524 | HITS-CLIP | 23142080 | Unspecified |
| mmu-let-7i-5p | MIMAT0000122 | Ptpn1 | 19246 | HITS-CLIP | 37986224\|23597149\|19536157 | Unspecified |
| mmu-let-7i-5p | MIMAT0000122 | Ptpn11 | 19247 | HITS-CLIP | 23597149\|23142080 | Unspecified |
| mmu-let-7i-5p | MIMAT0000122 | Ptpn12 | 19248 | HITS-CLIP | 37986224\|23597149\|29167373 | Unspecified |
| mmu-let-7i-5p | MIMAT0000122 | Ptpn2 | 19255 | HITS-CLIP | 23597149\|23142080 | Unspecified |
| mmu-let-7i-5p | MIMAT0000122 | Ptpn21 | 24000 | HITS-CLIP | 23597149\|23142080 | Unspecified |
| mmu-let-7i-5p | MIMAT0000122 | Ptpn22 | 19260 | HITS-CLIP | 23142080 | Unspecified |
| mmu-let-7i-5p | MIMAT0000122 | Ptpn23 | 104831 | HITS-CLIP | 23142080\|29167373 | Unspecified |
| mmu-let-7i-5p | MIMAT0000122 | Ptpn3 | 545622 | HITS-CLIP | 37986224\|19536157\|23597149\|23142 | Unspecified |
| mmu-let-7i-5p | MIMAT0000122 | Ptpn4 | 19258 | HITS-CLIP | 37986224\|19536157\|23142080 | Unspecified |
| mmu-let-7i-5p | MIMAT0000122 | Ptpn5 | 19259 | HITS-CLIP | 23142080 | Unspecified |
| mmu-let-7i-5p | MIMAT0000122 | Ptpn6 | 15170 | HITS-CLIP | 23142080 | Unspecified |
| mmu-let-7i-5p | MIMAT0000122 | Ptpn7 | 320139 | HITS-CLIP | 29167373 | Unspecified |
| mmu-let-7i-5p | MIMAT0000122 | Ptpn9 | 56294 | HITS-CLIP | 19536157 | Unspecified |
| mmu-let-7i-5p | MIMAT0000122 | Ptpra | 19262 | HITS-CLIP | 29167373 | Unspecified |
| mmu-let-7i-5p | MIMAT0000122 | Ptprb | 19263 | HITS-CLIP | 37986224\|23597149\|19536157 | Unspecified |
| mmu-let-7i-5p | MIMAT0000122 | Ptprc | 19264 | HITS-CLIP | 37986224\|23142080\|31433990\|29167 | Unspecified |
| mmu-let-7i-5p | MIMAT0000122 | Ptprd | 19266 | HITS-CLIP | 37986224\|19536157\|23597149 | Unspecified |
| mmu-let-7i-5p | MIMAT0000122 | Ptprf | 19268 | HITS-CLIP | 37986224 | Unspecified |
| mmu-let-7i-5p | MIMAT0000122 | Ptprj | 19271 | HITS-CLIP | 23142080\|19536157\|23597149\|29167 | Unspecified |
| mmu-let-7i-5p | MIMAT0000122 | Ptprk | 19272 | HITS-CLIP | 37986224\|23597149 | Unspecified |
| mmu-let-7i-5p | MIMAT0000122 | Ptprn | 19275 | HITS-CLIP | 37986224 | Unspecified |
| mmu-let-7i-5p | MIMAT0000122 | Ptpru | 19273 | HITS-CLIP | 37986224\|19536157 | Unspecified |
| mmu-let-7i-5p | MIMAT0000122 | Ptprz1 | 19283 | HITS-CLIP | 37986224\|19536157 | Unspecified |
| mmu-let-7i-5p | MIMAT0000122 | Ptrh2 | 217057 | HITS-CLIP | 23142080 | Unspecified |
| mmu-let-7i-5p | MIMAT0000122 | Pttg1ip | 108705 | HITS-CLIP | 37986224\|23597149\|23142080 | Unspecified |
| mmu-let-7i-5p | MIMAT0000122 | Puf60 | 67959 | HITS-CLIP | 23597149 | Unspecified |
| mmu-let-7i-5p | MIMAT0000122 | Pum1 | 80912 | HITS-CLIP | 37986224\|23597149\|23142080 | Unspecified |

| | | | | | | |
|---|---|---|---|---|---|---|
| mmu-let-7i-5p | MIMAT0000122 | Pum2 | 80913 | HITS-CLIP | 31433990\|23142080\|29167373 | Unspecified |
| mmu-let-7i-5p | MIMAT0000122 | Pura | 19290 | HITS-CLIP | 23142080 | Unspecified |
| mmu-let-7i-5p | MIMAT0000122 | Purb | 19291 | HITS-CLIP | 37986224\|23142080\|19536157\|23597 | Unspecified |
| mmu-let-7i-5p | MIMAT0000122 | Pus10 | 74467 | HITS-CLIP | 23142080 | Unspecified |
| mmu-let-7i-5p | MIMAT0000122 | Pvr | 52118 | HITS-CLIP | 37986224\|23597149\|23142080 | Unspecified |
| mmu-let-7i-5p | MIMAT0000122 | Pwp1 | 103136 | HITS-CLIP | 19536157\|23142080 | Unspecified |
| mmu-let-7i-5p | MIMAT0000122 | Pwwp2a | 70802 | HITS-CLIP | 37986224 | Unspecified |
| mmu-let-7i-5p | MIMAT0000122 | Pxdn | 69675 | HITS-CLIP | 37986224\|19536157\|23597149 | Unspecified |
| mmu-let-7i-5p | MIMAT0000122 | Pxk | 218699 | HITS-CLIP | 23597149 | Unspecified |
| mmu-let-7i-5p | MIMAT0000122 | Pxmp2 | 19301 | HITS-CLIP | 23597149 | Unspecified |
| mmu-let-7i-5p | MIMAT0000122 | Pxmp4 | 59038 | HITS-CLIP | 23597149\|23142080\|19536157 | Unspecified |
| mmu-let-7i-5p | MIMAT0000122 | Pygl | 110095 | HITS-CLIP | 23597149 | Unspecified |
| mmu-let-7i-5p | MIMAT0000122 | Pygm | 19309 | HITS-CLIP | 23597149 | Unspecified |
| mmu-let-7i-5p | MIMAT0000122 | Pygo1 | 72135 | HITS-CLIP | 19536157 | Unspecified |
| mmu-let-7i-5p | MIMAT0000122 | Pygo2 | 68911 | HITS-CLIP | 23597149\|23630078 | Unspecified |
| mmu-let-7i-5p | MIMAT0000122 | Pym1 | 84305 | HITS-CLIP | 23142080 | Unspecified |
| mmu-let-7i-5p | MIMAT0000122 | Pzp | 11287 | HITS-CLIP | 37986224\|23597149 | Unspecified |
| mmu-let-7i-5p | MIMAT0000122 | Qars | 97541 | HITS-CLIP//qCLASH | 37986224\|23597149\|23142080\|31393 | Unspecified |
| mmu-let-7i-5p | MIMAT0000122 | Qdpr | 110391 | HITS-CLIP | 23597149 | Unspecified |
| mmu-let-7i-5p | MIMAT0000122 | Qk | 19317 | HITS-CLIP | 37986224\|23142080\|29167373\|23597 | Unspecified |
| mmu-let-7i-5p | MIMAT0000122 | Qki | 19317 | HITS-CLIP | 26203562 | Unspecified |
| mmu-let-7i-5p | MIMAT0000122 | Qprt | 67375 | HITS-CLIP | 23142080 | Unspecified |
| mmu-let-7i-5p | MIMAT0000122 | Qrich1 | 69232 | HITS-CLIP | 23597149\|23142080\|29167373 | Unspecified |
| mmu-let-7i-5p | MIMAT0000122 | Qser1 | 99003 | HITS-CLIP | 37986224\|23142080\|23597149\|29167 | Unspecified |
| mmu-let-7i-5p | MIMAT0000122 | R3hdm1 | 226412 | HITS-CLIP | 29167373 | Unspecified |
| mmu-let-7i-5p | MIMAT0000122 | R3hdm2 | 71750 | HITS-CLIP | 23142080\|23597149\|19536157 | Unspecified |
| mmu-let-7i-5p | MIMAT0000122 | R3hdm4 | 109284 | HITS-CLIP | 23142080 | Unspecified |
| mmu-let-7i-5p | MIMAT0000122 | Rab10 | 19325 | HITS-CLIP | 23142080\|29167373 | Unspecified |
| mmu-let-7i-5p | MIMAT0000122 | Rab11a | 53869 | HITS-CLIP | 23142080 | Unspecified |
| mmu-let-7i-5p | MIMAT0000122 | Rab11fip1 | 75767 | HITS-CLIP | 23142080 | Unspecified |
| mmu-let-7i-5p | MIMAT0000122 | Rab11fip2 | 74998 | HITS-CLIP | 37986224\|23142080\|19536157\|23597 | Unspecified |
| mmu-let-7i-5p | MIMAT0000122 | Rab11fip3 | 215445 | HITS-CLIP | 23142080 | Unspecified |
| mmu-let-7i-5p | MIMAT0000122 | Rab11fip4 | 268451 | HITS-CLIP | 19536157\|23142080 | Unspecified |
| mmu-let-7i-5p | MIMAT0000122 | Rab12 | 19328 | HITS-CLIP | 31433990\|19536157\|23142080 | Unspecified |
| mmu-let-7i-5p | MIMAT0000122 | Rab14 | 68365 | HITS-CLIP | 37986224\|23597149\|23142080 | Unspecified |
| mmu-let-7i-5p | MIMAT0000122 | Rab18 | 19330 | HITS-CLIP | 23142080 | Unspecified |
| mmu-let-7i-5p | MIMAT0000122 | Rab1a | 5861 | HITS-CLIP | 23142080\|29167373 | Unspecified |
| mmu-let-7i-5p | MIMAT0000122 | Rab1b | 76308 | HITS-CLIP | 37986224\|23597149\|31433990\|23142 | Unspecified |
| mmu-let-7i-5p | MIMAT0000122 | Rab21 | 216344 | HITS-CLIP | 23142080 | Unspecified |

| | | | | | | |
|---|---|---|---|---|---|---|
| mmu-let-7i-5p | MIMAT0000122 | Rab22a | 19334 | HITS-CLIP | 31433990\|23142080 | Unspecified |
| mmu-let-7i-5p | MIMAT0000122 | Rab24 | 19336 | HITS-CLIP | 23142080 | Unspecified |
| mmu-let-7i-5p | MIMAT0000122 | Rab27a | 11891 | HITS-CLIP | 23142080 | Unspecified |
| mmu-let-7i-5p | MIMAT0000122 | Rab29 | 226422 | HITS-CLIP | 23142080 | Unspecified |
| mmu-let-7i-5p | MIMAT0000122 | Rab2a | 59021 | HITS-CLIP | 23597149 | Unspecified |
| mmu-let-7i-5p | MIMAT0000122 | Rab2b | 76338 | HITS-CLIP | 23597149 | Unspecified |
| mmu-let-7i-5p | MIMAT0000122 | Rab31 | 106572 | HITS-CLIP | 29167373 | Unspecified |
| mmu-let-7i-5p | MIMAT0000122 | Rab32 | 67844 | HITS-CLIP | 23597149 | Unspecified |
| mmu-let-7i-5p | MIMAT0000122 | Rab33b | 19338 | HITS-CLIP | 23142080 | Unspecified |
| mmu-let-7i-5p | MIMAT0000122 | Rab35 | 77407 | qCLASH//HITS-CLIP | 31393953\|29167373 | Unspecified |
| mmu-let-7i-5p | MIMAT0000122 | Rab39 | 270160 | HITS-CLIP | 23142080\|29167373 | Unspecified |
| mmu-let-7i-5p | MIMAT0000122 | Rab3a | 19339 | HITS-CLIP | 23142080\|29167373 | Unspecified |
| mmu-let-7i-5p | MIMAT0000122 | Rab3c | 67295 | HITS-CLIP | 37986224 | Unspecified |
| mmu-let-7i-5p | MIMAT0000122 | Rab3gap1 | 226407 | qCLASH//HITS-CLIP | 31393953\|23142080 | Unspecified |
| mmu-let-7i-5p | MIMAT0000122 | Rab3gap2 | 98732 | HITS-CLIP | 37986224\|19536157\|23142080\|29167 | Unspecified |
| mmu-let-7i-5p | MIMAT0000122 | Rab40c | 224624 | HITS-CLIP | 23142080\|23597149\|29167373 | Unspecified |
| mmu-let-7i-5p | MIMAT0000122 | Rab5b | 19344 | HITS-CLIP | 23142080 | Unspecified |
| mmu-let-7i-5p | MIMAT0000122 | Rab5if | 55969 | HITS-CLIP | 23597149\|23142080\|29167373 | Unspecified |
| mmu-let-7i-5p | MIMAT0000122 | Rab6b | 270192 | HITS-CLIP | 37986224 | Unspecified |
| mmu-let-7i-5p | MIMAT0000122 | Rab7b | 338382 | HITS-CLIP | 29167373 | Unspecified |
| mmu-let-7i-5p | MIMAT0000122 | Rab8b | 235442 | HITS-CLIP | 37986224\|23597149\|29167373\|23142 | Unspecified |
| mmu-let-7i-5p | MIMAT0000122 | Rabep1 | 54189 | HITS-CLIP | 19536157\|23142080\|29167373 | Unspecified |
| mmu-let-7i-5p | MIMAT0000122 | Rabepk | 227746 | HITS-CLIP | 23142080 | Unspecified |
| mmu-let-7i-5p | MIMAT0000122 | Rabgap1 | 227800 | HITS-CLIP | 23597149\|29167373\|23142080 | Unspecified |
| mmu-let-7i-5p | MIMAT0000122 | Rabgap1l | 29809 | HITS-CLIP | 37986224\|23597149\|23142080 | Unspecified |
| mmu-let-7i-5p | MIMAT0000122 | Rabggta | 56187 | HITS-CLIP | 23597149 | Unspecified |
| mmu-let-7i-5p | MIMAT0000122 | Rabggtb | 19352 | HITS-CLIP | 23142080 | Unspecified |
| mmu-let-7i-5p | MIMAT0000122 | Rabl3 | 67657 | HITS-CLIP//qCLASH | 37986224\|31393953 | Unspecified |
| mmu-let-7i-5p | MIMAT0000122 | Rac1 | 19353 | HITS-CLIP | 31433990\|23597149\|23142080 | Unspecified |
| mmu-let-7i-5p | MIMAT0000122 | Rac2 | 19354 | HITS-CLIP | 37986224\|23142080\|29167373 | Unspecified |
| mmu-let-7i-5p | MIMAT0000122 | Racgap1 | 26934 | HITS-CLIP | 23142080 | Unspecified |
| mmu-let-7i-5p | MIMAT0000122 | Rack1 | 10399 | HITS-CLIP//qCLASH | 23597149\|31393953\|23142080 | Unspecified |
| mmu-let-7i-5p | MIMAT0000122 | Rad18 | 58186 | HITS-CLIP | 23142080 | Unspecified |
| mmu-let-7i-5p | MIMAT0000122 | Rad21 | 19357 | HITS-CLIP | 37986224\|23142080\|29167373 | Unspecified |
| mmu-let-7i-5p | MIMAT0000122 | Rad23b | 19359 | HITS-CLIP | 23597149\|31433990\|23142080\|29167 | Unspecified |
| mmu-let-7i-5p | MIMAT0000122 | Rad51c | 114714 | HITS-CLIP | 23142080 | Unspecified |
| mmu-let-7i-5p | MIMAT0000122 | Rad54l2 | 81000 | HITS-CLIP | 37986224\|23597149\|29167373\|23142 | Unspecified |
| mmu-let-7i-5p | MIMAT0000122 | Rae1 | 66679 | HITS-CLIP | 37986224\|23142080 | Unspecified |
| mmu-let-7i-5p | MIMAT0000122 | Raet1e | 135250 | HITS-CLIP | 29167373 | Unspecified |

| | | | | | | |
|---|---|---|---|---|---|---|
| mmu-let-7i-5p | MIMAT0000122 | Rai1 | 19377 | HITS-CLIP | 37986224\|23597149\|23142080 | Unspecified |
| mmu-let-7i-5p | MIMAT0000122 | Ralb | 64143 | HITS-CLIP | 29167373 | Unspecified |
| mmu-let-7i-5p | MIMAT0000122 | Ralgapa1 | 56784 | HITS-CLIP | 23142080 | Unspecified |
| mmu-let-7i-5p | MIMAT0000122 | Ralgapa2 | 241694 | HITS-CLIP | 23597149\|19536157\|23142080\|29167 | Unspecified |
| mmu-let-7i-5p | MIMAT0000122 | Ralgapb | 228850 | HITS-CLIP | 23597149\|23142080 | Unspecified |
| mmu-let-7i-5p | MIMAT0000122 | Ralgds | 19730 | HITS-CLIP | 19536157\|29167373 | Unspecified |
| mmu-let-7i-5p | MIMAT0000122 | Ralgps1 | 241308 | HITS-CLIP | 23142080 | Unspecified |
| mmu-let-7i-5p | MIMAT0000122 | Ralgps2 | 78255 | HITS-CLIP | 19536157 | Unspecified |
| mmu-let-7i-5p | MIMAT0000122 | Raly | 19383 | HITS-CLIP | 23597149\|23142080 | Unspecified |
| mmu-let-7i-5p | MIMAT0000122 | Ramp2 | 54409 | HITS-CLIP | 23597149 | Unspecified |
| mmu-let-7i-5p | MIMAT0000122 | Ranbp10 | 74334 | HITS-CLIP | 37986224\|23597149 | Unspecified |
| mmu-let-7i-5p | MIMAT0000122 | Ranbp2 | 19386 | HITS-CLIP | 23597149\|23142080\|29167373 | Unspecified |
| mmu-let-7i-5p | MIMAT0000122 | Ranbp9 | 56705 | HITS-CLIP | 23142080\|23597149\|29167373 | Unspecified |
| mmu-let-7i-5p | MIMAT0000122 | Rangap1 | 19387 | HITS-CLIP | 23597149 | Unspecified |
| mmu-let-7i-5p | MIMAT0000122 | Rap1b | 215449 | HITS-CLIP | 23142080 | Unspecified |
| mmu-let-7i-5p | MIMAT0000122 | Rap1gds1 | 229877 | HITS-CLIP | 37986224\|19536157\|23142080\|29167 | Unspecified |
| mmu-let-7i-5p | MIMAT0000122 | Rap2c | 72065 | HITS-CLIP | 29167373\|23142080 | Unspecified |
| mmu-let-7i-5p | MIMAT0000122 | Rapgef1 | 107746 | HITS-CLIP | 29167373 | Unspecified |
| mmu-let-7i-5p | MIMAT0000122 | Rapgef2 | 76089 | HITS-CLIP | 23597149 | Unspecified |
| mmu-let-7i-5p | MIMAT0000122 | Rapgef4 | 56508 | HITS-CLIP | 23597149 | Unspecified |
| mmu-let-7i-5p | MIMAT0000122 | Rapgef5 | 217944 | HITS-CLIP | 19536157 | Unspecified |
| mmu-let-7i-5p | MIMAT0000122 | Rapgef6 | 192786 | HITS-CLIP | 23142080\|29167373 | Unspecified |
| mmu-let-7i-5p | MIMAT0000122 | Raph1 | 77300 | HITS-CLIP | 23142080\|23597149\|29167373 | Unspecified |
| mmu-let-7i-5p | MIMAT0000122 | Rarb | 218772 | HITS-CLIP | 23597149 | Unspecified |
| mmu-let-7i-5p | MIMAT0000122 | Rarres2 | 71660 | HITS-CLIP | 23597149 | Unspecified |
| mmu-let-7i-5p | MIMAT0000122 | Rars2 | 109093 | HITS-CLIP//qCLASH | 23597149\|31393953\|23142080 | Unspecified |
| mmu-let-7i-5p | MIMAT0000122 | Rasa1 | 218397 | qCLASH//HITS-CLIP | 31393953\|23142080\|29167373 | Unspecified |
| mmu-let-7i-5p | MIMAT0000122 | Rasa2 | 114713 | HITS-CLIP | 23597149\|23142080 | Unspecified |
| mmu-let-7i-5p | MIMAT0000122 | Rasa4 | 54153 | HITS-CLIP | 29167373 | Unspecified |
| mmu-let-7i-5p | MIMAT0000122 | Rasal2 | 226525 | HITS-CLIP | 37986224\|19536157 | Unspecified |
| mmu-let-7i-5p | MIMAT0000122 | Rasal3 | 320484 | HITS-CLIP | 23142080 | Unspecified |
| mmu-let-7i-5p | MIMAT0000122 | Rasd2 | 75141 | HITS-CLIP | 37986224 | Unspecified |
| mmu-let-7i-5p | MIMAT0000122 | Rasgrf2 | 19418 | HITS-CLIP | 19536157 | Unspecified |
| mmu-let-7i-5p | MIMAT0000122 | Rasgrp1 | 19419 | HITS-CLIP | 31433990\|19536157\|23142080 | Unspecified |
| mmu-let-7i-5p | MIMAT0000122 | Rasgrp3 | 240168 | HITS-CLIP | 23597149 | Unspecified |
| mmu-let-7i-5p | MIMAT0000122 | Rasl10b | 276952 | HITS-CLIP | 37986224\|19536157 | Unspecified |
| mmu-let-7i-5p | MIMAT0000122 | Rassf2 | 215653 | HITS-CLIP | 23142080\|29167373 | Unspecified |
| mmu-let-7i-5p | MIMAT0000122 | Rassf3 | 192678 | HITS-CLIP | 37986224\|23597149\|23142080 | Unspecified |
| mmu-let-7i-5p | MIMAT0000122 | Rassf4 | 213391 | HITS-CLIP | 29167373 | Unspecified |

| | | | | | | |
|---|---|---|---|---|---|---|
| mmu-let-7i-5p | MIMAT0000122 | Rassf5 | 54354 | HITS-CLIP | 23142080 | Unspecified |
| mmu-let-7i-5p | MIMAT0000122 | Rb1 | 19645 | HITS-CLIP | 37986224\|23142080\|19536157\|29167 | Unspecified |
| mmu-let-7i-5p | MIMAT0000122 | Rb1cc1 | 12421 | HITS-CLIP | 37986224\|23142080\|19536157\|29167 | Unspecified |
| mmu-let-7i-5p | MIMAT0000122 | Rbak | 57782 | HITS-CLIP | 23142080 | Unspecified |
| mmu-let-7i-5p | MIMAT0000122 | Rbbp4 | 19646 | HITS-CLIP | 23597149\|23142080\|29167373 | Unspecified |
| mmu-let-7i-5p | MIMAT0000122 | Rbbp5 | 213464 | HITS-CLIP | 23597149\|23142080 | Unspecified |
| mmu-let-7i-5p | MIMAT0000122 | Rbbp6 | 19647 | HITS-CLIP | 23597149\|23142080 | Unspecified |
| mmu-let-7i-5p | MIMAT0000122 | Rbbp7 | 245688 | HITS-CLIP | 37986224\|23597149\|23142080 | Unspecified |
| mmu-let-7i-5p | MIMAT0000122 | Rbbp8 | 225182 | HITS-CLIP | 23142080\|29167373 | Unspecified |
| mmu-let-7i-5p | MIMAT0000122 | Rbfox2 | 93686 | HITS-CLIP | 23597149\|26203562 | Unspecified |
| mmu-let-7i-5p | MIMAT0000122 | Rbfox3 | 52897 | HITS-CLIP | 19536157 | Unspecified |
| mmu-let-7i-5p | MIMAT0000122 | Rbl1 | 19650 | HITS-CLIP | 23142080 | Unspecified |
| mmu-let-7i-5p | MIMAT0000122 | Rbm10 | 236732 | HITS-CLIP | 23142080 | Unspecified |
| mmu-let-7i-5p | MIMAT0000122 | Rbm12 | 75710 | HITS-CLIP | 29167373 | Unspecified |
| mmu-let-7i-5p | MIMAT0000122 | Rbm14 | 56275 | HITS-CLIP | 23142080\|29167373 | Unspecified |
| mmu-let-7i-5p | MIMAT0000122 | Rbm15b | 109095 | HITS-CLIP | 23142080 | Unspecified |
| mmu-let-7i-5p | MIMAT0000122 | Rbm26 | 74213 | HITS-CLIP | 23142080\|19536157\|23597149\|29167 | Unspecified |
| mmu-let-7i-5p | MIMAT0000122 | Rbm27 | 225432 | HITS-CLIP | 23142080\|23597149\|29167373 | Unspecified |
| mmu-let-7i-5p | MIMAT0000122 | Rbm28 | 68272 | HITS-CLIP | 23142080 | Unspecified |
| mmu-let-7i-5p | MIMAT0000122 | Rbm3 | 19652 | HITS-CLIP | 23142080 | Unspecified |
| mmu-let-7i-5p | MIMAT0000122 | Rbm33 | 381626 | HITS-CLIP | 23142080\|29167373 | Unspecified |
| mmu-let-7i-5p | MIMAT0000122 | Rbm38 | 56190 | HITS-CLIP | 23142080 | Unspecified |
| mmu-let-7i-5p | MIMAT0000122 | Rbm39 | 170791 | HITS-CLIP | 23142080\|29167373 | Unspecified |
| mmu-let-7i-5p | MIMAT0000122 | Rbm4 | 19653 | HITS-CLIP | 37986224\|23142080\|23597149 | Unspecified |
| mmu-let-7i-5p | MIMAT0000122 | Rbm41 | 237073 | HITS-CLIP | 37986224 | Unspecified |
| mmu-let-7i-5p | MIMAT0000122 | Rbm45 | 241490 | HITS-CLIP | 23142080 | Unspecified |
| mmu-let-7i-5p | MIMAT0000122 | Rbm47 | 245945 | HITS-CLIP | 29167373 | Unspecified |
| mmu-let-7i-5p | MIMAT0000122 | Rbm48 | 269623 | HITS-CLIP | 23597149\|23142080 | Unspecified |
| mmu-let-7i-5p | MIMAT0000122 | Rbm5 | 83486 | HITS-CLIP | 23142080 | Unspecified |
| mmu-let-7i-5p | MIMAT0000122 | Rbm6 | 19654 | HITS-CLIP | 23142080 | Unspecified |
| mmu-let-7i-5p | MIMAT0000122 | Rbm7 | 67010 | HITS-CLIP | 23142080\|29167373 | Unspecified |
| mmu-let-7i-5p | MIMAT0000122 | Rbms1 | 56878 | HITS-CLIP | 37986224\|23597149\|19536157\|23142 | Unspecified |
| mmu-let-7i-5p | MIMAT0000122 | Rbp4 | 19662 | HITS-CLIP | 23597149 | Unspecified |
| mmu-let-7i-5p | MIMAT0000122 | Rbpj | 19664 | HITS-CLIP | 29167373 | Unspecified |
| mmu-let-7i-5p | MIMAT0000122 | Rbpms2 | 71973 | HITS-CLIP | 23597149 | Unspecified |
| mmu-let-7i-5p | MIMAT0000122 | Rbsn | 64145 | HITS-CLIP | 23142080 | Unspecified |
| mmu-let-7i-5p | MIMAT0000122 | Rc3h1 | 381305 | HITS-CLIP | 23142080\|19536157\|23597149\|29167 | Unspecified |
| mmu-let-7i-5p | MIMAT0000122 | Rc3h2 | 319817 | HITS-CLIP | 37986224\|23597149\|19536157\|23142 | Unspecified |
| mmu-let-7i-5p | MIMAT0000122 | Rcan1 | 54720 | HITS-CLIP | 23597149 | Unspecified |

| | | | | | | | |
|---|---|---|---|---|---|---|---|
| mmu-let-7i-5p | MIMAT0000122 | Rcbtb1 | 71330 | HITS-CLIP | | 23597149 | Unspecified |
| mmu-let-7i-5p | MIMAT0000122 | Rcbtb2 | 105670 | HITS-CLIP | 23142080\|23597149 | | Unspecified |
| mmu-let-7i-5p | MIMAT0000122 | Rcc2 | 108911 | HITS-CLIP | | 23142080 | Unspecified |
| mmu-let-7i-5p | MIMAT0000122 | Rcl1 | 59028 | HITS-CLIP | | 23597149 | Unspecified |
| mmu-let-7i-5p | MIMAT0000122 | Rcn2 | 26611 | HITS-CLIP | | 37986224 | Unspecified |
| mmu-let-7i-5p | MIMAT0000122 | Rcor1 | 217864 | HITS-CLIP | | 23142080 | Unspecified |
| mmu-let-7i-5p | MIMAT0000122 | Rdh10 | 98711 | HITS-CLIP | 37986224\|23142080\|19536157 | | Unspecified |
| mmu-let-7i-5p | MIMAT0000122 | Rdh11 | 17252 | HITS-CLIP | 23597149\|29167373 | | Unspecified |
| mmu-let-7i-5p | MIMAT0000122 | Rdh16f1 | | HITS-CLIP | | 23597149 | Unspecified |
| mmu-let-7i-5p | MIMAT0000122 | Rdh5 | 19682 | HITS-CLIP | | 23597149 | Unspecified |
| mmu-let-7i-5p | MIMAT0000122 | Rdm1 | 66599 | HITS-CLIP | | 23142080 | Unspecified |
| mmu-let-7i-5p | MIMAT0000122 | Rdx | 19684 | HITS-CLIP | 37986224\|23597149\|19536157\|23142 | | Unspecified |
| mmu-let-7i-5p | MIMAT0000122 | Reck | 53614 | HITS-CLIP | | 23142080 | Unspecified |
| mmu-let-7i-5p | MIMAT0000122 | Reep3 | 28193 | HITS-CLIP | 30971389\|19536157 | | Unspecified |
| mmu-let-7i-5p | MIMAT0000122 | Reep4 | 72549 | HITS-CLIP | 23597149\|23142080 | | Unspecified |
| mmu-let-7i-5p | MIMAT0000122 | Reep5 | 13476 | HITS-CLIP | | 29167373 | Unspecified |
| mmu-let-7i-5p | MIMAT0000122 | Reep6 | 70335 | HITS-CLIP | | 23597149 | Unspecified |
| mmu-let-7i-5p | MIMAT0000122 | Rel | 19696 | HITS-CLIP | 23142080\|29167373 | | Unspecified |
| mmu-let-7i-5p | MIMAT0000122 | Rela | 19697 | HITS-CLIP | | 23142080 | Unspecified |
| mmu-let-7i-5p | MIMAT0000122 | Relch | 57614 | HITS-CLIP | | 23142080 | Unspecified |
| mmu-let-7i-5p | MIMAT0000122 | Rell1 | 100532 | HITS-CLIP | | 23597149 | Unspecified |
| mmu-let-7i-5p | MIMAT0000122 | Reln | 19699 | HITS-CLIP | 19536157\|23597149 | | Unspecified |
| mmu-let-7i-5p | MIMAT0000122 | Repin1 | 58887 | HITS-CLIP | | 23597149 | Unspecified |
| mmu-let-7i-5p | MIMAT0000122 | Reps1 | 19707 | HITS-CLIP | 23142080\|29167373 | | Unspecified |
| mmu-let-7i-5p | MIMAT0000122 | Reps2 | 194590 | HITS-CLIP | | 23597149 | Unspecified |
| mmu-let-7i-5p | MIMAT0000122 | Rere | 68703 | HITS-CLIP | 19536157\|23597149\|23142080\|29167 | | Unspecified |
| mmu-let-7i-5p | MIMAT0000122 | Resf1 | 55196 | HITS-CLIP | | 23142080 | Unspecified |
| mmu-let-7i-5p | MIMAT0000122 | Rest | 19712 | HITS-CLIP | | 23142080 | Unspecified |
| mmu-let-7i-5p | MIMAT0000122 | Retreg1 | 66270 | HITS-CLIP | 23597149\|23142080 | | Unspecified |
| mmu-let-7i-5p | MIMAT0000122 | Retreg2 | 79137 | HITS-CLIP | 23597149\|23142080 | | Unspecified |
| mmu-let-7i-5p | MIMAT0000122 | Retreg3 | 162427 | HITS-CLIP | | 23597149 | Unspecified |
| mmu-let-7i-5p | MIMAT0000122 | Rev1 | 56210 | HITS-CLIP | 23142080\|29167373 | | Unspecified |
| mmu-let-7i-5p | MIMAT0000122 | Rev3l | 19714 | HITS-CLIP | 23597149\|23142080\|19536157 | | Unspecified |
| mmu-let-7i-5p | MIMAT0000122 | Rexo4 | 227656 | HITS-CLIP | | 29167373 | Unspecified |
| mmu-let-7i-5p | MIMAT0000122 | Rfc1 | 19687 | HITS-CLIP | 23142080\|29167373 | | Unspecified |
| mmu-let-7i-5p | MIMAT0000122 | Rfc2 | 19718 | HITS-CLIP | | 19536157 | Unspecified |
| mmu-let-7i-5p | MIMAT0000122 | Rfc4 | 106344 | HITS-CLIP | | 23142080 | Unspecified |
| mmu-let-7i-5p | MIMAT0000122 | Rffl | 67338 | HITS-CLIP | 23597149\|23142080 | | Unspecified |
| mmu-let-7i-5p | MIMAT0000122 | Rfk | 54391 | HITS-CLIP | | 23597149 | Unspecified |

| | | | | | | |
|---|---|---|---|---|---|---|
| mmu-let-7i-5p | MIMAT0000122 | Rft1 | 328370 | HITS-CLIP | 23142080 | Unspecified |
| mmu-let-7i-5p | MIMAT0000122 | Rftn1 | 76438 | HITS-CLIP | 37986224\|23142080 | Unspecified |
| mmu-let-7i-5p | MIMAT0000122 | Rfx1 | 19724 | HITS-CLIP | 23142080 | Unspecified |
| mmu-let-7i-5p | MIMAT0000122 | Rfx3 | 19726 | HITS-CLIP | 23142080 | Unspecified |
| mmu-let-7i-5p | MIMAT0000122 | Rfx5 | 53970 | HITS-CLIP | 19536157\|23142080\|29167373 | Unspecified |
| mmu-let-7i-5p | MIMAT0000122 | Rfx7 | 319758 | HITS-CLIP//qCLASH | 37986224\|23597149\|31393953\|1953[illegible] | Unspecified |
| mmu-let-7i-5p | MIMAT0000122 | Rfxap | 170767 | HITS-CLIP | 23142080 | Unspecified |
| mmu-let-7i-5p | MIMAT0000122 | Rgl1 | 19731 | HITS-CLIP | 23597149 | Unspecified |
| mmu-let-7i-5p | MIMAT0000122 | Rgl2 | 19732 | HITS-CLIP | 23597149\|23142080 | Unspecified |
| mmu-let-7i-5p | MIMAT0000122 | Rgmb | 68799 | HITS-CLIP | 23142080 | Unspecified |
| mmu-let-7i-5p | MIMAT0000122 | Rgp1 | 242406 | HITS-CLIP | 37986224\|19536157\|23597149\|2314[illegible] | Unspecified |
| mmu-let-7i-5p | MIMAT0000122 | Rgs12 | 71729 | HITS-CLIP | 29167373 | Unspecified |
| mmu-let-7i-5p | MIMAT0000122 | Rgs16 | 19734 | HITS-CLIP | 37986224\|23142080 | Unspecified |
| mmu-let-7i-5p | MIMAT0000122 | Rgs3 | 50780 | HITS-CLIP | 37986224\|23597149\|23142080 | Unspecified |
| mmu-let-7i-5p | MIMAT0000122 | Rgs4 | 19736 | HITS-CLIP | 37986224 | Unspecified |
| mmu-let-7i-5p | MIMAT0000122 | Rgs5 | 19737 | HITS-CLIP | 19536157 | Unspecified |
| mmu-let-7i-5p | MIMAT0000122 | Rgs7bp | 52882 | HITS-CLIP | 37986224\|19536157 | Unspecified |
| mmu-let-7i-5p | MIMAT0000122 | Rgs8 | 67792 | HITS-CLIP | 37986224 | Unspecified |
| mmu-let-7i-5p | MIMAT0000122 | Rhbdd1 | 76867 | HITS-CLIP | 31433990 | Unspecified |
| mmu-let-7i-5p | MIMAT0000122 | Rhbdd2 | 215160 | HITS-CLIP | 37986224\|23597149\|29167373 | Unspecified |
| mmu-let-7i-5p | MIMAT0000122 | Rhbdf2 | 217344 | HITS-CLIP | 29167373 | Unspecified |
| mmu-let-7i-5p | MIMAT0000122 | Rhbdl3 | 246104 | HITS-CLIP | 19536157 | Unspecified |
| mmu-let-7i-5p | MIMAT0000122 | Rhno1 | 72440 | HITS-CLIP | 37986224 | Unspecified |
| mmu-let-7i-5p | MIMAT0000122 | Rhoa | 11848 | HITS-CLIP | 23142080 | Unspecified |
| mmu-let-7i-5p | MIMAT0000122 | Rhob | 11852 | HITS-CLIP | 37986224\|23597149\|23142080\|2916[illegible] | Unspecified |
| mmu-let-7i-5p | MIMAT0000122 | Rhobtb1 | 69288 | HITS-CLIP | 23597149\|23142080 | Unspecified |
| mmu-let-7i-5p | MIMAT0000122 | Rhobtb3 | 73296 | HITS-CLIP | 37986224\|19536157 | Unspecified |
| mmu-let-7i-5p | MIMAT0000122 | Rhof | 23912 | HITS-CLIP | 23142080 | Unspecified |
| mmu-let-7i-5p | MIMAT0000122 | Rhog | 56212 | HITS-CLIP | 37986224\|23142080\|29167373 | Unspecified |
| mmu-let-7i-5p | MIMAT0000122 | Rhot1 | 59040 | HITS-CLIP | 23142080 | Unspecified |
| mmu-let-7i-5p | MIMAT0000122 | Rhot2 | 214952 | HITS-CLIP | 23597149\|23142080 | Unspecified |
| mmu-let-7i-5p | MIMAT0000122 | Ric1 | 850728 | HITS-CLIP | 23597149\|23142080\|29167373 | Unspecified |
| mmu-let-7i-5p | MIMAT0000122 | Ric8b | 237422 | HITS-CLIP | 23142080 | Unspecified |
| mmu-let-7i-5p | MIMAT0000122 | Rictor | 78757 | HITS-CLIP | 37986224\|23597149\|23142080 | Unspecified |
| mmu-let-7i-5p | MIMAT0000122 | Rif1 | 51869 | HITS-CLIP | 23142080 | Unspecified |
| mmu-let-7i-5p | MIMAT0000122 | Rigi | 23586 | HITS-CLIP | 23630078 | Unspecified |
| mmu-let-7i-5p | MIMAT0000122 | Rin3 | 217835 | HITS-CLIP | 23597149\|19536157 | Unspecified |
| mmu-let-7i-5p | MIMAT0000122 | Rinl | 320435 | HITS-CLIP | 23142080 | Unspecified |
| mmu-let-7i-5p | MIMAT0000122 | Rint1 | 72772 | HITS-CLIP | 23597149\|29167373 | Unspecified |

| | | | | | | |
|---|---|---|---|---|---|---|
| mmu-let-7i-5p | MIMAT0000122 | Riok2 | 67045 | HITS-CLIP | 23597149 | Unspecified |
| mmu-let-7i-5p | MIMAT0000122 | Riok3 | 66878 | HITS-CLIP | 37986224\|23597149\|23142080 | Unspecified |
| mmu-let-7i-5p | MIMAT0000122 | Riox1 | 79697 | HITS-CLIP | 23597149\|23142080 | Unspecified |
| mmu-let-7i-5p | MIMAT0000122 | Ripk1 | 19766 | HITS-CLIP | 31433990 | Unspecified |
| mmu-let-7i-5p | MIMAT0000122 | Ripk2 | 192656 | HITS-CLIP | 23597149\|29167373 | Unspecified |
| mmu-let-7i-5p | MIMAT0000122 | Ripor2 | 193385 | HITS-CLIP | 23142080 | Unspecified |
| mmu-let-7i-5p | MIMAT0000122 | Rlf | 109263 | HITS-CLIP | 23142080 | Unspecified |
| mmu-let-7i-5p | MIMAT0000122 | Rlim | 19820 | HITS-CLIP | 31433990\|23142080\|29167373 | Unspecified |
| mmu-let-7i-5p | MIMAT0000122 | Rmdn1 | 66302 | HITS-CLIP | 23597149\|23142080 | Unspecified |
| mmu-let-7i-5p | MIMAT0000122 | Rmdn2 | 381110 | HITS-CLIP | 23597149 | Unspecified |
| mmu-let-7i-5p | MIMAT0000122 | Rmnd5a | 68477 | HITS-CLIP | 19536157 | Unspecified |
| mmu-let-7i-5p | MIMAT0000122 | Rmnd5b | 66089 | HITS-CLIP | 23142080 | Unspecified |
| mmu-let-7i-5p | MIMAT0000122 | Rnasek | 52898 | HITS-CLIP | 37986224\|23597149 | Unspecified |
| mmu-let-7i-5p | MIMAT0000122 | Rnasel | 24014 | HITS-CLIP | 23142080\|29167373 | Unspecified |
| mmu-let-7i-5p | MIMAT0000122 | Rnf10 | 50849 | HITS-CLIP | 37986224\|23597149 | Unspecified |
| mmu-let-7i-5p | MIMAT0000122 | Rnf103 | 22644 | HITS-CLIP | 23597149 | Unspecified |
| mmu-let-7i-5p | MIMAT0000122 | Rnf111 | 93836 | HITS-CLIP | 23142080\|29167373\|19536157\|23597 | Unspecified |
| mmu-let-7i-5p | MIMAT0000122 | Rnf113a1 | 69942 | HITS-CLIP | 23142080 | Unspecified |
| mmu-let-7i-5p | MIMAT0000122 | Rnf113a2 | 66381 | HITS-CLIP | 37986224 | Unspecified |
| mmu-let-7i-5p | MIMAT0000122 | Rnf114 | 81018 | HITS-CLIP | 23597149 | Unspecified |
| mmu-let-7i-5p | MIMAT0000122 | Rnf115 | 67845 | HITS-CLIP | 37986224 | Unspecified |
| mmu-let-7i-5p | MIMAT0000122 | Rnf123 | 84585 | HITS-CLIP | 23142080\|23597149 | Unspecified |
| mmu-let-7i-5p | MIMAT0000122 | Rnf125 | 67664 | HITS-CLIP | 31433990 | Unspecified |
| mmu-let-7i-5p | MIMAT0000122 | Rnf128 | 66889 | HITS-CLIP | 37986224\|23597149 | Unspecified |
| mmu-let-7i-5p | MIMAT0000122 | Rnf138 | 56515 | HITS-CLIP | 23142080\|29167373 | Unspecified |
| mmu-let-7i-5p | MIMAT0000122 | Rnf139 | 75841 | HITS-CLIP | 23142080 | Unspecified |
| mmu-let-7i-5p | MIMAT0000122 | Rnf145 | 74315 | HITS-CLIP | 23142080 | Unspecified |
| mmu-let-7i-5p | MIMAT0000122 | Rnf146 | 68031 | HITS-CLIP | 23597149 | Unspecified |
| mmu-let-7i-5p | MIMAT0000122 | Rnf149 | 67702 | HITS-CLIP | 19536157 | Unspecified |
| mmu-let-7i-5p | MIMAT0000122 | Rnf150 | 330812 | HITS-CLIP | 29167373 | Unspecified |
| mmu-let-7i-5p | MIMAT0000122 | Rnf152 | 320311 | HITS-CLIP | 23142080 | Unspecified |
| mmu-let-7i-5p | MIMAT0000122 | Rnf165 | 225743 | HITS-CLIP | 19536157 | Unspecified |
| mmu-let-7i-5p | MIMAT0000122 | Rnf167 | 70510 | HITS-CLIP | 37986224\|31433990\|19536157\|23142 | Unspecified |
| mmu-let-7i-5p | MIMAT0000122 | Rnf168 | 70238 | HITS-CLIP | 23142080 | Unspecified |
| mmu-let-7i-5p | MIMAT0000122 | Rnf169 | 108937 | HITS-CLIP | 23142080 | Unspecified |
| mmu-let-7i-5p | MIMAT0000122 | Rnf181 | 66510 | HITS-CLIP | 23597149\|23142080 | Unspecified |
| mmu-let-7i-5p | MIMAT0000122 | Rnf185 | 193670 | HITS-CLIP | 23597149\|23142080 | Unspecified |
| mmu-let-7i-5p | MIMAT0000122 | Rnf19a | 30945 | HITS-CLIP | 23142080\|29167373 | Unspecified |
| mmu-let-7i-5p | MIMAT0000122 | Rnf19b | 75234 | HITS-CLIP | 23142080\|29167373 | Unspecified |

| | | | | | | |
|---|---|---|---|---|---|---|
| mmu-let-7i-5p | MIMAT0000122 | Rnf2 | 19821 | HITS-CLIP | 23142080 | Unspecified |
| mmu-let-7i-5p | MIMAT0000122 | Rnf20 | 109331 | HITS-CLIP | 23630078 | Unspecified |
| mmu-let-7i-5p | MIMAT0000122 | Rnf213 | 672511 | HITS-CLIP | 37986224\|23142080\|23597149\|29167 | Unspecified |
| mmu-let-7i-5p | MIMAT0000122 | Rnf214 | 235315 | HITS-CLIP | 23597149 | Unspecified |
| mmu-let-7i-5p | MIMAT0000122 | Rnf216 | 108086 | HITS-CLIP | 23142080 | Unspecified |
| mmu-let-7i-5p | MIMAT0000122 | Rnf217 | 268291 | HITS-CLIP | 37986224\|23597149\|19536157 | Unspecified |
| mmu-let-7i-5p | MIMAT0000122 | Rnf219 | 72486 | HITS-CLIP | 23142080 | Unspecified |
| mmu-let-7i-5p | MIMAT0000122 | Rnf220 | 66743 | HITS-CLIP | 23597149\|31433990 | Unspecified |
| mmu-let-7i-5p | MIMAT0000122 | Rnf225 | 381845 | qCLASH | 31393953 | Unspecified |
| mmu-let-7i-5p | MIMAT0000122 | Rnf25 | 57751 | HITS-CLIP | 23142080 | Unspecified |
| mmu-let-7i-5p | MIMAT0000122 | Rnf26 | 213211 | HITS-CLIP | 37986224 | Unspecified |
| mmu-let-7i-5p | MIMAT0000122 | Rnf31 | 268749 | HITS-CLIP | 37986224\|23142080 | Unspecified |
| mmu-let-7i-5p | MIMAT0000122 | Rnf38 | 73469 | HITS-CLIP | 37986224\|19536157\|23597149\|23142 | Unspecified |
| mmu-let-7i-5p | MIMAT0000122 | Rnf4 | 19822 | HITS-CLIP | 23142080\|23597149 | Unspecified |
| mmu-let-7i-5p | MIMAT0000122 | Rnf41 | 67588 | HITS-CLIP | 23597149 | Unspecified |
| mmu-let-7i-5p | MIMAT0000122 | Rnf44 | 105239 | HITS-CLIP | 37986224\|23597149\|19536157\|23142 | Unspecified |
| mmu-let-7i-5p | MIMAT0000122 | Rnf5 | 54197 | HITS-CLIP | 37986224\|23142080 | Unspecified |
| mmu-let-7i-5p | MIMAT0000122 | Rnf6 | 74132 | HITS-CLIP | 23597149 | Unspecified |
| mmu-let-7i-5p | MIMAT0000122 | Rnf7 | 19823 | HITS-CLIP | 37986224\|23597149\|19536157\|23142 | Unspecified |
| mmu-let-7i-5p | MIMAT0000122 | Rnft1 | 76892 | HITS-CLIP | 37986224\|23597149\|23142080\|19536 | Unspecified |
| mmu-let-7i-5p | MIMAT0000122 | Rnh1 | 107702 | HITS-CLIP | 23142080\|29167373 | Unspecified |
| mmu-let-7i-5p | MIMAT0000122 | Rnmt | 67897 | HITS-CLIP | 23597149\|23142080 | Unspecified |
| mmu-let-7i-5p | MIMAT0000122 | Rnpepl1 | 108657 | HITS-CLIP | 23597149\|19536157\|31433990\|23142 | Unspecified |
| mmu-let-7i-5p | MIMAT0000122 | Robo2 | 268902 | HITS-CLIP | 19536157 | Unspecified |
| mmu-let-7i-5p | MIMAT0000122 | Rock2 | 19878 | HITS-CLIP | 23142080\|29167373 | Unspecified |
| mmu-let-7i-5p | MIMAT0000122 | Ror1 | 26563 | HITS-CLIP | 23597149 | Unspecified |
| mmu-let-7i-5p | MIMAT0000122 | Rora | 19883 | HITS-CLIP | 23597149\|23142080 | Unspecified |
| mmu-let-7i-5p | MIMAT0000122 | Rorc | 19885 | HITS-CLIP | 23597149 | Unspecified |
| mmu-let-7i-5p | MIMAT0000122 | Rp2 | 19889 | HITS-CLIP | 23142080\|29167373 | Unspecified |
| mmu-let-7i-5p | MIMAT0000122 | Rp9 | 55934 | HITS-CLIP | 23142080 | Unspecified |
| mmu-let-7i-5p | MIMAT0000122 | Rpa1 | 68275 | HITS-CLIP | 23597149\|23142080 | Unspecified |
| mmu-let-7i-5p | MIMAT0000122 | Rpa3 | 68240 | HITS-CLIP | 23142080 | Unspecified |
| mmu-let-7i-5p | MIMAT0000122 | Rpap1 | 68925 | HITS-CLIP | 23142080 | Unspecified |
| mmu-let-7i-5p | MIMAT0000122 | Rpap2 | 231571 | HITS-CLIP | 23142080 | Unspecified |
| mmu-let-7i-5p | MIMAT0000122 | Rpf1 | 70285 | HITS-CLIP | 37986224 | Unspecified |
| mmu-let-7i-5p | MIMAT0000122 | Rpl13a | 22121 | HITS-CLIP | 31433990 | Unspecified |
| mmu-let-7i-5p | MIMAT0000122 | Rpl22 | 19934 | HITS-CLIP | 23142080 | Unspecified |
| mmu-let-7i-5p | MIMAT0000122 | Rpl31 | 114641 | HITS-CLIP | 23142080 | Unspecified |
| mmu-let-7i-5p | MIMAT0000122 | Rpl32 | 19951 | HITS-CLIP | 23142080 | Unspecified |

| | | | | | | |
|---|---|---|---|---|---|---|
| mmu-let-7i-5p | MIMAT0000122 | Rpl4 | 67891 | qCLASH//HITS-CLIP | 31393953\|23597149 | Unspecified |
| mmu-let-7i-5p | MIMAT0000122 | Rpl7 | 19989 | HITS-CLIP | 23142080 | Unspecified |
| mmu-let-7i-5p | MIMAT0000122 | Rpl8 | 26961 | HITS-CLIP | 23142080 | Unspecified |
| mmu-let-7i-5p | MIMAT0000122 | Rpn2 | 20014 | HITS-CLIP | 37986224\|23142080 | Unspecified |
| mmu-let-7i-5p | MIMAT0000122 | Rpp21 | 67676 | HITS-CLIP | 23142080 | Unspecified |
| mmu-let-7i-5p | MIMAT0000122 | Rprd1a | 225283 | HITS-CLIP | 29167373 | Unspecified |
| mmu-let-7i-5p | MIMAT0000122 | Rprd1b | 70470 | HITS-CLIP | 23142080 | Unspecified |
| mmu-let-7i-5p | MIMAT0000122 | Rprd2 | 75137 | HITS-CLIP | 23142080\|29167373 | Unspecified |
| mmu-let-7i-5p | MIMAT0000122 | Rps11 | 27207 | HITS-CLIP | 37986224\|23142080\|29167373 | Unspecified |
| mmu-let-7i-5p | MIMAT0000122 | Rps15a | 6210 | HITS-CLIP | 31433990 | Unspecified |
| mmu-let-7i-5p | MIMAT0000122 | Rps19bp1 | 66538 | HITS-CLIP | 23142080 | Unspecified |
| mmu-let-7i-5p | MIMAT0000122 | Rps2 | 16898 | HITS-CLIP | 23597149\|23142080\|29167373 | Unspecified |
| mmu-let-7i-5p | MIMAT0000122 | Rps3 | 27050 | HITS-CLIP | 23597149\|23142080 | Unspecified |
| mmu-let-7i-5p | MIMAT0000122 | Rps6ka1 | 20111 | HITS-CLIP | 29167373 | Unspecified |
| mmu-let-7i-5p | MIMAT0000122 | Rps6ka3 | 110651 | HITS-CLIP | 37986224\|23597149\|23142080\|29167 | Unspecified |
| mmu-let-7i-5p | MIMAT0000122 | Rps6ka4 | 56613 | HITS-CLIP | 23142080 | Unspecified |
| mmu-let-7i-5p | MIMAT0000122 | Rps6ka5 | 73086 | HITS-CLIP | 23142080 | Unspecified |
| mmu-let-7i-5p | MIMAT0000122 | Rps6kb1 | 72508 | HITS-CLIP | 37986224\|23142080\|23597149 | Unspecified |
| mmu-let-7i-5p | MIMAT0000122 | Rps6kb2 | 58988 | HITS-CLIP | 37986224\|23597149\|23142080\|26203 | Unspecified |
| mmu-let-7i-5p | MIMAT0000122 | Rps6kc1 | 320119 | HITS-CLIP | 23597149\|23142080 | Unspecified |
| mmu-let-7i-5p | MIMAT0000122 | Rptor | 74370 | HITS-CLIP | 29167373 | Unspecified |
| mmu-let-7i-5p | MIMAT0000122 | Rpusd2 | 271842 | HITS-CLIP | 37986224\|23597149 | Unspecified |
| mmu-let-7i-5p | MIMAT0000122 | Rpusd3 | 101122 | HITS-CLIP | 23597149 | Unspecified |
| mmu-let-7i-5p | MIMAT0000122 | Rraga | 68441 | HITS-CLIP | 23142080 | Unspecified |
| mmu-let-7i-5p | MIMAT0000122 | Rragd | 52187 | HITS-CLIP | 23597149\|19536157\|23142080 | Unspecified |
| mmu-let-7i-5p | MIMAT0000122 | Rras | 20130 | HITS-CLIP | 37986224 | Unspecified |
| mmu-let-7i-5p | MIMAT0000122 | Rrbp1 | 81910 | HITS-CLIP | 23597149\|29167373 | Unspecified |
| mmu-let-7i-5p | MIMAT0000122 | Rreb1 | 68750 | HITS-CLIP | 23142080\|23597149 | Unspecified |
| mmu-let-7i-5p | MIMAT0000122 | Rrm1 | 20133 | qCLASH//HITS-CLIP | 31393953\|23142080 | Unspecified |
| mmu-let-7i-5p | MIMAT0000122 | Rrm2 | 20135 | HITS-CLIP | 23142080\|31433990 | Unspecified |
| mmu-let-7i-5p | MIMAT0000122 | Rrn3 | 106298 | HITS-CLIP | 23597149\|23142080 | Unspecified |
| mmu-let-7i-5p | MIMAT0000122 | Rrnad1 | 229503 | HITS-CLIP | 29167373 | Unspecified |
| mmu-let-7i-5p | MIMAT0000122 | Rrp1 | 18114 | HITS-CLIP | 31433990 | Unspecified |
| mmu-let-7i-5p | MIMAT0000122 | Rrp1b | 72462 | HITS-CLIP | 37986224\|23142080\|26203562 | Unspecified |
| mmu-let-7i-5p | MIMAT0000122 | Rrp36 | 224823 | HITS-CLIP | 23142080 | Unspecified |
| mmu-let-7i-5p | MIMAT0000122 | Rsad2 | 58185 | HITS-CLIP | 23597149\|29167373 | Unspecified |
| mmu-let-7i-5p | MIMAT0000122 | Rsbn1 | 229675 | HITS-CLIP | 37986224\|19536157\|23597149\|23142 | Unspecified |
| mmu-let-7i-5p | MIMAT0000122 | Rsbn1l | 242860 | HITS-CLIP | 37986224\|23142080\|19536157\|23597 | Unspecified |
| mmu-let-7i-5p | MIMAT0000122 | Rsf1 | 233532 | HITS-CLIP | 23597149\|29167373 | Unspecified |

| | | | | | | |
|---|---|---|---|---|---|---|
| mmu-let-7i-5p | MIMAT0000122 | Rsl1d1 | 66409 | HITS-CLIP | 37986224\|23142080 | Unspecified |
| mmu-let-7i-5p | MIMAT0000122 | Rslcan18 | 432770 | HITS-CLIP | 23142080 | Unspecified |
| mmu-let-7i-5p | MIMAT0000122 | Rspry1 | 67610 | HITS-CLIP | 23597149\|23142080\|29167373 | Unspecified |
| mmu-let-7i-5p | MIMAT0000122 | Rsrp1 | 27981 | HITS-CLIP | 23597149\|29167373 | Unspecified |
| mmu-let-7i-5p | MIMAT0000122 | Rsu1 | 20163 | HITS-CLIP | 19536157\|23142080 | Unspecified |
| mmu-let-7i-5p | MIMAT0000122 | Rtca | 66368 | HITS-CLIP | 23597149\|23142080 | Unspecified |
| mmu-let-7i-5p | MIMAT0000122 | Rtf1 | 76246 | HITS-CLIP | 29167373\|23142080 | Unspecified |
| mmu-let-7i-5p | MIMAT0000122 | Rtf2 | 51507 | HITS-CLIP | 23597149 | Unspecified |
| mmu-let-7i-5p | MIMAT0000122 | Rtl6 | 223732 | HITS-CLIP | 19536157 | Unspecified |
| mmu-let-7i-5p | MIMAT0000122 | Rtn1 | 104001 | HITS-CLIP | 19536157 | Unspecified |
| mmu-let-7i-5p | MIMAT0000122 | Rtn3 | 20168 | HITS-CLIP | 37986224\|19536157\|23597149\|23142 | Unspecified |
| mmu-let-7i-5p | MIMAT0000122 | Rtn4 | 68585 | HITS-CLIP | 19536157\|23142080\|29167373 | Unspecified |
| mmu-let-7i-5p | MIMAT0000122 | Rtn4rl1 | 237847 | HITS-CLIP | 37986224\|23597149 | Unspecified |
| mmu-let-7i-5p | MIMAT0000122 | Rtp3 | 235636 | HITS-CLIP | 37986224\|23597149 | Unspecified |
| mmu-let-7i-5p | MIMAT0000122 | Rtraf | 51637 | HITS-CLIP | 23142080 | Unspecified |
| mmu-let-7i-5p | MIMAT0000122 | Rubcn | 9711 | HITS-CLIP | 23597149 | Unspecified |
| mmu-let-7i-5p | MIMAT0000122 | Rufy1 | 216724 | HITS-CLIP | 23142080 | Unspecified |
| mmu-let-7i-5p | MIMAT0000122 | Rufy2 | 70432 | HITS-CLIP | 23142080\|29167373 | Unspecified |
| mmu-let-7i-5p | MIMAT0000122 | Rufy3 | 52822 | qCLASH//HITS-CLIP | 31393953\|29167373 | Unspecified |
| mmu-let-7i-5p | MIMAT0000122 | Rundc1 | 217201 | HITS-CLIP | 23142080 | Unspecified |
| mmu-let-7i-5p | MIMAT0000122 | Runx1 | 12394 | HITS-CLIP | 29167373 | Unspecified |
| mmu-let-7i-5p | MIMAT0000122 | Runx1t1 | 12395 | HITS-CLIP | 37986224 | Unspecified |
| mmu-let-7i-5p | MIMAT0000122 | Runx2 | 12393 | HITS-CLIP | 23142080 | Unspecified |
| mmu-let-7i-5p | MIMAT0000122 | Runx3 | 12399 | HITS-CLIP | 23142080 | Unspecified |
| mmu-let-7i-5p | MIMAT0000122 | Rusc2 | 100213 | HITS-CLIP | 23597149\|29167373 | Unspecified |
| mmu-let-7i-5p | MIMAT0000122 | Rwdd2b | 53858 | HITS-CLIP | 23597149 | Unspecified |
| mmu-let-7i-5p | MIMAT0000122 | Rwdd4a | 192174 | HITS-CLIP | 29167373 | Unspecified |
| mmu-let-7i-5p | MIMAT0000122 | Rxra | 20181 | HITS-CLIP | 37986224\|23597149 | Unspecified |
| mmu-let-7i-5p | MIMAT0000122 | Rxrb | 20182 | HITS-CLIP | 19536157\|23142080 | Unspecified |
| mmu-let-7i-5p | MIMAT0000122 | Ryk | 20187 | HITS-CLIP | 37986224 | Unspecified |
| mmu-let-7i-5p | MIMAT0000122 | Ryr2 | 20191 | HITS-CLIP | 19536157 | Unspecified |
| mmu-let-7i-5p | MIMAT0000122 | S100a1 | 20193 | HITS-CLIP | 23597149\|23142080 | Unspecified |
| mmu-let-7i-5p | MIMAT0000122 | S100a11 | 20195 | HITS-CLIP | 23142080 | Unspecified |
| mmu-let-7i-5p | MIMAT0000122 | S100a13 | 20196 | HITS-CLIP | 23597149\|23142080 | Unspecified |
| mmu-let-7i-5p | MIMAT0000122 | S100a4 | 20198 | HITS-CLIP | 29167373 | Unspecified |
| mmu-let-7i-5p | MIMAT0000122 | S100pbp | 74648 | HITS-CLIP | 37986224\|23597149\|19536157\|23142 | Unspecified |
| mmu-let-7i-5p | MIMAT0000122 | S1pr1 | 13609 | HITS-CLIP | 23597149\|23142080 | Unspecified |
| mmu-let-7i-5p | MIMAT0000122 | S1pr2 | 14739 | HITS-CLIP | 37986224\|23597149\|23142080\|29167 | Unspecified |
| mmu-let-7i-5p | MIMAT0000122 | Saa2 | 20209 | HITS-CLIP | 23597149 | Unspecified |

| | | | | | | |
|---|---|---|---|---|---|---|
| mmu-let-7i-5p | MIMAT0000122 | Sacm1l | 83493 | HITS-CLIP | 37986224\|19536157\|23142080 | Unspecified |
| mmu-let-7i-5p | MIMAT0000122 | Sacs | 50720 | HITS-CLIP | 37986224\|19536157\|23142080\|29167 | Unspecified |
| mmu-let-7i-5p | MIMAT0000122 | Sall1 | 58198 | HITS-CLIP | 37986224\|23597149 | Unspecified |
| mmu-let-7i-5p | MIMAT0000122 | Sall2 | 50524 | HITS-CLIP | 23597149\|23142080 | Unspecified |
| mmu-let-7i-5p | MIMAT0000122 | Samd4b | 233033 | HITS-CLIP | 23142080\|29167373\|19536157\|23597 | Unspecified |
| mmu-let-7i-5p | MIMAT0000122 | Samd8 | 67630 | HITS-CLIP | 23597149\|23142080 | Unspecified |
| mmu-let-7i-5p | MIMAT0000122 | Samhd1 | 56045 | HITS-CLIP | 23597149\|23142080\|29167373 | Unspecified |
| mmu-let-7i-5p | MIMAT0000122 | Samm50 | 68653 | HITS-CLIP | 23142080 | Unspecified |
| mmu-let-7i-5p | MIMAT0000122 | Samsn1 | 67742 | HITS-CLIP | 23142080 | Unspecified |
| mmu-let-7i-5p | MIMAT0000122 | Sap130 | 269003 | HITS-CLIP | 23142080 | Unspecified |
| mmu-let-7i-5p | MIMAT0000122 | Sap30 | 60406 | HITS-CLIP | 23597149\|23142080 | Unspecified |
| mmu-let-7i-5p | MIMAT0000122 | Sar1a | 20224 | HITS-CLIP | 31433990\|23142080 | Unspecified |
| mmu-let-7i-5p | MIMAT0000122 | Sar1b | 66397 | HITS-CLIP | 23597149\|23142080 | Unspecified |
| mmu-let-7i-5p | MIMAT0000122 | Sardh | 192166 | HITS-CLIP | 23597149 | Unspecified |
| mmu-let-7i-5p | MIMAT0000122 | Sarnp | 66118 | HITS-CLIP | 23142080 | Unspecified |
| mmu-let-7i-5p | MIMAT0000122 | Sars | 20226 | HITS-CLIP | 23597149\|23142080 | Unspecified |
| mmu-let-7i-5p | MIMAT0000122 | Sash1 | 70097 | HITS-CLIP | 29167373 | Unspecified |
| mmu-let-7i-5p | MIMAT0000122 | Sash3 | 74131 | HITS-CLIP | 23142080 | Unspecified |
| mmu-let-7i-5p | MIMAT0000122 | Sat1 | 20229 | HITS-CLIP | 23142080 | Unspecified |
| mmu-let-7i-5p | MIMAT0000122 | Satb1 | 20230 | HITS-CLIP | 23142080\|31433990 | Unspecified |
| mmu-let-7i-5p | MIMAT0000122 | Sav1 | 64010 | HITS-CLIP | 29167373 | Unspecified |
| mmu-let-7i-5p | MIMAT0000122 | Sbf1 | 77980 | HITS-CLIP | 23597149\|19536157\|23142080\|29167 | Unspecified |
| mmu-let-7i-5p | MIMAT0000122 | Sbf2 | 319934 | HITS-CLIP | 23142080\|29167373 | Unspecified |
| mmu-let-7i-5p | MIMAT0000122 | Sbk1 | 104175 | HITS-CLIP | 37986224\|19536157\|23597149 | Unspecified |
| mmu-let-7i-5p | MIMAT0000122 | Sbno1 | 243272 | HITS-CLIP | 23142080\|19536157\|29167373 | Unspecified |
| mmu-let-7i-5p | MIMAT0000122 | Sbno2 | 216161 | HITS-CLIP | 23142080\|23597149 | Unspecified |
| mmu-let-7i-5p | MIMAT0000122 | Sc5d | 235293 | HITS-CLIP | 23597149\|23142080 | Unspecified |
| mmu-let-7i-5p | MIMAT0000122 | Scaf1 | 233208 | HITS-CLIP | 23597149 | Unspecified |
| mmu-let-7i-5p | MIMAT0000122 | Scaf11 | 72193 | HITS-CLIP | 23142080\|29167373 | Unspecified |
| mmu-let-7i-5p | MIMAT0000122 | Scaf4 | 224432 | HITS-CLIP | 23597149\|31433990\|29167373 | Unspecified |
| mmu-let-7i-5p | MIMAT0000122 | Scaf8 | 106583 | HITS-CLIP | 23142080 | Unspecified |
| mmu-let-7i-5p | MIMAT0000122 | Scai | 320271 | HITS-CLIP | 23142080 | Unspecified |
| mmu-let-7i-5p | MIMAT0000122 | Scamp1 | 107767 | HITS-CLIP | 23597149 | Unspecified |
| mmu-let-7i-5p | MIMAT0000122 | Scamp2 | 24044 | HITS-CLIP | 23142080 | Unspecified |
| mmu-let-7i-5p | MIMAT0000122 | Scamp3 | 24045 | HITS-CLIP | 23597149 | Unspecified |
| mmu-let-7i-5p | MIMAT0000122 | Scamp5 | 56807 | HITS-CLIP | 37986224\|19536157 | Unspecified |
| mmu-let-7i-5p | MIMAT0000122 | Scap | 235623 | HITS-CLIP | 23597149\|23142080 | Unspecified |
| mmu-let-7i-5p | MIMAT0000122 | Scaper | 244891 | HITS-CLIP | 23142080 | Unspecified |
| mmu-let-7i-5p | MIMAT0000122 | Scara3 | 219151 | HITS-CLIP | 37986224 | Unspecified |

| | | | | | | |
|---|---|---|---|---|---|---|
| mmu-let-7i-5p | MIMAT0000122 | Scara5 | 71145 | HITS-CLIP | 23597149 | Unspecified |
| mmu-let-7i-5p | MIMAT0000122 | Scarb1 | 20778 | HITS-CLIP | 23597149 | Unspecified |
| mmu-let-7i-5p | MIMAT0000122 | Scarb2 | 12492 | HITS-CLIP | 23142080\|29167373 | Unspecified |
| mmu-let-7i-5p | MIMAT0000122 | Scd1 | 20249 | HITS-CLIP | 37986224\|23597149 | Unspecified |
| mmu-let-7i-5p | MIMAT0000122 | Scd2 | 20250 | HITS-CLIP | 37986224\|23142080\|19536157\|29167 | Unspecified |
| mmu-let-7i-5p | MIMAT0000122 | Scfd1 | 76983 | HITS-CLIP | 29167373\|23142080 | Unspecified |
| mmu-let-7i-5p | MIMAT0000122 | Scfd2 | 212986 | HITS-CLIP | 23142080\|29167373 | Unspecified |
| mmu-let-7i-5p | MIMAT0000122 | Scimp | 327957 | HITS-CLIP | 23597149 | Unspecified |
| mmu-let-7i-5p | MIMAT0000122 | Scmh1 | 29871 | HITS-CLIP | 31433990 | Unspecified |
| mmu-let-7i-5p | MIMAT0000122 | Scml4 | 268297 | HITS-CLIP | 23142080 | Unspecified |
| mmu-let-7i-5p | MIMAT0000122 | Scn2a | 110876 | HITS-CLIP | 19536157 | Unspecified |
| mmu-let-7i-5p | MIMAT0000122 | Scn3b | 235281 | HITS-CLIP | 37986224 | Unspecified |
| mmu-let-7i-5p | MIMAT0000122 | Scn8a | 20273 | HITS-CLIP | 19536157 | Unspecified |
| mmu-let-7i-5p | MIMAT0000122 | Scn9a | 6335 | HITS-CLIP | 23142080 | Unspecified |
| mmu-let-7i-5p | MIMAT0000122 | Scp2 | 20280 | HITS-CLIP | 23597149 | Unspecified |
| mmu-let-7i-5p | MIMAT0000122 | Scrib | 105782 | HITS-CLIP | 23142080 | Unspecified |
| mmu-let-7i-5p | MIMAT0000122 | Scrt1 | 170729 | HITS-CLIP | 23142080 | Unspecified |
| mmu-let-7i-5p | MIMAT0000122 | Scyl1 | 78891 | HITS-CLIP | 37986224\|23597149 | Unspecified |
| mmu-let-7i-5p | MIMAT0000122 | Scyl2 | 213326 | HITS-CLIP | 37986224\|23142080\|29167373 | Unspecified |
| mmu-let-7i-5p | MIMAT0000122 | Sdad1 | 231452 | HITS-CLIP | 23630078 | Unspecified |
| mmu-let-7i-5p | MIMAT0000122 | Sdc1 | 20969 | HITS-CLIP | 26203562 | Unspecified |
| mmu-let-7i-5p | MIMAT0000122 | Sdc2 | 15529 | HITS-CLIP | 23597149 | Unspecified |
| mmu-let-7i-5p | MIMAT0000122 | Sdc3 | 20970 | HITS-CLIP | 29167373 | Unspecified |
| mmu-let-7i-5p | MIMAT0000122 | Sdcbp | 53378 | HITS-CLIP | 29167373\|23142080 | Unspecified |
| mmu-let-7i-5p | MIMAT0000122 | Sdf2l1 | 64136 | HITS-CLIP | 23142080 | Unspecified |
| mmu-let-7i-5p | MIMAT0000122 | Sdf4 | 20318 | HITS-CLIP | 23142080 | Unspecified |
| mmu-let-7i-5p | MIMAT0000122 | Sdha | 66945 | HITS-CLIP | 37986224\|23597149\|23142080\|1953( | Unspecified |
| mmu-let-7i-5p | MIMAT0000122 | Sdhaf2 | 66072 | HITS-CLIP | 23142080\|23597149 | Unspecified |
| mmu-let-7i-5p | MIMAT0000122 | Sdhaf3 | 57001 | HITS-CLIP | 23597149\|23142080 | Unspecified |
| mmu-let-7i-5p | MIMAT0000122 | Sdhb | 67680 | HITS-CLIP | 23142080 | Unspecified |
| mmu-let-7i-5p | MIMAT0000122 | Sdhc | 66052 | HITS-CLIP | 23597149\|23142080 | Unspecified |
| mmu-let-7i-5p | MIMAT0000122 | Sdhd | 66925 | HITS-CLIP | 23142080 | Unspecified |
| mmu-let-7i-5p | MIMAT0000122 | Sdr39u1 | 654795 | HITS-CLIP | 23142080 | Unspecified |
| mmu-let-7i-5p | MIMAT0000122 | Sdr42e1 | 74032 | HITS-CLIP | 23597149 | Unspecified |
| mmu-let-7i-5p | MIMAT0000122 | Sec13 | 110379 | HITS-CLIP | 37986224\|23597149\|23142080\|29167 | Unspecified |
| mmu-let-7i-5p | MIMAT0000122 | Sec14l1 | 74136 | HITS-CLIP | 37986224 | Unspecified |
| mmu-let-7i-5p | MIMAT0000122 | Sec14l2 | 67815 | HITS-CLIP | 23597149 | Unspecified |
| mmu-let-7i-5p | MIMAT0000122 | Sec14l4 | 103655 | HITS-CLIP | 23597149 | Unspecified |
| mmu-let-7i-5p | MIMAT0000122 | Sec16a | 227648 | HITS-CLIP | 37986224\|23597149\|23142080\|29167 | Unspecified |

| | | | | | | |
|---|---|---|---|---|---|---|
| mmu-let-7i-5p | MIMAT0000122 | Sec16b | 89867 | HITS-CLIP | 37986224\|23597149 | Unspecified |
| mmu-let-7i-5p | MIMAT0000122 | Sec22b | 20333 | HITS-CLIP | 19536157 | Unspecified |
| mmu-let-7i-5p | MIMAT0000122 | Sec23a | 20334 | HITS-CLIP | 23597149\|23142080 | Unspecified |
| mmu-let-7i-5p | MIMAT0000122 | Sec23b | 27054 | HITS-CLIP | 23597149\|29167373 | Unspecified |
| mmu-let-7i-5p | MIMAT0000122 | Sec23ip | 207352 | HITS-CLIP | 23142080 | Unspecified |
| mmu-let-7i-5p | MIMAT0000122 | Sec24a | 77371 | HITS-CLIP | 31433990\|23142080 | Unspecified |
| mmu-let-7i-5p | MIMAT0000122 | Sec24b | 99683 | HITS-CLIP | 37986224\|23142080\|29167373 | Unspecified |
| mmu-let-7i-5p | MIMAT0000122 | Sec24c | 218811 | HITS-CLIP//qCLASH | 37986224\|23597149\|31393953\|23142 | Unspecified |
| mmu-let-7i-5p | MIMAT0000122 | Sec24d | 69608 | HITS-CLIP | 23142080 | Unspecified |
| mmu-let-7i-5p | MIMAT0000122 | Sec31a | 69162 | HITS-CLIP | 23597149 | Unspecified |
| mmu-let-7i-5p | MIMAT0000122 | Sec61a1 | 53421 | HITS-CLIP | 29167373 | Unspecified |
| mmu-let-7i-5p | MIMAT0000122 | Sec62 | 69276 | HITS-CLIP | 37986224\|23597149\|23142080 | Unspecified |
| mmu-let-7i-5p | MIMAT0000122 | Sec63 | 140740 | HITS-CLIP | 37986224\|23597149\|23142080 | Unspecified |
| mmu-let-7i-5p | MIMAT0000122 | Secisbp2 | 75420 | HITS-CLIP | 23142080 | Unspecified |
| mmu-let-7i-5p | MIMAT0000122 | Secisbp2l | 70354 | HITS-CLIP | 37986224\|19536157\|23597149\|31433 | Unspecified |
| mmu-let-7i-5p | MIMAT0000122 | Seh1l | 72124 | HITS-CLIP | 23142080 | Unspecified |
| mmu-let-7i-5p | MIMAT0000122 | Sel1l | 20338 | HITS-CLIP | 37986224\|23597149\|31433990\|23142 | Unspecified |
| mmu-let-7i-5p | MIMAT0000122 | Sel1l3 | 231238 | HITS-CLIP | 37986224\|23597149\|19536157 | Unspecified |
| mmu-let-7i-5p | MIMAT0000122 | Selenbp2 | 20342 | HITS-CLIP | 23597149 | Unspecified |
| mmu-let-7i-5p | MIMAT0000122 | Selenoi | 28042 | HITS-CLIP | 23142080\|19536157 | Unspecified |
| mmu-let-7i-5p | MIMAT0000122 | Selenop | 6414 | HITS-CLIP | 23142080\|19536157\|23597149\|29167 | Unspecified |
| mmu-let-7i-5p | MIMAT0000122 | Selenot | 69227 | HITS-CLIP | 23142080\|29167373 | Unspecified |
| mmu-let-7i-5p | MIMAT0000122 | Sell | 20343 | HITS-CLIP | 23142080 | Unspecified |
| mmu-let-7i-5p | MIMAT0000122 | Sema3f | 20350 | HITS-CLIP | 23597149 | Unspecified |
| mmu-let-7i-5p | MIMAT0000122 | Sema4a | 20351 | HITS-CLIP | 23597149 | Unspecified |
| mmu-let-7i-5p | MIMAT0000122 | Sema4b | 20352 | HITS-CLIP | 19536157\|29167373\|23142080 | Unspecified |
| mmu-let-7i-5p | MIMAT0000122 | Sema4c | 20353 | HITS-CLIP | 37986224\|23142080\|19536157\|23597 | Unspecified |
| mmu-let-7i-5p | MIMAT0000122 | Sema4d | 20354 | HITS-CLIP | 37986224\|23142080\|29167373 | Unspecified |
| mmu-let-7i-5p | MIMAT0000122 | Sema4f | 20355 | HITS-CLIP | 37986224\|23142080\|19536157 | Unspecified |
| mmu-let-7i-5p | MIMAT0000122 | Sema4g | 26456 | HITS-CLIP | 37986224\|23597149\|19536157 | Unspecified |
| mmu-let-7i-5p | MIMAT0000122 | Sema6a | 20358 | HITS-CLIP | 23597149 | Unspecified |
| mmu-let-7i-5p | MIMAT0000122 | Sema6d | 214968 | HITS-CLIP | 19536157 | Unspecified |
| mmu-let-7i-5p | MIMAT0000122 | Sema7a | 20361 | HITS-CLIP | 23142080 | Unspecified |
| mmu-let-7i-5p | MIMAT0000122 | Senp1 | 223870 | HITS-CLIP | 23142080\|29167373 | Unspecified |
| mmu-let-7i-5p | MIMAT0000122 | Senp2 | 75826 | HITS-CLIP | 19536157\|23142080\|29167373 | Unspecified |
| mmu-let-7i-5p | MIMAT0000122 | Senp3 | 80886 | HITS-CLIP | 23597149\|23142080 | Unspecified |
| mmu-let-7i-5p | MIMAT0000122 | Senp5 | 320213 | HITS-CLIP | 37986224\|23142080\|23597149 | Unspecified |
| mmu-let-7i-5p | MIMAT0000122 | Senp6 | 215351 | HITS-CLIP | 23142080 | Unspecified |
| mmu-let-7i-5p | MIMAT0000122 | Senp7 | 66315 | HITS-CLIP | 23142080 | Unspecified |

| | | | | | | |
|---|---|---|---|---|---|---|
| mmu-let-7i-5p | MIMAT0000122 | Sephs1 | 109079 | HITS-CLIP | 23597149\|23142080 | Unspecified |
| mmu-let-7i-5p | MIMAT0000122 | Sephs2 | 20768 | HITS-CLIP | 37986224\|23142080 | Unspecified |
| mmu-let-7i-5p | MIMAT0000122 | Sepsecs | 211006 | HITS-CLIP | 23597149 | Unspecified |
| mmu-let-7i-5p | MIMAT0000122 | Sept1 | 54204 | HITS-CLIP | 23142080 | Unspecified |
| mmu-let-7i-5p | MIMAT0000122 | Sept11 | 52398 | HITS-CLIP | 37986224\|19536157 | Unspecified |
| mmu-let-7i-5p | MIMAT0000122 | Sept2 | 4735 | HITS-CLIP | 23597149\|29167373\|23142080 | Unspecified |
| mmu-let-7i-5p | MIMAT0000122 | Sept5 | 18951 | HITS-CLIP | 37986224 | Unspecified |
| mmu-let-7i-5p | MIMAT0000122 | Sept6 | 23157 | HITS-CLIP | 23142080 | Unspecified |
| mmu-let-7i-5p | MIMAT0000122 | Sept7 | 235072 | HITS-CLIP | 23597149\|23142080 | Unspecified |
| mmu-let-7i-5p | MIMAT0000122 | Sept9 | 53860 | HITS-CLIP | 23142080 | Unspecified |
| mmu-let-7i-5p | MIMAT0000122 | Serf2 | 378702 | HITS-CLIP | 29167373 | Unspecified |
| mmu-let-7i-5p | MIMAT0000122 | Serinc1 | 56442 | HITS-CLIP | 23142080\|19536157 | Unspecified |
| mmu-let-7i-5p | MIMAT0000122 | Serinc3 | 26943 | HITS-CLIP | 37986224\|23597149\|23142080\|19536 | Unspecified |
| mmu-let-7i-5p | MIMAT0000122 | Serinc5 | 218442 | HITS-CLIP | 37986224\|19536157\|23597149\|23142 | Unspecified |
| mmu-let-7i-5p | MIMAT0000122 | Serp1 | 28146 | HITS-CLIP | 37986224\|23597149\|23142080\|29167 | Unspecified |
| mmu-let-7i-5p | MIMAT0000122 | Serpina10 | 217847 | HITS-CLIP | 23597149 | Unspecified |
| mmu-let-7i-5p | MIMAT0000122 | Serpina1a | 20700 | HITS-CLIP | 23597149 | Unspecified |
| mmu-let-7i-5p | MIMAT0000122 | Serpina1b | 20701 | HITS-CLIP | 23597149 | Unspecified |
| mmu-let-7i-5p | MIMAT0000122 | Serpina3n | 20716 | HITS-CLIP | 23597149 | Unspecified |
| mmu-let-7i-5p | MIMAT0000122 | Serpina6 | 12401 | HITS-CLIP | 23597149 | Unspecified |
| mmu-let-7i-5p | MIMAT0000122 | Serpinb6a | 20719 | HITS-CLIP | 23142080\|23597149 | Unspecified |
| mmu-let-7i-5p | MIMAT0000122 | Serpinb9 | 20723 | HITS-CLIP | 23142080 | Unspecified |
| mmu-let-7i-5p | MIMAT0000122 | Serpinc1 | 11905 | HITS-CLIP | 23597149 | Unspecified |
| mmu-let-7i-5p | MIMAT0000122 | Serpind1 | 15160 | HITS-CLIP | 23597149 | Unspecified |
| mmu-let-7i-5p | MIMAT0000122 | Serpine1 | 18787 | HITS-CLIP | 23142080 | Unspecified |
| mmu-let-7i-5p | MIMAT0000122 | Serpinf1 | 20317 | HITS-CLIP | 23142080 | Unspecified |
| mmu-let-7i-5p | MIMAT0000122 | Serpinf2 | 18816 | HITS-CLIP | 23597149 | Unspecified |
| mmu-let-7i-5p | MIMAT0000122 | Serping1 | 12258 | HITS-CLIP | 23597149 | Unspecified |
| mmu-let-7i-5p | MIMAT0000122 | Sertad1 | 55942 | HITS-CLIP | 37986224\|23142080 | Unspecified |
| mmu-let-7i-5p | MIMAT0000122 | Sertad2 | 58172 | HITS-CLIP | 23597149\|23142080 | Unspecified |
| mmu-let-7i-5p | MIMAT0000122 | Sertad3 | 170742 | HITS-CLIP | 23142080 | Unspecified |
| mmu-let-7i-5p | MIMAT0000122 | Sesn1 | 140742 | HITS-CLIP | 37986224\|23597149\|19536157\|23142 | Unspecified |
| mmu-let-7i-5p | MIMAT0000122 | Sesn3 | 75747 | HITS-CLIP | 23142080\|19536157\|23597149 | Unspecified |
| mmu-let-7i-5p | MIMAT0000122 | Set | 56086 | HITS-CLIP | 23142080 | Unspecified |
| mmu-let-7i-5p | MIMAT0000122 | Setbp1 | 240427 | HITS-CLIP | 23142080 | Unspecified |
| mmu-let-7i-5p | MIMAT0000122 | Setd1a | 233904 | HITS-CLIP | 23597149 | Unspecified |
| mmu-let-7i-5p | MIMAT0000122 | Setd1b | 208043 | HITS-CLIP | 23597149\|29167373 | Unspecified |
| mmu-let-7i-5p | MIMAT0000122 | Setd2 | 235626 | HITS-CLIP | 23142080\|23597149\|29167373 | Unspecified |
| mmu-let-7i-5p | MIMAT0000122 | Setd3 | 52690 | HITS-CLIP | 23142080 | Unspecified |

| | | | | | | |
|---|---|---|---|---|---|---|
| mmu-let-7i-5p | MIMAT0000122 | Setd4 | 224440 | HITS-CLIP | 23597149 | Unspecified |
| mmu-let-7i-5p | MIMAT0000122 | Setd5 | 72895 | HITS-CLIP | 23597149\|29167373\|23142080 | Unspecified |
| mmu-let-7i-5p | MIMAT0000122 | Setd7 | 73251 | HITS-CLIP | 37986224\|23142080 | Unspecified |
| mmu-let-7i-5p | MIMAT0000122 | Setdb1 | 84505 | HITS-CLIP//qCLASH | 37986224\|23597149\|23142080\|3139 | Unspecified |
| mmu-let-7i-5p | MIMAT0000122 | Setx | 269254 | HITS-CLIP | 23142080 | Unspecified |
| mmu-let-7i-5p | MIMAT0000122 | Sez6l | 56747 | HITS-CLIP | 19536157 | Unspecified |
| mmu-let-7i-5p | MIMAT0000122 | Sf1 | 22668 | HITS-CLIP | 23597149 | Unspecified |
| mmu-let-7i-5p | MIMAT0000122 | Sf3a1 | 67465 | HITS-CLIP | 23142080 | Unspecified |
| mmu-let-7i-5p | MIMAT0000122 | Sf3a3 | 75062 | HITS-CLIP//qCLASH | 23597149\|31393953\|23142080 | Unspecified |
| mmu-let-7i-5p | MIMAT0000122 | Sf3b1 | 81898 | HITS-CLIP | 37986224\|23597149\|23142080 | Unspecified |
| mmu-let-7i-5p | MIMAT0000122 | Sf3b2 | 319322 | HITS-CLIP | 23597149 | Unspecified |
| mmu-let-7i-5p | MIMAT0000122 | Sf3b3 | 101943 | HITS-CLIP | 23142080 | Unspecified |
| mmu-let-7i-5p | MIMAT0000122 | Sf3b4 | 107701 | HITS-CLIP | 26203562 | Unspecified |
| mmu-let-7i-5p | MIMAT0000122 | Sfmbt1 | 54650 | HITS-CLIP | 37986224\|19536157\|23597149\|2314 | Unspecified |
| mmu-let-7i-5p | MIMAT0000122 | Sfpq | 71514 | HITS-CLIP | 29167373\|23142080 | Unspecified |
| mmu-let-7i-5p | MIMAT0000122 | Sfswap | 231769 | HITS-CLIP | 23142080 | Unspecified |
| mmu-let-7i-5p | MIMAT0000122 | Sft2d3 | 67158 | HITS-CLIP | 37986224\|19536157\|23142080 | Unspecified |
| mmu-let-7i-5p | MIMAT0000122 | Sfxn5 | 94282 | HITS-CLIP | 23597149 | Unspecified |
| mmu-let-7i-5p | MIMAT0000122 | Sgcb | 24051 | HITS-CLIP | 37986224 | Unspecified |
| mmu-let-7i-5p | MIMAT0000122 | Sgip1 | 73094 | HITS-CLIP | 37986224 | Unspecified |
| mmu-let-7i-5p | MIMAT0000122 | Sgk1 | 20393 | HITS-CLIP | 23597149\|23630078\|31433990 | Unspecified |
| mmu-let-7i-5p | MIMAT0000122 | Sgk3 | 170755 | HITS-CLIP | 23142080\|19536157\|29167373 | Unspecified |
| mmu-let-7i-5p | MIMAT0000122 | Sgms1 | 208449 | HITS-CLIP | 23142080\|29167373 | Unspecified |
| mmu-let-7i-5p | MIMAT0000122 | Sgms2 | 74442 | HITS-CLIP | 23597149 | Unspecified |
| mmu-let-7i-5p | MIMAT0000122 | Sgo2a | 68549 | HITS-CLIP | 23142080 | Unspecified |
| mmu-let-7i-5p | MIMAT0000122 | Sgpl1 | 20397 | HITS-CLIP | 19536157\|23597149\|29167373 | Unspecified |
| mmu-let-7i-5p | MIMAT0000122 | Sgta | 52551 | HITS-CLIP | 23597149 | Unspecified |
| mmu-let-7i-5p | MIMAT0000122 | Sh2b1 | 20399 | HITS-CLIP | 23142080 | Unspecified |
| mmu-let-7i-5p | MIMAT0000122 | Sh2b3 | 16923 | HITS-CLIP | 37986224\|23597149\|23142080\|2916 | Unspecified |
| mmu-let-7i-5p | MIMAT0000122 | Sh2d2a | 27371 | HITS-CLIP | 23142080 | Unspecified |
| mmu-let-7i-5p | MIMAT0000122 | Sh2d3c | 27387 | HITS-CLIP | 23142080 | Unspecified |
| mmu-let-7i-5p | MIMAT0000122 | Sh3bgrl | 56726 | HITS-CLIP | 23142080 | Unspecified |
| mmu-let-7i-5p | MIMAT0000122 | Sh3bgrl3 | 73723 | HITS-CLIP | 23597149\|23142080 | Unspecified |
| mmu-let-7i-5p | MIMAT0000122 | Sh3bp2 | 24055 | HITS-CLIP | 37986224\|23597149\|23142080\|2916 | Unspecified |
| mmu-let-7i-5p | MIMAT0000122 | Sh3d19 | 27059 | HITS-CLIP | 23597149 | Unspecified |
| mmu-let-7i-5p | MIMAT0000122 | Sh3gl1 | 20405 | HITS-CLIP | 37986224\|23597149\|23142080\|2916 | Unspecified |
| mmu-let-7i-5p | MIMAT0000122 | Sh3gl3 | 20408 | HITS-CLIP | 19536157\|23142080 | Unspecified |
| mmu-let-7i-5p | MIMAT0000122 | Sh3glb1 | 54673 | HITS-CLIP | 23142080\|23597149\|19536157\|2916 | Unspecified |
| mmu-let-7i-5p | MIMAT0000122 | Sh3kbp1 | 58194 | HITS-CLIP | 23142080 | Unspecified |

| | | | | | | |
|---|---|---|---|---|---|---|
| mmu-let-7i-5p | MIMAT0000122 | Sh3pxd2a | 14218 | HITS-CLIP | 37986224\|19536157\|23597149 | Unspecified |
| mmu-let-7i-5p | MIMAT0000122 | Sh3pxd2b | 268396 | HITS-CLIP | 29167373 | Unspecified |
| mmu-let-7i-5p | MIMAT0000122 | Sh3rf1 | 59009 | HITS-CLIP | 37986224\|23142080 | Unspecified |
| mmu-let-7i-5p | MIMAT0000122 | Shank1 | 243961 | HITS-CLIP | 19536157 | Unspecified |
| mmu-let-7i-5p | MIMAT0000122 | Shank3 | 58234 | HITS-CLIP | 23597149 | Unspecified |
| mmu-let-7i-5p | MIMAT0000122 | Sharpin | 106025 | HITS-CLIP | 23597149 | Unspecified |
| mmu-let-7i-5p | MIMAT0000122 | Shb | 230126 | HITS-CLIP | 23597149 | Unspecified |
| mmu-let-7i-5p | MIMAT0000122 | Shc1 | 20416 | HITS-CLIP | 23142080 | Unspecified |
| mmu-let-7i-5p | MIMAT0000122 | Shc4 | 271849 | HITS-CLIP | 23142080 | Unspecified |
| mmu-let-7i-5p | MIMAT0000122 | Shcbp1 | 20419 | HITS-CLIP | 23142080 | Unspecified |
| mmu-let-7i-5p | MIMAT0000122 | Shisa5 | 66940 | HITS-CLIP | 37986224\|23597149\|23142080\|29167 | Unspecified |
| mmu-let-7i-5p | MIMAT0000122 | Shisa6 | 380702 | HITS-CLIP | 19536157 | Unspecified |
| mmu-let-7i-5p | MIMAT0000122 | Shkbp1 | 192192 | HITS-CLIP | 37986224 | Unspecified |
| mmu-let-7i-5p | MIMAT0000122 | Shld2 | 54537 | HITS-CLIP | 23142080\|23597149 | Unspecified |
| mmu-let-7i-5p | MIMAT0000122 | Shmt2 | 108037 | HITS-CLIP | 23142080 | Unspecified |
| mmu-let-7i-5p | MIMAT0000122 | Shoc2 | 56392 | HITS-CLIP | 23142080 | Unspecified |
| mmu-let-7i-5p | MIMAT0000122 | Shpk | 74637 | HITS-CLIP | 23597149 | Unspecified |
| mmu-let-7i-5p | MIMAT0000122 | Shprh | 268281 | HITS-CLIP | 23597149\|23142080 | Unspecified |
| mmu-let-7i-5p | MIMAT0000122 | Shq1 | 72171 | HITS-CLIP | 23142080 | Unspecified |
| mmu-let-7i-5p | MIMAT0000122 | Shroom1 | 71774 | HITS-CLIP | 23597149 | Unspecified |
| mmu-let-7i-5p | MIMAT0000122 | Shroom2 | 110380 | HITS-CLIP | 23597149 | Unspecified |
| mmu-let-7i-5p | MIMAT0000122 | Shroom3 | 27428 | HITS-CLIP | 23597149 | Unspecified |
| mmu-let-7i-5p | MIMAT0000122 | Shtn1 | 57698 | HITS-CLIP | 23597149\|29167373 | Unspecified |
| mmu-let-7i-5p | MIMAT0000122 | Siae | 22619 | HITS-CLIP | 23142080 | Unspecified |
| mmu-let-7i-5p | MIMAT0000122 | Sidt2 | 214597 | HITS-CLIP | 31433990\|23142080 | Unspecified |
| mmu-let-7i-5p | MIMAT0000122 | Sigirr | 24058 | HITS-CLIP | 23142080 | Unspecified |
| mmu-let-7i-5p | MIMAT0000122 | Siglec1 | 20612 | HITS-CLIP | 23597149\|29167373 | Unspecified |
| mmu-let-7i-5p | MIMAT0000122 | Sigmar1 | 18391 | HITS-CLIP | 37986224\|23597149\|19536157 | Unspecified |
| mmu-let-7i-5p | MIMAT0000122 | Sik1 | 17691 | HITS-CLIP | 37986224\|23597149\|23142080 | Unspecified |
| mmu-let-7i-5p | MIMAT0000122 | Sik2 | 235344 | HITS-CLIP | 23142080 | Unspecified |
| mmu-let-7i-5p | MIMAT0000122 | Sik3 | 70661 | HITS-CLIP//qCLASH | 23597149\|31393953\|23142080 | Unspecified |
| mmu-let-7i-5p | MIMAT0000122 | Sike1 | 66641 | HITS-CLIP | 23597149\|23142080 | Unspecified |
| mmu-let-7i-5p | MIMAT0000122 | Sil1 | 81500 | HITS-CLIP | 23142080 | Unspecified |
| mmu-let-7i-5p | MIMAT0000122 | Simc1 | 319719 | HITS-CLIP | 23142080\|29167373 | Unspecified |
| mmu-let-7i-5p | MIMAT0000122 | Sin3a | 20466 | HITS-CLIP | 23597149 | Unspecified |
| mmu-let-7i-5p | MIMAT0000122 | Sin3b | 20467 | HITS-CLIP | 37986224\|23597149 | Unspecified |
| mmu-let-7i-5p | MIMAT0000122 | Sipa1 | 20469 | HITS-CLIP | 23142080 | Unspecified |
| mmu-let-7i-5p | MIMAT0000122 | Sipa1l1 | 217692 | HITS-CLIP | 23142080 | Unspecified |
| mmu-let-7i-5p | MIMAT0000122 | Sipa1l3 | 74206 | HITS-CLIP | 23142080\|23597149 | Unspecified |

| | | | | | | |
|---|---|---|---|---|---|---|
| mmu-let-7i-5p | MIMAT0000122 | Sirpa | 19261 | HITS-CLIP | 29167373 | Unspecified |
| mmu-let-7i-5p | MIMAT0000122 | Sirt1 | 93759 | HITS-CLIP | 19536157\|29167373 | Unspecified |
| mmu-let-7i-5p | MIMAT0000122 | Sirt7 | 209011 | HITS-CLIP | 23142080 | Unspecified |
| mmu-let-7i-5p | MIMAT0000122 | Ska2 | 66140 | HITS-CLIP | 23142080 | Unspecified |
| mmu-let-7i-5p | MIMAT0000122 | Skap2 | 54353 | HITS-CLIP | 29167373 | Unspecified |
| mmu-let-7i-5p | MIMAT0000122 | Ski | 20481 | HITS-CLIP | 31433990\|23142080 | Unspecified |
| mmu-let-7i-5p | MIMAT0000122 | Skil | 20482 | HITS-CLIP | 37986224\|19536157\|23142080\|29167 | Unspecified |
| mmu-let-7i-5p | MIMAT0000122 | Skiv2l | 108077 | HITS-CLIP | 23597149 | Unspecified |
| mmu-let-7i-5p | MIMAT0000122 | Skp2 | 27401 | HITS-CLIP | 23142080\|29167373 | Unspecified |
| mmu-let-7i-5p | MIMAT0000122 | Sla | 20491 | HITS-CLIP | 23142080 | Unspecified |
| mmu-let-7i-5p | MIMAT0000122 | Sla2 | 77799 | HITS-CLIP | 23142080 | Unspecified |
| mmu-let-7i-5p | MIMAT0000122 | Slain1 | 105439 | HITS-CLIP | 37986224\|23142080 | Unspecified |
| mmu-let-7i-5p | MIMAT0000122 | Slain2 | 75991 | HITS-CLIP | 23597149 | Unspecified |
| mmu-let-7i-5p | MIMAT0000122 | Slamf1 | 27218 | HITS-CLIP | 37986224 | Unspecified |
| mmu-let-7i-5p | MIMAT0000122 | Slamf6 | 30925 | HITS-CLIP | 23142080 | Unspecified |
| mmu-let-7i-5p | MIMAT0000122 | Slamf7 | 75345 | HITS-CLIP | 23142080\|29167373 | Unspecified |
| mmu-let-7i-5p | MIMAT0000122 | Slc10a1 | 20493 | HITS-CLIP | 23597149 | Unspecified |
| mmu-let-7i-5p | MIMAT0000122 | Slc10a3 | 214601 | HITS-CLIP | 23597149\|23142080 | Unspecified |
| mmu-let-7i-5p | MIMAT0000122 | Slc10a6 | 75750 | HITS-CLIP | 37986224\|23597149 | Unspecified |
| mmu-let-7i-5p | MIMAT0000122 | Slc10a7 | 76775 | HITS-CLIP | 37986224\|23597149\|31433990\|23142 | Unspecified |
| mmu-let-7i-5p | MIMAT0000122 | Slc11a2 | 18174 | HITS-CLIP | 23142080\|29167373 | Unspecified |
| mmu-let-7i-5p | MIMAT0000122 | Slc12a2 | 20496 | HITS-CLIP | 19536157 | Unspecified |
| mmu-let-7i-5p | MIMAT0000122 | Slc12a4 | 20498 | HITS-CLIP | 23142080 | Unspecified |
| mmu-let-7i-5p | MIMAT0000122 | Slc12a6 | 107723 | HITS-CLIP | 29167373\|23142080 | Unspecified |
| mmu-let-7i-5p | MIMAT0000122 | Slc12a9 | 83704 | HITS-CLIP | 23142080 | Unspecified |
| mmu-let-7i-5p | MIMAT0000122 | Slc15a3 | 65221 | HITS-CLIP | 29167373 | Unspecified |
| mmu-let-7i-5p | MIMAT0000122 | Slc16a1 | 20501 | HITS-CLIP | 19536157 | Unspecified |
| mmu-let-7i-5p | MIMAT0000122 | Slc16a10 | 72472 | HITS-CLIP | 37986224\|23597149\|23142080\|29167 | Unspecified |
| mmu-let-7i-5p | MIMAT0000122 | Slc16a12 | 240638 | HITS-CLIP | 37986224\|23597149 | Unspecified |
| mmu-let-7i-5p | MIMAT0000122 | Slc16a2 | 20502 | HITS-CLIP | 37986224\|23597149 | Unspecified |
| mmu-let-7i-5p | MIMAT0000122 | Slc16a3 | 80879 | HITS-CLIP | 37986224\|23142080\|29167373 | Unspecified |
| mmu-let-7i-5p | MIMAT0000122 | Slc16a5 | 217316 | HITS-CLIP | 37986224\|23597149 | Unspecified |
| mmu-let-7i-5p | MIMAT0000122 | Slc16a6 | 104681 | HITS-CLIP | 37986224\|23597149\|23142080 | Unspecified |
| mmu-let-7i-5p | MIMAT0000122 | Slc16a7 | 20503 | HITS-CLIP | 23597149 | Unspecified |
| mmu-let-7i-5p | MIMAT0000122 | Slc17a3 | 105355 | HITS-CLIP | 23597149 | Unspecified |
| mmu-let-7i-5p | MIMAT0000122 | Slc17a4 | 319848 | HITS-CLIP | 23597149 | Unspecified |
| mmu-let-7i-5p | MIMAT0000122 | Slc17a6 | 140919 | HITS-CLIP | 23142080 | Unspecified |
| mmu-let-7i-5p | MIMAT0000122 | Slc17a7 | 72961 | HITS-CLIP | 19536157 | Unspecified |
| mmu-let-7i-5p | MIMAT0000122 | Slc18b1 | 76306 | HITS-CLIP | 23597149 | Unspecified |

| | | | | | | |
|---|---|---|---|---|---|---|
| mmu-let-7i-5p | MIMAT0000122 | Slc19a1 | 20509 | HITS-CLIP | 23142080 | Unspecified |
| mmu-let-7i-5p | MIMAT0000122 | Slc19a2 | 116914 | HITS-CLIP | 37986224\|23597149\|23142080 | Unspecified |
| mmu-let-7i-5p | MIMAT0000122 | Slc1a2 | 20511 | HITS-CLIP | 37986224\|23597149\|19536157 | Unspecified |
| mmu-let-7i-5p | MIMAT0000122 | Slc20a1 | 20515 | HITS-CLIP | 37986224\|19536157\|29167373\|23142 | Unspecified |
| mmu-let-7i-5p | MIMAT0000122 | Slc20a2 | 20516 | HITS-CLIP | 23142080 | Unspecified |
| mmu-let-7i-5p | MIMAT0000122 | Slc22a1 | 20517 | HITS-CLIP | 23597149 | Unspecified |
| mmu-let-7i-5p | MIMAT0000122 | Slc22a23 | 73102 | HITS-CLIP | 37986224\|23597149\|19536157\|26203 | Unspecified |
| mmu-let-7i-5p | MIMAT0000122 | Slc22a7 | 108114 | HITS-CLIP | 23597149 | Unspecified |
| mmu-let-7i-5p | MIMAT0000122 | Slc23a1 | 20522 | HITS-CLIP | 23597149 | Unspecified |
| mmu-let-7i-5p | MIMAT0000122 | Slc23a2 | 54338 | HITS-CLIP | 37986224\|23597149\|29167373 | Unspecified |
| mmu-let-7i-5p | MIMAT0000122 | Slc25a1 | 13358 | HITS-CLIP | 23597149 | Unspecified |
| mmu-let-7i-5p | MIMAT0000122 | Slc25a11 | 67863 | HITS-CLIP | 23142080 | Unspecified |
| mmu-let-7i-5p | MIMAT0000122 | Slc25a12 | 78830 | HITS-CLIP | 37986224\|23142080\|19536157\|23597 | Unspecified |
| mmu-let-7i-5p | MIMAT0000122 | Slc25a13 | 50799 | HITS-CLIP | 37986224\|23142080 | Unspecified |
| mmu-let-7i-5p | MIMAT0000122 | Slc25a15 | 18408 | HITS-CLIP | 23597149 | Unspecified |
| mmu-let-7i-5p | MIMAT0000122 | Slc25a16 | 73132 | HITS-CLIP | 23597149 | Unspecified |
| mmu-let-7i-5p | MIMAT0000122 | Slc25a17 | 20524 | HITS-CLIP | 23597149\|23142080\|29167373 | Unspecified |
| mmu-let-7i-5p | MIMAT0000122 | Slc25a18 | 71803 | HITS-CLIP | 19536157 | Unspecified |
| mmu-let-7i-5p | MIMAT0000122 | Slc25a22 | 68267 | HITS-CLIP | 23597149 | Unspecified |
| mmu-let-7i-5p | MIMAT0000122 | Slc25a23 | 66972 | HITS-CLIP | 37986224\|23142080 | Unspecified |
| mmu-let-7i-5p | MIMAT0000122 | Slc25a24 | 229731 | HITS-CLIP | 23142080 | Unspecified |
| mmu-let-7i-5p | MIMAT0000122 | Slc25a25 | 227731 | HITS-CLIP | 37986224\|23597149\|23142080\|29167 | Unspecified |
| mmu-let-7i-5p | MIMAT0000122 | Slc25a3 | 18674 | HITS-CLIP | 23597149\|23142080 | Unspecified |
| mmu-let-7i-5p | MIMAT0000122 | Slc25a36 | 192287 | HITS-CLIP | 23142080 | Unspecified |
| mmu-let-7i-5p | MIMAT0000122 | Slc25a39 | 68066 | HITS-CLIP | 37986224\|23597149\|23142080 | Unspecified |
| mmu-let-7i-5p | MIMAT0000122 | Slc25a4 | 11739 | HITS-CLIP | 37986224\|29167373\|23142080 | Unspecified |
| mmu-let-7i-5p | MIMAT0000122 | Slc25a40 | 319653 | HITS-CLIP | 23142080 | Unspecified |
| mmu-let-7i-5p | MIMAT0000122 | Slc25a42 | 73095 | HITS-CLIP | 23597149 | Unspecified |
| mmu-let-7i-5p | MIMAT0000122 | Slc25a43 | 194744 | HITS-CLIP | 23597149 | Unspecified |
| mmu-let-7i-5p | MIMAT0000122 | Slc25a44 | 229517 | HITS-CLIP | 29167373 | Unspecified |
| mmu-let-7i-5p | MIMAT0000122 | Slc25a45 | 107375 | HITS-CLIP | 37986224\|23597149\|23142080\|29167 | Unspecified |
| mmu-let-7i-5p | MIMAT0000122 | Slc25a46 | 67453 | HITS-CLIP | 23142080 | Unspecified |
| mmu-let-7i-5p | MIMAT0000122 | Slc26a1 | 231583 | HITS-CLIP | 37986224\|23597149 | Unspecified |
| mmu-let-7i-5p | MIMAT0000122 | Slc26a11 | 268512 | HITS-CLIP | 37986224 | Unspecified |
| mmu-let-7i-5p | MIMAT0000122 | Slc26a2 | 13521 | HITS-CLIP | 23142080\|23597149\|19536157\|29167 | Unspecified |
| mmu-let-7i-5p | MIMAT0000122 | Slc26a6 | 65010 | HITS-CLIP | 23597149\|29167373 | Unspecified |
| mmu-let-7i-5p | MIMAT0000122 | Slc27a2 | 26458 | HITS-CLIP | 23597149 | Unspecified |
| mmu-let-7i-5p | MIMAT0000122 | Slc27a4 | 26569 | HITS-CLIP | 37986224 | Unspecified |
| mmu-let-7i-5p | MIMAT0000122 | Slc27a5 | 26459 | HITS-CLIP | 23597149 | Unspecified |

| | | | | | | |
|---|---|---|---|---|---|---|
| mmu-let-7i-5p | MIMAT0000122 | Slc29a1 | 63959 | HITS-CLIP | 37986224\|23142080 | Unspecified |
| mmu-let-7i-5p | MIMAT0000122 | Slc29a3 | 71279 | HITS-CLIP | 23142080 | Unspecified |
| mmu-let-7i-5p | MIMAT0000122 | Slc2a1 | 20525 | HITS-CLIP | 23142080 | Unspecified |
| mmu-let-7i-5p | MIMAT0000122 | Slc2a3 | 20527 | HITS-CLIP | 37986224\|23142080 | Unspecified |
| mmu-let-7i-5p | MIMAT0000122 | Slc2a5 | 56485 | HITS-CLIP | 37986224\|23597149 | Unspecified |
| mmu-let-7i-5p | MIMAT0000122 | Slc2a6 | 227659 | HITS-CLIP | 23142080 | Unspecified |
| mmu-let-7i-5p | MIMAT0000122 | Slc30a1 | 22782 | HITS-CLIP | 37986224\|19536157\|23597149\|23142 | Unspecified |
| mmu-let-7i-5p | MIMAT0000122 | Slc30a4 | 22785 | HITS-CLIP | 37986224\|23597149\|23142080 | Unspecified |
| mmu-let-7i-5p | MIMAT0000122 | Slc30a5 | 69048 | HITS-CLIP | 23142080 | Unspecified |
| mmu-let-7i-5p | MIMAT0000122 | Slc30a6 | 210148 | HITS-CLIP | 23597149\|23142080 | Unspecified |
| mmu-let-7i-5p | MIMAT0000122 | Slc30a7 | 66500 | HITS-CLIP | 23142080 | Unspecified |
| mmu-let-7i-5p | MIMAT0000122 | Slc30a9 | 109108 | HITS-CLIP | 29167373 | Unspecified |
| mmu-let-7i-5p | MIMAT0000122 | Slc31a1 | 20529 | HITS-CLIP | 37986224\|19536157\|23597149\|23142 | Unspecified |
| mmu-let-7i-5p | MIMAT0000122 | Slc31a2 | 20530 | HITS-CLIP | 37986224 | Unspecified |
| mmu-let-7i-5p | MIMAT0000122 | Slc35a1 | 24060 | HITS-CLIP | 23142080 | Unspecified |
| mmu-let-7i-5p | MIMAT0000122 | Slc35a3 | 229782 | HITS-CLIP | 29167373\|23142080 | Unspecified |
| mmu-let-7i-5p | MIMAT0000122 | Slc35a4 | 67843 | HITS-CLIP | 37986224\|23142080\|23597149\|19536 | Unspecified |
| mmu-let-7i-5p | MIMAT0000122 | Slc35a5 | 74102 | HITS-CLIP | 23597149\|23142080\|19536157 | Unspecified |
| mmu-let-7i-5p | MIMAT0000122 | Slc35b2 | 73836 | HITS-CLIP | 37986224\|23597149 | Unspecified |
| mmu-let-7i-5p | MIMAT0000122 | Slc35b4 | 58246 | HITS-CLIP | 37986224\|23597149\|23142080\|2916 | Unspecified |
| mmu-let-7i-5p | MIMAT0000122 | Slc35c1 | 228368 | HITS-CLIP | 23597149\|23142080 | Unspecified |
| mmu-let-7i-5p | MIMAT0000122 | Slc35d1 | 242585 | HITS-CLIP | 23597149\|23142080 | Unspecified |
| mmu-let-7i-5p | MIMAT0000122 | Slc35d2 | 70484 | HITS-CLIP | 23597149\|23142080\|29167373 | Unspecified |
| mmu-let-7i-5p | MIMAT0000122 | Slc35e1 | 270066 | HITS-CLIP | 23142080\|29167373 | Unspecified |
| mmu-let-7i-5p | MIMAT0000122 | Slc35e2 | 320541 | HITS-CLIP | 23597149 | Unspecified |
| mmu-let-7i-5p | MIMAT0000122 | Slc35f6 | 74919 | HITS-CLIP | 23142080 | Unspecified |
| mmu-let-7i-5p | MIMAT0000122 | Slc35g1 | 240660 | HITS-CLIP | 23597149\|23142080 | Unspecified |
| mmu-let-7i-5p | MIMAT0000122 | Slc36a1 | 215335 | HITS-CLIP | 23597149 | Unspecified |
| mmu-let-7i-5p | MIMAT0000122 | Slc37a2 | 56857 | HITS-CLIP | 29167373 | Unspecified |
| mmu-let-7i-5p | MIMAT0000122 | Slc37a4 | 14385 | HITS-CLIP | 37986224\|23142080\|23597149 | Unspecified |
| mmu-let-7i-5p | MIMAT0000122 | Slc38a1 | 105727 | HITS-CLIP | 23142080 | Unspecified |
| mmu-let-7i-5p | MIMAT0000122 | Slc38a2 | 67760 | HITS-CLIP | 37986224\|23597149\|23142080 | Unspecified |
| mmu-let-7i-5p | MIMAT0000122 | Slc38a3 | 76257 | HITS-CLIP | 37986224\|23597149 | Unspecified |
| mmu-let-7i-5p | MIMAT0000122 | Slc38a4 | 69354 | HITS-CLIP | 23597149 | Unspecified |
| mmu-let-7i-5p | MIMAT0000122 | Slc38a6 | 625098 | HITS-CLIP | 23142080 | Unspecified |
| mmu-let-7i-5p | MIMAT0000122 | Slc38a9 | 268706 | HITS-CLIP | 19536157\|23142080\|29167373 | Unspecified |
| mmu-let-7i-5p | MIMAT0000122 | Slc39a1 | 30791 | HITS-CLIP | 37986224 | Unspecified |
| mmu-let-7i-5p | MIMAT0000122 | Slc39a10 | 227059 | HITS-CLIP | 23142080 | Unspecified |
| mmu-let-7i-5p | MIMAT0000122 | Slc39a14 | 213053 | HITS-CLIP | 29167373 | Unspecified |

| | | | | | | |
|---|---|---|---|---|---|---|
| mmu-let-7i-5p | MIMAT0000122 | Slc39a4 | 72027 | HITS-CLIP | 23142080 | Unspecified |
| mmu-let-7i-5p | MIMAT0000122 | Slc39a6 | 106957 | HITS-CLIP | 23142080 | Unspecified |
| mmu-let-7i-5p | MIMAT0000122 | Slc39a7 | 14977 | HITS-CLIP | 23142080 | Unspecified |
| mmu-let-7i-5p | MIMAT0000122 | Slc39a9 | 328133 | HITS-CLIP | 23142080 | Unspecified |
| mmu-let-7i-5p | MIMAT0000122 | Slc3a1 | 20532 | HITS-CLIP | 23597149 | Unspecified |
| mmu-let-7i-5p | MIMAT0000122 | Slc3a2 | 17254 | HITS-CLIP | 23142080\|29167373 | Unspecified |
| mmu-let-7i-5p | MIMAT0000122 | Slc40a1 | 53945 | HITS-CLIP | 37986224\|23597149 | Unspecified |
| mmu-let-7i-5p | MIMAT0000122 | Slc41a1 | 98396 | HITS-CLIP | 23142080 | Unspecified |
| mmu-let-7i-5p | MIMAT0000122 | Slc41a2 | 338365 | HITS-CLIP | 37986224\|23597149\|29167373\|23142 | Unspecified |
| mmu-let-7i-5p | MIMAT0000122 | Slc43a2 | 215113 | HITS-CLIP | 37986224\|23142080\|23597149\|19536 | Unspecified |
| mmu-let-7i-5p | MIMAT0000122 | Slc43a3 | 58207 | HITS-CLIP | 23142080 | Unspecified |
| mmu-let-7i-5p | MIMAT0000122 | Slc44a1 | 100434 | HITS-CLIP | 23597149\|23142080\|29167373 | Unspecified |
| mmu-let-7i-5p | MIMAT0000122 | Slc45a3 | 212980 | HITS-CLIP | 23597149 | Unspecified |
| mmu-let-7i-5p | MIMAT0000122 | Slc46a1 | 52466 | HITS-CLIP | 23597149 | Unspecified |
| mmu-let-7i-5p | MIMAT0000122 | Slc48a1 | 67739 | HITS-CLIP | 37986224\|23597149\|23142080\|29167 | Unspecified |
| mmu-let-7i-5p | MIMAT0000122 | Slc4a10 | 94229 | HITS-CLIP | 19536157 | Unspecified |
| mmu-let-7i-5p | MIMAT0000122 | Slc4a2 | 20535 | HITS-CLIP | 23597149 | Unspecified |
| mmu-let-7i-5p | MIMAT0000122 | Slc4a4 | 54403 | HITS-CLIP | 37986224\|23142080\|23597149\|19536 | Unspecified |
| mmu-let-7i-5p | MIMAT0000122 | Slc4a7 | 218756 | HITS-CLIP | 37986224\|19536157\|23597149\|23142 | Unspecified |
| mmu-let-7i-5p | MIMAT0000122 | Slc50a1 | 19729 | HITS-CLIP | 23142080 | Unspecified |
| mmu-let-7i-5p | MIMAT0000122 | Slc5a3 | 53881 | HITS-CLIP | 23142080\|29167373 | Unspecified |
| mmu-let-7i-5p | MIMAT0000122 | Slc6a1 | 232333 | HITS-CLIP | 19536157 | Unspecified |
| mmu-let-7i-5p | MIMAT0000122 | Slc6a11 | 243616 | HITS-CLIP | 37986224\|19536157 | Unspecified |
| mmu-let-7i-5p | MIMAT0000122 | Slc6a13 | 14412 | HITS-CLIP | 23597149 | Unspecified |
| mmu-let-7i-5p | MIMAT0000122 | Slc6a17 | 229706 | HITS-CLIP | 19536157 | Unspecified |
| mmu-let-7i-5p | MIMAT0000122 | Slc6a6 | 21366 | HITS-CLIP | 37986224\|23597149\|19536157\|29167 | Unspecified |
| mmu-let-7i-5p | MIMAT0000122 | Slc6a8 | 102857 | HITS-CLIP | 29167373 | Unspecified |
| mmu-let-7i-5p | MIMAT0000122 | Slc6a9 | 14664 | HITS-CLIP | 23597149 | Unspecified |
| mmu-let-7i-5p | MIMAT0000122 | Slc7a1 | 11987 | HITS-CLIP | 23142080 | Unspecified |
| mmu-let-7i-5p | MIMAT0000122 | Slc7a10 | 53896 | HITS-CLIP | 37986224\|19536157 | Unspecified |
| mmu-let-7i-5p | MIMAT0000122 | Slc7a2 | 11988 | HITS-CLIP | 37986224\|23597149 | Unspecified |
| mmu-let-7i-5p | MIMAT0000122 | Slc7a4 | 224022 | HITS-CLIP | 19536157 | Unspecified |
| mmu-let-7i-5p | MIMAT0000122 | Slc7a5 | 20539 | HITS-CLIP | 23142080 | Unspecified |
| mmu-let-7i-5p | MIMAT0000122 | Slc7a6 | 330836 | HITS-CLIP | 23142080 | Unspecified |
| mmu-let-7i-5p | MIMAT0000122 | Slc7a8 | 50934 | HITS-CLIP | 23597149\|29167373 | Unspecified |
| mmu-let-7i-5p | MIMAT0000122 | Slc8b1 | 170756 | HITS-CLIP | 37986224\|23597149\|29167373 | Unspecified |
| mmu-let-7i-5p | MIMAT0000122 | Slc9a1 | 20544 | HITS-CLIP | 23142080 | Unspecified |
| mmu-let-7i-5p | MIMAT0000122 | Slc9a2 | 226999 | HITS-CLIP | 19536157 | Unspecified |
| mmu-let-7i-5p | MIMAT0000122 | Slc9a3r1 | 26941 | HITS-CLIP//qCLASH | 37986224\|23597149\|31393953 | Unspecified |

| | | | | | | |
|---|---|---|---|---|---|---|
| mmu-let-7i-5p | MIMAT0000122 | Slc9a6 | 236794 | HITS-CLIP | 23142080 | Unspecified |
| mmu-let-7i-5p | MIMAT0000122 | Slc9a7 | 236727 | HITS-CLIP | 37986224 | Unspecified |
| mmu-let-7i-5p | MIMAT0000122 | Slc9a9 | 331004 | HITS-CLIP | 37986224\|29167373 | Unspecified |
| mmu-let-7i-5p | MIMAT0000122 | Slco1a4 | 28250 | HITS-CLIP | 23597149 | Unspecified |
| mmu-let-7i-5p | MIMAT0000122 | Slco1c1 | 58807 | HITS-CLIP | 37986224 | Unspecified |
| mmu-let-7i-5p | MIMAT0000122 | Slco2a1 | 24059 | HITS-CLIP | 23597149 | Unspecified |
| mmu-let-7i-5p | MIMAT0000122 | Slco2b1 | 101488 | HITS-CLIP | 37986224\|23597149 | Unspecified |
| mmu-let-7i-5p | MIMAT0000122 | Slco4a1 | 108115 | HITS-CLIP | 29167373\|23142080 | Unspecified |
| mmu-let-7i-5p | MIMAT0000122 | Slf1 | 852127 | HITS-CLIP | 23142080 | Unspecified |
| mmu-let-7i-5p | MIMAT0000122 | Slf2 | 226151 | HITS-CLIP | 23597149\|23142080\|29167373 | Unspecified |
| mmu-let-7i-5p | MIMAT0000122 | Slfn1 | 20555 | HITS-CLIP | 23142080 | Unspecified |
| mmu-let-7i-5p | MIMAT0000122 | Slfn5 | 327978 | HITS-CLIP | 29167373 | Unspecified |
| mmu-let-7i-5p | MIMAT0000122 | Slitrk1 | 76965 | HITS-CLIP | 37986224 | Unspecified |
| mmu-let-7i-5p | MIMAT0000122 | Slitrk3 | 386750 | HITS-CLIP | 19536157 | Unspecified |
| mmu-let-7i-5p | MIMAT0000122 | Slitrk4 | 245446 | HITS-CLIP | 19536157 | Unspecified |
| mmu-let-7i-5p | MIMAT0000122 | Slitrk5 | 75409 | HITS-CLIP | 37986224\|19536157 | Unspecified |
| mmu-let-7i-5p | MIMAT0000122 | Slk | 20874 | HITS-CLIP | 23597149\|23142080\|29167373 | Unspecified |
| mmu-let-7i-5p | MIMAT0000122 | Slmap | 83997 | HITS-CLIP | 23597149\|23142080\|29167373 | Unspecified |
| mmu-let-7i-5p | MIMAT0000122 | Sltm | 66660 | HITS-CLIP | 23142080 | Unspecified |
| mmu-let-7i-5p | MIMAT0000122 | Slu7 | 193116 | HITS-CLIP | 23142080 | Unspecified |
| mmu-let-7i-5p | MIMAT0000122 | Slx1b | 75764 | HITS-CLIP | 23142080 | Unspecified |
| mmu-let-7i-5p | MIMAT0000122 | Smad1 | 17125 | HITS-CLIP | 23597149\|19536157 | Unspecified |
| mmu-let-7i-5p | MIMAT0000122 | Smad3 | 17127 | HITS-CLIP | 23142080 | Unspecified |
| mmu-let-7i-5p | MIMAT0000122 | Smad4 | 17128 | HITS-CLIP | 23597149\|23142080\|29167373 | Unspecified |
| mmu-let-7i-5p | MIMAT0000122 | Smad5 | 17129 | HITS-CLIP | 23597149\|23142080 | Unspecified |
| mmu-let-7i-5p | MIMAT0000122 | Smad6 | 17130 | HITS-CLIP | 23597149 | Unspecified |
| mmu-let-7i-5p | MIMAT0000122 | Smad7 | 17131 | HITS-CLIP | 23597149\|23142080 | Unspecified |
| mmu-let-7i-5p | MIMAT0000122 | Smad9 | 55994 | HITS-CLIP | 23597149 | Unspecified |
| mmu-let-7i-5p | MIMAT0000122 | Smap1 | 98366 | HITS-CLIP | 23597149\|23142080 | Unspecified |
| mmu-let-7i-5p | MIMAT0000122 | Smap2 | 69780 | HITS-CLIP | 23142080 | Unspecified |
| mmu-let-7i-5p | MIMAT0000122 | Smarca2 | 67155 | HITS-CLIP | 23597149 | Unspecified |
| mmu-let-7i-5p | MIMAT0000122 | Smarca4 | 20586 | HITS-CLIP | 37986224\|23597149\|23142080\|29167 | Unspecified |
| mmu-let-7i-5p | MIMAT0000122 | Smarcad1 | 13990 | HITS-CLIP | 37986224\|19536157\|23142080 | Unspecified |
| mmu-let-7i-5p | MIMAT0000122 | Smarcal1 | 54380 | HITS-CLIP | 23142080 | Unspecified |
| mmu-let-7i-5p | MIMAT0000122 | Smarcb1 | 20587 | HITS-CLIP | 23597149\|23142080 | Unspecified |
| mmu-let-7i-5p | MIMAT0000122 | Smarcc1 | 20588 | HITS-CLIP//qCLASH | 23142080\|31393953\|19536157\|23597 | Unspecified |
| mmu-let-7i-5p | MIMAT0000122 | Smarcc2 | 68094 | HITS-CLIP | 23597149\|23142080\|29167373 | Unspecified |
| mmu-let-7i-5p | MIMAT0000122 | Smarcd1 | 83797 | HITS-CLIP | 37986224\|23597149\|19536157\|23142 | Unspecified |
| mmu-let-7i-5p | MIMAT0000122 | Smarcd2 | 83796 | HITS-CLIP | 23597149\|23142080\|30971389\|1953( | Unspecified |

| | | | | | | | |
|---|---|---|---|---|---|---|---|
| mmu-let-7i-5p | MIMAT0000122 | Smarce1 | 57376 | HITS-CLIP | | 23142080 | Unspecified |
| mmu-let-7i-5p | MIMAT0000122 | Smc1a | 24061 | HITS-CLIP | 23597149\|23142080 | | Unspecified |
| mmu-let-7i-5p | MIMAT0000122 | Smc2 | 14211 | HITS-CLIP | | 23142080 | Unspecified |
| mmu-let-7i-5p | MIMAT0000122 | Smc4 | 70099 | HITS-CLIP | 37986224\|31433990\|23142080\|29167 | | Unspecified |
| mmu-let-7i-5p | MIMAT0000122 | Smc5 | 226026 | HITS-CLIP | 23142080\|29167373 | | Unspecified |
| mmu-let-7i-5p | MIMAT0000122 | Smc6 | 67241 | HITS-CLIP | | 23142080 | Unspecified |
| mmu-let-7i-5p | MIMAT0000122 | Smchd1 | 74355 | HITS-CLIP | 37986224\|23597149\|23142080\|29167 | | Unspecified |
| mmu-let-7i-5p | MIMAT0000122 | Smcr8 | 237782 | HITS-CLIP | 23597149\|23142080 | | Unspecified |
| mmu-let-7i-5p | MIMAT0000122 | Smg1 | 233789 | HITS-CLIP | 29167373\|23142080 | | Unspecified |
| mmu-let-7i-5p | MIMAT0000122 | Smg5 | 229512 | HITS-CLIP | | 23597149 | Unspecified |
| mmu-let-7i-5p | MIMAT0000122 | Smg7 | 226517 | HITS-CLIP | 37986224\|23597149\|29167373\|23142 | | Unspecified |
| mmu-let-7i-5p | MIMAT0000122 | Smg8 | 74133 | HITS-CLIP | | 23142080 | Unspecified |
| mmu-let-7i-5p | MIMAT0000122 | Smim13 | 108934 | HITS-CLIP | 37986224\|23142080\|19536157 | | Unspecified |
| mmu-let-7i-5p | MIMAT0000122 | Smim15 | 75616 | HITS-CLIP | | 37986224 | Unspecified |
| mmu-let-7i-5p | MIMAT0000122 | Smim8 | 66291 | HITS-CLIP | | 23142080 | Unspecified |
| mmu-let-7i-5p | MIMAT0000122 | Smpd1 | 20597 | HITS-CLIP | | 23142080 | Unspecified |
| mmu-let-7i-5p | MIMAT0000122 | Smpd2 | 20598 | HITS-CLIP | 23597149\|23142080 | | Unspecified |
| mmu-let-7i-5p | MIMAT0000122 | Smpd3 | 58994 | HITS-CLIP | 37986224\|19536157 | | Unspecified |
| mmu-let-7i-5p | MIMAT0000122 | Smpd4 | 77626 | HITS-CLIP | | 23142080 | Unspecified |
| mmu-let-7i-5p | MIMAT0000122 | Smpdl3a | 57319 | HITS-CLIP | 23597149\|23142080\|29167373 | | Unspecified |
| mmu-let-7i-5p | MIMAT0000122 | Smu1 | 74255 | HITS-CLIP | | 23142080 | Unspecified |
| mmu-let-7i-5p | MIMAT0000122 | Smug1 | 71726 | HITS-CLIP | 37986224\|23142080\|23597149\|31433 | | Unspecified |
| mmu-let-7i-5p | MIMAT0000122 | Smurf2 | 66313 | HITS-CLIP | 31433990\|23142080 | | Unspecified |
| mmu-let-7i-5p | MIMAT0000122 | Smyd2 | 226830 | HITS-CLIP | | 37986224 | Unspecified |
| mmu-let-7i-5p | MIMAT0000122 | Snai2 | 20583 | HITS-CLIP | | 23597149 | Unspecified |
| mmu-let-7i-5p | MIMAT0000122 | Snap23 | 20619 | HITS-CLIP | 37986224\|23142080 | | Unspecified |
| mmu-let-7i-5p | MIMAT0000122 | Snap29 | 67474 | HITS-CLIP | | 23142080 | Unspecified |
| mmu-let-7i-5p | MIMAT0000122 | Snap91 | 20616 | HITS-CLIP | | 19536157 | Unspecified |
| mmu-let-7i-5p | MIMAT0000122 | Snd1 | 56463 | HITS-CLIP | 23142080\|23597149\|29167373 | | Unspecified |
| mmu-let-7i-5p | MIMAT0000122 | Snrk | 20623 | HITS-CLIP | | 23597149 | Unspecified |
| mmu-let-7i-5p | MIMAT0000122 | Snrnp200 | 320632 | HITS-CLIP | 37986224\|23597149\|19536157\|23142 | | Unspecified |
| mmu-let-7i-5p | MIMAT0000122 | Snrnp70 | 20637 | HITS-CLIP | 23142080\|29167373 | | Unspecified |
| mmu-let-7i-5p | MIMAT0000122 | Snrpb | 20638 | HITS-CLIP | | 23142080 | Unspecified |
| mmu-let-7i-5p | MIMAT0000122 | Snrpd1 | 20641 | HITS-CLIP | 23142080\|29167373 | | Unspecified |
| mmu-let-7i-5p | MIMAT0000122 | Snupn | 66069 | HITS-CLIP | | 23142080 | Unspecified |
| mmu-let-7i-5p | MIMAT0000122 | Snx1 | 56440 | HITS-CLIP | | 23142080 | Unspecified |
| mmu-let-7i-5p | MIMAT0000122 | Snx10 | 71982 | HITS-CLIP | 23597149\|29167373 | | Unspecified |
| mmu-let-7i-5p | MIMAT0000122 | Snx12 | 55988 | HITS-CLIP | | 23142080 | Unspecified |
| mmu-let-7i-5p | MIMAT0000122 | Snx13 | 217463 | HITS-CLIP | | 23142080 | Unspecified |

| | | | | | | |
|---|---|---|---|---|---|---|
| mmu-let-7i-5p | MIMAT0000122 | Snx14 | 244962 | HITS-CLIP | 23597149\|19536157 | Unspecified |
| mmu-let-7i-5p | MIMAT0000122 | Snx15 | 69024 | HITS-CLIP | 23597149\|23142080 | Unspecified |
| mmu-let-7i-5p | MIMAT0000122 | Snx16 | 74718 | HITS-CLIP | 23142080 | Unspecified |
| mmu-let-7i-5p | MIMAT0000122 | Snx17 | 266781 | HITS-CLIP | 23597149\|23142080 | Unspecified |
| mmu-let-7i-5p | MIMAT0000122 | Snx18 | 170625 | HITS-CLIP | 37986224\|31433990\|23142080\|29167 | Unspecified |
| mmu-let-7i-5p | MIMAT0000122 | Snx19 | 102607 | HITS-CLIP | 37986224\|23597149 | Unspecified |
| mmu-let-7i-5p | MIMAT0000122 | Snx25 | 102141 | HITS-CLIP | 37986224\|23142080\|29167373 | Unspecified |
| mmu-let-7i-5p | MIMAT0000122 | Snx27 | 76742 | HITS-CLIP//qCLASH | 37986224\|31393953 | Unspecified |
| mmu-let-7i-5p | MIMAT0000122 | Snx29 | 74478 | HITS-CLIP | 29167373 | Unspecified |
| mmu-let-7i-5p | MIMAT0000122 | Snx30 | 209131 | HITS-CLIP | 23630078\|23142080\|29167373 | Unspecified |
| mmu-let-7i-5p | MIMAT0000122 | Snx33 | 235406 | HITS-CLIP | 23142080 | Unspecified |
| mmu-let-7i-5p | MIMAT0000122 | Snx4 | 69150 | HITS-CLIP | 23597149\|23142080\|19536157 | Unspecified |
| mmu-let-7i-5p | MIMAT0000122 | Snx5 | 69178 | HITS-CLIP | 37986224\|23597149\|19536157 | Unspecified |
| mmu-let-7i-5p | MIMAT0000122 | Snx9 | 66616 | HITS-CLIP | 23597149\|23142080 | Unspecified |
| mmu-let-7i-5p | MIMAT0000122 | Soat1 | 20652 | HITS-CLIP | 23142080\|26203562\|29167373 | Unspecified |
| mmu-let-7i-5p | MIMAT0000122 | Soat2 | 223920 | HITS-CLIP | 23597149 | Unspecified |
| mmu-let-7i-5p | MIMAT0000122 | Socs1 | 12703 | HITS-CLIP | 37986224\|23142080\|19536157\|23597 | Unspecified |
| mmu-let-7i-5p | MIMAT0000122 | Socs2 | 216233 | HITS-CLIP | 23142080 | Unspecified |
| mmu-let-7i-5p | MIMAT0000122 | Socs3 | 12702 | HITS-CLIP | 31433990 | Unspecified |
| mmu-let-7i-5p | MIMAT0000122 | Socs4 | 67296 | HITS-CLIP | 29167373 | Unspecified |
| mmu-let-7i-5p | MIMAT0000122 | Socs5 | 56468 | HITS-CLIP | 23142080 | Unspecified |
| mmu-let-7i-5p | MIMAT0000122 | Socs6 | 54607 | HITS-CLIP | 23142080 | Unspecified |
| mmu-let-7i-5p | MIMAT0000122 | Sod1 | 20655 | HITS-CLIP | 23597149\|23142080 | Unspecified |
| mmu-let-7i-5p | MIMAT0000122 | Sod2 | 20656 | HITS-CLIP | 23597149 | Unspecified |
| mmu-let-7i-5p | MIMAT0000122 | Son | 20658 | HITS-CLIP | 37986224\|19536157\|23597149\|23142 | Unspecified |
| mmu-let-7i-5p | MIMAT0000122 | Sorbs2 | 234214 | HITS-CLIP | 23597149\|19536157 | Unspecified |
| mmu-let-7i-5p | MIMAT0000122 | Sord | 20322 | HITS-CLIP | 37986224\|23597149 | Unspecified |
| mmu-let-7i-5p | MIMAT0000122 | Sorl1 | 20660 | HITS-CLIP | 19536157\|23142080 | Unspecified |
| mmu-let-7i-5p | MIMAT0000122 | Sort1 | 20661 | HITS-CLIP | 37986224\|23597149 | Unspecified |
| mmu-let-7i-5p | MIMAT0000122 | Sos1 | 20662 | HITS-CLIP | 37986224\|23597149\|23142080\|29167 | Unspecified |
| mmu-let-7i-5p | MIMAT0000122 | Sos2 | 20663 | HITS-CLIP | 23597149\|23142080 | Unspecified |
| mmu-let-7i-5p | MIMAT0000122 | Sowaha | 237761 | HITS-CLIP | 37986224\|19536157 | Unspecified |
| mmu-let-7i-5p | MIMAT0000122 | Sox11 | 20666 | HITS-CLIP | 37986224 | Unspecified |
| mmu-let-7i-5p | MIMAT0000122 | Sox5 | 20678 | HITS-CLIP | 37986224\|23597149\|19536157 | Unspecified |
| mmu-let-7i-5p | MIMAT0000122 | Sox6 | 20679 | HITS-CLIP | 30971389 | Unspecified |
| mmu-let-7i-5p | MIMAT0000122 | Sox7 | 20680 | HITS-CLIP | 37986224 | Unspecified |
| mmu-let-7i-5p | MIMAT0000122 | Sp1 | 20683 | HITS-CLIP//qCLASH | 37986224\|23597149\|31393953\|23142 | Unspecified |
| mmu-let-7i-5p | MIMAT0000122 | Sp100 | 20684 | HITS-CLIP | 23142080\|29167373 | Unspecified |
| mmu-let-7i-5p | MIMAT0000122 | Sp2 | 78912 | HITS-CLIP | 23142080 | Unspecified |

| | | | | | | |
|---|---|---|---|---|---|---|
| mmu-let-7i-5p | MIMAT0000122 | Sp3 | 20687 | HITS-CLIP | 37986224\|23597149\|23142080 | Unspecified |
| mmu-let-7i-5p | MIMAT0000122 | Sp4 | 20688 | HITS-CLIP | 23597149 | Unspecified |
| mmu-let-7i-5p | MIMAT0000122 | Spaca6 | 75202 | HITS-CLIP | 29167373 | Unspecified |
| mmu-let-7i-5p | MIMAT0000122 | Spag7 | 216873 | HITS-CLIP | 23142080 | Unspecified |
| mmu-let-7i-5p | MIMAT0000122 | Spag9 | 70834 | HITS-CLIP | 19536157\|23142080\|29167373 | Unspecified |
| mmu-let-7i-5p | MIMAT0000122 | Sparcl1 | 13602 | HITS-CLIP | 37986224\|19536157 | Unspecified |
| mmu-let-7i-5p | MIMAT0000122 | Spast | 50850 | HITS-CLIP//qCLASH | 23597149\|23142080\|31393953 | Unspecified |
| mmu-let-7i-5p | MIMAT0000122 | Spata13 | 219140 | HITS-CLIP | 37986224\|23142080 | Unspecified |
| mmu-let-7i-5p | MIMAT0000122 | Spata2 | 263876 | HITS-CLIP | 23142080 | Unspecified |
| mmu-let-7i-5p | MIMAT0000122 | Spata2l | 78779 | HITS-CLIP | 23597149 | Unspecified |
| mmu-let-7i-5p | MIMAT0000122 | Spata6 | 67946 | HITS-CLIP | 23142080 | Unspecified |
| mmu-let-7i-5p | MIMAT0000122 | Spc24 | 67629 | HITS-CLIP | 23142080 | Unspecified |
| mmu-let-7i-5p | MIMAT0000122 | Spcs3 | 76687 | HITS-CLIP | 37986224\|23597149\|31433990\|1953€ | Unspecified |
| mmu-let-7i-5p | MIMAT0000122 | Specc1 | 432572 | HITS-CLIP | 29167373 | Unspecified |
| mmu-let-7i-5p | MIMAT0000122 | Specc1l | 74392 | HITS-CLIP | 23597149 | Unspecified |
| mmu-let-7i-5p | MIMAT0000122 | Speg | 11790 | HITS-CLIP | 29167373 | Unspecified |
| mmu-let-7i-5p | MIMAT0000122 | Spen | 56381 | HITS-CLIP//qCLASH | 30971389\|31393953\|23597149\|23142 | Unspecified |
| mmu-let-7i-5p | MIMAT0000122 | Spg11 | 214585 | HITS-CLIP | 29167373 | Unspecified |
| mmu-let-7i-5p | MIMAT0000122 | Spg21 | 27965 | HITS-CLIP | 23142080 | Unspecified |
| mmu-let-7i-5p | MIMAT0000122 | Sphkap | 77629 | HITS-CLIP | 37986224 | Unspecified |
| mmu-let-7i-5p | MIMAT0000122 | Spin4 | 270624 | HITS-CLIP | 23142080 | Unspecified |
| mmu-let-7i-5p | MIMAT0000122 | Spindoc | 144097 | HITS-CLIP | 23142080 | Unspecified |
| mmu-let-7i-5p | MIMAT0000122 | Spint2 | 20733 | HITS-CLIP | 23597149\|23142080 | Unspecified |
| mmu-let-7i-5p | MIMAT0000122 | Spn | 20737 | HITS-CLIP | 23142080 | Unspecified |
| mmu-let-7i-5p | MIMAT0000122 | Spns1 | 73658 | HITS-CLIP | 37986224\|23597149\|23142080\|29167 | Unspecified |
| mmu-let-7i-5p | MIMAT0000122 | Spns2 | 216892 | HITS-CLIP | 23597149 | Unspecified |
| mmu-let-7i-5p | MIMAT0000122 | Spop | 20747 | HITS-CLIP | 29167373 | Unspecified |
| mmu-let-7i-5p | MIMAT0000122 | Sppl2a | 66552 | HITS-CLIP | 23597149\|23142080 | Unspecified |
| mmu-let-7i-5p | MIMAT0000122 | Sppl3 | 74585 | HITS-CLIP | 23142080 | Unspecified |
| mmu-let-7i-5p | MIMAT0000122 | Spred1 | 114715 | HITS-CLIP | 37986224\|23142080 | Unspecified |
| mmu-let-7i-5p | MIMAT0000122 | Spred2 | 114716 | HITS-CLIP | 37986224\|23142080 | Unspecified |
| mmu-let-7i-5p | MIMAT0000122 | Spry1 | 24063 | HITS-CLIP | 23142080 | Unspecified |
| mmu-let-7i-5p | MIMAT0000122 | Spry4 | 24066 | HITS-CLIP | 23597149 | Unspecified |
| mmu-let-7i-5p | MIMAT0000122 | Spryd3 | 223918 | HITS-CLIP | 23597149 | Unspecified |
| mmu-let-7i-5p | MIMAT0000122 | Spryd4 | 66701 | HITS-CLIP | 23597149 | Unspecified |
| mmu-let-7i-5p | MIMAT0000122 | Spsb3 | 90864 | HITS-CLIP | 23142080 | Unspecified |
| mmu-let-7i-5p | MIMAT0000122 | Sptan1 | 20740 | HITS-CLIP | 37986224\|19536157\|23597149 | Unspecified |
| mmu-let-7i-5p | MIMAT0000122 | Sptb | 20741 | HITS-CLIP | 19536157 | Unspecified |
| mmu-let-7i-5p | MIMAT0000122 | Sptbn1 | 20742 | HITS-CLIP | 37986224\|23597149\|19536157\|23142 | Unspecified |

| | | | | | | |
|---|---|---|---|---|---|---|
| mmu-let-7i-5p | MIMAT0000122 | Sptbn2 | 20743 | HITS-CLIP | 37986224\|23597149\|19536157 | Unspecified |
| mmu-let-7i-5p | MIMAT0000122 | Sptlc1 | 268656 | HITS-CLIP | 37986224\|23142080 | Unspecified |
| mmu-let-7i-5p | MIMAT0000122 | Sptlc2 | 20773 | HITS-CLIP | 23597149\|23142080\|19536157\|29167 | Unspecified |
| mmu-let-7i-5p | MIMAT0000122 | Sptssa | 104725 | HITS-CLIP | 23142080 | Unspecified |
| mmu-let-7i-5p | MIMAT0000122 | Spty2d1 | 101685 | HITS-CLIP | 37986224 | Unspecified |
| mmu-let-7i-5p | MIMAT0000122 | Sqle | 20775 | HITS-CLIP | 23597149\|23142080 | Unspecified |
| mmu-let-7i-5p | MIMAT0000122 | Sqor | 58472 | HITS-CLIP | 23597149\|23142080 | Unspecified |
| mmu-let-7i-5p | MIMAT0000122 | Sqstm1 | 18412 | HITS-CLIP | 23597149 | Unspecified |
| mmu-let-7i-5p | MIMAT0000122 | Sra1 | 24068 | HITS-CLIP | 37986224\|23142080 | Unspecified |
| mmu-let-7i-5p | MIMAT0000122 | Srcap | 100043597 | HITS-CLIP | 37986224\|23597149\|29167373 | Unspecified |
| mmu-let-7i-5p | MIMAT0000122 | Srd5a1 | 78925 | HITS-CLIP | 23597149 | Unspecified |
| mmu-let-7i-5p | MIMAT0000122 | Srd5a3 | 57357 | HITS-CLIP | 23597149 | Unspecified |
| mmu-let-7i-5p | MIMAT0000122 | Srebf1 | 20787 | HITS-CLIP | 23597149\|23142080\|19536157 | Unspecified |
| mmu-let-7i-5p | MIMAT0000122 | Srebf2 | 20788 | HITS-CLIP | 23597149\|23142080 | Unspecified |
| mmu-let-7i-5p | MIMAT0000122 | Srek1 | 218543 | HITS-CLIP | 37986224\|23142080\|23597149 | Unspecified |
| mmu-let-7i-5p | MIMAT0000122 | Srek1ip1 | 67288 | HITS-CLIP | 23142080\|29167373 | Unspecified |
| mmu-let-7i-5p | MIMAT0000122 | Srf | 20807 | HITS-CLIP | 37986224 | Unspecified |
| mmu-let-7i-5p | MIMAT0000122 | Srgap2 | 14270 | HITS-CLIP | 29167373 | Unspecified |
| mmu-let-7i-5p | MIMAT0000122 | Srgap3 | 259302 | HITS-CLIP | 19536157\|23142080 | Unspecified |
| mmu-let-7i-5p | MIMAT0000122 | Sri | 109552 | HITS-CLIP | 23597149\|23142080 | Unspecified |
| mmu-let-7i-5p | MIMAT0000122 | Srm | 20810 | HITS-CLIP | 37986224\|23597149 | Unspecified |
| mmu-let-7i-5p | MIMAT0000122 | Srp14 | 20813 | HITS-CLIP | 23142080 | Unspecified |
| mmu-let-7i-5p | MIMAT0000122 | Srp68 | 217337 | HITS-CLIP | 23142080 | Unspecified |
| mmu-let-7i-5p | MIMAT0000122 | Srp72 | 66661 | HITS-CLIP | 23597149\|23142080 | Unspecified |
| mmu-let-7i-5p | MIMAT0000122 | Srpk2 | 20817 | HITS-CLIP | 29167373 | Unspecified |
| mmu-let-7i-5p | MIMAT0000122 | Srpr | 67398 | HITS-CLIP | 23597149\|23142080 | Unspecified |
| mmu-let-7i-5p | MIMAT0000122 | Srprb | 20818 | HITS-CLIP | 23142080 | Unspecified |
| mmu-let-7i-5p | MIMAT0000122 | Srr | 27364 | HITS-CLIP | 37986224 | Unspecified |
| mmu-let-7i-5p | MIMAT0000122 | Srrm1 | 51796 | HITS-CLIP | 23142080\|29167373 | Unspecified |
| mmu-let-7i-5p | MIMAT0000122 | Srrm2 | 75956 | HITS-CLIP | 37986224\|23142080\|23597149\|29167 | Unspecified |
| mmu-let-7i-5p | MIMAT0000122 | Srrt | 83701 | HITS-CLIP | 23142080 | Unspecified |
| mmu-let-7i-5p | MIMAT0000122 | Srsf1 | 110809 | HITS-CLIP | 23142080\|23597149 | Unspecified |
| mmu-let-7i-5p | MIMAT0000122 | Srsf10 | 14105 | HITS-CLIP | 23597149 | Unspecified |
| mmu-let-7i-5p | MIMAT0000122 | Srsf2 | 20382 | HITS-CLIP | 23142080\|29167373 | Unspecified |
| mmu-let-7i-5p | MIMAT0000122 | Srsf6 | 67996 | HITS-CLIP | 23142080\|23597149\|29167373 | Unspecified |
| mmu-let-7i-5p | MIMAT0000122 | Srxn1 | 76650 | HITS-CLIP | 23597149 | Unspecified |
| mmu-let-7i-5p | MIMAT0000122 | Ss18 | 268996 | HITS-CLIP | 31433990\|30971389\|23142080 | Unspecified |
| mmu-let-7i-5p | MIMAT0000122 | Ss18l2 | 51188 | HITS-CLIP | 23142080 | Unspecified |
| mmu-let-7i-5p | MIMAT0000122 | Ssb | 20823 | HITS-CLIP | 23142080\|19536157\|29167373 | Unspecified |

| | | | | | | |
|---|---|---|---|---|---|---|
| mmu-let-7i-5p | MIMAT0000122 | Ssbp2 | 66970 | HITS-CLIP | 23142080 | Unspecified |
| mmu-let-7i-5p | MIMAT0000122 | Ssh1 | 231637 | HITS-CLIP | 23142080\|23597149\|19536157\|29167 | Unspecified |
| mmu-let-7i-5p | MIMAT0000122 | Ssh2 | 237860 | HITS-CLIP | 23597149\|23142080 | Unspecified |
| mmu-let-7i-5p | MIMAT0000122 | Ssr1 | 107513 | HITS-CLIP | 23597149\|23142080 | Unspecified |
| mmu-let-7i-5p | MIMAT0000122 | Ssr3 | 67437 | HITS-CLIP | 23597149 | Unspecified |
| mmu-let-7i-5p | MIMAT0000122 | Ssr4 | 20832 | HITS-CLIP | 23142080 | Unspecified |
| mmu-let-7i-5p | MIMAT0000122 | Ssrp1 | 20833 | qCLASH//HITS-CLIP | 31393953\|23142080 | Unspecified |
| mmu-let-7i-5p | MIMAT0000122 | Ssx2ip | 99167 | HITS-CLIP | 23597149 | Unspecified |
| mmu-let-7i-5p | MIMAT0000122 | St13 | 70356 | HITS-CLIP | 37986224\|23142080\|23597149\|19536 | Unspecified |
| mmu-let-7i-5p | MIMAT0000122 | St18 | 240690 | HITS-CLIP | 29167373 | Unspecified |
| mmu-let-7i-5p | MIMAT0000122 | St3gal1 | 20442 | HITS-CLIP | 37986224\|23142080\|23597149\|31433 | Unspecified |
| mmu-let-7i-5p | MIMAT0000122 | St3gal4 | 20443 | HITS-CLIP | 23142080 | Unspecified |
| mmu-let-7i-5p | MIMAT0000122 | St3gal5 | 20454 | HITS-CLIP | 23597149\|23142080 | Unspecified |
| mmu-let-7i-5p | MIMAT0000122 | St3gal6 | 54613 | HITS-CLIP | 23142080 | Unspecified |
| mmu-let-7i-5p | MIMAT0000122 | St6gal1 | 20440 | HITS-CLIP | 23597149\|23142080 | Unspecified |
| mmu-let-7i-5p | MIMAT0000122 | St6galnac5 | 26938 | HITS-CLIP | 37986224 | Unspecified |
| mmu-let-7i-5p | MIMAT0000122 | St7l | 229681 | HITS-CLIP | 23597149 | Unspecified |
| mmu-let-7i-5p | MIMAT0000122 | St8sia1 | 20449 | HITS-CLIP | 23142080 | Unspecified |
| mmu-let-7i-5p | MIMAT0000122 | St8sia4 | 20452 | HITS-CLIP | 23142080\|29167373 | Unspecified |
| mmu-let-7i-5p | MIMAT0000122 | Stab1 | 192187 | HITS-CLIP | 19536157 | Unspecified |
| mmu-let-7i-5p | MIMAT0000122 | Stab2 | 192188 | HITS-CLIP | 23597149 | Unspecified |
| mmu-let-7i-5p | MIMAT0000122 | Stag1 | 20842 | HITS-CLIP | 23142080\|23597149\|29167373 | Unspecified |
| mmu-let-7i-5p | MIMAT0000122 | Stag3 | 50878 | HITS-CLIP | 23142080 | Unspecified |
| mmu-let-7i-5p | MIMAT0000122 | Stam | 20844 | HITS-CLIP | 23597149\|19536157\|23142080 | Unspecified |
| mmu-let-7i-5p | MIMAT0000122 | Stam2 | 56324 | qCLASH//HITS-CLIP | 31393953\|23597149\|23142080\|29167 | Unspecified |
| mmu-let-7i-5p | MIMAT0000122 | Stambp | 70527 | HITS-CLIP | 31433990\|23142080 | Unspecified |
| mmu-let-7i-5p | MIMAT0000122 | Stap2 | 106766 | HITS-CLIP | 37986224 | Unspecified |
| mmu-let-7i-5p | MIMAT0000122 | Stard10 | 56018 | HITS-CLIP | 23597149 | Unspecified |
| mmu-let-7i-5p | MIMAT0000122 | Stard13 | 243362 | HITS-CLIP | 23597149\|19536157\|23142080 | Unspecified |
| mmu-let-7i-5p | MIMAT0000122 | Stard3nl | 76205 | HITS-CLIP | 37986224\|23142080 | Unspecified |
| mmu-let-7i-5p | MIMAT0000122 | Stard4 | 170459 | HITS-CLIP | 23142080 | Unspecified |
| mmu-let-7i-5p | MIMAT0000122 | Stard5 | 170460 | HITS-CLIP | 23597149 | Unspecified |
| mmu-let-7i-5p | MIMAT0000122 | Stard8 | 236920 | HITS-CLIP | 23597149 | Unspecified |
| mmu-let-7i-5p | MIMAT0000122 | Stard9 | 668880 | HITS-CLIP | 29167373 | Unspecified |
| mmu-let-7i-5p | MIMAT0000122 | Stat1 | 20846 | HITS-CLIP | 23142080 | Unspecified |
| mmu-let-7i-5p | MIMAT0000122 | Stat2 | 20847 | HITS-CLIP | 23597149\|23142080\|29167373 | Unspecified |
| mmu-let-7i-5p | MIMAT0000122 | Stat3 | 20848 | HITS-CLIP | 37986224\|23597149\|19536157\|23142 | Unspecified |
| mmu-let-7i-5p | MIMAT0000122 | Stat4 | 20849 | HITS-CLIP | 23142080\|29167373 | Unspecified |
| mmu-let-7i-5p | MIMAT0000122 | Stat5a | 20850 | HITS-CLIP | 23597149\|23142080 | Unspecified |

| | | | | | | |
|---|---|---|---|---|---|---|
| mmu-let-7i-5p | MIMAT0000122 | Stat5b | 20851 | HITS-CLIP | 23142080 | Unspecified |
| mmu-let-7i-5p | MIMAT0000122 | Stat6 | 20852 | HITS-CLIP | 23597149\|23142080 | Unspecified |
| mmu-let-7i-5p | MIMAT0000122 | Stau1 | 20853 | HITS-CLIP | 19536157\|23142080 | Unspecified |
| mmu-let-7i-5p | MIMAT0000122 | Stbd1 | 52331 | HITS-CLIP | 23597149 | Unspecified |
| mmu-let-7i-5p | MIMAT0000122 | Stc1 | 20855 | HITS-CLIP | 23142080 | Unspecified |
| mmu-let-7i-5p | MIMAT0000122 | Steap2 | 74051 | HITS-CLIP | 19536157 | Unspecified |
| mmu-let-7i-5p | MIMAT0000122 | Steap3 | 68428 | HITS-CLIP | 29167373 | Unspecified |
| mmu-let-7i-5p | MIMAT0000122 | Steap4 | 117167 | HITS-CLIP | 37986224\|23597149 | Unspecified |
| mmu-let-7i-5p | MIMAT0000122 | Stil | 20460 | HITS-CLIP | 23142080 | Unspecified |
| mmu-let-7i-5p | MIMAT0000122 | Stim1 | 20866 | HITS-CLIP | 37986224\|23142080 | Unspecified |
| mmu-let-7i-5p | MIMAT0000122 | Stim2 | 116873 | HITS-CLIP | 23142080 | Unspecified |
| mmu-let-7i-5p | MIMAT0000122 | Stip1 | 20867 | HITS-CLIP | 23597149\|23142080 | Unspecified |
| mmu-let-7i-5p | MIMAT0000122 | Stk10 | 20868 | HITS-CLIP | 37986224\|23597149\|23142080 | Unspecified |
| mmu-let-7i-5p | MIMAT0000122 | Stk11 | 20869 | HITS-CLIP | 23597149 | Unspecified |
| mmu-let-7i-5p | MIMAT0000122 | Stk11ip | 71728 | HITS-CLIP | 23597149\|23142080 | Unspecified |
| mmu-let-7i-5p | MIMAT0000122 | Stk17b | 98267 | HITS-CLIP | 23142080\|29167373 | Unspecified |
| mmu-let-7i-5p | MIMAT0000122 | Stk24 | 223255 | HITS-CLIP | 23597149\|19536157\|23142080\|29167 | Unspecified |
| mmu-let-7i-5p | MIMAT0000122 | Stk26 | 51765 | HITS-CLIP | 23142080 | Unspecified |
| mmu-let-7i-5p | MIMAT0000122 | Stk35 | 67333 | HITS-CLIP | 23142080\|23597149\|29167373 | Unspecified |
| mmu-let-7i-5p | MIMAT0000122 | Stk38 | 106504 | HITS-CLIP | 23142080\|29167373 | Unspecified |
| mmu-let-7i-5p | MIMAT0000122 | Stk39 | 53416 | HITS-CLIP | 37986224\|23142080 | Unspecified |
| mmu-let-7i-5p | MIMAT0000122 | Stk4 | 58231 | qCLASH | 31393953 | Unspecified |
| mmu-let-7i-5p | MIMAT0000122 | Stk40 | 74178 | HITS-CLIP | 37986224\|23142080\|26203562\|29167 | Unspecified |
| mmu-let-7i-5p | MIMAT0000122 | Stmn3 | 20262 | HITS-CLIP | 19536157 | Unspecified |
| mmu-let-7i-5p | MIMAT0000122 | Stn1 | 79991 | HITS-CLIP | 23142080 | Unspecified |
| mmu-let-7i-5p | MIMAT0000122 | Stox2 | 71069 | HITS-CLIP | 19536157 | Unspecified |
| mmu-let-7i-5p | MIMAT0000122 | Stra6 | 20897 | HITS-CLIP | 19536157 | Unspecified |
| mmu-let-7i-5p | MIMAT0000122 | Stra6l | 74152 | HITS-CLIP | 23597149 | Unspecified |
| mmu-let-7i-5p | MIMAT0000122 | Strada | 72149 | HITS-CLIP | 23142080 | Unspecified |
| mmu-let-7i-5p | MIMAT0000122 | Stradb | 227154 | HITS-CLIP | 23597149\|23142080 | Unspecified |
| mmu-let-7i-5p | MIMAT0000122 | Strip1 | 229707 | HITS-CLIP | 23142080 | Unspecified |
| mmu-let-7i-5p | MIMAT0000122 | Strn | 268980 | HITS-CLIP | 37986224\|23142080\|23597149\|19536 | Unspecified |
| mmu-let-7i-5p | MIMAT0000122 | Strn3 | 94186 | HITS-CLIP | 23142080 | Unspecified |
| mmu-let-7i-5p | MIMAT0000122 | Strn4 | 97387 | HITS-CLIP | 23597149\|19536157\|23142080\|29167 | Unspecified |
| mmu-let-7i-5p | MIMAT0000122 | Stt3a | 16430 | HITS-CLIP | 19536157\|31433990\|23142080 | Unspecified |
| mmu-let-7i-5p | MIMAT0000122 | Stt3b | 68292 | qCLASH//HITS-CLIP | 31393953\|23142080\|29167373 | Unspecified |
| mmu-let-7i-5p | MIMAT0000122 | Stx12 | 100226 | HITS-CLIP | 23597149\|23142080 | Unspecified |
| mmu-let-7i-5p | MIMAT0000122 | Stx17 | 67727 | HITS-CLIP | 23597149\|23630078\|23142080 | Unspecified |
| mmu-let-7i-5p | MIMAT0000122 | Stx3 | 20908 | HITS-CLIP | 37986224\|29167373 | Unspecified |

| | | | | | | |
|---|---|---|---|---|---|---|
| mmu-let-7i-5p | MIMAT0000122 | Stx6 | 58244 | HITS-CLIP | 23597149\|23142080 | Unspecified |
| mmu-let-7i-5p | MIMAT0000122 | Stx7 | 53331 | HITS-CLIP | 29167373\|19536157\|23597149 | Unspecified |
| mmu-let-7i-5p | MIMAT0000122 | Stx8 | 55943 | HITS-CLIP | 23142080 | Unspecified |
| mmu-let-7i-5p | MIMAT0000122 | Stxbp1 | 20910 | HITS-CLIP | 19536157 | Unspecified |
| mmu-let-7i-5p | MIMAT0000122 | Stxbp5 | 78808 | HITS-CLIP | 19536157\|23142080\|29167373 | Unspecified |
| mmu-let-7i-5p | MIMAT0000122 | Stxbp5l | 207227 | HITS-CLIP | 19536157 | Unspecified |
| mmu-let-7i-5p | MIMAT0000122 | Sub1 | 20024 | HITS-CLIP | 23142080 | Unspecified |
| mmu-let-7i-5p | MIMAT0000122 | Sucla2 | 20916 | HITS-CLIP | 23142080\|23597149\|19536157 | Unspecified |
| mmu-let-7i-5p | MIMAT0000122 | Sucnr1 | 84112 | HITS-CLIP | 23597149 | Unspecified |
| mmu-let-7i-5p | MIMAT0000122 | Suco | 226551 | HITS-CLIP | 37986224\|23597149\|19536157\|23142 | Unspecified |
| mmu-let-7i-5p | MIMAT0000122 | Sufu | 24069 | HITS-CLIP | 23142080 | Unspecified |
| mmu-let-7i-5p | MIMAT0000122 | Sulf2 | 72043 | HITS-CLIP | 29167373 | Unspecified |
| mmu-let-7i-5p | MIMAT0000122 | Sumf2 | 67902 | HITS-CLIP | 37986224\|23597149\|29167373 | Unspecified |
| mmu-let-7i-5p | MIMAT0000122 | Sun1 | 77053 | HITS-CLIP | 23142080 | Unspecified |
| mmu-let-7i-5p | MIMAT0000122 | Sun2 | 223697 | HITS-CLIP | 23597149\|23142080 | Unspecified |
| mmu-let-7i-5p | MIMAT0000122 | Suox | 211389 | HITS-CLIP | 37986224\|23597149 | Unspecified |
| mmu-let-7i-5p | MIMAT0000122 | Supt16 | 114741 | HITS-CLIP | 23142080\|29167373 | Unspecified |
| mmu-let-7i-5p | MIMAT0000122 | Supt20 | 56790 | HITS-CLIP | 23142080 | Unspecified |
| mmu-let-7i-5p | MIMAT0000122 | Supt5 | 20924 | HITS-CLIP | 29167373 | Unspecified |
| mmu-let-7i-5p | MIMAT0000122 | Supt6 | 20926 | HITS-CLIP | 23597149\|19536157\|29167373 | Unspecified |
| mmu-let-7i-5p | MIMAT0000122 | Supt7l | 72195 | HITS-CLIP | 23142080\|23597149 | Unspecified |
| mmu-let-7i-5p | MIMAT0000122 | Surf4 | 20932 | HITS-CLIP | 37986224\|23597149\|23142080\|29167 | Unspecified |
| mmu-let-7i-5p | MIMAT0000122 | Susd6 | 9766 | HITS-CLIP | 23597149\|23142080\|26203562\|29167 | Unspecified |
| mmu-let-7i-5p | MIMAT0000122 | Suv39h1 | 20937 | HITS-CLIP | 23597149 | Unspecified |
| mmu-let-7i-5p | MIMAT0000122 | Suv39h2 | 64707 | HITS-CLIP | 31433990\|23142080 | Unspecified |
| mmu-let-7i-5p | MIMAT0000122 | Suz12 | 52615 | HITS-CLIP | 37986224\|23142080 | Unspecified |
| mmu-let-7i-5p | MIMAT0000122 | Sv2a | 64051 | HITS-CLIP | 37986224 | Unspecified |
| mmu-let-7i-5p | MIMAT0000122 | Sv2b | 64176 | HITS-CLIP | 19536157 | Unspecified |
| mmu-let-7i-5p | MIMAT0000122 | Sv2c | 75209 | HITS-CLIP | 19536157 | Unspecified |
| mmu-let-7i-5p | MIMAT0000122 | Svil | 225115 | HITS-CLIP | 19536157\|23142080\|29167373 | Unspecified |
| mmu-let-7i-5p | MIMAT0000122 | Swap70 | 20947 | HITS-CLIP | 29167373 | Unspecified |
| mmu-let-7i-5p | MIMAT0000122 | Swsap1 | 66962 | HITS-CLIP | 23142080 | Unspecified |
| mmu-let-7i-5p | MIMAT0000122 | Swt1 | 66875 | HITS-CLIP | 23142080 | Unspecified |
| mmu-let-7i-5p | MIMAT0000122 | Syk | 20963 | HITS-CLIP | 29167373 | Unspecified |
| mmu-let-7i-5p | MIMAT0000122 | Sympk | 68188 | HITS-CLIP | 23597149\|23142080\|29167373 | Unspecified |
| mmu-let-7i-5p | MIMAT0000122 | Syn1 | 20964 | HITS-CLIP | 19536157 | Unspecified |
| mmu-let-7i-5p | MIMAT0000122 | Syncrip | 56403 | HITS-CLIP | 37986224\|23597149\|19536157\|23142 | Unspecified |
| mmu-let-7i-5p | MIMAT0000122 | Syne1 | 64009 | HITS-CLIP | 37986224\|23142080\|19536157\|23597 | Unspecified |
| mmu-let-7i-5p | MIMAT0000122 | Syne2 | 319565 | HITS-CLIP | 37986224\|23597149\|19536157 | Unspecified |

| | | | | | | |
|---|---|---|---|---|---|---|
| mmu-let-7i-5p | MIMAT0000122 | Syngap1 | 8831 | HITS-CLIP | 19536157 | Unspecified |
| mmu-let-7i-5p | MIMAT0000122 | Syngr1 | 20972 | HITS-CLIP | 19536157 | Unspecified |
| mmu-let-7i-5p | MIMAT0000122 | Synj1 | 104015 | HITS-CLIP | 19536157\|23142080\|29167373 | Unspecified |
| mmu-let-7i-5p | MIMAT0000122 | Synj2 | 20975 | HITS-CLIP | 29167373 | Unspecified |
| mmu-let-7i-5p | MIMAT0000122 | Synj2bp | 24071 | HITS-CLIP | 19536157 | Unspecified |
| mmu-let-7i-5p | MIMAT0000122 | Synrg | 217030 | HITS-CLIP | 37986224\|19536157\|23142080\|29167 | Unspecified |
| mmu-let-7i-5p | MIMAT0000122 | Sypl | 19027 | HITS-CLIP | 23142080 | Unspecified |
| mmu-let-7i-5p | MIMAT0000122 | Syt1 | 20979 | HITS-CLIP | 37986224\|19536157 | Unspecified |
| mmu-let-7i-5p | MIMAT0000122 | Syt11 | 229521 | HITS-CLIP | 29167373 | Unspecified |
| mmu-let-7i-5p | MIMAT0000122 | Syt2 | 20980 | HITS-CLIP | 19536157 | Unspecified |
| mmu-let-7i-5p | MIMAT0000122 | Syt3 | 20981 | HITS-CLIP | 19536157 | Unspecified |
| mmu-let-7i-5p | MIMAT0000122 | Syt5 | 53420 | HITS-CLIP | 37986224 | Unspecified |
| mmu-let-7i-5p | MIMAT0000122 | Syvn1 | 74126 | HITS-CLIP | 37986224\|23142080\|23597149\|1953€ | Unspecified |
| mmu-let-7i-5p | MIMAT0000122 | Szrd1 | 213491 | HITS-CLIP | 23597149\|19536157 | Unspecified |
| mmu-let-7i-5p | MIMAT0000122 | Szt2 | 230676 | HITS-CLIP | 23597149\|29167373 | Unspecified |
| mmu-let-7i-5p | MIMAT0000122 | Tab2 | 68652 | HITS-CLIP | 37986224\|23142080\|23597149\|29167 | Unspecified |
| mmu-let-7i-5p | MIMAT0000122 | Tab3 | 66724 | HITS-CLIP | 23597149\|23142080 | Unspecified |
| mmu-let-7i-5p | MIMAT0000122 | Tacc2 | 57752 | HITS-CLIP | 37986224\|19536157\|23597149 | Unspecified |
| mmu-let-7i-5p | MIMAT0000122 | Tacc3 | 21335 | HITS-CLIP | 23142080\|29167373 | Unspecified |
| mmu-let-7i-5p | MIMAT0000122 | Tada1 | 27878 | HITS-CLIP | 23142080 | Unspecified |
| mmu-let-7i-5p | MIMAT0000122 | Tada2a | 217031 | HITS-CLIP | 23597149 | Unspecified |
| mmu-let-7i-5p | MIMAT0000122 | Taf1 | 270627 | HITS-CLIP | 23142080 | Unspecified |
| mmu-let-7i-5p | MIMAT0000122 | Taf10 | 24075 | HITS-CLIP | 23142080 | Unspecified |
| mmu-let-7i-5p | MIMAT0000122 | Taf2 | 319944 | HITS-CLIP | 29167373\|23142080 | Unspecified |
| mmu-let-7i-5p | MIMAT0000122 | Taf4 | 6874 | HITS-CLIP | 23142080 | Unspecified |
| mmu-let-7i-5p | MIMAT0000122 | Taf4b | 72504 | HITS-CLIP | 23142080 | Unspecified |
| mmu-let-7i-5p | MIMAT0000122 | Taf5 | 226182 | HITS-CLIP | 37986224\|23142080 | Unspecified |
| mmu-let-7i-5p | MIMAT0000122 | Taf5l | 102162 | HITS-CLIP | 37986224\|23142080\|23597149\|1953€ | Unspecified |
| mmu-let-7i-5p | MIMAT0000122 | Taf6 | 21343 | HITS-CLIP | 37986224\|19536157\|23597149\|23142 | Unspecified |
| mmu-let-7i-5p | MIMAT0000122 | Taf7 | 24074 | HITS-CLIP | 37986224\|23142080\|29167373 | Unspecified |
| mmu-let-7i-5p | MIMAT0000122 | Taf8 | 63856 | HITS-CLIP | 23142080\|23597149 | Unspecified |
| mmu-let-7i-5p | MIMAT0000122 | Tagap | 72536 | HITS-CLIP | 23142080 | Unspecified |
| mmu-let-7i-5p | MIMAT0000122 | Tagln3 | 56370 | HITS-CLIP | 19536157 | Unspecified |
| mmu-let-7i-5p | MIMAT0000122 | Taldo1 | 21351 | HITS-CLIP | 23142080\|29167373 | Unspecified |
| mmu-let-7i-5p | MIMAT0000122 | Tanc1 | 66860 | HITS-CLIP | 23597149\|29167373\|23142080 | Unspecified |
| mmu-let-7i-5p | MIMAT0000122 | Tanc2 | 77097 | HITS-CLIP | 37986224\|19536157\|30971389\|23142 | Unspecified |
| mmu-let-7i-5p | MIMAT0000122 | Tank | 21353 | HITS-CLIP | 23142080\|29167373 | Unspecified |
| mmu-let-7i-5p | MIMAT0000122 | Taok1 | 216965 | HITS-CLIP | 37986224\|31433990\|19536157\|23142 | Unspecified |
| mmu-let-7i-5p | MIMAT0000122 | Taok2 | 381921 | HITS-CLIP | 23597149 | Unspecified |

| | | | | | | |
|---|---|---|---|---|---|---|
| mmu-let-7i-5p | MIMAT0000122 | Taok3 | 330177 | HITS-CLIP | 23597149\|29167373 | Unspecified |
| mmu-let-7i-5p | MIMAT0000122 | Tap1 | 21354 | HITS-CLIP | 23142080 | Unspecified |
| mmu-let-7i-5p | MIMAT0000122 | Tap2 | 21355 | HITS-CLIP | 23142080 | Unspecified |
| mmu-let-7i-5p | MIMAT0000122 | Tapbp | 21356 | HITS-CLIP | 23597149\|23142080\|29167373 | Unspecified |
| mmu-let-7i-5p | MIMAT0000122 | Tapbpl | 213233 | HITS-CLIP | 23597149\|23142080 | Unspecified |
| mmu-let-7i-5p | MIMAT0000122 | Tapt1 | 231225 | HITS-CLIP | 23142080 | Unspecified |
| mmu-let-7i-5p | MIMAT0000122 | Tardbp | 230908 | HITS-CLIP | 23597149\|23142080\|29167373 | Unspecified |
| mmu-let-7i-5p | MIMAT0000122 | Tars | 110960 | HITS-CLIP | 23142080 | Unspecified |
| mmu-let-7i-5p | MIMAT0000122 | Tarsl2 | 272396 | HITS-CLIP | 23142080 | Unspecified |
| mmu-let-7i-5p | MIMAT0000122 | Tasp1 | 75812 | HITS-CLIP | 23142080 | Unspecified |
| mmu-let-7i-5p | MIMAT0000122 | Tat | 234724 | HITS-CLIP | 37986224\|23597149 | Unspecified |
| mmu-let-7i-5p | MIMAT0000122 | Tax1bp1 | 52440 | HITS-CLIP | 37986224\|23597149\|23142080 | Unspecified |
| mmu-let-7i-5p | MIMAT0000122 | Tbc1d1 | 57915 | HITS-CLIP | 19536157\|23597149\|29167373\|23142 | Unspecified |
| mmu-let-7i-5p | MIMAT0000122 | Tbc1d10b | 68449 | HITS-CLIP | 29167373 | Unspecified |
| mmu-let-7i-5p | MIMAT0000122 | Tbc1d10c | 108995 | HITS-CLIP | 23142080 | Unspecified |
| mmu-let-7i-5p | MIMAT0000122 | Tbc1d14 | 100855 | HITS-CLIP | 29167373 | Unspecified |
| mmu-let-7i-5p | MIMAT0000122 | Tbc1d15 | 66687 | HITS-CLIP | 23142080 | Unspecified |
| mmu-let-7i-5p | MIMAT0000122 | Tbc1d16 | 207592 | HITS-CLIP | 23597149 | Unspecified |
| mmu-let-7i-5p | MIMAT0000122 | Tbc1d20 | 67231 | HITS-CLIP | 23142080 | Unspecified |
| mmu-let-7i-5p | MIMAT0000122 | Tbc1d23 | 67581 | HITS-CLIP | 23142080 | Unspecified |
| mmu-let-7i-5p | MIMAT0000122 | Tbc1d24 | 224617 | HITS-CLIP | 37986224\|19536157 | Unspecified |
| mmu-let-7i-5p | MIMAT0000122 | Tbc1d2b | 67016 | HITS-CLIP | 37986224\|23597149\|23142080\|29167 | Unspecified |
| mmu-let-7i-5p | MIMAT0000122 | Tbc1d30 | 74694 | HITS-CLIP | 23142080 | Unspecified |
| mmu-let-7i-5p | MIMAT0000122 | Tbc1d5 | 72238 | HITS-CLIP | 29167373 | Unspecified |
| mmu-let-7i-5p | MIMAT0000122 | Tbc1d7 | 67046 | HITS-CLIP | 23142080\|23597149 | Unspecified |
| mmu-let-7i-5p | MIMAT0000122 | Tbc1d8b | 245638 | HITS-CLIP | 31433990\|23142080 | Unspecified |
| mmu-let-7i-5p | MIMAT0000122 | Tbce | 70430 | HITS-CLIP | 23142080 | Unspecified |
| mmu-let-7i-5p | MIMAT0000122 | Tbcel | 272589 | HITS-CLIP | 23597149\|29167373 | Unspecified |
| mmu-let-7i-5p | MIMAT0000122 | Tbck | 271981 | HITS-CLIP | 23142080 | Unspecified |
| mmu-let-7i-5p | MIMAT0000122 | Tbk1 | 56480 | HITS-CLIP | 37986224\|23142080 | Unspecified |
| mmu-let-7i-5p | MIMAT0000122 | Tbkbp1 | 73174 | HITS-CLIP | 19536157\|23597149\|23142080\|29167 | Unspecified |
| mmu-let-7i-5p | MIMAT0000122 | Tbl1x | 21372 | HITS-CLIP | 23142080 | Unspecified |
| mmu-let-7i-5p | MIMAT0000122 | Tbl1xr1 | 81004 | HITS-CLIP | 37986224\|23142080\|23597149\|26203 | Unspecified |
| mmu-let-7i-5p | MIMAT0000122 | Tbl2 | 27368 | HITS-CLIP | 23597149 | Unspecified |
| mmu-let-7i-5p | MIMAT0000122 | Tbl3 | 213773 | HITS-CLIP | 19536157 | Unspecified |
| mmu-let-7i-5p | MIMAT0000122 | Tbp | 21374 | HITS-CLIP | 37986224\|23142080 | Unspecified |
| mmu-let-7i-5p | MIMAT0000122 | Tbpl1 | 237336 | HITS-CLIP | 23142080 | Unspecified |
| mmu-let-7i-5p | MIMAT0000122 | Tbrg4 | 21379 | HITS-CLIP | 23142080 | Unspecified |
| mmu-let-7i-5p | MIMAT0000122 | Tbx20 | 57246 | HITS-CLIP | 23597149 | Unspecified |

| | | | | | | |
|---|---|---|---|---|---|---|
| mmu-let-7i-5p | MIMAT0000122 | Tbx3 | 21386 | HITS-CLIP | 37986224\|23597149 | Unspecified |
| mmu-let-7i-5p | MIMAT0000122 | Tcaim | 382117 | HITS-CLIP | 30971389 | Unspecified |
| mmu-let-7i-5p | MIMAT0000122 | Tceal6 | 66104 | HITS-CLIP | 19536157 | Unspecified |
| mmu-let-7i-5p | MIMAT0000122 | Tceanc2 | 66526 | HITS-CLIP | 29167373 | Unspecified |
| mmu-let-7i-5p | MIMAT0000122 | Tcerg1 | 56070 | HITS-CLIP | 37986224\|23142080\|29167373 | Unspecified |
| mmu-let-7i-5p | MIMAT0000122 | Tcf12 | 21406 | HITS-CLIP | 23142080\|23597149\|29167373 | Unspecified |
| mmu-let-7i-5p | MIMAT0000122 | Tcf20 | 21411 | HITS-CLIP | 19536157\|23597149\|23142080\|29167 | Unspecified |
| mmu-let-7i-5p | MIMAT0000122 | Tcf25 | 66855 | HITS-CLIP | 23142080 | Unspecified |
| mmu-let-7i-5p | MIMAT0000122 | Tcf3 | 21423 | HITS-CLIP | 23142080\|29167373 | Unspecified |
| mmu-let-7i-5p | MIMAT0000122 | Tcf7 | 21414 | HITS-CLIP | 23142080 | Unspecified |
| mmu-let-7i-5p | MIMAT0000122 | Tcf7l1 | 21415 | HITS-CLIP | 23597149 | Unspecified |
| mmu-let-7i-5p | MIMAT0000122 | Tcf7l2 | 21416 | HITS-CLIP | 23597149 | Unspecified |
| mmu-let-7i-5p | MIMAT0000122 | Tcirg1 | 27060 | HITS-CLIP | 29167373 | Unspecified |
| mmu-let-7i-5p | MIMAT0000122 | Tcn2 | 21452 | HITS-CLIP | 23597149 | Unspecified |
| mmu-let-7i-5p | MIMAT0000122 | Tcof1 | 21453 | HITS-CLIP | 23142080\|29167373 | Unspecified |
| mmu-let-7i-5p | MIMAT0000122 | Tcp1 | 21454 | HITS-CLIP | 23142080\|23597149 | Unspecified |
| mmu-let-7i-5p | MIMAT0000122 | Tcp11l1 | 320554 | HITS-CLIP | 29167373 | Unspecified |
| mmu-let-7i-5p | MIMAT0000122 | Tcp11l2 | 216198 | HITS-CLIP | 23142080 | Unspecified |
| mmu-let-7i-5p | MIMAT0000122 | Tcta | 102791 | HITS-CLIP | 23597149 | Unspecified |
| mmu-let-7i-5p | MIMAT0000122 | Tdo2 | 56720 | HITS-CLIP | 23597149 | Unspecified |
| mmu-let-7i-5p | MIMAT0000122 | Tdp2 | 56196 | HITS-CLIP | 23142080 | Unspecified |
| mmu-let-7i-5p | MIMAT0000122 | Tdrd7 | 100121 | HITS-CLIP | 29167373 | Unspecified |
| mmu-let-7i-5p | MIMAT0000122 | Tecpr1 | 70381 | HITS-CLIP | 23597149\|23142080 | Unspecified |
| mmu-let-7i-5p | MIMAT0000122 | Tecr | 106529 | HITS-CLIP | 37986224\|23142080\|23597149 | Unspecified |
| mmu-let-7i-5p | MIMAT0000122 | Tef | 21685 | HITS-CLIP | 23597149 | Unspecified |
| mmu-let-7i-5p | MIMAT0000122 | Tek | 21687 | HITS-CLIP | 23597149 | Unspecified |
| mmu-let-7i-5p | MIMAT0000122 | Tent2 | 167153 | HITS-CLIP | 23142080\|31433990 | Unspecified |
| mmu-let-7i-5p | MIMAT0000122 | Tent4b | 64282 | HITS-CLIP | 23597149 | Unspecified |
| mmu-let-7i-5p | MIMAT0000122 | Tent5a | 55603 | HITS-CLIP | 23142080\|29167373 | Unspecified |
| mmu-let-7i-5p | MIMAT0000122 | Tent5c | 54855 | HITS-CLIP | 23597149\|23142080 | Unspecified |
| mmu-let-7i-5p | MIMAT0000122 | Tep1 | 21745 | HITS-CLIP | 29167373 | Unspecified |
| mmu-let-7i-5p | MIMAT0000122 | Terf2 | 21750 | HITS-CLIP | 31433990 | Unspecified |
| mmu-let-7i-5p | MIMAT0000122 | Tes | 21753 | HITS-CLIP | 37986224\|31433990\|23142080 | Unspecified |
| mmu-let-7i-5p | MIMAT0000122 | Tet1 | 52463 | HITS-CLIP | 23142080 | Unspecified |
| mmu-let-7i-5p | MIMAT0000122 | Tet2 | 214133 | HITS-CLIP | 37986224\|23597149\|23142080\|29167 | Unspecified |
| mmu-let-7i-5p | MIMAT0000122 | Tet3 | 194388 | HITS-CLIP | 37986224\|23597149\|19536157\|2314 | Unspecified |
| mmu-let-7i-5p | MIMAT0000122 | Tex10 | 269536 | HITS-CLIP | 23142080 | Unspecified |
| mmu-let-7i-5p | MIMAT0000122 | Tex15 | 104271 | HITS-CLIP | 23142080 | Unspecified |
| mmu-let-7i-5p | MIMAT0000122 | Tex2 | 21763 | HITS-CLIP | 29167373 | Unspecified |

| | | | | | | |
|---|---|---|---|---|---|---|
| mmu-let-7i-5p | MIMAT0000122 | Tex261 | 21766 | HITS-CLIP | 23597149 | Unspecified |
| mmu-let-7i-5p | MIMAT0000122 | Tfam | 21780 | HITS-CLIP | 23142080 | Unspecified |
| mmu-let-7i-5p | MIMAT0000122 | Tfap4 | 83383 | HITS-CLIP | 23142080 | Unspecified |
| mmu-let-7i-5p | MIMAT0000122 | Tfb1m | 224481 | HITS-CLIP | 37986224\|19536157\|23142080 | Unspecified |
| mmu-let-7i-5p | MIMAT0000122 | Tfcp2 | 21422 | HITS-CLIP | 23597149 | Unspecified |
| mmu-let-7i-5p | MIMAT0000122 | Tfdp1 | 21781 | HITS-CLIP | 23142080 | Unspecified |
| mmu-let-7i-5p | MIMAT0000122 | Tfdp2 | 211586 | HITS-CLIP | 23142080 | Unspecified |
| mmu-let-7i-5p | MIMAT0000122 | Tfe3 | 209446 | HITS-CLIP | 23597149 | Unspecified |
| mmu-let-7i-5p | MIMAT0000122 | Tfg | 21787 | HITS-CLIP | 23142080 | Unspecified |
| mmu-let-7i-5p | MIMAT0000122 | Tfip11 | 54723 | HITS-CLIP | 23597149\|23142080 | Unspecified |
| mmu-let-7i-5p | MIMAT0000122 | Tfr2 | 50765 | HITS-CLIP | 23597149 | Unspecified |
| mmu-let-7i-5p | MIMAT0000122 | Tfrc | 22042 | HITS-CLIP | 37986224\|23142080 | Unspecified |
| mmu-let-7i-5p | MIMAT0000122 | Tgds | 76355 | HITS-CLIP | 37986224\|23597149\|23142080\|29167 | Unspecified |
| mmu-let-7i-5p | MIMAT0000122 | Tgfb1 | 21803 | HITS-CLIP | 37986224\|23597149\|23142080\|29167 | Unspecified |
| mmu-let-7i-5p | MIMAT0000122 | Tgfbi | 21810 | HITS-CLIP | 23597149 | Unspecified |
| mmu-let-7i-5p | MIMAT0000122 | Tgfbr1 | 21812 | HITS-CLIP | 37986224\|23597149\|23142080\|23630 | Unspecified |
| mmu-let-7i-5p | MIMAT0000122 | Tgfbr3 | 21814 | HITS-CLIP | 37986224\|23597149\|19536157\|23142 | Unspecified |
| mmu-let-7i-5p | MIMAT0000122 | Tgif2 | 228839 | HITS-CLIP | 23142080 | Unspecified |
| mmu-let-7i-5p | MIMAT0000122 | Tgm2 | 21817 | HITS-CLIP | 23597149\|23142080 | Unspecified |
| mmu-let-7i-5p | MIMAT0000122 | Tgoln1 | 22134 | HITS-CLIP | 19536157\|23597149\|23142080\|29167 | Unspecified |
| mmu-let-7i-5p | MIMAT0000122 | Tgs1 | 116940 | HITS-CLIP | 37986224\|23630078\|19536157\|31433 | Unspecified |
| mmu-let-7i-5p | MIMAT0000122 | Thada | 240174 | HITS-CLIP | 23142080 | Unspecified |
| mmu-let-7i-5p | MIMAT0000122 | Thap2 | 66816 | HITS-CLIP | 19536157 | Unspecified |
| mmu-let-7i-5p | MIMAT0000122 | Thap4 | 67026 | HITS-CLIP | 23142080 | Unspecified |
| mmu-let-7i-5p | MIMAT0000122 | Thbd | 21824 | HITS-CLIP | 23597149\|29167373 | Unspecified |
| mmu-let-7i-5p | MIMAT0000122 | Thbs1 | 21825 | HITS-CLIP | 37986224\|19536157\|23142080 | Unspecified |
| mmu-let-7i-5p | MIMAT0000122 | Them6 | 223626 | HITS-CLIP | 37986224 | Unspecified |
| mmu-let-7i-5p | MIMAT0000122 | Themis | 210757 | HITS-CLIP | 23142080 | Unspecified |
| mmu-let-7i-5p | MIMAT0000122 | Thg1l | 66628 | HITS-CLIP | 23142080 | Unspecified |
| mmu-let-7i-5p | MIMAT0000122 | Thoc2 | 331401 | HITS-CLIP | 19536157\|23142080 | Unspecified |
| mmu-let-7i-5p | MIMAT0000122 | Thoc3 | 73666 | HITS-CLIP | 23142080 | Unspecified |
| mmu-let-7i-5p | MIMAT0000122 | Thoc5 | 8563 | HITS-CLIP | 29167373 | Unspecified |
| mmu-let-7i-5p | MIMAT0000122 | Thop1 | 50492 | HITS-CLIP | 23142080 | Unspecified |
| mmu-let-7i-5p | MIMAT0000122 | Thrap3 | 230753 | HITS-CLIP | 23597149 | Unspecified |
| mmu-let-7i-5p | MIMAT0000122 | Thrsp | 21835 | HITS-CLIP | 37986224\|23597149\|19536157 | Unspecified |
| mmu-let-7i-5p | MIMAT0000122 | Thsd7a | 330267 | HITS-CLIP | 19536157 | Unspecified |
| mmu-let-7i-5p | MIMAT0000122 | Thtpa | 105663 | HITS-CLIP | 23142080 | Unspecified |
| mmu-let-7i-5p | MIMAT0000122 | Thumpd3 | 14911 | HITS-CLIP | 23142080 | Unspecified |
| mmu-let-7i-5p | MIMAT0000122 | Thy1 | 21838 | HITS-CLIP | 31433990\|23142080 | Unspecified |

| | | | | | | |
|---|---|---|---|---|---|---|
| mmu-let-7i-5p | MIMAT0000122 | Tia1 | 21841 | HITS-CLIP | 19536157\|23142080 | Unspecified |
| mmu-let-7i-5p | MIMAT0000122 | Tial1 | 21843 | HITS-CLIP | 19536157 | Unspecified |
| mmu-let-7i-5p | MIMAT0000122 | Tiam1 | 21844 | HITS-CLIP | 23142080 | Unspecified |
| mmu-let-7i-5p | MIMAT0000122 | Ticrr | 77011 | HITS-CLIP | 23142080 | Unspecified |
| mmu-let-7i-5p | MIMAT0000122 | Timd2 | 171284 | HITS-CLIP | 23597149 | Unspecified |
| mmu-let-7i-5p | MIMAT0000122 | Timm10b | 14356 | HITS-CLIP | 37986224\|23142080 | Unspecified |
| mmu-let-7i-5p | MIMAT0000122 | Timm8b | 30057 | HITS-CLIP | 23142080 | Unspecified |
| mmu-let-7i-5p | MIMAT0000122 | Tinf2 | 28113 | HITS-CLIP | 23142080 | Unspecified |
| mmu-let-7i-5p | MIMAT0000122 | Tiparp | 99929 | HITS-CLIP | 23597149\|23142080 | Unspecified |
| mmu-let-7i-5p | MIMAT0000122 | Tipin | 66131 | HITS-CLIP | 23142080 | Unspecified |
| mmu-let-7i-5p | MIMAT0000122 | Tirap | 117149 | HITS-CLIP | 23597149\|29167373 | Unspecified |
| mmu-let-7i-5p | MIMAT0000122 | Tjap1 | 74094 | HITS-CLIP | 23597149 | Unspecified |
| mmu-let-7i-5p | MIMAT0000122 | Tjp1 | 21872 | HITS-CLIP | 37986224\|23597149 | Unspecified |
| mmu-let-7i-5p | MIMAT0000122 | Tjp2 | 21873 | HITS-CLIP | 23142080 | Unspecified |
| mmu-let-7i-5p | MIMAT0000122 | Tkfc | 26007 | HITS-CLIP | 23597149 | Unspecified |
| mmu-let-7i-5p | MIMAT0000122 | Tkt | 21881 | HITS-CLIP | 31433990 | Unspecified |
| mmu-let-7i-5p | MIMAT0000122 | Tle1 | 21885 | HITS-CLIP | 29167373 | Unspecified |
| mmu-let-7i-5p | MIMAT0000122 | Tle3 | 21887 | HITS-CLIP | 23142080 | Unspecified |
| mmu-let-7i-5p | MIMAT0000122 | Tlk1 | 228012 | HITS-CLIP | 23597149\|29167373\|23142080 | Unspecified |
| mmu-let-7i-5p | MIMAT0000122 | Tln1 | 21894 | HITS-CLIP | 23597149\|23142080\|29167373 | Unspecified |
| mmu-let-7i-5p | MIMAT0000122 | Tln2 | 70549 | HITS-CLIP | 26203562 | Unspecified |
| mmu-let-7i-5p | MIMAT0000122 | Tlnrd1 | 59274 | HITS-CLIP | 19536157\|23142080 | Unspecified |
| mmu-let-7i-5p | MIMAT0000122 | Tlr12 | 384059 | HITS-CLIP | 37986224\|19536157\|23597149 | Unspecified |
| mmu-let-7i-5p | MIMAT0000122 | Tlr13 | 279572 | HITS-CLIP | 29167373 | Unspecified |
| mmu-let-7i-5p | MIMAT0000122 | Tlr3 | 142980 | HITS-CLIP | 23597149 | Unspecified |
| mmu-let-7i-5p | MIMAT0000122 | Tlr5 | 53791 | HITS-CLIP | 23597149 | Unspecified |
| mmu-let-7i-5p | MIMAT0000122 | Tlr7 | 170743 | HITS-CLIP | 23142080\|29167373 | Unspecified |
| mmu-let-7i-5p | MIMAT0000122 | Tlr8 | 170744 | HITS-CLIP | 23597149 | Unspecified |
| mmu-let-7i-5p | MIMAT0000122 | Tm2d2 | 69742 | HITS-CLIP | 23597149\|23142080\|29167373 | Unspecified |
| mmu-let-7i-5p | MIMAT0000122 | Tm2d3 | 68634 | HITS-CLIP | 23142080 | Unspecified |
| mmu-let-7i-5p | MIMAT0000122 | Tm6sf1 | 107769 | HITS-CLIP | 23142080 | Unspecified |
| mmu-let-7i-5p | MIMAT0000122 | Tm6sf2 | 107770 | HITS-CLIP | 23597149 | Unspecified |
| mmu-let-7i-5p | MIMAT0000122 | Tm7sf2 | 73166 | HITS-CLIP | 23597149 | Unspecified |
| mmu-let-7i-5p | MIMAT0000122 | Tm9sf2 | 68059 | HITS-CLIP//qCLASH | 37986224\|23142080\|31393953\|29167 | Unspecified |
| mmu-let-7i-5p | MIMAT0000122 | Tm9sf3 | 107358 | HITS-CLIP | 37986224\|23597149\|23142080\|19536 | Unspecified |
| mmu-let-7i-5p | MIMAT0000122 | Tm9sf4 | 99237 | HITS-CLIP | 19536157\|23597149\|23142080\|29167 | Unspecified |
| mmu-let-7i-5p | MIMAT0000122 | Tma7 | 66167 | HITS-CLIP | 29167373 | Unspecified |
| mmu-let-7i-5p | MIMAT0000122 | Tmbim4 | 68212 | HITS-CLIP | 23142080 | Unspecified |
| mmu-let-7i-5p | MIMAT0000122 | Tmbim6 | 110213 | HITS-CLIP | 23597149\|31433990\|23142080\|19536 | Unspecified |

| | | | | | | |
|---|---|---|---|---|---|---|
| mmu-let-7i-5p | MIMAT0000122 | Tmc7 | 209760 | HITS-CLIP | 37986224\|23597149 | Unspecified |
| mmu-let-7i-5p | MIMAT0000122 | Tmc8 | 217356 | HITS-CLIP | 37986224\|23142080 | Unspecified |
| mmu-let-7i-5p | MIMAT0000122 | Tmcc1 | 330401 | HITS-CLIP | 37986224 | Unspecified |
| mmu-let-7i-5p | MIMAT0000122 | Tmco1 | 68944 | HITS-CLIP | 23597149\|29167373 | Unspecified |
| mmu-let-7i-5p | MIMAT0000122 | Tmco6 | 71983 | HITS-CLIP | 23597149\|23142080 | Unspecified |
| mmu-let-7i-5p | MIMAT0000122 | Tmed3 | 66111 | HITS-CLIP | 23597149 | Unspecified |
| mmu-let-7i-5p | MIMAT0000122 | Tmed4 | 103694 | HITS-CLIP | 23597149\|23142080 | Unspecified |
| mmu-let-7i-5p | MIMAT0000122 | Tmed5 | 73130 | HITS-CLIP | 37986224\|23597149\|19536157\|23142 | Unspecified |
| mmu-let-7i-5p | MIMAT0000122 | Tmed7 | 66676 | HITS-CLIP | 23142080\|29167373 | Unspecified |
| mmu-let-7i-5p | MIMAT0000122 | Tmed8 | 382620 | HITS-CLIP | 23142080 | Unspecified |
| mmu-let-7i-5p | MIMAT0000122 | Tmed9 | 67511 | HITS-CLIP | 23597149 | Unspecified |
| mmu-let-7i-5p | MIMAT0000122 | Tmem101 | 76547 | HITS-CLIP | 37986224\|23142080 | Unspecified |
| mmu-let-7i-5p | MIMAT0000122 | Tmem104 | 320534 | HITS-CLIP | 29167373 | Unspecified |
| mmu-let-7i-5p | MIMAT0000122 | Tmem106a | 217203 | HITS-CLIP | 29167373 | Unspecified |
| mmu-let-7i-5p | MIMAT0000122 | Tmem11 | 216821 | HITS-CLIP | 23142080 | Unspecified |
| mmu-let-7i-5p | MIMAT0000122 | Tmem120a | 215210 | qCLASH//HITS-CLIP | 31393953\|23597149\|23142080 | Unspecified |
| mmu-let-7i-5p | MIMAT0000122 | Tmem123 | 71929 | HITS-CLIP | 23597149\|29167373\|23142080 | Unspecified |
| mmu-let-7i-5p | MIMAT0000122 | Tmem127 | 69470 | HITS-CLIP | 29167373\|23142080 | Unspecified |
| mmu-let-7i-5p | MIMAT0000122 | Tmem129 | 68366 | HITS-CLIP | 23142080 | Unspecified |
| mmu-let-7i-5p | MIMAT0000122 | Tmem131 | 56030 | HITS-CLIP | 23142080\|19536157\|23597149 | Unspecified |
| mmu-let-7i-5p | MIMAT0000122 | Tmem131l | 23240 | HITS-CLIP | 23142080\|23597149 | Unspecified |
| mmu-let-7i-5p | MIMAT0000122 | Tmem132a | 98170 | HITS-CLIP | 19536157 | Unspecified |
| mmu-let-7i-5p | MIMAT0000122 | Tmem135 | 72759 | HITS-CLIP | 23142080 | Unspecified |
| mmu-let-7i-5p | MIMAT0000122 | Tmem143 | 70209 | HITS-CLIP | 23142080 | Unspecified |
| mmu-let-7i-5p | MIMAT0000122 | Tmem147 | 69804 | HITS-CLIP | 37986224\|23597149 | Unspecified |
| mmu-let-7i-5p | MIMAT0000122 | Tmem14c | 66154 | HITS-CLIP | 23597149\|23142080 | Unspecified |
| mmu-let-7i-5p | MIMAT0000122 | Tmem150a | 232086 | HITS-CLIP | 23597149 | Unspecified |
| mmu-let-7i-5p | MIMAT0000122 | Tmem151a | 381199 | HITS-CLIP | 37986224 | Unspecified |
| mmu-let-7i-5p | MIMAT0000122 | Tmem151b | 210573 | HITS-CLIP | 37986224 | Unspecified |
| mmu-let-7i-5p | MIMAT0000122 | Tmem154 | 320782 | HITS-CLIP | 23142080 | Unspecified |
| mmu-let-7i-5p | MIMAT0000122 | Tmem156 | 243025 | HITS-CLIP | 29167373\|23142080 | Unspecified |
| mmu-let-7i-5p | MIMAT0000122 | Tmem161b | 72745 | HITS-CLIP | 23142080 | Unspecified |
| mmu-let-7i-5p | MIMAT0000122 | Tmem165 | 21982 | HITS-CLIP//qCLASH | 37986224\|31393953\|19536157\|23142 | Unspecified |
| mmu-let-7i-5p | MIMAT0000122 | Tmem167 | 66074 | HITS-CLIP | 37986224\|23597149 | Unspecified |
| mmu-let-7i-5p | MIMAT0000122 | Tmem167b | 67495 | HITS-CLIP | 23142080 | Unspecified |
| mmu-let-7i-5p | MIMAT0000122 | Tmem168 | 101118 | HITS-CLIP | 37986224\|23142080 | Unspecified |
| mmu-let-7i-5p | MIMAT0000122 | Tmem170 | 66817 | HITS-CLIP | 23142080 | Unspecified |
| mmu-let-7i-5p | MIMAT0000122 | Tmem170b | 621976 | HITS-CLIP | 23142080\|19536157 | Unspecified |
| mmu-let-7i-5p | MIMAT0000122 | Tmem173 | 72512 | HITS-CLIP | 37986224\|23142080 | Unspecified |

| | | | | | | |
|---|---|---|---|---|---|---|
| mmu-let-7i-5p | MIMAT0000122 | Tmem175 | 72392 | HITS-CLIP | 37986224\|19536157\|23597149\|23142 | Unspecified |
| mmu-let-7i-5p | MIMAT0000122 | Tmem176b | 65963 | HITS-CLIP | 23597149 | Unspecified |
| mmu-let-7i-5p | MIMAT0000122 | Tmem178b | 434008 | HITS-CLIP | 19536157 | Unspecified |
| mmu-let-7i-5p | MIMAT0000122 | Tmem179b | 374395 | HITS-CLIP | 23142080 | Unspecified |
| mmu-let-7i-5p | MIMAT0000122 | Tmem181a | 77106 | HITS-CLIP | 19536157 | Unspecified |
| mmu-let-7i-5p | MIMAT0000122 | Tmem184b | 223693 | HITS-CLIP | 23142080\|29167373 | Unspecified |
| mmu-let-7i-5p | MIMAT0000122 | Tmem184c | 234463 | HITS-CLIP | 23142080 | Unspecified |
| mmu-let-7i-5p | MIMAT0000122 | Tmem185b | 226351 | HITS-CLIP | 23142080 | Unspecified |
| mmu-let-7i-5p | MIMAT0000122 | Tmem186 | 66690 | HITS-CLIP | 37986224\|19536157\|23597149\|23142 | Unspecified |
| mmu-let-7i-5p | MIMAT0000122 | Tmem189 | 407243 | HITS-CLIP | 37986224 | Unspecified |
| mmu-let-7i-5p | MIMAT0000122 | Tmem192 | 73067 | HITS-CLIP | 23142080\|29167373 | Unspecified |
| mmu-let-7i-5p | MIMAT0000122 | Tmem201 | 230917 | HITS-CLIP | 23597149\|23142080 | Unspecified |
| mmu-let-7i-5p | MIMAT0000122 | Tmem205 | 235043 | HITS-CLIP | 23597149 | Unspecified |
| mmu-let-7i-5p | MIMAT0000122 | Tmem209 | 72649 | HITS-CLIP | 23142080 | Unspecified |
| mmu-let-7i-5p | MIMAT0000122 | Tmem214 | 68796 | HITS-CLIP | 23597149 | Unspecified |
| mmu-let-7i-5p | MIMAT0000122 | Tmem223 | 79064 | HITS-CLIP | 23597149 | Unspecified |
| mmu-let-7i-5p | MIMAT0000122 | Tmem229b | 268567 | HITS-CLIP | 23142080 | Unspecified |
| mmu-let-7i-5p | MIMAT0000122 | Tmem237 | 381259 | HITS-CLIP | 23142080 | Unspecified |
| mmu-let-7i-5p | MIMAT0000122 | Tmem245 | 242474 | HITS-CLIP | 37986224 | Unspecified |
| mmu-let-7i-5p | MIMAT0000122 | Tmem256 | 254863 | HITS-CLIP | 23142080 | Unspecified |
| mmu-let-7i-5p | MIMAT0000122 | Tmem258 | 69038 | HITS-CLIP | 23597149 | Unspecified |
| mmu-let-7i-5p | MIMAT0000122 | Tmem260 | 218989 | HITS-CLIP | 23142080 | Unspecified |
| mmu-let-7i-5p | MIMAT0000122 | Tmem263 | 103266 | HITS-CLIP | 19536157\|23142080 | Unspecified |
| mmu-let-7i-5p | MIMAT0000122 | Tmem30a | 69981 | HITS-CLIP | 37986224\|23597149\|23142080 | Unspecified |
| mmu-let-7i-5p | MIMAT0000122 | Tmem33 | 67878 | HITS-CLIP | 37986224\|19536157\|23597149\|23142 | Unspecified |
| mmu-let-7i-5p | MIMAT0000122 | Tmem37 | 170706 | HITS-CLIP | 37986224 | Unspecified |
| mmu-let-7i-5p | MIMAT0000122 | Tmem38b | 52076 | HITS-CLIP | 37986224\|23597149 | Unspecified |
| mmu-let-7i-5p | MIMAT0000122 | Tmem39a | 67846 | HITS-CLIP | 23142080 | Unspecified |
| mmu-let-7i-5p | MIMAT0000122 | Tmem39b | 230770 | HITS-CLIP | 37986224 | Unspecified |
| mmu-let-7i-5p | MIMAT0000122 | Tmem41a | 66664 | HITS-CLIP | 37986224\|23597149\|23142080 | Unspecified |
| mmu-let-7i-5p | MIMAT0000122 | Tmem53 | 68777 | HITS-CLIP | 23597149 | Unspecified |
| mmu-let-7i-5p | MIMAT0000122 | Tmem56 | 99887 | HITS-CLIP | 37986224\|23597149\|19536157 | Unspecified |
| mmu-let-7i-5p | MIMAT0000122 | Tmem59 | 56374 | HITS-CLIP | 23597149\|23142080 | Unspecified |
| mmu-let-7i-5p | MIMAT0000122 | Tmem60 | 212090 | HITS-CLIP | 37986224\|19536157\|23142080 | Unspecified |
| mmu-let-7i-5p | MIMAT0000122 | Tmem64 | 100201 | HITS-CLIP | 31433990\|23142080 | Unspecified |
| mmu-let-7i-5p | MIMAT0000122 | Tmem65 | 74868 | HITS-CLIP | 37986224\|19536157\|23597149\|23142 | Unspecified |
| mmu-let-7i-5p | MIMAT0000122 | Tmem69 | 230657 | HITS-CLIP | 23142080 | Unspecified |
| mmu-let-7i-5p | MIMAT0000122 | Tmem8 | 60455 | HITS-CLIP | 23597149 | Unspecified |
| mmu-let-7i-5p | MIMAT0000122 | Tmem82 | 213989 | HITS-CLIP | 23597149 | Unspecified |

| | | | | | | |
|---|---|---|---|---|---|---|
| mmu-let-7i-5p | MIMAT0000122 | Tmem86a | 67893 | HITS-CLIP | 23142080 | Unspecified |
| mmu-let-7i-5p | MIMAT0000122 | Tmem86b | 68255 | HITS-CLIP | 23597149 | Unspecified |
| mmu-let-7i-5p | MIMAT0000122 | Tmem87a | 211499 | HITS-CLIP | 23142080 | Unspecified |
| mmu-let-7i-5p | MIMAT0000122 | Tmem87b | 72477 | HITS-CLIP | 23142080\|29167373 | Unspecified |
| mmu-let-7i-5p | MIMAT0000122 | Tmem9 | 66241 | HITS-CLIP | 23597149 | Unspecified |
| mmu-let-7i-5p | MIMAT0000122 | Tmem94 | 9772 | HITS-CLIP | 23142080 | Unspecified |
| mmu-let-7i-5p | MIMAT0000122 | Tmod2 | 50876 | HITS-CLIP | 19536157 | Unspecified |
| mmu-let-7i-5p | MIMAT0000122 | Tmod3 | 50875 | HITS-CLIP | 23142080 | Unspecified |
| mmu-let-7i-5p | MIMAT0000122 | Tmpo | 21917 | HITS-CLIP | 37986224\|23142080 | Unspecified |
| mmu-let-7i-5p | MIMAT0000122 | Tmppe | 100504715 | HITS-CLIP | 37986224 | Unspecified |
| mmu-let-7i-5p | MIMAT0000122 | Tmprss2 | 50528 | HITS-CLIP | 37986224\|23597149 | Unspecified |
| mmu-let-7i-5p | MIMAT0000122 | Tmsb10 | 19240 | HITS-CLIP | 23142080 | Unspecified |
| mmu-let-7i-5p | MIMAT0000122 | Tmsb4x | 19241 | HITS-CLIP | 19536157 | Unspecified |
| mmu-let-7i-5p | MIMAT0000122 | Tmtc2 | 278279 | HITS-CLIP | 23142080 | Unspecified |
| mmu-let-7i-5p | MIMAT0000122 | Tmtc3 | 237500 | HITS-CLIP | 23142080 | Unspecified |
| mmu-let-7i-5p | MIMAT0000122 | Tmx3 | 67988 | HITS-CLIP | 23142080 | Unspecified |
| mmu-let-7i-5p | MIMAT0000122 | Tmx4 | 52837 | HITS-CLIP | 37986224\|23597149\|29167373\|23142 | Unspecified |
| mmu-let-7i-5p | MIMAT0000122 | Tnc | 21923 | HITS-CLIP | 19536157 | Unspecified |
| mmu-let-7i-5p | MIMAT0000122 | Tnf | 21926 | HITS-CLIP | 23142080 | Unspecified |
| mmu-let-7i-5p | MIMAT0000122 | Tnfaip1 | 21927 | HITS-CLIP | 23597149\|23142080\|29167373 | Unspecified |
| mmu-let-7i-5p | MIMAT0000122 | Tnfaip3 | 21929 | HITS-CLIP | 23597149\|23142080\|29167373 | Unspecified |
| mmu-let-7i-5p | MIMAT0000122 | Tnfaip8 | 106869 | HITS-CLIP | 23142080 | Unspecified |
| mmu-let-7i-5p | MIMAT0000122 | Tnfaip8l2 | 69769 | HITS-CLIP | 23142080 | Unspecified |
| mmu-let-7i-5p | MIMAT0000122 | Tnfrsf10b | 21933 | HITS-CLIP | 23142080 | Unspecified |
| mmu-let-7i-5p | MIMAT0000122 | Tnfrsf11a | 21934 | HITS-CLIP | 29167373 | Unspecified |
| mmu-let-7i-5p | MIMAT0000122 | Tnfrsf1a | 21937 | HITS-CLIP | 37986224\|23597149\|29167373\|23142 | Unspecified |
| mmu-let-7i-5p | MIMAT0000122 | Tnfrsf1b | 21938 | HITS-CLIP | 37986224\|23597149\|29167373 | Unspecified |
| mmu-let-7i-5p | MIMAT0000122 | Tnfrsf26 | 244237 | HITS-CLIP | 21258322\|23142080 | Unspecified |
| mmu-let-7i-5p | MIMAT0000122 | Tnfrsf4 | 22163 | HITS-CLIP | 23142080 | Unspecified |
| mmu-let-7i-5p | MIMAT0000122 | Tnfrsf9 | 21942 | HITS-CLIP | 23142080 | Unspecified |
| mmu-let-7i-5p | MIMAT0000122 | Tnfsf11 | 21943 | HITS-CLIP | 23142080 | Unspecified |
| mmu-let-7i-5p | MIMAT0000122 | Tnfsf12 | 21944 | HITS-CLIP | 23597149\|29167373 | Unspecified |
| mmu-let-7i-5p | MIMAT0000122 | Tnfsf8 | 21949 | HITS-CLIP | 37986224\|23142080 | Unspecified |
| mmu-let-7i-5p | MIMAT0000122 | Tnfsf9 | 21950 | HITS-CLIP | 29167373 | Unspecified |
| mmu-let-7i-5p | MIMAT0000122 | Tnik | 665113 | HITS-CLIP | 23142080 | Unspecified |
| mmu-let-7i-5p | MIMAT0000122 | Tnk2 | 51789 | HITS-CLIP | 23597149 | Unspecified |
| mmu-let-7i-5p | MIMAT0000122 | Tnks | 21951 | HITS-CLIP | 23142080 | Unspecified |
| mmu-let-7i-5p | MIMAT0000122 | Tnks2 | 74493 | HITS-CLIP//qCLASH | 37986224\|23597149\|23142080\|31433 | Unspecified |
| mmu-let-7i-5p | MIMAT0000122 | Tnpo1 | 238799 | HITS-CLIP | 37986224\|19536157\|23142080\|29167 | Unspecified |

| | | | | | | |
|---|---|---|---|---|---|---|
| mmu-let-7i-5p | MIMAT0000122 | Tnpo2 | 212999 | HITS-CLIP | 23142080\|23597149\|29167373 | Unspecified |
| mmu-let-7i-5p | MIMAT0000122 | Tnrc18 | 231861 | HITS-CLIP | 23597149\|29167373 | Unspecified |
| mmu-let-7i-5p | MIMAT0000122 | Tnrc6a | 233833 | HITS-CLIP | 37986224\|23597149\|31433990\|19536 | Unspecified |
| mmu-let-7i-5p | MIMAT0000122 | Tnrc6b | 213988 | HITS-CLIP | 37986224\|19536157\|23597149\|23142 | Unspecified |
| mmu-let-7i-5p | MIMAT0000122 | Tnrc6c | 217351 | HITS-CLIP | 37986224\|31433990\|19536157\|23597 | Unspecified |
| mmu-let-7i-5p | MIMAT0000122 | Tns2 | 209039 | HITS-CLIP | 23597149 | Unspecified |
| mmu-let-7i-5p | MIMAT0000122 | Tns3 | 319939 | HITS-CLIP | 23597149\|19536157\|29167373 | Unspecified |
| mmu-let-7i-5p | MIMAT0000122 | Tob1 | 22057 | HITS-CLIP | 37986224\|23597149\|23142080 | Unspecified |
| mmu-let-7i-5p | MIMAT0000122 | Tob2 | 57259 | HITS-CLIP | 37986224\|23142080\|29167373 | Unspecified |
| mmu-let-7i-5p | MIMAT0000122 | Togaram1 | 23116 | HITS-CLIP | 23142080 | Unspecified |
| mmu-let-7i-5p | MIMAT0000122 | Tollip | 54473 | HITS-CLIP | 29167373 | Unspecified |
| mmu-let-7i-5p | MIMAT0000122 | Tom1l2 | 216810 | HITS-CLIP | 23597149 | Unspecified |
| mmu-let-7i-5p | MIMAT0000122 | Tomm70a | 28185 | HITS-CLIP | 23142080 | Unspecified |
| mmu-let-7i-5p | MIMAT0000122 | Tonsl | 72749 | HITS-CLIP | 37986224\|23142080 | Unspecified |
| mmu-let-7i-5p | MIMAT0000122 | Top1 | 21969 | HITS-CLIP | 23597149\|23142080\|29167373 | Unspecified |
| mmu-let-7i-5p | MIMAT0000122 | Top2a | 21973 | HITS-CLIP | 23597149\|23142080 | Unspecified |
| mmu-let-7i-5p | MIMAT0000122 | Top2b | 21974 | HITS-CLIP | 23597149\|23142080\|29167373 | Unspecified |
| mmu-let-7i-5p | MIMAT0000122 | Top3a | 21975 | HITS-CLIP | 29167373 | Unspecified |
| mmu-let-7i-5p | MIMAT0000122 | Top3b | 21976 | HITS-CLIP | 23142080 | Unspecified |
| mmu-let-7i-5p | MIMAT0000122 | Topbp1 | 235559 | HITS-CLIP | 29167373\|23142080 | Unspecified |
| mmu-let-7i-5p | MIMAT0000122 | Topors | 106021 | HITS-CLIP | 23142080 | Unspecified |
| mmu-let-7i-5p | MIMAT0000122 | Tor1aip1 | 208263 | HITS-CLIP | 37986224\|23142080\|29167373 | Unspecified |
| mmu-let-7i-5p | MIMAT0000122 | Tor1aip2 | 240832 | HITS-CLIP | 37986224\|23597149\|19536157\|23142 | Unspecified |
| mmu-let-7i-5p | MIMAT0000122 | Tor2a | 30933 | HITS-CLIP | 37986224\|23142080 | Unspecified |
| mmu-let-7i-5p | MIMAT0000122 | Tor4a | 227612 | HITS-CLIP | 23142080 | Unspecified |
| mmu-let-7i-5p | MIMAT0000122 | Tox | 252838 | HITS-CLIP | 23597149\|23142080 | Unspecified |
| mmu-let-7i-5p | MIMAT0000122 | Tox4 | 268741 | HITS-CLIP | 37986224\|23142080 | Unspecified |
| mmu-let-7i-5p | MIMAT0000122 | Tpcn1 | 252972 | HITS-CLIP | 37986224\|23597149\|23142080 | Unspecified |
| mmu-let-7i-5p | MIMAT0000122 | Tpd52l2 | 66314 | HITS-CLIP | 23142080 | Unspecified |
| mmu-let-7i-5p | MIMAT0000122 | Tpgs2 | 66648 | HITS-CLIP | 37986224\|19536157 | Unspecified |
| mmu-let-7i-5p | MIMAT0000122 | Tpi1 | 21991 | HITS-CLIP | 23142080\|23597149 | Unspecified |
| mmu-let-7i-5p | MIMAT0000122 | Tpk1 | 29807 | HITS-CLIP | 23597149 | Unspecified |
| mmu-let-7i-5p | MIMAT0000122 | Tpmt | 22017 | HITS-CLIP | 23597149 | Unspecified |
| mmu-let-7i-5p | MIMAT0000122 | Tpp1 | 12751 | HITS-CLIP | 23142080\|29167373 | Unspecified |
| mmu-let-7i-5p | MIMAT0000122 | Tpp2 | 22019 | HITS-CLIP | 37986224\|23597149\|19536157\|23142 | Unspecified |
| mmu-let-7i-5p | MIMAT0000122 | Tppp3 | 67971 | HITS-CLIP | 19536157 | Unspecified |
| mmu-let-7i-5p | MIMAT0000122 | Tpr | 108989 | HITS-CLIP | 23142080\|29167373 | Unspecified |
| mmu-let-7i-5p | MIMAT0000122 | Tprkb | 69786 | HITS-CLIP | 23597149 | Unspecified |
| mmu-let-7i-5p | MIMAT0000122 | Tprn | 97031 | HITS-CLIP | 19536157 | Unspecified |

| | | | | | | |
|---|---|---|---|---|---|---|
| mmu-let-7i-5p | MIMAT0000122 | Tpst2 | 22022 | HITS-CLIP | 37986224\|23597149 | Unspecified |
| mmu-let-7i-5p | MIMAT0000122 | Tpx2 | 72119 | HITS-CLIP | 23597149 | Unspecified |
| mmu-let-7i-5p | MIMAT0000122 | Tra2a | 101214 | HITS-CLIP | 23142080 | Unspecified |
| mmu-let-7i-5p | MIMAT0000122 | Trabd | 67976 | HITS-CLIP | 23597149\|23142080 | Unspecified |
| mmu-let-7i-5p | MIMAT0000122 | Traf1 | 22029 | HITS-CLIP | 23142080 | Unspecified |
| mmu-let-7i-5p | MIMAT0000122 | Traf3 | 22031 | HITS-CLIP | 23142080 | Unspecified |
| mmu-let-7i-5p | MIMAT0000122 | Traf3ip2 | 103213 | HITS-CLIP | 23142080 | Unspecified |
| mmu-let-7i-5p | MIMAT0000122 | Traf3ip3 | 215243 | HITS-CLIP | 23142080 | Unspecified |
| mmu-let-7i-5p | MIMAT0000122 | Traf6 | 22034 | HITS-CLIP | 23142080 | Unspecified |
| mmu-let-7i-5p | MIMAT0000122 | Traf7 | 224619 | HITS-CLIP | 23142080 | Unspecified |
| mmu-let-7i-5p | MIMAT0000122 | Trak1 | 67095 | HITS-CLIP | 23142080 | Unspecified |
| mmu-let-7i-5p | MIMAT0000122 | Trak2 | 70827 | HITS-CLIP | 23142080 | Unspecified |
| mmu-let-7i-5p | MIMAT0000122 | Tram1 | 72265 | HITS-CLIP | 23597149\|19536157\|23142080\|29167 | Unspecified |
| mmu-let-7i-5p | MIMAT0000122 | Tram1l1 | 229801 | HITS-CLIP | 37986224 | Unspecified |
| mmu-let-7i-5p | MIMAT0000122 | Tram2 | 170829 | HITS-CLIP | 37986224\|23597149\|23142080 | Unspecified |
| mmu-let-7i-5p | MIMAT0000122 | Trap1 | 68015 | HITS-CLIP | 23142080 | Unspecified |
| mmu-let-7i-5p | MIMAT0000122 | Trappc1 | 245828 | HITS-CLIP | 37986224\|23597149\|19536157\|23142 | Unspecified |
| mmu-let-7i-5p | MIMAT0000122 | Trappc10 | 216131 | HITS-CLIP | 37986224\|23597149 | Unspecified |
| mmu-let-7i-5p | MIMAT0000122 | Trappc11 | 320714 | HITS-CLIP | 37986224\|23597149\|23142080 | Unspecified |
| mmu-let-7i-5p | MIMAT0000122 | Trappc12 | 217449 | HITS-CLIP | 23597149\|29167373 | Unspecified |
| mmu-let-7i-5p | MIMAT0000122 | Trappc13 | 66975 | HITS-CLIP | 23142080 | Unspecified |
| mmu-let-7i-5p | MIMAT0000122 | Trappc3 | 27096 | HITS-CLIP | 23142080 | Unspecified |
| mmu-let-7i-5p | MIMAT0000122 | Trappc4 | 60409 | HITS-CLIP | 23142080 | Unspecified |
| mmu-let-7i-5p | MIMAT0000122 | Trappc5 | 66682 | HITS-CLIP | 37986224\|23142080 | Unspecified |
| mmu-let-7i-5p | MIMAT0000122 | Trappc8 | 75964 | HITS-CLIP | 23142080\|29167373 | Unspecified |
| mmu-let-7i-5p | MIMAT0000122 | Trappc9 | 76510 | HITS-CLIP | 23597149\|23142080 | Unspecified |
| mmu-let-7i-5p | MIMAT0000122 | Treml4 | 224840 | HITS-CLIP | 23597149 | Unspecified |
| mmu-let-7i-5p | MIMAT0000122 | Trerf1 | 224829 | HITS-CLIP | 31433990 | Unspecified |
| mmu-let-7i-5p | MIMAT0000122 | Trhde | 237553 | HITS-CLIP | 37986224\|19536157 | Unspecified |
| mmu-let-7i-5p | MIMAT0000122 | Triap1 | 69076 | HITS-CLIP | 23597149\|23142080 | Unspecified |
| mmu-let-7i-5p | MIMAT0000122 | Trib1 | 211770 | HITS-CLIP | 37986224\|23597149\|19536157\|23142 | Unspecified |
| mmu-let-7i-5p | MIMAT0000122 | Trib2 | 217410 | HITS-CLIP | 37986224\|23597149\|23142080\|19536 | Unspecified |
| mmu-let-7i-5p | MIMAT0000122 | Trib3 | 228775 | HITS-CLIP | 23597149 | Unspecified |
| mmu-let-7i-5p | MIMAT0000122 | Trim12c | 319236 | HITS-CLIP | 23142080 | Unspecified |
| mmu-let-7i-5p | MIMAT0000122 | Trim23 | 81003 | HITS-CLIP | 19536157\|23597149 | Unspecified |
| mmu-let-7i-5p | MIMAT0000122 | Trim25 | 217069 | HITS-CLIP | 37986224\|23142080\|29167373 | Unspecified |
| mmu-let-7i-5p | MIMAT0000122 | Trim26 | 22670 | HITS-CLIP | 23142080\|29167373 | Unspecified |
| mmu-let-7i-5p | MIMAT0000122 | Trim27 | 19720 | HITS-CLIP | 37986224\|23142080\|23597149 | Unspecified |
| mmu-let-7i-5p | MIMAT0000122 | Trim28 | 21849 | HITS-CLIP | 37986224\|23142080 | Unspecified |

| | | | | | | |
|---|---|---|---|---|---|---|
| mmu-let-7i-5p | MIMAT0000122 | Trim32 | 69807 | HITS-CLIP | 37986224\|23630078\|23597149\|26203 | Unspecified |
| mmu-let-7i-5p | MIMAT0000122 | Trim33 | 94093 | HITS-CLIP | 23597149\|23142080\|29167373 | Unspecified |
| mmu-let-7i-5p | MIMAT0000122 | Trim34a | 94094 | HITS-CLIP | 23142080 | Unspecified |
| mmu-let-7i-5p | MIMAT0000122 | Trim35 | 66854 | HITS-CLIP | 23142080\|29167373 | Unspecified |
| mmu-let-7i-5p | MIMAT0000122 | Trim41 | 211007 | HITS-CLIP | 37986224\|23142080\|29167373 | Unspecified |
| mmu-let-7i-5p | MIMAT0000122 | Trim56 | 384309 | HITS-CLIP | 23142080\|29167373 | Unspecified |
| mmu-let-7i-5p | MIMAT0000122 | Trim59 | 66949 | HITS-CLIP | 23142080 | Unspecified |
| mmu-let-7i-5p | MIMAT0000122 | Trim6 | 94088 | HITS-CLIP | 23142080 | Unspecified |
| mmu-let-7i-5p | MIMAT0000122 | Trim66 | 330627 | HITS-CLIP | 37986224 | Unspecified |
| mmu-let-7i-5p | MIMAT0000122 | Trim68 | 101700 | HITS-CLIP | 23597149\|23142080 | Unspecified |
| mmu-let-7i-5p | MIMAT0000122 | Trim71 | 636931 | HITS-CLIP | 23142080 | Unspecified |
| mmu-let-7i-5p | MIMAT0000122 | Trio | 223435 | HITS-CLIP | 19536157\|23597149\|23142080\|29167 | Unspecified |
| mmu-let-7i-5p | MIMAT0000122 | Triobp | 110253 | HITS-CLIP | 23142080 | Unspecified |
| mmu-let-7i-5p | MIMAT0000122 | Trip11 | 109181 | HITS-CLIP | 23597149\|23142080 | Unspecified |
| mmu-let-7i-5p | MIMAT0000122 | Trip12 | 14897 | HITS-CLIP | 37986224\|23597149\|19536157\|23142 | Unspecified |
| mmu-let-7i-5p | MIMAT0000122 | Trip13 | 69716 | HITS-CLIP | 37986224\|23142080 | Unspecified |
| mmu-let-7i-5p | MIMAT0000122 | Trip4 | 56404 | HITS-CLIP | 23597149 | Unspecified |
| mmu-let-7i-5p | MIMAT0000122 | Trmt10a | 108943 | HITS-CLIP | 23142080 | Unspecified |
| mmu-let-7i-5p | MIMAT0000122 | Trmt10c | 52575 | HITS-CLIP | 23597149\|23142080 | Unspecified |
| mmu-let-7i-5p | MIMAT0000122 | Trmt1l | 98685 | HITS-CLIP | 37986224\|29167373\|23142080 | Unspecified |
| mmu-let-7i-5p | MIMAT0000122 | Trmt5 | 76357 | HITS-CLIP | 37986224\|23142080 | Unspecified |
| mmu-let-7i-5p | MIMAT0000122 | Trmt6 | 66926 | HITS-CLIP | 23142080 | Unspecified |
| mmu-let-7i-5p | MIMAT0000122 | Trnp1 | 69539 | HITS-CLIP | 19536157 | Unspecified |
| mmu-let-7i-5p | MIMAT0000122 | Trnt1 | 70047 | HITS-CLIP | 23597149 | Unspecified |
| mmu-let-7i-5p | MIMAT0000122 | Trove2 | 20822 | HITS-CLIP | 23142080 | Unspecified |
| mmu-let-7i-5p | MIMAT0000122 | Trp53 | 22059 | HITS-CLIP | 23142080 | Unspecified |
| mmu-let-7i-5p | MIMAT0000122 | Trp53bp1 | 27223 | HITS-CLIP | 23142080 | Unspecified |
| mmu-let-7i-5p | MIMAT0000122 | Trp53bp2 | 209456 | HITS-CLIP | 23597149\|23142080 | Unspecified |
| mmu-let-7i-5p | MIMAT0000122 | Trp53inp1 | 60599 | HITS-CLIP | 37986224\|31433990\|23142080 | Unspecified |
| mmu-let-7i-5p | MIMAT0000122 | Trp53inp2 | 68728 | HITS-CLIP | 19536157 | Unspecified |
| mmu-let-7i-5p | MIMAT0000122 | Trp53rkb | 76367 | HITS-CLIP | 19536157 | Unspecified |
| mmu-let-7i-5p | MIMAT0000122 | Trpc4ap | 56407 | HITS-CLIP | 23597149\|23142080 | Unspecified |
| mmu-let-7i-5p | MIMAT0000122 | Trpm3 | 226025 | HITS-CLIP | 19536157 | Unspecified |
| mmu-let-7i-5p | MIMAT0000122 | Trpm7 | 58800 | HITS-CLIP | 23142080 | Unspecified |
| mmu-let-7i-5p | MIMAT0000122 | Trps1 | 83925 | HITS-CLIP | 37986224\|23142080 | Unspecified |
| mmu-let-7i-5p | MIMAT0000122 | Trrap | 100683 | HITS-CLIP | 37986224\|23597149\|19536157\|23142 | Unspecified |
| mmu-let-7i-5p | MIMAT0000122 | Tsc1 | 64930 | HITS-CLIP | 23142080 | Unspecified |
| mmu-let-7i-5p | MIMAT0000122 | Tsc22d2 | 72033 | HITS-CLIP | 37986224\|19536157\|23597149\|23142 | Unspecified |
| mmu-let-7i-5p | MIMAT0000122 | Tsc22d3 | 14605 | HITS-CLIP | 23597149\|23142080 | Unspecified |

| | | | | | | |
|---|---|---|---|---|---|---|
| mmu-let-7i-5p | MIMAT0000122 | Tsc22d4 | 78829 | HITS-CLIP | 23142080 | Unspecified |
| mmu-let-7i-5p | MIMAT0000122 | Tsen34 | 66078 | HITS-CLIP | 23142080 | Unspecified |
| mmu-let-7i-5p | MIMAT0000122 | Tshz1 | 110796 | HITS-CLIP | 23597149\|23142080 | Unspecified |
| mmu-let-7i-5p | MIMAT0000122 | Tshz2 | 228911 | HITS-CLIP | 23597149 | Unspecified |
| mmu-let-7i-5p | MIMAT0000122 | Tsn | 22099 | HITS-CLIP | 30971389\|23142080 | Unspecified |
| mmu-let-7i-5p | MIMAT0000122 | Tspan13 | 66109 | HITS-CLIP | 23142080 | Unspecified |
| mmu-let-7i-5p | MIMAT0000122 | Tspan14 | 52588 | HITS-CLIP | 23142080\|29167373\|19536157 | Unspecified |
| mmu-let-7i-5p | MIMAT0000122 | Tspan17 | 74257 | HITS-CLIP | 37986224\|19536157 | Unspecified |
| mmu-let-7i-5p | MIMAT0000122 | Tspan3 | 56434 | HITS-CLIP | 37986224\|19536157\|23142080 | Unspecified |
| mmu-let-7i-5p | MIMAT0000122 | Tspan31 | 67125 | HITS-CLIP | 19536157 | Unspecified |
| mmu-let-7i-5p | MIMAT0000122 | Tspan32 | 27027 | HITS-CLIP | 23142080 | Unspecified |
| mmu-let-7i-5p | MIMAT0000122 | Tspan4 | 64540 | HITS-CLIP | 23597149\|29167373 | Unspecified |
| mmu-let-7i-5p | MIMAT0000122 | Tspan5 | 56224 | HITS-CLIP | 37986224\|23142080 | Unspecified |
| mmu-let-7i-5p | MIMAT0000122 | Tspoap1 | 207777 | HITS-CLIP | 19536157 | Unspecified |
| mmu-let-7i-5p | MIMAT0000122 | Tspyl1 | 22110 | HITS-CLIP | 23142080 | Unspecified |
| mmu-let-7i-5p | MIMAT0000122 | Tspyl2 | 52808 | HITS-CLIP | 23142080 | Unspecified |
| mmu-let-7i-5p | MIMAT0000122 | Tspyl3 | 241732 | HITS-CLIP | 23142080 | Unspecified |
| mmu-let-7i-5p | MIMAT0000122 | Tsr1 | 104662 | HITS-CLIP | 26203562\|23142080 | Unspecified |
| mmu-let-7i-5p | MIMAT0000122 | Tsr3 | 68327 | HITS-CLIP | 23142080 | Unspecified |
| mmu-let-7i-5p | MIMAT0000122 | Tsta3 | 22122 | HITS-CLIP | 23142080 | Unspecified |
| mmu-let-7i-5p | MIMAT0000122 | Ttc13 | 234875 | HITS-CLIP | 29167373 | Unspecified |
| mmu-let-7i-5p | MIMAT0000122 | Ttc14 | 67120 | HITS-CLIP | 37986224\|31433990\|19536157\|23142 | Unspecified |
| mmu-let-7i-5p | MIMAT0000122 | Ttc17 | 74569 | HITS-CLIP | 23597149\|19536157\|23142080\|29167 | Unspecified |
| mmu-let-7i-5p | MIMAT0000122 | Ttc19 | 72795 | HITS-CLIP | 23142080 | Unspecified |
| mmu-let-7i-5p | MIMAT0000122 | Ttc23 | 67009 | HITS-CLIP | 23142080 | Unspecified |
| mmu-let-7i-5p | MIMAT0000122 | Ttc26 | 264134 | HITS-CLIP | 23142080 | Unspecified |
| mmu-let-7i-5p | MIMAT0000122 | Ttc27 | 74196 | HITS-CLIP | 19536157 | Unspecified |
| mmu-let-7i-5p | MIMAT0000122 | Ttc3 | 22129 | HITS-CLIP | 23142080\|19536157 | Unspecified |
| mmu-let-7i-5p | MIMAT0000122 | Ttc33 | 67515 | HITS-CLIP | 23142080 | Unspecified |
| mmu-let-7i-5p | MIMAT0000122 | Ttc36 | 192653 | HITS-CLIP | 23597149 | Unspecified |
| mmu-let-7i-5p | MIMAT0000122 | Ttc37 | 218343 | HITS-CLIP | 23142080 | Unspecified |
| mmu-let-7i-5p | MIMAT0000122 | Ttc39a | 230603 | HITS-CLIP | 23597149 | Unspecified |
| mmu-let-7i-5p | MIMAT0000122 | Ttc39b | 69863 | HITS-CLIP | 37986224\|23597149\|23142080 | Unspecified |
| mmu-let-7i-5p | MIMAT0000122 | Ttc39c | 72747 | HITS-CLIP | 23597149 | Unspecified |
| mmu-let-7i-5p | MIMAT0000122 | Ttc4 | 72354 | HITS-CLIP | 23142080 | Unspecified |
| mmu-let-7i-5p | MIMAT0000122 | Ttc7 | 225049 | HITS-CLIP | 23597149\|29167373\|23142080 | Unspecified |
| mmu-let-7i-5p | MIMAT0000122 | Ttc8 | 76260 | qCLASH | 31393953 | Unspecified |
| mmu-let-7i-5p | MIMAT0000122 | Ttc9c | 70387 | HITS-CLIP | 23142080 | Unspecified |
| mmu-let-7i-5p | MIMAT0000122 | Ttf2 | 74044 | HITS-CLIP | 23142080 | Unspecified |

| | | | | | | |
|---|---|---|---|---|---|---|
| mmu-let-7i-5p | MIMAT0000122 | Tti1 | 75425 | HITS-CLIP | 37986224\|23597149\|23142080\|29167 | Unspecified |
| mmu-let-7i-5p | MIMAT0000122 | Tti2 | 234138 | HITS-CLIP | 37986224\|23597149 | Unspecified |
| mmu-let-7i-5p | MIMAT0000122 | Ttk | 22137 | HITS-CLIP | 23142080 | Unspecified |
| mmu-let-7i-5p | MIMAT0000122 | Ttll4 | 67534 | HITS-CLIP | 37986224\|23597149\|23142080 | Unspecified |
| mmu-let-7i-5p | MIMAT0000122 | Ttpa | 50500 | HITS-CLIP | 23597149 | Unspecified |
| mmu-let-7i-5p | MIMAT0000122 | Ttpal | 76080 | HITS-CLIP | 37986224 | Unspecified |
| mmu-let-7i-5p | MIMAT0000122 | Ttyh1 | 57776 | HITS-CLIP | 37986224 | Unspecified |
| mmu-let-7i-5p | MIMAT0000122 | Ttyh2 | 117160 | HITS-CLIP | 19536157\|23597149\|29167373 | Unspecified |
| mmu-let-7i-5p | MIMAT0000122 | Ttyh3 | 78339 | HITS-CLIP | 19536157\|29167373\|23142080 | Unspecified |
| mmu-let-7i-5p | MIMAT0000122 | Tub | 22141 | HITS-CLIP | 37986224 | Unspecified |
| mmu-let-7i-5p | MIMAT0000122 | Tuba1a | 22142 | HITS-CLIP | 23142080 | Unspecified |
| mmu-let-7i-5p | MIMAT0000122 | Tuba4a | 22145 | HITS-CLIP | 37986224\|19536157 | Unspecified |
| mmu-let-7i-5p | MIMAT0000122 | Tubb4a | 22153 | HITS-CLIP | 37986224 | Unspecified |
| mmu-let-7i-5p | MIMAT0000122 | Tubb4b | 227613 | HITS-CLIP | 23597149 | Unspecified |
| mmu-let-7i-5p | MIMAT0000122 | Tubb5 | 22154 | HITS-CLIP | 37986224\|23597149\|19536157\|23142 | Unspecified |
| mmu-let-7i-5p | MIMAT0000122 | Tubd1 | 56427 | HITS-CLIP | 23142080 | Unspecified |
| mmu-let-7i-5p | MIMAT0000122 | Tube1 | 71924 | HITS-CLIP | 23142080 | Unspecified |
| mmu-let-7i-5p | MIMAT0000122 | Tubg1 | 103733 | HITS-CLIP | 23597149 | Unspecified |
| mmu-let-7i-5p | MIMAT0000122 | Tubgcp3 | 259279 | HITS-CLIP | 23142080 | Unspecified |
| mmu-let-7i-5p | MIMAT0000122 | Tubgcp4 | 51885 | qCLASH//HITS-CLIP | 31393953\|23142080 | Unspecified |
| mmu-let-7i-5p | MIMAT0000122 | Tubgcp5 | 233276 | HITS-CLIP | 23142080 | Unspecified |
| mmu-let-7i-5p | MIMAT0000122 | Tusc2 | 80385 | HITS-CLIP | 37986224\|19536157\|29167373 | Unspecified |
| mmu-let-7i-5p | MIMAT0000122 | Tut1 | 70044 | HITS-CLIP | 37986224\|23142080 | Unspecified |
| mmu-let-7i-5p | MIMAT0000122 | Tut4 | 23318 | HITS-CLIP | 19536157\|23597149\|23142080 | Unspecified |
| mmu-let-7i-5p | MIMAT0000122 | Tut7 | 79670 | HITS-CLIP | 23142080\|23597149 | Unspecified |
| mmu-let-7i-5p | MIMAT0000122 | Twf1 | 19230 | HITS-CLIP | 23597149\|23142080 | Unspecified |
| mmu-let-7i-5p | MIMAT0000122 | Twf2 | 23999 | HITS-CLIP | 23142080 | Unspecified |
| mmu-let-7i-5p | MIMAT0000122 | Txlna | 109658 | HITS-CLIP | 23597149\|23142080 | Unspecified |
| mmu-let-7i-5p | MIMAT0000122 | Txn2 | 56551 | HITS-CLIP | 37986224\|23597149\|19536157\|23142 | Unspecified |
| mmu-let-7i-5p | MIMAT0000122 | Txndc11 | 106200 | HITS-CLIP | 37986224\|30971389\|23597149\|29167 | Unspecified |
| mmu-let-7i-5p | MIMAT0000122 | Txndc12 | 66073 | HITS-CLIP | 23142080 | Unspecified |
| mmu-let-7i-5p | MIMAT0000122 | Txndc16 | 70561 | HITS-CLIP | 23142080 | Unspecified |
| mmu-let-7i-5p | MIMAT0000122 | Txndc5 | 105245 | HITS-CLIP | 23142080 | Unspecified |
| mmu-let-7i-5p | MIMAT0000122 | Txnip | 56338 | HITS-CLIP | 23597149\|23142080\|29167373 | Unspecified |
| mmu-let-7i-5p | MIMAT0000122 | Txnl1 | 53382 | HITS-CLIP | 23142080\|29167373 | Unspecified |
| mmu-let-7i-5p | MIMAT0000122 | Txnrd1 | 50493 | HITS-CLIP | 23597149 | Unspecified |
| mmu-let-7i-5p | MIMAT0000122 | Txnrd2 | 26462 | HITS-CLIP | 23597149 | Unspecified |
| mmu-let-7i-5p | MIMAT0000122 | Txnrd3 | 232223 | HITS-CLIP | 23597149 | Unspecified |
| mmu-let-7i-5p | MIMAT0000122 | Tyk2 | 54721 | HITS-CLIP | 37986224\|23142080\|23597149\|29167 | Unspecified |

| | | | | | | | |
|---|---|---|---|---|---|---|---|
| mmu-let-7i-5p | MIMAT0000122 | Tyro3 | 22174 | HITS-CLIP | | 19536157 | Unspecified |
| mmu-let-7i-5p | MIMAT0000122 | Tysnd1 | 71767 | HITS-CLIP | | 23142080 | Unspecified |
| mmu-let-7i-5p | MIMAT0000122 | Tyw1 | 100929 | HITS-CLIP | 37986224\|23597149 | | Unspecified |
| mmu-let-7i-5p | MIMAT0000122 | U2surp | 67958 | HITS-CLIP | | 23142080 | Unspecified |
| mmu-let-7i-5p | MIMAT0000122 | Uap1 | 107652 | HITS-CLIP | | 37986224 | Unspecified |
| mmu-let-7i-5p | MIMAT0000122 | Uba1 | 22201 | HITS-CLIP | 23597149\|23142080 | | Unspecified |
| mmu-let-7i-5p | MIMAT0000122 | Uba2 | 50995 | HITS-CLIP | | 23142080 | Unspecified |
| mmu-let-7i-5p | MIMAT0000122 | Uba3 | 22200 | HITS-CLIP | 23142080\|23597149 | | Unspecified |
| mmu-let-7i-5p | MIMAT0000122 | Uba6 | 231380 | HITS-CLIP | 23597149\|23142080 | | Unspecified |
| mmu-let-7i-5p | MIMAT0000122 | Ubac1 | 98766 | HITS-CLIP | 37986224\|23597149 | | Unspecified |
| mmu-let-7i-5p | MIMAT0000122 | Ubac2 | 68889 | HITS-CLIP | 37986224\|23597149\|23142080 | | Unspecified |
| mmu-let-7i-5p | MIMAT0000122 | Ubald1 | 207740 | HITS-CLIP | 23597149\|29167373 | | Unspecified |
| mmu-let-7i-5p | MIMAT0000122 | Ubap1 | 67123 | HITS-CLIP | 19536157\|23142080\|29167373 | | Unspecified |
| mmu-let-7i-5p | MIMAT0000122 | Ubap2 | 68926 | HITS-CLIP | 37986224\|23597149\|23142080\|29167 | | Unspecified |
| mmu-let-7i-5p | MIMAT0000122 | Ubap2l | 74383 | HITS-CLIP | 31433990\|29167373\|23142080 | | Unspecified |
| mmu-let-7i-5p | MIMAT0000122 | Ubash3a | 328795 | HITS-CLIP | | 23142080 | Unspecified |
| mmu-let-7i-5p | MIMAT0000122 | Ubash3b | 72828 | HITS-CLIP | | 23142080 | Unspecified |
| mmu-let-7i-5p | MIMAT0000122 | Ubc | 22190 | HITS-CLIP | 23142080\|23597149\|19536157\|29167 | | Unspecified |
| mmu-let-7i-5p | MIMAT0000122 | Ube2a | 22209 | HITS-CLIP | | 23142080 | Unspecified |
| mmu-let-7i-5p | MIMAT0000122 | Ube2b | 22210 | HITS-CLIP | 37986224\|23142080 | | Unspecified |
| mmu-let-7i-5p | MIMAT0000122 | Ube2c | 68612 | HITS-CLIP | | 23142080 | Unspecified |
| mmu-let-7i-5p | MIMAT0000122 | Ube2d2a | 56550 | HITS-CLIP | | 37986224 | Unspecified |
| mmu-let-7i-5p | MIMAT0000122 | Ube2d3 | 66105 | HITS-CLIP | 23142080\|23597149 | | Unspecified |
| mmu-let-7i-5p | MIMAT0000122 | Ube2e1 | 22194 | HITS-CLIP | | 23142080 | Unspecified |
| mmu-let-7i-5p | MIMAT0000122 | Ube2e3 | 22193 | HITS-CLIP | | 23142080 | Unspecified |
| mmu-let-7i-5p | MIMAT0000122 | Ube2g1 | 67128 | HITS-CLIP | | 23142080 | Unspecified |
| mmu-let-7i-5p | MIMAT0000122 | Ube2g2 | 22213 | HITS-CLIP | | 23142080 | Unspecified |
| mmu-let-7i-5p | MIMAT0000122 | Ube2h | 22214 | HITS-CLIP | 30971389\|23142080 | | Unspecified |
| mmu-let-7i-5p | MIMAT0000122 | Ube2j1 | 56228 | HITS-CLIP | 23597149\|29167373 | | Unspecified |
| mmu-let-7i-5p | MIMAT0000122 | Ube2k | 53323 | HITS-CLIP | | 23142080 | Unspecified |
| mmu-let-7i-5p | MIMAT0000122 | Ube2l3 | 22195 | HITS-CLIP | | 23142080 | Unspecified |
| mmu-let-7i-5p | MIMAT0000122 | Ube2l6 | 56791 | HITS-CLIP | 23597149\|23142080 | | Unspecified |
| mmu-let-7i-5p | MIMAT0000122 | Ube2m | 22192 | HITS-CLIP | 23597149\|23142080 | | Unspecified |
| mmu-let-7i-5p | MIMAT0000122 | Ube2o | 217342 | HITS-CLIP | | 19536157 | Unspecified |
| mmu-let-7i-5p | MIMAT0000122 | Ube2q1 | 70093 | HITS-CLIP | 19536157\|23142080\|29167373 | | Unspecified |
| mmu-let-7i-5p | MIMAT0000122 | Ube2q2 | 109161 | HITS-CLIP | 29167373\|23142080 | | Unspecified |
| mmu-let-7i-5p | MIMAT0000122 | Ube2r2 | 67615 | HITS-CLIP//qCLASH | 37986224\|31393953\|23142080 | | Unspecified |
| mmu-let-7i-5p | MIMAT0000122 | Ube2t | 67196 | HITS-CLIP | | 23142080 | Unspecified |
| mmu-let-7i-5p | MIMAT0000122 | Ube2v2 | 70620 | HITS-CLIP | | 23142080 | Unspecified |

| | | | | | | |
|---|---|---|---|---|---|---|
| mmu-let-7i-5p | MIMAT0000122 | Ube2w | 66799 | HITS-CLIP | 23597149\|23142080\|29167373 | Unspecified |
| mmu-let-7i-5p | MIMAT0000122 | Ube3a | 22215 | HITS-CLIP | 23142080\|29167373 | Unspecified |
| mmu-let-7i-5p | MIMAT0000122 | Ube3b | 117146 | HITS-CLIP | 23597149\|23142080 | Unspecified |
| mmu-let-7i-5p | MIMAT0000122 | Ube3c | 100763 | HITS-CLIP | 23142080 | Unspecified |
| mmu-let-7i-5p | MIMAT0000122 | Ube4a | 140630 | HITS-CLIP | 23142080 | Unspecified |
| mmu-let-7i-5p | MIMAT0000122 | Ube4b | 63958 | HITS-CLIP | 37986224\|23597149\|23142080 | Unspecified |
| mmu-let-7i-5p | MIMAT0000122 | Ubfd1 | 28018 | HITS-CLIP | 23597149\|19536157\|23142080 | Unspecified |
| mmu-let-7i-5p | MIMAT0000122 | Ubl3 | 24109 | HITS-CLIP | 23142080 | Unspecified |
| mmu-let-7i-5p | MIMAT0000122 | Ubn1 | 170644 | HITS-CLIP | 19536157\|23142080 | Unspecified |
| mmu-let-7i-5p | MIMAT0000122 | Ubn2 | 320538 | HITS-CLIP | 37986224\|23142080\|19536157\|23597 | Unspecified |
| mmu-let-7i-5p | MIMAT0000122 | Ubox5 | 140629 | HITS-CLIP | 37986224\|19536157\|29167373\|23142 | Unspecified |
| mmu-let-7i-5p | MIMAT0000122 | Ubqln1 | 56085 | HITS-CLIP | 23597149\|23142080 | Unspecified |
| mmu-let-7i-5p | MIMAT0000122 | Ubr1 | 22222 | HITS-CLIP | 19536157\|23142080 | Unspecified |
| mmu-let-7i-5p | MIMAT0000122 | Ubr2 | 224826 | HITS-CLIP//qCLASH | 37986224\|23597149\|31393953\|23142 | Unspecified |
| mmu-let-7i-5p | MIMAT0000122 | Ubr3 | 68795 | HITS-CLIP | 37986224\|23597149\|19536157\|23142 | Unspecified |
| mmu-let-7i-5p | MIMAT0000122 | Ubr4 | 69116 | HITS-CLIP | 37986224\|23597149\|23142080 | Unspecified |
| mmu-let-7i-5p | MIMAT0000122 | Ubr5 | 70790 | HITS-CLIP | 37986224\|23597149\|19536157\|23142 | Unspecified |
| mmu-let-7i-5p | MIMAT0000122 | Ubr7 | 66622 | HITS-CLIP | 23597149\|23142080 | Unspecified |
| mmu-let-7i-5p | MIMAT0000122 | Ubtd2 | 327900 | HITS-CLIP | 23142080 | Unspecified |
| mmu-let-7i-5p | MIMAT0000122 | Ubxn4 | 67812 | HITS-CLIP | 37986224\|23597149\|19536157\|23142 | Unspecified |
| mmu-let-7i-5p | MIMAT0000122 | Ubxn8 | 108159 | HITS-CLIP | 23142080 | Unspecified |
| mmu-let-7i-5p | MIMAT0000122 | Uchl1 | 22223 | HITS-CLIP | 19536157 | Unspecified |
| mmu-let-7i-5p | MIMAT0000122 | Uchl5 | 56207 | HITS-CLIP | 23142080 | Unspecified |
| mmu-let-7i-5p | MIMAT0000122 | Uck2 | 80914 | HITS-CLIP | 19536157 | Unspecified |
| mmu-let-7i-5p | MIMAT0000122 | Ucp2 | 22228 | HITS-CLIP | 37986224\|23142080\|29167373 | Unspecified |
| mmu-let-7i-5p | MIMAT0000122 | Ufd1 | 22230 | HITS-CLIP | 23597149 | Unspecified |
| mmu-let-7i-5p | MIMAT0000122 | Ufl1 | 67490 | HITS-CLIP | 37986224\|19536157\|23630078\|23597 | Unspecified |
| mmu-let-7i-5p | MIMAT0000122 | Ufsp2 | 192169 | HITS-CLIP | 23142080 | Unspecified |
| mmu-let-7i-5p | MIMAT0000122 | Ugcg | 22234 | HITS-CLIP | 37986224\|23597149\|19536157\|23142 | Unspecified |
| mmu-let-7i-5p | MIMAT0000122 | Ugdh | 22235 | HITS-CLIP | 23142080 | Unspecified |
| mmu-let-7i-5p | MIMAT0000122 | Uggt1 | 320011 | HITS-CLIP | 23597149 | Unspecified |
| mmu-let-7i-5p | MIMAT0000122 | Ugp2 | 216558 | HITS-CLIP | 23597149\|23142080 | Unspecified |
| mmu-let-7i-5p | MIMAT0000122 | Ugt1a7c | 394432 | HITS-CLIP | 23597149\|29167373 | Unspecified |
| mmu-let-7i-5p | MIMAT0000122 | Ugt2b34 | 100727 | HITS-CLIP | 23597149 | Unspecified |
| mmu-let-7i-5p | MIMAT0000122 | Ugt2b35 | 243085 | HITS-CLIP | 23597149 | Unspecified |
| mmu-let-7i-5p | MIMAT0000122 | Ugt2b36 | 231396 | HITS-CLIP | 23597149 | Unspecified |
| mmu-let-7i-5p | MIMAT0000122 | Uhmk1 | 16589 | HITS-CLIP | 37986224\|23597149\|19536157\|23142 | Unspecified |
| mmu-let-7i-5p | MIMAT0000122 | Uhrf1 | 18140 | HITS-CLIP | 29167373 | Unspecified |
| mmu-let-7i-5p | MIMAT0000122 | Uhrf1bp1l | 75089 | HITS-CLIP | 37986224\|23597149\|23142080\|29167 | Unspecified |

| | | | | | | |
|---|---|---|---|---|---|---|
| mmu-let-7i-5p | MIMAT0000122 | Uhrf2 | 109113 | HITS-CLIP | 37986224\|23597149\|19536157\|23142 | Unspecified |
| mmu-let-7i-5p | MIMAT0000122 | Uimc1 | 20184 | HITS-CLIP | 23142080 | Unspecified |
| mmu-let-7i-5p | MIMAT0000122 | Ulk1 | 22241 | HITS-CLIP | 23597149\|23142080 | Unspecified |
| mmu-let-7i-5p | MIMAT0000122 | Ulk2 | 29869 | HITS-CLIP | 37986224\|23597149\|23142080\|29167 | Unspecified |
| mmu-let-7i-5p | MIMAT0000122 | Ulk3 | 71742 | HITS-CLIP | 19536157\|23142080 | Unspecified |
| mmu-let-7i-5p | MIMAT0000122 | Unc119b | 106840 | HITS-CLIP | 23142080 | Unspecified |
| mmu-let-7i-5p | MIMAT0000122 | Unc13a | 382018 | HITS-CLIP | 19536157 | Unspecified |
| mmu-let-7i-5p | MIMAT0000122 | Unc45a | 101869 | HITS-CLIP | 23597149 | Unspecified |
| mmu-let-7i-5p | MIMAT0000122 | Unc5b | 107449 | HITS-CLIP | 19536157 | Unspecified |
| mmu-let-7i-5p | MIMAT0000122 | Unc5cl | 76589 | HITS-CLIP | 23142080 | Unspecified |
| mmu-let-7i-5p | MIMAT0000122 | Unc80 | 329178 | HITS-CLIP | 37986224\|19536157 | Unspecified |
| mmu-let-7i-5p | MIMAT0000122 | Unk | 217331 | HITS-CLIP | 23142080\|29167373 | Unspecified |
| mmu-let-7i-5p | MIMAT0000122 | Unkl | 74154 | HITS-CLIP | 23142080 | Unspecified |
| mmu-let-7i-5p | MIMAT0000122 | Uox | 22262 | HITS-CLIP | 23597149 | Unspecified |
| mmu-let-7i-5p | MIMAT0000122 | Upb1 | 103149 | HITS-CLIP | 23597149 | Unspecified |
| mmu-let-7i-5p | MIMAT0000122 | Upf1 | 19704 | HITS-CLIP//qCLASH | 37986224\|31393953 | Unspecified |
| mmu-let-7i-5p | MIMAT0000122 | Upf2 | 326622 | HITS-CLIP | 23142080 | Unspecified |
| mmu-let-7i-5p | MIMAT0000122 | Upp2 | 76654 | HITS-CLIP | 23597149 | Unspecified |
| mmu-let-7i-5p | MIMAT0000122 | Uprt | 331487 | HITS-CLIP | 23597149 | Unspecified |
| mmu-let-7i-5p | MIMAT0000122 | Uqcc1 | 56046 | HITS-CLIP | 23597149 | Unspecified |
| mmu-let-7i-5p | MIMAT0000122 | Uqcr10 | 66152 | HITS-CLIP | 23597149\|23142080 | Unspecified |
| mmu-let-7i-5p | MIMAT0000122 | Uqcrc1 | 22273 | HITS-CLIP//qCLASH | 37986224\|23597149\|31393953\|19536 | Unspecified |
| mmu-let-7i-5p | MIMAT0000122 | Uqcrc2 | 67003 | HITS-CLIP | 23142080 | Unspecified |
| mmu-let-7i-5p | MIMAT0000122 | Uqcrh | 66576 | HITS-CLIP | 37986224 | Unspecified |
| mmu-let-7i-5p | MIMAT0000122 | Urah | 76974 | HITS-CLIP | 23597149 | Unspecified |
| mmu-let-7i-5p | MIMAT0000122 | Urb2 | 382038 | HITS-CLIP | 37986224\|23142080 | Unspecified |
| mmu-let-7i-5p | MIMAT0000122 | Uri1 | 19777 | HITS-CLIP | 29167373 | Unspecified |
| mmu-let-7i-5p | MIMAT0000122 | Urm1 | 68205 | HITS-CLIP | 23142080 | Unspecified |
| mmu-let-7i-5p | MIMAT0000122 | Uroc1 | 243537 | HITS-CLIP | 23597149 | Unspecified |
| mmu-let-7i-5p | MIMAT0000122 | Usf3 | 205717 | HITS-CLIP | 23142080 | Unspecified |
| mmu-let-7i-5p | MIMAT0000122 | Ush1g | 16470 | HITS-CLIP | 23597149 | Unspecified |
| mmu-let-7i-5p | MIMAT0000122 | Uso1 | 56041 | HITS-CLIP | 23597149\|23142080\|29167373 | Unspecified |
| mmu-let-7i-5p | MIMAT0000122 | Usp1 | 230484 | HITS-CLIP | 23142080 | Unspecified |
| mmu-let-7i-5p | MIMAT0000122 | Usp10 | 22224 | HITS-CLIP | 23597149 | Unspecified |
| mmu-let-7i-5p | MIMAT0000122 | Usp11 | 236733 | HITS-CLIP | 37986224\|19536157 | Unspecified |
| mmu-let-7i-5p | MIMAT0000122 | Usp14 | 59025 | HITS-CLIP | 29167373 | Unspecified |
| mmu-let-7i-5p | MIMAT0000122 | Usp15 | 14479 | HITS-CLIP | 23142080\|19536157\|23597149\|29167 | Unspecified |
| mmu-let-7i-5p | MIMAT0000122 | Usp18 | 24110 | HITS-CLIP | 23142080 | Unspecified |
| mmu-let-7i-5p | MIMAT0000122 | Usp19 | 71472 | HITS-CLIP | 23142080 | Unspecified |

| | | | | | | |
|---|---|---|---|---|---|---|
| mmu-let-7i-5p | MIMAT0000122 | Usp2 | 53376 | HITS-CLIP | 23597149 | Unspecified |
| mmu-let-7i-5p | MIMAT0000122 | Usp21 | 30941 | HITS-CLIP | 23142080 | Unspecified |
| mmu-let-7i-5p | MIMAT0000122 | Usp24 | 329908 | HITS-CLIP | 37986224\|23142080\|19536157\|29167 | Unspecified |
| mmu-let-7i-5p | MIMAT0000122 | Usp25 | 30940 | HITS-CLIP | 23597149\|23142080\|29167373 | Unspecified |
| mmu-let-7i-5p | MIMAT0000122 | Usp28 | 235323 | HITS-CLIP | 19536157 | Unspecified |
| mmu-let-7i-5p | MIMAT0000122 | Usp3 | 235441 | HITS-CLIP | 23597149 | Unspecified |
| mmu-let-7i-5p | MIMAT0000122 | Usp31 | 76179 | HITS-CLIP | 37986224\|23142080\|29167373 | Unspecified |
| mmu-let-7i-5p | MIMAT0000122 | Usp32 | 237898 | HITS-CLIP | 37986224\|19536157\|23597149\|23142 | Unspecified |
| mmu-let-7i-5p | MIMAT0000122 | Usp33 | 170822 | HITS-CLIP | 23142080\|23597149\|19536157 | Unspecified |
| mmu-let-7i-5p | MIMAT0000122 | Usp34 | 17847 | HITS-CLIP | 37986224\|23142080\|23597149 | Unspecified |
| mmu-let-7i-5p | MIMAT0000122 | Usp36 | 72344 | HITS-CLIP | 29167373 | Unspecified |
| mmu-let-7i-5p | MIMAT0000122 | Usp37 | 319651 | HITS-CLIP | 23142080 | Unspecified |
| mmu-let-7i-5p | MIMAT0000122 | Usp38 | 74841 | HITS-CLIP | 37986224\|23597149\|29167373\|23142 | Unspecified |
| mmu-let-7i-5p | MIMAT0000122 | Usp4 | 22258 | HITS-CLIP | 37986224\|19536157\|23597149\|23142 | Unspecified |
| mmu-let-7i-5p | MIMAT0000122 | Usp42 | 76800 | HITS-CLIP | 37986224\|31433990\|23142080 | Unspecified |
| mmu-let-7i-5p | MIMAT0000122 | Usp45 | 77593 | HITS-CLIP | 23142080 | Unspecified |
| mmu-let-7i-5p | MIMAT0000122 | Usp46 | 69727 | HITS-CLIP | 23597149\|29167373 | Unspecified |
| mmu-let-7i-5p | MIMAT0000122 | Usp47 | 74996 | HITS-CLIP | 37986224\|23142080\|23597149\|29167 | Unspecified |
| mmu-let-7i-5p | MIMAT0000122 | Usp48 | 170707 | HITS-CLIP | 23142080 | Unspecified |
| mmu-let-7i-5p | MIMAT0000122 | Usp49 | 224836 | HITS-CLIP | 37986224\|23597149\|31433990\|23142 | Unspecified |
| mmu-let-7i-5p | MIMAT0000122 | Usp5 | 22225 | HITS-CLIP | 23597149\|19536157\|23142080\|29167 | Unspecified |
| mmu-let-7i-5p | MIMAT0000122 | Usp53 | 99526 | HITS-CLIP | 23142080 | Unspecified |
| mmu-let-7i-5p | MIMAT0000122 | Usp54 | 78787 | HITS-CLIP | 23597149\|19536157\|29167373 | Unspecified |
| mmu-let-7i-5p | MIMAT0000122 | Usp8 | 84092 | HITS-CLIP | 23597149\|19536157\|23142080\|29167 | Unspecified |
| mmu-let-7i-5p | MIMAT0000122 | Usp9x | 22284 | HITS-CLIP | 37986224\|23597149\|19536157\|23142 | Unspecified |
| mmu-let-7i-5p | MIMAT0000122 | Uspl1 | 231915 | HITS-CLIP | 23597149\|23142080 | Unspecified |
| mmu-let-7i-5p | MIMAT0000122 | Ust | 338362 | HITS-CLIP | 37986224\|19536157\|23142080 | Unspecified |
| mmu-let-7i-5p | MIMAT0000122 | Utp11 | 51118 | HITS-CLIP | 23597149 | Unspecified |
| mmu-let-7i-5p | MIMAT0000122 | Utp14b | 195434 | HITS-CLIP | 23630078\|19536157 | Unspecified |
| mmu-let-7i-5p | MIMAT0000122 | Utp18 | 217109 | HITS-CLIP | 23142080 | Unspecified |
| mmu-let-7i-5p | MIMAT0000122 | Utp20 | 70683 | HITS-CLIP | 23597149\|23142080 | Unspecified |
| mmu-let-7i-5p | MIMAT0000122 | Utp6 | 216987 | qCLASH | 31393953 | Unspecified |
| mmu-let-7i-5p | MIMAT0000122 | Utrn | 22288 | HITS-CLIP | 37986224\|23597149\|19536157\|23142 | Unspecified |
| mmu-let-7i-5p | MIMAT0000122 | Uty | 22290 | HITS-CLIP | 23142080 | Unspecified |
| mmu-let-7i-5p | MIMAT0000122 | Uvrag | 78610 | HITS-CLIP | 23142080 | Unspecified |
| mmu-let-7i-5p | MIMAT0000122 | Uxs1 | 67883 | HITS-CLIP | 23142080 | Unspecified |
| mmu-let-7i-5p | MIMAT0000122 | Vac14 | 234729 | HITS-CLIP | 37986224 | Unspecified |
| mmu-let-7i-5p | MIMAT0000122 | Vamp1 | 22317 | HITS-CLIP | 23142080 | Unspecified |
| mmu-let-7i-5p | MIMAT0000122 | Vamp3 | 22319 | HITS-CLIP | 23597149\|29167373\|23142080 | Unspecified |

| | | | | | | | |
|---|---|---|---|---|---|---|---|
| mmu-let-7i-5p | MIMAT0000122 | Vangl1 | 229658 | HITS-CLIP | | 19536157 | Unspecified |
| mmu-let-7i-5p | MIMAT0000122 | Vapa | 30960 | HITS-CLIP | 37986224\|23597149 | | Unspecified |
| mmu-let-7i-5p | MIMAT0000122 | Vapb | 56491 | HITS-CLIP | | 23142080 | Unspecified |
| mmu-let-7i-5p | MIMAT0000122 | Vars | 22321 | HITS-CLIP | 23597149\|23142080 | | Unspecified |
| mmu-let-7i-5p | MIMAT0000122 | Vasp | 22323 | HITS-CLIP | 23597149\|30971389\|23142080 | | Unspecified |
| mmu-let-7i-5p | MIMAT0000122 | Vav1 | 22324 | HITS-CLIP | | 23142080 | Unspecified |
| mmu-let-7i-5p | MIMAT0000122 | Vav3 | 57257 | HITS-CLIP | 19536157\|29167373\|23142080 | | Unspecified |
| mmu-let-7i-5p | MIMAT0000122 | Vbp1 | 22327 | HITS-CLIP | | 23142080 | Unspecified |
| mmu-let-7i-5p | MIMAT0000122 | Vcam1 | 22329 | HITS-CLIP | | 19536157 | Unspecified |
| mmu-let-7i-5p | MIMAT0000122 | Vcan | 13003 | HITS-CLIP | 37986224\|19536157\|29167373 | | Unspecified |
| mmu-let-7i-5p | MIMAT0000122 | Vcl | 22330 | HITS-CLIP | 37986224\|23597149 | | Unspecified |
| mmu-let-7i-5p | MIMAT0000122 | Vcpip1 | 70675 | HITS-CLIP | | 29167373 | Unspecified |
| mmu-let-7i-5p | MIMAT0000122 | Vcpkmt | 207965 | HITS-CLIP | | 23142080 | Unspecified |
| mmu-let-7i-5p | MIMAT0000122 | Vdac1 | 22333 | HITS-CLIP | 23597149\|23142080\|29167373 | | Unspecified |
| mmu-let-7i-5p | MIMAT0000122 | Vdac2 | 22334 | HITS-CLIP | 23597149\|23142080 | | Unspecified |
| mmu-let-7i-5p | MIMAT0000122 | Vegfa | 22339 | HITS-CLIP | 37986224\|23597149 | | Unspecified |
| mmu-let-7i-5p | MIMAT0000122 | Vegfc | 22341 | HITS-CLIP | | 23597149 | Unspecified |
| mmu-let-7i-5p | MIMAT0000122 | Vezf1 | 22344 | HITS-CLIP | 37986224\|23142080\|29167373 | | Unspecified |
| mmu-let-7i-5p | MIMAT0000122 | Vezt | 215008 | HITS-CLIP | 37986224\|19536157\|23142080 | | Unspecified |
| mmu-let-7i-5p | MIMAT0000122 | Vhl | 22346 | HITS-CLIP | | 23142080 | Unspecified |
| mmu-let-7i-5p | MIMAT0000122 | Vim | 22352 | HITS-CLIP | 37986224\|23597149\|29167373 | | Unspecified |
| mmu-let-7i-5p | MIMAT0000122 | Vipr1 | 22354 | HITS-CLIP | 37986224\|23597149 | | Unspecified |
| mmu-let-7i-5p | MIMAT0000122 | Virma | 25962 | qCLASH//HITS-CLIP | 31393953\|23597149\|23142080 | | Unspecified |
| mmu-let-7i-5p | MIMAT0000122 | Vkorc1 | 27973 | HITS-CLIP | 23597149\|19536157 | | Unspecified |
| mmu-let-7i-5p | MIMAT0000122 | Vkorc1l1 | 69568 | HITS-CLIP | | 19536157 | Unspecified |
| mmu-let-7i-5p | MIMAT0000122 | Vldlr | 22359 | HITS-CLIP | 23142080\|19536157 | | Unspecified |
| mmu-let-7i-5p | MIMAT0000122 | Vmac | 106639 | HITS-CLIP | | 23597149 | Unspecified |
| mmu-let-7i-5p | MIMAT0000122 | Vmp1 | 75909 | HITS-CLIP | 23597149\|23142080 | | Unspecified |
| mmu-let-7i-5p | MIMAT0000122 | Vps13a | 271564 | HITS-CLIP | 19536157\|23142080\|29167373 | | Unspecified |
| mmu-let-7i-5p | MIMAT0000122 | Vps13b | 666173 | HITS-CLIP | 23597149\|23142080\|29167373 | | Unspecified |
| mmu-let-7i-5p | MIMAT0000122 | Vps13c | 320528 | HITS-CLIP//qCLASH | 37986224\|23597149\|31393953\|23142 | | Unspecified |
| mmu-let-7i-5p | MIMAT0000122 | Vps13d | 230895 | HITS-CLIP | 23597149\|23142080\|29167373 | | Unspecified |
| mmu-let-7i-5p | MIMAT0000122 | Vps16 | 80743 | HITS-CLIP | 23142080\|29167373 | | Unspecified |
| mmu-let-7i-5p | MIMAT0000122 | Vps18 | 228545 | HITS-CLIP | | 29167373 | Unspecified |
| mmu-let-7i-5p | MIMAT0000122 | Vps26a | 30930 | HITS-CLIP | | 23142080 | Unspecified |
| mmu-let-7i-5p | MIMAT0000122 | Vps26b | 69091 | HITS-CLIP | 37986224\|23597149\|19536157\|23142 | | Unspecified |
| mmu-let-7i-5p | MIMAT0000122 | Vps26c | 135977289 | HITS-CLIP | | 23142080 | Unspecified |
| mmu-let-7i-5p | MIMAT0000122 | Vps28 | 51160 | HITS-CLIP | 23142080\|23597149 | | Unspecified |
| mmu-let-7i-5p | MIMAT0000122 | Vps35 | 65114 | HITS-CLIP | | 23597149 | Unspecified |

| | | | | | | |
|---|---|---|---|---|---|---|
| mmu-let-7i-5p | MIMAT0000122 | Vps35l | 57020 | HITS-CLIP | 19536157\|23142080 | Unspecified |
| mmu-let-7i-5p | MIMAT0000122 | Vps37a | 52348 | HITS-CLIP | 23597149\|23142080 | Unspecified |
| mmu-let-7i-5p | MIMAT0000122 | Vps37b | 330192 | HITS-CLIP | 23142080 | Unspecified |
| mmu-let-7i-5p | MIMAT0000122 | Vps37c | 107305 | HITS-CLIP | 23597149 | Unspecified |
| mmu-let-7i-5p | MIMAT0000122 | Vps39 | 269338 | HITS-CLIP | 23142080 | Unspecified |
| mmu-let-7i-5p | MIMAT0000122 | Vps4a | 116733 | HITS-CLIP | 23597149 | Unspecified |
| mmu-let-7i-5p | MIMAT0000122 | Vps4b | 20479 | HITS-CLIP | 23142080\|29167373 | Unspecified |
| mmu-let-7i-5p | MIMAT0000122 | Vps50 | 55610 | HITS-CLIP | 23142080 | Unspecified |
| mmu-let-7i-5p | MIMAT0000122 | Vps51 | 68505 | HITS-CLIP | 23142080 | Unspecified |
| mmu-let-7i-5p | MIMAT0000122 | Vps52 | 224705 | HITS-CLIP | 23142080 | Unspecified |
| mmu-let-7i-5p | MIMAT0000122 | Vps53 | 68299 | HITS-CLIP | 23597149 | Unspecified |
| mmu-let-7i-5p | MIMAT0000122 | Vps54 | 245944 | HITS-CLIP | 37986224\|23142080\|31433990\|29167 | Unspecified |
| mmu-let-7i-5p | MIMAT0000122 | Vps8 | 209018 | HITS-CLIP | 23142080 | Unspecified |
| mmu-let-7i-5p | MIMAT0000122 | Vrk3 | 101568 | HITS-CLIP | 23597149 | Unspecified |
| mmu-let-7i-5p | MIMAT0000122 | Vsir | 64115 | HITS-CLIP | 23597149\|23142080\|29167373 | Unspecified |
| mmu-let-7i-5p | MIMAT0000122 | Vsnl1 | 26950 | HITS-CLIP | 19536157 | Unspecified |
| mmu-let-7i-5p | MIMAT0000122 | Vstm5 | 69137 | HITS-CLIP | 23142080 | Unspecified |
| mmu-let-7i-5p | MIMAT0000122 | Vta1 | 66201 | HITS-CLIP | 23142080 | Unspecified |
| mmu-let-7i-5p | MIMAT0000122 | Vti1b | 53612 | HITS-CLIP | 23142080 | Unspecified |
| mmu-let-7i-5p | MIMAT0000122 | Vwa8 | 219189 | HITS-CLIP | 37986224\|23597149\|29167373 | Unspecified |
| mmu-let-7i-5p | MIMAT0000122 | Vwce | 71768 | HITS-CLIP | 37986224\|23597149 | Unspecified |
| mmu-let-7i-5p | MIMAT0000122 | Wac | 225131 | HITS-CLIP | 37986224\|23142080\|23597149\|29167 | Unspecified |
| mmu-let-7i-5p | MIMAT0000122 | Wapl | 23063 | HITS-CLIP | 23597149\|19536157\|23142080 | Unspecified |
| mmu-let-7i-5p | MIMAT0000122 | Wars | 22375 | HITS-CLIP | 23142080 | Unspecified |
| mmu-let-7i-5p | MIMAT0000122 | Wasf1 | 83767 | HITS-CLIP | 19536157 | Unspecified |
| mmu-let-7i-5p | MIMAT0000122 | Wasf2 | 242687 | HITS-CLIP | 19536157\|23142080\|29167373 | Unspecified |
| mmu-let-7i-5p | MIMAT0000122 | Washc2 | 28006 | HITS-CLIP | 23142080 | Unspecified |
| mmu-let-7i-5p | MIMAT0000122 | Washc3 | 51019 | HITS-CLIP | 23142080 | Unspecified |
| mmu-let-7i-5p | MIMAT0000122 | Washc5 | 9897 | HITS-CLIP | 23142080\|29167373 | Unspecified |
| mmu-let-7i-5p | MIMAT0000122 | Wasl | 73178 | HITS-CLIP | 37986224\|23597149\|31433990\|29167 | Unspecified |
| mmu-let-7i-5p | MIMAT0000122 | Wbp11 | 60321 | HITS-CLIP | 37986224\|23142080 | Unspecified |
| mmu-let-7i-5p | MIMAT0000122 | Wbp1l | 226178 | HITS-CLIP | 23597149\|23142080\|29167373 | Unspecified |
| mmu-let-7i-5p | MIMAT0000122 | Wbp2 | 22378 | HITS-CLIP | 23597149 | Unspecified |
| mmu-let-7i-5p | MIMAT0000122 | Wdfy3 | 72145 | HITS-CLIP | 37986224\|23597149\|29167373 | Unspecified |
| mmu-let-7i-5p | MIMAT0000122 | Wdfy4 | 545030 | HITS-CLIP | 29167373 | Unspecified |
| mmu-let-7i-5p | MIMAT0000122 | Wdhd1 | 218973 | HITS-CLIP | 23142080 | Unspecified |
| mmu-let-7i-5p | MIMAT0000122 | Wdr1 | 22388 | HITS-CLIP | 23597149\|23142080 | Unspecified |
| mmu-let-7i-5p | MIMAT0000122 | Wdr11 | 207425 | HITS-CLIP | 23597149\|23142080\|29167373 | Unspecified |
| mmu-let-7i-5p | MIMAT0000122 | Wdr18 | 216156 | HITS-CLIP | 37986224\|23142080\|29167373 | Unspecified |

| | | | | | | |
|---|---|---|---|---|---|---|
| mmu-let-7i-5p | MIMAT0000122 | Wdr26 | 226757 | HITS-CLIP | 37986224\|23142080\|31433990\|23597 | Unspecified |
| mmu-let-7i-5p | MIMAT0000122 | Wdr3 | 269470 | HITS-CLIP | 23142080 | Unspecified |
| mmu-let-7i-5p | MIMAT0000122 | Wdr33 | 74320 | HITS-CLIP | 23142080\|29167373 | Unspecified |
| mmu-let-7i-5p | MIMAT0000122 | Wdr37 | 207615 | HITS-CLIP | 37986224\|19536157\|23597149\|23142 | Unspecified |
| mmu-let-7i-5p | MIMAT0000122 | Wdr43 | 72515 | HITS-CLIP | 23142080 | Unspecified |
| mmu-let-7i-5p | MIMAT0000122 | Wdr44 | 72404 | HITS-CLIP | 23597149\|23142080 | Unspecified |
| mmu-let-7i-5p | MIMAT0000122 | Wdr45b | 66840 | HITS-CLIP | 31433990\|23142080 | Unspecified |
| mmu-let-7i-5p | MIMAT0000122 | Wdr48 | 67561 | HITS-CLIP | 37986224\|23597149\|23142080 | Unspecified |
| mmu-let-7i-5p | MIMAT0000122 | Wdr5 | 140858 | HITS-CLIP | 23142080 | Unspecified |
| mmu-let-7i-5p | MIMAT0000122 | Wdr59 | 319481 | HITS-CLIP | 19536157 | Unspecified |
| mmu-let-7i-5p | MIMAT0000122 | Wdr62 | 233064 | qCLASH | 31393953 | Unspecified |
| mmu-let-7i-5p | MIMAT0000122 | Wdr70 | 545085 | HITS-CLIP | 23142080 | Unspecified |
| mmu-let-7i-5p | MIMAT0000122 | Wdr73 | 71968 | HITS-CLIP | 23142080 | Unspecified |
| mmu-let-7i-5p | MIMAT0000122 | Wdr76 | 241627 | HITS-CLIP | 23142080 | Unspecified |
| mmu-let-7i-5p | MIMAT0000122 | Wdr77 | 70465 | HITS-CLIP | 31433990 | Unspecified |
| mmu-let-7i-5p | MIMAT0000122 | Wdr81 | 192652 | HITS-CLIP | 37986224\|23142080\|23597149\|29167 | Unspecified |
| mmu-let-7i-5p | MIMAT0000122 | Wdr89 | 72338 | HITS-CLIP | 23597149 | Unspecified |
| mmu-let-7i-5p | MIMAT0000122 | Wdtc1 | 230796 | HITS-CLIP | 23597149 | Unspecified |
| mmu-let-7i-5p | MIMAT0000122 | Wee1 | 22390 | HITS-CLIP | 19536157\|29167373\|23142080 | Unspecified |
| mmu-let-7i-5p | MIMAT0000122 | Wfs1 | 22393 | HITS-CLIP | 23597149 | Unspecified |
| mmu-let-7i-5p | MIMAT0000122 | Wipf1 | 215280 | HITS-CLIP | 31433990\|23142080 | Unspecified |
| mmu-let-7i-5p | MIMAT0000122 | Wipi1 | 52639 | HITS-CLIP | 37986224\|23597149\|23142080 | Unspecified |
| mmu-let-7i-5p | MIMAT0000122 | Wiz | 22404 | HITS-CLIP | 37986224\|23597149 | Unspecified |
| mmu-let-7i-5p | MIMAT0000122 | Wnk1 | 232341 | HITS-CLIP | 19536157\|23142080\|29167373 | Unspecified |
| mmu-let-7i-5p | MIMAT0000122 | Wnk3 | 279561 | HITS-CLIP | 19536157 | Unspecified |
| mmu-let-7i-5p | MIMAT0000122 | Wnt9a | 216795 | HITS-CLIP | 23597149\|19536157 | Unspecified |
| mmu-let-7i-5p | MIMAT0000122 | Wrb | 71446 | HITS-CLIP | 23142080 | Unspecified |
| mmu-let-7i-5p | MIMAT0000122 | Wrn | 22427 | HITS-CLIP | 31433990 | Unspecified |
| mmu-let-7i-5p | MIMAT0000122 | Wrnip1 | 78903 | HITS-CLIP | 23597149\|23142080 | Unspecified |
| mmu-let-7i-5p | MIMAT0000122 | Wsb1 | 78889 | HITS-CLIP | 23142080\|29167373 | Unspecified |
| mmu-let-7i-5p | MIMAT0000122 | Wtap | 60532 | HITS-CLIP | 23142080 | Unspecified |
| mmu-let-7i-5p | MIMAT0000122 | Wwc1 | 211652 | HITS-CLIP | 23597149\|23142080 | Unspecified |
| mmu-let-7i-5p | MIMAT0000122 | Wwp1 | 107568 | HITS-CLIP | 29167373 | Unspecified |
| mmu-let-7i-5p | MIMAT0000122 | Wwp2 | 66894 | HITS-CLIP | 23142080\|29167373 | Unspecified |
| mmu-let-7i-5p | MIMAT0000122 | Xab2 | 67439 | HITS-CLIP | 23597149 | Unspecified |
| mmu-let-7i-5p | MIMAT0000122 | Xbp1 | 22433 | HITS-CLIP | 23597149 | Unspecified |
| mmu-let-7i-5p | MIMAT0000122 | Xdh | 22436 | HITS-CLIP | 23597149\|29167373 | Unspecified |
| mmu-let-7i-5p | MIMAT0000122 | Xiap | 11798 | HITS-CLIP | 23597149\|23142080\|19536157\|29167 | Unspecified |
| mmu-let-7i-5p | MIMAT0000122 | Xkr6 | 219149 | HITS-CLIP | 19536157 | Unspecified |

| | | | | | | |
|---|---|---|---|---|---|---|
| mmu-let-7i-5p | MIMAT0000122 | Xkr8 | 381560 | HITS-CLIP | 23597149\|19536157\|23142080 | Unspecified |
| mmu-let-7i-5p | MIMAT0000122 | Xkr9 | 381246 | HITS-CLIP | 23597149 | Unspecified |
| mmu-let-7i-5p | MIMAT0000122 | Xpa | 22590 | HITS-CLIP | 23142080 | Unspecified |
| mmu-let-7i-5p | MIMAT0000122 | Xpc | 22591 | HITS-CLIP | 23597149 | Unspecified |
| mmu-let-7i-5p | MIMAT0000122 | Xpnpep1 | 170750 | HITS-CLIP | 23142080\|23597149 | Unspecified |
| mmu-let-7i-5p | MIMAT0000122 | Xpnpep2 | 170745 | HITS-CLIP | 23142080 | Unspecified |
| mmu-let-7i-5p | MIMAT0000122 | Xpnpep3 | 321003 | HITS-CLIP | 23142080\|23597149 | Unspecified |
| mmu-let-7i-5p | MIMAT0000122 | Xpo1 | 103573 | HITS-CLIP | 37986224\|19536157\|23142080 | Unspecified |
| mmu-let-7i-5p | MIMAT0000122 | Xpo4 | 57258 | HITS-CLIP | 23142080\|23597149 | Unspecified |
| mmu-let-7i-5p | MIMAT0000122 | Xpo5 | 72322 | HITS-CLIP | 37986224\|23142080\|29167373 | Unspecified |
| mmu-let-7i-5p | MIMAT0000122 | Xpo6 | 74204 | HITS-CLIP | 37986224\|23142080\|19536157\|23597 | Unspecified |
| mmu-let-7i-5p | MIMAT0000122 | Xpo7 | 65246 | HITS-CLIP | 23142080\|31433990\|23597149\|29167 | Unspecified |
| mmu-let-7i-5p | MIMAT0000122 | Xpot | 73192 | HITS-CLIP | 37986224\|23142080\|23597149\|2363( | Unspecified |
| mmu-let-7i-5p | MIMAT0000122 | Xpr1 | 19775 | HITS-CLIP | 37986224\|23597149\|19536157\|29167 | Unspecified |
| mmu-let-7i-5p | MIMAT0000122 | Xrcc2 | 57434 | HITS-CLIP | 31433990 | Unspecified |
| mmu-let-7i-5p | MIMAT0000122 | Xrcc5 | 22596 | HITS-CLIP | 23597149\|23142080 | Unspecified |
| mmu-let-7i-5p | MIMAT0000122 | Xrcc6 | 14375 | HITS-CLIP | 19536157\|23142080 | Unspecified |
| mmu-let-7i-5p | MIMAT0000122 | Xrn1 | 24127 | HITS-CLIP | 37986224\|23142080\|19536157\|29167 | Unspecified |
| mmu-let-7i-5p | MIMAT0000122 | Xrn2 | 24128 | HITS-CLIP | 37986224\|23597149\|19536157\|23142 | Unspecified |
| mmu-let-7i-5p | MIMAT0000122 | Xylb | 102448 | HITS-CLIP | 23597149 | Unspecified |
| mmu-let-7i-5p | MIMAT0000122 | Xylt1 | 233781 | HITS-CLIP | 19536157 | Unspecified |
| mmu-let-7i-5p | MIMAT0000122 | Xylt2 | 217119 | HITS-CLIP | 37986224\|23142080 | Unspecified |
| mmu-let-7i-5p | MIMAT0000122 | Yap1 | 22601 | HITS-CLIP | 23597149 | Unspecified |
| mmu-let-7i-5p | MIMAT0000122 | Yars | 107271 | HITS-CLIP | 29167373 | Unspecified |
| mmu-let-7i-5p | MIMAT0000122 | Ybey | 216119 | HITS-CLIP | 37986224 | Unspecified |
| mmu-let-7i-5p | MIMAT0000122 | Ybx1 | 22608 | HITS-CLIP | 23142080 | Unspecified |
| mmu-let-7i-5p | MIMAT0000122 | Ybx3 | 56449 | HITS-CLIP | 19536157 | Unspecified |
| mmu-let-7i-5p | MIMAT0000122 | Yeats2 | 208146 | HITS-CLIP | 23597149\|23142080\|29167373 | Unspecified |
| mmu-let-7i-5p | MIMAT0000122 | Yeats4 | 64050 | HITS-CLIP | 37986224\|23597149\|23142080 | Unspecified |
| mmu-let-7i-5p | MIMAT0000122 | Yes1 | 22612 | HITS-CLIP | 23597149 | Unspecified |
| mmu-let-7i-5p | MIMAT0000122 | Yipf1 | 230584 | HITS-CLIP | 31433990 | Unspecified |
| mmu-let-7i-5p | MIMAT0000122 | Yipf2 | 74766 | HITS-CLIP | 23142080 | Unspecified |
| mmu-let-7i-5p | MIMAT0000122 | Yipf3 | 28064 | HITS-CLIP | 23597149\|19536157\|23142080 | Unspecified |
| mmu-let-7i-5p | MIMAT0000122 | Yipf4 | 67864 | HITS-CLIP | 23142080 | Unspecified |
| mmu-let-7i-5p | MIMAT0000122 | Yipf5 | 67180 | HITS-CLIP | 23597149\|23142080 | Unspecified |
| mmu-let-7i-5p | MIMAT0000122 | Yipf6 | 77929 | HITS-CLIP | 29167373\|23142080 | Unspecified |
| mmu-let-7i-5p | MIMAT0000122 | Ylpm1 | 56531 | HITS-CLIP//qCLASH | 23597149\|23142080\|31393953\|1953( | Unspecified |
| mmu-let-7i-5p | MIMAT0000122 | Yme1l1 | 27377 | HITS-CLIP | 37986224\|23597149\|19536157\|23142 | Unspecified |
| mmu-let-7i-5p | MIMAT0000122 | Yod1 | 226418 | HITS-CLIP | 37986224\|23597149\|23142080 | Unspecified |

| | | | | | | |
|---|---|---|---|---|---|---|
| mmu-let-7i-5p | MIMAT0000122 | Ypel2 | 77864 | HITS-CLIP | 37986224\|23597149\|23142080 | Unspecified |
| mmu-let-7i-5p | MIMAT0000122 | Ypel3 | 66090 | HITS-CLIP | 31433990\|23142080 | Unspecified |
| mmu-let-7i-5p | MIMAT0000122 | Ypel4 | 241525 | HITS-CLIP | 37986224\|23142080 | Unspecified |
| mmu-let-7i-5p | MIMAT0000122 | Ypel5 | 383295 | HITS-CLIP | 23142080 | Unspecified |
| mmu-let-7i-5p | MIMAT0000122 | Yrdc | 230734 | qCLASH | 31393953 | Unspecified |
| mmu-let-7i-5p | MIMAT0000122 | Ythdf1 | 228994 | HITS-CLIP | 23142080 | Unspecified |
| mmu-let-7i-5p | MIMAT0000122 | Ythdf2 | 213541 | HITS-CLIP | 23142080 | Unspecified |
| mmu-let-7i-5p | MIMAT0000122 | Ythdf3 | 229096 | HITS-CLIP | 37986224\|23597149\|23142080\|2916 | Unspecified |
| mmu-let-7i-5p | MIMAT0000122 | Ywhab | 54401 | HITS-CLIP | 23142080\|19536157\|29167373 | Unspecified |
| mmu-let-7i-5p | MIMAT0000122 | Ywhae | 22627 | HITS-CLIP | 37986224\|23142080 | Unspecified |
| mmu-let-7i-5p | MIMAT0000122 | Ywhag | 22628 | HITS-CLIP | 37986224\|23597149\|19536157\|2314 | Unspecified |
| mmu-let-7i-5p | MIMAT0000122 | Ywhah | 22629 | HITS-CLIP | 23597149\|19536157 | Unspecified |
| mmu-let-7i-5p | MIMAT0000122 | Ywhaq | 22630 | qCLASH | 31393953 | Unspecified |
| mmu-let-7i-5p | MIMAT0000122 | Ywhaz | 22631 | HITS-CLIP | 23142080 | Unspecified |
| mmu-let-7i-5p | MIMAT0000122 | Yy1 | 22632 | HITS-CLIP | 23597149\|23142080 | Unspecified |
| mmu-let-7i-5p | MIMAT0000122 | Zadh2 | 225791 | HITS-CLIP | 23597149 | Unspecified |
| mmu-let-7i-5p | MIMAT0000122 | Zap70 | 22637 | HITS-CLIP | 23142080 | Unspecified |
| mmu-let-7i-5p | MIMAT0000122 | Zbed3 | 72114 | HITS-CLIP | 37986224\|23597149 | Unspecified |
| mmu-let-7i-5p | MIMAT0000122 | Zbp1 | 58203 | HITS-CLIP | 23142080 | Unspecified |
| mmu-let-7i-5p | MIMAT0000122 | Zbtb1 | 268564 | HITS-CLIP | 37986224\|23142080 | Unspecified |
| mmu-let-7i-5p | MIMAT0000122 | Zbtb11 | 271377 | HITS-CLIP | 29167373\|23142080 | Unspecified |
| mmu-let-7i-5p | MIMAT0000122 | Zbtb14 | 22666 | HITS-CLIP | 23142080 | Unspecified |
| mmu-let-7i-5p | MIMAT0000122 | Zbtb16 | 235320 | HITS-CLIP | 23597149 | Unspecified |
| mmu-let-7i-5p | MIMAT0000122 | Zbtb20 | 56490 | HITS-CLIP | 23597149\|23142080\|29167373 | Unspecified |
| mmu-let-7i-5p | MIMAT0000122 | Zbtb21 | 114565 | HITS-CLIP | 23597149\|23142080 | Unspecified |
| mmu-let-7i-5p | MIMAT0000122 | Zbtb22 | 81630 | HITS-CLIP | 23142080 | Unspecified |
| mmu-let-7i-5p | MIMAT0000122 | Zbtb25 | 109929 | HITS-CLIP | 23142080 | Unspecified |
| mmu-let-7i-5p | MIMAT0000122 | Zbtb33 | 56805 | HITS-CLIP | 29167373\|23142080 | Unspecified |
| mmu-let-7i-5p | MIMAT0000122 | Zbtb34 | 241311 | HITS-CLIP | 19536157\|23142080\|29167373 | Unspecified |
| mmu-let-7i-5p | MIMAT0000122 | Zbtb37 | 240869 | HITS-CLIP | 23142080 | Unspecified |
| mmu-let-7i-5p | MIMAT0000122 | Zbtb38 | 245007 | HITS-CLIP | 23142080\|23597149\|29167373 | Unspecified |
| mmu-let-7i-5p | MIMAT0000122 | Zbtb39 | 320080 | HITS-CLIP | 37986224\|23597149\|23142080 | Unspecified |
| mmu-let-7i-5p | MIMAT0000122 | Zbtb41 | 226470 | HITS-CLIP | 23597149\|23142080 | Unspecified |
| mmu-let-7i-5p | MIMAT0000122 | Zbtb42 | 382639 | HITS-CLIP | 23142080 | Unspecified |
| mmu-let-7i-5p | MIMAT0000122 | Zbtb43 | 71834 | HITS-CLIP | 23142080\|29167373 | Unspecified |
| mmu-let-7i-5p | MIMAT0000122 | Zbtb44 | 235132 | HITS-CLIP | 23142080 | Unspecified |
| mmu-let-7i-5p | MIMAT0000122 | Zbtb45 | 232879 | HITS-CLIP | 23142080 | Unspecified |
| mmu-let-7i-5p | MIMAT0000122 | Zbtb46 | 72147 | HITS-CLIP | 37986224 | Unspecified |
| mmu-let-7i-5p | MIMAT0000122 | Zbtb5 | 230119 | HITS-CLIP | 37986224 | Unspecified |

| | | | | | | | |
|---|---|---|---|---|---|---|---|
| mmu-let-7i-5p | MIMAT0000122 | Zbtb7a | 16969 | HITS-CLIP | | 23142080 | Unspecified |
| mmu-let-7i-5p | MIMAT0000122 | Zbtb7b | 22724 | HITS-CLIP | 23142080\|29167373 | | Unspecified |
| mmu-let-7i-5p | MIMAT0000122 | Zc3h10 | 103284 | HITS-CLIP | | 23142080 | Unspecified |
| mmu-let-7i-5p | MIMAT0000122 | Zc3h12d | 237256 | HITS-CLIP | | 23142080 | Unspecified |
| mmu-let-7i-5p | MIMAT0000122 | Zc3h13 | 67302 | HITS-CLIP | | 23142080 | Unspecified |
| mmu-let-7i-5p | MIMAT0000122 | Zc3h14 | 75553 | HITS-CLIP | | 23597149 | Unspecified |
| mmu-let-7i-5p | MIMAT0000122 | Zc3h4 | 330474 | HITS-CLIP | | 37986224 | Unspecified |
| mmu-let-7i-5p | MIMAT0000122 | Zc3h6 | 78751 | HITS-CLIP | | 23142080 | Unspecified |
| mmu-let-7i-5p | MIMAT0000122 | Zc3h7a | 106205 | HITS-CLIP | | 23142080 | Unspecified |
| mmu-let-7i-5p | MIMAT0000122 | Zc3h7b | 20286 | HITS-CLIP | | 29167373 | Unspecified |
| mmu-let-7i-5p | MIMAT0000122 | Zc3hav1 | 78781 | HITS-CLIP | 23142080\|29167373 | | Unspecified |
| mmu-let-7i-5p | MIMAT0000122 | Zc3hav1l | 209032 | HITS-CLIP | 37986224\|23142080 | | Unspecified |
| mmu-let-7i-5p | MIMAT0000122 | Zcchc10 | 67966 | HITS-CLIP | | 23597149 | Unspecified |
| mmu-let-7i-5p | MIMAT0000122 | Zcchc14 | 142682 | HITS-CLIP | 23597149\|29167373 | | Unspecified |
| mmu-let-7i-5p | MIMAT0000122 | Zcchc2 | 227449 | HITS-CLIP | 23597149\|31433990\|23142080\|29167 | | Unspecified |
| mmu-let-7i-5p | MIMAT0000122 | Zcchc24 | 71918 | HITS-CLIP | 19536157\|23597149\|23142080\|29167 | | Unspecified |
| mmu-let-7i-5p | MIMAT0000122 | Zcchc3 | 67917 | HITS-CLIP | | 23142080 | Unspecified |
| mmu-let-7i-5p | MIMAT0000122 | Zcchc4 | 78796 | HITS-CLIP | | 23142080 | Unspecified |
| mmu-let-7i-5p | MIMAT0000122 | Zcchc7 | 319885 | HITS-CLIP | | 29167373 | Unspecified |
| mmu-let-7i-5p | MIMAT0000122 | Zcchc8 | 70650 | HITS-CLIP | | 23142080 | Unspecified |
| mmu-let-7i-5p | MIMAT0000122 | Zcchc9 | 69085 | HITS-CLIP | 23597149\|23142080 | | Unspecified |
| mmu-let-7i-5p | MIMAT0000122 | Zcrb1 | 67197 | HITS-CLIP | | 23142080 | Unspecified |
| mmu-let-7i-5p | MIMAT0000122 | Zdhhc12 | 66220 | HITS-CLIP | 37986224\|23597149\|23142080 | | Unspecified |
| mmu-let-7i-5p | MIMAT0000122 | Zdhhc13 | 243983 | HITS-CLIP | 23597149\|19536157\|23142080\|29167 | | Unspecified |
| mmu-let-7i-5p | MIMAT0000122 | Zdhhc14 | 224454 | HITS-CLIP | | 29167373 | Unspecified |
| mmu-let-7i-5p | MIMAT0000122 | Zdhhc15 | 108672 | HITS-CLIP | | 23142080 | Unspecified |
| mmu-let-7i-5p | MIMAT0000122 | Zdhhc16 | 74168 | HITS-CLIP | 23597149\|23142080 | | Unspecified |
| mmu-let-7i-5p | MIMAT0000122 | Zdhhc18 | 503610 | HITS-CLIP | 37986224\|29167373 | | Unspecified |
| mmu-let-7i-5p | MIMAT0000122 | Zdhhc20 | 75965 | HITS-CLIP | | 23142080 | Unspecified |
| mmu-let-7i-5p | MIMAT0000122 | Zdhhc21 | 68268 | HITS-CLIP | 37986224\|23142080\|29167373 | | Unspecified |
| mmu-let-7i-5p | MIMAT0000122 | Zdhhc3 | 69035 | HITS-CLIP | 23142080\|23597149 | | Unspecified |
| mmu-let-7i-5p | MIMAT0000122 | Zdhhc7 | 102193 | HITS-CLIP | 19536157\|29167373 | | Unspecified |
| mmu-let-7i-5p | MIMAT0000122 | Zdhhc8 | 27801 | HITS-CLIP | | 37986224 | Unspecified |
| mmu-let-7i-5p | MIMAT0000122 | Zdhhc9 | 208884 | HITS-CLIP | | 23142080 | Unspecified |
| mmu-let-7i-5p | MIMAT0000122 | Zeb1 | 21417 | HITS-CLIP | 19536157\|23142080 | | Unspecified |
| mmu-let-7i-5p | MIMAT0000122 | Zeb2 | 24136 | HITS-CLIP | 23142080\|19536157\|29167373 | | Unspecified |
| mmu-let-7i-5p | MIMAT0000122 | Zer1 | 227693 | HITS-CLIP | 23597149\|23142080 | | Unspecified |
| mmu-let-7i-5p | MIMAT0000122 | Zfand3 | 21769 | HITS-CLIP | 37986224\|23142080\|19536157\|23597 | | Unspecified |
| mmu-let-7i-5p | MIMAT0000122 | Zfand4 | 67492 | HITS-CLIP | | 23597149 | Unspecified |

| | | | | | | |
|---|---|---|---|---|---|---|
| mmu-let-7i-5p | MIMAT0000122 | Zfand5 | 22682 | HITS-CLIP | 37986224\|23142080\|19536157\|29167 | Unspecified |
| mmu-let-7i-5p | MIMAT0000122 | Zfc3h1 | 216345 | HITS-CLIP | 37986224\|23597149\|23142080\|29167 | Unspecified |
| mmu-let-7i-5p | MIMAT0000122 | Zfhx2 | 239102 | HITS-CLIP | 37986224\|19536157\|23597149\|23142 | Unspecified |
| mmu-let-7i-5p | MIMAT0000122 | Zfhx3 | 11906 | HITS-CLIP | 37986224\|29167373\|23597149 | Unspecified |
| mmu-let-7i-5p | MIMAT0000122 | Zfhx4 | 80892 | HITS-CLIP | 37986224\|23597149 | Unspecified |
| mmu-let-7i-5p | MIMAT0000122 | Zfp1 | 22640 | HITS-CLIP | 29167373 | Unspecified |
| mmu-let-7i-5p | MIMAT0000122 | Zfp106 | 20402 | HITS-CLIP | 23597149\|23142080\|29167373 | Unspecified |
| mmu-let-7i-5p | MIMAT0000122 | Zfp12 | 231866 | HITS-CLIP | 37986224\|19536157\|23142080 | Unspecified |
| mmu-let-7i-5p | MIMAT0000122 | Zfp131 | 72465 | HITS-CLIP | 29167373 | Unspecified |
| mmu-let-7i-5p | MIMAT0000122 | Zfp141 | 434178 | HITS-CLIP | 23142080 | Unspecified |
| mmu-let-7i-5p | MIMAT0000122 | Zfp142 | 77264 | HITS-CLIP | 23142080 | Unspecified |
| mmu-let-7i-5p | MIMAT0000122 | Zfp143 | 20841 | HITS-CLIP | 23142080 | Unspecified |
| mmu-let-7i-5p | MIMAT0000122 | Zfp148 | 22661 | HITS-CLIP | 23142080\|29167373 | Unspecified |
| mmu-let-7i-5p | MIMAT0000122 | Zfp160 | 224585 | HITS-CLIP | 23142080 | Unspecified |
| mmu-let-7i-5p | MIMAT0000122 | Zfp169 | 67911 | HITS-CLIP | 23142080 | Unspecified |
| mmu-let-7i-5p | MIMAT0000122 | Zfp180 | 210135 | HITS-CLIP | 37986224\|29167373 | Unspecified |
| mmu-let-7i-5p | MIMAT0000122 | Zfp202 | 80902 | HITS-CLIP | 23142080 | Unspecified |
| mmu-let-7i-5p | MIMAT0000122 | Zfp207 | 22680 | HITS-CLIP | 37986224\|23142080\|23597149\|29167 | Unspecified |
| mmu-let-7i-5p | MIMAT0000122 | Zfp217 | 228913 | HITS-CLIP | 37986224\|23597149\|26203562\|23142 | Unspecified |
| mmu-let-7i-5p | MIMAT0000122 | Zfp236 | 329002 | HITS-CLIP | 23142080 | Unspecified |
| mmu-let-7i-5p | MIMAT0000122 | Zfp26 | 22688 | HITS-CLIP | 37986224\|23142080\|19536157\|29167 | Unspecified |
| mmu-let-7i-5p | MIMAT0000122 | Zfp260 | 26466 | HITS-CLIP | 37986224\|29167373\|23142080 | Unspecified |
| mmu-let-7i-5p | MIMAT0000122 | Zfp263 | 74120 | HITS-CLIP | 23142080\|29167373 | Unspecified |
| mmu-let-7i-5p | MIMAT0000122 | Zfp266 | 77519 | HITS-CLIP | 37986224\|23142080 | Unspecified |
| mmu-let-7i-5p | MIMAT0000122 | Zfp275 | 27081 | HITS-CLIP | 23142080\|29167373 | Unspecified |
| mmu-let-7i-5p | MIMAT0000122 | Zfp276 | 57247 | HITS-CLIP | 23142080 | Unspecified |
| mmu-let-7i-5p | MIMAT0000122 | Zfp277 | 246196 | HITS-CLIP | 23142080 | Unspecified |
| mmu-let-7i-5p | MIMAT0000122 | Zfp280b | 64453 | HITS-CLIP | 37986224\|19536157\|23597149\|23142 | Unspecified |
| mmu-let-7i-5p | MIMAT0000122 | Zfp280c | 208968 | HITS-CLIP | 23142080 | Unspecified |
| mmu-let-7i-5p | MIMAT0000122 | Zfp280d | 235469 | HITS-CLIP | 23142080 | Unspecified |
| mmu-let-7i-5p | MIMAT0000122 | Zfp281 | 226442 | HITS-CLIP | 37986224\|23597149\|23142080 | Unspecified |
| mmu-let-7i-5p | MIMAT0000122 | Zfp292 | 30046 | HITS-CLIP | 23142080\|29167373 | Unspecified |
| mmu-let-7i-5p | MIMAT0000122 | Zfp317 | 244713 | HITS-CLIP | 23142080 | Unspecified |
| mmu-let-7i-5p | MIMAT0000122 | Zfp318 | 57908 | HITS-CLIP | 37986224\|19536157\|23597149\|23142 | Unspecified |
| mmu-let-7i-5p | MIMAT0000122 | Zfp319 | 79233 | HITS-CLIP | 23142080 | Unspecified |
| mmu-let-7i-5p | MIMAT0000122 | Zfp322a | 218100 | HITS-CLIP | 23142080 | Unspecified |
| mmu-let-7i-5p | MIMAT0000122 | Zfp329 | 67230 | HITS-CLIP | 23142080 | Unspecified |
| mmu-let-7i-5p | MIMAT0000122 | Zfp335 | 329559 | HITS-CLIP | 23142080 | Unspecified |
| mmu-let-7i-5p | MIMAT0000122 | Zfp354b | 27274 | HITS-CLIP | 37986224 | Unspecified |

| | | | | | | |
|---|---|---|---|---|---|---|
| mmu-let-7i-5p | MIMAT0000122 | Zfp36 | 22695 | HITS-CLIP//qCLASH | 37986224\|23597149\|31393953\|23142 | Unspecified |
| mmu-let-7i-5p | MIMAT0000122 | Zfp367 | 238673 | HITS-CLIP | 23597149\|23142080 | Unspecified |
| mmu-let-7i-5p | MIMAT0000122 | Zfp36l1 | 12192 | HITS-CLIP | 37986224\|23597149\|23142080\|1953€ | Unspecified |
| mmu-let-7i-5p | MIMAT0000122 | Zfp384 | 269800 | HITS-CLIP | 23597149\|23142080 | Unspecified |
| mmu-let-7i-5p | MIMAT0000122 | Zfp386 | 56220 | HITS-CLIP | 37986224 | Unspecified |
| mmu-let-7i-5p | MIMAT0000122 | Zfp407 | 240476 | HITS-CLIP | 37986224\|23597149\|23142080 | Unspecified |
| mmu-let-7i-5p | MIMAT0000122 | Zfp414 | 328801 | HITS-CLIP | 37986224\|23142080 | Unspecified |
| mmu-let-7i-5p | MIMAT0000122 | Zfp426 | 235028 | HITS-CLIP | 23597149 | Unspecified |
| mmu-let-7i-5p | MIMAT0000122 | Zfp445 | 235682 | HITS-CLIP | 23142080\|23597149 | Unspecified |
| mmu-let-7i-5p | MIMAT0000122 | Zfp451 | 98403 | HITS-CLIP | 37986224\|23597149\|19536157\|23142 | Unspecified |
| mmu-let-7i-5p | MIMAT0000122 | Zfp455 | 218311 | HITS-CLIP | 23142080 | Unspecified |
| mmu-let-7i-5p | MIMAT0000122 | Zfp462 | 242466 | HITS-CLIP | 37986224\|19536157 | Unspecified |
| mmu-let-7i-5p | MIMAT0000122 | Zfp473 | 243963 | HITS-CLIP | 37986224\|23142080\|23597149 | Unspecified |
| mmu-let-7i-5p | MIMAT0000122 | Zfp51 | 22709 | HITS-CLIP | 23142080 | Unspecified |
| mmu-let-7i-5p | MIMAT0000122 | Zfp512 | 269639 | HITS-CLIP | 37986224\|23142080 | Unspecified |
| mmu-let-7i-5p | MIMAT0000122 | Zfp512b | 269401 | HITS-CLIP | 23597149\|23142080\|29167373 | Unspecified |
| mmu-let-7i-5p | MIMAT0000122 | Zfp513 | 101023 | HITS-CLIP | 29167373 | Unspecified |
| mmu-let-7i-5p | MIMAT0000122 | Zfp516 | 329003 | HITS-CLIP | 23142080\|29167373 | Unspecified |
| mmu-let-7i-5p | MIMAT0000122 | Zfp518a | 72672 | HITS-CLIP | 37986224\|19536157\|23597149\|23142 | Unspecified |
| mmu-let-7i-5p | MIMAT0000122 | Zfp521 | 225207 | HITS-CLIP | 37986224\|19536157 | Unspecified |
| mmu-let-7i-5p | MIMAT0000122 | Zfp523 | 7629 | HITS-CLIP | 19536157 | Unspecified |
| mmu-let-7i-5p | MIMAT0000122 | Zfp524 | 66056 | HITS-CLIP | 23142080 | Unspecified |
| mmu-let-7i-5p | MIMAT0000122 | Zfp526 | 210172 | HITS-CLIP | 29167373 | Unspecified |
| mmu-let-7i-5p | MIMAT0000122 | Zfp560 | 434377 | HITS-CLIP | 23142080 | Unspecified |
| mmu-let-7i-5p | MIMAT0000122 | Zfp568 | 243905 | HITS-CLIP | 37986224\|23142080\|23597149 | Unspecified |
| mmu-let-7i-5p | MIMAT0000122 | Zfp592 | 233410 | HITS-CLIP | 37986224\|23597149\|23142080 | Unspecified |
| mmu-let-7i-5p | MIMAT0000122 | Zfp597 | 71063 | HITS-CLIP | 37986224\|23142080 | Unspecified |
| mmu-let-7i-5p | MIMAT0000122 | Zfp598 | 213753 | HITS-CLIP | 23597149 | Unspecified |
| mmu-let-7i-5p | MIMAT0000122 | Zfp599 | 235048 | HITS-CLIP | 23142080 | Unspecified |
| mmu-let-7i-5p | MIMAT0000122 | Zfp60 | 22718 | HITS-CLIP | 23142080 | Unspecified |
| mmu-let-7i-5p | MIMAT0000122 | Zfp608 | 269023 | HITS-CLIP | 23142080 | Unspecified |
| mmu-let-7i-5p | MIMAT0000122 | Zfp609 | 214812 | HITS-CLIP | 23142080 | Unspecified |
| mmu-let-7i-5p | MIMAT0000122 | Zfp617 | 170938 | HITS-CLIP | 23142080 | Unspecified |
| mmu-let-7i-5p | MIMAT0000122 | Zfp618 | 72701 | HITS-CLIP | 29167373 | Unspecified |
| mmu-let-7i-5p | MIMAT0000122 | Zfp62 | 22720 | HITS-CLIP | 23142080\|29167373 | Unspecified |
| mmu-let-7i-5p | MIMAT0000122 | Zfp622 | 52521 | HITS-CLIP | 37986224\|23142080 | Unspecified |
| mmu-let-7i-5p | MIMAT0000122 | Zfp626 | 71163 | HITS-CLIP | 23142080 | Unspecified |
| mmu-let-7i-5p | MIMAT0000122 | Zfp629 | 320683 | HITS-CLIP | 31433990\|23142080 | Unspecified |
| mmu-let-7i-5p | MIMAT0000122 | Zfp638 | 27332 | HITS-CLIP | 29167373 | Unspecified |

| | | | | | | |
|---|---|---|---|---|---|---|
| mmu-let-7i-5p | MIMAT0000122 | Zfp641 | 239652 | HITS-CLIP | 37986224\|19536157 | Unspecified |
| mmu-let-7i-5p | MIMAT0000122 | Zfp644 | 52397 | HITS-CLIP | 37986224\|23597149\|31433990\|19536 | Unspecified |
| mmu-let-7i-5p | MIMAT0000122 | Zfp646 | 233905 | HITS-CLIP | 37986224\|23142080 | Unspecified |
| mmu-let-7i-5p | MIMAT0000122 | Zfp652 | 268469 | HITS-CLIP | 37986224\|23597149\|23142080\|29167 | Unspecified |
| mmu-let-7i-5p | MIMAT0000122 | Zfp654 | 72020 | HITS-CLIP | 37986224\|23142080\|23597149 | Unspecified |
| mmu-let-7i-5p | MIMAT0000122 | Zfp655 | 72611 | HITS-CLIP | 23142080\|29167373 | Unspecified |
| mmu-let-7i-5p | MIMAT0000122 | Zfp664 | 269704 | HITS-CLIP | 37986224\|23597149 | Unspecified |
| mmu-let-7i-5p | MIMAT0000122 | Zfp667 | 384763 | HITS-CLIP | 23142080 | Unspecified |
| mmu-let-7i-5p | MIMAT0000122 | Zfp683 | 100503878 | HITS-CLIP | 23142080 | Unspecified |
| mmu-let-7i-5p | MIMAT0000122 | Zfp687 | 78266 | HITS-CLIP | 23142080 | Unspecified |
| mmu-let-7i-5p | MIMAT0000122 | Zfp689 | 71131 | HITS-CLIP | 23142080 | Unspecified |
| mmu-let-7i-5p | MIMAT0000122 | Zfp691 | 195522 | HITS-CLIP | 23142080 | Unspecified |
| mmu-let-7i-5p | MIMAT0000122 | Zfp697 | 242109 | HITS-CLIP | 19536157 | Unspecified |
| mmu-let-7i-5p | MIMAT0000122 | Zfp7 | 223669 | HITS-CLIP | 23142080 | Unspecified |
| mmu-let-7i-5p | MIMAT0000122 | Zfp703 | 353310 | HITS-CLIP | 29167373 | Unspecified |
| mmu-let-7i-5p | MIMAT0000122 | Zfp704 | 170753 | HITS-CLIP | 37986224 | Unspecified |
| mmu-let-7i-5p | MIMAT0000122 | Zfp710 | 209225 | HITS-CLIP | 23142080\|23597149\|29167373 | Unspecified |
| mmu-let-7i-5p | MIMAT0000122 | Zfp712 | 78251 | HITS-CLIP | 23142080 | Unspecified |
| mmu-let-7i-5p | MIMAT0000122 | Zfp715 | 69930 | HITS-CLIP | 29167373 | Unspecified |
| mmu-let-7i-5p | MIMAT0000122 | Zfp719 | 210105 | HITS-CLIP | 23142080\|29167373 | Unspecified |
| mmu-let-7i-5p | MIMAT0000122 | Zfp729a | 212281 | HITS-CLIP | 23142080 | Unspecified |
| mmu-let-7i-5p | MIMAT0000122 | Zfp729b | 100416706 | HITS-CLIP | 23142080\|29167373 | Unspecified |
| mmu-let-7i-5p | MIMAT0000122 | Zfp738 | 408068 | HITS-CLIP | 23142080\|29167373 | Unspecified |
| mmu-let-7i-5p | MIMAT0000122 | Zfp740 | 68744 | HITS-CLIP | 37986224\|23142080 | Unspecified |
| mmu-let-7i-5p | MIMAT0000122 | Zfp746 | 69228 | HITS-CLIP | 23142080 | Unspecified |
| mmu-let-7i-5p | MIMAT0000122 | Zfp748 | 212276 | HITS-CLIP | 23142080 | Unspecified |
| mmu-let-7i-5p | MIMAT0000122 | Zfp760 | 240034 | HITS-CLIP | 23142080 | Unspecified |
| mmu-let-7i-5p | MIMAT0000122 | Zfp763 | 73451 | HITS-CLIP | 23597149\|23142080 | Unspecified |
| mmu-let-7i-5p | MIMAT0000122 | Zfp768 | 233890 | HITS-CLIP | 23597149 | Unspecified |
| mmu-let-7i-5p | MIMAT0000122 | Zfp770 | 228491 | HITS-CLIP | 19536157 | Unspecified |
| mmu-let-7i-5p | MIMAT0000122 | Zfp809 | 235047 | HITS-CLIP | 23142080\|23597149 | Unspecified |
| mmu-let-7i-5p | MIMAT0000122 | Zfp820 | 75424 | HITS-CLIP | 23142080 | Unspecified |
| mmu-let-7i-5p | MIMAT0000122 | Zfp827 | 622675 | HITS-CLIP | 19536157\|23142080 | Unspecified |
| mmu-let-7i-5p | MIMAT0000122 | Zfp830 | 66983 | HITS-CLIP | 23142080 | Unspecified |
| mmu-let-7i-5p | MIMAT0000122 | Zfp831 | 100043757 | HITS-CLIP | 37986224\|23142080 | Unspecified |
| mmu-let-7i-5p | MIMAT0000122 | Zfp839 | 72805 | HITS-CLIP | 29167373 | Unspecified |
| mmu-let-7i-5p | MIMAT0000122 | Zfp84 | 74352 | HITS-CLIP | 37986224\|23142080 | Unspecified |
| mmu-let-7i-5p | MIMAT0000122 | Zfp871 | 208292 | HITS-CLIP | 23142080\|19536157\|29167373 | Unspecified |
| mmu-let-7i-5p | MIMAT0000122 | Zfp91 | 109910 | qCLASH//HITS-CLIP | 31393953\|23142080\|29167373 | Unspecified |

| | | | | | | |
|---|---|---|---|---|---|---|
| mmu-let-7i-5p | MIMAT0000122 | Zfp930 | 234358 | qCLASH | 31393953 | Unspecified |
| mmu-let-7i-5p | MIMAT0000122 | Zfp933 | 242747 | HITS-CLIP | 23142080 | Unspecified |
| mmu-let-7i-5p | MIMAT0000122 | Zfp934 | 77117 | HITS-CLIP | 23142080 | Unspecified |
| mmu-let-7i-5p | MIMAT0000122 | Zfp942 | 73233 | HITS-CLIP | 23142080 | Unspecified |
| mmu-let-7i-5p | MIMAT0000122 | Zfp952 | 240067 | HITS-CLIP | 23142080 | Unspecified |
| mmu-let-7i-5p | MIMAT0000122 | Zfp955b | 100043468 | HITS-CLIP | 23597149\|23142080\|29167373 | Unspecified |
| mmu-let-7i-5p | MIMAT0000122 | Zfp97 | 22759 | qCLASH | 31393953 | Unspecified |
| mmu-let-7i-5p | MIMAT0000122 | Zfpl1 | 81909 | HITS-CLIP | 23142080 | Unspecified |
| mmu-let-7i-5p | MIMAT0000122 | Zfr | 22763 | HITS-CLIP | 19536157\|23597149\|23142080\|29167 | Unspecified |
| mmu-let-7i-5p | MIMAT0000122 | Zfx | 22764 | HITS-CLIP | 23597149\|23142080 | Unspecified |
| mmu-let-7i-5p | MIMAT0000122 | Zfyve16 | 218441 | HITS-CLIP | 37986224\|23142080 | Unspecified |
| mmu-let-7i-5p | MIMAT0000122 | Zfyve21 | 68520 | HITS-CLIP | 23142080 | Unspecified |
| mmu-let-7i-5p | MIMAT0000122 | Zfyve26 | 211978 | HITS-CLIP//qCLASH | 37986224\|23597149\|31393953\|23142 | Unspecified |
| mmu-let-7i-5p | MIMAT0000122 | Zfyve27 | 319740 | HITS-CLIP | 23597149 | Unspecified |
| mmu-let-7i-5p | MIMAT0000122 | Zfyve9 | 230597 | HITS-CLIP | 23597149 | Unspecified |
| mmu-let-7i-5p | MIMAT0000122 | Zhx1 | 22770 | HITS-CLIP | 23597149\|23142080 | Unspecified |
| mmu-let-7i-5p | MIMAT0000122 | Zhx3 | 320799 | HITS-CLIP | 23597149 | Unspecified |
| mmu-let-7i-5p | MIMAT0000122 | Zic2 | 22772 | HITS-CLIP | 19536157 | Unspecified |
| mmu-let-7i-5p | MIMAT0000122 | Zic5 | 65100 | HITS-CLIP | 37986224 | Unspecified |
| mmu-let-7i-5p | MIMAT0000122 | Zik1 | 22775 | HITS-CLIP | 37986224 | Unspecified |
| mmu-let-7i-5p | MIMAT0000122 | Zim1 | 22776 | HITS-CLIP | 23597149 | Unspecified |
| mmu-let-7i-5p | MIMAT0000122 | Zkscan1 | 74570 | HITS-CLIP | 37986224 | Unspecified |
| mmu-let-7i-5p | MIMAT0000122 | Zkscan14 | 67235 | HITS-CLIP | 23142080 | Unspecified |
| mmu-let-7i-5p | MIMAT0000122 | Zkscan2 | 210162 | HITS-CLIP | 19536157 | Unspecified |
| mmu-let-7i-5p | MIMAT0000122 | Zkscan7 | 382118 | HITS-CLIP | 19536157\|23597149 | Unspecified |
| mmu-let-7i-5p | MIMAT0000122 | Zmat1 | 215693 | HITS-CLIP | 23142080 | Unspecified |
| mmu-let-7i-5p | MIMAT0000122 | Zmat3 | 22401 | HITS-CLIP | 23597149\|23142080 | Unspecified |
| mmu-let-7i-5p | MIMAT0000122 | Zmat5 | 67178 | HITS-CLIP | 23142080 | Unspecified |
| mmu-let-7i-5p | MIMAT0000122 | Zmiz1 | 328365 | HITS-CLIP | 23142080\|23597149\|29167373 | Unspecified |
| mmu-let-7i-5p | MIMAT0000122 | Zmiz2 | 52915 | HITS-CLIP | 23142080 | Unspecified |
| mmu-let-7i-5p | MIMAT0000122 | Zmpste24 | 230709 | HITS-CLIP | 23597149\|23142080\|29167373 | Unspecified |
| mmu-let-7i-5p | MIMAT0000122 | Zmym1 | 68310 | HITS-CLIP | 23142080 | Unspecified |
| mmu-let-7i-5p | MIMAT0000122 | Zmym2 | 76007 | HITS-CLIP | 37986224\|31433990\|23142080\|19536 | Unspecified |
| mmu-let-7i-5p | MIMAT0000122 | Zmym3 | 56364 | HITS-CLIP | 23142080 | Unspecified |
| mmu-let-7i-5p | MIMAT0000122 | Zmym4 | 67785 | HITS-CLIP | 29167373 | Unspecified |
| mmu-let-7i-5p | MIMAT0000122 | Zmym6 | 100177 | HITS-CLIP | 23597149 | Unspecified |
| mmu-let-7i-5p | MIMAT0000122 | Zmynd11 | 66505 | HITS-CLIP | 19536157\|29167373 | Unspecified |
| mmu-let-7i-5p | MIMAT0000122 | Znfx1 | 98999 | HITS-CLIP | 23142080 | Unspecified |
| mmu-let-7i-5p | MIMAT0000122 | Znhit1 | 70103 | HITS-CLIP | 23142080 | Unspecified |

| | | | | | | |
|---|---|---|---|---|---|---|
| mmu-let-7i-5p | MIMAT0000122 | Znrd1as | 76416 | HITS-CLIP | 23142080 | Unspecified |
| mmu-let-7i-5p | MIMAT0000122 | Znrf2 | 387524 | HITS-CLIP | 31433990\|23142080\|29167373 | Unspecified |
| mmu-let-7i-5p | MIMAT0000122 | Znrf3 | 407821 | HITS-CLIP | 37986224\|23142080 | Unspecified |
| mmu-let-7i-5p | MIMAT0000122 | Zranb2 | 53861 | HITS-CLIP | 31433990\|23142080 | Unspecified |
| mmu-let-7i-5p | MIMAT0000122 | Zscan22 | 232878 | HITS-CLIP | 37986224 | Unspecified |
| mmu-let-7i-5p | MIMAT0000122 | Zscan29 | 99334 | HITS-CLIP | 23597149\|23142080 | Unspecified |
| mmu-let-7i-5p | MIMAT0000122 | Zswim4 | 212168 | qCLASH | 31393953 | Unspecified |
| mmu-let-7i-5p | MIMAT0000122 | Zswim5 | 74464 | HITS-CLIP | 37986224\|19536157\|23597149 | Unspecified |
| mmu-let-7i-5p | MIMAT0000122 | Zswim6 | 67263 | HITS-CLIP | 37986224 | Unspecified |
| mmu-let-7i-5p | MIMAT0000122 | Zswim8 | 268721 | HITS-CLIP | 23597149\|23142080 | Unspecified |
| mmu-let-7i-5p | MIMAT0000122 | Zup1 | 221302 | HITS-CLIP | 23142080 | Unspecified |
| mmu-let-7i-5p | MIMAT0000122 | Zwint | 52696 | HITS-CLIP | 37986224 | Unspecified |
| mmu-let-7i-5p | MIMAT0000122 | Zyg11b | 414872 | HITS-CLIP | 23142080 | Unspecified |
| mmu-let-7i-5p | MIMAT0000122 | Zzef1 | 195018 | HITS-CLIP | 37986224\|23142080\|23597149 | Unspecified |
| mmu-let-7i-5p | MIMAT0000122 | Zzz3 | 108946 | HITS-CLIP | 23597149\|23142080 | Unspecified |
| mmu-let-7i-5p | MIMAT0000122 | mt-Co1 | 4512 | HITS-CLIP | 23597149\|23142080\|29167373 | Unspecified |
| mmu-let-7i-5p | MIMAT0000122 | mt-Nd1 | 4535 | HITS-CLIP | 19536157\|23597149\|23142080\|29167 | Unspecified |
| mmu-let-7i-5p | MIMAT0000122 | mt-Nd2 | 4536 | HITS-CLIP//qCLASH | 23597149\|31393953\|19536157\|23142 | Unspecified |
| mmu-let-7i-5p | MIMAT0000122 | mt-Nd5 | 4540 | HITS-CLIP | 23597149\|19536157\|23142080\|29167 | Unspecified |
| mmu-miR-680 | MIMAT0003457 | Ap2b1 | 71770 | HITS-CLIP | 31433990 | Unspecified |
| mmu-miR-680 | MIMAT0003457 | C2cd5 | 74741 | HITS-CLIP | 31433990 | Unspecified |
| mmu-miR-680 | MIMAT0003457 | Runx1 | 12394 | HITS-CLIP | 31433990 | Unspecified |
| mmu-miR-3067-5p | MIMAT0014840 | Myb | 17863 | HITS-CLIP | 31433990 | Unspecified |
| mmu-miR-3067-5p | MIMAT0014840 | Shprh | 268281 | HITS-CLIP | 31433990 | Unspecified |
| mmu-miR-193a-3p | MIMAT0000223 | 1110038F14Rik | 117171 | HITS-CLIP | 31433990 | Unspecified |
| mmu-miR-193a-3p | MIMAT0000223 | 1300017J02Rik | 71775 | HITS-CLIP | 37986224\|23597149 | Unspecified |
| mmu-miR-193a-3p | MIMAT0000223 | 1700123O20Rik | 58248 | HITS-CLIP | 37986224\|23597149 | Unspecified |
| mmu-miR-193a-3p | MIMAT0000223 | 1810055G02Rik | 72056 | HITS-CLIP | 37986224 | Unspecified |
| mmu-miR-193a-3p | MIMAT0000223 | 2610507B11Rik | 72503 | HITS-CLIP//sRNA-Seq | 23597149\|33184237 | Unspecified |
| mmu-miR-193a-3p | MIMAT0000223 | 4833439L19Rik | 97820 | HITS-CLIP//sRNA-Seq | 23597149\|33184237 | Unspecified |
| mmu-miR-193a-3p | MIMAT0000223 | Aacs | 78894 | HITS-CLIP//sRNA-Seq | 23597149\|33184237 | Unspecified |
| mmu-miR-193a-3p | MIMAT0000223 | Aadac | 67758 | HITS-CLIP | 23597149 | Unspecified |
| mmu-miR-193a-3p | MIMAT0000223 | Abca1 | 11303 | HITS-CLIP | 23597149 | Unspecified |
| mmu-miR-193a-3p | MIMAT0000223 | Abcb7 | 11306 | HITS-CLIP | 23597149 | Unspecified |
| mmu-miR-193a-3p | MIMAT0000223 | Abcc2 | 12780 | HITS-CLIP | 37986224 | Unspecified |
| mmu-miR-193a-3p | MIMAT0000223 | Abcc3 | 76408 | HITS-CLIP | 37986224\|23597149 | Unspecified |
| mmu-miR-193a-3p | MIMAT0000223 | Abhd17b | 226016 | HITS-CLIP | 23597149 | Unspecified |
| mmu-miR-193a-3p | MIMAT0000223 | Abl2 | 11352 | HITS-CLIP | 23597149 | Unspecified |
| mmu-miR-193a-3p | MIMAT0000223 | Ablim1 | 226251 | sRNA-Seq//HITS-CLIP | 33184237\|23597149 | Unspecified |

| | | | | | | |
|---|---|---|---|---|---|---|
| mmu-miR-193a-3p | MIMAT0000223 | Acacb | 100705 | HITS-CLIP | 23597149 | Unspecified |
| mmu-miR-193a-3p | MIMAT0000223 | Acadvl | 11370 | HITS-CLIP | 37986224 | Unspecified |
| mmu-miR-193a-3p | MIMAT0000223 | Acmsd | 266645 | HITS-CLIP | 37986224\|23597149 | Unspecified |
| mmu-miR-193a-3p | MIMAT0000223 | Aco2 | 11429 | HITS-CLIP | 37986224 | Unspecified |
| mmu-miR-193a-3p | MIMAT0000223 | Acox3 | 80911 | HITS-CLIP | 23597149 | Unspecified |
| mmu-miR-193a-3p | MIMAT0000223 | Acp5 | 11433 | HITS-CLIP | 23597149 | Unspecified |
| mmu-miR-193a-3p | MIMAT0000223 | Acsl3 | 74205 | HITS-CLIP | 37986224 | Unspecified |
| mmu-miR-193a-3p | MIMAT0000223 | Acsm3 | 20216 | HITS-CLIP | 23597149 | Unspecified |
| mmu-miR-193a-3p | MIMAT0000223 | Actr2 | 66713 | HITS-CLIP//sRNA-Seq | 23597149\|33184237 | Unspecified |
| mmu-miR-193a-3p | MIMAT0000223 | Actr8 | 56249 | HITS-CLIP | 37986224\|23597149 | Unspecified |
| mmu-miR-193a-3p | MIMAT0000223 | Adcy9 | 11515 | HITS-CLIP | 23597149 | Unspecified |
| mmu-miR-193a-3p | MIMAT0000223 | Adhfe1 | 76187 | HITS-CLIP | 23597149 | Unspecified |
| mmu-miR-193a-3p | MIMAT0000223 | Adipor2 | 68465 | HITS-CLIP | 23597149 | Unspecified |
| mmu-miR-193a-3p | MIMAT0000223 | Adnp | 11538 | HITS-CLIP | 23597149 | Unspecified |
| mmu-miR-193a-3p | MIMAT0000223 | Adrm1 | 56436 | HITS-CLIP//sRNA-Seq | 37986224\|33184237\|23597149 | Unspecified |
| mmu-miR-193a-3p | MIMAT0000223 | Agpat1 | 55979 | HITS-CLIP//sRNA-Seq | 23597149\|33184237 | Unspecified |
| mmu-miR-193a-3p | MIMAT0000223 | Agt | 11606 | HITS-CLIP | 37986224\|23597149 | Unspecified |
| mmu-miR-193a-3p | MIMAT0000223 | Agxt | 11611 | HITS-CLIP | 23597149 | Unspecified |
| mmu-miR-193a-3p | MIMAT0000223 | Agxt2 | 268782 | HITS-CLIP | 23597149 | Unspecified |
| mmu-miR-193a-3p | MIMAT0000223 | Ahctf1 | 226747 | HITS-CLIP | 23597149 | Unspecified |
| mmu-miR-193a-3p | MIMAT0000223 | Ahdc1 | 230793 | HITS-CLIP | 19536157 | Unspecified |
| mmu-miR-193a-3p | MIMAT0000223 | Ahr | 11622 | HITS-CLIP | 23597149 | Unspecified |
| mmu-miR-193a-3p | MIMAT0000223 | Akap11 | 219181 | HITS-CLIP//sRNA-Seq | 19536157\|33184237 | Unspecified |
| mmu-miR-193a-3p | MIMAT0000223 | Akirin2 | 433693 | HITS-CLIP//sRNA-Seq | 19536157\|33184237 | Unspecified |
| mmu-miR-193a-3p | MIMAT0000223 | Akr1a1 | 58810 | HITS-CLIP//sRNA-Seq | 23597149\|33184237 | Unspecified |
| mmu-miR-193a-3p | MIMAT0000223 | Akr1c19 | 432720 | HITS-CLIP | 23597149 | Unspecified |
| mmu-miR-193a-3p | MIMAT0000223 | Akr1c6 | 83702 | HITS-CLIP | 37986224 | Unspecified |
| mmu-miR-193a-3p | MIMAT0000223 | Akr7a5 | 110198 | HITS-CLIP | 23597149 | Unspecified |
| mmu-miR-193a-3p | MIMAT0000223 | Alas1 | 11655 | HITS-CLIP//sRNA-Seq | 37986224\|33184237\|23597149 | Unspecified |
| mmu-miR-193a-3p | MIMAT0000223 | Aldh1a1 | 11668 | HITS-CLIP | 23597149 | Unspecified |
| mmu-miR-193a-3p | MIMAT0000223 | Aldh1l1 | 107747 | HITS-CLIP | 37986224 | Unspecified |
| mmu-miR-193a-3p | MIMAT0000223 | Aldh2 | 11669 | HITS-CLIP//sRNA-Seq | 23597149\|33184237 | Unspecified |
| mmu-miR-193a-3p | MIMAT0000223 | Aldh4a1 | 212647 | HITS-CLIP | 23597149 | Unspecified |
| mmu-miR-193a-3p | MIMAT0000223 | Aldoa | 11674 | HITS-CLIP//sRNA-Seq | 19536157\|33184237\|23597149 | Unspecified |
| mmu-miR-193a-3p | MIMAT0000223 | Aldob | 230163 | HITS-CLIP | 23597149 | Unspecified |
| mmu-miR-193a-3p | MIMAT0000223 | Alg14 | 66789 | HITS-CLIP | 23597149 | Unspecified |
| mmu-miR-193a-3p | MIMAT0000223 | Alkbh5 | 268420 | HITS-CLIP | 23597149 | Unspecified |
| mmu-miR-193a-3p | MIMAT0000223 | Ambra1 | 228361 | HITS-CLIP | 37986224\|23597149 | Unspecified |
| mmu-miR-193a-3p | MIMAT0000223 | Amigo3 | 320844 | HITS-CLIP | 23597149 | Unspecified |

| | | | | | | |
|---|---|---|---|---|---|---|
| mmu-miR-193a-3p | MIMAT0000223 | Ank3 | 11735 | HITS-CLIP//sRNA-Seq | 19536157\|33184237 | Unspecified |
| mmu-miR-193a-3p | MIMAT0000223 | Ankrd13a | 68420 | HITS-CLIP | 37986224 | Unspecified |
| mmu-miR-193a-3p | MIMAT0000223 | Anks1 | 224650 | HITS-CLIP | 23597149 | Unspecified |
| mmu-miR-193a-3p | MIMAT0000223 | Anxa2 | 12306 | HITS-CLIP//sRNA-Seq | 19536157\|33184237 | Unspecified |
| mmu-miR-193a-3p | MIMAT0000223 | Ap1g1 | 11765 | sRNA-Seq//HITS-CLIP | 33184237\|23597149 | Unspecified |
| mmu-miR-193a-3p | MIMAT0000223 | Ap2a1 | 11771 | HITS-CLIP//sRNA-Seq | 23597149\|33184237 | Unspecified |
| mmu-miR-193a-3p | MIMAT0000223 | Ap2b1 | 71770 | HITS-CLIP | 37986224 | Unspecified |
| mmu-miR-193a-3p | MIMAT0000223 | Ap2m1 | 11773 | HITS-CLIP//sRNA-Seq | 23597149\|33184237 | Unspecified |
| mmu-miR-193a-3p | MIMAT0000223 | Apex2 | 77622 | HITS-CLIP | 23597149 | Unspecified |
| mmu-miR-193a-3p | MIMAT0000223 | Apln | 30878 | HITS-CLIP | 19536157 | Unspecified |
| mmu-miR-193a-3p | MIMAT0000223 | Aplp2 | 11804 | HITS-CLIP | 37986224 | Unspecified |
| mmu-miR-193a-3p | MIMAT0000223 | Apoa5 | 66113 | HITS-CLIP | 23597149 | Unspecified |
| mmu-miR-193a-3p | MIMAT0000223 | Apob | 238055 | HITS-CLIP | 37986224\|23597149 | Unspecified |
| mmu-miR-193a-3p | MIMAT0000223 | Apoh | 350 | HITS-CLIP | 23597149 | Unspecified |
| mmu-miR-193a-3p | MIMAT0000223 | App | 11820 | sRNA-Seq//HITS-CLIP | 33184237\|26203562 | Unspecified |
| mmu-miR-193a-3p | MIMAT0000223 | Appl2 | 216190 | HITS-CLIP | 23597149 | Unspecified |
| mmu-miR-193a-3p | MIMAT0000223 | Aqp1 | 11826 | HITS-CLIP | 23597149 | Unspecified |
| mmu-miR-193a-3p | MIMAT0000223 | Arhgap29 | 214137 | HITS-CLIP | 23597149 | Unspecified |
| mmu-miR-193a-3p | MIMAT0000223 | Arhgap32 | 330914 | HITS-CLIP | 19536157 | Unspecified |
| mmu-miR-193a-3p | MIMAT0000223 | Arhgap42 | 71544 | HITS-CLIP | 37986224 | Unspecified |
| mmu-miR-193a-3p | MIMAT0000223 | Arhgap5 | 11855 | HITS-CLIP//sRNA-Seq | 37986224\|23597149\|33184237 | Unspecified |
| mmu-miR-193a-3p | MIMAT0000223 | Arhgef10 | 234094 | HITS-CLIP | 23597149 | Unspecified |
| mmu-miR-193a-3p | MIMAT0000223 | Arhgef12 | 69632 | HITS-CLIP | 37986224\|23597149 | Unspecified |
| mmu-miR-193a-3p | MIMAT0000223 | Arhgef26 | 622434 | HITS-CLIP | 37986224 | Unspecified |
| mmu-miR-193a-3p | MIMAT0000223 | Arhgef7 | 54126 | sRNA-Seq//HITS-CLIP | 33184237\|19536157 | Unspecified |
| mmu-miR-193a-3p | MIMAT0000223 | Arid1a | 93760 | HITS-CLIP | 37986224\|23597149 | Unspecified |
| mmu-miR-193a-3p | MIMAT0000223 | Armcx2 | 67416 | HITS-CLIP//sRNA-Seq | 23597149\|33184237 | Unspecified |
| mmu-miR-193a-3p | MIMAT0000223 | Arpc1b | 11867 | sRNA-Seq//HITS-CLIP | 33184237\|23597149 | Unspecified |
| mmu-miR-193a-3p | MIMAT0000223 | Arpp21 | 74100 | HITS-CLIP | 19536157 | Unspecified |
| mmu-miR-193a-3p | MIMAT0000223 | Asgr2 | 11890 | HITS-CLIP | 37986224 | Unspecified |
| mmu-miR-193a-3p | MIMAT0000223 | Ash1l | 192195 | HITS-CLIP | 19536157 | Unspecified |
| mmu-miR-193a-3p | MIMAT0000223 | Atad3a | 108888 | HITS-CLIP | 23597149 | Unspecified |
| mmu-miR-193a-3p | MIMAT0000223 | Atat1 | 73242 | HITS-CLIP | 23597149 | Unspecified |
| mmu-miR-193a-3p | MIMAT0000223 | Atf6 | 226641 | HITS-CLIP | 23597149 | Unspecified |
| mmu-miR-193a-3p | MIMAT0000223 | Atp13a3 | 224088 | HITS-CLIP | 23597149 | Unspecified |
| mmu-miR-193a-3p | MIMAT0000223 | Atp2a2 | 11938 | sRNA-Seq//HITS-CLIP | 33184237\|19536157 | Unspecified |
| mmu-miR-193a-3p | MIMAT0000223 | Atp2b2 | 11941 | HITS-CLIP//sRNA-Seq | 19536157\|33184237 | Unspecified |
| mmu-miR-193a-3p | MIMAT0000223 | Atp5a1 | 11946 | HITS-CLIP//sRNA-Seq | 23597149\|33184237 | Unspecified |
| mmu-miR-193a-3p | MIMAT0000223 | Atp5b | 11947 | HITS-CLIP//sRNA-Seq | 19536157\|33184237 | Unspecified |

| | | | | | | |
|---|---|---|---|---|---|---|
| mmu-miR-193a-3p | MIMAT0000223 | Atp6v0d1 | 11972 | HITS-CLIP//sRNA-Seq | 23597149\|33184237 | Unspecified |
| mmu-miR-193a-3p | MIMAT0000223 | Atp6v1c1 | 66335 | HITS-CLIP//sRNA-Seq | 23597149\|33184237 | Unspecified |
| mmu-miR-193a-3p | MIMAT0000223 | Atr | 245000 | HITS-CLIP | 23597149 | Unspecified |
| mmu-miR-193a-3p | MIMAT0000223 | Atrnl1 | 226255 | HITS-CLIP//sRNA-Seq | 23597149\|33184237 | Unspecified |
| mmu-miR-193a-3p | MIMAT0000223 | Atxn1l | 52335 | HITS-CLIP | 23597149 | Unspecified |
| mmu-miR-193a-3p | MIMAT0000223 | Atxn2 | 20239 | sRNA-Seq//HITS-CLIP | 33184237\|23597149 | Unspecified |
| mmu-miR-193a-3p | MIMAT0000223 | Atxn7 | 246103 | HITS-CLIP | 23597149 | Unspecified |
| mmu-miR-193a-3p | MIMAT0000223 | Aup1 | 11993 | HITS-CLIP | 23597149 | Unspecified |
| mmu-miR-193a-3p | MIMAT0000223 | Avl9 | 78937 | HITS-CLIP | 23597149 | Unspecified |
| mmu-miR-193a-3p | MIMAT0000223 | Axl | 26362 | HITS-CLIP | 23597149 | Unspecified |
| mmu-miR-193a-3p | MIMAT0000223 | B4galt1 | 14595 | HITS-CLIP | 37986224 | Unspecified |
| mmu-miR-193a-3p | MIMAT0000223 | B4galt5 | 56336 | HITS-CLIP | 37986224 | Unspecified |
| mmu-miR-193a-3p | MIMAT0000223 | BC004004 | 80748 | HITS-CLIP//sRNA-Seq | 37986224\|23597149\|33184237 | Unspecified |
| mmu-miR-193a-3p | MIMAT0000223 | Baat | 12012 | HITS-CLIP | 23597149 | Unspecified |
| mmu-miR-193a-3p | MIMAT0000223 | Banf1 | 23825 | HITS-CLIP//sRNA-Seq | 23597149\|33184237 | Unspecified |
| mmu-miR-193a-3p | MIMAT0000223 | Baz2a | 116848 | HITS-CLIP | 37986224\|23597149 | Unspecified |
| mmu-miR-193a-3p | MIMAT0000223 | Bbs7 | 71492 | HITS-CLIP | 23597149 | Unspecified |
| mmu-miR-193a-3p | MIMAT0000223 | Bcar1 | 12927 | HITS-CLIP | 37986224\|23597149 | Unspecified |
| mmu-miR-193a-3p | MIMAT0000223 | Bckdha | 12039 | HITS-CLIP | 37986224 | Unspecified |
| mmu-miR-193a-3p | MIMAT0000223 | Bcl11b | 58208 | HITS-CLIP | 19536157 | Unspecified |
| mmu-miR-193a-3p | MIMAT0000223 | Bcl6 | 12053 | HITS-CLIP | 37986224 | Unspecified |
| mmu-miR-193a-3p | MIMAT0000223 | Bclaf1 | 72567 | HITS-CLIP//sRNA-Seq | 19536157\|33184237 | Unspecified |
| mmu-miR-193a-3p | MIMAT0000223 | Bhlhe40 | 20893 | HITS-CLIP | 31433990 | Unspecified |
| mmu-miR-193a-3p | MIMAT0000223 | Bhlhe41 | 79362 | HITS-CLIP | 31433990 | Unspecified |
| mmu-miR-193a-3p | MIMAT0000223 | Bid | 12122 | HITS-CLIP | 23597149 | Unspecified |
| mmu-miR-193a-3p | MIMAT0000223 | Birc6 | 12211 | HITS-CLIP//sRNA-Seq | 23597149\|33184237 | Unspecified |
| mmu-miR-193a-3p | MIMAT0000223 | Bphl | 68021 | HITS-CLIP | 37986224 | Unspecified |
| mmu-miR-193a-3p | MIMAT0000223 | Brap | 72399 | HITS-CLIP | 23597149 | Unspecified |
| mmu-miR-193a-3p | MIMAT0000223 | Brd3 | 67382 | HITS-CLIP | 23597149 | Unspecified |
| mmu-miR-193a-3p | MIMAT0000223 | Bri3bp | 76809 | HITS-CLIP | 37986224 | Unspecified |
| mmu-miR-193a-3p | MIMAT0000223 | Brpf1 | 78783 | HITS-CLIP | 23597149 | Unspecified |
| mmu-miR-193a-3p | MIMAT0000223 | Brwd3 | 382236 | HITS-CLIP | 31433990 | Unspecified |
| mmu-miR-193a-3p | MIMAT0000223 | Bzw1 | 66882 | HITS-CLIP//sRNA-Seq | 37986224\|19536157\|33184237\|23597 | Unspecified |
| mmu-miR-193a-3p | MIMAT0000223 | C1galt1 | 94192 | HITS-CLIP | 23597149 | Unspecified |
| mmu-miR-193a-3p | MIMAT0000223 | C1ql3 | 227580 | HITS-CLIP | 19536157 | Unspecified |
| mmu-miR-193a-3p | MIMAT0000223 | C5ar1 | 12273 | HITS-CLIP | 23597149 | Unspecified |
| mmu-miR-193a-3p | MIMAT0000223 | C77080 | 97130 | HITS-CLIP | 23597149 | Unspecified |
| mmu-miR-193a-3p | MIMAT0000223 | Cabin1 | 104248 | HITS-CLIP | 23597149 | Unspecified |
| mmu-miR-193a-3p | MIMAT0000223 | Cables2 | 252966 | HITS-CLIP | 37986224 | Unspecified |

| | | | | | | |
|---|---|---|---|---|---|---|
| mmu-miR-193a-3p | MIMAT0000223 | Cacna1g | 12291 | HITS-CLIP | | 19536157 Unspecified |
| mmu-miR-193a-3p | MIMAT0000223 | Cadm1 | 54725 | HITS-CLIP//sRNA-Seq | 23597149\|33184237 | Unspecified |
| mmu-miR-193a-3p | MIMAT0000223 | Calm1 | 12313 | sRNA-Seq//HITS-CLIP | 33184237\|19536157 | Unspecified |
| mmu-miR-193a-3p | MIMAT0000223 | Calr | 12317 | HITS-CLIP//sRNA-Seq | 23597149\|19536157\|33184237 | Unspecified |
| mmu-miR-193a-3p | MIMAT0000223 | Calu | 12321 | HITS-CLIP//sRNA-Seq | 23597149\|33184237 | Unspecified |
| mmu-miR-193a-3p | MIMAT0000223 | Camta1 | 100072 | sRNA-Seq//HITS-CLIP | 33184237\|19536157 | Unspecified |
| mmu-miR-193a-3p | MIMAT0000223 | Canx | 12330 | sRNA-Seq//HITS-CLIP | 33184237\|23597149 | Unspecified |
| mmu-miR-193a-3p | MIMAT0000223 | Caprin1 | 53872 | HITS-CLIP//sRNA-Seq | 37986224\|23597149\|33184237 | Unspecified |
| mmu-miR-193a-3p | MIMAT0000223 | Carhsp1 | 52502 | HITS-CLIP//sRNA-Seq | 23597149\|33184237 | Unspecified |
| mmu-miR-193a-3p | MIMAT0000223 | Cavin2 | 8436 | HITS-CLIP | | 23597149 Unspecified |
| mmu-miR-193a-3p | MIMAT0000223 | Cbx6 | 494448 | HITS-CLIP | | 19536157 Unspecified |
| mmu-miR-193a-3p | MIMAT0000223 | Ccdc6 | 76551 | HITS-CLIP | 37986224\|23597149 | Unspecified |
| mmu-miR-193a-3p | MIMAT0000223 | Ccnd1 | 12443 | HITS-CLIP | | 23597149 Unspecified |
| mmu-miR-193a-3p | MIMAT0000223 | Cct2 | 12461 | HITS-CLIP//sRNA-Seq | 37986224\|33184237\|23597149 | Unspecified |
| mmu-miR-193a-3p | MIMAT0000223 | Cct3 | 12462 | HITS-CLIP//sRNA-Seq | 23597149\|33184237 | Unspecified |
| mmu-miR-193a-3p | MIMAT0000223 | Cct5 | 12465 | HITS-CLIP//sRNA-Seq | 23597149\|33184237 | Unspecified |
| mmu-miR-193a-3p | MIMAT0000223 | Cct7 | 12468 | HITS-CLIP//sRNA-Seq | 23597149\|33184237 | Unspecified |
| mmu-miR-193a-3p | MIMAT0000223 | Cd164 | 53599 | HITS-CLIP | | 23597149 Unspecified |
| mmu-miR-193a-3p | MIMAT0000223 | Cd47 | 16423 | sRNA-Seq//HITS-CLIP | 33184237\|19536157 | Unspecified |
| mmu-miR-193a-3p | MIMAT0000223 | Cdc25a | 12530 | HITS-CLIP | 37986224\|23597149 | Unspecified |
| mmu-miR-193a-3p | MIMAT0000223 | Cdh5 | 12562 | HITS-CLIP | | 23597149 Unspecified |
| mmu-miR-193a-3p | MIMAT0000223 | Cdk17 | 237459 | HITS-CLIP//sRNA-Seq | 23597149\|33184237 | Unspecified |
| mmu-miR-193a-3p | MIMAT0000223 | Cdk19 | 78334 | HITS-CLIP | | 19536157 Unspecified |
| mmu-miR-193a-3p | MIMAT0000223 | Cdkl2 | 53886 | HITS-CLIP | | 19536157 Unspecified |
| mmu-miR-193a-3p | MIMAT0000223 | Cds2 | 110911 | HITS-CLIP//sRNA-Seq | 23597149\|33184237 | Unspecified |
| mmu-miR-193a-3p | MIMAT0000223 | Cecr2 | 330409 | HITS-CLIP | | 23597149 Unspecified |
| mmu-miR-193a-3p | MIMAT0000223 | Celf2 | 14007 | HITS-CLIP | | 31433990 Unspecified |
| mmu-miR-193a-3p | MIMAT0000223 | Celsr2 | 53883 | HITS-CLIP | | 19536157 Unspecified |
| mmu-miR-193a-3p | MIMAT0000223 | Cenpb | 12616 | HITS-CLIP | | 19536157 Unspecified |
| mmu-miR-193a-3p | MIMAT0000223 | Cep170 | 545389 | HITS-CLIP//sRNA-Seq | 23597149\|33184237 | Unspecified |
| mmu-miR-193a-3p | MIMAT0000223 | Cep55 | 74107 | HITS-CLIP | | 23597149 Unspecified |
| mmu-miR-193a-3p | MIMAT0000223 | Cep85l | 100038725 | HITS-CLIP | | 23597149 Unspecified |
| mmu-miR-193a-3p | MIMAT0000223 | Ces1c | 13884 | HITS-CLIP | | 23597149 Unspecified |
| mmu-miR-193a-3p | MIMAT0000223 | Ces3b | 13909 | HITS-CLIP | | 23597149 Unspecified |
| mmu-miR-193a-3p | MIMAT0000223 | Cfdp1 | 23837 | HITS-CLIP//sRNA-Seq | 23597149\|33184237 | Unspecified |
| mmu-miR-193a-3p | MIMAT0000223 | Cfp | 18636 | HITS-CLIP | | 23597149 Unspecified |
| mmu-miR-193a-3p | MIMAT0000223 | Cgnl1 | 68178 | HITS-CLIP | | 23597149 Unspecified |
| mmu-miR-193a-3p | MIMAT0000223 | Chd7 | 320790 | HITS-CLIP | | 37986224 Unspecified |
| mmu-miR-193a-3p | MIMAT0000223 | Chd8 | 67772 | HITS-CLIP//sRNA-Seq | 23597149\|33184237 | Unspecified |

| | | | | | | |
|---|---|---|---|---|---|---|
| mmu-miR-193a-3p | MIMAT0000223 | Clec4f | 51811 | HITS-CLIP | 23597149 | Unspecified |
| mmu-miR-193a-3p | MIMAT0000223 | Clpb | 20480 | HITS-CLIP | 23597149 | Unspecified |
| mmu-miR-193a-3p | MIMAT0000223 | Clpx | 270166 | HITS-CLIP | 37986224\|23597149 | Unspecified |
| mmu-miR-193a-3p | MIMAT0000223 | Cltc | 67300 | HITS-CLIP//sRNA-Seq | 37986224\|19536157\|33184237 | Unspecified |
| mmu-miR-193a-3p | MIMAT0000223 | Clu | 12759 | HITS-CLIP//sRNA-Seq | 23597149\|33184237 | Unspecified |
| mmu-miR-193a-3p | MIMAT0000223 | Clybl | 69634 | HITS-CLIP | 23597149 | Unspecified |
| mmu-miR-193a-3p | MIMAT0000223 | Cmtm7 | 102545 | HITS-CLIP | 31433990 | Unspecified |
| mmu-miR-193a-3p | MIMAT0000223 | Cmtr1 | 74157 | HITS-CLIP | 23597149 | Unspecified |
| mmu-miR-193a-3p | MIMAT0000223 | Cnot6 | 104625 | HITS-CLIP//sRNA-Seq | 23597149\|33184237 | Unspecified |
| mmu-miR-193a-3p | MIMAT0000223 | Cnot8 | 69125 | HITS-CLIP | 23597149 | Unspecified |
| mmu-miR-193a-3p | MIMAT0000223 | Col18a1 | 12822 | HITS-CLIP | 37986224 | Unspecified |
| mmu-miR-193a-3p | MIMAT0000223 | Copa | 12847 | HITS-CLIP//sRNA-Seq | 23597149\|33184237 | Unspecified |
| mmu-miR-193a-3p | MIMAT0000223 | Copb2 | 50797 | HITS-CLIP | 37986224 | Unspecified |
| mmu-miR-193a-3p | MIMAT0000223 | Cp | 12870 | HITS-CLIP | 23597149 | Unspecified |
| mmu-miR-193a-3p | MIMAT0000223 | Cpn2 | 71756 | HITS-CLIP | 23597149 | Unspecified |
| mmu-miR-193a-3p | MIMAT0000223 | Cps1 | 227231 | HITS-CLIP | 23597149 | Unspecified |
| mmu-miR-193a-3p | MIMAT0000223 | Cpsf4l | 52670 | HITS-CLIP | 37986224 | Unspecified |
| mmu-miR-193a-3p | MIMAT0000223 | Cpsf7 | 269061 | HITS-CLIP | 37986224\|23597149 | Unspecified |
| mmu-miR-193a-3p | MIMAT0000223 | Cpt1a | 12894 | HITS-CLIP | 23597149 | Unspecified |
| mmu-miR-193a-3p | MIMAT0000223 | Cptp | 80772 | HITS-CLIP | 23597149 | Unspecified |
| mmu-miR-193a-3p | MIMAT0000223 | Creb3l3 | 208677 | HITS-CLIP | 23597149 | Unspecified |
| mmu-miR-193a-3p | MIMAT0000223 | Crebl2 | 232430 | HITS-CLIP | 23597149 | Unspecified |
| mmu-miR-193a-3p | MIMAT0000223 | Crk | 12928 | sRNA-Seq//HITS-CLIP | 33184237\|23597149 | Unspecified |
| mmu-miR-193a-3p | MIMAT0000223 | Crot | 74114 | HITS-CLIP | 23597149 | Unspecified |
| mmu-miR-193a-3p | MIMAT0000223 | Crtc1 | 382056 | HITS-CLIP | 37986224\|23597149 | Unspecified |
| mmu-miR-193a-3p | MIMAT0000223 | Csnk1g3 | 70425 | HITS-CLIP//sRNA-Seq | 19536157\|33184237 | Unspecified |
| mmu-miR-193a-3p | MIMAT0000223 | Ctdspl2 | 329506 | HITS-CLIP | 23597149 | Unspecified |
| mmu-miR-193a-3p | MIMAT0000223 | Cth | 107869 | HITS-CLIP | 23597149 | Unspecified |
| mmu-miR-193a-3p | MIMAT0000223 | Ctnnb1 | 12387 | HITS-CLIP//sRNA-Seq | 23597149\|19536157\|33184237 | Unspecified |
| mmu-miR-193a-3p | MIMAT0000223 | Ctsa | 19025 | HITS-CLIP | 23597149 | Unspecified |
| mmu-miR-193a-3p | MIMAT0000223 | Ctsl | 13039 | HITS-CLIP//sRNA-Seq | 23597149\|33184237 | Unspecified |
| mmu-miR-193a-3p | MIMAT0000223 | Ctso | 229445 | HITS-CLIP | 37986224 | Unspecified |
| mmu-miR-193a-3p | MIMAT0000223 | Cxadr | 13052 | HITS-CLIP | 37986224 | Unspecified |
| mmu-miR-193a-3p | MIMAT0000223 | Cxcl10 | 15945 | HITS-CLIP | 23597149 | Unspecified |
| mmu-miR-193a-3p | MIMAT0000223 | Cxcl12 | 20315 | HITS-CLIP | 23597149 | Unspecified |
| mmu-miR-193a-3p | MIMAT0000223 | Cyb5b | 66427 | HITS-CLIP | 37986224 | Unspecified |
| mmu-miR-193a-3p | MIMAT0000223 | Cyc1 | 66445 | HITS-CLIP//sRNA-Seq | 23597149\|33184237 | Unspecified |
| mmu-miR-193a-3p | MIMAT0000223 | Cyp26a1 | 13082 | HITS-CLIP | 23597149 | Unspecified |
| mmu-miR-193a-3p | MIMAT0000223 | Cyp27a1 | 104086 | HITS-CLIP | 23597149 | Unspecified |

| | | | | | | | |
|---|---|---|---|---|---|---|---|
| mmu-miR-193a-3p | MIMAT0000223 | Cyp2c67 | 545288 | HITS-CLIP | | 23597149 | Unspecified |
| mmu-miR-193a-3p | MIMAT0000223 | Cyp2c70 | 226105 | HITS-CLIP | | 37986224 | Unspecified |
| mmu-miR-193a-3p | MIMAT0000223 | Cyp2d26 | 76279 | HITS-CLIP | | 37986224 | Unspecified |
| mmu-miR-193a-3p | MIMAT0000223 | Cyp2e1 | 13106 | HITS-CLIP | | 23597149 | Unspecified |
| mmu-miR-193a-3p | MIMAT0000223 | D17H6S53E | 114585 | HITS-CLIP | | 37986224 | Unspecified |
| mmu-miR-193a-3p | MIMAT0000223 | D17Wsu92e | 224647 | HITS-CLIP | | 23597149 | Unspecified |
| mmu-miR-193a-3p | MIMAT0000223 | D430041D05Rik | 241589 | HITS-CLIP | | 19536157 | Unspecified |
| mmu-miR-193a-3p | MIMAT0000223 | Dact3 | 629378 | HITS-CLIP | | 19536157 | Unspecified |
| mmu-miR-193a-3p | MIMAT0000223 | Dazap2 | 23994 | HITS-CLIP | | 37986224 | Unspecified |
| mmu-miR-193a-3p | MIMAT0000223 | Dbp | 13170 | HITS-CLIP | | 23597149 | Unspecified |
| mmu-miR-193a-3p | MIMAT0000223 | Dbt | 13171 | HITS-CLIP | | 23597149 | Unspecified |
| mmu-miR-193a-3p | MIMAT0000223 | Dcaf10 | 242418 | HITS-CLIP | | 23597149 | Unspecified |
| mmu-miR-193a-3p | MIMAT0000223 | Dcaf5 | 320808 | HITS-CLIP | | 19536157 | Unspecified |
| mmu-miR-193a-3p | MIMAT0000223 | Dcaf7 | 71833 | HITS-CLIP//sRNA-Seq | 37986224\|23597149\|33184237 | | Unspecified |
| mmu-miR-193a-3p | MIMAT0000223 | Dcaf8 | 98193 | HITS-CLIP | 37986224\|23597149 | | Unspecified |
| mmu-miR-193a-3p | MIMAT0000223 | Dclk2 | 70762 | HITS-CLIP | | 19536157 | Unspecified |
| mmu-miR-193a-3p | MIMAT0000223 | Dcp2 | 70640 | HITS-CLIP | | 23597149 | Unspecified |
| mmu-miR-193a-3p | MIMAT0000223 | Dctn2 | 69654 | HITS-CLIP//sRNA-Seq | 23597149\|33184237 | | Unspecified |
| mmu-miR-193a-3p | MIMAT0000223 | Dctn3 | 53598 | HITS-CLIP//sRNA-Seq | 23597149\|33184237 | | Unspecified |
| mmu-miR-193a-3p | MIMAT0000223 | Ddx17 | 67040 | HITS-CLIP//sRNA-Seq | 23597149\|33184237 | | Unspecified |
| mmu-miR-193a-3p | MIMAT0000223 | Ddx19a | 13680 | qCLASH | | 31393953 | Unspecified |
| mmu-miR-193a-3p | MIMAT0000223 | Ddx19b | 234733 | HITS-CLIP | | 23597149 | Unspecified |
| mmu-miR-193a-3p | MIMAT0000223 | Ddx3x | 13205 | HITS-CLIP | | 37986224 | Unspecified |
| mmu-miR-193a-3p | MIMAT0000223 | Ddx6 | 13209 | sRNA-Seq//HITS-CLIP | 33184237\|31433990 | | Unspecified |
| mmu-miR-193a-3p | MIMAT0000223 | Deaf1 | 54006 | HITS-CLIP | | 23597149 | Unspecified |
| mmu-miR-193a-3p | MIMAT0000223 | Degs1 | 13244 | sRNA-Seq//HITS-CLIP | 33184237\|23597149 | | Unspecified |
| mmu-miR-193a-3p | MIMAT0000223 | Dele1 | 9812 | HITS-CLIP | | 23597149 | Unspecified |
| mmu-miR-193a-3p | MIMAT0000223 | Dennd4c | 329877 | HITS-CLIP | | 23597149 | Unspecified |
| mmu-miR-193a-3p | MIMAT0000223 | Dera | 232449 | HITS-CLIP | | 23597149 | Unspecified |
| mmu-miR-193a-3p | MIMAT0000223 | Dgcr2 | 13356 | HITS-CLIP//sRNA-Seq | 23597149\|33184237 | | Unspecified |
| mmu-miR-193a-3p | MIMAT0000223 | Dhrs4 | 28200 | HITS-CLIP | 37986224\|23597149 | | Unspecified |
| mmu-miR-193a-3p | MIMAT0000223 | Dhx9 | 13211 | HITS-CLIP//sRNA-Seq | 23597149\|33184237 | | Unspecified |
| mmu-miR-193a-3p | MIMAT0000223 | Dlc1 | 50768 | HITS-CLIP | 37986224\|23597149 | | Unspecified |
| mmu-miR-193a-3p | MIMAT0000223 | Dnaja3 | 83945 | HITS-CLIP | | 23597149 | Unspecified |
| mmu-miR-193a-3p | MIMAT0000223 | Dnajc10 | 66861 | HITS-CLIP//sRNA-Seq | 23597149\|33184237 | | Unspecified |
| mmu-miR-193a-3p | MIMAT0000223 | Dnajc13 | 235567 | sRNA-Seq//HITS-CLIP | 33184237\|23597149 | | Unspecified |
| mmu-miR-193a-3p | MIMAT0000223 | Dpp4 | 13482 | HITS-CLIP | 37986224\|23597149 | | Unspecified |
| mmu-miR-193a-3p | MIMAT0000223 | Dpt | 56429 | HITS-CLIP | | 23597149 | Unspecified |
| mmu-miR-193a-3p | MIMAT0000223 | Dpy19l1 | 244745 | HITS-CLIP | | 23597149 | Unspecified |

| | | | | | | |
|---|---|---|---|---|---|---|
| mmu-miR-193a-3p | MIMAT0000223 | Dr1 | 13486 | HITS-CLIP | 23597149 | Unspecified |
| mmu-miR-193a-3p | MIMAT0000223 | Drp2 | 13497 | HITS-CLIP | 19536157 | Unspecified |
| mmu-miR-193a-3p | MIMAT0000223 | Dsc2 | 13506 | HITS-CLIP | 23597149 | Unspecified |
| mmu-miR-193a-3p | MIMAT0000223 | Dst | 13518 | HITS-CLIP//sRNA-Seq | 37986224\|19536157\|33184237 | Unspecified |
| mmu-miR-193a-3p | MIMAT0000223 | Dstn | 56431 | sRNA-Seq//HITS-CLIP | 33184237\|23597149 | Unspecified |
| mmu-miR-193a-3p | MIMAT0000223 | Dtd1 | 66044 | HITS-CLIP | 23597149 | Unspecified |
| mmu-miR-193a-3p | MIMAT0000223 | Dtx2 | 74198 | HITS-CLIP | 23597149 | Unspecified |
| mmu-miR-193a-3p | MIMAT0000223 | Dusp6 | 67603 | HITS-CLIP//sRNA-Seq | 37986224\|23597149\|33184237 | Unspecified |
| mmu-miR-193a-3p | MIMAT0000223 | Dusp8 | 18218 | HITS-CLIP | 26203562 | Unspecified |
| mmu-miR-193a-3p | MIMAT0000223 | Dvl1 | 13542 | HITS-CLIP | 23597149 | Unspecified |
| mmu-miR-193a-3p | MIMAT0000223 | Dync1h1 | 13424 | HITS-CLIP | 37986224 | Unspecified |
| mmu-miR-193a-3p | MIMAT0000223 | Dync1li2 | 234663 | sRNA-Seq//HITS-CLIP | 33184237\|23597149 | Unspecified |
| mmu-miR-193a-3p | MIMAT0000223 | Ece1 | 230857 | HITS-CLIP//sRNA-Seq | 37986224\|33184237\|23597149 | Unspecified |
| mmu-miR-193a-3p | MIMAT0000223 | Echs1 | 93747 | HITS-CLIP//sRNA-Seq | 23597149\|33184237 | Unspecified |
| mmu-miR-193a-3p | MIMAT0000223 | Edc3 | 353190 | HITS-CLIP | 23597149 | Unspecified |
| mmu-miR-193a-3p | MIMAT0000223 | Eea1 | 216238 | HITS-CLIP | 23597149 | Unspecified |
| mmu-miR-193a-3p | MIMAT0000223 | Eef2 | 13629 | HITS-CLIP//sRNA-Seq | 23597149\|33184237 | Unspecified |
| mmu-miR-193a-3p | MIMAT0000223 | Egfl7 | 353156 | HITS-CLIP | 37986224\|23597149 | Unspecified |
| mmu-miR-193a-3p | MIMAT0000223 | Egfr | 13649 | HITS-CLIP | 23597149 | Unspecified |
| mmu-miR-193a-3p | MIMAT0000223 | Egln2 | 112406 | HITS-CLIP | 37986224 | Unspecified |
| mmu-miR-193a-3p | MIMAT0000223 | Ehbp1 | 216565 | HITS-CLIP | 23597149 | Unspecified |
| mmu-miR-193a-3p | MIMAT0000223 | Ehd3 | 57440 | HITS-CLIP | 23597149 | Unspecified |
| mmu-miR-193a-3p | MIMAT0000223 | Eif2ak1 | 15467 | HITS-CLIP | 37986224\|23597149 | Unspecified |
| mmu-miR-193a-3p | MIMAT0000223 | Eif3a | 13669 | sRNA-Seq//HITS-CLIP | 33184237\|19536157 | Unspecified |
| mmu-miR-193a-3p | MIMAT0000223 | Eif4ebp2 | 13688 | HITS-CLIP | 23597149 | Unspecified |
| mmu-miR-193a-3p | MIMAT0000223 | Eif4g2 | 13690 | HITS-CLIP//sRNA-Seq | 19536157\|33184237 | Unspecified |
| mmu-miR-193a-3p | MIMAT0000223 | Eif5 | 217869 | HITS-CLIP | 37986224 | Unspecified |
| mmu-miR-193a-3p | MIMAT0000223 | Elavl1 | 15568 | HITS-CLIP | 23597149 | Unspecified |
| mmu-miR-193a-3p | MIMAT0000223 | Ell2 | 192657 | HITS-CLIP | 23597149 | Unspecified |
| mmu-miR-193a-3p | MIMAT0000223 | Elovl2 | 54326 | HITS-CLIP | 23597149 | Unspecified |
| mmu-miR-193a-3p | MIMAT0000223 | Emc1 | 230866 | HITS-CLIP | 37986224 | Unspecified |
| mmu-miR-193a-3p | MIMAT0000223 | Emc4 | 68032 | sRNA-Seq//HITS-CLIP | 33184237\|19536157 | Unspecified |
| mmu-miR-193a-3p | MIMAT0000223 | En2 | 13799 | HITS-CLIP | 19536157 | Unspecified |
| mmu-miR-193a-3p | MIMAT0000223 | Epas1 | 13819 | HITS-CLIP | 23597149\|19536157\|31433990 | Unspecified |
| mmu-miR-193a-3p | MIMAT0000223 | Epha7 | 13841 | HITS-CLIP | 19536157 | Unspecified |
| mmu-miR-193a-3p | MIMAT0000223 | Erbin | 55914 | HITS-CLIP | 23597149 | Unspecified |
| mmu-miR-193a-3p | MIMAT0000223 | Erlin1 | 226144 | HITS-CLIP | 37986224 | Unspecified |
| mmu-miR-193a-3p | MIMAT0000223 | Errfi1 | 74155 | HITS-CLIP | 23597149 | Unspecified |
| mmu-miR-193a-3p | MIMAT0000223 | Esrp2 | 77411 | HITS-CLIP | 23597149 | Unspecified |

| | | | | | | |
|---|---|---|---|---|---|---|
| mmu-miR-193a-3p | MIMAT0000223 | Etnk1 | 75320 | HITS-CLIP//sRNA-Seq | 23597149\|33184237 | Unspecified |
| mmu-miR-193a-3p | MIMAT0000223 | Ewsr1 | 14030 | HITS-CLIP//sRNA-Seq | 23597149\|33184237 | Unspecified |
| mmu-miR-193a-3p | MIMAT0000223 | Exosc3 | 66362 | HITS-CLIP | 23597149 | Unspecified |
| mmu-miR-193a-3p | MIMAT0000223 | F2 | 14061 | HITS-CLIP | 37986224\|23597149 | Unspecified |
| mmu-miR-193a-3p | MIMAT0000223 | Faf2 | 76577 | HITS-CLIP//sRNA-Seq | 23597149\|33184237 | Unspecified |
| mmu-miR-193a-3p | MIMAT0000223 | Fah | 14085 | HITS-CLIP | 23597149 | Unspecified |
| mmu-miR-193a-3p | MIMAT0000223 | Fahd2a | 68126 | HITS-CLIP | 23597149 | Unspecified |
| mmu-miR-193a-3p | MIMAT0000223 | Fam114a1 | 68303 | HITS-CLIP | 23597149 | Unspecified |
| mmu-miR-193a-3p | MIMAT0000223 | Fam120a | 218236 | HITS-CLIP | 37986224\|23597149 | Unspecified |
| mmu-miR-193a-3p | MIMAT0000223 | Fam126b | 213056 | HITS-CLIP | 23597149 | Unspecified |
| mmu-miR-193a-3p | MIMAT0000223 | Fam160a1 | 229488 | HITS-CLIP | 23597149 | Unspecified |
| mmu-miR-193a-3p | MIMAT0000223 | Fam171a1 | 269233 | sRNA-Seq//HITS-CLIP | 33184237\|19536157 | Unspecified |
| mmu-miR-193a-3p | MIMAT0000223 | Fam210b | 67017 | HITS-CLIP | 23597149 | Unspecified |
| mmu-miR-193a-3p | MIMAT0000223 | Fam214a | 235493 | HITS-CLIP | 37986224\|23597149 | Unspecified |
| mmu-miR-193a-3p | MIMAT0000223 | Fasn | 14104 | sRNA-Seq//HITS-CLIP | 33184237\|23597149 | Unspecified |
| mmu-miR-193a-3p | MIMAT0000223 | Fbf1 | 217335 | HITS-CLIP | 31433990 | Unspecified |
| mmu-miR-193a-3p | MIMAT0000223 | Fbxl17 | 50758 | HITS-CLIP | 37986224\|23597149 | Unspecified |
| mmu-miR-193a-3p | MIMAT0000223 | Fbxo22 | 71999 | HITS-CLIP | 23597149 | Unspecified |
| mmu-miR-193a-3p | MIMAT0000223 | Fetub | 59083 | HITS-CLIP | 23597149 | Unspecified |
| mmu-miR-193a-3p | MIMAT0000223 | Fga | 14161 | HITS-CLIP | 23597149 | Unspecified |
| mmu-miR-193a-3p | MIMAT0000223 | Fgd6 | 13998 | HITS-CLIP | 23597149 | Unspecified |
| mmu-miR-193a-3p | MIMAT0000223 | Fgf1 | 14164 | HITS-CLIP | 23597149 | Unspecified |
| mmu-miR-193a-3p | MIMAT0000223 | Fkbp1b | 14226 | sRNA-Seq//HITS-CLIP | 33184237\|19536157 | Unspecified |
| mmu-miR-193a-3p | MIMAT0000223 | Flrt3 | 71436 | HITS-CLIP | 19536157 | Unspecified |
| mmu-miR-193a-3p | MIMAT0000223 | Flt4 | 14257 | HITS-CLIP | 37986224\|23597149 | Unspecified |
| mmu-miR-193a-3p | MIMAT0000223 | Fmo5 | 14263 | HITS-CLIP | 23597149 | Unspecified |
| mmu-miR-193a-3p | MIMAT0000223 | Fn1 | 14268 | HITS-CLIP | 23597149 | Unspecified |
| mmu-miR-193a-3p | MIMAT0000223 | Fndc3a | 319448 | HITS-CLIP | 23597149 | Unspecified |
| mmu-miR-193a-3p | MIMAT0000223 | Fndc3b | 72007 | HITS-CLIP | 23597149 | Unspecified |
| mmu-miR-193a-3p | MIMAT0000223 | Foxa3 | 15377 | HITS-CLIP | 23597149 | Unspecified |
| mmu-miR-193a-3p | MIMAT0000223 | Frmd4b | 232288 | HITS-CLIP | 23597149 | Unspecified |
| mmu-miR-193a-3p | MIMAT0000223 | Fxn | 14297 | HITS-CLIP | 23597149 | Unspecified |
| mmu-miR-193a-3p | MIMAT0000223 | Fxyd1 | 56188 | HITS-CLIP | 23597149 | Unspecified |
| mmu-miR-193a-3p | MIMAT0000223 | Fzr1 | 56371 | HITS-CLIP | 23597149 | Unspecified |
| mmu-miR-193a-3p | MIMAT0000223 | G3bp1 | 27041 | HITS-CLIP | 37986224 | Unspecified |
| mmu-miR-193a-3p | MIMAT0000223 | G6pc | 14377 | HITS-CLIP | 37986224 | Unspecified |
| mmu-miR-193a-3p | MIMAT0000223 | Galk2 | 69976 | HITS-CLIP | 23597149 | Unspecified |
| mmu-miR-193a-3p | MIMAT0000223 | Galm | 319625 | HITS-CLIP | 23597149 | Unspecified |
| mmu-miR-193a-3p | MIMAT0000223 | Galnt1 | 14423 | HITS-CLIP | 37986224\|23597149 | Unspecified |

| | | | | | | |
|---|---|---|---|---|---|---|
| mmu-miR-193a-3p | MIMAT0000223 | Gamt | 14431 | HITS-CLIP | 23597149 | Unspecified |
| mmu-miR-193a-3p | MIMAT0000223 | Gart | 14450 | HITS-CLIP | 23597149 | Unspecified |
| mmu-miR-193a-3p | MIMAT0000223 | Gatad2b | 229542 | HITS-CLIP | 23597149 | Unspecified |
| mmu-miR-193a-3p | MIMAT0000223 | Gba2 | 230101 | HITS-CLIP | 23597149 | Unspecified |
| mmu-miR-193a-3p | MIMAT0000223 | Gcc1 | 74375 | HITS-CLIP | 23597149 | Unspecified |
| mmu-miR-193a-3p | MIMAT0000223 | Gcgr | 14527 | HITS-CLIP | 37986224\|23597149 | Unspecified |
| mmu-miR-193a-3p | MIMAT0000223 | Gck | 103988 | HITS-CLIP | 23597149 | Unspecified |
| mmu-miR-193a-3p | MIMAT0000223 | Gclc | 14629 | HITS-CLIP | 23597149 | Unspecified |
| mmu-miR-193a-3p | MIMAT0000223 | Gda | 14544 | HITS-CLIP | 37986224\|23597149 | Unspecified |
| mmu-miR-193a-3p | MIMAT0000223 | Gemin5 | 216766 | HITS-CLIP | 23597149 | Unspecified |
| mmu-miR-193a-3p | MIMAT0000223 | Gfra1 | 14585 | HITS-CLIP | 23597149 | Unspecified |
| mmu-miR-193a-3p | MIMAT0000223 | Ggps1 | 14593 | HITS-CLIP//sRNA-Seq | 23597149\|33184237 | Unspecified |
| mmu-miR-193a-3p | MIMAT0000223 | Gigyf1 | 57330 | HITS-CLIP | 31433990\|19536157 | Unspecified |
| mmu-miR-193a-3p | MIMAT0000223 | Gjb2 | 14619 | HITS-CLIP | 23597149 | Unspecified |
| mmu-miR-193a-3p | MIMAT0000223 | Glce | 93683 | HITS-CLIP | 23597149 | Unspecified |
| mmu-miR-193a-3p | MIMAT0000223 | Glipr2 | 384009 | HITS-CLIP | 31433990 | Unspecified |
| mmu-miR-193a-3p | MIMAT0000223 | Glo1 | 109801 | sRNA-Seq//HITS-CLIP | 33184237\|23597149 | Unspecified |
| mmu-miR-193a-3p | MIMAT0000223 | Glrx | 93692 | HITS-CLIP | 37986224\|23597149 | Unspecified |
| mmu-miR-193a-3p | MIMAT0000223 | Gls2 | 216456 | HITS-CLIP | 37986224 | Unspecified |
| mmu-miR-193a-3p | MIMAT0000223 | Gm1673 | 381633 | sRNA-Seq//HITS-CLIP | 33184237\|19536157 | Unspecified |
| mmu-miR-193a-3p | MIMAT0000223 | Gm20425 | 6608681 | HITS-CLIP | 23597149 | Unspecified |
| mmu-miR-193a-3p | MIMAT0000223 | Gm38394 | 667118 | HITS-CLIP | 23597149 | Unspecified |
| mmu-miR-193a-3p | MIMAT0000223 | Gm45902 | 119065974 | HITS-CLIP | 31433990 | Unspecified |
| mmu-miR-193a-3p | MIMAT0000223 | Gm4788 | 214403 | HITS-CLIP | 23597149 | Unspecified |
| mmu-miR-193a-3p | MIMAT0000223 | Gm49336 | 127230315 | sRNA-Seq//HITS-CLIP | 33184237\|23597149 | Unspecified |
| mmu-miR-193a-3p | MIMAT0000223 | Gnai3 | 14679 | HITS-CLIP//sRNA-Seq | 37986224\|31433990\|33184237 | Unspecified |
| mmu-miR-193a-3p | MIMAT0000223 | Gne | 50798 | HITS-CLIP | 37986224\|23597149 | Unspecified |
| mmu-miR-193a-3p | MIMAT0000223 | Golga3 | 269682 | HITS-CLIP | 23597149 | Unspecified |
| mmu-miR-193a-3p | MIMAT0000223 | Gpam | 14732 | HITS-CLIP | 19536157\|23597149 | Unspecified |
| mmu-miR-193a-3p | MIMAT0000223 | Gpat4 | 137964 | HITS-CLIP | 23597149 | Unspecified |
| mmu-miR-193a-3p | MIMAT0000223 | Gpd1 | 14555 | HITS-CLIP | 37986224 | Unspecified |
| mmu-miR-193a-3p | MIMAT0000223 | Gphn | 268566 | HITS-CLIP | 23597149 | Unspecified |
| mmu-miR-193a-3p | MIMAT0000223 | Gpm6a | 234267 | HITS-CLIP | 19536157 | Unspecified |
| mmu-miR-193a-3p | MIMAT0000223 | Gpt2 | 108682 | HITS-CLIP | 23597149 | Unspecified |
| mmu-miR-193a-3p | MIMAT0000223 | Grb7 | 14786 | HITS-CLIP | 37986224\|23597149 | Unspecified |
| mmu-miR-193a-3p | MIMAT0000223 | Grem2 | 23893 | HITS-CLIP | 23597149 | Unspecified |
| mmu-miR-193a-3p | MIMAT0000223 | Gria4 | 14802 | HITS-CLIP//sRNA-Seq | 19536157\|33184237 | Unspecified |
| mmu-miR-193a-3p | MIMAT0000223 | Grwd1 | 101612 | HITS-CLIP | 37986224 | Unspecified |
| mmu-miR-193a-3p | MIMAT0000223 | Gstk1 | 76263 | HITS-CLIP | 23597149 | Unspecified |

| | | | | | | |
|---|---|---|---|---|---|---|
| mmu-miR-193a-3p | MIMAT0000223 | Gstm1 | 14862 | HITS-CLIP | 23597149 | Unspecified |
| mmu-miR-193a-3p | MIMAT0000223 | Gstm2 | 14863 | HITS-CLIP | 23597149 | Unspecified |
| mmu-miR-193a-3p | MIMAT0000223 | Gstm6 | 14867 | HITS-CLIP | 23597149 | Unspecified |
| mmu-miR-193a-3p | MIMAT0000223 | Gstm7 | 68312 | HITS-CLIP | 23597149 | Unspecified |
| mmu-miR-193a-3p | MIMAT0000223 | H2-Aa | 14960 | HITS-CLIP | 37986224\|23597149 | Unspecified |
| mmu-miR-193a-3p | MIMAT0000223 | H6pd | 100198 | HITS-CLIP | 37986224\|23597149 | Unspecified |
| mmu-miR-193a-3p | MIMAT0000223 | Hadh | 15107 | HITS-CLIP | 23597149 | Unspecified |
| mmu-miR-193a-3p | MIMAT0000223 | Hal | 15109 | HITS-CLIP | 37986224 | Unspecified |
| mmu-miR-193a-3p | MIMAT0000223 | Hars | 15115 | HITS-CLIP | 37986224 | Unspecified |
| mmu-miR-193a-3p | MIMAT0000223 | Hc | 15139 | HITS-CLIP | 23597149 | Unspecified |
| mmu-miR-193a-3p | MIMAT0000223 | Hectd1 | 207304 | HITS-CLIP//sRNA-Seq | 23597149\|33184237 | Unspecified |
| mmu-miR-193a-3p | MIMAT0000223 | Helz2 | 229003 | HITS-CLIP | 23597149 | Unspecified |
| mmu-miR-193a-3p | MIMAT0000223 | Herc1 | 235439 | HITS-CLIP//sRNA-Seq | 37986224\|23597149\|33184237 | Unspecified |
| mmu-miR-193a-3p | MIMAT0000223 | Herc2 | 15204 | HITS-CLIP//sRNA-Seq | 23597149\|33184237 | Unspecified |
| mmu-miR-193a-3p | MIMAT0000223 | Herpud1 | 64209 | HITS-CLIP | 23597149 | Unspecified |
| mmu-miR-193a-3p | MIMAT0000223 | Hfe2 | 69585 | HITS-CLIP | 23597149 | Unspecified |
| mmu-miR-193a-3p | MIMAT0000223 | Hipk2 | 15258 | HITS-CLIP | 23597149 | Unspecified |
| mmu-miR-193a-3p | MIMAT0000223 | Hira | 15260 | HITS-CLIP | 23597149 | Unspecified |
| mmu-miR-193a-3p | MIMAT0000223 | Hivep2 | 15273 | HITS-CLIP | 23597149 | Unspecified |
| mmu-miR-193a-3p | MIMAT0000223 | Hmgcr | 15357 | HITS-CLIP//sRNA-Seq | 19536157\|33184237 | Unspecified |
| mmu-miR-193a-3p | MIMAT0000223 | Hmgcs1 | 208715 | HITS-CLIP//sRNA-Seq | 23597149\|33184237 | Unspecified |
| mmu-miR-193a-3p | MIMAT0000223 | Hmgcs2 | 15360 | HITS-CLIP | 37986224\|23597149 | Unspecified |
| mmu-miR-193a-3p | MIMAT0000223 | Hp1bp3 | 15441 | HITS-CLIP//sRNA-Seq | 37986224\|23597149\|33184237 | Unspecified |
| mmu-miR-193a-3p | MIMAT0000223 | Hpd | 15445 | HITS-CLIP | 23597149 | Unspecified |
| mmu-miR-193a-3p | MIMAT0000223 | Hprt | 15452 | HITS-CLIP | 37986224 | Unspecified |
| mmu-miR-193a-3p | MIMAT0000223 | Hps6 | 20170 | HITS-CLIP | 31433990 | Unspecified |
| mmu-miR-193a-3p | MIMAT0000223 | Hr | 15460 | HITS-CLIP | 26203562 | Unspecified |
| mmu-miR-193a-3p | MIMAT0000223 | Hrg | 94175 | HITS-CLIP | 23597149 | Unspecified |
| mmu-miR-193a-3p | MIMAT0000223 | Hsd17b2 | 15486 | HITS-CLIP | 37986224 | Unspecified |
| mmu-miR-193a-3p | MIMAT0000223 | Hsd17b4 | 15488 | HITS-CLIP | 23597149 | Unspecified |
| mmu-miR-193a-3p | MIMAT0000223 | Hsd3b5 | 15496 | HITS-CLIP | 23597149 | Unspecified |
| mmu-miR-193a-3p | MIMAT0000223 | Hsd3b7 | 101502 | HITS-CLIP | 37986224\|23597149 | Unspecified |
| mmu-miR-193a-3p | MIMAT0000223 | Hsdl2 | 72479 | HITS-CLIP | 37986224 | Unspecified |
| mmu-miR-193a-3p | MIMAT0000223 | Hsp90ab1 | 15516 | HITS-CLIP//sRNA-Seq | 23597149\|33184237\|31433990 | Unspecified |
| mmu-miR-193a-3p | MIMAT0000223 | Hsp90b1 | 22027 | HITS-CLIP//sRNA-Seq | 37986224\|23597149\|33184237 | Unspecified |
| mmu-miR-193a-3p | MIMAT0000223 | Hspa4 | 15525 | HITS-CLIP//sRNA-Seq | 23597149\|33184237 | Unspecified |
| mmu-miR-193a-3p | MIMAT0000223 | Huwe1 | 59026 | HITS-CLIP//sRNA-Seq | 19536157\|33184237 | Unspecified |
| mmu-miR-193a-3p | MIMAT0000223 | Hyal1 | 15586 | HITS-CLIP | 23597149 | Unspecified |
| mmu-miR-193a-3p | MIMAT0000223 | Hyou1 | 12282 | HITS-CLIP | 23597149 | Unspecified |

| | | | | | | |
|---|---|---|---|---|---|---|
| mmu-miR-193a-3p | MIMAT0000223 | Idh3g | 15929 | sRNA-Seq//HITS-CLIP | 33184237\|23597149 | Unspecified |
| mmu-miR-193a-3p | MIMAT0000223 | Ifngr2 | 15980 | HITS-CLIP | 21258322\|31433990 | Unspecified |
| mmu-miR-193a-3p | MIMAT0000223 | Igf1r | 16001 | HITS-CLIP | 19536157 | Unspecified |
| mmu-miR-193a-3p | MIMAT0000223 | Ikbkg | 16151 | HITS-CLIP | 37986224 | Unspecified |
| mmu-miR-193a-3p | MIMAT0000223 | Il15ra | 16169 | HITS-CLIP | 31433990 | Unspecified |
| mmu-miR-193a-3p | MIMAT0000223 | Il2rb | 16185 | HITS-CLIP | 31433990 | Unspecified |
| mmu-miR-193a-3p | MIMAT0000223 | Il4ra | 16190 | HITS-CLIP | 37986224\|23597149 | Unspecified |
| mmu-miR-193a-3p | MIMAT0000223 | Il6st | 16195 | sRNA-Seq//HITS-CLIP | 33184237\|23597149 | Unspecified |
| mmu-miR-193a-3p | MIMAT0000223 | Imp4 | 27993 | HITS-CLIP | 23597149 | Unspecified |
| mmu-miR-193a-3p | MIMAT0000223 | Ing4 | 28019 | HITS-CLIP | 23597149 | Unspecified |
| mmu-miR-193a-3p | MIMAT0000223 | Inip | 66209 | HITS-CLIP | 23597149 | Unspecified |
| mmu-miR-193a-3p | MIMAT0000223 | Insr | 16337 | HITS-CLIP | 23597149\|19536157 | Unspecified |
| mmu-miR-193a-3p | MIMAT0000223 | Ipo11 | 76582 | HITS-CLIP | 23597149 | Unspecified |
| mmu-miR-193a-3p | MIMAT0000223 | Ipo7 | 233726 | HITS-CLIP//sRNA-Seq | 37986224\|23597149\|33184237 | Unspecified |
| mmu-miR-193a-3p | MIMAT0000223 | Ipp | 16351 | HITS-CLIP | 37986224 | Unspecified |
| mmu-miR-193a-3p | MIMAT0000223 | Irf2bp2 | 270110 | HITS-CLIP | 23597149 | Unspecified |
| mmu-miR-193a-3p | MIMAT0000223 | Irf4 | 16364 | HITS-CLIP | 31433990 | Unspecified |
| mmu-miR-193a-3p | MIMAT0000223 | Itch | 16396 | HITS-CLIP | 23597149 | Unspecified |
| mmu-miR-193a-3p | MIMAT0000223 | Itga5 | 16402 | HITS-CLIP | 23597149 | Unspecified |
| mmu-miR-193a-3p | MIMAT0000223 | Itih1 | 16424 | HITS-CLIP | 23597149 | Unspecified |
| mmu-miR-193a-3p | MIMAT0000223 | Itih2 | 16425 | HITS-CLIP | 23597149 | Unspecified |
| mmu-miR-193a-3p | MIMAT0000223 | Itih4 | 16427 | HITS-CLIP | 23597149 | Unspecified |
| mmu-miR-193a-3p | MIMAT0000223 | Itpr1 | 16438 | HITS-CLIP | 23597149 | Unspecified |
| mmu-miR-193a-3p | MIMAT0000223 | Iyd | 70337 | HITS-CLIP | 23597149 | Unspecified |
| mmu-miR-193a-3p | MIMAT0000223 | Jade1 | 269424 | HITS-CLIP | 19536157 | Unspecified |
| mmu-miR-193a-3p | MIMAT0000223 | Jarid2 | 16468 | HITS-CLIP//sRNA-Seq | 23597149\|33184237 | Unspecified |
| mmu-miR-193a-3p | MIMAT0000223 | Jcad | 57608 | HITS-CLIP | 19536157 | Unspecified |
| mmu-miR-193a-3p | MIMAT0000223 | Kansl3 | 226976 | HITS-CLIP | 26203562 | Unspecified |
| mmu-miR-193a-3p | MIMAT0000223 | Kat5 | 81601 | HITS-CLIP | 19536157 | Unspecified |
| mmu-miR-193a-3p | MIMAT0000223 | Kcnh1 | 16510 | HITS-CLIP | 19536157 | Unspecified |
| mmu-miR-193a-3p | MIMAT0000223 | Kcnk10 | 72258 | HITS-CLIP | 19536157 | Unspecified |
| mmu-miR-193a-3p | MIMAT0000223 | Kdm5a | 214899 | HITS-CLIP | 37986224 | Unspecified |
| mmu-miR-193a-3p | MIMAT0000223 | Kdr | 16542 | HITS-CLIP | 23597149 | Unspecified |
| mmu-miR-193a-3p | MIMAT0000223 | Khsrp | 16549 | HITS-CLIP | 31433990 | Unspecified |
| mmu-miR-193a-3p | MIMAT0000223 | Kif1c | 16562 | HITS-CLIP | 37986224\|23597149 | Unspecified |
| mmu-miR-193a-3p | MIMAT0000223 | Kif5b | 16573 | sRNA-Seq//HITS-CLIP | 33184237\|31433990 | Unspecified |
| mmu-miR-193a-3p | MIMAT0000223 | Klf3 | 16599 | HITS-CLIP | 19536157\|23597149 | Unspecified |
| mmu-miR-193a-3p | MIMAT0000223 | Klf5 | 12224 | HITS-CLIP | 26203562 | Unspecified |
| mmu-miR-193a-3p | MIMAT0000223 | Klf9 | 16601 | HITS-CLIP | 23597149 | Unspecified |

| | | | | | | |
|---|---|---|---|---|---|---|
| mmu-miR-193a-3p | MIMAT0000223 | Klhl22 | 224023 | HITS-CLIP | 37986224\|23597149 | Unspecified |
| mmu-miR-193a-3p | MIMAT0000223 | Klhl24 | 75785 | HITS-CLIP | 37986224 | Unspecified |
| mmu-miR-193a-3p | MIMAT0000223 | Klhl3 | 100503085 | HITS-CLIP | 23597149 | Unspecified |
| mmu-miR-193a-3p | MIMAT0000223 | Kmt2c | 231051 | HITS-CLIP | 19536157 | Unspecified |
| mmu-miR-193a-3p | MIMAT0000223 | Kmt2e | 69188 | HITS-CLIP | 23597149 | Unspecified |
| mmu-miR-193a-3p | MIMAT0000223 | Kras | 16653 | HITS-CLIP//sRNA-Seq | 23597149\|33184237 | Unspecified |
| mmu-miR-193a-3p | MIMAT0000223 | Kri1 | 215194 | HITS-CLIP | 23597149 | Unspecified |
| mmu-miR-193a-3p | MIMAT0000223 | Krt18 | 16668 | HITS-CLIP | 23597149 | Unspecified |
| mmu-miR-193a-3p | MIMAT0000223 | Kynu | 70789 | HITS-CLIP | 37986224 | Unspecified |
| mmu-miR-193a-3p | MIMAT0000223 | Lactb | 80907 | HITS-CLIP | 23597149 | Unspecified |
| mmu-miR-193a-3p | MIMAT0000223 | Lama5 | 16776 | HITS-CLIP | 23597149 | Unspecified |
| mmu-miR-193a-3p | MIMAT0000223 | Lamp2 | 16784 | sRNA-Seq//HITS-CLIP | 33184237\|19536157 | Unspecified |
| mmu-miR-193a-3p | MIMAT0000223 | Lap3 | 66988 | HITS-CLIP | 23597149 | Unspecified |
| mmu-miR-193a-3p | MIMAT0000223 | Larp1 | 73158 | sRNA-Seq//HITS-CLIP | 33184237\|26203562 | Unspecified |
| mmu-miR-193a-3p | MIMAT0000223 | Larp4b | 217980 | sRNA-Seq//HITS-CLIP | 33184237\|31433990 | Unspecified |
| mmu-miR-193a-3p | MIMAT0000223 | Lcn2 | 16819 | HITS-CLIP | 23597149 | Unspecified |
| mmu-miR-193a-3p | MIMAT0000223 | Ldlr | 16835 | sRNA-Seq//HITS-CLIP | 33184237\|31433990 | Unspecified |
| mmu-miR-193a-3p | MIMAT0000223 | Lgr4 | 107515 | HITS-CLIP | 37986224 | Unspecified |
| mmu-miR-193a-3p | MIMAT0000223 | Lhfp | 108927 | HITS-CLIP | 19536157\|23597149 | Unspecified |
| mmu-miR-193a-3p | MIMAT0000223 | Lig3 | 16882 | HITS-CLIP | 37986224 | Unspecified |
| mmu-miR-193a-3p | MIMAT0000223 | Lims2 | 225341 | HITS-CLIP | 23597149 | Unspecified |
| mmu-miR-193a-3p | MIMAT0000223 | Lin7c | 22343 | HITS-CLIP//sRNA-Seq | 37986224\|33184237\|19536157 | Unspecified |
| mmu-miR-193a-3p | MIMAT0000223 | Lmbr1 | 56873 | HITS-CLIP | 23597149 | Unspecified |
| mmu-miR-193a-3p | MIMAT0000223 | Lpin1 | 14245 | HITS-CLIP | 23597149 | Unspecified |
| mmu-miR-193a-3p | MIMAT0000223 | Lpin2 | 64898 | HITS-CLIP | 23597149 | Unspecified |
| mmu-miR-193a-3p | MIMAT0000223 | Lrba | 80877 | HITS-CLIP | 37986224\|23597149 | Unspecified |
| mmu-miR-193a-3p | MIMAT0000223 | Lrp1 | 16971 | HITS-CLIP | 23597149 | Unspecified |
| mmu-miR-193a-3p | MIMAT0000223 | Lrp10 | 65107 | HITS-CLIP | 23597149 | Unspecified |
| mmu-miR-193a-3p | MIMAT0000223 | Lrp5 | 16973 | HITS-CLIP | 23597149 | Unspecified |
| mmu-miR-193a-3p | MIMAT0000223 | Lrrc58 | 320184 | HITS-CLIP//sRNA-Seq | 23597149\|33184237 | Unspecified |
| mmu-miR-193a-3p | MIMAT0000223 | Lrrc8a | 241296 | HITS-CLIP | 37986224\|19536157\|23597149 | Unspecified |
| mmu-miR-193a-3p | MIMAT0000223 | Lsr | 54135 | HITS-CLIP | 23597149 | Unspecified |
| mmu-miR-193a-3p | MIMAT0000223 | Ltbr | 17000 | HITS-CLIP | 23597149 | Unspecified |
| mmu-miR-193a-3p | MIMAT0000223 | Ly6e | 17069 | HITS-CLIP//sRNA-Seq | 23597149\|19536157\|33184237 | Unspecified |
| mmu-miR-193a-3p | MIMAT0000223 | Lyar | 17089 | HITS-CLIP | 26203562 | Unspecified |
| mmu-miR-193a-3p | MIMAT0000223 | Macf1 | 11426 | HITS-CLIP//sRNA-Seq | 37986224\|23597149\|33184237\|1953( | Unspecified |
| mmu-miR-193a-3p | MIMAT0000223 | Maged1 | 94275 | HITS-CLIP//sRNA-Seq | 23597149\|33184237 | Unspecified |
| mmu-miR-193a-3p | MIMAT0000223 | Magee1 | 107528 | HITS-CLIP//sRNA-Seq | 19536157\|33184237 | Unspecified |
| mmu-miR-193a-3p | MIMAT0000223 | Magt1 | 67075 | HITS-CLIP | 37986224 | Unspecified |

| | | | | | | |
|---|---|---|---|---|---|---|
| mmu-miR-193a-3p | MIMAT0000223 | Maml1 | 103806 | HITS-CLIP | 23597149 | Unspecified |
| mmu-miR-193a-3p | MIMAT0000223 | Man2a1 | 17158 | HITS-CLIP | 23597149 | Unspecified |
| mmu-miR-193a-3p | MIMAT0000223 | Map1a | 17754 | sRNA-Seq//HITS-CLIP | 33184237\|19536157 | Unspecified |
| mmu-miR-193a-3p | MIMAT0000223 | Map1b | 17755 | HITS-CLIP//sRNA-Seq | 19536157\|33184237\|23597149 | Unspecified |
| mmu-miR-193a-3p | MIMAT0000223 | Map2k3 | 26397 | HITS-CLIP | 23597149 | Unspecified |
| mmu-miR-193a-3p | MIMAT0000223 | Map3k1 | 26401 | HITS-CLIP | 23597149 | Unspecified |
| mmu-miR-193a-3p | MIMAT0000223 | Map3k2 | 26405 | HITS-CLIP | 23597149 | Unspecified |
| mmu-miR-193a-3p | MIMAT0000223 | Map3k5 | 26408 | HITS-CLIP | 37986224 | Unspecified |
| mmu-miR-193a-3p | MIMAT0000223 | Mapk1 | 26413 | HITS-CLIP | 37986224 | Unspecified |
| mmu-miR-193a-3p | MIMAT0000223 | Mapk14 | 26416 | HITS-CLIP | 23597149 | Unspecified |
| mmu-miR-193a-3p | MIMAT0000223 | Mapkbp1 | 26390 | HITS-CLIP | 37986224 | Unspecified |
| mmu-miR-193a-3p | MIMAT0000223 | Mapre1 | 13589 | HITS-CLIP//sRNA-Seq | 23597149\|33184237 | Unspecified |
| mmu-miR-193a-3p | MIMAT0000223 | Marc1 | 66112 | HITS-CLIP | 23597149 | Unspecified |
| mmu-miR-193a-3p | MIMAT0000223 | March6 | 10299 | HITS-CLIP//sRNA-Seq | 23597149\|33184237 | Unspecified |
| mmu-miR-193a-3p | MIMAT0000223 | Mark2 | 13728 | HITS-CLIP | 37986224\|23597149 | Unspecified |
| mmu-miR-193a-3p | MIMAT0000223 | Masp1 | 17174 | HITS-CLIP | 23597149 | Unspecified |
| mmu-miR-193a-3p | MIMAT0000223 | Matr3 | 17184 | HITS-CLIP//sRNA-Seq | 23597149\|33184237 | Unspecified |
| mmu-miR-193a-3p | MIMAT0000223 | Mcm10 | 70024 | HITS-CLIP | 37986224 | Unspecified |
| mmu-miR-193a-3p | MIMAT0000223 | Mcm3 | 17215 | HITS-CLIP | 31433990 | Unspecified |
| mmu-miR-193a-3p | MIMAT0000223 | Mdm2 | 17246 | HITS-CLIP | 23597149 | Unspecified |
| mmu-miR-193a-3p | MIMAT0000223 | Med16 | 216154 | HITS-CLIP | 37986224\|23597149 | Unspecified |
| mmu-miR-193a-3p | MIMAT0000223 | Med20 | 56771 | HITS-CLIP | 23597149 | Unspecified |
| mmu-miR-193a-3p | MIMAT0000223 | Metap1 | 75624 | HITS-CLIP | 23597149 | Unspecified |
| mmu-miR-193a-3p | MIMAT0000223 | Mettl26 | 50438 | HITS-CLIP | 23597149 | Unspecified |
| mmu-miR-193a-3p | MIMAT0000223 | Mettl7a1 | 70152 | HITS-CLIP | 23597149 | Unspecified |
| mmu-miR-193a-3p | MIMAT0000223 | Mfn2 | 170731 | HITS-CLIP//sRNA-Seq | 23597149\|33184237 | Unspecified |
| mmu-miR-193a-3p | MIMAT0000223 | Mga | 29808 | HITS-CLIP//sRNA-Seq | 23597149\|33184237 | Unspecified |
| mmu-miR-193a-3p | MIMAT0000223 | Mgat1 | 17308 | HITS-CLIP | 23597149 | Unspecified |
| mmu-miR-193a-3p | MIMAT0000223 | Mgrn1 | 17237 | HITS-CLIP | 37986224\|23597149 | Unspecified |
| mmu-miR-193a-3p | MIMAT0000223 | Mia2 | 338320 | HITS-CLIP | 23597149 | Unspecified |
| mmu-miR-193a-3p | MIMAT0000223 | Mia3 | 338366 | HITS-CLIP//sRNA-Seq | 23597149\|33184237 | Unspecified |
| mmu-miR-193a-3p | MIMAT0000223 | Mib1 | 225164 | HITS-CLIP//sRNA-Seq | 37986224\|23597149\|33184237 | Unspecified |
| mmu-miR-193a-3p | MIMAT0000223 | Midn | 59090 | HITS-CLIP//sRNA-Seq | 31433990\|33184237\|23597149 | Unspecified |
| mmu-miR-193a-3p | MIMAT0000223 | Mlec | 109154 | HITS-CLIP | 19536157\|23597149 | Unspecified |
| mmu-miR-193a-3p | MIMAT0000223 | Mllt6 | 246198 | HITS-CLIP | 37986224 | Unspecified |
| mmu-miR-193a-3p | MIMAT0000223 | Mmp16 | 17389 | HITS-CLIP | 19536157 | Unspecified |
| mmu-miR-193a-3p | MIMAT0000223 | Mmp19 | 58223 | HITS-CLIP | 23597149 | Unspecified |
| mmu-miR-193a-3p | MIMAT0000223 | Mms19 | 72199 | HITS-CLIP | 23597149 | Unspecified |
| mmu-miR-193a-3p | MIMAT0000223 | Mob1a | 232157 | HITS-CLIP | 31433990 | Unspecified |

| | | | | | | | |
|---|---|---|---|---|---|---|---|
| mmu-miR-193a-3p | MIMAT0000223 | Mocs1 | 56738 | HITS-CLIP | | 23597149 | Unspecified |
| mmu-miR-193a-3p | MIMAT0000223 | Mosmo | 233812 | HITS-CLIP | | 23597149 | Unspecified |
| mmu-miR-193a-3p | MIMAT0000223 | Mpc2 | 70456 | HITS-CLIP//sRNA-Seq | 23597149\|33184237 | | Unspecified |
| mmu-miR-193a-3p | MIMAT0000223 | Mphosph8 | 75339 | HITS-CLIP | | 31433990 | Unspecified |
| mmu-miR-193a-3p | MIMAT0000223 | Mphosph9 | 269702 | HITS-CLIP | | 23597149 | Unspecified |
| mmu-miR-193a-3p | MIMAT0000223 | Mrpl39 | 27393 | HITS-CLIP | | 23597149 | Unspecified |
| mmu-miR-193a-3p | MIMAT0000223 | Mrps22 | 64655 | HITS-CLIP | 37986224\|23597149 | | Unspecified |
| mmu-miR-193a-3p | MIMAT0000223 | Mrtfb | 239719 | HITS-CLIP | | 23597149 | Unspecified |
| mmu-miR-193a-3p | MIMAT0000223 | Mtch2 | 56428 | HITS-CLIP//sRNA-Seq | 23597149\|33184237 | | Unspecified |
| mmu-miR-193a-3p | MIMAT0000223 | Mtf1 | 17764 | HITS-CLIP | | 23597149 | Unspecified |
| mmu-miR-193a-3p | MIMAT0000223 | Mtor | 56717 | HITS-CLIP | | 23597149 | Unspecified |
| mmu-miR-193a-3p | MIMAT0000223 | Mtpn | 14489 | sRNA-Seq//HITS-CLIP | 33184237\|19536157 | | Unspecified |
| mmu-miR-193a-3p | MIMAT0000223 | Mtus1 | 102103 | HITS-CLIP//sRNA-Seq | 23597149\|33184237 | | Unspecified |
| mmu-miR-193a-3p | MIMAT0000223 | Mycn | 18109 | HITS-CLIP | 37986224\|23597149 | | Unspecified |
| mmu-miR-193a-3p | MIMAT0000223 | Myo16 | 244281 | HITS-CLIP | | 19536157 | Unspecified |
| mmu-miR-193a-3p | MIMAT0000223 | Myo18a | 360013 | HITS-CLIP | | 23597149 | Unspecified |
| mmu-miR-193a-3p | MIMAT0000223 | Myo1d | 338367 | HITS-CLIP | | 23597149 | Unspecified |
| mmu-miR-193a-3p | MIMAT0000223 | Myo6 | 17920 | HITS-CLIP | | 23597149 | Unspecified |
| mmu-miR-193a-3p | MIMAT0000223 | Naa25 | 231713 | HITS-CLIP | | 23597149 | Unspecified |
| mmu-miR-193a-3p | MIMAT0000223 | Nbea | 26422 | HITS-CLIP | | 19536157 | Unspecified |
| mmu-miR-193a-3p | MIMAT0000223 | Ncln | 103425 | HITS-CLIP | | 30971389 | Unspecified |
| mmu-miR-193a-3p | MIMAT0000223 | Ncoa3 | 17979 | HITS-CLIP | | 23597149 | Unspecified |
| mmu-miR-193a-3p | MIMAT0000223 | Ndfip1 | 65113 | HITS-CLIP//sRNA-Seq | 23597149\|33184237 | | Unspecified |
| mmu-miR-193a-3p | MIMAT0000223 | Ndfip2 | 76273 | HITS-CLIP | | 37986224 | Unspecified |
| mmu-miR-193a-3p | MIMAT0000223 | Ndrg2 | 29811 | HITS-CLIP | | 23597149 | Unspecified |
| mmu-miR-193a-3p | MIMAT0000223 | Ndufs1 | 227197 | HITS-CLIP//sRNA-Seq | 23597149\|33184237 | | Unspecified |
| mmu-miR-193a-3p | MIMAT0000223 | Ndufs7 | 75406 | HITS-CLIP//sRNA-Seq | 23597149\|33184237 | | Unspecified |
| mmu-miR-193a-3p | MIMAT0000223 | Ndufv1 | 17995 | sRNA-Seq//HITS-CLIP | 33184237\|23597149 | | Unspecified |
| mmu-miR-193a-3p | MIMAT0000223 | Nedd4 | 17999 | HITS-CLIP//sRNA-Seq | 23597149\|33184237 | | Unspecified |
| mmu-miR-193a-3p | MIMAT0000223 | Nfasc | 269116 | HITS-CLIP//sRNA-Seq | 19536157\|33184237 | | Unspecified |
| mmu-miR-193a-3p | MIMAT0000223 | Nfatc1 | 18018 | HITS-CLIP | | 23597149 | Unspecified |
| mmu-miR-193a-3p | MIMAT0000223 | Nfe2l1 | 18023 | HITS-CLIP | | 37986224 | Unspecified |
| mmu-miR-193a-3p | MIMAT0000223 | Nfia | 18027 | HITS-CLIP | | 23597149 | Unspecified |
| mmu-miR-193a-3p | MIMAT0000223 | Nfib | 18028 | HITS-CLIP | | 23597149 | Unspecified |
| mmu-miR-193a-3p | MIMAT0000223 | Nfic | 18029 | HITS-CLIP | | 37986224 | Unspecified |
| mmu-miR-193a-3p | MIMAT0000223 | Nfkbia | 18035 | HITS-CLIP | 37986224\|23597149 | | Unspecified |
| mmu-miR-193a-3p | MIMAT0000223 | Nfx1 | 74164 | HITS-CLIP | | 37986224 | Unspecified |
| mmu-miR-193a-3p | MIMAT0000223 | Nisch | 64652 | HITS-CLIP//sRNA-Seq | 23597149\|33184237 | | Unspecified |
| mmu-miR-193a-3p | MIMAT0000223 | Nit1 | 27045 | HITS-CLIP | | 23597149 | Unspecified |

| | | | | | | |
|---|---|---|---|---|---|---|
| mmu-miR-193a-3p | MIMAT0000223 | Nme1 | 18102 | sRNA-Seq//HITS-CLIP | 33184237\|26203562 | Unspecified |
| mmu-miR-193a-3p | MIMAT0000223 | Nol6 | 230082 | HITS-CLIP | 23597149 | Unspecified |
| mmu-miR-193a-3p | MIMAT0000223 | Notch2 | 18129 | HITS-CLIP | 23597149 | Unspecified |
| mmu-miR-193a-3p | MIMAT0000223 | Npc2 | 67963 | HITS-CLIP | 23597149 | Unspecified |
| mmu-miR-193a-3p | MIMAT0000223 | Nr0b2 | 23957 | HITS-CLIP | 23597149 | Unspecified |
| mmu-miR-193a-3p | MIMAT0000223 | Nr2c2 | 22026 | HITS-CLIP | 37986224 | Unspecified |
| mmu-miR-193a-3p | MIMAT0000223 | Nr2f2 | 11819 | HITS-CLIP | 23597149 | Unspecified |
| mmu-miR-193a-3p | MIMAT0000223 | Nr4a1 | 15370 | HITS-CLIP | 31433990 | Unspecified |
| mmu-miR-193a-3p | MIMAT0000223 | Nrbp1 | 192292 | sRNA-Seq//HITS-CLIP | 33184237\|23597149 | Unspecified |
| mmu-miR-193a-3p | MIMAT0000223 | Nsf | 18195 | sRNA-Seq//HITS-CLIP | 33184237\|19536157 | Unspecified |
| mmu-miR-193a-3p | MIMAT0000223 | Nudt2 | 66401 | HITS-CLIP | 37986224 | Unspecified |
| mmu-miR-193a-3p | MIMAT0000223 | Numb | 18222 | HITS-CLIP | 37986224 | Unspecified |
| mmu-miR-193a-3p | MIMAT0000223 | Nup205 | 70699 | HITS-CLIP | 37986224 | Unspecified |
| mmu-miR-193a-3p | MIMAT0000223 | Nup98 | 269966 | HITS-CLIP | 23597149 | Unspecified |
| mmu-miR-193a-3p | MIMAT0000223 | Oat | 18242 | HITS-CLIP | 37986224 | Unspecified |
| mmu-miR-193a-3p | MIMAT0000223 | Ocln | 18260 | HITS-CLIP | 37986224 | Unspecified |
| mmu-miR-193a-3p | MIMAT0000223 | Ogt | 108155 | HITS-CLIP//sRNA-Seq | 37986224\|19536157\|33184237 | Unspecified |
| mmu-miR-193a-3p | MIMAT0000223 | Orai3 | 269999 | HITS-CLIP | 23597149 | Unspecified |
| mmu-miR-193a-3p | MIMAT0000223 | Osbp2 | 74309 | HITS-CLIP | 19536157 | Unspecified |
| mmu-miR-193a-3p | MIMAT0000223 | Osbpl1a | 64291 | HITS-CLIP | 37986224\|23597149 | Unspecified |
| mmu-miR-193a-3p | MIMAT0000223 | Osbpl8 | 237542 | HITS-CLIP//sRNA-Seq | 19536157\|33184237 | Unspecified |
| mmu-miR-193a-3p | MIMAT0000223 | Osgin1 | 71839 | HITS-CLIP | 37986224\|23597149 | Unspecified |
| mmu-miR-193a-3p | MIMAT0000223 | Ostc | 66357 | HITS-CLIP//sRNA-Seq//qCLASH | 37986224\|33184237\|23597149\|31393 | Unspecified |
| mmu-miR-193a-3p | MIMAT0000223 | Otud1 | 71198 | HITS-CLIP | 31433990 | Unspecified |
| mmu-miR-193a-3p | MIMAT0000223 | P2ry4 | 57385 | HITS-CLIP | 37986224\|23597149 | Unspecified |
| mmu-miR-193a-3p | MIMAT0000223 | Pak2 | 224105 | HITS-CLIP | 37986224 | Unspecified |
| mmu-miR-193a-3p | MIMAT0000223 | Pan3 | 72587 | HITS-CLIP | 23597149 | Unspecified |
| mmu-miR-193a-3p | MIMAT0000223 | Paqr9 | 75552 | HITS-CLIP | 23597149 | Unspecified |
| mmu-miR-193a-3p | MIMAT0000223 | Pbld1 | 68371 | HITS-CLIP | 23597149 | Unspecified |
| mmu-miR-193a-3p | MIMAT0000223 | Pcdh1 | 75599 | HITS-CLIP | 23597149\|19536157 | Unspecified |
| mmu-miR-193a-3p | MIMAT0000223 | Pcif1 | 228866 | HITS-CLIP | 19536157 | Unspecified |
| mmu-miR-193a-3p | MIMAT0000223 | Pck1 | 18534 | HITS-CLIP | 23597149 | Unspecified |
| mmu-miR-193a-3p | MIMAT0000223 | Pcnx3 | 399909 | HITS-CLIP | 23597149 | Unspecified |
| mmu-miR-193a-3p | MIMAT0000223 | Pcsk7 | 18554 | HITS-CLIP | 23597149 | Unspecified |
| mmu-miR-193a-3p | MIMAT0000223 | Pdcd4 | 18569 | HITS-CLIP | 23597149 | Unspecified |
| mmu-miR-193a-3p | MIMAT0000223 | Pdcd6ip | 18571 | HITS-CLIP | 23597149 | Unspecified |
| mmu-miR-193a-3p | MIMAT0000223 | Pde3b | 18576 | HITS-CLIP | 23597149 | Unspecified |
| mmu-miR-193a-3p | MIMAT0000223 | Pdia3 | 14827 | HITS-CLIP//sRNA-Seq | 23597149\|33184237 | Unspecified |
| mmu-miR-193a-3p | MIMAT0000223 | Pdzd8 | 107368 | HITS-CLIP | 37986224 | Unspecified |

| | | | | | | |
|---|---|---|---|---|---|---|
| mmu-miR-193a-3p | MIMAT0000223 | Pea15a | 18611 | sRNA-Seq//HITS-CLIP | 33184237\|19536157 | Unspecified |
| mmu-miR-193a-3p | MIMAT0000223 | Per2 | 18627 | HITS-CLIP | 37986224\|23597149 | Unspecified |
| mmu-miR-193a-3p | MIMAT0000223 | Pex3 | 56535 | HITS-CLIP | 23597149 | Unspecified |
| mmu-miR-193a-3p | MIMAT0000223 | Pfn1 | 18643 | HITS-CLIP//sRNA-Seq | 23597149\|19536157\|33184237 | Unspecified |
| mmu-miR-193a-3p | MIMAT0000223 | Pggt1b | 225467 | HITS-CLIP | 23597149 | Unspecified |
| mmu-miR-193a-3p | MIMAT0000223 | Pgpep1 | 66522 | HITS-CLIP | 23597149 | Unspecified |
| mmu-miR-193a-3p | MIMAT0000223 | Pgrmc1 | 53328 | HITS-CLIP//sRNA-Seq | 23597149\|33184237 | Unspecified |
| mmu-miR-193a-3p | MIMAT0000223 | Phka2 | 110094 | HITS-CLIP | 23597149 | Unspecified |
| mmu-miR-193a-3p | MIMAT0000223 | Phldb2 | 208177 | HITS-CLIP | 37986224\|23597149 | Unspecified |
| mmu-miR-193a-3p | MIMAT0000223 | Pi4ka | 224020 | HITS-CLIP//sRNA-Seq | 19536157\|33184237 | Unspecified |
| mmu-miR-193a-3p | MIMAT0000223 | Pi4kb | 107650 | HITS-CLIP | 23597149 | Unspecified |
| mmu-miR-193a-3p | MIMAT0000223 | Pias2 | 17344 | HITS-CLIP | 19536157 | Unspecified |
| mmu-miR-193a-3p | MIMAT0000223 | Pik3c2a | 18704 | HITS-CLIP | 23597149 | Unspecified |
| mmu-miR-193a-3p | MIMAT0000223 | Pik3r1 | 18708 | HITS-CLIP | 37986224 | Unspecified |
| mmu-miR-193a-3p | MIMAT0000223 | Pikfyve | 18711 | HITS-CLIP | 37986224\|23597149 | Unspecified |
| mmu-miR-193a-3p | MIMAT0000223 | Pink1 | 68943 | HITS-CLIP | 23597149 | Unspecified |
| mmu-miR-193a-3p | MIMAT0000223 | Pisd | 320951 | HITS-CLIP | 23597149 | Unspecified |
| mmu-miR-193a-3p | MIMAT0000223 | Pitpna | 18738 | HITS-CLIP//sRNA-Seq | 23597149\|33184237 | Unspecified |
| mmu-miR-193a-3p | MIMAT0000223 | Pklr | 18770 | HITS-CLIP | 23597149 | Unspecified |
| mmu-miR-193a-3p | MIMAT0000223 | Pknox1 | 18771 | HITS-CLIP | 37986224\|23597149 | Unspecified |
| mmu-miR-193a-3p | MIMAT0000223 | Pkp4 | 227937 | HITS-CLIP | 23597149 | Unspecified |
| mmu-miR-193a-3p | MIMAT0000223 | Plag1 | 56711 | HITS-CLIP | 26203562 | Unspecified |
| mmu-miR-193a-3p | MIMAT0000223 | Plcxd2 | 433022 | HITS-CLIP | 37986224\|23597149 | Unspecified |
| mmu-miR-193a-3p | MIMAT0000223 | Plec | 18810 | sRNA-Seq//HITS-CLIP | 33184237\|23597149 | Unspecified |
| mmu-miR-193a-3p | MIMAT0000223 | Plekha6 | 240753 | HITS-CLIP | 37986224 | Unspecified |
| mmu-miR-193a-3p | MIMAT0000223 | Plg | 18815 | HITS-CLIP | 37986224 | Unspecified |
| mmu-miR-193a-3p | MIMAT0000223 | Plppr4 | 229791 | HITS-CLIP | 19536157 | Unspecified |
| mmu-miR-193a-3p | MIMAT0000223 | Plxna1 | 18844 | HITS-CLIP | 37986224 | Unspecified |
| mmu-miR-193a-3p | MIMAT0000223 | Plxnc1 | 54712 | HITS-CLIP | 23597149 | Unspecified |
| mmu-miR-193a-3p | MIMAT0000223 | Pnisr | 66625 | HITS-CLIP | 23597149 | Unspecified |
| mmu-miR-193a-3p | MIMAT0000223 | Pnpla2 | 66853 | HITS-CLIP | 37986224 | Unspecified |
| mmu-miR-193a-3p | MIMAT0000223 | Pole3 | 59001 | HITS-CLIP | 23597149 | Unspecified |
| mmu-miR-193a-3p | MIMAT0000223 | Polr2a | 20020 | sRNA-Seq//HITS-CLIP | 33184237\|23597149 | Unspecified |
| mmu-miR-193a-3p | MIMAT0000223 | Ppara | 19013 | HITS-CLIP | 37986224\|23597149 | Unspecified |
| mmu-miR-193a-3p | MIMAT0000223 | Ppargc1a | 19017 | HITS-CLIP | 23597149 | Unspecified |
| mmu-miR-193a-3p | MIMAT0000223 | Ppif | 105675 | HITS-CLIP | 23597149 | Unspecified |
| mmu-miR-193a-3p | MIMAT0000223 | Ppl | 19041 | HITS-CLIP | 23597149 | Unspecified |
| mmu-miR-193a-3p | MIMAT0000223 | Ppm1k | 243382 | HITS-CLIP | 23597149 | Unspecified |
| mmu-miR-193a-3p | MIMAT0000223 | Ppp1r15b | 108954 | HITS-CLIP | 23597149 | Unspecified |

| | | | | | | |
|---|---|---|---|---|---|---|
| mmu-miR-193a-3p | MIMAT0000223 | Ppp2r2a | 71978 | sRNA-Seq//HITS-CLIP | 33184237\|23597149 | Unspecified |
| mmu-miR-193a-3p | MIMAT0000223 | Ppp2r3a | 235542 | HITS-CLIP | 37986224 | Unspecified |
| mmu-miR-193a-3p | MIMAT0000223 | Ppp2r5c | 26931 | sRNA-Seq//HITS-CLIP | 33184237\|23597149 | Unspecified |
| mmu-miR-193a-3p | MIMAT0000223 | Ppp5c | 19060 | HITS-CLIP//sRNA-Seq | 23597149\|33184237 | Unspecified |
| mmu-miR-193a-3p | MIMAT0000223 | Ppp6r1 | 243819 | HITS-CLIP//sRNA-Seq | 23597149\|33184237 | Unspecified |
| mmu-miR-193a-3p | MIMAT0000223 | Ppp6r3 | 52036 | HITS-CLIP//sRNA-Seq | 23597149\|33184237 | Unspecified |
| mmu-miR-193a-3p | MIMAT0000223 | Pqlc1 | 66943 | sRNA-Seq//HITS-CLIP | 33184237\|23597149 | Unspecified |
| mmu-miR-193a-3p | MIMAT0000223 | Prkaa2 | 108079 | HITS-CLIP | 37986224 | Unspecified |
| mmu-miR-193a-3p | MIMAT0000223 | Prkar1a | 19084 | HITS-CLIP//sRNA-Seq | 23597149\|33184237 | Unspecified |
| mmu-miR-193a-3p | MIMAT0000223 | Prkar2a | 19087 | sRNA-Seq//HITS-CLIP | 33184237\|23597149 | Unspecified |
| mmu-miR-193a-3p | MIMAT0000223 | Prkd3 | 75292 | HITS-CLIP | 23597149 | Unspecified |
| mmu-miR-193a-3p | MIMAT0000223 | Prlr | 19116 | HITS-CLIP | 23597149 | Unspecified |
| mmu-miR-193a-3p | MIMAT0000223 | Prnp | 19122 | HITS-CLIP//sRNA-Seq | 19536157\|33184237 | Unspecified |
| mmu-miR-193a-3p | MIMAT0000223 | Proc | 19123 | HITS-CLIP | 23597149 | Unspecified |
| mmu-miR-193a-3p | MIMAT0000223 | Prodh | 19125 | HITS-CLIP | 23597149 | Unspecified |
| mmu-miR-193a-3p | MIMAT0000223 | Prox1 | 19130 | HITS-CLIP | 37986224\|23597149 | Unspecified |
| mmu-miR-193a-3p | MIMAT0000223 | Prpf19 | 28000 | sRNA-Seq//HITS-CLIP | 33184237\|23597149 | Unspecified |
| mmu-miR-193a-3p | MIMAT0000223 | Prpf8 | 192159 | HITS-CLIP//sRNA-Seq | 23597149\|33184237 | Unspecified |
| mmu-miR-193a-3p | MIMAT0000223 | Prr14l | 215476 | HITS-CLIP | 23597149 | Unspecified |
| mmu-miR-193a-3p | MIMAT0000223 | Prrc2a | 53761 | HITS-CLIP//sRNA-Seq | 23597149\|33184237 | Unspecified |
| mmu-miR-193a-3p | MIMAT0000223 | Prrc2b | 227723 | sRNA-Seq//HITS-CLIP | 33184237\|19536157 | Unspecified |
| mmu-miR-193a-3p | MIMAT0000223 | Prrc2c | 226562 | sRNA-Seq//HITS-CLIP | 33184237\|23597149 | Unspecified |
| mmu-miR-193a-3p | MIMAT0000223 | Psap | 19156 | sRNA-Seq//HITS-CLIP | 33184237\|23597149 | Unspecified |
| mmu-miR-193a-3p | MIMAT0000223 | Psmd11 | 69077 | HITS-CLIP | 37986224 | Unspecified |
| mmu-miR-193a-3p | MIMAT0000223 | Ptcd2 | 68927 | HITS-CLIP | 23597149 | Unspecified |
| mmu-miR-193a-3p | MIMAT0000223 | Ptchd1 | 211612 | HITS-CLIP | 19536157 | Unspecified |
| mmu-miR-193a-3p | MIMAT0000223 | Ptpn23 | 104831 | HITS-CLIP | 23597149 | Unspecified |
| mmu-miR-193a-3p | MIMAT0000223 | Ptpn3 | 545622 | HITS-CLIP | 37986224 | Unspecified |
| mmu-miR-193a-3p | MIMAT0000223 | Ptprd | 19266 | HITS-CLIP//sRNA-Seq | 23597149\|33184237 | Unspecified |
| mmu-miR-193a-3p | MIMAT0000223 | Ptprs | 19280 | HITS-CLIP//sRNA-Seq | 19536157\|33184237 | Unspecified |
| mmu-miR-193a-3p | MIMAT0000223 | Purb | 19291 | HITS-CLIP | 19536157 | Unspecified |
| mmu-miR-193a-3p | MIMAT0000223 | Pxn | 19303 | HITS-CLIP | 23597149 | Unspecified |
| mmu-miR-193a-3p | MIMAT0000223 | Pycrl | 66194 | HITS-CLIP | 23597149 | Unspecified |
| mmu-miR-193a-3p | MIMAT0000223 | Pygl | 110095 | HITS-CLIP | 37986224 | Unspecified |
| mmu-miR-193a-3p | MIMAT0000223 | Pzp | 11287 | HITS-CLIP | 37986224\|23597149 | Unspecified |
| mmu-miR-193a-3p | MIMAT0000223 | Qdpr | 110391 | HITS-CLIP//sRNA-Seq | 23597149\|33184237 | Unspecified |
| mmu-miR-193a-3p | MIMAT0000223 | Qk | 19317 | HITS-CLIP | 19536157 | Unspecified |
| mmu-miR-193a-3p | MIMAT0000223 | R3hdm1 | 226412 | HITS-CLIP//sRNA-Seq | 19536157\|33184237 | Unspecified |
| mmu-miR-193a-3p | MIMAT0000223 | Rab11a | 53869 | HITS-CLIP | 37986224 | Unspecified |

| | | | | | | |
|---|---|---|---|---|---|---|
| mmu-miR-193a-3p | MIMAT0000223 | Rab5c | 19345 | HITS-CLIP//sRNA-Seq | 23597149\|33184237 | Unspecified |
| mmu-miR-193a-3p | MIMAT0000223 | Rab6b | 270192 | HITS-CLIP//sRNA-Seq | 19536157\|33184237 | Unspecified |
| mmu-miR-193a-3p | MIMAT0000223 | Rabep1 | 54189 | HITS-CLIP | 37986224 | Unspecified |
| mmu-miR-193a-3p | MIMAT0000223 | Rabl3 | 67657 | HITS-CLIP | 37986224\|23597149 | Unspecified |
| mmu-miR-193a-3p | MIMAT0000223 | Rack1 | 10399 | HITS-CLIP//sRNA-Seq | 23597149\|33184237 | Unspecified |
| mmu-miR-193a-3p | MIMAT0000223 | Ranbp2 | 19386 | HITS-CLIP//sRNA-Seq | 37986224\|23597149\|33184237\|1953( | Unspecified |
| mmu-miR-193a-3p | MIMAT0000223 | Rars2 | 109093 | HITS-CLIP | 37986224 | Unspecified |
| mmu-miR-193a-3p | MIMAT0000223 | Rassf1 | 56289 | qCLASH | 31393953 | Unspecified |
| mmu-miR-193a-3p | MIMAT0000223 | Raver1 | 71766 | HITS-CLIP | 37986224\|23597149 | Unspecified |
| mmu-miR-193a-3p | MIMAT0000223 | Rbbp4 | 19646 | HITS-CLIP | 37986224 | Unspecified |
| mmu-miR-193a-3p | MIMAT0000223 | Rbm47 | 245945 | HITS-CLIP | 23597149 | Unspecified |
| mmu-miR-193a-3p | MIMAT0000223 | Rbms1 | 56878 | HITS-CLIP//sRNA-Seq | 23597149\|33184237 | Unspecified |
| mmu-miR-193a-3p | MIMAT0000223 | Rbpms2 | 71973 | HITS-CLIP | 23597149 | Unspecified |
| mmu-miR-193a-3p | MIMAT0000223 | Rcor1 | 217864 | HITS-CLIP | 31433990 | Unspecified |
| mmu-miR-193a-3p | MIMAT0000223 | Reep1 | 52250 | sRNA-Seq//HITS-CLIP | 33184237\|19536157 | Unspecified |
| mmu-miR-193a-3p | MIMAT0000223 | Rfc1 | 19687 | HITS-CLIP | 23597149 | Unspecified |
| mmu-miR-193a-3p | MIMAT0000223 | Rgs16 | 19734 | HITS-CLIP | 23597149 | Unspecified |
| mmu-miR-193a-3p | MIMAT0000223 | Rhobtb2 | 246710 | HITS-CLIP | 19536157 | Unspecified |
| mmu-miR-193a-3p | MIMAT0000223 | Rhog | 56212 | HITS-CLIP | 23597149 | Unspecified |
| mmu-miR-193a-3p | MIMAT0000223 | Rif1 | 51869 | HITS-CLIP | 23597149 | Unspecified |
| mmu-miR-193a-3p | MIMAT0000223 | Rlf | 109263 | HITS-CLIP | 23597149 | Unspecified |
| mmu-miR-193a-3p | MIMAT0000223 | Rlim | 19820 | sRNA-Seq//HITS-CLIP | 33184237\|23597149\|31433990 | Unspecified |
| mmu-miR-193a-3p | MIMAT0000223 | Rnf111 | 93836 | HITS-CLIP | 23597149 | Unspecified |
| mmu-miR-193a-3p | MIMAT0000223 | Rnf149 | 67702 | HITS-CLIP | 23597149 | Unspecified |
| mmu-miR-193a-3p | MIMAT0000223 | Rnf34 | 80751 | HITS-CLIP | 23597149 | Unspecified |
| mmu-miR-193a-3p | MIMAT0000223 | Rnf43 | 207742 | HITS-CLIP | 23597149 | Unspecified |
| mmu-miR-193a-3p | MIMAT0000223 | Rnf44 | 105239 | HITS-CLIP | 23597149 | Unspecified |
| mmu-miR-193a-3p | MIMAT0000223 | Ror1 | 26563 | HITS-CLIP | 23597149 | Unspecified |
| mmu-miR-193a-3p | MIMAT0000223 | Rpn1 | 103963 | sRNA-Seq//HITS-CLIP | 33184237\|26203562 | Unspecified |
| mmu-miR-193a-3p | MIMAT0000223 | Rps6ka1 | 20111 | HITS-CLIP | 23597149 | Unspecified |
| mmu-miR-193a-3p | MIMAT0000223 | Rps6ka3 | 110651 | HITS-CLIP | 37986224 | Unspecified |
| mmu-miR-193a-3p | MIMAT0000223 | Rrbp1 | 81910 | HITS-CLIP | 37986224\|23597149 | Unspecified |
| mmu-miR-193a-3p | MIMAT0000223 | Rreb1 | 68750 | HITS-CLIP | 23597149 | Unspecified |
| mmu-miR-193a-3p | MIMAT0000223 | Rrn3 | 106298 | HITS-CLIP//sRNA-Seq | 23597149\|33184237 | Unspecified |
| mmu-miR-193a-3p | MIMAT0000223 | Rsl1d1 | 66409 | sRNA-Seq//HITS-CLIP | 33184237\|23597149 | Unspecified |
| mmu-miR-193a-3p | MIMAT0000223 | Rxra | 20181 | HITS-CLIP | 23597149 | Unspecified |
| mmu-miR-193a-3p | MIMAT0000223 | S1pr1 | 13609 | HITS-CLIP | 23597149 | Unspecified |
| mmu-miR-193a-3p | MIMAT0000223 | Saa4 | 20211 | HITS-CLIP | 23597149 | Unspecified |
| mmu-miR-193a-3p | MIMAT0000223 | Sall1 | 58198 | HITS-CLIP | 23597149 | Unspecified |

| | | | | | | |
|---|---|---|---|---|---|---|
| mmu-miR-193a-3p | MIMAT0000223 | Sar1a | 20224 | HITS-CLIP//sRNA-Seq | 37986224\|23597149\|33184237 | Unspecified |
| mmu-miR-193a-3p | MIMAT0000223 | Sardh | 192166 | HITS-CLIP | 23597149 | Unspecified |
| mmu-miR-193a-3p | MIMAT0000223 | Sbds | 66711 | HITS-CLIP//sRNA-Seq | 23597149\|33184237 | Unspecified |
| mmu-miR-193a-3p | MIMAT0000223 | Sbk1 | 104175 | sRNA-Seq//HITS-CLIP | 33184237\|23597149 | Unspecified |
| mmu-miR-193a-3p | MIMAT0000223 | Scara5 | 71145 | HITS-CLIP | 23597149 | Unspecified |
| mmu-miR-193a-3p | MIMAT0000223 | Scarf1 | 380713 | HITS-CLIP | 23597149 | Unspecified |
| mmu-miR-193a-3p | MIMAT0000223 | Scd1 | 20249 | HITS-CLIP//sRNA-Seq | 37986224\|23597149\|33184237 | Unspecified |
| mmu-miR-193a-3p | MIMAT0000223 | Scn1a | 20265 | HITS-CLIP | 19536157 | Unspecified |
| mmu-miR-193a-3p | MIMAT0000223 | Scn2a | 110876 | HITS-CLIP | 19536157 | Unspecified |
| mmu-miR-193a-3p | MIMAT0000223 | Scrn3 | 74616 | HITS-CLIP | 23597149 | Unspecified |
| mmu-miR-193a-3p | MIMAT0000223 | Scyl3 | 240880 | HITS-CLIP | 37986224\|23597149 | Unspecified |
| mmu-miR-193a-3p | MIMAT0000223 | Sdc1 | 20969 | HITS-CLIP | 26203562 | Unspecified |
| mmu-miR-193a-3p | MIMAT0000223 | Sdhd | 66925 | HITS-CLIP//sRNA-Seq | 23597149\|33184237 | Unspecified |
| mmu-miR-193a-3p | MIMAT0000223 | Sec13 | 110379 | HITS-CLIP//sRNA-Seq | 23597149\|33184237 | Unspecified |
| mmu-miR-193a-3p | MIMAT0000223 | Sec16a | 227648 | HITS-CLIP | 37986224\|23597149 | Unspecified |
| mmu-miR-193a-3p | MIMAT0000223 | Sec16b | 89867 | HITS-CLIP | 23597149 | Unspecified |
| mmu-miR-193a-3p | MIMAT0000223 | Sec23a | 20334 | HITS-CLIP//sRNA-Seq | 19536157\|33184237 | Unspecified |
| mmu-miR-193a-3p | MIMAT0000223 | Sec23b | 27054 | HITS-CLIP | 23597149 | Unspecified |
| mmu-miR-193a-3p | MIMAT0000223 | Sec24c | 218811 | HITS-CLIP//sRNA-Seq | 37986224\|33184237\|23597149 | Unspecified |
| mmu-miR-193a-3p | MIMAT0000223 | Sec31a | 69162 | HITS-CLIP//sRNA-Seq | 23597149\|33184237 | Unspecified |
| mmu-miR-193a-3p | MIMAT0000223 | Sec61a1 | 53421 | HITS-CLIP | 23597149 | Unspecified |
| mmu-miR-193a-3p | MIMAT0000223 | Sec63 | 140740 | HITS-CLIP | 23597149 | Unspecified |
| mmu-miR-193a-3p | MIMAT0000223 | Seh1l | 72124 | HITS-CLIP | 23597149 | Unspecified |
| mmu-miR-193a-3p | MIMAT0000223 | Selenbp2 | 20342 | HITS-CLIP | 23597149 | Unspecified |
| mmu-miR-193a-3p | MIMAT0000223 | Selenow | 135964823 | HITS-CLIP//sRNA-Seq | 23597149\|33184237 | Unspecified |
| mmu-miR-193a-3p | MIMAT0000223 | Sema4g | 26456 | HITS-CLIP | 23597149 | Unspecified |
| mmu-miR-193a-3p | MIMAT0000223 | Senp6 | 215351 | sRNA-Seq//HITS-CLIP | 33184237\|31433990 | Unspecified |
| mmu-miR-193a-3p | MIMAT0000223 | Sepsecs | 211006 | HITS-CLIP | 23597149 | Unspecified |
| mmu-miR-193a-3p | MIMAT0000223 | Sept7 | 235072 | HITS-CLIP | 37986224 | Unspecified |
| mmu-miR-193a-3p | MIMAT0000223 | Serinc3 | 26943 | HITS-CLIP//sRNA-Seq | 23597149\|33184237 | Unspecified |
| mmu-miR-193a-3p | MIMAT0000223 | Serpina10 | 217847 | HITS-CLIP | 23597149 | Unspecified |
| mmu-miR-193a-3p | MIMAT0000223 | Serpina11 | 327657 | HITS-CLIP | 23597149 | Unspecified |
| mmu-miR-193a-3p | MIMAT0000223 | Serpina1b | 20701 | HITS-CLIP | 23597149 | Unspecified |
| mmu-miR-193a-3p | MIMAT0000223 | Serpinc1 | 11905 | HITS-CLIP | 37986224\|23597149 | Unspecified |
| mmu-miR-193a-3p | MIMAT0000223 | Serpine1 | 18787 | HITS-CLIP | 23597149 | Unspecified |
| mmu-miR-193a-3p | MIMAT0000223 | Serpinf2 | 18816 | HITS-CLIP | 23597149 | Unspecified |
| mmu-miR-193a-3p | MIMAT0000223 | Sesn3 | 75747 | HITS-CLIP | 19536157 | Unspecified |
| mmu-miR-193a-3p | MIMAT0000223 | Setd1a | 233904 | HITS-CLIP | 37986224 | Unspecified |
| mmu-miR-193a-3p | MIMAT0000223 | Sf1 | 22668 | sRNA-Seq//HITS-CLIP | 33184237\|23597149 | Unspecified |

| | | | | | | |
|---|---|---|---|---|---|---|
| mmu-miR-193a-3p | MIMAT0000223 | Sf3b2 | 319322 | HITS-CLIP//sRNA-Seq | 23597149\|33184237 | Unspecified |
| mmu-miR-193a-3p | MIMAT0000223 | Sgpp1 | 81535 | HITS-CLIP | 23597149 | Unspecified |
| mmu-miR-193a-3p | MIMAT0000223 | She | 214547 | HITS-CLIP | 23597149 | Unspecified |
| mmu-miR-193a-3p | MIMAT0000223 | Shkbp1 | 192192 | HITS-CLIP | 23597149 | Unspecified |
| mmu-miR-193a-3p | MIMAT0000223 | Shroom2 | 110380 | HITS-CLIP | 19536157 | Unspecified |
| mmu-miR-193a-3p | MIMAT0000223 | Sigmar1 | 18391 | HITS-CLIP//sRNA-Seq | 23597149\|33184237 | Unspecified |
| mmu-miR-193a-3p | MIMAT0000223 | Sik1 | 17691 | HITS-CLIP | 37986224\|23597149 | Unspecified |
| mmu-miR-193a-3p | MIMAT0000223 | Sipa1l2 | 244668 | HITS-CLIP | 23597149 | Unspecified |
| mmu-miR-193a-3p | MIMAT0000223 | Slc19a2 | 116914 | HITS-CLIP | 23597149 | Unspecified |
| mmu-miR-193a-3p | MIMAT0000223 | Slc22a17 | 59049 | HITS-CLIP//sRNA-Seq | 19536157\|33184237 | Unspecified |
| mmu-miR-193a-3p | MIMAT0000223 | Slc22a18 | 18400 | HITS-CLIP | 23597149 | Unspecified |
| mmu-miR-193a-3p | MIMAT0000223 | Slc22a23 | 73102 | HITS-CLIP | 37986224 | Unspecified |
| mmu-miR-193a-3p | MIMAT0000223 | Slc25a1 | 13358 | HITS-CLIP//sRNA-Seq | 23597149\|33184237 | Unspecified |
| mmu-miR-193a-3p | MIMAT0000223 | Slc25a15 | 18408 | HITS-CLIP | 37986224\|23597149 | Unspecified |
| mmu-miR-193a-3p | MIMAT0000223 | Slc25a22 | 68267 | HITS-CLIP | 37986224\|23597149 | Unspecified |
| mmu-miR-193a-3p | MIMAT0000223 | Slc25a23 | 66972 | HITS-CLIP | 23597149 | Unspecified |
| mmu-miR-193a-3p | MIMAT0000223 | Slc25a3 | 18674 | sRNA-Seq//HITS-CLIP | 33184237\|19536157\|23597149 | Unspecified |
| mmu-miR-193a-3p | MIMAT0000223 | Slc25a39 | 68066 | HITS-CLIP//sRNA-Seq | 23597149\|33184237 | Unspecified |
| mmu-miR-193a-3p | MIMAT0000223 | Slc25a47 | 104910 | HITS-CLIP | 23597149 | Unspecified |
| mmu-miR-193a-3p | MIMAT0000223 | Slc27a5 | 26459 | HITS-CLIP | 23597149 | Unspecified |
| mmu-miR-193a-3p | MIMAT0000223 | Slc31a1 | 20529 | HITS-CLIP | 23597149 | Unspecified |
| mmu-miR-193a-3p | MIMAT0000223 | Slc35d1 | 242585 | HITS-CLIP | 23597149 | Unspecified |
| mmu-miR-193a-3p | MIMAT0000223 | Slc36a1 | 215335 | HITS-CLIP | 37986224\|23597149 | Unspecified |
| mmu-miR-193a-3p | MIMAT0000223 | Slc38a2 | 67760 | HITS-CLIP//sRNA-Seq | 23597149\|33184237 | Unspecified |
| mmu-miR-193a-3p | MIMAT0000223 | Slc39a14 | 213053 | HITS-CLIP | 23597149 | Unspecified |
| mmu-miR-193a-3p | MIMAT0000223 | Slc39a7 | 14977 | HITS-CLIP | 37986224 | Unspecified |
| mmu-miR-193a-3p | MIMAT0000223 | Slc40a1 | 53945 | HITS-CLIP | 37986224\|23597149 | Unspecified |
| mmu-miR-193a-3p | MIMAT0000223 | Slc41a2 | 338365 | HITS-CLIP | 23597149 | Unspecified |
| mmu-miR-193a-3p | MIMAT0000223 | Slc44a2 | 68682 | sRNA-Seq//HITS-CLIP | 33184237\|19536157 | Unspecified |
| mmu-miR-193a-3p | MIMAT0000223 | Slc44a5 | 242259 | HITS-CLIP | 19536157 | Unspecified |
| mmu-miR-193a-3p | MIMAT0000223 | Slc4a7 | 218756 | HITS-CLIP | 23597149 | Unspecified |
| mmu-miR-193a-3p | MIMAT0000223 | Slc7a2 | 11988 | HITS-CLIP | 23597149 | Unspecified |
| mmu-miR-193a-3p | MIMAT0000223 | Slc7a5 | 20539 | HITS-CLIP//sRNA-Seq | 19536157\|33184237 | Unspecified |
| mmu-miR-193a-3p | MIMAT0000223 | Slmap | 83997 | HITS-CLIP | 37986224 | Unspecified |
| mmu-miR-193a-3p | MIMAT0000223 | Smcr8 | 237782 | HITS-CLIP | 23597149 | Unspecified |
| mmu-miR-193a-3p | MIMAT0000223 | Smg6 | 103677 | HITS-CLIP | 23597149 | Unspecified |
| mmu-miR-193a-3p | MIMAT0000223 | Smpd1 | 20597 | HITS-CLIP//sRNA-Seq | 23597149\|33184237 | Unspecified |
| mmu-miR-193a-3p | MIMAT0000223 | Smyd3 | 69726 | HITS-CLIP | 26203562 | Unspecified |
| mmu-miR-193a-3p | MIMAT0000223 | Snap29 | 67474 | HITS-CLIP | 31433990 | Unspecified |

| | | | | | | |
|---|---|---|---|---|---|---|
| mmu-miR-193a-3p | MIMAT0000223 | Snca | 20617 | sRNA-Seq//HITS-CLIP | 33184237\|19536157 | Unspecified |
| mmu-miR-193a-3p | MIMAT0000223 | Snrnp200 | 320632 | HITS-CLIP//sRNA-Seq | 23597149\|33184237 | Unspecified |
| mmu-miR-193a-3p | MIMAT0000223 | Sntb2 | 20650 | HITS-CLIP | 19536157 | Unspecified |
| mmu-miR-193a-3p | MIMAT0000223 | Snx16 | 74718 | HITS-CLIP | 37986224 | Unspecified |
| mmu-miR-193a-3p | MIMAT0000223 | Snx18 | 170625 | HITS-CLIP | 23597149 | Unspecified |
| mmu-miR-193a-3p | MIMAT0000223 | Snx3 | 54198 | HITS-CLIP//sRNA-Seq | 31433990\|33184237 | Unspecified |
| mmu-miR-193a-3p | MIMAT0000223 | Sord | 20322 | HITS-CLIP | 23597149 | Unspecified |
| mmu-miR-193a-3p | MIMAT0000223 | Sorl1 | 20660 | HITS-CLIP | 19536157 | Unspecified |
| mmu-miR-193a-3p | MIMAT0000223 | Sparc | 20692 | HITS-CLIP | 23597149 | Unspecified |
| mmu-miR-193a-3p | MIMAT0000223 | Spock1 | 20745 | sRNA-Seq//HITS-CLIP | 33184237\|19536157 | Unspecified |
| mmu-miR-193a-3p | MIMAT0000223 | Spryd4 | 66701 | HITS-CLIP | 37986224 | Unspecified |
| mmu-miR-193a-3p | MIMAT0000223 | Spryd7 | 66674 | HITS-CLIP//sRNA-Seq | 23597149\|33184237 | Unspecified |
| mmu-miR-193a-3p | MIMAT0000223 | Spsb2 | 14794 | HITS-CLIP | 23597149 | Unspecified |
| mmu-miR-193a-3p | MIMAT0000223 | Sptan1 | 20740 | sRNA-Seq//HITS-CLIP | 33184237\|23597149 | Unspecified |
| mmu-miR-193a-3p | MIMAT0000223 | Sra1 | 24068 | sRNA-Seq//HITS-CLIP | 33184237\|23597149 | Unspecified |
| mmu-miR-193a-3p | MIMAT0000223 | Srebf2 | 20788 | HITS-CLIP//sRNA-Seq | 23597149\|33184237 | Unspecified |
| mmu-miR-193a-3p | MIMAT0000223 | Srrm4 | 68955 | HITS-CLIP | 23597149 | Unspecified |
| mmu-miR-193a-3p | MIMAT0000223 | Srsf9 | 108014 | HITS-CLIP | 23597149 | Unspecified |
| mmu-miR-193a-3p | MIMAT0000223 | St3gal5 | 20454 | HITS-CLIP | 23597149 | Unspecified |
| mmu-miR-193a-3p | MIMAT0000223 | St5 | 76954 | HITS-CLIP | 23597149 | Unspecified |
| mmu-miR-193a-3p | MIMAT0000223 | Stab2 | 192188 | HITS-CLIP | 23597149 | Unspecified |
| mmu-miR-193a-3p | MIMAT0000223 | Stag2 | 20843 | HITS-CLIP | 23597149 | Unspecified |
| mmu-miR-193a-3p | MIMAT0000223 | Stard3 | 59045 | HITS-CLIP | 23597149 | Unspecified |
| mmu-miR-193a-3p | MIMAT0000223 | Stard4 | 170459 | HITS-CLIP//sRNA-Seq | 23597149\|33184237 | Unspecified |
| mmu-miR-193a-3p | MIMAT0000223 | Stard5 | 170460 | HITS-CLIP | 37986224\|23597149 | Unspecified |
| mmu-miR-193a-3p | MIMAT0000223 | Stat6 | 20852 | HITS-CLIP | 23597149 | Unspecified |
| mmu-miR-193a-3p | MIMAT0000223 | Steap4 | 117167 | HITS-CLIP | 23597149 | Unspecified |
| mmu-miR-193a-3p | MIMAT0000223 | Stk17b | 98267 | HITS-CLIP | 37986224 | Unspecified |
| mmu-miR-193a-3p | MIMAT0000223 | Suclg2 | 20917 | HITS-CLIP | 23597149 | Unspecified |
| mmu-miR-193a-3p | MIMAT0000223 | Sulf2 | 72043 | HITS-CLIP | 23597149 | Unspecified |
| mmu-miR-193a-3p | MIMAT0000223 | Sult1d1 | 53315 | HITS-CLIP | 23597149 | Unspecified |
| mmu-miR-193a-3p | MIMAT0000223 | Sun1 | 77053 | HITS-CLIP | 23597149 | Unspecified |
| mmu-miR-193a-3p | MIMAT0000223 | Suox | 211389 | HITS-CLIP | 23597149 | Unspecified |
| mmu-miR-193a-3p | MIMAT0000223 | Supt5 | 20924 | HITS-CLIP//sRNA-Seq | 23597149\|33184237 | Unspecified |
| mmu-miR-193a-3p | MIMAT0000223 | Surf1 | 20930 | HITS-CLIP | 23597149 | Unspecified |
| mmu-miR-193a-3p | MIMAT0000223 | Susd6 | 9766 | HITS-CLIP | 23597149 | Unspecified |
| mmu-miR-193a-3p | MIMAT0000223 | Svil | 225115 | HITS-CLIP | 23597149 | Unspecified |
| mmu-miR-193a-3p | MIMAT0000223 | Syne2 | 319565 | HITS-CLIP | 23597149 | Unspecified |
| mmu-miR-193a-3p | MIMAT0000223 | Syp | 20977 | HITS-CLIP//sRNA-Seq | 19536157\|33184237 | Unspecified |

| | | | | | | | |
|---|---|---|---|---|---|---|---|
| mmu-miR-193a-3p | MIMAT0000223 | Sytl4 | 27359 | HITS-CLIP | | 23597149 | Unspecified |
| mmu-miR-193a-3p | MIMAT0000223 | Tab3 | 66724 | HITS-CLIP | | 23597149 | Unspecified |
| mmu-miR-193a-3p | MIMAT0000223 | Taok3 | 330177 | HITS-CLIP | | 37986224 | Unspecified |
| mmu-miR-193a-3p | MIMAT0000223 | Tasp1 | 75812 | HITS-CLIP | | 23597149 | Unspecified |
| mmu-miR-193a-3p | MIMAT0000223 | Tax1bp1 | 52440 | HITS-CLIP | | 37986224 | Unspecified |
| mmu-miR-193a-3p | MIMAT0000223 | Tbc1d14 | 100855 | sRNA-Seq//HITS-CLIP | 33184237\|23597149 | | Unspecified |
| mmu-miR-193a-3p | MIMAT0000223 | Tcea3 | 21401 | HITS-CLIP | | 23597149 | Unspecified |
| mmu-miR-193a-3p | MIMAT0000223 | Tet3 | 194388 | HITS-CLIP | | 23597149 | Unspecified |
| mmu-miR-193a-3p | MIMAT0000223 | Tfdp2 | 211586 | HITS-CLIP | | 23597149 | Unspecified |
| mmu-miR-193a-3p | MIMAT0000223 | Tgfbr1 | 21812 | HITS-CLIP | | 23597149 | Unspecified |
| mmu-miR-193a-3p | MIMAT0000223 | Tgoln1 | 22134 | HITS-CLIP | 37986224\|23597149 | | Unspecified |
| mmu-miR-193a-3p | MIMAT0000223 | Timm8b | 30057 | HITS-CLIP | | 37986224 | Unspecified |
| mmu-miR-193a-3p | MIMAT0000223 | Timp2 | 21858 | HITS-CLIP | | 19536157 | Unspecified |
| mmu-miR-193a-3p | MIMAT0000223 | Tirap | 117149 | HITS-CLIP | | 23597149 | Unspecified |
| mmu-miR-193a-3p | MIMAT0000223 | Tjp1 | 21872 | HITS-CLIP | | 37986224 | Unspecified |
| mmu-miR-193a-3p | MIMAT0000223 | Tkfc | 26007 | HITS-CLIP | | 23597149 | Unspecified |
| mmu-miR-193a-3p | MIMAT0000223 | Tmbim6 | 110213 | HITS-CLIP//sRNA-Seq | 23597149\|33184237 | | Unspecified |
| mmu-miR-193a-3p | MIMAT0000223 | Tmed1 | 17083 | HITS-CLIP | | 23597149 | Unspecified |
| mmu-miR-193a-3p | MIMAT0000223 | Tmed3 | 66111 | HITS-CLIP | 37986224\|23597149 | | Unspecified |
| mmu-miR-193a-3p | MIMAT0000223 | Tmed4 | 103694 | HITS-CLIP//sRNA-Seq | 37986224\|23597149\|33184237 | | Unspecified |
| mmu-miR-193a-3p | MIMAT0000223 | Tmem131 | 56030 | HITS-CLIP | | 31433990 | Unspecified |
| mmu-miR-193a-3p | MIMAT0000223 | Tmem131l | 23240 | HITS-CLIP | | 23597149 | Unspecified |
| mmu-miR-193a-3p | MIMAT0000223 | Tmem147 | 69804 | HITS-CLIP//sRNA-Seq | 23597149\|33184237 | | Unspecified |
| mmu-miR-193a-3p | MIMAT0000223 | Tmem170 | 66817 | HITS-CLIP | | 23597149 | Unspecified |
| mmu-miR-193a-3p | MIMAT0000223 | Tmem18 | 211986 | HITS-CLIP | | 23597149 | Unspecified |
| mmu-miR-193a-3p | MIMAT0000223 | Tmem30a | 69981 | HITS-CLIP//sRNA-Seq | 37986224\|23597149\|33184237\|1953( | | Unspecified |
| mmu-miR-193a-3p | MIMAT0000223 | Tmem33 | 67878 | HITS-CLIP | | 37986224 | Unspecified |
| mmu-miR-193a-3p | MIMAT0000223 | Tmem38b | 52076 | qCLASH | | 31393953 | Unspecified |
| mmu-miR-193a-3p | MIMAT0000223 | Tmem56 | 99887 | HITS-CLIP | | 23597149 | Unspecified |
| mmu-miR-193a-3p | MIMAT0000223 | Tmem86a | 67893 | HITS-CLIP | | 23597149 | Unspecified |
| mmu-miR-193a-3p | MIMAT0000223 | Tmprss6 | 71753 | HITS-CLIP | 37986224\|23597149 | | Unspecified |
| mmu-miR-193a-3p | MIMAT0000223 | Tnks1bp1 | 228140 | HITS-CLIP | | 23597149 | Unspecified |
| mmu-miR-193a-3p | MIMAT0000223 | Tnpo1 | 238799 | HITS-CLIP | | 23597149 | Unspecified |
| mmu-miR-193a-3p | MIMAT0000223 | Tnrc6a | 233833 | HITS-CLIP | 23597149\|19536157 | | Unspecified |
| mmu-miR-193a-3p | MIMAT0000223 | Tnrc6b | 213988 | HITS-CLIP | | 37986224 | Unspecified |
| mmu-miR-193a-3p | MIMAT0000223 | Tnrc6c | 217351 | HITS-CLIP | 23597149\|19536157\|31433990 | | Unspecified |
| mmu-miR-193a-3p | MIMAT0000223 | Tns1 | 21961 | HITS-CLIP | 37986224\|23597149 | | Unspecified |
| mmu-miR-193a-3p | MIMAT0000223 | Tollip | 54473 | HITS-CLIP | 37986224\|23597149 | | Unspecified |
| mmu-miR-193a-3p | MIMAT0000223 | Tomm6 | 66119 | sRNA-Seq//HITS-CLIP | 33184237\|23597149 | | Unspecified |

| | | | | | | |
|---|---|---|---|---|---|---|
| mmu-miR-193a-3p | MIMAT0000223 | Tomm70a | 28185 | qCLASH | 31393953 | Unspecified |
| mmu-miR-193a-3p | MIMAT0000223 | Tpr | 108989 | HITS-CLIP//sRNA-Seq | 37986224\|23597149\|33184237 | Unspecified |
| mmu-miR-193a-3p | MIMAT0000223 | Tpra1 | 24100 | HITS-CLIP | 30971389 | Unspecified |
| mmu-miR-193a-3p | MIMAT0000223 | Trib2 | 217410 | sRNA-Seq//HITS-CLIP | 33184237\|19536157 | Unspecified |
| mmu-miR-193a-3p | MIMAT0000223 | Trim33 | 94093 | HITS-CLIP | 19536157 | Unspecified |
| mmu-miR-193a-3p | MIMAT0000223 | Trir | 79002 | sRNA-Seq//HITS-CLIP | 33184237\|31433990 | Unspecified |
| mmu-miR-193a-3p | MIMAT0000223 | Trmt6 | 66926 | HITS-CLIP | 23597149 | Unspecified |
| mmu-miR-193a-3p | MIMAT0000223 | Trp53inp1 | 60599 | HITS-CLIP | 37986224\|23597149\|31433990 | Unspecified |
| mmu-miR-193a-3p | MIMAT0000223 | Trp53inp2 | 68728 | sRNA-Seq//HITS-CLIP | 33184237\|23597149 | Unspecified |
| mmu-miR-193a-3p | MIMAT0000223 | Trps1 | 83925 | HITS-CLIP | 19536157 | Unspecified |
| mmu-miR-193a-3p | MIMAT0000223 | Trrap | 100683 | HITS-CLIP//sRNA-Seq | 23597149\|33184237 | Unspecified |
| mmu-miR-193a-3p | MIMAT0000223 | Tsc22d1 | 21807 | HITS-CLIP//sRNA-Seq | 37986224\|33184237\|23597149 | Unspecified |
| mmu-miR-193a-3p | MIMAT0000223 | Ttc38 | 239570 | HITS-CLIP | 23597149 | Unspecified |
| mmu-miR-193a-3p | MIMAT0000223 | Tti1 | 75425 | HITS-CLIP | 37986224 | Unspecified |
| mmu-miR-193a-3p | MIMAT0000223 | Ttpa | 50500 | HITS-CLIP | 23597149 | Unspecified |
| mmu-miR-193a-3p | MIMAT0000223 | Tuba1b | 22143 | HITS-CLIP//sRNA-Seq | 31433990\|33184237 | Unspecified |
| mmu-miR-193a-3p | MIMAT0000223 | Tubb2a | 22151 | sRNA-Seq//HITS-CLIP | 33184237\|23597149\|19536157 | Unspecified |
| mmu-miR-193a-3p | MIMAT0000223 | Uba6 | 231380 | qCLASH | 31393953 | Unspecified |
| mmu-miR-193a-3p | MIMAT0000223 | Ubap1 | 67123 | HITS-CLIP | 37986224 | Unspecified |
| mmu-miR-193a-3p | MIMAT0000223 | Ubb | 22187 | HITS-CLIP//sRNA-Seq | 23597149\|33184237 | Unspecified |
| mmu-miR-193a-3p | MIMAT0000223 | Ube2m | 22192 | HITS-CLIP//sRNA-Seq | 23597149\|33184237 | Unspecified |
| mmu-miR-193a-3p | MIMAT0000223 | Ube3a | 22215 | HITS-CLIP//sRNA-Seq | 19536157\|33184237 | Unspecified |
| mmu-miR-193a-3p | MIMAT0000223 | Ube3c | 100763 | HITS-CLIP//sRNA-Seq | 23597149\|33184237 | Unspecified |
| mmu-miR-193a-3p | MIMAT0000223 | Ubfd1 | 28018 | HITS-CLIP//sRNA-Seq | 23597149\|33184237 | Unspecified |
| mmu-miR-193a-3p | MIMAT0000223 | Ubn2 | 320538 | HITS-CLIP | 23597149 | Unspecified |
| mmu-miR-193a-3p | MIMAT0000223 | Ubp1 | 22221 | HITS-CLIP | 37986224\|31433990\|23597149 | Unspecified |
| mmu-miR-193a-3p | MIMAT0000223 | Ubr4 | 69116 | HITS-CLIP//sRNA-Seq | 19536157\|23597149\|33184237 | Unspecified |
| mmu-miR-193a-3p | MIMAT0000223 | Ubr5 | 70790 | HITS-CLIP//sRNA-Seq | 23597149\|33184237 | Unspecified |
| mmu-miR-193a-3p | MIMAT0000223 | Ufc1 | 66155 | HITS-CLIP//sRNA-Seq | 23597149\|33184237 | Unspecified |
| mmu-miR-193a-3p | MIMAT0000223 | Ugp2 | 216558 | HITS-CLIP | 23597149 | Unspecified |
| mmu-miR-193a-3p | MIMAT0000223 | Ugt2b35 | 243085 | HITS-CLIP | 23597149 | Unspecified |
| mmu-miR-193a-3p | MIMAT0000223 | Unc119b | 106840 | HITS-CLIP | 37986224\|23597149 | Unspecified |
| mmu-miR-193a-3p | MIMAT0000223 | Unc5b | 107449 | HITS-CLIP | 19536157 | Unspecified |
| mmu-miR-193a-3p | MIMAT0000223 | Unc5c | 22253 | sRNA-Seq//HITS-CLIP | 33184237\|19536157 | Unspecified |
| mmu-miR-193a-3p | MIMAT0000223 | Upp2 | 76654 | HITS-CLIP | 37986224 | Unspecified |
| mmu-miR-193a-3p | MIMAT0000223 | Uqcc2 | 67267 | HITS-CLIP//sRNA-Seq | 23597149\|33184237 | Unspecified |
| mmu-miR-193a-3p | MIMAT0000223 | Urgcp | 72046 | HITS-CLIP | 19536157 | Unspecified |
| mmu-miR-193a-3p | MIMAT0000223 | Usf2 | 22282 | HITS-CLIP | 23597149 | Unspecified |
| mmu-miR-193a-3p | MIMAT0000223 | Usp10 | 22224 | HITS-CLIP | 37986224 | Unspecified |

| | | | | | | |
|---|---|---|---|---|---|---|
| mmu-miR-193a-3p | MIMAT0000223 | Usp19 | 71472 | HITS-CLIP//sRNA-Seq | 23597149\|33184237 | Unspecified |
| mmu-miR-193a-3p | MIMAT0000223 | Usp24 | 329908 | HITS-CLIP | 23597149 | Unspecified |
| mmu-miR-193a-3p | MIMAT0000223 | Usp25 | 30940 | HITS-CLIP | 23597149 | Unspecified |
| mmu-miR-193a-3p | MIMAT0000223 | Usp34 | 17847 | HITS-CLIP | 37986224 | Unspecified |
| mmu-miR-193a-3p | MIMAT0000223 | Usp49 | 224836 | HITS-CLIP | 23597149 | Unspecified |
| mmu-miR-193a-3p | MIMAT0000223 | Usp9x | 22284 | HITS-CLIP//sRNA-Seq | 31433990\|33184237 | Unspecified |
| mmu-miR-193a-3p | MIMAT0000223 | Vapb | 56491 | HITS-CLIP//sRNA-Seq | 23597149\|33184237 | Unspecified |
| mmu-miR-193a-3p | MIMAT0000223 | Vezf1 | 22344 | HITS-CLIP | 23597149 | Unspecified |
| mmu-miR-193a-3p | MIMAT0000223 | Vipas39 | 104799 | HITS-CLIP | 37986224\|23597149 | Unspecified |
| mmu-miR-193a-3p | MIMAT0000223 | Virma | 25962 | HITS-CLIP | 19536157\|23597149 | Unspecified |
| mmu-miR-193a-3p | MIMAT0000223 | Vmp1 | 75909 | sRNA-Seq//HITS-CLIP | 33184237\|31433990 | Unspecified |
| mmu-miR-193a-3p | MIMAT0000223 | Vps13a | 271564 | sRNA-Seq//HITS-CLIP | 33184237\|19536157 | Unspecified |
| mmu-miR-193a-3p | MIMAT0000223 | Vps13c | 320528 | HITS-CLIP//sRNA-Seq | 23597149\|33184237 | Unspecified |
| mmu-miR-193a-3p | MIMAT0000223 | Vps37b | 330192 | HITS-CLIP | 37986224\|23597149 | Unspecified |
| mmu-miR-193a-3p | MIMAT0000223 | Vps4b | 20479 | HITS-CLIP//sRNA-Seq | 23597149\|33184237 | Unspecified |
| mmu-miR-193a-3p | MIMAT0000223 | Vta1 | 66201 | HITS-CLIP//sRNA-Seq | 23597149\|33184237 | Unspecified |
| mmu-miR-193a-3p | MIMAT0000223 | Vwa8 | 219189 | HITS-CLIP | 23597149 | Unspecified |
| mmu-miR-193a-3p | MIMAT0000223 | Wac | 225131 | HITS-CLIP//sRNA-Seq | 37986224\|23597149\|33184237 | Unspecified |
| mmu-miR-193a-3p | MIMAT0000223 | Wbp1l | 226178 | HITS-CLIP | 37986224\|23597149 | Unspecified |
| mmu-miR-193a-3p | MIMAT0000223 | Wdfy3 | 72145 | sRNA-Seq//HITS-CLIP | 33184237\|19536157 | Unspecified |
| mmu-miR-193a-3p | MIMAT0000223 | Wdr11 | 207425 | HITS-CLIP | 23597149 | Unspecified |
| mmu-miR-193a-3p | MIMAT0000223 | Wdr36 | 225348 | HITS-CLIP | 37986224 | Unspecified |
| mmu-miR-193a-3p | MIMAT0000223 | Wdr37 | 207615 | sRNA-Seq//HITS-CLIP | 33184237\|23597149 | Unspecified |
| mmu-miR-193a-3p | MIMAT0000223 | Wdr61 | 66317 | HITS-CLIP | 23597149 | Unspecified |
| mmu-miR-193a-3p | MIMAT0000223 | Wdr7 | 104082 | HITS-CLIP | 23597149 | Unspecified |
| mmu-miR-193a-3p | MIMAT0000223 | Wdr82 | 77305 | HITS-CLIP//sRNA-Seq | 23597149\|33184237 | Unspecified |
| mmu-miR-193a-3p | MIMAT0000223 | Wdr89 | 72338 | HITS-CLIP | 23597149 | Unspecified |
| mmu-miR-193a-3p | MIMAT0000223 | Wdr91 | 101240 | HITS-CLIP | 23597149 | Unspecified |
| mmu-miR-193a-3p | MIMAT0000223 | Wfdc21 | 66107 | HITS-CLIP | 23597149 | Unspecified |
| mmu-miR-193a-3p | MIMAT0000223 | Wnk1 | 232341 | sRNA-Seq//HITS-CLIP | 33184237\|31433990 | Unspecified |
| mmu-miR-193a-3p | MIMAT0000223 | Wwc1 | 211652 | HITS-CLIP | 23597149 | Unspecified |
| mmu-miR-193a-3p | MIMAT0000223 | Xdh | 22436 | HITS-CLIP | 37986224\|23597149 | Unspecified |
| mmu-miR-193a-3p | MIMAT0000223 | Yipf5 | 67180 | HITS-CLIP | 23597149 | Unspecified |
| mmu-miR-193a-3p | MIMAT0000223 | Yrdc | 230734 | HITS-CLIP | 23597149 | Unspecified |
| mmu-miR-193a-3p | MIMAT0000223 | Ythdf3 | 229096 | HITS-CLIP//sRNA-Seq | 23597149\|33184237 | Unspecified |
| mmu-miR-193a-3p | MIMAT0000223 | Ywhaz | 22631 | sRNA-Seq//HITS-CLIP | 33184237\|31433990 | Unspecified |
| mmu-miR-193a-3p | MIMAT0000223 | Zadh2 | 225791 | HITS-CLIP | 23597149 | Unspecified |
| mmu-miR-193a-3p | MIMAT0000223 | Zbtb10 | 229055 | HITS-CLIP | 23597149 | Unspecified |
| mmu-miR-193a-3p | MIMAT0000223 | Zbtb11 | 271377 | HITS-CLIP | 37986224\|23597149 | Unspecified |

| | | | | | | | |
|---|---|---|---|---|---|---|---|
| mmu-miR-193a-3p | MIMAT0000223 | Zbtb18 | 30928 | HITS-CLIP | | 19536157 | Unspecified |
| mmu-miR-193a-3p | MIMAT0000223 | Zbtb20 | 56490 | HITS-CLIP | | 37986224 | Unspecified |
| mmu-miR-193a-3p | MIMAT0000223 | Zbtb33 | 56805 | HITS-CLIP | | 23597149 | Unspecified |
| mmu-miR-193a-3p | MIMAT0000223 | Zcchc24 | 71918 | HITS-CLIP | | 23597149 | Unspecified |
| mmu-miR-193a-3p | MIMAT0000223 | Zdhhc12 | 66220 | HITS-CLIP | | 23597149 | Unspecified |
| mmu-miR-193a-3p | MIMAT0000223 | Zfp106 | 20402 | HITS-CLIP//sRNA-Seq | 23597149\|33184237 | | Unspecified |
| mmu-miR-193a-3p | MIMAT0000223 | Zfp131 | 72465 | HITS-CLIP | | 31433990 | Unspecified |
| mmu-miR-193a-3p | MIMAT0000223 | Zfp148 | 22661 | HITS-CLIP | | 19536157 | Unspecified |
| mmu-miR-193a-3p | MIMAT0000223 | Zfp207 | 22680 | HITS-CLIP//sRNA-Seq | 31433990\|33184237 | | Unspecified |
| mmu-miR-193a-3p | MIMAT0000223 | Zfp281 | 226442 | HITS-CLIP | | 23597149 | Unspecified |
| mmu-miR-193a-3p | MIMAT0000223 | Zfp36 | 22695 | HITS-CLIP | 37986224\|23597149 | | Unspecified |
| mmu-miR-193a-3p | MIMAT0000223 | Zfp36l1 | 12192 | HITS-CLIP | 37986224\|23597149 | | Unspecified |
| mmu-miR-193a-3p | MIMAT0000223 | Zfp512b | 269401 | HITS-CLIP//sRNA-Seq | 23597149\|33184237 | | Unspecified |
| mmu-miR-193a-3p | MIMAT0000223 | Zfp609 | 214812 | HITS-CLIP | | 23597149 | Unspecified |
| mmu-miR-193a-3p | MIMAT0000223 | Zfp628 | 232816 | HITS-CLIP | | 23597149 | Unspecified |
| mmu-miR-193a-3p | MIMAT0000223 | Zfp704 | 170753 | HITS-CLIP | | 23597149 | Unspecified |
| mmu-miR-193a-3p | MIMAT0000223 | Zfp740 | 68744 | HITS-CLIP | | 23597149 | Unspecified |
| mmu-miR-193a-3p | MIMAT0000223 | Zfpl1 | 81909 | HITS-CLIP | | 23597149 | Unspecified |
| mmu-miR-193a-3p | MIMAT0000223 | Zhx3 | 320799 | HITS-CLIP | 37986224\|23597149 | | Unspecified |
| mmu-miR-193a-3p | MIMAT0000223 | Zmiz1 | 328365 | HITS-CLIP//sRNA-Seq | 23597149\|33184237 | | Unspecified |
| mmu-miR-193a-3p | MIMAT0000223 | Znhit2 | 29805 | HITS-CLIP | | 23597149 | Unspecified |
| mmu-miR-193a-3p | MIMAT0000223 | Znrf3 | 407821 | HITS-CLIP | | 23597149 | Unspecified |
| mmu-miR-193a-3p | MIMAT0000223 | Zzef1 | 195018 | HITS-CLIP | | 23597149 | Unspecified |
| mmu-miR-378a-3p | MIMAT0003151 | 1110059G10Rik | 66202 | HITS-CLIP | | 23142080 | Unspecified |
| mmu-miR-378a-3p | MIMAT0003151 | 1700017B05Rik | 74211 | HITS-CLIP | | 37986224 | Unspecified |
| mmu-miR-378a-3p | MIMAT0003151 | 1810013L24Rik | 69053 | HITS-CLIP | 37986224\|23142080\|25083871 | | Unspecified |
| mmu-miR-378a-3p | MIMAT0003151 | 2310061I04Rik | 69662 | HITS-CLIP | 23142080\|25083871 | | Unspecified |
| mmu-miR-378a-3p | MIMAT0003151 | 2610507B11Rik | 72503 | HITS-CLIP | 25083871\|29167373 | | Unspecified |
| mmu-miR-378a-3p | MIMAT0003151 | 2700081O15Rik | 108899 | HITS-CLIP | | 25083871 | Unspecified |
| mmu-miR-378a-3p | MIMAT0003151 | 4833439L19Rik | 97820 | HITS-CLIP | | 37986224 | Unspecified |
| mmu-miR-378a-3p | MIMAT0003151 | 4931406C07Rik | 70984 | HITS-CLIP | | 23597149 | Unspecified |
| mmu-miR-378a-3p | MIMAT0003151 | 4931406P16Rik | 233103 | HITS-CLIP | | 37986224 | Unspecified |
| mmu-miR-378a-3p | MIMAT0003151 | 4932438A13Rik | 229227 | HITS-CLIP | | 25083871 | Unspecified |
| mmu-miR-378a-3p | MIMAT0003151 | 5031439G07Rik | 223739 | HITS-CLIP | | 37986224 | Unspecified |
| mmu-miR-378a-3p | MIMAT0003151 | 8030462N17Rik | 212163 | HITS-CLIP | | 25083871 | Unspecified |
| mmu-miR-378a-3p | MIMAT0003151 | 9930021J03Rik | 240613 | HITS-CLIP | 37986224\|25083871 | | Unspecified |
| mmu-miR-378a-3p | MIMAT0003151 | AI182371 | 98870 | HITS-CLIP | | 23597149 | Unspecified |
| mmu-miR-378a-3p | MIMAT0003151 | AI597479 | 98404 | HITS-CLIP | | 25083871 | Unspecified |
| mmu-miR-378a-3p | MIMAT0003151 | Aacs | 78894 | HITS-CLIP | | 29167373 | Unspecified |

| | | | | | | |
|---|---|---|---|---|---|---|
| mmu-miR-378a-3p | MIMAT0003151 | Aass | 30956 | HITS-CLIP | 37986224\|23597149 | Unspecified |
| mmu-miR-378a-3p | MIMAT0003151 | Abcb1b | 18669 | Microarrays | 37986224\|22855601 | Unspecified |
| mmu-miR-378a-3p | MIMAT0003151 | Abcb6 | 74104 | HITS-CLIP | 37986224 | Unspecified |
| mmu-miR-378a-3p | MIMAT0003151 | Abcc1 | 17250 | Microarrays | 37986224\|22855601 | Unspecified |
| mmu-miR-378a-3p | MIMAT0003151 | Abcc5 | 27416 | HITS-CLIP | 37986224 | Unspecified |
| mmu-miR-378a-3p | MIMAT0003151 | Abcc6 | 27421 | HITS-CLIP | 23597149 | Unspecified |
| mmu-miR-378a-3p | MIMAT0003151 | Abce1 | 24015 | Microarrays | 37986224\|22855601 | Unspecified |
| mmu-miR-378a-3p | MIMAT0003151 | Abcf1 | 224742 | HITS-CLIP | 23142080 | Unspecified |
| mmu-miR-378a-3p | MIMAT0003151 | Abhd12 | 76192 | HITS-CLIP | 37986224\|25083871\|29167373 | Unspecified |
| mmu-miR-378a-3p | MIMAT0003151 | Abhd13 | 68904 | HITS-CLIP | 37986224 | Unspecified |
| mmu-miR-378a-3p | MIMAT0003151 | Abi1 | 11308 | HITS-CLIP | 37986224 | Unspecified |
| mmu-miR-378a-3p | MIMAT0003151 | Abl1 | 11350 | HITS-CLIP | 37986224\|23142080 | Unspecified |
| mmu-miR-378a-3p | MIMAT0003151 | Abl2 | 11352 | Microarrays | 37986224\|22855601 | Unspecified |
| mmu-miR-378a-3p | MIMAT0003151 | Abr | 109934 | HITS-CLIP | 25083871\|29167373 | Unspecified |
| mmu-miR-378a-3p | MIMAT0003151 | Abraxas2 | 23172 | HITS-CLIP | 25083871\|23597149 | Unspecified |
| mmu-miR-378a-3p | MIMAT0003151 | Acbd3 | 170760 | HITS-CLIP | 37986224 | Unspecified |
| mmu-miR-378a-3p | MIMAT0003151 | Acin1 | 56215 | HITS-CLIP | 37986224 | Unspecified |
| mmu-miR-378a-3p | MIMAT0003151 | Acsl4 | 50790 | Microarrays//HITS-CLIP | 37986224\|22855601\|23142080 | Unspecified |
| mmu-miR-378a-3p | MIMAT0003151 | Acsl5 | 433256 | HITS-CLIP | 37986224 | Unspecified |
| mmu-miR-378a-3p | MIMAT0003151 | Actn4 | 60595 | HITS-CLIP, Microarrays//Microarrays | 37986224\|22855601 | Unspecified |
| mmu-miR-378a-3p | MIMAT0003151 | Acvr1b | 11479 | HITS-CLIP | 37986224\|26203562 | Unspecified |
| mmu-miR-378a-3p | MIMAT0003151 | Acvr2a | 11480 | HITS-CLIP | 37986224 | Unspecified |
| mmu-miR-378a-3p | MIMAT0003151 | Adamts2 | 216725 | HITS-CLIP | 37986224 | Unspecified |
| mmu-miR-378a-3p | MIMAT0003151 | Adamts6 | 108154 | HITS-CLIP | 37986224\|25083871 | Unspecified |
| mmu-miR-378a-3p | MIMAT0003151 | Adarb1 | 110532 | HITS-CLIP | 37986224 | Unspecified |
| mmu-miR-378a-3p | MIMAT0003151 | Adgrf3 | 165082 | qCLASH | 31393953 | Unspecified |
| mmu-miR-378a-3p | MIMAT0003151 | Adgrg1 | 9289 | HITS-CLIP | 25083871 | Unspecified |
| mmu-miR-378a-3p | MIMAT0003151 | Adgrg6 | 57211 | HITS-CLIP | 25083871 | Unspecified |
| mmu-miR-378a-3p | MIMAT0003151 | Adipor2 | 68465 | HITS-CLIP | 37986224 | Unspecified |
| mmu-miR-378a-3p | MIMAT0003151 | Adnp | 11538 | HITS-CLIP | 37986224\|23142080 | Unspecified |
| mmu-miR-378a-3p | MIMAT0003151 | Adnp2 | 240442 | HITS-CLIP | 37986224 | Unspecified |
| mmu-miR-378a-3p | MIMAT0003151 | Adprhl1 | 113622 | HITS-CLIP | 25753418 | Unspecified |
| mmu-miR-378a-3p | MIMAT0003151 | Aebp1 | 11568 | HITS-CLIP | 37986224 | Unspecified |
| mmu-miR-378a-3p | MIMAT0003151 | Aebp2 | 11569 | HITS-CLIP | 37986224\|23142080 | Unspecified |
| mmu-miR-378a-3p | MIMAT0003151 | Aen | 68048 | Microarrays | 37986224\|22855601 | Unspecified |
| mmu-miR-378a-3p | MIMAT0003151 | Aff1 | 17355 | HITS-CLIP | 37986224 | Unspecified |
| mmu-miR-378a-3p | MIMAT0003151 | Aff4 | 93736 | HITS-CLIP | 37986224 | Unspecified |
| mmu-miR-378a-3p | MIMAT0003151 | Afg3l2 | 69597 | HITS-CLIP | 23597149 | Unspecified |
| mmu-miR-378a-3p | MIMAT0003151 | Aftph | 216549 | HITS-CLIP | 37986224 | Unspecified |

| | | | | | | |
|---|---|---|---|---|---|---|
| mmu-miR-378a-3p | MIMAT0003151 | Agfg1 | 15463 | HITS-CLIP, Microarrays//HITS-CLIP//Microa | 37986224\|23142080\|22855601\|25083 | Unspecified |
| mmu-miR-378a-3p | MIMAT0003151 | Ago1 | 236511 | HITS-CLIP | 37986224\|23142080 | Unspecified |
| mmu-miR-378a-3p | MIMAT0003151 | Ago3 | 214150 | HITS-CLIP | 23142080 | Unspecified |
| mmu-miR-378a-3p | MIMAT0003151 | Agpat3 | 28169 | HITS-CLIP | 37986224 | Unspecified |
| mmu-miR-378a-3p | MIMAT0003151 | Agtpbp1 | 67269 | HITS-CLIP | 25083871 | Unspecified |
| mmu-miR-378a-3p | MIMAT0003151 | Agtrap | 11610 | HITS-CLIP | 25753418 | Unspecified |
| mmu-miR-378a-3p | MIMAT0003151 | Agxt2 | 268782 | HITS-CLIP | 37986224 | Unspecified |
| mmu-miR-378a-3p | MIMAT0003151 | Ahctf1 | 226747 | HITS-CLIP | 37986224\|25083871\|23597149 | Unspecified |
| mmu-miR-378a-3p | MIMAT0003151 | Ahdc1 | 230793 | HITS-CLIP | 37986224 | Unspecified |
| mmu-miR-378a-3p | MIMAT0003151 | Ahnak | 66395 | HITS-CLIP | 37986224\|29167373 | Unspecified |
| mmu-miR-378a-3p | MIMAT0003151 | Ak4 | 11639 | HITS-CLIP | 37986224 | Unspecified |
| mmu-miR-378a-3p | MIMAT0003151 | Akap10 | 56697 | HITS-CLIP | 23142080\|29167373 | Unspecified |
| mmu-miR-378a-3p | MIMAT0003151 | Akap11 | 219181 | HITS-CLIP | 37986224 | Unspecified |
| mmu-miR-378a-3p | MIMAT0003151 | Akap13 | 75547 | HITS-CLIP | 37986224 | Unspecified |
| mmu-miR-378a-3p | MIMAT0003151 | Akap6 | 238161 | HITS-CLIP | 25083871\|25753418 | Unspecified |
| mmu-miR-378a-3p | MIMAT0003151 | Akap9 | 100986 | HITS-CLIP | 37986224\|25083871 | Unspecified |
| mmu-miR-378a-3p | MIMAT0003151 | Akt1 | 11651 | Microarrays | 37986224\|22855601 | Unspecified |
| mmu-miR-378a-3p | MIMAT0003151 | Alcam | 11658 | Microarrays | 37986224\|22855601 | Unspecified |
| mmu-miR-378a-3p | MIMAT0003151 | Alg10b | 380959 | HITS-CLIP | 37986224 | Unspecified |
| mmu-miR-378a-3p | MIMAT0003151 | Alg14 | 66789 | HITS-CLIP | 25753418 | Unspecified |
| mmu-miR-378a-3p | MIMAT0003151 | Alg2 | 56737 | HITS-CLIP | 37986224 | Unspecified |
| mmu-miR-378a-3p | MIMAT0003151 | Alg3 | 208624 | Microarrays | 37986224\|22855601 | Unspecified |
| mmu-miR-378a-3p | MIMAT0003151 | Alg5 | 66248 | HITS-CLIP | 23142080 | Unspecified |
| mmu-miR-378a-3p | MIMAT0003151 | Alg9 | 102580 | HITS-CLIP | 25083871 | Unspecified |
| mmu-miR-378a-3p | MIMAT0003151 | Alpk3 | 116904 | HITS-CLIP | 25753418 | Unspecified |
| mmu-miR-378a-3p | MIMAT0003151 | Amot | 27494 | HITS-CLIP | 37986224 | Unspecified |
| mmu-miR-378a-3p | MIMAT0003151 | Anapc2 | 99152 | HITS-CLIP | 37986224 | Unspecified |
| mmu-miR-378a-3p | MIMAT0003151 | Angel2 | 52477 | HITS-CLIP | 37986224 | Unspecified |
| mmu-miR-378a-3p | MIMAT0003151 | Ank | 11732 | Microarrays | 37986224\|22855601 | Unspecified |
| mmu-miR-378a-3p | MIMAT0003151 | Ank2 | 109676 | HITS-CLIP | 37986224 | Unspecified |
| mmu-miR-378a-3p | MIMAT0003151 | Ankfy1 | 11736 | HITS-CLIP | 37986224 | Unspecified |
| mmu-miR-378a-3p | MIMAT0003151 | Ankhd1 | 108857 | HITS-CLIP | 37986224 | Unspecified |
| mmu-miR-378a-3p | MIMAT0003151 | Ankib1 | 70797 | HITS-CLIP | 25083871\|23142080 | Unspecified |
| mmu-miR-378a-3p | MIMAT0003151 | Ankra2 | 68558 | HITS-CLIP | 37986224\|25083871 | Unspecified |
| mmu-miR-378a-3p | MIMAT0003151 | Ankrd10 | 102334 | HITS-CLIP | 37986224 | Unspecified |
| mmu-miR-378a-3p | MIMAT0003151 | Ankrd11 | 77087 | HITS-CLIP | 37986224\|25083871 | Unspecified |
| mmu-miR-378a-3p | MIMAT0003151 | Ankrd12 | 106585 | HITS-CLIP | 37986224 | Unspecified |
| mmu-miR-378a-3p | MIMAT0003151 | Ankrd13c | 433667 | HITS-CLIP | 23597149 | Unspecified |
| mmu-miR-378a-3p | MIMAT0003151 | Ankrd17 | 81702 | HITS-CLIP | 37986224\|25083871\|23142080\|29167 | Unspecified |

| | | | | | | |
|---|---|---|---|---|---|---|
| mmu-miR-378a-3p | MIMAT0003151 | Ankrd28 | 105522 | Microarrays | 37986224\|22855601 | Unspecified |
| mmu-miR-378a-3p | MIMAT0003151 | Ankrd37 | 654824 | HITS-CLIP | 23142080 | Unspecified |
| mmu-miR-378a-3p | MIMAT0003151 | Ankrd52 | 237615 | HITS-CLIP | 37986224\|23142080\|25083871 | Unspecified |
| mmu-miR-378a-3p | MIMAT0003151 | Anxa2 | 12306 | HITS-CLIP | 25083871 | Unspecified |
| mmu-miR-378a-3p | MIMAT0003151 | Anxa3 | 11745 | Microarrays | 37986224\|22855601 | Unspecified |
| mmu-miR-378a-3p | MIMAT0003151 | Ap1ar | 211556 | HITS-CLIP | 37986224 | Unspecified |
| mmu-miR-378a-3p | MIMAT0003151 | Ap1g1 | 11765 | HITS-CLIP | 37986224 | Unspecified |
| mmu-miR-378a-3p | MIMAT0003151 | Ap2b1 | 71770 | HITS-CLIP | 37986224 | Unspecified |
| mmu-miR-378a-3p | MIMAT0003151 | Ap3b1 | 11774 | HITS-CLIP | 23597149\|23142080 | Unspecified |
| mmu-miR-378a-3p | MIMAT0003151 | Ap3d1 | 11776 | HITS-CLIP | 37986224\|25083871 | Unspecified |
| mmu-miR-378a-3p | MIMAT0003151 | Ap4b1 | 67489 | HITS-CLIP | 37986224 | Unspecified |
| mmu-miR-378a-3p | MIMAT0003151 | Ap4e1 | 108011 | HITS-CLIP | 23597149\|25753418 | Unspecified |
| mmu-miR-378a-3p | MIMAT0003151 | Apbb2 | 11787 | HITS-CLIP | 25083871 | Unspecified |
| mmu-miR-378a-3p | MIMAT0003151 | Apc | 11789 | HITS-CLIP | 37986224\|25083871 | Unspecified |
| mmu-miR-378a-3p | MIMAT0003151 | Api5 | 11800 | HITS-CLIP | 37986224 | Unspecified |
| mmu-miR-378a-3p | MIMAT0003151 | Aplf | 72103 | Microarrays | 37986224 | Unspecified |
| mmu-miR-378a-3p | MIMAT0003151 | Apob | 238055 | HITS-CLIP | 37986224\|23597149 | Unspecified |
| mmu-miR-378a-3p | MIMAT0003151 | Apod | 11815 | Microarrays | 37986224 | Unspecified |
| mmu-miR-378a-3p | MIMAT0003151 | Aqp1 | 11826 | Microarrays | 37986224\|22855601 | Unspecified |
| mmu-miR-378a-3p | MIMAT0003151 | Arap3 | 106952 | HITS-CLIP | 25753418 | Unspecified |
| mmu-miR-378a-3p | MIMAT0003151 | Arcn1 | 213827 | HITS-CLIP | 37986224 | Unspecified |
| mmu-miR-378a-3p | MIMAT0003151 | Arf2 | 11841 | HITS-CLIP | 37986224 | Unspecified |
| mmu-miR-378a-3p | MIMAT0003151 | Arf6 | 11845 | HITS-CLIP | 37986224 | Unspecified |
| mmu-miR-378a-3p | MIMAT0003151 | Arfgap1 | 228998 | HITS-CLIP | 37986224\|25083871\|23142080\|29167 | Unspecified |
| mmu-miR-378a-3p | MIMAT0003151 | Arfgef1 | 211673 | Microarrays//HITS-CLIP | 37986224\|25083871\|22855601\|23142 | Unspecified |
| mmu-miR-378a-3p | MIMAT0003151 | Arglu1 | 234023 | HITS-CLIP | 37986224 | Unspecified |
| mmu-miR-378a-3p | MIMAT0003151 | Arhgap10 | 78514 | HITS-CLIP | 23142080 | Unspecified |
| mmu-miR-378a-3p | MIMAT0003151 | Arhgap31 | 12549 | HITS-CLIP | 37986224 | Unspecified |
| mmu-miR-378a-3p | MIMAT0003151 | Arhgap32 | 330914 | HITS-CLIP | 37986224 | Unspecified |
| mmu-miR-378a-3p | MIMAT0003151 | Arhgap35 | 232906 | HITS-CLIP | 37986224 | Unspecified |
| mmu-miR-378a-3p | MIMAT0003151 | Arhgap42 | 71544 | HITS-CLIP | 37986224\|25083871 | Unspecified |
| mmu-miR-378a-3p | MIMAT0003151 | Arhgap5 | 11855 | HITS-CLIP | 37986224 | Unspecified |
| mmu-miR-378a-3p | MIMAT0003151 | Arhgef10 | 234094 | HITS-CLIP | 37986224 | Unspecified |
| mmu-miR-378a-3p | MIMAT0003151 | Arhgef11 | 213498 | HITS-CLIP | 37986224\|25083871 | Unspecified |
| mmu-miR-378a-3p | MIMAT0003151 | Arhgef12 | 69632 | HITS-CLIP | 37986224\|25083871\|23597149 | Unspecified |
| mmu-miR-378a-3p | MIMAT0003151 | Arhgef2 | 16800 | HITS-CLIP | 25083871\|29167373 | Unspecified |
| mmu-miR-378a-3p | MIMAT0003151 | Arid1a | 93760 | HITS-CLIP | 25753418 | Unspecified |
| mmu-miR-378a-3p | MIMAT0003151 | Arid2 | 77044 | HITS-CLIP | 37986224\|25083871 | Unspecified |
| mmu-miR-378a-3p | MIMAT0003151 | Arid4b | 94246 | HITS-CLIP | 37986224\|25083871 | Unspecified |

| | | | | | | |
|---|---|---|---|---|---|---|
| mmu-miR-378a-3p | MIMAT0003151 | Arih1 | 23806 | HITS-CLIP | 23142080 | Unspecified |
| mmu-miR-378a-3p | MIMAT0003151 | Arl14ep | 212772 | Microarrays | 37986224\|22855601 | Unspecified |
| mmu-miR-378a-3p | MIMAT0003151 | Arl4c | 320982 | HITS-CLIP | 37986224 | Unspecified |
| mmu-miR-378a-3p | MIMAT0003151 | Arl5a | 75423 | HITS-CLIP | 37986224 | Unspecified |
| mmu-miR-378a-3p | MIMAT0003151 | Arl8b | 67166 | HITS-CLIP | 25083871 | Unspecified |
| mmu-miR-378a-3p | MIMAT0003151 | Arnt | 11863 | HITS-CLIP | 37986224 | Unspecified |
| mmu-miR-378a-3p | MIMAT0003151 | Arrb1 | 109689 | HITS-CLIP | 23142080\|29167373 | Unspecified |
| mmu-miR-378a-3p | MIMAT0003151 | Arrdc3 | 105171 | HITS-CLIP | 25083871 | Unspecified |
| mmu-miR-378a-3p | MIMAT0003151 | Asah1 | 11886 | Microarrays | 37986224\|22855601 | Unspecified |
| mmu-miR-378a-3p | MIMAT0003151 | Asap1 | 13196 | HITS-CLIP | 25083871 | Unspecified |
| mmu-miR-378a-3p | MIMAT0003151 | Asb2 | 65256 | HITS-CLIP | 25753418 | Unspecified |
| mmu-miR-378a-3p | MIMAT0003151 | Ash1l | 192195 | HITS-CLIP | 37986224\|25083871\|29167373 | Unspecified |
| mmu-miR-378a-3p | MIMAT0003151 | Aspm | 12316 | HITS-CLIP | 37986224 | Unspecified |
| mmu-miR-378a-3p | MIMAT0003151 | Aspn | 66695 | Microarrays | 37986224 | Unspecified |
| mmu-miR-378a-3p | MIMAT0003151 | Asxl1 | 228790 | HITS-CLIP | 37986224 | Unspecified |
| mmu-miR-378a-3p | MIMAT0003151 | Atad2 | 70472 | Microarrays | 37986224\|22855601 | Unspecified |
| mmu-miR-378a-3p | MIMAT0003151 | Atad5 | 237877 | HITS-CLIP | 25083871\|23142080 | Unspecified |
| mmu-miR-378a-3p | MIMAT0003151 | Atf2 | 11909 | HITS-CLIP | 25083871\|23142080 | Unspecified |
| mmu-miR-378a-3p | MIMAT0003151 | Atf7ip | 54343 | HITS-CLIP | 25083871\|25753418\|23142080 | Unspecified |
| mmu-miR-378a-3p | MIMAT0003151 | Atg2b | 76559 | HITS-CLIP | 37986224 | Unspecified |
| mmu-miR-378a-3p | MIMAT0003151 | Atg9a | 245860 | HITS-CLIP | 37986224 | Unspecified |
| mmu-miR-378a-3p | MIMAT0003151 | Atl2 | 56298 | HITS-CLIP | 23597149 | Unspecified |
| mmu-miR-378a-3p | MIMAT0003151 | Atl3 | 109168 | HITS-CLIP | 25083871 | Unspecified |
| mmu-miR-378a-3p | MIMAT0003151 | Atox1 | 11927 | HITS-CLIP | 23597149 | Unspecified |
| mmu-miR-378a-3p | MIMAT0003151 | Atp10a | 11982 | Microarrays | 37986224\|22855601 | Unspecified |
| mmu-miR-378a-3p | MIMAT0003151 | Atp11b | 76295 | HITS-CLIP | 37986224\|23142080 | Unspecified |
| mmu-miR-378a-3p | MIMAT0003151 | Atp1a2 | 98660 | HITS-CLIP | 37986224 | Unspecified |
| mmu-miR-378a-3p | MIMAT0003151 | Atp2b1 | 67972 | HITS-CLIP | 25083871\|23142080 | Unspecified |
| mmu-miR-378a-3p | MIMAT0003151 | Atp5a1 | 11946 | HITS-CLIP | 25753418 | Unspecified |
| mmu-miR-378a-3p | MIMAT0003151 | Atp6v0a1 | 11975 | HITS-CLIP | 37986224 | Unspecified |
| mmu-miR-378a-3p | MIMAT0003151 | Atp6v0b | 114143 | HITS-CLIP | 37986224 | Unspecified |
| mmu-miR-378a-3p | MIMAT0003151 | Atp6v1a | 11964 | HITS-CLIP | 37986224 | Unspecified |
| mmu-miR-378a-3p | MIMAT0003151 | Atp8b2 | 54667 | HITS-CLIP | 37986224 | Unspecified |
| mmu-miR-378a-3p | MIMAT0003151 | Atpaf1 | 230649 | qCLASH | 31393953 | Unspecified |
| mmu-miR-378a-3p | MIMAT0003151 | Atrnl1 | 226255 | HITS-CLIP | 37986224 | Unspecified |
| mmu-miR-378a-3p | MIMAT0003151 | Atrx | 22589 | HITS-CLIP | 25083871\|23142080 | Unspecified |
| mmu-miR-378a-3p | MIMAT0003151 | Atxn1 | 20238 | HITS-CLIP | 37986224\|23142080 | Unspecified |
| mmu-miR-378a-3p | MIMAT0003151 | Atxn1l | 52335 | HITS-CLIP | 37986224 | Unspecified |
| mmu-miR-378a-3p | MIMAT0003151 | Atxn2 | 20239 | HITS-CLIP | 25083871 | Unspecified |

| | | | | | | | |
|---|---|---|---|---|---|---|---|
| mmu-miR-378a-3p | MIMAT0003151 | Atxn3 | 110616 | qCLASH | | 31393953 | Unspecified |
| mmu-miR-378a-3p | MIMAT0003151 | Atxn7 | 246103 | HITS-CLIP | 37986224\|25083871 | | Unspecified |
| mmu-miR-378a-3p | MIMAT0003151 | Aup1 | 11993 | HITS-CLIP | 25083871\|29167373 | | Unspecified |
| mmu-miR-378a-3p | MIMAT0003151 | Avl9 | 78937 | HITS-CLIP | | 37986224 | Unspecified |
| mmu-miR-378a-3p | MIMAT0003151 | Azi2 | 27215 | HITS-CLIP | | 37986224 | Unspecified |
| mmu-miR-378a-3p | MIMAT0003151 | Azin1 | 54375 | Microarrays | 37986224\|22855601 | | Unspecified |
| mmu-miR-378a-3p | MIMAT0003151 | B3galnt2 | 97884 | Microarrays | 37986224\|22855601 | | Unspecified |
| mmu-miR-378a-3p | MIMAT0003151 | B3glct | 381694 | qCLASH | | 31393953 | Unspecified |
| mmu-miR-378a-3p | MIMAT0003151 | B4galnt1 | 14421 | HITS-CLIP | | 29167373 | Unspecified |
| mmu-miR-378a-3p | MIMAT0003151 | BC048403 | 270802 | HITS-CLIP | | 37986224 | Unspecified |
| mmu-miR-378a-3p | MIMAT0003151 | Bach1 | 12013 | HITS-CLIP | | 37986224 | Unspecified |
| mmu-miR-378a-3p | MIMAT0003151 | Bach2 | 12014 | HITS-CLIP | 37986224\|23142080 | | Unspecified |
| mmu-miR-378a-3p | MIMAT0003151 | Bag3 | 29810 | HITS-CLIP | | 25753418 | Unspecified |
| mmu-miR-378a-3p | MIMAT0003151 | Baz1b | 22385 | HITS-CLIP | 37986224\|25083871 | | Unspecified |
| mmu-miR-378a-3p | MIMAT0003151 | Bbs12 | 241950 | HITS-CLIP | | 37986224 | Unspecified |
| mmu-miR-378a-3p | MIMAT0003151 | Bcar1 | 12927 | HITS-CLIP | | 37986224 | Unspecified |
| mmu-miR-378a-3p | MIMAT0003151 | Bcl2 | 12043 | HITS-CLIP | | 37986224 | Unspecified |
| mmu-miR-378a-3p | MIMAT0003151 | Bcl2l1 | 12048 | HITS-CLIP, Microarrays//Microarrays | 37986224\|22855601 | | Unspecified |
| mmu-miR-378a-3p | MIMAT0003151 | Bcl2l2 | 12050 | HITS-CLIP | | 25083871 | Unspecified |
| mmu-miR-378a-3p | MIMAT0003151 | Bcl7a | 77045 | HITS-CLIP | | 37986224 | Unspecified |
| mmu-miR-378a-3p | MIMAT0003151 | Bcl9 | 77578 | HITS-CLIP | | 37986224 | Unspecified |
| mmu-miR-378a-3p | MIMAT0003151 | Bcl9l | 80288 | HITS-CLIP | 23597149\|25083871 | | Unspecified |
| mmu-miR-378a-3p | MIMAT0003151 | Bclaf1 | 72567 | HITS-CLIP | 25083871\|23142080 | | Unspecified |
| mmu-miR-378a-3p | MIMAT0003151 | Bcr | 110279 | HITS-CLIP | | 37986224 | Unspecified |
| mmu-miR-378a-3p | MIMAT0003151 | Bfar | 67118 | HITS-CLIP | | 37986224 | Unspecified |
| mmu-miR-378a-3p | MIMAT0003151 | Bhlhe40 | 20893 | HITS-CLIP | | 37986224 | Unspecified |
| mmu-miR-378a-3p | MIMAT0003151 | Bicc1 | 83675 | HITS-CLIP | | 25083871 | Unspecified |
| mmu-miR-378a-3p | MIMAT0003151 | Bicra | 29998 | HITS-CLIP | | 25083871 | Unspecified |
| mmu-miR-378a-3p | MIMAT0003151 | Birc6 | 12211 | HITS-CLIP | | 25083871 | Unspecified |
| mmu-miR-378a-3p | MIMAT0003151 | Bmf | 171543 | HITS-CLIP, Microarrays | | 37986224 | Unspecified |
| mmu-miR-378a-3p | MIMAT0003151 | Bmi1 | 12151 | HITS-CLIP | | 37986224 | Unspecified |
| mmu-miR-378a-3p | MIMAT0003151 | Bmpr2 | 12168 | HITS-CLIP | 37986224\|23597149 | | Unspecified |
| mmu-miR-378a-3p | MIMAT0003151 | Bnc2 | 242509 | HITS-CLIP | | 37986224 | Unspecified |
| mmu-miR-378a-3p | MIMAT0003151 | Bnip2 | 12175 | HITS-CLIP | | 23142080 | Unspecified |
| mmu-miR-378a-3p | MIMAT0003151 | Boc | 117606 | Microarrays | | 37986224 | Unspecified |
| mmu-miR-378a-3p | MIMAT0003151 | Bod1l | 665775 | HITS-CLIP | 37986224\|25083871 | | Unspecified |
| mmu-miR-378a-3p | MIMAT0003151 | Bora | 77744 | HITS-CLIP | | 23142080 | Unspecified |
| mmu-miR-378a-3p | MIMAT0003151 | Bpnt2 | 54928 | HITS-CLIP | | 25753418 | Unspecified |
| mmu-miR-378a-3p | MIMAT0003151 | Bptf | 207165 | HITS-CLIP | 37986224\|25083871\|23142080\|29167 | | Unspecified |

| | | | | | | |
|---|---|---|---|---|---|---|
| mmu-miR-378a-3p | MIMAT0003151 | Braf | 109880 | HITS-CLIP | 25083871 | Unspecified |
| mmu-miR-378a-3p | MIMAT0003151 | Brd3 | 67382 | HITS-CLIP | 37986224\|23597149 | Unspecified |
| mmu-miR-378a-3p | MIMAT0003151 | Brd4 | 57261 | HITS-CLIP | 37986224 | Unspecified |
| mmu-miR-378a-3p | MIMAT0003151 | Brinp3 | 215378 | HITS-CLIP | 25083871 | Unspecified |
| mmu-miR-378a-3p | MIMAT0003151 | Brk1 | 101314 | HITS-CLIP | 37986224 | Unspecified |
| mmu-miR-378a-3p | MIMAT0003151 | Brme1 | 74666 | qCLASH | 31393953 | Unspecified |
| mmu-miR-378a-3p | MIMAT0003151 | Brms1l | 52592 | HITS-CLIP | 23597149\|23142080 | Unspecified |
| mmu-miR-378a-3p | MIMAT0003151 | Brpf1 | 78783 | HITS-CLIP | 37986224 | Unspecified |
| mmu-miR-378a-3p | MIMAT0003151 | Brwd1 | 93871 | HITS-CLIP | 23142080 | Unspecified |
| mmu-miR-378a-3p | MIMAT0003151 | Brwd3 | 382236 | HITS-CLIP | 37986224 | Unspecified |
| mmu-miR-378a-3p | MIMAT0003151 | Btaf1 | 107182 | Microarrays | 37986224\|22855601 | Unspecified |
| mmu-miR-378a-3p | MIMAT0003151 | Btg2 | 12227 | Microarrays | 37986224\|22855601 | Unspecified |
| mmu-miR-378a-3p | MIMAT0003151 | Bub1 | 12235 | HITS-CLIP | 23142080 | Unspecified |
| mmu-miR-378a-3p | MIMAT0003151 | Bzw1 | 66882 | HITS-CLIP | 37986224\|25753418 | Unspecified |
| mmu-miR-378a-3p | MIMAT0003151 | C1galt1 | 94192 | HITS-CLIP, Microarrays//Microarrays | 37986224\|22855601 | Unspecified |
| mmu-miR-378a-3p | MIMAT0003151 | C2cd5 | 74741 | HITS-CLIP | 37986224 | Unspecified |
| mmu-miR-378a-3p | MIMAT0003151 | C3 | 12266 | HITS-CLIP | 23597149 | Unspecified |
| mmu-miR-378a-3p | MIMAT0003151 | C4b | 12268 | HITS-CLIP | 37986224 | Unspecified |
| mmu-miR-378a-3p | MIMAT0003151 | C8a | 230558 | HITS-CLIP | 37986224 | Unspecified |
| mmu-miR-378a-3p | MIMAT0003151 | Cab39 | 12283 | HITS-CLIP | 37986224 | Unspecified |
| mmu-miR-378a-3p | MIMAT0003151 | Cables2 | 252966 | HITS-CLIP | 37986224 | Unspecified |
| mmu-miR-378a-3p | MIMAT0003151 | Cacna2d1 | 12293 | HITS-CLIP | 25083871 | Unspecified |
| mmu-miR-378a-3p | MIMAT0003151 | Calcrl | 54598 | Microarrays | 37986224 | Unspecified |
| mmu-miR-378a-3p | MIMAT0003151 | Camk2d | 108058 | HITS-CLIP | 37986224\|25083871 | Unspecified |
| mmu-miR-378a-3p | MIMAT0003151 | Camk2n1 | 66259 | HITS-CLIP | 37986224 | Unspecified |
| mmu-miR-378a-3p | MIMAT0003151 | Camsap1 | 227634 | HITS-CLIP | 37986224 | Unspecified |
| mmu-miR-378a-3p | MIMAT0003151 | Camsap2 | 67886 | Microarrays | 37986224\|22855601 | Unspecified |
| mmu-miR-378a-3p | MIMAT0003151 | Cand1 | 71902 | HITS-CLIP | 37986224\|25083871\|23142080 | Unspecified |
| mmu-miR-378a-3p | MIMAT0003151 | Cand2 | 67088 | qCLASH | 31393953 | Unspecified |
| mmu-miR-378a-3p | MIMAT0003151 | Canx | 12330 | HITS-CLIP | 23142080 | Unspecified |
| mmu-miR-378a-3p | MIMAT0003151 | Caprin1 | 53872 | HITS-CLIP | 25083871\|23142080 | Unspecified |
| mmu-miR-378a-3p | MIMAT0003151 | Casp3 | 12367 | HITS-CLIP | 37986224 | Unspecified |
| mmu-miR-378a-3p | MIMAT0003151 | Casp8 | 12370 | HITS-CLIP | 37986224 | Unspecified |
| mmu-miR-378a-3p | MIMAT0003151 | Casp8ap2 | 26885 | HITS-CLIP | 25083871 | Unspecified |
| mmu-miR-378a-3p | MIMAT0003151 | Cavin2 | 8436 | Microarrays | 22855601 | Unspecified |
| mmu-miR-378a-3p | MIMAT0003151 | Cbfa2t2 | 12396 | HITS-CLIP | 25753418 | Unspecified |
| mmu-miR-378a-3p | MIMAT0003151 | Cbl | 12402 | HITS-CLIP | 37986224\|23142080\|29167373 | Unspecified |
| mmu-miR-378a-3p | MIMAT0003151 | Cbr1 | 12408 | HITS-CLIP | 37986224 | Unspecified |
| mmu-miR-378a-3p | MIMAT0003151 | Cbs | 12411 | HITS-CLIP | 37986224 | Unspecified |

| | | | | | | |
|---|---|---|---|---|---|---|
| mmu-miR-378a-3p | MIMAT0003151 | Cbx5 | 12419 | HITS-CLIP | 23142080 | Unspecified |
| mmu-miR-378a-3p | MIMAT0003151 | Cbx6 | 494448 | HITS-CLIP | 37986224 | Unspecified |
| mmu-miR-378a-3p | MIMAT0003151 | Ccar1 | 67500 | HITS-CLIP | 23142080 | Unspecified |
| mmu-miR-378a-3p | MIMAT0003151 | Ccdc141 | 545428 | HITS-CLIP | 25083871 | Unspecified |
| mmu-miR-378a-3p | MIMAT0003151 | Ccdc32 | 269336 | HITS-CLIP | 37986224 | Unspecified |
| mmu-miR-378a-3p | MIMAT0003151 | Ccl2 | 20296 | Microarrays | 37986224\|22855601 | Unspecified |
| mmu-miR-378a-3p | MIMAT0003151 | Ccl5 | 20304 | Microarrays | 37986224 | Unspecified |
| mmu-miR-378a-3p | MIMAT0003151 | Ccn2 | 1490 | Microarrays//HITS-CLIP | 22855601\|25083871 | Unspecified |
| mmu-miR-378a-3p | MIMAT0003151 | Ccnd1 | 12443 | Microarrays | 37986224\|22855601 | Unspecified |
| mmu-miR-378a-3p | MIMAT0003151 | Ccnd2 | 12444 | HITS-CLIP | 37986224 | Unspecified |
| mmu-miR-378a-3p | MIMAT0003151 | Ccne1 | 12447 | Microarrays | 37986224\|22855601 | Unspecified |
| mmu-miR-378a-3p | MIMAT0003151 | Ccng1 | 12450 | Microarrays | 37986224\|22855601 | Unspecified |
| mmu-miR-378a-3p | MIMAT0003151 | Ccnj | 240665 | HITS-CLIP | 23142080 | Unspecified |
| mmu-miR-378a-3p | MIMAT0003151 | Ccnk | 12454 | HITS-CLIP | 25083871 | Unspecified |
| mmu-miR-378a-3p | MIMAT0003151 | Ccnl2 | 56036 | HITS-CLIP | 25083871\|25753418 | Unspecified |
| mmu-miR-378a-3p | MIMAT0003151 | Ccnt2 | 72949 | HITS-CLIP | 23142080 | Unspecified |
| mmu-miR-378a-3p | MIMAT0003151 | Cct7 | 12468 | HITS-CLIP | 25753418 | Unspecified |
| mmu-miR-378a-3p | MIMAT0003151 | Cd109 | 235505 | Microarrays | 37986224\|22855601 | Unspecified |
| mmu-miR-378a-3p | MIMAT0003151 | Cd302 | 66205 | HITS-CLIP | 23597149 | Unspecified |
| mmu-miR-378a-3p | MIMAT0003151 | Cd36 | 12491 | HITS-CLIP | 29167373 | Unspecified |
| mmu-miR-378a-3p | MIMAT0003151 | Cd44 | 12505 | Microarrays | 37986224\|22855601 | Unspecified |
| mmu-miR-378a-3p | MIMAT0003151 | Cdc123 | 98828 | HITS-CLIP | 23142080 | Unspecified |
| mmu-miR-378a-3p | MIMAT0003151 | Cdc20 | 107995 | HITS-CLIP | 37986224 | Unspecified |
| mmu-miR-378a-3p | MIMAT0003151 | Cdc27 | 217232 | HITS-CLIP | 23142080 | Unspecified |
| mmu-miR-378a-3p | MIMAT0003151 | Cdc37l1 | 67072 | HITS-CLIP | 25083871 | Unspecified |
| mmu-miR-378a-3p | MIMAT0003151 | Cdc73 | 214498 | HITS-CLIP | 37986224 | Unspecified |
| mmu-miR-378a-3p | MIMAT0003151 | Cdca4 | 71963 | HITS-CLIP | 23142080 | Unspecified |
| mmu-miR-378a-3p | MIMAT0003151 | Cdca7l | 217946 | Microarrays | 37986224\|22855601 | Unspecified |
| mmu-miR-378a-3p | MIMAT0003151 | Cdh13 | 12554 | HITS-CLIP | 37986224 | Unspecified |
| mmu-miR-378a-3p | MIMAT0003151 | Cdip1 | 66626 | HITS-CLIP | 37986224 | Unspecified |
| mmu-miR-378a-3p | MIMAT0003151 | Cdk12 | 69131 | HITS-CLIP | 37986224\|25083871\|23597149 | Unspecified |
| mmu-miR-378a-3p | MIMAT0003151 | Cdk13 | 69562 | qCLASH | 31393953 | Unspecified |
| mmu-miR-378a-3p | MIMAT0003151 | Cdk6 | 12571 | Microarrays//HITS-CLIP | 37986224\|23142080\|22855601\|29167 | Unspecified |
| mmu-miR-378a-3p | MIMAT0003151 | Cdk7 | 12572 | HITS-CLIP | 37986224 | Unspecified |
| mmu-miR-378a-3p | MIMAT0003151 | Cdk8 | 264064 | HITS-CLIP | 25083871\|23142080 | Unspecified |
| mmu-miR-378a-3p | MIMAT0003151 | Cdkn2aip | 70925 | HITS-CLIP | 29167373 | Unspecified |
| mmu-miR-378a-3p | MIMAT0003151 | Cdo1 | 12583 | Microarrays//HITS-CLIP | 37986224\|23597149 | Unspecified |
| mmu-miR-378a-3p | MIMAT0003151 | Cds2 | 110911 | qCLASH | 31393953 | Unspecified |
| mmu-miR-378a-3p | MIMAT0003151 | Cdt1 | 67177 | Microarrays | 37986224\|22855601 | Unspecified |

| | | | | | | | |
|---|---|---|---|---|---|---|---|
| mmu-miR-378a-3p | MIMAT0003151 | Celf1 | 13046 | HITS-CLIP | | 37986224 | Unspecified |
| mmu-miR-378a-3p | MIMAT0003151 | Celsr1 | 12614 | HITS-CLIP | | 23142080 | Unspecified |
| mmu-miR-378a-3p | MIMAT0003151 | Cemip | 80982 | Microarrays | 37986224\|22855601 | | Unspecified |
| mmu-miR-378a-3p | MIMAT0003151 | Cemip2 | 23670 | HITS-CLIP | | 25083871 | Unspecified |
| mmu-miR-378a-3p | MIMAT0003151 | Cenpf | 108000 | HITS-CLIP | | 37986224 | Unspecified |
| mmu-miR-378a-3p | MIMAT0003151 | Cep104 | 230967 | HITS-CLIP | | 37986224 | Unspecified |
| mmu-miR-378a-3p | MIMAT0003151 | Cep170 | 545389 | HITS-CLIP | | 29167373 | Unspecified |
| mmu-miR-378a-3p | MIMAT0003151 | Cep350 | 74081 | HITS-CLIP | | 25083871 | Unspecified |
| mmu-miR-378a-3p | MIMAT0003151 | Cep55 | 74107 | Microarrays | 37986224\|22855601 | | Unspecified |
| mmu-miR-378a-3p | MIMAT0003151 | Cep85 | 70012 | HITS-CLIP | | 23597149 | Unspecified |
| mmu-miR-378a-3p | MIMAT0003151 | Cers5 | 71949 | HITS-CLIP | 25083871\|23142080 | | Unspecified |
| mmu-miR-378a-3p | MIMAT0003151 | Cers6 | 241447 | HITS-CLIP | | 25083871 | Unspecified |
| mmu-miR-378a-3p | MIMAT0003151 | Ces3a | 382053 | HITS-CLIP | | 37986224 | Unspecified |
| mmu-miR-378a-3p | MIMAT0003151 | Ces3b | 13909 | HITS-CLIP | | 37986224 | Unspecified |
| mmu-miR-378a-3p | MIMAT0003151 | Cetn3 | 12626 | HITS-CLIP | | 37986224 | Unspecified |
| mmu-miR-378a-3p | MIMAT0003151 | Cfap36 | 112942 | HITS-CLIP | | 25083871 | Unspecified |
| mmu-miR-378a-3p | MIMAT0003151 | Cfh | 12628 | HITS-CLIP | | 37986224 | Unspecified |
| mmu-miR-378a-3p | MIMAT0003151 | Cflar | 12633 | HITS-CLIP | | 37986224 | Unspecified |
| mmu-miR-378a-3p | MIMAT0003151 | Chaf1b | 110749 | Microarrays | 37986224\|22855601 | | Unspecified |
| mmu-miR-378a-3p | MIMAT0003151 | Chchd3 | 66075 | HITS-CLIP | | 23142080 | Unspecified |
| mmu-miR-378a-3p | MIMAT0003151 | Chd1 | 12648 | HITS-CLIP | | 25083871 | Unspecified |
| mmu-miR-378a-3p | MIMAT0003151 | Chd2 | 244059 | HITS-CLIP | | 25083871 | Unspecified |
| mmu-miR-378a-3p | MIMAT0003151 | Chd3 | 216848 | HITS-CLIP | | 25083871 | Unspecified |
| mmu-miR-378a-3p | MIMAT0003151 | Chd7 | 320790 | HITS-CLIP | | 23597149 | Unspecified |
| mmu-miR-378a-3p | MIMAT0003151 | Chd9 | 109151 | HITS-CLIP | | 25083871 | Unspecified |
| mmu-miR-378a-3p | MIMAT0003151 | Chek1 | 12649 | Microarrays | 37986224\|22855601 | | Unspecified |
| mmu-miR-378a-3p | MIMAT0003151 | Chic1 | 12212 | HITS-CLIP | | 37986224 | Unspecified |
| mmu-miR-378a-3p | MIMAT0003151 | Chka | 12660 | Microarrays | 37986224\|22855601 | | Unspecified |
| mmu-miR-378a-3p | MIMAT0003151 | Chkb | 12651 | HITS-CLIP | | 37986224 | Unspecified |
| mmu-miR-378a-3p | MIMAT0003151 | Chst2 | 54371 | HITS-CLIP | | 37986224 | Unspecified |
| mmu-miR-378a-3p | MIMAT0003151 | Chtf8 | 214987 | HITS-CLIP | 37986224\|23142080 | | Unspecified |
| mmu-miR-378a-3p | MIMAT0003151 | Ckap4 | 216197 | HITS-CLIP | | 37986224 | Unspecified |
| mmu-miR-378a-3p | MIMAT0003151 | Clasp1 | 76707 | HITS-CLIP | 23142080\|25083871 | | Unspecified |
| mmu-miR-378a-3p | MIMAT0003151 | Clasp2 | 76499 | HITS-CLIP | | 23142080 | Unspecified |
| mmu-miR-378a-3p | MIMAT0003151 | Clcn2 | 12724 | HITS-CLIP | | 37986224 | Unspecified |
| mmu-miR-378a-3p | MIMAT0003151 | Clcn3 | 12725 | HITS-CLIP | | 25083871 | Unspecified |
| mmu-miR-378a-3p | MIMAT0003151 | Cldn12 | 64945 | Microarrays | 37986224\|22855601 | | Unspecified |
| mmu-miR-378a-3p | MIMAT0003151 | Cldnd1 | 224250 | HITS-CLIP | | 25083871 | Unspecified |
| mmu-miR-378a-3p | MIMAT0003151 | Clint1 | 216705 | HITS-CLIP | 25083871\|23142080 | | Unspecified |

| | | | | | | | |
|---|---|---|---|---|---|---|---|
| mmu-miR-378a-3p | MIMAT0003151 | Clk4 | 12750 | HITS-CLIP | | 37986224 | Unspecified |
| mmu-miR-378a-3p | MIMAT0003151 | Clstn1 | 65945 | HITS-CLIP | 37986224\|25083871 | | Unspecified |
| mmu-miR-378a-3p | MIMAT0003151 | Cltc | 67300 | HITS-CLIP | | 23142080 | Unspecified |
| mmu-miR-378a-3p | MIMAT0003151 | Cmbl | 69574 | HITS-CLIP | | 37986224 | Unspecified |
| mmu-miR-378a-3p | MIMAT0003151 | Cmtr1 | 74157 | HITS-CLIP | | 37986224 | Unspecified |
| mmu-miR-378a-3p | MIMAT0003151 | Cnot1 | 234594 | HITS-CLIP | | 23597149 | Unspecified |
| mmu-miR-378a-3p | MIMAT0003151 | Cnot11 | 52846 | HITS-CLIP | | 37986224 | Unspecified |
| mmu-miR-378a-3p | MIMAT0003151 | Cnot4 | 53621 | HITS-CLIP | | 23142080 | Unspecified |
| mmu-miR-378a-3p | MIMAT0003151 | Cnot6 | 104625 | HITS-CLIP | 37986224\|23142080 | | Unspecified |
| mmu-miR-378a-3p | MIMAT0003151 | Cntln | 338349 | HITS-CLIP | | 37986224 | Unspecified |
| mmu-miR-378a-3p | MIMAT0003151 | Cntrl | 26920 | HITS-CLIP | | 23142080 | Unspecified |
| mmu-miR-378a-3p | MIMAT0003151 | Coasy | 71743 | HITS-CLIP | | 37986224 | Unspecified |
| mmu-miR-378a-3p | MIMAT0003151 | Cobll1 | 319876 | Microarrays | 37986224\|22855601 | | Unspecified |
| mmu-miR-378a-3p | MIMAT0003151 | Cog3 | 338337 | HITS-CLIP | | 37986224 | Unspecified |
| mmu-miR-378a-3p | MIMAT0003151 | Col12a1 | 12816 | HITS-CLIP | | 25083871 | Unspecified |
| mmu-miR-378a-3p | MIMAT0003151 | Col18a1 | 12822 | HITS-CLIP | | 37986224 | Unspecified |
| mmu-miR-378a-3p | MIMAT0003151 | Col1a2 | 12843 | HITS-CLIP | | 25083871 | Unspecified |
| mmu-miR-378a-3p | MIMAT0003151 | Col5a3 | 53867 | HITS-CLIP | | 37986224 | Unspecified |
| mmu-miR-378a-3p | MIMAT0003151 | Col6a3 | 12835 | HITS-CLIP | 37986224\|25083871 | | Unspecified |
| mmu-miR-378a-3p | MIMAT0003151 | Commd9 | 76501 | HITS-CLIP | | 37986224 | Unspecified |
| mmu-miR-378a-3p | MIMAT0003151 | Copa | 12847 | HITS-CLIP | | 23597149 | Unspecified |
| mmu-miR-378a-3p | MIMAT0003151 | Coq10a | 210582 | HITS-CLIP | | 25753418 | Unspecified |
| mmu-miR-378a-3p | MIMAT0003151 | Coq10b | 67876 | HITS-CLIP | 37986224\|25083871 | | Unspecified |
| mmu-miR-378a-3p | MIMAT0003151 | Coq8a | 56997 | HITS-CLIP | | 23597149 | Unspecified |
| mmu-miR-378a-3p | MIMAT0003151 | Cox10 | 70383 | HITS-CLIP | | 37986224 | Unspecified |
| mmu-miR-378a-3p | MIMAT0003151 | Cp | 12870 | HITS-CLIP | | 23597149 | Unspecified |
| mmu-miR-378a-3p | MIMAT0003151 | Cpeb2 | 231207 | HITS-CLIP | | 37986224 | Unspecified |
| mmu-miR-378a-3p | MIMAT0003151 | Cpeb4 | 67579 | HITS-CLIP | 25083871\|29167373 | | Unspecified |
| mmu-miR-378a-3p | MIMAT0003151 | Cps1 | 227231 | HITS-CLIP | | 23597149 | Unspecified |
| mmu-miR-378a-3p | MIMAT0003151 | Cpsf3 | 54451 | HITS-CLIP | | 29167373 | Unspecified |
| mmu-miR-378a-3p | MIMAT0003151 | Cpsf4 | 54188 | HITS-CLIP | | 23597149 | Unspecified |
| mmu-miR-378a-3p | MIMAT0003151 | Cramp1l | 57354 | HITS-CLIP | | 25083871 | Unspecified |
| mmu-miR-378a-3p | MIMAT0003151 | Crebl2 | 232430 | HITS-CLIP | 37986224\|23142080 | | Unspecified |
| mmu-miR-378a-3p | MIMAT0003151 | Crebrf | 77128 | HITS-CLIP | | 37986224 | Unspecified |
| mmu-miR-378a-3p | MIMAT0003151 | Crim1 | 50766 | Microarrays | 37986224\|22855601 | | Unspecified |
| mmu-miR-378a-3p | MIMAT0003151 | Crk | 12928 | qCLASH | | 31393953 | Unspecified |
| mmu-miR-378a-3p | MIMAT0003151 | Crkl | 12929 | HITS-CLIP | 23142080\|25083871 | | Unspecified |
| mmu-miR-378a-3p | MIMAT0003151 | Crnkl1 | 66877 | HITS-CLIP | | 37986224 | Unspecified |
| mmu-miR-378a-3p | MIMAT0003151 | Cs | 12974 | HITS-CLIP | | 23142080 | Unspecified |

| | | | | | | |
|---|---|---|---|---|---|---|
| mmu-miR-378a-3p | MIMAT0003151 | Csf1 | 12977 | Microarrays | 37986224\|22855601 | Unspecified |
| mmu-miR-378a-3p | MIMAT0003151 | Csgalnact2 | 78752 | HITS-CLIP | 37986224 | Unspecified |
| mmu-miR-378a-3p | MIMAT0003151 | Csk | 12988 | HITS-CLIP | 37986224\|23597149 | Unspecified |
| mmu-miR-378a-3p | MIMAT0003151 | Csnk1a1 | 93687 | HITS-CLIP | 37986224 | Unspecified |
| mmu-miR-378a-3p | MIMAT0003151 | Csnk1g1 | 214897 | HITS-CLIP | 37986224 | Unspecified |
| mmu-miR-378a-3p | MIMAT0003151 | Csnk1g3 | 70425 | HITS-CLIP | 37986224\|25083871\|23142080 | Unspecified |
| mmu-miR-378a-3p | MIMAT0003151 | Cspg4 | 121021 | Microarrays | 37986224\|22855601 | Unspecified |
| mmu-miR-378a-3p | MIMAT0003151 | Ctcf | 13018 | HITS-CLIP | 37986224 | Unspecified |
| mmu-miR-378a-3p | MIMAT0003151 | Ctdnep1 | 67181 | HITS-CLIP | 25083871 | Unspecified |
| mmu-miR-378a-3p | MIMAT0003151 | Ctdspl2 | 329506 | HITS-CLIP | 25083871\|23142080 | Unspecified |
| mmu-miR-378a-3p | MIMAT0003151 | Ctnna1 | 12385 | HITS-CLIP | 37986224 | Unspecified |
| mmu-miR-378a-3p | MIMAT0003151 | Ctnnb1 | 12387 | HITS-CLIP | 37986224\|23142080 | Unspecified |
| mmu-miR-378a-3p | MIMAT0003151 | Ctnnd1 | 12388 | HITS-CLIP | 25083871 | Unspecified |
| mmu-miR-378a-3p | MIMAT0003151 | Ctps | 51797 | Microarrays | 37986224\|22855601 | Unspecified |
| mmu-miR-378a-3p | MIMAT0003151 | Ctr9 | 22083 | HITS-CLIP | 37986224 | Unspecified |
| mmu-miR-378a-3p | MIMAT0003151 | Ctsb | 13030 | HITS-CLIP | 37986224 | Unspecified |
| mmu-miR-378a-3p | MIMAT0003151 | Ctsd | 13033 | HITS-CLIP | 37986224 | Unspecified |
| mmu-miR-378a-3p | MIMAT0003151 | Ctsz | 64138 | HITS-CLIP | 37986224 | Unspecified |
| mmu-miR-378a-3p | MIMAT0003151 | Cul1 | 26965 | HITS-CLIP | 25083871\|23142080 | Unspecified |
| mmu-miR-378a-3p | MIMAT0003151 | Cul2 | 71745 | HITS-CLIP | 23142080 | Unspecified |
| mmu-miR-378a-3p | MIMAT0003151 | Cul3 | 26554 | HITS-CLIP | 23142080 | Unspecified |
| mmu-miR-378a-3p | MIMAT0003151 | Cux1 | 13047 | HITS-CLIP | 37986224 | Unspecified |
| mmu-miR-378a-3p | MIMAT0003151 | Cwc22 | 80744 | HITS-CLIP | 23142080 | Unspecified |
| mmu-miR-378a-3p | MIMAT0003151 | Cwf19l2 | 244672 | HITS-CLIP | 37986224 | Unspecified |
| mmu-miR-378a-3p | MIMAT0003151 | Cyb5a | 109672 | Microarrays | 37986224 | Unspecified |
| mmu-miR-378a-3p | MIMAT0003151 | Cyb5b | 66427 | HITS-CLIP | 25083871\|23142080 | Unspecified |
| mmu-miR-378a-3p | MIMAT0003151 | Cyc1 | 66445 | HITS-CLIP | 37986224 | Unspecified |
| mmu-miR-378a-3p | MIMAT0003151 | Cyp2j5 | 13109 | HITS-CLIP | 23597149 | Unspecified |
| mmu-miR-378a-3p | MIMAT0003151 | Cyp4f14 | 64385 | HITS-CLIP | 23597149 | Unspecified |
| mmu-miR-378a-3p | MIMAT0003151 | Cyp4v3 | 102294 | HITS-CLIP | 37986224 | Unspecified |
| mmu-miR-378a-3p | MIMAT0003151 | Cysltr1 | 58861 | HITS-CLIP | 23142080 | Unspecified |
| mmu-miR-378a-3p | MIMAT0003151 | Cystm1 | 66060 | HITS-CLIP | 23142080 | Unspecified |
| mmu-miR-378a-3p | MIMAT0003151 | Cyth3 | 19159 | HITS-CLIP | 25083871 | Unspecified |
| mmu-miR-378a-3p | MIMAT0003151 | D17Wsu92e | 224647 | HITS-CLIP | 29167373 | Unspecified |
| mmu-miR-378a-3p | MIMAT0003151 | D5Ertd579e | 320661 | HITS-CLIP | 25083871 | Unspecified |
| mmu-miR-378a-3p | MIMAT0003151 | Daam2 | 76441 | HITS-CLIP | 37986224 | Unspecified |
| mmu-miR-378a-3p | MIMAT0003151 | Dab2 | 13132 | HITS-CLIP | 37986224\|25083871\|23597149\|29167 | Unspecified |
| mmu-miR-378a-3p | MIMAT0003151 | Dag1 | 13138 | HITS-CLIP | 25753418 | Unspecified |
| mmu-miR-378a-3p | MIMAT0003151 | Dars | 226414 | HITS-CLIP | 25083871 | Unspecified |

| | | | | | | |
|---|---|---|---|---|---|---|
| mmu-miR-378a-3p | MIMAT0003151 | Dazap2 | 23994 | HITS-CLIP | 37986224\|23142080\|25083871 | Unspecified |
| mmu-miR-378a-3p | MIMAT0003151 | Dbf4 | 27214 | HITS-CLIP | 25083871 | Unspecified |
| mmu-miR-378a-3p | MIMAT0003151 | Dcaf12 | 68970 | HITS-CLIP | 23142080 | Unspecified |
| mmu-miR-378a-3p | MIMAT0003151 | Dcaf5 | 320808 | HITS-CLIP | 37986224 | Unspecified |
| mmu-miR-378a-3p | MIMAT0003151 | Dcaf6 | 74106 | HITS-CLIP | 25083871 | Unspecified |
| mmu-miR-378a-3p | MIMAT0003151 | Dcaf7 | 71833 | HITS-CLIP | 37986224 | Unspecified |
| mmu-miR-378a-3p | MIMAT0003151 | Dcbld2 | 73379 | HITS-CLIP | 37986224\|25083871 | Unspecified |
| mmu-miR-378a-3p | MIMAT0003151 | Dclk1 | 13175 | HITS-CLIP | 37986224 | Unspecified |
| mmu-miR-378a-3p | MIMAT0003151 | Dclre1b | 140917 | HITS-CLIP | 37986224\|23597149 | Unspecified |
| mmu-miR-378a-3p | MIMAT0003151 | Dcp2 | 70640 | HITS-CLIP | 37986224\|23597149\|25083871\|29167 | Unspecified |
| mmu-miR-378a-3p | MIMAT0003151 | Dctn4 | 67665 | HITS-CLIP | 25753418 | Unspecified |
| mmu-miR-378a-3p | MIMAT0003151 | Dcun1d3 | 233805 | HITS-CLIP | 37986224 | Unspecified |
| mmu-miR-378a-3p | MIMAT0003151 | Ddah1 | 69219 | HITS-CLIP | 37986224 | Unspecified |
| mmu-miR-378a-3p | MIMAT0003151 | Ddhd1 | 114874 | qCLASH | 31393953 | Unspecified |
| mmu-miR-378a-3p | MIMAT0003151 | Ddias | 220042 | Microarrays | 22855601 | Unspecified |
| mmu-miR-378a-3p | MIMAT0003151 | Ddr2 | 18214 | HITS-CLIP | 37986224\|25083871 | Unspecified |
| mmu-miR-378a-3p | MIMAT0003151 | Ddx17 | 67040 | HITS-CLIP | 23142080 | Unspecified |
| mmu-miR-378a-3p | MIMAT0003151 | Ddx19b | 234733 | HITS-CLIP | 37986224 | Unspecified |
| mmu-miR-378a-3p | MIMAT0003151 | Ddx21 | 56200 | Microarrays | 37986224\|22855601 | Unspecified |
| mmu-miR-378a-3p | MIMAT0003151 | Ddx27 | 228889 | Microarrays | 37986224\|22855601 | Unspecified |
| mmu-miR-378a-3p | MIMAT0003151 | Ddx3x | 13205 | HITS-CLIP | 37986224\|23142080\|25083871 | Unspecified |
| mmu-miR-378a-3p | MIMAT0003151 | Ddx3y | 26900 | HITS-CLIP | 37986224\|25753418\|23142080 | Unspecified |
| mmu-miR-378a-3p | MIMAT0003151 | Ddx5 | 13207 | HITS-CLIP | 37986224 | Unspecified |
| mmu-miR-378a-3p | MIMAT0003151 | Ddx6 | 13209 | HITS-CLIP | 23142080 | Unspecified |
| mmu-miR-378a-3p | MIMAT0003151 | Dek | 110052 | HITS-CLIP//qCLASH | 23597149\|31393953\|23142080 | Unspecified |
| mmu-miR-378a-3p | MIMAT0003151 | Dele1 | 9812 | HITS-CLIP | 25753418 | Unspecified |
| mmu-miR-378a-3p | MIMAT0003151 | Dennd1b | 329260 | HITS-CLIP | 37986224\|25083871 | Unspecified |
| mmu-miR-378a-3p | MIMAT0003151 | Dennd4b | 229541 | HITS-CLIP | 37986224 | Unspecified |
| mmu-miR-378a-3p | MIMAT0003151 | Derl2 | 116891 | HITS-CLIP | 25083871 | Unspecified |
| mmu-miR-378a-3p | MIMAT0003151 | Des | 13346 | HITS-CLIP | 25753418 | Unspecified |
| mmu-miR-378a-3p | MIMAT0003151 | Desi2 | 78825 | HITS-CLIP | 25083871\|29167373 | Unspecified |
| mmu-miR-378a-3p | MIMAT0003151 | Dhps | 330817 | HITS-CLIP | 25083871 | Unspecified |
| mmu-miR-378a-3p | MIMAT0003151 | Dhx36 | 72162 | HITS-CLIP | 25083871 | Unspecified |
| mmu-miR-378a-3p | MIMAT0003151 | Dhx9 | 13211 | HITS-CLIP | 23597149\|23142080 | Unspecified |
| mmu-miR-378a-3p | MIMAT0003151 | Diaph1 | 1729 | HITS-CLIP | 23597149 | Unspecified |
| mmu-miR-378a-3p | MIMAT0003151 | Dido1 | 23856 | HITS-CLIP | 25083871 | Unspecified |
| mmu-miR-378a-3p | MIMAT0003151 | Dip2c | 208440 | HITS-CLIP | 37986224 | Unspecified |
| mmu-miR-378a-3p | MIMAT0003151 | Dkc1 | 245474 | HITS-CLIP//Microarrays | 21258322\|22855601 | Unspecified |
| mmu-miR-378a-3p | MIMAT0003151 | Dlat | 235339 | HITS-CLIP | 37986224 | Unspecified |

| | | | | | | |
|---|---|---|---|---|---|---|
| mmu-miR-378a-3p | MIMAT0003151 | Dlgap4 | 228836 | HITS-CLIP | 37986224 | Unspecified |
| mmu-miR-378a-3p | MIMAT0003151 | Dlst | 78920 | HITS-CLIP | 37986224 | Unspecified |
| mmu-miR-378a-3p | MIMAT0003151 | Dmgdh | 74129 | HITS-CLIP | 37986224\|23597149 | Unspecified |
| mmu-miR-378a-3p | MIMAT0003151 | Dnaja2 | 56445 | HITS-CLIP | 23597149\|25083871\|23142080 | Unspecified |
| mmu-miR-378a-3p | MIMAT0003151 | Dnajb14 | 70604 | HITS-CLIP | 37986224 | Unspecified |
| mmu-miR-378a-3p | MIMAT0003151 | Dnajb5 | 56323 | HITS-CLIP | 25753418 | Unspecified |
| mmu-miR-378a-3p | MIMAT0003151 | Dnajc1 | 13418 | HITS-CLIP | 37986224 | Unspecified |
| mmu-miR-378a-3p | MIMAT0003151 | Dnajc5 | 13002 | HITS-CLIP | 37986224\|25083871\|23597149\|23142 | Unspecified |
| mmu-miR-378a-3p | MIMAT0003151 | Dnal1 | 105000 | HITS-CLIP | 37986224 | Unspecified |
| mmu-miR-378a-3p | MIMAT0003151 | Dnm2 | 13430 | HITS-CLIP | 25083871\|23142080 | Unspecified |
| mmu-miR-378a-3p | MIMAT0003151 | Dnmbp | 71972 | HITS-CLIP | 37986224 | Unspecified |
| mmu-miR-378a-3p | MIMAT0003151 | Dock10 | 210293 | HITS-CLIP | 29167373 | Unspecified |
| mmu-miR-378a-3p | MIMAT0003151 | Dock8 | 76088 | HITS-CLIP | 29167373 | Unspecified |
| mmu-miR-378a-3p | MIMAT0003151 | Dot1l | 208266 | HITS-CLIP | 37986224 | Unspecified |
| mmu-miR-378a-3p | MIMAT0003151 | Dpp8 | 74388 | HITS-CLIP | 25083871\|29167373 | Unspecified |
| mmu-miR-378a-3p | MIMAT0003151 | Dpt | 56429 | Microarrays | 37986224 | Unspecified |
| mmu-miR-378a-3p | MIMAT0003151 | Dpy19l3 | 233115 | HITS-CLIP | 37986224 | Unspecified |
| mmu-miR-378a-3p | MIMAT0003151 | Dpyd | 99586 | HITS-CLIP | 37986224 | Unspecified |
| mmu-miR-378a-3p | MIMAT0003151 | Dr1 | 13486 | HITS-CLIP | 23142080 | Unspecified |
| mmu-miR-378a-3p | MIMAT0003151 | Dsc2 | 13506 | HITS-CLIP | 37986224 | Unspecified |
| mmu-miR-378a-3p | MIMAT0003151 | Dsel | 319901 | HITS-CLIP | 37986224 | Unspecified |
| mmu-miR-378a-3p | MIMAT0003151 | Dst | 13518 | HITS-CLIP | 25083871 | Unspecified |
| mmu-miR-378a-3p | MIMAT0003151 | Dstn | 56431 | HITS-CLIP | 23597149 | Unspecified |
| mmu-miR-378a-3p | MIMAT0003151 | Dtl | 76843 | Microarrays | 37986224\|22855601 | Unspecified |
| mmu-miR-378a-3p | MIMAT0003151 | Dusp22 | 105352 | HITS-CLIP | 37986224\|23142080 | Unspecified |
| mmu-miR-378a-3p | MIMAT0003151 | Dusp4 | 319520 | Microarrays | 37986224\|22855601 | Unspecified |
| mmu-miR-378a-3p | MIMAT0003151 | Dusp6 | 67603 | Microarrays | 37986224\|22855601 | Unspecified |
| mmu-miR-378a-3p | MIMAT0003151 | Dync1h1 | 13424 | HITS-CLIP | 37986224\|23597149\|23142080\|25083 | Unspecified |
| mmu-miR-378a-3p | MIMAT0003151 | Dync1li2 | 234663 | HITS-CLIP | 23142080 | Unspecified |
| mmu-miR-378a-3p | MIMAT0003151 | Dyrk1a | 13548 | HITS-CLIP | 37986224 | Unspecified |
| mmu-miR-378a-3p | MIMAT0003151 | Dyrk2 | 69181 | HITS-CLIP | 37986224\|25083871 | Unspecified |
| mmu-miR-378a-3p | MIMAT0003151 | E2f2 | 242705 | HITS-CLIP | 37986224\|23142080 | Unspecified |
| mmu-miR-378a-3p | MIMAT0003151 | E2f3 | 13557 | HITS-CLIP | 23142080 | Unspecified |
| mmu-miR-378a-3p | MIMAT0003151 | E2f4 | 104394 | Microarrays | 37986224\|22855601 | Unspecified |
| mmu-miR-378a-3p | MIMAT0003151 | Ect2 | 13605 | HITS-CLIP | 37986224 | Unspecified |
| mmu-miR-378a-3p | MIMAT0003151 | Eda2r | 245527 | Microarrays//HITS-CLIP | 37986224\|25083871\|22855601 | Unspecified |
| mmu-miR-378a-3p | MIMAT0003151 | Edem1 | 192193 | HITS-CLIP | 37986224 | Unspecified |
| mmu-miR-378a-3p | MIMAT0003151 | Edem3 | 66967 | HITS-CLIP | 25083871 | Unspecified |
| mmu-miR-378a-3p | MIMAT0003151 | Edil3 | 13612 | HITS-CLIP | 37986224 | Unspecified |

| | | | | | | |
|---|---|---|---|---|---|---|
| mmu-miR-378a-3p | MIMAT0003151 | Eea1 | 216238 | Microarrays//HITS-CLIP | 37986224\|25083871\|22855601 | Unspecified |
| mmu-miR-378a-3p | MIMAT0003151 | Eef1a1 | 13627 | HITS-CLIP | 37986224 | Unspecified |
| mmu-miR-378a-3p | MIMAT0003151 | Eef2 | 13629 | HITS-CLIP | 37986224\|25083871\|23597149\|23142 | Unspecified |
| mmu-miR-378a-3p | MIMAT0003151 | Eef2k | 13631 | HITS-CLIP | 37986224 | Unspecified |
| mmu-miR-378a-3p | MIMAT0003151 | Eepd1 | 67484 | HITS-CLIP | 29167373 | Unspecified |
| mmu-miR-378a-3p | MIMAT0003151 | Efcab14 | 230648 | qCLASH | 31393953 | Unspecified |
| mmu-miR-378a-3p | MIMAT0003151 | Efl1 | 1942 | qCLASH | 31393953 | Unspecified |
| mmu-miR-378a-3p | MIMAT0003151 | Efnb2 | 13642 | Microarrays | 37986224\|22855601 | Unspecified |
| mmu-miR-378a-3p | MIMAT0003151 | Efr3a | 76740 | HITS-CLIP | 25083871 | Unspecified |
| mmu-miR-378a-3p | MIMAT0003151 | Eftud2 | 20624 | HITS-CLIP | 23597149 | Unspecified |
| mmu-miR-378a-3p | MIMAT0003151 | Egr1 | 13653 | HITS-CLIP | 37986224 | Unspecified |
| mmu-miR-378a-3p | MIMAT0003151 | Egr2 | 13654 | HITS-CLIP | 23142080 | Unspecified |
| mmu-miR-378a-3p | MIMAT0003151 | Eif1ad | 69860 | Microarrays | 37986224\|22855601 | Unspecified |
| mmu-miR-378a-3p | MIMAT0003151 | Eif2ak2 | 19106 | HITS-CLIP | 37986224 | Unspecified |
| mmu-miR-378a-3p | MIMAT0003151 | Eif3a | 13669 | HITS-CLIP | 25083871\|25753418 | Unspecified |
| mmu-miR-378a-3p | MIMAT0003151 | Eif3d | 55944 | HITS-CLIP | 37986224 | Unspecified |
| mmu-miR-378a-3p | MIMAT0003151 | Eif3l | 223691 | qCLASH | 31393953 | Unspecified |
| mmu-miR-378a-3p | MIMAT0003151 | Eif4a1 | 13681 | qCLASH | 31393953 | Unspecified |
| mmu-miR-378a-3p | MIMAT0003151 | Eif4b | 75705 | HITS-CLIP | 23597149\|29167373 | Unspecified |
| mmu-miR-378a-3p | MIMAT0003151 | Eif4e2 | 26987 | HITS-CLIP | 37986224 | Unspecified |
| mmu-miR-378a-3p | MIMAT0003151 | Eif4enif1 | 74203 | qCLASH | 31393953 | Unspecified |
| mmu-miR-378a-3p | MIMAT0003151 | Eif4g1 | 208643 | HITS-CLIP, Microarrays//Microarrays | 37986224\|22855601 | Unspecified |
| mmu-miR-378a-3p | MIMAT0003151 | Eif4g3 | 230861 | HITS-CLIP | 37986224\|25083871\|23142080 | Unspecified |
| mmu-miR-378a-3p | MIMAT0003151 | Eif4h | 22384 | HITS-CLIP | 23142080\|23597149\|25083871 | Unspecified |
| mmu-miR-378a-3p | MIMAT0003151 | Eif5b | 226982 | HITS-CLIP | 37986224 | Unspecified |
| mmu-miR-378a-3p | MIMAT0003151 | Elavl1 | 15568 | HITS-CLIP | 23142080 | Unspecified |
| mmu-miR-378a-3p | MIMAT0003151 | Elk3 | 13713 | HITS-CLIP | 23142080 | Unspecified |
| mmu-miR-378a-3p | MIMAT0003151 | Elk4 | 13714 | HITS-CLIP | 37986224 | Unspecified |
| mmu-miR-378a-3p | MIMAT0003151 | Ell2 | 192657 | HITS-CLIP | 29167373 | Unspecified |
| mmu-miR-378a-3p | MIMAT0003151 | Elmsan1 | 238317 | HITS-CLIP | 29167373 | Unspecified |
| mmu-miR-378a-3p | MIMAT0003151 | Elovl2 | 54326 | HITS-CLIP | 37986224 | Unspecified |
| mmu-miR-378a-3p | MIMAT0003151 | Emc1 | 230866 | HITS-CLIP | 37986224 | Unspecified |
| mmu-miR-378a-3p | MIMAT0003151 | Emc3 | 66087 | HITS-CLIP | 37986224\|25083871 | Unspecified |
| mmu-miR-378a-3p | MIMAT0003151 | Eml4 | 78798 | HITS-CLIP | 37986224\|23142080\|29167373 | Unspecified |
| mmu-miR-378a-3p | MIMAT0003151 | Enah | 13800 | HITS-CLIP | 37986224 | Unspecified |
| mmu-miR-378a-3p | MIMAT0003151 | Enc1 | 13803 | HITS-CLIP | 37986224 | Unspecified |
| mmu-miR-378a-3p | MIMAT0003151 | Eng | 13805 | HITS-CLIP | 23597149 | Unspecified |
| mmu-miR-378a-3p | MIMAT0003151 | Enpp4 | 224794 | HITS-CLIP | 37986224 | Unspecified |
| mmu-miR-378a-3p | MIMAT0003151 | Ep300 | 328572 | HITS-CLIP | 37986224\|25083871 | Unspecified |

| | | | | | | | |
|---|---|---|---|---|---|---|---|
| mmu-miR-378a-3p | MIMAT0003151 | Epc2 | 227867 | HITS-CLIP | | 25083871 | Unspecified |
| mmu-miR-378a-3p | MIMAT0003151 | Epha2 | 13836 | Microarrays | 37986224\|22855601 | | Unspecified |
| mmu-miR-378a-3p | MIMAT0003151 | Epha4 | 13838 | HITS-CLIP | | 25753418 | Unspecified |
| mmu-miR-378a-3p | MIMAT0003151 | Ephb3 | 13845 | HITS-CLIP | | 37986224 | Unspecified |
| mmu-miR-378a-3p | MIMAT0003151 | Ephx1 | 13849 | Microarrays | 37986224\|22855601 | | Unspecified |
| mmu-miR-378a-3p | MIMAT0003151 | Ephx2 | 13850 | HITS-CLIP | | 23597149 | Unspecified |
| mmu-miR-378a-3p | MIMAT0003151 | Epn1 | 13854 | HITS-CLIP | | 37986224 | Unspecified |
| mmu-miR-378a-3p | MIMAT0003151 | Eprs | 107508 | HITS-CLIP | | 25083871 | Unspecified |
| mmu-miR-378a-3p | MIMAT0003151 | Ercc6l | 236930 | qCLASH | | 31393953 | Unspecified |
| mmu-miR-378a-3p | MIMAT0003151 | Ereg | 13874 | HITS-CLIP, Microarrays//Microarrays | 37986224\|22855601 | | Unspecified |
| mmu-miR-378a-3p | MIMAT0003151 | Ergic2 | 67456 | HITS-CLIP | | 37986224 | Unspecified |
| mmu-miR-378a-3p | MIMAT0003151 | Ero1a | 30001 | Microarrays | | 22855601 | Unspecified |
| mmu-miR-378a-3p | MIMAT0003151 | Ero1l | 50527 | Microarrays | | 37986224 | Unspecified |
| mmu-miR-378a-3p | MIMAT0003151 | Errfi1 | 74155 | Microarrays | 37986224\|22855601 | | Unspecified |
| mmu-miR-378a-3p | MIMAT0003151 | Esco2 | 71988 | HITS-CLIP | | 37986224 | Unspecified |
| mmu-miR-378a-3p | MIMAT0003151 | Esd | 13885 | HITS-CLIP | | 29167373 | Unspecified |
| mmu-miR-378a-3p | MIMAT0003151 | Esf1 | 66580 | Microarrays | 37986224\|22855601 | | Unspecified |
| mmu-miR-378a-3p | MIMAT0003151 | Etfdh | 66841 | HITS-CLIP | | 37986224 | Unspecified |
| mmu-miR-378a-3p | MIMAT0003151 | Etv5 | 104156 | Microarrays | 37986224\|22855601 | | Unspecified |
| mmu-miR-378a-3p | MIMAT0003151 | Etv6 | 14011 | HITS-CLIP | | 37986224 | Unspecified |
| mmu-miR-378a-3p | MIMAT0003151 | Ewsr1 | 14030 | qCLASH | | 31393953 | Unspecified |
| mmu-miR-378a-3p | MIMAT0003151 | Exo1 | 26909 | HITS-CLIP | | 37986224 | Unspecified |
| mmu-miR-378a-3p | MIMAT0003151 | Exoc5 | 105504 | HITS-CLIP | | 37986224 | Unspecified |
| mmu-miR-378a-3p | MIMAT0003151 | Extl3 | 54616 | HITS-CLIP | | 25083871 | Unspecified |
| mmu-miR-378a-3p | MIMAT0003151 | Eya3 | 14050 | HITS-CLIP | | 37986224 | Unspecified |
| mmu-miR-378a-3p | MIMAT0003151 | Ezh1 | 14055 | HITS-CLIP | | 37986224 | Unspecified |
| mmu-miR-378a-3p | MIMAT0003151 | Ezh2 | 14056 | HITS-CLIP | 25083871\|23142080 | | Unspecified |
| mmu-miR-378a-3p | MIMAT0003151 | Faah | 14073 | HITS-CLIP | | 37986224 | Unspecified |
| mmu-miR-378a-3p | MIMAT0003151 | Fads3 | 60527 | HITS-CLIP | | 25083871 | Unspecified |
| mmu-miR-378a-3p | MIMAT0003151 | Faf2 | 76577 | HITS-CLIP | | 25753418 | Unspecified |
| mmu-miR-378a-3p | MIMAT0003151 | Fam107b | 66540 | HITS-CLIP | | 37986224 | Unspecified |
| mmu-miR-378a-3p | MIMAT0003151 | Fam117a | 215512 | HITS-CLIP | | 25083871 | Unspecified |
| mmu-miR-378a-3p | MIMAT0003151 | Fam118a | 73225 | HITS-CLIP | | 37986224 | Unspecified |
| mmu-miR-378a-3p | MIMAT0003151 | Fam122b | 78755 | Microarrays | | 37986224 | Unspecified |
| mmu-miR-378a-3p | MIMAT0003151 | Fam129b | 227737 | HITS-CLIP | | 37986224 | Unspecified |
| mmu-miR-378a-3p | MIMAT0003151 | Fam135a | 68187 | HITS-CLIP | | 37986224 | Unspecified |
| mmu-miR-378a-3p | MIMAT0003151 | Fam13b | 225358 | HITS-CLIP | | 37986224 | Unspecified |
| mmu-miR-378a-3p | MIMAT0003151 | Fam168a | 319604 | HITS-CLIP | | 37986224 | Unspecified |
| mmu-miR-378a-3p | MIMAT0003151 | Fam168b | 214469 | HITS-CLIP | | 37986224 | Unspecified |

| | | | | | | |
|---|---|---|---|---|---|---|
| mmu-miR-378a-3p | MIMAT0003151 | Fam222b | 216971 | HITS-CLIP | 37986224\|25083871 | Unspecified |
| mmu-miR-378a-3p | MIMAT0003151 | Fam50a | 108160 | HITS-CLIP | 37986224\|23597149 | Unspecified |
| mmu-miR-378a-3p | MIMAT0003151 | Fam91a1 | 210998 | HITS-CLIP | 25083871 | Unspecified |
| mmu-miR-378a-3p | MIMAT0003151 | Fam98a | 72722 | HITS-CLIP | 25083871 | Unspecified |
| mmu-miR-378a-3p | MIMAT0003151 | Fam98b | 68215 | HITS-CLIP | 25083871\|23142080 | Unspecified |
| mmu-miR-378a-3p | MIMAT0003151 | Farp2 | 227377 | HITS-CLIP | 37986224 | Unspecified |
| mmu-miR-378a-3p | MIMAT0003151 | Fat1 | 14107 | HITS-CLIP, Microarrays//Microarrays//HITS | 37986224\|22855601\|25083871 | Unspecified |
| mmu-miR-378a-3p | MIMAT0003151 | Fbn1 | 14118 | HITS-CLIP | 37986224 | Unspecified |
| mmu-miR-378a-3p | MIMAT0003151 | Fbrs | 14123 | HITS-CLIP | 23142080\|25083871 | Unspecified |
| mmu-miR-378a-3p | MIMAT0003151 | Fbxl17 | 50758 | HITS-CLIP | 37986224 | Unspecified |
| mmu-miR-378a-3p | MIMAT0003151 | Fbxl20 | 72194 | HITS-CLIP | 25083871 | Unspecified |
| mmu-miR-378a-3p | MIMAT0003151 | Fbxl6 | 30840 | HITS-CLIP | 37986224 | Unspecified |
| mmu-miR-378a-3p | MIMAT0003151 | Fbxo10 | 269529 | HITS-CLIP | 37986224 | Unspecified |
| mmu-miR-378a-3p | MIMAT0003151 | Fbxo11 | 225055 | HITS-CLIP | 25083871 | Unspecified |
| mmu-miR-378a-3p | MIMAT0003151 | Fbxo28 | 67948 | HITS-CLIP//qCLASH | 37986224\|31393953 | Unspecified |
| mmu-miR-378a-3p | MIMAT0003151 | Fbxo30 | 71865 | HITS-CLIP | 37986224 | Unspecified |
| mmu-miR-378a-3p | MIMAT0003151 | Fbxo33 | 70611 | HITS-CLIP | 37986224 | Unspecified |
| mmu-miR-378a-3p | MIMAT0003151 | Fbxw8 | 231672 | HITS-CLIP | 23597149 | Unspecified |
| mmu-miR-378a-3p | MIMAT0003151 | Fcho2 | 218503 | qCLASH | 31393953 | Unspecified |
| mmu-miR-378a-3p | MIMAT0003151 | Fchsd2 | 207278 | HITS-CLIP | 23597149 | Unspecified |
| mmu-miR-378a-3p | MIMAT0003151 | Fermt2 | 218952 | HITS-CLIP | 25083871\|23597149 | Unspecified |
| mmu-miR-378a-3p | MIMAT0003151 | Fgd3 | 30938 | HITS-CLIP | 23142080 | Unspecified |
| mmu-miR-378a-3p | MIMAT0003151 | Fgfrl1 | 116701 | HITS-CLIP | 37986224 | Unspecified |
| mmu-miR-378a-3p | MIMAT0003151 | Fgl1 | 234199 | HITS-CLIP | 37986224\|23597149 | Unspecified |
| mmu-miR-378a-3p | MIMAT0003151 | Fh1 | 14194 | HITS-CLIP | 37986224\|25083871\|23142080 | Unspecified |
| mmu-miR-378a-3p | MIMAT0003151 | Fhdc1 | 229474 | Microarrays | 37986224\|22855601 | Unspecified |
| mmu-miR-378a-3p | MIMAT0003151 | Fhl2 | 14200 | Microarrays | 37986224\|22855601 | Unspecified |
| mmu-miR-378a-3p | MIMAT0003151 | Fignl1 | 60530 | HITS-CLIP | 37986224 | Unspecified |
| mmu-miR-378a-3p | MIMAT0003151 | Filip1 | 70598 | HITS-CLIP | 25083871 | Unspecified |
| mmu-miR-378a-3p | MIMAT0003151 | Filip1l | 78749 | HITS-CLIP | 25083871\|23142080 | Unspecified |
| mmu-miR-378a-3p | MIMAT0003151 | Fis1 | 66437 | HITS-CLIP | 25083871 | Unspecified |
| mmu-miR-378a-3p | MIMAT0003151 | Fitm2 | 228859 | HITS-CLIP | 37986224 | Unspecified |
| mmu-miR-378a-3p | MIMAT0003151 | Fkbp14 | 231997 | HITS-CLIP | 37986224\|25083871 | Unspecified |
| mmu-miR-378a-3p | MIMAT0003151 | Fkbp2 | 14227 | HITS-CLIP | 23142080 | Unspecified |
| mmu-miR-378a-3p | MIMAT0003151 | Fkbp4 | 14228 | HITS-CLIP | 37986224 | Unspecified |
| mmu-miR-378a-3p | MIMAT0003151 | Flcn | 216805 | HITS-CLIP | 37986224 | Unspecified |
| mmu-miR-378a-3p | MIMAT0003151 | Flnb | 286940 | HITS-CLIP, Microarrays//Microarrays | 37986224\|22855601 | Unspecified |
| mmu-miR-378a-3p | MIMAT0003151 | Flrt2 | 399558 | Microarrays//HITS-CLIP | 37986224\|25083871\|22855601 | Unspecified |
| mmu-miR-378a-3p | MIMAT0003151 | Fmnl2 | 71409 | HITS-CLIP | 23597149 | Unspecified |

| | | | | | | |
|---|---|---|---|---|---|---|
| mmu-miR-378a-3p | MIMAT0003151 | Fmo5 | 14263 | HITS-CLIP | 37986224\|23597149 | Unspecified |
| mmu-miR-378a-3p | MIMAT0003151 | Fn1 | 14268 | HITS-CLIP | 25083871 | Unspecified |
| mmu-miR-378a-3p | MIMAT0003151 | Fnbp4 | 55935 | HITS-CLIP | 37986224 | Unspecified |
| mmu-miR-378a-3p | MIMAT0003151 | Fndc5 | 384061 | HITS-CLIP | 25083871 | Unspecified |
| mmu-miR-378a-3p | MIMAT0003151 | Fnta | 14272 | HITS-CLIP | 37986224\|23142080 | Unspecified |
| mmu-miR-378a-3p | MIMAT0003151 | Fosl1 | 14283 | Microarrays | 37986224\|22855601 | Unspecified |
| mmu-miR-378a-3p | MIMAT0003151 | Fosl2 | 14284 | HITS-CLIP | 25083871 | Unspecified |
| mmu-miR-378a-3p | MIMAT0003151 | Foxj2 | 60611 | HITS-CLIP | 37986224\|25083871\|23142080 | Unspecified |
| mmu-miR-378a-3p | MIMAT0003151 | Foxn2 | 14236 | Microarrays | 37986224\|22855601 | Unspecified |
| mmu-miR-378a-3p | MIMAT0003151 | Foxo1 | 56458 | HITS-CLIP | 25083871\|23142080 | Unspecified |
| mmu-miR-378a-3p | MIMAT0003151 | Foxo3 | 56484 | HITS-CLIP | 37986224\|23597149\|23142080\|29167 | Unspecified |
| mmu-miR-378a-3p | MIMAT0003151 | Foxo6 | 329934 | HITS-CLIP | 25083871 | Unspecified |
| mmu-miR-378a-3p | MIMAT0003151 | Frrs1 | 20321 | Microarrays | 37986224\|22855601 | Unspecified |
| mmu-miR-378a-3p | MIMAT0003151 | Frs2 | 327826 | HITS-CLIP | 25083871\|23142080 | Unspecified |
| mmu-miR-378a-3p | MIMAT0003151 | Fry | 320365 | HITS-CLIP | 25753418 | Unspecified |
| mmu-miR-378a-3p | MIMAT0003151 | Ftsj3 | 56095 | Microarrays//HITS-CLIP | 37986224\|22855601\|25083871 | Unspecified |
| mmu-miR-378a-3p | MIMAT0003151 | Fubp1 | 51886 | HITS-CLIP | 37986224\|23142080\|25083871\|29167 | Unspecified |
| mmu-miR-378a-3p | MIMAT0003151 | Fyttd1 | 69823 | HITS-CLIP | 37986224 | Unspecified |
| mmu-miR-378a-3p | MIMAT0003151 | Fzd5 | 14367 | HITS-CLIP | 37986224 | Unspecified |
| mmu-miR-378a-3p | MIMAT0003151 | Gabbr1 | 54393 | HITS-CLIP | 37986224 | Unspecified |
| mmu-miR-378a-3p | MIMAT0003151 | Gabpb2 | 213054 | HITS-CLIP | 37986224 | Unspecified |
| mmu-miR-378a-3p | MIMAT0003151 | Gadd45a | 13197 | Microarrays | 37986224\|22855601 | Unspecified |
| mmu-miR-378a-3p | MIMAT0003151 | Galc | 14420 | HITS-CLIP | 37986224 | Unspecified |
| mmu-miR-378a-3p | MIMAT0003151 | Galnt11 | 231050 | HITS-CLIP | 29167373 | Unspecified |
| mmu-miR-378a-3p | MIMAT0003151 | Gamt | 14431 | HITS-CLIP | 23597149 | Unspecified |
| mmu-miR-378a-3p | MIMAT0003151 | Ganab | 14376 | HITS-CLIP | 23597149\|23142080 | Unspecified |
| mmu-miR-378a-3p | MIMAT0003151 | Gapdh | 14433 | qCLASH | 31393953 | Unspecified |
| mmu-miR-378a-3p | MIMAT0003151 | Gars | 353172 | HITS-CLIP | 23597149\|23142080 | Unspecified |
| mmu-miR-378a-3p | MIMAT0003151 | Gart | 14450 | HITS-CLIP, Microarrays//Microarrays | 37986224\|22855601 | Unspecified |
| mmu-miR-378a-3p | MIMAT0003151 | Gas2l3 | 237436 | Microarrays | 37986224\|22855601 | Unspecified |
| mmu-miR-378a-3p | MIMAT0003151 | Gask1b | 68659 | Microarrays | 22855601 | Unspecified |
| mmu-miR-378a-3p | MIMAT0003151 | Gbe1 | 74185 | HITS-CLIP | 37986224 | Unspecified |
| mmu-miR-378a-3p | MIMAT0003151 | Gbf1 | 107338 | HITS-CLIP | 23597149\|25083871\|25753418 | Unspecified |
| mmu-miR-378a-3p | MIMAT0003151 | Gcc2 | 70297 | HITS-CLIP | 37986224\|25083871 | Unspecified |
| mmu-miR-378a-3p | MIMAT0003151 | Gclc | 14629 | Microarrays | 37986224\|22855601 | Unspecified |
| mmu-miR-378a-3p | MIMAT0003151 | Gcnt4 | 218476 | HITS-CLIP, Microarrays//Microarrays | 37986224\|22855601 | Unspecified |
| mmu-miR-378a-3p | MIMAT0003151 | Gdi1 | 14567 | HITS-CLIP | 23142080 | Unspecified |
| mmu-miR-378a-3p | MIMAT0003151 | Get4 | 67604 | HITS-CLIP | 37986224 | Unspecified |
| mmu-miR-378a-3p | MIMAT0003151 | Gfm1 | 28030 | HITS-CLIP | 25083871\|23142080 | Unspecified |

| | | | | | | |
|---|---|---|---|---|---|---|
| mmu-miR-378a-3p | MIMAT0003151 | Ggta1 | 14594 | HITS-CLIP | 37986224 | Unspecified |
| mmu-miR-378a-3p | MIMAT0003151 | Gimap8 | 243374 | HITS-CLIP | 23142080 | Unspecified |
| mmu-miR-378a-3p | MIMAT0003151 | Git2 | 26431 | HITS-CLIP | 37986224 | Unspecified |
| mmu-miR-378a-3p | MIMAT0003151 | Gja1 | 14609 | HITS-CLIP, Microarrays//Microarrays | 37986224\|22855601 | Unspecified |
| mmu-miR-378a-3p | MIMAT0003151 | Gja4 | 14612 | HITS-CLIP | 37986224 | Unspecified |
| mmu-miR-378a-3p | MIMAT0003151 | Glipr2 | 384009 | HITS-CLIP | 25083871\|23142080 | Unspecified |
| mmu-miR-378a-3p | MIMAT0003151 | Gls | 14660 | HITS-CLIP | 25083871 | Unspecified |
| mmu-miR-378a-3p | MIMAT0003151 | Gls2 | 216456 | HITS-CLIP | 37986224\|23597149 | Unspecified |
| mmu-miR-378a-3p | MIMAT0003151 | Glul | 14645 | HITS-CLIP | 25753418 | Unspecified |
| mmu-miR-378a-3p | MIMAT0003151 | Gm1840 | 669429 | HITS-CLIP | 37986224 | Unspecified |
| mmu-miR-378a-3p | MIMAT0003151 | Gm20498 | 105940408 | HITS-CLIP | 37986224 | Unspecified |
| mmu-miR-378a-3p | MIMAT0003151 | Gm21988 | 6610016 | HITS-CLIP | 25753418 | Unspecified |
| mmu-miR-378a-3p | MIMAT0003151 | Gm38394 | 667118 | HITS-CLIP | 25083871 | Unspecified |
| mmu-miR-378a-3p | MIMAT0003151 | Gm49333 | 110599584 | HITS-CLIP | 23142080 | Unspecified |
| mmu-miR-378a-3p | MIMAT0003151 | Gm7694 | 665574 | HITS-CLIP | 23597149 | Unspecified |
| mmu-miR-378a-3p | MIMAT0003151 | Gm9803 | 100042179 | HITS-CLIP | 23142080 | Unspecified |
| mmu-miR-378a-3p | MIMAT0003151 | Gna12 | 14673 | HITS-CLIP | 37986224 | Unspecified |
| mmu-miR-378a-3p | MIMAT0003151 | Gnai2 | 14678 | HITS-CLIP | 23142080\|29167373 | Unspecified |
| mmu-miR-378a-3p | MIMAT0003151 | Gnai3 | 14679 | HITS-CLIP | 23142080\|25083871 | Unspecified |
| mmu-miR-378a-3p | MIMAT0003151 | Gnaq | 14682 | HITS-CLIP | 37986224 | Unspecified |
| mmu-miR-378a-3p | MIMAT0003151 | Gnas | 14683 | HITS-CLIP | 25083871\|23142080 | Unspecified |
| mmu-miR-378a-3p | MIMAT0003151 | Gnl3 | 30877 | Microarrays | 37986224\|22855601 | Unspecified |
| mmu-miR-378a-3p | MIMAT0003151 | Gnpat | 14712 | HITS-CLIP | 25083871 | Unspecified |
| mmu-miR-378a-3p | MIMAT0003151 | Gnptab | 432486 | Microarrays//HITS-CLIP | 37986224\|22855601\|23142080 | Unspecified |
| mmu-miR-378a-3p | MIMAT0003151 | Gns | 75612 | HITS-CLIP | 37986224\|25083871\|23597149\|29167 | Unspecified |
| mmu-miR-378a-3p | MIMAT0003151 | Golt1b | 66964 | HITS-CLIP | 29167373\|23142080 | Unspecified |
| mmu-miR-378a-3p | MIMAT0003151 | Gpatch8 | 237943 | HITS-CLIP | 37986224 | Unspecified |
| mmu-miR-378a-3p | MIMAT0003151 | Gpbp1l1 | 77110 | HITS-CLIP | 37986224 | Unspecified |
| mmu-miR-378a-3p | MIMAT0003151 | Gpi1 | 14751 | HITS-CLIP | 23597149\|29167373 | Unspecified |
| mmu-miR-378a-3p | MIMAT0003151 | Gpr137b | 83924 | Microarrays | 37986224\|22855601 | Unspecified |
| mmu-miR-378a-3p | MIMAT0003151 | Gpr176 | 381413 | Microarrays | 37986224\|22855601 | Unspecified |
| mmu-miR-378a-3p | MIMAT0003151 | Gprc5b | 64297 | HITS-CLIP | 37986224 | Unspecified |
| mmu-miR-378a-3p | MIMAT0003151 | Gprin3 | 243385 | HITS-CLIP | 37986224\|23597149 | Unspecified |
| mmu-miR-378a-3p | MIMAT0003151 | Gpsm1 | 67839 | HITS-CLIP | 37986224 | Unspecified |
| mmu-miR-378a-3p | MIMAT0003151 | Gpt | 76282 | HITS-CLIP | 23597149 | Unspecified |
| mmu-miR-378a-3p | MIMAT0003151 | Gpx8 | 69590 | HITS-CLIP | 25083871 | Unspecified |
| mmu-miR-378a-3p | MIMAT0003151 | Gramd1b | 235283 | HITS-CLIP | 37986224\|23142080\|29167373 | Unspecified |
| mmu-miR-378a-3p | MIMAT0003151 | Grb10 | 14783 | HITS-CLIP | 25753418 | Unspecified |
| mmu-miR-378a-3p | MIMAT0003151 | Grb14 | 50915 | Microarrays | 37986224\|22855601 | Unspecified |

| | | | | | | |
|---|---|---|---|---|---|---|
| mmu-miR-378a-3p | MIMAT0003151 | Grb2 | 14784 | HITS-CLIP | 37986224\|23142080 | Unspecified |
| mmu-miR-378a-3p | MIMAT0003151 | Grem1 | 23892 | HITS-CLIP | 37986224 | Unspecified |
| mmu-miR-378a-3p | MIMAT0003151 | Grem2 | 23893 | Microarrays | 37986224\|22855601 | Unspecified |
| mmu-miR-378a-3p | MIMAT0003151 | Gria3 | 53623 | Microarrays | 37986224\|22855601 | Unspecified |
| mmu-miR-378a-3p | MIMAT0003151 | Grik2 | 14806 | Microarrays | 37986224\|22855601 | Unspecified |
| mmu-miR-378a-3p | MIMAT0003151 | Grk1 | 24013 | qCLASH | 31393953 | Unspecified |
| mmu-miR-378a-3p | MIMAT0003151 | Grpel1 | 17713 | HITS-CLIP | 37986224\|23597149\|25083871 | Unspecified |
| mmu-miR-378a-3p | MIMAT0003151 | Grsf1 | 231413 | HITS-CLIP | 37986224 | Unspecified |
| mmu-miR-378a-3p | MIMAT0003151 | Grwd1 | 101612 | Microarrays | 37986224\|22855601 | Unspecified |
| mmu-miR-378a-3p | MIMAT0003151 | Gstm1 | 14862 | Microarrays | 37986224 | Unspecified |
| mmu-miR-378a-3p | MIMAT0003151 | Gstm2 | 14863 | Microarrays | 37986224 | Unspecified |
| mmu-miR-378a-3p | MIMAT0003151 | Gstm5 | 14866 | Microarrays | 37986224 | Unspecified |
| mmu-miR-378a-3p | MIMAT0003151 | Gsto1 | 14873 | HITS-CLIP, Microarrays//Microarrays | 37986224\|22855601 | Unspecified |
| mmu-miR-378a-3p | MIMAT0003151 | Gstt1 | 14871 | HITS-CLIP | 37986224 | Unspecified |
| mmu-miR-378a-3p | MIMAT0003151 | Gtf2i | 14886 | HITS-CLIP | 37986224 | Unspecified |
| mmu-miR-378a-3p | MIMAT0003151 | Gtf3c4 | 269252 | HITS-CLIP | 37986224\|23142080 | Unspecified |
| mmu-miR-378a-3p | MIMAT0003151 | Gtpbp4 | 69237 | Microarrays | 37986224\|22855601 | Unspecified |
| mmu-miR-378a-3p | MIMAT0003151 | Gucd1 | 68778 | HITS-CLIP | 37986224 | Unspecified |
| mmu-miR-378a-3p | MIMAT0003151 | Gzmb | 14939 | HITS-CLIP | 37986224 | Unspecified |
| mmu-miR-378a-3p | MIMAT0003151 | H1f0 | 14958 | HITS-CLIP | 37986224 | Unspecified |
| mmu-miR-378a-3p | MIMAT0003151 | H3f3b | 15081 | HITS-CLIP | 37986224\|23142080 | Unspecified |
| mmu-miR-378a-3p | MIMAT0003151 | Habp4 | 56541 | HITS-CLIP | 23597149\|25083871 | Unspecified |
| mmu-miR-378a-3p | MIMAT0003151 | Has2 | 15117 | HITS-CLIP, Microarrays//Microarrays | 37986224\|22855601 | Unspecified |
| mmu-miR-378a-3p | MIMAT0003151 | Hbegf | 15200 | Microarrays | 37986224\|22855601 | Unspecified |
| mmu-miR-378a-3p | MIMAT0003151 | Hbp1 | 73389 | HITS-CLIP | 37986224\|25083871 | Unspecified |
| mmu-miR-378a-3p | MIMAT0003151 | Hcfc1 | 15161 | HITS-CLIP | 25753418 | Unspecified |
| mmu-miR-378a-3p | MIMAT0003151 | Hcfc2 | 67933 | HITS-CLIP | 37986224 | Unspecified |
| mmu-miR-378a-3p | MIMAT0003151 | Hdac4 | 208727 | HITS-CLIP | 23142080 | Unspecified |
| mmu-miR-378a-3p | MIMAT0003151 | Hddc2 | 69692 | HITS-CLIP | 23142080 | Unspecified |
| mmu-miR-378a-3p | MIMAT0003151 | Hddc3 | 68695 | Microarrays | 37986224 | Unspecified |
| mmu-miR-378a-3p | MIMAT0003151 | Heatr1 | 217995 | Microarrays | 37986224\|22855601 | Unspecified |
| mmu-miR-378a-3p | MIMAT0003151 | Heatr5a | 320487 | HITS-CLIP | 29167373 | Unspecified |
| mmu-miR-378a-3p | MIMAT0003151 | Hectd1 | 207304 | HITS-CLIP | 23142080 | Unspecified |
| mmu-miR-378a-3p | MIMAT0003151 | Herc1 | 235439 | HITS-CLIP | 25083871 | Unspecified |
| mmu-miR-378a-3p | MIMAT0003151 | Herc2 | 15204 | HITS-CLIP | 25083871 | Unspecified |
| mmu-miR-378a-3p | MIMAT0003151 | Herc4 | 67345 | HITS-CLIP | 25083871 | Unspecified |
| mmu-miR-378a-3p | MIMAT0003151 | Hibadh | 58875 | HITS-CLIP | 37986224 | Unspecified |
| mmu-miR-378a-3p | MIMAT0003151 | Hif1a | 15251 | HITS-CLIP | 37986224\|23597149\|23142080\|2575[illegible] | Unspecified |
| mmu-miR-378a-3p | MIMAT0003151 | Hif1an | 319594 | HITS-CLIP | 37986224 | Unspecified |

| | | | | | | |
|---|---|---|---|---|---|---|
| mmu-miR-378a-3p | MIMAT0003151 | Hilpda | 69573 | HITS-CLIP | 37986224 | Unspecified |
| mmu-miR-378a-3p | MIMAT0003151 | Hip1 | 215114 | HITS-CLIP | 25083871 | Unspecified |
| mmu-miR-378a-3p | MIMAT0003151 | Hipk1 | 15257 | HITS-CLIP | 29167373 | Unspecified |
| mmu-miR-378a-3p | MIMAT0003151 | Hipk2 | 15258 | HITS-CLIP, Microarrays//Microarrays | 37986224\|22855601 | Unspecified |
| mmu-miR-378a-3p | MIMAT0003151 | Hivep2 | 15273 | HITS-CLIP | 25083871 | Unspecified |
| mmu-miR-378a-3p | MIMAT0003151 | Hk2 | 15277 | Microarrays | 37986224\|22855601 | Unspecified |
| mmu-miR-378a-3p | MIMAT0003151 | Hmga2 | 15364 | Microarrays | 37986224\|22855601 | Unspecified |
| mmu-miR-378a-3p | MIMAT0003151 | Hmgcr | 15357 | HITS-CLIP | 37986224 | Unspecified |
| mmu-miR-378a-3p | MIMAT0003151 | Hmgcs2 | 15360 | HITS-CLIP | 23597149 | Unspecified |
| mmu-miR-378a-3p | MIMAT0003151 | Hmox1 | 15368 | Microarrays | 37986224\|22855601 | Unspecified |
| mmu-miR-378a-3p | MIMAT0003151 | Hnrnpd | 11991 | HITS-CLIP | 37986224 | Unspecified |
| mmu-miR-378a-3p | MIMAT0003151 | Hnrnpl | 15388 | HITS-CLIP | 37986224 | Unspecified |
| mmu-miR-378a-3p | MIMAT0003151 | Hnrnpm | 76936 | HITS-CLIP | 23142080\|29167373 | Unspecified |
| mmu-miR-378a-3p | MIMAT0003151 | Hnrnpr | 74326 | HITS-CLIP | 37986224 | Unspecified |
| mmu-miR-378a-3p | MIMAT0003151 | Hnrnpu | 51810 | HITS-CLIP | 37986224\|25083871\|29167373 | Unspecified |
| mmu-miR-378a-3p | MIMAT0003151 | Hnrnpul2 | 68693 | HITS-CLIP | 37986224\|26203562\|25753418 | Unspecified |
| mmu-miR-378a-3p | MIMAT0003151 | Hp1bp3 | 15441 | HITS-CLIP | 37986224 | Unspecified |
| mmu-miR-378a-3p | MIMAT0003151 | Hpn | 15451 | HITS-CLIP | 23597149 | Unspecified |
| mmu-miR-378a-3p | MIMAT0003151 | Hspa12a | 73442 | HITS-CLIP | 37986224 | Unspecified |
| mmu-miR-378a-3p | MIMAT0003151 | Hspa4l | 18415 | Microarrays | 37986224\|22855601 | Unspecified |
| mmu-miR-378a-3p | MIMAT0003151 | Hspbap1 | 66667 | HITS-CLIP | 23142080 | Unspecified |
| mmu-miR-378a-3p | MIMAT0003151 | Hsph1 | 15505 | Microarrays//qCLASH | 37986224\|22855601\|31393953 | Unspecified |
| mmu-miR-378a-3p | MIMAT0003151 | Huwe1 | 59026 | HITS-CLIP | 23142080 | Unspecified |
| mmu-miR-378a-3p | MIMAT0003151 | Hyou1 | 12282 | HITS-CLIP | 37986224 | Unspecified |
| mmu-miR-378a-3p | MIMAT0003151 | Iars | 105148 | HITS-CLIP | 23142080\|23597149 | Unspecified |
| mmu-miR-378a-3p | MIMAT0003151 | Icmt | 57295 | HITS-CLIP | 37986224 | Unspecified |
| mmu-miR-378a-3p | MIMAT0003151 | Id2 | 15902 | Microarrays | 37986224 | Unspecified |
| mmu-miR-378a-3p | MIMAT0003151 | Ide | 15925 | HITS-CLIP | 23597149 | Unspecified |
| mmu-miR-378a-3p | MIMAT0003151 | Idh1 | 15926 | HITS-CLIP | 23597149\|23142080 | Unspecified |
| mmu-miR-378a-3p | MIMAT0003151 | Idh2 | 269951 | HITS-CLIP | 37986224 | Unspecified |
| mmu-miR-378a-3p | MIMAT0003151 | Idh3b | 170718 | HITS-CLIP | 37986224 | Unspecified |
| mmu-miR-378a-3p | MIMAT0003151 | Ier2 | 15936 | Microarrays | 37986224\|22855601 | Unspecified |
| mmu-miR-378a-3p | MIMAT0003151 | Ier5 | 15939 | HITS-CLIP | 29167373 | Unspecified |
| mmu-miR-378a-3p | MIMAT0003151 | Iffo1 | 320678 | HITS-CLIP | 37986224 | Unspecified |
| mmu-miR-378a-3p | MIMAT0003151 | Ifi27 | 52668 | Microarrays | 37986224 | Unspecified |
| mmu-miR-378a-3p | MIMAT0003151 | Ifrd1 | 15982 | Microarrays | 37986224\|22855601 | Unspecified |
| mmu-miR-378a-3p | MIMAT0003151 | Ifrd2 | 15983 | Microarrays | 37986224\|22855601 | Unspecified |
| mmu-miR-378a-3p | MIMAT0003151 | Ift122 | 81896 | HITS-CLIP | 25753418 | Unspecified |
| mmu-miR-378a-3p | MIMAT0003151 | Igf1 | 16000 | HITS-CLIP | 23597149 | Unspecified |

| | | | | | | |
|---|---|---|---|---|---|---|
| mmu-miR-378a-3p | MIMAT0003151 | Igf1r | 16001 | HITS-CLIP | 25083871\|23142080 | Unspecified |
| mmu-miR-378a-3p | MIMAT0003151 | Igfbp7 | 29817 | Microarrays | 37986224 | Unspecified |
| mmu-miR-378a-3p | MIMAT0003151 | Igsf3 | 78908 | HITS-CLIP | 37986224 | Unspecified |
| mmu-miR-378a-3p | MIMAT0003151 | Igsf8 | 140559 | HITS-CLIP | 37986224 | Unspecified |
| mmu-miR-378a-3p | MIMAT0003151 | Iigp1 | 60440 | HITS-CLIP | 37986224\|23597149 | Unspecified |
| mmu-miR-378a-3p | MIMAT0003151 | Ik | 24010 | HITS-CLIP | 23142080 | Unspecified |
| mmu-miR-378a-3p | MIMAT0003151 | Il13ra1 | 16164 | HITS-CLIP | 25083871 | Unspecified |
| mmu-miR-378a-3p | MIMAT0003151 | Il1rap | 16180 | Microarrays | 37986224\|22855601 | Unspecified |
| mmu-miR-378a-3p | MIMAT0003151 | Il1rl1 | 17082 | Microarrays | 37986224\|22855601 | Unspecified |
| mmu-miR-378a-3p | MIMAT0003151 | Il4ra | 16190 | Microarrays | 37986224\|22855601 | Unspecified |
| mmu-miR-378a-3p | MIMAT0003151 | Il6st | 16195 | HITS-CLIP | 37986224 | Unspecified |
| mmu-miR-378a-3p | MIMAT0003151 | Il9r | 3581 | qCLASH | 31393953 | Unspecified |
| mmu-miR-378a-3p | MIMAT0003151 | Immt | 76614 | HITS-CLIP | 25083871 | Unspecified |
| mmu-miR-378a-3p | MIMAT0003151 | Impad1 | 242291 | HITS-CLIP | 37986224 | Unspecified |
| mmu-miR-378a-3p | MIMAT0003151 | Inf2 | 70435 | HITS-CLIP | 37986224 | Unspecified |
| mmu-miR-378a-3p | MIMAT0003151 | Inhba | 16323 | Microarrays | 37986224\|22855601 | Unspecified |
| mmu-miR-378a-3p | MIMAT0003151 | Ino80 | 68142 | HITS-CLIP | 25083871 | Unspecified |
| mmu-miR-378a-3p | MIMAT0003151 | Ino80d | 227195 | HITS-CLIP | 37986224\|25083871 | Unspecified |
| mmu-miR-378a-3p | MIMAT0003151 | Inpp4b | 234515 | HITS-CLIP | 37986224\|23142080 | Unspecified |
| mmu-miR-378a-3p | MIMAT0003151 | Inpp5a | 212111 | HITS-CLIP | 37986224 | Unspecified |
| mmu-miR-378a-3p | MIMAT0003151 | Inpp5d | 16331 | HITS-CLIP | 29167373 | Unspecified |
| mmu-miR-378a-3p | MIMAT0003151 | Inppl1 | 16332 | HITS-CLIP | 25083871 | Unspecified |
| mmu-miR-378a-3p | MIMAT0003151 | Insig1 | 231070 | HITS-CLIP | 25083871\|23597149\|23142080 | Unspecified |
| mmu-miR-378a-3p | MIMAT0003151 | Ints8 | 72656 | HITS-CLIP | 25083871\|25753418\|29167373\|2314 | Unspecified |
| mmu-miR-378a-3p | MIMAT0003151 | Ip6k1 | 27399 | HITS-CLIP | 37986224 | Unspecified |
| mmu-miR-378a-3p | MIMAT0003151 | Ipo5 | 70572 | HITS-CLIP | 37986224\|23597149\|25083871\|2314 | Unspecified |
| mmu-miR-378a-3p | MIMAT0003151 | Ipo7 | 233726 | Microarrays | 37986224\|22855601 | Unspecified |
| mmu-miR-378a-3p | MIMAT0003151 | Iqcb1 | 320299 | HITS-CLIP | 37986224 | Unspecified |
| mmu-miR-378a-3p | MIMAT0003151 | Iqgap1 | 29875 | HITS-CLIP | 29167373 | Unspecified |
| mmu-miR-378a-3p | MIMAT0003151 | Iqgap2 | 544963 | HITS-CLIP | 23597149 | Unspecified |
| mmu-miR-378a-3p | MIMAT0003151 | Iqsec1 | 232227 | HITS-CLIP | 37986224 | Unspecified |
| mmu-miR-378a-3p | MIMAT0003151 | Irak1 | 16179 | HITS-CLIP | 37986224 | Unspecified |
| mmu-miR-378a-3p | MIMAT0003151 | Ireb2 | 64602 | HITS-CLIP | 23142080 | Unspecified |
| mmu-miR-378a-3p | MIMAT0003151 | Irf2bp2 | 270110 | Microarrays | 37986224\|22855601 | Unspecified |
| mmu-miR-378a-3p | MIMAT0003151 | Irf2bpl | 238330 | HITS-CLIP | 37986224 | Unspecified |
| mmu-miR-378a-3p | MIMAT0003151 | Irf4 | 16364 | HITS-CLIP | 37986224 | Unspecified |
| mmu-miR-378a-3p | MIMAT0003151 | Isoc1 | 66307 | HITS-CLIP | 37986224 | Unspecified |
| mmu-miR-378a-3p | MIMAT0003151 | Ist1 | 71955 | HITS-CLIP | 37986224 | Unspecified |
| mmu-miR-378a-3p | MIMAT0003151 | Itga5 | 16402 | Microarrays | 37986224\|22855601 | Unspecified |

| | | | | | | |
|---|---|---|---|---|---|---|
| mmu-miR-378a-3p | MIMAT0003151 | Itga6 | 16403 | Microarrays | 37986224\|22855601 | Unspecified |
| mmu-miR-378a-3p | MIMAT0003151 | Itga7 | 16404 | HITS-CLIP | 23142080\|25083871 | Unspecified |
| mmu-miR-378a-3p | MIMAT0003151 | Itgav | 16410 | Microarrays//HITS-CLIP | 37986224\|22855601\|23142080\|29167 | Unspecified |
| mmu-miR-378a-3p | MIMAT0003151 | Itgb3 | 16416 | HITS-CLIP | 37986224 | Unspecified |
| mmu-miR-378a-3p | MIMAT0003151 | Itih2 | 16425 | HITS-CLIP | 23597149 | Unspecified |
| mmu-miR-378a-3p | MIMAT0003151 | Itih3 | 16426 | HITS-CLIP | 23597149 | Unspecified |
| mmu-miR-378a-3p | MIMAT0003151 | Itpripl2 | 319622 | HITS-CLIP, Microarrays//Microarrays | 37986224\|22855601 | Unspecified |
| mmu-miR-378a-3p | MIMAT0003151 | Itsn1 | 16443 | HITS-CLIP | 37986224 | Unspecified |
| mmu-miR-378a-3p | MIMAT0003151 | Ivns1abp | 117198 | HITS-CLIP | 25083871 | Unspecified |
| mmu-miR-378a-3p | MIMAT0003151 | Iws1 | 73473 | HITS-CLIP | 37986224 | Unspecified |
| mmu-miR-378a-3p | MIMAT0003151 | Jade1 | 269424 | HITS-CLIP, Microarrays//Microarrays//HITS | 37986224\|22855601\|25083871\|23142 | Unspecified |
| mmu-miR-378a-3p | MIMAT0003151 | Jag1 | 16449 | HITS-CLIP, Microarrays//Microarrays | 37986224\|22855601 | Unspecified |
| mmu-miR-378a-3p | MIMAT0003151 | Jarid2 | 16468 | HITS-CLIP | 37986224\|23597149 | Unspecified |
| mmu-miR-378a-3p | MIMAT0003151 | Jkamp | 104771 | HITS-CLIP | 23597149\|23142080 | Unspecified |
| mmu-miR-378a-3p | MIMAT0003151 | Jmy | 57748 | HITS-CLIP | 37986224\|23142080 | Unspecified |
| mmu-miR-378a-3p | MIMAT0003151 | Jph2 | 59091 | HITS-CLIP | 37986224 | Unspecified |
| mmu-miR-378a-3p | MIMAT0003151 | Jun | 16476 | HITS-CLIP | 37986224 | Unspecified |
| mmu-miR-378a-3p | MIMAT0003151 | Jund | 16478 | HITS-CLIP | 37986224 | Unspecified |
| mmu-miR-378a-3p | MIMAT0003151 | Jup | 16480 | HITS-CLIP | 25753418 | Unspecified |
| mmu-miR-378a-3p | MIMAT0003151 | Kalrn | 545156 | HITS-CLIP | 37986224 | Unspecified |
| mmu-miR-378a-3p | MIMAT0003151 | Kansl1 | 76719 | HITS-CLIP | 37986224 | Unspecified |
| mmu-miR-378a-3p | MIMAT0003151 | Kat7 | 217127 | HITS-CLIP | 25083871 | Unspecified |
| mmu-miR-378a-3p | MIMAT0003151 | Kbtbd2 | 210973 | HITS-CLIP | 37986224\|23142080 | Unspecified |
| mmu-miR-378a-3p | MIMAT0003151 | Kbtbd7 | 211255 | HITS-CLIP | 37986224 | Unspecified |
| mmu-miR-378a-3p | MIMAT0003151 | Kcnk5 | 16529 | HITS-CLIP | 37986224 | Unspecified |
| mmu-miR-378a-3p | MIMAT0003151 | Kctd9 | 105440 | HITS-CLIP | 37986224\|25083871 | Unspecified |
| mmu-miR-378a-3p | MIMAT0003151 | Kdelr2 | 66913 | HITS-CLIP | 37986224 | Unspecified |
| mmu-miR-378a-3p | MIMAT0003151 | Kdelr3 | 105785 | Microarrays | 37986224 | Unspecified |
| mmu-miR-378a-3p | MIMAT0003151 | Kdm1a | 99982 | HITS-CLIP | 25083871 | Unspecified |
| mmu-miR-378a-3p | MIMAT0003151 | Kdm5a | 214899 | HITS-CLIP | 37986224\|25083871 | Unspecified |
| mmu-miR-378a-3p | MIMAT0003151 | Kdm5c | 20591 | qCLASH | 31393953 | Unspecified |
| mmu-miR-378a-3p | MIMAT0003151 | Kdm6a | 22289 | HITS-CLIP | 23597149\|29167373\|23142080 | Unspecified |
| mmu-miR-378a-3p | MIMAT0003151 | Kdm6b | 216850 | HITS-CLIP | 25083871\|25753418 | Unspecified |
| mmu-miR-378a-3p | MIMAT0003151 | Kdm7a | 338523 | HITS-CLIP | 25083871\|23142080 | Unspecified |
| mmu-miR-378a-3p | MIMAT0003151 | Kif15 | 209737 | HITS-CLIP | 21258322\|23142080 | Unspecified |
| mmu-miR-378a-3p | MIMAT0003151 | Kif16b | 16558 | HITS-CLIP | 25083871 | Unspecified |
| mmu-miR-378a-3p | MIMAT0003151 | Kif1b | 16561 | HITS-CLIP | 37986224 | Unspecified |
| mmu-miR-378a-3p | MIMAT0003151 | Kif20a | 19348 | Microarrays | 37986224\|22855601 | Unspecified |
| mmu-miR-378a-3p | MIMAT0003151 | Kif21a | 16564 | HITS-CLIP | 37986224 | Unspecified |

| | | | | | | |
|---|---|---|---|---|---|---|
| mmu-miR-378a-3p | MIMAT0003151 | Kif24 | 109242 | HITS-CLIP | 37986224 | Unspecified |
| mmu-miR-378a-3p | MIMAT0003151 | Kif26a | 668303 | HITS-CLIP | 37986224 | Unspecified |
| mmu-miR-378a-3p | MIMAT0003151 | Kif2a | 16563 | HITS-CLIP | 37986224 | Unspecified |
| mmu-miR-378a-3p | MIMAT0003151 | Kif5b | 16573 | Microarrays//HITS-CLIP | 37986224\|22855601\|25083871 | Unspecified |
| mmu-miR-378a-3p | MIMAT0003151 | Kifap3 | 16579 | HITS-CLIP | 37986224 | Unspecified |
| mmu-miR-378a-3p | MIMAT0003151 | Kitl | 17311 | Microarrays | 37986224\|22855601 | Unspecified |
| mmu-miR-378a-3p | MIMAT0003151 | Klf10 | 21847 | Microarrays | 37986224\|22855601 | Unspecified |
| mmu-miR-378a-3p | MIMAT0003151 | Klf13 | 50794 | HITS-CLIP | 37986224 | Unspecified |
| mmu-miR-378a-3p | MIMAT0003151 | Klf3 | 16599 | HITS-CLIP | 23142080 | Unspecified |
| mmu-miR-378a-3p | MIMAT0003151 | Klf4 | 16600 | Microarrays | 37986224\|22855601 | Unspecified |
| mmu-miR-378a-3p | MIMAT0003151 | Klf6 | 23849 | HITS-CLIP | 25083871 | Unspecified |
| mmu-miR-378a-3p | MIMAT0003151 | Klf7 | 93691 | Microarrays | 37986224\|22855601 | Unspecified |
| mmu-miR-378a-3p | MIMAT0003151 | Klf9 | 16601 | Microarrays | 37986224\|22855601 | Unspecified |
| mmu-miR-378a-3p | MIMAT0003151 | Klhdc1 | 271005 | HITS-CLIP | 37986224 | Unspecified |
| mmu-miR-378a-3p | MIMAT0003151 | Klhdc10 | 76788 | HITS-CLIP | 37986224 | Unspecified |
| mmu-miR-378a-3p | MIMAT0003151 | Klhdc8a | 213417 | Microarrays | 37986224\|22855601 | Unspecified |
| mmu-miR-378a-3p | MIMAT0003151 | Kmt2a | 214162 | HITS-CLIP | 37986224\|29167373 | Unspecified |
| mmu-miR-378a-3p | MIMAT0003151 | Kmt2c | 231051 | HITS-CLIP | 37986224\|25083871\|29167373 | Unspecified |
| mmu-miR-378a-3p | MIMAT0003151 | Kmt2d | 381022 | HITS-CLIP | 29167373 | Unspecified |
| mmu-miR-378a-3p | MIMAT0003151 | Kpna3 | 16648 | HITS-CLIP | 25083871 | Unspecified |
| mmu-miR-378a-3p | MIMAT0003151 | Kpna6 | 16650 | HITS-CLIP | 37986224 | Unspecified |
| mmu-miR-378a-3p | MIMAT0003151 | Kpnb1 | 16211 | HITS-CLIP | 37986224\|23597149\|23142080 | Unspecified |
| mmu-miR-378a-3p | MIMAT0003151 | Kras | 16653 | HITS-CLIP | 23142080 | Unspecified |
| mmu-miR-378a-3p | MIMAT0003151 | Ktn1 | 16709 | HITS-CLIP | 25083871 | Unspecified |
| mmu-miR-378a-3p | MIMAT0003151 | L1cam | 16728 | HITS-CLIP | 29167373 | Unspecified |
| mmu-miR-378a-3p | MIMAT0003151 | Lactb2 | 212442 | HITS-CLIP | 37986224 | Unspecified |
| mmu-miR-378a-3p | MIMAT0003151 | Lamp2 | 16784 | qCLASH | 31393953 | Unspecified |
| mmu-miR-378a-3p | MIMAT0003151 | Larp1 | 73158 | Microarrays | 37986224\|22855601 | Unspecified |
| mmu-miR-378a-3p | MIMAT0003151 | Larp4b | 217980 | HITS-CLIP, Microarrays//Microarrays | 37986224\|22855601 | Unspecified |
| mmu-miR-378a-3p | MIMAT0003151 | Lars | 107045 | qCLASH | 31393953 | Unspecified |
| mmu-miR-378a-3p | MIMAT0003151 | Lbh | 77889 | HITS-CLIP, Microarrays//Microarrays | 37986224\|22855601 | Unspecified |
| mmu-miR-378a-3p | MIMAT0003151 | Lclat1 | 225010 | Microarrays | 37986224\|22855601 | Unspecified |
| mmu-miR-378a-3p | MIMAT0003151 | Lcorl | 209707 | HITS-CLIP | 25083871\|23142080 | Unspecified |
| mmu-miR-378a-3p | MIMAT0003151 | Lcp1 | 18826 | HITS-CLIP | 23142080\|29167373 | Unspecified |
| mmu-miR-378a-3p | MIMAT0003151 | Lcp2 | 16822 | HITS-CLIP | 37986224 | Unspecified |
| mmu-miR-378a-3p | MIMAT0003151 | Ldha | 16828 | HITS-CLIP | 37986224 | Unspecified |
| mmu-miR-378a-3p | MIMAT0003151 | Ldhd | 52815 | HITS-CLIP | 23597149 | Unspecified |
| mmu-miR-378a-3p | MIMAT0003151 | Leng8 | 232798 | HITS-CLIP | 25083871 | Unspecified |
| mmu-miR-378a-3p | MIMAT0003151 | Lgals8 | 56048 | HITS-CLIP | 37986224 | Unspecified |

| | | | | | | |
|---|---|---|---|---|---|---|
| mmu-miR-378a-3p | MIMAT0003151 | Lgalsl | 216551 | HITS-CLIP | 37986224\|23597149\|23142080 | Unspecified |
| mmu-miR-378a-3p | MIMAT0003151 | Lgmn | 19141 | HITS-CLIP | 29167373 | Unspecified |
| mmu-miR-378a-3p | MIMAT0003151 | Lhfpl2 | 218454 | HITS-CLIP | 37986224\|25083871 | Unspecified |
| mmu-miR-378a-3p | MIMAT0003151 | Lima1 | 65970 | HITS-CLIP | 25083871 | Unspecified |
| mmu-miR-378a-3p | MIMAT0003151 | Lman1 | 70361 | HITS-CLIP | 37986224 | Unspecified |
| mmu-miR-378a-3p | MIMAT0003151 | Lmo7 | 380928 | HITS-CLIP | 37986224 | Unspecified |
| mmu-miR-378a-3p | MIMAT0003151 | Lnpk | 69605 | HITS-CLIP | 23142080\|29167373 | Unspecified |
| mmu-miR-378a-3p | MIMAT0003151 | Lnx2 | 140887 | HITS-CLIP | 37986224 | Unspecified |
| mmu-miR-378a-3p | MIMAT0003151 | Lonp2 | 66887 | HITS-CLIP | 23597149 | Unspecified |
| mmu-miR-378a-3p | MIMAT0003151 | Lpar6 | 67168 | HITS-CLIP | 37986224 | Unspecified |
| mmu-miR-378a-3p | MIMAT0003151 | Lpcat3 | 14792 | qCLASH | 31393953 | Unspecified |
| mmu-miR-378a-3p | MIMAT0003151 | Lpin1 | 14245 | HITS-CLIP | 37986224\|23142080 | Unspecified |
| mmu-miR-378a-3p | MIMAT0003151 | Lpin2 | 64898 | HITS-CLIP | 23597149\|29167373 | Unspecified |
| mmu-miR-378a-3p | MIMAT0003151 | Lrch3 | 70144 | HITS-CLIP | 37986224 | Unspecified |
| mmu-miR-378a-3p | MIMAT0003151 | Lrif1 | 321000 | HITS-CLIP | 37986224\|23142080 | Unspecified |
| mmu-miR-378a-3p | MIMAT0003151 | Lrig1 | 16206 | Microarrays | 37986224\|22855601 | Unspecified |
| mmu-miR-378a-3p | MIMAT0003151 | Lrig2 | 269473 | HITS-CLIP | 37986224 | Unspecified |
| mmu-miR-378a-3p | MIMAT0003151 | Lrp1 | 16971 | HITS-CLIP | 23597149 | Unspecified |
| mmu-miR-378a-3p | MIMAT0003151 | Lrp12 | 239393 | Microarrays | 37986224\|22855601 | Unspecified |
| mmu-miR-378a-3p | MIMAT0003151 | Lrp4 | 228357 | HITS-CLIP | 37986224 | Unspecified |
| mmu-miR-378a-3p | MIMAT0003151 | Lrp5 | 16973 | HITS-CLIP | 37986224 | Unspecified |
| mmu-miR-378a-3p | MIMAT0003151 | Lrp8 | 16975 | Microarrays//HITS-CLIP | 37986224\|22855601\|23142080 | Unspecified |
| mmu-miR-378a-3p | MIMAT0003151 | Lrrc17 | 74511 | Microarrays | 37986224 | Unspecified |
| mmu-miR-378a-3p | MIMAT0003151 | Lrrc57 | 66606 | HITS-CLIP | 23142080 | Unspecified |
| mmu-miR-378a-3p | MIMAT0003151 | Lrrc58 | 320184 | HITS-CLIP | 37986224 | Unspecified |
| mmu-miR-378a-3p | MIMAT0003151 | Lrrc59 | 98238 | HITS-CLIP | 37986224 | Unspecified |
| mmu-miR-378a-3p | MIMAT0003151 | Lrrc8a | 241296 | HITS-CLIP | 25083871\|23142080 | Unspecified |
| mmu-miR-378a-3p | MIMAT0003151 | Lrrc8c | 100604 | Microarrays | 37986224\|22855601 | Unspecified |
| mmu-miR-378a-3p | MIMAT0003151 | Lrrc8d | 231549 | HITS-CLIP | 37986224\|25753418 | Unspecified |
| mmu-miR-378a-3p | MIMAT0003151 | Lsm14b | 241846 | HITS-CLIP | 37986224 | Unspecified |
| mmu-miR-378a-3p | MIMAT0003151 | Lss | 16987 | HITS-CLIP | 37986224\|23142080 | Unspecified |
| mmu-miR-378a-3p | MIMAT0003151 | Ltbp2 | 16997 | Microarrays | 37986224\|22855601 | Unspecified |
| mmu-miR-378a-3p | MIMAT0003151 | Luc7l2 | 192196 | HITS-CLIP | 23142080\|29167373 | Unspecified |
| mmu-miR-378a-3p | MIMAT0003151 | Luc7l3 | 67684 | HITS-CLIP | 37986224\|23142080 | Unspecified |
| mmu-miR-378a-3p | MIMAT0003151 | Lum | 17022 | Microarrays | 37986224 | Unspecified |
| mmu-miR-378a-3p | MIMAT0003151 | Lurap1l | 52829 | HITS-CLIP | 23597149 | Unspecified |
| mmu-miR-378a-3p | MIMAT0003151 | Luzp1 | 269593 | HITS-CLIP | 37986224 | Unspecified |
| mmu-miR-378a-3p | MIMAT0003151 | Ly75 | 17076 | Microarrays | 37986224\|22855601 | Unspecified |
| mmu-miR-378a-3p | MIMAT0003151 | Lyst | 17101 | HITS-CLIP | 37986224\|25083871\|23142080\|29167 | Unspecified |

| | | | | | | |
|---|---|---|---|---|---|---|
| mmu-miR-378a-3p | MIMAT0003151 | Macf1 | 11426 | HITS-CLIP | 25083871\|25753418\|29167373 | Unspecified |
| mmu-miR-378a-3p | MIMAT0003151 | Maea | 59003 | HITS-CLIP | 25083871\|23142080\|29167373 | Unspecified |
| mmu-miR-378a-3p | MIMAT0003151 | Maff | 17133 | HITS-CLIP | 37986224\|25083871\|23142080 | Unspecified |
| mmu-miR-378a-3p | MIMAT0003151 | Magi3 | 99470 | HITS-CLIP | 37986224\|25753418 | Unspecified |
| mmu-miR-378a-3p | MIMAT0003151 | Map1b | 17755 | Microarrays | 37986224\|22855601 | Unspecified |
| mmu-miR-378a-3p | MIMAT0003151 | Map1lc3b | 67443 | HITS-CLIP | 37986224 | Unspecified |
| mmu-miR-378a-3p | MIMAT0003151 | Map2k4 | 26398 | HITS-CLIP | 37986224\|25083871 | Unspecified |
| mmu-miR-378a-3p | MIMAT0003151 | Map3k11 | 26403 | HITS-CLIP | 37986224 | Unspecified |
| mmu-miR-378a-3p | MIMAT0003151 | Map3k2 | 26405 | HITS-CLIP | 23142080 | Unspecified |
| mmu-miR-378a-3p | MIMAT0003151 | Map3k20 | 65964 | Microarrays | 22855601 | Unspecified |
| mmu-miR-378a-3p | MIMAT0003151 | Map3k7 | 26409 | HITS-CLIP | 37986224 | Unspecified |
| mmu-miR-378a-3p | MIMAT0003151 | Map4k4 | 26921 | Microarrays | 37986224\|22855601 | Unspecified |
| mmu-miR-378a-3p | MIMAT0003151 | Mapk1 | 26413 | HITS-CLIP//Luciferase reporter assay | 25083871\|23142080\|25529446 | Unspecified |
| mmu-miR-378a-3p | MIMAT0003151 | Mapk6 | 50772 | Microarrays | 37986224\|22855601 | Unspecified |
| mmu-miR-378a-3p | MIMAT0003151 | Mapk8ip3 | 30957 | HITS-CLIP | 37986224 | Unspecified |
| mmu-miR-378a-3p | MIMAT0003151 | March6 | 10299 | HITS-CLIP | 25083871\|23142080 | Unspecified |
| mmu-miR-378a-3p | MIMAT0003151 | March8 | 71779 | HITS-CLIP | 37986224 | Unspecified |
| mmu-miR-378a-3p | MIMAT0003151 | Mast2 | 17776 | Microarrays//HITS-CLIP | 37986224\|22855601\|25083871\|23142 | Unspecified |
| mmu-miR-378a-3p | MIMAT0003151 | Matr3 | 17184 | HITS-CLIP | 25083871\|23142080\|23597149\|29167 | Unspecified |
| mmu-miR-378a-3p | MIMAT0003151 | Mau2 | 74549 | HITS-CLIP | 37986224 | Unspecified |
| mmu-miR-378a-3p | MIMAT0003151 | Mbd5 | 109241 | HITS-CLIP | 37986224\|23142080 | Unspecified |
| mmu-miR-378a-3p | MIMAT0003151 | Mbnl2 | 105559 | HITS-CLIP | 37986224\|29167373 | Unspecified |
| mmu-miR-378a-3p | MIMAT0003151 | Mcc | 328949 | HITS-CLIP | 37986224 | Unspecified |
| mmu-miR-378a-3p | MIMAT0003151 | Mcl1 | 17210 | HITS-CLIP | 37986224\|23142080 | Unspecified |
| mmu-miR-378a-3p | MIMAT0003151 | Mcm2 | 17216 | HITS-CLIP | 29167373 | Unspecified |
| mmu-miR-378a-3p | MIMAT0003151 | Mcm3 | 17215 | Microarrays | 37986224\|22855601 | Unspecified |
| mmu-miR-378a-3p | MIMAT0003151 | Mcm3ap | 54387 | HITS-CLIP | 25083871 | Unspecified |
| mmu-miR-378a-3p | MIMAT0003151 | Mdfic | 16543 | HITS-CLIP | 37986224\|25083871\|23597149\|23142 | Unspecified |
| mmu-miR-378a-3p | MIMAT0003151 | Mdm2 | 17246 | Microarrays | 37986224\|22855601 | Unspecified |
| mmu-miR-378a-3p | MIMAT0003151 | Mdm4 | 17248 | HITS-CLIP | 37986224 | Unspecified |
| mmu-miR-378a-3p | MIMAT0003151 | Mdn1 | 100019 | HITS-CLIP, Microarrays//Microarrays | 37986224\|22855601 | Unspecified |
| mmu-miR-378a-3p | MIMAT0003151 | Mecom | 14013 | HITS-CLIP | 25083871 | Unspecified |
| mmu-miR-378a-3p | MIMAT0003151 | Mecp2 | 17257 | HITS-CLIP | 25083871 | Unspecified |
| mmu-miR-378a-3p | MIMAT0003151 | Med13 | 327987 | HITS-CLIP | 37986224\|25083871\|29167373 | Unspecified |
| mmu-miR-378a-3p | MIMAT0003151 | Med14 | 26896 | HITS-CLIP | 37986224 | Unspecified |
| mmu-miR-378a-3p | MIMAT0003151 | Med22 | 20933 | HITS-CLIP | 37986224 | Unspecified |
| mmu-miR-378a-3p | MIMAT0003151 | Med26 | 70625 | HITS-CLIP | 23142080 | Unspecified |
| mmu-miR-378a-3p | MIMAT0003151 | Med7 | 66213 | HITS-CLIP | 23597149 | Unspecified |
| mmu-miR-378a-3p | MIMAT0003151 | Mef2d | 17261 | HITS-CLIP | 37986224 | Unspecified |

| | | | | | | |
|---|---|---|---|---|---|---|
| mmu-miR-378a-3p | MIMAT0003151 | Megf9 | 230316 | HITS-CLIP | 23597149 | Unspecified |
| mmu-miR-378a-3p | MIMAT0003151 | Meis2 | 17536 | HITS-CLIP | 37986224 | Unspecified |
| mmu-miR-378a-3p | MIMAT0003151 | Mest | 17294 | HITS-CLIP, Microarrays | 37986224 | Unspecified |
| mmu-miR-378a-3p | MIMAT0003151 | Metap2 | 56307 | HITS-CLIP | 25083871 | Unspecified |
| mmu-miR-378a-3p | MIMAT0003151 | Mettl7b | 71664 | HITS-CLIP | 37986224 | Unspecified |
| mmu-miR-378a-3p | MIMAT0003151 | Mex3c | 240396 | Microarrays | 37986224\|22855601 | Unspecified |
| mmu-miR-378a-3p | MIMAT0003151 | Mff | 75734 | HITS-CLIP | 23597149\|29167373 | Unspecified |
| mmu-miR-378a-3p | MIMAT0003151 | Mga | 29808 | HITS-CLIP | 23142080 | Unspecified |
| mmu-miR-378a-3p | MIMAT0003151 | Mgat3 | 17309 | HITS-CLIP | 37986224 | Unspecified |
| mmu-miR-378a-3p | MIMAT0003151 | Mgrn1 | 17237 | HITS-CLIP | 37986224 | Unspecified |
| mmu-miR-378a-3p | MIMAT0003151 | Mib1 | 225164 | HITS-CLIP | 37986224 | Unspecified |
| mmu-miR-378a-3p | MIMAT0003151 | Mid1ip1 | 68041 | HITS-CLIP | 25083871 | Unspecified |
| mmu-miR-378a-3p | MIMAT0003151 | Mid2 | 23947 | HITS-CLIP | 23597149 | Unspecified |
| mmu-miR-378a-3p | MIMAT0003151 | Mief1 | 239555 | HITS-CLIP | 37986224\|23142080\|25083871 | Unspecified |
| mmu-miR-378a-3p | MIMAT0003151 | Mindy3 | 80013 | qCLASH | 31393953 | Unspecified |
| mmu-miR-378a-3p | MIMAT0003151 | Mki67 | 17345 | HITS-CLIP | 37986224 | Unspecified |
| mmu-miR-378a-3p | MIMAT0003151 | Mknk1 | 17346 | HITS-CLIP | 37986224 | Unspecified |
| mmu-miR-378a-3p | MIMAT0003151 | Mkrn1 | 54484 | HITS-CLIP | 25083871\|23142080 | Unspecified |
| mmu-miR-378a-3p | MIMAT0003151 | Mllt3 | 70122 | HITS-CLIP | 37986224\|23142080 | Unspecified |
| mmu-miR-378a-3p | MIMAT0003151 | Mmadhc | 109129 | HITS-CLIP | 37986224 | Unspecified |
| mmu-miR-378a-3p | MIMAT0003151 | Mmgt2 | 216829 | HITS-CLIP | 37986224 | Unspecified |
| mmu-miR-378a-3p | MIMAT0003151 | Mms19 | 72199 | HITS-CLIP | 23142080\|25083871 | Unspecified |
| mmu-miR-378a-3p | MIMAT0003151 | Mnt | 17428 | HITS-CLIP//qCLASH | 37986224\|31393953 | Unspecified |
| mmu-miR-378a-3p | MIMAT0003151 | Mob1a | 232157 | HITS-CLIP | 37986224 | Unspecified |
| mmu-miR-378a-3p | MIMAT0003151 | Mob3b | 214944 | HITS-CLIP | 37986224 | Unspecified |
| mmu-miR-378a-3p | MIMAT0003151 | Morc2a | 74522 | HITS-CLIP | 37986224\|25753418 | Unspecified |
| mmu-miR-378a-3p | MIMAT0003151 | Morf4l2 | 56397 | HITS-CLIP | 25083871\|23142080 | Unspecified |
| mmu-miR-378a-3p | MIMAT0003151 | Mosmo | 233812 | Microarrays | 22855601 | Unspecified |
| mmu-miR-378a-3p | MIMAT0003151 | Mphosph10 | 67973 | HITS-CLIP, Microarrays//Microarrays | 37986224\|22855601 | Unspecified |
| mmu-miR-378a-3p | MIMAT0003151 | Mprip | 26936 | HITS-CLIP | 37986224 | Unspecified |
| mmu-miR-378a-3p | MIMAT0003151 | Mrgbp | 73247 | HITS-CLIP | 37986224 | Unspecified |
| mmu-miR-378a-3p | MIMAT0003151 | Mrm3 | 67390 | HITS-CLIP | 23597149 | Unspecified |
| mmu-miR-378a-3p | MIMAT0003151 | Mroh1 | 223658 | HITS-CLIP | 29167373 | Unspecified |
| mmu-miR-378a-3p | MIMAT0003151 | Mrpl15 | 27395 | qCLASH | 31393953 | Unspecified |
| mmu-miR-378a-3p | MIMAT0003151 | Mrpl38 | 60441 | HITS-CLIP | 25753418 | Unspecified |
| mmu-miR-378a-3p | MIMAT0003151 | Mrpl39 | 27393 | HITS-CLIP | 23597149 | Unspecified |
| mmu-miR-378a-3p | MIMAT0003151 | Mrps10 | 64657 | HITS-CLIP | 37986224\|23142080 | Unspecified |
| mmu-miR-378a-3p | MIMAT0003151 | Msi2 | 76626 | HITS-CLIP | 25083871\|23142080 | Unspecified |
| mmu-miR-378a-3p | MIMAT0003151 | Msl2 | 77853 | HITS-CLIP | 37986224 | Unspecified |

| | | | | | | |
|---|---|---|---|---|---|---|
| mmu-miR-378a-3p | MIMAT0003151 | Msn | 17698 | HITS-CLIP | 37986224\|23142080 | Unspecified |
| mmu-miR-378a-3p | MIMAT0003151 | Msr1 | 20288 | HITS-CLIP | 23597149 | Unspecified |
| mmu-miR-378a-3p | MIMAT0003151 | Mtbp | 105837 | Microarrays | 37986224\|22855601 | Unspecified |
| mmu-miR-378a-3p | MIMAT0003151 | Mtcl1 | 68617 | HITS-CLIP | 37986224 | Unspecified |
| mmu-miR-378a-3p | MIMAT0003151 | Mtf1 | 17764 | HITS-CLIP | 37986224 | Unspecified |
| mmu-miR-378a-3p | MIMAT0003151 | Mthfd1 | 108156 | Microarrays | 37986224\|22855601 | Unspecified |
| mmu-miR-378a-3p | MIMAT0003151 | Mthfd1l | 270685 | Microarrays | 37986224\|22855601 | Unspecified |
| mmu-miR-378a-3p | MIMAT0003151 | Mtmr3 | 74302 | HITS-CLIP | 37986224\|23597149 | Unspecified |
| mmu-miR-378a-3p | MIMAT0003151 | Mtss1 | 211401 | HITS-CLIP | 37986224\|25083871 | Unspecified |
| mmu-miR-378a-3p | MIMAT0003151 | Mtx3 | 382793 | HITS-CLIP | 37986224 | Unspecified |
| mmu-miR-378a-3p | MIMAT0003151 | Mybbp1a | 18432 | Microarrays | 37986224\|22855601 | Unspecified |
| mmu-miR-378a-3p | MIMAT0003151 | Myc | 17869 | Microarrays | 37986224\|22855601 | Unspecified |
| mmu-miR-378a-3p | MIMAT0003151 | Mycbp2 | 105689 | HITS-CLIP | 23142080 | Unspecified |
| mmu-miR-378a-3p | MIMAT0003151 | Myh1 | 17879 | HITS-CLIP | 25083871 | Unspecified |
| mmu-miR-378a-3p | MIMAT0003151 | Myh10 | 77579 | HITS-CLIP | 37986224\|25083871 | Unspecified |
| mmu-miR-378a-3p | MIMAT0003151 | Myh3 | 17883 | HITS-CLIP | 25083871 | Unspecified |
| mmu-miR-378a-3p | MIMAT0003151 | Myh9 | 17886 | HITS-CLIP | 29167373 | Unspecified |
| mmu-miR-378a-3p | MIMAT0003151 | Mylip | 218203 | HITS-CLIP | 25753418 | Unspecified |
| mmu-miR-378a-3p | MIMAT0003151 | Mylk | 107589 | HITS-CLIP | 37986224 | Unspecified |
| mmu-miR-378a-3p | MIMAT0003151 | Myo1e | 71602 | HITS-CLIP | 25083871 | Unspecified |
| mmu-miR-378a-3p | MIMAT0003151 | Myo5a | 17918 | HITS-CLIP | 25083871\|29167373 | Unspecified |
| mmu-miR-378a-3p | MIMAT0003151 | Myo6 | 17920 | HITS-CLIP | 23597149\|25753418 | Unspecified |
| mmu-miR-378a-3p | MIMAT0003151 | Myo9a | 270163 | HITS-CLIP | 23142080 | Unspecified |
| mmu-miR-378a-3p | MIMAT0003151 | Myo9b | 17925 | HITS-CLIP | 25083871 | Unspecified |
| mmu-miR-378a-3p | MIMAT0003151 | Myog | 17928 | HITS-CLIP | 37986224 | Unspecified |
| mmu-miR-378a-3p | MIMAT0003151 | N4bp2l2 | 381695 | HITS-CLIP | 37986224 | Unspecified |
| mmu-miR-378a-3p | MIMAT0003151 | Naa10 | 56292 | HITS-CLIP | 23142080 | Unspecified |
| mmu-miR-378a-3p | MIMAT0003151 | Naa15 | 74838 | Microarrays | 37986224\|22855601 | Unspecified |
| mmu-miR-378a-3p | MIMAT0003151 | Naa16 | 66897 | HITS-CLIP | 23142080 | Unspecified |
| mmu-miR-378a-3p | MIMAT0003151 | Naa25 | 231713 | Microarrays//HITS-CLIP | 37986224\|22855601\|25083871\|23142 | Unspecified |
| mmu-miR-378a-3p | MIMAT0003151 | Naca | 17938 | HITS-CLIP | 37986224 | Unspecified |
| mmu-miR-378a-3p | MIMAT0003151 | Nacc1 | 66830 | HITS-CLIP | 37986224 | Unspecified |
| mmu-miR-378a-3p | MIMAT0003151 | Nadk | 192185 | HITS-CLIP | 37986224 | Unspecified |
| mmu-miR-378a-3p | MIMAT0003151 | Nadk2 | 68646 | HITS-CLIP | 37986224\|23597149 | Unspecified |
| mmu-miR-378a-3p | MIMAT0003151 | Nap1l1 | 53605 | HITS-CLIP | 37986224 | Unspecified |
| mmu-miR-378a-3p | MIMAT0003151 | Nars | 70223 | qCLASH | 31393953 | Unspecified |
| mmu-miR-378a-3p | MIMAT0003151 | Nav1 | 215690 | HITS-CLIP | 37986224 | Unspecified |
| mmu-miR-378a-3p | MIMAT0003151 | Nav2 | 78286 | HITS-CLIP | 25083871 | Unspecified |
| mmu-miR-378a-3p | MIMAT0003151 | Nav3 | 260315 | HITS-CLIP | 25083871 | Unspecified |

| | | | | | | |
|---|---|---|---|---|---|---|
| mmu-miR-378a-3p | MIMAT0003151 | Nbeal1 | 269198 | HITS-CLIP | 37986224 | Unspecified |
| mmu-miR-378a-3p | MIMAT0003151 | Nbeal2 | 235627 | HITS-CLIP | 23142080 | Unspecified |
| mmu-miR-378a-3p | MIMAT0003151 | Ncam1 | 17967 | HITS-CLIP | 37986224\|25083871 | Unspecified |
| mmu-miR-378a-3p | MIMAT0003151 | Ncapd2 | 68298 | HITS-CLIP | 23142080 | Unspecified |
| mmu-miR-378a-3p | MIMAT0003151 | Nceh1 | 320024 | Microarrays | 37986224\|22855601 | Unspecified |
| mmu-miR-378a-3p | MIMAT0003151 | Nckap1l | 105855 | HITS-CLIP | 23142080 | Unspecified |
| mmu-miR-378a-3p | MIMAT0003151 | Ncoa3 | 17979 | HITS-CLIP | 23142080\|29167373 | Unspecified |
| mmu-miR-378a-3p | MIMAT0003151 | Ncoa6 | 56406 | HITS-CLIP | 37986224\|23142080 | Unspecified |
| mmu-miR-378a-3p | MIMAT0003151 | Ncoa7 | 211329 | HITS-CLIP | 25083871\|23142080 | Unspecified |
| mmu-miR-378a-3p | MIMAT0003151 | Ncor1 | 20185 | HITS-CLIP | 37986224\|23597149\|25083871 | Unspecified |
| mmu-miR-378a-3p | MIMAT0003151 | Ncs1 | 14299 | Microarrays | 37986224\|22855601 | Unspecified |
| mmu-miR-378a-3p | MIMAT0003151 | Ndel1 | 83431 | HITS-CLIP | 25083871 | Unspecified |
| mmu-miR-378a-3p | MIMAT0003151 | Ndrg4 | 234593 | HITS-CLIP | 25083871 | Unspecified |
| mmu-miR-378a-3p | MIMAT0003151 | Ndufa9 | 66108 | Microarrays | 37986224\|22855601 | Unspecified |
| mmu-miR-378a-3p | MIMAT0003151 | Ndufaf7 | 73694 | HITS-CLIP | 23142080 | Unspecified |
| mmu-miR-378a-3p | MIMAT0003151 | Ndufc1 | 66377 | HITS-CLIP | 25753418 | Unspecified |
| mmu-miR-378a-3p | MIMAT0003151 | Ndufs3 | 68349 | HITS-CLIP | 25753418 | Unspecified |
| mmu-miR-378a-3p | MIMAT0003151 | Ndufs4 | 17993 | HITS-CLIP | 25753418 | Unspecified |
| mmu-miR-378a-3p | MIMAT0003151 | Necap1 | 67602 | HITS-CLIP | 25083871\|23597149\|23142080 | Unspecified |
| mmu-miR-378a-3p | MIMAT0003151 | Nedd1 | 17997 | HITS-CLIP | 25083871 | Unspecified |
| mmu-miR-378a-3p | MIMAT0003151 | Nek1 | 18004 | HITS-CLIP | 37986224 | Unspecified |
| mmu-miR-378a-3p | MIMAT0003151 | Nelfa | 24116 | HITS-CLIP | 25083871 | Unspecified |
| mmu-miR-378a-3p | MIMAT0003151 | Net1 | 56349 | HITS-CLIP | 37986224 | Unspecified |
| mmu-miR-378a-3p | MIMAT0003151 | Nfat5 | 54446 | HITS-CLIP | 23142080\|25083871 | Unspecified |
| mmu-miR-378a-3p | MIMAT0003151 | Nfatc3 | 18021 | HITS-CLIP | 37986224\|23142080\|29167373 | Unspecified |
| mmu-miR-378a-3p | MIMAT0003151 | Nfe2l1 | 18023 | HITS-CLIP | 37986224\|29167373 | Unspecified |
| mmu-miR-378a-3p | MIMAT0003151 | Nfia | 18027 | HITS-CLIP | 23597149 | Unspecified |
| mmu-miR-378a-3p | MIMAT0003151 | Nfib | 18028 | HITS-CLIP | 37986224 | Unspecified |
| mmu-miR-378a-3p | MIMAT0003151 | Nfil3 | 18030 | Microarrays | 37986224\|22855601 | Unspecified |
| mmu-miR-378a-3p | MIMAT0003151 | Nhlrc2 | 66866 | HITS-CLIP, Microarrays//Microarrays | 37986224\|22855601 | Unspecified |
| mmu-miR-378a-3p | MIMAT0003151 | Nid1 | 18073 | Microarrays | 37986224\|22855601 | Unspecified |
| mmu-miR-378a-3p | MIMAT0003151 | Nid2 | 18074 | Microarrays | 37986224 | Unspecified |
| mmu-miR-378a-3p | MIMAT0003151 | Nin | 18080 | HITS-CLIP | 37986224\|29167373\|25083871 | Unspecified |
| mmu-miR-378a-3p | MIMAT0003151 | Nipal1 | 70701 | Microarrays | 37986224\|22855601 | Unspecified |
| mmu-miR-378a-3p | MIMAT0003151 | Nipbl | 71175 | HITS-CLIP | 23142080 | Unspecified |
| mmu-miR-378a-3p | MIMAT0003151 | Nisch | 64652 | HITS-CLIP | 37986224 | Unspecified |
| mmu-miR-378a-3p | MIMAT0003151 | Nktr | 18087 | HITS-CLIP | 37986224 | Unspecified |
| mmu-miR-378a-3p | MIMAT0003151 | Nlrp6 | 101613 | HITS-CLIP | 37986224 | Unspecified |
| mmu-miR-378a-3p | MIMAT0003151 | Nmt2 | 18108 | HITS-CLIP, Microarrays//Microarrays | 37986224\|22855601 | Unspecified |

| | | | | | | |
|---|---|---|---|---|---|---|
| mmu-miR-378a-3p | MIMAT0003151 | Noa1 | 56412 | HITS-CLIP | 23597149 | Unspecified |
| mmu-miR-378a-3p | MIMAT0003151 | Noc3l | 57753 | Microarrays | 37986224\|22855601 | Unspecified |
| mmu-miR-378a-3p | MIMAT0003151 | Nol10 | 217431 | Microarrays | 37986224\|22855601 | Unspecified |
| mmu-miR-378a-3p | MIMAT0003151 | Nolc1 | 70769 | HITS-CLIP | 37986224\|25753418 | Unspecified |
| mmu-miR-378a-3p | MIMAT0003151 | Nomo1 | 211548 | Microarrays | 37986224\|22855601 | Unspecified |
| mmu-miR-378a-3p | MIMAT0003151 | Nop14 | 75416 | HITS-CLIP | 25083871\|23597149 | Unspecified |
| mmu-miR-378a-3p | MIMAT0003151 | Nop2 | 110109 | Microarrays | 37986224\|22855601 | Unspecified |
| mmu-miR-378a-3p | MIMAT0003151 | Nop58 | 55989 | Microarrays | 37986224\|22855601 | Unspecified |
| mmu-miR-378a-3p | MIMAT0003151 | Npdc1 | 18146 | HITS-CLIP | 37986224 | Unspecified |
| mmu-miR-378a-3p | MIMAT0003151 | Nploc4 | 217365 | HITS-CLIP | 23597149 | Unspecified |
| mmu-miR-378a-3p | MIMAT0003151 | Npm1 | 18148 | HITS-CLIP | 37986224 | Unspecified |
| mmu-miR-378a-3p | MIMAT0003151 | Nr1d1 | 217166 | HITS-CLIP, Microarrays//Microarrays | 37986224\|22855601 | Unspecified |
| mmu-miR-378a-3p | MIMAT0003151 | Nr2c2 | 22026 | HITS-CLIP | 37986224 | Unspecified |
| mmu-miR-378a-3p | MIMAT0003151 | Nr3c1 | 14815 | HITS-CLIP | 37986224\|23142080 | Unspecified |
| mmu-miR-378a-3p | MIMAT0003151 | Nr4a1 | 15370 | Microarrays | 37986224\|22855601 | Unspecified |
| mmu-miR-378a-3p | MIMAT0003151 | Nras | 18176 | HITS-CLIP | 37986224 | Unspecified |
| mmu-miR-378a-3p | MIMAT0003151 | Nrd1 | 230598 | HITS-CLIP | 25083871\|23142080\|29167373 | Unspecified |
| mmu-miR-378a-3p | MIMAT0003151 | Nrip1 | 268903 | HITS-CLIP | 23142080 | Unspecified |
| mmu-miR-378a-3p | MIMAT0003151 | Nrn1 | 68404 | Microarrays | 37986224\|22855601 | Unspecified |
| mmu-miR-378a-3p | MIMAT0003151 | Nrp2 | 18187 | HITS-CLIP | 25083871 | Unspecified |
| mmu-miR-378a-3p | MIMAT0003151 | Nsd1 | 18193 | HITS-CLIP | 37986224 | Unspecified |
| mmu-miR-378a-3p | MIMAT0003151 | Nsd2 | 107823 | HITS-CLIP | 23142080 | Unspecified |
| mmu-miR-378a-3p | MIMAT0003151 | Nsd3 | 234135 | HITS-CLIP | 25083871\|23142080\|29167373 | Unspecified |
| mmu-miR-378a-3p | MIMAT0003151 | Nt5c1a | 84618 | HITS-CLIP | 25753418 | Unspecified |
| mmu-miR-378a-3p | MIMAT0003151 | Nuak1 | 77976 | HITS-CLIP | 37986224 | Unspecified |
| mmu-miR-378a-3p | MIMAT0003151 | Nucks1 | 98415 | HITS-CLIP | 25083871\|23142080\|29167373 | Unspecified |
| mmu-miR-378a-3p | MIMAT0003151 | Nudt18 | 213484 | HITS-CLIP | 23597149 | Unspecified |
| mmu-miR-378a-3p | MIMAT0003151 | Nufip2 | 68564 | HITS-CLIP | 37986224\|25083871\|23142080 | Unspecified |
| mmu-miR-378a-3p | MIMAT0003151 | Numa1 | 101706 | HITS-CLIP | 37986224 | Unspecified |
| mmu-miR-378a-3p | MIMAT0003151 | Nup153 | 218210 | Microarrays//HITS-CLIP | 37986224\|22855601\|29167373 | Unspecified |
| mmu-miR-378a-3p | MIMAT0003151 | Nup155 | 170762 | Microarrays | 37986224\|22855601 | Unspecified |
| mmu-miR-378a-3p | MIMAT0003151 | Nusap1 | 108907 | HITS-CLIP | 25083871 | Unspecified |
| mmu-miR-378a-3p | MIMAT0003151 | Nxf1 | 53319 | HITS-CLIP | 37986224\|25083871\|23142080\|29167 | Unspecified |
| mmu-miR-378a-3p | MIMAT0003151 | Oaf | 102644 | HITS-CLIP | 37986224 | Unspecified |
| mmu-miR-378a-3p | MIMAT0003151 | Ociad1 | 68095 | HITS-CLIP | 37986224\|23142080\|23597149 | Unspecified |
| mmu-miR-378a-3p | MIMAT0003151 | Olfm1 | 56177 | HITS-CLIP | 25083871 | Unspecified |
| mmu-miR-378a-3p | MIMAT0003151 | Orc2 | 18393 | HITS-CLIP | 25083871\|23597149 | Unspecified |
| mmu-miR-378a-3p | MIMAT0003151 | Orc3 | 50793 | HITS-CLIP | 37986224 | Unspecified |
| mmu-miR-378a-3p | MIMAT0003151 | Orc6 | 56452 | HITS-CLIP | 25083871 | Unspecified |

| | | | | | | |
|---|---|---|---|---|---|---|
| mmu-miR-378a-3p | MIMAT0003151 | Ormdl1 | 227102 | HITS-CLIP | 37986224 | Unspecified |
| mmu-miR-378a-3p | MIMAT0003151 | Os9 | 216440 | HITS-CLIP | 37986224 | Unspecified |
| mmu-miR-378a-3p | MIMAT0003151 | Osbpl3 | 71720 | Microarrays | 37986224\|22855601 | Unspecified |
| mmu-miR-378a-3p | MIMAT0003151 | Otub2 | 68149 | HITS-CLIP | 37986224 | Unspecified |
| mmu-miR-378a-3p | MIMAT0003151 | Otud4 | 73945 | Microarrays | 37986224\|22855601 | Unspecified |
| mmu-miR-378a-3p | MIMAT0003151 | Oxa1l | 69089 | HITS-CLIP | 23597149 | Unspecified |
| mmu-miR-378a-3p | MIMAT0003151 | Pacsin2 | 23970 | HITS-CLIP | 37986224 | Unspecified |
| mmu-miR-378a-3p | MIMAT0003151 | Pafah1b1 | 18472 | HITS-CLIP | 23597149\|23142080 | Unspecified |
| mmu-miR-378a-3p | MIMAT0003151 | Pafah1b2 | 18475 | HITS-CLIP | 37986224 | Unspecified |
| mmu-miR-378a-3p | MIMAT0003151 | Paip1 | 218693 | HITS-CLIP | 25083871\|23142080\|29167373 | Unspecified |
| mmu-miR-378a-3p | MIMAT0003151 | Pak4 | 70584 | HITS-CLIP | 37986224 | Unspecified |
| mmu-miR-378a-3p | MIMAT0003151 | Papola | 18789 | HITS-CLIP | 37986224\|25083871\|23142080\|29167 | Unspecified |
| mmu-miR-378a-3p | MIMAT0003151 | Parg | 26430 | HITS-CLIP | 37986224\|25083871 | Unspecified |
| mmu-miR-378a-3p | MIMAT0003151 | Parl | 381038 | HITS-CLIP | 23142080 | Unspecified |
| mmu-miR-378a-3p | MIMAT0003151 | Parp14 | 547253 | HITS-CLIP | 37986224\|29167373 | Unspecified |
| mmu-miR-378a-3p | MIMAT0003151 | Pars2 | 230577 | HITS-CLIP | 23142080 | Unspecified |
| mmu-miR-378a-3p | MIMAT0003151 | Parva | 57342 | HITS-CLIP | 23597149\|25083871 | Unspecified |
| mmu-miR-378a-3p | MIMAT0003151 | Patl1 | 225929 | HITS-CLIP | 25083871 | Unspecified |
| mmu-miR-378a-3p | MIMAT0003151 | Pbrm1 | 66923 | HITS-CLIP | 25083871 | Unspecified |
| mmu-miR-378a-3p | MIMAT0003151 | Pbx3 | 18516 | HITS-CLIP | 25083871 | Unspecified |
| mmu-miR-378a-3p | MIMAT0003151 | Pbxip1 | 229534 | HITS-CLIP | 37986224 | Unspecified |
| mmu-miR-378a-3p | MIMAT0003151 | Pcbp2 | 18521 | HITS-CLIP | 37986224 | Unspecified |
| mmu-miR-378a-3p | MIMAT0003151 | Pcdh7 | 54216 | Microarrays//HITS-CLIP | 37986224\|25083871\|22855601 | Unspecified |
| mmu-miR-378a-3p | MIMAT0003151 | Pcdhga12 | 93724 | HITS-CLIP | 37986224 | Unspecified |
| mmu-miR-378a-3p | MIMAT0003151 | Pcm1 | 18536 | HITS-CLIP | 37986224 | Unspecified |
| mmu-miR-378a-3p | MIMAT0003151 | Pcmtd1 | 319263 | HITS-CLIP | 37986224 | Unspecified |
| mmu-miR-378a-3p | MIMAT0003151 | Pcnx | 54604 | HITS-CLIP | 23142080 | Unspecified |
| mmu-miR-378a-3p | MIMAT0003151 | Pcolce | 18542 | HITS-CLIP | 37986224 | Unspecified |
| mmu-miR-378a-3p | MIMAT0003151 | Pdcd6ip | 18571 | HITS-CLIP | 23597149 | Unspecified |
| mmu-miR-378a-3p | MIMAT0003151 | Pdcl | 67466 | HITS-CLIP | 37986224 | Unspecified |
| mmu-miR-378a-3p | MIMAT0003151 | Pde3b | 18576 | HITS-CLIP | 23597149 | Unspecified |
| mmu-miR-378a-3p | MIMAT0003151 | Pde4d | 238871 | HITS-CLIP | 37986224 | Unspecified |
| mmu-miR-378a-3p | MIMAT0003151 | Pde8a | 18584 | HITS-CLIP | 29167373 | Unspecified |
| mmu-miR-378a-3p | MIMAT0003151 | Pdgfa | 18590 | Microarrays | 37986224\|22855601 | Unspecified |
| mmu-miR-378a-3p | MIMAT0003151 | Pdgfc | 54635 | Microarrays | 37986224\|22855601 | Unspecified |
| mmu-miR-378a-3p | MIMAT0003151 | Pdia3 | 14827 | HITS-CLIP | 25083871 | Unspecified |
| mmu-miR-378a-3p | MIMAT0003151 | Pdia4 | 12304 | HITS-CLIP | 37986224 | Unspecified |
| mmu-miR-378a-3p | MIMAT0003151 | Pdlim5 | 56376 | Microarrays | 37986224\|22855601 | Unspecified |
| mmu-miR-378a-3p | MIMAT0003151 | Pdpk1 | 18607 | HITS-CLIP | 23142080 | Unspecified |

| | | | | | | |
|---|---|---|---|---|---|---|
| mmu-miR-378a-3p | MIMAT0003151 | Pdzd8 | 107368 | HITS-CLIP, Microarrays//Microarrays | 37986224\|22855601 | Unspecified |
| mmu-miR-378a-3p | MIMAT0003151 | Pdzk1 | 59020 | HITS-CLIP | 37986224 | Unspecified |
| mmu-miR-378a-3p | MIMAT0003151 | Peak1 | 244895 | HITS-CLIP | 37986224 | Unspecified |
| mmu-miR-378a-3p | MIMAT0003151 | Pef1 | 67898 | HITS-CLIP | 37986224\|23142080 | Unspecified |
| mmu-miR-378a-3p | MIMAT0003151 | Peli1 | 67245 | HITS-CLIP, Microarrays//Microarrays | 37986224\|22855601 | Unspecified |
| mmu-miR-378a-3p | MIMAT0003151 | Per2 | 18627 | HITS-CLIP | 37986224 | Unspecified |
| mmu-miR-378a-3p | MIMAT0003151 | Pex13 | 72129 | HITS-CLIP | 37986224 | Unspecified |
| mmu-miR-378a-3p | MIMAT0003151 | Pex5 | 19305 | HITS-CLIP | 37986224 | Unspecified |
| mmu-miR-378a-3p | MIMAT0003151 | Pfas | 237823 | Microarrays | 37986224\|22855601 | Unspecified |
| mmu-miR-378a-3p | MIMAT0003151 | Pgm2l1 | 70974 | HITS-CLIP | 29167373 | Unspecified |
| mmu-miR-378a-3p | MIMAT0003151 | Pgrmc1 | 53328 | HITS-CLIP | 37986224\|25083871\|23597149\|2314: | Unspecified |
| mmu-miR-378a-3p | MIMAT0003151 | Pgs1 | 74451 | Microarrays | 37986224\|22855601 | Unspecified |
| mmu-miR-378a-3p | MIMAT0003151 | Phc3 | 241915 | HITS-CLIP | 37986224\|23142080 | Unspecified |
| mmu-miR-378a-3p | MIMAT0003151 | Phf10 | 72057 | HITS-CLIP | 23142080 | Unspecified |
| mmu-miR-378a-3p | MIMAT0003151 | Phf12 | 268448 | HITS-CLIP | 37986224 | Unspecified |
| mmu-miR-378a-3p | MIMAT0003151 | Phf13 | 230936 | HITS-CLIP | 37986224 | Unspecified |
| mmu-miR-378a-3p | MIMAT0003151 | Phf23 | 78246 | HITS-CLIP | 23142080 | Unspecified |
| mmu-miR-378a-3p | MIMAT0003151 | Phldb2 | 208177 | HITS-CLIP | 25083871 | Unspecified |
| mmu-miR-378a-3p | MIMAT0003151 | Phrf1 | 101471 | HITS-CLIP | 37986224 | Unspecified |
| mmu-miR-378a-3p | MIMAT0003151 | Pi4k2b | 67073 | HITS-CLIP | 37986224 | Unspecified |
| mmu-miR-378a-3p | MIMAT0003151 | Pi4kb | 107650 | HITS-CLIP | 37986224 | Unspecified |
| mmu-miR-378a-3p | MIMAT0003151 | Pias1 | 56469 | HITS-CLIP | 29167373 | Unspecified |
| mmu-miR-378a-3p | MIMAT0003151 | Pias2 | 17344 | HITS-CLIP | 29167373 | Unspecified |
| mmu-miR-378a-3p | MIMAT0003151 | Piezo1 | 234839 | Microarrays | 37986224\|22855601 | Unspecified |
| mmu-miR-378a-3p | MIMAT0003151 | Piezo2 | 667742 | HITS-CLIP | 37986224 | Unspecified |
| mmu-miR-378a-3p | MIMAT0003151 | Pigc | 67292 | HITS-CLIP | 37986224 | Unspecified |
| mmu-miR-378a-3p | MIMAT0003151 | Pign | 27392 | HITS-CLIP | 25083871\|23142080 | Unspecified |
| mmu-miR-378a-3p | MIMAT0003151 | Pigu | 228812 | HITS-CLIP | 37986224 | Unspecified |
| mmu-miR-378a-3p | MIMAT0003151 | Pik3c2a | 18704 | HITS-CLIP | 29167373 | Unspecified |
| mmu-miR-378a-3p | MIMAT0003151 | Pik3c2b | 240752 | HITS-CLIP | 23142080 | Unspecified |
| mmu-miR-378a-3p | MIMAT0003151 | Pik3c3 | 225326 | HITS-CLIP | 23142080 | Unspecified |
| mmu-miR-378a-3p | MIMAT0003151 | Pik3cb | 74769 | Microarrays | 37986224\|22855601 | Unspecified |
| mmu-miR-378a-3p | MIMAT0003151 | Pik3cd | 18707 | HITS-CLIP | 29167373 | Unspecified |
| mmu-miR-378a-3p | MIMAT0003151 | Pik3r1 | 18708 | HITS-CLIP | 37986224 | Unspecified |
| mmu-miR-378a-3p | MIMAT0003151 | Pikfyve | 18711 | HITS-CLIP | 37986224\|29167373 | Unspecified |
| mmu-miR-378a-3p | MIMAT0003151 | Pimreg | 54478 | Microarrays | 22855601 | Unspecified |
| mmu-miR-378a-3p | MIMAT0003151 | Pin1 | 23988 | HITS-CLIP | 25083871 | Unspecified |
| mmu-miR-378a-3p | MIMAT0003151 | Pkd1 | 18763 | HITS-CLIP | 37986224 | Unspecified |
| mmu-miR-378a-3p | MIMAT0003151 | Pkp1 | 18772 | HITS-CLIP | 25083871 | Unspecified |

| | | | | | | | |
|---|---|---|---|---|---|---|---|
| mmu-miR-378a-3p | MIMAT0003151 | Pkp4 | 227937 | HITS-CLIP | 23597149\|25753418 | | Unspecified |
| mmu-miR-378a-3p | MIMAT0003151 | Pla2g4a | 18783 | Microarrays | 37986224\|22855601 | | Unspecified |
| mmu-miR-378a-3p | MIMAT0003151 | Pla2g7 | 27226 | Microarrays | 37986224\|22855601 | | Unspecified |
| mmu-miR-378a-3p | MIMAT0003151 | Plagl2 | 54711 | HITS-CLIP | | 37986224 | Unspecified |
| mmu-miR-378a-3p | MIMAT0003151 | Plau | 18792 | HITS-CLIP, Microarrays//Microarrays | 37986224\|22855601 | | Unspecified |
| mmu-miR-378a-3p | MIMAT0003151 | Plaur | 18793 | Microarrays | 37986224\|22855601 | | Unspecified |
| mmu-miR-378a-3p | MIMAT0003151 | Plbd2 | 71772 | HITS-CLIP | 37986224\|25753418 | | Unspecified |
| mmu-miR-378a-3p | MIMAT0003151 | Plcg1 | 18803 | HITS-CLIP | | 37986224 | Unspecified |
| mmu-miR-378a-3p | MIMAT0003151 | Plec | 18810 | Microarrays | 37986224\|22855601 | | Unspecified |
| mmu-miR-378a-3p | MIMAT0003151 | Plekha2 | 83436 | HITS-CLIP | | 37986224 | Unspecified |
| mmu-miR-378a-3p | MIMAT0003151 | Plekha5 | 109135 | HITS-CLIP | | 37986224 | Unspecified |
| mmu-miR-378a-3p | MIMAT0003151 | Plekha8 | 231999 | HITS-CLIP | | 37986224 | Unspecified |
| mmu-miR-378a-3p | MIMAT0003151 | Plekhb2 | 226971 | HITS-CLIP | | 37986224 | Unspecified |
| mmu-miR-378a-3p | MIMAT0003151 | Plekhm2 | 69582 | HITS-CLIP | | 25083871 | Unspecified |
| mmu-miR-378a-3p | MIMAT0003151 | Plekhm3 | 241075 | HITS-CLIP | | 37986224 | Unspecified |
| mmu-miR-378a-3p | MIMAT0003151 | Plk3 | 12795 | Microarrays | 37986224\|22855601 | | Unspecified |
| mmu-miR-378a-3p | MIMAT0003151 | Plk4 | 20873 | HITS-CLIP | | 23142080 | Unspecified |
| mmu-miR-378a-3p | MIMAT0003151 | Plod2 | 26432 | Microarrays | 37986224\|22855601 | | Unspecified |
| mmu-miR-378a-3p | MIMAT0003151 | Pls3 | 102866 | HITS-CLIP | 25083871\|29167373 | | Unspecified |
| mmu-miR-378a-3p | MIMAT0003151 | Plxna2 | 18845 | Microarrays | 37986224\|22855601 | | Unspecified |
| mmu-miR-378a-3p | MIMAT0003151 | Plxnb2 | 140570 | HITS-CLIP | | 25083871 | Unspecified |
| mmu-miR-378a-3p | MIMAT0003151 | Pmepa1 | 65112 | Microarrays | 37986224\|22855601 | | Unspecified |
| mmu-miR-378a-3p | MIMAT0003151 | Pml | 18854 | HITS-CLIP | | 23142080 | Unspecified |
| mmu-miR-378a-3p | MIMAT0003151 | Pnn | 18949 | HITS-CLIP | | 37986224 | Unspecified |
| mmu-miR-378a-3p | MIMAT0003151 | Pnpt1 | 71701 | HITS-CLIP | | 23142080 | Unspecified |
| mmu-miR-378a-3p | MIMAT0003151 | Pofut2 | 80294 | HITS-CLIP | | 37986224 | Unspecified |
| mmu-miR-378a-3p | MIMAT0003151 | Pola1 | 18968 | Microarrays | 37986224\|22855601 | | Unspecified |
| mmu-miR-378a-3p | MIMAT0003151 | Poldip3 | 73826 | HITS-CLIP | | 23597149 | Unspecified |
| mmu-miR-378a-3p | MIMAT0003151 | Pole | 18973 | HITS-CLIP | | 23597149 | Unspecified |
| mmu-miR-378a-3p | MIMAT0003151 | Polk | 27015 | Microarrays | 37986224\|22855601 | | Unspecified |
| mmu-miR-378a-3p | MIMAT0003151 | Polr1b | 20017 | Microarrays | 37986224\|22855601 | | Unspecified |
| mmu-miR-378a-3p | MIMAT0003151 | Polr2a | 20020 | HITS-CLIP | 25083871\|23142080 | | Unspecified |
| mmu-miR-378a-3p | MIMAT0003151 | Polr3f | 70408 | HITS-CLIP | | 37986224 | Unspecified |
| mmu-miR-378a-3p | MIMAT0003151 | Pom121 | 107939 | HITS-CLIP | 37986224\|25083871 | | Unspecified |
| mmu-miR-378a-3p | MIMAT0003151 | Pomgnt1 | 68273 | HITS-CLIP | | 37986224 | Unspecified |
| mmu-miR-378a-3p | MIMAT0003151 | Pon1 | 18979 | HITS-CLIP | | 37986224 | Unspecified |
| mmu-miR-378a-3p | MIMAT0003151 | Postn | 50706 | Microarrays//HITS-CLIP | 37986224\|25083871 | | Unspecified |
| mmu-miR-378a-3p | MIMAT0003151 | Ppat | 231327 | Microarrays | 37986224\|22855601 | | Unspecified |
| mmu-miR-378a-3p | MIMAT0003151 | Pphln1 | 223828 | HITS-CLIP | | 37986224 | Unspecified |

| | | | | | | |
|---|---|---|---|---|---|---|
| mmu-miR-378a-3p | MIMAT0003151 | Ppig | 228005 | HITS-CLIP | 37986224\|23142080\|29167373 | Unspecified |
| mmu-miR-378a-3p | MIMAT0003151 | Ppm1k | 243382 | HITS-CLIP | 37986224 | Unspecified |
| mmu-miR-378a-3p | MIMAT0003151 | Ppme1 | 72590 | HITS-CLIP | 25083871\|23142080 | Unspecified |
| mmu-miR-378a-3p | MIMAT0003151 | Ppp1r3a | 140491 | HITS-CLIP | 25753418 | Unspecified |
| mmu-miR-378a-3p | MIMAT0003151 | Ppp1r8 | 100336 | HITS-CLIP | 25083871 | Unspecified |
| mmu-miR-378a-3p | MIMAT0003151 | Ppp1r9a | 243725 | HITS-CLIP | 25753418 | Unspecified |
| mmu-miR-378a-3p | MIMAT0003151 | Ppp2ca | 19052 | HITS-CLIP | 37986224\|29167373 | Unspecified |
| mmu-miR-378a-3p | MIMAT0003151 | Ppp2r1b | 73699 | HITS-CLIP | 23597149\|23142080 | Unspecified |
| mmu-miR-378a-3p | MIMAT0003151 | Ppp2r2d | 52432 | HITS-CLIP | 23142080 | Unspecified |
| mmu-miR-378a-3p | MIMAT0003151 | Ppp3cb | 19056 | HITS-CLIP | 37986224\|23142080 | Unspecified |
| mmu-miR-378a-3p | MIMAT0003151 | Ppp3r1 | 19058 | HITS-CLIP | 25083871\|23142080 | Unspecified |
| mmu-miR-378a-3p | MIMAT0003151 | Ppp4r3a | 55671 | HITS-CLIP | 25083871\|23142080 | Unspecified |
| mmu-miR-378a-3p | MIMAT0003151 | Ppp6c | 67857 | HITS-CLIP | 37986224 | Unspecified |
| mmu-miR-378a-3p | MIMAT0003151 | Pptc7 | 320717 | HITS-CLIP | 37986224 | Unspecified |
| mmu-miR-378a-3p | MIMAT0003151 | Prdm4 | 72843 | HITS-CLIP | 25753418 | Unspecified |
| mmu-miR-378a-3p | MIMAT0003151 | Prdm9 | 213389 | HITS-CLIP | 23597149 | Unspecified |
| mmu-miR-378a-3p | MIMAT0003151 | Prdx3 | 11757 | HITS-CLIP | 37986224 | Unspecified |
| mmu-miR-378a-3p | MIMAT0003151 | Prdx6 | 11758 | HITS-CLIP | 25753418 | Unspecified |
| mmu-miR-378a-3p | MIMAT0003151 | Prep | 19072 | HITS-CLIP | 37986224 | Unspecified |
| mmu-miR-378a-3p | MIMAT0003151 | Prkaa1 | 105787 | Microarrays//HITS-CLIP | 37986224\|22855601\|29167373 | Unspecified |
| mmu-miR-378a-3p | MIMAT0003151 | Prkaca | 18747 | HITS-CLIP | 25753418 | Unspecified |
| mmu-miR-378a-3p | MIMAT0003151 | Prkacb | 18749 | HITS-CLIP | 23142080 | Unspecified |
| mmu-miR-378a-3p | MIMAT0003151 | Prkar1a | 19084 | HITS-CLIP | 23597149 | Unspecified |
| mmu-miR-378a-3p | MIMAT0003151 | Prkar2a | 19087 | HITS-CLIP, Microarrays//Microarrays//HITS | 37986224\|22855601\|25753418 | Unspecified |
| mmu-miR-378a-3p | MIMAT0003151 | Prkca | 18750 | HITS-CLIP, Microarrays//Microarrays | 37986224\|22855601 | Unspecified |
| mmu-miR-378a-3p | MIMAT0003151 | Prkci | 18759 | HITS-CLIP | 25083871\|23597149 | Unspecified |
| mmu-miR-378a-3p | MIMAT0003151 | Prkdc | 19090 | HITS-CLIP | 23142080 | Unspecified |
| mmu-miR-378a-3p | MIMAT0003151 | Prmt1 | 15469 | HITS-CLIP | 25083871 | Unspecified |
| mmu-miR-378a-3p | MIMAT0003151 | Prmt3 | 71974 | HITS-CLIP | 37986224 | Unspecified |
| mmu-miR-378a-3p | MIMAT0003151 | Prokr1 | 58182 | Microarrays | 37986224 | Unspecified |
| mmu-miR-378a-3p | MIMAT0003151 | Prpf3 | 70767 | HITS-CLIP | 23142080 | Unspecified |
| mmu-miR-378a-3p | MIMAT0003151 | Prpf38b | 66921 | HITS-CLIP | 23142080 | Unspecified |
| mmu-miR-378a-3p | MIMAT0003151 | Prpf40a | 56194 | HITS-CLIP | 25753418 | Unspecified |
| mmu-miR-378a-3p | MIMAT0003151 | Prr14 | 233895 | HITS-CLIP | 23597149 | Unspecified |
| mmu-miR-378a-3p | MIMAT0003151 | Prrc2a | 53761 | HITS-CLIP | 25083871 | Unspecified |
| mmu-miR-378a-3p | MIMAT0003151 | Prrc2b | 227723 | HITS-CLIP | 25083871\|23142080 | Unspecified |
| mmu-miR-378a-3p | MIMAT0003151 | Prune2 | 353211 | HITS-CLIP | 37986224 | Unspecified |
| mmu-miR-378a-3p | MIMAT0003151 | Psma1 | 26440 | HITS-CLIP | 37986224 | Unspecified |
| mmu-miR-378a-3p | MIMAT0003151 | Psma7 | 26444 | HITS-CLIP | 25083871 | Unspecified |

| | | | | | | |
|---|---|---|---|---|---|---|
| mmu-miR-378a-3p | MIMAT0003151 | Psmc6 | 67089 | HITS-CLIP | 23142080\|25083871 | Unspecified |
| mmu-miR-378a-3p | MIMAT0003151 | Psmd1 | 70247 | HITS-CLIP | 25083871 | Unspecified |
| mmu-miR-378a-3p | MIMAT0003151 | Psmd11 | 69077 | HITS-CLIP | 37986224 | Unspecified |
| mmu-miR-378a-3p | MIMAT0003151 | Psmd14 | 59029 | HITS-CLIP | 25083871 | Unspecified |
| mmu-miR-378a-3p | MIMAT0003151 | Psmd9 | 67151 | HITS-CLIP | 37986224 | Unspecified |
| mmu-miR-378a-3p | MIMAT0003151 | Psme4 | 103554 | HITS-CLIP | 25083871 | Unspecified |
| mmu-miR-378a-3p | MIMAT0003151 | Psrc1 | 56742 | Microarrays | 37986224\|22855601 | Unspecified |
| mmu-miR-378a-3p | MIMAT0003151 | Ptbp1 | 19205 | HITS-CLIP | 25083871 | Unspecified |
| mmu-miR-378a-3p | MIMAT0003151 | Ptbp2 | 56195 | Microarrays | 37986224\|22855601 | Unspecified |
| mmu-miR-378a-3p | MIMAT0003151 | Ptbp3 | 230257 | HITS-CLIP | 37986224\|23597149\|23142080 | Unspecified |
| mmu-miR-378a-3p | MIMAT0003151 | Pten | 19211 | HITS-CLIP | 37986224\|25083871\|23142080\|29167 | Unspecified |
| mmu-miR-378a-3p | MIMAT0003151 | Ptger2 | 19217 | HITS-CLIP | 23142080 | Unspecified |
| mmu-miR-378a-3p | MIMAT0003151 | Ptgfrn | 19221 | HITS-CLIP | 37986224 | Unspecified |
| mmu-miR-378a-3p | MIMAT0003151 | Ptgs2 | 19225 | Microarrays | 37986224\|22855601 | Unspecified |
| mmu-miR-378a-3p | MIMAT0003151 | Ptk2b | 19229 | HITS-CLIP | 29167373 | Unspecified |
| mmu-miR-378a-3p | MIMAT0003151 | Ptp4a2 | 19244 | HITS-CLIP | 37986224 | Unspecified |
| mmu-miR-378a-3p | MIMAT0003151 | Ptpn14 | 19250 | HITS-CLIP | 37986224 | Unspecified |
| mmu-miR-378a-3p | MIMAT0003151 | Ptpn4 | 19258 | HITS-CLIP | 37986224 | Unspecified |
| mmu-miR-378a-3p | MIMAT0003151 | Ptpn5 | 19259 | HITS-CLIP | 23142080 | Unspecified |
| mmu-miR-378a-3p | MIMAT0003151 | Ptprb | 19263 | HITS-CLIP | 37986224 | Unspecified |
| mmu-miR-378a-3p | MIMAT0003151 | Ptpre | 19267 | HITS-CLIP | 23142080 | Unspecified |
| mmu-miR-378a-3p | MIMAT0003151 | Ptprj | 19271 | HITS-CLIP | 37986224\|29167373 | Unspecified |
| mmu-miR-378a-3p | MIMAT0003151 | Ptprs | 19280 | HITS-CLIP | 25083871 | Unspecified |
| mmu-miR-378a-3p | MIMAT0003151 | Pum2 | 80913 | HITS-CLIP | 37986224\|25083871\|23597149\|23142 | Unspecified |
| mmu-miR-378a-3p | MIMAT0003151 | Pura | 19290 | HITS-CLIP | 37986224 | Unspecified |
| mmu-miR-378a-3p | MIMAT0003151 | Purb | 19291 | HITS-CLIP | 37986224\|23142080 | Unspecified |
| mmu-miR-378a-3p | MIMAT0003151 | Pvr | 52118 | HITS-CLIP, Microarrays//Microarrays | 37986224\|22855601 | Unspecified |
| mmu-miR-378a-3p | MIMAT0003151 | Pwwp2a | 70802 | HITS-CLIP | 37986224 | Unspecified |
| mmu-miR-378a-3p | MIMAT0003151 | Pxmp4 | 59038 | HITS-CLIP, Microarrays | 37986224 | Unspecified |
| mmu-miR-378a-3p | MIMAT0003151 | Pycr1 | 209027 | Microarrays | 37986224 | Unspecified |
| mmu-miR-378a-3p | MIMAT0003151 | Pygo2 | 68911 | HITS-CLIP | 25083871 | Unspecified |
| mmu-miR-378a-3p | MIMAT0003151 | Qser1 | 99003 | HITS-CLIP | 37986224 | Unspecified |
| mmu-miR-378a-3p | MIMAT0003151 | Qsox1 | 104009 | HITS-CLIP | 37986224 | Unspecified |
| mmu-miR-378a-3p | MIMAT0003151 | R3hdm1 | 226412 | HITS-CLIP | 25083871\|23142080 | Unspecified |
| mmu-miR-378a-3p | MIMAT0003151 | Rab10 | 19325 | HITS-CLIP | 37986224 | Unspecified |
| mmu-miR-378a-3p | MIMAT0003151 | Rab12 | 19328 | HITS-CLIP | 23142080 | Unspecified |
| mmu-miR-378a-3p | MIMAT0003151 | Rab1a | 5861 | HITS-CLIP | 23597149\|25083871\|23142080\|29167 | Unspecified |
| mmu-miR-378a-3p | MIMAT0003151 | Rab1b | 76308 | HITS-CLIP | 37986224 | Unspecified |
| mmu-miR-378a-3p | MIMAT0003151 | Rab22a | 19334 | HITS-CLIP | 25083871 | Unspecified |

| | | | | | | |
|---|---|---|---|---|---|---|
| mmu-miR-378a-3p | MIMAT0003151 | Rab34 | 19376 | HITS-CLIP | 25083871 | Unspecified |
| mmu-miR-378a-3p | MIMAT0003151 | Rab5c | 19345 | HITS-CLIP | 23142080 | Unspecified |
| mmu-miR-378a-3p | MIMAT0003151 | Rab7 | 19349 | HITS-CLIP | 37986224 | Unspecified |
| mmu-miR-378a-3p | MIMAT0003151 | Rab8b | 235442 | HITS-CLIP | 25083871 | Unspecified |
| mmu-miR-378a-3p | MIMAT0003151 | Rabgap1l | 29809 | HITS-CLIP | 23142080 | Unspecified |
| mmu-miR-378a-3p | MIMAT0003151 | Rabl6 | 227624 | HITS-CLIP | 25083871 | Unspecified |
| mmu-miR-378a-3p | MIMAT0003151 | Racgap1 | 26934 | HITS-CLIP | 37986224 | Unspecified |
| mmu-miR-378a-3p | MIMAT0003151 | Rack1 | 10399 | qCLASH | 31393953 | Unspecified |
| mmu-miR-378a-3p | MIMAT0003151 | Rad21 | 19357 | HITS-CLIP | 23142080 | Unspecified |
| mmu-miR-378a-3p | MIMAT0003151 | Rad50 | 19360 | HITS-CLIP | 37986224 | Unspecified |
| mmu-miR-378a-3p | MIMAT0003151 | Rad54l2 | 81000 | HITS-CLIP | 37986224 | Unspecified |
| mmu-miR-378a-3p | MIMAT0003151 | Raf1 | 110157 | HITS-CLIP | 37986224 | Unspecified |
| mmu-miR-378a-3p | MIMAT0003151 | Rai14 | 75646 | HITS-CLIP, Microarrays//Microarrays | 37986224\|22855601 | Unspecified |
| mmu-miR-378a-3p | MIMAT0003151 | Raly | 19383 | HITS-CLIP | 23597149\|23142080 | Unspecified |
| mmu-miR-378a-3p | MIMAT0003151 | Ranbp10 | 74334 | HITS-CLIP | 37986224\|25083871 | Unspecified |
| mmu-miR-378a-3p | MIMAT0003151 | Ranbp2 | 19386 | HITS-CLIP | 37986224\|23597149\|23142080\|25083 | Unspecified |
| mmu-miR-378a-3p | MIMAT0003151 | Ranbp9 | 56705 | HITS-CLIP | 23142080 | Unspecified |
| mmu-miR-378a-3p | MIMAT0003151 | Rap1b | 215449 | HITS-CLIP | 37986224\|23142080\|29167373 | Unspecified |
| mmu-miR-378a-3p | MIMAT0003151 | Rap1gds1 | 229877 | HITS-CLIP | 37986224 | Unspecified |
| mmu-miR-378a-3p | MIMAT0003151 | Rapgef2 | 76089 | HITS-CLIP | 37986224 | Unspecified |
| mmu-miR-378a-3p | MIMAT0003151 | Raph1 | 77300 | HITS-CLIP | 37986224 | Unspecified |
| mmu-miR-378a-3p | MIMAT0003151 | Rars2 | 109093 | HITS-CLIP | 37986224\|25753418 | Unspecified |
| mmu-miR-378a-3p | MIMAT0003151 | Rasal2 | 226525 | HITS-CLIP | 37986224\|23597149 | Unspecified |
| mmu-miR-378a-3p | MIMAT0003151 | Rassf8 | 71323 | Microarrays | 37986224\|22855601 | Unspecified |
| mmu-miR-378a-3p | MIMAT0003151 | Rb1cc1 | 12421 | HITS-CLIP | 25083871\|23142080 | Unspecified |
| mmu-miR-378a-3p | MIMAT0003151 | Rbm14 | 56275 | HITS-CLIP | 37986224\|25083871 | Unspecified |
| mmu-miR-378a-3p | MIMAT0003151 | Rbm15 | 229700 | HITS-CLIP//qCLASH | 37986224\|25083871\|31393953\|23142 | Unspecified |
| mmu-miR-378a-3p | MIMAT0003151 | Rbm20 | 73713 | HITS-CLIP | 37986224 | Unspecified |
| mmu-miR-378a-3p | MIMAT0003151 | Rbm26 | 74213 | HITS-CLIP | 37986224 | Unspecified |
| mmu-miR-378a-3p | MIMAT0003151 | Rbm4 | 19653 | HITS-CLIP | 37986224\|23597149 | Unspecified |
| mmu-miR-378a-3p | MIMAT0003151 | Rbm4b | 66704 | HITS-CLIP | 23597149\|23142080 | Unspecified |
| mmu-miR-378a-3p | MIMAT0003151 | Rbms1 | 56878 | HITS-CLIP | 23142080 | Unspecified |
| mmu-miR-378a-3p | MIMAT0003151 | Rc3h2 | 319817 | HITS-CLIP | 23142080 | Unspecified |
| mmu-miR-378a-3p | MIMAT0003151 | Rcc2 | 108911 | HITS-CLIP | 37986224\|25083871\|23142080 | Unspecified |
| mmu-miR-378a-3p | MIMAT0003151 | Rchy1 | 68098 | HITS-CLIP | 37986224\|23142080 | Unspecified |
| mmu-miR-378a-3p | MIMAT0003151 | Rcn2 | 26611 | HITS-CLIP | 37986224 | Unspecified |
| mmu-miR-378a-3p | MIMAT0003151 | Rcor1 | 217864 | HITS-CLIP | 25083871\|23142080 | Unspecified |
| mmu-miR-378a-3p | MIMAT0003151 | Rdh10 | 98711 | Microarrays | 37986224\|22855601 | Unspecified |
| mmu-miR-378a-3p | MIMAT0003151 | Rdh14 | 105014 | HITS-CLIP | 37986224 | Unspecified |

| | | | | | | |
|---|---|---|---|---|---|---|
| mmu-miR-378a-3p | MIMAT0003151 | Reep3 | 28193 | qCLASH | 31393953 | Unspecified |
| mmu-miR-378a-3p | MIMAT0003151 | Resf1 | 55196 | Microarrays | 22855601 | Unspecified |
| mmu-miR-378a-3p | MIMAT0003151 | Rest | 19712 | HITS-CLIP | 37986224 | Unspecified |
| mmu-miR-378a-3p | MIMAT0003151 | Rfc2 | 19718 | HITS-CLIP | 23142080 | Unspecified |
| mmu-miR-378a-3p | MIMAT0003151 | Rfx3 | 19726 | HITS-CLIP | 37986224 | Unspecified |
| mmu-miR-378a-3p | MIMAT0003151 | Rfx7 | 319758 | HITS-CLIP | 37986224\|25083871\|23142080\|29167 | Unspecified |
| mmu-miR-378a-3p | MIMAT0003151 | Rhbdd1 | 76867 | HITS-CLIP | 37986224 | Unspecified |
| mmu-miR-378a-3p | MIMAT0003151 | Rhobtb1 | 69288 | HITS-CLIP | 37986224\|23142080 | Unspecified |
| mmu-miR-378a-3p | MIMAT0003151 | Rhobtb3 | 73296 | HITS-CLIP | 37986224 | Unspecified |
| mmu-miR-378a-3p | MIMAT0003151 | Rhoq | 104215 | HITS-CLIP | 37986224 | Unspecified |
| mmu-miR-378a-3p | MIMAT0003151 | Ric8a | 60626 | HITS-CLIP | 23142080 | Unspecified |
| mmu-miR-378a-3p | MIMAT0003151 | Rictor | 78757 | HITS-CLIP | 37986224\|25083871\|23597149\|23142 | Unspecified |
| mmu-miR-378a-3p | MIMAT0003151 | Rif1 | 51869 | HITS-CLIP | 37986224 | Unspecified |
| mmu-miR-378a-3p | MIMAT0003151 | Rin1 | 225870 | Microarrays | 37986224\|22855601 | Unspecified |
| mmu-miR-378a-3p | MIMAT0003151 | Ripk2 | 192656 | qCLASH | 31393953 | Unspecified |
| mmu-miR-378a-3p | MIMAT0003151 | Rlf | 109263 | HITS-CLIP | 37986224 | Unspecified |
| mmu-miR-378a-3p | MIMAT0003151 | Rnasek | 52898 | HITS-CLIP | 25753418 | Unspecified |
| mmu-miR-378a-3p | MIMAT0003151 | Rnf10 | 50849 | HITS-CLIP | 25083871\|23142080 | Unspecified |
| mmu-miR-378a-3p | MIMAT0003151 | Rnf103 | 22644 | HITS-CLIP | 25083871 | Unspecified |
| mmu-miR-378a-3p | MIMAT0003151 | Rnf113a1 | 69942 | HITS-CLIP | 23142080 | Unspecified |
| mmu-miR-378a-3p | MIMAT0003151 | Rnf115 | 67845 | HITS-CLIP | 37986224 | Unspecified |
| mmu-miR-378a-3p | MIMAT0003151 | Rnf13 | 24017 | HITS-CLIP | 37986224 | Unspecified |
| mmu-miR-378a-3p | MIMAT0003151 | Rnf144b | 218215 | HITS-CLIP | 25753418 | Unspecified |
| mmu-miR-378a-3p | MIMAT0003151 | Rnf149 | 67702 | Microarrays | 37986224\|22855601 | Unspecified |
| mmu-miR-378a-3p | MIMAT0003151 | Rnf167 | 70510 | HITS-CLIP | 37986224 | Unspecified |
| mmu-miR-378a-3p | MIMAT0003151 | Rnf181 | 66510 | HITS-CLIP | 25753418 | Unspecified |
| mmu-miR-378a-3p | MIMAT0003151 | Rnf19a | 30945 | HITS-CLIP | 37986224 | Unspecified |
| mmu-miR-378a-3p | MIMAT0003151 | Rnf213 | 672511 | HITS-CLIP | 37986224 | Unspecified |
| mmu-miR-378a-3p | MIMAT0003151 | Rnf217 | 268291 | HITS-CLIP | 37986224 | Unspecified |
| mmu-miR-378a-3p | MIMAT0003151 | Rnf24 | 51902 | HITS-CLIP | 23597149 | Unspecified |
| mmu-miR-378a-3p | MIMAT0003151 | Rock1 | 19877 | HITS-CLIP | 37986224 | Unspecified |
| mmu-miR-378a-3p | MIMAT0003151 | Rock2 | 19878 | Microarrays | 37986224\|22855601 | Unspecified |
| mmu-miR-378a-3p | MIMAT0003151 | Rpa1 | 68275 | HITS-CLIP | 23597149\|23142080 | Unspecified |
| mmu-miR-378a-3p | MIMAT0003151 | Rpgrip1l | 244585 | qCLASH | 31393953 | Unspecified |
| mmu-miR-378a-3p | MIMAT0003151 | Rpl12 | 269261 | Microarrays | 37986224\|22855601 | Unspecified |
| mmu-miR-378a-3p | MIMAT0003151 | Rpl14 | 67115 | HITS-CLIP | 25753418 | Unspecified |
| mmu-miR-378a-3p | MIMAT0003151 | Rpl39 | 67248 | HITS-CLIP | 37986224 | Unspecified |
| mmu-miR-378a-3p | MIMAT0003151 | Rpn2 | 20014 | HITS-CLIP | 25083871 | Unspecified |
| mmu-miR-378a-3p | MIMAT0003151 | Rrad | 56437 | HITS-CLIP | 25083871 | Unspecified |

| | | | | | | |
|---|---|---|---|---|---|---|
| mmu-miR-378a-3p | MIMAT0003151 | Rras2 | 66922 | Microarrays | 37986224\|22855601 | Unspecified |
| mmu-miR-378a-3p | MIMAT0003151 | Rrbp1 | 81910 | HITS-CLIP | 37986224 | Unspecified |
| mmu-miR-378a-3p | MIMAT0003151 | Rreb1 | 68750 | HITS-CLIP | 37986224 | Unspecified |
| mmu-miR-378a-3p | MIMAT0003151 | Rrm1 | 20133 | HITS-CLIP | 37986224 | Unspecified |
| mmu-miR-378a-3p | MIMAT0003151 | Rrn3 | 106298 | Microarrays | 37986224\|22855601 | Unspecified |
| mmu-miR-378a-3p | MIMAT0003151 | Rrp12 | 107094 | Microarrays | 37986224\|22855601 | Unspecified |
| mmu-miR-378a-3p | MIMAT0003151 | Rrp1b | 72462 | Microarrays | 37986224\|22855601 | Unspecified |
| mmu-miR-378a-3p | MIMAT0003151 | Rrs1 | 59014 | Microarrays | 37986224\|22855601 | Unspecified |
| mmu-miR-378a-3p | MIMAT0003151 | Rsbn1l | 242860 | HITS-CLIP | 25753418 | Unspecified |
| mmu-miR-378a-3p | MIMAT0003151 | Rsf1 | 233532 | HITS-CLIP | 37986224\|23142080 | Unspecified |
| mmu-miR-378a-3p | MIMAT0003151 | Rspry1 | 67610 | HITS-CLIP | 23597149 | Unspecified |
| mmu-miR-378a-3p | MIMAT0003151 | Rtkn | 20166 | HITS-CLIP | 37986224 | Unspecified |
| mmu-miR-378a-3p | MIMAT0003151 | Rtn4 | 68585 | HITS-CLIP | 25083871\|29167373 | Unspecified |
| mmu-miR-378a-3p | MIMAT0003151 | Rundc1 | 217201 | HITS-CLIP | 31433990 | Unspecified |
| mmu-miR-378a-3p | MIMAT0003151 | Runx1 | 12394 | HITS-CLIP, Microarrays//Microarrays | 37986224\|22855601 | Unspecified |
| mmu-miR-378a-3p | MIMAT0003151 | Runx1t1 | 12395 | HITS-CLIP | 23597149 | Unspecified |
| mmu-miR-378a-3p | MIMAT0003151 | Runx3 | 12399 | HITS-CLIP | 37986224 | Unspecified |
| mmu-miR-378a-3p | MIMAT0003151 | Ryk | 20187 | HITS-CLIP | 25083871\|23142080 | Unspecified |
| mmu-miR-378a-3p | MIMAT0003151 | Ryr1 | 20190 | HITS-CLIP | 37986224\|25083871 | Unspecified |
| mmu-miR-378a-3p | MIMAT0003151 | S100pbp | 74648 | HITS-CLIP | 37986224 | Unspecified |
| mmu-miR-378a-3p | MIMAT0003151 | Sacm1l | 83493 | HITS-CLIP | 37986224 | Unspecified |
| mmu-miR-378a-3p | MIMAT0003151 | Samd1 | 666704 | HITS-CLIP | 37986224 | Unspecified |
| mmu-miR-378a-3p | MIMAT0003151 | Sap18b | 100041953 | HITS-CLIP | 23142080 | Unspecified |
| mmu-miR-378a-3p | MIMAT0003151 | Sar1a | 20224 | HITS-CLIP | 37986224 | Unspecified |
| mmu-miR-378a-3p | MIMAT0003151 | Sbds | 66711 | HITS-CLIP | 37986224 | Unspecified |
| mmu-miR-378a-3p | MIMAT0003151 | Sbk1 | 104175 | HITS-CLIP | 37986224 | Unspecified |
| mmu-miR-378a-3p | MIMAT0003151 | Sbno1 | 243272 | HITS-CLIP | 23142080 | Unspecified |
| mmu-miR-378a-3p | MIMAT0003151 | Sc5d | 235293 | HITS-CLIP | 23142080 | Unspecified |
| mmu-miR-378a-3p | MIMAT0003151 | Scarf2 | 224024 | HITS-CLIP | 37986224 | Unspecified |
| mmu-miR-378a-3p | MIMAT0003151 | Scd2 | 20250 | qCLASH | 31393953 | Unspecified |
| mmu-miR-378a-3p | MIMAT0003151 | Scn5a | 20271 | HITS-CLIP | 25083871 | Unspecified |
| mmu-miR-378a-3p | MIMAT0003151 | Scyl2 | 213326 | HITS-CLIP | 23142080 | Unspecified |
| mmu-miR-378a-3p | MIMAT0003151 | Sdad1 | 231452 | Microarrays | 37986224\|22855601 | Unspecified |
| mmu-miR-378a-3p | MIMAT0003151 | Sdcbp | 53378 | HITS-CLIP | 37986224 | Unspecified |
| mmu-miR-378a-3p | MIMAT0003151 | Sdf2 | 20316 | HITS-CLIP | 37986224 | Unspecified |
| mmu-miR-378a-3p | MIMAT0003151 | Sdf4 | 20318 | HITS-CLIP | 37986224 | Unspecified |
| mmu-miR-378a-3p | MIMAT0003151 | Sec16a | 227648 | HITS-CLIP | 37986224 | Unspecified |
| mmu-miR-378a-3p | MIMAT0003151 | Sec22b | 20333 | HITS-CLIP | 37986224 | Unspecified |
| mmu-miR-378a-3p | MIMAT0003151 | Sec23ip | 207352 | HITS-CLIP | 25083871\|23142080 | Unspecified |

| | | | | | | |
|---|---|---|---|---|---|---|
| mmu-miR-378a-3p | MIMAT0003151 | Sec24a | 77371 | HITS-CLIP | 25083871\|23597149\|23142080 | Unspecified |
| mmu-miR-378a-3p | MIMAT0003151 | Sec24b | 99683 | qCLASH//HITS-CLIP | 31393953\|25083871 | Unspecified |
| mmu-miR-378a-3p | MIMAT0003151 | Sec62 | 69276 | HITS-CLIP | 37986224 | Unspecified |
| mmu-miR-378a-3p | MIMAT0003151 | Secisbp2l | 70354 | HITS-CLIP | 37986224 | Unspecified |
| mmu-miR-378a-3p | MIMAT0003151 | Seh1l | 72124 | HITS-CLIP | 23142080 | Unspecified |
| mmu-miR-378a-3p | MIMAT0003151 | Selenoi | 28042 | Microarrays | 22855601 | Unspecified |
| mmu-miR-378a-3p | MIMAT0003151 | Senp1 | 223870 | Microarrays | 37986224\|22855601 | Unspecified |
| mmu-miR-378a-3p | MIMAT0003151 | Senp5 | 320213 | HITS-CLIP | 37986224 | Unspecified |
| mmu-miR-378a-3p | MIMAT0003151 | Sergef | 27414 | HITS-CLIP | 25753418 | Unspecified |
| mmu-miR-378a-3p | MIMAT0003151 | Serinc3 | 26943 | HITS-CLIP | 23142080 | Unspecified |
| mmu-miR-378a-3p | MIMAT0003151 | Serpina3n | 20716 | Microarrays | 37986224\|22855601 | Unspecified |
| mmu-miR-378a-3p | MIMAT0003151 | Serpinb8 | 20725 | Microarrays | 37986224 | Unspecified |
| mmu-miR-378a-3p | MIMAT0003151 | Serpinb9 | 20723 | HITS-CLIP | 37986224 | Unspecified |
| mmu-miR-378a-3p | MIMAT0003151 | Serpind1 | 15160 | HITS-CLIP | 23597149 | Unspecified |
| mmu-miR-378a-3p | MIMAT0003151 | Serpine1 | 18787 | Microarrays | 37986224\|22855601 | Unspecified |
| mmu-miR-378a-3p | MIMAT0003151 | Serpine2 | 20720 | HITS-CLIP | 37986224 | Unspecified |
| mmu-miR-378a-3p | MIMAT0003151 | Sertad3 | 170742 | Microarrays | 37986224 | Unspecified |
| mmu-miR-378a-3p | MIMAT0003151 | Sesn2 | 230784 | HITS-CLIP | 37986224 | Unspecified |
| mmu-miR-378a-3p | MIMAT0003151 | Sesn3 | 75747 | HITS-CLIP | 23142080\|25083871 | Unspecified |
| mmu-miR-378a-3p | MIMAT0003151 | Setbp1 | 240427 | HITS-CLIP | 37986224 | Unspecified |
| mmu-miR-378a-3p | MIMAT0003151 | Setx | 269254 | HITS-CLIP | 37986224\|25083871 | Unspecified |
| mmu-miR-378a-3p | MIMAT0003151 | Sf3b1 | 81898 | HITS-CLIP | 37986224 | Unspecified |
| mmu-miR-378a-3p | MIMAT0003151 | Sf3b3 | 101943 | qCLASH | 31393953 | Unspecified |
| mmu-miR-378a-3p | MIMAT0003151 | Sfrp2 | 20319 | Microarrays | 37986224 | Unspecified |
| mmu-miR-378a-3p | MIMAT0003151 | Sgk1 | 20393 | Microarrays//HITS-CLIP | 37986224\|22855601\|25083871\|29167 | Unspecified |
| mmu-miR-378a-3p | MIMAT0003151 | Sgk2 | 27219 | HITS-CLIP | 23597149 | Unspecified |
| mmu-miR-378a-3p | MIMAT0003151 | Sgo2a | 68549 | HITS-CLIP | 23142080 | Unspecified |
| mmu-miR-378a-3p | MIMAT0003151 | Sh3bp2 | 24055 | HITS-CLIP | 37986224 | Unspecified |
| mmu-miR-378a-3p | MIMAT0003151 | Sh3glb1 | 54673 | HITS-CLIP | 25083871 | Unspecified |
| mmu-miR-378a-3p | MIMAT0003151 | Sh3pxd2a | 14218 | HITS-CLIP | 37986224 | Unspecified |
| mmu-miR-378a-3p | MIMAT0003151 | Sh3pxd2b | 268396 | HITS-CLIP | 37986224 | Unspecified |
| mmu-miR-378a-3p | MIMAT0003151 | Shisa5 | 66940 | HITS-CLIP//qCLASH | 37986224\|25083871\|31393953\|23142 | Unspecified |
| mmu-miR-378a-3p | MIMAT0003151 | Shoc2 | 56392 | HITS-CLIP | 25083871\|23142080 | Unspecified |
| mmu-miR-378a-3p | MIMAT0003151 | Sidt2 | 214597 | HITS-CLIP | 37986224 | Unspecified |
| mmu-miR-378a-3p | MIMAT0003151 | Sik1 | 17691 | HITS-CLIP | 37986224 | Unspecified |
| mmu-miR-378a-3p | MIMAT0003151 | Sinhcaf | 58516 | Microarrays | 22855601 | Unspecified |
| mmu-miR-378a-3p | MIMAT0003151 | Six1 | 20471 | Microarrays | 37986224 | Unspecified |
| mmu-miR-378a-3p | MIMAT0003151 | Six4 | 20474 | HITS-CLIP | 25083871 | Unspecified |
| mmu-miR-378a-3p | MIMAT0003151 | Ski | 20481 | HITS-CLIP | 25083871 | Unspecified |

| | | | | | | |
|---|---|---|---|---|---|---|
| mmu-miR-378a-3p | MIMAT0003151 | Skil | 20482 | HITS-CLIP | 37986224 | Unspecified |
| mmu-miR-378a-3p | MIMAT0003151 | Skp2 | 27401 | HITS-CLIP, Microarrays//Microarrays//HITS | 37986224\|22855601\|23142080 | Unspecified |
| mmu-miR-378a-3p | MIMAT0003151 | Slbp | 20492 | HITS-CLIP | 23142080 | Unspecified |
| mmu-miR-378a-3p | MIMAT0003151 | Slc12a8 | 171286 | HITS-CLIP | 23142080 | Unspecified |
| mmu-miR-378a-3p | MIMAT0003151 | Slc16a1 | 20501 | Microarrays | 37986224\|22855601 | Unspecified |
| mmu-miR-378a-3p | MIMAT0003151 | Slc16a13 | 69309 | Microarrays | 37986224\|22855601 | Unspecified |
| mmu-miR-378a-3p | MIMAT0003151 | Slc19a2 | 116914 | Microarrays | 37986224\|22855601 | Unspecified |
| mmu-miR-378a-3p | MIMAT0003151 | Slc20a1 | 20515 | Microarrays//HITS-CLIP | 37986224\|22855601\|23142080 | Unspecified |
| mmu-miR-378a-3p | MIMAT0003151 | Slc20a2 | 20516 | Microarrays | 37986224\|22855601 | Unspecified |
| mmu-miR-378a-3p | MIMAT0003151 | Slc22a23 | 73102 | Microarrays | 37986224\|22855601 | Unspecified |
| mmu-miR-378a-3p | MIMAT0003151 | Slc25a12 | 78830 | qCLASH | 31393953 | Unspecified |
| mmu-miR-378a-3p | MIMAT0003151 | Slc25a24 | 229731 | HITS-CLIP | 37986224\|25083871 | Unspecified |
| mmu-miR-378a-3p | MIMAT0003151 | Slc25a32 | 69906 | Microarrays | 37986224\|22855601 | Unspecified |
| mmu-miR-378a-3p | MIMAT0003151 | Slc2a1 | 20525 | HITS-CLIP | 37986224 | Unspecified |
| mmu-miR-378a-3p | MIMAT0003151 | Slc2a3 | 20527 | HITS-CLIP | 37986224\|23142080 | Unspecified |
| mmu-miR-378a-3p | MIMAT0003151 | Slc30a4 | 22785 | Microarrays | 37986224\|22855601 | Unspecified |
| mmu-miR-378a-3p | MIMAT0003151 | Slc30a7 | 66500 | HITS-CLIP | 37986224\|25083871 | Unspecified |
| mmu-miR-378a-3p | MIMAT0003151 | Slc30a9 | 109108 | HITS-CLIP | 37986224 | Unspecified |
| mmu-miR-378a-3p | MIMAT0003151 | Slc31a1 | 20529 | HITS-CLIP | 25083871 | Unspecified |
| mmu-miR-378a-3p | MIMAT0003151 | Slc35a2 | 22232 | HITS-CLIP | 37986224\|25083871 | Unspecified |
| mmu-miR-378a-3p | MIMAT0003151 | Slc35a3 | 229782 | Microarrays | 37986224\|22855601 | Unspecified |
| mmu-miR-378a-3p | MIMAT0003151 | Slc35f5 | 74150 | HITS-CLIP | 25083871\|25753418 | Unspecified |
| mmu-miR-378a-3p | MIMAT0003151 | Slc38a1 | 105727 | HITS-CLIP | 37986224 | Unspecified |
| mmu-miR-378a-3p | MIMAT0003151 | Slc38a2 | 67760 | HITS-CLIP | 23142080 | Unspecified |
| mmu-miR-378a-3p | MIMAT0003151 | Slc38a6 | 625098 | HITS-CLIP | 25083871 | Unspecified |
| mmu-miR-378a-3p | MIMAT0003151 | Slc39a10 | 227059 | HITS-CLIP | 29167373 | Unspecified |
| mmu-miR-378a-3p | MIMAT0003151 | Slc39a13 | 68427 | HITS-CLIP | 37986224 | Unspecified |
| mmu-miR-378a-3p | MIMAT0003151 | Slc39a6 | 106957 | Microarrays | 37986224\|22855601 | Unspecified |
| mmu-miR-378a-3p | MIMAT0003151 | Slc41a2 | 338365 | HITS-CLIP | 37986224\|23597149 | Unspecified |
| mmu-miR-378a-3p | MIMAT0003151 | Slc4a7 | 218756 | HITS-CLIP | 25083871\|23142080 | Unspecified |
| mmu-miR-378a-3p | MIMAT0003151 | Slc6a6 | 21366 | Microarrays | 37986224\|22855601 | Unspecified |
| mmu-miR-378a-3p | MIMAT0003151 | Slc6a9 | 14664 | HITS-CLIP | 37986224 | Unspecified |
| mmu-miR-378a-3p | MIMAT0003151 | Slc7a1 | 11987 | Microarrays | 37986224\|22855601 | Unspecified |
| mmu-miR-378a-3p | MIMAT0003151 | Slc7a11 | 26570 | Microarrays | 37986224\|22855601 | Unspecified |
| mmu-miR-378a-3p | MIMAT0003151 | Slc7a2 | 11988 | HITS-CLIP | 25083871 | Unspecified |
| mmu-miR-378a-3p | MIMAT0003151 | Slc7a5 | 20539 | Microarrays | 37986224\|22855601 | Unspecified |
| mmu-miR-378a-3p | MIMAT0003151 | Slc9a6 | 236794 | HITS-CLIP | 37986224 | Unspecified |
| mmu-miR-378a-3p | MIMAT0003151 | Slc9a8 | 77031 | HITS-CLIP | 37986224\|25753418 | Unspecified |
| mmu-miR-378a-3p | MIMAT0003151 | Slco2a1 | 24059 | Microarrays | 37986224\|22855601 | Unspecified |

| | | | | | | |
|---|---|---|---|---|---|---|
| mmu-miR-378a-3p | MIMAT0003151 | Slf2 | 226151 | HITS-CLIP | 25083871 | Unspecified |
| mmu-miR-378a-3p | MIMAT0003151 | Slit2 | 20563 | Microarrays | 37986224\|22855601 | Unspecified |
| mmu-miR-378a-3p | MIMAT0003151 | Slk | 20874 | Microarrays//HITS-CLIP | 37986224\|25083871\|22855601 | Unspecified |
| mmu-miR-378a-3p | MIMAT0003151 | Slmap | 83997 | HITS-CLIP | 37986224 | Unspecified |
| mmu-miR-378a-3p | MIMAT0003151 | Sltm | 66660 | HITS-CLIP | 25083871 | Unspecified |
| mmu-miR-378a-3p | MIMAT0003151 | Smad1 | 17125 | HITS-CLIP | 37986224 | Unspecified |
| mmu-miR-378a-3p | MIMAT0003151 | Smad5 | 17129 | HITS-CLIP | 37986224\|25083871 | Unspecified |
| mmu-miR-378a-3p | MIMAT0003151 | Smad7 | 17131 | Microarrays | 37986224\|22855601 | Unspecified |
| mmu-miR-378a-3p | MIMAT0003151 | Smarca2 | 67155 | HITS-CLIP | 37986224\|25753418 | Unspecified |
| mmu-miR-378a-3p | MIMAT0003151 | Smarca5 | 93762 | HITS-CLIP | 25083871\|23142080 | Unspecified |
| mmu-miR-378a-3p | MIMAT0003151 | Smarcad1 | 13990 | HITS-CLIP | 25083871 | Unspecified |
| mmu-miR-378a-3p | MIMAT0003151 | Smarcc1 | 20588 | HITS-CLIP | 37986224 | Unspecified |
| mmu-miR-378a-3p | MIMAT0003151 | Smarcc2 | 68094 | HITS-CLIP | 37986224 | Unspecified |
| mmu-miR-378a-3p | MIMAT0003151 | Smchd1 | 74355 | HITS-CLIP | 23142080 | Unspecified |
| mmu-miR-378a-3p | MIMAT0003151 | Smcr8 | 237782 | HITS-CLIP | 37986224\|25083871\|23142080 | Unspecified |
| mmu-miR-378a-3p | MIMAT0003151 | Smg1 | 233789 | HITS-CLIP | 37986224\|23142080\|29167373 | Unspecified |
| mmu-miR-378a-3p | MIMAT0003151 | Smg8 | 74133 | HITS-CLIP | 23597149 | Unspecified |
| mmu-miR-378a-3p | MIMAT0003151 | Smim11 | 68936 | Microarrays | 37986224 | Unspecified |
| mmu-miR-378a-3p | MIMAT0003151 | Smoc2 | 64074 | Microarrays | 37986224 | Unspecified |
| mmu-miR-378a-3p | MIMAT0003151 | Smurf1 | 75788 | HITS-CLIP | 37986224 | Unspecified |
| mmu-miR-378a-3p | MIMAT0003151 | Snap47 | 67826 | HITS-CLIP | 37986224 | Unspecified |
| mmu-miR-378a-3p | MIMAT0003151 | Snn | 20621 | HITS-CLIP | 37986224 | Unspecified |
| mmu-miR-378a-3p | MIMAT0003151 | Snrpb | 20638 | HITS-CLIP | 23597149\|23142080 | Unspecified |
| mmu-miR-378a-3p | MIMAT0003151 | Snw1 | 66354 | HITS-CLIP | 25083871 | Unspecified |
| mmu-miR-378a-3p | MIMAT0003151 | Snx12 | 55988 | HITS-CLIP | 23142080 | Unspecified |
| mmu-miR-378a-3p | MIMAT0003151 | Snx13 | 217463 | HITS-CLIP | 37986224 | Unspecified |
| mmu-miR-378a-3p | MIMAT0003151 | Snx17 | 266781 | HITS-CLIP | 25083871\|23142080 | Unspecified |
| mmu-miR-378a-3p | MIMAT0003151 | Snx18 | 170625 | HITS-CLIP | 37986224 | Unspecified |
| mmu-miR-378a-3p | MIMAT0003151 | Snx20 | 71607 | HITS-CLIP | 29167373 | Unspecified |
| mmu-miR-378a-3p | MIMAT0003151 | Snx33 | 235406 | HITS-CLIP | 37986224 | Unspecified |
| mmu-miR-378a-3p | MIMAT0003151 | Soat1 | 20652 | HITS-CLIP | 37986224\|29167373 | Unspecified |
| mmu-miR-378a-3p | MIMAT0003151 | Sobp | 109205 | HITS-CLIP | 37986224 | Unspecified |
| mmu-miR-378a-3p | MIMAT0003151 | Son | 20658 | HITS-CLIP | 37986224\|25083871\|23142080 | Unspecified |
| mmu-miR-378a-3p | MIMAT0003151 | Sorbs2 | 234214 | HITS-CLIP | 25753418 | Unspecified |
| mmu-miR-378a-3p | MIMAT0003151 | Sord | 20322 | HITS-CLIP | 37986224 | Unspecified |
| mmu-miR-378a-3p | MIMAT0003151 | Sox4 | 20677 | HITS-CLIP, Microarrays//Microarrays | 37986224\|22855601 | Unspecified |
| mmu-miR-378a-3p | MIMAT0003151 | Sp1 | 20683 | HITS-CLIP | 37986224 | Unspecified |
| mmu-miR-378a-3p | MIMAT0003151 | Sp100 | 20684 | HITS-CLIP | 23142080 | Unspecified |
| mmu-miR-378a-3p | MIMAT0003151 | Sp2 | 78912 | HITS-CLIP | 37986224 | Unspecified |

| | | | | | | |
|---|---|---|---|---|---|---|
| mmu-miR-378a-3p | MIMAT0003151 | Sp3 | 20687 | HITS-CLIP | 37986224\|25083871\|23142080 | Unspecified |
| mmu-miR-378a-3p | MIMAT0003151 | Spag9 | 70834 | HITS-CLIP | 37986224 | Unspecified |
| mmu-miR-378a-3p | MIMAT0003151 | Sparc | 20692 | HITS-CLIP | 23597149\|25753418 | Unspecified |
| mmu-miR-378a-3p | MIMAT0003151 | Spast | 50850 | HITS-CLIP | 25083871 | Unspecified |
| mmu-miR-378a-3p | MIMAT0003151 | Spata2 | 263876 | HITS-CLIP | 23142080 | Unspecified |
| mmu-miR-378a-3p | MIMAT0003151 | Spata2l | 78779 | HITS-CLIP | 37986224 | Unspecified |
| mmu-miR-378a-3p | MIMAT0003151 | Spcs3 | 76687 | HITS-CLIP | 31433990 | Unspecified |
| mmu-miR-378a-3p | MIMAT0003151 | Specc1 | 432572 | HITS-CLIP | 37986224 | Unspecified |
| mmu-miR-378a-3p | MIMAT0003151 | Specc1l | 74392 | HITS-CLIP | 25083871 | Unspecified |
| mmu-miR-378a-3p | MIMAT0003151 | Speg | 11790 | HITS-CLIP | 37986224\|25753418 | Unspecified |
| mmu-miR-378a-3p | MIMAT0003151 | Spen | 56381 | HITS-CLIP | 25083871 | Unspecified |
| mmu-miR-378a-3p | MIMAT0003151 | Spg20 | 229285 | HITS-CLIP | 25083871\|23142080 | Unspecified |
| mmu-miR-378a-3p | MIMAT0003151 | Spin1 | 20729 | HITS-CLIP | 37986224 | Unspecified |
| mmu-miR-378a-3p | MIMAT0003151 | Spop | 20747 | HITS-CLIP | 37986224 | Unspecified |
| mmu-miR-378a-3p | MIMAT0003151 | Spp1 | 20750 | HITS-CLIP | 23142080 | Unspecified |
| mmu-miR-378a-3p | MIMAT0003151 | Sppl3 | 74585 | HITS-CLIP | 37986224\|25083871\|23597149 | Unspecified |
| mmu-miR-378a-3p | MIMAT0003151 | Spred1 | 114715 | HITS-CLIP, Microarrays//Microarrays | 37986224\|22855601 | Unspecified |
| mmu-miR-378a-3p | MIMAT0003151 | Spred2 | 114716 | HITS-CLIP | 37986224 | Unspecified |
| mmu-miR-378a-3p | MIMAT0003151 | Spred3 | 101809 | HITS-CLIP | 29167373 | Unspecified |
| mmu-miR-378a-3p | MIMAT0003151 | Spry4 | 24066 | HITS-CLIP | 37986224 | Unspecified |
| mmu-miR-378a-3p | MIMAT0003151 | Spryd7 | 66674 | HITS-CLIP | 23142080 | Unspecified |
| mmu-miR-378a-3p | MIMAT0003151 | Sptan1 | 20740 | HITS-CLIP | 25083871 | Unspecified |
| mmu-miR-378a-3p | MIMAT0003151 | Sptbn1 | 20742 | HITS-CLIP | 37986224\|23142080 | Unspecified |
| mmu-miR-378a-3p | MIMAT0003151 | Sqstm1 | 18412 | HITS-CLIP | 37986224 | Unspecified |
| mmu-miR-378a-3p | MIMAT0003151 | Srcap | 100043597 | HITS-CLIP | 37986224\|23142080 | Unspecified |
| mmu-miR-378a-3p | MIMAT0003151 | Srebf1 | 20787 | HITS-CLIP | 37986224 | Unspecified |
| mmu-miR-378a-3p | MIMAT0003151 | Srek1 | 218543 | HITS-CLIP | 37986224\|23142080 | Unspecified |
| mmu-miR-378a-3p | MIMAT0003151 | Srm | 20810 | HITS-CLIP | 37986224\|23597149 | Unspecified |
| mmu-miR-378a-3p | MIMAT0003151 | Srrm2 | 75956 | HITS-CLIP | 37986224\|25083871\|29167373 | Unspecified |
| mmu-miR-378a-3p | MIMAT0003151 | Srsf10 | 14105 | HITS-CLIP | 23142080 | Unspecified |
| mmu-miR-378a-3p | MIMAT0003151 | Srsf2 | 20382 | HITS-CLIP//qCLASH | 37986224\|31393953 | Unspecified |
| mmu-miR-378a-3p | MIMAT0003151 | Srsf3 | 20383 | HITS-CLIP | 37986224\|25083871 | Unspecified |
| mmu-miR-378a-3p | MIMAT0003151 | Srsf6 | 67996 | HITS-CLIP | 37986224\|23142080\|29167373 | Unspecified |
| mmu-miR-378a-3p | MIMAT0003151 | Ss18l1 | 269397 | HITS-CLIP | 23142080 | Unspecified |
| mmu-miR-378a-3p | MIMAT0003151 | Sspn | 16651 | HITS-CLIP | 37986224 | Unspecified |
| mmu-miR-378a-3p | MIMAT0003151 | Ssrp1 | 20833 | HITS-CLIP | 37986224 | Unspecified |
| mmu-miR-378a-3p | MIMAT0003151 | St3gal1 | 20442 | HITS-CLIP | 29167373\|23597149 | Unspecified |
| mmu-miR-378a-3p | MIMAT0003151 | St3gal5 | 20454 | HITS-CLIP | 37986224 | Unspecified |
| mmu-miR-378a-3p | MIMAT0003151 | St6gal1 | 20440 | HITS-CLIP | 37986224 | Unspecified |

| | | | | | | |
|---|---|---|---|---|---|---|
| mmu-miR-378a-3p | MIMAT0003151 | Stac2 | 217154 | HITS-CLIP | 25083871 | Unspecified |
| mmu-miR-378a-3p | MIMAT0003151 | Stam2 | 56324 | HITS-CLIP//qCLASH | 37986224\|25083871\|31393953 | Unspecified |
| mmu-miR-378a-3p | MIMAT0003151 | Stat6 | 20852 | HITS-CLIP | 23597149 | Unspecified |
| mmu-miR-378a-3p | MIMAT0003151 | Stau2 | 29819 | HITS-CLIP | 37986224 | Unspecified |
| mmu-miR-378a-3p | MIMAT0003151 | Steap4 | 117167 | HITS-CLIP | 37986224 | Unspecified |
| mmu-miR-378a-3p | MIMAT0003151 | Stk35 | 67333 | HITS-CLIP | 37986224 | Unspecified |
| mmu-miR-378a-3p | MIMAT0003151 | Stk39 | 53416 | Microarrays | 37986224\|22855601 | Unspecified |
| mmu-miR-378a-3p | MIMAT0003151 | Stk4 | 58231 | HITS-CLIP | 37986224 | Unspecified |
| mmu-miR-378a-3p | MIMAT0003151 | Stk40 | 74178 | Microarrays | 37986224\|22855601 | Unspecified |
| mmu-miR-378a-3p | MIMAT0003151 | Stmn1 | 16765 | Microarrays | 37986224\|22855601 | Unspecified |
| mmu-miR-378a-3p | MIMAT0003151 | Stom | 13830 | HITS-CLIP | 37986224 | Unspecified |
| mmu-miR-378a-3p | MIMAT0003151 | Ston1 | 77057 | HITS-CLIP | 37986224 | Unspecified |
| mmu-miR-378a-3p | MIMAT0003151 | Stradb | 227154 | HITS-CLIP | 37986224 | Unspecified |
| mmu-miR-378a-3p | MIMAT0003151 | Stt3b | 68292 | HITS-CLIP | 37986224 | Unspecified |
| mmu-miR-378a-3p | MIMAT0003151 | Stx11 | 74732 | HITS-CLIP | 37986224\|25083871\|23142080 | Unspecified |
| mmu-miR-378a-3p | MIMAT0003151 | Stx17 | 67727 | HITS-CLIP | 37986224\|25083871 | Unspecified |
| mmu-miR-378a-3p | MIMAT0003151 | Suco | 226551 | HITS-CLIP | 37986224 | Unspecified |
| mmu-miR-378a-3p | MIMAT0003151 | Sulf1 | 240725 | HITS-CLIP | 25083871 | Unspecified |
| mmu-miR-378a-3p | MIMAT0003151 | Supt20 | 56790 | HITS-CLIP | 23142080\|25083871 | Unspecified |
| mmu-miR-378a-3p | MIMAT0003151 | Supt6 | 20926 | HITS-CLIP | 25083871\|29167373 | Unspecified |
| mmu-miR-378a-3p | MIMAT0003151 | Surf4 | 20932 | HITS-CLIP | 37986224 | Unspecified |
| mmu-miR-378a-3p | MIMAT0003151 | Surf6 | 20935 | HITS-CLIP | 37986224 | Unspecified |
| mmu-miR-378a-3p | MIMAT0003151 | Susd6 | 9766 | Microarrays | 22855601 | Unspecified |
| mmu-miR-378a-3p | MIMAT0003151 | Svil | 225115 | HITS-CLIP | 37986224\|25083871\|25753418 | Unspecified |
| mmu-miR-378a-3p | MIMAT0003151 | Syncrip | 56403 | HITS-CLIP | 23142080 | Unspecified |
| mmu-miR-378a-3p | MIMAT0003151 | Synj1 | 104015 | HITS-CLIP | 37986224\|29167373 | Unspecified |
| mmu-miR-378a-3p | MIMAT0003151 | Synpo | 104027 | HITS-CLIP | 37986224 | Unspecified |
| mmu-miR-378a-3p | MIMAT0003151 | Synpo2 | 118449 | HITS-CLIP | 37986224 | Unspecified |
| mmu-miR-378a-3p | MIMAT0003151 | Synrg | 217030 | HITS-CLIP | 37986224 | Unspecified |
| mmu-miR-378a-3p | MIMAT0003151 | Szt2 | 230676 | HITS-CLIP | 37986224 | Unspecified |
| mmu-miR-378a-3p | MIMAT0003151 | Tab2 | 68652 | HITS-CLIP | 23142080\|25083871\|23597149\|29167 | Unspecified |
| mmu-miR-378a-3p | MIMAT0003151 | Taf1 | 270627 | HITS-CLIP | 25083871 | Unspecified |
| mmu-miR-378a-3p | MIMAT0003151 | Taf1d | 75316 | HITS-CLIP | 29167373 | Unspecified |
| mmu-miR-378a-3p | MIMAT0003151 | Taf2 | 319944 | HITS-CLIP | 37986224 | Unspecified |
| mmu-miR-378a-3p | MIMAT0003151 | Taf4b | 72504 | HITS-CLIP | 37986224 | Unspecified |
| mmu-miR-378a-3p | MIMAT0003151 | Tanc1 | 66860 | HITS-CLIP | 37986224 | Unspecified |
| mmu-miR-378a-3p | MIMAT0003151 | Tanc2 | 77097 | HITS-CLIP | 37986224 | Unspecified |
| mmu-miR-378a-3p | MIMAT0003151 | Taok1 | 216965 | HITS-CLIP | 23142080 | Unspecified |
| mmu-miR-378a-3p | MIMAT0003151 | Taok2 | 381921 | HITS-CLIP | 23142080 | Unspecified |

| | | | | | | |
|---|---|---|---|---|---|---|
| mmu-miR-378a-3p | MIMAT0003151 | Tapt1 | 231225 | HITS-CLIP | 37986224 | Unspecified |
| mmu-miR-378a-3p | MIMAT0003151 | Tardbp | 230908 | HITS-CLIP | 37986224 | Unspecified |
| mmu-miR-378a-3p | MIMAT0003151 | Tax1bp1 | 52440 | HITS-CLIP | 37986224\|25083871 | Unspecified |
| mmu-miR-378a-3p | MIMAT0003151 | Tbc1d10b | 68449 | HITS-CLIP | 37986224\|23142080 | Unspecified |
| mmu-miR-378a-3p | MIMAT0003151 | Tbc1d14 | 100855 | HITS-CLIP | 23597149 | Unspecified |
| mmu-miR-378a-3p | MIMAT0003151 | Tbc1d22a | 223754 | HITS-CLIP | 37986224 | Unspecified |
| mmu-miR-378a-3p | MIMAT0003151 | Tbcd | 108903 | qCLASH | 31393953 | Unspecified |
| mmu-miR-378a-3p | MIMAT0003151 | Tbkbp1 | 73174 | HITS-CLIP | 23142080 | Unspecified |
| mmu-miR-378a-3p | MIMAT0003151 | Tbl1x | 21372 | HITS-CLIP | 37986224 | Unspecified |
| mmu-miR-378a-3p | MIMAT0003151 | Tbl1xr1 | 81004 | HITS-CLIP | 23142080\|25083871 | Unspecified |
| mmu-miR-378a-3p | MIMAT0003151 | Tceanc2 | 66526 | HITS-CLIP | 29167373 | Unspecified |
| mmu-miR-378a-3p | MIMAT0003151 | Tcerg1 | 56070 | HITS-CLIP | 37986224 | Unspecified |
| mmu-miR-378a-3p | MIMAT0003151 | Tcf12 | 21406 | HITS-CLIP | 37986224\|25083871 | Unspecified |
| mmu-miR-378a-3p | MIMAT0003151 | Tcf20 | 21411 | HITS-CLIP | 37986224 | Unspecified |
| mmu-miR-378a-3p | MIMAT0003151 | Tcf25 | 66855 | HITS-CLIP | 37986224 | Unspecified |
| mmu-miR-378a-3p | MIMAT0003151 | Tcf4 | 21413 | HITS-CLIP | 25083871 | Unspecified |
| mmu-miR-378a-3p | MIMAT0003151 | Tdp2 | 56196 | HITS-CLIP | 37986224 | Unspecified |
| mmu-miR-378a-3p | MIMAT0003151 | Tdrd7 | 100121 | HITS-CLIP | 29167373 | Unspecified |
| mmu-miR-378a-3p | MIMAT0003151 | Tead1 | 21676 | HITS-CLIP | 37986224 | Unspecified |
| mmu-miR-378a-3p | MIMAT0003151 | Tead4 | 21679 | Microarrays | 37986224\|22855601 | Unspecified |
| mmu-miR-378a-3p | MIMAT0003151 | Tenm3 | 23965 | HITS-CLIP | 37986224 | Unspecified |
| mmu-miR-378a-3p | MIMAT0003151 | Tent4b | 64282 | Microarrays//HITS-CLIP | 22855601\|25083871\|23142080 | Unspecified |
| mmu-miR-378a-3p | MIMAT0003151 | Tent5a | 55603 | HITS-CLIP | 25083871 | Unspecified |
| mmu-miR-378a-3p | MIMAT0003151 | Tes | 21753 | Microarrays | 37986224\|22855601 | Unspecified |
| mmu-miR-378a-3p | MIMAT0003151 | Tesk1 | 21754 | HITS-CLIP | 37986224 | Unspecified |
| mmu-miR-378a-3p | MIMAT0003151 | Tet3 | 194388 | HITS-CLIP | 23142080\|29167373 | Unspecified |
| mmu-miR-378a-3p | MIMAT0003151 | Tex261 | 21766 | HITS-CLIP | 37986224 | Unspecified |
| mmu-miR-378a-3p | MIMAT0003151 | Tfdp1 | 21781 | HITS-CLIP | 29167373 | Unspecified |
| mmu-miR-378a-3p | MIMAT0003151 | Tfdp2 | 211586 | HITS-CLIP | 37986224 | Unspecified |
| mmu-miR-378a-3p | MIMAT0003151 | Tfrc | 22042 | Microarrays | 37986224\|22855601 | Unspecified |
| mmu-miR-378a-3p | MIMAT0003151 | Tgfb2 | 21808 | HITS-CLIP | 37986224\|25083871 | Unspecified |
| mmu-miR-378a-3p | MIMAT0003151 | Tgoln1 | 22134 | HITS-CLIP | 37986224 | Unspecified |
| mmu-miR-378a-3p | MIMAT0003151 | Tgs1 | 116940 | HITS-CLIP//qCLASH | 37986224\|31393953 | Unspecified |
| mmu-miR-378a-3p | MIMAT0003151 | Thbs1 | 21825 | Microarrays//HITS-CLIP | 37986224\|25083871\|22855601 | Unspecified |
| mmu-miR-378a-3p | MIMAT0003151 | Thoc2 | 331401 | HITS-CLIP | 37986224\|25083871 | Unspecified |
| mmu-miR-378a-3p | MIMAT0003151 | Thrap3 | 230753 | HITS-CLIP | 23142080 | Unspecified |
| mmu-miR-378a-3p | MIMAT0003151 | Tiam2 | 24001 | HITS-CLIP | 37986224 | Unspecified |
| mmu-miR-378a-3p | MIMAT0003151 | Timm8a1 | 30058 | Microarrays//qCLASH | 37986224\|22855601\|31393953 | Unspecified |
| mmu-miR-378a-3p | MIMAT0003151 | Timp2 | 21858 | HITS-CLIP | 29167373 | Unspecified |

| | | | | | | |
|---|---|---|---|---|---|---|
| mmu-miR-378a-3p | MIMAT0003151 | Tiparp | 99929 | Microarrays | 37986224\|22855601 | Unspecified |
| mmu-miR-378a-3p | MIMAT0003151 | Tjp2 | 21873 | Microarrays | 37986224\|22855601 | Unspecified |
| mmu-miR-378a-3p | MIMAT0003151 | Tkt | 21881 | Microarrays | 37986224\|22855601 | Unspecified |
| mmu-miR-378a-3p | MIMAT0003151 | Tle3 | 21887 | HITS-CLIP | 25083871 | Unspecified |
| mmu-miR-378a-3p | MIMAT0003151 | Tle4 | 21888 | HITS-CLIP | 23142080 | Unspecified |
| mmu-miR-378a-3p | MIMAT0003151 | Tlk1 | 228012 | HITS-CLIP | 25083871 | Unspecified |
| mmu-miR-378a-3p | MIMAT0003151 | Tlk2 | 24086 | HITS-CLIP | 37986224 | Unspecified |
| mmu-miR-378a-3p | MIMAT0003151 | Tm2d2 | 69742 | HITS-CLIP | 23597149\|25753418 | Unspecified |
| mmu-miR-378a-3p | MIMAT0003151 | Tm9sf2 | 68059 | HITS-CLIP | 37986224 | Unspecified |
| mmu-miR-378a-3p | MIMAT0003151 | Tmbim1 | 69660 | HITS-CLIP | 37986224 | Unspecified |
| mmu-miR-378a-3p | MIMAT0003151 | Tmcc1 | 330401 | HITS-CLIP | 37986224 | Unspecified |
| mmu-miR-378a-3p | MIMAT0003151 | Tmcc3 | 319880 | HITS-CLIP | 37986224 | Unspecified |
| mmu-miR-378a-3p | MIMAT0003151 | Tmed3 | 66111 | Microarrays | 37986224 | Unspecified |
| mmu-miR-378a-3p | MIMAT0003151 | Tmed5 | 73130 | HITS-CLIP | 37986224\|25083871\|23142080\|29167 | Unspecified |
| mmu-miR-378a-3p | MIMAT0003151 | Tmem101 | 76547 | HITS-CLIP | 25083871 | Unspecified |
| mmu-miR-378a-3p | MIMAT0003151 | Tmem106b | 71900 | HITS-CLIP | 37986224\|25083871 | Unspecified |
| mmu-miR-378a-3p | MIMAT0003151 | Tmem109 | 68539 | HITS-CLIP | 37986224 | Unspecified |
| mmu-miR-378a-3p | MIMAT0003151 | Tmem127 | 69470 | HITS-CLIP | 37986224 | Unspecified |
| mmu-miR-378a-3p | MIMAT0003151 | Tmem129 | 68366 | HITS-CLIP | 37986224 | Unspecified |
| mmu-miR-378a-3p | MIMAT0003151 | Tmem131l | 23240 | HITS-CLIP | 23597149 | Unspecified |
| mmu-miR-378a-3p | MIMAT0003151 | Tmem135 | 72759 | HITS-CLIP | 37986224 | Unspecified |
| mmu-miR-378a-3p | MIMAT0003151 | Tmem184b | 223693 | HITS-CLIP | 37986224 | Unspecified |
| mmu-miR-378a-3p | MIMAT0003151 | Tmem198b | 73827 | HITS-CLIP | 37986224 | Unspecified |
| mmu-miR-378a-3p | MIMAT0003151 | Tmem218 | 66279 | Microarrays | 37986224 | Unspecified |
| mmu-miR-378a-3p | MIMAT0003151 | Tmem245 | 242474 | HITS-CLIP | 23597149\|25753418 | Unspecified |
| mmu-miR-378a-3p | MIMAT0003151 | Tmem259 | 216157 | HITS-CLIP | 37986224 | Unspecified |
| mmu-miR-378a-3p | MIMAT0003151 | Tmem263 | 103266 | HITS-CLIP | 37986224 | Unspecified |
| mmu-miR-378a-3p | MIMAT0003151 | Tmem30a | 69981 | HITS-CLIP | 37986224 | Unspecified |
| mmu-miR-378a-3p | MIMAT0003151 | Tmem33 | 67878 | HITS-CLIP | 37986224 | Unspecified |
| mmu-miR-378a-3p | MIMAT0003151 | Tmem47 | 192216 | Microarrays | 37986224\|22855601 | Unspecified |
| mmu-miR-378a-3p | MIMAT0003151 | Tmem50a | 71817 | HITS-CLIP | 23142080 | Unspecified |
| mmu-miR-378a-3p | MIMAT0003151 | Tmem62 | 96957 | HITS-CLIP | 37986224\|25083871\|23597149 | Unspecified |
| mmu-miR-378a-3p | MIMAT0003151 | Tmem64 | 100201 | HITS-CLIP | 25753418\|23142080 | Unspecified |
| mmu-miR-378a-3p | MIMAT0003151 | Tmem68 | 72098 | HITS-CLIP | 37986224 | Unspecified |
| mmu-miR-378a-3p | MIMAT0003151 | Tmem94 | 9772 | HITS-CLIP | 25083871 | Unspecified |
| mmu-miR-378a-3p | MIMAT0003151 | Tmod1 | 21916 | HITS-CLIP | 25753418 | Unspecified |
| mmu-miR-378a-3p | MIMAT0003151 | Tmppe | 100504715 | HITS-CLIP | 37986224 | Unspecified |
| mmu-miR-378a-3p | MIMAT0003151 | Tnc | 21923 | Microarrays | 37986224\|22855601 | Unspecified |
| mmu-miR-378a-3p | MIMAT0003151 | Tnfaip8l2 | 69769 | HITS-CLIP | 23142080 | Unspecified |

| | | | | | | |
|---|---|---|---|---|---|---|
| mmu-miR-378a-3p | MIMAT0003151 | Tnfrsf10b | 21933 | HITS-CLIP | 37986224 | Unspecified |
| mmu-miR-378a-3p | MIMAT0003151 | Tnfrsf11b | 18383 | Microarrays | 37986224\|22855601 | Unspecified |
| mmu-miR-378a-3p | MIMAT0003151 | Tnfrsf12a | 27279 | Microarrays | 37986224\|22855601 | Unspecified |
| mmu-miR-378a-3p | MIMAT0003151 | Tnfsf8 | 21949 | HITS-CLIP | 37986224 | Unspecified |
| mmu-miR-378a-3p | MIMAT0003151 | Tnks | 21951 | HITS-CLIP | 23142080 | Unspecified |
| mmu-miR-378a-3p | MIMAT0003151 | Tnks2 | 74493 | HITS-CLIP | 37986224 | Unspecified |
| mmu-miR-378a-3p | MIMAT0003151 | Tnpo1 | 238799 | HITS-CLIP | 37986224\|25083871 | Unspecified |
| mmu-miR-378a-3p | MIMAT0003151 | Tnrc6a | 233833 | HITS-CLIP | 25083871\|23142080 | Unspecified |
| mmu-miR-378a-3p | MIMAT0003151 | Tnrc6b | 213988 | HITS-CLIP | 37986224 | Unspecified |
| mmu-miR-378a-3p | MIMAT0003151 | Tnrc6c | 217351 | HITS-CLIP | 37986224\|23142080 | Unspecified |
| mmu-miR-378a-3p | MIMAT0003151 | Tns1 | 21961 | HITS-CLIP | 37986224 | Unspecified |
| mmu-miR-378a-3p | MIMAT0003151 | Tns2 | 209039 | HITS-CLIP | 25083871 | Unspecified |
| mmu-miR-378a-3p | MIMAT0003151 | Tns3 | 319939 | HITS-CLIP | 29167373 | Unspecified |
| mmu-miR-378a-3p | MIMAT0003151 | Tob1 | 22057 | HITS-CLIP | 37986224 | Unspecified |
| mmu-miR-378a-3p | MIMAT0003151 | Tob2 | 57259 | HITS-CLIP | 37986224 | Unspecified |
| mmu-miR-378a-3p | MIMAT0003151 | Tomm70a | 28185 | Microarrays//HITS-CLIP | 37986224\|23142080\|22855601\|25083 | Unspecified |
| mmu-miR-378a-3p | MIMAT0003151 | Topors | 106021 | HITS-CLIP | 37986224\|25083871\|23597149\|23142 | Unspecified |
| mmu-miR-378a-3p | MIMAT0003151 | Tor1aip1 | 208263 | HITS-CLIP | 37986224 | Unspecified |
| mmu-miR-378a-3p | MIMAT0003151 | Tor1aip2 | 240832 | HITS-CLIP | 37986224 | Unspecified |
| mmu-miR-378a-3p | MIMAT0003151 | Tox4 | 268741 | HITS-CLIP | 25083871\|23142080 | Unspecified |
| mmu-miR-378a-3p | MIMAT0003151 | Traf6 | 22034 | HITS-CLIP | 37986224 | Unspecified |
| mmu-miR-378a-3p | MIMAT0003151 | Trak2 | 70827 | HITS-CLIP | 25083871 | Unspecified |
| mmu-miR-378a-3p | MIMAT0003151 | Tram2 | 170829 | HITS-CLIP | 37986224 | Unspecified |
| mmu-miR-378a-3p | MIMAT0003151 | Trap1 | 68015 | HITS-CLIP | 37986224\|23597149 | Unspecified |
| mmu-miR-378a-3p | MIMAT0003151 | Trappc8 | 75964 | HITS-CLIP | 37986224 | Unspecified |
| mmu-miR-378a-3p | MIMAT0003151 | Trib2 | 217410 | Microarrays | 37986224\|22855601 | Unspecified |
| mmu-miR-378a-3p | MIMAT0003151 | Trib3 | 228775 | Microarrays | 37986224\|22855601 | Unspecified |
| mmu-miR-378a-3p | MIMAT0003151 | Trim24 | 21848 | HITS-CLIP | 23142080 | Unspecified |
| mmu-miR-378a-3p | MIMAT0003151 | Trim26 | 22670 | HITS-CLIP | 37986224 | Unspecified |
| mmu-miR-378a-3p | MIMAT0003151 | Trim30a | 20128 | Microarrays | 37986224 | Unspecified |
| mmu-miR-378a-3p | MIMAT0003151 | Trim35 | 66854 | HITS-CLIP | 37986224 | Unspecified |
| mmu-miR-378a-3p | MIMAT0003151 | Trim37 | 68729 | HITS-CLIP | 23142080 | Unspecified |
| mmu-miR-378a-3p | MIMAT0003151 | Trim41 | 211007 | qCLASH | 31393953 | Unspecified |
| mmu-miR-378a-3p | MIMAT0003151 | Trim44 | 80985 | HITS-CLIP | 37986224 | Unspecified |
| mmu-miR-378a-3p | MIMAT0003151 | Trim56 | 384309 | HITS-CLIP | 37986224\|25083871 | Unspecified |
| mmu-miR-378a-3p | MIMAT0003151 | Trim59 | 66949 | HITS-CLIP | 37986224 | Unspecified |
| mmu-miR-378a-3p | MIMAT0003151 | Trip11 | 109181 | HITS-CLIP | 37986224 | Unspecified |
| mmu-miR-378a-3p | MIMAT0003151 | Trmt6 | 66926 | HITS-CLIP | 37986224\|23142080 | Unspecified |
| mmu-miR-378a-3p | MIMAT0003151 | Trmt61a | 328162 | Microarrays | 37986224\|22855601 | Unspecified |

| | | | | | | |
|---|---|---|---|---|---|---|
| mmu-miR-378a-3p | MIMAT0003151 | Trp53bp1 | 27223 | HITS-CLIP | 37986224 | Unspecified |
| mmu-miR-378a-3p | MIMAT0003151 | Trp53inp1 | 60599 | HITS-CLIP, Microarrays//Microarrays | 37986224\|22855601 | Unspecified |
| mmu-miR-378a-3p | MIMAT0003151 | Trps1 | 83925 | HITS-CLIP | 37986224 | Unspecified |
| mmu-miR-378a-3p | MIMAT0003151 | Trrap | 100683 | HITS-CLIP | 37986224\|29167373 | Unspecified |
| mmu-miR-378a-3p | MIMAT0003151 | Tsc1 | 64930 | HITS-CLIP | 37986224 | Unspecified |
| mmu-miR-378a-3p | MIMAT0003151 | Tsc2 | 22084 | HITS-CLIP | 37986224 | Unspecified |
| mmu-miR-378a-3p | MIMAT0003151 | Tsc22d1 | 21807 | HITS-CLIP | 37986224 | Unspecified |
| mmu-miR-378a-3p | MIMAT0003151 | Tsc22d3 | 14605 | HITS-CLIP | 37986224 | Unspecified |
| mmu-miR-378a-3p | MIMAT0003151 | Tsen2 | 381802 | HITS-CLIP | 23142080 | Unspecified |
| mmu-miR-378a-3p | MIMAT0003151 | Tsnax | 53424 | HITS-CLIP | 37986224 | Unspecified |
| mmu-miR-378a-3p | MIMAT0003151 | Tspan5 | 56224 | HITS-CLIP | 37986224 | Unspecified |
| mmu-miR-378a-3p | MIMAT0003151 | Tsr1 | 104662 | Microarrays | 37986224\|22855601 | Unspecified |
| mmu-miR-378a-3p | MIMAT0003151 | Ttbk2 | 140810 | HITS-CLIP | 37986224 | Unspecified |
| mmu-miR-378a-3p | MIMAT0003151 | Ttc14 | 67120 | HITS-CLIP | 25083871 | Unspecified |
| mmu-miR-378a-3p | MIMAT0003151 | Ttc17 | 74569 | HITS-CLIP | 37986224 | Unspecified |
| mmu-miR-378a-3p | MIMAT0003151 | Ttc27 | 74196 | HITS-CLIP | 23142080 | Unspecified |
| mmu-miR-378a-3p | MIMAT0003151 | Ttc28 | 209683 | HITS-CLIP | 37986224 | Unspecified |
| mmu-miR-378a-3p | MIMAT0003151 | Ttc39b | 69863 | HITS-CLIP | 37986224\|25083871 | Unspecified |
| mmu-miR-378a-3p | MIMAT0003151 | Ttl | 69737 | HITS-CLIP | 25083871\|29167373 | Unspecified |
| mmu-miR-378a-3p | MIMAT0003151 | Ttll4 | 67534 | HITS-CLIP | 37986224 | Unspecified |
| mmu-miR-378a-3p | MIMAT0003151 | Ttn | 22138 | HITS-CLIP | 37986224\|25083871 | Unspecified |
| mmu-miR-378a-3p | MIMAT0003151 | Txnl1 | 53382 | HITS-CLIP | 37986224\|23142080\|23597149\|25083 | Unspecified |
| mmu-miR-378a-3p | MIMAT0003151 | Txnrd1 | 50493 | Microarrays | 37986224\|22855601 | Unspecified |
| mmu-miR-378a-3p | MIMAT0003151 | U2surp | 67958 | HITS-CLIP | 37986224 | Unspecified |
| mmu-miR-378a-3p | MIMAT0003151 | Ubap2l | 74383 | HITS-CLIP | 25083871 | Unspecified |
| mmu-miR-378a-3p | MIMAT0003151 | Ubash3b | 72828 | HITS-CLIP | 29167373 | Unspecified |
| mmu-miR-378a-3p | MIMAT0003151 | Ube2a | 22209 | HITS-CLIP | 23142080 | Unspecified |
| mmu-miR-378a-3p | MIMAT0003151 | Ube2d2a | 56550 | HITS-CLIP | 37986224 | Unspecified |
| mmu-miR-378a-3p | MIMAT0003151 | Ube2h | 22214 | HITS-CLIP | 37986224 | Unspecified |
| mmu-miR-378a-3p | MIMAT0003151 | Ube2l3 | 22195 | HITS-CLIP | 37986224 | Unspecified |
| mmu-miR-378a-3p | MIMAT0003151 | Ube2l6 | 56791 | HITS-CLIP | 23597149 | Unspecified |
| mmu-miR-378a-3p | MIMAT0003151 | Ube2r2 | 67615 | HITS-CLIP//qCLASH | 25083871\|31393953\|23142080 | Unspecified |
| mmu-miR-378a-3p | MIMAT0003151 | Ube3a | 22215 | Microarrays | 37986224\|22855601 | Unspecified |
| mmu-miR-378a-3p | MIMAT0003151 | Ube3c | 100763 | HITS-CLIP | 37986224 | Unspecified |
| mmu-miR-378a-3p | MIMAT0003151 | Ube4a | 140630 | HITS-CLIP | 37986224 | Unspecified |
| mmu-miR-378a-3p | MIMAT0003151 | Ubfd1 | 28018 | HITS-CLIP | 25083871 | Unspecified |
| mmu-miR-378a-3p | MIMAT0003151 | Ubn1 | 170644 | HITS-CLIP | 37986224 | Unspecified |
| mmu-miR-378a-3p | MIMAT0003151 | Ubn2 | 320538 | HITS-CLIP | 37986224 | Unspecified |
| mmu-miR-378a-3p | MIMAT0003151 | Ubp1 | 22221 | HITS-CLIP | 37986224 | Unspecified |

| | | | | | | |
|---|---|---|---|---|---|---|
| mmu-miR-378a-3p | MIMAT0003151 | Ubqln1 | 56085 | HITS-CLIP | 37986224\|23142080 | Unspecified |
| mmu-miR-378a-3p | MIMAT0003151 | Ubqln2 | 54609 | HITS-CLIP | 23597149\|23142080 | Unspecified |
| mmu-miR-378a-3p | MIMAT0003151 | Ubr2 | 224826 | HITS-CLIP | 37986224 | Unspecified |
| mmu-miR-378a-3p | MIMAT0003151 | Ubr3 | 68795 | HITS-CLIP | 25083871\|23597149\|29167373 | Unspecified |
| mmu-miR-378a-3p | MIMAT0003151 | Ubr4 | 69116 | HITS-CLIP | 23142080 | Unspecified |
| mmu-miR-378a-3p | MIMAT0003151 | Ubr5 | 70790 | HITS-CLIP | 37986224\|25753418\|29167373 | Unspecified |
| mmu-miR-378a-3p | MIMAT0003151 | Ubr7 | 66622 | HITS-CLIP | 25083871 | Unspecified |
| mmu-miR-378a-3p | MIMAT0003151 | Ubtf | 21429 | HITS-CLIP | 37986224 | Unspecified |
| mmu-miR-378a-3p | MIMAT0003151 | Ubxn4 | 67812 | HITS-CLIP | 37986224 | Unspecified |
| mmu-miR-378a-3p | MIMAT0003151 | Ubxn7 | 224111 | HITS-CLIP | 25083871 | Unspecified |
| mmu-miR-378a-3p | MIMAT0003151 | Uchl3 | 50933 | HITS-CLIP | 37986224 | Unspecified |
| mmu-miR-378a-3p | MIMAT0003151 | Uck2 | 80914 | Microarrays//qCLASH | 37986224\|22855601\|31393953 | Unspecified |
| mmu-miR-378a-3p | MIMAT0003151 | Ufd1 | 22230 | HITS-CLIP | 23142080 | Unspecified |
| mmu-miR-378a-3p | MIMAT0003151 | Ufl1 | 67490 | HITS-CLIP | 37986224\|25083871\|25753418 | Unspecified |
| mmu-miR-378a-3p | MIMAT0003151 | Ugcg | 22234 | HITS-CLIP | 25083871\|23142080 | Unspecified |
| mmu-miR-378a-3p | MIMAT0003151 | Uhmk1 | 16589 | HITS-CLIP | 37986224 | Unspecified |
| mmu-miR-378a-3p | MIMAT0003151 | Uhrf1bp1l | 75089 | Microarrays | 37986224\|22855601 | Unspecified |
| mmu-miR-378a-3p | MIMAT0003151 | Umps | 22247 | Microarrays | 37986224\|22855601 | Unspecified |
| mmu-miR-378a-3p | MIMAT0003151 | Unk | 217331 | HITS-CLIP | 23142080 | Unspecified |
| mmu-miR-378a-3p | MIMAT0003151 | Uox | 22262 | HITS-CLIP | 23597149 | Unspecified |
| mmu-miR-378a-3p | MIMAT0003151 | Urb1 | 207932 | Microarrays | 37986224\|22855601 | Unspecified |
| mmu-miR-378a-3p | MIMAT0003151 | Urgcp | 72046 | HITS-CLIP | 37986224 | Unspecified |
| mmu-miR-378a-3p | MIMAT0003151 | Uri1 | 19777 | HITS-CLIP | 25083871 | Unspecified |
| mmu-miR-378a-3p | MIMAT0003151 | Uroc1 | 243537 | HITS-CLIP | 37986224 | Unspecified |
| mmu-miR-378a-3p | MIMAT0003151 | Usb1 | 101985 | HITS-CLIP | 37986224 | Unspecified |
| mmu-miR-378a-3p | MIMAT0003151 | Usf2 | 22282 | HITS-CLIP | 25083871 | Unspecified |
| mmu-miR-378a-3p | MIMAT0003151 | Usf3 | 205717 | HITS-CLIP | 25083871 | Unspecified |
| mmu-miR-378a-3p | MIMAT0003151 | Uso1 | 56041 | HITS-CLIP | 25083871 | Unspecified |
| mmu-miR-378a-3p | MIMAT0003151 | Usp10 | 22224 | HITS-CLIP | 37986224 | Unspecified |
| mmu-miR-378a-3p | MIMAT0003151 | Usp11 | 236733 | HITS-CLIP | 25083871 | Unspecified |
| mmu-miR-378a-3p | MIMAT0003151 | Usp14 | 59025 | HITS-CLIP | 37986224\|23142080 | Unspecified |
| mmu-miR-378a-3p | MIMAT0003151 | Usp19 | 71472 | HITS-CLIP | 23597149 | Unspecified |
| mmu-miR-378a-3p | MIMAT0003151 | Usp2 | 53376 | HITS-CLIP | 25083871 | Unspecified |
| mmu-miR-378a-3p | MIMAT0003151 | Usp24 | 329908 | HITS-CLIP | 23142080 | Unspecified |
| mmu-miR-378a-3p | MIMAT0003151 | Usp28 | 235323 | HITS-CLIP | 37986224 | Unspecified |
| mmu-miR-378a-3p | MIMAT0003151 | Usp3 | 235441 | HITS-CLIP | 23142080 | Unspecified |
| mmu-miR-378a-3p | MIMAT0003151 | Usp33 | 170822 | HITS-CLIP | 23597149 | Unspecified |
| mmu-miR-378a-3p | MIMAT0003151 | Usp34 | 17847 | HITS-CLIP | 23142080\|25083871 | Unspecified |
| mmu-miR-378a-3p | MIMAT0003151 | Usp38 | 74841 | HITS-CLIP | 25083871 | Unspecified |

| | | | | | | |
|---|---|---|---|---|---|---|
| mmu-miR-378a-3p | MIMAT0003151 | Usp45 | 77593 | HITS-CLIP | 25083871\|25753418\|23142080 | Unspecified |
| mmu-miR-378a-3p | MIMAT0003151 | Usp47 | 74996 | HITS-CLIP | 25083871 | Unspecified |
| mmu-miR-378a-3p | MIMAT0003151 | Usp5 | 22225 | HITS-CLIP | 25083871 | Unspecified |
| mmu-miR-378a-3p | MIMAT0003151 | Usp6nl | 98910 | HITS-CLIP | 37986224 | Unspecified |
| mmu-miR-378a-3p | MIMAT0003151 | Usp7 | 252870 | Microarrays | 37986224\|22855601 | Unspecified |
| mmu-miR-378a-3p | MIMAT0003151 | Usp9x | 22284 | HITS-CLIP | 25083871\|23142080 | Unspecified |
| mmu-miR-378a-3p | MIMAT0003151 | Utp14b | 195434 | HITS-CLIP | 37986224 | Unspecified |
| mmu-miR-378a-3p | MIMAT0003151 | Utp15 | 105372 | Microarrays | 37986224\|22855601 | Unspecified |
| mmu-miR-378a-3p | MIMAT0003151 | Utp20 | 70683 | Microarrays//qCLASH | 37986224\|22855601\|31393953 | Unspecified |
| mmu-miR-378a-3p | MIMAT0003151 | Utp4 | 21771 | Microarrays | 22855601 | Unspecified |
| mmu-miR-378a-3p | MIMAT0003151 | Utp6 | 216987 | HITS-CLIP | 25083871\|23597149\|23142080 | Unspecified |
| mmu-miR-378a-3p | MIMAT0003151 | Utrn | 22288 | HITS-CLIP | 23142080 | Unspecified |
| mmu-miR-378a-3p | MIMAT0003151 | Vcan | 13003 | HITS-CLIP | 37986224 | Unspecified |
| mmu-miR-378a-3p | MIMAT0003151 | Vcl | 22330 | Microarrays | 37986224\|22855601 | Unspecified |
| mmu-miR-378a-3p | MIMAT0003151 | Vegfa | 22339 | Microarrays | 37986224\|22855601 | Unspecified |
| mmu-miR-378a-3p | MIMAT0003151 | Vgll3 | 73569 | HITS-CLIP | 37986224 | Unspecified |
| mmu-miR-378a-3p | MIMAT0003151 | Vopp1 | 232023 | Microarrays | 37986224\|22855601 | Unspecified |
| mmu-miR-378a-3p | MIMAT0003151 | Vps13b | 666173 | HITS-CLIP | 29167373 | Unspecified |
| mmu-miR-378a-3p | MIMAT0003151 | Vps13c | 320528 | HITS-CLIP | 37986224 | Unspecified |
| mmu-miR-378a-3p | MIMAT0003151 | Vps13d | 230895 | HITS-CLIP | 37986224\|25753418 | Unspecified |
| mmu-miR-378a-3p | MIMAT0003151 | Vps16 | 80743 | HITS-CLIP | 29167373 | Unspecified |
| mmu-miR-378a-3p | MIMAT0003151 | Vps26c | 135977289 | Microarrays | 22855601 | Unspecified |
| mmu-miR-378a-3p | MIMAT0003151 | Vps28 | 51160 | HITS-CLIP | 23597149 | Unspecified |
| mmu-miR-378a-3p | MIMAT0003151 | Vps35 | 65114 | HITS-CLIP | 37986224\|25083871\|23142080\|29167 | Unspecified |
| mmu-miR-378a-3p | MIMAT0003151 | Vps37a | 52348 | HITS-CLIP | 23142080 | Unspecified |
| mmu-miR-378a-3p | MIMAT0003151 | Vps4b | 20479 | HITS-CLIP | 37986224\|25083871\|23142080 | Unspecified |
| mmu-miR-378a-3p | MIMAT0003151 | Vps54 | 245944 | HITS-CLIP | 25083871\|23142080 | Unspecified |
| mmu-miR-378a-3p | MIMAT0003151 | Wac | 225131 | HITS-CLIP | 23142080\|25083871 | Unspecified |
| mmu-miR-378a-3p | MIMAT0003151 | Washc2 | 28006 | HITS-CLIP | 29167373 | Unspecified |
| mmu-miR-378a-3p | MIMAT0003151 | Washc4 | 319277 | HITS-CLIP | 25083871 | Unspecified |
| mmu-miR-378a-3p | MIMAT0003151 | Wbp11 | 60321 | HITS-CLIP | 25083871\|23142080 | Unspecified |
| mmu-miR-378a-3p | MIMAT0003151 | Wdfy3 | 72145 | HITS-CLIP | 37986224\|25083871 | Unspecified |
| mmu-miR-378a-3p | MIMAT0003151 | Wdr18 | 216156 | HITS-CLIP | 37986224 | Unspecified |
| mmu-miR-378a-3p | MIMAT0003151 | Wdr26 | 226757 | HITS-CLIP | 23142080\|23597149\|25083871 | Unspecified |
| mmu-miR-378a-3p | MIMAT0003151 | Wdr3 | 269470 | Microarrays | 37986224\|22855601 | Unspecified |
| mmu-miR-378a-3p | MIMAT0003151 | Wdr36 | 225348 | Microarrays | 37986224\|22855601 | Unspecified |
| mmu-miR-378a-3p | MIMAT0003151 | Wdr37 | 207615 | HITS-CLIP | 37986224 | Unspecified |
| mmu-miR-378a-3p | MIMAT0003151 | Wdr43 | 72515 | Microarrays | 37986224\|22855601 | Unspecified |
| mmu-miR-378a-3p | MIMAT0003151 | Wdr45 | 54636 | HITS-CLIP | 23597149\|23142080 | Unspecified |

| | | | | | | | |
|---|---|---|---|---|---|---|---|
| mmu-miR-378a-3p | MIMAT0003151 | Wdr48 | 67561 | HITS-CLIP | | 37986224 | Unspecified |
| mmu-miR-378a-3p | MIMAT0003151 | Wdr6 | 83669 | HITS-CLIP | | 37986224 | Unspecified |
| mmu-miR-378a-3p | MIMAT0003151 | Wdr75 | 73674 | Microarrays | 37986224\|22855601 | | Unspecified |
| mmu-miR-378a-3p | MIMAT0003151 | Wdr91 | 101240 | HITS-CLIP | | 29167373 | Unspecified |
| mmu-miR-378a-3p | MIMAT0003151 | Wipf1 | 215280 | HITS-CLIP | | 37986224 | Unspecified |
| mmu-miR-378a-3p | MIMAT0003151 | Wipi1 | 52639 | HITS-CLIP | | 37986224 | Unspecified |
| mmu-miR-378a-3p | MIMAT0003151 | Wnt5a | 22418 | Microarrays | 37986224\|22855601 | | Unspecified |
| mmu-miR-378a-3p | MIMAT0003151 | Xpr1 | 19775 | HITS-CLIP | | 23142080 | Unspecified |
| mmu-miR-378a-3p | MIMAT0003151 | Xrcc6 | 14375 | HITS-CLIP | | 37986224 | Unspecified |
| mmu-miR-378a-3p | MIMAT0003151 | Xylt1 | 233781 | HITS-CLIP | | 29167373 | Unspecified |
| mmu-miR-378a-3p | MIMAT0003151 | Ylpm1 | 56531 | HITS-CLIP | | 37986224 | Unspecified |
| mmu-miR-378a-3p | MIMAT0003151 | Ypel2 | 77864 | HITS-CLIP | | 37986224 | Unspecified |
| mmu-miR-378a-3p | MIMAT0003151 | Ypel3 | 66090 | Microarrays | | 37986224 | Unspecified |
| mmu-miR-378a-3p | MIMAT0003151 | Ythdf1 | 228994 | HITS-CLIP | | 37986224 | Unspecified |
| mmu-miR-378a-3p | MIMAT0003151 | Ythdf2 | 213541 | HITS-CLIP | | 29167373 | Unspecified |
| mmu-miR-378a-3p | MIMAT0003151 | Zbtb20 | 56490 | HITS-CLIP | 37986224\|25083871 | | Unspecified |
| mmu-miR-378a-3p | MIMAT0003151 | Zbtb33 | 56805 | HITS-CLIP | | 37986224 | Unspecified |
| mmu-miR-378a-3p | MIMAT0003151 | Zbtb34 | 241311 | HITS-CLIP | | 37986224 | Unspecified |
| mmu-miR-378a-3p | MIMAT0003151 | Zbtb39 | 320080 | HITS-CLIP | | 37986224 | Unspecified |
| mmu-miR-378a-3p | MIMAT0003151 | Zbtb44 | 235132 | HITS-CLIP | | 37986224 | Unspecified |
| mmu-miR-378a-3p | MIMAT0003151 | Zc2hc1a | 67306 | HITS-CLIP | | 37986224 | Unspecified |
| mmu-miR-378a-3p | MIMAT0003151 | Zc3h12c | 244871 | HITS-CLIP | | 25083871 | Unspecified |
| mmu-miR-378a-3p | MIMAT0003151 | Zc3h15 | 69082 | HITS-CLIP | 25083871\|23142080 | | Unspecified |
| mmu-miR-378a-3p | MIMAT0003151 | Zc3h6 | 78751 | HITS-CLIP | 37986224\|25083871\|23142080 | | Unspecified |
| mmu-miR-378a-3p | MIMAT0003151 | Zc3h7a | 106205 | HITS-CLIP | | 37986224 | Unspecified |
| mmu-miR-378a-3p | MIMAT0003151 | Zc3hc1 | 51530 | HITS-CLIP | | 25753418 | Unspecified |
| mmu-miR-378a-3p | MIMAT0003151 | Zcchc14 | 142682 | HITS-CLIP | 37986224\|25083871 | | Unspecified |
| mmu-miR-378a-3p | MIMAT0003151 | Zcchc18 | 66995 | HITS-CLIP | | 23142080 | Unspecified |
| mmu-miR-378a-3p | MIMAT0003151 | Zcchc2 | 227449 | HITS-CLIP | 37986224\|25083871\|23597149 | | Unspecified |
| mmu-miR-378a-3p | MIMAT0003151 | Zdhhc12 | 66220 | HITS-CLIP | | 37986224 | Unspecified |
| mmu-miR-378a-3p | MIMAT0003151 | Zdhhc14 | 224454 | HITS-CLIP | | 29167373 | Unspecified |
| mmu-miR-378a-3p | MIMAT0003151 | Zdhhc17 | 320150 | HITS-CLIP | | 25083871 | Unspecified |
| mmu-miR-378a-3p | MIMAT0003151 | Zdhhc21 | 68268 | HITS-CLIP | | 37986224 | Unspecified |
| mmu-miR-378a-3p | MIMAT0003151 | Zdhhc24 | 70605 | HITS-CLIP | | 37986224 | Unspecified |
| mmu-miR-378a-3p | MIMAT0003151 | Zeb2 | 24136 | HITS-CLIP | | 25083871 | Unspecified |
| mmu-miR-378a-3p | MIMAT0003151 | Zfand2a | 100494 | HITS-CLIP | | 37986224 | Unspecified |
| mmu-miR-378a-3p | MIMAT0003151 | Zfand6 | 65098 | HITS-CLIP | | 23597149 | Unspecified |
| mmu-miR-378a-3p | MIMAT0003151 | Zfc3h1 | 216345 | HITS-CLIP | | 23142080 | Unspecified |
| mmu-miR-378a-3p | MIMAT0003151 | Zfhx3 | 11906 | HITS-CLIP | | 37986224 | Unspecified |

| | | | | | | |
|---|---|---|---|---|---|---|
| mmu-miR-378a-3p | MIMAT0003151 | Zfhx4 | 80892 | HITS-CLIP | 37986224\|25083871 | Unspecified |
| mmu-miR-378a-3p | MIMAT0003151 | Zfp106 | 20402 | HITS-CLIP | 25083871\|23142080 | Unspecified |
| mmu-miR-378a-3p | MIMAT0003151 | Zfp148 | 22661 | HITS-CLIP | 25083871\|29167373 | Unspecified |
| mmu-miR-378a-3p | MIMAT0003151 | Zfp217 | 228913 | HITS-CLIP | 37986224\|23142080 | Unspecified |
| mmu-miR-378a-3p | MIMAT0003151 | Zfp260 | 26466 | HITS-CLIP | 37986224 | Unspecified |
| mmu-miR-378a-3p | MIMAT0003151 | Zfp27 | 22689 | HITS-CLIP | 25083871 | Unspecified |
| mmu-miR-378a-3p | MIMAT0003151 | Zfp275 | 27081 | HITS-CLIP | 25083871 | Unspecified |
| mmu-miR-378a-3p | MIMAT0003151 | Zfp280c | 208968 | HITS-CLIP | 25083871 | Unspecified |
| mmu-miR-378a-3p | MIMAT0003151 | Zfp281 | 226442 | HITS-CLIP | 37986224 | Unspecified |
| mmu-miR-378a-3p | MIMAT0003151 | Zfp292 | 30046 | HITS-CLIP | 23142080 | Unspecified |
| mmu-miR-378a-3p | MIMAT0003151 | Zfp318 | 57908 | HITS-CLIP | 37986224 | Unspecified |
| mmu-miR-378a-3p | MIMAT0003151 | Zfp319 | 79233 | HITS-CLIP | 37986224 | Unspecified |
| mmu-miR-378a-3p | MIMAT0003151 | Zfp330 | 30932 | HITS-CLIP | 23142080 | Unspecified |
| mmu-miR-378a-3p | MIMAT0003151 | Zfp335 | 329559 | HITS-CLIP | 23142080 | Unspecified |
| mmu-miR-378a-3p | MIMAT0003151 | Zfp36 | 22695 | Microarrays | 37986224\|22855601 | Unspecified |
| mmu-miR-378a-3p | MIMAT0003151 | Zfp36l2 | 12193 | HITS-CLIP | 37986224\|23142080 | Unspecified |
| mmu-miR-378a-3p | MIMAT0003151 | Zfp410 | 52708 | HITS-CLIP | 25083871 | Unspecified |
| mmu-miR-378a-3p | MIMAT0003151 | Zfp451 | 98403 | HITS-CLIP | 37986224\|23142080 | Unspecified |
| mmu-miR-378a-3p | MIMAT0003151 | Zfp512 | 269639 | HITS-CLIP | 37986224 | Unspecified |
| mmu-miR-378a-3p | MIMAT0003151 | Zfp60 | 22718 | HITS-CLIP | 37986224 | Unspecified |
| mmu-miR-378a-3p | MIMAT0003151 | Zfp638 | 27332 | HITS-CLIP | 23597149 | Unspecified |
| mmu-miR-378a-3p | MIMAT0003151 | Zfp652 | 268469 | HITS-CLIP | 37986224\|25083871\|23142080 | Unspecified |
| mmu-miR-378a-3p | MIMAT0003151 | Zfp68 | 24135 | HITS-CLIP | 37986224 | Unspecified |
| mmu-miR-378a-3p | MIMAT0003151 | Zfp770 | 228491 | HITS-CLIP | 37986224 | Unspecified |
| mmu-miR-378a-3p | MIMAT0003151 | Zfp799 | 240064 | HITS-CLIP | 37986224 | Unspecified |
| mmu-miR-378a-3p | MIMAT0003151 | Zfp800 | 627049 | HITS-CLIP | 37986224\|25083871\|23142080 | Unspecified |
| mmu-miR-378a-3p | MIMAT0003151 | Zfp831 | 100043757 | HITS-CLIP | 37986224 | Unspecified |
| mmu-miR-378a-3p | MIMAT0003151 | Zfpl1 | 81909 | HITS-CLIP | 37986224\|25083871 | Unspecified |
| mmu-miR-378a-3p | MIMAT0003151 | Zfr | 22763 | HITS-CLIP | 37986224\|25083871\|23142080\|2575 | Unspecified |
| mmu-miR-378a-3p | MIMAT0003151 | Zfyve9 | 230597 | HITS-CLIP | 37986224 | Unspecified |
| mmu-miR-378a-3p | MIMAT0003151 | Zhx1 | 22770 | HITS-CLIP | 25083871\|23142080 | Unspecified |
| mmu-miR-378a-3p | MIMAT0003151 | Zkscan1 | 74570 | HITS-CLIP | 37986224\|25083871\|23142080 | Unspecified |
| mmu-miR-378a-3p | MIMAT0003151 | Zkscan3 | 72739 | HITS-CLIP | 37986224 | Unspecified |
| mmu-miR-378a-3p | MIMAT0003151 | Zmat3 | 22401 | HITS-CLIP, Microarrays//Microarrays | 37986224\|22855601 | Unspecified |
| mmu-miR-378a-3p | MIMAT0003151 | Zmpste24 | 230709 | HITS-CLIP | 37986224 | Unspecified |
| mmu-miR-378a-3p | MIMAT0003151 | Zmym2 | 76007 | HITS-CLIP | 37986224\|23142080 | Unspecified |
| mmu-miR-378a-3p | MIMAT0003151 | Zmym4 | 67785 | HITS-CLIP | 37986224 | Unspecified |
| mmu-miR-378a-3p | MIMAT0003151 | Znrf2 | 387524 | Microarrays | 37986224\|22855601 | Unspecified |
| mmu-miR-378a-3p | MIMAT0003151 | Znrf3 | 407821 | HITS-CLIP | 23142080 | Unspecified |

| | | | | | | |
|---|---|---|---|---|---|---|
| mmu-miR-378a-3p | MIMAT0003151 | Zranb2 | 53861 | HITS-CLIP | 37986224 | Unspecified |
| mmu-miR-378a-3p | MIMAT0003151 | Zrsr2 | 22184 | HITS-CLIP | 23142080 | Unspecified |
| mmu-miR-378a-3p | MIMAT0003151 | Zscan22 | 232878 | HITS-CLIP | 37986224 | Unspecified |
| mmu-miR-378a-3p | MIMAT0003151 | Zswim4 | 212168 | Microarrays | 37986224\|22855601 | Unspecified |
| mmu-miR-378a-3p | MIMAT0003151 | Zswim8 | 268721 | HITS-CLIP | 23597149\|25083871\|29167373 | Unspecified |
| mmu-miR-378a-3p | MIMAT0003151 | Zup1 | 221302 | HITS-CLIP | 23142080 | Unspecified |
| mmu-miR-378a-3p | MIMAT0003151 | Zxdb | 668166 | HITS-CLIP | 37986224 | Unspecified |
| mmu-miR-378a-3p | MIMAT0003151 | Zxdc | 80292 | Microarrays//HITS-CLIP | 37986224\|22855601\|23142080 | Unspecified |
| mmu-miR-378a-3p | MIMAT0003151 | Zyg11b | 414872 | HITS-CLIP | 37986224 | Unspecified |
| mmu-miR-378a-3p | MIMAT0003151 | Zyx | 22793 | HITS-CLIP | 37986224 | Unspecified |
| mmu-miR-378a-3p | MIMAT0003151 | Zzef1 | 195018 | HITS-CLIP | 25753418 | Unspecified |
| mmu-miR-378a-3p | MIMAT0003151 | mt-Cytb | 19893556 | HITS-CLIP | 23142080 | Unspecified |
| mmu-miR-1306-5p | MIMAT0019136 | Atp6v0a1 | 11975 | qCLASH | 31393953 | Unspecified |
| mmu-miR-1306-5p | MIMAT0019136 | Sez6l | 56747 | qCLASH | 31393953 | Unspecified |
| mmu-miR-125b-5p | MIMAT0000136 | 0610010F05Rik | 71675 | HITS-CLIP | 37986224\|25083871 | Unspecified |
| mmu-miR-125b-5p | MIMAT0000136 | 0610030E20Rik | 68364 | HITS-CLIP | 37986224 | Unspecified |
| mmu-miR-125b-5p | MIMAT0000136 | 1110012L19Rik | 68618 | Microarrays | 37986224 | Unspecified |
| mmu-miR-125b-5p | MIMAT0000136 | 1110032F04Rik | 68725 | Microarrays | 37986224\|22469780 | Unspecified |
| mmu-miR-125b-5p | MIMAT0000136 | 1600002K03Rik | 69770 | Microarrays | 37986224 | Unspecified |
| mmu-miR-125b-5p | MIMAT0000136 | 1600012H06Rik | 67912 | Microarrays | 37986224\|22469780 | Unspecified |
| mmu-miR-125b-5p | MIMAT0000136 | 1700013G24Rik | 69380 | Microarrays | 37986224 | Unspecified |
| mmu-miR-125b-5p | MIMAT0000136 | 1700017B05Rik | 74211 | qCLASH | 31393953 | Unspecified |
| mmu-miR-125b-5p | MIMAT0000136 | 1700019D03Rik | 67080 | Microarrays | 37986224 | Unspecified |
| mmu-miR-125b-5p | MIMAT0000136 | 1700020D05Rik | 75555 | Microarrays | 37986224 | Unspecified |
| mmu-miR-125b-5p | MIMAT0000136 | 1700020L24Rik | 66330 | Microarrays | 37986224\|22469780 | Unspecified |
| mmu-miR-125b-5p | MIMAT0000136 | 1700028P14Rik | 67483 | Microarrays | 37986224 | Unspecified |
| mmu-miR-125b-5p | MIMAT0000136 | 1700037H04Rik | 67326 | Microarrays | 37986224 | Unspecified |
| mmu-miR-125b-5p | MIMAT0000136 | 1700066M21Rik | 73467 | Microarrays | 37986224\|22469780 | Unspecified |
| mmu-miR-125b-5p | MIMAT0000136 | 1700088E04Rik | 27660 | Microarrays | 37986224 | Unspecified |
| mmu-miR-125b-5p | MIMAT0000136 | 1810062G17Rik | 72282 | Microarrays | 37986224 | Unspecified |
| mmu-miR-125b-5p | MIMAT0000136 | 2310022A10Rik | 66367 | HITS-CLIP | 25083871 | Unspecified |
| mmu-miR-125b-5p | MIMAT0000136 | 2310022B05Rik | 69551 | HITS-CLIP | 37986224 | Unspecified |
| mmu-miR-125b-5p | MIMAT0000136 | 2310057J18Rik | 67719 | Microarrays | 37986224\|22469780 | Unspecified |
| mmu-miR-125b-5p | MIMAT0000136 | 2310061I04Rik | 69662 | Microarrays | 37986224 | Unspecified |
| mmu-miR-125b-5p | MIMAT0000136 | 2410002F23Rik | 668661 | HITS-CLIP | 37986224 | Unspecified |
| mmu-miR-125b-5p | MIMAT0000136 | 2610318N02Rik | 70458 | Microarrays | 37986224\|22469780 | Unspecified |
| mmu-miR-125b-5p | MIMAT0000136 | 2700081O15Rik | 108899 | HITS-CLIP | 25083871 | Unspecified |
| mmu-miR-125b-5p | MIMAT0000136 | 4930451I11Rik | 78118 | Microarrays | 37986224 | Unspecified |
| mmu-miR-125b-5p | MIMAT0000136 | 4930522L14Rik | 100041734 | Microarrays | 37986224\|22469780 | Unspecified |

| | | | | | | |
|---|---|---|---|---|---|---|
| mmu-miR-125b-5p | MIMAT0000136 | 4930579F01Rik | 67741 | Microarrays | 37986224 | Unspecified |
| mmu-miR-125b-5p | MIMAT0000136 | 4931408C20Rik | 210940 | Microarrays | 37986224 | Unspecified |
| mmu-miR-125b-5p | MIMAT0000136 | 4932438A13Rik | 229227 | HITS-CLIP | 37986224 | Unspecified |
| mmu-miR-125b-5p | MIMAT0000136 | 5031439G07Rik | 223739 | HITS-CLIP | 37986224 | Unspecified |
| mmu-miR-125b-5p | MIMAT0000136 | 5730480H06Rik | 70592 | Microarrays | 22469780 | Unspecified |
| mmu-miR-125b-5p | MIMAT0000136 | 6030458C11Rik | 77877 | HITS-CLIP, Microarrays//HITS-CLIP//Microa | 37986224\|25083871\|22469780 | Unspecified |
| mmu-miR-125b-5p | MIMAT0000136 | 6430548M08Rik | 234797 | HITS-CLIP | 37986224 | Unspecified |
| mmu-miR-125b-5p | MIMAT0000136 | 8030462N17Rik | 212163 | HITS-CLIP | 37986224 | Unspecified |
| mmu-miR-125b-5p | MIMAT0000136 | 9230102O04Rik | 77682 | Microarrays | 37986224 | Unspecified |
| mmu-miR-125b-5p | MIMAT0000136 | 9430015G10Rik | 230996 | Microarrays | 37986224 | Unspecified |
| mmu-miR-125b-5p | MIMAT0000136 | 9530068E07Rik | 213673 | HITS-CLIP | 37986224 | Unspecified |
| mmu-miR-125b-5p | MIMAT0000136 | 9930012K11Rik | 268759 | Microarrays | 37986224\|22469780 | Unspecified |
| mmu-miR-125b-5p | MIMAT0000136 | 9930021J03Rik | 240613 | HITS-CLIP | 37986224\|25083871 | Unspecified |
| mmu-miR-125b-5p | MIMAT0000136 | 9930111J21Rik2 | 245240 | Microarrays | 37986224\|22469780 | Unspecified |
| mmu-miR-125b-5p | MIMAT0000136 | A130010J15Rik | 319266 | HITS-CLIP, Microarrays//Microarrays//HITS | 37986224\|22469780\|25753418 | Unspecified |
| mmu-miR-125b-5p | MIMAT0000136 | AA467197 | 433470 | Microarrays | 37986224 | Unspecified |
| mmu-miR-125b-5p | MIMAT0000136 | AB124611 | 382062 | Microarrays | 37986224\|22469780 | Unspecified |
| mmu-miR-125b-5p | MIMAT0000136 | AI606181 | 226118 | HITS-CLIP | 37986224 | Unspecified |
| mmu-miR-125b-5p | MIMAT0000136 | AU040320 | 100317 | HITS-CLIP | 37986224\|19536157 | Unspecified |
| mmu-miR-125b-5p | MIMAT0000136 | Aacs | 78894 | HITS-CLIP | 37986224 | Unspecified |
| mmu-miR-125b-5p | MIMAT0000136 | Aagab | 66939 | HITS-CLIP | 37986224 | Unspecified |
| mmu-miR-125b-5p | MIMAT0000136 | Aass | 30956 | HITS-CLIP | 23597149 | Unspecified |
| mmu-miR-125b-5p | MIMAT0000136 | Abca2 | 11305 | HITS-CLIP, Microarrays | 37986224 | Unspecified |
| mmu-miR-125b-5p | MIMAT0000136 | Abcb11 | 27413 | HITS-CLIP | 37986224 | Unspecified |
| mmu-miR-125b-5p | MIMAT0000136 | Abcb1a | 18671 | Microarrays | 37986224 | Unspecified |
| mmu-miR-125b-5p | MIMAT0000136 | Abcb4 | 18670 | Microarrays | 37986224 | Unspecified |
| mmu-miR-125b-5p | MIMAT0000136 | Abcb6 | 74104 | Microarrays | 37986224\|22469780 | Unspecified |
| mmu-miR-125b-5p | MIMAT0000136 | Abcb7 | 11306 | HITS-CLIP, Microarrays//Microarrays | 37986224\|22469780 | Unspecified |
| mmu-miR-125b-5p | MIMAT0000136 | Abcb9 | 56325 | Microarrays | 37986224 | Unspecified |
| mmu-miR-125b-5p | MIMAT0000136 | Abcc5 | 27416 | HITS-CLIP | 37986224\|25083871\|29167373 | Unspecified |
| mmu-miR-125b-5p | MIMAT0000136 | Abcd1 | 11666 | HITS-CLIP | 37986224 | Unspecified |
| mmu-miR-125b-5p | MIMAT0000136 | Abcg2 | 26357 | Microarrays | 37986224\|22469780 | Unspecified |
| mmu-miR-125b-5p | MIMAT0000136 | Abhd13 | 68904 | HITS-CLIP | 37986224 | Unspecified |
| mmu-miR-125b-5p | MIMAT0000136 | Abhd2 | 54608 | HITS-CLIP, Microarrays//Microarrays | 37986224\|22469780 | Unspecified |
| mmu-miR-125b-5p | MIMAT0000136 | Abhd3 | 106861 | HITS-CLIP | 37986224 | Unspecified |
| mmu-miR-125b-5p | MIMAT0000136 | Abhd6 | 66082 | HITS-CLIP | 37986224\|25083871 | Unspecified |
| mmu-miR-125b-5p | MIMAT0000136 | Abi2 | 329165 | HITS-CLIP | 37986224 | Unspecified |
| mmu-miR-125b-5p | MIMAT0000136 | Abl2 | 11352 | HITS-CLIP | 37986224\|25083871 | Unspecified |
| mmu-miR-125b-5p | MIMAT0000136 | Ablim1 | 226251 | HITS-CLIP | 37986224 | Unspecified |

| | | | | | | | |
|---|---|---|---|---|---|---|---|
| mmu-miR-125b-5p | MIMAT0000136 | Abr | 109934 | HITS-CLIP | | 37986224 | Unspecified |
| mmu-miR-125b-5p | MIMAT0000136 | Abra | 223513 | HITS-CLIP | | 25753418 | Unspecified |
| mmu-miR-125b-5p | MIMAT0000136 | Abtb1 | 80283 | Microarray//Microarrays//Other | 16495412\|22469780 | | Unspecified |
| mmu-miR-125b-5p | MIMAT0000136 | Acacb | 100705 | Microarrays | | 37986224 | Unspecified |
| mmu-miR-125b-5p | MIMAT0000136 | Acad12 | 338350 | HITS-CLIP | | 37986224 | Unspecified |
| mmu-miR-125b-5p | MIMAT0000136 | Acads | 11409 | HITS-CLIP | | 37986224 | Unspecified |
| mmu-miR-125b-5p | MIMAT0000136 | Acadvl | 11370 | HITS-CLIP | | 23597149 | Unspecified |
| mmu-miR-125b-5p | MIMAT0000136 | Acap3 | 140500 | HITS-CLIP | | 37986224 | Unspecified |
| mmu-miR-125b-5p | MIMAT0000136 | Aco2 | 11429 | Microarrays | | 37986224 | Unspecified |
| mmu-miR-125b-5p | MIMAT0000136 | Acot7 | 70025 | Microarrays | | 37986224 | Unspecified |
| mmu-miR-125b-5p | MIMAT0000136 | Acox3 | 80911 | HITS-CLIP | | 37986224 | Unspecified |
| mmu-miR-125b-5p | MIMAT0000136 | Acss2 | 60525 | Microarrays | | 37986224 | Unspecified |
| mmu-miR-125b-5p | MIMAT0000136 | Actb | 11461 | qCLASH | | 31393953 | Unspecified |
| mmu-miR-125b-5p | MIMAT0000136 | Actn1 | 109711 | HITS-CLIP | 23597149\|29167373 | | Unspecified |
| mmu-miR-125b-5p | MIMAT0000136 | Actn4 | 60595 | HITS-CLIP | | 37986224 | Unspecified |
| mmu-miR-125b-5p | MIMAT0000136 | Actr2 | 66713 | HITS-CLIP | 25083871\|29167373 | | Unspecified |
| mmu-miR-125b-5p | MIMAT0000136 | Actr8 | 56249 | HITS-CLIP | | 37986224 | Unspecified |
| mmu-miR-125b-5p | MIMAT0000136 | Acvr1 | 11477 | HITS-CLIP | | 25083871 | Unspecified |
| mmu-miR-125b-5p | MIMAT0000136 | Acvr1b | 11479 | HITS-CLIP | | 37986224 | Unspecified |
| mmu-miR-125b-5p | MIMAT0000136 | Acy3 | 71670 | Microarrays | | 37986224 | Unspecified |
| mmu-miR-125b-5p | MIMAT0000136 | Adal | 75894 | Microarrays | 37986224\|22469780 | | Unspecified |
| mmu-miR-125b-5p | MIMAT0000136 | Adam11 | 11488 | HITS-CLIP | | 37986224 | Unspecified |
| mmu-miR-125b-5p | MIMAT0000136 | Adam28 | 13522 | Microarrays | 37986224\|22469780 | | Unspecified |
| mmu-miR-125b-5p | MIMAT0000136 | Adam9 | 11502 | HITS-CLIP | | 25083871 | Unspecified |
| mmu-miR-125b-5p | MIMAT0000136 | Adamts10 | 224697 | Microarrays | | 37986224 | Unspecified |
| mmu-miR-125b-5p | MIMAT0000136 | Adamts12 | 239337 | HITS-CLIP | | 25083871 | Unspecified |
| mmu-miR-125b-5p | MIMAT0000136 | Adamts15 | 235130 | HITS-CLIP | | 37986224 | Unspecified |
| mmu-miR-125b-5p | MIMAT0000136 | Adar | 56417 | Microarrays | | 37986224 | Unspecified |
| mmu-miR-125b-5p | MIMAT0000136 | Adat3 | 100113398 | Microarrays | | 37986224 | Unspecified |
| mmu-miR-125b-5p | MIMAT0000136 | Adck1 | 72113 | HITS-CLIP | | 37986224 | Unspecified |
| mmu-miR-125b-5p | MIMAT0000136 | Adck2 | 57869 | HITS-CLIP | | 37986224 | Unspecified |
| mmu-miR-125b-5p | MIMAT0000136 | Adcy1 | 432530 | HITS-CLIP | 37986224\|19536157 | | Unspecified |
| mmu-miR-125b-5p | MIMAT0000136 | Adcy6 | 11512 | HITS-CLIP | | 37986224 | Unspecified |
| mmu-miR-125b-5p | MIMAT0000136 | Adcy9 | 11515 | Microarrays | | 37986224 | Unspecified |
| mmu-miR-125b-5p | MIMAT0000136 | Add3 | 27360 | Microarrays | | 37986224 | Unspecified |
| mmu-miR-125b-5p | MIMAT0000136 | Adgre5 | 976 | Microarrays//HITS-CLIP | 22469780\|25753418 | | Unspecified |
| mmu-miR-125b-5p | MIMAT0000136 | Adm2 | 223780 | HITS-CLIP | | 37986224 | Unspecified |
| mmu-miR-125b-5p | MIMAT0000136 | Adnp | 11538 | HITS-CLIP | | 29167373 | Unspecified |
| mmu-miR-125b-5p | MIMAT0000136 | Adra2a | 11551 | HITS-CLIP | | 37986224 | Unspecified |

| | | | | | | | |
|---|---|---|---|---|---|---|---|
| mmu-miR-125b-5p | MIMAT0000136 | Adrb2 | 11555 | Microarrays | | 37986224 | Unspecified |
| mmu-miR-125b-5p | MIMAT0000136 | Adssl1 | 11565 | Microarrays | | 37986224 | Unspecified |
| mmu-miR-125b-5p | MIMAT0000136 | Aebp1 | 11568 | HITS-CLIP | | 25083871 | Unspecified |
| mmu-miR-125b-5p | MIMAT0000136 | Aebp2 | 11569 | HITS-CLIP | | 37986224 | Unspecified |
| mmu-miR-125b-5p | MIMAT0000136 | Aen | 68048 | HITS-CLIP | | 37986224 | Unspecified |
| mmu-miR-125b-5p | MIMAT0000136 | Afap1l2 | 226250 | HITS-CLIP | | 37986224 | Unspecified |
| mmu-miR-125b-5p | MIMAT0000136 | Aff1 | 17355 | HITS-CLIP | 37986224\|25083871 | | Unspecified |
| mmu-miR-125b-5p | MIMAT0000136 | Afmid | 71562 | Microarrays | | 37986224 | Unspecified |
| mmu-miR-125b-5p | MIMAT0000136 | Aftph | 216549 | HITS-CLIP | | 37986224 | Unspecified |
| mmu-miR-125b-5p | MIMAT0000136 | Agap1 | 347722 | HITS-CLIP | | 25083871 | Unspecified |
| mmu-miR-125b-5p | MIMAT0000136 | Agap2 | 216439 | Microarrays | 37986224\|22469780 | | Unspecified |
| mmu-miR-125b-5p | MIMAT0000136 | Agfg1 | 15463 | HITS-CLIP | | 37986224 | Unspecified |
| mmu-miR-125b-5p | MIMAT0000136 | Ago1 | 236511 | HITS-CLIP | | 37986224 | Unspecified |
| mmu-miR-125b-5p | MIMAT0000136 | Ago2 | 239528 | HITS-CLIP, Microarrays//HITS-CLIP | 37986224\|25083871 | | Unspecified |
| mmu-miR-125b-5p | MIMAT0000136 | Ago3 | 214150 | HITS-CLIP | | 37986224 | Unspecified |
| mmu-miR-125b-5p | MIMAT0000136 | Agpat2 | 67512 | HITS-CLIP | | 23597149 | Unspecified |
| mmu-miR-125b-5p | MIMAT0000136 | Agpat3 | 28169 | HITS-CLIP | | 37986224 | Unspecified |
| mmu-miR-125b-5p | MIMAT0000136 | Agps | 228061 | Microarrays | 37986224\|22469780 | | Unspecified |
| mmu-miR-125b-5p | MIMAT0000136 | Agtrap | 11610 | HITS-CLIP | | 37986224 | Unspecified |
| mmu-miR-125b-5p | MIMAT0000136 | Ahcyl1 | 229709 | HITS-CLIP | 37986224\|23597149 | | Unspecified |
| mmu-miR-125b-5p | MIMAT0000136 | Ahcyl2 | 74340 | HITS-CLIP | | 37986224 | Unspecified |
| mmu-miR-125b-5p | MIMAT0000136 | Ahi1 | 52906 | Microarrays | | 37986224 | Unspecified |
| mmu-miR-125b-5p | MIMAT0000136 | Ahnak | 66395 | HITS-CLIP | 37986224\|25083871 | | Unspecified |
| mmu-miR-125b-5p | MIMAT0000136 | Ahsa2 | 268390 | HITS-CLIP | 37986224\|23597149 | | Unspecified |
| mmu-miR-125b-5p | MIMAT0000136 | Aif1 | 11629 | Microarrays | | 37986224 | Unspecified |
| mmu-miR-125b-5p | MIMAT0000136 | Aifm1 | 26926 | HITS-CLIP | 37986224\|25753418 | | Unspecified |
| mmu-miR-125b-5p | MIMAT0000136 | Ak1 | 11636 | Microarrays | | 37986224 | Unspecified |
| mmu-miR-125b-5p | MIMAT0000136 | Ak3 | 56248 | Microarrays | | 37986224 | Unspecified |
| mmu-miR-125b-5p | MIMAT0000136 | Ak8 | 68870 | Microarrays | | 37986224 | Unspecified |
| mmu-miR-125b-5p | MIMAT0000136 | Akap10 | 56697 | HITS-CLIP | 37986224\|25083871 | | Unspecified |
| mmu-miR-125b-5p | MIMAT0000136 | Akap13 | 75547 | HITS-CLIP | | 23597149 | Unspecified |
| mmu-miR-125b-5p | MIMAT0000136 | Akap17b | 338351 | HITS-CLIP | | 37986224 | Unspecified |
| mmu-miR-125b-5p | MIMAT0000136 | Akap2 | 11641 | HITS-CLIP | | 25083871 | Unspecified |
| mmu-miR-125b-5p | MIMAT0000136 | Akap7 | 432442 | Microarrays | | 37986224 | Unspecified |
| mmu-miR-125b-5p | MIMAT0000136 | Akap9 | 100986 | HITS-CLIP | | 25083871 | Unspecified |
| mmu-miR-125b-5p | MIMAT0000136 | Akna | 100182 | Microarrays//HITS-CLIP | 37986224\|25753418 | | Unspecified |
| mmu-miR-125b-5p | MIMAT0000136 | Akr1d1 | 208665 | HITS-CLIP | | 23597149 | Unspecified |
| mmu-miR-125b-5p | MIMAT0000136 | Akt3 | 23797 | HITS-CLIP | | 19536157 | Unspecified |
| mmu-miR-125b-5p | MIMAT0000136 | Alas1 | 11655 | HITS-CLIP | | 37986224 | Unspecified |

| | | | | | | | |
|---|---|---|---|---|---|---|---|
| mmu-miR-125b-5p | MIMAT0000136 | Aldh18a1 | 56454 | Microarrays | | 37986224 | Unspecified |
| mmu-miR-125b-5p | MIMAT0000136 | Aldh1l1 | 107747 | HITS-CLIP | 37986224\|23597149 | | Unspecified |
| mmu-miR-125b-5p | MIMAT0000136 | Aldh2 | 11669 | HITS-CLIP, Microarrays | | 37986224 | Unspecified |
| mmu-miR-125b-5p | MIMAT0000136 | Aldh3a1 | 11670 | Microarrays | | 37986224 | Unspecified |
| mmu-miR-125b-5p | MIMAT0000136 | Aldh3b1 | 67689 | Microarrays | 37986224\|22469780 | | Unspecified |
| mmu-miR-125b-5p | MIMAT0000136 | Aldh4a1 | 212647 | HITS-CLIP | | 23597149 | Unspecified |
| mmu-miR-125b-5p | MIMAT0000136 | Aldh6a1 | 104776 | HITS-CLIP | | 23597149 | Unspecified |
| mmu-miR-125b-5p | MIMAT0000136 | Aldob | 230163 | Microarrays | 37986224\|22469780 | | Unspecified |
| mmu-miR-125b-5p | MIMAT0000136 | Alg1 | 208211 | Microarrays | 37986224\|22469780 | | Unspecified |
| mmu-miR-125b-5p | MIMAT0000136 | Alg10b | 380959 | HITS-CLIP | | 37986224 | Unspecified |
| mmu-miR-125b-5p | MIMAT0000136 | Alg6 | 320438 | HITS-CLIP, Microarrays//HITS-CLIP//Microa | 37986224\|25083871\|22469780 | | Unspecified |
| mmu-miR-125b-5p | MIMAT0000136 | Alkbh1 | 211064 | HITS-CLIP | | 37986224 | Unspecified |
| mmu-miR-125b-5p | MIMAT0000136 | Alkbh5 | 268420 | HITS-CLIP | | 25083871 | Unspecified |
| mmu-miR-125b-5p | MIMAT0000136 | Alpk3 | 116904 | HITS-CLIP | | 37986224 | Unspecified |
| mmu-miR-125b-5p | MIMAT0000136 | Als2 | 74018 | Microarrays | | 37986224 | Unspecified |
| mmu-miR-125b-5p | MIMAT0000136 | Amacr | 17117 | HITS-CLIP | 37986224\|23597149 | | Unspecified |
| mmu-miR-125b-5p | MIMAT0000136 | Amer1 | 72345 | HITS-CLIP | | 37986224 | Unspecified |
| mmu-miR-125b-5p | MIMAT0000136 | Ammecr1 | 56068 | Microarrays//HITS-CLIP | 37986224\|25083871\|22469780 | | Unspecified |
| mmu-miR-125b-5p | MIMAT0000136 | Amn1 | 232566 | HITS-CLIP | | 37986224 | Unspecified |
| mmu-miR-125b-5p | MIMAT0000136 | Amot | 27494 | HITS-CLIP | | 37986224 | Unspecified |
| mmu-miR-125b-5p | MIMAT0000136 | Amotl1 | 75723 | HITS-CLIP | | 37986224 | Unspecified |
| mmu-miR-125b-5p | MIMAT0000136 | Amotl2 | 56332 | HITS-CLIP, Microarrays//HITS-CLIP//Microa | 37986224\|25083871\|22469780 | | Unspecified |
| mmu-miR-125b-5p | MIMAT0000136 | Anapc2 | 99152 | Microarrays | 37986224\|22469780 | | Unspecified |
| mmu-miR-125b-5p | MIMAT0000136 | Ang | 11727 | Microarrays | | 37986224 | Unspecified |
| mmu-miR-125b-5p | MIMAT0000136 | Angptl6 | 70726 | Microarrays | | 37986224 | Unspecified |
| mmu-miR-125b-5p | MIMAT0000136 | Ank | 11732 | HITS-CLIP, Microarrays | | 37986224 | Unspecified |
| mmu-miR-125b-5p | MIMAT0000136 | Ank2 | 109676 | HITS-CLIP | | 25083871 | Unspecified |
| mmu-miR-125b-5p | MIMAT0000136 | Ankfy1 | 11736 | HITS-CLIP | | 37986224 | Unspecified |
| mmu-miR-125b-5p | MIMAT0000136 | Ankhd1 | 108857 | HITS-CLIP | | 37986224 | Unspecified |
| mmu-miR-125b-5p | MIMAT0000136 | Ankib1 | 70797 | HITS-CLIP | | 37986224 | Unspecified |
| mmu-miR-125b-5p | MIMAT0000136 | Ankrd11 | 77087 | Microarrays | | 37986224 | Unspecified |
| mmu-miR-125b-5p | MIMAT0000136 | Ankrd13c | 433667 | HITS-CLIP | 37986224\|25083871\|23597149 | | Unspecified |
| mmu-miR-125b-5p | MIMAT0000136 | Ankrd33b | 67434 | HITS-CLIP | 37986224\|25083871\|23597149\|25753 | | Unspecified |
| mmu-miR-125b-5p | MIMAT0000136 | Ankrd37 | 654824 | Microarrays | | 37986224 | Unspecified |
| mmu-miR-125b-5p | MIMAT0000136 | Ankrd44 | 329154 | HITS-CLIP | | 37986224 | Unspecified |
| mmu-miR-125b-5p | MIMAT0000136 | Ankrd50 | 99696 | HITS-CLIP | 37986224\|25083871\|29167373 | | Unspecified |
| mmu-miR-125b-5p | MIMAT0000136 | Anks1 | 224650 | Microarrays | | 37986224 | Unspecified |
| mmu-miR-125b-5p | MIMAT0000136 | Anks4b | 72074 | HITS-CLIP | | 37986224 | Unspecified |
| mmu-miR-125b-5p | MIMAT0000136 | Ano5 | 233246 | Microarrays | | 37986224 | Unspecified |

| | | | | | | |
|---|---|---|---|---|---|---|
| mmu-miR-125b-5p | MIMAT0000136 | Anp32a | 11737 | HITS-CLIP | 37986224 | Unspecified |
| mmu-miR-125b-5p | MIMAT0000136 | Antxr2 | 71914 | HITS-CLIP | 25083871 | Unspecified |
| mmu-miR-125b-5p | MIMAT0000136 | Anxa5 | 11747 | Microarrays | 37986224 | Unspecified |
| mmu-miR-125b-5p | MIMAT0000136 | Anxa6 | 11749 | Microarrays | 37986224 | Unspecified |
| mmu-miR-125b-5p | MIMAT0000136 | Anxa9 | 71790 | Microarrays | 37986224\|22469780 | Unspecified |
| mmu-miR-125b-5p | MIMAT0000136 | Aoc2 | 237940 | Microarrays | 37986224\|22469780 | Unspecified |
| mmu-miR-125b-5p | MIMAT0000136 | Ap1s1 | 11769 | HITS-CLIP | 37986224 | Unspecified |
| mmu-miR-125b-5p | MIMAT0000136 | Ap2a1 | 11771 | Microarrays | 37986224 | Unspecified |
| mmu-miR-125b-5p | MIMAT0000136 | Ap2b1 | 71770 | HITS-CLIP, Microarrays | 37986224 | Unspecified |
| mmu-miR-125b-5p | MIMAT0000136 | Ap3d1 | 11776 | HITS-CLIP | 31433990\|25083871 | Unspecified |
| mmu-miR-125b-5p | MIMAT0000136 | Ap3m1 | 55946 | HITS-CLIP | 25083871 | Unspecified |
| mmu-miR-125b-5p | MIMAT0000136 | Ap3m2 | 64933 | HITS-CLIP | 37986224\|19536157 | Unspecified |
| mmu-miR-125b-5p | MIMAT0000136 | Ap4s1 | 11782 | Microarrays | 37986224\|22469780 | Unspecified |
| mmu-miR-125b-5p | MIMAT0000136 | Ap5m1 | 74385 | HITS-CLIP, Microarrays//Microarrays | 37986224\|22469780 | Unspecified |
| mmu-miR-125b-5p | MIMAT0000136 | Apba1 | 319924 | HITS-CLIP, Microarrays | 37986224 | Unspecified |
| mmu-miR-125b-5p | MIMAT0000136 | Apbb2 | 11787 | Microarrays | 37986224 | Unspecified |
| mmu-miR-125b-5p | MIMAT0000136 | Apc | 11789 | HITS-CLIP | 37986224\|25083871\|19536157 | Unspecified |
| mmu-miR-125b-5p | MIMAT0000136 | Apmap | 71881 | HITS-CLIP, Microarrays//Microarrays | 37986224\|22469780 | Unspecified |
| mmu-miR-125b-5p | MIMAT0000136 | Apob | 238055 | HITS-CLIP | 37986224\|23597149 | Unspecified |
| mmu-miR-125b-5p | MIMAT0000136 | Apobec1 | 11810 | Microarrays | 37986224 | Unspecified |
| mmu-miR-125b-5p | MIMAT0000136 | Apoc4 | 11425 | HITS-CLIP | 37986224 | Unspecified |
| mmu-miR-125b-5p | MIMAT0000136 | Apold1 | 381823 | Microarrays | 37986224 | Unspecified |
| mmu-miR-125b-5p | MIMAT0000136 | App | 11820 | HITS-CLIP | 37986224 | Unspecified |
| mmu-miR-125b-5p | MIMAT0000136 | Appbp2 | 66884 | HITS-CLIP | 37986224 | Unspecified |
| mmu-miR-125b-5p | MIMAT0000136 | Aqp4 | 11829 | HITS-CLIP | 37986224 | Unspecified |
| mmu-miR-125b-5p | MIMAT0000136 | Arcn1 | 213827 | HITS-CLIP | 37986224 | Unspecified |
| mmu-miR-125b-5p | MIMAT0000136 | Arel1 | 68497 | HITS-CLIP | 25083871 | Unspecified |
| mmu-miR-125b-5p | MIMAT0000136 | Arf6 | 11845 | HITS-CLIP | 37986224 | Unspecified |
| mmu-miR-125b-5p | MIMAT0000136 | Arfgef1 | 211673 | HITS-CLIP | 37986224\|25083871\|23597149\|29167 | Unspecified |
| mmu-miR-125b-5p | MIMAT0000136 | Arhgap10 | 78514 | Microarrays | 37986224 | Unspecified |
| mmu-miR-125b-5p | MIMAT0000136 | Arhgap18 | 73910 | Microarrays | 37986224 | Unspecified |
| mmu-miR-125b-5p | MIMAT0000136 | Arhgap21 | 71435 | HITS-CLIP | 37986224 | Unspecified |
| mmu-miR-125b-5p | MIMAT0000136 | Arhgap25 | 232201 | Microarrays | 37986224\|22469780 | Unspecified |
| mmu-miR-125b-5p | MIMAT0000136 | Arhgap32 | 330914 | HITS-CLIP | 37986224 | Unspecified |
| mmu-miR-125b-5p | MIMAT0000136 | Arhgap39 | 223666 | Microarrays | 37986224 | Unspecified |
| mmu-miR-125b-5p | MIMAT0000136 | Arhgap5 | 11855 | HITS-CLIP | 25083871 | Unspecified |
| mmu-miR-125b-5p | MIMAT0000136 | Arhgef1 | 16801 | Microarrays | 37986224 | Unspecified |
| mmu-miR-125b-5p | MIMAT0000136 | Arhgef10l | 72754 | HITS-CLIP | 37986224 | Unspecified |
| mmu-miR-125b-5p | MIMAT0000136 | Arhgef11 | 213498 | Microarrays | 37986224\|22469780 | Unspecified |

| | | | | | | |
|---|---|---|---|---|---|---|
| mmu-miR-125b-5p | MIMAT0000136 | Arhgef12 | 69632 | HITS-CLIP | 25083871\|23597149 | Unspecified |
| mmu-miR-125b-5p | MIMAT0000136 | Arhgef3 | 71704 | Microarrays | 37986224 | Unspecified |
| mmu-miR-125b-5p | MIMAT0000136 | Arhgef39 | 230098 | Microarrays | 37986224\|22469780 | Unspecified |
| mmu-miR-125b-5p | MIMAT0000136 | Arhgef4 | 226970 | Microarrays | 37986224 | Unspecified |
| mmu-miR-125b-5p | MIMAT0000136 | Arhgef40 | 268739 | HITS-CLIP | 37986224 | Unspecified |
| mmu-miR-125b-5p | MIMAT0000136 | Arid1a | 93760 | HITS-CLIP, Microarrays//HITS-CLIP | 37986224\|25083871 | Unspecified |
| mmu-miR-125b-5p | MIMAT0000136 | Arid2 | 77044 | HITS-CLIP | 37986224\|25083871\|23597149 | Unspecified |
| mmu-miR-125b-5p | MIMAT0000136 | Arid3a | 13496 | Microarray//Microarrays//Other | 16495412\|22469780 | Unspecified |
| mmu-miR-125b-5p | MIMAT0000136 | Arid4b | 94246 | HITS-CLIP | 37986224 | Unspecified |
| mmu-miR-125b-5p | MIMAT0000136 | Arih1 | 23806 | HITS-CLIP | 37986224\|25083871 | Unspecified |
| mmu-miR-125b-5p | MIMAT0000136 | Arl2 | 56327 | Microarrays | 37986224 | Unspecified |
| mmu-miR-125b-5p | MIMAT0000136 | Arl4d | 80981 | Microarrays | 37986224\|22469780 | Unspecified |
| mmu-miR-125b-5p | MIMAT0000136 | Arl5b | 75869 | HITS-CLIP | 37986224 | Unspecified |
| mmu-miR-125b-5p | MIMAT0000136 | Armc6 | 76813 | HITS-CLIP | 37986224 | Unspecified |
| mmu-miR-125b-5p | MIMAT0000136 | Armc8 | 74125 | HITS-CLIP | 25083871 | Unspecified |
| mmu-miR-125b-5p | MIMAT0000136 | Armcx2 | 67416 | HITS-CLIP | 37986224\|19536157 | Unspecified |
| mmu-miR-125b-5p | MIMAT0000136 | Armcx6 | 278097 | Microarrays | 37986224 | Unspecified |
| mmu-miR-125b-5p | MIMAT0000136 | Armt1 | 79624 | Microarrays | 22469780 | Unspecified |
| mmu-miR-125b-5p | MIMAT0000136 | Arpc1b | 11867 | Microarrays | 37986224 | Unspecified |
| mmu-miR-125b-5p | MIMAT0000136 | Arrb1 | 109689 | HITS-CLIP | 37986224 | Unspecified |
| mmu-miR-125b-5p | MIMAT0000136 | Arrb2 | 216869 | Microarrays | 37986224 | Unspecified |
| mmu-miR-125b-5p | MIMAT0000136 | Arrdc3 | 105171 | HITS-CLIP | 25083871\|23597149 | Unspecified |
| mmu-miR-125b-5p | MIMAT0000136 | Arrdc4 | 66412 | HITS-CLIP | 37986224 | Unspecified |
| mmu-miR-125b-5p | MIMAT0000136 | Arsg | 74008 | Microarrays | 37986224 | Unspecified |
| mmu-miR-125b-5p | MIMAT0000136 | Art4 | 109978 | Microarrays | 37986224\|22469780 | Unspecified |
| mmu-miR-125b-5p | MIMAT0000136 | Asah1 | 11886 | Microarrays | 37986224 | Unspecified |
| mmu-miR-125b-5p | MIMAT0000136 | Asah2 | 54447 | Microarrays | 37986224 | Unspecified |
| mmu-miR-125b-5p | MIMAT0000136 | Asb2 | 65256 | HITS-CLIP | 37986224 | Unspecified |
| mmu-miR-125b-5p | MIMAT0000136 | Asic1 | 11419 | Microarrays | 37986224\|22469780 | Unspecified |
| mmu-miR-125b-5p | MIMAT0000136 | Asl | 109900 | HITS-CLIP | 23597149 | Unspecified |
| mmu-miR-125b-5p | MIMAT0000136 | Asna1 | 56495 | HITS-CLIP | 25083871 | Unspecified |
| mmu-miR-125b-5p | MIMAT0000136 | Asph | 65973 | HITS-CLIP | 29167373 | Unspecified |
| mmu-miR-125b-5p | MIMAT0000136 | Aspm | 12316 | HITS-CLIP | 37986224 | Unspecified |
| mmu-miR-125b-5p | MIMAT0000136 | Aspn | 66695 | Microarrays | 37986224\|22469780 | Unspecified |
| mmu-miR-125b-5p | MIMAT0000136 | Asrgl1 | 66514 | HITS-CLIP | 37986224 | Unspecified |
| mmu-miR-125b-5p | MIMAT0000136 | Astn1 | 11899 | HITS-CLIP | 37986224 | Unspecified |
| mmu-miR-125b-5p | MIMAT0000136 | Astn2 | 56079 | Microarrays | 37986224\|22469780 | Unspecified |
| mmu-miR-125b-5p | MIMAT0000136 | Asxl1 | 228790 | HITS-CLIP | 37986224 | Unspecified |
| mmu-miR-125b-5p | MIMAT0000136 | Atad2 | 70472 | Microarrays | 37986224\|22469780 | Unspecified |

| | | | | | | |
|---|---|---|---|---|---|---|
| mmu-miR-125b-5p | MIMAT0000136 | Atad2b | 320817 | HITS-CLIP | 25083871 | Unspecified |
| mmu-miR-125b-5p | MIMAT0000136 | Atf5 | 107503 | HITS-CLIP | 37986224 | Unspecified |
| mmu-miR-125b-5p | MIMAT0000136 | Atf6 | 226641 | HITS-CLIP | 37986224 | Unspecified |
| mmu-miR-125b-5p | MIMAT0000136 | Atf7ip | 54343 | HITS-CLIP | 29167373 | Unspecified |
| mmu-miR-125b-5p | MIMAT0000136 | Atg101 | 68118 | HITS-CLIP | 37986224 | Unspecified |
| mmu-miR-125b-5p | MIMAT0000136 | Atg2b | 76559 | HITS-CLIP | 37986224 | Unspecified |
| mmu-miR-125b-5p | MIMAT0000136 | Atg4c | 242557 | Microarrays | 37986224 | Unspecified |
| mmu-miR-125b-5p | MIMAT0000136 | Atg4d | 235040 | Microarrays | 37986224\|22469780 | Unspecified |
| mmu-miR-125b-5p | MIMAT0000136 | Atg5 | 11793 | Microarrays | 37986224\|22469780 | Unspecified |
| mmu-miR-125b-5p | MIMAT0000136 | Atg9a | 245860 | Microarrays | 37986224 | Unspecified |
| mmu-miR-125b-5p | MIMAT0000136 | Atl2 | 56298 | HITS-CLIP | 37986224 | Unspecified |
| mmu-miR-125b-5p | MIMAT0000136 | Atm | 11920 | HITS-CLIP | 37986224 | Unspecified |
| mmu-miR-125b-5p | MIMAT0000136 | Atn1 | 13498 | Microarrays | 37986224 | Unspecified |
| mmu-miR-125b-5p | MIMAT0000136 | Atoh8 | 71093 | HITS-CLIP | 37986224 | Unspecified |
| mmu-miR-125b-5p | MIMAT0000136 | Atp10a | 11982 | Microarrays | 37986224 | Unspecified |
| mmu-miR-125b-5p | MIMAT0000136 | Atp11a | 50770 | HITS-CLIP | 25083871 | Unspecified |
| mmu-miR-125b-5p | MIMAT0000136 | Atp11b | 76295 | HITS-CLIP | 37986224\|25083871 | Unspecified |
| mmu-miR-125b-5p | MIMAT0000136 | Atp11c | 320940 | Microarrays | 37986224\|22469780 | Unspecified |
| mmu-miR-125b-5p | MIMAT0000136 | Atp1a1 | 11928 | Microarrays//qCLASH | 37986224\|22469780\|31393953 | Unspecified |
| mmu-miR-125b-5p | MIMAT0000136 | Atp1a2 | 98660 | HITS-CLIP | 19536157 | Unspecified |
| mmu-miR-125b-5p | MIMAT0000136 | Atp1b1 | 11931 | HITS-CLIP, Microarrays//HITS-CLIP//Microa | 37986224\|23597149\|22469780\|1953 | Unspecified |
| mmu-miR-125b-5p | MIMAT0000136 | Atp23 | 855754 | Microarrays | 22469780 | Unspecified |
| mmu-miR-125b-5p | MIMAT0000136 | Atp2b1 | 67972 | HITS-CLIP | 25083871\|29167373 | Unspecified |
| mmu-miR-125b-5p | MIMAT0000136 | Atp2b4 | 381290 | Microarrays | 37986224 | Unspecified |
| mmu-miR-125b-5p | MIMAT0000136 | Atp2c1 | 235574 | Microarrays | 37986224\|22469780 | Unspecified |
| mmu-miR-125b-5p | MIMAT0000136 | Atp5b | 11947 | HITS-CLIP | 37986224\|25083871 | Unspecified |
| mmu-miR-125b-5p | MIMAT0000136 | Atp5j2 | 57423 | HITS-CLIP | 25083871\|23597149 | Unspecified |
| mmu-miR-125b-5p | MIMAT0000136 | Atp5s | 27109 | HITS-CLIP | 23597149 | Unspecified |
| mmu-miR-125b-5p | MIMAT0000136 | Atp6v0a2 | 21871 | Microarrays | 37986224\|22469780 | Unspecified |
| mmu-miR-125b-5p | MIMAT0000136 | Atp6v0d1 | 11972 | Microarrays | 37986224 | Unspecified |
| mmu-miR-125b-5p | MIMAT0000136 | Atp6v1b2 | 11966 | HITS-CLIP | 37986224 | Unspecified |
| mmu-miR-125b-5p | MIMAT0000136 | Atp6v1c1 | 66335 | HITS-CLIP | 23597149\|25083871\|25753418 | Unspecified |
| mmu-miR-125b-5p | MIMAT0000136 | Atp7a | 11977 | HITS-CLIP | 37986224 | Unspecified |
| mmu-miR-125b-5p | MIMAT0000136 | Atp7b | 11979 | HITS-CLIP | 37986224 | Unspecified |
| mmu-miR-125b-5p | MIMAT0000136 | Atp8b4 | 241633 | HITS-CLIP | 29167373 | Unspecified |
| mmu-miR-125b-5p | MIMAT0000136 | Atxn1 | 20238 | HITS-CLIP | 37986224\|19536157 | Unspecified |
| mmu-miR-125b-5p | MIMAT0000136 | Atxn10 | 54138 | Microarrays | 37986224 | Unspecified |
| mmu-miR-125b-5p | MIMAT0000136 | Atxn1l | 52335 | HITS-CLIP//qCLASH | 37986224\|25083871\|31393953 | Unspecified |
| mmu-miR-125b-5p | MIMAT0000136 | Atxn2l | 233871 | Microarrays | 37986224 | Unspecified |

| | | | | | | | |
|---|---|---|---|---|---|---|---|
| mmu-miR-125b-5p | MIMAT0000136 | Atxn3 | 110616 | HITS-CLIP | | 37986224 | Unspecified |
| mmu-miR-125b-5p | MIMAT0000136 | Atxn7 | 246103 | HITS-CLIP | 37986224\|25083871 | | Unspecified |
| mmu-miR-125b-5p | MIMAT0000136 | Atxn7l1 | 380753 | HITS-CLIP | | 37986224 | Unspecified |
| mmu-miR-125b-5p | MIMAT0000136 | Atxn7l3 | 217218 | HITS-CLIP, Microarrays | | 37986224 | Unspecified |
| mmu-miR-125b-5p | MIMAT0000136 | Atxn7l3b | 382423 | HITS-CLIP | | 37986224 | Unspecified |
| mmu-miR-125b-5p | MIMAT0000136 | Aup1 | 11993 | HITS-CLIP | 37986224\|25083871 | | Unspecified |
| mmu-miR-125b-5p | MIMAT0000136 | Auts2 | 319974 | HITS-CLIP | | 37986224 | Unspecified |
| mmu-miR-125b-5p | MIMAT0000136 | Avl9 | 78937 | HITS-CLIP | | 37986224 | Unspecified |
| mmu-miR-125b-5p | MIMAT0000136 | Axl | 26362 | HITS-CLIP | | 37986224 | Unspecified |
| mmu-miR-125b-5p | MIMAT0000136 | Aym1 | 503692 | Microarrays | 37986224\|22469780 | | Unspecified |
| mmu-miR-125b-5p | MIMAT0000136 | Azi2 | 27215 | HITS-CLIP | | 37986224 | Unspecified |
| mmu-miR-125b-5p | MIMAT0000136 | B020031M17Rik | 333467 | Microarrays | | 37986224 | Unspecified |
| mmu-miR-125b-5p | MIMAT0000136 | B2m | 12010 | Microarrays | | 37986224 | Unspecified |
| mmu-miR-125b-5p | MIMAT0000136 | B3galnt2 | 97884 | HITS-CLIP | | 37986224 | Unspecified |
| mmu-miR-125b-5p | MIMAT0000136 | B3galt6 | 117592 | Microarrays | 37986224\|22469780 | | Unspecified |
| mmu-miR-125b-5p | MIMAT0000136 | B3gnt5 | 108105 | Microarrays | 37986224\|22469780 | | Unspecified |
| mmu-miR-125b-5p | MIMAT0000136 | B4galnt1 | 14421 | HITS-CLIP | | 37986224 | Unspecified |
| mmu-miR-125b-5p | MIMAT0000136 | B4galt5 | 56336 | HITS-CLIP | | 37986224 | Unspecified |
| mmu-miR-125b-5p | MIMAT0000136 | B4galt6 | 56386 | HITS-CLIP | | 37986224 | Unspecified |
| mmu-miR-125b-5p | MIMAT0000136 | BC005537 | 79555 | HITS-CLIP | | 37986224 | Unspecified |
| mmu-miR-125b-5p | MIMAT0000136 | BC005561 | 100042165 | Microarrays | | 37986224 | Unspecified |
| mmu-miR-125b-5p | MIMAT0000136 | BC005624 | 227707 | Microarrays | | 37986224 | Unspecified |
| mmu-miR-125b-5p | MIMAT0000136 | BC048507 | 408058 | Microarrays | 37986224\|22469780 | | Unspecified |
| mmu-miR-125b-5p | MIMAT0000136 | Bach1 | 12013 | HITS-CLIP | | 25083871 | Unspecified |
| mmu-miR-125b-5p | MIMAT0000136 | Bach2 | 12014 | HITS-CLIP | 37986224\|25753418 | | Unspecified |
| mmu-miR-125b-5p | MIMAT0000136 | Bag2 | 213539 | Microarrays | | 37986224 | Unspecified |
| mmu-miR-125b-5p | MIMAT0000136 | Bag4 | 67384 | HITS-CLIP | 37986224\|25083871 | | Unspecified |
| mmu-miR-125b-5p | MIMAT0000136 | Bahd1 | 228536 | HITS-CLIP | | 37986224 | Unspecified |
| mmu-miR-125b-5p | MIMAT0000136 | Bak1 | 12018 | Luciferase assay//qRT-PCR//Western blot/ | 21368288\|21935352\|25083871\|26024 | | Unspecified |
| mmu-miR-125b-5p | MIMAT0000136 | Banf2 | 403171 | Microarrays | 37986224\|22469780 | | Unspecified |
| mmu-miR-125b-5p | MIMAT0000136 | Bap1 | 104416 | HITS-CLIP | | 37986224 | Unspecified |
| mmu-miR-125b-5p | MIMAT0000136 | Batf2 | 74481 | Microarrays | | 37986224 | Unspecified |
| mmu-miR-125b-5p | MIMAT0000136 | Baz1b | 22385 | HITS-CLIP | | 25083871 | Unspecified |
| mmu-miR-125b-5p | MIMAT0000136 | Baz2a | 116848 | HITS-CLIP | 37986224\|25083871 | | Unspecified |
| mmu-miR-125b-5p | MIMAT0000136 | Bbc3 | 170770 | HITS-CLIP, Microarrays, Luciferase Reporte | 37986224\|21935352 | | Unspecified |
| mmu-miR-125b-5p | MIMAT0000136 | Bbs2 | 67378 | Microarrays | | 37986224 | Unspecified |
| mmu-miR-125b-5p | MIMAT0000136 | Bcar1 | 12927 | HITS-CLIP | | 37986224 | Unspecified |
| mmu-miR-125b-5p | MIMAT0000136 | Bcar3 | 29815 | HITS-CLIP | | 29167373 | Unspecified |
| mmu-miR-125b-5p | MIMAT0000136 | Bcat2 | 12036 | Microarrays | | 37986224 | Unspecified |

| | | | | | | |
|---|---|---|---|---|---|---|
| mmu-miR-125b-5p | MIMAT0000136 | Bcdin3d | 75284 | Microarrays | 37986224\|22469780 | Unspecified |
| mmu-miR-125b-5p | MIMAT0000136 | Bckdha | 12039 | HITS-CLIP | 37986224 | Unspecified |
| mmu-miR-125b-5p | MIMAT0000136 | Bcl2 | 12043 | HITS-CLIP | 25753418 | Unspecified |
| mmu-miR-125b-5p | MIMAT0000136 | Bcl2a1c | 12046 | Microarrays | 37986224 | Unspecified |
| mmu-miR-125b-5p | MIMAT0000136 | Bcl2a1d | 12047 | Microarrays | 37986224 | Unspecified |
| mmu-miR-125b-5p | MIMAT0000136 | Bcl2l1 | 12048 | Microarrays | 37986224 | Unspecified |
| mmu-miR-125b-5p | MIMAT0000136 | Bcl2l12 | 75736 | Microarrays | 37986224\|22469780 | Unspecified |
| mmu-miR-125b-5p | MIMAT0000136 | Bcl2l13 | 94044 | HITS-CLIP, Microarrays//Microarrays//HIT | 37986224\|22469780\|25083871 | Unspecified |
| mmu-miR-125b-5p | MIMAT0000136 | Bcl2l2 | 12050 | HITS-CLIP | 37986224 | Unspecified |
| mmu-miR-125b-5p | MIMAT0000136 | Bcl9 | 77578 | HITS-CLIP | 37986224 | Unspecified |
| mmu-miR-125b-5p | MIMAT0000136 | Bclaf1 | 72567 | HITS-CLIP | 37986224\|25083871 | Unspecified |
| mmu-miR-125b-5p | MIMAT0000136 | Bcr | 110279 | Microarrays | 37986224 | Unspecified |
| mmu-miR-125b-5p | MIMAT0000136 | Bdh2 | 69772 | Microarrays | 37986224 | Unspecified |
| mmu-miR-125b-5p | MIMAT0000136 | Becn1 | 56208 | Microarrays | 37986224 | Unspecified |
| mmu-miR-125b-5p | MIMAT0000136 | Begain | 380785 | Microarrays | 37986224 | Unspecified |
| mmu-miR-125b-5p | MIMAT0000136 | Bfar | 67118 | Microarrays | 22469780 | Unspecified |
| mmu-miR-125b-5p | MIMAT0000136 | Bfsp2 | 107993 | Microarrays | 37986224\|22469780 | Unspecified |
| mmu-miR-125b-5p | MIMAT0000136 | Bhlhe40 | 20893 | Microarrays | 37986224 | Unspecified |
| mmu-miR-125b-5p | MIMAT0000136 | Bhmt2 | 64918 | HITS-CLIP | 37986224 | Unspecified |
| mmu-miR-125b-5p | MIMAT0000136 | Birc5 | 11799 | Microarrays | 37986224\|22469780 | Unspecified |
| mmu-miR-125b-5p | MIMAT0000136 | Birc6 | 12211 | HITS-CLIP | 37986224\|25083871\|29167373 | Unspecified |
| mmu-miR-125b-5p | MIMAT0000136 | Bloc1s2 | 73689 | Microarrays | 37986224\|22469780 | Unspecified |
| mmu-miR-125b-5p | MIMAT0000136 | Bltp2 | 9703 | HITS-CLIP | 25753418 | Unspecified |
| mmu-miR-125b-5p | MIMAT0000136 | Blzf1 | 66352 | HITS-CLIP, Microarrays//Microarrays | 37986224\|22469780 | Unspecified |
| mmu-miR-125b-5p | MIMAT0000136 | Bmp3 | 110075 | Microarrays | 37986224\|22469780 | Unspecified |
| mmu-miR-125b-5p | MIMAT0000136 | Bmpr2 | 12168 | HITS-CLIP | 37986224 | Unspecified |
| mmu-miR-125b-5p | MIMAT0000136 | Bod1l | 665775 | HITS-CLIP | 25083871 | Unspecified |
| mmu-miR-125b-5p | MIMAT0000136 | Bpifa5 | 67135 | Microarrays | 37986224 | Unspecified |
| mmu-miR-125b-5p | MIMAT0000136 | Braf | 109880 | HITS-CLIP, Microarrays | 37986224 | Unspecified |
| mmu-miR-125b-5p | MIMAT0000136 | Brap | 72399 | HITS-CLIP | 23597149 | Unspecified |
| mmu-miR-125b-5p | MIMAT0000136 | Brd1 | 223770 | HITS-CLIP | 37986224 | Unspecified |
| mmu-miR-125b-5p | MIMAT0000136 | Brd2 | 14312 | HITS-CLIP | 37986224 | Unspecified |
| mmu-miR-125b-5p | MIMAT0000136 | Brd7 | 26992 | HITS-CLIP | 25083871 | Unspecified |
| mmu-miR-125b-5p | MIMAT0000136 | Brinp1 | 56710 | HITS-CLIP | 37986224 | Unspecified |
| mmu-miR-125b-5p | MIMAT0000136 | Brinp3 | 215378 | HITS-CLIP | 37986224\|25083871 | Unspecified |
| mmu-miR-125b-5p | MIMAT0000136 | Brix1 | 67832 | HITS-CLIP | 25083871 | Unspecified |
| mmu-miR-125b-5p | MIMAT0000136 | Brms1 | 107392 | Microarrays | 37986224\|22469780 | Unspecified |
| mmu-miR-125b-5p | MIMAT0000136 | Brpf1 | 78783 | HITS-CLIP | 37986224 | Unspecified |
| mmu-miR-125b-5p | MIMAT0000136 | Brwd1 | 93871 | HITS-CLIP | 37986224 | Unspecified |

| | | | | | | |
|---|---|---|---|---|---|---|
| mmu-miR-125b-5p | MIMAT0000136 | Bsn | 12217 | HITS-CLIP | 37986224 | Unspecified |
| mmu-miR-125b-5p | MIMAT0000136 | Bsph1 | 330470 | Microarrays | 37986224\|22469780 | Unspecified |
| mmu-miR-125b-5p | MIMAT0000136 | Bst2 | 69550 | Microarrays | 37986224 | Unspecified |
| mmu-miR-125b-5p | MIMAT0000136 | Btbd2 | 208198 | Microarrays | 37986224 | Unspecified |
| mmu-miR-125b-5p | MIMAT0000136 | Btg2 | 12227 | HITS-CLIP//Microarray//qRT-PCR | 37986224\|25753418\|33330617 | Unspecified |
| mmu-miR-125b-5p | MIMAT0000136 | Btg3 | 12228 | Microarrays | 37986224 | Unspecified |
| mmu-miR-125b-5p | MIMAT0000136 | Bud23 | 114049 | Microarrays | 22469780 | Unspecified |
| mmu-miR-125b-5p | MIMAT0000136 | Bud31 | 231889 | Microarrays | 37986224\|22469780 | Unspecified |
| mmu-miR-125b-5p | MIMAT0000136 | Bzw2 | 66912 | HITS-CLIP | 25083871 | Unspecified |
| mmu-miR-125b-5p | MIMAT0000136 | C1galt1 | 94192 | Microarrays//HITS-CLIP | 37986224\|25083871\|22469780 | Unspecified |
| mmu-miR-125b-5p | MIMAT0000136 | C1galt1c1 | 59048 | Microarrays | 37986224\|22469780 | Unspecified |
| mmu-miR-125b-5p | MIMAT0000136 | C1qa | 12259 | HITS-CLIP | 23597149\|29167373 | Unspecified |
| mmu-miR-125b-5p | MIMAT0000136 | C1qtnf1 | 56745 | HITS-CLIP | 37986224 | Unspecified |
| mmu-miR-125b-5p | MIMAT0000136 | C1qtnf6 | 72709 | HITS-CLIP | 37986224 | Unspecified |
| mmu-miR-125b-5p | MIMAT0000136 | C2cd4c | 237397 | Microarrays | 37986224 | Unspecified |
| mmu-miR-125b-5p | MIMAT0000136 | C3 | 12266 | HITS-CLIP | 23597149 | Unspecified |
| mmu-miR-125b-5p | MIMAT0000136 | C330007P06Rik | 77644 | Microarrays | 37986224 | Unspecified |
| mmu-miR-125b-5p | MIMAT0000136 | C330018D20Rik | 77422 | Microarrays | 37986224\|22469780 | Unspecified |
| mmu-miR-125b-5p | MIMAT0000136 | C4bp | 12269 | Microarrays | 37986224 | Unspecified |
| mmu-miR-125b-5p | MIMAT0000136 | C5ar2 | 319430 | Microarrays | 37986224\|22469780 | Unspecified |
| mmu-miR-125b-5p | MIMAT0000136 | C77080 | 97130 | HITS-CLIP | 37986224 | Unspecified |
| mmu-miR-125b-5p | MIMAT0000136 | Cabp7 | 192650 | HITS-CLIP | 37986224 | Unspecified |
| mmu-miR-125b-5p | MIMAT0000136 | Cacna1b | 12287 | HITS-CLIP | 19536157 | Unspecified |
| mmu-miR-125b-5p | MIMAT0000136 | Cacna1c | 12288 | HITS-CLIP | 37986224 | Unspecified |
| mmu-miR-125b-5p | MIMAT0000136 | Cacna2d1 | 12293 | HITS-CLIP | 25083871 | Unspecified |
| mmu-miR-125b-5p | MIMAT0000136 | Cacnb1 | 12295 | HITS-CLIP | 37986224 | Unspecified |
| mmu-miR-125b-5p | MIMAT0000136 | Cacul1 | 78832 | Microarrays | 37986224\|22469780 | Unspecified |
| mmu-miR-125b-5p | MIMAT0000136 | Cadm1 | 54725 | HITS-CLIP | 37986224\|25083871 | Unspecified |
| mmu-miR-125b-5p | MIMAT0000136 | Cadm4 | 260299 | HITS-CLIP | 37986224 | Unspecified |
| mmu-miR-125b-5p | MIMAT0000136 | Cald1 | 109624 | HITS-CLIP | 25083871 | Unspecified |
| mmu-miR-125b-5p | MIMAT0000136 | Calm1 | 12313 | HITS-CLIP | 37986224 | Unspecified |
| mmu-miR-125b-5p | MIMAT0000136 | Caln1 | 140904 | HITS-CLIP | 37986224 | Unspecified |
| mmu-miR-125b-5p | MIMAT0000136 | Calu | 12321 | HITS-CLIP | 25083871 | Unspecified |
| mmu-miR-125b-5p | MIMAT0000136 | Caly | 68566 | Microarrays | 37986224\|22469780 | Unspecified |
| mmu-miR-125b-5p | MIMAT0000136 | Camk1g | 215303 | Microarrays | 37986224 | Unspecified |
| mmu-miR-125b-5p | MIMAT0000136 | Camk2a | 12322 | HITS-CLIP | 37986224 | Unspecified |
| mmu-miR-125b-5p | MIMAT0000136 | Camk4 | 12326 | HITS-CLIP | 37986224 | Unspecified |
| mmu-miR-125b-5p | MIMAT0000136 | Cand1 | 71902 | HITS-CLIP | 37986224 | Unspecified |
| mmu-miR-125b-5p | MIMAT0000136 | Cant1 | 76025 | HITS-CLIP | 25083871 | Unspecified |

| | | | | | | |
|---|---|---|---|---|---|---|
| mmu-miR-125b-5p | MIMAT0000136 | Cap1 | 12331 | Microarrays | 37986224 | Unspecified |
| mmu-miR-125b-5p | MIMAT0000136 | Capg | 12332 | Microarrays | 37986224 | Unspecified |
| mmu-miR-125b-5p | MIMAT0000136 | Capn11 | 268958 | Microarrays | 37986224\|22469780 | Unspecified |
| mmu-miR-125b-5p | MIMAT0000136 | Capn2 | 12334 | HITS-CLIP | 37986224\|25083871 | Unspecified |
| mmu-miR-125b-5p | MIMAT0000136 | Capn5 | 12337 | Microarrays | 37986224 | Unspecified |
| mmu-miR-125b-5p | MIMAT0000136 | Capns1 | 12336 | Microarrays | 37986224 | Unspecified |
| mmu-miR-125b-5p | MIMAT0000136 | Caprin1 | 53872 | HITS-CLIP | 25083871\|19536157 | Unspecified |
| mmu-miR-125b-5p | MIMAT0000136 | Capza2 | 12343 | HITS-CLIP | 23597149\|25083871\|25753418 | Unspecified |
| mmu-miR-125b-5p | MIMAT0000136 | Car12 | 76459 | HITS-CLIP | 37986224 | Unspecified |
| mmu-miR-125b-5p | MIMAT0000136 | Car5a | 12352 | Microarrays | 37986224 | Unspecified |
| mmu-miR-125b-5p | MIMAT0000136 | Card11 | 108723 | Microarrays | 37986224 | Unspecified |
| mmu-miR-125b-5p | MIMAT0000136 | Carm1 | 59035 | HITS-CLIP, Microarrays | 37986224 | Unspecified |
| mmu-miR-125b-5p | MIMAT0000136 | Carmil1 | 55604 | HITS-CLIP | 19536157\|29167373 | Unspecified |
| mmu-miR-125b-5p | MIMAT0000136 | Casp2 | 12366 | HITS-CLIP | 37986224\|25083871 | Unspecified |
| mmu-miR-125b-5p | MIMAT0000136 | Casp3 | 12367 | HITS-CLIP | 37986224 | Unspecified |
| mmu-miR-125b-5p | MIMAT0000136 | Casp7 | 12369 | Microarrays | 37986224\|22469780 | Unspecified |
| mmu-miR-125b-5p | MIMAT0000136 | Casp8 | 12370 | Microarrays | 37986224 | Unspecified |
| mmu-miR-125b-5p | MIMAT0000136 | Casp9 | 12371 | Microarrays | 37986224\|22469780 | Unspecified |
| mmu-miR-125b-5p | MIMAT0000136 | Cbll1 | 104836 | HITS-CLIP | 37986224\|25083871 | Unspecified |
| mmu-miR-125b-5p | MIMAT0000136 | Cbr1 | 12408 | HITS-CLIP | 37986224 | Unspecified |
| mmu-miR-125b-5p | MIMAT0000136 | Cbr4 | 234309 | HITS-CLIP | 37986224 | Unspecified |
| mmu-miR-125b-5p | MIMAT0000136 | Cbx2 | 12416 | HITS-CLIP | 37986224 | Unspecified |
| mmu-miR-125b-5p | MIMAT0000136 | Cbx5 | 12419 | HITS-CLIP | 37986224 | Unspecified |
| mmu-miR-125b-5p | MIMAT0000136 | Cbx6 | 494448 | HITS-CLIP | 37986224 | Unspecified |
| mmu-miR-125b-5p | MIMAT0000136 | Cbx7 | 52609 | HITS-CLIP | 37986224 | Unspecified |
| mmu-miR-125b-5p | MIMAT0000136 | Cc2d1a | 212139 | Microarrays | 37986224 | Unspecified |
| mmu-miR-125b-5p | MIMAT0000136 | Ccdc102a | 234582 | HITS-CLIP, Microarrays | 37986224 | Unspecified |
| mmu-miR-125b-5p | MIMAT0000136 | Ccdc115 | 69668 | Microarrays | 37986224\|22469780 | Unspecified |
| mmu-miR-125b-5p | MIMAT0000136 | Ccdc117 | 104479 | HITS-CLIP | 37986224 | Unspecified |
| mmu-miR-125b-5p | MIMAT0000136 | Ccdc125 | 76041 | Microarrays | 37986224\|22469780 | Unspecified |
| mmu-miR-125b-5p | MIMAT0000136 | Ccdc126 | 57895 | HITS-CLIP | 37986224 | Unspecified |
| mmu-miR-125b-5p | MIMAT0000136 | Ccdc13 | 100502861 | Microarrays | 37986224\|22469780 | Unspecified |
| mmu-miR-125b-5p | MIMAT0000136 | Ccdc130 | 67736 | Microarrays | 37986224 | Unspecified |
| mmu-miR-125b-5p | MIMAT0000136 | Ccdc137 | 67291 | HITS-CLIP | 37986224 | Unspecified |
| mmu-miR-125b-5p | MIMAT0000136 | Ccdc141 | 545428 | HITS-CLIP | 25753418 | Unspecified |
| mmu-miR-125b-5p | MIMAT0000136 | Ccdc155 | 384619 | Microarrays | 37986224 | Unspecified |
| mmu-miR-125b-5p | MIMAT0000136 | Ccdc174 | 232236 | Microarrays | 37986224\|22469780 | Unspecified |
| mmu-miR-125b-5p | MIMAT0000136 | Ccdc18 | 73254 | Microarrays | 37986224 | Unspecified |
| mmu-miR-125b-5p | MIMAT0000136 | Ccdc71l | 72123 | HITS-CLIP, Microarrays//HITS-CLIP//Microa | 37986224\|25083871\|22469780\|29167 | Unspecified |

| | | | | | | |
|---|---|---|---|---|---|---|
| mmu-miR-125b-5p | MIMAT0000136 | Ccdc80 | 67896 | HITS-CLIP | 37986224 | Unspecified |
| mmu-miR-125b-5p | MIMAT0000136 | Ccdc82 | 66396 | HITS-CLIP | 37986224 | Unspecified |
| mmu-miR-125b-5p | MIMAT0000136 | Ccdc90b | 66365 | HITS-CLIP | 37986224 | Unspecified |
| mmu-miR-125b-5p | MIMAT0000136 | Ccl2 | 20296 | Microarrays | 37986224 | Unspecified |
| mmu-miR-125b-5p | MIMAT0000136 | Ccl3 | 20302 | Microarrays | 37986224 | Unspecified |
| mmu-miR-125b-5p | MIMAT0000136 | Ccl5 | 20304 | Microarrays | 37986224 | Unspecified |
| mmu-miR-125b-5p | MIMAT0000136 | Ccl9 | 20308 | Microarrays | 37986224 | Unspecified |
| mmu-miR-125b-5p | MIMAT0000136 | Ccm2 | 216527 | HITS-CLIP | 37986224 | Unspecified |
| mmu-miR-125b-5p | MIMAT0000136 | Ccna2 | 12428 | HITS-CLIP | 37986224 | Unspecified |
| mmu-miR-125b-5p | MIMAT0000136 | Ccnc | 51813 | HITS-CLIP, Microarrays//Microarrays | 37986224\|22469780 | Unspecified |
| mmu-miR-125b-5p | MIMAT0000136 | Ccnd1 | 12443 | Microarrays | 37986224 | Unspecified |
| mmu-miR-125b-5p | MIMAT0000136 | Ccnd2 | 12444 | Microarrays | 37986224 | Unspecified |
| mmu-miR-125b-5p | MIMAT0000136 | Ccnd3 | 12445 | HITS-CLIP | 37986224 | Unspecified |
| mmu-miR-125b-5p | MIMAT0000136 | Ccne1 | 12447 | HITS-CLIP | 23597149 | Unspecified |
| mmu-miR-125b-5p | MIMAT0000136 | Ccng1 | 12450 | HITS-CLIP | 37986224 | Unspecified |
| mmu-miR-125b-5p | MIMAT0000136 | Ccnj | 240665 | HITS-CLIP | 37986224 | Unspecified |
| mmu-miR-125b-5p | MIMAT0000136 | Ccnjl | 380694 | HITS-CLIP | 37986224\|19536157 | Unspecified |
| mmu-miR-125b-5p | MIMAT0000136 | Ccnq | 92002 | Microarrays | 22469780 | Unspecified |
| mmu-miR-125b-5p | MIMAT0000136 | Ccnt1 | 12455 | Microarrays//HITS-CLIP | 37986224\|25083871\|29167373 | Unspecified |
| mmu-miR-125b-5p | MIMAT0000136 | Ccpg1 | 72278 | HITS-CLIP | 25083871\|19536157 | Unspecified |
| mmu-miR-125b-5p | MIMAT0000136 | Ccr2 | 12772 | Microarrays | 37986224 | Unspecified |
| mmu-miR-125b-5p | MIMAT0000136 | Ccr5 | 12774 | Microarrays | 37986224 | Unspecified |
| mmu-miR-125b-5p | MIMAT0000136 | Ccr7 | 12775 | Microarrays | 37986224\|22469780 | Unspecified |
| mmu-miR-125b-5p | MIMAT0000136 | Ccrl2 | 54199 | Microarrays | 37986224 | Unspecified |
| mmu-miR-125b-5p | MIMAT0000136 | Cct4 | 12464 | HITS-CLIP | 25083871 | Unspecified |
| mmu-miR-125b-5p | MIMAT0000136 | Cct6a | 12466 | qCLASH | 31393953 | Unspecified |
| mmu-miR-125b-5p | MIMAT0000136 | Cd200 | 17470 | Microarrays | 37986224 | Unspecified |
| mmu-miR-125b-5p | MIMAT0000136 | Cd24a | 12484 | Microarrays | 37986224\|22469780 | Unspecified |
| mmu-miR-125b-5p | MIMAT0000136 | Cd274 | 60533 | Microarrays | 37986224 | Unspecified |
| mmu-miR-125b-5p | MIMAT0000136 | Cd276 | 102657 | HITS-CLIP | 37986224 | Unspecified |
| mmu-miR-125b-5p | MIMAT0000136 | Cd28 | 12487 | Microarrays | 37986224 | Unspecified |
| mmu-miR-125b-5p | MIMAT0000136 | Cd33 | 12489 | Microarrays | 37986224 | Unspecified |
| mmu-miR-125b-5p | MIMAT0000136 | Cd37 | 12493 | Microarrays | 37986224\|22469780 | Unspecified |
| mmu-miR-125b-5p | MIMAT0000136 | Cd3d | 12500 | Microarrays | 37986224\|22469780 | Unspecified |
| mmu-miR-125b-5p | MIMAT0000136 | Cd3e | 12501 | Microarrays | 37986224\|22469780 | Unspecified |
| mmu-miR-125b-5p | MIMAT0000136 | Cd69 | 12515 | Microarrays | 37986224 | Unspecified |
| mmu-miR-125b-5p | MIMAT0000136 | Cd79a | 12518 | Microarrays | 37986224 | Unspecified |
| mmu-miR-125b-5p | MIMAT0000136 | Cdc25b | 12531 | HITS-CLIP | 25083871\|29167373 | Unspecified |
| mmu-miR-125b-5p | MIMAT0000136 | Cdc37 | 12539 | Microarrays | 37986224\|22469780 | Unspecified |

| | | | | | | |
|---|---|---|---|---|---|---|
| mmu-miR-125b-5p | MIMAT0000136 | Cdc37l1 | 67072 | HITS-CLIP | 37986224\|25083871\|23597149 | Unspecified |
| mmu-miR-125b-5p | MIMAT0000136 | Cdc42 | 12540 | HITS-CLIP | 37986224 | Unspecified |
| mmu-miR-125b-5p | MIMAT0000136 | Cdc42ep3 | 260409 | Microarrays | 37986224\|22469780 | Unspecified |
| mmu-miR-125b-5p | MIMAT0000136 | Cdc42ep4 | 56699 | HITS-CLIP, Microarrays | 37986224 | Unspecified |
| mmu-miR-125b-5p | MIMAT0000136 | Cdc42se1 | 57912 | HITS-CLIP | 37986224\|23597149 | Unspecified |
| mmu-miR-125b-5p | MIMAT0000136 | Cdc6 | 23834 | Microarrays | 37986224\|22469780 | Unspecified |
| mmu-miR-125b-5p | MIMAT0000136 | Cdca7l | 217946 | HITS-CLIP | 37986224 | Unspecified |
| mmu-miR-125b-5p | MIMAT0000136 | Cdh13 | 12554 | HITS-CLIP | 25083871 | Unspecified |
| mmu-miR-125b-5p | MIMAT0000136 | Cdh5 | 12562 | Microarrays | 37986224 | Unspecified |
| mmu-miR-125b-5p | MIMAT0000136 | Cdip1 | 66626 | HITS-CLIP | 37986224 | Unspecified |
| mmu-miR-125b-5p | MIMAT0000136 | Cdk12 | 69131 | HITS-CLIP | 37986224\|25083871 | Unspecified |
| mmu-miR-125b-5p | MIMAT0000136 | Cdk13 | 69562 | HITS-CLIP | 37986224 | Unspecified |
| mmu-miR-125b-5p | MIMAT0000136 | Cdk16 | 18555 | HITS-CLIP, Microarrays | 37986224 | Unspecified |
| mmu-miR-125b-5p | MIMAT0000136 | Cdk17 | 237459 | HITS-CLIP | 25083871 | Unspecified |
| mmu-miR-125b-5p | MIMAT0000136 | Cdk18 | 18557 | HITS-CLIP | 37986224 | Unspecified |
| mmu-miR-125b-5p | MIMAT0000136 | Cdk19 | 78334 | HITS-CLIP | 37986224\|25083871 | Unspecified |
| mmu-miR-125b-5p | MIMAT0000136 | Cdk7 | 12572 | HITS-CLIP | 25083871 | Unspecified |
| mmu-miR-125b-5p | MIMAT0000136 | Cdk8 | 264064 | HITS-CLIP | 25083871 | Unspecified |
| mmu-miR-125b-5p | MIMAT0000136 | Cdkl1 | 71091 | Microarrays | 37986224\|22469780 | Unspecified |
| mmu-miR-125b-5p | MIMAT0000136 | Cdkn1a | 12575 | Microarrays | 37986224 | Unspecified |
| mmu-miR-125b-5p | MIMAT0000136 | Cdkn2a | 12578 | Microarrays | 37986224 | Unspecified |
| mmu-miR-125b-5p | MIMAT0000136 | Cdkn2aip | 70925 | HITS-CLIP, Microarrays//Microarrays | 37986224\|22469780 | Unspecified |
| mmu-miR-125b-5p | MIMAT0000136 | Cdkn2b | 12579 | Microarrays | 37986224 | Unspecified |
| mmu-miR-125b-5p | MIMAT0000136 | Cdon | 57810 | HITS-CLIP | 25083871 | Unspecified |
| mmu-miR-125b-5p | MIMAT0000136 | Cdr2l | 237988 | HITS-CLIP | 37986224\|25083871 | Unspecified |
| mmu-miR-125b-5p | MIMAT0000136 | Cdyl | 12593 | Microarrays | 37986224 | Unspecified |
| mmu-miR-125b-5p | MIMAT0000136 | Ceacam16 | 330483 | Microarrays | 37986224 | Unspecified |
| mmu-miR-125b-5p | MIMAT0000136 | Celf2 | 14007 | HITS-CLIP | 37986224 | Unspecified |
| mmu-miR-125b-5p | MIMAT0000136 | Celsr2 | 53883 | HITS-CLIP | 37986224 | Unspecified |
| mmu-miR-125b-5p | MIMAT0000136 | Cemip2 | 23670 | HITS-CLIP | 23597149 | Unspecified |
| mmu-miR-125b-5p | MIMAT0000136 | Cenpe | 229841 | HITS-CLIP | 37986224 | Unspecified |
| mmu-miR-125b-5p | MIMAT0000136 | Cenpf | 108000 | HITS-CLIP | 37986224 | Unspecified |
| mmu-miR-125b-5p | MIMAT0000136 | Cenpj | 219103 | HITS-CLIP | 37986224 | Unspecified |
| mmu-miR-125b-5p | MIMAT0000136 | Cenps | 378708 | Microarrays | 22469780 | Unspecified |
| mmu-miR-125b-5p | MIMAT0000136 | Cenpv | 73139 | Microarrays | 37986224 | Unspecified |
| mmu-miR-125b-5p | MIMAT0000136 | Cep162 | 382090 | Microarrays | 37986224 | Unspecified |
| mmu-miR-125b-5p | MIMAT0000136 | Cep170 | 545389 | Microarrays//HITS-CLIP | 37986224\|19536157 | Unspecified |
| mmu-miR-125b-5p | MIMAT0000136 | Cep19 | 66994 | Microarrays | 37986224\|22469780 | Unspecified |
| mmu-miR-125b-5p | MIMAT0000136 | Cep350 | 74081 | HITS-CLIP | 37986224 | Unspecified |

| | | | | | | | |
|---|---|---|---|---|---|---|---|
| mmu-miR-125b-5p | MIMAT0000136 | Cep68 | 216543 | HITS-CLIP | | 37986224 | Unspecified |
| mmu-miR-125b-5p | MIMAT0000136 | Cep83 | 77048 | Microarrays | | 37986224 | Unspecified |
| mmu-miR-125b-5p | MIMAT0000136 | Cep85 | 70012 | HITS-CLIP | | 23597149 | Unspecified |
| mmu-miR-125b-5p | MIMAT0000136 | Cep97 | 74201 | HITS-CLIP | | 37986224 | Unspecified |
| mmu-miR-125b-5p | MIMAT0000136 | Cers2 | 76893 | Microarrays | | 37986224 | Unspecified |
| mmu-miR-125b-5p | MIMAT0000136 | Ces1d | 104158 | HITS-CLIP | | 37986224 | Unspecified |
| mmu-miR-125b-5p | MIMAT0000136 | Ces1e | 13897 | Microarrays | | 37986224 | Unspecified |
| mmu-miR-125b-5p | MIMAT0000136 | Ces2a | 102022 | HITS-CLIP | | 37986224 | Unspecified |
| mmu-miR-125b-5p | MIMAT0000136 | Cetn3 | 12626 | Microarrays | 37986224\|22469780 | | Unspecified |
| mmu-miR-125b-5p | MIMAT0000136 | Cfap300 | 85016 | Microarrays | | 22469780 | Unspecified |
| mmu-miR-125b-5p | MIMAT0000136 | Cfap95 | 138255 | Microarrays | | 22469780 | Unspecified |
| mmu-miR-125b-5p | MIMAT0000136 | Cfl2 | 12632 | Microarrays | | 37986224 | Unspecified |
| mmu-miR-125b-5p | MIMAT0000136 | Cflar | 12633 | HITS-CLIP | 37986224\|25083871 | | Unspecified |
| mmu-miR-125b-5p | MIMAT0000136 | Cgnl1 | 68178 | HITS-CLIP | | 29167373 | Unspecified |
| mmu-miR-125b-5p | MIMAT0000136 | Cgref1 | 68567 | HITS-CLIP | | 37986224 | Unspecified |
| mmu-miR-125b-5p | MIMAT0000136 | Chchd10 | 103172 | HITS-CLIP | | 37986224 | Unspecified |
| mmu-miR-125b-5p | MIMAT0000136 | Chchd4 | 72170 | Microarrays | 37986224\|22469780 | | Unspecified |
| mmu-miR-125b-5p | MIMAT0000136 | Chd1 | 12648 | HITS-CLIP | 37986224\|25083871 | | Unspecified |
| mmu-miR-125b-5p | MIMAT0000136 | Chd2 | 244059 | HITS-CLIP | | 25083871 | Unspecified |
| mmu-miR-125b-5p | MIMAT0000136 | Chd3 | 216848 | Microarrays//HITS-CLIP | 37986224\|25083871\|29167373 | | Unspecified |
| mmu-miR-125b-5p | MIMAT0000136 | Chd5 | 269610 | Microarrays | | 37986224 | Unspecified |
| mmu-miR-125b-5p | MIMAT0000136 | Chd6 | 71389 | HITS-CLIP | | 37986224 | Unspecified |
| mmu-miR-125b-5p | MIMAT0000136 | Chd7 | 320790 | HITS-CLIP | 37986224\|25083871\|19536157\|2575[illegible] | | Unspecified |
| mmu-miR-125b-5p | MIMAT0000136 | Chd8 | 67772 | HITS-CLIP | | 25083871 | Unspecified |
| mmu-miR-125b-5p | MIMAT0000136 | Chd9 | 109151 | HITS-CLIP | 37986224\|25083871\|19536157 | | Unspecified |
| mmu-miR-125b-5p | MIMAT0000136 | Chdh | 218865 | HITS-CLIP | 37986224\|23597149 | | Unspecified |
| mmu-miR-125b-5p | MIMAT0000136 | Chgb | 12653 | HITS-CLIP | | 37986224 | Unspecified |
| mmu-miR-125b-5p | MIMAT0000136 | Chic1 | 12212 | HITS-CLIP | | 37986224 | Unspecified |
| mmu-miR-125b-5p | MIMAT0000136 | Chpf2 | 100910 | qCLASH | | 31393953 | Unspecified |
| mmu-miR-125b-5p | MIMAT0000136 | Chrnb2 | 11444 | Microarrays | | 37986224 | Unspecified |
| mmu-miR-125b-5p | MIMAT0000136 | Chrnd | 11447 | Microarrays | 37986224\|22469780 | | Unspecified |
| mmu-miR-125b-5p | MIMAT0000136 | Chst10 | 98388 | Microarrays | | 37986224 | Unspecified |
| mmu-miR-125b-5p | MIMAT0000136 | Chst15 | 77590 | Microarrays//HITS-CLIP | 37986224\|19536157 | | Unspecified |
| mmu-miR-125b-5p | MIMAT0000136 | Chst3 | 53374 | HITS-CLIP | 37986224\|25083871 | | Unspecified |
| mmu-miR-125b-5p | MIMAT0000136 | Chsy1 | 269941 | HITS-CLIP | | 37986224 | Unspecified |
| mmu-miR-125b-5p | MIMAT0000136 | Chsy3 | 78923 | HITS-CLIP | | 23597149 | Unspecified |
| mmu-miR-125b-5p | MIMAT0000136 | Chtf8 | 214987 | HITS-CLIP, Microarrays//Microarrays | 37986224\|22469780 | | Unspecified |
| mmu-miR-125b-5p | MIMAT0000136 | Cisd1 | 52637 | Microarrays | 37986224\|22469780 | | Unspecified |
| mmu-miR-125b-5p | MIMAT0000136 | Cited2 | 17684 | Microarrays | | 37986224 | Unspecified |

| | | | | | | |
|---|---|---|---|---|---|---|
| mmu-miR-125b-5p | MIMAT0000136 | Ckap2l | 70466 | HITS-CLIP | 37986224 | Unspecified |
| mmu-miR-125b-5p | MIMAT0000136 | Ckap5 | 75786 | HITS-CLIP | 25753418 | Unspecified |
| mmu-miR-125b-5p | MIMAT0000136 | Cks1b | 54124 | HITS-CLIP | 37986224 | Unspecified |
| mmu-miR-125b-5p | MIMAT0000136 | Clasp2 | 76499 | HITS-CLIP | 37986224\|25083871 | Unspecified |
| mmu-miR-125b-5p | MIMAT0000136 | Clcc1 | 229725 | HITS-CLIP | 37986224 | Unspecified |
| mmu-miR-125b-5p | MIMAT0000136 | Clcf1 | 56708 | Microarrays | 37986224 | Unspecified |
| mmu-miR-125b-5p | MIMAT0000136 | Clcn2 | 12724 | Microarrays | 37986224 | Unspecified |
| mmu-miR-125b-5p | MIMAT0000136 | Clcn3 | 12725 | HITS-CLIP | 25753418 | Unspecified |
| mmu-miR-125b-5p | MIMAT0000136 | Clcn4 | 12727 | Microarrays | 22469780 | Unspecified |
| mmu-miR-125b-5p | MIMAT0000136 | Cldn10 | 58187 | Microarrays | 37986224 | Unspecified |
| mmu-miR-125b-5p | MIMAT0000136 | Cldn18 | 56492 | Microarrays | 37986224\|22469780 | Unspecified |
| mmu-miR-125b-5p | MIMAT0000136 | Cldn34c1 | 73061 | Microarrays | 22469780 | Unspecified |
| mmu-miR-125b-5p | MIMAT0000136 | Clec1b | 56760 | Microarrays | 37986224 | Unspecified |
| mmu-miR-125b-5p | MIMAT0000136 | Clec4a4 | 474145 | Microarrays | 37986224\|22469780 | Unspecified |
| mmu-miR-125b-5p | MIMAT0000136 | Clec7a | 56644 | Microarrays | 37986224 | Unspecified |
| mmu-miR-125b-5p | MIMAT0000136 | Clic5 | 224796 | Microarrays | 37986224 | Unspecified |
| mmu-miR-125b-5p | MIMAT0000136 | Clip1 | 56430 | Microarrays | 37986224 | Unspecified |
| mmu-miR-125b-5p | MIMAT0000136 | Clip2 | 269713 | Microarrays | 37986224 | Unspecified |
| mmu-miR-125b-5p | MIMAT0000136 | Clip4 | 78785 | HITS-CLIP | 25753418 | Unspecified |
| mmu-miR-125b-5p | MIMAT0000136 | Clk1 | 12747 | HITS-CLIP | 37986224 | Unspecified |
| mmu-miR-125b-5p | MIMAT0000136 | Clk4 | 12750 | HITS-CLIP | 37986224 | Unspecified |
| mmu-miR-125b-5p | MIMAT0000136 | Clmp | 71566 | HITS-CLIP | 37986224 | Unspecified |
| mmu-miR-125b-5p | MIMAT0000136 | Cln6 | 76524 | HITS-CLIP | 37986224 | Unspecified |
| mmu-miR-125b-5p | MIMAT0000136 | Clock | 12753 | HITS-CLIP | 37986224 | Unspecified |
| mmu-miR-125b-5p | MIMAT0000136 | Clpb | 20480 | Microarrays | 37986224 | Unspecified |
| mmu-miR-125b-5p | MIMAT0000136 | Clpsl2 | 328788 | Microarrays | 37986224\|22469780 | Unspecified |
| mmu-miR-125b-5p | MIMAT0000136 | Clstn2 | 64085 | Microarrays | 37986224\|22469780 | Unspecified |
| mmu-miR-125b-5p | MIMAT0000136 | Cltc | 67300 | HITS-CLIP | 25083871 | Unspecified |
| mmu-miR-125b-5p | MIMAT0000136 | Clu | 12759 | HITS-CLIP | 23597149 | Unspecified |
| mmu-miR-125b-5p | MIMAT0000136 | Cluh | 74148 | HITS-CLIP | 23597149\|26203562 | Unspecified |
| mmu-miR-125b-5p | MIMAT0000136 | Cma2 | 545055 | Microarrays | 37986224\|22469780 | Unspecified |
| mmu-miR-125b-5p | MIMAT0000136 | Cmpk2 | 22169 | Microarrays | 37986224 | Unspecified |
| mmu-miR-125b-5p | MIMAT0000136 | Cmtm7 | 102545 | Microarrays | 37986224\|22469780 | Unspecified |
| mmu-miR-125b-5p | MIMAT0000136 | Cmtr2 | 234728 | Microarrays | 37986224\|22469780 | Unspecified |
| mmu-miR-125b-5p | MIMAT0000136 | Cmya5 | 76469 | Microarrays | 37986224 | Unspecified |
| mmu-miR-125b-5p | MIMAT0000136 | Cndp1 | 338403 | Microarrays | 37986224 | Unspecified |
| mmu-miR-125b-5p | MIMAT0000136 | Cnga1 | 12788 | Microarrays | 37986224\|22469780 | Unspecified |
| mmu-miR-125b-5p | MIMAT0000136 | Cnih2 | 12794 | HITS-CLIP | 37986224 | Unspecified |
| mmu-miR-125b-5p | MIMAT0000136 | Cnksr2 | 245684 | HITS-CLIP | 37986224 | Unspecified |

| | | | | | | | |
|---|---|---|---|---|---|---|---|
| mmu-miR-125b-5p | MIMAT0000136 | Cnn2 | 12798 | Microarrays | | 37986224 | Unspecified |
| mmu-miR-125b-5p | MIMAT0000136 | Cnot1 | 234594 | HITS-CLIP | 37986224\|25083871\|29167373 | | Unspecified |
| mmu-miR-125b-5p | MIMAT0000136 | Cnot2 | 72068 | HITS-CLIP | 37986224\|25083871\|23597149 | | Unspecified |
| mmu-miR-125b-5p | MIMAT0000136 | Cnot3 | 232791 | Microarrays | | 37986224 | Unspecified |
| mmu-miR-125b-5p | MIMAT0000136 | Cnot6 | 104625 | HITS-CLIP, Microarrays//Microarrays | 37986224\|22469780 | | Unspecified |
| mmu-miR-125b-5p | MIMAT0000136 | Cnot6l | 231464 | HITS-CLIP | | 29167373 | Unspecified |
| mmu-miR-125b-5p | MIMAT0000136 | Cnot7 | 18983 | HITS-CLIP | | 37986224 | Unspecified |
| mmu-miR-125b-5p | MIMAT0000136 | Cntnap5a | 636808 | HITS-CLIP | | 37986224 | Unspecified |
| mmu-miR-125b-5p | MIMAT0000136 | Cntrob | 216846 | Microarrays | | 37986224 | Unspecified |
| mmu-miR-125b-5p | MIMAT0000136 | Coa6 | 67892 | Microarrays | 37986224\|22469780 | | Unspecified |
| mmu-miR-125b-5p | MIMAT0000136 | Cog1 | 16834 | HITS-CLIP//qCLASH | 37986224\|31393953\|25083871 | | Unspecified |
| mmu-miR-125b-5p | MIMAT0000136 | Col12a1 | 12816 | HITS-CLIP | | 25083871 | Unspecified |
| mmu-miR-125b-5p | MIMAT0000136 | Col18a1 | 12822 | HITS-CLIP | | 37986224 | Unspecified |
| mmu-miR-125b-5p | MIMAT0000136 | Col1a1 | 12842 | qCLASH//HITS-CLIP | 31393953\|25083871 | | Unspecified |
| mmu-miR-125b-5p | MIMAT0000136 | Col4a1 | 12826 | HITS-CLIP | | 37986224 | Unspecified |
| mmu-miR-125b-5p | MIMAT0000136 | Col5a1 | 12831 | HITS-CLIP | 25083871\|23597149 | | Unspecified |
| mmu-miR-125b-5p | MIMAT0000136 | Col5a2 | 12832 | HITS-CLIP | | 25083871 | Unspecified |
| mmu-miR-125b-5p | MIMAT0000136 | Col5a3 | 53867 | Microarrays | | 37986224 | Unspecified |
| mmu-miR-125b-5p | MIMAT0000136 | Commd2 | 52245 | Microarrays | 37986224\|22469780 | | Unspecified |
| mmu-miR-125b-5p | MIMAT0000136 | Commd5 | 66398 | Microarrays | 37986224\|22469780 | | Unspecified |
| mmu-miR-125b-5p | MIMAT0000136 | Copg1 | 54161 | HITS-CLIP | | 37986224 | Unspecified |
| mmu-miR-125b-5p | MIMAT0000136 | Cops2 | 12848 | HITS-CLIP | | 25753418 | Unspecified |
| mmu-miR-125b-5p | MIMAT0000136 | Cops7b | 26895 | HITS-CLIP, Microarrays//Microarrays | 37986224\|22469780 | | Unspecified |
| mmu-miR-125b-5p | MIMAT0000136 | Copz2 | 56358 | Microarrays | | 37986224 | Unspecified |
| mmu-miR-125b-5p | MIMAT0000136 | Coq2 | 71883 | HITS-CLIP | | 37986224 | Unspecified |
| mmu-miR-125b-5p | MIMAT0000136 | Coro2b | 235431 | HITS-CLIP | | 37986224 | Unspecified |
| mmu-miR-125b-5p | MIMAT0000136 | Cort | 12854 | Microarrays | 37986224\|22469780 | | Unspecified |
| mmu-miR-125b-5p | MIMAT0000136 | Cox6a2 | 12862 | Microarrays | | 37986224 | Unspecified |
| mmu-miR-125b-5p | MIMAT0000136 | Cpd | 12874 | HITS-CLIP | | 37986224 | Unspecified |
| mmu-miR-125b-5p | MIMAT0000136 | Cpe | 12876 | HITS-CLIP | | 37986224 | Unspecified |
| mmu-miR-125b-5p | MIMAT0000136 | Cpeb1 | 12877 | HITS-CLIP | | 25083871 | Unspecified |
| mmu-miR-125b-5p | MIMAT0000136 | Cplx1 | 12889 | HITS-CLIP | | 19536157 | Unspecified |
| mmu-miR-125b-5p | MIMAT0000136 | Cplx2 | 12890 | HITS-CLIP | | 37986224 | Unspecified |
| mmu-miR-125b-5p | MIMAT0000136 | Cplx3 | 235415 | Microarrays | 37986224\|22469780 | | Unspecified |
| mmu-miR-125b-5p | MIMAT0000136 | Cpm | 70574 | Microarrays | 37986224\|22469780 | | Unspecified |
| mmu-miR-125b-5p | MIMAT0000136 | Cpn2 | 71756 | HITS-CLIP | | 37986224 | Unspecified |
| mmu-miR-125b-5p | MIMAT0000136 | Cpne2 | 234577 | HITS-CLIP, Microarrays//Microarrays | 37986224\|22469780 | | Unspecified |
| mmu-miR-125b-5p | MIMAT0000136 | Cpne7 | 102278 | Microarrays | 37986224\|22469780 | | Unspecified |
| mmu-miR-125b-5p | MIMAT0000136 | Cpox | 12892 | HITS-CLIP | | 37986224 | Unspecified |

| | | | | | | |
|---|---|---|---|---|---|---|
| mmu-miR-125b-5p | MIMAT0000136 | Cpsf6 | 432508 | HITS-CLIP | 37986224 | Unspecified |
| mmu-miR-125b-5p | MIMAT0000136 | Cpt2 | 12896 | HITS-CLIP | 25083871 | Unspecified |
| mmu-miR-125b-5p | MIMAT0000136 | Cr2 | 12902 | Microarrays | 37986224 | Unspecified |
| mmu-miR-125b-5p | MIMAT0000136 | Cramp1l | 57354 | HITS-CLIP | 37986224\|25083871 | Unspecified |
| mmu-miR-125b-5p | MIMAT0000136 | Creb3 | 12913 | Microarrays | 37986224\|22469780 | Unspecified |
| mmu-miR-125b-5p | MIMAT0000136 | Crebrf | 77128 | HITS-CLIP | 37986224\|25083871 | Unspecified |
| mmu-miR-125b-5p | MIMAT0000136 | Creg1 | 433375 | HITS-CLIP | 37986224 | Unspecified |
| mmu-miR-125b-5p | MIMAT0000136 | Creld1 | 171508 | HITS-CLIP | 37986224 | Unspecified |
| mmu-miR-125b-5p | MIMAT0000136 | Crip1 | 12925 | Microarrays | 37986224\|22469780 | Unspecified |
| mmu-miR-125b-5p | MIMAT0000136 | Crk | 12928 | HITS-CLIP | 37986224\|25083871 | Unspecified |
| mmu-miR-125b-5p | MIMAT0000136 | Crls1 | 66586 | HITS-CLIP, Microarrays//Microarrays | 37986224\|22469780 | Unspecified |
| mmu-miR-125b-5p | MIMAT0000136 | Crtc3 | 70461 | Microarrays | 37986224 | Unspecified |
| mmu-miR-125b-5p | MIMAT0000136 | Csde1 | 229663 | HITS-CLIP, Microarrays//Microarrays | 37986224\|22469780 | Unspecified |
| mmu-miR-125b-5p | MIMAT0000136 | Csnk1a1 | 93687 | HITS-CLIP | 25083871\|29167373 | Unspecified |
| mmu-miR-125b-5p | MIMAT0000136 | Csnk1e | 27373 | HITS-CLIP | 37986224 | Unspecified |
| mmu-miR-125b-5p | MIMAT0000136 | Csnk1g1 | 214897 | HITS-CLIP | 37986224 | Unspecified |
| mmu-miR-125b-5p | MIMAT0000136 | Csnk2a1 | 12995 | HITS-CLIP | 37986224 | Unspecified |
| mmu-miR-125b-5p | MIMAT0000136 | Csprs | 114564 | Microarrays | 37986224 | Unspecified |
| mmu-miR-125b-5p | MIMAT0000136 | Csrnp1 | 215418 | Microarrays | 37986224\|22469780 | Unspecified |
| mmu-miR-125b-5p | MIMAT0000136 | Csrnp3 | 77771 | Microarrays | 37986224\|22469780 | Unspecified |
| mmu-miR-125b-5p | MIMAT0000136 | Cstad | 78617 | Microarrays | 37986224 | Unspecified |
| mmu-miR-125b-5p | MIMAT0000136 | Cstf2t | 83410 | Microarrays | 37986224\|22469780 | Unspecified |
| mmu-miR-125b-5p | MIMAT0000136 | Ctbs | 74245 | Microarrays | 37986224\|22469780 | Unspecified |
| mmu-miR-125b-5p | MIMAT0000136 | Ctdsp2 | 52468 | HITS-CLIP | 37986224\|25083871\|19536157 | Unspecified |
| mmu-miR-125b-5p | MIMAT0000136 | Cth | 107869 | Microarrays | 37986224 | Unspecified |
| mmu-miR-125b-5p | MIMAT0000136 | Ctif | 269037 | HITS-CLIP | 37986224\|25083871 | Unspecified |
| mmu-miR-125b-5p | MIMAT0000136 | Ctla2a | 13024 | Microarrays | 37986224 | Unspecified |
| mmu-miR-125b-5p | MIMAT0000136 | Ctla2b | 13025 | Microarrays | 37986224 | Unspecified |
| mmu-miR-125b-5p | MIMAT0000136 | Ctnna1 | 12385 | HITS-CLIP | 25083871 | Unspecified |
| mmu-miR-125b-5p | MIMAT0000136 | Ctnnal1 | 54366 | HITS-CLIP | 25753418 | Unspecified |
| mmu-miR-125b-5p | MIMAT0000136 | Ctnnd1 | 12388 | HITS-CLIP | 29167373 | Unspecified |
| mmu-miR-125b-5p | MIMAT0000136 | Ctr9 | 22083 | Microarrays | 37986224 | Unspecified |
| mmu-miR-125b-5p | MIMAT0000136 | Ctsb | 13030 | HITS-CLIP | 25753418 | Unspecified |
| mmu-miR-125b-5p | MIMAT0000136 | Ctsd | 13033 | HITS-CLIP | 37986224 | Unspecified |
| mmu-miR-125b-5p | MIMAT0000136 | Ctsk | 13038 | Microarrays | 37986224\|22469780 | Unspecified |
| mmu-miR-125b-5p | MIMAT0000136 | Ctsw | 13041 | Microarrays | 37986224\|22469780 | Unspecified |
| mmu-miR-125b-5p | MIMAT0000136 | Ctxn1 | 330695 | Microarrays | 37986224 | Unspecified |
| mmu-miR-125b-5p | MIMAT0000136 | Cuedc1 | 103841 | HITS-CLIP | 25083871 | Unspecified |
| mmu-miR-125b-5p | MIMAT0000136 | Cul5 | 75717 | HITS-CLIP | 37986224\|25083871 | Unspecified |

| | | | | | | |
|---|---|---|---|---|---|---|
| mmu-miR-125b-5p | MIMAT0000136 | Cux1 | 13047 | HITS-CLIP, Microarrays | 37986224 | Unspecified |
| mmu-miR-125b-5p | MIMAT0000136 | Cux2 | 13048 | Microarrays | 37986224 | Unspecified |
| mmu-miR-125b-5p | MIMAT0000136 | Cxcl10 | 15945 | Microarrays | 37986224 | Unspecified |
| mmu-miR-125b-5p | MIMAT0000136 | Cxcl12 | 20315 | HITS-CLIP | 37986224 | Unspecified |
| mmu-miR-125b-5p | MIMAT0000136 | Cxcr4 | 12767 | Microarrays | 37986224\|22469780 | Unspecified |
| mmu-miR-125b-5p | MIMAT0000136 | Cxxc1 | 74322 | Microarrays | 37986224 | Unspecified |
| mmu-miR-125b-5p | MIMAT0000136 | Cxxc5 | 67393 | Microarrays | 37986224 | Unspecified |
| mmu-miR-125b-5p | MIMAT0000136 | Cyb561d1 | 72023 | HITS-CLIP, Microarrays//Microarrays | 37986224\|22469780 | Unspecified |
| mmu-miR-125b-5p | MIMAT0000136 | Cyb561d2 | 56368 | Microarrays | 37986224\|22469780 | Unspecified |
| mmu-miR-125b-5p | MIMAT0000136 | Cyb5r4 | 266690 | Microarrays | 37986224\|22469780 | Unspecified |
| mmu-miR-125b-5p | MIMAT0000136 | Cyc1 | 66445 | HITS-CLIP | 37986224 | Unspecified |
| mmu-miR-125b-5p | MIMAT0000136 | Cyfip1 | 20430 | HITS-CLIP | 37986224\|29167373 | Unspecified |
| mmu-miR-125b-5p | MIMAT0000136 | Cyp11a1 | 13070 | Microarrays | 37986224 | Unspecified |
| mmu-miR-125b-5p | MIMAT0000136 | Cyp17a1 | 13074 | Microarrays | 37986224\|22469780 | Unspecified |
| mmu-miR-125b-5p | MIMAT0000136 | Cyp1a2 | 13077 | HITS-CLIP | 37986224 | Unspecified |
| mmu-miR-125b-5p | MIMAT0000136 | Cyp2a12 | 13085 | HITS-CLIP | 37986224 | Unspecified |
| mmu-miR-125b-5p | MIMAT0000136 | Cyp2d12 | 380997 | Microarrays | 37986224 | Unspecified |
| mmu-miR-125b-5p | MIMAT0000136 | Cyp2f2 | 13107 | Microarrays | 37986224 | Unspecified |
| mmu-miR-125b-5p | MIMAT0000136 | Cyp2t4 | 384724 | Microarrays | 37986224\|22469780 | Unspecified |
| mmu-miR-125b-5p | MIMAT0000136 | Cyp4f13 | 170716 | Microarrays | 37986224 | Unspecified |
| mmu-miR-125b-5p | MIMAT0000136 | Cyp4f14 | 64385 | Microarrays | 37986224\|22469780 | Unspecified |
| mmu-miR-125b-5p | MIMAT0000136 | Cyrib | 51571 | Microarrays | 22469780 | Unspecified |
| mmu-miR-125b-5p | MIMAT0000136 | Cyth1 | 19157 | HITS-CLIP | 37986224\|25083871 | Unspecified |
| mmu-miR-125b-5p | MIMAT0000136 | Cyth3 | 19159 | HITS-CLIP | 37986224 | Unspecified |
| mmu-miR-125b-5p | MIMAT0000136 | Cyth4 | 72318 | Microarrays | 37986224 | Unspecified |
| mmu-miR-125b-5p | MIMAT0000136 | D10Wsu102e | 28109 | HITS-CLIP | 37986224 | Unspecified |
| mmu-miR-125b-5p | MIMAT0000136 | D17H6S53E | 114585 | HITS-CLIP | 37986224 | Unspecified |
| mmu-miR-125b-5p | MIMAT0000136 | Daam1 | 208846 | HITS-CLIP | 37986224\|25083871\|19536157\|29167 | Unspecified |
| mmu-miR-125b-5p | MIMAT0000136 | Dab2 | 13132 | HITS-CLIP | 37986224\|25083871 | Unspecified |
| mmu-miR-125b-5p | MIMAT0000136 | Dag1 | 13138 | HITS-CLIP, Microarrays | 37986224 | Unspecified |
| mmu-miR-125b-5p | MIMAT0000136 | Dazap2 | 23994 | HITS-CLIP | 37986224\|25083871 | Unspecified |
| mmu-miR-125b-5p | MIMAT0000136 | Dbn1 | 56320 | Microarrays | 37986224 | Unspecified |
| mmu-miR-125b-5p | MIMAT0000136 | Dcaf6 | 74106 | HITS-CLIP, Microarrays//HITS-CLIP | 37986224\|25753418 | Unspecified |
| mmu-miR-125b-5p | MIMAT0000136 | Dcakd | 68087 | Microarrays | 37986224 | Unspecified |
| mmu-miR-125b-5p | MIMAT0000136 | Dclk2 | 70762 | Microarrays | 37986224 | Unspecified |
| mmu-miR-125b-5p | MIMAT0000136 | Dcp2 | 70640 | HITS-CLIP, Microarrays//Microarrays | 37986224\|22469780 | Unspecified |
| mmu-miR-125b-5p | MIMAT0000136 | Dcun1d2 | 102323 | HITS-CLIP | 25083871 | Unspecified |
| mmu-miR-125b-5p | MIMAT0000136 | Dcxr | 67880 | HITS-CLIP | 37986224 | Unspecified |
| mmu-miR-125b-5p | MIMAT0000136 | Ddb1 | 13194 | HITS-CLIP//qCLASH | 25083871\|31393953 | Unspecified |

| | | | | | | | |
|---|---|---|---|---|---|---|---|
| mmu-miR-125b-5p | MIMAT0000136 | Ddit3 | 13198 | Microarrays | | 37986224 | Unspecified |
| mmu-miR-125b-5p | MIMAT0000136 | Ddit4 | 74747 | Microarrays | | 37986224 | Unspecified |
| mmu-miR-125b-5p | MIMAT0000136 | Ddr2 | 18214 | HITS-CLIP | | 37986224 | Unspecified |
| mmu-miR-125b-5p | MIMAT0000136 | Ddx17 | 67040 | HITS-CLIP | | 37986224 | Unspecified |
| mmu-miR-125b-5p | MIMAT0000136 | Ddx23 | 74351 | Microarrays | 37986224\|22469780 | | Unspecified |
| mmu-miR-125b-5p | MIMAT0000136 | Ddx3x | 13205 | HITS-CLIP | | 37986224 | Unspecified |
| mmu-miR-125b-5p | MIMAT0000136 | Ddx3y | 26900 | HITS-CLIP | | 25753418 | Unspecified |
| mmu-miR-125b-5p | MIMAT0000136 | Ddx46 | 212880 | qCLASH | | 31393953 | Unspecified |
| mmu-miR-125b-5p | MIMAT0000136 | Ddx5 | 13207 | HITS-CLIP | | 37986224 | Unspecified |
| mmu-miR-125b-5p | MIMAT0000136 | Ddx50 | 94213 | Microarrays | | 37986224 | Unspecified |
| mmu-miR-125b-5p | MIMAT0000136 | Ddx52 | 78394 | HITS-CLIP | | 37986224 | Unspecified |
| mmu-miR-125b-5p | MIMAT0000136 | Ddx56 | 52513 | Microarrays | 37986224\|22469780 | | Unspecified |
| mmu-miR-125b-5p | MIMAT0000136 | Ddx58 | 230073 | Microarrays | | 37986224 | Unspecified |
| mmu-miR-125b-5p | MIMAT0000136 | Dedd2 | 67379 | Microarrays | 37986224\|22469780 | | Unspecified |
| mmu-miR-125b-5p | MIMAT0000136 | Dennd1a | 227801 | Microarrays | 37986224\|22469780 | | Unspecified |
| mmu-miR-125b-5p | MIMAT0000136 | Dennd1b | 329260 | HITS-CLIP | | 37986224 | Unspecified |
| mmu-miR-125b-5p | MIMAT0000136 | Dennd4a | 102442 | HITS-CLIP | | 37986224 | Unspecified |
| mmu-miR-125b-5p | MIMAT0000136 | Dennd4c | 329877 | HITS-CLIP | | 37986224 | Unspecified |
| mmu-miR-125b-5p | MIMAT0000136 | Dennd5b | 320560 | HITS-CLIP | | 37986224 | Unspecified |
| mmu-miR-125b-5p | MIMAT0000136 | Dennd6a | 211922 | HITS-CLIP, Microarrays//HITS-CLIP//Microa | 37986224\|25083871\|22469780 | | Unspecified |
| mmu-miR-125b-5p | MIMAT0000136 | Deptor | 97998 | HITS-CLIP | | 37986224 | Unspecified |
| mmu-miR-125b-5p | MIMAT0000136 | Derl3 | 70377 | Microarrays | 37986224\|22469780 | | Unspecified |
| mmu-miR-125b-5p | MIMAT0000136 | Des | 13346 | HITS-CLIP | | 25083871 | Unspecified |
| mmu-miR-125b-5p | MIMAT0000136 | Dffa | 13347 | HITS-CLIP | | 37986224 | Unspecified |
| mmu-miR-125b-5p | MIMAT0000136 | Dgcr2 | 13356 | Microarrays | 37986224\|22469780 | | Unspecified |
| mmu-miR-125b-5p | MIMAT0000136 | Dgka | 13139 | HITS-CLIP | | 25083871 | Unspecified |
| mmu-miR-125b-5p | MIMAT0000136 | Dgkd | 227333 | Microarrays//HITS-CLIP | 37986224\|25083871 | | Unspecified |
| mmu-miR-125b-5p | MIMAT0000136 | Dgke | 56077 | HITS-CLIP | | 37986224 | Unspecified |
| mmu-miR-125b-5p | MIMAT0000136 | Dgkh | 380921 | HITS-CLIP, Microarrays | | 37986224 | Unspecified |
| mmu-miR-125b-5p | MIMAT0000136 | Dhcr24 | 74754 | HITS-CLIP | | 37986224 | Unspecified |
| mmu-miR-125b-5p | MIMAT0000136 | Dhcr7 | 13360 | HITS-CLIP | | 37986224 | Unspecified |
| mmu-miR-125b-5p | MIMAT0000136 | Dhrs13 | 70451 | HITS-CLIP | | 37986224 | Unspecified |
| mmu-miR-125b-5p | MIMAT0000136 | Dhx30 | 72831 | HITS-CLIP | | 37986224 | Unspecified |
| mmu-miR-125b-5p | MIMAT0000136 | Dhx33 | 216877 | HITS-CLIP, Microarrays//HITS-CLIP//Microa | 37986224\|25083871\|22469780 | | Unspecified |
| mmu-miR-125b-5p | MIMAT0000136 | Dhx58 | 80861 | Microarrays | | 37986224 | Unspecified |
| mmu-miR-125b-5p | MIMAT0000136 | Dhx9 | 13211 | Microarrays//HITS-CLIP | 37986224\|22469780\|25753418 | | Unspecified |
| mmu-miR-125b-5p | MIMAT0000136 | Dicer1 | 192119 | HITS-CLIP | | 37986224 | Unspecified |
| mmu-miR-125b-5p | MIMAT0000136 | Dido1 | 23856 | HITS-CLIP | | 37986224 | Unspecified |
| mmu-miR-125b-5p | MIMAT0000136 | Dio2 | 13371 | HITS-CLIP | | 37986224 | Unspecified |

| | | | | | | | |
|---|---|---|---|---|---|---|---|
| mmu-miR-125b-5p | MIMAT0000136 | Dip2b | 239667 | HITS-CLIP | | 37986224 | Unspecified |
| mmu-miR-125b-5p | MIMAT0000136 | Dis3 | 72662 | HITS-CLIP | | 25083871 | Unspecified |
| mmu-miR-125b-5p | MIMAT0000136 | Disp1 | 68897 | HITS-CLIP, Microarrays//Microarrays | 37986224\|22469780 | | Unspecified |
| mmu-miR-125b-5p | MIMAT0000136 | Dkc1 | 245474 | HITS-CLIP | | 25083871 | Unspecified |
| mmu-miR-125b-5p | MIMAT0000136 | Dlat | 235339 | HITS-CLIP | | 25083871 | Unspecified |
| mmu-miR-125b-5p | MIMAT0000136 | Dlc1 | 50768 | HITS-CLIP | | 23597149 | Unspecified |
| mmu-miR-125b-5p | MIMAT0000136 | Dld | 13382 | HITS-CLIP | | 25753418 | Unspecified |
| mmu-miR-125b-5p | MIMAT0000136 | Dlg3 | 53310 | Microarrays | | 37986224 | Unspecified |
| mmu-miR-125b-5p | MIMAT0000136 | Dlg4 | 13385 | HITS-CLIP | | 37986224 | Unspecified |
| mmu-miR-125b-5p | MIMAT0000136 | Dlgap1 | 224997 | HITS-CLIP | | 37986224 | Unspecified |
| mmu-miR-125b-5p | MIMAT0000136 | Dlx3 | 13393 | Microarrays | 37986224\|22469780 | | Unspecified |
| mmu-miR-125b-5p | MIMAT0000136 | Dlx6 | 13396 | qCLASH | | 31393953 | Unspecified |
| mmu-miR-125b-5p | MIMAT0000136 | Dmgdh | 74129 | HITS-CLIP | | 37986224 | Unspecified |
| mmu-miR-125b-5p | MIMAT0000136 | Dmpk | 13400 | HITS-CLIP | | 25753418 | Unspecified |
| mmu-miR-125b-5p | MIMAT0000136 | Dmrtc1a | 70887 | Microarrays | | 37986224 | Unspecified |
| mmu-miR-125b-5p | MIMAT0000136 | Dmtn | 13829 | Microarrays | 37986224\|22469780 | | Unspecified |
| mmu-miR-125b-5p | MIMAT0000136 | Dmxl1 | 240283 | HITS-CLIP | 37986224\|25083871 | | Unspecified |
| mmu-miR-125b-5p | MIMAT0000136 | Dnajb1 | 81489 | Microarrays | 37986224\|22469780 | | Unspecified |
| mmu-miR-125b-5p | MIMAT0000136 | Dnajb14 | 70604 | HITS-CLIP | | 37986224 | Unspecified |
| mmu-miR-125b-5p | MIMAT0000136 | Dnajb3 | 15504 | Microarrays | 37986224\|22469780 | | Unspecified |
| mmu-miR-125b-5p | MIMAT0000136 | Dnajb5 | 56323 | HITS-CLIP | 37986224\|25083871 | | Unspecified |
| mmu-miR-125b-5p | MIMAT0000136 | Dnajc1 | 13418 | Microarrays | | 37986224 | Unspecified |
| mmu-miR-125b-5p | MIMAT0000136 | Dnajc10 | 66861 | HITS-CLIP, Microarrays//HITS-CLIP//Microa | 37986224\|25083871\|19536157\|2246 | | Unspecified |
| mmu-miR-125b-5p | MIMAT0000136 | Dnajc13 | 235567 | Microarrays | | 37986224 | Unspecified |
| mmu-miR-125b-5p | MIMAT0000136 | Dnajc14 | 74330 | HITS-CLIP | 37986224\|25083871\|19536157 | | Unspecified |
| mmu-miR-125b-5p | MIMAT0000136 | Dnajc22 | 72778 | HITS-CLIP | | 23597149 | Unspecified |
| mmu-miR-125b-5p | MIMAT0000136 | Dnajc25 | 72429 | Microarrays | 37986224\|22469780 | | Unspecified |
| mmu-miR-125b-5p | MIMAT0000136 | Dnajc5 | 13002 | HITS-CLIP | | 37986224 | Unspecified |
| mmu-miR-125b-5p | MIMAT0000136 | Dnajc7 | 56354 | qCLASH | | 31393953 | Unspecified |
| mmu-miR-125b-5p | MIMAT0000136 | Dnmt3b | 13436 | HITS-CLIP | | 37986224 | Unspecified |
| mmu-miR-125b-5p | MIMAT0000136 | Dnph1 | 381101 | Microarrays | 37986224\|22469780 | | Unspecified |
| mmu-miR-125b-5p | MIMAT0000136 | Dntt | 21673 | Microarrays | | 37986224 | Unspecified |
| mmu-miR-125b-5p | MIMAT0000136 | Dnttip2 | 99480 | HITS-CLIP | | 37986224 | Unspecified |
| mmu-miR-125b-5p | MIMAT0000136 | Dock11 | 75974 | HITS-CLIP | | 37986224 | Unspecified |
| mmu-miR-125b-5p | MIMAT0000136 | Dok1 | 13448 | HITS-CLIP | 37986224\|25083871 | | Unspecified |
| mmu-miR-125b-5p | MIMAT0000136 | Dok2 | 13449 | Microarrays | | 37986224 | Unspecified |
| mmu-miR-125b-5p | MIMAT0000136 | Dok4 | 114255 | Microarrays | | 37986224 | Unspecified |
| mmu-miR-125b-5p | MIMAT0000136 | Dok6 | 623279 | Microarrays | 37986224\|22469780 | | Unspecified |
| mmu-miR-125b-5p | MIMAT0000136 | Dpp9 | 224897 | HITS-CLIP | | 37986224 | Unspecified |

| | | | | | | | |
|---|---|---|---|---|---|---|---|
| mmu-miR-125b-5p | MIMAT0000136 | Dpy19l1 | 244745 | HITS-CLIP | | 37986224 | Unspecified |
| mmu-miR-125b-5p | MIMAT0000136 | Dpysl2 | 12934 | HITS-CLIP | | 37986224 | Unspecified |
| mmu-miR-125b-5p | MIMAT0000136 | Dram2 | 67171 | HITS-CLIP | 37986224\|25083871\|23597149\|1953( | | Unspecified |
| mmu-miR-125b-5p | MIMAT0000136 | Dscc1 | 72107 | Microarrays | 37986224\|22469780 | | Unspecified |
| mmu-miR-125b-5p | MIMAT0000136 | Dse | 212898 | HITS-CLIP | | 25083871 | Unspecified |
| mmu-miR-125b-5p | MIMAT0000136 | Dsg2 | 13511 | HITS-CLIP | | 37986224 | Unspecified |
| mmu-miR-125b-5p | MIMAT0000136 | Dsp | 109620 | HITS-CLIP | | 23597149 | Unspecified |
| mmu-miR-125b-5p | MIMAT0000136 | Dst | 13518 | HITS-CLIP | 37986224\|25083871 | | Unspecified |
| mmu-miR-125b-5p | MIMAT0000136 | Dstyk | 213452 | HITS-CLIP | | 37986224 | Unspecified |
| mmu-miR-125b-5p | MIMAT0000136 | Dtna | 13527 | HITS-CLIP | | 25753418 | Unspecified |
| mmu-miR-125b-5p | MIMAT0000136 | Dtx3 | 80904 | Microarrays | | 37986224 | Unspecified |
| mmu-miR-125b-5p | MIMAT0000136 | Dtx4 | 207521 | Microarrays | | 37986224 | Unspecified |
| mmu-miR-125b-5p | MIMAT0000136 | Dtymk | 21915 | Microarrays | 37986224\|22469780 | | Unspecified |
| mmu-miR-125b-5p | MIMAT0000136 | Duoxa1 | 213696 | Microarrays | | 37986224 | Unspecified |
| mmu-miR-125b-5p | MIMAT0000136 | Dus1l | 68730 | Microarray//HITS-CLIP//qCLASH//Microarr | 16495412\|23597149\|25083871\|31393 | | Unspecified |
| mmu-miR-125b-5p | MIMAT0000136 | Dusp10 | 63953 | HITS-CLIP, Microarrays//Microarrays | 37986224\|22469780 | | Unspecified |
| mmu-miR-125b-5p | MIMAT0000136 | Dusp11 | 72102 | HITS-CLIP, Microarrays//Microarrays | 37986224\|22469780 | | Unspecified |
| mmu-miR-125b-5p | MIMAT0000136 | Dusp21 | 73547 | Microarrays | 37986224\|22469780 | | Unspecified |
| mmu-miR-125b-5p | MIMAT0000136 | Dusp23 | 68440 | Microarrays | 37986224\|22469780 | | Unspecified |
| mmu-miR-125b-5p | MIMAT0000136 | Dusp28 | 67446 | Microarrays | 37986224\|22469780 | | Unspecified |
| mmu-miR-125b-5p | MIMAT0000136 | Dusp3 | 72349 | HITS-CLIP | | 37986224 | Unspecified |
| mmu-miR-125b-5p | MIMAT0000136 | Dusp5 | 240672 | Microarrays | | 37986224 | Unspecified |
| mmu-miR-125b-5p | MIMAT0000136 | Dusp6 | 67603 | HITS-CLIP | 37986224\|25083871 | | Unspecified |
| mmu-miR-125b-5p | MIMAT0000136 | Dut | 110074 | Microarrays | 37986224\|22469780 | | Unspecified |
| mmu-miR-125b-5p | MIMAT0000136 | Dvl3 | 13544 | HITS-CLIP, Microarrays//Microarrays | 37986224\|22469780 | | Unspecified |
| mmu-miR-125b-5p | MIMAT0000136 | Dym | 69190 | HITS-CLIP | | 37986224 | Unspecified |
| mmu-miR-125b-5p | MIMAT0000136 | Dync1h1 | 13424 | HITS-CLIP, Microarrays//HITS-CLIP | 37986224\|23597149\|25083871 | | Unspecified |
| mmu-miR-125b-5p | MIMAT0000136 | Dync1li2 | 234663 | HITS-CLIP | 37986224\|29167373 | | Unspecified |
| mmu-miR-125b-5p | MIMAT0000136 | Dync2h1 | 110350 | Microarrays | | 37986224 | Unspecified |
| mmu-miR-125b-5p | MIMAT0000136 | Dynlt3 | 67117 | HITS-CLIP | 37986224\|25083871 | | Unspecified |
| mmu-miR-125b-5p | MIMAT0000136 | Dyrk1b | 13549 | Microarrays | 37986224\|22469780 | | Unspecified |
| mmu-miR-125b-5p | MIMAT0000136 | E130309D02Rik | 231868 | HITS-CLIP | | 37986224 | Unspecified |
| mmu-miR-125b-5p | MIMAT0000136 | E130311K13Rik | 329659 | HITS-CLIP | | 37986224 | Unspecified |
| mmu-miR-125b-5p | MIMAT0000136 | E2f2 | 242705 | HITS-CLIP, Microarrays//Microarrays | 37986224\|22469780 | | Unspecified |
| mmu-miR-125b-5p | MIMAT0000136 | E2f3 | 13557 | HITS-CLIP | | 37986224 | Unspecified |
| mmu-miR-125b-5p | MIMAT0000136 | E2f5 | 13559 | Microarrays | | 37986224 | Unspecified |
| mmu-miR-125b-5p | MIMAT0000136 | E2f8 | 108961 | Microarrays | 37986224\|22469780 | | Unspecified |
| mmu-miR-125b-5p | MIMAT0000136 | Eaf1 | 74427 | HITS-CLIP//qCLASH | 37986224\|23597149\|31393953 | | Unspecified |
| mmu-miR-125b-5p | MIMAT0000136 | Ebpl | 68177 | HITS-CLIP | | 37986224 | Unspecified |

| | | | | | | |
|---|---|---|---|---|---|---|
| mmu-miR-125b-5p | MIMAT0000136 | Ece1 | 230857 | HITS-CLIP | 37986224\|25083871 | Unspecified |
| mmu-miR-125b-5p | MIMAT0000136 | Echdc3 | 67856 | Microarrays | 37986224 | Unspecified |
| mmu-miR-125b-5p | MIMAT0000136 | Ecm1 | 13601 | HITS-CLIP | 37986224 | Unspecified |
| mmu-miR-125b-5p | MIMAT0000136 | Edc3 | 353190 | HITS-CLIP | 37986224 | Unspecified |
| mmu-miR-125b-5p | MIMAT0000136 | Edem1 | 192193 | HITS-CLIP | 37986224 | Unspecified |
| mmu-miR-125b-5p | MIMAT0000136 | Edem2 | 108687 | HITS-CLIP | 25083871 | Unspecified |
| mmu-miR-125b-5p | MIMAT0000136 | Edem3 | 66967 | HITS-CLIP | 37986224\|25083871\|23597149\|1953( | Unspecified |
| mmu-miR-125b-5p | MIMAT0000136 | Edrf1 | 214764 | HITS-CLIP | 37986224 | Unspecified |
| mmu-miR-125b-5p | MIMAT0000136 | Eea1 | 216238 | HITS-CLIP | 25083871 | Unspecified |
| mmu-miR-125b-5p | MIMAT0000136 | Eed | 13626 | Microarrays | 37986224\|22469780 | Unspecified |
| mmu-miR-125b-5p | MIMAT0000136 | Eef2 | 13629 | qCLASH | 31393953 | Unspecified |
| mmu-miR-125b-5p | MIMAT0000136 | Eef2k | 13631 | HITS-CLIP | 37986224\|25083871 | Unspecified |
| mmu-miR-125b-5p | MIMAT0000136 | Eefsec | 65967 | Microarrays | 37986224 | Unspecified |
| mmu-miR-125b-5p | MIMAT0000136 | Efemp1 | 216616 | Microarrays | 37986224\|22469780 | Unspecified |
| mmu-miR-125b-5p | MIMAT0000136 | Efhd1 | 98363 | Microarrays | 37986224 | Unspecified |
| mmu-miR-125b-5p | MIMAT0000136 | Efr3a | 76740 | HITS-CLIP | 37986224 | Unspecified |
| mmu-miR-125b-5p | MIMAT0000136 | Egfr | 13649 | HITS-CLIP | 37986224\|23597149 | Unspecified |
| mmu-miR-125b-5p | MIMAT0000136 | Egr1 | 13653 | Microarrays | 37986224 | Unspecified |
| mmu-miR-125b-5p | MIMAT0000136 | Egr3 | 13655 | HITS-CLIP | 37986224 | Unspecified |
| mmu-miR-125b-5p | MIMAT0000136 | Ehbp1 | 216565 | HITS-CLIP | 25083871 | Unspecified |
| mmu-miR-125b-5p | MIMAT0000136 | Ehbp1l1 | 114601 | HITS-CLIP | 37986224 | Unspecified |
| mmu-miR-125b-5p | MIMAT0000136 | Ehd2 | 259300 | HITS-CLIP, Microarrays//Microarrays | 37986224\|22469780 | Unspecified |
| mmu-miR-125b-5p | MIMAT0000136 | Ehd4 | 98878 | HITS-CLIP | 37986224 | Unspecified |
| mmu-miR-125b-5p | MIMAT0000136 | Ehmt1 | 77683 | HITS-CLIP, Microarrays//Microarrays | 37986224\|22469780 | Unspecified |
| mmu-miR-125b-5p | MIMAT0000136 | Eid2 | 386655 | Microarrays | 37986224\|22469780 | Unspecified |
| mmu-miR-125b-5p | MIMAT0000136 | Eif1ad | 69860 | HITS-CLIP | 25083871 | Unspecified |
| mmu-miR-125b-5p | MIMAT0000136 | Eif2ak2 | 19106 | Microarrays | 37986224 | Unspecified |
| mmu-miR-125b-5p | MIMAT0000136 | Eif2ak3 | 13666 | HITS-CLIP | 37986224 | Unspecified |
| mmu-miR-125b-5p | MIMAT0000136 | Eif2ak4 | 27103 | HITS-CLIP | 37986224 | Unspecified |
| mmu-miR-125b-5p | MIMAT0000136 | Eif2b4 | 13667 | HITS-CLIP | 23597149 | Unspecified |
| mmu-miR-125b-5p | MIMAT0000136 | Eif2s3x | 26905 | HITS-CLIP | 37986224 | Unspecified |
| mmu-miR-125b-5p | MIMAT0000136 | Eif3a | 13669 | qCLASH | 31393953 | Unspecified |
| mmu-miR-125b-5p | MIMAT0000136 | Eif4a2 | 13682 | HITS-CLIP | 25083871\|23597149\|29167373 | Unspecified |
| mmu-miR-125b-5p | MIMAT0000136 | Eif4a3 | 192170 | HITS-CLIP | 25083871\|19536157 | Unspecified |
| mmu-miR-125b-5p | MIMAT0000136 | Eif4e2 | 26987 | HITS-CLIP | 25083871 | Unspecified |
| mmu-miR-125b-5p | MIMAT0000136 | Eif4ebp3 | 108112 | Microarrays | 37986224 | Unspecified |
| mmu-miR-125b-5p | MIMAT0000136 | Eif4g1 | 208643 | HITS-CLIP | 37986224\|25083871\|23597149 | Unspecified |
| mmu-miR-125b-5p | MIMAT0000136 | Eif5a | 276770 | HITS-CLIP | 37986224 | Unspecified |
| mmu-miR-125b-5p | MIMAT0000136 | Eif5a2 | 208691 | HITS-CLIP | 37986224\|25083871\|29167373 | Unspecified |

| | | | | | | |
|---|---|---|---|---|---|---|
| mmu-miR-125b-5p | MIMAT0000136 | Eif5b | 226982 | HITS-CLIP | 25083871 | Unspecified |
| mmu-miR-125b-5p | MIMAT0000136 | Elane | 50701 | Microarrays | 37986224 | Unspecified |
| mmu-miR-125b-5p | MIMAT0000136 | Elavl1 | 15568 | HITS-CLIP | 37986224 | Unspecified |
| mmu-miR-125b-5p | MIMAT0000136 | Elavl3 | 15571 | HITS-CLIP | 37986224 | Unspecified |
| mmu-miR-125b-5p | MIMAT0000136 | Elf1 | 13709 | HITS-CLIP | 37986224 | Unspecified |
| mmu-miR-125b-5p | MIMAT0000136 | Elfn1 | 243312 | HITS-CLIP | 37986224 | Unspecified |
| mmu-miR-125b-5p | MIMAT0000136 | Elk1 | 13712 | HITS-CLIP | 37986224 | Unspecified |
| mmu-miR-125b-5p | MIMAT0000136 | Elk4 | 13714 | HITS-CLIP | 37986224\|25083871 | Unspecified |
| mmu-miR-125b-5p | MIMAT0000136 | Ell | 13716 | HITS-CLIP | 25083871 | Unspecified |
| mmu-miR-125b-5p | MIMAT0000136 | Ell3 | 269344 | Microarrays | 37986224\|22469780 | Unspecified |
| mmu-miR-125b-5p | MIMAT0000136 | Elmo1 | 140580 | Microarrays | 37986224 | Unspecified |
| mmu-miR-125b-5p | MIMAT0000136 | Elmod2 | 244548 | Microarrays | 37986224\|22469780 | Unspecified |
| mmu-miR-125b-5p | MIMAT0000136 | Elmsan1 | 238317 | HITS-CLIP | 37986224\|29167373 | Unspecified |
| mmu-miR-125b-5p | MIMAT0000136 | Elovl2 | 54326 | HITS-CLIP | 37986224 | Unspecified |
| mmu-miR-125b-5p | MIMAT0000136 | Elovl6 | 170439 | HITS-CLIP | 37986224 | Unspecified |
| mmu-miR-125b-5p | MIMAT0000136 | Elp6 | 72341 | Microarrays | 37986224\|22469780 | Unspecified |
| mmu-miR-125b-5p | MIMAT0000136 | Emc2 | 66736 | HITS-CLIP | 37986224 | Unspecified |
| mmu-miR-125b-5p | MIMAT0000136 | Emc3 | 66087 | HITS-CLIP | 25083871 | Unspecified |
| mmu-miR-125b-5p | MIMAT0000136 | Eml2 | 72205 | Microarrays | 37986224 | Unspecified |
| mmu-miR-125b-5p | MIMAT0000136 | Eml4 | 78798 | HITS-CLIP | 37986224 | Unspecified |
| mmu-miR-125b-5p | MIMAT0000136 | Eml5 | 319670 | HITS-CLIP | 37986224 | Unspecified |
| mmu-miR-125b-5p | MIMAT0000136 | Emp1 | 13730 | HITS-CLIP | 25083871\|29167373 | Unspecified |
| mmu-miR-125b-5p | MIMAT0000136 | En1 | 13798 | HITS-CLIP | 37986224 | Unspecified |
| mmu-miR-125b-5p | MIMAT0000136 | Enc1 | 13803 | HITS-CLIP | 37986224 | Unspecified |
| mmu-miR-125b-5p | MIMAT0000136 | Endou | 19011 | Microarrays | 37986224\|22469780 | Unspecified |
| mmu-miR-125b-5p | MIMAT0000136 | Eno1 | 13806 | Microarrays | 37986224 | Unspecified |
| mmu-miR-125b-5p | MIMAT0000136 | Eno1b | 433182 | Microarrays | 37986224 | Unspecified |
| mmu-miR-125b-5p | MIMAT0000136 | Enpep | 13809 | Microarrays | 37986224\|22469780 | Unspecified |
| mmu-miR-125b-5p | MIMAT0000136 | Entpd4 | 67464 | Microarray//Microarrays | 16495412\|22469780 | Unspecified |
| mmu-miR-125b-5p | MIMAT0000136 | Entpd5 | 12499 | Microarrays | 37986224\|22469780 | Unspecified |
| mmu-miR-125b-5p | MIMAT0000136 | Entpd6 | 12497 | HITS-CLIP | 29167373 | Unspecified |
| mmu-miR-125b-5p | MIMAT0000136 | Entpd7 | 93685 | HITS-CLIP | 37986224\|25083871 | Unspecified |
| mmu-miR-125b-5p | MIMAT0000136 | Eola1 | 91966 | Microarrays | 22469780 | Unspecified |
| mmu-miR-125b-5p | MIMAT0000136 | Ep400 | 75560 | HITS-CLIP | 37986224\|25083871 | Unspecified |
| mmu-miR-125b-5p | MIMAT0000136 | Epdr1 | 105298 | HITS-CLIP | 37986224 | Unspecified |
| mmu-miR-125b-5p | MIMAT0000136 | Epg5 | 100502841 | HITS-CLIP | 29167373 | Unspecified |
| mmu-miR-125b-5p | MIMAT0000136 | Epha2 | 13836 | Microarrays | 37986224\|22469780 | Unspecified |
| mmu-miR-125b-5p | MIMAT0000136 | Epha7 | 13841 | HITS-CLIP | 37986224 | Unspecified |
| mmu-miR-125b-5p | MIMAT0000136 | Eprs | 107508 | Microarrays | 37986224 | Unspecified |

| | | | | | | |
|---|---|---|---|---|---|---|
| mmu-miR-125b-5p | MIMAT0000136 | Erbin | 55914 | HITS-CLIP | 19536157\|29167373 | Unspecified |
| mmu-miR-125b-5p | MIMAT0000136 | Ercc6 | 319955 | HITS-CLIP | 37986224 | Unspecified |
| mmu-miR-125b-5p | MIMAT0000136 | Erdr1 | 170942 | Microarrays | 37986224 | Unspecified |
| mmu-miR-125b-5p | MIMAT0000136 | Ergic2 | 67456 | HITS-CLIP | 37986224 | Unspecified |
| mmu-miR-125b-5p | MIMAT0000136 | Ergic3 | 66366 | HITS-CLIP | 37986224\|25083871 | Unspecified |
| mmu-miR-125b-5p | MIMAT0000136 | Erlin1 | 226144 | HITS-CLIP | 37986224\|25083871 | Unspecified |
| mmu-miR-125b-5p | MIMAT0000136 | Erlin2 | 244373 | HITS-CLIP | 37986224 | Unspecified |
| mmu-miR-125b-5p | MIMAT0000136 | Ermn | 77767 | Microarrays | 37986224 | Unspecified |
| mmu-miR-125b-5p | MIMAT0000136 | Ermp1 | 226090 | HITS-CLIP | 37986224 | Unspecified |
| mmu-miR-125b-5p | MIMAT0000136 | Ero1l | 50527 | Microarrays | 37986224 | Unspecified |
| mmu-miR-125b-5p | MIMAT0000136 | Ero1lb | 67475 | HITS-CLIP | 37986224 | Unspecified |
| mmu-miR-125b-5p | MIMAT0000136 | Esco1 | 77805 | Microarrays | 37986224\|22469780 | Unspecified |
| mmu-miR-125b-5p | MIMAT0000136 | Esrra | 26379 | HITS-CLIP | 19536157 | Unspecified |
| mmu-miR-125b-5p | MIMAT0000136 | Esyt2 | 52635 | HITS-CLIP | 37986224\|25083871 | Unspecified |
| mmu-miR-125b-5p | MIMAT0000136 | Etd | 69501 | Microarrays | 37986224\|22469780 | Unspecified |
| mmu-miR-125b-5p | MIMAT0000136 | Etfdh | 66841 | Microarrays//HITS-CLIP | 37986224\|22469780\|25753418 | Unspecified |
| mmu-miR-125b-5p | MIMAT0000136 | Ethe1 | 66071 | Microarrays | 37986224 | Unspecified |
| mmu-miR-125b-5p | MIMAT0000136 | Ets1 | 23871 | HITS-CLIP | 37986224\|25083871 | Unspecified |
| mmu-miR-125b-5p | MIMAT0000136 | Etv3 | 27049 | HITS-CLIP | 37986224 | Unspecified |
| mmu-miR-125b-5p | MIMAT0000136 | Etv6 | 14011 | HITS-CLIP | 37986224 | Unspecified |
| mmu-miR-125b-5p | MIMAT0000136 | Eva1a | 232146 | HITS-CLIP | 37986224\|25083871 | Unspecified |
| mmu-miR-125b-5p | MIMAT0000136 | Exo1 | 26909 | Microarrays | 37986224\|22469780 | Unspecified |
| mmu-miR-125b-5p | MIMAT0000136 | Exoc6 | 107371 | HITS-CLIP | 29167373 | Unspecified |
| mmu-miR-125b-5p | MIMAT0000136 | Exoc7 | 53413 | HITS-CLIP | 37986224 | Unspecified |
| mmu-miR-125b-5p | MIMAT0000136 | Ext1 | 14042 | HITS-CLIP | 37986224 | Unspecified |
| mmu-miR-125b-5p | MIMAT0000136 | Extl1 | 56219 | Microarrays | 37986224 | Unspecified |
| mmu-miR-125b-5p | MIMAT0000136 | Extl3 | 54616 | HITS-CLIP | 37986224 | Unspecified |
| mmu-miR-125b-5p | MIMAT0000136 | Ezh1 | 14055 | HITS-CLIP | 37986224 | Unspecified |
| mmu-miR-125b-5p | MIMAT0000136 | Ezr | 22350 | HITS-CLIP | 25753418 | Unspecified |
| mmu-miR-125b-5p | MIMAT0000136 | F5 | 14067 | HITS-CLIP | 37986224 | Unspecified |
| mmu-miR-125b-5p | MIMAT0000136 | Fads2 | 56473 | HITS-CLIP | 37986224 | Unspecified |
| mmu-miR-125b-5p | MIMAT0000136 | Fah | 14085 | Microarrays | 37986224 | Unspecified |
| mmu-miR-125b-5p | MIMAT0000136 | Fam102a | 98952 | HITS-CLIP | 37986224 | Unspecified |
| mmu-miR-125b-5p | MIMAT0000136 | Fam107b | 66540 | HITS-CLIP | 37986224 | Unspecified |
| mmu-miR-125b-5p | MIMAT0000136 | Fam114a1 | 68303 | Microarrays | 37986224 | Unspecified |
| mmu-miR-125b-5p | MIMAT0000136 | Fam117a | 215512 | Microarrays | 37986224 | Unspecified |
| mmu-miR-125b-5p | MIMAT0000136 | Fam118a | 73225 | HITS-CLIP | 37986224 | Unspecified |
| mmu-miR-125b-5p | MIMAT0000136 | Fam120a | 218236 | HITS-CLIP | 37986224 | Unspecified |
| mmu-miR-125b-5p | MIMAT0000136 | Fam126b | 213056 | HITS-CLIP | 37986224 | Unspecified |

| | | | | | | |
|---|---|---|---|---|---|---|
| mmu-miR-125b-5p | MIMAT0000136 | Fam129b | 227737 | HITS-CLIP | 37986224\|25083871\|29167373 | Unspecified |
| mmu-miR-125b-5p | MIMAT0000136 | Fam131b | 76156 | HITS-CLIP | 37986224 | Unspecified |
| mmu-miR-125b-5p | MIMAT0000136 | Fam133b | 68152 | HITS-CLIP | 25083871 | Unspecified |
| mmu-miR-125b-5p | MIMAT0000136 | Fam13c | 71721 | Microarrays | 37986224\|22469780 | Unspecified |
| mmu-miR-125b-5p | MIMAT0000136 | Fam160b1 | 226252 | HITS-CLIP | 37986224 | Unspecified |
| mmu-miR-125b-5p | MIMAT0000136 | Fam161b | 217705 | Microarrays | 37986224\|22469780 | Unspecified |
| mmu-miR-125b-5p | MIMAT0000136 | Fam167b | 230766 | Microarrays | 37986224\|22469780 | Unspecified |
| mmu-miR-125b-5p | MIMAT0000136 | Fam168a | 319604 | HITS-CLIP, Microarrays | 37986224 | Unspecified |
| mmu-miR-125b-5p | MIMAT0000136 | Fam168b | 214469 | HITS-CLIP | 37986224 | Unspecified |
| mmu-miR-125b-5p | MIMAT0000136 | Fam171b | 241520 | Microarrays | 37986224 | Unspecified |
| mmu-miR-125b-5p | MIMAT0000136 | Fam174c | 55009 | Microarrays | 22469780 | Unspecified |
| mmu-miR-125b-5p | MIMAT0000136 | Fam193a | 231128 | HITS-CLIP | 37986224 | Unspecified |
| mmu-miR-125b-5p | MIMAT0000136 | Fam20b | 215015 | HITS-CLIP | 37986224\|25083871 | Unspecified |
| mmu-miR-125b-5p | MIMAT0000136 | Fam210a | 108654 | HITS-CLIP | 37986224 | Unspecified |
| mmu-miR-125b-5p | MIMAT0000136 | Fam217a | 71864 | Microarrays | 37986224\|22469780 | Unspecified |
| mmu-miR-125b-5p | MIMAT0000136 | Fam222b | 216971 | HITS-CLIP | 37986224 | Unspecified |
| mmu-miR-125b-5p | MIMAT0000136 | Fam3b | 52793 | Microarrays | 37986224 | Unspecified |
| mmu-miR-125b-5p | MIMAT0000136 | Fam43a | 224093 | HITS-CLIP | 37986224 | Unspecified |
| mmu-miR-125b-5p | MIMAT0000136 | Fam49b | 223601 | Microarrays | 37986224 | Unspecified |
| mmu-miR-125b-5p | MIMAT0000136 | Fam53b | 77938 | HITS-CLIP | 37986224 | Unspecified |
| mmu-miR-125b-5p | MIMAT0000136 | Fam53c | 66306 | HITS-CLIP | 37986224\|25083871\|29167373 | Unspecified |
| mmu-miR-125b-5p | MIMAT0000136 | Fam57a | 116972 | HITS-CLIP | 37986224 | Unspecified |
| mmu-miR-125b-5p | MIMAT0000136 | Fam71f2 | 245884 | Microarrays | 37986224 | Unspecified |
| mmu-miR-125b-5p | MIMAT0000136 | Fam78a | 241303 | HITS-CLIP | 37986224 | Unspecified |
| mmu-miR-125b-5p | MIMAT0000136 | Fam78b | 226610 | Microarrays | 37986224 | Unspecified |
| mmu-miR-125b-5p | MIMAT0000136 | Fam83h | 105732 | HITS-CLIP | 37986224 | Unspecified |
| mmu-miR-125b-5p | MIMAT0000136 | Fam98b | 68215 | HITS-CLIP | 25083871 | Unspecified |
| mmu-miR-125b-5p | MIMAT0000136 | Fam98c | 73833 | Microarrays | 37986224\|22469780 | Unspecified |
| mmu-miR-125b-5p | MIMAT0000136 | Fancb | 237211 | Microarrays | 37986224\|22469780 | Unspecified |
| mmu-miR-125b-5p | MIMAT0000136 | Fancf | 100040608 | Microarrays | 37986224\|22469780 | Unspecified |
| mmu-miR-125b-5p | MIMAT0000136 | Farsb | 23874 | Microarrays | 37986224 | Unspecified |
| mmu-miR-125b-5p | MIMAT0000136 | Fasn | 14104 | HITS-CLIP | 23597149 | Unspecified |
| mmu-miR-125b-5p | MIMAT0000136 | Fastkd1 | 320720 | HITS-CLIP | 37986224 | Unspecified |
| mmu-miR-125b-5p | MIMAT0000136 | Fastkd3 | 69577 | Microarrays | 37986224\|22469780 | Unspecified |
| mmu-miR-125b-5p | MIMAT0000136 | Fastkd5 | 380601 | Microarrays | 37986224\|22469780 | Unspecified |
| mmu-miR-125b-5p | MIMAT0000136 | Fat1 | 14107 | HITS-CLIP | 37986224 | Unspecified |
| mmu-miR-125b-5p | MIMAT0000136 | Fat4 | 329628 | HITS-CLIP | 37986224 | Unspecified |
| mmu-miR-125b-5p | MIMAT0000136 | Fbll1 | 237730 | HITS-CLIP | 37986224 | Unspecified |
| mmu-miR-125b-5p | MIMAT0000136 | Fbln2 | 14115 | HITS-CLIP | 25083871 | Unspecified |

| | | | | | | | |
|---|---|---|---|---|---|---|---|
| mmu-miR-125b-5p | MIMAT0000136 | Fbp1 | 14121 | Microarrays//HITS-CLIP | 37986224\|23597149 | | Unspecified |
| mmu-miR-125b-5p | MIMAT0000136 | Fbxl12 | 30843 | Microarrays | | 37986224 | Unspecified |
| mmu-miR-125b-5p | MIMAT0000136 | Fbxl17 | 50758 | Microarrays | 37986224\|22469780 | | Unspecified |
| mmu-miR-125b-5p | MIMAT0000136 | Fbxl20 | 72194 | HITS-CLIP | | 37986224 | Unspecified |
| mmu-miR-125b-5p | MIMAT0000136 | Fbxo10 | 269529 | HITS-CLIP | | 37986224 | Unspecified |
| mmu-miR-125b-5p | MIMAT0000136 | Fbxo11 | 225055 | HITS-CLIP | | 25083871 | Unspecified |
| mmu-miR-125b-5p | MIMAT0000136 | Fbxo28 | 67948 | Microarrays | | 37986224 | Unspecified |
| mmu-miR-125b-5p | MIMAT0000136 | Fbxo3 | 57443 | HITS-CLIP | | 37986224 | Unspecified |
| mmu-miR-125b-5p | MIMAT0000136 | Fbxo32 | 67731 | HITS-CLIP | | 37986224 | Unspecified |
| mmu-miR-125b-5p | MIMAT0000136 | Fbxo33 | 70611 | HITS-CLIP, Microarrays//Microarrays//HITS | 37986224\|22469780\|19536157 | | Unspecified |
| mmu-miR-125b-5p | MIMAT0000136 | Fbxo34 | 78938 | HITS-CLIP | | 37986224 | Unspecified |
| mmu-miR-125b-5p | MIMAT0000136 | Fbxo36 | 66153 | Microarrays | | 37986224 | Unspecified |
| mmu-miR-125b-5p | MIMAT0000136 | Fbxo38 | 107035 | Microarrays | 37986224\|22469780 | | Unspecified |
| mmu-miR-125b-5p | MIMAT0000136 | Fbxw7 | 50754 | HITS-CLIP | | 25083871 | Unspecified |
| mmu-miR-125b-5p | MIMAT0000136 | Fbxw9 | 68628 | Microarrays//HITS-CLIP | 37986224\|25083871 | | Unspecified |
| mmu-miR-125b-5p | MIMAT0000136 | Fchsd1 | 319262 | Microarrays | | 37986224 | Unspecified |
| mmu-miR-125b-5p | MIMAT0000136 | Fdps | 110196 | Microarrays | | 37986224 | Unspecified |
| mmu-miR-125b-5p | MIMAT0000136 | Fdxacb1 | 382137 | Microarrays | 37986224\|22469780 | | Unspecified |
| mmu-miR-125b-5p | MIMAT0000136 | Fech | 14151 | HITS-CLIP | | 37986224 | Unspecified |
| mmu-miR-125b-5p | MIMAT0000136 | Fem1b | 14155 | HITS-CLIP | | 37986224 | Unspecified |
| mmu-miR-125b-5p | MIMAT0000136 | Fem1c | 240263 | Microarrays | 37986224\|22469780 | | Unspecified |
| mmu-miR-125b-5p | MIMAT0000136 | Fen1 | 14156 | Microarrays | 37986224\|22469780 | | Unspecified |
| mmu-miR-125b-5p | MIMAT0000136 | Fermt2 | 218952 | HITS-CLIP | | 37986224 | Unspecified |
| mmu-miR-125b-5p | MIMAT0000136 | Fga | 14161 | HITS-CLIP | | 37986224 | Unspecified |
| mmu-miR-125b-5p | MIMAT0000136 | Fgd2 | 26382 | Microarrays | | 37986224 | Unspecified |
| mmu-miR-125b-5p | MIMAT0000136 | Fgd4 | 224014 | HITS-CLIP | | 37986224 | Unspecified |
| mmu-miR-125b-5p | MIMAT0000136 | Fgd6 | 13998 | HITS-CLIP | | 37986224 | Unspecified |
| mmu-miR-125b-5p | MIMAT0000136 | Fgf1 | 14164 | HITS-CLIP | | 25753418 | Unspecified |
| mmu-miR-125b-5p | MIMAT0000136 | Fgf3 | 14174 | Microarrays | | 37986224 | Unspecified |
| mmu-miR-125b-5p | MIMAT0000136 | Fgf9 | 14180 | Microarrays | | 37986224 | Unspecified |
| mmu-miR-125b-5p | MIMAT0000136 | Fgl1 | 234199 | HITS-CLIP | | 23597149 | Unspecified |
| mmu-miR-125b-5p | MIMAT0000136 | Fhl1 | 14199 | HITS-CLIP | 37986224\|25083871 | | Unspecified |
| mmu-miR-125b-5p | MIMAT0000136 | Fhod3 | 225288 | HITS-CLIP | | 37986224 | Unspecified |
| mmu-miR-125b-5p | MIMAT0000136 | Fignl1 | 60530 | Microarrays | 37986224\|22469780 | | Unspecified |
| mmu-miR-125b-5p | MIMAT0000136 | Filip1 | 70598 | Microarrays | | 37986224 | Unspecified |
| mmu-miR-125b-5p | MIMAT0000136 | Fip1l1 | 66899 | HITS-CLIP, Microarrays | | 37986224 | Unspecified |
| mmu-miR-125b-5p | MIMAT0000136 | Fkbp4 | 14228 | Microarrays | | 37986224 | Unspecified |
| mmu-miR-125b-5p | MIMAT0000136 | Fkbpl | 56299 | HITS-CLIP, Microarrays//Microarrays | 37986224\|22469780 | | Unspecified |
| mmu-miR-125b-5p | MIMAT0000136 | Flcn | 216805 | Microarrays | 37986224\|22469780 | | Unspecified |

| | | | | | | |
|---|---|---|---|---|---|---|
| mmu-miR-125b-5p | MIMAT0000136 | Flii | 14248 | HITS-CLIP | 37986224 | Unspecified |
| mmu-miR-125b-5p | MIMAT0000136 | Flna | 192176 | HITS-CLIP | 25083871 | Unspecified |
| mmu-miR-125b-5p | MIMAT0000136 | Flot1 | 14251 | Microarrays | 37986224 | Unspecified |
| mmu-miR-125b-5p | MIMAT0000136 | Flywch2 | 76917 | Microarrays | 37986224\|22469780 | Unspecified |
| mmu-miR-125b-5p | MIMAT0000136 | Fmnl1 | 57778 | Microarrays | 37986224 | Unspecified |
| mmu-miR-125b-5p | MIMAT0000136 | Fmr1 | 14265 | Microarrays | 37986224\|22469780 | Unspecified |
| mmu-miR-125b-5p | MIMAT0000136 | Fn1 | 14268 | HITS-CLIP | 37986224\|23597149 | Unspecified |
| mmu-miR-125b-5p | MIMAT0000136 | Fndc1 | 68655 | HITS-CLIP | 37986224 | Unspecified |
| mmu-miR-125b-5p | MIMAT0000136 | Fndc3a | 319448 | Microarrays | 37986224 | Unspecified |
| mmu-miR-125b-5p | MIMAT0000136 | Fndc3b | 72007 | HITS-CLIP | 37986224 | Unspecified |
| mmu-miR-125b-5p | MIMAT0000136 | Fnip2 | 329679 | Microarrays | 37986224 | Unspecified |
| mmu-miR-125b-5p | MIMAT0000136 | Fopnl | 66086 | HITS-CLIP | 37986224 | Unspecified |
| mmu-miR-125b-5p | MIMAT0000136 | Fosb | 14282 | Microarrays | 37986224 | Unspecified |
| mmu-miR-125b-5p | MIMAT0000136 | Foxf1 | 15227 | Microarrays | 37986224\|22469780 | Unspecified |
| mmu-miR-125b-5p | MIMAT0000136 | Foxn3 | 71375 | Microarrays//HITS-CLIP | 37986224\|25083871\|22469780 | Unspecified |
| mmu-miR-125b-5p | MIMAT0000136 | Foxp1 | 108655 | Microarrays//HITS-CLIP | 37986224\|25753418 | Unspecified |
| mmu-miR-125b-5p | MIMAT0000136 | Foxp4 | 74123 | Microarrays | 37986224 | Unspecified |
| mmu-miR-125b-5p | MIMAT0000136 | Foxq1 | 15220 | HITS-CLIP | 37986224 | Unspecified |
| mmu-miR-125b-5p | MIMAT0000136 | Foxs1 | 14239 | HITS-CLIP | 37986224 | Unspecified |
| mmu-miR-125b-5p | MIMAT0000136 | Fpr-rs4 | 14291 | Microarrays | 37986224\|22469780 | Unspecified |
| mmu-miR-125b-5p | MIMAT0000136 | Frmd4a | 209630 | HITS-CLIP | 25083871 | Unspecified |
| mmu-miR-125b-5p | MIMAT0000136 | Frrs1 | 20321 | HITS-CLIP | 37986224 | Unspecified |
| mmu-miR-125b-5p | MIMAT0000136 | Fryl | 72313 | HITS-CLIP | 37986224 | Unspecified |
| mmu-miR-125b-5p | MIMAT0000136 | Fscn3 | 56223 | Microarrays | 37986224\|22469780 | Unspecified |
| mmu-miR-125b-5p | MIMAT0000136 | Fsd1 | 240121 | HITS-CLIP | 37986224 | Unspecified |
| mmu-miR-125b-5p | MIMAT0000136 | Fto | 26383 | Microarrays//HITS-CLIP | 37986224\|25083871 | Unspecified |
| mmu-miR-125b-5p | MIMAT0000136 | Fubp1 | 51886 | HITS-CLIP | 25083871 | Unspecified |
| mmu-miR-125b-5p | MIMAT0000136 | Fus | 233908 | HITS-CLIP | 37986224 | Unspecified |
| mmu-miR-125b-5p | MIMAT0000136 | Fut1 | 14343 | HITS-CLIP | 37986224 | Unspecified |
| mmu-miR-125b-5p | MIMAT0000136 | Fut11 | 73068 | Microarrays | 37986224\|22469780 | Unspecified |
| mmu-miR-125b-5p | MIMAT0000136 | Fut9 | 14348 | HITS-CLIP | 37986224 | Unspecified |
| mmu-miR-125b-5p | MIMAT0000136 | Fxr1 | 14359 | qCLASH//HITS-CLIP | 31393953\|25083871\|19536157\|29167 | Unspecified |
| mmu-miR-125b-5p | MIMAT0000136 | Fxr2 | 23879 | HITS-CLIP | 25083871 | Unspecified |
| mmu-miR-125b-5p | MIMAT0000136 | Fxyd1 | 56188 | Microarrays | 37986224\|22469780 | Unspecified |
| mmu-miR-125b-5p | MIMAT0000136 | Fxyd7 | 57780 | Microarrays | 37986224 | Unspecified |
| mmu-miR-125b-5p | MIMAT0000136 | Fyn | 14360 | Microarrays | 37986224\|22469780 | Unspecified |
| mmu-miR-125b-5p | MIMAT0000136 | Fzd1 | 14362 | HITS-CLIP | 37986224 | Unspecified |
| mmu-miR-125b-5p | MIMAT0000136 | G6pc | 14377 | Microarrays | 37986224\|22469780 | Unspecified |
| mmu-miR-125b-5p | MIMAT0000136 | Gab3 | 210710 | Microarrays | 37986224 | Unspecified |

| | | | | | | | |
|---|---|---|---|---|---|---|---|
| mmu-miR-125b-5p | MIMAT0000136 | Gabarapl1 | 57436 | HITS-CLIP | | 37986224 | Unspecified |
| mmu-miR-125b-5p | MIMAT0000136 | Gabbr2 | 242425 | HITS-CLIP | | 37986224 | Unspecified |
| mmu-miR-125b-5p | MIMAT0000136 | Gabpb1 | 14391 | HITS-CLIP | | 25083871 | Unspecified |
| mmu-miR-125b-5p | MIMAT0000136 | Gabpb2 | 213054 | HITS-CLIP, Microarrays//Microarrays | 37986224\|22469780 | | Unspecified |
| mmu-miR-125b-5p | MIMAT0000136 | Gabrb1 | 14400 | Microarrays | 37986224\|22469780 | | Unspecified |
| mmu-miR-125b-5p | MIMAT0000136 | Gabrr2 | 14409 | Microarrays | 37986224\|22469780 | | Unspecified |
| mmu-miR-125b-5p | MIMAT0000136 | Gadd45g | 23882 | Microarrays | | 37986224 | Unspecified |
| mmu-miR-125b-5p | MIMAT0000136 | Galns | 50917 | Microarrays | 37986224\|22469780 | | Unspecified |
| mmu-miR-125b-5p | MIMAT0000136 | Galnt12 | 230145 | Microarrays | 37986224\|22469780 | | Unspecified |
| mmu-miR-125b-5p | MIMAT0000136 | Galnt7 | 108150 | HITS-CLIP | | 37986224 | Unspecified |
| mmu-miR-125b-5p | MIMAT0000136 | Ganc | 76051 | Microarrays | | 37986224 | Unspecified |
| mmu-miR-125b-5p | MIMAT0000136 | Gapdh | 14433 | Microarrays | | 37986224 | Unspecified |
| mmu-miR-125b-5p | MIMAT0000136 | Gapvd1 | 66691 | HITS-CLIP | | 25083871 | Unspecified |
| mmu-miR-125b-5p | MIMAT0000136 | Garnl3 | 99326 | Microarrays | | 37986224 | Unspecified |
| mmu-miR-125b-5p | MIMAT0000136 | Gas1 | 14451 | HITS-CLIP | | 37986224 | Unspecified |
| mmu-miR-125b-5p | MIMAT0000136 | Gas2l1 | 78926 | HITS-CLIP | | 37986224 | Unspecified |
| mmu-miR-125b-5p | MIMAT0000136 | Gas2l3 | 237436 | HITS-CLIP | | 37986224 | Unspecified |
| mmu-miR-125b-5p | MIMAT0000136 | Gata5 | 14464 | Microarrays | 37986224\|22469780 | | Unspecified |
| mmu-miR-125b-5p | MIMAT0000136 | Gata6 | 14465 | HITS-CLIP | | 25753418 | Unspecified |
| mmu-miR-125b-5p | MIMAT0000136 | Gatad1 | 67210 | Microarrays | 37986224\|22469780 | | Unspecified |
| mmu-miR-125b-5p | MIMAT0000136 | Gatad2a | 234366 | HITS-CLIP | | 23597149 | Unspecified |
| mmu-miR-125b-5p | MIMAT0000136 | Gbe1 | 74185 | HITS-CLIP | | 37986224 | Unspecified |
| mmu-miR-125b-5p | MIMAT0000136 | Gbp3 | 55932 | Microarrays | | 37986224 | Unspecified |
| mmu-miR-125b-5p | MIMAT0000136 | Gbp7 | 229900 | Microarrays | | 37986224 | Unspecified |
| mmu-miR-125b-5p | MIMAT0000136 | Gcc2 | 70297 | HITS-CLIP | | 37986224 | Unspecified |
| mmu-miR-125b-5p | MIMAT0000136 | Gcdh | 270076 | Microarrays | 37986224\|22469780 | | Unspecified |
| mmu-miR-125b-5p | MIMAT0000136 | Gch1 | 14528 | Microarrays | | 37986224 | Unspecified |
| mmu-miR-125b-5p | MIMAT0000136 | Gcm1 | 14531 | Microarrays | | 37986224 | Unspecified |
| mmu-miR-125b-5p | MIMAT0000136 | Gcn1 | 10985 | HITS-CLIP | | 25753418 | Unspecified |
| mmu-miR-125b-5p | MIMAT0000136 | Gcnt1 | 14537 | HITS-CLIP | | 37986224 | Unspecified |
| mmu-miR-125b-5p | MIMAT0000136 | Gdf11 | 14561 | HITS-CLIP | | 37986224 | Unspecified |
| mmu-miR-125b-5p | MIMAT0000136 | Gem | 14579 | Microarrays | 37986224\|22469780 | | Unspecified |
| mmu-miR-125b-5p | MIMAT0000136 | Gfi1 | 14581 | Microarrays | 37986224\|22469780 | | Unspecified |
| mmu-miR-125b-5p | MIMAT0000136 | Gfi1b | 14582 | Microarrays | | 37986224 | Unspecified |
| mmu-miR-125b-5p | MIMAT0000136 | Gfm2 | 320806 | HITS-CLIP | | 37986224 | Unspecified |
| mmu-miR-125b-5p | MIMAT0000136 | Gfpt1 | 14583 | HITS-CLIP | | 37986224 | Unspecified |
| mmu-miR-125b-5p | MIMAT0000136 | Gga2 | 74105 | HITS-CLIP | 37986224\|25083871 | | Unspecified |
| mmu-miR-125b-5p | MIMAT0000136 | Ggnbp2 | 217039 | HITS-CLIP | | 37986224 | Unspecified |
| mmu-miR-125b-5p | MIMAT0000136 | Gh | 14599 | Microarrays | 37986224\|22469780 | | Unspecified |

| | | | | | | | |
|---|---|---|---|---|---|---|---|
| mmu-miR-125b-5p | MIMAT0000136 | Ghitm | 66092 | HITS-CLIP | | 23597149 | Unspecified |
| mmu-miR-125b-5p | MIMAT0000136 | Gid4 | 66771 | HITS-CLIP | | 37986224 | Unspecified |
| mmu-miR-125b-5p | MIMAT0000136 | Gigyf1 | 57330 | HITS-CLIP | 37986224\|25083871 | | Unspecified |
| mmu-miR-125b-5p | MIMAT0000136 | Gimap5 | 317757 | Microarrays | 37986224\|22469780 | | Unspecified |
| mmu-miR-125b-5p | MIMAT0000136 | Gimap9 | 317758 | Microarrays | 37986224\|22469780 | | Unspecified |
| mmu-miR-125b-5p | MIMAT0000136 | Gin1 | 252876 | Microarrays | 37986224\|22469780 | | Unspecified |
| mmu-miR-125b-5p | MIMAT0000136 | Gipr | 381853 | Microarrays | | 37986224 | Unspecified |
| mmu-miR-125b-5p | MIMAT0000136 | Gja3 | 14611 | Microarrays | 37986224\|22469780 | | Unspecified |
| mmu-miR-125b-5p | MIMAT0000136 | Gjb1 | 14618 | HITS-CLIP | | 37986224 | Unspecified |
| mmu-miR-125b-5p | MIMAT0000136 | Gjc1 | 14615 | HITS-CLIP | 37986224\|25083871 | | Unspecified |
| mmu-miR-125b-5p | MIMAT0000136 | Gla | 11605 | HITS-CLIP | | 37986224 | Unspecified |
| mmu-miR-125b-5p | MIMAT0000136 | Glb1 | 12091 | Microarrays | | 37986224 | Unspecified |
| mmu-miR-125b-5p | MIMAT0000136 | Glcci1 | 170772 | HITS-CLIP | | 37986224 | Unspecified |
| mmu-miR-125b-5p | MIMAT0000136 | Glce | 93683 | HITS-CLIP | | 37986224 | Unspecified |
| mmu-miR-125b-5p | MIMAT0000136 | Glipr2 | 384009 | Microarrays | | 37986224 | Unspecified |
| mmu-miR-125b-5p | MIMAT0000136 | Gls | 14660 | HITS-CLIP, Microarrays//HITS-CLIP//Microa | 37986224\|25083871\|22469780\|19536 | | Unspecified |
| mmu-miR-125b-5p | MIMAT0000136 | Glud1 | 14661 | HITS-CLIP | | 23597149 | Unspecified |
| mmu-miR-125b-5p | MIMAT0000136 | Glul | 14645 | HITS-CLIP | | 37986224 | Unspecified |
| mmu-miR-125b-5p | MIMAT0000136 | Glyr1 | 74022 | HITS-CLIP//qCLASH | 23597149\|31393953 | | Unspecified |
| mmu-miR-125b-5p | MIMAT0000136 | Gm20498 | 105940408 | HITS-CLIP | | 37986224 | Unspecified |
| mmu-miR-125b-5p | MIMAT0000136 | Gm21833 | 6609710 | qCLASH | | 31393953 | Unspecified |
| mmu-miR-125b-5p | MIMAT0000136 | Gm266 | 212539 | Microarrays | 37986224\|22469780 | | Unspecified |
| mmu-miR-125b-5p | MIMAT0000136 | Gm3336 | 100502950 | Microarrays | | 37986224 | Unspecified |
| mmu-miR-125b-5p | MIMAT0000136 | Gm340 | 381224 | HITS-CLIP | | 37986224 | Unspecified |
| mmu-miR-125b-5p | MIMAT0000136 | Gm49322 | | Microarrays | | 22469780 | Unspecified |
| mmu-miR-125b-5p | MIMAT0000136 | Gm5168 | 382275 | Microarrays | | 37986224 | Unspecified |
| mmu-miR-125b-5p | MIMAT0000136 | Gm6729 | 627035 | Microarrays | 37986224\|22469780 | | Unspecified |
| mmu-miR-125b-5p | MIMAT0000136 | Gm8369 | 666926 | Microarrays | | 37986224 | Unspecified |
| mmu-miR-125b-5p | MIMAT0000136 | Gm9938 | 791290 | HITS-CLIP | | 37986224 | Unspecified |
| mmu-miR-125b-5p | MIMAT0000136 | Gmcl1 | 23885 | HITS-CLIP | | 37986224 | Unspecified |
| mmu-miR-125b-5p | MIMAT0000136 | Gmeb2 | 229004 | HITS-CLIP | 37986224\|25083871 | | Unspecified |
| mmu-miR-125b-5p | MIMAT0000136 | Gmps | 229363 | HITS-CLIP | | 29167373 | Unspecified |
| mmu-miR-125b-5p | MIMAT0000136 | Gna11 | 14672 | HITS-CLIP | | 37986224 | Unspecified |
| mmu-miR-125b-5p | MIMAT0000136 | Gna12 | 14673 | HITS-CLIP | 37986224\|23597149 | | Unspecified |
| mmu-miR-125b-5p | MIMAT0000136 | Gnai2 | 14678 | HITS-CLIP | | 37986224 | Unspecified |
| mmu-miR-125b-5p | MIMAT0000136 | Gnaq | 14682 | Microarrays | | 37986224 | Unspecified |
| mmu-miR-125b-5p | MIMAT0000136 | Gnb1 | 14688 | Microarrays | | 37986224 | Unspecified |
| mmu-miR-125b-5p | MIMAT0000136 | Gnb2 | 14693 | HITS-CLIP | | 23597149 | Unspecified |
| mmu-miR-125b-5p | MIMAT0000136 | Gnb5 | 14697 | Microarrays | | 37986224 | Unspecified |

| | | | | | | | |
|---|---|---|---|---|---|---|---|
| mmu-miR-125b-5p | MIMAT0000136 | Gng12 | 14701 | Microarrays | | 37986224 | Unspecified |
| mmu-miR-125b-5p | MIMAT0000136 | Gnpat | 14712 | HITS-CLIP | | 25083871 | Unspecified |
| mmu-miR-125b-5p | MIMAT0000136 | Golga3 | 269682 | HITS-CLIP | | 37986224 | Unspecified |
| mmu-miR-125b-5p | MIMAT0000136 | Golga4 | 54214 | qCLASH | | 31393953 | Unspecified |
| mmu-miR-125b-5p | MIMAT0000136 | Golgb1 | 224139 | HITS-CLIP | | 25083871 | Unspecified |
| mmu-miR-125b-5p | MIMAT0000136 | Golim4 | 73124 | HITS-CLIP | | 37986224 | Unspecified |
| mmu-miR-125b-5p | MIMAT0000136 | Gopc | 94221 | HITS-CLIP | | 37986224 | Unspecified |
| mmu-miR-125b-5p | MIMAT0000136 | Gosr2 | 56494 | HITS-CLIP | | 31433990 | Unspecified |
| mmu-miR-125b-5p | MIMAT0000136 | Got2 | 14719 | HITS-CLIP | 37986224\|23597149 | | Unspecified |
| mmu-miR-125b-5p | MIMAT0000136 | Gp9 | 54368 | Microarrays | | 37986224 | Unspecified |
| mmu-miR-125b-5p | MIMAT0000136 | Gpam | 14732 | HITS-CLIP | | 37986224 | Unspecified |
| mmu-miR-125b-5p | MIMAT0000136 | Gpat3 | 84803 | Microarrays//HITS-CLIP | 22469780\|25753418 | | Unspecified |
| mmu-miR-125b-5p | MIMAT0000136 | Gpatch8 | 237943 | HITS-CLIP | | 37986224 | Unspecified |
| mmu-miR-125b-5p | MIMAT0000136 | Gpc1 | 14733 | HITS-CLIP | | 37986224 | Unspecified |
| mmu-miR-125b-5p | MIMAT0000136 | Gpcpd1 | 74182 | Microarrays | 37986224\|22469780 | | Unspecified |
| mmu-miR-125b-5p | MIMAT0000136 | Gpd1 | 14555 | HITS-CLIP | | 37986224 | Unspecified |
| mmu-miR-125b-5p | MIMAT0000136 | Gpm6a | 234267 | HITS-CLIP | | 37986224 | Unspecified |
| mmu-miR-125b-5p | MIMAT0000136 | Gpr108 | 78308 | qCLASH | | 31393953 | Unspecified |
| mmu-miR-125b-5p | MIMAT0000136 | Gpr19 | 14760 | Microarrays | | 37986224 | Unspecified |
| mmu-miR-125b-5p | MIMAT0000136 | Gpr88 | 64378 | HITS-CLIP, Microarrays//Microarrays | 37986224\|22469780 | | Unspecified |
| mmu-miR-125b-5p | MIMAT0000136 | Gprasp2 | 245607 | HITS-CLIP | | 37986224 | Unspecified |
| mmu-miR-125b-5p | MIMAT0000136 | Gprc5c | 70355 | HITS-CLIP | | 37986224 | Unspecified |
| mmu-miR-125b-5p | MIMAT0000136 | Gpsm2 | 76123 | HITS-CLIP | | 37986224 | Unspecified |
| mmu-miR-125b-5p | MIMAT0000136 | Gpt2 | 108682 | Microarrays | | 37986224 | Unspecified |
| mmu-miR-125b-5p | MIMAT0000136 | Gramd1b | 235283 | HITS-CLIP | | 37986224 | Unspecified |
| mmu-miR-125b-5p | MIMAT0000136 | Gramd2 | 546134 | Microarrays | 37986224\|22469780 | | Unspecified |
| mmu-miR-125b-5p | MIMAT0000136 | Gramd3 | 107022 | Microarrays | | 37986224 | Unspecified |
| mmu-miR-125b-5p | MIMAT0000136 | Gramd4 | 223752 | HITS-CLIP | | 37986224 | Unspecified |
| mmu-miR-125b-5p | MIMAT0000136 | Grap2 | 17444 | Microarrays | | 37986224 | Unspecified |
| mmu-miR-125b-5p | MIMAT0000136 | Gria2 | 14800 | HITS-CLIP | | 19536157 | Unspecified |
| mmu-miR-125b-5p | MIMAT0000136 | Grik5 | 14809 | Microarrays | | 37986224 | Unspecified |
| mmu-miR-125b-5p | MIMAT0000136 | Grin2b | 14812 | HITS-CLIP | | 37986224 | Unspecified |
| mmu-miR-125b-5p | MIMAT0000136 | Grina | 66168 | HITS-CLIP | | 37986224 | Unspecified |
| mmu-miR-125b-5p | MIMAT0000136 | Grk1 | 24013 | qCLASH | | 31393953 | Unspecified |
| mmu-miR-125b-5p | MIMAT0000136 | Grk4 | 14772 | HITS-CLIP | | 37986224 | Unspecified |
| mmu-miR-125b-5p | MIMAT0000136 | Grm3 | 108069 | HITS-CLIP | | 37986224 | Unspecified |
| mmu-miR-125b-5p | MIMAT0000136 | Grn | 14824 | HITS-CLIP, Microarrays//HITS-CLIP | 37986224\|23597149 | | Unspecified |
| mmu-miR-125b-5p | MIMAT0000136 | Grsf1 | 231413 | HITS-CLIP | | 37986224 | Unspecified |
| mmu-miR-125b-5p | MIMAT0000136 | Gsap | 212167 | Microarrays | | 37986224 | Unspecified |

| | | | | | | |
|---|---|---|---|---|---|---|
| mmu-miR-125b-5p | MIMAT0000136 | Gsk3b | 56637 | HITS-CLIP | 37986224\|25083871\|29167373 | Unspecified |
| mmu-miR-125b-5p | MIMAT0000136 | Gsn | 227753 | HITS-CLIP | 37986224 | Unspecified |
| mmu-miR-125b-5p | MIMAT0000136 | Gstm1 | 14862 | Microarrays | 37986224 | Unspecified |
| mmu-miR-125b-5p | MIMAT0000136 | Gstm4 | 14865 | Microarrays | 37986224 | Unspecified |
| mmu-miR-125b-5p | MIMAT0000136 | Gstp3 | 225884 | Microarrays | 22469780 | Unspecified |
| mmu-miR-125b-5p | MIMAT0000136 | Gstt1 | 14871 | Microarrays | 37986224 | Unspecified |
| mmu-miR-125b-5p | MIMAT0000136 | Gtf2h3 | 209357 | HITS-CLIP | 37986224 | Unspecified |
| mmu-miR-125b-5p | MIMAT0000136 | Gtf3c2 | 71752 | HITS-CLIP | 37986224 | Unspecified |
| mmu-miR-125b-5p | MIMAT0000136 | Gtpbp2 | 56055 | HITS-CLIP | 37986224\|25083871\|29167373 | Unspecified |
| mmu-miR-125b-5p | MIMAT0000136 | Guca2b | 14916 | Microarrays | 37986224\|22469780 | Unspecified |
| mmu-miR-125b-5p | MIMAT0000136 | Gucd1 | 68778 | HITS-CLIP | 37986224 | Unspecified |
| mmu-miR-125b-5p | MIMAT0000136 | Guk1 | 14923 | HITS-CLIP | 25083871 | Unspecified |
| mmu-miR-125b-5p | MIMAT0000136 | Gvin1 | 74558 | Microarrays | 37986224 | Unspecified |
| mmu-miR-125b-5p | MIMAT0000136 | Gxylt1 | 223827 | HITS-CLIP | 37986224 | Unspecified |
| mmu-miR-125b-5p | MIMAT0000136 | Gys1 | 14936 | Microarrays | 37986224 | Unspecified |
| mmu-miR-125b-5p | MIMAT0000136 | Gzf1 | 74533 | HITS-CLIP | 37986224 | Unspecified |
| mmu-miR-125b-5p | MIMAT0000136 | H1f1 | 3024 | Microarrays | 22469780 | Unspecified |
| mmu-miR-125b-5p | MIMAT0000136 | H1f4 | 3008 | Microarrays | 22469780 | Unspecified |
| mmu-miR-125b-5p | MIMAT0000136 | H1f5 | 3009 | Microarrays | 22469780 | Unspecified |
| mmu-miR-125b-5p | MIMAT0000136 | H2-D1 | 14964 | Microarrays | 37986224 | Unspecified |
| mmu-miR-125b-5p | MIMAT0000136 | H2-DMa | 14998 | Microarrays | 37986224 | Unspecified |
| mmu-miR-125b-5p | MIMAT0000136 | H2-DMb1 | 14999 | Microarrays | 37986224 | Unspecified |
| mmu-miR-125b-5p | MIMAT0000136 | H2-Eb1 | 14969 | Microarrays | 37986224 | Unspecified |
| mmu-miR-125b-5p | MIMAT0000136 | H2-K1 | 14972 | HITS-CLIP | 37986224 | Unspecified |
| mmu-miR-125b-5p | MIMAT0000136 | H2-Ke6 | 14979 | Microarrays | 37986224 | Unspecified |
| mmu-miR-125b-5p | MIMAT0000136 | H2-Q4 | 15015 | Microarrays | 37986224 | Unspecified |
| mmu-miR-125b-5p | MIMAT0000136 | H2-T23 | 15040 | Microarrays | 37986224 | Unspecified |
| mmu-miR-125b-5p | MIMAT0000136 | H2-T24 | 15042 | Microarrays | 37986224 | Unspecified |
| mmu-miR-125b-5p | MIMAT0000136 | H2ac1 | 221613 | Microarrays | 22469780 | Unspecified |
| mmu-miR-125b-5p | MIMAT0000136 | H2ac21 | 317772 | Microarrays | 22469780 | Unspecified |
| mmu-miR-125b-5p | MIMAT0000136 | H2ac4 | 8335 | Microarrays | 22469780 | Unspecified |
| mmu-miR-125b-5p | MIMAT0000136 | H2al1e | 547160 | Microarrays | 22469780 | Unspecified |
| mmu-miR-125b-5p | MIMAT0000136 | H2bc3 | 3018 | Microarrays | 22469780 | Unspecified |
| mmu-miR-125b-5p | MIMAT0000136 | H3c1 | 8350 | Microarrays | 22469780 | Unspecified |
| mmu-miR-125b-5p | MIMAT0000136 | H3c11 | 8350 | Microarrays | 22469780 | Unspecified |
| mmu-miR-125b-5p | MIMAT0000136 | H3c14 | 126961 | Microarrays | 22469780 | Unspecified |
| mmu-miR-125b-5p | MIMAT0000136 | H3c4 | 8350 | Microarrays | 22469780 | Unspecified |
| mmu-miR-125b-5p | MIMAT0000136 | H3c7 | 8350 | Microarrays | 22469780 | Unspecified |
| mmu-miR-125b-5p | MIMAT0000136 | H3f3b | 15081 | HITS-CLIP | 25083871 | Unspecified |

| | | | | | | |
|---|---|---|---|---|---|---|
| mmu-miR-125b-5p | MIMAT0000136 | H4c11 | 8359 | Microarrays | 22469780 | Unspecified |
| mmu-miR-125b-5p | MIMAT0000136 | H4c12 | 8359 | Microarrays | 22469780 | Unspecified |
| mmu-miR-125b-5p | MIMAT0000136 | H4c14 | 8359 | Microarrays | 22469780 | Unspecified |
| mmu-miR-125b-5p | MIMAT0000136 | H4c16 | 8359 | Microarrays//qCLASH | 22469780\|31393953 | Unspecified |
| mmu-miR-125b-5p | MIMAT0000136 | H4c17 | 100041230 | Microarrays | 22469780 | Unspecified |
| mmu-miR-125b-5p | MIMAT0000136 | H4c3 | 8359 | Microarrays | 22469780 | Unspecified |
| mmu-miR-125b-5p | MIMAT0000136 | H4c4 | 8359 | Microarrays | 22469780 | Unspecified |
| mmu-miR-125b-5p | MIMAT0000136 | H4c9 | 8359 | Microarrays | 22469780 | Unspecified |
| mmu-miR-125b-5p | MIMAT0000136 | H6pd | 100198 | HITS-CLIP, Microarrays//Microarrays | 37986224\|22469780 | Unspecified |
| mmu-miR-125b-5p | MIMAT0000136 | Haao | 107766 | HITS-CLIP | 37986224\|23597149 | Unspecified |
| mmu-miR-125b-5p | MIMAT0000136 | Hacd3 | 57874 | Microarrays | 22469780 | Unspecified |
| mmu-miR-125b-5p | MIMAT0000136 | Haghl | 68977 | Microarrays | 37986224 | Unspecified |
| mmu-miR-125b-5p | MIMAT0000136 | Hal | 15109 | HITS-CLIP | 37986224 | Unspecified |
| mmu-miR-125b-5p | MIMAT0000136 | Hap1 | 15114 | Microarrays | 37986224\|22469780 | Unspecified |
| mmu-miR-125b-5p | MIMAT0000136 | Hapln1 | 12950 | HITS-CLIP | 37986224\|19536157 | Unspecified |
| mmu-miR-125b-5p | MIMAT0000136 | Haus3 | 231123 | HITS-CLIP | 25083871 | Unspecified |
| mmu-miR-125b-5p | MIMAT0000136 | Hbb-bt | 101488143 | Microarrays | 37986224 | Unspecified |
| mmu-miR-125b-5p | MIMAT0000136 | Hbp1 | 73389 | HITS-CLIP | 25083871 | Unspecified |
| mmu-miR-125b-5p | MIMAT0000136 | Hbs1l | 56422 | Microarrays | 37986224\|22469780 | Unspecified |
| mmu-miR-125b-5p | MIMAT0000136 | Hcfc1r1 | 353502 | Microarrays | 37986224 | Unspecified |
| mmu-miR-125b-5p | MIMAT0000136 | Hcn3 | 15168 | Microarrays | 37986224 | Unspecified |
| mmu-miR-125b-5p | MIMAT0000136 | Hdac1 | 433759 | Microarrays | 37986224\|22469780 | Unspecified |
| mmu-miR-125b-5p | MIMAT0000136 | Hdac3 | 15183 | HITS-CLIP, Microarrays//Microarrays | 37986224\|22469780 | Unspecified |
| mmu-miR-125b-5p | MIMAT0000136 | Hdac4 | 208727 | HITS-CLIP, Microarrays | 37986224 | Unspecified |
| mmu-miR-125b-5p | MIMAT0000136 | Hddc3 | 68695 | HITS-CLIP | 37986224 | Unspecified |
| mmu-miR-125b-5p | MIMAT0000136 | Hdgf | 15191 | HITS-CLIP | 37986224 | Unspecified |
| mmu-miR-125b-5p | MIMAT0000136 | Hdhd3 | 72748 | Microarrays | 37986224\|22469780 | Unspecified |
| mmu-miR-125b-5p | MIMAT0000136 | Hdlbp | 110611 | HITS-CLIP, Microarrays//HITS-CLIP | 37986224\|23597149 | Unspecified |
| mmu-miR-125b-5p | MIMAT0000136 | Heatr5b | 320473 | HITS-CLIP | 29167373 | Unspecified |
| mmu-miR-125b-5p | MIMAT0000136 | Hectd1 | 207304 | Microarrays | 37986224 | Unspecified |
| mmu-miR-125b-5p | MIMAT0000136 | Hectd2 | 226098 | HITS-CLIP | 25083871\|19536157 | Unspecified |
| mmu-miR-125b-5p | MIMAT0000136 | Heg1 | 77446 | HITS-CLIP | 37986224\|19536157 | Unspecified |
| mmu-miR-125b-5p | MIMAT0000136 | Hells | 15201 | Microarrays | 37986224\|22469780 | Unspecified |
| mmu-miR-125b-5p | MIMAT0000136 | Helz | 78455 | HITS-CLIP | 29167373 | Unspecified |
| mmu-miR-125b-5p | MIMAT0000136 | Helz2 | 229003 | Microarrays | 37986224 | Unspecified |
| mmu-miR-125b-5p | MIMAT0000136 | Hephl1 | 244698 | HITS-CLIP | 37986224 | Unspecified |
| mmu-miR-125b-5p | MIMAT0000136 | Herc1 | 235439 | HITS-CLIP | 37986224 | Unspecified |
| mmu-miR-125b-5p | MIMAT0000136 | Herc2 | 15204 | HITS-CLIP | 37986224\|23597149 | Unspecified |
| mmu-miR-125b-5p | MIMAT0000136 | Herc4 | 67345 | Microarrays | 37986224\|22469780 | Unspecified |

| | | | | | | | |
|---|---|---|---|---|---|---|---|
| mmu-miR-125b-5p | MIMAT0000136 | Herpud2 | 80517 | Microarrays | | 37986224 | Unspecified |
| mmu-miR-125b-5p | MIMAT0000136 | Hes1 | 15205 | Microarrays | 37986224\|22469780 | | Unspecified |
| mmu-miR-125b-5p | MIMAT0000136 | Hes2 | 15206 | Microarrays | 37986224\|22469780 | | Unspecified |
| mmu-miR-125b-5p | MIMAT0000136 | Hes5 | 15208 | Microarrays | 37986224\|22469780 | | Unspecified |
| mmu-miR-125b-5p | MIMAT0000136 | Hexim1 | 192231 | Microarrays | 37986224\|22469780 | | Unspecified |
| mmu-miR-125b-5p | MIMAT0000136 | Hgsnat | 52120 | HITS-CLIP | | 23597149 | Unspecified |
| mmu-miR-125b-5p | MIMAT0000136 | Hic2 | 58180 | HITS-CLIP | | 37986224 | Unspecified |
| mmu-miR-125b-5p | MIMAT0000136 | Hid1 | 217310 | Microarrays//HITS-CLIP | 37986224\|25083871 | | Unspecified |
| mmu-miR-125b-5p | MIMAT0000136 | Hilpda | 69573 | Microarrays | | 37986224 | Unspecified |
| mmu-miR-125b-5p | MIMAT0000136 | Hip1 | 215114 | Microarrays | | 37986224 | Unspecified |
| mmu-miR-125b-5p | MIMAT0000136 | Hipk1 | 15257 | HITS-CLIP | | 25083871 | Unspecified |
| mmu-miR-125b-5p | MIMAT0000136 | Hipk2 | 15258 | HITS-CLIP | 37986224\|23597149 | | Unspecified |
| mmu-miR-125b-5p | MIMAT0000136 | Hipk3 | 15259 | HITS-CLIP | | 37986224 | Unspecified |
| mmu-miR-125b-5p | MIMAT0000136 | Hipk4 | 233020 | Microarrays | 37986224\|22469780 | | Unspecified |
| mmu-miR-125b-5p | MIMAT0000136 | Hist1h4m | 100041230 | Microarrays | | 37986224 | Unspecified |
| mmu-miR-125b-5p | MIMAT0000136 | Hivep1 | 110521 | HITS-CLIP, Microarrays | | 37986224 | Unspecified |
| mmu-miR-125b-5p | MIMAT0000136 | Hivep2 | 15273 | HITS-CLIP | | 37986224 | Unspecified |
| mmu-miR-125b-5p | MIMAT0000136 | Hjv | 69606 | Microarrays | | 22469780 | Unspecified |
| mmu-miR-125b-5p | MIMAT0000136 | Hk2 | 15277 | Microarrays | | 37986224 | Unspecified |
| mmu-miR-125b-5p | MIMAT0000136 | Hlx | 15284 | Microarrays | | 37986224 | Unspecified |
| mmu-miR-125b-5p | MIMAT0000136 | Hmgb2 | 97165 | Microarrays | 37986224\|22469780 | | Unspecified |
| mmu-miR-125b-5p | MIMAT0000136 | Hmgcr | 15357 | HITS-CLIP, Microarrays//HITS-CLIP | 37986224\|25083871 | | Unspecified |
| mmu-miR-125b-5p | MIMAT0000136 | Hmmr | 15366 | Microarrays | | 37986224 | Unspecified |
| mmu-miR-125b-5p | MIMAT0000136 | Hmox1 | 15368 | HITS-CLIP, Microarrays | | 37986224 | Unspecified |
| mmu-miR-125b-5p | MIMAT0000136 | Hnf4a | 15378 | HITS-CLIP | | 23597149 | Unspecified |
| mmu-miR-125b-5p | MIMAT0000136 | Hnrnpa2b1 | 53379 | HITS-CLIP | | 37986224 | Unspecified |
| mmu-miR-125b-5p | MIMAT0000136 | Hnrnpdl | 50926 | Microarrays | 37986224\|22469780 | | Unspecified |
| mmu-miR-125b-5p | MIMAT0000136 | Hnrnph1 | 59013 | HITS-CLIP | | 37986224 | Unspecified |
| mmu-miR-125b-5p | MIMAT0000136 | Hnrnph2 | 56258 | HITS-CLIP | | 37986224 | Unspecified |
| mmu-miR-125b-5p | MIMAT0000136 | Hnrnpl | 15388 | HITS-CLIP | | 25083871 | Unspecified |
| mmu-miR-125b-5p | MIMAT0000136 | Hnrnpr | 74326 | HITS-CLIP | 37986224\|25083871 | | Unspecified |
| mmu-miR-125b-5p | MIMAT0000136 | Hnrnpu | 51810 | HITS-CLIP | 37986224\|23597149\|19536157\|29167 | | Unspecified |
| mmu-miR-125b-5p | MIMAT0000136 | Hnrnpul2 | 68693 | HITS-CLIP | | 37986224 | Unspecified |
| mmu-miR-125b-5p | MIMAT0000136 | Homez | 239099 | HITS-CLIP | 37986224\|25083871 | | Unspecified |
| mmu-miR-125b-5p | MIMAT0000136 | Hook1 | 77963 | HITS-CLIP | | 25753418 | Unspecified |
| mmu-miR-125b-5p | MIMAT0000136 | Hook3 | 320191 | HITS-CLIP | | 37986224 | Unspecified |
| mmu-miR-125b-5p | MIMAT0000136 | Hoxa3 | 15400 | HITS-CLIP | 37986224\|25083871 | | Unspecified |
| mmu-miR-125b-5p | MIMAT0000136 | Hoxb3 | 15410 | Microarrays | | 37986224 | Unspecified |
| mmu-miR-125b-5p | MIMAT0000136 | Hoxb4 | 15412 | Microarrays | 37986224\|22469780 | | Unspecified |

| | | | | | | |
|---|---|---|---|---|---|---|
| mmu-miR-125b-5p | MIMAT0000136 | Hoxb5 | 15413 | Microarrays | 37986224 | Unspecified |
| mmu-miR-125b-5p | MIMAT0000136 | Hoxb6 | 15414 | Microarrays | 37986224 | Unspecified |
| mmu-miR-125b-5p | MIMAT0000136 | Hoxb8 | 15416 | Microarrays | 37986224 | Unspecified |
| mmu-miR-125b-5p | MIMAT0000136 | Hoxc4 | 15423 | HITS-CLIP | 37986224 | Unspecified |
| mmu-miR-125b-5p | MIMAT0000136 | Hoxc6 | 15425 | HITS-CLIP | 37986224\|25083871 | Unspecified |
| mmu-miR-125b-5p | MIMAT0000136 | Hpdl | 242642 | Microarrays | 37986224\|22469780 | Unspecified |
| mmu-miR-125b-5p | MIMAT0000136 | Hprt | 15452 | HITS-CLIP | 37986224 | Unspecified |
| mmu-miR-125b-5p | MIMAT0000136 | Hpse | 15442 | Microarrays | 37986224 | Unspecified |
| mmu-miR-125b-5p | MIMAT0000136 | Hs2st1 | 23908 | Microarrays | 37986224\|22469780 | Unspecified |
| mmu-miR-125b-5p | MIMAT0000136 | Hs6st1 | 50785 | HITS-CLIP | 37986224 | Unspecified |
| mmu-miR-125b-5p | MIMAT0000136 | Hsd11b1 | 15483 | Microarrays | 37986224 | Unspecified |
| mmu-miR-125b-5p | MIMAT0000136 | Hsdl1 | 72552 | HITS-CLIP | 37986224 | Unspecified |
| mmu-miR-125b-5p | MIMAT0000136 | Hsdl2 | 72479 | HITS-CLIP | 37986224\|25753418 | Unspecified |
| mmu-miR-125b-5p | MIMAT0000136 | Hsf2 | 15500 | Microarrays | 37986224\|22469780 | Unspecified |
| mmu-miR-125b-5p | MIMAT0000136 | Hsp90aa1 | 15519 | HITS-CLIP | 37986224 | Unspecified |
| mmu-miR-125b-5p | MIMAT0000136 | Hsp90b1 | 22027 | qCLASH | 31393953 | Unspecified |
| mmu-miR-125b-5p | MIMAT0000136 | Hspa12a | 73442 | HITS-CLIP | 37986224 | Unspecified |
| mmu-miR-125b-5p | MIMAT0000136 | Hspa5 | 14828 | HITS-CLIP | 25753418 | Unspecified |
| mmu-miR-125b-5p | MIMAT0000136 | Hspa9 | 15526 | HITS-CLIP | 25753418 | Unspecified |
| mmu-miR-125b-5p | MIMAT0000136 | Hspb7 | 29818 | HITS-CLIP | 25083871 | Unspecified |
| mmu-miR-125b-5p | MIMAT0000136 | Hspb9 | 75482 | Microarrays | 37986224 | Unspecified |
| mmu-miR-125b-5p | MIMAT0000136 | Hsph1 | 15505 | HITS-CLIP | 23597149 | Unspecified |
| mmu-miR-125b-5p | MIMAT0000136 | Iars | 105148 | Microarrays | 37986224 | Unspecified |
| mmu-miR-125b-5p | MIMAT0000136 | Iba57 | 216792 | Microarrays | 37986224\|22469780 | Unspecified |
| mmu-miR-125b-5p | MIMAT0000136 | Icam1 | 15894 | Microarrays | 37986224 | Unspecified |
| mmu-miR-125b-5p | MIMAT0000136 | Icam2 | 15896 | Microarrays | 37986224\|22469780 | Unspecified |
| mmu-miR-125b-5p | MIMAT0000136 | Ice1 | 822287 | qCLASH | 31393953 | Unspecified |
| mmu-miR-125b-5p | MIMAT0000136 | Ick | 56542 | Microarrays | 37986224 | Unspecified |
| mmu-miR-125b-5p | MIMAT0000136 | Icmt | 57295 | HITS-CLIP | 37986224\|25083871 | Unspecified |
| mmu-miR-125b-5p | MIMAT0000136 | Id1 | 15901 | HITS-CLIP | 37986224 | Unspecified |
| mmu-miR-125b-5p | MIMAT0000136 | Id2 | 15902 | Microarrays | 37986224 | Unspecified |
| mmu-miR-125b-5p | MIMAT0000136 | Id3 | 15903 | Microarrays | 37986224\|22469780 | Unspecified |
| mmu-miR-125b-5p | MIMAT0000136 | Idh1 | 15926 | HITS-CLIP | 25083871\|25753418 | Unspecified |
| mmu-miR-125b-5p | MIMAT0000136 | Idh3b | 170718 | HITS-CLIP | 25753418 | Unspecified |
| mmu-miR-125b-5p | MIMAT0000136 | Ido2 | 209176 | HITS-CLIP | 37986224 | Unspecified |
| mmu-miR-125b-5p | MIMAT0000136 | Idua | 15932 | Microarrays | 37986224 | Unspecified |
| mmu-miR-125b-5p | MIMAT0000136 | Ier2 | 15936 | HITS-CLIP, Microarrays | 37986224 | Unspecified |
| mmu-miR-125b-5p | MIMAT0000136 | Ier3 | 15937 | HITS-CLIP | 37986224 | Unspecified |
| mmu-miR-125b-5p | MIMAT0000136 | Ifi204 | 15951 | Microarrays | 37986224 | Unspecified |

| | | | | | | |
|---|---|---|---|---|---|---|
| mmu-miR-125b-5p | MIMAT0000136 | Ifi27 | 52668 | Microarrays | 37986224 | Unspecified |
| mmu-miR-125b-5p | MIMAT0000136 | Ifi27l2a | 76933 | Microarrays | 37986224 | Unspecified |
| mmu-miR-125b-5p | MIMAT0000136 | Ifi35 | 70110 | Microarrays | 37986224 | Unspecified |
| mmu-miR-125b-5p | MIMAT0000136 | Ifih1 | 71586 | Microarrays | 37986224 | Unspecified |
| mmu-miR-125b-5p | MIMAT0000136 | Ifit1 | 15957 | Microarrays | 37986224 | Unspecified |
| mmu-miR-125b-5p | MIMAT0000136 | Ifit3 | 15959 | Microarrays | 37986224 | Unspecified |
| mmu-miR-125b-5p | MIMAT0000136 | Ifitm2 | 80876 | Microarrays | 37986224 | Unspecified |
| mmu-miR-125b-5p | MIMAT0000136 | Ifitm3 | 66141 | Microarrays | 37986224 | Unspecified |
| mmu-miR-125b-5p | MIMAT0000136 | Ifrd1 | 15982 | HITS-CLIP | 37986224 | Unspecified |
| mmu-miR-125b-5p | MIMAT0000136 | Ift172 | 67661 | Microarrays | 37986224 | Unspecified |
| mmu-miR-125b-5p | MIMAT0000136 | Ift81 | 12589 | Microarrays | 37986224 | Unspecified |
| mmu-miR-125b-5p | MIMAT0000136 | Igdcc4 | 56741 | Microarrays | 37986224 | Unspecified |
| mmu-miR-125b-5p | MIMAT0000136 | Igf1r | 16001 | HITS-CLIP | 37986224\|25083871 | Unspecified |
| mmu-miR-125b-5p | MIMAT0000136 | Igf2bp1 | 140486 | HITS-CLIP | 37986224 | Unspecified |
| mmu-miR-125b-5p | MIMAT0000136 | Igfbp6 | 16012 | HITS-CLIP | 37986224 | Unspecified |
| mmu-miR-125b-5p | MIMAT0000136 | Igfn1 | 226438 | HITS-CLIP | 37986224 | Unspecified |
| mmu-miR-125b-5p | MIMAT0000136 | Igll1 | 16136 | Microarrays | 37986224\|22469780 | Unspecified |
| mmu-miR-125b-5p | MIMAT0000136 | Igsf3 | 78908 | HITS-CLIP | 37986224 | Unspecified |
| mmu-miR-125b-5p | MIMAT0000136 | Igsf8 | 140559 | Microarrays | 37986224 | Unspecified |
| mmu-miR-125b-5p | MIMAT0000136 | Igtp | 16145 | Microarrays | 37986224 | Unspecified |
| mmu-miR-125b-5p | MIMAT0000136 | Ihh | 16147 | Microarrays | 37986224\|22469780 | Unspecified |
| mmu-miR-125b-5p | MIMAT0000136 | Ikbip | 67454 | HITS-CLIP | 37986224 | Unspecified |
| mmu-miR-125b-5p | MIMAT0000136 | Ikbke | 56489 | HITS-CLIP | 29167373 | Unspecified |
| mmu-miR-125b-5p | MIMAT0000136 | Ikzf2 | 22779 | HITS-CLIP | 37986224 | Unspecified |
| mmu-miR-125b-5p | MIMAT0000136 | Ikzf4 | 22781 | HITS-CLIP | 37986224 | Unspecified |
| mmu-miR-125b-5p | MIMAT0000136 | Ikzf5 | 67143 | Microarrays//HITS-CLIP | 37986224\|25083871\|22469780 | Unspecified |
| mmu-miR-125b-5p | MIMAT0000136 | Il10rb | 16155 | HITS-CLIP | 37986224 | Unspecified |
| mmu-miR-125b-5p | MIMAT0000136 | Il12a | 16159 | Microarrays | 37986224 | Unspecified |
| mmu-miR-125b-5p | MIMAT0000136 | Il12rb1 | 16161 | Microarrays | 37986224 | Unspecified |
| mmu-miR-125b-5p | MIMAT0000136 | Il13ra1 | 16164 | HITS-CLIP | 37986224 | Unspecified |
| mmu-miR-125b-5p | MIMAT0000136 | Il16 | 16170 | Microarrays | 37986224\|22469780 | Unspecified |
| mmu-miR-125b-5p | MIMAT0000136 | Il1r1 | 16177 | HITS-CLIP | 37986224 | Unspecified |
| mmu-miR-125b-5p | MIMAT0000136 | Il1rap | 16180 | HITS-CLIP | 37986224\|23597149 | Unspecified |
| mmu-miR-125b-5p | MIMAT0000136 | Il21r | 60504 | Microarrays | 37986224 | Unspecified |
| mmu-miR-125b-5p | MIMAT0000136 | Il27ra | 50931 | Microarrays | 37986224 | Unspecified |
| mmu-miR-125b-5p | MIMAT0000136 | Il2ra | 16184 | Microarrays | 37986224 | Unspecified |
| mmu-miR-125b-5p | MIMAT0000136 | Il3ra | 16188 | Microarrays | 37986224 | Unspecified |
| mmu-miR-125b-5p | MIMAT0000136 | Il4i1 | 14204 | Microarrays | 37986224 | Unspecified |
| mmu-miR-125b-5p | MIMAT0000136 | Il6ra | 16194 | HITS-CLIP | 37986224 | Unspecified |

| | | | | | | |
|---|---|---|---|---|---|---|
| mmu-miR-125b-5p | MIMAT0000136 | Il6st | 16195 | HITS-CLIP, Microarrays | 37986224 | Unspecified |
| mmu-miR-125b-5p | MIMAT0000136 | Immp1l | 66541 | Microarrays | 37986224\|22469780 | Unspecified |
| mmu-miR-125b-5p | MIMAT0000136 | Impad1 | 242291 | HITS-CLIP | 37986224 | Unspecified |
| mmu-miR-125b-5p | MIMAT0000136 | Inafm2 | 100505573 | HITS-CLIP | 25083871 | Unspecified |
| mmu-miR-125b-5p | MIMAT0000136 | Inf2 | 70435 | Microarrays | 37986224 | Unspecified |
| mmu-miR-125b-5p | MIMAT0000136 | Ing1 | 26356 | Microarrays | 37986224\|22469780 | Unspecified |
| mmu-miR-125b-5p | MIMAT0000136 | Ing3 | 71777 | HITS-CLIP | 25083871 | Unspecified |
| mmu-miR-125b-5p | MIMAT0000136 | Inip | 66209 | HITS-CLIP | 37986224 | Unspecified |
| mmu-miR-125b-5p | MIMAT0000136 | Ino80d | 227195 | HITS-CLIP | 37986224\|25083871 | Unspecified |
| mmu-miR-125b-5p | MIMAT0000136 | Inpp1 | 16329 | Microarrays | 37986224 | Unspecified |
| mmu-miR-125b-5p | MIMAT0000136 | Inpp4a | 269180 | HITS-CLIP | 37986224 | Unspecified |
| mmu-miR-125b-5p | MIMAT0000136 | Inppl1 | 16332 | HITS-CLIP | 25083871 | Unspecified |
| mmu-miR-125b-5p | MIMAT0000136 | Insig2 | 72999 | HITS-CLIP | 37986224 | Unspecified |
| mmu-miR-125b-5p | MIMAT0000136 | Insr | 16337 | HITS-CLIP | 37986224 | Unspecified |
| mmu-miR-125b-5p | MIMAT0000136 | Insrr | 23920 | Microarrays | 37986224 | Unspecified |
| mmu-miR-125b-5p | MIMAT0000136 | Ints7 | 77065 | HITS-CLIP | 37986224 | Unspecified |
| mmu-miR-125b-5p | MIMAT0000136 | Ip6k1 | 27399 | HITS-CLIP | 37986224 | Unspecified |
| mmu-miR-125b-5p | MIMAT0000136 | Ipmk | 69718 | HITS-CLIP | 25083871 | Unspecified |
| mmu-miR-125b-5p | MIMAT0000136 | Ipo8 | 320727 | HITS-CLIP | 37986224 | Unspecified |
| mmu-miR-125b-5p | MIMAT0000136 | Ipo9 | 226432 | HITS-CLIP | 37986224\|25083871\|29167373 | Unspecified |
| mmu-miR-125b-5p | MIMAT0000136 | Iqgap1 | 29875 | HITS-CLIP | 31433990\|25753418 | Unspecified |
| mmu-miR-125b-5p | MIMAT0000136 | Iqgap2 | 544963 | Microarrays | 37986224 | Unspecified |
| mmu-miR-125b-5p | MIMAT0000136 | Iqgap3 | 404710 | Microarrays | 37986224 | Unspecified |
| mmu-miR-125b-5p | MIMAT0000136 | Irf1 | 16362 | HITS-CLIP | 37986224 | Unspecified |
| mmu-miR-125b-5p | MIMAT0000136 | Irf2 | 16363 | HITS-CLIP | 37986224 | Unspecified |
| mmu-miR-125b-5p | MIMAT0000136 | Irf4 | 16364 | Microarrays | 37986224\|22469780 | Unspecified |
| mmu-miR-125b-5p | MIMAT0000136 | Irf5 | 27056 | HITS-CLIP | 37986224 | Unspecified |
| mmu-miR-125b-5p | MIMAT0000136 | Irf7 | 54123 | Microarrays | 37986224 | Unspecified |
| mmu-miR-125b-5p | MIMAT0000136 | Irf8 | 15900 | Microarrays | 37986224 | Unspecified |
| mmu-miR-125b-5p | MIMAT0000136 | Irf9 | 16391 | Microarrays | 37986224 | Unspecified |
| mmu-miR-125b-5p | MIMAT0000136 | Irgm1 | 15944 | Microarrays | 37986224 | Unspecified |
| mmu-miR-125b-5p | MIMAT0000136 | Irs1 | 16367 | HITS-CLIP | 37986224 | Unspecified |
| mmu-miR-125b-5p | MIMAT0000136 | Irx2 | 16372 | HITS-CLIP | 25083871 | Unspecified |
| mmu-miR-125b-5p | MIMAT0000136 | Isg15 | 100038882 | Microarrays | 37986224 | Unspecified |
| mmu-miR-125b-5p | MIMAT0000136 | Isg20 | 57444 | Microarrays | 37986224 | Unspecified |
| mmu-miR-125b-5p | MIMAT0000136 | Isg20l2 | 229504 | HITS-CLIP | 25083871 | Unspecified |
| mmu-miR-125b-5p | MIMAT0000136 | Ist1 | 71955 | HITS-CLIP | 37986224\|19536157 | Unspecified |
| mmu-miR-125b-5p | MIMAT0000136 | Itch | 16396 | HITS-CLIP, Microarrays, Luciferase Reporte | 37986224\|25083871\|21935352\|2246 | Unspecified |
| mmu-miR-125b-5p | MIMAT0000136 | Itga5 | 16402 | HITS-CLIP, Microarrays | 37986224 | Unspecified |

| | | | | | | | |
|---|---|---|---|---|---|---|---|
| mmu-miR-125b-5p | MIMAT0000136 | Itgb1 | 16412 | HITS-CLIP | | 37986224 | Unspecified |
| mmu-miR-125b-5p | MIMAT0000136 | Itgb3 | 16416 | Microarrays | | 37986224 | Unspecified |
| mmu-miR-125b-5p | MIMAT0000136 | Itgb6 | 16420 | Microarrays | 37986224\|22469780 | | Unspecified |
| mmu-miR-125b-5p | MIMAT0000136 | Itgb7 | 16421 | Microarrays | 37986224\|22469780 | | Unspecified |
| mmu-miR-125b-5p | MIMAT0000136 | Itih3 | 16426 | HITS-CLIP | | 23597149 | Unspecified |
| mmu-miR-125b-5p | MIMAT0000136 | Itih4 | 16427 | HITS-CLIP | | 23597149 | Unspecified |
| mmu-miR-125b-5p | MIMAT0000136 | Itih5 | 209378 | HITS-CLIP | | 37986224 | Unspecified |
| mmu-miR-125b-5p | MIMAT0000136 | Itln1 | 16429 | Microarrays | | 37986224 | Unspecified |
| mmu-miR-125b-5p | MIMAT0000136 | Itm2c | 64294 | Microarrays | | 37986224 | Unspecified |
| mmu-miR-125b-5p | MIMAT0000136 | Itpk1 | 217837 | Microarrays | | 37986224 | Unspecified |
| mmu-miR-125b-5p | MIMAT0000136 | Itpkb | 320404 | HITS-CLIP | | 37986224 | Unspecified |
| mmu-miR-125b-5p | MIMAT0000136 | Itpkc | 233011 | Microarrays | 37986224\|22469780 | | Unspecified |
| mmu-miR-125b-5p | MIMAT0000136 | Itpr1 | 16438 | HITS-CLIP | | 37986224 | Unspecified |
| mmu-miR-125b-5p | MIMAT0000136 | Itpripl2 | 319622 | HITS-CLIP | | 37986224 | Unspecified |
| mmu-miR-125b-5p | MIMAT0000136 | Itsn2 | 20403 | HITS-CLIP | | 37986224 | Unspecified |
| mmu-miR-125b-5p | MIMAT0000136 | Ivl | 16447 | Microarrays | | 37986224 | Unspecified |
| mmu-miR-125b-5p | MIMAT0000136 | Jade1 | 269424 | HITS-CLIP | | 37986224 | Unspecified |
| mmu-miR-125b-5p | MIMAT0000136 | Jade2 | 76901 | HITS-CLIP | | 37986224 | Unspecified |
| mmu-miR-125b-5p | MIMAT0000136 | Jak1 | 16451 | HITS-CLIP, Microarrays | | 37986224 | Unspecified |
| mmu-miR-125b-5p | MIMAT0000136 | Jam2 | 67374 | HITS-CLIP | | 25083871 | Unspecified |
| mmu-miR-125b-5p | MIMAT0000136 | Jam3 | 83964 | Microarrays | | 37986224 | Unspecified |
| mmu-miR-125b-5p | MIMAT0000136 | Jarid2 | 16468 | HITS-CLIP//qCLASH | 37986224\|25083871\|31393953\|1953€ | | Unspecified |
| mmu-miR-125b-5p | MIMAT0000136 | Jdp2 | 81703 | Microarrays | | 37986224 | Unspecified |
| mmu-miR-125b-5p | MIMAT0000136 | Jkamp | 104771 | HITS-CLIP | | 37986224 | Unspecified |
| mmu-miR-125b-5p | MIMAT0000136 | Jmjd1c | 108829 | HITS-CLIP | | 25083871 | Unspecified |
| mmu-miR-125b-5p | MIMAT0000136 | Josd1 | 74158 | HITS-CLIP | | 37986224 | Unspecified |
| mmu-miR-125b-5p | MIMAT0000136 | Jph1 | 57339 | HITS-CLIP | | 37986224 | Unspecified |
| mmu-miR-125b-5p | MIMAT0000136 | Junb | 16477 | Microarrays | | 37986224 | Unspecified |
| mmu-miR-125b-5p | MIMAT0000136 | Jup | 16480 | HITS-CLIP | | 37986224 | Unspecified |
| mmu-miR-125b-5p | MIMAT0000136 | Kank2 | 235041 | HITS-CLIP | | 37986224 | Unspecified |
| mmu-miR-125b-5p | MIMAT0000136 | Kansl3 | 226976 | HITS-CLIP | | 37986224 | Unspecified |
| mmu-miR-125b-5p | MIMAT0000136 | Kat6b | 54169 | Microarrays | | 37986224 | Unspecified |
| mmu-miR-125b-5p | MIMAT0000136 | Katna1 | 23924 | HITS-CLIP | | 37986224 | Unspecified |
| mmu-miR-125b-5p | MIMAT0000136 | Kbtbd3 | 69149 | Microarrays | 37986224\|22469780 | | Unspecified |
| mmu-miR-125b-5p | MIMAT0000136 | Kbtbd4 | 67136 | Microarrays | 37986224\|22469780 | | Unspecified |
| mmu-miR-125b-5p | MIMAT0000136 | Kbtbd8 | 243574 | Microarrays | 37986224\|22469780 | | Unspecified |
| mmu-miR-125b-5p | MIMAT0000136 | Kcmf1 | 74287 | HITS-CLIP | | 37986224 | Unspecified |
| mmu-miR-125b-5p | MIMAT0000136 | Kcnc1 | 16502 | HITS-CLIP | | 37986224 | Unspecified |
| mmu-miR-125b-5p | MIMAT0000136 | Kcng1 | 241794 | Microarrays | 37986224\|22469780 | | Unspecified |

| | | | | | | | |
|---|---|---|---|---|---|---|---|
| mmu-miR-125b-5p | MIMAT0000136 | Kcnh1 | 16510 | HITS-CLIP | | 37986224 | Unspecified |
| mmu-miR-125b-5p | MIMAT0000136 | Kcnh7 | 170738 | HITS-CLIP | | 37986224 | Unspecified |
| mmu-miR-125b-5p | MIMAT0000136 | Kcnj10 | 16513 | HITS-CLIP | | 37986224 | Unspecified |
| mmu-miR-125b-5p | MIMAT0000136 | Kcnmb1 | 16533 | Microarrays | | 37986224 | Unspecified |
| mmu-miR-125b-5p | MIMAT0000136 | Kcnq4 | 60613 | HITS-CLIP | | 37986224 | Unspecified |
| mmu-miR-125b-5p | MIMAT0000136 | Kcnt1 | 227632 | HITS-CLIP | | 37986224 | Unspecified |
| mmu-miR-125b-5p | MIMAT0000136 | Kcnt2 | 240776 | HITS-CLIP | | 37986224 | Unspecified |
| mmu-miR-125b-5p | MIMAT0000136 | Kctd11 | 216858 | HITS-CLIP | | 37986224 | Unspecified |
| mmu-miR-125b-5p | MIMAT0000136 | Kctd13 | 233877 | Microarrays | | 37986224 | Unspecified |
| mmu-miR-125b-5p | MIMAT0000136 | Kctd16 | 383348 | HITS-CLIP | 37986224\|19536157 | | Unspecified |
| mmu-miR-125b-5p | MIMAT0000136 | Kctd20 | 66989 | Microarrays | 37986224\|22469780 | | Unspecified |
| mmu-miR-125b-5p | MIMAT0000136 | Kctd21 | 622320 | HITS-CLIP | | 25083871 | Unspecified |
| mmu-miR-125b-5p | MIMAT0000136 | Kctd5 | 69259 | HITS-CLIP, Microarrays | | 37986224 | Unspecified |
| mmu-miR-125b-5p | MIMAT0000136 | Kdelr1 | 68137 | HITS-CLIP | | 25083871 | Unspecified |
| mmu-miR-125b-5p | MIMAT0000136 | Kdelr2 | 66913 | HITS-CLIP | | 25083871 | Unspecified |
| mmu-miR-125b-5p | MIMAT0000136 | Kdelr3 | 105785 | HITS-CLIP | | 25083871 | Unspecified |
| mmu-miR-125b-5p | MIMAT0000136 | Kdm2a | 225876 | HITS-CLIP | | 25083871 | Unspecified |
| mmu-miR-125b-5p | MIMAT0000136 | Kdm3b | 277250 | HITS-CLIP | 37986224\|23597149 | | Unspecified |
| mmu-miR-125b-5p | MIMAT0000136 | Kdm4b | 193796 | HITS-CLIP | | 23597149 | Unspecified |
| mmu-miR-125b-5p | MIMAT0000136 | Kdm4c | 76804 | HITS-CLIP | 37986224\|25083871 | | Unspecified |
| mmu-miR-125b-5p | MIMAT0000136 | Kdm5b | 75605 | HITS-CLIP | | 37986224 | Unspecified |
| mmu-miR-125b-5p | MIMAT0000136 | Kdm6a | 22289 | HITS-CLIP | 25083871\|23597149 | | Unspecified |
| mmu-miR-125b-5p | MIMAT0000136 | Kdm6b | 216850 | HITS-CLIP | | 37986224 | Unspecified |
| mmu-miR-125b-5p | MIMAT0000136 | Kdm7a | 338523 | HITS-CLIP | | 37986224 | Unspecified |
| mmu-miR-125b-5p | MIMAT0000136 | Kdsr | 70750 | Microarrays | 37986224\|22469780 | | Unspecified |
| mmu-miR-125b-5p | MIMAT0000136 | Khnyn | 219094 | HITS-CLIP | | 37986224 | Unspecified |
| mmu-miR-125b-5p | MIMAT0000136 | Kidins220 | 77480 | HITS-CLIP | | 25083871 | Unspecified |
| mmu-miR-125b-5p | MIMAT0000136 | Kif19a | 286942 | Microarrays | 37986224\|22469780 | | Unspecified |
| mmu-miR-125b-5p | MIMAT0000136 | Kif1a | 16560 | HITS-CLIP | 37986224\|19536157 | | Unspecified |
| mmu-miR-125b-5p | MIMAT0000136 | Kif1b | 16561 | HITS-CLIP | | 37986224 | Unspecified |
| mmu-miR-125b-5p | MIMAT0000136 | Kif1bp | 26128 | HITS-CLIP | | 25083871 | Unspecified |
| mmu-miR-125b-5p | MIMAT0000136 | Kif1c | 16562 | HITS-CLIP | | 37986224 | Unspecified |
| mmu-miR-125b-5p | MIMAT0000136 | Kif21b | 16565 | Microarrays | | 37986224 | Unspecified |
| mmu-miR-125b-5p | MIMAT0000136 | Kif24 | 109242 | HITS-CLIP | | 37986224 | Unspecified |
| mmu-miR-125b-5p | MIMAT0000136 | Kif26b | 269152 | HITS-CLIP | | 37986224 | Unspecified |
| mmu-miR-125b-5p | MIMAT0000136 | Kif3a | 16568 | Microarrays | | 37986224 | Unspecified |
| mmu-miR-125b-5p | MIMAT0000136 | Kif3b | 16569 | HITS-CLIP | | 25083871 | Unspecified |
| mmu-miR-125b-5p | MIMAT0000136 | Kif5a | 16572 | HITS-CLIP, Microarrays//HITS-CLIP | 37986224\|19536157 | | Unspecified |
| mmu-miR-125b-5p | MIMAT0000136 | Kif5c | 16574 | HITS-CLIP | | 25083871 | Unspecified |

| | | | | | | |
|---|---|---|---|---|---|---|
| mmu-miR-125b-5p | MIMAT0000136 | Kifc1 | 100502766 | Microarrays | 37986224 | Unspecified |
| mmu-miR-125b-5p | MIMAT0000136 | Kifc3 | 16582 | Microarrays | 37986224 | Unspecified |
| mmu-miR-125b-5p | MIMAT0000136 | Kirrel | 170643 | HITS-CLIP | 37986224 | Unspecified |
| mmu-miR-125b-5p | MIMAT0000136 | Klc2 | 16594 | HITS-CLIP | 37986224 | Unspecified |
| mmu-miR-125b-5p | MIMAT0000136 | Klc3 | 232943 | Microarrays | 37986224 | Unspecified |
| mmu-miR-125b-5p | MIMAT0000136 | Klf11 | 194655 | HITS-CLIP | 37986224 | Unspecified |
| mmu-miR-125b-5p | MIMAT0000136 | Klf14 | 619665 | HITS-CLIP | 37986224 | Unspecified |
| mmu-miR-125b-5p | MIMAT0000136 | Klf3 | 16599 | Microarrays//HITS-CLIP | 37986224\|25083871 | Unspecified |
| mmu-miR-125b-5p | MIMAT0000136 | Klf7 | 93691 | HITS-CLIP | 37986224\|25083871 | Unspecified |
| mmu-miR-125b-5p | MIMAT0000136 | Klf9 | 16601 | Microarrays | 37986224 | Unspecified |
| mmu-miR-125b-5p | MIMAT0000136 | Klhdc10 | 76788 | HITS-CLIP | 37986224 | Unspecified |
| mmu-miR-125b-5p | MIMAT0000136 | Klhdc3 | 71765 | HITS-CLIP | 25083871 | Unspecified |
| mmu-miR-125b-5p | MIMAT0000136 | Klhdc9 | 68874 | Microarrays | 37986224 | Unspecified |
| mmu-miR-125b-5p | MIMAT0000136 | Klhl20 | 226541 | Microarrays | 37986224 | Unspecified |
| mmu-miR-125b-5p | MIMAT0000136 | Klhl21 | 242785 | Microarrays | 37986224\|22469780 | Unspecified |
| mmu-miR-125b-5p | MIMAT0000136 | Klhl23 | 277396 | HITS-CLIP | 37986224 | Unspecified |
| mmu-miR-125b-5p | MIMAT0000136 | Klhl24 | 75785 | HITS-CLIP | 37986224 | Unspecified |
| mmu-miR-125b-5p | MIMAT0000136 | Klhl25 | 207952 | Microarrays | 37986224\|22469780 | Unspecified |
| mmu-miR-125b-5p | MIMAT0000136 | Klhl29 | 208439 | Microarrays | 37986224 | Unspecified |
| mmu-miR-125b-5p | MIMAT0000136 | Klhl32 | 212390 | Microarrays | 37986224\|22469780 | Unspecified |
| mmu-miR-125b-5p | MIMAT0000136 | Klhl36 | 234796 | HITS-CLIP | 37986224 | Unspecified |
| mmu-miR-125b-5p | MIMAT0000136 | Klhl40 | 72330 | HITS-CLIP | 25083871 | Unspecified |
| mmu-miR-125b-5p | MIMAT0000136 | Klhl9 | 242521 | Microarrays | 37986224\|22469780 | Unspecified |
| mmu-miR-125b-5p | MIMAT0000136 | Klra1 | 16627 | Microarrays | 37986224\|22469780 | Unspecified |
| mmu-miR-125b-5p | MIMAT0000136 | Klra4 | 16635 | Microarrays | 37986224\|22469780 | Unspecified |
| mmu-miR-125b-5p | MIMAT0000136 | Klra7 | 16638 | Microarrays | 37986224\|22469780 | Unspecified |
| mmu-miR-125b-5p | MIMAT0000136 | Klra9 | 16640 | Microarrays | 37986224\|22469780 | Unspecified |
| mmu-miR-125b-5p | MIMAT0000136 | Klrg2 | 74253 | qCLASH | 31393953 | Unspecified |
| mmu-miR-125b-5p | MIMAT0000136 | Kmt2a | 214162 | HITS-CLIP | 37986224\|25083871 | Unspecified |
| mmu-miR-125b-5p | MIMAT0000136 | Kmt2b | 75410 | HITS-CLIP | 37986224\|23597149\|19536157\|25083 | Unspecified |
| mmu-miR-125b-5p | MIMAT0000136 | Kmt2c | 231051 | HITS-CLIP | 37986224\|19536157\|25083871 | Unspecified |
| mmu-miR-125b-5p | MIMAT0000136 | Kmt2d | 381022 | HITS-CLIP | 37986224\|25083871 | Unspecified |
| mmu-miR-125b-5p | MIMAT0000136 | Kmt5b | 225888 | HITS-CLIP | 25083871 | Unspecified |
| mmu-miR-125b-5p | MIMAT0000136 | Kmt5c | 84787 | HITS-CLIP | 25083871 | Unspecified |
| mmu-miR-125b-5p | MIMAT0000136 | Knop1 | 66356 | HITS-CLIP, Microarrays//Microarrays | 37986224\|22469780 | Unspecified |
| mmu-miR-125b-5p | MIMAT0000136 | Kpna1 | 16646 | HITS-CLIP | 25083871 | Unspecified |
| mmu-miR-125b-5p | MIMAT0000136 | Kpna3 | 16648 | HITS-CLIP | 25083871\|25753418 | Unspecified |
| mmu-miR-125b-5p | MIMAT0000136 | Kpna4 | 16649 | HITS-CLIP, Microarrays//Microarrays | 37986224\|22469780 | Unspecified |
| mmu-miR-125b-5p | MIMAT0000136 | Kpna6 | 16650 | HITS-CLIP | 37986224\|25083871 | Unspecified |

| | | | | | | |
|---|---|---|---|---|---|---|
| mmu-miR-125b-5p | MIMAT0000136 | Krtap19-4 | 170654 | Microarrays | 37986224\|22469780 | Unspecified |
| mmu-miR-125b-5p | MIMAT0000136 | Krtap19-5 | 16704 | Microarrays | 37986224\|22469780 | Unspecified |
| mmu-miR-125b-5p | MIMAT0000136 | Kynu | 70789 | Microarrays | 37986224\|22469780 | Unspecified |
| mmu-miR-125b-5p | MIMAT0000136 | L1td1 | 381591 | Microarrays | 37986224 | Unspecified |
| mmu-miR-125b-5p | MIMAT0000136 | Lacc1 | 210808 | Microarrays | 37986224 | Unspecified |
| mmu-miR-125b-5p | MIMAT0000136 | Lactb | 80907 | HITS-CLIP, Microarrays//HITS-CLIP//Microa | 37986224\|25083871\|22469780\|19536 | Unspecified |
| mmu-miR-125b-5p | MIMAT0000136 | Lamb1 | 16777 | Microarrays | 37986224 | Unspecified |
| mmu-miR-125b-5p | MIMAT0000136 | Lamb2 | 16779 | Microarrays | 37986224 | Unspecified |
| mmu-miR-125b-5p | MIMAT0000136 | Lamc1 | 226519 | Microarrays//HITS-CLIP | 37986224\|25083871 | Unspecified |
| mmu-miR-125b-5p | MIMAT0000136 | Lamp2 | 16784 | HITS-CLIP | 37986224 | Unspecified |
| mmu-miR-125b-5p | MIMAT0000136 | Larp1 | 73158 | HITS-CLIP | 37986224\|25083871\|29167373 | Unspecified |
| mmu-miR-125b-5p | MIMAT0000136 | Larp4 | 207214 | HITS-CLIP | 37986224 | Unspecified |
| mmu-miR-125b-5p | MIMAT0000136 | Lars | 107045 | HITS-CLIP, Microarrays | 37986224 | Unspecified |
| mmu-miR-125b-5p | MIMAT0000136 | Lat | 16797 | Microarrays | 37986224 | Unspecified |
| mmu-miR-125b-5p | MIMAT0000136 | Lat2 | 56743 | Microarrays | 37986224 | Unspecified |
| mmu-miR-125b-5p | MIMAT0000136 | Lats1 | 16798 | Microarrays | 37986224\|22469780 | Unspecified |
| mmu-miR-125b-5p | MIMAT0000136 | Lats2 | 50523 | HITS-CLIP | 37986224 | Unspecified |
| mmu-miR-125b-5p | MIMAT0000136 | Lbh | 77889 | HITS-CLIP | 37986224 | Unspecified |
| mmu-miR-125b-5p | MIMAT0000136 | Lbx2 | 16815 | Microarrays | 37986224\|22469780 | Unspecified |
| mmu-miR-125b-5p | MIMAT0000136 | Lck | 16818 | Microarrays | 37986224\|22469780 | Unspecified |
| mmu-miR-125b-5p | MIMAT0000136 | Lcn10 | 332578 | Microarrays | 37986224\|22469780 | Unspecified |
| mmu-miR-125b-5p | MIMAT0000136 | Lcorl | 209707 | HITS-CLIP | 37986224 | Unspecified |
| mmu-miR-125b-5p | MIMAT0000136 | Ldb1 | 16825 | HITS-CLIP | 37986224 | Unspecified |
| mmu-miR-125b-5p | MIMAT0000136 | Ldb2 | 16826 | HITS-CLIP | 37986224 | Unspecified |
| mmu-miR-125b-5p | MIMAT0000136 | Ldha | 16828 | Microarrays | 37986224\|22469780 | Unspecified |
| mmu-miR-125b-5p | MIMAT0000136 | Ldlrad3 | 241576 | HITS-CLIP | 37986224\|25083871 | Unspecified |
| mmu-miR-125b-5p | MIMAT0000136 | Leng8 | 232798 | HITS-CLIP | 37986224 | Unspecified |
| mmu-miR-125b-5p | MIMAT0000136 | Leo1 | 235497 | Microarrays | 37986224\|22469780 | Unspecified |
| mmu-miR-125b-5p | MIMAT0000136 | Leprotl1 | 68192 | HITS-CLIP | 37986224\|25083871 | Unspecified |
| mmu-miR-125b-5p | MIMAT0000136 | Lfng | 16848 | HITS-CLIP | 37986224\|25083871\|23597149 | Unspecified |
| mmu-miR-125b-5p | MIMAT0000136 | Lgals1 | 16852 | Microarrays | 37986224 | Unspecified |
| mmu-miR-125b-5p | MIMAT0000136 | Lgals3bp | 19039 | Microarrays | 37986224 | Unspecified |
| mmu-miR-125b-5p | MIMAT0000136 | Lgals9 | 16859 | Microarrays | 37986224 | Unspecified |
| mmu-miR-125b-5p | MIMAT0000136 | Lgi3 | 213469 | HITS-CLIP | 19536157 | Unspecified |
| mmu-miR-125b-5p | MIMAT0000136 | Lgmn | 19141 | HITS-CLIP | 29167373 | Unspecified |
| mmu-miR-125b-5p | MIMAT0000136 | Lgr5 | 14160 | Microarrays | 37986224\|22469780 | Unspecified |
| mmu-miR-125b-5p | MIMAT0000136 | Lifr | 16880 | HITS-CLIP | 37986224\|23597149\|19536157 | Unspecified |
| mmu-miR-125b-5p | MIMAT0000136 | Limch1 | 77569 | Microarrays | 37986224\|22469780 | Unspecified |
| mmu-miR-125b-5p | MIMAT0000136 | Limd2 | 67803 | HITS-CLIP | 37986224 | Unspecified |

| | | | | | | |
|---|---|---|---|---|---|---|
| mmu-miR-125b-5p | MIMAT0000136 | Lin54 | 231506 | HITS-CLIP | 23597149 | Unspecified |
| mmu-miR-125b-5p | MIMAT0000136 | Lin7c | 22343 | HITS-CLIP | 25083871 | Unspecified |
| mmu-miR-125b-5p | MIMAT0000136 | Lipa | 16889 | Microarrays | 37986224\|22469780 | Unspecified |
| mmu-miR-125b-5p | MIMAT0000136 | Lipc | 15450 | HITS-CLIP, Microarrays//Microarrays | 37986224\|22469780 | Unspecified |
| mmu-miR-125b-5p | MIMAT0000136 | Lipt2 | 67164 | Microarrays | 37986224\|22469780 | Unspecified |
| mmu-miR-125b-5p | MIMAT0000136 | Lmbr1l | 74775 | Microarrays | 37986224\|22469780 | Unspecified |
| mmu-miR-125b-5p | MIMAT0000136 | Lmna | 16905 | HITS-CLIP, Microarrays | 37986224 | Unspecified |
| mmu-miR-125b-5p | MIMAT0000136 | Lmnb1 | 16906 | HITS-CLIP//qCLASH | 37986224\|31393953 | Unspecified |
| mmu-miR-125b-5p | MIMAT0000136 | Lmo2 | 16909 | Microarrays | 37986224 | Unspecified |
| mmu-miR-125b-5p | MIMAT0000136 | Lmo7 | 380928 | HITS-CLIP | 25083871 | Unspecified |
| mmu-miR-125b-5p | MIMAT0000136 | Lmtk2 | 231876 | HITS-CLIP | 37986224\|25753418 | Unspecified |
| mmu-miR-125b-5p | MIMAT0000136 | Lmtk3 | 381983 | Microarrays | 37986224 | Unspecified |
| mmu-miR-125b-5p | MIMAT0000136 | Lnpep | 240028 | HITS-CLIP, Microarrays//Microarrays | 37986224\|22469780 | Unspecified |
| mmu-miR-125b-5p | MIMAT0000136 | Lonp1 | 74142 | HITS-CLIP | 25083871 | Unspecified |
| mmu-miR-125b-5p | MIMAT0000136 | Lpar4 | 78134 | HITS-CLIP | 37986224 | Unspecified |
| mmu-miR-125b-5p | MIMAT0000136 | Lpar6 | 67168 | Microarrays//HITS-CLIP | 37986224\|23597149\|22469780 | Unspecified |
| mmu-miR-125b-5p | MIMAT0000136 | Lpcat1 | 210992 | HITS-CLIP | 37986224 | Unspecified |
| mmu-miR-125b-5p | MIMAT0000136 | Lpcat3 | 14792 | Microarrays | 37986224 | Unspecified |
| mmu-miR-125b-5p | MIMAT0000136 | Lpgat1 | 226856 | HITS-CLIP | 37986224 | Unspecified |
| mmu-miR-125b-5p | MIMAT0000136 | Lpin2 | 64898 | HITS-CLIP, Microarrays//HITS-CLIP | 37986224\|25083871\|29167373 | Unspecified |
| mmu-miR-125b-5p | MIMAT0000136 | Lpin3 | 64899 | HITS-CLIP | 37986224 | Unspecified |
| mmu-miR-125b-5p | MIMAT0000136 | Lpl | 16956 | HITS-CLIP | 25083871 | Unspecified |
| mmu-miR-125b-5p | MIMAT0000136 | Lpp | 210126 | HITS-CLIP, Microarrays//HITS-CLIP | 37986224\|25083871 | Unspecified |
| mmu-miR-125b-5p | MIMAT0000136 | Lrba | 80877 | HITS-CLIP | 37986224 | Unspecified |
| mmu-miR-125b-5p | MIMAT0000136 | Lrig1 | 16206 | HITS-CLIP//qCLASH | 37986224\|31393953\|23597149 | Unspecified |
| mmu-miR-125b-5p | MIMAT0000136 | Lrig3 | 320398 | HITS-CLIP | 25083871 | Unspecified |
| mmu-miR-125b-5p | MIMAT0000136 | Lrp4 | 228357 | HITS-CLIP | 37986224\|25083871 | Unspecified |
| mmu-miR-125b-5p | MIMAT0000136 | Lrr1 | 69706 | Microarrays | 37986224\|22469780 | Unspecified |
| mmu-miR-125b-5p | MIMAT0000136 | Lrrc20 | 216011 | Microarrays | 37986224 | Unspecified |
| mmu-miR-125b-5p | MIMAT0000136 | Lrrc30 | 240131 | HITS-CLIP | 37986224 | Unspecified |
| mmu-miR-125b-5p | MIMAT0000136 | Lrrc34 | 71827 | Microarrays | 37986224\|22469780 | Unspecified |
| mmu-miR-125b-5p | MIMAT0000136 | Lrrc41 | 230654 | HITS-CLIP | 25083871 | Unspecified |
| mmu-miR-125b-5p | MIMAT0000136 | Lrrc47 | 72946 | HITS-CLIP | 23597149 | Unspecified |
| mmu-miR-125b-5p | MIMAT0000136 | Lrrc59 | 98238 | HITS-CLIP | 29167373 | Unspecified |
| mmu-miR-125b-5p | MIMAT0000136 | Lrrc8a | 241296 | HITS-CLIP | 37986224\|25083871\|19536157\|23597 | Unspecified |
| mmu-miR-125b-5p | MIMAT0000136 | Lrrc8b | 433926 | HITS-CLIP | 37986224 | Unspecified |
| mmu-miR-125b-5p | MIMAT0000136 | Lrrc8c | 100604 | Microarrays | 37986224 | Unspecified |
| mmu-miR-125b-5p | MIMAT0000136 | Lrrc8d | 231549 | HITS-CLIP, Microarrays | 37986224 | Unspecified |
| mmu-miR-125b-5p | MIMAT0000136 | Lrrfip1 | 16978 | HITS-CLIP, Microarrays | 37986224 | Unspecified |

| | | | | | | |
|---|---|---|---|---|---|---|
| mmu-miR-125b-5p | MIMAT0000136 | Lrrk1 | 233328 | Microarrays | 37986224\|22469780 | Unspecified |
| mmu-miR-125b-5p | MIMAT0000136 | Lrrn3 | 16981 | Microarrays | 37986224\|22469780 | Unspecified |
| mmu-miR-125b-5p | MIMAT0000136 | Lrrtm1 | 74342 | Microarrays | 37986224 | Unspecified |
| mmu-miR-125b-5p | MIMAT0000136 | Lsm14b | 241846 | HITS-CLIP | 37986224 | Unspecified |
| mmu-miR-125b-5p | MIMAT0000136 | Lsm3 | 67678 | Microarrays | 37986224\|22469780 | Unspecified |
| mmu-miR-125b-5p | MIMAT0000136 | Lsm5 | 66373 | Microarrays | 37986224\|22469780 | Unspecified |
| mmu-miR-125b-5p | MIMAT0000136 | Lsm6 | 78651 | Microarrays | 37986224\|22469780 | Unspecified |
| mmu-miR-125b-5p | MIMAT0000136 | Lsm8 | 76522 | Microarrays | 37986224\|22469780 | Unspecified |
| mmu-miR-125b-5p | MIMAT0000136 | Lsp1 | 16985 | Microarrays | 37986224 | Unspecified |
| mmu-miR-125b-5p | MIMAT0000136 | Lss | 16987 | HITS-CLIP, Microarrays//HITS-CLIP | 37986224\|25083871 | Unspecified |
| mmu-miR-125b-5p | MIMAT0000136 | Ltb4r1 | 16995 | Microarrays | 37986224 | Unspecified |
| mmu-miR-125b-5p | MIMAT0000136 | Ltc4s | 17001 | Microarrays | 37986224\|22469780 | Unspecified |
| mmu-miR-125b-5p | MIMAT0000136 | Lurap1l | 52829 | HITS-CLIP | 37986224 | Unspecified |
| mmu-miR-125b-5p | MIMAT0000136 | Luzp1 | 269593 | HITS-CLIP | 37986224 | Unspecified |
| mmu-miR-125b-5p | MIMAT0000136 | Ly6e | 17069 | HITS-CLIP | 37986224 | Unspecified |
| mmu-miR-125b-5p | MIMAT0000136 | Ly6g6d | 114654 | Microarrays | 37986224\|22469780 | Unspecified |
| mmu-miR-125b-5p | MIMAT0000136 | Ly86 | 17084 | HITS-CLIP | 37986224 | Unspecified |
| mmu-miR-125b-5p | MIMAT0000136 | Ly96 | 17087 | Microarrays | 37986224\|22469780 | Unspecified |
| mmu-miR-125b-5p | MIMAT0000136 | Lyn | 17096 | Microarrays | 37986224\|22469780 | Unspecified |
| mmu-miR-125b-5p | MIMAT0000136 | Lypla2 | 26394 | HITS-CLIP | 37986224 | Unspecified |
| mmu-miR-125b-5p | MIMAT0000136 | Lysmd3 | 80289 | Microarrays | 37986224\|22469780 | Unspecified |
| mmu-miR-125b-5p | MIMAT0000136 | Lyst | 17101 | HITS-CLIP | 37986224\|25083871\|29167373 | Unspecified |
| mmu-miR-125b-5p | MIMAT0000136 | Lyz1 | 17110 | Microarrays | 37986224 | Unspecified |
| mmu-miR-125b-5p | MIMAT0000136 | Lyz2 | 17105 | Microarrays | 37986224 | Unspecified |
| mmu-miR-125b-5p | MIMAT0000136 | Lzts2 | 226154 | Microarrays | 37986224 | Unspecified |
| mmu-miR-125b-5p | MIMAT0000136 | M6pr | 17113 | Microarrays | 37986224\|22469780 | Unspecified |
| mmu-miR-125b-5p | MIMAT0000136 | Macf1 | 11426 | HITS-CLIP | 37986224\|19536157\|29167373 | Unspecified |
| mmu-miR-125b-5p | MIMAT0000136 | Macrod1 | 107227 | Microarrays | 37986224 | Unspecified |
| mmu-miR-125b-5p | MIMAT0000136 | Maf | 17132 | HITS-CLIP | 37986224 | Unspecified |
| mmu-miR-125b-5p | MIMAT0000136 | Maff | 17133 | Microarrays | 37986224 | Unspecified |
| mmu-miR-125b-5p | MIMAT0000136 | Maged1 | 94275 | HITS-CLIP | 37986224 | Unspecified |
| mmu-miR-125b-5p | MIMAT0000136 | Magee1 | 107528 | HITS-CLIP | 37986224 | Unspecified |
| mmu-miR-125b-5p | MIMAT0000136 | Magee2 | 272790 | Microarrays | 37986224\|22469780 | Unspecified |
| mmu-miR-125b-5p | MIMAT0000136 | Magoh | 17149 | Microarrays | 37986224\|22469780 | Unspecified |
| mmu-miR-125b-5p | MIMAT0000136 | Mamdc2 | 71738 | HITS-CLIP | 37986224 | Unspecified |
| mmu-miR-125b-5p | MIMAT0000136 | Man1a | 17155 | HITS-CLIP | 37986224 | Unspecified |
| mmu-miR-125b-5p | MIMAT0000136 | Man1a2 | 17156 | HITS-CLIP | 37986224 | Unspecified |
| mmu-miR-125b-5p | MIMAT0000136 | Man1b1 | 227619 | HITS-CLIP | 37986224\|25083871 | Unspecified |
| mmu-miR-125b-5p | MIMAT0000136 | Man1c1 | 230815 | HITS-CLIP | 25083871\|29167373 | Unspecified |

| | | | | | | |
|---|---|---|---|---|---|---|
| mmu-miR-125b-5p | MIMAT0000136 | Man2b2 | 17160 | Microarrays | 37986224 | Unspecified |
| mmu-miR-125b-5p | MIMAT0000136 | Maneal | 215090 | HITS-CLIP | 37986224 | Unspecified |
| mmu-miR-125b-5p | MIMAT0000136 | Map1a | 17754 | HITS-CLIP | 37986224 | Unspecified |
| mmu-miR-125b-5p | MIMAT0000136 | Map2k1 | 26395 | Microarrays | 37986224 | Unspecified |
| mmu-miR-125b-5p | MIMAT0000136 | Map2k3 | 26397 | HITS-CLIP | 37986224 | Unspecified |
| mmu-miR-125b-5p | MIMAT0000136 | Map2k4 | 26398 | HITS-CLIP | 37986224 | Unspecified |
| mmu-miR-125b-5p | MIMAT0000136 | Map2k7 | 26400 | B-globin reporter assay//Microarray//HITS | 16495412\|23597149 | Unspecified |
| mmu-miR-125b-5p | MIMAT0000136 | Map3k1 | 26401 | HITS-CLIP, Luciferase Reporter Assay//HITS | 37986224\|25083871\|23597149 | Unspecified |
| mmu-miR-125b-5p | MIMAT0000136 | Map3k11 | 26403 | HITS-CLIP, Luciferase Reporter Assay//HITS | 37986224\|23597149 | Unspecified |
| mmu-miR-125b-5p | MIMAT0000136 | Map3k2 | 26405 | HITS-CLIP | 37986224 | Unspecified |
| mmu-miR-125b-5p | MIMAT0000136 | Map4 | 17758 | HITS-CLIP | 37986224\|29167373 | Unspecified |
| mmu-miR-125b-5p | MIMAT0000136 | Map9 | 213582 | Microarrays | 37986224 | Unspecified |
| mmu-miR-125b-5p | MIMAT0000136 | Mapk1 | 26413 | HITS-CLIP | 25083871 | Unspecified |
| mmu-miR-125b-5p | MIMAT0000136 | Mapk14 | 26416 | HITS-CLIP, Luciferase Reporter Assay | 37986224 | Unspecified |
| mmu-miR-125b-5p | MIMAT0000136 | Mapk1ip1l | 218975 | HITS-CLIP | 37986224 | Unspecified |
| mmu-miR-125b-5p | MIMAT0000136 | Mapk3 | 26417 | Microarrays | 37986224 | Unspecified |
| mmu-miR-125b-5p | MIMAT0000136 | Mapk8 | 26419 | HITS-CLIP | 25083871 | Unspecified |
| mmu-miR-125b-5p | MIMAT0000136 | Mapk8ip2 | 60597 | HITS-CLIP | 37986224 | Unspecified |
| mmu-miR-125b-5p | MIMAT0000136 | Mapk8ip3 | 30957 | Microarrays | 37986224 | Unspecified |
| mmu-miR-125b-5p | MIMAT0000136 | Mapkap1 | 227743 | HITS-CLIP | 37986224 | Unspecified |
| mmu-miR-125b-5p | MIMAT0000136 | Mapt | 17762 | HITS-CLIP | 37986224\|19536157 | Unspecified |
| mmu-miR-125b-5p | MIMAT0000136 | March6 | 10299 | HITS-CLIP | 25083871 | Unspecified |
| mmu-miR-125b-5p | MIMAT0000136 | March7 | 57438 | HITS-CLIP | 37986224\|25083871 | Unspecified |
| mmu-miR-125b-5p | MIMAT0000136 | March8 | 71779 | HITS-CLIP | 37986224 | Unspecified |
| mmu-miR-125b-5p | MIMAT0000136 | March9 | 216438 | HITS-CLIP | 37986224 | Unspecified |
| mmu-miR-125b-5p | MIMAT0000136 | Marchf1 | 72925 | Microarrays | 37986224 | Unspecified |
| mmu-miR-125b-5p | MIMAT0000136 | Marcks | 17118 | HITS-CLIP, Microarrays//Microarrays | 37986224\|22469780 | Unspecified |
| mmu-miR-125b-5p | MIMAT0000136 | Marf1 | 223989 | HITS-CLIP | 37986224 | Unspecified |
| mmu-miR-125b-5p | MIMAT0000136 | Mark4 | 232944 | Microarrays | 37986224 | Unspecified |
| mmu-miR-125b-5p | MIMAT0000136 | Mars2 | 212679 | Microarrays | 37986224\|22469780 | Unspecified |
| mmu-miR-125b-5p | MIMAT0000136 | Marveld1 | 277010 | HITS-CLIP | 23597149 | Unspecified |
| mmu-miR-125b-5p | MIMAT0000136 | Masp2 | 17175 | HITS-CLIP | 23597149 | Unspecified |
| mmu-miR-125b-5p | MIMAT0000136 | Mast2 | 17776 | Microarrays | 37986224 | Unspecified |
| mmu-miR-125b-5p | MIMAT0000136 | Mat1a | 11720 | HITS-CLIP, Microarrays//Microarrays | 37986224\|22469780 | Unspecified |
| mmu-miR-125b-5p | MIMAT0000136 | Mat2a | 232087 | Microarrays//HITS-CLIP | 37986224\|22469780\|19536157 | Unspecified |
| mmu-miR-125b-5p | MIMAT0000136 | Mat2b | 108645 | Microarrays | 37986224\|22469780 | Unspecified |
| mmu-miR-125b-5p | MIMAT0000136 | Matn2 | 17181 | HITS-CLIP | 37986224 | Unspecified |
| mmu-miR-125b-5p | MIMAT0000136 | Mau2 | 74549 | HITS-CLIP | 37986224 | Unspecified |
| mmu-miR-125b-5p | MIMAT0000136 | Mbd1 | 17190 | HITS-CLIP | 25083871 | Unspecified |

| | | | | | | |
|---|---|---|---|---|---|---|
| mmu-miR-125b-5p | MIMAT0000136 | Mblac1 | 330216 | Microarrays | 37986224\|22469780 | Unspecified |
| mmu-miR-125b-5p | MIMAT0000136 | Mbnl3 | 171170 | Microarrays | 37986224\|22469780 | Unspecified |
| mmu-miR-125b-5p | MIMAT0000136 | Mboat7 | 77582 | HITS-CLIP | 37986224 | Unspecified |
| mmu-miR-125b-5p | MIMAT0000136 | Mbp | 17196 | HITS-CLIP | 37986224 | Unspecified |
| mmu-miR-125b-5p | MIMAT0000136 | Mbtps1 | 56453 | HITS-CLIP | 37986224 | Unspecified |
| mmu-miR-125b-5p | MIMAT0000136 | Mcidas | 622408 | Microarrays | 37986224 | Unspecified |
| mmu-miR-125b-5p | MIMAT0000136 | Mcl1 | 17210 | HITS-CLIP//qCLASH | 37986224\|25083871\|31393953\|23597 | Unspecified |
| mmu-miR-125b-5p | MIMAT0000136 | Mdn1 | 100019 | HITS-CLIP | 29167373 | Unspecified |
| mmu-miR-125b-5p | MIMAT0000136 | Mecp2 | 17257 | HITS-CLIP, Microarrays//HITS-CLIP | 37986224\|25083871 | Unspecified |
| mmu-miR-125b-5p | MIMAT0000136 | Med1 | 19014 | HITS-CLIP | 25083871 | Unspecified |
| mmu-miR-125b-5p | MIMAT0000136 | Med12 | 59024 | HITS-CLIP | 25083871 | Unspecified |
| mmu-miR-125b-5p | MIMAT0000136 | Med13 | 327987 | HITS-CLIP | 37986224\|25083871 | Unspecified |
| mmu-miR-125b-5p | MIMAT0000136 | Med13l | 76199 | Microarrays//HITS-CLIP | 37986224\|25083871 | Unspecified |
| mmu-miR-125b-5p | MIMAT0000136 | Med14 | 26896 | HITS-CLIP | 37986224\|25083871 | Unspecified |
| mmu-miR-125b-5p | MIMAT0000136 | Med15 | 94112 | HITS-CLIP | 37986224 | Unspecified |
| mmu-miR-125b-5p | MIMAT0000136 | Med17 | 234959 | HITS-CLIP | 25083871 | Unspecified |
| mmu-miR-125b-5p | MIMAT0000136 | Med25 | 75613 | Microarrays | 37986224 | Unspecified |
| mmu-miR-125b-5p | MIMAT0000136 | Med9 | 192191 | HITS-CLIP | 37986224 | Unspecified |
| mmu-miR-125b-5p | MIMAT0000136 | Mef2a | 17258 | HITS-CLIP | 37986224 | Unspecified |
| mmu-miR-125b-5p | MIMAT0000136 | Mef2c | 17260 | Microarrays | 37986224 | Unspecified |
| mmu-miR-125b-5p | MIMAT0000136 | Mef2d | 17261 | HITS-CLIP | 37986224 | Unspecified |
| mmu-miR-125b-5p | MIMAT0000136 | Megf10 | 70417 | Microarrays | 37986224\|22469780 | Unspecified |
| mmu-miR-125b-5p | MIMAT0000136 | Megf9 | 230316 | HITS-CLIP | 37986224\|19536157 | Unspecified |
| mmu-miR-125b-5p | MIMAT0000136 | Memo1 | 76890 | HITS-CLIP | 37986224 | Unspecified |
| mmu-miR-125b-5p | MIMAT0000136 | Mertk | 17289 | HITS-CLIP | 37986224 | Unspecified |
| mmu-miR-125b-5p | MIMAT0000136 | Mettl14 | 210529 | HITS-CLIP | 23597149 | Unspecified |
| mmu-miR-125b-5p | MIMAT0000136 | Mettl18 | 69962 | Microarrays | 37986224\|22469780 | Unspecified |
| mmu-miR-125b-5p | MIMAT0000136 | Mettl22 | 239706 | Microarrays | 37986224 | Unspecified |
| mmu-miR-125b-5p | MIMAT0000136 | Mettl25 | 216292 | Microarrays | 37986224\|22469780 | Unspecified |
| mmu-miR-125b-5p | MIMAT0000136 | Mettl5 | 75422 | Microarrays | 37986224\|22469780 | Unspecified |
| mmu-miR-125b-5p | MIMAT0000136 | Mettl7a1 | 70152 | HITS-CLIP | 37986224 | Unspecified |
| mmu-miR-125b-5p | MIMAT0000136 | Mettl7a2 | 393082 | Microarrays | 37986224 | Unspecified |
| mmu-miR-125b-5p | MIMAT0000136 | Mex3a | 72640 | HITS-CLIP | 37986224 | Unspecified |
| mmu-miR-125b-5p | MIMAT0000136 | Mfap1b | 100034361 | HITS-CLIP | 37986224 | Unspecified |
| mmu-miR-125b-5p | MIMAT0000136 | Mfap3l | 71306 | HITS-CLIP | 37986224 | Unspecified |
| mmu-miR-125b-5p | MIMAT0000136 | Mfhas1 | 52065 | HITS-CLIP | 37986224\|25083871\|19536157 | Unspecified |
| mmu-miR-125b-5p | MIMAT0000136 | Mfn1 | 67414 | HITS-CLIP | 25753418 | Unspecified |
| mmu-miR-125b-5p | MIMAT0000136 | Mfsd12 | 73822 | qCLASH | 31393953 | Unspecified |
| mmu-miR-125b-5p | MIMAT0000136 | Mfsd2a | 76574 | HITS-CLIP | 23597149\|25083871 | Unspecified |

| | | | | | | |
|---|---|---|---|---|---|---|
| mmu-miR-125b-5p | MIMAT0000136 | Mfsd9 | 211798 | HITS-CLIP//qCLASH | 37986224\|31393953 | Unspecified |
| mmu-miR-125b-5p | MIMAT0000136 | Mga | 29808 | HITS-CLIP | 37986224\|25083871\|19536157 | Unspecified |
| mmu-miR-125b-5p | MIMAT0000136 | Mgat3 | 17309 | HITS-CLIP | 37986224 | Unspecified |
| mmu-miR-125b-5p | MIMAT0000136 | Mgat4b | 103534 | Microarrays | 37986224 | Unspecified |
| mmu-miR-125b-5p | MIMAT0000136 | Mgst1 | 56615 | HITS-CLIP | 37986224 | Unspecified |
| mmu-miR-125b-5p | MIMAT0000136 | Mia3 | 338366 | HITS-CLIP | 37986224\|23597149 | Unspecified |
| mmu-miR-125b-5p | MIMAT0000136 | Mib1 | 225164 | HITS-CLIP | 37986224\|25083871 | Unspecified |
| mmu-miR-125b-5p | MIMAT0000136 | Mib2 | 76580 | Microarrays | 37986224 | Unspecified |
| mmu-miR-125b-5p | MIMAT0000136 | Mical2 | 320878 | HITS-CLIP | 37986224 | Unspecified |
| mmu-miR-125b-5p | MIMAT0000136 | Mical3 | 194401 | HITS-CLIP | 37986224 | Unspecified |
| mmu-miR-125b-5p | MIMAT0000136 | Micu1 | 216001 | Microarrays | 37986224 | Unspecified |
| mmu-miR-125b-5p | MIMAT0000136 | Micu2 | 68514 | HITS-CLIP | 37986224 | Unspecified |
| mmu-miR-125b-5p | MIMAT0000136 | Midn | 59090 | HITS-CLIP | 25083871\|23597149 | Unspecified |
| mmu-miR-125b-5p | MIMAT0000136 | Mink1 | 50932 | HITS-CLIP, Microarrays//HITS-CLIP | 37986224\|23597149 | Unspecified |
| mmu-miR-125b-5p | MIMAT0000136 | Mis12 | 67139 | HITS-CLIP | 37986224 | Unspecified |
| mmu-miR-125b-5p | MIMAT0000136 | Mitf | 17342 | HITS-CLIP | 37986224 | Unspecified |
| mmu-miR-125b-5p | MIMAT0000136 | Mixl1 | 27217 | Microarrays | 37986224 | Unspecified |
| mmu-miR-125b-5p | MIMAT0000136 | Mki67 | 17345 | HITS-CLIP | 37986224 | Unspecified |
| mmu-miR-125b-5p | MIMAT0000136 | Mknk2 | 17347 | HITS-CLIP | 37986224 | Unspecified |
| mmu-miR-125b-5p | MIMAT0000136 | Mlf2 | 30853 | HITS-CLIP | 37986224 | Unspecified |
| mmu-miR-125b-5p | MIMAT0000136 | Mlip | 69642 | HITS-CLIP | 37986224 | Unspecified |
| mmu-miR-125b-5p | MIMAT0000136 | Mllt6 | 246198 | HITS-CLIP | 25083871 | Unspecified |
| mmu-miR-125b-5p | MIMAT0000136 | Mlst8 | 56716 | HITS-CLIP | 37986224 | Unspecified |
| mmu-miR-125b-5p | MIMAT0000136 | Mlxip | 208104 | HITS-CLIP | 37986224 | Unspecified |
| mmu-miR-125b-5p | MIMAT0000136 | Mlycd | 56690 | HITS-CLIP | 37986224 | Unspecified |
| mmu-miR-125b-5p | MIMAT0000136 | Mmp13 | 17386 | Microarrays | 37986224 | Unspecified |
| mmu-miR-125b-5p | MIMAT0000136 | Mmp14 | 17387 | Microarrays | 37986224 | Unspecified |
| mmu-miR-125b-5p | MIMAT0000136 | Mmp15 | 17388 | HITS-CLIP | 25083871 | Unspecified |
| mmu-miR-125b-5p | MIMAT0000136 | Mmrn2 | 105450 | Microarrays | 37986224\|22469780 | Unspecified |
| mmu-miR-125b-5p | MIMAT0000136 | Mndal | 100040462 | Microarrays | 37986224 | Unspecified |
| mmu-miR-125b-5p | MIMAT0000136 | Mob1a | 232157 | HITS-CLIP | 37986224 | Unspecified |
| mmu-miR-125b-5p | MIMAT0000136 | Mogs | 57377 | Microarrays | 37986224\|22469780 | Unspecified |
| mmu-miR-125b-5p | MIMAT0000136 | Mon2 | 67074 | HITS-CLIP | 37986224\|23597149 | Unspecified |
| mmu-miR-125b-5p | MIMAT0000136 | Morc2a | 74522 | HITS-CLIP | 37986224 | Unspecified |
| mmu-miR-125b-5p | MIMAT0000136 | Morf4l2 | 56397 | HITS-CLIP | 37986224 | Unspecified |
| mmu-miR-125b-5p | MIMAT0000136 | Morn1 | 76866 | Microarrays | 37986224 | Unspecified |
| mmu-miR-125b-5p | MIMAT0000136 | Mov10 | 17454 | Microarrays | 37986224 | Unspecified |
| mmu-miR-125b-5p | MIMAT0000136 | Mpdu1 | 24070 | Microarrays | 37986224\|22469780 | Unspecified |
| mmu-miR-125b-5p | MIMAT0000136 | Mpeg1 | 17476 | Microarrays//HITS-CLIP | 37986224\|29167373 | Unspecified |

| | | | | | | |
|---|---|---|---|---|---|---|
| mmu-miR-125b-5p | MIMAT0000136 | Mphosph6 | 68533 | HITS-CLIP, Microarrays//Microarrays | 37986224\|22469780 | Unspecified |
| mmu-miR-125b-5p | MIMAT0000136 | Mprip | 26936 | HITS-CLIP | 25083871 | Unspecified |
| mmu-miR-125b-5p | MIMAT0000136 | Mpz | 17528 | Microarrays | 37986224\|22469780 | Unspecified |
| mmu-miR-125b-5p | MIMAT0000136 | Mr1 | 15064 | Microarrays | 37986224 | Unspecified |
| mmu-miR-125b-5p | MIMAT0000136 | Mrc1 | 17533 | Microarrays | 37986224 | Unspecified |
| mmu-miR-125b-5p | MIMAT0000136 | Mreg | 381269 | Microarrays | 37986224\|22469780 | Unspecified |
| mmu-miR-125b-5p | MIMAT0000136 | Mrgprb3 | 404238 | Microarrays | 37986224\|22469780 | Unspecified |
| mmu-miR-125b-5p | MIMAT0000136 | Mrpl16 | 94063 | Microarrays | 37986224\|22469780 | Unspecified |
| mmu-miR-125b-5p | MIMAT0000136 | Mrpl35 | 66223 | HITS-CLIP | 25753418 | Unspecified |
| mmu-miR-125b-5p | MIMAT0000136 | Mrpl37 | 56280 | HITS-CLIP | 25753418 | Unspecified |
| mmu-miR-125b-5p | MIMAT0000136 | Mrpl46 | 67308 | Microarrays | 37986224\|22469780 | Unspecified |
| mmu-miR-125b-5p | MIMAT0000136 | Mrps14 | 64659 | Microarrays | 37986224\|22469780 | Unspecified |
| mmu-miR-125b-5p | MIMAT0000136 | Mrps18c | 68735 | Microarrays | 37986224\|22469780 | Unspecified |
| mmu-miR-125b-5p | MIMAT0000136 | Mrtfa | 57591 | HITS-CLIP | 25083871 | Unspecified |
| mmu-miR-125b-5p | MIMAT0000136 | Ms4a4b | 60361 | Microarrays | 37986224\|22469780 | Unspecified |
| mmu-miR-125b-5p | MIMAT0000136 | Ms4a6c | 73656 | Microarrays | 37986224 | Unspecified |
| mmu-miR-125b-5p | MIMAT0000136 | Msantd2 | 235184 | Microarrays | 37986224\|22469780 | Unspecified |
| mmu-miR-125b-5p | MIMAT0000136 | Msl1 | 74026 | HITS-CLIP | 37986224 | Unspecified |
| mmu-miR-125b-5p | MIMAT0000136 | Msl2 | 77853 | HITS-CLIP, Microarrays//Microarrays | 37986224\|22469780 | Unspecified |
| mmu-miR-125b-5p | MIMAT0000136 | Msrb1 | 27361 | HITS-CLIP | 37986224 | Unspecified |
| mmu-miR-125b-5p | MIMAT0000136 | Msrb3 | 320183 | HITS-CLIP | 37986224\|25083871 | Unspecified |
| mmu-miR-125b-5p | MIMAT0000136 | Mt3 | 17751 | Microarrays | 37986224\|22469780 | Unspecified |
| mmu-miR-125b-5p | MIMAT0000136 | Mta1 | 116870 | Microarrays | 37986224 | Unspecified |
| mmu-miR-125b-5p | MIMAT0000136 | Mtch2 | 56428 | HITS-CLIP | 37986224 | Unspecified |
| mmu-miR-125b-5p | MIMAT0000136 | Mtdh | 67154 | Microarrays | 37986224 | Unspecified |
| mmu-miR-125b-5p | MIMAT0000136 | Mtfp1 | 67900 | Microarrays | 37986224\|22469780 | Unspecified |
| mmu-miR-125b-5p | MIMAT0000136 | Mtfr1l | 76824 | HITS-CLIP | 25083871 | Unspecified |
| mmu-miR-125b-5p | MIMAT0000136 | Mthfd2 | 17768 | Microarrays | 37986224 | Unspecified |
| mmu-miR-125b-5p | MIMAT0000136 | Mtmr3 | 74302 | HITS-CLIP | 37986224 | Unspecified |
| mmu-miR-125b-5p | MIMAT0000136 | Mtr | 238505 | HITS-CLIP | 37986224\|25753418 | Unspecified |
| mmu-miR-125b-5p | MIMAT0000136 | Mtrex | 23517 | HITS-CLIP | 25753418 | Unspecified |
| mmu-miR-125b-5p | MIMAT0000136 | Mtss1 | 211401 | HITS-CLIP | 37986224 | Unspecified |
| mmu-miR-125b-5p | MIMAT0000136 | Mtus1 | 102103 | HITS-CLIP | 37986224 | Unspecified |
| mmu-miR-125b-5p | MIMAT0000136 | Mtx3 | 382793 | HITS-CLIP | 37986224 | Unspecified |
| mmu-miR-125b-5p | MIMAT0000136 | Mug1 | 17836 | HITS-CLIP | 37986224 | Unspecified |
| mmu-miR-125b-5p | MIMAT0000136 | Mvd | 192156 | Microarrays | 37986224 | Unspecified |
| mmu-miR-125b-5p | MIMAT0000136 | Mx1 | 17857 | Microarrays | 37986224 | Unspecified |
| mmu-miR-125b-5p | MIMAT0000136 | Mxd4 | 17122 | HITS-CLIP, Microarrays | 37986224 | Unspecified |
| mmu-miR-125b-5p | MIMAT0000136 | Mxi1 | 17859 | Microarrays | 37986224\|22469780 | Unspecified |

| | | | | | | |
|---|---|---|---|---|---|---|
| mmu-miR-125b-5p | MIMAT0000136 | Mxra8 | 74761 | Microarrays | 37986224 | Unspecified |
| mmu-miR-125b-5p | MIMAT0000136 | Mybbp1a | 18432 | HITS-CLIP | 37986224 | Unspecified |
| mmu-miR-125b-5p | MIMAT0000136 | Mybl1 | 17864 | HITS-CLIP | 25083871 | Unspecified |
| mmu-miR-125b-5p | MIMAT0000136 | Mybl2 | 17865 | Microarrays | 37986224\|22469780 | Unspecified |
| mmu-miR-125b-5p | MIMAT0000136 | Myc | 17869 | Microarrays | 37986224 | Unspecified |
| mmu-miR-125b-5p | MIMAT0000136 | Mycbp2 | 105689 | HITS-CLIP | 37986224\|25083871\|29167373 | Unspecified |
| mmu-miR-125b-5p | MIMAT0000136 | Myh1 | 17879 | HITS-CLIP | 25083871 | Unspecified |
| mmu-miR-125b-5p | MIMAT0000136 | Myh10 | 77579 | Microarrays | 37986224\|22469780 | Unspecified |
| mmu-miR-125b-5p | MIMAT0000136 | Myl4 | 17896 | Microarrays | 37986224\|22469780 | Unspecified |
| mmu-miR-125b-5p | MIMAT0000136 | Myl6 | 17904 | Microarrays | 37986224\|22469780 | Unspecified |
| mmu-miR-125b-5p | MIMAT0000136 | Myl6b | 216459 | Microarrays | 37986224 | Unspecified |
| mmu-miR-125b-5p | MIMAT0000136 | Myo10 | 17909 | HITS-CLIP | 37986224\|25083871 | Unspecified |
| mmu-miR-125b-5p | MIMAT0000136 | Myo18a | 360013 | HITS-CLIP, Microarrays | 37986224 | Unspecified |
| mmu-miR-125b-5p | MIMAT0000136 | Myo18b | 74376 | HITS-CLIP | 25083871 | Unspecified |
| mmu-miR-125b-5p | MIMAT0000136 | Myo1c | 17913 | HITS-CLIP | 37986224\|25753418 | Unspecified |
| mmu-miR-125b-5p | MIMAT0000136 | Myo1e | 71602 | HITS-CLIP | 25753418\|29167373 | Unspecified |
| mmu-miR-125b-5p | MIMAT0000136 | Myo1f | 17916 | Microarrays | 37986224 | Unspecified |
| mmu-miR-125b-5p | MIMAT0000136 | Myo9a | 270163 | HITS-CLIP | 37986224\|25083871\|23597149 | Unspecified |
| mmu-miR-125b-5p | MIMAT0000136 | Myom1 | 17929 | Microarrays | 37986224 | Unspecified |
| mmu-miR-125b-5p | MIMAT0000136 | Myom3 | 242702 | Microarrays | 37986224 | Unspecified |
| mmu-miR-125b-5p | MIMAT0000136 | Mzb1 | 69816 | Microarrays | 37986224\|22469780 | Unspecified |
| mmu-miR-125b-5p | MIMAT0000136 | Naa15 | 74838 | HITS-CLIP | 37986224 | Unspecified |
| mmu-miR-125b-5p | MIMAT0000136 | Naa40 | 70999 | HITS-CLIP, Microarrays//Microarrays | 37986224\|22469780 | Unspecified |
| mmu-miR-125b-5p | MIMAT0000136 | Naalad2 | 72560 | Microarrays | 37986224\|22469780 | Unspecified |
| mmu-miR-125b-5p | MIMAT0000136 | Nab1 | 17936 | Microarrays | 37986224 | Unspecified |
| mmu-miR-125b-5p | MIMAT0000136 | Naca | 17938 | HITS-CLIP | 37986224 | Unspecified |
| mmu-miR-125b-5p | MIMAT0000136 | Nacc1 | 66830 | HITS-CLIP | 29167373 | Unspecified |
| mmu-miR-125b-5p | MIMAT0000136 | Naga | 17939 | Microarrays | 37986224 | Unspecified |
| mmu-miR-125b-5p | MIMAT0000136 | Naif1 | 71254 | HITS-CLIP | 25083871 | Unspecified |
| mmu-miR-125b-5p | MIMAT0000136 | Nampt | 59027 | HITS-CLIP, Microarrays | 37986224 | Unspecified |
| mmu-miR-125b-5p | MIMAT0000136 | Nanp | 67311 | Microarrays | 37986224\|22469780 | Unspecified |
| mmu-miR-125b-5p | MIMAT0000136 | Napg | 108123 | HITS-CLIP | 37986224 | Unspecified |
| mmu-miR-125b-5p | MIMAT0000136 | Narf | 67608 | HITS-CLIP, Microarrays//Microarrays | 37986224\|22469780 | Unspecified |
| mmu-miR-125b-5p | MIMAT0000136 | Nars | 70223 | Microarrays//qCLASH | 37986224\|31393953 | Unspecified |
| mmu-miR-125b-5p | MIMAT0000136 | Nat8l | 269642 | HITS-CLIP | 37986224 | Unspecified |
| mmu-miR-125b-5p | MIMAT0000136 | Natd1 | 24083 | Microarrays | 22469780 | Unspecified |
| mmu-miR-125b-5p | MIMAT0000136 | Nav1 | 215690 | HITS-CLIP | 37986224 | Unspecified |
| mmu-miR-125b-5p | MIMAT0000136 | Nbeal1 | 269198 | HITS-CLIP | 37986224\|23597149 | Unspecified |
| mmu-miR-125b-5p | MIMAT0000136 | Ncan | 13004 | HITS-CLIP | 37986224\|19536157 | Unspecified |

| | | | | | | |
|---|---|---|---|---|---|---|
| mmu-miR-125b-5p | MIMAT0000136 | Ncapg2 | 76044 | HITS-CLIP | 37986224 | Unspecified |
| mmu-miR-125b-5p | MIMAT0000136 | Ncbp1 | 433702 | HITS-CLIP | 37986224 | Unspecified |
| mmu-miR-125b-5p | MIMAT0000136 | Ncbp3 | 66874 | HITS-CLIP | 25083871 | Unspecified |
| mmu-miR-125b-5p | MIMAT0000136 | Ncf1 | 17969 | Microarrays | 37986224 | Unspecified |
| mmu-miR-125b-5p | MIMAT0000136 | Nck2 | 17974 | HITS-CLIP, Microarrays//HITS-CLIP//Microa | 37986224\|25083871\|22469780 | Unspecified |
| mmu-miR-125b-5p | MIMAT0000136 | Nckap5l | 380969 | HITS-CLIP | 37986224\|25083871 | Unspecified |
| mmu-miR-125b-5p | MIMAT0000136 | Ncl | 17975 | HITS-CLIP | 37986224 | Unspecified |
| mmu-miR-125b-5p | MIMAT0000136 | Ncln | 103425 | HITS-CLIP | 37986224\|25083871 | Unspecified |
| mmu-miR-125b-5p | MIMAT0000136 | Ncoa1 | 17977 | HITS-CLIP, Microarrays | 37986224 | Unspecified |
| mmu-miR-125b-5p | MIMAT0000136 | Ncoa3 | 17979 | HITS-CLIP | 37986224\|23597149 | Unspecified |
| mmu-miR-125b-5p | MIMAT0000136 | Ncor2 | 20602 | HITS-CLIP | 37986224 | Unspecified |
| mmu-miR-125b-5p | MIMAT0000136 | Ncstn | 59287 | HITS-CLIP | 37986224 | Unspecified |
| mmu-miR-125b-5p | MIMAT0000136 | Ndel1 | 83431 | HITS-CLIP | 37986224 | Unspecified |
| mmu-miR-125b-5p | MIMAT0000136 | Ndn | 17984 | HITS-CLIP | 37986224 | Unspecified |
| mmu-miR-125b-5p | MIMAT0000136 | Ndor1 | 78797 | HITS-CLIP | 37986224 | Unspecified |
| mmu-miR-125b-5p | MIMAT0000136 | Ndrg1 | 17988 | Microarrays | 37986224 | Unspecified |
| mmu-miR-125b-5p | MIMAT0000136 | Ndst1 | 15531 | HITS-CLIP | 37986224 | Unspecified |
| mmu-miR-125b-5p | MIMAT0000136 | Ndufaf8 | 284184 | Microarrays | 22469780 | Unspecified |
| mmu-miR-125b-5p | MIMAT0000136 | Ndufb2 | 68198 | Microarrays | 37986224\|22469780 | Unspecified |
| mmu-miR-125b-5p | MIMAT0000136 | Ndufc1 | 66377 | Microarrays | 37986224\|22469780 | Unspecified |
| mmu-miR-125b-5p | MIMAT0000136 | Ndufs3 | 68349 | qCLASH//HITS-CLIP | 31393953\|25753418 | Unspecified |
| mmu-miR-125b-5p | MIMAT0000136 | Ndufs4 | 17993 | HITS-CLIP | 37986224\|25083871 | Unspecified |
| mmu-miR-125b-5p | MIMAT0000136 | Nectin3 | 58998 | HITS-CLIP | 23597149 | Unspecified |
| mmu-miR-125b-5p | MIMAT0000136 | Nectin4 | 71740 | Microarrays | 22469780 | Unspecified |
| mmu-miR-125b-5p | MIMAT0000136 | Nedd8 | 18002 | HITS-CLIP | 37986224 | Unspecified |
| mmu-miR-125b-5p | MIMAT0000136 | Nek2 | 18005 | HITS-CLIP | 37986224 | Unspecified |
| mmu-miR-125b-5p | MIMAT0000136 | Nek6 | 59126 | Microarrays | 37986224 | Unspecified |
| mmu-miR-125b-5p | MIMAT0000136 | Nek7 | 59125 | HITS-CLIP | 37986224 | Unspecified |
| mmu-miR-125b-5p | MIMAT0000136 | Nek8 | 140859 | Microarrays | 37986224 | Unspecified |
| mmu-miR-125b-5p | MIMAT0000136 | Nemp1 | 23306 | HITS-CLIP//Microarrays | 25083871\|22469780 | Unspecified |
| mmu-miR-125b-5p | MIMAT0000136 | Neo1 | 18007 | HITS-CLIP | 37986224 | Unspecified |
| mmu-miR-125b-5p | MIMAT0000136 | Nes | 18008 | HITS-CLIP | 37986224 | Unspecified |
| mmu-miR-125b-5p | MIMAT0000136 | Neto2 | 74513 | HITS-CLIP | 37986224 | Unspecified |
| mmu-miR-125b-5p | MIMAT0000136 | Neu1 | 18010 | Microarrays | 37986224\|22469780 | Unspecified |
| mmu-miR-125b-5p | MIMAT0000136 | Neurl1b | 240055 | HITS-CLIP | 37986224 | Unspecified |
| mmu-miR-125b-5p | MIMAT0000136 | Nexn | 68810 | Microarrays | 37986224 | Unspecified |
| mmu-miR-125b-5p | MIMAT0000136 | Nf1 | 18015 | HITS-CLIP | 37986224\|25083871 | Unspecified |
| mmu-miR-125b-5p | MIMAT0000136 | Nfat5 | 54446 | HITS-CLIP | 25753418 | Unspecified |
| mmu-miR-125b-5p | MIMAT0000136 | Nfe2 | 18022 | Microarrays | 37986224 | Unspecified |

| | | | | | | |
|---|---|---|---|---|---|---|
| mmu-miR-125b-5p | MIMAT0000136 | Nfe2l1 | 18023 | HITS-CLIP | 37986224 | Unspecified |
| mmu-miR-125b-5p | MIMAT0000136 | Nfib | 18028 | HITS-CLIP | 37986224 | Unspecified |
| mmu-miR-125b-5p | MIMAT0000136 | Nfic | 18029 | Microarrays//HITS-CLIP | 37986224\|25083871\|29167373 | Unspecified |
| mmu-miR-125b-5p | MIMAT0000136 | Nfix | 18032 | HITS-CLIP | 37986224 | Unspecified |
| mmu-miR-125b-5p | MIMAT0000136 | Nfkb2 | 18034 | Microarrays | 37986224 | Unspecified |
| mmu-miR-125b-5p | MIMAT0000136 | Nfkbid | 243910 | Microarrays | 37986224 | Unspecified |
| mmu-miR-125b-5p | MIMAT0000136 | Nfx1 | 74164 | HITS-CLIP | 25083871\|25753418 | Unspecified |
| mmu-miR-125b-5p | MIMAT0000136 | Ngly1 | 59007 | HITS-CLIP | 37986224 | Unspecified |
| mmu-miR-125b-5p | MIMAT0000136 | Nhsl1 | 215819 | HITS-CLIP | 25083871 | Unspecified |
| mmu-miR-125b-5p | MIMAT0000136 | Nicn1 | 66257 | HITS-CLIP | 37986224 | Unspecified |
| mmu-miR-125b-5p | MIMAT0000136 | Nifk | 67949 | Microarrays | 37986224\|22469780 | Unspecified |
| mmu-miR-125b-5p | MIMAT0000136 | Nin | 18080 | HITS-CLIP | 37986224 | Unspecified |
| mmu-miR-125b-5p | MIMAT0000136 | Ninl | 78177 | HITS-CLIP | 37986224\|25083871 | Unspecified |
| mmu-miR-125b-5p | MIMAT0000136 | Nipa2 | 93790 | Microarrays | 37986224\|22469780 | Unspecified |
| mmu-miR-125b-5p | MIMAT0000136 | Nipal3 | 74552 | HITS-CLIP | 37986224 | Unspecified |
| mmu-miR-125b-5p | MIMAT0000136 | Nisch | 64652 | HITS-CLIP | 37986224\|25753418 | Unspecified |
| mmu-miR-125b-5p | MIMAT0000136 | Nkap | 67050 | HITS-CLIP | 37986224\|25083871 | Unspecified |
| mmu-miR-125b-5p | MIMAT0000136 | Nlgn2 | 216856 | HITS-CLIP, Microarrays | 37986224 | Unspecified |
| mmu-miR-125b-5p | MIMAT0000136 | Nmd3 | 97112 | HITS-CLIP | 25083871 | Unspecified |
| mmu-miR-125b-5p | MIMAT0000136 | Nmt1 | 18107 | HITS-CLIP | 37986224 | Unspecified |
| mmu-miR-125b-5p | MIMAT0000136 | Nnat | 18111 | HITS-CLIP | 37986224 | Unspecified |
| mmu-miR-125b-5p | MIMAT0000136 | Noc3l | 57753 | HITS-CLIP | 37986224 | Unspecified |
| mmu-miR-125b-5p | MIMAT0000136 | Nod1 | 107607 | Microarrays | 37986224 | Unspecified |
| mmu-miR-125b-5p | MIMAT0000136 | Nog | 18121 | Microarrays | 37986224\|22469780 | Unspecified |
| mmu-miR-125b-5p | MIMAT0000136 | Nol9 | 74035 | HITS-CLIP | 37986224 | Unspecified |
| mmu-miR-125b-5p | MIMAT0000136 | Nono | 53610 | HITS-CLIP | 25083871 | Unspecified |
| mmu-miR-125b-5p | MIMAT0000136 | Nop14 | 75416 | Microarrays | 37986224 | Unspecified |
| mmu-miR-125b-5p | MIMAT0000136 | Nos1 | 18125 | HITS-CLIP | 37986224 | Unspecified |
| mmu-miR-125b-5p | MIMAT0000136 | Notch2 | 18129 | HITS-CLIP | 25083871 | Unspecified |
| mmu-miR-125b-5p | MIMAT0000136 | Noxo1 | 71893 | Microarrays | 37986224 | Unspecified |
| mmu-miR-125b-5p | MIMAT0000136 | Noxred1 | 71275 | Microarrays | 37986224 | Unspecified |
| mmu-miR-125b-5p | MIMAT0000136 | Npat | 244879 | HITS-CLIP | 37986224 | Unspecified |
| mmu-miR-125b-5p | MIMAT0000136 | Npc1 | 18145 | HITS-CLIP | 25083871 | Unspecified |
| mmu-miR-125b-5p | MIMAT0000136 | Npl | 74091 | Microarrays | 37986224\|22469780 | Unspecified |
| mmu-miR-125b-5p | MIMAT0000136 | Nprl3 | 17168 | Microarrays | 37986224 | Unspecified |
| mmu-miR-125b-5p | MIMAT0000136 | Nptx1 | 18164 | HITS-CLIP | 37986224 | Unspecified |
| mmu-miR-125b-5p | MIMAT0000136 | Npvf | 60531 | Microarrays | 37986224\|22469780 | Unspecified |
| mmu-miR-125b-5p | MIMAT0000136 | Npy4r | 19065 | Microarrays | 37986224\|22469780 | Unspecified |
| mmu-miR-125b-5p | MIMAT0000136 | Nr1d1 | 217166 | HITS-CLIP | 37986224\|25083871 | Unspecified |

| | | | | | | |
|---|---|---|---|---|---|---|
| mmu-miR-125b-5p | MIMAT0000136 | Nr1h5 | 381463 | Microarrays | 37986224 | Unspecified |
| mmu-miR-125b-5p | MIMAT0000136 | Nr4a1 | 15370 | Microarrays | 37986224 | Unspecified |
| mmu-miR-125b-5p | MIMAT0000136 | Nr6a1 | 14536 | HITS-CLIP | 37986224 | Unspecified |
| mmu-miR-125b-5p | MIMAT0000136 | Nrde2 | 217827 | HITS-CLIP | 25083871 | Unspecified |
| mmu-miR-125b-5p | MIMAT0000136 | Nrm | 106582 | HITS-CLIP, Microarrays//Microarrays | 37986224\|22469780 | Unspecified |
| mmu-miR-125b-5p | MIMAT0000136 | Nrn1 | 68404 | HITS-CLIP | 37986224 | Unspecified |
| mmu-miR-125b-5p | MIMAT0000136 | Nrp1 | 18186 | HITS-CLIP, Microarrays | 37986224 | Unspecified |
| mmu-miR-125b-5p | MIMAT0000136 | Nrsn2 | 228777 | HITS-CLIP | 37986224 | Unspecified |
| mmu-miR-125b-5p | MIMAT0000136 | Nrxn2 | 18190 | Microarrays | 37986224 | Unspecified |
| mmu-miR-125b-5p | MIMAT0000136 | Nscme3l | 75555 | Microarrays | 22469780 | Unspecified |
| mmu-miR-125b-5p | MIMAT0000136 | Nsd1 | 18193 | HITS-CLIP | 37986224\|29167373 | Unspecified |
| mmu-miR-125b-5p | MIMAT0000136 | Nsd2 | 107823 | HITS-CLIP | 25083871\|23597149 | Unspecified |
| mmu-miR-125b-5p | MIMAT0000136 | Nsg2 | 18197 | Microarrays | 37986224 | Unspecified |
| mmu-miR-125b-5p | MIMAT0000136 | Nsmaf | 18201 | HITS-CLIP | 37986224\|25083871 | Unspecified |
| mmu-miR-125b-5p | MIMAT0000136 | Nt5dc1 | 319638 | HITS-CLIP, Microarrays//Microarrays | 37986224\|22469780 | Unspecified |
| mmu-miR-125b-5p | MIMAT0000136 | Nt5dc3 | 103466 | HITS-CLIP | 37986224 | Unspecified |
| mmu-miR-125b-5p | MIMAT0000136 | Nub1 | 53312 | HITS-CLIP | 37986224 | Unspecified |
| mmu-miR-125b-5p | MIMAT0000136 | Nucb1 | 18220 | HITS-CLIP | 37986224 | Unspecified |
| mmu-miR-125b-5p | MIMAT0000136 | Nudcd2 | 52653 | HITS-CLIP//qCLASH | 37986224\|31393953\|19536157 | Unspecified |
| mmu-miR-125b-5p | MIMAT0000136 | Nudt18 | 213484 | HITS-CLIP | 37986224 | Unspecified |
| mmu-miR-125b-5p | MIMAT0000136 | Nudt3 | 56409 | Microarrays | 37986224\|22469780 | Unspecified |
| mmu-miR-125b-5p | MIMAT0000136 | Nudt5 | 53893 | Microarrays | 37986224\|22469780 | Unspecified |
| mmu-miR-125b-5p | MIMAT0000136 | Nufip2 | 68564 | HITS-CLIP | 25083871 | Unspecified |
| mmu-miR-125b-5p | MIMAT0000136 | Numbl | 18223 | HITS-CLIP | 37986224 | Unspecified |
| mmu-miR-125b-5p | MIMAT0000136 | Nup153 | 218210 | HITS-CLIP | 37986224\|25083871 | Unspecified |
| mmu-miR-125b-5p | MIMAT0000136 | Nup160 | 59015 | HITS-CLIP | 37986224 | Unspecified |
| mmu-miR-125b-5p | MIMAT0000136 | Nup210 | 54563 | Microarrays | 37986224\|22469780 | Unspecified |
| mmu-miR-125b-5p | MIMAT0000136 | Nup214 | 227720 | HITS-CLIP | 37986224\|25083871\|19536157\|29167 | Unspecified |
| mmu-miR-125b-5p | MIMAT0000136 | Nup50 | 18141 | HITS-CLIP | 37986224 | Unspecified |
| mmu-miR-125b-5p | MIMAT0000136 | Nupr1 | 56312 | Microarrays | 37986224 | Unspecified |
| mmu-miR-125b-5p | MIMAT0000136 | Nxpe4 | 244853 | Microarrays | 37986224\|22469780 | Unspecified |
| mmu-miR-125b-5p | MIMAT0000136 | Nxph4 | 104080 | Microarrays | 37986224 | Unspecified |
| mmu-miR-125b-5p | MIMAT0000136 | Nxt1 | 56488 | Microarrays | 37986224\|22469780 | Unspecified |
| mmu-miR-125b-5p | MIMAT0000136 | Nxt2 | 237082 | HITS-CLIP, Microarrays//Microarrays | 37986224\|22469780 | Unspecified |
| mmu-miR-125b-5p | MIMAT0000136 | Oaf | 102644 | HITS-CLIP | 23597149 | Unspecified |
| mmu-miR-125b-5p | MIMAT0000136 | Oas2 | 246728 | Microarrays | 37986224 | Unspecified |
| mmu-miR-125b-5p | MIMAT0000136 | Oasl1 | 231655 | Microarrays | 37986224 | Unspecified |
| mmu-miR-125b-5p | MIMAT0000136 | Oasl2 | 23962 | Microarrays | 37986224 | Unspecified |
| mmu-miR-125b-5p | MIMAT0000136 | Oaz2 | 18247 | HITS-CLIP | 37986224 | Unspecified |

| | | | | | | | |
|---|---|---|---|---|---|---|---|
| mmu-miR-125b-5p | MIMAT0000136 | Obp2a | 227627 | Microarrays | | 37986224 | Unspecified |
| mmu-miR-125b-5p | MIMAT0000136 | Obsl1 | 98733 | Microarrays | | 37986224 | Unspecified |
| mmu-miR-125b-5p | MIMAT0000136 | Odf2 | 18286 | HITS-CLIP | | 37986224 | Unspecified |
| mmu-miR-125b-5p | MIMAT0000136 | Odf4 | 252868 | Microarrays | 37986224\|22469780 | | Unspecified |
| mmu-miR-125b-5p | MIMAT0000136 | Ogg1 | 18294 | Microarrays | 37986224\|22469780 | | Unspecified |
| mmu-miR-125b-5p | MIMAT0000136 | Ogt | 108155 | HITS-CLIP | | 25753418 | Unspecified |
| mmu-miR-125b-5p | MIMAT0000136 | Olfr1016 | 257915 | Microarrays | | 37986224 | Unspecified |
| mmu-miR-125b-5p | MIMAT0000136 | Olfr1033 | 258571 | Microarrays | | 37986224 | Unspecified |
| mmu-miR-125b-5p | MIMAT0000136 | Olfr1094 | 258362 | Microarrays | | 37986224 | Unspecified |
| mmu-miR-125b-5p | MIMAT0000136 | Olfr1121 | 258345 | Microarrays | | 37986224 | Unspecified |
| mmu-miR-125b-5p | MIMAT0000136 | Olfr1178 | 258203 | Microarrays | | 37986224 | Unspecified |
| mmu-miR-125b-5p | MIMAT0000136 | Olfr1188 | 258921 | Microarrays | | 37986224 | Unspecified |
| mmu-miR-125b-5p | MIMAT0000136 | Olfr1239 | 258972 | Microarrays | | 37986224 | Unspecified |
| mmu-miR-125b-5p | MIMAT0000136 | Olfr1305 | 258396 | Microarrays | | 37986224 | Unspecified |
| mmu-miR-125b-5p | MIMAT0000136 | Olfr1443 | 258693 | Microarrays | | 37986224 | Unspecified |
| mmu-miR-125b-5p | MIMAT0000136 | Olfr18 | 18315 | Microarrays | | 37986224 | Unspecified |
| mmu-miR-125b-5p | MIMAT0000136 | Olfr187 | 258319 | Microarrays | | 37986224 | Unspecified |
| mmu-miR-125b-5p | MIMAT0000136 | Olfr198 | 258036 | Microarrays | | 37986224 | Unspecified |
| mmu-miR-125b-5p | MIMAT0000136 | Olfr221 | 258420 | Microarrays | | 37986224 | Unspecified |
| mmu-miR-125b-5p | MIMAT0000136 | Olfr235 | 258681 | Microarrays | | 37986224 | Unspecified |
| mmu-miR-125b-5p | MIMAT0000136 | Olfr294 | 257904 | Microarrays | | 37986224 | Unspecified |
| mmu-miR-125b-5p | MIMAT0000136 | Olfr299 | 257929 | Microarrays | | 37986224 | Unspecified |
| mmu-miR-125b-5p | MIMAT0000136 | Olfr361 | 258365 | Microarrays | | 37986224 | Unspecified |
| mmu-miR-125b-5p | MIMAT0000136 | Olfr455 | 546896 | Microarrays | | 37986224 | Unspecified |
| mmu-miR-125b-5p | MIMAT0000136 | Olfr544 | 257926 | Microarrays | | 37986224 | Unspecified |
| mmu-miR-125b-5p | MIMAT0000136 | Olfr593 | 258378 | Microarrays | | 37986224 | Unspecified |
| mmu-miR-125b-5p | MIMAT0000136 | Olfr726 | 258313 | Microarrays | | 37986224 | Unspecified |
| mmu-miR-125b-5p | MIMAT0000136 | Olfr763 | 258861 | Microarrays | | 37986224 | Unspecified |
| mmu-miR-125b-5p | MIMAT0000136 | Olfr781 | 258723 | Microarrays | | 37986224 | Unspecified |
| mmu-miR-125b-5p | MIMAT0000136 | Olfr827 | 258297 | Microarrays | | 37986224 | Unspecified |
| mmu-miR-125b-5p | MIMAT0000136 | Olfr862 | 258555 | Microarrays | | 37986224 | Unspecified |
| mmu-miR-125b-5p | MIMAT0000136 | Olfr870 | 57251 | Microarrays | | 37986224 | Unspecified |
| mmu-miR-125b-5p | MIMAT0000136 | Olfr875 | 258744 | Microarrays | | 37986224 | Unspecified |
| mmu-miR-125b-5p | MIMAT0000136 | Olfr969 | 258823 | Microarrays | | 37986224 | Unspecified |
| mmu-miR-125b-5p | MIMAT0000136 | Omd | 27047 | Microarrays | | 37986224 | Unspecified |
| mmu-miR-125b-5p | MIMAT0000136 | Opa1 | 74143 | HITS-CLIP | | 37986224 | Unspecified |
| mmu-miR-125b-5p | MIMAT0000136 | Or10ac1 | 546896 | Microarrays | | 22469780 | Unspecified |
| mmu-miR-125b-5p | MIMAT0000136 | Or10p21 | 258861 | Microarrays | | 22469780 | Unspecified |
| mmu-miR-125b-5p | MIMAT0000136 | Or12k8 | 258365 | Microarrays | | 22469780 | Unspecified |

| | | | | | | | |
|---|---|---|---|---|---|---|---|
| mmu-miR-125b-5p | MIMAT0000136 | Or14a256 | 257904 | Microarrays | | 22469780 | Unspecified |
| mmu-miR-125b-5p | MIMAT0000136 | Or4a2 | 258972 | Microarrays | | 22469780 | Unspecified |
| mmu-miR-125b-5p | MIMAT0000136 | Or4c101 | 258921 | Microarrays | | 22469780 | Unspecified |
| mmu-miR-125b-5p | MIMAT0000136 | Or4f14b | 257956 | qCLASH | | 31393953 | Unspecified |
| mmu-miR-125b-5p | MIMAT0000136 | Or4p7 | 258203 | Microarrays | | 22469780 | Unspecified |
| mmu-miR-125b-5p | MIMAT0000136 | Or52s1 | 258378 | Microarrays | | 22469780 | Unspecified |
| mmu-miR-125b-5p | MIMAT0000136 | Or55b4 | 257926 | Microarrays | | 22469780 | Unspecified |
| mmu-miR-125b-5p | MIMAT0000136 | Or5ac16 | 258036 | Microarrays | | 22469780 | Unspecified |
| mmu-miR-125b-5p | MIMAT0000136 | Or5b95 | 258693 | Microarrays | | 22469780 | Unspecified |
| mmu-miR-125b-5p | MIMAT0000136 | Or5h19 | 258319 | Microarrays | | 22469780 | Unspecified |
| mmu-miR-125b-5p | MIMAT0000136 | Or5m3b | 258571 | Microarrays | | 22469780 | Unspecified |
| mmu-miR-125b-5p | MIMAT0000136 | Or5t9 | 258362 | Microarrays | | 22469780 | Unspecified |
| mmu-miR-125b-5p | MIMAT0000136 | Or6c35 | 258723 | Microarrays | | 22469780 | Unspecified |
| mmu-miR-125b-5p | MIMAT0000136 | Or8g54 | 258823 | Microarrays | | 22469780 | Unspecified |
| mmu-miR-125b-5p | MIMAT0000136 | Orc2 | 18393 | HITS-CLIP | 37986224\|25083871 | | Unspecified |
| mmu-miR-125b-5p | MIMAT0000136 | Orc5 | 26429 | HITS-CLIP | | 29167373 | Unspecified |
| mmu-miR-125b-5p | MIMAT0000136 | Osbpl2 | 228983 | Microarrays | | 37986224 | Unspecified |
| mmu-miR-125b-5p | MIMAT0000136 | Osbpl9 | 100273 | HITS-CLIP | 37986224\|23597149 | | Unspecified |
| mmu-miR-125b-5p | MIMAT0000136 | Oscar | 232790 | Microarrays | 37986224\|22469780 | | Unspecified |
| mmu-miR-125b-5p | MIMAT0000136 | Osgin2 | 209212 | Microarrays | 37986224\|22469780 | | Unspecified |
| mmu-miR-125b-5p | MIMAT0000136 | Osr1 | 23967 | Microarrays | 37986224\|22469780 | | Unspecified |
| mmu-miR-125b-5p | MIMAT0000136 | Ostm1 | 14628 | HITS-CLIP | | 37986224 | Unspecified |
| mmu-miR-125b-5p | MIMAT0000136 | Otub2 | 68149 | Microarrays | | 37986224 | Unspecified |
| mmu-miR-125b-5p | MIMAT0000136 | Otx1 | 18423 | Microarrays | | 37986224 | Unspecified |
| mmu-miR-125b-5p | MIMAT0000136 | Ovca2 | 246257 | Microarrays | 37986224\|22469780 | | Unspecified |
| mmu-miR-125b-5p | MIMAT0000136 | Oxct1 | 67041 | HITS-CLIP | | 25753418 | Unspecified |
| mmu-miR-125b-5p | MIMAT0000136 | P2rx3 | 228139 | Microarrays | | 37986224 | Unspecified |
| mmu-miR-125b-5p | MIMAT0000136 | P2rx4 | 18438 | Microarrays | 37986224\|22469780 | | Unspecified |
| mmu-miR-125b-5p | MIMAT0000136 | P2rx7 | 18439 | HITS-CLIP | | 37986224 | Unspecified |
| mmu-miR-125b-5p | MIMAT0000136 | P2ry10b | 213438 | Microarrays | | 22469780 | Unspecified |
| mmu-miR-125b-5p | MIMAT0000136 | P2ry6 | 233571 | Microarrays | | 37986224 | Unspecified |
| mmu-miR-125b-5p | MIMAT0000136 | P3h1 | 64175 | Microarrays | | 22469780 | Unspecified |
| mmu-miR-125b-5p | MIMAT0000136 | Pa2g4 | 18813 | qCLASH | | 31393953 | Unspecified |
| mmu-miR-125b-5p | MIMAT0000136 | Pabpc1 | 18458 | HITS-CLIP, Microarrays//HITS-CLIP | 37986224\|23597149\|25083871\|29167 | | Unspecified |
| mmu-miR-125b-5p | MIMAT0000136 | Pacrgl | 66768 | Microarrays | | 37986224 | Unspecified |
| mmu-miR-125b-5p | MIMAT0000136 | Pacs1 | 107975 | Microarrays | | 37986224 | Unspecified |
| mmu-miR-125b-5p | MIMAT0000136 | Pacs2 | 217893 | HITS-CLIP | | 37986224 | Unspecified |
| mmu-miR-125b-5p | MIMAT0000136 | Pafah1b1 | 18472 | HITS-CLIP | 37986224\|19536157\|29167373 | | Unspecified |
| mmu-miR-125b-5p | MIMAT0000136 | Pah | 18478 | HITS-CLIP | | 23597149 | Unspecified |

| | | | | | | |
|---|---|---|---|---|---|---|
| mmu-miR-125b-5p | MIMAT0000136 | Paip2b | 232164 | HITS-CLIP | 37986224 | Unspecified |
| mmu-miR-125b-5p | MIMAT0000136 | Pak3 | 18481 | HITS-CLIP | 37986224 | Unspecified |
| mmu-miR-125b-5p | MIMAT0000136 | Pak6 | 214230 | HITS-CLIP | 37986224 | Unspecified |
| mmu-miR-125b-5p | MIMAT0000136 | Pakap | 677884 | HITS-CLIP | 25753418 | Unspecified |
| mmu-miR-125b-5p | MIMAT0000136 | Pald1 | 27355 | HITS-CLIP | 37986224 | Unspecified |
| mmu-miR-125b-5p | MIMAT0000136 | Palm | 18483 | Microarrays | 37986224 | Unspecified |
| mmu-miR-125b-5p | MIMAT0000136 | Pam | 18484 | Microarrays | 37986224 | Unspecified |
| mmu-miR-125b-5p | MIMAT0000136 | Pamr1 | 210622 | Microarrays | 37986224 | Unspecified |
| mmu-miR-125b-5p | MIMAT0000136 | Pan2 | 103135 | HITS-CLIP | 37986224\|25083871 | Unspecified |
| mmu-miR-125b-5p | MIMAT0000136 | Pank2 | 74450 | HITS-CLIP, Microarrays//Microarrays | 37986224\|22469780 | Unspecified |
| mmu-miR-125b-5p | MIMAT0000136 | Papola | 18789 | HITS-CLIP | 37986224 | Unspecified |
| mmu-miR-125b-5p | MIMAT0000136 | Parp1 | 11545 | HITS-CLIP | 37986224 | Unspecified |
| mmu-miR-125b-5p | MIMAT0000136 | Parp14 | 547253 | Microarrays | 37986224 | Unspecified |
| mmu-miR-125b-5p | MIMAT0000136 | Parp16 | 214424 | HITS-CLIP | 23597149 | Unspecified |
| mmu-miR-125b-5p | MIMAT0000136 | Parvg | 64099 | Microarrays | 37986224 | Unspecified |
| mmu-miR-125b-5p | MIMAT0000136 | Patl1 | 225929 | Microarrays | 37986224 | Unspecified |
| mmu-miR-125b-5p | MIMAT0000136 | Pbrm1 | 66923 | HITS-CLIP | 25083871 | Unspecified |
| mmu-miR-125b-5p | MIMAT0000136 | Pbx1 | 18514 | HITS-CLIP | 37986224 | Unspecified |
| mmu-miR-125b-5p | MIMAT0000136 | Pbxip1 | 229534 | HITS-CLIP | 25083871 | Unspecified |
| mmu-miR-125b-5p | MIMAT0000136 | Pcbp2 | 18521 | HITS-CLIP | 37986224 | Unspecified |
| mmu-miR-125b-5p | MIMAT0000136 | Pcbp4 | 59092 | HITS-CLIP | 37986224 | Unspecified |
| mmu-miR-125b-5p | MIMAT0000136 | Pcdh1 | 75599 | HITS-CLIP | 37986224\|19536157 | Unspecified |
| mmu-miR-125b-5p | MIMAT0000136 | Pcdh17 | 219228 | HITS-CLIP | 37986224 | Unspecified |
| mmu-miR-125b-5p | MIMAT0000136 | Pcdh7 | 54216 | HITS-CLIP | 37986224 | Unspecified |
| mmu-miR-125b-5p | MIMAT0000136 | Pcdhb22 | 93893 | HITS-CLIP, Microarrays//Microarrays | 37986224\|22469780 | Unspecified |
| mmu-miR-125b-5p | MIMAT0000136 | Pcdhb7 | 93878 | HITS-CLIP | 37986224 | Unspecified |
| mmu-miR-125b-5p | MIMAT0000136 | Pcdhga12 | 93724 | HITS-CLIP | 37986224\|25083871 | Unspecified |
| mmu-miR-125b-5p | MIMAT0000136 | Pcf11 | 74737 | HITS-CLIP | 37986224 | Unspecified |
| mmu-miR-125b-5p | MIMAT0000136 | Pcgf2 | 22658 | Microarrays | 37986224 | Unspecified |
| mmu-miR-125b-5p | MIMAT0000136 | Pcgf6 | 71041 | HITS-CLIP | 37986224\|25083871 | Unspecified |
| mmu-miR-125b-5p | MIMAT0000136 | Pck1 | 18534 | Microarrays | 37986224\|22469780 | Unspecified |
| mmu-miR-125b-5p | MIMAT0000136 | Pck2 | 74551 | Microarrays | 37986224 | Unspecified |
| mmu-miR-125b-5p | MIMAT0000136 | Pclaf | 68026 | Microarrays | 22469780 | Unspecified |
| mmu-miR-125b-5p | MIMAT0000136 | Pcmtd2 | 245867 | HITS-CLIP | 37986224 | Unspecified |
| mmu-miR-125b-5p | MIMAT0000136 | Pcna | 18538 | Microarrays | 37986224\|22469780 | Unspecified |
| mmu-miR-125b-5p | MIMAT0000136 | Pcnx | 54604 | HITS-CLIP | 25083871 | Unspecified |
| mmu-miR-125b-5p | MIMAT0000136 | Pcsk7 | 18554 | HITS-CLIP | 37986224 | Unspecified |
| mmu-miR-125b-5p | MIMAT0000136 | Pctp | 18559 | HITS-CLIP | 37986224\|23597149\|29167373 | Unspecified |
| mmu-miR-125b-5p | MIMAT0000136 | Pcx | 18563 | Microarrays | 37986224 | Unspecified |

| | | | | | | |
|---|---|---|---|---|---|---|
| mmu-miR-125b-5p | MIMAT0000136 | Pcyox1 | 66881 | HITS-CLIP | 37986224 | Unspecified |
| mmu-miR-125b-5p | MIMAT0000136 | Pcyt1a | 13026 | Microarrays | 37986224 | Unspecified |
| mmu-miR-125b-5p | MIMAT0000136 | Pcyt1b | 236899 | Microarrays | 37986224 | Unspecified |
| mmu-miR-125b-5p | MIMAT0000136 | Pdcd4 | 18569 | Microarrays | 37986224 | Unspecified |
| mmu-miR-125b-5p | MIMAT0000136 | Pdcd6ip | 18571 | Microarrays//HITS-CLIP | 37986224\|25083871 | Unspecified |
| mmu-miR-125b-5p | MIMAT0000136 | Pde1b | 18574 | Microarrays | 37986224 | Unspecified |
| mmu-miR-125b-5p | MIMAT0000136 | Pde4a | 18577 | Microarrays | 37986224 | Unspecified |
| mmu-miR-125b-5p | MIMAT0000136 | Pde4b | 18578 | Microarrays | 37986224 | Unspecified |
| mmu-miR-125b-5p | MIMAT0000136 | Pde4dip | 83679 | HITS-CLIP | 25083871 | Unspecified |
| mmu-miR-125b-5p | MIMAT0000136 | Pde5a | 242202 | HITS-CLIP | 25753418 | Unspecified |
| mmu-miR-125b-5p | MIMAT0000136 | Pde6d | 18582 | Microarrays | 37986224 | Unspecified |
| mmu-miR-125b-5p | MIMAT0000136 | Pde6g | 18588 | Microarrays | 37986224 | Unspecified |
| mmu-miR-125b-5p | MIMAT0000136 | Pde8a | 18584 | HITS-CLIP | 37986224 | Unspecified |
| mmu-miR-125b-5p | MIMAT0000136 | Pde9a | 18585 | HITS-CLIP | 37986224\|23597149 | Unspecified |
| mmu-miR-125b-5p | MIMAT0000136 | Pdf | 68023 | Microarrays | 37986224\|22469780 | Unspecified |
| mmu-miR-125b-5p | MIMAT0000136 | Pdgfrb | 18596 | HITS-CLIP | 37986224 | Unspecified |
| mmu-miR-125b-5p | MIMAT0000136 | Pdhx | 27402 | Microarrays | 37986224\|22469780 | Unspecified |
| mmu-miR-125b-5p | MIMAT0000136 | Pdk2 | 18604 | HITS-CLIP | 37986224 | Unspecified |
| mmu-miR-125b-5p | MIMAT0000136 | Pds5a | 71521 | HITS-CLIP | 37986224 | Unspecified |
| mmu-miR-125b-5p | MIMAT0000136 | Pdxk | 216134 | HITS-CLIP | 37986224 | Unspecified |
| mmu-miR-125b-5p | MIMAT0000136 | Pdzd4 | 245469 | Microarrays | 37986224 | Unspecified |
| mmu-miR-125b-5p | MIMAT0000136 | Pea15a | 18611 | Microarrays | 37986224 | Unspecified |
| mmu-miR-125b-5p | MIMAT0000136 | Peak1 | 244895 | HITS-CLIP | 37986224 | Unspecified |
| mmu-miR-125b-5p | MIMAT0000136 | Peli2 | 93834 | HITS-CLIP | 37986224\|25083871 | Unspecified |
| mmu-miR-125b-5p | MIMAT0000136 | Pes1 | 64934 | Microarrays | 37986224\|22469780 | Unspecified |
| mmu-miR-125b-5p | MIMAT0000136 | Pex14 | 56273 | Microarrays | 37986224\|22469780 | Unspecified |
| mmu-miR-125b-5p | MIMAT0000136 | Pf4 | 56744 | Microarrays | 37986224\|22469780 | Unspecified |
| mmu-miR-125b-5p | MIMAT0000136 | Pfkl | 18641 | Microarrays//HITS-CLIP | 37986224\|25083871 | Unspecified |
| mmu-miR-125b-5p | MIMAT0000136 | Pfkm | 18642 | Microarrays//HITS-CLIP | 37986224\|25753418 | Unspecified |
| mmu-miR-125b-5p | MIMAT0000136 | Pfn1 | 18643 | Microarrays | 37986224 | Unspecified |
| mmu-miR-125b-5p | MIMAT0000136 | Pgap2 | 233575 | Microarrays | 37986224 | Unspecified |
| mmu-miR-125b-5p | MIMAT0000136 | Pgap3 | 320655 | Microarrays | 37986224\|22469780 | Unspecified |
| mmu-miR-125b-5p | MIMAT0000136 | Pgrmc2 | 70804 | HITS-CLIP | 37986224 | Unspecified |
| mmu-miR-125b-5p | MIMAT0000136 | Phactr1 | 218194 | HITS-CLIP | 37986224 | Unspecified |
| mmu-miR-125b-5p | MIMAT0000136 | Phc3 | 241915 | HITS-CLIP | 37986224 | Unspecified |
| mmu-miR-125b-5p | MIMAT0000136 | Pheta1 | 144717 | Microarrays | 22469780 | Unspecified |
| mmu-miR-125b-5p | MIMAT0000136 | Phf20 | 228829 | HITS-CLIP | 37986224\|25083871\|23597149 | Unspecified |
| mmu-miR-125b-5p | MIMAT0000136 | Phf3 | 213109 | HITS-CLIP | 37986224 | Unspecified |
| mmu-miR-125b-5p | MIMAT0000136 | Phf8 | 320595 | HITS-CLIP | 37986224 | Unspecified |

| | | | | | | | |
|---|---|---|---|---|---|---|---|
| mmu-miR-125b-5p | MIMAT0000136 | Phip | 83946 | HITS-CLIP | | 25083871 | Unspecified |
| mmu-miR-125b-5p | MIMAT0000136 | Phlda1 | 21664 | Microarrays | | 37986224 | Unspecified |
| mmu-miR-125b-5p | MIMAT0000136 | Phldb2 | 208177 | HITS-CLIP | | 37986224 | Unspecified |
| mmu-miR-125b-5p | MIMAT0000136 | Phlpp1 | 98432 | Microarrays | | 37986224 | Unspecified |
| mmu-miR-125b-5p | MIMAT0000136 | Phospho2 | 73373 | HITS-CLIP | | 37986224 | Unspecified |
| mmu-miR-125b-5p | MIMAT0000136 | Pi4k2b | 67073 | HITS-CLIP | | 25083871 | Unspecified |
| mmu-miR-125b-5p | MIMAT0000136 | Pi4kb | 107650 | HITS-CLIP, Microarrays//Microarrays | 37986224\|22469780 | | Unspecified |
| mmu-miR-125b-5p | MIMAT0000136 | Pid1 | 98496 | HITS-CLIP | | 37986224 | Unspecified |
| mmu-miR-125b-5p | MIMAT0000136 | Piezo1 | 234839 | Microarrays | | 37986224 | Unspecified |
| mmu-miR-125b-5p | MIMAT0000136 | Piezo2 | 667742 | HITS-CLIP | | 37986224 | Unspecified |
| mmu-miR-125b-5p | MIMAT0000136 | Pigg | 433931 | Microarrays | | 37986224 | Unspecified |
| mmu-miR-125b-5p | MIMAT0000136 | Pigm | 67556 | HITS-CLIP | | 37986224 | Unspecified |
| mmu-miR-125b-5p | MIMAT0000136 | Pigo | 56703 | Microarrays | 37986224\|22469780 | | Unspecified |
| mmu-miR-125b-5p | MIMAT0000136 | Pigr | 18703 | Microarrays | | 37986224 | Unspecified |
| mmu-miR-125b-5p | MIMAT0000136 | Pigt | 78928 | Microarrays | 37986224\|22469780 | | Unspecified |
| mmu-miR-125b-5p | MIMAT0000136 | Pigw | 70325 | Microarrays | 37986224\|22469780 | | Unspecified |
| mmu-miR-125b-5p | MIMAT0000136 | Pik3c2b | 240752 | HITS-CLIP | | 37986224 | Unspecified |
| mmu-miR-125b-5p | MIMAT0000136 | Pik3ca | 18706 | HITS-CLIP | | 37986224 | Unspecified |
| mmu-miR-125b-5p | MIMAT0000136 | Pik3cb | 74769 | HITS-CLIP | | 23597149 | Unspecified |
| mmu-miR-125b-5p | MIMAT0000136 | Pik3cg | 30955 | Microarrays | | 37986224 | Unspecified |
| mmu-miR-125b-5p | MIMAT0000136 | Pik3ip1 | 216505 | HITS-CLIP | | 25753418 | Unspecified |
| mmu-miR-125b-5p | MIMAT0000136 | Pik3r2 | 18709 | HITS-CLIP, Microarrays | | 37986224 | Unspecified |
| mmu-miR-125b-5p | MIMAT0000136 | Pik3r3 | 18710 | HITS-CLIP | | 37986224 | Unspecified |
| mmu-miR-125b-5p | MIMAT0000136 | Pikfyve | 18711 | HITS-CLIP | | 37986224 | Unspecified |
| mmu-miR-125b-5p | MIMAT0000136 | Pinx1 | 72400 | Microarrays | | 37986224 | Unspecified |
| mmu-miR-125b-5p | MIMAT0000136 | Pip4k2a | 18718 | Microarrays | | 37986224 | Unspecified |
| mmu-miR-125b-5p | MIMAT0000136 | Pip4k2b | 108083 | HITS-CLIP, Microarrays | | 37986224 | Unspecified |
| mmu-miR-125b-5p | MIMAT0000136 | Pipox | 19193 | Microarrays | 37986224\|22469780 | | Unspecified |
| mmu-miR-125b-5p | MIMAT0000136 | Pitpna | 18738 | HITS-CLIP | 37986224\|29167373 | | Unspecified |
| mmu-miR-125b-5p | MIMAT0000136 | Pitpnm1 | 18739 | Microarrays | | 37986224 | Unspecified |
| mmu-miR-125b-5p | MIMAT0000136 | Pja2 | 224938 | HITS-CLIP | | 37986224 | Unspecified |
| mmu-miR-125b-5p | MIMAT0000136 | Pkd1 | 18763 | HITS-CLIP, Microarrays | | 37986224 | Unspecified |
| mmu-miR-125b-5p | MIMAT0000136 | Pkd1l3 | 244646 | Microarrays | | 37986224 | Unspecified |
| mmu-miR-125b-5p | MIMAT0000136 | Pkhd1 | 241035 | HITS-CLIP | | 37986224 | Unspecified |
| mmu-miR-125b-5p | MIMAT0000136 | Pklr | 18770 | Microarrays | | 37986224 | Unspecified |
| mmu-miR-125b-5p | MIMAT0000136 | Pkm | 18746 | HITS-CLIP, Microarrays | | 37986224 | Unspecified |
| mmu-miR-125b-5p | MIMAT0000136 | Pkn1 | 320795 | HITS-CLIP | | 25083871 | Unspecified |
| mmu-miR-125b-5p | MIMAT0000136 | Pkp4 | 227937 | Microarrays | | 37986224 | Unspecified |
| mmu-miR-125b-5p | MIMAT0000136 | Pla2g15 | 192654 | HITS-CLIP, Microarrays | | 37986224 | Unspecified |

| | | | | | | |
|---|---|---|---|---|---|---|
| mmu-miR-125b-5p | MIMAT0000136 | Pla2g2e | 26970 | Microarrays | 37986224\|22469780 | Unspecified |
| mmu-miR-125b-5p | MIMAT0000136 | Pla2g4b | 211429 | Microarrays | 37986224\|22469780 | Unspecified |
| mmu-miR-125b-5p | MIMAT0000136 | Pla2g6 | 53357 | Microarrays | 37986224 | Unspecified |
| mmu-miR-125b-5p | MIMAT0000136 | Plagl2 | 54711 | HITS-CLIP | 25083871\|19536157 | Unspecified |
| mmu-miR-125b-5p | MIMAT0000136 | Plat | 18791 | HITS-CLIP | 25753418 | Unspecified |
| mmu-miR-125b-5p | MIMAT0000136 | Plau | 18792 | Microarrays | 37986224 | Unspecified |
| mmu-miR-125b-5p | MIMAT0000136 | Plbd2 | 71772 | HITS-CLIP | 37986224 | Unspecified |
| mmu-miR-125b-5p | MIMAT0000136 | Plcb4 | 18798 | HITS-CLIP | 37986224 | Unspecified |
| mmu-miR-125b-5p | MIMAT0000136 | Plcd1 | 18799 | HITS-CLIP | 37986224 | Unspecified |
| mmu-miR-125b-5p | MIMAT0000136 | Plcxd2 | 433022 | HITS-CLIP | 37986224\|25083871 | Unspecified |
| mmu-miR-125b-5p | MIMAT0000136 | Pld4 | 104759 | Microarrays | 37986224\|22469780 | Unspecified |
| mmu-miR-125b-5p | MIMAT0000136 | Plec | 18810 | HITS-CLIP | 37986224\|29167373 | Unspecified |
| mmu-miR-125b-5p | MIMAT0000136 | Plek | 56193 | Microarrays | 37986224 | Unspecified |
| mmu-miR-125b-5p | MIMAT0000136 | Plekha5 | 109135 | Microarrays | 37986224 | Unspecified |
| mmu-miR-125b-5p | MIMAT0000136 | Plekhb2 | 226971 | HITS-CLIP | 37986224 | Unspecified |
| mmu-miR-125b-5p | MIMAT0000136 | Plekhm2 | 69582 | HITS-CLIP | 37986224 | Unspecified |
| mmu-miR-125b-5p | MIMAT0000136 | Plekhm3 | 241075 | HITS-CLIP | 37986224\|29167373 | Unspecified |
| mmu-miR-125b-5p | MIMAT0000136 | Plekho2 | 102595 | Microarrays | 37986224 | Unspecified |
| mmu-miR-125b-5p | MIMAT0000136 | Plin3 | 66905 | Microarrays | 37986224 | Unspecified |
| mmu-miR-125b-5p | MIMAT0000136 | Plk2 | 20620 | HITS-CLIP | 37986224 | Unspecified |
| mmu-miR-125b-5p | MIMAT0000136 | Plk3 | 12795 | HITS-CLIP | 37986224 | Unspecified |
| mmu-miR-125b-5p | MIMAT0000136 | Plod2 | 26432 | HITS-CLIP | 37986224\|25083871 | Unspecified |
| mmu-miR-125b-5p | MIMAT0000136 | Plpp1 | 8611 | HITS-CLIP | 25083871 | Unspecified |
| mmu-miR-125b-5p | MIMAT0000136 | Plpp7 | 227721 | HITS-CLIP | 25083871 | Unspecified |
| mmu-miR-125b-5p | MIMAT0000136 | Plscr3 | 70310 | Microarrays | 37986224 | Unspecified |
| mmu-miR-125b-5p | MIMAT0000136 | Plscr4 | 235527 | Microarrays | 37986224 | Unspecified |
| mmu-miR-125b-5p | MIMAT0000136 | Pltp | 18830 | Microarrays | 37986224 | Unspecified |
| mmu-miR-125b-5p | MIMAT0000136 | Plxna1 | 18844 | HITS-CLIP, Microarrays//HITS-CLIP | 37986224\|25083871 | Unspecified |
| mmu-miR-125b-5p | MIMAT0000136 | Plxnb3 | 140571 | Microarrays | 37986224 | Unspecified |
| mmu-miR-125b-5p | MIMAT0000136 | Plxnc1 | 54712 | HITS-CLIP | 23597149 | Unspecified |
| mmu-miR-125b-5p | MIMAT0000136 | Plxnd1 | 67784 | Microarrays | 37986224 | Unspecified |
| mmu-miR-125b-5p | MIMAT0000136 | Pml | 18854 | Microarrays | 37986224 | Unspecified |
| mmu-miR-125b-5p | MIMAT0000136 | Pmvk | 68603 | Microarrays | 37986224 | Unspecified |
| mmu-miR-125b-5p | MIMAT0000136 | Pnma2 | 239157 | Microarrays | 37986224 | Unspecified |
| mmu-miR-125b-5p | MIMAT0000136 | Pnn | 18949 | Microarrays | 37986224\|22469780 | Unspecified |
| mmu-miR-125b-5p | MIMAT0000136 | Pno1 | 66249 | HITS-CLIP, Microarrays//Microarrays | 37986224\|22469780 | Unspecified |
| mmu-miR-125b-5p | MIMAT0000136 | Pnpla2 | 66853 | HITS-CLIP | 25753418 | Unspecified |
| mmu-miR-125b-5p | MIMAT0000136 | Pnpo | 103711 | Microarrays | 37986224\|22469780 | Unspecified |
| mmu-miR-125b-5p | MIMAT0000136 | Polk | 27015 | Microarrays | 37986224 | Unspecified |

| | | | | | | | |
|---|---|---|---|---|---|---|---|
| mmu-miR-125b-5p | MIMAT0000136 | Polm | 54125 | Microarrays | 37986224\|22469780 | | Unspecified |
| mmu-miR-125b-5p | MIMAT0000136 | Polr2a | 20020 | HITS-CLIP | | 29167373 | Unspecified |
| mmu-miR-125b-5p | MIMAT0000136 | Polr2g | 67710 | HITS-CLIP | | 37986224 | Unspecified |
| mmu-miR-125b-5p | MIMAT0000136 | Polr2h | 245841 | Microarrays | 37986224\|22469780 | | Unspecified |
| mmu-miR-125b-5p | MIMAT0000136 | Polr2k | 17749 | Microarrays | 37986224\|22469780 | | Unspecified |
| mmu-miR-125b-5p | MIMAT0000136 | Polr3k | 67005 | Microarrays | 37986224\|22469780 | | Unspecified |
| mmu-miR-125b-5p | MIMAT0000136 | Pomgnt1 | 68273 | Microarrays | 37986224\|22469780 | | Unspecified |
| mmu-miR-125b-5p | MIMAT0000136 | Pop5 | 117109 | Microarrays | 37986224\|22469780 | | Unspecified |
| mmu-miR-125b-5p | MIMAT0000136 | Pop7 | 74097 | Microarrays | 37986224\|22469780 | | Unspecified |
| mmu-miR-125b-5p | MIMAT0000136 | Pot1a | 101185 | Microarrays | 37986224\|22469780 | | Unspecified |
| mmu-miR-125b-5p | MIMAT0000136 | Pou2f2 | 18987 | Microarrays | | 37986224 | Unspecified |
| mmu-miR-125b-5p | MIMAT0000136 | Pou3f1 | 18991 | Microarrays | 37986224\|22469780 | | Unspecified |
| mmu-miR-125b-5p | MIMAT0000136 | Pou3f3 | 18993 | HITS-CLIP | | 37986224 | Unspecified |
| mmu-miR-125b-5p | MIMAT0000136 | Pou5f2 | 75507 | HITS-CLIP, Microarrays | | 37986224 | Unspecified |
| mmu-miR-125b-5p | MIMAT0000136 | Ppara | 19013 | HITS-CLIP | | 37986224 | Unspecified |
| mmu-miR-125b-5p | MIMAT0000136 | Ppargc1b | 170826 | HITS-CLIP, Microarrays | | 37986224 | Unspecified |
| mmu-miR-125b-5p | MIMAT0000136 | Ppat | 231327 | HITS-CLIP, Microarrays//HITS-CLIP//Microa | 37986224\|25083871\|22469780 | | Unspecified |
| mmu-miR-125b-5p | MIMAT0000136 | Ppdpf | 66496 | Microarrays | | 37986224 | Unspecified |
| mmu-miR-125b-5p | MIMAT0000136 | Ppfia2 | 327814 | HITS-CLIP | | 37986224 | Unspecified |
| mmu-miR-125b-5p | MIMAT0000136 | Ppfia4 | 68507 | HITS-CLIP | | 37986224 | Unspecified |
| mmu-miR-125b-5p | MIMAT0000136 | Ppfibp1 | 67533 | HITS-CLIP | | 25083871 | Unspecified |
| mmu-miR-125b-5p | MIMAT0000136 | Ppif | 105675 | HITS-CLIP | | 37986224 | Unspecified |
| mmu-miR-125b-5p | MIMAT0000136 | Ppig | 228005 | HITS-CLIP | | 37986224 | Unspecified |
| mmu-miR-125b-5p | MIMAT0000136 | Ppih | 66101 | Microarrays | 37986224\|22469780 | | Unspecified |
| mmu-miR-125b-5p | MIMAT0000136 | Ppil4 | 67418 | HITS-CLIP, Microarrays//Microarrays | 37986224\|22469780 | | Unspecified |
| mmu-miR-125b-5p | MIMAT0000136 | Ppm1b | 19043 | HITS-CLIP | | 37986224 | Unspecified |
| mmu-miR-125b-5p | MIMAT0000136 | Ppm1f | 68606 | HITS-CLIP | | 37986224 | Unspecified |
| mmu-miR-125b-5p | MIMAT0000136 | Ppm1j | 71887 | Microarrays | | 37986224 | Unspecified |
| mmu-miR-125b-5p | MIMAT0000136 | Ppme1 | 72590 | HITS-CLIP | | 37986224 | Unspecified |
| mmu-miR-125b-5p | MIMAT0000136 | Ppp1ca | 19045 | Luciferase assay//qRT-PCR//Western blot/ | | 21935352 | Unspecified |
| mmu-miR-125b-5p | MIMAT0000136 | Ppp1r12a | 17931 | HITS-CLIP | | 37986224 | Unspecified |
| mmu-miR-125b-5p | MIMAT0000136 | Ppp1r12b | 329251 | HITS-CLIP | | 37986224 | Unspecified |
| mmu-miR-125b-5p | MIMAT0000136 | Ppp1r12c | 232807 | Microarrays | | 37986224 | Unspecified |
| mmu-miR-125b-5p | MIMAT0000136 | Ppp1r14c | 76142 | HITS-CLIP | | 25753418 | Unspecified |
| mmu-miR-125b-5p | MIMAT0000136 | Ppp1r16a | 73062 | HITS-CLIP | | 37986224 | Unspecified |
| mmu-miR-125b-5p | MIMAT0000136 | Ppp1r32 | 67752 | Microarrays | | 37986224 | Unspecified |
| mmu-miR-125b-5p | MIMAT0000136 | Ppp1r37 | 232947 | HITS-CLIP | | 37986224 | Unspecified |
| mmu-miR-125b-5p | MIMAT0000136 | Ppp1r3d | 228966 | HITS-CLIP | | 37986224 | Unspecified |
| mmu-miR-125b-5p | MIMAT0000136 | Ppp1r7 | 66385 | HITS-CLIP | | 37986224 | Unspecified |

| | | | | | | |
|---|---|---|---|---|---|---|
| mmu-miR-125b-5p | MIMAT0000136 | Ppp1r9a | 243725 | HITS-CLIP | 37986224 | Unspecified |
| mmu-miR-125b-5p | MIMAT0000136 | Ppp1r9b | 217124 | Microarrays//HITS-CLIP | 37986224\|25083871 | Unspecified |
| mmu-miR-125b-5p | MIMAT0000136 | Ppp2ca | 19052 | HITS-CLIP, Luciferase Reporter Assay//Luci | 37986224\|21935352\|25083871 | Unspecified |
| mmu-miR-125b-5p | MIMAT0000136 | Ppp2r1a | 51792 | Microarrays//HITS-CLIP | 37986224\|23597149 | Unspecified |
| mmu-miR-125b-5p | MIMAT0000136 | Ppp2r1b | 73699 | HITS-CLIP | 23597149 | Unspecified |
| mmu-miR-125b-5p | MIMAT0000136 | Ppp2r3a | 235542 | HITS-CLIP | 25083871 | Unspecified |
| mmu-miR-125b-5p | MIMAT0000136 | Ppp2r5c | 26931 | HITS-CLIP | 37986224 | Unspecified |
| mmu-miR-125b-5p | MIMAT0000136 | Ppp2r5e | 26932 | HITS-CLIP | 37986224 | Unspecified |
| mmu-miR-125b-5p | MIMAT0000136 | Ppp3ca | 19055 | HITS-CLIP | 25083871 | Unspecified |
| mmu-miR-125b-5p | MIMAT0000136 | Ppp3cb | 19056 | HITS-CLIP | 37986224 | Unspecified |
| mmu-miR-125b-5p | MIMAT0000136 | Ppp4r3a | 55671 | Microarrays//HITS-CLIP | 22469780\|29167373 | Unspecified |
| mmu-miR-125b-5p | MIMAT0000136 | Ppp6r1 | 243819 | HITS-CLIP | 37986224 | Unspecified |
| mmu-miR-125b-5p | MIMAT0000136 | Ppp6r2 | 71474 | HITS-CLIP | 23597149 | Unspecified |
| mmu-miR-125b-5p | MIMAT0000136 | Ppp6r3 | 52036 | HITS-CLIP | 37986224 | Unspecified |
| mmu-miR-125b-5p | MIMAT0000136 | Ppt1 | 19063 | HITS-CLIP | 37986224 | Unspecified |
| mmu-miR-125b-5p | MIMAT0000136 | Ppwd1 | 238831 | HITS-CLIP, Microarrays//Microarrays | 37986224\|22469780 | Unspecified |
| mmu-miR-125b-5p | MIMAT0000136 | Praf2 | 54637 | Microarrays | 37986224 | Unspecified |
| mmu-miR-125b-5p | MIMAT0000136 | Prdm1 | 12142 | HITS-CLIP | 37986224\|25083871\|29167373 | Unspecified |
| mmu-miR-125b-5p | MIMAT0000136 | Prdm2 | 110593 | HITS-CLIP | 37986224 | Unspecified |
| mmu-miR-125b-5p | MIMAT0000136 | Prelid2 | 77619 | Microarrays | 37986224 | Unspecified |
| mmu-miR-125b-5p | MIMAT0000136 | Prkaa2 | 108079 | HITS-CLIP | 37986224 | Unspecified |
| mmu-miR-125b-5p | MIMAT0000136 | Prkaca | 18747 | Microarrays | 37986224 | Unspecified |
| mmu-miR-125b-5p | MIMAT0000136 | Prkag3 | 241113 | Microarrays | 37986224 | Unspecified |
| mmu-miR-125b-5p | MIMAT0000136 | Prkca | 18750 | HITS-CLIP | 37986224 | Unspecified |
| mmu-miR-125b-5p | MIMAT0000136 | Prkdc | 19090 | Microarrays | 37986224 | Unspecified |
| mmu-miR-125b-5p | MIMAT0000136 | Prkra | 23992 | Luciferase assay//qRT-PCR//Western blot/ | 21935352 | Unspecified |
| mmu-miR-125b-5p | MIMAT0000136 | Prmt5 | 27374 | Microarrays | 37986224 | Unspecified |
| mmu-miR-125b-5p | MIMAT0000136 | Prmt9 | 80204 | Microarrays | 22469780 | Unspecified |
| mmu-miR-125b-5p | MIMAT0000136 | Proser1 | 212127 | HITS-CLIP | 37986224 | Unspecified |
| mmu-miR-125b-5p | MIMAT0000136 | Prox1 | 19130 | HITS-CLIP | 23597149 | Unspecified |
| mmu-miR-125b-5p | MIMAT0000136 | Prpf18 | 67229 | Microarrays | 37986224\|22469780 | Unspecified |
| mmu-miR-125b-5p | MIMAT0000136 | Prpf38b | 66921 | HITS-CLIP | 37986224 | Unspecified |
| mmu-miR-125b-5p | MIMAT0000136 | Prpf39 | 328110 | HITS-CLIP | 37986224 | Unspecified |
| mmu-miR-125b-5p | MIMAT0000136 | Prpf4 | 70052 | Microarrays | 37986224\|22469780 | Unspecified |
| mmu-miR-125b-5p | MIMAT0000136 | Prpf40b | 54614 | Microarrays | 37986224 | Unspecified |
| mmu-miR-125b-5p | MIMAT0000136 | Prpf4b | 19134 | HITS-CLIP | 37986224 | Unspecified |
| mmu-miR-125b-5p | MIMAT0000136 | Prpf6 | 68879 | HITS-CLIP | 23597149 | Unspecified |
| mmu-miR-125b-5p | MIMAT0000136 | Prps1l3 | 328099 | Microarrays | 37986224\|22469780 | Unspecified |
| mmu-miR-125b-5p | MIMAT0000136 | Prr12 | 233210 | Microarrays | 37986224 | Unspecified |

| | | | | | | |
|---|---|---|---|---|---|---|
| mmu-miR-125b-5p | MIMAT0000136 | Prr14l | 215476 | HITS-CLIP | 25083871 | Unspecified |
| mmu-miR-125b-5p | MIMAT0000136 | Prrc1 | 73137 | HITS-CLIP | 37986224\|25083871 | Unspecified |
| mmu-miR-125b-5p | MIMAT0000136 | Prrc2a | 53761 | HITS-CLIP, Microarrays//HITS-CLIP//Microa | 37986224\|25083871\|22469780 | Unspecified |
| mmu-miR-125b-5p | MIMAT0000136 | Prrc2b | 227723 | HITS-CLIP, Microarrays//HITS-CLIP | 37986224\|25083871\|19536157 | Unspecified |
| mmu-miR-125b-5p | MIMAT0000136 | Prrc2c | 226562 | HITS-CLIP | 25083871 | Unspecified |
| mmu-miR-125b-5p | MIMAT0000136 | Prrx1 | 18933 | HITS-CLIP | 25083871 | Unspecified |
| mmu-miR-125b-5p | MIMAT0000136 | Prss16 | 54373 | Microarrays | 37986224 | Unspecified |
| mmu-miR-125b-5p | MIMAT0000136 | Prss57 | 73106 | Microarrays | 37986224\|22469780 | Unspecified |
| mmu-miR-125b-5p | MIMAT0000136 | Prtg | 235472 | HITS-CLIP, Microarrays//Microarrays | 37986224\|22469780 | Unspecified |
| mmu-miR-125b-5p | MIMAT0000136 | Prune2 | 353211 | HITS-CLIP | 25083871 | Unspecified |
| mmu-miR-125b-5p | MIMAT0000136 | Prx | 19153 | Microarrays | 37986224 | Unspecified |
| mmu-miR-125b-5p | MIMAT0000136 | Pskh1 | 244631 | HITS-CLIP | 25083871 | Unspecified |
| mmu-miR-125b-5p | MIMAT0000136 | Psma1 | 26440 | HITS-CLIP | 37986224 | Unspecified |
| mmu-miR-125b-5p | MIMAT0000136 | Psmc1 | 19179 | HITS-CLIP | 37986224\|25083871 | Unspecified |
| mmu-miR-125b-5p | MIMAT0000136 | Psmd6 | 66413 | HITS-CLIP//qCLASH | 37986224\|31393953 | Unspecified |
| mmu-miR-125b-5p | MIMAT0000136 | Psmd7 | 17463 | HITS-CLIP | 37986224 | Unspecified |
| mmu-miR-125b-5p | MIMAT0000136 | Psme1 | 19186 | Microarrays | 37986224 | Unspecified |
| mmu-miR-125b-5p | MIMAT0000136 | Psme4 | 103554 | HITS-CLIP | 37986224\|25083871 | Unspecified |
| mmu-miR-125b-5p | MIMAT0000136 | Psmg3 | 66506 | HITS-CLIP, Microarrays//Microarrays | 37986224\|22469780 | Unspecified |
| mmu-miR-125b-5p | MIMAT0000136 | Ptar1 | 72351 | HITS-CLIP | 19536157\|25083871 | Unspecified |
| mmu-miR-125b-5p | MIMAT0000136 | Ptgir | 19222 | Microarrays | 37986224 | Unspecified |
| mmu-miR-125b-5p | MIMAT0000136 | Ptgr2 | 77219 | HITS-CLIP | 23597149 | Unspecified |
| mmu-miR-125b-5p | MIMAT0000136 | Ptk2 | 14083 | HITS-CLIP | 37986224 | Unspecified |
| mmu-miR-125b-5p | MIMAT0000136 | Ptk2b | 19229 | Microarrays | 37986224 | Unspecified |
| mmu-miR-125b-5p | MIMAT0000136 | Ptk7 | 71461 | HITS-CLIP, Microarrays//Microarrays | 37986224\|22469780 | Unspecified |
| mmu-miR-125b-5p | MIMAT0000136 | Ptms | 69202 | Microarrays | 37986224\|22469780 | Unspecified |
| mmu-miR-125b-5p | MIMAT0000136 | Ptp4a3 | 19245 | Microarrays | 37986224 | Unspecified |
| mmu-miR-125b-5p | MIMAT0000136 | Ptpa | 5524 | HITS-CLIP | 23597149 | Unspecified |
| mmu-miR-125b-5p | MIMAT0000136 | Ptpn1 | 19246 | HITS-CLIP | 37986224\|25083871\|19536157\|29167 | Unspecified |
| mmu-miR-125b-5p | MIMAT0000136 | Ptpn11 | 19247 | HITS-CLIP | 37986224 | Unspecified |
| mmu-miR-125b-5p | MIMAT0000136 | Ptpn14 | 19250 | HITS-CLIP | 37986224 | Unspecified |
| mmu-miR-125b-5p | MIMAT0000136 | Ptpn18 | 19253 | Microarrays | 37986224\|22469780 | Unspecified |
| mmu-miR-125b-5p | MIMAT0000136 | Ptpn21 | 24000 | HITS-CLIP | 37986224 | Unspecified |
| mmu-miR-125b-5p | MIMAT0000136 | Ptpn23 | 104831 | Microarrays | 37986224 | Unspecified |
| mmu-miR-125b-5p | MIMAT0000136 | Ptpn3 | 545622 | HITS-CLIP | 37986224\|25083871\|25753418 | Unspecified |
| mmu-miR-125b-5p | MIMAT0000136 | Ptpn9 | 56294 | Microarrays | 37986224 | Unspecified |
| mmu-miR-125b-5p | MIMAT0000136 | Ptprc | 19264 | Microarrays | 37986224 | Unspecified |
| mmu-miR-125b-5p | MIMAT0000136 | Ptprd | 19266 | HITS-CLIP | 37986224 | Unspecified |
| mmu-miR-125b-5p | MIMAT0000136 | Ptprf | 19268 | HITS-CLIP | 37986224 | Unspecified |

| | | | | | | |
|---|---|---|---|---|---|---|
| mmu-miR-125b-5p | MIMAT0000136 | Ptprg | 19270 | HITS-CLIP | 37986224 | Unspecified |
| mmu-miR-125b-5p | MIMAT0000136 | Ptprr | 19279 | Microarrays | 37986224 | Unspecified |
| mmu-miR-125b-5p | MIMAT0000136 | Ptprz1 | 19283 | HITS-CLIP | 37986224 | Unspecified |
| mmu-miR-125b-5p | MIMAT0000136 | Pum1 | 80912 | HITS-CLIP | 37986224\|25083871\|29167373 | Unspecified |
| mmu-miR-125b-5p | MIMAT0000136 | Pum2 | 80913 | HITS-CLIP | 25083871 | Unspecified |
| mmu-miR-125b-5p | MIMAT0000136 | Purb | 19291 | HITS-CLIP | 37986224\|19536157 | Unspecified |
| mmu-miR-125b-5p | MIMAT0000136 | Pvr | 52118 | HITS-CLIP | 37986224 | Unspecified |
| mmu-miR-125b-5p | MIMAT0000136 | Pwwp2a | 70802 | HITS-CLIP | 37986224 | Unspecified |
| mmu-miR-125b-5p | MIMAT0000136 | Pycr1 | 209027 | HITS-CLIP, Microarrays | 37986224 | Unspecified |
| mmu-miR-125b-5p | MIMAT0000136 | Pygb | 110078 | Microarrays | 37986224 | Unspecified |
| mmu-miR-125b-5p | MIMAT0000136 | Qars | 97541 | HITS-CLIP | 37986224 | Unspecified |
| mmu-miR-125b-5p | MIMAT0000136 | Qk | 19317 | HITS-CLIP | 37986224\|25083871\|29167373 | Unspecified |
| mmu-miR-125b-5p | MIMAT0000136 | Qrfp | 227717 | Microarrays | 37986224 | Unspecified |
| mmu-miR-125b-5p | MIMAT0000136 | Qrich1 | 69232 | HITS-CLIP | 37986224 | Unspecified |
| mmu-miR-125b-5p | MIMAT0000136 | Qser1 | 99003 | HITS-CLIP | 37986224\|19536157\|29167373 | Unspecified |
| mmu-miR-125b-5p | MIMAT0000136 | Qsox1 | 104009 | HITS-CLIP | 37986224 | Unspecified |
| mmu-miR-125b-5p | MIMAT0000136 | Qsox2 | 227638 | HITS-CLIP | 37986224 | Unspecified |
| mmu-miR-125b-5p | MIMAT0000136 | R3hdm4 | 109284 | Microarrays | 37986224 | Unspecified |
| mmu-miR-125b-5p | MIMAT0000136 | Rab11a | 53869 | HITS-CLIP | 37986224 | Unspecified |
| mmu-miR-125b-5p | MIMAT0000136 | Rab11fip2 | 74998 | HITS-CLIP | 25083871 | Unspecified |
| mmu-miR-125b-5p | MIMAT0000136 | Rab11fip3 | 215445 | qCLASH | 31393953 | Unspecified |
| mmu-miR-125b-5p | MIMAT0000136 | Rab11fip4 | 268451 | HITS-CLIP | 37986224 | Unspecified |
| mmu-miR-125b-5p | MIMAT0000136 | Rab11fip5 | 52055 | Microarrays | 37986224 | Unspecified |
| mmu-miR-125b-5p | MIMAT0000136 | Rab14 | 68365 | HITS-CLIP | 37986224 | Unspecified |
| mmu-miR-125b-5p | MIMAT0000136 | Rab19 | 19331 | Microarrays | 37986224\|22469780 | Unspecified |
| mmu-miR-125b-5p | MIMAT0000136 | Rab31 | 106572 | HITS-CLIP | 37986224 | Unspecified |
| mmu-miR-125b-5p | MIMAT0000136 | Rab34 | 19376 | HITS-CLIP | 37986224 | Unspecified |
| mmu-miR-125b-5p | MIMAT0000136 | Rab3a | 19339 | Microarrays | 37986224\|22469780 | Unspecified |
| mmu-miR-125b-5p | MIMAT0000136 | Rab3gap1 | 226407 | HITS-CLIP | 25083871 | Unspecified |
| mmu-miR-125b-5p | MIMAT0000136 | Rab44 | 442827 | Microarrays | 37986224 | Unspecified |
| mmu-miR-125b-5p | MIMAT0000136 | Rab5b | 19344 | HITS-CLIP | 37986224 | Unspecified |
| mmu-miR-125b-5p | MIMAT0000136 | Rab6b | 270192 | HITS-CLIP | 37986224 | Unspecified |
| mmu-miR-125b-5p | MIMAT0000136 | Rab8b | 235442 | HITS-CLIP | 37986224\|25083871 | Unspecified |
| mmu-miR-125b-5p | MIMAT0000136 | Rabl2 | 68708 | HITS-CLIP | 37986224 | Unspecified |
| mmu-miR-125b-5p | MIMAT0000136 | Rabl6 | 227624 | Microarray//HITS-CLIP//Other | 16495412\|25083871 | Unspecified |
| mmu-miR-125b-5p | MIMAT0000136 | Rac2 | 19354 | Microarrays | 37986224 | Unspecified |
| mmu-miR-125b-5p | MIMAT0000136 | Rad17 | 19356 | HITS-CLIP | 25083871 | Unspecified |
| mmu-miR-125b-5p | MIMAT0000136 | Rad51ap2 | 209550 | Microarrays | 37986224\|22469780 | Unspecified |
| mmu-miR-125b-5p | MIMAT0000136 | Rad54l2 | 81000 | HITS-CLIP | 37986224 | Unspecified |

| | | | | | | |
|---|---|---|---|---|---|---|
| mmu-miR-125b-5p | MIMAT0000136 | Rai1 | 19377 | Microarrays | 37986224 | Unspecified |
| mmu-miR-125b-5p | MIMAT0000136 | Rala | 56044 | HITS-CLIP | 37986224 | Unspecified |
| mmu-miR-125b-5p | MIMAT0000136 | Ralgapa1 | 56784 | Microarrays | 37986224 | Unspecified |
| mmu-miR-125b-5p | MIMAT0000136 | Ralgapa2 | 241694 | HITS-CLIP | 25753418 | Unspecified |
| mmu-miR-125b-5p | MIMAT0000136 | Ralgapb | 228850 | HITS-CLIP | 37986224 | Unspecified |
| mmu-miR-125b-5p | MIMAT0000136 | Ralgds | 19730 | HITS-CLIP | 37986224\|25083871 | Unspecified |
| mmu-miR-125b-5p | MIMAT0000136 | Ranbp10 | 74334 | HITS-CLIP | 37986224\|25083871 | Unspecified |
| mmu-miR-125b-5p | MIMAT0000136 | Ranbp2 | 19386 | HITS-CLIP | 37986224\|25083871\|29167373 | Unspecified |
| mmu-miR-125b-5p | MIMAT0000136 | Rap1a | 109905 | HITS-CLIP | 37986224 | Unspecified |
| mmu-miR-125b-5p | MIMAT0000136 | Rap1gap2 | 380711 | Microarrays | 37986224\|22469780 | Unspecified |
| mmu-miR-125b-5p | MIMAT0000136 | Rap2b | 74012 | HITS-CLIP | 37986224 | Unspecified |
| mmu-miR-125b-5p | MIMAT0000136 | Rapgefl1 | 268480 | Microarrays | 37986224 | Unspecified |
| mmu-miR-125b-5p | MIMAT0000136 | Raph1 | 77300 | HITS-CLIP | 37986224 | Unspecified |
| mmu-miR-125b-5p | MIMAT0000136 | Rarg | 19411 | Microarrays | 37986224 | Unspecified |
| mmu-miR-125b-5p | MIMAT0000136 | Rasa2 | 114713 | HITS-CLIP | 37986224 | Unspecified |
| mmu-miR-125b-5p | MIMAT0000136 | Rasa4 | 54153 | Microarrays | 37986224 | Unspecified |
| mmu-miR-125b-5p | MIMAT0000136 | Rasal1 | 19415 | Microarrays | 37986224 | Unspecified |
| mmu-miR-125b-5p | MIMAT0000136 | Rasal2 | 226525 | HITS-CLIP | 37986224 | Unspecified |
| mmu-miR-125b-5p | MIMAT0000136 | Rasgrp2 | 19395 | Microarrays | 37986224\|22469780 | Unspecified |
| mmu-miR-125b-5p | MIMAT0000136 | Rasgrp3 | 240168 | Microarrays | 37986224 | Unspecified |
| mmu-miR-125b-5p | MIMAT0000136 | Rassf3 | 192678 | HITS-CLIP | 37986224 | Unspecified |
| mmu-miR-125b-5p | MIMAT0000136 | Rassf4 | 213391 | Microarrays | 37986224 | Unspecified |
| mmu-miR-125b-5p | MIMAT0000136 | Rassf5 | 54354 | HITS-CLIP | 37986224 | Unspecified |
| mmu-miR-125b-5p | MIMAT0000136 | Rassf8 | 71323 | HITS-CLIP | 37986224 | Unspecified |
| mmu-miR-125b-5p | MIMAT0000136 | Rb1 | 19645 | HITS-CLIP | 23597149 | Unspecified |
| mmu-miR-125b-5p | MIMAT0000136 | Rb1cc1 | 12421 | HITS-CLIP | 25083871 | Unspecified |
| mmu-miR-125b-5p | MIMAT0000136 | Rbbp6 | 19647 | HITS-CLIP | 37986224 | Unspecified |
| mmu-miR-125b-5p | MIMAT0000136 | Rbfox2 | 93686 | HITS-CLIP | 37986224 | Unspecified |
| mmu-miR-125b-5p | MIMAT0000136 | Rbis | 401466 | Microarrays | 22469780 | Unspecified |
| mmu-miR-125b-5p | MIMAT0000136 | Rbm12 | 75710 | HITS-CLIP, Microarrays//Microarrays | 37986224\|22469780 | Unspecified |
| mmu-miR-125b-5p | MIMAT0000136 | Rbm12b1 | 72397 | Microarrays | 37986224\|22469780 | Unspecified |
| mmu-miR-125b-5p | MIMAT0000136 | Rbm12b2 | 77604 | HITS-CLIP | 37986224 | Unspecified |
| mmu-miR-125b-5p | MIMAT0000136 | Rbm14 | 56275 | Microarrays | 37986224\|22469780 | Unspecified |
| mmu-miR-125b-5p | MIMAT0000136 | Rbm15b | 109095 | HITS-CLIP | 37986224 | Unspecified |
| mmu-miR-125b-5p | MIMAT0000136 | Rbm17 | 76938 | HITS-CLIP | 25083871 | Unspecified |
| mmu-miR-125b-5p | MIMAT0000136 | Rbm18 | 67889 | HITS-CLIP | 37986224\|25083871 | Unspecified |
| mmu-miR-125b-5p | MIMAT0000136 | Rbm19 | 74111 | HITS-CLIP | 37986224 | Unspecified |
| mmu-miR-125b-5p | MIMAT0000136 | Rbm20 | 73713 | HITS-CLIP | 37986224\|25083871 | Unspecified |
| mmu-miR-125b-5p | MIMAT0000136 | Rbm22 | 66810 | Microarrays | 37986224 | Unspecified |

| | | | | | | |
|---|---|---|---|---|---|---|
| mmu-miR-125b-5p | MIMAT0000136 | Rbm24 | 666794 | HITS-CLIP | 37986224 | Unspecified |
| mmu-miR-125b-5p | MIMAT0000136 | Rbm25 | 67039 | HITS-CLIP | 25083871 | Unspecified |
| mmu-miR-125b-5p | MIMAT0000136 | Rbm38 | 56190 | HITS-CLIP | 37986224 | Unspecified |
| mmu-miR-125b-5p | MIMAT0000136 | Rbm39 | 170791 | HITS-CLIP | 37986224 | Unspecified |
| mmu-miR-125b-5p | MIMAT0000136 | Rbm4 | 19653 | Microarrays | 37986224\|22469780 | Unspecified |
| mmu-miR-125b-5p | MIMAT0000136 | Rbm45 | 241490 | Microarrays | 37986224\|22469780 | Unspecified |
| mmu-miR-125b-5p | MIMAT0000136 | Rbm6 | 19654 | HITS-CLIP | 37986224 | Unspecified |
| mmu-miR-125b-5p | MIMAT0000136 | Rbms2 | 56516 | Microarrays | 37986224 | Unspecified |
| mmu-miR-125b-5p | MIMAT0000136 | Rc3h1 | 381305 | HITS-CLIP | 37986224\|25083871\|23597149 | Unspecified |
| mmu-miR-125b-5p | MIMAT0000136 | Rc3h2 | 319817 | HITS-CLIP | 37986224 | Unspecified |
| mmu-miR-125b-5p | MIMAT0000136 | Rce1 | 19671 | HITS-CLIP, Microarrays//Microarrays | 37986224\|22469780 | Unspecified |
| mmu-miR-125b-5p | MIMAT0000136 | Rcor1 | 217864 | HITS-CLIP | 25083871 | Unspecified |
| mmu-miR-125b-5p | MIMAT0000136 | Rdh10 | 98711 | Microarrays | 37986224\|22469780 | Unspecified |
| mmu-miR-125b-5p | MIMAT0000136 | Rdh7 | 54150 | Microarrays | 37986224\|22469780 | Unspecified |
| mmu-miR-125b-5p | MIMAT0000136 | Reep3 | 28193 | HITS-CLIP | 37986224 | Unspecified |
| mmu-miR-125b-5p | MIMAT0000136 | Reep4 | 72549 | HITS-CLIP | 37986224 | Unspecified |
| mmu-miR-125b-5p | MIMAT0000136 | Reep6 | 70335 | Microarrays | 37986224 | Unspecified |
| mmu-miR-125b-5p | MIMAT0000136 | Rela | 19697 | HITS-CLIP | 25083871 | Unspecified |
| mmu-miR-125b-5p | MIMAT0000136 | Relb | 19698 | Microarrays | 37986224 | Unspecified |
| mmu-miR-125b-5p | MIMAT0000136 | Rell1 | 100532 | Microarrays | 37986224 | Unspecified |
| mmu-miR-125b-5p | MIMAT0000136 | Reps1 | 19707 | Microarrays | 37986224 | Unspecified |
| mmu-miR-125b-5p | MIMAT0000136 | Resp18 | 19711 | Microarrays | 37986224\|22469780 | Unspecified |
| mmu-miR-125b-5p | MIMAT0000136 | Rest | 19712 | HITS-CLIP | 37986224\|25083871\|29167373 | Unspecified |
| mmu-miR-125b-5p | MIMAT0000136 | Ret | 19713 | HITS-CLIP | 37986224 | Unspecified |
| mmu-miR-125b-5p | MIMAT0000136 | Retnlb | 57263 | Microarrays | 37986224\|22469780 | Unspecified |
| mmu-miR-125b-5p | MIMAT0000136 | Retreg1 | 66270 | HITS-CLIP | 23597149\|25083871\|29167373 | Unspecified |
| mmu-miR-125b-5p | MIMAT0000136 | Retreg2 | 79137 | HITS-CLIP | 25083871 | Unspecified |
| mmu-miR-125b-5p | MIMAT0000136 | Retreg3 | 162427 | HITS-CLIP//Microarrays | 23597149\|22469780 | Unspecified |
| mmu-miR-125b-5p | MIMAT0000136 | Rev1 | 56210 | HITS-CLIP | 37986224 | Unspecified |
| mmu-miR-125b-5p | MIMAT0000136 | Rev3l | 19714 | HITS-CLIP, Microarrays | 37986224 | Unspecified |
| mmu-miR-125b-5p | MIMAT0000136 | Rex2 | 100043034 | Microarrays | 37986224\|22469780 | Unspecified |
| mmu-miR-125b-5p | MIMAT0000136 | Rfc5 | 72151 | Microarrays | 37986224\|22469780 | Unspecified |
| mmu-miR-125b-5p | MIMAT0000136 | Rffl | 67338 | Microarrays | 37986224 | Unspecified |
| mmu-miR-125b-5p | MIMAT0000136 | Rfx3 | 19726 | HITS-CLIP | 37986224 | Unspecified |
| mmu-miR-125b-5p | MIMAT0000136 | Rfx7 | 319758 | HITS-CLIP | 37986224\|19536157\|29167373 | Unspecified |
| mmu-miR-125b-5p | MIMAT0000136 | Rgcc | 66214 | Microarrays | 37986224\|22469780 | Unspecified |
| mmu-miR-125b-5p | MIMAT0000136 | Rgp1 | 242406 | HITS-CLIP | 37986224 | Unspecified |
| mmu-miR-125b-5p | MIMAT0000136 | Rgs1 | 50778 | Microarrays | 37986224 | Unspecified |
| mmu-miR-125b-5p | MIMAT0000136 | Rgs16 | 19734 | Microarrays | 37986224 | Unspecified |

| | | | | | | | |
|---|---|---|---|---|---|---|---|
| mmu-miR-125b-5p | MIMAT0000136 | Rgs2 | 19735 | Microarrays | | 37986224 | Unspecified |
| mmu-miR-125b-5p | MIMAT0000136 | Rgs7bp | 52882 | HITS-CLIP | 37986224\|19536157 | | Unspecified |
| mmu-miR-125b-5p | MIMAT0000136 | Rgs8 | 67792 | HITS-CLIP | | 37986224 | Unspecified |
| mmu-miR-125b-5p | MIMAT0000136 | Rgsl1 | 240816 | Microarrays | 37986224\|22469780 | | Unspecified |
| mmu-miR-125b-5p | MIMAT0000136 | Rhebl1 | 69159 | Microarray//Microarrays//HITS-CLIP//Oth | 16495412\|22469780\|25083871 | | Unspecified |
| mmu-miR-125b-5p | MIMAT0000136 | Rhobtb2 | 246710 | HITS-CLIP | | 37986224 | Unspecified |
| mmu-miR-125b-5p | MIMAT0000136 | Rhoq | 104215 | HITS-CLIP | 37986224\|19536157 | | Unspecified |
| mmu-miR-125b-5p | MIMAT0000136 | Rhot2 | 214952 | HITS-CLIP, Microarrays//Microarrays | 37986224\|22469780 | | Unspecified |
| mmu-miR-125b-5p | MIMAT0000136 | Rhou | 69581 | HITS-CLIP | | 37986224 | Unspecified |
| mmu-miR-125b-5p | MIMAT0000136 | Rhox5 | 18617 | Microarrays | | 37986224 | Unspecified |
| mmu-miR-125b-5p | MIMAT0000136 | Rhox8 | 434768 | Microarrays//qCLASH | 37986224\|31393953 | | Unspecified |
| mmu-miR-125b-5p | MIMAT0000136 | Rif1 | 51869 | HITS-CLIP | | 37986224 | Unspecified |
| mmu-miR-125b-5p | MIMAT0000136 | Rilpl1 | 75695 | HITS-CLIP, Microarrays | | 37986224 | Unspecified |
| mmu-miR-125b-5p | MIMAT0000136 | Rin1 | 225870 | HITS-CLIP | | 37986224 | Unspecified |
| mmu-miR-125b-5p | MIMAT0000136 | Riok1 | 71340 | HITS-CLIP | | 37986224 | Unspecified |
| mmu-miR-125b-5p | MIMAT0000136 | Ripk2 | 192656 | Microarrays | 37986224\|22469780 | | Unspecified |
| mmu-miR-125b-5p | MIMAT0000136 | Ripor1 | 75687 | HITS-CLIP | 25083871\|23597149 | | Unspecified |
| mmu-miR-125b-5p | MIMAT0000136 | Rlf | 109263 | HITS-CLIP | 37986224\|25083871 | | Unspecified |
| mmu-miR-125b-5p | MIMAT0000136 | Rlim | 19820 | HITS-CLIP | | 37986224 | Unspecified |
| mmu-miR-125b-5p | MIMAT0000136 | Rnase12 | 497106 | Microarrays | | 37986224 | Unspecified |
| mmu-miR-125b-5p | MIMAT0000136 | Rnase4 | 58809 | Microarrays | | 37986224 | Unspecified |
| mmu-miR-125b-5p | MIMAT0000136 | Rnase6 | 78416 | Microarrays | | 37986224 | Unspecified |
| mmu-miR-125b-5p | MIMAT0000136 | Rnaseh1 | 19819 | HITS-CLIP | | 37986224 | Unspecified |
| mmu-miR-125b-5p | MIMAT0000136 | Rnaseh2b | 67153 | Microarrays | 37986224\|22469780 | | Unspecified |
| mmu-miR-125b-5p | MIMAT0000136 | Rnasek | 52898 | Microarrays | | 37986224 | Unspecified |
| mmu-miR-125b-5p | MIMAT0000136 | Rnd1 | 223881 | Microarrays | | 37986224 | Unspecified |
| mmu-miR-125b-5p | MIMAT0000136 | Rnf103 | 22644 | HITS-CLIP | | 25083871 | Unspecified |
| mmu-miR-125b-5p | MIMAT0000136 | Rnf11 | 29864 | HITS-CLIP | | 37986224 | Unspecified |
| mmu-miR-125b-5p | MIMAT0000136 | Rnf111 | 93836 | HITS-CLIP | 37986224\|25083871 | | Unspecified |
| mmu-miR-125b-5p | MIMAT0000136 | Rnf123 | 84585 | Microarrays | | 37986224 | Unspecified |
| mmu-miR-125b-5p | MIMAT0000136 | Rnf149 | 67702 | Microarrays | 37986224\|22469780 | | Unspecified |
| mmu-miR-125b-5p | MIMAT0000136 | Rnf157 | 217340 | HITS-CLIP, Microarrays | | 37986224 | Unspecified |
| mmu-miR-125b-5p | MIMAT0000136 | Rnf166 | 68718 | Microarrays | | 37986224 | Unspecified |
| mmu-miR-125b-5p | MIMAT0000136 | Rnf169 | 108937 | HITS-CLIP | | 23597149 | Unspecified |
| mmu-miR-125b-5p | MIMAT0000136 | Rnf181 | 66510 | Microarrays | | 37986224 | Unspecified |
| mmu-miR-125b-5p | MIMAT0000136 | Rnf19a | 30945 | HITS-CLIP | | 29167373 | Unspecified |
| mmu-miR-125b-5p | MIMAT0000136 | Rnf20 | 109331 | HITS-CLIP | 25083871\|25753418 | | Unspecified |
| mmu-miR-125b-5p | MIMAT0000136 | Rnf217 | 268291 | HITS-CLIP | | 37986224 | Unspecified |
| mmu-miR-125b-5p | MIMAT0000136 | Rnf24 | 51902 | Microarrays | | 37986224 | Unspecified |

| | | | | | | |
|---|---|---|---|---|---|---|
| mmu-miR-125b-5p | MIMAT0000136 | Rnf31 | 268749 | Microarrays | 37986224 | Unspecified |
| mmu-miR-125b-5p | MIMAT0000136 | Rnf38 | 73469 | HITS-CLIP | 37986224\|25083871\|19536157 | Unspecified |
| mmu-miR-125b-5p | MIMAT0000136 | Rnf4 | 19822 | HITS-CLIP | 37986224 | Unspecified |
| mmu-miR-125b-5p | MIMAT0000136 | Rnf41 | 67588 | HITS-CLIP | 37986224 | Unspecified |
| mmu-miR-125b-5p | MIMAT0000136 | Rnf44 | 105239 | HITS-CLIP | 37986224\|25083871 | Unspecified |
| mmu-miR-125b-5p | MIMAT0000136 | Rnmt | 67897 | HITS-CLIP | 37986224 | Unspecified |
| mmu-miR-125b-5p | MIMAT0000136 | Rnpepl1 | 108657 | HITS-CLIP | 25083871 | Unspecified |
| mmu-miR-125b-5p | MIMAT0000136 | Robo1 | 19876 | HITS-CLIP | 37986224 | Unspecified |
| mmu-miR-125b-5p | MIMAT0000136 | Rom1 | 19881 | Microarrays | 37986224\|22469780 | Unspecified |
| mmu-miR-125b-5p | MIMAT0000136 | Ropn1l | 252967 | Microarrays | 37986224 | Unspecified |
| mmu-miR-125b-5p | MIMAT0000136 | Rora | 19883 | HITS-CLIP | 37986224\|19536157\|25753418 | Unspecified |
| mmu-miR-125b-5p | MIMAT0000136 | Ros1 | 19886 | Microarrays | 37986224 | Unspecified |
| mmu-miR-125b-5p | MIMAT0000136 | Rpa1 | 68275 | HITS-CLIP | 37986224 | Unspecified |
| mmu-miR-125b-5p | MIMAT0000136 | Rpgrip1l | 244585 | HITS-CLIP | 37986224 | Unspecified |
| mmu-miR-125b-5p | MIMAT0000136 | Rpl35 | 66489 | qCLASH | 31393953 | Unspecified |
| mmu-miR-125b-5p | MIMAT0000136 | Rpl4 | 67891 | qCLASH | 31393953 | Unspecified |
| mmu-miR-125b-5p | MIMAT0000136 | Rprd1a | 225283 | HITS-CLIP | 25753418 | Unspecified |
| mmu-miR-125b-5p | MIMAT0000136 | Rprd2 | 75137 | HITS-CLIP | 37986224 | Unspecified |
| mmu-miR-125b-5p | MIMAT0000136 | Rps19 | 20085 | HITS-CLIP | 25083871 | Unspecified |
| mmu-miR-125b-5p | MIMAT0000136 | Rps6ka3 | 110651 | HITS-CLIP | 25083871 | Unspecified |
| mmu-miR-125b-5p | MIMAT0000136 | Rps7 | 20115 | Microarrays | 37986224\|22469780 | Unspecified |
| mmu-miR-125b-5p | MIMAT0000136 | Rragc | 54170 | HITS-CLIP | 25083871 | Unspecified |
| mmu-miR-125b-5p | MIMAT0000136 | Rras | 20130 | Microarrays | 37986224 | Unspecified |
| mmu-miR-125b-5p | MIMAT0000136 | Rrbp1 | 81910 | HITS-CLIP | 37986224 | Unspecified |
| mmu-miR-125b-5p | MIMAT0000136 | Rreb1 | 68750 | HITS-CLIP | 29167373 | Unspecified |
| mmu-miR-125b-5p | MIMAT0000136 | Rrm1 | 20133 | HITS-CLIP | 37986224 | Unspecified |
| mmu-miR-125b-5p | MIMAT0000136 | Rrp36 | 224823 | HITS-CLIP | 37986224 | Unspecified |
| mmu-miR-125b-5p | MIMAT0000136 | Rsad2 | 58185 | Microarrays | 37986224 | Unspecified |
| mmu-miR-125b-5p | MIMAT0000136 | Rsbn1 | 229675 | HITS-CLIP | 37986224\|25753418 | Unspecified |
| mmu-miR-125b-5p | MIMAT0000136 | Rsl1d1 | 66409 | HITS-CLIP | 23597149 | Unspecified |
| mmu-miR-125b-5p | MIMAT0000136 | Rsrc1 | 66880 | HITS-CLIP | 37986224 | Unspecified |
| mmu-miR-125b-5p | MIMAT0000136 | Rsu1 | 20163 | HITS-CLIP | 29167373 | Unspecified |
| mmu-miR-125b-5p | MIMAT0000136 | Rtkn2 | 170799 | Microarrays | 37986224 | Unspecified |
| mmu-miR-125b-5p | MIMAT0000136 | Rtn1 | 104001 | Microarrays | 37986224 | Unspecified |
| mmu-miR-125b-5p | MIMAT0000136 | Rtp4 | 67775 | Microarrays | 37986224 | Unspecified |
| mmu-miR-125b-5p | MIMAT0000136 | Rufy3 | 52822 | HITS-CLIP | 25083871 | Unspecified |
| mmu-miR-125b-5p | MIMAT0000136 | Runx1t1 | 12395 | HITS-CLIP | 37986224 | Unspecified |
| mmu-miR-125b-5p | MIMAT0000136 | Runx3 | 12399 | HITS-CLIP | 37986224 | Unspecified |
| mmu-miR-125b-5p | MIMAT0000136 | Rusc2 | 100213 | HITS-CLIP | 25753418 | Unspecified |

| | | | | | | |
|---|---|---|---|---|---|---|
| mmu-miR-125b-5p | MIMAT0000136 | Ruvbl2 | 20174 | Microarrays | 37986224 | Unspecified |
| mmu-miR-125b-5p | MIMAT0000136 | Rxra | 20181 | HITS-CLIP | 37986224 | Unspecified |
| mmu-miR-125b-5p | MIMAT0000136 | Ryr2 | 20191 | HITS-CLIP | 19536157 | Unspecified |
| mmu-miR-125b-5p | MIMAT0000136 | Ryr3 | 20192 | HITS-CLIP | 37986224 | Unspecified |
| mmu-miR-125b-5p | MIMAT0000136 | S100a13 | 20196 | Microarrays | 37986224\|22469780 | Unspecified |
| mmu-miR-125b-5p | MIMAT0000136 | S100a4 | 20198 | Microarrays | 37986224 | Unspecified |
| mmu-miR-125b-5p | MIMAT0000136 | S100a6 | 20200 | Microarrays | 37986224 | Unspecified |
| mmu-miR-125b-5p | MIMAT0000136 | S1pr2 | 14739 | Microarrays | 37986224 | Unspecified |
| mmu-miR-125b-5p | MIMAT0000136 | Sacs | 50720 | HITS-CLIP | 37986224 | Unspecified |
| mmu-miR-125b-5p | MIMAT0000136 | Samd4 | 74480 | HITS-CLIP | 37986224 | Unspecified |
| mmu-miR-125b-5p | MIMAT0000136 | Samd9l | 209086 | Microarrays | 37986224 | Unspecified |
| mmu-miR-125b-5p | MIMAT0000136 | Sap130 | 269003 | HITS-CLIP | 37986224\|25083871 | Unspecified |
| mmu-miR-125b-5p | MIMAT0000136 | Sap25 | 751865 | Microarrays | 37986224 | Unspecified |
| mmu-miR-125b-5p | MIMAT0000136 | Sap30bp | 57230 | Microarrays | 37986224\|22469780 | Unspecified |
| mmu-miR-125b-5p | MIMAT0000136 | Sapcd1 | 78376 | Microarrays | 37986224 | Unspecified |
| mmu-miR-125b-5p | MIMAT0000136 | Sars | 20226 | Microarrays | 37986224 | Unspecified |
| mmu-miR-125b-5p | MIMAT0000136 | Sash1 | 70097 | HITS-CLIP | 37986224 | Unspecified |
| mmu-miR-125b-5p | MIMAT0000136 | Satb1 | 20230 | Microarrays | 37986224 | Unspecified |
| mmu-miR-125b-5p | MIMAT0000136 | Sav1 | 64010 | HITS-CLIP | 37986224 | Unspecified |
| mmu-miR-125b-5p | MIMAT0000136 | Sbno1 | 243272 | HITS-CLIP | 37986224 | Unspecified |
| mmu-miR-125b-5p | MIMAT0000136 | Sbno2 | 216161 | HITS-CLIP | 37986224\|25083871 | Unspecified |
| mmu-miR-125b-5p | MIMAT0000136 | Scaf11 | 72193 | HITS-CLIP | 37986224\|25083871\|29167373 | Unspecified |
| mmu-miR-125b-5p | MIMAT0000136 | Scamp2 | 24044 | HITS-CLIP | 37986224 | Unspecified |
| mmu-miR-125b-5p | MIMAT0000136 | Scamp3 | 24045 | HITS-CLIP | 37986224 | Unspecified |
| mmu-miR-125b-5p | MIMAT0000136 | Scara5 | 71145 | HITS-CLIP | 37986224 | Unspecified |
| mmu-miR-125b-5p | MIMAT0000136 | Scarb1 | 20778 | HITS-CLIP | 37986224\|23597149 | Unspecified |
| mmu-miR-125b-5p | MIMAT0000136 | Scd1 | 20249 | Microarrays | 37986224 | Unspecified |
| mmu-miR-125b-5p | MIMAT0000136 | Scg3 | 20255 | HITS-CLIP | 37986224 | Unspecified |
| mmu-miR-125b-5p | MIMAT0000136 | Scml4 | 268297 | Microarrays | 37986224\|22469780 | Unspecified |
| mmu-miR-125b-5p | MIMAT0000136 | Scn2b | 72821 | HITS-CLIP | 37986224 | Unspecified |
| mmu-miR-125b-5p | MIMAT0000136 | Scn3b | 235281 | HITS-CLIP | 37986224 | Unspecified |
| mmu-miR-125b-5p | MIMAT0000136 | Scn4a | 110880 | HITS-CLIP | 37986224 | Unspecified |
| mmu-miR-125b-5p | MIMAT0000136 | Scn5a | 20271 | HITS-CLIP, Microarrays//Microarrays | 37986224\|22469780 | Unspecified |
| mmu-miR-125b-5p | MIMAT0000136 | Scoc | 56367 | Microarrays | 37986224 | Unspecified |
| mmu-miR-125b-5p | MIMAT0000136 | Scpep1 | 74617 | Microarrays | 37986224 | Unspecified |
| mmu-miR-125b-5p | MIMAT0000136 | Scrt2 | 545474 | HITS-CLIP | 37986224 | Unspecified |
| mmu-miR-125b-5p | MIMAT0000136 | Scyl2 | 213326 | HITS-CLIP | 37986224 | Unspecified |
| mmu-miR-125b-5p | MIMAT0000136 | Sdc1 | 20969 | HITS-CLIP | 37986224 | Unspecified |
| mmu-miR-125b-5p | MIMAT0000136 | Sdc3 | 20970 | HITS-CLIP | 37986224 | Unspecified |

| | | | | | | |
|---|---|---|---|---|---|---|
| mmu-miR-125b-5p | MIMAT0000136 | Sdc4 | 20971 | Microarrays | 37986224\|22469780 | Unspecified |
| mmu-miR-125b-5p | MIMAT0000136 | Sdcbp2 | 228765 | Microarrays | 37986224 | Unspecified |
| mmu-miR-125b-5p | MIMAT0000136 | Sdhc | 66052 | HITS-CLIP | 37986224 | Unspecified |
| mmu-miR-125b-5p | MIMAT0000136 | Sec14l2 | 67815 | HITS-CLIP, Microarrays//HITS-CLIP//Microa | 37986224\|25083871\|22469780 | Unspecified |
| mmu-miR-125b-5p | MIMAT0000136 | Sec16a | 227648 | HITS-CLIP | 37986224 | Unspecified |
| mmu-miR-125b-5p | MIMAT0000136 | Sec16b | 89867 | HITS-CLIP | 37986224\|23597149 | Unspecified |
| mmu-miR-125b-5p | MIMAT0000136 | Sec22b | 20333 | HITS-CLIP | 37986224 | Unspecified |
| mmu-miR-125b-5p | MIMAT0000136 | Sec23a | 20334 | HITS-CLIP | 25083871 | Unspecified |
| mmu-miR-125b-5p | MIMAT0000136 | Sec23b | 27054 | HITS-CLIP | 23597149\|25083871 | Unspecified |
| mmu-miR-125b-5p | MIMAT0000136 | Sec23ip | 207352 | HITS-CLIP | 37986224\|25083871 | Unspecified |
| mmu-miR-125b-5p | MIMAT0000136 | Sec24a | 77371 | HITS-CLIP | 37986224 | Unspecified |
| mmu-miR-125b-5p | MIMAT0000136 | Sec24b | 99683 | HITS-CLIP | 37986224\|25083871 | Unspecified |
| mmu-miR-125b-5p | MIMAT0000136 | Sec63 | 140740 | HITS-CLIP | 25083871 | Unspecified |
| mmu-miR-125b-5p | MIMAT0000136 | Secisbp2l | 70354 | HITS-CLIP | 37986224 | Unspecified |
| mmu-miR-125b-5p | MIMAT0000136 | Sel1l | 20338 | HITS-CLIP, Luciferase Reporter Assay//Luci | 37986224\|21935352 | Unspecified |
| mmu-miR-125b-5p | MIMAT0000136 | Sel1l3 | 231238 | HITS-CLIP | 37986224 | Unspecified |
| mmu-miR-125b-5p | MIMAT0000136 | Sell | 20343 | Microarrays | 37986224 | Unspecified |
| mmu-miR-125b-5p | MIMAT0000136 | Selplg | 20345 | Microarrays | 37986224 | Unspecified |
| mmu-miR-125b-5p | MIMAT0000136 | Sema3e | 20349 | HITS-CLIP, Microarrays | 37986224 | Unspecified |
| mmu-miR-125b-5p | MIMAT0000136 | Sema4b | 20352 | HITS-CLIP, Microarrays//Microarrays | 37986224\|22469780 | Unspecified |
| mmu-miR-125b-5p | MIMAT0000136 | Sema4c | 20353 | HITS-CLIP | 37986224 | Unspecified |
| mmu-miR-125b-5p | MIMAT0000136 | Sema4d | 20354 | HITS-CLIP | 37986224 | Unspecified |
| mmu-miR-125b-5p | MIMAT0000136 | Sema4f | 20355 | HITS-CLIP | 37986224 | Unspecified |
| mmu-miR-125b-5p | MIMAT0000136 | Sema4g | 26456 | HITS-CLIP | 23597149 | Unspecified |
| mmu-miR-125b-5p | MIMAT0000136 | Sema6a | 20358 | HITS-CLIP | 25083871 | Unspecified |
| mmu-miR-125b-5p | MIMAT0000136 | Sema6b | 20359 | Microarrays//HITS-CLIP | 37986224\|19536157 | Unspecified |
| mmu-miR-125b-5p | MIMAT0000136 | Senp5 | 320213 | HITS-CLIP | 37986224 | Unspecified |
| mmu-miR-125b-5p | MIMAT0000136 | Senp8 | 71599 | HITS-CLIP | 37986224 | Unspecified |
| mmu-miR-125b-5p | MIMAT0000136 | Sephs2 | 20768 | Microarrays//HITS-CLIP | 37986224\|25083871\|22469780 | Unspecified |
| mmu-miR-125b-5p | MIMAT0000136 | Sept2 | 4735 | HITS-CLIP | 25083871 | Unspecified |
| mmu-miR-125b-5p | MIMAT0000136 | Sept8 | 20362 | HITS-CLIP | 37986224 | Unspecified |
| mmu-miR-125b-5p | MIMAT0000136 | Septin6 | 56526 | Microarrays | 37986224 | Unspecified |
| mmu-miR-125b-5p | MIMAT0000136 | Serinc5 | 218442 | HITS-CLIP | 37986224 | Unspecified |
| mmu-miR-125b-5p | MIMAT0000136 | Serpina3k | 20714 | Microarrays | 37986224\|22469780 | Unspecified |
| mmu-miR-125b-5p | MIMAT0000136 | Serpinb6e | 435350 | Microarrays | 37986224 | Unspecified |
| mmu-miR-125b-5p | MIMAT0000136 | Serpinb9e | 20710 | Microarrays | 37986224 | Unspecified |
| mmu-miR-125b-5p | MIMAT0000136 | Serpind1 | 15160 | HITS-CLIP | 23597149 | Unspecified |
| mmu-miR-125b-5p | MIMAT0000136 | Sertad2 | 58172 | Microarrays | 37986224 | Unspecified |
| mmu-miR-125b-5p | MIMAT0000136 | Sertad3 | 170742 | HITS-CLIP, Microarrays//Microarrays | 37986224\|22469780 | Unspecified |

| | | | | | | | |
|---|---|---|---|---|---|---|---|
| mmu-miR-125b-5p | MIMAT0000136 | Sesn1 | 140742 | Microarrays | | 37986224 | Unspecified |
| mmu-miR-125b-5p | MIMAT0000136 | Sesn2 | 230784 | Microarrays | | 37986224 | Unspecified |
| mmu-miR-125b-5p | MIMAT0000136 | Sesn3 | 75747 | Microarrays | 37986224\|22469780 | | Unspecified |
| mmu-miR-125b-5p | MIMAT0000136 | Sestd1 | 228071 | Microarrays | | 37986224 | Unspecified |
| mmu-miR-125b-5p | MIMAT0000136 | Setbp1 | 240427 | HITS-CLIP | | 37986224 | Unspecified |
| mmu-miR-125b-5p | MIMAT0000136 | Setd2 | 235626 | HITS-CLIP | | 37986224 | Unspecified |
| mmu-miR-125b-5p | MIMAT0000136 | Setd5 | 72895 | HITS-CLIP | | 29167373 | Unspecified |
| mmu-miR-125b-5p | MIMAT0000136 | Setd7 | 73251 | HITS-CLIP | | 37986224 | Unspecified |
| mmu-miR-125b-5p | MIMAT0000136 | Setx | 269254 | HITS-CLIP | | 37986224 | Unspecified |
| mmu-miR-125b-5p | MIMAT0000136 | Sez6l | 56747 | qCLASH | | 31393953 | Unspecified |
| mmu-miR-125b-5p | MIMAT0000136 | Sez6l2 | 233878 | Microarrays | | 37986224 | Unspecified |
| mmu-miR-125b-5p | MIMAT0000136 | Sf3a2 | 20222 | Microarrays | | 37986224 | Unspecified |
| mmu-miR-125b-5p | MIMAT0000136 | Sfmbt1 | 54650 | HITS-CLIP | | 25083871 | Unspecified |
| mmu-miR-125b-5p | MIMAT0000136 | Sfpq | 71514 | Microarrays//HITS-CLIP | 37986224\|25083871\|22469780 | | Unspecified |
| mmu-miR-125b-5p | MIMAT0000136 | Sft2d3 | 67158 | HITS-CLIP | | 37986224 | Unspecified |
| mmu-miR-125b-5p | MIMAT0000136 | Sgms2 | 74442 | HITS-CLIP | | 37986224 | Unspecified |
| mmu-miR-125b-5p | MIMAT0000136 | Sgpl1 | 20397 | HITS-CLIP | | 37986224 | Unspecified |
| mmu-miR-125b-5p | MIMAT0000136 | Sgtb | 218544 | Microarrays | | 37986224 | Unspecified |
| mmu-miR-125b-5p | MIMAT0000136 | Sh2b2 | 23921 | Microarrays | | 37986224 | Unspecified |
| mmu-miR-125b-5p | MIMAT0000136 | Sh2b3 | 16923 | HITS-CLIP | | 37986224 | Unspecified |
| mmu-miR-125b-5p | MIMAT0000136 | Sh2d3c | 27387 | Microarrays | | 37986224 | Unspecified |
| mmu-miR-125b-5p | MIMAT0000136 | Sh3bgrl3 | 73723 | Microarrays | | 37986224 | Unspecified |
| mmu-miR-125b-5p | MIMAT0000136 | Sh3bp2 | 24055 | HITS-CLIP | | 37986224 | Unspecified |
| mmu-miR-125b-5p | MIMAT0000136 | Sh3bp4 | 98402 | HITS-CLIP | | 37986224 | Unspecified |
| mmu-miR-125b-5p | MIMAT0000136 | Sh3bp5l | 79566 | HITS-CLIP, Microarrays//Microarrays | 37986224\|22469780 | | Unspecified |
| mmu-miR-125b-5p | MIMAT0000136 | Sh3d19 | 27059 | HITS-CLIP | | 23597149 | Unspecified |
| mmu-miR-125b-5p | MIMAT0000136 | Sh3pxd2b | 268396 | HITS-CLIP | | 37986224 | Unspecified |
| mmu-miR-125b-5p | MIMAT0000136 | Shc1 | 20416 | HITS-CLIP | | 25083871 | Unspecified |
| mmu-miR-125b-5p | MIMAT0000136 | Shmt1 | 20425 | HITS-CLIP | | 37986224 | Unspecified |
| mmu-miR-125b-5p | MIMAT0000136 | Shoc2 | 56392 | Microarrays | | 37986224 | Unspecified |
| mmu-miR-125b-5p | MIMAT0000136 | Siah1b | 20438 | Microarrays | 37986224\|22469780 | | Unspecified |
| mmu-miR-125b-5p | MIMAT0000136 | Sirt1 | 93759 | Microarrays | 37986224\|22469780 | | Unspecified |
| mmu-miR-125b-5p | MIMAT0000136 | Sirt2 | 64383 | Microarrays | | 37986224 | Unspecified |
| mmu-miR-125b-5p | MIMAT0000136 | Sirt6 | 50721 | HITS-CLIP | | 37986224 | Unspecified |
| mmu-miR-125b-5p | MIMAT0000136 | Sirt7 | 209011 | HITS-CLIP | | 37986224 | Unspecified |
| mmu-miR-125b-5p | MIMAT0000136 | Six4 | 20474 | HITS-CLIP | | 25083871 | Unspecified |
| mmu-miR-125b-5p | MIMAT0000136 | Skil | 20482 | HITS-CLIP | 25083871\|29167373 | | Unspecified |
| mmu-miR-125b-5p | MIMAT0000136 | Sla | 20491 | Microarrays | | 37986224 | Unspecified |
| mmu-miR-125b-5p | MIMAT0000136 | Sla2 | 77799 | Microarrays | | 37986224 | Unspecified |

| | | | | | | |
|---|---|---|---|---|---|---|
| mmu-miR-125b-5p | MIMAT0000136 | Slamf6 | 30925 | Microarrays | 37986224\|22469780 | Unspecified |
| mmu-miR-125b-5p | MIMAT0000136 | Slamf8 | 74748 | Microarrays | 37986224\|22469780 | Unspecified |
| mmu-miR-125b-5p | MIMAT0000136 | Slamf9 | 98365 | Microarrays | 37986224 | Unspecified |
| mmu-miR-125b-5p | MIMAT0000136 | Slc10a3 | 214601 | Microarrays | 37986224\|22469780 | Unspecified |
| mmu-miR-125b-5p | MIMAT0000136 | Slc11a1 | 18173 | Microarrays | 37986224 | Unspecified |
| mmu-miR-125b-5p | MIMAT0000136 | Slc12a2 | 20496 | HITS-CLIP | 25083871 | Unspecified |
| mmu-miR-125b-5p | MIMAT0000136 | Slc12a7 | 20499 | HITS-CLIP | 37986224 | Unspecified |
| mmu-miR-125b-5p | MIMAT0000136 | Slc13a2 | 20500 | Microarrays | 37986224 | Unspecified |
| mmu-miR-125b-5p | MIMAT0000136 | Slc14a1 | 108052 | Microarrays | 37986224 | Unspecified |
| mmu-miR-125b-5p | MIMAT0000136 | Slc16a1 | 20501 | HITS-CLIP | 37986224\|25083871 | Unspecified |
| mmu-miR-125b-5p | MIMAT0000136 | Slc16a12 | 240638 | Microarrays | 37986224\|22469780 | Unspecified |
| mmu-miR-125b-5p | MIMAT0000136 | Slc16a6 | 104681 | HITS-CLIP | 37986224 | Unspecified |
| mmu-miR-125b-5p | MIMAT0000136 | Slc17a7 | 72961 | HITS-CLIP | 37986224\|25083871 | Unspecified |
| mmu-miR-125b-5p | MIMAT0000136 | Slc1a2 | 20511 | HITS-CLIP | 37986224 | Unspecified |
| mmu-miR-125b-5p | MIMAT0000136 | Slc1a4 | 55963 | HITS-CLIP | 25083871 | Unspecified |
| mmu-miR-125b-5p | MIMAT0000136 | Slc1a5 | 20514 | HITS-CLIP, Microarrays | 37986224 | Unspecified |
| mmu-miR-125b-5p | MIMAT0000136 | Slc22a18 | 18400 | HITS-CLIP | 37986224 | Unspecified |
| mmu-miR-125b-5p | MIMAT0000136 | Slc23a2 | 54338 | Microarrays | 37986224\|22469780 | Unspecified |
| mmu-miR-125b-5p | MIMAT0000136 | Slc25a1 | 13358 | Microarrays | 37986224 | Unspecified |
| mmu-miR-125b-5p | MIMAT0000136 | Slc25a15 | 18408 | HITS-CLIP | 37986224 | Unspecified |
| mmu-miR-125b-5p | MIMAT0000136 | Slc25a16 | 73132 | HITS-CLIP | 37986224 | Unspecified |
| mmu-miR-125b-5p | MIMAT0000136 | Slc25a23 | 66972 | Microarrays | 37986224 | Unspecified |
| mmu-miR-125b-5p | MIMAT0000136 | Slc25a30 | 67554 | Microarrays | 37986224 | Unspecified |
| mmu-miR-125b-5p | MIMAT0000136 | Slc25a32 | 69906 | HITS-CLIP | 25083871 | Unspecified |
| mmu-miR-125b-5p | MIMAT0000136 | Slc25a35 | 71998 | Microarrays//HITS-CLIP | 37986224\|23597149 | Unspecified |
| mmu-miR-125b-5p | MIMAT0000136 | Slc25a42 | 73095 | Microarrays | 37986224 | Unspecified |
| mmu-miR-125b-5p | MIMAT0000136 | Slc25a45 | 107375 | HITS-CLIP | 37986224 | Unspecified |
| mmu-miR-125b-5p | MIMAT0000136 | Slc25a46 | 67453 | HITS-CLIP | 37986224 | Unspecified |
| mmu-miR-125b-5p | MIMAT0000136 | Slc25a51 | 230125 | HITS-CLIP | 37986224\|25083871 | Unspecified |
| mmu-miR-125b-5p | MIMAT0000136 | Slc29a1 | 63959 | HITS-CLIP | 23597149 | Unspecified |
| mmu-miR-125b-5p | MIMAT0000136 | Slc2a3 | 20527 | Microarrays | 37986224 | Unspecified |
| mmu-miR-125b-5p | MIMAT0000136 | Slc2a5 | 56485 | Microarrays | 37986224 | Unspecified |
| mmu-miR-125b-5p | MIMAT0000136 | Slc30a4 | 22785 | HITS-CLIP | 25083871 | Unspecified |
| mmu-miR-125b-5p | MIMAT0000136 | Slc30a6 | 210148 | Microarrays | 37986224\|22469780 | Unspecified |
| mmu-miR-125b-5p | MIMAT0000136 | Slc35a1 | 24060 | HITS-CLIP | 25083871 | Unspecified |
| mmu-miR-125b-5p | MIMAT0000136 | Slc35b2 | 73836 | Microarrays | 37986224\|22469780 | Unspecified |
| mmu-miR-125b-5p | MIMAT0000136 | Slc35c1 | 228368 | HITS-CLIP | 37986224 | Unspecified |
| mmu-miR-125b-5p | MIMAT0000136 | Slc35e1 | 270066 | HITS-CLIP | 37986224 | Unspecified |
| mmu-miR-125b-5p | MIMAT0000136 | Slc35e4 | 103710 | HITS-CLIP | 37986224 | Unspecified |

| | | | | | | |
|---|---|---|---|---|---|---|
| mmu-miR-125b-5p | MIMAT0000136 | Slc35g1 | 240660 | Microarrays | 37986224\|22469780 | Unspecified |
| mmu-miR-125b-5p | MIMAT0000136 | Slc37a3 | 72144 | Microarrays | 37986224\|22469780 | Unspecified |
| mmu-miR-125b-5p | MIMAT0000136 | Slc38a1 | 105727 | HITS-CLIP | 37986224 | Unspecified |
| mmu-miR-125b-5p | MIMAT0000136 | Slc38a10 | 72055 | Microarrays | 37986224\|22469780 | Unspecified |
| mmu-miR-125b-5p | MIMAT0000136 | Slc38a5 | 209837 | Microarrays | 37986224\|22469780 | Unspecified |
| mmu-miR-125b-5p | MIMAT0000136 | Slc38a7 | 234595 | Microarrays | 37986224 | Unspecified |
| mmu-miR-125b-5p | MIMAT0000136 | Slc38a9 | 268706 | HITS-CLIP | 37986224 | Unspecified |
| mmu-miR-125b-5p | MIMAT0000136 | Slc39a10 | 227059 | Microarrays | 37986224 | Unspecified |
| mmu-miR-125b-5p | MIMAT0000136 | Slc39a14 | 213053 | HITS-CLIP | 25083871\|23597149 | Unspecified |
| mmu-miR-125b-5p | MIMAT0000136 | Slc39a7 | 14977 | HITS-CLIP | 37986224 | Unspecified |
| mmu-miR-125b-5p | MIMAT0000136 | Slc39a8 | 67547 | Microarrays | 37986224\|22469780 | Unspecified |
| mmu-miR-125b-5p | MIMAT0000136 | Slc39a9 | 328133 | HITS-CLIP | 37986224\|25083871 | Unspecified |
| mmu-miR-125b-5p | MIMAT0000136 | Slc3a2 | 17254 | Microarrays//HITS-CLIP | 37986224\|25083871 | Unspecified |
| mmu-miR-125b-5p | MIMAT0000136 | Slc41a1 | 98396 | Microarrays | 37986224 | Unspecified |
| mmu-miR-125b-5p | MIMAT0000136 | Slc43a2 | 215113 | HITS-CLIP | 37986224 | Unspecified |
| mmu-miR-125b-5p | MIMAT0000136 | Slc44a2 | 68682 | HITS-CLIP | 25083871 | Unspecified |
| mmu-miR-125b-5p | MIMAT0000136 | Slc44a3 | 213603 | Microarrays | 37986224 | Unspecified |
| mmu-miR-125b-5p | MIMAT0000136 | Slc45a4 | 106068 | HITS-CLIP | 37986224\|19536157\|29167373 | Unspecified |
| mmu-miR-125b-5p | MIMAT0000136 | Slc46a3 | 71706 | Microarrays | 37986224\|22469780 | Unspecified |
| mmu-miR-125b-5p | MIMAT0000136 | Slc48a1 | 67739 | HITS-CLIP | 37986224 | Unspecified |
| mmu-miR-125b-5p | MIMAT0000136 | Slc4a4 | 54403 | HITS-CLIP | 37986224 | Unspecified |
| mmu-miR-125b-5p | MIMAT0000136 | Slc4a7 | 218756 | HITS-CLIP, Microarrays//HITS-CLIP | 37986224\|25083871\|23597149 | Unspecified |
| mmu-miR-125b-5p | MIMAT0000136 | Slc50a1 | 19729 | Microarrays | 37986224 | Unspecified |
| mmu-miR-125b-5p | MIMAT0000136 | Slc5a6 | 330064 | Microarrays | 37986224\|22469780 | Unspecified |
| mmu-miR-125b-5p | MIMAT0000136 | Slc6a2 | 20538 | Microarrays | 37986224\|22469780 | Unspecified |
| mmu-miR-125b-5p | MIMAT0000136 | Slc6a8 | 102857 | HITS-CLIP | 25753418 | Unspecified |
| mmu-miR-125b-5p | MIMAT0000136 | Slc6a9 | 14664 | Microarrays | 37986224 | Unspecified |
| mmu-miR-125b-5p | MIMAT0000136 | Slc7a1 | 11987 | HITS-CLIP | 37986224 | Unspecified |
| mmu-miR-125b-5p | MIMAT0000136 | Slc7a2 | 11988 | HITS-CLIP | 37986224 | Unspecified |
| mmu-miR-125b-5p | MIMAT0000136 | Slc7a5 | 20539 | HITS-CLIP | 37986224 | Unspecified |
| mmu-miR-125b-5p | MIMAT0000136 | Slc7a6 | 330836 | HITS-CLIP | 37986224 | Unspecified |
| mmu-miR-125b-5p | MIMAT0000136 | Slc8b1 | 170756 | HITS-CLIP | 37986224 | Unspecified |
| mmu-miR-125b-5p | MIMAT0000136 | Slc9a1 | 20544 | HITS-CLIP | 25083871 | Unspecified |
| mmu-miR-125b-5p | MIMAT0000136 | Slc9a6 | 236794 | HITS-CLIP | 37986224 | Unspecified |
| mmu-miR-125b-5p | MIMAT0000136 | Slc9a7 | 236727 | Microarrays | 37986224\|22469780 | Unspecified |
| mmu-miR-125b-5p | MIMAT0000136 | Slc9a9 | 331004 | HITS-CLIP, Microarrays | 37986224 | Unspecified |
| mmu-miR-125b-5p | MIMAT0000136 | Slco3a1 | 108116 | HITS-CLIP | 37986224 | Unspecified |
| mmu-miR-125b-5p | MIMAT0000136 | Slco4a1 | 108115 | Microarrays | 37986224\|22469780 | Unspecified |
| mmu-miR-125b-5p | MIMAT0000136 | Slit1 | 20562 | Microarrays | 37986224 | Unspecified |

| | | | | | | |
|---|---|---|---|---|---|---|
| mmu-miR-125b-5p | MIMAT0000136 | Slitrk6 | 239250 | HITS-CLIP | 37986224\|19536157 | Unspecified |
| mmu-miR-125b-5p | MIMAT0000136 | Slk | 20874 | HITS-CLIP | 25083871 | Unspecified |
| mmu-miR-125b-5p | MIMAT0000136 | Smad1 | 17125 | HITS-CLIP | 25753418 | Unspecified |
| mmu-miR-125b-5p | MIMAT0000136 | Smad2 | 17126 | HITS-CLIP | 37986224 | Unspecified |
| mmu-miR-125b-5p | MIMAT0000136 | Smad4 | 17128 | HITS-CLIP | 25083871\|29167373 | Unspecified |
| mmu-miR-125b-5p | MIMAT0000136 | Smad5 | 17129 | HITS-CLIP | 37986224 | Unspecified |
| mmu-miR-125b-5p | MIMAT0000136 | Smad6 | 17130 | HITS-CLIP | 37986224 | Unspecified |
| mmu-miR-125b-5p | MIMAT0000136 | Smap1 | 98366 | HITS-CLIP | 37986224\|25083871 | Unspecified |
| mmu-miR-125b-5p | MIMAT0000136 | Smap2 | 69780 | HITS-CLIP, Microarrays | 37986224 | Unspecified |
| mmu-miR-125b-5p | MIMAT0000136 | Smarcd1 | 83797 | HITS-CLIP | 37986224\|25083871 | Unspecified |
| mmu-miR-125b-5p | MIMAT0000136 | Smc1a | 24061 | HITS-CLIP | 37986224 | Unspecified |
| mmu-miR-125b-5p | MIMAT0000136 | Smg1 | 233789 | HITS-CLIP | 37986224\|25083871\|29167373 | Unspecified |
| mmu-miR-125b-5p | MIMAT0000136 | Smg6 | 103677 | Microarrays | 37986224\|22469780 | Unspecified |
| mmu-miR-125b-5p | MIMAT0000136 | Smim12 | 80284 | HITS-CLIP, Microarrays//Microarrays | 37986224\|22469780 | Unspecified |
| mmu-miR-125b-5p | MIMAT0000136 | Smim19 | 102032 | Microarrays | 37986224\|22469780 | Unspecified |
| mmu-miR-125b-5p | MIMAT0000136 | Smim4 | 66487 | HITS-CLIP | 37986224 | Unspecified |
| mmu-miR-125b-5p | MIMAT0000136 | Smim7 | 66818 | Microarrays | 37986224\|22469780 | Unspecified |
| mmu-miR-125b-5p | MIMAT0000136 | Smoc1 | 64075 | HITS-CLIP | 37986224\|23597149 | Unspecified |
| mmu-miR-125b-5p | MIMAT0000136 | Smurf1 | 75788 | Microarrays | 37986224 | Unspecified |
| mmu-miR-125b-5p | MIMAT0000136 | Snap23 | 20619 | HITS-CLIP | 37986224 | Unspecified |
| mmu-miR-125b-5p | MIMAT0000136 | Snap25 | 20614 | Microarrays | 37986224 | Unspecified |
| mmu-miR-125b-5p | MIMAT0000136 | Snap47 | 67826 | HITS-CLIP | 37986224 | Unspecified |
| mmu-miR-125b-5p | MIMAT0000136 | Snd1 | 56463 | Microarrays//HITS-CLIP | 37986224\|23597149\|25753418 | Unspecified |
| mmu-miR-125b-5p | MIMAT0000136 | Snrnp40 | 66585 | HITS-CLIP//Microarrays | 25083871\|22469780 | Unspecified |
| mmu-miR-125b-5p | MIMAT0000136 | Snrnp70 | 20637 | HITS-CLIP | 37986224\|25083871\|29167373 | Unspecified |
| mmu-miR-125b-5p | MIMAT0000136 | Snrpa | 53607 | Microarrays | 37986224 | Unspecified |
| mmu-miR-125b-5p | MIMAT0000136 | Snrpd1 | 20641 | HITS-CLIP | 25083871 | Unspecified |
| mmu-miR-125b-5p | MIMAT0000136 | Snta1 | 20648 | Microarrays | 37986224 | Unspecified |
| mmu-miR-125b-5p | MIMAT0000136 | Sntb2 | 20650 | HITS-CLIP | 37986224 | Unspecified |
| mmu-miR-125b-5p | MIMAT0000136 | Snx27 | 76742 | HITS-CLIP | 37986224 | Unspecified |
| mmu-miR-125b-5p | MIMAT0000136 | Snx4 | 69150 | HITS-CLIP | 37986224 | Unspecified |
| mmu-miR-125b-5p | MIMAT0000136 | Snx8 | 231834 | Microarrays | 37986224\|22469780 | Unspecified |
| mmu-miR-125b-5p | MIMAT0000136 | Snx9 | 66616 | Microarrays | 37986224 | Unspecified |
| mmu-miR-125b-5p | MIMAT0000136 | Sobp | 109205 | HITS-CLIP | 37986224 | Unspecified |
| mmu-miR-125b-5p | MIMAT0000136 | Socs4 | 67296 | Microarrays//HITS-CLIP | 37986224\|22469780\|19536157 | Unspecified |
| mmu-miR-125b-5p | MIMAT0000136 | Socs5 | 56468 | HITS-CLIP | 37986224\|25083871 | Unspecified |
| mmu-miR-125b-5p | MIMAT0000136 | Soga3 | 67412 | Microarrays | 37986224 | Unspecified |
| mmu-miR-125b-5p | MIMAT0000136 | Sohlh2 | 74434 | Microarrays | 37986224\|22469780 | Unspecified |
| mmu-miR-125b-5p | MIMAT0000136 | Sox11 | 20666 | HITS-CLIP//Western blotting//qRT-PCR | 37986224\|25083871\|33584192 | Unspecified |

| | | | | | | |
|---|---|---|---|---|---|---|
| mmu-miR-125b-5p | MIMAT0000136 | Sox12 | 20667 | qCLASH | 31393953 | Unspecified |
| mmu-miR-125b-5p | MIMAT0000136 | Sox4 | 20677 | HITS-CLIP | 37986224 | Unspecified |
| mmu-miR-125b-5p | MIMAT0000136 | Sox5 | 20678 | HITS-CLIP | 23597149 | Unspecified |
| mmu-miR-125b-5p | MIMAT0000136 | Sp1 | 20683 | HITS-CLIP | 37986224\|25083871\|29167373 | Unspecified |
| mmu-miR-125b-5p | MIMAT0000136 | Sp100 | 20684 | Microarrays | 37986224 | Unspecified |
| mmu-miR-125b-5p | MIMAT0000136 | Sp110 | 109032 | Microarrays | 37986224 | Unspecified |
| mmu-miR-125b-5p | MIMAT0000136 | Sp140 | 434484 | Microarrays | 37986224 | Unspecified |
| mmu-miR-125b-5p | MIMAT0000136 | Sp3 | 20687 | HITS-CLIP | 37986224\|25083871\|29167373 | Unspecified |
| mmu-miR-125b-5p | MIMAT0000136 | Spag9 | 70834 | HITS-CLIP | 37986224\|25083871\|19536157 | Unspecified |
| mmu-miR-125b-5p | MIMAT0000136 | Sparc | 20692 | Microarrays | 37986224 | Unspecified |
| mmu-miR-125b-5p | MIMAT0000136 | Spata6 | 67946 | HITS-CLIP | 37986224 | Unspecified |
| mmu-miR-125b-5p | MIMAT0000136 | Spcs3 | 76687 | HITS-CLIP | 37986224 | Unspecified |
| mmu-miR-125b-5p | MIMAT0000136 | Speer4f1 | 70935 | Microarrays | 22469780 | Unspecified |
| mmu-miR-125b-5p | MIMAT0000136 | Speg | 11790 | HITS-CLIP | 37986224\|25083871 | Unspecified |
| mmu-miR-125b-5p | MIMAT0000136 | Spg20 | 229285 | HITS-CLIP | 37986224 | Unspecified |
| mmu-miR-125b-5p | MIMAT0000136 | Spin1 | 20729 | HITS-CLIP | 37986224 | Unspecified |
| mmu-miR-125b-5p | MIMAT0000136 | Spink13 | 100038417 | Microarrays | 37986224\|22469780 | Unspecified |
| mmu-miR-125b-5p | MIMAT0000136 | Spint2 | 20733 | Microarrays | 37986224\|22469780 | Unspecified |
| mmu-miR-125b-5p | MIMAT0000136 | Spn | 20737 | Microarrays | 37986224\|22469780 | Unspecified |
| mmu-miR-125b-5p | MIMAT0000136 | Spock2 | 94214 | HITS-CLIP | 37986224 | Unspecified |
| mmu-miR-125b-5p | MIMAT0000136 | Spop | 20747 | HITS-CLIP | 37986224 | Unspecified |
| mmu-miR-125b-5p | MIMAT0000136 | Spred3 | 101809 | HITS-CLIP | 37986224 | Unspecified |
| mmu-miR-125b-5p | MIMAT0000136 | Sprr2a1 | 20755 | Microarrays | 37986224\|22469780 | Unspecified |
| mmu-miR-125b-5p | MIMAT0000136 | Sprr4 | 229562 | Microarrays | 37986224 | Unspecified |
| mmu-miR-125b-5p | MIMAT0000136 | Sptan1 | 20740 | HITS-CLIP | 25083871 | Unspecified |
| mmu-miR-125b-5p | MIMAT0000136 | Sptb | 20741 | HITS-CLIP | 37986224 | Unspecified |
| mmu-miR-125b-5p | MIMAT0000136 | Sptbn1 | 20742 | HITS-CLIP | 19536157 | Unspecified |
| mmu-miR-125b-5p | MIMAT0000136 | Sptbn2 | 20743 | HITS-CLIP | 37986224 | Unspecified |
| mmu-miR-125b-5p | MIMAT0000136 | Spty2d1 | 101685 | HITS-CLIP | 25083871 | Unspecified |
| mmu-miR-125b-5p | MIMAT0000136 | Srebf2 | 20788 | HITS-CLIP | 37986224 | Unspecified |
| mmu-miR-125b-5p | MIMAT0000136 | Srf | 20807 | HITS-CLIP | 37986224 | Unspecified |
| mmu-miR-125b-5p | MIMAT0000136 | Srfbp1 | 67222 | Microarrays | 37986224\|22469780 | Unspecified |
| mmu-miR-125b-5p | MIMAT0000136 | Srgap3 | 259302 | Microarrays | 37986224 | Unspecified |
| mmu-miR-125b-5p | MIMAT0000136 | Srl | 106393 | HITS-CLIP | 37986224 | Unspecified |
| mmu-miR-125b-5p | MIMAT0000136 | Srp19 | 66384 | Microarrays | 37986224\|22469780 | Unspecified |
| mmu-miR-125b-5p | MIMAT0000136 | Srpk1 | 20815 | HITS-CLIP | 37986224 | Unspecified |
| mmu-miR-125b-5p | MIMAT0000136 | Srpk2 | 20817 | HITS-CLIP | 37986224 | Unspecified |
| mmu-miR-125b-5p | MIMAT0000136 | Srpr | 67398 | Microarrays | 37986224 | Unspecified |
| mmu-miR-125b-5p | MIMAT0000136 | Srpx2 | 68792 | Microarrays | 37986224\|22469780 | Unspecified |

| | | | | | | |
|---|---|---|---|---|---|---|
| mmu-miR-125b-5p | MIMAT0000136 | Srrm2 | 75956 | HITS-CLIP | 23597149 | Unspecified |
| mmu-miR-125b-5p | MIMAT0000136 | Srsf1 | 110809 | Microarrays | 37986224\|22469780 | Unspecified |
| mmu-miR-125b-5p | MIMAT0000136 | Srsf10 | 14105 | Microarrays | 37986224\|22469780 | Unspecified |
| mmu-miR-125b-5p | MIMAT0000136 | Srsf3 | 20383 | Microarrays | 37986224\|22469780 | Unspecified |
| mmu-miR-125b-5p | MIMAT0000136 | Srsf6 | 67996 | HITS-CLIP | 37986224\|25083871 | Unspecified |
| mmu-miR-125b-5p | MIMAT0000136 | Srsf7 | 225027 | Microarrays//HITS-CLIP | 37986224\|22469780\|19536157 | Unspecified |
| mmu-miR-125b-5p | MIMAT0000136 | Ssh1 | 231637 | HITS-CLIP | 37986224 | Unspecified |
| mmu-miR-125b-5p | MIMAT0000136 | Ssh2 | 237860 | Microarrays//HITS-CLIP | 37986224\|29167373 | Unspecified |
| mmu-miR-125b-5p | MIMAT0000136 | Ssh3 | 245857 | Microarrays | 37986224 | Unspecified |
| mmu-miR-125b-5p | MIMAT0000136 | Ssr1 | 107513 | HITS-CLIP | 25083871 | Unspecified |
| mmu-miR-125b-5p | MIMAT0000136 | Ssr3 | 67437 | HITS-CLIP | 37986224 | Unspecified |
| mmu-miR-125b-5p | MIMAT0000136 | Ssxb8 | 631002 | Microarrays | 37986224\|22469780 | Unspecified |
| mmu-miR-125b-5p | MIMAT0000136 | St3gal1 | 20442 | HITS-CLIP | 37986224\|29167373 | Unspecified |
| mmu-miR-125b-5p | MIMAT0000136 | St5 | 76954 | Microarrays | 37986224 | Unspecified |
| mmu-miR-125b-5p | MIMAT0000136 | St6galnac4 | 20448 | HITS-CLIP | 37986224 | Unspecified |
| mmu-miR-125b-5p | MIMAT0000136 | St8sia4 | 20452 | Microarrays//HITS-CLIP | 37986224\|22469780\|29167373 | Unspecified |
| mmu-miR-125b-5p | MIMAT0000136 | Stab1 | 192187 | HITS-CLIP | 29167373 | Unspecified |
| mmu-miR-125b-5p | MIMAT0000136 | Stard13 | 243362 | HITS-CLIP | 37986224 | Unspecified |
| mmu-miR-125b-5p | MIMAT0000136 | Stard9 | 668880 | HITS-CLIP | 37986224 | Unspecified |
| mmu-miR-125b-5p | MIMAT0000136 | Stat2 | 20847 | Microarrays | 37986224 | Unspecified |
| mmu-miR-125b-5p | MIMAT0000136 | Stat4 | 20849 | Microarrays | 37986224 | Unspecified |
| mmu-miR-125b-5p | MIMAT0000136 | Stbd1 | 52331 | Microarrays | 37986224 | Unspecified |
| mmu-miR-125b-5p | MIMAT0000136 | Steep1 | 63932 | Microarrays | 22469780 | Unspecified |
| mmu-miR-125b-5p | MIMAT0000136 | Stim1 | 20866 | HITS-CLIP | 37986224 | Unspecified |
| mmu-miR-125b-5p | MIMAT0000136 | Stim2 | 116873 | Microarrays | 37986224 | Unspecified |
| mmu-miR-125b-5p | MIMAT0000136 | Stk10 | 20868 | HITS-CLIP | 25083871 | Unspecified |
| mmu-miR-125b-5p | MIMAT0000136 | Stk11ip | 71728 | HITS-CLIP, Microarrays//Microarrays | 37986224\|22469780 | Unspecified |
| mmu-miR-125b-5p | MIMAT0000136 | Stk25 | 59041 | HITS-CLIP | 37986224 | Unspecified |
| mmu-miR-125b-5p | MIMAT0000136 | Stk35 | 67333 | HITS-CLIP | 37986224 | Unspecified |
| mmu-miR-125b-5p | MIMAT0000136 | Stk38l | 232533 | HITS-CLIP | 37986224\|25083871 | Unspecified |
| mmu-miR-125b-5p | MIMAT0000136 | Stk40 | 74178 | Microarrays | 37986224 | Unspecified |
| mmu-miR-125b-5p | MIMAT0000136 | Stoml1 | 69106 | Microarrays | 37986224 | Unspecified |
| mmu-miR-125b-5p | MIMAT0000136 | Stradb | 227154 | Microarrays | 37986224 | Unspecified |
| mmu-miR-125b-5p | MIMAT0000136 | Strn | 268980 | HITS-CLIP | 37986224\|25083871 | Unspecified |
| mmu-miR-125b-5p | MIMAT0000136 | Stt3a | 16430 | HITS-CLIP | 37986224 | Unspecified |
| mmu-miR-125b-5p | MIMAT0000136 | Stx11 | 74732 | Microarrays | 37986224 | Unspecified |
| mmu-miR-125b-5p | MIMAT0000136 | Stx1b | 56216 | HITS-CLIP | 37986224 | Unspecified |
| mmu-miR-125b-5p | MIMAT0000136 | Stx3 | 20908 | HITS-CLIP | 37986224 | Unspecified |
| mmu-miR-125b-5p | MIMAT0000136 | Stx4a | 20909 | Microarrays | 37986224 | Unspecified |

| | | | | | | |
|---|---|---|---|---|---|---|
| mmu-miR-125b-5p | MIMAT0000136 | Stxbp1 | 20910 | Microarrays | 37986224 | Unspecified |
| mmu-miR-125b-5p | MIMAT0000136 | Sumo3 | 20610 | HITS-CLIP | 37986224 | Unspecified |
| mmu-miR-125b-5p | MIMAT0000136 | Sun1 | 77053 | HITS-CLIP, Microarrays//Microarrays | 37986224\|22469780 | Unspecified |
| mmu-miR-125b-5p | MIMAT0000136 | Supt16 | 114741 | HITS-CLIP | 37986224 | Unspecified |
| mmu-miR-125b-5p | MIMAT0000136 | Supt4a | 20922 | Microarrays | 37986224\|22469780 | Unspecified |
| mmu-miR-125b-5p | MIMAT0000136 | Susd6 | 9766 | HITS-CLIP | 25083871 | Unspecified |
| mmu-miR-125b-5p | MIMAT0000136 | Suv39h1 | 20937 | Luciferase assay//Western blot//HITS-CLIP | 20699419\|25083871\|22469780 | Unspecified |
| mmu-miR-125b-5p | MIMAT0000136 | Suz12 | 52615 | HITS-CLIP | 37986224\|25083871 | Unspecified |
| mmu-miR-125b-5p | MIMAT0000136 | Sv2a | 64051 | HITS-CLIP | 37986224 | Unspecified |
| mmu-miR-125b-5p | MIMAT0000136 | Svip | 75744 | HITS-CLIP | 37986224 | Unspecified |
| mmu-miR-125b-5p | MIMAT0000136 | Svopl | 320590 | Microarrays | 37986224 | Unspecified |
| mmu-miR-125b-5p | MIMAT0000136 | Swap70 | 20947 | HITS-CLIP | 37986224 | Unspecified |
| mmu-miR-125b-5p | MIMAT0000136 | Syk | 20963 | Microarrays | 37986224\|22469780 | Unspecified |
| mmu-miR-125b-5p | MIMAT0000136 | Sympk | 68188 | Microarrays | 37986224 | Unspecified |
| mmu-miR-125b-5p | MIMAT0000136 | Syn2 | 20965 | HITS-CLIP | 37986224\|19536157 | Unspecified |
| mmu-miR-125b-5p | MIMAT0000136 | Sync | 68828 | HITS-CLIP | 37986224 | Unspecified |
| mmu-miR-125b-5p | MIMAT0000136 | Syncrip | 56403 | Microarrays | 37986224\|22469780 | Unspecified |
| mmu-miR-125b-5p | MIMAT0000136 | Syne1 | 64009 | HITS-CLIP, Microarrays//qCLASH | 37986224\|31393953 | Unspecified |
| mmu-miR-125b-5p | MIMAT0000136 | Syne3 | 212073 | HITS-CLIP | 37986224 | Unspecified |
| mmu-miR-125b-5p | MIMAT0000136 | Syngr2 | 20973 | HITS-CLIP | 37986224 | Unspecified |
| mmu-miR-125b-5p | MIMAT0000136 | Synj2 | 20975 | HITS-CLIP | 37986224 | Unspecified |
| mmu-miR-125b-5p | MIMAT0000136 | Synpo | 104027 | HITS-CLIP | 37986224\|25083871 | Unspecified |
| mmu-miR-125b-5p | MIMAT0000136 | Synpo2l | 68760 | HITS-CLIP | 25753418 | Unspecified |
| mmu-miR-125b-5p | MIMAT0000136 | Syt10 | 54526 | Microarrays | 37986224 | Unspecified |
| mmu-miR-125b-5p | MIMAT0000136 | Syvn1 | 74126 | HITS-CLIP | 25083871 | Unspecified |
| mmu-miR-125b-5p | MIMAT0000136 | Szrd1 | 213491 | HITS-CLIP | 37986224\|25083871 | Unspecified |
| mmu-miR-125b-5p | MIMAT0000136 | Tab2 | 68652 | Microarrays | 37986224 | Unspecified |
| mmu-miR-125b-5p | MIMAT0000136 | Tab3 | 66724 | HITS-CLIP | 37986224 | Unspecified |
| mmu-miR-125b-5p | MIMAT0000136 | Tacc2 | 57752 | HITS-CLIP | 37986224\|25083871 | Unspecified |
| mmu-miR-125b-5p | MIMAT0000136 | Tada3 | 101206 | HITS-CLIP | 37986224 | Unspecified |
| mmu-miR-125b-5p | MIMAT0000136 | Taf1 | 270627 | HITS-CLIP | 23597149 | Unspecified |
| mmu-miR-125b-5p | MIMAT0000136 | Taf11 | 68776 | Microarrays | 37986224\|22469780 | Unspecified |
| mmu-miR-125b-5p | MIMAT0000136 | Taf1a | 21339 | Microarrays | 37986224\|22469780 | Unspecified |
| mmu-miR-125b-5p | MIMAT0000136 | Taf4 | 6874 | Microarrays | 22469780 | Unspecified |
| mmu-miR-125b-5p | MIMAT0000136 | Taf4b | 72504 | HITS-CLIP | 37986224 | Unspecified |
| mmu-miR-125b-5p | MIMAT0000136 | Taf8 | 63856 | Microarrays | 37986224\|22469780 | Unspecified |
| mmu-miR-125b-5p | MIMAT0000136 | Taok1 | 216965 | HITS-CLIP | 29167373 | Unspecified |
| mmu-miR-125b-5p | MIMAT0000136 | Taok2 | 381921 | HITS-CLIP | 37986224 | Unspecified |
| mmu-miR-125b-5p | MIMAT0000136 | Tarbp2 | 21357 | Microarrays | 37986224\|22469780 | Unspecified |

| | | | | | | |
|---|---|---|---|---|---|---|
| mmu-miR-125b-5p | MIMAT0000136 | Tas2r109 | 387343 | Microarrays | 37986224\|22469780 | Unspecified |
| mmu-miR-125b-5p | MIMAT0000136 | Tas2r137 | 574417 | Microarrays | 37986224 | Unspecified |
| mmu-miR-125b-5p | MIMAT0000136 | Tatdn2 | 381801 | Microarrays | 37986224\|22469780 | Unspecified |
| mmu-miR-125b-5p | MIMAT0000136 | Tax1bp1 | 52440 | Microarrays | 37986224 | Unspecified |
| mmu-miR-125b-5p | MIMAT0000136 | Taz | 66826 | HITS-CLIP | 37986224 | Unspecified |
| mmu-miR-125b-5p | MIMAT0000136 | Tbc1d1 | 57915 | HITS-CLIP | 37986224 | Unspecified |
| mmu-miR-125b-5p | MIMAT0000136 | Tbc1d16 | 207592 | HITS-CLIP | 37986224 | Unspecified |
| mmu-miR-125b-5p | MIMAT0000136 | Tbc1d22b | 381085 | HITS-CLIP | 37986224 | Unspecified |
| mmu-miR-125b-5p | MIMAT0000136 | Tbc1d25 | 209815 | HITS-CLIP | 37986224 | Unspecified |
| mmu-miR-125b-5p | MIMAT0000136 | Tbc1d30 | 74694 | HITS-CLIP | 37986224 | Unspecified |
| mmu-miR-125b-5p | MIMAT0000136 | Tbc1d31 | 210544 | HITS-CLIP | 37986224 | Unspecified |
| mmu-miR-125b-5p | MIMAT0000136 | Tbc1d8 | 54610 | HITS-CLIP | 37986224 | Unspecified |
| mmu-miR-125b-5p | MIMAT0000136 | Tbc1d8b | 245638 | HITS-CLIP | 37986224 | Unspecified |
| mmu-miR-125b-5p | MIMAT0000136 | Tbc1d9b | 76795 | Microarrays | 37986224 | Unspecified |
| mmu-miR-125b-5p | MIMAT0000136 | Tbce | 70430 | HITS-CLIP | 37986224 | Unspecified |
| mmu-miR-125b-5p | MIMAT0000136 | Tbcel | 272589 | HITS-CLIP | 37986224 | Unspecified |
| mmu-miR-125b-5p | MIMAT0000136 | Tbl1xr1 | 81004 | HITS-CLIP | 25083871\|25753418 | Unspecified |
| mmu-miR-125b-5p | MIMAT0000136 | Tbl2 | 27368 | HITS-CLIP | 37986224 | Unspecified |
| mmu-miR-125b-5p | MIMAT0000136 | Tbl3 | 213773 | HITS-CLIP | 37986224 | Unspecified |
| mmu-miR-125b-5p | MIMAT0000136 | Tbp | 21374 | HITS-CLIP | 37986224 | Unspecified |
| mmu-miR-125b-5p | MIMAT0000136 | Tbr1 | 21375 | HITS-CLIP | 37986224 | Unspecified |
| mmu-miR-125b-5p | MIMAT0000136 | Tbrg1 | 21376 | qCLASH | 31393953 | Unspecified |
| mmu-miR-125b-5p | MIMAT0000136 | Tbx15 | 21384 | HITS-CLIP | 37986224 | Unspecified |
| mmu-miR-125b-5p | MIMAT0000136 | Tbxa2r | 21390 | Microarrays | 37986224 | Unspecified |
| mmu-miR-125b-5p | MIMAT0000136 | Tceal8 | 66684 | HITS-CLIP | 25083871 | Unspecified |
| mmu-miR-125b-5p | MIMAT0000136 | Tcf20 | 21411 | HITS-CLIP, Microarrays//HITS-CLIP | 37986224\|25083871 | Unspecified |
| mmu-miR-125b-5p | MIMAT0000136 | Tcf25 | 66855 | HITS-CLIP | 37986224 | Unspecified |
| mmu-miR-125b-5p | MIMAT0000136 | Tcf7 | 21414 | HITS-CLIP | 37986224\|25083871 | Unspecified |
| mmu-miR-125b-5p | MIMAT0000136 | Tcf7l1 | 21415 | HITS-CLIP | 37986224 | Unspecified |
| mmu-miR-125b-5p | MIMAT0000136 | Tcf7l2 | 21416 | HITS-CLIP | 37986224 | Unspecified |
| mmu-miR-125b-5p | MIMAT0000136 | Tchp | 77832 | Microarrays | 37986224 | Unspecified |
| mmu-miR-125b-5p | MIMAT0000136 | Tcirg1 | 27060 | HITS-CLIP | 37986224 | Unspecified |
| mmu-miR-125b-5p | MIMAT0000136 | Tcstv3 | 236219 | Microarrays | 37986224 | Unspecified |
| mmu-miR-125b-5p | MIMAT0000136 | Tdg | 21665 | Luciferase Reporter Assay | 37986224\|21935352 | Unspecified |
| mmu-miR-125b-5p | MIMAT0000136 | Tdh | 58865 | Microarrays | 37986224 | Unspecified |
| mmu-miR-125b-5p | MIMAT0000136 | Tdp2 | 56196 | HITS-CLIP | 37986224 | Unspecified |
| mmu-miR-125b-5p | MIMAT0000136 | Tdrd7 | 100121 | Microarrays | 37986224 | Unspecified |
| mmu-miR-125b-5p | MIMAT0000136 | Tead1 | 21676 | HITS-CLIP | 37986224 | Unspecified |
| mmu-miR-125b-5p | MIMAT0000136 | Tecr | 106529 | Microarrays | 37986224 | Unspecified |

| | | | | | | | |
|---|---|---|---|---|---|---|---|
| mmu-miR-125b-5p | MIMAT0000136 | Tedc1 | 104732 | Microarrays | | 22469780 | Unspecified |
| mmu-miR-125b-5p | MIMAT0000136 | Tedc2 | 72016 | qCLASH | | 31393953 | Unspecified |
| mmu-miR-125b-5p | MIMAT0000136 | Tef | 21685 | HITS-CLIP | | 37986224 | Unspecified |
| mmu-miR-125b-5p | MIMAT0000136 | Tekt4 | 71840 | Microarrays | 37986224\|22469780 | | Unspecified |
| mmu-miR-125b-5p | MIMAT0000136 | Tent4b | 64282 | Microarrays | | 22469780 | Unspecified |
| mmu-miR-125b-5p | MIMAT0000136 | Tep1 | 21745 | Microarrays | | 37986224 | Unspecified |
| mmu-miR-125b-5p | MIMAT0000136 | Tet2 | 214133 | HITS-CLIP | | 25083871 | Unspecified |
| mmu-miR-125b-5p | MIMAT0000136 | Tet3 | 194388 | HITS-CLIP | | 37986224 | Unspecified |
| mmu-miR-125b-5p | MIMAT0000136 | Tfap4 | 83383 | Microarrays | 37986224\|22469780 | | Unspecified |
| mmu-miR-125b-5p | MIMAT0000136 | Tfcp2 | 21422 | HITS-CLIP | | 25083871 | Unspecified |
| mmu-miR-125b-5p | MIMAT0000136 | Tfcp2l1 | 81879 | HITS-CLIP | | 37986224 | Unspecified |
| mmu-miR-125b-5p | MIMAT0000136 | Tfdp2 | 211586 | HITS-CLIP | 37986224\|25753418 | | Unspecified |
| mmu-miR-125b-5p | MIMAT0000136 | Tfec | 21426 | Microarrays | | 37986224 | Unspecified |
| mmu-miR-125b-5p | MIMAT0000136 | Tfrc | 22042 | HITS-CLIP | 37986224\|19536157 | | Unspecified |
| mmu-miR-125b-5p | MIMAT0000136 | Tgfb1 | 21803 | Microarrays | | 37986224 | Unspecified |
| mmu-miR-125b-5p | MIMAT0000136 | Tgfb3 | 21809 | Microarrays | | 37986224 | Unspecified |
| mmu-miR-125b-5p | MIMAT0000136 | Tgfbr1 | 21812 | HITS-CLIP | 37986224\|25083871 | | Unspecified |
| mmu-miR-125b-5p | MIMAT0000136 | Tgfbr2 | 21813 | HITS-CLIP | | 37986224 | Unspecified |
| mmu-miR-125b-5p | MIMAT0000136 | Tgfbr3 | 21814 | HITS-CLIP | | 37986224 | Unspecified |
| mmu-miR-125b-5p | MIMAT0000136 | Tgif1 | 21815 | HITS-CLIP | | 37986224 | Unspecified |
| mmu-miR-125b-5p | MIMAT0000136 | Tgoln1 | 22134 | HITS-CLIP | | 37986224 | Unspecified |
| mmu-miR-125b-5p | MIMAT0000136 | Tgs1 | 116940 | HITS-CLIP | 37986224\|25083871 | | Unspecified |
| mmu-miR-125b-5p | MIMAT0000136 | Tgtp2 | 100039796 | Microarrays | | 37986224 | Unspecified |
| mmu-miR-125b-5p | MIMAT0000136 | Thap3 | 69876 | HITS-CLIP | | 37986224 | Unspecified |
| mmu-miR-125b-5p | MIMAT0000136 | Thbs1 | 21825 | HITS-CLIP | | 25083871 | Unspecified |
| mmu-miR-125b-5p | MIMAT0000136 | Them6 | 223626 | HITS-CLIP | | 37986224 | Unspecified |
| mmu-miR-125b-5p | MIMAT0000136 | Thg1l | 66628 | Microarrays | 37986224\|22469780 | | Unspecified |
| mmu-miR-125b-5p | MIMAT0000136 | Thoc1 | 225160 | Microarrays | 37986224\|22469780 | | Unspecified |
| mmu-miR-125b-5p | MIMAT0000136 | Thra | 21833 | Microarrays | | 37986224 | Unspecified |
| mmu-miR-125b-5p | MIMAT0000136 | Thrap3 | 230753 | HITS-CLIP | | 37986224 | Unspecified |
| mmu-miR-125b-5p | MIMAT0000136 | Thumpd3 | 14911 | Microarrays | 37986224\|22469780 | | Unspecified |
| mmu-miR-125b-5p | MIMAT0000136 | Tiam1 | 21844 | Microarrays | | 37986224 | Unspecified |
| mmu-miR-125b-5p | MIMAT0000136 | Tifa | 211550 | Microarrays | 37986224\|22469780 | | Unspecified |
| mmu-miR-125b-5p | MIMAT0000136 | Tifab | 212937 | Microarrays | | 37986224 | Unspecified |
| mmu-miR-125b-5p | MIMAT0000136 | Tigd2 | 68140 | HITS-CLIP | | 37986224 | Unspecified |
| mmu-miR-125b-5p | MIMAT0000136 | Tigd3 | 332359 | Microarrays | 37986224\|22469780 | | Unspecified |
| mmu-miR-125b-5p | MIMAT0000136 | Tiparp | 99929 | HITS-CLIP | | 37986224 | Unspecified |
| mmu-miR-125b-5p | MIMAT0000136 | Tjp1 | 21872 | HITS-CLIP | | 19536157 | Unspecified |
| mmu-miR-125b-5p | MIMAT0000136 | Tjp3 | 27375 | Microarrays | | 37986224 | Unspecified |

| | | | | | | |
|---|---|---|---|---|---|---|
| mmu-miR-125b-5p | MIMAT0000136 | Tlcd1 | 68385 | HITS-CLIP | 37986224 | Unspecified |
| mmu-miR-125b-5p | MIMAT0000136 | Tlcd4 | 99887 | Microarrays | 22469780 | Unspecified |
| mmu-miR-125b-5p | MIMAT0000136 | Tle1 | 21885 | HITS-CLIP | 37986224 | Unspecified |
| mmu-miR-125b-5p | MIMAT0000136 | Tle3 | 21887 | HITS-CLIP | 37986224\|25083871 | Unspecified |
| mmu-miR-125b-5p | MIMAT0000136 | Tle5 | 166 | Microarrays | 22469780 | Unspecified |
| mmu-miR-125b-5p | MIMAT0000136 | Tlk1 | 228012 | HITS-CLIP | 25083871 | Unspecified |
| mmu-miR-125b-5p | MIMAT0000136 | Tlk2 | 24086 | HITS-CLIP | 25083871 | Unspecified |
| mmu-miR-125b-5p | MIMAT0000136 | Tln2 | 70549 | HITS-CLIP | 37986224 | Unspecified |
| mmu-miR-125b-5p | MIMAT0000136 | Tlr4 | 21898 | HITS-CLIP | 25083871 | Unspecified |
| mmu-miR-125b-5p | MIMAT0000136 | Tlr7 | 170743 | Microarrays | 37986224 | Unspecified |
| mmu-miR-125b-5p | MIMAT0000136 | Tm6sf1 | 107769 | Microarrays | 37986224 | Unspecified |
| mmu-miR-125b-5p | MIMAT0000136 | Tm6sf2 | 107770 | Microarrays | 37986224\|22469780 | Unspecified |
| mmu-miR-125b-5p | MIMAT0000136 | Tm9sf2 | 68059 | HITS-CLIP | 25083871 | Unspecified |
| mmu-miR-125b-5p | MIMAT0000136 | Tmbim6 | 110213 | HITS-CLIP | 37986224 | Unspecified |
| mmu-miR-125b-5p | MIMAT0000136 | Tmcc1 | 330401 | HITS-CLIP | 37986224 | Unspecified |
| mmu-miR-125b-5p | MIMAT0000136 | Tmed4 | 103694 | HITS-CLIP | 37986224 | Unspecified |
| mmu-miR-125b-5p | MIMAT0000136 | Tmed7 | 66676 | HITS-CLIP | 37986224 | Unspecified |
| mmu-miR-125b-5p | MIMAT0000136 | Tmed9 | 67511 | Microarrays//HITS-CLIP | 37986224\|22469780\|25083871 | Unspecified |
| mmu-miR-125b-5p | MIMAT0000136 | Tmem100 | 67888 | HITS-CLIP | 37986224 | Unspecified |
| mmu-miR-125b-5p | MIMAT0000136 | Tmem106a | 217203 | Microarrays | 37986224 | Unspecified |
| mmu-miR-125b-5p | MIMAT0000136 | Tmem106b | 71900 | HITS-CLIP | 37986224 | Unspecified |
| mmu-miR-125b-5p | MIMAT0000136 | Tmem106c | 380967 | Microarrays | 37986224 | Unspecified |
| mmu-miR-125b-5p | MIMAT0000136 | Tmem120b | 330189 | HITS-CLIP | 37986224 | Unspecified |
| mmu-miR-125b-5p | MIMAT0000136 | Tmem121 | 69195 | HITS-CLIP | 37986224 | Unspecified |
| mmu-miR-125b-5p | MIMAT0000136 | Tmem123 | 71929 | Microarrays | 37986224\|22469780 | Unspecified |
| mmu-miR-125b-5p | MIMAT0000136 | Tmem126a | 66271 | Microarrays | 37986224\|22469780 | Unspecified |
| mmu-miR-125b-5p | MIMAT0000136 | Tmem128 | 66309 | Microarrays | 37986224\|22469780 | Unspecified |
| mmu-miR-125b-5p | MIMAT0000136 | Tmem135 | 72759 | HITS-CLIP | 37986224 | Unspecified |
| mmu-miR-125b-5p | MIMAT0000136 | Tmem139 | 109218 | Microarrays | 37986224 | Unspecified |
| mmu-miR-125b-5p | MIMAT0000136 | Tmem161b | 72745 | HITS-CLIP | 37986224 | Unspecified |
| mmu-miR-125b-5p | MIMAT0000136 | Tmem168 | 101118 | Microarrays//HITS-CLIP | 37986224\|25083871\|22469780\|25753 | Unspecified |
| mmu-miR-125b-5p | MIMAT0000136 | Tmem170b | 621976 | HITS-CLIP | 37986224 | Unspecified |
| mmu-miR-125b-5p | MIMAT0000136 | Tmem178b | 434008 | HITS-CLIP | 37986224 | Unspecified |
| mmu-miR-125b-5p | MIMAT0000136 | Tmem184b | 223693 | HITS-CLIP | 25083871 | Unspecified |
| mmu-miR-125b-5p | MIMAT0000136 | Tmem185a | 84548 | Microarrays | 22469780 | Unspecified |
| mmu-miR-125b-5p | MIMAT0000136 | Tmem186 | 66690 | HITS-CLIP | 37986224 | Unspecified |
| mmu-miR-125b-5p | MIMAT0000136 | Tmem19 | 67226 | HITS-CLIP, Microarrays//Microarrays | 37986224\|22469780 | Unspecified |
| mmu-miR-125b-5p | MIMAT0000136 | Tmem214 | 68796 | qCLASH | 31393953 | Unspecified |
| mmu-miR-125b-5p | MIMAT0000136 | Tmem231 | 234740 | HITS-CLIP | 37986224 | Unspecified |

| | | | | | | |
|---|---|---|---|---|---|---|
| mmu-miR-125b-5p | MIMAT0000136 | Tmem234 | 76799 | HITS-CLIP | 37986224\|25083871\|23597149 | Unspecified |
| mmu-miR-125b-5p | MIMAT0000136 | Tmem243 | 652925 | Microarrays | 37986224\|22469780 | Unspecified |
| mmu-miR-125b-5p | MIMAT0000136 | Tmem245 | 242474 | HITS-CLIP//qCLASH | 37986224\|31393953 | Unspecified |
| mmu-miR-125b-5p | MIMAT0000136 | Tmem253 | 619301 | Microarrays | 37986224\|22469780 | Unspecified |
| mmu-miR-125b-5p | MIMAT0000136 | Tmem255a | 245386 | Microarrays | 37986224\|22469780 | Unspecified |
| mmu-miR-125b-5p | MIMAT0000136 | Tmem38b | 52076 | qCLASH | 31393953 | Unspecified |
| mmu-miR-125b-5p | MIMAT0000136 | Tmem42 | 66079 | Microarrays | 37986224\|22469780 | Unspecified |
| mmu-miR-125b-5p | MIMAT0000136 | Tmem50b | 77975 | Microarrays | 37986224\|22469780 | Unspecified |
| mmu-miR-125b-5p | MIMAT0000136 | Tmem56 | 99887 | Microarrays | 37986224 | Unspecified |
| mmu-miR-125b-5p | MIMAT0000136 | Tmem64 | 100201 | Microarrays | 37986224\|22469780 | Unspecified |
| mmu-miR-125b-5p | MIMAT0000136 | Tmem69 | 230657 | Microarrays | 37986224\|22469780 | Unspecified |
| mmu-miR-125b-5p | MIMAT0000136 | Tmem87a | 211499 | HITS-CLIP | 37986224 | Unspecified |
| mmu-miR-125b-5p | MIMAT0000136 | Tmem87b | 72477 | Microarrays | 37986224 | Unspecified |
| mmu-miR-125b-5p | MIMAT0000136 | Tmf1 | 232286 | Microarrays | 37986224 | Unspecified |
| mmu-miR-125b-5p | MIMAT0000136 | Tmod2 | 50876 | HITS-CLIP | 37986224 | Unspecified |
| mmu-miR-125b-5p | MIMAT0000136 | Tmpo | 21917 | HITS-CLIP | 23597149 | Unspecified |
| mmu-miR-125b-5p | MIMAT0000136 | Tmppe | 100504715 | HITS-CLIP | 37986224 | Unspecified |
| mmu-miR-125b-5p | MIMAT0000136 | Tmsb10 | 19240 | Microarrays | 37986224 | Unspecified |
| mmu-miR-125b-5p | MIMAT0000136 | Tmtc1 | 387314 | Microarrays | 37986224\|22469780 | Unspecified |
| mmu-miR-125b-5p | MIMAT0000136 | Tmtc2 | 278279 | HITS-CLIP | 37986224\|25083871 | Unspecified |
| mmu-miR-125b-5p | MIMAT0000136 | Tmx4 | 52837 | Microarrays | 37986224 | Unspecified |
| mmu-miR-125b-5p | MIMAT0000136 | Tnfrsf19 | 29820 | Microarrays | 37986224\|22469780 | Unspecified |
| mmu-miR-125b-5p | MIMAT0000136 | Tnfrsf1a | 21937 | HITS-CLIP | 37986224 | Unspecified |
| mmu-miR-125b-5p | MIMAT0000136 | Tnfrsf1b | 21938 | HITS-CLIP//qCLASH | 37986224\|31393953 | Unspecified |
| mmu-miR-125b-5p | MIMAT0000136 | Tnfsf4 | 22164 | Microarrays | 37986224\|22469780 | Unspecified |
| mmu-miR-125b-5p | MIMAT0000136 | Tnk2 | 51789 | Microarrays | 37986224 | Unspecified |
| mmu-miR-125b-5p | MIMAT0000136 | Tnks | 21951 | HITS-CLIP | 37986224\|19536157 | Unspecified |
| mmu-miR-125b-5p | MIMAT0000136 | Tnks1bp1 | 228140 | HITS-CLIP | 37986224\|25083871 | Unspecified |
| mmu-miR-125b-5p | MIMAT0000136 | Tnni2 | 21953 | Microarrays | 37986224 | Unspecified |
| mmu-miR-125b-5p | MIMAT0000136 | Tnnt2 | 21956 | Microarrays | 37986224 | Unspecified |
| mmu-miR-125b-5p | MIMAT0000136 | Tnpo1 | 238799 | HITS-CLIP | 23597149 | Unspecified |
| mmu-miR-125b-5p | MIMAT0000136 | Tnr | 21960 | HITS-CLIP | 37986224 | Unspecified |
| mmu-miR-125b-5p | MIMAT0000136 | Tnrc6a | 233833 | HITS-CLIP | 37986224\|25083871\|29167373 | Unspecified |
| mmu-miR-125b-5p | MIMAT0000136 | Tnrc6b | 213988 | HITS-CLIP | 37986224\|25083871 | Unspecified |
| mmu-miR-125b-5p | MIMAT0000136 | Tnrc6c | 217351 | HITS-CLIP | 37986224 | Unspecified |
| mmu-miR-125b-5p | MIMAT0000136 | Tns3 | 319939 | HITS-CLIP | 25753418 | Unspecified |
| mmu-miR-125b-5p | MIMAT0000136 | Tob1 | 22057 | HITS-CLIP | 37986224 | Unspecified |
| mmu-miR-125b-5p | MIMAT0000136 | Tomm70a | 28185 | HITS-CLIP | 37986224 | Unspecified |
| mmu-miR-125b-5p | MIMAT0000136 | Top3b | 21976 | Microarrays | 37986224 | Unspecified |

| | | | | | | |
|---|---|---|---|---|---|---|
| mmu-miR-125b-5p | MIMAT0000136 | Tor2a | 30933 | Microarray//Microarrays//qCLASH//Other | 16495412\|22469780\|31393953 | Unspecified |
| mmu-miR-125b-5p | MIMAT0000136 | Tor3a | 30935 | HITS-CLIP | 37986224\|29167373 | Unspecified |
| mmu-miR-125b-5p | MIMAT0000136 | Tpgs1 | 110012 | Microarrays | 37986224\|22469780 | Unspecified |
| mmu-miR-125b-5p | MIMAT0000136 | Tph1 | 21990 | Microarrays | 37986224\|22469780 | Unspecified |
| mmu-miR-125b-5p | MIMAT0000136 | Tpm1 | 22003 | HITS-CLIP | 37986224 | Unspecified |
| mmu-miR-125b-5p | MIMAT0000136 | Tpm2 | 22004 | HITS-CLIP | 25753418 | Unspecified |
| mmu-miR-125b-5p | MIMAT0000136 | Tpm4 | 326618 | HITS-CLIP | 37986224\|29167373 | Unspecified |
| mmu-miR-125b-5p | MIMAT0000136 | Tpp2 | 22019 | HITS-CLIP | 25083871 | Unspecified |
| mmu-miR-125b-5p | MIMAT0000136 | Tppp | 72948 | HITS-CLIP | 37986224 | Unspecified |
| mmu-miR-125b-5p | MIMAT0000136 | Tpr | 108989 | HITS-CLIP | 25083871\|29167373 | Unspecified |
| mmu-miR-125b-5p | MIMAT0000136 | Tra2a | 101214 | Microarrays | 37986224\|22469780 | Unspecified |
| mmu-miR-125b-5p | MIMAT0000136 | Traf3 | 22031 | Microarrays | 37986224\|22469780 | Unspecified |
| mmu-miR-125b-5p | MIMAT0000136 | Traf4 | 22032 | HITS-CLIP, Microarrays//Microarrays | 37986224\|22469780 | Unspecified |
| mmu-miR-125b-5p | MIMAT0000136 | Traf6 | 22034 | HITS-CLIP | 37986224 | Unspecified |
| mmu-miR-125b-5p | MIMAT0000136 | Tram1 | 72265 | HITS-CLIP | 37986224\|19536157 | Unspecified |
| mmu-miR-125b-5p | MIMAT0000136 | Trap1 | 68015 | HITS-CLIP | 25083871 | Unspecified |
| mmu-miR-125b-5p | MIMAT0000136 | Trappc11 | 320714 | HITS-CLIP | 25083871 | Unspecified |
| mmu-miR-125b-5p | MIMAT0000136 | Trappc5 | 66682 | HITS-CLIP | 37986224 | Unspecified |
| mmu-miR-125b-5p | MIMAT0000136 | Trappc6b | 78232 | HITS-CLIP | 37986224 | Unspecified |
| mmu-miR-125b-5p | MIMAT0000136 | Trappc8 | 75964 | HITS-CLIP | 25083871 | Unspecified |
| mmu-miR-125b-5p | MIMAT0000136 | Trat1 | 77647 | Microarrays | 37986224 | Unspecified |
| mmu-miR-125b-5p | MIMAT0000136 | Trib2 | 217410 | HITS-CLIP | 37986224 | Unspecified |
| mmu-miR-125b-5p | MIMAT0000136 | Trib3 | 228775 | Microarrays | 37986224 | Unspecified |
| mmu-miR-125b-5p | MIMAT0000136 | Tril | 66873 | HITS-CLIP | 37986224 | Unspecified |
| mmu-miR-125b-5p | MIMAT0000136 | Trim11 | 94091 | HITS-CLIP | 37986224 | Unspecified |
| mmu-miR-125b-5p | MIMAT0000136 | Trim12c | 319236 | Microarrays | 37986224 | Unspecified |
| mmu-miR-125b-5p | MIMAT0000136 | Trim23 | 81003 | HITS-CLIP | 37986224 | Unspecified |
| mmu-miR-125b-5p | MIMAT0000136 | Trim30a | 20128 | Microarrays | 37986224 | Unspecified |
| mmu-miR-125b-5p | MIMAT0000136 | Trim30d | 209387 | Microarrays | 37986224 | Unspecified |
| mmu-miR-125b-5p | MIMAT0000136 | Trim35 | 66854 | HITS-CLIP | 37986224 | Unspecified |
| mmu-miR-125b-5p | MIMAT0000136 | Trim46 | 360213 | Microarrays | 37986224 | Unspecified |
| mmu-miR-125b-5p | MIMAT0000136 | Trim65 | 338364 | Microarrays | 37986224\|22469780 | Unspecified |
| mmu-miR-125b-5p | MIMAT0000136 | Trio | 223435 | HITS-CLIP | 25083871 | Unspecified |
| mmu-miR-125b-5p | MIMAT0000136 | Trip10 | 106628 | Microarrays | 37986224 | Unspecified |
| mmu-miR-125b-5p | MIMAT0000136 | Trip6 | 22051 | Microarrays | 37986224 | Unspecified |
| mmu-miR-125b-5p | MIMAT0000136 | Trit1 | 66966 | Microarrays | 37986224 | Unspecified |
| mmu-miR-125b-5p | MIMAT0000136 | Trmt12 | 68260 | Microarrays | 37986224\|22469780 | Unspecified |
| mmu-miR-125b-5p | MIMAT0000136 | Trmt5 | 76357 | HITS-CLIP | 25753418 | Unspecified |
| mmu-miR-125b-5p | MIMAT0000136 | Trp53 | 22059 | Microarrays | 37986224\|22469780 | Unspecified |

| | | | | | | | |
|---|---|---|---|---|---|---|---|
| mmu-miR-125b-5p | MIMAT0000136 | Trp53i11 | 277414 | Microarrays | | 37986224 | Unspecified |
| mmu-miR-125b-5p | MIMAT0000136 | Trp53inp1 | 60599 | HITS-CLIP, Microarrays//HITS-CLIP//Microa | 37986224\|23597149\|25083871\|22469 | | Unspecified |
| mmu-miR-125b-5p | MIMAT0000136 | Trp53rkb | 76367 | Microarrays | | 22469780 | Unspecified |
| mmu-miR-125b-5p | MIMAT0000136 | Trpc3 | 22065 | Microarrays | | 37986224 | Unspecified |
| mmu-miR-125b-5p | MIMAT0000136 | Trps1 | 83925 | HITS-CLIP | | 37986224 | Unspecified |
| mmu-miR-125b-5p | MIMAT0000136 | Trub1 | 72133 | HITS-CLIP | 37986224\|25083871 | | Unspecified |
| mmu-miR-125b-5p | MIMAT0000136 | Tsacc | 76927 | Microarrays | 37986224\|22469780 | | Unspecified |
| mmu-miR-125b-5p | MIMAT0000136 | Tsc22d1 | 21807 | HITS-CLIP | | 25083871 | Unspecified |
| mmu-miR-125b-5p | MIMAT0000136 | Tsc22d3 | 14605 | Microarrays | | 37986224 | Unspecified |
| mmu-miR-125b-5p | MIMAT0000136 | Tsc22d4 | 78829 | HITS-CLIP | | 37986224 | Unspecified |
| mmu-miR-125b-5p | MIMAT0000136 | Tspan14 | 52588 | HITS-CLIP | | 37986224 | Unspecified |
| mmu-miR-125b-5p | MIMAT0000136 | Tspan2 | 70747 | Microarrays | 37986224\|22469780 | | Unspecified |
| mmu-miR-125b-5p | MIMAT0000136 | Tspan5 | 56224 | HITS-CLIP | | 37986224 | Unspecified |
| mmu-miR-125b-5p | MIMAT0000136 | Tspoap1 | 207777 | Microarrays | | 22469780 | Unspecified |
| mmu-miR-125b-5p | MIMAT0000136 | Tspyl1 | 22110 | HITS-CLIP | | 37986224 | Unspecified |
| mmu-miR-125b-5p | MIMAT0000136 | Tstd2 | 272027 | HITS-CLIP | | 37986224 | Unspecified |
| mmu-miR-125b-5p | MIMAT0000136 | Ttbk1 | 106763 | HITS-CLIP | | 37986224 | Unspecified |
| mmu-miR-125b-5p | MIMAT0000136 | Ttbk2 | 140810 | Microarrays | | 37986224 | Unspecified |
| mmu-miR-125b-5p | MIMAT0000136 | Ttc13 | 234875 | Microarrays | 37986224\|22469780 | | Unspecified |
| mmu-miR-125b-5p | MIMAT0000136 | Ttc17 | 74569 | HITS-CLIP | | 25083871 | Unspecified |
| mmu-miR-125b-5p | MIMAT0000136 | Ttc19 | 72795 | HITS-CLIP | | 37986224 | Unspecified |
| mmu-miR-125b-5p | MIMAT0000136 | Ttc26 | 264134 | Microarrays | | 37986224 | Unspecified |
| mmu-miR-125b-5p | MIMAT0000136 | Ttc33 | 67515 | Microarrays | 37986224\|22469780 | | Unspecified |
| mmu-miR-125b-5p | MIMAT0000136 | Ttc7 | 225049 | HITS-CLIP | 37986224\|23597149 | | Unspecified |
| mmu-miR-125b-5p | MIMAT0000136 | Ttl | 69737 | HITS-CLIP | | 37986224 | Unspecified |
| mmu-miR-125b-5p | MIMAT0000136 | Ttll4 | 67534 | HITS-CLIP | | 37986224 | Unspecified |
| mmu-miR-125b-5p | MIMAT0000136 | Ttll9 | 74711 | Microarrays | 37986224\|22469780 | | Unspecified |
| mmu-miR-125b-5p | MIMAT0000136 | Ttn | 22138 | HITS-CLIP | 37986224\|25083871\|25753418 | | Unspecified |
| mmu-miR-125b-5p | MIMAT0000136 | Ttpal | 76080 | HITS-CLIP | | 37986224 | Unspecified |
| mmu-miR-125b-5p | MIMAT0000136 | Ttyh1 | 57776 | HITS-CLIP | | 37986224 | Unspecified |
| mmu-miR-125b-5p | MIMAT0000136 | Ttyh3 | 78339 | HITS-CLIP | | 37986224 | Unspecified |
| mmu-miR-125b-5p | MIMAT0000136 | Tuba8 | 53857 | Microarrays | | 37986224 | Unspecified |
| mmu-miR-125b-5p | MIMAT0000136 | Tubb2b | 73710 | HITS-CLIP | | 37986224 | Unspecified |
| mmu-miR-125b-5p | MIMAT0000136 | Tubb6 | 67951 | Microarrays | | 37986224 | Unspecified |
| mmu-miR-125b-5p | MIMAT0000136 | Tube1 | 71924 | Microarrays | 37986224\|22469780 | | Unspecified |
| mmu-miR-125b-5p | MIMAT0000136 | Tubg2 | 103768 | Microarrays | | 37986224 | Unspecified |
| mmu-miR-125b-5p | MIMAT0000136 | Tulp3 | 22158 | Microarrays | | 37986224 | Unspecified |
| mmu-miR-125b-5p | MIMAT0000136 | Txn1 | 22166 | Microarrays | 37986224\|22469780 | | Unspecified |
| mmu-miR-125b-5p | MIMAT0000136 | Txn2 | 56551 | HITS-CLIP | | 19536157 | Unspecified |

| | | | | | | |
|---|---|---|---|---|---|---|
| mmu-miR-125b-5p | MIMAT0000136 | Txnip | 56338 | HITS-CLIP, Microarrays//qCLASH//HITS-CLI | 37986224\|31393953\|23597149\|29167 | Unspecified |
| mmu-miR-125b-5p | MIMAT0000136 | Txnrd1 | 50493 | HITS-CLIP | 37986224 | Unspecified |
| mmu-miR-125b-5p | MIMAT0000136 | Tyrobp | 22177 | Microarrays | 37986224 | Unspecified |
| mmu-miR-125b-5p | MIMAT0000136 | U2surp | 67958 | HITS-CLIP | 37986224 | Unspecified |
| mmu-miR-125b-5p | MIMAT0000136 | Uba1 | 22201 | HITS-CLIP | 25083871 | Unspecified |
| mmu-miR-125b-5p | MIMAT0000136 | Ubap2 | 68926 | HITS-CLIP | 37986224\|29167373 | Unspecified |
| mmu-miR-125b-5p | MIMAT0000136 | Ubap2l | 74383 | HITS-CLIP | 25083871\|29167373 | Unspecified |
| mmu-miR-125b-5p | MIMAT0000136 | Ube2d1 | 216080 | Microarrays | 37986224 | Unspecified |
| mmu-miR-125b-5p | MIMAT0000136 | Ube2g1 | 67128 | Microarrays | 37986224\|22469780 | Unspecified |
| mmu-miR-125b-5p | MIMAT0000136 | Ube2g2 | 22213 | HITS-CLIP | 37986224 | Unspecified |
| mmu-miR-125b-5p | MIMAT0000136 | Ube2k | 53323 | HITS-CLIP | 25083871 | Unspecified |
| mmu-miR-125b-5p | MIMAT0000136 | Ube2l6 | 56791 | Microarrays | 37986224 | Unspecified |
| mmu-miR-125b-5p | MIMAT0000136 | Ube2m | 22192 | Microarrays | 37986224 | Unspecified |
| mmu-miR-125b-5p | MIMAT0000136 | Ube2q1 | 70093 | HITS-CLIP//qCLASH | 37986224\|31393953 | Unspecified |
| mmu-miR-125b-5p | MIMAT0000136 | Ube2r2 | 67615 | HITS-CLIP | 37986224 | Unspecified |
| mmu-miR-125b-5p | MIMAT0000136 | Ube2z | 268470 | HITS-CLIP | 37986224 | Unspecified |
| mmu-miR-125b-5p | MIMAT0000136 | Ube4a | 140630 | Microarrays//HITS-CLIP | 37986224\|22469780\|25753418 | Unspecified |
| mmu-miR-125b-5p | MIMAT0000136 | Ubl7 | 69459 | Microarrays | 37986224 | Unspecified |
| mmu-miR-125b-5p | MIMAT0000136 | Ubn1 | 170644 | HITS-CLIP, Microarrays//HITS-CLIP//Microa | 37986224\|25083871\|22469780\|29167 | Unspecified |
| mmu-miR-125b-5p | MIMAT0000136 | Ubn2 | 320538 | HITS-CLIP | 37986224\|25083871 | Unspecified |
| mmu-miR-125b-5p | MIMAT0000136 | Ubqln2 | 54609 | HITS-CLIP | 37986224 | Unspecified |
| mmu-miR-125b-5p | MIMAT0000136 | Ubr3 | 68795 | HITS-CLIP | 25083871\|25753418 | Unspecified |
| mmu-miR-125b-5p | MIMAT0000136 | Ubr4 | 69116 | Microarrays | 37986224 | Unspecified |
| mmu-miR-125b-5p | MIMAT0000136 | Ubr7 | 66622 | HITS-CLIP | 37986224 | Unspecified |
| mmu-miR-125b-5p | MIMAT0000136 | Ubtd1 | 226122 | HITS-CLIP | 37986224\|25083871 | Unspecified |
| mmu-miR-125b-5p | MIMAT0000136 | Ubxn11 | 67586 | Microarrays | 37986224 | Unspecified |
| mmu-miR-125b-5p | MIMAT0000136 | Uck2 | 80914 | qCLASH//HITS-CLIP | 31393953\|19536157 | Unspecified |
| mmu-miR-125b-5p | MIMAT0000136 | Ucp2 | 22228 | Microarrays | 37986224 | Unspecified |
| mmu-miR-125b-5p | MIMAT0000136 | Uevld | 54122 | Microarrays | 37986224 | Unspecified |
| mmu-miR-125b-5p | MIMAT0000136 | Uggt1 | 320011 | HITS-CLIP | 25083871 | Unspecified |
| mmu-miR-125b-5p | MIMAT0000136 | Uggt2 | 66435 | HITS-CLIP | 37986224 | Unspecified |
| mmu-miR-125b-5p | MIMAT0000136 | Ugp2 | 216558 | HITS-CLIP | 37986224 | Unspecified |
| mmu-miR-125b-5p | MIMAT0000136 | Ugt1a6a | 94284 | Microarrays | 37986224\|22469780 | Unspecified |
| mmu-miR-125b-5p | MIMAT0000136 | Ugt1a7c | 394432 | HITS-CLIP | 37986224\|23597149 | Unspecified |
| mmu-miR-125b-5p | MIMAT0000136 | Ugt2b35 | 243085 | HITS-CLIP | 23597149 | Unspecified |
| mmu-miR-125b-5p | MIMAT0000136 | Uhmk1 | 16589 | HITS-CLIP | 37986224 | Unspecified |
| mmu-miR-125b-5p | MIMAT0000136 | Ulk3 | 71742 | HITS-CLIP | 37986224\|25083871\|23597149\|1953€ | Unspecified |
| mmu-miR-125b-5p | MIMAT0000136 | Unc5b | 107449 | HITS-CLIP | 37986224\|25083871 | Unspecified |
| mmu-miR-125b-5p | MIMAT0000136 | Unc5c | 22253 | HITS-CLIP | 37986224 | Unspecified |

| | | | | | | |
|---|---|---|---|---|---|---|
| mmu-miR-125b-5p | MIMAT0000136 | Unc80 | 329178 | HITS-CLIP | 19536157 | Unspecified |
| mmu-miR-125b-5p | MIMAT0000136 | Upf1 | 19704 | HITS-CLIP | 25083871\|23597149 | Unspecified |
| mmu-miR-125b-5p | MIMAT0000136 | Upf3b | 68134 | HITS-CLIP | 37986224\|25083871 | Unspecified |
| mmu-miR-125b-5p | MIMAT0000136 | Upp1 | 22271 | Microarrays | 37986224\|22469780 | Unspecified |
| mmu-miR-125b-5p | MIMAT0000136 | Upp2 | 76654 | HITS-CLIP | 37986224 | Unspecified |
| mmu-miR-125b-5p | MIMAT0000136 | Uqcc1 | 56046 | HITS-CLIP | 37986224 | Unspecified |
| mmu-miR-125b-5p | MIMAT0000136 | Uros | 22276 | Microarrays | 37986224\|22469780 | Unspecified |
| mmu-miR-125b-5p | MIMAT0000136 | Usf3 | 205717 | HITS-CLIP | 25083871\|19536157 | Unspecified |
| mmu-miR-125b-5p | MIMAT0000136 | Ushbp1 | 234395 | Microarrays | 37986224 | Unspecified |
| mmu-miR-125b-5p | MIMAT0000136 | Uso1 | 56041 | HITS-CLIP | 25083871 | Unspecified |
| mmu-miR-125b-5p | MIMAT0000136 | Usp11 | 236733 | Microarrays | 37986224 | Unspecified |
| mmu-miR-125b-5p | MIMAT0000136 | Usp12 | 22217 | HITS-CLIP, Microarrays//HITS-CLIP//Microa | 37986224\|25083871\|22469780 | Unspecified |
| mmu-miR-125b-5p | MIMAT0000136 | Usp18 | 24110 | Microarrays | 37986224 | Unspecified |
| mmu-miR-125b-5p | MIMAT0000136 | Usp2 | 53376 | HITS-CLIP | 37986224 | Unspecified |
| mmu-miR-125b-5p | MIMAT0000136 | Usp22 | 216825 | HITS-CLIP | 25083871 | Unspecified |
| mmu-miR-125b-5p | MIMAT0000136 | Usp32 | 237898 | HITS-CLIP | 25083871 | Unspecified |
| mmu-miR-125b-5p | MIMAT0000136 | Usp34 | 17847 | HITS-CLIP | 37986224\|25083871 | Unspecified |
| mmu-miR-125b-5p | MIMAT0000136 | Usp36 | 72344 | HITS-CLIP | 37986224 | Unspecified |
| mmu-miR-125b-5p | MIMAT0000136 | Usp40 | 227334 | Microarrays | 37986224 | Unspecified |
| mmu-miR-125b-5p | MIMAT0000136 | Usp45 | 77593 | HITS-CLIP | 37986224\|23597149 | Unspecified |
| mmu-miR-125b-5p | MIMAT0000136 | Usp46 | 69727 | HITS-CLIP | 37986224 | Unspecified |
| mmu-miR-125b-5p | MIMAT0000136 | Usp47 | 74996 | HITS-CLIP | 37986224\|23597149\|19536157\|2508 | Unspecified |
| mmu-miR-125b-5p | MIMAT0000136 | Usp53 | 99526 | HITS-CLIP | 37986224 | Unspecified |
| mmu-miR-125b-5p | MIMAT0000136 | Usp7 | 252870 | HITS-CLIP | 37986224\|25083871 | Unspecified |
| mmu-miR-125b-5p | MIMAT0000136 | Usp8 | 84092 | Microarrays | 37986224 | Unspecified |
| mmu-miR-125b-5p | MIMAT0000136 | Usp9x | 22284 | Microarrays//HITS-CLIP | 37986224\|23597149 | Unspecified |
| mmu-miR-125b-5p | MIMAT0000136 | Ust | 338362 | Microarrays | 37986224\|22469780 | Unspecified |
| mmu-miR-125b-5p | MIMAT0000136 | Utp14b | 195434 | HITS-CLIP | 37986224\|25083871 | Unspecified |
| mmu-miR-125b-5p | MIMAT0000136 | Utrn | 22288 | Microarrays | 37986224 | Unspecified |
| mmu-miR-125b-5p | MIMAT0000136 | Uts2b | 224065 | Microarrays | 37986224 | Unspecified |
| mmu-miR-125b-5p | MIMAT0000136 | Uts2r | 217369 | Microarrays | 37986224\|22469780 | Unspecified |
| mmu-miR-125b-5p | MIMAT0000136 | Uvrag | 78610 | HITS-CLIP | 37986224 | Unspecified |
| mmu-miR-125b-5p | MIMAT0000136 | Vamp1 | 22317 | HITS-CLIP | 37986224 | Unspecified |
| mmu-miR-125b-5p | MIMAT0000136 | Vamp2 | 22318 | Microarrays | 37986224 | Unspecified |
| mmu-miR-125b-5p | MIMAT0000136 | Vamp7 | 20955 | Microarrays | 37986224 | Unspecified |
| mmu-miR-125b-5p | MIMAT0000136 | Vangl1 | 229658 | HITS-CLIP | 37986224 | Unspecified |
| mmu-miR-125b-5p | MIMAT0000136 | Vars | 22321 | Microarrays | 37986224 | Unspecified |
| mmu-miR-125b-5p | MIMAT0000136 | Vash1 | 238328 | HITS-CLIP | 37986224 | Unspecified |
| mmu-miR-125b-5p | MIMAT0000136 | Vash2 | 226841 | HITS-CLIP | 37986224 | Unspecified |

| | | | | | | |
|---|---|---|---|---|---|---|
| mmu-miR-125b-5p | MIMAT0000136 | Vasn | 246154 | HITS-CLIP | 37986224 | Unspecified |
| mmu-miR-125b-5p | MIMAT0000136 | Vasp | 22323 | Microarrays | 37986224 | Unspecified |
| mmu-miR-125b-5p | MIMAT0000136 | Vat1 | 26949 | HITS-CLIP, Microarrays | 37986224 | Unspecified |
| mmu-miR-125b-5p | MIMAT0000136 | Vat1l | 270097 | HITS-CLIP | 37986224 | Unspecified |
| mmu-miR-125b-5p | MIMAT0000136 | Vav2 | 22325 | HITS-CLIP | 37986224 | Unspecified |
| mmu-miR-125b-5p | MIMAT0000136 | Vcan | 13003 | HITS-CLIP | 37986224 | Unspecified |
| mmu-miR-125b-5p | MIMAT0000136 | Vcl | 22330 | Microarrays//HITS-CLIP | 37986224\|25083871 | Unspecified |
| mmu-miR-125b-5p | MIMAT0000136 | Vcpip1 | 70675 | HITS-CLIP | 37986224\|25083871 | Unspecified |
| mmu-miR-125b-5p | MIMAT0000136 | Vegfb | 22340 | Microarrays | 37986224 | Unspecified |
| mmu-miR-125b-5p | MIMAT0000136 | Vezf1 | 22344 | HITS-CLIP | 37986224 | Unspecified |
| mmu-miR-125b-5p | MIMAT0000136 | Vim | 22352 | Microarrays | 37986224\|22469780 | Unspecified |
| mmu-miR-125b-5p | MIMAT0000136 | Vipr1 | 22354 | Microarrays | 37986224\|22469780 | Unspecified |
| mmu-miR-125b-5p | MIMAT0000136 | Vmn1r52 | 113849 | Microarrays | 37986224 | Unspecified |
| mmu-miR-125b-5p | MIMAT0000136 | Vmn1r56 | 81015 | Microarrays | 37986224 | Unspecified |
| mmu-miR-125b-5p | MIMAT0000136 | Vmn1r6 | 171193 | Microarrays | 37986224 | Unspecified |
| mmu-miR-125b-5p | MIMAT0000136 | Vmn1r87 | 171261 | Microarrays | 37986224\|22469780 | Unspecified |
| mmu-miR-125b-5p | MIMAT0000136 | Vmn2r110 | 224582 | Microarrays | 37986224 | Unspecified |
| mmu-miR-125b-5p | MIMAT0000136 | Vmn2r26 | 56552 | Microarrays | 37986224\|22469780 | Unspecified |
| mmu-miR-125b-5p | MIMAT0000136 | Vmn2r79 | 621430 | qCLASH | 31393953 | Unspecified |
| mmu-miR-125b-5p | MIMAT0000136 | Vmp1 | 75909 | HITS-CLIP | 25753418 | Unspecified |
| mmu-miR-125b-5p | MIMAT0000136 | Vps13a | 271564 | HITS-CLIP | 37986224\|25083871\|29167373 | Unspecified |
| mmu-miR-125b-5p | MIMAT0000136 | Vps13b | 666173 | HITS-CLIP | 37986224\|23597149 | Unspecified |
| mmu-miR-125b-5p | MIMAT0000136 | Vps13c | 320528 | HITS-CLIP | 37986224\|25083871 | Unspecified |
| mmu-miR-125b-5p | MIMAT0000136 | Vps13d | 230895 | HITS-CLIP | 37986224 | Unspecified |
| mmu-miR-125b-5p | MIMAT0000136 | Vps25 | 28084 | HITS-CLIP | 37986224 | Unspecified |
| mmu-miR-125b-5p | MIMAT0000136 | Vps36 | 70160 | HITS-CLIP | 37986224\|25083871 | Unspecified |
| mmu-miR-125b-5p | MIMAT0000136 | Vps37a | 52348 | HITS-CLIP | 37986224 | Unspecified |
| mmu-miR-125b-5p | MIMAT0000136 | Vps37b | 330192 | HITS-CLIP | 37986224\|25083871 | Unspecified |
| mmu-miR-125b-5p | MIMAT0000136 | Vps4b | 20479 | Luciferase assay//HITS-CLIP//Luciferase re | 25403182\|25083871 | Unspecified |
| mmu-miR-125b-5p | MIMAT0000136 | Vsig8 | 240916 | qCLASH | 31393953 | Unspecified |
| mmu-miR-125b-5p | MIMAT0000136 | Vsir | 64115 | HITS-CLIP | 19536157 | Unspecified |
| mmu-miR-125b-5p | MIMAT0000136 | Vwa8 | 219189 | HITS-CLIP | 37986224\|25083871\|23597149 | Unspecified |
| mmu-miR-125b-5p | MIMAT0000136 | Wars | 22375 | HITS-CLIP | 37986224 | Unspecified |
| mmu-miR-125b-5p | MIMAT0000136 | Wasf2 | 242687 | HITS-CLIP | 37986224 | Unspecified |
| mmu-miR-125b-5p | MIMAT0000136 | Wbp2 | 22378 | Microarrays | 37986224 | Unspecified |
| mmu-miR-125b-5p | MIMAT0000136 | Wdr18 | 216156 | HITS-CLIP | 37986224 | Unspecified |
| mmu-miR-125b-5p | MIMAT0000136 | Wdr43 | 72515 | HITS-CLIP | 37986224 | Unspecified |
| mmu-miR-125b-5p | MIMAT0000136 | Wdr5 | 140858 | HITS-CLIP | 37986224 | Unspecified |
| mmu-miR-125b-5p | MIMAT0000136 | Wdr55 | 67936 | Microarrays | 37986224\|22469780 | Unspecified |

| | | | | | | |
|---|---|---|---|---|---|---|
| mmu-miR-125b-5p | MIMAT0000136 | Wdr5b | 69544 | Microarrays | 37986224\|22469780 | Unspecified |
| mmu-miR-125b-5p | MIMAT0000136 | Wdr6 | 83669 | HITS-CLIP | 37986224\|25753418 | Unspecified |
| mmu-miR-125b-5p | MIMAT0000136 | Wdr60 | 217935 | Microarrays | 37986224 | Unspecified |
| mmu-miR-125b-5p | MIMAT0000136 | Wdr7 | 104082 | HITS-CLIP | 25083871\|25753418 | Unspecified |
| mmu-miR-125b-5p | MIMAT0000136 | Wdr73 | 71968 | Microarrays | 37986224\|22469780 | Unspecified |
| mmu-miR-125b-5p | MIMAT0000136 | Wdr76 | 241627 | HITS-CLIP, Microarrays//Microarrays | 37986224\|22469780 | Unspecified |
| mmu-miR-125b-5p | MIMAT0000136 | Wdr81 | 192652 | Microarrays//HITS-CLIP | 37986224\|22469780\|23597149 | Unspecified |
| mmu-miR-125b-5p | MIMAT0000136 | Wdr82 | 77305 | HITS-CLIP | 25083871 | Unspecified |
| mmu-miR-125b-5p | MIMAT0000136 | Wdr91 | 101240 | Microarrays | 37986224 | Unspecified |
| mmu-miR-125b-5p | MIMAT0000136 | Wfdc17 | 100034251 | Microarrays | 37986224 | Unspecified |
| mmu-miR-125b-5p | MIMAT0000136 | Wfs1 | 22393 | Microarrays | 37986224 | Unspecified |
| mmu-miR-125b-5p | MIMAT0000136 | Whamm | 434204 | Microarrays | 37986224 | Unspecified |
| mmu-miR-125b-5p | MIMAT0000136 | Wif1 | 24117 | Microarrays | 37986224 | Unspecified |
| mmu-miR-125b-5p | MIMAT0000136 | Wipf1 | 215280 | HITS-CLIP | 37986224 | Unspecified |
| mmu-miR-125b-5p | MIMAT0000136 | Wipf2 | 68524 | HITS-CLIP | 37986224\|25083871 | Unspecified |
| mmu-miR-125b-5p | MIMAT0000136 | Wiz | 22404 | HITS-CLIP | 37986224\|25083871 | Unspecified |
| mmu-miR-125b-5p | MIMAT0000136 | Wnk1 | 232341 | HITS-CLIP | 25083871\|25753418\|29167373 | Unspecified |
| mmu-miR-125b-5p | MIMAT0000136 | Wnt2 | 22413 | HITS-CLIP | 37986224\|23597149 | Unspecified |
| mmu-miR-125b-5p | MIMAT0000136 | Wsb2 | 59043 | Microarrays | 37986224\|22469780 | Unspecified |
| mmu-miR-125b-5p | MIMAT0000136 | Wscd2 | 320916 | HITS-CLIP, Microarrays | 37986224 | Unspecified |
| mmu-miR-125b-5p | MIMAT0000136 | Wwc2 | 52357 | HITS-CLIP | 37986224 | Unspecified |
| mmu-miR-125b-5p | MIMAT0000136 | Wwox | 80707 | Microarrays | 37986224 | Unspecified |
| mmu-miR-125b-5p | MIMAT0000136 | Wwp1 | 107568 | HITS-CLIP | 25083871\|23597149 | Unspecified |
| mmu-miR-125b-5p | MIMAT0000136 | Xaf1 | 327959 | Microarrays | 37986224 | Unspecified |
| mmu-miR-125b-5p | MIMAT0000136 | Xbp1 | 22433 | HITS-CLIP | 37986224 | Unspecified |
| mmu-miR-125b-5p | MIMAT0000136 | Xdh | 22436 | HITS-CLIP | 37986224 | Unspecified |
| mmu-miR-125b-5p | MIMAT0000136 | Xirp1 | 22437 | HITS-CLIP | 37986224 | Unspecified |
| mmu-miR-125b-5p | MIMAT0000136 | Xpo1 | 103573 | HITS-CLIP | 25083871 | Unspecified |
| mmu-miR-125b-5p | MIMAT0000136 | Xpo4 | 57258 | HITS-CLIP | 37986224 | Unspecified |
| mmu-miR-125b-5p | MIMAT0000136 | Xpo6 | 74204 | Microarrays | 37986224 | Unspecified |
| mmu-miR-125b-5p | MIMAT0000136 | Xrcc6 | 14375 | HITS-CLIP | 37986224 | Unspecified |
| mmu-miR-125b-5p | MIMAT0000136 | Xrn1 | 24127 | Microarrays//HITS-CLIP | 37986224\|25083871\|22469780 | Unspecified |
| mmu-miR-125b-5p | MIMAT0000136 | Xxylt1 | 268880 | Microarrays | 37986224 | Unspecified |
| mmu-miR-125b-5p | MIMAT0000136 | Ybx1 | 22608 | HITS-CLIP | 25753418 | Unspecified |
| mmu-miR-125b-5p | MIMAT0000136 | Yeats2 | 208146 | Microarrays | 37986224 | Unspecified |
| mmu-miR-125b-5p | MIMAT0000136 | Yeats4 | 64050 | Microarrays | 37986224\|22469780 | Unspecified |
| mmu-miR-125b-5p | MIMAT0000136 | Yes1 | 22612 | HITS-CLIP | 37986224 | Unspecified |
| mmu-miR-125b-5p | MIMAT0000136 | Yod1 | 226418 | HITS-CLIP | 37986224\|23597149\|29167373 | Unspecified |
| mmu-miR-125b-5p | MIMAT0000136 | Ypel2 | 77864 | HITS-CLIP | 37986224 | Unspecified |

| | | | | | | |
|---|---|---|---|---|---|---|
| mmu-miR-125b-5p | MIMAT0000136 | Ypel3 | 66090 | Microarrays | 37986224 | Unspecified |
| mmu-miR-125b-5p | MIMAT0000136 | Ypel5 | 383295 | Microarrays | 37986224 | Unspecified |
| mmu-miR-125b-5p | MIMAT0000136 | Ythdf2 | 213541 | HITS-CLIP | 37986224 | Unspecified |
| mmu-miR-125b-5p | MIMAT0000136 | Ythdf3 | 229096 | HITS-CLIP//qCLASH | 37986224\|31393953\|29167373 | Unspecified |
| mmu-miR-125b-5p | MIMAT0000136 | Ywhab | 54401 | HITS-CLIP | 25083871 | Unspecified |
| mmu-miR-125b-5p | MIMAT0000136 | Ywhag | 22628 | HITS-CLIP | 25083871 | Unspecified |
| mmu-miR-125b-5p | MIMAT0000136 | Ywhaz | 22631 | HITS-CLIP | 37986224 | Unspecified |
| mmu-miR-125b-5p | MIMAT0000136 | Zadh2 | 225791 | Microarrays | 37986224\|22469780 | Unspecified |
| mmu-miR-125b-5p | MIMAT0000136 | Zap70 | 22637 | Microarrays | 37986224\|22469780 | Unspecified |
| mmu-miR-125b-5p | MIMAT0000136 | Zbed4 | 223773 | Microarrays | 37986224\|22469780 | Unspecified |
| mmu-miR-125b-5p | MIMAT0000136 | Zbed6 | 667118 | HITS-CLIP | 37986224 | Unspecified |
| mmu-miR-125b-5p | MIMAT0000136 | Zbtb12 | 193736 | HITS-CLIP | 37986224 | Unspecified |
| mmu-miR-125b-5p | MIMAT0000136 | Zbtb18 | 30928 | Microarrays | 37986224 | Unspecified |
| mmu-miR-125b-5p | MIMAT0000136 | Zbtb20 | 56490 | HITS-CLIP, Microarrays//HITS-CLIP | 37986224\|25753418 | Unspecified |
| mmu-miR-125b-5p | MIMAT0000136 | Zbtb24 | 268294 | HITS-CLIP | 29167373 | Unspecified |
| mmu-miR-125b-5p | MIMAT0000136 | Zbtb32 | 58206 | Microarrays | 37986224 | Unspecified |
| mmu-miR-125b-5p | MIMAT0000136 | Zbtb33 | 56805 | Microarrays | 37986224\|22469780 | Unspecified |
| mmu-miR-125b-5p | MIMAT0000136 | Zbtb34 | 241311 | HITS-CLIP | 37986224 | Unspecified |
| mmu-miR-125b-5p | MIMAT0000136 | Zbtb37 | 240869 | HITS-CLIP | 37986224 | Unspecified |
| mmu-miR-125b-5p | MIMAT0000136 | Zbtb39 | 320080 | HITS-CLIP | 37986224\|25083871 | Unspecified |
| mmu-miR-125b-5p | MIMAT0000136 | Zbtb4 | 75580 | HITS-CLIP, Microarrays | 37986224 | Unspecified |
| mmu-miR-125b-5p | MIMAT0000136 | Zbtb7a | 16969 | HITS-CLIP | 37986224 | Unspecified |
| mmu-miR-125b-5p | MIMAT0000136 | Zbtb9 | 474156 | Microarrays | 37986224\|22469780 | Unspecified |
| mmu-miR-125b-5p | MIMAT0000136 | Zc3h10 | 103284 | Microarrays | 37986224 | Unspecified |
| mmu-miR-125b-5p | MIMAT0000136 | Zc3h12c | 244871 | HITS-CLIP | 37986224 | Unspecified |
| mmu-miR-125b-5p | MIMAT0000136 | Zc3h14 | 75553 | Microarrays//HITS-CLIP | 37986224\|25083871 | Unspecified |
| mmu-miR-125b-5p | MIMAT0000136 | Zc3h6 | 78751 | Microarrays | 37986224 | Unspecified |
| mmu-miR-125b-5p | MIMAT0000136 | Zc3h7b | 20286 | HITS-CLIP | 37986224 | Unspecified |
| mmu-miR-125b-5p | MIMAT0000136 | Zcchc14 | 142682 | HITS-CLIP | 37986224 | Unspecified |
| mmu-miR-125b-5p | MIMAT0000136 | Zcchc2 | 227449 | HITS-CLIP, Microarrays | 37986224 | Unspecified |
| mmu-miR-125b-5p | MIMAT0000136 | Zcchc24 | 71918 | Microarrays | 37986224 | Unspecified |
| mmu-miR-125b-5p | MIMAT0000136 | Zcchc7 | 319885 | Microarrays | 37986224 | Unspecified |
| mmu-miR-125b-5p | MIMAT0000136 | Zdbf2 | 73884 | HITS-CLIP | 37986224 | Unspecified |
| mmu-miR-125b-5p | MIMAT0000136 | Zdhhc1 | 70796 | Microarrays | 37986224 | Unspecified |
| mmu-miR-125b-5p | MIMAT0000136 | Zdhhc14 | 224454 | Microarrays | 37986224 | Unspecified |
| mmu-miR-125b-5p | MIMAT0000136 | Zdhhc18 | 503610 | HITS-CLIP | 37986224 | Unspecified |
| mmu-miR-125b-5p | MIMAT0000136 | Zdhhc20 | 75965 | HITS-CLIP | 37986224\|29167373 | Unspecified |
| mmu-miR-125b-5p | MIMAT0000136 | Zdhhc21 | 68268 | HITS-CLIP | 25753418 | Unspecified |
| mmu-miR-125b-5p | MIMAT0000136 | Zdhhc22 | 238331 | Microarrays | 37986224 | Unspecified |

| | | | | | | |
|---|---|---|---|---|---|---|
| mmu-miR-125b-5p | MIMAT0000136 | Zdhhc24 | 70605 | Microarrays | 37986224\|22469780 | Unspecified |
| mmu-miR-125b-5p | MIMAT0000136 | Zdhhc5 | 228136 | HITS-CLIP | 37986224 | Unspecified |
| mmu-miR-125b-5p | MIMAT0000136 | Zdhhc7 | 102193 | HITS-CLIP | 37986224\|19536157 | Unspecified |
| mmu-miR-125b-5p | MIMAT0000136 | Zdhhc9 | 208884 | HITS-CLIP | 37986224 | Unspecified |
| mmu-miR-125b-5p | MIMAT0000136 | Zeb1 | 21417 | HITS-CLIP, Microarrays//HITS-CLIP | 37986224\|25753418 | Unspecified |
| mmu-miR-125b-5p | MIMAT0000136 | Zeb2 | 24136 | HITS-CLIP | 37986224 | Unspecified |
| mmu-miR-125b-5p | MIMAT0000136 | Zer1 | 227693 | Microarrays | 37986224 | Unspecified |
| mmu-miR-125b-5p | MIMAT0000136 | Zfand2a | 100494 | Microarrays | 37986224\|22469780 | Unspecified |
| mmu-miR-125b-5p | MIMAT0000136 | Zfand3 | 21769 | HITS-CLIP | 25083871 | Unspecified |
| mmu-miR-125b-5p | MIMAT0000136 | Zfhx2 | 239102 | HITS-CLIP | 37986224\|25753418 | Unspecified |
| mmu-miR-125b-5p | MIMAT0000136 | Zfhx3 | 11906 | HITS-CLIP | 25083871 | Unspecified |
| mmu-miR-125b-5p | MIMAT0000136 | Zfhx4 | 80892 | HITS-CLIP | 37986224\|25083871 | Unspecified |
| mmu-miR-125b-5p | MIMAT0000136 | Zfp106 | 20402 | Microarrays | 37986224 | Unspecified |
| mmu-miR-125b-5p | MIMAT0000136 | Zfp113 | 56314 | HITS-CLIP | 37986224 | Unspecified |
| mmu-miR-125b-5p | MIMAT0000136 | Zfp120 | 104348 | HITS-CLIP | 37986224 | Unspecified |
| mmu-miR-125b-5p | MIMAT0000136 | Zfp148 | 22661 | HITS-CLIP | 37986224 | Unspecified |
| mmu-miR-125b-5p | MIMAT0000136 | Zfp24 | 59057 | Microarrays | 22469780 | Unspecified |
| mmu-miR-125b-5p | MIMAT0000136 | Zfp248 | 72720 | HITS-CLIP | 37986224 | Unspecified |
| mmu-miR-125b-5p | MIMAT0000136 | Zfp26 | 22688 | HITS-CLIP | 37986224 | Unspecified |
| mmu-miR-125b-5p | MIMAT0000136 | Zfp260 | 26466 | Microarrays | 37986224\|22469780 | Unspecified |
| mmu-miR-125b-5p | MIMAT0000136 | Zfp281 | 226442 | HITS-CLIP, Microarrays//HITS-CLIP//qCLASI | 37986224\|25083871\|31393953\|23597 | Unspecified |
| mmu-miR-125b-5p | MIMAT0000136 | Zfp282 | 101095 | HITS-CLIP, Microarrays//Microarrays | 37986224\|22469780 | Unspecified |
| mmu-miR-125b-5p | MIMAT0000136 | Zfp292 | 30046 | HITS-CLIP | 37986224\|25083871\|25753418 | Unspecified |
| mmu-miR-125b-5p | MIMAT0000136 | Zfp319 | 79233 | HITS-CLIP | 37986224 | Unspecified |
| mmu-miR-125b-5p | MIMAT0000136 | Zfp322a | 218100 | HITS-CLIP, Microarrays//HITS-CLIP//Microa | 37986224\|25083871\|22469780 | Unspecified |
| mmu-miR-125b-5p | MIMAT0000136 | Zfp329 | 67230 | HITS-CLIP | 37986224 | Unspecified |
| mmu-miR-125b-5p | MIMAT0000136 | Zfp346 | 26919 | HITS-CLIP, Microarrays | 37986224 | Unspecified |
| mmu-miR-125b-5p | MIMAT0000136 | Zfp352 | 236537 | Microarrays | 37986224\|22469780 | Unspecified |
| mmu-miR-125b-5p | MIMAT0000136 | Zfp362 | 230761 | HITS-CLIP | 37986224 | Unspecified |
| mmu-miR-125b-5p | MIMAT0000136 | Zfp36l2 | 12193 | Microarrays | 37986224 | Unspecified |
| mmu-miR-125b-5p | MIMAT0000136 | Zfp36l3 | 333473 | Microarrays | 37986224\|22469780 | Unspecified |
| mmu-miR-125b-5p | MIMAT0000136 | Zfp385a | 29813 | Microarray//HITS-CLIP//Other | 16495412\|25083871 | Unspecified |
| mmu-miR-125b-5p | MIMAT0000136 | Zfp395 | 380912 | Microarrays | 37986224 | Unspecified |
| mmu-miR-125b-5p | MIMAT0000136 | Zfp397 | 69256 | HITS-CLIP | 37986224 | Unspecified |
| mmu-miR-125b-5p | MIMAT0000136 | Zfp408 | 381410 | HITS-CLIP | 37986224 | Unspecified |
| mmu-miR-125b-5p | MIMAT0000136 | Zfp41 | 22701 | qCLASH | 31393953 | Unspecified |
| mmu-miR-125b-5p | MIMAT0000136 | Zfp410 | 52708 | HITS-CLIP | 37986224 | Unspecified |
| mmu-miR-125b-5p | MIMAT0000136 | Zfp414 | 328801 | HITS-CLIP | 37986224 | Unspecified |
| mmu-miR-125b-5p | MIMAT0000136 | Zfp451 | 98403 | HITS-CLIP | 37986224 | Unspecified |

| | | | | | | |
|---|---|---|---|---|---|---|
| mmu-miR-125b-5p | MIMAT0000136 | Zfp507 | 668501 | Microarrays | 37986224\|22469780 | Unspecified |
| mmu-miR-125b-5p | MIMAT0000136 | Zfp516 | 329003 | HITS-CLIP | 37986224 | Unspecified |
| mmu-miR-125b-5p | MIMAT0000136 | Zfp518a | 72672 | HITS-CLIP | 37986224 | Unspecified |
| mmu-miR-125b-5p | MIMAT0000136 | Zfp523 | 7629 | HITS-CLIP | 25753418 | Unspecified |
| mmu-miR-125b-5p | MIMAT0000136 | Zfp536 | 243937 | Microarrays | 37986224 | Unspecified |
| mmu-miR-125b-5p | MIMAT0000136 | Zfp568 | 243905 | Microarrays | 37986224\|22469780 | Unspecified |
| mmu-miR-125b-5p | MIMAT0000136 | Zfp57 | 22715 | HITS-CLIP | 37986224 | Unspecified |
| mmu-miR-125b-5p | MIMAT0000136 | Zfp597 | 71063 | HITS-CLIP | 37986224 | Unspecified |
| mmu-miR-125b-5p | MIMAT0000136 | Zfp608 | 269023 | HITS-CLIP | 37986224 | Unspecified |
| mmu-miR-125b-5p | MIMAT0000136 | Zfp609 | 214812 | HITS-CLIP, Microarrays | 37986224 | Unspecified |
| mmu-miR-125b-5p | MIMAT0000136 | Zfp62 | 22720 | HITS-CLIP, Microarrays//Microarrays | 37986224\|22469780 | Unspecified |
| mmu-miR-125b-5p | MIMAT0000136 | Zfp652 | 268469 | HITS-CLIP | 37986224\|23597149 | Unspecified |
| mmu-miR-125b-5p | MIMAT0000136 | Zfp653 | 319601 | Microarrays | 37986224\|22469780 | Unspecified |
| mmu-miR-125b-5p | MIMAT0000136 | Zfp654 | 72020 | qCLASH | 31393953 | Unspecified |
| mmu-miR-125b-5p | MIMAT0000136 | Zfp664 | 269704 | HITS-CLIP | 37986224 | Unspecified |
| mmu-miR-125b-5p | MIMAT0000136 | Zfp672 | 319475 | Microarrays | 37986224\|22469780 | Unspecified |
| mmu-miR-125b-5p | MIMAT0000136 | Zfp697 | 242109 | HITS-CLIP | 37986224 | Unspecified |
| mmu-miR-125b-5p | MIMAT0000136 | Zfp703 | 353310 | HITS-CLIP | 37986224 | Unspecified |
| mmu-miR-125b-5p | MIMAT0000136 | Zfp747 | 269997 | Microarrays | 37986224\|22469780 | Unspecified |
| mmu-miR-125b-5p | MIMAT0000136 | Zfp758 | 224598 | HITS-CLIP | 37986224 | Unspecified |
| mmu-miR-125b-5p | MIMAT0000136 | Zfp799 | 240064 | Microarrays | 37986224\|22469780 | Unspecified |
| mmu-miR-125b-5p | MIMAT0000136 | Zfp820 | 75424 | HITS-CLIP | 37986224\|25083871 | Unspecified |
| mmu-miR-125b-5p | MIMAT0000136 | Zfp827 | 622675 | Microarrays | 37986224\|22469780 | Unspecified |
| mmu-miR-125b-5p | MIMAT0000136 | Zfp830 | 66983 | Microarrays | 37986224\|22469780 | Unspecified |
| mmu-miR-125b-5p | MIMAT0000136 | Zfp84 | 74352 | Microarrays | 37986224\|22469780 | Unspecified |
| mmu-miR-125b-5p | MIMAT0000136 | Zfp879 | 214779 | Microarrays | 37986224\|22469780 | Unspecified |
| mmu-miR-125b-5p | MIMAT0000136 | Zfp91 | 109910 | HITS-CLIP | 37986224 | Unspecified |
| mmu-miR-125b-5p | MIMAT0000136 | Zfp938 | 237411 | Microarrays | 37986224\|22469780 | Unspecified |
| mmu-miR-125b-5p | MIMAT0000136 | Zfp943 | 74670 | Microarrays | 37986224\|22469780 | Unspecified |
| mmu-miR-125b-5p | MIMAT0000136 | Zfp947 | 210853 | Microarrays | 37986224\|22469780 | Unspecified |
| mmu-miR-125b-5p | MIMAT0000136 | Zfp949 | 71640 | HITS-CLIP | 37986224 | Unspecified |
| mmu-miR-125b-5p | MIMAT0000136 | Zfp953 | 629016 | Microarrays | 37986224\|22469780 | Unspecified |
| mmu-miR-125b-5p | MIMAT0000136 | Zfp982 | 195531 | Microarrays | 22469780 | Unspecified |
| mmu-miR-125b-5p | MIMAT0000136 | Zfr | 22763 | Microarrays | 37986224\|22469780 | Unspecified |
| mmu-miR-125b-5p | MIMAT0000136 | Zfr2 | 103406 | Microarrays | 37986224 | Unspecified |
| mmu-miR-125b-5p | MIMAT0000136 | Zfyve1 | 217695 | HITS-CLIP | 37986224\|25083871 | Unspecified |
| mmu-miR-125b-5p | MIMAT0000136 | Zfyve21 | 68520 | HITS-CLIP | 37986224 | Unspecified |
| mmu-miR-125b-5p | MIMAT0000136 | Zfyve26 | 211978 | HITS-CLIP | 37986224 | Unspecified |
| mmu-miR-125b-5p | MIMAT0000136 | Zfyve9 | 230597 | HITS-CLIP | 37986224 | Unspecified |

| | | | | | | |
|---|---|---|---|---|---|---|
| mmu-miR-125b-5p | MIMAT0000136 | Zgrf1 | 71643 | Microarrays | 37986224\|22469780 | Unspecified |
| mmu-miR-125b-5p | MIMAT0000136 | Zhx3 | 320799 | HITS-CLIP | 37986224 | Unspecified |
| mmu-miR-125b-5p | MIMAT0000136 | Zim1 | 22776 | HITS-CLIP | 37986224 | Unspecified |
| mmu-miR-125b-5p | MIMAT0000136 | Zkscan17 | 268417 | HITS-CLIP | 37986224 | Unspecified |
| mmu-miR-125b-5p | MIMAT0000136 | Zkscan3 | 72739 | HITS-CLIP | 37986224 | Unspecified |
| mmu-miR-125b-5p | MIMAT0000136 | Zkscan8 | 93681 | HITS-CLIP, Microarrays//HITS-CLIP//Microa | 37986224\|25083871\|22469780 | Unspecified |
| mmu-miR-125b-5p | MIMAT0000136 | Zmiz2 | 52915 | Microarrays | 37986224 | Unspecified |
| mmu-miR-125b-5p | MIMAT0000136 | Zmym1 | 68310 | Microarrays | 37986224\|22469780 | Unspecified |
| mmu-miR-125b-5p | MIMAT0000136 | Zmym2 | 76007 | HITS-CLIP | 37986224\|25083871\|29167373 | Unspecified |
| mmu-miR-125b-5p | MIMAT0000136 | Zmynd12 | 332934 | Microarrays | 37986224 | Unspecified |
| mmu-miR-125b-5p | MIMAT0000136 | Zmynd8 | 228880 | HITS-CLIP | 37986224 | Unspecified |
| mmu-miR-125b-5p | MIMAT0000136 | Znfx1 | 98999 | HITS-CLIP | 37986224 | Unspecified |
| mmu-miR-125b-5p | MIMAT0000136 | Znhit3 | 448850 | Microarrays | 37986224\|22469780 | Unspecified |
| mmu-miR-125b-5p | MIMAT0000136 | Znrf1 | 170737 | HITS-CLIP | 37986224 | Unspecified |
| mmu-miR-125b-5p | MIMAT0000136 | Znrf2 | 387524 | HITS-CLIP | 37986224 | Unspecified |
| mmu-miR-125b-5p | MIMAT0000136 | Znrf3 | 407821 | HITS-CLIP, Microarrays//HITS-CLIP | 37986224\|25083871 | Unspecified |
| mmu-miR-125b-5p | MIMAT0000136 | Zscan2 | 22691 | Microarrays | 37986224 | Unspecified |
| mmu-miR-125b-5p | MIMAT0000136 | Zscan21 | 22697 | Microarrays | 37986224\|22469780 | Unspecified |
| mmu-miR-125b-5p | MIMAT0000136 | Zscan26 | 432731 | HITS-CLIP | 37986224 | Unspecified |
| mmu-miR-125b-5p | MIMAT0000136 | Zscan29 | 99334 | HITS-CLIP | 37986224\|25083871 | Unspecified |
| mmu-miR-125b-5p | MIMAT0000136 | Zswim3 | 67538 | HITS-CLIP | 29167373 | Unspecified |
| mmu-miR-125b-5p | MIMAT0000136 | Zswim5 | 74464 | HITS-CLIP | 37986224 | Unspecified |
| mmu-miR-125b-5p | MIMAT0000136 | Zswim6 | 67263 | HITS-CLIP | 37986224 | Unspecified |
| mmu-miR-125b-5p | MIMAT0000136 | Zswim8 | 268721 | HITS-CLIP | 25083871 | Unspecified |
| mmu-miR-125b-5p | MIMAT0000136 | Zxdc | 80292 | HITS-CLIP | 37986224 | Unspecified |
| mmu-miR-125b-5p | MIMAT0000136 | mt-Co1 | 4512 | qCLASH | 31393953 | Unspecified |
| mmu-miR-125b-5p | MIMAT0000136 | mt-Nd1 | 4535 | HITS-CLIP//qCLASH | 23597149\|31393953 | Unspecified |
| mmu-miR-125b-5p | MIMAT0000136 | mt-Nd3 | 4537 | Microarrays | 22469780 | Unspecified |
| mmu-miR-125b-5p | MIMAT0000136 | mt-Nd6 | 4541 | qCLASH | 31393953 | Unspecified |
| mmu-miR-3095-5p | MIMAT0014911 | Antxr2 | 71914 | HITS-CLIP | 31433990 | Unspecified |
| mmu-miR-3095-5p | MIMAT0014911 | Stk39 | 53416 | HITS-CLIP | 31433990 | Unspecified |
| mmu-miR-3079-3p | MIMAT0014867 | B4galt1 | 14595 | HITS-CLIP | 26203562 | Unspecified |
| mmu-miR-3079-3p | MIMAT0014867 | Srsf6 | 67996 | HITS-CLIP | 31433990 | Unspecified |
| mmu-miR-3079-3p | MIMAT0014867 | Zfp292 | 30046 | HITS-CLIP | 31433990 | Unspecified |
| mmu-miR-509-3p | MIMAT0004891 | AW146154 | 101835 | HITS-CLIP | 26203562 | Unspecified |
| mmu-miR-509-3p | MIMAT0004891 | Ahcyl2 | 74340 | HITS-CLIP | 31433990 | Unspecified |
| mmu-miR-509-3p | MIMAT0004891 | Brms1l | 52592 | HITS-CLIP | 26203562 | Unspecified |
| mmu-miR-509-3p | MIMAT0004891 | Ccar1 | 67500 | HITS-CLIP | 26203562 | Unspecified |
| mmu-miR-509-3p | MIMAT0004891 | Dcp2 | 70640 | HITS-CLIP | 31433990 | Unspecified |

| | | | | | | |
|---|---|---|---|---|---|---|
| mmu-miR-509-3p | MIMAT0004891 | Dgkd | 227333 | HITS-CLIP | 31433990 | Unspecified |
| mmu-miR-509-3p | MIMAT0004891 | Dglucy | 217830 | HITS-CLIP | 26203562 | Unspecified |
| mmu-miR-509-3p | MIMAT0004891 | Dnajc3 | 100037258 | HITS-CLIP | 26203562 | Unspecified |
| mmu-miR-509-3p | MIMAT0004891 | Epc1 | 13831 | HITS-CLIP | 31433990 | Unspecified |
| mmu-miR-509-3p | MIMAT0004891 | Foxo3 | 56484 | HITS-CLIP | 31433990 | Unspecified |
| mmu-miR-509-3p | MIMAT0004891 | Gimap6 | 231931 | HITS-CLIP | 31433990 | Unspecified |
| mmu-miR-509-3p | MIMAT0004891 | Gm45234 | | HITS-CLIP | 31433990 | Unspecified |
| mmu-miR-509-3p | MIMAT0004891 | Grcc10 | 14790 | HITS-CLIP | 31433990 | Unspecified |
| mmu-miR-509-3p | MIMAT0004891 | Il2ra | 16184 | HITS-CLIP | 31433990 | Unspecified |
| mmu-miR-509-3p | MIMAT0004891 | Kif5b | 16573 | HITS-CLIP | 31433990 | Unspecified |
| mmu-miR-509-3p | MIMAT0004891 | Midn | 59090 | HITS-CLIP | 31433990 | Unspecified |
| mmu-miR-509-3p | MIMAT0004891 | Ndufs6 | 407785 | HITS-CLIP | 31433990 | Unspecified |
| mmu-miR-509-3p | MIMAT0004891 | Neurl4 | 216860 | HITS-CLIP | 26203562 | Unspecified |
| mmu-miR-509-3p | MIMAT0004891 | Otud1 | 71198 | HITS-CLIP | 31433990 | Unspecified |
| mmu-miR-509-3p | MIMAT0004891 | Plcxd2 | 433022 | HITS-CLIP | 26203562 | Unspecified |
| mmu-miR-509-3p | MIMAT0004891 | Plk3 | 12795 | HITS-CLIP | 26203562 | Unspecified |
| mmu-miR-509-3p | MIMAT0004891 | S100a10 | 20194 | HITS-CLIP | 31433990 | Unspecified |
| mmu-miR-509-3p | MIMAT0004891 | Tsnax | 53424 | HITS-CLIP | 31433990 | Unspecified |
| mmu-miR-509-3p | MIMAT0004891 | Tubgcp3 | 259279 | HITS-CLIP | 31433990 | Unspecified |
| mmu-miR-509-3p | MIMAT0004891 | Zfp367 | 238673 | HITS-CLIP | 26203562 | Unspecified |
| mmu-miR-3068-5p | MIMAT0014842 | Cct6a | 12466 | HITS-CLIP | 37986224 | Unspecified |
| mmu-miR-3068-5p | MIMAT0014842 | Chp1 | 56398 | qCLASH | 31393953 | Unspecified |
| mmu-miR-3068-5p | MIMAT0014842 | Dcaf10 | 242418 | HITS-CLIP | 37986224 | Unspecified |
| mmu-miR-3068-5p | MIMAT0014842 | Kdm5a | 214899 | qCLASH | 31393953 | Unspecified |
| mmu-miR-3068-5p | MIMAT0014842 | Sfpq | 71514 | HITS-CLIP | 37986224 | Unspecified |
| mmu-miR-3068-5p | MIMAT0014842 | Trak1 | 67095 | HITS-CLIP | 37986224 | Unspecified |
| mmu-miR-3068-5p | MIMAT0014842 | Usp31 | 76179 | HITS-CLIP | 37986224 | Unspecified |
| mmu-miR-384-3p | MIMAT0001076 | Chmp1b | 67078 | HITS-CLIP | 26203562 | Unspecified |
| mmu-miR-384-3p | MIMAT0001076 | Rad18 | 58186 | HITS-CLIP | 31433990 | Unspecified |
| mmu-miR-384-3p | MIMAT0001076 | Spcs3 | 76687 | HITS-CLIP | 31433990 | Unspecified |
| mmu-miR-28c | MIMAT0019339 | Creb1 | 12912 | HITS-CLIP | 31433990 | Unspecified |
| mmu-miR-28c | MIMAT0019339 | Foxk1 | 17425 | HITS-CLIP | 31433990 | Unspecified |
| mmu-miR-28c | MIMAT0019339 | Hnrnpa0 | 77134 | HITS-CLIP | 31433990 | Unspecified |
| mmu-miR-28c | MIMAT0019339 | Srsf6 | 67996 | HITS-CLIP | 31433990 | Unspecified |
| mmu-miR-147-3p | MIMAT0004857 | Aco1 | 11428 | HITS-CLIP | 26203562 | Unspecified |
| mmu-miR-147-3p | MIMAT0004857 | Actb | 11461 | HITS-CLIP | 31433990 | Unspecified |
| mmu-miR-147-3p | MIMAT0004857 | Actg1 | 11465 | HITS-CLIP | 31433990 | Unspecified |
| mmu-miR-147-3p | MIMAT0004857 | Acvr1b | 11479 | HITS-CLIP | 31433990\|26203562 | Unspecified |
| mmu-miR-147-3p | MIMAT0004857 | Aplp2 | 11804 | HITS-CLIP | 26203562 | Unspecified |

| | | | | | | |
|---|---|---|---|---|---|---|
| mmu-miR-147-3p | MIMAT0004857 | Arl6ip1 | 54208 | HITS-CLIP | 31433990 | Unspecified |
| mmu-miR-147-3p | MIMAT0004857 | Atad2b | 320817 | HITS-CLIP | 26203562 | Unspecified |
| mmu-miR-147-3p | MIMAT0004857 | Atp2a2 | 11938 | HITS-CLIP | 31433990 | Unspecified |
| mmu-miR-147-3p | MIMAT0004857 | Bach2 | 12014 | HITS-CLIP | 31433990 | Unspecified |
| mmu-miR-147-3p | MIMAT0004857 | Bcl11b | 58208 | HITS-CLIP | 31433990 | Unspecified |
| mmu-miR-147-3p | MIMAT0004857 | Cep120 | 225523 | HITS-CLIP | 31433990 | Unspecified |
| mmu-miR-147-3p | MIMAT0004857 | Cibar1 | 137392 | HITS-CLIP | 31433990 | Unspecified |
| mmu-miR-147-3p | MIMAT0004857 | Cnot6l | 231464 | HITS-CLIP | 31433990 | Unspecified |
| mmu-miR-147-3p | MIMAT0004857 | Dcun1d3 | 233805 | HITS-CLIP | 31433990 | Unspecified |
| mmu-miR-147-3p | MIMAT0004857 | Dennd1b | 329260 | HITS-CLIP | 31433990 | Unspecified |
| mmu-miR-147-3p | MIMAT0004857 | Ercc1 | 13870 | HITS-CLIP | 31433990 | Unspecified |
| mmu-miR-147-3p | MIMAT0004857 | Fkbp8 | 14232 | HITS-CLIP | 31433990 | Unspecified |
| mmu-miR-147-3p | MIMAT0004857 | Gimap6 | 231931 | HITS-CLIP | 31433990 | Unspecified |
| mmu-miR-147-3p | MIMAT0004857 | Gnas | 14683 | HITS-CLIP | 31433990 | Unspecified |
| mmu-miR-147-3p | MIMAT0004857 | Igfbp2 | 16008 | HITS-CLIP | 26203562 | Unspecified |
| mmu-miR-147-3p | MIMAT0004857 | Insig1 | 231070 | HITS-CLIP | 31433990 | Unspecified |
| mmu-miR-147-3p | MIMAT0004857 | Kank2 | 235041 | HITS-CLIP | 26203562 | Unspecified |
| mmu-miR-147-3p | MIMAT0004857 | Klhdc3 | 71765 | HITS-CLIP | 31433990 | Unspecified |
| mmu-miR-147-3p | MIMAT0004857 | Kmt2a | 214162 | HITS-CLIP | 31433990\|26203562 | Unspecified |
| mmu-miR-147-3p | MIMAT0004857 | Larp4b | 217980 | HITS-CLIP | 31433990 | Unspecified |
| mmu-miR-147-3p | MIMAT0004857 | Mob1a | 232157 | HITS-CLIP | 31433990 | Unspecified |
| mmu-miR-147-3p | MIMAT0004857 | Myc | 17869 | HITS-CLIP | 31433990 | Unspecified |
| mmu-miR-147-3p | MIMAT0004857 | Ndufa4 | 17992 | qCLASH | 31393953 | Unspecified |
| mmu-miR-147-3p | MIMAT0004857 | Nedd4 | 17999 | HITS-CLIP | 26203562 | Unspecified |
| mmu-miR-147-3p | MIMAT0004857 | Nkiras2 | 71966 | HITS-CLIP | 31433990 | Unspecified |
| mmu-miR-147-3p | MIMAT0004857 | Nsmce1 | 67711 | HITS-CLIP | 31433990 | Unspecified |
| mmu-miR-147-3p | MIMAT0004857 | Ovol1 | 18426 | HITS-CLIP | 26203562 | Unspecified |
| mmu-miR-147-3p | MIMAT0004857 | Pde3b | 18576 | HITS-CLIP | 31433990 | Unspecified |
| mmu-miR-147-3p | MIMAT0004857 | Peg3 | 18616 | HITS-CLIP | 26203562 | Unspecified |
| mmu-miR-147-3p | MIMAT0004857 | Polr2a | 20020 | HITS-CLIP | 31433990 | Unspecified |
| mmu-miR-147-3p | MIMAT0004857 | Prkaca | 18747 | HITS-CLIP | 31433990 | Unspecified |
| mmu-miR-147-3p | MIMAT0004857 | Prpf40a | 56194 | HITS-CLIP | 31433990 | Unspecified |
| mmu-miR-147-3p | MIMAT0004857 | Rnf2 | 19821 | HITS-CLIP | 31433990 | Unspecified |
| mmu-miR-147-3p | MIMAT0004857 | Ski | 20481 | HITS-CLIP | 31433990 | Unspecified |
| mmu-miR-147-3p | MIMAT0004857 | Socs4 | 67296 | HITS-CLIP | 31433990 | Unspecified |
| mmu-miR-147-3p | MIMAT0004857 | Sptbn1 | 20742 | HITS-CLIP | 31433990 | Unspecified |
| mmu-miR-147-3p | MIMAT0004857 | St7 | 7982 | HITS-CLIP | 31433990 | Unspecified |
| mmu-miR-147-3p | MIMAT0004857 | Taf4 | 6874 | HITS-CLIP | 26203562 | Unspecified |
| mmu-miR-147-3p | MIMAT0004857 | Ttc7 | 225049 | HITS-CLIP | 31433990 | Unspecified |

| | | | | | | |
|---|---|---|---|---|---|---|
| mmu-miR-147-3p | MIMAT0004857 | Ube2s | 77891 | HITS-CLIP | 31433990 | Unspecified |
| mmu-miR-147-3p | MIMAT0004857 | Ulbp1 | 77777 | HITS-CLIP | 31433990 | Unspecified |
| mmu-miR-147-3p | MIMAT0004857 | Usp14 | 59025 | HITS-CLIP | 31433990 | Unspecified |
| mmu-miR-147-3p | MIMAT0004857 | Wnk1 | 232341 | HITS-CLIP | 26203562 | Unspecified |
| mmu-miR-721 | MIMAT0003515 | Ago2 | 239528 | HITS-CLIP | 31433990 | Unspecified |
| mmu-miR-721 | MIMAT0003515 | Fam107b | 66540 | HITS-CLIP | 26203562 | Unspecified |
| mmu-miR-721 | MIMAT0003515 | Hdac7 | 56233 | HITS-CLIP | 31433990 | Unspecified |
| mmu-miR-721 | MIMAT0003515 | Kidins220 | 77480 | HITS-CLIP | 31433990 | Unspecified |
| mmu-miR-721 | MIMAT0003515 | Lats1 | 16798 | HITS-CLIP | 31433990 | Unspecified |
| mmu-miR-721 | MIMAT0003515 | Parp3 | 235587 | HITS-CLIP | 31433990 | Unspecified |
| mmu-miR-721 | MIMAT0003515 | Plpbp | 11212 | HITS-CLIP | 31433990 | Unspecified |
| mmu-miR-721 | MIMAT0003515 | Smad7 | 17131 | HITS-CLIP | 31433990 | Unspecified |
| mmu-miR-721 | MIMAT0003515 | Smarcd2 | 83796 | HITS-CLIP | 26203562 | Unspecified |
| mmu-miR-721 | MIMAT0003515 | Sub1 | 20024 | HITS-CLIP | 31433990 | Unspecified |
| mmu-miR-721 | MIMAT0003515 | Suz12 | 52615 | HITS-CLIP | 31433990 | Unspecified |
| mmu-miR-721 | MIMAT0003515 | Tbc1d10c | 108995 | HITS-CLIP | 31433990 | Unspecified |
| mmu-miR-721 | MIMAT0003515 | Tgfbr1 | 21812 | HITS-CLIP | 31433990 | Unspecified |
| mmu-miR-721 | MIMAT0003515 | Zbtb39 | 320080 | HITS-CLIP | 31433990 | Unspecified |
| mmu-miR-721 | MIMAT0003515 | Zfp800 | 627049 | HITS-CLIP | 31433990 | Unspecified |
| mmu-miR-383-5p | MIMAT0000748 | Adrb2 | 11555 | HITS-CLIP | 26203562 | Unspecified |
| mmu-miR-383-5p | MIMAT0000748 | B4galt1 | 14595 | HITS-CLIP | 26203562 | Unspecified |
| mmu-miR-383-5p | MIMAT0000748 | Bcl7c | 12055 | HITS-CLIP | 26203562 | Unspecified |
| mmu-miR-383-5p | MIMAT0000748 | Clptm1l | 218335 | HITS-CLIP | 31433990 | Unspecified |
| mmu-miR-383-5p | MIMAT0000748 | Dgke | 56077 | HITS-CLIP | 26203562 | Unspecified |
| mmu-miR-383-5p | MIMAT0000748 | Eme2 | 193838 | HITS-CLIP | 26203562 | Unspecified |
| mmu-miR-383-5p | MIMAT0000748 | Epha2 | 13836 | HITS-CLIP | 26203562 | Unspecified |
| mmu-miR-383-5p | MIMAT0000748 | Fas | 14102 | qPCR | 37986224 | Unspecified |
| mmu-miR-383-5p | MIMAT0000748 | Mapk8ip3 | 30957 | HITS-CLIP | 26203562 | Unspecified |
| mmu-miR-383-5p | MIMAT0000748 | Marchf8 | 220972 | HITS-CLIP | 26203562 | Unspecified |
| mmu-miR-383-5p | MIMAT0000748 | Myc | 17869 | Luciferase Reporter Assay, qPCR, Western | 37986224\|22593182 | Unspecified |
| mmu-miR-383-5p | MIMAT0000748 | Nfe2l1 | 18023 | HITS-CLIP | 26203562 | Unspecified |
| mmu-miR-383-5p | MIMAT0000748 | Nfil3 | 18030 | HITS-CLIP | 26203562 | Unspecified |
| mmu-miR-383-5p | MIMAT0000748 | Polr2l | 66491 | HITS-CLIP | 31433990 | Unspecified |
| mmu-miR-383-5p | MIMAT0000748 | Prr13 | 66151 | HITS-CLIP | 31433990 | Unspecified |
| mmu-miR-383-5p | MIMAT0000748 | Psme4 | 103554 | HITS-CLIP | 26203562 | Unspecified |
| mmu-miR-383-5p | MIMAT0000748 | Seh1l | 72124 | HITS-CLIP | 31433990 | Unspecified |
| mmu-miR-383-5p | MIMAT0000748 | Selenot | 69227 | HITS-CLIP | 31433990 | Unspecified |
| mmu-miR-383-5p | MIMAT0000748 | Sf1 | 22668 | Western Blot | 37986224 | Unspecified |
| mmu-miR-383-5p | MIMAT0000748 | Ube2z | 268470 | HITS-CLIP | 31433990 | Unspecified |

| | | | | | | |
|---|---|---|---|---|---|---|
| mmu-miR-3079-5p | MIMAT0014866 | Ehbp1l1 | 114601 | HITS-CLIP | 31433990 | Unspecified |
| mmu-miR-3079-5p | MIMAT0014866 | Hipk3 | 15259 | HITS-CLIP | 31433990 | Unspecified |
| mmu-miR-3079-5p | MIMAT0014866 | Lck | 16818 | HITS-CLIP | 31433990 | Unspecified |
| mmu-miR-125b-2-3p | MIMAT0004529 | 0610012G03Rik | 106264 | HITS-CLIP | 25753418 | Unspecified |
| mmu-miR-125b-2-3p | MIMAT0004529 | 1190002N15Rik | 68861 | HITS-CLIP | 23597149 | Unspecified |
| mmu-miR-125b-2-3p | MIMAT0004529 | 2610507B11Rik | 72503 | HITS-CLIP | 23597149\|29167373 | Unspecified |
| mmu-miR-125b-2-3p | MIMAT0004529 | 9530068E07Rik | 213673 | HITS-CLIP | 23597149 | Unspecified |
| mmu-miR-125b-2-3p | MIMAT0004529 | AC115631.1 | | HITS-CLIP | 23597149 | Unspecified |
| mmu-miR-125b-2-3p | MIMAT0004529 | Abat | 268860 | HITS-CLIP | 23597149 | Unspecified |
| mmu-miR-125b-2-3p | MIMAT0004529 | Abca2 | 11305 | HITS-CLIP | 25753418 | Unspecified |
| mmu-miR-125b-2-3p | MIMAT0004529 | Abca8a | 217258 | HITS-CLIP | 37986224\|23597149 | Unspecified |
| mmu-miR-125b-2-3p | MIMAT0004529 | Abcb6 | 74104 | HITS-CLIP | 23597149 | Unspecified |
| mmu-miR-125b-2-3p | MIMAT0004529 | Abcd3 | 19299 | HITS-CLIP | 23597149 | Unspecified |
| mmu-miR-125b-2-3p | MIMAT0004529 | Abhd17b | 226016 | HITS-CLIP | 23597149 | Unspecified |
| mmu-miR-125b-2-3p | MIMAT0004529 | Acaca | 107476 | HITS-CLIP | 29167373 | Unspecified |
| mmu-miR-125b-2-3p | MIMAT0004529 | Acacb | 100705 | HITS-CLIP | 25753418 | Unspecified |
| mmu-miR-125b-2-3p | MIMAT0004529 | Acbd5 | 74159 | HITS-CLIP | 23597149 | Unspecified |
| mmu-miR-125b-2-3p | MIMAT0004529 | Acly | 104112 | HITS-CLIP | 37986224 | Unspecified |
| mmu-miR-125b-2-3p | MIMAT0004529 | Acox3 | 80911 | HITS-CLIP | 37986224 | Unspecified |
| mmu-miR-125b-2-3p | MIMAT0004529 | Acsf2 | 264895 | HITS-CLIP | 37986224 | Unspecified |
| mmu-miR-125b-2-3p | MIMAT0004529 | Acsl1 | 14081 | HITS-CLIP | 23597149 | Unspecified |
| mmu-miR-125b-2-3p | MIMAT0004529 | Acsl5 | 433256 | HITS-CLIP | 37986224 | Unspecified |
| mmu-miR-125b-2-3p | MIMAT0004529 | Acsm1 | 117147 | HITS-CLIP | 23597149 | Unspecified |
| mmu-miR-125b-2-3p | MIMAT0004529 | Acsm5 | 272428 | HITS-CLIP | 23597149 | Unspecified |
| mmu-miR-125b-2-3p | MIMAT0004529 | Actr2 | 66713 | HITS-CLIP | 29167373 | Unspecified |
| mmu-miR-125b-2-3p | MIMAT0004529 | Adam17 | 11491 | HITS-CLIP | 31433990 | Unspecified |
| mmu-miR-125b-2-3p | MIMAT0004529 | Adar | 56417 | HITS-CLIP | 37986224\|23597149 | Unspecified |
| mmu-miR-125b-2-3p | MIMAT0004529 | Add3 | 27360 | HITS-CLIP | 23597149 | Unspecified |
| mmu-miR-125b-2-3p | MIMAT0004529 | Adgrl4 | 64123 | HITS-CLIP | 23597149 | Unspecified |
| mmu-miR-125b-2-3p | MIMAT0004529 | Adh1 | 11522 | HITS-CLIP | 23597149 | Unspecified |
| mmu-miR-125b-2-3p | MIMAT0004529 | Adipor1 | 72674 | HITS-CLIP | 23597149 | Unspecified |
| mmu-miR-125b-2-3p | MIMAT0004529 | Agl | 77559 | HITS-CLIP | 25753418 | Unspecified |
| mmu-miR-125b-2-3p | MIMAT0004529 | Akap9 | 100986 | HITS-CLIP | 37986224\|23597149\|29167373 | Unspecified |
| mmu-miR-125b-2-3p | MIMAT0004529 | Akr1d1 | 208665 | HITS-CLIP | 23597149 | Unspecified |
| mmu-miR-125b-2-3p | MIMAT0004529 | Alb | 213 | HITS-CLIP | 23597149 | Unspecified |
| mmu-miR-125b-2-3p | MIMAT0004529 | Aldh1l1 | 107747 | HITS-CLIP | 23597149 | Unspecified |
| mmu-miR-125b-2-3p | MIMAT0004529 | Aldh6a1 | 104776 | HITS-CLIP | 23597149 | Unspecified |
| mmu-miR-125b-2-3p | MIMAT0004529 | Aldh7a1 | 110695 | HITS-CLIP | 37986224\|23597149 | Unspecified |
| mmu-miR-125b-2-3p | MIMAT0004529 | Aldoa | 11674 | HITS-CLIP | 23597149 | Unspecified |

| | | | | | | | |
|---|---|---|---|---|---|---|---|
| mmu-miR-125b-2-3p | MIMAT0004529 | Aldob | 230163 | HITS-CLIP | | 37986224 | Unspecified |
| mmu-miR-125b-2-3p | MIMAT0004529 | Anapc1 | 17222 | HITS-CLIP | 37986224\|23597149 | | Unspecified |
| mmu-miR-125b-2-3p | MIMAT0004529 | Ang | 11727 | HITS-CLIP | | 37986224 | Unspecified |
| mmu-miR-125b-2-3p | MIMAT0004529 | Ank | 11732 | HITS-CLIP | | 29167373 | Unspecified |
| mmu-miR-125b-2-3p | MIMAT0004529 | Ankfy1 | 11736 | HITS-CLIP | | 29167373 | Unspecified |
| mmu-miR-125b-2-3p | MIMAT0004529 | Ankrd28 | 105522 | HITS-CLIP | 37986224\|23597149 | | Unspecified |
| mmu-miR-125b-2-3p | MIMAT0004529 | Ankrd50 | 99696 | HITS-CLIP | | 29167373 | Unspecified |
| mmu-miR-125b-2-3p | MIMAT0004529 | Anks4b | 72074 | HITS-CLIP | 37986224\|23597149 | | Unspecified |
| mmu-miR-125b-2-3p | MIMAT0004529 | Ap1b1 | 11764 | HITS-CLIP | | 37986224 | Unspecified |
| mmu-miR-125b-2-3p | MIMAT0004529 | Api5 | 11800 | HITS-CLIP | | 29167373 | Unspecified |
| mmu-miR-125b-2-3p | MIMAT0004529 | Apob | 238055 | HITS-CLIP | 37986224\|23597149 | | Unspecified |
| mmu-miR-125b-2-3p | MIMAT0004529 | Aqp11 | 66333 | HITS-CLIP | | 23597149 | Unspecified |
| mmu-miR-125b-2-3p | MIMAT0004529 | Arf3 | 11842 | HITS-CLIP | | 25753418 | Unspecified |
| mmu-miR-125b-2-3p | MIMAT0004529 | Arfgap2 | 77038 | HITS-CLIP | 37986224\|23597149 | | Unspecified |
| mmu-miR-125b-2-3p | MIMAT0004529 | Arhgef12 | 69632 | HITS-CLIP | | 37986224 | Unspecified |
| mmu-miR-125b-2-3p | MIMAT0004529 | Arih1 | 23806 | HITS-CLIP | | 29167373 | Unspecified |
| mmu-miR-125b-2-3p | MIMAT0004529 | Arrdc3 | 105171 | HITS-CLIP | | 37986224 | Unspecified |
| mmu-miR-125b-2-3p | MIMAT0004529 | Art4 | 109978 | HITS-CLIP | | 23597149 | Unspecified |
| mmu-miR-125b-2-3p | MIMAT0004529 | Asap1 | 13196 | HITS-CLIP | 23597149\|29167373 | | Unspecified |
| mmu-miR-125b-2-3p | MIMAT0004529 | Ascc3 | 77987 | HITS-CLIP | | 25753418 | Unspecified |
| mmu-miR-125b-2-3p | MIMAT0004529 | Atg4a | 666468 | HITS-CLIP | | 25753418 | Unspecified |
| mmu-miR-125b-2-3p | MIMAT0004529 | Atl3 | 109168 | HITS-CLIP | | 23597149 | Unspecified |
| mmu-miR-125b-2-3p | MIMAT0004529 | Atp11b | 76295 | HITS-CLIP | | 25753418 | Unspecified |
| mmu-miR-125b-2-3p | MIMAT0004529 | Atp1b1 | 11931 | HITS-CLIP | | 23597149 | Unspecified |
| mmu-miR-125b-2-3p | MIMAT0004529 | Atp6v1b2 | 11966 | HITS-CLIP | 23597149\|29167373 | | Unspecified |
| mmu-miR-125b-2-3p | MIMAT0004529 | Atp6v1c1 | 66335 | HITS-CLIP | 23597149\|29167373 | | Unspecified |
| mmu-miR-125b-2-3p | MIMAT0004529 | Atrx | 22589 | HITS-CLIP | | 29167373 | Unspecified |
| mmu-miR-125b-2-3p | MIMAT0004529 | Atxn7l3b | 382423 | HITS-CLIP | | 37986224 | Unspecified |
| mmu-miR-125b-2-3p | MIMAT0004529 | Azin1 | 54375 | HITS-CLIP | | 37986224 | Unspecified |
| mmu-miR-125b-2-3p | MIMAT0004529 | B4galt1 | 14595 | HITS-CLIP | 25753418\|29167373 | | Unspecified |
| mmu-miR-125b-2-3p | MIMAT0004529 | B4galt6 | 56386 | HITS-CLIP | | 29167373 | Unspecified |
| mmu-miR-125b-2-3p | MIMAT0004529 | Baat | 12012 | HITS-CLIP | | 23597149 | Unspecified |
| mmu-miR-125b-2-3p | MIMAT0004529 | Bach1 | 12013 | HITS-CLIP | | 37986224 | Unspecified |
| mmu-miR-125b-2-3p | MIMAT0004529 | Baz2b | 407823 | HITS-CLIP | | 29167373 | Unspecified |
| mmu-miR-125b-2-3p | MIMAT0004529 | Bckdha | 12039 | HITS-CLIP | | 25753418 | Unspecified |
| mmu-miR-125b-2-3p | MIMAT0004529 | Bhlhe40 | 20893 | HITS-CLIP | | 23597149 | Unspecified |
| mmu-miR-125b-2-3p | MIMAT0004529 | Birc6 | 12211 | HITS-CLIP | 37986224\|23597149 | | Unspecified |
| mmu-miR-125b-2-3p | MIMAT0004529 | Bmpr1a | 12166 | HITS-CLIP | | 37986224 | Unspecified |
| mmu-miR-125b-2-3p | MIMAT0004529 | Brd2 | 14312 | HITS-CLIP | | 29167373 | Unspecified |

| | | | | | | |
|---|---|---|---|---|---|---|
| mmu-miR-125b-2-3p | MIMAT0004529 | Btf3 | 218490 | HITS-CLIP | 23597149 | Unspecified |
| mmu-miR-125b-2-3p | MIMAT0004529 | Bzw1 | 66882 | HITS-CLIP | 37986224 | Unspecified |
| mmu-miR-125b-2-3p | MIMAT0004529 | C1galt1 | 94192 | HITS-CLIP | 23597149 | Unspecified |
| mmu-miR-125b-2-3p | MIMAT0004529 | C5ar2 | 319430 | HITS-CLIP | 29167373 | Unspecified |
| mmu-miR-125b-2-3p | MIMAT0004529 | C6 | 12274 | HITS-CLIP | 23597149 | Unspecified |
| mmu-miR-125b-2-3p | MIMAT0004529 | Cand1 | 71902 | HITS-CLIP | 23597149 | Unspecified |
| mmu-miR-125b-2-3p | MIMAT0004529 | Canx | 12330 | HITS-CLIP | 37986224 | Unspecified |
| mmu-miR-125b-2-3p | MIMAT0004529 | Car8 | 12319 | HITS-CLIP | 23597149\|25753418 | Unspecified |
| mmu-miR-125b-2-3p | MIMAT0004529 | Casd1 | 213819 | HITS-CLIP | 25753418 | Unspecified |
| mmu-miR-125b-2-3p | MIMAT0004529 | Cat | 12359 | HITS-CLIP | 37986224 | Unspecified |
| mmu-miR-125b-2-3p | MIMAT0004529 | Cbr4 | 234309 | HITS-CLIP | 37986224 | Unspecified |
| mmu-miR-125b-2-3p | MIMAT0004529 | Ccdc71l | 72123 | HITS-CLIP | 29167373 | Unspecified |
| mmu-miR-125b-2-3p | MIMAT0004529 | Ccna2 | 12428 | HITS-CLIP//Luciferase reporter assay | 31433990\|27020049 | Unspecified |
| mmu-miR-125b-2-3p | MIMAT0004529 | Ccnl2 | 56036 | HITS-CLIP | 23597149 | Unspecified |
| mmu-miR-125b-2-3p | MIMAT0004529 | Cd164 | 53599 | HITS-CLIP | 23597149 | Unspecified |
| mmu-miR-125b-2-3p | MIMAT0004529 | Cd47 | 16423 | HITS-CLIP | 23597149 | Unspecified |
| mmu-miR-125b-2-3p | MIMAT0004529 | Cd99l2 | 171486 | HITS-CLIP | 25753418 | Unspecified |
| mmu-miR-125b-2-3p | MIMAT0004529 | Cdc42se2 | 72729 | HITS-CLIP | 29167373 | Unspecified |
| mmu-miR-125b-2-3p | MIMAT0004529 | Cdh2 | 12558 | HITS-CLIP | 25753418 | Unspecified |
| mmu-miR-125b-2-3p | MIMAT0004529 | Cdk13 | 69562 | HITS-CLIP | 29167373 | Unspecified |
| mmu-miR-125b-2-3p | MIMAT0004529 | Cdk18 | 18557 | HITS-CLIP | 25753418 | Unspecified |
| mmu-miR-125b-2-3p | MIMAT0004529 | Cds2 | 110911 | HITS-CLIP | 37986224 | Unspecified |
| mmu-miR-125b-2-3p | MIMAT0004529 | Celf2 | 14007 | HITS-CLIP | 29167373 | Unspecified |
| mmu-miR-125b-2-3p | MIMAT0004529 | Ces3a | 382053 | HITS-CLIP | 37986224 | Unspecified |
| mmu-miR-125b-2-3p | MIMAT0004529 | Cfl2 | 12632 | HITS-CLIP | 29167373 | Unspecified |
| mmu-miR-125b-2-3p | MIMAT0004529 | Chd1 | 12648 | HITS-CLIP | 37986224\|23597149\|29167373 | Unspecified |
| mmu-miR-125b-2-3p | MIMAT0004529 | Chd4 | 107932 | HITS-CLIP | 29167373 | Unspecified |
| mmu-miR-125b-2-3p | MIMAT0004529 | Chd7 | 320790 | HITS-CLIP | 23597149\|25753418 | Unspecified |
| mmu-miR-125b-2-3p | MIMAT0004529 | Cipc | 217732 | HITS-CLIP | 23597149 | Unspecified |
| mmu-miR-125b-2-3p | MIMAT0004529 | Clcn2 | 12724 | HITS-CLIP | 23597149 | Unspecified |
| mmu-miR-125b-2-3p | MIMAT0004529 | Clcn3 | 12725 | HITS-CLIP | 23597149 | Unspecified |
| mmu-miR-125b-2-3p | MIMAT0004529 | Clcn5 | 12728 | HITS-CLIP | 29167373 | Unspecified |
| mmu-miR-125b-2-3p | MIMAT0004529 | Clint1 | 216705 | HITS-CLIP | 29167373 | Unspecified |
| mmu-miR-125b-2-3p | MIMAT0004529 | Clk3 | 102414 | HITS-CLIP | 23597149 | Unspecified |
| mmu-miR-125b-2-3p | MIMAT0004529 | Cltc | 67300 | HITS-CLIP | 37986224\|23597149\|29167373 | Unspecified |
| mmu-miR-125b-2-3p | MIMAT0004529 | Cnnm3 | 94218 | HITS-CLIP | 37986224 | Unspecified |
| mmu-miR-125b-2-3p | MIMAT0004529 | Cnot1 | 234594 | HITS-CLIP | 37986224\|23597149 | Unspecified |
| mmu-miR-125b-2-3p | MIMAT0004529 | Cobll1 | 319876 | HITS-CLIP | 23597149 | Unspecified |
| mmu-miR-125b-2-3p | MIMAT0004529 | Col4a1 | 12826 | HITS-CLIP | 37986224 | Unspecified |

| | | | | | | | |
|---|---|---|---|---|---|---|---|
| mmu-miR-125b-2-3p | MIMAT0004529 | Comt | 12846 | HITS-CLIP | | 23597149 | Unspecified |
| mmu-miR-125b-2-3p | MIMAT0004529 | Cops7b | 26895 | HITS-CLIP | | 37986224 | Unspecified |
| mmu-miR-125b-2-3p | MIMAT0004529 | Cox7a2 | 12866 | HITS-CLIP | | 25753418 | Unspecified |
| mmu-miR-125b-2-3p | MIMAT0004529 | Cpd | 12874 | HITS-CLIP | | 29167373 | Unspecified |
| mmu-miR-125b-2-3p | MIMAT0004529 | Cpeb2 | 231207 | HITS-CLIP | | 37986224 | Unspecified |
| mmu-miR-125b-2-3p | MIMAT0004529 | Cpeb4 | 67579 | HITS-CLIP | | 37986224 | Unspecified |
| mmu-miR-125b-2-3p | MIMAT0004529 | Cpped1 | 223978 | HITS-CLIP | | 23597149 | Unspecified |
| mmu-miR-125b-2-3p | MIMAT0004529 | Cps1 | 227231 | HITS-CLIP | | 23597149 | Unspecified |
| mmu-miR-125b-2-3p | MIMAT0004529 | Cpt1a | 12894 | HITS-CLIP | | 37986224 | Unspecified |
| mmu-miR-125b-2-3p | MIMAT0004529 | Crebrf | 77128 | HITS-CLIP | | 29167373 | Unspecified |
| mmu-miR-125b-2-3p | MIMAT0004529 | Crk | 12928 | HITS-CLIP | | 29167373 | Unspecified |
| mmu-miR-125b-2-3p | MIMAT0004529 | Ctnnb1 | 12387 | HITS-CLIP | 37986224\|23597149 | | Unspecified |
| mmu-miR-125b-2-3p | MIMAT0004529 | Ctss | 13040 | HITS-CLIP | | 29167373 | Unspecified |
| mmu-miR-125b-2-3p | MIMAT0004529 | Cul3 | 26554 | HITS-CLIP | | 25753418 | Unspecified |
| mmu-miR-125b-2-3p | MIMAT0004529 | Cyp3a13 | 13113 | HITS-CLIP | | 37986224 | Unspecified |
| mmu-miR-125b-2-3p | MIMAT0004529 | Cyp4v3 | 102294 | HITS-CLIP | | 23597149 | Unspecified |
| mmu-miR-125b-2-3p | MIMAT0004529 | Daam1 | 208846 | HITS-CLIP | | 29167373 | Unspecified |
| mmu-miR-125b-2-3p | MIMAT0004529 | Dcbld2 | 73379 | HITS-CLIP | | 29167373 | Unspecified |
| mmu-miR-125b-2-3p | MIMAT0004529 | Ddx17 | 67040 | HITS-CLIP | | 29167373 | Unspecified |
| mmu-miR-125b-2-3p | MIMAT0004529 | Ddx21 | 56200 | HITS-CLIP | | 29167373 | Unspecified |
| mmu-miR-125b-2-3p | MIMAT0004529 | Ddx3x | 13205 | HITS-CLIP | 23597149\|29167373 | | Unspecified |
| mmu-miR-125b-2-3p | MIMAT0004529 | Ddx3y | 26900 | HITS-CLIP | | 25753418 | Unspecified |
| mmu-miR-125b-2-3p | MIMAT0004529 | Ddx60 | 234311 | HITS-CLIP | | 29167373 | Unspecified |
| mmu-miR-125b-2-3p | MIMAT0004529 | Dennd1b | 329260 | HITS-CLIP | | 23597149 | Unspecified |
| mmu-miR-125b-2-3p | MIMAT0004529 | Depdc7 | 211896 | HITS-CLIP | | 37986224 | Unspecified |
| mmu-miR-125b-2-3p | MIMAT0004529 | Det1 | 76375 | HITS-CLIP | | 29167373 | Unspecified |
| mmu-miR-125b-2-3p | MIMAT0004529 | Dhrs3 | 20148 | HITS-CLIP | | 29167373 | Unspecified |
| mmu-miR-125b-2-3p | MIMAT0004529 | Dhx36 | 72162 | HITS-CLIP | | 23597149 | Unspecified |
| mmu-miR-125b-2-3p | MIMAT0004529 | Dhx9 | 13211 | HITS-CLIP | | 23597149 | Unspecified |
| mmu-miR-125b-2-3p | MIMAT0004529 | Diaph2 | 54004 | HITS-CLIP | | 29167373 | Unspecified |
| mmu-miR-125b-2-3p | MIMAT0004529 | Dmxl1 | 240283 | HITS-CLIP | | 29167373 | Unspecified |
| mmu-miR-125b-2-3p | MIMAT0004529 | Dnajb14 | 70604 | HITS-CLIP | 23597149\|29167373 | | Unspecified |
| mmu-miR-125b-2-3p | MIMAT0004529 | Dnajc13 | 235567 | HITS-CLIP | 37986224\|23597149\|29167373 | | Unspecified |
| mmu-miR-125b-2-3p | MIMAT0004529 | Dnmt3a | 13435 | HITS-CLIP | | 23597149 | Unspecified |
| mmu-miR-125b-2-3p | MIMAT0004529 | Dock4 | 238130 | HITS-CLIP | | 25753418 | Unspecified |
| mmu-miR-125b-2-3p | MIMAT0004529 | Dot1l | 208266 | HITS-CLIP | | 29167373 | Unspecified |
| mmu-miR-125b-2-3p | MIMAT0004529 | Dpep2 | 319446 | HITS-CLIP | | 29167373 | Unspecified |
| mmu-miR-125b-2-3p | MIMAT0004529 | Dpysl2 | 12934 | HITS-CLIP | | 29167373 | Unspecified |
| mmu-miR-125b-2-3p | MIMAT0004529 | Dpysl3 | 22240 | HITS-CLIP | | 37986224 | Unspecified |

| | | | | | | |
|---|---|---|---|---|---|---|
| mmu-miR-125b-2-3p | MIMAT0004529 | Dr1 | 13486 | HITS-CLIP | 31433990 | Unspecified |
| mmu-miR-125b-2-3p | MIMAT0004529 | Dram2 | 67171 | HITS-CLIP | 37986224\|23597149 | Unspecified |
| mmu-miR-125b-2-3p | MIMAT0004529 | Dsp | 109620 | HITS-CLIP | 37986224\|23597149\|25753418 | Unspecified |
| mmu-miR-125b-2-3p | MIMAT0004529 | Dst | 13518 | HITS-CLIP | 25753418 | Unspecified |
| mmu-miR-125b-2-3p | MIMAT0004529 | Dstn | 56431 | HITS-CLIP | 23597149\|29167373 | Unspecified |
| mmu-miR-125b-2-3p | MIMAT0004529 | Dusp1 | 19252 | HITS-CLIP | 37986224 | Unspecified |
| mmu-miR-125b-2-3p | MIMAT0004529 | Dusp5 | 240672 | HITS-CLIP | 25753418 | Unspecified |
| mmu-miR-125b-2-3p | MIMAT0004529 | Dusp7 | 235584 | HITS-CLIP | 29167373 | Unspecified |
| mmu-miR-125b-2-3p | MIMAT0004529 | Dync1h1 | 13424 | HITS-CLIP | 23597149\|29167373 | Unspecified |
| mmu-miR-125b-2-3p | MIMAT0004529 | E130308A19Rik | 230259 | HITS-CLIP | 25753418 | Unspecified |
| mmu-miR-125b-2-3p | MIMAT0004529 | E130309D02Rik | 231868 | HITS-CLIP | 37986224 | Unspecified |
| mmu-miR-125b-2-3p | MIMAT0004529 | E2f8 | 108961 | HITS-CLIP | 29167373 | Unspecified |
| mmu-miR-125b-2-3p | MIMAT0004529 | Eef1b2 | 55949 | HITS-CLIP | 25753418 | Unspecified |
| mmu-miR-125b-2-3p | MIMAT0004529 | Ehd3 | 57440 | HITS-CLIP | 37986224 | Unspecified |
| mmu-miR-125b-2-3p | MIMAT0004529 | Eif4g1 | 208643 | HITS-CLIP | 23597149 | Unspecified |
| mmu-miR-125b-2-3p | MIMAT0004529 | Eif5 | 217869 | HITS-CLIP | 23597149\|29167373 | Unspecified |
| mmu-miR-125b-2-3p | MIMAT0004529 | Elmod2 | 244548 | HITS-CLIP | 37986224\|23597149 | Unspecified |
| mmu-miR-125b-2-3p | MIMAT0004529 | Elovl5 | 68801 | HITS-CLIP | 37986224 | Unspecified |
| mmu-miR-125b-2-3p | MIMAT0004529 | Emc2 | 66736 | HITS-CLIP | 23597149 | Unspecified |
| mmu-miR-125b-2-3p | MIMAT0004529 | Entpd5 | 12499 | HITS-CLIP | 23597149 | Unspecified |
| mmu-miR-125b-2-3p | MIMAT0004529 | Eny2 | 223527 | HITS-CLIP | 23597149 | Unspecified |
| mmu-miR-125b-2-3p | MIMAT0004529 | Eprs | 107508 | HITS-CLIP | 29167373 | Unspecified |
| mmu-miR-125b-2-3p | MIMAT0004529 | Erbb2 | 13866 | HITS-CLIP | 23597149 | Unspecified |
| mmu-miR-125b-2-3p | MIMAT0004529 | Erbb3 | 13867 | HITS-CLIP | 37986224 | Unspecified |
| mmu-miR-125b-2-3p | MIMAT0004529 | Etnppl | 71760 | HITS-CLIP | 23597149 | Unspecified |
| mmu-miR-125b-2-3p | MIMAT0004529 | Ets1 | 23871 | HITS-CLIP | 25753418 | Unspecified |
| mmu-miR-125b-2-3p | MIMAT0004529 | Fam149a | 212326 | HITS-CLIP | 23597149 | Unspecified |
| mmu-miR-125b-2-3p | MIMAT0004529 | Fam162a | 70186 | HITS-CLIP | 23597149 | Unspecified |
| mmu-miR-125b-2-3p | MIMAT0004529 | Fam3a | 66294 | HITS-CLIP | 25753418 | Unspecified |
| mmu-miR-125b-2-3p | MIMAT0004529 | Fam69a | 388650 | HITS-CLIP | 29167373 | Unspecified |
| mmu-miR-125b-2-3p | MIMAT0004529 | Fasn | 14104 | HITS-CLIP | 23597149 | Unspecified |
| mmu-miR-125b-2-3p | MIMAT0004529 | Fbxw11 | 103583 | HITS-CLIP | 29167373 | Unspecified |
| mmu-miR-125b-2-3p | MIMAT0004529 | Fbxw7 | 50754 | HITS-CLIP | 23597149 | Unspecified |
| mmu-miR-125b-2-3p | MIMAT0004529 | Fcer1g | 14127 | HITS-CLIP | 29167373 | Unspecified |
| mmu-miR-125b-2-3p | MIMAT0004529 | Fem1a | 14154 | HITS-CLIP | 25753418 | Unspecified |
| mmu-miR-125b-2-3p | MIMAT0004529 | Fetub | 59083 | HITS-CLIP | 23597149 | Unspecified |
| mmu-miR-125b-2-3p | MIMAT0004529 | Filip1l | 78749 | HITS-CLIP | 25753418 | Unspecified |
| mmu-miR-125b-2-3p | MIMAT0004529 | Fkbp8 | 14232 | HITS-CLIP | 23597149 | Unspecified |
| mmu-miR-125b-2-3p | MIMAT0004529 | Flnc | 68794 | HITS-CLIP | 25753418 | Unspecified |

| | | | | | | |
|---|---|---|---|---|---|---|
| mmu-miR-125b-2-3p | MIMAT0004529 | Flt4 | 14257 | HITS-CLIP | 37986224 | Unspecified |
| mmu-miR-125b-2-3p | MIMAT0004529 | Fmo5 | 14263 | HITS-CLIP | 23597149 | Unspecified |
| mmu-miR-125b-2-3p | MIMAT0004529 | Foxo1 | 56458 | HITS-CLIP | 37986224 | Unspecified |
| mmu-miR-125b-2-3p | MIMAT0004529 | Foxo3 | 56484 | HITS-CLIP | 23597149 | Unspecified |
| mmu-miR-125b-2-3p | MIMAT0004529 | Foxp2 | 114142 | HITS-CLIP | 23597149 | Unspecified |
| mmu-miR-125b-2-3p | MIMAT0004529 | Frk | 14302 | HITS-CLIP | 23597149 | Unspecified |
| mmu-miR-125b-2-3p | MIMAT0004529 | G3bp1 | 27041 | HITS-CLIP | 29167373 | Unspecified |
| mmu-miR-125b-2-3p | MIMAT0004529 | Gas7 | 14457 | HITS-CLIP | 29167373 | Unspecified |
| mmu-miR-125b-2-3p | MIMAT0004529 | Gba | 14466 | HITS-CLIP | 23597149 | Unspecified |
| mmu-miR-125b-2-3p | MIMAT0004529 | Gga3 | 260302 | HITS-CLIP | 37986224\|25753418 | Unspecified |
| mmu-miR-125b-2-3p | MIMAT0004529 | Ghitm | 66092 | HITS-CLIP | 23597149 | Unspecified |
| mmu-miR-125b-2-3p | MIMAT0004529 | Gid4 | 66771 | HITS-CLIP | 23597149 | Unspecified |
| mmu-miR-125b-2-3p | MIMAT0004529 | Glg1 | 20340 | HITS-CLIP | 23597149 | Unspecified |
| mmu-miR-125b-2-3p | MIMAT0004529 | Glt8d1 | 76485 | HITS-CLIP | 37986224 | Unspecified |
| mmu-miR-125b-2-3p | MIMAT0004529 | Glul | 14645 | HITS-CLIP | 37986224 | Unspecified |
| mmu-miR-125b-2-3p | MIMAT0004529 | Gm45837 | | HITS-CLIP | 25753418 | Unspecified |
| mmu-miR-125b-2-3p | MIMAT0004529 | Gm5878 | 545861 | HITS-CLIP | 37986224 | Unspecified |
| mmu-miR-125b-2-3p | MIMAT0004529 | Gnai2 | 14678 | HITS-CLIP | 23597149 | Unspecified |
| mmu-miR-125b-2-3p | MIMAT0004529 | Gnai3 | 14679 | HITS-CLIP | 23597149 | Unspecified |
| mmu-miR-125b-2-3p | MIMAT0004529 | Gnaq | 14682 | HITS-CLIP | 23597149 | Unspecified |
| mmu-miR-125b-2-3p | MIMAT0004529 | Gpd2 | 14571 | HITS-CLIP | 23597149 | Unspecified |
| mmu-miR-125b-2-3p | MIMAT0004529 | Gpr21 | 338346 | HITS-CLIP | 29167373 | Unspecified |
| mmu-miR-125b-2-3p | MIMAT0004529 | Gsk3b | 56637 | HITS-CLIP | 29167373 | Unspecified |
| mmu-miR-125b-2-3p | MIMAT0004529 | Gskip | 66787 | HITS-CLIP | 25753418 | Unspecified |
| mmu-miR-125b-2-3p | MIMAT0004529 | Gspt1 | 14852 | HITS-CLIP | 37986224\|23597149 | Unspecified |
| mmu-miR-125b-2-3p | MIMAT0004529 | Gtpbp2 | 56055 | HITS-CLIP | 29167373 | Unspecified |
| mmu-miR-125b-2-3p | MIMAT0004529 | H2afx | 15270 | HITS-CLIP | 37986224 | Unspecified |
| mmu-miR-125b-2-3p | MIMAT0004529 | Hadhb | 231086 | HITS-CLIP | 23597149 | Unspecified |
| mmu-miR-125b-2-3p | MIMAT0004529 | Haus8 | 76478 | HITS-CLIP | 25753418 | Unspecified |
| mmu-miR-125b-2-3p | MIMAT0004529 | Hebp1 | 15199 | HITS-CLIP | 23597149 | Unspecified |
| mmu-miR-125b-2-3p | MIMAT0004529 | Hectd1 | 207304 | HITS-CLIP | 23597149 | Unspecified |
| mmu-miR-125b-2-3p | MIMAT0004529 | Helz2 | 229003 | HITS-CLIP | 37986224 | Unspecified |
| mmu-miR-125b-2-3p | MIMAT0004529 | Herc1 | 235439 | HITS-CLIP | 23597149 | Unspecified |
| mmu-miR-125b-2-3p | MIMAT0004529 | Hgd | 15233 | HITS-CLIP | 23597149 | Unspecified |
| mmu-miR-125b-2-3p | MIMAT0004529 | Hif1a | 15251 | HITS-CLIP | 23597149 | Unspecified |
| mmu-miR-125b-2-3p | MIMAT0004529 | Hipk3 | 15259 | HITS-CLIP | 29167373 | Unspecified |
| mmu-miR-125b-2-3p | MIMAT0004529 | Hnrnpc | 15381 | HITS-CLIP | 23597149 | Unspecified |
| mmu-miR-125b-2-3p | MIMAT0004529 | Hnrnpu | 51810 | HITS-CLIP | 23597149 | Unspecified |
| mmu-miR-125b-2-3p | MIMAT0004529 | Hp1bp3 | 15441 | HITS-CLIP | 37986224 | Unspecified |

| | | | | | | |
|---|---|---|---|---|---|---|
| mmu-miR-125b-2-3p | MIMAT0004529 | Hpgds | 54486 | HITS-CLIP | 29167373 | Unspecified |
| mmu-miR-125b-2-3p | MIMAT0004529 | Hrc | 15464 | HITS-CLIP | 25753418 | Unspecified |
| mmu-miR-125b-2-3p | MIMAT0004529 | Hspb8 | 80888 | HITS-CLIP | 37986224 | Unspecified |
| mmu-miR-125b-2-3p | MIMAT0004529 | Huwe1 | 59026 | HITS-CLIP | 37986224 | Unspecified |
| mmu-miR-125b-2-3p | MIMAT0004529 | Hyou1 | 12282 | HITS-CLIP | 37986224 | Unspecified |
| mmu-miR-125b-2-3p | MIMAT0004529 | Idh1 | 15926 | HITS-CLIP | 23597149 | Unspecified |
| mmu-miR-125b-2-3p | MIMAT0004529 | Ier5 | 15939 | HITS-CLIP | 29167373 | Unspecified |
| mmu-miR-125b-2-3p | MIMAT0004529 | Igf2r | 16004 | HITS-CLIP | 29167373 | Unspecified |
| mmu-miR-125b-2-3p | MIMAT0004529 | Igfbp4 | 16010 | HITS-CLIP | 37986224\|23597149 | Unspecified |
| mmu-miR-125b-2-3p | MIMAT0004529 | Ikbkg | 16151 | HITS-CLIP | 37986224 | Unspecified |
| mmu-miR-125b-2-3p | MIMAT0004529 | Il12rb2 | 16162 | HITS-CLIP | 31433990 | Unspecified |
| mmu-miR-125b-2-3p | MIMAT0004529 | Inpp5d | 16331 | HITS-CLIP | 29167373 | Unspecified |
| mmu-miR-125b-2-3p | MIMAT0004529 | Ipo8 | 320727 | HITS-CLIP | 29167373 | Unspecified |
| mmu-miR-125b-2-3p | MIMAT0004529 | Iqgap2 | 544963 | HITS-CLIP | 23597149 | Unspecified |
| mmu-miR-125b-2-3p | MIMAT0004529 | Itgb1 | 16412 | HITS-CLIP | 29167373 | Unspecified |
| mmu-miR-125b-2-3p | MIMAT0004529 | Itih2 | 16425 | HITS-CLIP | 37986224\|23597149 | Unspecified |
| mmu-miR-125b-2-3p | MIMAT0004529 | Itpr3 | 16440 | HITS-CLIP | 29167373 | Unspecified |
| mmu-miR-125b-2-3p | MIMAT0004529 | Ivd | 56357 | HITS-CLIP | 23597149 | Unspecified |
| mmu-miR-125b-2-3p | MIMAT0004529 | Ivns1abp | 117198 | HITS-CLIP | 25753418 | Unspecified |
| mmu-miR-125b-2-3p | MIMAT0004529 | Jam2 | 67374 | HITS-CLIP | 23597149 | Unspecified |
| mmu-miR-125b-2-3p | MIMAT0004529 | Kat6a | 244349 | HITS-CLIP | 37986224\|23597149\|29167373 | Unspecified |
| mmu-miR-125b-2-3p | MIMAT0004529 | Kbtbd2 | 210973 | HITS-CLIP | 29167373 | Unspecified |
| mmu-miR-125b-2-3p | MIMAT0004529 | Kbtbd3 | 69149 | HITS-CLIP | 25753418 | Unspecified |
| mmu-miR-125b-2-3p | MIMAT0004529 | Kbtbd4 | 67136 | HITS-CLIP | 25753418 | Unspecified |
| mmu-miR-125b-2-3p | MIMAT0004529 | Kctd12 | 239217 | HITS-CLIP | 29167373 | Unspecified |
| mmu-miR-125b-2-3p | MIMAT0004529 | Kdm2a | 225876 | HITS-CLIP | 29167373 | Unspecified |
| mmu-miR-125b-2-3p | MIMAT0004529 | Kif13b | 16554 | HITS-CLIP | 29167373 | Unspecified |
| mmu-miR-125b-2-3p | MIMAT0004529 | Kif1b | 16561 | HITS-CLIP | 37986224 | Unspecified |
| mmu-miR-125b-2-3p | MIMAT0004529 | Kif21a | 16564 | HITS-CLIP | 23597149 | Unspecified |
| mmu-miR-125b-2-3p | MIMAT0004529 | Kif5b | 16573 | HITS-CLIP | 37986224 | Unspecified |
| mmu-miR-125b-2-3p | MIMAT0004529 | Klf7 | 93691 | HITS-CLIP | 29167373 | Unspecified |
| mmu-miR-125b-2-3p | MIMAT0004529 | Klf9 | 16601 | HITS-CLIP | 23597149 | Unspecified |
| mmu-miR-125b-2-3p | MIMAT0004529 | Klhdc10 | 76788 | HITS-CLIP | 37986224 | Unspecified |
| mmu-miR-125b-2-3p | MIMAT0004529 | Klhl3 | 100503085 | HITS-CLIP | 37986224\|23597149 | Unspecified |
| mmu-miR-125b-2-3p | MIMAT0004529 | Kyat3 | 56267 | HITS-CLIP | 23597149 | Unspecified |
| mmu-miR-125b-2-3p | MIMAT0004529 | Lamtor1 | 66508 | HITS-CLIP | 37986224\|23597149 | Unspecified |
| mmu-miR-125b-2-3p | MIMAT0004529 | Laptm4a | 17775 | HITS-CLIP | 37986224 | Unspecified |
| mmu-miR-125b-2-3p | MIMAT0004529 | Laptm5 | 16792 | HITS-CLIP | 37986224 | Unspecified |
| mmu-miR-125b-2-3p | MIMAT0004529 | Larp4b | 217980 | HITS-CLIP | 23597149 | Unspecified |

| | | | | | | |
|---|---|---|---|---|---|---|
| mmu-miR-125b-2-3p | MIMAT0004529 | Lmna | 16905 | HITS-CLIP | 23597149 | Unspecified |
| mmu-miR-125b-2-3p | MIMAT0004529 | Lonp2 | 66887 | HITS-CLIP | 23597149 | Unspecified |
| mmu-miR-125b-2-3p | MIMAT0004529 | Lrch4 | 231798 | HITS-CLIP | 29167373 | Unspecified |
| mmu-miR-125b-2-3p | MIMAT0004529 | Lrp6 | 16974 | HITS-CLIP | 25753418 | Unspecified |
| mmu-miR-125b-2-3p | MIMAT0004529 | Lrpap1 | 16976 | HITS-CLIP | 25753418 | Unspecified |
| mmu-miR-125b-2-3p | MIMAT0004529 | Lrpprc | 72416 | HITS-CLIP | 23597149 | Unspecified |
| mmu-miR-125b-2-3p | MIMAT0004529 | Lrrc8a | 241296 | HITS-CLIP | 37986224\|23597149\|29167373 | Unspecified |
| mmu-miR-125b-2-3p | MIMAT0004529 | Lss | 16987 | HITS-CLIP | 23597149 | Unspecified |
| mmu-miR-125b-2-3p | MIMAT0004529 | Luc7l2 | 192196 | HITS-CLIP | 29167373 | Unspecified |
| mmu-miR-125b-2-3p | MIMAT0004529 | Macf1 | 11426 | HITS-CLIP | 29167373 | Unspecified |
| mmu-miR-125b-2-3p | MIMAT0004529 | Maged1 | 94275 | HITS-CLIP | 37986224 | Unspecified |
| mmu-miR-125b-2-3p | MIMAT0004529 | Magi3 | 99470 | HITS-CLIP | 23597149 | Unspecified |
| mmu-miR-125b-2-3p | MIMAT0004529 | Marchf6 | 223455 | HITS-CLIP | 37986224 | Unspecified |
| mmu-miR-125b-2-3p | MIMAT0004529 | Mat1a | 11720 | HITS-CLIP | 23597149 | Unspecified |
| mmu-miR-125b-2-3p | MIMAT0004529 | Mavs | 228607 | HITS-CLIP | 25753418 | Unspecified |
| mmu-miR-125b-2-3p | MIMAT0004529 | Mccc1 | 72039 | HITS-CLIP | 25753418\|23597149 | Unspecified |
| mmu-miR-125b-2-3p | MIMAT0004529 | Mcm5 | 17218 | HITS-CLIP | 23597149 | Unspecified |
| mmu-miR-125b-2-3p | MIMAT0004529 | Mdh1 | 17449 | HITS-CLIP | 23597149 | Unspecified |
| mmu-miR-125b-2-3p | MIMAT0004529 | Med13 | 327987 | HITS-CLIP | 23597149 | Unspecified |
| mmu-miR-125b-2-3p | MIMAT0004529 | Med13l | 76199 | HITS-CLIP | 37986224 | Unspecified |
| mmu-miR-125b-2-3p | MIMAT0004529 | Mef2a | 17258 | HITS-CLIP | 23597149 | Unspecified |
| mmu-miR-125b-2-3p | MIMAT0004529 | Mef2d | 17261 | HITS-CLIP | 29167373 | Unspecified |
| mmu-miR-125b-2-3p | MIMAT0004529 | Memo1 | 76890 | HITS-CLIP | 29167373 | Unspecified |
| mmu-miR-125b-2-3p | MIMAT0004529 | Metap1 | 75624 | HITS-CLIP | 23597149 | Unspecified |
| mmu-miR-125b-2-3p | MIMAT0004529 | Metap2 | 56307 | HITS-CLIP | 37986224 | Unspecified |
| mmu-miR-125b-2-3p | MIMAT0004529 | Mga | 29808 | HITS-CLIP | 23597149 | Unspecified |
| mmu-miR-125b-2-3p | MIMAT0004529 | Mgea5 | 76055 | HITS-CLIP | 23597149\|29167373 | Unspecified |
| mmu-miR-125b-2-3p | MIMAT0004529 | Mms19 | 72199 | HITS-CLIP | 23597149 | Unspecified |
| mmu-miR-125b-2-3p | MIMAT0004529 | Mob1b | 68473 | HITS-CLIP | 29167373 | Unspecified |
| mmu-miR-125b-2-3p | MIMAT0004529 | Mpp6 | 56524 | HITS-CLIP | 23597149 | Unspecified |
| mmu-miR-125b-2-3p | MIMAT0004529 | Mpzl2 | 14012 | HITS-CLIP | 23597149 | Unspecified |
| mmu-miR-125b-2-3p | MIMAT0004529 | Msn | 17698 | HITS-CLIP | 37986224 | Unspecified |
| mmu-miR-125b-2-3p | MIMAT0004529 | Mtdh | 67154 | HITS-CLIP | 29167373 | Unspecified |
| mmu-miR-125b-2-3p | MIMAT0004529 | Mtmr10 | 233315 | HITS-CLIP | 23597149 | Unspecified |
| mmu-miR-125b-2-3p | MIMAT0004529 | Mtpap | 67440 | HITS-CLIP | 25753418 | Unspecified |
| mmu-miR-125b-2-3p | MIMAT0004529 | Mug1 | 17836 | HITS-CLIP | 37986224 | Unspecified |
| mmu-miR-125b-2-3p | MIMAT0004529 | Myo18a | 360013 | HITS-CLIP | 37986224 | Unspecified |
| mmu-miR-125b-2-3p | MIMAT0004529 | Naa25 | 231713 | HITS-CLIP | 29167373 | Unspecified |
| mmu-miR-125b-2-3p | MIMAT0004529 | Nbeal1 | 269198 | HITS-CLIP | 23597149 | Unspecified |

| | | | | | | |
|---|---|---|---|---|---|---|
| mmu-miR-125b-2-3p | MIMAT0004529 | Nckap1 | 50884 | HITS-CLIP | 23597149 | Unspecified |
| mmu-miR-125b-2-3p | MIMAT0004529 | Nckap1l | 105855 | HITS-CLIP | 29167373 | Unspecified |
| mmu-miR-125b-2-3p | MIMAT0004529 | Ncln | 103425 | HITS-CLIP | 23597149\|29167373 | Unspecified |
| mmu-miR-125b-2-3p | MIMAT0004529 | Ndufb11 | 104130 | HITS-CLIP | 23597149 | Unspecified |
| mmu-miR-125b-2-3p | MIMAT0004529 | Ndufb9 | 66218 | HITS-CLIP | 23597149 | Unspecified |
| mmu-miR-125b-2-3p | MIMAT0004529 | Necap1 | 67602 | HITS-CLIP | 29167373 | Unspecified |
| mmu-miR-125b-2-3p | MIMAT0004529 | Nedd4 | 17999 | HITS-CLIP | 23597149 | Unspecified |
| mmu-miR-125b-2-3p | MIMAT0004529 | Nek9 | 217718 | HITS-CLIP | 25753418 | Unspecified |
| mmu-miR-125b-2-3p | MIMAT0004529 | Nemf | 66244 | HITS-CLIP | 23597149 | Unspecified |
| mmu-miR-125b-2-3p | MIMAT0004529 | Nemp1 | 23306 | HITS-CLIP | 23597149 | Unspecified |
| mmu-miR-125b-2-3p | MIMAT0004529 | Neo1 | 18007 | HITS-CLIP | 23597149 | Unspecified |
| mmu-miR-125b-2-3p | MIMAT0004529 | Neurl4 | 216860 | HITS-CLIP | 23597149 | Unspecified |
| mmu-miR-125b-2-3p | MIMAT0004529 | Nfkbid | 243910 | HITS-CLIP | 29167373 | Unspecified |
| mmu-miR-125b-2-3p | MIMAT0004529 | Ngly1 | 59007 | HITS-CLIP | 37986224\|23597149 | Unspecified |
| mmu-miR-125b-2-3p | MIMAT0004529 | Notch2 | 18129 | HITS-CLIP | 37986224 | Unspecified |
| mmu-miR-125b-2-3p | MIMAT0004529 | Nploc4 | 217365 | HITS-CLIP | 23597149 | Unspecified |
| mmu-miR-125b-2-3p | MIMAT0004529 | Nr4a1 | 15370 | HITS-CLIP | 23597149 | Unspecified |
| mmu-miR-125b-2-3p | MIMAT0004529 | Nras | 18176 | HITS-CLIP | 23597149\|29167373 | Unspecified |
| mmu-miR-125b-2-3p | MIMAT0004529 | Nrep | 27528 | HITS-CLIP | 23597149 | Unspecified |
| mmu-miR-125b-2-3p | MIMAT0004529 | Nrp2 | 18187 | HITS-CLIP | 29167373 | Unspecified |
| mmu-miR-125b-2-3p | MIMAT0004529 | Nsd3 | 234135 | HITS-CLIP | 29167373 | Unspecified |
| mmu-miR-125b-2-3p | MIMAT0004529 | Nsmce4a | 67872 | HITS-CLIP | 29167373 | Unspecified |
| mmu-miR-125b-2-3p | MIMAT0004529 | Nup50 | 18141 | HITS-CLIP | 29167373 | Unspecified |
| mmu-miR-125b-2-3p | MIMAT0004529 | Ogfrl1 | 70155 | HITS-CLIP | 29167373 | Unspecified |
| mmu-miR-125b-2-3p | MIMAT0004529 | Opa1 | 74143 | HITS-CLIP | 37986224 | Unspecified |
| mmu-miR-125b-2-3p | MIMAT0004529 | Ophn1 | 94190 | HITS-CLIP | 29167373 | Unspecified |
| mmu-miR-125b-2-3p | MIMAT0004529 | Os9 | 216440 | HITS-CLIP | 37986224 | Unspecified |
| mmu-miR-125b-2-3p | MIMAT0004529 | Osbpl8 | 237542 | HITS-CLIP | 29167373 | Unspecified |
| mmu-miR-125b-2-3p | MIMAT0004529 | Osgin1 | 71839 | HITS-CLIP | 23597149 | Unspecified |
| mmu-miR-125b-2-3p | MIMAT0004529 | Otud4 | 73945 | HITS-CLIP | 37986224\|23597149 | Unspecified |
| mmu-miR-125b-2-3p | MIMAT0004529 | P4ha1 | 18451 | HITS-CLIP | 29167373 | Unspecified |
| mmu-miR-125b-2-3p | MIMAT0004529 | Pa2g4 | 18813 | HITS-CLIP | 23597149 | Unspecified |
| mmu-miR-125b-2-3p | MIMAT0004529 | Pabpc1 | 18458 | HITS-CLIP | 23597149 | Unspecified |
| mmu-miR-125b-2-3p | MIMAT0004529 | Pafah1b1 | 18472 | HITS-CLIP | 29167373 | Unspecified |
| mmu-miR-125b-2-3p | MIMAT0004529 | Paip2 | 67869 | HITS-CLIP | 23597149 | Unspecified |
| mmu-miR-125b-2-3p | MIMAT0004529 | Palmd | 114301 | HITS-CLIP | 37986224 | Unspecified |
| mmu-miR-125b-2-3p | MIMAT0004529 | Papola | 18789 | HITS-CLIP | 29167373 | Unspecified |
| mmu-miR-125b-2-3p | MIMAT0004529 | Papss2 | 23972 | HITS-CLIP | 23597149 | Unspecified |
| mmu-miR-125b-2-3p | MIMAT0004529 | Parp14 | 547253 | HITS-CLIP | 29167373 | Unspecified |

| | | | | | | | |
|---|---|---|---|---|---|---|---|
| mmu-miR-125b-2-3p | MIMAT0004529 | Paxbp1 | 67367 | HITS-CLIP | | 29167373 | Unspecified |
| mmu-miR-125b-2-3p | MIMAT0004529 | Paxip1 | 55982 | HITS-CLIP | | 23597149 | Unspecified |
| mmu-miR-125b-2-3p | MIMAT0004529 | Pbrm1 | 66923 | HITS-CLIP | | 37986224 | Unspecified |
| mmu-miR-125b-2-3p | MIMAT0004529 | Pcca | 110821 | HITS-CLIP | | 37986224 | Unspecified |
| mmu-miR-125b-2-3p | MIMAT0004529 | Pcdh7 | 54216 | HITS-CLIP | | 29167373 | Unspecified |
| mmu-miR-125b-2-3p | MIMAT0004529 | Pdcd6ip | 18571 | HITS-CLIP | | 29167373 | Unspecified |
| mmu-miR-125b-2-3p | MIMAT0004529 | Pde2a | 207728 | HITS-CLIP | | 25753418 | Unspecified |
| mmu-miR-125b-2-3p | MIMAT0004529 | Pde4b | 18578 | HITS-CLIP | | 29167373 | Unspecified |
| mmu-miR-125b-2-3p | MIMAT0004529 | Pde4dip | 83679 | HITS-CLIP | | 23597149 | Unspecified |
| mmu-miR-125b-2-3p | MIMAT0004529 | Pdlim5 | 56376 | HITS-CLIP | | 23597149 | Unspecified |
| mmu-miR-125b-2-3p | MIMAT0004529 | Pdzd8 | 107368 | HITS-CLIP | | 29167373 | Unspecified |
| mmu-miR-125b-2-3p | MIMAT0004529 | Pgm1 | 66681 | HITS-CLIP | | 23597149 | Unspecified |
| mmu-miR-125b-2-3p | MIMAT0004529 | Phf20l1 | 239510 | HITS-CLIP | 29167373\|23597149 | | Unspecified |
| mmu-miR-125b-2-3p | MIMAT0004529 | Phf3 | 213109 | HITS-CLIP | | 23597149 | Unspecified |
| mmu-miR-125b-2-3p | MIMAT0004529 | Phip | 83946 | HITS-CLIP | | 29167373 | Unspecified |
| mmu-miR-125b-2-3p | MIMAT0004529 | Pik3ca | 18706 | HITS-CLIP | | 25753418 | Unspecified |
| mmu-miR-125b-2-3p | MIMAT0004529 | Pik3cb | 74769 | HITS-CLIP | | 23597149 | Unspecified |
| mmu-miR-125b-2-3p | MIMAT0004529 | Pik3r1 | 18708 | HITS-CLIP | | 23597149 | Unspecified |
| mmu-miR-125b-2-3p | MIMAT0004529 | Pip4k2b | 108083 | HITS-CLIP | | 29167373 | Unspecified |
| mmu-miR-125b-2-3p | MIMAT0004529 | Pitpnb | 56305 | HITS-CLIP | | 23597149 | Unspecified |
| mmu-miR-125b-2-3p | MIMAT0004529 | Pkp4 | 227937 | HITS-CLIP | 37986224\|23597149 | | Unspecified |
| mmu-miR-125b-2-3p | MIMAT0004529 | Plcb1 | 18795 | HITS-CLIP | | 23597149 | Unspecified |
| mmu-miR-125b-2-3p | MIMAT0004529 | Plcxd2 | 433022 | HITS-CLIP | | 23597149 | Unspecified |
| mmu-miR-125b-2-3p | MIMAT0004529 | Plek | 56193 | HITS-CLIP | | 23597149 | Unspecified |
| mmu-miR-125b-2-3p | MIMAT0004529 | Plxna1 | 18844 | HITS-CLIP | | 29167373 | Unspecified |
| mmu-miR-125b-2-3p | MIMAT0004529 | Polr2m | 28015 | HITS-CLIP | | 37986224 | Unspecified |
| mmu-miR-125b-2-3p | MIMAT0004529 | Polr3f | 70408 | HITS-CLIP | | 29167373 | Unspecified |
| mmu-miR-125b-2-3p | MIMAT0004529 | Ppara | 19013 | HITS-CLIP | | 23597149 | Unspecified |
| mmu-miR-125b-2-3p | MIMAT0004529 | Ppip5k2 | 227399 | HITS-CLIP | | 25753418 | Unspecified |
| mmu-miR-125b-2-3p | MIMAT0004529 | Ppp1r3a | 140491 | HITS-CLIP | | 25753418 | Unspecified |
| mmu-miR-125b-2-3p | MIMAT0004529 | Ppp2r1b | 73699 | HITS-CLIP | | 37986224 | Unspecified |
| mmu-miR-125b-2-3p | MIMAT0004529 | Ppp3ca | 19055 | HITS-CLIP | | 23597149 | Unspecified |
| mmu-miR-125b-2-3p | MIMAT0004529 | Ppp6c | 67857 | HITS-CLIP | 37986224\|23597149 | | Unspecified |
| mmu-miR-125b-2-3p | MIMAT0004529 | Pptc7 | 320717 | HITS-CLIP | | 25753418 | Unspecified |
| mmu-miR-125b-2-3p | MIMAT0004529 | Prex2 | 109294 | HITS-CLIP | | 37986224 | Unspecified |
| mmu-miR-125b-2-3p | MIMAT0004529 | Prkaca | 18747 | HITS-CLIP | | 23597149 | Unspecified |
| mmu-miR-125b-2-3p | MIMAT0004529 | Prnp | 19122 | HITS-CLIP | | 25753418 | Unspecified |
| mmu-miR-125b-2-3p | MIMAT0004529 | Prpf8 | 192159 | HITS-CLIP | | 37986224 | Unspecified |
| mmu-miR-125b-2-3p | MIMAT0004529 | Psma1 | 26440 | HITS-CLIP | | 37986224 | Unspecified |

| | | | | | | |
|---|---|---|---|---|---|---|
| mmu-miR-125b-2-3p | MIMAT0004529 | Psmb3 | 26446 | HITS-CLIP | 29167373 | Unspecified |
| mmu-miR-125b-2-3p | MIMAT0004529 | Psmb9 | 16912 | HITS-CLIP | 23597149 | Unspecified |
| mmu-miR-125b-2-3p | MIMAT0004529 | Psme4 | 103554 | HITS-CLIP | 29167373 | Unspecified |
| mmu-miR-125b-2-3p | MIMAT0004529 | Pten | 19211 | HITS-CLIP | 29167373 | Unspecified |
| mmu-miR-125b-2-3p | MIMAT0004529 | Ptpn2 | 19255 | HITS-CLIP | 23597149 | Unspecified |
| mmu-miR-125b-2-3p | MIMAT0004529 | Pum2 | 80913 | HITS-CLIP | 29167373 | Unspecified |
| mmu-miR-125b-2-3p | MIMAT0004529 | Pura | 19290 | HITS-CLIP | 29167373 | Unspecified |
| mmu-miR-125b-2-3p | MIMAT0004529 | Pycard | 66824 | HITS-CLIP | 29167373 | Unspecified |
| mmu-miR-125b-2-3p | MIMAT0004529 | Qk | 19317 | HITS-CLIP | 29167373\|23597149 | Unspecified |
| mmu-miR-125b-2-3p | MIMAT0004529 | Rab14 | 68365 | HITS-CLIP | 23597149 | Unspecified |
| mmu-miR-125b-2-3p | MIMAT0004529 | Rab7 | 19349 | HITS-CLIP | 23597149 | Unspecified |
| mmu-miR-125b-2-3p | MIMAT0004529 | Ralgapb | 228850 | HITS-CLIP | 23597149\|29167373 | Unspecified |
| mmu-miR-125b-2-3p | MIMAT0004529 | Ranbp2 | 19386 | HITS-CLIP | 37986224\|23597149\|25753418\|29167 | Unspecified |
| mmu-miR-125b-2-3p | MIMAT0004529 | Rapgef5 | 217944 | HITS-CLIP | 29167373 | Unspecified |
| mmu-miR-125b-2-3p | MIMAT0004529 | Rasgrp3 | 240168 | HITS-CLIP | 37986224 | Unspecified |
| mmu-miR-125b-2-3p | MIMAT0004529 | Rbbp4 | 19646 | HITS-CLIP | 29167373 | Unspecified |
| mmu-miR-125b-2-3p | MIMAT0004529 | Rbm47 | 245945 | HITS-CLIP | 37986224 | Unspecified |
| mmu-miR-125b-2-3p | MIMAT0004529 | Rbm6 | 19654 | HITS-CLIP | 23597149 | Unspecified |
| mmu-miR-125b-2-3p | MIMAT0004529 | Rcl1 | 59028 | HITS-CLIP | 37986224 | Unspecified |
| mmu-miR-125b-2-3p | MIMAT0004529 | Rgs10 | 67865 | HITS-CLIP | 29167373 | Unspecified |
| mmu-miR-125b-2-3p | MIMAT0004529 | Rhbdd2 | 215160 | HITS-CLIP | 25753418 | Unspecified |
| mmu-miR-125b-2-3p | MIMAT0004529 | Rhob | 11852 | HITS-CLIP | 29167373 | Unspecified |
| mmu-miR-125b-2-3p | MIMAT0004529 | Rictor | 78757 | HITS-CLIP | 29167373 | Unspecified |
| mmu-miR-125b-2-3p | MIMAT0004529 | Riok1 | 71340 | HITS-CLIP | 29167373 | Unspecified |
| mmu-miR-125b-2-3p | MIMAT0004529 | Ripor1 | 75687 | HITS-CLIP | 25753418 | Unspecified |
| mmu-miR-125b-2-3p | MIMAT0004529 | Rlf | 109263 | HITS-CLIP | 37986224 | Unspecified |
| mmu-miR-125b-2-3p | MIMAT0004529 | Rmnd5a | 68477 | HITS-CLIP | 37986224 | Unspecified |
| mmu-miR-125b-2-3p | MIMAT0004529 | Rnd3 | 74194 | HITS-CLIP | 25753418 | Unspecified |
| mmu-miR-125b-2-3p | MIMAT0004529 | Rnf103 | 22644 | HITS-CLIP | 37986224 | Unspecified |
| mmu-miR-125b-2-3p | MIMAT0004529 | Rnf111 | 93836 | HITS-CLIP | 23597149 | Unspecified |
| mmu-miR-125b-2-3p | MIMAT0004529 | Rnf123 | 84585 | HITS-CLIP | 25753418 | Unspecified |
| mmu-miR-125b-2-3p | MIMAT0004529 | Rnf149 | 67702 | HITS-CLIP | 29167373 | Unspecified |
| mmu-miR-125b-2-3p | MIMAT0004529 | Rnf213 | 672511 | HITS-CLIP | 23597149\|29167373 | Unspecified |
| mmu-miR-125b-2-3p | MIMAT0004529 | Rnf214 | 235315 | HITS-CLIP | 23597149 | Unspecified |
| mmu-miR-125b-2-3p | MIMAT0004529 | Rnf217 | 268291 | HITS-CLIP | 23597149 | Unspecified |
| mmu-miR-125b-2-3p | MIMAT0004529 | Rps3a1 | 20091 | HITS-CLIP | 37986224\|29167373 | Unspecified |
| mmu-miR-125b-2-3p | MIMAT0004529 | Rsbn1l | 242860 | HITS-CLIP | 29167373 | Unspecified |
| mmu-miR-125b-2-3p | MIMAT0004529 | Rtn3 | 20168 | HITS-CLIP | 29167373 | Unspecified |
| mmu-miR-125b-2-3p | MIMAT0004529 | S100a1 | 20193 | HITS-CLIP | 23597149 | Unspecified |

| | | | | | | |
|---|---|---|---|---|---|---|
| mmu-miR-125b-2-3p | MIMAT0004529 | Sap30 | 60406 | HITS-CLIP | 23597149 | Unspecified |
| mmu-miR-125b-2-3p | MIMAT0004529 | Sbno1 | 243272 | HITS-CLIP | 29167373 | Unspecified |
| mmu-miR-125b-2-3p | MIMAT0004529 | Scaf8 | 106583 | HITS-CLIP | 29167373 | Unspecified |
| mmu-miR-125b-2-3p | MIMAT0004529 | Scara5 | 71145 | HITS-CLIP | 37986224 | Unspecified |
| mmu-miR-125b-2-3p | MIMAT0004529 | Scn1b | 20266 | HITS-CLIP | 23597149 | Unspecified |
| mmu-miR-125b-2-3p | MIMAT0004529 | Scn4a | 110880 | HITS-CLIP | 25753418 | Unspecified |
| mmu-miR-125b-2-3p | MIMAT0004529 | Scrib | 105782 | HITS-CLIP | 23597149 | Unspecified |
| mmu-miR-125b-2-3p | MIMAT0004529 | Sdc4 | 20971 | HITS-CLIP | 29167373 | Unspecified |
| mmu-miR-125b-2-3p | MIMAT0004529 | Sec24a | 77371 | HITS-CLIP | 37986224 | Unspecified |
| mmu-miR-125b-2-3p | MIMAT0004529 | Sec24c | 218811 | HITS-CLIP | 37986224 | Unspecified |
| mmu-miR-125b-2-3p | MIMAT0004529 | Sec31a | 69162 | HITS-CLIP | 37986224 | Unspecified |
| mmu-miR-125b-2-3p | MIMAT0004529 | Senp5 | 320213 | HITS-CLIP | 23597149\|29167373 | Unspecified |
| mmu-miR-125b-2-3p | MIMAT0004529 | Senp6 | 215351 | HITS-CLIP | 29167373 | Unspecified |
| mmu-miR-125b-2-3p | MIMAT0004529 | Septin2 | 18000 | HITS-CLIP | 25753418 | Unspecified |
| mmu-miR-125b-2-3p | MIMAT0004529 | Serpinc1 | 11905 | HITS-CLIP | 23597149 | Unspecified |
| mmu-miR-125b-2-3p | MIMAT0004529 | Serpine1 | 18787 | HITS-CLIP | 23142080\|23597149 | Unspecified |
| mmu-miR-125b-2-3p | MIMAT0004529 | Serpinf2 | 18816 | HITS-CLIP | 37986224 | Unspecified |
| mmu-miR-125b-2-3p | MIMAT0004529 | Setd2 | 235626 | HITS-CLIP | 23597149 | Unspecified |
| mmu-miR-125b-2-3p | MIMAT0004529 | Setd5 | 72895 | HITS-CLIP | 23597149\|29167373 | Unspecified |
| mmu-miR-125b-2-3p | MIMAT0004529 | Sf1 | 22668 | HITS-CLIP | 29167373 | Unspecified |
| mmu-miR-125b-2-3p | MIMAT0004529 | Sfpq | 71514 | HITS-CLIP | 29167373 | Unspecified |
| mmu-miR-125b-2-3p | MIMAT0004529 | Sfxn2 | 94279 | HITS-CLIP | 23597149 | Unspecified |
| mmu-miR-125b-2-3p | MIMAT0004529 | Sgk1 | 20393 | HITS-CLIP | 25753418 | Unspecified |
| mmu-miR-125b-2-3p | MIMAT0004529 | Shroom2 | 110380 | HITS-CLIP | 23597149 | Unspecified |
| mmu-miR-125b-2-3p | MIMAT0004529 | Siah2 | 20439 | HITS-CLIP | 23597149 | Unspecified |
| mmu-miR-125b-2-3p | MIMAT0004529 | Sirpa | 19261 | HITS-CLIP | 29167373 | Unspecified |
| mmu-miR-125b-2-3p | MIMAT0004529 | Sirt1 | 93759 | HITS-CLIP | 23597149 | Unspecified |
| mmu-miR-125b-2-3p | MIMAT0004529 | Slc16a1 | 20501 | HITS-CLIP | 23597149 | Unspecified |
| mmu-miR-125b-2-3p | MIMAT0004529 | Slc17a3 | 105355 | HITS-CLIP | 23597149 | Unspecified |
| mmu-miR-125b-2-3p | MIMAT0004529 | Slc25a20 | 57279 | HITS-CLIP | 37986224 | Unspecified |
| mmu-miR-125b-2-3p | MIMAT0004529 | Slc25a36 | 192287 | HITS-CLIP | 29167373 | Unspecified |
| mmu-miR-125b-2-3p | MIMAT0004529 | Slc31a2 | 20530 | HITS-CLIP | 25753418 | Unspecified |
| mmu-miR-125b-2-3p | MIMAT0004529 | Slc37a4 | 14385 | HITS-CLIP | 37986224\|23597149 | Unspecified |
| mmu-miR-125b-2-3p | MIMAT0004529 | Slc39a3 | 106947 | HITS-CLIP | 29167373 | Unspecified |
| mmu-miR-125b-2-3p | MIMAT0004529 | Slc4a7 | 218756 | HITS-CLIP | 29167373 | Unspecified |
| mmu-miR-125b-2-3p | MIMAT0004529 | Slc7a2 | 11988 | HITS-CLIP | 37986224\|23597149 | Unspecified |
| mmu-miR-125b-2-3p | MIMAT0004529 | Slmap | 83997 | HITS-CLIP | 37986224\|29167373 | Unspecified |
| mmu-miR-125b-2-3p | MIMAT0004529 | Sln | 66402 | HITS-CLIP | 25753418 | Unspecified |
| mmu-miR-125b-2-3p | MIMAT0004529 | Smarca2 | 67155 | HITS-CLIP | 23597149 | Unspecified |

| | | | | | | |
|---|---|---|---|---|---|---|
| mmu-miR-125b-2-3p | MIMAT0004529 | Smg1 | 233789 | HITS-CLIP | 29167373 | Unspecified |
| mmu-miR-125b-2-3p | MIMAT0004529 | Snx13 | 217463 | HITS-CLIP | 37986224 | Unspecified |
| mmu-miR-125b-2-3p | MIMAT0004529 | Son | 20658 | HITS-CLIP | 23597149\|29167373 | Unspecified |
| mmu-miR-125b-2-3p | MIMAT0004529 | Spcs3 | 76687 | HITS-CLIP | 23597149 | Unspecified |
| mmu-miR-125b-2-3p | MIMAT0004529 | Sqstm1 | 18412 | HITS-CLIP | 37986224\|23597149 | Unspecified |
| mmu-miR-125b-2-3p | MIMAT0004529 | Srcap | 100043597 | HITS-CLIP | 29167373 | Unspecified |
| mmu-miR-125b-2-3p | MIMAT0004529 | Srgap2 | 14270 | HITS-CLIP | 29167373 | Unspecified |
| mmu-miR-125b-2-3p | MIMAT0004529 | Ssb | 20823 | HITS-CLIP | 37986224 | Unspecified |
| mmu-miR-125b-2-3p | MIMAT0004529 | St13 | 70356 | HITS-CLIP | 23597149 | Unspecified |
| mmu-miR-125b-2-3p | MIMAT0004529 | St3gal4 | 20443 | HITS-CLIP | 37986224 | Unspecified |
| mmu-miR-125b-2-3p | MIMAT0004529 | Stard5 | 170460 | HITS-CLIP | 37986224 | Unspecified |
| mmu-miR-125b-2-3p | MIMAT0004529 | Strap | 20901 | HITS-CLIP | 23597149 | Unspecified |
| mmu-miR-125b-2-3p | MIMAT0004529 | Sucla2 | 20916 | HITS-CLIP | 23597149 | Unspecified |
| mmu-miR-125b-2-3p | MIMAT0004529 | Surf1 | 20930 | HITS-CLIP | 37986224 | Unspecified |
| mmu-miR-125b-2-3p | MIMAT0004529 | Susd6 | 9766 | HITS-CLIP | 29167373 | Unspecified |
| mmu-miR-125b-2-3p | MIMAT0004529 | Svil | 225115 | HITS-CLIP | 23597149\|25753418 | Unspecified |
| mmu-miR-125b-2-3p | MIMAT0004529 | Syngr1 | 20972 | HITS-CLIP | 29167373 | Unspecified |
| mmu-miR-125b-2-3p | MIMAT0004529 | Syt11 | 229521 | HITS-CLIP | 31433990 | Unspecified |
| mmu-miR-125b-2-3p | MIMAT0004529 | Taf1 | 270627 | HITS-CLIP | 23597149 | Unspecified |
| mmu-miR-125b-2-3p | MIMAT0004529 | Taok1 | 216965 | HITS-CLIP | 23597149 | Unspecified |
| mmu-miR-125b-2-3p | MIMAT0004529 | Tat | 234724 | HITS-CLIP | 23597149 | Unspecified |
| mmu-miR-125b-2-3p | MIMAT0004529 | Tbc1d2b | 67016 | HITS-CLIP | 37986224 | Unspecified |
| mmu-miR-125b-2-3p | MIMAT0004529 | Tead1 | 21676 | HITS-CLIP | 23597149 | Unspecified |
| mmu-miR-125b-2-3p | MIMAT0004529 | Tent4a | 11044 | HITS-CLIP | 23597149\|29167373 | Unspecified |
| mmu-miR-125b-2-3p | MIMAT0004529 | Thoc2 | 331401 | HITS-CLIP | 37986224 | Unspecified |
| mmu-miR-125b-2-3p | MIMAT0004529 | Thrb | 21834 | HITS-CLIP | 23597149 | Unspecified |
| mmu-miR-125b-2-3p | MIMAT0004529 | Tigar | 319801 | HITS-CLIP | 25753418 | Unspecified |
| mmu-miR-125b-2-3p | MIMAT0004529 | Tle4 | 21888 | HITS-CLIP | 23597149 | Unspecified |
| mmu-miR-125b-2-3p | MIMAT0004529 | Tmem106a | 217203 | HITS-CLIP | 37986224\|23597149 | Unspecified |
| mmu-miR-125b-2-3p | MIMAT0004529 | Tmem18 | 211986 | HITS-CLIP | 37986224 | Unspecified |
| mmu-miR-125b-2-3p | MIMAT0004529 | Tmem248 | 71667 | HITS-CLIP | 23597149 | Unspecified |
| mmu-miR-125b-2-3p | MIMAT0004529 | Tmem30a | 69981 | HITS-CLIP | 37986224 | Unspecified |
| mmu-miR-125b-2-3p | MIMAT0004529 | Tmem87a | 211499 | HITS-CLIP | 23597149\|29167373 | Unspecified |
| mmu-miR-125b-2-3p | MIMAT0004529 | Tnf | 21926 | HITS-CLIP | 29167373 | Unspecified |
| mmu-miR-125b-2-3p | MIMAT0004529 | Tnks2 | 74493 | HITS-CLIP | 37986224\|23597149 | Unspecified |
| mmu-miR-125b-2-3p | MIMAT0004529 | Tnpo3 | 320938 | HITS-CLIP | 37986224\|25753418\|29167373 | Unspecified |
| mmu-miR-125b-2-3p | MIMAT0004529 | Toe1 | 68276 | HITS-CLIP | 37986224 | Unspecified |
| mmu-miR-125b-2-3p | MIMAT0004529 | Tollip | 54473 | HITS-CLIP | 29167373 | Unspecified |
| mmu-miR-125b-2-3p | MIMAT0004529 | Tor1aip2 | 240832 | HITS-CLIP | 23597149 | Unspecified |

| | | | | | | |
|---|---|---|---|---|---|---|
| mmu-miR-125b-2-3p | MIMAT0004529 | Tpp1 | 12751 | HITS-CLIP | 23597149 | Unspecified |
| mmu-miR-125b-2-3p | MIMAT0004529 | Trappc1 | 245828 | HITS-CLIP | 29167373 | Unspecified |
| mmu-miR-125b-2-3p | MIMAT0004529 | Trib1 | 211770 | HITS-CLIP | 37986224 | Unspecified |
| mmu-miR-125b-2-3p | MIMAT0004529 | Trim44 | 80985 | HITS-CLIP | 29167373 | Unspecified |
| mmu-miR-125b-2-3p | MIMAT0004529 | Trip12 | 14897 | HITS-CLIP | 37986224\|23597149\|29167373 | Unspecified |
| mmu-miR-125b-2-3p | MIMAT0004529 | Trp53inp2 | 68728 | HITS-CLIP | 23597149 | Unspecified |
| mmu-miR-125b-2-3p | MIMAT0004529 | Tsc22d3 | 14605 | HITS-CLIP | 37986224 | Unspecified |
| mmu-miR-125b-2-3p | MIMAT0004529 | Ttc17 | 74569 | HITS-CLIP | 29167373 | Unspecified |
| mmu-miR-125b-2-3p | MIMAT0004529 | Ttc7 | 225049 | HITS-CLIP | 23597149 | Unspecified |
| mmu-miR-125b-2-3p | MIMAT0004529 | Ttn | 22138 | HITS-CLIP | 25753418 | Unspecified |
| mmu-miR-125b-2-3p | MIMAT0004529 | Tuba1b | 22143 | HITS-CLIP | 23597149 | Unspecified |
| mmu-miR-125b-2-3p | MIMAT0004529 | Tuba4a | 22145 | HITS-CLIP | 23597149 | Unspecified |
| mmu-miR-125b-2-3p | MIMAT0004529 | Tubb2a | 22151 | HITS-CLIP | 23597149 | Unspecified |
| mmu-miR-125b-2-3p | MIMAT0004529 | Tut4 | 23318 | HITS-CLIP | 23597149 | Unspecified |
| mmu-miR-125b-2-3p | MIMAT0004529 | Txnip | 56338 | HITS-CLIP | 23597149\|29167373 | Unspecified |
| mmu-miR-125b-2-3p | MIMAT0004529 | Ube3a | 22215 | HITS-CLIP | 25753418 | Unspecified |
| mmu-miR-125b-2-3p | MIMAT0004529 | Ube4a | 140630 | HITS-CLIP | 23597149 | Unspecified |
| mmu-miR-125b-2-3p | MIMAT0004529 | Ubfd1 | 28018 | HITS-CLIP | 23597149 | Unspecified |
| mmu-miR-125b-2-3p | MIMAT0004529 | Ugcg | 22234 | HITS-CLIP | 25753418 | Unspecified |
| mmu-miR-125b-2-3p | MIMAT0004529 | Ugt2b1 | 71773 | HITS-CLIP | 23597149 | Unspecified |
| mmu-miR-125b-2-3p | MIMAT0004529 | Unc119b | 106840 | HITS-CLIP | 31433990 | Unspecified |
| mmu-miR-125b-2-3p | MIMAT0004529 | Upf2 | 326622 | HITS-CLIP | 29167373 | Unspecified |
| mmu-miR-125b-2-3p | MIMAT0004529 | Upp2 | 76654 | HITS-CLIP | 37986224 | Unspecified |
| mmu-miR-125b-2-3p | MIMAT0004529 | Uqcrc2 | 67003 | HITS-CLIP | 25753418 | Unspecified |
| mmu-miR-125b-2-3p | MIMAT0004529 | Usp15 | 14479 | HITS-CLIP | 37986224 | Unspecified |
| mmu-miR-125b-2-3p | MIMAT0004529 | Usp2 | 53376 | HITS-CLIP | 23597149 | Unspecified |
| mmu-miR-125b-2-3p | MIMAT0004529 | Usp7 | 252870 | HITS-CLIP | 29167373 | Unspecified |
| mmu-miR-125b-2-3p | MIMAT0004529 | Usp9x | 22284 | HITS-CLIP | 23597149\|29167373 | Unspecified |
| mmu-miR-125b-2-3p | MIMAT0004529 | Uty | 22290 | HITS-CLIP | 25753418 | Unspecified |
| mmu-miR-125b-2-3p | MIMAT0004529 | Vav3 | 57257 | HITS-CLIP | 29167373 | Unspecified |
| mmu-miR-125b-2-3p | MIMAT0004529 | Vcl | 22330 | HITS-CLIP | 25753418 | Unspecified |
| mmu-miR-125b-2-3p | MIMAT0004529 | Vdac2 | 22334 | HITS-CLIP | 37986224 | Unspecified |
| mmu-miR-125b-2-3p | MIMAT0004529 | Vil1 | 22349 | HITS-CLIP | 25753418 | Unspecified |
| mmu-miR-125b-2-3p | MIMAT0004529 | Vps13a | 271564 | HITS-CLIP | 29167373 | Unspecified |
| mmu-miR-125b-2-3p | MIMAT0004529 | Vps13b | 666173 | HITS-CLIP | 37986224 | Unspecified |
| mmu-miR-125b-2-3p | MIMAT0004529 | Vps50 | 55610 | HITS-CLIP | 23597149 | Unspecified |
| mmu-miR-125b-2-3p | MIMAT0004529 | Wdr26 | 226757 | HITS-CLIP | 23597149 | Unspecified |
| mmu-miR-125b-2-3p | MIMAT0004529 | Wnk2 | 75607 | HITS-CLIP | 25753418 | Unspecified |
| mmu-miR-125b-2-3p | MIMAT0004529 | Wtap | 60532 | HITS-CLIP | 29167373 | Unspecified |

| | | | | | | |
|---|---|---|---|---|---|---|
| mmu-miR-125b-2-3p | MIMAT0004529 | Xiap | 11798 | HITS-CLIP | 29167373 | Unspecified |
| mmu-miR-125b-2-3p | MIMAT0004529 | Xpo1 | 103573 | HITS-CLIP | 23597149 | Unspecified |
| mmu-miR-125b-2-3p | MIMAT0004529 | Yod1 | 226418 | HITS-CLIP | 29167373 | Unspecified |
| mmu-miR-125b-2-3p | MIMAT0004529 | Ythdc1 | 231386 | HITS-CLIP | 29167373 | Unspecified |
| mmu-miR-125b-2-3p | MIMAT0004529 | Zbtb5 | 230119 | HITS-CLIP | 23597149\|25753418 | Unspecified |
| mmu-miR-125b-2-3p | MIMAT0004529 | Zbtb7a | 16969 | HITS-CLIP | 29167373 | Unspecified |
| mmu-miR-125b-2-3p | MIMAT0004529 | Zc3h12c | 244871 | HITS-CLIP | 37986224 | Unspecified |
| mmu-miR-125b-2-3p | MIMAT0004529 | Zfhx3 | 11906 | HITS-CLIP | 25753418 | Unspecified |
| mmu-miR-125b-2-3p | MIMAT0004529 | Zfp532 | 328977 | HITS-CLIP | 25753418 | Unspecified |
| mmu-miR-125b-2-3p | MIMAT0004529 | Zfp568 | 243905 | HITS-CLIP | 37986224 | Unspecified |
| mmu-miR-125b-2-3p | MIMAT0004529 | Zfp592 | 233410 | HITS-CLIP | 29167373 | Unspecified |
| mmu-miR-125b-2-3p | MIMAT0004529 | Zfp672 | 319475 | HITS-CLIP | 37986224 | Unspecified |
| mmu-miR-125b-2-3p | MIMAT0004529 | Zfp706 | 68036 | HITS-CLIP | 37986224 | Unspecified |
| mmu-miR-125b-2-3p | MIMAT0004529 | Zmat2 | 66492 | HITS-CLIP | 37986224 | Unspecified |
| mmu-miR-125b-2-3p | MIMAT0004529 | Zmpste24 | 230709 | HITS-CLIP | 37986224 | Unspecified |
| mmu-miR-125b-2-3p | MIMAT0004529 | Zmym4 | 67785 | HITS-CLIP | 29167373 | Unspecified |
| mmu-miR-125b-2-3p | MIMAT0004529 | mt-Co1 | 4512 | HITS-CLIP | 29167373\|23597149 | Unspecified |
| mmu-miR-1900 | MIMAT0007870 | Mknk2 | 17347 | HITS-CLIP | 31433990 | Unspecified |
| mmu-miR-100-5p | MIMAT0000655 | 0610030E20Rik | 68364 | HITS-CLIP | 25753418 | Unspecified |
| mmu-miR-100-5p | MIMAT0000655 | Abcc9 | 20928 | HITS-CLIP | 25753418 | Unspecified |
| mmu-miR-100-5p | MIMAT0000655 | Agl | 77559 | HITS-CLIP | 25753418 | Unspecified |
| mmu-miR-100-5p | MIMAT0000655 | Cap2 | 67252 | HITS-CLIP | 25753418 | Unspecified |
| mmu-miR-100-5p | MIMAT0000655 | Cdk8 | 264064 | HITS-CLIP | 25753418 | Unspecified |
| mmu-miR-100-5p | MIMAT0000655 | Ctnna1 | 12385 | HITS-CLIP | 25753418 | Unspecified |
| mmu-miR-100-5p | MIMAT0000655 | Dag1 | 13138 | HITS-CLIP | 25753418 | Unspecified |
| mmu-miR-100-5p | MIMAT0000655 | Dcaf12 | 68970 | HITS-CLIP | 25753418 | Unspecified |
| mmu-miR-100-5p | MIMAT0000655 | Dnajb5 | 56323 | HITS-CLIP | 25753418 | Unspecified |
| mmu-miR-100-5p | MIMAT0000655 | Dnmt3a | 13435 | HITS-CLIP | 25753418 | Unspecified |
| mmu-miR-100-5p | MIMAT0000655 | Fam3a | 66294 | HITS-CLIP | 25753418 | Unspecified |
| mmu-miR-100-5p | MIMAT0000655 | Fzd8 | 14370 | HITS-CLIP | 25753418 | Unspecified |
| mmu-miR-100-5p | MIMAT0000655 | Gigyf1 | 57330 | HITS-CLIP | 25753418\|26203562 | Unspecified |
| mmu-miR-100-5p | MIMAT0000655 | Gm28035 | 118451630 | HITS-CLIP | 31433990 | Unspecified |
| mmu-miR-100-5p | MIMAT0000655 | Gpc4 | 14735 | HITS-CLIP | 25753418 | Unspecified |
| mmu-miR-100-5p | MIMAT0000655 | Impdh1 | 23917 | HITS-CLIP | 25753418 | Unspecified |
| mmu-miR-100-5p | MIMAT0000655 | Ing4 | 28019 | HITS-CLIP | 25753418 | Unspecified |
| mmu-miR-100-5p | MIMAT0000655 | Ivd | 56357 | HITS-CLIP | 25753418 | Unspecified |
| mmu-miR-100-5p | MIMAT0000655 | Lrrc8a | 241296 | HITS-CLIP | 31433990 | Unspecified |
| mmu-miR-100-5p | MIMAT0000655 | Man2c1 | 73744 | HITS-CLIP | 25753418 | Unspecified |
| mmu-miR-100-5p | MIMAT0000655 | Mical2 | 320878 | HITS-CLIP | 25753418 | Unspecified |

| | | | | | | |
|---|---|---|---|---|---|---|
| mmu-miR-100-5p | MIMAT0000655 | Nars | 70223 | HITS-CLIP | 37986224 | Unspecified |
| mmu-miR-100-5p | MIMAT0000655 | Ndufs6 | 407785 | HITS-CLIP | 31433990\|25753418 | Unspecified |
| mmu-miR-100-5p | MIMAT0000655 | Obsl1 | 98733 | HITS-CLIP | 25753418 | Unspecified |
| mmu-miR-100-5p | MIMAT0000655 | Parp2 | 11546 | HITS-CLIP | 25753418 | Unspecified |
| mmu-miR-100-5p | MIMAT0000655 | Pde3a | 54611 | HITS-CLIP | 25753418 | Unspecified |
| mmu-miR-100-5p | MIMAT0000655 | Ppm1j | 71887 | HITS-CLIP | 25753418 | Unspecified |
| mmu-miR-100-5p | MIMAT0000655 | Prdm4 | 72843 | HITS-CLIP | 25753418 | Unspecified |
| mmu-miR-100-5p | MIMAT0000655 | Prkce | 18754 | HITS-CLIP | 25753418 | Unspecified |
| mmu-miR-100-5p | MIMAT0000655 | Prpf8 | 192159 | HITS-CLIP | 25753418 | Unspecified |
| mmu-miR-100-5p | MIMAT0000655 | Pten | 19211 | HITS-CLIP | 25753418 | Unspecified |
| mmu-miR-100-5p | MIMAT0000655 | Pygb | 110078 | HITS-CLIP | 25753418 | Unspecified |
| mmu-miR-100-5p | MIMAT0000655 | Rasa1 | 218397 | HITS-CLIP | 25753418 | Unspecified |
| mmu-miR-100-5p | MIMAT0000655 | Rdx | 19684 | HITS-CLIP | 31433990\|25753418 | Unspecified |
| mmu-miR-100-5p | MIMAT0000655 | Rmdn1 | 66302 | HITS-CLIP | 25753418 | Unspecified |
| mmu-miR-100-5p | MIMAT0000655 | Rnf139 | 75841 | HITS-CLIP | 25753418 | Unspecified |
| mmu-miR-100-5p | MIMAT0000655 | Runx1t1 | 12395 | HITS-CLIP | 25753418 | Unspecified |
| mmu-miR-100-5p | MIMAT0000655 | Sik3 | 70661 | HITS-CLIP | 25753418 | Unspecified |
| mmu-miR-100-5p | MIMAT0000655 | Slc8a1 | 20541 | HITS-CLIP | 25753418 | Unspecified |
| mmu-miR-100-5p | MIMAT0000655 | Sptb | 20741 | HITS-CLIP | 25753418 | Unspecified |
| mmu-miR-100-5p | MIMAT0000655 | Sptbn1 | 20742 | HITS-CLIP | 25753418 | Unspecified |
| mmu-miR-100-5p | MIMAT0000655 | Stat2 | 20847 | HITS-CLIP | 25753418 | Unspecified |
| mmu-miR-100-5p | MIMAT0000655 | Steap3 | 68428 | HITS-CLIP | 25753418 | Unspecified |
| mmu-miR-100-5p | MIMAT0000655 | Supt6 | 20926 | HITS-CLIP | 31433990\|25753418 | Unspecified |
| mmu-miR-100-5p | MIMAT0000655 | Timm44 | 21856 | HITS-CLIP | 25753418 | Unspecified |
| mmu-miR-100-5p | MIMAT0000655 | Tmem38b | 52076 | qCLASH | 31393953 | Unspecified |
| mmu-miR-100-5p | MIMAT0000655 | Tnrc18 | 231861 | HITS-CLIP | 25753418 | Unspecified |
| mmu-miR-100-5p | MIMAT0000655 | Trim71 | 636931 | HITS-CLIP | 37986224 | Unspecified |
| mmu-miR-100-5p | MIMAT0000655 | Uqcrc2 | 67003 | HITS-CLIP | 25753418 | Unspecified |
| mmu-miR-100-5p | MIMAT0000655 | Zfp839 | 72805 | HITS-CLIP | 25753418 | Unspecified |
| mmu-miR-3086-5p | MIMAT0014880 | Ckap5 | 75786 | HITS-CLIP | 31433990 | Unspecified |
| mmu-miR-3086-5p | MIMAT0014880 | Endou | 19011 | HITS-CLIP | 26203562 | Unspecified |
| mmu-miR-3086-5p | MIMAT0014880 | Fam219b | 78323 | HITS-CLIP | 31433990 | Unspecified |
| mmu-miR-3086-5p | MIMAT0014880 | Fosl2 | 14284 | HITS-CLIP | 31433990 | Unspecified |
| mmu-miR-3086-5p | MIMAT0014880 | Rbm12 | 75710 | HITS-CLIP | 37986224 | Unspecified |
| mmu-miR-3086-5p | MIMAT0014880 | Trim28 | 21849 | HITS-CLIP | 37986224 | Unspecified |
| mmu-miR-3086-5p | MIMAT0014880 | Trps1 | 83925 | HITS-CLIP | 37986224 | Unspecified |
| mmu-miR-3086-5p | MIMAT0014880 | Yars | 107271 | HITS-CLIP | 37986224 | Unspecified |
| mmu-miR-3086-5p | MIMAT0014880 | Yeats2 | 208146 | HITS-CLIP | 31433990 | Unspecified |
| mmu-miR-3086-5p | MIMAT0014880 | Zfp532 | 328977 | HITS-CLIP | 37986224 | Unspecified |

| | | | | | | |
|---|---|---|---|---|---|---|
| mmu-miR-378b | MIMAT0019348 | 0610030E20Rik | 68364 | HITS-CLIP | 37986224 | Unspecified |
| mmu-miR-378b | MIMAT0019348 | 4833439L19Rik | 97820 | HITS-CLIP | 37986224 | Unspecified |
| mmu-miR-378b | MIMAT0019348 | AI182371 | 98870 | HITS-CLIP | 23597149 | Unspecified |
| mmu-miR-378b | MIMAT0019348 | Aacs | 78894 | HITS-CLIP | 37986224 | Unspecified |
| mmu-miR-378b | MIMAT0019348 | Aass | 30956 | HITS-CLIP | 23597149 | Unspecified |
| mmu-miR-378b | MIMAT0019348 | Abcb6 | 74104 | HITS-CLIP | 23597149 | Unspecified |
| mmu-miR-378b | MIMAT0019348 | Abcc6 | 27421 | HITS-CLIP | 23597149 | Unspecified |
| mmu-miR-378b | MIMAT0019348 | Abcf3 | 27406 | HITS-CLIP | 23597149 | Unspecified |
| mmu-miR-378b | MIMAT0019348 | Acaa2 | 52538 | HITS-CLIP | 37986224 | Unspecified |
| mmu-miR-378b | MIMAT0019348 | Acacb | 100705 | HITS-CLIP | 37986224 | Unspecified |
| mmu-miR-378b | MIMAT0019348 | Acox1 | 11430 | HITS-CLIP | 23597149 | Unspecified |
| mmu-miR-378b | MIMAT0019348 | Acsl1 | 14081 | HITS-CLIP | 23597149 | Unspecified |
| mmu-miR-378b | MIMAT0019348 | Acsl4 | 50790 | HITS-CLIP | 37986224 | Unspecified |
| mmu-miR-378b | MIMAT0019348 | Acsm1 | 117147 | HITS-CLIP | 23597149 | Unspecified |
| mmu-miR-378b | MIMAT0019348 | Acsm5 | 272428 | HITS-CLIP | 23597149 | Unspecified |
| mmu-miR-378b | MIMAT0019348 | Adck5 | 268822 | HITS-CLIP | 23597149 | Unspecified |
| mmu-miR-378b | MIMAT0019348 | Ahctf1 | 226747 | HITS-CLIP | 23597149 | Unspecified |
| mmu-miR-378b | MIMAT0019348 | Ahdc1 | 230793 | HITS-CLIP | 37986224 | Unspecified |
| mmu-miR-378b | MIMAT0019348 | Ahnak | 66395 | HITS-CLIP | 37986224 | Unspecified |
| mmu-miR-378b | MIMAT0019348 | Ak3 | 56248 | HITS-CLIP | 23597149 | Unspecified |
| mmu-miR-378b | MIMAT0019348 | Akr1c6 | 83702 | HITS-CLIP | 23597149 | Unspecified |
| mmu-miR-378b | MIMAT0019348 | Akt2 | 11652 | HITS-CLIP | 37986224 | Unspecified |
| mmu-miR-378b | MIMAT0019348 | Aktip | 14339 | HITS-CLIP | 23597149 | Unspecified |
| mmu-miR-378b | MIMAT0019348 | Alas2 | 11656 | HITS-CLIP | 23597149 | Unspecified |
| mmu-miR-378b | MIMAT0019348 | Aldh4a1 | 212647 | HITS-CLIP | 37986224 | Unspecified |
| mmu-miR-378b | MIMAT0019348 | Aldh6a1 | 104776 | HITS-CLIP | 37986224 | Unspecified |
| mmu-miR-378b | MIMAT0019348 | Aldoa | 11674 | HITS-CLIP | 23597149 | Unspecified |
| mmu-miR-378b | MIMAT0019348 | Alg11 | 207958 | HITS-CLIP | 37986224 | Unspecified |
| mmu-miR-378b | MIMAT0019348 | Ankrd13c | 433667 | HITS-CLIP | 23597149 | Unspecified |
| mmu-miR-378b | MIMAT0019348 | Ap2a1 | 11771 | HITS-CLIP | 37986224\|23597149 | Unspecified |
| mmu-miR-378b | MIMAT0019348 | Ap3d1 | 11776 | HITS-CLIP | 37986224 | Unspecified |
| mmu-miR-378b | MIMAT0019348 | Ap4e1 | 108011 | HITS-CLIP | 23597149 | Unspecified |
| mmu-miR-378b | MIMAT0019348 | Aplp2 | 11804 | HITS-CLIP | 37986224 | Unspecified |
| mmu-miR-378b | MIMAT0019348 | Apob | 238055 | HITS-CLIP | 23597149 | Unspecified |
| mmu-miR-378b | MIMAT0019348 | Arhgap32 | 330914 | HITS-CLIP | 37986224 | Unspecified |
| mmu-miR-378b | MIMAT0019348 | Arhgef12 | 69632 | HITS-CLIP | 37986224 | Unspecified |
| mmu-miR-378b | MIMAT0019348 | Atox1 | 11927 | HITS-CLIP | 23597149 | Unspecified |
| mmu-miR-378b | MIMAT0019348 | Atp5c1 | 11949 | HITS-CLIP | 23597149 | Unspecified |
| mmu-miR-378b | MIMAT0019348 | Atrn | 11990 | HITS-CLIP | 37986224\|23597149 | Unspecified |

| | | | | | | |
|---|---|---|---|---|---|---|
| mmu-miR-378b | MIMAT0019348 | Azin1 | 54375 | HITS-CLIP | 23597149 | Unspecified |
| mmu-miR-378b | MIMAT0019348 | Baat | 12012 | HITS-CLIP | 37986224 | Unspecified |
| mmu-miR-378b | MIMAT0019348 | Baz1b | 22385 | HITS-CLIP | 23597149 | Unspecified |
| mmu-miR-378b | MIMAT0019348 | Bcr | 110279 | HITS-CLIP | 37986224 | Unspecified |
| mmu-miR-378b | MIMAT0019348 | Becn1 | 56208 | HITS-CLIP | 23597149 | Unspecified |
| mmu-miR-378b | MIMAT0019348 | Bhlhe40 | 20893 | HITS-CLIP | 23597149 | Unspecified |
| mmu-miR-378b | MIMAT0019348 | Brd3 | 67382 | HITS-CLIP | 23597149 | Unspecified |
| mmu-miR-378b | MIMAT0019348 | Brms1l | 52592 | HITS-CLIP | 23597149 | Unspecified |
| mmu-miR-378b | MIMAT0019348 | C3 | 12266 | HITS-CLIP | 23597149 | Unspecified |
| mmu-miR-378b | MIMAT0019348 | C4b | 12268 | HITS-CLIP | 37986224\|23597149 | Unspecified |
| mmu-miR-378b | MIMAT0019348 | C77080 | 97130 | HITS-CLIP | 37986224 | Unspecified |
| mmu-miR-378b | MIMAT0019348 | Camk2d | 108058 | HITS-CLIP | 23597149 | Unspecified |
| mmu-miR-378b | MIMAT0019348 | Cand1 | 71902 | HITS-CLIP | 23597149 | Unspecified |
| mmu-miR-378b | MIMAT0019348 | Casp9 | 12371 | HITS-CLIP | 23597149 | Unspecified |
| mmu-miR-378b | MIMAT0019348 | Cct4 | 12464 | HITS-CLIP | 23597149 | Unspecified |
| mmu-miR-378b | MIMAT0019348 | Cd1d1 | 12479 | HITS-CLIP | 37986224 | Unspecified |
| mmu-miR-378b | MIMAT0019348 | Cdc20 | 107995 | HITS-CLIP | 23597149 | Unspecified |
| mmu-miR-378b | MIMAT0019348 | Cdc37 | 12539 | HITS-CLIP | 37986224 | Unspecified |
| mmu-miR-378b | MIMAT0019348 | Cdc42 | 12540 | HITS-CLIP | 23597149 | Unspecified |
| mmu-miR-378b | MIMAT0019348 | Cdh5 | 12562 | HITS-CLIP | 37986224 | Unspecified |
| mmu-miR-378b | MIMAT0019348 | Cdip1 | 66626 | HITS-CLIP | 23597149 | Unspecified |
| mmu-miR-378b | MIMAT0019348 | Cdo1 | 12583 | HITS-CLIP | 37986224 | Unspecified |
| mmu-miR-378b | MIMAT0019348 | Cecr2 | 330409 | HITS-CLIP | 23597149 | Unspecified |
| mmu-miR-378b | MIMAT0019348 | Cep350 | 74081 | HITS-CLIP | 23597149 | Unspecified |
| mmu-miR-378b | MIMAT0019348 | Cep85 | 70012 | HITS-CLIP | 23597149 | Unspecified |
| mmu-miR-378b | MIMAT0019348 | Ces3a | 382053 | HITS-CLIP | 37986224 | Unspecified |
| mmu-miR-378b | MIMAT0019348 | Cfh | 12628 | HITS-CLIP | 37986224 | Unspecified |
| mmu-miR-378b | MIMAT0019348 | Cgnl1 | 68178 | HITS-CLIP | 37986224 | Unspecified |
| mmu-miR-378b | MIMAT0019348 | Chd7 | 320790 | HITS-CLIP | 37986224 | Unspecified |
| mmu-miR-378b | MIMAT0019348 | Chd9 | 109151 | HITS-CLIP | 23597149 | Unspecified |
| mmu-miR-378b | MIMAT0019348 | Cipc | 217732 | HITS-CLIP | 23597149 | Unspecified |
| mmu-miR-378b | MIMAT0019348 | Clasp2 | 76499 | HITS-CLIP | 23597149 | Unspecified |
| mmu-miR-378b | MIMAT0019348 | Cluh | 74148 | HITS-CLIP | 37986224 | Unspecified |
| mmu-miR-378b | MIMAT0019348 | Col18a1 | 12822 | HITS-CLIP | 37986224 | Unspecified |
| mmu-miR-378b | MIMAT0019348 | Copb1 | 70349 | HITS-CLIP | 37986224 | Unspecified |
| mmu-miR-378b | MIMAT0019348 | Copg2 | 54160 | HITS-CLIP | 23597149 | Unspecified |
| mmu-miR-378b | MIMAT0019348 | Coq8a | 56997 | HITS-CLIP | 23597149 | Unspecified |
| mmu-miR-378b | MIMAT0019348 | Cps1 | 227231 | HITS-CLIP | 23597149 | Unspecified |
| mmu-miR-378b | MIMAT0019348 | Cpsf4l | 52670 | HITS-CLIP | 23597149 | Unspecified |

| | | | | | | |
|---|---|---|---|---|---|---|
| mmu-miR-378b | MIMAT0019348 | Creg1 | 433375 | HITS-CLIP | 23597149 | Unspecified |
| mmu-miR-378b | MIMAT0019348 | Crot | 74114 | HITS-CLIP | 37986224 | Unspecified |
| mmu-miR-378b | MIMAT0019348 | Cryz | 12972 | HITS-CLIP | 23597149 | Unspecified |
| mmu-miR-378b | MIMAT0019348 | Csnk1a1 | 93687 | HITS-CLIP | 37986224 | Unspecified |
| mmu-miR-378b | MIMAT0019348 | Ctnnd1 | 12388 | HITS-CLIP | 23597149 | Unspecified |
| mmu-miR-378b | MIMAT0019348 | Ctr9 | 22083 | HITS-CLIP | 37986224 | Unspecified |
| mmu-miR-378b | MIMAT0019348 | Ctsd | 13033 | HITS-CLIP | 37986224 | Unspecified |
| mmu-miR-378b | MIMAT0019348 | Ctsl | 13039 | HITS-CLIP | 23597149 | Unspecified |
| mmu-miR-378b | MIMAT0019348 | Cyld | 74256 | HITS-CLIP | 37986224 | Unspecified |
| mmu-miR-378b | MIMAT0019348 | Cyp1a2 | 13077 | HITS-CLIP | 37986224 | Unspecified |
| mmu-miR-378b | MIMAT0019348 | Cyp2c29 | 13095 | HITS-CLIP | 23597149 | Unspecified |
| mmu-miR-378b | MIMAT0019348 | Cyp2j5 | 13109 | HITS-CLIP | 23597149 | Unspecified |
| mmu-miR-378b | MIMAT0019348 | Cyp4a10 | 13117 | HITS-CLIP | 37986224\|23597149 | Unspecified |
| mmu-miR-378b | MIMAT0019348 | Cyp4a14 | 13119 | HITS-CLIP | 37986224\|23597149 | Unspecified |
| mmu-miR-378b | MIMAT0019348 | Cyp4f14 | 64385 | HITS-CLIP | 23597149 | Unspecified |
| mmu-miR-378b | MIMAT0019348 | Cyp4f15 | 106648 | HITS-CLIP | 37986224 | Unspecified |
| mmu-miR-378b | MIMAT0019348 | Cyp4v3 | 102294 | HITS-CLIP | 23597149 | Unspecified |
| mmu-miR-378b | MIMAT0019348 | Dapk1 | 69635 | HITS-CLIP | 23597149 | Unspecified |
| mmu-miR-378b | MIMAT0019348 | Dclre1b | 140917 | HITS-CLIP | 23597149 | Unspecified |
| mmu-miR-378b | MIMAT0019348 | Dcp2 | 70640 | HITS-CLIP | 23597149 | Unspecified |
| mmu-miR-378b | MIMAT0019348 | Ddrgk1 | 77006 | HITS-CLIP | 23597149 | Unspecified |
| mmu-miR-378b | MIMAT0019348 | Ddx1 | 104721 | HITS-CLIP | 23597149 | Unspecified |
| mmu-miR-378b | MIMAT0019348 | Ddx6 | 13209 | HITS-CLIP | 37986224\|23597149 | Unspecified |
| mmu-miR-378b | MIMAT0019348 | Dek | 110052 | HITS-CLIP | 23597149 | Unspecified |
| mmu-miR-378b | MIMAT0019348 | Dglucy | 217830 | HITS-CLIP | 23597149 | Unspecified |
| mmu-miR-378b | MIMAT0019348 | Dhx38 | 64340 | HITS-CLIP | 23597149 | Unspecified |
| mmu-miR-378b | MIMAT0019348 | Dhx9 | 13211 | HITS-CLIP | 37986224 | Unspecified |
| mmu-miR-378b | MIMAT0019348 | Dnaja2 | 56445 | HITS-CLIP | 37986224 | Unspecified |
| mmu-miR-378b | MIMAT0019348 | Dnmt1 | 13433 | HITS-CLIP | 23597149 | Unspecified |
| mmu-miR-378b | MIMAT0019348 | Dop1b | 9980 | HITS-CLIP | 23597149 | Unspecified |
| mmu-miR-378b | MIMAT0019348 | Dsg2 | 13511 | HITS-CLIP | 23597149 | Unspecified |
| mmu-miR-378b | MIMAT0019348 | Dstn | 56431 | HITS-CLIP | 23597149 | Unspecified |
| mmu-miR-378b | MIMAT0019348 | Dync1h1 | 13424 | HITS-CLIP | 23597149 | Unspecified |
| mmu-miR-378b | MIMAT0019348 | Eef2 | 13629 | HITS-CLIP | 23597149 | Unspecified |
| mmu-miR-378b | MIMAT0019348 | Eftud2 | 20624 | HITS-CLIP | 23597149 | Unspecified |
| mmu-miR-378b | MIMAT0019348 | Ehhadh | 74147 | HITS-CLIP | 37986224 | Unspecified |
| mmu-miR-378b | MIMAT0019348 | Eif3a | 13669 | HITS-CLIP | 37986224 | Unspecified |
| mmu-miR-378b | MIMAT0019348 | Eif3d | 55944 | HITS-CLIP | 37986224 | Unspecified |
| mmu-miR-378b | MIMAT0019348 | Eif3l | 223691 | HITS-CLIP | 37986224 | Unspecified |

| | | | | | | |
|---|---|---|---|---|---|---|
| mmu-miR-378b | MIMAT0019348 | Eif4b | 75705 | HITS-CLIP | 23597149 | Unspecified |
| mmu-miR-378b | MIMAT0019348 | Emsy | 56946 | HITS-CLIP | 23597149 | Unspecified |
| mmu-miR-378b | MIMAT0019348 | Eng | 13805 | HITS-CLIP | 37986224\|23597149 | Unspecified |
| mmu-miR-378b | MIMAT0019348 | Ep300 | 328572 | HITS-CLIP | 23597149 | Unspecified |
| mmu-miR-378b | MIMAT0019348 | Ep400 | 75560 | HITS-CLIP | 23597149 | Unspecified |
| mmu-miR-378b | MIMAT0019348 | Epb41l5 | 226352 | HITS-CLIP | 23597149 | Unspecified |
| mmu-miR-378b | MIMAT0019348 | Ephx2 | 13850 | HITS-CLIP | 23597149 | Unspecified |
| mmu-miR-378b | MIMAT0019348 | Epn2 | 13855 | HITS-CLIP | 23597149 | Unspecified |
| mmu-miR-378b | MIMAT0019348 | Etf1 | 225363 | HITS-CLIP | 37986224 | Unspecified |
| mmu-miR-378b | MIMAT0019348 | Etfa | 110842 | HITS-CLIP | 23597149 | Unspecified |
| mmu-miR-378b | MIMAT0019348 | F5 | 14067 | HITS-CLIP | 37986224\|23597149 | Unspecified |
| mmu-miR-378b | MIMAT0019348 | Faah | 14073 | HITS-CLIP | 37986224 | Unspecified |
| mmu-miR-378b | MIMAT0019348 | Fads1 | 76267 | HITS-CLIP | 37986224 | Unspecified |
| mmu-miR-378b | MIMAT0019348 | Fads6 | 328035 | HITS-CLIP | 23597149 | Unspecified |
| mmu-miR-378b | MIMAT0019348 | Fam50a | 108160 | HITS-CLIP | 23597149 | Unspecified |
| mmu-miR-378b | MIMAT0019348 | Fasn | 14104 | HITS-CLIP | 23597149 | Unspecified |
| mmu-miR-378b | MIMAT0019348 | Fbxl4 | 269514 | HITS-CLIP | 23597149 | Unspecified |
| mmu-miR-378b | MIMAT0019348 | Fcho2 | 218503 | HITS-CLIP | 37986224 | Unspecified |
| mmu-miR-378b | MIMAT0019348 | Fchsd2 | 207278 | HITS-CLIP | 23597149 | Unspecified |
| mmu-miR-378b | MIMAT0019348 | Fermt2 | 218952 | HITS-CLIP | 23597149 | Unspecified |
| mmu-miR-378b | MIMAT0019348 | Fgl1 | 234199 | HITS-CLIP | 23597149 | Unspecified |
| mmu-miR-378b | MIMAT0019348 | Fkbp4 | 14228 | HITS-CLIP | 23597149 | Unspecified |
| mmu-miR-378b | MIMAT0019348 | Foxk1 | 17425 | HITS-CLIP | 23597149 | Unspecified |
| mmu-miR-378b | MIMAT0019348 | Gamt | 14431 | HITS-CLIP | 23597149 | Unspecified |
| mmu-miR-378b | MIMAT0019348 | Gbf1 | 107338 | HITS-CLIP | 23597149 | Unspecified |
| mmu-miR-378b | MIMAT0019348 | Gclc | 14629 | HITS-CLIP | 23597149 | Unspecified |
| mmu-miR-378b | MIMAT0019348 | Gigyf1 | 57330 | HITS-CLIP | 23597149 | Unspecified |
| mmu-miR-378b | MIMAT0019348 | Gjb2 | 14619 | HITS-CLIP | 37986224 | Unspecified |
| mmu-miR-378b | MIMAT0019348 | Gls2 | 216456 | HITS-CLIP | 23597149 | Unspecified |
| mmu-miR-378b | MIMAT0019348 | Glul | 14645 | HITS-CLIP | 37986224 | Unspecified |
| mmu-miR-378b | MIMAT0019348 | Gm4737 | 11615 | HITS-CLIP | 23597149 | Unspecified |
| mmu-miR-378b | MIMAT0019348 | Gnai2 | 14678 | HITS-CLIP | 37986224 | Unspecified |
| mmu-miR-378b | MIMAT0019348 | Gns | 75612 | HITS-CLIP | 23597149 | Unspecified |
| mmu-miR-378b | MIMAT0019348 | Gopc | 94221 | HITS-CLIP | 37986224\|23597149 | Unspecified |
| mmu-miR-378b | MIMAT0019348 | Gphn | 268566 | HITS-CLIP | 23597149 | Unspecified |
| mmu-miR-378b | MIMAT0019348 | Gpi1 | 14751 | HITS-CLIP | 23597149 | Unspecified |
| mmu-miR-378b | MIMAT0019348 | Gprin3 | 243385 | HITS-CLIP | 23597149 | Unspecified |
| mmu-miR-378b | MIMAT0019348 | Gpt | 76282 | HITS-CLIP | 23597149 | Unspecified |
| mmu-miR-378b | MIMAT0019348 | Gpt2 | 108682 | HITS-CLIP | 37986224 | Unspecified |

| | | | | | | |
|---|---|---|---|---|---|---|
| mmu-miR-378b | MIMAT0019348 | Grb7 | 14786 | HITS-CLIP | 37986224 | Unspecified |
| mmu-miR-378b | MIMAT0019348 | Grpel1 | 17713 | HITS-CLIP | 23597149 | Unspecified |
| mmu-miR-378b | MIMAT0019348 | Gsk3b | 56637 | HITS-CLIP | 23597149 | Unspecified |
| mmu-miR-378b | MIMAT0019348 | Gstp3 | 225884 | HITS-CLIP | 31433990 | Unspecified |
| mmu-miR-378b | MIMAT0019348 | Gtf2h4 | 14885 | HITS-CLIP | 23597149 | Unspecified |
| mmu-miR-378b | MIMAT0019348 | Gtpbp8 | 66067 | HITS-CLIP | 23597149 | Unspecified |
| mmu-miR-378b | MIMAT0019348 | H1f0 | 14958 | HITS-CLIP | 37986224 | Unspecified |
| mmu-miR-378b | MIMAT0019348 | Habp4 | 56541 | HITS-CLIP | 23597149 | Unspecified |
| mmu-miR-378b | MIMAT0019348 | Hacd3 | 57874 | HITS-CLIP | 23597149 | Unspecified |
| mmu-miR-378b | MIMAT0019348 | Hal | 15109 | HITS-CLIP | 37986224 | Unspecified |
| mmu-miR-378b | MIMAT0019348 | Hc | 15139 | HITS-CLIP | 23597149 | Unspecified |
| mmu-miR-378b | MIMAT0019348 | Hectd1 | 207304 | HITS-CLIP | 37986224 | Unspecified |
| mmu-miR-378b | MIMAT0019348 | Hectd3 | 76608 | HITS-CLIP | 37986224 | Unspecified |
| mmu-miR-378b | MIMAT0019348 | Hjurp | 381280 | HITS-CLIP | 37986224 | Unspecified |
| mmu-miR-378b | MIMAT0019348 | Hnrnpl | 15388 | HITS-CLIP | 37986224 | Unspecified |
| mmu-miR-378b | MIMAT0019348 | Hrg | 94175 | HITS-CLIP | 37986224 | Unspecified |
| mmu-miR-378b | MIMAT0019348 | Hspa4l | 18415 | HITS-CLIP | 23597149 | Unspecified |
| mmu-miR-378b | MIMAT0019348 | Hyou1 | 12282 | HITS-CLIP | 23597149 | Unspecified |
| mmu-miR-378b | MIMAT0019348 | Igf1 | 16000 | HITS-CLIP | 23597149 | Unspecified |
| mmu-miR-378b | MIMAT0019348 | Iigp1 | 60440 | HITS-CLIP | 23597149 | Unspecified |
| mmu-miR-378b | MIMAT0019348 | Impad1 | 242291 | HITS-CLIP | 37986224 | Unspecified |
| mmu-miR-378b | MIMAT0019348 | Iqgap2 | 544963 | HITS-CLIP | 23597149 | Unspecified |
| mmu-miR-378b | MIMAT0019348 | Ireb2 | 64602 | HITS-CLIP | 37986224 | Unspecified |
| mmu-miR-378b | MIMAT0019348 | Irf2 | 16363 | HITS-CLIP | 37986224\|23597149 | Unspecified |
| mmu-miR-378b | MIMAT0019348 | Itih2 | 16425 | HITS-CLIP | 23597149 | Unspecified |
| mmu-miR-378b | MIMAT0019348 | Itih3 | 16426 | HITS-CLIP | 23597149 | Unspecified |
| mmu-miR-378b | MIMAT0019348 | Jak1 | 16451 | HITS-CLIP | 23597149 | Unspecified |
| mmu-miR-378b | MIMAT0019348 | Jarid2 | 16468 | HITS-CLIP | 37986224 | Unspecified |
| mmu-miR-378b | MIMAT0019348 | Jkamp | 104771 | HITS-CLIP | 23597149 | Unspecified |
| mmu-miR-378b | MIMAT0019348 | Jund | 16478 | HITS-CLIP | 37986224 | Unspecified |
| mmu-miR-378b | MIMAT0019348 | Kank1 | 107351 | HITS-CLIP | 23597149 | Unspecified |
| mmu-miR-378b | MIMAT0019348 | Kcnk10 | 72258 | HITS-CLIP | 37986224 | Unspecified |
| mmu-miR-378b | MIMAT0019348 | Kdm5a | 214899 | HITS-CLIP | 37986224 | Unspecified |
| mmu-miR-378b | MIMAT0019348 | Kdm6a | 22289 | HITS-CLIP | 23597149 | Unspecified |
| mmu-miR-378b | MIMAT0019348 | Klf9 | 16601 | HITS-CLIP | 37986224\|23597149 | Unspecified |
| mmu-miR-378b | MIMAT0019348 | Klhl3 | 100503085 | HITS-CLIP | 23597149 | Unspecified |
| mmu-miR-378b | MIMAT0019348 | Kmt2d | 381022 | HITS-CLIP | 23597149 | Unspecified |
| mmu-miR-378b | MIMAT0019348 | Krit1 | 79264 | HITS-CLIP | 37986224\|23597149 | Unspecified |
| mmu-miR-378b | MIMAT0019348 | Kyat3 | 56267 | HITS-CLIP | 23597149 | Unspecified |

| | | | | | | |
|---|---|---|---|---|---|---|
| mmu-miR-378b | MIMAT0019348 | Lars | 107045 | qCLASH | 31393953 | Unspecified |
| mmu-miR-378b | MIMAT0019348 | Ldha | 16828 | HITS-CLIP | 37986224 | Unspecified |
| mmu-miR-378b | MIMAT0019348 | Ldhd | 52815 | HITS-CLIP | 23597149 | Unspecified |
| mmu-miR-378b | MIMAT0019348 | Lgals9 | 16859 | HITS-CLIP | 23597149 | Unspecified |
| mmu-miR-378b | MIMAT0019348 | Lgalsl | 216551 | HITS-CLIP | 37986224\|23597149 | Unspecified |
| mmu-miR-378b | MIMAT0019348 | Lima1 | 65970 | HITS-CLIP | 23597149 | Unspecified |
| mmu-miR-378b | MIMAT0019348 | Lipc | 15450 | HITS-CLIP | 23597149 | Unspecified |
| mmu-miR-378b | MIMAT0019348 | Lnpep | 240028 | HITS-CLIP | 23597149 | Unspecified |
| mmu-miR-378b | MIMAT0019348 | Lrp1 | 16971 | HITS-CLIP | 23597149 | Unspecified |
| mmu-miR-378b | MIMAT0019348 | Lrp10 | 65107 | HITS-CLIP | 23597149 | Unspecified |
| mmu-miR-378b | MIMAT0019348 | Lsm14b | 241846 | HITS-CLIP | 37986224 | Unspecified |
| mmu-miR-378b | MIMAT0019348 | Macf1 | 11426 | HITS-CLIP | 23597149 | Unspecified |
| mmu-miR-378b | MIMAT0019348 | Map3k1 | 26401 | HITS-CLIP | 23597149 | Unspecified |
| mmu-miR-378b | MIMAT0019348 | Mapk14 | 26416 | HITS-CLIP | 37986224 | Unspecified |
| mmu-miR-378b | MIMAT0019348 | Mapkap1 | 227743 | HITS-CLIP | 23597149 | Unspecified |
| mmu-miR-378b | MIMAT0019348 | Marc1 | 66112 | HITS-CLIP | 23597149 | Unspecified |
| mmu-miR-378b | MIMAT0019348 | Marc2 | 67247 | HITS-CLIP | 37986224 | Unspecified |
| mmu-miR-378b | MIMAT0019348 | Matr3 | 17184 | HITS-CLIP | 23597149 | Unspecified |
| mmu-miR-378b | MIMAT0019348 | Mavs | 228607 | HITS-CLIP | 37986224 | Unspecified |
| mmu-miR-378b | MIMAT0019348 | Mcc | 328949 | HITS-CLIP | 37986224 | Unspecified |
| mmu-miR-378b | MIMAT0019348 | Mdn1 | 100019 | HITS-CLIP | 37986224 | Unspecified |
| mmu-miR-378b | MIMAT0019348 | Med13 | 327987 | HITS-CLIP | 37986224 | Unspecified |
| mmu-miR-378b | MIMAT0019348 | Med18 | 67219 | HITS-CLIP | 31433990 | Unspecified |
| mmu-miR-378b | MIMAT0019348 | Med7 | 66213 | HITS-CLIP | 23597149 | Unspecified |
| mmu-miR-378b | MIMAT0019348 | Megf9 | 230316 | HITS-CLIP | 37986224 | Unspecified |
| mmu-miR-378b | MIMAT0019348 | Mettl7b | 71664 | HITS-CLIP | 23597149 | Unspecified |
| mmu-miR-378b | MIMAT0019348 | Micu1 | 216001 | HITS-CLIP | 37986224 | Unspecified |
| mmu-miR-378b | MIMAT0019348 | Mknk1 | 17346 | HITS-CLIP | 23597149 | Unspecified |
| mmu-miR-378b | MIMAT0019348 | Mmadhc | 109129 | HITS-CLIP | 37986224 | Unspecified |
| mmu-miR-378b | MIMAT0019348 | Mob3b | 214944 | HITS-CLIP | 23597149 | Unspecified |
| mmu-miR-378b | MIMAT0019348 | Mrm3 | 67390 | HITS-CLIP | 23597149 | Unspecified |
| mmu-miR-378b | MIMAT0019348 | Mrpl39 | 27393 | HITS-CLIP | 23597149 | Unspecified |
| mmu-miR-378b | MIMAT0019348 | Mtmr3 | 74302 | HITS-CLIP | 23597149 | Unspecified |
| mmu-miR-378b | MIMAT0019348 | Mtss1 | 211401 | HITS-CLIP | 37986224 | Unspecified |
| mmu-miR-378b | MIMAT0019348 | Myh9 | 17886 | HITS-CLIP | 37986224 | Unspecified |
| mmu-miR-378b | MIMAT0019348 | Mylk | 107589 | HITS-CLIP | 23597149 | Unspecified |
| mmu-miR-378b | MIMAT0019348 | Myo1b | 17912 | HITS-CLIP | 37986224 | Unspecified |
| mmu-miR-378b | MIMAT0019348 | Naa60 | 74763 | HITS-CLIP | 23597149 | Unspecified |
| mmu-miR-378b | MIMAT0019348 | Nadk2 | 68646 | HITS-CLIP | 23597149 | Unspecified |

| | | | | | | |
|---|---|---|---|---|---|---|
| mmu-miR-378b | MIMAT0019348 | Nbr1 | 17966 | HITS-CLIP | 37986224\|23597149 | Unspecified |
| mmu-miR-378b | MIMAT0019348 | Ncor1 | 20185 | HITS-CLIP | 23597149 | Unspecified |
| mmu-miR-378b | MIMAT0019348 | Ndufa8 | 68375 | HITS-CLIP | 37986224 | Unspecified |
| mmu-miR-378b | MIMAT0019348 | Ndufa9 | 66108 | HITS-CLIP | 23597149 | Unspecified |
| mmu-miR-378b | MIMAT0019348 | Nfe2l2 | 18024 | HITS-CLIP | 37986224 | Unspecified |
| mmu-miR-378b | MIMAT0019348 | Nfia | 18027 | HITS-CLIP | 23597149 | Unspecified |
| mmu-miR-378b | MIMAT0019348 | Nfix | 18032 | HITS-CLIP | 37986224 | Unspecified |
| mmu-miR-378b | MIMAT0019348 | Noa1 | 56412 | HITS-CLIP | 23597149 | Unspecified |
| mmu-miR-378b | MIMAT0019348 | Nploc4 | 217365 | HITS-CLIP | 23597149 | Unspecified |
| mmu-miR-378b | MIMAT0019348 | Ociad1 | 68095 | HITS-CLIP | 23597149 | Unspecified |
| mmu-miR-378b | MIMAT0019348 | Onecut2 | 225631 | HITS-CLIP | 37986224 | Unspecified |
| mmu-miR-378b | MIMAT0019348 | Ormdl3 | 66612 | HITS-CLIP | 23597149 | Unspecified |
| mmu-miR-378b | MIMAT0019348 | Oxsr1 | 108737 | HITS-CLIP | 23597149 | Unspecified |
| mmu-miR-378b | MIMAT0019348 | Pacsin2 | 23970 | HITS-CLIP | 37986224 | Unspecified |
| mmu-miR-378b | MIMAT0019348 | Pafah1b1 | 18472 | HITS-CLIP | 23597149 | Unspecified |
| mmu-miR-378b | MIMAT0019348 | Paics | 67054 | HITS-CLIP | 37986224 | Unspecified |
| mmu-miR-378b | MIMAT0019348 | Parp14 | 547253 | HITS-CLIP | 37986224 | Unspecified |
| mmu-miR-378b | MIMAT0019348 | Parva | 57342 | HITS-CLIP | 23597149 | Unspecified |
| mmu-miR-378b | MIMAT0019348 | Patl1 | 225929 | HITS-CLIP | 31433990 | Unspecified |
| mmu-miR-378b | MIMAT0019348 | Pcyt1b | 236899 | HITS-CLIP | 31433990 | Unspecified |
| mmu-miR-378b | MIMAT0019348 | Pdcd4 | 18569 | HITS-CLIP | 37986224 | Unspecified |
| mmu-miR-378b | MIMAT0019348 | Pdcd6ip | 18571 | HITS-CLIP | 37986224\|23597149 | Unspecified |
| mmu-miR-378b | MIMAT0019348 | Pdia4 | 12304 | HITS-CLIP | 23597149 | Unspecified |
| mmu-miR-378b | MIMAT0019348 | Pdk2 | 18604 | HITS-CLIP | 23597149 | Unspecified |
| mmu-miR-378b | MIMAT0019348 | Peak1 | 244895 | HITS-CLIP | 23597149 | Unspecified |
| mmu-miR-378b | MIMAT0019348 | Pgm1 | 66681 | HITS-CLIP | 37986224 | Unspecified |
| mmu-miR-378b | MIMAT0019348 | Pgpep1 | 66522 | HITS-CLIP | 37986224 | Unspecified |
| mmu-miR-378b | MIMAT0019348 | Pip4k2c | 117150 | HITS-CLIP | 23597149 | Unspecified |
| mmu-miR-378b | MIMAT0019348 | Pip4p1 | 90809 | HITS-CLIP | 23597149 | Unspecified |
| mmu-miR-378b | MIMAT0019348 | Pole | 18973 | HITS-CLIP | 23597149 | Unspecified |
| mmu-miR-378b | MIMAT0019348 | Por | 18984 | HITS-CLIP | 23597149 | Unspecified |
| mmu-miR-378b | MIMAT0019348 | Pou2f1 | 18986 | HITS-CLIP | 23597149 | Unspecified |
| mmu-miR-378b | MIMAT0019348 | Prep | 19072 | HITS-CLIP | 23597149 | Unspecified |
| mmu-miR-378b | MIMAT0019348 | Prkci | 18759 | HITS-CLIP | 23597149 | Unspecified |
| mmu-miR-378b | MIMAT0019348 | Prrc2c | 226562 | HITS-CLIP | 23597149 | Unspecified |
| mmu-miR-378b | MIMAT0019348 | Ptbp3 | 230257 | HITS-CLIP | 23597149 | Unspecified |
| mmu-miR-378b | MIMAT0019348 | Ptges2 | 96979 | HITS-CLIP | 23597149 | Unspecified |
| mmu-miR-378b | MIMAT0019348 | Ptprj | 19271 | HITS-CLIP | 23597149 | Unspecified |
| mmu-miR-378b | MIMAT0019348 | Pum2 | 80913 | HITS-CLIP | 23597149 | Unspecified |

| | | | | | | | |
|---|---|---|---|---|---|---|---|
| mmu-miR-378b | MIMAT0019348 | Rab11fip1 | 75767 | HITS-CLIP | | 37986224 | Unspecified |
| mmu-miR-378b | MIMAT0019348 | Rab1a | 5861 | HITS-CLIP | | 23597149 | Unspecified |
| mmu-miR-378b | MIMAT0019348 | Rab5c | 19345 | HITS-CLIP | | 23597149 | Unspecified |
| mmu-miR-378b | MIMAT0019348 | Rab7 | 19349 | HITS-CLIP | | 23597149 | Unspecified |
| mmu-miR-378b | MIMAT0019348 | Ralgapa2 | 241694 | HITS-CLIP | | 37986224 | Unspecified |
| mmu-miR-378b | MIMAT0019348 | Raly | 19383 | HITS-CLIP | | 23597149 | Unspecified |
| mmu-miR-378b | MIMAT0019348 | Ranbp2 | 19386 | HITS-CLIP | | 23597149 | Unspecified |
| mmu-miR-378b | MIMAT0019348 | Rbm15 | 229700 | HITS-CLIP | | 23597149 | Unspecified |
| mmu-miR-378b | MIMAT0019348 | Rbm4 | 19653 | HITS-CLIP | | 23597149 | Unspecified |
| mmu-miR-378b | MIMAT0019348 | Rbm4b | 66704 | HITS-CLIP | | 23597149 | Unspecified |
| mmu-miR-378b | MIMAT0019348 | Repin1 | 58887 | HITS-CLIP | | 23597149 | Unspecified |
| mmu-miR-378b | MIMAT0019348 | Rnf13 | 24017 | HITS-CLIP | | 37986224 | Unspecified |
| mmu-miR-378b | MIMAT0019348 | Rnf213 | 672511 | HITS-CLIP | | 37986224 | Unspecified |
| mmu-miR-378b | MIMAT0019348 | Rpp14 | 67053 | HITS-CLIP | | 23597149 | Unspecified |
| mmu-miR-378b | MIMAT0019348 | Rtf2 | 51507 | HITS-CLIP | | 23597149 | Unspecified |
| mmu-miR-378b | MIMAT0019348 | Rtp3 | 235636 | HITS-CLIP | | 37986224 | Unspecified |
| mmu-miR-378b | MIMAT0019348 | Runx1t1 | 12395 | HITS-CLIP | | 23597149 | Unspecified |
| mmu-miR-378b | MIMAT0019348 | Sbk1 | 104175 | HITS-CLIP | | 37986224 | Unspecified |
| mmu-miR-378b | MIMAT0019348 | Sec16a | 227648 | HITS-CLIP | | 23597149 | Unspecified |
| mmu-miR-378b | MIMAT0019348 | Sec24a | 77371 | HITS-CLIP | | 23597149 | Unspecified |
| mmu-miR-378b | MIMAT0019348 | Sec24d | 69608 | HITS-CLIP | | 23597149 | Unspecified |
| mmu-miR-378b | MIMAT0019348 | Sema4g | 26456 | HITS-CLIP | | 23597149 | Unspecified |
| mmu-miR-378b | MIMAT0019348 | Serpina10 | 217847 | HITS-CLIP | | 23597149 | Unspecified |
| mmu-miR-378b | MIMAT0019348 | Serpina1a | 20700 | HITS-CLIP | | 37986224 | Unspecified |
| mmu-miR-378b | MIMAT0019348 | Serpina1d | 20703 | HITS-CLIP | | 37986224 | Unspecified |
| mmu-miR-378b | MIMAT0019348 | Serpind1 | 15160 | HITS-CLIP | | 23597149 | Unspecified |
| mmu-miR-378b | MIMAT0019348 | Sf1 | 22668 | HITS-CLIP | | 37986224 | Unspecified |
| mmu-miR-378b | MIMAT0019348 | Sfxn5 | 94282 | HITS-CLIP | 37986224\|23597149 | | Unspecified |
| mmu-miR-378b | MIMAT0019348 | Sgk2 | 27219 | HITS-CLIP | 37986224\|23597149 | | Unspecified |
| mmu-miR-378b | MIMAT0019348 | Shank2 | 210274 | HITS-CLIP | | 23597149 | Unspecified |
| mmu-miR-378b | MIMAT0019348 | Slain2 | 75991 | HITS-CLIP | | 37986224 | Unspecified |
| mmu-miR-378b | MIMAT0019348 | Slc18b1 | 76306 | HITS-CLIP | | 23597149 | Unspecified |
| mmu-miR-378b | MIMAT0019348 | Slc25a45 | 107375 | HITS-CLIP | | 37986224 | Unspecified |
| mmu-miR-378b | MIMAT0019348 | Slc26a1 | 231583 | HITS-CLIP | | 37986224 | Unspecified |
| mmu-miR-378b | MIMAT0019348 | Slc30a1 | 22782 | HITS-CLIP | | 23597149 | Unspecified |
| mmu-miR-378b | MIMAT0019348 | Slc35a5 | 74102 | HITS-CLIP | | 23597149 | Unspecified |
| mmu-miR-378b | MIMAT0019348 | Slc38a7 | 234595 | HITS-CLIP | | 23597149 | Unspecified |
| mmu-miR-378b | MIMAT0019348 | Slc41a2 | 338365 | HITS-CLIP | | 37986224 | Unspecified |
| mmu-miR-378b | MIMAT0019348 | Slc7a2 | 11988 | HITS-CLIP | | 23597149 | Unspecified |

| | | | | | | | |
|---|---|---|---|---|---|---|---|
| mmu-miR-378b | MIMAT0019348 | Slc9a3r1 | 26941 | HITS-CLIP | | 37986224 | Unspecified |
| mmu-miR-378b | MIMAT0019348 | Smarca4 | 20586 | HITS-CLIP | | 23597149 | Unspecified |
| mmu-miR-378b | MIMAT0019348 | Smc6 | 67241 | HITS-CLIP | | 37986224 | Unspecified |
| mmu-miR-378b | MIMAT0019348 | Smg7 | 226517 | HITS-CLIP | | 23597149 | Unspecified |
| mmu-miR-378b | MIMAT0019348 | Smg9 | 71997 | HITS-CLIP | | 23597149 | Unspecified |
| mmu-miR-378b | MIMAT0019348 | Snai2 | 20583 | HITS-CLIP | | 37986224 | Unspecified |
| mmu-miR-378b | MIMAT0019348 | Snrpb | 20638 | HITS-CLIP | | 23597149 | Unspecified |
| mmu-miR-378b | MIMAT0019348 | Snx17 | 266781 | HITS-CLIP | | 23597149 | Unspecified |
| mmu-miR-378b | MIMAT0019348 | Sparc | 20692 | HITS-CLIP | | 23597149 | Unspecified |
| mmu-miR-378b | MIMAT0019348 | Spata2 | 263876 | HITS-CLIP | | 37986224 | Unspecified |
| mmu-miR-378b | MIMAT0019348 | Spata2l | 78779 | HITS-CLIP | | 37986224 | Unspecified |
| mmu-miR-378b | MIMAT0019348 | Spcs3 | 76687 | HITS-CLIP | | 31433990 | Unspecified |
| mmu-miR-378b | MIMAT0019348 | Sppl3 | 74585 | HITS-CLIP | | 23597149 | Unspecified |
| mmu-miR-378b | MIMAT0019348 | Srebf1 | 20787 | HITS-CLIP | | 23597149 | Unspecified |
| mmu-miR-378b | MIMAT0019348 | Srrm2 | 75956 | HITS-CLIP | | 23597149 | Unspecified |
| mmu-miR-378b | MIMAT0019348 | Srsf2 | 20382 | HITS-CLIP | 37986224\|23597149 | | Unspecified |
| mmu-miR-378b | MIMAT0019348 | Srsf6 | 67996 | HITS-CLIP | | 23597149 | Unspecified |
| mmu-miR-378b | MIMAT0019348 | St13 | 70356 | HITS-CLIP | | 37986224 | Unspecified |
| mmu-miR-378b | MIMAT0019348 | Stard5 | 170460 | HITS-CLIP | | 37986224 | Unspecified |
| mmu-miR-378b | MIMAT0019348 | Stat3 | 20848 | HITS-CLIP | 23597149\|31433990 | | Unspecified |
| mmu-miR-378b | MIMAT0019348 | Stat6 | 20852 | HITS-CLIP | | 23597149 | Unspecified |
| mmu-miR-378b | MIMAT0019348 | Stt3b | 68292 | HITS-CLIP | | 23597149 | Unspecified |
| mmu-miR-378b | MIMAT0019348 | Suz12 | 52615 | HITS-CLIP | | 31433990 | Unspecified |
| mmu-miR-378b | MIMAT0019348 | Taok1 | 216965 | HITS-CLIP | | 23597149 | Unspecified |
| mmu-miR-378b | MIMAT0019348 | Tfip11 | 54723 | HITS-CLIP | | 23597149 | Unspecified |
| mmu-miR-378b | MIMAT0019348 | Tm4sf4 | 229302 | HITS-CLIP | | 37986224 | Unspecified |
| mmu-miR-378b | MIMAT0019348 | Tm9sf1 | 74140 | HITS-CLIP | | 23597149 | Unspecified |
| mmu-miR-378b | MIMAT0019348 | Tmem131l | 23240 | HITS-CLIP | | 23597149 | Unspecified |
| mmu-miR-378b | MIMAT0019348 | Tmem135 | 72759 | HITS-CLIP | | 23597149 | Unspecified |
| mmu-miR-378b | MIMAT0019348 | Tmem170b | 621976 | HITS-CLIP | | 37986224 | Unspecified |
| mmu-miR-378b | MIMAT0019348 | Tmem62 | 96957 | HITS-CLIP | | 23597149 | Unspecified |
| mmu-miR-378b | MIMAT0019348 | Tnks2 | 74493 | HITS-CLIP | | 23597149 | Unspecified |
| mmu-miR-378b | MIMAT0019348 | Tnrc6a | 233833 | HITS-CLIP | | 23597149 | Unspecified |
| mmu-miR-378b | MIMAT0019348 | Tns1 | 21961 | HITS-CLIP | | 37986224 | Unspecified |
| mmu-miR-378b | MIMAT0019348 | Tob1 | 22057 | HITS-CLIP | | 37986224 | Unspecified |
| mmu-miR-378b | MIMAT0019348 | Tomm70a | 28185 | HITS-CLIP | | 23597149 | Unspecified |
| mmu-miR-378b | MIMAT0019348 | Top2b | 21974 | HITS-CLIP | | 23597149 | Unspecified |
| mmu-miR-378b | MIMAT0019348 | Topors | 106021 | HITS-CLIP | | 23597149 | Unspecified |
| mmu-miR-378b | MIMAT0019348 | Tor1aip2 | 240832 | HITS-CLIP | 37986224\|23597149 | | Unspecified |

| | | | | | | |
|---|---|---|---|---|---|---|
| mmu-miR-378b | MIMAT0019348 | Trappc13 | 66975 | HITS-CLIP | 23597149 | Unspecified |
| mmu-miR-378b | MIMAT0019348 | Trim30a | 20128 | HITS-CLIP | 37986224 | Unspecified |
| mmu-miR-378b | MIMAT0019348 | Tuba4a | 22145 | HITS-CLIP | 37986224 | Unspecified |
| mmu-miR-378b | MIMAT0019348 | Txnl1 | 53382 | HITS-CLIP | 23597149 | Unspecified |
| mmu-miR-378b | MIMAT0019348 | Txnrd1 | 50493 | HITS-CLIP | 23597149 | Unspecified |
| mmu-miR-378b | MIMAT0019348 | Ubb | 22187 | HITS-CLIP | 23597149 | Unspecified |
| mmu-miR-378b | MIMAT0019348 | Ube3b | 117146 | HITS-CLIP | 23597149 | Unspecified |
| mmu-miR-378b | MIMAT0019348 | Ubqln1 | 56085 | HITS-CLIP | 37986224 | Unspecified |
| mmu-miR-378b | MIMAT0019348 | Ubr4 | 69116 | HITS-CLIP | 37986224 | Unspecified |
| mmu-miR-378b | MIMAT0019348 | Ubr7 | 66622 | HITS-CLIP | 23597149 | Unspecified |
| mmu-miR-378b | MIMAT0019348 | Uhmk1 | 16589 | HITS-CLIP | 37986224 | Unspecified |
| mmu-miR-378b | MIMAT0019348 | Unk | 217331 | HITS-CLIP | 23597149 | Unspecified |
| mmu-miR-378b | MIMAT0019348 | Uroc1 | 243537 | HITS-CLIP | 37986224 | Unspecified |
| mmu-miR-378b | MIMAT0019348 | Usp10 | 22224 | HITS-CLIP | 23597149 | Unspecified |
| mmu-miR-378b | MIMAT0019348 | Vps41 | 218035 | HITS-CLIP | 37986224 | Unspecified |
| mmu-miR-378b | MIMAT0019348 | Vwa8 | 219189 | HITS-CLIP | 23597149 | Unspecified |
| mmu-miR-378b | MIMAT0019348 | Wdr26 | 226757 | HITS-CLIP | 23597149 | Unspecified |
| mmu-miR-378b | MIMAT0019348 | Whamm | 434204 | HITS-CLIP | 23597149 | Unspecified |
| mmu-miR-378b | MIMAT0019348 | Ythdf3 | 229096 | HITS-CLIP | 37986224 | Unspecified |
| mmu-miR-378b | MIMAT0019348 | Zc3h15 | 69082 | HITS-CLIP | 23597149 | Unspecified |
| mmu-miR-378b | MIMAT0019348 | Zcchc2 | 227449 | HITS-CLIP | 37986224 | Unspecified |
| mmu-miR-378b | MIMAT0019348 | Zcchc24 | 71918 | HITS-CLIP | 37986224 | Unspecified |
| mmu-miR-378b | MIMAT0019348 | Zdhhc13 | 243983 | HITS-CLIP | 23597149 | Unspecified |
| mmu-miR-378b | MIMAT0019348 | Zfp24 | 59057 | HITS-CLIP | 23597149 | Unspecified |
| mmu-miR-378b | MIMAT0019348 | Zfp36 | 22695 | HITS-CLIP | 23597149 | Unspecified |
| mmu-miR-378b | MIMAT0019348 | Zfp36l1 | 12192 | HITS-CLIP | 37986224 | Unspecified |
| mmu-miR-378b | MIMAT0019348 | Zfp638 | 27332 | HITS-CLIP | 23597149 | Unspecified |
| mmu-miR-378b | MIMAT0019348 | Zmat3 | 22401 | HITS-CLIP | 37986224 | Unspecified |
| mmu-miR-708-5p | MIMAT0004828 | 1600014C10Rik | 72244 | HITS-CLIP | 37986224 | Unspecified |
| mmu-miR-708-5p | MIMAT0004828 | 2310022A10Rik | 66367 | HITS-CLIP | 37986224 | Unspecified |
| mmu-miR-708-5p | MIMAT0004828 | 2610507B11Rik | 72503 | HITS-CLIP | 37986224\|23597149 | Unspecified |
| mmu-miR-708-5p | MIMAT0004828 | 2900026A02Rik | 243219 | HITS-CLIP | 37986224 | Unspecified |
| mmu-miR-708-5p | MIMAT0004828 | 4932438A13Rik | 229227 | HITS-CLIP | 19536157 | Unspecified |
| mmu-miR-708-5p | MIMAT0004828 | 6430548M08Rik | 234797 | HITS-CLIP | 37986224 | Unspecified |
| mmu-miR-708-5p | MIMAT0004828 | Aadac | 67758 | HITS-CLIP | 23597149 | Unspecified |
| mmu-miR-708-5p | MIMAT0004828 | Aak1 | 269774 | HITS-CLIP | 19536157 | Unspecified |
| mmu-miR-708-5p | MIMAT0004828 | Aars | 234734 | HITS-CLIP | 23597149 | Unspecified |
| mmu-miR-708-5p | MIMAT0004828 | Abat | 268860 | HITS-CLIP | 37986224 | Unspecified |
| mmu-miR-708-5p | MIMAT0004828 | Abca1 | 11303 | HITS-CLIP | 37986224\|23597149 | Unspecified |

| | | | | | | | |
|---|---|---|---|---|---|---|---|
| mmu-miR-708-5p | MIMAT0004828 | Abca6 | 76184 | HITS-CLIP | | 37986224 | Unspecified |
| mmu-miR-708-5p | MIMAT0004828 | Abcb11 | 27413 | HITS-CLIP | | 37986224\|23597149 | Unspecified |
| mmu-miR-708-5p | MIMAT0004828 | Abcb6 | 74104 | HITS-CLIP | | 23597149 | Unspecified |
| mmu-miR-708-5p | MIMAT0004828 | Abcd3 | 19299 | HITS-CLIP | | 37986224 | Unspecified |
| mmu-miR-708-5p | MIMAT0004828 | Abcg8 | 67470 | HITS-CLIP | | 23597149 | Unspecified |
| mmu-miR-708-5p | MIMAT0004828 | Abhd14b | 76491 | HITS-CLIP | | 23597149 | Unspecified |
| mmu-miR-708-5p | MIMAT0004828 | Abhd17b | 226016 | HITS-CLIP | | 19536157 | Unspecified |
| mmu-miR-708-5p | MIMAT0004828 | Abhd2 | 54608 | HITS-CLIP | | 23597149 | Unspecified |
| mmu-miR-708-5p | MIMAT0004828 | Ablim3 | 319713 | HITS-CLIP | | 23597149 | Unspecified |
| mmu-miR-708-5p | MIMAT0004828 | Acacb | 100705 | HITS-CLIP | | 37986224 | Unspecified |
| mmu-miR-708-5p | MIMAT0004828 | Acap3 | 140500 | HITS-CLIP | | 37986224 | Unspecified |
| mmu-miR-708-5p | MIMAT0004828 | Acbd3 | 170760 | HITS-CLIP | | 23597149 | Unspecified |
| mmu-miR-708-5p | MIMAT0004828 | Acbd5 | 74159 | HITS-CLIP | | 23597149 | Unspecified |
| mmu-miR-708-5p | MIMAT0004828 | Acly | 104112 | HITS-CLIP | | 23597149\|19536157 | Unspecified |
| mmu-miR-708-5p | MIMAT0004828 | Acmsd | 266645 | HITS-CLIP | | 23597149 | Unspecified |
| mmu-miR-708-5p | MIMAT0004828 | Acp2 | 11432 | HITS-CLIP | | 37986224 | Unspecified |
| mmu-miR-708-5p | MIMAT0004828 | Acsf3 | 257633 | HITS-CLIP | | 23597149 | Unspecified |
| mmu-miR-708-5p | MIMAT0004828 | Acsl4 | 50790 | HITS-CLIP | | 23597149 | Unspecified |
| mmu-miR-708-5p | MIMAT0004828 | Acsm5 | 272428 | HITS-CLIP | | 37986224 | Unspecified |
| mmu-miR-708-5p | MIMAT0004828 | Acss1 | 68738 | HITS-CLIP | | 37986224 | Unspecified |
| mmu-miR-708-5p | MIMAT0004828 | Actb | 11461 | HITS-CLIP | | 37986224\|23597149 | Unspecified |
| mmu-miR-708-5p | MIMAT0004828 | Adam10 | 11487 | HITS-CLIP | | 37986224\|23597149 | Unspecified |
| mmu-miR-708-5p | MIMAT0004828 | Adcy1 | 432530 | HITS-CLIP | | 37986224 | Unspecified |
| mmu-miR-708-5p | MIMAT0004828 | Adcy5 | 224129 | HITS-CLIP | | 37986224 | Unspecified |
| mmu-miR-708-5p | MIMAT0004828 | Adcyap1r1 | 11517 | HITS-CLIP | | 37986224 | Unspecified |
| mmu-miR-708-5p | MIMAT0004828 | Add2 | 11519 | HITS-CLIP | | 37986224 | Unspecified |
| mmu-miR-708-5p | MIMAT0004828 | Adgre4 | 52614 | HITS-CLIP | | 23597149 | Unspecified |
| mmu-miR-708-5p | MIMAT0004828 | Adhfe1 | 76187 | HITS-CLIP | | 23597149 | Unspecified |
| mmu-miR-708-5p | MIMAT0004828 | Adi1 | 104923 | HITS-CLIP | | 37986224 | Unspecified |
| mmu-miR-708-5p | MIMAT0004828 | Adipor1 | 72674 | HITS-CLIP | | 37986224 | Unspecified |
| mmu-miR-708-5p | MIMAT0004828 | Adipor2 | 68465 | HITS-CLIP | | 23597149 | Unspecified |
| mmu-miR-708-5p | MIMAT0004828 | Adora2a | 11540 | HITS-CLIP | | 37986224 | Unspecified |
| mmu-miR-708-5p | MIMAT0004828 | Adrb2 | 11555 | HITS-CLIP | | 37986224 | Unspecified |
| mmu-miR-708-5p | MIMAT0004828 | Aebp2 | 11569 | HITS-CLIP | | 37986224 | Unspecified |
| mmu-miR-708-5p | MIMAT0004828 | Aff3 | 16764 | HITS-CLIP | | 19536157 | Unspecified |
| mmu-miR-708-5p | MIMAT0004828 | Aff4 | 93736 | HITS-CLIP | | 37986224 | Unspecified |
| mmu-miR-708-5p | MIMAT0004828 | Ago3 | 214150 | HITS-CLIP | | 37986224 | Unspecified |
| mmu-miR-708-5p | MIMAT0004828 | Agpat3 | 28169 | HITS-CLIP | | 37986224 | Unspecified |
| mmu-miR-708-5p | MIMAT0004828 | Agpat5 | 52123 | HITS-CLIP | | 37986224 | Unspecified |

| | | | | | | |
|---|---|---|---|---|---|---|
| mmu-miR-708-5p | MIMAT0004828 | Agtpbp1 | 67269 | HITS-CLIP | 19536157 | Unspecified |
| mmu-miR-708-5p | MIMAT0004828 | Ahcyl2 | 74340 | HITS-CLIP | 19536157 | Unspecified |
| mmu-miR-708-5p | MIMAT0004828 | Ahdc1 | 230793 | HITS-CLIP | 37986224\|19536157 | Unspecified |
| mmu-miR-708-5p | MIMAT0004828 | Ahr | 11622 | HITS-CLIP | 37986224 | Unspecified |
| mmu-miR-708-5p | MIMAT0004828 | Ak2 | 11637 | HITS-CLIP | 23597149 | Unspecified |
| mmu-miR-708-5p | MIMAT0004828 | Akap1 | 11640 | HITS-CLIP | 37986224 | Unspecified |
| mmu-miR-708-5p | MIMAT0004828 | Akap17b | 338351 | HITS-CLIP | 37986224 | Unspecified |
| mmu-miR-708-5p | MIMAT0004828 | Alas1 | 11655 | HITS-CLIP | 37986224\|23597149 | Unspecified |
| mmu-miR-708-5p | MIMAT0004828 | Alcam | 11658 | HITS-CLIP | 19536157 | Unspecified |
| mmu-miR-708-5p | MIMAT0004828 | Aldh1l1 | 107747 | HITS-CLIP | 23597149 | Unspecified |
| mmu-miR-708-5p | MIMAT0004828 | Aldh3a2 | 11671 | HITS-CLIP | 37986224 | Unspecified |
| mmu-miR-708-5p | MIMAT0004828 | Aldh4a1 | 212647 | HITS-CLIP | 37986224 | Unspecified |
| mmu-miR-708-5p | MIMAT0004828 | Alg14 | 66789 | HITS-CLIP | 37986224 | Unspecified |
| mmu-miR-708-5p | MIMAT0004828 | Alg6 | 320438 | HITS-CLIP | 37986224 | Unspecified |
| mmu-miR-708-5p | MIMAT0004828 | Alms1 | 236266 | HITS-CLIP | 37986224 | Unspecified |
| mmu-miR-708-5p | MIMAT0004828 | Ammecr1 | 56068 | HITS-CLIP | 37986224 | Unspecified |
| mmu-miR-708-5p | MIMAT0004828 | Anapc2 | 99152 | HITS-CLIP | 37986224 | Unspecified |
| mmu-miR-708-5p | MIMAT0004828 | Ank2 | 109676 | HITS-CLIP | 19536157 | Unspecified |
| mmu-miR-708-5p | MIMAT0004828 | Ank3 | 11735 | HITS-CLIP | 37986224\|19536157 | Unspecified |
| mmu-miR-708-5p | MIMAT0004828 | Ankfy1 | 11736 | HITS-CLIP | 23597149 | Unspecified |
| mmu-miR-708-5p | MIMAT0004828 | Ankrd17 | 81702 | HITS-CLIP | 19536157 | Unspecified |
| mmu-miR-708-5p | MIMAT0004828 | Ankrd28 | 105522 | HITS-CLIP | 37986224 | Unspecified |
| mmu-miR-708-5p | MIMAT0004828 | Ankrd44 | 329154 | HITS-CLIP | 23597149 | Unspecified |
| mmu-miR-708-5p | MIMAT0004828 | Ankrd52 | 237615 | HITS-CLIP | 37986224 | Unspecified |
| mmu-miR-708-5p | MIMAT0004828 | Anks1b | 77531 | HITS-CLIP | 37986224 | Unspecified |
| mmu-miR-708-5p | MIMAT0004828 | Anp32a | 11737 | HITS-CLIP | 37986224 | Unspecified |
| mmu-miR-708-5p | MIMAT0004828 | Anp32e | 66471 | HITS-CLIP | 23597149 | Unspecified |
| mmu-miR-708-5p | MIMAT0004828 | Antxr1 | 69538 | HITS-CLIP | 37986224 | Unspecified |
| mmu-miR-708-5p | MIMAT0004828 | Anxa7 | 11750 | HITS-CLIP | 37986224 | Unspecified |
| mmu-miR-708-5p | MIMAT0004828 | Ap1b1 | 11764 | HITS-CLIP | 37986224 | Unspecified |
| mmu-miR-708-5p | MIMAT0004828 | Ap2a2 | 11772 | HITS-CLIP | 23597149 | Unspecified |
| mmu-miR-708-5p | MIMAT0004828 | Ap2b1 | 71770 | HITS-CLIP | 37986224 | Unspecified |
| mmu-miR-708-5p | MIMAT0004828 | Ap3s2 | 11778 | HITS-CLIP | 37986224 | Unspecified |
| mmu-miR-708-5p | MIMAT0004828 | Apba1 | 319924 | HITS-CLIP | 37986224 | Unspecified |
| mmu-miR-708-5p | MIMAT0004828 | Apc | 11789 | HITS-CLIP | 37986224 | Unspecified |
| mmu-miR-708-5p | MIMAT0004828 | Apcdd1 | 494504 | HITS-CLIP | 37986224 | Unspecified |
| mmu-miR-708-5p | MIMAT0004828 | Apob | 238055 | HITS-CLIP | 23597149 | Unspecified |
| mmu-miR-708-5p | MIMAT0004828 | Apoc2 | 11813 | HITS-CLIP | 37986224 | Unspecified |
| mmu-miR-708-5p | MIMAT0004828 | App | 11820 | HITS-CLIP | 37986224 | Unspecified |

| | | | | | | |
|---|---|---|---|---|---|---|
| mmu-miR-708-5p | MIMAT0004828 | Aqp11 | 66333 | HITS-CLIP | 37986224\|23597149 | Unspecified |
| mmu-miR-708-5p | MIMAT0004828 | Aqp9 | 64008 | HITS-CLIP | 23597149 | Unspecified |
| mmu-miR-708-5p | MIMAT0004828 | Arf3 | 11842 | HITS-CLIP | 37986224 | Unspecified |
| mmu-miR-708-5p | MIMAT0004828 | Arg1 | 11846 | HITS-CLIP | 23597149 | Unspecified |
| mmu-miR-708-5p | MIMAT0004828 | Arhgap11a | 228482 | HITS-CLIP | 37986224\|23597149 | Unspecified |
| mmu-miR-708-5p | MIMAT0004828 | Arhgap23 | 58996 | HITS-CLIP | 19536157 | Unspecified |
| mmu-miR-708-5p | MIMAT0004828 | Arhgap32 | 330914 | HITS-CLIP | 37986224 | Unspecified |
| mmu-miR-708-5p | MIMAT0004828 | Arhgap35 | 232906 | HITS-CLIP | 23597149\|19536157 | Unspecified |
| mmu-miR-708-5p | MIMAT0004828 | Arhgef10l | 72754 | HITS-CLIP | 23597149 | Unspecified |
| mmu-miR-708-5p | MIMAT0004828 | Arhgef11 | 213498 | HITS-CLIP | 37986224 | Unspecified |
| mmu-miR-708-5p | MIMAT0004828 | Arhgef12 | 69632 | HITS-CLIP | 23597149 | Unspecified |
| mmu-miR-708-5p | MIMAT0004828 | Arhgef19 | 213649 | HITS-CLIP | 23597149 | Unspecified |
| mmu-miR-708-5p | MIMAT0004828 | Arid2 | 77044 | HITS-CLIP | 37986224 | Unspecified |
| mmu-miR-708-5p | MIMAT0004828 | Arih2 | 23807 | HITS-CLIP | 37986224 | Unspecified |
| mmu-miR-708-5p | MIMAT0004828 | Arl5b | 75869 | HITS-CLIP | 23597149 | Unspecified |
| mmu-miR-708-5p | MIMAT0004828 | Arrb1 | 109689 | HITS-CLIP | 37986224 | Unspecified |
| mmu-miR-708-5p | MIMAT0004828 | Arrdc3 | 105171 | HITS-CLIP | 23597149 | Unspecified |
| mmu-miR-708-5p | MIMAT0004828 | Ascc3 | 77987 | HITS-CLIP | 37986224\|23597149 | Unspecified |
| mmu-miR-708-5p | MIMAT0004828 | Ash1l | 192195 | HITS-CLIP | 37986224 | Unspecified |
| mmu-miR-708-5p | MIMAT0004828 | Asph | 65973 | HITS-CLIP | 37986224 | Unspecified |
| mmu-miR-708-5p | MIMAT0004828 | Ate1 | 11907 | HITS-CLIP | 37986224 | Unspecified |
| mmu-miR-708-5p | MIMAT0004828 | Atg9a | 245860 | HITS-CLIP | 19536157 | Unspecified |
| mmu-miR-708-5p | MIMAT0004828 | Atp1a1 | 11928 | HITS-CLIP | 23597149 | Unspecified |
| mmu-miR-708-5p | MIMAT0004828 | Atp1a3 | 232975 | HITS-CLIP | 37986224 | Unspecified |
| mmu-miR-708-5p | MIMAT0004828 | Atp1b1 | 11931 | HITS-CLIP | 37986224 | Unspecified |
| mmu-miR-708-5p | MIMAT0004828 | Atp1b2 | 11932 | HITS-CLIP | 37986224 | Unspecified |
| mmu-miR-708-5p | MIMAT0004828 | Atp2a2 | 11938 | HITS-CLIP | 19536157 | Unspecified |
| mmu-miR-708-5p | MIMAT0004828 | Atp2b4 | 381290 | HITS-CLIP | 19536157\|23597149 | Unspecified |
| mmu-miR-708-5p | MIMAT0004828 | Atp6ap1 | 54411 | HITS-CLIP | 37986224 | Unspecified |
| mmu-miR-708-5p | MIMAT0004828 | Atp6v1a | 11964 | HITS-CLIP | 19536157 | Unspecified |
| mmu-miR-708-5p | MIMAT0004828 | Atp6v1b2 | 11966 | HITS-CLIP | 37986224 | Unspecified |
| mmu-miR-708-5p | MIMAT0004828 | Atxn1 | 20238 | HITS-CLIP | 23597149\|19536157 | Unspecified |
| mmu-miR-708-5p | MIMAT0004828 | Atxn1l | 52335 | HITS-CLIP | 37986224 | Unspecified |
| mmu-miR-708-5p | MIMAT0004828 | Atxn2 | 20239 | HITS-CLIP | 23597149 | Unspecified |
| mmu-miR-708-5p | MIMAT0004828 | Avl9 | 78937 | HITS-CLIP | 37986224 | Unspecified |
| mmu-miR-708-5p | MIMAT0004828 | B3gat1 | 76898 | HITS-CLIP | 37986224 | Unspecified |
| mmu-miR-708-5p | MIMAT0004828 | B4galt3 | 57370 | HITS-CLIP | 19536157 | Unspecified |
| mmu-miR-708-5p | MIMAT0004828 | BC005537 | 79555 | HITS-CLIP | 31433990 | Unspecified |
| mmu-miR-708-5p | MIMAT0004828 | Bag6 | 224727 | HITS-CLIP | 37986224\|23597149 | Unspecified |

| | | | | | | | |
|---|---|---|---|---|---|---|---|
| mmu-miR-708-5p | MIMAT0004828 | Baiap2l1 | 66898 | HITS-CLIP | | 37986224 | Unspecified |
| mmu-miR-708-5p | MIMAT0004828 | Baz1b | 22385 | HITS-CLIP | | 23597149 | Unspecified |
| mmu-miR-708-5p | MIMAT0004828 | Bbip1 | 100503572 | HITS-CLIP | | 23597149 | Unspecified |
| mmu-miR-708-5p | MIMAT0004828 | Bcl6 | 12053 | HITS-CLIP | | 23597149 | Unspecified |
| mmu-miR-708-5p | MIMAT0004828 | Bet1l | 54399 | HITS-CLIP | | 37986224 | Unspecified |
| mmu-miR-708-5p | MIMAT0004828 | Bhlhe40 | 20893 | HITS-CLIP | | 23597149 | Unspecified |
| mmu-miR-708-5p | MIMAT0004828 | Bmp2k | 140780 | HITS-CLIP | | 37986224 | Unspecified |
| mmu-miR-708-5p | MIMAT0004828 | Bmpr1a | 12166 | HITS-CLIP | | 23597149 | Unspecified |
| mmu-miR-708-5p | MIMAT0004828 | Bptf | 207165 | HITS-CLIP | | 19536157 | Unspecified |
| mmu-miR-708-5p | MIMAT0004828 | Brd2 | 14312 | HITS-CLIP | | 23597149 | Unspecified |
| mmu-miR-708-5p | MIMAT0004828 | Brinp1 | 56710 | HITS-CLIP | | 37986224 | Unspecified |
| mmu-miR-708-5p | MIMAT0004828 | Bscl2 | 14705 | HITS-CLIP | | 37986224 | Unspecified |
| mmu-miR-708-5p | MIMAT0004828 | Bsn | 12217 | HITS-CLIP | | 37986224 | Unspecified |
| mmu-miR-708-5p | MIMAT0004828 | Btbd3 | 228662 | HITS-CLIP | | 19536157 | Unspecified |
| mmu-miR-708-5p | MIMAT0004828 | Btf3 | 218490 | HITS-CLIP | | 31433990 | Unspecified |
| mmu-miR-708-5p | MIMAT0004828 | Btg2 | 12227 | HITS-CLIP | 37986224\|23597149 | | Unspecified |
| mmu-miR-708-5p | MIMAT0004828 | C1galt1 | 94192 | HITS-CLIP | | 23597149 | Unspecified |
| mmu-miR-708-5p | MIMAT0004828 | C1galt1c1 | 59048 | HITS-CLIP | | 23597149 | Unspecified |
| mmu-miR-708-5p | MIMAT0004828 | C1qtnf5 | 235312 | HITS-CLIP | | 37986224 | Unspecified |
| mmu-miR-708-5p | MIMAT0004828 | C2cd2l | 71764 | HITS-CLIP | | 23597149 | Unspecified |
| mmu-miR-708-5p | MIMAT0004828 | C4b | 12268 | HITS-CLIP | | 37986224 | Unspecified |
| mmu-miR-708-5p | MIMAT0004828 | C77080 | 97130 | HITS-CLIP | | 37986224 | Unspecified |
| mmu-miR-708-5p | MIMAT0004828 | C8b | 110382 | HITS-CLIP | 37986224\|23597149 | | Unspecified |
| mmu-miR-708-5p | MIMAT0004828 | C9orf72 | 203228 | HITS-CLIP | | 23597149 | Unspecified |
| mmu-miR-708-5p | MIMAT0004828 | Cald1 | 109624 | HITS-CLIP | | 23597149 | Unspecified |
| mmu-miR-708-5p | MIMAT0004828 | Calm1 | 12313 | HITS-CLIP | | 37986224 | Unspecified |
| mmu-miR-708-5p | MIMAT0004828 | Calm3 | 12315 | HITS-CLIP | | 37986224 | Unspecified |
| mmu-miR-708-5p | MIMAT0004828 | Caln1 | 140904 | HITS-CLIP | | 37986224 | Unspecified |
| mmu-miR-708-5p | MIMAT0004828 | Calr | 12317 | HITS-CLIP | 37986224\|23597149 | | Unspecified |
| mmu-miR-708-5p | MIMAT0004828 | Calu | 12321 | HITS-CLIP | 37986224\|23597149 | | Unspecified |
| mmu-miR-708-5p | MIMAT0004828 | Caly | 68566 | HITS-CLIP | | 37986224 | Unspecified |
| mmu-miR-708-5p | MIMAT0004828 | Canx | 12330 | HITS-CLIP | | 37986224 | Unspecified |
| mmu-miR-708-5p | MIMAT0004828 | Capn1 | 12333 | HITS-CLIP | | 23597149 | Unspecified |
| mmu-miR-708-5p | MIMAT0004828 | Cars | 27267 | HITS-CLIP | | 23597149 | Unspecified |
| mmu-miR-708-5p | MIMAT0004828 | Casp8 | 12370 | HITS-CLIP | | 37986224 | Unspecified |
| mmu-miR-708-5p | MIMAT0004828 | Cbs | 12411 | HITS-CLIP | 37986224\|23597149 | | Unspecified |
| mmu-miR-708-5p | MIMAT0004828 | Cbx5 | 12419 | HITS-CLIP | | 37986224 | Unspecified |
| mmu-miR-708-5p | MIMAT0004828 | Cbx6 | 494448 | HITS-CLIP | | 37986224 | Unspecified |
| mmu-miR-708-5p | MIMAT0004828 | Ccdc117 | 104479 | HITS-CLIP | | 37986224 | Unspecified |

| | | | | | | |
|---|---|---|---|---|---|---|
| mmu-miR-708-5p | MIMAT0004828 | Ccdc6 | 76551 | HITS-CLIP | 37986224 | Unspecified |
| mmu-miR-708-5p | MIMAT0004828 | Ccdc88a | 108686 | HITS-CLIP | 37986224 | Unspecified |
| mmu-miR-708-5p | MIMAT0004828 | Ccdc92 | 215707 | HITS-CLIP | 37986224 | Unspecified |
| mmu-miR-708-5p | MIMAT0004828 | Ccnt1 | 12455 | HITS-CLIP | 23597149 | Unspecified |
| mmu-miR-708-5p | MIMAT0004828 | Ccnt2 | 72949 | HITS-CLIP | 23597149 | Unspecified |
| mmu-miR-708-5p | MIMAT0004828 | Cct2 | 12461 | HITS-CLIP | 37986224 | Unspecified |
| mmu-miR-708-5p | MIMAT0004828 | Cd151 | 12476 | HITS-CLIP | 37986224 | Unspecified |
| mmu-miR-708-5p | MIMAT0004828 | Cd164 | 53599 | HITS-CLIP | 37986224 | Unspecified |
| mmu-miR-708-5p | MIMAT0004828 | Cd2ap | 12488 | HITS-CLIP | 23597149 | Unspecified |
| mmu-miR-708-5p | MIMAT0004828 | Cd93 | 17064 | HITS-CLIP | 37986224 | Unspecified |
| mmu-miR-708-5p | MIMAT0004828 | Cdc37l1 | 67072 | HITS-CLIP | 19536157 | Unspecified |
| mmu-miR-708-5p | MIMAT0004828 | Cdc40 | 71713 | HITS-CLIP | 23597149 | Unspecified |
| mmu-miR-708-5p | MIMAT0004828 | Cdc42 | 12540 | HITS-CLIP | 37986224 | Unspecified |
| mmu-miR-708-5p | MIMAT0004828 | Cdca5 | 67849 | HITS-CLIP | 37986224 | Unspecified |
| mmu-miR-708-5p | MIMAT0004828 | Cdh1 | 12550 | HITS-CLIP | 37986224 | Unspecified |
| mmu-miR-708-5p | MIMAT0004828 | Cdh8 | 12564 | HITS-CLIP | 19536157 | Unspecified |
| mmu-miR-708-5p | MIMAT0004828 | Cdip1 | 66626 | HITS-CLIP | 23597149 | Unspecified |
| mmu-miR-708-5p | MIMAT0004828 | Cdk4 | 12567 | HITS-CLIP | 37986224 | Unspecified |
| mmu-miR-708-5p | MIMAT0004828 | Cdkal1 | 68916 | HITS-CLIP | 23597149 | Unspecified |
| mmu-miR-708-5p | MIMAT0004828 | Cebpa | 12606 | HITS-CLIP | 37986224\|23597149 | Unspecified |
| mmu-miR-708-5p | MIMAT0004828 | Cebpg | 12611 | HITS-CLIP | 37986224 | Unspecified |
| mmu-miR-708-5p | MIMAT0004828 | Celf1 | 13046 | HITS-CLIP | 23597149 | Unspecified |
| mmu-miR-708-5p | MIMAT0004828 | Celf2 | 14007 | HITS-CLIP | 37986224 | Unspecified |
| mmu-miR-708-5p | MIMAT0004828 | Celf3 | 78784 | HITS-CLIP | 37986224 | Unspecified |
| mmu-miR-708-5p | MIMAT0004828 | Cep350 | 74081 | HITS-CLIP | 37986224\|23597149 | Unspecified |
| mmu-miR-708-5p | MIMAT0004828 | Cep85l | 100038725 | HITS-CLIP | 23597149 | Unspecified |
| mmu-miR-708-5p | MIMAT0004828 | Ces1f | 234564 | HITS-CLIP | 37986224 | Unspecified |
| mmu-miR-708-5p | MIMAT0004828 | Ces1g | 12623 | HITS-CLIP | 23597149 | Unspecified |
| mmu-miR-708-5p | MIMAT0004828 | Ces3a | 382053 | HITS-CLIP | 37986224 | Unspecified |
| mmu-miR-708-5p | MIMAT0004828 | Ces3b | 13909 | HITS-CLIP | 37986224 | Unspecified |
| mmu-miR-708-5p | MIMAT0004828 | Cfdp1 | 23837 | HITS-CLIP | 37986224 | Unspecified |
| mmu-miR-708-5p | MIMAT0004828 | Cgnl1 | 68178 | HITS-CLIP | 23597149 | Unspecified |
| mmu-miR-708-5p | MIMAT0004828 | Chd3 | 216848 | HITS-CLIP | 19536157 | Unspecified |
| mmu-miR-708-5p | MIMAT0004828 | Chd4 | 107932 | HITS-CLIP | 37986224\|23597149 | Unspecified |
| mmu-miR-708-5p | MIMAT0004828 | Chd8 | 67772 | HITS-CLIP | 26203562 | Unspecified |
| mmu-miR-708-5p | MIMAT0004828 | Cherp | 27967 | HITS-CLIP | 37986224 | Unspecified |
| mmu-miR-708-5p | MIMAT0004828 | Cideb | 12684 | HITS-CLIP | 23597149 | Unspecified |
| mmu-miR-708-5p | MIMAT0004828 | Ckap5 | 75786 | HITS-CLIP | 19536157 | Unspecified |
| mmu-miR-708-5p | MIMAT0004828 | Clic4 | 29876 | HITS-CLIP | 26203562 | Unspecified |

| | | | | | | | |
|---|---|---|---|---|---|---|---|
| mmu-miR-708-5p | MIMAT0004828 | Clptm1 | 56457 | HITS-CLIP | | 23597149 | Unspecified |
| mmu-miR-708-5p | MIMAT0004828 | Clpx | 270166 | HITS-CLIP | 23597149\|19536157 | | Unspecified |
| mmu-miR-708-5p | MIMAT0004828 | Clstn3 | 232370 | HITS-CLIP | | 37986224 | Unspecified |
| mmu-miR-708-5p | MIMAT0004828 | Cltc | 67300 | HITS-CLIP | 37986224\|23597149 | | Unspecified |
| mmu-miR-708-5p | MIMAT0004828 | Cmah | 12763 | HITS-CLIP | | 23597149 | Unspecified |
| mmu-miR-708-5p | MIMAT0004828 | Cmip | 74440 | HITS-CLIP | | 37986224 | Unspecified |
| mmu-miR-708-5p | MIMAT0004828 | Cmtm8 | 70031 | HITS-CLIP | 37986224\|23597149 | | Unspecified |
| mmu-miR-708-5p | MIMAT0004828 | Cnep1r1 | 382030 | HITS-CLIP | | 19536157 | Unspecified |
| mmu-miR-708-5p | MIMAT0004828 | Cntn2 | 21367 | HITS-CLIP | | 37986224 | Unspecified |
| mmu-miR-708-5p | MIMAT0004828 | Coa5 | 76178 | HITS-CLIP | | 37986224 | Unspecified |
| mmu-miR-708-5p | MIMAT0004828 | Col18a1 | 12822 | HITS-CLIP | 37986224\|23597149 | | Unspecified |
| mmu-miR-708-5p | MIMAT0004828 | Col1a2 | 12843 | HITS-CLIP | | 37986224 | Unspecified |
| mmu-miR-708-5p | MIMAT0004828 | Col3a1 | 12825 | HITS-CLIP | | 23597149 | Unspecified |
| mmu-miR-708-5p | MIMAT0004828 | Col4a1 | 12826 | HITS-CLIP | | 37986224 | Unspecified |
| mmu-miR-708-5p | MIMAT0004828 | Col4a2 | 12827 | HITS-CLIP | | 37986224 | Unspecified |
| mmu-miR-708-5p | MIMAT0004828 | Col8a2 | 329941 | HITS-CLIP | | 37986224 | Unspecified |
| mmu-miR-708-5p | MIMAT0004828 | Copb2 | 50797 | HITS-CLIP | | 23597149 | Unspecified |
| mmu-miR-708-5p | MIMAT0004828 | Copg1 | 54161 | HITS-CLIP | | 23597149 | Unspecified |
| mmu-miR-708-5p | MIMAT0004828 | Coq8a | 56997 | HITS-CLIP | | 23597149 | Unspecified |
| mmu-miR-708-5p | MIMAT0004828 | Cox14 | 66379 | HITS-CLIP | | 37986224 | Unspecified |
| mmu-miR-708-5p | MIMAT0004828 | Cox15 | 226139 | HITS-CLIP | | 23597149 | Unspecified |
| mmu-miR-708-5p | MIMAT0004828 | Cox6a1 | 12861 | HITS-CLIP | | 37986224 | Unspecified |
| mmu-miR-708-5p | MIMAT0004828 | Cp | 12870 | HITS-CLIP | | 23597149 | Unspecified |
| mmu-miR-708-5p | MIMAT0004828 | Cpe | 12876 | HITS-CLIP | 37986224\|19536157 | | Unspecified |
| mmu-miR-708-5p | MIMAT0004828 | Cpeb2 | 231207 | HITS-CLIP | | 19536157 | Unspecified |
| mmu-miR-708-5p | MIMAT0004828 | Cplx1 | 12889 | HITS-CLIP | | 19536157 | Unspecified |
| mmu-miR-708-5p | MIMAT0004828 | Cpn1 | 93721 | HITS-CLIP | | 23597149 | Unspecified |
| mmu-miR-708-5p | MIMAT0004828 | Cpne3 | 70568 | HITS-CLIP | | 37986224 | Unspecified |
| mmu-miR-708-5p | MIMAT0004828 | Cps1 | 227231 | HITS-CLIP | 37986224\|23597149 | | Unspecified |
| mmu-miR-708-5p | MIMAT0004828 | Cpt1a | 12894 | HITS-CLIP | | 37986224 | Unspecified |
| mmu-miR-708-5p | MIMAT0004828 | Crat | 12908 | HITS-CLIP | | 23597149 | Unspecified |
| mmu-miR-708-5p | MIMAT0004828 | Crebrf | 77128 | HITS-CLIP | | 37986224 | Unspecified |
| mmu-miR-708-5p | MIMAT0004828 | Crem | 12916 | HITS-CLIP | | 23597149 | Unspecified |
| mmu-miR-708-5p | MIMAT0004828 | Crls1 | 66586 | HITS-CLIP | | 23597149 | Unspecified |
| mmu-miR-708-5p | MIMAT0004828 | Crnkl1 | 66877 | HITS-CLIP | 37986224\|23597149 | | Unspecified |
| mmu-miR-708-5p | MIMAT0004828 | Crot | 74114 | HITS-CLIP | | 37986224 | Unspecified |
| mmu-miR-708-5p | MIMAT0004828 | Csdc2 | 105859 | HITS-CLIP | | 37986224 | Unspecified |
| mmu-miR-708-5p | MIMAT0004828 | Csde1 | 229663 | HITS-CLIP | | 23597149 | Unspecified |
| mmu-miR-708-5p | MIMAT0004828 | Cse1l | 110750 | HITS-CLIP | | 37986224 | Unspecified |

| | | | | | | |
|---|---|---|---|---|---|---|
| mmu-miR-708-5p | MIMAT0004828 | Csmd2 | 329942 | HITS-CLIP | 19536157 | Unspecified |
| mmu-miR-708-5p | MIMAT0004828 | Csnk1a1 | 93687 | HITS-CLIP | 23597149 | Unspecified |
| mmu-miR-708-5p | MIMAT0004828 | Csnk1d | 104318 | HITS-CLIP | 37986224 | Unspecified |
| mmu-miR-708-5p | MIMAT0004828 | Csnk2a2 | 13000 | HITS-CLIP | 23597149 | Unspecified |
| mmu-miR-708-5p | MIMAT0004828 | Csrnp2 | 207785 | HITS-CLIP | 37986224 | Unspecified |
| mmu-miR-708-5p | MIMAT0004828 | Cst3 | 13010 | HITS-CLIP | 37986224 | Unspecified |
| mmu-miR-708-5p | MIMAT0004828 | Cstf3 | 228410 | HITS-CLIP | 23597149 | Unspecified |
| mmu-miR-708-5p | MIMAT0004828 | Ctcf | 13018 | HITS-CLIP | 23597149 | Unspecified |
| mmu-miR-708-5p | MIMAT0004828 | Ctnnb1 | 12387 | HITS-CLIP | 37986224 | Unspecified |
| mmu-miR-708-5p | MIMAT0004828 | Ctnnd1 | 12388 | HITS-CLIP | 23597149 | Unspecified |
| mmu-miR-708-5p | MIMAT0004828 | Ctsb | 13030 | HITS-CLIP | 37986224\|23597149 | Unspecified |
| mmu-miR-708-5p | MIMAT0004828 | Cx3cl1 | 20312 | HITS-CLIP | 37986224 | Unspecified |
| mmu-miR-708-5p | MIMAT0004828 | Cyb5b | 66427 | HITS-CLIP | 37986224 | Unspecified |
| mmu-miR-708-5p | MIMAT0004828 | Cyfip2 | 76884 | HITS-CLIP | 19536157 | Unspecified |
| mmu-miR-708-5p | MIMAT0004828 | Cyp2c29 | 13095 | HITS-CLIP | 37986224 | Unspecified |
| mmu-miR-708-5p | MIMAT0004828 | Cyp2e1 | 13106 | HITS-CLIP | 23597149 | Unspecified |
| mmu-miR-708-5p | MIMAT0004828 | Cyp4a14 | 13119 | HITS-CLIP | 37986224 | Unspecified |
| mmu-miR-708-5p | MIMAT0004828 | Cyp4b1 | 13120 | HITS-CLIP | 37986224 | Unspecified |
| mmu-miR-708-5p | MIMAT0004828 | Cyp7b1 | 13123 | HITS-CLIP | 23597149 | Unspecified |
| mmu-miR-708-5p | MIMAT0004828 | D230025D16Rik | 234678 | HITS-CLIP | 37986224\|23597149 | Unspecified |
| mmu-miR-708-5p | MIMAT0004828 | D430041D05Rik | 241589 | HITS-CLIP | 37986224 | Unspecified |
| mmu-miR-708-5p | MIMAT0004828 | Dag1 | 13138 | HITS-CLIP | 37986224\|23597149 | Unspecified |
| mmu-miR-708-5p | MIMAT0004828 | Dalrd3 | 67789 | HITS-CLIP | 23597149 | Unspecified |
| mmu-miR-708-5p | MIMAT0004828 | Dap | 223453 | HITS-CLIP | 37986224 | Unspecified |
| mmu-miR-708-5p | MIMAT0004828 | Dap3 | 65111 | HITS-CLIP | 23597149 | Unspecified |
| mmu-miR-708-5p | MIMAT0004828 | Dazap2 | 23994 | HITS-CLIP | 23597149 | Unspecified |
| mmu-miR-708-5p | MIMAT0004828 | Dcaf7 | 71833 | HITS-CLIP | 37986224 | Unspecified |
| mmu-miR-708-5p | MIMAT0004828 | Dcaf8 | 98193 | HITS-CLIP | 23597149 | Unspecified |
| mmu-miR-708-5p | MIMAT0004828 | Dctn1 | 13191 | HITS-CLIP | 23597149 | Unspecified |
| mmu-miR-708-5p | MIMAT0004828 | Ddah1 | 69219 | HITS-CLIP | 37986224 | Unspecified |
| mmu-miR-708-5p | MIMAT0004828 | Ddb1 | 13194 | HITS-CLIP | 23597149 | Unspecified |
| mmu-miR-708-5p | MIMAT0004828 | Ddx17 | 67040 | HITS-CLIP | 37986224 | Unspecified |
| mmu-miR-708-5p | MIMAT0004828 | Ddx3x | 13205 | HITS-CLIP | 37986224 | Unspecified |
| mmu-miR-708-5p | MIMAT0004828 | Ddx5 | 13207 | HITS-CLIP | 37986224\|23597149\|19536157 | Unspecified |
| mmu-miR-708-5p | MIMAT0004828 | Degs1 | 13244 | HITS-CLIP | 37986224 | Unspecified |
| mmu-miR-708-5p | MIMAT0004828 | Depdc7 | 211896 | HITS-CLIP | 37986224\|23597149 | Unspecified |
| mmu-miR-708-5p | MIMAT0004828 | Derl2 | 116891 | HITS-CLIP | 23597149 | Unspecified |
| mmu-miR-708-5p | MIMAT0004828 | Dgat2 | 67800 | HITS-CLIP | 23597149 | Unspecified |
| mmu-miR-708-5p | MIMAT0004828 | Dgkd | 227333 | HITS-CLIP | 37986224 | Unspecified |

| | | | | | | |
|---|---|---|---|---|---|---|
| mmu-miR-708-5p | MIMAT0004828 | Dgkh | 380921 | HITS-CLIP | 37986224 | Unspecified |
| mmu-miR-708-5p | MIMAT0004828 | Dhcr24 | 74754 | HITS-CLIP | 37986224 | Unspecified |
| mmu-miR-708-5p | MIMAT0004828 | Dhx29 | 218629 | HITS-CLIP | 23597149 | Unspecified |
| mmu-miR-708-5p | MIMAT0004828 | Dicer1 | 192119 | HITS-CLIP | 37986224\|23597149 | Unspecified |
| mmu-miR-708-5p | MIMAT0004828 | Dip2b | 239667 | HITS-CLIP | 37986224\|19536157 | Unspecified |
| mmu-miR-708-5p | MIMAT0004828 | Dip2c | 208440 | HITS-CLIP | 23597149 | Unspecified |
| mmu-miR-708-5p | MIMAT0004828 | Diras1 | 208666 | HITS-CLIP | 37986224 | Unspecified |
| mmu-miR-708-5p | MIMAT0004828 | Disp2 | 214240 | HITS-CLIP | 37986224 | Unspecified |
| mmu-miR-708-5p | MIMAT0004828 | Dlat | 235339 | HITS-CLIP | 37986224 | Unspecified |
| mmu-miR-708-5p | MIMAT0004828 | Dlc1 | 50768 | HITS-CLIP | 37986224 | Unspecified |
| mmu-miR-708-5p | MIMAT0004828 | Dld | 13382 | HITS-CLIP | 37986224 | Unspecified |
| mmu-miR-708-5p | MIMAT0004828 | Dmgdh | 74129 | HITS-CLIP | 37986224 | Unspecified |
| mmu-miR-708-5p | MIMAT0004828 | Dnajb12 | 56709 | HITS-CLIP | 37986224\|23597149 | Unspecified |
| mmu-miR-708-5p | MIMAT0004828 | Dnajb5 | 56323 | HITS-CLIP | 37986224 | Unspecified |
| mmu-miR-708-5p | MIMAT0004828 | Dnajc1 | 13418 | HITS-CLIP | 37986224 | Unspecified |
| mmu-miR-708-5p | MIMAT0004828 | Dnajc10 | 66861 | HITS-CLIP | 23597149 | Unspecified |
| mmu-miR-708-5p | MIMAT0004828 | Dnajc14 | 74330 | HITS-CLIP | 37986224 | Unspecified |
| mmu-miR-708-5p | MIMAT0004828 | Dnajc5 | 13002 | HITS-CLIP | 37986224 | Unspecified |
| mmu-miR-708-5p | MIMAT0004828 | Dnm1 | 13429 | HITS-CLIP | 37986224 | Unspecified |
| mmu-miR-708-5p | MIMAT0004828 | Dnmbp | 71972 | HITS-CLIP | 23597149 | Unspecified |
| mmu-miR-708-5p | MIMAT0004828 | Dnmt3a | 13435 | HITS-CLIP | 23597149\|19536157 | Unspecified |
| mmu-miR-708-5p | MIMAT0004828 | Dot1l | 208266 | HITS-CLIP | 37986224 | Unspecified |
| mmu-miR-708-5p | MIMAT0004828 | Dpagt1 | 13478 | HITS-CLIP | 37986224 | Unspecified |
| mmu-miR-708-5p | MIMAT0004828 | Dpyd | 99586 | HITS-CLIP | 37986224\|23597149 | Unspecified |
| mmu-miR-708-5p | MIMAT0004828 | Dram2 | 67171 | HITS-CLIP | 23597149 | Unspecified |
| mmu-miR-708-5p | MIMAT0004828 | Dstn | 56431 | HITS-CLIP | 37986224 | Unspecified |
| mmu-miR-708-5p | MIMAT0004828 | Dtymk | 21915 | HITS-CLIP | 23597149 | Unspecified |
| mmu-miR-708-5p | MIMAT0004828 | Dusp1 | 19252 | HITS-CLIP | 37986224\|23597149 | Unspecified |
| mmu-miR-708-5p | MIMAT0004828 | Dusp8 | 18218 | HITS-CLIP | 37986224 | Unspecified |
| mmu-miR-708-5p | MIMAT0004828 | Dync1h1 | 13424 | HITS-CLIP | 37986224\|23597149\|19536157 | Unspecified |
| mmu-miR-708-5p | MIMAT0004828 | Dync1i2 | 13427 | HITS-CLIP | 37986224 | Unspecified |
| mmu-miR-708-5p | MIMAT0004828 | Dync1li2 | 234663 | HITS-CLIP | 37986224 | Unspecified |
| mmu-miR-708-5p | MIMAT0004828 | Dynll2 | 68097 | HITS-CLIP | 23597149\|19536157 | Unspecified |
| mmu-miR-708-5p | MIMAT0004828 | Dyrk1a | 13548 | HITS-CLIP | 23597149 | Unspecified |
| mmu-miR-708-5p | MIMAT0004828 | E130308A19Rik | 230259 | HITS-CLIP | 37986224 | Unspecified |
| mmu-miR-708-5p | MIMAT0004828 | Ece1 | 230857 | HITS-CLIP | 37986224 | Unspecified |
| mmu-miR-708-5p | MIMAT0004828 | Ecsit | 26940 | HITS-CLIP | 23597149 | Unspecified |
| mmu-miR-708-5p | MIMAT0004828 | Edem1 | 192193 | HITS-CLIP | 37986224\|23597149 | Unspecified |
| mmu-miR-708-5p | MIMAT0004828 | Eea1 | 216238 | HITS-CLIP | 19536157 | Unspecified |

| | | | | | | |
|---|---|---|---|---|---|---|
| mmu-miR-708-5p | MIMAT0004828 | Eef1a1 | 13627 | HITS-CLIP | 37986224\|23597149 | Unspecified |
| mmu-miR-708-5p | MIMAT0004828 | Eef2 | 13629 | HITS-CLIP | 37986224 | Unspecified |
| mmu-miR-708-5p | MIMAT0004828 | Efcab14 | 230648 | HITS-CLIP | 19536157 | Unspecified |
| mmu-miR-708-5p | MIMAT0004828 | Efemp2 | 58859 | HITS-CLIP | 23597149 | Unspecified |
| mmu-miR-708-5p | MIMAT0004828 | Efr3a | 76740 | HITS-CLIP | 37986224 | Unspecified |
| mmu-miR-708-5p | MIMAT0004828 | Egln1 | 112405 | HITS-CLIP | 23597149 | Unspecified |
| mmu-miR-708-5p | MIMAT0004828 | Ehd1 | 13660 | HITS-CLIP | 37986224\|23597149 | Unspecified |
| mmu-miR-708-5p | MIMAT0004828 | Ehd3 | 57440 | HITS-CLIP | 23597149 | Unspecified |
| mmu-miR-708-5p | MIMAT0004828 | Ehf | 13661 | HITS-CLIP | 37986224 | Unspecified |
| mmu-miR-708-5p | MIMAT0004828 | Eif2s3y | 26908 | HITS-CLIP | 23597149 | Unspecified |
| mmu-miR-708-5p | MIMAT0004828 | Eif3c | 56347 | HITS-CLIP | 37986224 | Unspecified |
| mmu-miR-708-5p | MIMAT0004828 | Eif3f | 66085 | HITS-CLIP | 19536157 | Unspecified |
| mmu-miR-708-5p | MIMAT0004828 | Eif4ebp1 | 13685 | HITS-CLIP | 37986224 | Unspecified |
| mmu-miR-708-5p | MIMAT0004828 | Eif4ebp2 | 13688 | HITS-CLIP | 37986224\|23597149 | Unspecified |
| mmu-miR-708-5p | MIMAT0004828 | Eif4g1 | 208643 | HITS-CLIP | 37986224\|23597149 | Unspecified |
| mmu-miR-708-5p | MIMAT0004828 | Eif4g2 | 13690 | HITS-CLIP | 19536157\|23597149 | Unspecified |
| mmu-miR-708-5p | MIMAT0004828 | Eif5 | 217869 | HITS-CLIP | 23597149 | Unspecified |
| mmu-miR-708-5p | MIMAT0004828 | Ell | 13716 | HITS-CLIP | 26203562 | Unspecified |
| mmu-miR-708-5p | MIMAT0004828 | Ell2 | 192657 | HITS-CLIP | 37986224 | Unspecified |
| mmu-miR-708-5p | MIMAT0004828 | Elmo1 | 140580 | HITS-CLIP | 37986224 | Unspecified |
| mmu-miR-708-5p | MIMAT0004828 | Eloa | 6924 | HITS-CLIP | 23597149 | Unspecified |
| mmu-miR-708-5p | MIMAT0004828 | Elovl6 | 170439 | HITS-CLIP | 37986224 | Unspecified |
| mmu-miR-708-5p | MIMAT0004828 | En2 | 13799 | HITS-CLIP | 37986224 | Unspecified |
| mmu-miR-708-5p | MIMAT0004828 | Eno2 | 13807 | HITS-CLIP | 19536157 | Unspecified |
| mmu-miR-708-5p | MIMAT0004828 | Enpp1 | 18605 | HITS-CLIP | 23597149 | Unspecified |
| mmu-miR-708-5p | MIMAT0004828 | Enpp2 | 18606 | HITS-CLIP | 23597149 | Unspecified |
| mmu-miR-708-5p | MIMAT0004828 | Entpd5 | 12499 | HITS-CLIP | 23597149 | Unspecified |
| mmu-miR-708-5p | MIMAT0004828 | Eogt | 101351 | HITS-CLIP | 37986224 | Unspecified |
| mmu-miR-708-5p | MIMAT0004828 | Ep300 | 328572 | HITS-CLIP | 37986224 | Unspecified |
| mmu-miR-708-5p | MIMAT0004828 | Epc2 | 227867 | HITS-CLIP | 37986224 | Unspecified |
| mmu-miR-708-5p | MIMAT0004828 | Ephx1 | 13849 | HITS-CLIP | 23597149 | Unspecified |
| mmu-miR-708-5p | MIMAT0004828 | Eps15 | 13858 | HITS-CLIP | 23597149 | Unspecified |
| mmu-miR-708-5p | MIMAT0004828 | Erc1 | 111173 | HITS-CLIP | 23597149 | Unspecified |
| mmu-miR-708-5p | MIMAT0004828 | Ercc3 | 13872 | HITS-CLIP | 23597149 | Unspecified |
| mmu-miR-708-5p | MIMAT0004828 | Ergic3 | 66366 | HITS-CLIP | 23597149 | Unspecified |
| mmu-miR-708-5p | MIMAT0004828 | Erlin2 | 244373 | HITS-CLIP | 37986224 | Unspecified |
| mmu-miR-708-5p | MIMAT0004828 | Ermp1 | 226090 | HITS-CLIP | 37986224 | Unspecified |
| mmu-miR-708-5p | MIMAT0004828 | Ero1lb | 67475 | HITS-CLIP | 37986224 | Unspecified |
| mmu-miR-708-5p | MIMAT0004828 | Errfi1 | 74155 | HITS-CLIP | 23597149 | Unspecified |

| | | | | | | | |
|---|---|---|---|---|---|---|---|
| mmu-miR-708-5p | MIMAT0004828 | Esr1 | 13982 | HITS-CLIP | | 37986224 | Unspecified |
| mmu-miR-708-5p | MIMAT0004828 | Esyt2 | 52635 | HITS-CLIP | | 37986224 | Unspecified |
| mmu-miR-708-5p | MIMAT0004828 | Etf1 | 225363 | HITS-CLIP | | 37986224 | Unspecified |
| mmu-miR-708-5p | MIMAT0004828 | Etfdh | 66841 | HITS-CLIP | | 23597149 | Unspecified |
| mmu-miR-708-5p | MIMAT0004828 | Etl4 | 208618 | HITS-CLIP | | 37986224 | Unspecified |
| mmu-miR-708-5p | MIMAT0004828 | Etnk2 | 214253 | HITS-CLIP | | 37986224 | Unspecified |
| mmu-miR-708-5p | MIMAT0004828 | Exoc4 | 20336 | HITS-CLIP | | 23597149 | Unspecified |
| mmu-miR-708-5p | MIMAT0004828 | Exoc6b | 75914 | HITS-CLIP | | 37986224 | Unspecified |
| mmu-miR-708-5p | MIMAT0004828 | Exph5 | 320051 | HITS-CLIP | | 23597149 | Unspecified |
| mmu-miR-708-5p | MIMAT0004828 | Ext1 | 14042 | HITS-CLIP | | 37986224 | Unspecified |
| mmu-miR-708-5p | MIMAT0004828 | Extl2 | 58193 | HITS-CLIP | | 37986224 | Unspecified |
| mmu-miR-708-5p | MIMAT0004828 | Ezh2 | 14056 | HITS-CLIP | | 23597149 | Unspecified |
| mmu-miR-708-5p | MIMAT0004828 | F12 | 58992 | HITS-CLIP | | 37986224 | Unspecified |
| mmu-miR-708-5p | MIMAT0004828 | F5 | 14067 | HITS-CLIP | 37986224\|23597149 | | Unspecified |
| mmu-miR-708-5p | MIMAT0004828 | F8 | 14069 | HITS-CLIP | | 23597149 | Unspecified |
| mmu-miR-708-5p | MIMAT0004828 | F9 | 14071 | HITS-CLIP | | 37986224 | Unspecified |
| mmu-miR-708-5p | MIMAT0004828 | Fads1 | 76267 | HITS-CLIP | | 37986224 | Unspecified |
| mmu-miR-708-5p | MIMAT0004828 | Faf2 | 76577 | HITS-CLIP | | 37986224 | Unspecified |
| mmu-miR-708-5p | MIMAT0004828 | Fam124a | 629059 | HITS-CLIP | | 37986224 | Unspecified |
| mmu-miR-708-5p | MIMAT0004828 | Fam126b | 213056 | HITS-CLIP | | 19536157 | Unspecified |
| mmu-miR-708-5p | MIMAT0004828 | Fam193a | 231128 | HITS-CLIP | | 23597149 | Unspecified |
| mmu-miR-708-5p | MIMAT0004828 | Fam214a | 235493 | HITS-CLIP | 37986224\|23597149 | | Unspecified |
| mmu-miR-708-5p | MIMAT0004828 | Fam43a | 224093 | HITS-CLIP | 37986224\|19536157 | | Unspecified |
| mmu-miR-708-5p | MIMAT0004828 | Fam53b | 77938 | HITS-CLIP | | 37986224 | Unspecified |
| mmu-miR-708-5p | MIMAT0004828 | Fam76a | 230789 | HITS-CLIP | | 37986224 | Unspecified |
| mmu-miR-708-5p | MIMAT0004828 | Fastkd2 | 75619 | HITS-CLIP | | 37986224 | Unspecified |
| mmu-miR-708-5p | MIMAT0004828 | Fat1 | 14107 | HITS-CLIP | 37986224\|26203562 | | Unspecified |
| mmu-miR-708-5p | MIMAT0004828 | Fat3 | 270120 | HITS-CLIP | | 19536157 | Unspecified |
| mmu-miR-708-5p | MIMAT0004828 | Fbxl5 | 242960 | HITS-CLIP | | 23597149 | Unspecified |
| mmu-miR-708-5p | MIMAT0004828 | Fbxo21 | 231670 | HITS-CLIP | | 37986224 | Unspecified |
| mmu-miR-708-5p | MIMAT0004828 | Fchsd2 | 207278 | HITS-CLIP | 37986224\|23597149 | | Unspecified |
| mmu-miR-708-5p | MIMAT0004828 | Fga | 14161 | HITS-CLIP | | 23597149 | Unspecified |
| mmu-miR-708-5p | MIMAT0004828 | Fgd4 | 224014 | HITS-CLIP | | 37986224 | Unspecified |
| mmu-miR-708-5p | MIMAT0004828 | Fgd6 | 13998 | HITS-CLIP | | 37986224 | Unspecified |
| mmu-miR-708-5p | MIMAT0004828 | Fgfr3 | 14184 | HITS-CLIP | | 37986224 | Unspecified |
| mmu-miR-708-5p | MIMAT0004828 | Fgfrl1 | 116701 | HITS-CLIP | | 37986224 | Unspecified |
| mmu-miR-708-5p | MIMAT0004828 | Fgl1 | 234199 | HITS-CLIP | | 23597149 | Unspecified |
| mmu-miR-708-5p | MIMAT0004828 | Fkbp15 | 338355 | HITS-CLIP | | 23597149 | Unspecified |
| mmu-miR-708-5p | MIMAT0004828 | Fkbp1a | 14225 | HITS-CLIP | | 37986224 | Unspecified |

| | | | | | | | |
|---|---|---|---|---|---|---|---|
| mmu-miR-708-5p | MIMAT0004828 | Flnb | 286940 | HITS-CLIP | | 23597149 | Unspecified |
| mmu-miR-708-5p | MIMAT0004828 | Flot2 | 14252 | HITS-CLIP | | 37986224 | Unspecified |
| mmu-miR-708-5p | MIMAT0004828 | Flt1 | 14254 | HITS-CLIP | | 37986224 | Unspecified |
| mmu-miR-708-5p | MIMAT0004828 | Fmnl2 | 71409 | HITS-CLIP | | 23597149 | Unspecified |
| mmu-miR-708-5p | MIMAT0004828 | Fn3krp | 238024 | HITS-CLIP | | 23597149 | Unspecified |
| mmu-miR-708-5p | MIMAT0004828 | Fnbp1 | 14269 | HITS-CLIP | | 37986224 | Unspecified |
| mmu-miR-708-5p | MIMAT0004828 | Fndc3b | 72007 | HITS-CLIP | | 37986224 | Unspecified |
| mmu-miR-708-5p | MIMAT0004828 | Fos | 14281 | HITS-CLIP | | 37986224 | Unspecified |
| mmu-miR-708-5p | MIMAT0004828 | Fosb | 14282 | HITS-CLIP | | 23597149 | Unspecified |
| mmu-miR-708-5p | MIMAT0004828 | Foxj3 | 230700 | HITS-CLIP | 37986224\|23597149 | | Unspecified |
| mmu-miR-708-5p | MIMAT0004828 | Foxk1 | 17425 | HITS-CLIP | 23597149\|31433990 | | Unspecified |
| mmu-miR-708-5p | MIMAT0004828 | Foxk2 | 68837 | HITS-CLIP | | 37986224 | Unspecified |
| mmu-miR-708-5p | MIMAT0004828 | Foxp2 | 114142 | HITS-CLIP | | 37986224 | Unspecified |
| mmu-miR-708-5p | MIMAT0004828 | Fpgs | 14287 | HITS-CLIP | 37986224\|23597149 | | Unspecified |
| mmu-miR-708-5p | MIMAT0004828 | Frmd4b | 232288 | HITS-CLIP | | 37986224 | Unspecified |
| mmu-miR-708-5p | MIMAT0004828 | Fstl1 | 14314 | HITS-CLIP | 37986224\|19536157 | | Unspecified |
| mmu-miR-708-5p | MIMAT0004828 | Ftcd | 14317 | HITS-CLIP | | 23597149 | Unspecified |
| mmu-miR-708-5p | MIMAT0004828 | Ftl1 | 14325 | HITS-CLIP | | 37986224 | Unspecified |
| mmu-miR-708-5p | MIMAT0004828 | Fut9 | 14348 | HITS-CLIP | | 37986224 | Unspecified |
| mmu-miR-708-5p | MIMAT0004828 | Fxyd7 | 57780 | HITS-CLIP | | 37986224 | Unspecified |
| mmu-miR-708-5p | MIMAT0004828 | Fyn | 14360 | HITS-CLIP | | 37986224 | Unspecified |
| mmu-miR-708-5p | MIMAT0004828 | Fyttd1 | 69823 | HITS-CLIP | | 23597149 | Unspecified |
| mmu-miR-708-5p | MIMAT0004828 | G3bp2 | 23881 | HITS-CLIP | | 23597149 | Unspecified |
| mmu-miR-708-5p | MIMAT0004828 | Gab1 | 14388 | HITS-CLIP | | 37986224 | Unspecified |
| mmu-miR-708-5p | MIMAT0004828 | Gabarapl1 | 57436 | HITS-CLIP | | 37986224 | Unspecified |
| mmu-miR-708-5p | MIMAT0004828 | Gabra1 | 14394 | HITS-CLIP | 37986224\|19536157 | | Unspecified |
| mmu-miR-708-5p | MIMAT0004828 | Gabrg2 | 14406 | HITS-CLIP | | 37986224 | Unspecified |
| mmu-miR-708-5p | MIMAT0004828 | Gal3st3 | 545276 | HITS-CLIP | | 37986224 | Unspecified |
| mmu-miR-708-5p | MIMAT0004828 | Ganab | 14376 | HITS-CLIP | | 23597149 | Unspecified |
| mmu-miR-708-5p | MIMAT0004828 | Gapvd1 | 66691 | HITS-CLIP | 37986224\|23597149 | | Unspecified |
| mmu-miR-708-5p | MIMAT0004828 | Gatd1 | 347862 | HITS-CLIP | | 23597149 | Unspecified |
| mmu-miR-708-5p | MIMAT0004828 | Gcgr | 14527 | HITS-CLIP | | 23597149 | Unspecified |
| mmu-miR-708-5p | MIMAT0004828 | Gckr | 231103 | HITS-CLIP | | 23597149 | Unspecified |
| mmu-miR-708-5p | MIMAT0004828 | Gcsh | 68133 | HITS-CLIP | | 37986224 | Unspecified |
| mmu-miR-708-5p | MIMAT0004828 | Gda | 14544 | HITS-CLIP | | 37986224 | Unspecified |
| mmu-miR-708-5p | MIMAT0004828 | Gde1 | 56209 | HITS-CLIP | | 23597149 | Unspecified |
| mmu-miR-708-5p | MIMAT0004828 | Gid4 | 66771 | HITS-CLIP | | 23597149 | Unspecified |
| mmu-miR-708-5p | MIMAT0004828 | Gigyf2 | 227331 | HITS-CLIP | 37986224\|23597149 | | Unspecified |
| mmu-miR-708-5p | MIMAT0004828 | Gja1 | 14609 | HITS-CLIP | | 37986224 | Unspecified |

| | | | | | | |
|---|---|---|---|---|---|---|
| mmu-miR-708-5p | MIMAT0004828 | Gk | 14933 | HITS-CLIP | 23597149 | Unspecified |
| mmu-miR-708-5p | MIMAT0004828 | Glcci1 | 170772 | HITS-CLIP | 37986224 | Unspecified |
| mmu-miR-708-5p | MIMAT0004828 | Glce | 93683 | HITS-CLIP | 19536157 | Unspecified |
| mmu-miR-708-5p | MIMAT0004828 | Gldc | 104174 | HITS-CLIP | 37986224\|23597149 | Unspecified |
| mmu-miR-708-5p | MIMAT0004828 | Glg1 | 20340 | HITS-CLIP | 37986224 | Unspecified |
| mmu-miR-708-5p | MIMAT0004828 | Glis2 | 83396 | HITS-CLIP | 19536157 | Unspecified |
| mmu-miR-708-5p | MIMAT0004828 | Gm20708 | 6608141 | HITS-CLIP | 23597149 | Unspecified |
| mmu-miR-708-5p | MIMAT0004828 | Gm2a | 14667 | HITS-CLIP | 37986224 | Unspecified |
| mmu-miR-708-5p | MIMAT0004828 | Gm38394 | 667118 | HITS-CLIP | 23597149 | Unspecified |
| mmu-miR-708-5p | MIMAT0004828 | Gmfb | 63985 | HITS-CLIP | 23597149 | Unspecified |
| mmu-miR-708-5p | MIMAT0004828 | Gmps | 229363 | HITS-CLIP | 23597149 | Unspecified |
| mmu-miR-708-5p | MIMAT0004828 | Gna13 | 14674 | HITS-CLIP | 37986224 | Unspecified |
| mmu-miR-708-5p | MIMAT0004828 | Gnao1 | 14681 | HITS-CLIP | 37986224 | Unspecified |
| mmu-miR-708-5p | MIMAT0004828 | Gnas | 14683 | HITS-CLIP | 31433990 | Unspecified |
| mmu-miR-708-5p | MIMAT0004828 | Gnb1 | 14688 | HITS-CLIP | 37986224\|23597149 | Unspecified |
| mmu-miR-708-5p | MIMAT0004828 | Gnb5 | 14697 | HITS-CLIP | 37986224 | Unspecified |
| mmu-miR-708-5p | MIMAT0004828 | Gne | 50798 | HITS-CLIP | 23597149 | Unspecified |
| mmu-miR-708-5p | MIMAT0004828 | Gnl1 | 14670 | HITS-CLIP | 37986224 | Unspecified |
| mmu-miR-708-5p | MIMAT0004828 | Gnpat | 14712 | HITS-CLIP | 19536157 | Unspecified |
| mmu-miR-708-5p | MIMAT0004828 | Gns | 75612 | HITS-CLIP | 37986224 | Unspecified |
| mmu-miR-708-5p | MIMAT0004828 | Golgb1 | 224139 | HITS-CLIP | 23597149 | Unspecified |
| mmu-miR-708-5p | MIMAT0004828 | Golph3 | 66629 | HITS-CLIP | 37986224 | Unspecified |
| mmu-miR-708-5p | MIMAT0004828 | Got1 | 14718 | HITS-CLIP | 23597149 | Unspecified |
| mmu-miR-708-5p | MIMAT0004828 | Got2 | 14719 | HITS-CLIP | 23597149 | Unspecified |
| mmu-miR-708-5p | MIMAT0004828 | Gpatch2 | 67769 | HITS-CLIP | 37986224 | Unspecified |
| mmu-miR-708-5p | MIMAT0004828 | Gpatch8 | 237943 | HITS-CLIP | 37986224 | Unspecified |
| mmu-miR-708-5p | MIMAT0004828 | Gphn | 268566 | HITS-CLIP | 37986224\|23597149 | Unspecified |
| mmu-miR-708-5p | MIMAT0004828 | Gpld1 | 14756 | HITS-CLIP | 23597149 | Unspecified |
| mmu-miR-708-5p | MIMAT0004828 | Gpm6a | 234267 | HITS-CLIP | 37986224 | Unspecified |
| mmu-miR-708-5p | MIMAT0004828 | Gpr146 | 80290 | HITS-CLIP | 37986224 | Unspecified |
| mmu-miR-708-5p | MIMAT0004828 | Gpr34 | 23890 | HITS-CLIP | 37986224 | Unspecified |
| mmu-miR-708-5p | MIMAT0004828 | Gpr85 | 64450 | HITS-CLIP | 37986224 | Unspecified |
| mmu-miR-708-5p | MIMAT0004828 | Gpx1 | 14775 | HITS-CLIP | 37986224 | Unspecified |
| mmu-miR-708-5p | MIMAT0004828 | Gpx7 | 67305 | HITS-CLIP | 37986224 | Unspecified |
| mmu-miR-708-5p | MIMAT0004828 | Gria3 | 53623 | HITS-CLIP | 37986224 | Unspecified |
| mmu-miR-708-5p | MIMAT0004828 | Grid1 | 14803 | HITS-CLIP | 37986224 | Unspecified |
| mmu-miR-708-5p | MIMAT0004828 | Grik2 | 14806 | HITS-CLIP | 19536157 | Unspecified |
| mmu-miR-708-5p | MIMAT0004828 | Grin1 | 14810 | HITS-CLIP | 37986224 | Unspecified |
| mmu-miR-708-5p | MIMAT0004828 | Grin2b | 14812 | HITS-CLIP | 37986224\|19536157 | Unspecified |

| | | | | | | |
|---|---|---|---|---|---|---|
| mmu-miR-708-5p | MIMAT0004828 | Grm2 | 108068 | HITS-CLIP | 37986224 | Unspecified |
| mmu-miR-708-5p | MIMAT0004828 | Grm4 | 268934 | HITS-CLIP | 37986224 | Unspecified |
| mmu-miR-708-5p | MIMAT0004828 | Grm5 | 108071 | HITS-CLIP | 19536157 | Unspecified |
| mmu-miR-708-5p | MIMAT0004828 | Gsk3a | 606496 | HITS-CLIP | 19536157 | Unspecified |
| mmu-miR-708-5p | MIMAT0004828 | Gsk3b | 56637 | HITS-CLIP | 37986224 | Unspecified |
| mmu-miR-708-5p | MIMAT0004828 | Gtf2a1 | 83602 | HITS-CLIP | 37986224 | Unspecified |
| mmu-miR-708-5p | MIMAT0004828 | Gtf3c1 | 233863 | HITS-CLIP | 37986224\|23597149 | Unspecified |
| mmu-miR-708-5p | MIMAT0004828 | Gtf3c2 | 71752 | HITS-CLIP | 23597149 | Unspecified |
| mmu-miR-708-5p | MIMAT0004828 | Gtf3c4 | 269252 | HITS-CLIP | 23597149 | Unspecified |
| mmu-miR-708-5p | MIMAT0004828 | H13 | 14950 | HITS-CLIP | 31433990\|23597149 | Unspecified |
| mmu-miR-708-5p | MIMAT0004828 | H1f0 | 14958 | HITS-CLIP | 37986224 | Unspecified |
| mmu-miR-708-5p | MIMAT0004828 | Hadha | 97212 | HITS-CLIP | 37986224 | Unspecified |
| mmu-miR-708-5p | MIMAT0004828 | Hcfc2 | 67933 | HITS-CLIP | 37986224\|23597149 | Unspecified |
| mmu-miR-708-5p | MIMAT0004828 | Hdlbp | 110611 | HITS-CLIP | 23597149 | Unspecified |
| mmu-miR-708-5p | MIMAT0004828 | Heatr5b | 320473 | HITS-CLIP | 37986224 | Unspecified |
| mmu-miR-708-5p | MIMAT0004828 | Hectd1 | 207304 | HITS-CLIP | 19536157\|23597149 | Unspecified |
| mmu-miR-708-5p | MIMAT0004828 | Herc1 | 235439 | HITS-CLIP | 37986224 | Unspecified |
| mmu-miR-708-5p | MIMAT0004828 | Herc2 | 15204 | HITS-CLIP | 23597149 | Unspecified |
| mmu-miR-708-5p | MIMAT0004828 | Herpud1 | 64209 | HITS-CLIP | 23597149 | Unspecified |
| mmu-miR-708-5p | MIMAT0004828 | Hes1 | 15205 | HITS-CLIP | 23597149 | Unspecified |
| mmu-miR-708-5p | MIMAT0004828 | Hexim1 | 192231 | HITS-CLIP | 23597149 | Unspecified |
| mmu-miR-708-5p | MIMAT0004828 | Hgfac | 54426 | HITS-CLIP | 23597149 | Unspecified |
| mmu-miR-708-5p | MIMAT0004828 | Hic1 | 15248 | HITS-CLIP | 23597149 | Unspecified |
| mmu-miR-708-5p | MIMAT0004828 | Hif1an | 319594 | HITS-CLIP | 37986224 | Unspecified |
| mmu-miR-708-5p | MIMAT0004828 | Hip1 | 215114 | HITS-CLIP | 37986224 | Unspecified |
| mmu-miR-708-5p | MIMAT0004828 | Hipk1 | 15257 | HITS-CLIP | 19536157 | Unspecified |
| mmu-miR-708-5p | MIMAT0004828 | Hipk3 | 15259 | HITS-CLIP | 23597149 | Unspecified |
| mmu-miR-708-5p | MIMAT0004828 | Hira | 15260 | HITS-CLIP | 37986224 | Unspecified |
| mmu-miR-708-5p | MIMAT0004828 | Hist1h1c | 50708 | HITS-CLIP | 23597149 | Unspecified |
| mmu-miR-708-5p | MIMAT0004828 | Hivep1 | 110521 | HITS-CLIP | 37986224 | Unspecified |
| mmu-miR-708-5p | MIMAT0004828 | Hivep2 | 15273 | HITS-CLIP | 37986224\|23597149 | Unspecified |
| mmu-miR-708-5p | MIMAT0004828 | Hmgcr | 15357 | HITS-CLIP | 37986224\|19536157 | Unspecified |
| mmu-miR-708-5p | MIMAT0004828 | Hmgcs2 | 15360 | HITS-CLIP | 23597149 | Unspecified |
| mmu-miR-708-5p | MIMAT0004828 | Hmgxb3 | 106894 | HITS-CLIP | 37986224 | Unspecified |
| mmu-miR-708-5p | MIMAT0004828 | Hmmr | 15366 | HITS-CLIP | 37986224 | Unspecified |
| mmu-miR-708-5p | MIMAT0004828 | Hnf1b | 21410 | HITS-CLIP | 23597149 | Unspecified |
| mmu-miR-708-5p | MIMAT0004828 | Hnf4a | 15378 | HITS-CLIP | 23597149 | Unspecified |
| mmu-miR-708-5p | MIMAT0004828 | Hnrnpd | 11991 | HITS-CLIP | 19536157 | Unspecified |
| mmu-miR-708-5p | MIMAT0004828 | Hp1bp3 | 15441 | HITS-CLIP | 37986224\|23597149 | Unspecified |

| | | | | | | | |
|---|---|---|---|---|---|---|---|
| mmu-miR-708-5p | MIMAT0004828 | Hpgd | 15446 | HITS-CLIP | | 37986224 | Unspecified |
| mmu-miR-708-5p | MIMAT0004828 | Hs6st1 | 50785 | HITS-CLIP | | 37986224 | Unspecified |
| mmu-miR-708-5p | MIMAT0004828 | Hsd17b14 | 66065 | HITS-CLIP | | 37986224 | Unspecified |
| mmu-miR-708-5p | MIMAT0004828 | Hsd17b4 | 15488 | HITS-CLIP | | 23597149 | Unspecified |
| mmu-miR-708-5p | MIMAT0004828 | Hsf2 | 15500 | HITS-CLIP | | 23597149 | Unspecified |
| mmu-miR-708-5p | MIMAT0004828 | Hsp90ab1 | 15516 | HITS-CLIP | | 23597149 | Unspecified |
| mmu-miR-708-5p | MIMAT0004828 | Hspa4 | 15525 | HITS-CLIP | | 37986224 | Unspecified |
| mmu-miR-708-5p | MIMAT0004828 | Hspa4l | 18415 | HITS-CLIP | | 37986224 | Unspecified |
| mmu-miR-708-5p | MIMAT0004828 | Hspa9 | 15526 | HITS-CLIP | 37986224\|23597149 | | Unspecified |
| mmu-miR-708-5p | MIMAT0004828 | Htr2c | 15560 | HITS-CLIP | | 37986224 | Unspecified |
| mmu-miR-708-5p | MIMAT0004828 | Huwe1 | 59026 | HITS-CLIP | 37986224\|23597149 | | Unspecified |
| mmu-miR-708-5p | MIMAT0004828 | Iars | 105148 | HITS-CLIP | | 37986224 | Unspecified |
| mmu-miR-708-5p | MIMAT0004828 | Ice2 | 93697 | HITS-CLIP | | 23597149 | Unspecified |
| mmu-miR-708-5p | MIMAT0004828 | Idh1 | 15926 | HITS-CLIP | | 23597149 | Unspecified |
| mmu-miR-708-5p | MIMAT0004828 | Idnk | 75731 | HITS-CLIP | | 37986224 | Unspecified |
| mmu-miR-708-5p | MIMAT0004828 | Ido2 | 209176 | HITS-CLIP | 37986224\|23597149 | | Unspecified |
| mmu-miR-708-5p | MIMAT0004828 | Ids | 15931 | HITS-CLIP | | 37986224 | Unspecified |
| mmu-miR-708-5p | MIMAT0004828 | Ifnar1 | 15975 | HITS-CLIP | | 37986224 | Unspecified |
| mmu-miR-708-5p | MIMAT0004828 | Igbp1 | 18518 | HITS-CLIP | | 37986224 | Unspecified |
| mmu-miR-708-5p | MIMAT0004828 | Igf2 | 16002 | HITS-CLIP | | 37986224 | Unspecified |
| mmu-miR-708-5p | MIMAT0004828 | Igf2bp1 | 140486 | HITS-CLIP | | 37986224 | Unspecified |
| mmu-miR-708-5p | MIMAT0004828 | Igf2r | 16004 | HITS-CLIP | | 37986224 | Unspecified |
| mmu-miR-708-5p | MIMAT0004828 | Igfbp5 | 16011 | HITS-CLIP | | 37986224 | Unspecified |
| mmu-miR-708-5p | MIMAT0004828 | Iglon5 | 210094 | HITS-CLIP | | 37986224 | Unspecified |
| mmu-miR-708-5p | MIMAT0004828 | Igsf9b | 235086 | HITS-CLIP | | 37986224 | Unspecified |
| mmu-miR-708-5p | MIMAT0004828 | Ikbkb | 16150 | HITS-CLIP | | 31433990 | Unspecified |
| mmu-miR-708-5p | MIMAT0004828 | Ikzf4 | 22781 | HITS-CLIP | 37986224\|31433990 | | Unspecified |
| mmu-miR-708-5p | MIMAT0004828 | Il4ra | 16190 | HITS-CLIP | | 23597149 | Unspecified |
| mmu-miR-708-5p | MIMAT0004828 | Ildr2 | 100039795 | HITS-CLIP | | 37986224 | Unspecified |
| mmu-miR-708-5p | MIMAT0004828 | Ino80d | 227195 | HITS-CLIP | | 37986224 | Unspecified |
| mmu-miR-708-5p | MIMAT0004828 | Insr | 16337 | HITS-CLIP | | 37986224 | Unspecified |
| mmu-miR-708-5p | MIMAT0004828 | Ints3 | 229543 | HITS-CLIP | | 23597149 | Unspecified |
| mmu-miR-708-5p | MIMAT0004828 | Ints8 | 72656 | HITS-CLIP | | 23597149 | Unspecified |
| mmu-miR-708-5p | MIMAT0004828 | Ip6k1 | 27399 | HITS-CLIP | 19536157\|23597149 | | Unspecified |
| mmu-miR-708-5p | MIMAT0004828 | Ipo5 | 70572 | HITS-CLIP | | 23597149 | Unspecified |
| mmu-miR-708-5p | MIMAT0004828 | Iqgap3 | 404710 | HITS-CLIP | | 23597149 | Unspecified |
| mmu-miR-708-5p | MIMAT0004828 | Irf2 | 16363 | HITS-CLIP | 37986224\|23597149 | | Unspecified |
| mmu-miR-708-5p | MIMAT0004828 | Irgq | 210146 | HITS-CLIP | | 37986224 | Unspecified |
| mmu-miR-708-5p | MIMAT0004828 | Irs1 | 16367 | HITS-CLIP | | 37986224 | Unspecified |

| | | | | | | |
|---|---|---|---|---|---|---|
| mmu-miR-708-5p | MIMAT0004828 | Isca2 | 74316 | HITS-CLIP | 37986224 | Unspecified |
| mmu-miR-708-5p | MIMAT0004828 | Isg20l2 | 229504 | HITS-CLIP | 37986224 | Unspecified |
| mmu-miR-708-5p | MIMAT0004828 | Isoc1 | 66307 | HITS-CLIP | 37986224\|23597149 | Unspecified |
| mmu-miR-708-5p | MIMAT0004828 | Ist1 | 71955 | HITS-CLIP | 37986224 | Unspecified |
| mmu-miR-708-5p | MIMAT0004828 | Itga9 | 104099 | HITS-CLIP | 23597149 | Unspecified |
| mmu-miR-708-5p | MIMAT0004828 | Itpkb | 320404 | HITS-CLIP | 23597149 | Unspecified |
| mmu-miR-708-5p | MIMAT0004828 | Jun | 16476 | HITS-CLIP | 37986224 | Unspecified |
| mmu-miR-708-5p | MIMAT0004828 | Jund | 16478 | HITS-CLIP | 23597149 | Unspecified |
| mmu-miR-708-5p | MIMAT0004828 | Jup | 16480 | HITS-CLIP | 37986224\|23597149 | Unspecified |
| mmu-miR-708-5p | MIMAT0004828 | Kantr | 102723508 | HITS-CLIP | 19536157 | Unspecified |
| mmu-miR-708-5p | MIMAT0004828 | Kat5 | 81601 | HITS-CLIP | 23597149 | Unspecified |
| mmu-miR-708-5p | MIMAT0004828 | Kat7 | 217127 | HITS-CLIP | 37986224 | Unspecified |
| mmu-miR-708-5p | MIMAT0004828 | Kcmf1 | 74287 | HITS-CLIP | 23597149 | Unspecified |
| mmu-miR-708-5p | MIMAT0004828 | Kcna1 | 16485 | HITS-CLIP | 37986224\|19536157 | Unspecified |
| mmu-miR-708-5p | MIMAT0004828 | Kcna2 | 16490 | HITS-CLIP | 19536157 | Unspecified |
| mmu-miR-708-5p | MIMAT0004828 | Kcnc1 | 16502 | HITS-CLIP | 37986224 | Unspecified |
| mmu-miR-708-5p | MIMAT0004828 | Kcnc4 | 99738 | HITS-CLIP | 37986224 | Unspecified |
| mmu-miR-708-5p | MIMAT0004828 | Kcnj10 | 16513 | HITS-CLIP | 37986224 | Unspecified |
| mmu-miR-708-5p | MIMAT0004828 | Kcnk3 | 16527 | HITS-CLIP | 37986224\|19536157 | Unspecified |
| mmu-miR-708-5p | MIMAT0004828 | Kcnma1 | 16531 | HITS-CLIP | 19536157 | Unspecified |
| mmu-miR-708-5p | MIMAT0004828 | Kcnv1 | 67498 | HITS-CLIP | 37986224 | Unspecified |
| mmu-miR-708-5p | MIMAT0004828 | Kctd15 | 233107 | HITS-CLIP | 37986224 | Unspecified |
| mmu-miR-708-5p | MIMAT0004828 | Kctd5 | 69259 | HITS-CLIP | 23597149 | Unspecified |
| mmu-miR-708-5p | MIMAT0004828 | Kdm3b | 277250 | HITS-CLIP | 23597149 | Unspecified |
| mmu-miR-708-5p | MIMAT0004828 | Kdm4b | 193796 | HITS-CLIP | 37986224 | Unspecified |
| mmu-miR-708-5p | MIMAT0004828 | Kdm4c | 76804 | HITS-CLIP | 23597149 | Unspecified |
| mmu-miR-708-5p | MIMAT0004828 | Kdm5c | 20591 | HITS-CLIP | 37986224 | Unspecified |
| mmu-miR-708-5p | MIMAT0004828 | Kdr | 16542 | HITS-CLIP | 37986224 | Unspecified |
| mmu-miR-708-5p | MIMAT0004828 | Kif1a | 16560 | HITS-CLIP | 37986224\|19536157 | Unspecified |
| mmu-miR-708-5p | MIMAT0004828 | Kif1b | 16561 | HITS-CLIP | 37986224\|23597149 | Unspecified |
| mmu-miR-708-5p | MIMAT0004828 | Kif1c | 16562 | HITS-CLIP | 37986224 | Unspecified |
| mmu-miR-708-5p | MIMAT0004828 | Kif21b | 16565 | HITS-CLIP | 37986224 | Unspecified |
| mmu-miR-708-5p | MIMAT0004828 | Kif22 | 110033 | HITS-CLIP | 37986224 | Unspecified |
| mmu-miR-708-5p | MIMAT0004828 | Kif3b | 16569 | HITS-CLIP | 37986224 | Unspecified |
| mmu-miR-708-5p | MIMAT0004828 | Klf3 | 16599 | HITS-CLIP | 37986224 | Unspecified |
| mmu-miR-708-5p | MIMAT0004828 | Klf7 | 93691 | HITS-CLIP | 37986224 | Unspecified |
| mmu-miR-708-5p | MIMAT0004828 | Klhl32 | 212390 | HITS-CLIP | 19536157 | Unspecified |
| mmu-miR-708-5p | MIMAT0004828 | Klhl7 | 52323 | HITS-CLIP | 23597149 | Unspecified |
| mmu-miR-708-5p | MIMAT0004828 | Klhl9 | 242521 | HITS-CLIP | 23597149 | Unspecified |

| | | | | | | |
|---|---|---|---|---|---|---|
| mmu-miR-708-5p | MIMAT0004828 | Kmo | 98256 | HITS-CLIP | 37986224\|23597149 | Unspecified |
| mmu-miR-708-5p | MIMAT0004828 | Kmt5b | 225888 | HITS-CLIP | 19536157 | Unspecified |
| mmu-miR-708-5p | MIMAT0004828 | Kpna4 | 16649 | HITS-CLIP | 23597149 | Unspecified |
| mmu-miR-708-5p | MIMAT0004828 | Kpnb1 | 16211 | HITS-CLIP | 37986224 | Unspecified |
| mmu-miR-708-5p | MIMAT0004828 | Krit1 | 79264 | HITS-CLIP | 23597149 | Unspecified |
| mmu-miR-708-5p | MIMAT0004828 | L1cam | 16728 | HITS-CLIP | 37986224 | Unspecified |
| mmu-miR-708-5p | MIMAT0004828 | Lamp2 | 16784 | HITS-CLIP | 37986224\|23597149\|19536157 | Unspecified |
| mmu-miR-708-5p | MIMAT0004828 | Laptm4a | 17775 | HITS-CLIP | 37986224 | Unspecified |
| mmu-miR-708-5p | MIMAT0004828 | Larp1 | 73158 | HITS-CLIP | 37986224 | Unspecified |
| mmu-miR-708-5p | MIMAT0004828 | Lbp | 16803 | HITS-CLIP | 23597149 | Unspecified |
| mmu-miR-708-5p | MIMAT0004828 | Ldlr | 16835 | HITS-CLIP | 37986224 | Unspecified |
| mmu-miR-708-5p | MIMAT0004828 | Letm1 | 56384 | HITS-CLIP | 37986224 | Unspecified |
| mmu-miR-708-5p | MIMAT0004828 | Lifr | 16880 | HITS-CLIP | 37986224 | Unspecified |
| mmu-miR-708-5p | MIMAT0004828 | Limch1 | 77569 | HITS-CLIP | 37986224 | Unspecified |
| mmu-miR-708-5p | MIMAT0004828 | Lims2 | 225341 | HITS-CLIP | 23597149 | Unspecified |
| mmu-miR-708-5p | MIMAT0004828 | Lin7c | 22343 | HITS-CLIP | 37986224 | Unspecified |
| mmu-miR-708-5p | MIMAT0004828 | Lingo1 | 235402 | HITS-CLIP | 37986224 | Unspecified |
| mmu-miR-708-5p | MIMAT0004828 | Lipc | 15450 | HITS-CLIP | 37986224 | Unspecified |
| mmu-miR-708-5p | MIMAT0004828 | Lmf1 | 76483 | HITS-CLIP | 37986224 | Unspecified |
| mmu-miR-708-5p | MIMAT0004828 | Lmtk2 | 231876 | HITS-CLIP | 37986224 | Unspecified |
| mmu-miR-708-5p | MIMAT0004828 | Lnpep | 240028 | HITS-CLIP | 19536157 | Unspecified |
| mmu-miR-708-5p | MIMAT0004828 | Lonp1 | 74142 | HITS-CLIP | 23597149 | Unspecified |
| mmu-miR-708-5p | MIMAT0004828 | Lpar6 | 67168 | HITS-CLIP | 37986224 | Unspecified |
| mmu-miR-708-5p | MIMAT0004828 | Lpcat1 | 210992 | HITS-CLIP | 23597149 | Unspecified |
| mmu-miR-708-5p | MIMAT0004828 | Lpcat3 | 14792 | HITS-CLIP | 37986224 | Unspecified |
| mmu-miR-708-5p | MIMAT0004828 | Lpin1 | 14245 | HITS-CLIP | 37986224 | Unspecified |
| mmu-miR-708-5p | MIMAT0004828 | Lpin2 | 64898 | HITS-CLIP | 37986224\|23597149 | Unspecified |
| mmu-miR-708-5p | MIMAT0004828 | Lrig2 | 269473 | HITS-CLIP | 37986224 | Unspecified |
| mmu-miR-708-5p | MIMAT0004828 | Lrp1 | 16971 | HITS-CLIP | 23597149 | Unspecified |
| mmu-miR-708-5p | MIMAT0004828 | Lrrc28 | 67867 | HITS-CLIP | 23597149 | Unspecified |
| mmu-miR-708-5p | MIMAT0004828 | Lrrc58 | 320184 | HITS-CLIP | 37986224\|23597149 | Unspecified |
| mmu-miR-708-5p | MIMAT0004828 | Lrrfip1 | 16978 | HITS-CLIP | 37986224 | Unspecified |
| mmu-miR-708-5p | MIMAT0004828 | Lrrn1 | 16979 | HITS-CLIP | 37986224 | Unspecified |
| mmu-miR-708-5p | MIMAT0004828 | Luc7l | 66978 | HITS-CLIP | 37986224 | Unspecified |
| mmu-miR-708-5p | MIMAT0004828 | Luc7l3 | 67684 | HITS-CLIP | 37986224 | Unspecified |
| mmu-miR-708-5p | MIMAT0004828 | Luzp1 | 269593 | HITS-CLIP | 37986224\|19536157 | Unspecified |
| mmu-miR-708-5p | MIMAT0004828 | Ly6e | 17069 | HITS-CLIP | 23597149 | Unspecified |
| mmu-miR-708-5p | MIMAT0004828 | Lypla1 | 18777 | HITS-CLIP | 23597149\|19536157 | Unspecified |
| mmu-miR-708-5p | MIMAT0004828 | Man1a | 17155 | HITS-CLIP | 23597149 | Unspecified |

| | | | | | | | |
|---|---|---|---|---|---|---|---|
| mmu-miR-708-5p | MIMAT0004828 | Man2a1 | 17158 | HITS-CLIP | | 23597149 | Unspecified |
| mmu-miR-708-5p | MIMAT0004828 | Maneal | 215090 | HITS-CLIP | | 37986224 | Unspecified |
| mmu-miR-708-5p | MIMAT0004828 | Map1b | 17755 | HITS-CLIP | 37986224\|19536157 | | Unspecified |
| mmu-miR-708-5p | MIMAT0004828 | Map2 | 17756 | HITS-CLIP | 37986224\|19536157 | | Unspecified |
| mmu-miR-708-5p | MIMAT0004828 | Map3k12 | 26404 | HITS-CLIP | | 37986224 | Unspecified |
| mmu-miR-708-5p | MIMAT0004828 | Map3k9 | 338372 | HITS-CLIP | | 37986224 | Unspecified |
| mmu-miR-708-5p | MIMAT0004828 | Map6 | 17760 | HITS-CLIP | | 37986224 | Unspecified |
| mmu-miR-708-5p | MIMAT0004828 | Mapk10 | 26414 | HITS-CLIP | | 37986224 | Unspecified |
| mmu-miR-708-5p | MIMAT0004828 | Mapk14 | 26416 | HITS-CLIP | 37986224\|23597149 | | Unspecified |
| mmu-miR-708-5p | MIMAT0004828 | Mapk3 | 26417 | HITS-CLIP | | 23597149 | Unspecified |
| mmu-miR-708-5p | MIMAT0004828 | Mapk4 | 225724 | HITS-CLIP | | 37986224 | Unspecified |
| mmu-miR-708-5p | MIMAT0004828 | Mapk9 | 26420 | HITS-CLIP | 37986224\|19536157 | | Unspecified |
| mmu-miR-708-5p | MIMAT0004828 | Marc2 | 67247 | HITS-CLIP | | 23597149 | Unspecified |
| mmu-miR-708-5p | MIMAT0004828 | Marchf6 | 223455 | HITS-CLIP | 37986224\|19536157\|23597149 | | Unspecified |
| mmu-miR-708-5p | MIMAT0004828 | Marf1 | 223989 | HITS-CLIP | 37986224\|23597149 | | Unspecified |
| mmu-miR-708-5p | MIMAT0004828 | Mars2 | 212679 | HITS-CLIP | | 37986224 | Unspecified |
| mmu-miR-708-5p | MIMAT0004828 | Matr3 | 17184 | HITS-CLIP | | 37986224 | Unspecified |
| mmu-miR-708-5p | MIMAT0004828 | Maz | 17188 | HITS-CLIP | | 37986224 | Unspecified |
| mmu-miR-708-5p | MIMAT0004828 | Mbd6 | 110962 | HITS-CLIP | | 37986224 | Unspecified |
| mmu-miR-708-5p | MIMAT0004828 | Mbp | 17196 | HITS-CLIP | | 37986224 | Unspecified |
| mmu-miR-708-5p | MIMAT0004828 | Mcfd2 | 193813 | HITS-CLIP | 23597149\|19536157 | | Unspecified |
| mmu-miR-708-5p | MIMAT0004828 | Mdh2 | 17448 | HITS-CLIP | | 37986224 | Unspecified |
| mmu-miR-708-5p | MIMAT0004828 | Med1 | 19014 | HITS-CLIP | 37986224\|19536157 | | Unspecified |
| mmu-miR-708-5p | MIMAT0004828 | Med13 | 327987 | HITS-CLIP | | 19536157 | Unspecified |
| mmu-miR-708-5p | MIMAT0004828 | Med20 | 56771 | HITS-CLIP | | 37986224 | Unspecified |
| mmu-miR-708-5p | MIMAT0004828 | Med28 | 66999 | HITS-CLIP | | 37986224 | Unspecified |
| mmu-miR-708-5p | MIMAT0004828 | Mepce | 231803 | HITS-CLIP | | 37986224 | Unspecified |
| mmu-miR-708-5p | MIMAT0004828 | Mertk | 17289 | HITS-CLIP | | 37986224 | Unspecified |
| mmu-miR-708-5p | MIMAT0004828 | Mettl7b | 71664 | HITS-CLIP | 37986224\|23597149 | | Unspecified |
| mmu-miR-708-5p | MIMAT0004828 | Mfap3l | 71306 | HITS-CLIP | | 37986224 | Unspecified |
| mmu-miR-708-5p | MIMAT0004828 | Mfrp | 259172 | HITS-CLIP | | 37986224 | Unspecified |
| mmu-miR-708-5p | MIMAT0004828 | Mfsd11 | 69900 | HITS-CLIP | | 37986224 | Unspecified |
| mmu-miR-708-5p | MIMAT0004828 | Mgam | 232714 | HITS-CLIP | | 37986224 | Unspecified |
| mmu-miR-708-5p | MIMAT0004828 | Mgat3 | 17309 | HITS-CLIP | | 37986224 | Unspecified |
| mmu-miR-708-5p | MIMAT0004828 | Mgp | 17313 | HITS-CLIP | | 37986224 | Unspecified |
| mmu-miR-708-5p | MIMAT0004828 | Mia2 | 338320 | HITS-CLIP | | 23597149 | Unspecified |
| mmu-miR-708-5p | MIMAT0004828 | Mib1 | 225164 | HITS-CLIP | | 37986224 | Unspecified |
| mmu-miR-708-5p | MIMAT0004828 | Mical3 | 194401 | HITS-CLIP | 37986224\|23597149 | | Unspecified |
| mmu-miR-708-5p | MIMAT0004828 | Mid2 | 23947 | HITS-CLIP | | 37986224 | Unspecified |

| | | | | | | | |
|---|---|---|---|---|---|---|---|
| mmu-miR-708-5p | MIMAT0004828 | Mier3 | 218613 | HITS-CLIP | | 37986224 | Unspecified |
| mmu-miR-708-5p | MIMAT0004828 | Mindy3 | 80013 | HITS-CLIP | | 23597149 | Unspecified |
| mmu-miR-708-5p | MIMAT0004828 | Minpp1 | 17330 | HITS-CLIP | | 19536157 | Unspecified |
| mmu-miR-708-5p | MIMAT0004828 | Mki67 | 17345 | HITS-CLIP | | 37986224 | Unspecified |
| mmu-miR-708-5p | MIMAT0004828 | Mknk2 | 17347 | HITS-CLIP | | 37986224 | Unspecified |
| mmu-miR-708-5p | MIMAT0004828 | Mlxip | 208104 | HITS-CLIP | | 37986224 | Unspecified |
| mmu-miR-708-5p | MIMAT0004828 | Mmaa | 109136 | HITS-CLIP | | 23597149 | Unspecified |
| mmu-miR-708-5p | MIMAT0004828 | Mmadhc | 109129 | HITS-CLIP | | 23597149 | Unspecified |
| mmu-miR-708-5p | MIMAT0004828 | Morf4l2 | 56397 | HITS-CLIP | | 23597149 | Unspecified |
| mmu-miR-708-5p | MIMAT0004828 | Mpp1 | 17524 | HITS-CLIP | | 37986224 | Unspecified |
| mmu-miR-708-5p | MIMAT0004828 | Mpst | 246221 | HITS-CLIP | 37986224\|23597149 | | Unspecified |
| mmu-miR-708-5p | MIMAT0004828 | Msl1 | 74026 | HITS-CLIP | | 37986224 | Unspecified |
| mmu-miR-708-5p | MIMAT0004828 | Msn | 17698 | HITS-CLIP | | 23597149 | Unspecified |
| mmu-miR-708-5p | MIMAT0004828 | Msr1 | 20288 | HITS-CLIP | | 37986224 | Unspecified |
| mmu-miR-708-5p | MIMAT0004828 | Msra | 110265 | HITS-CLIP | | 37986224 | Unspecified |
| mmu-miR-708-5p | MIMAT0004828 | Msrb1 | 27361 | HITS-CLIP | | 37986224 | Unspecified |
| mmu-miR-708-5p | MIMAT0004828 | Mtch2 | 56428 | HITS-CLIP | | 37986224 | Unspecified |
| mmu-miR-708-5p | MIMAT0004828 | Mtdh | 67154 | HITS-CLIP | | 19536157 | Unspecified |
| mmu-miR-708-5p | MIMAT0004828 | Mtmr3 | 74302 | HITS-CLIP | 37986224\|23597149 | | Unspecified |
| mmu-miR-708-5p | MIMAT0004828 | Mtpap | 67440 | HITS-CLIP | | 23597149 | Unspecified |
| mmu-miR-708-5p | MIMAT0004828 | Mtpn | 14489 | HITS-CLIP | | 37986224 | Unspecified |
| mmu-miR-708-5p | MIMAT0004828 | Mycbp2 | 105689 | HITS-CLIP | | 19536157 | Unspecified |
| mmu-miR-708-5p | MIMAT0004828 | Myo18a | 360013 | HITS-CLIP | | 23597149 | Unspecified |
| mmu-miR-708-5p | MIMAT0004828 | Myo1e | 71602 | HITS-CLIP | | 19536157 | Unspecified |
| mmu-miR-708-5p | MIMAT0004828 | Myt1 | 17932 | HITS-CLIP | | 37986224 | Unspecified |
| mmu-miR-708-5p | MIMAT0004828 | Myt1l | 17933 | HITS-CLIP | | 19536157 | Unspecified |
| mmu-miR-708-5p | MIMAT0004828 | N4bp2l2 | 381695 | HITS-CLIP | | 37986224 | Unspecified |
| mmu-miR-708-5p | MIMAT0004828 | Naa25 | 231713 | HITS-CLIP | | 37986224 | Unspecified |
| mmu-miR-708-5p | MIMAT0004828 | Nampt | 59027 | HITS-CLIP | 37986224\|23597149 | | Unspecified |
| mmu-miR-708-5p | MIMAT0004828 | Nbea | 26422 | HITS-CLIP | | 37986224 | Unspecified |
| mmu-miR-708-5p | MIMAT0004828 | Nbeal1 | 269198 | HITS-CLIP | | 37986224 | Unspecified |
| mmu-miR-708-5p | MIMAT0004828 | Nbr1 | 17966 | HITS-CLIP | | 23597149 | Unspecified |
| mmu-miR-708-5p | MIMAT0004828 | Ncam1 | 17967 | HITS-CLIP | | 37986224 | Unspecified |
| mmu-miR-708-5p | MIMAT0004828 | Ncapd3 | 78658 | HITS-CLIP | | 37986224 | Unspecified |
| mmu-miR-708-5p | MIMAT0004828 | Ncbp1 | 433702 | HITS-CLIP | | 37986224 | Unspecified |
| mmu-miR-708-5p | MIMAT0004828 | Nckap1 | 50884 | HITS-CLIP | | 23597149 | Unspecified |
| mmu-miR-708-5p | MIMAT0004828 | Ncl | 17975 | HITS-CLIP | | 23597149 | Unspecified |
| mmu-miR-708-5p | MIMAT0004828 | Ncstn | 59287 | HITS-CLIP | | 37986224 | Unspecified |
| mmu-miR-708-5p | MIMAT0004828 | Ndfip1 | 65113 | HITS-CLIP | | 37986224 | Unspecified |

| | | | | | | |
|---|---|---|---|---|---|---|
| mmu-miR-708-5p | MIMAT0004828 | Ndrg2 | 29811 | HITS-CLIP | 37986224\|19536157 | Unspecified |
| mmu-miR-708-5p | MIMAT0004828 | Ndufs2 | 226646 | HITS-CLIP | 37986224 | Unspecified |
| mmu-miR-708-5p | MIMAT0004828 | Nedd4 | 17999 | HITS-CLIP | 37986224 | Unspecified |
| mmu-miR-708-5p | MIMAT0004828 | Nelfa | 24116 | HITS-CLIP | 23597149 | Unspecified |
| mmu-miR-708-5p | MIMAT0004828 | Neo1 | 18007 | HITS-CLIP | 37986224 | Unspecified |
| mmu-miR-708-5p | MIMAT0004828 | Net1 | 56349 | HITS-CLIP | 23597149 | Unspecified |
| mmu-miR-708-5p | MIMAT0004828 | Neto1 | 246317 | HITS-CLIP | 19536157 | Unspecified |
| mmu-miR-708-5p | MIMAT0004828 | Neurod1 | 18012 | HITS-CLIP | 37986224 | Unspecified |
| mmu-miR-708-5p | MIMAT0004828 | Nfasc | 269116 | HITS-CLIP | 19536157 | Unspecified |
| mmu-miR-708-5p | MIMAT0004828 | Nfat5 | 54446 | HITS-CLIP | 37986224 | Unspecified |
| mmu-miR-708-5p | MIMAT0004828 | Nfia | 18027 | HITS-CLIP | 37986224 | Unspecified |
| mmu-miR-708-5p | MIMAT0004828 | Nfib | 18028 | HITS-CLIP | 19536157 | Unspecified |
| mmu-miR-708-5p | MIMAT0004828 | Nfix | 18032 | HITS-CLIP | 37986224 | Unspecified |
| mmu-miR-708-5p | MIMAT0004828 | Nipa1 | 233280 | HITS-CLIP | 37986224\|19536157 | Unspecified |
| mmu-miR-708-5p | MIMAT0004828 | Nisch | 64652 | HITS-CLIP | 37986224 | Unspecified |
| mmu-miR-708-5p | MIMAT0004828 | Nkiras1 | 69721 | HITS-CLIP | 23597149 | Unspecified |
| mmu-miR-708-5p | MIMAT0004828 | Nln | 75805 | HITS-CLIP | 23597149 | Unspecified |
| mmu-miR-708-5p | MIMAT0004828 | Nmi | 64685 | HITS-CLIP | 37986224 | Unspecified |
| mmu-miR-708-5p | MIMAT0004828 | Nnat | 18111 | HITS-CLIP | 37986224\|19536157 | Unspecified |
| mmu-miR-708-5p | MIMAT0004828 | Noct | 12457 | HITS-CLIP | 19536157 | Unspecified |
| mmu-miR-708-5p | MIMAT0004828 | Nol7 | 70078 | HITS-CLIP | 23597149 | Unspecified |
| mmu-miR-708-5p | MIMAT0004828 | Nos1ap | 70729 | HITS-CLIP | 37986224\|23597149 | Unspecified |
| mmu-miR-708-5p | MIMAT0004828 | Nos3 | 18127 | HITS-CLIP | 23597149 | Unspecified |
| mmu-miR-708-5p | MIMAT0004828 | Nostrin | 329416 | HITS-CLIP | 37986224 | Unspecified |
| mmu-miR-708-5p | MIMAT0004828 | Notch2 | 18129 | HITS-CLIP | 37986224\|23597149 | Unspecified |
| mmu-miR-708-5p | MIMAT0004828 | Nova1 | 664883 | HITS-CLIP | 37986224 | Unspecified |
| mmu-miR-708-5p | MIMAT0004828 | Nova2 | 384569 | HITS-CLIP | 37986224 | Unspecified |
| mmu-miR-708-5p | MIMAT0004828 | Npat | 244879 | HITS-CLIP | 37986224 | Unspecified |
| mmu-miR-708-5p | MIMAT0004828 | Nptx1 | 18164 | HITS-CLIP | 37986224 | Unspecified |
| mmu-miR-708-5p | MIMAT0004828 | Nr1d1 | 217166 | HITS-CLIP | 37986224 | Unspecified |
| mmu-miR-708-5p | MIMAT0004828 | Nr1h3 | 22259 | HITS-CLIP | 23597149 | Unspecified |
| mmu-miR-708-5p | MIMAT0004828 | Nr3c2 | 110784 | HITS-CLIP | 23597149 | Unspecified |
| mmu-miR-708-5p | MIMAT0004828 | Nr4a1 | 15370 | HITS-CLIP | 37986224 | Unspecified |
| mmu-miR-708-5p | MIMAT0004828 | Nr6a1 | 14536 | HITS-CLIP | 26203562 | Unspecified |
| mmu-miR-708-5p | MIMAT0004828 | Nras | 18176 | HITS-CLIP | 37986224 | Unspecified |
| mmu-miR-708-5p | MIMAT0004828 | Nrbp1 | 192292 | HITS-CLIP | 19536157 | Unspecified |
| mmu-miR-708-5p | MIMAT0004828 | Nrgn | 64011 | HITS-CLIP | 37986224 | Unspecified |
| mmu-miR-708-5p | MIMAT0004828 | Nrxn1 | 18189 | HITS-CLIP | 37986224 | Unspecified |
| mmu-miR-708-5p | MIMAT0004828 | Nrxn3 | 18191 | HITS-CLIP | 37986224 | Unspecified |

| | | | | | | |
|---|---|---|---|---|---|---|
| mmu-miR-708-5p | MIMAT0004828 | Nsd1 | 18193 | HITS-CLIP | 37986224 | Unspecified |
| mmu-miR-708-5p | MIMAT0004828 | Nsg1 | 18196 | HITS-CLIP | 37986224 | Unspecified |
| mmu-miR-708-5p | MIMAT0004828 | Nsrp1 | 237859 | HITS-CLIP | 23597149 | Unspecified |
| mmu-miR-708-5p | MIMAT0004828 | Nt5dc3 | 103466 | HITS-CLIP | 37986224 | Unspecified |
| mmu-miR-708-5p | MIMAT0004828 | Ntrk2 | 18212 | HITS-CLIP | 19536157 | Unspecified |
| mmu-miR-708-5p | MIMAT0004828 | Nucks1 | 98415 | HITS-CLIP | 37986224 | Unspecified |
| mmu-miR-708-5p | MIMAT0004828 | Nup85 | 445007 | HITS-CLIP | 37986224 | Unspecified |
| mmu-miR-708-5p | MIMAT0004828 | Nus1 | 52014 | HITS-CLIP | 37986224 | Unspecified |
| mmu-miR-708-5p | MIMAT0004828 | Nxpe4 | 244853 | HITS-CLIP | 37986224 | Unspecified |
| mmu-miR-708-5p | MIMAT0004828 | Oaz2 | 18247 | HITS-CLIP | 37986224 | Unspecified |
| mmu-miR-708-5p | MIMAT0004828 | Onecut1 | 15379 | HITS-CLIP | 37986224 | Unspecified |
| mmu-miR-708-5p | MIMAT0004828 | Opcml | 330908 | HITS-CLIP | 37986224 | Unspecified |
| mmu-miR-708-5p | MIMAT0004828 | Oprl1 | 18389 | HITS-CLIP | 37986224 | Unspecified |
| mmu-miR-708-5p | MIMAT0004828 | Os9 | 216440 | HITS-CLIP | 37986224 | Unspecified |
| mmu-miR-708-5p | MIMAT0004828 | Osbpl11 | 106326 | HITS-CLIP | 37986224 | Unspecified |
| mmu-miR-708-5p | MIMAT0004828 | Osbpl6 | 99031 | HITS-CLIP | 19536157 | Unspecified |
| mmu-miR-708-5p | MIMAT0004828 | Osgin1 | 71839 | HITS-CLIP | 23597149 | Unspecified |
| mmu-miR-708-5p | MIMAT0004828 | Pabpc1 | 18458 | HITS-CLIP | 23597149 | Unspecified |
| mmu-miR-708-5p | MIMAT0004828 | Pag1 | 94212 | HITS-CLIP | 37986224 | Unspecified |
| mmu-miR-708-5p | MIMAT0004828 | Paip1 | 218693 | HITS-CLIP | 37986224\|23597149 | Unspecified |
| mmu-miR-708-5p | MIMAT0004828 | Pak2 | 224105 | HITS-CLIP | 37986224 | Unspecified |
| mmu-miR-708-5p | MIMAT0004828 | Paqr9 | 75552 | HITS-CLIP | 19536157 | Unspecified |
| mmu-miR-708-5p | MIMAT0004828 | Pbrm1 | 66923 | HITS-CLIP | 19536157 | Unspecified |
| mmu-miR-708-5p | MIMAT0004828 | Pccb | 66904 | HITS-CLIP | 23597149 | Unspecified |
| mmu-miR-708-5p | MIMAT0004828 | Pcdh15 | 11994 | HITS-CLIP | 37986224 | Unspecified |
| mmu-miR-708-5p | MIMAT0004828 | Pcnx | 54604 | HITS-CLIP | 37986224 | Unspecified |
| mmu-miR-708-5p | MIMAT0004828 | Pcnx3 | 399909 | HITS-CLIP | 23597149 | Unspecified |
| mmu-miR-708-5p | MIMAT0004828 | Pctp | 18559 | HITS-CLIP | 23597149 | Unspecified |
| mmu-miR-708-5p | MIMAT0004828 | Pcx | 18563 | HITS-CLIP | 23597149 | Unspecified |
| mmu-miR-708-5p | MIMAT0004828 | Pdcd4 | 18569 | HITS-CLIP | 37986224\|26203562 | Unspecified |
| mmu-miR-708-5p | MIMAT0004828 | Pdgfb | 18591 | HITS-CLIP | 37986224 | Unspecified |
| mmu-miR-708-5p | MIMAT0004828 | Pdia3 | 14827 | HITS-CLIP | 23597149 | Unspecified |
| mmu-miR-708-5p | MIMAT0004828 | Pdia5 | 72599 | HITS-CLIP | 23597149 | Unspecified |
| mmu-miR-708-5p | MIMAT0004828 | Pdia6 | 71853 | HITS-CLIP | 23597149 | Unspecified |
| mmu-miR-708-5p | MIMAT0004828 | Pdk2 | 18604 | HITS-CLIP | 23597149 | Unspecified |
| mmu-miR-708-5p | MIMAT0004828 | Pdlim5 | 56376 | HITS-CLIP | 37986224\|23597149 | Unspecified |
| mmu-miR-708-5p | MIMAT0004828 | Pdyn | 18610 | HITS-CLIP | 37986224 | Unspecified |
| mmu-miR-708-5p | MIMAT0004828 | Pdzd8 | 107368 | HITS-CLIP | 37986224 | Unspecified |
| mmu-miR-708-5p | MIMAT0004828 | Pef1 | 67898 | HITS-CLIP | 23597149 | Unspecified |

| | | | | | | |
|---|---|---|---|---|---|---|
| mmu-miR-708-5p | MIMAT0004828 | Peg3 | 18616 | HITS-CLIP | 37986224 | Unspecified |
| mmu-miR-708-5p | MIMAT0004828 | Pelp1 | 75273 | HITS-CLIP | 37986224 | Unspecified |
| mmu-miR-708-5p | MIMAT0004828 | Pepd | 18624 | HITS-CLIP | 23597149 | Unspecified |
| mmu-miR-708-5p | MIMAT0004828 | Pex12 | 103737 | HITS-CLIP | 23597149 | Unspecified |
| mmu-miR-708-5p | MIMAT0004828 | Pex5 | 19305 | HITS-CLIP | 23597149 | Unspecified |
| mmu-miR-708-5p | MIMAT0004828 | Pex6 | 224824 | HITS-CLIP | 23597149 | Unspecified |
| mmu-miR-708-5p | MIMAT0004828 | Pgrmc1 | 53328 | HITS-CLIP | 23597149 | Unspecified |
| mmu-miR-708-5p | MIMAT0004828 | Pgrmc2 | 70804 | HITS-CLIP | 23597149 | Unspecified |
| mmu-miR-708-5p | MIMAT0004828 | Pgs1 | 74451 | HITS-CLIP | 31433990 | Unspecified |
| mmu-miR-708-5p | MIMAT0004828 | Phactr4 | 100169 | HITS-CLIP | 37986224 | Unspecified |
| mmu-miR-708-5p | MIMAT0004828 | Phf3 | 213109 | HITS-CLIP | 37986224 | Unspecified |
| mmu-miR-708-5p | MIMAT0004828 | Phldb2 | 208177 | HITS-CLIP | 23597149 | Unspecified |
| mmu-miR-708-5p | MIMAT0004828 | Phlpp1 | 98432 | HITS-CLIP | 23597149 | Unspecified |
| mmu-miR-708-5p | MIMAT0004828 | Phtf2 | 68770 | HITS-CLIP | 37986224 | Unspecified |
| mmu-miR-708-5p | MIMAT0004828 | Phyhip | 105653 | HITS-CLIP | 37986224 | Unspecified |
| mmu-miR-708-5p | MIMAT0004828 | Pias2 | 17344 | HITS-CLIP | 37986224 | Unspecified |
| mmu-miR-708-5p | MIMAT0004828 | Pias4 | 59004 | HITS-CLIP | 23597149 | Unspecified |
| mmu-miR-708-5p | MIMAT0004828 | Pigp | 56176 | HITS-CLIP | 23597149 | Unspecified |
| mmu-miR-708-5p | MIMAT0004828 | Pigs | 276846 | HITS-CLIP | 23597149 | Unspecified |
| mmu-miR-708-5p | MIMAT0004828 | Pik3r1 | 18708 | HITS-CLIP | 19536157 | Unspecified |
| mmu-miR-708-5p | MIMAT0004828 | Pithd1 | 66193 | HITS-CLIP | 37986224 | Unspecified |
| mmu-miR-708-5p | MIMAT0004828 | Pitpna | 18738 | HITS-CLIP | 37986224 | Unspecified |
| mmu-miR-708-5p | MIMAT0004828 | Plbd2 | 71772 | HITS-CLIP | 37986224 | Unspecified |
| mmu-miR-708-5p | MIMAT0004828 | Plec | 18810 | HITS-CLIP | 23597149 | Unspecified |
| mmu-miR-708-5p | MIMAT0004828 | Plp1 | 18823 | HITS-CLIP | 19536157 | Unspecified |
| mmu-miR-708-5p | MIMAT0004828 | Pls3 | 102866 | HITS-CLIP | 23597149\|19536157 | Unspecified |
| mmu-miR-708-5p | MIMAT0004828 | Plxna4 | 243743 | HITS-CLIP | 37986224 | Unspecified |
| mmu-miR-708-5p | MIMAT0004828 | Plxnb1 | 235611 | HITS-CLIP | 23597149 | Unspecified |
| mmu-miR-708-5p | MIMAT0004828 | Pnkd | 56695 | HITS-CLIP | 37986224 | Unspecified |
| mmu-miR-708-5p | MIMAT0004828 | Pnmal2 | 434128 | HITS-CLIP | 37986224 | Unspecified |
| mmu-miR-708-5p | MIMAT0004828 | Pnpla6 | 50767 | HITS-CLIP | 19536157 | Unspecified |
| mmu-miR-708-5p | MIMAT0004828 | Podxl | 27205 | HITS-CLIP | 37986224 | Unspecified |
| mmu-miR-708-5p | MIMAT0004828 | Pofut1 | 140484 | HITS-CLIP | 37986224 | Unspecified |
| mmu-miR-708-5p | MIMAT0004828 | Poldip2 | 67811 | HITS-CLIP | 23597149 | Unspecified |
| mmu-miR-708-5p | MIMAT0004828 | Postn | 50706 | HITS-CLIP | 23597149 | Unspecified |
| mmu-miR-708-5p | MIMAT0004828 | Ppara | 19013 | HITS-CLIP | 37986224\|23597149 | Unspecified |
| mmu-miR-708-5p | MIMAT0004828 | Ppargc1b | 170826 | HITS-CLIP | 23597149 | Unspecified |
| mmu-miR-708-5p | MIMAT0004828 | Ppm1k | 243382 | HITS-CLIP | 37986224 | Unspecified |
| mmu-miR-708-5p | MIMAT0004828 | Ppm1l | 242083 | HITS-CLIP | 37986224 | Unspecified |

| | | | | | | | |
|---|---|---|---|---|---|---|---|
| mmu-miR-708-5p | MIMAT0004828 | Ppp1r12a | 17931 | HITS-CLIP | | 37986224 | Unspecified |
| mmu-miR-708-5p | MIMAT0004828 | Ppp1r3d | 228966 | HITS-CLIP | | 37986224 | Unspecified |
| mmu-miR-708-5p | MIMAT0004828 | Ppp1r7 | 66385 | HITS-CLIP | | 37986224 | Unspecified |
| mmu-miR-708-5p | MIMAT0004828 | Ppp2ca | 19052 | HITS-CLIP | | 23597149 | Unspecified |
| mmu-miR-708-5p | MIMAT0004828 | Ppp2r2a | 71978 | HITS-CLIP | | 23597149 | Unspecified |
| mmu-miR-708-5p | MIMAT0004828 | Ppp3ca | 19055 | HITS-CLIP | | 19536157 | Unspecified |
| mmu-miR-708-5p | MIMAT0004828 | Ppp3cb | 19056 | HITS-CLIP | | 19536157 | Unspecified |
| mmu-miR-708-5p | MIMAT0004828 | Ppp6c | 67857 | HITS-CLIP | | 37986224 | Unspecified |
| mmu-miR-708-5p | MIMAT0004828 | Ppp6r1 | 243819 | HITS-CLIP | | 23597149 | Unspecified |
| mmu-miR-708-5p | MIMAT0004828 | Ppp6r2 | 71474 | HITS-CLIP | | 37986224 | Unspecified |
| mmu-miR-708-5p | MIMAT0004828 | Pqlc1 | 66943 | HITS-CLIP | | 37986224 | Unspecified |
| mmu-miR-708-5p | MIMAT0004828 | Pramef8 | 242736 | HITS-CLIP | | 37986224 | Unspecified |
| mmu-miR-708-5p | MIMAT0004828 | Prkaa1 | 105787 | HITS-CLIP | | 37986224 | Unspecified |
| mmu-miR-708-5p | MIMAT0004828 | Prkag2 | 108099 | HITS-CLIP | 37986224\|23597149 | | Unspecified |
| mmu-miR-708-5p | MIMAT0004828 | Prkar2a | 19087 | HITS-CLIP | | 37986224 | Unspecified |
| mmu-miR-708-5p | MIMAT0004828 | Prkcb | 18751 | HITS-CLIP | | 19536157 | Unspecified |
| mmu-miR-708-5p | MIMAT0004828 | Prmt3 | 71974 | HITS-CLIP | | 37986224 | Unspecified |
| mmu-miR-708-5p | MIMAT0004828 | Pros1 | 19128 | HITS-CLIP | | 23597149 | Unspecified |
| mmu-miR-708-5p | MIMAT0004828 | Prox1 | 19130 | HITS-CLIP | | 23597149 | Unspecified |
| mmu-miR-708-5p | MIMAT0004828 | Prpf8 | 192159 | HITS-CLIP | | 23597149 | Unspecified |
| mmu-miR-708-5p | MIMAT0004828 | Prps1 | 19139 | HITS-CLIP | | 37986224 | Unspecified |
| mmu-miR-708-5p | MIMAT0004828 | Prrc2b | 227723 | HITS-CLIP | | 37986224 | Unspecified |
| mmu-miR-708-5p | MIMAT0004828 | Prrc2c | 226562 | HITS-CLIP | | 19536157 | Unspecified |
| mmu-miR-708-5p | MIMAT0004828 | Pskh1 | 244631 | HITS-CLIP | | 37986224 | Unspecified |
| mmu-miR-708-5p | MIMAT0004828 | Psma3 | 19167 | HITS-CLIP | | 37986224 | Unspecified |
| mmu-miR-708-5p | MIMAT0004828 | Psma5 | 26442 | HITS-CLIP | 37986224\|23597149 | | Unspecified |
| mmu-miR-708-5p | MIMAT0004828 | Psmb2 | 26445 | HITS-CLIP | 37986224\|23597149 | | Unspecified |
| mmu-miR-708-5p | MIMAT0004828 | Psmc5 | 19184 | HITS-CLIP | | 23597149 | Unspecified |
| mmu-miR-708-5p | MIMAT0004828 | Psmd13 | 23997 | HITS-CLIP | | 19536157 | Unspecified |
| mmu-miR-708-5p | MIMAT0004828 | Psme3 | 19192 | HITS-CLIP | | 31433990 | Unspecified |
| mmu-miR-708-5p | MIMAT0004828 | Psme4 | 103554 | HITS-CLIP | | 37986224 | Unspecified |
| mmu-miR-708-5p | MIMAT0004828 | Ptcd3 | 69956 | HITS-CLIP | | 37986224 | Unspecified |
| mmu-miR-708-5p | MIMAT0004828 | Pten | 19211 | HITS-CLIP | 37986224\|23597149 | | Unspecified |
| mmu-miR-708-5p | MIMAT0004828 | Ptk2 | 14083 | HITS-CLIP | | 23597149 | Unspecified |
| mmu-miR-708-5p | MIMAT0004828 | Ptpn1 | 19246 | HITS-CLIP | | 37986224 | Unspecified |
| mmu-miR-708-5p | MIMAT0004828 | Ptpn4 | 19258 | HITS-CLIP | | 37986224 | Unspecified |
| mmu-miR-708-5p | MIMAT0004828 | Ptprb | 19263 | HITS-CLIP | | 23597149 | Unspecified |
| mmu-miR-708-5p | MIMAT0004828 | Ptprf | 19268 | HITS-CLIP | | 37986224 | Unspecified |
| mmu-miR-708-5p | MIMAT0004828 | Ptprk | 19272 | HITS-CLIP | | 19536157 | Unspecified |

| | | | | | | |
|---|---|---|---|---|---|---|
| mmu-miR-708-5p | MIMAT0004828 | Ptprz1 | 19283 | HITS-CLIP | 37986224\|19536157 | Unspecified |
| mmu-miR-708-5p | MIMAT0004828 | Pum1 | 80912 | HITS-CLIP | 23597149 | Unspecified |
| mmu-miR-708-5p | MIMAT0004828 | Pum2 | 80913 | HITS-CLIP | 23597149 | Unspecified |
| mmu-miR-708-5p | MIMAT0004828 | Pura | 19290 | HITS-CLIP | 23597149 | Unspecified |
| mmu-miR-708-5p | MIMAT0004828 | Purb | 19291 | HITS-CLIP | 37986224 | Unspecified |
| mmu-miR-708-5p | MIMAT0004828 | Pygb | 110078 | HITS-CLIP | 37986224\|19536157 | Unspecified |
| mmu-miR-708-5p | MIMAT0004828 | Pygl | 110095 | HITS-CLIP | 23597149 | Unspecified |
| mmu-miR-708-5p | MIMAT0004828 | Pyurf | 66459 | HITS-CLIP | 23597149 | Unspecified |
| mmu-miR-708-5p | MIMAT0004828 | Qdpr | 110391 | HITS-CLIP | 23597149 | Unspecified |
| mmu-miR-708-5p | MIMAT0004828 | Qk | 19317 | HITS-CLIP | 37986224\|19536157 | Unspecified |
| mmu-miR-708-5p | MIMAT0004828 | R3hdm2 | 71750 | HITS-CLIP | 37986224 | Unspecified |
| mmu-miR-708-5p | MIMAT0004828 | R3hdm4 | 109284 | HITS-CLIP | 37986224 | Unspecified |
| mmu-miR-708-5p | MIMAT0004828 | Rab11a | 53869 | HITS-CLIP | 23597149 | Unspecified |
| mmu-miR-708-5p | MIMAT0004828 | Rab11fip1 | 75767 | HITS-CLIP | 23597149 | Unspecified |
| mmu-miR-708-5p | MIMAT0004828 | Rab30 | 75985 | HITS-CLIP | 37986224 | Unspecified |
| mmu-miR-708-5p | MIMAT0004828 | Rab35 | 77407 | HITS-CLIP | 37986224\|23597149 | Unspecified |
| mmu-miR-708-5p | MIMAT0004828 | Rabl3 | 67657 | HITS-CLIP | 23597149 | Unspecified |
| mmu-miR-708-5p | MIMAT0004828 | Rai1 | 19377 | HITS-CLIP | 23597149 | Unspecified |
| mmu-miR-708-5p | MIMAT0004828 | Ralgapb | 228850 | HITS-CLIP | 37986224 | Unspecified |
| mmu-miR-708-5p | MIMAT0004828 | Ranbp10 | 74334 | HITS-CLIP | 23597149 | Unspecified |
| mmu-miR-708-5p | MIMAT0004828 | Ranbp2 | 19386 | HITS-CLIP | 37986224 | Unspecified |
| mmu-miR-708-5p | MIMAT0004828 | Rap1b | 215449 | HITS-CLIP | 37986224\|19536157 | Unspecified |
| mmu-miR-708-5p | MIMAT0004828 | Rap1gds1 | 229877 | HITS-CLIP | 23597149 | Unspecified |
| mmu-miR-708-5p | MIMAT0004828 | Rap2c | 72065 | HITS-CLIP | 37986224 | Unspecified |
| mmu-miR-708-5p | MIMAT0004828 | Rapgef4 | 56508 | HITS-CLIP | 23597149 | Unspecified |
| mmu-miR-708-5p | MIMAT0004828 | Rbm25 | 67039 | HITS-CLIP | 23597149 | Unspecified |
| mmu-miR-708-5p | MIMAT0004828 | Rc3h2 | 319817 | HITS-CLIP | 23597149 | Unspecified |
| mmu-miR-708-5p | MIMAT0004828 | Rce1 | 19671 | HITS-CLIP | 37986224 | Unspecified |
| mmu-miR-708-5p | MIMAT0004828 | Rdh7 | 54150 | HITS-CLIP | 37986224 | Unspecified |
| mmu-miR-708-5p | MIMAT0004828 | Reep2 | 225362 | HITS-CLIP | 37986224 | Unspecified |
| mmu-miR-708-5p | MIMAT0004828 | Reep3 | 28193 | HITS-CLIP | 37986224 | Unspecified |
| mmu-miR-708-5p | MIMAT0004828 | Rela | 19697 | HITS-CLIP | 37986224\|23597149 | Unspecified |
| mmu-miR-708-5p | MIMAT0004828 | Rgs5 | 19737 | HITS-CLIP | 37986224 | Unspecified |
| mmu-miR-708-5p | MIMAT0004828 | Rhobtb1 | 69288 | HITS-CLIP | 37986224 | Unspecified |
| mmu-miR-708-5p | MIMAT0004828 | Rit1 | 19769 | HITS-CLIP | 37986224 | Unspecified |
| mmu-miR-708-5p | MIMAT0004828 | Rmnd5a | 68477 | HITS-CLIP | 37986224 | Unspecified |
| mmu-miR-708-5p | MIMAT0004828 | Rnasek | 52898 | HITS-CLIP | 37986224 | Unspecified |
| mmu-miR-708-5p | MIMAT0004828 | Rnd3 | 74194 | HITS-CLIP | 23597149 | Unspecified |
| mmu-miR-708-5p | MIMAT0004828 | Rnf13 | 24017 | HITS-CLIP | 19536157 | Unspecified |

| | | | | | | |
|---|---|---|---|---|---|---|
| mmu-miR-708-5p | MIMAT0004828 | Rnf130 | 59044 | HITS-CLIP | 37986224 | Unspecified |
| mmu-miR-708-5p | MIMAT0004828 | Rnf185 | 193670 | HITS-CLIP | 23597149 | Unspecified |
| mmu-miR-708-5p | MIMAT0004828 | Rnf216 | 108086 | HITS-CLIP | 37986224 | Unspecified |
| mmu-miR-708-5p | MIMAT0004828 | Rnf38 | 73469 | HITS-CLIP | 23597149 | Unspecified |
| mmu-miR-708-5p | MIMAT0004828 | Rnf44 | 105239 | HITS-CLIP | 37986224 | Unspecified |
| mmu-miR-708-5p | MIMAT0004828 | Rnpepl1 | 108657 | HITS-CLIP | 37986224 | Unspecified |
| mmu-miR-708-5p | MIMAT0004828 | Robo2 | 268902 | HITS-CLIP | 19536157 | Unspecified |
| mmu-miR-708-5p | MIMAT0004828 | Rogdi | 66049 | HITS-CLIP | 37986224 | Unspecified |
| mmu-miR-708-5p | MIMAT0004828 | Rph3al | 380714 | HITS-CLIP | 37986224 | Unspecified |
| mmu-miR-708-5p | MIMAT0004828 | Rplp0 | 11837 | HITS-CLIP | 37986224 | Unspecified |
| mmu-miR-708-5p | MIMAT0004828 | Rpn1 | 103963 | HITS-CLIP | 23597149 | Unspecified |
| mmu-miR-708-5p | MIMAT0004828 | Rps5 | 20103 | HITS-CLIP | 23597149 | Unspecified |
| mmu-miR-708-5p | MIMAT0004828 | Rps6ka3 | 110651 | HITS-CLIP | 37986224 | Unspecified |
| mmu-miR-708-5p | MIMAT0004828 | Rraga | 68441 | HITS-CLIP | 37986224 | Unspecified |
| mmu-miR-708-5p | MIMAT0004828 | Rras | 20130 | HITS-CLIP | 37986224 | Unspecified |
| mmu-miR-708-5p | MIMAT0004828 | Rrbp1 | 81910 | HITS-CLIP | 23597149 | Unspecified |
| mmu-miR-708-5p | MIMAT0004828 | Rrn3 | 106298 | HITS-CLIP | 23597149 | Unspecified |
| mmu-miR-708-5p | MIMAT0004828 | Rsl1d1 | 66409 | HITS-CLIP | 23597149 | Unspecified |
| mmu-miR-708-5p | MIMAT0004828 | Rtcb | 28088 | HITS-CLIP | 37986224 | Unspecified |
| mmu-miR-708-5p | MIMAT0004828 | Rtn3 | 20168 | HITS-CLIP | 19536157 | Unspecified |
| mmu-miR-708-5p | MIMAT0004828 | Rusc1 | 72296 | HITS-CLIP | 19536157 | Unspecified |
| mmu-miR-708-5p | MIMAT0004828 | Rxra | 20181 | HITS-CLIP | 37986224 | Unspecified |
| mmu-miR-708-5p | MIMAT0004828 | Ryr2 | 20191 | HITS-CLIP | 19536157 | Unspecified |
| mmu-miR-708-5p | MIMAT0004828 | Sacs | 50720 | HITS-CLIP | 37986224 | Unspecified |
| mmu-miR-708-5p | MIMAT0004828 | Sbno1 | 243272 | HITS-CLIP | 19536157 | Unspecified |
| mmu-miR-708-5p | MIMAT0004828 | Scamp3 | 24045 | HITS-CLIP | 37986224 | Unspecified |
| mmu-miR-708-5p | MIMAT0004828 | Scamp5 | 56807 | HITS-CLIP | 37986224 | Unspecified |
| mmu-miR-708-5p | MIMAT0004828 | Scd1 | 20249 | HITS-CLIP | 37986224 | Unspecified |
| mmu-miR-708-5p | MIMAT0004828 | Scd2 | 20250 | HITS-CLIP | 37986224 | Unspecified |
| mmu-miR-708-5p | MIMAT0004828 | Scmh1 | 29871 | HITS-CLIP | 37986224 | Unspecified |
| mmu-miR-708-5p | MIMAT0004828 | Scn3a | 20269 | HITS-CLIP | 19536157 | Unspecified |
| mmu-miR-708-5p | MIMAT0004828 | Scrg1 | 20284 | HITS-CLIP | 37986224 | Unspecified |
| mmu-miR-708-5p | MIMAT0004828 | Scyl3 | 240880 | HITS-CLIP | 37986224 | Unspecified |
| mmu-miR-708-5p | MIMAT0004828 | Sdha | 66945 | HITS-CLIP | 37986224\|23597149\|19536157 | Unspecified |
| mmu-miR-708-5p | MIMAT0004828 | Sds | 231691 | HITS-CLIP | 23597149 | Unspecified |
| mmu-miR-708-5p | MIMAT0004828 | Sec14l1 | 74136 | HITS-CLIP | 37986224 | Unspecified |
| mmu-miR-708-5p | MIMAT0004828 | Sec14l2 | 67815 | HITS-CLIP | 37986224\|23597149 | Unspecified |
| mmu-miR-708-5p | MIMAT0004828 | Sec16a | 227648 | HITS-CLIP | 37986224\|23597149 | Unspecified |
| mmu-miR-708-5p | MIMAT0004828 | Sec16b | 89867 | HITS-CLIP | 23597149 | Unspecified |

| | | | | | | | |
|---|---|---|---|---|---|---|---|
| mmu-miR-708-5p | MIMAT0004828 | Sec23b | 27054 | HITS-CLIP | | 37986224 | Unspecified |
| mmu-miR-708-5p | MIMAT0004828 | Sec24b | 99683 | HITS-CLIP | | 37986224 | Unspecified |
| mmu-miR-708-5p | MIMAT0004828 | Sec24c | 218811 | HITS-CLIP | | 23597149 | Unspecified |
| mmu-miR-708-5p | MIMAT0004828 | Sec61a1 | 53421 | HITS-CLIP | | 37986224 | Unspecified |
| mmu-miR-708-5p | MIMAT0004828 | Sec62 | 69276 | HITS-CLIP | | 37986224 | Unspecified |
| mmu-miR-708-5p | MIMAT0004828 | Sel1l | 20338 | HITS-CLIP | | 37986224\|23597149 | Unspecified |
| mmu-miR-708-5p | MIMAT0004828 | Sema4g | 26456 | HITS-CLIP | | 37986224\|23597149 | Unspecified |
| mmu-miR-708-5p | MIMAT0004828 | Sema5a | 20356 | HITS-CLIP | | 37986224 | Unspecified |
| mmu-miR-708-5p | MIMAT0004828 | Sepsecs | 211006 | HITS-CLIP | | 23597149 | Unspecified |
| mmu-miR-708-5p | MIMAT0004828 | Serpina3n | 20716 | HITS-CLIP | | 37986224\|23597149 | Unspecified |
| mmu-miR-708-5p | MIMAT0004828 | Serpinf1 | 20317 | HITS-CLIP | | 23597149 | Unspecified |
| mmu-miR-708-5p | MIMAT0004828 | Serpinf2 | 18816 | HITS-CLIP | | 23597149 | Unspecified |
| mmu-miR-708-5p | MIMAT0004828 | Sertad2 | 58172 | HITS-CLIP | | 23597149 | Unspecified |
| mmu-miR-708-5p | MIMAT0004828 | Sestd1 | 228071 | HITS-CLIP | | 37986224 | Unspecified |
| mmu-miR-708-5p | MIMAT0004828 | Setx | 269254 | HITS-CLIP | | 23597149 | Unspecified |
| mmu-miR-708-5p | MIMAT0004828 | Sf1 | 22668 | HITS-CLIP | | 23597149 | Unspecified |
| mmu-miR-708-5p | MIMAT0004828 | Sft2d2 | 108735 | HITS-CLIP | | 37986224 | Unspecified |
| mmu-miR-708-5p | MIMAT0004828 | Sgms2 | 74442 | HITS-CLIP | | 37986224 | Unspecified |
| mmu-miR-708-5p | MIMAT0004828 | Sgpl1 | 20397 | HITS-CLIP | | 37986224 | Unspecified |
| mmu-miR-708-5p | MIMAT0004828 | Sh3gl2 | 20404 | HITS-CLIP | | 37986224 | Unspecified |
| mmu-miR-708-5p | MIMAT0004828 | Sh3pxd2b | 268396 | HITS-CLIP | | 23597149 | Unspecified |
| mmu-miR-708-5p | MIMAT0004828 | Shank1 | 243961 | HITS-CLIP | | 37986224 | Unspecified |
| mmu-miR-708-5p | MIMAT0004828 | Shmt2 | 108037 | HITS-CLIP | | 23597149 | Unspecified |
| mmu-miR-708-5p | MIMAT0004828 | Sik1 | 17691 | HITS-CLIP | | 37986224 | Unspecified |
| mmu-miR-708-5p | MIMAT0004828 | Sil1 | 81500 | HITS-CLIP | | 23597149 | Unspecified |
| mmu-miR-708-5p | MIMAT0004828 | Sipa1l1 | 217692 | HITS-CLIP | | 19536157 | Unspecified |
| mmu-miR-708-5p | MIMAT0004828 | Sirt3 | 64384 | HITS-CLIP | | 37986224 | Unspecified |
| mmu-miR-708-5p | MIMAT0004828 | Slc10a5 | 241877 | HITS-CLIP | | 37986224 | Unspecified |
| mmu-miR-708-5p | MIMAT0004828 | Slc12a5 | 57138 | HITS-CLIP | | 37986224 | Unspecified |
| mmu-miR-708-5p | MIMAT0004828 | Slc17a7 | 72961 | HITS-CLIP | | 37986224 | Unspecified |
| mmu-miR-708-5p | MIMAT0004828 | Slc1a2 | 20511 | HITS-CLIP | | 37986224 | Unspecified |
| mmu-miR-708-5p | MIMAT0004828 | Slc20a2 | 20516 | HITS-CLIP | | 37986224 | Unspecified |
| mmu-miR-708-5p | MIMAT0004828 | Slc22a1 | 20517 | HITS-CLIP | | 23597149 | Unspecified |
| mmu-miR-708-5p | MIMAT0004828 | Slc22a17 | 59049 | HITS-CLIP | | 19536157 | Unspecified |
| mmu-miR-708-5p | MIMAT0004828 | Slc25a1 | 13358 | HITS-CLIP | | 23597149 | Unspecified |
| mmu-miR-708-5p | MIMAT0004828 | Slc25a12 | 78830 | HITS-CLIP | | 23597149\|19536157 | Unspecified |
| mmu-miR-708-5p | MIMAT0004828 | Slc25a13 | 50799 | HITS-CLIP | | 23597149 | Unspecified |
| mmu-miR-708-5p | MIMAT0004828 | Slc25a20 | 57279 | HITS-CLIP | | 37986224\|23597149 | Unspecified |
| mmu-miR-708-5p | MIMAT0004828 | Slc25a39 | 68066 | HITS-CLIP | | 23597149 | Unspecified |

| | | | | | | |
|---|---|---|---|---|---|---|
| mmu-miR-708-5p | MIMAT0004828 | Slc25a42 | 73095 | HITS-CLIP | 23597149 | Unspecified |
| mmu-miR-708-5p | MIMAT0004828 | Slc27a5 | 26459 | HITS-CLIP | 23597149 | Unspecified |
| mmu-miR-708-5p | MIMAT0004828 | Slc2a8 | 56017 | HITS-CLIP | 37986224 | Unspecified |
| mmu-miR-708-5p | MIMAT0004828 | Slc30a9 | 109108 | HITS-CLIP | 23597149 | Unspecified |
| mmu-miR-708-5p | MIMAT0004828 | Slc31a2 | 20530 | HITS-CLIP | 37986224\|19536157 | Unspecified |
| mmu-miR-708-5p | MIMAT0004828 | Slc35e2 | 320541 | HITS-CLIP | 37986224 | Unspecified |
| mmu-miR-708-5p | MIMAT0004828 | Slc36a1 | 215335 | HITS-CLIP | 37986224 | Unspecified |
| mmu-miR-708-5p | MIMAT0004828 | Slc37a4 | 14385 | HITS-CLIP | 37986224 | Unspecified |
| mmu-miR-708-5p | MIMAT0004828 | Slc38a4 | 69354 | HITS-CLIP | 23597149 | Unspecified |
| mmu-miR-708-5p | MIMAT0004828 | Slc39a14 | 213053 | HITS-CLIP | 37986224\|19536157 | Unspecified |
| mmu-miR-708-5p | MIMAT0004828 | Slc40a1 | 53945 | HITS-CLIP | 37986224\|23597149 | Unspecified |
| mmu-miR-708-5p | MIMAT0004828 | Slc4a4 | 54403 | HITS-CLIP | 23597149 | Unspecified |
| mmu-miR-708-5p | MIMAT0004828 | Slc6a1 | 232333 | HITS-CLIP | 37986224 | Unspecified |
| mmu-miR-708-5p | MIMAT0004828 | Slc6a13 | 14412 | HITS-CLIP | 37986224\|23597149 | Unspecified |
| mmu-miR-708-5p | MIMAT0004828 | Slc6a6 | 21366 | HITS-CLIP | 37986224 | Unspecified |
| mmu-miR-708-5p | MIMAT0004828 | Slc7a1 | 11987 | HITS-CLIP | 37986224 | Unspecified |
| mmu-miR-708-5p | MIMAT0004828 | Slitrk2 | 245450 | HITS-CLIP | 37986224 | Unspecified |
| mmu-miR-708-5p | MIMAT0004828 | Smad5 | 17129 | HITS-CLIP | 37986224\|23597149 | Unspecified |
| mmu-miR-708-5p | MIMAT0004828 | Smap2 | 69780 | HITS-CLIP | 23597149 | Unspecified |
| mmu-miR-708-5p | MIMAT0004828 | Smarca2 | 67155 | HITS-CLIP | 37986224\|23597149 | Unspecified |
| mmu-miR-708-5p | MIMAT0004828 | Smcr8 | 237782 | HITS-CLIP | 37986224 | Unspecified |
| mmu-miR-708-5p | MIMAT0004828 | Smg1 | 233789 | HITS-CLIP | 31433990 | Unspecified |
| mmu-miR-708-5p | MIMAT0004828 | Smg7 | 226517 | HITS-CLIP | 37986224 | Unspecified |
| mmu-miR-708-5p | MIMAT0004828 | Smg8 | 74133 | HITS-CLIP | 37986224 | Unspecified |
| mmu-miR-708-5p | MIMAT0004828 | Smpdl3a | 57319 | HITS-CLIP | 37986224 | Unspecified |
| mmu-miR-708-5p | MIMAT0004828 | Smurf2 | 66313 | HITS-CLIP | 37986224 | Unspecified |
| mmu-miR-708-5p | MIMAT0004828 | Snap25 | 20614 | HITS-CLIP | 37986224\|19536157 | Unspecified |
| mmu-miR-708-5p | MIMAT0004828 | Snap91 | 20616 | HITS-CLIP | 19536157 | Unspecified |
| mmu-miR-708-5p | MIMAT0004828 | Snf8 | 27681 | HITS-CLIP | 23597149 | Unspecified |
| mmu-miR-708-5p | MIMAT0004828 | Snhg11 | 319317 | HITS-CLIP | 37986224 | Unspecified |
| mmu-miR-708-5p | MIMAT0004828 | Snrnp200 | 320632 | HITS-CLIP | 37986224\|23597149 | Unspecified |
| mmu-miR-708-5p | MIMAT0004828 | Snw1 | 66354 | HITS-CLIP | 37986224 | Unspecified |
| mmu-miR-708-5p | MIMAT0004828 | Snx13 | 217463 | HITS-CLIP | 37986224 | Unspecified |
| mmu-miR-708-5p | MIMAT0004828 | Snx6 | 72183 | HITS-CLIP | 37986224 | Unspecified |
| mmu-miR-708-5p | MIMAT0004828 | Socs3 | 12702 | HITS-CLIP | 37986224\|23597149 | Unspecified |
| mmu-miR-708-5p | MIMAT0004828 | Socs4 | 67296 | HITS-CLIP | 37986224 | Unspecified |
| mmu-miR-708-5p | MIMAT0004828 | Socs7 | 192157 | HITS-CLIP | 37986224 | Unspecified |
| mmu-miR-708-5p | MIMAT0004828 | Soga1 | 320706 | HITS-CLIP | 37986224 | Unspecified |
| mmu-miR-708-5p | MIMAT0004828 | Son | 20658 | HITS-CLIP | 37986224\|19536157\|23597149 | Unspecified |

| | | | | | | |
|---|---|---|---|---|---|---|
| mmu-miR-708-5p | MIMAT0004828 | Sorl1 | 20660 | HITS-CLIP | 37986224\|19536157 | Unspecified |
| mmu-miR-708-5p | MIMAT0004828 | Sox10 | 20665 | HITS-CLIP | 37986224 | Unspecified |
| mmu-miR-708-5p | MIMAT0004828 | Sox11 | 20666 | HITS-CLIP | 37986224 | Unspecified |
| mmu-miR-708-5p | MIMAT0004828 | Sparc | 20692 | HITS-CLIP | 37986224 | Unspecified |
| mmu-miR-708-5p | MIMAT0004828 | Sparcl1 | 13602 | HITS-CLIP | 37986224\|19536157 | Unspecified |
| mmu-miR-708-5p | MIMAT0004828 | Spcs2 | 66624 | HITS-CLIP | 37986224 | Unspecified |
| mmu-miR-708-5p | MIMAT0004828 | Spg20 | 229285 | HITS-CLIP | 23597149 | Unspecified |
| mmu-miR-708-5p | MIMAT0004828 | Spns2 | 216892 | HITS-CLIP | 37986224 | Unspecified |
| mmu-miR-708-5p | MIMAT0004828 | Spock2 | 94214 | HITS-CLIP | 37986224 | Unspecified |
| mmu-miR-708-5p | MIMAT0004828 | Sppl3 | 74585 | HITS-CLIP | 37986224 | Unspecified |
| mmu-miR-708-5p | MIMAT0004828 | Spred1 | 114715 | HITS-CLIP | 37986224 | Unspecified |
| mmu-miR-708-5p | MIMAT0004828 | Spry4 | 24066 | HITS-CLIP | 23597149 | Unspecified |
| mmu-miR-708-5p | MIMAT0004828 | Sptan1 | 20740 | HITS-CLIP | 19536157 | Unspecified |
| mmu-miR-708-5p | MIMAT0004828 | Sptbn1 | 20742 | HITS-CLIP | 19536157 | Unspecified |
| mmu-miR-708-5p | MIMAT0004828 | Sptbn2 | 20743 | HITS-CLIP | 37986224 | Unspecified |
| mmu-miR-708-5p | MIMAT0004828 | Srebf2 | 20788 | HITS-CLIP | 37986224 | Unspecified |
| mmu-miR-708-5p | MIMAT0004828 | Srek1 | 218543 | HITS-CLIP | 37986224 | Unspecified |
| mmu-miR-708-5p | MIMAT0004828 | Srpr | 67398 | HITS-CLIP | 37986224 | Unspecified |
| mmu-miR-708-5p | MIMAT0004828 | Srrm2 | 75956 | HITS-CLIP | 19536157\|23597149 | Unspecified |
| mmu-miR-708-5p | MIMAT0004828 | Ssbp3 | 72475 | HITS-CLIP | 23597149 | Unspecified |
| mmu-miR-708-5p | MIMAT0004828 | St3gal5 | 20454 | HITS-CLIP | 37986224 | Unspecified |
| mmu-miR-708-5p | MIMAT0004828 | Stab2 | 192188 | HITS-CLIP | 37986224\|23597149 | Unspecified |
| mmu-miR-708-5p | MIMAT0004828 | Stag2 | 20843 | HITS-CLIP | 37986224\|19536157 | Unspecified |
| mmu-miR-708-5p | MIMAT0004828 | Stard10 | 56018 | HITS-CLIP | 23597149 | Unspecified |
| mmu-miR-708-5p | MIMAT0004828 | Stat5b | 20851 | HITS-CLIP | 37986224 | Unspecified |
| mmu-miR-708-5p | MIMAT0004828 | Stbd1 | 52331 | HITS-CLIP | 37986224 | Unspecified |
| mmu-miR-708-5p | MIMAT0004828 | Stim2 | 116873 | HITS-CLIP | 23597149 | Unspecified |
| mmu-miR-708-5p | MIMAT0004828 | Stk25 | 59041 | HITS-CLIP | 37986224\|23597149 | Unspecified |
| mmu-miR-708-5p | MIMAT0004828 | Stk35 | 67333 | HITS-CLIP | 37986224\|23597149 | Unspecified |
| mmu-miR-708-5p | MIMAT0004828 | Stmn2 | 20257 | HITS-CLIP | 37986224\|19536157 | Unspecified |
| mmu-miR-708-5p | MIMAT0004828 | Stmn3 | 20262 | HITS-CLIP | 37986224 | Unspecified |
| mmu-miR-708-5p | MIMAT0004828 | Stra6 | 20897 | HITS-CLIP | 37986224 | Unspecified |
| mmu-miR-708-5p | MIMAT0004828 | Stradb | 227154 | HITS-CLIP | 37986224 | Unspecified |
| mmu-miR-708-5p | MIMAT0004828 | Strn | 268980 | HITS-CLIP | 37986224 | Unspecified |
| mmu-miR-708-5p | MIMAT0004828 | Stt3b | 68292 | HITS-CLIP | 37986224 | Unspecified |
| mmu-miR-708-5p | MIMAT0004828 | Stub1 | 56424 | HITS-CLIP | 23597149 | Unspecified |
| mmu-miR-708-5p | MIMAT0004828 | Stx1a | 20907 | HITS-CLIP | 37986224 | Unspecified |
| mmu-miR-708-5p | MIMAT0004828 | Stxbp1 | 20910 | HITS-CLIP | 19536157 | Unspecified |
| mmu-miR-708-5p | MIMAT0004828 | Suds3 | 71954 | HITS-CLIP | 23597149 | Unspecified |

| | | | | | | |
|---|---|---|---|---|---|---|
| mmu-miR-708-5p | MIMAT0004828 | Sult1d1 | 53315 | HITS-CLIP | 23597149 | Unspecified |
| mmu-miR-708-5p | MIMAT0004828 | Sun1 | 77053 | HITS-CLIP | 37986224 | Unspecified |
| mmu-miR-708-5p | MIMAT0004828 | Syn2 | 20965 | HITS-CLIP | 19536157 | Unspecified |
| mmu-miR-708-5p | MIMAT0004828 | Synrg | 217030 | HITS-CLIP | 23597149 | Unspecified |
| mmu-miR-708-5p | MIMAT0004828 | Syvn1 | 74126 | HITS-CLIP | 37986224 | Unspecified |
| mmu-miR-708-5p | MIMAT0004828 | Tab2 | 68652 | HITS-CLIP | 37986224\|23597149\|19536157 | Unspecified |
| mmu-miR-708-5p | MIMAT0004828 | Tab3 | 66724 | HITS-CLIP | 37986224 | Unspecified |
| mmu-miR-708-5p | MIMAT0004828 | Tacc2 | 57752 | HITS-CLIP | 23597149 | Unspecified |
| mmu-miR-708-5p | MIMAT0004828 | Tagln3 | 56370 | HITS-CLIP | 37986224 | Unspecified |
| mmu-miR-708-5p | MIMAT0004828 | Tanc2 | 77097 | HITS-CLIP | 37986224 | Unspecified |
| mmu-miR-708-5p | MIMAT0004828 | Tango2 | 27883 | HITS-CLIP | 37986224 | Unspecified |
| mmu-miR-708-5p | MIMAT0004828 | Taok1 | 216965 | HITS-CLIP | 37986224 | Unspecified |
| mmu-miR-708-5p | MIMAT0004828 | Tardbp | 230908 | HITS-CLIP | 37986224 | Unspecified |
| mmu-miR-708-5p | MIMAT0004828 | Tbc1d24 | 224617 | HITS-CLIP | 37986224 | Unspecified |
| mmu-miR-708-5p | MIMAT0004828 | Tbpl1 | 237336 | HITS-CLIP | 23597149 | Unspecified |
| mmu-miR-708-5p | MIMAT0004828 | Tbrg1 | 21376 | HITS-CLIP | 23597149 | Unspecified |
| mmu-miR-708-5p | MIMAT0004828 | Tcf12 | 21406 | HITS-CLIP | 37986224\|23597149 | Unspecified |
| mmu-miR-708-5p | MIMAT0004828 | Tcn2 | 21452 | HITS-CLIP | 23597149 | Unspecified |
| mmu-miR-708-5p | MIMAT0004828 | Tdo2 | 56720 | HITS-CLIP | 23597149 | Unspecified |
| mmu-miR-708-5p | MIMAT0004828 | Tedc2 | 72016 | HITS-CLIP | 23597149 | Unspecified |
| mmu-miR-708-5p | MIMAT0004828 | Tef | 21685 | HITS-CLIP | 37986224 | Unspecified |
| mmu-miR-708-5p | MIMAT0004828 | Tent5c | 54855 | HITS-CLIP | 23597149 | Unspecified |
| mmu-miR-708-5p | MIMAT0004828 | Tet2 | 214133 | HITS-CLIP | 37986224 | Unspecified |
| mmu-miR-708-5p | MIMAT0004828 | Tet3 | 194388 | HITS-CLIP | 37986224 | Unspecified |
| mmu-miR-708-5p | MIMAT0004828 | Tex2 | 21763 | HITS-CLIP | 37986224 | Unspecified |
| mmu-miR-708-5p | MIMAT0004828 | Tex261 | 21766 | HITS-CLIP | 37986224 | Unspecified |
| mmu-miR-708-5p | MIMAT0004828 | Tfap4 | 83383 | HITS-CLIP | 23597149 | Unspecified |
| mmu-miR-708-5p | MIMAT0004828 | Tfr2 | 50765 | HITS-CLIP | 37986224 | Unspecified |
| mmu-miR-708-5p | MIMAT0004828 | Tgfb1i1 | 21804 | HITS-CLIP | 19536157 | Unspecified |
| mmu-miR-708-5p | MIMAT0004828 | Tgfbr3 | 21814 | HITS-CLIP | 37986224 | Unspecified |
| mmu-miR-708-5p | MIMAT0004828 | Tgfbrap1 | 73122 | HITS-CLIP | 37986224 | Unspecified |
| mmu-miR-708-5p | MIMAT0004828 | Tgs1 | 116940 | HITS-CLIP | 23597149 | Unspecified |
| mmu-miR-708-5p | MIMAT0004828 | Thoc1 | 225160 | HITS-CLIP | 31433990 | Unspecified |
| mmu-miR-708-5p | MIMAT0004828 | Thrap3 | 230753 | HITS-CLIP | 37986224 | Unspecified |
| mmu-miR-708-5p | MIMAT0004828 | Tiprl | 226591 | HITS-CLIP | 23597149 | Unspecified |
| mmu-miR-708-5p | MIMAT0004828 | Tjp1 | 21872 | HITS-CLIP | 37986224\|23597149 | Unspecified |
| mmu-miR-708-5p | MIMAT0004828 | Tk1 | 21877 | HITS-CLIP | 23597149 | Unspecified |
| mmu-miR-708-5p | MIMAT0004828 | Tkfc | 26007 | HITS-CLIP | 23597149 | Unspecified |
| mmu-miR-708-5p | MIMAT0004828 | Tlk2 | 24086 | HITS-CLIP | 37986224 | Unspecified |

| | | | | | | |
|---|---|---|---|---|---|---|
| mmu-miR-708-5p | MIMAT0004828 | Tln1 | 21894 | HITS-CLIP | 37986224\|23597149 | Unspecified |
| mmu-miR-708-5p | MIMAT0004828 | Tmed3 | 66111 | HITS-CLIP | 37986224 | Unspecified |
| mmu-miR-708-5p | MIMAT0004828 | Tmed9 | 67511 | HITS-CLIP | 37986224 | Unspecified |
| mmu-miR-708-5p | MIMAT0004828 | Tmem151b | 210573 | HITS-CLIP | 37986224 | Unspecified |
| mmu-miR-708-5p | MIMAT0004828 | Tmem178 | 68027 | HITS-CLIP | 37986224 | Unspecified |
| mmu-miR-708-5p | MIMAT0004828 | Tmem256 | 254863 | HITS-CLIP | 23597149 | Unspecified |
| mmu-miR-708-5p | MIMAT0004828 | Tmem258 | 69038 | HITS-CLIP | 23597149 | Unspecified |
| mmu-miR-708-5p | MIMAT0004828 | Tmem41a | 66664 | HITS-CLIP | 37986224 | Unspecified |
| mmu-miR-708-5p | MIMAT0004828 | Tmem50a | 71817 | HITS-CLIP | 37986224 | Unspecified |
| mmu-miR-708-5p | MIMAT0004828 | Tmem56 | 99887 | HITS-CLIP | 37986224 | Unspecified |
| mmu-miR-708-5p | MIMAT0004828 | Tmem59 | 56374 | HITS-CLIP | 23597149 | Unspecified |
| mmu-miR-708-5p | MIMAT0004828 | Tmem62 | 96957 | HITS-CLIP | 37986224 | Unspecified |
| mmu-miR-708-5p | MIMAT0004828 | Tmem86b | 68255 | HITS-CLIP | 37986224 | Unspecified |
| mmu-miR-708-5p | MIMAT0004828 | Tmem9 | 66241 | HITS-CLIP | 37986224 | Unspecified |
| mmu-miR-708-5p | MIMAT0004828 | Tmod2 | 50876 | HITS-CLIP | 19536157 | Unspecified |
| mmu-miR-708-5p | MIMAT0004828 | Tmpo | 21917 | HITS-CLIP | 23597149 | Unspecified |
| mmu-miR-708-5p | MIMAT0004828 | Tmtc2 | 278279 | HITS-CLIP | 37986224 | Unspecified |
| mmu-miR-708-5p | MIMAT0004828 | Tnfrsf1b | 21938 | HITS-CLIP | 37986224 | Unspecified |
| mmu-miR-708-5p | MIMAT0004828 | Tnfrsf21 | 94185 | HITS-CLIP | 37986224 | Unspecified |
| mmu-miR-708-5p | MIMAT0004828 | Tnik | 665113 | HITS-CLIP | 37986224 | Unspecified |
| mmu-miR-708-5p | MIMAT0004828 | Tnks | 21951 | HITS-CLIP | 37986224 | Unspecified |
| mmu-miR-708-5p | MIMAT0004828 | Tnks2 | 74493 | HITS-CLIP | 37986224 | Unspecified |
| mmu-miR-708-5p | MIMAT0004828 | Tnrc6b | 213988 | HITS-CLIP | 37986224\|19536157\|23597149 | Unspecified |
| mmu-miR-708-5p | MIMAT0004828 | Tns1 | 21961 | HITS-CLIP | 23597149 | Unspecified |
| mmu-miR-708-5p | MIMAT0004828 | Tomm70a | 28185 | HITS-CLIP | 19536157 | Unspecified |
| mmu-miR-708-5p | MIMAT0004828 | Top3a | 21975 | HITS-CLIP | 23597149 | Unspecified |
| mmu-miR-708-5p | MIMAT0004828 | Topors | 106021 | HITS-CLIP | 37986224 | Unspecified |
| mmu-miR-708-5p | MIMAT0004828 | Tor1aip1 | 208263 | HITS-CLIP | 23597149 | Unspecified |
| mmu-miR-708-5p | MIMAT0004828 | Tor2a | 30933 | HITS-CLIP | 23597149 | Unspecified |
| mmu-miR-708-5p | MIMAT0004828 | Tpcn1 | 252972 | HITS-CLIP | 37986224 | Unspecified |
| mmu-miR-708-5p | MIMAT0004828 | Tpr | 108989 | HITS-CLIP | 19536157\|23597149 | Unspecified |
| mmu-miR-708-5p | MIMAT0004828 | Tpra1 | 24100 | HITS-CLIP | 19536157 | Unspecified |
| mmu-miR-708-5p | MIMAT0004828 | Tprgl | 67808 | HITS-CLIP | 37986224 | Unspecified |
| mmu-miR-708-5p | MIMAT0004828 | Trak1 | 67095 | HITS-CLIP | 23597149 | Unspecified |
| mmu-miR-708-5p | MIMAT0004828 | Trak2 | 70827 | HITS-CLIP | 37986224 | Unspecified |
| mmu-miR-708-5p | MIMAT0004828 | Tram1l1 | 229801 | HITS-CLIP | 37986224 | Unspecified |
| mmu-miR-708-5p | MIMAT0004828 | Trappc4 | 60409 | HITS-CLIP | 37986224 | Unspecified |
| mmu-miR-708-5p | MIMAT0004828 | Trib1 | 211770 | HITS-CLIP | 37986224\|23597149 | Unspecified |
| mmu-miR-708-5p | MIMAT0004828 | Trim28 | 21849 | HITS-CLIP | 37986224 | Unspecified |

| | | | | | | |
|---|---|---|---|---|---|---|
| mmu-miR-708-5p | MIMAT0004828 | Trim33 | 94093 | HITS-CLIP | 37986224\|23597149 | Unspecified |
| mmu-miR-708-5p | MIMAT0004828 | Trip12 | 14897 | HITS-CLIP | 23597149 | Unspecified |
| mmu-miR-708-5p | MIMAT0004828 | Trit1 | 66966 | HITS-CLIP | 37986224 | Unspecified |
| mmu-miR-708-5p | MIMAT0004828 | Trp53 | 22059 | HITS-CLIP | 23597149 | Unspecified |
| mmu-miR-708-5p | MIMAT0004828 | Trp53bp2 | 209456 | HITS-CLIP | 23597149 | Unspecified |
| mmu-miR-708-5p | MIMAT0004828 | Trp53i11 | 277414 | HITS-CLIP | 37986224 | Unspecified |
| mmu-miR-708-5p | MIMAT0004828 | Trp53inp1 | 60599 | HITS-CLIP | 37986224 | Unspecified |
| mmu-miR-708-5p | MIMAT0004828 | Trp53inp2 | 68728 | HITS-CLIP | 37986224\|23597149\|19536157 | Unspecified |
| mmu-miR-708-5p | MIMAT0004828 | Trpc4ap | 56407 | HITS-CLIP | 37986224 | Unspecified |
| mmu-miR-708-5p | MIMAT0004828 | Trps1 | 83925 | HITS-CLIP | 19536157 | Unspecified |
| mmu-miR-708-5p | MIMAT0004828 | Trrap | 100683 | HITS-CLIP | 23597149 | Unspecified |
| mmu-miR-708-5p | MIMAT0004828 | Tsc1 | 64930 | HITS-CLIP | 19536157 | Unspecified |
| mmu-miR-708-5p | MIMAT0004828 | Tsc22d3 | 14605 | HITS-CLIP | 37986224 | Unspecified |
| mmu-miR-708-5p | MIMAT0004828 | Tshz1 | 110796 | HITS-CLIP | 37986224 | Unspecified |
| mmu-miR-708-5p | MIMAT0004828 | Tspan13 | 66109 | HITS-CLIP | 37986224 | Unspecified |
| mmu-miR-708-5p | MIMAT0004828 | Tspan3 | 56434 | HITS-CLIP | 37986224 | Unspecified |
| mmu-miR-708-5p | MIMAT0004828 | Tspyl1 | 22110 | HITS-CLIP | 23597149\|19536157 | Unspecified |
| mmu-miR-708-5p | MIMAT0004828 | Tst | 22117 | HITS-CLIP | 37986224 | Unspecified |
| mmu-miR-708-5p | MIMAT0004828 | Ttc3 | 22129 | HITS-CLIP | 37986224 | Unspecified |
| mmu-miR-708-5p | MIMAT0004828 | Ttc39b | 69863 | HITS-CLIP | 37986224\|23597149 | Unspecified |
| mmu-miR-708-5p | MIMAT0004828 | Ttc7 | 225049 | HITS-CLIP | 37986224 | Unspecified |
| mmu-miR-708-5p | MIMAT0004828 | Tti1 | 75425 | HITS-CLIP | 31433990 | Unspecified |
| mmu-miR-708-5p | MIMAT0004828 | Ttr | 22139 | HITS-CLIP | 37986224 | Unspecified |
| mmu-miR-708-5p | MIMAT0004828 | Tuba4a | 22145 | HITS-CLIP | 37986224\|23597149 | Unspecified |
| mmu-miR-708-5p | MIMAT0004828 | Tubb5 | 22154 | HITS-CLIP | 37986224 | Unspecified |
| mmu-miR-708-5p | MIMAT0004828 | Tut4 | 23318 | HITS-CLIP | 23597149 | Unspecified |
| mmu-miR-708-5p | MIMAT0004828 | Tut7 | 79670 | HITS-CLIP | 23597149 | Unspecified |
| mmu-miR-708-5p | MIMAT0004828 | Twf1 | 19230 | HITS-CLIP | 23597149 | Unspecified |
| mmu-miR-708-5p | MIMAT0004828 | Txndc11 | 106200 | HITS-CLIP | 23597149\|19536157 | Unspecified |
| mmu-miR-708-5p | MIMAT0004828 | Txnl1 | 53382 | HITS-CLIP | 37986224\|19536157\|23597149 | Unspecified |
| mmu-miR-708-5p | MIMAT0004828 | Txnrd1 | 50493 | HITS-CLIP | 37986224 | Unspecified |
| mmu-miR-708-5p | MIMAT0004828 | Uba6 | 231380 | HITS-CLIP | 19536157 | Unspecified |
| mmu-miR-708-5p | MIMAT0004828 | Ube2h | 22214 | HITS-CLIP | 37986224 | Unspecified |
| mmu-miR-708-5p | MIMAT0004828 | Ube2j1 | 56228 | HITS-CLIP | 23597149 | Unspecified |
| mmu-miR-708-5p | MIMAT0004828 | Ube2n | 93765 | HITS-CLIP | 37986224 | Unspecified |
| mmu-miR-708-5p | MIMAT0004828 | Ube3b | 117146 | HITS-CLIP | 37986224 | Unspecified |
| mmu-miR-708-5p | MIMAT0004828 | Ube3c | 100763 | HITS-CLIP | 37986224\|19536157 | Unspecified |
| mmu-miR-708-5p | MIMAT0004828 | Ube4a | 140630 | HITS-CLIP | 23597149 | Unspecified |
| mmu-miR-708-5p | MIMAT0004828 | Ubn1 | 170644 | HITS-CLIP | 37986224 | Unspecified |

| | | | | | | |
|---|---|---|---|---|---|---|
| mmu-miR-708-5p | MIMAT0004828 | Ubn2 | 320538 | HITS-CLIP | 37986224 | Unspecified |
| mmu-miR-708-5p | MIMAT0004828 | Ubr1 | 22222 | HITS-CLIP | 37986224 | Unspecified |
| mmu-miR-708-5p | MIMAT0004828 | Ubr4 | 69116 | HITS-CLIP | 23597149 | Unspecified |
| mmu-miR-708-5p | MIMAT0004828 | Ubxn4 | 67812 | HITS-CLIP | 37986224 | Unspecified |
| mmu-miR-708-5p | MIMAT0004828 | Ugcg | 22234 | HITS-CLIP | 19536157 | Unspecified |
| mmu-miR-708-5p | MIMAT0004828 | Ugt8a | 22239 | HITS-CLIP | 37986224 | Unspecified |
| mmu-miR-708-5p | MIMAT0004828 | Uhmk1 | 16589 | HITS-CLIP | 19536157 | Unspecified |
| mmu-miR-708-5p | MIMAT0004828 | Uhrf1bp1l | 75089 | HITS-CLIP | 23597149 | Unspecified |
| mmu-miR-708-5p | MIMAT0004828 | Ulk1 | 22241 | HITS-CLIP | 37986224 | Unspecified |
| mmu-miR-708-5p | MIMAT0004828 | Umps | 22247 | HITS-CLIP | 23597149 | Unspecified |
| mmu-miR-708-5p | MIMAT0004828 | Unc13c | 208898 | HITS-CLIP | 19536157 | Unspecified |
| mmu-miR-708-5p | MIMAT0004828 | Unc5b | 107449 | HITS-CLIP | 37986224 | Unspecified |
| mmu-miR-708-5p | MIMAT0004828 | Unc5d | 210801 | HITS-CLIP | 37986224 | Unspecified |
| mmu-miR-708-5p | MIMAT0004828 | Upp2 | 76654 | HITS-CLIP | 23597149 | Unspecified |
| mmu-miR-708-5p | MIMAT0004828 | Uqcrc1 | 22273 | HITS-CLIP | 23597149\|26203562 | Unspecified |
| mmu-miR-708-5p | MIMAT0004828 | Uqcrc2 | 67003 | HITS-CLIP | 23597149 | Unspecified |
| mmu-miR-708-5p | MIMAT0004828 | Usp24 | 329908 | HITS-CLIP | 23597149 | Unspecified |
| mmu-miR-708-5p | MIMAT0004828 | Usp32 | 237898 | HITS-CLIP | 37986224 | Unspecified |
| mmu-miR-708-5p | MIMAT0004828 | Usp4 | 22258 | HITS-CLIP | 23597149 | Unspecified |
| mmu-miR-708-5p | MIMAT0004828 | Usp48 | 170707 | HITS-CLIP | 23597149 | Unspecified |
| mmu-miR-708-5p | MIMAT0004828 | Usp8 | 84092 | HITS-CLIP | 37986224\|23597149 | Unspecified |
| mmu-miR-708-5p | MIMAT0004828 | Utp3 | 65961 | HITS-CLIP | 23597149 | Unspecified |
| mmu-miR-708-5p | MIMAT0004828 | Vamp2 | 22318 | HITS-CLIP | 37986224 | Unspecified |
| mmu-miR-708-5p | MIMAT0004828 | Vcl | 22330 | HITS-CLIP | 37986224 | Unspecified |
| mmu-miR-708-5p | MIMAT0004828 | Vcpip1 | 70675 | HITS-CLIP | 37986224 | Unspecified |
| mmu-miR-708-5p | MIMAT0004828 | Vldlr | 22359 | HITS-CLIP | 37986224 | Unspecified |
| mmu-miR-708-5p | MIMAT0004828 | Vps13b | 666173 | HITS-CLIP | 23597149 | Unspecified |
| mmu-miR-708-5p | MIMAT0004828 | Vps13c | 320528 | HITS-CLIP | 23597149 | Unspecified |
| mmu-miR-708-5p | MIMAT0004828 | Vps28 | 51160 | HITS-CLIP | 23597149 | Unspecified |
| mmu-miR-708-5p | MIMAT0004828 | Vps54 | 245944 | HITS-CLIP | 19536157 | Unspecified |
| mmu-miR-708-5p | MIMAT0004828 | Vrk3 | 101568 | HITS-CLIP | 37986224 | Unspecified |
| mmu-miR-708-5p | MIMAT0004828 | Vtn | 22370 | HITS-CLIP | 23597149 | Unspecified |
| mmu-miR-708-5p | MIMAT0004828 | Vwf | 22371 | HITS-CLIP | 23597149 | Unspecified |
| mmu-miR-708-5p | MIMAT0004828 | Wasf2 | 242687 | HITS-CLIP | 37986224 | Unspecified |
| mmu-miR-708-5p | MIMAT0004828 | Wbp11 | 60321 | HITS-CLIP | 37986224 | Unspecified |
| mmu-miR-708-5p | MIMAT0004828 | Wbp1l | 226178 | HITS-CLIP | 37986224 | Unspecified |
| mmu-miR-708-5p | MIMAT0004828 | Wdfy3 | 72145 | HITS-CLIP | 23597149 | Unspecified |
| mmu-miR-708-5p | MIMAT0004828 | Wdr20 | 69641 | HITS-CLIP | 37986224 | Unspecified |
| mmu-miR-708-5p | MIMAT0004828 | Wdr45b | 66840 | HITS-CLIP | 23597149 | Unspecified |

| | | | | | | |
|---|---|---|---|---|---|---|
| mmu-miR-708-5p | MIMAT0004828 | Wdr83os | 414077 | HITS-CLIP | 37986224 | Unspecified |
| mmu-miR-708-5p | MIMAT0004828 | Wipf2 | 68524 | HITS-CLIP | 37986224 | Unspecified |
| mmu-miR-708-5p | MIMAT0004828 | Wls | 68151 | HITS-CLIP | 37986224 | Unspecified |
| mmu-miR-708-5p | MIMAT0004828 | Wnk1 | 232341 | HITS-CLIP | 23597149 | Unspecified |
| mmu-miR-708-5p | MIMAT0004828 | Xbp1 | 22433 | HITS-CLIP | 23597149 | Unspecified |
| mmu-miR-708-5p | MIMAT0004828 | Xkr6 | 219149 | HITS-CLIP | 19536157 | Unspecified |
| mmu-miR-708-5p | MIMAT0004828 | Xpo5 | 72322 | HITS-CLIP | 23597149 | Unspecified |
| mmu-miR-708-5p | MIMAT0004828 | Yeats2 | 208146 | HITS-CLIP | 23597149 | Unspecified |
| mmu-miR-708-5p | MIMAT0004828 | Yeats4 | 64050 | HITS-CLIP | 37986224 | Unspecified |
| mmu-miR-708-5p | MIMAT0004828 | Yipf6 | 77929 | HITS-CLIP | 19536157 | Unspecified |
| mmu-miR-708-5p | MIMAT0004828 | Ylpm1 | 56531 | HITS-CLIP | 23597149 | Unspecified |
| mmu-miR-708-5p | MIMAT0004828 | Ythdf2 | 213541 | HITS-CLIP | 37986224\|23597149 | Unspecified |
| mmu-miR-708-5p | MIMAT0004828 | Ythdf3 | 229096 | HITS-CLIP | 23597149 | Unspecified |
| mmu-miR-708-5p | MIMAT0004828 | Ywhag | 22628 | HITS-CLIP | 37986224 | Unspecified |
| mmu-miR-708-5p | MIMAT0004828 | Zadh2 | 225791 | HITS-CLIP | 23597149 | Unspecified |
| mmu-miR-708-5p | MIMAT0004828 | Zbed6 | 667118 | HITS-CLIP | 37986224 | Unspecified |
| mmu-miR-708-5p | MIMAT0004828 | Zbtb17 | 22642 | HITS-CLIP | 37986224 | Unspecified |
| mmu-miR-708-5p | MIMAT0004828 | Zbtb20 | 56490 | HITS-CLIP | 37986224 | Unspecified |
| mmu-miR-708-5p | MIMAT0004828 | Zbtb24 | 268294 | HITS-CLIP | 23597149 | Unspecified |
| mmu-miR-708-5p | MIMAT0004828 | Zbtb38 | 245007 | HITS-CLIP | 37986224 | Unspecified |
| mmu-miR-708-5p | MIMAT0004828 | Zc3h10 | 103284 | HITS-CLIP | 37986224\|23597149 | Unspecified |
| mmu-miR-708-5p | MIMAT0004828 | Zc3h12c | 244871 | HITS-CLIP | 37986224 | Unspecified |
| mmu-miR-708-5p | MIMAT0004828 | Zdhhc5 | 228136 | HITS-CLIP | 37986224 | Unspecified |
| mmu-miR-708-5p | MIMAT0004828 | Zdhhc7 | 102193 | HITS-CLIP | 37986224 | Unspecified |
| mmu-miR-708-5p | MIMAT0004828 | Zfc3h1 | 216345 | HITS-CLIP | 37986224 | Unspecified |
| mmu-miR-708-5p | MIMAT0004828 | Zfhx4 | 80892 | HITS-CLIP | 19536157 | Unspecified |
| mmu-miR-708-5p | MIMAT0004828 | Zfp148 | 22661 | HITS-CLIP | 37986224\|19536157 | Unspecified |
| mmu-miR-708-5p | MIMAT0004828 | Zfp174 | 385674 | HITS-CLIP | 37986224 | Unspecified |
| mmu-miR-708-5p | MIMAT0004828 | Zfp217 | 228913 | HITS-CLIP | 31433990 | Unspecified |
| mmu-miR-708-5p | MIMAT0004828 | Zfp236 | 329002 | HITS-CLIP | 19536157 | Unspecified |
| mmu-miR-708-5p | MIMAT0004828 | Zfp281 | 226442 | HITS-CLIP | 23597149 | Unspecified |
| mmu-miR-708-5p | MIMAT0004828 | Zfp568 | 243905 | HITS-CLIP | 37986224 | Unspecified |
| mmu-miR-708-5p | MIMAT0004828 | Zfp592 | 233410 | HITS-CLIP | 37986224 | Unspecified |
| mmu-miR-708-5p | MIMAT0004828 | Zfp598 | 213753 | HITS-CLIP | 23597149 | Unspecified |
| mmu-miR-708-5p | MIMAT0004828 | Zfp622 | 52521 | HITS-CLIP | 37986224 | Unspecified |
| mmu-miR-708-5p | MIMAT0004828 | Zfp638 | 27332 | HITS-CLIP | 19536157 | Unspecified |
| mmu-miR-708-5p | MIMAT0004828 | Zfp644 | 52397 | HITS-CLIP | 37986224 | Unspecified |
| mmu-miR-708-5p | MIMAT0004828 | Zfp687 | 78266 | HITS-CLIP | 37986224 | Unspecified |
| mmu-miR-708-5p | MIMAT0004828 | Zfp704 | 170753 | HITS-CLIP | 37986224 | Unspecified |

| | | | | | | |
|---|---|---|---|---|---|---|
| mmu-miR-708-5p | MIMAT0004828 | Zfp91 | 109910 | HITS-CLIP | 37986224\|23597149 | Unspecified |
| mmu-miR-708-5p | MIMAT0004828 | Zfyve26 | 211978 | HITS-CLIP | 23597149 | Unspecified |
| mmu-miR-708-5p | MIMAT0004828 | Zmiz1 | 328365 | HITS-CLIP | 37986224\|23597149 | Unspecified |
| mmu-miR-708-5p | MIMAT0004828 | Zranb1 | 360216 | HITS-CLIP | 23597149 | Unspecified |
| mmu-miR-708-5p | MIMAT0004828 | Zranb2 | 53861 | HITS-CLIP | 37986224\|23597149 | Unspecified |
| mmu-miR-708-5p | MIMAT0004828 | Zswim6 | 67263 | HITS-CLIP | 37986224 | Unspecified |
| mmu-miR-708-5p | MIMAT0004828 | Zswim8 | 268721 | HITS-CLIP | 37986224\|19536157 | Unspecified |
| mmu-miR-708-5p | MIMAT0004828 | Zyg11b | 414872 | HITS-CLIP | 37986224 | Unspecified |
| mmu-miR-708-5p | MIMAT0004828 | Zyx | 22793 | HITS-CLIP | 37986224 | Unspecified |
| mmu-miR-29b-1-5p | MIMAT0004523 | 1110002E22Rik | 102634333 | HITS-CLIP | 25083871 | Unspecified |
| mmu-miR-29b-1-5p | MIMAT0004523 | 2610002M06Rik | 67028 | HITS-CLIP | 25083871 | Unspecified |
| mmu-miR-29b-1-5p | MIMAT0004523 | 4930402H24Rik | 228602 | HITS-CLIP | 37986224\|25083871 | Unspecified |
| mmu-miR-29b-1-5p | MIMAT0004523 | 4930453N24Rik | 67609 | HITS-CLIP | 37986224 | Unspecified |
| mmu-miR-29b-1-5p | MIMAT0004523 | 4933434E20Rik | 99650 | HITS-CLIP | 37986224 | Unspecified |
| mmu-miR-29b-1-5p | MIMAT0004523 | 9930021J03Rik | 240613 | HITS-CLIP | 37986224\|25083871 | Unspecified |
| mmu-miR-29b-1-5p | MIMAT0004523 | AI597479 | 98404 | HITS-CLIP | 25083871 | Unspecified |
| mmu-miR-29b-1-5p | MIMAT0004523 | AI606181 | 226118 | HITS-CLIP | 37986224 | Unspecified |
| mmu-miR-29b-1-5p | MIMAT0004523 | Aak1 | 269774 | HITS-CLIP | 37986224 | Unspecified |
| mmu-miR-29b-1-5p | MIMAT0004523 | Abca5 | 217265 | HITS-CLIP | 25083871 | Unspecified |
| mmu-miR-29b-1-5p | MIMAT0004523 | Abcc5 | 27416 | HITS-CLIP | 25083871 | Unspecified |
| mmu-miR-29b-1-5p | MIMAT0004523 | Abcg1 | 11307 | HITS-CLIP | 37986224 | Unspecified |
| mmu-miR-29b-1-5p | MIMAT0004523 | Abhd4 | 105501 | HITS-CLIP | 25083871 | Unspecified |
| mmu-miR-29b-1-5p | MIMAT0004523 | Abl2 | 11352 | HITS-CLIP | 37986224 | Unspecified |
| mmu-miR-29b-1-5p | MIMAT0004523 | Acap3 | 140500 | HITS-CLIP | 25083871 | Unspecified |
| mmu-miR-29b-1-5p | MIMAT0004523 | Acly | 104112 | HITS-CLIP | 37986224\|25083871 | Unspecified |
| mmu-miR-29b-1-5p | MIMAT0004523 | Acsl3 | 74205 | HITS-CLIP | 25083871 | Unspecified |
| mmu-miR-29b-1-5p | MIMAT0004523 | Actc1 | 11464 | HITS-CLIP | 37986224 | Unspecified |
| mmu-miR-29b-1-5p | MIMAT0004523 | Actn3 | 11474 | HITS-CLIP | 25083871 | Unspecified |
| mmu-miR-29b-1-5p | MIMAT0004523 | Acvr2a | 11480 | HITS-CLIP | 37986224\|25083871 | Unspecified |
| mmu-miR-29b-1-5p | MIMAT0004523 | Acvr2b | 11481 | HITS-CLIP | 25083871 | Unspecified |
| mmu-miR-29b-1-5p | MIMAT0004523 | Adam19 | 11492 | HITS-CLIP | 37986224 | Unspecified |
| mmu-miR-29b-1-5p | MIMAT0004523 | Adamts5 | 23794 | HITS-CLIP | 37986224 | Unspecified |
| mmu-miR-29b-1-5p | MIMAT0004523 | Adamtsl5 | 66548 | HITS-CLIP | 25083871 | Unspecified |
| mmu-miR-29b-1-5p | MIMAT0004523 | Adcy7 | 11513 | HITS-CLIP | 37986224 | Unspecified |
| mmu-miR-29b-1-5p | MIMAT0004523 | Adgrg1 | 9289 | HITS-CLIP | 25083871 | Unspecified |
| mmu-miR-29b-1-5p | MIMAT0004523 | Adi1 | 104923 | HITS-CLIP | 37986224 | Unspecified |
| mmu-miR-29b-1-5p | MIMAT0004523 | Adipor1 | 72674 | HITS-CLIP | 37986224 | Unspecified |
| mmu-miR-29b-1-5p | MIMAT0004523 | Adm | 11535 | HITS-CLIP | 37986224 | Unspecified |
| mmu-miR-29b-1-5p | MIMAT0004523 | Adnp | 11538 | HITS-CLIP | 25083871 | Unspecified |

| | | | | | | |
|---|---|---|---|---|---|---|
| mmu-miR-29b-1-5p | MIMAT0004523 | Adprh | 11544 | HITS-CLIP | 37986224 | Unspecified |
| mmu-miR-29b-1-5p | MIMAT0004523 | Aebp1 | 11568 | HITS-CLIP | 37986224 | Unspecified |
| mmu-miR-29b-1-5p | MIMAT0004523 | Agap1 | 347722 | HITS-CLIP | 25083871 | Unspecified |
| mmu-miR-29b-1-5p | MIMAT0004523 | Ago1 | 236511 | HITS-CLIP | 25083871 | Unspecified |
| mmu-miR-29b-1-5p | MIMAT0004523 | Agpat3 | 28169 | HITS-CLIP | 37986224 | Unspecified |
| mmu-miR-29b-1-5p | MIMAT0004523 | Ahdc1 | 230793 | HITS-CLIP | 37986224\|25083871 | Unspecified |
| mmu-miR-29b-1-5p | MIMAT0004523 | Ak3 | 56248 | HITS-CLIP | 37986224 | Unspecified |
| mmu-miR-29b-1-5p | MIMAT0004523 | Akap13 | 75547 | HITS-CLIP | 25083871 | Unspecified |
| mmu-miR-29b-1-5p | MIMAT0004523 | Akap6 | 238161 | HITS-CLIP | 37986224 | Unspecified |
| mmu-miR-29b-1-5p | MIMAT0004523 | Akirin2 | 433693 | HITS-CLIP | 25083871 | Unspecified |
| mmu-miR-29b-1-5p | MIMAT0004523 | Akr1a1 | 58810 | HITS-CLIP | 25083871 | Unspecified |
| mmu-miR-29b-1-5p | MIMAT0004523 | Aldh18a1 | 56454 | HITS-CLIP | 25083871 | Unspecified |
| mmu-miR-29b-1-5p | MIMAT0004523 | Alg10b | 380959 | HITS-CLIP | 37986224 | Unspecified |
| mmu-miR-29b-1-5p | MIMAT0004523 | Ampd2 | 109674 | HITS-CLIP | 37986224 | Unspecified |
| mmu-miR-29b-1-5p | MIMAT0004523 | Anapc1 | 17222 | HITS-CLIP | 37986224\|25083871 | Unspecified |
| mmu-miR-29b-1-5p | MIMAT0004523 | Ankrd12 | 106585 | HITS-CLIP | 37986224 | Unspecified |
| mmu-miR-29b-1-5p | MIMAT0004523 | Ankrd13a | 68420 | HITS-CLIP | 37986224 | Unspecified |
| mmu-miR-29b-1-5p | MIMAT0004523 | Ankrd16 | 320816 | HITS-CLIP | 37986224 | Unspecified |
| mmu-miR-29b-1-5p | MIMAT0004523 | Ankrd17 | 81702 | HITS-CLIP | 37986224\|25083871 | Unspecified |
| mmu-miR-29b-1-5p | MIMAT0004523 | Ankrd33b | 67434 | HITS-CLIP | 37986224 | Unspecified |
| mmu-miR-29b-1-5p | MIMAT0004523 | Ankrd50 | 99696 | HITS-CLIP | 25083871 | Unspecified |
| mmu-miR-29b-1-5p | MIMAT0004523 | Ano10 | 102566 | HITS-CLIP | 37986224 | Unspecified |
| mmu-miR-29b-1-5p | MIMAT0004523 | Antxr2 | 71914 | HITS-CLIP | 25083871 | Unspecified |
| mmu-miR-29b-1-5p | MIMAT0004523 | Anxa8 | 11752 | HITS-CLIP | 25083871 | Unspecified |
| mmu-miR-29b-1-5p | MIMAT0004523 | Ap2a2 | 11772 | HITS-CLIP | 25083871 | Unspecified |
| mmu-miR-29b-1-5p | MIMAT0004523 | Ap2b1 | 71770 | HITS-CLIP | 25083871 | Unspecified |
| mmu-miR-29b-1-5p | MIMAT0004523 | Apbb2 | 11787 | HITS-CLIP | 37986224 | Unspecified |
| mmu-miR-29b-1-5p | MIMAT0004523 | Apc | 11789 | HITS-CLIP | 37986224\|25083871 | Unspecified |
| mmu-miR-29b-1-5p | MIMAT0004523 | Appbp2 | 66884 | HITS-CLIP | 37986224 | Unspecified |
| mmu-miR-29b-1-5p | MIMAT0004523 | Appl1 | 72993 | HITS-CLIP | 37986224 | Unspecified |
| mmu-miR-29b-1-5p | MIMAT0004523 | Aqp5 | 11830 | HITS-CLIP | 37986224 | Unspecified |
| mmu-miR-29b-1-5p | MIMAT0004523 | Arel1 | 68497 | HITS-CLIP | 37986224\|25083871 | Unspecified |
| mmu-miR-29b-1-5p | MIMAT0004523 | Arf2 | 11841 | HITS-CLIP | 37986224 | Unspecified |
| mmu-miR-29b-1-5p | MIMAT0004523 | Arfgap1 | 228998 | HITS-CLIP | 37986224 | Unspecified |
| mmu-miR-29b-1-5p | MIMAT0004523 | Arhgap1 | 228359 | HITS-CLIP | 37986224 | Unspecified |
| mmu-miR-29b-1-5p | MIMAT0004523 | Arhgap5 | 11855 | HITS-CLIP | 25083871 | Unspecified |
| mmu-miR-29b-1-5p | MIMAT0004523 | Arhgef12 | 69632 | HITS-CLIP | 37986224\|25083871 | Unspecified |
| mmu-miR-29b-1-5p | MIMAT0004523 | Arid1b | 239985 | HITS-CLIP | 37986224 | Unspecified |
| mmu-miR-29b-1-5p | MIMAT0004523 | Arid4a | 238247 | HITS-CLIP | 37986224 | Unspecified |

| | | | | | | |
|---|---|---|---|---|---|---|
| mmu-miR-29b-1-5p | MIMAT0004523 | Arid5b | 71371 | HITS-CLIP | 37986224 | Unspecified |
| mmu-miR-29b-1-5p | MIMAT0004523 | Arih1 | 23806 | HITS-CLIP | 37986224\|25083871 | Unspecified |
| mmu-miR-29b-1-5p | MIMAT0004523 | Arl15 | 218639 | HITS-CLIP | 25083871 | Unspecified |
| mmu-miR-29b-1-5p | MIMAT0004523 | Arl4c | 320982 | HITS-CLIP | 37986224 | Unspecified |
| mmu-miR-29b-1-5p | MIMAT0004523 | Arl6ip1 | 54208 | HITS-CLIP | 37986224 | Unspecified |
| mmu-miR-29b-1-5p | MIMAT0004523 | Arl8b | 67166 | HITS-CLIP | 25083871 | Unspecified |
| mmu-miR-29b-1-5p | MIMAT0004523 | Arnt | 11863 | HITS-CLIP | 37986224 | Unspecified |
| mmu-miR-29b-1-5p | MIMAT0004523 | Arrdc3 | 105171 | HITS-CLIP | 37986224 | Unspecified |
| mmu-miR-29b-1-5p | MIMAT0004523 | Asb1 | 65247 | HITS-CLIP | 37986224 | Unspecified |
| mmu-miR-29b-1-5p | MIMAT0004523 | Asf1a | 66403 | HITS-CLIP | 37986224 | Unspecified |
| mmu-miR-29b-1-5p | MIMAT0004523 | Ash1l | 192195 | HITS-CLIP | 37986224\|25083871 | Unspecified |
| mmu-miR-29b-1-5p | MIMAT0004523 | Asxl1 | 228790 | HITS-CLIP | 37986224 | Unspecified |
| mmu-miR-29b-1-5p | MIMAT0004523 | Atad2b | 320817 | HITS-CLIP | 37986224\|25083871 | Unspecified |
| mmu-miR-29b-1-5p | MIMAT0004523 | Atf4 | 11911 | HITS-CLIP | 25083871 | Unspecified |
| mmu-miR-29b-1-5p | MIMAT0004523 | Atf6 | 226641 | HITS-CLIP | 37986224 | Unspecified |
| mmu-miR-29b-1-5p | MIMAT0004523 | Atf7 | 223922 | HITS-CLIP | 37986224 | Unspecified |
| mmu-miR-29b-1-5p | MIMAT0004523 | Atg2a | 329015 | HITS-CLIP | 37986224 | Unspecified |
| mmu-miR-29b-1-5p | MIMAT0004523 | Atic | 108147 | HITS-CLIP | 25083871 | Unspecified |
| mmu-miR-29b-1-5p | MIMAT0004523 | Atp10a | 11982 | HITS-CLIP | 25083871 | Unspecified |
| mmu-miR-29b-1-5p | MIMAT0004523 | Atp2a1 | 11937 | HITS-CLIP | 25083871 | Unspecified |
| mmu-miR-29b-1-5p | MIMAT0004523 | Atp5a1 | 11946 | HITS-CLIP | 37986224 | Unspecified |
| mmu-miR-29b-1-5p | MIMAT0004523 | Atp5h | 71679 | HITS-CLIP | 37986224\|25083871 | Unspecified |
| mmu-miR-29b-1-5p | MIMAT0004523 | Atxn1l | 52335 | HITS-CLIP | 37986224 | Unspecified |
| mmu-miR-29b-1-5p | MIMAT0004523 | Atxn2 | 20239 | HITS-CLIP | 37986224 | Unspecified |
| mmu-miR-29b-1-5p | MIMAT0004523 | Atxn2l | 233871 | HITS-CLIP | 25083871 | Unspecified |
| mmu-miR-29b-1-5p | MIMAT0004523 | Atxn3 | 110616 | HITS-CLIP | 37986224 | Unspecified |
| mmu-miR-29b-1-5p | MIMAT0004523 | Atxn7l1 | 380753 | HITS-CLIP | 37986224 | Unspecified |
| mmu-miR-29b-1-5p | MIMAT0004523 | Avl9 | 78937 | HITS-CLIP | 37986224 | Unspecified |
| mmu-miR-29b-1-5p | MIMAT0004523 | BC005537 | 79555 | HITS-CLIP | 37986224 | Unspecified |
| mmu-miR-29b-1-5p | MIMAT0004523 | BC005561 | 100042165 | HITS-CLIP | 37986224 | Unspecified |
| mmu-miR-29b-1-5p | MIMAT0004523 | BC031181 | 407819 | HITS-CLIP | 37986224 | Unspecified |
| mmu-miR-29b-1-5p | MIMAT0004523 | Bag6 | 224727 | HITS-CLIP | 25083871 | Unspecified |
| mmu-miR-29b-1-5p | MIMAT0004523 | Baiap2l1 | 66898 | HITS-CLIP | 37986224 | Unspecified |
| mmu-miR-29b-1-5p | MIMAT0004523 | Baz1a | 217578 | HITS-CLIP | 25083871 | Unspecified |
| mmu-miR-29b-1-5p | MIMAT0004523 | Baz1b | 22385 | HITS-CLIP | 37986224 | Unspecified |
| mmu-miR-29b-1-5p | MIMAT0004523 | Baz2a | 116848 | HITS-CLIP | 25083871 | Unspecified |
| mmu-miR-29b-1-5p | MIMAT0004523 | Bbx | 70508 | HITS-CLIP | 37986224 | Unspecified |
| mmu-miR-29b-1-5p | MIMAT0004523 | Bcl9l | 80288 | HITS-CLIP | 25083871 | Unspecified |
| mmu-miR-29b-1-5p | MIMAT0004523 | Bcorl1 | 320376 | HITS-CLIP | 37986224 | Unspecified |

| | | | | | | | |
|---|---|---|---|---|---|---|---|
| mmu-miR-29b-1-5p | MIMAT0004523 | Bdp1 | 544971 | HITS-CLIP | | 25083871 | Unspecified |
| mmu-miR-29b-1-5p | MIMAT0004523 | Bicd2 | 76895 | HITS-CLIP | | 25083871 | Unspecified |
| mmu-miR-29b-1-5p | MIMAT0004523 | Birc2 | 11797 | HITS-CLIP | | 37986224 | Unspecified |
| mmu-miR-29b-1-5p | MIMAT0004523 | Bivm | 246229 | HITS-CLIP | | 37986224 | Unspecified |
| mmu-miR-29b-1-5p | MIMAT0004523 | Bmi1 | 12151 | HITS-CLIP | 37986224\|25083871 | | Unspecified |
| mmu-miR-29b-1-5p | MIMAT0004523 | Bmpr2 | 12168 | HITS-CLIP | | 37986224 | Unspecified |
| mmu-miR-29b-1-5p | MIMAT0004523 | Bms1 | 213895 | HITS-CLIP | | 25083871 | Unspecified |
| mmu-miR-29b-1-5p | MIMAT0004523 | Bnc2 | 242509 | HITS-CLIP | | 25083871 | Unspecified |
| mmu-miR-29b-1-5p | MIMAT0004523 | Bnip3l | 12177 | HITS-CLIP | | 37986224 | Unspecified |
| mmu-miR-29b-1-5p | MIMAT0004523 | Bod1l | 665775 | HITS-CLIP | | 37986224 | Unspecified |
| mmu-miR-29b-1-5p | MIMAT0004523 | Bptf | 207165 | HITS-CLIP | | 25083871 | Unspecified |
| mmu-miR-29b-1-5p | MIMAT0004523 | Brap | 72399 | HITS-CLIP | | 25083871 | Unspecified |
| mmu-miR-29b-1-5p | MIMAT0004523 | Brca2 | 12190 | HITS-CLIP | | 25083871 | Unspecified |
| mmu-miR-29b-1-5p | MIMAT0004523 | Brd1 | 223770 | HITS-CLIP | | 37986224 | Unspecified |
| mmu-miR-29b-1-5p | MIMAT0004523 | Brd2 | 14312 | HITS-CLIP | | 25083871 | Unspecified |
| mmu-miR-29b-1-5p | MIMAT0004523 | Brd4 | 57261 | HITS-CLIP | | 37986224 | Unspecified |
| mmu-miR-29b-1-5p | MIMAT0004523 | Brf1 | 72308 | HITS-CLIP | | 25083871 | Unspecified |
| mmu-miR-29b-1-5p | MIMAT0004523 | Brpf3 | 268936 | HITS-CLIP | | 25083871 | Unspecified |
| mmu-miR-29b-1-5p | MIMAT0004523 | Brwd3 | 382236 | HITS-CLIP | | 37986224 | Unspecified |
| mmu-miR-29b-1-5p | MIMAT0004523 | Btaf1 | 107182 | HITS-CLIP | | 37986224 | Unspecified |
| mmu-miR-29b-1-5p | MIMAT0004523 | Btbd2 | 208198 | HITS-CLIP | | 25083871 | Unspecified |
| mmu-miR-29b-1-5p | MIMAT0004523 | Btf3 | 218490 | HITS-CLIP | | 37986224 | Unspecified |
| mmu-miR-29b-1-5p | MIMAT0004523 | Bzw1 | 66882 | HITS-CLIP | | 37986224 | Unspecified |
| mmu-miR-29b-1-5p | MIMAT0004523 | Bzw2 | 66912 | HITS-CLIP | | 37986224 | Unspecified |
| mmu-miR-29b-1-5p | MIMAT0004523 | C1galt1 | 94192 | HITS-CLIP | 37986224\|25083871 | | Unspecified |
| mmu-miR-29b-1-5p | MIMAT0004523 | C1qtnf1 | 56745 | HITS-CLIP | | 25083871 | Unspecified |
| mmu-miR-29b-1-5p | MIMAT0004523 | C77080 | 97130 | HITS-CLIP | | 37986224 | Unspecified |
| mmu-miR-29b-1-5p | MIMAT0004523 | Cab39 | 12283 | HITS-CLIP | | 25083871 | Unspecified |
| mmu-miR-29b-1-5p | MIMAT0004523 | Cacna1s | 12292 | HITS-CLIP | | 37986224 | Unspecified |
| mmu-miR-29b-1-5p | MIMAT0004523 | Cadm1 | 54725 | HITS-CLIP | | 37986224 | Unspecified |
| mmu-miR-29b-1-5p | MIMAT0004523 | Cald1 | 109624 | HITS-CLIP | | 25083871 | Unspecified |
| mmu-miR-29b-1-5p | MIMAT0004523 | Calm1 | 12313 | HITS-CLIP | | 37986224 | Unspecified |
| mmu-miR-29b-1-5p | MIMAT0004523 | Calu | 12321 | HITS-CLIP | 37986224\|25083871 | | Unspecified |
| mmu-miR-29b-1-5p | MIMAT0004523 | Camk2n1 | 66259 | HITS-CLIP | | 37986224 | Unspecified |
| mmu-miR-29b-1-5p | MIMAT0004523 | Camsap1 | 227634 | HITS-CLIP | 37986224\|25083871 | | Unspecified |
| mmu-miR-29b-1-5p | MIMAT0004523 | Camta1 | 100072 | HITS-CLIP | | 25083871 | Unspecified |
| mmu-miR-29b-1-5p | MIMAT0004523 | Cand1 | 71902 | HITS-CLIP | | 25083871 | Unspecified |
| mmu-miR-29b-1-5p | MIMAT0004523 | Cand2 | 67088 | HITS-CLIP | | 25083871 | Unspecified |
| mmu-miR-29b-1-5p | MIMAT0004523 | Capn3 | 12335 | HITS-CLIP | | 25083871 | Unspecified |

| | | | | | | |
|---|---|---|---|---|---|---|
| mmu-miR-29b-1-5p | MIMAT0004523 | Capn6 | 12338 | HITS-CLIP | 37986224 | Unspecified |
| mmu-miR-29b-1-5p | MIMAT0004523 | Capzb | 12345 | HITS-CLIP | 37986224 | Unspecified |
| mmu-miR-29b-1-5p | MIMAT0004523 | Car13 | 71934 | HITS-CLIP | 37986224 | Unspecified |
| mmu-miR-29b-1-5p | MIMAT0004523 | Cask | 12361 | HITS-CLIP | 37986224 | Unspecified |
| mmu-miR-29b-1-5p | MIMAT0004523 | Cast | 12380 | HITS-CLIP | 25083871 | Unspecified |
| mmu-miR-29b-1-5p | MIMAT0004523 | Cav1 | 12389 | HITS-CLIP | 25083871 | Unspecified |
| mmu-miR-29b-1-5p | MIMAT0004523 | Cav2 | 12390 | HITS-CLIP | 25083871 | Unspecified |
| mmu-miR-29b-1-5p | MIMAT0004523 | Cavin4 | 347273 | HITS-CLIP | 25083871 | Unspecified |
| mmu-miR-29b-1-5p | MIMAT0004523 | Cbfa2t2 | 12396 | HITS-CLIP | 37986224 | Unspecified |
| mmu-miR-29b-1-5p | MIMAT0004523 | Cblb | 208650 | HITS-CLIP | 25083871 | Unspecified |
| mmu-miR-29b-1-5p | MIMAT0004523 | Cbx1 | 12412 | HITS-CLIP | 37986224 | Unspecified |
| mmu-miR-29b-1-5p | MIMAT0004523 | Cbx7 | 52609 | HITS-CLIP | 25083871 | Unspecified |
| mmu-miR-29b-1-5p | MIMAT0004523 | Ccdc117 | 104479 | HITS-CLIP | 37986224\|25083871 | Unspecified |
| mmu-miR-29b-1-5p | MIMAT0004523 | Ccdc25 | 67179 | HITS-CLIP | 37986224 | Unspecified |
| mmu-miR-29b-1-5p | MIMAT0004523 | Ccdc50 | 67501 | HITS-CLIP | 25083871 | Unspecified |
| mmu-miR-29b-1-5p | MIMAT0004523 | Ccdc6 | 76551 | HITS-CLIP | 25083871 | Unspecified |
| mmu-miR-29b-1-5p | MIMAT0004523 | Ccl7 | 20306 | HITS-CLIP | 37986224 | Unspecified |
| mmu-miR-29b-1-5p | MIMAT0004523 | Ccnd1 | 12443 | HITS-CLIP | 25083871 | Unspecified |
| mmu-miR-29b-1-5p | MIMAT0004523 | Ccnl2 | 56036 | HITS-CLIP | 37986224 | Unspecified |
| mmu-miR-29b-1-5p | MIMAT0004523 | Ccpg1 | 72278 | HITS-CLIP | 37986224 | Unspecified |
| mmu-miR-29b-1-5p | MIMAT0004523 | Ccser2 | 72972 | HITS-CLIP | 37986224 | Unspecified |
| mmu-miR-29b-1-5p | MIMAT0004523 | Cct4 | 12464 | HITS-CLIP | 25083871 | Unspecified |
| mmu-miR-29b-1-5p | MIMAT0004523 | Cd2ap | 12488 | HITS-CLIP | 37986224 | Unspecified |
| mmu-miR-29b-1-5p | MIMAT0004523 | Cd47 | 16423 | HITS-CLIP | 37986224 | Unspecified |
| mmu-miR-29b-1-5p | MIMAT0004523 | Cd99l2 | 171486 | HITS-CLIP | 37986224 | Unspecified |
| mmu-miR-29b-1-5p | MIMAT0004523 | Cdc14b | 218294 | HITS-CLIP | 25083871 | Unspecified |
| mmu-miR-29b-1-5p | MIMAT0004523 | Cdc40 | 71713 | HITS-CLIP | 25083871 | Unspecified |
| mmu-miR-29b-1-5p | MIMAT0004523 | Cdc42bpa | 226751 | HITS-CLIP | 37986224 | Unspecified |
| mmu-miR-29b-1-5p | MIMAT0004523 | Cdc42se1 | 57912 | HITS-CLIP | 25083871 | Unspecified |
| mmu-miR-29b-1-5p | MIMAT0004523 | Cdip1 | 66626 | HITS-CLIP | 37986224 | Unspecified |
| mmu-miR-29b-1-5p | MIMAT0004523 | Cdk14 | 18647 | HITS-CLIP | 25083871 | Unspecified |
| mmu-miR-29b-1-5p | MIMAT0004523 | Cdk4 | 12567 | HITS-CLIP | 25083871 | Unspecified |
| mmu-miR-29b-1-5p | MIMAT0004523 | Cdk5r1 | 12569 | HITS-CLIP | 25083871 | Unspecified |
| mmu-miR-29b-1-5p | MIMAT0004523 | Cdk6 | 12571 | HITS-CLIP | 25083871 | Unspecified |
| mmu-miR-29b-1-5p | MIMAT0004523 | Cdk7 | 12572 | HITS-CLIP | 37986224 | Unspecified |
| mmu-miR-29b-1-5p | MIMAT0004523 | Cdk8 | 264064 | HITS-CLIP | 37986224 | Unspecified |
| mmu-miR-29b-1-5p | MIMAT0004523 | Cdkn1b | 12576 | HITS-CLIP | 37986224 | Unspecified |
| mmu-miR-29b-1-5p | MIMAT0004523 | Cdon | 57810 | HITS-CLIP | 25083871 | Unspecified |
| mmu-miR-29b-1-5p | MIMAT0004523 | Cdr2l | 237988 | HITS-CLIP | 37986224 | Unspecified |

| | | | | | | |
|---|---|---|---|---|---|---|
| mmu-miR-29b-1-5p | MIMAT0004523 | Cebpg | 12611 | HITS-CLIP | 25083871 | Unspecified |
| mmu-miR-29b-1-5p | MIMAT0004523 | Celf1 | 13046 | HITS-CLIP | 37986224\|25083871 | Unspecified |
| mmu-miR-29b-1-5p | MIMAT0004523 | Celf2 | 14007 | HITS-CLIP | 37986224 | Unspecified |
| mmu-miR-29b-1-5p | MIMAT0004523 | Cenpb | 12616 | HITS-CLIP | 25083871 | Unspecified |
| mmu-miR-29b-1-5p | MIMAT0004523 | Cenpe | 229841 | HITS-CLIP | 37986224\|25083871 | Unspecified |
| mmu-miR-29b-1-5p | MIMAT0004523 | Cenpf | 108000 | HITS-CLIP | 37986224 | Unspecified |
| mmu-miR-29b-1-5p | MIMAT0004523 | Cenpq | 83815 | HITS-CLIP | 25083871 | Unspecified |
| mmu-miR-29b-1-5p | MIMAT0004523 | Cep170b | 217882 | HITS-CLIP | 37986224 | Unspecified |
| mmu-miR-29b-1-5p | MIMAT0004523 | Cep350 | 74081 | HITS-CLIP | 25083871 | Unspecified |
| mmu-miR-29b-1-5p | MIMAT0004523 | Cep57l1 | 103268 | HITS-CLIP | 37986224 | Unspecified |
| mmu-miR-29b-1-5p | MIMAT0004523 | Cep68 | 216543 | HITS-CLIP | 37986224\|25083871 | Unspecified |
| mmu-miR-29b-1-5p | MIMAT0004523 | Cers6 | 241447 | HITS-CLIP | 37986224 | Unspecified |
| mmu-miR-29b-1-5p | MIMAT0004523 | Cggbp1 | 106143 | HITS-CLIP | 25083871 | Unspecified |
| mmu-miR-29b-1-5p | MIMAT0004523 | Cgnl1 | 68178 | HITS-CLIP | 37986224 | Unspecified |
| mmu-miR-29b-1-5p | MIMAT0004523 | Chd1 | 12648 | HITS-CLIP | 25083871 | Unspecified |
| mmu-miR-29b-1-5p | MIMAT0004523 | Chd8 | 67772 | HITS-CLIP | 25083871 | Unspecified |
| mmu-miR-29b-1-5p | MIMAT0004523 | Chd9 | 109151 | HITS-CLIP | 25083871 | Unspecified |
| mmu-miR-29b-1-5p | MIMAT0004523 | Cherp | 27967 | HITS-CLIP | 37986224 | Unspecified |
| mmu-miR-29b-1-5p | MIMAT0004523 | Chic1 | 12212 | HITS-CLIP | 25083871 | Unspecified |
| mmu-miR-29b-1-5p | MIMAT0004523 | Chmp4b | 75608 | HITS-CLIP | 37986224 | Unspecified |
| mmu-miR-29b-1-5p | MIMAT0004523 | Chordc1 | 66917 | HITS-CLIP | 25083871 | Unspecified |
| mmu-miR-29b-1-5p | MIMAT0004523 | Chrna1 | 11435 | HITS-CLIP | 37986224 | Unspecified |
| mmu-miR-29b-1-5p | MIMAT0004523 | Chrnb1 | 11443 | HITS-CLIP | 37986224 | Unspecified |
| mmu-miR-29b-1-5p | MIMAT0004523 | Chst11 | 58250 | HITS-CLIP | 37986224 | Unspecified |
| mmu-miR-29b-1-5p | MIMAT0004523 | Chsy1 | 269941 | HITS-CLIP | 37986224\|25083871 | Unspecified |
| mmu-miR-29b-1-5p | MIMAT0004523 | Cited2 | 17684 | HITS-CLIP | 37986224 | Unspecified |
| mmu-miR-29b-1-5p | MIMAT0004523 | Ckap2l | 70466 | HITS-CLIP | 37986224 | Unspecified |
| mmu-miR-29b-1-5p | MIMAT0004523 | Clasp1 | 76707 | HITS-CLIP | 25083871 | Unspecified |
| mmu-miR-29b-1-5p | MIMAT0004523 | Clasp2 | 76499 | HITS-CLIP | 25083871 | Unspecified |
| mmu-miR-29b-1-5p | MIMAT0004523 | Clip1 | 56430 | HITS-CLIP | 37986224 | Unspecified |
| mmu-miR-29b-1-5p | MIMAT0004523 | Clock | 12753 | HITS-CLIP | 37986224\|25083871 | Unspecified |
| mmu-miR-29b-1-5p | MIMAT0004523 | Clpx | 270166 | HITS-CLIP | 37986224\|25083871 | Unspecified |
| mmu-miR-29b-1-5p | MIMAT0004523 | Cltb | 74325 | HITS-CLIP | 25083871 | Unspecified |
| mmu-miR-29b-1-5p | MIMAT0004523 | Cnn3 | 71994 | HITS-CLIP | 25083871 | Unspecified |
| mmu-miR-29b-1-5p | MIMAT0004523 | Cnot11 | 52846 | HITS-CLIP | 25083871 | Unspecified |
| mmu-miR-29b-1-5p | MIMAT0004523 | Cnot4 | 53621 | HITS-CLIP | 25083871 | Unspecified |
| mmu-miR-29b-1-5p | MIMAT0004523 | Cnot7 | 18983 | HITS-CLIP | 37986224 | Unspecified |
| mmu-miR-29b-1-5p | MIMAT0004523 | Cntln | 338349 | HITS-CLIP | 37986224 | Unspecified |
| mmu-miR-29b-1-5p | MIMAT0004523 | Cog3 | 338337 | HITS-CLIP | 37986224 | Unspecified |

| | | | | | | |
|---|---|---|---|---|---|---|
| mmu-miR-29b-1-5p | MIMAT0004523 | Col1a2 | 12843 | HITS-CLIP//Immunohistochemistry//qRT-P | 25083871\|26524535 | Unspecified |
| mmu-miR-29b-1-5p | MIMAT0004523 | Col4a3bp | 68018 | HITS-CLIP | 37986224 | Unspecified |
| mmu-miR-29b-1-5p | MIMAT0004523 | Col5a1 | 12831 | HITS-CLIP | 25083871 | Unspecified |
| mmu-miR-29b-1-5p | MIMAT0004523 | Col6a3 | 12835 | HITS-CLIP | 25083871 | Unspecified |
| mmu-miR-29b-1-5p | MIMAT0004523 | Colec12 | 140792 | HITS-CLIP | 25083871 | Unspecified |
| mmu-miR-29b-1-5p | MIMAT0004523 | Comt | 12846 | HITS-CLIP | 37986224 | Unspecified |
| mmu-miR-29b-1-5p | MIMAT0004523 | Copb1 | 70349 | HITS-CLIP | 25083871 | Unspecified |
| mmu-miR-29b-1-5p | MIMAT0004523 | Copg2 | 54160 | HITS-CLIP | 25083871 | Unspecified |
| mmu-miR-29b-1-5p | MIMAT0004523 | Cops2 | 12848 | HITS-CLIP | 37986224 | Unspecified |
| mmu-miR-29b-1-5p | MIMAT0004523 | Coq9 | 67914 | HITS-CLIP | 25083871 | Unspecified |
| mmu-miR-29b-1-5p | MIMAT0004523 | Cox6b1 | 110323 | HITS-CLIP | 25083871 | Unspecified |
| mmu-miR-29b-1-5p | MIMAT0004523 | Cpeb4 | 67579 | HITS-CLIP | 37986224 | Unspecified |
| mmu-miR-29b-1-5p | MIMAT0004523 | Cpsf1 | 94230 | HITS-CLIP | 37986224\|25083871 | Unspecified |
| mmu-miR-29b-1-5p | MIMAT0004523 | Cpsf2 | 51786 | HITS-CLIP | 37986224\|25083871 | Unspecified |
| mmu-miR-29b-1-5p | MIMAT0004523 | Cpsf6 | 432508 | HITS-CLIP | 25083871 | Unspecified |
| mmu-miR-29b-1-5p | MIMAT0004523 | Cpsf7 | 269061 | HITS-CLIP | 37986224 | Unspecified |
| mmu-miR-29b-1-5p | MIMAT0004523 | Crbn | 58799 | HITS-CLIP | 37986224 | Unspecified |
| mmu-miR-29b-1-5p | MIMAT0004523 | Creb3l2 | 208647 | HITS-CLIP | 37986224 | Unspecified |
| mmu-miR-29b-1-5p | MIMAT0004523 | Crebbp | 12914 | HITS-CLIP | 37986224\|25083871 | Unspecified |
| mmu-miR-29b-1-5p | MIMAT0004523 | Crebzf | 233490 | HITS-CLIP | 37986224 | Unspecified |
| mmu-miR-29b-1-5p | MIMAT0004523 | Crim1 | 50766 | HITS-CLIP | 37986224 | Unspecified |
| mmu-miR-29b-1-5p | MIMAT0004523 | Cryab | 12955 | HITS-CLIP | 25083871 | Unspecified |
| mmu-miR-29b-1-5p | MIMAT0004523 | Csde1 | 229663 | HITS-CLIP | 25083871 | Unspecified |
| mmu-miR-29b-1-5p | MIMAT0004523 | Csnk1a1 | 93687 | HITS-CLIP | 37986224\|25083871 | Unspecified |
| mmu-miR-29b-1-5p | MIMAT0004523 | Cspg4 | 121021 | HITS-CLIP | 37986224\|25083871 | Unspecified |
| mmu-miR-29b-1-5p | MIMAT0004523 | Ctcf | 13018 | HITS-CLIP | 25083871 | Unspecified |
| mmu-miR-29b-1-5p | MIMAT0004523 | Ctnnd1 | 12388 | HITS-CLIP | 25083871 | Unspecified |
| mmu-miR-29b-1-5p | MIMAT0004523 | Ctns | 83429 | HITS-CLIP | 37986224 | Unspecified |
| mmu-miR-29b-1-5p | MIMAT0004523 | Cuedc1 | 103841 | HITS-CLIP | 37986224 | Unspecified |
| mmu-miR-29b-1-5p | MIMAT0004523 | Cul4b | 72584 | HITS-CLIP | 25083871 | Unspecified |
| mmu-miR-29b-1-5p | MIMAT0004523 | Cul5 | 75717 | HITS-CLIP | 37986224 | Unspecified |
| mmu-miR-29b-1-5p | MIMAT0004523 | Cux1 | 13047 | HITS-CLIP | 37986224\|25083871 | Unspecified |
| mmu-miR-29b-1-5p | MIMAT0004523 | Cxcl12 | 20315 | HITS-CLIP//Luciferase reporter assay//Wes | 37986224\|25083871\|32395570 | Unspecified |
| mmu-miR-29b-1-5p | MIMAT0004523 | Cyp1b1 | 13078 | HITS-CLIP | 37986224 | Unspecified |
| mmu-miR-29b-1-5p | MIMAT0004523 | Cyp51 | 13121 | HITS-CLIP | 25083871 | Unspecified |
| mmu-miR-29b-1-5p | MIMAT0004523 | Daam1 | 208846 | HITS-CLIP | 37986224 | Unspecified |
| mmu-miR-29b-1-5p | MIMAT0004523 | Daam2 | 76441 | HITS-CLIP | 37986224 | Unspecified |
| mmu-miR-29b-1-5p | MIMAT0004523 | Dab2 | 13132 | HITS-CLIP | 25083871 | Unspecified |
| mmu-miR-29b-1-5p | MIMAT0004523 | Dab2ip | 69601 | HITS-CLIP | 25083871 | Unspecified |

| | | | | | | |
|---|---|---|---|---|---|---|
| mmu-miR-29b-1-5p | MIMAT0004523 | Dag1 | 13138 | HITS-CLIP | 37986224 | Unspecified |
| mmu-miR-29b-1-5p | MIMAT0004523 | Dazap1 | 70248 | HITS-CLIP | 37986224 | Unspecified |
| mmu-miR-29b-1-5p | MIMAT0004523 | Dbr1 | 83703 | HITS-CLIP | 25083871 | Unspecified |
| mmu-miR-29b-1-5p | MIMAT0004523 | Dcaf1 | 41836 | HITS-CLIP | 25083871 | Unspecified |
| mmu-miR-29b-1-5p | MIMAT0004523 | Dcbld1 | 66686 | HITS-CLIP | 37986224 | Unspecified |
| mmu-miR-29b-1-5p | MIMAT0004523 | Dclk1 | 13175 | HITS-CLIP | 25083871 | Unspecified |
| mmu-miR-29b-1-5p | MIMAT0004523 | Dcp1a | 75901 | HITS-CLIP | 37986224 | Unspecified |
| mmu-miR-29b-1-5p | MIMAT0004523 | Dcun1d3 | 233805 | HITS-CLIP | 37986224 | Unspecified |
| mmu-miR-29b-1-5p | MIMAT0004523 | Ddost | 13200 | HITS-CLIP | 37986224 | Unspecified |
| mmu-miR-29b-1-5p | MIMAT0004523 | Ddr1 | 12305 | HITS-CLIP | 37986224 | Unspecified |
| mmu-miR-29b-1-5p | MIMAT0004523 | Ddr2 | 18214 | HITS-CLIP | 37986224 | Unspecified |
| mmu-miR-29b-1-5p | MIMAT0004523 | Ddx19a | 13680 | HITS-CLIP | 25083871 | Unspecified |
| mmu-miR-29b-1-5p | MIMAT0004523 | Ddx54 | 71990 | HITS-CLIP | 31433990 | Unspecified |
| mmu-miR-29b-1-5p | MIMAT0004523 | Dedd | 21945 | HITS-CLIP | 37986224 | Unspecified |
| mmu-miR-29b-1-5p | MIMAT0004523 | Dennd5a | 19347 | HITS-CLIP | 37986224\|25083871 | Unspecified |
| mmu-miR-29b-1-5p | MIMAT0004523 | Dennd5b | 320560 | HITS-CLIP | 37986224 | Unspecified |
| mmu-miR-29b-1-5p | MIMAT0004523 | Des | 13346 | HITS-CLIP | 25083871 | Unspecified |
| mmu-miR-29b-1-5p | MIMAT0004523 | Desi2 | 78825 | HITS-CLIP | 25083871 | Unspecified |
| mmu-miR-29b-1-5p | MIMAT0004523 | Dgkd | 227333 | HITS-CLIP | 37986224 | Unspecified |
| mmu-miR-29b-1-5p | MIMAT0004523 | Dgke | 56077 | HITS-CLIP | 25083871 | Unspecified |
| mmu-miR-29b-1-5p | MIMAT0004523 | Dhcr24 | 74754 | HITS-CLIP | 37986224 | Unspecified |
| mmu-miR-29b-1-5p | MIMAT0004523 | Dhx32 | 101437 | HITS-CLIP | 37986224 | Unspecified |
| mmu-miR-29b-1-5p | MIMAT0004523 | Dhx40 | 67487 | HITS-CLIP | 25083871 | Unspecified |
| mmu-miR-29b-1-5p | MIMAT0004523 | Dio2 | 13371 | HITS-CLIP | 37986224 | Unspecified |
| mmu-miR-29b-1-5p | MIMAT0004523 | Dlc1 | 50768 | HITS-CLIP | 37986224 | Unspecified |
| mmu-miR-29b-1-5p | MIMAT0004523 | Dlg1 | 13383 | HITS-CLIP | 37986224\|25083871 | Unspecified |
| mmu-miR-29b-1-5p | MIMAT0004523 | Dll1 | 13388 | HITS-CLIP | 37986224 | Unspecified |
| mmu-miR-29b-1-5p | MIMAT0004523 | Dlst | 78920 | HITS-CLIP | 37986224 | Unspecified |
| mmu-miR-29b-1-5p | MIMAT0004523 | Dmpk | 13400 | HITS-CLIP | 25083871 | Unspecified |
| mmu-miR-29b-1-5p | MIMAT0004523 | Dmtf1 | 23857 | HITS-CLIP | 25083871 | Unspecified |
| mmu-miR-29b-1-5p | MIMAT0004523 | Dmwd | 13401 | HITS-CLIP | 37986224 | Unspecified |
| mmu-miR-29b-1-5p | MIMAT0004523 | Dmxl1 | 240283 | HITS-CLIP | 37986224 | Unspecified |
| mmu-miR-29b-1-5p | MIMAT0004523 | Dnaja2 | 56445 | HITS-CLIP | 25083871 | Unspecified |
| mmu-miR-29b-1-5p | MIMAT0004523 | Dnajb11 | 67838 | HITS-CLIP | 37986224 | Unspecified |
| mmu-miR-29b-1-5p | MIMAT0004523 | Dnajc13 | 235567 | HITS-CLIP | 25083871 | Unspecified |
| mmu-miR-29b-1-5p | MIMAT0004523 | Dnajc3 | 100037258 | HITS-CLIP | 25083871 | Unspecified |
| mmu-miR-29b-1-5p | MIMAT0004523 | Dnttip2 | 99480 | HITS-CLIP | 25083871 | Unspecified |
| mmu-miR-29b-1-5p | MIMAT0004523 | Dot1l | 208266 | HITS-CLIP | 37986224 | Unspecified |
| mmu-miR-29b-1-5p | MIMAT0004523 | Dpy30 | 66310 | HITS-CLIP | 25083871 | Unspecified |

| | | | | | | | |
|---|---|---|---|---|---|---|---|
| mmu-miR-29b-1-5p | MIMAT0004523 | Dram1 | 71712 | HITS-CLIP | | 37986224 | Unspecified |
| mmu-miR-29b-1-5p | MIMAT0004523 | Dstn | 56431 | HITS-CLIP | | 25083871 | Unspecified |
| mmu-miR-29b-1-5p | MIMAT0004523 | Dtd2 | 328092 | HITS-CLIP | | 25083871 | Unspecified |
| mmu-miR-29b-1-5p | MIMAT0004523 | Dtna | 13527 | HITS-CLIP | | 25083871 | Unspecified |
| mmu-miR-29b-1-5p | MIMAT0004523 | Dus1l | 68730 | HITS-CLIP | | 25083871 | Unspecified |
| mmu-miR-29b-1-5p | MIMAT0004523 | Dusp7 | 235584 | HITS-CLIP | | 25083871 | Unspecified |
| mmu-miR-29b-1-5p | MIMAT0004523 | Dvl3 | 13544 | HITS-CLIP | | 37986224 | Unspecified |
| mmu-miR-29b-1-5p | MIMAT0004523 | Dync1h1 | 13424 | HITS-CLIP | 37986224\|25083871 | | Unspecified |
| mmu-miR-29b-1-5p | MIMAT0004523 | Dync1li2 | 234663 | HITS-CLIP | | 37986224 | Unspecified |
| mmu-miR-29b-1-5p | MIMAT0004523 | Dyrk1a | 13548 | HITS-CLIP | | 37986224 | Unspecified |
| mmu-miR-29b-1-5p | MIMAT0004523 | Dyrk2 | 69181 | HITS-CLIP | | 37986224 | Unspecified |
| mmu-miR-29b-1-5p | MIMAT0004523 | E2f8 | 108961 | HITS-CLIP | | 25083871 | Unspecified |
| mmu-miR-29b-1-5p | MIMAT0004523 | Eaf1 | 74427 | HITS-CLIP | | 25083871 | Unspecified |
| mmu-miR-29b-1-5p | MIMAT0004523 | Ebf1 | 13591 | HITS-CLIP | | 37986224 | Unspecified |
| mmu-miR-29b-1-5p | MIMAT0004523 | Ecm1 | 13601 | HITS-CLIP | | 37986224 | Unspecified |
| mmu-miR-29b-1-5p | MIMAT0004523 | Efcab14 | 230648 | HITS-CLIP | | 37986224 | Unspecified |
| mmu-miR-29b-1-5p | MIMAT0004523 | Efna2 | 13637 | HITS-CLIP | | 25083871 | Unspecified |
| mmu-miR-29b-1-5p | MIMAT0004523 | Efr3a | 76740 | HITS-CLIP | | 25083871 | Unspecified |
| mmu-miR-29b-1-5p | MIMAT0004523 | Ehbp1l1 | 114601 | HITS-CLIP | 37986224\|25083871 | | Unspecified |
| mmu-miR-29b-1-5p | MIMAT0004523 | Ehmt1 | 77683 | HITS-CLIP | | 37986224 | Unspecified |
| mmu-miR-29b-1-5p | MIMAT0004523 | Eid1 | 58521 | HITS-CLIP | | 37986224 | Unspecified |
| mmu-miR-29b-1-5p | MIMAT0004523 | Eif1ad | 69860 | HITS-CLIP | | 37986224 | Unspecified |
| mmu-miR-29b-1-5p | MIMAT0004523 | Eif3c | 56347 | HITS-CLIP | | 25083871 | Unspecified |
| mmu-miR-29b-1-5p | MIMAT0004523 | Eif3l | 223691 | HITS-CLIP | | 31433990 | Unspecified |
| mmu-miR-29b-1-5p | MIMAT0004523 | Eif4g1 | 208643 | HITS-CLIP | 37986224\|25083871 | | Unspecified |
| mmu-miR-29b-1-5p | MIMAT0004523 | Eif4g2 | 13690 | HITS-CLIP | 37986224\|25083871 | | Unspecified |
| mmu-miR-29b-1-5p | MIMAT0004523 | Eif4g3 | 230861 | HITS-CLIP | 37986224\|25083871 | | Unspecified |
| mmu-miR-29b-1-5p | MIMAT0004523 | Eif5b | 226982 | HITS-CLIP | | 25083871 | Unspecified |
| mmu-miR-29b-1-5p | MIMAT0004523 | Elk4 | 13714 | HITS-CLIP | | 37986224 | Unspecified |
| mmu-miR-29b-1-5p | MIMAT0004523 | Ell2 | 192657 | HITS-CLIP | | 25083871 | Unspecified |
| mmu-miR-29b-1-5p | MIMAT0004523 | Elmsan1 | 238317 | HITS-CLIP | | 37986224 | Unspecified |
| mmu-miR-29b-1-5p | MIMAT0004523 | Enah | 13800 | HITS-CLIP | | 37986224 | Unspecified |
| mmu-miR-29b-1-5p | MIMAT0004523 | Ep300 | 328572 | HITS-CLIP | 37986224\|25083871 | | Unspecified |
| mmu-miR-29b-1-5p | MIMAT0004523 | Epg5 | 100502841 | HITS-CLIP | | 37986224 | Unspecified |
| mmu-miR-29b-1-5p | MIMAT0004523 | Ephb3 | 13845 | HITS-CLIP | | 25083871 | Unspecified |
| mmu-miR-29b-1-5p | MIMAT0004523 | Epm2aip1 | 77781 | HITS-CLIP | 37986224\|25083871 | | Unspecified |
| mmu-miR-29b-1-5p | MIMAT0004523 | Eprs | 107508 | HITS-CLIP | | 25083871 | Unspecified |
| mmu-miR-29b-1-5p | MIMAT0004523 | Ercc6 | 319955 | HITS-CLIP | | 37986224 | Unspecified |
| mmu-miR-29b-1-5p | MIMAT0004523 | Ergic1 | 67458 | HITS-CLIP | | 37986224 | Unspecified |

| | | | | | | |
|---|---|---|---|---|---|---|
| mmu-miR-29b-1-5p | MIMAT0004523 | Erlin1 | 226144 | HITS-CLIP | 25083871 | Unspecified |
| mmu-miR-29b-1-5p | MIMAT0004523 | Errfi1 | 74155 | HITS-CLIP | 25083871 | Unspecified |
| mmu-miR-29b-1-5p | MIMAT0004523 | Esf1 | 66580 | HITS-CLIP | 37986224 | Unspecified |
| mmu-miR-29b-1-5p | MIMAT0004523 | Etf1 | 225363 | HITS-CLIP | 37986224\|25083871 | Unspecified |
| mmu-miR-29b-1-5p | MIMAT0004523 | Etfdh | 66841 | HITS-CLIP | 37986224 | Unspecified |
| mmu-miR-29b-1-5p | MIMAT0004523 | Ewsr1 | 14030 | HITS-CLIP | 37986224 | Unspecified |
| mmu-miR-29b-1-5p | MIMAT0004523 | Exoc5 | 105504 | HITS-CLIP | 37986224 | Unspecified |
| mmu-miR-29b-1-5p | MIMAT0004523 | Exosc2 | 227715 | HITS-CLIP | 37986224 | Unspecified |
| mmu-miR-29b-1-5p | MIMAT0004523 | Ext1 | 14042 | HITS-CLIP | 37986224 | Unspecified |
| mmu-miR-29b-1-5p | MIMAT0004523 | Eya3 | 14050 | HITS-CLIP | 25083871 | Unspecified |
| mmu-miR-29b-1-5p | MIMAT0004523 | Eya4 | 14051 | HITS-CLIP | 37986224 | Unspecified |
| mmu-miR-29b-1-5p | MIMAT0004523 | Fads3 | 60527 | HITS-CLIP | 37986224\|25083871 | Unspecified |
| mmu-miR-29b-1-5p | MIMAT0004523 | Fam110a | 73847 | HITS-CLIP | 25083871 | Unspecified |
| mmu-miR-29b-1-5p | MIMAT0004523 | Fam118a | 73225 | HITS-CLIP | 25083871 | Unspecified |
| mmu-miR-29b-1-5p | MIMAT0004523 | Fam126a | 84652 | HITS-CLIP | 37986224 | Unspecified |
| mmu-miR-29b-1-5p | MIMAT0004523 | Fam135a | 68187 | HITS-CLIP | 37986224 | Unspecified |
| mmu-miR-29b-1-5p | MIMAT0004523 | Fam160b1 | 226252 | HITS-CLIP | 25083871 | Unspecified |
| mmu-miR-29b-1-5p | MIMAT0004523 | Fam193a | 231128 | HITS-CLIP | 37986224 | Unspecified |
| mmu-miR-29b-1-5p | MIMAT0004523 | Fam208a | 218850 | HITS-CLIP | 25083871 | Unspecified |
| mmu-miR-29b-1-5p | MIMAT0004523 | Fam20b | 215015 | HITS-CLIP | 37986224 | Unspecified |
| mmu-miR-29b-1-5p | MIMAT0004523 | Fam210a | 108654 | HITS-CLIP | 37986224 | Unspecified |
| mmu-miR-29b-1-5p | MIMAT0004523 | Fam214a | 235493 | HITS-CLIP | 37986224 | Unspecified |
| mmu-miR-29b-1-5p | MIMAT0004523 | Fam222b | 216971 | HITS-CLIP | 37986224\|25083871 | Unspecified |
| mmu-miR-29b-1-5p | MIMAT0004523 | Fam89b | 17826 | HITS-CLIP | 25083871 | Unspecified |
| mmu-miR-29b-1-5p | MIMAT0004523 | Fam91a1 | 210998 | HITS-CLIP | 25083871 | Unspecified |
| mmu-miR-29b-1-5p | MIMAT0004523 | Fancf | 100040608 | HITS-CLIP | 37986224 | Unspecified |
| mmu-miR-29b-1-5p | MIMAT0004523 | Far1 | 67420 | HITS-CLIP | 37986224 | Unspecified |
| mmu-miR-29b-1-5p | MIMAT0004523 | Farp2 | 227377 | HITS-CLIP | 37986224 | Unspecified |
| mmu-miR-29b-1-5p | MIMAT0004523 | Fat1 | 14107 | HITS-CLIP | 25083871 | Unspecified |
| mmu-miR-29b-1-5p | MIMAT0004523 | Fat4 | 329628 | HITS-CLIP | 37986224 | Unspecified |
| mmu-miR-29b-1-5p | MIMAT0004523 | Fbrsl1 | 381668 | HITS-CLIP | 25083871 | Unspecified |
| mmu-miR-29b-1-5p | MIMAT0004523 | Fbxl17 | 50758 | HITS-CLIP | 37986224 | Unspecified |
| mmu-miR-29b-1-5p | MIMAT0004523 | Fbxl18 | 231863 | HITS-CLIP | 37986224 | Unspecified |
| mmu-miR-29b-1-5p | MIMAT0004523 | Fbxw5 | 30839 | HITS-CLIP | 37986224 | Unspecified |
| mmu-miR-29b-1-5p | MIMAT0004523 | Fgd6 | 13998 | HITS-CLIP | 25083871 | Unspecified |
| mmu-miR-29b-1-5p | MIMAT0004523 | Fgf7 | 14178 | HITS-CLIP | 37986224 | Unspecified |
| mmu-miR-29b-1-5p | MIMAT0004523 | Fgfr1op2 | 67529 | HITS-CLIP | 37986224 | Unspecified |
| mmu-miR-29b-1-5p | MIMAT0004523 | Fkbp10 | 14230 | HITS-CLIP | 37986224\|25083871 | Unspecified |
| mmu-miR-29b-1-5p | MIMAT0004523 | Fkbp15 | 338355 | HITS-CLIP | 25083871 | Unspecified |

| | | | | | | | |
|---|---|---|---|---|---|---|---|
| mmu-miR-29b-1-5p | MIMAT0004523 | Fkbp3 | 30795 | HITS-CLIP | | 37986224 | Unspecified |
| mmu-miR-29b-1-5p | MIMAT0004523 | Flcn | 216805 | HITS-CLIP | | 37986224 | Unspecified |
| mmu-miR-29b-1-5p | MIMAT0004523 | Flii | 14248 | HITS-CLIP | | 25083871 | Unspecified |
| mmu-miR-29b-1-5p | MIMAT0004523 | Flna | 192176 | HITS-CLIP | | 25083871 | Unspecified |
| mmu-miR-29b-1-5p | MIMAT0004523 | Flnb | 286940 | HITS-CLIP | | 37986224 | Unspecified |
| mmu-miR-29b-1-5p | MIMAT0004523 | Flrt2 | 399558 | HITS-CLIP | 37986224\|25083871 | | Unspecified |
| mmu-miR-29b-1-5p | MIMAT0004523 | Fmo1 | 14261 | HITS-CLIP | | 25083871 | Unspecified |
| mmu-miR-29b-1-5p | MIMAT0004523 | Fmod | 14264 | HITS-CLIP | | 37986224 | Unspecified |
| mmu-miR-29b-1-5p | MIMAT0004523 | Fndc3a | 319448 | HITS-CLIP | | 25083871 | Unspecified |
| mmu-miR-29b-1-5p | MIMAT0004523 | Fndc3b | 72007 | HITS-CLIP | | 25083871 | Unspecified |
| mmu-miR-29b-1-5p | MIMAT0004523 | Fndc5 | 384061 | HITS-CLIP | | 25083871 | Unspecified |
| mmu-miR-29b-1-5p | MIMAT0004523 | Fnip1 | 216742 | HITS-CLIP | | 37986224 | Unspecified |
| mmu-miR-29b-1-5p | MIMAT0004523 | Fosl2 | 14284 | HITS-CLIP | 37986224\|25083871 | | Unspecified |
| mmu-miR-29b-1-5p | MIMAT0004523 | Foxj3 | 230700 | HITS-CLIP | | 25083871 | Unspecified |
| mmu-miR-29b-1-5p | MIMAT0004523 | Foxo3 | 56484 | HITS-CLIP | 37986224\|25083871 | | Unspecified |
| mmu-miR-29b-1-5p | MIMAT0004523 | Foxo6 | 329934 | HITS-CLIP | | 25083871 | Unspecified |
| mmu-miR-29b-1-5p | MIMAT0004523 | Frk | 14302 | HITS-CLIP | | 25083871 | Unspecified |
| mmu-miR-29b-1-5p | MIMAT0004523 | Frmd4a | 209630 | HITS-CLIP | 37986224\|25083871 | | Unspecified |
| mmu-miR-29b-1-5p | MIMAT0004523 | Fstl3 | 83554 | HITS-CLIP | | 37986224 | Unspecified |
| mmu-miR-29b-1-5p | MIMAT0004523 | Fubp1 | 51886 | HITS-CLIP | | 37986224 | Unspecified |
| mmu-miR-29b-1-5p | MIMAT0004523 | Fzd6 | 14368 | HITS-CLIP | 37986224\|25083871 | | Unspecified |
| mmu-miR-29b-1-5p | MIMAT0004523 | G3bp1 | 27041 | HITS-CLIP | | 37986224 | Unspecified |
| mmu-miR-29b-1-5p | MIMAT0004523 | Gabarap | 56486 | HITS-CLIP | | 37986224 | Unspecified |
| mmu-miR-29b-1-5p | MIMAT0004523 | Gadd45gip1 | 102060 | HITS-CLIP | | 25083871 | Unspecified |
| mmu-miR-29b-1-5p | MIMAT0004523 | Galnt1 | 14423 | HITS-CLIP | | 25083871 | Unspecified |
| mmu-miR-29b-1-5p | MIMAT0004523 | Galnt2 | 108148 | HITS-CLIP | | 37986224 | Unspecified |
| mmu-miR-29b-1-5p | MIMAT0004523 | Gan | 209239 | HITS-CLIP | | 25083871 | Unspecified |
| mmu-miR-29b-1-5p | MIMAT0004523 | Ganab | 14376 | HITS-CLIP | | 37986224 | Unspecified |
| mmu-miR-29b-1-5p | MIMAT0004523 | Gas1 | 14451 | HITS-CLIP | | 25083871 | Unspecified |
| mmu-miR-29b-1-5p | MIMAT0004523 | Gcc2 | 70297 | HITS-CLIP | | 37986224 | Unspecified |
| mmu-miR-29b-1-5p | MIMAT0004523 | Gdap2 | 14547 | HITS-CLIP | | 25083871 | Unspecified |
| mmu-miR-29b-1-5p | MIMAT0004523 | Gdi2 | 14569 | HITS-CLIP | | 25083871 | Unspecified |
| mmu-miR-29b-1-5p | MIMAT0004523 | Ghr | 14600 | HITS-CLIP | | 37986224 | Unspecified |
| mmu-miR-29b-1-5p | MIMAT0004523 | Git1 | 216963 | HITS-CLIP | | 37986224 | Unspecified |
| mmu-miR-29b-1-5p | MIMAT0004523 | Glud1 | 14661 | HITS-CLIP | | 25083871 | Unspecified |
| mmu-miR-29b-1-5p | MIMAT0004523 | Gm49336 | 127230315 | HITS-CLIP | | 25083871 | Unspecified |
| mmu-miR-29b-1-5p | MIMAT0004523 | Gmppb | 331026 | HITS-CLIP | | 25083871 | Unspecified |
| mmu-miR-29b-1-5p | MIMAT0004523 | Gnas | 14683 | HITS-CLIP | | 25083871 | Unspecified |
| mmu-miR-29b-1-5p | MIMAT0004523 | Gnb1 | 14688 | HITS-CLIP | | 37986224 | Unspecified |

| | | | | | | | |
|---|---|---|---|---|---|---|---|
| mmu-miR-29b-1-5p | MIMAT0004523 | Gnl3l | 237107 | HITS-CLIP | | 37986224 | Unspecified |
| mmu-miR-29b-1-5p | MIMAT0004523 | Gnpat | 14712 | HITS-CLIP | 37986224\|25083871 | | Unspecified |
| mmu-miR-29b-1-5p | MIMAT0004523 | Golga1 | 76899 | HITS-CLIP | | 25083871 | Unspecified |
| mmu-miR-29b-1-5p | MIMAT0004523 | Golga4 | 54214 | HITS-CLIP | | 25083871 | Unspecified |
| mmu-miR-29b-1-5p | MIMAT0004523 | Golgb1 | 224139 | HITS-CLIP | | 37986224 | Unspecified |
| mmu-miR-29b-1-5p | MIMAT0004523 | Gon4l | 76022 | HITS-CLIP | | 25083871 | Unspecified |
| mmu-miR-29b-1-5p | MIMAT0004523 | Gorasp2 | 70231 | HITS-CLIP | | 25083871 | Unspecified |
| mmu-miR-29b-1-5p | MIMAT0004523 | Gpatch8 | 237943 | HITS-CLIP | | 37986224 | Unspecified |
| mmu-miR-29b-1-5p | MIMAT0004523 | Gpcpd1 | 74182 | HITS-CLIP | | 25083871 | Unspecified |
| mmu-miR-29b-1-5p | MIMAT0004523 | Gpd2 | 14571 | HITS-CLIP | 37986224\|25083871 | | Unspecified |
| mmu-miR-29b-1-5p | MIMAT0004523 | Gpr63 | 81006 | HITS-CLIP | | 37986224 | Unspecified |
| mmu-miR-29b-1-5p | MIMAT0004523 | Gpt2 | 108682 | HITS-CLIP | | 25083871 | Unspecified |
| mmu-miR-29b-1-5p | MIMAT0004523 | Gpx4 | 625249 | HITS-CLIP | | 25083871 | Unspecified |
| mmu-miR-29b-1-5p | MIMAT0004523 | Grem1 | 23892 | HITS-CLIP | 37986224\|25083871 | | Unspecified |
| mmu-miR-29b-1-5p | MIMAT0004523 | Grina | 66168 | HITS-CLIP | | 25083871 | Unspecified |
| mmu-miR-29b-1-5p | MIMAT0004523 | Grk5 | 14773 | HITS-CLIP | | 37986224 | Unspecified |
| mmu-miR-29b-1-5p | MIMAT0004523 | Grk6 | 26385 | HITS-CLIP | | 25083871 | Unspecified |
| mmu-miR-29b-1-5p | MIMAT0004523 | Grsf1 | 231413 | HITS-CLIP | | 37986224 | Unspecified |
| mmu-miR-29b-1-5p | MIMAT0004523 | Gsn | 227753 | HITS-CLIP | | 25083871 | Unspecified |
| mmu-miR-29b-1-5p | MIMAT0004523 | Gtf2a1 | 83602 | HITS-CLIP | | 25083871 | Unspecified |
| mmu-miR-29b-1-5p | MIMAT0004523 | Gtf2b | 229906 | HITS-CLIP | | 25083871 | Unspecified |
| mmu-miR-29b-1-5p | MIMAT0004523 | Gtf2e1 | 74197 | HITS-CLIP | | 37986224 | Unspecified |
| mmu-miR-29b-1-5p | MIMAT0004523 | Gtf2i | 14886 | HITS-CLIP | | 25083871 | Unspecified |
| mmu-miR-29b-1-5p | MIMAT0004523 | Gtf2ird1 | 57080 | HITS-CLIP | | 25083871 | Unspecified |
| mmu-miR-29b-1-5p | MIMAT0004523 | Gtpbp2 | 56055 | HITS-CLIP | | 25083871 | Unspecified |
| mmu-miR-29b-1-5p | MIMAT0004523 | Gtpbp4 | 69237 | HITS-CLIP | | 37986224 | Unspecified |
| mmu-miR-29b-1-5p | MIMAT0004523 | Gyg | 27357 | HITS-CLIP | | 25083871 | Unspecified |
| mmu-miR-29b-1-5p | MIMAT0004523 | Habp4 | 56541 | HITS-CLIP | | 25083871 | Unspecified |
| mmu-miR-29b-1-5p | MIMAT0004523 | Has2 | 15117 | HITS-CLIP | | 37986224 | Unspecified |
| mmu-miR-29b-1-5p | MIMAT0004523 | Hcfc1 | 15161 | HITS-CLIP | | 25083871 | Unspecified |
| mmu-miR-29b-1-5p | MIMAT0004523 | Hdac4 | 208727 | HITS-CLIP | | 25083871 | Unspecified |
| mmu-miR-29b-1-5p | MIMAT0004523 | Hdgf | 15191 | HITS-CLIP | | 25083871 | Unspecified |
| mmu-miR-29b-1-5p | MIMAT0004523 | Heatr3 | 234549 | HITS-CLIP | | 25083871 | Unspecified |
| mmu-miR-29b-1-5p | MIMAT0004523 | Hectd1 | 207304 | HITS-CLIP | | 37986224 | Unspecified |
| mmu-miR-29b-1-5p | MIMAT0004523 | Hectd2 | 226098 | HITS-CLIP | | 37986224 | Unspecified |
| mmu-miR-29b-1-5p | MIMAT0004523 | Herc1 | 235439 | HITS-CLIP | | 25083871 | Unspecified |
| mmu-miR-29b-1-5p | MIMAT0004523 | Hexim1 | 192231 | HITS-CLIP | 37986224\|25083871 | | Unspecified |
| mmu-miR-29b-1-5p | MIMAT0004523 | Hfe2 | 69585 | HITS-CLIP | | 25083871 | Unspecified |
| mmu-miR-29b-1-5p | MIMAT0004523 | Hic2 | 58180 | HITS-CLIP | | 37986224 | Unspecified |

| | | | | | | | |
|---|---|---|---|---|---|---|---|
| mmu-miR-29b-1-5p | MIMAT0004523 | Hif1a | 15251 | HITS-CLIP | | 25083871 | Unspecified |
| mmu-miR-29b-1-5p | MIMAT0004523 | Hif1an | 319594 | HITS-CLIP | | 37986224 | Unspecified |
| mmu-miR-29b-1-5p | MIMAT0004523 | Hipk1 | 15257 | HITS-CLIP | | 37986224 | Unspecified |
| mmu-miR-29b-1-5p | MIMAT0004523 | Hipk2 | 15258 | HITS-CLIP | 37986224\|25083871 | | Unspecified |
| mmu-miR-29b-1-5p | MIMAT0004523 | Hipk3 | 15259 | HITS-CLIP | | 37986224 | Unspecified |
| mmu-miR-29b-1-5p | MIMAT0004523 | Hivep1 | 110521 | HITS-CLIP | | 37986224 | Unspecified |
| mmu-miR-29b-1-5p | MIMAT0004523 | Hmga2 | 15364 | HITS-CLIP | | 37986224 | Unspecified |
| mmu-miR-29b-1-5p | MIMAT0004523 | Hmgcl | 15356 | HITS-CLIP | | 37986224 | Unspecified |
| mmu-miR-29b-1-5p | MIMAT0004523 | Hmgxb4 | 70823 | HITS-CLIP | | 37986224 | Unspecified |
| mmu-miR-29b-1-5p | MIMAT0004523 | Hnrnpa2b1 | 53379 | HITS-CLIP | | 37986224 | Unspecified |
| mmu-miR-29b-1-5p | MIMAT0004523 | Hnrnpab | 15384 | HITS-CLIP | 37986224\|25083871 | | Unspecified |
| mmu-miR-29b-1-5p | MIMAT0004523 | Hnrnpl | 15388 | HITS-CLIP | | 37986224 | Unspecified |
| mmu-miR-29b-1-5p | MIMAT0004523 | Hnrnpu | 51810 | HITS-CLIP | | 25083871 | Unspecified |
| mmu-miR-29b-1-5p | MIMAT0004523 | Hnrnpul1 | 232989 | HITS-CLIP | | 25083871 | Unspecified |
| mmu-miR-29b-1-5p | MIMAT0004523 | Hook3 | 320191 | HITS-CLIP | | 37986224 | Unspecified |
| mmu-miR-29b-1-5p | MIMAT0004523 | Hoxa6 | 15403 | HITS-CLIP | | 37986224 | Unspecified |
| mmu-miR-29b-1-5p | MIMAT0004523 | Hoxc4 | 15423 | HITS-CLIP | | 25083871 | Unspecified |
| mmu-miR-29b-1-5p | MIMAT0004523 | Hoxc9 | 15427 | HITS-CLIP | | 37986224 | Unspecified |
| mmu-miR-29b-1-5p | MIMAT0004523 | Hspa4 | 15525 | HITS-CLIP | | 25083871 | Unspecified |
| mmu-miR-29b-1-5p | MIMAT0004523 | Hspa4l | 18415 | HITS-CLIP | | 25083871 | Unspecified |
| mmu-miR-29b-1-5p | MIMAT0004523 | Htra1 | 56213 | HITS-CLIP | | 25083871 | Unspecified |
| mmu-miR-29b-1-5p | MIMAT0004523 | Huwe1 | 59026 | HITS-CLIP | | 25083871 | Unspecified |
| mmu-miR-29b-1-5p | MIMAT0004523 | Ibtk | 108837 | HITS-CLIP | | 25083871 | Unspecified |
| mmu-miR-29b-1-5p | MIMAT0004523 | Ier3 | 15937 | HITS-CLIP | | 26203562 | Unspecified |
| mmu-miR-29b-1-5p | MIMAT0004523 | Ifngr1 | 15979 | HITS-CLIP | | 37986224 | Unspecified |
| mmu-miR-29b-1-5p | MIMAT0004523 | Igf2r | 16004 | HITS-CLIP | | 25083871 | Unspecified |
| mmu-miR-29b-1-5p | MIMAT0004523 | Igsf3 | 78908 | HITS-CLIP | | 25083871 | Unspecified |
| mmu-miR-29b-1-5p | MIMAT0004523 | Ik | 24010 | HITS-CLIP | | 25083871 | Unspecified |
| mmu-miR-29b-1-5p | MIMAT0004523 | Ikzf2 | 22779 | HITS-CLIP | | 37986224 | Unspecified |
| mmu-miR-29b-1-5p | MIMAT0004523 | Il12a | 16159 | HITS-CLIP | 37986224\|25083871 | | Unspecified |
| mmu-miR-29b-1-5p | MIMAT0004523 | Il6st | 16195 | HITS-CLIP | 37986224\|25083871 | | Unspecified |
| mmu-miR-29b-1-5p | MIMAT0004523 | Ilf2 | 67781 | HITS-CLIP | | 25083871 | Unspecified |
| mmu-miR-29b-1-5p | MIMAT0004523 | Ilf3 | 16201 | HITS-CLIP | | 25083871 | Unspecified |
| mmu-miR-29b-1-5p | MIMAT0004523 | Immt | 76614 | HITS-CLIP | 37986224\|25083871 | | Unspecified |
| mmu-miR-29b-1-5p | MIMAT0004523 | Ing3 | 71777 | HITS-CLIP | | 25083871 | Unspecified |
| mmu-miR-29b-1-5p | MIMAT0004523 | Inhbb | 16324 | HITS-CLIP | | 37986224 | Unspecified |
| mmu-miR-29b-1-5p | MIMAT0004523 | Ino80c | 225280 | HITS-CLIP | | 25083871 | Unspecified |
| mmu-miR-29b-1-5p | MIMAT0004523 | Ino80d | 227195 | HITS-CLIP | 37986224\|25083871 | | Unspecified |
| mmu-miR-29b-1-5p | MIMAT0004523 | Inpp4b | 234515 | HITS-CLIP | | 25083871 | Unspecified |

| | | | | | | |
|---|---|---|---|---|---|---|
| mmu-miR-29b-1-5p | MIMAT0004523 | Ints2 | 70422 | HITS-CLIP | 37986224 | Unspecified |
| mmu-miR-29b-1-5p | MIMAT0004523 | Ints4 | 101861 | HITS-CLIP | 37986224 | Unspecified |
| mmu-miR-29b-1-5p | MIMAT0004523 | Ints6 | 18130 | HITS-CLIP | 37986224 | Unspecified |
| mmu-miR-29b-1-5p | MIMAT0004523 | Ip6k3 | 271424 | HITS-CLIP | 37986224 | Unspecified |
| mmu-miR-29b-1-5p | MIMAT0004523 | Iqgap1 | 29875 | HITS-CLIP | 37986224\|25083871 | Unspecified |
| mmu-miR-29b-1-5p | MIMAT0004523 | Iqsec2 | 245666 | HITS-CLIP | 25083871 | Unspecified |
| mmu-miR-29b-1-5p | MIMAT0004523 | Irak4 | 266632 | HITS-CLIP | 37986224 | Unspecified |
| mmu-miR-29b-1-5p | MIMAT0004523 | Ireb2 | 64602 | HITS-CLIP | 37986224 | Unspecified |
| mmu-miR-29b-1-5p | MIMAT0004523 | Irs1 | 16367 | HITS-CLIP | 37986224\|25083871 | Unspecified |
| mmu-miR-29b-1-5p | MIMAT0004523 | Ist1 | 71955 | HITS-CLIP | 37986224 | Unspecified |
| mmu-miR-29b-1-5p | MIMAT0004523 | Itsn1 | 16443 | HITS-CLIP | 37986224 | Unspecified |
| mmu-miR-29b-1-5p | MIMAT0004523 | Iws1 | 73473 | HITS-CLIP | 37986224 | Unspecified |
| mmu-miR-29b-1-5p | MIMAT0004523 | Jam2 | 67374 | HITS-CLIP | 37986224 | Unspecified |
| mmu-miR-29b-1-5p | MIMAT0004523 | Jdp2 | 81703 | HITS-CLIP | 37986224 | Unspecified |
| mmu-miR-29b-1-5p | MIMAT0004523 | Jmjd1c | 108829 | HITS-CLIP | 37986224\|25083871 | Unspecified |
| mmu-miR-29b-1-5p | MIMAT0004523 | Jmjd4 | 194952 | HITS-CLIP | 37986224 | Unspecified |
| mmu-miR-29b-1-5p | MIMAT0004523 | Kat6a | 244349 | HITS-CLIP | 37986224\|25083871 | Unspecified |
| mmu-miR-29b-1-5p | MIMAT0004523 | Kbtbd4 | 67136 | HITS-CLIP | 37986224 | Unspecified |
| mmu-miR-29b-1-5p | MIMAT0004523 | Kbtbd8 | 243574 | HITS-CLIP | 37986224 | Unspecified |
| mmu-miR-29b-1-5p | MIMAT0004523 | Kcnn3 | 140493 | HITS-CLIP | 37986224\|25083871 | Unspecified |
| mmu-miR-29b-1-5p | MIMAT0004523 | Kctd10 | 330171 | HITS-CLIP | 37986224 | Unspecified |
| mmu-miR-29b-1-5p | MIMAT0004523 | Kctd5 | 69259 | HITS-CLIP | 37986224 | Unspecified |
| mmu-miR-29b-1-5p | MIMAT0004523 | Kctd9 | 105440 | HITS-CLIP | 37986224 | Unspecified |
| mmu-miR-29b-1-5p | MIMAT0004523 | Kdm1b | 218214 | HITS-CLIP | 37986224 | Unspecified |
| mmu-miR-29b-1-5p | MIMAT0004523 | Khsrp | 16549 | HITS-CLIP | 25083871 | Unspecified |
| mmu-miR-29b-1-5p | MIMAT0004523 | Kif1bp | 26128 | HITS-CLIP | 25083871 | Unspecified |
| mmu-miR-29b-1-5p | MIMAT0004523 | Kif1c | 16562 | HITS-CLIP | 25083871 | Unspecified |
| mmu-miR-29b-1-5p | MIMAT0004523 | Kif20a | 19348 | HITS-CLIP | 37986224\|25083871 | Unspecified |
| mmu-miR-29b-1-5p | MIMAT0004523 | Kif21b | 16565 | HITS-CLIP | 37986224\|25083871 | Unspecified |
| mmu-miR-29b-1-5p | MIMAT0004523 | Kif2a | 16563 | HITS-CLIP | 25083871 | Unspecified |
| mmu-miR-29b-1-5p | MIMAT0004523 | Klc1 | 16593 | HITS-CLIP | 25083871 | Unspecified |
| mmu-miR-29b-1-5p | MIMAT0004523 | Klc3 | 232943 | HITS-CLIP | 31433990 | Unspecified |
| mmu-miR-29b-1-5p | MIMAT0004523 | Klf10 | 21847 | HITS-CLIP | 37986224 | Unspecified |
| mmu-miR-29b-1-5p | MIMAT0004523 | Klf13 | 50794 | HITS-CLIP | 37986224 | Unspecified |
| mmu-miR-29b-1-5p | MIMAT0004523 | Klf5 | 12224 | HITS-CLIP | 37986224\|25083871 | Unspecified |
| mmu-miR-29b-1-5p | MIMAT0004523 | Klf7 | 93691 | HITS-CLIP | 37986224 | Unspecified |
| mmu-miR-29b-1-5p | MIMAT0004523 | Klf9 | 16601 | HITS-CLIP | 25083871 | Unspecified |
| mmu-miR-29b-1-5p | MIMAT0004523 | Klhdc10 | 76788 | HITS-CLIP | 37986224 | Unspecified |
| mmu-miR-29b-1-5p | MIMAT0004523 | Klhl2 | 77113 | HITS-CLIP | 25083871 | Unspecified |

| | | | | | | |
|---|---|---|---|---|---|---|
| mmu-miR-29b-1-5p | MIMAT0004523 | Klhl24 | 75785 | HITS-CLIP | 37986224\|25083871 | Unspecified |
| mmu-miR-29b-1-5p | MIMAT0004523 | Kmt2a | 214162 | HITS-CLIP | 37986224 | Unspecified |
| mmu-miR-29b-1-5p | MIMAT0004523 | Kmt2b | 75410 | HITS-CLIP | 25083871 | Unspecified |
| mmu-miR-29b-1-5p | MIMAT0004523 | Kmt2c | 231051 | HITS-CLIP | 37986224\|25083871 | Unspecified |
| mmu-miR-29b-1-5p | MIMAT0004523 | Kmt2d | 381022 | HITS-CLIP | 37986224\|25083871 | Unspecified |
| mmu-miR-29b-1-5p | MIMAT0004523 | Kmt2e | 69188 | HITS-CLIP | 25083871 | Unspecified |
| mmu-miR-29b-1-5p | MIMAT0004523 | Knop1 | 66356 | HITS-CLIP | 37986224 | Unspecified |
| mmu-miR-29b-1-5p | MIMAT0004523 | Kpna1 | 16646 | HITS-CLIP | 37986224\|25083871 | Unspecified |
| mmu-miR-29b-1-5p | MIMAT0004523 | Kpna3 | 16648 | HITS-CLIP | 37986224 | Unspecified |
| mmu-miR-29b-1-5p | MIMAT0004523 | Krcc1 | 57896 | HITS-CLIP | 37986224 | Unspecified |
| mmu-miR-29b-1-5p | MIMAT0004523 | Lats1 | 16798 | HITS-CLIP | 37986224 | Unspecified |
| mmu-miR-29b-1-5p | MIMAT0004523 | Lbh | 77889 | HITS-CLIP | 25083871 | Unspecified |
| mmu-miR-29b-1-5p | MIMAT0004523 | Ldlr | 16835 | HITS-CLIP | 25083871 | Unspecified |
| mmu-miR-29b-1-5p | MIMAT0004523 | Ldlrap1 | 100017 | HITS-CLIP | 25083871 | Unspecified |
| mmu-miR-29b-1-5p | MIMAT0004523 | Lemd3 | 380664 | HITS-CLIP | 25083871 | Unspecified |
| mmu-miR-29b-1-5p | MIMAT0004523 | Leo1 | 235497 | HITS-CLIP | 25083871 | Unspecified |
| mmu-miR-29b-1-5p | MIMAT0004523 | Letmd1 | 68614 | HITS-CLIP | 37986224 | Unspecified |
| mmu-miR-29b-1-5p | MIMAT0004523 | Lgals1 | 16852 | HITS-CLIP | 25083871 | Unspecified |
| mmu-miR-29b-1-5p | MIMAT0004523 | Lhfpl2 | 218454 | HITS-CLIP | 37986224 | Unspecified |
| mmu-miR-29b-1-5p | MIMAT0004523 | Lims1 | 110829 | HITS-CLIP | 37986224 | Unspecified |
| mmu-miR-29b-1-5p | MIMAT0004523 | Lmod3 | 320502 | HITS-CLIP | 25083871 | Unspecified |
| mmu-miR-29b-1-5p | MIMAT0004523 | Lonrf1 | 244421 | HITS-CLIP | 25083871 | Unspecified |
| mmu-miR-29b-1-5p | MIMAT0004523 | Loxl1 | 16949 | HITS-CLIP | 37986224 | Unspecified |
| mmu-miR-29b-1-5p | MIMAT0004523 | Lpar1 | 14745 | HITS-CLIP | 37986224 | Unspecified |
| mmu-miR-29b-1-5p | MIMAT0004523 | Lpp | 210126 | HITS-CLIP | 37986224\|25083871 | Unspecified |
| mmu-miR-29b-1-5p | MIMAT0004523 | Lrch4 | 231798 | HITS-CLIP | 25083871 | Unspecified |
| mmu-miR-29b-1-5p | MIMAT0004523 | Lrpprc | 72416 | HITS-CLIP | 25083871 | Unspecified |
| mmu-miR-29b-1-5p | MIMAT0004523 | Lrrc58 | 320184 | HITS-CLIP | 25083871 | Unspecified |
| mmu-miR-29b-1-5p | MIMAT0004523 | Lrrc8c | 100604 | HITS-CLIP | 37986224 | Unspecified |
| mmu-miR-29b-1-5p | MIMAT0004523 | Lrrk1 | 233328 | HITS-CLIP | 37986224 | Unspecified |
| mmu-miR-29b-1-5p | MIMAT0004523 | Lsm6 | 78651 | HITS-CLIP | 25083871 | Unspecified |
| mmu-miR-29b-1-5p | MIMAT0004523 | Luc7l2 | 192196 | HITS-CLIP | 37986224 | Unspecified |
| mmu-miR-29b-1-5p | MIMAT0004523 | Lypla1 | 18777 | HITS-CLIP | 37986224\|25083871 | Unspecified |
| mmu-miR-29b-1-5p | MIMAT0004523 | Macf1 | 11426 | HITS-CLIP | 25083871 | Unspecified |
| mmu-miR-29b-1-5p | MIMAT0004523 | Maco1 | 55219 | HITS-CLIP | 25083871 | Unspecified |
| mmu-miR-29b-1-5p | MIMAT0004523 | Maf | 17132 | HITS-CLIP | 37986224 | Unspecified |
| mmu-miR-29b-1-5p | MIMAT0004523 | Maged1 | 94275 | HITS-CLIP | 25083871 | Unspecified |
| mmu-miR-29b-1-5p | MIMAT0004523 | Magi3 | 99470 | HITS-CLIP | 37986224 | Unspecified |
| mmu-miR-29b-1-5p | MIMAT0004523 | Maml2 | 270118 | HITS-CLIP | 37986224 | Unspecified |

| | | | | | | |
|---|---|---|---|---|---|---|
| mmu-miR-29b-1-5p | MIMAT0004523 | Maml3 | 433586 | HITS-CLIP | 37986224 | Unspecified |
| mmu-miR-29b-1-5p | MIMAT0004523 | Manea | 242362 | HITS-CLIP | 37986224 | Unspecified |
| mmu-miR-29b-1-5p | MIMAT0004523 | Map1a | 17754 | HITS-CLIP | 37986224\|25083871 | Unspecified |
| mmu-miR-29b-1-5p | MIMAT0004523 | Map3k1 | 26401 | HITS-CLIP | 37986224 | Unspecified |
| mmu-miR-29b-1-5p | MIMAT0004523 | Map3k3 | 26406 | HITS-CLIP | 37986224 | Unspecified |
| mmu-miR-29b-1-5p | MIMAT0004523 | Map4 | 17758 | HITS-CLIP | 37986224 | Unspecified |
| mmu-miR-29b-1-5p | MIMAT0004523 | Map7d1 | 245877 | HITS-CLIP | 37986224\|25083871 | Unspecified |
| mmu-miR-29b-1-5p | MIMAT0004523 | Mapk14 | 26416 | HITS-CLIP | 31433990 | Unspecified |
| mmu-miR-29b-1-5p | MIMAT0004523 | Mapk3 | 26417 | HITS-CLIP | 25083871 | Unspecified |
| mmu-miR-29b-1-5p | MIMAT0004523 | Mapk8 | 26419 | HITS-CLIP | 37986224 | Unspecified |
| mmu-miR-29b-1-5p | MIMAT0004523 | Mapk8ip3 | 30957 | HITS-CLIP | 37986224 | Unspecified |
| mmu-miR-29b-1-5p | MIMAT0004523 | Mapkapk2 | 17164 | HITS-CLIP | 37986224 | Unspecified |
| mmu-miR-29b-1-5p | MIMAT0004523 | Marcks | 17118 | HITS-CLIP | 37986224 | Unspecified |
| mmu-miR-29b-1-5p | MIMAT0004523 | Mast2 | 17776 | HITS-CLIP | 25083871 | Unspecified |
| mmu-miR-29b-1-5p | MIMAT0004523 | Mast4 | 328329 | HITS-CLIP | 37986224\|25083871 | Unspecified |
| mmu-miR-29b-1-5p | MIMAT0004523 | Matr3 | 17184 | HITS-CLIP | 25083871 | Unspecified |
| mmu-miR-29b-1-5p | MIMAT0004523 | Mbtd1 | 103537 | HITS-CLIP | 37986224 | Unspecified |
| mmu-miR-29b-1-5p | MIMAT0004523 | Mdn1 | 100019 | HITS-CLIP | 37986224 | Unspecified |
| mmu-miR-29b-1-5p | MIMAT0004523 | Mecom | 14013 | HITS-CLIP | 37986224 | Unspecified |
| mmu-miR-29b-1-5p | MIMAT0004523 | Mecp2 | 17257 | HITS-CLIP | 37986224 | Unspecified |
| mmu-miR-29b-1-5p | MIMAT0004523 | Med15 | 94112 | HITS-CLIP | 25083871 | Unspecified |
| mmu-miR-29b-1-5p | MIMAT0004523 | Med22 | 20933 | HITS-CLIP | 37986224 | Unspecified |
| mmu-miR-29b-1-5p | MIMAT0004523 | Med26 | 70625 | HITS-CLIP | 37986224 | Unspecified |
| mmu-miR-29b-1-5p | MIMAT0004523 | Mef2d | 17261 | HITS-CLIP | 37986224 | Unspecified |
| mmu-miR-29b-1-5p | MIMAT0004523 | Men1 | 17283 | HITS-CLIP | 25083871 | Unspecified |
| mmu-miR-29b-1-5p | MIMAT0004523 | Mepce | 231803 | HITS-CLIP | 37986224 | Unspecified |
| mmu-miR-29b-1-5p | MIMAT0004523 | Met | 17295 | HITS-CLIP | 25083871 | Unspecified |
| mmu-miR-29b-1-5p | MIMAT0004523 | Mettl16 | 67493 | HITS-CLIP | 37986224 | Unspecified |
| mmu-miR-29b-1-5p | MIMAT0004523 | Mfap3l | 71306 | HITS-CLIP | 37986224 | Unspecified |
| mmu-miR-29b-1-5p | MIMAT0004523 | Mfsd14a | 15247 | HITS-CLIP | 25083871 | Unspecified |
| mmu-miR-29b-1-5p | MIMAT0004523 | Mga | 29808 | HITS-CLIP | 37986224\|25083871 | Unspecified |
| mmu-miR-29b-1-5p | MIMAT0004523 | Mgat1 | 17308 | HITS-CLIP | 37986224 | Unspecified |
| mmu-miR-29b-1-5p | MIMAT0004523 | Micall1 | 27008 | HITS-CLIP | 37986224 | Unspecified |
| mmu-miR-29b-1-5p | MIMAT0004523 | Mid1ip1 | 68041 | HITS-CLIP | 37986224 | Unspecified |
| mmu-miR-29b-1-5p | MIMAT0004523 | Midn | 59090 | HITS-CLIP | 25083871 | Unspecified |
| mmu-miR-29b-1-5p | MIMAT0004523 | Mitf | 17342 | HITS-CLIP | 37986224 | Unspecified |
| mmu-miR-29b-1-5p | MIMAT0004523 | Mki67 | 17345 | HITS-CLIP | 37986224 | Unspecified |
| mmu-miR-29b-1-5p | MIMAT0004523 | Mkks | 59030 | HITS-CLIP | 37986224 | Unspecified |
| mmu-miR-29b-1-5p | MIMAT0004523 | Mknk2 | 17347 | HITS-CLIP | 25083871 | Unspecified |

| | | | | | | |
|---|---|---|---|---|---|---|
| mmu-miR-29b-1-5p | MIMAT0004523 | Mllt6 | 246198 | HITS-CLIP | 37986224\|25083871 | Unspecified |
| mmu-miR-29b-1-5p | MIMAT0004523 | Mlxip | 208104 | HITS-CLIP | 25083871 | Unspecified |
| mmu-miR-29b-1-5p | MIMAT0004523 | Mmgt1 | 236792 | HITS-CLIP | 37986224 | Unspecified |
| mmu-miR-29b-1-5p | MIMAT0004523 | Mon1b | 270096 | HITS-CLIP | 37986224 | Unspecified |
| mmu-miR-29b-1-5p | MIMAT0004523 | Mpdz | 17475 | HITS-CLIP | 25083871 | Unspecified |
| mmu-miR-29b-1-5p | MIMAT0004523 | Mprip | 26936 | HITS-CLIP | 37986224 | Unspecified |
| mmu-miR-29b-1-5p | MIMAT0004523 | Mras | 17532 | HITS-CLIP | 25083871 | Unspecified |
| mmu-miR-29b-1-5p | MIMAT0004523 | Mrpl1 | 94061 | HITS-CLIP | 37986224 | Unspecified |
| mmu-miR-29b-1-5p | MIMAT0004523 | Mrpl15 | 27395 | HITS-CLIP | 25083871 | Unspecified |
| mmu-miR-29b-1-5p | MIMAT0004523 | Mrpl34 | 94065 | HITS-CLIP | 37986224 | Unspecified |
| mmu-miR-29b-1-5p | MIMAT0004523 | Mrtfb | 239719 | HITS-CLIP | 25083871 | Unspecified |
| mmu-miR-29b-1-5p | MIMAT0004523 | Msi2 | 76626 | HITS-CLIP | 37986224 | Unspecified |
| mmu-miR-29b-1-5p | MIMAT0004523 | Msl2 | 77853 | HITS-CLIP | 25083871 | Unspecified |
| mmu-miR-29b-1-5p | MIMAT0004523 | Msl3 | 17692 | HITS-CLIP | 25083871 | Unspecified |
| mmu-miR-29b-1-5p | MIMAT0004523 | Msn | 17698 | HITS-CLIP | 25083871 | Unspecified |
| mmu-miR-29b-1-5p | MIMAT0004523 | Mthfd2 | 17768 | HITS-CLIP | 25083871 | Unspecified |
| mmu-miR-29b-1-5p | MIMAT0004523 | Mthfsd | 234814 | HITS-CLIP | 25083871 | Unspecified |
| mmu-miR-29b-1-5p | MIMAT0004523 | Mtss1 | 211401 | HITS-CLIP | 37986224 | Unspecified |
| mmu-miR-29b-1-5p | MIMAT0004523 | Mtus1 | 102103 | HITS-CLIP | 37986224 | Unspecified |
| mmu-miR-29b-1-5p | MIMAT0004523 | Myadm | 50918 | HITS-CLIP | 25083871 | Unspecified |
| mmu-miR-29b-1-5p | MIMAT0004523 | Myc | 17869 | HITS-CLIP | 37986224 | Unspecified |
| mmu-miR-29b-1-5p | MIMAT0004523 | Myh3 | 17883 | HITS-CLIP | 25083871 | Unspecified |
| mmu-miR-29b-1-5p | MIMAT0004523 | Myh8 | 17885 | HITS-CLIP | 25083871 | Unspecified |
| mmu-miR-29b-1-5p | MIMAT0004523 | Myh9 | 17886 | HITS-CLIP | 37986224 | Unspecified |
| mmu-miR-29b-1-5p | MIMAT0004523 | Myo18a | 360013 | HITS-CLIP | 25083871 | Unspecified |
| mmu-miR-29b-1-5p | MIMAT0004523 | Myo5a | 17918 | HITS-CLIP | 25083871 | Unspecified |
| mmu-miR-29b-1-5p | MIMAT0004523 | Myo9a | 270163 | HITS-CLIP | 25083871 | Unspecified |
| mmu-miR-29b-1-5p | MIMAT0004523 | Myom2 | 17930 | HITS-CLIP | 25083871 | Unspecified |
| mmu-miR-29b-1-5p | MIMAT0004523 | N4bp2l2 | 381695 | HITS-CLIP | 37986224 | Unspecified |
| mmu-miR-29b-1-5p | MIMAT0004523 | Naa25 | 231713 | HITS-CLIP | 37986224 | Unspecified |
| mmu-miR-29b-1-5p | MIMAT0004523 | Nacc2 | 67991 | HITS-CLIP | 37986224\|25083871 | Unspecified |
| mmu-miR-29b-1-5p | MIMAT0004523 | Napg | 108123 | HITS-CLIP | 37986224 | Unspecified |
| mmu-miR-29b-1-5p | MIMAT0004523 | Narf | 67608 | HITS-CLIP | 37986224 | Unspecified |
| mmu-miR-29b-1-5p | MIMAT0004523 | Nav2 | 78286 | HITS-CLIP | 37986224 | Unspecified |
| mmu-miR-29b-1-5p | MIMAT0004523 | Nav3 | 260315 | HITS-CLIP | 25083871 | Unspecified |
| mmu-miR-29b-1-5p | MIMAT0004523 | Nbeal1 | 269198 | HITS-CLIP | 37986224 | Unspecified |
| mmu-miR-29b-1-5p | MIMAT0004523 | Nbr1 | 17966 | HITS-CLIP | 37986224 | Unspecified |
| mmu-miR-29b-1-5p | MIMAT0004523 | Ncam1 | 17967 | HITS-CLIP | 37986224\|25083871 | Unspecified |
| mmu-miR-29b-1-5p | MIMAT0004523 | Ncapd3 | 78658 | HITS-CLIP | 37986224 | Unspecified |

| | | | | | | |
|---|---|---|---|---|---|---|
| mmu-miR-29b-1-5p | MIMAT0004523 | Ncaph | 215387 | HITS-CLIP | 37986224 | Unspecified |
| mmu-miR-29b-1-5p | MIMAT0004523 | Nckap1 | 50884 | HITS-CLIP | 25083871 | Unspecified |
| mmu-miR-29b-1-5p | MIMAT0004523 | Ncl | 17975 | HITS-CLIP | 37986224 | Unspecified |
| mmu-miR-29b-1-5p | MIMAT0004523 | Ncln | 103425 | HITS-CLIP | 37986224 | Unspecified |
| mmu-miR-29b-1-5p | MIMAT0004523 | Ncoa1 | 17977 | HITS-CLIP | 37986224 | Unspecified |
| mmu-miR-29b-1-5p | MIMAT0004523 | Ncoa2 | 17978 | HITS-CLIP | 37986224 | Unspecified |
| mmu-miR-29b-1-5p | MIMAT0004523 | Ncoa5 | 228869 | HITS-CLIP | 37986224 | Unspecified |
| mmu-miR-29b-1-5p | MIMAT0004523 | Ncoa6 | 56406 | HITS-CLIP | 25083871 | Unspecified |
| mmu-miR-29b-1-5p | MIMAT0004523 | Ncor1 | 20185 | HITS-CLIP | 25083871 | Unspecified |
| mmu-miR-29b-1-5p | MIMAT0004523 | Ncor2 | 20602 | HITS-CLIP | 25083871 | Unspecified |
| mmu-miR-29b-1-5p | MIMAT0004523 | Ncs1 | 14299 | HITS-CLIP | 25083871 | Unspecified |
| mmu-miR-29b-1-5p | MIMAT0004523 | Ndel1 | 83431 | HITS-CLIP | 25083871 | Unspecified |
| mmu-miR-29b-1-5p | MIMAT0004523 | Ndfip1 | 65113 | HITS-CLIP | 37986224 | Unspecified |
| mmu-miR-29b-1-5p | MIMAT0004523 | Ndrg2 | 29811 | HITS-CLIP | 25083871 | Unspecified |
| mmu-miR-29b-1-5p | MIMAT0004523 | Ndst1 | 15531 | HITS-CLIP | 25083871 | Unspecified |
| mmu-miR-29b-1-5p | MIMAT0004523 | Neb | 17996 | HITS-CLIP | 37986224\|25083871 | Unspecified |
| mmu-miR-29b-1-5p | MIMAT0004523 | Nedd4 | 17999 | HITS-CLIP | 25083871 | Unspecified |
| mmu-miR-29b-1-5p | MIMAT0004523 | Nek7 | 59125 | HITS-CLIP | 25083871 | Unspecified |
| mmu-miR-29b-1-5p | MIMAT0004523 | Nemf | 66244 | HITS-CLIP | 37986224 | Unspecified |
| mmu-miR-29b-1-5p | MIMAT0004523 | Nes | 18008 | HITS-CLIP | 25083871 | Unspecified |
| mmu-miR-29b-1-5p | MIMAT0004523 | Nf1 | 18015 | HITS-CLIP | 25083871 | Unspecified |
| mmu-miR-29b-1-5p | MIMAT0004523 | Nfat5 | 54446 | HITS-CLIP | 25083871 | Unspecified |
| mmu-miR-29b-1-5p | MIMAT0004523 | Nfatc3 | 18021 | HITS-CLIP | 37986224 | Unspecified |
| mmu-miR-29b-1-5p | MIMAT0004523 | Nfe2l1 | 18023 | HITS-CLIP | 37986224 | Unspecified |
| mmu-miR-29b-1-5p | MIMAT0004523 | Nfib | 18028 | HITS-CLIP | 25083871 | Unspecified |
| mmu-miR-29b-1-5p | MIMAT0004523 | Nfkb1 | 18033 | HITS-CLIP | 37986224 | Unspecified |
| mmu-miR-29b-1-5p | MIMAT0004523 | Nfrkb | 235134 | HITS-CLIP | 37986224 | Unspecified |
| mmu-miR-29b-1-5p | MIMAT0004523 | Nid2 | 18074 | HITS-CLIP | 25083871 | Unspecified |
| mmu-miR-29b-1-5p | MIMAT0004523 | Nipbl | 71175 | HITS-CLIP | 25083871 | Unspecified |
| mmu-miR-29b-1-5p | MIMAT0004523 | Nisch | 64652 | HITS-CLIP | 37986224\|25083871 | Unspecified |
| mmu-miR-29b-1-5p | MIMAT0004523 | Nkap | 67050 | HITS-CLIP | 37986224 | Unspecified |
| mmu-miR-29b-1-5p | MIMAT0004523 | Nktr | 18087 | HITS-CLIP | 37986224 | Unspecified |
| mmu-miR-29b-1-5p | MIMAT0004523 | Nlgn2 | 216856 | HITS-CLIP | 37986224 | Unspecified |
| mmu-miR-29b-1-5p | MIMAT0004523 | Nln | 75805 | HITS-CLIP | 37986224 | Unspecified |
| mmu-miR-29b-1-5p | MIMAT0004523 | Notch1 | 18128 | HITS-CLIP | 25083871 | Unspecified |
| mmu-miR-29b-1-5p | MIMAT0004523 | Notch2 | 18129 | HITS-CLIP | 25083871 | Unspecified |
| mmu-miR-29b-1-5p | MIMAT0004523 | Notch3 | 18131 | HITS-CLIP | 25083871 | Unspecified |
| mmu-miR-29b-1-5p | MIMAT0004523 | Npc1 | 18145 | HITS-CLIP | 37986224 | Unspecified |
| mmu-miR-29b-1-5p | MIMAT0004523 | Nploc4 | 217365 | HITS-CLIP | 25083871 | Unspecified |

| | | | | | | |
|---|---|---|---|---|---|---|
| mmu-miR-29b-1-5p | MIMAT0004523 | Npnt | 114249 | HITS-CLIP | 37986224 | Unspecified |
| mmu-miR-29b-1-5p | MIMAT0004523 | Nptn | 20320 | HITS-CLIP | 25083871 | Unspecified |
| mmu-miR-29b-1-5p | MIMAT0004523 | Nr1d2 | 353187 | HITS-CLIP | 37986224\|25083871 | Unspecified |
| mmu-miR-29b-1-5p | MIMAT0004523 | Nr2f2 | 11819 | HITS-CLIP | 37986224\|25083871 | Unspecified |
| mmu-miR-29b-1-5p | MIMAT0004523 | Nr2f6 | 13864 | HITS-CLIP | 25083871 | Unspecified |
| mmu-miR-29b-1-5p | MIMAT0004523 | Nr3c1 | 14815 | HITS-CLIP | 37986224 | Unspecified |
| mmu-miR-29b-1-5p | MIMAT0004523 | Nr6a1 | 14536 | HITS-CLIP | 37986224\|25083871 | Unspecified |
| mmu-miR-29b-1-5p | MIMAT0004523 | Nrbp1 | 192292 | HITS-CLIP | 25083871 | Unspecified |
| mmu-miR-29b-1-5p | MIMAT0004523 | Nrd1 | 230598 | HITS-CLIP | 25083871 | Unspecified |
| mmu-miR-29b-1-5p | MIMAT0004523 | Nrf1 | 18181 | HITS-CLIP | 37986224 | Unspecified |
| mmu-miR-29b-1-5p | MIMAT0004523 | Nrp1 | 18186 | HITS-CLIP | 37986224 | Unspecified |
| mmu-miR-29b-1-5p | MIMAT0004523 | Nsd1 | 18193 | HITS-CLIP | 37986224 | Unspecified |
| mmu-miR-29b-1-5p | MIMAT0004523 | Nsd3 | 234135 | HITS-CLIP | 25083871 | Unspecified |
| mmu-miR-29b-1-5p | MIMAT0004523 | Nsmce3 | 56160 | HITS-CLIP | 25083871 | Unspecified |
| mmu-miR-29b-1-5p | MIMAT0004523 | Nt5dc3 | 103466 | HITS-CLIP | 37986224 | Unspecified |
| mmu-miR-29b-1-5p | MIMAT0004523 | Ntn4 | 57764 | HITS-CLIP | 25083871 | Unspecified |
| mmu-miR-29b-1-5p | MIMAT0004523 | Nup107 | 103468 | HITS-CLIP | 25083871 | Unspecified |
| mmu-miR-29b-1-5p | MIMAT0004523 | Nup153 | 218210 | HITS-CLIP | 25083871 | Unspecified |
| mmu-miR-29b-1-5p | MIMAT0004523 | Nup214 | 227720 | HITS-CLIP | 37986224 | Unspecified |
| mmu-miR-29b-1-5p | MIMAT0004523 | Nup62 | 18226 | HITS-CLIP | 37986224 | Unspecified |
| mmu-miR-29b-1-5p | MIMAT0004523 | Nup93 | 71805 | HITS-CLIP | 37986224 | Unspecified |
| mmu-miR-29b-1-5p | MIMAT0004523 | Nxn | 18230 | HITS-CLIP | 37986224 | Unspecified |
| mmu-miR-29b-1-5p | MIMAT0004523 | Obscn | 380698 | HITS-CLIP | 25083871 | Unspecified |
| mmu-miR-29b-1-5p | MIMAT0004523 | Ociad1 | 68095 | HITS-CLIP | 37986224 | Unspecified |
| mmu-miR-29b-1-5p | MIMAT0004523 | Ogn | 18295 | HITS-CLIP | 25083871 | Unspecified |
| mmu-miR-29b-1-5p | MIMAT0004523 | Ogt | 108155 | HITS-CLIP | 37986224 | Unspecified |
| mmu-miR-29b-1-5p | MIMAT0004523 | Ola1 | 67059 | HITS-CLIP | 25083871 | Unspecified |
| mmu-miR-29b-1-5p | MIMAT0004523 | Olfm1 | 56177 | HITS-CLIP | 25083871 | Unspecified |
| mmu-miR-29b-1-5p | MIMAT0004523 | Olfml2b | 320078 | HITS-CLIP | 37986224 | Unspecified |
| mmu-miR-29b-1-5p | MIMAT0004523 | Orai1 | 109305 | HITS-CLIP | 25083871 | Unspecified |
| mmu-miR-29b-1-5p | MIMAT0004523 | Orc2 | 18393 | HITS-CLIP | 37986224 | Unspecified |
| mmu-miR-29b-1-5p | MIMAT0004523 | Orc6 | 56452 | HITS-CLIP | 37986224 | Unspecified |
| mmu-miR-29b-1-5p | MIMAT0004523 | Otud4 | 73945 | HITS-CLIP | 37986224 | Unspecified |
| mmu-miR-29b-1-5p | MIMAT0004523 | Oxsr1 | 108737 | HITS-CLIP | 37986224 | Unspecified |
| mmu-miR-29b-1-5p | MIMAT0004523 | P4hb | 18453 | HITS-CLIP | 25083871 | Unspecified |
| mmu-miR-29b-1-5p | MIMAT0004523 | Pa2g4 | 18813 | HITS-CLIP | 25083871 | Unspecified |
| mmu-miR-29b-1-5p | MIMAT0004523 | Pabpc4 | 230721 | HITS-CLIP | 37986224 | Unspecified |
| mmu-miR-29b-1-5p | MIMAT0004523 | Pacs2 | 217893 | HITS-CLIP | 25083871 | Unspecified |
| mmu-miR-29b-1-5p | MIMAT0004523 | Pafah1b2 | 18475 | HITS-CLIP | 37986224\|25083871 | Unspecified |

| | | | | | | | |
|---|---|---|---|---|---|---|---|
| mmu-miR-29b-1-5p | MIMAT0004523 | Paip2 | 67869 | HITS-CLIP | | 25083871 | Unspecified |
| mmu-miR-29b-1-5p | MIMAT0004523 | Pak2 | 224105 | HITS-CLIP | | 37986224 | Unspecified |
| mmu-miR-29b-1-5p | MIMAT0004523 | Pan3 | 72587 | HITS-CLIP | | 25083871 | Unspecified |
| mmu-miR-29b-1-5p | MIMAT0004523 | Pank3 | 211347 | HITS-CLIP | | 37986224 | Unspecified |
| mmu-miR-29b-1-5p | MIMAT0004523 | Papola | 18789 | HITS-CLIP | | 37986224 | Unspecified |
| mmu-miR-29b-1-5p | MIMAT0004523 | Papss1 | 23971 | HITS-CLIP | | 37986224 | Unspecified |
| mmu-miR-29b-1-5p | MIMAT0004523 | Pard3 | 93742 | HITS-CLIP | | 37986224 | Unspecified |
| mmu-miR-29b-1-5p | MIMAT0004523 | Parg | 26430 | HITS-CLIP | | 37986224 | Unspecified |
| mmu-miR-29b-1-5p | MIMAT0004523 | Parp3 | 235587 | HITS-CLIP | | 37986224 | Unspecified |
| mmu-miR-29b-1-5p | MIMAT0004523 | Parva | 57342 | HITS-CLIP | | 37986224 | Unspecified |
| mmu-miR-29b-1-5p | MIMAT0004523 | Patl1 | 225929 | HITS-CLIP | | 37986224 | Unspecified |
| mmu-miR-29b-1-5p | MIMAT0004523 | Pbrm1 | 66923 | HITS-CLIP | | 25083871 | Unspecified |
| mmu-miR-29b-1-5p | MIMAT0004523 | Pcdhb21 | 93892 | HITS-CLIP | | 37986224 | Unspecified |
| mmu-miR-29b-1-5p | MIMAT0004523 | Pcdhga12 | 93724 | HITS-CLIP | 37986224\|25083871 | | Unspecified |
| mmu-miR-29b-1-5p | MIMAT0004523 | Pcgf3 | 69587 | HITS-CLIP | | 25083871 | Unspecified |
| mmu-miR-29b-1-5p | MIMAT0004523 | Pcm1 | 18536 | HITS-CLIP | | 25083871 | Unspecified |
| mmu-miR-29b-1-5p | MIMAT0004523 | Pcnx | 54604 | HITS-CLIP | | 25083871 | Unspecified |
| mmu-miR-29b-1-5p | MIMAT0004523 | Pdcl | 67466 | HITS-CLIP | | 37986224 | Unspecified |
| mmu-miR-29b-1-5p | MIMAT0004523 | Pde4d | 238871 | HITS-CLIP | | 37986224 | Unspecified |
| mmu-miR-29b-1-5p | MIMAT0004523 | Pde4dip | 83679 | HITS-CLIP | | 25083871 | Unspecified |
| mmu-miR-29b-1-5p | MIMAT0004523 | Pdhb | 68263 | HITS-CLIP | | 25083871 | Unspecified |
| mmu-miR-29b-1-5p | MIMAT0004523 | Pdik1l | 230809 | HITS-CLIP | | 37986224 | Unspecified |
| mmu-miR-29b-1-5p | MIMAT0004523 | Pdk2 | 18604 | HITS-CLIP | | 25083871 | Unspecified |
| mmu-miR-29b-1-5p | MIMAT0004523 | Pdlim7 | 67399 | HITS-CLIP | | 25083871 | Unspecified |
| mmu-miR-29b-1-5p | MIMAT0004523 | Pdzd8 | 107368 | HITS-CLIP | | 25083871 | Unspecified |
| mmu-miR-29b-1-5p | MIMAT0004523 | Pea15a | 18611 | HITS-CLIP | | 37986224 | Unspecified |
| mmu-miR-29b-1-5p | MIMAT0004523 | Peak1 | 244895 | HITS-CLIP | | 25083871 | Unspecified |
| mmu-miR-29b-1-5p | MIMAT0004523 | Peli1 | 67245 | HITS-CLIP | | 37986224 | Unspecified |
| mmu-miR-29b-1-5p | MIMAT0004523 | Pex13 | 72129 | HITS-CLIP | | 25083871 | Unspecified |
| mmu-miR-29b-1-5p | MIMAT0004523 | Pgrmc2 | 70804 | HITS-CLIP | | 37986224 | Unspecified |
| mmu-miR-29b-1-5p | MIMAT0004523 | Phax | 56698 | HITS-CLIP | | 37986224 | Unspecified |
| mmu-miR-29b-1-5p | MIMAT0004523 | Phf10 | 72057 | HITS-CLIP | | 25083871 | Unspecified |
| mmu-miR-29b-1-5p | MIMAT0004523 | Phf20l1 | 239510 | HITS-CLIP | 37986224\|25083871 | | Unspecified |
| mmu-miR-29b-1-5p | MIMAT0004523 | Phip | 83946 | HITS-CLIP | | 37986224 | Unspecified |
| mmu-miR-29b-1-5p | MIMAT0004523 | Phlpp1 | 98432 | HITS-CLIP | 37986224\|25083871 | | Unspecified |
| mmu-miR-29b-1-5p | MIMAT0004523 | Phtf2 | 68770 | HITS-CLIP | | 25083871 | Unspecified |
| mmu-miR-29b-1-5p | MIMAT0004523 | Pi4ka | 224020 | HITS-CLIP | | 37986224 | Unspecified |
| mmu-miR-29b-1-5p | MIMAT0004523 | Pias1 | 56469 | HITS-CLIP | | 37986224 | Unspecified |
| mmu-miR-29b-1-5p | MIMAT0004523 | Piezo2 | 667742 | HITS-CLIP | | 25083871 | Unspecified |

| | | | | | | |
|---|---|---|---|---|---|---|
| mmu-miR-29b-1-5p | MIMAT0004523 | Pigm | 67556 | HITS-CLIP | 37986224 | Unspecified |
| mmu-miR-29b-1-5p | MIMAT0004523 | Pign | 27392 | HITS-CLIP | 37986224 | Unspecified |
| mmu-miR-29b-1-5p | MIMAT0004523 | Pigp | 56176 | HITS-CLIP | 37986224 | Unspecified |
| mmu-miR-29b-1-5p | MIMAT0004523 | Pik3ca | 18706 | HITS-CLIP | 37986224 | Unspecified |
| mmu-miR-29b-1-5p | MIMAT0004523 | Pip4k2c | 117150 | HITS-CLIP | 37986224 | Unspecified |
| mmu-miR-29b-1-5p | MIMAT0004523 | Pkd1 | 18763 | HITS-CLIP | 37986224 | Unspecified |
| mmu-miR-29b-1-5p | MIMAT0004523 | Plaa | 18786 | HITS-CLIP | 25083871 | Unspecified |
| mmu-miR-29b-1-5p | MIMAT0004523 | Plbd2 | 71772 | HITS-CLIP | 25083871 | Unspecified |
| mmu-miR-29b-1-5p | MIMAT0004523 | Plekha8 | 231999 | HITS-CLIP | 37986224 | Unspecified |
| mmu-miR-29b-1-5p | MIMAT0004523 | Plekhg5 | 269608 | HITS-CLIP | 25083871 | Unspecified |
| mmu-miR-29b-1-5p | MIMAT0004523 | Plin2 | 11520 | HITS-CLIP | 25083871 | Unspecified |
| mmu-miR-29b-1-5p | MIMAT0004523 | Plin3 | 66905 | HITS-CLIP | 37986224 | Unspecified |
| mmu-miR-29b-1-5p | MIMAT0004523 | Plk2 | 20620 | HITS-CLIP | 25083871 | Unspecified |
| mmu-miR-29b-1-5p | MIMAT0004523 | Pls3 | 102866 | HITS-CLIP | 37986224 | Unspecified |
| mmu-miR-29b-1-5p | MIMAT0004523 | Plscr4 | 235527 | HITS-CLIP | 37986224 | Unspecified |
| mmu-miR-29b-1-5p | MIMAT0004523 | Plxna2 | 18845 | HITS-CLIP | 37986224 | Unspecified |
| mmu-miR-29b-1-5p | MIMAT0004523 | Plxnb1 | 235611 | HITS-CLIP | 37986224 | Unspecified |
| mmu-miR-29b-1-5p | MIMAT0004523 | Plxnd1 | 67784 | HITS-CLIP | 37986224 | Unspecified |
| mmu-miR-29b-1-5p | MIMAT0004523 | Pmepa1 | 65112 | HITS-CLIP | 37986224 | Unspecified |
| mmu-miR-29b-1-5p | MIMAT0004523 | Pnn | 18949 | HITS-CLIP | 25083871 | Unspecified |
| mmu-miR-29b-1-5p | MIMAT0004523 | Pnrc1 | 108767 | HITS-CLIP | 25083871 | Unspecified |
| mmu-miR-29b-1-5p | MIMAT0004523 | Pofut2 | 80294 | HITS-CLIP | 37986224 | Unspecified |
| mmu-miR-29b-1-5p | MIMAT0004523 | Pogz | 229584 | HITS-CLIP | 37986224 | Unspecified |
| mmu-miR-29b-1-5p | MIMAT0004523 | Poldip2 | 67811 | HITS-CLIP | 37986224 | Unspecified |
| mmu-miR-29b-1-5p | MIMAT0004523 | Pop5 | 117109 | HITS-CLIP | 37986224 | Unspecified |
| mmu-miR-29b-1-5p | MIMAT0004523 | Pou2f1 | 18986 | HITS-CLIP | 25083871 | Unspecified |
| mmu-miR-29b-1-5p | MIMAT0004523 | Ppargc1b | 170826 | HITS-CLIP | 25083871 | Unspecified |
| mmu-miR-29b-1-5p | MIMAT0004523 | Ppm1g | 14208 | HITS-CLIP | 25083871 | Unspecified |
| mmu-miR-29b-1-5p | MIMAT0004523 | Ppp1r10 | 52040 | HITS-CLIP | 37986224 | Unspecified |
| mmu-miR-29b-1-5p | MIMAT0004523 | Ppp1r12a | 17931 | HITS-CLIP | 25083871 | Unspecified |
| mmu-miR-29b-1-5p | MIMAT0004523 | Ppp1r15b | 108954 | HITS-CLIP | 37986224 | Unspecified |
| mmu-miR-29b-1-5p | MIMAT0004523 | Ppp2cb | 19053 | HITS-CLIP | 25083871 | Unspecified |
| mmu-miR-29b-1-5p | MIMAT0004523 | Ppp3ca | 19055 | HITS-CLIP | 37986224 | Unspecified |
| mmu-miR-29b-1-5p | MIMAT0004523 | Ppp3r1 | 19058 | HITS-CLIP | 25083871 | Unspecified |
| mmu-miR-29b-1-5p | MIMAT0004523 | Ppp6r3 | 52036 | HITS-CLIP | 37986224 | Unspecified |
| mmu-miR-29b-1-5p | MIMAT0004523 | Prag1 | 157285 | HITS-CLIP | 25083871 | Unspecified |
| mmu-miR-29b-1-5p | MIMAT0004523 | Prdm2 | 110593 | HITS-CLIP | 37986224 | Unspecified |
| mmu-miR-29b-1-5p | MIMAT0004523 | Prelid3b | 51012 | HITS-CLIP | 25083871 | Unspecified |
| mmu-miR-29b-1-5p | MIMAT0004523 | Prelp | 116847 | HITS-CLIP | 37986224 | Unspecified |

| | | | | | | |
|---|---|---|---|---|---|---|
| mmu-miR-29b-1-5p | MIMAT0004523 | Prkacb | 18749 | HITS-CLIP | 37986224 | Unspecified |
| mmu-miR-29b-1-5p | MIMAT0004523 | Prkar1a | 19084 | HITS-CLIP | 25083871 | Unspecified |
| mmu-miR-29b-1-5p | MIMAT0004523 | Prkx | 19108 | HITS-CLIP | 37986224 | Unspecified |
| mmu-miR-29b-1-5p | MIMAT0004523 | Prpf40a | 56194 | HITS-CLIP | 37986224\|25083871 | Unspecified |
| mmu-miR-29b-1-5p | MIMAT0004523 | Prpf8 | 192159 | HITS-CLIP | 25083871 | Unspecified |
| mmu-miR-29b-1-5p | MIMAT0004523 | Prr14l | 215476 | HITS-CLIP | 37986224 | Unspecified |
| mmu-miR-29b-1-5p | MIMAT0004523 | Prrc1 | 73137 | HITS-CLIP | 37986224 | Unspecified |
| mmu-miR-29b-1-5p | MIMAT0004523 | Prrc2a | 53761 | HITS-CLIP | 37986224 | Unspecified |
| mmu-miR-29b-1-5p | MIMAT0004523 | Prrc2b | 227723 | HITS-CLIP | 25083871 | Unspecified |
| mmu-miR-29b-1-5p | MIMAT0004523 | Prrx1 | 18933 | HITS-CLIP | 37986224 | Unspecified |
| mmu-miR-29b-1-5p | MIMAT0004523 | Prune2 | 353211 | HITS-CLIP | 37986224\|25083871 | Unspecified |
| mmu-miR-29b-1-5p | MIMAT0004523 | Psap | 19156 | HITS-CLIP | 25083871 | Unspecified |
| mmu-miR-29b-1-5p | MIMAT0004523 | Psmc3 | 19182 | HITS-CLIP | 25083871 | Unspecified |
| mmu-miR-29b-1-5p | MIMAT0004523 | Psmc6 | 67089 | HITS-CLIP | 25083871 | Unspecified |
| mmu-miR-29b-1-5p | MIMAT0004523 | Pstpip2 | 19201 | HITS-CLIP | 25083871 | Unspecified |
| mmu-miR-29b-1-5p | MIMAT0004523 | Ptar1 | 72351 | HITS-CLIP | 25083871 | Unspecified |
| mmu-miR-29b-1-5p | MIMAT0004523 | Ptbp1 | 19205 | HITS-CLIP | 37986224 | Unspecified |
| mmu-miR-29b-1-5p | MIMAT0004523 | Ptbp3 | 230257 | HITS-CLIP | 37986224 | Unspecified |
| mmu-miR-29b-1-5p | MIMAT0004523 | Ptdss1 | 19210 | HITS-CLIP | 25083871 | Unspecified |
| mmu-miR-29b-1-5p | MIMAT0004523 | Ptger4 | 19219 | HITS-CLIP | 37986224 | Unspecified |
| mmu-miR-29b-1-5p | MIMAT0004523 | Ptk2 | 14083 | HITS-CLIP | 37986224\|25083871 | Unspecified |
| mmu-miR-29b-1-5p | MIMAT0004523 | Ptp4a2 | 19244 | HITS-CLIP | 37986224\|25083871 | Unspecified |
| mmu-miR-29b-1-5p | MIMAT0004523 | Ptpa | 5524 | HITS-CLIP | 25083871 | Unspecified |
| mmu-miR-29b-1-5p | MIMAT0004523 | Ptpn11 | 19247 | HITS-CLIP | 37986224\|25083871 | Unspecified |
| mmu-miR-29b-1-5p | MIMAT0004523 | Ptpn3 | 545622 | HITS-CLIP | 37986224 | Unspecified |
| mmu-miR-29b-1-5p | MIMAT0004523 | Ptpn4 | 19258 | HITS-CLIP | 25083871 | Unspecified |
| mmu-miR-29b-1-5p | MIMAT0004523 | Pum1 | 80912 | HITS-CLIP | 37986224 | Unspecified |
| mmu-miR-29b-1-5p | MIMAT0004523 | Pum2 | 80913 | HITS-CLIP | 25083871 | Unspecified |
| mmu-miR-29b-1-5p | MIMAT0004523 | Pura | 19290 | HITS-CLIP | 25083871 | Unspecified |
| mmu-miR-29b-1-5p | MIMAT0004523 | Purb | 19291 | HITS-CLIP | 37986224\|25083871 | Unspecified |
| mmu-miR-29b-1-5p | MIMAT0004523 | Pwwp2a | 70802 | HITS-CLIP | 37986224 | Unspecified |
| mmu-miR-29b-1-5p | MIMAT0004523 | Pxn | 19303 | HITS-CLIP | 25083871 | Unspecified |
| mmu-miR-29b-1-5p | MIMAT0004523 | Pxylp1 | 235534 | HITS-CLIP | 37986224\|25083871 | Unspecified |
| mmu-miR-29b-1-5p | MIMAT0004523 | Qser1 | 99003 | HITS-CLIP | 37986224 | Unspecified |
| mmu-miR-29b-1-5p | MIMAT0004523 | R3hcc1l | 52013 | HITS-CLIP | 37986224 | Unspecified |
| mmu-miR-29b-1-5p | MIMAT0004523 | R3hdm1 | 226412 | HITS-CLIP | 37986224 | Unspecified |
| mmu-miR-29b-1-5p | MIMAT0004523 | R3hdm2 | 71750 | HITS-CLIP | 25083871 | Unspecified |
| mmu-miR-29b-1-5p | MIMAT0004523 | Rab1a | 5861 | HITS-CLIP | 25083871 | Unspecified |
| mmu-miR-29b-1-5p | MIMAT0004523 | Rab31 | 106572 | HITS-CLIP | 37986224 | Unspecified |

| | | | | | | |
|---|---|---|---|---|---|---|
| mmu-miR-29b-1-5p | MIMAT0004523 | Rab35 | 77407 | HITS-CLIP | 37986224 | Unspecified |
| mmu-miR-29b-1-5p | MIMAT0004523 | Rab6a | 19346 | HITS-CLIP | 25083871 | Unspecified |
| mmu-miR-29b-1-5p | MIMAT0004523 | Rabgap1 | 227800 | HITS-CLIP | 37986224 | Unspecified |
| mmu-miR-29b-1-5p | MIMAT0004523 | Rabl6 | 227624 | HITS-CLIP | 37986224 | Unspecified |
| mmu-miR-29b-1-5p | MIMAT0004523 | Rack1 | 10399 | HITS-CLIP | 25083871 | Unspecified |
| mmu-miR-29b-1-5p | MIMAT0004523 | Rad21 | 19357 | HITS-CLIP | 37986224\|25083871 | Unspecified |
| mmu-miR-29b-1-5p | MIMAT0004523 | Rad54l2 | 81000 | HITS-CLIP | 37986224 | Unspecified |
| mmu-miR-29b-1-5p | MIMAT0004523 | Rai14 | 75646 | HITS-CLIP | 25083871 | Unspecified |
| mmu-miR-29b-1-5p | MIMAT0004523 | Ralbp1 | 19765 | HITS-CLIP | 37986224 | Unspecified |
| mmu-miR-29b-1-5p | MIMAT0004523 | Ranbp2 | 19386 | HITS-CLIP | 37986224\|25083871 | Unspecified |
| mmu-miR-29b-1-5p | MIMAT0004523 | Rap1gds1 | 229877 | HITS-CLIP | 37986224 | Unspecified |
| mmu-miR-29b-1-5p | MIMAT0004523 | Rapgef1 | 107746 | HITS-CLIP | 25083871 | Unspecified |
| mmu-miR-29b-1-5p | MIMAT0004523 | Rapgef2 | 76089 | HITS-CLIP | 37986224\|25083871 | Unspecified |
| mmu-miR-29b-1-5p | MIMAT0004523 | Rapsn | 19400 | HITS-CLIP | 37986224 | Unspecified |
| mmu-miR-29b-1-5p | MIMAT0004523 | Rara | 19401 | HITS-CLIP | 37986224 | Unspecified |
| mmu-miR-29b-1-5p | MIMAT0004523 | Rasal2 | 226525 | HITS-CLIP | 37986224 | Unspecified |
| mmu-miR-29b-1-5p | MIMAT0004523 | Rassf8 | 71323 | HITS-CLIP | 37986224 | Unspecified |
| mmu-miR-29b-1-5p | MIMAT0004523 | Raver1 | 71766 | HITS-CLIP | 37986224 | Unspecified |
| mmu-miR-29b-1-5p | MIMAT0004523 | Rb1cc1 | 12421 | HITS-CLIP | 37986224 | Unspecified |
| mmu-miR-29b-1-5p | MIMAT0004523 | Rbbp6 | 19647 | HITS-CLIP | 37986224 | Unspecified |
| mmu-miR-29b-1-5p | MIMAT0004523 | Rbm12 | 75710 | HITS-CLIP | 37986224 | Unspecified |
| mmu-miR-29b-1-5p | MIMAT0004523 | Rbm14 | 56275 | HITS-CLIP | 37986224 | Unspecified |
| mmu-miR-29b-1-5p | MIMAT0004523 | Rbm20 | 73713 | HITS-CLIP | 37986224 | Unspecified |
| mmu-miR-29b-1-5p | MIMAT0004523 | Rbm24 | 666794 | HITS-CLIP | 25083871 | Unspecified |
| mmu-miR-29b-1-5p | MIMAT0004523 | Rbm26 | 74213 | HITS-CLIP | 25083871 | Unspecified |
| mmu-miR-29b-1-5p | MIMAT0004523 | Rbm27 | 225432 | HITS-CLIP | 37986224\|25083871 | Unspecified |
| mmu-miR-29b-1-5p | MIMAT0004523 | Rbm33 | 381626 | HITS-CLIP | 37986224 | Unspecified |
| mmu-miR-29b-1-5p | MIMAT0004523 | Rbms3 | 207181 | HITS-CLIP | 37986224 | Unspecified |
| mmu-miR-29b-1-5p | MIMAT0004523 | Rc3h1 | 381305 | HITS-CLIP | 37986224 | Unspecified |
| mmu-miR-29b-1-5p | MIMAT0004523 | Rc3h2 | 319817 | HITS-CLIP | 25083871 | Unspecified |
| mmu-miR-29b-1-5p | MIMAT0004523 | Rcbtb1 | 71330 | HITS-CLIP | 37986224 | Unspecified |
| mmu-miR-29b-1-5p | MIMAT0004523 | Rcor3 | 214742 | HITS-CLIP | 37986224 | Unspecified |
| mmu-miR-29b-1-5p | MIMAT0004523 | Reep3 | 28193 | HITS-CLIP | 25083871 | Unspecified |
| mmu-miR-29b-1-5p | MIMAT0004523 | Reep4 | 72549 | HITS-CLIP | 25083871 | Unspecified |
| mmu-miR-29b-1-5p | MIMAT0004523 | Resf1 | 55196 | HITS-CLIP | 25083871 | Unspecified |
| mmu-miR-29b-1-5p | MIMAT0004523 | Rexo4 | 227656 | HITS-CLIP | 37986224 | Unspecified |
| mmu-miR-29b-1-5p | MIMAT0004523 | Rfc1 | 19687 | HITS-CLIP | 37986224 | Unspecified |
| mmu-miR-29b-1-5p | MIMAT0004523 | Rfx3 | 19726 | HITS-CLIP | 37986224 | Unspecified |
| mmu-miR-29b-1-5p | MIMAT0004523 | Rfx7 | 319758 | HITS-CLIP | 25083871 | Unspecified |

| | | | | | | | | |
|---|---|---|---|---|---|---|---|---|
| mmu-miR-29b-1-5p | MIMAT0004523 | Rgs19 | 56470 | HITS-CLIP | | | 37986224 | Unspecified |
| mmu-miR-29b-1-5p | MIMAT0004523 | Rhobtb1 | 69288 | HITS-CLIP | | | 25083871 | Unspecified |
| mmu-miR-29b-1-5p | MIMAT0004523 | Rhobtb2 | 246710 | HITS-CLIP | | | 37986224 | Unspecified |
| mmu-miR-29b-1-5p | MIMAT0004523 | Rictor | 78757 | HITS-CLIP | | | 25083871 | Unspecified |
| mmu-miR-29b-1-5p | MIMAT0004523 | Rlf | 109263 | HITS-CLIP | | | 25083871 | Unspecified |
| mmu-miR-29b-1-5p | MIMAT0004523 | Rlim | 19820 | HITS-CLIP | | | 25083871 | Unspecified |
| mmu-miR-29b-1-5p | MIMAT0004523 | Rmnd5b | 66089 | HITS-CLIP | | | 37986224 | Unspecified |
| mmu-miR-29b-1-5p | MIMAT0004523 | Rnf111 | 93836 | HITS-CLIP | | | 25083871 | Unspecified |
| mmu-miR-29b-1-5p | MIMAT0004523 | Rnf145 | 74315 | HITS-CLIP | | | 25083871 | Unspecified |
| mmu-miR-29b-1-5p | MIMAT0004523 | Rnf150 | 330812 | HITS-CLIP | | | 37986224 | Unspecified |
| mmu-miR-29b-1-5p | MIMAT0004523 | Rnf213 | 672511 | HITS-CLIP | | | 37986224 | Unspecified |
| mmu-miR-29b-1-5p | MIMAT0004523 | Rpgrip1l | 244585 | HITS-CLIP | | 37986224\|25083871 | | Unspecified |
| mmu-miR-29b-1-5p | MIMAT0004523 | Rpl4 | 67891 | HITS-CLIP | | | 25083871 | Unspecified |
| mmu-miR-29b-1-5p | MIMAT0004523 | Rps6ka3 | 110651 | HITS-CLIP | | | 37986224 | Unspecified |
| mmu-miR-29b-1-5p | MIMAT0004523 | Rps6kb1 | 72508 | HITS-CLIP | | | 37986224 | Unspecified |
| mmu-miR-29b-1-5p | MIMAT0004523 | Rreb1 | 68750 | HITS-CLIP | | | 25083871 | Unspecified |
| mmu-miR-29b-1-5p | MIMAT0004523 | Rrm2 | 20135 | HITS-CLIP | | | 37986224 | Unspecified |
| mmu-miR-29b-1-5p | MIMAT0004523 | Rrp12 | 107094 | HITS-CLIP | | | 25083871 | Unspecified |
| mmu-miR-29b-1-5p | MIMAT0004523 | Rrp1b | 72462 | HITS-CLIP | | | 37986224 | Unspecified |
| mmu-miR-29b-1-5p | MIMAT0004523 | Rrs1 | 59014 | HITS-CLIP | | | 37986224 | Unspecified |
| mmu-miR-29b-1-5p | MIMAT0004523 | Rsbn1 | 229675 | HITS-CLIP | | | 25083871 | Unspecified |
| mmu-miR-29b-1-5p | MIMAT0004523 | Rsbn1l | 242860 | HITS-CLIP | | | 25083871 | Unspecified |
| mmu-miR-29b-1-5p | MIMAT0004523 | Rspry1 | 67610 | HITS-CLIP | | | 25083871 | Unspecified |
| mmu-miR-29b-1-5p | MIMAT0004523 | Rtf1 | 76246 | HITS-CLIP | | 37986224\|25083871 | | Unspecified |
| mmu-miR-29b-1-5p | MIMAT0004523 | Rtn4 | 68585 | HITS-CLIP | | | 25083871 | Unspecified |
| mmu-miR-29b-1-5p | MIMAT0004523 | Rufy2 | 70432 | HITS-CLIP | | | 37986224 | Unspecified |
| mmu-miR-29b-1-5p | MIMAT0004523 | Rxra | 20181 | HITS-CLIP | | | 37986224 | Unspecified |
| mmu-miR-29b-1-5p | MIMAT0004523 | Ryr1 | 20190 | HITS-CLIP | | | 37986224 | Unspecified |
| mmu-miR-29b-1-5p | MIMAT0004523 | Sacm1l | 83493 | HITS-CLIP | | | 25083871 | Unspecified |
| mmu-miR-29b-1-5p | MIMAT0004523 | Sacs | 50720 | HITS-CLIP | | 37986224\|25083871 | | Unspecified |
| mmu-miR-29b-1-5p | MIMAT0004523 | Samd4 | 74480 | HITS-CLIP | | | 37986224 | Unspecified |
| mmu-miR-29b-1-5p | MIMAT0004523 | Samd8 | 67630 | HITS-CLIP | | 37986224\|25083871 | | Unspecified |
| mmu-miR-29b-1-5p | MIMAT0004523 | Sars | 20226 | HITS-CLIP | | | 37986224 | Unspecified |
| mmu-miR-29b-1-5p | MIMAT0004523 | Satb1 | 20230 | HITS-CLIP | | | 37986224 | Unspecified |
| mmu-miR-29b-1-5p | MIMAT0004523 | Sav1 | 64010 | HITS-CLIP | | | 25083871 | Unspecified |
| mmu-miR-29b-1-5p | MIMAT0004523 | Sbds | 66711 | HITS-CLIP | | | 37986224 | Unspecified |
| mmu-miR-29b-1-5p | MIMAT0004523 | Sbf2 | 319934 | HITS-CLIP | | | 37986224 | Unspecified |
| mmu-miR-29b-1-5p | MIMAT0004523 | Sbno1 | 243272 | HITS-CLIP | | | 25083871 | Unspecified |
| mmu-miR-29b-1-5p | MIMAT0004523 | Sbno2 | 216161 | HITS-CLIP | | | 25083871 | Unspecified |

| | | | | | | | |
|---|---|---|---|---|---|---|---|
| mmu-miR-29b-1-5p | MIMAT0004523 | Scaf11 | 72193 | HITS-CLIP | | 37986224 | Unspecified |
| mmu-miR-29b-1-5p | MIMAT0004523 | Scpep1 | 74617 | HITS-CLIP | | 37986224 | Unspecified |
| mmu-miR-29b-1-5p | MIMAT0004523 | Scrib | 105782 | HITS-CLIP | | 37986224 | Unspecified |
| mmu-miR-29b-1-5p | MIMAT0004523 | Sdf4 | 20318 | HITS-CLIP | | 37986224 | Unspecified |
| mmu-miR-29b-1-5p | MIMAT0004523 | Sec14l2 | 67815 | HITS-CLIP | | 25083871 | Unspecified |
| mmu-miR-29b-1-5p | MIMAT0004523 | Sec16a | 227648 | HITS-CLIP | 37986224\|25083871 | | Unspecified |
| mmu-miR-29b-1-5p | MIMAT0004523 | Sec24a | 77371 | HITS-CLIP | | 25083871 | Unspecified |
| mmu-miR-29b-1-5p | MIMAT0004523 | Sec24c | 218811 | HITS-CLIP | | 25083871 | Unspecified |
| mmu-miR-29b-1-5p | MIMAT0004523 | Sec62 | 69276 | HITS-CLIP | | 25083871 | Unspecified |
| mmu-miR-29b-1-5p | MIMAT0004523 | Secisbp2l | 70354 | HITS-CLIP | | 37986224 | Unspecified |
| mmu-miR-29b-1-5p | MIMAT0004523 | Sema3d | 108151 | HITS-CLIP | | 37986224 | Unspecified |
| mmu-miR-29b-1-5p | MIMAT0004523 | Sema7a | 20361 | HITS-CLIP | | 25083871 | Unspecified |
| mmu-miR-29b-1-5p | MIMAT0004523 | Senp6 | 215351 | HITS-CLIP | | 25083871 | Unspecified |
| mmu-miR-29b-1-5p | MIMAT0004523 | Senp7 | 66315 | HITS-CLIP | | 25083871 | Unspecified |
| mmu-miR-29b-1-5p | MIMAT0004523 | Sept11 | 52398 | HITS-CLIP | | 25083871 | Unspecified |
| mmu-miR-29b-1-5p | MIMAT0004523 | Serbp1 | 66870 | HITS-CLIP | 37986224\|25083871 | | Unspecified |
| mmu-miR-29b-1-5p | MIMAT0004523 | Serinc1 | 56442 | HITS-CLIP | | 25083871 | Unspecified |
| mmu-miR-29b-1-5p | MIMAT0004523 | Serinc3 | 26943 | HITS-CLIP | | 37986224 | Unspecified |
| mmu-miR-29b-1-5p | MIMAT0004523 | Sertad1 | 55942 | HITS-CLIP | | 37986224 | Unspecified |
| mmu-miR-29b-1-5p | MIMAT0004523 | Sertad4 | 214791 | HITS-CLIP | | 37986224 | Unspecified |
| mmu-miR-29b-1-5p | MIMAT0004523 | Sesn2 | 230784 | HITS-CLIP | | 25083871 | Unspecified |
| mmu-miR-29b-1-5p | MIMAT0004523 | Set | 56086 | HITS-CLIP | | 25083871 | Unspecified |
| mmu-miR-29b-1-5p | MIMAT0004523 | Setbp1 | 240427 | HITS-CLIP | | 37986224 | Unspecified |
| mmu-miR-29b-1-5p | MIMAT0004523 | Setd1b | 208043 | HITS-CLIP | | 25083871 | Unspecified |
| mmu-miR-29b-1-5p | MIMAT0004523 | Setd5 | 72895 | HITS-CLIP | | 25083871 | Unspecified |
| mmu-miR-29b-1-5p | MIMAT0004523 | Setdb1 | 84505 | HITS-CLIP | | 25083871 | Unspecified |
| mmu-miR-29b-1-5p | MIMAT0004523 | Setx | 269254 | HITS-CLIP | | 37986224 | Unspecified |
| mmu-miR-29b-1-5p | MIMAT0004523 | Sf3b5 | 66125 | HITS-CLIP | | 37986224 | Unspecified |
| mmu-miR-29b-1-5p | MIMAT0004523 | Sf3b6 | 66055 | HITS-CLIP | | 25083871 | Unspecified |
| mmu-miR-29b-1-5p | MIMAT0004523 | Sfpq | 71514 | HITS-CLIP | | 37986224 | Unspecified |
| mmu-miR-29b-1-5p | MIMAT0004523 | Sfswap | 231769 | HITS-CLIP | | 37986224 | Unspecified |
| mmu-miR-29b-1-5p | MIMAT0004523 | Sft2d2 | 108735 | HITS-CLIP | | 37986224 | Unspecified |
| mmu-miR-29b-1-5p | MIMAT0004523 | Sh2b3 | 16923 | HITS-CLIP | | 37986224 | Unspecified |
| mmu-miR-29b-1-5p | MIMAT0004523 | Sh3bp2 | 24055 | HITS-CLIP | | 25083871 | Unspecified |
| mmu-miR-29b-1-5p | MIMAT0004523 | Sh3d19 | 27059 | HITS-CLIP | 37986224\|25083871 | | Unspecified |
| mmu-miR-29b-1-5p | MIMAT0004523 | Sh3pxd2a | 14218 | HITS-CLIP | | 25083871 | Unspecified |
| mmu-miR-29b-1-5p | MIMAT0004523 | Siah2 | 20439 | HITS-CLIP | | 37986224 | Unspecified |
| mmu-miR-29b-1-5p | MIMAT0004523 | Sidt2 | 214597 | HITS-CLIP | | 37986224 | Unspecified |
| mmu-miR-29b-1-5p | MIMAT0004523 | Sik3 | 70661 | HITS-CLIP | | 37986224 | Unspecified |

| | | | | | | |
|---|---|---|---|---|---|---|
| mmu-miR-29b-1-5p | MIMAT0004523 | Sike1 | 66641 | HITS-CLIP | 37986224 | Unspecified |
| mmu-miR-29b-1-5p | MIMAT0004523 | Sin3a | 20466 | HITS-CLIP | 37986224 | Unspecified |
| mmu-miR-29b-1-5p | MIMAT0004523 | Sirt1 | 93759 | HITS-CLIP | 37986224 | Unspecified |
| mmu-miR-29b-1-5p | MIMAT0004523 | Six1 | 20471 | HITS-CLIP | 37986224 | Unspecified |
| mmu-miR-29b-1-5p | MIMAT0004523 | Six4 | 20474 | HITS-CLIP | 37986224 | Unspecified |
| mmu-miR-29b-1-5p | MIMAT0004523 | Ska2 | 66140 | HITS-CLIP | 37986224 | Unspecified |
| mmu-miR-29b-1-5p | MIMAT0004523 | Ski | 20481 | HITS-CLIP | 25083871 | Unspecified |
| mmu-miR-29b-1-5p | MIMAT0004523 | Slc12a6 | 107723 | HITS-CLIP | 37986224 | Unspecified |
| mmu-miR-29b-1-5p | MIMAT0004523 | Slc1a4 | 55963 | HITS-CLIP | 37986224 | Unspecified |
| mmu-miR-29b-1-5p | MIMAT0004523 | Slc23a2 | 54338 | HITS-CLIP | 37986224 | Unspecified |
| mmu-miR-29b-1-5p | MIMAT0004523 | Slc24a3 | 94249 | HITS-CLIP | 37986224 | Unspecified |
| mmu-miR-29b-1-5p | MIMAT0004523 | Slc25a36 | 192287 | HITS-CLIP | 37986224 | Unspecified |
| mmu-miR-29b-1-5p | MIMAT0004523 | Slc27a4 | 26569 | HITS-CLIP | 37986224 | Unspecified |
| mmu-miR-29b-1-5p | MIMAT0004523 | Slc29a1 | 63959 | HITS-CLIP | 37986224 | Unspecified |
| mmu-miR-29b-1-5p | MIMAT0004523 | Slc2a3 | 20527 | HITS-CLIP | 37986224 | Unspecified |
| mmu-miR-29b-1-5p | MIMAT0004523 | Slc38a1 | 105727 | HITS-CLIP | 37986224 | Unspecified |
| mmu-miR-29b-1-5p | MIMAT0004523 | Slc38a2 | 67760 | HITS-CLIP | 25083871 | Unspecified |
| mmu-miR-29b-1-5p | MIMAT0004523 | Slc38a7 | 234595 | HITS-CLIP | 25083871 | Unspecified |
| mmu-miR-29b-1-5p | MIMAT0004523 | Slc39a10 | 227059 | HITS-CLIP | 37986224\|25083871 | Unspecified |
| mmu-miR-29b-1-5p | MIMAT0004523 | Slc39a11 | 69806 | HITS-CLIP | 37986224 | Unspecified |
| mmu-miR-29b-1-5p | MIMAT0004523 | Slc41a2 | 338365 | HITS-CLIP | 37986224 | Unspecified |
| mmu-miR-29b-1-5p | MIMAT0004523 | Slc5a3 | 53881 | HITS-CLIP | 25083871 | Unspecified |
| mmu-miR-29b-1-5p | MIMAT0004523 | Slc7a1 | 11987 | HITS-CLIP | 37986224 | Unspecified |
| mmu-miR-29b-1-5p | MIMAT0004523 | Slco5a1 | 240726 | HITS-CLIP | 37986224 | Unspecified |
| mmu-miR-29b-1-5p | MIMAT0004523 | Slk | 20874 | HITS-CLIP | 37986224 | Unspecified |
| mmu-miR-29b-1-5p | MIMAT0004523 | Smad3 | 17127 | HITS-CLIP | 25083871 | Unspecified |
| mmu-miR-29b-1-5p | MIMAT0004523 | Smad5 | 17129 | HITS-CLIP | 37986224 | Unspecified |
| mmu-miR-29b-1-5p | MIMAT0004523 | Smarca2 | 67155 | HITS-CLIP | 25083871 | Unspecified |
| mmu-miR-29b-1-5p | MIMAT0004523 | Smarca4 | 20586 | HITS-CLIP | 25083871 | Unspecified |
| mmu-miR-29b-1-5p | MIMAT0004523 | Smarcad1 | 13990 | HITS-CLIP | 25083871 | Unspecified |
| mmu-miR-29b-1-5p | MIMAT0004523 | Smc1a | 24061 | HITS-CLIP | 37986224\|25083871 | Unspecified |
| mmu-miR-29b-1-5p | MIMAT0004523 | Smc3 | 13006 | HITS-CLIP | 25083871 | Unspecified |
| mmu-miR-29b-1-5p | MIMAT0004523 | Smcr8 | 237782 | HITS-CLIP | 37986224\|25083871 | Unspecified |
| mmu-miR-29b-1-5p | MIMAT0004523 | Smg6 | 103677 | HITS-CLIP | 37986224 | Unspecified |
| mmu-miR-29b-1-5p | MIMAT0004523 | Smg7 | 226517 | HITS-CLIP | 37986224\|25083871 | Unspecified |
| mmu-miR-29b-1-5p | MIMAT0004523 | Smg8 | 74133 | HITS-CLIP | 37986224 | Unspecified |
| mmu-miR-29b-1-5p | MIMAT0004523 | Smurf1 | 75788 | HITS-CLIP | 37986224 | Unspecified |
| mmu-miR-29b-1-5p | MIMAT0004523 | Snai2 | 20583 | HITS-CLIP | 25083871 | Unspecified |
| mmu-miR-29b-1-5p | MIMAT0004523 | Snap47 | 67826 | HITS-CLIP | 37986224 | Unspecified |

| | | | | | | | |
|---|---|---|---|---|---|---|---|
| mmu-miR-29b-1-5p | MIMAT0004523 | Snrnp200 | 320632 | HITS-CLIP | | 25083871 | Unspecified |
| mmu-miR-29b-1-5p | MIMAT0004523 | Sntb2 | 20650 | HITS-CLIP | | 37986224 | Unspecified |
| mmu-miR-29b-1-5p | MIMAT0004523 | Snx2 | 67804 | HITS-CLIP | | 25083871 | Unspecified |
| mmu-miR-29b-1-5p | MIMAT0004523 | Snx33 | 235406 | HITS-CLIP | | 37986224 | Unspecified |
| mmu-miR-29b-1-5p | MIMAT0004523 | Sobp | 109205 | HITS-CLIP | 37986224\|25083871 | | Unspecified |
| mmu-miR-29b-1-5p | MIMAT0004523 | Socs1 | 12703 | HITS-CLIP | | 25083871 | Unspecified |
| mmu-miR-29b-1-5p | MIMAT0004523 | Socs2 | 216233 | HITS-CLIP | | 37986224 | Unspecified |
| mmu-miR-29b-1-5p | MIMAT0004523 | Socs3 | 12702 | HITS-CLIP | | 25083871 | Unspecified |
| mmu-miR-29b-1-5p | MIMAT0004523 | Socs4 | 67296 | HITS-CLIP | | 37986224 | Unspecified |
| mmu-miR-29b-1-5p | MIMAT0004523 | Son | 20658 | HITS-CLIP | | 25083871 | Unspecified |
| mmu-miR-29b-1-5p | MIMAT0004523 | Sos1 | 20662 | HITS-CLIP | | 25083871 | Unspecified |
| mmu-miR-29b-1-5p | MIMAT0004523 | Sox12 | 20667 | HITS-CLIP | 37986224\|25083871 | | Unspecified |
| mmu-miR-29b-1-5p | MIMAT0004523 | Sp1 | 20683 | HITS-CLIP | 37986224\|25083871 | | Unspecified |
| mmu-miR-29b-1-5p | MIMAT0004523 | Sp2 | 78912 | HITS-CLIP | 37986224\|25083871 | | Unspecified |
| mmu-miR-29b-1-5p | MIMAT0004523 | Specc1 | 432572 | HITS-CLIP | | 25083871 | Unspecified |
| mmu-miR-29b-1-5p | MIMAT0004523 | Specc1l | 74392 | HITS-CLIP | | 37986224 | Unspecified |
| mmu-miR-29b-1-5p | MIMAT0004523 | Speg | 11790 | HITS-CLIP | | 37986224 | Unspecified |
| mmu-miR-29b-1-5p | MIMAT0004523 | Spen | 56381 | HITS-CLIP | 37986224\|25083871 | | Unspecified |
| mmu-miR-29b-1-5p | MIMAT0004523 | Spop | 20747 | HITS-CLIP | | 37986224 | Unspecified |
| mmu-miR-29b-1-5p | MIMAT0004523 | Spp1 | 20750 | HITS-CLIP | | 25083871 | Unspecified |
| mmu-miR-29b-1-5p | MIMAT0004523 | Sppl3 | 74585 | HITS-CLIP | | 25083871 | Unspecified |
| mmu-miR-29b-1-5p | MIMAT0004523 | Spred1 | 114715 | HITS-CLIP | | 25083871 | Unspecified |
| mmu-miR-29b-1-5p | MIMAT0004523 | Spryd3 | 223918 | HITS-CLIP | | 25083871 | Unspecified |
| mmu-miR-29b-1-5p | MIMAT0004523 | Sptan1 | 20740 | HITS-CLIP | 37986224\|25083871 | | Unspecified |
| mmu-miR-29b-1-5p | MIMAT0004523 | Srcap | 100043597 | HITS-CLIP | | 25083871 | Unspecified |
| mmu-miR-29b-1-5p | MIMAT0004523 | Srebf2 | 20788 | HITS-CLIP | | 25083871 | Unspecified |
| mmu-miR-29b-1-5p | MIMAT0004523 | Srgap2 | 14270 | HITS-CLIP | | 37986224 | Unspecified |
| mmu-miR-29b-1-5p | MIMAT0004523 | Srgap3 | 259302 | HITS-CLIP | | 37986224 | Unspecified |
| mmu-miR-29b-1-5p | MIMAT0004523 | Srp72 | 66661 | HITS-CLIP | 37986224\|25083871 | | Unspecified |
| mmu-miR-29b-1-5p | MIMAT0004523 | Srrm2 | 75956 | HITS-CLIP | 37986224\|25083871 | | Unspecified |
| mmu-miR-29b-1-5p | MIMAT0004523 | Srsf2 | 20382 | HITS-CLIP | | 25083871 | Unspecified |
| mmu-miR-29b-1-5p | MIMAT0004523 | Srsf3 | 20383 | HITS-CLIP | | 37986224 | Unspecified |
| mmu-miR-29b-1-5p | MIMAT0004523 | Ssh2 | 237860 | HITS-CLIP | | 37986224 | Unspecified |
| mmu-miR-29b-1-5p | MIMAT0004523 | Sspn | 16651 | HITS-CLIP | | 37986224 | Unspecified |
| mmu-miR-29b-1-5p | MIMAT0004523 | Ssr2 | 66256 | HITS-CLIP | | 37986224 | Unspecified |
| mmu-miR-29b-1-5p | MIMAT0004523 | St13 | 70356 | HITS-CLIP | | 37986224 | Unspecified |
| mmu-miR-29b-1-5p | MIMAT0004523 | St3gal1 | 20442 | HITS-CLIP | | 37986224 | Unspecified |
| mmu-miR-29b-1-5p | MIMAT0004523 | St7l | 229681 | HITS-CLIP | | 37986224 | Unspecified |
| mmu-miR-29b-1-5p | MIMAT0004523 | Stac2 | 217154 | HITS-CLIP | | 37986224 | Unspecified |

| | | | | | | |
|---|---|---|---|---|---|---|
| mmu-miR-29b-1-5p | MIMAT0004523 | Stag2 | 20843 | HITS-CLIP | 25083871 | Unspecified |
| mmu-miR-29b-1-5p | MIMAT0004523 | Stam | 20844 | HITS-CLIP | 37986224\|25083871 | Unspecified |
| mmu-miR-29b-1-5p | MIMAT0004523 | Stambp | 70527 | HITS-CLIP | 25083871 | Unspecified |
| mmu-miR-29b-1-5p | MIMAT0004523 | Stard9 | 668880 | HITS-CLIP | 37986224 | Unspecified |
| mmu-miR-29b-1-5p | MIMAT0004523 | Stat3 | 20848 | HITS-CLIP | 37986224 | Unspecified |
| mmu-miR-29b-1-5p | MIMAT0004523 | Stau1 | 20853 | HITS-CLIP | 25083871 | Unspecified |
| mmu-miR-29b-1-5p | MIMAT0004523 | Steap2 | 74051 | HITS-CLIP | 37986224 | Unspecified |
| mmu-miR-29b-1-5p | MIMAT0004523 | Stip1 | 20867 | HITS-CLIP | 25083871 | Unspecified |
| mmu-miR-29b-1-5p | MIMAT0004523 | Stk10 | 20868 | HITS-CLIP | 25083871 | Unspecified |
| mmu-miR-29b-1-5p | MIMAT0004523 | Stk35 | 67333 | HITS-CLIP | 37986224 | Unspecified |
| mmu-miR-29b-1-5p | MIMAT0004523 | Stk4 | 58231 | HITS-CLIP | 37986224 | Unspecified |
| mmu-miR-29b-1-5p | MIMAT0004523 | Stx16 | 228960 | HITS-CLIP | 37986224\|25083871 | Unspecified |
| mmu-miR-29b-1-5p | MIMAT0004523 | Sun2 | 223697 | HITS-CLIP | 25083871 | Unspecified |
| mmu-miR-29b-1-5p | MIMAT0004523 | Supt6 | 20926 | HITS-CLIP | 25083871 | Unspecified |
| mmu-miR-29b-1-5p | MIMAT0004523 | Syde1 | 71709 | HITS-CLIP | 37986224 | Unspecified |
| mmu-miR-29b-1-5p | MIMAT0004523 | Syncrip | 56403 | HITS-CLIP | 25083871 | Unspecified |
| mmu-miR-29b-1-5p | MIMAT0004523 | Synj2 | 20975 | HITS-CLIP | 25083871 | Unspecified |
| mmu-miR-29b-1-5p | MIMAT0004523 | Synpo2 | 118449 | HITS-CLIP | 37986224 | Unspecified |
| mmu-miR-29b-1-5p | MIMAT0004523 | Synpo2l | 68760 | HITS-CLIP | 25083871 | Unspecified |
| mmu-miR-29b-1-5p | MIMAT0004523 | Synrg | 217030 | HITS-CLIP | 25083871 | Unspecified |
| mmu-miR-29b-1-5p | MIMAT0004523 | Szrd1 | 213491 | HITS-CLIP | 37986224 | Unspecified |
| mmu-miR-29b-1-5p | MIMAT0004523 | Tab1 | 66513 | HITS-CLIP | 37986224 | Unspecified |
| mmu-miR-29b-1-5p | MIMAT0004523 | Tab2 | 68652 | HITS-CLIP | 37986224\|25083871 | Unspecified |
| mmu-miR-29b-1-5p | MIMAT0004523 | Tacc1 | 320165 | HITS-CLIP | 37986224 | Unspecified |
| mmu-miR-29b-1-5p | MIMAT0004523 | Taf1 | 270627 | HITS-CLIP | 37986224\|25083871 | Unspecified |
| mmu-miR-29b-1-5p | MIMAT0004523 | Taf12 | 66464 | HITS-CLIP | 37986224 | Unspecified |
| mmu-miR-29b-1-5p | MIMAT0004523 | Taf3 | 209361 | HITS-CLIP | 37986224 | Unspecified |
| mmu-miR-29b-1-5p | MIMAT0004523 | Tanc1 | 66860 | HITS-CLIP | 37986224 | Unspecified |
| mmu-miR-29b-1-5p | MIMAT0004523 | Taok3 | 330177 | HITS-CLIP | 25083871 | Unspecified |
| mmu-miR-29b-1-5p | MIMAT0004523 | Tapt1 | 231225 | HITS-CLIP | 25083871 | Unspecified |
| mmu-miR-29b-1-5p | MIMAT0004523 | Tardbp | 230908 | HITS-CLIP | 37986224 | Unspecified |
| mmu-miR-29b-1-5p | MIMAT0004523 | Tbc1d15 | 66687 | HITS-CLIP | 37986224 | Unspecified |
| mmu-miR-29b-1-5p | MIMAT0004523 | Tbc1d16 | 207592 | HITS-CLIP | 37986224 | Unspecified |
| mmu-miR-29b-1-5p | MIMAT0004523 | Tbc1d5 | 72238 | HITS-CLIP | 37986224 | Unspecified |
| mmu-miR-29b-1-5p | MIMAT0004523 | Tbccd1 | 70573 | HITS-CLIP | 37986224 | Unspecified |
| mmu-miR-29b-1-5p | MIMAT0004523 | Tbl1x | 21372 | HITS-CLIP | 37986224\|25083871 | Unspecified |
| mmu-miR-29b-1-5p | MIMAT0004523 | Tcf12 | 21406 | HITS-CLIP | 25083871 | Unspecified |
| mmu-miR-29b-1-5p | MIMAT0004523 | Tcf20 | 21411 | HITS-CLIP | 25083871 | Unspecified |
| mmu-miR-29b-1-5p | MIMAT0004523 | Tcf4 | 21413 | HITS-CLIP | 25083871 | Unspecified |

| | | | | | | |
|---|---|---|---|---|---|---|
| mmu-miR-29b-1-5p | MIMAT0004523 | Tcf7 | 21414 | HITS-CLIP | 37986224 | Unspecified |
| mmu-miR-29b-1-5p | MIMAT0004523 | Tcp11l1 | 320554 | HITS-CLIP | 37986224 | Unspecified |
| mmu-miR-29b-1-5p | MIMAT0004523 | Tead1 | 21676 | HITS-CLIP | 37986224\|25083871 | Unspecified |
| mmu-miR-29b-1-5p | MIMAT0004523 | Tenm3 | 23965 | HITS-CLIP | 37986224 | Unspecified |
| mmu-miR-29b-1-5p | MIMAT0004523 | Tent4b | 64282 | HITS-CLIP | 25083871 | Unspecified |
| mmu-miR-29b-1-5p | MIMAT0004523 | Terf2 | 21750 | HITS-CLIP | 37986224 | Unspecified |
| mmu-miR-29b-1-5p | MIMAT0004523 | Tes | 21753 | HITS-CLIP | 37986224\|25083871 | Unspecified |
| mmu-miR-29b-1-5p | MIMAT0004523 | Tet1 | 52463 | HITS-CLIP | 37986224 | Unspecified |
| mmu-miR-29b-1-5p | MIMAT0004523 | Tet3 | 194388 | HITS-CLIP | 37986224 | Unspecified |
| mmu-miR-29b-1-5p | MIMAT0004523 | Tex10 | 269536 | HITS-CLIP | 37986224 | Unspecified |
| mmu-miR-29b-1-5p | MIMAT0004523 | Tex2 | 21763 | HITS-CLIP | 37986224 | Unspecified |
| mmu-miR-29b-1-5p | MIMAT0004523 | Tfam | 21780 | HITS-CLIP | 37986224 | Unspecified |
| mmu-miR-29b-1-5p | MIMAT0004523 | Tfdp1 | 21781 | HITS-CLIP | 25083871 | Unspecified |
| mmu-miR-29b-1-5p | MIMAT0004523 | Tfrc | 22042 | HITS-CLIP | 37986224 | Unspecified |
| mmu-miR-29b-1-5p | MIMAT0004523 | Tgfb1 | 21803 | HITS-CLIP | 25083871 | Unspecified |
| mmu-miR-29b-1-5p | MIMAT0004523 | Tgfb2 | 21808 | HITS-CLIP | 25083871 | Unspecified |
| mmu-miR-29b-1-5p | MIMAT0004523 | Thbs2 | 21826 | HITS-CLIP | 37986224 | Unspecified |
| mmu-miR-29b-1-5p | MIMAT0004523 | Tia1 | 21841 | HITS-CLIP | 25083871 | Unspecified |
| mmu-miR-29b-1-5p | MIMAT0004523 | Tiam2 | 24001 | HITS-CLIP | 37986224 | Unspecified |
| mmu-miR-29b-1-5p | MIMAT0004523 | Timp3 | 21859 | HITS-CLIP | 25083871 | Unspecified |
| mmu-miR-29b-1-5p | MIMAT0004523 | Tiparp | 99929 | HITS-CLIP | 37986224 | Unspecified |
| mmu-miR-29b-1-5p | MIMAT0004523 | Tjp1 | 21872 | HITS-CLIP | 25083871 | Unspecified |
| mmu-miR-29b-1-5p | MIMAT0004523 | Tle3 | 21887 | HITS-CLIP | 25083871 | Unspecified |
| mmu-miR-29b-1-5p | MIMAT0004523 | Tlk2 | 24086 | HITS-CLIP | 25083871 | Unspecified |
| mmu-miR-29b-1-5p | MIMAT0004523 | Tm9sf2 | 68059 | HITS-CLIP | 25083871 | Unspecified |
| mmu-miR-29b-1-5p | MIMAT0004523 | Tmbim1 | 69660 | HITS-CLIP | 37986224 | Unspecified |
| mmu-miR-29b-1-5p | MIMAT0004523 | Tmbim6 | 110213 | HITS-CLIP | 25083871 | Unspecified |
| mmu-miR-29b-1-5p | MIMAT0004523 | Tmed10 | 68581 | HITS-CLIP | 25083871 | Unspecified |
| mmu-miR-29b-1-5p | MIMAT0004523 | Tmem131 | 56030 | HITS-CLIP | 25083871 | Unspecified |
| mmu-miR-29b-1-5p | MIMAT0004523 | Tmem183a | 57439 | HITS-CLIP | 37986224 | Unspecified |
| mmu-miR-29b-1-5p | MIMAT0004523 | Tmem184c | 234463 | HITS-CLIP | 37986224 | Unspecified |
| mmu-miR-29b-1-5p | MIMAT0004523 | Tmem198b | 73827 | HITS-CLIP | 37986224 | Unspecified |
| mmu-miR-29b-1-5p | MIMAT0004523 | Tmem214 | 68796 | HITS-CLIP | 25083871 | Unspecified |
| mmu-miR-29b-1-5p | MIMAT0004523 | Tmem47 | 192216 | HITS-CLIP | 37986224 | Unspecified |
| mmu-miR-29b-1-5p | MIMAT0004523 | Tmem70 | 70397 | HITS-CLIP | 37986224\|25083871 | Unspecified |
| mmu-miR-29b-1-5p | MIMAT0004523 | Tmod1 | 21916 | HITS-CLIP | 37986224 | Unspecified |
| mmu-miR-29b-1-5p | MIMAT0004523 | Tnfrsf11b | 18383 | HITS-CLIP | 37986224 | Unspecified |
| mmu-miR-29b-1-5p | MIMAT0004523 | Tnrc6a | 233833 | HITS-CLIP | 25083871 | Unspecified |
| mmu-miR-29b-1-5p | MIMAT0004523 | Tnrc6b | 213988 | HITS-CLIP | 25083871 | Unspecified |

| | | | | | | |
|---|---|---|---|---|---|---|
| mmu-miR-29b-1-5p | MIMAT0004523 | Tnrc6c | 217351 | HITS-CLIP | 37986224\|25083871 | Unspecified |
| mmu-miR-29b-1-5p | MIMAT0004523 | Tns1 | 21961 | HITS-CLIP | 37986224 | Unspecified |
| mmu-miR-29b-1-5p | MIMAT0004523 | Tob1 | 22057 | HITS-CLIP | 37986224 | Unspecified |
| mmu-miR-29b-1-5p | MIMAT0004523 | Top2a | 21973 | HITS-CLIP | 37986224\|25083871 | Unspecified |
| mmu-miR-29b-1-5p | MIMAT0004523 | Top2b | 21974 | HITS-CLIP | 25083871 | Unspecified |
| mmu-miR-29b-1-5p | MIMAT0004523 | Tor1aip1 | 208263 | HITS-CLIP | 37986224\|25083871 | Unspecified |
| mmu-miR-29b-1-5p | MIMAT0004523 | Tox4 | 268741 | HITS-CLIP | 37986224 | Unspecified |
| mmu-miR-29b-1-5p | MIMAT0004523 | Tpp2 | 22019 | HITS-CLIP | 25083871 | Unspecified |
| mmu-miR-29b-1-5p | MIMAT0004523 | Traf6 | 22034 | HITS-CLIP | 37986224 | Unspecified |
| mmu-miR-29b-1-5p | MIMAT0004523 | Trappc8 | 75964 | HITS-CLIP | 37986224\|25083871 | Unspecified |
| mmu-miR-29b-1-5p | MIMAT0004523 | Trim44 | 80985 | HITS-CLIP | 37986224\|25083871 | Unspecified |
| mmu-miR-29b-1-5p | MIMAT0004523 | Trim56 | 384309 | HITS-CLIP | 37986224 | Unspecified |
| mmu-miR-29b-1-5p | MIMAT0004523 | Trio | 223435 | HITS-CLIP | 37986224\|25083871 | Unspecified |
| mmu-miR-29b-1-5p | MIMAT0004523 | Trip11 | 109181 | HITS-CLIP | 37986224 | Unspecified |
| mmu-miR-29b-1-5p | MIMAT0004523 | Trip12 | 14897 | HITS-CLIP | 37986224\|25083871 | Unspecified |
| mmu-miR-29b-1-5p | MIMAT0004523 | Trmt10a | 108943 | HITS-CLIP | 37986224 | Unspecified |
| mmu-miR-29b-1-5p | MIMAT0004523 | Trmt1l | 98685 | HITS-CLIP | 25083871 | Unspecified |
| mmu-miR-29b-1-5p | MIMAT0004523 | Trnt1 | 70047 | HITS-CLIP | 37986224 | Unspecified |
| mmu-miR-29b-1-5p | MIMAT0004523 | Trp53inp2 | 68728 | HITS-CLIP | 37986224 | Unspecified |
| mmu-miR-29b-1-5p | MIMAT0004523 | Tsc22d1 | 21807 | HITS-CLIP | 37986224 | Unspecified |
| mmu-miR-29b-1-5p | MIMAT0004523 | Tsc22d4 | 78829 | HITS-CLIP | 37986224 | Unspecified |
| mmu-miR-29b-1-5p | MIMAT0004523 | Tsku | 244152 | HITS-CLIP | 37986224 | Unspecified |
| mmu-miR-29b-1-5p | MIMAT0004523 | Tspan3 | 56434 | HITS-CLIP | 25083871 | Unspecified |
| mmu-miR-29b-1-5p | MIMAT0004523 | Tspan5 | 56224 | HITS-CLIP | 37986224 | Unspecified |
| mmu-miR-29b-1-5p | MIMAT0004523 | Tspyl1 | 22110 | HITS-CLIP | 37986224 | Unspecified |
| mmu-miR-29b-1-5p | MIMAT0004523 | Tspyl2 | 52808 | HITS-CLIP | 37986224 | Unspecified |
| mmu-miR-29b-1-5p | MIMAT0004523 | Tsr1 | 104662 | HITS-CLIP | 37986224 | Unspecified |
| mmu-miR-29b-1-5p | MIMAT0004523 | Ttc4 | 72354 | HITS-CLIP | 25083871 | Unspecified |
| mmu-miR-29b-1-5p | MIMAT0004523 | Ttc7b | 104718 | HITS-CLIP | 37986224 | Unspecified |
| mmu-miR-29b-1-5p | MIMAT0004523 | Ttll4 | 67534 | HITS-CLIP | 37986224 | Unspecified |
| mmu-miR-29b-1-5p | MIMAT0004523 | Ttn | 22138 | HITS-CLIP | 37986224\|25083871 | Unspecified |
| mmu-miR-29b-1-5p | MIMAT0004523 | Ttyh2 | 117160 | HITS-CLIP | 37986224 | Unspecified |
| mmu-miR-29b-1-5p | MIMAT0004523 | Tubb2a | 22151 | HITS-CLIP | 37986224 | Unspecified |
| mmu-miR-29b-1-5p | MIMAT0004523 | Tubb4b | 227613 | HITS-CLIP | 25083871 | Unspecified |
| mmu-miR-29b-1-5p | MIMAT0004523 | Tubb5 | 22154 | HITS-CLIP | 37986224\|25083871 | Unspecified |
| mmu-miR-29b-1-5p | MIMAT0004523 | Tut4 | 23318 | HITS-CLIP | 25083871 | Unspecified |
| mmu-miR-29b-1-5p | MIMAT0004523 | Txndc12 | 66073 | HITS-CLIP | 25083871 | Unspecified |
| mmu-miR-29b-1-5p | MIMAT0004523 | Uba6 | 231380 | HITS-CLIP | 37986224 | Unspecified |
| mmu-miR-29b-1-5p | MIMAT0004523 | Ubac1 | 98766 | HITS-CLIP | 37986224 | Unspecified |

| | | | | | | |
|---|---|---|---|---|---|---|
| mmu-miR-29b-1-5p | MIMAT0004523 | Ubap2l | 74383 | HITS-CLIP | 37986224 | Unspecified |
| mmu-miR-29b-1-5p | MIMAT0004523 | Ube2h | 22214 | HITS-CLIP | 37986224 | Unspecified |
| mmu-miR-29b-1-5p | MIMAT0004523 | Ube2k | 53323 | HITS-CLIP | 25083871 | Unspecified |
| mmu-miR-29b-1-5p | MIMAT0004523 | Ube4b | 63958 | HITS-CLIP | 37986224 | Unspecified |
| mmu-miR-29b-1-5p | MIMAT0004523 | Ubl3 | 24109 | HITS-CLIP | 37986224 | Unspecified |
| mmu-miR-29b-1-5p | MIMAT0004523 | Ubn1 | 170644 | HITS-CLIP | 37986224 | Unspecified |
| mmu-miR-29b-1-5p | MIMAT0004523 | Ubn2 | 320538 | HITS-CLIP | 37986224\|25083871 | Unspecified |
| mmu-miR-29b-1-5p | MIMAT0004523 | Ubqln2 | 54609 | HITS-CLIP | 37986224 | Unspecified |
| mmu-miR-29b-1-5p | MIMAT0004523 | Ubr3 | 68795 | HITS-CLIP | 25083871 | Unspecified |
| mmu-miR-29b-1-5p | MIMAT0004523 | Ubr4 | 69116 | HITS-CLIP | 37986224\|25083871 | Unspecified |
| mmu-miR-29b-1-5p | MIMAT0004523 | Ubr5 | 70790 | HITS-CLIP | 37986224\|25083871 | Unspecified |
| mmu-miR-29b-1-5p | MIMAT0004523 | Ubtd1 | 226122 | HITS-CLIP | 25083871 | Unspecified |
| mmu-miR-29b-1-5p | MIMAT0004523 | Ubxn4 | 67812 | HITS-CLIP | 25083871 | Unspecified |
| mmu-miR-29b-1-5p | MIMAT0004523 | Uimc1 | 20184 | HITS-CLIP | 37986224 | Unspecified |
| mmu-miR-29b-1-5p | MIMAT0004523 | Upf2 | 326622 | HITS-CLIP | 37986224 | Unspecified |
| mmu-miR-29b-1-5p | MIMAT0004523 | Usf3 | 205717 | HITS-CLIP | 25083871 | Unspecified |
| mmu-miR-29b-1-5p | MIMAT0004523 | Usp10 | 22224 | HITS-CLIP | 37986224 | Unspecified |
| mmu-miR-29b-1-5p | MIMAT0004523 | Usp24 | 329908 | HITS-CLIP | 37986224 | Unspecified |
| mmu-miR-29b-1-5p | MIMAT0004523 | Usp28 | 235323 | HITS-CLIP | 37986224 | Unspecified |
| mmu-miR-29b-1-5p | MIMAT0004523 | Usp32 | 237898 | HITS-CLIP | 37986224\|25083871 | Unspecified |
| mmu-miR-29b-1-5p | MIMAT0004523 | Usp34 | 17847 | HITS-CLIP | 25083871 | Unspecified |
| mmu-miR-29b-1-5p | MIMAT0004523 | Usp4 | 22258 | HITS-CLIP | 37986224 | Unspecified |
| mmu-miR-29b-1-5p | MIMAT0004523 | Usp47 | 74996 | HITS-CLIP | 25083871 | Unspecified |
| mmu-miR-29b-1-5p | MIMAT0004523 | Usp48 | 170707 | HITS-CLIP | 37986224 | Unspecified |
| mmu-miR-29b-1-5p | MIMAT0004523 | Usp49 | 224836 | HITS-CLIP | 37986224 | Unspecified |
| mmu-miR-29b-1-5p | MIMAT0004523 | Usp53 | 99526 | HITS-CLIP | 37986224 | Unspecified |
| mmu-miR-29b-1-5p | MIMAT0004523 | Usp7 | 252870 | HITS-CLIP | 37986224 | Unspecified |
| mmu-miR-29b-1-5p | MIMAT0004523 | Usp8 | 84092 | HITS-CLIP | 37986224 | Unspecified |
| mmu-miR-29b-1-5p | MIMAT0004523 | Vav2 | 22325 | HITS-CLIP | 37986224\|25083871 | Unspecified |
| mmu-miR-29b-1-5p | MIMAT0004523 | Vcan | 13003 | HITS-CLIP | 37986224\|25083871 | Unspecified |
| mmu-miR-29b-1-5p | MIMAT0004523 | Vezf1 | 22344 | HITS-CLIP | 37986224\|25083871 | Unspecified |
| mmu-miR-29b-1-5p | MIMAT0004523 | Vezt | 215008 | HITS-CLIP | 37986224 | Unspecified |
| mmu-miR-29b-1-5p | MIMAT0004523 | Virma | 25962 | HITS-CLIP | 25083871 | Unspecified |
| mmu-miR-29b-1-5p | MIMAT0004523 | Vps13a | 271564 | HITS-CLIP | 25083871 | Unspecified |
| mmu-miR-29b-1-5p | MIMAT0004523 | Vps13b | 666173 | HITS-CLIP | 37986224\|25083871 | Unspecified |
| mmu-miR-29b-1-5p | MIMAT0004523 | Vps13c | 320528 | HITS-CLIP | 37986224 | Unspecified |
| mmu-miR-29b-1-5p | MIMAT0004523 | Vps13d | 230895 | HITS-CLIP | 25083871 | Unspecified |
| mmu-miR-29b-1-5p | MIMAT0004523 | Vps26a | 30930 | HITS-CLIP | 37986224 | Unspecified |
| mmu-miR-29b-1-5p | MIMAT0004523 | Vps37a | 52348 | HITS-CLIP | 37986224 | Unspecified |

| | | | | | | | |
|---|---|---|---|---|---|---|---|
| mmu-miR-29b-1-5p | MIMAT0004523 | Vps41 | 218035 | HITS-CLIP | | 25083871 | Unspecified |
| mmu-miR-29b-1-5p | MIMAT0004523 | Vps54 | 245944 | HITS-CLIP | | 37986224 | Unspecified |
| mmu-miR-29b-1-5p | MIMAT0004523 | Vta1 | 66201 | HITS-CLIP | | 25083871 | Unspecified |
| mmu-miR-29b-1-5p | MIMAT0004523 | Wac | 225131 | HITS-CLIP | | 25083871 | Unspecified |
| mmu-miR-29b-1-5p | MIMAT0004523 | Wapl | 23063 | HITS-CLIP | | 25083871 | Unspecified |
| mmu-miR-29b-1-5p | MIMAT0004523 | Wasf2 | 242687 | HITS-CLIP | | 25083871 | Unspecified |
| mmu-miR-29b-1-5p | MIMAT0004523 | Washc4 | 319277 | HITS-CLIP | | 25083871 | Unspecified |
| mmu-miR-29b-1-5p | MIMAT0004523 | Wdr26 | 226757 | HITS-CLIP | | 37986224 | Unspecified |
| mmu-miR-29b-1-5p | MIMAT0004523 | Wdr48 | 67561 | HITS-CLIP | | 37986224 | Unspecified |
| mmu-miR-29b-1-5p | MIMAT0004523 | Whrn | 73750 | HITS-CLIP | 37986224\|25083871 | | Unspecified |
| mmu-miR-29b-1-5p | MIMAT0004523 | Wipf1 | 215280 | HITS-CLIP | | 37986224 | Unspecified |
| mmu-miR-29b-1-5p | MIMAT0004523 | Wipi2 | 74781 | HITS-CLIP | | 37986224 | Unspecified |
| mmu-miR-29b-1-5p | MIMAT0004523 | Wnt9a | 216795 | HITS-CLIP | 37986224\|25083871 | | Unspecified |
| mmu-miR-29b-1-5p | MIMAT0004523 | Wwtr1 | 97064 | HITS-CLIP | | 37986224 | Unspecified |
| mmu-miR-29b-1-5p | MIMAT0004523 | Xbp1 | 22433 | HITS-CLIP | | 37986224 | Unspecified |
| mmu-miR-29b-1-5p | MIMAT0004523 | Xiap | 11798 | HITS-CLIP | | 25083871 | Unspecified |
| mmu-miR-29b-1-5p | MIMAT0004523 | Xpo7 | 65246 | HITS-CLIP | 37986224\|25083871 | | Unspecified |
| mmu-miR-29b-1-5p | MIMAT0004523 | Xpot | 73192 | HITS-CLIP | | 37986224 | Unspecified |
| mmu-miR-29b-1-5p | MIMAT0004523 | Xrn2 | 24128 | HITS-CLIP | | 25083871 | Unspecified |
| mmu-miR-29b-1-5p | MIMAT0004523 | Xylt2 | 217119 | HITS-CLIP | | 25083871 | Unspecified |
| mmu-miR-29b-1-5p | MIMAT0004523 | Yipf5 | 67180 | HITS-CLIP | | 37986224 | Unspecified |
| mmu-miR-29b-1-5p | MIMAT0004523 | Yipf6 | 77929 | HITS-CLIP | | 25083871 | Unspecified |
| mmu-miR-29b-1-5p | MIMAT0004523 | Ylpm1 | 56531 | HITS-CLIP | | 37986224 | Unspecified |
| mmu-miR-29b-1-5p | MIMAT0004523 | Yod1 | 226418 | HITS-CLIP | 37986224\|25083871 | | Unspecified |
| mmu-miR-29b-1-5p | MIMAT0004523 | Ythdf1 | 228994 | HITS-CLIP | | 37986224 | Unspecified |
| mmu-miR-29b-1-5p | MIMAT0004523 | Ythdf3 | 229096 | HITS-CLIP | 37986224\|25083871 | | Unspecified |
| mmu-miR-29b-1-5p | MIMAT0004523 | Ywhae | 22627 | HITS-CLIP | | 37986224 | Unspecified |
| mmu-miR-29b-1-5p | MIMAT0004523 | Zbtb18 | 30928 | HITS-CLIP | | 25083871 | Unspecified |
| mmu-miR-29b-1-5p | MIMAT0004523 | Zbtb20 | 56490 | HITS-CLIP | 37986224\|25083871 | | Unspecified |
| mmu-miR-29b-1-5p | MIMAT0004523 | Zbtb25 | 109929 | HITS-CLIP | | 37986224 | Unspecified |
| mmu-miR-29b-1-5p | MIMAT0004523 | Zbtb34 | 241311 | HITS-CLIP | | 37986224 | Unspecified |
| mmu-miR-29b-1-5p | MIMAT0004523 | Zbtb37 | 240869 | HITS-CLIP | | 37986224 | Unspecified |
| mmu-miR-29b-1-5p | MIMAT0004523 | Zbtb4 | 75580 | HITS-CLIP | | 37986224 | Unspecified |
| mmu-miR-29b-1-5p | MIMAT0004523 | Zbtb41 | 226470 | HITS-CLIP | 37986224\|25083871 | | Unspecified |
| mmu-miR-29b-1-5p | MIMAT0004523 | Zbtb43 | 71834 | HITS-CLIP | 37986224\|25083871 | | Unspecified |
| mmu-miR-29b-1-5p | MIMAT0004523 | Zbtb44 | 235132 | HITS-CLIP | | 37986224 | Unspecified |
| mmu-miR-29b-1-5p | MIMAT0004523 | Zbtb49 | 75079 | HITS-CLIP | | 37986224 | Unspecified |
| mmu-miR-29b-1-5p | MIMAT0004523 | Zbtb7a | 16969 | HITS-CLIP | | 37986224 | Unspecified |
| mmu-miR-29b-1-5p | MIMAT0004523 | Zc3h12c | 244871 | HITS-CLIP | | 37986224 | Unspecified |

| | | | | | | |
|---|---|---|---|---|---|---|
| mmu-miR-29b-1-5p | MIMAT0004523 | Zc3h13 | 67302 | HITS-CLIP | 37986224 | Unspecified |
| mmu-miR-29b-1-5p | MIMAT0004523 | Zc3h7b | 20286 | HITS-CLIP | 25083871 | Unspecified |
| mmu-miR-29b-1-5p | MIMAT0004523 | Zc3hav1l | 209032 | HITS-CLIP | 37986224 | Unspecified |
| mmu-miR-29b-1-5p | MIMAT0004523 | Zdhhc13 | 243983 | HITS-CLIP | 37986224 | Unspecified |
| mmu-miR-29b-1-5p | MIMAT0004523 | Zer1 | 227693 | HITS-CLIP | 37986224 | Unspecified |
| mmu-miR-29b-1-5p | MIMAT0004523 | Zfand3 | 21769 | HITS-CLIP | 25083871 | Unspecified |
| mmu-miR-29b-1-5p | MIMAT0004523 | Zfc3h1 | 216345 | HITS-CLIP | 25083871 | Unspecified |
| mmu-miR-29b-1-5p | MIMAT0004523 | Zfhx3 | 11906 | HITS-CLIP | 37986224\|25083871 | Unspecified |
| mmu-miR-29b-1-5p | MIMAT0004523 | Zfhx4 | 80892 | HITS-CLIP | 37986224\|25083871 | Unspecified |
| mmu-miR-29b-1-5p | MIMAT0004523 | Zfp106 | 20402 | HITS-CLIP | 25083871 | Unspecified |
| mmu-miR-29b-1-5p | MIMAT0004523 | Zfp12 | 231866 | HITS-CLIP | 37986224 | Unspecified |
| mmu-miR-29b-1-5p | MIMAT0004523 | Zfp143 | 20841 | HITS-CLIP | 37986224 | Unspecified |
| mmu-miR-29b-1-5p | MIMAT0004523 | Zfp148 | 22661 | HITS-CLIP | 37986224\|25083871 | Unspecified |
| mmu-miR-29b-1-5p | MIMAT0004523 | Zfp157 | 72154 | HITS-CLIP | 25083871 | Unspecified |
| mmu-miR-29b-1-5p | MIMAT0004523 | Zfp207 | 22680 | HITS-CLIP | 25083871 | Unspecified |
| mmu-miR-29b-1-5p | MIMAT0004523 | Zfp275 | 27081 | HITS-CLIP | 37986224 | Unspecified |
| mmu-miR-29b-1-5p | MIMAT0004523 | Zfp292 | 30046 | HITS-CLIP | 37986224\|25083871 | Unspecified |
| mmu-miR-29b-1-5p | MIMAT0004523 | Zfp318 | 57908 | HITS-CLIP | 37986224\|25083871 | Unspecified |
| mmu-miR-29b-1-5p | MIMAT0004523 | Zfp319 | 79233 | HITS-CLIP | 37986224 | Unspecified |
| mmu-miR-29b-1-5p | MIMAT0004523 | Zfp326 | 54367 | HITS-CLIP | 37986224 | Unspecified |
| mmu-miR-29b-1-5p | MIMAT0004523 | Zfp36l1 | 12192 | HITS-CLIP | 37986224 | Unspecified |
| mmu-miR-29b-1-5p | MIMAT0004523 | Zfp385a | 29813 | HITS-CLIP | 25083871 | Unspecified |
| mmu-miR-29b-1-5p | MIMAT0004523 | Zfp451 | 98403 | HITS-CLIP | 37986224\|25083871 | Unspecified |
| mmu-miR-29b-1-5p | MIMAT0004523 | Zfp513 | 101023 | HITS-CLIP | 37986224 | Unspecified |
| mmu-miR-29b-1-5p | MIMAT0004523 | Zfp60 | 22718 | HITS-CLIP | 37986224 | Unspecified |
| mmu-miR-29b-1-5p | MIMAT0004523 | Zfp606 | 67370 | HITS-CLIP | 37986224 | Unspecified |
| mmu-miR-29b-1-5p | MIMAT0004523 | Zfp609 | 214812 | HITS-CLIP | 37986224\|25083871 | Unspecified |
| mmu-miR-29b-1-5p | MIMAT0004523 | Zfp652 | 268469 | HITS-CLIP | 37986224 | Unspecified |
| mmu-miR-29b-1-5p | MIMAT0004523 | Zfp655 | 72611 | HITS-CLIP | 37986224 | Unspecified |
| mmu-miR-29b-1-5p | MIMAT0004523 | Zfp687 | 78266 | HITS-CLIP | 37986224 | Unspecified |
| mmu-miR-29b-1-5p | MIMAT0004523 | Zfp715 | 69930 | HITS-CLIP | 37986224 | Unspecified |
| mmu-miR-29b-1-5p | MIMAT0004523 | Zfp760 | 240034 | HITS-CLIP | 37986224 | Unspecified |
| mmu-miR-29b-1-5p | MIMAT0004523 | Zfp768 | 233890 | HITS-CLIP | 37986224 | Unspecified |
| mmu-miR-29b-1-5p | MIMAT0004523 | Zfp800 | 627049 | HITS-CLIP | 37986224 | Unspecified |
| mmu-miR-29b-1-5p | MIMAT0004523 | Zfr | 22763 | HITS-CLIP | 25083871 | Unspecified |
| mmu-miR-29b-1-5p | MIMAT0004523 | Zhx1 | 22770 | HITS-CLIP | 37986224 | Unspecified |
| mmu-miR-29b-1-5p | MIMAT0004523 | Zim1 | 22776 | HITS-CLIP | 37986224 | Unspecified |
| mmu-miR-29b-1-5p | MIMAT0004523 | Zkscan1 | 74570 | HITS-CLIP | 25083871 | Unspecified |
| mmu-miR-29b-1-5p | MIMAT0004523 | Zmat3 | 22401 | HITS-CLIP | 25083871 | Unspecified |

| | | | | | | |
|---|---|---|---|---|---|---|
| mmu-miR-29b-1-5p | MIMAT0004523 | Zmiz1 | 328365 | HITS-CLIP | 37986224 | Unspecified |
| mmu-miR-29b-1-5p | MIMAT0004523 | Zmiz2 | 52915 | HITS-CLIP | 25083871 | Unspecified |
| mmu-miR-29b-1-5p | MIMAT0004523 | Znfx1 | 98999 | HITS-CLIP | 37986224 | Unspecified |
| mmu-miR-29b-1-5p | MIMAT0004523 | Znrf3 | 407821 | HITS-CLIP | 25083871 | Unspecified |
| mmu-miR-29b-1-5p | MIMAT0004523 | Zwint | 52696 | HITS-CLIP | 25083871 | Unspecified |
| mmu-miR-29b-1-5p | MIMAT0004523 | Zzz3 | 108946 | HITS-CLIP | 37986224 | Unspecified |
| mmu-miR-378a-5p | MIMAT0000742 | 0610030E20Rik | 68364 | HITS-CLIP | 37986224 | Unspecified |
| mmu-miR-378a-5p | MIMAT0000742 | 0610037L13Rik | 74098 | HITS-CLIP | 23597149 | Unspecified |
| mmu-miR-378a-5p | MIMAT0000742 | 1110051M20Rik | 228356 | HITS-CLIP | 37986224 | Unspecified |
| mmu-miR-378a-5p | MIMAT0000742 | 1700123O20Rik | 58248 | HITS-CLIP | 37986224 | Unspecified |
| mmu-miR-378a-5p | MIMAT0000742 | 2310057M21Rik | 68277 | HITS-CLIP | 37986224 | Unspecified |
| mmu-miR-378a-5p | MIMAT0000742 | 2510009E07Rik | 72190 | HITS-CLIP | 25083871\|29167373 | Unspecified |
| mmu-miR-378a-5p | MIMAT0000742 | 3425401B19Rik | 100504518 | HITS-CLIP | 37986224 | Unspecified |
| mmu-miR-378a-5p | MIMAT0000742 | 4921524J17Rik | 66714 | HITS-CLIP | 37986224 | Unspecified |
| mmu-miR-378a-5p | MIMAT0000742 | 4933434E20Rik | 99650 | HITS-CLIP | 37986224 | Unspecified |
| mmu-miR-378a-5p | MIMAT0000742 | AI987944 | 233168 | HITS-CLIP | 23597149 | Unspecified |
| mmu-miR-378a-5p | MIMAT0000742 | Aars | 234734 | HITS-CLIP | 37986224 | Unspecified |
| mmu-miR-378a-5p | MIMAT0000742 | Abat | 268860 | HITS-CLIP | 37986224 | Unspecified |
| mmu-miR-378a-5p | MIMAT0000742 | Abca1 | 11303 | HITS-CLIP | 37986224 | Unspecified |
| mmu-miR-378a-5p | MIMAT0000742 | Abcb11 | 27413 | HITS-CLIP | 23597149 | Unspecified |
| mmu-miR-378a-5p | MIMAT0000742 | Abcb7 | 11306 | HITS-CLIP | 37986224\|25753418 | Unspecified |
| mmu-miR-378a-5p | MIMAT0000742 | Abcd3 | 19299 | HITS-CLIP | 25753418 | Unspecified |
| mmu-miR-378a-5p | MIMAT0000742 | Abi2 | 329165 | HITS-CLIP | 37986224 | Unspecified |
| mmu-miR-378a-5p | MIMAT0000742 | Abr | 109934 | HITS-CLIP | 37986224\|29167373 | Unspecified |
| mmu-miR-378a-5p | MIMAT0000742 | Abtb1 | 80283 | HITS-CLIP | 37986224 | Unspecified |
| mmu-miR-378a-5p | MIMAT0000742 | Acacb | 100705 | HITS-CLIP | 25753418 | Unspecified |
| mmu-miR-378a-5p | MIMAT0000742 | Acad11 | 102632 | HITS-CLIP | 37986224 | Unspecified |
| mmu-miR-378a-5p | MIMAT0000742 | Acads | 11409 | HITS-CLIP | 37986224 | Unspecified |
| mmu-miR-378a-5p | MIMAT0000742 | Acbd3 | 170760 | HITS-CLIP | 37986224 | Unspecified |
| mmu-miR-378a-5p | MIMAT0000742 | Acly | 104112 | HITS-CLIP | 37986224 | Unspecified |
| mmu-miR-378a-5p | MIMAT0000742 | Aco1 | 11428 | HITS-CLIP | 37986224\|23597149 | Unspecified |
| mmu-miR-378a-5p | MIMAT0000742 | Aco2 | 11429 | HITS-CLIP | 37986224 | Unspecified |
| mmu-miR-378a-5p | MIMAT0000742 | Acox2 | 93732 | HITS-CLIP | 37986224 | Unspecified |
| mmu-miR-378a-5p | MIMAT0000742 | Acsl3 | 74205 | HITS-CLIP | 25083871 | Unspecified |
| mmu-miR-378a-5p | MIMAT0000742 | Acss2 | 60525 | HITS-CLIP | 37986224 | Unspecified |
| mmu-miR-378a-5p | MIMAT0000742 | Actn1 | 109711 | HITS-CLIP | 37986224\|23597149\|29167373 | Unspecified |
| mmu-miR-378a-5p | MIMAT0000742 | Actn3 | 11474 | HITS-CLIP | 25083871 | Unspecified |
| mmu-miR-378a-5p | MIMAT0000742 | Actn4 | 60595 | HITS-CLIP | 37986224\|23597149\|25753418 | Unspecified |
| mmu-miR-378a-5p | MIMAT0000742 | Acvr2a | 11480 | HITS-CLIP | 37986224 | Unspecified |

| | | | | | | |
|---|---|---|---|---|---|---|
| mmu-miR-378a-5p | MIMAT0000742 | Adam19 | 11492 | HITS-CLIP | 37986224\|25753418 | Unspecified |
| mmu-miR-378a-5p | MIMAT0000742 | Add1 | 11518 | HITS-CLIP | 25083871 | Unspecified |
| mmu-miR-378a-5p | MIMAT0000742 | Adipor1 | 72674 | HITS-CLIP | 37986224 | Unspecified |
| mmu-miR-378a-5p | MIMAT0000742 | Adnp | 11538 | HITS-CLIP | 29167373 | Unspecified |
| mmu-miR-378a-5p | MIMAT0000742 | Aes | 14797 | HITS-CLIP | 25083871 | Unspecified |
| mmu-miR-378a-5p | MIMAT0000742 | Afap1 | 70292 | HITS-CLIP | 37986224 | Unspecified |
| mmu-miR-378a-5p | MIMAT0000742 | Aff1 | 17355 | HITS-CLIP | 23597149 | Unspecified |
| mmu-miR-378a-5p | MIMAT0000742 | Aftph | 216549 | HITS-CLIP | 37986224 | Unspecified |
| mmu-miR-378a-5p | MIMAT0000742 | Agl | 77559 | HITS-CLIP | 37986224 | Unspecified |
| mmu-miR-378a-5p | MIMAT0000742 | Agmat | 75986 | HITS-CLIP | 23597149 | Unspecified |
| mmu-miR-378a-5p | MIMAT0000742 | Ago1 | 236511 | HITS-CLIP | 37986224 | Unspecified |
| mmu-miR-378a-5p | MIMAT0000742 | Ago2 | 239528 | qCLASH | 31393953 | Unspecified |
| mmu-miR-378a-5p | MIMAT0000742 | Ago3 | 214150 | HITS-CLIP | 29167373 | Unspecified |
| mmu-miR-378a-5p | MIMAT0000742 | Agpat5 | 52123 | HITS-CLIP | 37986224 | Unspecified |
| mmu-miR-378a-5p | MIMAT0000742 | Ahctf1 | 226747 | HITS-CLIP | 37986224 | Unspecified |
| mmu-miR-378a-5p | MIMAT0000742 | Ahnak | 66395 | HITS-CLIP | 25083871 | Unspecified |
| mmu-miR-378a-5p | MIMAT0000742 | Ahsa2 | 268390 | HITS-CLIP | 23597149 | Unspecified |
| mmu-miR-378a-5p | MIMAT0000742 | Akap13 | 75547 | HITS-CLIP | 37986224\|25083871 | Unspecified |
| mmu-miR-378a-5p | MIMAT0000742 | Akap6 | 238161 | HITS-CLIP | 37986224 | Unspecified |
| mmu-miR-378a-5p | MIMAT0000742 | Akap9 | 100986 | HITS-CLIP | 37986224\|25083871 | Unspecified |
| mmu-miR-378a-5p | MIMAT0000742 | Akirin1 | 68050 | HITS-CLIP | 37986224 | Unspecified |
| mmu-miR-378a-5p | MIMAT0000742 | Alg10b | 380959 | HITS-CLIP | 37986224\|25083871\|29167373 | Unspecified |
| mmu-miR-378a-5p | MIMAT0000742 | Alg11 | 207958 | HITS-CLIP | 37986224 | Unspecified |
| mmu-miR-378a-5p | MIMAT0000742 | Alg2 | 56737 | HITS-CLIP | 37986224 | Unspecified |
| mmu-miR-378a-5p | MIMAT0000742 | Alkbh5 | 268420 | HITS-CLIP | 25083871 | Unspecified |
| mmu-miR-378a-5p | MIMAT0000742 | Alms1 | 236266 | HITS-CLIP | 37986224 | Unspecified |
| mmu-miR-378a-5p | MIMAT0000742 | Amfr | 23802 | HITS-CLIP | 25083871 | Unspecified |
| mmu-miR-378a-5p | MIMAT0000742 | Ammecr1l | 225339 | HITS-CLIP | 37986224\|29167373 | Unspecified |
| mmu-miR-378a-5p | MIMAT0000742 | Amotl1 | 75723 | HITS-CLIP | 37986224 | Unspecified |
| mmu-miR-378a-5p | MIMAT0000742 | Anapc1 | 17222 | HITS-CLIP | 37986224 | Unspecified |
| mmu-miR-378a-5p | MIMAT0000742 | Anapc5 | 59008 | HITS-CLIP | 37986224 | Unspecified |
| mmu-miR-378a-5p | MIMAT0000742 | Anapc7 | 56317 | HITS-CLIP | 37986224 | Unspecified |
| mmu-miR-378a-5p | MIMAT0000742 | Angel2 | 52477 | HITS-CLIP | 37986224 | Unspecified |
| mmu-miR-378a-5p | MIMAT0000742 | Ank | 11732 | HITS-CLIP | 37986224 | Unspecified |
| mmu-miR-378a-5p | MIMAT0000742 | Ankhd1 | 108857 | HITS-CLIP | 37986224 | Unspecified |
| mmu-miR-378a-5p | MIMAT0000742 | Ankle2 | 71782 | HITS-CLIP | 37986224 | Unspecified |
| mmu-miR-378a-5p | MIMAT0000742 | Ankrd11 | 77087 | HITS-CLIP | 25083871 | Unspecified |
| mmu-miR-378a-5p | MIMAT0000742 | Ankrd12 | 106585 | HITS-CLIP | 37986224 | Unspecified |
| mmu-miR-378a-5p | MIMAT0000742 | Ankrd17 | 81702 | HITS-CLIP | 37986224\|25083871 | Unspecified |

| | | | | | | |
|---|---|---|---|---|---|---|
| mmu-miR-378a-5p | MIMAT0000742 | Ankrd28 | 105522 | HITS-CLIP | 37986224\|23597149\|25083871 | Unspecified |
| mmu-miR-378a-5p | MIMAT0000742 | Ankrd46 | 68839 | HITS-CLIP | 37986224 | Unspecified |
| mmu-miR-378a-5p | MIMAT0000742 | Ankrd49 | 56503 | HITS-CLIP | 37986224 | Unspecified |
| mmu-miR-378a-5p | MIMAT0000742 | Ankrd50 | 99696 | HITS-CLIP | 29167373 | Unspecified |
| mmu-miR-378a-5p | MIMAT0000742 | Anks1 | 224650 | HITS-CLIP | 25753418 | Unspecified |
| mmu-miR-378a-5p | MIMAT0000742 | Anxa4 | 11746 | HITS-CLIP | 29167373 | Unspecified |
| mmu-miR-378a-5p | MIMAT0000742 | Ap1s1 | 11769 | HITS-CLIP | 25753418 | Unspecified |
| mmu-miR-378a-5p | MIMAT0000742 | Ap3d1 | 11776 | HITS-CLIP | 25083871 | Unspecified |
| mmu-miR-378a-5p | MIMAT0000742 | Ap3m1 | 55946 | HITS-CLIP | 25083871\|23597149 | Unspecified |
| mmu-miR-378a-5p | MIMAT0000742 | Apbb2 | 11787 | HITS-CLIP | 37986224\|25753418 | Unspecified |
| mmu-miR-378a-5p | MIMAT0000742 | Apc | 11789 | HITS-CLIP | 37986224\|25083871 | Unspecified |
| mmu-miR-378a-5p | MIMAT0000742 | Apoa5 | 66113 | HITS-CLIP | 37986224 | Unspecified |
| mmu-miR-378a-5p | MIMAT0000742 | Apob | 238055 | HITS-CLIP | 37986224 | Unspecified |
| mmu-miR-378a-5p | MIMAT0000742 | Appbp2 | 66884 | HITS-CLIP | 25083871 | Unspecified |
| mmu-miR-378a-5p | MIMAT0000742 | Araf | 11836 | HITS-CLIP | 37986224 | Unspecified |
| mmu-miR-378a-5p | MIMAT0000742 | Arel1 | 68497 | HITS-CLIP | 37986224 | Unspecified |
| mmu-miR-378a-5p | MIMAT0000742 | Arf3 | 11842 | HITS-CLIP | 25753418 | Unspecified |
| mmu-miR-378a-5p | MIMAT0000742 | Arfgef1 | 211673 | HITS-CLIP | 23597149 | Unspecified |
| mmu-miR-378a-5p | MIMAT0000742 | Arhgap11a | 228482 | HITS-CLIP | 37986224 | Unspecified |
| mmu-miR-378a-5p | MIMAT0000742 | Arhgap28 | 268970 | HITS-CLIP | 37986224 | Unspecified |
| mmu-miR-378a-5p | MIMAT0000742 | Arhgap29 | 214137 | HITS-CLIP | 37986224\|23597149 | Unspecified |
| mmu-miR-378a-5p | MIMAT0000742 | Arhgap35 | 232906 | HITS-CLIP | 37986224 | Unspecified |
| mmu-miR-378a-5p | MIMAT0000742 | Arhgap42 | 71544 | HITS-CLIP | 37986224\|25083871\|23597149 | Unspecified |
| mmu-miR-378a-5p | MIMAT0000742 | Arhgef12 | 69632 | HITS-CLIP | 37986224\|25083871\|25753418 | Unspecified |
| mmu-miR-378a-5p | MIMAT0000742 | Arhgef2 | 16800 | HITS-CLIP | 25083871 | Unspecified |
| mmu-miR-378a-5p | MIMAT0000742 | Arhgef7 | 54126 | HITS-CLIP | 37986224\|25083871 | Unspecified |
| mmu-miR-378a-5p | MIMAT0000742 | Arid1b | 239985 | HITS-CLIP | 37986224\|25083871 | Unspecified |
| mmu-miR-378a-5p | MIMAT0000742 | Arid2 | 77044 | HITS-CLIP | 37986224 | Unspecified |
| mmu-miR-378a-5p | MIMAT0000742 | Arid3b | 56380 | HITS-CLIP | 37986224 | Unspecified |
| mmu-miR-378a-5p | MIMAT0000742 | Arid4b | 94246 | HITS-CLIP | 37986224 | Unspecified |
| mmu-miR-378a-5p | MIMAT0000742 | Arid5b | 71371 | HITS-CLIP | 37986224 | Unspecified |
| mmu-miR-378a-5p | MIMAT0000742 | Arl6 | 56297 | HITS-CLIP | 25083871 | Unspecified |
| mmu-miR-378a-5p | MIMAT0000742 | Armc1 | 74252 | HITS-CLIP | 25083871 | Unspecified |
| mmu-miR-378a-5p | MIMAT0000742 | Armcx2 | 67416 | HITS-CLIP | 37986224 | Unspecified |
| mmu-miR-378a-5p | MIMAT0000742 | Armcx4 | 100503043 | HITS-CLIP | 37986224 | Unspecified |
| mmu-miR-378a-5p | MIMAT0000742 | Asah1 | 11886 | HITS-CLIP | 37986224 | Unspecified |
| mmu-miR-378a-5p | MIMAT0000742 | Ascc3 | 77987 | HITS-CLIP | 37986224\|25083871 | Unspecified |
| mmu-miR-378a-5p | MIMAT0000742 | Ash1l | 192195 | HITS-CLIP | 37986224 | Unspecified |
| mmu-miR-378a-5p | MIMAT0000742 | Asnsd1 | 70396 | HITS-CLIP | 37986224 | Unspecified |

| | | | | | | |
|---|---|---|---|---|---|---|
| mmu-miR-378a-5p | MIMAT0000742 | Asph | 65973 | HITS-CLIP | 37986224 | Unspecified |
| mmu-miR-378a-5p | MIMAT0000742 | Ass1 | 11898 | HITS-CLIP | 37986224 | Unspecified |
| mmu-miR-378a-5p | MIMAT0000742 | Asxl1 | 228790 | HITS-CLIP | 37986224 | Unspecified |
| mmu-miR-378a-5p | MIMAT0000742 | Atad1 | 67979 | HITS-CLIP | 37986224\|25083871 | Unspecified |
| mmu-miR-378a-5p | MIMAT0000742 | Atad2b | 320817 | HITS-CLIP | 25083871 | Unspecified |
| mmu-miR-378a-5p | MIMAT0000742 | Ate1 | 11907 | HITS-CLIP | 25083871 | Unspecified |
| mmu-miR-378a-5p | MIMAT0000742 | Atf4 | 11911 | HITS-CLIP | 37986224 | Unspecified |
| mmu-miR-378a-5p | MIMAT0000742 | Atf7 | 223922 | HITS-CLIP | 37986224 | Unspecified |
| mmu-miR-378a-5p | MIMAT0000742 | Atf7ip | 54343 | HITS-CLIP | 25753418 | Unspecified |
| mmu-miR-378a-5p | MIMAT0000742 | Atic | 108147 | HITS-CLIP | 37986224 | Unspecified |
| mmu-miR-378a-5p | MIMAT0000742 | Atl2 | 56298 | HITS-CLIP | 25083871 | Unspecified |
| mmu-miR-378a-5p | MIMAT0000742 | Atl3 | 109168 | HITS-CLIP | 37986224 | Unspecified |
| mmu-miR-378a-5p | MIMAT0000742 | Atp11a | 50770 | HITS-CLIP | 37986224 | Unspecified |
| mmu-miR-378a-5p | MIMAT0000742 | Atp13a3 | 224088 | HITS-CLIP | 37986224 | Unspecified |
| mmu-miR-378a-5p | MIMAT0000742 | Atp1a2 | 98660 | HITS-CLIP | 37986224 | Unspecified |
| mmu-miR-378a-5p | MIMAT0000742 | Atp2a2 | 11938 | HITS-CLIP | 37986224 | Unspecified |
| mmu-miR-378a-5p | MIMAT0000742 | Atp2b4 | 381290 | HITS-CLIP | 37986224 | Unspecified |
| mmu-miR-378a-5p | MIMAT0000742 | Atp2c1 | 235574 | HITS-CLIP | 37986224 | Unspecified |
| mmu-miR-378a-5p | MIMAT0000742 | Atp6v0a1 | 11975 | HITS-CLIP | 37986224 | Unspecified |
| mmu-miR-378a-5p | MIMAT0000742 | Atp9b | 50771 | HITS-CLIP | 37986224 | Unspecified |
| mmu-miR-378a-5p | MIMAT0000742 | Atxn1l | 52335 | HITS-CLIP | 37986224\|25083871 | Unspecified |
| mmu-miR-378a-5p | MIMAT0000742 | Atxn2 | 20239 | HITS-CLIP | 25083871 | Unspecified |
| mmu-miR-378a-5p | MIMAT0000742 | Atxn3 | 110616 | HITS-CLIP | 37986224 | Unspecified |
| mmu-miR-378a-5p | MIMAT0000742 | Atxn7 | 246103 | HITS-CLIP | 37986224 | Unspecified |
| mmu-miR-378a-5p | MIMAT0000742 | Atxn7l3 | 217218 | HITS-CLIP | 37986224 | Unspecified |
| mmu-miR-378a-5p | MIMAT0000742 | Atxn7l3b | 382423 | HITS-CLIP | 37986224 | Unspecified |
| mmu-miR-378a-5p | MIMAT0000742 | Auh | 11992 | HITS-CLIP | 25753418 | Unspecified |
| mmu-miR-378a-5p | MIMAT0000742 | Axl | 26362 | HITS-CLIP | 37986224 | Unspecified |
| mmu-miR-378a-5p | MIMAT0000742 | B4galt5 | 56336 | HITS-CLIP | 37986224 | Unspecified |
| mmu-miR-378a-5p | MIMAT0000742 | BC052040 | 399568 | HITS-CLIP | 37986224 | Unspecified |
| mmu-miR-378a-5p | MIMAT0000742 | Bace1 | 23821 | HITS-CLIP | 37986224 | Unspecified |
| mmu-miR-378a-5p | MIMAT0000742 | Bag3 | 29810 | HITS-CLIP | 37986224 | Unspecified |
| mmu-miR-378a-5p | MIMAT0000742 | Bard1 | 12021 | HITS-CLIP | 37986224 | Unspecified |
| mmu-miR-378a-5p | MIMAT0000742 | Baz1a | 217578 | HITS-CLIP | 31433990 | Unspecified |
| mmu-miR-378a-5p | MIMAT0000742 | Baz1b | 22385 | HITS-CLIP | 37986224 | Unspecified |
| mmu-miR-378a-5p | MIMAT0000742 | Baz2a | 116848 | HITS-CLIP | 37986224 | Unspecified |
| mmu-miR-378a-5p | MIMAT0000742 | Baz2b | 407823 | HITS-CLIP | 37986224\|25083871 | Unspecified |
| mmu-miR-378a-5p | MIMAT0000742 | Bckdha | 12039 | HITS-CLIP | 37986224 | Unspecified |
| mmu-miR-378a-5p | MIMAT0000742 | Bcl2l13 | 94044 | HITS-CLIP | 25083871 | Unspecified |

| | | | | | | | |
|---|---|---|---|---|---|---|---|
| mmu-miR-378a-5p | MIMAT0000742 | Bcl7a | 77045 | HITS-CLIP | | 25083871 | Unspecified |
| mmu-miR-378a-5p | MIMAT0000742 | Bclaf1 | 72567 | HITS-CLIP | | 29167373 | Unspecified |
| mmu-miR-378a-5p | MIMAT0000742 | Bend5 | 67621 | HITS-CLIP | | 23597149 | Unspecified |
| mmu-miR-378a-5p | MIMAT0000742 | Bet1l | 54399 | HITS-CLIP | | 37986224 | Unspecified |
| mmu-miR-378a-5p | MIMAT0000742 | Bicd2 | 76895 | HITS-CLIP | | 37986224 | Unspecified |
| mmu-miR-378a-5p | MIMAT0000742 | Bin1 | 30948 | HITS-CLIP | | 37986224 | Unspecified |
| mmu-miR-378a-5p | MIMAT0000742 | Birc6 | 12211 | HITS-CLIP | | 37986224 | Unspecified |
| mmu-miR-378a-5p | MIMAT0000742 | Blzf1 | 66352 | HITS-CLIP | | 37986224 | Unspecified |
| mmu-miR-378a-5p | MIMAT0000742 | Bmp4 | 12159 | HITS-CLIP | | 37986224 | Unspecified |
| mmu-miR-378a-5p | MIMAT0000742 | Bmper | 73230 | HITS-CLIP | | 37986224 | Unspecified |
| mmu-miR-378a-5p | MIMAT0000742 | Bnc2 | 242509 | HITS-CLIP | | 37986224 | Unspecified |
| mmu-miR-378a-5p | MIMAT0000742 | Bnip2 | 12175 | HITS-CLIP | 37986224\|25083871 | | Unspecified |
| mmu-miR-378a-5p | MIMAT0000742 | Brd1 | 223770 | HITS-CLIP | | 37986224 | Unspecified |
| mmu-miR-378a-5p | MIMAT0000742 | Brd3 | 67382 | HITS-CLIP | | 25083871 | Unspecified |
| mmu-miR-378a-5p | MIMAT0000742 | Brd7 | 26992 | HITS-CLIP | | 29167373 | Unspecified |
| mmu-miR-378a-5p | MIMAT0000742 | Brox | 71678 | HITS-CLIP | | 37986224 | Unspecified |
| mmu-miR-378a-5p | MIMAT0000742 | Brpf1 | 78783 | HITS-CLIP | | 37986224 | Unspecified |
| mmu-miR-378a-5p | MIMAT0000742 | Brpf3 | 268936 | HITS-CLIP | | 37986224 | Unspecified |
| mmu-miR-378a-5p | MIMAT0000742 | Brwd1 | 93871 | HITS-CLIP | | 37986224 | Unspecified |
| mmu-miR-378a-5p | MIMAT0000742 | Bsg | 12215 | HITS-CLIP | 25083871\|23597149 | | Unspecified |
| mmu-miR-378a-5p | MIMAT0000742 | Btrc | 12234 | HITS-CLIP | 25083871\|23597149 | | Unspecified |
| mmu-miR-378a-5p | MIMAT0000742 | C1qtnf6 | 72709 | HITS-CLIP | | 37986224 | Unspecified |
| mmu-miR-378a-5p | MIMAT0000742 | C1ra | 50909 | HITS-CLIP | | 37986224 | Unspecified |
| mmu-miR-378a-5p | MIMAT0000742 | C2cd2 | 207781 | HITS-CLIP | | 37986224 | Unspecified |
| mmu-miR-378a-5p | MIMAT0000742 | C9orf72 | 203228 | HITS-CLIP | 25083871\|25753418 | | Unspecified |
| mmu-miR-378a-5p | MIMAT0000742 | Cab39 | 12283 | HITS-CLIP | | 37986224 | Unspecified |
| mmu-miR-378a-5p | MIMAT0000742 | Cacna2d1 | 12293 | HITS-CLIP | | 25083871 | Unspecified |
| mmu-miR-378a-5p | MIMAT0000742 | Cald1 | 109624 | HITS-CLIP | | 25083871 | Unspecified |
| mmu-miR-378a-5p | MIMAT0000742 | Camk2g | 12325 | HITS-CLIP | | 37986224 | Unspecified |
| mmu-miR-378a-5p | MIMAT0000742 | Camkk2 | 207565 | HITS-CLIP | | 37986224 | Unspecified |
| mmu-miR-378a-5p | MIMAT0000742 | Camsap2 | 67886 | HITS-CLIP | | 37986224 | Unspecified |
| mmu-miR-378a-5p | MIMAT0000742 | Canx | 12330 | HITS-CLIP | 25753418\|25083871 | | Unspecified |
| mmu-miR-378a-5p | MIMAT0000742 | Capn6 | 12338 | HITS-CLIP | | 37986224 | Unspecified |
| mmu-miR-378a-5p | MIMAT0000742 | Capn7 | 12339 | HITS-CLIP | | 37986224 | Unspecified |
| mmu-miR-378a-5p | MIMAT0000742 | Car14 | 23831 | HITS-CLIP | | 23597149 | Unspecified |
| mmu-miR-378a-5p | MIMAT0000742 | Carnmt1 | 67383 | qCLASH | | 31393953 | Unspecified |
| mmu-miR-378a-5p | MIMAT0000742 | Casc4 | 319996 | HITS-CLIP | 25083871\|23597149 | | Unspecified |
| mmu-miR-378a-5p | MIMAT0000742 | Casp6 | 12368 | HITS-CLIP | | 37986224 | Unspecified |
| mmu-miR-378a-5p | MIMAT0000742 | Casq1 | 12372 | HITS-CLIP | | 37986224 | Unspecified |

| | | | | | | |
|---|---|---|---|---|---|---|
| mmu-miR-378a-5p | MIMAT0000742 | Casz1 | 69743 | HITS-CLIP | 37986224 | Unspecified |
| mmu-miR-378a-5p | MIMAT0000742 | Cblb | 208650 | HITS-CLIP | 37986224 | Unspecified |
| mmu-miR-378a-5p | MIMAT0000742 | Cbll1 | 104836 | HITS-CLIP | 37986224 | Unspecified |
| mmu-miR-378a-5p | MIMAT0000742 | Cbx2 | 12416 | HITS-CLIP | 37986224 | Unspecified |
| mmu-miR-378a-5p | MIMAT0000742 | Cbx6 | 494448 | HITS-CLIP | 37986224 | Unspecified |
| mmu-miR-378a-5p | MIMAT0000742 | Ccar1 | 67500 | HITS-CLIP | 25753418 | Unspecified |
| mmu-miR-378a-5p | MIMAT0000742 | Ccdc117 | 104479 | HITS-CLIP | 37986224 | Unspecified |
| mmu-miR-378a-5p | MIMAT0000742 | Ccdc32 | 269336 | HITS-CLIP | 37986224 | Unspecified |
| mmu-miR-378a-5p | MIMAT0000742 | Ccdc82 | 66396 | HITS-CLIP | 37986224 | Unspecified |
| mmu-miR-378a-5p | MIMAT0000742 | Ccdc88a | 108686 | HITS-CLIP | 37986224 | Unspecified |
| mmu-miR-378a-5p | MIMAT0000742 | Ccdc88c | 68339 | HITS-CLIP | 37986224 | Unspecified |
| mmu-miR-378a-5p | MIMAT0000742 | Ccnd1 | 12443 | HITS-CLIP | 25753418 | Unspecified |
| mmu-miR-378a-5p | MIMAT0000742 | Ccnl1 | 56706 | HITS-CLIP | 37986224 | Unspecified |
| mmu-miR-378a-5p | MIMAT0000742 | Ccnl2 | 56036 | HITS-CLIP | 37986224 | Unspecified |
| mmu-miR-378a-5p | MIMAT0000742 | Ccnt2 | 72949 | HITS-CLIP | 25083871\|23597149 | Unspecified |
| mmu-miR-378a-5p | MIMAT0000742 | Ccpg1 | 72278 | HITS-CLIP | 25083871 | Unspecified |
| mmu-miR-378a-5p | MIMAT0000742 | Ccs | 12460 | HITS-CLIP | 37986224 | Unspecified |
| mmu-miR-378a-5p | MIMAT0000742 | Cct5 | 12465 | qCLASH//HITS-CLIP | 31393953\|29167373 | Unspecified |
| mmu-miR-378a-5p | MIMAT0000742 | Cct8 | 12469 | HITS-CLIP | 25753418 | Unspecified |
| mmu-miR-378a-5p | MIMAT0000742 | Cd44 | 12505 | HITS-CLIP | 25083871\|25753418 | Unspecified |
| mmu-miR-378a-5p | MIMAT0000742 | Cd47 | 16423 | HITS-CLIP | 37986224 | Unspecified |
| mmu-miR-378a-5p | MIMAT0000742 | Cd82 | 12521 | HITS-CLIP | 23597149 | Unspecified |
| mmu-miR-378a-5p | MIMAT0000742 | Cd99l2 | 171486 | HITS-CLIP | 37986224 | Unspecified |
| mmu-miR-378a-5p | MIMAT0000742 | Cdc25a | 12530 | HITS-CLIP | 37986224 | Unspecified |
| mmu-miR-378a-5p | MIMAT0000742 | Cdc42bpg | 240505 | HITS-CLIP | 37986224 | Unspecified |
| mmu-miR-378a-5p | MIMAT0000742 | Cdc42ep3 | 260409 | HITS-CLIP | 37986224 | Unspecified |
| mmu-miR-378a-5p | MIMAT0000742 | Cdc5l | 71702 | HITS-CLIP | 25083871 | Unspecified |
| mmu-miR-378a-5p | MIMAT0000742 | Cdc73 | 214498 | HITS-CLIP | 25083871 | Unspecified |
| mmu-miR-378a-5p | MIMAT0000742 | Cdh2 | 12558 | HITS-CLIP | 25083871 | Unspecified |
| mmu-miR-378a-5p | MIMAT0000742 | Cdk5r1 | 12569 | HITS-CLIP | 37986224 | Unspecified |
| mmu-miR-378a-5p | MIMAT0000742 | Cdk6 | 12571 | HITS-CLIP | 25083871 | Unspecified |
| mmu-miR-378a-5p | MIMAT0000742 | Cdkn2c | 12580 | HITS-CLIP | 37986224 | Unspecified |
| mmu-miR-378a-5p | MIMAT0000742 | Cdo1 | 12583 | HITS-CLIP | 23597149 | Unspecified |
| mmu-miR-378a-5p | MIMAT0000742 | Cdr2 | 12585 | HITS-CLIP | 37986224 | Unspecified |
| mmu-miR-378a-5p | MIMAT0000742 | Cds2 | 110911 | HITS-CLIP | 37986224\|25083871 | Unspecified |
| mmu-miR-378a-5p | MIMAT0000742 | Cebpg | 12611 | HITS-CLIP | 37986224 | Unspecified |
| mmu-miR-378a-5p | MIMAT0000742 | Cecr2 | 330409 | HITS-CLIP | 37986224 | Unspecified |
| mmu-miR-378a-5p | MIMAT0000742 | Celf1 | 13046 | HITS-CLIP | 25083871\|23597149 | Unspecified |
| mmu-miR-378a-5p | MIMAT0000742 | Celf2 | 14007 | HITS-CLIP | 37986224 | Unspecified |

| | | | | | | |
|---|---|---|---|---|---|---|
| mmu-miR-378a-5p | MIMAT0000742 | Cemip2 | 23670 | HITS-CLIP | 25083871 | Unspecified |
| mmu-miR-378a-5p | MIMAT0000742 | Cenpf | 108000 | HITS-CLIP | 25083871 | Unspecified |
| mmu-miR-378a-5p | MIMAT0000742 | Cep120 | 225523 | HITS-CLIP | 29167373 | Unspecified |
| mmu-miR-378a-5p | MIMAT0000742 | Cep152 | 99100 | HITS-CLIP | 25753418 | Unspecified |
| mmu-miR-378a-5p | MIMAT0000742 | Cep162 | 382090 | HITS-CLIP | 37986224 | Unspecified |
| mmu-miR-378a-5p | MIMAT0000742 | Cep350 | 74081 | HITS-CLIP | 37986224\|25083871 | Unspecified |
| mmu-miR-378a-5p | MIMAT0000742 | Cep85l | 100038725 | HITS-CLIP | 37986224 | Unspecified |
| mmu-miR-378a-5p | MIMAT0000742 | Cers5 | 71949 | HITS-CLIP | 25753418 | Unspecified |
| mmu-miR-378a-5p | MIMAT0000742 | Cers6 | 241447 | HITS-CLIP | 37986224 | Unspecified |
| mmu-miR-378a-5p | MIMAT0000742 | Cfh | 12628 | HITS-CLIP | 37986224 | Unspecified |
| mmu-miR-378a-5p | MIMAT0000742 | Chd3 | 216848 | HITS-CLIP | 37986224 | Unspecified |
| mmu-miR-378a-5p | MIMAT0000742 | Chd4 | 107932 | HITS-CLIP | 25083871\|29167373 | Unspecified |
| mmu-miR-378a-5p | MIMAT0000742 | Chd6 | 71389 | HITS-CLIP | 37986224 | Unspecified |
| mmu-miR-378a-5p | MIMAT0000742 | Chd7 | 320790 | HITS-CLIP | 25083871\|25753418 | Unspecified |
| mmu-miR-378a-5p | MIMAT0000742 | Chd8 | 67772 | HITS-CLIP | 37986224 | Unspecified |
| mmu-miR-378a-5p | MIMAT0000742 | Chd9 | 109151 | HITS-CLIP | 37986224 | Unspecified |
| mmu-miR-378a-5p | MIMAT0000742 | Chdh | 218865 | HITS-CLIP | 23597149 | Unspecified |
| mmu-miR-378a-5p | MIMAT0000742 | Cherp | 27967 | HITS-CLIP | 37986224 | Unspecified |
| mmu-miR-378a-5p | MIMAT0000742 | Chmp4b | 75608 | HITS-CLIP | 37986224 | Unspecified |
| mmu-miR-378a-5p | MIMAT0000742 | Chordc1 | 66917 | HITS-CLIP | 37986224 | Unspecified |
| mmu-miR-378a-5p | MIMAT0000742 | Chrnd | 11447 | HITS-CLIP | 37986224 | Unspecified |
| mmu-miR-378a-5p | MIMAT0000742 | Chsy1 | 269941 | HITS-CLIP | 37986224 | Unspecified |
| mmu-miR-378a-5p | MIMAT0000742 | Cic | 71722 | HITS-CLIP | 37986224 | Unspecified |
| mmu-miR-378a-5p | MIMAT0000742 | Cipc | 217732 | HITS-CLIP | 37986224 | Unspecified |
| mmu-miR-378a-5p | MIMAT0000742 | Ckap4 | 216197 | HITS-CLIP | 25083871 | Unspecified |
| mmu-miR-378a-5p | MIMAT0000742 | Clasp1 | 76707 | HITS-CLIP | 25083871 | Unspecified |
| mmu-miR-378a-5p | MIMAT0000742 | Clcn2 | 12724 | HITS-CLIP | 23597149 | Unspecified |
| mmu-miR-378a-5p | MIMAT0000742 | Clcn3 | 12725 | HITS-CLIP | 37986224 | Unspecified |
| mmu-miR-378a-5p | MIMAT0000742 | Clcn5 | 12728 | HITS-CLIP | 25083871 | Unspecified |
| mmu-miR-378a-5p | MIMAT0000742 | Cldn12 | 64945 | HITS-CLIP | 37986224 | Unspecified |
| mmu-miR-378a-5p | MIMAT0000742 | Clec2d | 93694 | HITS-CLIP | 37986224 | Unspecified |
| mmu-miR-378a-5p | MIMAT0000742 | Clint1 | 216705 | HITS-CLIP | 37986224\|25083871\|23597149 | Unspecified |
| mmu-miR-378a-5p | MIMAT0000742 | Clk1 | 12747 | HITS-CLIP | 37986224 | Unspecified |
| mmu-miR-378a-5p | MIMAT0000742 | Clk2 | 12748 | HITS-CLIP | 37986224 | Unspecified |
| mmu-miR-378a-5p | MIMAT0000742 | Clk4 | 12750 | HITS-CLIP | 37986224 | Unspecified |
| mmu-miR-378a-5p | MIMAT0000742 | Cln5 | 211286 | HITS-CLIP | 25083871 | Unspecified |
| mmu-miR-378a-5p | MIMAT0000742 | Clock | 12753 | HITS-CLIP | 37986224\|25083871 | Unspecified |
| mmu-miR-378a-5p | MIMAT0000742 | Cltc | 67300 | HITS-CLIP | 25083871\|25753418 | Unspecified |
| mmu-miR-378a-5p | MIMAT0000742 | Clu | 12759 | HITS-CLIP | 23597149 | Unspecified |

| | | | | | | | |
|---|---|---|---|---|---|---|---|
| mmu-miR-378a-5p | MIMAT0000742 | Cluh | 74148 | HITS-CLIP | | 37986224 | Unspecified |
| mmu-miR-378a-5p | MIMAT0000742 | Cmip | 74440 | HITS-CLIP | | 29167373 | Unspecified |
| mmu-miR-378a-5p | MIMAT0000742 | Cmtr1 | 74157 | HITS-CLIP | 37986224\|23597149 | | Unspecified |
| mmu-miR-378a-5p | MIMAT0000742 | Cnksr3 | 215748 | HITS-CLIP | | 37986224 | Unspecified |
| mmu-miR-378a-5p | MIMAT0000742 | Cnn2 | 12798 | HITS-CLIP | | 37986224 | Unspecified |
| mmu-miR-378a-5p | MIMAT0000742 | Cnnm4 | 94220 | HITS-CLIP | 37986224\|23597149 | | Unspecified |
| mmu-miR-378a-5p | MIMAT0000742 | Cnot1 | 234594 | HITS-CLIP | | 29167373 | Unspecified |
| mmu-miR-378a-5p | MIMAT0000742 | Cnot10 | 78893 | HITS-CLIP | | 23597149 | Unspecified |
| mmu-miR-378a-5p | MIMAT0000742 | Cnot7 | 18983 | HITS-CLIP | | 37986224 | Unspecified |
| mmu-miR-378a-5p | MIMAT0000742 | Cnp | 12799 | HITS-CLIP | | 37986224 | Unspecified |
| mmu-miR-378a-5p | MIMAT0000742 | Cnpy3 | 72029 | HITS-CLIP | | 37986224 | Unspecified |
| mmu-miR-378a-5p | MIMAT0000742 | Coa5 | 76178 | HITS-CLIP | | 25083871 | Unspecified |
| mmu-miR-378a-5p | MIMAT0000742 | Coasy | 71743 | HITS-CLIP | | 37986224 | Unspecified |
| mmu-miR-378a-5p | MIMAT0000742 | Cobll1 | 319876 | HITS-CLIP | | 37986224 | Unspecified |
| mmu-miR-378a-5p | MIMAT0000742 | Col6a2 | 12834 | HITS-CLIP | | 37986224 | Unspecified |
| mmu-miR-378a-5p | MIMAT0000742 | Col6a3 | 12835 | HITS-CLIP | 37986224\|25083871 | | Unspecified |
| mmu-miR-378a-5p | MIMAT0000742 | Col8a1 | 12837 | HITS-CLIP | | 37986224 | Unspecified |
| mmu-miR-378a-5p | MIMAT0000742 | Comt | 12846 | HITS-CLIP | | 37986224 | Unspecified |
| mmu-miR-378a-5p | MIMAT0000742 | Cops5 | 26754 | HITS-CLIP | 37986224\|25083871 | | Unspecified |
| mmu-miR-378a-5p | MIMAT0000742 | Coq2 | 71883 | HITS-CLIP | | 37986224 | Unspecified |
| mmu-miR-378a-5p | MIMAT0000742 | Coq9 | 67914 | HITS-CLIP | | 37986224 | Unspecified |
| mmu-miR-378a-5p | MIMAT0000742 | Coro6 | 216961 | HITS-CLIP | | 37986224 | Unspecified |
| mmu-miR-378a-5p | MIMAT0000742 | Cox10 | 70383 | HITS-CLIP | | 37986224 | Unspecified |
| mmu-miR-378a-5p | MIMAT0000742 | Cpd | 12874 | HITS-CLIP | | 25083871 | Unspecified |
| mmu-miR-378a-5p | MIMAT0000742 | Cpeb4 | 67579 | HITS-CLIP | | 25083871 | Unspecified |
| mmu-miR-378a-5p | MIMAT0000742 | Cplx2 | 12890 | HITS-CLIP | | 37986224 | Unspecified |
| mmu-miR-378a-5p | MIMAT0000742 | Cpn2 | 71756 | HITS-CLIP | | 37986224 | Unspecified |
| mmu-miR-378a-5p | MIMAT0000742 | Cpne1 | 266692 | HITS-CLIP | | 37986224 | Unspecified |
| mmu-miR-378a-5p | MIMAT0000742 | Cpne2 | 234577 | HITS-CLIP | | 37986224 | Unspecified |
| mmu-miR-378a-5p | MIMAT0000742 | Cpox | 12892 | HITS-CLIP | | 37986224 | Unspecified |
| mmu-miR-378a-5p | MIMAT0000742 | Cpsf2 | 51786 | HITS-CLIP | | 25083871 | Unspecified |
| mmu-miR-378a-5p | MIMAT0000742 | Cpsf6 | 432508 | HITS-CLIP | | 37986224 | Unspecified |
| mmu-miR-378a-5p | MIMAT0000742 | Cpt1a | 12894 | HITS-CLIP | | 23597149 | Unspecified |
| mmu-miR-378a-5p | MIMAT0000742 | Creb1 | 12912 | HITS-CLIP | 37986224\|25083871 | | Unspecified |
| mmu-miR-378a-5p | MIMAT0000742 | Crebrf | 77128 | HITS-CLIP | 25083871\|23597149 | | Unspecified |
| mmu-miR-378a-5p | MIMAT0000742 | Creg1 | 433375 | HITS-CLIP | | 25753418 | Unspecified |
| mmu-miR-378a-5p | MIMAT0000742 | Crem | 12916 | HITS-CLIP | | 37986224 | Unspecified |
| mmu-miR-378a-5p | MIMAT0000742 | Crk | 12928 | HITS-CLIP | | 37986224 | Unspecified |
| mmu-miR-378a-5p | MIMAT0000742 | Crlf1 | 12931 | HITS-CLIP | | 25083871 | Unspecified |

| | | | | | | |
|---|---|---|---|---|---|---|
| mmu-miR-378a-5p | MIMAT0000742 | Crls1 | 66586 | HITS-CLIP | 37986224 | Unspecified |
| mmu-miR-378a-5p | MIMAT0000742 | Crp | 12944 | HITS-CLIP | 37986224 | Unspecified |
| mmu-miR-378a-5p | MIMAT0000742 | Csnk1g1 | 214897 | HITS-CLIP | 37986224 | Unspecified |
| mmu-miR-378a-5p | MIMAT0000742 | Csnk1g3 | 70425 | HITS-CLIP | 37986224\|25083871 | Unspecified |
| mmu-miR-378a-5p | MIMAT0000742 | Ctbp1 | 13016 | HITS-CLIP | 37986224 | Unspecified |
| mmu-miR-378a-5p | MIMAT0000742 | Ctdsp2 | 52468 | HITS-CLIP | 37986224 | Unspecified |
| mmu-miR-378a-5p | MIMAT0000742 | Ctdspl2 | 329506 | HITS-CLIP | 37986224\|25083871 | Unspecified |
| mmu-miR-378a-5p | MIMAT0000742 | Ctnna1 | 12385 | HITS-CLIP | 25083871 | Unspecified |
| mmu-miR-378a-5p | MIMAT0000742 | Ctnnd1 | 12388 | HITS-CLIP | 25083871 | Unspecified |
| mmu-miR-378a-5p | MIMAT0000742 | Cttnbp2nl | 80281 | HITS-CLIP | 25083871 | Unspecified |
| mmu-miR-378a-5p | MIMAT0000742 | Cul4a | 99375 | HITS-CLIP | 25083871 | Unspecified |
| mmu-miR-378a-5p | MIMAT0000742 | Cux1 | 13047 | HITS-CLIP | 37986224\|25083871 | Unspecified |
| mmu-miR-378a-5p | MIMAT0000742 | Cxcl12 | 20315 | HITS-CLIP | 23597149 | Unspecified |
| mmu-miR-378a-5p | MIMAT0000742 | Cyp2j5 | 13109 | HITS-CLIP | 37986224 | Unspecified |
| mmu-miR-378a-5p | MIMAT0000742 | Cyp2u1 | 71519 | HITS-CLIP | 37986224 | Unspecified |
| mmu-miR-378a-5p | MIMAT0000742 | Cyp51 | 13121 | HITS-CLIP | 37986224 | Unspecified |
| mmu-miR-378a-5p | MIMAT0000742 | Cyth3 | 19159 | HITS-CLIP | 37986224 | Unspecified |
| mmu-miR-378a-5p | MIMAT0000742 | D17H6S53E | 114585 | HITS-CLIP | 37986224 | Unspecified |
| mmu-miR-378a-5p | MIMAT0000742 | D17Wsu92e | 224647 | HITS-CLIP | 23597149 | Unspecified |
| mmu-miR-378a-5p | MIMAT0000742 | Dab2 | 13132 | HITS-CLIP | 37986224 | Unspecified |
| mmu-miR-378a-5p | MIMAT0000742 | Dab2ip | 69601 | HITS-CLIP | 37986224\|25083871 | Unspecified |
| mmu-miR-378a-5p | MIMAT0000742 | Dbt | 13171 | HITS-CLIP | 37986224 | Unspecified |
| mmu-miR-378a-5p | MIMAT0000742 | Dcaf12 | 68970 | HITS-CLIP | 23597149\|25083871 | Unspecified |
| mmu-miR-378a-5p | MIMAT0000742 | Dcaf17 | 75763 | HITS-CLIP | 37986224\|23597149 | Unspecified |
| mmu-miR-378a-5p | MIMAT0000742 | Dcaf8 | 98193 | HITS-CLIP | 25083871 | Unspecified |
| mmu-miR-378a-5p | MIMAT0000742 | Ddb1 | 13194 | HITS-CLIP | 23597149\|25083871 | Unspecified |
| mmu-miR-378a-5p | MIMAT0000742 | Ddi2 | 68817 | HITS-CLIP | 37986224 | Unspecified |
| mmu-miR-378a-5p | MIMAT0000742 | Ddr2 | 18214 | HITS-CLIP | 37986224\|25083871 | Unspecified |
| mmu-miR-378a-5p | MIMAT0000742 | Ddx1 | 104721 | HITS-CLIP | 37986224\|25083871 | Unspecified |
| mmu-miR-378a-5p | MIMAT0000742 | Ddx17 | 67040 | HITS-CLIP | 37986224\|25083871 | Unspecified |
| mmu-miR-378a-5p | MIMAT0000742 | Ddx19a | 13680 | HITS-CLIP | 37986224 | Unspecified |
| mmu-miR-378a-5p | MIMAT0000742 | Ddx39b | 53817 | HITS-CLIP | 37986224 | Unspecified |
| mmu-miR-378a-5p | MIMAT0000742 | Ddx5 | 13207 | HITS-CLIP | 29167373 | Unspecified |
| mmu-miR-378a-5p | MIMAT0000742 | Ddx54 | 71990 | HITS-CLIP | 37986224 | Unspecified |
| mmu-miR-378a-5p | MIMAT0000742 | Ddx6 | 13209 | HITS-CLIP | 37986224\|25083871\|29167373 | Unspecified |
| mmu-miR-378a-5p | MIMAT0000742 | Dedd2 | 67379 | HITS-CLIP | 37986224 | Unspecified |
| mmu-miR-378a-5p | MIMAT0000742 | Degs1 | 13244 | HITS-CLIP | 37986224 | Unspecified |
| mmu-miR-378a-5p | MIMAT0000742 | Dennd5a | 19347 | HITS-CLIP | 37986224\|25753418 | Unspecified |
| mmu-miR-378a-5p | MIMAT0000742 | Dennd5b | 320560 | HITS-CLIP | 37986224 | Unspecified |

| | | | | | | |
|---|---|---|---|---|---|---|
| mmu-miR-378a-5p | MIMAT0000742 | Dennd6a | 211922 | HITS-CLIP | 25083871 | Unspecified |
| mmu-miR-378a-5p | MIMAT0000742 | Derl1 | 67819 | qCLASH | 31393953 | Unspecified |
| mmu-miR-378a-5p | MIMAT0000742 | Des | 13346 | HITS-CLIP | 37986224 | Unspecified |
| mmu-miR-378a-5p | MIMAT0000742 | Desi2 | 78825 | HITS-CLIP | 37986224 | Unspecified |
| mmu-miR-378a-5p | MIMAT0000742 | Dgkd | 227333 | HITS-CLIP | 29167373 | Unspecified |
| mmu-miR-378a-5p | MIMAT0000742 | Dguok | 27369 | HITS-CLIP | 23597149 | Unspecified |
| mmu-miR-378a-5p | MIMAT0000742 | Dhdh | 71755 | HITS-CLIP | 23597149 | Unspecified |
| mmu-miR-378a-5p | MIMAT0000742 | Dhrs1 | 52585 | HITS-CLIP | 37986224 | Unspecified |
| mmu-miR-378a-5p | MIMAT0000742 | Dhx35 | 71715 | HITS-CLIP | 37986224 | Unspecified |
| mmu-miR-378a-5p | MIMAT0000742 | Dhx40 | 67487 | HITS-CLIP | 37986224 | Unspecified |
| mmu-miR-378a-5p | MIMAT0000742 | Dhx9 | 13211 | HITS-CLIP | 23597149 | Unspecified |
| mmu-miR-378a-5p | MIMAT0000742 | Dido1 | 23856 | HITS-CLIP | 37986224 | Unspecified |
| mmu-miR-378a-5p | MIMAT0000742 | Dio2 | 13371 | HITS-CLIP | 37986224 | Unspecified |
| mmu-miR-378a-5p | MIMAT0000742 | Dlc1 | 50768 | HITS-CLIP | 37986224 | Unspecified |
| mmu-miR-378a-5p | MIMAT0000742 | Dlg1 | 13383 | HITS-CLIP | 37986224 | Unspecified |
| mmu-miR-378a-5p | MIMAT0000742 | Dlg4 | 13385 | HITS-CLIP | 25083871 | Unspecified |
| mmu-miR-378a-5p | MIMAT0000742 | Dlst | 78920 | HITS-CLIP | 37986224 | Unspecified |
| mmu-miR-378a-5p | MIMAT0000742 | Dlx3 | 13393 | HITS-CLIP | 37986224\|25083871 | Unspecified |
| mmu-miR-378a-5p | MIMAT0000742 | Dmxl1 | 240283 | HITS-CLIP | 37986224 | Unspecified |
| mmu-miR-378a-5p | MIMAT0000742 | Dnaja4 | 58233 | HITS-CLIP | 37986224 | Unspecified |
| mmu-miR-378a-5p | MIMAT0000742 | Dnajb2 | 56812 | HITS-CLIP | 37986224 | Unspecified |
| mmu-miR-378a-5p | MIMAT0000742 | Dnajc10 | 66861 | HITS-CLIP | 25083871 | Unspecified |
| mmu-miR-378a-5p | MIMAT0000742 | Dnajc14 | 74330 | HITS-CLIP | 37986224 | Unspecified |
| mmu-miR-378a-5p | MIMAT0000742 | Dnajc21 | 78244 | HITS-CLIP | 25083871 | Unspecified |
| mmu-miR-378a-5p | MIMAT0000742 | Dnajc27 | 217378 | HITS-CLIP | 37986224 | Unspecified |
| mmu-miR-378a-5p | MIMAT0000742 | Dnajc5 | 13002 | HITS-CLIP | 37986224 | Unspecified |
| mmu-miR-378a-5p | MIMAT0000742 | Dnal1 | 105000 | HITS-CLIP | 37986224 | Unspecified |
| mmu-miR-378a-5p | MIMAT0000742 | Dnmt1 | 13433 | HITS-CLIP | 29167373 | Unspecified |
| mmu-miR-378a-5p | MIMAT0000742 | Dnpep | 13437 | HITS-CLIP | 23597149 | Unspecified |
| mmu-miR-378a-5p | MIMAT0000742 | Dnttip2 | 99480 | HITS-CLIP | 25753418 | Unspecified |
| mmu-miR-378a-5p | MIMAT0000742 | Dock5 | 68813 | HITS-CLIP | 37986224 | Unspecified |
| mmu-miR-378a-5p | MIMAT0000742 | Dpm1 | 13480 | HITS-CLIP | 25753418 | Unspecified |
| mmu-miR-378a-5p | MIMAT0000742 | Dpp8 | 74388 | HITS-CLIP | 25083871\|23597149\|25753418 | Unspecified |
| mmu-miR-378a-5p | MIMAT0000742 | Dpy19l1 | 244745 | HITS-CLIP | 23597149 | Unspecified |
| mmu-miR-378a-5p | MIMAT0000742 | Dpy19l4 | 381510 | HITS-CLIP | 29167373 | Unspecified |
| mmu-miR-378a-5p | MIMAT0000742 | Dpysl2 | 12934 | HITS-CLIP | 37986224 | Unspecified |
| mmu-miR-378a-5p | MIMAT0000742 | Drosha | 14000 | HITS-CLIP | 25083871 | Unspecified |
| mmu-miR-378a-5p | MIMAT0000742 | Drp2 | 13497 | HITS-CLIP | 37986224 | Unspecified |
| mmu-miR-378a-5p | MIMAT0000742 | Dse | 212898 | HITS-CLIP | 37986224\|25083871\|23597149 | Unspecified |

| | | | | | | |
|---|---|---|---|---|---|---|
| mmu-miR-378a-5p | MIMAT0000742 | Dst | 13518 | HITS-CLIP | 25083871 | Unspecified |
| mmu-miR-378a-5p | MIMAT0000742 | Dstn | 56431 | HITS-CLIP | 37986224 | Unspecified |
| mmu-miR-378a-5p | MIMAT0000742 | Dtwd2 | 68857 | HITS-CLIP | 37986224 | Unspecified |
| mmu-miR-378a-5p | MIMAT0000742 | Dusp3 | 72349 | HITS-CLIP | 37986224 | Unspecified |
| mmu-miR-378a-5p | MIMAT0000742 | Dusp5 | 240672 | HITS-CLIP | 37986224 | Unspecified |
| mmu-miR-378a-5p | MIMAT0000742 | Dvl3 | 13544 | HITS-CLIP | 25083871 | Unspecified |
| mmu-miR-378a-5p | MIMAT0000742 | Dync1h1 | 13424 | HITS-CLIP | 37986224\|23597149 | Unspecified |
| mmu-miR-378a-5p | MIMAT0000742 | Dyrk1a | 13548 | HITS-CLIP | 37986224 | Unspecified |
| mmu-miR-378a-5p | MIMAT0000742 | E130308A19Rik | 230259 | HITS-CLIP | 23597149 | Unspecified |
| mmu-miR-378a-5p | MIMAT0000742 | E130309D02Rik | 231868 | HITS-CLIP | 37986224 | Unspecified |
| mmu-miR-378a-5p | MIMAT0000742 | E2f5 | 13559 | HITS-CLIP | 37986224 | Unspecified |
| mmu-miR-378a-5p | MIMAT0000742 | E2f6 | 50496 | HITS-CLIP | 25083871 | Unspecified |
| mmu-miR-378a-5p | MIMAT0000742 | E2f8 | 108961 | HITS-CLIP | 25083871 | Unspecified |
| mmu-miR-378a-5p | MIMAT0000742 | Ebag9 | 55960 | HITS-CLIP | 37986224 | Unspecified |
| mmu-miR-378a-5p | MIMAT0000742 | Ebf3 | 13593 | HITS-CLIP | 37986224 | Unspecified |
| mmu-miR-378a-5p | MIMAT0000742 | Ece1 | 230857 | HITS-CLIP | 37986224 | Unspecified |
| mmu-miR-378a-5p | MIMAT0000742 | Ech1 | 51798 | HITS-CLIP | 25083871\|23597149 | Unspecified |
| mmu-miR-378a-5p | MIMAT0000742 | Echdc2 | 52430 | HITS-CLIP | 37986224 | Unspecified |
| mmu-miR-378a-5p | MIMAT0000742 | Edc3 | 353190 | HITS-CLIP | 23597149 | Unspecified |
| mmu-miR-378a-5p | MIMAT0000742 | Edem1 | 192193 | HITS-CLIP | 23597149 | Unspecified |
| mmu-miR-378a-5p | MIMAT0000742 | Eea1 | 216238 | HITS-CLIP | 37986224\|25083871 | Unspecified |
| mmu-miR-378a-5p | MIMAT0000742 | Eef2 | 13629 | qCLASH | 31393953 | Unspecified |
| mmu-miR-378a-5p | MIMAT0000742 | Efhd2 | 27984 | HITS-CLIP | 37986224 | Unspecified |
| mmu-miR-378a-5p | MIMAT0000742 | Efr3a | 76740 | HITS-CLIP | 25083871 | Unspecified |
| mmu-miR-378a-5p | MIMAT0000742 | Ehbp1l1 | 114601 | HITS-CLIP | 37986224\|25083871 | Unspecified |
| mmu-miR-378a-5p | MIMAT0000742 | Eif1a | 13664 | HITS-CLIP | 37986224\|25083871 | Unspecified |
| mmu-miR-378a-5p | MIMAT0000742 | Eif2b4 | 13667 | HITS-CLIP | 37986224 | Unspecified |
| mmu-miR-378a-5p | MIMAT0000742 | Eif4g1 | 208643 | HITS-CLIP | 37986224 | Unspecified |
| mmu-miR-378a-5p | MIMAT0000742 | Eif4g2 | 13690 | HITS-CLIP | 29167373 | Unspecified |
| mmu-miR-378a-5p | MIMAT0000742 | Eif4h | 22384 | HITS-CLIP | 37986224 | Unspecified |
| mmu-miR-378a-5p | MIMAT0000742 | Eif5b | 226982 | HITS-CLIP | 37986224\|25083871 | Unspecified |
| mmu-miR-378a-5p | MIMAT0000742 | Elavl1 | 15568 | HITS-CLIP | 37986224 | Unspecified |
| mmu-miR-378a-5p | MIMAT0000742 | Elk3 | 13713 | HITS-CLIP | 37986224\|29167373 | Unspecified |
| mmu-miR-378a-5p | MIMAT0000742 | Elmo1 | 140580 | HITS-CLIP | 29167373 | Unspecified |
| mmu-miR-378a-5p | MIMAT0000742 | Elovl1 | 54325 | HITS-CLIP | 37986224\|25083871 | Unspecified |
| mmu-miR-378a-5p | MIMAT0000742 | Elovl5 | 68801 | HITS-CLIP | 23597149 | Unspecified |
| mmu-miR-378a-5p | MIMAT0000742 | Eml1 | 68519 | HITS-CLIP | 37986224 | Unspecified |
| mmu-miR-378a-5p | MIMAT0000742 | Eml3 | 225898 | HITS-CLIP | 23597149 | Unspecified |
| mmu-miR-378a-5p | MIMAT0000742 | Eml4 | 78798 | HITS-CLIP | 25083871 | Unspecified |

| | | | | | | |
|---|---|---|---|---|---|---|
| mmu-miR-378a-5p | MIMAT0000742 | Enah | 13800 | HITS-CLIP | 25083871 | Unspecified |
| mmu-miR-378a-5p | MIMAT0000742 | Enpp1 | 18605 | HITS-CLIP | 37986224 | Unspecified |
| mmu-miR-378a-5p | MIMAT0000742 | Enpp4 | 224794 | HITS-CLIP | 37986224 | Unspecified |
| mmu-miR-378a-5p | MIMAT0000742 | Entpd5 | 12499 | HITS-CLIP | 37986224 | Unspecified |
| mmu-miR-378a-5p | MIMAT0000742 | Entpd7 | 93685 | HITS-CLIP | 37986224 | Unspecified |
| mmu-miR-378a-5p | MIMAT0000742 | Entr1 | 10807 | HITS-CLIP | 25083871 | Unspecified |
| mmu-miR-378a-5p | MIMAT0000742 | Ep300 | 328572 | HITS-CLIP | 25083871 | Unspecified |
| mmu-miR-378a-5p | MIMAT0000742 | Epg5 | 100502841 | HITS-CLIP | 37986224 | Unspecified |
| mmu-miR-378a-5p | MIMAT0000742 | Eps15 | 13858 | HITS-CLIP | 23597149 | Unspecified |
| mmu-miR-378a-5p | MIMAT0000742 | Erc1 | 111173 | HITS-CLIP | 37986224 | Unspecified |
| mmu-miR-378a-5p | MIMAT0000742 | Ermp1 | 226090 | HITS-CLIP | 37986224 | Unspecified |
| mmu-miR-378a-5p | MIMAT0000742 | Ern1 | 78943 | HITS-CLIP | 23597149 | Unspecified |
| mmu-miR-378a-5p | MIMAT0000742 | Erp44 | 76299 | HITS-CLIP | 37986224 | Unspecified |
| mmu-miR-378a-5p | MIMAT0000742 | Esco1 | 77805 | HITS-CLIP | 37986224 | Unspecified |
| mmu-miR-378a-5p | MIMAT0000742 | Ets1 | 23871 | HITS-CLIP | 37986224 | Unspecified |
| mmu-miR-378a-5p | MIMAT0000742 | Evi5 | 14020 | HITS-CLIP | 37986224 | Unspecified |
| mmu-miR-378a-5p | MIMAT0000742 | Ezh1 | 14055 | HITS-CLIP | 25753418 | Unspecified |
| mmu-miR-378a-5p | MIMAT0000742 | F12 | 58992 | HITS-CLIP | 37986224 | Unspecified |
| mmu-miR-378a-5p | MIMAT0000742 | Fah | 14085 | HITS-CLIP | 37986224 | Unspecified |
| mmu-miR-378a-5p | MIMAT0000742 | Fam107b | 66540 | HITS-CLIP | 37986224 | Unspecified |
| mmu-miR-378a-5p | MIMAT0000742 | Fam111a | 107373 | HITS-CLIP | 37986224 | Unspecified |
| mmu-miR-378a-5p | MIMAT0000742 | Fam114a2 | 67726 | HITS-CLIP | 37986224 | Unspecified |
| mmu-miR-378a-5p | MIMAT0000742 | Fam129b | 227737 | HITS-CLIP | 25083871 | Unspecified |
| mmu-miR-378a-5p | MIMAT0000742 | Fam131b | 76156 | HITS-CLIP | 37986224 | Unspecified |
| mmu-miR-378a-5p | MIMAT0000742 | Fam133b | 68152 | HITS-CLIP | 25083871 | Unspecified |
| mmu-miR-378a-5p | MIMAT0000742 | Fam135a | 68187 | HITS-CLIP | 25083871 | Unspecified |
| mmu-miR-378a-5p | MIMAT0000742 | Fam168a | 319604 | HITS-CLIP | 37986224 | Unspecified |
| mmu-miR-378a-5p | MIMAT0000742 | Fam171a1 | 269233 | HITS-CLIP | 37986224 | Unspecified |
| mmu-miR-378a-5p | MIMAT0000742 | Fam193a | 231128 | HITS-CLIP | 37986224 | Unspecified |
| mmu-miR-378a-5p | MIMAT0000742 | Fam204a | 76539 | HITS-CLIP | 37986224 | Unspecified |
| mmu-miR-378a-5p | MIMAT0000742 | Fam208b | 54906 | HITS-CLIP | 25083871 | Unspecified |
| mmu-miR-378a-5p | MIMAT0000742 | Fam20a | 208659 | HITS-CLIP | 37986224 | Unspecified |
| mmu-miR-378a-5p | MIMAT0000742 | Fam210a | 108654 | HITS-CLIP | 37986224 | Unspecified |
| mmu-miR-378a-5p | MIMAT0000742 | Fam234b | 57613 | HITS-CLIP | 23597149 | Unspecified |
| mmu-miR-378a-5p | MIMAT0000742 | Fam8a1 | 51439 | HITS-CLIP | 31433990 | Unspecified |
| mmu-miR-378a-5p | MIMAT0000742 | Fam91a1 | 210998 | HITS-CLIP | 25083871 | Unspecified |
| mmu-miR-378a-5p | MIMAT0000742 | Fasn | 14104 | HITS-CLIP | 23597149 | Unspecified |
| mmu-miR-378a-5p | MIMAT0000742 | Fastkd2 | 75619 | HITS-CLIP | 37986224 | Unspecified |
| mmu-miR-378a-5p | MIMAT0000742 | Fat1 | 14107 | HITS-CLIP | 37986224\|25083871 | Unspecified |

| | | | | | | | |
|---|---|---|---|---|---|---|---|
| mmu-miR-378a-5p | MIMAT0000742 | Fbln5 | 23876 | HITS-CLIP | | 25083871 | Unspecified |
| mmu-miR-378a-5p | MIMAT0000742 | Fbxl18 | 231863 | HITS-CLIP | | 37986224 | Unspecified |
| mmu-miR-378a-5p | MIMAT0000742 | Fbxl3 | 50789 | HITS-CLIP | | 37986224 | Unspecified |
| mmu-miR-378a-5p | MIMAT0000742 | Fbxo28 | 67948 | HITS-CLIP | 37986224\|25083871 | | Unspecified |
| mmu-miR-378a-5p | MIMAT0000742 | Fbxo42 | 213499 | HITS-CLIP | | 37986224 | Unspecified |
| mmu-miR-378a-5p | MIMAT0000742 | Fcho2 | 218503 | HITS-CLIP | | 37986224 | Unspecified |
| mmu-miR-378a-5p | MIMAT0000742 | Fchsd2 | 207278 | HITS-CLIP | | 23597149 | Unspecified |
| mmu-miR-378a-5p | MIMAT0000742 | Fem1c | 240263 | qCLASH | | 31393953 | Unspecified |
| mmu-miR-378a-5p | MIMAT0000742 | Fen1 | 14156 | HITS-CLIP | | 37986224 | Unspecified |
| mmu-miR-378a-5p | MIMAT0000742 | Fgd6 | 13998 | HITS-CLIP | | 29167373 | Unspecified |
| mmu-miR-378a-5p | MIMAT0000742 | Fgfr1 | 14182 | HITS-CLIP | | 37986224 | Unspecified |
| mmu-miR-378a-5p | MIMAT0000742 | Fignl2 | 668225 | HITS-CLIP | | 37986224 | Unspecified |
| mmu-miR-378a-5p | MIMAT0000742 | Filip1 | 70598 | HITS-CLIP | | 37986224 | Unspecified |
| mmu-miR-378a-5p | MIMAT0000742 | Fktn | 246179 | HITS-CLIP | | 37986224 | Unspecified |
| mmu-miR-378a-5p | MIMAT0000742 | Flii | 14248 | HITS-CLIP | | 37986224 | Unspecified |
| mmu-miR-378a-5p | MIMAT0000742 | Flnb | 286940 | HITS-CLIP | | 25083871 | Unspecified |
| mmu-miR-378a-5p | MIMAT0000742 | Flnc | 68794 | HITS-CLIP | 37986224\|25083871 | | Unspecified |
| mmu-miR-378a-5p | MIMAT0000742 | Fmnl3 | 22379 | HITS-CLIP | | 25083871 | Unspecified |
| mmu-miR-378a-5p | MIMAT0000742 | Fn1 | 14268 | HITS-CLIP | 37986224\|23597149 | | Unspecified |
| mmu-miR-378a-5p | MIMAT0000742 | Fnip1 | 216742 | HITS-CLIP | 37986224\|25083871 | | Unspecified |
| mmu-miR-378a-5p | MIMAT0000742 | Fos | 14281 | HITS-CLIP | | 23597149 | Unspecified |
| mmu-miR-378a-5p | MIMAT0000742 | Foxn3 | 71375 | HITS-CLIP | 37986224\|25083871 | | Unspecified |
| mmu-miR-378a-5p | MIMAT0000742 | Foxo3 | 56484 | HITS-CLIP | | 37986224 | Unspecified |
| mmu-miR-378a-5p | MIMAT0000742 | Frmd4a | 209630 | HITS-CLIP | 37986224\|25083871 | | Unspecified |
| mmu-miR-378a-5p | MIMAT0000742 | Frmd6 | 319710 | HITS-CLIP | | 37986224 | Unspecified |
| mmu-miR-378a-5p | MIMAT0000742 | Frs2 | 327826 | HITS-CLIP | 37986224\|25083871 | | Unspecified |
| mmu-miR-378a-5p | MIMAT0000742 | Fry | 320365 | HITS-CLIP | | 25753418 | Unspecified |
| mmu-miR-378a-5p | MIMAT0000742 | Fscn1 | 14086 | HITS-CLIP | | 25083871 | Unspecified |
| mmu-miR-378a-5p | MIMAT0000742 | Fst | 14313 | HITS-CLIP | | 37986224 | Unspecified |
| mmu-miR-378a-5p | MIMAT0000742 | Fstl1 | 14314 | HITS-CLIP | | 25083871 | Unspecified |
| mmu-miR-378a-5p | MIMAT0000742 | Fto | 26383 | HITS-CLIP | | 25083871 | Unspecified |
| mmu-miR-378a-5p | MIMAT0000742 | Ftsj3 | 56095 | HITS-CLIP | 37986224\|25083871 | | Unspecified |
| mmu-miR-378a-5p | MIMAT0000742 | Fxr1 | 14359 | HITS-CLIP | | 25083871 | Unspecified |
| mmu-miR-378a-5p | MIMAT0000742 | Fxr2 | 23879 | HITS-CLIP | | 25083871 | Unspecified |
| mmu-miR-378a-5p | MIMAT0000742 | Fyn | 14360 | HITS-CLIP | | 37986224 | Unspecified |
| mmu-miR-378a-5p | MIMAT0000742 | G3bp1 | 27041 | HITS-CLIP | | 37986224 | Unspecified |
| mmu-miR-378a-5p | MIMAT0000742 | Gabarapl1 | 57436 | HITS-CLIP | | 37986224 | Unspecified |
| mmu-miR-378a-5p | MIMAT0000742 | Gabpa | 14390 | HITS-CLIP | 37986224\|25083871 | | Unspecified |
| mmu-miR-378a-5p | MIMAT0000742 | Gabpb2 | 213054 | HITS-CLIP | 37986224\|25083871 | | Unspecified |

| | | | | | | |
|---|---|---|---|---|---|---|
| mmu-miR-378a-5p | MIMAT0000742 | Galnt4 | 14426 | HITS-CLIP | 37986224 | Unspecified |
| mmu-miR-378a-5p | MIMAT0000742 | Gamt | 14431 | HITS-CLIP | 37986224 | Unspecified |
| mmu-miR-378a-5p | MIMAT0000742 | Gata3 | 14462 | HITS-CLIP | 37986224 | Unspecified |
| mmu-miR-378a-5p | MIMAT0000742 | Gatad2a | 234366 | HITS-CLIP | 37986224\|25083871 | Unspecified |
| mmu-miR-378a-5p | MIMAT0000742 | Gbe1 | 74185 | HITS-CLIP | 37986224 | Unspecified |
| mmu-miR-378a-5p | MIMAT0000742 | Gbf1 | 107338 | HITS-CLIP | 37986224 | Unspecified |
| mmu-miR-378a-5p | MIMAT0000742 | Gc | 14473 | HITS-CLIP | 23597149 | Unspecified |
| mmu-miR-378a-5p | MIMAT0000742 | Gcsh | 68133 | HITS-CLIP | 37986224\|23597149 | Unspecified |
| mmu-miR-378a-5p | MIMAT0000742 | Gem | 14579 | HITS-CLIP | 25083871 | Unspecified |
| mmu-miR-378a-5p | MIMAT0000742 | Gfod1 | 328232 | HITS-CLIP | 37986224 | Unspecified |
| mmu-miR-378a-5p | MIMAT0000742 | Ghr | 14600 | HITS-CLIP | 25753418 | Unspecified |
| mmu-miR-378a-5p | MIMAT0000742 | Gid4 | 66771 | HITS-CLIP | 37986224 | Unspecified |
| mmu-miR-378a-5p | MIMAT0000742 | Gigyf2 | 227331 | HITS-CLIP | 37986224\|25083871 | Unspecified |
| mmu-miR-378a-5p | MIMAT0000742 | Glce | 93683 | HITS-CLIP | 37986224 | Unspecified |
| mmu-miR-378a-5p | MIMAT0000742 | Gls2 | 216456 | HITS-CLIP | 23597149 | Unspecified |
| mmu-miR-378a-5p | MIMAT0000742 | Glul | 14645 | HITS-CLIP | 37986224 | Unspecified |
| mmu-miR-378a-5p | MIMAT0000742 | Glyat | 107146 | HITS-CLIP | 37986224 | Unspecified |
| mmu-miR-378a-5p | MIMAT0000742 | Gm20716 | 105980075 | HITS-CLIP | 25753418 | Unspecified |
| mmu-miR-378a-5p | MIMAT0000742 | Gm2a | 14667 | HITS-CLIP | 37986224 | Unspecified |
| mmu-miR-378a-5p | MIMAT0000742 | Gm49336 | 127230315 | HITS-CLIP | 25083871 | Unspecified |
| mmu-miR-378a-5p | MIMAT0000742 | Gm5422 | 432448 | HITS-CLIP | 37986224 | Unspecified |
| mmu-miR-378a-5p | MIMAT0000742 | Gm5577 | 434064 | HITS-CLIP | 37986224 | Unspecified |
| mmu-miR-378a-5p | MIMAT0000742 | Gmcl1 | 23885 | HITS-CLIP | 37986224 | Unspecified |
| mmu-miR-378a-5p | MIMAT0000742 | Gmfb | 63985 | HITS-CLIP | 37986224 | Unspecified |
| mmu-miR-378a-5p | MIMAT0000742 | Gnai3 | 14679 | HITS-CLIP | 37986224 | Unspecified |
| mmu-miR-378a-5p | MIMAT0000742 | Gnaq | 14682 | HITS-CLIP | 37986224\|25083871\|29167373 | Unspecified |
| mmu-miR-378a-5p | MIMAT0000742 | Gnb1 | 14688 | HITS-CLIP | 37986224 | Unspecified |
| mmu-miR-378a-5p | MIMAT0000742 | Gnb2 | 14693 | HITS-CLIP | 37986224 | Unspecified |
| mmu-miR-378a-5p | MIMAT0000742 | Golga3 | 269682 | HITS-CLIP | 37986224 | Unspecified |
| mmu-miR-378a-5p | MIMAT0000742 | Golga4 | 54214 | HITS-CLIP | 25083871 | Unspecified |
| mmu-miR-378a-5p | MIMAT0000742 | Gon4l | 76022 | HITS-CLIP | 37986224 | Unspecified |
| mmu-miR-378a-5p | MIMAT0000742 | Gopc | 94221 | HITS-CLIP | 37986224 | Unspecified |
| mmu-miR-378a-5p | MIMAT0000742 | Got1 | 14718 | HITS-CLIP | 37986224 | Unspecified |
| mmu-miR-378a-5p | MIMAT0000742 | Gpalpp1 | 67467 | HITS-CLIP | 37986224 | Unspecified |
| mmu-miR-378a-5p | MIMAT0000742 | Gpam | 14732 | HITS-CLIP | 37986224\|25083871 | Unspecified |
| mmu-miR-378a-5p | MIMAT0000742 | Gpr146 | 80290 | HITS-CLIP | 37986224 | Unspecified |
| mmu-miR-378a-5p | MIMAT0000742 | Gpr161 | 240888 | HITS-CLIP | 37986224 | Unspecified |
| mmu-miR-378a-5p | MIMAT0000742 | Gprc5c | 70355 | HITS-CLIP | 25083871\|23597149 | Unspecified |
| mmu-miR-378a-5p | MIMAT0000742 | Gpt2 | 108682 | HITS-CLIP | 37986224 | Unspecified |

| | | | | | | |
|---|---|---|---|---|---|---|
| mmu-miR-378a-5p | MIMAT0000742 | Gramd1b | 235283 | HITS-CLIP | 37986224 | Unspecified |
| mmu-miR-378a-5p | MIMAT0000742 | Grasp | 56149 | HITS-CLIP | 23597149 | Unspecified |
| mmu-miR-378a-5p | MIMAT0000742 | Grk2 | 156 | HITS-CLIP | 25083871 | Unspecified |
| mmu-miR-378a-5p | MIMAT0000742 | Grsf1 | 231413 | HITS-CLIP | 25753418 | Unspecified |
| mmu-miR-378a-5p | MIMAT0000742 | Gspt1 | 14852 | HITS-CLIP | 37986224 | Unspecified |
| mmu-miR-378a-5p | MIMAT0000742 | Gtf2h1 | 14884 | HITS-CLIP | 37986224 | Unspecified |
| mmu-miR-378a-5p | MIMAT0000742 | Gtf2ird1 | 57080 | HITS-CLIP | 37986224\|25083871 | Unspecified |
| mmu-miR-378a-5p | MIMAT0000742 | Gucd1 | 68778 | HITS-CLIP | 37986224 | Unspecified |
| mmu-miR-378a-5p | MIMAT0000742 | Gxylt1 | 223827 | HITS-CLIP | 37986224 | Unspecified |
| mmu-miR-378a-5p | MIMAT0000742 | Habp2 | 226243 | HITS-CLIP | 37986224 | Unspecified |
| mmu-miR-378a-5p | MIMAT0000742 | Haus6 | 230376 | HITS-CLIP | 37986224 | Unspecified |
| mmu-miR-378a-5p | MIMAT0000742 | Hbs1l | 56422 | HITS-CLIP | 25083871 | Unspecified |
| mmu-miR-378a-5p | MIMAT0000742 | Hdac3 | 15183 | HITS-CLIP | 37986224 | Unspecified |
| mmu-miR-378a-5p | MIMAT0000742 | Hdac4 | 208727 | HITS-CLIP | 37986224 | Unspecified |
| mmu-miR-378a-5p | MIMAT0000742 | Hdac7 | 56233 | HITS-CLIP | 37986224 | Unspecified |
| mmu-miR-378a-5p | MIMAT0000742 | Hdgfl2 | 84717 | HITS-CLIP | 25753418 | Unspecified |
| mmu-miR-378a-5p | MIMAT0000742 | Hdlbp | 110611 | HITS-CLIP | 37986224 | Unspecified |
| mmu-miR-378a-5p | MIMAT0000742 | Heatr5a | 320487 | HITS-CLIP | 29167373 | Unspecified |
| mmu-miR-378a-5p | MIMAT0000742 | Hectd1 | 207304 | HITS-CLIP | 37986224\|25083871\|29167373 | Unspecified |
| mmu-miR-378a-5p | MIMAT0000742 | Heg1 | 77446 | HITS-CLIP | 37986224 | Unspecified |
| mmu-miR-378a-5p | MIMAT0000742 | Helz | 78455 | HITS-CLIP | 37986224\|25083871 | Unspecified |
| mmu-miR-378a-5p | MIMAT0000742 | Herc1 | 235439 | HITS-CLIP | 25083871 | Unspecified |
| mmu-miR-378a-5p | MIMAT0000742 | Herc4 | 67345 | HITS-CLIP | 25083871 | Unspecified |
| mmu-miR-378a-5p | MIMAT0000742 | Herpud2 | 80517 | HITS-CLIP | 37986224\|25083871 | Unspecified |
| mmu-miR-378a-5p | MIMAT0000742 | Hey1 | 15213 | HITS-CLIP | 37986224 | Unspecified |
| mmu-miR-378a-5p | MIMAT0000742 | Hira | 15260 | HITS-CLIP | 37986224 | Unspecified |
| mmu-miR-378a-5p | MIMAT0000742 | Hivep2 | 15273 | HITS-CLIP | 37986224\|25083871 | Unspecified |
| mmu-miR-378a-5p | MIMAT0000742 | Hjurp | 381280 | HITS-CLIP | 37986224 | Unspecified |
| mmu-miR-378a-5p | MIMAT0000742 | Hmgcr | 15357 | HITS-CLIP | 23597149\|25083871 | Unspecified |
| mmu-miR-378a-5p | MIMAT0000742 | Hnf4a | 15378 | HITS-CLIP | 37986224 | Unspecified |
| mmu-miR-378a-5p | MIMAT0000742 | Hnrnpa2b1 | 53379 | qCLASH | 31393953 | Unspecified |
| mmu-miR-378a-5p | MIMAT0000742 | Hnrnpab | 15384 | HITS-CLIP | 25083871 | Unspecified |
| mmu-miR-378a-5p | MIMAT0000742 | Hnrnpu | 51810 | HITS-CLIP | 37986224 | Unspecified |
| mmu-miR-378a-5p | MIMAT0000742 | Hnrnpul1 | 232989 | HITS-CLIP | 25083871 | Unspecified |
| mmu-miR-378a-5p | MIMAT0000742 | Hook1 | 77963 | HITS-CLIP | 23597149 | Unspecified |
| mmu-miR-378a-5p | MIMAT0000742 | Hoxa3 | 15400 | HITS-CLIP | 25083871 | Unspecified |
| mmu-miR-378a-5p | MIMAT0000742 | Hoxc4 | 15423 | HITS-CLIP | 37986224 | Unspecified |
| mmu-miR-378a-5p | MIMAT0000742 | Hprt | 15452 | HITS-CLIP | 37986224 | Unspecified |
| mmu-miR-378a-5p | MIMAT0000742 | Hpx | 15458 | HITS-CLIP | 37986224 | Unspecified |

| | | | | | | |
|---|---|---|---|---|---|---|
| mmu-miR-378a-5p | MIMAT0000742 | Hs6st1 | 50785 | HITS-CLIP | 37986224 | Unspecified |
| mmu-miR-378a-5p | MIMAT0000742 | Hsdl2 | 72479 | HITS-CLIP | 25753418 | Unspecified |
| mmu-miR-378a-5p | MIMAT0000742 | Hsp90ab1 | 15516 | HITS-CLIP | 37986224 | Unspecified |
| mmu-miR-378a-5p | MIMAT0000742 | Hspa4l | 18415 | HITS-CLIP | 25083871 | Unspecified |
| mmu-miR-378a-5p | MIMAT0000742 | Hspd1 | 15510 | HITS-CLIP | 37986224 | Unspecified |
| mmu-miR-378a-5p | MIMAT0000742 | Hspg2 | 15530 | HITS-CLIP | 37986224\|25083871 | Unspecified |
| mmu-miR-378a-5p | MIMAT0000742 | Huwe1 | 59026 | HITS-CLIP | 23597149 | Unspecified |
| mmu-miR-378a-5p | MIMAT0000742 | Hyal2 | 15587 | HITS-CLIP | 37986224 | Unspecified |
| mmu-miR-378a-5p | MIMAT0000742 | Ibtk | 108837 | HITS-CLIP | 37986224\|25083871 | Unspecified |
| mmu-miR-378a-5p | MIMAT0000742 | Icmt | 57295 | HITS-CLIP | 37986224 | Unspecified |
| mmu-miR-378a-5p | MIMAT0000742 | Ifi207 | 226691 | HITS-CLIP | 29167373 | Unspecified |
| mmu-miR-378a-5p | MIMAT0000742 | Ifnar1 | 15975 | HITS-CLIP | 37986224 | Unspecified |
| mmu-miR-378a-5p | MIMAT0000742 | Igf1 | 16000 | HITS-CLIP | 37986224 | Unspecified |
| mmu-miR-378a-5p | MIMAT0000742 | Igf1r | 16001 | HITS-CLIP | 37986224\|25083871 | Unspecified |
| mmu-miR-378a-5p | MIMAT0000742 | Ikbip | 67454 | HITS-CLIP | 37986224 | Unspecified |
| mmu-miR-378a-5p | MIMAT0000742 | Ikzf1 | 22778 | HITS-CLIP | 29167373 | Unspecified |
| mmu-miR-378a-5p | MIMAT0000742 | Ikzf2 | 22779 | HITS-CLIP | 37986224 | Unspecified |
| mmu-miR-378a-5p | MIMAT0000742 | Il1r1 | 16177 | HITS-CLIP | 37986224 | Unspecified |
| mmu-miR-378a-5p | MIMAT0000742 | Il6st | 16195 | HITS-CLIP | 37986224 | Unspecified |
| mmu-miR-378a-5p | MIMAT0000742 | Impad1 | 242291 | HITS-CLIP | 37986224 | Unspecified |
| mmu-miR-378a-5p | MIMAT0000742 | Ing1 | 26356 | HITS-CLIP | 37986224 | Unspecified |
| mmu-miR-378a-5p | MIMAT0000742 | Ing5 | 66262 | HITS-CLIP | 37986224 | Unspecified |
| mmu-miR-378a-5p | MIMAT0000742 | Ino80d | 227195 | HITS-CLIP | 29167373 | Unspecified |
| mmu-miR-378a-5p | MIMAT0000742 | Inpp4b | 234515 | HITS-CLIP | 25083871 | Unspecified |
| mmu-miR-378a-5p | MIMAT0000742 | Insr | 16337 | HITS-CLIP | 37986224 | Unspecified |
| mmu-miR-378a-5p | MIMAT0000742 | Ints8 | 72656 | HITS-CLIP | 25083871\|25753418 | Unspecified |
| mmu-miR-378a-5p | MIMAT0000742 | Ip6k2 | 76500 | HITS-CLIP | 25083871 | Unspecified |
| mmu-miR-378a-5p | MIMAT0000742 | Ipo4 | 75751 | HITS-CLIP | 37986224 | Unspecified |
| mmu-miR-378a-5p | MIMAT0000742 | Ipo7 | 233726 | HITS-CLIP | 25083871 | Unspecified |
| mmu-miR-378a-5p | MIMAT0000742 | Ipo8 | 320727 | HITS-CLIP | 37986224\|25083871 | Unspecified |
| mmu-miR-378a-5p | MIMAT0000742 | Ipo9 | 226432 | HITS-CLIP | 37986224\|25083871\|29167373 | Unspecified |
| mmu-miR-378a-5p | MIMAT0000742 | Ipp | 16351 | HITS-CLIP | 23597149 | Unspecified |
| mmu-miR-378a-5p | MIMAT0000742 | Iqgap1 | 29875 | HITS-CLIP | 37986224 | Unspecified |
| mmu-miR-378a-5p | MIMAT0000742 | Iqgap2 | 544963 | HITS-CLIP | 37986224 | Unspecified |
| mmu-miR-378a-5p | MIMAT0000742 | Iqgap3 | 404710 | HITS-CLIP | 23597149 | Unspecified |
| mmu-miR-378a-5p | MIMAT0000742 | Iqsec2 | 245666 | HITS-CLIP | 37986224 | Unspecified |
| mmu-miR-378a-5p | MIMAT0000742 | Irak4 | 266632 | HITS-CLIP | 37986224 | Unspecified |
| mmu-miR-378a-5p | MIMAT0000742 | Ireb2 | 64602 | HITS-CLIP | 37986224 | Unspecified |
| mmu-miR-378a-5p | MIMAT0000742 | Irf1 | 16362 | HITS-CLIP | 23597149 | Unspecified |

| | | | | | | | |
|---|---|---|---|---|---|---|---|
| mmu-miR-378a-5p | MIMAT0000742 | Irx2 | 16372 | HITS-CLIP | | 37986224 | Unspecified |
| mmu-miR-378a-5p | MIMAT0000742 | Isg20l2 | 229504 | HITS-CLIP | | 37986224 | Unspecified |
| mmu-miR-378a-5p | MIMAT0000742 | Itga5 | 16402 | HITS-CLIP | | 37986224 | Unspecified |
| mmu-miR-378a-5p | MIMAT0000742 | Itgb8 | 320910 | HITS-CLIP | | 37986224 | Unspecified |
| mmu-miR-378a-5p | MIMAT0000742 | Itih4 | 16427 | HITS-CLIP | | 23597149 | Unspecified |
| mmu-miR-378a-5p | MIMAT0000742 | Itpripl2 | 319622 | HITS-CLIP | | 37986224 | Unspecified |
| mmu-miR-378a-5p | MIMAT0000742 | Itsn1 | 16443 | HITS-CLIP | | 37986224 | Unspecified |
| mmu-miR-378a-5p | MIMAT0000742 | Ivns1abp | 117198 | HITS-CLIP | | 25083871 | Unspecified |
| mmu-miR-378a-5p | MIMAT0000742 | Iws1 | 73473 | HITS-CLIP | | 37986224 | Unspecified |
| mmu-miR-378a-5p | MIMAT0000742 | Jagn1 | 67767 | HITS-CLIP | | 37986224 | Unspecified |
| mmu-miR-378a-5p | MIMAT0000742 | Jarid2 | 16468 | HITS-CLIP | 37986224\|25083871 | | Unspecified |
| mmu-miR-378a-5p | MIMAT0000742 | Jchain | 16069 | qCLASH | | 31393953 | Unspecified |
| mmu-miR-378a-5p | MIMAT0000742 | Jdp2 | 81703 | HITS-CLIP | 25083871\|29167373 | | Unspecified |
| mmu-miR-378a-5p | MIMAT0000742 | Jmjd1c | 108829 | HITS-CLIP | | 25753418 | Unspecified |
| mmu-miR-378a-5p | MIMAT0000742 | Josd1 | 74158 | HITS-CLIP | | 37986224 | Unspecified |
| mmu-miR-378a-5p | MIMAT0000742 | Josd2 | 66124 | HITS-CLIP | | 37986224 | Unspecified |
| mmu-miR-378a-5p | MIMAT0000742 | Jph2 | 59091 | HITS-CLIP | | 37986224 | Unspecified |
| mmu-miR-378a-5p | MIMAT0000742 | Katnal1 | 231912 | HITS-CLIP | | 37986224 | Unspecified |
| mmu-miR-378a-5p | MIMAT0000742 | Kbtbd11 | 74901 | HITS-CLIP | | 37986224 | Unspecified |
| mmu-miR-378a-5p | MIMAT0000742 | Kbtbd4 | 67136 | HITS-CLIP | | 37986224 | Unspecified |
| mmu-miR-378a-5p | MIMAT0000742 | Kcnj2 | 16518 | HITS-CLIP | | 25083871 | Unspecified |
| mmu-miR-378a-5p | MIMAT0000742 | Kcnn3 | 140493 | HITS-CLIP | | 37986224 | Unspecified |
| mmu-miR-378a-5p | MIMAT0000742 | Kcnn4 | 16534 | HITS-CLIP | | 29167373 | Unspecified |
| mmu-miR-378a-5p | MIMAT0000742 | Kctd11 | 216858 | HITS-CLIP | | 37986224 | Unspecified |
| mmu-miR-378a-5p | MIMAT0000742 | Kdm3a | 104263 | HITS-CLIP | 37986224\|23597149 | | Unspecified |
| mmu-miR-378a-5p | MIMAT0000742 | Kdm3b | 277250 | HITS-CLIP | | 37986224 | Unspecified |
| mmu-miR-378a-5p | MIMAT0000742 | Kdm5a | 214899 | HITS-CLIP | | 37986224 | Unspecified |
| mmu-miR-378a-5p | MIMAT0000742 | Kdm7a | 338523 | HITS-CLIP | | 25083871 | Unspecified |
| mmu-miR-378a-5p | MIMAT0000742 | Keg1 | 64697 | HITS-CLIP | 37986224\|23597149 | | Unspecified |
| mmu-miR-378a-5p | MIMAT0000742 | Khdrbs1 | 20218 | HITS-CLIP | | 23597149 | Unspecified |
| mmu-miR-378a-5p | MIMAT0000742 | Khk | 16548 | HITS-CLIP | 37986224\|23597149 | | Unspecified |
| mmu-miR-378a-5p | MIMAT0000742 | Kif13b | 16554 | HITS-CLIP | | 37986224 | Unspecified |
| mmu-miR-378a-5p | MIMAT0000742 | Kif1b | 16561 | HITS-CLIP | | 37986224 | Unspecified |
| mmu-miR-378a-5p | MIMAT0000742 | Kif26b | 269152 | HITS-CLIP | | 37986224 | Unspecified |
| mmu-miR-378a-5p | MIMAT0000742 | Kif3b | 16569 | HITS-CLIP | | 37986224 | Unspecified |
| mmu-miR-378a-5p | MIMAT0000742 | Kif5b | 16573 | HITS-CLIP | 23597149\|25083871 | | Unspecified |
| mmu-miR-378a-5p | MIMAT0000742 | Klf9 | 16601 | HITS-CLIP | | 37986224 | Unspecified |
| mmu-miR-378a-5p | MIMAT0000742 | Klhdc3 | 71765 | HITS-CLIP | | 25083871 | Unspecified |
| mmu-miR-378a-5p | MIMAT0000742 | Klhl15 | 236904 | HITS-CLIP | | 37986224 | Unspecified |

| | | | | | | |
|---|---|---|---|---|---|---|
| mmu-miR-378a-5p | MIMAT0000742 | Klhl24 | 75785 | HITS-CLIP | 37986224\|25083871 | Unspecified |
| mmu-miR-378a-5p | MIMAT0000742 | Kmt2a | 214162 | HITS-CLIP | 25083871\|25753418 | Unspecified |
| mmu-miR-378a-5p | MIMAT0000742 | Kmt2b | 75410 | HITS-CLIP | 37986224 | Unspecified |
| mmu-miR-378a-5p | MIMAT0000742 | Kmt2c | 231051 | HITS-CLIP | 37986224\|25083871 | Unspecified |
| mmu-miR-378a-5p | MIMAT0000742 | Kng1 | 16644 | HITS-CLIP | 37986224 | Unspecified |
| mmu-miR-378a-5p | MIMAT0000742 | Knop1 | 66356 | HITS-CLIP | 37986224 | Unspecified |
| mmu-miR-378a-5p | MIMAT0000742 | Kpna1 | 16646 | HITS-CLIP | 37986224 | Unspecified |
| mmu-miR-378a-5p | MIMAT0000742 | Kpna3 | 16648 | HITS-CLIP | 25083871\|25753418 | Unspecified |
| mmu-miR-378a-5p | MIMAT0000742 | Kpna4 | 16649 | HITS-CLIP | 37986224 | Unspecified |
| mmu-miR-378a-5p | MIMAT0000742 | Kras | 16653 | HITS-CLIP | 37986224 | Unspecified |
| mmu-miR-378a-5p | MIMAT0000742 | Krr1 | 52705 | HITS-CLIP | 37986224 | Unspecified |
| mmu-miR-378a-5p | MIMAT0000742 | Krt8 | 16691 | HITS-CLIP | 37986224 | Unspecified |
| mmu-miR-378a-5p | MIMAT0000742 | Ktn1 | 16709 | HITS-CLIP | 25083871 | Unspecified |
| mmu-miR-378a-5p | MIMAT0000742 | Lamb1 | 16777 | HITS-CLIP | 37986224\|25083871 | Unspecified |
| mmu-miR-378a-5p | MIMAT0000742 | Lamc2 | 16782 | HITS-CLIP | 37986224 | Unspecified |
| mmu-miR-378a-5p | MIMAT0000742 | Lamtor3 | 56692 | qCLASH | 31393953 | Unspecified |
| mmu-miR-378a-5p | MIMAT0000742 | Larp1 | 73158 | HITS-CLIP | 25083871\|29167373 | Unspecified |
| mmu-miR-378a-5p | MIMAT0000742 | Larp4b | 217980 | HITS-CLIP | 25083871 | Unspecified |
| mmu-miR-378a-5p | MIMAT0000742 | Lats1 | 16798 | HITS-CLIP | 23597149 | Unspecified |
| mmu-miR-378a-5p | MIMAT0000742 | Lbh | 77889 | HITS-CLIP | 25753418 | Unspecified |
| mmu-miR-378a-5p | MIMAT0000742 | Lbr | 98386 | HITS-CLIP | 37986224 | Unspecified |
| mmu-miR-378a-5p | MIMAT0000742 | Lcor | 212391 | HITS-CLIP | 37986224\|25083871 | Unspecified |
| mmu-miR-378a-5p | MIMAT0000742 | Ldb1 | 16825 | HITS-CLIP | 37986224 | Unspecified |
| mmu-miR-378a-5p | MIMAT0000742 | Ldlr | 16835 | HITS-CLIP | 37986224 | Unspecified |
| mmu-miR-378a-5p | MIMAT0000742 | Ldlrad3 | 241576 | HITS-CLIP | 37986224 | Unspecified |
| mmu-miR-378a-5p | MIMAT0000742 | Leng8 | 232798 | HITS-CLIP | 37986224 | Unspecified |
| mmu-miR-378a-5p | MIMAT0000742 | Leprot | 230514 | HITS-CLIP | 23597149 | Unspecified |
| mmu-miR-378a-5p | MIMAT0000742 | Leprotl1 | 68192 | HITS-CLIP | 37986224 | Unspecified |
| mmu-miR-378a-5p | MIMAT0000742 | Lgals9 | 16859 | HITS-CLIP | 23597149 | Unspecified |
| mmu-miR-378a-5p | MIMAT0000742 | Lima1 | 65970 | HITS-CLIP | 37986224 | Unspecified |
| mmu-miR-378a-5p | MIMAT0000742 | Limd1 | 29806 | HITS-CLIP | 37986224 | Unspecified |
| mmu-miR-378a-5p | MIMAT0000742 | Lin54 | 231506 | HITS-CLIP | 37986224 | Unspecified |
| mmu-miR-378a-5p | MIMAT0000742 | Lman1 | 70361 | HITS-CLIP | 25083871 | Unspecified |
| mmu-miR-378a-5p | MIMAT0000742 | Lmbr1l | 74775 | HITS-CLIP | 37986224 | Unspecified |
| mmu-miR-378a-5p | MIMAT0000742 | Lmna | 16905 | HITS-CLIP | 37986224 | Unspecified |
| mmu-miR-378a-5p | MIMAT0000742 | Lnpep | 240028 | HITS-CLIP | 37986224 | Unspecified |
| mmu-miR-378a-5p | MIMAT0000742 | Loxl3 | 16950 | HITS-CLIP | 37986224 | Unspecified |
| mmu-miR-378a-5p | MIMAT0000742 | Loxl4 | 67573 | HITS-CLIP | 37986224 | Unspecified |
| mmu-miR-378a-5p | MIMAT0000742 | Lpar4 | 78134 | HITS-CLIP | 37986224 | Unspecified |

| | | | | | | |
|---|---|---|---|---|---|---|
| mmu-miR-378a-5p | MIMAT0000742 | Lpp | 210126 | HITS-CLIP | 37986224\|25083871 | Unspecified |
| mmu-miR-378a-5p | MIMAT0000742 | Lrch1 | 380916 | HITS-CLIP | 25083871 | Unspecified |
| mmu-miR-378a-5p | MIMAT0000742 | Lrch3 | 70144 | HITS-CLIP | 37986224 | Unspecified |
| mmu-miR-378a-5p | MIMAT0000742 | Lrch4 | 231798 | HITS-CLIP | 25083871 | Unspecified |
| mmu-miR-378a-5p | MIMAT0000742 | Lrif1 | 321000 | HITS-CLIP | 37986224 | Unspecified |
| mmu-miR-378a-5p | MIMAT0000742 | Lrig3 | 320398 | HITS-CLIP | 25083871 | Unspecified |
| mmu-miR-378a-5p | MIMAT0000742 | Lrp1 | 16971 | HITS-CLIP | 37986224\|23597149\|29167373 | Unspecified |
| mmu-miR-378a-5p | MIMAT0000742 | Lrrc58 | 320184 | HITS-CLIP | 37986224\|25083871 | Unspecified |
| mmu-miR-378a-5p | MIMAT0000742 | Lrrfip1 | 16978 | HITS-CLIP | 37986224 | Unspecified |
| mmu-miR-378a-5p | MIMAT0000742 | Lsm12 | 268490 | HITS-CLIP | 37986224 | Unspecified |
| mmu-miR-378a-5p | MIMAT0000742 | Lsm3 | 67678 | HITS-CLIP | 37986224 | Unspecified |
| mmu-miR-378a-5p | MIMAT0000742 | Lss | 16987 | HITS-CLIP | 37986224 | Unspecified |
| mmu-miR-378a-5p | MIMAT0000742 | Ltbp2 | 16997 | HITS-CLIP | 25753418 | Unspecified |
| mmu-miR-378a-5p | MIMAT0000742 | Luc7l | 66978 | HITS-CLIP | 37986224 | Unspecified |
| mmu-miR-378a-5p | MIMAT0000742 | Luc7l2 | 192196 | HITS-CLIP | 37986224 | Unspecified |
| mmu-miR-378a-5p | MIMAT0000742 | Luc7l3 | 67684 | HITS-CLIP | 37986224\|25083871 | Unspecified |
| mmu-miR-378a-5p | MIMAT0000742 | Lurap1l | 52829 | HITS-CLIP | 23597149 | Unspecified |
| mmu-miR-378a-5p | MIMAT0000742 | Luzp1 | 269593 | HITS-CLIP | 37986224 | Unspecified |
| mmu-miR-378a-5p | MIMAT0000742 | Lypla2 | 26394 | HITS-CLIP | 37986224 | Unspecified |
| mmu-miR-378a-5p | MIMAT0000742 | Lyst | 17101 | HITS-CLIP | 37986224\|25083871 | Unspecified |
| mmu-miR-378a-5p | MIMAT0000742 | Macf1 | 11426 | HITS-CLIP | 25083871\|25753418 | Unspecified |
| mmu-miR-378a-5p | MIMAT0000742 | Magi3 | 99470 | HITS-CLIP | 37986224 | Unspecified |
| mmu-miR-378a-5p | MIMAT0000742 | Mak16 | 67920 | HITS-CLIP | 37986224 | Unspecified |
| mmu-miR-378a-5p | MIMAT0000742 | Maml3 | 433586 | HITS-CLIP | 37986224 | Unspecified |
| mmu-miR-378a-5p | MIMAT0000742 | Map2k1 | 26395 | HITS-CLIP | 37986224\|25083871 | Unspecified |
| mmu-miR-378a-5p | MIMAT0000742 | Map2k3 | 26397 | HITS-CLIP | 37986224 | Unspecified |
| mmu-miR-378a-5p | MIMAT0000742 | Map3k11 | 26403 | HITS-CLIP | 25083871 | Unspecified |
| mmu-miR-378a-5p | MIMAT0000742 | Map3k2 | 26405 | HITS-CLIP | 37986224\|25083871 | Unspecified |
| mmu-miR-378a-5p | MIMAT0000742 | Map3k3 | 26406 | HITS-CLIP | 23597149 | Unspecified |
| mmu-miR-378a-5p | MIMAT0000742 | Map3k9 | 338372 | HITS-CLIP | 29167373 | Unspecified |
| mmu-miR-378a-5p | MIMAT0000742 | Map4 | 17758 | HITS-CLIP | 37986224\|25083871 | Unspecified |
| mmu-miR-378a-5p | MIMAT0000742 | Map4k4 | 26921 | HITS-CLIP | 25083871 | Unspecified |
| mmu-miR-378a-5p | MIMAT0000742 | Mapk8 | 26419 | HITS-CLIP | 37986224\|25083871 | Unspecified |
| mmu-miR-378a-5p | MIMAT0000742 | March8 | 71779 | HITS-CLIP | 37986224 | Unspecified |
| mmu-miR-378a-5p | MIMAT0000742 | Marf1 | 223989 | HITS-CLIP | 25083871 | Unspecified |
| mmu-miR-378a-5p | MIMAT0000742 | Marveld1 | 277010 | HITS-CLIP | 37986224\|25083871 | Unspecified |
| mmu-miR-378a-5p | MIMAT0000742 | Mbd1 | 17190 | HITS-CLIP | 37986224\|25083871 | Unspecified |
| mmu-miR-378a-5p | MIMAT0000742 | Mbip | 217588 | HITS-CLIP | 25753418 | Unspecified |
| mmu-miR-378a-5p | MIMAT0000742 | Mbtd1 | 103537 | HITS-CLIP | 37986224 | Unspecified |

| | | | | | | |
|---|---|---|---|---|---|---|
| mmu-miR-378a-5p | MIMAT0000742 | Mcl1 | 17210 | HITS-CLIP | 37986224\|23597149\|29167373 | Unspecified |
| mmu-miR-378a-5p | MIMAT0000742 | Mcm6 | 17219 | HITS-CLIP | 25083871 | Unspecified |
| mmu-miR-378a-5p | MIMAT0000742 | Mdm4 | 17248 | HITS-CLIP | 37986224 | Unspecified |
| mmu-miR-378a-5p | MIMAT0000742 | Me1 | 17436 | HITS-CLIP | 37986224 | Unspecified |
| mmu-miR-378a-5p | MIMAT0000742 | Mecp2 | 17257 | HITS-CLIP | 37986224\|25083871 | Unspecified |
| mmu-miR-378a-5p | MIMAT0000742 | Med1 | 19014 | HITS-CLIP | 37986224\|23597149 | Unspecified |
| mmu-miR-378a-5p | MIMAT0000742 | Med13l | 76199 | HITS-CLIP | 37986224 | Unspecified |
| mmu-miR-378a-5p | MIMAT0000742 | Med14 | 26896 | HITS-CLIP | 37986224 | Unspecified |
| mmu-miR-378a-5p | MIMAT0000742 | Med28 | 66999 | HITS-CLIP | 37986224 | Unspecified |
| mmu-miR-378a-5p | MIMAT0000742 | Medag | 70717 | HITS-CLIP | 37986224 | Unspecified |
| mmu-miR-378a-5p | MIMAT0000742 | Megf10 | 70417 | HITS-CLIP | 37986224 | Unspecified |
| mmu-miR-378a-5p | MIMAT0000742 | Mex3a | 72640 | HITS-CLIP | 37986224 | Unspecified |
| mmu-miR-378a-5p | MIMAT0000742 | Mex3b | 108797 | HITS-CLIP | 25083871 | Unspecified |
| mmu-miR-378a-5p | MIMAT0000742 | Mex3c | 240396 | HITS-CLIP | 25083871 | Unspecified |
| mmu-miR-378a-5p | MIMAT0000742 | Mfn2 | 170731 | HITS-CLIP | 37986224\|25083871 | Unspecified |
| mmu-miR-378a-5p | MIMAT0000742 | Mfsd1 | 66868 | HITS-CLIP | 37986224 | Unspecified |
| mmu-miR-378a-5p | MIMAT0000742 | Mfsd12 | 73822 | HITS-CLIP | 37986224 | Unspecified |
| mmu-miR-378a-5p | MIMAT0000742 | Mfsd8 | 72175 | HITS-CLIP | 25753418 | Unspecified |
| mmu-miR-378a-5p | MIMAT0000742 | Mga | 29808 | HITS-CLIP | 37986224 | Unspecified |
| mmu-miR-378a-5p | MIMAT0000742 | Mgat1 | 17308 | HITS-CLIP | 37986224 | Unspecified |
| mmu-miR-378a-5p | MIMAT0000742 | Mgll | 23945 | HITS-CLIP | 37986224\|23597149 | Unspecified |
| mmu-miR-378a-5p | MIMAT0000742 | Micu2 | 68514 | HITS-CLIP | 23597149 | Unspecified |
| mmu-miR-378a-5p | MIMAT0000742 | Micu3 | 78506 | HITS-CLIP | 37986224 | Unspecified |
| mmu-miR-378a-5p | MIMAT0000742 | Mis12 | 67139 | HITS-CLIP | 37986224 | Unspecified |
| mmu-miR-378a-5p | MIMAT0000742 | Mki67 | 17345 | HITS-CLIP | 37986224 | Unspecified |
| mmu-miR-378a-5p | MIMAT0000742 | Mknk2 | 17347 | HITS-CLIP | 37986224 | Unspecified |
| mmu-miR-378a-5p | MIMAT0000742 | Mlec | 109154 | HITS-CLIP | 37986224\|25083871\|23597149\|29167 | Unspecified |
| mmu-miR-378a-5p | MIMAT0000742 | Mllt6 | 246198 | HITS-CLIP | 37986224 | Unspecified |
| mmu-miR-378a-5p | MIMAT0000742 | Mlxip | 208104 | HITS-CLIP | 37986224 | Unspecified |
| mmu-miR-378a-5p | MIMAT0000742 | Mmd | 67468 | HITS-CLIP | 23597149\|29167373 | Unspecified |
| mmu-miR-378a-5p | MIMAT0000742 | Mob4 | 19070 | HITS-CLIP | 37986224 | Unspecified |
| mmu-miR-378a-5p | MIMAT0000742 | Morc2a | 74522 | HITS-CLIP | 29167373 | Unspecified |
| mmu-miR-378a-5p | MIMAT0000742 | Morc3 | 338467 | HITS-CLIP | 25083871 | Unspecified |
| mmu-miR-378a-5p | MIMAT0000742 | Mpp5 | 56217 | HITS-CLIP | 37986224 | Unspecified |
| mmu-miR-378a-5p | MIMAT0000742 | Mprip | 26936 | HITS-CLIP | 37986224\|25083871 | Unspecified |
| mmu-miR-378a-5p | MIMAT0000742 | Mre11a | 17535 | HITS-CLIP | 37986224 | Unspecified |
| mmu-miR-378a-5p | MIMAT0000742 | Mrgpre | 244238 | HITS-CLIP | 37986224 | Unspecified |
| mmu-miR-378a-5p | MIMAT0000742 | Mri1 | 67873 | HITS-CLIP | 37986224 | Unspecified |
| mmu-miR-378a-5p | MIMAT0000742 | Mrpl18 | 67681 | HITS-CLIP | 23597149 | Unspecified |

| | | | | | | | |
|---|---|---|---|---|---|---|---|
| mmu-miR-378a-5p | MIMAT0000742 | Mrpl32 | 75398 | HITS-CLIP | | 37986224 | Unspecified |
| mmu-miR-378a-5p | MIMAT0000742 | Mrpl52 | 68836 | HITS-CLIP | 23597149\|25083871\|25753418 | | Unspecified |
| mmu-miR-378a-5p | MIMAT0000742 | Mrps15 | 66407 | HITS-CLIP | | 25083871 | Unspecified |
| mmu-miR-378a-5p | MIMAT0000742 | Msantd3 | 66665 | HITS-CLIP | | 37986224 | Unspecified |
| mmu-miR-378a-5p | MIMAT0000742 | Msh6 | 17688 | HITS-CLIP | | 37986224 | Unspecified |
| mmu-miR-378a-5p | MIMAT0000742 | Msl2 | 77853 | HITS-CLIP | | 37986224 | Unspecified |
| mmu-miR-378a-5p | MIMAT0000742 | Msrb3 | 320183 | HITS-CLIP | | 23597149 | Unspecified |
| mmu-miR-378a-5p | MIMAT0000742 | Mtcl1 | 68617 | HITS-CLIP | | 37986224 | Unspecified |
| mmu-miR-378a-5p | MIMAT0000742 | Mtss1 | 211401 | HITS-CLIP | | 37986224 | Unspecified |
| mmu-miR-378a-5p | MIMAT0000742 | Mtus1 | 102103 | HITS-CLIP | 37986224\|25083871 | | Unspecified |
| mmu-miR-378a-5p | MIMAT0000742 | Myadm | 50918 | HITS-CLIP | | 25753418 | Unspecified |
| mmu-miR-378a-5p | MIMAT0000742 | Mycbp2 | 105689 | HITS-CLIP | | 29167373 | Unspecified |
| mmu-miR-378a-5p | MIMAT0000742 | Myh1 | 17879 | HITS-CLIP | | 25083871 | Unspecified |
| mmu-miR-378a-5p | MIMAT0000742 | Myh8 | 17885 | HITS-CLIP | | 25083871 | Unspecified |
| mmu-miR-378a-5p | MIMAT0000742 | Myo18a | 360013 | HITS-CLIP | 37986224\|25083871 | | Unspecified |
| mmu-miR-378a-5p | MIMAT0000742 | Myo1d | 338367 | HITS-CLIP | | 37986224 | Unspecified |
| mmu-miR-378a-5p | MIMAT0000742 | Myo1e | 71602 | HITS-CLIP | | 37986224 | Unspecified |
| mmu-miR-378a-5p | MIMAT0000742 | Myo5a | 17918 | HITS-CLIP | | 37986224 | Unspecified |
| mmu-miR-378a-5p | MIMAT0000742 | Myo9b | 17925 | HITS-CLIP | | 29167373 | Unspecified |
| mmu-miR-378a-5p | MIMAT0000742 | Myocd | 214384 | HITS-CLIP | | 25753418 | Unspecified |
| mmu-miR-378a-5p | MIMAT0000742 | Myof | 226101 | HITS-CLIP | | 25083871 | Unspecified |
| mmu-miR-378a-5p | MIMAT0000742 | Myog | 17928 | HITS-CLIP | 37986224\|25083871 | | Unspecified |
| mmu-miR-378a-5p | MIMAT0000742 | Myom2 | 17930 | HITS-CLIP | | 25083871 | Unspecified |
| mmu-miR-378a-5p | MIMAT0000742 | Mysm1 | 320713 | HITS-CLIP | | 37986224 | Unspecified |
| mmu-miR-378a-5p | MIMAT0000742 | Naa15 | 74838 | HITS-CLIP | 37986224\|25083871 | | Unspecified |
| mmu-miR-378a-5p | MIMAT0000742 | Naa25 | 231713 | HITS-CLIP | | 25083871 | Unspecified |
| mmu-miR-378a-5p | MIMAT0000742 | Naa40 | 70999 | HITS-CLIP | | 37986224 | Unspecified |
| mmu-miR-378a-5p | MIMAT0000742 | Naa50 | 72117 | HITS-CLIP | | 37986224 | Unspecified |
| mmu-miR-378a-5p | MIMAT0000742 | Naa60 | 74763 | HITS-CLIP | | 37986224 | Unspecified |
| mmu-miR-378a-5p | MIMAT0000742 | Nacc2 | 67991 | HITS-CLIP | | 37986224 | Unspecified |
| mmu-miR-378a-5p | MIMAT0000742 | Nadk | 192185 | HITS-CLIP | | 25083871 | Unspecified |
| mmu-miR-378a-5p | MIMAT0000742 | Nap1l4 | 17955 | HITS-CLIP | | 37986224 | Unspecified |
| mmu-miR-378a-5p | MIMAT0000742 | Napg | 108123 | HITS-CLIP | | 23597149 | Unspecified |
| mmu-miR-378a-5p | MIMAT0000742 | Nbea | 26422 | HITS-CLIP | | 37986224 | Unspecified |
| mmu-miR-378a-5p | MIMAT0000742 | Nbeal1 | 269198 | HITS-CLIP | | 37986224 | Unspecified |
| mmu-miR-378a-5p | MIMAT0000742 | Ncapd3 | 78658 | HITS-CLIP | | 37986224 | Unspecified |
| mmu-miR-378a-5p | MIMAT0000742 | Ncaph2 | 52683 | HITS-CLIP | | 29167373 | Unspecified |
| mmu-miR-378a-5p | MIMAT0000742 | Ncbp2 | 68092 | HITS-CLIP | | 37986224 | Unspecified |
| mmu-miR-378a-5p | MIMAT0000742 | Nceh1 | 320024 | HITS-CLIP | 37986224\|29167373 | | Unspecified |

| | | | | | | |
|---|---|---|---|---|---|---|
| mmu-miR-378a-5p | MIMAT0000742 | Nckap1 | 50884 | HITS-CLIP | 37986224 | Unspecified |
| mmu-miR-378a-5p | MIMAT0000742 | Nckap5l | 380969 | HITS-CLIP | 37986224 | Unspecified |
| mmu-miR-378a-5p | MIMAT0000742 | Ncl | 17975 | HITS-CLIP | 37986224\|25753418 | Unspecified |
| mmu-miR-378a-5p | MIMAT0000742 | Ncoa1 | 17977 | HITS-CLIP | 29167373 | Unspecified |
| mmu-miR-378a-5p | MIMAT0000742 | Ncoa2 | 17978 | HITS-CLIP | 37986224 | Unspecified |
| mmu-miR-378a-5p | MIMAT0000742 | Ncoa3 | 17979 | HITS-CLIP | 37986224 | Unspecified |
| mmu-miR-378a-5p | MIMAT0000742 | Ncoa6 | 56406 | HITS-CLIP | 37986224 | Unspecified |
| mmu-miR-378a-5p | MIMAT0000742 | Ncor1 | 20185 | HITS-CLIP | 37986224\|25083871 | Unspecified |
| mmu-miR-378a-5p | MIMAT0000742 | Ncor2 | 20602 | HITS-CLIP | 25083871 | Unspecified |
| mmu-miR-378a-5p | MIMAT0000742 | Ndel1 | 83431 | HITS-CLIP | 25083871 | Unspecified |
| mmu-miR-378a-5p | MIMAT0000742 | Ndrg3 | 29812 | HITS-CLIP | 37986224 | Unspecified |
| mmu-miR-378a-5p | MIMAT0000742 | Ndst1 | 15531 | HITS-CLIP | 37986224 | Unspecified |
| mmu-miR-378a-5p | MIMAT0000742 | Ndufb2 | 68198 | HITS-CLIP | 25753418 | Unspecified |
| mmu-miR-378a-5p | MIMAT0000742 | Necap1 | 67602 | HITS-CLIP | 37986224 | Unspecified |
| mmu-miR-378a-5p | MIMAT0000742 | Nedd4l | 83814 | HITS-CLIP | 37986224 | Unspecified |
| mmu-miR-378a-5p | MIMAT0000742 | Nedd9 | 18003 | HITS-CLIP | 37986224 | Unspecified |
| mmu-miR-378a-5p | MIMAT0000742 | Nek9 | 217718 | HITS-CLIP | 37986224 | Unspecified |
| mmu-miR-378a-5p | MIMAT0000742 | Nemf | 66244 | HITS-CLIP | 25083871 | Unspecified |
| mmu-miR-378a-5p | MIMAT0000742 | Nes | 18008 | HITS-CLIP | 37986224 | Unspecified |
| mmu-miR-378a-5p | MIMAT0000742 | Nfat5 | 54446 | HITS-CLIP | 37986224\|25083871 | Unspecified |
| mmu-miR-378a-5p | MIMAT0000742 | Nfatc3 | 18021 | HITS-CLIP | 25083871 | Unspecified |
| mmu-miR-378a-5p | MIMAT0000742 | Nfe2l1 | 18023 | HITS-CLIP | 37986224\|25083871\|23597149 | Unspecified |
| mmu-miR-378a-5p | MIMAT0000742 | Nfia | 18027 | HITS-CLIP | 37986224 | Unspecified |
| mmu-miR-378a-5p | MIMAT0000742 | Nfib | 18028 | HITS-CLIP | 37986224 | Unspecified |
| mmu-miR-378a-5p | MIMAT0000742 | Nfic | 18029 | HITS-CLIP | 25753418 | Unspecified |
| mmu-miR-378a-5p | MIMAT0000742 | Nfs1 | 18041 | HITS-CLIP | 37986224 | Unspecified |
| mmu-miR-378a-5p | MIMAT0000742 | Nfx1 | 74164 | HITS-CLIP | 37986224 | Unspecified |
| mmu-miR-378a-5p | MIMAT0000742 | Nhlrc2 | 66866 | HITS-CLIP | 37986224 | Unspecified |
| mmu-miR-378a-5p | MIMAT0000742 | Nhlrc3 | 212114 | HITS-CLIP | 37986224 | Unspecified |
| mmu-miR-378a-5p | MIMAT0000742 | Nin | 18080 | HITS-CLIP | 29167373 | Unspecified |
| mmu-miR-378a-5p | MIMAT0000742 | Ninj1 | 18081 | HITS-CLIP | 37986224 | Unspecified |
| mmu-miR-378a-5p | MIMAT0000742 | Nisch | 64652 | HITS-CLIP | 37986224 | Unspecified |
| mmu-miR-378a-5p | MIMAT0000742 | Nktr | 18087 | HITS-CLIP | 37986224 | Unspecified |
| mmu-miR-378a-5p | MIMAT0000742 | Nlrp12 | 378425 | HITS-CLIP | 23597149 | Unspecified |
| mmu-miR-378a-5p | MIMAT0000742 | Noct | 12457 | HITS-CLIP | 23597149 | Unspecified |
| mmu-miR-378a-5p | MIMAT0000742 | Nop2 | 110109 | HITS-CLIP | 25083871 | Unspecified |
| mmu-miR-378a-5p | MIMAT0000742 | Notch2 | 18129 | HITS-CLIP | 37986224 | Unspecified |
| mmu-miR-378a-5p | MIMAT0000742 | Nr0b2 | 23957 | HITS-CLIP | 23597149 | Unspecified |
| mmu-miR-378a-5p | MIMAT0000742 | Nr1d1 | 217166 | HITS-CLIP | 25083871 | Unspecified |

| | | | | | | |
|---|---|---|---|---|---|---|
| mmu-miR-378a-5p | MIMAT0000742 | Nr1h4 | 20186 | HITS-CLIP | 37986224 | Unspecified |
| mmu-miR-378a-5p | MIMAT0000742 | Nr2c1 | 22025 | HITS-CLIP | 37986224 | Unspecified |
| mmu-miR-378a-5p | MIMAT0000742 | Nr2f2 | 11819 | HITS-CLIP | 23597149 | Unspecified |
| mmu-miR-378a-5p | MIMAT0000742 | Nr3c1 | 14815 | HITS-CLIP | 37986224 | Unspecified |
| mmu-miR-378a-5p | MIMAT0000742 | Nr6a1 | 14536 | HITS-CLIP | 37986224 | Unspecified |
| mmu-miR-378a-5p | MIMAT0000742 | Nrap | 18175 | HITS-CLIP | 25753418 | Unspecified |
| mmu-miR-378a-5p | MIMAT0000742 | Nrn1 | 68404 | HITS-CLIP | 37986224 | Unspecified |
| mmu-miR-378a-5p | MIMAT0000742 | Nrp1 | 18186 | HITS-CLIP | 37986224 | Unspecified |
| mmu-miR-378a-5p | MIMAT0000742 | Nrp2 | 18187 | HITS-CLIP | 29167373 | Unspecified |
| mmu-miR-378a-5p | MIMAT0000742 | Nsd1 | 18193 | HITS-CLIP | 37986224 | Unspecified |
| mmu-miR-378a-5p | MIMAT0000742 | Nsd3 | 234135 | HITS-CLIP | 25083871 | Unspecified |
| mmu-miR-378a-5p | MIMAT0000742 | Nsf | 18195 | HITS-CLIP | 37986224 | Unspecified |
| mmu-miR-378a-5p | MIMAT0000742 | Nt5dc3 | 103466 | HITS-CLIP | 37986224 | Unspecified |
| mmu-miR-378a-5p | MIMAT0000742 | Nubp2 | 26426 | HITS-CLIP | 37986224 | Unspecified |
| mmu-miR-378a-5p | MIMAT0000742 | Nudcd3 | 209586 | HITS-CLIP | 25753418 | Unspecified |
| mmu-miR-378a-5p | MIMAT0000742 | Nudt4 | 71207 | HITS-CLIP | 37986224 | Unspecified |
| mmu-miR-378a-5p | MIMAT0000742 | Nudt9 | 74167 | HITS-CLIP | 25753418 | Unspecified |
| mmu-miR-378a-5p | MIMAT0000742 | Nup155 | 170762 | HITS-CLIP | 25753418 | Unspecified |
| mmu-miR-378a-5p | MIMAT0000742 | Nup205 | 70699 | HITS-CLIP | 37986224 | Unspecified |
| mmu-miR-378a-5p | MIMAT0000742 | Nup93 | 71805 | HITS-CLIP | 37986224 | Unspecified |
| mmu-miR-378a-5p | MIMAT0000742 | Nxpe3 | 385658 | HITS-CLIP | 37986224 | Unspecified |
| mmu-miR-378a-5p | MIMAT0000742 | Nynrin | 277154 | HITS-CLIP | 37986224 | Unspecified |
| mmu-miR-378a-5p | MIMAT0000742 | Oaf | 102644 | HITS-CLIP | 37986224 | Unspecified |
| mmu-miR-378a-5p | MIMAT0000742 | Oat | 18242 | HITS-CLIP | 25083871 | Unspecified |
| mmu-miR-378a-5p | MIMAT0000742 | Oaz1 | 18245 | HITS-CLIP | 25753418 | Unspecified |
| mmu-miR-378a-5p | MIMAT0000742 | Obscn | 380698 | HITS-CLIP | 25083871 | Unspecified |
| mmu-miR-378a-5p | MIMAT0000742 | Odf2 | 18286 | HITS-CLIP | 25753418 | Unspecified |
| mmu-miR-378a-5p | MIMAT0000742 | Ogt | 108155 | HITS-CLIP | 37986224 | Unspecified |
| mmu-miR-378a-5p | MIMAT0000742 | Ormdl2 | 66844 | HITS-CLIP | 37986224 | Unspecified |
| mmu-miR-378a-5p | MIMAT0000742 | Osbpl3 | 71720 | HITS-CLIP | 37986224 | Unspecified |
| mmu-miR-378a-5p | MIMAT0000742 | Osbpl7 | 71240 | HITS-CLIP | 25083871 | Unspecified |
| mmu-miR-378a-5p | MIMAT0000742 | Osbpl8 | 237542 | HITS-CLIP | 37986224 | Unspecified |
| mmu-miR-378a-5p | MIMAT0000742 | Osbpl9 | 100273 | HITS-CLIP | 23597149 | Unspecified |
| mmu-miR-378a-5p | MIMAT0000742 | Ostm1 | 14628 | HITS-CLIP | 37986224\|25083871 | Unspecified |
| mmu-miR-378a-5p | MIMAT0000742 | P4ha1 | 18451 | HITS-CLIP | 37986224 | Unspecified |
| mmu-miR-378a-5p | MIMAT0000742 | Pacsin3 | 80708 | HITS-CLIP | 37986224 | Unspecified |
| mmu-miR-378a-5p | MIMAT0000742 | Pafah2 | 100163 | HITS-CLIP | 23597149 | Unspecified |
| mmu-miR-378a-5p | MIMAT0000742 | Pakap | 677884 | HITS-CLIP | 25753418 | Unspecified |
| mmu-miR-378a-5p | MIMAT0000742 | Pan3 | 72587 | HITS-CLIP | 37986224 | Unspecified |

| | | | | | | |
|---|---|---|---|---|---|---|
| mmu-miR-378a-5p | MIMAT0000742 | Pank2 | 74450 | HITS-CLIP | 37986224 | Unspecified |
| mmu-miR-378a-5p | MIMAT0000742 | Papola | 18789 | HITS-CLIP | 37986224 | Unspecified |
| mmu-miR-378a-5p | MIMAT0000742 | Pax7 | 18509 | HITS-CLIP | 37986224 | Unspecified |
| mmu-miR-378a-5p | MIMAT0000742 | Pbdc1 | 67683 | HITS-CLIP | 37986224 | Unspecified |
| mmu-miR-378a-5p | MIMAT0000742 | Pbrm1 | 66923 | HITS-CLIP | 25083871 | Unspecified |
| mmu-miR-378a-5p | MIMAT0000742 | Pcdh12 | 53601 | HITS-CLIP | 23597149 | Unspecified |
| mmu-miR-378a-5p | MIMAT0000742 | Pcdh7 | 54216 | HITS-CLIP | 37986224 | Unspecified |
| mmu-miR-378a-5p | MIMAT0000742 | Pcgf3 | 69587 | HITS-CLIP | 37986224 | Unspecified |
| mmu-miR-378a-5p | MIMAT0000742 | Pcyt2 | 68671 | HITS-CLIP | 37986224 | Unspecified |
| mmu-miR-378a-5p | MIMAT0000742 | Pdap1 | 231887 | HITS-CLIP | 25083871 | Unspecified |
| mmu-miR-378a-5p | MIMAT0000742 | Pdcd6ip | 18571 | HITS-CLIP | 37986224\|29167373\|25083871 | Unspecified |
| mmu-miR-378a-5p | MIMAT0000742 | Pde4dip | 83679 | HITS-CLIP | 25083871 | Unspecified |
| mmu-miR-378a-5p | MIMAT0000742 | Pdgfd | 71785 | HITS-CLIP | 25753418 | Unspecified |
| mmu-miR-378a-5p | MIMAT0000742 | Pdia5 | 72599 | HITS-CLIP | 23597149 | Unspecified |
| mmu-miR-378a-5p | MIMAT0000742 | Pdk1 | 228026 | HITS-CLIP | 37986224 | Unspecified |
| mmu-miR-378a-5p | MIMAT0000742 | Pdlim4 | 30794 | HITS-CLIP | 37986224 | Unspecified |
| mmu-miR-378a-5p | MIMAT0000742 | Pdp1 | 381511 | HITS-CLIP | 37986224 | Unspecified |
| mmu-miR-378a-5p | MIMAT0000742 | Pdpk1 | 18607 | HITS-CLIP | 37986224 | Unspecified |
| mmu-miR-378a-5p | MIMAT0000742 | Pdxdc1 | 94184 | HITS-CLIP | 37986224 | Unspecified |
| mmu-miR-378a-5p | MIMAT0000742 | Pdzd8 | 107368 | HITS-CLIP | 37986224\|25083871 | Unspecified |
| mmu-miR-378a-5p | MIMAT0000742 | Pdzrn3 | 55983 | HITS-CLIP | 37986224 | Unspecified |
| mmu-miR-378a-5p | MIMAT0000742 | Peg10 | 170676 | HITS-CLIP | 37986224\|25083871 | Unspecified |
| mmu-miR-378a-5p | MIMAT0000742 | Peg3 | 18616 | HITS-CLIP | 37986224 | Unspecified |
| mmu-miR-378a-5p | MIMAT0000742 | Peli1 | 67245 | HITS-CLIP | 37986224 | Unspecified |
| mmu-miR-378a-5p | MIMAT0000742 | Per3 | 18628 | HITS-CLIP | 37986224 | Unspecified |
| mmu-miR-378a-5p | MIMAT0000742 | Pex19 | 19298 | HITS-CLIP | 37986224 | Unspecified |
| mmu-miR-378a-5p | MIMAT0000742 | Pex5 | 19305 | HITS-CLIP | 37986224 | Unspecified |
| mmu-miR-378a-5p | MIMAT0000742 | Pfkfb4 | 270198 | HITS-CLIP | 37986224 | Unspecified |
| mmu-miR-378a-5p | MIMAT0000742 | Pfn1 | 18643 | HITS-CLIP//qCLASH | 37986224\|23597149\|31393953 | Unspecified |
| mmu-miR-378a-5p | MIMAT0000742 | Pgm1 | 66681 | HITS-CLIP | 25083871 | Unspecified |
| mmu-miR-378a-5p | MIMAT0000742 | Pgm2 | 72157 | HITS-CLIP | 37986224 | Unspecified |
| mmu-miR-378a-5p | MIMAT0000742 | Pgp | 67078 | HITS-CLIP | 37986224 | Unspecified |
| mmu-miR-378a-5p | MIMAT0000742 | Phc3 | 241915 | HITS-CLIP | 37986224 | Unspecified |
| mmu-miR-378a-5p | MIMAT0000742 | Phf10 | 72057 | HITS-CLIP | 37986224\|25083871 | Unspecified |
| mmu-miR-378a-5p | MIMAT0000742 | Phf3 | 213109 | HITS-CLIP | 37986224 | Unspecified |
| mmu-miR-378a-5p | MIMAT0000742 | Phldb2 | 208177 | HITS-CLIP | 23597149 | Unspecified |
| mmu-miR-378a-5p | MIMAT0000742 | Phtf1 | 18685 | HITS-CLIP | 37986224 | Unspecified |
| mmu-miR-378a-5p | MIMAT0000742 | Pi4k2b | 67073 | HITS-CLIP | 37986224 | Unspecified |
| mmu-miR-378a-5p | MIMAT0000742 | Pi4ka | 224020 | HITS-CLIP | 29167373\|25083871 | Unspecified |

| | | | | | | |
|---|---|---|---|---|---|---|
| mmu-miR-378a-5p | MIMAT0000742 | Pi4kb | 107650 | HITS-CLIP | 37986224 | Unspecified |
| mmu-miR-378a-5p | MIMAT0000742 | Pias4 | 59004 | HITS-CLIP | 37986224 | Unspecified |
| mmu-miR-378a-5p | MIMAT0000742 | Pigm | 67556 | HITS-CLIP | 37986224 | Unspecified |
| mmu-miR-378a-5p | MIMAT0000742 | Pign | 27392 | HITS-CLIP | 37986224 | Unspecified |
| mmu-miR-378a-5p | MIMAT0000742 | Pip5k1a | 18720 | HITS-CLIP | 37986224 | Unspecified |
| mmu-miR-378a-5p | MIMAT0000742 | Pithd1 | 66193 | HITS-CLIP | 25083871 | Unspecified |
| mmu-miR-378a-5p | MIMAT0000742 | Pitrm1 | 69617 | HITS-CLIP | 25083871 | Unspecified |
| mmu-miR-378a-5p | MIMAT0000742 | Pitx2 | 18741 | HITS-CLIP | 25083871 | Unspecified |
| mmu-miR-378a-5p | MIMAT0000742 | Pkm | 18746 | HITS-CLIP | 37986224\|25083871 | Unspecified |
| mmu-miR-378a-5p | MIMAT0000742 | Pkn1 | 320795 | HITS-CLIP | 37986224 | Unspecified |
| mmu-miR-378a-5p | MIMAT0000742 | Plcb4 | 18798 | HITS-CLIP | 37986224 | Unspecified |
| mmu-miR-378a-5p | MIMAT0000742 | Plcl2 | 224860 | HITS-CLIP | 37986224 | Unspecified |
| mmu-miR-378a-5p | MIMAT0000742 | Plec | 18810 | HITS-CLIP | 37986224\|23597149 | Unspecified |
| mmu-miR-378a-5p | MIMAT0000742 | Plekhg5 | 269608 | HITS-CLIP | 37986224 | Unspecified |
| mmu-miR-378a-5p | MIMAT0000742 | Plekhh2 | 213556 | HITS-CLIP | 37986224 | Unspecified |
| mmu-miR-378a-5p | MIMAT0000742 | Plekhm2 | 69582 | HITS-CLIP | 37986224 | Unspecified |
| mmu-miR-378a-5p | MIMAT0000742 | Plin4 | 57435 | HITS-CLIP | 25753418 | Unspecified |
| mmu-miR-378a-5p | MIMAT0000742 | Plk2 | 20620 | HITS-CLIP | 37986224\|25083871\|29167373 | Unspecified |
| mmu-miR-378a-5p | MIMAT0000742 | Plod1 | 18822 | HITS-CLIP | 23597149 | Unspecified |
| mmu-miR-378a-5p | MIMAT0000742 | Plod2 | 26432 | HITS-CLIP | 25083871 | Unspecified |
| mmu-miR-378a-5p | MIMAT0000742 | Plxna2 | 18845 | HITS-CLIP | 25083871 | Unspecified |
| mmu-miR-378a-5p | MIMAT0000742 | Plxnb2 | 140570 | HITS-CLIP | 23597149 | Unspecified |
| mmu-miR-378a-5p | MIMAT0000742 | Plxnd1 | 67784 | HITS-CLIP | 25753418 | Unspecified |
| mmu-miR-378a-5p | MIMAT0000742 | Pm20d2 | 242377 | HITS-CLIP | 25753418 | Unspecified |
| mmu-miR-378a-5p | MIMAT0000742 | Pnisr | 66625 | HITS-CLIP | 37986224\|25083871\|25753418 | Unspecified |
| mmu-miR-378a-5p | MIMAT0000742 | Pnkd | 56695 | HITS-CLIP | 37986224 | Unspecified |
| mmu-miR-378a-5p | MIMAT0000742 | Pnrc1 | 108767 | HITS-CLIP | 29167373 | Unspecified |
| mmu-miR-378a-5p | MIMAT0000742 | Poglut1 | 224143 | HITS-CLIP | 37986224 | Unspecified |
| mmu-miR-378a-5p | MIMAT0000742 | Pogz | 229584 | HITS-CLIP | 37986224 | Unspecified |
| mmu-miR-378a-5p | MIMAT0000742 | Poldip3 | 73826 | HITS-CLIP | 37986224 | Unspecified |
| mmu-miR-378a-5p | MIMAT0000742 | Polr2a | 20020 | HITS-CLIP | 37986224 | Unspecified |
| mmu-miR-378a-5p | MIMAT0000742 | Polr2b | 231329 | HITS-CLIP | 37986224 | Unspecified |
| mmu-miR-378a-5p | MIMAT0000742 | Por | 18984 | HITS-CLIP | 37986224 | Unspecified |
| mmu-miR-378a-5p | MIMAT0000742 | Ppfia1 | 233977 | HITS-CLIP | 37986224 | Unspecified |
| mmu-miR-378a-5p | MIMAT0000742 | Ppfia4 | 68507 | HITS-CLIP | 37986224 | Unspecified |
| mmu-miR-378a-5p | MIMAT0000742 | Ppfibp1 | 67533 | HITS-CLIP | 25083871 | Unspecified |
| mmu-miR-378a-5p | MIMAT0000742 | Pphln1 | 223828 | HITS-CLIP | 37986224 | Unspecified |
| mmu-miR-378a-5p | MIMAT0000742 | Ppil4 | 67418 | HITS-CLIP | 37986224 | Unspecified |
| mmu-miR-378a-5p | MIMAT0000742 | Ppp1r12a | 17931 | HITS-CLIP | 25083871 | Unspecified |

| | | | | | | |
|---|---|---|---|---|---|---|
| mmu-miR-378a-5p | MIMAT0000742 | Ppp1r12b | 329251 | HITS-CLIP | 37986224\|25083871 | Unspecified |
| mmu-miR-378a-5p | MIMAT0000742 | Ppp1r15b | 108954 | HITS-CLIP | 37986224 | Unspecified |
| mmu-miR-378a-5p | MIMAT0000742 | Ppp1r16a | 73062 | HITS-CLIP | 37986224 | Unspecified |
| mmu-miR-378a-5p | MIMAT0000742 | Ppp1r3a | 140491 | HITS-CLIP | 25753418 | Unspecified |
| mmu-miR-378a-5p | MIMAT0000742 | Ppp1r7 | 66385 | HITS-CLIP | 37986224 | Unspecified |
| mmu-miR-378a-5p | MIMAT0000742 | Ppp2r1b | 73699 | HITS-CLIP | 37986224 | Unspecified |
| mmu-miR-378a-5p | MIMAT0000742 | Ppp2r2a | 71978 | HITS-CLIP | 37986224 | Unspecified |
| mmu-miR-378a-5p | MIMAT0000742 | Ppp2r3a | 235542 | HITS-CLIP | 37986224 | Unspecified |
| mmu-miR-378a-5p | MIMAT0000742 | Ppp2r5e | 26932 | HITS-CLIP | 37986224\|23597149 | Unspecified |
| mmu-miR-378a-5p | MIMAT0000742 | Ppp4r2 | 232314 | HITS-CLIP | 29167373 | Unspecified |
| mmu-miR-378a-5p | MIMAT0000742 | Pprc1 | 226169 | HITS-CLIP | 25083871 | Unspecified |
| mmu-miR-378a-5p | MIMAT0000742 | Ppt2 | 54397 | HITS-CLIP | 37986224 | Unspecified |
| mmu-miR-378a-5p | MIMAT0000742 | Prdm2 | 110593 | HITS-CLIP | 37986224 | Unspecified |
| mmu-miR-378a-5p | MIMAT0000742 | Prkaa1 | 105787 | HITS-CLIP | 29167373 | Unspecified |
| mmu-miR-378a-5p | MIMAT0000742 | Prkx | 19108 | HITS-CLIP | 37986224 | Unspecified |
| mmu-miR-378a-5p | MIMAT0000742 | Prpf39 | 328110 | HITS-CLIP | 23597149 | Unspecified |
| mmu-miR-378a-5p | MIMAT0000742 | Prpf40a | 56194 | HITS-CLIP | 37986224 | Unspecified |
| mmu-miR-378a-5p | MIMAT0000742 | Prpf4b | 19134 | HITS-CLIP | 37986224 | Unspecified |
| mmu-miR-378a-5p | MIMAT0000742 | Prpf8 | 192159 | HITS-CLIP | 25083871\|29167373 | Unspecified |
| mmu-miR-378a-5p | MIMAT0000742 | Prps1 | 19139 | HITS-CLIP | 37986224\|25083871 | Unspecified |
| mmu-miR-378a-5p | MIMAT0000742 | Prps1l3 | 328099 | HITS-CLIP | 37986224 | Unspecified |
| mmu-miR-378a-5p | MIMAT0000742 | Prpsap1 | 67763 | HITS-CLIP | 37986224 | Unspecified |
| mmu-miR-378a-5p | MIMAT0000742 | Prr14l | 215476 | HITS-CLIP | 37986224 | Unspecified |
| mmu-miR-378a-5p | MIMAT0000742 | Prrc2a | 53761 | HITS-CLIP | 25083871\|25753418 | Unspecified |
| mmu-miR-378a-5p | MIMAT0000742 | Prrc2b | 227723 | HITS-CLIP | 37986224\|25083871 | Unspecified |
| mmu-miR-378a-5p | MIMAT0000742 | Prrc2c | 226562 | HITS-CLIP | 23597149 | Unspecified |
| mmu-miR-378a-5p | MIMAT0000742 | Prrx1 | 18933 | HITS-CLIP | 37986224\|25083871 | Unspecified |
| mmu-miR-378a-5p | MIMAT0000742 | Psma6 | 26443 | HITS-CLIP | 23597149 | Unspecified |
| mmu-miR-378a-5p | MIMAT0000742 | Psmb4 | 19172 | HITS-CLIP | 25083871 | Unspecified |
| mmu-miR-378a-5p | MIMAT0000742 | Psmd1 | 70247 | HITS-CLIP | 37986224\|23597149 | Unspecified |
| mmu-miR-378a-5p | MIMAT0000742 | Psme3 | 19192 | HITS-CLIP | 37986224 | Unspecified |
| mmu-miR-378a-5p | MIMAT0000742 | Ptbp3 | 230257 | HITS-CLIP | 23597149\|25083871\|25753418 | Unspecified |
| mmu-miR-378a-5p | MIMAT0000742 | Ptch1 | 19206 | HITS-CLIP | 23597149 | Unspecified |
| mmu-miR-378a-5p | MIMAT0000742 | Ptgs2 | 19225 | HITS-CLIP | 37986224 | Unspecified |
| mmu-miR-378a-5p | MIMAT0000742 | Ptpa | 5524 | HITS-CLIP | 25083871\|25753418 | Unspecified |
| mmu-miR-378a-5p | MIMAT0000742 | Ptpn11 | 19247 | HITS-CLIP | 37986224 | Unspecified |
| mmu-miR-378a-5p | MIMAT0000742 | Ptpn12 | 19248 | HITS-CLIP | 25083871 | Unspecified |
| mmu-miR-378a-5p | MIMAT0000742 | Ptpn14 | 19250 | HITS-CLIP | 37986224 | Unspecified |
| mmu-miR-378a-5p | MIMAT0000742 | Ptprk | 19272 | HITS-CLIP | 23597149 | Unspecified |

| | | | | | | |
|---|---|---|---|---|---|---|
| mmu-miR-378a-5p | MIMAT0000742 | Ptprs | 19280 | HITS-CLIP | 37986224 | Unspecified |
| mmu-miR-378a-5p | MIMAT0000742 | Pttg1ip | 108705 | HITS-CLIP | 25083871 | Unspecified |
| mmu-miR-378a-5p | MIMAT0000742 | Pura | 19290 | HITS-CLIP | 37986224 | Unspecified |
| mmu-miR-378a-5p | MIMAT0000742 | Pus10 | 74467 | HITS-CLIP | 37986224 | Unspecified |
| mmu-miR-378a-5p | MIMAT0000742 | Pxdn | 69675 | HITS-CLIP | 37986224 | Unspecified |
| mmu-miR-378a-5p | MIMAT0000742 | Pxmp2 | 19301 | HITS-CLIP | 25753418 | Unspecified |
| mmu-miR-378a-5p | MIMAT0000742 | Pxn | 19303 | HITS-CLIP | 37986224 | Unspecified |
| mmu-miR-378a-5p | MIMAT0000742 | Pzp | 11287 | HITS-CLIP | 37986224 | Unspecified |
| mmu-miR-378a-5p | MIMAT0000742 | Qk | 19317 | HITS-CLIP | 25083871 | Unspecified |
| mmu-miR-378a-5p | MIMAT0000742 | Qser1 | 99003 | HITS-CLIP | 37986224\|29167373 | Unspecified |
| mmu-miR-378a-5p | MIMAT0000742 | Rab11fip2 | 74998 | HITS-CLIP | 37986224 | Unspecified |
| mmu-miR-378a-5p | MIMAT0000742 | Rab1b | 76308 | HITS-CLIP | 37986224 | Unspecified |
| mmu-miR-378a-5p | MIMAT0000742 | Rab22a | 19334 | HITS-CLIP | 25083871 | Unspecified |
| mmu-miR-378a-5p | MIMAT0000742 | Rab3gap2 | 98732 | HITS-CLIP | 37986224 | Unspecified |
| mmu-miR-378a-5p | MIMAT0000742 | Rab4b | 19342 | HITS-CLIP | 23597149 | Unspecified |
| mmu-miR-378a-5p | MIMAT0000742 | Rab5a | 271457 | HITS-CLIP | 25083871 | Unspecified |
| mmu-miR-378a-5p | MIMAT0000742 | Rab5c | 19345 | HITS-CLIP | 37986224 | Unspecified |
| mmu-miR-378a-5p | MIMAT0000742 | Rab6a | 19346 | HITS-CLIP | 23597149 | Unspecified |
| mmu-miR-378a-5p | MIMAT0000742 | Rabgap1 | 227800 | HITS-CLIP | 37986224 | Unspecified |
| mmu-miR-378a-5p | MIMAT0000742 | Rabgef1 | 56715 | HITS-CLIP | 37986224 | Unspecified |
| mmu-miR-378a-5p | MIMAT0000742 | Rad23a | 19358 | HITS-CLIP | 37986224 | Unspecified |
| mmu-miR-378a-5p | MIMAT0000742 | Rad54l2 | 81000 | HITS-CLIP | 37986224\|25083871\|23597149 | Unspecified |
| mmu-miR-378a-5p | MIMAT0000742 | Rala | 56044 | HITS-CLIP | 25083871 | Unspecified |
| mmu-miR-378a-5p | MIMAT0000742 | Ralbp1 | 19765 | HITS-CLIP | 37986224 | Unspecified |
| mmu-miR-378a-5p | MIMAT0000742 | Ralgapb | 228850 | HITS-CLIP | 37986224 | Unspecified |
| mmu-miR-378a-5p | MIMAT0000742 | Ralgps2 | 78255 | HITS-CLIP | 25083871 | Unspecified |
| mmu-miR-378a-5p | MIMAT0000742 | Ranbp10 | 74334 | HITS-CLIP | 37986224 | Unspecified |
| mmu-miR-378a-5p | MIMAT0000742 | Ranbp6 | 240614 | HITS-CLIP | 37986224 | Unspecified |
| mmu-miR-378a-5p | MIMAT0000742 | Ranbp9 | 56705 | HITS-CLIP | 37986224 | Unspecified |
| mmu-miR-378a-5p | MIMAT0000742 | Rap1b | 215449 | HITS-CLIP | 25083871 | Unspecified |
| mmu-miR-378a-5p | MIMAT0000742 | Rap2a | 76108 | HITS-CLIP | 25083871 | Unspecified |
| mmu-miR-378a-5p | MIMAT0000742 | Rapgef1 | 107746 | HITS-CLIP | 37986224\|25083871 | Unspecified |
| mmu-miR-378a-5p | MIMAT0000742 | Rb1cc1 | 12421 | HITS-CLIP | 37986224\|25083871 | Unspecified |
| mmu-miR-378a-5p | MIMAT0000742 | Rbbp6 | 19647 | HITS-CLIP | 25083871\|29167373 | Unspecified |
| mmu-miR-378a-5p | MIMAT0000742 | Rbm12 | 75710 | HITS-CLIP | 37986224 | Unspecified |
| mmu-miR-378a-5p | MIMAT0000742 | Rbm12b2 | 77604 | HITS-CLIP | 37986224 | Unspecified |
| mmu-miR-378a-5p | MIMAT0000742 | Rbm14 | 56275 | HITS-CLIP | 37986224 | Unspecified |
| mmu-miR-378a-5p | MIMAT0000742 | Rbm15b | 109095 | HITS-CLIP | 25083871 | Unspecified |
| mmu-miR-378a-5p | MIMAT0000742 | Rbm20 | 73713 | HITS-CLIP | 37986224 | Unspecified |

| | | | | | | | |
|---|---|---|---|---|---|---|---|
| mmu-miR-378a-5p | MIMAT0000742 | Rbm41 | 237073 | HITS-CLIP | | 37986224 | Unspecified |
| mmu-miR-378a-5p | MIMAT0000742 | Rbms1 | 56878 | HITS-CLIP | | 29167373 | Unspecified |
| mmu-miR-378a-5p | MIMAT0000742 | Rc3h1 | 381305 | HITS-CLIP | | 25083871 | Unspecified |
| mmu-miR-378a-5p | MIMAT0000742 | Rcan1 | 54720 | HITS-CLIP | | 25753418 | Unspecified |
| mmu-miR-378a-5p | MIMAT0000742 | Rcor1 | 217864 | HITS-CLIP | | 37986224 | Unspecified |
| mmu-miR-378a-5p | MIMAT0000742 | Rere | 68703 | HITS-CLIP | 25083871\|29167373 | | Unspecified |
| mmu-miR-378a-5p | MIMAT0000742 | Rexo1 | 66932 | HITS-CLIP | | 29167373 | Unspecified |
| mmu-miR-378a-5p | MIMAT0000742 | Rfc3 | 69263 | qCLASH | | 31393953 | Unspecified |
| mmu-miR-378a-5p | MIMAT0000742 | Rft1 | 328370 | HITS-CLIP | | 37986224 | Unspecified |
| mmu-miR-378a-5p | MIMAT0000742 | Rfwd3 | 234736 | HITS-CLIP | | 37986224 | Unspecified |
| mmu-miR-378a-5p | MIMAT0000742 | Rfx7 | 319758 | HITS-CLIP | 37986224\|25083871 | | Unspecified |
| mmu-miR-378a-5p | MIMAT0000742 | Rgp1 | 242406 | HITS-CLIP | | 37986224 | Unspecified |
| mmu-miR-378a-5p | MIMAT0000742 | Ric1 | 850728 | HITS-CLIP | | 29167373 | Unspecified |
| mmu-miR-378a-5p | MIMAT0000742 | Rif1 | 51869 | HITS-CLIP | | 37986224 | Unspecified |
| mmu-miR-378a-5p | MIMAT0000742 | Rilpl1 | 75695 | HITS-CLIP | | 37986224 | Unspecified |
| mmu-miR-378a-5p | MIMAT0000742 | Rin2 | 74030 | HITS-CLIP | 37986224\|25083871 | | Unspecified |
| mmu-miR-378a-5p | MIMAT0000742 | Rin3 | 217835 | HITS-CLIP | | 37986224 | Unspecified |
| mmu-miR-378a-5p | MIMAT0000742 | Ripk2 | 192656 | HITS-CLIP | | 25083871 | Unspecified |
| mmu-miR-378a-5p | MIMAT0000742 | Rmnd5a | 68477 | HITS-CLIP | 37986224\|23597149 | | Unspecified |
| mmu-miR-378a-5p | MIMAT0000742 | Rnf111 | 93836 | HITS-CLIP | | 37986224 | Unspecified |
| mmu-miR-378a-5p | MIMAT0000742 | Rnf13 | 24017 | HITS-CLIP | | 29167373 | Unspecified |
| mmu-miR-378a-5p | MIMAT0000742 | Rnf138 | 56515 | HITS-CLIP | | 37986224 | Unspecified |
| mmu-miR-378a-5p | MIMAT0000742 | Rnf150 | 330812 | HITS-CLIP | | 37986224 | Unspecified |
| mmu-miR-378a-5p | MIMAT0000742 | Rnf20 | 109331 | HITS-CLIP | | 37986224 | Unspecified |
| mmu-miR-378a-5p | MIMAT0000742 | Rnf213 | 672511 | HITS-CLIP | | 37986224 | Unspecified |
| mmu-miR-378a-5p | MIMAT0000742 | Rnf217 | 268291 | HITS-CLIP | | 23597149 | Unspecified |
| mmu-miR-378a-5p | MIMAT0000742 | Rnf24 | 51902 | HITS-CLIP | | 37986224 | Unspecified |
| mmu-miR-378a-5p | MIMAT0000742 | Rnf26 | 213211 | HITS-CLIP | | 25083871 | Unspecified |
| mmu-miR-378a-5p | MIMAT0000742 | Rnf38 | 73469 | HITS-CLIP | | 25083871 | Unspecified |
| mmu-miR-378a-5p | MIMAT0000742 | Rnf44 | 105239 | HITS-CLIP | | 37986224 | Unspecified |
| mmu-miR-378a-5p | MIMAT0000742 | Rpp25l | 69961 | HITS-CLIP | | 25753418 | Unspecified |
| mmu-miR-378a-5p | MIMAT0000742 | Rprd2 | 75137 | HITS-CLIP | 37986224\|25083871 | | Unspecified |
| mmu-miR-378a-5p | MIMAT0000742 | Rps20 | 67427 | HITS-CLIP | | 37986224 | Unspecified |
| mmu-miR-378a-5p | MIMAT0000742 | Rps6kb1 | 72508 | HITS-CLIP | | 37986224 | Unspecified |
| mmu-miR-378a-5p | MIMAT0000742 | Rrn3 | 106298 | HITS-CLIP | | 37986224 | Unspecified |
| mmu-miR-378a-5p | MIMAT0000742 | Rrp1 | 18114 | HITS-CLIP | | 37986224 | Unspecified |
| mmu-miR-378a-5p | MIMAT0000742 | Rrp1b | 72462 | HITS-CLIP | | 37986224 | Unspecified |
| mmu-miR-378a-5p | MIMAT0000742 | Rsbn1l | 242860 | HITS-CLIP | | 29167373 | Unspecified |
| mmu-miR-378a-5p | MIMAT0000742 | Rtn3 | 20168 | HITS-CLIP | | 37986224 | Unspecified |

| | | | | | | |
|---|---|---|---|---|---|---|
| mmu-miR-378a-5p | MIMAT0000742 | Rtn4 | 68585 | HITS-CLIP | 37986224 | Unspecified |
| mmu-miR-378a-5p | MIMAT0000742 | Ruvbl1 | 56505 | HITS-CLIP | 37986224 | Unspecified |
| mmu-miR-378a-5p | MIMAT0000742 | Rxra | 20181 | HITS-CLIP | 37986224 | Unspecified |
| mmu-miR-378a-5p | MIMAT0000742 | Ryr1 | 20190 | HITS-CLIP | 25083871 | Unspecified |
| mmu-miR-378a-5p | MIMAT0000742 | Ryr3 | 20192 | HITS-CLIP | 37986224 | Unspecified |
| mmu-miR-378a-5p | MIMAT0000742 | S100pbp | 74648 | HITS-CLIP | 25083871 | Unspecified |
| mmu-miR-378a-5p | MIMAT0000742 | Sacs | 50720 | HITS-CLIP | 37986224 | Unspecified |
| mmu-miR-378a-5p | MIMAT0000742 | Samd8 | 67630 | HITS-CLIP | 25083871 | Unspecified |
| mmu-miR-378a-5p | MIMAT0000742 | Sars | 20226 | HITS-CLIP | 37986224\|25083871\|29167373 | Unspecified |
| mmu-miR-378a-5p | MIMAT0000742 | Satb2 | 212712 | HITS-CLIP | 37986224 | Unspecified |
| mmu-miR-378a-5p | MIMAT0000742 | Sav1 | 64010 | HITS-CLIP | 37986224 | Unspecified |
| mmu-miR-378a-5p | MIMAT0000742 | Sbf1 | 77980 | HITS-CLIP | 25083871 | Unspecified |
| mmu-miR-378a-5p | MIMAT0000742 | Sbno1 | 243272 | HITS-CLIP | 37986224 | Unspecified |
| mmu-miR-378a-5p | MIMAT0000742 | Scaf11 | 72193 | HITS-CLIP | 37986224 | Unspecified |
| mmu-miR-378a-5p | MIMAT0000742 | Scmh1 | 29871 | HITS-CLIP | 37986224 | Unspecified |
| mmu-miR-378a-5p | MIMAT0000742 | Sdc2 | 15529 | HITS-CLIP | 37986224 | Unspecified |
| mmu-miR-378a-5p | MIMAT0000742 | Sdc3 | 20970 | HITS-CLIP | 25753418 | Unspecified |
| mmu-miR-378a-5p | MIMAT0000742 | Sdc4 | 20971 | HITS-CLIP | 37986224 | Unspecified |
| mmu-miR-378a-5p | MIMAT0000742 | Sde2 | 208768 | HITS-CLIP | 37986224 | Unspecified |
| mmu-miR-378a-5p | MIMAT0000742 | Sdhaf2 | 66072 | HITS-CLIP | 37986224 | Unspecified |
| mmu-miR-378a-5p | MIMAT0000742 | Sdhd | 66925 | HITS-CLIP | 37986224 | Unspecified |
| mmu-miR-378a-5p | MIMAT0000742 | Sec14l2 | 67815 | HITS-CLIP | 37986224\|25083871 | Unspecified |
| mmu-miR-378a-5p | MIMAT0000742 | Sec16a | 227648 | HITS-CLIP | 37986224 | Unspecified |
| mmu-miR-378a-5p | MIMAT0000742 | Sec22a | 317717 | HITS-CLIP | 37986224 | Unspecified |
| mmu-miR-378a-5p | MIMAT0000742 | Sec22b | 20333 | HITS-CLIP | 37986224\|23597149\|25083871 | Unspecified |
| mmu-miR-378a-5p | MIMAT0000742 | Sec24b | 99683 | HITS-CLIP | 37986224 | Unspecified |
| mmu-miR-378a-5p | MIMAT0000742 | Secisbp2l | 70354 | HITS-CLIP | 37986224 | Unspecified |
| mmu-miR-378a-5p | MIMAT0000742 | Sel1l | 20338 | HITS-CLIP | 37986224 | Unspecified |
| mmu-miR-378a-5p | MIMAT0000742 | Selenof | 9403 | HITS-CLIP | 25083871 | Unspecified |
| mmu-miR-378a-5p | MIMAT0000742 | Selenon | 57190 | HITS-CLIP | 25083871 | Unspecified |
| mmu-miR-378a-5p | MIMAT0000742 | Senp5 | 320213 | HITS-CLIP | 37986224 | Unspecified |
| mmu-miR-378a-5p | MIMAT0000742 | Sephs2 | 20768 | HITS-CLIP | 37986224 | Unspecified |
| mmu-miR-378a-5p | MIMAT0000742 | Sept10 | 103080 | HITS-CLIP | 37986224 | Unspecified |
| mmu-miR-378a-5p | MIMAT0000742 | Sept9 | 53860 | HITS-CLIP | 37986224 | Unspecified |
| mmu-miR-378a-5p | MIMAT0000742 | Serf2 | 378702 | HITS-CLIP | 37986224 | Unspecified |
| mmu-miR-378a-5p | MIMAT0000742 | Serinc5 | 218442 | HITS-CLIP | 37986224\|25083871 | Unspecified |
| mmu-miR-378a-5p | MIMAT0000742 | Serpina1b | 20701 | HITS-CLIP | 23597149 | Unspecified |
| mmu-miR-378a-5p | MIMAT0000742 | Serpind1 | 15160 | HITS-CLIP | 37986224 | Unspecified |
| mmu-miR-378a-5p | MIMAT0000742 | Serpine1 | 18787 | HITS-CLIP | 25083871 | Unspecified |

| | | | | | | |
|---|---|---|---|---|---|---|
| mmu-miR-378a-5p | MIMAT0000742 | Sertad3 | 170742 | HITS-CLIP | 37986224 | Unspecified |
| mmu-miR-378a-5p | MIMAT0000742 | Sesn2 | 230784 | HITS-CLIP | 25083871 | Unspecified |
| mmu-miR-378a-5p | MIMAT0000742 | Set | 56086 | HITS-CLIP | 37986224\|25083871 | Unspecified |
| mmu-miR-378a-5p | MIMAT0000742 | Setd1a | 233904 | HITS-CLIP | 37986224 | Unspecified |
| mmu-miR-378a-5p | MIMAT0000742 | Setd2 | 235626 | HITS-CLIP | 37986224 | Unspecified |
| mmu-miR-378a-5p | MIMAT0000742 | Setd5 | 72895 | HITS-CLIP | 37986224 | Unspecified |
| mmu-miR-378a-5p | MIMAT0000742 | Setd7 | 73251 | HITS-CLIP | 37986224 | Unspecified |
| mmu-miR-378a-5p | MIMAT0000742 | Setdb1 | 84505 | HITS-CLIP | 37986224 | Unspecified |
| mmu-miR-378a-5p | MIMAT0000742 | Sft2d2 | 108735 | HITS-CLIP | 37986224 | Unspecified |
| mmu-miR-378a-5p | MIMAT0000742 | Sft2d3 | 67158 | HITS-CLIP | 37986224 | Unspecified |
| mmu-miR-378a-5p | MIMAT0000742 | Sfxn1 | 14057 | HITS-CLIP | 37986224 | Unspecified |
| mmu-miR-378a-5p | MIMAT0000742 | Sgpl1 | 20397 | HITS-CLIP | 37986224\|23597149 | Unspecified |
| mmu-miR-378a-5p | MIMAT0000742 | Sh3bgrl3 | 73723 | HITS-CLIP | 25083871 | Unspecified |
| mmu-miR-378a-5p | MIMAT0000742 | Sh3glb1 | 54673 | HITS-CLIP | 25083871 | Unspecified |
| mmu-miR-378a-5p | MIMAT0000742 | Sh3kbp1 | 58194 | HITS-CLIP | 37986224 | Unspecified |
| mmu-miR-378a-5p | MIMAT0000742 | Sh3pxd2b | 268396 | HITS-CLIP | 37986224 | Unspecified |
| mmu-miR-378a-5p | MIMAT0000742 | Shank3 | 58234 | HITS-CLIP | 25753418 | Unspecified |
| mmu-miR-378a-5p | MIMAT0000742 | Shc1 | 20416 | HITS-CLIP | 37986224 | Unspecified |
| mmu-miR-378a-5p | MIMAT0000742 | Shisa5 | 66940 | HITS-CLIP | 29167373 | Unspecified |
| mmu-miR-378a-5p | MIMAT0000742 | Sik3 | 70661 | HITS-CLIP | 25753418 | Unspecified |
| mmu-miR-378a-5p | MIMAT0000742 | Sil1 | 81500 | HITS-CLIP | 37986224 | Unspecified |
| mmu-miR-378a-5p | MIMAT0000742 | Simc1 | 319719 | HITS-CLIP | 37986224 | Unspecified |
| mmu-miR-378a-5p | MIMAT0000742 | Sin3a | 20466 | HITS-CLIP | 25083871\|23597149 | Unspecified |
| mmu-miR-378a-5p | MIMAT0000742 | Sin3b | 20467 | HITS-CLIP | 23597149 | Unspecified |
| mmu-miR-378a-5p | MIMAT0000742 | Sipa1 | 20469 | HITS-CLIP | 37986224 | Unspecified |
| mmu-miR-378a-5p | MIMAT0000742 | Sipa1l3 | 74206 | HITS-CLIP | 37986224\|29167373 | Unspecified |
| mmu-miR-378a-5p | MIMAT0000742 | Sirt1 | 93759 | HITS-CLIP | 25083871 | Unspecified |
| mmu-miR-378a-5p | MIMAT0000742 | Sirt6 | 50721 | HITS-CLIP | 37986224 | Unspecified |
| mmu-miR-378a-5p | MIMAT0000742 | Six4 | 20474 | HITS-CLIP | 37986224 | Unspecified |
| mmu-miR-378a-5p | MIMAT0000742 | Slc11a2 | 18174 | HITS-CLIP | 37986224 | Unspecified |
| mmu-miR-378a-5p | MIMAT0000742 | Slc12a7 | 20499 | HITS-CLIP | 23597149\|25083871 | Unspecified |
| mmu-miR-378a-5p | MIMAT0000742 | Slc15a4 | 100561 | HITS-CLIP | 37986224 | Unspecified |
| mmu-miR-378a-5p | MIMAT0000742 | Slc16a1 | 20501 | HITS-CLIP | 25753418 | Unspecified |
| mmu-miR-378a-5p | MIMAT0000742 | Slc1a4 | 55963 | HITS-CLIP | 25083871 | Unspecified |
| mmu-miR-378a-5p | MIMAT0000742 | Slc23a2 | 54338 | HITS-CLIP | 23597149 | Unspecified |
| mmu-miR-378a-5p | MIMAT0000742 | Slc25a15 | 18408 | HITS-CLIP | 23597149 | Unspecified |
| mmu-miR-378a-5p | MIMAT0000742 | Slc25a37 | 67712 | HITS-CLIP | 37986224 | Unspecified |
| mmu-miR-378a-5p | MIMAT0000742 | Slc25a45 | 107375 | HITS-CLIP | 29167373 | Unspecified |
| mmu-miR-378a-5p | MIMAT0000742 | Slc25a47 | 104910 | HITS-CLIP | 37986224 | Unspecified |

| | | | | | | |
|---|---|---|---|---|---|---|
| mmu-miR-378a-5p | MIMAT0000742 | Slc25a51 | 230125 | HITS-CLIP | 37986224 | Unspecified |
| mmu-miR-378a-5p | MIMAT0000742 | Slc27a4 | 26569 | HITS-CLIP | 37986224 | Unspecified |
| mmu-miR-378a-5p | MIMAT0000742 | Slc29a1 | 63959 | HITS-CLIP | 25083871 | Unspecified |
| mmu-miR-378a-5p | MIMAT0000742 | Slc30a4 | 22785 | HITS-CLIP | 37986224 | Unspecified |
| mmu-miR-378a-5p | MIMAT0000742 | Slc30a5 | 69048 | HITS-CLIP | 25083871\|29167373 | Unspecified |
| mmu-miR-378a-5p | MIMAT0000742 | Slc35a2 | 22232 | HITS-CLIP | 25083871 | Unspecified |
| mmu-miR-378a-5p | MIMAT0000742 | Slc35a5 | 74102 | HITS-CLIP | 37986224 | Unspecified |
| mmu-miR-378a-5p | MIMAT0000742 | Slc39a13 | 68427 | HITS-CLIP | 37986224 | Unspecified |
| mmu-miR-378a-5p | MIMAT0000742 | Slc39a9 | 328133 | HITS-CLIP | 37986224 | Unspecified |
| mmu-miR-378a-5p | MIMAT0000742 | Slc44a1 | 100434 | HITS-CLIP | 37986224\|25083871 | Unspecified |
| mmu-miR-378a-5p | MIMAT0000742 | Slc5a3 | 53881 | HITS-CLIP | 37986224 | Unspecified |
| mmu-miR-378a-5p | MIMAT0000742 | Slc6a13 | 14412 | HITS-CLIP | 23597149 | Unspecified |
| mmu-miR-378a-5p | MIMAT0000742 | Slc6a6 | 21366 | HITS-CLIP | 25083871 | Unspecified |
| mmu-miR-378a-5p | MIMAT0000742 | Slc7a1 | 11987 | HITS-CLIP | 37986224 | Unspecified |
| mmu-miR-378a-5p | MIMAT0000742 | Slc7a2 | 11988 | HITS-CLIP | 37986224 | Unspecified |
| mmu-miR-378a-5p | MIMAT0000742 | Slc8b1 | 170756 | HITS-CLIP | 37986224 | Unspecified |
| mmu-miR-378a-5p | MIMAT0000742 | Slu7 | 193116 | HITS-CLIP | 37986224 | Unspecified |
| mmu-miR-378a-5p | MIMAT0000742 | Smad3 | 17127 | HITS-CLIP | 37986224 | Unspecified |
| mmu-miR-378a-5p | MIMAT0000742 | Smad4 | 17128 | HITS-CLIP | 37986224 | Unspecified |
| mmu-miR-378a-5p | MIMAT0000742 | Smad7 | 17131 | HITS-CLIP | 37986224 | Unspecified |
| mmu-miR-378a-5p | MIMAT0000742 | Smap2 | 69780 | qCLASH | 31393953 | Unspecified |
| mmu-miR-378a-5p | MIMAT0000742 | Smarcd1 | 83797 | HITS-CLIP | 37986224 | Unspecified |
| mmu-miR-378a-5p | MIMAT0000742 | Smc1a | 24061 | HITS-CLIP | 29167373 | Unspecified |
| mmu-miR-378a-5p | MIMAT0000742 | Smchd1 | 74355 | HITS-CLIP | 37986224 | Unspecified |
| mmu-miR-378a-5p | MIMAT0000742 | Smcr8 | 237782 | HITS-CLIP | 37986224 | Unspecified |
| mmu-miR-378a-5p | MIMAT0000742 | Smg1 | 233789 | HITS-CLIP | 37986224\|25083871\|29167373 | Unspecified |
| mmu-miR-378a-5p | MIMAT0000742 | Smg8 | 74133 | HITS-CLIP | 25083871 | Unspecified |
| mmu-miR-378a-5p | MIMAT0000742 | Snap23 | 20619 | HITS-CLIP | 37986224 | Unspecified |
| mmu-miR-378a-5p | MIMAT0000742 | Snd1 | 56463 | HITS-CLIP | 37986224 | Unspecified |
| mmu-miR-378a-5p | MIMAT0000742 | Sntb2 | 20650 | HITS-CLIP | 37986224 | Unspecified |
| mmu-miR-378a-5p | MIMAT0000742 | Snx18 | 170625 | HITS-CLIP | 37986224 | Unspecified |
| mmu-miR-378a-5p | MIMAT0000742 | Soat1 | 20652 | HITS-CLIP | 37986224\|25083871 | Unspecified |
| mmu-miR-378a-5p | MIMAT0000742 | Socs3 | 12702 | HITS-CLIP | 37986224 | Unspecified |
| mmu-miR-378a-5p | MIMAT0000742 | Sod1 | 20655 | HITS-CLIP | 37986224 | Unspecified |
| mmu-miR-378a-5p | MIMAT0000742 | Son | 20658 | HITS-CLIP | 37986224 | Unspecified |
| mmu-miR-378a-5p | MIMAT0000742 | Sorbs2 | 234214 | HITS-CLIP | 25083871 | Unspecified |
| mmu-miR-378a-5p | MIMAT0000742 | Sos2 | 20663 | HITS-CLIP | 25083871 | Unspecified |
| mmu-miR-378a-5p | MIMAT0000742 | Sox11 | 20666 | HITS-CLIP | 37986224\|25083871 | Unspecified |
| mmu-miR-378a-5p | MIMAT0000742 | Sox9 | 20682 | HITS-CLIP | 37986224 | Unspecified |

| | | | | | | |
|---|---|---|---|---|---|---|
| mmu-miR-378a-5p | MIMAT0000742 | Sp1 | 20683 | HITS-CLIP | 25083871 | Unspecified |
| mmu-miR-378a-5p | MIMAT0000742 | Sp3 | 20687 | HITS-CLIP | 25083871 | Unspecified |
| mmu-miR-378a-5p | MIMAT0000742 | Spaca6 | 75202 | HITS-CLIP | 37986224 | Unspecified |
| mmu-miR-378a-5p | MIMAT0000742 | Spata13 | 219140 | HITS-CLIP | 37986224 | Unspecified |
| mmu-miR-378a-5p | MIMAT0000742 | Spata2 | 263876 | HITS-CLIP | 37986224 | Unspecified |
| mmu-miR-378a-5p | MIMAT0000742 | Spc25 | 66442 | HITS-CLIP | 29167373 | Unspecified |
| mmu-miR-378a-5p | MIMAT0000742 | Spcs3 | 76687 | HITS-CLIP | 37986224 | Unspecified |
| mmu-miR-378a-5p | MIMAT0000742 | Specc1 | 432572 | HITS-CLIP | 37986224 | Unspecified |
| mmu-miR-378a-5p | MIMAT0000742 | Specc1l | 74392 | HITS-CLIP | 23597149 | Unspecified |
| mmu-miR-378a-5p | MIMAT0000742 | Spg20 | 229285 | HITS-CLIP | 25083871 | Unspecified |
| mmu-miR-378a-5p | MIMAT0000742 | Spryd7 | 66674 | HITS-CLIP | 25753418 | Unspecified |
| mmu-miR-378a-5p | MIMAT0000742 | Spsb1 | 74646 | HITS-CLIP | 37986224 | Unspecified |
| mmu-miR-378a-5p | MIMAT0000742 | Sptbn1 | 20742 | HITS-CLIP | 37986224\|25083871\|23597149 | Unspecified |
| mmu-miR-378a-5p | MIMAT0000742 | Sptssa | 104725 | HITS-CLIP | 37986224\|25083871\|23597149 | Unspecified |
| mmu-miR-378a-5p | MIMAT0000742 | Sqstm1 | 18412 | HITS-CLIP | 23597149 | Unspecified |
| mmu-miR-378a-5p | MIMAT0000742 | Srcap | 100043597 | HITS-CLIP | 37986224\|25083871 | Unspecified |
| mmu-miR-378a-5p | MIMAT0000742 | Srd5a3 | 57357 | HITS-CLIP | 37986224 | Unspecified |
| mmu-miR-378a-5p | MIMAT0000742 | Srebf2 | 20788 | HITS-CLIP | 25083871 | Unspecified |
| mmu-miR-378a-5p | MIMAT0000742 | Srp19 | 66384 | HITS-CLIP | 25083871 | Unspecified |
| mmu-miR-378a-5p | MIMAT0000742 | Srpx2 | 68792 | HITS-CLIP | 25083871 | Unspecified |
| mmu-miR-378a-5p | MIMAT0000742 | Srrd | 70118 | HITS-CLIP | 31433990 | Unspecified |
| mmu-miR-378a-5p | MIMAT0000742 | Srrm1 | 51796 | HITS-CLIP | 25083871\|23597149\|29167373 | Unspecified |
| mmu-miR-378a-5p | MIMAT0000742 | Srrm2 | 75956 | HITS-CLIP | 37986224\|25083871\|29167373 | Unspecified |
| mmu-miR-378a-5p | MIMAT0000742 | Ss18 | 268996 | HITS-CLIP | 25083871 | Unspecified |
| mmu-miR-378a-5p | MIMAT0000742 | Ssh1 | 231637 | HITS-CLIP | 37986224 | Unspecified |
| mmu-miR-378a-5p | MIMAT0000742 | Ssh2 | 237860 | HITS-CLIP | 37986224 | Unspecified |
| mmu-miR-378a-5p | MIMAT0000742 | Sspn | 16651 | HITS-CLIP | 37986224 | Unspecified |
| mmu-miR-378a-5p | MIMAT0000742 | Ssr1 | 107513 | HITS-CLIP | 25083871 | Unspecified |
| mmu-miR-378a-5p | MIMAT0000742 | Ssr2 | 66256 | HITS-CLIP | 25753418 | Unspecified |
| mmu-miR-378a-5p | MIMAT0000742 | St3gal1 | 20442 | HITS-CLIP | 37986224 | Unspecified |
| mmu-miR-378a-5p | MIMAT0000742 | St6galnac4 | 20448 | HITS-CLIP | 37986224 | Unspecified |
| mmu-miR-378a-5p | MIMAT0000742 | St8sia2 | 20450 | HITS-CLIP | 25083871 | Unspecified |
| mmu-miR-378a-5p | MIMAT0000742 | Stk10 | 20868 | HITS-CLIP | 37986224 | Unspecified |
| mmu-miR-378a-5p | MIMAT0000742 | Stk11 | 20869 | HITS-CLIP | 37986224 | Unspecified |
| mmu-miR-378a-5p | MIMAT0000742 | Stk4 | 58231 | HITS-CLIP | 37986224 | Unspecified |
| mmu-miR-378a-5p | MIMAT0000742 | Stradb | 227154 | HITS-CLIP | 37986224 | Unspecified |
| mmu-miR-378a-5p | MIMAT0000742 | Strn | 268980 | HITS-CLIP | 37986224 | Unspecified |
| mmu-miR-378a-5p | MIMAT0000742 | Strn3 | 94186 | HITS-CLIP | 25083871 | Unspecified |
| mmu-miR-378a-5p | MIMAT0000742 | Strn4 | 97387 | HITS-CLIP | 23597149 | Unspecified |

| | | | | | | |
|---|---|---|---|---|---|---|
| mmu-miR-378a-5p | MIMAT0000742 | Stt3a | 16430 | HITS-CLIP | 37986224\|25083871 | Unspecified |
| mmu-miR-378a-5p | MIMAT0000742 | Stx17 | 67727 | HITS-CLIP | 37986224 | Unspecified |
| mmu-miR-378a-5p | MIMAT0000742 | Stxbp1 | 20910 | HITS-CLIP | 37986224 | Unspecified |
| mmu-miR-378a-5p | MIMAT0000742 | Stxbp2 | 20911 | qCLASH | 31393953 | Unspecified |
| mmu-miR-378a-5p | MIMAT0000742 | Sucla2 | 20916 | HITS-CLIP | 25083871 | Unspecified |
| mmu-miR-378a-5p | MIMAT0000742 | Suco | 226551 | HITS-CLIP | 25083871 | Unspecified |
| mmu-miR-378a-5p | MIMAT0000742 | Supt20 | 56790 | HITS-CLIP | 37986224 | Unspecified |
| mmu-miR-378a-5p | MIMAT0000742 | Supt6 | 20926 | HITS-CLIP | 25083871 | Unspecified |
| mmu-miR-378a-5p | MIMAT0000742 | Surf4 | 20932 | HITS-CLIP | 37986224 | Unspecified |
| mmu-miR-378a-5p | MIMAT0000742 | Surf6 | 20935 | HITS-CLIP | 37986224 | Unspecified |
| mmu-miR-378a-5p | MIMAT0000742 | Svil | 225115 | HITS-CLIP | 23597149 | Unspecified |
| mmu-miR-378a-5p | MIMAT0000742 | Syne1 | 64009 | HITS-CLIP | 37986224 | Unspecified |
| mmu-miR-378a-5p | MIMAT0000742 | Synj1 | 104015 | HITS-CLIP | 25083871 | Unspecified |
| mmu-miR-378a-5p | MIMAT0000742 | Syt13 | 80976 | HITS-CLIP | 37986224 | Unspecified |
| mmu-miR-378a-5p | MIMAT0000742 | Syvn1 | 74126 | HITS-CLIP | 25083871 | Unspecified |
| mmu-miR-378a-5p | MIMAT0000742 | Szrd1 | 213491 | HITS-CLIP | 37986224 | Unspecified |
| mmu-miR-378a-5p | MIMAT0000742 | Tab3 | 66724 | HITS-CLIP | 37986224 | Unspecified |
| mmu-miR-378a-5p | MIMAT0000742 | Taf2 | 319944 | HITS-CLIP | 25083871 | Unspecified |
| mmu-miR-378a-5p | MIMAT0000742 | Taf5l | 102162 | HITS-CLIP | 37986224 | Unspecified |
| mmu-miR-378a-5p | MIMAT0000742 | Tanc1 | 66860 | HITS-CLIP | 37986224 | Unspecified |
| mmu-miR-378a-5p | MIMAT0000742 | Tanc2 | 77097 | HITS-CLIP | 37986224 | Unspecified |
| mmu-miR-378a-5p | MIMAT0000742 | Taok3 | 330177 | HITS-CLIP | 37986224 | Unspecified |
| mmu-miR-378a-5p | MIMAT0000742 | Tardbp | 230908 | HITS-CLIP | 25083871 | Unspecified |
| mmu-miR-378a-5p | MIMAT0000742 | Tat | 234724 | HITS-CLIP | 23597149 | Unspecified |
| mmu-miR-378a-5p | MIMAT0000742 | Tbc1d1 | 57915 | HITS-CLIP | 25083871 | Unspecified |
| mmu-miR-378a-5p | MIMAT0000742 | Tbc1d10a | 103724 | HITS-CLIP | 37986224 | Unspecified |
| mmu-miR-378a-5p | MIMAT0000742 | Tbc1d10b | 68449 | HITS-CLIP | 37986224 | Unspecified |
| mmu-miR-378a-5p | MIMAT0000742 | Tbc1d13 | 70296 | HITS-CLIP | 37986224 | Unspecified |
| mmu-miR-378a-5p | MIMAT0000742 | Tbc1d17 | 233204 | HITS-CLIP | 23597149 | Unspecified |
| mmu-miR-378a-5p | MIMAT0000742 | Tbc1d24 | 224617 | HITS-CLIP | 37986224 | Unspecified |
| mmu-miR-378a-5p | MIMAT0000742 | Tbcd | 108903 | HITS-CLIP | 25083871 | Unspecified |
| mmu-miR-378a-5p | MIMAT0000742 | Tbl1xr1 | 81004 | HITS-CLIP | 37986224 | Unspecified |
| mmu-miR-378a-5p | MIMAT0000742 | Tcf12 | 21406 | HITS-CLIP | 37986224 | Unspecified |
| mmu-miR-378a-5p | MIMAT0000742 | Tcf20 | 21411 | HITS-CLIP | 37986224 | Unspecified |
| mmu-miR-378a-5p | MIMAT0000742 | Tcf25 | 66855 | HITS-CLIP | 25083871 | Unspecified |
| mmu-miR-378a-5p | MIMAT0000742 | Tcf4 | 21413 | HITS-CLIP | 37986224\|25083871 | Unspecified |
| mmu-miR-378a-5p | MIMAT0000742 | Tcf7l2 | 21416 | HITS-CLIP | 37986224 | Unspecified |
| mmu-miR-378a-5p | MIMAT0000742 | Tdrd7 | 100121 | HITS-CLIP | 37986224 | Unspecified |
| mmu-miR-378a-5p | MIMAT0000742 | Tdrp | 72148 | HITS-CLIP | 37986224 | Unspecified |

| | | | | | | |
|---|---|---|---|---|---|---|
| mmu-miR-378a-5p | MIMAT0000742 | Tead1 | 21676 | HITS-CLIP | 37986224 | Unspecified |
| mmu-miR-378a-5p | MIMAT0000742 | Tet2 | 214133 | HITS-CLIP | 37986224 | Unspecified |
| mmu-miR-378a-5p | MIMAT0000742 | Tet3 | 194388 | HITS-CLIP | 37986224\|25083871\|29167373 | Unspecified |
| mmu-miR-378a-5p | MIMAT0000742 | Tex264 | 21767 | HITS-CLIP | 23597149 | Unspecified |
| mmu-miR-378a-5p | MIMAT0000742 | Tfdp1 | 21781 | HITS-CLIP | 37986224 | Unspecified |
| mmu-miR-378a-5p | MIMAT0000742 | Tfdp2 | 211586 | HITS-CLIP | 37986224 | Unspecified |
| mmu-miR-378a-5p | MIMAT0000742 | Tgfbr1 | 21812 | HITS-CLIP | 23597149\|25083871 | Unspecified |
| mmu-miR-378a-5p | MIMAT0000742 | Thbs1 | 21825 | HITS-CLIP | 25083871 | Unspecified |
| mmu-miR-378a-5p | MIMAT0000742 | Thra | 21833 | HITS-CLIP | 37986224 | Unspecified |
| mmu-miR-378a-5p | MIMAT0000742 | Thrap3 | 230753 | HITS-CLIP | 25083871 | Unspecified |
| mmu-miR-378a-5p | MIMAT0000742 | Tigar | 319801 | HITS-CLIP | 25753418 | Unspecified |
| mmu-miR-378a-5p | MIMAT0000742 | Tiparp | 99929 | HITS-CLIP | 37986224 | Unspecified |
| mmu-miR-378a-5p | MIMAT0000742 | Tjp2 | 21873 | HITS-CLIP | 29167373 | Unspecified |
| mmu-miR-378a-5p | MIMAT0000742 | Tk1 | 21877 | HITS-CLIP | 37986224 | Unspecified |
| mmu-miR-378a-5p | MIMAT0000742 | Tlk1 | 228012 | HITS-CLIP | 37986224 | Unspecified |
| mmu-miR-378a-5p | MIMAT0000742 | Tln1 | 21894 | HITS-CLIP | 37986224\|23597149\|25753418\|29167 | Unspecified |
| mmu-miR-378a-5p | MIMAT0000742 | Tln2 | 70549 | HITS-CLIP | 37986224 | Unspecified |
| mmu-miR-378a-5p | MIMAT0000742 | Tm9sf1 | 74140 | HITS-CLIP | 37986224 | Unspecified |
| mmu-miR-378a-5p | MIMAT0000742 | Tm9sf3 | 107358 | HITS-CLIP | 37986224 | Unspecified |
| mmu-miR-378a-5p | MIMAT0000742 | Tmed4 | 103694 | HITS-CLIP | 37986224 | Unspecified |
| mmu-miR-378a-5p | MIMAT0000742 | Tmed8 | 382620 | HITS-CLIP | 37986224 | Unspecified |
| mmu-miR-378a-5p | MIMAT0000742 | Tmem131 | 56030 | HITS-CLIP | 37986224 | Unspecified |
| mmu-miR-378a-5p | MIMAT0000742 | Tmem165 | 21982 | HITS-CLIP | 25083871 | Unspecified |
| mmu-miR-378a-5p | MIMAT0000742 | Tmem168 | 101118 | HITS-CLIP | 25083871\|25753418 | Unspecified |
| mmu-miR-378a-5p | MIMAT0000742 | Tmem170b | 621976 | HITS-CLIP | 37986224 | Unspecified |
| mmu-miR-378a-5p | MIMAT0000742 | Tmem176a | 66058 | HITS-CLIP | 37986224 | Unspecified |
| mmu-miR-378a-5p | MIMAT0000742 | Tmem192 | 73067 | HITS-CLIP | 25083871 | Unspecified |
| mmu-miR-378a-5p | MIMAT0000742 | Tmem209 | 72649 | HITS-CLIP | 25083871 | Unspecified |
| mmu-miR-378a-5p | MIMAT0000742 | Tmem219 | 68742 | HITS-CLIP | 25083871 | Unspecified |
| mmu-miR-378a-5p | MIMAT0000742 | Tmem234 | 76799 | HITS-CLIP | 37986224 | Unspecified |
| mmu-miR-378a-5p | MIMAT0000742 | Tmem245 | 242474 | HITS-CLIP | 37986224\|25083871 | Unspecified |
| mmu-miR-378a-5p | MIMAT0000742 | Tmem248 | 71667 | HITS-CLIP | 37986224\|25083871 | Unspecified |
| mmu-miR-378a-5p | MIMAT0000742 | Tmem263 | 103266 | HITS-CLIP | 37986224 | Unspecified |
| mmu-miR-378a-5p | MIMAT0000742 | Tmem38b | 52076 | qCLASH | 31393953 | Unspecified |
| mmu-miR-378a-5p | MIMAT0000742 | Tmem64 | 100201 | HITS-CLIP | 37986224 | Unspecified |
| mmu-miR-378a-5p | MIMAT0000742 | Tmem8b | 242409 | HITS-CLIP | 25753418 | Unspecified |
| mmu-miR-378a-5p | MIMAT0000742 | Tmub2 | 72053 | HITS-CLIP | 37986224 | Unspecified |
| mmu-miR-378a-5p | MIMAT0000742 | Tmx3 | 67988 | HITS-CLIP | 37986224 | Unspecified |
| mmu-miR-378a-5p | MIMAT0000742 | Tnfaip1 | 21927 | HITS-CLIP | 25753418 | Unspecified |

| | | | | | | | |
|---|---|---|---|---|---|---|---|
| mmu-miR-378a-5p | MIMAT0000742 | Tnfaip2 | 21928 | HITS-CLIP | | 25083871 | Unspecified |
| mmu-miR-378a-5p | MIMAT0000742 | Tnfaip3 | 21929 | HITS-CLIP | | 25083871 | Unspecified |
| mmu-miR-378a-5p | MIMAT0000742 | Tnfrsf1b | 21938 | HITS-CLIP | | 23597149 | Unspecified |
| mmu-miR-378a-5p | MIMAT0000742 | Tnnt1 | 21955 | HITS-CLIP | | 25083871 | Unspecified |
| mmu-miR-378a-5p | MIMAT0000742 | Tnnt3 | 21957 | HITS-CLIP | | 25083871 | Unspecified |
| mmu-miR-378a-5p | MIMAT0000742 | Tnpo1 | 238799 | HITS-CLIP | | 37986224 | Unspecified |
| mmu-miR-378a-5p | MIMAT0000742 | Tnrc18 | 231861 | HITS-CLIP | | 29167373 | Unspecified |
| mmu-miR-378a-5p | MIMAT0000742 | Tnrc6b | 213988 | HITS-CLIP | 37986224\|25083871 | | Unspecified |
| mmu-miR-378a-5p | MIMAT0000742 | Tns3 | 319939 | HITS-CLIP | 37986224\|29167373 | | Unspecified |
| mmu-miR-378a-5p | MIMAT0000742 | Tob1 | 22057 | HITS-CLIP | 37986224\|25753418 | | Unspecified |
| mmu-miR-378a-5p | MIMAT0000742 | Tob2 | 57259 | HITS-CLIP | | 37986224 | Unspecified |
| mmu-miR-378a-5p | MIMAT0000742 | Togaram1 | 23116 | HITS-CLIP | | 25753418 | Unspecified |
| mmu-miR-378a-5p | MIMAT0000742 | Tonsl | 72749 | HITS-CLIP | | 37986224 | Unspecified |
| mmu-miR-378a-5p | MIMAT0000742 | Top1 | 21969 | HITS-CLIP | | 25083871 | Unspecified |
| mmu-miR-378a-5p | MIMAT0000742 | Topors | 106021 | HITS-CLIP | | 25753418 | Unspecified |
| mmu-miR-378a-5p | MIMAT0000742 | Tox4 | 268741 | HITS-CLIP | | 25083871 | Unspecified |
| mmu-miR-378a-5p | MIMAT0000742 | Tpcn1 | 252972 | HITS-CLIP | | 37986224 | Unspecified |
| mmu-miR-378a-5p | MIMAT0000742 | Tpp2 | 22019 | HITS-CLIP | 37986224\|25083871 | | Unspecified |
| mmu-miR-378a-5p | MIMAT0000742 | Tpr | 108989 | HITS-CLIP | 25083871\|29167373 | | Unspecified |
| mmu-miR-378a-5p | MIMAT0000742 | Trabd | 67976 | HITS-CLIP | | 37986224 | Unspecified |
| mmu-miR-378a-5p | MIMAT0000742 | Trappc13 | 66975 | HITS-CLIP | | 37986224 | Unspecified |
| mmu-miR-378a-5p | MIMAT0000742 | Trappc2 | 66226 | HITS-CLIP | | 37986224 | Unspecified |
| mmu-miR-378a-5p | MIMAT0000742 | Trib1 | 211770 | HITS-CLIP | 37986224\|29167373 | | Unspecified |
| mmu-miR-378a-5p | MIMAT0000742 | Trib3 | 228775 | HITS-CLIP | | 37986224 | Unspecified |
| mmu-miR-378a-5p | MIMAT0000742 | Trim35 | 66854 | HITS-CLIP | | 37986224 | Unspecified |
| mmu-miR-378a-5p | MIMAT0000742 | Trim39 | 79263 | HITS-CLIP | | 37986224 | Unspecified |
| mmu-miR-378a-5p | MIMAT0000742 | Trim56 | 384309 | HITS-CLIP | | 37986224 | Unspecified |
| mmu-miR-378a-5p | MIMAT0000742 | Trim72 | 434246 | HITS-CLIP | | 25083871 | Unspecified |
| mmu-miR-378a-5p | MIMAT0000742 | Trim8 | 93679 | HITS-CLIP | | 37986224 | Unspecified |
| mmu-miR-378a-5p | MIMAT0000742 | Trip11 | 109181 | HITS-CLIP | | 37986224 | Unspecified |
| mmu-miR-378a-5p | MIMAT0000742 | Trip6 | 22051 | HITS-CLIP | | 23597149 | Unspecified |
| mmu-miR-378a-5p | MIMAT0000742 | Trp53 | 22059 | HITS-CLIP | | 37986224 | Unspecified |
| mmu-miR-378a-5p | MIMAT0000742 | Trp53bp2 | 209456 | HITS-CLIP | | 37986224 | Unspecified |
| mmu-miR-378a-5p | MIMAT0000742 | Trp53inp2 | 68728 | HITS-CLIP | | 23597149 | Unspecified |
| mmu-miR-378a-5p | MIMAT0000742 | Trpc4ap | 56407 | HITS-CLIP | | 37986224 | Unspecified |
| mmu-miR-378a-5p | MIMAT0000742 | Tsc22d2 | 72033 | HITS-CLIP | | 37986224 | Unspecified |
| mmu-miR-378a-5p | MIMAT0000742 | Tsen2 | 381802 | HITS-CLIP | | 37986224 | Unspecified |
| mmu-miR-378a-5p | MIMAT0000742 | Tshz3 | 243931 | HITS-CLIP | | 37986224 | Unspecified |
| mmu-miR-378a-5p | MIMAT0000742 | Tspan9 | 109246 | HITS-CLIP | | 37986224 | Unspecified |

| | | | | | | | |
|---|---|---|---|---|---|---|---|
| mmu-miR-378a-5p | MIMAT0000742 | Tstd2 | 272027 | HITS-CLIP | | 37986224 | Unspecified |
| mmu-miR-378a-5p | MIMAT0000742 | Ttbk2 | 140810 | HITS-CLIP | | 37986224 | Unspecified |
| mmu-miR-378a-5p | MIMAT0000742 | Ttc28 | 209683 | HITS-CLIP | | 37986224 | Unspecified |
| mmu-miR-378a-5p | MIMAT0000742 | Ttc3 | 22129 | HITS-CLIP | | 25083871 | Unspecified |
| mmu-miR-378a-5p | MIMAT0000742 | Ttc39c | 72747 | HITS-CLIP | | 37986224 | Unspecified |
| mmu-miR-378a-5p | MIMAT0000742 | Ttn | 22138 | HITS-CLIP | | 37986224 | Unspecified |
| mmu-miR-378a-5p | MIMAT0000742 | Ttr | 22139 | HITS-CLIP | | 23597149 | Unspecified |
| mmu-miR-378a-5p | MIMAT0000742 | Txlna | 109658 | HITS-CLIP | | 37986224 | Unspecified |
| mmu-miR-378a-5p | MIMAT0000742 | Txnip | 56338 | HITS-CLIP | | 37986224 | Unspecified |
| mmu-miR-378a-5p | MIMAT0000742 | Tyms | 22171 | HITS-CLIP | | 37986224 | Unspecified |
| mmu-miR-378a-5p | MIMAT0000742 | U2surp | 67958 | HITS-CLIP | | 29167373 | Unspecified |
| mmu-miR-378a-5p | MIMAT0000742 | Uaca | 72565 | HITS-CLIP | 37986224\|25083871 | | Unspecified |
| mmu-miR-378a-5p | MIMAT0000742 | Uap1l1 | 227620 | HITS-CLIP | | 37986224 | Unspecified |
| mmu-miR-378a-5p | MIMAT0000742 | Uba6 | 231380 | HITS-CLIP | | 37986224 | Unspecified |
| mmu-miR-378a-5p | MIMAT0000742 | Ubap1 | 67123 | HITS-CLIP | 25083871\|23597149 | | Unspecified |
| mmu-miR-378a-5p | MIMAT0000742 | Ubap2l | 74383 | HITS-CLIP | 23597149\|29167373 | | Unspecified |
| mmu-miR-378a-5p | MIMAT0000742 | Ube2d2a | 56550 | HITS-CLIP | | 26203562 | Unspecified |
| mmu-miR-378a-5p | MIMAT0000742 | Ube2e2 | 218793 | HITS-CLIP | | 37986224 | Unspecified |
| mmu-miR-378a-5p | MIMAT0000742 | Ube2e3 | 22193 | HITS-CLIP | | 29167373 | Unspecified |
| mmu-miR-378a-5p | MIMAT0000742 | Ube2i | 22196 | HITS-CLIP | | 37986224 | Unspecified |
| mmu-miR-378a-5p | MIMAT0000742 | Ube2r2 | 67615 | HITS-CLIP | | 37986224 | Unspecified |
| mmu-miR-378a-5p | MIMAT0000742 | Ube2w | 66799 | HITS-CLIP | | 37986224 | Unspecified |
| mmu-miR-378a-5p | MIMAT0000742 | Ube4b | 63958 | HITS-CLIP | 37986224\|25753418 | | Unspecified |
| mmu-miR-378a-5p | MIMAT0000742 | Ubfd1 | 28018 | HITS-CLIP | | 37986224 | Unspecified |
| mmu-miR-378a-5p | MIMAT0000742 | Ubn1 | 170644 | HITS-CLIP | | 37986224 | Unspecified |
| mmu-miR-378a-5p | MIMAT0000742 | Ubn2 | 320538 | HITS-CLIP | 37986224\|25083871 | | Unspecified |
| mmu-miR-378a-5p | MIMAT0000742 | Ubr1 | 22222 | HITS-CLIP | | 37986224 | Unspecified |
| mmu-miR-378a-5p | MIMAT0000742 | Ubr2 | 224826 | HITS-CLIP | | 37986224 | Unspecified |
| mmu-miR-378a-5p | MIMAT0000742 | Ubr3 | 68795 | HITS-CLIP | | 23597149 | Unspecified |
| mmu-miR-378a-5p | MIMAT0000742 | Ubr4 | 69116 | HITS-CLIP | | 25083871 | Unspecified |
| mmu-miR-378a-5p | MIMAT0000742 | Ubtf | 21429 | HITS-CLIP | | 37986224 | Unspecified |
| mmu-miR-378a-5p | MIMAT0000742 | Ubxn1 | 225896 | HITS-CLIP | | 37986224 | Unspecified |
| mmu-miR-378a-5p | MIMAT0000742 | Ubxn7 | 224111 | HITS-CLIP | | 37986224 | Unspecified |
| mmu-miR-378a-5p | MIMAT0000742 | Uevld | 54122 | HITS-CLIP | | 37986224 | Unspecified |
| mmu-miR-378a-5p | MIMAT0000742 | Uggt1 | 320011 | HITS-CLIP | | 37986224 | Unspecified |
| mmu-miR-378a-5p | MIMAT0000742 | Uhrf1bp1 | 224648 | HITS-CLIP | | 37986224 | Unspecified |
| mmu-miR-378a-5p | MIMAT0000742 | Uhrf1bp1l | 75089 | HITS-CLIP | | 25083871 | Unspecified |
| mmu-miR-378a-5p | MIMAT0000742 | Uhrf2 | 109113 | HITS-CLIP | | 37986224 | Unspecified |
| mmu-miR-378a-5p | MIMAT0000742 | Umps | 22247 | HITS-CLIP | | 37986224 | Unspecified |

| | | | | | | |
|---|---|---|---|---|---|---|
| mmu-miR-378a-5p | MIMAT0000742 | Unc5b | 107449 | HITS-CLIP | 37986224 | Unspecified |
| mmu-miR-378a-5p | MIMAT0000742 | Upf1 | 19704 | HITS-CLIP | 25083871 | Unspecified |
| mmu-miR-378a-5p | MIMAT0000742 | Upp2 | 76654 | HITS-CLIP | 23597149 | Unspecified |
| mmu-miR-378a-5p | MIMAT0000742 | Uqcrfs1 | 66694 | HITS-CLIP | 37986224 | Unspecified |
| mmu-miR-378a-5p | MIMAT0000742 | Uso1 | 56041 | HITS-CLIP | 37986224 | Unspecified |
| mmu-miR-378a-5p | MIMAT0000742 | Usp13 | 72607 | HITS-CLIP | 25753418 | Unspecified |
| mmu-miR-378a-5p | MIMAT0000742 | Usp15 | 14479 | HITS-CLIP | 25083871 | Unspecified |
| mmu-miR-378a-5p | MIMAT0000742 | Usp22 | 216825 | HITS-CLIP | 25083871 | Unspecified |
| mmu-miR-378a-5p | MIMAT0000742 | Usp24 | 329908 | HITS-CLIP | 37986224 | Unspecified |
| mmu-miR-378a-5p | MIMAT0000742 | Usp34 | 17847 | HITS-CLIP | 37986224\|25083871\|23597149 | Unspecified |
| mmu-miR-378a-5p | MIMAT0000742 | Usp38 | 74841 | HITS-CLIP | 25083871 | Unspecified |
| mmu-miR-378a-5p | MIMAT0000742 | Usp4 | 22258 | HITS-CLIP | 37986224 | Unspecified |
| mmu-miR-378a-5p | MIMAT0000742 | Usp42 | 76800 | HITS-CLIP | 37986224 | Unspecified |
| mmu-miR-378a-5p | MIMAT0000742 | Usp45 | 77593 | HITS-CLIP | 37986224 | Unspecified |
| mmu-miR-378a-5p | MIMAT0000742 | Usp48 | 170707 | HITS-CLIP | 37986224 | Unspecified |
| mmu-miR-378a-5p | MIMAT0000742 | Vars | 22321 | HITS-CLIP | 37986224 | Unspecified |
| mmu-miR-378a-5p | MIMAT0000742 | Vav3 | 57257 | HITS-CLIP | 37986224 | Unspecified |
| mmu-miR-378a-5p | MIMAT0000742 | Vcam1 | 22329 | HITS-CLIP | 37986224 | Unspecified |
| mmu-miR-378a-5p | MIMAT0000742 | Vcan | 13003 | HITS-CLIP | 37986224 | Unspecified |
| mmu-miR-378a-5p | MIMAT0000742 | Vegfa | 22339 | HITS-CLIP | 25083871 | Unspecified |
| mmu-miR-378a-5p | MIMAT0000742 | Vgll3 | 73569 | HITS-CLIP | 37986224 | Unspecified |
| mmu-miR-378a-5p | MIMAT0000742 | Vhl | 22346 | HITS-CLIP | 37986224 | Unspecified |
| mmu-miR-378a-5p | MIMAT0000742 | Vps26a | 30930 | HITS-CLIP | 37986224 | Unspecified |
| mmu-miR-378a-5p | MIMAT0000742 | Vps26b | 69091 | HITS-CLIP | 37986224 | Unspecified |
| mmu-miR-378a-5p | MIMAT0000742 | Vps33a | 77573 | HITS-CLIP | 37986224 | Unspecified |
| mmu-miR-378a-5p | MIMAT0000742 | Vps37a | 52348 | HITS-CLIP | 37986224 | Unspecified |
| mmu-miR-378a-5p | MIMAT0000742 | Vps37c | 107305 | HITS-CLIP | 37986224 | Unspecified |
| mmu-miR-378a-5p | MIMAT0000742 | Vps4b | 20479 | HITS-CLIP | 37986224 | Unspecified |
| mmu-miR-378a-5p | MIMAT0000742 | Vtn | 22370 | HITS-CLIP | 23597149 | Unspecified |
| mmu-miR-378a-5p | MIMAT0000742 | Wasf2 | 242687 | HITS-CLIP | 25083871 | Unspecified |
| mmu-miR-378a-5p | MIMAT0000742 | Washc4 | 319277 | HITS-CLIP | 25083871 | Unspecified |
| mmu-miR-378a-5p | MIMAT0000742 | Wasl | 73178 | HITS-CLIP | 37986224 | Unspecified |
| mmu-miR-378a-5p | MIMAT0000742 | Wbp11 | 60321 | HITS-CLIP | 25083871 | Unspecified |
| mmu-miR-378a-5p | MIMAT0000742 | Wdr26 | 226757 | HITS-CLIP | 29167373\|25083871 | Unspecified |
| mmu-miR-378a-5p | MIMAT0000742 | Wdr82 | 77305 | HITS-CLIP | 23597149 | Unspecified |
| mmu-miR-378a-5p | MIMAT0000742 | Wdr91 | 101240 | HITS-CLIP | 29167373 | Unspecified |
| mmu-miR-378a-5p | MIMAT0000742 | Wipi2 | 74781 | HITS-CLIP | 37986224 | Unspecified |
| mmu-miR-378a-5p | MIMAT0000742 | Wnk1 | 232341 | HITS-CLIP | 25083871 | Unspecified |
| mmu-miR-378a-5p | MIMAT0000742 | Wrb | 71446 | HITS-CLIP | 37986224 | Unspecified |

| | | | | | | |
|---|---|---|---|---|---|---|
| mmu-miR-378a-5p | MIMAT0000742 | Wsb1 | 78889 | HITS-CLIP | 37986224\|25083871 | Unspecified |
| mmu-miR-378a-5p | MIMAT0000742 | Wwc1 | 211652 | HITS-CLIP | 37986224 | Unspecified |
| mmu-miR-378a-5p | MIMAT0000742 | Xbp1 | 22433 | HITS-CLIP | 37986224 | Unspecified |
| mmu-miR-378a-5p | MIMAT0000742 | Ylpm1 | 56531 | HITS-CLIP | 37986224 | Unspecified |
| mmu-miR-378a-5p | MIMAT0000742 | Ythdc1 | 231386 | HITS-CLIP | 37986224 | Unspecified |
| mmu-miR-378a-5p | MIMAT0000742 | Zbed3 | 72114 | HITS-CLIP | 25083871 | Unspecified |
| mmu-miR-378a-5p | MIMAT0000742 | Zbtb10 | 229055 | HITS-CLIP | 25083871 | Unspecified |
| mmu-miR-378a-5p | MIMAT0000742 | Zbtb16 | 235320 | HITS-CLIP | 25753418 | Unspecified |
| mmu-miR-378a-5p | MIMAT0000742 | Zbtb18 | 30928 | HITS-CLIP | 23597149 | Unspecified |
| mmu-miR-378a-5p | MIMAT0000742 | Zbtb20 | 56490 | HITS-CLIP | 37986224\|25083871 | Unspecified |
| mmu-miR-378a-5p | MIMAT0000742 | Zbtb38 | 245007 | HITS-CLIP | 25083871 | Unspecified |
| mmu-miR-378a-5p | MIMAT0000742 | Zbtb4 | 75580 | HITS-CLIP | 25083871 | Unspecified |
| mmu-miR-378a-5p | MIMAT0000742 | Zbtb44 | 235132 | HITS-CLIP | 37986224 | Unspecified |
| mmu-miR-378a-5p | MIMAT0000742 | Zbtb6 | 241322 | HITS-CLIP | 37986224 | Unspecified |
| mmu-miR-378a-5p | MIMAT0000742 | Zbtb7a | 16969 | HITS-CLIP | 25083871 | Unspecified |
| mmu-miR-378a-5p | MIMAT0000742 | Zbtb7b | 22724 | HITS-CLIP | 37986224 | Unspecified |
| mmu-miR-378a-5p | MIMAT0000742 | Zc3h10 | 103284 | HITS-CLIP | 37986224 | Unspecified |
| mmu-miR-378a-5p | MIMAT0000742 | Zc3h12c | 244871 | HITS-CLIP | 37986224 | Unspecified |
| mmu-miR-378a-5p | MIMAT0000742 | Zc3h13 | 67302 | HITS-CLIP | 25083871 | Unspecified |
| mmu-miR-378a-5p | MIMAT0000742 | Zc3h14 | 75553 | HITS-CLIP | 25083871 | Unspecified |
| mmu-miR-378a-5p | MIMAT0000742 | Zdhhc21 | 68268 | HITS-CLIP | 37986224\|25083871 | Unspecified |
| mmu-miR-378a-5p | MIMAT0000742 | Zdhhc3 | 69035 | HITS-CLIP | 37986224 | Unspecified |
| mmu-miR-378a-5p | MIMAT0000742 | Zeb1 | 21417 | HITS-CLIP | 25083871\|25753418 | Unspecified |
| mmu-miR-378a-5p | MIMAT0000742 | Zfand3 | 21769 | HITS-CLIP | 37986224 | Unspecified |
| mmu-miR-378a-5p | MIMAT0000742 | Zfhx3 | 11906 | HITS-CLIP | 25083871 | Unspecified |
| mmu-miR-378a-5p | MIMAT0000742 | Zfhx4 | 80892 | HITS-CLIP | 25083871 | Unspecified |
| mmu-miR-378a-5p | MIMAT0000742 | Zfp106 | 20402 | HITS-CLIP | 37986224\|25083871 | Unspecified |
| mmu-miR-378a-5p | MIMAT0000742 | Zfp12 | 231866 | HITS-CLIP | 37986224 | Unspecified |
| mmu-miR-378a-5p | MIMAT0000742 | Zfp142 | 77264 | HITS-CLIP | 37986224 | Unspecified |
| mmu-miR-378a-5p | MIMAT0000742 | Zfp180 | 210135 | HITS-CLIP | 37986224 | Unspecified |
| mmu-miR-378a-5p | MIMAT0000742 | Zfp236 | 329002 | HITS-CLIP | 37986224\|25083871 | Unspecified |
| mmu-miR-378a-5p | MIMAT0000742 | Zfp266 | 77519 | HITS-CLIP | 37986224 | Unspecified |
| mmu-miR-378a-5p | MIMAT0000742 | Zfp280c | 208968 | HITS-CLIP | 37986224 | Unspecified |
| mmu-miR-378a-5p | MIMAT0000742 | Zfp280d | 235469 | HITS-CLIP | 37986224 | Unspecified |
| mmu-miR-378a-5p | MIMAT0000742 | Zfp281 | 226442 | HITS-CLIP | 25083871 | Unspecified |
| mmu-miR-378a-5p | MIMAT0000742 | Zfp292 | 30046 | HITS-CLIP | 37986224 | Unspecified |
| mmu-miR-378a-5p | MIMAT0000742 | Zfp318 | 57908 | HITS-CLIP | 37986224 | Unspecified |
| mmu-miR-378a-5p | MIMAT0000742 | Zfp319 | 79233 | HITS-CLIP | 37986224 | Unspecified |
| mmu-miR-378a-5p | MIMAT0000742 | Zfp322a | 218100 | HITS-CLIP | 37986224 | Unspecified |

| | | | | | | |
|---|---|---|---|---|---|---|
| mmu-miR-378a-5p | MIMAT0000742 | Zfp334 | 228876 | HITS-CLIP | 37986224 | Unspecified |
| mmu-miR-378a-5p | MIMAT0000742 | Zfp346 | 26919 | HITS-CLIP | 37986224 | Unspecified |
| mmu-miR-378a-5p | MIMAT0000742 | Zfp367 | 238673 | HITS-CLIP | 37986224 | Unspecified |
| mmu-miR-378a-5p | MIMAT0000742 | Zfp36l1 | 12192 | HITS-CLIP | 37986224 | Unspecified |
| mmu-miR-378a-5p | MIMAT0000742 | Zfp397 | 69256 | HITS-CLIP | 37986224 | Unspecified |
| mmu-miR-378a-5p | MIMAT0000742 | Zfp40 | 22700 | HITS-CLIP | 37986224 | Unspecified |
| mmu-miR-378a-5p | MIMAT0000742 | Zfp451 | 98403 | HITS-CLIP | 37986224 | Unspecified |
| mmu-miR-378a-5p | MIMAT0000742 | Zfp458 | 238690 | HITS-CLIP | 37986224 | Unspecified |
| mmu-miR-378a-5p | MIMAT0000742 | Zfp507 | 668501 | HITS-CLIP | 37986224 | Unspecified |
| mmu-miR-378a-5p | MIMAT0000742 | Zfp516 | 329003 | HITS-CLIP | 37986224 | Unspecified |
| mmu-miR-378a-5p | MIMAT0000742 | Zfp553 | 233887 | HITS-CLIP | 37986224 | Unspecified |
| mmu-miR-378a-5p | MIMAT0000742 | Zfp57 | 22715 | HITS-CLIP | 37986224 | Unspecified |
| mmu-miR-378a-5p | MIMAT0000742 | Zfp606 | 67370 | HITS-CLIP | 37986224 | Unspecified |
| mmu-miR-378a-5p | MIMAT0000742 | Zfp608 | 269023 | HITS-CLIP | 37986224 | Unspecified |
| mmu-miR-378a-5p | MIMAT0000742 | Zfp609 | 214812 | HITS-CLIP | 37986224 | Unspecified |
| mmu-miR-378a-5p | MIMAT0000742 | Zfp617 | 170938 | HITS-CLIP | 37986224 | Unspecified |
| mmu-miR-378a-5p | MIMAT0000742 | Zfp652 | 268469 | HITS-CLIP | 37986224\|25083871 | Unspecified |
| mmu-miR-378a-5p | MIMAT0000742 | Zfp760 | 240034 | HITS-CLIP | 37986224 | Unspecified |
| mmu-miR-378a-5p | MIMAT0000742 | Zfp770 | 228491 | HITS-CLIP | 37986224 | Unspecified |
| mmu-miR-378a-5p | MIMAT0000742 | Zfp821 | 75871 | HITS-CLIP | 25083871 | Unspecified |
| mmu-miR-378a-5p | MIMAT0000742 | Zfpl1 | 81909 | HITS-CLIP | 23597149 | Unspecified |
| mmu-miR-378a-5p | MIMAT0000742 | Zhx1 | 22770 | HITS-CLIP | 25083871 | Unspecified |
| mmu-miR-378a-5p | MIMAT0000742 | Zim1 | 22776 | HITS-CLIP | 37986224 | Unspecified |
| mmu-miR-378a-5p | MIMAT0000742 | Zkscan1 | 74570 | HITS-CLIP | 37986224\|25083871 | Unspecified |
| mmu-miR-378a-5p | MIMAT0000742 | Zkscan3 | 72739 | HITS-CLIP | 37986224 | Unspecified |
| mmu-miR-378a-5p | MIMAT0000742 | Zkscan7 | 382118 | HITS-CLIP | 37986224 | Unspecified |
| mmu-miR-378a-5p | MIMAT0000742 | Zkscan8 | 93681 | HITS-CLIP | 37986224 | Unspecified |
| mmu-miR-378a-5p | MIMAT0000742 | Zmynd8 | 228880 | HITS-CLIP | 37986224 | Unspecified |
| mmu-miR-378a-5p | MIMAT0000742 | Zranb1 | 360216 | HITS-CLIP | 37986224 | Unspecified |
| mmu-miR-378a-5p | MIMAT0000742 | Zscan22 | 232878 | HITS-CLIP | 37986224 | Unspecified |
| mmu-miR-378a-5p | MIMAT0000742 | Zscan29 | 99334 | HITS-CLIP | 37986224 | Unspecified |
| mmu-miR-378a-5p | MIMAT0000742 | Zswim6 | 67263 | HITS-CLIP | 37986224 | Unspecified |
| mmu-miR-378a-5p | MIMAT0000742 | Zyg11b | 414872 | HITS-CLIP | 37986224 | Unspecified |
| mmu-miR-5620-3p | MIMAT0022368 | 2310033P09Rik | 67862 | HITS-CLIP | 37986224 | Unspecified |
| mmu-miR-5620-3p | MIMAT0022368 | Aacs | 78894 | HITS-CLIP | 37986224 | Unspecified |
| mmu-miR-5620-3p | MIMAT0022368 | Aadat | 23923 | HITS-CLIP | 23597149 | Unspecified |
| mmu-miR-5620-3p | MIMAT0022368 | Abat | 268860 | HITS-CLIP | 37986224 | Unspecified |
| mmu-miR-5620-3p | MIMAT0022368 | Abca2 | 11305 | HITS-CLIP | 23597149 | Unspecified |
| mmu-miR-5620-3p | MIMAT0022368 | Abhd15 | 67477 | HITS-CLIP | 37986224 | Unspecified |

| | | | | | | | |
|---|---|---|---|---|---|---|---|
| mmu-miR-5620-3p | MIMAT0022368 | Abi1 | 11308 | HITS-CLIP | | 37986224 | Unspecified |
| mmu-miR-5620-3p | MIMAT0022368 | Acaa2 | 52538 | HITS-CLIP | | 37986224 | Unspecified |
| mmu-miR-5620-3p | MIMAT0022368 | Acat1 | 110446 | HITS-CLIP | 37986224\|23597149 | | Unspecified |
| mmu-miR-5620-3p | MIMAT0022368 | Acat2 | 110460 | HITS-CLIP | | 37986224 | Unspecified |
| mmu-miR-5620-3p | MIMAT0022368 | Acbd5 | 74159 | HITS-CLIP | | 37986224 | Unspecified |
| mmu-miR-5620-3p | MIMAT0022368 | Acox1 | 11430 | HITS-CLIP | | 37986224 | Unspecified |
| mmu-miR-5620-3p | MIMAT0022368 | Adcy6 | 11512 | HITS-CLIP | | 23597149 | Unspecified |
| mmu-miR-5620-3p | MIMAT0022368 | Adipor1 | 72674 | HITS-CLIP | | 37986224 | Unspecified |
| mmu-miR-5620-3p | MIMAT0022368 | Agtr1a | 11607 | HITS-CLIP | | 37986224 | Unspecified |
| mmu-miR-5620-3p | MIMAT0022368 | Ahr | 11622 | HITS-CLIP | 37986224\|23597149 | | Unspecified |
| mmu-miR-5620-3p | MIMAT0022368 | Akirin2 | 433693 | HITS-CLIP | | 23597149 | Unspecified |
| mmu-miR-5620-3p | MIMAT0022368 | Aldh1a7 | 26358 | HITS-CLIP | | 37986224 | Unspecified |
| mmu-miR-5620-3p | MIMAT0022368 | Aldh4a1 | 212647 | HITS-CLIP | | 37986224 | Unspecified |
| mmu-miR-5620-3p | MIMAT0022368 | Amfr | 23802 | HITS-CLIP | | 23597149 | Unspecified |
| mmu-miR-5620-3p | MIMAT0022368 | Ankrd44 | 329154 | HITS-CLIP | | 23597149 | Unspecified |
| mmu-miR-5620-3p | MIMAT0022368 | Anpep | 16790 | HITS-CLIP | | 23597149 | Unspecified |
| mmu-miR-5620-3p | MIMAT0022368 | Ap1b1 | 11764 | HITS-CLIP | 37986224\|23597149 | | Unspecified |
| mmu-miR-5620-3p | MIMAT0022368 | Apob | 238055 | HITS-CLIP | 37986224\|23597149 | | Unspecified |
| mmu-miR-5620-3p | MIMAT0022368 | Arf3 | 11842 | HITS-CLIP | | 23597149 | Unspecified |
| mmu-miR-5620-3p | MIMAT0022368 | Arg1 | 11846 | HITS-CLIP | | 37986224 | Unspecified |
| mmu-miR-5620-3p | MIMAT0022368 | Arl4a | 11861 | HITS-CLIP | | 37986224 | Unspecified |
| mmu-miR-5620-3p | MIMAT0022368 | Armc1 | 74252 | HITS-CLIP | | 23597149 | Unspecified |
| mmu-miR-5620-3p | MIMAT0022368 | Arrb2 | 216869 | HITS-CLIP | | 37986224 | Unspecified |
| mmu-miR-5620-3p | MIMAT0022368 | Atf5 | 107503 | HITS-CLIP | | 23597149 | Unspecified |
| mmu-miR-5620-3p | MIMAT0022368 | Atg5 | 11793 | HITS-CLIP | | 37986224 | Unspecified |
| mmu-miR-5620-3p | MIMAT0022368 | Atp13a3 | 224088 | HITS-CLIP | | 23597149 | Unspecified |
| mmu-miR-5620-3p | MIMAT0022368 | Atp1a1 | 11928 | HITS-CLIP | | 37986224 | Unspecified |
| mmu-miR-5620-3p | MIMAT0022368 | Atp5j2 | 57423 | HITS-CLIP | | 37986224 | Unspecified |
| mmu-miR-5620-3p | MIMAT0022368 | Bcar3 | 29815 | HITS-CLIP | | 23597149 | Unspecified |
| mmu-miR-5620-3p | MIMAT0022368 | Bcl2l1 | 12048 | HITS-CLIP | | 37986224 | Unspecified |
| mmu-miR-5620-3p | MIMAT0022368 | Bcr | 110279 | HITS-CLIP | | 37986224 | Unspecified |
| mmu-miR-5620-3p | MIMAT0022368 | Birc6 | 12211 | HITS-CLIP | | 23597149 | Unspecified |
| mmu-miR-5620-3p | MIMAT0022368 | Brd2 | 14312 | HITS-CLIP | | 37986224 | Unspecified |
| mmu-miR-5620-3p | MIMAT0022368 | C4bp | 12269 | HITS-CLIP | | 37986224 | Unspecified |
| mmu-miR-5620-3p | MIMAT0022368 | Calm3 | 12315 | HITS-CLIP | | 37986224 | Unspecified |
| mmu-miR-5620-3p | MIMAT0022368 | Canx | 12330 | HITS-CLIP | | 37986224 | Unspecified |
| mmu-miR-5620-3p | MIMAT0022368 | Cct2 | 12461 | HITS-CLIP | | 37986224 | Unspecified |
| mmu-miR-5620-3p | MIMAT0022368 | Cct5 | 12465 | HITS-CLIP | 37986224\|23597149 | | Unspecified |
| mmu-miR-5620-3p | MIMAT0022368 | Cdc34 | 216150 | HITS-CLIP | | 37986224 | Unspecified |

| | | | | | | |
|---|---|---|---|---|---|---|
| mmu-miR-5620-3p | MIMAT0022368 | Cdc42 | 12540 | HITS-CLIP | 37986224 | Unspecified |
| mmu-miR-5620-3p | MIMAT0022368 | Cdk16 | 18555 | HITS-CLIP | 37986224\|23597149 | Unspecified |
| mmu-miR-5620-3p | MIMAT0022368 | Cep350 | 74081 | HITS-CLIP | 37986224 | Unspecified |
| mmu-miR-5620-3p | MIMAT0022368 | Ces1f | 234564 | HITS-CLIP | 37986224 | Unspecified |
| mmu-miR-5620-3p | MIMAT0022368 | Chdh | 218865 | HITS-CLIP | 23597149 | Unspecified |
| mmu-miR-5620-3p | MIMAT0022368 | Chuk | 12675 | HITS-CLIP | 23597149 | Unspecified |
| mmu-miR-5620-3p | MIMAT0022368 | Clcn3 | 12725 | HITS-CLIP | 37986224 | Unspecified |
| mmu-miR-5620-3p | MIMAT0022368 | Cldn2 | 12738 | HITS-CLIP | 37986224 | Unspecified |
| mmu-miR-5620-3p | MIMAT0022368 | Clpx | 270166 | HITS-CLIP | 23597149 | Unspecified |
| mmu-miR-5620-3p | MIMAT0022368 | Cmbl | 69574 | HITS-CLIP | 37986224 | Unspecified |
| mmu-miR-5620-3p | MIMAT0022368 | Cnbp | 12785 | HITS-CLIP | 37986224 | Unspecified |
| mmu-miR-5620-3p | MIMAT0022368 | Coq9 | 67914 | HITS-CLIP | 23597149 | Unspecified |
| mmu-miR-5620-3p | MIMAT0022368 | Coro1b | 23789 | HITS-CLIP | 23597149 | Unspecified |
| mmu-miR-5620-3p | MIMAT0022368 | Creld1 | 171508 | HITS-CLIP | 23597149 | Unspecified |
| mmu-miR-5620-3p | MIMAT0022368 | Crk | 12928 | HITS-CLIP | 37986224 | Unspecified |
| mmu-miR-5620-3p | MIMAT0022368 | Crlf3 | 54394 | HITS-CLIP | 37986224 | Unspecified |
| mmu-miR-5620-3p | MIMAT0022368 | Csnk1d | 104318 | HITS-CLIP | 23597149 | Unspecified |
| mmu-miR-5620-3p | MIMAT0022368 | Ctsd | 13033 | HITS-CLIP | 37986224 | Unspecified |
| mmu-miR-5620-3p | MIMAT0022368 | Cux1 | 13047 | HITS-CLIP | 23597149 | Unspecified |
| mmu-miR-5620-3p | MIMAT0022368 | Cxcl12 | 20315 | HITS-CLIP | 37986224 | Unspecified |
| mmu-miR-5620-3p | MIMAT0022368 | Cyb561a3 | 225912 | HITS-CLIP | 37986224 | Unspecified |
| mmu-miR-5620-3p | MIMAT0022368 | Cyp1a2 | 13077 | HITS-CLIP | 23597149 | Unspecified |
| mmu-miR-5620-3p | MIMAT0022368 | Cyp2a5 | 13087 | HITS-CLIP | 37986224 | Unspecified |
| mmu-miR-5620-3p | MIMAT0022368 | Cyp2e1 | 13106 | HITS-CLIP | 23597149 | Unspecified |
| mmu-miR-5620-3p | MIMAT0022368 | Cyp4a12a | 277753 | HITS-CLIP | 37986224 | Unspecified |
| mmu-miR-5620-3p | MIMAT0022368 | Dars | 226414 | HITS-CLIP | 37986224 | Unspecified |
| mmu-miR-5620-3p | MIMAT0022368 | Dcaf7 | 71833 | HITS-CLIP | 37986224 | Unspecified |
| mmu-miR-5620-3p | MIMAT0022368 | Ddx3x | 13205 | HITS-CLIP | 23597149 | Unspecified |
| mmu-miR-5620-3p | MIMAT0022368 | Ddx3y | 26900 | HITS-CLIP | 37986224 | Unspecified |
| mmu-miR-5620-3p | MIMAT0022368 | Ddx5 | 13207 | HITS-CLIP | 23597149 | Unspecified |
| mmu-miR-5620-3p | MIMAT0022368 | Dhcr24 | 74754 | HITS-CLIP | 37986224 | Unspecified |
| mmu-miR-5620-3p | MIMAT0022368 | Dhx37 | 208144 | HITS-CLIP | 37986224 | Unspecified |
| mmu-miR-5620-3p | MIMAT0022368 | Dmd | 13405 | HITS-CLIP | 37986224 | Unspecified |
| mmu-miR-5620-3p | MIMAT0022368 | Dnajc16 | 214063 | HITS-CLIP | 37986224 | Unspecified |
| mmu-miR-5620-3p | MIMAT0022368 | Dynll2 | 68097 | HITS-CLIP | 37986224\|23597149 | Unspecified |
| mmu-miR-5620-3p | MIMAT0022368 | Edc3 | 353190 | HITS-CLIP | 37986224\|23597149 | Unspecified |
| mmu-miR-5620-3p | MIMAT0022368 | Efna1 | 13636 | HITS-CLIP | 37986224 | Unspecified |
| mmu-miR-5620-3p | MIMAT0022368 | Ehmt2 | 110147 | HITS-CLIP | 37986224 | Unspecified |
| mmu-miR-5620-3p | MIMAT0022368 | Eif2b2 | 217715 | HITS-CLIP | 37986224 | Unspecified |

| | | | | | | |
|---|---|---|---|---|---|---|
| mmu-miR-5620-3p | MIMAT0022368 | Eif4g1 | 208643 | HITS-CLIP | 37986224\|23597149 | Unspecified |
| mmu-miR-5620-3p | MIMAT0022368 | Elp2 | 58523 | HITS-CLIP | 23597149 | Unspecified |
| mmu-miR-5620-3p | MIMAT0022368 | Epas1 | 13819 | HITS-CLIP | 23597149 | Unspecified |
| mmu-miR-5620-3p | MIMAT0022368 | Ephx2 | 13850 | HITS-CLIP | 23597149 | Unspecified |
| mmu-miR-5620-3p | MIMAT0022368 | Erp44 | 76299 | HITS-CLIP | 37986224\|23597149 | Unspecified |
| mmu-miR-5620-3p | MIMAT0022368 | Esrp2 | 77411 | HITS-CLIP | 37986224\|23597149 | Unspecified |
| mmu-miR-5620-3p | MIMAT0022368 | Etfb | 110826 | HITS-CLIP | 23597149 | Unspecified |
| mmu-miR-5620-3p | MIMAT0022368 | Exoc6 | 107371 | HITS-CLIP | 23597149 | Unspecified |
| mmu-miR-5620-3p | MIMAT0022368 | Exosc7 | 66446 | HITS-CLIP | 37986224 | Unspecified |
| mmu-miR-5620-3p | MIMAT0022368 | Faah | 14073 | HITS-CLIP | 23597149 | Unspecified |
| mmu-miR-5620-3p | MIMAT0022368 | Fam76b | 72826 | HITS-CLIP | 37986224 | Unspecified |
| mmu-miR-5620-3p | MIMAT0022368 | Fbf1 | 217335 | HITS-CLIP | 37986224 | Unspecified |
| mmu-miR-5620-3p | MIMAT0022368 | Flad1 | 319945 | HITS-CLIP | 37986224 | Unspecified |
| mmu-miR-5620-3p | MIMAT0022368 | Flt4 | 14257 | HITS-CLIP | 23597149 | Unspecified |
| mmu-miR-5620-3p | MIMAT0022368 | Fn1 | 14268 | HITS-CLIP | 37986224 | Unspecified |
| mmu-miR-5620-3p | MIMAT0022368 | Fndc3a | 319448 | HITS-CLIP | 37986224 | Unspecified |
| mmu-miR-5620-3p | MIMAT0022368 | Foxj3 | 230700 | HITS-CLIP | 37986224 | Unspecified |
| mmu-miR-5620-3p | MIMAT0022368 | Foxn3 | 71375 | HITS-CLIP | 23597149 | Unspecified |
| mmu-miR-5620-3p | MIMAT0022368 | Fuom | 69064 | HITS-CLIP | 37986224 | Unspecified |
| mmu-miR-5620-3p | MIMAT0022368 | Gbf1 | 107338 | HITS-CLIP | 23597149 | Unspecified |
| mmu-miR-5620-3p | MIMAT0022368 | Gcc2 | 70297 | HITS-CLIP | 37986224 | Unspecified |
| mmu-miR-5620-3p | MIMAT0022368 | Gcsh | 68133 | HITS-CLIP | 37986224 | Unspecified |
| mmu-miR-5620-3p | MIMAT0022368 | Ghr | 14600 | HITS-CLIP | 37986224 | Unspecified |
| mmu-miR-5620-3p | MIMAT0022368 | Glyctk | 235582 | HITS-CLIP | 37986224 | Unspecified |
| mmu-miR-5620-3p | MIMAT0022368 | Gnas | 14683 | HITS-CLIP | 37986224 | Unspecified |
| mmu-miR-5620-3p | MIMAT0022368 | Gnb1 | 14688 | HITS-CLIP | 37986224 | Unspecified |
| mmu-miR-5620-3p | MIMAT0022368 | Gorasp2 | 70231 | HITS-CLIP | 23597149 | Unspecified |
| mmu-miR-5620-3p | MIMAT0022368 | Got1 | 14718 | HITS-CLIP | 37986224 | Unspecified |
| mmu-miR-5620-3p | MIMAT0022368 | Gpd1l | 333433 | HITS-CLIP | 37986224 | Unspecified |
| mmu-miR-5620-3p | MIMAT0022368 | Grpel1 | 17713 | HITS-CLIP | 23597149 | Unspecified |
| mmu-miR-5620-3p | MIMAT0022368 | Gtpbp1 | 14904 | HITS-CLIP | 23597149 | Unspecified |
| mmu-miR-5620-3p | MIMAT0022368 | H2-Aa | 14960 | HITS-CLIP | 37986224 | Unspecified |
| mmu-miR-5620-3p | MIMAT0022368 | H2-Ab1 | 14961 | HITS-CLIP | 37986224 | Unspecified |
| mmu-miR-5620-3p | MIMAT0022368 | H2bc13 | 8340 | HITS-CLIP | 31433990 | Unspecified |
| mmu-miR-5620-3p | MIMAT0022368 | Habp2 | 226243 | HITS-CLIP | 37986224 | Unspecified |
| mmu-miR-5620-3p | MIMAT0022368 | Hacl1 | 56794 | HITS-CLIP | 23597149 | Unspecified |
| mmu-miR-5620-3p | MIMAT0022368 | Hadha | 97212 | HITS-CLIP | 23597149 | Unspecified |
| mmu-miR-5620-3p | MIMAT0022368 | Hagh | 14651 | HITS-CLIP | 23597149 | Unspecified |
| mmu-miR-5620-3p | MIMAT0022368 | Hbs1l | 56422 | HITS-CLIP | 23597149 | Unspecified |

| | | | | | | | |
|---|---|---|---|---|---|---|---|
| mmu-miR-5620-3p | MIMAT0022368 | Hcfc1 | 15161 | HITS-CLIP | | 37986224 | Unspecified |
| mmu-miR-5620-3p | MIMAT0022368 | Hdgfl2 | 84717 | HITS-CLIP | | 23597149 | Unspecified |
| mmu-miR-5620-3p | MIMAT0022368 | Helz | 78455 | HITS-CLIP | | 23597149 | Unspecified |
| mmu-miR-5620-3p | MIMAT0022368 | Hes1 | 15205 | HITS-CLIP | | 23597149 | Unspecified |
| mmu-miR-5620-3p | MIMAT0022368 | Hibadh | 58875 | HITS-CLIP | | 37986224 | Unspecified |
| mmu-miR-5620-3p | MIMAT0022368 | Hif1an | 319594 | HITS-CLIP | | 37986224 | Unspecified |
| mmu-miR-5620-3p | MIMAT0022368 | Hnrnpl | 15388 | HITS-CLIP | | 23597149 | Unspecified |
| mmu-miR-5620-3p | MIMAT0022368 | Hoga1 | 67432 | HITS-CLIP | | 23597149 | Unspecified |
| mmu-miR-5620-3p | MIMAT0022368 | Hsp90ab1 | 15516 | HITS-CLIP | | 23597149 | Unspecified |
| mmu-miR-5620-3p | MIMAT0022368 | Hspa5 | 14828 | HITS-CLIP | | 23597149 | Unspecified |
| mmu-miR-5620-3p | MIMAT0022368 | Hspa9 | 15526 | HITS-CLIP | | 37986224 | Unspecified |
| mmu-miR-5620-3p | MIMAT0022368 | Huwe1 | 59026 | HITS-CLIP | | 37986224 | Unspecified |
| mmu-miR-5620-3p | MIMAT0022368 | Hyal2 | 15587 | HITS-CLIP | | 37986224 | Unspecified |
| mmu-miR-5620-3p | MIMAT0022368 | Hyou1 | 12282 | HITS-CLIP | | 37986224 | Unspecified |
| mmu-miR-5620-3p | MIMAT0022368 | Igfbp2 | 16008 | HITS-CLIP | | 23597149 | Unspecified |
| mmu-miR-5620-3p | MIMAT0022368 | Iigp1 | 60440 | HITS-CLIP | | 37986224 | Unspecified |
| mmu-miR-5620-3p | MIMAT0022368 | Il1rap | 16180 | HITS-CLIP | | 23597149 | Unspecified |
| mmu-miR-5620-3p | MIMAT0022368 | Il4ra | 16190 | HITS-CLIP | | 37986224 | Unspecified |
| mmu-miR-5620-3p | MIMAT0022368 | Irf2bp2 | 270110 | HITS-CLIP | | 23597149 | Unspecified |
| mmu-miR-5620-3p | MIMAT0022368 | Itgb1 | 16412 | HITS-CLIP | | 23597149 | Unspecified |
| mmu-miR-5620-3p | MIMAT0022368 | Itsn1 | 16443 | HITS-CLIP | | 37986224 | Unspecified |
| mmu-miR-5620-3p | MIMAT0022368 | Jak1 | 16451 | HITS-CLIP | | 37986224 | Unspecified |
| mmu-miR-5620-3p | MIMAT0022368 | Jmjd1c | 108829 | HITS-CLIP | | 37986224 | Unspecified |
| mmu-miR-5620-3p | MIMAT0022368 | Jmy | 57748 | HITS-CLIP | | 37986224 | Unspecified |
| mmu-miR-5620-3p | MIMAT0022368 | Kdm5c | 20591 | HITS-CLIP | | 23597149 | Unspecified |
| mmu-miR-5620-3p | MIMAT0022368 | Kidins220 | 77480 | HITS-CLIP | | 37986224 | Unspecified |
| mmu-miR-5620-3p | MIMAT0022368 | Kynu | 70789 | HITS-CLIP | | 37986224 | Unspecified |
| mmu-miR-5620-3p | MIMAT0022368 | Lactb2 | 212442 | HITS-CLIP | | 37986224 | Unspecified |
| mmu-miR-5620-3p | MIMAT0022368 | Lamp2 | 16784 | HITS-CLIP | | 37986224 | Unspecified |
| mmu-miR-5620-3p | MIMAT0022368 | Laptm4a | 17775 | HITS-CLIP | | 23597149 | Unspecified |
| mmu-miR-5620-3p | MIMAT0022368 | Larp1 | 73158 | HITS-CLIP | | 23597149 | Unspecified |
| mmu-miR-5620-3p | MIMAT0022368 | Lemd3 | 380664 | HITS-CLIP | | 37986224 | Unspecified |
| mmu-miR-5620-3p | MIMAT0022368 | Lif | 16878 | HITS-CLIP | | 31433990 | Unspecified |
| mmu-miR-5620-3p | MIMAT0022368 | Lifr | 16880 | HITS-CLIP | | 37986224 | Unspecified |
| mmu-miR-5620-3p | MIMAT0022368 | Lmnb1 | 16906 | HITS-CLIP | | 37986224 | Unspecified |
| mmu-miR-5620-3p | MIMAT0022368 | Lmtk2 | 231876 | HITS-CLIP | | 37986224 | Unspecified |
| mmu-miR-5620-3p | MIMAT0022368 | Lpin2 | 64898 | HITS-CLIP | | 37986224 | Unspecified |
| mmu-miR-5620-3p | MIMAT0022368 | Lrat | 79235 | HITS-CLIP | | 37986224 | Unspecified |
| mmu-miR-5620-3p | MIMAT0022368 | Lrp1 | 16971 | HITS-CLIP | 37986224\|23597149 | | Unspecified |

| | | | | | | |
|---|---|---|---|---|---|---|
| mmu-miR-5620-3p | MIMAT0022368 | Lrp4 | 228357 | HITS-CLIP | 23597149 | Unspecified |
| mmu-miR-5620-3p | MIMAT0022368 | Lrrc8a | 241296 | HITS-CLIP | 37986224 | Unspecified |
| mmu-miR-5620-3p | MIMAT0022368 | Lypla1 | 18777 | HITS-CLIP | 37986224 | Unspecified |
| mmu-miR-5620-3p | MIMAT0022368 | Maged1 | 94275 | HITS-CLIP | 37986224 | Unspecified |
| mmu-miR-5620-3p | MIMAT0022368 | Magi3 | 99470 | HITS-CLIP | 37986224 | Unspecified |
| mmu-miR-5620-3p | MIMAT0022368 | Mat2a | 232087 | HITS-CLIP | 37986224 | Unspecified |
| mmu-miR-5620-3p | MIMAT0022368 | Mau2 | 74549 | HITS-CLIP | 37986224 | Unspecified |
| mmu-miR-5620-3p | MIMAT0022368 | Mcl1 | 17210 | HITS-CLIP | 37986224 | Unspecified |
| mmu-miR-5620-3p | MIMAT0022368 | Mff | 75734 | HITS-CLIP | 23597149 | Unspecified |
| mmu-miR-5620-3p | MIMAT0022368 | Mia3 | 338366 | HITS-CLIP | 37986224 | Unspecified |
| mmu-miR-5620-3p | MIMAT0022368 | Mlec | 109154 | HITS-CLIP | 37986224 | Unspecified |
| mmu-miR-5620-3p | MIMAT0022368 | Mocos | 68591 | HITS-CLIP | 37986224 | Unspecified |
| mmu-miR-5620-3p | MIMAT0022368 | Mrpl28 | 68611 | HITS-CLIP | 37986224 | Unspecified |
| mmu-miR-5620-3p | MIMAT0022368 | Msl2 | 77853 | HITS-CLIP | 37986224 | Unspecified |
| mmu-miR-5620-3p | MIMAT0022368 | Msl3 | 17692 | HITS-CLIP | 23597149 | Unspecified |
| mmu-miR-5620-3p | MIMAT0022368 | Mthfd1 | 108156 | HITS-CLIP | 23597149 | Unspecified |
| mmu-miR-5620-3p | MIMAT0022368 | Mtus1 | 102103 | HITS-CLIP | 37986224 | Unspecified |
| mmu-miR-5620-3p | MIMAT0022368 | Mup3 | 17842 | HITS-CLIP | 23597149 | Unspecified |
| mmu-miR-5620-3p | MIMAT0022368 | Mybl1 | 17864 | HITS-CLIP | 23597149 | Unspecified |
| mmu-miR-5620-3p | MIMAT0022368 | Naa38 | 78304 | HITS-CLIP | 23597149 | Unspecified |
| mmu-miR-5620-3p | MIMAT0022368 | Narf | 67608 | HITS-CLIP | 37986224 | Unspecified |
| mmu-miR-5620-3p | MIMAT0022368 | Ncbp3 | 66874 | HITS-CLIP | 23597149 | Unspecified |
| mmu-miR-5620-3p | MIMAT0022368 | Ndufb11 | 104130 | HITS-CLIP | 23597149 | Unspecified |
| mmu-miR-5620-3p | MIMAT0022368 | Ndufb9 | 66218 | HITS-CLIP | 37986224 | Unspecified |
| mmu-miR-5620-3p | MIMAT0022368 | Ndufs1 | 227197 | HITS-CLIP | 23597149 | Unspecified |
| mmu-miR-5620-3p | MIMAT0022368 | Ndufv1 | 17995 | HITS-CLIP | 37986224 | Unspecified |
| mmu-miR-5620-3p | MIMAT0022368 | Nfkbia | 18035 | HITS-CLIP | 23597149 | Unspecified |
| mmu-miR-5620-3p | MIMAT0022368 | Nid2 | 18074 | HITS-CLIP | 37986224 | Unspecified |
| mmu-miR-5620-3p | MIMAT0022368 | Nisch | 64652 | HITS-CLIP | 37986224\|23597149 | Unspecified |
| mmu-miR-5620-3p | MIMAT0022368 | Npepl1 | 228961 | HITS-CLIP | 37986224 | Unspecified |
| mmu-miR-5620-3p | MIMAT0022368 | Nr3c1 | 14815 | HITS-CLIP | 37986224 | Unspecified |
| mmu-miR-5620-3p | MIMAT0022368 | Nrde2 | 217827 | HITS-CLIP | 37986224 | Unspecified |
| mmu-miR-5620-3p | MIMAT0022368 | Nrn1 | 68404 | HITS-CLIP | 37986224 | Unspecified |
| mmu-miR-5620-3p | MIMAT0022368 | Nup155 | 170762 | HITS-CLIP | 23597149 | Unspecified |
| mmu-miR-5620-3p | MIMAT0022368 | Oat | 18242 | HITS-CLIP | 37986224 | Unspecified |
| mmu-miR-5620-3p | MIMAT0022368 | Ormdl3 | 66612 | HITS-CLIP | 23597149 | Unspecified |
| mmu-miR-5620-3p | MIMAT0022368 | Oxsr1 | 108737 | HITS-CLIP | 37986224 | Unspecified |
| mmu-miR-5620-3p | MIMAT0022368 | Pafah1b1 | 18472 | HITS-CLIP | 37986224 | Unspecified |
| mmu-miR-5620-3p | MIMAT0022368 | Papola | 18789 | HITS-CLIP | 37986224 | Unspecified |

| | | | | | | |
|---|---|---|---|---|---|---|
| mmu-miR-5620-3p | MIMAT0022368 | Pdcd4 | 18569 | HITS-CLIP | 23597149 | Unspecified |
| mmu-miR-5620-3p | MIMAT0022368 | Pdcd6 | 18570 | HITS-CLIP | 37986224 | Unspecified |
| mmu-miR-5620-3p | MIMAT0022368 | Pde4b | 18578 | HITS-CLIP | 37986224 | Unspecified |
| mmu-miR-5620-3p | MIMAT0022368 | Phf20 | 228829 | HITS-CLIP | 37986224 | Unspecified |
| mmu-miR-5620-3p | MIMAT0022368 | Plau | 18792 | HITS-CLIP | 37986224 | Unspecified |
| mmu-miR-5620-3p | MIMAT0022368 | Plekha6 | 240753 | HITS-CLIP | 37986224 | Unspecified |
| mmu-miR-5620-3p | MIMAT0022368 | Polr2a | 20020 | HITS-CLIP | 23597149 | Unspecified |
| mmu-miR-5620-3p | MIMAT0022368 | Ppil4 | 67418 | HITS-CLIP | 31433990 | Unspecified |
| mmu-miR-5620-3p | MIMAT0022368 | Ppip5k2 | 227399 | HITS-CLIP | 37986224\|23597149 | Unspecified |
| mmu-miR-5620-3p | MIMAT0022368 | Ppp1ca | 19045 | HITS-CLIP | 37986224 | Unspecified |
| mmu-miR-5620-3p | MIMAT0022368 | Ppp1cb | 19046 | HITS-CLIP | 23597149 | Unspecified |
| mmu-miR-5620-3p | MIMAT0022368 | Ppp4c | 56420 | HITS-CLIP | 37986224 | Unspecified |
| mmu-miR-5620-3p | MIMAT0022368 | Prdm1 | 12142 | HITS-CLIP | 23597149 | Unspecified |
| mmu-miR-5620-3p | MIMAT0022368 | Prg4 | 96875 | HITS-CLIP | 23597149 | Unspecified |
| mmu-miR-5620-3p | MIMAT0022368 | Prpf8 | 192159 | HITS-CLIP | 37986224 | Unspecified |
| mmu-miR-5620-3p | MIMAT0022368 | Psen1 | 19164 | HITS-CLIP | 23597149 | Unspecified |
| mmu-miR-5620-3p | MIMAT0022368 | Psma1 | 26440 | HITS-CLIP | 37986224 | Unspecified |
| mmu-miR-5620-3p | MIMAT0022368 | Psma7 | 26444 | HITS-CLIP | 23597149 | Unspecified |
| mmu-miR-5620-3p | MIMAT0022368 | Psmb2 | 26445 | HITS-CLIP | 37986224 | Unspecified |
| mmu-miR-5620-3p | MIMAT0022368 | Psmb4 | 19172 | HITS-CLIP | 37986224 | Unspecified |
| mmu-miR-5620-3p | MIMAT0022368 | Psmd2 | 21762 | HITS-CLIP | 37986224 | Unspecified |
| mmu-miR-5620-3p | MIMAT0022368 | Pum1 | 80912 | HITS-CLIP | 37986224 | Unspecified |
| mmu-miR-5620-3p | MIMAT0022368 | Pus10 | 74467 | HITS-CLIP | 37986224 | Unspecified |
| mmu-miR-5620-3p | MIMAT0022368 | Pzp | 11287 | HITS-CLIP | 23597149 | Unspecified |
| mmu-miR-5620-3p | MIMAT0022368 | Qrich1 | 69232 | HITS-CLIP | 37986224 | Unspecified |
| mmu-miR-5620-3p | MIMAT0022368 | Qsox1 | 104009 | HITS-CLIP | 23597149 | Unspecified |
| mmu-miR-5620-3p | MIMAT0022368 | Rab11a | 53869 | HITS-CLIP | 23597149 | Unspecified |
| mmu-miR-5620-3p | MIMAT0022368 | Rab5if | 55969 | HITS-CLIP | 23597149 | Unspecified |
| mmu-miR-5620-3p | MIMAT0022368 | Rabep2 | 70314 | HITS-CLIP | 23597149 | Unspecified |
| mmu-miR-5620-3p | MIMAT0022368 | Rad21 | 19357 | HITS-CLIP | 37986224 | Unspecified |
| mmu-miR-5620-3p | MIMAT0022368 | Rbm45 | 241490 | HITS-CLIP | 37986224 | Unspecified |
| mmu-miR-5620-3p | MIMAT0022368 | Rdx | 19684 | HITS-CLIP | 37986224 | Unspecified |
| mmu-miR-5620-3p | MIMAT0022368 | Rhot1 | 59040 | HITS-CLIP | 23597149 | Unspecified |
| mmu-miR-5620-3p | MIMAT0022368 | Ric1 | 850728 | HITS-CLIP | 23597149 | Unspecified |
| mmu-miR-5620-3p | MIMAT0022368 | Rlf | 109263 | HITS-CLIP | 37986224\|23597149 | Unspecified |
| mmu-miR-5620-3p | MIMAT0022368 | Rlim | 19820 | HITS-CLIP | 37986224\|23597149 | Unspecified |
| mmu-miR-5620-3p | MIMAT0022368 | Rnd3 | 74194 | HITS-CLIP | 37986224 | Unspecified |
| mmu-miR-5620-3p | MIMAT0022368 | Rnf11 | 29864 | HITS-CLIP | 37986224 | Unspecified |
| mmu-miR-5620-3p | MIMAT0022368 | Rnf13 | 24017 | HITS-CLIP | 37986224 | Unspecified |

| | | | | | | |
|---|---|---|---|---|---|---|
| mmu-miR-5620-3p | MIMAT0022368 | Rnf6 | 74132 | HITS-CLIP | 23597149 | Unspecified |
| mmu-miR-5620-3p | MIMAT0022368 | Rngtt | 24018 | HITS-CLIP | 37986224 | Unspecified |
| mmu-miR-5620-3p | MIMAT0022368 | Rprd1b | 70470 | HITS-CLIP | 37986224\|23597149 | Unspecified |
| mmu-miR-5620-3p | MIMAT0022368 | Rps5 | 20103 | HITS-CLIP | 23597149 | Unspecified |
| mmu-miR-5620-3p | MIMAT0022368 | Rxra | 20181 | HITS-CLIP | 23597149 | Unspecified |
| mmu-miR-5620-3p | MIMAT0022368 | Sardh | 192166 | HITS-CLIP | 37986224 | Unspecified |
| mmu-miR-5620-3p | MIMAT0022368 | Sars | 20226 | HITS-CLIP | 37986224 | Unspecified |
| mmu-miR-5620-3p | MIMAT0022368 | Scamp1 | 107767 | HITS-CLIP | 37986224 | Unspecified |
| mmu-miR-5620-3p | MIMAT0022368 | Sdc2 | 15529 | HITS-CLIP | 37986224 | Unspecified |
| mmu-miR-5620-3p | MIMAT0022368 | Sdr42e1 | 74032 | HITS-CLIP | 37986224 | Unspecified |
| mmu-miR-5620-3p | MIMAT0022368 | Sec14l2 | 67815 | HITS-CLIP | 37986224 | Unspecified |
| mmu-miR-5620-3p | MIMAT0022368 | Sec24a | 77371 | HITS-CLIP | 23597149 | Unspecified |
| mmu-miR-5620-3p | MIMAT0022368 | Sema6d | 214968 | HITS-CLIP | 37986224 | Unspecified |
| mmu-miR-5620-3p | MIMAT0022368 | Sept10 | 103080 | HITS-CLIP | 37986224 | Unspecified |
| mmu-miR-5620-3p | MIMAT0022368 | Serpina3n | 20716 | HITS-CLIP | 37986224 | Unspecified |
| mmu-miR-5620-3p | MIMAT0022368 | Serpind1 | 15160 | HITS-CLIP | 23597149 | Unspecified |
| mmu-miR-5620-3p | MIMAT0022368 | Serpine1 | 18787 | HITS-CLIP | 23597149 | Unspecified |
| mmu-miR-5620-3p | MIMAT0022368 | Sgpl1 | 20397 | HITS-CLIP | 37986224 | Unspecified |
| mmu-miR-5620-3p | MIMAT0022368 | Sgpp1 | 81535 | HITS-CLIP | 37986224 | Unspecified |
| mmu-miR-5620-3p | MIMAT0022368 | Slc10a5 | 241877 | HITS-CLIP | 37986224 | Unspecified |
| mmu-miR-5620-3p | MIMAT0022368 | Slc25a10 | 27376 | HITS-CLIP | 37986224 | Unspecified |
| mmu-miR-5620-3p | MIMAT0022368 | Slc25a3 | 18674 | HITS-CLIP | 37986224 | Unspecified |
| mmu-miR-5620-3p | MIMAT0022368 | Slc27a2 | 26458 | HITS-CLIP | 37986224 | Unspecified |
| mmu-miR-5620-3p | MIMAT0022368 | Slc7a2 | 11988 | HITS-CLIP | 37986224\|23597149 | Unspecified |
| mmu-miR-5620-3p | MIMAT0022368 | Slco1b2 | 28253 | HITS-CLIP | 23597149 | Unspecified |
| mmu-miR-5620-3p | MIMAT0022368 | Slfn5 | 327978 | HITS-CLIP | 37986224 | Unspecified |
| mmu-miR-5620-3p | MIMAT0022368 | Son | 20658 | HITS-CLIP | 37986224\|23597149 | Unspecified |
| mmu-miR-5620-3p | MIMAT0022368 | Spata2l | 78779 | HITS-CLIP | 37986224 | Unspecified |
| mmu-miR-5620-3p | MIMAT0022368 | Sppl3 | 74585 | HITS-CLIP | 23597149 | Unspecified |
| mmu-miR-5620-3p | MIMAT0022368 | Sqstm1 | 18412 | HITS-CLIP | 37986224\|23597149 | Unspecified |
| mmu-miR-5620-3p | MIMAT0022368 | Srebf2 | 20788 | HITS-CLIP | 37986224\|23597149 | Unspecified |
| mmu-miR-5620-3p | MIMAT0022368 | Srp19 | 66384 | HITS-CLIP | 23597149 | Unspecified |
| mmu-miR-5620-3p | MIMAT0022368 | Srrt | 83701 | HITS-CLIP | 37986224 | Unspecified |
| mmu-miR-5620-3p | MIMAT0022368 | Stk11 | 20869 | HITS-CLIP | 23597149 | Unspecified |
| mmu-miR-5620-3p | MIMAT0022368 | Sucla2 | 20916 | HITS-CLIP | 37986224 | Unspecified |
| mmu-miR-5620-3p | MIMAT0022368 | Syne1 | 64009 | HITS-CLIP | 23597149 | Unspecified |
| mmu-miR-5620-3p | MIMAT0022368 | Taf1 | 270627 | HITS-CLIP | 23597149 | Unspecified |
| mmu-miR-5620-3p | MIMAT0022368 | Tbc1d31 | 210544 | HITS-CLIP | 23597149 | Unspecified |
| mmu-miR-5620-3p | MIMAT0022368 | Tcp11l2 | 216198 | HITS-CLIP | 23597149 | Unspecified |

| | | | | | | |
|---|---|---|---|---|---|---|
| mmu-miR-5620-3p | MIMAT0022368 | Tek | 21687 | HITS-CLIP | 37986224\|23597149 | Unspecified |
| mmu-miR-5620-3p | MIMAT0022368 | Tex2 | 21763 | HITS-CLIP | 37986224 | Unspecified |
| mmu-miR-5620-3p | MIMAT0022368 | Tgfbrap1 | 73122 | HITS-CLIP | 23597149 | Unspecified |
| mmu-miR-5620-3p | MIMAT0022368 | Tgoln1 | 22134 | HITS-CLIP | 37986224 | Unspecified |
| mmu-miR-5620-3p | MIMAT0022368 | Thap11 | 59016 | HITS-CLIP | 23597149 | Unspecified |
| mmu-miR-5620-3p | MIMAT0022368 | Tln1 | 21894 | HITS-CLIP | 37986224 | Unspecified |
| mmu-miR-5620-3p | MIMAT0022368 | Tm2d2 | 69742 | HITS-CLIP | 37986224\|23597149 | Unspecified |
| mmu-miR-5620-3p | MIMAT0022368 | Tm9sf1 | 74140 | HITS-CLIP | 37986224 | Unspecified |
| mmu-miR-5620-3p | MIMAT0022368 | Tmem189 | 407243 | HITS-CLIP | 37986224 | Unspecified |
| mmu-miR-5620-3p | MIMAT0022368 | Tmem220 | 338369 | HITS-CLIP | 23597149 | Unspecified |
| mmu-miR-5620-3p | MIMAT0022368 | Tnpo3 | 320938 | HITS-CLIP | 31433990 | Unspecified |
| mmu-miR-5620-3p | MIMAT0022368 | Tpr | 108989 | HITS-CLIP | 23597149 | Unspecified |
| mmu-miR-5620-3p | MIMAT0022368 | Tpx2 | 72119 | HITS-CLIP | 37986224 | Unspecified |
| mmu-miR-5620-3p | MIMAT0022368 | Trim26 | 22670 | HITS-CLIP | 37986224 | Unspecified |
| mmu-miR-5620-3p | MIMAT0022368 | Trip12 | 14897 | HITS-CLIP | 37986224 | Unspecified |
| mmu-miR-5620-3p | MIMAT0022368 | Trmt6 | 66926 | HITS-CLIP | 23597149 | Unspecified |
| mmu-miR-5620-3p | MIMAT0022368 | Tsc22d3 | 14605 | HITS-CLIP | 37986224\|23597149 | Unspecified |
| mmu-miR-5620-3p | MIMAT0022368 | Tuba4a | 22145 | HITS-CLIP | 37986224 | Unspecified |
| mmu-miR-5620-3p | MIMAT0022368 | Tubb2a | 22151 | HITS-CLIP | 37986224 | Unspecified |
| mmu-miR-5620-3p | MIMAT0022368 | Twsg1 | 65960 | HITS-CLIP | 37986224 | Unspecified |
| mmu-miR-5620-3p | MIMAT0022368 | Txn1 | 22166 | HITS-CLIP | 23597149 | Unspecified |
| mmu-miR-5620-3p | MIMAT0022368 | Txndc15 | 69672 | HITS-CLIP | 37986224 | Unspecified |
| mmu-miR-5620-3p | MIMAT0022368 | Ube2g2 | 22213 | HITS-CLIP | 37986224 | Unspecified |
| mmu-miR-5620-3p | MIMAT0022368 | Ube2r2 | 67615 | HITS-CLIP | 37986224 | Unspecified |
| mmu-miR-5620-3p | MIMAT0022368 | Ubr2 | 224826 | HITS-CLIP | 37986224 | Unspecified |
| mmu-miR-5620-3p | MIMAT0022368 | Ugt1a1 | 394436 | HITS-CLIP | 37986224 | Unspecified |
| mmu-miR-5620-3p | MIMAT0022368 | Ugt1a7c | 394432 | HITS-CLIP | 37986224 | Unspecified |
| mmu-miR-5620-3p | MIMAT0022368 | Upf3b | 68134 | HITS-CLIP | 37986224 | Unspecified |
| mmu-miR-5620-3p | MIMAT0022368 | Urod | 22275 | HITS-CLIP | 23597149 | Unspecified |
| mmu-miR-5620-3p | MIMAT0022368 | Ush1g | 16470 | HITS-CLIP | 37986224 | Unspecified |
| mmu-miR-5620-3p | MIMAT0022368 | Usp19 | 71472 | HITS-CLIP | 37986224\|23597149 | Unspecified |
| mmu-miR-5620-3p | MIMAT0022368 | Usp9x | 22284 | HITS-CLIP | 23597149 | Unspecified |
| mmu-miR-5620-3p | MIMAT0022368 | Uspl1 | 231915 | HITS-CLIP | 37986224 | Unspecified |
| mmu-miR-5620-3p | MIMAT0022368 | Vapa | 30960 | HITS-CLIP | 23597149 | Unspecified |
| mmu-miR-5620-3p | MIMAT0022368 | Vps13c | 320528 | HITS-CLIP | 37986224 | Unspecified |
| mmu-miR-5620-3p | MIMAT0022368 | Vps41 | 218035 | HITS-CLIP | 23597149 | Unspecified |
| mmu-miR-5620-3p | MIMAT0022368 | Wapl | 23063 | HITS-CLIP | 23597149 | Unspecified |
| mmu-miR-5620-3p | MIMAT0022368 | Wdr26 | 226757 | HITS-CLIP | 23597149 | Unspecified |
| mmu-miR-5620-3p | MIMAT0022368 | Wdr82 | 77305 | HITS-CLIP | 37986224 | Unspecified |

| | | | | | | |
|---|---|---|---|---|---|---|
| mmu-miR-5620-3p | MIMAT0022368 | Xkr9 | 381246 | HITS-CLIP | 37986224 | Unspecified |
| mmu-miR-5620-3p | MIMAT0022368 | Ythdf3 | 229096 | HITS-CLIP | 37986224 | Unspecified |
| mmu-miR-5620-3p | MIMAT0022368 | Zfhx4 | 80892 | HITS-CLIP | 37986224 | Unspecified |
| mmu-miR-5620-3p | MIMAT0022368 | Zfp655 | 72611 | HITS-CLIP | 37986224 | Unspecified |
| mmu-miR-5620-3p | MIMAT0022368 | Zmiz2 | 52915 | HITS-CLIP | 37986224 | Unspecified |
| mmu-miR-21a-3p | MIMAT0004628 | 1700017B05Rik | 74211 | HITS-CLIP | 23142080 | Unspecified |
| mmu-miR-21a-3p | MIMAT0004628 | 1700025G04Rik | 69399 | HITS-CLIP | 23142080 | Unspecified |
| mmu-miR-21a-3p | MIMAT0004628 | 1810013L24Rik | 69053 | HITS-CLIP | 23142080\|29167373 | Unspecified |
| mmu-miR-21a-3p | MIMAT0004628 | 2010111I01Rik | 72061 | HITS-CLIP | 29167373 | Unspecified |
| mmu-miR-21a-3p | MIMAT0004628 | 2310022B05Rik | 69551 | HITS-CLIP | 23142080 | Unspecified |
| mmu-miR-21a-3p | MIMAT0004628 | 2510039O18Rik | 77034 | qCLASH | 31393953 | Unspecified |
| mmu-miR-21a-3p | MIMAT0004628 | 2610001J05Rik | 66520 | HITS-CLIP | 23142080 | Unspecified |
| mmu-miR-21a-3p | MIMAT0004628 | 2610507B11Rik | 72503 | HITS-CLIP | 23142080 | Unspecified |
| mmu-miR-21a-3p | MIMAT0004628 | 4921524J17Rik | 66714 | HITS-CLIP | 23142080 | Unspecified |
| mmu-miR-21a-3p | MIMAT0004628 | 4930402H24Rik | 228602 | HITS-CLIP | 23142080 | Unspecified |
| mmu-miR-21a-3p | MIMAT0004628 | 4930503L19Rik | 269033 | HITS-CLIP | 23142080 | Unspecified |
| mmu-miR-21a-3p | MIMAT0004628 | 4931406P16Rik | 233103 | HITS-CLIP | 23142080 | Unspecified |
| mmu-miR-21a-3p | MIMAT0004628 | 4932438A13Rik | 229227 | HITS-CLIP | 23142080\|29167373 | Unspecified |
| mmu-miR-21a-3p | MIMAT0004628 | 4933434E20Rik | 99650 | HITS-CLIP | 23142080 | Unspecified |
| mmu-miR-21a-3p | MIMAT0004628 | 6430548M08Rik | 234797 | HITS-CLIP | 29167373 | Unspecified |
| mmu-miR-21a-3p | MIMAT0004628 | 8030462N17Rik | 212163 | HITS-CLIP | 23142080\|29167373 | Unspecified |
| mmu-miR-21a-3p | MIMAT0004628 | 9530068E07Rik | 213673 | HITS-CLIP | 37986224\|23142080 | Unspecified |
| mmu-miR-21a-3p | MIMAT0004628 | 9930021J03Rik | 240613 | HITS-CLIP | 23142080 | Unspecified |
| mmu-miR-21a-3p | MIMAT0004628 | Aaas | 223921 | HITS-CLIP | 23142080 | Unspecified |
| mmu-miR-21a-3p | MIMAT0004628 | Aars | 234734 | HITS-CLIP | 23142080 | Unspecified |
| mmu-miR-21a-3p | MIMAT0004628 | Abca1 | 11303 | HITS-CLIP | 29167373 | Unspecified |
| mmu-miR-21a-3p | MIMAT0004628 | Abca5 | 217265 | HITS-CLIP | 23142080 | Unspecified |
| mmu-miR-21a-3p | MIMAT0004628 | Abcb1b | 18669 | HITS-CLIP | 23142080 | Unspecified |
| mmu-miR-21a-3p | MIMAT0004628 | Abcb7 | 11306 | HITS-CLIP | 23142080 | Unspecified |
| mmu-miR-21a-3p | MIMAT0004628 | Abcc1 | 17250 | HITS-CLIP | 29167373 | Unspecified |
| mmu-miR-21a-3p | MIMAT0004628 | Abce1 | 24015 | HITS-CLIP | 23142080 | Unspecified |
| mmu-miR-21a-3p | MIMAT0004628 | Abcg1 | 11307 | HITS-CLIP | 23142080 | Unspecified |
| mmu-miR-21a-3p | MIMAT0004628 | Abhd17b | 226016 | HITS-CLIP | 23142080 | Unspecified |
| mmu-miR-21a-3p | MIMAT0004628 | Abhd17c | 70178 | HITS-CLIP | 23142080 | Unspecified |
| mmu-miR-21a-3p | MIMAT0004628 | Abhd18 | 269423 | HITS-CLIP | 23142080 | Unspecified |
| mmu-miR-21a-3p | MIMAT0004628 | Abhd2 | 54608 | HITS-CLIP | 23142080 | Unspecified |
| mmu-miR-21a-3p | MIMAT0004628 | Abi2 | 329165 | HITS-CLIP | 37986224\|23142080 | Unspecified |
| mmu-miR-21a-3p | MIMAT0004628 | Abr | 109934 | HITS-CLIP | 29167373 | Unspecified |
| mmu-miR-21a-3p | MIMAT0004628 | Abraxas2 | 23172 | HITS-CLIP | 23142080 | Unspecified |

| | | | | | | |
|---|---|---|---|---|---|---|
| mmu-miR-21a-3p | MIMAT0004628 | Abtb2 | 99382 | HITS-CLIP | 23142080 | Unspecified |
| mmu-miR-21a-3p | MIMAT0004628 | Acadsb | 66885 | HITS-CLIP | 23142080 | Unspecified |
| mmu-miR-21a-3p | MIMAT0004628 | Acap1 | 216859 | HITS-CLIP | 37986224\|23142080 | Unspecified |
| mmu-miR-21a-3p | MIMAT0004628 | Acap2 | 78618 | HITS-CLIP | 29167373 | Unspecified |
| mmu-miR-21a-3p | MIMAT0004628 | Acat1 | 110446 | HITS-CLIP | 23142080 | Unspecified |
| mmu-miR-21a-3p | MIMAT0004628 | Acbd3 | 170760 | HITS-CLIP | 37986224\|23142080\|29167373 | Unspecified |
| mmu-miR-21a-3p | MIMAT0004628 | Acin1 | 56215 | HITS-CLIP | 29167373 | Unspecified |
| mmu-miR-21a-3p | MIMAT0004628 | Acly | 104112 | HITS-CLIP | 23142080 | Unspecified |
| mmu-miR-21a-3p | MIMAT0004628 | Acsbg1 | 94180 | HITS-CLIP | 37986224\|23142080 | Unspecified |
| mmu-miR-21a-3p | MIMAT0004628 | Acsl3 | 74205 | HITS-CLIP | 23142080 | Unspecified |
| mmu-miR-21a-3p | MIMAT0004628 | Acsl4 | 50790 | HITS-CLIP | 23142080 | Unspecified |
| mmu-miR-21a-3p | MIMAT0004628 | Actb | 11461 | HITS-CLIP | 23142080 | Unspecified |
| mmu-miR-21a-3p | MIMAT0004628 | Actr10 | 56444 | HITS-CLIP | 23142080 | Unspecified |
| mmu-miR-21a-3p | MIMAT0004628 | Actr2 | 66713 | HITS-CLIP | 23142080 | Unspecified |
| mmu-miR-21a-3p | MIMAT0004628 | Acvr1b | 11479 | HITS-CLIP | 23142080 | Unspecified |
| mmu-miR-21a-3p | MIMAT0004628 | Acvr2a | 11480 | HITS-CLIP | 23142080\|29167373 | Unspecified |
| mmu-miR-21a-3p | MIMAT0004628 | Acvr2b | 11481 | HITS-CLIP | 23142080 | Unspecified |
| mmu-miR-21a-3p | MIMAT0004628 | Adam17 | 11491 | HITS-CLIP | 23142080 | Unspecified |
| mmu-miR-21a-3p | MIMAT0004628 | Adam19 | 11492 | HITS-CLIP | 23142080 | Unspecified |
| mmu-miR-21a-3p | MIMAT0004628 | Adap1 | 231821 | HITS-CLIP | 23142080 | Unspecified |
| mmu-miR-21a-3p | MIMAT0004628 | Adarb1 | 110532 | HITS-CLIP | 23142080 | Unspecified |
| mmu-miR-21a-3p | MIMAT0004628 | Add1 | 11518 | HITS-CLIP | 23142080 | Unspecified |
| mmu-miR-21a-3p | MIMAT0004628 | Add3 | 27360 | HITS-CLIP | 23142080 | Unspecified |
| mmu-miR-21a-3p | MIMAT0004628 | Adgrl2 | 23266 | HITS-CLIP | 29167373 | Unspecified |
| mmu-miR-21a-3p | MIMAT0004628 | Adh5 | 11532 | HITS-CLIP | 23142080 | Unspecified |
| mmu-miR-21a-3p | MIMAT0004628 | Adipor2 | 68465 | HITS-CLIP | 23142080 | Unspecified |
| mmu-miR-21a-3p | MIMAT0004628 | Adk | 11534 | HITS-CLIP | 23142080 | Unspecified |
| mmu-miR-21a-3p | MIMAT0004628 | Adnp | 11538 | HITS-CLIP | 37986224\|23142080\|29167373 | Unspecified |
| mmu-miR-21a-3p | MIMAT0004628 | Ado | 211488 | HITS-CLIP | 37986224\|23142080 | Unspecified |
| mmu-miR-21a-3p | MIMAT0004628 | Adprh | 11544 | HITS-CLIP | 23142080 | Unspecified |
| mmu-miR-21a-3p | MIMAT0004628 | Adss | 11566 | HITS-CLIP | 37986224 | Unspecified |
| mmu-miR-21a-3p | MIMAT0004628 | Aebp2 | 11569 | HITS-CLIP | 37986224\|23142080 | Unspecified |
| mmu-miR-21a-3p | MIMAT0004628 | Afdn | 4301 | HITS-CLIP | 29167373 | Unspecified |
| mmu-miR-21a-3p | MIMAT0004628 | Aff4 | 93736 | HITS-CLIP | 37986224\|23142080 | Unspecified |
| mmu-miR-21a-3p | MIMAT0004628 | Afg3l2 | 69597 | HITS-CLIP | 23142080 | Unspecified |
| mmu-miR-21a-3p | MIMAT0004628 | Agfg1 | 15463 | HITS-CLIP | 23142080 | Unspecified |
| mmu-miR-21a-3p | MIMAT0004628 | Aggf1 | 66549 | HITS-CLIP | 29167373 | Unspecified |
| mmu-miR-21a-3p | MIMAT0004628 | Ago1 | 236511 | HITS-CLIP | 37986224 | Unspecified |
| mmu-miR-21a-3p | MIMAT0004628 | Ago2 | 239528 | HITS-CLIP | 37986224\|23142080 | Unspecified |

| | | | | | | |
|---|---|---|---|---|---|---|
| mmu-miR-21a-3p | MIMAT0004628 | Ago3 | 214150 | HITS-CLIP | 37986224\|23142080\|29167373 | Unspecified |
| mmu-miR-21a-3p | MIMAT0004628 | Agpat1 | 55979 | HITS-CLIP | 23142080 | Unspecified |
| mmu-miR-21a-3p | MIMAT0004628 | Agpat4 | 68262 | HITS-CLIP | 23142080 | Unspecified |
| mmu-miR-21a-3p | MIMAT0004628 | Agps | 228061 | HITS-CLIP | 23142080 | Unspecified |
| mmu-miR-21a-3p | MIMAT0004628 | Ahctf1 | 226747 | HITS-CLIP | 37986224\|23142080 | Unspecified |
| mmu-miR-21a-3p | MIMAT0004628 | Ahcyl2 | 74340 | HITS-CLIP | 37986224\|23142080 | Unspecified |
| mmu-miR-21a-3p | MIMAT0004628 | Ahdc1 | 230793 | HITS-CLIP | 23142080 | Unspecified |
| mmu-miR-21a-3p | MIMAT0004628 | Ahnak | 66395 | HITS-CLIP | 23142080\|29167373 | Unspecified |
| mmu-miR-21a-3p | MIMAT0004628 | Ahr | 11622 | HITS-CLIP | 23142080 | Unspecified |
| mmu-miR-21a-3p | MIMAT0004628 | Aimp1 | 13722 | HITS-CLIP | 23142080 | Unspecified |
| mmu-miR-21a-3p | MIMAT0004628 | Ajuba | 16475 | HITS-CLIP | 23142080 | Unspecified |
| mmu-miR-21a-3p | MIMAT0004628 | Ak4 | 11639 | HITS-CLIP | 23142080 | Unspecified |
| mmu-miR-21a-3p | MIMAT0004628 | Akap11 | 219181 | HITS-CLIP | 37986224\|23142080 | Unspecified |
| mmu-miR-21a-3p | MIMAT0004628 | Akap13 | 75547 | HITS-CLIP | 23142080\|29167373 | Unspecified |
| mmu-miR-21a-3p | MIMAT0004628 | Akap9 | 100986 | HITS-CLIP | 29167373 | Unspecified |
| mmu-miR-21a-3p | MIMAT0004628 | Akirin1 | 68050 | HITS-CLIP | 23142080 | Unspecified |
| mmu-miR-21a-3p | MIMAT0004628 | Akirin2 | 433693 | HITS-CLIP | 23142080 | Unspecified |
| mmu-miR-21a-3p | MIMAT0004628 | Akt3 | 23797 | HITS-CLIP | 23142080 | Unspecified |
| mmu-miR-21a-3p | MIMAT0004628 | Alcam | 11658 | HITS-CLIP | 23142080 | Unspecified |
| mmu-miR-21a-3p | MIMAT0004628 | Aldoa | 11674 | HITS-CLIP | 37986224\|23142080 | Unspecified |
| mmu-miR-21a-3p | MIMAT0004628 | Aldoc | 11676 | HITS-CLIP | 23142080 | Unspecified |
| mmu-miR-21a-3p | MIMAT0004628 | Alg10b | 380959 | HITS-CLIP | 23142080 | Unspecified |
| mmu-miR-21a-3p | MIMAT0004628 | Alg2 | 56737 | HITS-CLIP | 23142080 | Unspecified |
| mmu-miR-21a-3p | MIMAT0004628 | Alg8 | 381903 | HITS-CLIP | 29167373 | Unspecified |
| mmu-miR-21a-3p | MIMAT0004628 | Alg9 | 102580 | HITS-CLIP | 23142080 | Unspecified |
| mmu-miR-21a-3p | MIMAT0004628 | Alkbh1 | 211064 | HITS-CLIP | 23142080 | Unspecified |
| mmu-miR-21a-3p | MIMAT0004628 | Alkbh5 | 268420 | HITS-CLIP | 23142080\|29167373 | Unspecified |
| mmu-miR-21a-3p | MIMAT0004628 | Als2cl | 235633 | HITS-CLIP | 23142080 | Unspecified |
| mmu-miR-21a-3p | MIMAT0004628 | Alyref2 | 56009 | HITS-CLIP | 23142080 | Unspecified |
| mmu-miR-21a-3p | MIMAT0004628 | Ambra1 | 228361 | HITS-CLIP | 23142080\|29167373 | Unspecified |
| mmu-miR-21a-3p | MIMAT0004628 | Amer1 | 72345 | HITS-CLIP | 23142080 | Unspecified |
| mmu-miR-21a-3p | MIMAT0004628 | Amfr | 23802 | HITS-CLIP | 23142080 | Unspecified |
| mmu-miR-21a-3p | MIMAT0004628 | Ammecr1 | 56068 | HITS-CLIP | 23142080 | Unspecified |
| mmu-miR-21a-3p | MIMAT0004628 | Amz2 | 13929 | HITS-CLIP | 37986224\|23142080 | Unspecified |
| mmu-miR-21a-3p | MIMAT0004628 | Anapc1 | 17222 | HITS-CLIP | 23142080\|29167373 | Unspecified |
| mmu-miR-21a-3p | MIMAT0004628 | Anapc2 | 99152 | HITS-CLIP | 23142080 | Unspecified |
| mmu-miR-21a-3p | MIMAT0004628 | Anapc4 | 52206 | HITS-CLIP | 23142080 | Unspecified |
| mmu-miR-21a-3p | MIMAT0004628 | Angel2 | 52477 | HITS-CLIP | 23142080 | Unspecified |
| mmu-miR-21a-3p | MIMAT0004628 | Ankfy1 | 11736 | HITS-CLIP | 23142080\|29167373 | Unspecified |

| | | | | | | |
|---|---|---|---|---|---|---|
| mmu-miR-21a-3p | MIMAT0004628 | Ankhd1 | 108857 | HITS-CLIP | 23142080 | Unspecified |
| mmu-miR-21a-3p | MIMAT0004628 | Ankib1 | 70797 | HITS-CLIP | 37986224\|23142080 | Unspecified |
| mmu-miR-21a-3p | MIMAT0004628 | Ankrd10 | 102334 | HITS-CLIP | 23142080 | Unspecified |
| mmu-miR-21a-3p | MIMAT0004628 | Ankrd11 | 77087 | HITS-CLIP | 23142080 | Unspecified |
| mmu-miR-21a-3p | MIMAT0004628 | Ankrd13c | 433667 | HITS-CLIP | 23142080 | Unspecified |
| mmu-miR-21a-3p | MIMAT0004628 | Ankrd17 | 81702 | HITS-CLIP | 37986224\|23142080\|29167373 | Unspecified |
| mmu-miR-21a-3p | MIMAT0004628 | Ankrd28 | 105522 | HITS-CLIP | 23142080\|29167373 | Unspecified |
| mmu-miR-21a-3p | MIMAT0004628 | Ankrd44 | 329154 | HITS-CLIP | 23142080 | Unspecified |
| mmu-miR-21a-3p | MIMAT0004628 | Ankrd46 | 68839 | HITS-CLIP | 37986224\|23142080 | Unspecified |
| mmu-miR-21a-3p | MIMAT0004628 | Ankrd50 | 99696 | HITS-CLIP | 23142080 | Unspecified |
| mmu-miR-21a-3p | MIMAT0004628 | Ankrd52 | 237615 | HITS-CLIP | 23142080\|29167373 | Unspecified |
| mmu-miR-21a-3p | MIMAT0004628 | Ankrd9 | 74251 | HITS-CLIP | 23142080 | Unspecified |
| mmu-miR-21a-3p | MIMAT0004628 | Anks1 | 224650 | HITS-CLIP | 23142080 | Unspecified |
| mmu-miR-21a-3p | MIMAT0004628 | Anks3 | 72615 | HITS-CLIP | 23142080 | Unspecified |
| mmu-miR-21a-3p | MIMAT0004628 | Anp32a | 11737 | HITS-CLIP | 23142080 | Unspecified |
| mmu-miR-21a-3p | MIMAT0004628 | Anp32e | 66471 | HITS-CLIP | 23142080 | Unspecified |
| mmu-miR-21a-3p | MIMAT0004628 | Antxr2 | 71914 | HITS-CLIP | 23142080 | Unspecified |
| mmu-miR-21a-3p | MIMAT0004628 | Anxa2 | 12306 | HITS-CLIP | 23142080 | Unspecified |
| mmu-miR-21a-3p | MIMAT0004628 | Anxa7 | 11750 | HITS-CLIP | 23142080 | Unspecified |
| mmu-miR-21a-3p | MIMAT0004628 | Ap1b1 | 11764 | HITS-CLIP | 37986224\|23142080 | Unspecified |
| mmu-miR-21a-3p | MIMAT0004628 | Ap1g1 | 11765 | HITS-CLIP | 23142080\|29167373 | Unspecified |
| mmu-miR-21a-3p | MIMAT0004628 | Ap1s2 | 108012 | HITS-CLIP | 23142080 | Unspecified |
| mmu-miR-21a-3p | MIMAT0004628 | Ap1s3 | 252903 | HITS-CLIP | 23142080 | Unspecified |
| mmu-miR-21a-3p | MIMAT0004628 | Ap2a1 | 11771 | HITS-CLIP | 23142080 | Unspecified |
| mmu-miR-21a-3p | MIMAT0004628 | Ap2b1 | 71770 | HITS-CLIP | 37986224 | Unspecified |
| mmu-miR-21a-3p | MIMAT0004628 | Ap3b1 | 11774 | HITS-CLIP | 23142080 | Unspecified |
| mmu-miR-21a-3p | MIMAT0004628 | Ap3d1 | 11776 | HITS-CLIP | 23142080 | Unspecified |
| mmu-miR-21a-3p | MIMAT0004628 | Ap4b1 | 67489 | HITS-CLIP | 23142080 | Unspecified |
| mmu-miR-21a-3p | MIMAT0004628 | Ap5b1 | 381201 | HITS-CLIP | 29167373 | Unspecified |
| mmu-miR-21a-3p | MIMAT0004628 | Ap5m1 | 74385 | HITS-CLIP | 23142080 | Unspecified |
| mmu-miR-21a-3p | MIMAT0004628 | Apaf1 | 11783 | HITS-CLIP | 23142080 | Unspecified |
| mmu-miR-21a-3p | MIMAT0004628 | Apbb1ip | 54519 | HITS-CLIP | 29167373 | Unspecified |
| mmu-miR-21a-3p | MIMAT0004628 | Apbb3 | 225372 | HITS-CLIP | 23142080 | Unspecified |
| mmu-miR-21a-3p | MIMAT0004628 | Apc | 11789 | HITS-CLIP | 23142080 | Unspecified |
| mmu-miR-21a-3p | MIMAT0004628 | Apeh | 235606 | HITS-CLIP | 23142080 | Unspecified |
| mmu-miR-21a-3p | MIMAT0004628 | Api5 | 11800 | HITS-CLIP | 23142080 | Unspecified |
| mmu-miR-21a-3p | MIMAT0004628 | Apobec3 | 80287 | HITS-CLIP | 23142080 | Unspecified |
| mmu-miR-21a-3p | MIMAT0004628 | Appbp2 | 66884 | HITS-CLIP | 23142080 | Unspecified |
| mmu-miR-21a-3p | MIMAT0004628 | Appl1 | 72993 | HITS-CLIP | 37986224\|23142080 | Unspecified |

| | | | | | | |
|---|---|---|---|---|---|---|
| mmu-miR-21a-3p | MIMAT0004628 | Aptx | 66408 | HITS-CLIP | 23142080 | Unspecified |
| mmu-miR-21a-3p | MIMAT0004628 | Aqp11 | 66333 | HITS-CLIP | 23142080 | Unspecified |
| mmu-miR-21a-3p | MIMAT0004628 | Ar | 11835 | HITS-CLIP | 23142080 | Unspecified |
| mmu-miR-21a-3p | MIMAT0004628 | Arel1 | 68497 | HITS-CLIP | 23142080 | Unspecified |
| mmu-miR-21a-3p | MIMAT0004628 | Arf1 | 11840 | HITS-CLIP | 23142080 | Unspecified |
| mmu-miR-21a-3p | MIMAT0004628 | Arf3 | 11842 | HITS-CLIP | 37986224\|29167373\|23142080 | Unspecified |
| mmu-miR-21a-3p | MIMAT0004628 | Arf4 | 11843 | HITS-CLIP | 23142080 | Unspecified |
| mmu-miR-21a-3p | MIMAT0004628 | Arfgef1 | 211673 | HITS-CLIP | 23142080\|29167373 | Unspecified |
| mmu-miR-21a-3p | MIMAT0004628 | Arfgef2 | 99371 | HITS-CLIP | 29167373 | Unspecified |
| mmu-miR-21a-3p | MIMAT0004628 | Arglu1 | 234023 | HITS-CLIP | 23142080 | Unspecified |
| mmu-miR-21a-3p | MIMAT0004628 | Arhgap11a | 228482 | HITS-CLIP | 37986224\|23142080 | Unspecified |
| mmu-miR-21a-3p | MIMAT0004628 | Arhgap12 | 75415 | HITS-CLIP | 23142080 | Unspecified |
| mmu-miR-21a-3p | MIMAT0004628 | Arhgap15 | 76117 | HITS-CLIP | 37986224\|23142080 | Unspecified |
| mmu-miR-21a-3p | MIMAT0004628 | Arhgap19 | 71085 | HITS-CLIP | 23142080 | Unspecified |
| mmu-miR-21a-3p | MIMAT0004628 | Arhgap26 | 71302 | HITS-CLIP | 23142080 | Unspecified |
| mmu-miR-21a-3p | MIMAT0004628 | Arhgap35 | 232906 | HITS-CLIP | 23142080 | Unspecified |
| mmu-miR-21a-3p | MIMAT0004628 | Arhgef11 | 213498 | HITS-CLIP | 29167373 | Unspecified |
| mmu-miR-21a-3p | MIMAT0004628 | Arhgef2 | 16800 | HITS-CLIP | 23142080 | Unspecified |
| mmu-miR-21a-3p | MIMAT0004628 | Arhgef3 | 71704 | HITS-CLIP | 23142080 | Unspecified |
| mmu-miR-21a-3p | MIMAT0004628 | Arhgef6 | 73341 | HITS-CLIP | 23142080 | Unspecified |
| mmu-miR-21a-3p | MIMAT0004628 | Arid1a | 93760 | HITS-CLIP | 23142080 | Unspecified |
| mmu-miR-21a-3p | MIMAT0004628 | Arid1b | 239985 | HITS-CLIP | 23142080\|29167373 | Unspecified |
| mmu-miR-21a-3p | MIMAT0004628 | Arid2 | 77044 | HITS-CLIP | 23142080 | Unspecified |
| mmu-miR-21a-3p | MIMAT0004628 | Arid3b | 56380 | HITS-CLIP | 37986224\|23142080 | Unspecified |
| mmu-miR-21a-3p | MIMAT0004628 | Arid4a | 238247 | HITS-CLIP | 23142080\|29167373 | Unspecified |
| mmu-miR-21a-3p | MIMAT0004628 | Arid4b | 94246 | HITS-CLIP | 37986224\|23142080\|29167373 | Unspecified |
| mmu-miR-21a-3p | MIMAT0004628 | Arid5b | 71371 | HITS-CLIP | 37986224\|23142080 | Unspecified |
| mmu-miR-21a-3p | MIMAT0004628 | Arih2 | 23807 | HITS-CLIP | 23142080 | Unspecified |
| mmu-miR-21a-3p | MIMAT0004628 | Arl2bp | 107566 | HITS-CLIP | 23142080 | Unspecified |
| mmu-miR-21a-3p | MIMAT0004628 | Arl4c | 320982 | HITS-CLIP | 23142080 | Unspecified |
| mmu-miR-21a-3p | MIMAT0004628 | Arl5b | 75869 | HITS-CLIP | 23142080 | Unspecified |
| mmu-miR-21a-3p | MIMAT0004628 | Arl6ip1 | 54208 | HITS-CLIP | 37986224\|23142080\|29167373 | Unspecified |
| mmu-miR-21a-3p | MIMAT0004628 | Arl6ip4 | 65105 | HITS-CLIP | 23142080 | Unspecified |
| mmu-miR-21a-3p | MIMAT0004628 | Arl6ip5 | 65106 | HITS-CLIP | 37986224\|23142080 | Unspecified |
| mmu-miR-21a-3p | MIMAT0004628 | Arl6ip6 | 65103 | HITS-CLIP | 23142080 | Unspecified |
| mmu-miR-21a-3p | MIMAT0004628 | Arl8b | 67166 | HITS-CLIP | 23142080 | Unspecified |
| mmu-miR-21a-3p | MIMAT0004628 | Armc8 | 74125 | HITS-CLIP | 23142080 | Unspecified |
| mmu-miR-21a-3p | MIMAT0004628 | Armh3 | 79591 | HITS-CLIP | 23142080 | Unspecified |
| mmu-miR-21a-3p | MIMAT0004628 | Arntl | 11865 | HITS-CLIP | 37986224 | Unspecified |

| | | | | | | |
|---|---|---|---|---|---|---|
| mmu-miR-21a-3p | MIMAT0004628 | Arpc1a | 56443 | HITS-CLIP | 23142080 | Unspecified |
| mmu-miR-21a-3p | MIMAT0004628 | Arpc2 | 76709 | HITS-CLIP | 23142080 | Unspecified |
| mmu-miR-21a-3p | MIMAT0004628 | Arpc5 | 67771 | HITS-CLIP | 23142080 | Unspecified |
| mmu-miR-21a-3p | MIMAT0004628 | Arrdc3 | 105171 | HITS-CLIP | 23142080 | Unspecified |
| mmu-miR-21a-3p | MIMAT0004628 | Asah1 | 11886 | HITS-CLIP | 23142080 | Unspecified |
| mmu-miR-21a-3p | MIMAT0004628 | Asap1 | 13196 | HITS-CLIP | 37986224\|23142080\|29167373 | Unspecified |
| mmu-miR-21a-3p | MIMAT0004628 | Asb3 | 65257 | HITS-CLIP | 23142080 | Unspecified |
| mmu-miR-21a-3p | MIMAT0004628 | Asb7 | 117589 | HITS-CLIP | 23142080 | Unspecified |
| mmu-miR-21a-3p | MIMAT0004628 | Asb8 | 78541 | HITS-CLIP | 23142080 | Unspecified |
| mmu-miR-21a-3p | MIMAT0004628 | Asf1a | 66403 | HITS-CLIP | 23142080 | Unspecified |
| mmu-miR-21a-3p | MIMAT0004628 | Ash1l | 192195 | HITS-CLIP | 37986224 | Unspecified |
| mmu-miR-21a-3p | MIMAT0004628 | Asnsd1 | 70396 | HITS-CLIP | 23142080 | Unspecified |
| mmu-miR-21a-3p | MIMAT0004628 | Aspm | 12316 | HITS-CLIP | 37986224\|23142080 | Unspecified |
| mmu-miR-21a-3p | MIMAT0004628 | Atad1 | 67979 | HITS-CLIP | 37986224\|23142080 | Unspecified |
| mmu-miR-21a-3p | MIMAT0004628 | Atad2 | 70472 | HITS-CLIP | 23142080 | Unspecified |
| mmu-miR-21a-3p | MIMAT0004628 | Atad2b | 320817 | HITS-CLIP | 29167373\|23142080 | Unspecified |
| mmu-miR-21a-3p | MIMAT0004628 | Atad5 | 237877 | HITS-CLIP | 23142080 | Unspecified |
| mmu-miR-21a-3p | MIMAT0004628 | Ate1 | 11907 | HITS-CLIP | 29167373 | Unspecified |
| mmu-miR-21a-3p | MIMAT0004628 | Atf1 | 11908 | HITS-CLIP | 23142080 | Unspecified |
| mmu-miR-21a-3p | MIMAT0004628 | Atf3 | 11910 | HITS-CLIP | 23142080 | Unspecified |
| mmu-miR-21a-3p | MIMAT0004628 | Atf6 | 226641 | HITS-CLIP | 23142080 | Unspecified |
| mmu-miR-21a-3p | MIMAT0004628 | Atf7 | 223922 | HITS-CLIP | 23142080\|29167373 | Unspecified |
| mmu-miR-21a-3p | MIMAT0004628 | Atf7ip | 54343 | HITS-CLIP | 37986224\|23142080 | Unspecified |
| mmu-miR-21a-3p | MIMAT0004628 | Atg16l2 | 73683 | HITS-CLIP | 23142080 | Unspecified |
| mmu-miR-21a-3p | MIMAT0004628 | Atg4b | 66615 | HITS-CLIP | 29167373 | Unspecified |
| mmu-miR-21a-3p | MIMAT0004628 | Atg5 | 11793 | HITS-CLIP | 29167373 | Unspecified |
| mmu-miR-21a-3p | MIMAT0004628 | Atg9a | 245860 | HITS-CLIP | 23142080 | Unspecified |
| mmu-miR-21a-3p | MIMAT0004628 | Atl3 | 109168 | HITS-CLIP | 37986224\|23142080 | Unspecified |
| mmu-miR-21a-3p | MIMAT0004628 | Atm | 11920 | HITS-CLIP | 23142080 | Unspecified |
| mmu-miR-21a-3p | MIMAT0004628 | Atmin | 234776 | HITS-CLIP | 23142080 | Unspecified |
| mmu-miR-21a-3p | MIMAT0004628 | Atp10a | 11982 | HITS-CLIP | 23142080 | Unspecified |
| mmu-miR-21a-3p | MIMAT0004628 | Atp11a | 50770 | HITS-CLIP | 37986224\|23142080 | Unspecified |
| mmu-miR-21a-3p | MIMAT0004628 | Atp11b | 76295 | HITS-CLIP | 37986224\|23142080 | Unspecified |
| mmu-miR-21a-3p | MIMAT0004628 | Atp11c | 320940 | HITS-CLIP | 23142080 | Unspecified |
| mmu-miR-21a-3p | MIMAT0004628 | Atp13a2 | 23400 | HITS-CLIP | 23142080 | Unspecified |
| mmu-miR-21a-3p | MIMAT0004628 | Atp13a3 | 224088 | HITS-CLIP | 23142080\|29167373 | Unspecified |
| mmu-miR-21a-3p | MIMAT0004628 | Atp2a2 | 11938 | HITS-CLIP | 29167373 | Unspecified |
| mmu-miR-21a-3p | MIMAT0004628 | Atp2a3 | 53313 | HITS-CLIP | 23142080 | Unspecified |
| mmu-miR-21a-3p | MIMAT0004628 | Atp2b1 | 67972 | HITS-CLIP | 23142080\|29167373 | Unspecified |

| | | | | | | |
|---|---|---|---|---|---|---|
| mmu-miR-21a-3p | MIMAT0004628 | Atp5b | 11947 | HITS-CLIP | 37986224\|23142080 | Unspecified |
| mmu-miR-21a-3p | MIMAT0004628 | Atp6ap1 | 54411 | HITS-CLIP | 23142080 | Unspecified |
| mmu-miR-21a-3p | MIMAT0004628 | Atp6v0a2 | 21871 | HITS-CLIP | 23142080 | Unspecified |
| mmu-miR-21a-3p | MIMAT0004628 | Atp6v1a | 11964 | HITS-CLIP | 23142080\|29167373 | Unspecified |
| mmu-miR-21a-3p | MIMAT0004628 | Atp6v1b2 | 11966 | HITS-CLIP | 37986224\|23142080\|29167373 | Unspecified |
| mmu-miR-21a-3p | MIMAT0004628 | Atp6v1f | 66144 | HITS-CLIP | 23142080 | Unspecified |
| mmu-miR-21a-3p | MIMAT0004628 | Atp6v1h | 108664 | HITS-CLIP | 37986224\|29167373\|23142080 | Unspecified |
| mmu-miR-21a-3p | MIMAT0004628 | Atp7a | 11977 | HITS-CLIP | 23142080\|26203562\|29167373 | Unspecified |
| mmu-miR-21a-3p | MIMAT0004628 | Atp9a | 11981 | HITS-CLIP | 23142080 | Unspecified |
| mmu-miR-21a-3p | MIMAT0004628 | Atr | 245000 | HITS-CLIP | 23142080 | Unspecified |
| mmu-miR-21a-3p | MIMAT0004628 | Atrn | 11990 | HITS-CLIP | 23142080 | Unspecified |
| mmu-miR-21a-3p | MIMAT0004628 | Atrx | 22589 | HITS-CLIP | 23142080 | Unspecified |
| mmu-miR-21a-3p | MIMAT0004628 | Atxn1 | 20238 | HITS-CLIP | 23142080 | Unspecified |
| mmu-miR-21a-3p | MIMAT0004628 | Atxn10 | 54138 | HITS-CLIP | 23142080 | Unspecified |
| mmu-miR-21a-3p | MIMAT0004628 | Atxn1l | 52335 | HITS-CLIP | 37986224\|23142080\|29167373 | Unspecified |
| mmu-miR-21a-3p | MIMAT0004628 | Atxn2l | 233871 | HITS-CLIP | 23142080 | Unspecified |
| mmu-miR-21a-3p | MIMAT0004628 | Atxn3 | 110616 | HITS-CLIP | 23142080\|29167373 | Unspecified |
| mmu-miR-21a-3p | MIMAT0004628 | Atxn7 | 246103 | HITS-CLIP | 29167373 | Unspecified |
| mmu-miR-21a-3p | MIMAT0004628 | Atxn7l3 | 217218 | HITS-CLIP | 37986224\|23142080 | Unspecified |
| mmu-miR-21a-3p | MIMAT0004628 | Atxn7l3b | 382423 | HITS-CLIP//qCLASH | 37986224\|31393953\|23142080\|29167 | Unspecified |
| mmu-miR-21a-3p | MIMAT0004628 | Avl9 | 78937 | HITS-CLIP | 29167373\|23142080 | Unspecified |
| mmu-miR-21a-3p | MIMAT0004628 | B230219D22Rik | 78521 | HITS-CLIP | 29167373\|23142080 | Unspecified |
| mmu-miR-21a-3p | MIMAT0004628 | B3galnt2 | 97884 | HITS-CLIP | 23142080 | Unspecified |
| mmu-miR-21a-3p | MIMAT0004628 | B3gnt2 | 53625 | HITS-CLIP | 23142080 | Unspecified |
| mmu-miR-21a-3p | MIMAT0004628 | B4galt1 | 14595 | HITS-CLIP | 23142080 | Unspecified |
| mmu-miR-21a-3p | MIMAT0004628 | B4galt7 | 218271 | HITS-CLIP | 23142080 | Unspecified |
| mmu-miR-21a-3p | MIMAT0004628 | B4gat1 | 135972575 | HITS-CLIP | 23142080 | Unspecified |
| mmu-miR-21a-3p | MIMAT0004628 | BC005537 | 79555 | HITS-CLIP | 23142080 | Unspecified |
| mmu-miR-21a-3p | MIMAT0004628 | Babam2 | 9577 | HITS-CLIP | 23142080 | Unspecified |
| mmu-miR-21a-3p | MIMAT0004628 | Bach2 | 12014 | HITS-CLIP | 37986224\|23142080 | Unspecified |
| mmu-miR-21a-3p | MIMAT0004628 | Bag6 | 224727 | HITS-CLIP | 23142080 | Unspecified |
| mmu-miR-21a-3p | MIMAT0004628 | Baz1a | 217578 | HITS-CLIP | 37986224\|23142080 | Unspecified |
| mmu-miR-21a-3p | MIMAT0004628 | Baz1b | 22385 | HITS-CLIP | 23142080 | Unspecified |
| mmu-miR-21a-3p | MIMAT0004628 | Bbc3 | 170770 | HITS-CLIP | 37986224\|23142080 | Unspecified |
| mmu-miR-21a-3p | MIMAT0004628 | Bbs12 | 241950 | HITS-CLIP | 23142080 | Unspecified |
| mmu-miR-21a-3p | MIMAT0004628 | Bbs5 | 72569 | HITS-CLIP | 23142080 | Unspecified |
| mmu-miR-21a-3p | MIMAT0004628 | Bbx | 70508 | HITS-CLIP | 23142080 | Unspecified |
| mmu-miR-21a-3p | MIMAT0004628 | Bcar3 | 29815 | HITS-CLIP | 29167373 | Unspecified |
| mmu-miR-21a-3p | MIMAT0004628 | Bcas2 | 68183 | HITS-CLIP | 37986224\|23142080 | Unspecified |

| | | | | | | |
|---|---|---|---|---|---|---|
| mmu-miR-21a-3p | MIMAT0004628 | Bcl11b | 58208 | HITS-CLIP | 37986224\|23142080 | Unspecified |
| mmu-miR-21a-3p | MIMAT0004628 | Bcl2l1 | 12048 | HITS-CLIP | 29167373 | Unspecified |
| mmu-miR-21a-3p | MIMAT0004628 | Bcl2l11 | 12125 | HITS-CLIP | 23142080 | Unspecified |
| mmu-miR-21a-3p | MIMAT0004628 | Bcl6 | 12053 | HITS-CLIP | 23142080\|29167373 | Unspecified |
| mmu-miR-21a-3p | MIMAT0004628 | Bclaf1 | 72567 | HITS-CLIP | 29167373 | Unspecified |
| mmu-miR-21a-3p | MIMAT0004628 | Bclaf3 | 382252 | HITS-CLIP | 23142080 | Unspecified |
| mmu-miR-21a-3p | MIMAT0004628 | Bcor | 71458 | HITS-CLIP | 23142080 | Unspecified |
| mmu-miR-21a-3p | MIMAT0004628 | Bcorl1 | 320376 | HITS-CLIP | 23142080 | Unspecified |
| mmu-miR-21a-3p | MIMAT0004628 | Bcr | 110279 | HITS-CLIP | 37986224 | Unspecified |
| mmu-miR-21a-3p | MIMAT0004628 | Bdh1 | 71911 | HITS-CLIP | 23142080 | Unspecified |
| mmu-miR-21a-3p | MIMAT0004628 | Becn1 | 56208 | HITS-CLIP | 23142080 | Unspecified |
| mmu-miR-21a-3p | MIMAT0004628 | Best3 | 382427 | HITS-CLIP | 23142080 | Unspecified |
| mmu-miR-21a-3p | MIMAT0004628 | Bet1 | 12068 | HITS-CLIP | 37986224\|23142080 | Unspecified |
| mmu-miR-21a-3p | MIMAT0004628 | Bhlhb9 | 70237 | HITS-CLIP | 23142080 | Unspecified |
| mmu-miR-21a-3p | MIMAT0004628 | Bhlhe40 | 20893 | HITS-CLIP | 37986224 | Unspecified |
| mmu-miR-21a-3p | MIMAT0004628 | Bhlhe41 | 79362 | HITS-CLIP | 37986224\|29167373\|23142080 | Unspecified |
| mmu-miR-21a-3p | MIMAT0004628 | Bicd2 | 76895 | HITS-CLIP | 23142080 | Unspecified |
| mmu-miR-21a-3p | MIMAT0004628 | Bicdl1 | 92558 | HITS-CLIP | 23142080 | Unspecified |
| mmu-miR-21a-3p | MIMAT0004628 | Bicral | 23506 | HITS-CLIP | 23142080 | Unspecified |
| mmu-miR-21a-3p | MIMAT0004628 | Birc2 | 11797 | HITS-CLIP | 37986224 | Unspecified |
| mmu-miR-21a-3p | MIMAT0004628 | Birc3 | 11796 | HITS-CLIP | 23142080 | Unspecified |
| mmu-miR-21a-3p | MIMAT0004628 | Birc6 | 12211 | HITS-CLIP | 37986224\|23142080\|29167373 | Unspecified |
| mmu-miR-21a-3p | MIMAT0004628 | Blcap | 53619 | HITS-CLIP | 29167373 | Unspecified |
| mmu-miR-21a-3p | MIMAT0004628 | Blm | 12144 | HITS-CLIP | 37986224 | Unspecified |
| mmu-miR-21a-3p | MIMAT0004628 | Bloc1s6 | 18457 | HITS-CLIP | 37986224\|23142080 | Unspecified |
| mmu-miR-21a-3p | MIMAT0004628 | Bmpr1a | 12166 | HITS-CLIP | 23142080 | Unspecified |
| mmu-miR-21a-3p | MIMAT0004628 | Bmpr2 | 12168 | HITS-CLIP | 23142080 | Unspecified |
| mmu-miR-21a-3p | MIMAT0004628 | Bms1 | 213895 | HITS-CLIP | 23142080 | Unspecified |
| mmu-miR-21a-3p | MIMAT0004628 | Bmt2 | 101148 | HITS-CLIP | 23142080 | Unspecified |
| mmu-miR-21a-3p | MIMAT0004628 | Bnip2 | 12175 | HITS-CLIP | 23142080 | Unspecified |
| mmu-miR-21a-3p | MIMAT0004628 | Bnip3l | 12177 | HITS-CLIP | 23142080 | Unspecified |
| mmu-miR-21a-3p | MIMAT0004628 | Bphl | 68021 | HITS-CLIP | 23142080 | Unspecified |
| mmu-miR-21a-3p | MIMAT0004628 | Bptf | 207165 | HITS-CLIP | 37986224 | Unspecified |
| mmu-miR-21a-3p | MIMAT0004628 | Braf | 109880 | HITS-CLIP | 23142080 | Unspecified |
| mmu-miR-21a-3p | MIMAT0004628 | Brat1 | 231841 | HITS-CLIP | 23142080 | Unspecified |
| mmu-miR-21a-3p | MIMAT0004628 | Brca1 | 12189 | HITS-CLIP | 23142080 | Unspecified |
| mmu-miR-21a-3p | MIMAT0004628 | Brd2 | 14312 | HITS-CLIP | 23142080 | Unspecified |
| mmu-miR-21a-3p | MIMAT0004628 | Brd4 | 57261 | HITS-CLIP | 29167373 | Unspecified |
| mmu-miR-21a-3p | MIMAT0004628 | Brd7 | 26992 | HITS-CLIP | 23142080 | Unspecified |

| | | | | | | |
|---|---|---|---|---|---|---|
| mmu-miR-21a-3p | MIMAT0004628 | Brd8 | 78656 | HITS-CLIP | 23142080 | Unspecified |
| mmu-miR-21a-3p | MIMAT0004628 | Bri3bp | 76809 | HITS-CLIP | 23142080 | Unspecified |
| mmu-miR-21a-3p | MIMAT0004628 | Brip1 | 237911 | HITS-CLIP | 23142080 | Unspecified |
| mmu-miR-21a-3p | MIMAT0004628 | Brox | 71678 | HITS-CLIP | 23142080 | Unspecified |
| mmu-miR-21a-3p | MIMAT0004628 | Brwd3 | 382236 | HITS-CLIP | 23142080 | Unspecified |
| mmu-miR-21a-3p | MIMAT0004628 | Bsg | 12215 | HITS-CLIP | 23142080 | Unspecified |
| mmu-miR-21a-3p | MIMAT0004628 | Btaf1 | 107182 | HITS-CLIP | 23142080\|29167373 | Unspecified |
| mmu-miR-21a-3p | MIMAT0004628 | Btbd7 | 238386 | HITS-CLIP | 23142080 | Unspecified |
| mmu-miR-21a-3p | MIMAT0004628 | Btbd8 | 100503185 | HITS-CLIP | 23142080 | Unspecified |
| mmu-miR-21a-3p | MIMAT0004628 | Btg1 | 12226 | HITS-CLIP | 29167373 | Unspecified |
| mmu-miR-21a-3p | MIMAT0004628 | Btla | 208154 | HITS-CLIP | 37986224\|23142080 | Unspecified |
| mmu-miR-21a-3p | MIMAT0004628 | Bub1b | 12236 | HITS-CLIP | 23142080 | Unspecified |
| mmu-miR-21a-3p | MIMAT0004628 | C1d | 57316 | HITS-CLIP | 23142080 | Unspecified |
| mmu-miR-21a-3p | MIMAT0004628 | C1galt1 | 94192 | HITS-CLIP | 23142080 | Unspecified |
| mmu-miR-21a-3p | MIMAT0004628 | C1galt1c1 | 59048 | HITS-CLIP | 23142080 | Unspecified |
| mmu-miR-21a-3p | MIMAT0004628 | C1qc | 12262 | HITS-CLIP | 29167373 | Unspecified |
| mmu-miR-21a-3p | MIMAT0004628 | C2cd5 | 74741 | HITS-CLIP | 23142080 | Unspecified |
| mmu-miR-21a-3p | MIMAT0004628 | C9orf72 | 203228 | HITS-CLIP | 23142080 | Unspecified |
| mmu-miR-21a-3p | MIMAT0004628 | Caap1 | 67770 | HITS-CLIP | 23142080 | Unspecified |
| mmu-miR-21a-3p | MIMAT0004628 | Cab39 | 12283 | HITS-CLIP | 37986224\|23142080 | Unspecified |
| mmu-miR-21a-3p | MIMAT0004628 | Cacna1d | 12289 | HITS-CLIP | 29167373 | Unspecified |
| mmu-miR-21a-3p | MIMAT0004628 | Cacul1 | 78832 | HITS-CLIP | 23142080\|29167373 | Unspecified |
| mmu-miR-21a-3p | MIMAT0004628 | Cacybp | 12301 | HITS-CLIP | 23142080 | Unspecified |
| mmu-miR-21a-3p | MIMAT0004628 | Cadm1 | 54725 | HITS-CLIP | 29167373 | Unspecified |
| mmu-miR-21a-3p | MIMAT0004628 | Calcrl | 54598 | HITS-CLIP | 23142080 | Unspecified |
| mmu-miR-21a-3p | MIMAT0004628 | Calm1 | 12313 | HITS-CLIP | 23142080\|29167373 | Unspecified |
| mmu-miR-21a-3p | MIMAT0004628 | Calm2 | 12314 | HITS-CLIP | 23142080\|29167373 | Unspecified |
| mmu-miR-21a-3p | MIMAT0004628 | Camk2g | 12325 | HITS-CLIP | 23142080 | Unspecified |
| mmu-miR-21a-3p | MIMAT0004628 | Camkk2 | 207565 | HITS-CLIP | 29167373 | Unspecified |
| mmu-miR-21a-3p | MIMAT0004628 | Caml | 12328 | HITS-CLIP | 23142080 | Unspecified |
| mmu-miR-21a-3p | MIMAT0004628 | Camsap2 | 67886 | HITS-CLIP | 23142080 | Unspecified |
| mmu-miR-21a-3p | MIMAT0004628 | Canx | 12330 | HITS-CLIP | 37986224\|23142080 | Unspecified |
| mmu-miR-21a-3p | MIMAT0004628 | Capn7 | 12339 | HITS-CLIP | 37986224 | Unspecified |
| mmu-miR-21a-3p | MIMAT0004628 | Caprin1 | 53872 | HITS-CLIP | 23142080 | Unspecified |
| mmu-miR-21a-3p | MIMAT0004628 | Capza2 | 12343 | HITS-CLIP | 29167373 | Unspecified |
| mmu-miR-21a-3p | MIMAT0004628 | Casp2 | 12366 | HITS-CLIP | 23142080 | Unspecified |
| mmu-miR-21a-3p | MIMAT0004628 | Casp8ap2 | 26885 | HITS-CLIP | 29167373 | Unspecified |
| mmu-miR-21a-3p | MIMAT0004628 | Cast | 12380 | HITS-CLIP | 23142080 | Unspecified |
| mmu-miR-21a-3p | MIMAT0004628 | Castor1 | 652968 | HITS-CLIP | 23142080 | Unspecified |

| | | | | | | |
|---|---|---|---|---|---|---|
| mmu-miR-21a-3p | MIMAT0004628 | Cbarp | 100503659 | HITS-CLIP | 23142080 | Unspecified |
| mmu-miR-21a-3p | MIMAT0004628 | Cbl | 12402 | HITS-CLIP | 29167373 | Unspecified |
| mmu-miR-21a-3p | MIMAT0004628 | Cblb | 208650 | HITS-CLIP | 23142080 | Unspecified |
| mmu-miR-21a-3p | MIMAT0004628 | Cbx5 | 12419 | HITS-CLIP | 23142080 | Unspecified |
| mmu-miR-21a-3p | MIMAT0004628 | Ccar1 | 67500 | HITS-CLIP | 23142080\|29167373 | Unspecified |
| mmu-miR-21a-3p | MIMAT0004628 | Ccar2 | 219158 | HITS-CLIP | 23142080 | Unspecified |
| mmu-miR-21a-3p | MIMAT0004628 | Ccdc126 | 57895 | HITS-CLIP | 23142080 | Unspecified |
| mmu-miR-21a-3p | MIMAT0004628 | Ccdc130 | 67736 | HITS-CLIP | 37986224\|23142080 | Unspecified |
| mmu-miR-21a-3p | MIMAT0004628 | Ccdc141 | 545428 | HITS-CLIP | 37986224\|23142080 | Unspecified |
| mmu-miR-21a-3p | MIMAT0004628 | Ccdc18 | 73254 | HITS-CLIP | 23142080 | Unspecified |
| mmu-miR-21a-3p | MIMAT0004628 | Ccdc186 | 55088 | HITS-CLIP | 29167373 | Unspecified |
| mmu-miR-21a-3p | MIMAT0004628 | Ccdc22 | 54638 | HITS-CLIP | 23142080 | Unspecified |
| mmu-miR-21a-3p | MIMAT0004628 | Ccdc28a | 215814 | HITS-CLIP | 23142080 | Unspecified |
| mmu-miR-21a-3p | MIMAT0004628 | Ccdc47 | 67163 | HITS-CLIP | 37986224\|23142080 | Unspecified |
| mmu-miR-21a-3p | MIMAT0004628 | Ccdc50 | 67501 | HITS-CLIP | 23142080\|29167373 | Unspecified |
| mmu-miR-21a-3p | MIMAT0004628 | Ccdc6 | 76551 | HITS-CLIP | 23142080 | Unspecified |
| mmu-miR-21a-3p | MIMAT0004628 | Ccdc71 | 72454 | HITS-CLIP | 23142080 | Unspecified |
| mmu-miR-21a-3p | MIMAT0004628 | Ccdc71l | 72123 | HITS-CLIP | 23142080 | Unspecified |
| mmu-miR-21a-3p | MIMAT0004628 | Ccl2 | 20296 | HITS-CLIP | 29167373 | Unspecified |
| mmu-miR-21a-3p | MIMAT0004628 | Ccl3 | 20302 | HITS-CLIP | 23142080 | Unspecified |
| mmu-miR-21a-3p | MIMAT0004628 | Ccl4 | 20303 | HITS-CLIP | 37986224\|23142080 | Unspecified |
| mmu-miR-21a-3p | MIMAT0004628 | Ccl9 | 20308 | HITS-CLIP | 29167373 | Unspecified |
| mmu-miR-21a-3p | MIMAT0004628 | Ccna2 | 12428 | HITS-CLIP | 23142080 | Unspecified |
| mmu-miR-21a-3p | MIMAT0004628 | Ccnb1 | 268697 | HITS-CLIP | 23142080 | Unspecified |
| mmu-miR-21a-3p | MIMAT0004628 | Ccnc | 51813 | HITS-CLIP | 23142080 | Unspecified |
| mmu-miR-21a-3p | MIMAT0004628 | Ccnd2 | 12444 | HITS-CLIP | 23142080 | Unspecified |
| mmu-miR-21a-3p | MIMAT0004628 | Ccne2 | 12448 | HITS-CLIP | 23142080 | Unspecified |
| mmu-miR-21a-3p | MIMAT0004628 | Ccnf | 12449 | HITS-CLIP | 23142080 | Unspecified |
| mmu-miR-21a-3p | MIMAT0004628 | Ccni | 12453 | HITS-CLIP | 23142080 | Unspecified |
| mmu-miR-21a-3p | MIMAT0004628 | Ccnj | 240665 | HITS-CLIP | 23142080 | Unspecified |
| mmu-miR-21a-3p | MIMAT0004628 | Ccnl2 | 56036 | HITS-CLIP | 23142080 | Unspecified |
| mmu-miR-21a-3p | MIMAT0004628 | Ccnt1 | 12455 | HITS-CLIP | 37986224\|23142080 | Unspecified |
| mmu-miR-21a-3p | MIMAT0004628 | Ccnt2 | 72949 | HITS-CLIP | 37986224\|23142080\|29167373 | Unspecified |
| mmu-miR-21a-3p | MIMAT0004628 | Ccp110 | 101565 | HITS-CLIP | 29167373 | Unspecified |
| mmu-miR-21a-3p | MIMAT0004628 | Ccr7 | 12775 | HITS-CLIP | 23142080 | Unspecified |
| mmu-miR-21a-3p | MIMAT0004628 | Ccs | 12460 | HITS-CLIP | 23142080 | Unspecified |
| mmu-miR-21a-3p | MIMAT0004628 | Ccser2 | 72972 | HITS-CLIP | 23142080 | Unspecified |
| mmu-miR-21a-3p | MIMAT0004628 | Cct5 | 12465 | HITS-CLIP | 23142080 | Unspecified |
| mmu-miR-21a-3p | MIMAT0004628 | Cct7 | 12468 | HITS-CLIP | 23142080 | Unspecified |

| | | | | | | |
|---|---|---|---|---|---|---|
| mmu-miR-21a-3p | MIMAT0004628 | Cd101 | 630146 | HITS-CLIP | 23142080 | Unspecified |
| mmu-miR-21a-3p | MIMAT0004628 | Cd14 | 12475 | HITS-CLIP | 29167373 | Unspecified |
| mmu-miR-21a-3p | MIMAT0004628 | Cd151 | 12476 | HITS-CLIP | 23142080 | Unspecified |
| mmu-miR-21a-3p | MIMAT0004628 | Cd164 | 53599 | HITS-CLIP | 37986224\|23142080 | Unspecified |
| mmu-miR-21a-3p | MIMAT0004628 | Cd200 | 17470 | HITS-CLIP | 37986224\|23142080 | Unspecified |
| mmu-miR-21a-3p | MIMAT0004628 | Cd247 | 12503 | HITS-CLIP | 37986224 | Unspecified |
| mmu-miR-21a-3p | MIMAT0004628 | Cd28 | 12487 | HITS-CLIP | 37986224\|23142080 | Unspecified |
| mmu-miR-21a-3p | MIMAT0004628 | Cd2ap | 12488 | HITS-CLIP | 23142080 | Unspecified |
| mmu-miR-21a-3p | MIMAT0004628 | Cd300a | 217303 | HITS-CLIP | 29167373 | Unspecified |
| mmu-miR-21a-3p | MIMAT0004628 | Cd36 | 12491 | HITS-CLIP | 29167373 | Unspecified |
| mmu-miR-21a-3p | MIMAT0004628 | Cd3d | 12500 | HITS-CLIP | 23142080 | Unspecified |
| mmu-miR-21a-3p | MIMAT0004628 | Cd40lg | 959 | HITS-CLIP | 23142080 | Unspecified |
| mmu-miR-21a-3p | MIMAT0004628 | Cd47 | 16423 | HITS-CLIP | 23142080 | Unspecified |
| mmu-miR-21a-3p | MIMAT0004628 | Cd53 | 12508 | HITS-CLIP | 29167373\|23142080 | Unspecified |
| mmu-miR-21a-3p | MIMAT0004628 | Cd63 | 12512 | HITS-CLIP | 29167373 | Unspecified |
| mmu-miR-21a-3p | MIMAT0004628 | Cd79a | 12518 | HITS-CLIP | 23142080 | Unspecified |
| mmu-miR-21a-3p | MIMAT0004628 | Cd81 | 12520 | HITS-CLIP | 23142080 | Unspecified |
| mmu-miR-21a-3p | MIMAT0004628 | Cd82 | 12521 | HITS-CLIP | 23142080 | Unspecified |
| mmu-miR-21a-3p | MIMAT0004628 | Cd8a | 12525 | HITS-CLIP | 37986224 | Unspecified |
| mmu-miR-21a-3p | MIMAT0004628 | Cd93 | 17064 | HITS-CLIP | 29167373 | Unspecified |
| mmu-miR-21a-3p | MIMAT0004628 | Cdan1 | 68968 | HITS-CLIP | 23142080 | Unspecified |
| mmu-miR-21a-3p | MIMAT0004628 | Cdc14a | 229776 | HITS-CLIP | 23142080 | Unspecified |
| mmu-miR-21a-3p | MIMAT0004628 | Cdc14b | 218294 | HITS-CLIP | 23142080 | Unspecified |
| mmu-miR-21a-3p | MIMAT0004628 | Cdc16 | 69957 | HITS-CLIP | 23142080 | Unspecified |
| mmu-miR-21a-3p | MIMAT0004628 | Cdc20 | 107995 | HITS-CLIP | 23142080 | Unspecified |
| mmu-miR-21a-3p | MIMAT0004628 | Cdc23 | 52563 | HITS-CLIP | 23142080 | Unspecified |
| mmu-miR-21a-3p | MIMAT0004628 | Cdc27 | 217232 | HITS-CLIP | 23142080 | Unspecified |
| mmu-miR-21a-3p | MIMAT0004628 | Cdc34 | 216150 | HITS-CLIP | 23142080 | Unspecified |
| mmu-miR-21a-3p | MIMAT0004628 | Cdc37l1 | 67072 | HITS-CLIP | 37986224\|23142080 | Unspecified |
| mmu-miR-21a-3p | MIMAT0004628 | Cdc40 | 71713 | HITS-CLIP | 23142080 | Unspecified |
| mmu-miR-21a-3p | MIMAT0004628 | Cdc42 | 12540 | HITS-CLIP | 29167373\|23142080 | Unspecified |
| mmu-miR-21a-3p | MIMAT0004628 | Cdc7 | 12545 | HITS-CLIP | 23142080 | Unspecified |
| mmu-miR-21a-3p | MIMAT0004628 | Cdca8 | 52276 | HITS-CLIP | 23142080 | Unspecified |
| mmu-miR-21a-3p | MIMAT0004628 | Cdk12 | 69131 | HITS-CLIP | 29167373 | Unspecified |
| mmu-miR-21a-3p | MIMAT0004628 | Cdk13 | 69562 | HITS-CLIP | 23142080 | Unspecified |
| mmu-miR-21a-3p | MIMAT0004628 | Cdk16 | 18555 | HITS-CLIP | 23142080 | Unspecified |
| mmu-miR-21a-3p | MIMAT0004628 | Cdk17 | 237459 | HITS-CLIP | 37986224\|23142080\|29167373 | Unspecified |
| mmu-miR-21a-3p | MIMAT0004628 | Cdk19 | 78334 | HITS-CLIP | 37986224\|23142080 | Unspecified |
| mmu-miR-21a-3p | MIMAT0004628 | Cdk2ap2 | 52004 | HITS-CLIP | 23142080 | Unspecified |

| | | | | | | |
|---|---|---|---|---|---|---|
| mmu-miR-21a-3p | MIMAT0004628 | Cdk5rap2 | 214444 | HITS-CLIP | 37986224 | Unspecified |
| mmu-miR-21a-3p | MIMAT0004628 | Cdk6 | 12571 | HITS-CLIP | 29167373 | Unspecified |
| mmu-miR-21a-3p | MIMAT0004628 | Cdk7 | 12572 | HITS-CLIP | 23142080 | Unspecified |
| mmu-miR-21a-3p | MIMAT0004628 | Cdk8 | 264064 | HITS-CLIP | 23142080 | Unspecified |
| mmu-miR-21a-3p | MIMAT0004628 | Cdk9 | 107951 | HITS-CLIP | 37986224\|23142080 | Unspecified |
| mmu-miR-21a-3p | MIMAT0004628 | Cdkn1b | 12576 | HITS-CLIP | 23142080 | Unspecified |
| mmu-miR-21a-3p | MIMAT0004628 | Cdkn2d | 12581 | HITS-CLIP | 23142080 | Unspecified |
| mmu-miR-21a-3p | MIMAT0004628 | Cdon | 57810 | HITS-CLIP | 37986224\|23142080 | Unspecified |
| mmu-miR-21a-3p | MIMAT0004628 | Cds2 | 110911 | HITS-CLIP | 23142080 | Unspecified |
| mmu-miR-21a-3p | MIMAT0004628 | Cdt1 | 67177 | HITS-CLIP | 23142080 | Unspecified |
| mmu-miR-21a-3p | MIMAT0004628 | Cdyl | 12593 | HITS-CLIP | 23142080 | Unspecified |
| mmu-miR-21a-3p | MIMAT0004628 | Celf1 | 13046 | HITS-CLIP | 37986224\|23142080\|29167373 | Unspecified |
| mmu-miR-21a-3p | MIMAT0004628 | Celf2 | 14007 | HITS-CLIP | 37986224\|23142080 | Unspecified |
| mmu-miR-21a-3p | MIMAT0004628 | Cemip2 | 23670 | HITS-CLIP | 23142080 | Unspecified |
| mmu-miR-21a-3p | MIMAT0004628 | Cenpa | 12615 | HITS-CLIP | 37986224 | Unspecified |
| mmu-miR-21a-3p | MIMAT0004628 | Cenpi | 102920 | HITS-CLIP | 23142080 | Unspecified |
| mmu-miR-21a-3p | MIMAT0004628 | Cep104 | 230967 | HITS-CLIP | 23142080 | Unspecified |
| mmu-miR-21a-3p | MIMAT0004628 | Cep120 | 225523 | HITS-CLIP | 37986224\|29167373\|23142080 | Unspecified |
| mmu-miR-21a-3p | MIMAT0004628 | Cep170 | 545389 | HITS-CLIP | 37986224\|23142080 | Unspecified |
| mmu-miR-21a-3p | MIMAT0004628 | Cep192 | 70799 | HITS-CLIP | 23142080 | Unspecified |
| mmu-miR-21a-3p | MIMAT0004628 | Cep350 | 74081 | HITS-CLIP | 23142080 | Unspecified |
| mmu-miR-21a-3p | MIMAT0004628 | Cep55 | 74107 | HITS-CLIP | 37986224\|23142080 | Unspecified |
| mmu-miR-21a-3p | MIMAT0004628 | Cep97 | 74201 | HITS-CLIP | 37986224\|23142080 | Unspecified |
| mmu-miR-21a-3p | MIMAT0004628 | Cept1 | 99712 | HITS-CLIP | 23142080 | Unspecified |
| mmu-miR-21a-3p | MIMAT0004628 | Cfap97 | 57587 | HITS-CLIP | 23142080 | Unspecified |
| mmu-miR-21a-3p | MIMAT0004628 | Cfl1 | 12631 | HITS-CLIP | 23142080\|29167373 | Unspecified |
| mmu-miR-21a-3p | MIMAT0004628 | Ch25h | 12642 | HITS-CLIP | 29167373 | Unspecified |
| mmu-miR-21a-3p | MIMAT0004628 | Chd1 | 12648 | HITS-CLIP | 23142080 | Unspecified |
| mmu-miR-21a-3p | MIMAT0004628 | Chd2 | 244059 | HITS-CLIP | 37986224\|23142080\|29167373 | Unspecified |
| mmu-miR-21a-3p | MIMAT0004628 | Chd3 | 216848 | HITS-CLIP | 29167373 | Unspecified |
| mmu-miR-21a-3p | MIMAT0004628 | Chd7 | 320790 | HITS-CLIP | 37986224\|23142080 | Unspecified |
| mmu-miR-21a-3p | MIMAT0004628 | Chd9 | 109151 | HITS-CLIP | 23142080 | Unspecified |
| mmu-miR-21a-3p | MIMAT0004628 | Chek1 | 12649 | HITS-CLIP | 23142080 | Unspecified |
| mmu-miR-21a-3p | MIMAT0004628 | Chic2 | 74277 | HITS-CLIP | 37986224 | Unspecified |
| mmu-miR-21a-3p | MIMAT0004628 | Chm | 12662 | HITS-CLIP | 23142080 | Unspecified |
| mmu-miR-21a-3p | MIMAT0004628 | Chmp4b | 75608 | HITS-CLIP | 37986224\|23142080 | Unspecified |
| mmu-miR-21a-3p | MIMAT0004628 | Chp1 | 56398 | HITS-CLIP | 23142080 | Unspecified |
| mmu-miR-21a-3p | MIMAT0004628 | Chpt1 | 212862 | HITS-CLIP | 23142080 | Unspecified |
| mmu-miR-21a-3p | MIMAT0004628 | Chrna6 | 8973 | HITS-CLIP | 23142080 | Unspecified |

| | | | | | | |
|---|---|---|---|---|---|---|
| mmu-miR-21a-3p | MIMAT0004628 | Churc1 | 211151 | HITS-CLIP | 23142080 | Unspecified |
| mmu-miR-21a-3p | MIMAT0004628 | Cib1 | 23991 | HITS-CLIP | 23142080 | Unspecified |
| mmu-miR-21a-3p | MIMAT0004628 | Cic | 71722 | HITS-CLIP | 23142080\|29167373 | Unspecified |
| mmu-miR-21a-3p | MIMAT0004628 | Cip2a | 57650 | HITS-CLIP | 23142080 | Unspecified |
| mmu-miR-21a-3p | MIMAT0004628 | Cit | 12704 | HITS-CLIP | 23142080 | Unspecified |
| mmu-miR-21a-3p | MIMAT0004628 | Cited2 | 17684 | HITS-CLIP | 23142080 | Unspecified |
| mmu-miR-21a-3p | MIMAT0004628 | Ckap5 | 75786 | HITS-CLIP | 23142080 | Unspecified |
| mmu-miR-21a-3p | MIMAT0004628 | Clasp1 | 76707 | HITS-CLIP | 23142080 | Unspecified |
| mmu-miR-21a-3p | MIMAT0004628 | Clcc1 | 229725 | HITS-CLIP | 29167373 | Unspecified |
| mmu-miR-21a-3p | MIMAT0004628 | Clcf1 | 56708 | HITS-CLIP | 23142080 | Unspecified |
| mmu-miR-21a-3p | MIMAT0004628 | Clcn3 | 12725 | HITS-CLIP | 37986224\|23142080 | Unspecified |
| mmu-miR-21a-3p | MIMAT0004628 | Cldn10 | 58187 | HITS-CLIP | 23142080 | Unspecified |
| mmu-miR-21a-3p | MIMAT0004628 | Cldn12 | 64945 | HITS-CLIP | 23142080 | Unspecified |
| mmu-miR-21a-3p | MIMAT0004628 | Clec12a | 160364 | HITS-CLIP | 29167373 | Unspecified |
| mmu-miR-21a-3p | MIMAT0004628 | Clec4a3 | 73149 | HITS-CLIP | 29167373 | Unspecified |
| mmu-miR-21a-3p | MIMAT0004628 | Clec4n | 56620 | HITS-CLIP | 29167373 | Unspecified |
| mmu-miR-21a-3p | MIMAT0004628 | Clint1 | 216705 | HITS-CLIP | 23142080 | Unspecified |
| mmu-miR-21a-3p | MIMAT0004628 | Clip1 | 56430 | HITS-CLIP | 23142080\|29167373 | Unspecified |
| mmu-miR-21a-3p | MIMAT0004628 | Cln6 | 76524 | HITS-CLIP | 23142080 | Unspecified |
| mmu-miR-21a-3p | MIMAT0004628 | Clock | 12753 | HITS-CLIP | 29167373 | Unspecified |
| mmu-miR-21a-3p | MIMAT0004628 | Clpx | 270166 | HITS-CLIP | 23142080\|29167373 | Unspecified |
| mmu-miR-21a-3p | MIMAT0004628 | Clta | 12757 | HITS-CLIP | 23142080\|29167373 | Unspecified |
| mmu-miR-21a-3p | MIMAT0004628 | Cltc | 67300 | HITS-CLIP | 37986224\|23142080\|29167373 | Unspecified |
| mmu-miR-21a-3p | MIMAT0004628 | Cluh | 74148 | HITS-CLIP | 23142080 | Unspecified |
| mmu-miR-21a-3p | MIMAT0004628 | Cmtm6 | 67213 | HITS-CLIP | 23142080 | Unspecified |
| mmu-miR-21a-3p | MIMAT0004628 | Cnih1 | 12793 | HITS-CLIP | 37986224\|23142080 | Unspecified |
| mmu-miR-21a-3p | MIMAT0004628 | Cnn3 | 71994 | HITS-CLIP | 23142080 | Unspecified |
| mmu-miR-21a-3p | MIMAT0004628 | Cnnm4 | 94220 | HITS-CLIP | 23142080 | Unspecified |
| mmu-miR-21a-3p | MIMAT0004628 | Cnot1 | 234594 | HITS-CLIP | 37986224\|23142080\|29167373 | Unspecified |
| mmu-miR-21a-3p | MIMAT0004628 | Cnot10 | 78893 | HITS-CLIP | 23142080 | Unspecified |
| mmu-miR-21a-3p | MIMAT0004628 | Cnot11 | 52846 | HITS-CLIP | 23142080 | Unspecified |
| mmu-miR-21a-3p | MIMAT0004628 | Cnot4 | 53621 | HITS-CLIP | 29167373\|23142080 | Unspecified |
| mmu-miR-21a-3p | MIMAT0004628 | Cnot6 | 104625 | HITS-CLIP | 37986224\|23142080 | Unspecified |
| mmu-miR-21a-3p | MIMAT0004628 | Cnot6l | 231464 | HITS-CLIP | 37986224\|23142080 | Unspecified |
| mmu-miR-21a-3p | MIMAT0004628 | Cnot7 | 18983 | HITS-CLIP | 23142080 | Unspecified |
| mmu-miR-21a-3p | MIMAT0004628 | Cnot8 | 69125 | HITS-CLIP | 23142080 | Unspecified |
| mmu-miR-21a-3p | MIMAT0004628 | Cntrl | 26920 | HITS-CLIP | 23142080 | Unspecified |
| mmu-miR-21a-3p | MIMAT0004628 | Cobll1 | 319876 | HITS-CLIP | 23142080 | Unspecified |
| mmu-miR-21a-3p | MIMAT0004628 | Cog3 | 338337 | HITS-CLIP | 23142080\|29167373 | Unspecified |

| | | | | | | |
|---|---|---|---|---|---|---|
| mmu-miR-21a-3p | MIMAT0004628 | Cog5 | 238123 | HITS-CLIP | 23142080 | Unspecified |
| mmu-miR-21a-3p | MIMAT0004628 | Col18a1 | 12822 | HITS-CLIP | 29167373 | Unspecified |
| mmu-miR-21a-3p | MIMAT0004628 | Col4a3bp | 68018 | HITS-CLIP | 23142080\|29167373 | Unspecified |
| mmu-miR-21a-3p | MIMAT0004628 | Commd10 | 69456 | HITS-CLIP | 29167373 | Unspecified |
| mmu-miR-21a-3p | MIMAT0004628 | Copa | 12847 | HITS-CLIP | 23142080 | Unspecified |
| mmu-miR-21a-3p | MIMAT0004628 | Copb1 | 70349 | HITS-CLIP | 23142080\|29167373 | Unspecified |
| mmu-miR-21a-3p | MIMAT0004628 | Copb2 | 50797 | HITS-CLIP | 23142080 | Unspecified |
| mmu-miR-21a-3p | MIMAT0004628 | Cope | 59042 | HITS-CLIP | 23142080 | Unspecified |
| mmu-miR-21a-3p | MIMAT0004628 | Copg1 | 54161 | HITS-CLIP | 23142080 | Unspecified |
| mmu-miR-21a-3p | MIMAT0004628 | Copg2 | 54160 | HITS-CLIP | 23142080 | Unspecified |
| mmu-miR-21a-3p | MIMAT0004628 | Cops2 | 12848 | HITS-CLIP | 23142080 | Unspecified |
| mmu-miR-21a-3p | MIMAT0004628 | Cops3 | 26572 | HITS-CLIP | 23142080 | Unspecified |
| mmu-miR-21a-3p | MIMAT0004628 | Cops5 | 26754 | HITS-CLIP | 23142080 | Unspecified |
| mmu-miR-21a-3p | MIMAT0004628 | Cops7a | 26894 | HITS-CLIP | 23142080 | Unspecified |
| mmu-miR-21a-3p | MIMAT0004628 | Coq10b | 67876 | HITS-CLIP | 23142080 | Unspecified |
| mmu-miR-21a-3p | MIMAT0004628 | Cox17 | 12856 | HITS-CLIP | 23142080 | Unspecified |
| mmu-miR-21a-3p | MIMAT0004628 | Cox4i1 | 12857 | HITS-CLIP | 23142080 | Unspecified |
| mmu-miR-21a-3p | MIMAT0004628 | Cox5a | 12858 | HITS-CLIP | 23142080 | Unspecified |
| mmu-miR-21a-3p | MIMAT0004628 | Cox7a2l | 20463 | HITS-CLIP | 23142080 | Unspecified |
| mmu-miR-21a-3p | MIMAT0004628 | Cox8a | 12868 | HITS-CLIP | 37986224 | Unspecified |
| mmu-miR-21a-3p | MIMAT0004628 | Cpd | 12874 | HITS-CLIP | 31433990\|23142080 | Unspecified |
| mmu-miR-21a-3p | MIMAT0004628 | Cpeb2 | 231207 | HITS-CLIP | 37986224\|23142080\|29167373 | Unspecified |
| mmu-miR-21a-3p | MIMAT0004628 | Cpeb4 | 67579 | HITS-CLIP | 37986224\|23142080\|29167373 | Unspecified |
| mmu-miR-21a-3p | MIMAT0004628 | Cplane1 | 65250 | HITS-CLIP | 23142080 | Unspecified |
| mmu-miR-21a-3p | MIMAT0004628 | Cpne1 | 266692 | HITS-CLIP | 23142080 | Unspecified |
| mmu-miR-21a-3p | MIMAT0004628 | Cpne3 | 70568 | HITS-CLIP | 23142080 | Unspecified |
| mmu-miR-21a-3p | MIMAT0004628 | Cpq | 54381 | HITS-CLIP | 23142080 | Unspecified |
| mmu-miR-21a-3p | MIMAT0004628 | Cpsf2 | 51786 | HITS-CLIP | 29167373 | Unspecified |
| mmu-miR-21a-3p | MIMAT0004628 | Cpsf6 | 432508 | HITS-CLIP | 37986224\|23142080 | Unspecified |
| mmu-miR-21a-3p | MIMAT0004628 | Cpsf7 | 269061 | HITS-CLIP | 29167373 | Unspecified |
| mmu-miR-21a-3p | MIMAT0004628 | Cr1l | 12946 | HITS-CLIP | 23142080 | Unspecified |
| mmu-miR-21a-3p | MIMAT0004628 | Cramp1l | 57354 | HITS-CLIP | 23142080 | Unspecified |
| mmu-miR-21a-3p | MIMAT0004628 | Creb3l2 | 208647 | HITS-CLIP | 23142080 | Unspecified |
| mmu-miR-21a-3p | MIMAT0004628 | Crebbp | 12914 | HITS-CLIP | 23142080\|29167373 | Unspecified |
| mmu-miR-21a-3p | MIMAT0004628 | Crebrf | 77128 | HITS-CLIP | 37986224\|23142080 | Unspecified |
| mmu-miR-21a-3p | MIMAT0004628 | Crebzf | 233490 | HITS-CLIP | 23142080 | Unspecified |
| mmu-miR-21a-3p | MIMAT0004628 | Crem | 12916 | HITS-CLIP | 37986224\|23142080 | Unspecified |
| mmu-miR-21a-3p | MIMAT0004628 | Crim1 | 50766 | HITS-CLIP | 23142080 | Unspecified |
| mmu-miR-21a-3p | MIMAT0004628 | Cript | 56724 | HITS-CLIP | 37986224\|23142080\|29167373 | Unspecified |

| | | | | | | |
|---|---|---|---|---|---|---|
| mmu-miR-21a-3p | MIMAT0004628 | Crk | 12928 | HITS-CLIP | 23142080\|29167373 | Unspecified |
| mmu-miR-21a-3p | MIMAT0004628 | Crtc3 | 70461 | HITS-CLIP | 23142080 | Unspecified |
| mmu-miR-21a-3p | MIMAT0004628 | Crybg2 | 55057 | HITS-CLIP | 23142080 | Unspecified |
| mmu-miR-21a-3p | MIMAT0004628 | Csde1 | 229663 | HITS-CLIP | 23142080 | Unspecified |
| mmu-miR-21a-3p | MIMAT0004628 | Cse1l | 110750 | HITS-CLIP | 23142080 | Unspecified |
| mmu-miR-21a-3p | MIMAT0004628 | Csf1r | 12978 | HITS-CLIP | 29167373 | Unspecified |
| mmu-miR-21a-3p | MIMAT0004628 | Csgalnact2 | 78752 | HITS-CLIP | 23142080 | Unspecified |
| mmu-miR-21a-3p | MIMAT0004628 | Csk | 12988 | HITS-CLIP | 23142080\|29167373 | Unspecified |
| mmu-miR-21a-3p | MIMAT0004628 | Csmd1 | 94109 | HITS-CLIP | 23142080 | Unspecified |
| mmu-miR-21a-3p | MIMAT0004628 | Csnk1a1 | 93687 | HITS-CLIP | 37986224\|23142080\|29167373 | Unspecified |
| mmu-miR-21a-3p | MIMAT0004628 | Csnk1d | 104318 | HITS-CLIP | 23142080 | Unspecified |
| mmu-miR-21a-3p | MIMAT0004628 | Csnk1g1 | 214897 | HITS-CLIP | 23142080 | Unspecified |
| mmu-miR-21a-3p | MIMAT0004628 | Csnk1g3 | 70425 | HITS-CLIP | 23142080 | Unspecified |
| mmu-miR-21a-3p | MIMAT0004628 | Cspp1 | 211660 | HITS-CLIP | 23142080 | Unspecified |
| mmu-miR-21a-3p | MIMAT0004628 | Cst3 | 13010 | HITS-CLIP | 23142080\|29167373 | Unspecified |
| mmu-miR-21a-3p | MIMAT0004628 | Cstf1 | 67337 | HITS-CLIP | 23142080 | Unspecified |
| mmu-miR-21a-3p | MIMAT0004628 | Cstf2 | 108062 | HITS-CLIP | 23142080 | Unspecified |
| mmu-miR-21a-3p | MIMAT0004628 | Ctbp2 | 13017 | HITS-CLIP | 37986224 | Unspecified |
| mmu-miR-21a-3p | MIMAT0004628 | Ctc1 | 68964 | HITS-CLIP | 23142080 | Unspecified |
| mmu-miR-21a-3p | MIMAT0004628 | Ctcf | 13018 | HITS-CLIP | 23142080 | Unspecified |
| mmu-miR-21a-3p | MIMAT0004628 | Ctdspl | 69274 | HITS-CLIP | 37986224 | Unspecified |
| mmu-miR-21a-3p | MIMAT0004628 | Ctdspl2 | 329506 | HITS-CLIP | 37986224\|23142080 | Unspecified |
| mmu-miR-21a-3p | MIMAT0004628 | Ctla4 | 12477 | HITS-CLIP | 37986224\|23142080 | Unspecified |
| mmu-miR-21a-3p | MIMAT0004628 | Ctnna1 | 12385 | HITS-CLIP | 23142080 | Unspecified |
| mmu-miR-21a-3p | MIMAT0004628 | Ctnnb1 | 12387 | HITS-CLIP | 37986224\|31433990\|23142080 | Unspecified |
| mmu-miR-21a-3p | MIMAT0004628 | Ctnnd1 | 12388 | HITS-CLIP | 29167373 | Unspecified |
| mmu-miR-21a-3p | MIMAT0004628 | Ctr9 | 22083 | HITS-CLIP | 23142080 | Unspecified |
| mmu-miR-21a-3p | MIMAT0004628 | Ctsb | 13030 | HITS-CLIP | 23142080 | Unspecified |
| mmu-miR-21a-3p | MIMAT0004628 | Ctsc | 13032 | HITS-CLIP | 29167373\|23142080 | Unspecified |
| mmu-miR-21a-3p | MIMAT0004628 | Ctsd | 13033 | HITS-CLIP | 29167373 | Unspecified |
| mmu-miR-21a-3p | MIMAT0004628 | Ctso | 229445 | HITS-CLIP | 23142080 | Unspecified |
| mmu-miR-21a-3p | MIMAT0004628 | Ctsz | 64138 | HITS-CLIP | 23142080 | Unspecified |
| mmu-miR-21a-3p | MIMAT0004628 | Cul1 | 26965 | HITS-CLIP | 23142080 | Unspecified |
| mmu-miR-21a-3p | MIMAT0004628 | Cul2 | 71745 | HITS-CLIP | 23142080 | Unspecified |
| mmu-miR-21a-3p | MIMAT0004628 | Cul3 | 26554 | HITS-CLIP | 23142080 | Unspecified |
| mmu-miR-21a-3p | MIMAT0004628 | Cul4a | 99375 | HITS-CLIP | 23142080 | Unspecified |
| mmu-miR-21a-3p | MIMAT0004628 | Cul5 | 75717 | HITS-CLIP | 23142080 | Unspecified |
| mmu-miR-21a-3p | MIMAT0004628 | Cul7 | 66515 | HITS-CLIP | 23142080 | Unspecified |
| mmu-miR-21a-3p | MIMAT0004628 | Cul9 | 78309 | HITS-CLIP | 29167373 | Unspecified |

| | | | | | | |
|---|---|---|---|---|---|---|
| mmu-miR-21a-3p | MIMAT0004628 | Cux1 | 13047 | HITS-CLIP | 23142080 | Unspecified |
| mmu-miR-21a-3p | MIMAT0004628 | Cxcr4 | 12767 | HITS-CLIP | 37986224\|23142080\|29167373 | Unspecified |
| mmu-miR-21a-3p | MIMAT0004628 | Cyb561 | 13056 | HITS-CLIP | 23142080 | Unspecified |
| mmu-miR-21a-3p | MIMAT0004628 | Cyb5b | 66427 | HITS-CLIP | 37986224\|23142080 | Unspecified |
| mmu-miR-21a-3p | MIMAT0004628 | Cyb5r3 | 109754 | HITS-CLIP | 23142080\|29167373 | Unspecified |
| mmu-miR-21a-3p | MIMAT0004628 | Cyld | 74256 | HITS-CLIP | 37986224\|23142080 | Unspecified |
| mmu-miR-21a-3p | MIMAT0004628 | Cyp51 | 13121 | HITS-CLIP | 37986224\|23142080 | Unspecified |
| mmu-miR-21a-3p | MIMAT0004628 | Cyth3 | 19159 | HITS-CLIP | 23142080 | Unspecified |
| mmu-miR-21a-3p | MIMAT0004628 | D16Ertd472e | 67102 | HITS-CLIP | 23142080 | Unspecified |
| mmu-miR-21a-3p | MIMAT0004628 | D1Ertd622e | 52392 | HITS-CLIP | 29167373 | Unspecified |
| mmu-miR-21a-3p | MIMAT0004628 | D2hgdh | 98314 | HITS-CLIP | 37986224 | Unspecified |
| mmu-miR-21a-3p | MIMAT0004628 | D5Ertd579e | 320661 | HITS-CLIP | 23142080 | Unspecified |
| mmu-miR-21a-3p | MIMAT0004628 | Daam1 | 208846 | HITS-CLIP | 29167373 | Unspecified |
| mmu-miR-21a-3p | MIMAT0004628 | Dag1 | 13138 | HITS-CLIP | 37986224\|23142080 | Unspecified |
| mmu-miR-21a-3p | MIMAT0004628 | Dap3 | 65111 | HITS-CLIP | 23142080 | Unspecified |
| mmu-miR-21a-3p | MIMAT0004628 | Dapk2 | 13143 | HITS-CLIP | 23142080 | Unspecified |
| mmu-miR-21a-3p | MIMAT0004628 | Dars | 226414 | HITS-CLIP | 23142080 | Unspecified |
| mmu-miR-21a-3p | MIMAT0004628 | Dbf4 | 27214 | HITS-CLIP | 23142080 | Unspecified |
| mmu-miR-21a-3p | MIMAT0004628 | Dbi | 13167 | HITS-CLIP | 23142080 | Unspecified |
| mmu-miR-21a-3p | MIMAT0004628 | Dcaf1 | 41836 | HITS-CLIP | 23142080 | Unspecified |
| mmu-miR-21a-3p | MIMAT0004628 | Dcaf10 | 242418 | HITS-CLIP | 23142080 | Unspecified |
| mmu-miR-21a-3p | MIMAT0004628 | Dcaf12 | 68970 | qCLASH//HITS-CLIP | 31393953\|23142080 | Unspecified |
| mmu-miR-21a-3p | MIMAT0004628 | Dcaf15 | 212123 | HITS-CLIP | 23142080 | Unspecified |
| mmu-miR-21a-3p | MIMAT0004628 | Dcaf7 | 71833 | HITS-CLIP | 37986224 | Unspecified |
| mmu-miR-21a-3p | MIMAT0004628 | Dcp1a | 75901 | HITS-CLIP | 23142080 | Unspecified |
| mmu-miR-21a-3p | MIMAT0004628 | Dcp2 | 70640 | HITS-CLIP | 37986224\|23142080 | Unspecified |
| mmu-miR-21a-3p | MIMAT0004628 | Dctn4 | 67665 | HITS-CLIP | 23142080 | Unspecified |
| mmu-miR-21a-3p | MIMAT0004628 | Dcun1d2 | 102323 | HITS-CLIP | 29167373 | Unspecified |
| mmu-miR-21a-3p | MIMAT0004628 | Dcun1d3 | 233805 | HITS-CLIP | 23142080 | Unspecified |
| mmu-miR-21a-3p | MIMAT0004628 | Dcun1d4 | 100737 | HITS-CLIP | 23142080 | Unspecified |
| mmu-miR-21a-3p | MIMAT0004628 | Dcun1d5 | 76863 | HITS-CLIP | 23142080 | Unspecified |
| mmu-miR-21a-3p | MIMAT0004628 | Dda1 | 66498 | HITS-CLIP | 23142080 | Unspecified |
| mmu-miR-21a-3p | MIMAT0004628 | Ddb1 | 13194 | HITS-CLIP | 23142080 | Unspecified |
| mmu-miR-21a-3p | MIMAT0004628 | Ddhd1 | 114874 | HITS-CLIP | 23142080 | Unspecified |
| mmu-miR-21a-3p | MIMAT0004628 | Ddi2 | 68817 | HITS-CLIP | 23142080 | Unspecified |
| mmu-miR-21a-3p | MIMAT0004628 | Ddit4 | 74747 | HITS-CLIP | 23142080 | Unspecified |
| mmu-miR-21a-3p | MIMAT0004628 | Ddx10 | 77591 | HITS-CLIP | 23142080 | Unspecified |
| mmu-miR-21a-3p | MIMAT0004628 | Ddx17 | 67040 | HITS-CLIP | 23142080 | Unspecified |
| mmu-miR-21a-3p | MIMAT0004628 | Ddx19a | 13680 | HITS-CLIP | 23142080 | Unspecified |

| | | | | | | |
|---|---|---|---|---|---|---|
| mmu-miR-21a-3p | MIMAT0004628 | Ddx20 | 53975 | HITS-CLIP | 23142080 | Unspecified |
| mmu-miR-21a-3p | MIMAT0004628 | Ddx21 | 56200 | HITS-CLIP | 23142080 | Unspecified |
| mmu-miR-21a-3p | MIMAT0004628 | Ddx23 | 74351 | HITS-CLIP | 29167373 | Unspecified |
| mmu-miR-21a-3p | MIMAT0004628 | Ddx24 | 27225 | HITS-CLIP | 23142080 | Unspecified |
| mmu-miR-21a-3p | MIMAT0004628 | Ddx3x | 13205 | HITS-CLIP | 23142080 | Unspecified |
| mmu-miR-21a-3p | MIMAT0004628 | Ddx3y | 26900 | HITS-CLIP | 23142080 | Unspecified |
| mmu-miR-21a-3p | MIMAT0004628 | Ddx46 | 212880 | HITS-CLIP | 23142080 | Unspecified |
| mmu-miR-21a-3p | MIMAT0004628 | Ddx5 | 13207 | HITS-CLIP | 23142080\|29167373 | Unspecified |
| mmu-miR-21a-3p | MIMAT0004628 | Ddx6 | 13209 | HITS-CLIP | 23142080 | Unspecified |
| mmu-miR-21a-3p | MIMAT0004628 | Deaf1 | 54006 | HITS-CLIP | 23142080 | Unspecified |
| mmu-miR-21a-3p | MIMAT0004628 | Decr2 | 26378 | HITS-CLIP | 23142080 | Unspecified |
| mmu-miR-21a-3p | MIMAT0004628 | Dennd1b | 329260 | HITS-CLIP | 37986224\|23142080 | Unspecified |
| mmu-miR-21a-3p | MIMAT0004628 | Dennd4b | 229541 | HITS-CLIP | 23142080 | Unspecified |
| mmu-miR-21a-3p | MIMAT0004628 | Depdc1b | 218581 | HITS-CLIP | 37986224\|23142080 | Unspecified |
| mmu-miR-21a-3p | MIMAT0004628 | Depdc5 | 277854 | HITS-CLIP | 23142080 | Unspecified |
| mmu-miR-21a-3p | MIMAT0004628 | Desi2 | 78825 | HITS-CLIP | 23142080 | Unspecified |
| mmu-miR-21a-3p | MIMAT0004628 | Dgkd | 227333 | HITS-CLIP | 29167373 | Unspecified |
| mmu-miR-21a-3p | MIMAT0004628 | Dgkh | 380921 | HITS-CLIP | 23142080 | Unspecified |
| mmu-miR-21a-3p | MIMAT0004628 | Dgkz | 104418 | HITS-CLIP | 23142080\|29167373 | Unspecified |
| mmu-miR-21a-3p | MIMAT0004628 | Dhcr24 | 74754 | HITS-CLIP | 23142080 | Unspecified |
| mmu-miR-21a-3p | MIMAT0004628 | Dhdh | 71755 | HITS-CLIP | 29167373 | Unspecified |
| mmu-miR-21a-3p | MIMAT0004628 | Dhx15 | 13204 | HITS-CLIP | 23142080 | Unspecified |
| mmu-miR-21a-3p | MIMAT0004628 | Dhx29 | 218629 | HITS-CLIP | 23142080 | Unspecified |
| mmu-miR-21a-3p | MIMAT0004628 | Dhx34 | 71723 | HITS-CLIP | 23142080 | Unspecified |
| mmu-miR-21a-3p | MIMAT0004628 | Dhx36 | 72162 | HITS-CLIP | 23142080 | Unspecified |
| mmu-miR-21a-3p | MIMAT0004628 | Dhx40 | 67487 | HITS-CLIP | 23142080 | Unspecified |
| mmu-miR-21a-3p | MIMAT0004628 | Dhx58 | 80861 | HITS-CLIP | 23142080\|29167373 | Unspecified |
| mmu-miR-21a-3p | MIMAT0004628 | Dhx9 | 13211 | HITS-CLIP | 23142080\|29167373 | Unspecified |
| mmu-miR-21a-3p | MIMAT0004628 | Diaph1 | 1729 | HITS-CLIP | 23142080\|29167373 | Unspecified |
| mmu-miR-21a-3p | MIMAT0004628 | Dicer1 | 192119 | HITS-CLIP | 23142080 | Unspecified |
| mmu-miR-21a-3p | MIMAT0004628 | Dido1 | 23856 | HITS-CLIP | 37986224 | Unspecified |
| mmu-miR-21a-3p | MIMAT0004628 | Dip2b | 239667 | HITS-CLIP | 23142080\|29167373 | Unspecified |
| mmu-miR-21a-3p | MIMAT0004628 | Dixdc1 | 330938 | HITS-CLIP | 23142080 | Unspecified |
| mmu-miR-21a-3p | MIMAT0004628 | Dkc1 | 245474 | HITS-CLIP | 23142080 | Unspecified |
| mmu-miR-21a-3p | MIMAT0004628 | Dlat | 235339 | HITS-CLIP | 23142080 | Unspecified |
| mmu-miR-21a-3p | MIMAT0004628 | Dlg3 | 53310 | HITS-CLIP | 23142080 | Unspecified |
| mmu-miR-21a-3p | MIMAT0004628 | Dlgap5 | 218977 | HITS-CLIP | 23142080 | Unspecified |
| mmu-miR-21a-3p | MIMAT0004628 | Dmtf1 | 23857 | HITS-CLIP | 37986224\|23142080 | Unspecified |
| mmu-miR-21a-3p | MIMAT0004628 | Dmxl1 | 240283 | HITS-CLIP | 37986224\|23142080\|29167373 | Unspecified |

| | | | | | | |
|---|---|---|---|---|---|---|
| mmu-miR-21a-3p | MIMAT0004628 | Dnah7a | 627872 | HITS-CLIP | 23142080 | Unspecified |
| mmu-miR-21a-3p | MIMAT0004628 | Dnaja2 | 56445 | HITS-CLIP | 37986224\|23142080 | Unspecified |
| mmu-miR-21a-3p | MIMAT0004628 | Dnajb14 | 70604 | HITS-CLIP | 23142080\|29167373 | Unspecified |
| mmu-miR-21a-3p | MIMAT0004628 | Dnajb4 | 67035 | HITS-CLIP | 23142080 | Unspecified |
| mmu-miR-21a-3p | MIMAT0004628 | Dnajc11 | 230935 | HITS-CLIP | 23142080 | Unspecified |
| mmu-miR-21a-3p | MIMAT0004628 | Dnajc14 | 74330 | HITS-CLIP | 23142080 | Unspecified |
| mmu-miR-21a-3p | MIMAT0004628 | Dnajc18 | 76594 | HITS-CLIP | 29167373 | Unspecified |
| mmu-miR-21a-3p | MIMAT0004628 | Dnajc2 | 22791 | HITS-CLIP | 23142080 | Unspecified |
| mmu-miR-21a-3p | MIMAT0004628 | Dnajc5 | 13002 | HITS-CLIP | 37986224\|23142080 | Unspecified |
| mmu-miR-21a-3p | MIMAT0004628 | Dnajc7 | 56354 | HITS-CLIP | 23142080 | Unspecified |
| mmu-miR-21a-3p | MIMAT0004628 | Dnajc8 | 68598 | HITS-CLIP | 23142080 | Unspecified |
| mmu-miR-21a-3p | MIMAT0004628 | Dnase1l1 | 69537 | HITS-CLIP | 29167373 | Unspecified |
| mmu-miR-21a-3p | MIMAT0004628 | Dnlz | 52838 | HITS-CLIP | 23142080 | Unspecified |
| mmu-miR-21a-3p | MIMAT0004628 | Dnmt1 | 13433 | HITS-CLIP | 37986224\|23142080\|29167373 | Unspecified |
| mmu-miR-21a-3p | MIMAT0004628 | Dnmt3a | 13435 | HITS-CLIP | 37986224\|23142080 | Unspecified |
| mmu-miR-21a-3p | MIMAT0004628 | Dnpep | 13437 | HITS-CLIP | 23142080 | Unspecified |
| mmu-miR-21a-3p | MIMAT0004628 | Dock10 | 210293 | HITS-CLIP | 37986224\|23142080 | Unspecified |
| mmu-miR-21a-3p | MIMAT0004628 | Dock11 | 75974 | HITS-CLIP | 23142080 | Unspecified |
| mmu-miR-21a-3p | MIMAT0004628 | Dock4 | 238130 | HITS-CLIP | 29167373 | Unspecified |
| mmu-miR-21a-3p | MIMAT0004628 | Dock8 | 76088 | HITS-CLIP | 23142080\|29167373 | Unspecified |
| mmu-miR-21a-3p | MIMAT0004628 | Dock9 | 105445 | HITS-CLIP | 23142080 | Unspecified |
| mmu-miR-21a-3p | MIMAT0004628 | Dop1b | 9980 | HITS-CLIP | 23142080 | Unspecified |
| mmu-miR-21a-3p | MIMAT0004628 | Dpy19l3 | 233115 | HITS-CLIP | 23142080 | Unspecified |
| mmu-miR-21a-3p | MIMAT0004628 | Dpy19l4 | 381510 | HITS-CLIP | 23142080 | Unspecified |
| mmu-miR-21a-3p | MIMAT0004628 | Dpysl2 | 12934 | HITS-CLIP | 23142080 | Unspecified |
| mmu-miR-21a-3p | MIMAT0004628 | Dram2 | 67171 | HITS-CLIP | 37986224\|23142080 | Unspecified |
| mmu-miR-21a-3p | MIMAT0004628 | Drg1 | 13494 | HITS-CLIP | 23142080 | Unspecified |
| mmu-miR-21a-3p | MIMAT0004628 | Drg2 | 13495 | HITS-CLIP | 23142080 | Unspecified |
| mmu-miR-21a-3p | MIMAT0004628 | Dsel | 319901 | HITS-CLIP | 23142080 | Unspecified |
| mmu-miR-21a-3p | MIMAT0004628 | Dtd2 | 328092 | HITS-CLIP | 23142080 | Unspecified |
| mmu-miR-21a-3p | MIMAT0004628 | Dtl | 76843 | HITS-CLIP | 23142080 | Unspecified |
| mmu-miR-21a-3p | MIMAT0004628 | Dus2 | 66369 | HITS-CLIP | 23142080 | Unspecified |
| mmu-miR-21a-3p | MIMAT0004628 | Dusp4 | 319520 | HITS-CLIP | 23142080 | Unspecified |
| mmu-miR-21a-3p | MIMAT0004628 | Dut | 110074 | HITS-CLIP | 23142080 | Unspecified |
| mmu-miR-21a-3p | MIMAT0004628 | Dvl3 | 13544 | HITS-CLIP | 23142080 | Unspecified |
| mmu-miR-21a-3p | MIMAT0004628 | Dym | 69190 | HITS-CLIP | 37986224\|23142080 | Unspecified |
| mmu-miR-21a-3p | MIMAT0004628 | Dync1h1 | 13424 | HITS-CLIP | 23142080\|29167373 | Unspecified |
| mmu-miR-21a-3p | MIMAT0004628 | Dync1i2 | 13427 | HITS-CLIP | 23142080\|29167373 | Unspecified |
| mmu-miR-21a-3p | MIMAT0004628 | Dync1li2 | 234663 | HITS-CLIP | 23142080 | Unspecified |

| | | | | | | |
|---|---|---|---|---|---|---|
| mmu-miR-21a-3p | MIMAT0004628 | Dynlrb1 | 67068 | HITS-CLIP | 23142080 | Unspecified |
| mmu-miR-21a-3p | MIMAT0004628 | Dynlt3 | 67117 | HITS-CLIP | 23142080 | Unspecified |
| mmu-miR-21a-3p | MIMAT0004628 | Dyrk1a | 13548 | HITS-CLIP | 37986224\|23142080\|29167373 | Unspecified |
| mmu-miR-21a-3p | MIMAT0004628 | Dyrk2 | 69181 | HITS-CLIP | 23142080\|29167373 | Unspecified |
| mmu-miR-21a-3p | MIMAT0004628 | E2f2 | 242705 | HITS-CLIP | 23142080 | Unspecified |
| mmu-miR-21a-3p | MIMAT0004628 | E2f3 | 13557 | HITS-CLIP | 37986224\|23142080 | Unspecified |
| mmu-miR-21a-3p | MIMAT0004628 | E2f7 | 52679 | HITS-CLIP | 23142080 | Unspecified |
| mmu-miR-21a-3p | MIMAT0004628 | E2f8 | 108961 | HITS-CLIP | 37986224\|23142080 | Unspecified |
| mmu-miR-21a-3p | MIMAT0004628 | E330009J07Rik | 243780 | HITS-CLIP | 23142080 | Unspecified |
| mmu-miR-21a-3p | MIMAT0004628 | Eaf1 | 74427 | HITS-CLIP | 23142080 | Unspecified |
| mmu-miR-21a-3p | MIMAT0004628 | Eapp | 66266 | HITS-CLIP | 23142080 | Unspecified |
| mmu-miR-21a-3p | MIMAT0004628 | Ebna1bp2 | 69072 | HITS-CLIP | 23142080 | Unspecified |
| mmu-miR-21a-3p | MIMAT0004628 | Ecd | 70601 | HITS-CLIP | 23142080 | Unspecified |
| mmu-miR-21a-3p | MIMAT0004628 | Ecpas | 23392 | HITS-CLIP | 23142080 | Unspecified |
| mmu-miR-21a-3p | MIMAT0004628 | Ect2 | 13605 | HITS-CLIP | 23142080 | Unspecified |
| mmu-miR-21a-3p | MIMAT0004628 | Edc3 | 353190 | HITS-CLIP | 23142080 | Unspecified |
| mmu-miR-21a-3p | MIMAT0004628 | Edc4 | 234699 | HITS-CLIP | 23142080 | Unspecified |
| mmu-miR-21a-3p | MIMAT0004628 | Edem1 | 192193 | HITS-CLIP | 37986224 | Unspecified |
| mmu-miR-21a-3p | MIMAT0004628 | Edf1 | 59022 | HITS-CLIP | 23142080\|29167373 | Unspecified |
| mmu-miR-21a-3p | MIMAT0004628 | Eea1 | 216238 | HITS-CLIP | 23142080\|29167373 | Unspecified |
| mmu-miR-21a-3p | MIMAT0004628 | Eed | 13626 | HITS-CLIP | 23142080 | Unspecified |
| mmu-miR-21a-3p | MIMAT0004628 | Eef1a1 | 13627 | HITS-CLIP | 23142080 | Unspecified |
| mmu-miR-21a-3p | MIMAT0004628 | Eef1b2 | 55949 | HITS-CLIP | 23142080 | Unspecified |
| mmu-miR-21a-3p | MIMAT0004628 | Efcab14 | 230648 | HITS-CLIP | 29167373 | Unspecified |
| mmu-miR-21a-3p | MIMAT0004628 | Efl1 | 1942 | HITS-CLIP | 23142080 | Unspecified |
| mmu-miR-21a-3p | MIMAT0004628 | Efr3a | 76740 | HITS-CLIP | 23142080\|29167373 | Unspecified |
| mmu-miR-21a-3p | MIMAT0004628 | Egln3 | 112407 | HITS-CLIP | 23142080 | Unspecified |
| mmu-miR-21a-3p | MIMAT0004628 | Egr1 | 13653 | HITS-CLIP | 23142080 | Unspecified |
| mmu-miR-21a-3p | MIMAT0004628 | Egr3 | 13655 | HITS-CLIP | 23142080 | Unspecified |
| mmu-miR-21a-3p | MIMAT0004628 | Ehhadh | 74147 | HITS-CLIP | 37986224\|23142080 | Unspecified |
| mmu-miR-21a-3p | MIMAT0004628 | Ehmt1 | 77683 | HITS-CLIP | 23142080 | Unspecified |
| mmu-miR-21a-3p | MIMAT0004628 | Eif1a | 13664 | HITS-CLIP | 23142080 | Unspecified |
| mmu-miR-21a-3p | MIMAT0004628 | Eif1b | 68969 | HITS-CLIP | 23142080 | Unspecified |
| mmu-miR-21a-3p | MIMAT0004628 | Eif2a | 229317 | HITS-CLIP | 23142080 | Unspecified |
| mmu-miR-21a-3p | MIMAT0004628 | Eif2ak4 | 27103 | HITS-CLIP | 23142080 | Unspecified |
| mmu-miR-21a-3p | MIMAT0004628 | Eif2s2 | 67204 | HITS-CLIP | 23142080 | Unspecified |
| mmu-miR-21a-3p | MIMAT0004628 | Eif2s3y | 26908 | HITS-CLIP | 23142080 | Unspecified |
| mmu-miR-21a-3p | MIMAT0004628 | Eif3a | 13669 | HITS-CLIP | 23142080\|29167373 | Unspecified |
| mmu-miR-21a-3p | MIMAT0004628 | Eif3c | 56347 | HITS-CLIP | 37986224\|23142080 | Unspecified |

| | | | | | | |
|---|---|---|---|---|---|---|
| mmu-miR-21a-3p | MIMAT0004628 | Eif3e | 16341 | HITS-CLIP | 23142080 | Unspecified |
| mmu-miR-21a-3p | MIMAT0004628 | Eif3g | 8666 | HITS-CLIP | 23142080 | Unspecified |
| mmu-miR-21a-3p | MIMAT0004628 | Eif3h | 68135 | HITS-CLIP | 23142080 | Unspecified |
| mmu-miR-21a-3p | MIMAT0004628 | Eif4a2 | 13682 | HITS-CLIP | 37986224\|23142080\|29167373 | Unspecified |
| mmu-miR-21a-3p | MIMAT0004628 | Eif4b | 75705 | HITS-CLIP | 23142080 | Unspecified |
| mmu-miR-21a-3p | MIMAT0004628 | Eif4e | 13684 | HITS-CLIP | 29167373\|23142080 | Unspecified |
| mmu-miR-21a-3p | MIMAT0004628 | Eif4e3 | 66892 | HITS-CLIP | 23142080 | Unspecified |
| mmu-miR-21a-3p | MIMAT0004628 | Eif4ebp2 | 13688 | HITS-CLIP | 23142080 | Unspecified |
| mmu-miR-21a-3p | MIMAT0004628 | Eif4g1 | 208643 | HITS-CLIP | 37986224\|23142080\|29167373 | Unspecified |
| mmu-miR-21a-3p | MIMAT0004628 | Eif4h | 22384 | HITS-CLIP | 37986224\|23142080 | Unspecified |
| mmu-miR-21a-3p | MIMAT0004628 | Eif5a | 276770 | HITS-CLIP | 37986224\|23142080 | Unspecified |
| mmu-miR-21a-3p | MIMAT0004628 | Eif5b | 226982 | HITS-CLIP | 23142080 | Unspecified |
| mmu-miR-21a-3p | MIMAT0004628 | Eif6 | 16418 | HITS-CLIP | 23142080 | Unspecified |
| mmu-miR-21a-3p | MIMAT0004628 | Elac2 | 68626 | HITS-CLIP | 23142080 | Unspecified |
| mmu-miR-21a-3p | MIMAT0004628 | Elf1 | 13709 | HITS-CLIP | 23142080\|29167373 | Unspecified |
| mmu-miR-21a-3p | MIMAT0004628 | Elf2 | 69257 | HITS-CLIP | 37986224\|23142080 | Unspecified |
| mmu-miR-21a-3p | MIMAT0004628 | Elf4 | 56501 | HITS-CLIP | 23142080 | Unspecified |
| mmu-miR-21a-3p | MIMAT0004628 | Elk3 | 13713 | HITS-CLIP | 23142080 | Unspecified |
| mmu-miR-21a-3p | MIMAT0004628 | Elmo1 | 140580 | HITS-CLIP | 23142080\|29167373 | Unspecified |
| mmu-miR-21a-3p | MIMAT0004628 | Elmsan1 | 238317 | HITS-CLIP | 23142080 | Unspecified |
| mmu-miR-21a-3p | MIMAT0004628 | Eloc | 6921 | HITS-CLIP | 23142080 | Unspecified |
| mmu-miR-21a-3p | MIMAT0004628 | Elovl5 | 68801 | HITS-CLIP | 37986224\|29167373\|23142080 | Unspecified |
| mmu-miR-21a-3p | MIMAT0004628 | Elovl6 | 170439 | HITS-CLIP | 23142080 | Unspecified |
| mmu-miR-21a-3p | MIMAT0004628 | Elp1 | 37455 | HITS-CLIP | 28735896\|23142080 | Unspecified |
| mmu-miR-21a-3p | MIMAT0004628 | Emc2 | 66736 | HITS-CLIP | 23142080 | Unspecified |
| mmu-miR-21a-3p | MIMAT0004628 | Emg1 | 14791 | HITS-CLIP | 23142080 | Unspecified |
| mmu-miR-21a-3p | MIMAT0004628 | Eml4 | 78798 | HITS-CLIP | 23142080 | Unspecified |
| mmu-miR-21a-3p | MIMAT0004628 | Emp1 | 13730 | HITS-CLIP | 29167373 | Unspecified |
| mmu-miR-21a-3p | MIMAT0004628 | Emsy | 56946 | HITS-CLIP | 23142080 | Unspecified |
| mmu-miR-21a-3p | MIMAT0004628 | Enah | 13800 | HITS-CLIP | 37986224 | Unspecified |
| mmu-miR-21a-3p | MIMAT0004628 | Enc1 | 13803 | HITS-CLIP | 37986224\|23142080 | Unspecified |
| mmu-miR-21a-3p | MIMAT0004628 | Ensa | 56205 | HITS-CLIP | 23142080 | Unspecified |
| mmu-miR-21a-3p | MIMAT0004628 | Entpd5 | 12499 | HITS-CLIP | 23142080 | Unspecified |
| mmu-miR-21a-3p | MIMAT0004628 | Eomes | 13813 | HITS-CLIP | 23142080 | Unspecified |
| mmu-miR-21a-3p | MIMAT0004628 | Ep300 | 328572 | HITS-CLIP | 23142080 | Unspecified |
| mmu-miR-21a-3p | MIMAT0004628 | Epas1 | 13819 | HITS-CLIP | 23142080 | Unspecified |
| mmu-miR-21a-3p | MIMAT0004628 | Epc1 | 13831 | HITS-CLIP | 29167373 | Unspecified |
| mmu-miR-21a-3p | MIMAT0004628 | Epc2 | 227867 | HITS-CLIP | 23142080\|29167373 | Unspecified |
| mmu-miR-21a-3p | MIMAT0004628 | Epm2aip1 | 77781 | HITS-CLIP | 23142080 | Unspecified |

| | | | | | | |
|---|---|---|---|---|---|---|
| mmu-miR-21a-3p | MIMAT0004628 | Eps15 | 13858 | HITS-CLIP | 31433990\|23142080 | Unspecified |
| mmu-miR-21a-3p | MIMAT0004628 | Erbin | 55914 | HITS-CLIP | 23142080\|29167373 | Unspecified |
| mmu-miR-21a-3p | MIMAT0004628 | Ercc6 | 319955 | HITS-CLIP | 23142080 | Unspecified |
| mmu-miR-21a-3p | MIMAT0004628 | Ercc6l | 236930 | HITS-CLIP | 23142080 | Unspecified |
| mmu-miR-21a-3p | MIMAT0004628 | Ergic2 | 67456 | HITS-CLIP | 29167373 | Unspecified |
| mmu-miR-21a-3p | MIMAT0004628 | Eri3 | 140546 | HITS-CLIP | 37986224 | Unspecified |
| mmu-miR-21a-3p | MIMAT0004628 | Erlin1 | 226144 | HITS-CLIP | 23142080 | Unspecified |
| mmu-miR-21a-3p | MIMAT0004628 | Ero1l | 50527 | HITS-CLIP | 37986224\|23142080 | Unspecified |
| mmu-miR-21a-3p | MIMAT0004628 | Esd | 13885 | HITS-CLIP | 23142080 | Unspecified |
| mmu-miR-21a-3p | MIMAT0004628 | Esf1 | 66580 | HITS-CLIP | 31433990\|23142080 | Unspecified |
| mmu-miR-21a-3p | MIMAT0004628 | Esyt2 | 52635 | HITS-CLIP | 37986224\|23142080 | Unspecified |
| mmu-miR-21a-3p | MIMAT0004628 | Etf1 | 225363 | HITS-CLIP | 37986224\|23142080\|29167373 | Unspecified |
| mmu-miR-21a-3p | MIMAT0004628 | Etfa | 110842 | HITS-CLIP | 23142080 | Unspecified |
| mmu-miR-21a-3p | MIMAT0004628 | Etfrf1 | 67636 | HITS-CLIP | 23142080 | Unspecified |
| mmu-miR-21a-3p | MIMAT0004628 | Etnk1 | 75320 | HITS-CLIP | 37986224\|29167373\|23142080 | Unspecified |
| mmu-miR-21a-3p | MIMAT0004628 | Ets1 | 23871 | HITS-CLIP | 37986224\|23142080 | Unspecified |
| mmu-miR-21a-3p | MIMAT0004628 | Evi5 | 14020 | HITS-CLIP | 23142080 | Unspecified |
| mmu-miR-21a-3p | MIMAT0004628 | Ewsr1 | 14030 | HITS-CLIP | 23142080 | Unspecified |
| mmu-miR-21a-3p | MIMAT0004628 | Exoc2 | 66482 | HITS-CLIP | 37986224 | Unspecified |
| mmu-miR-21a-3p | MIMAT0004628 | Ezh1 | 14055 | HITS-CLIP | 23142080 | Unspecified |
| mmu-miR-21a-3p | MIMAT0004628 | Ezh2 | 14056 | HITS-CLIP | 23142080 | Unspecified |
| mmu-miR-21a-3p | MIMAT0004628 | F2rl2 | 14064 | HITS-CLIP | 23142080 | Unspecified |
| mmu-miR-21a-3p | MIMAT0004628 | F830016B08Rik | 240328 | HITS-CLIP | 23142080 | Unspecified |
| mmu-miR-21a-3p | MIMAT0004628 | F8a | 14070 | HITS-CLIP | 23142080 | Unspecified |
| mmu-miR-21a-3p | MIMAT0004628 | Fads1 | 76267 | HITS-CLIP | 37986224\|23142080 | Unspecified |
| mmu-miR-21a-3p | MIMAT0004628 | Faf1 | 14084 | HITS-CLIP | 23142080 | Unspecified |
| mmu-miR-21a-3p | MIMAT0004628 | Fam107b | 66540 | HITS-CLIP | 37986224\|23142080 | Unspecified |
| mmu-miR-21a-3p | MIMAT0004628 | Fam117b | 72750 | HITS-CLIP | 37986224\|29167373\|23142080 | Unspecified |
| mmu-miR-21a-3p | MIMAT0004628 | Fam120a | 218236 | HITS-CLIP | 23142080\|29167373 | Unspecified |
| mmu-miR-21a-3p | MIMAT0004628 | Fam126a | 84652 | HITS-CLIP | 29167373 | Unspecified |
| mmu-miR-21a-3p | MIMAT0004628 | Fam126b | 213056 | HITS-CLIP | 37986224\|23142080 | Unspecified |
| mmu-miR-21a-3p | MIMAT0004628 | Fam129b | 227737 | HITS-CLIP | 29167373 | Unspecified |
| mmu-miR-21a-3p | MIMAT0004628 | Fam133b | 68152 | HITS-CLIP | 23142080 | Unspecified |
| mmu-miR-21a-3p | MIMAT0004628 | Fam13b | 225358 | HITS-CLIP | 29167373 | Unspecified |
| mmu-miR-21a-3p | MIMAT0004628 | Fam168b | 214469 | HITS-CLIP | 23142080 | Unspecified |
| mmu-miR-21a-3p | MIMAT0004628 | Fam172a | 68675 | HITS-CLIP | 23142080 | Unspecified |
| mmu-miR-21a-3p | MIMAT0004628 | Fam193b | 212483 | HITS-CLIP | 23142080 | Unspecified |
| mmu-miR-21a-3p | MIMAT0004628 | Fam208a | 218850 | HITS-CLIP | 23142080 | Unspecified |
| mmu-miR-21a-3p | MIMAT0004628 | Fam208b | 54906 | HITS-CLIP | 23142080 | Unspecified |

| | | | | | | | |
|---|---|---|---|---|---|---|---|
| mmu-miR-21a-3p | MIMAT0004628 | Fam210a | 108654 | HITS-CLIP | | 23142080 | Unspecified |
| mmu-miR-21a-3p | MIMAT0004628 | Fam234b | 57613 | HITS-CLIP | | 23142080 | Unspecified |
| mmu-miR-21a-3p | MIMAT0004628 | Fam43a | 224093 | HITS-CLIP | | 37986224 | Unspecified |
| mmu-miR-21a-3p | MIMAT0004628 | Fam49b | 223601 | HITS-CLIP | | 37986224 | Unspecified |
| mmu-miR-21a-3p | MIMAT0004628 | Fam53b | 77938 | HITS-CLIP | | 23142080 | Unspecified |
| mmu-miR-21a-3p | MIMAT0004628 | Fam69a | 388650 | HITS-CLIP | | 23142080 | Unspecified |
| mmu-miR-21a-3p | MIMAT0004628 | Fam72a | 108900 | HITS-CLIP | | 23142080 | Unspecified |
| mmu-miR-21a-3p | MIMAT0004628 | Fam76b | 72826 | HITS-CLIP | 37986224\|23142080 | | Unspecified |
| mmu-miR-21a-3p | MIMAT0004628 | Fam8a1 | 51439 | HITS-CLIP | | 23142080 | Unspecified |
| mmu-miR-21a-3p | MIMAT0004628 | Fam91a1 | 210998 | HITS-CLIP | | 23142080 | Unspecified |
| mmu-miR-21a-3p | MIMAT0004628 | Fancb | 237211 | HITS-CLIP | | 23142080 | Unspecified |
| mmu-miR-21a-3p | MIMAT0004628 | Fancd2 | 211651 | HITS-CLIP | | 23142080 | Unspecified |
| mmu-miR-21a-3p | MIMAT0004628 | Fanci | 208836 | HITS-CLIP | | 23142080 | Unspecified |
| mmu-miR-21a-3p | MIMAT0004628 | Far1 | 67420 | HITS-CLIP | | 23142080 | Unspecified |
| mmu-miR-21a-3p | MIMAT0004628 | Farsb | 23874 | HITS-CLIP | | 23142080 | Unspecified |
| mmu-miR-21a-3p | MIMAT0004628 | Fasl | 14103 | HITS-CLIP | | 23142080 | Unspecified |
| mmu-miR-21a-3p | MIMAT0004628 | Fasn | 14104 | HITS-CLIP | | 23142080 | Unspecified |
| mmu-miR-21a-3p | MIMAT0004628 | Fbxl3 | 50789 | HITS-CLIP | 37986224\|23142080 | | Unspecified |
| mmu-miR-21a-3p | MIMAT0004628 | Fbxl5 | 242960 | HITS-CLIP | 29167373\|23142080 | | Unspecified |
| mmu-miR-21a-3p | MIMAT0004628 | Fbxo11 | 225055 | HITS-CLIP | | 23142080 | Unspecified |
| mmu-miR-21a-3p | MIMAT0004628 | Fbxo21 | 231670 | HITS-CLIP | | 37986224 | Unspecified |
| mmu-miR-21a-3p | MIMAT0004628 | Fbxo22 | 71999 | HITS-CLIP | | 29167373 | Unspecified |
| mmu-miR-21a-3p | MIMAT0004628 | Fbxo3 | 57443 | HITS-CLIP | | 23142080 | Unspecified |
| mmu-miR-21a-3p | MIMAT0004628 | Fbxo32 | 67731 | HITS-CLIP | | 23142080 | Unspecified |
| mmu-miR-21a-3p | MIMAT0004628 | Fbxw11 | 103583 | HITS-CLIP | | 23142080 | Unspecified |
| mmu-miR-21a-3p | MIMAT0004628 | Fbxw2 | 30050 | HITS-CLIP | | 23142080 | Unspecified |
| mmu-miR-21a-3p | MIMAT0004628 | Fbxw7 | 50754 | HITS-CLIP | | 23142080 | Unspecified |
| mmu-miR-21a-3p | MIMAT0004628 | Fdx1 | 14148 | HITS-CLIP | | 23142080 | Unspecified |
| mmu-miR-21a-3p | MIMAT0004628 | Fem1b | 14155 | HITS-CLIP | 23142080\|29167373 | | Unspecified |
| mmu-miR-21a-3p | MIMAT0004628 | Fem1c | 240263 | HITS-CLIP | 37986224\|23142080 | | Unspecified |
| mmu-miR-21a-3p | MIMAT0004628 | Fen1 | 14156 | HITS-CLIP | | 37986224 | Unspecified |
| mmu-miR-21a-3p | MIMAT0004628 | Fgf2 | 14173 | HITS-CLIP | | 23142080 | Unspecified |
| mmu-miR-21a-3p | MIMAT0004628 | Fignl1 | 60530 | HITS-CLIP | | 37986224 | Unspecified |
| mmu-miR-21a-3p | MIMAT0004628 | Fis1 | 66437 | HITS-CLIP | | 23142080 | Unspecified |
| mmu-miR-21a-3p | MIMAT0004628 | Fkbp1a | 14225 | HITS-CLIP | 37986224\|23142080 | | Unspecified |
| mmu-miR-21a-3p | MIMAT0004628 | Fkbp4 | 14228 | HITS-CLIP | | 23142080 | Unspecified |
| mmu-miR-21a-3p | MIMAT0004628 | Fktn | 246179 | HITS-CLIP | | 29167373 | Unspecified |
| mmu-miR-21a-3p | MIMAT0004628 | Fli1 | 14247 | HITS-CLIP | | 23142080 | Unspecified |
| mmu-miR-21a-3p | MIMAT0004628 | Flna | 192176 | HITS-CLIP | 23142080\|29167373 | | Unspecified |

| | | | | | | |
|---|---|---|---|---|---|---|
| mmu-miR-21a-3p | MIMAT0004628 | Flt3l | 14256 | HITS-CLIP | 23142080 | Unspecified |
| mmu-miR-21a-3p | MIMAT0004628 | Fmnl1 | 57778 | HITS-CLIP | 23142080 | Unspecified |
| mmu-miR-21a-3p | MIMAT0004628 | Fmr1 | 14265 | HITS-CLIP | 23142080 | Unspecified |
| mmu-miR-21a-3p | MIMAT0004628 | Fnbp1 | 14269 | HITS-CLIP | 29167373 | Unspecified |
| mmu-miR-21a-3p | MIMAT0004628 | Fnbp4 | 55935 | HITS-CLIP | 37986224\|23142080 | Unspecified |
| mmu-miR-21a-3p | MIMAT0004628 | Fndc3a | 319448 | HITS-CLIP | 37986224\|23142080\|29167373 | Unspecified |
| mmu-miR-21a-3p | MIMAT0004628 | Fndc3b | 72007 | HITS-CLIP | 37986224\|23142080\|29167373 | Unspecified |
| mmu-miR-21a-3p | MIMAT0004628 | Fnip1 | 216742 | HITS-CLIP | 37986224\|23142080 | Unspecified |
| mmu-miR-21a-3p | MIMAT0004628 | Fnip2 | 329679 | HITS-CLIP | 29167373 | Unspecified |
| mmu-miR-21a-3p | MIMAT0004628 | Focad | 230393 | HITS-CLIP | 23142080 | Unspecified |
| mmu-miR-21a-3p | MIMAT0004628 | Fopnl | 66086 | HITS-CLIP | 23142080 | Unspecified |
| mmu-miR-21a-3p | MIMAT0004628 | Fos | 14281 | HITS-CLIP | 23142080 | Unspecified |
| mmu-miR-21a-3p | MIMAT0004628 | Fosl2 | 14284 | HITS-CLIP | 23142080 | Unspecified |
| mmu-miR-21a-3p | MIMAT0004628 | Foxj2 | 60611 | HITS-CLIP | 37986224 | Unspecified |
| mmu-miR-21a-3p | MIMAT0004628 | Foxj3 | 230700 | HITS-CLIP | 23142080 | Unspecified |
| mmu-miR-21a-3p | MIMAT0004628 | Foxk2 | 68837 | HITS-CLIP | 23142080\|29167373 | Unspecified |
| mmu-miR-21a-3p | MIMAT0004628 | Foxn3 | 71375 | HITS-CLIP | 23142080 | Unspecified |
| mmu-miR-21a-3p | MIMAT0004628 | Foxo1 | 56458 | HITS-CLIP | 23142080 | Unspecified |
| mmu-miR-21a-3p | MIMAT0004628 | Foxo3 | 56484 | HITS-CLIP | 37986224 | Unspecified |
| mmu-miR-21a-3p | MIMAT0004628 | Foxp1 | 108655 | HITS-CLIP | 23142080 | Unspecified |
| mmu-miR-21a-3p | MIMAT0004628 | Frg1 | 14300 | HITS-CLIP | 23142080 | Unspecified |
| mmu-miR-21a-3p | MIMAT0004628 | Frmd6 | 319710 | HITS-CLIP | 23142080 | Unspecified |
| mmu-miR-21a-3p | MIMAT0004628 | Frmd8 | 67457 | HITS-CLIP | 31433990 | Unspecified |
| mmu-miR-21a-3p | MIMAT0004628 | Frs2 | 327826 | HITS-CLIP | 37986224\|23142080 | Unspecified |
| mmu-miR-21a-3p | MIMAT0004628 | Fryl | 72313 | HITS-CLIP | 23142080 | Unspecified |
| mmu-miR-21a-3p | MIMAT0004628 | Fsd1l | 319636 | HITS-CLIP | 23142080 | Unspecified |
| mmu-miR-21a-3p | MIMAT0004628 | Fubp1 | 51886 | HITS-CLIP | 37986224\|23142080\|29167373 | Unspecified |
| mmu-miR-21a-3p | MIMAT0004628 | Fxr1 | 14359 | HITS-CLIP | 23142080\|29167373 | Unspecified |
| mmu-miR-21a-3p | MIMAT0004628 | Fxyd5 | 18301 | HITS-CLIP | 23142080 | Unspecified |
| mmu-miR-21a-3p | MIMAT0004628 | Fyn | 14360 | HITS-CLIP | 23142080 | Unspecified |
| mmu-miR-21a-3p | MIMAT0004628 | Fyttd1 | 69823 | HITS-CLIP | 23142080 | Unspecified |
| mmu-miR-21a-3p | MIMAT0004628 | G2e3 | 217558 | HITS-CLIP | 37986224\|23142080 | Unspecified |
| mmu-miR-21a-3p | MIMAT0004628 | G3bp2 | 23881 | HITS-CLIP | 23142080 | Unspecified |
| mmu-miR-21a-3p | MIMAT0004628 | Gabarap | 56486 | HITS-CLIP | 23142080 | Unspecified |
| mmu-miR-21a-3p | MIMAT0004628 | Gadd45a | 13197 | HITS-CLIP | 23142080 | Unspecified |
| mmu-miR-21a-3p | MIMAT0004628 | Gadd45g | 23882 | HITS-CLIP | 37986224 | Unspecified |
| mmu-miR-21a-3p | MIMAT0004628 | Galk2 | 69976 | HITS-CLIP | 23142080 | Unspecified |
| mmu-miR-21a-3p | MIMAT0004628 | Galnt1 | 14423 | HITS-CLIP | 23142080 | Unspecified |
| mmu-miR-21a-3p | MIMAT0004628 | Galnt2 | 108148 | HITS-CLIP | 29167373 | Unspecified |

| | | | | | | |
|---|---|---|---|---|---|---|
| mmu-miR-21a-3p | MIMAT0004628 | Galnt7 | 108150 | HITS-CLIP | 29167373 | Unspecified |
| mmu-miR-21a-3p | MIMAT0004628 | Gan | 209239 | HITS-CLIP | 23142080 | Unspecified |
| mmu-miR-21a-3p | MIMAT0004628 | Gars | 353172 | HITS-CLIP | 23142080 | Unspecified |
| mmu-miR-21a-3p | MIMAT0004628 | Gas2l3 | 237436 | HITS-CLIP | 37986224\|23142080 | Unspecified |
| mmu-miR-21a-3p | MIMAT0004628 | Gas7 | 14457 | HITS-CLIP | 29167373 | Unspecified |
| mmu-miR-21a-3p | MIMAT0004628 | Gata3 | 14462 | HITS-CLIP | 37986224\|23142080 | Unspecified |
| mmu-miR-21a-3p | MIMAT0004628 | Gatd1 | 347862 | HITS-CLIP | 23142080 | Unspecified |
| mmu-miR-21a-3p | MIMAT0004628 | Gbp5 | 229898 | HITS-CLIP | 23142080 | Unspecified |
| mmu-miR-21a-3p | MIMAT0004628 | Gcfc2 | 330361 | HITS-CLIP | 23142080 | Unspecified |
| mmu-miR-21a-3p | MIMAT0004628 | Gcsh | 68133 | HITS-CLIP | 23142080 | Unspecified |
| mmu-miR-21a-3p | MIMAT0004628 | Gdap2 | 14547 | HITS-CLIP | 23142080 | Unspecified |
| mmu-miR-21a-3p | MIMAT0004628 | Gdi1 | 14567 | HITS-CLIP | 23142080 | Unspecified |
| mmu-miR-21a-3p | MIMAT0004628 | Gem | 14579 | HITS-CLIP | 37986224\|23142080 | Unspecified |
| mmu-miR-21a-3p | MIMAT0004628 | Gfm2 | 320806 | HITS-CLIP | 23142080 | Unspecified |
| mmu-miR-21a-3p | MIMAT0004628 | Ggnbp1 | 70772 | HITS-CLIP | 37986224 | Unspecified |
| mmu-miR-21a-3p | MIMAT0004628 | Ggnbp2 | 217039 | HITS-CLIP | 23142080 | Unspecified |
| mmu-miR-21a-3p | MIMAT0004628 | Ggta1 | 14594 | HITS-CLIP | 23142080\|29167373 | Unspecified |
| mmu-miR-21a-3p | MIMAT0004628 | Ghitm | 66092 | HITS-CLIP | 23142080 | Unspecified |
| mmu-miR-21a-3p | MIMAT0004628 | Gid4 | 66771 | HITS-CLIP | 23142080 | Unspecified |
| mmu-miR-21a-3p | MIMAT0004628 | Gigyf1 | 57330 | HITS-CLIP | 37986224 | Unspecified |
| mmu-miR-21a-3p | MIMAT0004628 | Gimap4 | 107526 | HITS-CLIP | 37986224 | Unspecified |
| mmu-miR-21a-3p | MIMAT0004628 | Gimap6 | 231931 | HITS-CLIP | 23142080 | Unspecified |
| mmu-miR-21a-3p | MIMAT0004628 | Gimap8 | 243374 | HITS-CLIP | 37986224 | Unspecified |
| mmu-miR-21a-3p | MIMAT0004628 | Gimap9 | 317758 | HITS-CLIP | 37986224\|23142080 | Unspecified |
| mmu-miR-21a-3p | MIMAT0004628 | Glmn | 170823 | HITS-CLIP | 23142080 | Unspecified |
| mmu-miR-21a-3p | MIMAT0004628 | Glrx | 93692 | HITS-CLIP | 37986224 | Unspecified |
| mmu-miR-21a-3p | MIMAT0004628 | Glrx3 | 30926 | HITS-CLIP | 29167373 | Unspecified |
| mmu-miR-21a-3p | MIMAT0004628 | Gls | 14660 | HITS-CLIP | 37986224\|23142080 | Unspecified |
| mmu-miR-21a-3p | MIMAT0004628 | Glud1 | 14661 | HITS-CLIP | 23142080 | Unspecified |
| mmu-miR-21a-3p | MIMAT0004628 | Glyr1 | 74022 | HITS-CLIP | 23142080\|29167373 | Unspecified |
| mmu-miR-21a-3p | MIMAT0004628 | Gm10320 | 6617047 | HITS-CLIP | 23142080 | Unspecified |
| mmu-miR-21a-3p | MIMAT0004628 | Gm16286 | 117903916 | HITS-CLIP | 23142080 | Unspecified |
| mmu-miR-21a-3p | MIMAT0004628 | Gm20498 | 105940408 | HITS-CLIP | 23142080 | Unspecified |
| mmu-miR-21a-3p | MIMAT0004628 | Gm38394 | 667118 | HITS-CLIP | 23142080 | Unspecified |
| mmu-miR-21a-3p | MIMAT0004628 | Gmfb | 63985 | HITS-CLIP | 23142080\|29167373 | Unspecified |
| mmu-miR-21a-3p | MIMAT0004628 | Gmnn | 57441 | HITS-CLIP | 37986224 | Unspecified |
| mmu-miR-21a-3p | MIMAT0004628 | Gmps | 229363 | HITS-CLIP | 23142080 | Unspecified |
| mmu-miR-21a-3p | MIMAT0004628 | Gna13 | 14674 | HITS-CLIP | 23142080 | Unspecified |
| mmu-miR-21a-3p | MIMAT0004628 | Gnai3 | 14679 | HITS-CLIP | 23142080 | Unspecified |

| | | | | | | |
|---|---|---|---|---|---|---|
| mmu-miR-21a-3p | MIMAT0004628 | Gnaq | 14682 | HITS-CLIP | 23142080 | Unspecified |
| mmu-miR-21a-3p | MIMAT0004628 | Gnb1 | 14688 | HITS-CLIP | 29167373 | Unspecified |
| mmu-miR-21a-3p | MIMAT0004628 | Gng12 | 14701 | HITS-CLIP | 23142080 | Unspecified |
| mmu-miR-21a-3p | MIMAT0004628 | Gnl2 | 230737 | HITS-CLIP | 23142080 | Unspecified |
| mmu-miR-21a-3p | MIMAT0004628 | Golga1 | 76899 | HITS-CLIP | 23142080 | Unspecified |
| mmu-miR-21a-3p | MIMAT0004628 | Golga3 | 269682 | HITS-CLIP | 37986224 | Unspecified |
| mmu-miR-21a-3p | MIMAT0004628 | Golga4 | 54214 | HITS-CLIP | 29167373 | Unspecified |
| mmu-miR-21a-3p | MIMAT0004628 | Golph3 | 66629 | HITS-CLIP | 23142080 | Unspecified |
| mmu-miR-21a-3p | MIMAT0004628 | Gorasp2 | 70231 | HITS-CLIP | 23142080 | Unspecified |
| mmu-miR-21a-3p | MIMAT0004628 | Gosr1 | 53334 | HITS-CLIP | 23142080 | Unspecified |
| mmu-miR-21a-3p | MIMAT0004628 | Gpam | 14732 | HITS-CLIP | 23142080 | Unspecified |
| mmu-miR-21a-3p | MIMAT0004628 | Gpat3 | 84803 | HITS-CLIP | 23142080 | Unspecified |
| mmu-miR-21a-3p | MIMAT0004628 | Gpatch8 | 237943 | HITS-CLIP | 23142080 | Unspecified |
| mmu-miR-21a-3p | MIMAT0004628 | Gpbp1l1 | 77110 | HITS-CLIP | 23142080 | Unspecified |
| mmu-miR-21a-3p | MIMAT0004628 | Gpcpd1 | 74182 | HITS-CLIP | 23142080 | Unspecified |
| mmu-miR-21a-3p | MIMAT0004628 | Gpd1l | 333433 | HITS-CLIP | 23142080 | Unspecified |
| mmu-miR-21a-3p | MIMAT0004628 | Gpld1 | 14756 | HITS-CLIP | 23142080 | Unspecified |
| mmu-miR-21a-3p | MIMAT0004628 | Gpm6b | 14758 | HITS-CLIP | 23142080 | Unspecified |
| mmu-miR-21a-3p | MIMAT0004628 | Gpr107 | 277463 | HITS-CLIP | 29167373 | Unspecified |
| mmu-miR-21a-3p | MIMAT0004628 | Gpr146 | 80290 | HITS-CLIP | 23142080 | Unspecified |
| mmu-miR-21a-3p | MIMAT0004628 | Gpr171 | 229323 | HITS-CLIP | 23142080 | Unspecified |
| mmu-miR-21a-3p | MIMAT0004628 | Gpr18 | 110168 | HITS-CLIP | 23142080 | Unspecified |
| mmu-miR-21a-3p | MIMAT0004628 | Gpr68 | 238377 | HITS-CLIP | 37986224 | Unspecified |
| mmu-miR-21a-3p | MIMAT0004628 | Gprasp1 | 67298 | HITS-CLIP | 23142080 | Unspecified |
| mmu-miR-21a-3p | MIMAT0004628 | Gprin3 | 243385 | HITS-CLIP | 37986224 | Unspecified |
| mmu-miR-21a-3p | MIMAT0004628 | Gpx1 | 14775 | HITS-CLIP | 23142080 | Unspecified |
| mmu-miR-21a-3p | MIMAT0004628 | Gramd1a | 52857 | HITS-CLIP | 29167373 | Unspecified |
| mmu-miR-21a-3p | MIMAT0004628 | Gramd1b | 235283 | HITS-CLIP | 23142080 | Unspecified |
| mmu-miR-21a-3p | MIMAT0004628 | Gramd3 | 107022 | HITS-CLIP | 37986224\|23142080 | Unspecified |
| mmu-miR-21a-3p | MIMAT0004628 | Grpel1 | 17713 | HITS-CLIP | 23142080 | Unspecified |
| mmu-miR-21a-3p | MIMAT0004628 | Grwd1 | 101612 | HITS-CLIP | 23142080 | Unspecified |
| mmu-miR-21a-3p | MIMAT0004628 | Gse1 | 382034 | HITS-CLIP | 37986224\|23142080 | Unspecified |
| mmu-miR-21a-3p | MIMAT0004628 | Gsk3b | 56637 | HITS-CLIP | 23142080\|29167373 | Unspecified |
| mmu-miR-21a-3p | MIMAT0004628 | Gstcd | 67553 | HITS-CLIP | 29167373 | Unspecified |
| mmu-miR-21a-3p | MIMAT0004628 | Gsto1 | 14873 | HITS-CLIP | 23142080 | Unspecified |
| mmu-miR-21a-3p | MIMAT0004628 | Gstt1 | 14871 | HITS-CLIP | 23142080 | Unspecified |
| mmu-miR-21a-3p | MIMAT0004628 | Gtf2a1 | 83602 | HITS-CLIP | 29167373 | Unspecified |
| mmu-miR-21a-3p | MIMAT0004628 | Gtf2e1 | 74197 | HITS-CLIP | 23142080 | Unspecified |
| mmu-miR-21a-3p | MIMAT0004628 | Gtf2f2 | 68705 | HITS-CLIP | 23142080 | Unspecified |

| | | | | | | |
|---|---|---|---|---|---|---|
| mmu-miR-21a-3p | MIMAT0004628 | Gtf2i | 14886 | HITS-CLIP | 23142080 | Unspecified |
| mmu-miR-21a-3p | MIMAT0004628 | Gtf3c1 | 233863 | HITS-CLIP | 23142080\|29167373 | Unspecified |
| mmu-miR-21a-3p | MIMAT0004628 | Gtf3c3 | 98488 | HITS-CLIP | 23142080 | Unspecified |
| mmu-miR-21a-3p | MIMAT0004628 | Gtpbp1 | 14904 | HITS-CLIP | 23142080 | Unspecified |
| mmu-miR-21a-3p | MIMAT0004628 | Gtpbp2 | 56055 | HITS-CLIP | 23142080 | Unspecified |
| mmu-miR-21a-3p | MIMAT0004628 | Gucd1 | 68778 | HITS-CLIP | 23142080 | Unspecified |
| mmu-miR-21a-3p | MIMAT0004628 | Gucy1a1 | 60596 | HITS-CLIP | 23142080 | Unspecified |
| mmu-miR-21a-3p | MIMAT0004628 | Guf1 | 231279 | HITS-CLIP | 23142080 | Unspecified |
| mmu-miR-21a-3p | MIMAT0004628 | Gxylt1 | 223827 | HITS-CLIP | 23142080 | Unspecified |
| mmu-miR-21a-3p | MIMAT0004628 | Gzmb | 14939 | HITS-CLIP | 23142080 | Unspecified |
| mmu-miR-21a-3p | MIMAT0004628 | H13 | 14950 | HITS-CLIP | 23142080\|29167373 | Unspecified |
| mmu-miR-21a-3p | MIMAT0004628 | H2-D1 | 14964 | HITS-CLIP | 37986224\|23142080 | Unspecified |
| mmu-miR-21a-3p | MIMAT0004628 | H2-Eb1 | 14969 | HITS-CLIP | 23142080 | Unspecified |
| mmu-miR-21a-3p | MIMAT0004628 | H2-K1 | 14972 | HITS-CLIP | 23142080\|29167373 | Unspecified |
| mmu-miR-21a-3p | MIMAT0004628 | H2-M3 | 14991 | HITS-CLIP | 37986224\|23142080 | Unspecified |
| mmu-miR-21a-3p | MIMAT0004628 | H2-Q4 | 15015 | HITS-CLIP | 37986224 | Unspecified |
| mmu-miR-21a-3p | MIMAT0004628 | H2afv | 77605 | HITS-CLIP | 23142080 | Unspecified |
| mmu-miR-21a-3p | MIMAT0004628 | H2afy | 9555 | HITS-CLIP | 23142080 | Unspecified |
| mmu-miR-21a-3p | MIMAT0004628 | H3f3a | 15078 | HITS-CLIP | 23142080 | Unspecified |
| mmu-miR-21a-3p | MIMAT0004628 | H3f3b | 15081 | HITS-CLIP | 23142080\|29167373 | Unspecified |
| mmu-miR-21a-3p | MIMAT0004628 | Hacd3 | 57874 | HITS-CLIP | 23142080 | Unspecified |
| mmu-miR-21a-3p | MIMAT0004628 | Hat1 | 107435 | HITS-CLIP | 23142080 | Unspecified |
| mmu-miR-21a-3p | MIMAT0004628 | Haus6 | 230376 | HITS-CLIP | 37986224 | Unspecified |
| mmu-miR-21a-3p | MIMAT0004628 | Havcr2 | 171285 | HITS-CLIP | 37986224 | Unspecified |
| mmu-miR-21a-3p | MIMAT0004628 | Hbegf | 15200 | HITS-CLIP | 23142080 | Unspecified |
| mmu-miR-21a-3p | MIMAT0004628 | Hbp1 | 73389 | HITS-CLIP | 37986224\|23142080 | Unspecified |
| mmu-miR-21a-3p | MIMAT0004628 | Hbs1l | 56422 | HITS-CLIP | 23142080\|29167373 | Unspecified |
| mmu-miR-21a-3p | MIMAT0004628 | Hccs | 15159 | HITS-CLIP | 23142080 | Unspecified |
| mmu-miR-21a-3p | MIMAT0004628 | Hcfc1 | 15161 | HITS-CLIP | 23142080\|29167373 | Unspecified |
| mmu-miR-21a-3p | MIMAT0004628 | Hcfc2 | 67933 | HITS-CLIP | 37986224\|23142080 | Unspecified |
| mmu-miR-21a-3p | MIMAT0004628 | Hck | 15162 | HITS-CLIP | 29167373 | Unspecified |
| mmu-miR-21a-3p | MIMAT0004628 | Hdac4 | 208727 | HITS-CLIP | 23142080 | Unspecified |
| mmu-miR-21a-3p | MIMAT0004628 | Hdac7 | 56233 | HITS-CLIP | 23142080 | Unspecified |
| mmu-miR-21a-3p | MIMAT0004628 | Hdc | 15186 | HITS-CLIP | 23142080 | Unspecified |
| mmu-miR-21a-3p | MIMAT0004628 | Hdlbp | 110611 | HITS-CLIP | 23142080 | Unspecified |
| mmu-miR-21a-3p | MIMAT0004628 | Heatr1 | 217995 | HITS-CLIP | 23142080 | Unspecified |
| mmu-miR-21a-3p | MIMAT0004628 | Heatr5a | 320487 | HITS-CLIP | 29167373 | Unspecified |
| mmu-miR-21a-3p | MIMAT0004628 | Heatr5b | 320473 | HITS-CLIP | 23142080 | Unspecified |
| mmu-miR-21a-3p | MIMAT0004628 | Heca | 380629 | HITS-CLIP | 29167373 | Unspecified |

| | | | | | | |
|---|---|---|---|---|---|---|
| mmu-miR-21a-3p | MIMAT0004628 | Hectd1 | 207304 | HITS-CLIP | 23142080 | Unspecified |
| mmu-miR-21a-3p | MIMAT0004628 | Hectd4 | 283450 | HITS-CLIP | 23142080 | Unspecified |
| mmu-miR-21a-3p | MIMAT0004628 | Heg1 | 77446 | HITS-CLIP | 23142080 | Unspecified |
| mmu-miR-21a-3p | MIMAT0004628 | Helb | 117599 | HITS-CLIP | 23142080 | Unspecified |
| mmu-miR-21a-3p | MIMAT0004628 | Hells | 15201 | HITS-CLIP | 37986224 | Unspecified |
| mmu-miR-21a-3p | MIMAT0004628 | Helz | 78455 | HITS-CLIP | 23142080 | Unspecified |
| mmu-miR-21a-3p | MIMAT0004628 | Helz2 | 229003 | HITS-CLIP | 23142080\|29167373 | Unspecified |
| mmu-miR-21a-3p | MIMAT0004628 | Hemk1 | 69536 | HITS-CLIP | 23142080 | Unspecified |
| mmu-miR-21a-3p | MIMAT0004628 | Herc1 | 235439 | HITS-CLIP | 23142080 | Unspecified |
| mmu-miR-21a-3p | MIMAT0004628 | Herc2 | 15204 | HITS-CLIP | 23142080 | Unspecified |
| mmu-miR-21a-3p | MIMAT0004628 | Herc3 | 73998 | HITS-CLIP | 29167373\|23142080 | Unspecified |
| mmu-miR-21a-3p | MIMAT0004628 | Herc4 | 67345 | HITS-CLIP | 23142080 | Unspecified |
| mmu-miR-21a-3p | MIMAT0004628 | Herpud1 | 64209 | HITS-CLIP | 37986224\|23142080 | Unspecified |
| mmu-miR-21a-3p | MIMAT0004628 | Herpud2 | 80517 | HITS-CLIP | 23142080 | Unspecified |
| mmu-miR-21a-3p | MIMAT0004628 | Hexim1 | 192231 | HITS-CLIP | 23142080 | Unspecified |
| mmu-miR-21a-3p | MIMAT0004628 | Hibadh | 58875 | HITS-CLIP | 23142080 | Unspecified |
| mmu-miR-21a-3p | MIMAT0004628 | Hibch | 227095 | HITS-CLIP | 23142080 | Unspecified |
| mmu-miR-21a-3p | MIMAT0004628 | Hic2 | 58180 | HITS-CLIP | 23142080 | Unspecified |
| mmu-miR-21a-3p | MIMAT0004628 | Hif1a | 15251 | HITS-CLIP | 37986224\|23142080\|29167373 | Unspecified |
| mmu-miR-21a-3p | MIMAT0004628 | Higd1a | 56295 | HITS-CLIP | 23142080 | Unspecified |
| mmu-miR-21a-3p | MIMAT0004628 | Hipk1 | 15257 | HITS-CLIP | 37986224\|23142080 | Unspecified |
| mmu-miR-21a-3p | MIMAT0004628 | Hipk2 | 15258 | HITS-CLIP | 23142080 | Unspecified |
| mmu-miR-21a-3p | MIMAT0004628 | Hipk3 | 15259 | HITS-CLIP | 23142080\|29167373 | Unspecified |
| mmu-miR-21a-3p | MIMAT0004628 | Hist1h1b | 3009 | HITS-CLIP | 23142080 | Unspecified |
| mmu-miR-21a-3p | MIMAT0004628 | Hist1h2af | 319173 | HITS-CLIP | 23142080 | Unspecified |
| mmu-miR-21a-3p | MIMAT0004628 | Hist2h2bb | 319189 | HITS-CLIP | 23142080 | Unspecified |
| mmu-miR-21a-3p | MIMAT0004628 | Hivep1 | 110521 | HITS-CLIP | 23142080 | Unspecified |
| mmu-miR-21a-3p | MIMAT0004628 | Hk2 | 15277 | HITS-CLIP | 23142080 | Unspecified |
| mmu-miR-21a-3p | MIMAT0004628 | Hltf | 20585 | HITS-CLIP | 23142080 | Unspecified |
| mmu-miR-21a-3p | MIMAT0004628 | Hmbs | 15288 | HITS-CLIP | 23142080 | Unspecified |
| mmu-miR-21a-3p | MIMAT0004628 | Hmga1 | 15361 | HITS-CLIP | 23142080 | Unspecified |
| mmu-miR-21a-3p | MIMAT0004628 | Hmga2 | 15364 | HITS-CLIP | 29167373 | Unspecified |
| mmu-miR-21a-3p | MIMAT0004628 | Hmgcr | 15357 | HITS-CLIP | 37986224 | Unspecified |
| mmu-miR-21a-3p | MIMAT0004628 | Hmgcs1 | 208715 | HITS-CLIP | 23142080 | Unspecified |
| mmu-miR-21a-3p | MIMAT0004628 | Hmgxb4 | 70823 | HITS-CLIP | 23142080 | Unspecified |
| mmu-miR-21a-3p | MIMAT0004628 | Hmmr | 15366 | HITS-CLIP | 23142080 | Unspecified |
| mmu-miR-21a-3p | MIMAT0004628 | Hmox2 | 15369 | HITS-CLIP | 23142080 | Unspecified |
| mmu-miR-21a-3p | MIMAT0004628 | Hnrnpa2b1 | 53379 | HITS-CLIP | 37986224\|23142080 | Unspecified |
| mmu-miR-21a-3p | MIMAT0004628 | Hnrnpc | 15381 | HITS-CLIP | 37986224\|23142080 | Unspecified |

| | | | | | | |
|---|---|---|---|---|---|---|
| mmu-miR-21a-3p | MIMAT0004628 | Hnrnpd | 11991 | HITS-CLIP | 23142080 | Unspecified |
| mmu-miR-21a-3p | MIMAT0004628 | Hnrnph1 | 59013 | HITS-CLIP | 23142080 | Unspecified |
| mmu-miR-21a-3p | MIMAT0004628 | Hnrnpk | 15387 | HITS-CLIP | 23142080\|29167373 | Unspecified |
| mmu-miR-21a-3p | MIMAT0004628 | Hnrnpl | 15388 | HITS-CLIP | 23142080 | Unspecified |
| mmu-miR-21a-3p | MIMAT0004628 | Hnrnpu | 51810 | HITS-CLIP | 37986224\|23142080\|29167373 | Unspecified |
| mmu-miR-21a-3p | MIMAT0004628 | Hnrnpul2 | 68693 | HITS-CLIP | 37986224\|23142080 | Unspecified |
| mmu-miR-21a-3p | MIMAT0004628 | Hook1 | 77963 | HITS-CLIP | 23142080 | Unspecified |
| mmu-miR-21a-3p | MIMAT0004628 | Hook3 | 320191 | HITS-CLIP | 29167373\|23142080 | Unspecified |
| mmu-miR-21a-3p | MIMAT0004628 | Hopx | 74318 | HITS-CLIP | 37986224 | Unspecified |
| mmu-miR-21a-3p | MIMAT0004628 | Hp1bp3 | 15441 | HITS-CLIP | 37986224\|23142080\|29167373 | Unspecified |
| mmu-miR-21a-3p | MIMAT0004628 | Hpf1 | 54969 | HITS-CLIP | 23142080 | Unspecified |
| mmu-miR-21a-3p | MIMAT0004628 | Hprt | 15452 | HITS-CLIP | 23142080 | Unspecified |
| mmu-miR-21a-3p | MIMAT0004628 | Hps6 | 20170 | HITS-CLIP | 23142080 | Unspecified |
| mmu-miR-21a-3p | MIMAT0004628 | Hs3st3b1 | 54710 | HITS-CLIP | 23142080 | Unspecified |
| mmu-miR-21a-3p | MIMAT0004628 | Hsbp1 | 68196 | HITS-CLIP | 23142080 | Unspecified |
| mmu-miR-21a-3p | MIMAT0004628 | Hsd17b4 | 15488 | HITS-CLIP | 23142080 | Unspecified |
| mmu-miR-21a-3p | MIMAT0004628 | Hsp90aa1 | 15519 | HITS-CLIP | 37986224 | Unspecified |
| mmu-miR-21a-3p | MIMAT0004628 | Hsp90b1 | 22027 | HITS-CLIP | 23142080 | Unspecified |
| mmu-miR-21a-3p | MIMAT0004628 | Hspa13 | 110920 | HITS-CLIP | 23142080 | Unspecified |
| mmu-miR-21a-3p | MIMAT0004628 | Hspa14 | 50497 | HITS-CLIP | 23142080 | Unspecified |
| mmu-miR-21a-3p | MIMAT0004628 | Hspa4 | 15525 | HITS-CLIP | 23142080 | Unspecified |
| mmu-miR-21a-3p | MIMAT0004628 | Hspa4l | 18415 | HITS-CLIP | 37986224\|23142080 | Unspecified |
| mmu-miR-21a-3p | MIMAT0004628 | Hspa8 | 15481 | HITS-CLIP | 37986224 | Unspecified |
| mmu-miR-21a-3p | MIMAT0004628 | Hspa9 | 15526 | HITS-CLIP | 23142080 | Unspecified |
| mmu-miR-21a-3p | MIMAT0004628 | Hspd1 | 15510 | HITS-CLIP | 37986224\|23142080 | Unspecified |
| mmu-miR-21a-3p | MIMAT0004628 | Htra2 | 64704 | HITS-CLIP | 37986224\|29167373 | Unspecified |
| mmu-miR-21a-3p | MIMAT0004628 | Htt | 15194 | HITS-CLIP | 23142080 | Unspecified |
| mmu-miR-21a-3p | MIMAT0004628 | Huwe1 | 59026 | HITS-CLIP | 23142080 | Unspecified |
| mmu-miR-21a-3p | MIMAT0004628 | Iah1 | 67732 | HITS-CLIP | 23142080 | Unspecified |
| mmu-miR-21a-3p | MIMAT0004628 | Iars | 105148 | HITS-CLIP | 23142080 | Unspecified |
| mmu-miR-21a-3p | MIMAT0004628 | Ibtk | 108837 | HITS-CLIP | 29167373\|23142080 | Unspecified |
| mmu-miR-21a-3p | MIMAT0004628 | Ick | 56542 | HITS-CLIP | 23142080 | Unspecified |
| mmu-miR-21a-3p | MIMAT0004628 | Icos | 54167 | HITS-CLIP | 37986224 | Unspecified |
| mmu-miR-21a-3p | MIMAT0004628 | Id2 | 15902 | HITS-CLIP | 23142080 | Unspecified |
| mmu-miR-21a-3p | MIMAT0004628 | Id3 | 15903 | HITS-CLIP | 26203562 | Unspecified |
| mmu-miR-21a-3p | MIMAT0004628 | Idh1 | 15926 | HITS-CLIP | 23142080 | Unspecified |
| mmu-miR-21a-3p | MIMAT0004628 | Idh3b | 170718 | HITS-CLIP | 23142080 | Unspecified |
| mmu-miR-21a-3p | MIMAT0004628 | Idh3g | 15929 | HITS-CLIP | 23142080 | Unspecified |
| mmu-miR-21a-3p | MIMAT0004628 | Idnk | 75731 | HITS-CLIP | 23142080 | Unspecified |

| | | | | | | |
|---|---|---|---|---|---|---|
| mmu-miR-21a-3p | MIMAT0004628 | Iffo2 | 212632 | HITS-CLIP | 23142080\|29167373 | Unspecified |
| mmu-miR-21a-3p | MIMAT0004628 | Ifrd1 | 15982 | HITS-CLIP | 23142080 | Unspecified |
| mmu-miR-21a-3p | MIMAT0004628 | Ift46 | 76568 | HITS-CLIP | 23142080 | Unspecified |
| mmu-miR-21a-3p | MIMAT0004628 | Igdcc4 | 56741 | HITS-CLIP | 23142080 | Unspecified |
| mmu-miR-21a-3p | MIMAT0004628 | Igf1 | 16000 | HITS-CLIP | 29167373 | Unspecified |
| mmu-miR-21a-3p | MIMAT0004628 | Igf1r | 16001 | HITS-CLIP | 23142080 | Unspecified |
| mmu-miR-21a-3p | MIMAT0004628 | Igf2bp3 | 140488 | HITS-CLIP | 23142080 | Unspecified |
| mmu-miR-21a-3p | MIMAT0004628 | Igf2r | 16004 | HITS-CLIP | 23142080 | Unspecified |
| mmu-miR-21a-3p | MIMAT0004628 | Ikbke | 56489 | HITS-CLIP | 23142080 | Unspecified |
| mmu-miR-21a-3p | MIMAT0004628 | Ikzf1 | 22778 | HITS-CLIP | 37986224 | Unspecified |
| mmu-miR-21a-3p | MIMAT0004628 | Ikzf2 | 22779 | HITS-CLIP | 23142080 | Unspecified |
| mmu-miR-21a-3p | MIMAT0004628 | Il10 | 16153 | HITS-CLIP | 37986224 | Unspecified |
| mmu-miR-21a-3p | MIMAT0004628 | Il10ra | 16154 | HITS-CLIP | 37986224 | Unspecified |
| mmu-miR-21a-3p | MIMAT0004628 | Il10rb | 16155 | HITS-CLIP | 37986224\|23142080\|29167373 | Unspecified |
| mmu-miR-21a-3p | MIMAT0004628 | Il12rb2 | 16162 | HITS-CLIP | 23142080 | Unspecified |
| mmu-miR-21a-3p | MIMAT0004628 | Il16 | 16170 | HITS-CLIP | 23142080 | Unspecified |
| mmu-miR-21a-3p | MIMAT0004628 | Il4ra | 16190 | HITS-CLIP | 37986224\|23142080 | Unspecified |
| mmu-miR-21a-3p | MIMAT0004628 | Il6ra | 16194 | HITS-CLIP | 23142080 | Unspecified |
| mmu-miR-21a-3p | MIMAT0004628 | Il6st | 16195 | HITS-CLIP | 37986224\|23142080 | Unspecified |
| mmu-miR-21a-3p | MIMAT0004628 | Il7r | 16197 | HITS-CLIP | 23142080 | Unspecified |
| mmu-miR-21a-3p | MIMAT0004628 | Immt | 76614 | HITS-CLIP | 23142080 | Unspecified |
| mmu-miR-21a-3p | MIMAT0004628 | Imp3 | 102462 | HITS-CLIP | 23142080 | Unspecified |
| mmu-miR-21a-3p | MIMAT0004628 | Impa1 | 55980 | HITS-CLIP | 23142080 | Unspecified |
| mmu-miR-21a-3p | MIMAT0004628 | Impa2 | 114663 | HITS-CLIP | 23142080 | Unspecified |
| mmu-miR-21a-3p | MIMAT0004628 | Impact | 16210 | HITS-CLIP | 29167373 | Unspecified |
| mmu-miR-21a-3p | MIMAT0004628 | Impad1 | 242291 | HITS-CLIP | 37986224 | Unspecified |
| mmu-miR-21a-3p | MIMAT0004628 | Inafm2 | 100505573 | HITS-CLIP | 23142080\|29167373 | Unspecified |
| mmu-miR-21a-3p | MIMAT0004628 | Incenp | 16319 | HITS-CLIP | 23142080 | Unspecified |
| mmu-miR-21a-3p | MIMAT0004628 | Inf2 | 70435 | HITS-CLIP | 29167373 | Unspecified |
| mmu-miR-21a-3p | MIMAT0004628 | Ing1 | 26356 | HITS-CLIP | 23142080 | Unspecified |
| mmu-miR-21a-3p | MIMAT0004628 | Inhba | 16323 | HITS-CLIP | 23142080 | Unspecified |
| mmu-miR-21a-3p | MIMAT0004628 | Ino80d | 227195 | HITS-CLIP | 23142080 | Unspecified |
| mmu-miR-21a-3p | MIMAT0004628 | Inpp1 | 16329 | HITS-CLIP | 37986224 | Unspecified |
| mmu-miR-21a-3p | MIMAT0004628 | Inpp4a | 269180 | HITS-CLIP | 37986224\|23142080 | Unspecified |
| mmu-miR-21a-3p | MIMAT0004628 | Inpp5d | 16331 | HITS-CLIP | 29167373 | Unspecified |
| mmu-miR-21a-3p | MIMAT0004628 | Insig2 | 72999 | HITS-CLIP | 37986224 | Unspecified |
| mmu-miR-21a-3p | MIMAT0004628 | Insr | 16337 | HITS-CLIP | 23142080 | Unspecified |
| mmu-miR-21a-3p | MIMAT0004628 | Ints11 | 54973 | HITS-CLIP | 23142080 | Unspecified |
| mmu-miR-21a-3p | MIMAT0004628 | Ints5 | 80789 | HITS-CLIP | 23142080 | Unspecified |

| | | | | | | |
|---|---|---|---|---|---|---|
| mmu-miR-21a-3p | MIMAT0004628 | Ints6 | 18130 | HITS-CLIP | 23142080|29167373 | Unspecified |
| mmu-miR-21a-3p | MIMAT0004628 | Ints7 | 77065 | HITS-CLIP | 23142080 | Unspecified |
| mmu-miR-21a-3p | MIMAT0004628 | Ints8 | 72656 | HITS-CLIP | 23142080 | Unspecified |
| mmu-miR-21a-3p | MIMAT0004628 | Ip6k1 | 27399 | HITS-CLIP | 37986224 | Unspecified |
| mmu-miR-21a-3p | MIMAT0004628 | Ipmk | 69718 | HITS-CLIP | 23142080|29167373 | Unspecified |
| mmu-miR-21a-3p | MIMAT0004628 | Ipo11 | 76582 | HITS-CLIP | 37986224|23142080 | Unspecified |
| mmu-miR-21a-3p | MIMAT0004628 | Ipo7 | 233726 | HITS-CLIP | 23142080 | Unspecified |
| mmu-miR-21a-3p | MIMAT0004628 | Ipo9 | 226432 | HITS-CLIP | 29167373|23142080 | Unspecified |
| mmu-miR-21a-3p | MIMAT0004628 | Ippk | 75678 | HITS-CLIP | 37986224 | Unspecified |
| mmu-miR-21a-3p | MIMAT0004628 | Iqgap1 | 29875 | HITS-CLIP | 23142080|29167373 | Unspecified |
| mmu-miR-21a-3p | MIMAT0004628 | Iqsec1 | 232227 | HITS-CLIP | 23142080|29167373 | Unspecified |
| mmu-miR-21a-3p | MIMAT0004628 | Irak4 | 266632 | HITS-CLIP | 37986224 | Unspecified |
| mmu-miR-21a-3p | MIMAT0004628 | Ireb2 | 64602 | HITS-CLIP | 23142080|29167373 | Unspecified |
| mmu-miR-21a-3p | MIMAT0004628 | Irf2 | 16363 | HITS-CLIP | 23142080 | Unspecified |
| mmu-miR-21a-3p | MIMAT0004628 | Irf2bp2 | 270110 | HITS-CLIP | 37986224|23142080 | Unspecified |
| mmu-miR-21a-3p | MIMAT0004628 | Irf2bpl | 238330 | HITS-CLIP | 29167373 | Unspecified |
| mmu-miR-21a-3p | MIMAT0004628 | Irf4 | 16364 | HITS-CLIP | 37986224|23142080 | Unspecified |
| mmu-miR-21a-3p | MIMAT0004628 | Irgm1 | 15944 | HITS-CLIP | 37986224 | Unspecified |
| mmu-miR-21a-3p | MIMAT0004628 | Irgm2 | 54396 | HITS-CLIP | 23142080|31433990 | Unspecified |
| mmu-miR-21a-3p | MIMAT0004628 | Irs2 | 384783 | HITS-CLIP | 23142080 | Unspecified |
| mmu-miR-21a-3p | MIMAT0004628 | Isoc2a | 664994 | HITS-CLIP | 23142080 | Unspecified |
| mmu-miR-21a-3p | MIMAT0004628 | Ist1 | 71955 | HITS-CLIP | 37986224|23142080 | Unspecified |
| mmu-miR-21a-3p | MIMAT0004628 | Isyna1 | 71780 | HITS-CLIP | 23142080 | Unspecified |
| mmu-miR-21a-3p | MIMAT0004628 | Itch | 16396 | HITS-CLIP | 23142080 | Unspecified |
| mmu-miR-21a-3p | MIMAT0004628 | Itga4 | 16401 | HITS-CLIP | 37986224|23142080 | Unspecified |
| mmu-miR-21a-3p | MIMAT0004628 | Itga6 | 16403 | HITS-CLIP | 23142080|29167373 | Unspecified |
| mmu-miR-21a-3p | MIMAT0004628 | Itgal | 16408 | HITS-CLIP | 23142080 | Unspecified |
| mmu-miR-21a-3p | MIMAT0004628 | Itgav | 16410 | HITS-CLIP | 23142080 | Unspecified |
| mmu-miR-21a-3p | MIMAT0004628 | Itgb2 | 16414 | HITS-CLIP | 37986224 | Unspecified |
| mmu-miR-21a-3p | MIMAT0004628 | Itgb5 | 16419 | HITS-CLIP | 29167373 | Unspecified |
| mmu-miR-21a-3p | MIMAT0004628 | Itk | 16428 | HITS-CLIP | 23142080 | Unspecified |
| mmu-miR-21a-3p | MIMAT0004628 | Itm2a | 16431 | HITS-CLIP | 23142080 | Unspecified |
| mmu-miR-21a-3p | MIMAT0004628 | Itm2b | 16432 | HITS-CLIP | 23142080 | Unspecified |
| mmu-miR-21a-3p | MIMAT0004628 | Itpk1 | 217837 | HITS-CLIP | 23142080 | Unspecified |
| mmu-miR-21a-3p | MIMAT0004628 | Itpkb | 320404 | HITS-CLIP | 37986224 | Unspecified |
| mmu-miR-21a-3p | MIMAT0004628 | Itpr1 | 16438 | HITS-CLIP | 23142080 | Unspecified |
| mmu-miR-21a-3p | MIMAT0004628 | Itpr2 | 16439 | HITS-CLIP | 23142080 | Unspecified |
| mmu-miR-21a-3p | MIMAT0004628 | Itsn1 | 16443 | HITS-CLIP | 29167373 | Unspecified |
| mmu-miR-21a-3p | MIMAT0004628 | Itsn2 | 20403 | HITS-CLIP | 23142080|29167373 | Unspecified |

| | | | | | | |
|---|---|---|---|---|---|---|
| mmu-miR-21a-3p | MIMAT0004628 | Iws1 | 73473 | HITS-CLIP | 23142080 | Unspecified |
| mmu-miR-21a-3p | MIMAT0004628 | Jade1 | 269424 | HITS-CLIP | 23142080 | Unspecified |
| mmu-miR-21a-3p | MIMAT0004628 | Jag1 | 16449 | HITS-CLIP | 29167373 | Unspecified |
| mmu-miR-21a-3p | MIMAT0004628 | Jag2 | 16450 | HITS-CLIP | 23142080 | Unspecified |
| mmu-miR-21a-3p | MIMAT0004628 | Jak1 | 16451 | HITS-CLIP | 37986224\|23142080 | Unspecified |
| mmu-miR-21a-3p | MIMAT0004628 | Jak2 | 16452 | HITS-CLIP | 23142080 | Unspecified |
| mmu-miR-21a-3p | MIMAT0004628 | Jarid2 | 16468 | HITS-CLIP | 37986224\|23142080\|29167373 | Unspecified |
| mmu-miR-21a-3p | MIMAT0004628 | Jazf1 | 231986 | HITS-CLIP | 23142080 | Unspecified |
| mmu-miR-21a-3p | MIMAT0004628 | Jmjd1c | 108829 | HITS-CLIP | 37986224\|23142080 | Unspecified |
| mmu-miR-21a-3p | MIMAT0004628 | Jmy | 57748 | HITS-CLIP | 23142080 | Unspecified |
| mmu-miR-21a-3p | MIMAT0004628 | Josd1 | 74158 | HITS-CLIP | 31433990 | Unspecified |
| mmu-miR-21a-3p | MIMAT0004628 | Jund | 16478 | HITS-CLIP | 23142080 | Unspecified |
| mmu-miR-21a-3p | MIMAT0004628 | Kansl1 | 76719 | HITS-CLIP | 23142080 | Unspecified |
| mmu-miR-21a-3p | MIMAT0004628 | Kat2b | 18519 | HITS-CLIP | 23142080 | Unspecified |
| mmu-miR-21a-3p | MIMAT0004628 | Kat6b | 54169 | HITS-CLIP | 23142080\|29167373 | Unspecified |
| mmu-miR-21a-3p | MIMAT0004628 | Katnbl1 | 72425 | HITS-CLIP | 29167373\|23142080 | Unspecified |
| mmu-miR-21a-3p | MIMAT0004628 | Kazn | 71529 | HITS-CLIP | 23142080 | Unspecified |
| mmu-miR-21a-3p | MIMAT0004628 | Kbtbd11 | 74901 | HITS-CLIP | 23142080 | Unspecified |
| mmu-miR-21a-3p | MIMAT0004628 | Kbtbd2 | 210973 | HITS-CLIP | 37986224\|23142080\|29167373 | Unspecified |
| mmu-miR-21a-3p | MIMAT0004628 | Kcmf1 | 74287 | HITS-CLIP | 23142080 | Unspecified |
| mmu-miR-21a-3p | MIMAT0004628 | Kcnab2 | 16498 | HITS-CLIP | 29167373 | Unspecified |
| mmu-miR-21a-3p | MIMAT0004628 | Kcnn4 | 16534 | HITS-CLIP | 37986224\|23142080 | Unspecified |
| mmu-miR-21a-3p | MIMAT0004628 | Kctd10 | 330171 | HITS-CLIP | 23142080 | Unspecified |
| mmu-miR-21a-3p | MIMAT0004628 | Kctd5 | 69259 | HITS-CLIP | 23142080\|29167373 | Unspecified |
| mmu-miR-21a-3p | MIMAT0004628 | Kctd9 | 105440 | HITS-CLIP | 23142080 | Unspecified |
| mmu-miR-21a-3p | MIMAT0004628 | Kdelr1 | 68137 | HITS-CLIP | 23142080 | Unspecified |
| mmu-miR-21a-3p | MIMAT0004628 | Kdm1a | 99982 | HITS-CLIP | 23142080 | Unspecified |
| mmu-miR-21a-3p | MIMAT0004628 | Kdm2a | 225876 | HITS-CLIP | 23142080\|29167373 | Unspecified |
| mmu-miR-21a-3p | MIMAT0004628 | Kdm2b | 30841 | HITS-CLIP | 37986224 | Unspecified |
| mmu-miR-21a-3p | MIMAT0004628 | Kdm4c | 76804 | HITS-CLIP | 23142080 | Unspecified |
| mmu-miR-21a-3p | MIMAT0004628 | Kdm5a | 214899 | HITS-CLIP | 23142080 | Unspecified |
| mmu-miR-21a-3p | MIMAT0004628 | Kdm5b | 75605 | HITS-CLIP | 37986224\|23142080 | Unspecified |
| mmu-miR-21a-3p | MIMAT0004628 | Kdm5c | 20591 | HITS-CLIP | 29167373 | Unspecified |
| mmu-miR-21a-3p | MIMAT0004628 | Kdm6a | 22289 | HITS-CLIP | 23142080 | Unspecified |
| mmu-miR-21a-3p | MIMAT0004628 | Kdm7a | 338523 | HITS-CLIP | 37986224\|23142080 | Unspecified |
| mmu-miR-21a-3p | MIMAT0004628 | Kdsr | 70750 | HITS-CLIP | 23142080 | Unspecified |
| mmu-miR-21a-3p | MIMAT0004628 | Khdc4 | 22889 | HITS-CLIP | 23142080 | Unspecified |
| mmu-miR-21a-3p | MIMAT0004628 | Khdrbs1 | 20218 | HITS-CLIP | 23142080 | Unspecified |
| mmu-miR-21a-3p | MIMAT0004628 | Khnyn | 219094 | HITS-CLIP | 37986224 | Unspecified |

| | | | | | | |
|---|---|---|---|---|---|---|
| mmu-miR-21a-3p | MIMAT0004628 | Khsrp | 16549 | HITS-CLIP | 23142080 | Unspecified |
| mmu-miR-21a-3p | MIMAT0004628 | Kidins220 | 77480 | HITS-CLIP | 23142080 | Unspecified |
| mmu-miR-21a-3p | MIMAT0004628 | Kif11 | 16551 | HITS-CLIP | 23142080 | Unspecified |
| mmu-miR-21a-3p | MIMAT0004628 | Kif18a | 228421 | HITS-CLIP | 23142080 | Unspecified |
| mmu-miR-21a-3p | MIMAT0004628 | Kif1b | 16561 | HITS-CLIP | 29167373\|23142080 | Unspecified |
| mmu-miR-21a-3p | MIMAT0004628 | Kif1bp | 26128 | HITS-CLIP | 23142080 | Unspecified |
| mmu-miR-21a-3p | MIMAT0004628 | Kif21b | 16565 | HITS-CLIP | 23142080 | Unspecified |
| mmu-miR-21a-3p | MIMAT0004628 | Kif23 | 71819 | HITS-CLIP | 23142080 | Unspecified |
| mmu-miR-21a-3p | MIMAT0004628 | Kif4 | 16571 | HITS-CLIP | 23142080 | Unspecified |
| mmu-miR-21a-3p | MIMAT0004628 | Kif5b | 16573 | HITS-CLIP | 37986224\|23142080\|29167373 | Unspecified |
| mmu-miR-21a-3p | MIMAT0004628 | Klc1 | 16593 | HITS-CLIP | 29167373 | Unspecified |
| mmu-miR-21a-3p | MIMAT0004628 | Klf5 | 12224 | HITS-CLIP | 26203562 | Unspecified |
| mmu-miR-21a-3p | MIMAT0004628 | Klf6 | 23849 | HITS-CLIP | 37986224\|23142080\|29167373 | Unspecified |
| mmu-miR-21a-3p | MIMAT0004628 | Klf7 | 93691 | HITS-CLIP | 23142080 | Unspecified |
| mmu-miR-21a-3p | MIMAT0004628 | Klhdc1 | 271005 | HITS-CLIP | 23142080 | Unspecified |
| mmu-miR-21a-3p | MIMAT0004628 | Klhdc10 | 76788 | HITS-CLIP | 31433990 | Unspecified |
| mmu-miR-21a-3p | MIMAT0004628 | Klhdc3 | 71765 | HITS-CLIP | 23142080 | Unspecified |
| mmu-miR-21a-3p | MIMAT0004628 | Klhl2 | 77113 | HITS-CLIP | 23142080 | Unspecified |
| mmu-miR-21a-3p | MIMAT0004628 | Klhl24 | 75785 | HITS-CLIP | 23142080 | Unspecified |
| mmu-miR-21a-3p | MIMAT0004628 | Klhl28 | 66689 | HITS-CLIP | 23142080 | Unspecified |
| mmu-miR-21a-3p | MIMAT0004628 | Klhl9 | 242521 | HITS-CLIP | 23142080 | Unspecified |
| mmu-miR-21a-3p | MIMAT0004628 | Kmt2a | 214162 | HITS-CLIP | 23142080 | Unspecified |
| mmu-miR-21a-3p | MIMAT0004628 | Kmt2c | 231051 | HITS-CLIP | 37986224\|23142080\|29167373 | Unspecified |
| mmu-miR-21a-3p | MIMAT0004628 | Kmt2d | 381022 | HITS-CLIP | 37986224\|23142080\|29167373 | Unspecified |
| mmu-miR-21a-3p | MIMAT0004628 | Kmt2e | 69188 | HITS-CLIP | 23142080\|29167373 | Unspecified |
| mmu-miR-21a-3p | MIMAT0004628 | Knl1 | 57082 | HITS-CLIP | 23142080 | Unspecified |
| mmu-miR-21a-3p | MIMAT0004628 | Knop1 | 66356 | HITS-CLIP | 37986224\|23142080 | Unspecified |
| mmu-miR-21a-3p | MIMAT0004628 | Kntc1 | 208628 | HITS-CLIP | 23142080 | Unspecified |
| mmu-miR-21a-3p | MIMAT0004628 | Kpna1 | 16646 | HITS-CLIP | 23142080 | Unspecified |
| mmu-miR-21a-3p | MIMAT0004628 | Kpna3 | 16648 | HITS-CLIP | 23142080\|29167373 | Unspecified |
| mmu-miR-21a-3p | MIMAT0004628 | Kpna4 | 16649 | HITS-CLIP | 37986224\|29167373\|23142080 | Unspecified |
| mmu-miR-21a-3p | MIMAT0004628 | Kpna6 | 16650 | HITS-CLIP | 23142080 | Unspecified |
| mmu-miR-21a-3p | MIMAT0004628 | Kpnb1 | 16211 | HITS-CLIP | 29167373 | Unspecified |
| mmu-miR-21a-3p | MIMAT0004628 | Kras | 16653 | HITS-CLIP | 23142080 | Unspecified |
| mmu-miR-21a-3p | MIMAT0004628 | Kremen2 | 73016 | HITS-CLIP | 23142080 | Unspecified |
| mmu-miR-21a-3p | MIMAT0004628 | Krit1 | 79264 | HITS-CLIP | 23142080 | Unspecified |
| mmu-miR-21a-3p | MIMAT0004628 | Ktn1 | 16709 | HITS-CLIP | 23142080\|29167373 | Unspecified |
| mmu-miR-21a-3p | MIMAT0004628 | Lamc1 | 226519 | HITS-CLIP | 23142080 | Unspecified |
| mmu-miR-21a-3p | MIMAT0004628 | Lamp2 | 16784 | HITS-CLIP | 23142080 | Unspecified |

| | | | | | | |
|---|---|---|---|---|---|---|
| mmu-miR-21a-3p | MIMAT0004628 | Laptm5 | 16792 | HITS-CLIP | 23142080\|29167373 | Unspecified |
| mmu-miR-21a-3p | MIMAT0004628 | Larp4b | 217980 | HITS-CLIP | 37986224\|23142080 | Unspecified |
| mmu-miR-21a-3p | MIMAT0004628 | Lasp1 | 16796 | HITS-CLIP | 23142080 | Unspecified |
| mmu-miR-21a-3p | MIMAT0004628 | Lat2 | 56743 | HITS-CLIP | 29167373 | Unspecified |
| mmu-miR-21a-3p | MIMAT0004628 | Lats1 | 16798 | HITS-CLIP | 37986224\|23142080\|29167373 | Unspecified |
| mmu-miR-21a-3p | MIMAT0004628 | Lax1 | 240754 | HITS-CLIP | 37986224 | Unspecified |
| mmu-miR-21a-3p | MIMAT0004628 | Lbh | 77889 | HITS-CLIP | 23142080 | Unspecified |
| mmu-miR-21a-3p | MIMAT0004628 | Lbr | 98386 | HITS-CLIP | 37986224 | Unspecified |
| mmu-miR-21a-3p | MIMAT0004628 | Lck | 16818 | HITS-CLIP | 23142080 | Unspecified |
| mmu-miR-21a-3p | MIMAT0004628 | Lcor | 212391 | HITS-CLIP | 23142080 | Unspecified |
| mmu-miR-21a-3p | MIMAT0004628 | Lcorl | 209707 | HITS-CLIP | 23142080 | Unspecified |
| mmu-miR-21a-3p | MIMAT0004628 | Lcp1 | 18826 | HITS-CLIP | 23142080 | Unspecified |
| mmu-miR-21a-3p | MIMAT0004628 | Lcp2 | 16822 | HITS-CLIP | 37986224\|23142080 | Unspecified |
| mmu-miR-21a-3p | MIMAT0004628 | Lemd3 | 380664 | HITS-CLIP | 23142080 | Unspecified |
| mmu-miR-21a-3p | MIMAT0004628 | Leprotl1 | 68192 | HITS-CLIP | 23142080 | Unspecified |
| mmu-miR-21a-3p | MIMAT0004628 | Lfng | 16848 | HITS-CLIP | 23142080\|29167373 | Unspecified |
| mmu-miR-21a-3p | MIMAT0004628 | Lgals9 | 16859 | HITS-CLIP | 23142080 | Unspecified |
| mmu-miR-21a-3p | MIMAT0004628 | Lgalsl | 216551 | HITS-CLIP | 23142080 | Unspecified |
| mmu-miR-21a-3p | MIMAT0004628 | Lif | 16878 | HITS-CLIP | 23142080 | Unspecified |
| mmu-miR-21a-3p | MIMAT0004628 | Limd2 | 67803 | HITS-CLIP | 37986224\|23142080 | Unspecified |
| mmu-miR-21a-3p | MIMAT0004628 | Limk2 | 16886 | HITS-CLIP | 23142080 | Unspecified |
| mmu-miR-21a-3p | MIMAT0004628 | Lims1 | 110829 | HITS-CLIP | 37986224\|23142080\|29167373 | Unspecified |
| mmu-miR-21a-3p | MIMAT0004628 | Lin7c | 22343 | HITS-CLIP | 23142080 | Unspecified |
| mmu-miR-21a-3p | MIMAT0004628 | Lin9 | 72568 | HITS-CLIP | 23142080 | Unspecified |
| mmu-miR-21a-3p | MIMAT0004628 | Lman2 | 66890 | HITS-CLIP | 37986224\|23142080 | Unspecified |
| mmu-miR-21a-3p | MIMAT0004628 | Lmbrd1 | 68421 | HITS-CLIP | 37986224\|23142080 | Unspecified |
| mmu-miR-21a-3p | MIMAT0004628 | Lmbrd2 | 320506 | HITS-CLIP | 23142080 | Unspecified |
| mmu-miR-21a-3p | MIMAT0004628 | Lmnb1 | 16906 | HITS-CLIP | 23142080 | Unspecified |
| mmu-miR-21a-3p | MIMAT0004628 | Lnpep | 240028 | HITS-CLIP | 37986224\|23142080\|29167373 | Unspecified |
| mmu-miR-21a-3p | MIMAT0004628 | Lonp1 | 74142 | HITS-CLIP | 23142080 | Unspecified |
| mmu-miR-21a-3p | MIMAT0004628 | Lonp2 | 66887 | HITS-CLIP | 23142080 | Unspecified |
| mmu-miR-21a-3p | MIMAT0004628 | Lonrf1 | 244421 | HITS-CLIP | 37986224 | Unspecified |
| mmu-miR-21a-3p | MIMAT0004628 | Lpcat3 | 14792 | HITS-CLIP | 23142080 | Unspecified |
| mmu-miR-21a-3p | MIMAT0004628 | Lpin1 | 14245 | HITS-CLIP | 37986224\|23142080 | Unspecified |
| mmu-miR-21a-3p | MIMAT0004628 | Lpin2 | 64898 | HITS-CLIP | 23142080\|29167373 | Unspecified |
| mmu-miR-21a-3p | MIMAT0004628 | Lpp | 210126 | HITS-CLIP | 37986224\|23142080 | Unspecified |
| mmu-miR-21a-3p | MIMAT0004628 | Lrba | 80877 | HITS-CLIP | 23142080 | Unspecified |
| mmu-miR-21a-3p | MIMAT0004628 | Lrch3 | 70144 | HITS-CLIP | 23142080 | Unspecified |
| mmu-miR-21a-3p | MIMAT0004628 | Lrig1 | 16206 | HITS-CLIP | 23142080 | Unspecified |

| | | | | | | |
|---|---|---|---|---|---|---|
| mmu-miR-21a-3p | MIMAT0004628 | Lrig2 | 269473 | HITS-CLIP | 29167373\|23142080 | Unspecified |
| mmu-miR-21a-3p | MIMAT0004628 | Lrp12 | 239393 | HITS-CLIP | 23142080\|29167373 | Unspecified |
| mmu-miR-21a-3p | MIMAT0004628 | Lrp6 | 16974 | HITS-CLIP | 37986224\|29167373\|23142080 | Unspecified |
| mmu-miR-21a-3p | MIMAT0004628 | Lrpprc | 72416 | HITS-CLIP | 23142080 | Unspecified |
| mmu-miR-21a-3p | MIMAT0004628 | Lrr1 | 69706 | HITS-CLIP | 23142080 | Unspecified |
| mmu-miR-21a-3p | MIMAT0004628 | Lrrc40 | 67144 | HITS-CLIP | 23142080 | Unspecified |
| mmu-miR-21a-3p | MIMAT0004628 | Lrrc41 | 230654 | HITS-CLIP | 23142080 | Unspecified |
| mmu-miR-21a-3p | MIMAT0004628 | Lrrc42 | 77809 | HITS-CLIP | 23142080 | Unspecified |
| mmu-miR-21a-3p | MIMAT0004628 | Lrrc47 | 72946 | HITS-CLIP | 23142080 | Unspecified |
| mmu-miR-21a-3p | MIMAT0004628 | Lrrc57 | 66606 | HITS-CLIP | 23142080 | Unspecified |
| mmu-miR-21a-3p | MIMAT0004628 | Lrrc58 | 320184 | HITS-CLIP | 23142080 | Unspecified |
| mmu-miR-21a-3p | MIMAT0004628 | Lrrc59 | 98238 | HITS-CLIP | 23142080 | Unspecified |
| mmu-miR-21a-3p | MIMAT0004628 | Lrrc8c | 100604 | HITS-CLIP | 29167373 | Unspecified |
| mmu-miR-21a-3p | MIMAT0004628 | Lrrcc1 | 71710 | HITS-CLIP | 23142080 | Unspecified |
| mmu-miR-21a-3p | MIMAT0004628 | Lrrfip1 | 16978 | HITS-CLIP | 37986224\|23142080\|29167373 | Unspecified |
| mmu-miR-21a-3p | MIMAT0004628 | Lsm14a | 67070 | HITS-CLIP | 23142080\|29167373 | Unspecified |
| mmu-miR-21a-3p | MIMAT0004628 | Lsm14b | 241846 | HITS-CLIP | 37986224 | Unspecified |
| mmu-miR-21a-3p | MIMAT0004628 | Lss | 16987 | HITS-CLIP | 23142080 | Unspecified |
| mmu-miR-21a-3p | MIMAT0004628 | Ltn1 | 78913 | HITS-CLIP | 23142080 | Unspecified |
| mmu-miR-21a-3p | MIMAT0004628 | Luc7l2 | 192196 | HITS-CLIP | 23142080\|29167373 | Unspecified |
| mmu-miR-21a-3p | MIMAT0004628 | Luzp1 | 269593 | HITS-CLIP | 29167373 | Unspecified |
| mmu-miR-21a-3p | MIMAT0004628 | Ly6e | 17069 | HITS-CLIP | 23142080 | Unspecified |
| mmu-miR-21a-3p | MIMAT0004628 | Lyar | 17089 | HITS-CLIP | 26203562\|23142080 | Unspecified |
| mmu-miR-21a-3p | MIMAT0004628 | Lyn | 17096 | HITS-CLIP | 29167373 | Unspecified |
| mmu-miR-21a-3p | MIMAT0004628 | Lypla1 | 18777 | HITS-CLIP | 23142080 | Unspecified |
| mmu-miR-21a-3p | MIMAT0004628 | Lyst | 17101 | HITS-CLIP | 23142080\|29167373 | Unspecified |
| mmu-miR-21a-3p | MIMAT0004628 | M6pr | 17113 | HITS-CLIP | 29167373 | Unspecified |
| mmu-miR-21a-3p | MIMAT0004628 | Macf1 | 11426 | HITS-CLIP | 23142080\|29167373 | Unspecified |
| mmu-miR-21a-3p | MIMAT0004628 | Mad2l1 | 56150 | HITS-CLIP | 37986224 | Unspecified |
| mmu-miR-21a-3p | MIMAT0004628 | Madd | 228355 | HITS-CLIP | 37986224 | Unspecified |
| mmu-miR-21a-3p | MIMAT0004628 | Maea | 59003 | HITS-CLIP | 23142080 | Unspecified |
| mmu-miR-21a-3p | MIMAT0004628 | Maf | 17132 | HITS-CLIP | 23142080 | Unspecified |
| mmu-miR-21a-3p | MIMAT0004628 | Mafg | 17134 | HITS-CLIP | 23142080 | Unspecified |
| mmu-miR-21a-3p | MIMAT0004628 | Mafk | 17135 | HITS-CLIP | 23142080 | Unspecified |
| mmu-miR-21a-3p | MIMAT0004628 | Maip1 | 79568 | HITS-CLIP | 23142080 | Unspecified |
| mmu-miR-21a-3p | MIMAT0004628 | Malt1 | 240354 | HITS-CLIP | 37986224 | Unspecified |
| mmu-miR-21a-3p | MIMAT0004628 | Maml2 | 270118 | HITS-CLIP | 23142080 | Unspecified |
| mmu-miR-21a-3p | MIMAT0004628 | Man1a | 17155 | HITS-CLIP | 23142080 | Unspecified |
| mmu-miR-21a-3p | MIMAT0004628 | Man1a2 | 17156 | HITS-CLIP | 23142080\|29167373 | Unspecified |

| | | | | | | |
|---|---|---|---|---|---|---|
| mmu-miR-21a-3p | MIMAT0004628 | Man2a1 | 17158 | HITS-CLIP | 23142080\|29167373 | Unspecified |
| mmu-miR-21a-3p | MIMAT0004628 | Man2b1 | 17159 | HITS-CLIP | 29167373 | Unspecified |
| mmu-miR-21a-3p | MIMAT0004628 | Map2k1 | 26395 | HITS-CLIP | 37986224\|23142080 | Unspecified |
| mmu-miR-21a-3p | MIMAT0004628 | Map2k4 | 26398 | HITS-CLIP | 23142080 | Unspecified |
| mmu-miR-21a-3p | MIMAT0004628 | Map3k1 | 26401 | HITS-CLIP | 37986224\|23142080\|29167373 | Unspecified |
| mmu-miR-21a-3p | MIMAT0004628 | Map3k12 | 26404 | HITS-CLIP | 23142080 | Unspecified |
| mmu-miR-21a-3p | MIMAT0004628 | Map3k2 | 26405 | HITS-CLIP | 23142080 | Unspecified |
| mmu-miR-21a-3p | MIMAT0004628 | Map3k3 | 26406 | HITS-CLIP | 23142080 | Unspecified |
| mmu-miR-21a-3p | MIMAT0004628 | Map3k4 | 26407 | HITS-CLIP | 23142080 | Unspecified |
| mmu-miR-21a-3p | MIMAT0004628 | Map3k5 | 26408 | HITS-CLIP | 23142080 | Unspecified |
| mmu-miR-21a-3p | MIMAT0004628 | Map3k7 | 26409 | HITS-CLIP | 23142080 | Unspecified |
| mmu-miR-21a-3p | MIMAT0004628 | Map4k4 | 26921 | HITS-CLIP | 23142080 | Unspecified |
| mmu-miR-21a-3p | MIMAT0004628 | Mapk14 | 26416 | HITS-CLIP | 23142080 | Unspecified |
| mmu-miR-21a-3p | MIMAT0004628 | Mapk1ip1l | 218975 | HITS-CLIP | 37986224\|23142080 | Unspecified |
| mmu-miR-21a-3p | MIMAT0004628 | Mapk6 | 50772 | HITS-CLIP | 23142080 | Unspecified |
| mmu-miR-21a-3p | MIMAT0004628 | Mapk8 | 26419 | HITS-CLIP | 29167373 | Unspecified |
| mmu-miR-21a-3p | MIMAT0004628 | Mapk9 | 26420 | HITS-CLIP | 23142080 | Unspecified |
| mmu-miR-21a-3p | MIMAT0004628 | Mapkapk2 | 17164 | HITS-CLIP | 23142080 | Unspecified |
| mmu-miR-21a-3p | MIMAT0004628 | March7 | 57438 | HITS-CLIP | 23142080 | Unspecified |
| mmu-miR-21a-3p | MIMAT0004628 | March9 | 216438 | HITS-CLIP | 23142080 | Unspecified |
| mmu-miR-21a-3p | MIMAT0004628 | Marchf6 | 223455 | HITS-CLIP | 37986224\|23142080\|29167373 | Unspecified |
| mmu-miR-21a-3p | MIMAT0004628 | Marcks | 17118 | HITS-CLIP | 23142080\|29167373 | Unspecified |
| mmu-miR-21a-3p | MIMAT0004628 | Marf1 | 223989 | HITS-CLIP | 37986224\|23142080 | Unspecified |
| mmu-miR-21a-3p | MIMAT0004628 | Mars2 | 212679 | HITS-CLIP | 23142080 | Unspecified |
| mmu-miR-21a-3p | MIMAT0004628 | Mastl | 67121 | HITS-CLIP | 37986224\|23142080 | Unspecified |
| mmu-miR-21a-3p | MIMAT0004628 | Mat2a | 232087 | HITS-CLIP | 23142080 | Unspecified |
| mmu-miR-21a-3p | MIMAT0004628 | Mat2b | 108645 | HITS-CLIP | 37986224\|23142080 | Unspecified |
| mmu-miR-21a-3p | MIMAT0004628 | Matr3 | 17184 | HITS-CLIP | 23142080 | Unspecified |
| mmu-miR-21a-3p | MIMAT0004628 | Mbd2 | 17191 | HITS-CLIP | 23142080 | Unspecified |
| mmu-miR-21a-3p | MIMAT0004628 | Mbd4 | 17193 | HITS-CLIP | 23142080 | Unspecified |
| mmu-miR-21a-3p | MIMAT0004628 | Mbd5 | 109241 | HITS-CLIP | 29167373 | Unspecified |
| mmu-miR-21a-3p | MIMAT0004628 | Mbnl1 | 56758 | HITS-CLIP | 37986224\|23142080 | Unspecified |
| mmu-miR-21a-3p | MIMAT0004628 | Mbnl3 | 171170 | HITS-CLIP | 23142080 | Unspecified |
| mmu-miR-21a-3p | MIMAT0004628 | Mboat1 | 218121 | HITS-CLIP | 23142080 | Unspecified |
| mmu-miR-21a-3p | MIMAT0004628 | Mboat2 | 67216 | HITS-CLIP | 23142080 | Unspecified |
| mmu-miR-21a-3p | MIMAT0004628 | Mboat7 | 77582 | HITS-CLIP | 23142080 | Unspecified |
| mmu-miR-21a-3p | MIMAT0004628 | Mcl1 | 17210 | HITS-CLIP | 23142080\|29167373 | Unspecified |
| mmu-miR-21a-3p | MIMAT0004628 | Mcm2 | 17216 | HITS-CLIP | 37986224\|23142080 | Unspecified |
| mmu-miR-21a-3p | MIMAT0004628 | Mcm3 | 17215 | HITS-CLIP | 23142080 | Unspecified |

| | | | | | | |
|---|---|---|---|---|---|---|
| mmu-miR-21a-3p | MIMAT0004628 | Mcm3ap | 54387 | HITS-CLIP | 23142080 | Unspecified |
| mmu-miR-21a-3p | MIMAT0004628 | Mcm4 | 17217 | HITS-CLIP | 23142080 | Unspecified |
| mmu-miR-21a-3p | MIMAT0004628 | Mcm6 | 17219 | HITS-CLIP | 23142080 | Unspecified |
| mmu-miR-21a-3p | MIMAT0004628 | Mctp1 | 78771 | HITS-CLIP | 23142080 | Unspecified |
| mmu-miR-21a-3p | MIMAT0004628 | Mctp2 | 244049 | HITS-CLIP | 23142080 | Unspecified |
| mmu-miR-21a-3p | MIMAT0004628 | Mcts1 | 68995 | HITS-CLIP | 23142080 | Unspecified |
| mmu-miR-21a-3p | MIMAT0004628 | Mdfic | 16543 | HITS-CLIP | 37986224\|23142080\|29167373 | Unspecified |
| mmu-miR-21a-3p | MIMAT0004628 | Mdh1 | 17449 | HITS-CLIP | 23142080 | Unspecified |
| mmu-miR-21a-3p | MIMAT0004628 | Mdn1 | 100019 | HITS-CLIP | 23142080\|29167373 | Unspecified |
| mmu-miR-21a-3p | MIMAT0004628 | Med1 | 19014 | HITS-CLIP | 23142080 | Unspecified |
| mmu-miR-21a-3p | MIMAT0004628 | Med12 | 59024 | HITS-CLIP | 23142080 | Unspecified |
| mmu-miR-21a-3p | MIMAT0004628 | Med13 | 327987 | HITS-CLIP | 31433990\|23142080 | Unspecified |
| mmu-miR-21a-3p | MIMAT0004628 | Med14 | 26896 | HITS-CLIP | 23142080 | Unspecified |
| mmu-miR-21a-3p | MIMAT0004628 | Med20 | 56771 | HITS-CLIP | 37986224\|23142080\|29167373 | Unspecified |
| mmu-miR-21a-3p | MIMAT0004628 | Med21 | 108098 | HITS-CLIP | 23142080 | Unspecified |
| mmu-miR-21a-3p | MIMAT0004628 | Med23 | 70208 | HITS-CLIP | 23142080 | Unspecified |
| mmu-miR-21a-3p | MIMAT0004628 | Mef2a | 17258 | HITS-CLIP | 23142080 | Unspecified |
| mmu-miR-21a-3p | MIMAT0004628 | Mef2d | 17261 | HITS-CLIP | 37986224\|23142080 | Unspecified |
| mmu-miR-21a-3p | MIMAT0004628 | Mepce | 231803 | HITS-CLIP | 23142080 | Unspecified |
| mmu-miR-21a-3p | MIMAT0004628 | Metrnl | 210029 | HITS-CLIP | 23142080\|29167373 | Unspecified |
| mmu-miR-21a-3p | MIMAT0004628 | Mettl14 | 210529 | HITS-CLIP | 23142080 | Unspecified |
| mmu-miR-21a-3p | MIMAT0004628 | Mettl15 | 76894 | HITS-CLIP | 23142080 | Unspecified |
| mmu-miR-21a-3p | MIMAT0004628 | Mettl26 | 50438 | HITS-CLIP | 23142080 | Unspecified |
| mmu-miR-21a-3p | MIMAT0004628 | Mettl9 | 59052 | HITS-CLIP | 23142080 | Unspecified |
| mmu-miR-21a-3p | MIMAT0004628 | Mex3c | 240396 | HITS-CLIP | 23142080 | Unspecified |
| mmu-miR-21a-3p | MIMAT0004628 | Mff | 75734 | HITS-CLIP | 37986224\|23142080 | Unspecified |
| mmu-miR-21a-3p | MIMAT0004628 | Mfn1 | 67414 | HITS-CLIP | 23142080 | Unspecified |
| mmu-miR-21a-3p | MIMAT0004628 | Mfsd1 | 66868 | HITS-CLIP | 23142080 | Unspecified |
| mmu-miR-21a-3p | MIMAT0004628 | Mfsd10 | 68294 | HITS-CLIP | 23142080 | Unspecified |
| mmu-miR-21a-3p | MIMAT0004628 | Mfsd13a | 79847 | HITS-CLIP | 29167373 | Unspecified |
| mmu-miR-21a-3p | MIMAT0004628 | Mfsd14a | 15247 | HITS-CLIP | 23142080 | Unspecified |
| mmu-miR-21a-3p | MIMAT0004628 | Mfsd14b | 84641 | HITS-CLIP | 23142080 | Unspecified |
| mmu-miR-21a-3p | MIMAT0004628 | Mga | 29808 | HITS-CLIP | 23142080 | Unspecified |
| mmu-miR-21a-3p | MIMAT0004628 | Mgat1 | 17308 | HITS-CLIP | 23142080 | Unspecified |
| mmu-miR-21a-3p | MIMAT0004628 | Mgat4a | 269181 | HITS-CLIP | 37986224\|23142080 | Unspecified |
| mmu-miR-21a-3p | MIMAT0004628 | Mgea5 | 76055 | HITS-CLIP | 23142080 | Unspecified |
| mmu-miR-21a-3p | MIMAT0004628 | Mia2 | 338320 | HITS-CLIP | 23142080 | Unspecified |
| mmu-miR-21a-3p | MIMAT0004628 | Mib1 | 225164 | HITS-CLIP | 29167373 | Unspecified |
| mmu-miR-21a-3p | MIMAT0004628 | Mid1ip1 | 68041 | HITS-CLIP | 23142080 | Unspecified |

| | | | | | | | |
|---|---|---|---|---|---|---|---|
| mmu-miR-21a-3p | MIMAT0004628 | Mid2 | 23947 | HITS-CLIP | | 23142080 | Unspecified |
| mmu-miR-21a-3p | MIMAT0004628 | Mier3 | 218613 | HITS-CLIP | | 23142080\|29167373 | Unspecified |
| mmu-miR-21a-3p | MIMAT0004628 | Mindy2 | 235461 | HITS-CLIP | | 23142080\|29167373 | Unspecified |
| mmu-miR-21a-3p | MIMAT0004628 | Mindy3 | 80013 | HITS-CLIP | | 23142080 | Unspecified |
| mmu-miR-21a-3p | MIMAT0004628 | Mios | 252875 | HITS-CLIP | | 23142080 | Unspecified |
| mmu-miR-21a-3p | MIMAT0004628 | Mlec | 109154 | HITS-CLIP | | 23142080 | Unspecified |
| mmu-miR-21a-3p | MIMAT0004628 | Mllt3 | 70122 | HITS-CLIP | | 23142080 | Unspecified |
| mmu-miR-21a-3p | MIMAT0004628 | Mlxip | 208104 | HITS-CLIP | | 23142080 | Unspecified |
| mmu-miR-21a-3p | MIMAT0004628 | Mmadhc | 109129 | HITS-CLIP | | 23142080 | Unspecified |
| mmu-miR-21a-3p | MIMAT0004628 | Mmgt1 | 236792 | HITS-CLIP | | 23142080 | Unspecified |
| mmu-miR-21a-3p | MIMAT0004628 | Mms22l | 212377 | HITS-CLIP | | 23142080 | Unspecified |
| mmu-miR-21a-3p | MIMAT0004628 | Mnt | 17428 | HITS-CLIP | | 23142080 | Unspecified |
| mmu-miR-21a-3p | MIMAT0004628 | Mob1a | 232157 | HITS-CLIP | | 37986224\|29167373\|23142080 | Unspecified |
| mmu-miR-21a-3p | MIMAT0004628 | Mob1b | 68473 | HITS-CLIP | | 37986224\|23142080 | Unspecified |
| mmu-miR-21a-3p | MIMAT0004628 | Mob3a | 208228 | HITS-CLIP | | 23142080 | Unspecified |
| mmu-miR-21a-3p | MIMAT0004628 | Mob3c | 100465 | HITS-CLIP | | 23142080 | Unspecified |
| mmu-miR-21a-3p | MIMAT0004628 | Mob4 | 19070 | HITS-CLIP | | 23142080 | Unspecified |
| mmu-miR-21a-3p | MIMAT0004628 | Mocs2 | 17434 | HITS-CLIP | | 23142080 | Unspecified |
| mmu-miR-21a-3p | MIMAT0004628 | Mon2 | 67074 | HITS-CLIP | | 23142080\|29167373 | Unspecified |
| mmu-miR-21a-3p | MIMAT0004628 | Morc2a | 74522 | HITS-CLIP | | 23142080 | Unspecified |
| mmu-miR-21a-3p | MIMAT0004628 | Morc3 | 338467 | HITS-CLIP | | 37986224\|23142080 | Unspecified |
| mmu-miR-21a-3p | MIMAT0004628 | Mosmo | 233812 | HITS-CLIP | | 23142080\|29167373 | Unspecified |
| mmu-miR-21a-3p | MIMAT0004628 | Mospd1 | 70380 | HITS-CLIP | | 23142080 | Unspecified |
| mmu-miR-21a-3p | MIMAT0004628 | Mospd2 | 76763 | HITS-CLIP | | 23142080 | Unspecified |
| mmu-miR-21a-3p | MIMAT0004628 | Mov10 | 17454 | HITS-CLIP | | 37986224 | Unspecified |
| mmu-miR-21a-3p | MIMAT0004628 | Mpdu1 | 24070 | HITS-CLIP | | 23142080 | Unspecified |
| mmu-miR-21a-3p | MIMAT0004628 | Mphosph8 | 75339 | HITS-CLIP | | 23142080 | Unspecified |
| mmu-miR-21a-3p | MIMAT0004628 | Mphosph9 | 269702 | HITS-CLIP | | 37986224 | Unspecified |
| mmu-miR-21a-3p | MIMAT0004628 | Mpp5 | 56217 | HITS-CLIP | | 23142080 | Unspecified |
| mmu-miR-21a-3p | MIMAT0004628 | Mppe1 | 225651 | HITS-CLIP | | 23142080 | Unspecified |
| mmu-miR-21a-3p | MIMAT0004628 | Mpzl3 | 319742 | HITS-CLIP | | 23142080 | Unspecified |
| mmu-miR-21a-3p | MIMAT0004628 | Mrpl28 | 68611 | HITS-CLIP | | 23142080 | Unspecified |
| mmu-miR-21a-3p | MIMAT0004628 | Mrpl32 | 75398 | HITS-CLIP | | 23142080 | Unspecified |
| mmu-miR-21a-3p | MIMAT0004628 | Mrpl50 | 28028 | HITS-CLIP | | 23142080 | Unspecified |
| mmu-miR-21a-3p | MIMAT0004628 | Mrps18b | 66973 | HITS-CLIP | | 23142080 | Unspecified |
| mmu-miR-21a-3p | MIMAT0004628 | Mrps22 | 64655 | HITS-CLIP | | 23142080 | Unspecified |
| mmu-miR-21a-3p | MIMAT0004628 | Mrps24 | 64660 | HITS-CLIP | | 29167373 | Unspecified |
| mmu-miR-21a-3p | MIMAT0004628 | Mrtfb | 239719 | HITS-CLIP | | 23142080 | Unspecified |
| mmu-miR-21a-3p | MIMAT0004628 | Ms4a4b | 60361 | HITS-CLIP | | 23142080 | Unspecified |

| | | | | | | |
|---|---|---|---|---|---|---|
| mmu-miR-21a-3p | MIMAT0004628 | Ms4a4c | 64380 | HITS-CLIP | 23142080 | Unspecified |
| mmu-miR-21a-3p | MIMAT0004628 | Ms4a6b | 69774 | HITS-CLIP | 23142080 | Unspecified |
| mmu-miR-21a-3p | MIMAT0004628 | Msantd2 | 235184 | HITS-CLIP | 23142080 | Unspecified |
| mmu-miR-21a-3p | MIMAT0004628 | Msh6 | 17688 | HITS-CLIP | 23142080 | Unspecified |
| mmu-miR-21a-3p | MIMAT0004628 | Msi2 | 76626 | HITS-CLIP | 23142080\|29167373 | Unspecified |
| mmu-miR-21a-3p | MIMAT0004628 | Msl1 | 74026 | HITS-CLIP | 23142080 | Unspecified |
| mmu-miR-21a-3p | MIMAT0004628 | Msl2 | 77853 | HITS-CLIP | 23142080\|29167373 | Unspecified |
| mmu-miR-21a-3p | MIMAT0004628 | Msn | 17698 | HITS-CLIP | 23142080 | Unspecified |
| mmu-miR-21a-3p | MIMAT0004628 | Mt2 | 17750 | HITS-CLIP | 23142080 | Unspecified |
| mmu-miR-21a-3p | MIMAT0004628 | Mta2 | 23942 | HITS-CLIP | 23142080 | Unspecified |
| mmu-miR-21a-3p | MIMAT0004628 | Mta3 | 116871 | HITS-CLIP | 29167373 | Unspecified |
| mmu-miR-21a-3p | MIMAT0004628 | Mtbp | 105837 | HITS-CLIP | 23142080 | Unspecified |
| mmu-miR-21a-3p | MIMAT0004628 | Mtch2 | 56428 | HITS-CLIP | 23142080 | Unspecified |
| mmu-miR-21a-3p | MIMAT0004628 | Mtdh | 67154 | HITS-CLIP | 37986224\|23142080\|29167373 | Unspecified |
| mmu-miR-21a-3p | MIMAT0004628 | Mtf2 | 17765 | HITS-CLIP | 37986224\|23142080 | Unspecified |
| mmu-miR-21a-3p | MIMAT0004628 | Mthfd1l | 270685 | HITS-CLIP | 37986224 | Unspecified |
| mmu-miR-21a-3p | MIMAT0004628 | Mtmr3 | 74302 | HITS-CLIP | 23142080 | Unspecified |
| mmu-miR-21a-3p | MIMAT0004628 | Mtmr4 | 170749 | HITS-CLIP | 23142080 | Unspecified |
| mmu-miR-21a-3p | MIMAT0004628 | Mtmr6 | 219135 | HITS-CLIP | 23142080\|29167373 | Unspecified |
| mmu-miR-21a-3p | MIMAT0004628 | Mtmr9 | 210376 | HITS-CLIP | 37986224\|23142080\|29167373 | Unspecified |
| mmu-miR-21a-3p | MIMAT0004628 | Mtor | 56717 | HITS-CLIP | 23142080\|29167373 | Unspecified |
| mmu-miR-21a-3p | MIMAT0004628 | Mtpn | 14489 | HITS-CLIP | 37986224\|29167373\|23142080 | Unspecified |
| mmu-miR-21a-3p | MIMAT0004628 | Mtr | 238505 | HITS-CLIP | 23142080 | Unspecified |
| mmu-miR-21a-3p | MIMAT0004628 | Mtss1 | 211401 | HITS-CLIP | 23142080\|29167373 | Unspecified |
| mmu-miR-21a-3p | MIMAT0004628 | Mtx2 | 53375 | HITS-CLIP | 23142080 | Unspecified |
| mmu-miR-21a-3p | MIMAT0004628 | Mxi1 | 17859 | HITS-CLIP | 37986224 | Unspecified |
| mmu-miR-21a-3p | MIMAT0004628 | Myadm | 50918 | HITS-CLIP | 29167373 | Unspecified |
| mmu-miR-21a-3p | MIMAT0004628 | Myb | 17863 | HITS-CLIP | 23142080 | Unspecified |
| mmu-miR-21a-3p | MIMAT0004628 | Mybbp1a | 18432 | HITS-CLIP | 29167373 | Unspecified |
| mmu-miR-21a-3p | MIMAT0004628 | Mybl1 | 17864 | HITS-CLIP | 23142080 | Unspecified |
| mmu-miR-21a-3p | MIMAT0004628 | Mycbp2 | 105689 | HITS-CLIP | 23142080\|29167373 | Unspecified |
| mmu-miR-21a-3p | MIMAT0004628 | Myh9 | 17886 | HITS-CLIP | 23142080 | Unspecified |
| mmu-miR-21a-3p | MIMAT0004628 | Myo5a | 17918 | HITS-CLIP | 23142080\|29167373 | Unspecified |
| mmu-miR-21a-3p | MIMAT0004628 | Myo6 | 17920 | HITS-CLIP | 23142080 | Unspecified |
| mmu-miR-21a-3p | MIMAT0004628 | Myo9a | 270163 | HITS-CLIP | 23142080\|29167373 | Unspecified |
| mmu-miR-21a-3p | MIMAT0004628 | Myof | 226101 | HITS-CLIP | 29167373 | Unspecified |
| mmu-miR-21a-3p | MIMAT0004628 | Mysm1 | 320713 | HITS-CLIP | 37986224\|23142080 | Unspecified |
| mmu-miR-21a-3p | MIMAT0004628 | Mzt1 | 76789 | HITS-CLIP | 23142080 | Unspecified |
| mmu-miR-21a-3p | MIMAT0004628 | N4bp2 | 333789 | HITS-CLIP | 37986224\|23142080 | Unspecified |

| | | | | | | |
|---|---|---|---|---|---|---|
| mmu-miR-21a-3p | MIMAT0004628 | N4bp2l2 | 381695 | HITS-CLIP | 37986224 | Unspecified |
| mmu-miR-21a-3p | MIMAT0004628 | Naa15 | 74838 | HITS-CLIP | 37986224\|23142080 | Unspecified |
| mmu-miR-21a-3p | MIMAT0004628 | Naa16 | 66897 | HITS-CLIP | 23142080 | Unspecified |
| mmu-miR-21a-3p | MIMAT0004628 | Naa25 | 231713 | qCLASH//HITS-CLIP | 31393953\|29167373\|23142080 | Unspecified |
| mmu-miR-21a-3p | MIMAT0004628 | Naa35 | 78689 | HITS-CLIP | 23142080 | Unspecified |
| mmu-miR-21a-3p | MIMAT0004628 | Naa40 | 70999 | HITS-CLIP | 23142080 | Unspecified |
| mmu-miR-21a-3p | MIMAT0004628 | Naa50 | 72117 | HITS-CLIP | 23142080 | Unspecified |
| mmu-miR-21a-3p | MIMAT0004628 | Nab1 | 17936 | HITS-CLIP | 23142080 | Unspecified |
| mmu-miR-21a-3p | MIMAT0004628 | Nabp1 | 109019 | HITS-CLIP | 23142080\|29167373 | Unspecified |
| mmu-miR-21a-3p | MIMAT0004628 | Nampt | 59027 | HITS-CLIP | 23142080 | Unspecified |
| mmu-miR-21a-3p | MIMAT0004628 | Nanos1 | 332397 | HITS-CLIP | 23142080 | Unspecified |
| mmu-miR-21a-3p | MIMAT0004628 | Nap1l1 | 53605 | HITS-CLIP | 23142080 | Unspecified |
| mmu-miR-21a-3p | MIMAT0004628 | Nasp | 50927 | HITS-CLIP | 37986224\|23142080 | Unspecified |
| mmu-miR-21a-3p | MIMAT0004628 | Nbeal1 | 269198 | HITS-CLIP | 23142080 | Unspecified |
| mmu-miR-21a-3p | MIMAT0004628 | Nbeal2 | 235627 | HITS-CLIP | 37986224 | Unspecified |
| mmu-miR-21a-3p | MIMAT0004628 | Nbn | 27354 | HITS-CLIP | 23142080 | Unspecified |
| mmu-miR-21a-3p | MIMAT0004628 | Nbr1 | 17966 | HITS-CLIP | 37986224 | Unspecified |
| mmu-miR-21a-3p | MIMAT0004628 | Ncapd2 | 68298 | HITS-CLIP | 37986224\|23142080\|29167373 | Unspecified |
| mmu-miR-21a-3p | MIMAT0004628 | Ncapd3 | 78658 | HITS-CLIP | 37986224\|23142080 | Unspecified |
| mmu-miR-21a-3p | MIMAT0004628 | Ncapg | 54392 | HITS-CLIP | 23142080 | Unspecified |
| mmu-miR-21a-3p | MIMAT0004628 | Ncapg2 | 76044 | HITS-CLIP | 23142080 | Unspecified |
| mmu-miR-21a-3p | MIMAT0004628 | Ncaph | 215387 | HITS-CLIP | 23142080 | Unspecified |
| mmu-miR-21a-3p | MIMAT0004628 | Ncbp1 | 433702 | HITS-CLIP | 23142080 | Unspecified |
| mmu-miR-21a-3p | MIMAT0004628 | Ncf1 | 17969 | HITS-CLIP | 23142080\|29167373 | Unspecified |
| mmu-miR-21a-3p | MIMAT0004628 | Nck1 | 17973 | HITS-CLIP | 23142080\|29167373 | Unspecified |
| mmu-miR-21a-3p | MIMAT0004628 | Nckap1 | 50884 | HITS-CLIP | 23142080 | Unspecified |
| mmu-miR-21a-3p | MIMAT0004628 | Ncl | 17975 | HITS-CLIP | 23142080\|29167373 | Unspecified |
| mmu-miR-21a-3p | MIMAT0004628 | Ncoa2 | 17978 | HITS-CLIP | 37986224 | Unspecified |
| mmu-miR-21a-3p | MIMAT0004628 | Ncoa3 | 17979 | HITS-CLIP | 37986224\|23142080 | Unspecified |
| mmu-miR-21a-3p | MIMAT0004628 | Ncoa7 | 211329 | HITS-CLIP | 23142080 | Unspecified |
| mmu-miR-21a-3p | MIMAT0004628 | Ncor1 | 20185 | HITS-CLIP | 23142080\|29167373 | Unspecified |
| mmu-miR-21a-3p | MIMAT0004628 | Ncor2 | 20602 | HITS-CLIP | 29167373 | Unspecified |
| mmu-miR-21a-3p | MIMAT0004628 | Ndfip1 | 65113 | HITS-CLIP | 23142080 | Unspecified |
| mmu-miR-21a-3p | MIMAT0004628 | Ndfip2 | 76273 | HITS-CLIP | 23142080 | Unspecified |
| mmu-miR-21a-3p | MIMAT0004628 | Ndrg1 | 17988 | HITS-CLIP | 37986224\|23142080 | Unspecified |
| mmu-miR-21a-3p | MIMAT0004628 | Ndrg2 | 29811 | HITS-CLIP | 23142080 | Unspecified |
| mmu-miR-21a-3p | MIMAT0004628 | Ndst1 | 15531 | HITS-CLIP | 23142080 | Unspecified |
| mmu-miR-21a-3p | MIMAT0004628 | Ndufaf4 | 68493 | HITS-CLIP | 31433990\|23142080 | Unspecified |
| mmu-miR-21a-3p | MIMAT0004628 | Ndufs1 | 227197 | HITS-CLIP | 23142080\|29167373 | Unspecified |

| | | | | | | |
|---|---|---|---|---|---|---|
| mmu-miR-21a-3p | MIMAT0004628 | Necap1 | 67602 | HITS-CLIP | 37986224\|23142080\|29167373 | Unspecified |
| mmu-miR-21a-3p | MIMAT0004628 | Nedd1 | 17997 | HITS-CLIP | 23142080 | Unspecified |
| mmu-miR-21a-3p | MIMAT0004628 | Nek7 | 59125 | HITS-CLIP | 23142080 | Unspecified |
| mmu-miR-21a-3p | MIMAT0004628 | Nelfe | 27632 | HITS-CLIP | 23142080 | Unspecified |
| mmu-miR-21a-3p | MIMAT0004628 | Nemf | 66244 | HITS-CLIP | 23142080 | Unspecified |
| mmu-miR-21a-3p | MIMAT0004628 | Nemp1 | 23306 | HITS-CLIP | 23142080 | Unspecified |
| mmu-miR-21a-3p | MIMAT0004628 | Nemp2 | 227094 | HITS-CLIP | 23142080 | Unspecified |
| mmu-miR-21a-3p | MIMAT0004628 | Nf1 | 18015 | HITS-CLIP | 23142080\|29167373 | Unspecified |
| mmu-miR-21a-3p | MIMAT0004628 | Nfat5 | 54446 | HITS-CLIP | 23142080\|29167373 | Unspecified |
| mmu-miR-21a-3p | MIMAT0004628 | Nfatc3 | 18021 | HITS-CLIP | 23142080 | Unspecified |
| mmu-miR-21a-3p | MIMAT0004628 | Nfe2l2 | 18024 | HITS-CLIP | 37986224\|29167373 | Unspecified |
| mmu-miR-21a-3p | MIMAT0004628 | Nfil3 | 18030 | HITS-CLIP | 23142080 | Unspecified |
| mmu-miR-21a-3p | MIMAT0004628 | Nfkb1 | 18033 | HITS-CLIP | 37986224 | Unspecified |
| mmu-miR-21a-3p | MIMAT0004628 | Nfx1 | 74164 | HITS-CLIP | 23142080 | Unspecified |
| mmu-miR-21a-3p | MIMAT0004628 | Nfxl1 | 100978 | HITS-CLIP | 29167373 | Unspecified |
| mmu-miR-21a-3p | MIMAT0004628 | Nfya | 18044 | HITS-CLIP | 23142080 | Unspecified |
| mmu-miR-21a-3p | MIMAT0004628 | Nfyb | 18045 | HITS-CLIP | 23142080 | Unspecified |
| mmu-miR-21a-3p | MIMAT0004628 | Ngly1 | 59007 | HITS-CLIP | 23142080 | Unspecified |
| mmu-miR-21a-3p | MIMAT0004628 | Nhlrc2 | 66866 | HITS-CLIP | 29167373 | Unspecified |
| mmu-miR-21a-3p | MIMAT0004628 | Nifk | 67949 | HITS-CLIP | 23142080 | Unspecified |
| mmu-miR-21a-3p | MIMAT0004628 | Nipal1 | 70701 | HITS-CLIP | 23142080 | Unspecified |
| mmu-miR-21a-3p | MIMAT0004628 | Nipsnap1 | 18082 | HITS-CLIP | 23142080 | Unspecified |
| mmu-miR-21a-3p | MIMAT0004628 | Nipsnap3b | 66536 | HITS-CLIP | 23142080 | Unspecified |
| mmu-miR-21a-3p | MIMAT0004628 | Nisch | 64652 | HITS-CLIP | 23142080 | Unspecified |
| mmu-miR-21a-3p | MIMAT0004628 | Nkiras1 | 69721 | HITS-CLIP | 23142080 | Unspecified |
| mmu-miR-21a-3p | MIMAT0004628 | Nktr | 18087 | HITS-CLIP | 23142080 | Unspecified |
| mmu-miR-21a-3p | MIMAT0004628 | Nlrc5 | 434341 | HITS-CLIP | 23142080 | Unspecified |
| mmu-miR-21a-3p | MIMAT0004628 | Nme7 | 171567 | HITS-CLIP | 23142080 | Unspecified |
| mmu-miR-21a-3p | MIMAT0004628 | Noc2l | 57741 | HITS-CLIP | 23142080 | Unspecified |
| mmu-miR-21a-3p | MIMAT0004628 | Nol4l | 329540 | HITS-CLIP | 23142080\|29167373 | Unspecified |
| mmu-miR-21a-3p | MIMAT0004628 | Nol6 | 230082 | HITS-CLIP | 23142080 | Unspecified |
| mmu-miR-21a-3p | MIMAT0004628 | Nop56 | 67134 | HITS-CLIP | 23142080 | Unspecified |
| mmu-miR-21a-3p | MIMAT0004628 | Nop58 | 55989 | HITS-CLIP | 23142080 | Unspecified |
| mmu-miR-21a-3p | MIMAT0004628 | Nop9 | 67842 | HITS-CLIP | 23142080 | Unspecified |
| mmu-miR-21a-3p | MIMAT0004628 | Notch1 | 18128 | HITS-CLIP | 37986224\|23142080 | Unspecified |
| mmu-miR-21a-3p | MIMAT0004628 | Notch2 | 18129 | HITS-CLIP | 23142080 | Unspecified |
| mmu-miR-21a-3p | MIMAT0004628 | Npas2 | 18143 | HITS-CLIP | 37986224\|23142080 | Unspecified |
| mmu-miR-21a-3p | MIMAT0004628 | Npat | 244879 | HITS-CLIP | 37986224\|23142080 | Unspecified |
| mmu-miR-21a-3p | MIMAT0004628 | Npc2 | 67963 | HITS-CLIP | 37986224\|23142080 | Unspecified |

| | | | | | | |
|---|---|---|---|---|---|---|
| mmu-miR-21a-3p | MIMAT0004628 | Npepps | 19155 | HITS-CLIP | 37986224\|23142080 | Unspecified |
| mmu-miR-21a-3p | MIMAT0004628 | Nploc4 | 217365 | HITS-CLIP | 23142080 | Unspecified |
| mmu-miR-21a-3p | MIMAT0004628 | Nptn | 20320 | HITS-CLIP | 37986224 | Unspecified |
| mmu-miR-21a-3p | MIMAT0004628 | Nr2c2 | 22026 | HITS-CLIP | 23142080 | Unspecified |
| mmu-miR-21a-3p | MIMAT0004628 | Nr3c1 | 14815 | HITS-CLIP | 37986224\|23142080\|29167373 | Unspecified |
| mmu-miR-21a-3p | MIMAT0004628 | Nr4a2 | 18227 | HITS-CLIP | 23142080 | Unspecified |
| mmu-miR-21a-3p | MIMAT0004628 | Nr4a3 | 18124 | HITS-CLIP | 23142080 | Unspecified |
| mmu-miR-21a-3p | MIMAT0004628 | Nras | 18176 | HITS-CLIP | 37986224\|23142080\|29167373 | Unspecified |
| mmu-miR-21a-3p | MIMAT0004628 | Nrip1 | 268903 | HITS-CLIP | 23142080 | Unspecified |
| mmu-miR-21a-3p | MIMAT0004628 | Nrn1 | 68404 | HITS-CLIP | 37986224\|23142080 | Unspecified |
| mmu-miR-21a-3p | MIMAT0004628 | Nrp2 | 18187 | HITS-CLIP | 29167373 | Unspecified |
| mmu-miR-21a-3p | MIMAT0004628 | Nsd1 | 18193 | HITS-CLIP | 23142080\|29167373 | Unspecified |
| mmu-miR-21a-3p | MIMAT0004628 | Nsd2 | 107823 | HITS-CLIP | 23142080\|29167373 | Unspecified |
| mmu-miR-21a-3p | MIMAT0004628 | Nsd3 | 234135 | HITS-CLIP | 23142080 | Unspecified |
| mmu-miR-21a-3p | MIMAT0004628 | Nsmaf | 18201 | HITS-CLIP | 23142080 | Unspecified |
| mmu-miR-21a-3p | MIMAT0004628 | Nsun2 | 28114 | HITS-CLIP | 23142080 | Unspecified |
| mmu-miR-21a-3p | MIMAT0004628 | Nsun4 | 72181 | HITS-CLIP | 23142080 | Unspecified |
| mmu-miR-21a-3p | MIMAT0004628 | Nucks1 | 98415 | HITS-CLIP | 23142080 | Unspecified |
| mmu-miR-21a-3p | MIMAT0004628 | Nudcd3 | 209586 | HITS-CLIP | 23142080 | Unspecified |
| mmu-miR-21a-3p | MIMAT0004628 | Nudt3 | 56409 | HITS-CLIP | 29167373 | Unspecified |
| mmu-miR-21a-3p | MIMAT0004628 | Nudt4 | 71207 | HITS-CLIP | 29167373 | Unspecified |
| mmu-miR-21a-3p | MIMAT0004628 | Nufip2 | 68564 | HITS-CLIP | 37986224\|23142080\|29167373 | Unspecified |
| mmu-miR-21a-3p | MIMAT0004628 | Numa1 | 101706 | HITS-CLIP | 23142080 | Unspecified |
| mmu-miR-21a-3p | MIMAT0004628 | Nup107 | 103468 | HITS-CLIP | 37986224\|23142080 | Unspecified |
| mmu-miR-21a-3p | MIMAT0004628 | Nup133 | 234865 | HITS-CLIP | 23142080\|29167373 | Unspecified |
| mmu-miR-21a-3p | MIMAT0004628 | Nup153 | 218210 | HITS-CLIP | 37986224\|23142080\|29167373 | Unspecified |
| mmu-miR-21a-3p | MIMAT0004628 | Nup160 | 59015 | HITS-CLIP | 37986224\|23142080 | Unspecified |
| mmu-miR-21a-3p | MIMAT0004628 | Nup188 | 227699 | HITS-CLIP | 23142080 | Unspecified |
| mmu-miR-21a-3p | MIMAT0004628 | Nup210 | 54563 | HITS-CLIP | 23142080 | Unspecified |
| mmu-miR-21a-3p | MIMAT0004628 | Nup50 | 18141 | HITS-CLIP | 23142080 | Unspecified |
| mmu-miR-21a-3p | MIMAT0004628 | Nup88 | 19069 | HITS-CLIP | 23142080 | Unspecified |
| mmu-miR-21a-3p | MIMAT0004628 | Nup93 | 71805 | HITS-CLIP | 23142080 | Unspecified |
| mmu-miR-21a-3p | MIMAT0004628 | Nup98 | 269966 | HITS-CLIP | 23142080 | Unspecified |
| mmu-miR-21a-3p | MIMAT0004628 | Nxf1 | 53319 | HITS-CLIP | 23142080 | Unspecified |
| mmu-miR-21a-3p | MIMAT0004628 | Nxt1 | 56488 | HITS-CLIP | 23142080 | Unspecified |
| mmu-miR-21a-3p | MIMAT0004628 | Oat | 18242 | HITS-CLIP | 23142080 | Unspecified |
| mmu-miR-21a-3p | MIMAT0004628 | Ocrl | 320634 | HITS-CLIP | 23142080 | Unspecified |
| mmu-miR-21a-3p | MIMAT0004628 | Odc1 | 18263 | HITS-CLIP | 23142080 | Unspecified |
| mmu-miR-21a-3p | MIMAT0004628 | Odf2 | 18286 | HITS-CLIP | 23142080 | Unspecified |

| | | | | | | | |
|---|---|---|---|---|---|---|---|
| mmu-miR-21a-3p | MIMAT0004628 | Odf2l | 52184 | HITS-CLIP | | 23142080 | Unspecified |
| mmu-miR-21a-3p | MIMAT0004628 | Ogt | 108155 | HITS-CLIP | | 23142080 | Unspecified |
| mmu-miR-21a-3p | MIMAT0004628 | Opa1 | 74143 | HITS-CLIP | | 23142080 | Unspecified |
| mmu-miR-21a-3p | MIMAT0004628 | Orai1 | 109305 | HITS-CLIP | | 23142080 | Unspecified |
| mmu-miR-21a-3p | MIMAT0004628 | Orc2 | 18393 | HITS-CLIP | | 23142080 | Unspecified |
| mmu-miR-21a-3p | MIMAT0004628 | Orc3 | 50793 | HITS-CLIP | | 23142080 | Unspecified |
| mmu-miR-21a-3p | MIMAT0004628 | Os9 | 216440 | HITS-CLIP | | 23142080 | Unspecified |
| mmu-miR-21a-3p | MIMAT0004628 | Osbp | 76303 | HITS-CLIP | 37986224\|23142080 | | Unspecified |
| mmu-miR-21a-3p | MIMAT0004628 | Osbpl8 | 237542 | HITS-CLIP | | 29167373 | Unspecified |
| mmu-miR-21a-3p | MIMAT0004628 | Osbpl9 | 100273 | HITS-CLIP | | 23142080 | Unspecified |
| mmu-miR-21a-3p | MIMAT0004628 | Oser1 | 51526 | HITS-CLIP | | 29167373 | Unspecified |
| mmu-miR-21a-3p | MIMAT0004628 | Ostm1 | 14628 | HITS-CLIP | | 23142080 | Unspecified |
| mmu-miR-21a-3p | MIMAT0004628 | Otud1 | 71198 | HITS-CLIP | | 23142080 | Unspecified |
| mmu-miR-21a-3p | MIMAT0004628 | Otud4 | 73945 | HITS-CLIP | 37986224\|23142080 | | Unspecified |
| mmu-miR-21a-3p | MIMAT0004628 | Otud7b | 229603 | HITS-CLIP | 37986224\|23142080 | | Unspecified |
| mmu-miR-21a-3p | MIMAT0004628 | Oxct1 | 67041 | HITS-CLIP | | 23142080 | Unspecified |
| mmu-miR-21a-3p | MIMAT0004628 | Oxr1 | 170719 | HITS-CLIP | 37986224\|23142080\|29167373 | | Unspecified |
| mmu-miR-21a-3p | MIMAT0004628 | Oxsr1 | 108737 | HITS-CLIP | 37986224\|23142080 | | Unspecified |
| mmu-miR-21a-3p | MIMAT0004628 | P2rx4 | 18438 | HITS-CLIP | | 37986224 | Unspecified |
| mmu-miR-21a-3p | MIMAT0004628 | P4ha1 | 18451 | HITS-CLIP | 23142080\|29167373 | | Unspecified |
| mmu-miR-21a-3p | MIMAT0004628 | P4ha2 | 18452 | HITS-CLIP | 37986224\|23142080 | | Unspecified |
| mmu-miR-21a-3p | MIMAT0004628 | P4hb | 18453 | HITS-CLIP | 23142080\|29167373 | | Unspecified |
| mmu-miR-21a-3p | MIMAT0004628 | Pa2g4 | 18813 | HITS-CLIP | | 23142080 | Unspecified |
| mmu-miR-21a-3p | MIMAT0004628 | Pabpc1 | 18458 | HITS-CLIP | 37986224\|23142080 | | Unspecified |
| mmu-miR-21a-3p | MIMAT0004628 | Paf1 | 54624 | HITS-CLIP | | 37986224 | Unspecified |
| mmu-miR-21a-3p | MIMAT0004628 | Pafah1b1 | 18472 | HITS-CLIP | | 23142080 | Unspecified |
| mmu-miR-21a-3p | MIMAT0004628 | Pafah1b2 | 18475 | HITS-CLIP | | 23142080 | Unspecified |
| mmu-miR-21a-3p | MIMAT0004628 | Paip2b | 232164 | HITS-CLIP | | 23142080 | Unspecified |
| mmu-miR-21a-3p | MIMAT0004628 | Pak2 | 224105 | HITS-CLIP | | 23142080 | Unspecified |
| mmu-miR-21a-3p | MIMAT0004628 | Pank3 | 211347 | HITS-CLIP | 37986224\|23142080 | | Unspecified |
| mmu-miR-21a-3p | MIMAT0004628 | Panx1 | 55991 | HITS-CLIP | | 23142080 | Unspecified |
| mmu-miR-21a-3p | MIMAT0004628 | Papola | 18789 | HITS-CLIP | | 29167373 | Unspecified |
| mmu-miR-21a-3p | MIMAT0004628 | Pard6b | 58220 | HITS-CLIP | 37986224\|23142080 | | Unspecified |
| mmu-miR-21a-3p | MIMAT0004628 | Parg | 26430 | HITS-CLIP | | 23142080 | Unspecified |
| mmu-miR-21a-3p | MIMAT0004628 | Parp14 | 547253 | HITS-CLIP | 29167373\|23142080 | | Unspecified |
| mmu-miR-21a-3p | MIMAT0004628 | Parp16 | 214424 | HITS-CLIP | | 37986224 | Unspecified |
| mmu-miR-21a-3p | MIMAT0004628 | Parp6 | 67287 | HITS-CLIP | | 23142080 | Unspecified |
| mmu-miR-21a-3p | MIMAT0004628 | Pask | 269224 | HITS-CLIP | | 23142080 | Unspecified |
| mmu-miR-21a-3p | MIMAT0004628 | Patl1 | 225929 | HITS-CLIP | 29167373\|23142080 | | Unspecified |

| | | | | | | |
|---|---|---|---|---|---|---|
| mmu-miR-21a-3p | MIMAT0004628 | Pbrm1 | 66923 | HITS-CLIP | 23142080 | Unspecified |
| mmu-miR-21a-3p | MIMAT0004628 | Pcbp1 | 23983 | HITS-CLIP | 23142080 | Unspecified |
| mmu-miR-21a-3p | MIMAT0004628 | Pcbp2 | 18521 | HITS-CLIP | 23142080 | Unspecified |
| mmu-miR-21a-3p | MIMAT0004628 | Pcdh7 | 54216 | HITS-CLIP | 29167373 | Unspecified |
| mmu-miR-21a-3p | MIMAT0004628 | Pcf11 | 74737 | HITS-CLIP | 23142080\|29167373 | Unspecified |
| mmu-miR-21a-3p | MIMAT0004628 | Pcgf3 | 69587 | HITS-CLIP | 23142080 | Unspecified |
| mmu-miR-21a-3p | MIMAT0004628 | Pcgf5 | 76073 | HITS-CLIP | 23142080\|29167373 | Unspecified |
| mmu-miR-21a-3p | MIMAT0004628 | Pcid2 | 234069 | HITS-CLIP | 23142080 | Unspecified |
| mmu-miR-21a-3p | MIMAT0004628 | Pcm1 | 18536 | HITS-CLIP | 23142080 | Unspecified |
| mmu-miR-21a-3p | MIMAT0004628 | Pcmt1 | 18537 | HITS-CLIP | 23142080 | Unspecified |
| mmu-miR-21a-3p | MIMAT0004628 | Pcnp | 76302 | HITS-CLIP | 23142080 | Unspecified |
| mmu-miR-21a-3p | MIMAT0004628 | Pcnx | 54604 | HITS-CLIP | 23142080 | Unspecified |
| mmu-miR-21a-3p | MIMAT0004628 | Pcx | 18563 | HITS-CLIP | 23142080 | Unspecified |
| mmu-miR-21a-3p | MIMAT0004628 | Pcyt2 | 68671 | HITS-CLIP | 23142080 | Unspecified |
| mmu-miR-21a-3p | MIMAT0004628 | Pdcd2l | 68079 | HITS-CLIP | 23142080 | Unspecified |
| mmu-miR-21a-3p | MIMAT0004628 | Pdcd6ip | 18571 | HITS-CLIP | 37986224\|29167373\|23142080 | Unspecified |
| mmu-miR-21a-3p | MIMAT0004628 | Pdcd7 | 50996 | HITS-CLIP | 29167373\|23142080 | Unspecified |
| mmu-miR-21a-3p | MIMAT0004628 | Pdcl | 67466 | HITS-CLIP | 37986224 | Unspecified |
| mmu-miR-21a-3p | MIMAT0004628 | Pde4b | 18578 | HITS-CLIP | 23142080 | Unspecified |
| mmu-miR-21a-3p | MIMAT0004628 | Pde7a | 18583 | HITS-CLIP | 23142080 | Unspecified |
| mmu-miR-21a-3p | MIMAT0004628 | Pdgfb | 18591 | HITS-CLIP | 23142080\|29167373 | Unspecified |
| mmu-miR-21a-3p | MIMAT0004628 | Pdha1 | 18597 | HITS-CLIP | 23142080 | Unspecified |
| mmu-miR-21a-3p | MIMAT0004628 | Pdhb | 68263 | HITS-CLIP | 37986224\|23142080 | Unspecified |
| mmu-miR-21a-3p | MIMAT0004628 | Pdia3 | 14827 | HITS-CLIP | 37986224\|23142080 | Unspecified |
| mmu-miR-21a-3p | MIMAT0004628 | Pdk1 | 228026 | HITS-CLIP | 37986224\|23142080 | Unspecified |
| mmu-miR-21a-3p | MIMAT0004628 | Pdp1 | 381511 | HITS-CLIP | 23142080 | Unspecified |
| mmu-miR-21a-3p | MIMAT0004628 | Pdpk1 | 18607 | HITS-CLIP | 23142080 | Unspecified |
| mmu-miR-21a-3p | MIMAT0004628 | Pds5a | 71521 | HITS-CLIP | 23142080 | Unspecified |
| mmu-miR-21a-3p | MIMAT0004628 | Pdzd8 | 107368 | HITS-CLIP | 23142080 | Unspecified |
| mmu-miR-21a-3p | MIMAT0004628 | Peak1 | 244895 | HITS-CLIP | 23142080\|29167373 | Unspecified |
| mmu-miR-21a-3p | MIMAT0004628 | Peg10 | 170676 | HITS-CLIP | 37986224 | Unspecified |
| mmu-miR-21a-3p | MIMAT0004628 | Peli1 | 67245 | HITS-CLIP | 37986224\|23142080\|29167373 | Unspecified |
| mmu-miR-21a-3p | MIMAT0004628 | Per1 | 18626 | HITS-CLIP | 23142080 | Unspecified |
| mmu-miR-21a-3p | MIMAT0004628 | Pfdn2 | 18637 | HITS-CLIP | 37986224\|23142080 | Unspecified |
| mmu-miR-21a-3p | MIMAT0004628 | Pfkp | 56421 | HITS-CLIP | 23142080 | Unspecified |
| mmu-miR-21a-3p | MIMAT0004628 | Pgm2l1 | 70974 | HITS-CLIP | 23142080 | Unspecified |
| mmu-miR-21a-3p | MIMAT0004628 | Pgrmc1 | 53328 | HITS-CLIP | 23142080 | Unspecified |
| mmu-miR-21a-3p | MIMAT0004628 | Pgrmc2 | 70804 | HITS-CLIP | 23142080 | Unspecified |
| mmu-miR-21a-3p | MIMAT0004628 | Pgs1 | 74451 | HITS-CLIP | 37986224 | Unspecified |

| | | | | | | |
|---|---|---|---|---|---|---|
| mmu-miR-21a-3p | MIMAT0004628 | Phactr2 | 215789 | HITS-CLIP | 23142080 | Unspecified |
| mmu-miR-21a-3p | MIMAT0004628 | Phc3 | 241915 | HITS-CLIP | 37986224\|23142080 | Unspecified |
| mmu-miR-21a-3p | MIMAT0004628 | Phf11b | 236451 | HITS-CLIP | 23142080 | Unspecified |
| mmu-miR-21a-3p | MIMAT0004628 | Phf12 | 268448 | HITS-CLIP | 29167373 | Unspecified |
| mmu-miR-21a-3p | MIMAT0004628 | Phf13 | 230936 | HITS-CLIP | 37986224 | Unspecified |
| mmu-miR-21a-3p | MIMAT0004628 | Phf14 | 75725 | HITS-CLIP | 23142080 | Unspecified |
| mmu-miR-21a-3p | MIMAT0004628 | Phf20 | 228829 | HITS-CLIP | 23142080 | Unspecified |
| mmu-miR-21a-3p | MIMAT0004628 | Phf20l1 | 239510 | HITS-CLIP | 23142080 | Unspecified |
| mmu-miR-21a-3p | MIMAT0004628 | Phip | 83946 | HITS-CLIP | 23142080 | Unspecified |
| mmu-miR-21a-3p | MIMAT0004628 | Phka2 | 110094 | HITS-CLIP | 23142080 | Unspecified |
| mmu-miR-21a-3p | MIMAT0004628 | Phlda3 | 27280 | HITS-CLIP | 23142080 | Unspecified |
| mmu-miR-21a-3p | MIMAT0004628 | Phlpp2 | 244650 | HITS-CLIP | 23142080 | Unspecified |
| mmu-miR-21a-3p | MIMAT0004628 | Phtf2 | 68770 | HITS-CLIP | 37986224\|23142080 | Unspecified |
| mmu-miR-21a-3p | MIMAT0004628 | Pi4k2b | 67073 | HITS-CLIP | 37986224\|23142080 | Unspecified |
| mmu-miR-21a-3p | MIMAT0004628 | Pi4ka | 224020 | HITS-CLIP | 23142080\|29167373 | Unspecified |
| mmu-miR-21a-3p | MIMAT0004628 | Pi4kb | 107650 | HITS-CLIP | 29167373 | Unspecified |
| mmu-miR-21a-3p | MIMAT0004628 | Pias1 | 56469 | HITS-CLIP | 23142080\|29167373 | Unspecified |
| mmu-miR-21a-3p | MIMAT0004628 | Pias4 | 59004 | HITS-CLIP | 23142080 | Unspecified |
| mmu-miR-21a-3p | MIMAT0004628 | Picalm | 233489 | HITS-CLIP | 23142080\|29167373 | Unspecified |
| mmu-miR-21a-3p | MIMAT0004628 | Piezo1 | 234839 | HITS-CLIP | 23142080 | Unspecified |
| mmu-miR-21a-3p | MIMAT0004628 | Piga | 18700 | HITS-CLIP | 23142080 | Unspecified |
| mmu-miR-21a-3p | MIMAT0004628 | Pigc | 67292 | HITS-CLIP | 37986224\|23142080 | Unspecified |
| mmu-miR-21a-3p | MIMAT0004628 | Pigm | 67556 | HITS-CLIP | 23142080 | Unspecified |
| mmu-miR-21a-3p | MIMAT0004628 | Pigo | 56703 | HITS-CLIP | 23142080 | Unspecified |
| mmu-miR-21a-3p | MIMAT0004628 | Pigq | 14755 | HITS-CLIP | 23142080 | Unspecified |
| mmu-miR-21a-3p | MIMAT0004628 | Pigx | 72084 | HITS-CLIP | 37986224 | Unspecified |
| mmu-miR-21a-3p | MIMAT0004628 | Pik3c2a | 18704 | HITS-CLIP | 23142080\|29167373 | Unspecified |
| mmu-miR-21a-3p | MIMAT0004628 | Pik3ca | 18706 | HITS-CLIP | 23142080 | Unspecified |
| mmu-miR-21a-3p | MIMAT0004628 | Pik3r1 | 18708 | HITS-CLIP | 23142080 | Unspecified |
| mmu-miR-21a-3p | MIMAT0004628 | Pik3r5 | 320207 | HITS-CLIP | 29167373 | Unspecified |
| mmu-miR-21a-3p | MIMAT0004628 | Pikfyve | 18711 | HITS-CLIP | 37986224\|23142080\|29167373 | Unspecified |
| mmu-miR-21a-3p | MIMAT0004628 | Pim1 | 18712 | HITS-CLIP | 23142080 | Unspecified |
| mmu-miR-21a-3p | MIMAT0004628 | Pip4k2a | 18718 | HITS-CLIP | 23142080\|29167373 | Unspecified |
| mmu-miR-21a-3p | MIMAT0004628 | Pitpna | 18738 | HITS-CLIP | 37986224 | Unspecified |
| mmu-miR-21a-3p | MIMAT0004628 | Pitpnb | 56305 | HITS-CLIP | 23142080 | Unspecified |
| mmu-miR-21a-3p | MIMAT0004628 | Pitpnc1 | 71795 | HITS-CLIP | 23142080 | Unspecified |
| mmu-miR-21a-3p | MIMAT0004628 | Pja1 | 18744 | HITS-CLIP | 23142080 | Unspecified |
| mmu-miR-21a-3p | MIMAT0004628 | Pja2 | 224938 | HITS-CLIP | 37986224\|23142080 | Unspecified |
| mmu-miR-21a-3p | MIMAT0004628 | Pkd1 | 18763 | HITS-CLIP | 29167373 | Unspecified |

| | | | | | | |
|---|---|---|---|---|---|---|
| mmu-miR-21a-3p | MIMAT0004628 | Pkm | 18746 | HITS-CLIP | 23142080\|29167373 | Unspecified |
| mmu-miR-21a-3p | MIMAT0004628 | Pkmyt1 | 268930 | HITS-CLIP | 23142080 | Unspecified |
| mmu-miR-21a-3p | MIMAT0004628 | Pkn2 | 109333 | HITS-CLIP | 23142080 | Unspecified |
| mmu-miR-21a-3p | MIMAT0004628 | Pla2g15 | 192654 | HITS-CLIP | 37986224\|26203562 | Unspecified |
| mmu-miR-21a-3p | MIMAT0004628 | Plaa | 18786 | HITS-CLIP | 23142080 | Unspecified |
| mmu-miR-21a-3p | MIMAT0004628 | Plagl1 | 22634 | HITS-CLIP | 23142080 | Unspecified |
| mmu-miR-21a-3p | MIMAT0004628 | Plagl2 | 54711 | HITS-CLIP | 37986224\|23142080 | Unspecified |
| mmu-miR-21a-3p | MIMAT0004628 | Plcb4 | 18798 | HITS-CLIP | 23142080 | Unspecified |
| mmu-miR-21a-3p | MIMAT0004628 | Plcl2 | 224860 | HITS-CLIP | 29167373\|23142080 | Unspecified |
| mmu-miR-21a-3p | MIMAT0004628 | Pld3 | 18807 | HITS-CLIP | 23142080 | Unspecified |
| mmu-miR-21a-3p | MIMAT0004628 | Plec | 18810 | HITS-CLIP | 23142080\|29167373 | Unspecified |
| mmu-miR-21a-3p | MIMAT0004628 | Plekha5 | 109135 | HITS-CLIP | 23142080 | Unspecified |
| mmu-miR-21a-3p | MIMAT0004628 | Plekhb2 | 226971 | HITS-CLIP | 37986224\|23142080\|29167373 | Unspecified |
| mmu-miR-21a-3p | MIMAT0004628 | Plk2 | 20620 | HITS-CLIP | 23142080\|29167373 | Unspecified |
| mmu-miR-21a-3p | MIMAT0004628 | Plk3 | 12795 | HITS-CLIP | 23142080 | Unspecified |
| mmu-miR-21a-3p | MIMAT0004628 | Plod1 | 18822 | HITS-CLIP | 29167373 | Unspecified |
| mmu-miR-21a-3p | MIMAT0004628 | Plod2 | 26432 | HITS-CLIP | 23142080 | Unspecified |
| mmu-miR-21a-3p | MIMAT0004628 | Plrg1 | 53317 | HITS-CLIP | 23142080 | Unspecified |
| mmu-miR-21a-3p | MIMAT0004628 | Plxna1 | 18844 | HITS-CLIP | 29167373 | Unspecified |
| mmu-miR-21a-3p | MIMAT0004628 | Plxnc1 | 54712 | HITS-CLIP | 23142080 | Unspecified |
| mmu-miR-21a-3p | MIMAT0004628 | Pmp22 | 18858 | HITS-CLIP | 29167373 | Unspecified |
| mmu-miR-21a-3p | MIMAT0004628 | Pmpcb | 73078 | HITS-CLIP | 23142080 | Unspecified |
| mmu-miR-21a-3p | MIMAT0004628 | Pms2 | 18861 | HITS-CLIP | 23142080 | Unspecified |
| mmu-miR-21a-3p | MIMAT0004628 | Pmvk | 68603 | HITS-CLIP | 23142080 | Unspecified |
| mmu-miR-21a-3p | MIMAT0004628 | Pnisr | 66625 | HITS-CLIP | 23142080\|29167373 | Unspecified |
| mmu-miR-21a-3p | MIMAT0004628 | Pnrc1 | 108767 | HITS-CLIP | 23142080 | Unspecified |
| mmu-miR-21a-3p | MIMAT0004628 | Pofut2 | 80294 | HITS-CLIP | 29167373 | Unspecified |
| mmu-miR-21a-3p | MIMAT0004628 | Pola1 | 18968 | HITS-CLIP | 23142080 | Unspecified |
| mmu-miR-21a-3p | MIMAT0004628 | Pold2 | 18972 | HITS-CLIP | 23142080 | Unspecified |
| mmu-miR-21a-3p | MIMAT0004628 | Pold3 | 67967 | HITS-CLIP | 31433990 | Unspecified |
| mmu-miR-21a-3p | MIMAT0004628 | Polh | 80905 | HITS-CLIP | 23142080 | Unspecified |
| mmu-miR-21a-3p | MIMAT0004628 | Polq | 77782 | HITS-CLIP | 23142080 | Unspecified |
| mmu-miR-21a-3p | MIMAT0004628 | Polr1d | 20018 | HITS-CLIP | 37986224\|23142080 | Unspecified |
| mmu-miR-21a-3p | MIMAT0004628 | Polr2a | 20020 | HITS-CLIP | 23142080 | Unspecified |
| mmu-miR-21a-3p | MIMAT0004628 | Polr2b | 231329 | HITS-CLIP | 23142080 | Unspecified |
| mmu-miR-21a-3p | MIMAT0004628 | Polr2g | 67710 | HITS-CLIP | 37986224\|23142080 | Unspecified |
| mmu-miR-21a-3p | MIMAT0004628 | Polr3c | 74414 | HITS-CLIP | 23142080 | Unspecified |
| mmu-miR-21a-3p | MIMAT0004628 | Polr3k | 67005 | HITS-CLIP | 23142080 | Unspecified |
| mmu-miR-21a-3p | MIMAT0004628 | Pop1 | 67724 | HITS-CLIP | 23142080 | Unspecified |

| | | | | | | | |
|---|---|---|---|---|---|---|---|
| mmu-miR-21a-3p | MIMAT0004628 | Pou2f1 | 18986 | HITS-CLIP | | 23142080 | Unspecified |
| mmu-miR-21a-3p | MIMAT0004628 | Pou2f2 | 18987 | HITS-CLIP | | 29167373 | Unspecified |
| mmu-miR-21a-3p | MIMAT0004628 | Ppa2 | 74776 | HITS-CLIP | | 23142080 | Unspecified |
| mmu-miR-21a-3p | MIMAT0004628 | Ppard | 19015 | HITS-CLIP | | 23142080 | Unspecified |
| mmu-miR-21a-3p | MIMAT0004628 | Ppig | 228005 | HITS-CLIP | 29167373\|23142080 | | Unspecified |
| mmu-miR-21a-3p | MIMAT0004628 | Ppil4 | 67418 | HITS-CLIP | 37986224\|23142080 | | Unspecified |
| mmu-miR-21a-3p | MIMAT0004628 | Ppip5k2 | 227399 | HITS-CLIP | | 23142080 | Unspecified |
| mmu-miR-21a-3p | MIMAT0004628 | Ppm1a | 19042 | HITS-CLIP | | 23142080 | Unspecified |
| mmu-miR-21a-3p | MIMAT0004628 | Ppm1b | 19043 | HITS-CLIP | | 23142080 | Unspecified |
| mmu-miR-21a-3p | MIMAT0004628 | Ppp1cb | 19046 | HITS-CLIP | | 23142080 | Unspecified |
| mmu-miR-21a-3p | MIMAT0004628 | Ppp1cc | 19047 | HITS-CLIP | | 29167373 | Unspecified |
| mmu-miR-21a-3p | MIMAT0004628 | Ppp1r11 | 76497 | HITS-CLIP | | 23142080 | Unspecified |
| mmu-miR-21a-3p | MIMAT0004628 | Ppp1r12a | 17931 | HITS-CLIP | | 23142080 | Unspecified |
| mmu-miR-21a-3p | MIMAT0004628 | Ppp1r16b | 228852 | HITS-CLIP | | 23142080 | Unspecified |
| mmu-miR-21a-3p | MIMAT0004628 | Ppp2ca | 19052 | HITS-CLIP | | 23142080 | Unspecified |
| mmu-miR-21a-3p | MIMAT0004628 | Ppp2r1b | 73699 | HITS-CLIP | | 23142080 | Unspecified |
| mmu-miR-21a-3p | MIMAT0004628 | Ppp2r2a | 71978 | HITS-CLIP | | 23142080 | Unspecified |
| mmu-miR-21a-3p | MIMAT0004628 | Ppp2r5e | 26932 | HITS-CLIP | | 23142080 | Unspecified |
| mmu-miR-21a-3p | MIMAT0004628 | Ppp3cb | 19056 | HITS-CLIP | | 23142080 | Unspecified |
| mmu-miR-21a-3p | MIMAT0004628 | Ppp3cc | 19057 | HITS-CLIP | | 23142080 | Unspecified |
| mmu-miR-21a-3p | MIMAT0004628 | Ppp4r1 | 70351 | HITS-CLIP | 37986224\|23142080 | | Unspecified |
| mmu-miR-21a-3p | MIMAT0004628 | Ppp4r2 | 232314 | HITS-CLIP | | 23142080 | Unspecified |
| mmu-miR-21a-3p | MIMAT0004628 | Ppp4r3b | 57223 | HITS-CLIP | 23142080\|29167373 | | Unspecified |
| mmu-miR-21a-3p | MIMAT0004628 | Ppp6c | 67857 | HITS-CLIP | | 23142080 | Unspecified |
| mmu-miR-21a-3p | MIMAT0004628 | Ppp6r3 | 52036 | HITS-CLIP | 37986224\|23142080 | | Unspecified |
| mmu-miR-21a-3p | MIMAT0004628 | Pprc1 | 226169 | HITS-CLIP | | 29167373 | Unspecified |
| mmu-miR-21a-3p | MIMAT0004628 | Pptc7 | 320717 | HITS-CLIP | | 23142080 | Unspecified |
| mmu-miR-21a-3p | MIMAT0004628 | Pqbp1 | 10084 | HITS-CLIP | | 23142080 | Unspecified |
| mmu-miR-21a-3p | MIMAT0004628 | Prc1 | 233406 | HITS-CLIP | | 37986224 | Unspecified |
| mmu-miR-21a-3p | MIMAT0004628 | Prcc | 94315 | HITS-CLIP | | 37986224 | Unspecified |
| mmu-miR-21a-3p | MIMAT0004628 | Prcp | 72461 | HITS-CLIP | | 23142080 | Unspecified |
| mmu-miR-21a-3p | MIMAT0004628 | Prdm1 | 12142 | HITS-CLIP | 37986224\|23142080 | | Unspecified |
| mmu-miR-21a-3p | MIMAT0004628 | Prdm16 | 70673 | HITS-CLIP | | 23142080 | Unspecified |
| mmu-miR-21a-3p | MIMAT0004628 | Prdx1 | 18477 | HITS-CLIP | 37986224\|23142080\|29167373 | | Unspecified |
| mmu-miR-21a-3p | MIMAT0004628 | Prickle3 | 54630 | HITS-CLIP | | 23142080 | Unspecified |
| mmu-miR-21a-3p | MIMAT0004628 | Prkab2 | 108097 | HITS-CLIP | | 23142080 | Unspecified |
| mmu-miR-21a-3p | MIMAT0004628 | Prkaca | 18747 | HITS-CLIP | 23142080\|29167373 | | Unspecified |
| mmu-miR-21a-3p | MIMAT0004628 | Prkacb | 18749 | HITS-CLIP | | 23142080 | Unspecified |
| mmu-miR-21a-3p | MIMAT0004628 | Prkar1a | 19084 | HITS-CLIP | 37986224\|23142080\|29167373 | | Unspecified |

| | | | | | | |
|---|---|---|---|---|---|---|
| mmu-miR-21a-3p | MIMAT0004628 | Prkar2a | 19087 | HITS-CLIP | 23142080 | Unspecified |
| mmu-miR-21a-3p | MIMAT0004628 | Prkca | 18750 | HITS-CLIP | 23142080 | Unspecified |
| mmu-miR-21a-3p | MIMAT0004628 | Prkci | 18759 | HITS-CLIP | 23142080 | Unspecified |
| mmu-miR-21a-3p | MIMAT0004628 | Prkcq | 18761 | HITS-CLIP | 23142080 | Unspecified |
| mmu-miR-21a-3p | MIMAT0004628 | Prkd2 | 101540 | HITS-CLIP | 23142080 | Unspecified |
| mmu-miR-21a-3p | MIMAT0004628 | Prkdc | 19090 | HITS-CLIP | 31433990 | Unspecified |
| mmu-miR-21a-3p | MIMAT0004628 | Prkx | 19108 | HITS-CLIP | 23142080 | Unspecified |
| mmu-miR-21a-3p | MIMAT0004628 | Prmt7 | 214572 | HITS-CLIP | 23142080 | Unspecified |
| mmu-miR-21a-3p | MIMAT0004628 | Pros1 | 19128 | HITS-CLIP | 23142080 | Unspecified |
| mmu-miR-21a-3p | MIMAT0004628 | Proser1 | 212127 | HITS-CLIP | 29167373 | Unspecified |
| mmu-miR-21a-3p | MIMAT0004628 | Prpf19 | 28000 | HITS-CLIP | 23142080 | Unspecified |
| mmu-miR-21a-3p | MIMAT0004628 | Prpf3 | 70767 | HITS-CLIP | 29167373 | Unspecified |
| mmu-miR-21a-3p | MIMAT0004628 | Prpf38b | 66921 | HITS-CLIP | 23142080 | Unspecified |
| mmu-miR-21a-3p | MIMAT0004628 | Prpf39 | 328110 | HITS-CLIP | 23142080\|29167373 | Unspecified |
| mmu-miR-21a-3p | MIMAT0004628 | Prpf40a | 56194 | HITS-CLIP | 31433990\|23142080 | Unspecified |
| mmu-miR-21a-3p | MIMAT0004628 | Prpf8 | 192159 | HITS-CLIP | 23142080\|29167373 | Unspecified |
| mmu-miR-21a-3p | MIMAT0004628 | Prps2 | 110639 | HITS-CLIP | 23142080 | Unspecified |
| mmu-miR-21a-3p | MIMAT0004628 | Prr11 | 270906 | HITS-CLIP | 37986224 | Unspecified |
| mmu-miR-21a-3p | MIMAT0004628 | Prr14l | 215476 | HITS-CLIP | 23142080\|29167373 | Unspecified |
| mmu-miR-21a-3p | MIMAT0004628 | Prr3 | 75210 | HITS-CLIP | 23142080 | Unspecified |
| mmu-miR-21a-3p | MIMAT0004628 | Prr5 | 109270 | HITS-CLIP | 37986224 | Unspecified |
| mmu-miR-21a-3p | MIMAT0004628 | Prrc2a | 53761 | HITS-CLIP | 23142080 | Unspecified |
| mmu-miR-21a-3p | MIMAT0004628 | Prrc2b | 227723 | HITS-CLIP | 37986224\|23142080 | Unspecified |
| mmu-miR-21a-3p | MIMAT0004628 | Prrc2c | 226562 | HITS-CLIP | 37986224\|23142080\|29167373 | Unspecified |
| mmu-miR-21a-3p | MIMAT0004628 | Psap | 19156 | HITS-CLIP | 23142080\|29167373 | Unspecified |
| mmu-miR-21a-3p | MIMAT0004628 | Psd3 | 234353 | HITS-CLIP | 29167373 | Unspecified |
| mmu-miR-21a-3p | MIMAT0004628 | Psen1 | 19164 | HITS-CLIP | 29167373 | Unspecified |
| mmu-miR-21a-3p | MIMAT0004628 | Psip1 | 101739 | HITS-CLIP | 23142080 | Unspecified |
| mmu-miR-21a-3p | MIMAT0004628 | Psma1 | 26440 | HITS-CLIP | 37986224\|23142080 | Unspecified |
| mmu-miR-21a-3p | MIMAT0004628 | Psma4 | 26441 | HITS-CLIP | 23142080 | Unspecified |
| mmu-miR-21a-3p | MIMAT0004628 | Psma6 | 26443 | HITS-CLIP | 23142080 | Unspecified |
| mmu-miR-21a-3p | MIMAT0004628 | Psmb1 | 19170 | HITS-CLIP | 23142080 | Unspecified |
| mmu-miR-21a-3p | MIMAT0004628 | Psmb4 | 19172 | HITS-CLIP | 23142080 | Unspecified |
| mmu-miR-21a-3p | MIMAT0004628 | Psmb8 | 16913 | HITS-CLIP | 23142080 | Unspecified |
| mmu-miR-21a-3p | MIMAT0004628 | Psmb9 | 16912 | HITS-CLIP | 37986224 | Unspecified |
| mmu-miR-21a-3p | MIMAT0004628 | Psmc1 | 19179 | HITS-CLIP | 23142080 | Unspecified |
| mmu-miR-21a-3p | MIMAT0004628 | Psmc2 | 19181 | HITS-CLIP | 23142080 | Unspecified |
| mmu-miR-21a-3p | MIMAT0004628 | Psmd1 | 70247 | HITS-CLIP | 23142080\|29167373 | Unspecified |
| mmu-miR-21a-3p | MIMAT0004628 | Psmd10 | 53380 | HITS-CLIP | 23142080 | Unspecified |

| | | | | | | |
|---|---|---|---|---|---|---|
| mmu-miR-21a-3p | MIMAT0004628 | Psmd11 | 69077 | HITS-CLIP | 23142080 | Unspecified |
| mmu-miR-21a-3p | MIMAT0004628 | Psmd12 | 66997 | HITS-CLIP | 23142080 | Unspecified |
| mmu-miR-21a-3p | MIMAT0004628 | Psmd14 | 59029 | HITS-CLIP | 23142080 | Unspecified |
| mmu-miR-21a-3p | MIMAT0004628 | Psmd5 | 66998 | HITS-CLIP | 23142080 | Unspecified |
| mmu-miR-21a-3p | MIMAT0004628 | Psme4 | 103554 | HITS-CLIP | 23142080 | Unspecified |
| mmu-miR-21a-3p | MIMAT0004628 | Pspc1 | 66645 | HITS-CLIP | 23142080 | Unspecified |
| mmu-miR-21a-3p | MIMAT0004628 | Ptbp2 | 56195 | HITS-CLIP | 29167373 | Unspecified |
| mmu-miR-21a-3p | MIMAT0004628 | Ptbp3 | 230257 | HITS-CLIP | 37986224\|29167373\|23142080 | Unspecified |
| mmu-miR-21a-3p | MIMAT0004628 | Ptcd3 | 69956 | HITS-CLIP | 23142080 | Unspecified |
| mmu-miR-21a-3p | MIMAT0004628 | Pten | 19211 | HITS-CLIP | 23142080 | Unspecified |
| mmu-miR-21a-3p | MIMAT0004628 | Ptger2 | 19217 | HITS-CLIP | 23142080 | Unspecified |
| mmu-miR-21a-3p | MIMAT0004628 | Ptger4 | 19219 | HITS-CLIP | 37986224\|23142080\|29167373 | Unspecified |
| mmu-miR-21a-3p | MIMAT0004628 | Ptgfrn | 19221 | HITS-CLIP | 23142080 | Unspecified |
| mmu-miR-21a-3p | MIMAT0004628 | Ptgir | 19222 | HITS-CLIP | 23142080 | Unspecified |
| mmu-miR-21a-3p | MIMAT0004628 | Ptk2b | 19229 | HITS-CLIP | 29167373 | Unspecified |
| mmu-miR-21a-3p | MIMAT0004628 | Ptma | 19231 | HITS-CLIP | 23142080 | Unspecified |
| mmu-miR-21a-3p | MIMAT0004628 | Ptms | 69202 | HITS-CLIP | 37986224 | Unspecified |
| mmu-miR-21a-3p | MIMAT0004628 | Ptp4a1 | 19243 | HITS-CLIP | 23142080 | Unspecified |
| mmu-miR-21a-3p | MIMAT0004628 | Ptp4a2 | 19244 | HITS-CLIP | 37986224\|23142080 | Unspecified |
| mmu-miR-21a-3p | MIMAT0004628 | Ptpn1 | 19246 | HITS-CLIP | 23142080 | Unspecified |
| mmu-miR-21a-3p | MIMAT0004628 | Ptpn12 | 19248 | HITS-CLIP | 23142080 | Unspecified |
| mmu-miR-21a-3p | MIMAT0004628 | Ptpra | 19262 | HITS-CLIP | 23142080 | Unspecified |
| mmu-miR-21a-3p | MIMAT0004628 | Ptprc | 19264 | HITS-CLIP | 23142080 | Unspecified |
| mmu-miR-21a-3p | MIMAT0004628 | Ptprcap | 19265 | HITS-CLIP | 37986224 | Unspecified |
| mmu-miR-21a-3p | MIMAT0004628 | Pttg1ip | 108705 | HITS-CLIP | 23142080 | Unspecified |
| mmu-miR-21a-3p | MIMAT0004628 | Pum1 | 80912 | HITS-CLIP | 23142080 | Unspecified |
| mmu-miR-21a-3p | MIMAT0004628 | Pum3 | 52874 | HITS-CLIP | 29167373 | Unspecified |
| mmu-miR-21a-3p | MIMAT0004628 | Pura | 19290 | HITS-CLIP | 37986224\|23142080\|29167373 | Unspecified |
| mmu-miR-21a-3p | MIMAT0004628 | Purb | 19291 | HITS-CLIP | 37986224\|23142080\|29167373 | Unspecified |
| mmu-miR-21a-3p | MIMAT0004628 | Pus10 | 74467 | HITS-CLIP | 23142080 | Unspecified |
| mmu-miR-21a-3p | MIMAT0004628 | Pus3 | 67049 | HITS-CLIP | 23142080 | Unspecified |
| mmu-miR-21a-3p | MIMAT0004628 | Pwwp2a | 70802 | HITS-CLIP | 23142080 | Unspecified |
| mmu-miR-21a-3p | MIMAT0004628 | Pxk | 218699 | HITS-CLIP | 23142080 | Unspecified |
| mmu-miR-21a-3p | MIMAT0004628 | Pxmp4 | 59038 | HITS-CLIP | 23142080 | Unspecified |
| mmu-miR-21a-3p | MIMAT0004628 | Pygb | 110078 | HITS-CLIP | 29167373\|23142080 | Unspecified |
| mmu-miR-21a-3p | MIMAT0004628 | Qars | 97541 | HITS-CLIP | 37986224 | Unspecified |
| mmu-miR-21a-3p | MIMAT0004628 | Qk | 19317 | HITS-CLIP | 23142080\|29167373 | Unspecified |
| mmu-miR-21a-3p | MIMAT0004628 | Qrich1 | 69232 | HITS-CLIP | 23142080 | Unspecified |
| mmu-miR-21a-3p | MIMAT0004628 | R3hdm1 | 226412 | HITS-CLIP | 23142080\|29167373 | Unspecified |

| | | | | | | |
|---|---|---|---|---|---|---|
| mmu-miR-21a-3p | MIMAT0004628 | RECK | 8434 | HITS-CLIP//Luciferase reporter assay//qRT- | 23142080\|25893734 | Unspecified |
| mmu-miR-21a-3p | MIMAT0004628 | Rab11a | 53869 | HITS-CLIP | 23142080\|29167373 | Unspecified |
| mmu-miR-21a-3p | MIMAT0004628 | Rab12 | 19328 | HITS-CLIP | 37986224\|23142080 | Unspecified |
| mmu-miR-21a-3p | MIMAT0004628 | Rab14 | 68365 | HITS-CLIP | 37986224\|23142080 | Unspecified |
| mmu-miR-21a-3p | MIMAT0004628 | Rab1a | 5861 | HITS-CLIP | 23142080 | Unspecified |
| mmu-miR-21a-3p | MIMAT0004628 | Rab1b | 76308 | HITS-CLIP | 23142080 | Unspecified |
| mmu-miR-21a-3p | MIMAT0004628 | Rab21 | 216344 | HITS-CLIP | 23142080\|29167373 | Unspecified |
| mmu-miR-21a-3p | MIMAT0004628 | Rab24 | 19336 | HITS-CLIP | 23142080 | Unspecified |
| mmu-miR-21a-3p | MIMAT0004628 | Rab27a | 11891 | HITS-CLIP | 23142080 | Unspecified |
| mmu-miR-21a-3p | MIMAT0004628 | Rab2a | 59021 | HITS-CLIP | 37986224\|23142080 | Unspecified |
| mmu-miR-21a-3p | MIMAT0004628 | Rab32 | 67844 | HITS-CLIP | 29167373 | Unspecified |
| mmu-miR-21a-3p | MIMAT0004628 | Rab33b | 19338 | HITS-CLIP | 23142080 | Unspecified |
| mmu-miR-21a-3p | MIMAT0004628 | Rab3a | 19339 | HITS-CLIP | 23142080 | Unspecified |
| mmu-miR-21a-3p | MIMAT0004628 | Rab3gap1 | 226407 | HITS-CLIP | 23142080 | Unspecified |
| mmu-miR-21a-3p | MIMAT0004628 | Rab3il1 | 74760 | HITS-CLIP | 29167373 | Unspecified |
| mmu-miR-21a-3p | MIMAT0004628 | Rab40c | 224624 | HITS-CLIP | 37986224\|23142080 | Unspecified |
| mmu-miR-21a-3p | MIMAT0004628 | Rab5a | 271457 | HITS-CLIP | 23142080\|29167373 | Unspecified |
| mmu-miR-21a-3p | MIMAT0004628 | Rab6a | 19346 | HITS-CLIP | 29167373 | Unspecified |
| mmu-miR-21a-3p | MIMAT0004628 | Rab7 | 19349 | HITS-CLIP | 23142080\|29167373 | Unspecified |
| mmu-miR-21a-3p | MIMAT0004628 | Rab8a | 17274 | HITS-CLIP | 23142080 | Unspecified |
| mmu-miR-21a-3p | MIMAT0004628 | Rab8b | 235442 | HITS-CLIP | 37986224\|23142080 | Unspecified |
| mmu-miR-21a-3p | MIMAT0004628 | Rabac1 | 14470 | HITS-CLIP | 37986224\|23142080 | Unspecified |
| mmu-miR-21a-3p | MIMAT0004628 | Rabgap1 | 227800 | HITS-CLIP | 23142080 | Unspecified |
| mmu-miR-21a-3p | MIMAT0004628 | Rabgap1l | 29809 | HITS-CLIP | 23142080 | Unspecified |
| mmu-miR-21a-3p | MIMAT0004628 | Rabl3 | 67657 | HITS-CLIP | 23142080 | Unspecified |
| mmu-miR-21a-3p | MIMAT0004628 | Rabl6 | 227624 | HITS-CLIP | 29167373 | Unspecified |
| mmu-miR-21a-3p | MIMAT0004628 | Rac1 | 19353 | HITS-CLIP | 23142080 | Unspecified |
| mmu-miR-21a-3p | MIMAT0004628 | Rac2 | 19354 | HITS-CLIP | 37986224\|23142080\|29167373 | Unspecified |
| mmu-miR-21a-3p | MIMAT0004628 | Rack1 | 10399 | HITS-CLIP | 23142080 | Unspecified |
| mmu-miR-21a-3p | MIMAT0004628 | Rad23b | 19359 | HITS-CLIP | 23142080 | Unspecified |
| mmu-miR-21a-3p | MIMAT0004628 | Rad50 | 19360 | HITS-CLIP | 23142080 | Unspecified |
| mmu-miR-21a-3p | MIMAT0004628 | Rad51b | 5890 | HITS-CLIP | 23142080 | Unspecified |
| mmu-miR-21a-3p | MIMAT0004628 | Rae1 | 66679 | HITS-CLIP | 23142080 | Unspecified |
| mmu-miR-21a-3p | MIMAT0004628 | Rai1 | 19377 | HITS-CLIP | 23142080 | Unspecified |
| mmu-miR-21a-3p | MIMAT0004628 | Ralbp1 | 19765 | HITS-CLIP | 37986224 | Unspecified |
| mmu-miR-21a-3p | MIMAT0004628 | Ralgapa1 | 56784 | HITS-CLIP | 23142080 | Unspecified |
| mmu-miR-21a-3p | MIMAT0004628 | Ralgps1 | 241308 | HITS-CLIP | 37986224 | Unspecified |
| mmu-miR-21a-3p | MIMAT0004628 | Ramp3 | 56089 | HITS-CLIP | 23142080 | Unspecified |
| mmu-miR-21a-3p | MIMAT0004628 | Ran | 19384 | HITS-CLIP | 23142080\|29167373 | Unspecified |

| | | | | | | |
|---|---|---|---|---|---|---|
| mmu-miR-21a-3p | MIMAT0004628 | Ranbp1 | 19385 | HITS-CLIP | 23142080 | Unspecified |
| mmu-miR-21a-3p | MIMAT0004628 | Ranbp2 | 19386 | HITS-CLIP | 37986224\|23142080 | Unspecified |
| mmu-miR-21a-3p | MIMAT0004628 | Ranbp9 | 56705 | HITS-CLIP | 23142080 | Unspecified |
| mmu-miR-21a-3p | MIMAT0004628 | Rangap1 | 19387 | HITS-CLIP | 23142080 | Unspecified |
| mmu-miR-21a-3p | MIMAT0004628 | Rap1b | 215449 | HITS-CLIP | 37986224\|23142080\|29167373 | Unspecified |
| mmu-miR-21a-3p | MIMAT0004628 | Rap1gds1 | 229877 | HITS-CLIP | 23142080 | Unspecified |
| mmu-miR-21a-3p | MIMAT0004628 | Rap2a | 76108 | HITS-CLIP | 23142080 | Unspecified |
| mmu-miR-21a-3p | MIMAT0004628 | Rap2b | 74012 | HITS-CLIP | 23142080\|29167373 | Unspecified |
| mmu-miR-21a-3p | MIMAT0004628 | Rap2c | 72065 | HITS-CLIP | 23142080 | Unspecified |
| mmu-miR-21a-3p | MIMAT0004628 | Rapgef1 | 107746 | HITS-CLIP | 23142080 | Unspecified |
| mmu-miR-21a-3p | MIMAT0004628 | Rapgef2 | 76089 | HITS-CLIP | 23142080 | Unspecified |
| mmu-miR-21a-3p | MIMAT0004628 | Rapgef6 | 192786 | HITS-CLIP | 31433990\|23142080\|29167373 | Unspecified |
| mmu-miR-21a-3p | MIMAT0004628 | Rapgefl1 | 268480 | HITS-CLIP | 23142080 | Unspecified |
| mmu-miR-21a-3p | MIMAT0004628 | Rarg | 19411 | HITS-CLIP | 23142080 | Unspecified |
| mmu-miR-21a-3p | MIMAT0004628 | Rars | 104458 | HITS-CLIP | 23142080 | Unspecified |
| mmu-miR-21a-3p | MIMAT0004628 | Rars2 | 109093 | HITS-CLIP | 23142080 | Unspecified |
| mmu-miR-21a-3p | MIMAT0004628 | Rasa1 | 218397 | HITS-CLIP | 37986224\|23142080\|29167373 | Unspecified |
| mmu-miR-21a-3p | MIMAT0004628 | Rasa2 | 114713 | HITS-CLIP | 37986224\|23142080 | Unspecified |
| mmu-miR-21a-3p | MIMAT0004628 | Rasgrp1 | 19419 | HITS-CLIP | 23142080 | Unspecified |
| mmu-miR-21a-3p | MIMAT0004628 | Rassf2 | 215653 | HITS-CLIP | 23142080 | Unspecified |
| mmu-miR-21a-3p | MIMAT0004628 | Rassf3 | 192678 | HITS-CLIP | 29167373 | Unspecified |
| mmu-miR-21a-3p | MIMAT0004628 | Raver1 | 71766 | HITS-CLIP | 23142080\|29167373 | Unspecified |
| mmu-miR-21a-3p | MIMAT0004628 | Rb1 | 19645 | HITS-CLIP | 23142080 | Unspecified |
| mmu-miR-21a-3p | MIMAT0004628 | Rb1cc1 | 12421 | HITS-CLIP | 23142080 | Unspecified |
| mmu-miR-21a-3p | MIMAT0004628 | Rbbp5 | 213464 | HITS-CLIP | 23142080 | Unspecified |
| mmu-miR-21a-3p | MIMAT0004628 | Rbbp6 | 19647 | HITS-CLIP | 23142080 | Unspecified |
| mmu-miR-21a-3p | MIMAT0004628 | Rbbp7 | 245688 | HITS-CLIP | 23142080 | Unspecified |
| mmu-miR-21a-3p | MIMAT0004628 | Rbbp8 | 225182 | HITS-CLIP | 23142080 | Unspecified |
| mmu-miR-21a-3p | MIMAT0004628 | Rbl2 | 19651 | HITS-CLIP | 23142080 | Unspecified |
| mmu-miR-21a-3p | MIMAT0004628 | Rbm12 | 75710 | HITS-CLIP | 23142080 | Unspecified |
| mmu-miR-21a-3p | MIMAT0004628 | Rbm15 | 229700 | HITS-CLIP | 23142080 | Unspecified |
| mmu-miR-21a-3p | MIMAT0004628 | Rbm15b | 109095 | HITS-CLIP | 23142080 | Unspecified |
| mmu-miR-21a-3p | MIMAT0004628 | Rbm27 | 225432 | HITS-CLIP | 23142080 | Unspecified |
| mmu-miR-21a-3p | MIMAT0004628 | Rbm33 | 381626 | HITS-CLIP | 23142080\|29167373 | Unspecified |
| mmu-miR-21a-3p | MIMAT0004628 | Rbm38 | 56190 | HITS-CLIP | 23142080 | Unspecified |
| mmu-miR-21a-3p | MIMAT0004628 | Rbm39 | 170791 | HITS-CLIP | 23142080 | Unspecified |
| mmu-miR-21a-3p | MIMAT0004628 | Rbm45 | 241490 | HITS-CLIP | 23142080 | Unspecified |
| mmu-miR-21a-3p | MIMAT0004628 | Rbm5 | 83486 | HITS-CLIP | 23142080 | Unspecified |
| mmu-miR-21a-3p | MIMAT0004628 | Rbms1 | 56878 | HITS-CLIP | 37986224\|23142080 | Unspecified |

| | | | | | | |
|---|---|---|---|---|---|---|
| mmu-miR-21a-3p | MIMAT0004628 | Rc3h1 | 381305 | HITS-CLIP | 37986224\|23142080 | Unspecified |
| mmu-miR-21a-3p | MIMAT0004628 | Rc3h2 | 319817 | HITS-CLIP | 23142080 | Unspecified |
| mmu-miR-21a-3p | MIMAT0004628 | Rcbtb1 | 71330 | HITS-CLIP | 23142080 | Unspecified |
| mmu-miR-21a-3p | MIMAT0004628 | Rcc2 | 108911 | HITS-CLIP | 37986224\|23142080 | Unspecified |
| mmu-miR-21a-3p | MIMAT0004628 | Rcor1 | 217864 | HITS-CLIP | 37986224\|23142080 | Unspecified |
| mmu-miR-21a-3p | MIMAT0004628 | Rcor3 | 214742 | HITS-CLIP | 23142080 | Unspecified |
| mmu-miR-21a-3p | MIMAT0004628 | Rdh10 | 98711 | HITS-CLIP | 23142080 | Unspecified |
| mmu-miR-21a-3p | MIMAT0004628 | Rdh14 | 105014 | HITS-CLIP | 23142080 | Unspecified |
| mmu-miR-21a-3p | MIMAT0004628 | Rdx | 19684 | HITS-CLIP | 37986224\|23142080\|29167373 | Unspecified |
| mmu-miR-21a-3p | MIMAT0004628 | Reep1 | 52250 | HITS-CLIP | 23142080 | Unspecified |
| mmu-miR-21a-3p | MIMAT0004628 | Reep5 | 13476 | HITS-CLIP | 23142080 | Unspecified |
| mmu-miR-21a-3p | MIMAT0004628 | Rela | 19697 | HITS-CLIP | 23142080\|29167373 | Unspecified |
| mmu-miR-21a-3p | MIMAT0004628 | Relch | 57614 | HITS-CLIP | 23142080\|29167373 | Unspecified |
| mmu-miR-21a-3p | MIMAT0004628 | Rer1 | 67830 | HITS-CLIP | 29167373\|23142080 | Unspecified |
| mmu-miR-21a-3p | MIMAT0004628 | Retreg2 | 79137 | HITS-CLIP | 23142080 | Unspecified |
| mmu-miR-21a-3p | MIMAT0004628 | Retreg3 | 162427 | HITS-CLIP | 23142080 | Unspecified |
| mmu-miR-21a-3p | MIMAT0004628 | Rev3l | 19714 | HITS-CLIP | 23142080 | Unspecified |
| mmu-miR-21a-3p | MIMAT0004628 | Rexo2 | 104444 | HITS-CLIP | 23142080 | Unspecified |
| mmu-miR-21a-3p | MIMAT0004628 | Rffl | 67338 | HITS-CLIP | 23142080 | Unspecified |
| mmu-miR-21a-3p | MIMAT0004628 | Rftn1 | 76438 | HITS-CLIP | 37986224\|23142080 | Unspecified |
| mmu-miR-21a-3p | MIMAT0004628 | Rfwd3 | 234736 | HITS-CLIP | 23142080 | Unspecified |
| mmu-miR-21a-3p | MIMAT0004628 | Rfx3 | 19726 | HITS-CLIP | 23142080 | Unspecified |
| mmu-miR-21a-3p | MIMAT0004628 | Rfx7 | 319758 | HITS-CLIP | 23142080 | Unspecified |
| mmu-miR-21a-3p | MIMAT0004628 | Rgs1 | 50778 | HITS-CLIP | 37986224 | Unspecified |
| mmu-miR-21a-3p | MIMAT0004628 | Rgs3 | 50780 | HITS-CLIP | 23142080 | Unspecified |
| mmu-miR-21a-3p | MIMAT0004628 | Rhbdf2 | 217344 | HITS-CLIP | 29167373 | Unspecified |
| mmu-miR-21a-3p | MIMAT0004628 | Rheb | 19744 | HITS-CLIP | 23142080 | Unspecified |
| mmu-miR-21a-3p | MIMAT0004628 | Rhoq | 104215 | HITS-CLIP | 37986224\|23142080 | Unspecified |
| mmu-miR-21a-3p | MIMAT0004628 | Rhot1 | 59040 | HITS-CLIP | 23142080 | Unspecified |
| mmu-miR-21a-3p | MIMAT0004628 | Ric1 | 850728 | HITS-CLIP | 23142080 | Unspecified |
| mmu-miR-21a-3p | MIMAT0004628 | Rictor | 78757 | HITS-CLIP | 23142080 | Unspecified |
| mmu-miR-21a-3p | MIMAT0004628 | Rif1 | 51869 | HITS-CLIP | 23142080 | Unspecified |
| mmu-miR-21a-3p | MIMAT0004628 | Rin2 | 74030 | HITS-CLIP | 29167373 | Unspecified |
| mmu-miR-21a-3p | MIMAT0004628 | Riok3 | 66878 | HITS-CLIP | 23142080 | Unspecified |
| mmu-miR-21a-3p | MIMAT0004628 | Ripk2 | 192656 | HITS-CLIP | 23142080 | Unspecified |
| mmu-miR-21a-3p | MIMAT0004628 | Rlf | 109263 | HITS-CLIP | 23142080 | Unspecified |
| mmu-miR-21a-3p | MIMAT0004628 | Rlim | 19820 | HITS-CLIP | 23142080\|29167373 | Unspecified |
| mmu-miR-21a-3p | MIMAT0004628 | Rmdn1 | 66302 | HITS-CLIP | 23142080 | Unspecified |
| mmu-miR-21a-3p | MIMAT0004628 | Rmi1 | 74386 | HITS-CLIP | 29167373 | Unspecified |

| | | | | | | |
|---|---|---|---|---|---|---|
| mmu-miR-21a-3p | MIMAT0004628 | Rmnd5a | 68477 | HITS-CLIP | 37986224\|23142080 | Unspecified |
| mmu-miR-21a-3p | MIMAT0004628 | Rmnd5b | 66089 | HITS-CLIP | 23142080\|29167373 | Unspecified |
| mmu-miR-21a-3p | MIMAT0004628 | Rnf10 | 50849 | HITS-CLIP | 23142080 | Unspecified |
| mmu-miR-21a-3p | MIMAT0004628 | Rnf111 | 93836 | HITS-CLIP | 37986224\|23142080\|29167373 | Unspecified |
| mmu-miR-21a-3p | MIMAT0004628 | Rnf123 | 84585 | HITS-CLIP | 23142080 | Unspecified |
| mmu-miR-21a-3p | MIMAT0004628 | Rnf125 | 67664 | HITS-CLIP | 23142080 | Unspecified |
| mmu-miR-21a-3p | MIMAT0004628 | Rnf128 | 66889 | HITS-CLIP | 29167373 | Unspecified |
| mmu-miR-21a-3p | MIMAT0004628 | Rnf138 | 56515 | HITS-CLIP | 37986224\|23142080 | Unspecified |
| mmu-miR-21a-3p | MIMAT0004628 | Rnf139 | 75841 | HITS-CLIP | 23142080 | Unspecified |
| mmu-miR-21a-3p | MIMAT0004628 | Rnf141 | 67150 | HITS-CLIP | 23142080 | Unspecified |
| mmu-miR-21a-3p | MIMAT0004628 | Rnf145 | 74315 | HITS-CLIP | 23142080 | Unspecified |
| mmu-miR-21a-3p | MIMAT0004628 | Rnf152 | 320311 | HITS-CLIP | 23142080 | Unspecified |
| mmu-miR-21a-3p | MIMAT0004628 | Rnf167 | 70510 | HITS-CLIP | 23142080 | Unspecified |
| mmu-miR-21a-3p | MIMAT0004628 | Rnf19a | 30945 | HITS-CLIP | 23142080 | Unspecified |
| mmu-miR-21a-3p | MIMAT0004628 | Rnf19b | 75234 | HITS-CLIP | 23142080\|29167373 | Unspecified |
| mmu-miR-21a-3p | MIMAT0004628 | Rnf2 | 19821 | HITS-CLIP | 23142080 | Unspecified |
| mmu-miR-21a-3p | MIMAT0004628 | Rnf213 | 672511 | HITS-CLIP | 29167373 | Unspecified |
| mmu-miR-21a-3p | MIMAT0004628 | Rnf220 | 66743 | HITS-CLIP | 23142080 | Unspecified |
| mmu-miR-21a-3p | MIMAT0004628 | Rnf38 | 73469 | HITS-CLIP | 37986224\|23142080\|29167373 | Unspecified |
| mmu-miR-21a-3p | MIMAT0004628 | Rnf4 | 19822 | HITS-CLIP | 23142080 | Unspecified |
| mmu-miR-21a-3p | MIMAT0004628 | Rnf41 | 67588 | HITS-CLIP | 23142080\|29167373 | Unspecified |
| mmu-miR-21a-3p | MIMAT0004628 | Rock1 | 19877 | HITS-CLIP | 37986224\|31433990\|23142080 | Unspecified |
| mmu-miR-21a-3p | MIMAT0004628 | Rock2 | 19878 | HITS-CLIP | 23142080\|29167373 | Unspecified |
| mmu-miR-21a-3p | MIMAT0004628 | Rora | 19883 | HITS-CLIP | 37986224\|23142080 | Unspecified |
| mmu-miR-21a-3p | MIMAT0004628 | Rpap1 | 68925 | HITS-CLIP | 23142080 | Unspecified |
| mmu-miR-21a-3p | MIMAT0004628 | Rpap2 | 231571 | HITS-CLIP | 23142080 | Unspecified |
| mmu-miR-21a-3p | MIMAT0004628 | Rpap3 | 71919 | HITS-CLIP | 23142080 | Unspecified |
| mmu-miR-21a-3p | MIMAT0004628 | Rpia | 19895 | HITS-CLIP | 37986224\|23142080 | Unspecified |
| mmu-miR-21a-3p | MIMAT0004628 | Rpl13a | 22121 | HITS-CLIP | 37986224 | Unspecified |
| mmu-miR-21a-3p | MIMAT0004628 | Rpl4 | 67891 | HITS-CLIP | 23142080\|29167373 | Unspecified |
| mmu-miR-21a-3p | MIMAT0004628 | Rplp0 | 11837 | HITS-CLIP | 23142080 | Unspecified |
| mmu-miR-21a-3p | MIMAT0004628 | Rpn1 | 103963 | HITS-CLIP | 23142080 | Unspecified |
| mmu-miR-21a-3p | MIMAT0004628 | Rpn2 | 20014 | HITS-CLIP | 37986224 | Unspecified |
| mmu-miR-21a-3p | MIMAT0004628 | Rpp14 | 67053 | HITS-CLIP | 23142080 | Unspecified |
| mmu-miR-21a-3p | MIMAT0004628 | Rpp30 | 54364 | HITS-CLIP | 23142080 | Unspecified |
| mmu-miR-21a-3p | MIMAT0004628 | Rpp40 | 208366 | HITS-CLIP | 23142080 | Unspecified |
| mmu-miR-21a-3p | MIMAT0004628 | Rprd2 | 75137 | HITS-CLIP | 29167373 | Unspecified |
| mmu-miR-21a-3p | MIMAT0004628 | Rps13 | 68052 | HITS-CLIP | 37986224\|23142080 | Unspecified |
| mmu-miR-21a-3p | MIMAT0004628 | Rps5 | 20103 | HITS-CLIP | 23142080\|29167373 | Unspecified |

| | | | | | | |
|---|---|---|---|---|---|---|
| mmu-miR-21a-3p | MIMAT0004628 | Rps8 | 20116 | HITS-CLIP | 37986224\|23142080 | Unspecified |
| mmu-miR-21a-3p | MIMAT0004628 | Rps9 | 76846 | HITS-CLIP | 23142080 | Unspecified |
| mmu-miR-21a-3p | MIMAT0004628 | Rrad | 56437 | HITS-CLIP | 23142080 | Unspecified |
| mmu-miR-21a-3p | MIMAT0004628 | Rraga | 68441 | HITS-CLIP | 37986224\|23142080 | Unspecified |
| mmu-miR-21a-3p | MIMAT0004628 | Rragc | 54170 | HITS-CLIP | 23142080\|29167373 | Unspecified |
| mmu-miR-21a-3p | MIMAT0004628 | Rras2 | 66922 | HITS-CLIP | 37986224\|23142080 | Unspecified |
| mmu-miR-21a-3p | MIMAT0004628 | Rreb1 | 68750 | HITS-CLIP | 37986224 | Unspecified |
| mmu-miR-21a-3p | MIMAT0004628 | Rrm2 | 20135 | HITS-CLIP | 37986224\|23142080 | Unspecified |
| mmu-miR-21a-3p | MIMAT0004628 | Rrp15 | 67223 | HITS-CLIP | 23142080 | Unspecified |
| mmu-miR-21a-3p | MIMAT0004628 | Rrp36 | 224823 | HITS-CLIP | 23142080 | Unspecified |
| mmu-miR-21a-3p | MIMAT0004628 | Rsad2 | 58185 | HITS-CLIP | 29167373 | Unspecified |
| mmu-miR-21a-3p | MIMAT0004628 | Rsbn1 | 229675 | HITS-CLIP | 23142080\|29167373 | Unspecified |
| mmu-miR-21a-3p | MIMAT0004628 | Rsbn1l | 242860 | HITS-CLIP | 23142080\|29167373 | Unspecified |
| mmu-miR-21a-3p | MIMAT0004628 | Rsf1 | 233532 | HITS-CLIP | 23142080 | Unspecified |
| mmu-miR-21a-3p | MIMAT0004628 | Rsl1d1 | 66409 | HITS-CLIP | 23142080 | Unspecified |
| mmu-miR-21a-3p | MIMAT0004628 | Rsrc2 | 208606 | HITS-CLIP | 23142080 | Unspecified |
| mmu-miR-21a-3p | MIMAT0004628 | Rsrp1 | 27981 | HITS-CLIP | 23142080\|29167373 | Unspecified |
| mmu-miR-21a-3p | MIMAT0004628 | Rtcb | 28088 | HITS-CLIP | 23142080 | Unspecified |
| mmu-miR-21a-3p | MIMAT0004628 | Rtf1 | 76246 | HITS-CLIP | 37986224\|23142080 | Unspecified |
| mmu-miR-21a-3p | MIMAT0004628 | Rtn3 | 20168 | HITS-CLIP | 23142080 | Unspecified |
| mmu-miR-21a-3p | MIMAT0004628 | Rtn4 | 68585 | HITS-CLIP | 37986224\|23142080\|29167373 | Unspecified |
| mmu-miR-21a-3p | MIMAT0004628 | Rtraf | 51637 | HITS-CLIP | 23142080 | Unspecified |
| mmu-miR-21a-3p | MIMAT0004628 | Rufy3 | 52822 | HITS-CLIP | 29167373 | Unspecified |
| mmu-miR-21a-3p | MIMAT0004628 | Runx1 | 12394 | HITS-CLIP | 23142080 | Unspecified |
| mmu-miR-21a-3p | MIMAT0004628 | Rusc2 | 100213 | HITS-CLIP | 29167373 | Unspecified |
| mmu-miR-21a-3p | MIMAT0004628 | Rwdd2a | 69519 | HITS-CLIP | 23142080 | Unspecified |
| mmu-miR-21a-3p | MIMAT0004628 | Rxylt1 | 10329 | HITS-CLIP | 29167373 | Unspecified |
| mmu-miR-21a-3p | MIMAT0004628 | S100pbp | 74648 | HITS-CLIP | 37986224\|23142080 | Unspecified |
| mmu-miR-21a-3p | MIMAT0004628 | S1pr1 | 13609 | HITS-CLIP | 23142080 | Unspecified |
| mmu-miR-21a-3p | MIMAT0004628 | Sacs | 50720 | HITS-CLIP | 37986224\|23142080 | Unspecified |
| mmu-miR-21a-3p | MIMAT0004628 | Safb2 | 224902 | HITS-CLIP | 23142080\|29167373 | Unspecified |
| mmu-miR-21a-3p | MIMAT0004628 | Sall2 | 50524 | HITS-CLIP | 23142080 | Unspecified |
| mmu-miR-21a-3p | MIMAT0004628 | Samd4b | 233033 | HITS-CLIP | 23142080 | Unspecified |
| mmu-miR-21a-3p | MIMAT0004628 | Samd8 | 67630 | HITS-CLIP | 37986224 | Unspecified |
| mmu-miR-21a-3p | MIMAT0004628 | Samhd1 | 56045 | HITS-CLIP | 37986224\|23142080 | Unspecified |
| mmu-miR-21a-3p | MIMAT0004628 | Samsn1 | 67742 | HITS-CLIP | 23142080 | Unspecified |
| mmu-miR-21a-3p | MIMAT0004628 | Sap130 | 269003 | HITS-CLIP | 23142080\|29167373 | Unspecified |
| mmu-miR-21a-3p | MIMAT0004628 | Sap30 | 60406 | HITS-CLIP | 37986224\|23142080 | Unspecified |
| mmu-miR-21a-3p | MIMAT0004628 | Sar1b | 66397 | HITS-CLIP | 23142080 | Unspecified |

| | | | | | | |
|---|---|---|---|---|---|---|
| mmu-miR-21a-3p | MIMAT0004628 | Saraf | 51669 | HITS-CLIP | 23142080\|29167373 | Unspecified |
| mmu-miR-21a-3p | MIMAT0004628 | Sart1 | 20227 | HITS-CLIP | 23142080 | Unspecified |
| mmu-miR-21a-3p | MIMAT0004628 | Sart3 | 53890 | HITS-CLIP | 23142080 | Unspecified |
| mmu-miR-21a-3p | MIMAT0004628 | Sash1 | 70097 | HITS-CLIP | 29167373 | Unspecified |
| mmu-miR-21a-3p | MIMAT0004628 | Satb1 | 20230 | HITS-CLIP | 37986224\|23142080 | Unspecified |
| mmu-miR-21a-3p | MIMAT0004628 | Sbf2 | 319934 | HITS-CLIP | 29167373\|23142080 | Unspecified |
| mmu-miR-21a-3p | MIMAT0004628 | Sbno1 | 243272 | HITS-CLIP | 23142080\|29167373 | Unspecified |
| mmu-miR-21a-3p | MIMAT0004628 | Scaf11 | 72193 | HITS-CLIP | 23142080 | Unspecified |
| mmu-miR-21a-3p | MIMAT0004628 | Scaf4 | 224432 | HITS-CLIP | 23142080 | Unspecified |
| mmu-miR-21a-3p | MIMAT0004628 | Scai | 320271 | HITS-CLIP | 23142080 | Unspecified |
| mmu-miR-21a-3p | MIMAT0004628 | Scarb2 | 12492 | HITS-CLIP | 23142080 | Unspecified |
| mmu-miR-21a-3p | MIMAT0004628 | Scd2 | 20250 | HITS-CLIP | 37986224\|23142080 | Unspecified |
| mmu-miR-21a-3p | MIMAT0004628 | Scfd1 | 76983 | HITS-CLIP | 23142080 | Unspecified |
| mmu-miR-21a-3p | MIMAT0004628 | Scn1b | 20266 | HITS-CLIP | 23142080 | Unspecified |
| mmu-miR-21a-3p | MIMAT0004628 | Scrn3 | 74616 | HITS-CLIP | 23142080 | Unspecified |
| mmu-miR-21a-3p | MIMAT0004628 | Sdad1 | 231452 | HITS-CLIP | 23142080 | Unspecified |
| mmu-miR-21a-3p | MIMAT0004628 | Sdc3 | 20970 | HITS-CLIP | 23142080\|29167373 | Unspecified |
| mmu-miR-21a-3p | MIMAT0004628 | Sdc4 | 20971 | HITS-CLIP | 37986224 | Unspecified |
| mmu-miR-21a-3p | MIMAT0004628 | Sdcbp | 53378 | HITS-CLIP | 23142080 | Unspecified |
| mmu-miR-21a-3p | MIMAT0004628 | Sdf4 | 20318 | HITS-CLIP | 37986224\|23142080 | Unspecified |
| mmu-miR-21a-3p | MIMAT0004628 | Sdha | 66945 | HITS-CLIP | 23142080 | Unspecified |
| mmu-miR-21a-3p | MIMAT0004628 | Sec11a | 56529 | HITS-CLIP | 37986224\|23142080 | Unspecified |
| mmu-miR-21a-3p | MIMAT0004628 | Sec11c | 66286 | HITS-CLIP | 23142080 | Unspecified |
| mmu-miR-21a-3p | MIMAT0004628 | Sec22a | 317717 | HITS-CLIP | 23142080 | Unspecified |
| mmu-miR-21a-3p | MIMAT0004628 | Sec23a | 20334 | HITS-CLIP | 23142080 | Unspecified |
| mmu-miR-21a-3p | MIMAT0004628 | Sec23ip | 207352 | HITS-CLIP | 23142080 | Unspecified |
| mmu-miR-21a-3p | MIMAT0004628 | Sec24a | 77371 | HITS-CLIP | 37986224\|23142080 | Unspecified |
| mmu-miR-21a-3p | MIMAT0004628 | Sec24b | 99683 | HITS-CLIP | 23142080 | Unspecified |
| mmu-miR-21a-3p | MIMAT0004628 | Sec24c | 218811 | HITS-CLIP | 23142080 | Unspecified |
| mmu-miR-21a-3p | MIMAT0004628 | Sec31a | 69162 | HITS-CLIP | 23142080 | Unspecified |
| mmu-miR-21a-3p | MIMAT0004628 | Sec61b | 66212 | HITS-CLIP | 23142080 | Unspecified |
| mmu-miR-21a-3p | MIMAT0004628 | Secisbp2l | 70354 | HITS-CLIP | 37986224\|23142080 | Unspecified |
| mmu-miR-21a-3p | MIMAT0004628 | Seh1l | 72124 | HITS-CLIP | 37986224\|23142080 | Unspecified |
| mmu-miR-21a-3p | MIMAT0004628 | Selenop | 6414 | HITS-CLIP | 23142080\|29167373 | Unspecified |
| mmu-miR-21a-3p | MIMAT0004628 | Sell | 20343 | HITS-CLIP | 23142080 | Unspecified |
| mmu-miR-21a-3p | MIMAT0004628 | Sema4d | 20354 | HITS-CLIP | 37986224\|23142080 | Unspecified |
| mmu-miR-21a-3p | MIMAT0004628 | Sema6d | 214968 | HITS-CLIP | 23142080 | Unspecified |
| mmu-miR-21a-3p | MIMAT0004628 | Senp1 | 223870 | HITS-CLIP | 29167373 | Unspecified |
| mmu-miR-21a-3p | MIMAT0004628 | Senp5 | 320213 | HITS-CLIP | 23142080\|29167373 | Unspecified |

| | | | | | | |
|---|---|---|---|---|---|---|
| mmu-miR-21a-3p | MIMAT0004628 | Senp7 | 66315 | HITS-CLIP | 23142080 | Unspecified |
| mmu-miR-21a-3p | MIMAT0004628 | Sephs2 | 20768 | HITS-CLIP | 37986224\|23142080 | Unspecified |
| mmu-miR-21a-3p | MIMAT0004628 | Sept11 | 52398 | HITS-CLIP | 37986224\|23142080\|29167373 | Unspecified |
| mmu-miR-21a-3p | MIMAT0004628 | Sept6 | 23157 | HITS-CLIP | 23142080 | Unspecified |
| mmu-miR-21a-3p | MIMAT0004628 | Sept7 | 235072 | HITS-CLIP | 23142080\|29167373 | Unspecified |
| mmu-miR-21a-3p | MIMAT0004628 | Septin7 | 989 | HITS-CLIP | 31433990 | Unspecified |
| mmu-miR-21a-3p | MIMAT0004628 | Serbp1 | 66870 | HITS-CLIP | 23142080 | Unspecified |
| mmu-miR-21a-3p | MIMAT0004628 | Serinc1 | 56442 | HITS-CLIP | 37986224\|23142080\|29167373 | Unspecified |
| mmu-miR-21a-3p | MIMAT0004628 | Serinc3 | 26943 | HITS-CLIP | 23142080 | Unspecified |
| mmu-miR-21a-3p | MIMAT0004628 | Serinc5 | 218442 | HITS-CLIP | 23142080 | Unspecified |
| mmu-miR-21a-3p | MIMAT0004628 | Serp1 | 28146 | HITS-CLIP | 37986224\|23142080 | Unspecified |
| mmu-miR-21a-3p | MIMAT0004628 | Serpinb9 | 20723 | HITS-CLIP | 37986224 | Unspecified |
| mmu-miR-21a-3p | MIMAT0004628 | Serpine1 | 18787 | HITS-CLIP | 23142080 | Unspecified |
| mmu-miR-21a-3p | MIMAT0004628 | Sesn1 | 140742 | HITS-CLIP | 23142080\|29167373 | Unspecified |
| mmu-miR-21a-3p | MIMAT0004628 | Sesn3 | 75747 | HITS-CLIP | 23142080 | Unspecified |
| mmu-miR-21a-3p | MIMAT0004628 | Setbp1 | 240427 | HITS-CLIP | 23142080 | Unspecified |
| mmu-miR-21a-3p | MIMAT0004628 | Setd2 | 235626 | HITS-CLIP | 37986224\|29167373 | Unspecified |
| mmu-miR-21a-3p | MIMAT0004628 | Setd5 | 72895 | HITS-CLIP | 23142080\|29167373 | Unspecified |
| mmu-miR-21a-3p | MIMAT0004628 | Setd7 | 73251 | HITS-CLIP | 23142080 | Unspecified |
| mmu-miR-21a-3p | MIMAT0004628 | Setx | 269254 | HITS-CLIP | 37986224\|23142080\|29167373 | Unspecified |
| mmu-miR-21a-3p | MIMAT0004628 | Sf3b1 | 81898 | HITS-CLIP | 37986224\|23142080\|29167373 | Unspecified |
| mmu-miR-21a-3p | MIMAT0004628 | Sf3b2 | 319322 | HITS-CLIP | 23142080 | Unspecified |
| mmu-miR-21a-3p | MIMAT0004628 | Sf3b3 | 101943 | HITS-CLIP | 23142080 | Unspecified |
| mmu-miR-21a-3p | MIMAT0004628 | Sfmbt1 | 54650 | HITS-CLIP | 29167373\|23142080 | Unspecified |
| mmu-miR-21a-3p | MIMAT0004628 | Sfpq | 71514 | HITS-CLIP | 23142080 | Unspecified |
| mmu-miR-21a-3p | MIMAT0004628 | Sfxn3 | 94280 | HITS-CLIP | 23142080 | Unspecified |
| mmu-miR-21a-3p | MIMAT0004628 | Sgpp1 | 81535 | HITS-CLIP | 23142080 | Unspecified |
| mmu-miR-21a-3p | MIMAT0004628 | Sh2b1 | 20399 | HITS-CLIP | 37986224\|29167373\|23142080 | Unspecified |
| mmu-miR-21a-3p | MIMAT0004628 | Sh2d2a | 27371 | HITS-CLIP | 23142080 | Unspecified |
| mmu-miR-21a-3p | MIMAT0004628 | Sh2d4b | 328381 | HITS-CLIP | 29167373 | Unspecified |
| mmu-miR-21a-3p | MIMAT0004628 | Sh3bp5 | 24056 | HITS-CLIP | 23142080 | Unspecified |
| mmu-miR-21a-3p | MIMAT0004628 | Sh3gl3 | 20408 | HITS-CLIP | 23142080 | Unspecified |
| mmu-miR-21a-3p | MIMAT0004628 | Sh3glb1 | 54673 | HITS-CLIP//qCLASH | 23142080\|31393953\|29167373 | Unspecified |
| mmu-miR-21a-3p | MIMAT0004628 | Sh3kbp1 | 58194 | HITS-CLIP | 23142080 | Unspecified |
| mmu-miR-21a-3p | MIMAT0004628 | Shisa5 | 66940 | HITS-CLIP | 37986224\|23142080\|29167373 | Unspecified |
| mmu-miR-21a-3p | MIMAT0004628 | Siah1a | 20437 | HITS-CLIP | 37986224\|23142080 | Unspecified |
| mmu-miR-21a-3p | MIMAT0004628 | Siah1b | 20438 | HITS-CLIP | 23142080 | Unspecified |
| mmu-miR-21a-3p | MIMAT0004628 | Sik1 | 17691 | HITS-CLIP | 23142080 | Unspecified |
| mmu-miR-21a-3p | MIMAT0004628 | Sik3 | 70661 | HITS-CLIP | 23142080 | Unspecified |

| | | | | | | |
|---|---|---|---|---|---|---|
| mmu-miR-21a-3p | MIMAT0004628 | Sike1 | 66641 | HITS-CLIP | 23142080 | Unspecified |
| mmu-miR-21a-3p | MIMAT0004628 | Sil1 | 81500 | HITS-CLIP | 23142080 | Unspecified |
| mmu-miR-21a-3p | MIMAT0004628 | Sin3a | 20466 | HITS-CLIP | 23142080 | Unspecified |
| mmu-miR-21a-3p | MIMAT0004628 | Sin3b | 20467 | HITS-CLIP | 37986224\|23142080 | Unspecified |
| mmu-miR-21a-3p | MIMAT0004628 | Sipa1l2 | 244668 | HITS-CLIP | 37986224\|23142080 | Unspecified |
| mmu-miR-21a-3p | MIMAT0004628 | Sirpb1c | 100038947 | HITS-CLIP | 29167373 | Unspecified |
| mmu-miR-21a-3p | MIMAT0004628 | Sirt1 | 93759 | HITS-CLIP | 23142080 | Unspecified |
| mmu-miR-21a-3p | MIMAT0004628 | Ska2 | 66140 | HITS-CLIP | 37986224 | Unspecified |
| mmu-miR-21a-3p | MIMAT0004628 | Skil | 20482 | HITS-CLIP | 37986224\|23142080\|29167373 | Unspecified |
| mmu-miR-21a-3p | MIMAT0004628 | Skp1a | 21402 | HITS-CLIP | 23142080 | Unspecified |
| mmu-miR-21a-3p | MIMAT0004628 | Sla | 20491 | HITS-CLIP | 37986224 | Unspecified |
| mmu-miR-21a-3p | MIMAT0004628 | Slain1 | 105439 | HITS-CLIP | 37986224\|23142080 | Unspecified |
| mmu-miR-21a-3p | MIMAT0004628 | Slain2 | 75991 | HITS-CLIP | 23142080 | Unspecified |
| mmu-miR-21a-3p | MIMAT0004628 | Slc11a2 | 18174 | HITS-CLIP | 29167373 | Unspecified |
| mmu-miR-21a-3p | MIMAT0004628 | Slc12a2 | 20496 | HITS-CLIP | 23142080 | Unspecified |
| mmu-miR-21a-3p | MIMAT0004628 | Slc12a4 | 20498 | HITS-CLIP | 23142080 | Unspecified |
| mmu-miR-21a-3p | MIMAT0004628 | Slc15a4 | 100561 | HITS-CLIP | 23142080 | Unspecified |
| mmu-miR-21a-3p | MIMAT0004628 | Slc16a1 | 20501 | HITS-CLIP | 37986224\|23142080 | Unspecified |
| mmu-miR-21a-3p | MIMAT0004628 | Slc16a10 | 72472 | HITS-CLIP | 29167373 | Unspecified |
| mmu-miR-21a-3p | MIMAT0004628 | Slc16a3 | 80879 | HITS-CLIP | 23142080 | Unspecified |
| mmu-miR-21a-3p | MIMAT0004628 | Slc16a6 | 104681 | HITS-CLIP | 37986224\|23142080 | Unspecified |
| mmu-miR-21a-3p | MIMAT0004628 | Slc17a6 | 140919 | HITS-CLIP | 37986224\|23142080 | Unspecified |
| mmu-miR-21a-3p | MIMAT0004628 | Slc19a2 | 116914 | HITS-CLIP | 37986224\|23142080 | Unspecified |
| mmu-miR-21a-3p | MIMAT0004628 | Slc20a1 | 20515 | HITS-CLIP | 37986224\|23142080 | Unspecified |
| mmu-miR-21a-3p | MIMAT0004628 | Slc20a2 | 20516 | HITS-CLIP | 29167373 | Unspecified |
| mmu-miR-21a-3p | MIMAT0004628 | Slc23a2 | 54338 | HITS-CLIP | 37986224\|23142080 | Unspecified |
| mmu-miR-21a-3p | MIMAT0004628 | Slc25a11 | 67863 | HITS-CLIP | 23142080 | Unspecified |
| mmu-miR-21a-3p | MIMAT0004628 | Slc25a13 | 50799 | HITS-CLIP | 29167373 | Unspecified |
| mmu-miR-21a-3p | MIMAT0004628 | Slc25a17 | 20524 | HITS-CLIP | 23142080 | Unspecified |
| mmu-miR-21a-3p | MIMAT0004628 | Slc25a3 | 18674 | HITS-CLIP | 37986224\|23142080 | Unspecified |
| mmu-miR-21a-3p | MIMAT0004628 | Slc25a36 | 192287 | HITS-CLIP | 23142080\|29167373 | Unspecified |
| mmu-miR-21a-3p | MIMAT0004628 | Slc25a46 | 67453 | HITS-CLIP | 37986224\|23142080 | Unspecified |
| mmu-miR-21a-3p | MIMAT0004628 | Slc25a5 | 11740 | HITS-CLIP | 23142080 | Unspecified |
| mmu-miR-21a-3p | MIMAT0004628 | Slc25a51 | 230125 | HITS-CLIP | 37986224\|23142080 | Unspecified |
| mmu-miR-21a-3p | MIMAT0004628 | Slc2a1 | 20525 | HITS-CLIP | 23142080 | Unspecified |
| mmu-miR-21a-3p | MIMAT0004628 | Slc2a3 | 20527 | HITS-CLIP | 23142080 | Unspecified |
| mmu-miR-21a-3p | MIMAT0004628 | Slc30a4 | 22785 | HITS-CLIP | 23142080 | Unspecified |
| mmu-miR-21a-3p | MIMAT0004628 | Slc30a5 | 69048 | HITS-CLIP | 23142080 | Unspecified |
| mmu-miR-21a-3p | MIMAT0004628 | Slc30a6 | 210148 | HITS-CLIP | 23142080 | Unspecified |

| | | | | | | |
|---|---|---|---|---|---|---|
| mmu-miR-21a-3p | MIMAT0004628 | Slc33a1 | 11416 | HITS-CLIP | 23142080 | Unspecified |
| mmu-miR-21a-3p | MIMAT0004628 | Slc35a1 | 24060 | HITS-CLIP | 23142080 | Unspecified |
| mmu-miR-21a-3p | MIMAT0004628 | Slc35a3 | 229782 | HITS-CLIP | 23142080 | Unspecified |
| mmu-miR-21a-3p | MIMAT0004628 | Slc35e2 | 320541 | HITS-CLIP | 23142080 | Unspecified |
| mmu-miR-21a-3p | MIMAT0004628 | Slc37a1 | 224674 | HITS-CLIP | 23142080 | Unspecified |
| mmu-miR-21a-3p | MIMAT0004628 | Slc37a3 | 72144 | HITS-CLIP | 23142080 | Unspecified |
| mmu-miR-21a-3p | MIMAT0004628 | Slc37a4 | 14385 | HITS-CLIP | 23142080 | Unspecified |
| mmu-miR-21a-3p | MIMAT0004628 | Slc38a2 | 67760 | HITS-CLIP | 23142080 | Unspecified |
| mmu-miR-21a-3p | MIMAT0004628 | Slc39a1 | 30791 | HITS-CLIP | 37986224\|23142080 | Unspecified |
| mmu-miR-21a-3p | MIMAT0004628 | Slc39a10 | 227059 | HITS-CLIP | 23142080 | Unspecified |
| mmu-miR-21a-3p | MIMAT0004628 | Slc3a2 | 17254 | HITS-CLIP | 23142080 | Unspecified |
| mmu-miR-21a-3p | MIMAT0004628 | Slc41a1 | 98396 | HITS-CLIP | 23142080 | Unspecified |
| mmu-miR-21a-3p | MIMAT0004628 | Slc43a2 | 215113 | HITS-CLIP | 23142080\|29167373 | Unspecified |
| mmu-miR-21a-3p | MIMAT0004628 | Slc43a3 | 58207 | HITS-CLIP | 23142080 | Unspecified |
| mmu-miR-21a-3p | MIMAT0004628 | Slc44a2 | 68682 | HITS-CLIP | 23142080 | Unspecified |
| mmu-miR-21a-3p | MIMAT0004628 | Slc45a4 | 106068 | HITS-CLIP | 29167373 | Unspecified |
| mmu-miR-21a-3p | MIMAT0004628 | Slc4a2 | 20535 | HITS-CLIP | 23142080\|29167373 | Unspecified |
| mmu-miR-21a-3p | MIMAT0004628 | Slc4a7 | 218756 | HITS-CLIP | 29167373\|23142080 | Unspecified |
| mmu-miR-21a-3p | MIMAT0004628 | Slc5a3 | 53881 | HITS-CLIP | 37986224\|23142080\|29167373 | Unspecified |
| mmu-miR-21a-3p | MIMAT0004628 | Slc6a6 | 21366 | HITS-CLIP | 23142080 | Unspecified |
| mmu-miR-21a-3p | MIMAT0004628 | Slc7a1 | 11987 | HITS-CLIP | 37986224 | Unspecified |
| mmu-miR-21a-3p | MIMAT0004628 | Slc9a1 | 20544 | HITS-CLIP | 23142080 | Unspecified |
| mmu-miR-21a-3p | MIMAT0004628 | Slc9a3r1 | 26941 | HITS-CLIP | 23142080 | Unspecified |
| mmu-miR-21a-3p | MIMAT0004628 | Slc9a6 | 236794 | HITS-CLIP | 23142080 | Unspecified |
| mmu-miR-21a-3p | MIMAT0004628 | Slf2 | 226151 | HITS-CLIP | 23142080 | Unspecified |
| mmu-miR-21a-3p | MIMAT0004628 | Sltm | 66660 | HITS-CLIP | 23142080 | Unspecified |
| mmu-miR-21a-3p | MIMAT0004628 | Smad2 | 17126 | HITS-CLIP | 23142080 | Unspecified |
| mmu-miR-21a-3p | MIMAT0004628 | Smad4 | 17128 | HITS-CLIP | 23142080 | Unspecified |
| mmu-miR-21a-3p | MIMAT0004628 | Smad5 | 17129 | HITS-CLIP | 37986224\|23142080\|29167373 | Unspecified |
| mmu-miR-21a-3p | MIMAT0004628 | Smap1 | 98366 | HITS-CLIP | 23142080 | Unspecified |
| mmu-miR-21a-3p | MIMAT0004628 | Smarca5 | 93762 | HITS-CLIP | 23142080 | Unspecified |
| mmu-miR-21a-3p | MIMAT0004628 | Smarcad1 | 13990 | HITS-CLIP | 37986224\|23142080 | Unspecified |
| mmu-miR-21a-3p | MIMAT0004628 | Smarcal1 | 54380 | HITS-CLIP | 23142080 | Unspecified |
| mmu-miR-21a-3p | MIMAT0004628 | Smarcc2 | 68094 | HITS-CLIP | 37986224\|23142080\|29167373 | Unspecified |
| mmu-miR-21a-3p | MIMAT0004628 | Smarcd1 | 83797 | HITS-CLIP | 29167373 | Unspecified |
| mmu-miR-21a-3p | MIMAT0004628 | Smc2 | 14211 | HITS-CLIP | 23142080 | Unspecified |
| mmu-miR-21a-3p | MIMAT0004628 | Smc3 | 13006 | HITS-CLIP | 37986224\|23142080 | Unspecified |
| mmu-miR-21a-3p | MIMAT0004628 | Smc4 | 70099 | HITS-CLIP | 23142080 | Unspecified |
| mmu-miR-21a-3p | MIMAT0004628 | Smc6 | 67241 | HITS-CLIP | 23142080\|29167373 | Unspecified |

| | | | | | | |
|---|---|---|---|---|---|---|
| mmu-miR-21a-3p | MIMAT0004628 | Smchd1 | 74355 | HITS-CLIP | 23142080 | Unspecified |
| mmu-miR-21a-3p | MIMAT0004628 | Smcr8 | 237782 | HITS-CLIP | 37986224\|23142080 | Unspecified |
| mmu-miR-21a-3p | MIMAT0004628 | Smg1 | 233789 | HITS-CLIP | 29167373\|23142080 | Unspecified |
| mmu-miR-21a-3p | MIMAT0004628 | Smg5 | 229512 | HITS-CLIP | 23142080 | Unspecified |
| mmu-miR-21a-3p | MIMAT0004628 | Smg7 | 226517 | HITS-CLIP | 23142080 | Unspecified |
| mmu-miR-21a-3p | MIMAT0004628 | Smg9 | 71997 | HITS-CLIP | 29167373 | Unspecified |
| mmu-miR-21a-3p | MIMAT0004628 | Smndc1 | 76479 | HITS-CLIP | 23142080 | Unspecified |
| mmu-miR-21a-3p | MIMAT0004628 | Smpd1 | 20597 | HITS-CLIP | 37986224 | Unspecified |
| mmu-miR-21a-3p | MIMAT0004628 | Smpdl3a | 57319 | HITS-CLIP | 29167373 | Unspecified |
| mmu-miR-21a-3p | MIMAT0004628 | Smurf2 | 66313 | HITS-CLIP | 37986224\|23142080\|29167373 | Unspecified |
| mmu-miR-21a-3p | MIMAT0004628 | Smyd5 | 232187 | HITS-CLIP | 23142080 | Unspecified |
| mmu-miR-21a-3p | MIMAT0004628 | Snip1 | 76793 | HITS-CLIP | 23142080 | Unspecified |
| mmu-miR-21a-3p | MIMAT0004628 | Snn | 20621 | HITS-CLIP | 23142080 | Unspecified |
| mmu-miR-21a-3p | MIMAT0004628 | Snrnp200 | 320632 | HITS-CLIP | 37986224\|23142080 | Unspecified |
| mmu-miR-21a-3p | MIMAT0004628 | Snrpb | 20638 | HITS-CLIP | 23142080 | Unspecified |
| mmu-miR-21a-3p | MIMAT0004628 | Snrpb2 | 20639 | HITS-CLIP | 23142080 | Unspecified |
| mmu-miR-21a-3p | MIMAT0004628 | Snw1 | 66354 | HITS-CLIP | 23142080 | Unspecified |
| mmu-miR-21a-3p | MIMAT0004628 | Snx18 | 170625 | HITS-CLIP | 37986224\|23142080 | Unspecified |
| mmu-miR-21a-3p | MIMAT0004628 | Snx27 | 76742 | HITS-CLIP | 37986224\|23142080 | Unspecified |
| mmu-miR-21a-3p | MIMAT0004628 | Snx30 | 209131 | HITS-CLIP | 29167373\|23142080 | Unspecified |
| mmu-miR-21a-3p | MIMAT0004628 | Snx4 | 69150 | HITS-CLIP | 23142080 | Unspecified |
| mmu-miR-21a-3p | MIMAT0004628 | Snx5 | 69178 | HITS-CLIP | 29167373 | Unspecified |
| mmu-miR-21a-3p | MIMAT0004628 | Snx6 | 72183 | HITS-CLIP | 23142080 | Unspecified |
| mmu-miR-21a-3p | MIMAT0004628 | Soat1 | 20652 | HITS-CLIP | 37986224\|23142080 | Unspecified |
| mmu-miR-21a-3p | MIMAT0004628 | Socs1 | 12703 | HITS-CLIP | 23142080 | Unspecified |
| mmu-miR-21a-3p | MIMAT0004628 | Socs4 | 67296 | HITS-CLIP | 23142080 | Unspecified |
| mmu-miR-21a-3p | MIMAT0004628 | Socs5 | 56468 | HITS-CLIP | 23142080 | Unspecified |
| mmu-miR-21a-3p | MIMAT0004628 | Socs7 | 192157 | HITS-CLIP | 29167373 | Unspecified |
| mmu-miR-21a-3p | MIMAT0004628 | Son | 20658 | HITS-CLIP | 23142080\|29167373 | Unspecified |
| mmu-miR-21a-3p | MIMAT0004628 | Sorl1 | 20660 | HITS-CLIP | 23142080 | Unspecified |
| mmu-miR-21a-3p | MIMAT0004628 | Sos1 | 20662 | HITS-CLIP | 23142080 | Unspecified |
| mmu-miR-21a-3p | MIMAT0004628 | Sox5 | 20678 | HITS-CLIP | 23142080 | Unspecified |
| mmu-miR-21a-3p | MIMAT0004628 | Sp3 | 20687 | HITS-CLIP | 23142080\|29167373 | Unspecified |
| mmu-miR-21a-3p | MIMAT0004628 | Sp4 | 20688 | HITS-CLIP | 23142080 | Unspecified |
| mmu-miR-21a-3p | MIMAT0004628 | Spag9 | 70834 | HITS-CLIP | 23142080\|29167373 | Unspecified |
| mmu-miR-21a-3p | MIMAT0004628 | Spast | 50850 | HITS-CLIP | 23142080\|29167373 | Unspecified |
| mmu-miR-21a-3p | MIMAT0004628 | Spata13 | 219140 | HITS-CLIP | 23142080 | Unspecified |
| mmu-miR-21a-3p | MIMAT0004628 | Spata2 | 263876 | HITS-CLIP | 26203562 | Unspecified |
| mmu-miR-21a-3p | MIMAT0004628 | Spcs2 | 66624 | HITS-CLIP | 37986224\|23142080 | Unspecified |

| | | | | | | |
|---|---|---|---|---|---|---|
| mmu-miR-21a-3p | MIMAT0004628 | Spg11 | 214585 | HITS-CLIP | 29167373\|23142080 | Unspecified |
| mmu-miR-21a-3p | MIMAT0004628 | Spg21 | 27965 | HITS-CLIP | 37986224 | Unspecified |
| mmu-miR-21a-3p | MIMAT0004628 | Spidr | 224008 | HITS-CLIP | 23142080 | Unspecified |
| mmu-miR-21a-3p | MIMAT0004628 | Spin1 | 20729 | HITS-CLIP | 23142080 | Unspecified |
| mmu-miR-21a-3p | MIMAT0004628 | Spn | 20737 | HITS-CLIP | 37986224 | Unspecified |
| mmu-miR-21a-3p | MIMAT0004628 | Spop | 20747 | HITS-CLIP | 23142080 | Unspecified |
| mmu-miR-21a-3p | MIMAT0004628 | Spp1 | 20750 | HITS-CLIP | 37986224\|29167373 | Unspecified |
| mmu-miR-21a-3p | MIMAT0004628 | Spred1 | 114715 | HITS-CLIP | 29167373\|23142080 | Unspecified |
| mmu-miR-21a-3p | MIMAT0004628 | Spred2 | 114716 | HITS-CLIP | 23142080 | Unspecified |
| mmu-miR-21a-3p | MIMAT0004628 | Spry2 | 24064 | HITS-CLIP | 23142080 | Unspecified |
| mmu-miR-21a-3p | MIMAT0004628 | Spry4 | 24066 | HITS-CLIP | 23142080 | Unspecified |
| mmu-miR-21a-3p | MIMAT0004628 | Spryd3 | 223918 | HITS-CLIP | 23142080\|29167373 | Unspecified |
| mmu-miR-21a-3p | MIMAT0004628 | Sptan1 | 20740 | HITS-CLIP | 23142080 | Unspecified |
| mmu-miR-21a-3p | MIMAT0004628 | Sptbn1 | 20742 | HITS-CLIP | 37986224\|23142080\|29167373 | Unspecified |
| mmu-miR-21a-3p | MIMAT0004628 | Sptlc1 | 268656 | HITS-CLIP | 37986224\|23142080 | Unspecified |
| mmu-miR-21a-3p | MIMAT0004628 | Sptlc2 | 20773 | HITS-CLIP | 29167373 | Unspecified |
| mmu-miR-21a-3p | MIMAT0004628 | Spty2d1 | 101685 | HITS-CLIP | 37986224\|23142080 | Unspecified |
| mmu-miR-21a-3p | MIMAT0004628 | Sqle | 20775 | HITS-CLIP | 23142080 | Unspecified |
| mmu-miR-21a-3p | MIMAT0004628 | Sqstm1 | 18412 | HITS-CLIP | 23142080 | Unspecified |
| mmu-miR-21a-3p | MIMAT0004628 | Srcap | 100043597 | HITS-CLIP | 23142080\|29167373 | Unspecified |
| mmu-miR-21a-3p | MIMAT0004628 | Srebf1 | 20787 | HITS-CLIP | 23142080 | Unspecified |
| mmu-miR-21a-3p | MIMAT0004628 | Srebf2 | 20788 | HITS-CLIP | 37986224\|23142080 | Unspecified |
| mmu-miR-21a-3p | MIMAT0004628 | Srgap2 | 14270 | HITS-CLIP | 29167373 | Unspecified |
| mmu-miR-21a-3p | MIMAT0004628 | Srgap3 | 259302 | HITS-CLIP | 23142080 | Unspecified |
| mmu-miR-21a-3p | MIMAT0004628 | Srgn | 19073 | HITS-CLIP | 37986224\|23142080 | Unspecified |
| mmu-miR-21a-3p | MIMAT0004628 | Srp14 | 20813 | HITS-CLIP | 23142080 | Unspecified |
| mmu-miR-21a-3p | MIMAT0004628 | Srpk1 | 20815 | HITS-CLIP | 37986224\|23142080 | Unspecified |
| mmu-miR-21a-3p | MIMAT0004628 | Srrm1 | 51796 | HITS-CLIP | 37986224\|23142080\|29167373 | Unspecified |
| mmu-miR-21a-3p | MIMAT0004628 | Srrm2 | 75956 | HITS-CLIP | 37986224\|23142080 | Unspecified |
| mmu-miR-21a-3p | MIMAT0004628 | Srsf1 | 110809 | HITS-CLIP | 23142080 | Unspecified |
| mmu-miR-21a-3p | MIMAT0004628 | Srsf11 | 69207 | HITS-CLIP | 23142080 | Unspecified |
| mmu-miR-21a-3p | MIMAT0004628 | Srsf2 | 20382 | HITS-CLIP | 37986224\|23142080 | Unspecified |
| mmu-miR-21a-3p | MIMAT0004628 | Srsf4 | 57317 | HITS-CLIP | 23142080 | Unspecified |
| mmu-miR-21a-3p | MIMAT0004628 | Srsf6 | 67996 | HITS-CLIP | 23142080 | Unspecified |
| mmu-miR-21a-3p | MIMAT0004628 | Ssb | 20823 | HITS-CLIP | 23142080 | Unspecified |
| mmu-miR-21a-3p | MIMAT0004628 | Ssbp2 | 66970 | HITS-CLIP | 23142080 | Unspecified |
| mmu-miR-21a-3p | MIMAT0004628 | Ssh1 | 231637 | HITS-CLIP | 37986224 | Unspecified |
| mmu-miR-21a-3p | MIMAT0004628 | Ssh2 | 237860 | HITS-CLIP | 37986224\|29167373\|23142080 | Unspecified |
| mmu-miR-21a-3p | MIMAT0004628 | Ssr1 | 107513 | HITS-CLIP | 37986224\|23142080\|29167373 | Unspecified |

| | | | | | | |
|---|---|---|---|---|---|---|
| mmu-miR-21a-3p | MIMAT0004628 | Ssr2 | 66256 | HITS-CLIP | 23142080 | Unspecified |
| mmu-miR-21a-3p | MIMAT0004628 | Ssr3 | 67437 | HITS-CLIP | 37986224\|23142080 | Unspecified |
| mmu-miR-21a-3p | MIMAT0004628 | St8sia1 | 20449 | HITS-CLIP | 23142080 | Unspecified |
| mmu-miR-21a-3p | MIMAT0004628 | St8sia4 | 20452 | HITS-CLIP | 23142080 | Unspecified |
| mmu-miR-21a-3p | MIMAT0004628 | Stag1 | 20842 | HITS-CLIP | 23142080\|29167373 | Unspecified |
| mmu-miR-21a-3p | MIMAT0004628 | Stag2 | 20843 | HITS-CLIP | 23142080\|29167373 | Unspecified |
| mmu-miR-21a-3p | MIMAT0004628 | Stam | 20844 | HITS-CLIP | 23142080\|29167373 | Unspecified |
| mmu-miR-21a-3p | MIMAT0004628 | Stambp | 70527 | HITS-CLIP | 23142080 | Unspecified |
| mmu-miR-21a-3p | MIMAT0004628 | Stambpl1 | 76630 | HITS-CLIP | 37986224 | Unspecified |
| mmu-miR-21a-3p | MIMAT0004628 | Stap1 | 56792 | HITS-CLIP | 23142080 | Unspecified |
| mmu-miR-21a-3p | MIMAT0004628 | Stard4 | 170459 | HITS-CLIP | 23142080 | Unspecified |
| mmu-miR-21a-3p | MIMAT0004628 | Stard7 | 99138 | HITS-CLIP | 23142080 | Unspecified |
| mmu-miR-21a-3p | MIMAT0004628 | Stat1 | 20846 | HITS-CLIP | 37986224 | Unspecified |
| mmu-miR-21a-3p | MIMAT0004628 | Stat5a | 20850 | HITS-CLIP | 23142080 | Unspecified |
| mmu-miR-21a-3p | MIMAT0004628 | Stat5b | 20851 | HITS-CLIP | 37986224\|23142080\|29167373 | Unspecified |
| mmu-miR-21a-3p | MIMAT0004628 | Stat6 | 20852 | HITS-CLIP | 23142080 | Unspecified |
| mmu-miR-21a-3p | MIMAT0004628 | Stau1 | 20853 | HITS-CLIP | 23142080 | Unspecified |
| mmu-miR-21a-3p | MIMAT0004628 | Stil | 20460 | HITS-CLIP | 23142080 | Unspecified |
| mmu-miR-21a-3p | MIMAT0004628 | Stim1 | 20866 | HITS-CLIP | 23142080 | Unspecified |
| mmu-miR-21a-3p | MIMAT0004628 | Stim2 | 116873 | HITS-CLIP | 37986224\|23142080 | Unspecified |
| mmu-miR-21a-3p | MIMAT0004628 | Stk10 | 20868 | HITS-CLIP | 23142080 | Unspecified |
| mmu-miR-21a-3p | MIMAT0004628 | Stk11 | 20869 | HITS-CLIP | 23142080 | Unspecified |
| mmu-miR-21a-3p | MIMAT0004628 | Stk17b | 98267 | HITS-CLIP | 23142080 | Unspecified |
| mmu-miR-21a-3p | MIMAT0004628 | Stk26 | 51765 | HITS-CLIP | 23142080 | Unspecified |
| mmu-miR-21a-3p | MIMAT0004628 | Stk35 | 67333 | HITS-CLIP | 29167373 | Unspecified |
| mmu-miR-21a-3p | MIMAT0004628 | Stk38 | 106504 | HITS-CLIP | 23142080\|29167373 | Unspecified |
| mmu-miR-21a-3p | MIMAT0004628 | Stk39 | 53416 | HITS-CLIP | 23142080 | Unspecified |
| mmu-miR-21a-3p | MIMAT0004628 | Stk4 | 58231 | HITS-CLIP | 37986224\|23142080 | Unspecified |
| mmu-miR-21a-3p | MIMAT0004628 | Stk40 | 74178 | HITS-CLIP | 37986224\|23142080 | Unspecified |
| mmu-miR-21a-3p | MIMAT0004628 | Strap | 20901 | HITS-CLIP | 23142080 | Unspecified |
| mmu-miR-21a-3p | MIMAT0004628 | Strbp | 20744 | HITS-CLIP | 37986224 | Unspecified |
| mmu-miR-21a-3p | MIMAT0004628 | Strn | 268980 | HITS-CLIP | 23142080 | Unspecified |
| mmu-miR-21a-3p | MIMAT0004628 | Strn3 | 94186 | HITS-CLIP | 37986224 | Unspecified |
| mmu-miR-21a-3p | MIMAT0004628 | Stt3a | 16430 | HITS-CLIP | 23142080 | Unspecified |
| mmu-miR-21a-3p | MIMAT0004628 | Stt3b | 68292 | HITS-CLIP | 23142080 | Unspecified |
| mmu-miR-21a-3p | MIMAT0004628 | Stxbp3 | 20912 | HITS-CLIP | 23142080 | Unspecified |
| mmu-miR-21a-3p | MIMAT0004628 | Stxbp5 | 78808 | HITS-CLIP | 29167373 | Unspecified |
| mmu-miR-21a-3p | MIMAT0004628 | Sucla2 | 20916 | HITS-CLIP | 37986224\|23142080 | Unspecified |
| mmu-miR-21a-3p | MIMAT0004628 | Sun1 | 77053 | HITS-CLIP | 23142080 | Unspecified |

| | | | | | | |
|---|---|---|---|---|---|---|
| mmu-miR-21a-3p | MIMAT0004628 | Supt20 | 56790 | HITS-CLIP | 31433990 | Unspecified |
| mmu-miR-21a-3p | MIMAT0004628 | Supt6 | 20926 | HITS-CLIP | 23142080 | Unspecified |
| mmu-miR-21a-3p | MIMAT0004628 | Surf4 | 20932 | HITS-CLIP | 23142080 | Unspecified |
| mmu-miR-21a-3p | MIMAT0004628 | Susd6 | 9766 | HITS-CLIP | 29167373 | Unspecified |
| mmu-miR-21a-3p | MIMAT0004628 | Suv39h1 | 20937 | HITS-CLIP | 23142080 | Unspecified |
| mmu-miR-21a-3p | MIMAT0004628 | Suz12 | 52615 | HITS-CLIP | 37986224\|23142080 | Unspecified |
| mmu-miR-21a-3p | MIMAT0004628 | Swt1 | 66875 | HITS-CLIP | 23142080 | Unspecified |
| mmu-miR-21a-3p | MIMAT0004628 | Syne2 | 319565 | HITS-CLIP | 23142080 | Unspecified |
| mmu-miR-21a-3p | MIMAT0004628 | Syne3 | 212073 | HITS-CLIP | 23142080 | Unspecified |
| mmu-miR-21a-3p | MIMAT0004628 | Synj1 | 104015 | HITS-CLIP | 37986224\|23142080\|29167373 | Unspecified |
| mmu-miR-21a-3p | MIMAT0004628 | Sypl | 19027 | HITS-CLIP | 23142080 | Unspecified |
| mmu-miR-21a-3p | MIMAT0004628 | Szrd1 | 213491 | HITS-CLIP | 23142080 | Unspecified |
| mmu-miR-21a-3p | MIMAT0004628 | Tab2 | 68652 | HITS-CLIP | 23142080\|31433990 | Unspecified |
| mmu-miR-21a-3p | MIMAT0004628 | Tacc1 | 320165 | HITS-CLIP | 23142080 | Unspecified |
| mmu-miR-21a-3p | MIMAT0004628 | Taf1 | 270627 | HITS-CLIP | 23142080 | Unspecified |
| mmu-miR-21a-3p | MIMAT0004628 | Taf15 | 70439 | HITS-CLIP | 23142080 | Unspecified |
| mmu-miR-21a-3p | MIMAT0004628 | Taf4b | 72504 | HITS-CLIP | 23142080 | Unspecified |
| mmu-miR-21a-3p | MIMAT0004628 | Tagap | 72536 | HITS-CLIP | 23142080 | Unspecified |
| mmu-miR-21a-3p | MIMAT0004628 | Taldo1 | 21351 | HITS-CLIP | 23142080 | Unspecified |
| mmu-miR-21a-3p | MIMAT0004628 | Taok1 | 216965 | HITS-CLIP | 37986224\|23142080 | Unspecified |
| mmu-miR-21a-3p | MIMAT0004628 | Tap2 | 21355 | HITS-CLIP | 37986224 | Unspecified |
| mmu-miR-21a-3p | MIMAT0004628 | Tapbp | 21356 | HITS-CLIP | 23142080 | Unspecified |
| mmu-miR-21a-3p | MIMAT0004628 | Tardbp | 230908 | HITS-CLIP | 37986224\|23142080\|29167373 | Unspecified |
| mmu-miR-21a-3p | MIMAT0004628 | Tbc1d1 | 57915 | HITS-CLIP | 23142080 | Unspecified |
| mmu-miR-21a-3p | MIMAT0004628 | Tbc1d10b | 68449 | HITS-CLIP | 37986224\|23142080\|29167373 | Unspecified |
| mmu-miR-21a-3p | MIMAT0004628 | Tbc1d14 | 100855 | HITS-CLIP | 23142080 | Unspecified |
| mmu-miR-21a-3p | MIMAT0004628 | Tbc1d15 | 66687 | HITS-CLIP | 23142080 | Unspecified |
| mmu-miR-21a-3p | MIMAT0004628 | Tbc1d22b | 381085 | HITS-CLIP | 23142080 | Unspecified |
| mmu-miR-21a-3p | MIMAT0004628 | Tbc1d4 | 210789 | HITS-CLIP | 37986224\|23142080 | Unspecified |
| mmu-miR-21a-3p | MIMAT0004628 | Tbc1d8b | 245638 | HITS-CLIP | 23142080 | Unspecified |
| mmu-miR-21a-3p | MIMAT0004628 | Tbca | 21371 | HITS-CLIP | 23142080 | Unspecified |
| mmu-miR-21a-3p | MIMAT0004628 | Tbcel | 272589 | HITS-CLIP | 23142080\|29167373 | Unspecified |
| mmu-miR-21a-3p | MIMAT0004628 | Tbk1 | 56480 | HITS-CLIP | 23142080 | Unspecified |
| mmu-miR-21a-3p | MIMAT0004628 | Tbl1xr1 | 81004 | HITS-CLIP | 37986224\|23142080\|29167373 | Unspecified |
| mmu-miR-21a-3p | MIMAT0004628 | Tbp | 21374 | HITS-CLIP | 37986224\|23142080 | Unspecified |
| mmu-miR-21a-3p | MIMAT0004628 | Tcea1 | 21399 | HITS-CLIP | 29167373 | Unspecified |
| mmu-miR-21a-3p | MIMAT0004628 | Tcerg1 | 56070 | HITS-CLIP | 37986224\|23142080 | Unspecified |
| mmu-miR-21a-3p | MIMAT0004628 | Tcf20 | 21411 | HITS-CLIP | 23142080 | Unspecified |
| mmu-miR-21a-3p | MIMAT0004628 | Tcf7 | 21414 | HITS-CLIP | 37986224 | Unspecified |

| | | | | | | |
|---|---|---|---|---|---|---|
| mmu-miR-21a-3p | MIMAT0004628 | Tcp1 | 21454 | HITS-CLIP | 37986224\|23142080\|29167373 | Unspecified |
| mmu-miR-21a-3p | MIMAT0004628 | Tcp11l2 | 216198 | HITS-CLIP | 23142080 | Unspecified |
| mmu-miR-21a-3p | MIMAT0004628 | Tecr | 106529 | HITS-CLIP | 23142080 | Unspecified |
| mmu-miR-21a-3p | MIMAT0004628 | Tent4a | 11044 | HITS-CLIP | 23142080 | Unspecified |
| mmu-miR-21a-3p | MIMAT0004628 | Tent4b | 64282 | HITS-CLIP | 23142080 | Unspecified |
| mmu-miR-21a-3p | MIMAT0004628 | Tent5a | 55603 | HITS-CLIP | 29167373 | Unspecified |
| mmu-miR-21a-3p | MIMAT0004628 | Tent5c | 54855 | HITS-CLIP | 23142080 | Unspecified |
| mmu-miR-21a-3p | MIMAT0004628 | Tep1 | 21745 | HITS-CLIP | 23142080\|29167373 | Unspecified |
| mmu-miR-21a-3p | MIMAT0004628 | Tet1 | 52463 | HITS-CLIP | 37986224\|23142080 | Unspecified |
| mmu-miR-21a-3p | MIMAT0004628 | Tet2 | 214133 | HITS-CLIP | 37986224\|23142080\|29167373 | Unspecified |
| mmu-miR-21a-3p | MIMAT0004628 | Tet3 | 194388 | HITS-CLIP | 23142080\|29167373 | Unspecified |
| mmu-miR-21a-3p | MIMAT0004628 | Tex264 | 21767 | HITS-CLIP | 23142080 | Unspecified |
| mmu-miR-21a-3p | MIMAT0004628 | Tfdp1 | 21781 | HITS-CLIP | 23142080 | Unspecified |
| mmu-miR-21a-3p | MIMAT0004628 | Tfg | 21787 | HITS-CLIP | 23142080 | Unspecified |
| mmu-miR-21a-3p | MIMAT0004628 | Tg | 21819 | HITS-CLIP | 37986224\|23142080 | Unspecified |
| mmu-miR-21a-3p | MIMAT0004628 | Tgds | 76355 | HITS-CLIP | 23142080 | Unspecified |
| mmu-miR-21a-3p | MIMAT0004628 | Tgfbr1 | 21812 | HITS-CLIP | 23142080 | Unspecified |
| mmu-miR-21a-3p | MIMAT0004628 | Tgfbr2 | 21813 | HITS-CLIP | 29167373 | Unspecified |
| mmu-miR-21a-3p | MIMAT0004628 | Tgif1 | 21815 | HITS-CLIP | 23142080 | Unspecified |
| mmu-miR-21a-3p | MIMAT0004628 | Tgoln1 | 22134 | HITS-CLIP | 37986224\|23142080\|29167373 | Unspecified |
| mmu-miR-21a-3p | MIMAT0004628 | Tgs1 | 116940 | HITS-CLIP | 29167373\|23142080 | Unspecified |
| mmu-miR-21a-3p | MIMAT0004628 | Thada | 240174 | HITS-CLIP | 23142080 | Unspecified |
| mmu-miR-21a-3p | MIMAT0004628 | Thap12 | 72981 | HITS-CLIP | 23142080 | Unspecified |
| mmu-miR-21a-3p | MIMAT0004628 | Themis | 210757 | HITS-CLIP | 23142080 | Unspecified |
| mmu-miR-21a-3p | MIMAT0004628 | Thoc1 | 225160 | HITS-CLIP | 23142080 | Unspecified |
| mmu-miR-21a-3p | MIMAT0004628 | Thoc2 | 331401 | HITS-CLIP | 23142080 | Unspecified |
| mmu-miR-21a-3p | MIMAT0004628 | Thy1 | 21838 | HITS-CLIP | 23142080 | Unspecified |
| mmu-miR-21a-3p | MIMAT0004628 | Tial1 | 21843 | HITS-CLIP | 23142080 | Unspecified |
| mmu-miR-21a-3p | MIMAT0004628 | Tiam1 | 21844 | HITS-CLIP | 23142080 | Unspecified |
| mmu-miR-21a-3p | MIMAT0004628 | Tigit | 100043314 | HITS-CLIP | 23142080 | Unspecified |
| mmu-miR-21a-3p | MIMAT0004628 | Timm10b | 14356 | HITS-CLIP | 23142080 | Unspecified |
| mmu-miR-21a-3p | MIMAT0004628 | Timp2 | 21858 | HITS-CLIP | 23142080\|29167373 | Unspecified |
| mmu-miR-21a-3p | MIMAT0004628 | Tiparp | 99929 | HITS-CLIP | 37986224\|23142080 | Unspecified |
| mmu-miR-21a-3p | MIMAT0004628 | Tiprl | 226591 | HITS-CLIP | 37986224 | Unspecified |
| mmu-miR-21a-3p | MIMAT0004628 | Tle3 | 21887 | HITS-CLIP | 23142080 | Unspecified |
| mmu-miR-21a-3p | MIMAT0004628 | Tle4 | 21888 | HITS-CLIP | 37986224\|23142080 | Unspecified |
| mmu-miR-21a-3p | MIMAT0004628 | Tlk1 | 228012 | HITS-CLIP | 23142080 | Unspecified |
| mmu-miR-21a-3p | MIMAT0004628 | Tlk2 | 24086 | HITS-CLIP | 23142080 | Unspecified |
| mmu-miR-21a-3p | MIMAT0004628 | Tlr13 | 279572 | HITS-CLIP | 29167373 | Unspecified |

| | | | | | | | |
|---|---|---|---|---|---|---|---|
| mmu-miR-21a-3p | MIMAT0004628 | Tm9sf2 | 68059 | HITS-CLIP | | 23142080 | Unspecified |
| mmu-miR-21a-3p | MIMAT0004628 | Tm9sf3 | 107358 | HITS-CLIP | | 37986224\|23142080 | Unspecified |
| mmu-miR-21a-3p | MIMAT0004628 | Tmc8 | 217356 | HITS-CLIP | | 37986224 | Unspecified |
| mmu-miR-21a-3p | MIMAT0004628 | Tmcc1 | 330401 | HITS-CLIP | | 23142080 | Unspecified |
| mmu-miR-21a-3p | MIMAT0004628 | Tmed8 | 382620 | HITS-CLIP | | 23142080 | Unspecified |
| mmu-miR-21a-3p | MIMAT0004628 | Tmem101 | 76547 | HITS-CLIP | | 23142080 | Unspecified |
| mmu-miR-21a-3p | MIMAT0004628 | Tmem115 | 56395 | HITS-CLIP | | 23142080 | Unspecified |
| mmu-miR-21a-3p | MIMAT0004628 | Tmem123 | 71929 | HITS-CLIP | | 23142080 | Unspecified |
| mmu-miR-21a-3p | MIMAT0004628 | Tmem126b | 68472 | HITS-CLIP | | 23142080 | Unspecified |
| mmu-miR-21a-3p | MIMAT0004628 | Tmem127 | 69470 | HITS-CLIP | | 23142080 | Unspecified |
| mmu-miR-21a-3p | MIMAT0004628 | Tmem128 | 66309 | HITS-CLIP | | 23142080 | Unspecified |
| mmu-miR-21a-3p | MIMAT0004628 | Tmem131 | 56030 | HITS-CLIP | | 23142080 | Unspecified |
| mmu-miR-21a-3p | MIMAT0004628 | Tmem147 | 69804 | HITS-CLIP | | 23142080 | Unspecified |
| mmu-miR-21a-3p | MIMAT0004628 | Tmem164 | 209497 | HITS-CLIP | | 23142080 | Unspecified |
| mmu-miR-21a-3p | MIMAT0004628 | Tmem167 | 66074 | HITS-CLIP | | 23142080 | Unspecified |
| mmu-miR-21a-3p | MIMAT0004628 | Tmem167b | 67495 | HITS-CLIP | | 23142080 | Unspecified |
| mmu-miR-21a-3p | MIMAT0004628 | Tmem168 | 101118 | HITS-CLIP | | 23142080 | Unspecified |
| mmu-miR-21a-3p | MIMAT0004628 | Tmem170b | 621976 | HITS-CLIP | | 23142080 | Unspecified |
| mmu-miR-21a-3p | MIMAT0004628 | Tmem175 | 72392 | HITS-CLIP | | 23142080 | Unspecified |
| mmu-miR-21a-3p | MIMAT0004628 | Tmem179b | 374395 | HITS-CLIP | | 23142080 | Unspecified |
| mmu-miR-21a-3p | MIMAT0004628 | Tmem184c | 234463 | HITS-CLIP | | 29167373 | Unspecified |
| mmu-miR-21a-3p | MIMAT0004628 | Tmem185a | 84548 | HITS-CLIP | | 23142080 | Unspecified |
| mmu-miR-21a-3p | MIMAT0004628 | Tmem185b | 226351 | HITS-CLIP | | 29167373 | Unspecified |
| mmu-miR-21a-3p | MIMAT0004628 | Tmem201 | 230917 | HITS-CLIP | | 23142080 | Unspecified |
| mmu-miR-21a-3p | MIMAT0004628 | Tmem209 | 72649 | HITS-CLIP | | 29167373 | Unspecified |
| mmu-miR-21a-3p | MIMAT0004628 | Tmem230 | 70612 | HITS-CLIP | | 23142080 | Unspecified |
| mmu-miR-21a-3p | MIMAT0004628 | Tmem258 | 69038 | HITS-CLIP | | 37986224\|23142080 | Unspecified |
| mmu-miR-21a-3p | MIMAT0004628 | Tmem30a | 69981 | HITS-CLIP | | 37986224\|23142080 | Unspecified |
| mmu-miR-21a-3p | MIMAT0004628 | Tmem33 | 67878 | HITS-CLIP | | 23142080 | Unspecified |
| mmu-miR-21a-3p | MIMAT0004628 | Tmem38b | 52076 | HITS-CLIP | | 23142080 | Unspecified |
| mmu-miR-21a-3p | MIMAT0004628 | Tmem40 | 94346 | HITS-CLIP | | 23142080 | Unspecified |
| mmu-miR-21a-3p | MIMAT0004628 | Tmem45a | 56277 | HITS-CLIP | | 23142080 | Unspecified |
| mmu-miR-21a-3p | MIMAT0004628 | Tmem59 | 56374 | HITS-CLIP | | 23142080 | Unspecified |
| mmu-miR-21a-3p | MIMAT0004628 | Tmem60 | 212090 | HITS-CLIP | | 23142080 | Unspecified |
| mmu-miR-21a-3p | MIMAT0004628 | Tmem63b | 224807 | HITS-CLIP | | 23142080 | Unspecified |
| mmu-miR-21a-3p | MIMAT0004628 | Tmem64 | 100201 | HITS-CLIP | | 37986224\|23142080 | Unspecified |
| mmu-miR-21a-3p | MIMAT0004628 | Tmem67 | 329795 | HITS-CLIP | | 23142080 | Unspecified |
| mmu-miR-21a-3p | MIMAT0004628 | Tmem87a | 211499 | HITS-CLIP | | 23142080 | Unspecified |
| mmu-miR-21a-3p | MIMAT0004628 | Tmem94 | 9772 | HITS-CLIP | | 23142080 | Unspecified |

| | | | | | | |
|---|---|---|---|---|---|---|
| mmu-miR-21a-3p | MIMAT0004628 | Tmem97 | 69071 | qCLASH | 31393953 | Unspecified |
| mmu-miR-21a-3p | MIMAT0004628 | Tmpo | 21917 | HITS-CLIP | 37986224\|23142080 | Unspecified |
| mmu-miR-21a-3p | MIMAT0004628 | Tmx3 | 67988 | HITS-CLIP | 23142080 | Unspecified |
| mmu-miR-21a-3p | MIMAT0004628 | Tmx4 | 52837 | HITS-CLIP | 29167373 | Unspecified |
| mmu-miR-21a-3p | MIMAT0004628 | Tnfaip3 | 21929 | HITS-CLIP | 23142080 | Unspecified |
| mmu-miR-21a-3p | MIMAT0004628 | Tnfsf8 | 21949 | HITS-CLIP | 23142080 | Unspecified |
| mmu-miR-21a-3p | MIMAT0004628 | Tnks | 21951 | HITS-CLIP | 29167373\|23142080 | Unspecified |
| mmu-miR-21a-3p | MIMAT0004628 | Tnpo1 | 238799 | HITS-CLIP | 37986224\|23142080 | Unspecified |
| mmu-miR-21a-3p | MIMAT0004628 | Tnpo2 | 212999 | HITS-CLIP | 23142080 | Unspecified |
| mmu-miR-21a-3p | MIMAT0004628 | Tnrc18 | 231861 | HITS-CLIP | 23142080\|29167373 | Unspecified |
| mmu-miR-21a-3p | MIMAT0004628 | Tnrc6a | 233833 | HITS-CLIP | 37986224\|23142080\|29167373 | Unspecified |
| mmu-miR-21a-3p | MIMAT0004628 | Tnrc6b | 213988 | HITS-CLIP | 37986224\|23142080 | Unspecified |
| mmu-miR-21a-3p | MIMAT0004628 | Tnrc6c | 217351 | HITS-CLIP | 37986224\|23142080\|29167373 | Unspecified |
| mmu-miR-21a-3p | MIMAT0004628 | Togaram1 | 23116 | HITS-CLIP | 23142080 | Unspecified |
| mmu-miR-21a-3p | MIMAT0004628 | Tollip | 54473 | HITS-CLIP | 37986224 | Unspecified |
| mmu-miR-21a-3p | MIMAT0004628 | Tomm34 | 67145 | HITS-CLIP | 23142080 | Unspecified |
| mmu-miR-21a-3p | MIMAT0004628 | Tomm70a | 28185 | HITS-CLIP | 37986224\|23142080 | Unspecified |
| mmu-miR-21a-3p | MIMAT0004628 | Top1 | 21969 | HITS-CLIP | 23142080 | Unspecified |
| mmu-miR-21a-3p | MIMAT0004628 | Top2a | 21973 | HITS-CLIP | 37986224\|23142080\|29167373 | Unspecified |
| mmu-miR-21a-3p | MIMAT0004628 | Top2b | 21974 | HITS-CLIP | 37986224\|23142080 | Unspecified |
| mmu-miR-21a-3p | MIMAT0004628 | Topbp1 | 235559 | HITS-CLIP | 23142080 | Unspecified |
| mmu-miR-21a-3p | MIMAT0004628 | Tor1aip1 | 208263 | HITS-CLIP | 29167373 | Unspecified |
| mmu-miR-21a-3p | MIMAT0004628 | Tor1aip2 | 240832 | HITS-CLIP | 37986224\|23142080\|29167373 | Unspecified |
| mmu-miR-21a-3p | MIMAT0004628 | Tpcn1 | 252972 | HITS-CLIP | 37986224\|29167373\|23142080 | Unspecified |
| mmu-miR-21a-3p | MIMAT0004628 | Tpd52l2 | 66314 | HITS-CLIP | 23142080 | Unspecified |
| mmu-miR-21a-3p | MIMAT0004628 | Tpi1 | 21991 | HITS-CLIP | 23142080 | Unspecified |
| mmu-miR-21a-3p | MIMAT0004628 | Tpm4 | 326618 | HITS-CLIP | 37986224 | Unspecified |
| mmu-miR-21a-3p | MIMAT0004628 | Tpp2 | 22019 | HITS-CLIP | 23142080 | Unspecified |
| mmu-miR-21a-3p | MIMAT0004628 | Tpr | 108989 | HITS-CLIP | 37986224\|23142080\|29167373 | Unspecified |
| mmu-miR-21a-3p | MIMAT0004628 | Tra2a | 101214 | HITS-CLIP | 37986224\|23142080 | Unspecified |
| mmu-miR-21a-3p | MIMAT0004628 | Traf1 | 22029 | HITS-CLIP | 37986224 | Unspecified |
| mmu-miR-21a-3p | MIMAT0004628 | Traf7 | 224619 | HITS-CLIP | 37986224\|23142080 | Unspecified |
| mmu-miR-21a-3p | MIMAT0004628 | Tram1 | 72265 | HITS-CLIP | 37986224 | Unspecified |
| mmu-miR-21a-3p | MIMAT0004628 | Trappc11 | 320714 | HITS-CLIP | 23142080 | Unspecified |
| mmu-miR-21a-3p | MIMAT0004628 | Trappc13 | 66975 | HITS-CLIP | 23142080 | Unspecified |
| mmu-miR-21a-3p | MIMAT0004628 | Trdmt1 | 13434 | HITS-CLIP | 23142080 | Unspecified |
| mmu-miR-21a-3p | MIMAT0004628 | Treml2 | 328833 | HITS-CLIP | 37986224\|23142080 | Unspecified |
| mmu-miR-21a-3p | MIMAT0004628 | Triap1 | 69076 | HITS-CLIP | 23142080 | Unspecified |
| mmu-miR-21a-3p | MIMAT0004628 | Trib1 | 211770 | HITS-CLIP | 23142080\|29167373 | Unspecified |

| | | | | | | | |
|---|---|---|---|---|---|---|---|
| mmu-miR-21a-3p | MIMAT0004628 | Trib2 | 217410 | HITS-CLIP | | 23142080 | Unspecified |
| mmu-miR-21a-3p | MIMAT0004628 | Trim23 | 81003 | HITS-CLIP | | 23142080 | Unspecified |
| mmu-miR-21a-3p | MIMAT0004628 | Trim24 | 21848 | HITS-CLIP | | 23142080 | Unspecified |
| mmu-miR-21a-3p | MIMAT0004628 | Trim25 | 217069 | HITS-CLIP | | 37986224 | Unspecified |
| mmu-miR-21a-3p | MIMAT0004628 | Trim28 | 21849 | HITS-CLIP | 37986224\|23142080 | | Unspecified |
| mmu-miR-21a-3p | MIMAT0004628 | Trim32 | 69807 | HITS-CLIP | | 23142080 | Unspecified |
| mmu-miR-21a-3p | MIMAT0004628 | Trim33 | 94093 | HITS-CLIP | 37986224\|23142080 | | Unspecified |
| mmu-miR-21a-3p | MIMAT0004628 | Trim37 | 68729 | HITS-CLIP | | 23142080 | Unspecified |
| mmu-miR-21a-3p | MIMAT0004628 | Trim41 | 211007 | HITS-CLIP | | 23142080 | Unspecified |
| mmu-miR-21a-3p | MIMAT0004628 | Trip11 | 109181 | HITS-CLIP | 37986224\|23142080\|29167373 | | Unspecified |
| mmu-miR-21a-3p | MIMAT0004628 | Trip12 | 14897 | HITS-CLIP | 31433990\|23142080\|29167373 | | Unspecified |
| mmu-miR-21a-3p | MIMAT0004628 | Trmt5 | 76357 | HITS-CLIP | | 23142080 | Unspecified |
| mmu-miR-21a-3p | MIMAT0004628 | Trmt6 | 66926 | HITS-CLIP | | 23142080 | Unspecified |
| mmu-miR-21a-3p | MIMAT0004628 | Trmu | 72026 | HITS-CLIP | | 23142080 | Unspecified |
| mmu-miR-21a-3p | MIMAT0004628 | Trp53bp1 | 27223 | HITS-CLIP | | 23142080 | Unspecified |
| mmu-miR-21a-3p | MIMAT0004628 | Trp53inp1 | 60599 | HITS-CLIP | 23142080\|29167373 | | Unspecified |
| mmu-miR-21a-3p | MIMAT0004628 | Trpm4 | 68667 | HITS-CLIP | | 23142080 | Unspecified |
| mmu-miR-21a-3p | MIMAT0004628 | Trpm7 | 58800 | HITS-CLIP | 23142080\|29167373 | | Unspecified |
| mmu-miR-21a-3p | MIMAT0004628 | Trps1 | 83925 | HITS-CLIP | 23142080\|29167373 | | Unspecified |
| mmu-miR-21a-3p | MIMAT0004628 | Trrap | 100683 | HITS-CLIP | 23142080\|29167373 | | Unspecified |
| mmu-miR-21a-3p | MIMAT0004628 | Tsc1 | 64930 | HITS-CLIP | | 29167373 | Unspecified |
| mmu-miR-21a-3p | MIMAT0004628 | Tsc22d2 | 72033 | HITS-CLIP | | 23142080 | Unspecified |
| mmu-miR-21a-3p | MIMAT0004628 | Tsc22d3 | 14605 | HITS-CLIP | | 23142080 | Unspecified |
| mmu-miR-21a-3p | MIMAT0004628 | Tsfm | 10102 | HITS-CLIP | | 23142080 | Unspecified |
| mmu-miR-21a-3p | MIMAT0004628 | Tsg101 | 22088 | HITS-CLIP | | 23142080 | Unspecified |
| mmu-miR-21a-3p | MIMAT0004628 | Tshz1 | 110796 | HITS-CLIP | | 23142080 | Unspecified |
| mmu-miR-21a-3p | MIMAT0004628 | Tsn | 22099 | HITS-CLIP | | 23142080 | Unspecified |
| mmu-miR-21a-3p | MIMAT0004628 | Tspan13 | 66109 | HITS-CLIP | 37986224\|23142080 | | Unspecified |
| mmu-miR-21a-3p | MIMAT0004628 | Tspan14 | 52588 | HITS-CLIP | 37986224\|23142080 | | Unspecified |
| mmu-miR-21a-3p | MIMAT0004628 | Tspan4 | 64540 | HITS-CLIP | | 23142080 | Unspecified |
| mmu-miR-21a-3p | MIMAT0004628 | Tspan9 | 109246 | HITS-CLIP | | 23142080 | Unspecified |
| mmu-miR-21a-3p | MIMAT0004628 | Tspyl4 | 72480 | HITS-CLIP | | 23142080 | Unspecified |
| mmu-miR-21a-3p | MIMAT0004628 | Ttbk2 | 140810 | HITS-CLIP | | 29167373 | Unspecified |
| mmu-miR-21a-3p | MIMAT0004628 | Ttc27 | 74196 | HITS-CLIP | | 23142080 | Unspecified |
| mmu-miR-21a-3p | MIMAT0004628 | Ttc3 | 22129 | HITS-CLIP | | 23142080 | Unspecified |
| mmu-miR-21a-3p | MIMAT0004628 | Ttc33 | 67515 | HITS-CLIP | | 23142080 | Unspecified |
| mmu-miR-21a-3p | MIMAT0004628 | Ttc37 | 218343 | HITS-CLIP | | 23142080 | Unspecified |
| mmu-miR-21a-3p | MIMAT0004628 | Ttc39b | 69863 | HITS-CLIP | 37986224\|23142080 | | Unspecified |
| mmu-miR-21a-3p | MIMAT0004628 | Ttc7b | 104718 | HITS-CLIP | 29167373\|23142080 | | Unspecified |

| | | | | | | |
|---|---|---|---|---|---|---|
| mmu-miR-21a-3p | MIMAT0004628 | Ttf2 | 74044 | HITS-CLIP | 23142080 | Unspecified |
| mmu-miR-21a-3p | MIMAT0004628 | Ttll12 | 223723 | HITS-CLIP | 23142080 | Unspecified |
| mmu-miR-21a-3p | MIMAT0004628 | Ttyh3 | 78339 | HITS-CLIP | 23142080 | Unspecified |
| mmu-miR-21a-3p | MIMAT0004628 | Tubgcp3 | 259279 | HITS-CLIP | 37986224\|23142080\|29167373 | Unspecified |
| mmu-miR-21a-3p | MIMAT0004628 | Tubgcp5 | 233276 | HITS-CLIP | 29167373\|23142080 | Unspecified |
| mmu-miR-21a-3p | MIMAT0004628 | Tubgcp6 | 328580 | HITS-CLIP | 23142080 | Unspecified |
| mmu-miR-21a-3p | MIMAT0004628 | Tut4 | 23318 | HITS-CLIP | 23142080 | Unspecified |
| mmu-miR-21a-3p | MIMAT0004628 | Txk | 22165 | HITS-CLIP | 23142080 | Unspecified |
| mmu-miR-21a-3p | MIMAT0004628 | Txndc11 | 106200 | HITS-CLIP | 23142080\|29167373 | Unspecified |
| mmu-miR-21a-3p | MIMAT0004628 | Txndc12 | 66073 | HITS-CLIP | 23142080 | Unspecified |
| mmu-miR-21a-3p | MIMAT0004628 | Txndc15 | 69672 | HITS-CLIP | 23142080 | Unspecified |
| mmu-miR-21a-3p | MIMAT0004628 | Txndc17 | 52700 | HITS-CLIP | 23142080 | Unspecified |
| mmu-miR-21a-3p | MIMAT0004628 | Txndc9 | 98258 | HITS-CLIP | 23142080 | Unspecified |
| mmu-miR-21a-3p | MIMAT0004628 | Txnl1 | 53382 | HITS-CLIP | 23142080 | Unspecified |
| mmu-miR-21a-3p | MIMAT0004628 | U2af2 | 22185 | qCLASH | 31393953 | Unspecified |
| mmu-miR-21a-3p | MIMAT0004628 | U2surp | 67958 | qCLASH//HITS-CLIP | 31393953\|23142080\|29167373 | Unspecified |
| mmu-miR-21a-3p | MIMAT0004628 | Uap1 | 107652 | HITS-CLIP | 31433990 | Unspecified |
| mmu-miR-21a-3p | MIMAT0004628 | Uba1 | 22201 | HITS-CLIP | 23142080 | Unspecified |
| mmu-miR-21a-3p | MIMAT0004628 | Uba3 | 22200 | HITS-CLIP | 37986224\|23142080 | Unspecified |
| mmu-miR-21a-3p | MIMAT0004628 | Uba6 | 231380 | HITS-CLIP | 37986224 | Unspecified |
| mmu-miR-21a-3p | MIMAT0004628 | Ubash3b | 72828 | HITS-CLIP | 23142080 | Unspecified |
| mmu-miR-21a-3p | MIMAT0004628 | Ubc | 22190 | qCLASH | 31393953 | Unspecified |
| mmu-miR-21a-3p | MIMAT0004628 | Ube2b | 22210 | HITS-CLIP | 37986224\|23142080\|29167373 | Unspecified |
| mmu-miR-21a-3p | MIMAT0004628 | Ube2d1 | 216080 | HITS-CLIP | 23142080 | Unspecified |
| mmu-miR-21a-3p | MIMAT0004628 | Ube2d2a | 56550 | HITS-CLIP | 23142080\|29167373 | Unspecified |
| mmu-miR-21a-3p | MIMAT0004628 | Ube2g1 | 67128 | HITS-CLIP | 23142080 | Unspecified |
| mmu-miR-21a-3p | MIMAT0004628 | Ube2g2 | 22213 | HITS-CLIP | 23142080 | Unspecified |
| mmu-miR-21a-3p | MIMAT0004628 | Ube2j1 | 56228 | HITS-CLIP | 37986224\|23142080 | Unspecified |
| mmu-miR-21a-3p | MIMAT0004628 | Ube2l3 | 22195 | HITS-CLIP | 23142080 | Unspecified |
| mmu-miR-21a-3p | MIMAT0004628 | Ube2w | 66799 | HITS-CLIP | 23142080 | Unspecified |
| mmu-miR-21a-3p | MIMAT0004628 | Ube2z | 268470 | HITS-CLIP | 23142080 | Unspecified |
| mmu-miR-21a-3p | MIMAT0004628 | Ube3a | 22215 | HITS-CLIP | 23142080 | Unspecified |
| mmu-miR-21a-3p | MIMAT0004628 | Ube3c | 100763 | HITS-CLIP | 23142080 | Unspecified |
| mmu-miR-21a-3p | MIMAT0004628 | Ube4a | 140630 | HITS-CLIP | 23142080\|29167373 | Unspecified |
| mmu-miR-21a-3p | MIMAT0004628 | Ube4b | 63958 | HITS-CLIP | 37986224\|23142080 | Unspecified |
| mmu-miR-21a-3p | MIMAT0004628 | Ubfd1 | 28018 | HITS-CLIP | 37986224\|23142080 | Unspecified |
| mmu-miR-21a-3p | MIMAT0004628 | Ubn1 | 170644 | HITS-CLIP | 23142080 | Unspecified |
| mmu-miR-21a-3p | MIMAT0004628 | Ubn2 | 320538 | HITS-CLIP | 23142080 | Unspecified |
| mmu-miR-21a-3p | MIMAT0004628 | Ubp1 | 22221 | HITS-CLIP | 23142080 | Unspecified |

| | | | | | | |
|---|---|---|---|---|---|---|
| mmu-miR-21a-3p | MIMAT0004628 | Ubqln1 | 56085 | HITS-CLIP | 23142080 | Unspecified |
| mmu-miR-21a-3p | MIMAT0004628 | Ubr2 | 224826 | HITS-CLIP | 23142080 | Unspecified |
| mmu-miR-21a-3p | MIMAT0004628 | Ubr3 | 68795 | HITS-CLIP | 23142080\|29167373 | Unspecified |
| mmu-miR-21a-3p | MIMAT0004628 | Ubr4 | 69116 | HITS-CLIP | 23142080\|29167373 | Unspecified |
| mmu-miR-21a-3p | MIMAT0004628 | Ubr5 | 70790 | HITS-CLIP | 37986224\|23142080 | Unspecified |
| mmu-miR-21a-3p | MIMAT0004628 | Ubxn4 | 67812 | HITS-CLIP | 23142080 | Unspecified |
| mmu-miR-21a-3p | MIMAT0004628 | Uck2 | 80914 | HITS-CLIP | 23142080 | Unspecified |
| mmu-miR-21a-3p | MIMAT0004628 | Ucp2 | 22228 | HITS-CLIP | 23142080\|29167373 | Unspecified |
| mmu-miR-21a-3p | MIMAT0004628 | Ufd1 | 22230 | HITS-CLIP | 23142080\|29167373 | Unspecified |
| mmu-miR-21a-3p | MIMAT0004628 | Ufl1 | 67490 | HITS-CLIP | 23142080 | Unspecified |
| mmu-miR-21a-3p | MIMAT0004628 | Ufm1 | 67890 | HITS-CLIP | 23142080 | Unspecified |
| mmu-miR-21a-3p | MIMAT0004628 | Ugcg | 22234 | HITS-CLIP | 23142080 | Unspecified |
| mmu-miR-21a-3p | MIMAT0004628 | Uggt1 | 320011 | HITS-CLIP | 29167373\|23142080 | Unspecified |
| mmu-miR-21a-3p | MIMAT0004628 | Ugp2 | 216558 | HITS-CLIP | 23142080 | Unspecified |
| mmu-miR-21a-3p | MIMAT0004628 | Ugt1a5 | 394433 | HITS-CLIP | 29167373 | Unspecified |
| mmu-miR-21a-3p | MIMAT0004628 | Uhmk1 | 16589 | HITS-CLIP | 37986224\|23142080\|29167373 | Unspecified |
| mmu-miR-21a-3p | MIMAT0004628 | Uhrf1 | 18140 | HITS-CLIP | 23142080 | Unspecified |
| mmu-miR-21a-3p | MIMAT0004628 | Uhrf1bp1 | 224648 | HITS-CLIP | 23142080 | Unspecified |
| mmu-miR-21a-3p | MIMAT0004628 | Uhrf1bp1l | 75089 | HITS-CLIP | 23142080\|29167373 | Unspecified |
| mmu-miR-21a-3p | MIMAT0004628 | Uhrf2 | 109113 | HITS-CLIP | 23142080\|29167373 | Unspecified |
| mmu-miR-21a-3p | MIMAT0004628 | Ulk1 | 22241 | HITS-CLIP | 23142080 | Unspecified |
| mmu-miR-21a-3p | MIMAT0004628 | Umps | 22247 | HITS-CLIP | 23142080 | Unspecified |
| mmu-miR-21a-3p | MIMAT0004628 | Unc50 | 67387 | HITS-CLIP | 37986224\|23142080 | Unspecified |
| mmu-miR-21a-3p | MIMAT0004628 | Unc93b1 | 54445 | HITS-CLIP | 29167373 | Unspecified |
| mmu-miR-21a-3p | MIMAT0004628 | Upf1 | 19704 | HITS-CLIP | 37986224\|23142080 | Unspecified |
| mmu-miR-21a-3p | MIMAT0004628 | Uqcc1 | 56046 | HITS-CLIP | 23142080\|29167373 | Unspecified |
| mmu-miR-21a-3p | MIMAT0004628 | Uqcr10 | 66152 | HITS-CLIP | 23142080 | Unspecified |
| mmu-miR-21a-3p | MIMAT0004628 | Uqcrc2 | 67003 | HITS-CLIP | 23142080 | Unspecified |
| mmu-miR-21a-3p | MIMAT0004628 | Uri1 | 19777 | HITS-CLIP | 23142080 | Unspecified |
| mmu-miR-21a-3p | MIMAT0004628 | Usf3 | 205717 | HITS-CLIP | 29167373 | Unspecified |
| mmu-miR-21a-3p | MIMAT0004628 | Uso1 | 56041 | HITS-CLIP | 23142080 | Unspecified |
| mmu-miR-21a-3p | MIMAT0004628 | Usp12 | 22217 | HITS-CLIP | 23142080 | Unspecified |
| mmu-miR-21a-3p | MIMAT0004628 | Usp14 | 59025 | HITS-CLIP | 37986224\|23142080 | Unspecified |
| mmu-miR-21a-3p | MIMAT0004628 | Usp15 | 14479 | HITS-CLIP | 37986224\|23142080\|29167373 | Unspecified |
| mmu-miR-21a-3p | MIMAT0004628 | Usp16 | 74112 | HITS-CLIP | 37986224\|23142080 | Unspecified |
| mmu-miR-21a-3p | MIMAT0004628 | Usp19 | 71472 | HITS-CLIP | 37986224\|23142080 | Unspecified |
| mmu-miR-21a-3p | MIMAT0004628 | Usp22 | 216825 | HITS-CLIP | 23142080 | Unspecified |
| mmu-miR-21a-3p | MIMAT0004628 | Usp25 | 30940 | HITS-CLIP | 23142080 | Unspecified |
| mmu-miR-21a-3p | MIMAT0004628 | Usp28 | 235323 | HITS-CLIP | 26203562\|23142080 | Unspecified |

| | | | | | | |
|---|---|---|---|---|---|---|
| mmu-miR-21a-3p | MIMAT0004628 | Usp3 | 235441 | HITS-CLIP | 23142080 | Unspecified |
| mmu-miR-21a-3p | MIMAT0004628 | Usp31 | 76179 | HITS-CLIP | 23142080 | Unspecified |
| mmu-miR-21a-3p | MIMAT0004628 | Usp33 | 170822 | HITS-CLIP | 37986224\|23142080 | Unspecified |
| mmu-miR-21a-3p | MIMAT0004628 | Usp34 | 17847 | HITS-CLIP | 37986224\|23142080 | Unspecified |
| mmu-miR-21a-3p | MIMAT0004628 | Usp37 | 319651 | HITS-CLIP | 37986224\|23142080 | Unspecified |
| mmu-miR-21a-3p | MIMAT0004628 | Usp38 | 74841 | HITS-CLIP | 23142080\|29167373 | Unspecified |
| mmu-miR-21a-3p | MIMAT0004628 | Usp45 | 77593 | HITS-CLIP | 23142080 | Unspecified |
| mmu-miR-21a-3p | MIMAT0004628 | Usp46 | 69727 | HITS-CLIP | 23142080 | Unspecified |
| mmu-miR-21a-3p | MIMAT0004628 | Usp47 | 74996 | HITS-CLIP | 37986224\|23142080 | Unspecified |
| mmu-miR-21a-3p | MIMAT0004628 | Usp7 | 252870 | HITS-CLIP | 23142080 | Unspecified |
| mmu-miR-21a-3p | MIMAT0004628 | Usp8 | 84092 | HITS-CLIP | 37986224\|23142080 | Unspecified |
| mmu-miR-21a-3p | MIMAT0004628 | Usp9x | 22284 | HITS-CLIP | 23142080 | Unspecified |
| mmu-miR-21a-3p | MIMAT0004628 | Utp3 | 65961 | HITS-CLIP | 23142080 | Unspecified |
| mmu-miR-21a-3p | MIMAT0004628 | Utp4 | 21771 | HITS-CLIP | 23142080 | Unspecified |
| mmu-miR-21a-3p | MIMAT0004628 | Utrn | 22288 | HITS-CLIP | 37986224\|23142080 | Unspecified |
| mmu-miR-21a-3p | MIMAT0004628 | Vapa | 30960 | HITS-CLIP | 37986224\|23142080 | Unspecified |
| mmu-miR-21a-3p | MIMAT0004628 | Vapb | 56491 | HITS-CLIP | 29167373 | Unspecified |
| mmu-miR-21a-3p | MIMAT0004628 | Vars | 22321 | HITS-CLIP | 23142080 | Unspecified |
| mmu-miR-21a-3p | MIMAT0004628 | Vasp | 22323 | HITS-CLIP | 23142080 | Unspecified |
| mmu-miR-21a-3p | MIMAT0004628 | Vbp1 | 22327 | HITS-CLIP | 23142080 | Unspecified |
| mmu-miR-21a-3p | MIMAT0004628 | Vcpip1 | 70675 | HITS-CLIP | 23142080\|29167373 | Unspecified |
| mmu-miR-21a-3p | MIMAT0004628 | Vdac2 | 22334 | HITS-CLIP | 23142080 | Unspecified |
| mmu-miR-21a-3p | MIMAT0004628 | Vegfa | 22339 | HITS-CLIP | 37986224\|23142080 | Unspecified |
| mmu-miR-21a-3p | MIMAT0004628 | Vgll4 | 232334 | HITS-CLIP | 37986224\|23142080 | Unspecified |
| mmu-miR-21a-3p | MIMAT0004628 | Vim | 22352 | HITS-CLIP | 29167373\|23142080 | Unspecified |
| mmu-miR-21a-3p | MIMAT0004628 | Virma | 25962 | HITS-CLIP | 23142080\|29167373 | Unspecified |
| mmu-miR-21a-3p | MIMAT0004628 | Vkorc1l1 | 69568 | HITS-CLIP | 37986224\|23142080 | Unspecified |
| mmu-miR-21a-3p | MIMAT0004628 | Vldlr | 22359 | HITS-CLIP | 23142080 | Unspecified |
| mmu-miR-21a-3p | MIMAT0004628 | Vmp1 | 75909 | HITS-CLIP | 23142080 | Unspecified |
| mmu-miR-21a-3p | MIMAT0004628 | Vopp1 | 232023 | HITS-CLIP | 23142080\|29167373 | Unspecified |
| mmu-miR-21a-3p | MIMAT0004628 | Vps13a | 271564 | HITS-CLIP | 23142080\|29167373 | Unspecified |
| mmu-miR-21a-3p | MIMAT0004628 | Vps13c | 320528 | HITS-CLIP | 23142080 | Unspecified |
| mmu-miR-21a-3p | MIMAT0004628 | Vps13d | 230895 | HITS-CLIP | 23142080 | Unspecified |
| mmu-miR-21a-3p | MIMAT0004628 | Vps18 | 228545 | HITS-CLIP | 23142080 | Unspecified |
| mmu-miR-21a-3p | MIMAT0004628 | Vps26b | 69091 | HITS-CLIP | 23142080 | Unspecified |
| mmu-miR-21a-3p | MIMAT0004628 | Vps33b | 233405 | HITS-CLIP | 23142080 | Unspecified |
| mmu-miR-21a-3p | MIMAT0004628 | Vps35 | 65114 | HITS-CLIP | 37986224\|23142080\|29167373 | Unspecified |
| mmu-miR-21a-3p | MIMAT0004628 | Vps36 | 70160 | HITS-CLIP | 23142080 | Unspecified |
| mmu-miR-21a-3p | MIMAT0004628 | Vps37c | 107305 | HITS-CLIP | 23142080 | Unspecified |

| | | | | | | |
|---|---|---|---|---|---|---|
| mmu-miR-21a-3p | MIMAT0004628 | Vps4b | 20479 | HITS-CLIP | 23142080 | Unspecified |
| mmu-miR-21a-3p | MIMAT0004628 | Vps51 | 68505 | HITS-CLIP | 23142080 | Unspecified |
| mmu-miR-21a-3p | MIMAT0004628 | Vps54 | 245944 | HITS-CLIP | 37986224\|23142080 | Unspecified |
| mmu-miR-21a-3p | MIMAT0004628 | Vps8 | 209018 | HITS-CLIP | 23142080 | Unspecified |
| mmu-miR-21a-3p | MIMAT0004628 | Vrk1 | 22367 | HITS-CLIP | 23142080 | Unspecified |
| mmu-miR-21a-3p | MIMAT0004628 | Vsir | 64115 | HITS-CLIP | 23142080\|29167373 | Unspecified |
| mmu-miR-21a-3p | MIMAT0004628 | Vta1 | 66201 | HITS-CLIP | 23142080\|29167373 | Unspecified |
| mmu-miR-21a-3p | MIMAT0004628 | Vti1b | 53612 | HITS-CLIP | 37986224\|23142080\|29167373 | Unspecified |
| mmu-miR-21a-3p | MIMAT0004628 | Wac | 225131 | HITS-CLIP | 23142080 | Unspecified |
| mmu-miR-21a-3p | MIMAT0004628 | Wapl | 23063 | HITS-CLIP | 29167373\|23142080 | Unspecified |
| mmu-miR-21a-3p | MIMAT0004628 | Wars | 22375 | HITS-CLIP | 23142080 | Unspecified |
| mmu-miR-21a-3p | MIMAT0004628 | Washc4 | 319277 | HITS-CLIP | 23142080\|29167373 | Unspecified |
| mmu-miR-21a-3p | MIMAT0004628 | Wasl | 73178 | HITS-CLIP | 23142080 | Unspecified |
| mmu-miR-21a-3p | MIMAT0004628 | Wbp11 | 60321 | HITS-CLIP | 23142080 | Unspecified |
| mmu-miR-21a-3p | MIMAT0004628 | Wbp1l | 226178 | HITS-CLIP | 23142080\|29167373 | Unspecified |
| mmu-miR-21a-3p | MIMAT0004628 | Wdr11 | 207425 | HITS-CLIP | 23142080 | Unspecified |
| mmu-miR-21a-3p | MIMAT0004628 | Wdr20 | 69641 | HITS-CLIP | 23142080 | Unspecified |
| mmu-miR-21a-3p | MIMAT0004628 | Wdr24 | 268933 | HITS-CLIP | 23142080 | Unspecified |
| mmu-miR-21a-3p | MIMAT0004628 | Wdr26 | 226757 | HITS-CLIP | 23142080 | Unspecified |
| mmu-miR-21a-3p | MIMAT0004628 | Wdr3 | 269470 | HITS-CLIP | 23142080 | Unspecified |
| mmu-miR-21a-3p | MIMAT0004628 | Wdr33 | 74320 | HITS-CLIP | 29167373 | Unspecified |
| mmu-miR-21a-3p | MIMAT0004628 | Wdr35 | 74682 | HITS-CLIP | 23142080 | Unspecified |
| mmu-miR-21a-3p | MIMAT0004628 | Wdr36 | 225348 | HITS-CLIP | 29167373\|23142080 | Unspecified |
| mmu-miR-21a-3p | MIMAT0004628 | Wdr37 | 207615 | HITS-CLIP | 37986224\|23142080 | Unspecified |
| mmu-miR-21a-3p | MIMAT0004628 | Wdr44 | 72404 | HITS-CLIP | 23142080 | Unspecified |
| mmu-miR-21a-3p | MIMAT0004628 | Wdr45 | 54636 | HITS-CLIP | 23142080 | Unspecified |
| mmu-miR-21a-3p | MIMAT0004628 | Wdr45b | 66840 | HITS-CLIP | 23142080 | Unspecified |
| mmu-miR-21a-3p | MIMAT0004628 | Wdr47 | 99512 | HITS-CLIP | 23142080 | Unspecified |
| mmu-miR-21a-3p | MIMAT0004628 | Wdr55 | 67936 | HITS-CLIP | 23142080\|29167373 | Unspecified |
| mmu-miR-21a-3p | MIMAT0004628 | Wdr82 | 77305 | HITS-CLIP | 23142080 | Unspecified |
| mmu-miR-21a-3p | MIMAT0004628 | Wdsub1 | 72137 | HITS-CLIP | 23142080 | Unspecified |
| mmu-miR-21a-3p | MIMAT0004628 | Wee1 | 22390 | HITS-CLIP | 29167373\|23142080 | Unspecified |
| mmu-miR-21a-3p | MIMAT0004628 | Wipf1 | 215280 | HITS-CLIP | 37986224\|23142080 | Unspecified |
| mmu-miR-21a-3p | MIMAT0004628 | Wipf2 | 68524 | HITS-CLIP | 23142080\|29167373 | Unspecified |
| mmu-miR-21a-3p | MIMAT0004628 | Wipi2 | 74781 | HITS-CLIP | 23142080 | Unspecified |
| mmu-miR-21a-3p | MIMAT0004628 | Wls | 68151 | HITS-CLIP | 23142080 | Unspecified |
| mmu-miR-21a-3p | MIMAT0004628 | Wnk1 | 232341 | HITS-CLIP | 37986224\|23142080\|29167373 | Unspecified |
| mmu-miR-21a-3p | MIMAT0004628 | Wnt10b | 22410 | HITS-CLIP | 23142080 | Unspecified |
| mmu-miR-21a-3p | MIMAT0004628 | Wsb1 | 78889 | HITS-CLIP | 37986224\|23142080\|29167373 | Unspecified |

| | | | | | | |
|---|---|---|---|---|---|---|
| mmu-miR-21a-3p | MIMAT0004628 | Wtap | 60532 | HITS-CLIP | 37986224\|23142080 | Unspecified |
| mmu-miR-21a-3p | MIMAT0004628 | Xiap | 11798 | HITS-CLIP | 23142080 | Unspecified |
| mmu-miR-21a-3p | MIMAT0004628 | Xkr6 | 219149 | HITS-CLIP | 37986224 | Unspecified |
| mmu-miR-21a-3p | MIMAT0004628 | Xpo1 | 103573 | HITS-CLIP | 23142080\|29167373 | Unspecified |
| mmu-miR-21a-3p | MIMAT0004628 | Xpo4 | 57258 | HITS-CLIP | 23142080\|29167373 | Unspecified |
| mmu-miR-21a-3p | MIMAT0004628 | Xpo5 | 72322 | HITS-CLIP | 23142080 | Unspecified |
| mmu-miR-21a-3p | MIMAT0004628 | Xpo6 | 74204 | HITS-CLIP | 23142080 | Unspecified |
| mmu-miR-21a-3p | MIMAT0004628 | Xpo7 | 65246 | HITS-CLIP | 23142080 | Unspecified |
| mmu-miR-21a-3p | MIMAT0004628 | Xpot | 73192 | HITS-CLIP | 23142080 | Unspecified |
| mmu-miR-21a-3p | MIMAT0004628 | Xxylt1 | 268880 | HITS-CLIP | 23142080 | Unspecified |
| mmu-miR-21a-3p | MIMAT0004628 | Xylt1 | 233781 | HITS-CLIP | 23142080 | Unspecified |
| mmu-miR-21a-3p | MIMAT0004628 | Yars | 107271 | HITS-CLIP | 23142080 | Unspecified |
| mmu-miR-21a-3p | MIMAT0004628 | Yju2 | 853767 | HITS-CLIP | 23142080 | Unspecified |
| mmu-miR-21a-3p | MIMAT0004628 | Ylpm1 | 56531 | HITS-CLIP | 23142080 | Unspecified |
| mmu-miR-21a-3p | MIMAT0004628 | Ypel3 | 66090 | HITS-CLIP | 23142080 | Unspecified |
| mmu-miR-21a-3p | MIMAT0004628 | Ythdc1 | 231386 | HITS-CLIP | 23142080 | Unspecified |
| mmu-miR-21a-3p | MIMAT0004628 | Ythdf1 | 228994 | HITS-CLIP | 23142080 | Unspecified |
| mmu-miR-21a-3p | MIMAT0004628 | Ythdf3 | 229096 | HITS-CLIP | 37986224\|23142080 | Unspecified |
| mmu-miR-21a-3p | MIMAT0004628 | Ywhae | 22627 | HITS-CLIP | 23142080 | Unspecified |
| mmu-miR-21a-3p | MIMAT0004628 | Ywhaz | 22631 | HITS-CLIP | 37986224 | Unspecified |
| mmu-miR-21a-3p | MIMAT0004628 | Yy1 | 22632 | HITS-CLIP | 23142080 | Unspecified |
| mmu-miR-21a-3p | MIMAT0004628 | Zadh2 | 225791 | HITS-CLIP | 23142080 | Unspecified |
| mmu-miR-21a-3p | MIMAT0004628 | Zbed6 | 667118 | HITS-CLIP | 37986224 | Unspecified |
| mmu-miR-21a-3p | MIMAT0004628 | Zbtb1 | 268564 | HITS-CLIP | 23142080 | Unspecified |
| mmu-miR-21a-3p | MIMAT0004628 | Zbtb11 | 271377 | HITS-CLIP | 37986224\|23142080 | Unspecified |
| mmu-miR-21a-3p | MIMAT0004628 | Zbtb14 | 22666 | HITS-CLIP | 23142080 | Unspecified |
| mmu-miR-21a-3p | MIMAT0004628 | Zbtb18 | 30928 | HITS-CLIP | 37986224\|23142080 | Unspecified |
| mmu-miR-21a-3p | MIMAT0004628 | Zbtb20 | 56490 | HITS-CLIP | 31433990\|23142080\|29167373 | Unspecified |
| mmu-miR-21a-3p | MIMAT0004628 | Zbtb33 | 56805 | HITS-CLIP | 23142080 | Unspecified |
| mmu-miR-21a-3p | MIMAT0004628 | Zbtb37 | 240869 | HITS-CLIP | 23142080 | Unspecified |
| mmu-miR-21a-3p | MIMAT0004628 | Zbtb38 | 245007 | HITS-CLIP | 37986224\|23142080 | Unspecified |
| mmu-miR-21a-3p | MIMAT0004628 | Zbtb41 | 226470 | HITS-CLIP | 23142080 | Unspecified |
| mmu-miR-21a-3p | MIMAT0004628 | Zbtb44 | 235132 | HITS-CLIP | 37986224\|23142080 | Unspecified |
| mmu-miR-21a-3p | MIMAT0004628 | Zbtb5 | 230119 | HITS-CLIP | 23142080 | Unspecified |
| mmu-miR-21a-3p | MIMAT0004628 | Zbtb6 | 241322 | HITS-CLIP | 23142080 | Unspecified |
| mmu-miR-21a-3p | MIMAT0004628 | Zbtb8a | 73680 | HITS-CLIP | 23142080 | Unspecified |
| mmu-miR-21a-3p | MIMAT0004628 | Zbtb8os | 67106 | HITS-CLIP | 23142080 | Unspecified |
| mmu-miR-21a-3p | MIMAT0004628 | Zbtb9 | 474156 | HITS-CLIP | 23142080 | Unspecified |
| mmu-miR-21a-3p | MIMAT0004628 | Zc3h15 | 69082 | HITS-CLIP | 23142080 | Unspecified |

| | | | | | | |
|---|---|---|---|---|---|---|
| mmu-miR-21a-3p | MIMAT0004628 | Zc3h6 | 78751 | HITS-CLIP | 37986224\|23142080 | Unspecified |
| mmu-miR-21a-3p | MIMAT0004628 | Zcchc10 | 67966 | HITS-CLIP | 23142080 | Unspecified |
| mmu-miR-21a-3p | MIMAT0004628 | Zdhhc12 | 66220 | HITS-CLIP | 23142080 | Unspecified |
| mmu-miR-21a-3p | MIMAT0004628 | Zdhhc15 | 108672 | HITS-CLIP | 23142080 | Unspecified |
| mmu-miR-21a-3p | MIMAT0004628 | Zdhhc20 | 75965 | HITS-CLIP | 37986224\|23142080\|29167373 | Unspecified |
| mmu-miR-21a-3p | MIMAT0004628 | Zdhhc21 | 68268 | HITS-CLIP | 37986224\|29167373\|23142080 | Unspecified |
| mmu-miR-21a-3p | MIMAT0004628 | Zeb2 | 24136 | HITS-CLIP | 23142080 | Unspecified |
| mmu-miR-21a-3p | MIMAT0004628 | Zfc3h1 | 216345 | HITS-CLIP | 23142080\|29167373 | Unspecified |
| mmu-miR-21a-3p | MIMAT0004628 | Zfp101 | 22643 | HITS-CLIP | 29167373 | Unspecified |
| mmu-miR-21a-3p | MIMAT0004628 | Zfp148 | 22661 | HITS-CLIP | 23142080\|29167373 | Unspecified |
| mmu-miR-21a-3p | MIMAT0004628 | Zfp207 | 22680 | HITS-CLIP | 23142080\|29167373 | Unspecified |
| mmu-miR-21a-3p | MIMAT0004628 | Zfp217 | 228913 | HITS-CLIP | 37986224 | Unspecified |
| mmu-miR-21a-3p | MIMAT0004628 | Zfp236 | 329002 | HITS-CLIP | 23142080 | Unspecified |
| mmu-miR-21a-3p | MIMAT0004628 | Zfp248 | 72720 | HITS-CLIP | 23142080 | Unspecified |
| mmu-miR-21a-3p | MIMAT0004628 | Zfp26 | 22688 | HITS-CLIP | 37986224\|23142080 | Unspecified |
| mmu-miR-21a-3p | MIMAT0004628 | Zfp275 | 27081 | HITS-CLIP | 23142080 | Unspecified |
| mmu-miR-21a-3p | MIMAT0004628 | Zfp280b | 64453 | HITS-CLIP | 37986224 | Unspecified |
| mmu-miR-21a-3p | MIMAT0004628 | Zfp280c | 208968 | HITS-CLIP | 23142080 | Unspecified |
| mmu-miR-21a-3p | MIMAT0004628 | Zfp280d | 235469 | HITS-CLIP | 23142080 | Unspecified |
| mmu-miR-21a-3p | MIMAT0004628 | Zfp281 | 226442 | HITS-CLIP | 23142080 | Unspecified |
| mmu-miR-21a-3p | MIMAT0004628 | Zfp292 | 30046 | HITS-CLIP | 29167373\|23142080 | Unspecified |
| mmu-miR-21a-3p | MIMAT0004628 | Zfp318 | 57908 | HITS-CLIP | 23142080 | Unspecified |
| mmu-miR-21a-3p | MIMAT0004628 | Zfp319 | 79233 | HITS-CLIP | 23142080 | Unspecified |
| mmu-miR-21a-3p | MIMAT0004628 | Zfp322a | 218100 | HITS-CLIP | 23142080 | Unspecified |
| mmu-miR-21a-3p | MIMAT0004628 | Zfp329 | 67230 | HITS-CLIP | 23142080 | Unspecified |
| mmu-miR-21a-3p | MIMAT0004628 | Zfp36 | 22695 | HITS-CLIP | 23142080 | Unspecified |
| mmu-miR-21a-3p | MIMAT0004628 | Zfp367 | 238673 | HITS-CLIP | 23142080 | Unspecified |
| mmu-miR-21a-3p | MIMAT0004628 | Zfp36l2 | 12193 | HITS-CLIP | 23142080 | Unspecified |
| mmu-miR-21a-3p | MIMAT0004628 | Zfp407 | 240476 | HITS-CLIP | 23142080 | Unspecified |
| mmu-miR-21a-3p | MIMAT0004628 | Zfp445 | 235682 | HITS-CLIP | 23142080 | Unspecified |
| mmu-miR-21a-3p | MIMAT0004628 | Zfp451 | 98403 | HITS-CLIP | 23142080 | Unspecified |
| mmu-miR-21a-3p | MIMAT0004628 | Zfp507 | 668501 | HITS-CLIP | 23142080 | Unspecified |
| mmu-miR-21a-3p | MIMAT0004628 | Zfp512 | 269639 | HITS-CLIP | 23142080 | Unspecified |
| mmu-miR-21a-3p | MIMAT0004628 | Zfp512b | 269401 | HITS-CLIP | 23142080 | Unspecified |
| mmu-miR-21a-3p | MIMAT0004628 | Zfp516 | 329003 | HITS-CLIP | 23142080 | Unspecified |
| mmu-miR-21a-3p | MIMAT0004628 | Zfp518a | 72672 | HITS-CLIP | 23142080 | Unspecified |
| mmu-miR-21a-3p | MIMAT0004628 | Zfp592 | 233410 | HITS-CLIP | 23142080 | Unspecified |
| mmu-miR-21a-3p | MIMAT0004628 | Zfp609 | 214812 | HITS-CLIP | 23142080 | Unspecified |
| mmu-miR-21a-3p | MIMAT0004628 | Zfp622 | 52521 | HITS-CLIP | 23142080 | Unspecified |

| | | | | | | |
|---|---|---|---|---|---|---|
| mmu-miR-21a-3p | MIMAT0004628 | Zfp638 | 27332 | HITS-CLIP | 23142080 | Unspecified |
| mmu-miR-21a-3p | MIMAT0004628 | Zfp644 | 52397 | HITS-CLIP | 23142080 | Unspecified |
| mmu-miR-21a-3p | MIMAT0004628 | Zfp646 | 233905 | HITS-CLIP | 23142080\|29167373 | Unspecified |
| mmu-miR-21a-3p | MIMAT0004628 | Zfp652 | 268469 | HITS-CLIP | 23142080 | Unspecified |
| mmu-miR-21a-3p | MIMAT0004628 | Zfp655 | 72611 | HITS-CLIP | 23142080\|29167373 | Unspecified |
| mmu-miR-21a-3p | MIMAT0004628 | Zfp706 | 68036 | HITS-CLIP | 37986224\|23142080 | Unspecified |
| mmu-miR-21a-3p | MIMAT0004628 | Zfp715 | 69930 | HITS-CLIP | 29167373\|23142080 | Unspecified |
| mmu-miR-21a-3p | MIMAT0004628 | Zfp740 | 68744 | HITS-CLIP | 23142080 | Unspecified |
| mmu-miR-21a-3p | MIMAT0004628 | Zfp746 | 69228 | HITS-CLIP | 23142080 | Unspecified |
| mmu-miR-21a-3p | MIMAT0004628 | Zfp770 | 228491 | HITS-CLIP | 29167373 | Unspecified |
| mmu-miR-21a-3p | MIMAT0004628 | Zfp773 | 76373 | HITS-CLIP | 23142080 | Unspecified |
| mmu-miR-21a-3p | MIMAT0004628 | Zfp827 | 622675 | HITS-CLIP | 23142080 | Unspecified |
| mmu-miR-21a-3p | MIMAT0004628 | Zfp871 | 208292 | HITS-CLIP | 37986224\|23142080 | Unspecified |
| mmu-miR-21a-3p | MIMAT0004628 | Zfp91 | 109910 | HITS-CLIP | 29167373\|23142080 | Unspecified |
| mmu-miR-21a-3p | MIMAT0004628 | Zfp983 | 73229 | HITS-CLIP | 23142080 | Unspecified |
| mmu-miR-21a-3p | MIMAT0004628 | Zfr | 22763 | HITS-CLIP | 37986224\|23142080\|29167373 | Unspecified |
| mmu-miR-21a-3p | MIMAT0004628 | Zfx | 22764 | HITS-CLIP | 23142080 | Unspecified |
| mmu-miR-21a-3p | MIMAT0004628 | Zfyve16 | 218441 | HITS-CLIP | 23142080 | Unspecified |
| mmu-miR-21a-3p | MIMAT0004628 | Zhx1 | 22770 | HITS-CLIP | 23142080\|29167373 | Unspecified |
| mmu-miR-21a-3p | MIMAT0004628 | Zkscan1 | 74570 | HITS-CLIP | 23142080 | Unspecified |
| mmu-miR-21a-3p | MIMAT0004628 | Zmiz1 | 328365 | HITS-CLIP | 23142080 | Unspecified |
| mmu-miR-21a-3p | MIMAT0004628 | Zmym1 | 68310 | HITS-CLIP | 23142080 | Unspecified |
| mmu-miR-21a-3p | MIMAT0004628 | Zmym2 | 76007 | HITS-CLIP | 23142080 | Unspecified |
| mmu-miR-21a-3p | MIMAT0004628 | Zmym3 | 56364 | HITS-CLIP | 23142080 | Unspecified |
| mmu-miR-21a-3p | MIMAT0004628 | Zmym4 | 67785 | HITS-CLIP | 23142080 | Unspecified |
| mmu-miR-21a-3p | MIMAT0004628 | Zmym6 | 100177 | HITS-CLIP | 29167373 | Unspecified |
| mmu-miR-21a-3p | MIMAT0004628 | Zmynd11 | 66505 | HITS-CLIP | 23142080 | Unspecified |
| mmu-miR-21a-3p | MIMAT0004628 | Zmynd8 | 228880 | HITS-CLIP | 23142080 | Unspecified |
| mmu-miR-21a-3p | MIMAT0004628 | Znrf1 | 170737 | HITS-CLIP | 23142080 | Unspecified |
| mmu-miR-21a-3p | MIMAT0004628 | Znrf2 | 387524 | HITS-CLIP | 23142080 | Unspecified |
| mmu-miR-21a-3p | MIMAT0004628 | Znrf3 | 407821 | HITS-CLIP | 23142080 | Unspecified |
| mmu-miR-21a-3p | MIMAT0004628 | Zranb1 | 360216 | HITS-CLIP | 23142080 | Unspecified |
| mmu-miR-21a-3p | MIMAT0004628 | Zwint | 52696 | HITS-CLIP | 23142080 | Unspecified |
| mmu-miR-21a-3p | MIMAT0004628 | Zxdb | 668166 | HITS-CLIP | 23142080 | Unspecified |
| mmu-miR-21a-3p | MIMAT0004628 | Zyg11b | 414872 | HITS-CLIP | 23142080 | Unspecified |
| mmu-miR-21a-3p | MIMAT0004628 | Zzef1 | 195018 | HITS-CLIP | 23142080 | Unspecified |
| mmu-miR-193a-5p | MIMAT0004544 | Actn4 | 60595 | HITS-CLIP | 37986224 | Unspecified |
| mmu-miR-193a-5p | MIMAT0004544 | Anln | 68743 | HITS-CLIP | 37986224 | Unspecified |
| mmu-miR-193a-5p | MIMAT0004544 | Anp32b | 67628 | HITS-CLIP | 37986224 | Unspecified |

| | | | | | | |
|---|---|---|---|---|---|---|
| mmu-miR-193a-5p | MIMAT0004544 | Brat1 | 231841 | HITS-CLIP | 37986224 | Unspecified |
| mmu-miR-193a-5p | MIMAT0004544 | Cct7 | 12468 | HITS-CLIP | 37986224 | Unspecified |
| mmu-miR-193a-5p | MIMAT0004544 | Dtymk | 21915 | HITS-CLIP | 31433990 | Unspecified |
| mmu-miR-193a-5p | MIMAT0004544 | Ercc6l | 236930 | HITS-CLIP | 37986224 | Unspecified |
| mmu-miR-193a-5p | MIMAT0004544 | Fxyd5 | 18301 | HITS-CLIP | 31433990 | Unspecified |
| mmu-miR-193a-5p | MIMAT0004544 | Homer2 | 26557 | HITS-CLIP | 37986224 | Unspecified |
| mmu-miR-193a-5p | MIMAT0004544 | Il6st | 16195 | HITS-CLIP | 31433990 | Unspecified |
| mmu-miR-193a-5p | MIMAT0004544 | Kpna4 | 16649 | HITS-CLIP | 37986224 | Unspecified |
| mmu-miR-193a-5p | MIMAT0004544 | Lifr | 16880 | HITS-CLIP | 37986224 | Unspecified |
| mmu-miR-193a-5p | MIMAT0004544 | Nop10 | 66181 | HITS-CLIP | 37986224 | Unspecified |
| mmu-miR-193a-5p | MIMAT0004544 | Nsd1 | 18193 | HITS-CLIP | 37986224 | Unspecified |
| mmu-miR-193a-5p | MIMAT0004544 | Oxsr1 | 108737 | HITS-CLIP | 37986224 | Unspecified |
| mmu-miR-193a-5p | MIMAT0004544 | Pdik1l | 230809 | HITS-CLIP | 31433990 | Unspecified |
| mmu-miR-193a-5p | MIMAT0004544 | Plekha7 | 233765 | HITS-CLIP | 37986224 | Unspecified |
| mmu-miR-193a-5p | MIMAT0004544 | Sema3f | 20350 | HITS-CLIP | 37986224 | Unspecified |
| mmu-miR-193a-5p | MIMAT0004544 | Sf3b3 | 101943 | HITS-CLIP | 31433990 | Unspecified |
| mmu-miR-193a-5p | MIMAT0004544 | Srrm2 | 75956 | HITS-CLIP | 37986224 | Unspecified |
| mmu-miR-193a-5p | MIMAT0004544 | Top2a | 21973 | HITS-CLIP | 31433990 | Unspecified |
| mmu-miR-193a-5p | MIMAT0004544 | Zfp280b | 64453 | HITS-CLIP | 37986224 | Unspecified |
| mmu-miR-30c-2-3p | MIMAT0005438 | Abi2 | 329165 | HITS-CLIP | 37986224 | Unspecified |
| mmu-miR-30c-2-3p | MIMAT0005438 | Cbx7 | 52609 | HITS-CLIP | 37986224 | Unspecified |
| mmu-miR-30c-2-3p | MIMAT0005438 | Csnk1a1 | 93687 | HITS-CLIP | 37986224 | Unspecified |
| mmu-miR-30c-2-3p | MIMAT0005438 | Ctbp2 | 13017 | HITS-CLIP | 37986224 | Unspecified |
| mmu-miR-30c-2-3p | MIMAT0005438 | Ddx3x | 13205 | HITS-CLIP | 37986224 | Unspecified |
| mmu-miR-30c-2-3p | MIMAT0005438 | Dlgap4 | 228836 | HITS-CLIP | 37986224 | Unspecified |
| mmu-miR-30c-2-3p | MIMAT0005438 | Dnajc1 | 13418 | HITS-CLIP | 37986224 | Unspecified |
| mmu-miR-30c-2-3p | MIMAT0005438 | Enc1 | 13803 | HITS-CLIP | 31433990 | Unspecified |
| mmu-miR-30c-2-3p | MIMAT0005438 | Ikzf1 | 22778 | HITS-CLIP | 31433990 | Unspecified |
| mmu-miR-30c-2-3p | MIMAT0005438 | Mafg | 17134 | HITS-CLIP | 37986224 | Unspecified |
| mmu-miR-30c-2-3p | MIMAT0005438 | Mtrex | 23517 | HITS-CLIP | 31433990 | Unspecified |
| mmu-miR-30c-2-3p | MIMAT0005438 | Papola | 18789 | HITS-CLIP | 31433990 | Unspecified |
| mmu-miR-30c-2-3p | MIMAT0005438 | Pbrm1 | 66923 | HITS-CLIP | 31433990 | Unspecified |
| mmu-miR-30c-2-3p | MIMAT0005438 | Psmd14 | 59029 | HITS-CLIP | 31433990 | Unspecified |
| mmu-miR-30c-2-3p | MIMAT0005438 | Rai1 | 19377 | HITS-CLIP | 37986224 | Unspecified |
| mmu-miR-30c-2-3p | MIMAT0005438 | Rpl32 | 19951 | HITS-CLIP | 26203562 | Unspecified |
| mmu-miR-30c-2-3p | MIMAT0005438 | Sin3a | 20466 | HITS-CLIP | 37986224 | Unspecified |
| mmu-miR-30c-2-3p | MIMAT0005438 | Tnpo3 | 320938 | HITS-CLIP | 37986224 | Unspecified |
| mmu-let-7i-3p | MIMAT0004520 | 1810013L24Rik | 69053 | HITS-CLIP | 29167373 | Unspecified |
| mmu-let-7i-3p | MIMAT0004520 | 2510009E07Rik | 72190 | HITS-CLIP | 23597149 | Unspecified |

| | | | | | | |
|---|---|---|---|---|---|---|
| mmu-let-7i-3p | MIMAT0004520 | 9930021J03Rik | 240613 | HITS-CLIP | 23597149 | Unspecified |
| mmu-let-7i-3p | MIMAT0004520 | Abhd2 | 54608 | HITS-CLIP | 23597149 | Unspecified |
| mmu-let-7i-3p | MIMAT0004520 | Abi1 | 11308 | HITS-CLIP | 29167373 | Unspecified |
| mmu-let-7i-3p | MIMAT0004520 | Acadl | 11363 | HITS-CLIP | 29167373 | Unspecified |
| mmu-let-7i-3p | MIMAT0004520 | Acat1 | 110446 | HITS-CLIP | 23597149 | Unspecified |
| mmu-let-7i-3p | MIMAT0004520 | Acly | 104112 | HITS-CLIP | 23597149 | Unspecified |
| mmu-let-7i-3p | MIMAT0004520 | Aco1 | 11428 | HITS-CLIP | 23597149 | Unspecified |
| mmu-let-7i-3p | MIMAT0004520 | Acox1 | 11430 | HITS-CLIP | 23597149 | Unspecified |
| mmu-let-7i-3p | MIMAT0004520 | Acss2 | 60525 | HITS-CLIP | 23597149 | Unspecified |
| mmu-let-7i-3p | MIMAT0004520 | Actb | 11461 | HITS-CLIP | 37986224\|23597149 | Unspecified |
| mmu-let-7i-3p | MIMAT0004520 | Actn4 | 60595 | HITS-CLIP | 23597149 | Unspecified |
| mmu-let-7i-3p | MIMAT0004520 | Actr1a | 54130 | HITS-CLIP | 37986224 | Unspecified |
| mmu-let-7i-3p | MIMAT0004520 | Actr1b | 226977 | HITS-CLIP | 29167373 | Unspecified |
| mmu-let-7i-3p | MIMAT0004520 | Adi1 | 104923 | HITS-CLIP | 23597149 | Unspecified |
| mmu-let-7i-3p | MIMAT0004520 | Adsl | 11564 | HITS-CLIP | 23597149 | Unspecified |
| mmu-let-7i-3p | MIMAT0004520 | Aff4 | 93736 | HITS-CLIP | 29167373 | Unspecified |
| mmu-let-7i-3p | MIMAT0004520 | Afm | 280662 | HITS-CLIP | 23597149 | Unspecified |
| mmu-let-7i-3p | MIMAT0004520 | Ago2 | 239528 | HITS-CLIP | 29167373 | Unspecified |
| mmu-let-7i-3p | MIMAT0004520 | Ahctf1 | 226747 | HITS-CLIP | 29167373 | Unspecified |
| mmu-let-7i-3p | MIMAT0004520 | Akap13 | 75547 | HITS-CLIP | 23597149 | Unspecified |
| mmu-let-7i-3p | MIMAT0004520 | Akirin2 | 433693 | HITS-CLIP | 23597149 | Unspecified |
| mmu-let-7i-3p | MIMAT0004520 | Alas2 | 11656 | HITS-CLIP | 23597149 | Unspecified |
| mmu-let-7i-3p | MIMAT0004520 | Alb | 213 | HITS-CLIP | 23597149 | Unspecified |
| mmu-let-7i-3p | MIMAT0004520 | Aldh1a1 | 11668 | HITS-CLIP | 23597149 | Unspecified |
| mmu-let-7i-3p | MIMAT0004520 | Aldh1a7 | 26358 | HITS-CLIP | 23597149 | Unspecified |
| mmu-let-7i-3p | MIMAT0004520 | Aldh1l1 | 107747 | HITS-CLIP | 23597149 | Unspecified |
| mmu-let-7i-3p | MIMAT0004520 | Aldh2 | 11669 | HITS-CLIP | 23597149 | Unspecified |
| mmu-let-7i-3p | MIMAT0004520 | Alg9 | 102580 | HITS-CLIP | 23597149 | Unspecified |
| mmu-let-7i-3p | MIMAT0004520 | Ankrd17 | 81702 | HITS-CLIP | 23597149 | Unspecified |
| mmu-let-7i-3p | MIMAT0004520 | Anxa2 | 12306 | HITS-CLIP | 29167373 | Unspecified |
| mmu-let-7i-3p | MIMAT0004520 | Ap2a2 | 11772 | HITS-CLIP | 37986224\|23597149 | Unspecified |
| mmu-let-7i-3p | MIMAT0004520 | Ap2s1 | 232910 | HITS-CLIP | 23597149 | Unspecified |
| mmu-let-7i-3p | MIMAT0004520 | Apc | 11789 | HITS-CLIP | 29167373 | Unspecified |
| mmu-let-7i-3p | MIMAT0004520 | Appl1 | 72993 | HITS-CLIP | 29167373 | Unspecified |
| mmu-let-7i-3p | MIMAT0004520 | Arf4 | 11843 | HITS-CLIP | 29167373 | Unspecified |
| mmu-let-7i-3p | MIMAT0004520 | Arfgef2 | 99371 | HITS-CLIP | 29167373 | Unspecified |
| mmu-let-7i-3p | MIMAT0004520 | Arhgap35 | 232906 | HITS-CLIP | 23597149 | Unspecified |
| mmu-let-7i-3p | MIMAT0004520 | Arhgdia | 192662 | HITS-CLIP | 29167373 | Unspecified |
| mmu-let-7i-3p | MIMAT0004520 | Arhgef26 | 622434 | HITS-CLIP | 37986224 | Unspecified |

| | | | | | | |
|---|---|---|---|---|---|---|
| mmu-let-7i-3p | MIMAT0004520 | Arl4c | 320982 | HITS-CLIP | 29167373 | Unspecified |
| mmu-let-7i-3p | MIMAT0004520 | Arpc1a | 56443 | HITS-CLIP | 23597149 | Unspecified |
| mmu-let-7i-3p | MIMAT0004520 | Arpc2 | 76709 | HITS-CLIP | 23597149 | Unspecified |
| mmu-let-7i-3p | MIMAT0004520 | Asah2 | 54447 | HITS-CLIP | 29167373 | Unspecified |
| mmu-let-7i-3p | MIMAT0004520 | Atf7ip | 54343 | HITS-CLIP | 23597149\|29167373 | Unspecified |
| mmu-let-7i-3p | MIMAT0004520 | Atp1b1 | 11931 | HITS-CLIP | 23597149 | Unspecified |
| mmu-let-7i-3p | MIMAT0004520 | Atp2a2 | 11938 | HITS-CLIP | 23597149\|29167373 | Unspecified |
| mmu-let-7i-3p | MIMAT0004520 | Atp2b1 | 67972 | HITS-CLIP | 29167373 | Unspecified |
| mmu-let-7i-3p | MIMAT0004520 | Atp5a1 | 11946 | HITS-CLIP | 23597149 | Unspecified |
| mmu-let-7i-3p | MIMAT0004520 | Atp5b | 11947 | HITS-CLIP | 37986224 | Unspecified |
| mmu-let-7i-3p | MIMAT0004520 | Atp5d | 66043 | HITS-CLIP | 23597149 | Unspecified |
| mmu-let-7i-3p | MIMAT0004520 | Atp6v1b2 | 11966 | HITS-CLIP | 23597149 | Unspecified |
| mmu-let-7i-3p | MIMAT0004520 | Atp6v1c1 | 66335 | HITS-CLIP | 23597149 | Unspecified |
| mmu-let-7i-3p | MIMAT0004520 | Atp6v1e1 | 11973 | HITS-CLIP | 23597149 | Unspecified |
| mmu-let-7i-3p | MIMAT0004520 | Atp6v1h | 108664 | HITS-CLIP | 23597149 | Unspecified |
| mmu-let-7i-3p | MIMAT0004520 | Atp8a1 | 11980 | HITS-CLIP | 23597149\|29167373 | Unspecified |
| mmu-let-7i-3p | MIMAT0004520 | BC005537 | 79555 | HITS-CLIP | 29167373 | Unspecified |
| mmu-let-7i-3p | MIMAT0004520 | Bahcc1 | 268515 | HITS-CLIP | 23597149 | Unspecified |
| mmu-let-7i-3p | MIMAT0004520 | Baiap2 | 108100 | HITS-CLIP | 37986224 | Unspecified |
| mmu-let-7i-3p | MIMAT0004520 | Bcl6 | 12053 | HITS-CLIP | 29167373 | Unspecified |
| mmu-let-7i-3p | MIMAT0004520 | Bcorl1 | 320376 | HITS-CLIP | 23597149 | Unspecified |
| mmu-let-7i-3p | MIMAT0004520 | Bhlhe40 | 20893 | HITS-CLIP | 37986224\|29167373 | Unspecified |
| mmu-let-7i-3p | MIMAT0004520 | Btf3 | 218490 | HITS-CLIP | 23597149 | Unspecified |
| mmu-let-7i-3p | MIMAT0004520 | C9 | 12279 | HITS-CLIP | 23597149 | Unspecified |
| mmu-let-7i-3p | MIMAT0004520 | Cacul1 | 78832 | HITS-CLIP | 29167373 | Unspecified |
| mmu-let-7i-3p | MIMAT0004520 | Camk1d | 227541 | HITS-CLIP | 29167373 | Unspecified |
| mmu-let-7i-3p | MIMAT0004520 | Caprin1 | 53872 | HITS-CLIP | 23597149 | Unspecified |
| mmu-let-7i-3p | MIMAT0004520 | Cat | 12359 | HITS-CLIP | 23597149 | Unspecified |
| mmu-let-7i-3p | MIMAT0004520 | Cav2 | 12390 | HITS-CLIP | 29167373 | Unspecified |
| mmu-let-7i-3p | MIMAT0004520 | Cblc | 80794 | HITS-CLIP | 23597149 | Unspecified |
| mmu-let-7i-3p | MIMAT0004520 | Ccdc25 | 67179 | HITS-CLIP | 23597149\|29167373 | Unspecified |
| mmu-let-7i-3p | MIMAT0004520 | Ccdc6 | 76551 | HITS-CLIP | 29167373 | Unspecified |
| mmu-let-7i-3p | MIMAT0004520 | Ccnt1 | 12455 | HITS-CLIP | 29167373 | Unspecified |
| mmu-let-7i-3p | MIMAT0004520 | Cct2 | 12461 | HITS-CLIP | 37986224 | Unspecified |
| mmu-let-7i-3p | MIMAT0004520 | Cct6a | 12466 | qCLASH | 31393953 | Unspecified |
| mmu-let-7i-3p | MIMAT0004520 | Cd164 | 53599 | HITS-CLIP//qCLASH | 23597149\|31393953 | Unspecified |
| mmu-let-7i-3p | MIMAT0004520 | Cd84 | 12523 | HITS-CLIP | 29167373 | Unspecified |
| mmu-let-7i-3p | MIMAT0004520 | Cdk14 | 18647 | HITS-CLIP | 29167373 | Unspecified |
| mmu-let-7i-3p | MIMAT0004520 | Cdk16 | 18555 | HITS-CLIP | 23597149 | Unspecified |

| | | | | | | | |
|---|---|---|---|---|---|---|---|
| mmu-let-7i-3p | MIMAT0004520 | Cdk17 | 237459 | HITS-CLIP | | 29167373 | Unspecified |
| mmu-let-7i-3p | MIMAT0004520 | Cdk5rap3 | 80280 | HITS-CLIP | | 23597149 | Unspecified |
| mmu-let-7i-3p | MIMAT0004520 | Cdkn1b | 12576 | HITS-CLIP | | 29167373 | Unspecified |
| mmu-let-7i-3p | MIMAT0004520 | Cdr2l | 237988 | HITS-CLIP | | 23597149 | Unspecified |
| mmu-let-7i-3p | MIMAT0004520 | Cdv3 | 321022 | HITS-CLIP | 23597149\|29167373 | | Unspecified |
| mmu-let-7i-3p | MIMAT0004520 | Cebpa | 12606 | HITS-CLIP | | 23597149 | Unspecified |
| mmu-let-7i-3p | MIMAT0004520 | Cebpb | 12608 | HITS-CLIP | | 23597149 | Unspecified |
| mmu-let-7i-3p | MIMAT0004520 | Cebpz | 12607 | HITS-CLIP | | 23597149 | Unspecified |
| mmu-let-7i-3p | MIMAT0004520 | Celf1 | 13046 | HITS-CLIP | | 29167373 | Unspecified |
| mmu-let-7i-3p | MIMAT0004520 | Celf2 | 14007 | HITS-CLIP | | 29167373 | Unspecified |
| mmu-let-7i-3p | MIMAT0004520 | Cept1 | 99712 | HITS-CLIP | | 29167373 | Unspecified |
| mmu-let-7i-3p | MIMAT0004520 | Cggbp1 | 106143 | HITS-CLIP | | 29167373 | Unspecified |
| mmu-let-7i-3p | MIMAT0004520 | Cgnl1 | 68178 | HITS-CLIP | 23597149\|29167373 | | Unspecified |
| mmu-let-7i-3p | MIMAT0004520 | Chd7 | 320790 | HITS-CLIP | | 29167373 | Unspecified |
| mmu-let-7i-3p | MIMAT0004520 | Cherp | 27967 | HITS-CLIP | | 23597149 | Unspecified |
| mmu-let-7i-3p | MIMAT0004520 | Chmp1a | 234852 | HITS-CLIP | | 23597149 | Unspecified |
| mmu-let-7i-3p | MIMAT0004520 | Chmp2b | 68942 | HITS-CLIP | | 29167373 | Unspecified |
| mmu-let-7i-3p | MIMAT0004520 | Ciapin1 | 109006 | HITS-CLIP | | 23597149 | Unspecified |
| mmu-let-7i-3p | MIMAT0004520 | Cic | 71722 | HITS-CLIP | | 29167373 | Unspecified |
| mmu-let-7i-3p | MIMAT0004520 | Clasp2 | 76499 | HITS-CLIP | | 29167373 | Unspecified |
| mmu-let-7i-3p | MIMAT0004520 | Clec5a | 23845 | HITS-CLIP | | 29167373 | Unspecified |
| mmu-let-7i-3p | MIMAT0004520 | Clpx | 270166 | HITS-CLIP | 37986224\|23597149 | | Unspecified |
| mmu-let-7i-3p | MIMAT0004520 | Cluh | 74148 | HITS-CLIP | | 23597149 | Unspecified |
| mmu-let-7i-3p | MIMAT0004520 | Cnot1 | 234594 | HITS-CLIP | | 23597149 | Unspecified |
| mmu-let-7i-3p | MIMAT0004520 | Copg1 | 54161 | HITS-CLIP | | 23597149 | Unspecified |
| mmu-let-7i-3p | MIMAT0004520 | Cops2 | 12848 | HITS-CLIP | | 29167373 | Unspecified |
| mmu-let-7i-3p | MIMAT0004520 | Coq8a | 56997 | HITS-CLIP | | 23597149 | Unspecified |
| mmu-let-7i-3p | MIMAT0004520 | Cox14 | 66379 | HITS-CLIP | | 37986224 | Unspecified |
| mmu-let-7i-3p | MIMAT0004520 | Cox5a | 12858 | HITS-CLIP | | 23597149 | Unspecified |
| mmu-let-7i-3p | MIMAT0004520 | Cp | 12870 | HITS-CLIP | | 23597149 | Unspecified |
| mmu-let-7i-3p | MIMAT0004520 | Cpne2 | 234577 | HITS-CLIP | | 23597149 | Unspecified |
| mmu-let-7i-3p | MIMAT0004520 | Cpt1a | 12894 | HITS-CLIP | | 23597149 | Unspecified |
| mmu-let-7i-3p | MIMAT0004520 | Crat | 12908 | HITS-CLIP | | 23597149 | Unspecified |
| mmu-let-7i-3p | MIMAT0004520 | Creb5 | 231991 | HITS-CLIP | | 29167373 | Unspecified |
| mmu-let-7i-3p | MIMAT0004520 | Crk | 12928 | HITS-CLIP | | 23597149 | Unspecified |
| mmu-let-7i-3p | MIMAT0004520 | Crot | 74114 | HITS-CLIP | | 23597149 | Unspecified |
| mmu-let-7i-3p | MIMAT0004520 | Cry1 | 12952 | HITS-CLIP | | 23597149 | Unspecified |
| mmu-let-7i-3p | MIMAT0004520 | Csrp1 | 13007 | HITS-CLIP | | 23597149 | Unspecified |
| mmu-let-7i-3p | MIMAT0004520 | Ctnna1 | 12385 | HITS-CLIP | | 29167373 | Unspecified |

| | | | | | | |
|---|---|---|---|---|---|---|
| mmu-let-7i-3p | MIMAT0004520 | Ctsa | 19025 | HITS-CLIP | 23597149 | Unspecified |
| mmu-let-7i-3p | MIMAT0004520 | Cttnbp2nl | 80281 | HITS-CLIP | 29167373 | Unspecified |
| mmu-let-7i-3p | MIMAT0004520 | Cul4a | 99375 | HITS-CLIP | 23597149 | Unspecified |
| mmu-let-7i-3p | MIMAT0004520 | Daam1 | 208846 | HITS-CLIP | 23597149 | Unspecified |
| mmu-let-7i-3p | MIMAT0004520 | Dap | 223453 | HITS-CLIP | 23597149 | Unspecified |
| mmu-let-7i-3p | MIMAT0004520 | Dapk1 | 69635 | HITS-CLIP | 23597149 | Unspecified |
| mmu-let-7i-3p | MIMAT0004520 | Dctn1 | 13191 | HITS-CLIP | 23597149 | Unspecified |
| mmu-let-7i-3p | MIMAT0004520 | Dctn3 | 53598 | HITS-CLIP | 29167373 | Unspecified |
| mmu-let-7i-3p | MIMAT0004520 | Ddi2 | 68817 | HITS-CLIP | 23597149 | Unspecified |
| mmu-let-7i-3p | MIMAT0004520 | Ddx17 | 67040 | HITS-CLIP | 23597149 | Unspecified |
| mmu-let-7i-3p | MIMAT0004520 | Ddx39b | 53817 | HITS-CLIP | 37986224 | Unspecified |
| mmu-let-7i-3p | MIMAT0004520 | Decr1 | 67460 | HITS-CLIP | 23597149 | Unspecified |
| mmu-let-7i-3p | MIMAT0004520 | Depdc5 | 277854 | HITS-CLIP | 23597149 | Unspecified |
| mmu-let-7i-3p | MIMAT0004520 | Dhcr24 | 74754 | HITS-CLIP | 23597149 | Unspecified |
| mmu-let-7i-3p | MIMAT0004520 | Dkc1 | 245474 | HITS-CLIP | 23597149 | Unspecified |
| mmu-let-7i-3p | MIMAT0004520 | Dlg3 | 53310 | HITS-CLIP | 37986224 | Unspecified |
| mmu-let-7i-3p | MIMAT0004520 | Dmxl1 | 240283 | HITS-CLIP | 29167373 | Unspecified |
| mmu-let-7i-3p | MIMAT0004520 | Dnajb12 | 56709 | HITS-CLIP | 23597149 | Unspecified |
| mmu-let-7i-3p | MIMAT0004520 | Dock7 | 67299 | HITS-CLIP | 29167373 | Unspecified |
| mmu-let-7i-3p | MIMAT0004520 | Dpy19l1 | 244745 | HITS-CLIP | 23597149 | Unspecified |
| mmu-let-7i-3p | MIMAT0004520 | Dsg2 | 13511 | HITS-CLIP | 23597149 | Unspecified |
| mmu-let-7i-3p | MIMAT0004520 | E2f4 | 104394 | HITS-CLIP | 23597149 | Unspecified |
| mmu-let-7i-3p | MIMAT0004520 | Eci2 | 23986 | HITS-CLIP | 23597149 | Unspecified |
| mmu-let-7i-3p | MIMAT0004520 | Ecpas | 23392 | HITS-CLIP | 23597149 | Unspecified |
| mmu-let-7i-3p | MIMAT0004520 | Eea1 | 216238 | HITS-CLIP | 29167373 | Unspecified |
| mmu-let-7i-3p | MIMAT0004520 | Eef2 | 13629 | HITS-CLIP | 23597149 | Unspecified |
| mmu-let-7i-3p | MIMAT0004520 | Efr3a | 76740 | HITS-CLIP | 23597149 | Unspecified |
| mmu-let-7i-3p | MIMAT0004520 | Egfr | 13649 | HITS-CLIP | 23597149 | Unspecified |
| mmu-let-7i-3p | MIMAT0004520 | Egr1 | 13653 | HITS-CLIP | 37986224 | Unspecified |
| mmu-let-7i-3p | MIMAT0004520 | Eif3e | 16341 | HITS-CLIP | 23597149 | Unspecified |
| mmu-let-7i-3p | MIMAT0004520 | Eif4g1 | 208643 | HITS-CLIP | 23597149\|29167373 | Unspecified |
| mmu-let-7i-3p | MIMAT0004520 | Eif5a2 | 208691 | HITS-CLIP | 29167373 | Unspecified |
| mmu-let-7i-3p | MIMAT0004520 | Ell2 | 192657 | HITS-CLIP | 23597149\|29167373 | Unspecified |
| mmu-let-7i-3p | MIMAT0004520 | Eloa | 6924 | HITS-CLIP | 23597149 | Unspecified |
| mmu-let-7i-3p | MIMAT0004520 | Emc1 | 230866 | HITS-CLIP | 23597149 | Unspecified |
| mmu-let-7i-3p | MIMAT0004520 | Ep400 | 75560 | HITS-CLIP | 29167373 | Unspecified |
| mmu-let-7i-3p | MIMAT0004520 | Eps8 | 13860 | HITS-CLIP | 23597149 | Unspecified |
| mmu-let-7i-3p | MIMAT0004520 | Erap1 | 80898 | HITS-CLIP | 37986224 | Unspecified |
| mmu-let-7i-3p | MIMAT0004520 | Erbb3 | 13867 | HITS-CLIP | 23597149 | Unspecified |

| | | | | | | |
|---|---|---|---|---|---|---|
| mmu-let-7i-3p | MIMAT0004520 | Erp44 | 76299 | HITS-CLIP | 23597149 | Unspecified |
| mmu-let-7i-3p | MIMAT0004520 | Etnppl | 71760 | HITS-CLIP | 23597149 | Unspecified |
| mmu-let-7i-3p | MIMAT0004520 | Etv3 | 27049 | HITS-CLIP | 23597149 | Unspecified |
| mmu-let-7i-3p | MIMAT0004520 | F2 | 14061 | HITS-CLIP | 23597149 | Unspecified |
| mmu-let-7i-3p | MIMAT0004520 | Faah | 14073 | HITS-CLIP | 23597149 | Unspecified |
| mmu-let-7i-3p | MIMAT0004520 | Fam193a | 231128 | HITS-CLIP | 29167373 | Unspecified |
| mmu-let-7i-3p | MIMAT0004520 | Far1 | 67420 | HITS-CLIP | 29167373 | Unspecified |
| mmu-let-7i-3p | MIMAT0004520 | Fbp1 | 14121 | HITS-CLIP | 23597149 | Unspecified |
| mmu-let-7i-3p | MIMAT0004520 | Fbxo21 | 231670 | HITS-CLIP | 23597149 | Unspecified |
| mmu-let-7i-3p | MIMAT0004520 | Fbxo45 | 268882 | HITS-CLIP | 23597149 | Unspecified |
| mmu-let-7i-3p | MIMAT0004520 | Fh1 | 14194 | HITS-CLIP | 23597149 | Unspecified |
| mmu-let-7i-3p | MIMAT0004520 | Flna | 192176 | HITS-CLIP | 29167373 | Unspecified |
| mmu-let-7i-3p | MIMAT0004520 | Fmnl2 | 71409 | HITS-CLIP | 23597149 | Unspecified |
| mmu-let-7i-3p | MIMAT0004520 | Fn1 | 14268 | HITS-CLIP | 23597149 | Unspecified |
| mmu-let-7i-3p | MIMAT0004520 | Fndc3b | 72007 | HITS-CLIP | 23597149 | Unspecified |
| mmu-let-7i-3p | MIMAT0004520 | Fnip1 | 216742 | HITS-CLIP | 29167373 | Unspecified |
| mmu-let-7i-3p | MIMAT0004520 | Fos | 14281 | HITS-CLIP | 29167373 | Unspecified |
| mmu-let-7i-3p | MIMAT0004520 | Foxk1 | 17425 | HITS-CLIP | 37986224\|23597149 | Unspecified |
| mmu-let-7i-3p | MIMAT0004520 | Foxk2 | 68837 | HITS-CLIP | 29167373 | Unspecified |
| mmu-let-7i-3p | MIMAT0004520 | Foxo3 | 56484 | HITS-CLIP | 23597149 | Unspecified |
| mmu-let-7i-3p | MIMAT0004520 | Foxp1 | 108655 | HITS-CLIP | 29167373 | Unspecified |
| mmu-let-7i-3p | MIMAT0004520 | Foxp2 | 114142 | HITS-CLIP | 23597149 | Unspecified |
| mmu-let-7i-3p | MIMAT0004520 | G3bp1 | 27041 | HITS-CLIP | 29167373 | Unspecified |
| mmu-let-7i-3p | MIMAT0004520 | G3bp2 | 23881 | HITS-CLIP | 23597149 | Unspecified |
| mmu-let-7i-3p | MIMAT0004520 | Gan | 209239 | HITS-CLIP | 29167373 | Unspecified |
| mmu-let-7i-3p | MIMAT0004520 | Gc | 14473 | HITS-CLIP | 23597149 | Unspecified |
| mmu-let-7i-3p | MIMAT0004520 | Gcn1l1 | 10985 | HITS-CLIP | 23597149 | Unspecified |
| mmu-let-7i-3p | MIMAT0004520 | Gcnt2 | 14538 | HITS-CLIP | 23597149 | Unspecified |
| mmu-let-7i-3p | MIMAT0004520 | Gfpt1 | 14583 | HITS-CLIP | 23597149\|29167373 | Unspecified |
| mmu-let-7i-3p | MIMAT0004520 | Ggnbp2 | 217039 | HITS-CLIP | 23597149 | Unspecified |
| mmu-let-7i-3p | MIMAT0004520 | Ghitm | 66092 | HITS-CLIP | 23597149\|29167373 | Unspecified |
| mmu-let-7i-3p | MIMAT0004520 | Git1 | 216963 | HITS-CLIP | 23597149\|29167373 | Unspecified |
| mmu-let-7i-3p | MIMAT0004520 | Gja4 | 14612 | HITS-CLIP | 23597149 | Unspecified |
| mmu-let-7i-3p | MIMAT0004520 | Gls | 14660 | HITS-CLIP | 29167373 | Unspecified |
| mmu-let-7i-3p | MIMAT0004520 | Gmfb | 63985 | HITS-CLIP | 29167373 | Unspecified |
| mmu-let-7i-3p | MIMAT0004520 | Gne | 50798 | HITS-CLIP | 23597149 | Unspecified |
| mmu-let-7i-3p | MIMAT0004520 | Gopc | 94221 | HITS-CLIP | 23597149\|29167373 | Unspecified |
| mmu-let-7i-3p | MIMAT0004520 | Gpam | 14732 | HITS-CLIP | 23597149 | Unspecified |
| mmu-let-7i-3p | MIMAT0004520 | Gpcpd1 | 74182 | HITS-CLIP | 23597149 | Unspecified |

| | | | | | | | |
|---|---|---|---|---|---|---|---|
| mmu-let-7i-3p | MIMAT0004520 | Gpnmb | 93695 | HITS-CLIP | | 29167373 | Unspecified |
| mmu-let-7i-3p | MIMAT0004520 | Gpr39 | 71111 | HITS-CLIP | | 23597149 | Unspecified |
| mmu-let-7i-3p | MIMAT0004520 | Grhpr | 76238 | HITS-CLIP | | 23597149 | Unspecified |
| mmu-let-7i-3p | MIMAT0004520 | Gsk3b | 56637 | HITS-CLIP | | 29167373 | Unspecified |
| mmu-let-7i-3p | MIMAT0004520 | Gspt1 | 14852 | HITS-CLIP | | 29167373 | Unspecified |
| mmu-let-7i-3p | MIMAT0004520 | Gstz1 | 14874 | HITS-CLIP | | 23597149 | Unspecified |
| mmu-let-7i-3p | MIMAT0004520 | Gtf2h4 | 14885 | HITS-CLIP | | 23597149 | Unspecified |
| mmu-let-7i-3p | MIMAT0004520 | Gtf3c3 | 98488 | HITS-CLIP | | 29167373 | Unspecified |
| mmu-let-7i-3p | MIMAT0004520 | Gys2 | 232493 | HITS-CLIP | | 23597149 | Unspecified |
| mmu-let-7i-3p | MIMAT0004520 | Gzf1 | 74533 | HITS-CLIP | | 23597149 | Unspecified |
| mmu-let-7i-3p | MIMAT0004520 | Habp2 | 226243 | HITS-CLIP | | 23597149 | Unspecified |
| mmu-let-7i-3p | MIMAT0004520 | Hacd3 | 57874 | HITS-CLIP | | 23597149 | Unspecified |
| mmu-let-7i-3p | MIMAT0004520 | Hagh | 14651 | HITS-CLIP | | 23597149 | Unspecified |
| mmu-let-7i-3p | MIMAT0004520 | Hc | 15139 | HITS-CLIP | | 23597149 | Unspecified |
| mmu-let-7i-3p | MIMAT0004520 | Hcfc1 | 15161 | HITS-CLIP | | 23597149 | Unspecified |
| mmu-let-7i-3p | MIMAT0004520 | Hectd1 | 207304 | HITS-CLIP | | 23597149 | Unspecified |
| mmu-let-7i-3p | MIMAT0004520 | Herpud1 | 64209 | qCLASH | | 31393953 | Unspecified |
| mmu-let-7i-3p | MIMAT0004520 | Hhex | 15242 | HITS-CLIP | | 23597149 | Unspecified |
| mmu-let-7i-3p | MIMAT0004520 | Hmg20a | 66867 | HITS-CLIP | | 23597149 | Unspecified |
| mmu-let-7i-3p | MIMAT0004520 | Hnf4a | 15378 | HITS-CLIP | | 23597149 | Unspecified |
| mmu-let-7i-3p | MIMAT0004520 | Hnrnpdl | 50926 | HITS-CLIP | | 29167373 | Unspecified |
| mmu-let-7i-3p | MIMAT0004520 | Hnrnph1 | 59013 | HITS-CLIP | | 29167373 | Unspecified |
| mmu-let-7i-3p | MIMAT0004520 | Hnrnph3 | 432467 | HITS-CLIP | | 23597149 | Unspecified |
| mmu-let-7i-3p | MIMAT0004520 | Hnrnpk | 15387 | HITS-CLIP | | 29167373 | Unspecified |
| mmu-let-7i-3p | MIMAT0004520 | Hnrnpu | 51810 | HITS-CLIP | 23597149\|29167373 | | Unspecified |
| mmu-let-7i-3p | MIMAT0004520 | Hook3 | 320191 | HITS-CLIP | 23597149\|29167373 | | Unspecified |
| mmu-let-7i-3p | MIMAT0004520 | Hpd | 15445 | HITS-CLIP | | 23597149 | Unspecified |
| mmu-let-7i-3p | MIMAT0004520 | Hsd17b12 | 56348 | HITS-CLIP | | 23597149 | Unspecified |
| mmu-let-7i-3p | MIMAT0004520 | Iah1 | 67732 | HITS-CLIP | | 23597149 | Unspecified |
| mmu-let-7i-3p | MIMAT0004520 | Iars | 105148 | HITS-CLIP | | 23597149 | Unspecified |
| mmu-let-7i-3p | MIMAT0004520 | Idh3g | 15929 | HITS-CLIP | | 23597149 | Unspecified |
| mmu-let-7i-3p | MIMAT0004520 | Inhba | 16323 | HITS-CLIP | | 23597149 | Unspecified |
| mmu-let-7i-3p | MIMAT0004520 | Inmt | 21743 | HITS-CLIP | | 23597149 | Unspecified |
| mmu-let-7i-3p | MIMAT0004520 | Ints6l | 236790 | HITS-CLIP | | 29167373 | Unspecified |
| mmu-let-7i-3p | MIMAT0004520 | Ip6k2 | 76500 | HITS-CLIP | | 23597149 | Unspecified |
| mmu-let-7i-3p | MIMAT0004520 | Ipo7 | 233726 | HITS-CLIP | 23597149\|29167373 | | Unspecified |
| mmu-let-7i-3p | MIMAT0004520 | Ist1 | 71955 | HITS-CLIP | | 23597149 | Unspecified |
| mmu-let-7i-3p | MIMAT0004520 | Itgb1 | 16412 | HITS-CLIP | 23597149\|29167373 | | Unspecified |
| mmu-let-7i-3p | MIMAT0004520 | Itih4 | 16427 | HITS-CLIP | | 23597149 | Unspecified |

| | | | | | | |
|---|---|---|---|---|---|---|
| mmu-let-7i-3p | MIMAT0004520 | Itsn1 | 16443 | HITS-CLIP | 29167373 | Unspecified |
| mmu-let-7i-3p | MIMAT0004520 | Ivns1abp | 117198 | HITS-CLIP | 29167373 | Unspecified |
| mmu-let-7i-3p | MIMAT0004520 | Jmjd6 | 107817 | HITS-CLIP | 23597149 | Unspecified |
| mmu-let-7i-3p | MIMAT0004520 | Jun | 16476 | HITS-CLIP | 29167373 | Unspecified |
| mmu-let-7i-3p | MIMAT0004520 | Kbtbd2 | 210973 | HITS-CLIP | 23597149 | Unspecified |
| mmu-let-7i-3p | MIMAT0004520 | Kcmf1 | 74287 | HITS-CLIP | 23597149 | Unspecified |
| mmu-let-7i-3p | MIMAT0004520 | Kdelr2 | 66913 | HITS-CLIP | 29167373 | Unspecified |
| mmu-let-7i-3p | MIMAT0004520 | Kdr | 16542 | HITS-CLIP | 23597149 | Unspecified |
| mmu-let-7i-3p | MIMAT0004520 | Kidins220 | 77480 | HITS-CLIP | 29167373 | Unspecified |
| mmu-let-7i-3p | MIMAT0004520 | Klf10 | 21847 | HITS-CLIP | 29167373 | Unspecified |
| mmu-let-7i-3p | MIMAT0004520 | Klf2 | 16598 | HITS-CLIP | 23597149 | Unspecified |
| mmu-let-7i-3p | MIMAT0004520 | Klf6 | 23849 | HITS-CLIP | 29167373 | Unspecified |
| mmu-let-7i-3p | MIMAT0004520 | Kmt2c | 231051 | HITS-CLIP | 29167373 | Unspecified |
| mmu-let-7i-3p | MIMAT0004520 | Kmt2d | 381022 | HITS-CLIP | 23597149 | Unspecified |
| mmu-let-7i-3p | MIMAT0004520 | Kng1 | 16644 | HITS-CLIP | 23597149 | Unspecified |
| mmu-let-7i-3p | MIMAT0004520 | Kpna4 | 16649 | HITS-CLIP | 23597149 | Unspecified |
| mmu-let-7i-3p | MIMAT0004520 | Kpnb1 | 16211 | HITS-CLIP | 23597149 | Unspecified |
| mmu-let-7i-3p | MIMAT0004520 | Krt8 | 16691 | HITS-CLIP | 23597149 | Unspecified |
| mmu-let-7i-3p | MIMAT0004520 | Lamb1 | 16777 | HITS-CLIP | 23597149 | Unspecified |
| mmu-let-7i-3p | MIMAT0004520 | Lats1 | 16798 | HITS-CLIP | 23597149\|29167373 | Unspecified |
| mmu-let-7i-3p | MIMAT0004520 | Lats2 | 50523 | HITS-CLIP | 29167373 | Unspecified |
| mmu-let-7i-3p | MIMAT0004520 | Lcor | 212391 | HITS-CLIP | 23597149 | Unspecified |
| mmu-let-7i-3p | MIMAT0004520 | Ldha | 16828 | HITS-CLIP | 29167373 | Unspecified |
| mmu-let-7i-3p | MIMAT0004520 | Lin7c | 22343 | HITS-CLIP | 29167373 | Unspecified |
| mmu-let-7i-3p | MIMAT0004520 | Lipa | 16889 | HITS-CLIP | 29167373 | Unspecified |
| mmu-let-7i-3p | MIMAT0004520 | Litaf | 56722 | HITS-CLIP | 23597149 | Unspecified |
| mmu-let-7i-3p | MIMAT0004520 | Lpin2 | 64898 | HITS-CLIP | 23597149 | Unspecified |
| mmu-let-7i-3p | MIMAT0004520 | Lrch3 | 70144 | HITS-CLIP | 29167373 | Unspecified |
| mmu-let-7i-3p | MIMAT0004520 | Lrif1 | 321000 | HITS-CLIP | 29167373 | Unspecified |
| mmu-let-7i-3p | MIMAT0004520 | Lrp1 | 16971 | HITS-CLIP | 23597149\|29167373 | Unspecified |
| mmu-let-7i-3p | MIMAT0004520 | Lrrc58 | 320184 | HITS-CLIP | 29167373 | Unspecified |
| mmu-let-7i-3p | MIMAT0004520 | Lsm8 | 76522 | HITS-CLIP | 29167373 | Unspecified |
| mmu-let-7i-3p | MIMAT0004520 | Lss | 16987 | HITS-CLIP | 23597149 | Unspecified |
| mmu-let-7i-3p | MIMAT0004520 | Lyn | 17096 | HITS-CLIP | 29167373 | Unspecified |
| mmu-let-7i-3p | MIMAT0004520 | Macf1 | 11426 | HITS-CLIP | 29167373 | Unspecified |
| mmu-let-7i-3p | MIMAT0004520 | Maip1 | 79568 | HITS-CLIP | 23597149 | Unspecified |
| mmu-let-7i-3p | MIMAT0004520 | Mal2 | 105853 | HITS-CLIP | 23597149 | Unspecified |
| mmu-let-7i-3p | MIMAT0004520 | Map2k4 | 26398 | HITS-CLIP | 23597149 | Unspecified |
| mmu-let-7i-3p | MIMAT0004520 | Map3k11 | 26403 | HITS-CLIP | 29167373 | Unspecified |

| | | | | | | | |
|---|---|---|---|---|---|---|---|
| mmu-let-7i-3p | MIMAT0004520 | Mapk1 | 26413 | HITS-CLIP | | 29167373 | Unspecified |
| mmu-let-7i-3p | MIMAT0004520 | Marcks | 17118 | HITS-CLIP | | 29167373 | Unspecified |
| mmu-let-7i-3p | MIMAT0004520 | Marf1 | 223989 | HITS-CLIP | | 29167373 | Unspecified |
| mmu-let-7i-3p | MIMAT0004520 | Matr3 | 17184 | HITS-CLIP | | 23597149 | Unspecified |
| mmu-let-7i-3p | MIMAT0004520 | Mbnl2 | 105559 | HITS-CLIP | | 29167373 | Unspecified |
| mmu-let-7i-3p | MIMAT0004520 | Mcfd2 | 193813 | HITS-CLIP | | 23597149 | Unspecified |
| mmu-let-7i-3p | MIMAT0004520 | Mcm3 | 17215 | HITS-CLIP | | 37986224 | Unspecified |
| mmu-let-7i-3p | MIMAT0004520 | Med13 | 327987 | HITS-CLIP | | 23597149 | Unspecified |
| mmu-let-7i-3p | MIMAT0004520 | Med25 | 75613 | HITS-CLIP | | 23597149 | Unspecified |
| mmu-let-7i-3p | MIMAT0004520 | Met | 17295 | HITS-CLIP | | 23597149 | Unspecified |
| mmu-let-7i-3p | MIMAT0004520 | Mettl7b | 71664 | HITS-CLIP | | 23597149 | Unspecified |
| mmu-let-7i-3p | MIMAT0004520 | Mfn1 | 67414 | HITS-CLIP | | 23597149 | Unspecified |
| mmu-let-7i-3p | MIMAT0004520 | Mfsd5 | 106073 | HITS-CLIP | | 29167373 | Unspecified |
| mmu-let-7i-3p | MIMAT0004520 | Mfsd6 | 98682 | HITS-CLIP | | 26203562 | Unspecified |
| mmu-let-7i-3p | MIMAT0004520 | Mgrn1 | 17237 | HITS-CLIP | 23597149\|29167373 | | Unspecified |
| mmu-let-7i-3p | MIMAT0004520 | Mgst1 | 56615 | HITS-CLIP | | 23597149 | Unspecified |
| mmu-let-7i-3p | MIMAT0004520 | Midn | 59090 | HITS-CLIP | 23597149\|29167373 | | Unspecified |
| mmu-let-7i-3p | MIMAT0004520 | Mknk2 | 17347 | HITS-CLIP | | 29167373 | Unspecified |
| mmu-let-7i-3p | MIMAT0004520 | Mlec | 109154 | HITS-CLIP | | 23597149 | Unspecified |
| mmu-let-7i-3p | MIMAT0004520 | Mllt10 | 17354 | HITS-CLIP | | 29167373 | Unspecified |
| mmu-let-7i-3p | MIMAT0004520 | Mms19 | 72199 | HITS-CLIP | | 23597149 | Unspecified |
| mmu-let-7i-3p | MIMAT0004520 | Mob4 | 19070 | HITS-CLIP | | 23597149 | Unspecified |
| mmu-let-7i-3p | MIMAT0004520 | Mon1b | 270096 | HITS-CLIP | | 23597149 | Unspecified |
| mmu-let-7i-3p | MIMAT0004520 | Mosmo | 233812 | HITS-CLIP | | 29167373 | Unspecified |
| mmu-let-7i-3p | MIMAT0004520 | Mpv17l | 93734 | HITS-CLIP | | 23597149 | Unspecified |
| mmu-let-7i-3p | MIMAT0004520 | Mroh1 | 223658 | HITS-CLIP | | 23597149 | Unspecified |
| mmu-let-7i-3p | MIMAT0004520 | Mrps24 | 64660 | HITS-CLIP | | 23597149 | Unspecified |
| mmu-let-7i-3p | MIMAT0004520 | Mrps9 | 69527 | HITS-CLIP | | 23597149 | Unspecified |
| mmu-let-7i-3p | MIMAT0004520 | Mtmr12 | 268783 | HITS-CLIP | | 29167373 | Unspecified |
| mmu-let-7i-3p | MIMAT0004520 | Mug1 | 17836 | HITS-CLIP | | 23597149 | Unspecified |
| mmu-let-7i-3p | MIMAT0004520 | Myo18a | 360013 | HITS-CLIP | | 23597149 | Unspecified |
| mmu-let-7i-3p | MIMAT0004520 | Myo1c | 17913 | HITS-CLIP | | 29167373 | Unspecified |
| mmu-let-7i-3p | MIMAT0004520 | Myo1f | 17916 | HITS-CLIP | | 29167373 | Unspecified |
| mmu-let-7i-3p | MIMAT0004520 | Myo5a | 17918 | HITS-CLIP | | 29167373 | Unspecified |
| mmu-let-7i-3p | MIMAT0004520 | Nab1 | 17936 | HITS-CLIP | | 29167373 | Unspecified |
| mmu-let-7i-3p | MIMAT0004520 | Narf | 67608 | HITS-CLIP | | 23597149 | Unspecified |
| mmu-let-7i-3p | MIMAT0004520 | Nceh1 | 320024 | HITS-CLIP | | 29167373 | Unspecified |
| mmu-let-7i-3p | MIMAT0004520 | Ncoa3 | 17979 | HITS-CLIP | | 29167373 | Unspecified |
| mmu-let-7i-3p | MIMAT0004520 | Ndor1 | 78797 | HITS-CLIP | | 29167373 | Unspecified |

| | | | | | | | |
|---|---|---|---|---|---|---|---|
| mmu-let-7i-3p | MIMAT0004520 | Ndrg2 | 29811 | HITS-CLIP | | 23597149 | Unspecified |
| mmu-let-7i-3p | MIMAT0004520 | Necap1 | 67602 | HITS-CLIP | | 29167373 | Unspecified |
| mmu-let-7i-3p | MIMAT0004520 | Nek6 | 59126 | HITS-CLIP | | 23597149 | Unspecified |
| mmu-let-7i-3p | MIMAT0004520 | Nelfb | 58202 | HITS-CLIP | | 23597149 | Unspecified |
| mmu-let-7i-3p | MIMAT0004520 | Nfat5 | 54446 | HITS-CLIP | | 29167373 | Unspecified |
| mmu-let-7i-3p | MIMAT0004520 | Nfib | 18028 | HITS-CLIP | | 23597149 | Unspecified |
| mmu-let-7i-3p | MIMAT0004520 | Nfil3 | 18030 | HITS-CLIP | | 31433990 | Unspecified |
| mmu-let-7i-3p | MIMAT0004520 | Nfkbid | 243910 | HITS-CLIP | | 29167373 | Unspecified |
| mmu-let-7i-3p | MIMAT0004520 | Nfxl1 | 100978 | HITS-CLIP | | 29167373 | Unspecified |
| mmu-let-7i-3p | MIMAT0004520 | Nol4l | 329540 | HITS-CLIP | | 29167373 | Unspecified |
| mmu-let-7i-3p | MIMAT0004520 | Nr2f6 | 13864 | HITS-CLIP | | 37986224 | Unspecified |
| mmu-let-7i-3p | MIMAT0004520 | Nrros | 224109 | HITS-CLIP | | 31433990 | Unspecified |
| mmu-let-7i-3p | MIMAT0004520 | Nup98 | 269966 | HITS-CLIP | | 23597149 | Unspecified |
| mmu-let-7i-3p | MIMAT0004520 | Oaf | 102644 | HITS-CLIP | | 23597149 | Unspecified |
| mmu-let-7i-3p | MIMAT0004520 | Ociad2 | 433904 | HITS-CLIP | | 29167373 | Unspecified |
| mmu-let-7i-3p | MIMAT0004520 | Ola1 | 67059 | HITS-CLIP | 23597149\|29167373 | | Unspecified |
| mmu-let-7i-3p | MIMAT0004520 | Os9 | 216440 | HITS-CLIP | | 23597149 | Unspecified |
| mmu-let-7i-3p | MIMAT0004520 | Otulinl | 54491 | HITS-CLIP | | 29167373 | Unspecified |
| mmu-let-7i-3p | MIMAT0004520 | Pacs2 | 217893 | HITS-CLIP | | 29167373 | Unspecified |
| mmu-let-7i-3p | MIMAT0004520 | Pank1 | 75735 | HITS-CLIP | | 23597149 | Unspecified |
| mmu-let-7i-3p | MIMAT0004520 | Parg | 26430 | HITS-CLIP | | 29167373 | Unspecified |
| mmu-let-7i-3p | MIMAT0004520 | Pbld1 | 68371 | HITS-CLIP | | 23597149 | Unspecified |
| mmu-let-7i-3p | MIMAT0004520 | Pcca | 110821 | HITS-CLIP | | 23597149 | Unspecified |
| mmu-let-7i-3p | MIMAT0004520 | Pced1a | 319513 | HITS-CLIP | | 23597149 | Unspecified |
| mmu-let-7i-3p | MIMAT0004520 | Pcmt1 | 18537 | HITS-CLIP | | 29167373 | Unspecified |
| mmu-let-7i-3p | MIMAT0004520 | Pcx | 18563 | HITS-CLIP | | 23597149 | Unspecified |
| mmu-let-7i-3p | MIMAT0004520 | Pcyt2 | 68671 | HITS-CLIP | | 23597149 | Unspecified |
| mmu-let-7i-3p | MIMAT0004520 | Pdcd6ip | 18571 | HITS-CLIP | | 23597149 | Unspecified |
| mmu-let-7i-3p | MIMAT0004520 | Pde4dip | 83679 | HITS-CLIP | | 23597149 | Unspecified |
| mmu-let-7i-3p | MIMAT0004520 | Pdk4 | 27273 | HITS-CLIP | | 26203562 | Unspecified |
| mmu-let-7i-3p | MIMAT0004520 | Pdlim5 | 56376 | HITS-CLIP | | 29167373 | Unspecified |
| mmu-let-7i-3p | MIMAT0004520 | Pds5a | 71521 | HITS-CLIP | | 29167373 | Unspecified |
| mmu-let-7i-3p | MIMAT0004520 | Peli1 | 67245 | HITS-CLIP | | 31433990 | Unspecified |
| mmu-let-7i-3p | MIMAT0004520 | Phb2 | 12034 | HITS-CLIP | | 23597149 | Unspecified |
| mmu-let-7i-3p | MIMAT0004520 | Phc3 | 241915 | HITS-CLIP | | 29167373 | Unspecified |
| mmu-let-7i-3p | MIMAT0004520 | Phlpp2 | 244650 | HITS-CLIP | | 23597149 | Unspecified |
| mmu-let-7i-3p | MIMAT0004520 | Picalm | 233489 | HITS-CLIP | | 23597149 | Unspecified |
| mmu-let-7i-3p | MIMAT0004520 | Pik3cb | 74769 | HITS-CLIP | | 29167373 | Unspecified |
| mmu-let-7i-3p | MIMAT0004520 | Pip5k1c | 18717 | HITS-CLIP | 23597149\|29167373 | | Unspecified |

| | | | | | | |
|---|---|---|---|---|---|---|
| mmu-let-7i-3p | MIMAT0004520 | Pitpnm2 | 19679 | HITS-CLIP | 23597149 | Unspecified |
| mmu-let-7i-3p | MIMAT0004520 | Pklr | 18770 | HITS-CLIP | 23597149 | Unspecified |
| mmu-let-7i-3p | MIMAT0004520 | Plau | 18792 | HITS-CLIP | 29167373 | Unspecified |
| mmu-let-7i-3p | MIMAT0004520 | Plcxd2 | 433022 | HITS-CLIP | 23597149 | Unspecified |
| mmu-let-7i-3p | MIMAT0004520 | Plec | 18810 | HITS-CLIP | 29167373 | Unspecified |
| mmu-let-7i-3p | MIMAT0004520 | Plod1 | 18822 | HITS-CLIP | 23597149 | Unspecified |
| mmu-let-7i-3p | MIMAT0004520 | Plxnd1 | 67784 | HITS-CLIP | 23597149\|29167373 | Unspecified |
| mmu-let-7i-3p | MIMAT0004520 | Pnkp | 59047 | HITS-CLIP | 23597149 | Unspecified |
| mmu-let-7i-3p | MIMAT0004520 | Pnpla7 | 241274 | HITS-CLIP | 23597149 | Unspecified |
| mmu-let-7i-3p | MIMAT0004520 | Polr2b | 231329 | HITS-CLIP | 29167373 | Unspecified |
| mmu-let-7i-3p | MIMAT0004520 | Ppa1 | 67895 | HITS-CLIP | 23597149 | Unspecified |
| mmu-let-7i-3p | MIMAT0004520 | Ppara | 19013 | HITS-CLIP | 23597149 | Unspecified |
| mmu-let-7i-3p | MIMAT0004520 | Ppp1cb | 19046 | HITS-CLIP | 29167373 | Unspecified |
| mmu-let-7i-3p | MIMAT0004520 | Ppp1r12a | 17931 | HITS-CLIP | 29167373\|23597149 | Unspecified |
| mmu-let-7i-3p | MIMAT0004520 | Ppp6r1 | 243819 | HITS-CLIP | 29167373 | Unspecified |
| mmu-let-7i-3p | MIMAT0004520 | Ppp6r3 | 52036 | HITS-CLIP | 29167373 | Unspecified |
| mmu-let-7i-3p | MIMAT0004520 | Ppt1 | 19063 | HITS-CLIP | 29167373 | Unspecified |
| mmu-let-7i-3p | MIMAT0004520 | Pptc7 | 320717 | HITS-CLIP | 23597149 | Unspecified |
| mmu-let-7i-3p | MIMAT0004520 | Pqlc3 | 217430 | HITS-CLIP | 29167373 | Unspecified |
| mmu-let-7i-3p | MIMAT0004520 | Prep | 19072 | HITS-CLIP | 29167373 | Unspecified |
| mmu-let-7i-3p | MIMAT0004520 | Prkcd | 18753 | HITS-CLIP | 29167373 | Unspecified |
| mmu-let-7i-3p | MIMAT0004520 | Prox1 | 19130 | HITS-CLIP | 37986224 | Unspecified |
| mmu-let-7i-3p | MIMAT0004520 | Proz | 66901 | HITS-CLIP | 23597149 | Unspecified |
| mmu-let-7i-3p | MIMAT0004520 | Prpf40a | 56194 | HITS-CLIP | 37986224 | Unspecified |
| mmu-let-7i-3p | MIMAT0004520 | Prr11 | 270906 | HITS-CLIP | 23597149 | Unspecified |
| mmu-let-7i-3p | MIMAT0004520 | Psen1 | 19164 | HITS-CLIP | 29167373 | Unspecified |
| mmu-let-7i-3p | MIMAT0004520 | Psmb4 | 19172 | HITS-CLIP | 29167373 | Unspecified |
| mmu-let-7i-3p | MIMAT0004520 | Psmb8 | 16913 | HITS-CLIP | 23597149 | Unspecified |
| mmu-let-7i-3p | MIMAT0004520 | Psmd13 | 23997 | HITS-CLIP | 23597149 | Unspecified |
| mmu-let-7i-3p | MIMAT0004520 | Ptafr | 19204 | HITS-CLIP | 29167373 | Unspecified |
| mmu-let-7i-3p | MIMAT0004520 | Ptbp3 | 230257 | HITS-CLIP | 23597149\|29167373 | Unspecified |
| mmu-let-7i-3p | MIMAT0004520 | Ptpn12 | 19248 | HITS-CLIP | 23597149 | Unspecified |
| mmu-let-7i-3p | MIMAT0004520 | Ptpn6 | 15170 | HITS-CLIP | 29167373 | Unspecified |
| mmu-let-7i-3p | MIMAT0004520 | Puf60 | 67959 | HITS-CLIP | 29167373 | Unspecified |
| mmu-let-7i-3p | MIMAT0004520 | Pum2 | 80913 | HITS-CLIP | 29167373 | Unspecified |
| mmu-let-7i-3p | MIMAT0004520 | Qsox1 | 104009 | HITS-CLIP | 23597149 | Unspecified |
| mmu-let-7i-3p | MIMAT0004520 | Rab18 | 19330 | HITS-CLIP | 23597149 | Unspecified |
| mmu-let-7i-3p | MIMAT0004520 | Rab23 | 19335 | HITS-CLIP | 23597149 | Unspecified |
| mmu-let-7i-3p | MIMAT0004520 | Rab28 | 100972 | HITS-CLIP | 29167373 | Unspecified |

| | | | | | | |
|---|---|---|---|---|---|---|
| mmu-let-7i-3p | MIMAT0004520 | Rab43 | 69834 | HITS-CLIP | 23597149 | Unspecified |
| mmu-let-7i-3p | MIMAT0004520 | Rabac1 | 14470 | HITS-CLIP | 23597149 | Unspecified |
| mmu-let-7i-3p | MIMAT0004520 | Rabgap1 | 227800 | HITS-CLIP | 23597149 | Unspecified |
| mmu-let-7i-3p | MIMAT0004520 | Rad21 | 19357 | HITS-CLIP | 29167373 | Unspecified |
| mmu-let-7i-3p | MIMAT0004520 | Ralgapb | 228850 | HITS-CLIP | 23597149 | Unspecified |
| mmu-let-7i-3p | MIMAT0004520 | Ranbp2 | 19386 | HITS-CLIP | 23597149 | Unspecified |
| mmu-let-7i-3p | MIMAT0004520 | Rapgef5 | 217944 | HITS-CLIP | 29167373 | Unspecified |
| mmu-let-7i-3p | MIMAT0004520 | Raph1 | 77300 | HITS-CLIP | 29167373 | Unspecified |
| mmu-let-7i-3p | MIMAT0004520 | Rasa1 | 218397 | qCLASH | 31393953 | Unspecified |
| mmu-let-7i-3p | MIMAT0004520 | Rasal2 | 226525 | HITS-CLIP | 23597149 | Unspecified |
| mmu-let-7i-3p | MIMAT0004520 | Rasgef1b | 320292 | HITS-CLIP | 23597149 | Unspecified |
| mmu-let-7i-3p | MIMAT0004520 | Rbm12 | 75710 | HITS-CLIP | 29167373 | Unspecified |
| mmu-let-7i-3p | MIMAT0004520 | Rbm14 | 56275 | HITS-CLIP | 29167373 | Unspecified |
| mmu-let-7i-3p | MIMAT0004520 | Rdx | 19684 | HITS-CLIP | 29167373 | Unspecified |
| mmu-let-7i-3p | MIMAT0004520 | Rer1 | 67830 | HITS-CLIP | 29167373 | Unspecified |
| mmu-let-7i-3p | MIMAT0004520 | Retreg2 | 79137 | HITS-CLIP | 23597149 | Unspecified |
| mmu-let-7i-3p | MIMAT0004520 | Rexo1 | 66932 | HITS-CLIP | 29167373 | Unspecified |
| mmu-let-7i-3p | MIMAT0004520 | Rftn1 | 76438 | HITS-CLIP | 29167373 | Unspecified |
| mmu-let-7i-3p | MIMAT0004520 | Rfx1 | 19724 | HITS-CLIP | 23597149 | Unspecified |
| mmu-let-7i-3p | MIMAT0004520 | Rgs5 | 19737 | HITS-CLIP | 23597149 | Unspecified |
| mmu-let-7i-3p | MIMAT0004520 | Rhobtb1 | 69288 | HITS-CLIP | 23597149 | Unspecified |
| mmu-let-7i-3p | MIMAT0004520 | Rictor | 78757 | HITS-CLIP | 29167373 | Unspecified |
| mmu-let-7i-3p | MIMAT0004520 | Rlim | 19820 | HITS-CLIP | 29167373 | Unspecified |
| mmu-let-7i-3p | MIMAT0004520 | Rmdn3 | 67809 | HITS-CLIP | 23597149 | Unspecified |
| mmu-let-7i-3p | MIMAT0004520 | Rnf111 | 93836 | HITS-CLIP | 37986224\|29167373\|23597149 | Unspecified |
| mmu-let-7i-3p | MIMAT0004520 | Rnf213 | 672511 | HITS-CLIP | 23597149 | Unspecified |
| mmu-let-7i-3p | MIMAT0004520 | Rnf217 | 268291 | HITS-CLIP | 23597149\|29167373 | Unspecified |
| mmu-let-7i-3p | MIMAT0004520 | Rnf225 | 381845 | qCLASH | 31393953 | Unspecified |
| mmu-let-7i-3p | MIMAT0004520 | Rpl5 | 100503670 | qCLASH | 31393953 | Unspecified |
| mmu-let-7i-3p | MIMAT0004520 | Rps6ka4 | 56613 | HITS-CLIP | 23597149 | Unspecified |
| mmu-let-7i-3p | MIMAT0004520 | Rundc1 | 217201 | HITS-CLIP | 23597149 | Unspecified |
| mmu-let-7i-3p | MIMAT0004520 | Sae1 | 56459 | HITS-CLIP | 23597149 | Unspecified |
| mmu-let-7i-3p | MIMAT0004520 | Samd4b | 233033 | HITS-CLIP | 29167373 | Unspecified |
| mmu-let-7i-3p | MIMAT0004520 | Sar1b | 66397 | HITS-CLIP | 23597149\|29167373 | Unspecified |
| mmu-let-7i-3p | MIMAT0004520 | Sardh | 192166 | HITS-CLIP | 23597149 | Unspecified |
| mmu-let-7i-3p | MIMAT0004520 | Scly | 50880 | HITS-CLIP | 23597149 | Unspecified |
| mmu-let-7i-3p | MIMAT0004520 | Scp2 | 20280 | HITS-CLIP | 23597149 | Unspecified |
| mmu-let-7i-3p | MIMAT0004520 | Scyl2 | 213326 | HITS-CLIP | 23597149 | Unspecified |
| mmu-let-7i-3p | MIMAT0004520 | Sec22a | 317717 | HITS-CLIP | 29167373 | Unspecified |

| | | | | | | | |
|---|---|---|---|---|---|---|---|
| mmu-let-7i-3p | MIMAT0004520 | Secisbp2 | 75420 | HITS-CLIP | | 23597149 | Unspecified |
| mmu-let-7i-3p | MIMAT0004520 | Selenop | 6414 | HITS-CLIP | | 23597149 | Unspecified |
| mmu-let-7i-3p | MIMAT0004520 | Sema4g | 26456 | HITS-CLIP | | 23597149 | Unspecified |
| mmu-let-7i-3p | MIMAT0004520 | Sema6a | 20358 | HITS-CLIP | | 23597149 | Unspecified |
| mmu-let-7i-3p | MIMAT0004520 | Sephs1 | 109079 | HITS-CLIP | | 23597149 | Unspecified |
| mmu-let-7i-3p | MIMAT0004520 | Serpina1a | 20700 | HITS-CLIP | | 23597149 | Unspecified |
| mmu-let-7i-3p | MIMAT0004520 | Serpinc1 | 11905 | HITS-CLIP | | 23597149 | Unspecified |
| mmu-let-7i-3p | MIMAT0004520 | Setd1a | 233904 | HITS-CLIP | | 29167373 | Unspecified |
| mmu-let-7i-3p | MIMAT0004520 | Setd5 | 72895 | HITS-CLIP | | 29167373 | Unspecified |
| mmu-let-7i-3p | MIMAT0004520 | Sez6l | 56747 | qCLASH | | 31393953 | Unspecified |
| mmu-let-7i-3p | MIMAT0004520 | Sgk1 | 20393 | HITS-CLIP | | 29167373 | Unspecified |
| mmu-let-7i-3p | MIMAT0004520 | Sgsm3 | 105835 | HITS-CLIP | | 23597149 | Unspecified |
| mmu-let-7i-3p | MIMAT0004520 | Sh3bgrl2 | 212531 | HITS-CLIP | | 23597149 | Unspecified |
| mmu-let-7i-3p | MIMAT0004520 | Sh3glb1 | 54673 | HITS-CLIP | | 29167373 | Unspecified |
| mmu-let-7i-3p | MIMAT0004520 | Sh3kbp1 | 58194 | HITS-CLIP | | 29167373 | Unspecified |
| mmu-let-7i-3p | MIMAT0004520 | Sh3pxd2a | 14218 | HITS-CLIP | | 29167373 | Unspecified |
| mmu-let-7i-3p | MIMAT0004520 | Shb | 230126 | HITS-CLIP | | 23597149 | Unspecified |
| mmu-let-7i-3p | MIMAT0004520 | Shmt2 | 108037 | HITS-CLIP | | 23597149 | Unspecified |
| mmu-let-7i-3p | MIMAT0004520 | Sigmar1 | 18391 | HITS-CLIP | | 37986224 | Unspecified |
| mmu-let-7i-3p | MIMAT0004520 | Sik2 | 235344 | HITS-CLIP | | 31433990 | Unspecified |
| mmu-let-7i-3p | MIMAT0004520 | Sirpa | 19261 | HITS-CLIP | 23597149\|29167373 | | Unspecified |
| mmu-let-7i-3p | MIMAT0004520 | Skiv2l | 108077 | HITS-CLIP | | 37986224 | Unspecified |
| mmu-let-7i-3p | MIMAT0004520 | Slain2 | 75991 | HITS-CLIP | | 23597149 | Unspecified |
| mmu-let-7i-3p | MIMAT0004520 | Slc16a12 | 240638 | HITS-CLIP | | 23597149 | Unspecified |
| mmu-let-7i-3p | MIMAT0004520 | Slc25a12 | 78830 | HITS-CLIP | | 23597149 | Unspecified |
| mmu-let-7i-3p | MIMAT0004520 | Slc25a13 | 50799 | HITS-CLIP | | 23597149 | Unspecified |
| mmu-let-7i-3p | MIMAT0004520 | Slc27a2 | 26458 | HITS-CLIP | | 23597149 | Unspecified |
| mmu-let-7i-3p | MIMAT0004520 | Slc35e2 | 320541 | HITS-CLIP | | 23597149 | Unspecified |
| mmu-let-7i-3p | MIMAT0004520 | Slc38a4 | 69354 | HITS-CLIP | | 23597149 | Unspecified |
| mmu-let-7i-3p | MIMAT0004520 | Slc4a7 | 218756 | HITS-CLIP | | 29167373 | Unspecified |
| mmu-let-7i-3p | MIMAT0004520 | Slc5a3 | 53881 | HITS-CLIP | | 29167373 | Unspecified |
| mmu-let-7i-3p | MIMAT0004520 | Slc6a6 | 21366 | HITS-CLIP | 23597149\|29167373 | | Unspecified |
| mmu-let-7i-3p | MIMAT0004520 | Slc7a2 | 11988 | HITS-CLIP | | 23597149 | Unspecified |
| mmu-let-7i-3p | MIMAT0004520 | Slf1 | 852127 | HITS-CLIP | | 29167373 | Unspecified |
| mmu-let-7i-3p | MIMAT0004520 | Smad1 | 17125 | HITS-CLIP | | 23597149 | Unspecified |
| mmu-let-7i-3p | MIMAT0004520 | Smad2 | 17126 | HITS-CLIP | | 23597149 | Unspecified |
| mmu-let-7i-3p | MIMAT0004520 | Smap2 | 69780 | HITS-CLIP | | 23597149 | Unspecified |
| mmu-let-7i-3p | MIMAT0004520 | Smarca4 | 20586 | HITS-CLIP | | 29167373 | Unspecified |
| mmu-let-7i-3p | MIMAT0004520 | Smarcd2 | 83796 | HITS-CLIP | | 23597149 | Unspecified |

| | | | | | | |
|---|---|---|---|---|---|---|
| mmu-let-7i-3p | MIMAT0004520 | Smchd1 | 74355 | HITS-CLIP | 23597149 | Unspecified |
| mmu-let-7i-3p | MIMAT0004520 | Snd1 | 56463 | HITS-CLIP | 37986224 | Unspecified |
| mmu-let-7i-3p | MIMAT0004520 | Snx18 | 170625 | HITS-CLIP | 29167373 | Unspecified |
| mmu-let-7i-3p | MIMAT0004520 | Snx30 | 209131 | HITS-CLIP | 29167373 | Unspecified |
| mmu-let-7i-3p | MIMAT0004520 | Soga1 | 320706 | HITS-CLIP | 29167373 | Unspecified |
| mmu-let-7i-3p | MIMAT0004520 | Sp3 | 20687 | HITS-CLIP | 23597149 | Unspecified |
| mmu-let-7i-3p | MIMAT0004520 | Spred1 | 114715 | HITS-CLIP | 29167373 | Unspecified |
| mmu-let-7i-3p | MIMAT0004520 | Sptan1 | 20740 | HITS-CLIP | 37986224 | Unspecified |
| mmu-let-7i-3p | MIMAT0004520 | Sptssa | 104725 | HITS-CLIP | 29167373 | Unspecified |
| mmu-let-7i-3p | MIMAT0004520 | Srd5a3 | 57357 | HITS-CLIP | 23597149 | Unspecified |
| mmu-let-7i-3p | MIMAT0004520 | Srrm2 | 75956 | HITS-CLIP | 23597149 | Unspecified |
| mmu-let-7i-3p | MIMAT0004520 | Ssh2 | 237860 | HITS-CLIP | 23597149\|29167373 | Unspecified |
| mmu-let-7i-3p | MIMAT0004520 | Ssr1 | 107513 | HITS-CLIP | 29167373 | Unspecified |
| mmu-let-7i-3p | MIMAT0004520 | Stag2 | 20843 | HITS-CLIP | 23597149 | Unspecified |
| mmu-let-7i-3p | MIMAT0004520 | Stard10 | 56018 | HITS-CLIP | 37986224 | Unspecified |
| mmu-let-7i-3p | MIMAT0004520 | Stat2 | 20847 | HITS-CLIP | 29167373 | Unspecified |
| mmu-let-7i-3p | MIMAT0004520 | Stk38l | 232533 | HITS-CLIP | 29167373 | Unspecified |
| mmu-let-7i-3p | MIMAT0004520 | Sult1a1 | 20887 | HITS-CLIP | 23597149 | Unspecified |
| mmu-let-7i-3p | MIMAT0004520 | Sult2a8 | 76971 | HITS-CLIP | 23597149 | Unspecified |
| mmu-let-7i-3p | MIMAT0004520 | Suox | 211389 | HITS-CLIP | 23597149 | Unspecified |
| mmu-let-7i-3p | MIMAT0004520 | Taf1 | 270627 | HITS-CLIP | 37986224 | Unspecified |
| mmu-let-7i-3p | MIMAT0004520 | Taok1 | 216965 | HITS-CLIP | 23597149\|29167373 | Unspecified |
| mmu-let-7i-3p | MIMAT0004520 | Tars | 110960 | HITS-CLIP | 29167373 | Unspecified |
| mmu-let-7i-3p | MIMAT0004520 | Tat | 234724 | HITS-CLIP | 23597149 | Unspecified |
| mmu-let-7i-3p | MIMAT0004520 | Tbc1d1 | 57915 | HITS-CLIP | 29167373 | Unspecified |
| mmu-let-7i-3p | MIMAT0004520 | Tbc1d20 | 67231 | HITS-CLIP | 23597149 | Unspecified |
| mmu-let-7i-3p | MIMAT0004520 | Tbl1xr1 | 81004 | HITS-CLIP | 29167373 | Unspecified |
| mmu-let-7i-3p | MIMAT0004520 | Tcea3 | 21401 | HITS-CLIP | 23597149 | Unspecified |
| mmu-let-7i-3p | MIMAT0004520 | Tent4a | 11044 | HITS-CLIP | 23597149\|29167373 | Unspecified |
| mmu-let-7i-3p | MIMAT0004520 | Tex2 | 21763 | HITS-CLIP | 23597149 | Unspecified |
| mmu-let-7i-3p | MIMAT0004520 | Tgfbr1 | 21812 | HITS-CLIP | 23597149\|31433990 | Unspecified |
| mmu-let-7i-3p | MIMAT0004520 | Tgs1 | 116940 | HITS-CLIP | 29167373 | Unspecified |
| mmu-let-7i-3p | MIMAT0004520 | Timp2 | 21858 | HITS-CLIP | 23597149 | Unspecified |
| mmu-let-7i-3p | MIMAT0004520 | Tinagl1 | 94242 | HITS-CLIP | 23597149 | Unspecified |
| mmu-let-7i-3p | MIMAT0004520 | Tinf2 | 28113 | HITS-CLIP | 23597149 | Unspecified |
| mmu-let-7i-3p | MIMAT0004520 | Tiparp | 99929 | HITS-CLIP | 29167373 | Unspecified |
| mmu-let-7i-3p | MIMAT0004520 | Tkt | 21881 | HITS-CLIP | 29167373 | Unspecified |
| mmu-let-7i-3p | MIMAT0004520 | Tle4 | 21888 | HITS-CLIP | 23597149 | Unspecified |
| mmu-let-7i-3p | MIMAT0004520 | Tlk1 | 228012 | HITS-CLIP | 29167373 | Unspecified |

| | | | | | | |
|---|---|---|---|---|---|---|
| mmu-let-7i-3p | MIMAT0004520 | Tln1 | 21894 | HITS-CLIP | 23597149\|29167373 | Unspecified |
| mmu-let-7i-3p | MIMAT0004520 | Tlr7 | 170743 | HITS-CLIP | 29167373 | Unspecified |
| mmu-let-7i-3p | MIMAT0004520 | Tmem129 | 68366 | HITS-CLIP | 23597149 | Unspecified |
| mmu-let-7i-3p | MIMAT0004520 | Tmem176a | 66058 | HITS-CLIP | 23597149 | Unspecified |
| mmu-let-7i-3p | MIMAT0004520 | Tmem184c | 234463 | HITS-CLIP | 37986224\|23597149 | Unspecified |
| mmu-let-7i-3p | MIMAT0004520 | Tmem209 | 72649 | HITS-CLIP | 29167373 | Unspecified |
| mmu-let-7i-3p | MIMAT0004520 | Tmem8 | 60455 | HITS-CLIP | 29167373 | Unspecified |
| mmu-let-7i-3p | MIMAT0004520 | Tmx4 | 52837 | HITS-CLIP | 23597149 | Unspecified |
| mmu-let-7i-3p | MIMAT0004520 | Tnrc6a | 233833 | HITS-CLIP | 23597149 | Unspecified |
| mmu-let-7i-3p | MIMAT0004520 | Tob2 | 57259 | HITS-CLIP | 23597149 | Unspecified |
| mmu-let-7i-3p | MIMAT0004520 | Top1 | 21969 | HITS-CLIP | 37986224 | Unspecified |
| mmu-let-7i-3p | MIMAT0004520 | Tor1aip2 | 240832 | HITS-CLIP | 29167373 | Unspecified |
| mmu-let-7i-3p | MIMAT0004520 | Tpp2 | 22019 | HITS-CLIP | 29167373 | Unspecified |
| mmu-let-7i-3p | MIMAT0004520 | Tpr | 108989 | HITS-CLIP | 23597149 | Unspecified |
| mmu-let-7i-3p | MIMAT0004520 | Trappc10 | 216131 | HITS-CLIP | 23597149 | Unspecified |
| mmu-let-7i-3p | MIMAT0004520 | Trib3 | 228775 | HITS-CLIP | 23597149 | Unspecified |
| mmu-let-7i-3p | MIMAT0004520 | Trim11 | 94091 | qCLASH | 31393953 | Unspecified |
| mmu-let-7i-3p | MIMAT0004520 | Trmt13 | 229780 | HITS-CLIP | 29167373 | Unspecified |
| mmu-let-7i-3p | MIMAT0004520 | Tsc22d3 | 14605 | HITS-CLIP | 23597149 | Unspecified |
| mmu-let-7i-3p | MIMAT0004520 | Tsku | 244152 | HITS-CLIP | 23597149 | Unspecified |
| mmu-let-7i-3p | MIMAT0004520 | Ttc7 | 225049 | HITS-CLIP | 23597149 | Unspecified |
| mmu-let-7i-3p | MIMAT0004520 | Tut7 | 79670 | HITS-CLIP | 29167373 | Unspecified |
| mmu-let-7i-3p | MIMAT0004520 | Txndc5 | 105245 | HITS-CLIP | 23597149 | Unspecified |
| mmu-let-7i-3p | MIMAT0004520 | Txnip | 56338 | HITS-CLIP | 23597149 | Unspecified |
| mmu-let-7i-3p | MIMAT0004520 | Uap1l1 | 227620 | HITS-CLIP | 29167373 | Unspecified |
| mmu-let-7i-3p | MIMAT0004520 | Ubap2l | 74383 | HITS-CLIP | 23597149 | Unspecified |
| mmu-let-7i-3p | MIMAT0004520 | Ubr2 | 224826 | HITS-CLIP | 23597149\|29167373 | Unspecified |
| mmu-let-7i-3p | MIMAT0004520 | Ubr5 | 70790 | HITS-CLIP | 37986224 | Unspecified |
| mmu-let-7i-3p | MIMAT0004520 | Ubxn4 | 67812 | HITS-CLIP | 23597149 | Unspecified |
| mmu-let-7i-3p | MIMAT0004520 | Uchl3 | 50933 | HITS-CLIP | 37986224\|23597149 | Unspecified |
| mmu-let-7i-3p | MIMAT0004520 | Ugdh | 22235 | HITS-CLIP | 23597149 | Unspecified |
| mmu-let-7i-3p | MIMAT0004520 | Ugt2b1 | 71773 | HITS-CLIP | 37986224 | Unspecified |
| mmu-let-7i-3p | MIMAT0004520 | Unc93b1 | 54445 | HITS-CLIP | 37986224\|23597149 | Unspecified |
| mmu-let-7i-3p | MIMAT0004520 | Upp2 | 76654 | HITS-CLIP | 23597149 | Unspecified |
| mmu-let-7i-3p | MIMAT0004520 | Uqcrc1 | 22273 | HITS-CLIP | 23597149 | Unspecified |
| mmu-let-7i-3p | MIMAT0004520 | Uqcrfs1 | 66694 | HITS-CLIP | 23597149 | Unspecified |
| mmu-let-7i-3p | MIMAT0004520 | Usp15 | 14479 | HITS-CLIP | 23597149 | Unspecified |
| mmu-let-7i-3p | MIMAT0004520 | Usp34 | 17847 | HITS-CLIP | 23597149 | Unspecified |
| mmu-let-7i-3p | MIMAT0004520 | Usp47 | 74996 | HITS-CLIP | 23597149\|29167373 | Unspecified |

| | | | | | | |
|---|---|---|---|---|---|---|
| mmu-let-7i-3p | MIMAT0004520 | Usp8 | 84092 | HITS-CLIP | 29167373 | Unspecified |
| mmu-let-7i-3p | MIMAT0004520 | Usp9x | 22284 | HITS-CLIP | 29167373 | Unspecified |
| mmu-let-7i-3p | MIMAT0004520 | Uvrag | 78610 | HITS-CLIP | 23597149 | Unspecified |
| mmu-let-7i-3p | MIMAT0004520 | Vcl | 22330 | HITS-CLIP | 23597149 | Unspecified |
| mmu-let-7i-3p | MIMAT0004520 | Vcpip1 | 70675 | HITS-CLIP | 29167373 | Unspecified |
| mmu-let-7i-3p | MIMAT0004520 | Vps13d | 230895 | HITS-CLIP | 23597149 | Unspecified |
| mmu-let-7i-3p | MIMAT0004520 | Vps41 | 218035 | HITS-CLIP | 31433990 | Unspecified |
| mmu-let-7i-3p | MIMAT0004520 | Wdfy1 | 69368 | HITS-CLIP | 23597149 | Unspecified |
| mmu-let-7i-3p | MIMAT0004520 | Wdr26 | 226757 | HITS-CLIP | 29167373 | Unspecified |
| mmu-let-7i-3p | MIMAT0004520 | Wnk1 | 232341 | HITS-CLIP | 29167373 | Unspecified |
| mmu-let-7i-3p | MIMAT0004520 | Wsb1 | 78889 | HITS-CLIP | 23597149 | Unspecified |
| mmu-let-7i-3p | MIMAT0004520 | Wtap | 60532 | HITS-CLIP | 23597149\|29167373 | Unspecified |
| mmu-let-7i-3p | MIMAT0004520 | Wwp2 | 66894 | HITS-CLIP | 29167373 | Unspecified |
| mmu-let-7i-3p | MIMAT0004520 | Xpo5 | 72322 | HITS-CLIP | 29167373 | Unspecified |
| mmu-let-7i-3p | MIMAT0004520 | Ywhag | 22628 | HITS-CLIP | 23597149 | Unspecified |
| mmu-let-7i-3p | MIMAT0004520 | Ywhah | 22629 | HITS-CLIP | 29167373 | Unspecified |
| mmu-let-7i-3p | MIMAT0004520 | Ywhaq | 22630 | HITS-CLIP | 29167373 | Unspecified |
| mmu-let-7i-3p | MIMAT0004520 | Yy1 | 22632 | HITS-CLIP | 29167373 | Unspecified |
| mmu-let-7i-3p | MIMAT0004520 | Zbtb20 | 56490 | HITS-CLIP | 29167373 | Unspecified |
| mmu-let-7i-3p | MIMAT0004520 | Zbtb7a | 16969 | HITS-CLIP | 29167373 | Unspecified |
| mmu-let-7i-3p | MIMAT0004520 | Zdhhc5 | 228136 | HITS-CLIP | 23597149\|29167373 | Unspecified |
| mmu-let-7i-3p | MIMAT0004520 | Zfc3h1 | 216345 | HITS-CLIP | 29167373 | Unspecified |
| mmu-let-7i-3p | MIMAT0004520 | Zfp236 | 329002 | HITS-CLIP | 29167373 | Unspecified |
| mmu-let-7i-3p | MIMAT0004520 | Zfp292 | 30046 | HITS-CLIP | 29167373 | Unspecified |
| mmu-let-7i-3p | MIMAT0004520 | Zfp330 | 30932 | HITS-CLIP | 29167373 | Unspecified |
| mmu-let-7i-3p | MIMAT0004520 | Zfp36l1 | 12192 | HITS-CLIP | 23597149 | Unspecified |
| mmu-let-7i-3p | MIMAT0004520 | Zfp36l2 | 12193 | HITS-CLIP | 29167373 | Unspecified |
| mmu-let-7i-3p | MIMAT0004520 | Zfp609 | 214812 | HITS-CLIP | 23597149 | Unspecified |
| mmu-let-7i-3p | MIMAT0004520 | Zfp710 | 209225 | HITS-CLIP | 23597149 | Unspecified |
| mmu-let-7i-3p | MIMAT0004520 | Zfp91 | 109910 | HITS-CLIP | 23597149 | Unspecified |
| mmu-let-7i-3p | MIMAT0004520 | Zfr | 22763 | HITS-CLIP | 23597149\|29167373 | Unspecified |
| mmu-let-7i-3p | MIMAT0004520 | Zmiz1 | 328365 | HITS-CLIP | 31433990 | Unspecified |
| mmu-let-7i-3p | MIMAT0004520 | Zmiz2 | 52915 | HITS-CLIP | 37986224 | Unspecified |
| mmu-let-7i-3p | MIMAT0004520 | Zmym3 | 56364 | HITS-CLIP | 23597149 | Unspecified |
| mmu-let-7i-3p | MIMAT0004520 | Znfx1 | 98999 | HITS-CLIP | 23597149\|29167373 | Unspecified |
| mmu-let-7i-3p | MIMAT0004520 | Zranb3 | 226409 | HITS-CLIP | 23597149 | Unspecified |
| mmu-let-7i-3p | MIMAT0004520 | Zyx | 22793 | HITS-CLIP | 23597149 | Unspecified |
| mmu-miR-708-3p | MIMAT0003498 | Atxn3 | 110616 | HITS-CLIP | 26203562 | Unspecified |
| mmu-miR-708-3p | MIMAT0003498 | Ei24 | 13663 | HITS-CLIP | 26203562 | Unspecified |

| | | | | | | |
|---|---|---|---|---|---|---|
| mmu-miR-708-3p | MIMAT0003498 | Kmt2d | 381022 | HITS-CLIP | 37986224 | Unspecified |
| mmu-miR-708-3p | MIMAT0003498 | Lin28a | 83557 | HITS-CLIP | 37986224 | Unspecified |
| mmu-miR-708-3p | MIMAT0003498 | Peg3 | 18616 | HITS-CLIP | 26203562 | Unspecified |
| mmu-miR-708-3p | MIMAT0003498 | Rbms1 | 56878 | HITS-CLIP | 31433990 | Unspecified |
| mmu-miR-708-3p | MIMAT0003498 | Senp1 | 223870 | HITS-CLIP | 37986224 | Unspecified |
| mmu-miR-708-3p | MIMAT0003498 | Slc37a3 | 72144 | HITS-CLIP | 31433990 | Unspecified |
| mmu-miR-708-3p | MIMAT0003498 | Tmem184b | 223693 | HITS-CLIP | 37986224 | Unspecified |
| mmu-miR-24-2-5p | MIMAT0005440 | Adam23 | 23792 | HITS-CLIP | 37986224 | Unspecified |
| mmu-miR-24-2-5p | MIMAT0005440 | Adh5 | 11532 | HITS-CLIP | 25753418 | Unspecified |
| mmu-miR-24-2-5p | MIMAT0005440 | Aldh4a1 | 212647 | HITS-CLIP | 25753418 | Unspecified |
| mmu-miR-24-2-5p | MIMAT0005440 | Alpk3 | 116904 | HITS-CLIP | 25753418 | Unspecified |
| mmu-miR-24-2-5p | MIMAT0005440 | Ap2s1 | 232910 | HITS-CLIP | 25753418 | Unspecified |
| mmu-miR-24-2-5p | MIMAT0005440 | Arid1a | 93760 | HITS-CLIP | 25753418 | Unspecified |
| mmu-miR-24-2-5p | MIMAT0005440 | Atp2a2 | 11938 | HITS-CLIP | 25753418 | Unspecified |
| mmu-miR-24-2-5p | MIMAT0005440 | Brwd1 | 93871 | HITS-CLIP | 37986224 | Unspecified |
| mmu-miR-24-2-5p | MIMAT0005440 | Cct6a | 12466 | HITS-CLIP | 37986224 | Unspecified |
| mmu-miR-24-2-5p | MIMAT0005440 | Cdc73 | 214498 | HITS-CLIP | 37986224 | Unspecified |
| mmu-miR-24-2-5p | MIMAT0005440 | Cnksr1 | 194231 | HITS-CLIP | 25753418 | Unspecified |
| mmu-miR-24-2-5p | MIMAT0005440 | Cnot6l | 231464 | HITS-CLIP | 25753418 | Unspecified |
| mmu-miR-24-2-5p | MIMAT0005440 | Dcaf10 | 242418 | HITS-CLIP | 25753418 | Unspecified |
| mmu-miR-24-2-5p | MIMAT0005440 | Ddx17 | 67040 | HITS-CLIP | 37986224 | Unspecified |
| mmu-miR-24-2-5p | MIMAT0005440 | Ddx3y | 26900 | HITS-CLIP | 25753418 | Unspecified |
| mmu-miR-24-2-5p | MIMAT0005440 | Dnm2 | 13430 | HITS-CLIP | 25753418 | Unspecified |
| mmu-miR-24-2-5p | MIMAT0005440 | Dst | 13518 | HITS-CLIP | 25753418 | Unspecified |
| mmu-miR-24-2-5p | MIMAT0005440 | Egr1 | 13653 | HITS-CLIP | 25753418 | Unspecified |
| mmu-miR-24-2-5p | MIMAT0005440 | Eif3a | 13669 | HITS-CLIP | 37986224 | Unspecified |
| mmu-miR-24-2-5p | MIMAT0005440 | Flnc | 68794 | HITS-CLIP | 25753418 | Unspecified |
| mmu-miR-24-2-5p | MIMAT0005440 | Gng5 | 14707 | HITS-CLIP | 37986224 | Unspecified |
| mmu-miR-24-2-5p | MIMAT0005440 | Kdm3b | 277250 | HITS-CLIP | 37986224 | Unspecified |
| mmu-miR-24-2-5p | MIMAT0005440 | Kif1c | 16562 | HITS-CLIP | 25753418 | Unspecified |
| mmu-miR-24-2-5p | MIMAT0005440 | Klhl41 | 228003 | HITS-CLIP | 25753418 | Unspecified |
| mmu-miR-24-2-5p | MIMAT0005440 | Laptm4a | 17775 | HITS-CLIP | 25753418 | Unspecified |
| mmu-miR-24-2-5p | MIMAT0005440 | Lsm3 | 67678 | HITS-CLIP | 25753418 | Unspecified |
| mmu-miR-24-2-5p | MIMAT0005440 | Luc7l | 66978 | HITS-CLIP | 37986224 | Unspecified |
| mmu-miR-24-2-5p | MIMAT0005440 | Map2k2 | 26396 | HITS-CLIP | 25753418 | Unspecified |
| mmu-miR-24-2-5p | MIMAT0005440 | Mat2b | 108645 | HITS-CLIP | 25753418 | Unspecified |
| mmu-miR-24-2-5p | MIMAT0005440 | Matr3 | 17184 | HITS-CLIP | 37986224 | Unspecified |
| mmu-miR-24-2-5p | MIMAT0005440 | Me1 | 17436 | HITS-CLIP | 25753418 | Unspecified |
| mmu-miR-24-2-5p | MIMAT0005440 | Mmp15 | 17388 | HITS-CLIP | 25753418 | Unspecified |

| | | | | | | |
|---|---|---|---|---|---|---|
| mmu-miR-24-2-5p | MIMAT0005440 | Mtap | 66902 | HITS-CLIP | 25753418 | Unspecified |
| mmu-miR-24-2-5p | MIMAT0005440 | Myo18a | 360013 | HITS-CLIP | 25753418 | Unspecified |
| mmu-miR-24-2-5p | MIMAT0005440 | Myorg | 57462 | HITS-CLIP | 25753418 | Unspecified |
| mmu-miR-24-2-5p | MIMAT0005440 | Nae1 | 234664 | HITS-CLIP | 37986224 | Unspecified |
| mmu-miR-24-2-5p | MIMAT0005440 | Oard1 | 106821 | HITS-CLIP | 37986224 | Unspecified |
| mmu-miR-24-2-5p | MIMAT0005440 | Orc3 | 50793 | HITS-CLIP | 25753418 | Unspecified |
| mmu-miR-24-2-5p | MIMAT0005440 | Pa2g4 | 18813 | HITS-CLIP | 25753418 | Unspecified |
| mmu-miR-24-2-5p | MIMAT0005440 | Patl1 | 225929 | HITS-CLIP | 25753418 | Unspecified |
| mmu-miR-24-2-5p | MIMAT0005440 | Pgap4 | 84302 | HITS-CLIP | 25753418 | Unspecified |
| mmu-miR-24-2-5p | MIMAT0005440 | Pon2 | 330260 | HITS-CLIP | 25753418 | Unspecified |
| mmu-miR-24-2-5p | MIMAT0005440 | Prkaa2 | 108079 | HITS-CLIP | 25753418 | Unspecified |
| mmu-miR-24-2-5p | MIMAT0005440 | Prrc2a | 53761 | HITS-CLIP | 37986224 | Unspecified |
| mmu-miR-24-2-5p | MIMAT0005440 | R3hdm2 | 71750 | HITS-CLIP | 25753418 | Unspecified |
| mmu-miR-24-2-5p | MIMAT0005440 | Rasal2 | 226525 | HITS-CLIP | 25753418 | Unspecified |
| mmu-miR-24-2-5p | MIMAT0005440 | Rgs2 | 19735 | HITS-CLIP | 25753418 | Unspecified |
| mmu-miR-24-2-5p | MIMAT0005440 | Rhoq | 104215 | HITS-CLIP | 31433990 | Unspecified |
| mmu-miR-24-2-5p | MIMAT0005440 | Rxrb | 20182 | HITS-CLIP | 25753418 | Unspecified |
| mmu-miR-24-2-5p | MIMAT0005440 | Secisbp2l | 70354 | HITS-CLIP | 25753418 | Unspecified |
| mmu-miR-24-2-5p | MIMAT0005440 | Shank3 | 58234 | HITS-CLIP | 25753418 | Unspecified |
| mmu-miR-24-2-5p | MIMAT0005440 | Smcr8 | 237782 | HITS-CLIP | 25753418 | Unspecified |
| mmu-miR-24-2-5p | MIMAT0005440 | Sntb2 | 20650 | HITS-CLIP | 37986224 | Unspecified |
| mmu-miR-24-2-5p | MIMAT0005440 | Socs4 | 67296 | HITS-CLIP | 25753418 | Unspecified |
| mmu-miR-24-2-5p | MIMAT0005440 | Spag9 | 70834 | HITS-CLIP | 25753418 | Unspecified |
| mmu-miR-24-2-5p | MIMAT0005440 | Srsf2 | 20382 | HITS-CLIP | 31433990 | Unspecified |
| mmu-miR-24-2-5p | MIMAT0005440 | Tbc1d4 | 210789 | HITS-CLIP | 25753418 | Unspecified |
| mmu-miR-24-2-5p | MIMAT0005440 | Tmod1 | 21916 | HITS-CLIP | 25753418 | Unspecified |
| mmu-miR-24-2-5p | MIMAT0005440 | Tmtc1 | 387314 | HITS-CLIP | 25753418 | Unspecified |
| mmu-miR-24-2-5p | MIMAT0005440 | Tnk2 | 51789 | HITS-CLIP | 37986224 | Unspecified |
| mmu-miR-24-2-5p | MIMAT0005440 | Tns2 | 209039 | HITS-CLIP | 25753418 | Unspecified |
| mmu-miR-24-2-5p | MIMAT0005440 | Topors | 106021 | HITS-CLIP | 25753418 | Unspecified |
| mmu-miR-24-2-5p | MIMAT0005440 | Ubr5 | 70790 | HITS-CLIP | 25753418 | Unspecified |
| mmu-miR-24-2-5p | MIMAT0005440 | Yap1 | 22601 | HITS-CLIP | 37986224\|25753418 | Unspecified |
| mmu-miR-24-2-5p | MIMAT0005440 | Zfp110 | 65020 | HITS-CLIP | 25753418 | Unspecified |
| mmu-miR-24-2-5p | MIMAT0005440 | Zfx | 22764 | HITS-CLIP | 25753418 | Unspecified |
| mmu-miR-100-3p | MIMAT0017051 | Dennd1b | 329260 | HITS-CLIP | 31433990 | Unspecified |
| mmu-miR-100-3p | MIMAT0017051 | Kmt2d | 381022 | HITS-CLIP | 31433990 | Unspecified |
| mmu-miR-1231-5p | MIMAT0022357 | Cul2 | 71745 | HITS-CLIP | 26203562 | Unspecified |
| mmu-miR-1231-5p | MIMAT0022357 | Pard6b | 58220 | HITS-CLIP | 26203562 | Unspecified |
| mmu-miR-1231-5p | MIMAT0022357 | Zfp850 | 100043772 | HITS-CLIP | 31433990 | Unspecified |

| | | | | | | |
|---|---|---|---|---|---|---|
| mmu-miR-1249-5p | MIMAT0014804 | Nsd3 | 234135 | HITS-CLIP | 26203562 | Unspecified |
| mmu-miR-133c | MIMAT0025078 | Ksr1 | 16706 | HITS-CLIP | 31433990 | Unspecified |
| mmu-miR-133c | MIMAT0025078 | Tollip | 54473 | HITS-CLIP | 26203562 | Unspecified |
| mmu-miR-145b | MIMAT0025105 | Fasn | 14104 | HITS-CLIP | 31433990 | Unspecified |
| mmu-miR-145b | MIMAT0025105 | Paxip1 | 55982 | HITS-CLIP | 31433990 | Unspecified |
| mmu-miR-145b | MIMAT0025105 | Rpa3 | 68240 | HITS-CLIP | 31433990 | Unspecified |
| mmu-miR-145b | MIMAT0025105 | Smc3 | 13006 | HITS-CLIP | 31433990 | Unspecified |
| mmu-miR-145b | MIMAT0025105 | Tmem30a | 69981 | HITS-CLIP | 26203562 | Unspecified |
| mmu-miR-145b | MIMAT0025105 | Zswim6 | 67263 | HITS-CLIP | 31433990 | Unspecified |
| mmu-miR-147-5p | MIMAT0017269 | A630001G21Rik | 319997 | HITS-CLIP | 31433990 | Unspecified |
| mmu-miR-147-5p | MIMAT0017269 | Arl5b | 75869 | HITS-CLIP | 31433990 | Unspecified |
| mmu-miR-147-5p | MIMAT0017269 | Atxn1l | 52335 | HITS-CLIP | 31433990 | Unspecified |
| mmu-miR-147-5p | MIMAT0017269 | Bach2 | 12014 | HITS-CLIP | 31433990 | Unspecified |
| mmu-miR-147-5p | MIMAT0017269 | Birc6 | 12211 | HITS-CLIP | 31433990 | Unspecified |
| mmu-miR-147-5p | MIMAT0017269 | Bmt2 | 101148 | HITS-CLIP | 26203562 | Unspecified |
| mmu-miR-147-5p | MIMAT0017269 | Ccdc71l | 72123 | HITS-CLIP | 26203562 | Unspecified |
| mmu-miR-147-5p | MIMAT0017269 | Cntrob | 216846 | HITS-CLIP | 26203562 | Unspecified |
| mmu-miR-147-5p | MIMAT0017269 | Ctnnd1 | 12388 | HITS-CLIP | 26203562 | Unspecified |
| mmu-miR-147-5p | MIMAT0017269 | Fam107b | 66540 | HITS-CLIP | 31433990 | Unspecified |
| mmu-miR-147-5p | MIMAT0017269 | Foxp1 | 108655 | HITS-CLIP | 31433990 | Unspecified |
| mmu-miR-147-5p | MIMAT0017269 | Gm28635 | | HITS-CLIP | 26203562 | Unspecified |
| mmu-miR-147-5p | MIMAT0017269 | Hcls1 | 15163 | HITS-CLIP | 31433990 | Unspecified |
| mmu-miR-147-5p | MIMAT0017269 | Ifi47 | 15953 | HITS-CLIP | 31433990 | Unspecified |
| mmu-miR-147-5p | MIMAT0017269 | Krt6a | 3853 | HITS-CLIP | 26203562 | Unspecified |
| mmu-miR-147-5p | MIMAT0017269 | Limk2 | 16886 | HITS-CLIP | 31433990 | Unspecified |
| mmu-miR-147-5p | MIMAT0017269 | Mcl1 | 17210 | HITS-CLIP | 31433990 | Unspecified |
| mmu-miR-147-5p | MIMAT0017269 | Ncapd2 | 68298 | HITS-CLIP | 31433990 | Unspecified |
| mmu-miR-147-5p | MIMAT0017269 | Nup153 | 218210 | HITS-CLIP | 31433990 | Unspecified |
| mmu-miR-147-5p | MIMAT0017269 | Nup62 | 18226 | HITS-CLIP | 31433990 | Unspecified |
| mmu-miR-147-5p | MIMAT0017269 | Psmc2 | 19181 | HITS-CLIP | 31433990 | Unspecified |
| mmu-miR-147-5p | MIMAT0017269 | Ptgs2 | 19225 | HITS-CLIP | 26203562 | Unspecified |
| mmu-miR-147-5p | MIMAT0017269 | Sgpp1 | 81535 | HITS-CLIP | 26203562 | Unspecified |
| mmu-miR-147-5p | MIMAT0017269 | Tab2 | 68652 | HITS-CLIP | 31433990 | Unspecified |
| mmu-miR-147-5p | MIMAT0017269 | Tmsb10 | 19240 | HITS-CLIP | 31433990 | Unspecified |
| mmu-miR-147-5p | MIMAT0017269 | Trp53inp1 | 60599 | HITS-CLIP | 26203562 | Unspecified |
| mmu-miR-147-5p | MIMAT0017269 | Zbtb7b | 22724 | HITS-CLIP | 26203562 | Unspecified |
| mmu-miR-152-5p | MIMAT0016991 | 2900026A02Rik | 243219 | HITS-CLIP | 29167373 | Unspecified |
| mmu-miR-152-5p | MIMAT0016991 | Abcc1 | 17250 | HITS-CLIP | 29167373 | Unspecified |
| mmu-miR-152-5p | MIMAT0016991 | Acaa2 | 52538 | HITS-CLIP | 29167373 | Unspecified |

| | | | | | | |
|---|---|---|---|---|---|---|
| mmu-miR-152-5p | MIMAT0016991 | Ahnak | 66395 | HITS-CLIP | 29167373 | Unspecified |
| mmu-miR-152-5p | MIMAT0016991 | Apaf1 | 11783 | HITS-CLIP | 29167373 | Unspecified |
| mmu-miR-152-5p | MIMAT0016991 | Apbb2 | 11787 | HITS-CLIP | 29167373 | Unspecified |
| mmu-miR-152-5p | MIMAT0016991 | Axin1 | 12005 | HITS-CLIP | 29167373 | Unspecified |
| mmu-miR-152-5p | MIMAT0016991 | Cadm1 | 54725 | HITS-CLIP | 29167373 | Unspecified |
| mmu-miR-152-5p | MIMAT0016991 | Ccdc71l | 72123 | HITS-CLIP | 29167373 | Unspecified |
| mmu-miR-152-5p | MIMAT0016991 | Cd36 | 12491 | HITS-CLIP | 29167373 | Unspecified |
| mmu-miR-152-5p | MIMAT0016991 | Cebpb | 12608 | HITS-CLIP | 29167373 | Unspecified |
| mmu-miR-152-5p | MIMAT0016991 | Cntrl | 26920 | HITS-CLIP | 29167373 | Unspecified |
| mmu-miR-152-5p | MIMAT0016991 | Cwf19l2 | 244672 | HITS-CLIP | 31433990 | Unspecified |
| mmu-miR-152-5p | MIMAT0016991 | Dennd4c | 329877 | HITS-CLIP | 29167373 | Unspecified |
| mmu-miR-152-5p | MIMAT0016991 | Dgkd | 227333 | HITS-CLIP | 29167373 | Unspecified |
| mmu-miR-152-5p | MIMAT0016991 | Ehd4 | 98878 | HITS-CLIP | 29167373 | Unspecified |
| mmu-miR-152-5p | MIMAT0016991 | Elmo1 | 140580 | HITS-CLIP | 29167373 | Unspecified |
| mmu-miR-152-5p | MIMAT0016991 | Ggnbp2 | 217039 | HITS-CLIP | 29167373 | Unspecified |
| mmu-miR-152-5p | MIMAT0016991 | Glg1 | 20340 | HITS-CLIP | 29167373 | Unspecified |
| mmu-miR-152-5p | MIMAT0016991 | Glipr2 | 384009 | HITS-CLIP | 31433990 | Unspecified |
| mmu-miR-152-5p | MIMAT0016991 | H4c14 | 8359 | HITS-CLIP | 31433990 | Unspecified |
| mmu-miR-152-5p | MIMAT0016991 | Heatr5a | 320487 | HITS-CLIP | 29167373 | Unspecified |
| mmu-miR-152-5p | MIMAT0016991 | Iffo2 | 212632 | HITS-CLIP | 29167373 | Unspecified |
| mmu-miR-152-5p | MIMAT0016991 | Ipo7 | 233726 | HITS-CLIP | 29167373 | Unspecified |
| mmu-miR-152-5p | MIMAT0016991 | Kpnb1 | 16211 | HITS-CLIP | 29167373 | Unspecified |
| mmu-miR-152-5p | MIMAT0016991 | Lpcat2 | 270084 | HITS-CLIP | 29167373 | Unspecified |
| mmu-miR-152-5p | MIMAT0016991 | Mapkapk2 | 17164 | HITS-CLIP | 29167373 | Unspecified |
| mmu-miR-152-5p | MIMAT0016991 | Myo5a | 17918 | HITS-CLIP | 29167373 | Unspecified |
| mmu-miR-152-5p | MIMAT0016991 | Myo9b | 17925 | HITS-CLIP | 29167373 | Unspecified |
| mmu-miR-152-5p | MIMAT0016991 | Nfxl1 | 100978 | HITS-CLIP | 29167373 | Unspecified |
| mmu-miR-152-5p | MIMAT0016991 | Nipbl | 71175 | HITS-CLIP | 29167373 | Unspecified |
| mmu-miR-152-5p | MIMAT0016991 | Odf2l | 52184 | HITS-CLIP | 29167373 | Unspecified |
| mmu-miR-152-5p | MIMAT0016991 | Osbpl8 | 237542 | HITS-CLIP | 29167373 | Unspecified |
| mmu-miR-152-5p | MIMAT0016991 | Otud5 | 54644 | HITS-CLIP | 29167373 | Unspecified |
| mmu-miR-152-5p | MIMAT0016991 | Parp11 | 101187 | HITS-CLIP | 29167373 | Unspecified |
| mmu-miR-152-5p | MIMAT0016991 | Pea15a | 18611 | HITS-CLIP | 29167373 | Unspecified |
| mmu-miR-152-5p | MIMAT0016991 | Pip4k2a | 18718 | HITS-CLIP | 29167373 | Unspecified |
| mmu-miR-152-5p | MIMAT0016991 | Plec | 18810 | HITS-CLIP | 29167373 | Unspecified |
| mmu-miR-152-5p | MIMAT0016991 | Plekhm3 | 241075 | HITS-CLIP | 29167373 | Unspecified |
| mmu-miR-152-5p | MIMAT0016991 | Plk2 | 20620 | HITS-CLIP | 29167373 | Unspecified |
| mmu-miR-152-5p | MIMAT0016991 | Ppp1r3b | 244416 | HITS-CLIP | 29167373 | Unspecified |
| mmu-miR-152-5p | MIMAT0016991 | Ppp4r3a | 55671 | HITS-CLIP | 29167373 | Unspecified |

| | | | | | | |
|---|---|---|---|---|---|---|
| mmu-miR-152-5p | MIMAT0016991 | Ppp4r3b | 57223 | HITS-CLIP | 29167373 | Unspecified |
| mmu-miR-152-5p | MIMAT0016991 | Ptpn6 | 15170 | HITS-CLIP | 29167373 | Unspecified |
| mmu-miR-152-5p | MIMAT0016991 | Pum1 | 80912 | HITS-CLIP | 29167373 | Unspecified |
| mmu-miR-152-5p | MIMAT0016991 | Rapgef2 | 76089 | HITS-CLIP | 29167373 | Unspecified |
| mmu-miR-152-5p | MIMAT0016991 | Rbms1 | 56878 | HITS-CLIP | 29167373 | Unspecified |
| mmu-miR-152-5p | MIMAT0016991 | Rock2 | 19878 | HITS-CLIP | 29167373 | Unspecified |
| mmu-miR-152-5p | MIMAT0016991 | Seh1l | 72124 | HITS-CLIP | 29167373 | Unspecified |
| mmu-miR-152-5p | MIMAT0016991 | Sipa1l2 | 244668 | HITS-CLIP | 29167373 | Unspecified |
| mmu-miR-152-5p | MIMAT0016991 | Son | 20658 | HITS-CLIP | 29167373 | Unspecified |
| mmu-miR-152-5p | MIMAT0016991 | Stat2 | 20847 | HITS-CLIP | 29167373 | Unspecified |
| mmu-miR-152-5p | MIMAT0016991 | Syne1 | 64009 | HITS-CLIP | 29167373 | Unspecified |
| mmu-miR-152-5p | MIMAT0016991 | Tardbp | 230908 | HITS-CLIP | 29167373 | Unspecified |
| mmu-miR-152-5p | MIMAT0016991 | Tent2 | 167153 | HITS-CLIP | 29167373 | Unspecified |
| mmu-miR-152-5p | MIMAT0016991 | Tet3 | 194388 | HITS-CLIP | 29167373 | Unspecified |
| mmu-miR-152-5p | MIMAT0016991 | Tgif1 | 21815 | HITS-CLIP | 29167373 | Unspecified |
| mmu-miR-152-5p | MIMAT0016991 | Tle4 | 21888 | HITS-CLIP | 29167373 | Unspecified |
| mmu-miR-152-5p | MIMAT0016991 | Tm6sf1 | 107769 | HITS-CLIP | 29167373 | Unspecified |
| mmu-miR-152-5p | MIMAT0016991 | Tubb5 | 22154 | HITS-CLIP | 29167373 | Unspecified |
| mmu-miR-152-5p | MIMAT0016991 | Uchl5 | 56207 | HITS-CLIP | 29167373 | Unspecified |
| mmu-miR-152-5p | MIMAT0016991 | Uso1 | 56041 | HITS-CLIP | 29167373 | Unspecified |
| mmu-miR-152-5p | MIMAT0016991 | Xpo4 | 57258 | HITS-CLIP | 29167373 | Unspecified |
| mmu-miR-152-5p | MIMAT0016991 | Xrn2 | 24128 | HITS-CLIP | 29167373 | Unspecified |
| mmu-miR-152-5p | MIMAT0016991 | Zbtb10 | 229055 | HITS-CLIP | 31433990 | Unspecified |
| mmu-miR-152-5p | MIMAT0016991 | Zbtb41 | 226470 | HITS-CLIP | 29167373 | Unspecified |
| mmu-miR-1963 | MIMAT0009436 | Smc4 | 70099 | HITS-CLIP | 26203562 | Unspecified |
| mmu-miR-216b-3p | MIMAT0017233 | Clcf1 | 56708 | HITS-CLIP | 26203562 | Unspecified |
| mmu-miR-216b-3p | MIMAT0017233 | Gm45928 | 105948585 | HITS-CLIP | 26203562 | Unspecified |
| mmu-miR-216b-3p | MIMAT0017233 | Rragd | 52187 | HITS-CLIP | 31433990 | Unspecified |
| mmu-miR-21c | MIMAT0025148 | Bmt2 | 101148 | HITS-CLIP | 31433990 | Unspecified |
| mmu-miR-21c | MIMAT0025148 | C1qbp | 12261 | HITS-CLIP | 31433990 | Unspecified |
| mmu-miR-21c | MIMAT0025148 | Cd69 | 12515 | HITS-CLIP | 31433990 | Unspecified |
| mmu-miR-21c | MIMAT0025148 | Chd7 | 320790 | HITS-CLIP | 31433990 | Unspecified |
| mmu-miR-21c | MIMAT0025148 | Cul2 | 71745 | HITS-CLIP | 31433990 | Unspecified |
| mmu-miR-21c | MIMAT0025148 | Ddx28 | 71986 | HITS-CLIP | 31433990 | Unspecified |
| mmu-miR-21c | MIMAT0025148 | Derl1 | 67819 | HITS-CLIP | 31433990 | Unspecified |
| mmu-miR-21c | MIMAT0025148 | Dhrs7b | 216820 | HITS-CLIP | 26203562 | Unspecified |
| mmu-miR-21c | MIMAT0025148 | H3c11 | 8350 | HITS-CLIP | 31433990 | Unspecified |
| mmu-miR-21c | MIMAT0025148 | Hspa4 | 15525 | HITS-CLIP | 31433990 | Unspecified |
| mmu-miR-21c | MIMAT0025148 | Itgb3 | 16416 | HITS-CLIP | 31433990 | Unspecified |

| | | | | | | |
|---|---|---|---|---|---|---|
| mmu-miR-21c | MIMAT0025148 | Jak1 | 16451 | HITS-CLIP | 31433990 | Unspecified |
| mmu-miR-21c | MIMAT0025148 | Jcad | 57608 | HITS-CLIP | 31433990 | Unspecified |
| mmu-miR-21c | MIMAT0025148 | Josd1 | 74158 | HITS-CLIP | 31433990 | Unspecified |
| mmu-miR-21c | MIMAT0025148 | Mindy2 | 235461 | HITS-CLIP | 31433990 | Unspecified |
| mmu-miR-21c | MIMAT0025148 | Pgap1 | 241062 | HITS-CLIP | 26203562 | Unspecified |
| mmu-miR-21c | MIMAT0025148 | Plp2 | 18824 | HITS-CLIP | 31433990 | Unspecified |
| mmu-miR-21c | MIMAT0025148 | Prkci | 18759 | HITS-CLIP | 31433990 | Unspecified |
| mmu-miR-21c | MIMAT0025148 | Pvr | 52118 | HITS-CLIP | 31433990 | Unspecified |
| mmu-miR-21c | MIMAT0025148 | Reep3 | 28193 | HITS-CLIP | 26203562 | Unspecified |
| mmu-miR-21c | MIMAT0025148 | Relt | 320100 | HITS-CLIP | 31433990 | Unspecified |
| mmu-miR-21c | MIMAT0025148 | Sec11c | 66286 | HITS-CLIP | 31433990 | Unspecified |
| mmu-miR-21c | MIMAT0025148 | Sema4b | 20352 | HITS-CLIP | 26203562 | Unspecified |
| mmu-miR-21c | MIMAT0025148 | Strn4 | 97387 | HITS-CLIP | 26203562 | Unspecified |
| mmu-miR-21c | MIMAT0025148 | Tnrc6c | 217351 | HITS-CLIP | 31433990 | Unspecified |
| mmu-miR-21c | MIMAT0025148 | Trpv2 | 22368 | HITS-CLIP | 31433990 | Unspecified |
| mmu-miR-21c | MIMAT0025148 | Utp25 | 27042 | HITS-CLIP | 31433990 | Unspecified |
| mmu-miR-3068-3p | MIMAT0014843 | Abr | 109934 | HITS-CLIP | 29167373 | Unspecified |
| mmu-miR-3068-3p | MIMAT0014843 | Adnp | 11538 | HITS-CLIP | 29167373 | Unspecified |
| mmu-miR-3068-3p | MIMAT0014843 | Agl | 77559 | HITS-CLIP | 25753418 | Unspecified |
| mmu-miR-3068-3p | MIMAT0014843 | Ago1 | 236511 | HITS-CLIP | 29167373 | Unspecified |
| mmu-miR-3068-3p | MIMAT0014843 | Agrn | 11603 | HITS-CLIP | 25753418 | Unspecified |
| mmu-miR-3068-3p | MIMAT0014843 | Akirin2 | 433693 | HITS-CLIP | 25753418 | Unspecified |
| mmu-miR-3068-3p | MIMAT0014843 | Ankrd11 | 77087 | HITS-CLIP | 29167373 | Unspecified |
| mmu-miR-3068-3p | MIMAT0014843 | Aqr | 11834 | HITS-CLIP | 29167373 | Unspecified |
| mmu-miR-3068-3p | MIMAT0014843 | Arhgap21 | 71435 | HITS-CLIP | 25753418 | Unspecified |
| mmu-miR-3068-3p | MIMAT0014843 | Arhgap45 | 23526 | HITS-CLIP | 29167373 | Unspecified |
| mmu-miR-3068-3p | MIMAT0014843 | Asap1 | 13196 | HITS-CLIP | 29167373 | Unspecified |
| mmu-miR-3068-3p | MIMAT0014843 | Asxl1 | 228790 | HITS-CLIP | 25753418 | Unspecified |
| mmu-miR-3068-3p | MIMAT0014843 | Atosa | 56204 | HITS-CLIP | 25753418 | Unspecified |
| mmu-miR-3068-3p | MIMAT0014843 | Atp2a2 | 11938 | HITS-CLIP | 25753418 | Unspecified |
| mmu-miR-3068-3p | MIMAT0014843 | Atp6v1b2 | 11966 | HITS-CLIP | 29167373 | Unspecified |
| mmu-miR-3068-3p | MIMAT0014843 | B2m | 12010 | HITS-CLIP | 29167373 | Unspecified |
| mmu-miR-3068-3p | MIMAT0014843 | Bod1l | 665775 | HITS-CLIP | 29167373 | Unspecified |
| mmu-miR-3068-3p | MIMAT0014843 | Cd47 | 16423 | HITS-CLIP | 29167373 | Unspecified |
| mmu-miR-3068-3p | MIMAT0014843 | Chm | 12662 | HITS-CLIP | 25753418 | Unspecified |
| mmu-miR-3068-3p | MIMAT0014843 | Cited2 | 17684 | HITS-CLIP | 25753418 | Unspecified |
| mmu-miR-3068-3p | MIMAT0014843 | Clcn3 | 12725 | HITS-CLIP | 29167373 | Unspecified |
| mmu-miR-3068-3p | MIMAT0014843 | Cpeb4 | 67579 | HITS-CLIP | 29167373 | Unspecified |
| mmu-miR-3068-3p | MIMAT0014843 | Ctnna1 | 12385 | HITS-CLIP | 25753418 | Unspecified |

| | | | | | | |
|---|---|---|---|---|---|---|
| mmu-miR-3068-3p | MIMAT0014843 | Cul2 | 71745 | HITS-CLIP | 29167373 | Unspecified |
| mmu-miR-3068-3p | MIMAT0014843 | D5Ertd579e | 320661 | HITS-CLIP | 29167373 | Unspecified |
| mmu-miR-3068-3p | MIMAT0014843 | Dab2 | 13132 | HITS-CLIP | 29167373 | Unspecified |
| mmu-miR-3068-3p | MIMAT0014843 | Dctn3 | 53598 | HITS-CLIP | 25753418 | Unspecified |
| mmu-miR-3068-3p | MIMAT0014843 | Ddx3y | 26900 | HITS-CLIP | 25753418 | Unspecified |
| mmu-miR-3068-3p | MIMAT0014843 | Dicer1 | 192119 | HITS-CLIP | 29167373 | Unspecified |
| mmu-miR-3068-3p | MIMAT0014843 | Dld | 13382 | HITS-CLIP | 25753418 | Unspecified |
| mmu-miR-3068-3p | MIMAT0014843 | Dock2 | 94176 | HITS-CLIP | 29167373 | Unspecified |
| mmu-miR-3068-3p | MIMAT0014843 | Dusp7 | 235584 | HITS-CLIP | 29167373 | Unspecified |
| mmu-miR-3068-3p | MIMAT0014843 | Ecpas | 23392 | HITS-CLIP | 25753418 | Unspecified |
| mmu-miR-3068-3p | MIMAT0014843 | Egr1 | 13653 | HITS-CLIP | 25753418 | Unspecified |
| mmu-miR-3068-3p | MIMAT0014843 | Ep300 | 328572 | HITS-CLIP | 29167373 | Unspecified |
| mmu-miR-3068-3p | MIMAT0014843 | Erbin | 55914 | HITS-CLIP | 29167373 | Unspecified |
| mmu-miR-3068-3p | MIMAT0014843 | Ergic1 | 67458 | HITS-CLIP | 29167373 | Unspecified |
| mmu-miR-3068-3p | MIMAT0014843 | Esrrg | 26381 | HITS-CLIP | 25753418 | Unspecified |
| mmu-miR-3068-3p | MIMAT0014843 | Esyt1 | 23943 | HITS-CLIP | 29167373 | Unspecified |
| mmu-miR-3068-3p | MIMAT0014843 | Ezh1 | 14055 | HITS-CLIP | 25753418 | Unspecified |
| mmu-miR-3068-3p | MIMAT0014843 | Flna | 192176 | HITS-CLIP | 29167373 | Unspecified |
| mmu-miR-3068-3p | MIMAT0014843 | Flnc | 68794 | HITS-CLIP | 25753418 | Unspecified |
| mmu-miR-3068-3p | MIMAT0014843 | Foxk2 | 68837 | HITS-CLIP | 29167373 | Unspecified |
| mmu-miR-3068-3p | MIMAT0014843 | Ggnbp2 | 217039 | HITS-CLIP | 31433990 | Unspecified |
| mmu-miR-3068-3p | MIMAT0014843 | Ghr | 14600 | HITS-CLIP | 25753418 | Unspecified |
| mmu-miR-3068-3p | MIMAT0014843 | Gk | 14933 | HITS-CLIP | 29167373 | Unspecified |
| mmu-miR-3068-3p | MIMAT0014843 | Gm38431 | 100568458 | HITS-CLIP | 25753418 | Unspecified |
| mmu-miR-3068-3p | MIMAT0014843 | Gpcpd1 | 74182 | HITS-CLIP | 25753418 | Unspecified |
| mmu-miR-3068-3p | MIMAT0014843 | Heatr5b | 320473 | HITS-CLIP | 29167373 | Unspecified |
| mmu-miR-3068-3p | MIMAT0014843 | Hif1an | 319594 | HITS-CLIP | 25753418 | Unspecified |
| mmu-miR-3068-3p | MIMAT0014843 | Hrc | 15464 | HITS-CLIP | 25753418 | Unspecified |
| mmu-miR-3068-3p | MIMAT0014843 | Huwe1 | 59026 | HITS-CLIP | 29167373 | Unspecified |
| mmu-miR-3068-3p | MIMAT0014843 | Itpkb | 320404 | HITS-CLIP | 29167373 | Unspecified |
| mmu-miR-3068-3p | MIMAT0014843 | Junb | 16477 | HITS-CLIP | 29167373 | Unspecified |
| mmu-miR-3068-3p | MIMAT0014843 | Klf3 | 16599 | HITS-CLIP | 29167373 | Unspecified |
| mmu-miR-3068-3p | MIMAT0014843 | Klf4 | 16600 | HITS-CLIP | 25753418 | Unspecified |
| mmu-miR-3068-3p | MIMAT0014843 | Lemd3 | 380664 | HITS-CLIP | 29167373 | Unspecified |
| mmu-miR-3068-3p | MIMAT0014843 | Map3k1 | 26401 | HITS-CLIP | 25753418 | Unspecified |
| mmu-miR-3068-3p | MIMAT0014843 | Map3k12 | 26404 | HITS-CLIP | 31433990 | Unspecified |
| mmu-miR-3068-3p | MIMAT0014843 | Map4k4 | 26921 | HITS-CLIP | 29167373 | Unspecified |
| mmu-miR-3068-3p | MIMAT0014843 | Mcm4 | 17217 | HITS-CLIP | 29167373 | Unspecified |
| mmu-miR-3068-3p | MIMAT0014843 | Mdn1 | 100019 | HITS-CLIP | 25753418 | Unspecified |

| | | | | | | |
|---|---|---|---|---|---|---|
| mmu-miR-3068-3p | MIMAT0014843 | Mef2a | 17258 | HITS-CLIP | 29167373 | Unspecified |
| mmu-miR-3068-3p | MIMAT0014843 | Mplkip | 66308 | HITS-CLIP | 29167373 | Unspecified |
| mmu-miR-3068-3p | MIMAT0014843 | Mrc1 | 17533 | HITS-CLIP | 29167373 | Unspecified |
| mmu-miR-3068-3p | MIMAT0014843 | Msl1 | 74026 | HITS-CLIP | 29167373 | Unspecified |
| mmu-miR-3068-3p | MIMAT0014843 | Myh9 | 17886 | HITS-CLIP | 29167373 | Unspecified |
| mmu-miR-3068-3p | MIMAT0014843 | Naa15 | 74838 | HITS-CLIP | 29167373 | Unspecified |
| mmu-miR-3068-3p | MIMAT0014843 | Ndst2 | 17423 | HITS-CLIP | 29167373 | Unspecified |
| mmu-miR-3068-3p | MIMAT0014843 | Nf1 | 18015 | HITS-CLIP | 29167373 | Unspecified |
| mmu-miR-3068-3p | MIMAT0014843 | Nfib | 18028 | HITS-CLIP | 25753418 | Unspecified |
| mmu-miR-3068-3p | MIMAT0014843 | Osbpl8 | 237542 | HITS-CLIP | 29167373 | Unspecified |
| mmu-miR-3068-3p | MIMAT0014843 | Otulinl | 54491 | HITS-CLIP | 29167373 | Unspecified |
| mmu-miR-3068-3p | MIMAT0014843 | Oxa1l | 69089 | HITS-CLIP | 25753418 | Unspecified |
| mmu-miR-3068-3p | MIMAT0014843 | Pde8a | 18584 | HITS-CLIP | 29167373 | Unspecified |
| mmu-miR-3068-3p | MIMAT0014843 | Phactr4 | 100169 | HITS-CLIP | 25753418 | Unspecified |
| mmu-miR-3068-3p | MIMAT0014843 | Pip4k2b | 108083 | HITS-CLIP | 29167373 | Unspecified |
| mmu-miR-3068-3p | MIMAT0014843 | Pip4p1 | 90809 | HITS-CLIP | 29167373 | Unspecified |
| mmu-miR-3068-3p | MIMAT0014843 | Plcb2 | 18796 | HITS-CLIP | 29167373 | Unspecified |
| mmu-miR-3068-3p | MIMAT0014843 | Plin2 | 11520 | HITS-CLIP | 29167373 | Unspecified |
| mmu-miR-3068-3p | MIMAT0014843 | Pole3 | 59001 | HITS-CLIP | 25753418 | Unspecified |
| mmu-miR-3068-3p | MIMAT0014843 | Ppp1r2 | 66849 | HITS-CLIP | 29167373 | Unspecified |
| mmu-miR-3068-3p | MIMAT0014843 | Ppp1r3a | 140491 | HITS-CLIP | 25753418 | Unspecified |
| mmu-miR-3068-3p | MIMAT0014843 | Ptprc | 19264 | HITS-CLIP | 29167373 | Unspecified |
| mmu-miR-3068-3p | MIMAT0014843 | Rab14 | 68365 | HITS-CLIP | 29167373 | Unspecified |
| mmu-miR-3068-3p | MIMAT0014843 | Ranbp2 | 19386 | HITS-CLIP | 29167373 | Unspecified |
| mmu-miR-3068-3p | MIMAT0014843 | Raver1 | 71766 | HITS-CLIP | 25753418 | Unspecified |
| mmu-miR-3068-3p | MIMAT0014843 | Rc3h1 | 381305 | HITS-CLIP | 29167373 | Unspecified |
| mmu-miR-3068-3p | MIMAT0014843 | Rel | 19696 | HITS-CLIP | 29167373 | Unspecified |
| mmu-miR-3068-3p | MIMAT0014843 | Rin2 | 74030 | HITS-CLIP | 29167373 | Unspecified |
| mmu-miR-3068-3p | MIMAT0014843 | Rmdn1 | 66302 | HITS-CLIP | 25753418 | Unspecified |
| mmu-miR-3068-3p | MIMAT0014843 | Rock2 | 19878 | HITS-CLIP | 29167373 | Unspecified |
| mmu-miR-3068-3p | MIMAT0014843 | Sez6l | 56747 | qCLASH | 31393953 | Unspecified |
| mmu-miR-3068-3p | MIMAT0014843 | Sf3b2 | 319322 | HITS-CLIP | 29167373 | Unspecified |
| mmu-miR-3068-3p | MIMAT0014843 | Slain2 | 75991 | HITS-CLIP | 29167373 | Unspecified |
| mmu-miR-3068-3p | MIMAT0014843 | Slbp | 20492 | HITS-CLIP | 29167373 | Unspecified |
| mmu-miR-3068-3p | MIMAT0014843 | Slc6a8 | 102857 | HITS-CLIP | 25753418 | Unspecified |
| mmu-miR-3068-3p | MIMAT0014843 | Smad5 | 17129 | HITS-CLIP | 29167373 | Unspecified |
| mmu-miR-3068-3p | MIMAT0014843 | Snx18 | 170625 | HITS-CLIP | 29167373 | Unspecified |
| mmu-miR-3068-3p | MIMAT0014843 | Srek1 | 218543 | HITS-CLIP | 29167373 | Unspecified |
| mmu-miR-3068-3p | MIMAT0014843 | Srsf7 | 225027 | HITS-CLIP | 29167373 | Unspecified |

| | | | | | | |
|---|---|---|---|---|---|---|
| mmu-miR-3068-3p | MIMAT0014843 | Stk16 | 20872 | HITS-CLIP | 25753418 | Unspecified |
| mmu-miR-3068-3p | MIMAT0014843 | Strip2 | 320609 | HITS-CLIP | 25753418 | Unspecified |
| mmu-miR-3068-3p | MIMAT0014843 | Taldo1 | 21351 | HITS-CLIP | 29167373 | Unspecified |
| mmu-miR-3068-3p | MIMAT0014843 | Tgfbi | 21810 | HITS-CLIP | 29167373 | Unspecified |
| mmu-miR-3068-3p | MIMAT0014843 | Timp3 | 21859 | HITS-CLIP | 25753418 | Unspecified |
| mmu-miR-3068-3p | MIMAT0014843 | Tmem144 | 70652 | HITS-CLIP | 25753418 | Unspecified |
| mmu-miR-3068-3p | MIMAT0014843 | Tmem185b | 226351 | HITS-CLIP | 29167373 | Unspecified |
| mmu-miR-3068-3p | MIMAT0014843 | Tmem241 | 338363 | HITS-CLIP | 29167373 | Unspecified |
| mmu-miR-3068-3p | MIMAT0014843 | Tmem38b | 52076 | qCLASH | 31393953 | Unspecified |
| mmu-miR-3068-3p | MIMAT0014843 | Tomm70a | 28185 | HITS-CLIP | 29167373 | Unspecified |
| mmu-miR-3068-3p | MIMAT0014843 | Tor1aip2 | 240832 | HITS-CLIP | 29167373 | Unspecified |
| mmu-miR-3068-3p | MIMAT0014843 | Trpm7 | 58800 | HITS-CLIP | 29167373 | Unspecified |
| mmu-miR-3068-3p | MIMAT0014843 | Ttc17 | 74569 | HITS-CLIP | 25753418 | Unspecified |
| mmu-miR-3068-3p | MIMAT0014843 | Ttn | 22138 | HITS-CLIP | 25753418 | Unspecified |
| mmu-miR-3068-3p | MIMAT0014843 | Uba1 | 22201 | HITS-CLIP | 29167373 | Unspecified |
| mmu-miR-3068-3p | MIMAT0014843 | Ube2r2 | 67615 | HITS-CLIP | 25753418 | Unspecified |
| mmu-miR-3068-3p | MIMAT0014843 | Ubr3 | 68795 | HITS-CLIP | 29167373 | Unspecified |
| mmu-miR-3068-3p | MIMAT0014843 | Wasf2 | 242687 | HITS-CLIP | 29167373 | Unspecified |
| mmu-miR-3068-3p | MIMAT0014843 | Wdr5 | 140858 | HITS-CLIP | 29167373 | Unspecified |
| mmu-miR-3068-3p | MIMAT0014843 | Xiap | 11798 | HITS-CLIP | 29167373 | Unspecified |
| mmu-miR-3068-3p | MIMAT0014843 | Xrn2 | 24128 | HITS-CLIP | 29167373 | Unspecified |
| mmu-miR-3068-3p | MIMAT0014843 | Yme1l1 | 27377 | HITS-CLIP | 25753418 | Unspecified |
| mmu-miR-3068-3p | MIMAT0014843 | Yy1 | 22632 | HITS-CLIP | 29167373 | Unspecified |
| mmu-miR-3068-3p | MIMAT0014843 | Zeb2 | 24136 | HITS-CLIP | 29167373 | Unspecified |
| mmu-miR-3068-3p | MIMAT0014843 | Zfp36l1 | 12192 | HITS-CLIP | 29167373 | Unspecified |
| mmu-miR-3068-3p | MIMAT0014843 | Zfp800 | 627049 | HITS-CLIP | 29167373 | Unspecified |
| mmu-miR-3068-3p | MIMAT0014843 | Zmiz2 | 52915 | HITS-CLIP | 29167373 | Unspecified |
| mmu-miR-3092-5p | MIMAT0014905 | Dnajc16 | 214063 | HITS-CLIP | 26203562 | Unspecified |
| mmu-miR-3092-5p | MIMAT0014905 | Fbxw2 | 30050 | HITS-CLIP | 26203562 | Unspecified |
| mmu-miR-3092-5p | MIMAT0014905 | Ptdss1 | 19210 | HITS-CLIP | 31433990 | Unspecified |
| mmu-miR-3092-5p | MIMAT0014905 | Wbp1l | 226178 | HITS-CLIP | 26203562 | Unspecified |
| mmu-miR-3106-3p | MIMAT0014818 | Dot1l | 208266 | HITS-CLIP | 26203562 | Unspecified |
| mmu-miR-3106-3p | MIMAT0014818 | Fbxw7 | 50754 | HITS-CLIP | 31433990 | Unspecified |
| mmu-miR-3106-3p | MIMAT0014818 | H4c14 | 8359 | HITS-CLIP | 31433990 | Unspecified |
| mmu-miR-3106-3p | MIMAT0014818 | Lars2 | 102436 | HITS-CLIP | 31433990 | Unspecified |
| mmu-miR-3106-3p | MIMAT0014818 | Luc7l3 | 67684 | HITS-CLIP | 31433990 | Unspecified |
| mmu-miR-3108-3p | MIMAT0014948 | Actn1 | 109711 | HITS-CLIP | 26203562 | Unspecified |
| mmu-miR-3108-3p | MIMAT0014948 | Heca | 380629 | HITS-CLIP | 31433990 | Unspecified |
| mmu-miR-3108-3p | MIMAT0014948 | Krt16 | 16666 | HITS-CLIP | 26203562 | Unspecified |

| | | | | | | |
|---|---|---|---|---|---|---|
| mmu-miR-3108-3p | MIMAT0014948 | Nol11 | 68979 | HITS-CLIP | 31433990 | Unspecified |
| mmu-miR-3108-3p | MIMAT0014948 | Tiparp | 99929 | HITS-CLIP | 26203562 | Unspecified |
| mmu-miR-3108-3p | MIMAT0014948 | Zfp217 | 228913 | HITS-CLIP | 31433990 | Unspecified |
| mmu-miR-3109-5p | MIMAT0014949 | Acvr1b | 11479 | HITS-CLIP | 31433990\|26203562 | Unspecified |
| mmu-miR-3109-5p | MIMAT0014949 | Chd7 | 320790 | HITS-CLIP | 31433990 | Unspecified |
| mmu-miR-3109-5p | MIMAT0014949 | Purb | 19291 | HITS-CLIP | 31433990 | Unspecified |
| mmu-miR-383-3p | MIMAT0017082 | Chd7 | 320790 | HITS-CLIP | 26203562 | Unspecified |
| mmu-miR-496b | MIMAT0025158 | Ecpas | 23392 | HITS-CLIP | 31433990 | Unspecified |
| mmu-miR-5615-3p | MIMAT0022356 | Arid5b | 71371 | HITS-CLIP | 26203562 | Unspecified |
| mmu-miR-5615-3p | MIMAT0022356 | Bmpr1a | 12166 | HITS-CLIP | 26203562 | Unspecified |
| mmu-miR-5615-3p | MIMAT0022356 | Ctsd | 13033 | HITS-CLIP | 26203562 | Unspecified |
| mmu-miR-5615-3p | MIMAT0022356 | Dhx16 | 69192 | HITS-CLIP | 31433990 | Unspecified |
| mmu-miR-5615-3p | MIMAT0022356 | Dnajc13 | 235567 | HITS-CLIP | 26203562 | Unspecified |
| mmu-miR-5615-3p | MIMAT0022356 | Gata3 | 14462 | HITS-CLIP | 31433990 | Unspecified |
| mmu-miR-5615-3p | MIMAT0022356 | Kif3a | 16568 | HITS-CLIP | 31433990 | Unspecified |
| mmu-miR-5615-3p | MIMAT0022356 | Lrp6 | 16974 | HITS-CLIP | 26203562 | Unspecified |
| mmu-miR-5615-3p | MIMAT0022356 | Map3k1 | 26401 | HITS-CLIP | 31433990 | Unspecified |
| mmu-miR-5615-3p | MIMAT0022356 | Mtarc2 | 54996 | HITS-CLIP | 31433990 | Unspecified |
| mmu-miR-5615-3p | MIMAT0022356 | Myb | 17863 | HITS-CLIP | 31433990 | Unspecified |
| mmu-miR-5615-3p | MIMAT0022356 | Npepps | 19155 | HITS-CLIP | 31433990 | Unspecified |
| mmu-miR-5615-3p | MIMAT0022356 | Plk3 | 12795 | HITS-CLIP | 31433990 | Unspecified |
| mmu-miR-5615-3p | MIMAT0022356 | Ppp4r1 | 70351 | HITS-CLIP | 31433990 | Unspecified |
| mmu-miR-5615-3p | MIMAT0022356 | Prss8 | 76560 | HITS-CLIP | 26203562 | Unspecified |
| mmu-miR-5615-3p | MIMAT0022356 | Shcbp1 | 20419 | HITS-CLIP | 31433990 | Unspecified |
| mmu-miR-5615-3p | MIMAT0022356 | Sik1 | 17691 | HITS-CLIP | 26203562 | Unspecified |
| mmu-miR-5615-3p | MIMAT0022356 | Smc1a | 24061 | HITS-CLIP | 31433990 | Unspecified |
| mmu-miR-5615-3p | MIMAT0022356 | Tbl1xr1 | 81004 | HITS-CLIP | 26203562 | Unspecified |
| mmu-miR-5615-3p | MIMAT0022356 | Tmem184b | 223693 | HITS-CLIP | 26203562 | Unspecified |
| mmu-miR-5615-3p | MIMAT0022356 | Trio | 223435 | HITS-CLIP | 26203562 | Unspecified |
| mmu-miR-5615-3p | MIMAT0022356 | Wnt9a | 216795 | HITS-CLIP | 26203562 | Unspecified |
| mmu-miR-5615-3p | MIMAT0022356 | Zbtb7b | 22724 | HITS-CLIP | 26203562 | Unspecified |
| mmu-miR-5615-5p | MIMAT0022355 | Atp5a1 | 11946 | HITS-CLIP | 31433990 | Unspecified |
| mmu-miR-5615-5p | MIMAT0022355 | Bach2 | 12014 | HITS-CLIP | 31433990 | Unspecified |
| mmu-miR-5615-5p | MIMAT0022355 | C1qbp | 12261 | HITS-CLIP | 31433990 | Unspecified |
| mmu-miR-5615-5p | MIMAT0022355 | Grhl1 | 195733 | HITS-CLIP | 26203562 | Unspecified |
| mmu-miR-5615-5p | MIMAT0022355 | Iqgap2 | 544963 | HITS-CLIP | 31433990 | Unspecified |
| mmu-miR-5615-5p | MIMAT0022355 | Ppp6c | 67857 | HITS-CLIP | 31433990 | Unspecified |
| mmu-miR-5615-5p | MIMAT0022355 | Snx17 | 266781 | HITS-CLIP | 31433990 | Unspecified |
| mmu-miR-5615-5p | MIMAT0022355 | Vcl | 22330 | HITS-CLIP | 26203562 | Unspecified |

| | | | | | | |
|---|---|---|---|---|---|---|
| mmu-miR-5615-5p | MIMAT0022355 | Zbtb7b | 22724 | HITS-CLIP | 26203562 | Unspecified |
| mmu-miR-5623-3p | MIMAT0022374 | Ap3d1 | 11776 | HITS-CLIP | 26203562 | Unspecified |
| mmu-miR-5623-3p | MIMAT0022374 | Chaf1a | 27221 | HITS-CLIP | 31433990 | Unspecified |
| mmu-miR-5623-3p | MIMAT0022374 | Cwf19l2 | 244672 | HITS-CLIP | 31433990 | Unspecified |
| mmu-miR-5623-3p | MIMAT0022374 | Fbxo28 | 67948 | HITS-CLIP | 31433990 | Unspecified |
| mmu-miR-5623-3p | MIMAT0022374 | Itpr1 | 16438 | HITS-CLIP | 31433990 | Unspecified |
| mmu-miR-5623-3p | MIMAT0022374 | Larp1 | 73158 | HITS-CLIP | 26203562 | Unspecified |
| mmu-miR-5623-3p | MIMAT0022374 | Map3k1 | 26401 | HITS-CLIP | 31433990 | Unspecified |
| mmu-miR-5623-3p | MIMAT0022374 | Rnf31 | 268749 | HITS-CLIP | 31433990 | Unspecified |
| mmu-miR-5623-3p | MIMAT0022374 | Rock1 | 19877 | HITS-CLIP | 31433990 | Unspecified |
| mmu-miR-5623-3p | MIMAT0022374 | Setd2 | 235626 | HITS-CLIP | 31433990 | Unspecified |
| mmu-miR-5623-3p | MIMAT0022374 | Spen | 56381 | HITS-CLIP | 26203562 | Unspecified |
| mmu-miR-5623-3p | MIMAT0022374 | Stk40 | 74178 | HITS-CLIP | 26203562 | Unspecified |
| mmu-miR-5623-3p | MIMAT0022374 | Top1 | 21969 | HITS-CLIP | 26203562 | Unspecified |
| mmu-miR-5623-5p | MIMAT0022373 | Caprin1 | 53872 | HITS-CLIP | 31433990 | Unspecified |
| mmu-miR-5623-5p | MIMAT0022373 | Ccnd2 | 12444 | HITS-CLIP | 31433990 | Unspecified |
| mmu-miR-5623-5p | MIMAT0022373 | Cd44 | 12505 | HITS-CLIP | 31433990 | Unspecified |
| mmu-miR-5623-5p | MIMAT0022373 | Mcm2 | 17216 | HITS-CLIP | 31433990 | Unspecified |
| mmu-miR-5623-5p | MIMAT0022373 | Pbx2 | 18515 | HITS-CLIP | 31433990\|26203562 | Unspecified |
| mmu-miR-5623-5p | MIMAT0022373 | Pkm | 18746 | HITS-CLIP | 31433990 | Unspecified |
| mmu-miR-5623-5p | MIMAT0022373 | Wsb1 | 78889 | HITS-CLIP | 31433990 | Unspecified |
| mmu-miR-5626-3p | MIMAT0022382 | 0610030E20Rik | 68364 | HITS-CLIP | 31433990 | Unspecified |
| mmu-miR-5626-3p | MIMAT0022382 | Lrrc8c | 100604 | HITS-CLIP | 31433990 | Unspecified |
| mmu-miR-5626-3p | MIMAT0022382 | Nbeal2 | 235627 | HITS-CLIP | 31433990 | Unspecified |
| mmu-miR-5626-3p | MIMAT0022382 | Ndufaf4 | 68493 | HITS-CLIP | 31433990 | Unspecified |
| mmu-miR-5626-3p | MIMAT0022382 | Pank3 | 211347 | HITS-CLIP | 31433990 | Unspecified |
| mmu-miR-5626-3p | MIMAT0022382 | Pigs | 276846 | HITS-CLIP | 31433990 | Unspecified |
| mmu-miR-5626-3p | MIMAT0022382 | Pip4p2 | 55529 | HITS-CLIP | 31433990 | Unspecified |
| mmu-miR-5626-3p | MIMAT0022382 | Sik3 | 70661 | HITS-CLIP | 31433990 | Unspecified |
| mmu-miR-5626-5p | MIMAT0022381 | Wdr37 | 207615 | HITS-CLIP | 31433990 | Unspecified |
| mmu-miR-5709-3p | MIMAT0027104 | Trio | 223435 | HITS-CLIP | 26203562 | Unspecified |
| mmu-miR-5709-5p | MIMAT0022504 | Ccny | 67974 | HITS-CLIP | 26203562 | Unspecified |
| mmu-miR-592-3p | MIMAT0017234 | Cdc26 | 66440 | HITS-CLIP | 31433990 | Unspecified |
| mmu-miR-592-3p | MIMAT0017234 | Map3k1 | 26401 | HITS-CLIP | 31433990 | Unspecified |
| mmu-miR-6238 | MIMAT0024859 | Sox12 | 20667 | qCLASH | 31393953 | Unspecified |
| mmu-miR-6238 | MIMAT0024859 | Wasl | 73178 | HITS-CLIP | 31433990 | Unspecified |
| mmu-miR-6238 | MIMAT0024859 | Xndc1 | 102443350 | HITS-CLIP | 31433990 | Unspecified |
| mmu-miR-6239 | MIMAT0024860 | Chd7 | 320790 | HITS-CLIP | 31433990 | Unspecified |
| mmu-miR-6342 | MIMAT0025085 | Ahcyl2 | 74340 | HITS-CLIP | 31433990 | Unspecified |

| | | | | | | |
|---|---|---|---|---|---|---|
| mmu-miR-6342 | MIMAT0025085 | Casd1 | 213819 | HITS-CLIP | 31433990 | Unspecified |
| mmu-miR-6342 | MIMAT0025085 | Chd7 | 320790 | HITS-CLIP | 31433990 | Unspecified |
| mmu-miR-6342 | MIMAT0025085 | Ctdspl2 | 329506 | HITS-CLIP | 31433990 | Unspecified |
| mmu-miR-6342 | MIMAT0025085 | Prdx6 | 11758 | HITS-CLIP | 31433990 | Unspecified |
| mmu-miR-6342 | MIMAT0025085 | Sipa1l3 | 74206 | HITS-CLIP | 31433990 | Unspecified |
| mmu-miR-6342 | MIMAT0025085 | Slc39a3 | 106947 | HITS-CLIP | 26203562 | Unspecified |
| mmu-miR-6342 | MIMAT0025085 | Vcp | 269523 | HITS-CLIP | 31433990 | Unspecified |
| mmu-miR-6342 | MIMAT0025085 | Wasl | 73178 | HITS-CLIP | 31433990 | Unspecified |
| mmu-miR-6342 | MIMAT0025085 | Zfp292 | 30046 | HITS-CLIP | 26203562 | Unspecified |
| mmu-miR-6343 | MIMAT0025086 | Abcg1 | 11307 | HITS-CLIP | 31433990 | Unspecified |
| mmu-miR-6343 | MIMAT0025086 | Bmt2 | 101148 | HITS-CLIP | 31433990 | Unspecified |
| mmu-miR-6343 | MIMAT0025086 | Dcp2 | 70640 | HITS-CLIP | 31433990 | Unspecified |
| mmu-miR-6343 | MIMAT0025086 | Heatr3 | 234549 | HITS-CLIP | 31433990 | Unspecified |
| mmu-miR-6343 | MIMAT0025086 | Hprt | 15452 | HITS-CLIP | 31433990\|26203562 | Unspecified |
| mmu-miR-6343 | MIMAT0025086 | Hsp90ab1 | 15516 | HITS-CLIP | 26203562 | Unspecified |
| mmu-miR-6343 | MIMAT0025086 | Trp53inp1 | 60599 | HITS-CLIP | 31433990 | Unspecified |
| mmu-miR-6343 | MIMAT0025086 | Tubb5 | 22154 | HITS-CLIP | 31433990 | Unspecified |
| mmu-miR-6348 | MIMAT0025091 | Ankmy2 | 217473 | HITS-CLIP | 31433990 | Unspecified |
| mmu-miR-6348 | MIMAT0025091 | Bsg | 12215 | HITS-CLIP | 31433990 | Unspecified |
| mmu-miR-6348 | MIMAT0025091 | Mtmr3 | 74302 | HITS-CLIP | 31433990 | Unspecified |
| mmu-miR-6348 | MIMAT0025091 | Trim27 | 19720 | HITS-CLIP | 31433990 | Unspecified |
| mmu-miR-6348 | MIMAT0025091 | Zfp36 | 22695 | HITS-CLIP | 31433990 | Unspecified |
| mmu-miR-6354 | MIMAT0025097 | Cmtm4 | 97487 | HITS-CLIP | 31433990 | Unspecified |
| mmu-miR-6354 | MIMAT0025097 | Edem1 | 192193 | HITS-CLIP | 26203562 | Unspecified |
| mmu-miR-6354 | MIMAT0025097 | Eif3c | 56347 | HITS-CLIP | 31433990 | Unspecified |
| mmu-miR-6354 | MIMAT0025097 | Nup210 | 54563 | HITS-CLIP | 31433990 | Unspecified |
| mmu-miR-6354 | MIMAT0025097 | Plec | 18810 | HITS-CLIP | 26203562 | Unspecified |
| mmu-miR-6354 | MIMAT0025097 | Stk40 | 74178 | HITS-CLIP | 26203562 | Unspecified |
| mmu-miR-6354 | MIMAT0025097 | Tmsb4x | 19241 | HITS-CLIP | 31433990 | Unspecified |
| mmu-miR-6354 | MIMAT0025097 | Zbtb10 | 229055 | HITS-CLIP | 31433990 | Unspecified |
| mmu-miR-6360 | MIMAT0025103 | Arhgap10 | 78514 | HITS-CLIP | 31433990 | Unspecified |
| mmu-miR-6360 | MIMAT0025103 | Dpp8 | 74388 | HITS-CLIP | 31433990 | Unspecified |
| mmu-miR-6360 | MIMAT0025103 | Itpr1 | 16438 | HITS-CLIP | 31433990 | Unspecified |
| mmu-miR-6360 | MIMAT0025103 | Osm | 18413 | HITS-CLIP | 31433990 | Unspecified |
| mmu-miR-6360 | MIMAT0025103 | Rgl2 | 19732 | HITS-CLIP | 31433990 | Unspecified |
| mmu-miR-6360 | MIMAT0025103 | Sik2 | 235344 | HITS-CLIP | 31433990 | Unspecified |
| mmu-miR-6360 | MIMAT0025103 | Ube2k | 53323 | HITS-CLIP | 31433990 | Unspecified |
| mmu-miR-6360 | MIMAT0025103 | Vps54 | 245944 | HITS-CLIP | 31433990 | Unspecified |
| mmu-miR-6366 | MIMAT0025110 | Bcor | 71458 | HITS-CLIP | 31433990 | Unspecified |

| | | | | | | |
|---|---|---|---|---|---|---|
| mmu-miR-6366 | MIMAT0025110 | Ip6k1 | 27399 | HITS-CLIP | 31433990 | Unspecified |
| mmu-miR-6366 | MIMAT0025110 | Yy1 | 22632 | HITS-CLIP | 26203562 | Unspecified |
| mmu-miR-6378 | MIMAT0025124 | Ago2 | 239528 | HITS-CLIP | 31433990 | Unspecified |
| mmu-miR-6378 | MIMAT0025124 | Asxl1 | 228790 | HITS-CLIP | 26203562 | Unspecified |
| mmu-miR-6378 | MIMAT0025124 | Canx | 12330 | HITS-CLIP | 31433990 | Unspecified |
| mmu-miR-6378 | MIMAT0025124 | Hsp90ab1 | 15516 | HITS-CLIP | 31433990 | Unspecified |
| mmu-miR-6378 | MIMAT0025124 | Mdfic | 16543 | HITS-CLIP | 31433990 | Unspecified |
| mmu-miR-6378 | MIMAT0025124 | Midn | 59090 | HITS-CLIP | 31433990 | Unspecified |
| mmu-miR-6378 | MIMAT0025124 | Mlec | 109154 | HITS-CLIP | 31433990 | Unspecified |
| mmu-miR-6378 | MIMAT0025124 | Sf3b3 | 101943 | HITS-CLIP | 31433990 | Unspecified |
| mmu-miR-6378 | MIMAT0025124 | Tspyl1 | 22110 | HITS-CLIP | 31433990 | Unspecified |
| mmu-miR-6378 | MIMAT0025124 | Ugcg | 22234 | HITS-CLIP | 31433990 | Unspecified |
| mmu-miR-6384 | MIMAT0025130 | Pdik1l | 230809 | HITS-CLIP | 31433990 | Unspecified |
| mmu-miR-6403 | MIMAT0025155 | Carmil1 | 55604 | HITS-CLIP | 26203562 | Unspecified |
| mmu-miR-6403 | MIMAT0025155 | Srsf6 | 67996 | HITS-CLIP | 31433990 | Unspecified |
| mmu-miR-6405 | MIMAT0025157 | Flot2 | 14252 | HITS-CLIP | 26203562 | Unspecified |
| mmu-miR-6405 | MIMAT0025157 | Itsn2 | 20403 | HITS-CLIP | 31433990 | Unspecified |
| mmu-miR-6405 | MIMAT0025157 | Lamp1 | 16783 | HITS-CLIP | 31433990 | Unspecified |
| mmu-miR-6405 | MIMAT0025157 | Msn | 17698 | HITS-CLIP | 31433990 | Unspecified |
| mmu-miR-6405 | MIMAT0025157 | Rad54l | 19366 | HITS-CLIP | 31433990 | Unspecified |
| mmu-miR-6405 | MIMAT0025157 | Zfx | 22764 | HITS-CLIP | 31433990 | Unspecified |
| mmu-miR-6516-3p | MIMAT0027344 | 2510039O18Rik | 77034 | HITS-CLIP | 26203562 | Unspecified |
| mmu-miR-6516-3p | MIMAT0027344 | Agpat1 | 55979 | HITS-CLIP | 26203562 | Unspecified |
| mmu-miR-6516-3p | MIMAT0027344 | Atp2a2 | 11938 | HITS-CLIP | 31433990 | Unspecified |
| mmu-miR-6516-3p | MIMAT0027344 | Cdc6 | 23834 | HITS-CLIP | 31433990 | Unspecified |
| mmu-miR-6516-3p | MIMAT0027344 | Chaf1a | 27221 | HITS-CLIP | 31433990 | Unspecified |
| mmu-miR-6516-3p | MIMAT0027344 | Chd1 | 12648 | HITS-CLIP | 31433990 | Unspecified |
| mmu-miR-6516-3p | MIMAT0027344 | Dhx15 | 13204 | HITS-CLIP | 26203562 | Unspecified |
| mmu-miR-6516-3p | MIMAT0027344 | Dusp10 | 63953 | HITS-CLIP | 31433990 | Unspecified |
| mmu-miR-6516-3p | MIMAT0027344 | Eps15 | 13858 | HITS-CLIP | 31433990 | Unspecified |
| mmu-miR-6516-3p | MIMAT0027344 | Fam117b | 72750 | HITS-CLIP | 31433990 | Unspecified |
| mmu-miR-6516-3p | MIMAT0027344 | Focad | 230393 | HITS-CLIP | 31433990 | Unspecified |
| mmu-miR-6516-3p | MIMAT0027344 | Gna11 | 14672 | HITS-CLIP | 26203562 | Unspecified |
| mmu-miR-6516-3p | MIMAT0027344 | Grk2 | 156 | HITS-CLIP | 31433990 | Unspecified |
| mmu-miR-6516-3p | MIMAT0027344 | Gtf2f1 | 98053 | HITS-CLIP | 31433990 | Unspecified |
| mmu-miR-6516-3p | MIMAT0027344 | Inppl1 | 16332 | HITS-CLIP | 31433990 | Unspecified |
| mmu-miR-6516-3p | MIMAT0027344 | Insr | 16337 | HITS-CLIP | 31433990 | Unspecified |
| mmu-miR-6516-3p | MIMAT0027344 | Macroh2a1 | 9555 | HITS-CLIP | 31433990 | Unspecified |
| mmu-miR-6516-3p | MIMAT0027344 | Mark3 | 17169 | HITS-CLIP | 31433990 | Unspecified |

| | | | | | | |
|---|---|---|---|---|---|---|
| mmu-miR-6516-3p | MIMAT0027344 | Med31 | 67279 | HITS-CLIP | 26203562 | Unspecified |
| mmu-miR-6516-3p | MIMAT0027344 | Nbeal2 | 235627 | HITS-CLIP | 26203562 | Unspecified |
| mmu-miR-6516-3p | MIMAT0027344 | Ranbp2 | 19386 | HITS-CLIP | 26203562 | Unspecified |
| mmu-miR-6516-3p | MIMAT0027344 | Rreb1 | 68750 | HITS-CLIP | 31433990\|26203562 | Unspecified |
| mmu-miR-6516-3p | MIMAT0027344 | Slc9a1 | 20544 | HITS-CLIP | 26203562 | Unspecified |
| mmu-miR-6516-3p | MIMAT0027344 | Tfrc | 22042 | HITS-CLIP | 31433990 | Unspecified |
| mmu-miR-6516-3p | MIMAT0027344 | Tgfbr2 | 21813 | HITS-CLIP | 26203562 | Unspecified |
| mmu-miR-6516-3p | MIMAT0027344 | Tial1 | 21843 | HITS-CLIP | 31433990 | Unspecified |
| mmu-miR-6516-3p | MIMAT0027344 | Trio | 223435 | HITS-CLIP | 26203562 | Unspecified |
| mmu-miR-6516-3p | MIMAT0027344 | Ttll12 | 223723 | HITS-CLIP | 31433990 | Unspecified |
| mmu-miR-6516-3p | MIMAT0027344 | Ube2z | 268470 | HITS-CLIP | 31433990 | Unspecified |
| mmu-miR-6516-3p | MIMAT0027344 | Vezt | 215008 | HITS-CLIP | 26203562 | Unspecified |
| mmu-miR-6516-5p | MIMAT0027343 | Carmil1 | 55604 | HITS-CLIP | 26203562 | Unspecified |
| mmu-miR-6516-5p | MIMAT0027343 | Ctsd | 13033 | HITS-CLIP | 26203562 | Unspecified |
| mmu-miR-6516-5p | MIMAT0027343 | Cwf19l2 | 244672 | HITS-CLIP | 31433990 | Unspecified |
| mmu-miR-6516-5p | MIMAT0027343 | Plagl2 | 54711 | HITS-CLIP | 31433990 | Unspecified |
| mmu-miR-6516-5p | MIMAT0027343 | Plau | 18792 | HITS-CLIP | 26203562 | Unspecified |
| mmu-miR-6516-5p | MIMAT0027343 | Rrm1 | 20133 | HITS-CLIP | 31433990 | Unspecified |
| mmu-miR-6516-5p | MIMAT0027343 | Safb | 224903 | HITS-CLIP | 31433990 | Unspecified |
| mmu-miR-6516-5p | MIMAT0027343 | Srpk1 | 20815 | HITS-CLIP | 31433990 | Unspecified |
| mmu-miR-6516-5p | MIMAT0027343 | Tln1 | 21894 | HITS-CLIP | 31433990 | Unspecified |
| mmu-miR-6516-5p | MIMAT0027343 | Zfp511 | 69752 | HITS-CLIP | 26203562 | Unspecified |
| mmu-miR-6516-5p | MIMAT0027343 | Zfp710 | 209225 | HITS-CLIP | 31433990 | Unspecified |
| mmu-miR-6537-3p | MIMAT0025582 | Ankfy1 | 11736 | HITS-CLIP | 31433990 | Unspecified |
| mmu-miR-6537-3p | MIMAT0025582 | Cdc26 | 66440 | HITS-CLIP | 26203562\|31433990 | Unspecified |
| mmu-miR-6537-3p | MIMAT0025582 | Golph3 | 66629 | HITS-CLIP | 31433990 | Unspecified |
| mmu-miR-6537-3p | MIMAT0025582 | Il1r1 | 16177 | HITS-CLIP | 26203562 | Unspecified |
| mmu-miR-6537-3p | MIMAT0025582 | Rock1 | 19877 | HITS-CLIP | 26203562 | Unspecified |
| mmu-miR-6537-3p | MIMAT0025582 | Tnfrsf23 | 79201 | HITS-CLIP | 31433990 | Unspecified |
| mmu-miR-6537-3p | MIMAT0025582 | Ubc | 22190 | HITS-CLIP | 26203562 | Unspecified |
| mmu-miR-6537-5p | MIMAT0025581 | Cldn4 | 12740 | HITS-CLIP | 26203562 | Unspecified |
| mmu-miR-6537-5p | MIMAT0025581 | Ece1 | 230857 | HITS-CLIP | 26203562 | Unspecified |
| mmu-miR-6537-5p | MIMAT0025581 | Irf1 | 16362 | HITS-CLIP | 31433990 | Unspecified |
| mmu-miR-6537-5p | MIMAT0025581 | Krt14 | 16664 | HITS-CLIP | 26203562 | Unspecified |
| mmu-miR-6537-5p | MIMAT0025581 | Med28 | 66999 | HITS-CLIP | 26203562 | Unspecified |
| mmu-miR-6537-5p | MIMAT0025581 | Nrros | 224109 | HITS-CLIP | 31433990 | Unspecified |
| mmu-miR-6537-5p | MIMAT0025581 | Psap | 19156 | HITS-CLIP | 31433990 | Unspecified |
| mmu-miR-6537-5p | MIMAT0025581 | Rrm2 | 20135 | HITS-CLIP | 31433990 | Unspecified |
| mmu-miR-6539 | MIMAT0025584 | Atxn1l | 52335 | HITS-CLIP | 26203562 | Unspecified |

| | | | | | | |
|---|---|---|---|---|---|---|
| mmu-miR-6539 | MIMAT0025584 | Cds2 | 110911 | HITS-CLIP | 26203562 | Unspecified |
| mmu-miR-6539 | MIMAT0025584 | Clec2d | 93694 | HITS-CLIP | 31433990 | Unspecified |
| mmu-miR-6539 | MIMAT0025584 | Cpne3 | 70568 | HITS-CLIP | 26203562 | Unspecified |
| mmu-miR-6539 | MIMAT0025584 | Diaph1 | 1729 | HITS-CLIP | 26203562 | Unspecified |
| mmu-miR-6539 | MIMAT0025584 | Hectd4 | 283450 | HITS-CLIP | 31433990 | Unspecified |
| mmu-miR-6539 | MIMAT0025584 | Iffo2 | 212632 | HITS-CLIP | 26203562 | Unspecified |
| mmu-miR-6539 | MIMAT0025584 | Ints8 | 72656 | HITS-CLIP | 31433990 | Unspecified |
| mmu-miR-6539 | MIMAT0025584 | Klhdc3 | 71765 | HITS-CLIP | 31433990 | Unspecified |
| mmu-miR-6539 | MIMAT0025584 | Ppp1cb | 19046 | HITS-CLIP | 31433990 | Unspecified |
| mmu-miR-6539 | MIMAT0025584 | Prpf8 | 192159 | HITS-CLIP | 31433990 | Unspecified |
| mmu-miR-6539 | MIMAT0025584 | Rab2a | 59021 | HITS-CLIP | 31433990 | Unspecified |
| mmu-miR-6539 | MIMAT0025584 | Slfn2 | 20556 | HITS-CLIP | 31433990 | Unspecified |
| mmu-miR-6539 | MIMAT0025584 | Topors | 106021 | HITS-CLIP | 31433990 | Unspecified |
| mmu-miR-6541 | MIMAT0025588 | H1f3 | 3007 | HITS-CLIP | 31433990 | Unspecified |
| mmu-miR-6541 | MIMAT0025588 | H4c4 | 8359 | HITS-CLIP | 31433990 | Unspecified |
| mmu-miR-6541 | MIMAT0025588 | Hif1a | 15251 | HITS-CLIP | 31433990 | Unspecified |
| mmu-miR-6541 | MIMAT0025588 | Lrba | 80877 | HITS-CLIP | 31433990 | Unspecified |
| mmu-miR-6541 | MIMAT0025588 | Mdfic | 16543 | HITS-CLIP | 31433990 | Unspecified |
| mmu-miR-674-3p | MIMAT0003741 | Aco1 | 11428 | HITS-CLIP | 25753418 | Unspecified |
| mmu-miR-674-3p | MIMAT0003741 | Adgrl2 | 23266 | HITS-CLIP | 29167373 | Unspecified |
| mmu-miR-674-3p | MIMAT0003741 | Akap11 | 219181 | HITS-CLIP | 29167373 | Unspecified |
| mmu-miR-674-3p | MIMAT0003741 | Ankrd28 | 105522 | HITS-CLIP | 29167373 | Unspecified |
| mmu-miR-674-3p | MIMAT0003741 | Arid1a | 93760 | HITS-CLIP | 29167373 | Unspecified |
| mmu-miR-674-3p | MIMAT0003741 | Arpc2 | 76709 | HITS-CLIP | 29167373 | Unspecified |
| mmu-miR-674-3p | MIMAT0003741 | Atp6v1a | 11964 | HITS-CLIP | 29167373 | Unspecified |
| mmu-miR-674-3p | MIMAT0003741 | Bcl9 | 77578 | HITS-CLIP | 29167373 | Unspecified |
| mmu-miR-674-3p | MIMAT0003741 | C1galt1 | 94192 | HITS-CLIP | 25753418\|29167373 | Unspecified |
| mmu-miR-674-3p | MIMAT0003741 | Capg | 12332 | HITS-CLIP | 29167373 | Unspecified |
| mmu-miR-674-3p | MIMAT0003741 | Casp2 | 12366 | HITS-CLIP | 29167373 | Unspecified |
| mmu-miR-674-3p | MIMAT0003741 | Cct5 | 12465 | HITS-CLIP | 25753418 | Unspecified |
| mmu-miR-674-3p | MIMAT0003741 | Cdip1 | 66626 | HITS-CLIP | 25753418 | Unspecified |
| mmu-miR-674-3p | MIMAT0003741 | Cisd2 | 67006 | HITS-CLIP | 25753418 | Unspecified |
| mmu-miR-674-3p | MIMAT0003741 | Ckb | 12709 | HITS-CLIP | 29167373 | Unspecified |
| mmu-miR-674-3p | MIMAT0003741 | Clcn3 | 12725 | HITS-CLIP | 29167373 | Unspecified |
| mmu-miR-674-3p | MIMAT0003741 | Cnot6 | 104625 | HITS-CLIP | 29167373 | Unspecified |
| mmu-miR-674-3p | MIMAT0003741 | Cpsf6 | 432508 | HITS-CLIP | 29167373 | Unspecified |
| mmu-miR-674-3p | MIMAT0003741 | Crybg3 | 224273 | HITS-CLIP | 29167373 | Unspecified |
| mmu-miR-674-3p | MIMAT0003741 | Csnk1a1 | 93687 | HITS-CLIP | 29167373 | Unspecified |
| mmu-miR-674-3p | MIMAT0003741 | Ctnnal1 | 54366 | HITS-CLIP | 25753418 | Unspecified |

| | | | | | | |
|---|---|---|---|---|---|---|
| mmu-miR-674-3p | MIMAT0003741 | Ctsb | 13030 | HITS-CLIP | 29167373 | Unspecified |
| mmu-miR-674-3p | MIMAT0003741 | Cybb | 13058 | HITS-CLIP | 29167373 | Unspecified |
| mmu-miR-674-3p | MIMAT0003741 | D5Ertd579e | 320661 | HITS-CLIP | 29167373 | Unspecified |
| mmu-miR-674-3p | MIMAT0003741 | Dagla | 269060 | HITS-CLIP | 29167373 | Unspecified |
| mmu-miR-674-3p | MIMAT0003741 | Dbf4 | 27214 | HITS-CLIP | 29167373 | Unspecified |
| mmu-miR-674-3p | MIMAT0003741 | Ddx21 | 56200 | qCLASH | 31393953 | Unspecified |
| mmu-miR-674-3p | MIMAT0003741 | Dhx9 | 13211 | HITS-CLIP | 29167373 | Unspecified |
| mmu-miR-674-3p | MIMAT0003741 | Dmxl1 | 240283 | HITS-CLIP | 29167373 | Unspecified |
| mmu-miR-674-3p | MIMAT0003741 | Dram2 | 67171 | HITS-CLIP | 29167373 | Unspecified |
| mmu-miR-674-3p | MIMAT0003741 | Dync1h1 | 13424 | HITS-CLIP | 25753418 | Unspecified |
| mmu-miR-674-3p | MIMAT0003741 | Ecpas | 23392 | HITS-CLIP | 29167373 | Unspecified |
| mmu-miR-674-3p | MIMAT0003741 | Eif3e | 16341 | HITS-CLIP | 25753418 | Unspecified |
| mmu-miR-674-3p | MIMAT0003741 | Fbxo10 | 269529 | HITS-CLIP | 25753418 | Unspecified |
| mmu-miR-674-3p | MIMAT0003741 | Flnc | 68794 | HITS-CLIP | 25753418 | Unspecified |
| mmu-miR-674-3p | MIMAT0003741 | Foxk1 | 17425 | HITS-CLIP | 25753418 | Unspecified |
| mmu-miR-674-3p | MIMAT0003741 | Gclc | 14629 | HITS-CLIP | 29167373 | Unspecified |
| mmu-miR-674-3p | MIMAT0003741 | Gpam | 14732 | HITS-CLIP | 25753418 | Unspecified |
| mmu-miR-674-3p | MIMAT0003741 | Hnrnpc | 15381 | HITS-CLIP | 29167373 | Unspecified |
| mmu-miR-674-3p | MIMAT0003741 | Hspg2 | 15530 | HITS-CLIP | 25753418 | Unspecified |
| mmu-miR-674-3p | MIMAT0003741 | Ibtk | 108837 | HITS-CLIP | 29167373 | Unspecified |
| mmu-miR-674-3p | MIMAT0003741 | Iqgap2 | 544963 | HITS-CLIP | 29167373 | Unspecified |
| mmu-miR-674-3p | MIMAT0003741 | Itga6 | 16403 | HITS-CLIP | 29167373 | Unspecified |
| mmu-miR-674-3p | MIMAT0003741 | Itpripl2 | 319622 | HITS-CLIP | 29167373 | Unspecified |
| mmu-miR-674-3p | MIMAT0003741 | Kansl1 | 76719 | HITS-CLIP | 29167373 | Unspecified |
| mmu-miR-674-3p | MIMAT0003741 | Katnbl1 | 72425 | HITS-CLIP | 29167373 | Unspecified |
| mmu-miR-674-3p | MIMAT0003741 | Kbtbd2 | 210973 | HITS-CLIP | 29167373 | Unspecified |
| mmu-miR-674-3p | MIMAT0003741 | Kidins220 | 77480 | HITS-CLIP | 29167373 | Unspecified |
| mmu-miR-674-3p | MIMAT0003741 | Klhl24 | 75785 | HITS-CLIP | 25753418 | Unspecified |
| mmu-miR-674-3p | MIMAT0003741 | Kmt2c | 231051 | HITS-CLIP | 29167373 | Unspecified |
| mmu-miR-674-3p | MIMAT0003741 | Kmt2d | 381022 | HITS-CLIP | 25753418 | Unspecified |
| mmu-miR-674-3p | MIMAT0003741 | Kpna4 | 16649 | HITS-CLIP | 29167373 | Unspecified |
| mmu-miR-674-3p | MIMAT0003741 | Larp4 | 207214 | HITS-CLIP | 29167373 | Unspecified |
| mmu-miR-674-3p | MIMAT0003741 | Lmod2 | 93677 | HITS-CLIP | 25753418 | Unspecified |
| mmu-miR-674-3p | MIMAT0003741 | Maged1 | 94275 | HITS-CLIP | 29167373 | Unspecified |
| mmu-miR-674-3p | MIMAT0003741 | Maz | 17188 | HITS-CLIP | 29167373 | Unspecified |
| mmu-miR-674-3p | MIMAT0003741 | Mbnl1 | 56758 | HITS-CLIP | 29167373 | Unspecified |
| mmu-miR-674-3p | MIMAT0003741 | Mbnl2 | 105559 | HITS-CLIP | 29167373 | Unspecified |
| mmu-miR-674-3p | MIMAT0003741 | Mcm6 | 17219 | HITS-CLIP | 29167373 | Unspecified |
| mmu-miR-674-3p | MIMAT0003741 | Med1 | 19014 | HITS-CLIP | 29167373 | Unspecified |

| | | | | | | |
|---|---|---|---|---|---|---|
| mmu-miR-674-3p | MIMAT0003741 | Met | 17295 | HITS-CLIP | 25753418 | Unspecified |
| mmu-miR-674-3p | MIMAT0003741 | Mfhas1 | 52065 | HITS-CLIP | 29167373 | Unspecified |
| mmu-miR-674-3p | MIMAT0003741 | Mga | 29808 | HITS-CLIP | 29167373 | Unspecified |
| mmu-miR-674-3p | MIMAT0003741 | Mnt | 17428 | HITS-CLIP | 29167373 | Unspecified |
| mmu-miR-674-3p | MIMAT0003741 | Mybbp1a | 18432 | HITS-CLIP | 29167373 | Unspecified |
| mmu-miR-674-3p | MIMAT0003741 | Mybpc3 | 17868 | HITS-CLIP | 25753418 | Unspecified |
| mmu-miR-674-3p | MIMAT0003741 | Myo18a | 360013 | HITS-CLIP | 29167373 | Unspecified |
| mmu-miR-674-3p | MIMAT0003741 | Naa35 | 78689 | HITS-CLIP | 29167373 | Unspecified |
| mmu-miR-674-3p | MIMAT0003741 | Ndufa3 | 66091 | HITS-CLIP | 25753418 | Unspecified |
| mmu-miR-674-3p | MIMAT0003741 | Ndufaf6 | 76947 | HITS-CLIP | 25753418 | Unspecified |
| mmu-miR-674-3p | MIMAT0003741 | Nktr | 18087 | HITS-CLIP | 29167373 | Unspecified |
| mmu-miR-674-3p | MIMAT0003741 | Nom1 | 433864 | HITS-CLIP | 29167373 | Unspecified |
| mmu-miR-674-3p | MIMAT0003741 | Nup214 | 227720 | HITS-CLIP | 29167373 | Unspecified |
| mmu-miR-674-3p | MIMAT0003741 | Nup98 | 269966 | HITS-CLIP | 29167373 | Unspecified |
| mmu-miR-674-3p | MIMAT0003741 | Oas2 | 246728 | HITS-CLIP | 29167373 | Unspecified |
| mmu-miR-674-3p | MIMAT0003741 | Orc3 | 50793 | HITS-CLIP | 25753418 | Unspecified |
| mmu-miR-674-3p | MIMAT0003741 | Pabpc1 | 18458 | HITS-CLIP | 25753418\|29167373 | Unspecified |
| mmu-miR-674-3p | MIMAT0003741 | Pam | 18484 | HITS-CLIP | 25753418 | Unspecified |
| mmu-miR-674-3p | MIMAT0003741 | Pcyt1a | 13026 | HITS-CLIP | 29167373 | Unspecified |
| mmu-miR-674-3p | MIMAT0003741 | Pik3ip1 | 216505 | HITS-CLIP | 25753418 | Unspecified |
| mmu-miR-674-3p | MIMAT0003741 | Pkp2 | 67451 | HITS-CLIP | 25753418 | Unspecified |
| mmu-miR-674-3p | MIMAT0003741 | Plec | 18810 | HITS-CLIP | 25753418 | Unspecified |
| mmu-miR-674-3p | MIMAT0003741 | Prkacb | 18749 | HITS-CLIP | 29167373 | Unspecified |
| mmu-miR-674-3p | MIMAT0003741 | Prpf8 | 192159 | HITS-CLIP | 25753418 | Unspecified |
| mmu-miR-674-3p | MIMAT0003741 | Prrc2c | 226562 | HITS-CLIP | 25753418\|29167373 | Unspecified |
| mmu-miR-674-3p | MIMAT0003741 | Psmf1 | 9491 | HITS-CLIP | 25753418 | Unspecified |
| mmu-miR-674-3p | MIMAT0003741 | Qk | 19317 | HITS-CLIP | 29167373 | Unspecified |
| mmu-miR-674-3p | MIMAT0003741 | Rad23b | 19359 | HITS-CLIP | 25753418 | Unspecified |
| mmu-miR-674-3p | MIMAT0003741 | Rbbp6 | 19647 | HITS-CLIP | 29167373 | Unspecified |
| mmu-miR-674-3p | MIMAT0003741 | Rfc1 | 19687 | HITS-CLIP | 25753418 | Unspecified |
| mmu-miR-674-3p | MIMAT0003741 | Rnf144b | 218215 | HITS-CLIP | 29167373 | Unspecified |
| mmu-miR-674-3p | MIMAT0003741 | Rnf19b | 75234 | qCLASH | 31393953 | Unspecified |
| mmu-miR-674-3p | MIMAT0003741 | Samd8 | 67630 | HITS-CLIP | 25753418 | Unspecified |
| mmu-miR-674-3p | MIMAT0003741 | Saraf | 51669 | HITS-CLIP | 29167373 | Unspecified |
| mmu-miR-674-3p | MIMAT0003741 | Sash1 | 70097 | HITS-CLIP | 29167373 | Unspecified |
| mmu-miR-674-3p | MIMAT0003741 | Scyl2 | 213326 | HITS-CLIP | 29167373 | Unspecified |
| mmu-miR-674-3p | MIMAT0003741 | Sdc3 | 20970 | HITS-CLIP | 29167373 | Unspecified |
| mmu-miR-674-3p | MIMAT0003741 | Sez6l | 56747 | qCLASH | 31393953 | Unspecified |
| mmu-miR-674-3p | MIMAT0003741 | Skil | 20482 | HITS-CLIP | 29167373 | Unspecified |

| | | | | | | |
|---|---|---|---|---|---|---|
| mmu-miR-674-3p | MIMAT0003741 | Slc23a2 | 54338 | HITS-CLIP | 29167373 | Unspecified |
| mmu-miR-674-3p | MIMAT0003741 | Slc39a1 | 30791 | HITS-CLIP | 29167373 | Unspecified |
| mmu-miR-674-3p | MIMAT0003741 | Slc4a7 | 218756 | HITS-CLIP | 29167373 | Unspecified |
| mmu-miR-674-3p | MIMAT0003741 | Slc9a6 | 236794 | HITS-CLIP | 29167373 | Unspecified |
| mmu-miR-674-3p | MIMAT0003741 | Smim8 | 66291 | HITS-CLIP | 25753418 | Unspecified |
| mmu-miR-674-3p | MIMAT0003741 | Srek1 | 218543 | HITS-CLIP | 29167373 | Unspecified |
| mmu-miR-674-3p | MIMAT0003741 | Sri | 109552 | HITS-CLIP | 29167373 | Unspecified |
| mmu-miR-674-3p | MIMAT0003741 | Srpk1 | 20815 | HITS-CLIP | 25753418 | Unspecified |
| mmu-miR-674-3p | MIMAT0003741 | Ssh2 | 237860 | HITS-CLIP | 25753418 | Unspecified |
| mmu-miR-674-3p | MIMAT0003741 | St8sia4 | 20452 | HITS-CLIP | 29167373 | Unspecified |
| mmu-miR-674-3p | MIMAT0003741 | Stat2 | 20847 | HITS-CLIP | 25753418 | Unspecified |
| mmu-miR-674-3p | MIMAT0003741 | Synpo2l | 68760 | HITS-CLIP | 25753418 | Unspecified |
| mmu-miR-674-3p | MIMAT0003741 | Tcf20 | 21411 | HITS-CLIP | 29167373 | Unspecified |
| mmu-miR-674-3p | MIMAT0003741 | Tgfbr1 | 21812 | HITS-CLIP | 29167373 | Unspecified |
| mmu-miR-674-3p | MIMAT0003741 | Tkt | 21881 | HITS-CLIP | 29167373 | Unspecified |
| mmu-miR-674-3p | MIMAT0003741 | Tmbim6 | 110213 | HITS-CLIP | 31433990 | Unspecified |
| mmu-miR-674-3p | MIMAT0003741 | Tmem38b | 52076 | qCLASH | 31393953 | Unspecified |
| mmu-miR-674-3p | MIMAT0003741 | Tmem65 | 74868 | HITS-CLIP | 25753418 | Unspecified |
| mmu-miR-674-3p | MIMAT0003741 | Tnpo1 | 238799 | HITS-CLIP | 29167373 | Unspecified |
| mmu-miR-674-3p | MIMAT0003741 | Tnpo3 | 320938 | HITS-CLIP | 29167373 | Unspecified |
| mmu-miR-674-3p | MIMAT0003741 | Tnrc6c | 217351 | HITS-CLIP | 29167373 | Unspecified |
| mmu-miR-674-3p | MIMAT0003741 | Ttn | 22138 | HITS-CLIP | 25753418 | Unspecified |
| mmu-miR-674-3p | MIMAT0003741 | Ube2j1 | 56228 | HITS-CLIP | 25753418 | Unspecified |
| mmu-miR-674-3p | MIMAT0003741 | Ubr5 | 70790 | HITS-CLIP | 29167373 | Unspecified |
| mmu-miR-674-3p | MIMAT0003741 | Upf1 | 19704 | HITS-CLIP | 25753418 | Unspecified |
| mmu-miR-674-3p | MIMAT0003741 | Usf3 | 205717 | HITS-CLIP | 29167373 | Unspecified |
| mmu-miR-674-3p | MIMAT0003741 | Vps13b | 666173 | HITS-CLIP | 25753418\|29167373 | Unspecified |
| mmu-miR-674-3p | MIMAT0003741 | Vwa8 | 219189 | HITS-CLIP | 25753418 | Unspecified |
| mmu-miR-674-3p | MIMAT0003741 | Wdr81 | 192652 | HITS-CLIP | 29167373 | Unspecified |
| mmu-miR-674-3p | MIMAT0003741 | Wnk1 | 232341 | HITS-CLIP | 29167373 | Unspecified |
| mmu-miR-674-3p | MIMAT0003741 | Zbtb5 | 230119 | HITS-CLIP | 25753418 | Unspecified |
| mmu-miR-674-3p | MIMAT0003741 | Zeb2 | 24136 | HITS-CLIP | 29167373 | Unspecified |
| mmu-miR-674-3p | MIMAT0003741 | Zfp36l1 | 12192 | HITS-CLIP | 29167373 | Unspecified |
| mmu-miR-674-3p | MIMAT0003741 | Zfp36l2 | 12193 | HITS-CLIP | 29167373 | Unspecified |
| mmu-miR-674-3p | MIMAT0003741 | Zfp81 | 224694 | HITS-CLIP | 29167373 | Unspecified |
| mmu-miR-674-3p | MIMAT0003741 | Zzef1 | 195018 | HITS-CLIP | 29167373 | Unspecified |
| mmu-miR-6896-3p | MIMAT0027693 | Glipr2 | 384009 | HITS-CLIP | 31433990 | Unspecified |
| mmu-miR-6896-3p | MIMAT0027693 | Morc2a | 74522 | HITS-CLIP | 31433990 | Unspecified |
| mmu-miR-6896-3p | MIMAT0027693 | Ywhae | 22627 | HITS-CLIP | 31433990 | Unspecified |

| | | | | | | |
|---|---|---|---|---|---|---|
| mmu-miR-6896-5p | MIMAT0027692 | Chac1 | 69065 | HITS-CLIP | 31433990 | Unspecified |
| mmu-miR-6896-5p | MIMAT0027692 | Otud4 | 73945 | HITS-CLIP | 31433990 | Unspecified |
| mmu-miR-6896-5p | MIMAT0027692 | Rock1 | 19877 | HITS-CLIP | 31433990 | Unspecified |
| mmu-miR-6898-3p | MIMAT0027697 | 2310050C09Rik | 66533 | HITS-CLIP | 26203562 | Unspecified |
| mmu-miR-6898-3p | MIMAT0027697 | Cd2ap | 12488 | HITS-CLIP | 26203562 | Unspecified |
| mmu-miR-6898-3p | MIMAT0027697 | Klf6 | 23849 | HITS-CLIP | 26203562 | Unspecified |
| mmu-miR-6898-3p | MIMAT0027697 | Krt6a | 3853 | HITS-CLIP | 26203562 | Unspecified |
| mmu-miR-6898-3p | MIMAT0027697 | Ttc14 | 67120 | HITS-CLIP | 31433990 | Unspecified |
| mmu-miR-6898-5p | MIMAT0027696 | Cbll1 | 104836 | HITS-CLIP | 31433990 | Unspecified |
| mmu-miR-6898-5p | MIMAT0027696 | Ormdl3 | 66612 | HITS-CLIP | 31433990 | Unspecified |
| mmu-miR-6898-5p | MIMAT0027696 | Ubtf | 21429 | HITS-CLIP | 26203562 | Unspecified |
| mmu-miR-6906-3p | MIMAT0027713 | Tmem33 | 67878 | HITS-CLIP | 26203562 | Unspecified |
| mmu-miR-6906-5p | MIMAT0027712 | Khnyn | 219094 | HITS-CLIP | 31433990 | Unspecified |
| mmu-miR-6906-5p | MIMAT0027712 | Myc | 17869 | HITS-CLIP | 31433990 | Unspecified |
| mmu-miR-6906-5p | MIMAT0027712 | Ndufs6 | 407785 | HITS-CLIP | 31433990 | Unspecified |
| mmu-miR-6906-5p | MIMAT0027712 | Plagl2 | 54711 | HITS-CLIP | 31433990 | Unspecified |
| mmu-miR-6924-3p | MIMAT0027749 | Tnrc6a | 233833 | HITS-CLIP | 31433990 | Unspecified |
| mmu-miR-6924-5p | MIMAT0027748 | Aff4 | 93736 | HITS-CLIP | 31433990 | Unspecified |
| mmu-miR-6924-5p | MIMAT0027748 | Arpc3 | 56378 | HITS-CLIP | 31433990 | Unspecified |
| mmu-miR-6924-5p | MIMAT0027748 | Crybg1 | 202 | HITS-CLIP | 31433990 | Unspecified |
| mmu-miR-6924-5p | MIMAT0027748 | Eno1b | 433182 | HITS-CLIP | 31433990 | Unspecified |
| mmu-miR-6924-5p | MIMAT0027748 | Psme3 | 19192 | HITS-CLIP | 31433990 | Unspecified |
| mmu-miR-6924-5p | MIMAT0027748 | Rpap1 | 68925 | HITS-CLIP | 31433990 | Unspecified |
| mmu-miR-6924-5p | MIMAT0027748 | S100a11 | 20195 | HITS-CLIP | 26203562 | Unspecified |
| mmu-miR-6924-5p | MIMAT0027748 | Taf5 | 226182 | HITS-CLIP | 26203562 | Unspecified |
| mmu-miR-6924-5p | MIMAT0027748 | Yod1 | 226418 | HITS-CLIP | 26203562 | Unspecified |
| mmu-miR-6924-5p | MIMAT0027748 | Zfp598 | 213753 | HITS-CLIP | 26203562 | Unspecified |
| mmu-miR-6934-5p | MIMAT0027768 | Atad2b | 320817 | HITS-CLIP | 26203562 | Unspecified |
| mmu-miR-6934-5p | MIMAT0027768 | Cfl1 | 12631 | HITS-CLIP | 31433990 | Unspecified |
| mmu-miR-6934-5p | MIMAT0027768 | Dusp11 | 72102 | HITS-CLIP | 31433990 | Unspecified |
| mmu-miR-6934-5p | MIMAT0027768 | Ehbp1l1 | 114601 | HITS-CLIP | 31433990 | Unspecified |
| mmu-miR-6934-5p | MIMAT0027768 | Emc4 | 68032 | HITS-CLIP | 31433990 | Unspecified |
| mmu-miR-6934-5p | MIMAT0027768 | H3c7 | 8350 | HITS-CLIP | 31433990 | Unspecified |
| mmu-miR-6934-5p | MIMAT0027768 | Irgm2 | 54396 | HITS-CLIP | 31433990 | Unspecified |
| mmu-miR-6934-5p | MIMAT0027768 | Krt16 | 16666 | HITS-CLIP | 26203562 | Unspecified |
| mmu-miR-6934-5p | MIMAT0027768 | Nfe2l1 | 18023 | HITS-CLIP | 26203562 | Unspecified |
| mmu-miR-6934-5p | MIMAT0027768 | Socs1 | 12703 | HITS-CLIP | 31433990 | Unspecified |
| mmu-miR-6934-5p | MIMAT0027768 | Thoc1 | 225160 | HITS-CLIP | 31433990 | Unspecified |
| mmu-miR-6938-3p | MIMAT0027777 | Canx | 12330 | HITS-CLIP | 31433990 | Unspecified |

| | | | | | | |
|---|---|---|---|---|---|---|
| mmu-miR-6938-3p | MIMAT0027777 | Hps6 | 20170 | HITS-CLIP | 31433990 | Unspecified |
| mmu-miR-6938-3p | MIMAT0027777 | Hsd17b4 | 15488 | HITS-CLIP | 31433990 | Unspecified |
| mmu-miR-6938-3p | MIMAT0027777 | Man2b1 | 17159 | HITS-CLIP | 31433990 | Unspecified |
| mmu-miR-6938-3p | MIMAT0027777 | Palb2 | 233826 | HITS-CLIP | 31433990 | Unspecified |
| mmu-miR-6938-3p | MIMAT0027777 | Unk | 217331 | HITS-CLIP | 31433990 | Unspecified |
| mmu-miR-6938-5p | MIMAT0027776 | Atp1a1 | 11928 | HITS-CLIP | 31433990 | Unspecified |
| mmu-miR-6938-5p | MIMAT0027776 | Bcas2 | 68183 | HITS-CLIP | 31433990 | Unspecified |
| mmu-miR-6938-5p | MIMAT0027776 | Stat4 | 20849 | HITS-CLIP | 31433990 | Unspecified |
| mmu-miR-6962-3p | MIMAT0027825 | Myadm | 50918 | HITS-CLIP | 26203562 | Unspecified |
| mmu-miR-6962-3p | MIMAT0027825 | Plagl2 | 54711 | HITS-CLIP | 31433990 | Unspecified |
| mmu-miR-6962-5p | MIMAT0027824 | Pag1 | 94212 | HITS-CLIP | 31433990 | Unspecified |
| mmu-miR-6962-5p | MIMAT0027824 | Tbl1xr1 | 81004 | HITS-CLIP | 31433990 | Unspecified |
| mmu-miR-6962-5p | MIMAT0027824 | Tnrc6a | 233833 | HITS-CLIP | 31433990 | Unspecified |
| mmu-miR-6962-5p | MIMAT0027824 | Yeats2 | 208146 | HITS-CLIP | 26203562 | Unspecified |
| mmu-miR-6972-3p | MIMAT0027847 | Arhgef2 | 16800 | HITS-CLIP | 26203562 | Unspecified |
| mmu-miR-6972-5p | MIMAT0027846 | Itsn2 | 20403 | HITS-CLIP | 31433990 | Unspecified |
| mmu-miR-6972-5p | MIMAT0027846 | Prkar1a | 19084 | HITS-CLIP | 26203562 | Unspecified |
| mmu-miR-6972-5p | MIMAT0027846 | Srsf2 | 20382 | HITS-CLIP | 31433990 | Unspecified |
| mmu-miR-6988-3p | MIMAT0027879 | Agfg1 | 15463 | HITS-CLIP | 26203562 | Unspecified |
| mmu-miR-6988-3p | MIMAT0027879 | C2cd5 | 74741 | HITS-CLIP | 31433990 | Unspecified |
| mmu-miR-6988-3p | MIMAT0027879 | Fam8a1 | 51439 | HITS-CLIP | 31433990 | Unspecified |
| mmu-miR-6988-3p | MIMAT0027879 | Kmt2e | 69188 | HITS-CLIP | 31433990 | Unspecified |
| mmu-miR-6988-3p | MIMAT0027879 | Ncoa3 | 17979 | HITS-CLIP | 31433990 | Unspecified |
| mmu-miR-6988-3p | MIMAT0027879 | Notch1 | 18128 | HITS-CLIP | 26203562 | Unspecified |
| mmu-miR-6988-3p | MIMAT0027879 | Ppp1r8 | 100336 | HITS-CLIP | 31433990 | Unspecified |
| mmu-miR-6988-3p | MIMAT0027879 | Shprh | 268281 | HITS-CLIP | 31433990 | Unspecified |
| mmu-miR-6988-3p | MIMAT0027879 | Snd1 | 56463 | HITS-CLIP | 31433990 | Unspecified |
| mmu-miR-6988-3p | MIMAT0027879 | Ube2o | 217342 | HITS-CLIP | 26203562 | Unspecified |
| mmu-miR-6988-5p | MIMAT0027878 | Smc2 | 14211 | HITS-CLIP | 26203562 | Unspecified |
| mmu-miR-6995-3p | MIMAT0027893 | Cited2 | 17684 | HITS-CLIP | 31433990 | Unspecified |
| mmu-miR-6995-3p | MIMAT0027893 | Top2a | 21973 | HITS-CLIP | 31433990 | Unspecified |
| mmu-miR-6995-5p | MIMAT0027892 | Cdk2ap2 | 52004 | HITS-CLIP | 31433990 | Unspecified |
| mmu-miR-6995-5p | MIMAT0027892 | Kbtbd2 | 210973 | HITS-CLIP | 31433990 | Unspecified |
| mmu-miR-6995-5p | MIMAT0027892 | Myc | 17869 | HITS-CLIP | 31433990 | Unspecified |
| mmu-miR-7021-3p | MIMAT0027947 | Actn4 | 60595 | HITS-CLIP | 31433990 | Unspecified |
| mmu-miR-7021-3p | MIMAT0027947 | Dusp4 | 319520 | HITS-CLIP | 31433990 | Unspecified |
| mmu-miR-7021-3p | MIMAT0027947 | Ldlrap1 | 100017 | HITS-CLIP | 26203562 | Unspecified |
| mmu-miR-7021-3p | MIMAT0027947 | Rpl23 | 65019 | HITS-CLIP | 31433990 | Unspecified |
| mmu-miR-7021-3p | MIMAT0027947 | Srrm2 | 75956 | HITS-CLIP | 31433990 | Unspecified |

| | | | | | | |
|---|---|---|---|---|---|---|
| mmu-miR-7021-5p | MIMAT0027946 | B3gnt7 | 227327 | HITS-CLIP | 31433990 | Unspecified |
| mmu-miR-7021-5p | MIMAT0027946 | Hycc2 | 285172 | HITS-CLIP | 31433990 | Unspecified |
| mmu-miR-7021-5p | MIMAT0027946 | Net1 | 56349 | HITS-CLIP | 31433990 | Unspecified |
| mmu-miR-7021-5p | MIMAT0027946 | Rspry1 | 67610 | HITS-CLIP | 31433990 | Unspecified |
| mmu-miR-7021-5p | MIMAT0027946 | Trp53inp1 | 60599 | HITS-CLIP | 31433990 | Unspecified |
| mmu-miR-7025-3p | MIMAT0027955 | Dcps | 69305 | HITS-CLIP | 31433990 | Unspecified |
| mmu-miR-7025-3p | MIMAT0027955 | Kif21a | 16564 | HITS-CLIP | 26203562 | Unspecified |
| mmu-miR-7025-3p | MIMAT0027955 | Sdc1 | 20969 | HITS-CLIP | 26203562 | Unspecified |
| mmu-miR-7025-3p | MIMAT0027955 | Trp53inp1 | 60599 | HITS-CLIP | 26203562 | Unspecified |
| mmu-miR-7025-3p | MIMAT0027955 | Zwint | 52696 | HITS-CLIP | 31433990 | Unspecified |
| mmu-miR-7027-3p | MIMAT0027959 | Cry2 | 12953 | HITS-CLIP | 26203562 | Unspecified |
| mmu-miR-7027-3p | MIMAT0027959 | H4c14 | 8359 | HITS-CLIP | 31433990 | Unspecified |
| mmu-miR-7027-3p | MIMAT0027959 | Macroh2a1 | 9555 | HITS-CLIP | 31433990 | Unspecified |
| mmu-miR-7027-3p | MIMAT0027959 | Scd2 | 20250 | HITS-CLIP | 26203562 | Unspecified |
| mmu-miR-7027-3p | MIMAT0027959 | Trio | 223435 | HITS-CLIP | 26203562 | Unspecified |
| mmu-miR-7027-5p | MIMAT0027958 | Bend4 | 666938 | HITS-CLIP | 26203562 | Unspecified |
| mmu-miR-7027-5p | MIMAT0027958 | Cdc37 | 12539 | HITS-CLIP | 31433990 | Unspecified |
| mmu-miR-7027-5p | MIMAT0027958 | Msn | 17698 | HITS-CLIP | 31433990 | Unspecified |
| mmu-miR-7027-5p | MIMAT0027958 | Pon2 | 330260 | HITS-CLIP | 26203562 | Unspecified |
| mmu-miR-7027-5p | MIMAT0027958 | S100a10 | 20194 | HITS-CLIP | 31433990 | Unspecified |
| mmu-miR-7027-5p | MIMAT0027958 | Sdha | 66945 | HITS-CLIP | 26203562 | Unspecified |
| mmu-miR-7027-5p | MIMAT0027958 | Shprh | 268281 | HITS-CLIP | 31433990 | Unspecified |
| mmu-miR-7027-5p | MIMAT0027958 | Smyd5 | 232187 | HITS-CLIP | 31433990 | Unspecified |
| mmu-miR-7027-5p | MIMAT0027958 | Trim24 | 21848 | HITS-CLIP | 31433990 | Unspecified |
| mmu-miR-7036b-3p | MIMAT0029809 | Arhgef3 | 71704 | HITS-CLIP | 31433990 | Unspecified |
| mmu-miR-7036b-3p | MIMAT0029809 | Prkar1a | 19084 | HITS-CLIP | 26203562 | Unspecified |
| mmu-miR-7036b-3p | MIMAT0029809 | Supt16 | 114741 | HITS-CLIP | 31433990 | Unspecified |
| mmu-miR-7036b-3p | MIMAT0029809 | Tgtp1 | 21822 | HITS-CLIP | 31433990 | Unspecified |
| mmu-miR-7036b-5p | MIMAT0029808 | Larp4b | 217980 | HITS-CLIP | 31433990 | Unspecified |
| mmu-miR-7063-3p | MIMAT0028031 | Cdca4 | 71963 | HITS-CLIP | 31433990 | Unspecified |
| mmu-miR-7063-5p | MIMAT0028030 | Ago2 | 239528 | HITS-CLIP | 31433990 | Unspecified |
| mmu-miR-7063-5p | MIMAT0028030 | Anln | 68743 | HITS-CLIP | 31433990 | Unspecified |
| mmu-miR-7063-5p | MIMAT0028030 | Anxa7 | 11750 | HITS-CLIP | 31433990 | Unspecified |
| mmu-miR-7063-5p | MIMAT0028030 | Atg2b | 76559 | HITS-CLIP | 31433990 | Unspecified |
| mmu-miR-7063-5p | MIMAT0028030 | Cnot1 | 234594 | HITS-CLIP | 31433990 | Unspecified |
| mmu-miR-7063-5p | MIMAT0028030 | Col27a1 | 373864 | HITS-CLIP | 31433990 | Unspecified |
| mmu-miR-7063-5p | MIMAT0028030 | Cops2 | 12848 | HITS-CLIP | 31433990 | Unspecified |
| mmu-miR-7063-5p | MIMAT0028030 | Cse1l | 110750 | HITS-CLIP | 31433990 | Unspecified |
| mmu-miR-7063-5p | MIMAT0028030 | Dicer1 | 192119 | HITS-CLIP | 31433990 | Unspecified |

| | | | | | | | |
|---|---|---|---|---|---|---|---|
| mmu-miR-7063-5p | MIMAT0028030 | Dnajc11 | 230935 | HITS-CLIP | | 26203562 | Unspecified |
| mmu-miR-7063-5p | MIMAT0028030 | Dnmt1 | 13433 | HITS-CLIP | | 31433990 | Unspecified |
| mmu-miR-7063-5p | MIMAT0028030 | Edem1 | 192193 | HITS-CLIP | | 31433990 | Unspecified |
| mmu-miR-7063-5p | MIMAT0028030 | Ei24 | 13663 | HITS-CLIP | | 26203562 | Unspecified |
| mmu-miR-7063-5p | MIMAT0028030 | Errfi1 | 74155 | HITS-CLIP | | 26203562 | Unspecified |
| mmu-miR-7063-5p | MIMAT0028030 | Fam107b | 66540 | HITS-CLIP | | 26203562 | Unspecified |
| mmu-miR-7063-5p | MIMAT0028030 | Fat1 | 14107 | HITS-CLIP | | 26203562 | Unspecified |
| mmu-miR-7063-5p | MIMAT0028030 | Foxk1 | 17425 | HITS-CLIP | | 31433990 | Unspecified |
| mmu-miR-7063-5p | MIMAT0028030 | Gbp2b | 14468 | HITS-CLIP | | 31433990 | Unspecified |
| mmu-miR-7063-5p | MIMAT0028030 | Hnrnpdl | 50926 | HITS-CLIP | | 31433990 | Unspecified |
| mmu-miR-7063-5p | MIMAT0028030 | Hspa14 | 50497 | HITS-CLIP | | 31433990 | Unspecified |
| mmu-miR-7063-5p | MIMAT0028030 | Igtp | 16145 | HITS-CLIP | | 31433990 | Unspecified |
| mmu-miR-7063-5p | MIMAT0028030 | Il2rb | 16185 | HITS-CLIP | | 31433990 | Unspecified |
| mmu-miR-7063-5p | MIMAT0028030 | Lmtk2 | 231876 | HITS-CLIP | | 31433990 | Unspecified |
| mmu-miR-7063-5p | MIMAT0028030 | Mtmr3 | 74302 | HITS-CLIP | | 31433990 | Unspecified |
| mmu-miR-7063-5p | MIMAT0028030 | Myo1c | 17913 | HITS-CLIP | | 31433990 | Unspecified |
| mmu-miR-7063-5p | MIMAT0028030 | Numb | 18222 | HITS-CLIP | | 31433990 | Unspecified |
| mmu-miR-7063-5p | MIMAT0028030 | Ppp6r3 | 52036 | HITS-CLIP | | 31433990 | Unspecified |
| mmu-miR-7063-5p | MIMAT0028030 | Rps9 | 76846 | HITS-CLIP | | 31433990 | Unspecified |
| mmu-miR-7063-5p | MIMAT0028030 | Rrnad1 | 229503 | HITS-CLIP | | 31433990 | Unspecified |
| mmu-miR-7063-5p | MIMAT0028030 | Scd2 | 20250 | HITS-CLIP | | 31433990\|26203562 | Unspecified |
| mmu-miR-7063-5p | MIMAT0028030 | Senp3 | 80886 | HITS-CLIP | | 31433990 | Unspecified |
| mmu-miR-7063-5p | MIMAT0028030 | Siva1 | 30954 | HITS-CLIP | | 31433990 | Unspecified |
| mmu-miR-7063-5p | MIMAT0028030 | Slk | 20874 | HITS-CLIP | | 31433990 | Unspecified |
| mmu-miR-7063-5p | MIMAT0028030 | Stip1 | 20867 | HITS-CLIP | | 31433990 | Unspecified |
| mmu-miR-7063-5p | MIMAT0028030 | Tasor | 23272 | HITS-CLIP | | 31433990 | Unspecified |
| mmu-miR-7063-5p | MIMAT0028030 | Wdr82 | 77305 | HITS-CLIP | | 31433990 | Unspecified |
| mmu-miR-7063-5p | MIMAT0028030 | Zfp280b | 64453 | HITS-CLIP | | 31433990 | Unspecified |
| mmu-miR-7063-5p | MIMAT0028030 | Zkscan1 | 74570 | HITS-CLIP | | 31433990 | Unspecified |
| mmu-miR-7074-3p | MIMAT0028055 | Bend4 | 666938 | HITS-CLIP | | 26203562 | Unspecified |
| mmu-miR-7074-3p | MIMAT0028055 | Hif1a | 15251 | HITS-CLIP | | 31433990 | Unspecified |
| mmu-miR-7074-3p | MIMAT0028055 | Prss8 | 76560 | HITS-CLIP | | 26203562 | Unspecified |
| mmu-miR-7074-5p | MIMAT0028054 | Afg3l2 | 69597 | HITS-CLIP | | 31433990 | Unspecified |
| mmu-miR-7074-5p | MIMAT0028054 | Bmpr1a | 12166 | HITS-CLIP | | 26203562 | Unspecified |
| mmu-miR-7074-5p | MIMAT0028054 | Cct6a | 12466 | HITS-CLIP | | 31433990 | Unspecified |
| mmu-miR-7074-5p | MIMAT0028054 | Dennd1b | 329260 | HITS-CLIP | | 31433990 | Unspecified |
| mmu-miR-7074-5p | MIMAT0028054 | Grwd1 | 101612 | HITS-CLIP | | 31433990 | Unspecified |
| mmu-miR-7074-5p | MIMAT0028054 | Mob4 | 19070 | HITS-CLIP | | 31433990 | Unspecified |
| mmu-miR-7074-5p | MIMAT0028054 | Psmc1 | 19179 | HITS-CLIP | | 31433990 | Unspecified |

| | | | | | | |
|---|---|---|---|---|---|---|
| mmu-miR-7074-5p | MIMAT0028054 | Rad18 | 58186 | HITS-CLIP | 31433990 | Unspecified |
| mmu-miR-7074-5p | MIMAT0028054 | Rest | 19712 | HITS-CLIP | 26203562 | Unspecified |
| mmu-miR-7074-5p | MIMAT0028054 | Smc3 | 13006 | HITS-CLIP | 31433990 | Unspecified |
| mmu-miR-7074-5p | MIMAT0028054 | Trp53inp1 | 60599 | HITS-CLIP | 31433990 | Unspecified |
| mmu-miR-7074-5p | MIMAT0028054 | Tubb5 | 22154 | HITS-CLIP | 31433990 | Unspecified |
| mmu-miR-7074-5p | MIMAT0028054 | Xpr1 | 19775 | HITS-CLIP | 31433990 | Unspecified |
| mmu-miR-7074-5p | MIMAT0028054 | Zfp217 | 228913 | HITS-CLIP | 26203562 | Unspecified |
| mmu-miR-7074-5p | MIMAT0028054 | Zmiz1 | 328365 | HITS-CLIP | 31433990 | Unspecified |
| mmu-miR-7076-3p | MIMAT0028059 | Flnb | 286940 | HITS-CLIP | 26203562 | Unspecified |
| mmu-miR-7076-5p | MIMAT0028058 | Bach2 | 12014 | HITS-CLIP | 31433990 | Unspecified |
| mmu-miR-7076-5p | MIMAT0028058 | Chst11 | 58250 | HITS-CLIP | 31433990 | Unspecified |
| mmu-miR-7076-5p | MIMAT0028058 | Cul2 | 71745 | HITS-CLIP | 31433990 | Unspecified |
| mmu-miR-7076-5p | MIMAT0028058 | Dcun1d4 | 100737 | HITS-CLIP | 31433990 | Unspecified |
| mmu-miR-7076-5p | MIMAT0028058 | Zfx | 22764 | HITS-CLIP | 31433990 | Unspecified |
| mmu-miR-7091-3p | MIMAT0028089 | Dgkd | 227333 | HITS-CLIP | 31433990 | Unspecified |
| mmu-miR-7091-3p | MIMAT0028089 | Iqgap2 | 544963 | HITS-CLIP | 31433990 | Unspecified |
| mmu-miR-7091-3p | MIMAT0028089 | Ppard | 19015 | HITS-CLIP | 31433990 | Unspecified |
| mmu-miR-7091-3p | MIMAT0028089 | Sik2 | 235344 | HITS-CLIP | 31433990 | Unspecified |
| mmu-miR-7091-5p | MIMAT0028088 | Appbp2 | 66884 | HITS-CLIP | 31433990 | Unspecified |
| mmu-miR-7091-5p | MIMAT0028088 | Cbfb | 12400 | HITS-CLIP | 31433990 | Unspecified |
| mmu-miR-7091-5p | MIMAT0028088 | Epas1 | 13819 | HITS-CLIP | 31433990 | Unspecified |
| mmu-miR-7240-5p | MIMAT0028448 | Arid2 | 77044 | HITS-CLIP | 31433990 | Unspecified |
| mmu-miR-7240-5p | MIMAT0028448 | Atp13a3 | 224088 | HITS-CLIP | 31433990 | Unspecified |
| mmu-miR-7240-5p | MIMAT0028448 | Lgmn | 19141 | HITS-CLIP | 26203562 | Unspecified |
| mmu-miR-7240-5p | MIMAT0028448 | Pbx2 | 18515 | HITS-CLIP | 26203562 | Unspecified |
| mmu-miR-7240-5p | MIMAT0028448 | Ppfia1 | 233977 | HITS-CLIP | 31433990 | Unspecified |
| mmu-miR-7241-3p | MIMAT0028451 | 1700017B05Rik | 74211 | HITS-CLIP | 26203562 | Unspecified |
| mmu-miR-7241-3p | MIMAT0028451 | Adnp | 11538 | HITS-CLIP | 26203562 | Unspecified |
| mmu-miR-7241-3p | MIMAT0028451 | Aifm1 | 26926 | HITS-CLIP | 26203562 | Unspecified |
| mmu-miR-7241-3p | MIMAT0028451 | Arid4a | 238247 | HITS-CLIP | 26203562 | Unspecified |
| mmu-miR-7241-3p | MIMAT0028451 | Atl2 | 56298 | HITS-CLIP | 31433990 | Unspecified |
| mmu-miR-7241-3p | MIMAT0028451 | Atxn1l | 52335 | HITS-CLIP | 26203562 | Unspecified |
| mmu-miR-7241-3p | MIMAT0028451 | Bcl11b | 58208 | HITS-CLIP | 31433990 | Unspecified |
| mmu-miR-7241-3p | MIMAT0028451 | Bicd2 | 76895 | HITS-CLIP | 26203562 | Unspecified |
| mmu-miR-7241-3p | MIMAT0028451 | Camta2 | 216874 | HITS-CLIP | 31433990 | Unspecified |
| mmu-miR-7241-3p | MIMAT0028451 | Cfl2 | 12632 | HITS-CLIP | 26203562 | Unspecified |
| mmu-miR-7241-3p | MIMAT0028451 | Cmas | 12764 | HITS-CLIP | 26203562 | Unspecified |
| mmu-miR-7241-3p | MIMAT0028451 | Cops3 | 26572 | HITS-CLIP | 31433990 | Unspecified |
| mmu-miR-7241-3p | MIMAT0028451 | Crk | 12928 | HITS-CLIP | 31433990 | Unspecified |

| | | | | | | |
|---|---|---|---|---|---|---|
| mmu-miR-7241-3p | MIMAT0028451 | Dnajc3 | 100037258 | HITS-CLIP | 26203562 | Unspecified |
| mmu-miR-7241-3p | MIMAT0028451 | Eif3a | 13669 | HITS-CLIP | 26203562 | Unspecified |
| mmu-miR-7241-3p | MIMAT0028451 | Ep300 | 328572 | HITS-CLIP | 31433990 | Unspecified |
| mmu-miR-7241-3p | MIMAT0028451 | Gata3 | 14462 | HITS-CLIP | 31433990 | Unspecified |
| mmu-miR-7241-3p | MIMAT0028451 | Gm20716 | 105980075 | HITS-CLIP | 26203562 | Unspecified |
| mmu-miR-7241-3p | MIMAT0028451 | Grsf1 | 231413 | HITS-CLIP | 26203562 | Unspecified |
| mmu-miR-7241-3p | MIMAT0028451 | Hectd1 | 207304 | HITS-CLIP | 31433990 | Unspecified |
| mmu-miR-7241-3p | MIMAT0028451 | Hnrnpul2 | 68693 | HITS-CLIP | 26203562 | Unspecified |
| mmu-miR-7241-3p | MIMAT0028451 | Il2ra | 16184 | HITS-CLIP | 31433990 | Unspecified |
| mmu-miR-7241-3p | MIMAT0028451 | Ilkap | 67444 | HITS-CLIP | 26203562 | Unspecified |
| mmu-miR-7241-3p | MIMAT0028451 | Itga3 | 16400 | HITS-CLIP | 26203562 | Unspecified |
| mmu-miR-7241-3p | MIMAT0028451 | Kbtbd2 | 210973 | HITS-CLIP | 31433990 | Unspecified |
| mmu-miR-7241-3p | MIMAT0028451 | Klf6 | 23849 | HITS-CLIP | 26203562 | Unspecified |
| mmu-miR-7241-3p | MIMAT0028451 | Lamc1 | 226519 | HITS-CLIP | 31433990 | Unspecified |
| mmu-miR-7241-3p | MIMAT0028451 | Macf1 | 11426 | HITS-CLIP | 26203562 | Unspecified |
| mmu-miR-7241-3p | MIMAT0028451 | Marcks | 17118 | HITS-CLIP | 26203562 | Unspecified |
| mmu-miR-7241-3p | MIMAT0028451 | Mbnl3 | 171170 | HITS-CLIP | 26203562 | Unspecified |
| mmu-miR-7241-3p | MIMAT0028451 | Mfsd2a | 76574 | HITS-CLIP | 31433990 | Unspecified |
| mmu-miR-7241-3p | MIMAT0028451 | Mki67 | 17345 | HITS-CLIP | 26203562 | Unspecified |
| mmu-miR-7241-3p | MIMAT0028451 | Nfkbid | 243910 | HITS-CLIP | 31433990 | Unspecified |
| mmu-miR-7241-3p | MIMAT0028451 | Nr3c1 | 14815 | HITS-CLIP | 31433990 | Unspecified |
| mmu-miR-7241-3p | MIMAT0028451 | Oga | 10724 | HITS-CLIP | 31433990 | Unspecified |
| mmu-miR-7241-3p | MIMAT0028451 | Osm | 18413 | HITS-CLIP | 31433990 | Unspecified |
| mmu-miR-7241-3p | MIMAT0028451 | Pcyox1 | 66881 | HITS-CLIP | 26203562 | Unspecified |
| mmu-miR-7241-3p | MIMAT0028451 | Pip4p2 | 55529 | HITS-CLIP | 31433990 | Unspecified |
| mmu-miR-7241-3p | MIMAT0028451 | Rreb1 | 68750 | HITS-CLIP | 31433990\|26203562 | Unspecified |
| mmu-miR-7241-3p | MIMAT0028451 | Sema4d | 20354 | HITS-CLIP | 31433990 | Unspecified |
| mmu-miR-7241-3p | MIMAT0028451 | Senp5 | 320213 | HITS-CLIP | 31433990 | Unspecified |
| mmu-miR-7241-3p | MIMAT0028451 | Sh3pxd2a | 14218 | HITS-CLIP | 26203562 | Unspecified |
| mmu-miR-7241-3p | MIMAT0028451 | Slc30a5 | 69048 | HITS-CLIP | 31433990 | Unspecified |
| mmu-miR-7241-3p | MIMAT0028451 | Slc6a8 | 102857 | HITS-CLIP | 26203562 | Unspecified |
| mmu-miR-7241-3p | MIMAT0028451 | Srek1 | 218543 | HITS-CLIP | 31433990 | Unspecified |
| mmu-miR-7241-3p | MIMAT0028451 | Tbc1d17 | 233204 | HITS-CLIP | 26203562 | Unspecified |
| mmu-miR-7241-3p | MIMAT0028451 | Tmem245 | 242474 | HITS-CLIP | 26203562 | Unspecified |
| mmu-miR-7241-3p | MIMAT0028451 | Ubr4 | 69116 | HITS-CLIP | 31433990 | Unspecified |
| mmu-miR-7241-3p | MIMAT0028451 | Ufl1 | 67490 | HITS-CLIP | 31433990 | Unspecified |
| mmu-miR-7241-3p | MIMAT0028451 | Ywhag | 22628 | HITS-CLIP | 26203562 | Unspecified |
| mmu-miR-7241-3p | MIMAT0028451 | Zfp280b | 64453 | HITS-CLIP | 31433990 | Unspecified |
| mmu-miR-7241-5p | MIMAT0028450 | Acap1 | 216859 | HITS-CLIP | 31433990 | Unspecified |

| | | | | | | |
|---|---|---|---|---|---|---|
| mmu-miR-7241-5p | MIMAT0028450 | Gzmb | 14939 | HITS-CLIP | | 31433990 Unspecified |
| mmu-miR-7241-5p | MIMAT0028450 | Sec11c | 66286 | HITS-CLIP | | 31433990 Unspecified |
| mmu-miR-7646-3p | MIMAT0029795 | Capza2 | 12343 | HITS-CLIP | | 31433990 Unspecified |
| mmu-miR-7646-5p | MIMAT0029794 | Abl1 | 11350 | HITS-CLIP | | 31433990 Unspecified |
| mmu-miR-7646-5p | MIMAT0029794 | Adora2b | 11541 | HITS-CLIP | | 26203562 Unspecified |
| mmu-miR-7646-5p | MIMAT0029794 | Arhgap21 | 71435 | HITS-CLIP | | 31433990 Unspecified |
| mmu-miR-7646-5p | MIMAT0029794 | Cad | 69719 | HITS-CLIP | | 31433990 Unspecified |
| mmu-miR-7646-5p | MIMAT0029794 | Ccnd2 | 12444 | HITS-CLIP | | 31433990 Unspecified |
| mmu-miR-7646-5p | MIMAT0029794 | Cflar | 12633 | HITS-CLIP | | 31433990 Unspecified |
| mmu-miR-7646-5p | MIMAT0029794 | Chd8 | 67772 | HITS-CLIP | | 31433990 Unspecified |
| mmu-miR-7646-5p | MIMAT0029794 | Cldnd1 | 224250 | HITS-CLIP | | 31433990 Unspecified |
| mmu-miR-7646-5p | MIMAT0029794 | Dhx57 | 106794 | HITS-CLIP | | 26203562 Unspecified |
| mmu-miR-7646-5p | MIMAT0029794 | Dyrk1a | 13548 | HITS-CLIP | | 31433990 Unspecified |
| mmu-miR-7646-5p | MIMAT0029794 | Ergic2 | 67456 | HITS-CLIP | | 31433990 Unspecified |
| mmu-miR-7646-5p | MIMAT0029794 | Fbxl20 | 72194 | HITS-CLIP | | 31433990 Unspecified |
| mmu-miR-7646-5p | MIMAT0029794 | Fem1c | 240263 | HITS-CLIP | | 31433990 Unspecified |
| mmu-miR-7646-5p | MIMAT0029794 | Ikzf4 | 22781 | HITS-CLIP | | 31433990 Unspecified |
| mmu-miR-7646-5p | MIMAT0029794 | Ilkap | 67444 | HITS-CLIP | | 31433990 Unspecified |
| mmu-miR-7646-5p | MIMAT0029794 | Kdm3a | 104263 | HITS-CLIP | | 31433990 Unspecified |
| mmu-miR-7646-5p | MIMAT0029794 | Marchf8 | 220972 | HITS-CLIP | | 31433990 Unspecified |
| mmu-miR-7646-5p | MIMAT0029794 | Myh9 | 17886 | HITS-CLIP | | 26203562 Unspecified |
| mmu-miR-7646-5p | MIMAT0029794 | Nid1 | 18073 | HITS-CLIP | | 26203562 Unspecified |
| mmu-miR-7646-5p | MIMAT0029794 | Nipal4 | 214112 | HITS-CLIP | | 26203562 Unspecified |
| mmu-miR-7646-5p | MIMAT0029794 | Npepps | 19155 | HITS-CLIP | | 31433990 Unspecified |
| mmu-miR-7646-5p | MIMAT0029794 | Numbl | 18223 | HITS-CLIP | | 26203562 Unspecified |
| mmu-miR-7646-5p | MIMAT0029794 | Nup98 | 269966 | HITS-CLIP | | 26203562 Unspecified |
| mmu-miR-7646-5p | MIMAT0029794 | Pdcd4 | 18569 | HITS-CLIP | | 26203562 Unspecified |
| mmu-miR-7646-5p | MIMAT0029794 | Prrc2c | 226562 | HITS-CLIP | | 31433990 Unspecified |
| mmu-miR-7646-5p | MIMAT0029794 | Rngtt | 24018 | HITS-CLIP | | 31433990 Unspecified |
| mmu-miR-7646-5p | MIMAT0029794 | Satb1 | 20230 | HITS-CLIP | | 31433990 Unspecified |
| mmu-miR-7646-5p | MIMAT0029794 | Ssh2 | 237860 | HITS-CLIP | | 31433990 Unspecified |
| mmu-miR-7646-5p | MIMAT0029794 | Top2a | 21973 | HITS-CLIP | | 31433990 Unspecified |
| mmu-miR-7646-5p | MIMAT0029794 | Trabd | 67976 | HITS-CLIP | | 26203562 Unspecified |
| mmu-miR-7646-5p | MIMAT0029794 | Ube2o | 217342 | HITS-CLIP | | 26203562 Unspecified |
| mmu-miR-7646-5p | MIMAT0029794 | Zfp280b | 64453 | HITS-CLIP | | 31433990 Unspecified |
| mmu-miR-7646-5p | MIMAT0029794 | Zfp280c | 208968 | HITS-CLIP | | 26203562 Unspecified |
| mmu-miR-7650-3p | MIMAT0029803 | Alkbh5 | 268420 | HITS-CLIP | | 31433990 Unspecified |
| mmu-miR-7650-3p | MIMAT0029803 | Cenpc1 | 12617 | HITS-CLIP | | 31433990 Unspecified |
| mmu-miR-7650-3p | MIMAT0029803 | Ehbp1l1 | 114601 | HITS-CLIP | | 31433990 Unspecified |

| | | | | | | |
|---|---|---|---|---|---|---|
| mmu-miR-7650-3p | MIMAT0029803 | Hras | 15461 | HITS-CLIP | 31433990 | Unspecified |
| mmu-miR-7650-3p | MIMAT0029803 | Nfe2l1 | 18023 | HITS-CLIP | 26203562 | Unspecified |
| mmu-miR-7650-5p | MIMAT0029802 | Dnajc16 | 214063 | HITS-CLIP | 26203562 | Unspecified |
| mmu-miR-7650-5p | MIMAT0029802 | Srrm2 | 75956 | HITS-CLIP | 31433990 | Unspecified |
| mmu-miR-7650-5p | MIMAT0029802 | Tln1 | 21894 | HITS-CLIP | 31433990 | Unspecified |
| mmu-miR-7656-3p | MIMAT0029819 | Arhgap35 | 232906 | HITS-CLIP | 31433990 | Unspecified |
| mmu-miR-7656-3p | MIMAT0029819 | Atp5a1 | 11946 | HITS-CLIP | 31433990 | Unspecified |
| mmu-miR-7656-3p | MIMAT0029819 | Bach2 | 12014 | HITS-CLIP | 31433990 | Unspecified |
| mmu-miR-7656-3p | MIMAT0029819 | Elk4 | 13714 | HITS-CLIP | 31433990 | Unspecified |
| mmu-miR-7656-3p | MIMAT0029819 | H2bc14 | 8342 | HITS-CLIP | 31433990 | Unspecified |
| mmu-miR-7656-3p | MIMAT0029819 | Herc2 | 15204 | HITS-CLIP | 31433990 | Unspecified |
| mmu-miR-7656-3p | MIMAT0029819 | Ing5 | 66262 | HITS-CLIP | 31433990 | Unspecified |
| mmu-miR-7656-3p | MIMAT0029819 | Mdh2 | 17448 | HITS-CLIP | 31433990 | Unspecified |
| mmu-miR-7656-3p | MIMAT0029819 | Midn | 59090 | HITS-CLIP | 31433990 | Unspecified |
| mmu-miR-7656-3p | MIMAT0029819 | Pygo2 | 68911 | HITS-CLIP | 26203562 | Unspecified |
| mmu-miR-7656-3p | MIMAT0029819 | Spred1 | 114715 | HITS-CLIP | 31433990 | Unspecified |
| mmu-miR-7656-3p | MIMAT0029819 | Zmiz1 | 328365 | HITS-CLIP | 31433990 | Unspecified |
| mmu-miR-7656-5p | MIMAT0029818 | 1810013L24Rik | 69053 | HITS-CLIP | 31433990 | Unspecified |
| mmu-miR-7656-5p | MIMAT0029818 | Atp6v1a | 11964 | HITS-CLIP | 31433990 | Unspecified |
| mmu-miR-7656-5p | MIMAT0029818 | Cdc34 | 216150 | HITS-CLIP | 31433990 | Unspecified |
| mmu-miR-7656-5p | MIMAT0029818 | Clint1 | 216705 | HITS-CLIP | 31433990 | Unspecified |
| mmu-miR-7656-5p | MIMAT0029818 | Entpd6 | 12497 | HITS-CLIP | 31433990 | Unspecified |
| mmu-miR-7656-5p | MIMAT0029818 | Ep400 | 75560 | HITS-CLIP | 31433990 | Unspecified |
| mmu-miR-7656-5p | MIMAT0029818 | Fkbp8 | 14232 | HITS-CLIP | 31433990 | Unspecified |
| mmu-miR-7656-5p | MIMAT0029818 | Foxk1 | 17425 | HITS-CLIP | 31433990 | Unspecified |
| mmu-miR-7656-5p | MIMAT0029818 | Ggta1 | 14594 | HITS-CLIP | 26203562 | Unspecified |
| mmu-miR-7656-5p | MIMAT0029818 | Gpr137 | 107173 | HITS-CLIP | 26203562 | Unspecified |
| mmu-miR-7656-5p | MIMAT0029818 | Kpna3 | 16648 | HITS-CLIP | 31433990 | Unspecified |
| mmu-miR-7656-5p | MIMAT0029818 | Krt16 | 16666 | HITS-CLIP | 26203562 | Unspecified |
| mmu-miR-7656-5p | MIMAT0029818 | Lnpep | 240028 | HITS-CLIP | 31433990 | Unspecified |
| mmu-miR-7656-5p | MIMAT0029818 | Naif1 | 71254 | HITS-CLIP | 31433990 | Unspecified |
| mmu-miR-7656-5p | MIMAT0029818 | Notch2 | 18129 | HITS-CLIP | 26203562 | Unspecified |
| mmu-miR-7656-5p | MIMAT0029818 | Npepps | 19155 | HITS-CLIP | 31433990 | Unspecified |
| mmu-miR-7656-5p | MIMAT0029818 | Ppig | 228005 | HITS-CLIP | 31433990 | Unspecified |
| mmu-miR-7656-5p | MIMAT0029818 | Rnft1 | 76892 | HITS-CLIP | 31433990 | Unspecified |
| mmu-miR-7656-5p | MIMAT0029818 | Rps6ka1 | 20111 | HITS-CLIP | 31433990 | Unspecified |
| mmu-miR-7656-5p | MIMAT0029818 | Sufu | 24069 | HITS-CLIP | 26203562 | Unspecified |
| mmu-miR-7656-5p | MIMAT0029818 | Zfp516 | 329003 | HITS-CLIP | 26203562 | Unspecified |
| mmu-miR-7656-5p | MIMAT0029818 | Zfp955b | 100043468 | HITS-CLIP | 31433990 | Unspecified |

| | | | | | | |
|---|---|---|---|---|---|---|
| mmu-miR-7656-5p | MIMAT0029818 | Zmiz1 | 328365 | HITS-CLIP | 31433990 | Unspecified |
| mmu-miR-7657-3p | MIMAT0029821 | Adam9 | 11502 | HITS-CLIP | 26203562 | Unspecified |
| mmu-miR-7657-3p | MIMAT0029821 | Ash1l | 192195 | HITS-CLIP | 31433990 | Unspecified |
| mmu-miR-7657-3p | MIMAT0029821 | Brd2 | 14312 | HITS-CLIP | 26203562 | Unspecified |
| mmu-miR-7657-3p | MIMAT0029821 | Cdc42se1 | 57912 | HITS-CLIP | 26203562 | Unspecified |
| mmu-miR-7657-3p | MIMAT0029821 | Ece1 | 230857 | HITS-CLIP | 26203562 | Unspecified |
| mmu-miR-7657-3p | MIMAT0029821 | Errfi1 | 74155 | HITS-CLIP | 26203562 | Unspecified |
| mmu-miR-7657-3p | MIMAT0029821 | Foxo3 | 56484 | HITS-CLIP | 31433990 | Unspecified |
| mmu-miR-7657-3p | MIMAT0029821 | Golph3 | 66629 | HITS-CLIP | 31433990 | Unspecified |
| mmu-miR-7657-3p | MIMAT0029821 | Il12rb2 | 16162 | HITS-CLIP | 31433990 | Unspecified |
| mmu-miR-7657-3p | MIMAT0029821 | Iqgap1 | 29875 | HITS-CLIP | 31433990 | Unspecified |
| mmu-miR-7657-3p | MIMAT0029821 | Kmt2a | 214162 | HITS-CLIP | 26203562 | Unspecified |
| mmu-miR-7657-3p | MIMAT0029821 | Lyst | 17101 | HITS-CLIP | 31433990 | Unspecified |
| mmu-miR-7657-3p | MIMAT0029821 | Rgp1 | 242406 | HITS-CLIP | 31433990 | Unspecified |
| mmu-miR-7657-3p | MIMAT0029821 | Tnrc6a | 233833 | HITS-CLIP | 26203562 | Unspecified |
| mmu-miR-7657-3p | MIMAT0029821 | Top2a | 21973 | HITS-CLIP | 31433990 | Unspecified |
| mmu-miR-7660-3p | MIMAT0029827 | Itpr1 | 16438 | HITS-CLIP | 31433990 | Unspecified |
| mmu-miR-7660-3p | MIMAT0029827 | Nr6a1 | 14536 | HITS-CLIP | 26203562 | Unspecified |
| mmu-miR-7660-3p | MIMAT0029827 | Phf12 | 268448 | HITS-CLIP | 31433990 | Unspecified |
| mmu-miR-7660-3p | MIMAT0029827 | Rps9 | 76846 | HITS-CLIP | 31433990 | Unspecified |
| mmu-miR-7660-3p | MIMAT0029827 | Scamp1 | 107767 | HITS-CLIP | 26203562 | Unspecified |
| mmu-miR-7660-3p | MIMAT0029827 | Tnpo1 | 238799 | HITS-CLIP | 31433990 | Unspecified |
| mmu-miR-7660-3p | MIMAT0029827 | Ubtf | 21429 | HITS-CLIP | 26203562 | Unspecified |
| mmu-miR-7660-5p | MIMAT0029826 | Appbp2 | 66884 | HITS-CLIP | 31433990 | Unspecified |
| mmu-miR-7660-5p | MIMAT0029826 | Raly | 19383 | HITS-CLIP | 31433990 | Unspecified |
| mmu-miR-7660-5p | MIMAT0029826 | Setd1a | 233904 | HITS-CLIP | 31433990 | Unspecified |
| mmu-miR-7660-5p | MIMAT0029826 | Wdr37 | 207615 | HITS-CLIP | 31433990 | Unspecified |
| mmu-miR-7663-3p | MIMAT0029833 | 1810013L24Rik | 69053 | HITS-CLIP | 31433990 | Unspecified |
| mmu-miR-7663-3p | MIMAT0029833 | A630001G21Rik | 319997 | HITS-CLIP | 31433990 | Unspecified |
| mmu-miR-7663-3p | MIMAT0029833 | Actn1 | 109711 | HITS-CLIP | 26203562 | Unspecified |
| mmu-miR-7663-3p | MIMAT0029833 | Atad2 | 70472 | HITS-CLIP | 31433990 | Unspecified |
| mmu-miR-7663-3p | MIMAT0029833 | Btg2 | 12227 | HITS-CLIP | 31433990 | Unspecified |
| mmu-miR-7663-3p | MIMAT0029833 | Celf1 | 13046 | HITS-CLIP | 26203562 | Unspecified |
| mmu-miR-7663-3p | MIMAT0029833 | Dars | 226414 | HITS-CLIP | 31433990 | Unspecified |
| mmu-miR-7663-3p | MIMAT0029833 | Dcaf12 | 68970 | HITS-CLIP | 31433990 | Unspecified |
| mmu-miR-7663-3p | MIMAT0029833 | Ddx5 | 13207 | HITS-CLIP | 31433990 | Unspecified |
| mmu-miR-7663-3p | MIMAT0029833 | Dennd1b | 329260 | HITS-CLIP | 31433990 | Unspecified |
| mmu-miR-7663-3p | MIMAT0029833 | Dock8 | 76088 | HITS-CLIP | 31433990 | Unspecified |
| mmu-miR-7663-3p | MIMAT0029833 | Ezh2 | 14056 | HITS-CLIP | 31433990 | Unspecified |

| | | | | | | |
|---|---|---|---|---|---|---|
| mmu-miR-7663-3p | MIMAT0029833 | Gemin4 | 276919 | HITS-CLIP | 31433990 | Unspecified |
| mmu-miR-7663-3p | MIMAT0029833 | Gimap8 | 243374 | HITS-CLIP | 31433990 | Unspecified |
| mmu-miR-7663-3p | MIMAT0029833 | Gm42715 | | HITS-CLIP | 31433990 | Unspecified |
| mmu-miR-7663-3p | MIMAT0029833 | Gpr146 | 80290 | HITS-CLIP | 31433990 | Unspecified |
| mmu-miR-7663-3p | MIMAT0029833 | H13 | 14950 | HITS-CLIP | 31433990 | Unspecified |
| mmu-miR-7663-3p | MIMAT0029833 | H3f3b | 15081 | HITS-CLIP | 31433990 | Unspecified |
| mmu-miR-7663-3p | MIMAT0029833 | Igf1r | 16001 | HITS-CLIP | 26203562 | Unspecified |
| mmu-miR-7663-3p | MIMAT0029833 | Il2ra | 16184 | HITS-CLIP | 31433990 | Unspecified |
| mmu-miR-7663-3p | MIMAT0029833 | Itga5 | 16402 | HITS-CLIP | 26203562 | Unspecified |
| mmu-miR-7663-3p | MIMAT0029833 | Itgb3 | 16416 | HITS-CLIP | 31433990 | Unspecified |
| mmu-miR-7663-3p | MIMAT0029833 | Kmt2a | 214162 | HITS-CLIP | 31433990 | Unspecified |
| mmu-miR-7663-3p | MIMAT0029833 | Lamtor4 | 66096 | HITS-CLIP | 26203562 | Unspecified |
| mmu-miR-7663-3p | MIMAT0029833 | Macf1 | 11426 | HITS-CLIP | 26203562 | Unspecified |
| mmu-miR-7663-3p | MIMAT0029833 | Ncoa3 | 17979 | HITS-CLIP | 31433990 | Unspecified |
| mmu-miR-7663-3p | MIMAT0029833 | Ndufs6 | 407785 | HITS-CLIP | 31433990 | Unspecified |
| mmu-miR-7663-3p | MIMAT0029833 | Nfatc1 | 18018 | HITS-CLIP | 31433990 | Unspecified |
| mmu-miR-7663-3p | MIMAT0029833 | Nup98 | 269966 | HITS-CLIP | 26203562 | Unspecified |
| mmu-miR-7663-3p | MIMAT0029833 | Pacs2 | 217893 | HITS-CLIP | 26203562 | Unspecified |
| mmu-miR-7663-3p | MIMAT0029833 | Pdpr | 319518 | HITS-CLIP | 26203562 | Unspecified |
| mmu-miR-7663-3p | MIMAT0029833 | Peli1 | 67245 | HITS-CLIP | 31433990 | Unspecified |
| mmu-miR-7663-3p | MIMAT0029833 | Psmc1 | 19179 | HITS-CLIP | 31433990 | Unspecified |
| mmu-miR-7663-3p | MIMAT0029833 | Ptpn11 | 19247 | HITS-CLIP | 31433990 | Unspecified |
| mmu-miR-7663-3p | MIMAT0029833 | Rabgap1 | 227800 | HITS-CLIP | 26203562 | Unspecified |
| mmu-miR-7663-3p | MIMAT0029833 | Rad21 | 19357 | HITS-CLIP | 31433990 | Unspecified |
| mmu-miR-7663-3p | MIMAT0029833 | Rbm15b | 109095 | HITS-CLIP | 31433990 | Unspecified |
| mmu-miR-7663-3p | MIMAT0029833 | Rbm7 | 67010 | HITS-CLIP | 31433990 | Unspecified |
| mmu-miR-7663-3p | MIMAT0029833 | Rbms1 | 56878 | HITS-CLIP | 26203562 | Unspecified |
| mmu-miR-7663-3p | MIMAT0029833 | Rhbdf2 | 217344 | HITS-CLIP | 31433990 | Unspecified |
| mmu-miR-7663-3p | MIMAT0029833 | Riok3 | 66878 | HITS-CLIP | 31433990 | Unspecified |
| mmu-miR-7663-3p | MIMAT0029833 | Rnft1 | 76892 | HITS-CLIP | 31433990 | Unspecified |
| mmu-miR-7663-3p | MIMAT0029833 | Rpl23 | 65019 | HITS-CLIP | 31433990 | Unspecified |
| mmu-miR-7663-3p | MIMAT0029833 | Scd2 | 20250 | HITS-CLIP | 31433990 | Unspecified |
| mmu-miR-7663-3p | MIMAT0029833 | Sesn2 | 230784 | HITS-CLIP | 31433990 | Unspecified |
| mmu-miR-7663-3p | MIMAT0029833 | Slc43a2 | 215113 | HITS-CLIP | 31433990 | Unspecified |
| mmu-miR-7663-3p | MIMAT0029833 | Slco4a1 | 108115 | HITS-CLIP | 31433990 | Unspecified |
| mmu-miR-7663-3p | MIMAT0029833 | Smpd4 | 77626 | HITS-CLIP | 31433990 | Unspecified |
| mmu-miR-7663-3p | MIMAT0029833 | Smurf2 | 66313 | HITS-CLIP | 31433990 | Unspecified |
| mmu-miR-7663-3p | MIMAT0029833 | Snx25 | 102141 | HITS-CLIP | 26203562 | Unspecified |
| mmu-miR-7663-3p | MIMAT0029833 | Srcap | 100043597 | HITS-CLIP | 31433990 | Unspecified |

| | | | | | | |
|---|---|---|---|---|---|---|
| mmu-miR-7663-3p | MIMAT0029833 | Srsf1 | 110809 | HITS-CLIP | 31433990 | Unspecified |
| mmu-miR-7663-3p | MIMAT0029833 | Sufu | 24069 | HITS-CLIP | 26203562 | Unspecified |
| mmu-miR-7663-3p | MIMAT0029833 | Supt16 | 114741 | HITS-CLIP | 31433990 | Unspecified |
| mmu-miR-7663-3p | MIMAT0029833 | Syde1 | 71709 | HITS-CLIP | 26203562 | Unspecified |
| mmu-miR-7663-3p | MIMAT0029833 | Tigar | 319801 | HITS-CLIP | 26203562 | Unspecified |
| mmu-miR-7663-3p | MIMAT0029833 | Tkt | 21881 | HITS-CLIP | 31433990 | Unspecified |
| mmu-miR-7663-3p | MIMAT0029833 | Tmsb10 | 19240 | HITS-CLIP | 31433990 | Unspecified |
| mmu-miR-7663-3p | MIMAT0029833 | Trmt10c | 52575 | HITS-CLIP | 26203562 | Unspecified |
| mmu-miR-7663-3p | MIMAT0029833 | Trp53inp1 | 60599 | HITS-CLIP | 31433990\|26203562 | Unspecified |
| mmu-miR-7663-3p | MIMAT0029833 | Trpv3 | 246788 | HITS-CLIP | 26203562 | Unspecified |
| mmu-miR-7663-3p | MIMAT0029833 | Ucp2 | 22228 | HITS-CLIP | 31433990 | Unspecified |
| mmu-miR-7663-3p | MIMAT0029833 | Wasf2 | 242687 | HITS-CLIP | 31433990 | Unspecified |
| mmu-miR-7663-3p | MIMAT0029833 | Wdr26 | 226757 | HITS-CLIP | 31433990 | Unspecified |
| mmu-miR-7663-3p | MIMAT0029833 | Zfp710 | 209225 | HITS-CLIP | 31433990 | Unspecified |
| mmu-miR-7663-3p | MIMAT0029833 | Zkscan8 | 93681 | HITS-CLIP | 31433990 | Unspecified |
| mmu-miR-7663-5p | MIMAT0029832 | Ccne2 | 12448 | HITS-CLIP | 31433990 | Unspecified |
| mmu-miR-7663-5p | MIMAT0029832 | H4c14 | 8359 | HITS-CLIP | 31433990 | Unspecified |
| mmu-miR-7663-5p | MIMAT0029832 | Soat1 | 20652 | HITS-CLIP | 31433990 | Unspecified |
| mmu-miR-7663-5p | MIMAT0029832 | Sppl2a | 66552 | HITS-CLIP | 31433990 | Unspecified |
| mmu-miR-7663-5p | MIMAT0029832 | Thoc1 | 225160 | HITS-CLIP | 31433990 | Unspecified |
| mmu-miR-7663-5p | MIMAT0029832 | Tnrc6a | 233833 | HITS-CLIP | 31433990 | Unspecified |
| mmu-miR-8099 | MIMAT0031401 | Runx3 | 12399 | HITS-CLIP | 31433990 | Unspecified |
| mmu-miR-8108 | MIMAT0031413 | Cdk2ap2 | 52004 | HITS-CLIP | 31433990 | Unspecified |
| mmu-miR-8108 | MIMAT0031413 | Erbb3 | 13867 | HITS-CLIP | 26203562 | Unspecified |
| mmu-miR-8108 | MIMAT0031413 | Rrs1 | 59014 | HITS-CLIP | 31433990 | Unspecified |
| mmu-miR-8108 | MIMAT0031413 | Sbno1 | 243272 | HITS-CLIP | 26203562 | Unspecified |
| mmu-miR-8108 | MIMAT0031413 | Zfp36 | 22695 | HITS-CLIP | 31433990 | Unspecified |
| mmu-miR-8118 | MIMAT0031424 | Ankrd49 | 56503 | HITS-CLIP | 31433990 | Unspecified |
| mmu-miR-8118 | MIMAT0031424 | Anxa2 | 12306 | HITS-CLIP | 26203562 | Unspecified |
| mmu-miR-8118 | MIMAT0031424 | Bmt2 | 101148 | HITS-CLIP | 31433990 | Unspecified |
| mmu-miR-8118 | MIMAT0031424 | Echdc1 | 52665 | HITS-CLIP | 26203562 | Unspecified |
| mmu-miR-8118 | MIMAT0031424 | Mtdh | 67154 | HITS-CLIP | 31433990\|26203562 | Unspecified |
| mmu-miR-8118 | MIMAT0031424 | Sec22b | 20333 | HITS-CLIP | 31433990 | Unspecified |
| mmu-miR-8118 | MIMAT0031424 | Suco | 226551 | HITS-CLIP | 31433990 | Unspecified |
| mmu-miR-8118 | MIMAT0031424 | Sypl | 19027 | HITS-CLIP | 26203562 | Unspecified |
| mmu-miR-8118 | MIMAT0031424 | Trp53inp1 | 60599 | HITS-CLIP | 31433990 | Unspecified |
| mmu-miR-8118 | MIMAT0031424 | Zfp217 | 228913 | HITS-CLIP | 26203562 | Unspecified |
| mmu-miR-8120 | MIMAT0031426 | Asprv1 | 67855 | HITS-CLIP | 26203562 | Unspecified |
| mmu-miR-8120 | MIMAT0031426 | Dnmt1 | 13433 | HITS-CLIP | 31433990 | Unspecified |

| | | | | | | |
|---|---|---|---|---|---|---|
| mmu-miR-8120 | MIMAT0031426 | Etf1 | 225363 | HITS-CLIP | 31433990 | Unspecified |
| mmu-miR-8120 | MIMAT0031426 | Tgfbr1 | 21812 | HITS-CLIP | 31433990 | Unspecified |
| mmu-miR-9-3p | MIMAT0000143 | Acap3 | 140500 | HITS-CLIP | 31433990 | Unspecified |
| mmu-miR-9-3p | MIMAT0000143 | Ago2 | 239528 | qCLASH | 31393953 | Unspecified |
| mmu-miR-9-3p | MIMAT0000143 | Rcc2 | 108911 | qCLASH | 31393953 | Unspecified |
| mmu-miR-9-3p | MIMAT0000143 | Tmem38b | 52076 | qCLASH | 31393953 | Unspecified |
| mmu-miR-9-3p | MIMAT0000143 | Zbtb18 | 30928 | qCLASH | 31393953 | Unspecified |
